## Supplementary material for "Comparative pathogenesis of two lineages of Powassan virus reveals distinct clinical outcome, neuropathology, and inflammation": Table S1

| Gene | baseMean | log2FoldChange | lfcSE |
|---|---|---|---|
| 4933401J01Rik | 0 | NA | NA |
| Gm26206 | 0 | NA | NA |
| Xkr4 | 1236.723264 | -0.477716117 | 0.099301475 |
| Gm18956 | 0.257781492 | -0.724462757 | 2.517660746 |
| Gm37180 | 183.2012087 | -0.772553756 | 0.242241596 |
| Gm37363 | 156.7123429 | -0.910711259 | 0.241241069 |
| Gm37686 | 55.26816088 | -0.790974548 | 0.285629244 |
| Gm1992 | 0 | NA | NA |
| Gm37329 | 74.22829483 | -0.648670201 | 0.242432006 |
| Gm7341 | 0.472128458 | 1.377835395 | 1.537325731 |
| Gm38148 | 143.524156 | -0.877078077 | 0.270110165 |
| Gm19938 | 181.8288108 | -0.869873335 | 0.256367103 |
| Gm10568 | 17.47471811 | 1.331757259 | 0.351488906 |
| Gm38385 | 0 | NA | NA |
| Gm27396 | 0 | NA | NA |
| Gm37381 | 0.042716945 | -0.243571806 | 3.724945049 |
| Gm6101 | 0.038981395 | 0.718207094 | 3.724945049 |
| Rp1 | 1.19669966 | -0.599832503 | 1.123532023 |
| Gm37483 | 0 | NA | NA |
| Sox17 | 14.01852502 | -0.660819791 | 0.313878124 |
| Gm37587 | 0.590622753 | 1.151766375 | 1.428457232 |
| Gm7357 | 0.431020911 | 0.529440003 | 1.519925415 |
| Gm22307 | 0.205539566 | 1.078122579 | 2.601261389 |
| Gm38076 | 0 | NA | NA |
| Gm37323 | 0.424160584 | -1.834526795 | 1.403428972 |
| Gm7369 | 0.062426873 | 0.237312084 | 3.724945049 |
| Gm6085 | 7.026365196 | 0.187719625 | 0.420642279 |
| Gm6119 | 1.007284973 | -0.308876938 | 1.028389998 |
| Gm25493 | 0.04393175 | -0.243571806 | 3.724945049 |
| Gm2053 | 0.509840029 | -1.496486045 | 1.353081268 |
| Gm6123 | 6.523572364 | 0.407912357 | 0.37919244 |
| Mrpl15 | 339.6237885 | 0.149261462 | 0.098288241 |
| Gm37144 | 10.84448109 | -0.784434186 | 0.403882487 |
| Lypla1 | 301.4400782 | 0.076500393 | 0.104226113 |
| Gm37988 | 0.099106094 | 1.209894437 | 3.72418822 |
| Tcea1 | 660.8047814 | 0.404507978 | 0.098012463 |
| Gm6104 | 7.397713871 | -0.314087065 | 0.358800149 |
| Gm37277 | 41.2031627 | 0.580445019 | 0.182461145 |
| Rgs20 | 298.6651195 | 0.428385673 | 0.101984066 |
| Gm17100 | 2.600587843 | -0.293815888 | 0.635523205 |
| Gm37079 | 17.87741427 | -0.520229798 | 0.346106546 |
| Gm16041 | 0.448921586 | 0.877252522 | 1.371243987 |
| Gm17101 | 1.407567115 | -0.05829236 | 0.834036566 |
| Atp6v1h | 1710.406335 | 0.296576032 | 0.092177769 |
| Gm7182 | 0 | NA | NA |
| Gm37567 | 5.040767136 | -1.13186006 | 0.571857059 |

| | | | |
|---|---|---|---|
| Gm38264 | 0.54045421 | -1.152547281 | 1.35060343 |
| Oprk1 | 98.2787742 | -0.975174296 | 0.15392665 |
| Gm36965 | 0 | NA | NA |
| Gm22463 | 0 | NA | NA |
| Gm37429 | 0 | NA | NA |
| Npbwr1 | 11.64878932 | 0.087202255 | 0.324616464 |
| Gm19214 | 0.066050875 | 0.71820487 | 3.724945049 |
| Rb1cc1 | 2300.690646 | -0.169418183 | 0.053725881 |
| 4732440D04Rik | 117.8176336 | -0.335259135 | 0.165170711 |
| Fam150a | 0.409266314 | 0.229423084 | 1.475799672 |
| Gm2147 | 0.280898025 | 0.706377586 | 1.636673867 |
| Gm7417 | 0 | NA | NA |
| Gm19026 | 17.7062556 | -0.57448667 | 0.243558875 |
| St18 | 573.9728649 | -0.512335729 | 0.130294574 |
| Gm37405 | 0 | NA | NA |
| Gm7449 | 0.090074004 | -0.24357403 | 3.724945049 |
| Gm37108 | 10.32889666 | 0.56808921 | 0.306742483 |
| Gm37275 | 0 | NA | NA |
| Gm37225 | 0 | NA | NA |
| Gm37489 | 0 | NA | NA |
| Gm5694 | 0 | NA | NA |
| Pcmtd1 | 1844.737977 | 0.009123368 | 0.081330872 |
| Gm26983 | 0 | NA | NA |
| Gm38372 | 224.9946318 | -0.605977904 | 0.235251585 |
| Gm9826 | 6.912980464 | 0.05438992 | 0.398776051 |
| Nras-ps2 | 0 | NA | NA |
| Gm23274 | 0 | NA | NA |
| Gm19002 | 0.085327104 | 1.179809211 | 3.723500796 |
| Gm26901 | 31.62112565 | -1.129210283 | 0.304517195 |
| Gm18984 | 0 | NA | NA |
| Gm30414 | 0 | NA | NA |
| Gm37791 | 0 | NA | NA |
| Gm7470 | 0.066050875 | 0.71820487 | 3.724945049 |
| Rps2-ps2 | 0 | NA | NA |
| Gm38216 | 0 | NA | NA |
| Sntg1 | 622.888287 | -0.670901441 | 0.113345126 |
| Gm38024 | 20.53816256 | -0.710362834 | 0.399002201 |
| Gm16284 | 4.895741945 | -1.2074529 | 0.598639924 |
| Gm38259 | 46.04207795 | -1.145774805 | 0.258858116 |
| Gm23358 | 1.395067866 | -0.107302163 | 0.938164416 |
| Gm37005 | 8.035261982 | -0.445226237 | 0.392840858 |
| Gm15452 | 1.564680506 | -0.825300003 | 0.795246911 |
| Gm36964 | 0.112396584 | 1.192083721 | 3.724434897 |
| Gm24276 | 0.547110457 | -0.110399596 | 1.530403139 |
| Gm6152 | 0.180148697 | -0.243569188 | 2.705533632 |
| Gm7445 | 0.326379708 | 0.830132249 | 3.710140185 |
| Gm7493 | 10.60824579 | -0.807172723 | 0.489846224 |

| | | | |
|---|---|---|---|
| Gm37629 | 21.84260411 | -1.198841324 | 0.293928791 |
| Gm38008 | 8.876683074 | -0.755424257 | 0.407260997 |
| Gm37143 | 0 | NA | NA |
| Gm6187 | 0.1709406 | -0.724458201 | 3.38513154 |
| Gm18299 | 0 | NA | NA |
| Gm7512 | 0 | NA | NA |
| Gm24765 | 0 | NA | NA |
| Gm18300 | 1.926091363 | -0.574661014 | 0.679445829 |
| Rrs1 | 218.6880439 | 0.619929025 | 0.122220053 |
| Adhfe1 | 178.1879691 | 0.189596369 | 0.147171513 |
| 2610203C22Rik | 5.506898143 | -0.748836203 | 0.460441806 |
| Gm6161 | 3.14279713 | 0.214794725 | 0.651090869 |
| 3110035E14Rik | 1247.772309 | -1.190836492 | 0.182522992 |
| Gm29520 | 0 | NA | NA |
| Mybl1 | 100.6319816 | -0.003761593 | 0.162290948 |
| Vcpip1 | 2191.495749 | 0.455858879 | 0.07749525 |
| 1700034P13Rik | 2.716773814 | -0.824992282 | 0.614786266 |
| Sgk3 | 387.7590365 | 0.642437451 | 0.110343882 |
| Gm6195 | 2.037763675 | -0.111336398 | 0.886676605 |
| Gm22607 | 0 | NA | NA |
| Mcmdc2 | 2.919172438 | 0.685026961 | 0.599479292 |
| Snhg6 | 57.04786074 | 0.297000133 | 0.198311257 |
| Snord87 | 1.794878382 | -0.298149292 | 0.767663506 |
| Tcf24 | 4.573727287 | 0.988732883 | 0.492519507 |
| E330040D14Rik | 0.234971822 | 0.237329747 | 2.121545891 |
| Ppp1r42 | 4.644333044 | -0.711079784 | 0.481884515 |
| Gm15818 | 0 | NA | NA |
| Gm24674 | 0 | NA | NA |
| Cops5 | 641.9122966 | 0.35483567 | 0.098760933 |
| Cspp1 | 856.404923 | -0.803117982 | 0.169156264 |
| 1700047N06Rik | 0 | NA | NA |
| Arfgef1 | 2686.960645 | 0.094423873 | 0.050192454 |
| Gm26348 | 0 | NA | NA |
| Gm37569 | 39.97338864 | -0.517392548 | 0.305158477 |
| Cpa6 | 18.52223898 | -0.673828465 | 0.511881254 |
| Gm38005 | 2.757436252 | -1.802156669 | 0.846499177 |
| Gm15604 | 1.883799777 | -0.725891465 | 0.876203262 |
| Gm25253 | 0.347911031 | -0.756331046 | 3.220823089 |
| Gm15603 | 0.042716945 | -0.243571806 | 3.724945049 |
| Gm37133 | 0 | NA | NA |
| Gm5522 | 0.046345709 | 0.71820487 | 3.724945049 |
| Gm28659 | 0 | NA | NA |
| Gm22963 | 0 | NA | NA |
| Prex2 | 1191.400917 | -1.072717829 | 0.156742819 |
| Gm24173 | 0 | NA | NA |
| Gm28686 | 0 | NA | NA |
| Gm37410 | 0.060174537 | 0.237309859 | 3.724945049 |

| | | | |
|---|---|---|---|
| A830018L16Rik | 657.3086267 | -1.173793004 | 0.106772939 |
| Gm38178 | 28.55896715 | -1.128120639 | 0.312306998 |
| Gm29273 | 0 | NA | NA |
| Gm38069 | 9.696124784 | -0.969109796 | 0.472650927 |
| Gm7560 | 0.344328494 | 0.237330337 | 1.728125218 |
| Gm22633 | 0 | NA | NA |
| Mir6341 | 0 | NA | NA |
| Gm2383 | 0.481037375 | -1.249545905 | 1.411355751 |
| Gm6216 | 0 | NA | NA |
| Gm17644 | 0.118371171 | 0.718202646 | 3.724945049 |
| Gm29663 | 0 | NA | NA |
| Sulf1 | 425.5919252 | 0.864551473 | 0.128498565 |
| Slco5a1 | 118.4898984 | 0.270791308 | 0.145957054 |
| Gm37162 | 0 | NA | NA |
| Gm5250 | 0 | NA | NA |
| Gm29283 | 0 | NA | NA |
| Prdm14 | 0.178837887 | 0.685271736 | 2.696671491 |
| Ncoa2 | 2049.993035 | -0.089445254 | 0.046081091 |
| Gm38376 | 24.50757996 | -0.908866505 | 0.382400449 |
| Gm37409 | 22.73165489 | -0.850417322 | 0.324351705 |
| Gm27881 | 0 | NA | NA |
| Gm38223 | 5.5083264 | -1.382936985 | 0.432656628 |
| Gm37702 | 18.1938087 | -1.273626043 | 0.292866493 |
| Gm23169 | 4.580170235 | -0.948542657 | 0.574073669 |
| Gm38120 | 65.50301172 | -0.83005539 | 0.25159376 |
| Gm38380 | 51.36537569 | -0.60449664 | 0.300864158 |
| Gm29570 | 1.379700947 | -0.571036923 | 0.85660375 |
| Gm7593 | 0.038981395 | 0.718207094 | 3.724945049 |
| Tram1 | 601.6593407 | 0.777769333 | 0.1012804 |
| Lactb2 | 271.8818834 | 0.0124421 | 0.095141207 |
| Gm38319 | 9.470931737 | -0.281716037 | 0.527574981 |
| Xkr9 | 0.562487033 | 2.099216513 | 1.390930192 |
| Gm26273 | 0 | NA | NA |
| Gm5523 | 0.58821479 | 1.124588366 | 1.181993123 |
| Gm36947 | 0.09689823 | -0.724453471 | 3.724945049 |
| Gm25491 | 0 | NA | NA |
| Gm37400 | 0 | NA | NA |
| Eya1 | 126.1615464 | -0.584917762 | 0.130937106 |
| Gm22616 | 0 | NA | NA |
| Gm37444 | 2.632415511 | -2.606291748 | 0.947172531 |
| Gm9947 | 16.04594573 | -0.881005537 | 0.290387481 |
| Msc | 6.16192073 | 0.223253648 | 0.425052812 |
| Gm37412 | 7.68759231 | -1.083098624 | 0.403255378 |
| Smt3h2-ps4 | 2.907781834 | -0.210407084 | 0.602841782 |
| Rpl5-ps1 | 8.684837921 | 0.008158839 | 0.353460573 |
| Trpa1 | 3.284363746 | -0.776292489 | 0.57528164 |
| Gm26345 | 0 | NA | NA |

| | | | |
|---|---|---|---|
| Gm10566 | 6.613104402 | 0.336599266 | 0.402822929 |
| Kcnb2 | 762.7240551 | -0.737322119 | 0.105056722 |
| Gm38116 | 50.40214469 | -1.468695497 | 0.347879394 |
| Gm37138 | 26.46315223 | -1.023609146 | 0.278672482 |
| Gm25227 | 0 | NA | NA |
| Gm25168 | 0.155005853 | -1.181207458 | 2.822657929 |
| Gm6060 | 0.066050875 | 0.71820487 | 3.724945049 |
| Gm28669 | 0.095435502 | 0.718202646 | 3.724945049 |
| Terf1 | 191.9488994 | -0.225453819 | 0.12967766 |
| Gm37509 | 44.48996355 | -0.349777644 | 0.324783746 |
| Sbspon | 3.624262949 | -0.940398244 | 0.551922029 |
| Gm23259 | 0.061920613 | 0.718207094 | 3.724945049 |
| Gm7634 | 0 | NA | NA |
| 4930444P10Rik | 4.660125666 | -0.067600023 | 0.448755845 |
| Rpl7 | 1699.234633 | 0.75531637 | 0.113830889 |
| Rdh10 | 199.9939912 | 0.928754635 | 0.157413188 |
| Gm38249 | 4.612676316 | 0.14504782 | 0.459425396 |
| Gm28095 | 0 | NA | NA |
| Stau2 | 1742.3238 | -0.479919301 | 0.07588268 |
| Gm7568 | 0.218065207 | 0.718219249 | 2.537878383 |
| Ube2w | 820.6053904 | -0.099729577 | 0.058276316 |
| Gm38342 | 3.77249287 | 1.296284471 | 0.563425926 |
| Tceb1 | 826.0456009 | -0.134102588 | 0.083622386 |
| D030040B21Rik | 0.250367046 | 0.237326992 | 2.484698294 |
| Gm7654 | 0.171036444 | -0.63727414 | 2.766397645 |
| Tmem70 | 222.9577256 | 0.18306644 | 0.10553525 |
| Ly96 | 21.7470159 | 0.960166237 | 0.254409703 |
| Gm29649 | 0 | NA | NA |
| Gm5828 | 11.26054785 | 0.879782977 | 0.300915702 |
| Gm28376 | 0.229484813 | -0.724461516 | 3.114678745 |
| 4921511E07Rik | 0 | NA | NA |
| Jph1 | 403.3093643 | -1.026794623 | 0.0954734 |
| Gm28783 | 0.484663642 | 0.654841537 | 1.382238946 |
| Gdap1 | 1402.76127 | -0.372418066 | 0.06127229 |
| Gm28784 | 4.433742602 | -0.400565491 | 0.462423866 |
| Gm28785 | 0.049553047 | 0.71820487 | 3.724945049 |
| Gm5251 | 0.15956642 | 0.719166641 | 3.418731035 |
| Rbm6-ps1 | 6.916369844 | 0.140151884 | 0.410909581 |
| Gm6075 | 0.038981395 | 0.718207094 | 3.724945049 |
| Gm25166 | 0 | NA | NA |
| Pi15 | 14.69231755 | 0.396875798 | 0.278514082 |
| Gm28154 | 8.974735632 | -1.407822569 | 0.391459465 |
| Gm16070 | 24.47860644 | -1.428333843 | 0.250929671 |
| Crispld1 | 57.83831356 | -1.492461217 | 0.164707717 |
| Gm28153 | 1.598622982 | -0.781445498 | 0.911338713 |
| Gm7690 | 0 | NA | NA |
| Gm18775 | 13.00822407 | -0.899287121 | 0.283989562 |

| | | | |
|---|---|---|---|
| Gm17969 | 0 | NA | NA |
| Gm28755 | 1.336776882 | -0.307953336 | 0.866336686 |
| Gm28756 | 10.18614479 | -1.331505285 | 0.379966695 |
| Crisp4 | 0 | NA | NA |
| Defb44-ps | 0.047948698 | -0.243569582 | 3.724945049 |
| Defb18 | 0 | NA | NA |
| Gm15386 | 0 | NA | NA |
| Gm17837 | 0.172125443 | 0.237328282 | 2.321733346 |
| Gm29006 | 0 | NA | NA |
| Gm5252 | 0 | NA | NA |
| Gm24075 | 0 | NA | NA |
| Gm28343 | 0 | NA | NA |
| Gm28344 | 0 | NA | NA |
| Gm15825 | 1.225430203 | -0.038768914 | 0.979288088 |
| Tfap2d | 23.18306498 | -0.487671305 | 0.278816646 |
| Gm10075 | 1.913880918 | 0.296935149 | 0.73302359 |
| Tfap2b | 140.1785569 | -0.271759115 | 0.153933828 |
| Gm28340 | 0.106520246 | 0.718200421 | 3.724945049 |
| Gm28341 | 0 | NA | NA |
| Gm4849 | 4.195570756 | 1.361541398 | 0.504869663 |
| Gm24633 | 0 | NA | NA |
| Gm38382 | 0.088383099 | -0.72445792 | 3.724945049 |
| Pkhd1 | 2.807822083 | -0.100135529 | 0.67924981 |
| Gm37315 | 0 | NA | NA |
| 4930486I03Rik | 0.427990203 | -2.151776858 | 1.668316819 |
| Gm15795 | 0.244548831 | 0.235495601 | 2.108129304 |
| Gm24162 | 0 | NA | NA |
| Mir206 | 0 | NA | NA |
| Gm28653 | 1.730304082 | -1.301554281 | 0.748790566 |
| Mir133b | 0 | NA | NA |
| Il17a | 0 | NA | NA |
| Il17f | 0.328867834 | -0.151938115 | 2.038935859 |
| Mcm3 | 85.70957654 | 2.368407207 | 0.27113762 |
| Gm20587 | 0 | NA | NA |
| Gm28065 | 0.068097359 | -0.243569582 | 3.724945049 |
| Gm24723 | 0 | NA | NA |
| 6720483E21Rik | 3.174408629 | -1.179936929 | 0.688871415 |
| Paqr8 | 1990.654714 | -0.34324187 | 0.112513996 |
| Gm24171 | 0 | NA | NA |
| Gm28064 | 5.60213491 | -0.71715585 | 0.456059189 |
| Efhc1 | 14.45078694 | 0.194301597 | 0.277262833 |
| Tram2 | 100.5415102 | 2.398688402 | 0.186475459 |
| Gm28287 | 0.218494371 | -0.243563275 | 2.166806322 |
| Gm2693 | 2.478342139 | -0.945574722 | 0.714260968 |
| Tmem14a | 140.3427173 | -1.040154496 | 0.15601435 |
| Gm38224 | 3.730747895 | -0.360577515 | 0.496568626 |
| Gsta3 | 15.33756154 | -1.153235776 | 0.359262182 |

| | | | |
|---|---|---|---|
| Gm28836 | 1.863919813 | -1.142909578 | 0.763626284 |
| Gm7094 | 1.910334788 | 0.072570748 | 0.825555661 |
| Gm4956 | 0.357620736 | 0.967352264 | 1.524107044 |
| Gm21909 | 0.066050875 | 0.71820487 | 3.724945049 |
| Khdc1a | 0.075043205 | 0.237309859 | 3.724945049 |
| Gm16329 | 0 | NA | NA |
| Gm16327 | 0 | NA | NA |
| Khdc1c | 0.097139547 | 0.718207094 | 3.724945049 |
| Gm2709 | 0 | NA | NA |
| Gm16533 | 0 | NA | NA |
| Khdc1b | 0 | NA | NA |
| Gm16328 | 0 | NA | NA |
| Kcnq5 | 996.2770014 | -1.084006691 | 0.144566077 |
| Gm36949 | 26.09595903 | -1.361284139 | 0.330917564 |
| Gm7658 | 100.9699313 | 0.400708607 | 0.163275113 |
| Gm38243 | 62.35611887 | -1.541947234 | 0.346070903 |
| Gm26580 | 1.456296832 | -0.236606652 | 0.896485301 |
| Gm5696 | 0 | NA | NA |
| Gm28239 | 0 | NA | NA |
| Rims1 | 3306.597451 | -0.908918454 | 0.09509572 |
| Gm26866 | 0.048949532 | -0.243569582 | 3.724945049 |
| Gm7761 | 0.676990728 | -0.831253858 | 1.205524146 |
| Gm29506 | 0.481916195 | -0.684057482 | 1.476282644 |
| 4933415F23Rik | 0.213028513 | 0.237325393 | 2.157312989 |
| Gm24176 | 0 | NA | NA |
| Gm6420 | 0.044191549 | -0.243571806 | 3.724945049 |
| Mir30a | 2.056819932 | 0.422460205 | 0.693064426 |
| Gm27028 | 24.51280018 | -0.413584813 | 0.216913236 |
| Mir30c-2 | 0 | NA | NA |
| Gm29107 | 3.847956237 | -0.260746255 | 0.512081887 |
| Gm20954 | 0.386554245 | 0.804317854 | 3.259347973 |
| Ogfrl1 | 2085.851771 | -0.440263664 | 0.074595347 |
| Gm7784 | 0.046345709 | 0.71820487 | 3.724945049 |
| Gm28822 | 1.28195289 | 0.568584748 | 0.857010976 |
| B3gat2 | 201.421169 | -0.829306075 | 0.106212911 |
| Smap1 | 968.2258995 | -0.193949146 | 0.061060234 |
| Sdhaf4 | 66.50993585 | -0.373031583 | 0.171778778 |
| Gm26524 | 0.087261593 | -0.243576254 | 3.724945049 |
| Fam135a | 882.3529123 | -0.124809173 | 0.07489092 |
| Gm37580 | 2.220335296 | -0.592124891 | 0.653715737 |
| Gm26607 | 0 | NA | NA |
| Col9a1 | 27.58382149 | 0.956390989 | 0.315298206 |
| Col19a1 | 256.8225277 | -1.669896658 | 0.20319917 |
| Gm26418 | 0.607053417 | -2.576999365 | 1.297988907 |
| Tubb4b-ps2 | 1.899816927 | 0.353342129 | 0.691720204 |
| Gm28439 | 270.7744438 | -0.448311755 | 0.162642026 |
| Gm10222 | 81.6198627 | -0.28851025 | 0.184911575 |

| | | | |
|---|---|---|---|
| Gm28438 | 233.7213687 | 0.021639377 | 0.186047182 |
| Gm28437 | 654.6201061 | -0.128850454 | 0.18597131 |
| Gm10925 | 568.5256472 | -0.14165156 | 0.192118567 |
| Gm28661 | 361.8513258 | -0.250730856 | 0.140875606 |
| Gm29216 | 452.3609317 | 0.182133464 | 0.178922126 |
| Lmbrd1 | 1834.054128 | -0.412851539 | 0.058346673 |
| Gm28594 | 0 | NA | NA |
| Gm28237 | 0.181654796 | 1.679598083 | 3.311020364 |
| Gm9443 | 0.049553047 | 0.71820487 | 3.724945049 |
| Gm29414 | 0.167977605 | 0.718216306 | 2.75817218 |
| Adgrb3 | 2707.39029 | -0.852988246 | 0.121608385 |
| Gm15702 | 0.267089184 | 0.529400815 | 1.781032321 |
| Gm20172 | 56.45506021 | -1.142025075 | 0.288072061 |
| Gm22260 | 10.97169895 | -1.11839917 | 0.366234356 |
| Gm37543 | 46.77977661 | -1.179466486 | 0.337548531 |
| Gm37331 | 188.7440919 | -1.02884955 | 0.203903015 |
| Gm37211 | 123.4678815 | -1.093846013 | 0.243784097 |
| Gm37862 | 145.3838556 | -0.967048573 | 0.238037421 |
| Gm38286 | 155.9317435 | -1.209756364 | 0.232982078 |
| Gm37880 | 213.8342296 | -1.13282349 | 0.298940633 |
| Gm25294 | 0 | NA | NA |
| Gm37011 | 137.8405266 | -1.104508851 | 0.184438418 |
| Gm29214 | 43.40613233 | -1.060927355 | 0.228874588 |
| Gm37051 | 172.0940709 | -1.131357134 | 0.218590261 |
| Gm38260 | 342.1641184 | -1.118440467 | 0.176113819 |
| Gm37628 | 460.1944707 | -1.200952253 | 0.247156107 |
| 8030445P17Rik | 222.3518729 | -1.151949665 | 0.245959286 |
| Gm38198 | 365.9544784 | -1.193994608 | 0.295172488 |
| Gm37077 | 292.4722539 | -1.172699225 | 0.186837818 |
| Gm37895 | 566.8750103 | -1.109654778 | 0.182429523 |
| Gm36952 | 499.6234323 | -1.159187702 | 0.20916443 |
| Gm37602 | 362.6410622 | -0.970403335 | 0.150958371 |
| 2900022M07Rik | 83.257532 | -1.370113334 | 0.273488953 |
| 2900002M20Rik | 64.13250466 | -1.37187886 | 0.252311339 |
| Gm9884 | 9.752194563 | -1.12705866 | 0.35563167 |
| Gm18443 | 0 | NA | NA |
| Gm19123 | 0 | NA | NA |
| Gm24064 | 0 | NA | NA |
| 4931408C20Rik | 0 | NA | NA |
| n-R5s209 | 0 | NA | NA |
| Gm19641 | 0 | NA | NA |
| Gm5524 | 0.04393175 | -0.243571806 | 3.724945049 |
| Gm18776 | 0 | NA | NA |
| Gm19028 | 0.124853745 | 0.237309606 | 3.724945049 |
| Gm38089 | 0 | NA | NA |
| Gm7846 | 6.28322175 | 0.510495081 | 0.401645537 |
| Gm597 | 0.068097359 | -0.243569582 | 3.724945049 |

| | | | |
|---|---|---|---|
| Gm5697 | 0.151010504 | -0.243562199 | 2.854938236 |
| Gm6462 | 0 | NA | NA |
| Gm18033 | 0 | NA | NA |
| Gm25586 | 0 | NA | NA |
| Gm22679 | 0 | NA | NA |
| Gm23785 | 0 | NA | NA |
| Mir6342 | 0.042716945 | -0.243571806 | 3.724945049 |
| Gm5525 | 5.440920671 | 1.262793718 | 0.440050843 |
| Gm7858 | 0 | NA | NA |
| Gm23771 | 0 | NA | NA |
| Gm27430 | 0.117046891 | -0.724453471 | 3.724945049 |
| Gm9898 | 0.52260535 | -1.427828563 | 1.356865862 |
| Gm6473 | 7.058970522 | -0.510293983 | 0.396466893 |
| Gm2914 | 0.311886478 | 1.199122453 | 1.775331237 |
| Phf3 | 1888.771435 | -0.127242103 | 0.058761759 |
| Gm37522 | 62.34479475 | -1.047133284 | 0.234992659 |
| Gm37906 | 105.3489438 | -0.833067583 | 0.225688534 |
| Ptp4a1 | 625.4291984 | 0.70018233 | 0.115221383 |
| Gm5698 | 0 | NA | NA |
| Gm9458 | 0 | NA | NA |
| Gm5699 | 0.061920613 | 0.718207094 | 3.724945049 |
| Gm29669 | 0 | NA | NA |
| Gm28644 | 0 | NA | NA |
| Gm27362 | 0 | NA | NA |
| 4931428L18Rik | 0.048949532 | -0.243569582 | 3.724945049 |
| Gm29128 | 0 | NA | NA |
| Gm7893 | 0 | NA | NA |
| Lgsn | 0.247864911 | 1.577184326 | 2.505377127 |
| Pih1d3 | 0 | NA | NA |
| Gm26145 | 0 | NA | NA |
| Gm4850 | 0 | NA | NA |
| Gm28645 | 0.066050875 | 0.71820487 | 3.724945049 |
| Gm6489 | 0 | NA | NA |
| Gm10307 | 0 | NA | NA |
| Gm19680 | 8.167021386 | 0.907922218 | 0.377539764 |
| Gm23032 | 0 | NA | NA |
| Gm7910 | 0 | NA | NA |
| Khdrbs2 | 245.3494503 | -0.507096262 | 0.13857162 |
| Gm25814 | 0.068097359 | -0.243569582 | 3.724945049 |
| Gm37999 | 19.71845274 | -0.900517623 | 0.360758996 |
| Gm37110 | 1.003269596 | -1.710109047 | 1.064112476 |
| Gm37911 | 37.09896806 | -0.586045899 | 0.30136525 |
| Gm9839 | 0.136963818 | 0.718204821 | 3.59954433 |
| Gm37792 | 46.79344322 | -1.075110379 | 0.244559636 |
| Gm5415 | 0.486008019 | -0.254717907 | 1.347638049 |
| Gm24732 | 0 | NA | NA |
| Gm37426 | 6.59084447 | -0.755233808 | 0.457049246 |

| | | | |
|---|---|---|---|
| Gm23024 | 0.184258256 | -0.243569838 | 2.697073173 |
| Gm37151 | 1.344706689 | -0.376965954 | 1.148827234 |
| Gm28624 | 0.510604427 | -1.176520489 | 1.592165618 |
| Gm37724 | 1.002429578 | 0.429809386 | 1.099101059 |
| Gm7114 | 3.168220283 | -0.105277256 | 0.563047152 |
| Gm4785 | 0.341712212 | -0.230929107 | 1.594399959 |
| Gm24901 | 0 | NA | NA |
| Gm37591 | 0.710426184 | 0.718228122 | 1.494807356 |
| Gm2987 | 0 | NA | NA |
| Gm25792 | 0 | NA | NA |
| Gm18799 | 2.333919785 | -0.306350318 | 0.756512511 |
| Gm7933 | 0.088383099 | -0.72445792 | 3.724945049 |
| Gm23453 | 0 | NA | NA |
| Prim2 | 97.92590214 | -0.054400787 | 0.125615201 |
| Gm29228 | 0 | NA | NA |
| Gm37618 | 13.20070015 | -1.13075933 | 0.304428517 |
| Gm17813 | 0 | NA | NA |
| 1700001G17Rik | 1.688998503 | -0.544036412 | 0.720046497 |
| Gm38173 | 0.528487055 | 0.604585752 | 1.513222394 |
| Rab23 | 419.6998972 | -0.133969312 | 0.059034066 |
| Bag2 | 54.75618287 | -0.218394574 | 0.163883697 |
| Gm37905 | 0.392625049 | -1.344498049 | 1.46063845 |
| Zfp451 | 1940.361675 | -0.180915454 | 0.068904612 |
| Gm15455 | 3.94145278 | -0.622795633 | 0.629772772 |
| Bend6 | 1529.651517 | -0.609447672 | 0.12953275 |
| Gm37067 | 37.0527924 | -1.443682669 | 0.370024047 |
| Dst | 13187.49407 | -0.168768983 | 0.062135744 |
| Gm37354 | 31.03348255 | -1.26147564 | 0.326823516 |
| Gm28631 | 0 | NA | NA |
| Gm23792 | 0 | NA | NA |
| Gm38157 | 47.50224476 | -0.740384427 | 0.327366864 |
| mmu-mir-6896 | 1.465090628 | -1.66944572 | 1.002266708 |
| Gm26788 | 4.676338759 | 0.16961239 | 0.480918504 |
| D630036G22Rik | 69.39949552 | 0.044648184 | 0.200263216 |
| A930005N03Rik | 0.933986728 | -0.879964417 | 1.081216924 |
| Gm37958 | 0 | NA | NA |
| Gm37385 | 0.204111068 | 0.718218362 | 2.600299193 |
| Gm5266 | 0 | NA | NA |
| Gm3052 | 10.39265568 | -1.103334298 | 0.350213208 |
| Gm37233 | 84.28593999 | -0.562434866 | 0.272819128 |
| Mir5103 | 0 | NA | NA |
| Ccdc115 | 140.6358465 | 0.198247628 | 0.119458807 |
| Gm28306 | 0 | NA | NA |
| Imp4 | 379.5447932 | 0.445509097 | 0.084116442 |
| Ptpn18 | 38.27817853 | 2.249754256 | 0.26848323 |
| Gm28417 | 1.241452806 | 2.666945293 | 1.02498559 |
| 4930568A12Rik | 0.934086492 | 1.238125241 | 1.13363337 |

| | | | |
|---|---|---|---|
| Prss39 | 0.301806123 | 1.556824779 | 1.704208501 |
| Gm28416 | 0 | NA | NA |
| Cfc1 | 0.191490881 | -0.724462325 | 2.704974322 |
| Gm28419 | 0 | NA | NA |
| Prss40 | 0 | NA | NA |
| 1700101I19Rik | 0.559566189 | -1.398891658 | 3.608868373 |
| Amer3 | 415.894219 | -0.417974533 | 0.094302682 |
| Gm28418 | 0.045882455 | 0.237312084 | 3.724945049 |
| Gm36984 | 0.129408498 | 0.718212916 | 2.991811555 |
| Gm29730 | 0.628347421 | 1.19977852 | 1.372037131 |
| AA619741 | 4.444019085 | 3.762794857 | 1.066963429 |
| Arhgef4 | 1837.032545 | -0.508175871 | 0.050841059 |
| AC107766.1 | 0 | NA | NA |
| Gm37224 | 4.445654744 | -0.653835124 | 0.481688129 |
| Gm37845 | 4.660398994 | -0.627844276 | 0.63948808 |
| Gm38336 | 10.69144326 | -0.548135728 | 0.401321872 |
| Fam168b | 3287.598993 | -0.100758502 | 0.048100109 |
| Gm37127 | 1.263832448 | -0.531015327 | 1.117395009 |
| Plekhb2 | 2551.489122 | -0.064345418 | 0.072009189 |
| Gm19430 | 0.332159689 | 2.368701561 | 1.924907817 |
| Gm37670 | 0.517156391 | -0.535583685 | 1.364452887 |
| Gm8009 | 0.062426873 | 0.237312084 | 3.724945049 |
| Gm38027 | 0 | NA | NA |
| Gm25096 | 0 | NA | NA |
| Gm37068 | 0.62070299 | 0.481405279 | 1.200649632 |
| Gm25634 | 0 | NA | NA |
| Gm28415 | 0 | NA | NA |
| 1110002O04Rik | 0.877137428 | -0.547767586 | 0.947905822 |
| Gm8022 | 0 | NA | NA |
| Gm37146 | 0 | NA | NA |
| Gm33222 | 0.331946588 | -0.727855631 | 1.717255376 |
| 4930535G08Rik | 0 | NA | NA |
| Gm38049 | 0.068097359 | -0.243569582 | 3.724945049 |
| Hs6st1 | 777.2491163 | -0.014717516 | 0.07541613 |
| Gm37335 | 0.364365233 | -1.075841271 | 1.542152934 |
| Gm33280 | 0.253548292 | -0.236660215 | 1.831709371 |
| Uggt1 | 1213.9728 | 0.009631093 | 0.05921891 |
| Mir6897 | 0.539626406 | -0.421116531 | 1.394656329 |
| Neurl3 | 71.81858134 | 4.940571222 | 0.363539231 |
| Arid5a | 533.3445611 | 2.137257256 | 0.248081922 |
| 4930403P22Rik | 0.909075398 | 0.769458861 | 2.249672223 |
| Kansl3 | 1110.948051 | 0.240964998 | 0.070645205 |
| Mir6898 | 0.044191549 | -0.243571806 | 3.724945049 |
| Fer1l5 | 1.367163391 | 2.2583457 | 1.039054265 |
| Lman2l | 453.4304442 | -0.15209132 | 0.091106917 |
| Cnnm4 | 233.328149 | -0.100851304 | 0.107436533 |
| Gm37909 | 10.35866414 | -1.109460905 | 0.385703326 |

| | | | |
|---|---|---|---|
| AC084391.1 | 0 | NA | NA |
| Cnnm3 | 394.0709263 | -0.313495563 | 0.080652185 |
| Gm38033 | 0.17077006 | 0.718214265 | 2.740691057 |
| Ankrd23 | 35.9631227 | -0.043080939 | 0.184952667 |
| Ankrd39 | 86.73678357 | -0.085633554 | 0.139752539 |
| Sema4c | 276.9910831 | 0.646336085 | 0.11749613 |
| D430040D24Rik | 19.3189591 | 0.13827721 | 0.2625185 |
| Fam178b | 2.063825205 | 0.478953772 | 0.670870923 |
| Gm6428 | 1.366437448 | -0.20403202 | 0.804029803 |
| Gm24133 | 0 | NA | NA |
| Cox5b | 1330.17609 | -0.101722769 | 0.118479344 |
| Actr1b | 2273.379145 | -0.172188388 | 0.03998131 |
| 4933424G06Rik | 1.049948633 | 1.021873986 | 1.083250247 |
| Gm33533 | 12.17216712 | -1.829344302 | 0.463743523 |
| RP23-247A1.6 | 1.790960893 | -1.435292666 | 0.981992425 |
| Zap70 | 42.79799715 | 2.416105842 | 0.291503354 |
| Gm18828 | 0 | NA | NA |
| Tmem131 | 1521.534467 | -0.031778246 | 0.071153662 |
| Gm37506 | 4.051657186 | -1.652881915 | 0.579158547 |
| Gm38115 | 35.47402398 | -0.485738886 | 0.312648839 |
| F830112A20Rik | 59.60486576 | -0.327575452 | 0.257172753 |
| Rpl12-ps1 | 0.348012853 | 0.237329955 | 1.977104605 |
| n-R5s210 | 0 | NA | NA |
| Vwa3b | 17.13175727 | 0.22454779 | 0.314419249 |
| Gm37020 | 0.834730223 | -3.054054473 | 1.210539203 |
| Cnga3 | 3.017996072 | 1.37999974 | 0.731223608 |
| 1700074A21Rik | 0 | NA | NA |
| Inpp4a | 3160.431147 | -0.674475272 | 0.061043411 |
| 4930439A04Rik | 0.093831153 | -0.243571806 | 3.724945049 |
| Gm37135 | 0 | NA | NA |
| Coa5 | 899.7403321 | 0.23702237 | 0.106631647 |
| Unc50 | 302.5469516 | 0.214230998 | 0.101954781 |
| Mgat4a | 840.1121684 | -0.265646001 | 0.05083342 |
| 4930594C11Rik | 47.43194901 | -1.029064423 | 0.322941788 |
| 2010300C02Rik | 1765.094823 | -0.985270972 | 0.173840641 |
| 4930556I23Rik | 0.066050875 | 0.71820487 | 3.724945049 |
| 4930470B04Rik | 0.188205214 | 1.067748746 | 2.673953749 |
| Gm19863 | 38.19950049 | -2.0397683 | 0.422551661 |
| Tsga10 | 318.6599432 | -0.465262556 | 0.122591468 |
| Gm26805 | 21.25672272 | -2.046704474 | 0.336980385 |
| Lipt1 | 33.80092537 | -0.292129416 | 0.183378957 |
| Mitd1 | 202.445721 | 2.474609137 | 0.180722787 |
| Mrpl30 | 491.4665078 | 0.743755159 | 0.104605316 |
| Lyg2 | 0.741337718 | -3.057475973 | 1.249819513 |
| Gm15457 | 0.21650593 | 2.065999099 | 2.130658308 |
| Lyg1 | 0.136120942 | 1.15080397 | 3.595683408 |
| Gm23409 | 0 | NA | NA |

| | | | |
|---|---|---|---|
| Txndc9 | 544.4411505 | 0.658604945 | 0.088888427 |
| Eif5b | 954.1717507 | 0.33745292 | 0.116658469 |
| Rev1 | 696.4174186 | -0.377743951 | 0.112200398 |
| Gm37062 | 35.92510916 | -0.750857273 | 0.228438065 |
| Gm5099 | 0.141094079 | 0.718203391 | 3.563850454 |
| Aff3 | 992.3491355 | -0.557308066 | 0.074977857 |
| Gm16150 | 1.482000077 | 0.472032948 | 0.891339635 |
| Gm16151 | 0 | NA | NA |
| Gm16152 | 1.435447043 | 1.479838186 | 0.820349238 |
| Gm34727 | 0 | NA | NA |
| Gm38135 | 0 | NA | NA |
| 2610300A13Rik | 0 | NA | NA |
| Lonrf2 | 2520.190482 | -0.501531082 | 0.079429755 |
| Chst10 | 707.3698007 | -0.572143084 | 0.055931132 |
| Gm37707 | 0 | NA | NA |
| Gm6621 | 0.169398393 | 0.237310727 | 3.403281961 |
| Nms | 1.222583469 | 2.111396112 | 0.965139062 |
| Pdcl3 | 219.8699854 | 0.457624881 | 0.128738815 |
| Gm37309 | 20.51143474 | -0.930557216 | 0.344799224 |
| Gm37821 | 0.54509763 | 1.410745359 | 1.360632918 |
| Gm37091 | 0.205567437 | -0.243568543 | 2.594414786 |
| Mir6349 | 0 | NA | NA |
| Npas2 | 366.6556838 | -0.837279208 | 0.159681284 |
| Gm20428 | 0.263795272 | -0.268699157 | 1.804537573 |
| Gm3617 | 6.543142137 | -1.238592939 | 0.570200996 |
| Rpl31 | 1558.50621 | 0.658313669 | 0.125676232 |
| Tbc1d8 | 368.5922422 | -0.67461131 | 0.106445848 |
| Gm37265 | 6.813130323 | -1.41266323 | 0.480268582 |
| Gm24826 | 0 | NA | NA |
| ACEA_U3 | 3.059522033 | 0.662926718 | 0.531631097 |
| Cnot11 | 371.6623297 | 0.229228956 | 0.098782422 |
| Gm15832 | 37.25351844 | 0.247514854 | 0.301696357 |
| Snord89 | 3.528574751 | 0.402971302 | 0.518423635 |
| Rnf149 | 381.317406 | 1.305688346 | 0.175840115 |
| Gm23722 | 10.82063883 | 1.076597963 | 0.388747722 |
| Gm5100 | 0.671689889 | 0.677823808 | 1.206973085 |
| Creg2 | 1118.864952 | -0.341566863 | 0.150179538 |
| D930019O06Rik | 42.32280546 | -0.414996641 | 0.221193694 |
| Rfx8 | 1.59726488 | -0.545359501 | 0.887774061 |
| Gm3646 | 0.042716945 | -0.243571806 | 3.724945049 |
| 1700066B17Rik | 0.337370548 | -0.279628134 | 2.018292847 |
| Map4k4 | 1991.375243 | 0.132464228 | 0.047059702 |
| 4930456G14Rik | 5.719135214 | -0.958109502 | 0.522429244 |
| Il1r2 | 32.73985817 | 5.434364042 | 0.834842977 |
| Gm16894 | 21.14797858 | -1.049982133 | 0.26092252 |
| Gm37347 | 2.446674312 | 3.383841368 | 0.973454731 |
| Il1r1 | 110.9761096 | 1.348103954 | 0.159589879 |

| | | | |
|---|---|---|---|
| Gm38070 | 5.586992304 | 0.543512956 | 0.425078693 |
| Il1rl2 | 14.87794066 | 0.942325444 | 0.3205381 |
| Gm17970 | 0.509421185 | -0.407135207 | 1.347500019 |
| Il1rl1 | 3.917850035 | 0.919034514 | 0.516605523 |
| Il18r1 | 13.77975115 | 6.26411384 | 0.88025181 |
| Il18rap | 55.22165596 | 8.439306481 | 0.869774977 |
| Gm35801 | 0.072025462 | 0.237312084 | 3.724945049 |
| Slc9a4 | 29.46342029 | -0.276727734 | 0.224651909 |
| Gm37435 | 0.132101749 | 1.209894437 | 3.72418822 |
| Slc9a2 | 85.94852822 | -0.315616027 | 0.175418629 |
| Gm37623 | 0 | NA | NA |
| Gm37915 | 0.583248015 | -0.156791303 | 1.277521972 |
| Mfsd9 | 54.36389778 | 0.287474582 | 0.191748985 |
| Tmem182 | 1.853692403 | -0.668491179 | 0.911381857 |
| Gm5973 | 0 | NA | NA |
| 4930448I06Rik | 0.044544648 | 0.237309859 | 3.724945049 |
| Gm28634 | 0.143022002 | 0.719176055 | 3.547735711 |
| Gm29260 | 4.366569511 | 0.445700873 | 0.450262754 |
| Gm5267 | 0 | NA | NA |
| Gm8176 | 0 | NA | NA |
| Gm9915 | 0.24066848 | 0.712599087 | 2.104550697 |
| Gm29664 | 0 | NA | NA |
| Gm28175 | 3.438849202 | -0.850618331 | 0.552751779 |
| Gm28140 | 0.100902008 | 1.15059983 | 3.720931964 |
| Gm37047 | 0.333556256 | -1.846740268 | 1.51414344 |
| Pantr1 | 157.9407554 | -0.885217332 | 0.160680558 |
| 4930420N18Rik | 0 | NA | NA |
| Gm20646 | 38.88834011 | -1.337648805 | 0.198098462 |
| Pou3f3 | 1367.814843 | -0.340567583 | 0.089715636 |
| Pantr2 | 14.1555736 | -1.016802883 | 0.358858806 |
| Dalir | 16.94524304 | -1.776052471 | 0.374259152 |
| Mrps9 | 398.9447759 | -0.205105981 | 0.086176747 |
| Gm23126 | 0 | NA | NA |
| Gpr45 | 153.2010998 | -0.240783821 | 0.136013008 |
| Gm24001 | 0.318202806 | -0.806539375 | 2.004696787 |
| Tgfbrap1 | 851.4724595 | -0.011987961 | 0.056632661 |
| Gm28782 | 1.198752213 | -0.940626163 | 1.047315356 |
| 8430432A02Rik | 7.526368882 | -0.900375364 | 0.480830028 |
| AI597479 | 352.8821411 | -0.267314406 | 0.080488355 |
| Fhl2 | 255.1157235 | 0.504739751 | 0.120611596 |
| Gm8210 | 95.32441064 | 0.588931212 | 0.188820275 |
| Gm29040 | 0.048949532 | -0.243569582 | 3.724945049 |
| Gm29610 | 0 | NA | NA |
| Gm29041 | 0 | NA | NA |
| Nck2 | 373.3238872 | -0.169250879 | 0.081844929 |
| Gm37536 | 17.59062222 | -1.042513588 | 0.351706562 |
| Gm38228 | 18.03991603 | -1.161544575 | 0.37169011 |

| | | | |
|---|---|---|---|
| 1500015O10Rik | 35.22777033 | -0.532895149 | 0.278134348 |
| Gm29155 | 11.99474671 | -0.448833191 | 0.360261191 |
| Gm29157 | 0.378942021 | -2.122519219 | 1.500514049 |
| Gm29156 | 12.6916125 | -0.396868278 | 0.345229182 |
| Uxs1 | 508.6202783 | -0.13987556 | 0.082473862 |
| Gm16103 | 0 | NA | NA |
| Gm19587 | 1.101887088 | -0.342593346 | 1.005389245 |
| Gm8241 | 0 | NA | NA |
| Gm26352 | 0 | NA | NA |
| Gm24180 | 0.220091977 | -0.242833966 | 2.162314248 |
| Tpp2 | 1340.114203 | 0.087024001 | 0.083785301 |
| Mettl21c | 0.903763266 | 0.094389494 | 1.158784322 |
| Gm15838 | 0 | NA | NA |
| Gm8251 | 2.971797373 | 0.764397468 | 0.603143793 |
| Tex30 | 39.96396815 | 0.323725189 | 0.234203234 |
| Gm29247 | 0 | NA | NA |
| Kdelc1 | 74.7610229 | -0.226163713 | 0.128648174 |
| Bivm | 338.2242815 | -0.228105272 | 0.072797166 |
| Ercc5 | 350.2066079 | -0.279213837 | 0.060361046 |
| Mettl21e | 0.63532651 | -1.213765056 | 1.34870643 |
| Gm15833 | 0.607153663 | -0.898715715 | 1.317799698 |
| Gm5700 | 0 | NA | NA |
| Gm28893 | 0 | NA | NA |
| Txn-ps1 | 3.667605082 | 1.521637559 | 0.610610568 |
| Gulp1 | 83.35613432 | 0.063870658 | 0.152088076 |
| Gm37626 | 20.77771962 | -0.636059848 | 0.302742523 |
| Gm16102 | 1.145880322 | -2.586570008 | 1.02890491 |
| 4930521E06Rik | 0.181654796 | 1.679598083 | 3.311020364 |
| Gm38110 | 3.191642098 | -0.806060831 | 0.601022806 |
| Gm3489 | 0.12092566 | 0.237307635 | 3.724945049 |
| Gm19480 | 0 | NA | NA |
| Gm3496 | 0 | NA | NA |
| Gm5268 | 0 | NA | NA |
| Col3a1 | 232.7682847 | 1.948203253 | 0.188905725 |
| AC125167.1 | 0 | NA | NA |
| Col5a2 | 180.1321098 | 1.625690073 | 0.186523725 |
| Gm38211 | 9.579960985 | 1.072436971 | 0.38343299 |
| Gm8296 | 0.048949532 | -0.243569582 | 3.724945049 |
| Gm7063 | 0.241951302 | 0.787409104 | 2.11326256 |
| Gm8304 | 0 | NA | NA |
| Gm23216 | 0 | NA | NA |
| Gm23974 | 0 | NA | NA |
| Gm8307 | 0.148708979 | 1.198497773 | 2.868467391 |
| Gm23984 | 0 | NA | NA |
| Wdr75 | 417.7461979 | 0.350405471 | 0.066577521 |
| Gm28906 | 0 | NA | NA |
| Gm5526 | 49.19564311 | 0.298661644 | 0.215595977 |

| | | | |
|---|---|---|---|
| Gm5269 | 0.154938607 | 0.237326521 | 2.832095734 |
| Gm18303 | 0.136201742 | -0.231510441 | 2.944131489 |
| Slc40a1 | 53.73490417 | -1.197855612 | 0.259276272 |
| Gm9496 | 0 | NA | NA |
| Rpl23a-ps1 | 0 | NA | NA |
| Gm3852 | 0.218641893 | 0.789453977 | 2.17612268 |
| Stk-ps2 | 0 | NA | NA |
| Gm18849 | 0 | NA | NA |
| Dnah7b | 166.4217659 | -0.782491861 | 0.148814213 |
| n-R5s211 | 0.044191549 | -0.243571806 | 3.724945049 |
| Gm8326 | 1.175435645 | -1.050576595 | 1.069563352 |
| Dnah7c | 35.29196087 | -0.950289647 | 0.237003759 |
| Slc39a10 | 1977.950324 | -0.316639788 | 0.11685326 |
| Gm28151 | 62.07798039 | -0.217356173 | 0.23212895 |
| Gm37393 | 2.121468972 | -3.51304744 | 1.198111389 |
| Gm28527 | 0.844069991 | -1.332967813 | 1.252378122 |
| Gm28825 | 0.476041195 | 0.384261965 | 1.374631194 |
| Gm28826 | 0.77428951 | 2.669849043 | 1.198968608 |
| Gm4852 | 3.57865669 | 0.122075463 | 0.534380222 |
| Gm38090 | 0 | NA | NA |
| Mir6350 | 0 | NA | NA |
| Gm18117 | 0.885425998 | 2.958440019 | 1.483158466 |
| Gm37196 | 1.287564413 | 1.921454915 | 1.137952962 |
| Gm8337 | 21.49979967 | 0.298395405 | 0.276892718 |
| Gm22419 | 0 | NA | NA |
| Gm23829 | 0 | NA | NA |
| Gm19206 | 0 | NA | NA |
| Gm5974 | 0 | NA | NA |
| Gm23240 | 0 | NA | NA |
| Gm28830 | 0 | NA | NA |
| Gm8354 | 0 | NA | NA |
| Gm19124 | 0 | NA | NA |
| Gm8357 | 0 | NA | NA |
| Gm22693 | 0 | NA | NA |
| Gm28740 | 0 | NA | NA |
| Gm8367 | 0 | NA | NA |
| C230029F24Rik | 0 | NA | NA |
| Gm6757 | 0 | NA | NA |
| Gm3605 | 0 | NA | NA |
| Gm28388 | 0 | NA | NA |
| Gm8384 | 0 | NA | NA |
| 4933411E06Rik | 0.068097359 | -0.243569582 | 3.724945049 |
| Gm8391 | 0 | NA | NA |
| Gm27330 | 0 | NA | NA |
| Gm29665 | 0 | NA | NA |
| Gm20118 | 0 | NA | NA |
| Gm25843 | 0 | NA | NA |

| | | | |
|---|---|---|---|
| Gm28321 | 0.092691418 | 1.183079489 | 3.724560917 |
| Tmeff2 | 912.3017426 | -0.158904491 | 0.073578574 |
| Gm28322 | 2.545969209 | -0.917371176 | 0.709422569 |
| Gm25736 | 12.09809119 | -0.870401158 | 0.34397241 |
| Gm28319 | 0.231621402 | 0.237327852 | 2.52240785 |
| Gm28320 | 83.01456838 | -0.862749426 | 0.225084206 |
| Gm29325 | 37.76672809 | -0.729557622 | 0.267335996 |
| 9330175M20Rik | 4.588839173 | 0.348683997 | 0.535731467 |
| Sdpr | 92.2652614 | 0.657775486 | 0.147842395 |
| Gm5253 | 0.982135049 | -0.422910771 | 0.978030784 |
| 1700072G22Rik | 0.174149299 | 1.185086784 | 3.359371254 |
| Nabp1 | 240.5774001 | 0.956761666 | 0.193099383 |
| Gm17767 | 3.178614829 | 1.642971978 | 0.649966034 |
| Gm37535 | 0.140088945 | -1.149030578 | 3.601861891 |
| Gm28055 | 0.781209455 | 0.252970875 | 1.177088733 |
| Myo1b | 450.2003702 | -0.364321494 | 0.131038231 |
| Gm29453 | 3.059357988 | -0.032195809 | 0.662488849 |
| Gm28323 | 3.764028883 | -1.138115702 | 0.501015526 |
| Gm8419 | 0 | NA | NA |
| Gm27607 | 0 | NA | NA |
| Gm8420 | 0.200957741 | 1.97380807 | 3.229647234 |
| Stat4 | 19.2643103 | 2.149290592 | 0.308025405 |
| Gm28177 | 2.000698716 | 3.9519873 | 0.966455452 |
| Gm3940 | 0.598154365 | 0.18462713 | 1.284127788 |
| Stat1 | 2376.29098 | 5.019602675 | 0.114315694 |
| Gm18301 | 0.092228164 | 0.718202646 | 3.724945049 |
| Gls | 3738.856511 | -0.909043109 | 0.068440969 |
| Gm5975 | 0.042716945 | -0.243571806 | 3.724945049 |
| Gm23460 | 0 | NA | NA |
| Nab1 | 590.3308437 | 0.467810443 | 0.114468383 |
| Gm5976 | 0.038981395 | 0.718207094 | 3.724945049 |
| Rps27a-ps1 | 3.694411362 | -0.614455782 | 0.629119768 |
| Gm553 | 3.129775993 | -0.886250786 | 0.610668039 |
| Gm28178 | 0.452230046 | 2.34036879 | 1.296979963 |
| Gm5527 | 9.620821135 | 0.35519284 | 0.373293735 |
| Tmem194b | 84.55902287 | -0.162183973 | 0.184075378 |
| Gm37242 | 11.17186488 | -0.69442025 | 0.318390442 |
| Mfsd6 | 1706.135027 | -0.169063593 | 0.078658615 |
| Gm29670 | 25.59481537 | -1.181078264 | 0.305902916 |
| Gm28650 | 0.805100971 | -0.399769319 | 1.101722176 |
| Inpp1 | 600.5457346 | 0.677592465 | 0.106340565 |
| Hibch | 147.5834649 | -0.091116764 | 0.163751034 |
| Gm16107 | 1.228146034 | 1.064980244 | 1.178990142 |
| 1700019D03Rik | 237.2523422 | -0.096832264 | 0.088699037 |
| Mstn | 4.314567637 | 0.440704214 | 0.50200556 |
| Gm24349 | 0 | NA | NA |
| 1700019A02Rik | 5.126622909 | -0.768058157 | 0.54088677 |

| | | | |
|---|---|---|---|
| Pms1 | 234.6633514 | -0.283947907 | 0.093161904 |
| Gm28777 | 0 | NA | NA |
| Ormdl1 | 92.39055438 | 0.566791883 | 0.153985881 |
| Gm28778 | 0.044191549 | -0.243571806 | 3.724945049 |
| Osgepl1 | 175.8191109 | -0.207823678 | 0.108761557 |
| Gm28551 | 1.277925253 | -0.501549392 | 0.940310427 |
| Gm18800 | 2.110212028 | -1.044339053 | 0.694801362 |
| Asnsd1 | 39.89699009 | 0.193069096 | 0.172626153 |
| Gm28779 | 0.097139547 | 0.718207094 | 3.724945049 |
| Gm6799 | 0 | NA | NA |
| Gm6801 | 0 | NA | NA |
| Dnah7a | 93.9630857 | -0.330202918 | 0.172684433 |
| Gm25240 | 0.108123235 | -0.24357403 | 3.724945049 |
| Stk17b | 400.9618622 | 1.52838357 | 0.143813733 |
| Gm37863 | 0 | NA | NA |
| Hecw2 | 1282.188723 | -0.580703611 | 0.076668272 |
| Mir7681 | 0.068097359 | -0.243569582 | 3.724945049 |
| Gm24251 | 2.00647346 | -0.831715613 | 0.866662371 |
| Gm37633 | 69.5255469 | -1.528959831 | 0.351110391 |
| Gm25829 | 0.15586508 | 0.23732528 | 2.82680955 |
| Ccdc150 | 3.346209518 | -2.241762869 | 0.685964587 |
| Gtf3c3 | 606.9969703 | 0.004491021 | 0.065095034 |
| Pgap1 | 1020.253931 | -0.543421786 | 0.080178273 |
| Gm36955 | 30.15051617 | -1.148996124 | 0.244694383 |
| Gm19637 | 0.11197992 | 0.718202646 | 3.724945049 |
| Ankrd44 | 326.5543126 | 0.265194662 | 0.096311073 |
| 4930444A19Rik | 0.477440416 | 1.136738303 | 1.551367125 |
| Gm37370 | 15.65747994 | -0.213216055 | 0.30556559 |
| Gm24053 | 0 | NA | NA |
| A130048G24Rik | 23.27167967 | -0.129121081 | 0.320068231 |
| Sf3b1 | 4398.794146 | 0.155195526 | 0.072706194 |
| E330011M16Rik | 2.812432955 | 0.812823029 | 0.638265853 |
| Coq10b | 351.6495304 | 0.044406933 | 0.130643908 |
| Gm17971 | 11.17290908 | -1.494880556 | 0.520233828 |
| Hspd1 | 1608.718732 | 0.490170547 | 0.102066749 |
| Hspe1 | 313.2638214 | 0.566749112 | 0.149150078 |
| Gm5147 | 0.483827566 | -0.697538563 | 1.464866369 |
| Gm16581 | 13.32551551 | 0.065500953 | 0.354779113 |
| Mob4 | 696.0879921 | 0.144078784 | 0.090435045 |
| Gm6822 | 0.477893029 | 0.686912002 | 1.342723773 |
| Rftn2 | 338.1297183 | -0.377689144 | 0.127618612 |
| Gm10561 | 20.71569524 | -0.495814414 | 0.254085582 |
| Mars2 | 418.0161997 | -0.381679676 | 0.112308157 |
| Boll | 0.988924806 | 1.773181483 | 1.166204538 |
| Gm8292 | 51.67441755 | 0.636819643 | 0.191080977 |
| Gm29138 | 0 | NA | NA |
| Gm8531 | 0.047948698 | -0.243569582 | 3.724945049 |

| | | | |
|---|---|---|---|
| Plcl1 | 931.6952418 | 0.036545801 | 0.094030213 |
| Gm38056 | 155.7691939 | -0.667841599 | 0.367990376 |
| Gm6644 | 4.857481711 | 1.113299883 | 0.48689882 |
| Gm37382 | 65.4083701 | -0.557919423 | 0.265298554 |
| 9130227L01Rik | 1.675708655 | -0.774237866 | 0.81121098 |
| Gm36986 | 0 | NA | NA |
| 1700003I22Rik | 0 | NA | NA |
| Gm28900 | 0.298162442 | 2.260866236 | 1.69085046 |
| Gm28240 | 0 | NA | NA |
| Hsfy2 | 0 | NA | NA |
| Satb2 | 608.4974551 | -0.795448187 | 0.197906307 |
| 9130024F11Rik | 127.5850177 | -1.594993087 | 0.241781764 |
| Gm4081 | 9.66177043 | -2.828502154 | 0.616365017 |
| Gm23943 | 0 | NA | NA |
| 1700126A01Rik | 0.044544648 | 0.237309859 | 3.724945049 |
| BC055402 | 2.819616475 | 1.070952304 | 0.648603441 |
| Gm5254 | 0 | NA | NA |
| 4930558J18Rik | 2.344527321 | 0.120774608 | 0.678327497 |
| 1700066M21Rik | 269.4751219 | -0.278931353 | 0.101234578 |
| Tyw5 | 138.2526831 | 0.502253862 | 0.092923511 |
| 9430016H08Rik | 289.7250437 | 0.341486913 | 0.115043043 |
| Gm22371 | 0 | NA | NA |
| Gm24014 | 0 | NA | NA |
| Gm8581 | 0 | NA | NA |
| Gm37373 | 0.354472284 | -2.062455695 | 3.188679565 |
| Gm17929 | 0.044544648 | 0.237309859 | 3.724945049 |
| Spats2l | 531.7349419 | 1.339424194 | 0.087958118 |
| Gm17234 | 0.08486385 | 0.717551239 | 3.724898127 |
| Kctd18 | 231.8874669 | 0.066871403 | 0.125431343 |
| Sgol2a | 34.52874643 | 0.582541543 | 0.276582603 |
| Aox1 | 275.5358162 | 0.939974927 | 0.11862377 |
| Aox3 | 36.77314627 | 0.334769277 | 0.221639895 |
| Gm24548 | 0 | NA | NA |
| Aox4 | 64.4352454 | -0.030644702 | 0.223964971 |
| Gm15759 | 0.15100991 | 0.718202046 | 3.527597384 |
| Aox2 | 2.732211411 | -0.686000626 | 0.617087139 |
| Gm37607 | 38.38692177 | -0.31840831 | 0.25808239 |
| Bzw1 | 2353.558553 | 0.366957234 | 0.09446461 |
| Clk1 | 1464.941916 | -0.374183054 | 0.088383588 |
| Ppil3 | 142.411199 | 0.125481141 | 0.093626767 |
| Nif3l1 | 211.5522702 | 0.244459698 | 0.090983395 |
| Orc2 | 434.4142678 | 0.169716267 | 0.093235075 |
| Gm23966 | 4.595236341 | -0.545702782 | 0.639497271 |
| Gm15834 | 26.01032292 | 0.108393739 | 0.252106369 |
| Fam126b | 3027.102502 | -0.027556776 | 0.085810341 |
| Ndufb3 | 313.2837583 | -0.188835366 | 0.110778973 |
| Gm29389 | 0 | NA | NA |

| | | | |
|---|---|---|---|
| Gm18802 | 0 | NA | NA |
| Gm10068 | 3.8142007 | 0.14887428 | 0.490041869 |
| Als2cr12 | 0.590977814 | 3.022279646 | 1.197734252 |
| Gm20257 | 14.75086306 | 2.016449386 | 0.393674366 |
| Gm28803 | 0 | NA | NA |
| Cflar | 951.4956611 | 2.311331206 | 0.130935643 |
| Gm3841 | 0 | NA | NA |
| Casp8 | 157.9166945 | 4.286924759 | 0.235637677 |
| Gm37760 | 18.59560002 | 2.735455079 | 0.380917289 |
| Gm28802 | 1.046132847 | 1.462226053 | 0.949103995 |
| Trak2 | 2825.777811 | 0.018979002 | 0.069676051 |
| Stradb | 332.0964291 | -0.497894333 | 0.142830385 |
| Gm33589 | 0.118371171 | 0.718202646 | 3.724945049 |
| Gm29018 | 0.22549826 | 1.638241031 | 2.128349096 |
| Als2cr11 | 0.178692882 | 0.237326262 | 2.707499347 |
| Gm25502 | 0 | NA | NA |
| Tmem237 | 159.7002594 | -0.019190275 | 0.105192103 |
| G730003C15Rik | 13.69276046 | 0.041911005 | 0.29545652 |
| Mpp4 | 26.50791038 | -0.793984694 | 0.256958154 |
| Gm29017 | 0.060174537 | 0.237309859 | 3.724945049 |
| Gm19125 | 0 | NA | NA |
| Als2 | 1234.116172 | -0.237924781 | 0.107233346 |
| Cdk15 | 4.87154909 | -0.426036024 | 0.458337829 |
| Gm29016 | 0 | NA | NA |
| Gm37737 | 0.176324247 | -0.208510939 | 2.757307263 |
| Fzd7 | 304.579202 | 0.376058657 | 0.154321517 |
| Gm26813 | 0.044191549 | -0.243571806 | 3.724945049 |
| Gm973 | 103.013939 | 0.061373071 | 0.154141144 |
| Gm28411 | 0.118371171 | 0.718202646 | 3.724945049 |
| 1700122D07Rik | 0.136120942 | 1.15080397 | 3.595683408 |
| Sumo1 | 630.9605461 | 0.437445249 | 0.128430688 |
| Gm24545 | 0.226675736 | 0.23732959 | 2.143799133 |
| D430013B06Rik | 32.24759346 | -0.681313817 | 0.303592805 |
| Gm29596 | 1.33134391 | -0.993427772 | 0.867871392 |
| Nop58 | 820.0043818 | 0.313620981 | 0.106352503 |
| Snord70 | 4.644160981 | -0.516385551 | 0.488296113 |
| Gm26293 | 0 | NA | NA |
| Gm26287 | 2.814965633 | -1.022852089 | 0.638567107 |
| Snord11 | 1.304845187 | 1.068971528 | 1.024905271 |
| Bmpr2 | 5482.687723 | -0.161654999 | 0.055846313 |
| Gm38125 | 29.65508468 | -0.84843868 | 0.251796901 |
| Gm37977 | 0.18514425 | -1.201115342 | 3.292450568 |
| Fam117b | 1173.743163 | -0.15811529 | 0.069626544 |
| Gm37531 | 63.74214748 | -0.813276518 | 0.275555431 |
| Gm18829 | 0 | NA | NA |
| Ica1l | 384.5618481 | -0.639175985 | 0.076846265 |
| Wdr12 | 444.9492766 | -0.000836301 | 0.076669636 |

| | | | |
|---|---|---|---|
| Carf | 373.7108772 | -0.728231384 | 0.105386987 |
| Gm15464 | 5.596760458 | -0.254055157 | 0.474376137 |
| Nbeal1 | 1755.709698 | -0.16857163 | 0.083099902 |
| Gm29332 | 0 | NA | NA |
| Cyp20a1 | 130.1781194 | 0.910288577 | 0.155823856 |
| 6030460B20Rik | 3.025787973 | -0.471931365 | 0.602191458 |
| Abi2 | 3457.462493 | -0.62912246 | 0.075977858 |
| Raph1 | 1729.282084 | -0.00209925 | 0.104618352 |
| Gm11575 | 9.518548027 | -0.984063702 | 0.375250547 |
| Gm11576 | 0.448283844 | 2.117288945 | 1.496955839 |
| Gm11578 | 18.19495742 | -1.42498708 | 0.398957871 |
| Gm38388 | 0.262256756 | 0.237330204 | 2.055088433 |
| Gm23762 | 0 | NA | NA |
| Gm11577 | 0 | NA | NA |
| Eif4a-ps4 | 428.6153442 | 0.897091148 | 0.115790711 |
| Cd28 | 53.32452365 | 4.660487578 | 0.357543286 |
| Gm11579 | 0.085327104 | 1.179809211 | 3.723500796 |
| 2310016D23Rik | 0.256420178 | 1.655875485 | 2.450467423 |
| Gm37198 | 0.546045768 | 2.891424291 | 1.308256555 |
| Gm11609 | 0.066050875 | 0.71820487 | 3.724945049 |
| Gm11581 | 0.146451277 | -0.243562218 | 2.882358805 |
| Gm38137 | 1.155240215 | -1.699775839 | 1.024696628 |
| Ctla4 | 27.96364358 | 3.633669709 | 0.473449406 |
| Rpl18-ps1 | 12.04120041 | 0.626089799 | 0.364990454 |
| Gm11582 | 0.048949532 | -0.243569582 | 3.724945049 |
| Icos | 90.9935698 | 7.734517452 | 0.751790998 |
| Gm37039 | 2.679819182 | 5.06791897 | 1.067425611 |
| Rpl17-ps1 | 1.010478028 | 2.00061811 | 1.210466702 |
| Gm11590 | 0 | NA | NA |
| Gm11591 | 0 | NA | NA |
| Gm28083 | 0 | NA | NA |
| Gm11587 | 0.127971488 | 1.23029335 | 3.685533863 |
| 9530026F06Rik | 0 | NA | NA |
| Gm11588 | 0 | NA | NA |
| Gm25839 | 0 | NA | NA |
| Gm28449 | 0.11147366 | 1.230376831 | 3.722845817 |
| Gm36943 | 0 | NA | NA |
| Gm28094 | 0.046345709 | 0.71820487 | 3.724945049 |
| 4930587A21Rik | 0 | NA | NA |
| Pard3b | 166.061435 | 0.6329222 | 0.132194617 |
| Gm37205 | 11.98801516 | 0.785535548 | 0.350580863 |
| Gm38012 | 7.816280804 | -0.254295211 | 0.379147508 |
| Gm38278 | 3.411970267 | 0.287658505 | 0.549411049 |
| Pard3bos1 | 0 | NA | NA |
| Gm38075 | 5.595486472 | 0.554247497 | 0.440983557 |
| Gm29640 | 0 | NA | NA |
| Gm29641 | 0.044191549 | -0.243571806 | 3.724945049 |

| | | | |
|---|---|---|---|
| Pard3bos2 | 0.453587956 | 0.179317133 | 1.293915533 |
| Gm38351 | 3.154252535 | 0.303902007 | 0.794180979 |
| Pard3bos3 | 0.304629397 | 0.906998208 | 2.014741898 |
| Gm29084 | 2.918337729 | -0.009088437 | 0.609250989 |
| Gm11600 | 0 | NA | NA |
| Gm11599 | 0.770699074 | 0.337456348 | 1.272738912 |
| Nrp2 | 974.5911931 | 0.579320198 | 0.090401573 |
| Gm29083 | 5.840962313 | 0.451781997 | 0.444678267 |
| Gm37121 | 13.64577828 | -1.277385319 | 0.375364783 |
| Gm4208 | 0.515886961 | 0.665431895 | 1.417667265 |
| Ino80d | 1352.80268 | -0.036735177 | 0.065762698 |
| AL645685.1 | 0 | NA | NA |
| Gm24942 | 0 | NA | NA |
| Rps13-ps7 | 0 | NA | NA |
| Rps27-ps1 | 0 | NA | NA |
| Ino80dos | 70.0990274 | -0.473360948 | 0.140203336 |
| Gm20342 | 231.0873436 | -0.858133726 | 0.227864027 |
| Ndufs1 | 1507.181369 | -0.282784053 | 0.053115053 |
| Gm24557 | 0.222111871 | 0.237311956 | 3.129813646 |
| Eef1b2 | 871.8357536 | 0.504247674 | 0.133282687 |
| Gm27390 | 0 | NA | NA |
| Gm26457 | 0.157314084 | 0.718207559 | 3.435173412 |
| snoZ196 | 0 | NA | NA |
| Snora41 | 4.156744695 | 0.841360179 | 0.519589859 |
| Gpr1 | 4.89974049 | -0.905995726 | 0.456452673 |
| Gm27512 | 0.049553047 | 0.71820487 | 3.724945049 |
| Platr12 | 0.049553047 | 0.71820487 | 3.724945049 |
| 1700039I01Rik | 0.097899064 | -0.724453471 | 3.724945049 |
| Gm11608 | 0.112029109 | -0.724455696 | 3.724945049 |
| Gm24513 | 0.109124069 | -0.24357403 | 3.724945049 |
| Zdbf2 | 969.9858277 | -0.587805521 | 0.152017471 |
| Gm23448 | 0.180187022 | -0.243570275 | 2.70035346 |
| Gm11606 | 0 | NA | NA |
| Gm11605 | 3.581982469 | 0.493391695 | 0.545591807 |
| Adam23 | 3251.808784 | -0.756087666 | 0.059188156 |
| Gm22281 | 0 | NA | NA |
| Dytn | 1.029483559 | -0.466405942 | 0.988514487 |
| Gm12748 | 0 | NA | NA |
| Mdh1b | 8.408068065 | -0.689173474 | 0.366656452 |
| Fastkd2 | 291.2035401 | 0.148512142 | 0.125756408 |
| 4933402D24Rik | 1.548786521 | 0.437772902 | 0.786502708 |
| Gm13751 | 0.449839025 | 2.063821873 | 1.479552087 |
| Gm13754 | 0.863190071 | -1.976121083 | 1.222583281 |
| Gm13749 | 1.958865323 | -2.581722653 | 0.842386688 |
| Klf7 | 1205.650868 | -0.04425414 | 0.10312074 |
| Gm26649 | 0.52188035 | 2.642222141 | 1.462838754 |
| Mir6899 | 0.842388825 | -0.042373932 | 1.170657921 |

| | | | |
|---|---|---|---|
| Gm13748 | 0.622058045 | 0.399289782 | 1.381875646 |
| Gm13750 | 1.235703043 | 0.819696156 | 0.918919042 |
| Gm28981 | 0.770914299 | -0.14265494 | 1.095975831 |
| Gm25748 | 0 | NA | NA |
| Gm28980 | 0 | NA | NA |
| Creb1 | 1118.909764 | -0.165584417 | 0.084659684 |
| Mettl21a | 95.46113856 | -0.073763216 | 0.140364411 |
| Gm28982 | 1.624436933 | 0.917994512 | 0.813773817 |
| Gm29152 | 0 | NA | NA |
| 2810408I11Rik | 4.24223269 | 1.40388364 | 0.488619808 |
| Ccnyl1 | 180.7599231 | -0.000927369 | 0.132139736 |
| Fzd5 | 105.8313593 | 0.785059824 | 0.188291422 |
| Gm38058 | 2.181430594 | -0.428074199 | 0.698782037 |
| Plekhm3 | 907.4639202 | 0.063910149 | 0.057166489 |
| Rpl31-ps24 | 0 | NA | NA |
| Rpl10a-ps1 | 39.84853431 | 0.647779613 | 0.197799184 |
| Gm24788 | 0 | NA | NA |
| Akr1cl | 0.229212118 | -0.243570082 | 2.547047007 |
| Gm8550 | 0 | NA | NA |
| Cryge | 0 | NA | NA |
| Crygd | 0 | NA | NA |
| Crygc | 0 | NA | NA |
| Crygb | 0.150086409 | 0.237305146 | 3.724945049 |
| Gm7329 | 4.40840556 | 0.247984941 | 0.461025082 |
| Cryga | 0.97358459 | -1.654239305 | 1.29556966 |
| Gm28845 | 0.23654427 | -0.243569555 | 2.563195005 |
| D630023F18Rik | 33.29031821 | 0.054990595 | 0.209151542 |
| Idh1 | 461.6797512 | -0.085350181 | 0.126363179 |
| Pikfyve | 2046.792619 | 0.315856555 | 0.071132063 |
| Pth2r | 15.63521041 | -0.054850552 | 0.324097622 |
| Gm28927 | 0 | NA | NA |
| Gm28926 | 0 | NA | NA |
| Gm28925 | 0 | NA | NA |
| Gm28923 | 0 | NA | NA |
| Gm28922 | 0 | NA | NA |
| Gm28728 | 0 | NA | NA |
| Gm5255 | 0 | NA | NA |
| Gm8805 | 0 | NA | NA |
| Crygf | 0 | NA | NA |
| Gm8809 | 0 | NA | NA |
| Gm15659 | 0 | NA | NA |
| Gm8812 | 0 | NA | NA |
| Map2 | 21973.53413 | -0.235651432 | 0.096473416 |
| Gm24729 | 8.150369374 | -0.950939258 | 0.471622628 |
| Gm10558 | 1.106847758 | 0.867863483 | 1.036990037 |
| Gm25411 | 10.74259572 | -0.44808211 | 0.355541679 |
| Unc80 | 7728.076732 | -0.82496161 | 0.06595355 |

| | | | |
|---|---|---|---|
| Gm37181 | 0.701668669 | -2.192981787 | 1.347699607 |
| Rpe | 507.539684 | 0.493086013 | 0.083052053 |
| Kansl1l | 470.4693895 | 0.094027774 | 0.081086203 |
| Gm37570 | 61.3256198 | -0.690032007 | 0.244705057 |
| Gm38387 | 64.00509308 | -0.685148898 | 0.192294722 |
| Gm38162 | 96.75562574 | 0.317189384 | 0.192480364 |
| Gm15789 | 30.55906383 | 0.410329383 | 0.224037354 |
| 6820402A03Rik | 78.68479837 | -0.367858219 | 0.187194617 |
| Gm37285 | 31.02313678 | -0.059840725 | 0.224809072 |
| Gm37755 | 15.79173624 | 1.243087346 | 0.373098397 |
| Gm38218 | 8.540597391 | 0.512175704 | 0.373671317 |
| Acadl | 460.9663652 | 0.90859519 | 0.12666443 |
| Gm15793 | 6.435362979 | 1.824062317 | 0.442106312 |
| Gm27934 | 0.075043205 | 0.237309859 | 3.724945049 |
| Gm25832 | 0.12092566 | 0.237307635 | 3.724945049 |
| Myl1 | 8.248283288 | 0.109106654 | 0.433872194 |
| Rpl31-ps14 | 4.920430513 | 0.374055841 | 0.471328589 |
| Gm15826 | 0.731126313 | 1.025405946 | 1.533315052 |
| Lancl1 | 1237.017238 | -0.578440254 | 0.047319012 |
| Cps1 | 8.332986231 | -0.361594659 | 0.374030525 |
| Gm15668 | 0.088534442 | 1.179809211 | 3.723500796 |
| Gm26342 | 0 | NA | NA |
| Kif22-ps | 0 | NA | NA |
| Gm22548 | 0 | NA | NA |
| Gm15669 | 0 | NA | NA |
| Gm37957 | 0 | NA | NA |
| Gm23189 | 0 | NA | NA |
| Gm15671 | 0 | NA | NA |
| Erbb4 | 1732.331576 | -0.926451013 | 0.066521096 |
| Gm37100 | 8.451353278 | -1.137507663 | 0.404361969 |
| Gm15670 | 8.648476067 | -0.880911348 | 0.483899828 |
| Gm25542 | 1.041370757 | -0.668887888 | 1.154012292 |
| Gm19335 | 10.47773709 | -0.584733241 | 0.371462796 |
| Gm37735 | 8.967735426 | -1.118636711 | 0.343199137 |
| Gm37769 | 40.95679204 | -0.699607176 | 0.230852418 |
| Gm37061 | 10.97852267 | -0.949392742 | 0.453838762 |
| Gm16076 | 36.27563093 | -1.107519254 | 0.271213535 |
| Gm8840 | 0 | NA | NA |
| Pced1c-ps | 0 | NA | NA |
| Gm28497 | 0.047948698 | -0.243569582 | 3.724945049 |
| Gm29113 | 9.644466874 | -0.316965471 | 0.343646246 |
| Gm29114 | 0 | NA | NA |
| Ikzf2 | 269.8286907 | 0.531746787 | 0.10602418 |
| Gm29112 | 12.57536217 | 0.540152705 | 0.345118708 |
| Gm28112 | 19.23386575 | -0.111553247 | 0.307031742 |
| Gm37930 | 0.044191549 | -0.243571806 | 3.724945049 |
| Spag16 | 19.54478478 | 0.065813405 | 0.238432783 |

| | | | |
|---|---|---|---|
| Gm22665 | 0 | NA | NA |
| Gm23879 | 0 | NA | NA |
| Gm28580 | 0 | NA | NA |
| Gm38272 | 0.042716945 | -0.243571806 | 3.724945049 |
| Gm37456 | 0.272865106 | 0.237320973 | 2.458126347 |
| Gm23422 | 0 | NA | NA |
| Vwc2l | 194.8593581 | -0.282265526 | 0.091558524 |
| Bard1 | 62.05484921 | 1.781266914 | 0.226473097 |
| Gm16236 | 1.002654625 | 0.433912247 | 0.988720185 |
| Gm8870 | 0 | NA | NA |
| Abca12 | 5.342748304 | 0.215286271 | 0.570919089 |
| Gm6947 | 0 | NA | NA |
| Atic | 353.5973081 | 0.346045928 | 0.108742344 |
| Fn1 | 1063.525137 | 1.052811088 | 0.224245413 |
| Apol7d | 1.07675232 | 1.423705544 | 1.053385752 |
| Gm5829 | 1.23306525 | 1.182967212 | 0.923337736 |
| Gm5256 | 3.462762875 | -0.200156703 | 0.521626812 |
| Gm37217 | 0 | NA | NA |
| Gm8883 | 0 | NA | NA |
| Gm28818 | 0.213332449 | 0.237327985 | 2.174774587 |
| 4933417E11Rik | 0.198105924 | -0.576345274 | 2.625732037 |
| 4930556G22Rik | 0 | NA | NA |
| Gm5528 | 0 | NA | NA |
| Gm8885 | 0.131209559 | 1.165289508 | 2.972496041 |
| Gm4319 | 0.520182433 | 0.261177354 | 1.279132269 |
| Mreg | 141.6531737 | 0.108681646 | 0.131247744 |
| Gm25360 | 67.27497613 | 0.959734677 | 0.213041811 |
| D230017M19Rik | 46.21119991 | -0.7978843 | 0.162794658 |
| Gm23472 | 572.9451334 | -0.609364889 | 0.804439939 |
| Gm24497 | 54.97402124 | -0.572009153 | 0.372742004 |
| Gm25939 | 1027.212574 | 0.327616207 | 0.171693622 |
| Pecr | 29.32302085 | -0.490311203 | 0.196795216 |
| Tmem169 | 181.7981465 | -0.335266289 | 0.11264105 |
| Xrcc5 | 253.261693 | 0.281645459 | 0.078108349 |
| 4-Mar | 295.8236201 | -0.475822968 | 0.123996051 |
| Gm15843 | 0.209261529 | 1.185614193 | 2.617572548 |
| Smarcal1 | 689.8282747 | -0.185119834 | 0.054587619 |
| Ankar | 0.251200408 | 0.305727746 | 2.090476875 |
| Rpl37a | 999.4688314 | 0.343536998 | 0.127281123 |
| Igfbp2 | 490.8609578 | 0.983757868 | 0.190824897 |
| Igfbp5 | 3859.554287 | 0.251438634 | 0.14026055 |
| 1700027A15Rik | 1.147127115 | 0.941179044 | 0.882126854 |
| Tnp1 | 0 | NA | NA |
| Gm38106 | 0 | NA | NA |
| Gm29185 | 0.044544648 | 0.237309859 | 3.724945049 |
| Gm29184 | 0 | NA | NA |
| Pinc | 0.243249674 | -0.243566194 | 2.557831606 |

| | | | |
|---|---|---|---|
| Gm27315 | 0 | NA | NA |
| D530049I02Rik | 0 | NA | NA |
| Gm9553 | 0.046345709 | 0.71820487 | 3.724945049 |
| C530043A13Rik | 1.12921268 | 2.335698835 | 0.993865749 |
| Gm29186 | 0 | NA | NA |
| 6030407O03Rik | 6.071023221 | -1.314799657 | 0.429477167 |
| Gm29183 | 1.411578533 | -1.946433526 | 0.971673328 |
| Gm25329 | 0 | NA | NA |
| Mir6351 | 0.439338454 | -0.272873232 | 1.515409601 |
| Tns1 | 929.0676011 | 0.356816977 | 0.115928321 |
| Rufy4 | 2.528566672 | 2.393090608 | 0.738801109 |
| Cxcr2 | 2.628643724 | 3.249486322 | 0.924746137 |
| Cxcr1 | 0.182828849 | 1.122216213 | 2.253829118 |
| Mir1928 | 0 | NA | NA |
| Gm25524 | 0 | NA | NA |
| Arpc2 | 1724.327588 | 0.676705345 | 0.085813246 |
| Gpbar1 | 0.083526043 | 0.718202646 | 3.724945049 |
| Aamp | 1126.602311 | 0.162278283 | 0.081396932 |
| Gm28364 | 0.299075359 | 0.936683935 | 1.723164074 |
| Pnkd | 1381.963466 | -0.668367854 | 0.091840724 |
| Gm37503 | 2.04875246 | 0.192242308 | 0.754991446 |
| Tmbim1 | 824.4000618 | 1.441027958 | 0.152496033 |
| Gm29358 | 41.10390762 | -1.476437508 | 0.34071963 |
| Gm37931 | 25.45680512 | -1.563029553 | 0.452921006 |
| Catip | 16.47013026 | -0.021476012 | 0.242934595 |
| Slc11a1 | 156.1085199 | 4.157794051 | 0.243500283 |
| Ctdsp1 | 480.9035942 | 0.893890687 | 0.121325465 |
| Mir26b | 0.460912027 | -0.478031477 | 1.390925652 |
| Vil1 | 4.671085706 | 0.856228385 | 0.454106851 |
| Usp37 | 682.5650065 | 0.052637628 | 0.071955321 |
| Gm37733 | 34.22973971 | -0.523184616 | 0.291905341 |
| Rqcd1 | 401.5594137 | 0.274485267 | 0.061949005 |
| Plcd4 | 276.691242 | 0.649452585 | 0.094205752 |
| Zfp142 | 808.2705847 | 0.180201047 | 0.080813916 |
| Gm25035 | 2.10081791 | -0.847926191 | 0.74761681 |
| Bcs1l | 123.2340065 | -0.389206044 | 0.124032871 |
| Rnf25 | 329.113652 | 0.271344963 | 0.092375008 |
| Stk36 | 70.93351887 | 0.213132101 | 0.163334395 |
| Ttll4 | 457.8466587 | -0.239627062 | 0.110103568 |
| Cyp27a1 | 60.63009196 | -0.185652473 | 0.164064566 |
| Prkag3 | 1.196546384 | 0.193740494 | 0.742038793 |
| Gm27507 | 0 | NA | NA |
| Wnt6 | 20.88210294 | -0.74635607 | 0.302995082 |
| Gm29539 | 0.08169834 | 0.23732098 | 3.724945049 |
| Wnt10a | 51.88315531 | 0.332827335 | 0.235109344 |
| Gm16582 | 1.058309834 | -0.539395697 | 1.016261925 |
| Cdk5r2 | 2117.926955 | -0.807712608 | 0.114122538 |

| | | | |
|---|---|---|---|
| Fev | 2.243593674 | 1.133060864 | 0.926090749 |
| Cryba2 | 3.578913291 | 1.430256584 | 0.570308908 |
| Mir375 | 0 | NA | NA |
| Ccdc108 | 54.38524299 | -1.074634344 | 0.187021487 |
| Gm15841 | 0.048949532 | -0.243569582 | 3.724945049 |
| Ihh | 2.70016216 | -0.198766605 | 0.612818782 |
| Gm37744 | 0.044544648 | 0.237309859 | 3.724945049 |
| Nhej1 | 18.60744933 | 0.209987201 | 0.269424512 |
| Slc23a3 | 6.004310424 | 1.329603418 | 0.491316335 |
| Cnppd1 | 349.0250262 | 0.644596736 | 0.095006305 |
| Fam134a | 1705.406746 | -0.13684841 | 0.048365532 |
| Zfand2b | 100.1118125 | 0.264528829 | 0.139704348 |
| Abcb6 | 14.09945832 | -0.707758518 | 0.32137839 |
| Gm29253 | 0.323844909 | -0.10714251 | 2.022410361 |
| Atg9a | 756.6561104 | -0.405660442 | 0.056323437 |
| Ankzf1 | 160.4436551 | -0.488416319 | 0.144922455 |
| Glb1l | 98.7643794 | 0.105913667 | 0.128068038 |
| Gm27943 | 0 | NA | NA |
| Stk16 | 338.2768 | 0.307017149 | 0.066542246 |
| Tuba4a | 2899.871314 | -0.829394175 | 0.071069476 |
| A630095N17Rik | 0.826064167 | 1.437698783 | 1.141668905 |
| Dnajb2 | 784.3513935 | -0.008194217 | 0.080927949 |
| Ptprn | 4437.838423 | 0.013321506 | 0.069395596 |
| Gm28294 | 1.007042841 | -1.250860243 | 1.124408179 |
| Resp18 | 711.8094129 | 0.508903035 | 0.154423887 |
| Gm28902 | 0.705697236 | -0.61948123 | 1.067939374 |
| Dnpep | 296.9380932 | 0.484126646 | 0.111191564 |
| Des | 14.87844252 | 1.737954102 | 0.29398698 |
| Gm15179 | 4.05260616 | -0.50683558 | 0.535439056 |
| Speg | 1539.141981 | -0.814049715 | 0.141686088 |
| Gm15178 | 0 | NA | NA |
| Gmppa | 265.8934165 | 0.132337523 | 0.081625451 |
| Asic4 | 155.5102748 | -1.177572505 | 0.133429953 |
| Gm37264 | 4.269864839 | 0.719082234 | 0.509209828 |
| Chpf | 816.6335401 | 0.176958827 | 0.087251146 |
| Obsl1 | 195.5669334 | -0.106670484 | 0.091334451 |
| Tmem198 | 376.0167297 | -0.049760678 | 0.127673117 |
| Inha | 132.6233029 | -1.129160774 | 0.185369703 |
| Gm15181 | 0 | NA | NA |
| Stk11ip | 336.2431366 | -0.086317551 | 0.080813308 |
| Gm15183 | 0.113999573 | 0.23732098 | 3.724945049 |
| Slc4a3 | 1347.377089 | -0.961763304 | 0.082762176 |
| Gm15180 | 0.245985165 | 1.136059542 | 2.496946097 |
| Gm15182 | 0.115603922 | 1.209894437 | 3.72418822 |
| Gm5257 | 0.104637558 | 0.23732098 | 3.724945049 |
| Gm29065 | 0 | NA | NA |
| Gm29068 | 0 | NA | NA |

| | | | |
|---|---|---|---|
| Gm29069 | 0.135858026 | -1.180985464 | 2.941468349 |
| Gm17751 | 4.194728469 | 0.706235451 | 0.499190878 |
| Mir6343 | 0 | NA | NA |
| Gm816 | 0.297786538 | -1.922234716 | 1.531524621 |
| Gm38265 | 0 | NA | NA |
| Gm25974 | 0 | NA | NA |
| Epha4 | 2103.014444 | -0.936157635 | 0.128978877 |
| Mir6352 | 3.045583208 | -2.189670267 | 0.779740966 |
| Gm28386 | 2.146862605 | -1.467755735 | 0.866895479 |
| Gm28385 | 0.301967822 | -1.794287944 | 3.695942541 |
| Gm26263 | 0 | NA | NA |
| Gm28387 | 0.044544648 | 0.237309859 | 3.724945049 |
| Pax3 | 26.95793292 | -0.11012376 | 0.219356646 |
| Sgpp2 | 367.8063322 | -0.355645373 | 0.109075391 |
| Gm28410 | 0.413627519 | -0.243565967 | 1.549102166 |
| Gm37884 | 3.006274965 | -1.270904836 | 0.63830332 |
| Farsb | 522.3359989 | 0.283321908 | 0.080810447 |
| Gm28995 | 0 | NA | NA |
| 4833412K13Rik | 33.12266117 | -0.794946411 | 0.2719363 |
| BC035947 | 16.4961278 | -0.650087024 | 0.286857331 |
| Mogat1 | 1.455068488 | -0.399556102 | 0.910237861 |
| 5730419F03Rik | 0.372487069 | 0.921450266 | 1.668348854 |
| Gm29611 | 0 | NA | NA |
| Acsl3 | 1660.961095 | -0.206103559 | 0.077565353 |
| Utp14b | 706.5540839 | 0.027877124 | 0.088931905 |
| Gm6159 | 2.778261606 | 0.668059693 | 0.578647114 |
| Gm29536 | 0.417935612 | 2.510880266 | 1.689107383 |
| Kcne4 | 23.89478986 | 0.583852454 | 0.285865106 |
| Gm29187 | 0.218130886 | 0.237328552 | 2.17376931 |
| Gm5830 | 3.384143917 | -0.042942463 | 0.555003815 |
| Scg2 | 2880.675548 | 0.53107739 | 0.146197509 |
| Gm19027 | 0.061920613 | 0.718207094 | 3.724945049 |
| Gm24160 | 0 | NA | NA |
| Ap1s3 | 62.26140866 | 1.531890979 | 0.202755752 |
| Gm37886 | 2.695407072 | -0.1330423 | 0.711564473 |
| Wdfy1 | 946.0370216 | 0.626141133 | 0.064492193 |
| Gm10555 | 0.378505399 | 2.592817337 | 1.596592709 |
| Mrpl44 | 178.4373768 | 0.283794288 | 0.092142953 |
| Serpine2 | 1567.911986 | 0.175474473 | 0.124357169 |
| Gm5529 | 18.26108814 | -0.106022675 | 0.256943255 |
| Gm28071 | 2.944880665 | 1.173118106 | 0.725748287 |
| Gm28058 | 0 | NA | NA |
| Gm23934 | 0.310678561 | -0.28040752 | 1.589900785 |
| Gm38339 | 0.456870417 | 2.830970337 | 3.387090691 |
| 9830004L10Rik | 0 | NA | NA |
| Fam124b | 3.258857179 | 5.019640029 | 0.919931657 |
| 2310015K22Rik | 0 | NA | NA |

| | | | |
|---|---|---|---|
| Gm37932 | 4.562962678 | 0.130390782 | 0.444212583 |
| Cul3 | 2324.447608 | 0.118920724 | 0.080124668 |
| Gm37902 | 63.67772732 | -0.084361549 | 0.163906635 |
| Gm37262 | 28.6341495 | 0.021718457 | 0.226890093 |
| Gm29125 | 1.604780936 | 0.909370369 | 0.812256414 |
| Gm6189 | 0.828249129 | 3.67547539 | 1.278750434 |
| 1700016L21Rik | 0.083172944 | 0.23732098 | 3.724945049 |
| Dock10 | 1627.450511 | 0.504543531 | 0.092977125 |
| Gm37645 | 36.84305094 | 0.388500373 | 0.240285644 |
| Gm38062 | 55.87989744 | 0.329074056 | 0.337292427 |
| A630081D01Rik | 65.65902623 | 0.267619376 | 0.224974708 |
| Gm29124 | 0 | NA | NA |
| Gm28066 | 0 | NA | NA |
| Nyap2 | 954.1496905 | -0.706438476 | 0.094017627 |
| Gm37210 | 0 | NA | NA |
| Gm6198 | 0 | NA | NA |
| Gm5530 | 3.003564229 | 3.764933988 | 0.903959192 |
| Gm7516 | 0 | NA | NA |
| Gm37114 | 0 | NA | NA |
| Mir6344 | 0 | NA | NA |
| Irs1 | 784.393355 | -0.27129766 | 0.078223814 |
| Gm9747 | 0.131672813 | 1.557858112 | 3.660706558 |
| Rhbdd1 | 192.9071811 | 1.195394064 | 0.163584871 |
| Gm19552 | 8.274522055 | 0.766830216 | 0.387383263 |
| Gm28940 | 0.217873223 | -1.000641016 | 2.53644881 |
| Gm22112 | 0.107803068 | 0.719187033 | 3.724874737 |
| Col4a4 | 19.40116672 | -0.108984669 | 0.264998623 |
| Col4a3 | 9.983396163 | -0.894159435 | 0.337259991 |
| n-R5s213 | 0 | NA | NA |
| C430014B12Rik | 2.511728143 | 0.763006956 | 0.595380044 |
| Mff | 1852.754622 | -0.111131768 | 0.044963929 |
| Gm28941 | 40.41534512 | -0.564718686 | 0.262768557 |
| Tm4sf20 | 0.089062654 | 0.237318756 | 3.724945049 |
| Gm19257 | 0 | NA | NA |
| Gm36999 | 0.100902008 | 1.15059983 | 3.720931964 |
| Gm28942 | 0.253304709 | 1.614833275 | 1.813084145 |
| Gm17764 | 0 | NA | NA |
| Gm22396 | 0.22174029 | 0.33374547 | 3.147484704 |
| Agfg1 | 1873.69111 | 0.185402339 | 0.060313595 |
| Gm7553 | 0.943831312 | 0.039528102 | 1.026413606 |
| Slc19a3 | 12.62212474 | -0.535758611 | 0.329565119 |
| Gm7539 | 0 | NA | NA |
| Krtap28-10 | 0 | NA | NA |
| A030005K14Rik | 0 | NA | NA |
| Krtap28-13 | 0.044191549 | -0.243571806 | 3.724945049 |
| Gm29518 | 0.066050875 | 0.71820487 | 3.724945049 |
| Ccl20 | 0.166497232 | 1.729737333 | 3.388675328 |

| | | | |
|---|---|---|---|
| Gm29519 | 0 | NA | NA |
| Daw1 | 3.764882787 | -0.449600862 | 0.578421983 |
| Gm6244 | 0 | NA | NA |
| n-R5s214 | 0 | NA | NA |
| Sphkap | 3563.73903 | -0.51985519 | 0.101680099 |
| Gm6374 | 0.068097359 | -0.243569582 | 3.724945049 |
| Gm37517 | 48.29894096 | -1.603255489 | 0.363588662 |
| Gm37989 | 0 | NA | NA |
| 4933436I20Rik | 1.025359323 | 1.244617201 | 1.195099636 |
| Gm25754 | 0.150086409 | 0.237305146 | 3.724945049 |
| Gm24555 | 0 | NA | NA |
| Pid1 | 538.7136745 | 0.246660266 | 0.081400169 |
| Gm37058 | 25.37561462 | -0.303958975 | 0.316938567 |
| Gm18304 | 0 | NA | NA |
| Mir6353 | 3.815514299 | -0.133952836 | 0.664960703 |
| Dner | 2450.532823 | -0.283513645 | 0.108615722 |
| Gm37959 | 19.69556321 | -0.915532091 | 0.284578139 |
| Gm37925 | 43.42924116 | -1.175823839 | 0.272979044 |
| Mir5126 | 0.291915594 | -1.047256401 | 2.003117566 |
| Trip12 | 5110.806766 | 0.420121482 | 0.052446671 |
| Gm38374 | 71.20713703 | -0.140521781 | 0.190866977 |
| Gm37738 | 93.58139767 | -0.414729553 | 0.247735069 |
| Fbxo36 | 28.2317926 | -0.245071296 | 0.249540399 |
| Slc16a14 | 112.1372938 | -0.296692024 | 0.148270541 |
| Gm15433 | 3.38607774 | 5.721983907 | 0.91372043 |
| Gm29284 | 0.509711911 | 0.237331669 | 1.546512486 |
| C130026I21Rik | 27.88048967 | 6.060716049 | 0.663601693 |
| Rpl19-ps1 | 2.447118844 | -0.0297312 | 0.666010969 |
| Gm27907 | 0 | NA | NA |
| Gm7582 | 5.427132103 | 5.66088322 | 0.939965991 |
| Gm2389 | 0 | NA | NA |
| Gm18342 | 8.221653517 | 6.430341258 | 0.885527582 |
| A530032D15Rik | 14.11902256 | 7.626839898 | 0.871306942 |
| Gm10553 | 0.099106094 | 1.209894437 | 3.72418822 |
| A530040E14Rik | 19.7285084 | 7.630093742 | 0.88080328 |
| Gm16038 | 0.038981395 | 0.718207094 | 3.724945049 |
| Gm2635 | 0.088534442 | 1.179809211 | 3.723500796 |
| Gm6264 | 1.445224006 | 1.952838996 | 1.170797374 |
| Gm7609 | 9.276052769 | 7.032275321 | 0.872405676 |
| Gm16028 | 0.132101749 | 1.209894437 | 3.72418822 |
| Gm16026 | 7.688870986 | 6.120401441 | 0.882038064 |
| Gm2619 | 4.165531338 | 5.57240492 | 0.906364464 |
| Gm16025 | 3.376741903 | 0.719673151 | 0.511409584 |
| Gm7281 | 0.301876039 | 1.941911015 | 1.732504146 |
| Gm2666 | 0.742190307 | 3.539168451 | 1.188986468 |
| Gm2427 | 1.636398745 | 4.576632948 | 1.103011664 |
| Gm16092 | 29.37301029 | -0.467434852 | 0.27249506 |

| | | | |
|---|---|---|---|
| Gm7592 | 2.43285919 | 5.165790648 | 1.038559178 |
| G530012D18Rik | 0 | NA | NA |
| Sp110 | 246.5019786 | 6.564966825 | 0.286553477 |
| Gm16094 | 4.539762321 | 5.914551977 | 0.949796953 |
| Sp140 | 215.8469477 | 6.247326639 | 0.282235644 |
| Gm10552 | 1.59243937 | 4.639495636 | 1.004116828 |
| Gm17017 | 6.466014695 | 0.652082235 | 0.44680751 |
| Sp100 | 678.1539777 | 6.748293539 | 0.290108179 |
| n-R5s215 | 0 | NA | NA |
| A630001G21Rik | 22.19689395 | 2.841918803 | 0.322815217 |
| Mir8096 | 0 | NA | NA |
| Gm25472 | 0 | NA | NA |
| 9930111H07Rik | 0.389848623 | 2.067752346 | 1.651351578 |
| Gm22906 | 0 | NA | NA |
| Cab39 | 2445.880552 | 0.086934095 | 0.088881568 |
| Itm2c | 5468.611581 | -0.173635202 | 0.073716954 |
| 4933407L21Rik | 23.38036317 | -1.312912299 | 0.26581955 |
| Gm28884 | 0.751763609 | -1.451104614 | 1.287594271 |
| Gpr55 | 7.561829528 | 3.274663004 | 0.594042651 |
| Gm28100 | 0.240280111 | 0.237329113 | 2.463083575 |
| Gm18180 | 25.54930956 | 0.245793802 | 0.233252295 |
| Spata3 | 2.778035728 | -0.392472542 | 0.59007711 |
| 2810459M11Rik | 146.6634441 | 1.177206341 | 0.156188336 |
| Psmd1 | 1536.480455 | 0.334420966 | 0.063588729 |
| Gm22786 | 4.654196711 | -0.236516154 | 0.504509971 |
| Htr2b | 18.41870533 | 4.649758858 | 0.53312353 |
| Gm37914 | 41.69621495 | -0.166291929 | 0.263770818 |
| Gm21972 | 0.320475213 | -2.230003071 | 2.280147898 |
| Armc9 | 417.345602 | -0.14702635 | 0.177088934 |
| Gm16341 | 0.537632031 | 2.946467554 | 1.533946285 |
| B3gnt7 | 21.27627369 | 1.851590755 | 0.293386229 |
| Gm28626 | 1.81028421 | 2.052575052 | 1.121590238 |
| Gm5258 | 0 | NA | NA |
| Gm28627 | 0 | NA | NA |
| Ncl | 3170.074209 | 0.442471542 | 0.101053535 |
| Snora75 | 0 | NA | NA |
| mmu-mir-3535 | 0.451658936 | 0.209366165 | 1.335982742 |
| Gm24148 | 0 | NA | NA |
| Snord82 | 0.870629472 | -0.649183547 | 1.169873783 |
| C130036L24Rik | 32.9831117 | -0.377471699 | 0.237876531 |
| Nmur1 | 2.498311769 | 0.486294774 | 0.617844086 |
| 1700019O17Rik | 0.04393175 | -0.243571806 | 3.724945049 |
| Rpl30-ps6 | 0 | NA | NA |
| Ptma | 1852.709718 | 0.766008249 | 0.137081504 |
| Pde6d | 314.3105157 | -0.032099979 | 0.108047923 |
| Gm37152 | 0.163519603 | -0.243571882 | 2.778606197 |
| Cops7b | 225.9968041 | 0.012729506 | 0.081450974 |

| | | | |
|---|---|---|---|
| Gm22955 | 0 | NA | NA |
| Gm37022 | 4.10661082 | 0.421197871 | 0.62512408 |
| Nppc | 53.44822697 | 0.437074448 | 0.242374661 |
| Dis3l2 | 456.7534953 | -0.252071322 | 0.079917126 |
| Gm29097 | 0.04393175 | -0.243571806 | 3.724945049 |
| Gm29055 | 12.11508896 | -0.280037881 | 0.427954423 |
| Gm37485 | 23.25905464 | -0.985584512 | 0.309775155 |
| Gm38142 | 24.33645793 | -0.874908445 | 0.31125203 |
| Gm38010 | 20.30950254 | -0.539739574 | 0.273436906 |
| Gm37541 | 10.5940186 | -0.671836968 | 0.432644069 |
| Gm38021 | 12.72797742 | -1.228608587 | 0.328816382 |
| Gm6136 | 179.3671153 | 0.387325409 | 0.110894505 |
| Gm37967 | 11.04668228 | -1.106690396 | 0.346936238 |
| Gm28375 | 0.045882455 | 0.237312084 | 3.724945049 |
| Gm37017 | 0.873848832 | -1.590081252 | 1.247138089 |
| Gm37447 | 21.46227591 | -1.18085098 | 0.304179994 |
| Akp-ps1 | 0.752438969 | -0.243516094 | 1.132560034 |
| Gm29371 | 0 | NA | NA |
| Alppl2 | 0 | NA | NA |
| Alpi | 0 | NA | NA |
| Gm29372 | 0 | NA | NA |
| Akp3 | 0.332159689 | 2.368701561 | 1.924907817 |
| Gm29374 | 5.217580992 | 1.623738204 | 0.461166722 |
| Ecel1 | 298.3676729 | 0.97259774 | 0.140962412 |
| Prss56 | 20.4462505 | 2.565888012 | 0.78089528 |
| Chrnd | 0 | NA | NA |
| Chrng | 0.171131556 | 1.200054296 | 3.383981483 |
| Eif4e2 | 578.568997 | 0.112194049 | 0.061765719 |
| Gm26187 | 0.955307282 | 1.12950764 | 1.305340479 |
| Efhd1 | 197.7653228 | -0.962344709 | 0.150888652 |
| Efhd1os | 0.337798032 | -1.292916574 | 1.904396051 |
| Gigyf2 | 1430.757424 | -0.236811271 | 0.082343457 |
| Gm22549 | 0 | NA | NA |
| Kcnj13 | 41.11332588 | -1.214316696 | 0.265296009 |
| Gm23504 | 0 | NA | NA |
| 3110079O15Rik | 0.366579243 | -1.109469799 | 1.559531583 |
| Ngef | 1590.707428 | -0.912559499 | 0.129097068 |
| Neu2 | 18.78655172 | -0.58963223 | 0.262281315 |
| Gm17181 | 0.432611478 | -0.632793661 | 1.427671415 |
| Gm6153 | 1.379575143 | 0.814309043 | 0.892630348 |
| Inpp5d | 416.1094285 | 2.162707573 | 0.13138299 |
| Atg16l1 | 673.2924239 | -0.363825261 | 0.123119899 |
| Gm25395 | 742.8621918 | -0.131609399 | 0.184930018 |
| Scarna6 | 154.5847017 | 0.127898738 | 0.155973939 |
| Sag | 17.39870792 | 0.716554134 | 0.286440248 |
| Dgkd | 1519.408834 | -0.178205224 | 0.072053209 |
| Gm19582 | 6.959264896 | 0.664012771 | 0.479215723 |

| | | | |
|---|---|---|---|
| 4833421G17Rik | 11.9590053 | -0.601420881 | 0.332787207 |
| Gm38365 | 57.83653722 | -0.827336793 | 0.316756165 |
| Usp40 | 453.472211 | 0.071609093 | 0.114405954 |
| Gm25198 | 0 | NA | NA |
| Gm5832 | 0.105852363 | 0.23732098 | 3.724945049 |
| Gm15375 | 0.049553047 | 0.71820487 | 3.724945049 |
| Ugt1a10 | 0.161949631 | 1.583504458 | 2.7626009 |
| Gm20528 | 0.038981395 | 0.718207094 | 3.724945049 |
| Ugt1a9 | 0.038981395 | 0.718207094 | 3.724945049 |
| Ugt1a8 | 0 | NA | NA |
| Ugt1a7c | 12.7903233 | 2.277912352 | 0.36795777 |
| Gm15373 | 0.042716945 | -0.243571806 | 3.724945049 |
| Ugt1a6b | 1.167440347 | 2.604109555 | 1.111603798 |
| Gm15372 | 1.333048035 | 2.681344636 | 1.139564058 |
| Gm15370 | 0.204364325 | -0.24271053 | 2.645665856 |
| Ugt1a6a | 1.585662245 | 1.96562743 | 0.940958557 |
| Gm15369 | 0.593490712 | 2.28394426 | 1.252098314 |
| Gm15374 | 1.464328245 | 0.37977074 | 1.170473478 |
| Gm15371 | 0.512448221 | 1.620745718 | 1.357591756 |
| Ugt1a5 | 1.213945837 | -0.040904425 | 1.070978103 |
| Gm15368 | 0.624217637 | 1.451862923 | 1.422751407 |
| Gm15376 | 0.474048072 | 1.684889558 | 1.499090133 |
| Ugt1a2 | 0.875486306 | 0.881051262 | 1.177825845 |
| Dnajb3 | 7.809597006 | 0.848064531 | 0.373392187 |
| Ugt1a1 | 0.279519214 | -0.243563166 | 1.8014598 |
| Mroh2a | 3.442503238 | -0.069274187 | 0.603194516 |
| Hjurp | 549.4911097 | 0.031024828 | 0.083756812 |
| Trpm8 | 6.249743756 | -0.043066462 | 0.433123711 |
| 4930453O03Rik | 0.121388914 | 0.718200421 | 3.724945049 |
| Spp2 | 1.584204494 | 2.041240126 | 0.953193555 |
| Gm29538 | 0.330861516 | -0.955369174 | 1.719154108 |
| Gm28888 | 0 | NA | NA |
| Glrp1 | 2.778605519 | 4.635209237 | 0.925338691 |
| Gm19589 | 0.038981395 | 0.718207094 | 3.724945049 |
| Gm38130 | 0.038981395 | 0.718207094 | 3.724945049 |
| Gm29336 | 0.085433889 | -0.72445792 | 3.724945049 |
| Arl4c | 850.2294767 | 0.216351961 | 0.101024867 |
| Gm29335 | 0 | NA | NA |
| Platr5 | 0.048949532 | -0.243569582 | 3.724945049 |
| Gm29337 | 0 | NA | NA |
| Gm4753 | 0 | NA | NA |
| 1700067G17Rik | 0 | NA | NA |
| Gm18699 | 0 | NA | NA |
| Sh3bp4 | 304.2424635 | -0.002797862 | 0.140086713 |
| Gm38312 | 5.824867419 | -1.017688039 | 0.475881965 |
| Gm5259 | 0.045882455 | 0.237312084 | 3.724945049 |
| Gm28342 | 0.049553047 | 0.71820487 | 3.724945049 |

| | | | |
|---|---|---|---|
| C030007H22Rik | 0.466539735 | 0.668461678 | 3.679320886 |
| Agap1 | 3469.23624 | -0.161022067 | 0.065375518 |
| Gm37297 | 49.83626593 | -0.831341345 | 0.338739686 |
| Gm25180 | 3.378117529 | -1.037804393 | 0.769774857 |
| Gm28499 | 9.617358175 | -0.828124023 | 0.473825749 |
| 4933400F21Rik | 0.445470297 | 0.730169473 | 1.670526368 |
| Gm26037 | 7.202871343 | -1.457290007 | 0.521241277 |
| Gm37521 | 23.73742709 | -1.002588286 | 0.351643413 |
| Gbx2 | 22.87581777 | 0.312611097 | 0.272205568 |
| D130058E05Rik | 7.849306383 | -0.314942555 | 0.450656231 |
| Asb18 | 37.68452797 | -0.678915252 | 0.192681341 |
| Iqca | 7.094191665 | -0.726687411 | 0.399472249 |
| 4930434B07Rik | 0.106465261 | 0.718202646 | 3.724945049 |
| Ackr3 | 127.6655458 | 1.080354474 | 0.240984434 |
| Gm38277 | 0.04393175 | -0.243571806 | 3.724945049 |
| Gm28721 | 0.047948698 | -0.243569582 | 3.724945049 |
| Gm28723 | 1.104344843 | 1.307206901 | 1.189906975 |
| Gm27648 | 0 | NA | NA |
| Gm28722 | 0.261620403 | 1.517854941 | 2.050256886 |
| Gm35048 | 0.834783721 | 1.327607062 | 1.172097561 |
| Cops8 | 733.5162779 | 0.225317405 | 0.075337094 |
| Gm9991 | 0.061920613 | 0.718207094 | 3.724945049 |
| Col6a3 | 94.47715798 | -0.324445367 | 0.152706343 |
| Gm36970 | 0 | NA | NA |
| 4930474B08Rik | 0 | NA | NA |
| Mlph | 41.52328994 | -0.77552426 | 0.210282823 |
| Prlh | 0.989340548 | 1.086398395 | 1.175182664 |
| Rab17 | 0.340708601 | 0.678779797 | 1.77538034 |
| Gm28199 | 0 | NA | NA |
| Lrrfip1 | 605.1451411 | 0.292163909 | 0.118470578 |
| Rbm44 | 1.28530837 | 2.157132746 | 1.064769696 |
| Gm7776 | 0.372084114 | 1.689459333 | 1.971393409 |
| Ramp1 | 339.6276143 | -0.427069503 | 0.091664403 |
| Ube2f | 531.3694709 | 0.119269721 | 0.122723234 |
| Gm7785 | 19.25179799 | 0.660810865 | 0.324376394 |
| Gm26683 | 0.0885994 | -0.24357403 | 3.724945049 |
| Scly | 285.0654636 | 0.541873292 | 0.092733904 |
| Gm17090 | 0.087930927 | 0.236669296 | 3.724898107 |
| Espnl | 4.028718013 | 0.881230521 | 0.532164799 |
| Klhl30 | 7.180205943 | 0.443353708 | 0.555142684 |
| Fam132b | 9.846500243 | 2.460121683 | 0.537175449 |
| Ilkap | 381.4142211 | -0.033178678 | 0.105680246 |
| 1700020N18Rik | 1.184503545 | 0.285247592 | 1.141085518 |
| Hes6 | 86.06443169 | -0.164506538 | 0.139690327 |
| Per2 | 585.5330981 | -1.243477081 | 0.196359007 |
| Gm28381 | 0 | NA | NA |
| Gm28382 | 0 | NA | NA |

| | | | |
|---|---|---|---|
| Gm19085 | 0.292057199 | 0.771016847 | 2.434115916 |
| Traf3ip1 | 223.3191922 | -0.297038676 | 0.106572914 |
| Asb1 | 423.9897668 | -0.50278072 | 0.081534549 |
| Gm28380 | 0.180091664 | -0.243565765 | 2.750740754 |
| Gm29100 | 8.346887551 | -2.636391194 | 0.483078813 |
| Twist2 | 5.328893884 | -0.413007389 | 0.41250423 |
| Gm29101 | 0.685283607 | -1.473512469 | 1.233836894 |
| Gm37600 | 0 | NA | NA |
| Hdac4 | 1369.404079 | -0.181448136 | 0.080696284 |
| Gm37048 | 0.281888464 | -0.243566637 | 1.802651685 |
| Gm26720 | 0 | NA | NA |
| Gm37036 | 19.75259723 | -0.895468117 | 0.327384179 |
| Gm29099 | 0 | NA | NA |
| Gm7837 | 0.143650742 | 1.184932321 | 3.581766369 |
| Ndufa10 | 1126.271048 | -0.202983116 | 0.055076638 |
| Mir6900 | 0.79482971 | -1.17182572 | 1.128967815 |
| Olfr1416 | 0 | NA | NA |
| Olfr1415 | 0 | NA | NA |
| Olfr1414 | 0 | NA | NA |
| 1700054K02Rik | 0 | NA | NA |
| Olfr1413 | 0 | NA | NA |
| Gm28478 | 0 | NA | NA |
| Gm28479 | 0 | NA | NA |
| Olfr1412 | 0 | NA | NA |
| Olfr1411 | 0 | NA | NA |
| Olfr1410 | 0 | NA | NA |
| Olfr12 | 0 | NA | NA |
| Myeov2 | 377.7320121 | 0.126175973 | 0.113576377 |
| Otos | 0.964645659 | -0.886353956 | 1.046419214 |
| Gm29483 | 1.804989916 | -0.609252509 | 0.745150025 |
| Gm29482 | 0 | NA | NA |
| Gm29481 | 0.153668717 | -0.243571235 | 2.839403532 |
| Gpc1 | 1142.72074 | 0.287044244 | 0.079900965 |
| Gm29480 | 6.109138398 | 0.137018408 | 0.457572469 |
| Mir149 | 1.829161676 | -1.010442931 | 0.790786279 |
| Ankmy1 | 5.039844727 | 1.727748447 | 0.543625462 |
| Dusp28 | 267.5008543 | 1.099067599 | 0.167199752 |
| 5033417F24Rik | 38.62813476 | -0.362956347 | 0.180874314 |
| Rnpepl1 | 468.7456855 | 0.733265545 | 0.093301198 |
| 9430060I03Rik | 3.610602274 | 0.220056933 | 0.63924899 |
| Capn10 | 256.0341623 | -0.013475935 | 0.078724955 |
| Gpr35 | 139.3197398 | 5.435041159 | 0.389475942 |
| Gm18697 | 0.51839282 | 2.986657627 | 1.598680961 |
| Aqp12 | 0.175102337 | 1.440451655 | 3.306427423 |
| Kif1a | 18866.61453 | -0.314738135 | 0.054089003 |
| Gm28086 | 0.753737496 | 0.220906116 | 1.278359852 |
| Agxt | 0.11197992 | 0.718202646 | 3.724945049 |

| | | | |
|---|---|---|---|
| 2310007B03Rik | 0.800599448 | 1.137604607 | 1.197987442 |
| Crocc2 | 17.59480686 | -0.02010329 | 0.271745367 |
| Sned1 | 257.5400578 | -0.966470285 | 0.124209425 |
| Mir6901 | 0 | NA | NA |
| Mterf4 | 142.5738671 | -0.000356842 | 0.10761384 |
| Gm28535 | 1.935100933 | -0.702620052 | 0.711431501 |
| Gm7867 | 1.11968618 | -0.194319088 | 0.902830754 |
| Pask | 68.98058163 | -0.105365032 | 0.209625724 |
| Ppp1r7 | 1320.763185 | -0.290943864 | 0.062142211 |
| mmu-mir-6902 | 0.22467814 | -0.550980333 | 2.141189051 |
| Mir6345 | 0.94748184 | -1.451645783 | 1.04467613 |
| Gm26446 | 2.680156834 | -1.773363067 | 0.720201318 |
| Ano7 | 2.740637547 | -0.066156499 | 0.694258642 |
| Hdlbp | 3229.635038 | 0.262658533 | 0.048307855 |
| Gm17415 | 4.04170504 | 0.938518518 | 0.590597352 |
| 2-Sep | 1408.822543 | 0.725576439 | 0.084213324 |
| Farp2 | 211.4310669 | -0.294579208 | 0.115556288 |
| Gm37250 | 18.96035902 | -1.403584043 | 0.344490605 |
| Stk25 | 1506.774651 | -0.300117056 | 0.061068629 |
| Gm28536 | 1.8285836 | -1.088909618 | 0.700351075 |
| Bok | 205.8186137 | -0.39621242 | 0.09215782 |
| Thap4 | 221.3880966 | -0.098913638 | 0.07928103 |
| Gm10550 | 0.177745132 | -0.244115745 | 2.300135904 |
| Atg4b | 616.7307566 | -0.093126175 | 0.047728729 |
| Dtymk | 176.2881088 | -0.114633323 | 0.110288707 |
| Ing5 | 203.0410209 | 0.331286358 | 0.081982078 |
| Gm22089 | 0 | NA | NA |
| D2hgdh | 49.47669206 | -0.374629998 | 0.215140262 |
| Gal3st2 | 0.259214904 | 0.237330074 | 1.83327597 |
| Gm18666 | 2.840417843 | -0.417605181 | 0.757506897 |
| Gm9994 | 1.809180368 | -0.75222173 | 0.674755913 |
| Gm7889 | 3.175371776 | -0.022688658 | 0.552935949 |
| Gm6086 | 0.846372671 | -0.239561683 | 1.04162614 |
| Neu4 | 89.65460407 | -0.596256757 | 0.196081022 |
| Pdcd1 | 42.57420289 | 6.759576895 | 0.873375254 |
| Gm7891 | 0.255524584 | 1.28117929 | 2.490892268 |
| Gm7895 | 67.17479994 | -0.009327619 | 0.195434772 |
| Gm23389 | 0 | NA | NA |
| 4930440C22Rik | 0.220980774 | 1.68041995 | 3.10409395 |
| Gm23181 | 0 | NA | NA |
| Gm5264 | 0.086908494 | -0.72445792 | 3.724945049 |
| Fam174a | 321.8174294 | -0.135757514 | 0.140217083 |
| BC055308 | 22.09994013 | -0.44589987 | 0.26315363 |
| Gm15427 | 268.0268351 | 0.72192708 | 0.12823094 |
| St8sia4 | 341.5027806 | 0.039599544 | 0.15711606 |
| Gm37642 | 12.46481921 | 0.095644292 | 0.288292777 |
| 4930598F16Rik | 0 | NA | NA |

| | | | |
|---|---|---|---|
| Gm37761 | 0 | NA | NA |
| Gm22360 | 1.558628739 | -0.057527497 | 0.822313762 |
| 1700063A18Rik | 0 | NA | NA |
| Gm24586 | 0 | NA | NA |
| Gm22917 | 0 | NA | NA |
| Gm37076 | 0 | NA | NA |
| Gm29601 | 0.044544648 | 0.237309859 | 3.724945049 |
| 4930533P14Rik | 0 | NA | NA |
| Gm26245 | 0 | NA | NA |
| Gm1833 | 0.161949631 | 1.583504458 | 2.7626009 |
| Gm26078 | 0 | NA | NA |
| Slco4c1 | 49.5126318 | -0.383020198 | 0.191448081 |
| Panct2 | 37.25735819 | -0.162667906 | 0.228315435 |
| Slco6b1 | 0.161026707 | 1.711863256 | 3.451443504 |
| Gm9499 | 0 | NA | NA |
| Gm28901 | 0 | NA | NA |
| Gm6430 | 14.31094521 | 1.263605157 | 0.294894942 |
| Slco6c1 | 4.084609871 | -0.817303748 | 0.505309671 |
| Gm7133 | 0.216806889 | 0.237323933 | 2.60159912 |
| Gm7135 | 0.141094079 | 0.718203391 | 3.563850454 |
| Gm37171 | 2.848103185 | -0.120687309 | 0.625783306 |
| D1Ertd622e | 880.9560972 | -0.069978061 | 0.062172008 |
| Ppip5k2 | 911.1116556 | -0.254793485 | 0.096155688 |
| Gin1 | 94.83341611 | -0.042834312 | 0.121748846 |
| Pam | 2285.737651 | -0.342192221 | 0.062397383 |
| Gm37140 | 76.97524024 | -0.659866052 | 0.241491833 |
| Gm25413 | 0 | NA | NA |
| Gm3531 | 31.35057007 | 0.734017832 | 0.205138172 |
| B230216N24Rik | 130.1992663 | -1.182004968 | 0.232930378 |
| Gm24465 | 0.049553047 | 0.71820487 | 3.724945049 |
| 1810006J02Rik | 0.496068496 | 1.446538427 | 1.271630444 |
| Gm29461 | 0.196785726 | -0.243567967 | 2.22010112 |
| Gm7952 | 0 | NA | NA |
| Gm29462 | 0.477857232 | -0.920512415 | 1.380734951 |
| Slco6d1 | 0.368527898 | 0.265661978 | 1.589073013 |
| Gm3555 | 28.33472675 | 0.623590324 | 0.230699175 |
| Gm26134 | 0 | NA | NA |
| Gm7967 | 10.59292213 | 0.02896614 | 0.359183001 |
| Gm29460 | 1.272524668 | 0.672818338 | 1.050127244 |
| Gm23434 | 0.075043205 | 0.237309859 | 3.724945049 |
| Gm18248 | 0.075043205 | 0.237309859 | 3.724945049 |
| Cntnap5b | 377.3845069 | -0.689868208 | 0.094525102 |
| Tdpx-ps1 | 13.77002096 | 0.378537157 | 0.277658551 |
| Gm37877 | 23.5485973 | -0.828943117 | 0.28347372 |
| Gm19029 | 3.564592169 | -0.964957665 | 0.68510261 |
| Gm23602 | 0 | NA | NA |
| Gm29668 | 2.625920046 | -1.029033951 | 0.611919729 |

| | | | |
|---|---|---|---|
| Gm29667 | 14.43481637 | -0.716599095 | 0.334587967 |
| Gm18700 | 0 | NA | NA |
| Gm18185 | 0 | NA | NA |
| Gm29334 | 0 | NA | NA |
| Gm25537 | 0 | NA | NA |
| Gm20268 | 0 | NA | NA |
| Gm28801 | 0 | NA | NA |
| Psmb6-ps2 | 0.576768135 | 0.081753064 | 1.2722285 |
| Gm18969 | 0 | NA | NA |
| Gm28188 | 0.072025462 | 0.237312084 | 3.724945049 |
| Gm28187 | 1.345496586 | -0.04473324 | 0.854858802 |
| Gm24122 | 0 | NA | NA |
| Gm28623 | 0 | NA | NA |
| Gm23036 | 0 | NA | NA |
| Gm3116 | 0 | NA | NA |
| Gm20281 | 0 | NA | NA |
| Gm22034 | 0 | NA | NA |
| Gm18728 | 0 | NA | NA |
| Gm23965 | 0 | NA | NA |
| Gm29141 | 1.908747754 | 4.381282345 | 1.014036417 |
| Gm37789 | 0.149576918 | 1.19849824 | 2.863264735 |
| Gm18447 | 0 | NA | NA |
| Gm29140 | 0.04393175 | -0.243571806 | 3.724945049 |
| Gm23537 | 0 | NA | NA |
| Gm5260 | 0 | NA | NA |
| Gm29592 | 0 | NA | NA |
| Gm29011 | 0.220980774 | 1.68041995 | 3.10409395 |
| Cdh20 | 327.0013875 | 0.105995535 | 0.107141278 |
| Gm38031 | 85.93314083 | -0.101246346 | 0.193201762 |
| n-R5s216 | 0 | NA | NA |
| Gm36969 | 21.4981108 | -0.687343455 | 0.34795571 |
| Gm15699 | 0 | NA | NA |
| Gm29012 | 0.089089296 | 0.237305797 | 3.724945049 |
| Gm29013 | 0.046345709 | 0.71820487 | 3.724945049 |
| Rnf152 | 731.5992459 | -0.304063496 | 0.089356172 |
| Gm29014 | 0 | NA | NA |
| Gm17634 | 9.160375891 | -0.096505076 | 0.363985913 |
| Gm28403 | 1.125166341 | -0.537275166 | 1.03789594 |
| Gm3608 | 115.3994121 | 0.954997917 | 0.227208287 |
| Pign | 477.5995311 | 0.207138743 | 0.112657561 |
| Gm28404 | 2.874152792 | -0.385257967 | 0.593738566 |
| Gm20302 | 1.560084669 | 0.81738963 | 0.747756577 |
| Gm22426 | 0 | NA | NA |
| Gm8004 | 0 | NA | NA |
| 2310035C23Rik | 1589.964662 | -0.120644004 | 0.079338258 |
| Gm37566 | 68.8298285 | -0.360660215 | 0.188988839 |
| Gm10193 | 1.224645236 | 0.939728032 | 1.073369193 |

| | | | |
|---|---|---|---|
| Tnfrsf11a | 174.3773544 | 1.515480563 | 0.158762953 |
| Gm37779 | 0.176451472 | 0.7182167 | 2.72333919 |
| Gm26214 | 0 | NA | NA |
| Gm18801 | 2.226781727 | 5.091609208 | 0.978325288 |
| Gm7160 | 28.83074711 | 2.034555249 | 0.280649274 |
| Zcchc2 | 822.9552979 | 1.020637709 | 0.110203611 |
| Gm37372 | 0.112396584 | 1.192083721 | 3.724434897 |
| Gm26509 | 3.937004826 | -0.122725526 | 0.521822936 |
| Gm20753 | 4.418263327 | -1.26433133 | 0.544775875 |
| Phlpp1 | 1704.681168 | -0.311152195 | 0.075996671 |
| Gm38235 | 158.3680684 | -0.875628081 | 0.244298018 |
| A530053M12Rik | 0.224329914 | 1.721266092 | 2.570323433 |
| Gm6651 | 0.568438163 | 2.869673038 | 1.293236973 |
| Bcl2 | 493.9864849 | 0.217238175 | 0.113600293 |
| D630008O14Rik | 0.626527047 | 1.818707821 | 1.659442815 |
| Mir3473f | 0.393472992 | 0.890737583 | 1.784430073 |
| Gm37053 | 17.84686388 | -0.344515393 | 0.374106096 |
| Kdsr | 626.2639973 | -0.029275918 | 0.061075233 |
| Vps4b | 768.3375885 | 0.244013384 | 0.076733511 |
| Serpinb5 | 4.060285521 | 3.263330083 | 0.924618662 |
| Pou2f3-rs1 | 0 | NA | NA |
| Serpinb12 | 0.340899075 | 0.79627348 | 1.984870557 |
| Serpinb13 | 0 | NA | NA |
| Gm8080 | 0.048949532 | -0.243569582 | 3.724945049 |
| Serpinb3a | 0.112642008 | -0.24357403 | 3.724945049 |
| Serpinb3d | 0.044544648 | 0.237309859 | 3.724945049 |
| Gm15388 | 0 | NA | NA |
| Serpinb3b | 0.36088792 | -0.243588148 | 3.724945049 |
| Gm8084 | 0.045882455 | 0.237312084 | 3.724945049 |
| Gm15390 | 0 | NA | NA |
| Serpinb3c | 0.256447508 | 1.225217374 | 2.469852681 |
| Gm8089 | 0.10503227 | 1.179809211 | 3.723500796 |
| Serpinb11 | 0.313959609 | -0.641027915 | 1.541183901 |
| Gm15391 | 0 | NA | NA |
| Serpinb7 | 0.394543937 | 1.682292757 | 1.661165665 |
| Gm24553 | 0 | NA | NA |
| Serpinb2 | 8.118383424 | 4.385813481 | 0.76955513 |
| Serpinb10 | 0.745572437 | 2.769755536 | 1.246175041 |
| Serpinb8 | 106.9109276 | 0.672105064 | 0.17758355 |
| Gm15389 | 0 | NA | NA |
| Gm24267 | 0 | NA | NA |
| Gm22438 | 0.044544648 | 0.237309859 | 3.724945049 |
| D830032E09Rik | 0 | NA | NA |
| Gm18448 | 0.11147366 | 1.230376831 | 3.722845817 |
| Gm28282 | 0 | NA | NA |
| Gm5702 | 0 | NA | NA |
| Gm38212 | 0.044191549 | -0.243571806 | 3.724945049 |

| | | | |
|---|---|---|---|
| Gm28283 | 0 | NA | NA |
| Gm6679 | 0 | NA | NA |
| Gm17847 | 0 | NA | NA |
| Gm28281 | 0 | NA | NA |
| Gm22159 | 0.068097359 | -0.243569582 | 3.724945049 |
| Gm25293 | 0 | NA | NA |
| Gm8141 | 0.177100366 | -1.491187236 | 3.367753361 |
| Cdh7 | 599.3121521 | -0.502559168 | 0.119822755 |
| Gm18552 | 0 | NA | NA |
| Gm3332 | 0 | NA | NA |
| Gm6693 | 0 | NA | NA |
| Cdh19 | 348.6925766 | 0.26673618 | 0.131143629 |
| Gm38255 | 57.38585182 | -0.317657618 | 0.209631268 |
| Gm36944 | 56.47739328 | -0.616535793 | 0.246233587 |
| Gm29088 | 1.097863377 | 1.031558744 | 1.026782985 |
| Gm19927 | 0.068097359 | -0.243569582 | 3.724945049 |
| Gm8173 | 15.82173517 | -0.608136314 | 0.365758707 |
| Gm37934 | 0 | NA | NA |
| Dsel | 373.0186971 | -0.245766636 | 0.080296439 |
| Gm6028 | 0.150520013 | 0.237326327 | 2.857840489 |
| Gm17752 | 0 | NA | NA |
| Gm6717 | 0 | NA | NA |
| Gm19086 | 0 | NA | NA |
| Gm18406 | 9.163048501 | -0.243768766 | 0.329105813 |
| Gm22331 | 0 | NA | NA |
| Gm8204 | 0 | NA | NA |
| Gm28190 | 0 | NA | NA |
| Gm28189 | 0 | NA | NA |
| Gm17946 | 0 | NA | NA |
| Gm18181 | 0 | NA | NA |
| Gm24937 | 0 | NA | NA |
| 9330185C12Rik | 1.904432073 | 0.945023334 | 0.774688761 |
| Gm4854 | 0 | NA | NA |
| Gm17916 | 16.58061403 | -0.953504332 | 0.273802975 |
| Cntnap5a | 1043.802266 | -0.764074871 | 0.0848221 |
| Gm23393 | 0 | NA | NA |
| Gm28263 | 0.417031551 | -1.199453908 | 1.554389566 |
| Gm19965 | 1.062057355 | 0.57967775 | 1.14474647 |
| Gm17777 | 0 | NA | NA |
| Gm24030 | 0 | NA | NA |
| Gm27785 | 0 | NA | NA |
| Gm28361 | 0 | NA | NA |
| Zfp813-ps | 0 | NA | NA |
| Gm28362 | 0 | NA | NA |
| Gm28363 | 0 | NA | NA |
| Gm28359 | 0 | NA | NA |
| Gm18182 | 0 | NA | NA |

| | | | |
|---|---|---|---|
| B020011L13Rik | 0 | NA | NA |
| Gm28360 | 0 | NA | NA |
| Gm28169 | 0.218500166 | 1.192671525 | 2.170343423 |
| Gm7195 | 0 | NA | NA |
| Gm28168 | 0 | NA | NA |
| Gm7145 | 0 | NA | NA |
| Gm28358 | 0 | NA | NA |
| Gm29489 | 0 | NA | NA |
| Gm29490 | 0.044191549 | -0.243571806 | 3.724945049 |
| Gm29104 | 0 | NA | NA |
| Gm29106 | 0.046345709 | 0.71820487 | 3.724945049 |
| Gm7200 | 0 | NA | NA |
| Tsn | 868.3708414 | -0.043328543 | 0.069112187 |
| Nifk | 240.6864645 | 0.563856802 | 0.119739606 |
| Gm28867 | 0.180050447 | 0.719165083 | 3.327036943 |
| Clasp1 | 2304.76087 | -0.388771131 | 0.06369516 |
| Gm26080 | 0 | NA | NA |
| Gm22710 | 2.569247428 | -0.203195419 | 1.041868967 |
| Gm28466 | 0 | NA | NA |
| Tfcp2l1 | 115.5571692 | 0.431633231 | 0.154678529 |
| Gm22094 | 0 | NA | NA |
| Gm28882 | 0 | NA | NA |
| Gm23497 | 0 | NA | NA |
| Gm28467 | 0 | NA | NA |
| Gli2 | 130.8498884 | 0.061113055 | 0.170243804 |
| Mir6346 | 0 | NA | NA |
| Gm26831 | 0.564941401 | -2.096866313 | 1.207021302 |
| Gm8321 | 0 | NA | NA |
| Gm3508 | 0 | NA | NA |
| Gm29456 | 0 | NA | NA |
| Gm29455 | 2.475683411 | -0.956897562 | 0.686911097 |
| Inhbb | 109.178341 | 0.075910803 | 0.189239978 |
| Ralb | 386.9036816 | 1.064182336 | 0.096673531 |
| Gm29454 | 0 | NA | NA |
| Gm27184 | 4.248472554 | -0.116721785 | 0.55325418 |
| Tmem185b | 22.02441361 | 0.232423605 | 0.243659736 |
| AC132393.1 | 0 | NA | NA |
| 3830432H09Rik | 2.926677728 | 0.588487103 | 0.590069467 |
| Gm25322 | 0.15596086 | -1.187811386 | 3.483969583 |
| Epb4.1l5 | 218.5589968 | -0.010407723 | 0.089671417 |
| Gm37219 | 2.565042467 | -1.339810276 | 0.670454414 |
| Ptpn4 | 3441.616975 | -0.384312843 | 0.089458488 |
| Gm3851 | 0.583829782 | -0.782784483 | 1.210807789 |
| Gm37745 | 125.9235229 | -0.803496741 | 0.242259725 |
| Gm29657 | 0 | NA | NA |
| Gm8338 | 0 | NA | NA |
| Gm29658 | 0.060174537 | 0.237309859 | 3.724945049 |

| | | | |
|---|---|---|---|
| Gm3551 | 0.190043611 | 0.23732668 | 2.655002295 |
| Gm28209 | 0 | NA | NA |
| Tmem177 | 83.02492162 | -0.279757118 | 0.125788587 |
| Cfap221 | 5.416510947 | -0.212953854 | 0.464758213 |
| Gm17973 | 0 | NA | NA |
| Sctr | 5.377310654 | -0.94348459 | 0.450687038 |
| Tmem37 | 24.45334209 | 1.736801678 | 0.311331615 |
| Dbi | 1017.882369 | 0.416933972 | 0.144365234 |
| 3110009E18Rik | 15.78515723 | -0.36547272 | 0.271161201 |
| Steap3 | 67.01145874 | 0.426088856 | 0.244405517 |
| C1ql2 | 92.61537039 | 0.07994173 | 0.164279529 |
| 1700012E03Rik | 0.060174537 | 0.237309859 | 3.724945049 |
| Marco | 0.34718572 | 0.84142025 | 1.732470564 |
| En1 | 26.69343009 | 0.175672719 | 0.210963056 |
| RP24-358J4.2 | 0.343046485 | 0.237332353 | 1.583631074 |
| 2610027F03Rik | 0 | NA | NA |
| Gm29346 | 0.099155932 | 0.718202646 | 3.724945049 |
| Gm29345 | 0.060174537 | 0.237309859 | 3.724945049 |
| Gm29344 | 0 | NA | NA |
| Gm29348 | 0.333474911 | 1.126008713 | 1.724948368 |
| Gm29347 | 0.045882455 | 0.237312084 | 3.724945049 |
| Gm37043 | 0 | NA | NA |
| Celrr | 13.48815634 | -1.935130937 | 0.451041866 |
| Gm29359 | 0 | NA | NA |
| Gm38283 | 4.643660642 | -0.534903976 | 0.532577102 |
| Insig2 | 471.7823211 | 0.040293947 | 0.082951266 |
| Gm37174 | 21.08112134 | -0.3050107 | 0.259973746 |
| 2210011K15Rik | 0.099106094 | 1.209894437 | 3.72418822 |
| Ccdc93 | 456.8414215 | -0.121914292 | 0.103729615 |
| Gm15392 | 0 | NA | NA |
| Gm15393 | 0.061920613 | 0.718207094 | 3.724945049 |
| Htr5b | 20.85714652 | -0.536757115 | 0.319012594 |
| Gm8217 | 0 | NA | NA |
| Ddx18 | 253.9984606 | 0.658566019 | 0.080529487 |
| Gm28591 | 0 | NA | NA |
| Gm37920 | 0 | NA | NA |
| Gm15394 | 0 | NA | NA |
| Gm8392 | 0 | NA | NA |
| Gm26472 | 0 | NA | NA |
| Gm28590 | 0 | NA | NA |
| Gm24547 | 0 | NA | NA |
| Gm36982 | 0 | NA | NA |
| Gm25771 | 0 | NA | NA |
| Gm29192 | 0 | NA | NA |
| Gm37970 | 0.148650935 | -0.243572541 | 2.863172849 |
| Gm8403 | 0.085327104 | 1.179809211 | 3.723500796 |
| Gm28911 | 0.104106287 | -0.243576254 | 3.724945049 |

| | | | |
|---|---|---|---|
| Gm20058 | 0.094097695 | 0.718200421 | 3.724945049 |
| Gm28912 | 0 | NA | NA |
| Dpp10 | 1832.665455 | -0.777967962 | 0.087269022 |
| Gm37478 | 62.41200189 | -0.930158257 | 0.277325386 |
| Gm37221 | 43.7843784 | -0.988360104 | 0.282979983 |
| Gm24791 | 5.806203063 | -1.797106914 | 0.550253952 |
| Gm37066 | 38.05353759 | -1.398589071 | 0.295965577 |
| Gm37781 | 35.98403262 | -1.087676949 | 0.325519729 |
| Gm37396 | 18.40949471 | -1.269785446 | 0.464302897 |
| Gm38263 | 58.17025342 | -1.395555748 | 0.305657001 |
| Gm28929 | 0 | NA | NA |
| Gm18875 | 20.78588443 | -1.622777507 | 0.389961106 |
| Gm37551 | 88.74313094 | -1.424656542 | 0.296060986 |
| Gm38073 | 157.9840511 | -1.356034018 | 0.445878317 |
| Gm25578 | 0 | NA | NA |
| Gm28928 | 91.8867847 | -1.07348148 | 0.421566175 |
| Gm28299 | 17.41287975 | -1.048838772 | 0.39245177 |
| Gm37717 | 149.0700901 | -1.492749373 | 0.460826043 |
| 1700121L03Rik | 0.097139547 | 0.718207094 | 3.724945049 |
| Actr3 | 2282.827765 | 0.646296206 | 0.087057884 |
| Gm28706 | 0 | NA | NA |
| Slc35f5 | 263.8309605 | 0.613158936 | 0.093693104 |
| Gm18444 | 0 | NA | NA |
| Gpr39 | 2.657777857 | 1.162698795 | 0.752648308 |
| Lypd1 | 535.5413066 | -1.091893151 | 0.086362088 |
| Nckap5 | 192.1177816 | 0.233593141 | 0.148275738 |
| Gm37161 | 5.830223875 | -0.163183273 | 0.478332998 |
| Gm37936 | 6.414345313 | 0.042715175 | 0.510907154 |
| Gm38177 | 4.318687175 | -0.095976549 | 0.538292978 |
| 4930599A14Rik | 0 | NA | NA |
| Gm5261 | 0 | NA | NA |
| Mgat5 | 2217.417936 | -0.226755757 | 0.061016266 |
| Gm37772 | 11.00123342 | -0.539407149 | 0.400041583 |
| Gm37510 | 48.12361392 | -0.684648857 | 0.322528137 |
| Gm8451 | 18.99290832 | 0.667380856 | 0.319348927 |
| SCARNA4 | 0.095435502 | 0.718202646 | 3.724945049 |
| SCARNA15 | 0 | NA | NA |
| Gm23734 | 4.197363489 | -0.660119177 | 0.600253181 |
| Gm38301 | 6.512391526 | -0.984627792 | 0.465643772 |
| Tmem163 | 261.7619097 | -0.103242202 | 0.079513522 |
| Acmsd | 0.917997538 | 1.484168698 | 1.165316734 |
| 2900009J06Rik | 4.736596173 | 0.04125531 | 0.480825277 |
| Ccnt2 | 1690.484171 | -0.270208901 | 0.132350391 |
| Map3k19 | 75.93146868 | -0.227076543 | 0.18839706 |
| Rab3gap1 | 1284.44217 | -0.04763644 | 0.046906756 |
| Zranb3 | 101.8848498 | -0.142072214 | 0.136936449 |
| Gm28800 | 35.95127173 | -1.125942946 | 0.195555437 |

| | | | |
|---|---|---|---|
| Gm38248 | 15.05177351 | -0.348831969 | 0.409349535 |
| Rpl28-ps1 | 66.91960972 | 0.721965376 | 0.163153365 |
| Gm37625 | 7.67879382 | -0.722387594 | 0.387910239 |
| Gm38026 | 3.696876593 | -0.592443332 | 0.552182071 |
| R3hdm1 | 4415.135964 | -0.949110983 | 0.148195278 |
| Gm29383 | 0 | NA | NA |
| Gm37588 | 55.83216539 | -0.916980315 | 0.232302767 |
| Gm37868 | 81.33316158 | -1.076370612 | 0.280363378 |
| Gm37407 | 115.1470443 | -1.126765079 | 0.202207999 |
| Mir128-1 | 17.81336244 | -1.561789301 | 0.31392728 |
| Ubxn4 | 978.701728 | 0.498837625 | 0.09308967 |
| Lct | 31.88336858 | -1.488387461 | 0.316416013 |
| Mcm6 | 72.35290303 | 2.712432326 | 0.262354427 |
| Dars | 621.5308942 | 0.417735264 | 0.099695946 |
| Gm23902 | 0 | NA | NA |
| Gm29598 | 0 | NA | NA |
| Gm29597 | 0 | NA | NA |
| Gm25384 | 0 | NA | NA |
| Gm6170 | 19.62597736 | 0.712282274 | 0.276778548 |
| Cxcr4 | 127.9107672 | 3.493312862 | 0.282785173 |
| Gm29599 | 0.045882455 | 0.237312084 | 3.724945049 |
| Gm26686 | 0.389802664 | 0.606010867 | 1.433305784 |
| Gm23056 | 0 | NA | NA |
| Thsd7b | 229.954111 | -0.004831773 | 0.126138981 |
| Gm38240 | 12.58215362 | -0.778277733 | 0.314093038 |
| Gm38330 | 4.986765232 | -0.310017058 | 0.515798133 |
| Gm36951 | 10.68103066 | -0.268056106 | 0.362000284 |
| Gm15790 | 8.257872952 | 0.517852785 | 0.370036496 |
| Rpl31-ps9 | 4.393169526 | 0.445480906 | 0.5234412 |
| Gm37479 | 19.45303257 | -0.368230007 | 0.275002327 |
| Gm37064 | 10.99610456 | -0.452177111 | 0.325823088 |
| Gm37278 | 4.9996304 | -0.236621705 | 0.491029902 |
| Gm29428 | 1.653651842 | -2.246921047 | 0.965946475 |
| Gm16081 | 0 | NA | NA |
| Daf2 | 0.977993353 | 0.391971983 | 1.012612927 |
| Gm15674 | 0.068097359 | -0.243569582 | 3.724945049 |
| Gm15675 | 15.76810382 | -0.828044114 | 0.358800739 |
| Cd55 | 67.64943717 | -0.061652236 | 0.145387133 |
| Gm16083 | 14.70625333 | -0.830779982 | 0.286478274 |
| Gm37109 | 7.99913161 | -1.356206683 | 0.494262611 |
| Gm37283 | 5.739234123 | -1.865398802 | 0.507525151 |
| Zp3r | 2.754922463 | -0.294332512 | 0.654639374 |
| Gm23623 | 0.044544648 | 0.237309859 | 3.724945049 |
| C4bp | 2.027159789 | 2.135068344 | 0.751266026 |
| Gm29427 | 1.964410769 | -0.408937483 | 0.718736979 |
| C4bp-ps1 | 4.021019734 | -0.51622024 | 0.48043721 |
| Pfkfb2 | 323.8491531 | -0.524904335 | 0.120086717 |

| | | | |
|---|---|---|---|
| Yod1 | 565.3277476 | -0.048182793 | 0.109949862 |
| Gm37163 | 0.049553047 | 0.71820487 | 3.724945049 |
| AA986860 | 42.90179849 | 0.153626057 | 0.184827447 |
| Gm28857 | 0.066050875 | 0.71820487 | 3.724945049 |
| Gm28856 | 0.17254495 | 0.237326017 | 2.737704594 |
| Fcamr | 0.566848388 | 3.386419972 | 1.401950852 |
| Gm15848 | 0.042716945 | -0.243571806 | 3.724945049 |
| Pigr | 0.796384302 | 3.07466102 | 1.181156832 |
| Fcmr | 1.860507999 | 1.297122676 | 0.826877518 |
| Il24 | 0.119234754 | -0.243576254 | 3.724945049 |
| Il20 | 0.112396584 | 1.192083721 | 3.724434897 |
| Il19 | 1.495078259 | 2.300400192 | 1.223346412 |
| Il10 | 32.26081669 | 8.667698958 | 1.032372328 |
| Mapkapk2 | 535.196025 | 1.19536803 | 0.138276051 |
| Gm25549 | 2.426982663 | -0.20216933 | 0.649073259 |
| Gm37084 | 67.84258375 | 0.176176946 | 0.274407215 |
| Gm25717 | 0.038981395 | 0.718207094 | 3.724945049 |
| Dyrk3 | 126.7275624 | 0.97587082 | 0.183620829 |
| Eif2d | 340.1033352 | -0.304768141 | 0.12981848 |
| Rassf5 | 359.522077 | 0.467067668 | 0.087418303 |
| Gm17796 | 0.165470308 | 0.237318778 | 2.761559324 |
| Ikbke | 98.54742827 | 3.465182767 | 0.247058946 |
| Gm28914 | 1.174162655 | -1.348457933 | 1.021204617 |
| Gm28913 | 1.688430464 | -0.288974975 | 0.763061431 |
| Srgap2 | 1739.750525 | -0.269855712 | 0.079873105 |
| Gm38262 | 21.98513846 | -0.596830058 | 0.308394113 |
| Gm37954 | 37.75257877 | -0.850958087 | 0.296776854 |
| Gm29488 | 96.98125143 | -0.422567464 | 0.232148912 |
| Gm29487 | 5.722531327 | 0.36229752 | 0.600515657 |
| Gm24273 | 9.213624824 | -0.587799176 | 0.390038735 |
| Fam72a | 24.35166078 | 0.898618465 | 0.232804581 |
| Avpr1b | 0.995114343 | 1.050389988 | 1.31817369 |
| Gm8532 | 1.429318806 | 2.055885554 | 0.921618258 |
| Ctse | 45.61320862 | 5.360019827 | 0.459812131 |
| Rab7b | 58.99444633 | 2.076616796 | 0.310703319 |
| Slc26a9 | 1.868919118 | 0.315515563 | 0.713141331 |
| Pm20d1 | 45.2287229 | -0.250421091 | 0.263897286 |
| Slc41a1 | 1235.334159 | 0.493744458 | 0.11053708 |
| Rab29 | 79.93970694 | 2.062075574 | 0.219124674 |
| Gm29105 | 0 | NA | NA |
| Nucks1 | 3423.866732 | -0.058379021 | 0.076491092 |
| Gm38067 | 40.11111897 | -0.588385208 | 0.276557273 |
| Gm37676 | 0.045882455 | 0.237312084 | 3.724945049 |
| Gm29103 | 0 | NA | NA |
| Slc45a3 | 23.9080149 | -0.259538983 | 0.233966721 |
| Elk4 | 760.1203314 | 0.492088364 | 0.071685978 |
| Mfsd4 | 533.3513531 | -0.698830368 | 0.095556181 |

| | | | |
|---|---|---|---|
| Gm26892 | 3.660297346 | 1.021050946 | 0.668269613 |
| Gm26034 | 0.106465261 | 0.718202646 | 3.724945049 |
| Gm37120 | 6.573103098 | 0.849479443 | 0.52824234 |
| Cdk18 | 350.945801 | -0.088842501 | 0.127147578 |
| Gm29629 | 1.14102893 | 0.886568869 | 0.96956921 |
| Lemd1 | 70.07703057 | -0.838142948 | 0.217739831 |
| Mir135b | 0.432115038 | -0.585459293 | 1.562228799 |
| 1700037F24Rik | 0 | NA | NA |
| Gm10188 | 0.044544648 | 0.237309859 | 3.724945049 |
| Gm29630 | 0 | NA | NA |
| Gm7241 | 0.177688696 | -0.243572061 | 2.724664815 |
| Klhdc8a | 280.7923049 | 0.428289563 | 0.150186426 |
| Nuak2 | 109.3043444 | 3.623088778 | 0.266855462 |
| F730311O21Rik | 4.129176112 | 4.348547439 | 0.891966291 |
| Tmcc2 | 2513.213478 | -0.212901649 | 0.097366163 |
| Gm15849 | 0.048949532 | -0.243569582 | 3.724945049 |
| Dstyk | 977.2630414 | 0.028313465 | 0.05103275 |
| 6030442K20Rik | 15.5449471 | -0.727585885 | 0.326257679 |
| Rbbp5 | 612.646602 | 0.086887078 | 0.057950515 |
| Tmem81 | 45.06443145 | -0.149649197 | 0.167614946 |
| Cntn2 | 3239.191646 | 0.02089761 | 0.084193864 |
| Nfasc | 5772.548432 | -0.072404375 | 0.057385947 |
| Gm10538 | 0.168483864 | 0.237319907 | 2.75185196 |
| Gm22609 | 0.089089296 | 0.237305797 | 3.724945049 |
| Lrrn2 | 1233.035506 | -0.624368868 | 0.095621357 |
| AC124095.1 | 0.988133447 | -1.081936031 | 1.459012232 |
| Gm38308 | 6.071337244 | -0.462362856 | 0.634791683 |
| Mdm4 | 1609.398272 | -0.204367165 | 0.117626744 |
| Gm29257 | 3.367574844 | 0.902211899 | 0.536144262 |
| Pik3c2b | 418.6707706 | -0.935433586 | 0.13097228 |
| Gm28609 | 1.26125615 | 2.074012093 | 0.952640549 |
| Gm26616 | 7.818633818 | 0.902411116 | 0.413260031 |
| Ppp1r15b | 1586.837848 | 1.020873926 | 0.122111993 |
| Plekha6 | 1798.908558 | -0.493537013 | 0.087300393 |
| Gm19461 | 22.93070562 | 0.167136996 | 0.23965463 |
| Golt1a | 0.589467498 | -0.309161415 | 1.530964751 |
| Gm28040 | 0.066050875 | 0.71820487 | 3.724945049 |
| Kiss1 | 0.282483437 | 0.237331422 | 1.786847593 |
| Ren1 | 0.441022512 | -1.402863461 | 1.467792809 |
| Etnk2 | 45.75414376 | -1.151622367 | 0.209166825 |
| Sox13 | 89.13343506 | -0.11429429 | 0.140264714 |
| Gm8596 | 0 | NA | NA |
| Gm23939 | 0 | NA | NA |
| Snrpe | 146.4574117 | 0.562825384 | 0.145680577 |
| Gm26706 | 0.926160546 | -0.701916372 | 1.091063534 |
| Zc3h11a | 237.5869313 | -0.157984182 | 0.120621315 |
| Gm38394 | 135.9246581 | -0.273045736 | 0.160566589 |

| | | | |
|---|---|---|---|
| Zbed6 | 0 | NA | NA |
| Lax1 | 23.20725604 | 4.099024374 | 0.408589826 |
| Atp2b4 | 2855.857049 | -0.802104077 | 0.100272903 |
| Mir6903 | 0.148659141 | 1.513781604 | 3.716315176 |
| Gm37368 | 12.07769997 | -0.463765127 | 0.401296472 |
| Gm10537 | 0.578745683 | 1.598475336 | 1.21924862 |
| Gm28441 | 0.061920613 | 0.718207094 | 3.724945049 |
| Optc | 6.651424553 | 0.432019958 | 0.394140726 |
| Gm15851 | 0.236932206 | -0.724459918 | 2.538669821 |
| Prelp | 162.3998482 | -0.190378161 | 0.213895367 |
| Gm1627 | 0.141089779 | -1.1571752 | 2.901102929 |
| Gm18970 | 0.109124069 | -0.24357403 | 3.724945049 |
| Fmod | 152.7762662 | -2.175891075 | 0.366935389 |
| Btg2 | 283.4370204 | 0.621038702 | 0.29136738 |
| Gm38140 | 0.628069627 | 0.958277626 | 1.325675569 |
| Gm8618 | 0.104637558 | 0.23732098 | 3.724945049 |
| Chit1 | 1.035276703 | -0.360615122 | 1.092804603 |
| Chil1 | 168.3470407 | 1.347793881 | 0.25282483 |
| Mybph | 3.909958598 | 2.528722306 | 0.723607246 |
| Adora1 | 2485.958102 | -0.120023104 | 0.073327774 |
| Gm24517 | 0.075043205 | 0.237309859 | 3.724945049 |
| Myog | 0.249503106 | 1.141214347 | 2.112394226 |
| Ppfia4 | 1225.389552 | -0.933524965 | 0.113323744 |
| Tmem183a | 645.1063955 | 0.14255674 | 0.084679042 |
| 4933406M09Rik | 0 | NA | NA |
| Cyb5r1 | 155.8595938 | 1.054179929 | 0.112034002 |
| Adipor1 | 881.11371 | 0.488644542 | 0.074101004 |
| Platr1 | 1.832214876 | 2.273082053 | 1.064116151 |
| Gm37935 | 2.337573936 | -0.602525136 | 0.782741161 |
| Klhl12 | 493.9899022 | -0.320764422 | 0.080123831 |
| Rabif | 301.4519563 | 0.059027333 | 0.082083405 |
| Gm25612 | 0.720424393 | -0.290846176 | 1.188017374 |
| Gm15454 | 0.589661845 | 0.988136037 | 1.277531208 |
| Gm29376 | 0.461892809 | -2.367989515 | 1.425596206 |
| Mgat4e | 0.04393175 | -0.243571806 | 3.724945049 |
| Kdm5b | 1415.896103 | -0.232505304 | 0.081662747 |
| Gm26783 | 4.085067263 | -0.843218193 | 0.554969314 |
| Syt2 | 2912.71677 | -0.348746362 | 0.134311013 |
| Ppp1r12b | 1746.350506 | -0.703080272 | 0.095326319 |
| Gm37949 | 36.84559847 | -1.109887572 | 0.328880425 |
| Gm22232 | 0.044191549 | -0.243571806 | 3.724945049 |
| Gm37677 | 160.3400663 | -1.089607618 | 0.241599577 |
| Gm28892 | 1.029532974 | 0.575919238 | 1.437114782 |
| Ube2t | 14.99856726 | 0.590769242 | 0.278351189 |
| Lgr6 | 136.8876649 | 0.019248183 | 0.276993967 |
| Ptprv | 31.89205934 | -1.648975642 | 0.355316262 |
| Gm10535 | 0.133946075 | 0.7191812 | 3.626570343 |

| | | | |
|---|---|---|---|
| Ptpn7 | 31.7926034 | 2.406065995 | 0.311083953 |
| Arl8a | 2134.195625 | 0.209684242 | 0.060979316 |
| Gm26280 | 0.068097359 | -0.243569582 | 3.724945049 |
| Gm15445 | 11.41385022 | -0.218122799 | 0.433859606 |
| Gpr37l1 | 2282.91187 | 0.428202233 | 0.148701527 |
| Gm4204 | 333.7656317 | -0.061633274 | 0.105746185 |
| Gm26642 | 0.168999367 | 0.661544699 | 2.739468562 |
| Elf3 | 3.18304154 | 1.284931243 | 0.626497937 |
| Rnpep | 506.4393182 | 0.720933411 | 0.077799524 |
| Timm17a | 833.5513504 | -0.206682394 | 0.065417593 |
| Lmod1 | 6.027761192 | -0.29118594 | 0.464329235 |
| Gm22796 | 0.04393175 | -0.243571806 | 3.724945049 |
| Shisa4 | 891.8069629 | -0.130626106 | 0.08684164 |
| Ipo9 | 1823.820265 | -0.26329725 | 0.047271391 |
| Gm28277 | 10.85246338 | -0.97392284 | 0.449394886 |
| Nav1 | 1957.583854 | -0.738806166 | 0.101017216 |
| Mir1231 | 0 | NA | NA |
| 2610012C04Rik | 6.062032631 | -1.511369135 | 0.462619638 |
| Gm37333 | 28.22584607 | -1.319526899 | 0.403938346 |
| Gm4793 | 13.11457074 | -0.869242528 | 0.320825627 |
| Gm38399 | 26.61474392 | 1.448882185 | 0.225696239 |
| Csrp1 | 2908.739873 | 1.030598227 | 0.153098403 |
| Phlda3 | 300.6871226 | 1.011822155 | 0.156435998 |
| Tnni1 | 43.38678691 | 0.861581903 | 0.220654195 |
| Lad1 | 13.14689316 | -0.89193161 | 0.391658139 |
| Tnnt2 | 21.42389219 | -1.029349959 | 0.308505767 |
| Gm28883 | 1.230445023 | 0.064541061 | 0.936637926 |
| Pkp1 | 4.016384854 | -0.24642564 | 0.653065603 |
| Gm36938 | 0.07796279 | 1.072698098 | 3.721829099 |
| Igfn1 | 106.9666211 | -2.052707241 | 0.29387953 |
| Tmem9 | 526.3167638 | 0.569021424 | 0.110945053 |
| Ascl5 | 1.900208836 | 0.308105055 | 0.689112832 |
| Cacna1s | 20.1854872 | 0.449306005 | 0.273971502 |
| Gm37014 | 0 | NA | NA |
| Gm37759 | 0.707021654 | 0.369083272 | 1.346698793 |
| Gm15850 | 4.717623853 | -0.540219461 | 0.49360855 |
| Kif21b | 1190.983809 | -0.510567778 | 0.095336842 |
| Mroh3 | 7.987918312 | 1.246125591 | 0.391508027 |
| 5730559C18Rik | 56.42049381 | 1.350653992 | 0.18160685 |
| Gm26568 | 0.061920613 | 0.718207094 | 3.724945049 |
| A430034D21Rik | 0 | NA | NA |
| Gm37552 | 0.476395801 | 0.224427713 | 1.287284809 |
| Gm26781 | 4.326125698 | -0.093660535 | 0.446136238 |
| Gpr25 | 16.1131376 | -0.715571497 | 0.270515052 |
| Camsap2 | 3490.88954 | -0.179688754 | 0.051648217 |
| 9230116N13Rik | 6.573994958 | 0.550420281 | 0.523117466 |
| Ddx59 | 80.65548895 | 0.370778263 | 0.135699816 |

| | | | |
|---|---|---|---|
| Kif14 | 11.46902475 | 1.788446115 | 0.393097306 |
| Gm29486 | 0.097139547 | 0.718207094 | 3.724945049 |
| Gm29485 | 0.285872948 | -1.024226464 | 2.080135064 |
| Gm37799 | 0.092691418 | 1.183079489 | 3.724560917 |
| Gm33994 | 0.860407559 | 2.864167443 | 1.10975795 |
| 4933409D19Rik | 0.218406194 | 1.873529017 | 2.130879821 |
| Platr23 | 2.85951047 | 1.46389408 | 0.65638927 |
| Zfp281 | 1236.560802 | 0.749991875 | 0.109137939 |
| Gm19705 | 24.95245937 | 0.959227685 | 0.225444626 |
| Platr22 | 0.971167388 | 0.890335198 | 1.15278347 |
| Gm24086 | 0.066050875 | 0.71820487 | 3.724945049 |
| Gm22132 | 0 | NA | NA |
| Gm8762 | 0 | NA | NA |
| Gm29718 | 0.856212859 | -0.362369159 | 1.107975653 |
| Gm17781 | 0 | NA | NA |
| Nr5a2 | 2.338360347 | 0.661469012 | 0.669704703 |
| Gm23763 | 0 | NA | NA |
| Gm25609 | 0.046345709 | 0.71820487 | 3.724945049 |
| Gm37057 | 0.333318405 | -0.724461668 | 1.736697444 |
| 1700006P03Rik | 0 | NA | NA |
| Gm23534 | 0 | NA | NA |
| Gm37903 | 0.499362493 | -2.4326201 | 1.56586478 |
| Gm22727 | 0 | NA | NA |
| Gm37099 | 0.251787852 | 0.23732545 | 2.081282806 |
| Gm37293 | 0 | NA | NA |
| Gm4258 | 79.79539825 | -0.800585779 | 0.248763539 |
| Gm37298 | 205.364216 | -0.983815094 | 0.197357563 |
| Gm26979 | 0 | NA | NA |
| Gm37124 | 62.91602665 | -1.005620006 | 0.231967465 |
| A130050O07Rik | 109.5267668 | -0.89896547 | 0.259386137 |
| Gm26936 | 0 | NA | NA |
| Gm26930 | 92.36817408 | -0.952086169 | 0.274115706 |
| D130019J16Rik | 127.5229106 | -0.704712392 | 0.211688414 |
| Gm37534 | 242.1756396 | -0.668453627 | 0.198732065 |
| Mir181a-1 | 0.399769715 | -0.163963678 | 1.517709569 |
| A430106G13Rik | 56.85494254 | -0.623046527 | 0.273278752 |
| Mir181b-1 | 0 | NA | NA |
| Gm37632 | 17.06609017 | -0.490774014 | 0.252019142 |
| Gm23782 | 0 | NA | NA |
| Ptprc | 1013.820159 | 4.939741948 | 0.192312493 |
| A130071D04Rik | 74.31188421 | 5.494426173 | 0.421286585 |
| Gm18667 | 0 | NA | NA |
| Gm28556 | 0 | NA | NA |
| 1700113B19Rik | 0 | NA | NA |
| Atp6v1g3 | 0.18576184 | 1.601702808 | 3.71335388 |
| Gm28500 | 0.232206954 | -0.711958894 | 2.55780212 |
| Nek7 | 865.5502947 | 0.46961032 | 0.122197521 |

| | | | |
|---|---|---|---|
| Gm28501 | 0.215166026 | -0.243568332 | 2.556929146 |
| Gm37080 | 8.379318968 | 0.209127456 | 0.446969938 |
| Gm37101 | 16.00512593 | 0.913140711 | 0.364497472 |
| Gm5833 | 0.04393175 | -0.243571806 | 3.724945049 |
| Gm8790 | 0 | NA | NA |
| Lhx9 | 105.8300051 | -0.186054029 | 0.103920525 |
| 2310009B15Rik | 35.43042701 | -0.04620043 | 0.186663266 |
| 1700019P21Rik | 0 | NA | NA |
| n-R5s217 | 0.075043205 | 0.237309859 | 3.724945049 |
| Gm3933 | 0 | NA | NA |
| Gm37045 | 1.708528675 | 0.348551816 | 0.702055234 |
| Dennd1b | 1082.456846 | 0.510210508 | 0.075849131 |
| Gm37423 | 58.57700844 | -0.100679949 | 0.253742555 |
| AC117623.1 | 17.28746858 | 0.16693692 | 0.356103676 |
| Gm18553 | 0.072025462 | 0.237312084 | 3.724945049 |
| Gm16305 | 0 | NA | NA |
| Gm37470 | 63.54316749 | 0.177285933 | 0.19661718 |
| RP23-114L12.4 | 43.15415538 | -0.085189679 | 0.249266736 |
| Gm37655 | 0.107078754 | 0.236669296 | 3.724898107 |
| Gm16304 | 0 | NA | NA |
| Gm37718 | 31.22466683 | 0.059900398 | 0.254784026 |
| Crb1 | 30.89787103 | -0.315554964 | 0.259680036 |
| 4933436E23Rik | 0.112288909 | -0.724455696 | 3.724945049 |
| Gm34816 | 0 | NA | NA |
| Zbtb41 | 682.7308389 | -0.155591832 | 0.071891652 |
| Aspm | 33.81106085 | 1.155195309 | 0.344608052 |
| F13b | 0.541494639 | 1.941545005 | 1.408591273 |
| Cfhr1 | 0.094294407 | 0.23732098 | 3.724945049 |
| Cfhr3 | 0 | NA | NA |
| Gm26271 | 0 | NA | NA |
| Gm4788 | 0.298414574 | -0.214870003 | 1.809491374 |
| Gm25619 | 0 | NA | NA |
| Cfhr2 | 0.451902873 | -0.48406587 | 1.316707293 |
| Gm16332 | 4.277969054 | 1.072854003 | 0.479856933 |
| Gm5837 | 9.46935442 | 0.323196649 | 0.339120392 |
| Gm5834 | 1.243360777 | 0.251090762 | 1.040424077 |
| Gm38266 | 0 | NA | NA |
| Gm26048 | 0 | NA | NA |
| Gm38019 | 0.981599412 | -0.202304771 | 1.013959794 |
| Cfh | 496.518771 | 1.002989086 | 0.131300674 |
| Kcnt2 | 523.8644755 | -0.926998543 | 0.118651426 |
| Gm37777 | 49.81740225 | -1.691082312 | 0.248319065 |
| 4930590L20Rik | 0 | NA | NA |
| Gm37371 | 0.255648062 | -0.340187584 | 1.828685357 |
| Gm4845 | 0 | NA | NA |
| Gm25457 | 0 | NA | NA |
| Gm22016 | 0 | NA | NA |

| | | | |
|---|---|---|---|
| Gm37599 | 0.182775946 | 0.237309473 | 3.335217586 |
| Gm37142 | 0 | NA | NA |
| Gm37604 | 0 | NA | NA |
| Gm18183 | 0.075043205 | 0.237309859 | 3.724945049 |
| Gm28724 | 0 | NA | NA |
| Gm37796 | 0 | NA | NA |
| Gm5835 | 7.977887189 | 0.296148563 | 0.380135381 |
| Cdc73 | 1577.910083 | -0.059165248 | 0.054710868 |
| B3galt2 | 609.7109376 | -0.78313842 | 0.087566737 |
| Gm6397 | 3.497495822 | 0.331754663 | 0.574292368 |
| Glrx2 | 603.4570244 | -0.483571345 | 0.063122537 |
| Trove2 | 1714.422575 | 0.046085819 | 0.098197272 |
| Uchl5 | 265.6573748 | -0.267369541 | 0.140880839 |
| Gm29170 | 125.043805 | -0.372933987 | 0.117939186 |
| Gm25663 | 0 | NA | NA |
| Rgs2 | 585.4803157 | 0.496773313 | 0.203111943 |
| Rgs13 | 2.523201682 | 1.181596052 | 0.699751596 |
| Rbm6-ps2 | 0.220340683 | 0.730544822 | 2.190228816 |
| Rgs1 | 170.7604979 | 8.136123831 | 0.673600302 |
| Gm7266 | 0.253428164 | 1.210747498 | 2.095827001 |
| Rgs21 | 0 | NA | NA |
| Gm23672 | 0 | NA | NA |
| Rgs18 | 3.121267796 | 2.784099868 | 0.810058904 |
| Gm8856 | 0 | NA | NA |
| Gm22681 | 0 | NA | NA |
| Gm25708 | 0 | NA | NA |
| Gm6550 | 0 | NA | NA |
| Gm5262 | 0 | NA | NA |
| Gm25460 | 0 | NA | NA |
| Gm28392 | 0 | NA | NA |
| Gm38269 | 0.279407529 | -0.243563821 | 2.076631989 |
| Gm29514 | 42.66995015 | -0.779067861 | 0.217919188 |
| Gm5263 | 0.047948698 | -0.243569582 | 3.724945049 |
| Gm38187 | 45.19927449 | -1.066804337 | 0.287914711 |
| Brinp3 | 455.9540705 | -0.513282103 | 0.088370697 |
| Gm15583 | 12.60536537 | -0.456866514 | 0.39028232 |
| A230059L01Rik | 0.205212924 | 0.71821544 | 2.646361116 |
| Gm15584 | 0.127515837 | 1.482540432 | 3.685333214 |
| Gm37749 | 0.118371171 | 0.718202646 | 3.724945049 |
| Gm9931 | 0 | NA | NA |
| Gm17867 | 0.136963818 | 0.718204821 | 3.59954433 |
| Gm38284 | 0.044544648 | 0.237309859 | 3.724945049 |
| Gm29516 | 0 | NA | NA |
| Gm29515 | 0 | NA | NA |
| Gm22966 | 0 | NA | NA |
| Gm19891 | 0 | NA | NA |
| Gm18505 | 0.044191549 | -0.243571806 | 3.724945049 |

| | | | |
|---|---|---|---|
| Gm4034 | 0 | NA | NA |
| Gm29022 | 0 | NA | NA |
| Gm5703 | 0 | NA | NA |
| Gm15443 | 3.976562986 | 0.228989458 | 0.487533893 |
| Gm18751 | 0 | NA | NA |
| Gm29020 | 0 | NA | NA |
| Gm18701 | 0 | NA | NA |
| Gm37630 | 0.062426873 | 0.237312084 | 3.724945049 |
| Gm38189 | 0.200297358 | -0.243568666 | 2.615966688 |
| Gm29398 | 0.199606257 | 0.237329886 | 2.218693415 |
| Gm37096 | 0.086648695 | -0.72445792 | 3.724945049 |
| Gm37775 | 0.044544648 | 0.237309859 | 3.724945049 |
| Gm23535 | 0 | NA | NA |
| n-R5s218 | 0 | NA | NA |
| Pla2g4a | 121.1500737 | 1.967540252 | 0.214538341 |
| n-R5s219 | 0 | NA | NA |
| Gm19136 | 0.350167968 | 0.176055762 | 1.585463992 |
| AC114655.1 | 0.088736197 | -0.243576254 | 3.724945049 |
| Ptgs2os | 10.82272229 | 1.116157439 | 0.366508913 |
| Ptgs2 | 389.8698901 | 2.331081875 | 0.35268983 |
| Gm9687 | 0.529068976 | -1.630710541 | 1.253175268 |
| Ptgs2os2 | 1.911122854 | 0.962031701 | 0.793452285 |
| Gm15453 | 44.68224246 | 1.405867009 | 0.23850468 |
| Gm4322 | 0.586444394 | -0.871746583 | 1.245399855 |
| Pdc | 0.117760149 | -0.243576254 | 3.724945049 |
| 2310030A07Rik | 0.378962075 | 0.718227392 | 1.528537616 |
| Gm37764 | 0 | NA | NA |
| BC003331 | 544.608875 | 0.162984876 | 0.063768822 |
| Tpr | 2374.975998 | 0.147805771 | 0.081529759 |
| Prg4 | 160.3573172 | 2.569624497 | 0.388137494 |
| Ska2l-ps | 0.636218932 | -1.280063132 | 1.258297152 |
| Hmcn1 | 64.89396023 | -1.161770862 | 0.167189642 |
| Gm20631 | 0.097139547 | 0.718207094 | 3.724945049 |
| Gm29188 | 0 | NA | NA |
| Gm22756 | 0 | NA | NA |
| Gm37671 | 0.106358623 | -0.24357403 | 3.724945049 |
| Gm19087 | 0 | NA | NA |
| Gm8941 | 0.044544648 | 0.237309859 | 3.724945049 |
| C730036E19Rik | 0.469314954 | 1.199124328 | 1.553733539 |
| Gm24402 | 0 | NA | NA |
| Gm10138 | 35.28446059 | -1.294254626 | 0.200635004 |
| Ivns1abp | 1648.737111 | -0.295715688 | 0.074869035 |
| Swt1 | 331.9812924 | -0.261147032 | 0.113428404 |
| Trmt1l | 460.2408708 | 0.146292909 | 0.193881132 |
| mmu-mir-7682 | 1.074470705 | 0.606990689 | 0.926021448 |
| Gm37893 | 50.46302794 | -0.762030805 | 0.301866164 |
| Rnf2 | 494.0673276 | 0.510348508 | 0.097566421 |

| | | | |
|---|---|---|---|
| Gm36527 | 43.64518431 | -0.278558748 | 0.348592671 |
| Gm29072 | 0 | NA | NA |
| Gm19503 | 0.173853943 | 1.25450333 | 2.738478996 |
| Gm28181 | 0.049553047 | 0.71820487 | 3.724945049 |
| Fam129a | 103.521438 | 1.614202777 | 0.137653566 |
| 2810414N06Rik | 13.51786206 | -0.251966295 | 0.298309007 |
| Edem3 | 1232.170216 | 0.249050157 | 0.063284911 |
| 1700025G04Rik | 1157.852342 | -0.625654953 | 0.061956413 |
| Gm25279 | 0.423681387 | 0.806528963 | 1.722229482 |
| Gm28610 | 0.118371171 | 0.718202646 | 3.724945049 |
| Gm8964 | 0 | NA | NA |
| Gm38238 | 0.047948698 | -0.243569582 | 3.724945049 |
| Tsen15 | 205.9704903 | 0.06952222 | 0.090386963 |
| Colgalt2 | 66.2940545 | -0.463678259 | 0.144288674 |
| Gm15618 | 0.134880151 | 1.580366464 | 2.92708711 |
| Rgl1 | 1379.502362 | 0.228136352 | 0.08996518 |
| Gm15479 | 6.574668662 | -1.342043692 | 0.48301494 |
| Gm7278 | 19.06484076 | -1.348394243 | 0.318525312 |
| Gm28792 | 7.043371214 | -2.025893309 | 0.486817582 |
| Apobec4 | 0.396489138 | 0.68446566 | 1.555503935 |
| Arpc5 | 1009.34046 | 1.056928895 | 0.12005901 |
| Ncf2 | 128.1024071 | 2.924020672 | 0.199545768 |
| Smg7 | 2499.293702 | 0.654996906 | 0.085988859 |
| Gm28791 | 48.57147184 | -0.215832151 | 0.254384396 |
| 1700040K01Rik | 0.217189781 | 0.541998101 | 2.165401053 |
| Gm8976 | 0.43436012 | 0.756888051 | 1.740600579 |
| Nmnat2 | 3031.207894 | -0.066292336 | 0.076231022 |
| Gm28960 | 0 | NA | NA |
| Lamc2 | 154.9877848 | 0.708482856 | 0.13759271 |
| Lamc1 | 1030.31094 | 0.879320568 | 0.121824125 |
| Gm38009 | 27.75065565 | -0.622325484 | 0.28687019 |
| Gm8818 | 0.931703337 | 3.355384202 | 1.165196504 |
| E330020D12Rik | 5.955571085 | 0.780366828 | 0.397204161 |
| Shcbp1l | 22.81337923 | -1.475651967 | 0.279888646 |
| Gm25588 | 0.946605489 | -1.726480958 | 1.310866655 |
| Gm37669 | 3.285918002 | -1.854458611 | 0.62374498 |
| Dhx9 | 2116.613299 | 0.260762751 | 0.077458684 |
| Npl | 38.83715233 | 0.153443989 | 0.212782621 |
| Gm6652 | 5.116277746 | 0.992261499 | 0.601721074 |
| Rgs8 | 2498.85835 | -0.466319402 | 0.16004879 |
| Gm29529 | 0.238359071 | 0.260178449 | 2.11567261 |
| 5530400K19Rik | 2.012029913 | 2.200264527 | 0.84739925 |
| C230024C17Rik | 0.83423333 | -0.045762925 | 1.102729187 |
| Rgs16 | 197.3246427 | 0.149302726 | 0.132503609 |
| Gm28513 | 0.329560409 | 1.164192507 | 1.771919072 |
| Rnasel | 213.55156 | 1.434229541 | 0.146735703 |
| Gm28512 | 4.354440161 | -0.4665985 | 0.549247835 |

| | | | |
|---|---|---|---|
| Rgsl1 | 23.12760851 | -1.370859565 | 0.323272221 |
| Teddm2 | 33.79161557 | -1.101375383 | 0.300330019 |
| Teddm1b | 0.047948698 | -0.243569582 | 3.724945049 |
| Gm38352 | 5.642899575 | -2.123398249 | 0.546541238 |
| Teddm1a | 0.061920613 | 0.718207094 | 3.724945049 |
| Mir8114 | 0.170693195 | 1.254668918 | 2.754365917 |
| Glul | 15318.94443 | -0.43291682 | 0.158661352 |
| Gm29290 | 0.209920398 | 0.246671621 | 2.624885119 |
| Gm29291 | 0.447162105 | -0.101561958 | 1.31571575 |
| Gm28286 | 0.825983375 | -3.062594848 | 1.141840862 |
| A830008E24Rik | 2.805075798 | -0.730565386 | 0.64555058 |
| Gm25336 | 0 NA | | NA |
| Zfp648 | 4.036848141 | -0.031499065 | 0.541575008 |
| 4930452N14Rik | 0.785070532 | 1.403461796 | 1.253406265 |
| 4930532M18Rik | 0.278668908 | -0.243563663 | 1.798586118 |
| n-R5s220 | 0.221251082 | 1.181583011 | 2.57570221 |
| Gm18727 | 0.332553061 | 1.633244836 | 1.951456385 |
| Cacna1e | 5959.244093 | -1.015841946 | 0.15777827 |
| Gm29282 | 1.600249739 | 0.083041674 | 0.756279566 |
| Gm29441 | 0.044544648 | 0.237309859 | 3.724945049 |
| Gm9530 | 4.308798137 | -0.641746695 | 0.487362217 |
| Ier5 | 750.1768822 | 0.701456806 | 0.146329064 |
| Mr1 | 88.50137408 | 0.498440934 | 0.152849445 |
| Stx6 | 782.513869 | -0.157466601 | 0.052787036 |
| Gm8850 | 14.83397756 | -0.891533716 | 0.376976514 |
| BC034090 | 156.7399653 | 0.857841221 | 0.103814691 |
| Gm37622 | 0.256551671 | 0.71821729 | 2.467846442 |
| Xpr1 | 2184.107984 | -0.16729836 | 0.058844139 |
| Gm22569 | 0 NA | | NA |
| Gm5532 | 8.533642962 | 2.850522705 | 0.556793298 |
| Acbd6 | 499.0889261 | 0.124688063 | 0.082330448 |
| Lhx4 | 8.533691663 | -0.370796755 | 0.862143035 |
| Gm37089 | 0 NA | | NA |
| Qsox1 | 389.4008561 | 0.483071912 | 0.087364213 |
| Gm37539 | 0 NA | | NA |
| Gm37571 | 0.346370079 | 2.23498019 | 2.004836123 |
| Cep350 | 1964.651825 | 0.026203737 | 0.07536094 |
| Gm38038 | 0 NA | | NA |
| Gm36934 | 0.108309328 | 0.237310006 | 3.724945049 |
| Gm37383 | 6.027087886 | -0.618499793 | 0.444752114 |
| Gm23744 | 0.061920613 | 0.718207094 | 3.724945049 |
| Gm9694 | 3.918626087 | 0.477988491 | 0.688173478 |
| Tor1aip1 | 1314.956962 | 1.993991015 | 0.142086796 |
| 9430034N14Rik | 86.42678797 | 0.995236166 | 0.280715972 |
| Tor1aip2 | 1490.887513 | 1.827028647 | 0.131523302 |
| Fam163a | 99.41997446 | -0.008866579 | 0.144307984 |
| Gm38113 | 0 NA | | NA |

| | | | |
|---|---|---|---|
| Tdrd5 | 36.39050195 | -0.462271284 | 0.241664434 |
| A430050A11Rik | 1.164548605 | -0.639595428 | 1.084467106 |
| 4930518J20Rik | 0.382980858 | 0.723989016 | 1.573934475 |
| Nphs2 | 3.238706364 | 0.899128837 | 0.590794167 |
| Axdnd1 | 8.537287892 | -0.110712783 | 0.394305979 |
| Gm2000 | 140.1670121 | 0.744860351 | 0.146933029 |
| Soat1 | 740.7127753 | 1.053923287 | 0.14835447 |
| Gm37648 | 0.139856492 | 0.237322262 | 3.574411025 |
| Gm10031 | 2.652532074 | -0.17739592 | 0.634476269 |
| AC114588.1 | 0 | NA | NA |
| Abl2 | 1607.186212 | 0.254688006 | 0.107672513 |
| Gm38043 | 100.360574 | -0.636778301 | 0.30031299 |
| Tor3a | 2329.182894 | 5.116182992 | 0.248577357 |
| Fam20b | 1176.797779 | 0.04063323 | 0.050639806 |
| Gm19918 | 0 | NA | NA |
| Gm15428 | 0.152810971 | 1.199997388 | 2.844490072 |
| Ralgps2 | 727.6286885 | -0.504542416 | 0.088474905 |
| Gm28693 | 10.3212494 | -0.966189773 | 0.408259424 |
| Angptl1 | 9.659502383 | -0.525267192 | 0.323977007 |
| Gm37802 | 27.9942778 | -0.919774813 | 0.287062328 |
| Gm37943 | 22.83945072 | -1.569852415 | 0.394867564 |
| 4930439D14Rik | 2.895518095 | -0.895330369 | 0.633054199 |
| Gm28694 | 0 | NA | NA |
| Gm10531 | 0.066050875 | 0.71820487 | 3.724945049 |
| Tex35 | 6.225377881 | 1.24240552 | 0.956213288 |
| Rasal2 | 2052.323456 | -0.316018791 | 0.064314229 |
| Gm37788 | 31.13971489 | -0.637719411 | 0.237521759 |
| Gm37578 | 17.57312196 | -1.021978247 | 0.335147021 |
| Gm6058 | 12.1668676 | -0.747918716 | 0.37952401 |
| Gm37060 | 72.32838032 | -0.728228802 | 0.22664867 |
| 2810025M15Rik | 74.04682442 | 0.47933996 | 0.151023977 |
| BC026585 | 13.64787826 | 0.76274634 | 0.385455338 |
| Gm15486 | 0.137416547 | -0.240825018 | 3.63730774 |
| Sec16b | 13.26555679 | 0.588399691 | 0.295053906 |
| Gm37682 | 0 | NA | NA |
| Gm7412 | 0 | NA | NA |
| Gm37713 | 0 | NA | NA |
| Gm38213 | 0 | NA | NA |
| Gm38256 | 0.119234754 | -0.243576254 | 3.724945049 |
| Brinp2 | 1002.677425 | -0.701008458 | 0.124928343 |
| Astn1 | 3735.831764 | -0.66317555 | 0.063126609 |
| Gm37861 | 45.70823912 | -1.037226465 | 0.346181475 |
| Gm37679 | 1.033175386 | -2.444661521 | 1.126904702 |
| Mir488 | 2.924473085 | -1.125327614 | 0.77134365 |
| Gm37367 | 35.2279183 | -1.484343832 | 0.357633847 |
| Gm37848 | 2.466912353 | -3.595993827 | 0.871030121 |
| Pappa2 | 44.61674889 | 0.444750315 | 0.256262247 |

| | | | |
|---|---|---|---|
| 1600012P17Rik | 0.046345709 | 0.71820487 | 3.724945049 |
| 4930562F17Rik | 0 | NA | NA |
| Gm4953 | 0 | NA | NA |
| Gm7430 | 0.038981395 | 0.718207094 | 3.724945049 |
| Rfwd2 | 802.379122 | -0.138676446 | 0.103676231 |
| Gm37115 | 30.16231669 | -0.811523293 | 0.272929098 |
| Gm37088 | 0.075043205 | 0.237309859 | 3.724945049 |
| Scarna3a | 3.950984661 | -0.03509351 | 0.59587239 |
| mmu-mir-1843b | 0 | NA | NA |
| Gm31256 | 1.235093012 | -0.734557887 | 1.076005827 |
| Gm37757 | 0.066050875 | 0.71820487 | 3.724945049 |
| Tnr | 2666.999354 | -0.38286717 | 0.080008149 |
| Gm37731 | 39.71725266 | -1.428040196 | 0.292581574 |
| Gm10530 | 19.32941874 | -1.518248145 | 0.411855679 |
| Gm38039 | 0.270352934 | 1.172554502 | 1.788474954 |
| Gm24719 | 0 | NA | NA |
| Gm37294 | 0 | NA | NA |
| 4930523C07Rik | 65.68144704 | 0.856310087 | 0.192472846 |
| 4930562F07Rik | 0.26443487 | 1.096537527 | 1.783334947 |
| Tnn | 13.34929327 | 2.398226114 | 0.369784533 |
| Mrps14 | 388.8406474 | 0.105397428 | 0.092017596 |
| Cacybp | 851.1120869 | 0.763854912 | 0.132175441 |
| 4933417C20Rik | 5.899375495 | 0.132160748 | 0.451110252 |
| Rabgap1l | 3165.401473 | -0.443513202 | 0.064227724 |
| Gm36975 | 0.049553047 | 0.71820487 | 3.724945049 |
| Mir1927 | 6.385949591 | -0.789890104 | 0.414498593 |
| 9430087J23Rik | 22.80069575 | -1.082977209 | 0.261031992 |
| Gm37529 | 68.10051105 | -0.717350403 | 0.303606653 |
| Gm37212 | 65.71378451 | -0.686254914 | 0.250832258 |
| Gm37215 | 27.68718647 | -0.231810581 | 0.292913794 |
| Gm38302 | 37.68841078 | -0.597973361 | 0.356859572 |
| 6720464F23Rik | 73.41064915 | -0.580401763 | 0.252407781 |
| Gm23528 | 0 | NA | NA |
| Gm37873 | 11.00141775 | -0.621896125 | 0.364625299 |
| Gm37328 | 9.255702839 | -0.12807004 | 0.365873039 |
| Gm38329 | 12.82265238 | -0.224760055 | 0.278526263 |
| Gm38126 | 0.106465261 | 0.718202646 | 3.724945049 |
| Gm37083 | 0.417055923 | 0.093826953 | 1.49208459 |
| Gm38060 | 0.123841226 | 1.292225235 | 3.719439735 |
| Gm24988 | 0 | NA | NA |
| Gm6177 | 0.068097359 | -0.243569582 | 3.724945049 |
| Rc3h1 | 1747.242785 | -0.041890987 | 0.047010339 |
| Gm37052 | 30.04310741 | -0.265773083 | 0.24883268 |
| Gm37809 | 11.17045412 | -0.642105411 | 0.3690494 |
| Gm37653 | 63.56315506 | -0.511158332 | 0.222190001 |
| Serpinc1 | 4.738076569 | 0.068326054 | 0.466435191 |
| Gm37729 | 0.047948698 | -0.243569582 | 3.724945049 |

| | | | |
|---|---|---|---|
| Gm37072 | 1.390520144 | 0.14324893 | 0.950874311 |
| Zbtb37 | 638.7082528 | -0.43850183 | 0.09635271 |
| Gm37362 | 0.22549826 | 1.638241031 | 2.128349096 |
| Gas5 | 565.6631599 | 0.931029155 | 0.117603026 |
| Gm26224 | 0 | NA | NA |
| Gm22489 | 0 | NA | NA |
| Gm22357 | 0 | NA | NA |
| Gm23212 | 0 | NA | NA |
| Snord78 | 0 | NA | NA |
| Mir5117 | 0 | NA | NA |
| Snord80 | 0 | NA | NA |
| SNORD77 | 0 | NA | NA |
| Snord47 | 0 | NA | NA |
| Gm25789 | 0 | NA | NA |
| Dars2 | 123.0133515 | -0.208787069 | 0.129998094 |
| Gm37634 | 2.14016008 | -0.198358555 | 0.849210699 |
| Cenpl | 31.81213049 | 0.870341303 | 0.267815325 |
| Klhl20 | 300.3613963 | -0.003269054 | 0.114848802 |
| Ankrd45 | 441.8861501 | -0.253985116 | 0.121893602 |
| 4930469G21Rik | 0 | NA | NA |
| Gm6185 | 1.432525857 | 1.20103394 | 0.820969064 |
| Prdx6 | 2507.892268 | 1.362381361 | 0.222523738 |
| Gm19057 | 0 | NA | NA |
| Tnfsf4 | 0.470075119 | 2.747710125 | 1.474089603 |
| Tnfsf18 | 2.177622544 | 4.364577942 | 1.026884753 |
| Gm25488 | 0 | NA | NA |
| Gm37638 | 0 | NA | NA |
| Gm15429 | 9.641298891 | 0.266054055 | 0.326541172 |
| Gm5049 | 0 | NA | NA |
| Fasl | 19.90409191 | 3.645899478 | 0.38791097 |
| Gm37357 | 16.51548777 | -0.586021548 | 0.326076441 |
| Suco | 1095.050559 | 0.370227024 | 0.093348232 |
| Gm7496 | 0 | NA | NA |
| Gm31925 | 0.049553047 | 0.71820487 | 3.724945049 |
| 4930558K02Rik | 1.743960802 | 0.655490286 | 0.889605887 |
| Pigc | 130.768612 | -0.043759914 | 0.129206721 |
| Gm38304 | 0.16282664 | 1.267090096 | 3.411118614 |
| Dnm3 | 4093.372687 | -0.377286705 | 0.059195673 |
| 2810442N19Rik | 0.321884861 | -0.710270623 | 1.774869743 |
| Gm37767 | 5.981304473 | -1.591401149 | 0.51979793 |
| Dnm3os | 3.891078412 | 0.533430999 | 0.481030813 |
| Mir199a-2 | 0 | NA | NA |
| Mir214 | 0 | NA | NA |
| Gm16101 | 0 | NA | NA |
| Gm10176 | 1.890308503 | -0.275277827 | 0.999484557 |
| Mettl13 | 182.4821947 | 0.121732895 | 0.091903452 |
| Vamp4 | 850.2108809 | -0.132886558 | 0.055851281 |

| | | | |
|---|---|---|---|
| Gm5705 | 0.038981395 | 0.718207094 | 3.724945049 |
| Myoc | 87.94067266 | -1.564000306 | 0.452497336 |
| 7420461P10Rik | 0.321637403 | 2.242891068 | 1.994796695 |
| Prrc2c | 5776.579411 | 0.133426139 | 0.073428999 |
| Gm27616 | 4.734205595 | 0.388408669 | 0.438978545 |
| Fmo4 | 5.221071454 | 1.429628802 | 0.619844926 |
| Fmo1 | 94.46220945 | 0.144604259 | 0.177944834 |
| Fmo2 | 41.24708577 | -0.198868478 | 0.269304788 |
| Fmo6 | 0.536071372 | -0.132513698 | 1.354312104 |
| Fmo3 | 0.266623809 | 0.210789672 | 1.806782831 |
| Gm37273 | 0.04393175 | -0.243571806 | 3.724945049 |
| Mroh9 | 0.04393175 | -0.243571806 | 3.724945049 |
| Gm22671 | 0 | NA | NA |
| Gm37049 | 0.128271896 | -0.243573716 | 3.679089475 |
| Gm22434 | 0 | NA | NA |
| Prrx1 | 291.7242276 | 1.302318875 | 0.184212351 |
| Gm20471 | 0.226778023 | 0.237329592 | 2.143518328 |
| Gm26523 | 7.941957124 | 0.301690148 | 0.395026466 |
| Gm37644 | 11.16339375 | 0.395243626 | 0.325164863 |
| Gorab | 151.8566572 | 0.641428378 | 0.121118021 |
| Gm24940 | 0 | NA | NA |
| Gm29500 | 0.061920613 | 0.718207094 | 3.724945049 |
| Rpsa-ps1 | 2.716655785 | 0.666667729 | 0.597495152 |
| Mettl11b | 4.382455216 | -2.681158676 | 0.755143978 |
| Gm2343 | 0 | NA | NA |
| Kifap3 | 2725.982408 | -0.577177098 | 0.052046709 |
| Scyl3 | 425.2181155 | 0.259853362 | 0.103304163 |
| BC055324 | 25.44071188 | 0.000815196 | 0.221490924 |
| Mettl18 | 30.62632286 | 0.238001823 | 0.19982197 |
| Gm37119 | 0 | NA | NA |
| Sele | 15.49838402 | 7.419953303 | 1.157015238 |
| Sell | 313.8249993 | 11.70380262 | 0.969196061 |
| Selp | 27.22107536 | 8.448938566 | 0.917105284 |
| Gm16587 | 0.246813481 | 1.991878576 | 1.785371673 |
| Gm16548 | 0.29488824 | -0.768535452 | 1.810213875 |
| F5 | 60.57846656 | -0.533799104 | 0.234642984 |
| Gm16588 | 0.11197992 | 0.718202646 | 3.724945049 |
| Slc19a2 | 224.9421596 | 0.274642626 | 0.110991659 |
| Ccdc181 | 152.8998847 | -0.024148128 | 0.151304405 |
| Blzf1 | 283.9203218 | 0.20552809 | 0.083009316 |
| Nme7 | 157.2285977 | -0.406224498 | 0.106262308 |
| Gm37411 | 15.43722212 | -0.834505518 | 0.413266603 |
| Atp1b1 | 14543.60903 | -0.325249452 | 0.05868666 |
| Gm32391 | 15.06111228 | 0.122471193 | 0.287930948 |
| Gm37732 | 0 | NA | NA |
| Gm26685 | 0.956865499 | 2.421454061 | 1.148236823 |
| D630023O14Rik | 0.652111363 | 0.944259295 | 1.267499781 |

| | | | |
|---|---|---|---|
| Gm37754 | 0.289849351 | -0.712527053 | 1.777838034 |
| Gm37853 | 0 | NA | NA |
| Gm37184 | 2.721089389 | 1.226619638 | 0.779798901 |
| Gm36945 | 0 | NA | NA |
| Dpt | 2.400487258 | 1.400972696 | 0.760958216 |
| Gm32569 | 2.899492925 | -0.149323772 | 0.574126999 |
| Gm20743 | 18.05075087 | 0.040006246 | 0.281072684 |
| Gm37403 | 2.741653471 | -0.690760629 | 0.556606114 |
| Xcl1 | 1.167394941 | 3.844900712 | 1.0439555 |
| 1700029M03Rik | 0.062426873 | 0.237312084 | 3.724945049 |
| Gm26042 | 0 | NA | NA |
| 4930568G15Rik | 2.644077932 | 0.284204041 | 0.603915977 |
| Tbx19 | 0.755699022 | 0.184923051 | 1.27728826 |
| Gm16550 | 1.768574416 | 0.363884006 | 0.712625676 |
| Sft2d2 | 402.8459024 | 0.183766823 | 0.126203217 |
| Tiprl | 413.2944695 | 0.002540445 | 0.091798488 |
| Gm38190 | 112.7353609 | -0.908312169 | 0.237666515 |
| Gpr161 | 332.1613004 | -0.311159926 | 0.10581873 |
| Dcaf6 | 876.2034936 | -0.401123058 | 0.064593687 |
| Gm38368 | 0.038981395 | 0.718207094 | 3.724945049 |
| Gm24637 | 0.087930927 | 0.236669296 | 3.724898107 |
| Gm37864 | 59.27064257 | -0.760315782 | 0.275215239 |
| Mpc2 | 532.4916413 | -0.25703731 | 0.056782986 |
| Gm37860 | 11.90237977 | -1.463578277 | 0.422629887 |
| Adcy10 | 12.42222246 | 0.564546569 | 0.326211559 |
| Gm18186 | 0.674049174 | 0.820968592 | 1.439981674 |
| Gm38325 | 1.250018007 | -2.459081998 | 1.044606179 |
| Gm23599 | 0.39032937 | -1.465489021 | 3.26657777 |
| Gm23208 | 0.712615371 | -0.094328905 | 1.131141704 |
| Gm38210 | 0.496695103 | 1.793623132 | 1.34282774 |
| Mpzl1 | 254.7787603 | -0.140261245 | 0.109503418 |
| Gm37559 | 11.16705096 | -1.924258268 | 0.568428278 |
| Gm18407 | 0.233595724 | -1.120986794 | 2.531372763 |
| Rcsd1 | 77.36566827 | 0.495440031 | 0.13606981 |
| AC124587.1 | 0 | NA | NA |
| Gm37073 | 0.097139547 | 0.718207094 | 3.724945049 |
| Gm34767 | 0.288888109 | 0.237328504 | 2.080058444 |
| Creg1 | 384.0157734 | 0.502683479 | 0.156064433 |
| Gm36972 | 0 | NA | NA |
| Gm16565 | 4.973903582 | 1.741190703 | 0.496887086 |
| Gm24808 | 0 | NA | NA |
| Cd247 | 27.67774213 | 5.100492887 | 0.541772693 |
| Gm23402 | 0.088736197 | -0.243576254 | 3.724945049 |
| Gm32999 | 0.098502579 | 0.23732098 | 3.724945049 |
| Gm37856 | 0.167459142 | 1.165958262 | 2.753748546 |
| Gm37469 | 0.04393175 | -0.243571806 | 3.724945049 |
| Pou2f1 | 796.3877359 | -0.435594228 | 0.109580126 |

| | | | |
|---|---|---|---|
| Gm17976 | 0 | NA | NA |
| Gm26665 | 0 | NA | NA |
| Gm7626 | 0 | NA | NA |
| Gm37923 | 0.047948698 | -0.243569582 | 3.724945049 |
| Dusp27 | 9.458378533 | -0.712027024 | 0.356347141 |
| Gpa33 | 0.260737493 | 1.210352386 | 2.05630574 |
| Mael | 2.369035115 | 0.690212802 | 0.992870432 |
| Ildr2 | 1753.501267 | -0.293766494 | 0.097530775 |
| 5330438I03Rik | 1.619090465 | -0.437804577 | 0.723236405 |
| Gm15853 | 26.8966923 | 0.039932985 | 0.217598956 |
| Tada1 | 290.2844388 | -0.151354502 | 0.082712099 |
| Pogk | 1000.722577 | -0.124971098 | 0.069846733 |
| Gm16418 | 40.55579905 | -0.514066418 | 0.201491372 |
| Gm33354 | 0 | NA | NA |
| Gm4846 | 0 | NA | NA |
| Gm22992 | 0.042716945 | -0.243571806 | 3.724945049 |
| Gm4847 | 0.370767229 | 0.23733153 | 1.708795898 |
| Fmo9 | 0.364366871 | 1.625782864 | 1.521051526 |
| Gm4954 | 0 | NA | NA |
| Gm37904 | 0.072025462 | 0.237312084 | 3.724945049 |
| Gm6286 | 20.94810405 | 1.133879057 | 0.267327787 |
| Gm16701 | 90.39573473 | -0.417924093 | 0.12806797 |
| Fam78b | 369.0147211 | -0.239852273 | 0.085553443 |
| Gm17868 | 0.454564407 | 0.579409188 | 1.687620611 |
| Gm37994 | 0.075043205 | 0.237309859 | 3.724945049 |
| Uck2 | 476.1615417 | 0.367084514 | 0.112936752 |
| Gm37982 | 0.042716945 | -0.243571806 | 3.724945049 |
| Gm2453 | 1.234537134 | -0.51804803 | 0.874149139 |
| Mir6347 | 0 | NA | NA |
| Gm37232 | 0.061920613 | 0.718207094 | 3.724945049 |
| Tmco1 | 304.4300019 | 0.297990555 | 0.110199193 |
| Gm25911 | 23.94632856 | 1.346735879 | 1.062811049 |
| Gm24182 | 0.992449309 | -0.894597414 | 1.015814099 |
| Aldh9a1 | 305.1290703 | -0.011636019 | 0.115746003 |
| Mgst3 | 487.8742445 | -0.820716207 | 0.116876706 |
| Lrrc52 | 0.467496845 | 1.019062237 | 1.4001792 |
| Rxrg | 90.84925895 | -0.922395126 | 0.142076662 |
| Lmx1a | 12.68353485 | 0.225023888 | 0.341265843 |
| Gm20711 | 0.175176114 | -0.306860756 | 3.35577183 |
| Pbx1 | 3030.795283 | -0.374548561 | 0.082612855 |
| Gm37524 | 93.77681393 | -0.966008387 | 0.266208571 |
| Gm38381 | 118.8604427 | -1.053691345 | 0.251809652 |
| Mir6348 | 0.044544648 | 0.237309859 | 3.724945049 |
| 1700063I16Rik | 0.141317957 | 0.718202955 | 3.59624077 |
| Mir6354 | 0 | NA | NA |
| Gm37839 | 0 | NA | NA |
| Gm5265 | 0 | NA | NA |

| | | | |
|---|---|---|---|
| Nuf2 | 31.87590655 | 0.352858094 | 0.347045528 |
| Gm18900 | 0.075043205 | 0.237309859 | 3.724945049 |
| Gm36980 | 0 | NA | NA |
| Rgs5 | 507.967864 | -0.893710634 | 0.171426336 |
| Rgs4 | 2513.987794 | -0.351300957 | 0.151111996 |
| Gm22114 | 0 | NA | NA |
| Gm37586 | 0 | NA | NA |
| 1700084C01Rik | 205.568575 | 0.809118205 | 0.179434224 |
| Gm15852 | 1.264098847 | -0.111735059 | 0.864919123 |
| Hsd17b7 | 333.9167156 | 0.531643567 | 0.128773306 |
| 3110045C21Rik | 6.064642518 | 0.805429908 | 0.382511205 |
| Ddr2 | 402.5981112 | 1.520878139 | 0.136877116 |
| Uap1 | 505.8478976 | 0.232534972 | 0.069373555 |
| Gm37748 | 2.515876782 | -0.515247911 | 0.607672623 |
| Gm37502 | 0.166709681 | -0.724459871 | 2.767631179 |
| Uhmk1 | 4307.233663 | -0.367922225 | 0.065662769 |
| 4930500M09Rik | 0.668413162 | 0.887940034 | 1.169253068 |
| Sh2d1b2 | 3.139971286 | 5.061966392 | 0.974809776 |
| Gm6345 | 2.814645699 | -0.260362523 | 0.611027635 |
| Sh2d1b1 | 8.798894628 | 3.973567179 | 0.605270571 |
| Gm7694 | 71.28747091 | 0.140258586 | 0.143023238 |
| Nos1ap | 949.0976046 | -0.68076902 | 0.170872865 |
| 1700015E13Rik | 0 | NA | NA |
| Gm7299 | 0.686918141 | 0.32413693 | 1.085088263 |
| Olfml2b | 107.0665219 | -0.166216479 | 0.121796042 |
| Atf6 | 1426.693892 | 0.117234876 | 0.057379999 |
| Gm23523 | 2.532308763 | 0.119771739 | 0.813842315 |
| Gm25235 | 4.986884554 | -0.522764664 | 0.705186656 |
| Gm9929 | 5.148652476 | -0.470164927 | 0.563196476 |
| Dusp12 | 141.0307956 | -0.10749731 | 0.123653484 |
| Gm26620 | 3.756679203 | 0.532200247 | 0.595040746 |
| Fcrlb | 20.36960655 | 2.866282031 | 0.440308703 |
| Fcrla | 4.0864204 | 4.10788269 | 1.051579216 |
| Gm2962 | 27.33055226 | 0.611602974 | 0.247993004 |
| Gm37030 | 0.072025462 | 0.237312084 | 3.724945049 |
| Fcgr2b | 292.4194751 | 4.082472858 | 0.213363121 |
| Gm38204 | 0.062426873 | 0.237312084 | 3.724945049 |
| Gm10522 | 0.106465261 | 0.718202646 | 3.724945049 |
| Fcgr4 | 282.2570289 | 11.34980735 | 0.892807758 |
| Gm26110 | 0.545665175 | 3.266594514 | 1.267708001 |
| Fcgr3 | 124.3411756 | 1.944487732 | 0.151395429 |
| Mir6546 | 0.741170898 | 2.882268515 | 1.176055792 |
| Gm27552 | 0.046345709 | 0.71820487 | 3.724945049 |
| Gm27844 | 0 | NA | NA |
| Cfap126 | 16.16273733 | 0.608184472 | 0.354148718 |
| Sdhc | 664.894161 | 0.049466079 | 0.107247263 |
| Mpz | 3.16266539 | 0.458910836 | 0.601546005 |

| | | | |
|---|---|---|---|
| Pcp4l1 | 1008.326566 | -0.29868296 | 0.109351771 |
| Nr1i3 | 2.346028351 | -1.14540495 | 0.684926135 |
| Tomm40l | 561.1166494 | 0.287889652 | 0.092969748 |
| Apoa2 | 0.57971966 | -0.667978992 | 1.305731576 |
| Fcer1g | 255.0199519 | 4.525372114 | 0.250999332 |
| Ndufs2 | 1322.178398 | 0.084277363 | 0.073944708 |
| Adamts4 | 381.9510903 | 0.42610661 | 0.122921818 |
| B4galt3 | 352.3072303 | 0.151081761 | 0.086217449 |
| Ppox | 128.0565181 | -0.464071451 | 0.133238955 |
| Usp21 | 286.6155179 | -0.249964429 | 0.098152423 |
| Ufc1 | 373.6742995 | 0.188928749 | 0.093556379 |
| Rpl27-ps1 | 1.899360943 | 1.520520574 | 0.79745939 |
| Gm25151 | 0 | NA | NA |
| Gm20045 | 91.15361111 | -0.694668419 | 0.222751792 |
| Dedd | 384.4686255 | 0.393173948 | 0.086210072 |
| Nit1 | 175.9259044 | -0.24081218 | 0.098145791 |
| Pfdn2 | 372.1457708 | -0.061658742 | 0.170651762 |
| Gm37154 | 10.37578366 | -1.240816647 | 0.384842706 |
| Klhdc9 | 28.71586709 | -0.633536828 | 0.238164323 |
| Gm37097 | 0.580816694 | -1.641416686 | 1.364649111 |
| Pvrl4 | 89.21914686 | 1.26773246 | 0.133868071 |
| Arhgap30 | 261.1304874 | 3.700223813 | 0.184712217 |
| Usf1 | 598.6805458 | 0.488442311 | 0.080924132 |
| Tstd1 | 8.585507017 | 0.261142821 | 0.34510943 |
| Gm26641 | 0.661565226 | -0.857469587 | 1.179965518 |
| F11r | 61.91267919 | 1.188480784 | 0.160175365 |
| B930036N10Rik | 1.495946937 | 0.431693823 | 0.921759509 |
| Gm24871 | 0 | NA | NA |
| Alyref2 | 25.03101443 | 0.739299097 | 0.329296796 |
| Gm9765 | 0.094831987 | -0.243571806 | 3.724945049 |
| Itln1 | 0.674729533 | 1.199123858 | 1.555763706 |
| Gm37450 | 0 | NA | NA |
| Cd244 | 24.4635172 | 5.236492372 | 0.575717596 |
| Ly9 | 63.06084566 | 5.851377803 | 0.433741488 |
| Slamf7 | 242.896543 | 9.15677883 | 0.697212715 |
| Gm37787 | 0.62421388 | 1.20198773 | 1.4616355 |
| Mir7683 | 0.465770878 | 2.482311649 | 1.646954533 |
| A630035G10Rik | 0.725223345 | 2.218869468 | 1.299467367 |
| Gm37981 | 0.097139547 | 0.718207094 | 3.724945049 |
| Cd48 | 54.8175684 | 5.087792988 | 0.397925289 |
| Gm37010 | 3.46582388 | 2.243267257 | 0.737306921 |
| Tma7-ps | 109.4366652 | 0.365180936 | 0.185600377 |
| Gm37693 | 0 | NA | NA |
| Gm19030 | 0.512648943 | 2.928531923 | 3.239285742 |
| Gm26412 | 0 | NA | NA |
| Gm22950 | 0 | NA | NA |
| Slamf1 | 7.891299907 | 2.779744735 | 0.504976894 |

| | | | |
|---|---|---|---|
| Cd84 | 247.5967198 | 4.786374593 | 0.258149373 |
| Gm10521 | 6.960180967 | 6.559263292 | 0.963314738 |
| Gm37065 | 0.847774459 | 1.384392828 | 1.508845902 |
| Slamf6 | 33.56933231 | 4.422096771 | 0.393882677 |
| Vangl2 | 264.8998109 | -0.262237146 | 0.100035307 |
| Nhlh1 | 18.19327503 | 1.14500239 | 0.506119727 |
| Ncstn | 716.1302082 | 0.622558617 | 0.074064731 |
| Copa | 2652.049841 | 0.495525388 | 0.066815925 |
| Gm38164 | 3.312973942 | -0.292010815 | 0.586845847 |
| Gm37756 | 0 | NA | NA |
| Pex19 | 729.4998742 | 0.060115039 | 0.078526805 |
| Gm10171 | 8.347475085 | 0.64735599 | 0.368870295 |
| Dcaf8 | 1703.591968 | -0.138481504 | 0.051660531 |
| Pea15a | 5072.520127 | 0.315027968 | 0.129694292 |
| Casq1 | 10.6483708 | 0.798428269 | 0.369391637 |
| Gm17224 | 5.574451365 | -0.876299987 | 0.884701108 |
| Atp1a4 | 4.799918733 | -0.169873138 | 0.839655907 |
| Gm37950 | 1.138245853 | -0.191024419 | 1.478169728 |
| Igsf8 | 2030.429245 | 0.210118941 | 0.093137736 |
| Atp1a2 | 9906.868086 | -0.541800659 | 0.137386091 |
| Kcnj9 | 920.7093185 | -0.882770006 | 0.167484578 |
| Gm36937 | 1.071485715 | 0.394677263 | 1.02130681 |
| Kcnj10 | 4831.592137 | -0.129607309 | 0.155325041 |
| Pigm | 357.594723 | 0.315774245 | 0.126560994 |
| Gm37145 | 0.222528853 | 1.207457414 | 2.574761424 |
| Gm7804 | 0.35063021 | 2.108538442 | 1.909758361 |
| Slamf9 | 59.21569036 | 4.59640928 | 0.328955267 |
| Igsf9 | 57.64957815 | -0.009679018 | 0.190991236 |
| Tagln2 | 422.7865102 | 3.624667203 | 0.214672284 |
| Gm37125 | 0.235978602 | 0.741141459 | 2.121839885 |
| Cfap45 | 24.01669692 | 0.118068938 | 0.309168173 |
| 4933439K11Rik | 2.636750292 | 0.01578412 | 0.736562602 |
| Vsig8 | 4.194719996 | 0.204414073 | 0.45323471 |
| Slamf8 | 27.5563302 | 4.095487786 | 0.43716368 |
| Fcrl6 | 0.659227985 | 3.491970068 | 1.378260064 |
| Dusp23 | 20.89498152 | 0.150676276 | 0.276280738 |
| Gm2710 | 3.560626721 | 0.497781042 | 0.577247738 |
| Gm22383 | 0 | NA | NA |
| Crp | 0.134618652 | -0.243572095 | 2.958498134 |
| Gm38051 | 0 | NA | NA |
| Gm24817 | 0 | NA | NA |
| 9430083B18Rik | 0 | NA | NA |
| Apcs | 0.049553047 | 0.71820487 | 3.724945049 |
| Gm26077 | 0.04393175 | -0.243571806 | 3.724945049 |
| Olfr16 | 0 | NA | NA |
| RP23-127L7.9 | 0.046345709 | 0.71820487 | 3.724945049 |
| RP23-127L7.8 | 0 | NA | NA |

| | | | |
|---|---|---|---|
| Olfr1409-ps1 | 0 | NA | NA |
| Olfr1408 | 0.042716945 | -0.243571806 | 3.724945049 |
| Olfr1407-ps1 | 0 | NA | NA |
| Olfr1406 | 0.153801061 | -0.243562187 | 2.838638549 |
| Olfr218 | 0 | NA | NA |
| Olfr1404 | 0 | NA | NA |
| Fcer1a | 0.138573873 | 1.200070582 | 3.628829444 |
| Olfr418 | 0 | NA | NA |
| Mptx2 | 0.116944185 | 1.302646088 | 3.719336636 |
| Ackr1 | 337.5743771 | 0.359955772 | 0.115305748 |
| Cadm3 | 5210.19823 | -0.449550147 | 0.102472083 |
| Aim2 | 59.97870837 | 2.78158702 | 0.226931931 |
| Gm4955 | 118.0231463 | 10.64486787 | 0.919750957 |
| BC094916 | 233.0659656 | 10.76405816 | 0.950413616 |
| AC084073.1 | 0 | NA | NA |
| Pydc4 | 70.39675328 | 8.890528343 | 0.895605904 |
| Pyhin1 | 372.8526333 | 10.87360762 | 0.882983346 |
| AC117682.1 | 0 | NA | NA |
| Pydc3 | 163.3167361 | 10.13907065 | 0.906116295 |
| Gm18752 | 7.837557856 | 6.801171068 | 0.909854177 |
| Gm7871 | 0.290529814 | 1.127328445 | 2.016067982 |
| AI607873 | 591.904346 | 8.922068087 | 0.465289829 |
| Ifi204 | 992.1935414 | 11.62831724 | 0.741215942 |
| Gm16340 | 27.30976778 | 6.471723336 | 0.688359189 |
| Gm7897 | 1.063697798 | 4.258116353 | 1.100990483 |
| Gm38050 | 43.49727825 | 8.924006655 | 0.881056907 |
| Mndal | 289.3519551 | 7.157719999 | 0.384507802 |
| Mnda | 254.7490093 | 11.29839382 | 0.953831939 |
| Ifi203 | 735.2718426 | 6.902532264 | 0.340092946 |
| Ifi202b | 1532.515879 | 10.2553198 | 0.507960055 |
| Gm18445 | 3.120173769 | 5.333616018 | 1.07242837 |
| Ifi205 | 186.9498831 | 11.4044614 | 1.000743535 |
| Olfr433 | 1.889349206 | 4.198687053 | 1.028207716 |
| Olfr432 | 0.498029041 | 2.911470631 | 1.590912353 |
| Olfr431-ps1 | 0.481158361 | 2.888720054 | 1.372630695 |
| Olfr430 | 0 | NA | NA |
| Olfr429 | 0.127515837 | 1.482540432 | 3.685333214 |
| RP23-343E12.13 | 0.066050875 | 0.71820487 | 3.724945049 |
| Olfr427 | 0 | NA | NA |
| Gm37417 | 0 | NA | NA |
| Olfr231 | 0 | NA | NA |
| Olfr424 | 0 | NA | NA |
| Olfr258-ps1 | 0 | NA | NA |
| Olfr421-ps1 | 0.097139547 | 0.718207094 | 3.724945049 |
| Olfr420 | 0 | NA | NA |
| RP23-343E12.14 | 0 | NA | NA |
| Spta1 | 4.811948516 | 1.731615038 | 0.642215925 |

| | | | |
|---|---|---|---|
| Olfr419 | 0 | NA | NA |
| Mptx1 | 0.066050875 | 0.71820487 | 3.724945049 |
| Olfr417 | 0 | NA | NA |
| Olfr416-ps1 | 0 | NA | NA |
| Olfr248 | 0 | NA | NA |
| Olfr414 | 0.117046891 | -0.724453471 | 3.724945049 |
| Olfr220 | 0.244461041 | 0.316998841 | 2.51399217 |
| Fmn2 | 1375.015257 | -0.506296116 | 0.056921174 |
| Grem2 | 314.3592299 | 0.402634524 | 0.170031038 |
| Rgs7 | 979.1331399 | -0.056051684 | 0.094122689 |
| 5730408A14Rik | 66.38004792 | -0.60736478 | 0.237821277 |
| Gm36307 | 0.200008102 | 1.980545204 | 2.610701512 |
| Fh1 | 588.3356071 | -0.153637603 | 0.077644437 |
| Kmo | 33.53845944 | 4.228062549 | 0.403853722 |
| Opn3 | 46.18549766 | -0.039924173 | 0.200924043 |
| Chml | 277.8742486 | -0.167324491 | 0.079463701 |
| Wdr64 | 0.404309506 | -1.700955579 | 1.482711164 |
| Gm25560 | 0.044191549 | -0.243571806 | 3.724945049 |
| Exo1 | 11.02950793 | 1.175380941 | 0.343288167 |
| Gm23805 | 0 | NA | NA |
| Gm38100 | 0.094097695 | 0.718200421 | 3.724945049 |
| Gm37668 | 9.723507462 | -0.463231382 | 0.396329844 |
| Pld5 | 273.9533392 | -0.486790168 | 0.098412291 |
| B020018G12Rik | 7.849527246 | -1.309459253 | 0.432834857 |
| Gm38081 | 26.68633732 | -1.321771489 | 0.326104176 |
| Gm37916 | 0.048949532 | -0.243569582 | 3.724945049 |
| Gm38101 | 8.572675195 | -0.209830284 | 0.356988284 |
| Gm17965 | 0.042716945 | -0.243571806 | 3.724945049 |
| Gm38152 | 0.194279094 | 1.292225235 | 3.719439735 |
| Gm36536 | 0.394395078 | -0.670290039 | 1.563003353 |
| Gm17966 | 0.161076545 | 1.165842935 | 2.787893623 |
| Gm26104 | 0 | NA | NA |
| Cep170 | 2565.856711 | -0.196864197 | 0.060675707 |
| 3110062G12Rik | 32.76611244 | -0.455448089 | 0.21796112 |
| Gm38104 | 128.4296084 | -0.507853692 | 0.185420344 |
| Gm36992 | 0.232229192 | -1.854763361 | 3.096265346 |
| Mir350 | 3.5612593 | -0.425806243 | 0.539332593 |
| Gm25288 | 0 | NA | NA |
| Sdccag8 | 235.1494786 | -0.314798473 | 0.167887941 |
| Gm38158 | 1.079846962 | -0.61109636 | 0.94036072 |
| Gm38053 | 0.047948698 | -0.243569582 | 3.724945049 |
| Gm25993 | 2.439810687 | -0.382994248 | 0.658117242 |
| Gm37246 | 8.305594911 | -1.159040803 | 0.469659629 |
| Gm37490 | 6.173150445 | -1.032688667 | 0.405575599 |
| Gm15423 | 5.069600121 | -0.144761536 | 0.522651741 |
| Gm37519 | 17.14460392 | -0.796358671 | 0.443892001 |
| Akt3 | 2779.8936 | -0.113096519 | 0.061452165 |

| | | | |
|---|---|---|---|
| Gm37960 | 79.23742208 | -0.486426033 | 0.216196402 |
| Gm37106 | 32.57544906 | -0.743716024 | 0.257878069 |
| Gm37598 | 54.69327838 | -0.198565077 | 0.295270541 |
| Gm38146 | 68.83238682 | -0.452822541 | 0.200583089 |
| Gm37706 | 171.6006066 | -0.36719837 | 0.241642256 |
| Gm37463 | 61.45825268 | -0.580147936 | 0.274313741 |
| Gm38328 | 3.391924355 | -0.565337809 | 0.538599989 |
| Gm26801 | 7.020008498 | -0.047395288 | 0.356116195 |
| Zbtb18 | 3593.895032 | -0.52765285 | 0.103685795 |
| Gm29856 | 8.835626795 | -0.705071456 | 0.560531426 |
| Gm37306 | 1.917068816 | -0.89475596 | 1.020066462 |
| Gm37280 | 0.17486102 | 0.087909285 | 2.709259681 |
| Gm37858 | 0.209701617 | 1.556073118 | 2.585494606 |
| 2310043L19Rik | 0.252619382 | 0.239808777 | 2.088635892 |
| 1700016C15Rik | 0.143022002 | 0.719176055 | 3.547735711 |
| Adss | 882.0855769 | -0.131033696 | 0.070687659 |
| Gm16585 | 0 | NA | NA |
| Gm7068 | 2.009888605 | -2.041536547 | 0.914465996 |
| Gm16432 | 157.8944835 | -1.19029275 | 0.136494843 |
| Gm37313 | 0.130017973 | 0.238301601 | 3.662714184 |
| Gm25312 | 0 | NA | NA |
| Gm8010 | 62.50190762 | -0.182858794 | 0.140876618 |
| Desi2 | 449.7295651 | -0.134853988 | 0.081688394 |
| Gm8023 | 0.25372768 | 1.240871483 | 2.070088427 |
| 4930527J03Rik | 0.49872596 | -0.662161863 | 1.499443633 |
| Gm37486 | 42.26792344 | 1.452740502 | 0.202933165 |
| B230369F24Rik | 55.62345512 | -0.335462597 | 0.147954096 |
| Cox20 | 107.0345127 | 0.080943976 | 0.189700226 |
| Gm16586 | 574.5462637 | -0.378897882 | 0.104461762 |
| Gm28062 | 3.424742753 | -1.104319888 | 0.611733687 |
| Hnrnpu | 2062.938983 | -0.178144991 | 0.065163063 |
| Gm38300 | 25.20762262 | -0.153568306 | 0.240639248 |
| Gm27458 | 0.042716945 | -0.243571806 | 3.724945049 |
| Gm24919 | 0.075043205 | 0.237309859 | 3.724945049 |
| Efcab2 | 203.061846 | -0.320480225 | 0.094620872 |
| Kif26b | 843.0367765 | -0.604199371 | 0.110167673 |
| Gm24405 | 1.624132796 | -1.803074927 | 0.807909479 |
| Gm34176 | 4.619527546 | -1.781043482 | 0.624197482 |
| Gm16564 | 2.140983095 | -1.099747272 | 0.874994479 |
| Smyd3 | 427.0445891 | -0.549845456 | 0.078568148 |
| Gm5561 | 0.317629712 | -0.727267551 | 1.787835547 |
| Tfb2m | 166.5859778 | -0.187576305 | 0.093381684 |
| Gm15480 | 0 | NA | NA |
| Cnst | 707.9421141 | -0.358446638 | 0.056011795 |
| Sccpdh | 630.1321158 | 0.103738137 | 0.095396277 |
| Gm1305 | 0.735920534 | -0.243897125 | 1.065091644 |
| Ahctf1 | 1340.102284 | -0.257436613 | 0.081477201 |

| | | | |
|---|---|---|---|
| Gm10518 | 1.39994924 | -1.008463677 | 0.897827881 |
| Gm37070 | 7.2364301 | 0.770296928 | 0.419714006 |
| Cdc42bpa | 5990.889057 | -0.567387161 | 0.059578267 |
| Gm38169 | 41.0227843 | -1.260860933 | 0.276074684 |
| Gm38331 | 0.311669542 | 0.237878437 | 1.750282718 |
| Adck3 | 280.2963435 | -0.611135869 | 0.127571085 |
| Gm37336 | 1.602826546 | -0.67692674 | 0.94390445 |
| Psen2 | 248.7304506 | 0.691738337 | 0.125785905 |
| Gm31728 | 3.493469674 | -0.377884855 | 0.616802556 |
| Gm5069 | 31.42240588 | -0.401674097 | 0.217073318 |
| Itpkb | 1738.483568 | 1.313924837 | 0.135116181 |
| Gm37033 | 72.29879847 | 0.999574031 | 0.23365187 |
| Gm37390 | 9.932302052 | 0.214845794 | 0.378109991 |
| Gm36933 | 22.96547666 | 0.917153925 | 0.318044003 |
| 6330403A02Rik | 2006.529945 | -1.622342371 | 0.230286333 |
| Gm37984 | 25.25468169 | -2.672340129 | 0.381352081 |
| Gm37267 | 0.197212531 | -0.243571361 | 2.633261182 |
| Parp1 | 872.0659159 | -0.093941696 | 0.086911212 |
| Mir6904 | 1.534744651 | -2.598021342 | 0.937454268 |
| Lin9 | 91.26341556 | 0.123544131 | 0.154228364 |
| Mixl1 | 0.441773728 | 0.185874183 | 1.297918165 |
| Gm8101 | 0 | NA | NA |
| Acbd3 | 598.1054057 | 0.411073577 | 0.080072995 |
| Gm37768 | 0.122991902 | -0.24357403 | 3.724945049 |
| Gm38293 | 0.815026891 | -0.948523034 | 1.125864548 |
| H3f3a | 906.3881747 | 0.167412978 | 0.113496507 |
| Gm17275 | 0.368617932 | 1.345741296 | 1.728627706 |
| H3f3aos | 16.19441088 | -0.106709049 | 0.2887404 |
| Gm24836 | 0 | NA | NA |
| Sde2 | 325.6542778 | 0.419748566 | 0.100764688 |
| Gm20305 | 1.024514732 | 0.846769976 | 0.960870144 |
| Gm18036 | 0 | NA | NA |
| Lefty2 | 3.909599014 | -3.500543575 | 0.896374024 |
| Pycr2 | 201.3612007 | 0.1163456 | 0.107797434 |
| Lefty1 | 9.891254432 | -1.280770931 | 0.423568369 |
| Tmem63a | 404.5987899 | 0.113573486 | 0.123092046 |
| 2210411M09Rik | 0.048949532 | -0.243569582 | 3.724945049 |
| Gm36961 | 0 | NA | NA |
| Ephx1 | 210.2340185 | -0.232703734 | 0.131774009 |
| Gm37431 | 0.049553047 | 0.71820487 | 3.724945049 |
| Gm37086 | 0 | NA | NA |
| 9130409I23Rik | 0.044544648 | 0.237309859 | 3.724945049 |
| Gm17967 | 0.193566712 | 1.949803564 | 2.630756597 |
| Gm37406 | 0 | NA | NA |
| Gm6606 | 0.061920613 | 0.718207094 | 3.724945049 |
| Nvl | 960.5660003 | -0.318279733 | 0.119648211 |
| Fgfr3-ps | 0 | NA | NA |

| | | | |
|---|---|---|---|
| Cnih4 | 409.2563578 | 0.346891405 | 0.071824105 |
| Wdr26 | 2480.374956 | 0.266624789 | 0.07704938 |
| A430110L20Rik | 18.50512536 | -0.951587219 | 0.333323849 |
| Rpl35a-ps2 | 28.95887429 | 0.582642192 | 0.214548688 |
| Gm8146 | 0.858591824 | 0.384743463 | 1.112053547 |
| Cnih3 | 290.3343059 | -0.737143748 | 0.120951386 |
| 2900069G24Rik | 4.593070302 | -1.758904163 | 0.573177807 |
| Gm16547 | 0 | NA | NA |
| Gm16539 | 0 | NA | NA |
| Ccdc121 | 5.062754143 | 0.177102011 | 0.50340416 |
| Dnah14 | 10.40201644 | -0.118600931 | 0.35829479 |
| Gm37664 | 0.09176491 | 0.237310218 | 3.724945049 |
| Gm23690 | 1.324611736 | 0.068229778 | 1.058865359 |
| Lbr | 304.5066716 | 0.598670243 | 0.106212887 |
| Gm26004 | 0.608946563 | 1.611266411 | 1.137811249 |
| Gm5533 | 0.046345709 | 0.71820487 | 3.724945049 |
| Enah | 3997.842318 | -0.133247615 | 0.06446825 |
| 2700078F05Rik | 123.0838922 | -0.597998839 | 0.208683008 |
| Gm38359 | 39.36611411 | -0.734690439 | 0.250993508 |
| Gm37018 | 80.20486681 | -0.64356827 | 0.280611647 |
| Gm37258 | 42.3525549 | -0.70383433 | 0.235514081 |
| Gm37177 | 0.238685809 | -0.243571931 | 2.550065257 |
| Srp9 | 514.3250057 | 0.420840574 | 0.133500564 |
| Gm17968 | 0.075043205 | 0.237309859 | 3.724945049 |
| Vmn1r1 | 0 | NA | NA |
| Degs1 | 871.076032 | 0.379382369 | 0.092136931 |
| 1700047M11Rik | 71.18991338 | -1.949455947 | 0.24249605 |
| Fbxo28 | 708.639217 | 0.172663812 | 0.101753779 |
| Gm5706 | 0.042716945 | -0.243571806 | 3.724945049 |
| Gm10517 | 2.305351242 | -0.674260837 | 0.678015799 |
| Trp53bp2 | 449.2976652 | -0.337111423 | 0.080944485 |
| Capn2 | 1213.867559 | 0.954228029 | 0.104940603 |
| Gm37885 | 64.78973926 | -1.277004319 | 0.284510948 |
| Gm37069 | 19.58732916 | -2.219065447 | 0.47679279 |
| Gm8197 | 0 | NA | NA |
| Capn8 | 0.352854448 | -1.425272868 | 1.53903905 |
| Gm37516 | 0.044544648 | 0.237309859 | 3.724945049 |
| Gm37218 | 0 | NA | NA |
| Ccdc185 | 0.590978791 | 1.043865909 | 1.193548059 |
| Susd4 | 1048.966416 | -0.507805586 | 0.060549104 |
| Gm37542 | 0.158320866 | 0.733224281 | 2.813533472 |
| Tlr5 | 15.84910574 | 1.694470613 | 0.48042228 |
| Gm38079 | 0.535911413 | 0.961730273 | 1.293078528 |
| Gm24975 | 0.165856615 | -0.243568151 | 2.764544693 |
| Disp1 | 84.25718148 | -0.160762533 | 0.11672316 |
| BC085271 | 7.295759161 | 0.069167313 | 0.421952922 |
| Brox | 846.8803349 | 0.29241405 | 0.057293599 |

| | | | |
|---|---|---|---|
| Aida | 1155.608412 | 1.811160742 | 0.148893947 |
| Mia3 | 1173.748108 | -0.0739104 | 0.063027245 |
| Gm37339 | 0.824550338 | -0.092323575 | 1.104154423 |
| Taf1a | 110.8777169 | 0.207223198 | 0.14680686 |
| Gm37986 | 3.238027032 | 0.152279645 | 0.656351088 |
| Hhipl2 | 10.87507933 | 0.89843062 | 0.377645286 |
| Gm37214 | 0.761058687 | -0.018736111 | 1.056659976 |
| Gm33973 | 0.152667883 | -0.243571276 | 2.845214345 |
| Gm8214 | 0.087930927 | 0.236669296 | 3.724898107 |
| Gm34068 | 0.293108411 | 0.718195705 | 3.724945049 |
| Gm38332 | 0.075043205 | 0.237309859 | 3.724945049 |
| Gm38108 | 0.089089296 | 0.237305797 | 3.724945049 |
| Dusp10 | 189.900222 | 0.55158403 | 0.160707438 |
| 1700056E22Rik | 3.04764041 | 1.730792326 | 0.591417579 |
| Gm37223 | 0.046345709 | 0.71820487 | 3.724945049 |
| Gm37712 | 0 | NA | NA |
| 2900092O11Rik | 0.068097359 | -0.243569582 | 3.724945049 |
| 1700112H15Rik | 0.11193333 | 0.718202646 | 3.724945049 |
| Gm23349 | 0 | NA | NA |
| Gm37800 | 0.214735602 | 1.504082435 | 2.170026297 |
| Rpl21-ps1 | 1.040400475 | 0.127614668 | 1.151255706 |
| Gm38358 | 0.24875112 | -0.243563807 | 1.849989481 |
| 4930488B22Rik | 0 | NA | NA |
| Hlx | 28.3948279 | 3.627221509 | 0.321941701 |
| Gm34342 | 0.33098618 | 2.28446159 | 3.693032049 |
| 1-Mar | 1.626573545 | 0.497952829 | 0.757688625 |
| 2-Mar | 436.1785284 | 0.454911931 | 0.10917552 |
| Mir1981 | 0 | NA | NA |
| Gm23780 | 0 | NA | NA |
| C130074G19Rik | 294.2439593 | 0.068584698 | 0.175234778 |
| Mark1 | 1064.898702 | -0.415651863 | 0.078366787 |
| Gm38251 | 0.231207844 | 1.915286386 | 3.039939723 |
| Gm34732 | 0.152865955 | 1.145152106 | 3.499663468 |
| 4930433J02Rik | 0 | NA | NA |
| Gm37662 | 0 | NA | NA |
| Rab3gap2 | 1548.083402 | -0.075769261 | 0.056537348 |
| Snora36b | 0.393059373 | -0.679456963 | 1.536973798 |
| mmu-mir-664 | 0 | NA | NA |
| Gm34882 | 0 | NA | NA |
| Iars2 | 594.1626988 | -0.142406865 | 0.054681056 |
| 2010103J01Rik | 0 | NA | NA |
| Mir194-1 | 0 | NA | NA |
| Mir215 | 0 | NA | NA |
| Gm37785 | 5.812857154 | -0.072190913 | 0.435054886 |
| Bpnt1 | 456.4610138 | -0.176290537 | 0.080888528 |
| 4930532G15Rik | 1.624021655 | -1.748977825 | 0.976546362 |
| Eprs | 2033.228255 | 0.512199749 | 0.088031519 |

| | | | |
|---|---|---|---|
| 9630028B13Rik | 4.198644665 | -0.971345062 | 0.59254266 |
| Gm2061 | 20.60515688 | -1.626287439 | 0.257234906 |
| Slc30a10 | 131.5005642 | -1.603298099 | 0.182148959 |
| 5033404E19Rik | 1.621213957 | -2.279063605 | 0.95810858 |
| Gm38093 | 0 | NA | NA |
| Gm9722 | 0.233998759 | 0.23545975 | 2.135092051 |
| Lyplal1 | 31.22453688 | -0.10981064 | 0.192083766 |
| Gm38268 | 1.323601962 | -2.665878081 | 0.973243462 |
| Gm23821 | 0 | NA | NA |
| Gm37134 | 0 | NA | NA |
| Gm37272 | 0 | NA | NA |
| Gm37491 | 0 | NA | NA |
| A730004F24Rik | 0.094097695 | 0.718200421 | 3.724945049 |
| Tgfb2 | 267.8778118 | 0.501885438 | 0.119506484 |
| Gm26169 | 0.061920613 | 0.718207094 | 3.724945049 |
| Gm29491 | 2.076177089 | -0.126692953 | 0.670386036 |
| Gm37849 | 4.193292652 | -0.566941669 | 0.549301468 |
| Rrp15 | 96.16436869 | -0.004982107 | 0.176753183 |
| A430105J06Rik | 9.038545814 | -0.482126709 | 0.397524739 |
| 1700023G09Rik | 0 | NA | NA |
| Gm3809 | 0.939714841 | 0.438754819 | 0.978952425 |
| Gm37912 | 0.240869569 | 1.212829121 | 3.089662278 |
| D1Pas1 | 0.137647365 | 0.237309079 | 3.724945049 |
| Gm19058 | 0.109869311 | 0.238305784 | 3.724874707 |
| Spata17 | 10.75281961 | 0.077455609 | 0.288898343 |
| Gm15508 | 0.503467097 | -0.080268206 | 1.332768817 |
| RP23-438D3.4 | 1.299896474 | -1.090583979 | 0.916237264 |
| RP23-438D3.3 | 1.714015964 | -0.48821149 | 0.712355877 |
| Gm15509 | 0.816619922 | 0.122855026 | 0.992757499 |
| 1700007P06Rik | 4.898989025 | -0.970442357 | 0.53618955 |
| Gpatch2 | 371.7543322 | -0.080414131 | 0.083430755 |
| Gm21710 | 76.9518533 | -0.219132395 | 0.204998441 |
| Gm38155 | 0.312806312 | 1.134065129 | 1.756502848 |
| Gm37896 | 5.45108928 | -0.533265968 | 0.598289137 |
| Esrrg | 699.5845546 | -0.301321136 | 0.099350994 |
| Gm36388 | 5.428564864 | -0.578785642 | 0.499160804 |
| Gm37929 | 0 | NA | NA |
| 9330162B11Rik | 0.045882455 | 0.237312084 | 3.724945049 |
| Gm38315 | 9.475868802 | -1.55362518 | 0.481930158 |
| Gm38064 | 0.045882455 | 0.237312084 | 3.724945049 |
| Ush2a | 7.7862644 | -1.356204147 | 0.39855564 |
| Gm25095 | 0 | NA | NA |
| Gm25269 | 0 | NA | NA |
| Kctd3 | 958.3543934 | -0.222168353 | 0.062080884 |
| Gm37178 | 0.897014366 | -0.119699205 | 1.141166599 |
| Gm18698 | 0.26837777 | -0.243563818 | 2.092313189 |
| Kcnk2 | 433.3427616 | -0.617955342 | 0.102933104 |

| | | | |
|---|---|---|---|
| 2900042K21Rik | 2.348091134 | -3.333583341 | 1.030604581 |
| Gm3837 | 46.20088099 | 0.631267267 | 0.286464261 |
| A430027H14Rik | 10.33216563 | -1.333049481 | 0.325085124 |
| Gm24092 | 0.042716945 | -0.243571806 | 3.724945049 |
| Gm38122 | 0.072025462 | 0.237312084 | 3.724945049 |
| Cenpf | 46.80573964 | 1.345070649 | 0.382542907 |
| Gm37239 | 0.092881282 | -0.724455696 | 3.724945049 |
| Ptpn14 | 196.8365016 | -0.563761156 | 0.145262662 |
| Gm30280 | 0.99636849 | 0.598597384 | 1.053101255 |
| Gm26574 | 0.255031674 | 2.261512368 | 2.981745413 |
| Gm37422 | 1.509855825 | 0.225559208 | 0.858959625 |
| Smyd2 | 468.8475306 | -0.289406365 | 0.074400782 |
| Gm38245 | 23.42935459 | -1.668887033 | 0.38169796 |
| Prox1 | 753.5982863 | 0.119839928 | 0.072009864 |
| Gm28172 | 0.53361457 | -0.644475318 | 1.460471131 |
| Prox1os | 9.870830895 | 0.096268504 | 0.314920484 |
| Gm29678 | 0.175900428 | 0.265060088 | 2.724668472 |
| Gm30725 | 0.194279094 | 1.292225235 | 3.719439735 |
| Gm23153 | 0.062426873 | 0.237312084 | 3.724945049 |
| AC124613.1 | 0 | NA | NA |
| Gm30932 | 0 | NA | NA |
| Rps6kc1 | 776.7345912 | -0.198896502 | 0.061340428 |
| Angel2 | 626.1745066 | 0.007825927 | 0.05432057 |
| Vash2 | 94.15710728 | 0.35983318 | 0.173532496 |
| Gm31373 | 0.061920613 | 0.718207094 | 3.724945049 |
| Gm37095 | 4.062122998 | -1.161094444 | 0.571155641 |
| Gm8407 | 6.023452583 | -1.623323794 | 0.497831253 |
| Mfsd7b | 308.2483388 | 0.037557782 | 0.100721602 |
| A230020J21Rik | 0.378086136 | -1.166501804 | 1.764472797 |
| Spata45 | 0.216916081 | 0.237328524 | 2.17728567 |
| Tatdn3 | 79.86263131 | -0.462575356 | 0.139108572 |
| Nsl1 | 10.37462125 | 0.789205263 | 0.328204537 |
| Gm38188 | 0 | NA | NA |
| Batf3 | 9.56190983 | 1.18372927 | 0.370010538 |
| Gm38161 | 0 | NA | NA |
| Fam71a | 0.398364559 | 1.634313233 | 1.696876718 |
| Atf3 | 253.8533324 | 5.885313172 | 0.460482845 |
| Gm24748 | 0 | NA | NA |
| D730003I15Rik | 30.70567579 | -0.854400747 | 0.351033019 |
| Gm37074 | 9.818226126 | -1.155872229 | 0.532973327 |
| Gm37168 | 4.648304818 | -0.12426963 | 0.43317373 |
| Nenf | 240.7507074 | -0.090112233 | 0.106408657 |
| Tmem206 | 104.9785531 | -0.147175923 | 0.117733507 |
| Ppp2r5a | 813.5882951 | 0.122809822 | 0.08001811 |
| Gm2272 | 0 | NA | NA |
| Gm32200 | 9.706590373 | -0.713549829 | 0.388917528 |
| Gm37688 | 0.11193333 | 0.718202646 | 3.724945049 |

| | | | |
|---|---|---|---|
| Gm37432 | 0.52136618 | 1.094979604 | 1.503202567 |
| 9430037O13Rik | 2.32288591 | 1.70112001 | 0.869934863 |
| Dtl | 77.14293698 | 1.302409946 | 0.182043959 |
| Ints7 | 338.6046142 | 0.113368445 | 0.0730352 |
| Gm37349 | 5.323050413 | -0.052143622 | 0.530011653 |
| Lpgat1 | 3755.069215 | -0.299805875 | 0.058156137 |
| Gm22446 | 0 | NA | NA |
| Gm26203 | 0 | NA | NA |
| Gm20203 | 0 | NA | NA |
| Gm38037 | 1.052686423 | -1.593763672 | 1.081248815 |
| Nek2 | 70.81536854 | 0.80756373 | 0.318978104 |
| 1700034H15Rik | 43.54077039 | -0.911871182 | 0.187691094 |
| Slc30a1 | 668.252428 | 0.453582844 | 0.115421732 |
| Rd3 | 12.96523121 | 0.684724833 | 0.297273192 |
| Gm26670 | 1.595190335 | -0.069759847 | 0.728577351 |
| Gm32460 | 0.281516365 | -0.724460639 | 1.788251217 |
| Traf5 | 35.39043388 | 1.094942744 | 0.216885799 |
| Mir3473c | 0 | NA | NA |
| Gm3934 | 136.9920633 | -0.154878677 | 0.118928004 |
| Gm26879 | 0 | NA | NA |
| Rcor3 | 532.0108606 | -0.208076382 | 0.111241069 |
| Gm37055 | 0.109124069 | -0.24357403 | 3.724945049 |
| Gm10516 | 32.23956246 | -0.350706036 | 0.191338531 |
| Kcnh1 | 1278.299755 | -0.255517983 | 0.147609587 |
| Hhat | 36.35158037 | 0.392473999 | 0.240579993 |
| Gm22265 | 0.377885927 | 1.82567923 | 1.648581981 |
| Gm38360 | 0 | NA | NA |
| A730013G03Rik | 1.204108136 | -0.191258639 | 1.058975698 |
| 1700065J18Rik | 0.258344737 | 1.266663676 | 2.120230285 |
| Sertad4 | 214.5306265 | -0.390975865 | 0.120611796 |
| Gm15867 | 2.355781424 | 0.789735525 | 0.700104593 |
| Gm2367 | 0.23837135 | 0.156750474 | 2.545605147 |
| Syt14 | 711.6526152 | -0.666240016 | 0.110542375 |
| Gm23583 | 5.178783273 | -1.191714824 | 0.525331505 |
| Gm31406 | 1.718171857 | 0.628157116 | 0.795138794 |
| Gm37103 | 0 | NA | NA |
| Diexf | 261.3112737 | -0.137721805 | 0.095161451 |
| Irf6 | 4.847835626 | -0.472540463 | 0.438509773 |
| A130010J15Rik | 56.29500924 | -0.009377259 | 0.155198646 |
| Traf3ip3 | 31.36915684 | 3.390106427 | 0.313335409 |
| Gm15872 | 0.108123235 | -0.24357403 | 3.724945049 |
| 4930570N18Rik | 0.062426873 | 0.237312084 | 3.724945049 |
| Lamb3 | 38.67870685 | -0.004387379 | 0.212721822 |
| Hsd11b1 | 94.82910089 | -0.178044407 | 0.189171132 |
| Gm38118 | 1.886096619 | 0.274709337 | 0.797684291 |
| G0s2 | 30.35106119 | -0.184710046 | 0.193141041 |
| Gm37691 | 3.727288002 | -0.504576172 | 0.526276202 |

| | | | |
|---|---|---|---|
| Gm37650 | 0 | NA | NA |
| Camk1g | 290.0946269 | -1.055617193 | 0.149543643 |
| Gm37942 | 0 | NA | NA |
| Gm38022 | 0 | NA | NA |
| Gm38322 | 0.094097695 | 0.718200421 | 3.724945049 |
| 4631405K08Rik | 0 | NA | NA |
| Mir205 | 0 | NA | NA |
| Gm21362 | 0 | NA | NA |
| 4930503O07Rik | 0.33841178 | -0.986812756 | 3.706267102 |
| Plxna2 | 2388.321215 | -0.692692984 | 0.100854795 |
| Gm37783 | 41.75688494 | -1.018352571 | 0.294253998 |
| Gm37674 | 39.82514733 | -1.307651319 | 0.314685374 |
| Gm38036 | 64.68369131 | -1.157591531 | 0.334513008 |
| 2900035J10Rik | 6.043773575 | -1.322030399 | 0.480018782 |
| Gm19777 | 0.047948698 | -0.243569582 | 3.724945049 |
| Mir3962 | 0.624784964 | -0.615995136 | 2.726079717 |
| Gm23885 | 0.198152624 | 1.757323469 | 3.707396009 |
| Cd34 | 275.1221153 | -0.739159236 | 0.13608725 |
| Gm16897 | 7.560911453 | 0.079305966 | 0.373391747 |
| A330023F24Rik | 1658.07212 | -0.714200915 | 0.287243557 |
| Gm32250 | 0.045882455 | 0.237312084 | 3.724945049 |
| Gm37334 | 228.9475402 | -0.998423051 | 0.236144567 |
| 9630010A21Rik | 156.1316392 | -0.984040878 | 0.276020732 |
| Gm37027 | 0.870290022 | 1.867479255 | 1.290936679 |
| Cd46 | 147.5920642 | -0.753198169 | 0.201204666 |
| Mir29b-2 | 0 | NA | NA |
| Mir29c | 0 | NA | NA |
| Gm37366 | 35.61469171 | -0.603376594 | 0.250531865 |
| Gm37132 | 36.12461116 | -1.557768949 | 0.316204488 |
| Cr1l | 392.2221467 | 0.081743927 | 0.083341493 |
| Cr2 | 5.163701346 | -0.136476365 | 0.445149601 |
| Gm37887 | 0.066050875 | 0.71820487 | 3.724945049 |
| Gm24802 | 0.044191549 | -0.243571806 | 3.724945049 |
| Gm27940 | 0 | NA | NA |
| Gm38046 | 0 | NA | NA |
| Gm38194 | 0.659436564 | 0.605335844 | 1.342820669 |
| 9230019H11Rik | 0 | NA | NA |
| H60c | 0 | NA | NA |
| Gm26752 | 1.987374958 | -0.677486122 | 0.908370064 |
| Ppp1r14c | 150.8264208 | -0.302412495 | 0.098697118 |
| Gm24482 | 0.106358623 | -0.24357403 | 3.724945049 |
| Iyd | 3.589799332 | -1.544874866 | 0.711703096 |
| Plekhg1 | 783.4816878 | -0.005946644 | 0.078828449 |
| Gm16149 | 0.045882455 | 0.237312084 | 3.724945049 |
| Gm23023 | 0 | NA | NA |
| Gm16148 | 0.049553047 | 0.71820487 | 3.724945049 |
| Mthfd1l | 264.3319854 | -0.299064874 | 0.098041938 |

| | | | |
|---|---|---|---|
| Gm16074 | 3.696842725 | -1.411380475 | 0.762272129 |
| Gm25515 | 0.67351111 | 0.188466386 | 1.484353804 |
| Gm25369 | 0.211762239 | -0.724460933 | 2.616650133 |
| Akap12 | 1387.377891 | 1.328239294 | 0.272402261 |
| Zbtb2 | 206.2791963 | 0.480125642 | 0.165703398 |
| Gm21781 | 207.0306887 | -0.58450287 | 0.247297692 |
| Rmnd1 | 211.8870746 | -0.512579866 | 0.146603828 |
| Gm16153 | 6.393807032 | -0.082340682 | 0.427104023 |
| Armt1 | 185.8306212 | -0.157556935 | 0.095932765 |
| Ccdc170 | 10.84234273 | 0.65761593 | 0.339270801 |
| Gm22739 | 0.087261593 | -0.243576254 | 3.724945049 |
| Esr1 | 111.3424344 | 1.132296737 | 0.161672391 |
| Syne1 | 6019.024116 | -0.904430644 | 0.122919682 |
| Gm25694 | 9.729705334 | -1.62265591 | 0.437307598 |
| Gm10097 | 0.451409952 | 2.974289607 | 1.36028713 |
| Gm23573 | 0 | NA | NA |
| Myct1 | 1.954060925 | 1.627057754 | 0.770159607 |
| Vip | 156.4683625 | -0.332697268 | 0.164584925 |
| Fbxo5 | 28.97769736 | 1.112414716 | 0.201579091 |
| Mtrf1l | 83.2976707 | 0.071816874 | 0.133064194 |
| Rgs17 | 1666.499129 | -0.018914002 | 0.083872781 |
| Gm10945 | 0.044544648 | 0.237309859 | 3.724945049 |
| Oprm1 | 120.7739671 | -0.357959497 | 0.137171206 |
| Ipcef1 | 966.4889659 | -0.496290702 | 0.152851071 |
| Gm15536 | 9.233390511 | 0.497399788 | 0.351415609 |
| Cnksr3 | 268.184003 | -0.427818485 | 0.148229594 |
| Ulbp1 | 7.246076538 | 3.284841232 | 0.712082068 |
| Lrp11 | 1467.900515 | -0.274471299 | 0.052514823 |
| Gm16254 | 0.342362695 | 1.320070697 | 3.711827473 |
| Pcmt1 | 1204.251679 | -0.244084349 | 0.051329753 |
| Nup43 | 42.06222489 | 0.63337072 | 0.264956169 |
| Lats1 | 1258.401443 | 0.399558678 | 0.079279765 |
| Katna1 | 226.4998142 | 1.235430047 | 0.113794672 |
| Gm20470 | 0.150550172 | -0.724459893 | 2.857661711 |
| Ginm1 | 172.1070177 | 0.707980727 | 0.128276505 |
| B430219N15Rik | 0.972892922 | 1.614960635 | 1.090892705 |
| Ppil4 | 480.6494858 | -0.071394781 | 0.105674944 |
| Gm15560 | 0.152667883 | -0.243571276 | 2.845214345 |
| Zc3h12d | 64.81458113 | 6.998321255 | 0.612024036 |
| Mir5104 | 0.256698001 | 1.625097676 | 2.443902994 |
| Gm24726 | 0 | NA | NA |
| Tab2 | 1189.099294 | 0.260767266 | 0.076491981 |
| Ust | 495.4936296 | -0.332204257 | 0.093956771 |
| Gm24374 | 0 | NA | NA |
| Sash1 | 1877.282141 | 0.25639133 | 0.115851798 |
| Gm25410 | 1.528398176 | -0.099436437 | 0.774991205 |
| Gm26674 | 0.49731986 | 0.659293989 | 1.340028465 |

| | | | |
|---|---|---|---|
| Gm28904 | 9.097189116 | -1.262783203 | 0.382423547 |
| Gm9930 | 39.07755842 | -1.51452278 | 0.385400708 |
| Samd5 | 168.1922456 | -0.682340847 | 0.116886617 |
| Stxbp5 | 2460.054131 | -0.194070391 | 0.081507314 |
| Gm25635 | 3.189004039 | -0.658123726 | 0.692976318 |
| Gm28905 | 7.03479442 | -2.023019267 | 0.466330837 |
| Gm25682 | 1.985393048 | -1.902023079 | 0.927916427 |
| Adgb | 11.41173491 | 0.088195209 | 0.335222997 |
| Gm15568 | 0.044544648 | 0.237309859 | 3.724945049 |
| Rab32 | 75.72339442 | 3.579990217 | 0.243590344 |
| Gm5177 | 0.397592075 | 1.147977019 | 1.750412728 |
| Gm23601 | 0 | NA | NA |
| Gm10944 | 0 | NA | NA |
| Grm1 | 1592.406977 | -0.478360311 | 0.092220621 |
| Shprh | 1486.660042 | -0.362413702 | 0.088114509 |
| Gm16577 | 1.038701827 | 1.132089211 | 0.973764317 |
| 4930432B10Rik | 21.06911396 | -1.027048707 | 0.381470204 |
| Fbxo30 | 323.9844007 | 0.490473674 | 0.094242824 |
| Gm25979 | 0.044191549 | -0.243571806 | 3.724945049 |
| Epm2a | 429.8223933 | -0.594811028 | 0.10423921 |
| Rpl17-ps11 | 0 | NA | NA |
| Gm9797 | 3.625096175 | 0.205624749 | 0.563997218 |
| Gm22546 | 0.038981395 | 0.718207094 | 3.724945049 |
| Gm22194 | 0 | NA | NA |
| Gm25011 | 0 | NA | NA |
| Gm23680 | 0 | NA | NA |
| Gm25139 | 0 | NA | NA |
| Utrn | 1645.064442 | 0.351688221 | 0.056364798 |
| Gm23044 | 1.167901279 | 1.429707836 | 0.96757372 |
| B230208H11Rik | 2.934711433 | -0.02343617 | 0.595747606 |
| Stx11 | 71.27238607 | 7.675589503 | 0.77371461 |
| Sf3b5 | 177.9606754 | 0.414588178 | 0.135931269 |
| Plagl1 | 355.2404399 | 0.344794575 | 0.104969459 |
| Gm27741 | 0.062426873 | 0.237312084 | 3.724945049 |
| Zc2hc1b | 0.135956459 | -0.243575439 | 3.642411469 |
| Ltv1 | 143.8273288 | 0.451304914 | 0.133390456 |
| Phactr2 | 555.9939026 | -0.391198652 | 0.081100981 |
| Fuca2 | 561.995271 | 0.544164166 | 0.097496688 |
| Pex3 | 290.4366472 | 0.02648022 | 0.076760332 |
| Adat2 | 56.9225531 | -0.133874839 | 0.145070228 |
| Aig1 | 695.0458195 | -0.005624377 | 0.061183926 |
| Gm22995 | 0 | NA | NA |
| Gm8330 | 0.285805375 | 0.62998616 | 1.780482102 |
| Hivep2 | 5288.46622 | -0.55477791 | 0.092876165 |
| Gm26835 | 0.195722939 | 0.675788417 | 2.666087806 |
| Gm28289 | 0.0878635 | -0.72445792 | 3.724945049 |
| Gm24343 | 0.447929572 | -1.011757063 | 1.407295642 |

| | | | |
|---|---|---|---|
| Gm37299 | 0.353350569 | 1.094293812 | 1.696113045 |
| Adgrg6 | 79.60354276 | 1.04462197 | 0.179594333 |
| Gm10335 | 39.8430328 | 0.712047722 | 0.264919188 |
| Vta1 | 390.7247282 | 0.41836806 | 0.065844686 |
| Gm16234 | 0.304277014 | 0.730131765 | 2.033724514 |
| Nmbr | 18.58630088 | -0.767928089 | 0.268504062 |
| Gje1 | 0.044191549 | -0.243571806 | 3.724945049 |
| Gm8355 | 90.21878376 | 0.457891847 | 0.145468248 |
| B230364G03Rik | 0.277885724 | -1.203488933 | 1.779822051 |
| Gm23892 | 0 | NA | NA |
| Gm20125 | 6.549262826 | 0.731572136 | 0.468816158 |
| Gm25382 | 0 | NA | NA |
| Cited2 | 577.6609852 | 0.6917401 | 0.148213646 |
| Gm20655 | 8.589832509 | -0.446590268 | 0.393439312 |
| Txlnb | 15.59385469 | -1.280707558 | 0.308410635 |
| Gm7584 | 0.106056992 | 0.237307635 | 3.724945049 |
| Heca | 505.4761868 | 0.045596678 | 0.092492136 |
| Abracl | 74.37190115 | 1.022800586 | 0.18689384 |
| Gm17711 | 0.109982625 | 0.237318756 | 3.724945049 |
| Reps1 | 606.0807053 | -0.053240851 | 0.093312999 |
| Ect2l | 5.746393814 | 0.328056477 | 0.427704704 |
| Ccdc28a | 182.412726 | -0.426993399 | 0.141910183 |
| Gm10827 | 7.130661091 | 1.041821735 | 0.533872618 |
| Nhsl1 | 1061.864252 | 0.619391084 | 0.098043424 |
| Hebp2 | 41.95025653 | 0.922326954 | 0.23115981 |
| Arfgef3 | 2592.376606 | -0.58944283 | 0.091451196 |
| Gm4922 | 0 | NA | NA |
| Perp | 62.70509499 | 0.212888699 | 0.203517843 |
| Gm38175 | 0 | NA | NA |
| Tnfaip3 | 387.4893776 | 3.280636173 | 0.374828102 |
| Olig3 | 0.674822466 | -0.297776437 | 1.102733491 |
| Ifngr1 | 246.1757685 | 1.136529477 | 0.125940854 |
| Il22ra2 | 0.165580463 | 0.23732449 | 2.773571271 |
| 9230106D20Rik | 0 | NA | NA |
| Il20ra | 10.38114758 | 0.00812954 | 0.346111304 |
| Gm25256 | 0.617475372 | 1.122248232 | 1.334407953 |
| Slc35d3 | 65.99433708 | -1.105266782 | 0.161740594 |
| Pex7 | 193.8965027 | -0.393655225 | 0.082865248 |
| Map3k5 | 659.4939879 | -0.426232673 | 0.075120036 |
| Map7 | 996.5614196 | -0.554724172 | 0.086047648 |
| Mir6413 | 0 | NA | NA |
| 4933406P04Rik | 7.134388926 | -0.084021231 | 0.375796651 |
| Bclaf1 | 2858.72397 | 0.070394008 | 0.069565572 |
| Mtfr2 | 15.4157451 | 1.200522819 | 0.353355077 |
| Pde7b | 825.652375 | -0.214098653 | 0.111260365 |
| 4930403O15Rik | 14.60270139 | -0.675835097 | 0.282935832 |
| Gm17229 | 23.00707965 | -0.222664126 | 0.345441769 |

| | | | |
|---|---|---|---|
| Gm17230 | 1.594890904 | 3.623579651 | 0.999174952 |
| Ahi1 | 2789.699633 | -0.616896859 | 0.058913515 |
| Myb | 21.21849641 | 1.012143057 | 0.254537891 |
| Gm26577 | 5.818997596 | -0.529514402 | 0.409540234 |
| Gm2467 | 0 | NA | NA |
| Hbs1l | 606.1812586 | 0.154383169 | 0.089120013 |
| Aldh8a1 | 4.539391149 | 0.862837339 | 0.435809482 |
| 1700021A07Rik | 0.178864798 | 1.594141258 | 3.264221614 |
| 1700020N01Rik | 0.490798134 | 2.727605257 | 1.461364214 |
| Gm5420 | 0.09349418 | -0.24357403 | 3.724945049 |
| Gm29323 | 0 | NA | NA |
| Platr6 | 2.028201447 | -0.955153305 | 0.749087621 |
| Gm23839 | 0.264066224 | -0.243569711 | 2.448621796 |
| Sgk1 | 2023.552573 | 1.054308143 | 0.156168275 |
| Gm8540 | 0.066050875 | 0.71820487 | 3.724945049 |
| Gm7672 | 1.404341276 | 0.158824322 | 1.038276815 |
| Gm34267 | 0 | NA | NA |
| 4930444G20Rik | 0 | NA | NA |
| Gm2539 | 0.516867762 | 2.283206947 | 1.644432757 |
| Gm19791 | 0.170186936 | 1.71882767 | 3.39900844 |
| E030030I06Rik | 0.513032606 | 0.57178014 | 1.439475446 |
| Gm7678 | 0 | NA | NA |
| Gm4895 | 29.59563694 | -0.036405661 | 0.342594258 |
| Raet1e | 4.132199462 | 4.195933659 | 0.903072051 |
| H60b | 4.341397877 | 0.078937759 | 0.528435939 |
| Gm26740 | 3.074037044 | 2.294333743 | 0.755577523 |
| Gm26581 | 0.35312086 | 2.697693235 | 3.67922251 |
| Gm1972 | 0.209168331 | 1.916771144 | 2.554677273 |
| Hmgb1-ps8 | 0.251652017 | 0.237328285 | 2.094757334 |
| Gm26528 | 1.473604354 | 4.253458776 | 1.084711127 |
| C920009B18Rik | 12.28007475 | 5.04909346 | 0.707167801 |
| Gm5421 | 64.80517792 | -0.187933384 | 0.156157264 |
| Raet1d | 2.579584798 | 4.857781984 | 0.969735467 |
| Gm10825 | 0.745135914 | 3.577350403 | 1.248646852 |
| Gm16531 | 0.100902008 | 1.15059983 | 3.720931964 |
| Gm26585 | 0.066050875 | 0.71820487 | 3.724945049 |
| Slc2a12 | 91.64299392 | 0.209438715 | 0.169980713 |
| Tbpl1 | 649.6904834 | -0.076924705 | 0.049663702 |
| Mir7663 | 9.275244636 | -0.240512223 | 0.416380095 |
| Tcf21 | 0.038981395 | 0.718207094 | 3.724945049 |
| Eya4 | 83.33865471 | 0.355977364 | 0.123974938 |
| Gm24655 | 0.526083665 | 0.131386557 | 1.300738193 |
| Rps12 | 490.9795335 | 0.83073325 | 0.087081455 |
| Snora33 | 5.090834635 | -0.332498891 | 0.471970494 |
| Snord100 | 3.200779369 | 0.726529029 | 0.624898936 |
| Gm23130 | 6.373645778 | -0.176801562 | 0.547293568 |
| Slc18b1 | 157.4498406 | -0.4853363 | 0.117678531 |

| | | | |
|---|---|---|---|
| Vnn3 | 3.620663213 | 3.839914161 | 0.866288606 |
| Vnn1 | 16.91961705 | 1.50851124 | 0.305251111 |
| Taar1 | 0.366161499 | 1.396293037 | 1.712994641 |
| Taar2 | 0 | NA | NA |
| Taar3 | 0.061920613 | 0.718207094 | 3.724945049 |
| Taar4 | 0.07796279 | 1.072698098 | 3.721829099 |
| Taar5 | 0 | NA | NA |
| Taar6 | 0 | NA | NA |
| Taar7a | 0 | NA | NA |
| Taar7b | 0.046345709 | 0.71820487 | 3.724945049 |
| Taar7c-ps | 0 | NA | NA |
| Gm15136 | 0 | NA | NA |
| Taar7d | 0 | NA | NA |
| Taar7e | 0 | NA | NA |
| Gm6884 | 0 | NA | NA |
| Taar7f | 0 | NA | NA |
| Taar8a | 0 | NA | NA |
| Taar8b | 0 | NA | NA |
| Taar8c | 0 | NA | NA |
| Taar9 | 0 | NA | NA |
| Gm15137 | 0 | NA | NA |
| Stx7 | 1125.137819 | 0.171257185 | 0.061365791 |
| Moxd1 | 76.92434119 | 0.265211529 | 0.134424622 |
| Gm15271 | 0.245390278 | -0.731040768 | 2.144183146 |
| Gm15272 | 0.315650473 | -1.03666008 | 1.723946956 |
| Gm15270 | 6.965063545 | -0.352307972 | 0.400081422 |
| Gm6893 | 0.385058506 | -1.10241282 | 1.452470314 |
| Ctgf | 300.6738772 | 0.903803105 | 0.20637717 |
| Enpp1 | 114.2531846 | 0.134004329 | 0.125731721 |
| Enpp3 | 5.598650687 | 0.285169069 | 0.43141078 |
| Med23 | 495.4221233 | -0.200940725 | 0.08759123 |
| Gm23410 | 0 | NA | NA |
| Mir6905 | 0.793572527 | -0.261303107 | 1.212196154 |
| Arg1 | 19.82112316 | 4.167541541 | 0.440635377 |
| Gm22566 | 0.337983087 | -1.214930473 | 1.594539911 |
| Akap7 | 458.6354203 | -0.402073825 | 0.073212484 |
| Gm25016 | 0.821005853 | -1.245523113 | 1.279850065 |
| Gm26167 | 1.164232618 | -1.632089752 | 0.99153907 |
| Epb4.1l2 | 1228.23948 | 0.465633559 | 0.084007652 |
| Smlr1 | 0.237992309 | -0.243563174 | 2.113697579 |
| Gm29571 | 1.193153915 | 1.499200583 | 1.150999537 |
| Tmem200a | 188.3161123 | -0.032335678 | 0.145836678 |
| Gm9767 | 4.750744946 | 0.203117483 | 0.572315782 |
| Samd3 | 18.30979764 | -0.327352715 | 0.283102671 |
| L3mbtl3 | 344.0952442 | -0.033635999 | 0.084144737 |
| Gm27754 | 1.24360625 | -1.110026075 | 0.893869652 |
| Gm23757 | 0 | NA | NA |

| | | | |
|---|---|---|---|
| Gm8709 | 0 | NA | NA |
| Arhgap18 | 194.7190015 | 0.49640438 | 0.112567161 |
| Lama2 | 360.511942 | -0.454665265 | 0.15504657 |
| 4930579H20Rik | 0.409009542 | 1.240625149 | 1.71890835 |
| Gm23353 | 0.042716945 | -0.243571806 | 3.724945049 |
| Gm36962 | 9.987815949 | -0.797305378 | 0.341631221 |
| Gm10145 | 22.9043597 | 0.826986041 | 0.262824905 |
| Ptprk | 857.1284641 | -0.272934405 | 0.066665657 |
| Gm22370 | 0 | NA | NA |
| Themis | 33.46669582 | 6.197329093 | 0.683261233 |
| 4930519F09Rik | 0.493807599 | 0.192257644 | 1.37745337 |
| Gm22811 | 0 | NA | NA |
| 2310057J18Rik | 0.049553047 | 0.71820487 | 3.724945049 |
| Gm9824 | 52.77535247 | -0.259523003 | 0.191828188 |
| Gm9996 | 16.71528001 | -0.210736612 | 0.248766024 |
| Soga3 | 1394.424857 | 0.083470967 | 0.071416796 |
| 9330159F19Rik | 1284.714598 | -0.143199517 | 0.075921773 |
| Echdc1 | 288.0513829 | -0.144071397 | 0.11204704 |
| Rnf146 | 973.2977547 | 0.066726513 | 0.112701883 |
| Gm25596 | 5.415374794 | -0.477614704 | 0.457528681 |
| Gm24286 | 0 | NA | NA |
| Rspo3 | 164.9902427 | 0.68432376 | 0.147769862 |
| Gm10275 | 252.6440296 | 0.648491652 | 0.142390945 |
| Cenpw | 14.39510704 | -0.293588705 | 0.376775991 |
| Gm22623 | 0.372763955 | 1.534873015 | 1.427888839 |
| Trmt11 | 140.2023999 | -0.019191353 | 0.095383207 |
| Hint3 | 162.1893017 | -0.409467375 | 0.112737018 |
| Ncoa7 | 1617.001063 | 0.297802864 | 0.097393734 |
| Gm24189 | 3.823607117 | -0.513868885 | 0.537769006 |
| Hey2 | 73.53388155 | -0.178023178 | 0.253933045 |
| Gm24168 | 0 | NA | NA |
| Gm23790 | 0 | NA | NA |
| Gm5422 | 32.64956114 | 0.391477449 | 0.203901488 |
| Hddc2 | 80.54346223 | 0.016396343 | 0.156668967 |
| n-R5s76 | 0 | NA | NA |
| Tpd52l1 | 150.8148291 | 0.279111883 | 0.109755799 |
| Rnf217 | 787.7636301 | 0.226467636 | 0.087743714 |
| Gm7909 | 13.66660191 | -0.488195634 | 0.302670516 |
| Gm8793 | 0.447136588 | 0.913485677 | 1.374209043 |
| Nkain2 | 470.4928905 | -0.942960357 | 0.077803257 |
| Trdn | 7.453187596 | -1.388434551 | 0.718739144 |
| Clvs2 | 440.1642126 | -0.479621341 | 0.096605437 |
| Gm15939 | 3.414634292 | 0.413037585 | 0.556323094 |
| Gm4794 | 0 | NA | NA |
| Gm10327 | 7.81528087 | 0.031845611 | 0.36177313 |
| Sult3a1 | 0.245625509 | 0.237331042 | 1.849893626 |
| Rsph4a | 69.28623393 | 0.731260113 | 0.180977098 |

| | | | |
|---|---|---|---|
| Zufsp | 477.2138068 | 1.698825579 | 0.135247091 |
| Gm17567 | 17.61824852 | -1.23185549 | 0.280068323 |
| Rwdd1 | 291.9026565 | 0.213051406 | 0.098798975 |
| Trappc3l | 1.780324605 | -0.57100128 | 0.794407029 |
| Fam26d | 0.532386167 | 2.648621525 | 1.496992739 |
| Fam26e | 35.94992537 | 0.359701778 | 0.200242058 |
| Fam26f | 40.02314751 | 8.061549097 | 0.898986475 |
| Dse | 148.6253122 | 1.127063358 | 0.151443995 |
| Tspyl1 | 2089.883738 | -0.101105098 | 0.066425275 |
| Nt5dc1 | 71.24363843 | 0.876903747 | 0.162117781 |
| Gm26564 | 26.65952218 | -0.086776412 | 0.260680419 |
| Tspyl4 | 5615.800342 | -0.33107246 | 0.070540078 |
| Col10a1 | 2.713943931 | 0.453151934 | 0.630801 |
| Frk | 10.56880692 | 1.145062527 | 0.341893304 |
| Gm26341 | 0 | NA | NA |
| Amd2 | 40.7342373 | -0.123399269 | 0.161190728 |
| Hs3st5 | 126.4096297 | -0.535852051 | 0.152164873 |
| Gm22554 | 0 | NA | NA |
| Hdac2 | 735.3892852 | -0.140671293 | 0.059316961 |
| Gm38231 | 0 | NA | NA |
| Marcks | 2411.516162 | -0.088765793 | 0.115645554 |
| 5930403N24Rik | 7.127291511 | -1.01511185 | 0.416030962 |
| Gm26535 | 0.348734193 | 1.276440667 | 1.659864432 |
| Gm24710 | 0 | NA | NA |
| Gm24889 | 0 | NA | NA |
| Gm24297 | 0 | NA | NA |
| Gm22911 | 0 | NA | NA |
| Rfpl4b | 0.174199136 | 0.718215474 | 2.729447628 |
| Lama4 | 161.5765428 | 0.513996375 | 0.10766143 |
| Fam229b | 68.30184625 | -0.78245696 | 0.191266185 |
| Tube1 | 37.79269275 | -0.163332308 | 0.186610151 |
| Wisp3 | 1.507103499 | -0.493319637 | 0.828189428 |
| Fyn | 1361.385442 | -0.050630614 | 0.062233086 |
| Gm6963 | 0 | NA | NA |
| Gm16364 | 1.605310233 | 2.062898172 | 1.000276493 |
| Gm16365 | 6.794706781 | 1.06407747 | 0.457433395 |
| Traf3ip2 | 50.94917886 | 2.158706195 | 0.253115588 |
| E130307A14Rik | 318.1438882 | -0.75681432 | 0.120136692 |
| Rev3l | 1908.960953 | -0.209376464 | 0.096262544 |
| Gm22105 | 5.849095356 | -0.682055379 | 0.494525627 |
| Gm16210 | 0 | NA | NA |
| Gm26824 | 0.048949532 | -0.243569582 | 3.724945049 |
| AA474331 | 80.17615184 | -0.434074939 | 0.171228956 |
| 4930547M16Rik | 7.251366017 | -0.741098238 | 0.440694693 |
| 2010001E11Rik | 0.234609624 | 0.718219822 | 2.487251491 |
| G630090E17Rik | 3.289605183 | 5.226887106 | 0.994252265 |
| BC021785 | 2.350275864 | 1.834919077 | 0.769073881 |

| | | | |
|---|---|---|---|
| AI317395 | 5.519524843 | -0.032729528 | 0.409404032 |
| Gm8055 | 1.673936437 | -0.396623998 | 0.812670922 |
| Slc16a10 | 102.3548944 | 1.557518735 | 0.182258404 |
| Gm25613 | 0 | NA | NA |
| Gm37786 | 11.97329313 | -0.718852343 | 0.307702392 |
| Rpf2 | 140.9629893 | 0.338205869 | 0.12311284 |
| Gtf3c6 | 109.1455228 | 0.58884386 | 0.149295731 |
| U3 | 0.099155932 | 0.718202646 | 3.724945049 |
| Rnu3a | 0 | NA | NA |
| Amd1 | 757.3382757 | -0.214055955 | 0.068685249 |
| Cdk19os | 1.037245751 | 1.296179883 | 1.005277924 |
| Cdk19 | 1184.871889 | -0.26723275 | 0.095503323 |
| Slc22a16 | 0 | NA | NA |
| Ddo | 72.37407717 | -0.054101575 | 0.174584487 |
| Mettl24 | 13.24789061 | -0.727527814 | 0.31345026 |
| Cdc40 | 528.5442356 | -0.158286745 | 0.067028924 |
| Gm22948 | 2.318641632 | -2.212354228 | 0.786542353 |
| Wasf1 | 1683.342798 | -0.949031368 | 0.100770216 |
| Gpr6 | 59.52113351 | -1.432627617 | 0.264230741 |
| Fig4 | 411.9453483 | -0.187068916 | 0.078254213 |
| Gm25526 | 0 | NA | NA |
| Ak9 | 11.82815708 | -0.616683835 | 0.429208905 |
| Gm25626 | 0.159956389 | 0.219274127 | 2.798409695 |
| Gm27783 | 0 | NA | NA |
| Zbtb24 | 171.9864541 | 0.215544863 | 0.087991818 |
| Mical1 | 186.6857392 | 1.23578953 | 0.127804934 |
| Smpd2 | 91.56976239 | 0.009143316 | 0.146358271 |
| Ppil6 | 32.28546686 | -0.661407606 | 0.216474477 |
| Cd164 | 1390.831552 | 1.077651186 | 0.128744976 |
| Ccdc162 | 21.51099621 | 1.192755335 | 0.250173021 |
| Gm17196 | 0.189991899 | 0.118258099 | 2.666886151 |
| 5730435O14Rik | 0.634867864 | 1.139316209 | 1.115735807 |
| Gm29095 | 0.661465078 | 1.821689723 | 1.205701396 |
| Gm29096 | 5.25281502 | 0.558712591 | 0.434241839 |
| Gm6976 | 9.172692315 | -0.508429409 | 0.36648526 |
| Mir3473b | 0 | NA | NA |
| Cep57l1 | 78.55169042 | -0.235327244 | 0.193340397 |
| BC048559 | 0 | NA | NA |
| Sesn1 | 535.6668866 | -0.484736482 | 0.080139311 |
| Armc2 | 66.24966393 | -1.170369602 | 0.285087566 |
| Gm9855 | 93.44015898 | 0.376607772 | 0.148661713 |
| Foxo3 | 2059.889726 | -0.361164896 | 0.066401488 |
| Gm23533 | 0 | NA | NA |
| Lace1 | 140.8858273 | -0.064530547 | 0.110318674 |
| Gm15197 | 3.036843357 | -1.237262267 | 0.661869375 |
| Gm15198 | 0 | NA | NA |
| Gm25277 | 0 | NA | NA |

| | | | |
|---|---|---|---|
| Snx3 | 1059.401976 | 0.317348594 | 0.071569179 |
| Nr2e1 | 94.72567818 | -0.468647171 | 0.213797673 |
| Ostm1 | 722.7865687 | 0.417598916 | 0.081582048 |
| Gm15200 | 0.23382787 | 1.024375858 | 2.121198209 |
| Gm15199 | 0.066050875 | 0.71820487 | 3.724945049 |
| Sec63 | 1231.43905 | 0.041164866 | 0.064847766 |
| Gm26860 | 0.331010926 | 0.414158775 | 1.743941059 |
| Gm26464 | 0.049553047 | 0.71820487 | 3.724945049 |
| Scml4 | 87.95950352 | 0.883056572 | 0.16482706 |
| Sobp | 1030.323232 | -0.749675047 | 0.084778084 |
| Gm29245 | 0 | NA | NA |
| Gm29246 | 2.290242257 | -1.315240073 | 0.774561553 |
| 9030612E09Rik | 15.14098368 | -0.596567567 | 0.26424887 |
| Pdss2 | 173.4444041 | -0.291464031 | 0.093444308 |
| Gm16546 | 3.059482485 | -1.26441592 | 0.670872074 |
| Rps19-ps11 | 0 | NA | NA |
| Gm3699 | 19.13216094 | 0.999244679 | 0.285788974 |
| Bend3 | 116.2620847 | 0.477660937 | 0.178536685 |
| Gm9803 | 11.92917661 | 0.318101775 | 0.34045922 |
| 1700021F05Rik | 133.4447187 | 0.047391584 | 0.129963315 |
| Cd24a | 328.2481463 | 1.970000101 | 0.215732841 |
| F930017D23Rik | 0.149263833 | 0.23730699 | 3.49219411 |
| Gm9034 | 0.045882455 | 0.237312084 | 3.724945049 |
| Qrsl1 | 139.8918007 | -0.212256711 | 0.092683606 |
| Rtn4ip1 | 176.0176104 | 0.079279949 | 0.084571981 |
| Aim1 | 288.648516 | 4.358861536 | 0.308314366 |
| Gm10812 | 1.879239285 | 1.498942665 | 0.793979116 |
| Speer5-ps1 | 0.049553047 | 0.71820487 | 3.724945049 |
| Atg5 | 276.1062705 | 0.428060693 | 0.097535315 |
| Mir1929 | 2.194932386 | -0.499388719 | 0.727405564 |
| Prdm1 | 54.05971938 | 2.062568062 | 0.331445154 |
| Gm22087 | 0 | NA | NA |
| Gm27977 | 0 | NA | NA |
| Gm22503 | 0 | NA | NA |
| Gm24271 | 0 | NA | NA |
| Gm4795 | 0.55021313 | 0.355651914 | 1.212858365 |
| Prep | 326.8201584 | 0.174801579 | 0.067165935 |
| Gm23766 | 0 | NA | NA |
| Popdc3 | 11.87580344 | 0.274084211 | 0.289527113 |
| Gm27723 | 0 | NA | NA |
| Gm26016 | 0 | NA | NA |
| Gm15934 | 0.066050875 | 0.71820487 | 3.724945049 |
| Bves | 10.58168931 | -0.67853999 | 0.336705292 |
| Lin28b | 106.2737535 | -0.438634536 | 0.160397947 |
| Gm27731 | 0 | NA | NA |
| Hace1 | 690.7874818 | -0.804680348 | 0.103284508 |
| Gm27410 | 0 | NA | NA |

| | | | |
|---|---|---|---|
| Gm25740 | 0 | NA | NA |
| Gm23481 | 0 | NA | NA |
| Gm25650 | 0 | NA | NA |
| Gm23584 | 0 | NA | NA |
| Gm24256 | 0 | NA | NA |
| Grik2 | 2090.882152 | -0.320542992 | 0.09650603 |
| Gm26543 | 111.823433 | -0.309931506 | 0.193855967 |
| Ascc3 | 1896.390272 | 1.697148323 | 0.093659271 |
| Gm23883 | 0 | NA | NA |
| Sim1 | 14.16261422 | -0.715297027 | 0.301330289 |
| Gm26257 | 0 | NA | NA |
| Gp49a | 332.5075168 | 9.072562612 | 0.665756372 |
| Lilrb4 | 459.2260594 | 7.892734168 | 0.485082771 |
| Fam162b | 0 | NA | NA |
| Gprc6a | 0.291554621 | -1.15186453 | 1.769679715 |
| Rfx6 | 0.232111299 | 0.718217912 | 2.511643541 |
| 4933411G06Rik | 0 | NA | NA |
| Gm23906 | 0 | NA | NA |
| Vgll2 | 1.144635215 | 0.467626646 | 1.016749268 |
| Gm24084 | 0.042716945 | -0.243571806 | 3.724945049 |
| Ros1 | 0 | NA | NA |
| Gm20630 | 0 | NA | NA |
| Gm18513 | 0 | NA | NA |
| Dcbld1 | 120.8113824 | -0.219890958 | 0.128037141 |
| Gopc | 773.3070799 | -0.075531219 | 0.068485796 |
| Gm26741 | 0 | NA | NA |
| Rps19-ps10 | 0 | NA | NA |
| Nepn | 0.397314471 | -0.838291332 | 1.522863886 |
| Nus1 | 1151.136521 | 0.185053553 | 0.083952052 |
| Zfa-ps | 7.158863588 | 0.509144474 | 0.490637836 |
| Slc35f1 | 918.4030169 | -0.298796371 | 0.064793007 |
| Gm25664 | 1.948438448 | -1.740699328 | 0.88402009 |
| Cep85l | 359.2181035 | -0.271485991 | 0.123442755 |
| Pln | 12.74356708 | 0.531690695 | 0.355359669 |
| Gm5423 | 0.736769121 | 0.488847543 | 1.367976545 |
| Mcm9 | 170.9008201 | 0.935925253 | 0.108095193 |
| Asf1a | 203.4365805 | 0.400163984 | 0.108875469 |
| Fam184a | 493.2642014 | -0.444778615 | 0.091570737 |
| Man1a | 671.9857947 | 0.053862465 | 0.077921863 |
| Gm16998 | 3.462133719 | -0.995155352 | 0.597180745 |
| Gm26177 | 0 | NA | NA |
| Gm25602 | 0 | NA | NA |
| Tbc1d32 | 271.3183319 | -0.323693438 | 0.107064873 |
| Msl3l2 | 168.4746344 | 0.863694588 | 0.103682526 |
| Rps8-ps1 | 3.747707297 | 0.005533493 | 0.512961392 |
| Gja1 | 2426.848436 | -0.644765118 | 0.175415649 |
| Gm9795 | 5.035917182 | 0.051825076 | 0.443620815 |

| | | | |
|---|---|---|---|
| Gm9956 | 0.322113463 | -0.125376343 | 1.760740588 |
| Gm24285 | 0 | NA | NA |
| 4930467K11Rik | 0.534824616 | -0.595278305 | 1.32180863 |
| Hsf2 | 367.3738673 | 0.091169079 | 0.080619803 |
| Serinc1 | 9098.13279 | -0.222411993 | 0.070033218 |
| Rpl48-ps1 | 0 | NA | NA |
| Pkib | 225.7964083 | -0.215775555 | 0.157835216 |
| Gm5777 | 0.168032776 | 0.778740122 | 3.411704968 |
| Fabp7 | 502.0611919 | 0.795640363 | 0.333970109 |
| Smpdl3a | 299.0070279 | 1.361679118 | 0.17455332 |
| AW822073 | 0.119587853 | 0.237305575 | 3.724945049 |
| Gm10807 | 0.089089296 | 0.237305797 | 3.724945049 |
| Duxf3 | 0.097139547 | 0.718207094 | 3.724945049 |
| Gm4981 | 0.424461334 | 0.280491604 | 3.119829035 |
| Gm19459 | 0.218736334 | -0.243564953 | 2.218267398 |
| Gcc2 | 1409.09042 | -0.248346221 | 0.06337238 |
| Lims1 | 608.1636776 | 0.863582238 | 0.10552547 |
| Gm23058 | 0 | NA | NA |
| Ranbp2 | 4662.349267 | 0.325109075 | 0.075426016 |
| Ccdc138 | 45.03437437 | -0.360400238 | 0.198172341 |
| Edar | 4.213958645 | 0.043134265 | 0.495986009 |
| BB019430 | 0 | NA | NA |
| Gm25055 | 0 | NA | NA |
| Gm17542 | 0 | NA | NA |
| Gm9987 | 7.178240949 | -1.053574078 | 0.536177862 |
| Sh3rf3 | 355.2242412 | -0.584501417 | 0.175844751 |
| Gm27672 | 1.813029451 | -2.353266261 | 0.831280528 |
| 10-Sep | 84.1994719 | 0.591725459 | 0.179866712 |
| Gm23223 | 0.038981395 | 0.718207094 | 3.724945049 |
| AC153941.1 | 0 | NA | NA |
| Sowahc | 0 | NA | NA |
| P4ha1 | 725.4105145 | 0.327856646 | 0.100678197 |
| Gm10273 | 13.1323639 | 0.417831063 | 0.325407598 |
| Pla2g12b | 0.130524232 | -0.243571427 | 3.657871338 |
| Oit3 | 3.202472028 | 2.773500017 | 0.790491682 |
| Mcu | 514.9080926 | -0.237565775 | 0.063832012 |
| Gm10322 | 0.281194119 | -0.242888666 | 2.014260973 |
| Micu1 | 809.3638985 | -0.075147255 | 0.059006777 |
| Gm17148 | 2.409158253 | 0.269724596 | 0.749305426 |
| Gm17059 | 0.044544648 | 0.237309859 | 3.724945049 |
| Gm7399 | 5.948605025 | -1.011585753 | 0.459391942 |
| Gm23803 | 0 | NA | NA |
| Dnajb12 | 642.9386256 | -0.00333238 | 0.064963677 |
| Ddit4 | 632.8851748 | 0.196649518 | 0.175086085 |
| Anapc16 | 407.1600245 | 0.162979659 | 0.083912799 |
| Ascc1 | 125.3215762 | -0.176906934 | 0.103833595 |
| Spock2 | 9371.899727 | -0.543519988 | 0.061901584 |

| | | | |
|---|---|---|---|
| mmu-mir-6906 | 1.418442629 | -0.777525897 | 0.905999767 |
| Chst3 | 65.59010548 | 0.951334563 | 0.176012234 |
| Gm22117 | 0.042716945 | -0.243571806 | 3.724945049 |
| Psap | 17356.86525 | 0.672954779 | 0.103842392 |
| Cdh23 | 85.39716704 | 0.032704273 | 0.223829824 |
| 4632428N05Rik | 287.6489372 | 0.559743473 | 0.178554938 |
| Gm17455 | 4.08792871 | 1.33812776 | 0.497521371 |
| Slc29a3 | 540.1404158 | 1.556275982 | 0.13126667 |
| Unc5b | 525.6811087 | -0.187274916 | 0.098683951 |
| Mir6408 | 0 | NA | NA |
| Pldi | 0.297702773 | 2.461644078 | 1.882744355 |
| Gm26947 | 9.224316412 | 0.968555174 | 0.395013966 |
| Mir466j | 0.085433889 | -0.72445792 | 3.724945049 |
| Gm20611 | 0.272151178 | -1.31766086 | 2.460064227 |
| Gm20625 | 0.049553047 | 0.71820487 | 3.724945049 |
| Gm20610 | 0 | NA | NA |
| Pcbd1 | 47.59642594 | -0.049726826 | 0.183587461 |
| Sgpl1 | 896.1568652 | 1.371453288 | 0.132485026 |
| AC123530.1 | 1.014434794 | -0.822056769 | 1.172043155 |
| Tbata | 34.89983757 | -1.516181492 | 0.377319975 |
| Adamts14 | 36.59661118 | 3.859028884 | 0.362881886 |
| Prf1 | 187.0444178 | 10.90466853 | 1.021389876 |
| Rpl27a-ps1 | 0.285326127 | -0.707964549 | 1.775579644 |
| Pald1 | 146.0309485 | 0.060681463 | 0.140055661 |
| Nodal | 1.922318947 | -0.688325535 | 0.690675178 |
| Eif4ebp2 | 324.7426737 | -0.114053875 | 0.095064752 |
| D830039M14Rik | 3.979968375 | -0.57309734 | 0.552681476 |
| Lrrc20 | 191.831978 | -0.422151036 | 0.098063834 |
| Gm28447 | 0.497289099 | 0.206038459 | 1.45464695 |
| Npffr1 | 23.64047625 | 0.063870842 | 0.223801098 |
| Ppa1 | 944.9657682 | 1.122902018 | 0.137899411 |
| Sar1a | 1213.944438 | 0.557895802 | 0.103674192 |
| Tysnd1 | 151.8466627 | 0.102209194 | 0.097549636 |
| Aifm2 | 86.05177613 | 0.676501716 | 0.155687074 |
| H2afy2 | 209.0800445 | -0.07115207 | 0.113458924 |
| Mir5108 | 0 | NA | NA |
| Col13a1 | 11.90106664 | -0.6109616 | 0.338704693 |
| Gm5424 | 120.3562083 | 0.611952402 | 0.158883395 |
| 2010107G23Rik | 182.7260212 | -0.51364178 | 0.106805542 |
| Neurog3 | 0.108123235 | -0.24357403 | 3.724945049 |
| Tspan15 | 244.3145109 | 0.527908078 | 0.149414898 |
| Mir7662 | 1.416932329 | -0.801105164 | 0.809539864 |
| Tacr2 | 2.342138215 | 1.468303116 | 0.704818695 |
| Hk1 | 4224.69477 | 0.092295694 | 0.055975823 |
| Hk1os | 8.888663959 | 0.473810017 | 0.331366349 |
| Hkdc1 | 74.93408703 | -1.143452119 | 0.181223166 |
| Supv3l1 | 235.6531214 | 0.092687578 | 0.101924097 |

| | | | |
|---|---|---|---|
| 4930507D05Rik | 5.319148665 | 0.063022649 | 0.420677544 |
| Vps26a | 797.6808432 | 0.677562158 | 0.081471432 |
| Srgn | 508.9808035 | 5.330810906 | 0.391845658 |
| Kif1bp | 1369.8405 | -0.032273283 | 0.062402691 |
| RP24-255E8.1 | 2.889734049 | 0.516697699 | 0.672366829 |
| Ddx21 | 587.0568038 | 1.046594942 | 0.117242785 |
| Ddx50 | 804.8841463 | 0.072285819 | 0.07504889 |
| Stox1 | 37.04183788 | -0.826765256 | 0.24166301 |
| mmu-mir-7215 | 0 | NA | NA |
| Ccar1 | 875.9317859 | 0.212184631 | 0.090498411 |
| Snord98 | 8.868178155 | 0.352745976 | 0.432370101 |
| Tet1 | 553.5602397 | -0.274750821 | 0.120778916 |
| Gm26479 | 0 | NA | NA |
| Gm22803 | 0.09349418 | -0.24357403 | 3.724945049 |
| Slc25a16 | 401.0961555 | -0.346920735 | 0.092114573 |
| Gm25862 | 0 | NA | NA |
| Gm16128 | 5.788109619 | -0.932975738 | 0.593268004 |
| Dna2 | 37.65220541 | 0.42671558 | 0.246648238 |
| Rufy2 | 990.6841044 | -0.447223716 | 0.110401832 |
| Gm16135 | 8.108709463 | -0.688733826 | 0.498465554 |
| Hnrnph3 | 390.3370152 | -0.243674981 | 0.13587515 |
| Pbld2 | 14.50480443 | -0.483419975 | 0.275160801 |
| Gm16143 | 0.144050924 | 0.237309606 | 3.724945049 |
| Gm38063 | 0.592709862 | 0.222563672 | 1.20381418 |
| Atoh7 | 10.98456152 | 0.019837525 | 0.417780567 |
| Mypn | 24.12272005 | -1.350621392 | 0.228911623 |
| 1700120B22Rik | 2.541048356 | 0.15736104 | 0.586478208 |
| Herc4 | 532.4503219 | -0.018609766 | 0.064624092 |
| Gm26789 | 0.091666476 | -0.724455696 | 3.724945049 |
| Sirt1 | 314.9020831 | 0.026611574 | 0.082452254 |
| Gm16145 | 0.37202309 | -0.710575342 | 1.594164034 |
| Dnajc12 | 114.8880949 | 0.243811436 | 0.132906796 |
| Gm7075 | 0.075043205 | 0.237309859 | 3.724945049 |
| Ctnna3 | 57.98869911 | -1.288762342 | 0.174053522 |
| Gm10118 | 118.090575 | -1.469715794 | 0.217350164 |
| Lrrtm3 | 650.2187372 | -1.099071087 | 0.081248784 |
| AC116138.1 | 1.265617311 | -1.800524773 | 0.899462955 |
| Gm23854 | 0 | NA | NA |
| Gm28881 | 0 | NA | NA |
| 1700023F02Rik | 0.746473191 | -0.868763619 | 1.264593429 |
| Gm22829 | 0 | NA | NA |
| Gm22594 | 0 | NA | NA |
| Gm28139 | 0 | NA | NA |
| 1110002J07Rik | 9.655946214 | 1.118112741 | 0.418565482 |
| Gm26576 | 2.265170197 | 0.516042391 | 0.65771268 |
| Reep3 | 781.8290574 | 0.350341803 | 0.08549931 |
| Jmjd1c | 3049.812434 | 0.069040044 | 0.09304321 |

| | | | |
|---|---|---|---|
| Gm26359 | 0 | NA | NA |
| Nrbf2 | 127.8010227 | 0.213077747 | 0.09801702 |
| Egr2 | 185.0575354 | 0.620860395 | 0.527220149 |
| Ado | 961.6363525 | -0.116906739 | 0.082152729 |
| Gm10797 | 2.84885825 | -1.067901443 | 0.632611781 |
| Zfp365 | 3160.688214 | -0.594352244 | 0.126711773 |
| Rtkn2 | 179.8255072 | 1.963602083 | 0.154547308 |
| Gm16212 | 0.075043205 | 0.237309859 | 3.724945049 |
| Arid5b | 664.009088 | 0.552937064 | 0.105883894 |
| 1700040L02Rik | 9.415632204 | -1.717588006 | 0.389913638 |
| Tmem26 | 24.00413336 | 2.172989244 | 0.275718637 |
| Gm24172 | 0 | NA | NA |
| Rhobtb1 | 46.1805193 | -0.358283429 | 0.186900838 |
| A930033H14Rik | 22.60880403 | -0.402272546 | 0.233700224 |
| Cdk1 | 20.93667416 | 2.625558621 | 0.480432643 |
| Ank3 | 10782.92156 | -0.62113015 | 0.090930848 |
| Ccdc6 | 1500.046561 | 0.581067661 | 0.088723114 |
| Mrln | 8.103814522 | 1.293257845 | 0.452379858 |
| Slc16a9 | 87.74711258 | 1.75329295 | 0.189155444 |
| Gm7097 | 0.061920613 | 0.718207094 | 3.724945049 |
| Gm9877 | 0.095435502 | 0.718202646 | 3.724945049 |
| Fam13c | 629.0128233 | -0.385245304 | 0.096224831 |
| Gm22320 | 0.300192116 | 0.71820249 | 3.724945049 |
| Phyhipl | 1745.320192 | -0.039759923 | 0.05712465 |
| 4930533K18Rik | 0.42462922 | -0.879360189 | 1.485340574 |
| Gm24486 | 0.340478385 | -0.227649542 | 1.493527237 |
| Bicc1 | 210.8407525 | 0.968726757 | 0.131649149 |
| Gm15647 | 0 | NA | NA |
| Tfam | 397.4176102 | 0.320007533 | 0.104468601 |
| Ube2d1 | 468.1548561 | -0.229458317 | 0.057378691 |
| Gm20609 | 0.553453779 | -1.39764474 | 1.226957129 |
| Cisd1 | 406.1124699 | -0.22500996 | 0.072910426 |
| Ipmk | 440.7198033 | -0.295602434 | 0.071141658 |
| Gm22937 | 0 | NA | NA |
| 1700049L16Rik | 0 | NA | NA |
| Gm9923 | 40.21032448 | 0.285946876 | 0.174601188 |
| Gm24984 | 0 | NA | NA |
| Zwint | 7079.662798 | 0.041449971 | 0.07464549 |
| Pcdh15 | 611.2248178 | 0.058473036 | 0.189292771 |
| AC159477.1 | 0 | NA | NA |
| Gm37438 | 0 | NA | NA |
| Gm15398 | 0.158818843 | -0.196148845 | 2.807459993 |
| Gm5778 | 18.43137054 | -0.352976653 | 0.511946028 |
| Gm15397 | 18.38539889 | -0.138513752 | 0.32033553 |
| Gm6407 | 0 | NA | NA |
| Gm6419 | 9.498614017 | -0.138816089 | 0.340192757 |
| Rsph14 | 2.362931495 | -0.900889164 | 0.664836582 |

| | | | |
|---|---|---|---|
| Gnaz | 847.2906744 | -0.426198662 | 0.094382379 |
| Rab36 | 225.5149465 | -0.67875297 | 0.12769376 |
| Bcr | 2079.456346 | -0.204115324 | 0.072092141 |
| Specc1l | 984.3676726 | 0.032607089 | 0.049450225 |
| Adora2a | 245.9031525 | -1.748867692 | 0.177566679 |
| Gucd1 | 173.5614678 | 0.199625798 | 0.095837435 |
| Upb1 | 7.757773806 | 0.157687514 | 0.406250503 |
| Gm25545 | 0.075043205 | 0.237309859 | 3.724945049 |
| Snrpd3 | 401.6260286 | 0.118540895 | 0.089938007 |
| Lrrc75b | 242.2640389 | -0.662781703 | 0.102170393 |
| Ggt1 | 11.84378073 | 0.248562113 | 0.329381041 |
| Ggt5 | 50.67091169 | 2.139003238 | 0.268726268 |
| Gm9985 | 0.329446336 | 1.216669267 | 1.718738605 |
| Gm26402 | 0 | NA | NA |
| Susd2 | 127.9307858 | -0.629711701 | 0.117918311 |
| Cabin1 | 1380.689238 | -0.00355639 | 0.0713275 |
| Ddt | 134.0537612 | -0.094602482 | 0.109255333 |
| Gm20441 | 0 | NA | NA |
| Gstt3 | 21.78233339 | -0.172811245 | 0.254223911 |
| Gstt1 | 101.288934 | 0.757410947 | 0.188630539 |
| Gstt4 | 0.243832141 | 1.928591825 | 3.023648315 |
| Gstt2 | 19.40726938 | 0.455852831 | 0.232389677 |
| Mif | 1051.951317 | 0.372782631 | 0.103128021 |
| Derl3 | 13.80484017 | -0.279821623 | 0.291812751 |
| Smarcb1 | 629.9954016 | 0.148611606 | 0.062522521 |
| Mmp11 | 15.83720611 | 1.083131058 | 0.258687595 |
| Chchd10 | 1094.33119 | -0.134031904 | 0.091697251 |
| Gm867 | 15.27197981 | 2.363301289 | 0.31042749 |
| Vpreb3 | 1.181927862 | 2.366974531 | 1.067219621 |
| Gm16221 | 0.106112162 | 0.23732098 | 3.724945049 |
| Gm5134 | 1.323702316 | 0.626459732 | 0.92115323 |
| Gm16219 | 0 | NA | NA |
| Gm16220 | 0 | NA | NA |
| Mir6409 | 0 | NA | NA |
| Zfp280b | 318.8465618 | -0.439197593 | 0.095777158 |
| Slc5a4b | 0.19103638 | -0.243563445 | 2.252891137 |
| Gm16222 | 0.688368266 | 1.256323302 | 1.533068685 |
| Slc5a4a | 0.337090955 | 2.242783769 | 1.704524101 |
| Prmt2 | 399.9587037 | -0.070346972 | 0.10790117 |
| Mir678 | 0 | NA | NA |
| S100b | 1036.502615 | -0.658577169 | 0.147283818 |
| Dip2a | 1191.452903 | -0.647544233 | 0.086184739 |
| Gm16240 | 0.04393175 | -0.243571806 | 3.724945049 |
| Pcnt | 701.5738582 | -0.177055188 | 0.089566037 |
| 1700094J05Rik | 0.456503428 | -0.720072019 | 1.584762299 |
| 2610028H24Rik | 0.44299472 | -0.095165444 | 1.36637991 |
| Ybey | 77.35572085 | 0.191471363 | 0.126500964 |

| | | | |
|---|---|---|---|
| Mcm3ap | 786.2211096 | -0.199517848 | 0.089184206 |
| Lss | 497.5698186 | -0.230947528 | 0.072019317 |
| 4930483K19Rik | 0 | NA | NA |
| Spatc1l | 0.383664054 | -0.505518787 | 1.495604587 |
| Gm15343 | 0 | NA | NA |
| Ftcd | 0.710593519 | 0.223770508 | 1.149645922 |
| Col6a2 | 230.4261296 | 0.565675853 | 0.13920187 |
| Col6a1 | 323.7855034 | 0.223703201 | 0.185804626 |
| Pcbp3 | 627.0954025 | -0.072160569 | 0.11551717 |
| Gm37782 | 0.137011352 | -0.24356365 | 2.942485862 |
| Gm10787 | 0.044544648 | 0.237309859 | 3.724945049 |
| Slc19a1 | 146.9527464 | -0.264383569 | 0.10037672 |
| Col18a1 | 325.5916588 | 0.701941881 | 0.165363876 |
| Gm10941 | 3.197615884 | -0.740983098 | 0.617148013 |
| Pofut2 | 316.8806264 | 0.386705497 | 0.083753834 |
| Adarb1 | 3284.804805 | -0.58228059 | 0.073633376 |
| Fam207a | 347.3627384 | 0.373226204 | 0.098392546 |
| 2810425M01Rik | 0.856869739 | -1.199789418 | 1.111751341 |
| Itgb2 | 410.5844244 | 4.401730761 | 0.183867286 |
| Pttg1ip | 760.5419193 | 1.09720141 | 0.124563687 |
| Sumo3 | 1158.6635 | 0.405501693 | 0.08890671 |
| Ube2g2 | 330.4207087 | 0.503902033 | 0.088278571 |
| Mir1930 | 0 | NA | NA |
| Gm19402 | 0 | NA | NA |
| Gm10272 | 0.197138355 | 0.71821699 | 2.624156999 |
| Gm10024 | 0.182127695 | -0.573980991 | 3.345561342 |
| Gm10142 | 0 | NA | NA |
| Krtap12-1 | 0.19999695 | 1.253166828 | 2.619555179 |
| Gm10100 | 0.235931372 | -0.724459919 | 2.541968987 |
| Gm3233 | 0 | NA | NA |
| Gm3238 | 0 | NA | NA |
| Gm7138 | 0 | NA | NA |
| Gm3250 | 0.062426873 | 0.237312084 | 3.724945049 |
| Gm7137 | 0 | NA | NA |
| Gm19668 | 0 | NA | NA |
| Gm9507 | 0.060174537 | 0.237309859 | 3.724945049 |
| Gm2696 | 0 | NA | NA |
| Krtap10-4 | 0 | NA | NA |
| AC153874.1 | 0 | NA | NA |
| Tspear | 5.130515031 | -0.352150509 | 0.419019488 |
| Gm10318 | 0.127104694 | -0.243563766 | 3.010998535 |
| 1700009J07Rik | 0.532715572 | -0.149221904 | 1.518521208 |
| Lrrc3 | 123.618813 | -0.81286494 | 0.205707185 |
| Trpm2 | 446.088107 | -0.93086875 | 0.139596627 |
| Gm22394 | 0.066050875 | 0.71820487 | 3.724945049 |
| Gm22871 | 0.044544648 | 0.237309859 | 3.724945049 |
| 1810043G02Rik | 263.2113337 | -0.274124234 | 0.084332257 |

| | | | |
|---|---|---|---|
| Pfkl | 1143.605665 | -0.313693299 | 0.082361389 |
| Aire | 5.516484393 | -0.127557524 | 0.753791063 |
| Dnmt3l | 4.167386937 | 3.469565721 | 0.686281212 |
| Icosl | 151.2157367 | 2.262146933 | 0.170449065 |
| D10Jhu81e | 415.5710078 | -0.465650433 | 0.087376968 |
| Pwp2 | 271.1694499 | 0.148465625 | 0.068705251 |
| Trappc10 | 1524.4217 | 0.038817845 | 0.04244074 |
| Agpat3 | 3193.730264 | -0.173055779 | 0.063756575 |
| n-R5s77 | 0 | NA | NA |
| Mir7659 | 1.249419009 | -0.619782462 | 0.870775435 |
| Gm22514 | 0 | NA | NA |
| Gm10146 | 20.48723405 | 0.099730988 | 0.250665835 |
| Rrp1 | 2027.885291 | 0.043970171 | 0.089022751 |
| Mir6907 | 2.924483123 | -0.287228712 | 0.725740904 |
| Cstb | 362.581673 | 2.271861563 | 0.200407564 |
| Mir6410 | 0 | NA | NA |
| Pdxk | 2399.16992 | -0.280786674 | 0.081936745 |
| Mir6908 | 0.341053521 | 0.719843925 | 1.733511284 |
| Gm9978 | 0.176220594 | -0.724458674 | 2.719488399 |
| Gm25164 | 0 | NA | NA |
| Gm27541 | 0.044191549 | -0.243571806 | 3.724945049 |
| Ilvbl | 215.6243573 | -0.038043961 | 0.120541773 |
| Syde1 | 248.2945122 | 1.575886155 | 0.160906312 |
| Gm37514 | 0.458997541 | 1.544648928 | 1.484563164 |
| Olfr1357 | 0 | NA | NA |
| Casp14 | 0.09176491 | 0.237310218 | 3.724945049 |
| Ccdc105 | 3.747387982 | 0.259606407 | 0.514963646 |
| Slc1a6 | 295.8032966 | 0.102570383 | 0.167823339 |
| Olfr1356 | 0 | NA | NA |
| Olfr1355 | 0.089814205 | -0.24357403 | 3.724945049 |
| Olfr1354 | 0.11147366 | 1.230376831 | 3.722845817 |
| Olfr8 | 0 | NA | NA |
| Olfr1353 | 0.097139547 | 0.718207094 | 3.724945049 |
| Olfr1352 | 0 | NA | NA |
| Olfr1351 | 0 | NA | NA |
| Olfr57 | 0.139856492 | 0.237322262 | 3.574411025 |
| 2610008E11Rik | 487.1302078 | -0.273006833 | 0.07847338 |
| Vmn2r80 | 0 | NA | NA |
| Vmn2r81 | 0 | NA | NA |
| Vmn2r82 | 0.049553047 | 0.71820487 | 3.724945049 |
| Vmn2r83 | 0 | NA | NA |
| Ppap2c | 40.71318606 | 0.45276891 | 0.222499624 |
| Mir6909 | 0.048949532 | -0.243569582 | 3.724945049 |
| Mier2 | 291.2295781 | -0.042420779 | 0.091179912 |
| Theg | 1.989322217 | 0.967816401 | 0.807902499 |
| C2cd4c | 123.155375 | -0.633033522 | 0.16888065 |
| Shc2 | 358.7182919 | -0.710695979 | 0.162551519 |

| | | | |
|---|---|---|---|
| Odf3l2 | 1.872511713 | -1.153332695 | 0.956864042 |
| Madcam1 | 2.384226188 | 4.035620015 | 1.080256942 |
| Tpgs1 | 315.7362571 | 0.302951991 | 0.098746161 |
| Cdc34 | 422.4012987 | 0.709036007 | 0.100330764 |
| Gzmm | 21.34813532 | 0.676169732 | 0.332725179 |
| Gm25794 | 0 | NA | NA |
| Bsg | 2599.444032 | 0.069794397 | 0.081995182 |
| Hcn2 | 1794.794068 | -0.343021603 | 0.094777368 |
| Polrmt | 300.0243024 | -0.16398028 | 0.106073652 |
| Fgf22 | 7.710007984 | -1.448389799 | 0.394168105 |
| Rnf126 | 503.0038443 | 0.029694443 | 0.075708465 |
| Fstl3 | 34.70897125 | 2.7388172 | 0.327934401 |
| Prss57 | 5.311167766 | 0.339131911 | 0.426122922 |
| Palm | 1414.659549 | -0.54490094 | 0.082063584 |
| Misp | 27.35574496 | 3.06077079 | 0.341781236 |
| E130317F20Rik | 72.27215724 | -0.197487284 | 0.149719864 |
| Ptbp1 | 656.5387669 | 1.490793189 | 0.11983379 |
| Lppr3 | 374.0295548 | -1.258468226 | 0.113114419 |
| Mir6910 | 0 | NA | NA |
| Gm17134 | 0.135598227 | -1.1790103 | 2.942835517 |
| Prtn3 | 7.905640509 | -0.346026579 | 0.439684491 |
| Elane | 0.826815689 | 2.032231151 | 1.160570195 |
| Cfd | 0.08486385 | 0.717551239 | 3.724898127 |
| Med16 | 415.3832138 | -0.276936457 | 0.070260535 |
| Gm24019 | 2.36422992 | 0.890726784 | 0.672318749 |
| R3hdm4 | 1784.00259 | -0.549146272 | 0.080066828 |
| Gm26602 | 0.222471884 | -1.158650817 | 2.586134815 |
| Kiss1r | 45.63901687 | -0.726957856 | 0.168697171 |
| Arid3a | 140.4446512 | 0.129139414 | 0.126747371 |
| Wdr18 | 517.0323779 | 0.191544279 | 0.068178459 |
| Grin3b | 6.451345326 | -1.165625376 | 0.414208023 |
| Tmem259 | 829.796043 | 0.016580669 | 0.064283382 |
| Gm26243 | 1.926543226 | 0.312233091 | 0.712093456 |
| Cnn2 | 130.197291 | 2.331638386 | 0.188131447 |
| Abca7 | 370.3957148 | -0.338655978 | 0.097946156 |
| Hmha1 | 218.2921494 | 2.889840502 | 0.1621645 |
| Polr2e | 305.7771488 | -0.106958681 | 0.076205391 |
| Gpx4 | 1189.407151 | 0.304207733 | 0.116746855 |
| Sbno2 | 850.3215173 | 3.279113627 | 0.23802572 |
| Stk11 | 787.6366792 | 0.471855988 | 0.067915612 |
| Dos | 2722.989964 | -0.397673044 | 0.098508029 |
| Atp5d | 1392.897457 | -0.057538556 | 0.079562322 |
| Midn | 1117.501778 | 0.640896627 | 0.127814761 |
| Cirbp | 669.8346141 | -0.141318481 | 0.137737943 |
| 1600002K03Rik | 19.50566266 | 0.312384242 | 0.265379276 |
| Efna2 | 60.39084732 | -0.59293945 | 0.160956918 |
| Mum1 | 291.2372399 | -0.528492513 | 0.080497285 |

| | | | |
|---|---|---|---|
| Ndufs7 | 538.4769027 | 0.06613589 | 0.076013619 |
| Gamt | 97.19118593 | -1.195007021 | 0.140425918 |
| Dazap1 | 600.5946169 | 0.213838959 | 0.092587928 |
| Mir6911 | 0.644009716 | -2.162045274 | 1.248615535 |
| Gm15122 | 0 | NA | NA |
| Rps15 | 2187.489196 | 0.34524756 | 0.109653584 |
| Apc2 | 2618.790862 | -0.673102308 | 0.050129613 |
| 2310011J03Rik | 240.725416 | 0.816497962 | 0.096385401 |
| Pcsk4 | 31.63242481 | -1.109182549 | 0.260336237 |
| Reep6 | 48.59301236 | -1.080179292 | 0.16136176 |
| Adamtsl5 | 57.94529312 | 0.020030967 | 0.187722062 |
| Plk5 | 149.4888237 | -1.383248759 | 0.219036937 |
| Gm15123 | 0 | NA | NA |
| Gm15124 | 0.780980462 | -0.509356998 | 1.016546655 |
| Mex3d | 199.4244648 | -0.032183202 | 0.107910795 |
| Gm22721 | 0 | NA | NA |
| Mbd3 | 699.1099546 | 0.183337167 | 0.08330897 |
| Uqcr11 | 557.7849959 | -0.433756762 | 0.125086023 |
| Tcf3 | 302.852696 | 0.286277925 | 0.107945393 |
| Gm25044 | 0.104637558 | 0.23732098 | 3.724945049 |
| Onecut3 | 5.071639938 | -0.480568116 | 0.440199985 |
| 2610034O05Rik | 0.287784626 | -1.003101036 | 1.760443753 |
| Atp8b3 | 7.756145753 | 0.302087042 | 0.341967748 |
| Rexo1 | 665.8855935 | -0.051589686 | 0.094831321 |
| Klf16 | 435.0080941 | -0.572929682 | 0.122415402 |
| Gm26710 | 2.145254238 | -1.116260541 | 0.708059264 |
| Abhd17a | 822.7097596 | -0.235594052 | 0.079802974 |
| Gm29093 | 14.5563475 | -0.788666 | 0.352132148 |
| Adat3 | 37.40970625 | -0.484924812 | 0.27974158 |
| Scamp4 | 2.483253496 | 0.419631343 | 0.688622168 |
| Mir6912 | 0.975114657 | -0.947808244 | 0.992554297 |
| Csnk1g2 | 1099.862112 | -0.169774832 | 0.057105567 |
| Btbd2 | 650.9398297 | -0.345715677 | 0.088937523 |
| Mknk2 | 631.5413471 | 0.602967665 | 0.100082671 |
| Gm26541 | 0.467763849 | 0.54237417 | 1.45420688 |
| Mob3a | 177.9043315 | 1.094184917 | 0.167426696 |
| Izumo4 | 105.0430274 | -1.229432945 | 0.187859146 |
| Ap3d1 | 2190.778951 | 0.164727876 | 0.050593864 |
| Dot1l | 1006.566257 | 0.083684914 | 0.153810157 |
| Gm17151 | 0.089089296 | 0.237305797 | 3.724945049 |
| Plekhj1 | 280.4915415 | 0.407736054 | 0.08552939 |
| Sf3a2 | 462.5313837 | 0.450185944 | 0.108901734 |
| Amh | 11.52291177 | -0.791086667 | 0.411348018 |
| Jsrp1 | 14.91585505 | 1.410234583 | 0.424777073 |
| Oaz1 | 1026.280487 | 0.021263593 | 0.106360522 |
| Mir1982 | 0 | NA | NA |
| Lingo3 | 479.2193401 | -0.779252131 | 0.125688523 |

| | | | |
|---|---|---|---|
| Lsm7 | 91.89504733 | 0.135291539 | 0.142435212 |
| Sppl2b | 350.2345351 | -0.061995462 | 0.076492686 |
| Tmprss9 | 6.250140971 | 0.021753083 | 0.40341445 |
| Timm13 | 280.1181752 | 0.183624916 | 0.11127732 |
| Lmnb2 | 221.9066422 | -0.239197036 | 0.095188878 |
| Gadd45b | 364.2798403 | 2.83698559 | 0.311860619 |
| Gng7 | 1229.082418 | -1.54547811 | 0.241343794 |
| Gm3828 | 3.291353755 | -2.044791222 | 0.834848745 |
| Diras1 | 1155.444641 | -0.467421545 | 0.067613496 |
| Gm16099 | 2.516192357 | -0.86411937 | 0.62817162 |
| Slc39a3 | 692.1380104 | -0.232250821 | 0.073907443 |
| Sgta | 1016.513989 | 0.172513105 | 0.055245668 |
| Thop1 | 406.0563204 | 0.082629341 | 0.064670542 |
| Creb3l3 | 2.644382858 | 0.240526511 | 0.648572161 |
| mmu-mir-5615-1 | 0.100902008 | 1.15059983 | 3.720931964 |
| Mir5615-2 | 0 | NA | NA |
| Map2k2 | 675.7442005 | 0.446606023 | 0.086579032 |
| Zbtb7a | 2167.929022 | -0.043801676 | 0.070186631 |
| Pias4 | 256.0514789 | 0.196825585 | 0.100679337 |
| Eef2 | 12379.31089 | 0.280544905 | 0.074245585 |
| Snord37 | 3.244100523 | -0.220598063 | 0.510659172 |
| 4930442H23Rik | 0.124853745 | 0.237309606 | 3.724945049 |
| Dapk3 | 292.8022491 | 0.211203341 | 0.091548774 |
| Atcayos | 2.229235169 | 0.954381266 | 0.742805385 |
| Nmrk2 | 2.519283505 | 1.946025016 | 0.709252849 |
| Atcay | 1736.526661 | -0.697689989 | 0.074186631 |
| Zfr2 | 517.1457258 | -0.905022862 | 0.138935269 |
| Matk | 696.4187115 | -1.000804985 | 0.148222043 |
| Mrpl54 | 179.3062319 | 0.598989463 | 0.121560252 |
| Apba3 | 193.0437716 | 0.464854862 | 0.131194566 |
| Mir3057 | 0.237113133 | 0.21008858 | 1.860946881 |
| Tjp3 | 12.76057694 | 0.939924961 | 0.293197626 |
| Pip5k1c | 3939.793252 | -0.353655679 | 0.090197372 |
| Gm16315 | 2.415294132 | 0.718803826 | 0.65609284 |
| Cactin | 253.0737681 | 0.390674764 | 0.104366813 |
| Tbxa2r | 4.861979082 | 1.058178858 | 0.430756809 |
| Gipc3 | 3.688286577 | 1.618805048 | 0.559837832 |
| Hmg20b | 157.6554839 | 0.540672614 | 0.103065336 |
| Mfsd12 | 595.8008876 | 0.706054324 | 0.110945491 |
| 4930404N11Rik | 5.09738268 | 0.047379513 | 0.648449506 |
| Fzr1 | 541.0904011 | -0.151082693 | 0.059714401 |
| Dohh | 443.5414654 | 0.108631532 | 0.075568133 |
| Mir6913 | 0.75335248 | 0.28375002 | 1.097743835 |
| Smim24 | 5.77545002 | 0.153555813 | 0.471264125 |
| Nfic | 2277.670415 | 0.083499122 | 0.064739172 |
| Gm16104 | 0.970068437 | 1.668509593 | 0.985743966 |
| mmu-mir-1191b | 3.498764646 | -0.885748122 | 0.692255895 |

| | | | |
|---|---|---|---|
| Celf5 | 1725.007547 | -1.049150998 | 0.172696942 |
| Gm16105 | 0.342577373 | 0.678589637 | 1.77306583 |
| Ncln | 601.4543055 | 0.605021648 | 0.068488715 |
| S1pr4 | 11.27433504 | 3.123422036 | 0.445693273 |
| Gna15 | 25.30164811 | 1.413701924 | 0.316864675 |
| Gm16106 | 0.086908494 | -0.72445792 | 3.724945049 |
| Gm25595 | 0 | NA | NA |
| Gna11 | 1114.577472 | -0.215439496 | 0.064328262 |
| Aes | 3534.40744 | -0.177103227 | 0.073519876 |
| Gm15917 | 0.222899254 | -0.735469674 | 2.149106675 |
| Tle2 | 226.3530908 | -1.455668481 | 0.220229401 |
| Tle6 | 20.29999438 | -0.542731165 | 0.233202933 |
| BC025920 | 34.61278247 | -0.583420997 | 0.212852142 |
| Gm26896 | 0.097501745 | 0.23732098 | 3.724945049 |
| Sirt6 | 169.5015077 | 0.073263929 | 0.094194535 |
| Ankrd24 | 225.4576987 | -1.036414856 | 0.167312031 |
| Gm10778 | 8.434511479 | 0.090215435 | 0.365113955 |
| Gm8112 | 0.038981395 | 0.718207094 | 3.724945049 |
| Zfp433 | 19.00695932 | -0.133078258 | 0.232276681 |
| Gm3055 | 1.030732002 | -0.029102157 | 0.968214559 |
| Gm4767 | 1.089198015 | -1.422010247 | 1.188703128 |
| Zfp781 | 1.228365715 | -0.144595138 | 0.91838631 |
| Zfp873 | 74.79504174 | -0.206203152 | 0.159075295 |
| AU041133 | 155.8068101 | -0.019733968 | 0.108806407 |
| Gm15608 | 10.40952602 | -0.284134608 | 0.364480291 |
| Zfp938 | 255.4037447 | -0.099245897 | 0.103406971 |
| 4932415D10Rik | 0.046345709 | 0.71820487 | 3.724945049 |
| Gm4924 | 47.95002397 | -0.107999337 | 0.170319111 |
| Gm1553 | 0 | NA | NA |
| Ap3m1-ps | 10.60277313 | 0.521271485 | 0.350676862 |
| 1190007I07Rik | 27.31729701 | -0.483397999 | 0.293605637 |
| Tdg | 118.6799162 | 0.050313281 | 0.147637123 |
| Glt8d2 | 72.76759065 | -0.829849813 | 0.281868429 |
| Hcfc2 | 483.7093496 | -0.26133413 | 0.069499746 |
| Nfyb | 378.4988816 | -0.192608476 | 0.074919418 |
| 1700028I16Rik | 0.19184735 | 1.55431524 | 2.659048956 |
| Txnrd1 | 1189.720619 | 1.200319454 | 0.133100909 |
| Gm4799 | 4.084400745 | 1.702465135 | 0.583173929 |
| Chst11 | 1110.164625 | 0.945858385 | 0.106349622 |
| 1700025N21Rik | 0.075043205 | 0.237309859 | 3.724945049 |
| Gm23122 | 0 | NA | NA |
| Slc41a2 | 340.536907 | -0.085294798 | 0.063535002 |
| D10Wsu102e | 409.3325013 | -0.181626432 | 0.078977278 |
| Aldh1l2 | 203.3414108 | 0.876877179 | 0.131095767 |
| A230046K03Rik | 1351.145369 | 0.841835695 | 0.079418562 |
| Appl2 | 956.0063235 | -1.072803428 | 0.183406421 |
| 1500009L16Rik | 86.40011601 | -0.424423514 | 0.133017815 |

| | | | |
|---|---|---|---|
| Gm37908 | 0 | NA | NA |
| Nuak1 | 1233.243768 | -0.320585136 | 0.106251459 |
| 4930463O16Rik | 0.038981395 | 0.718207094 | 3.724945049 |
| Ckap4 | 392.0257556 | 0.840524999 | 0.116175266 |
| Platr7 | 0.182269687 | 0.718205987 | 3.313323614 |
| Tcp11l2 | 249.9597353 | -0.108658206 | 0.114176524 |
| Gm25657 | 0.602022574 | 1.230570406 | 1.327560395 |
| Gm17249 | 4.368992961 | -0.749068106 | 0.467989139 |
| Polr3b | 386.4218719 | 0.555539508 | 0.070139305 |
| Rfx4 | 218.129252 | -0.334470036 | 0.148211417 |
| Gm24226 | 0.075043205 | 0.237309859 | 3.724945049 |
| Ric8b | 669.9951763 | -0.355727573 | 0.052060685 |
| Fhl4 | 12.95936291 | 0.349858462 | 0.320629452 |
| Tmem263 | 569.4293162 | -0.049563401 | 0.081253085 |
| Mterf2 | 117.1077398 | -0.332013336 | 0.151661444 |
| Cry1 | 350.2670687 | -0.006300335 | 0.152600251 |
| Gm27392 | 0 | NA | NA |
| Gm8394 | 52.17199378 | 0.742658736 | 0.204959775 |
| Gm25281 | 0 | NA | NA |
| Btbd11 | 507.0576605 | -0.320562852 | 0.098857439 |
| Pwp1 | 247.3605164 | 0.473700695 | 0.089788699 |
| Prdm4 | 408.5061188 | 0.355608132 | 0.070679577 |
| Ascl4 | 2.415104612 | -1.235234067 | 0.739521533 |
| Rtcb | 1117.008586 | 0.445496922 | 0.082384519 |
| Bpifc | 3.577274519 | 1.552783449 | 0.619291671 |
| A230060F14Rik | 5.145365424 | 0.70877209 | 0.472097102 |
| Fbxo7 | 420.5312043 | 0.519665451 | 0.08641196 |
| Gm24430 | 0.537185458 | 1.532752009 | 1.407310175 |
| Syn3 | 338.3405326 | -0.768670802 | 0.169467671 |
| 4930486F22Rik | 0.066050875 | 0.71820487 | 3.724945049 |
| Gm15990 | 3.87042643 | 5.850307584 | 0.912720492 |
| Timp3 | 2834.917203 | 1.62252741 | 0.13732858 |
| Gm6729 | 0 | NA | NA |
| Gm5174 | 1.40433107 | -0.364811535 | 1.022863855 |
| 1810014B01Rik | 16.79727865 | -0.468686019 | 0.271909083 |
| Hsp90b1 | 4072.803122 | 0.638216816 | 0.106141139 |
| Gm15344 | 13.86957958 | -1.817364419 | 0.327974704 |
| Gm25745 | 0 | NA | NA |
| Ttc41 | 22.08483189 | 0.046729438 | 0.285454807 |
| Fabp3-ps1 | 8.356713702 | -0.547508341 | 0.419091217 |
| Nt5dc3 | 1819.324601 | 0.021881548 | 0.085394555 |
| Gm16268 | 9.292864951 | -0.225960904 | 0.36454004 |
| Stab2 | 7.172886002 | -0.060389864 | 0.469378 |
| Gm16280 | 0 | NA | NA |
| Gm16270 | 0.987182834 | 3.526778622 | 1.10841096 |
| Gm16271 | 0.570993603 | -0.450995333 | 1.462656734 |
| Gm16269 | 0 | NA | NA |

| | | | |
|---|---|---|---|
| 1700113H08Rik | 0.362813327 | -0.813032014 | 1.533591682 |
| Gm10764 | 0.093831153 | -0.243571806 | 3.724945049 |
| Gm23191 | 0 | NA | NA |
| Ascl1 | 36.60879247 | -1.064660983 | 0.21976964 |
| Pah | 2.972707099 | -1.064400261 | 0.624106452 |
| Igf1os | 0.184770788 | 0.718214619 | 2.691102959 |
| Igf1 | 193.7112454 | -0.215170822 | 0.115449432 |
| Pmch | 109.1827471 | -0.715281151 | 0.169213537 |
| Parpbp | 16.45101383 | 0.132449958 | 0.337070512 |
| Nup37 | 44.56539038 | 0.64164432 | 0.190609521 |
| Gm26357 | 0 | NA | NA |
| Ccdc53 | 129.2902162 | 0.275862363 | 0.142203401 |
| Dram1 | 46.36865287 | 2.094786694 | 0.256770156 |
| Gnptab | 1221.700717 | 0.393833402 | 0.054148902 |
| Gm26546 | 4.34695503 | 0.779407673 | 0.455191792 |
| Gm16235 | 0.918591783 | 0.840769493 | 1.023026733 |
| Chpt1 | 1036.030604 | -0.624552314 | 0.103312104 |
| Sycp3 | 7.536531091 | 0.169539534 | 0.45059956 |
| Mybpc1 | 139.6857782 | -0.591490696 | 0.228286663 |
| Spic | 2.940423805 | 4.320233236 | 0.932287974 |
| n-R5s79 | 0 | NA | NA |
| Gm4925 | 6.36087602 | -0.224971074 | 0.384363638 |
| Arl1 | 1132.198676 | 0.323509797 | 0.083055778 |
| Utp20 | 565.1990458 | 0.334624351 | 0.072777897 |
| Slc5a8 | 3.475576094 | 1.219445651 | 0.709823849 |
| Ano4 | 411.0563822 | -0.290693813 | 0.101854923 |
| Gas2l3 | 106.5758252 | 1.398160064 | 0.250910357 |
| Nr1h4 | 10.81026118 | 1.117357075 | 0.608888767 |
| Slc17a8 | 21.16662692 | -0.152837837 | 0.319774843 |
| Scyl2 | 725.9700778 | 0.116033219 | 0.07738054 |
| 1500026H17Rik | 20.55184199 | -0.926734208 | 0.220321722 |
| Actr6 | 166.1310719 | 0.161209164 | 0.111592351 |
| Uhrf1bp1l | 1814.744137 | 0.248712509 | 0.06572477 |
| Gm26180 | 0.045882455 | 0.237312084 | 3.724945049 |
| Anks1b | 4400.438249 | -0.546345369 | 0.093473942 |
| Gm22602 | 2.708289425 | -1.139445301 | 0.703539865 |
| Apaf1 | 344.5692301 | 0.922029733 | 0.11511977 |
| Gm25390 | 0 | NA | NA |
| Ikbip | 137.6747652 | -0.059684468 | 0.135340146 |
| Slc25a3 | 3200.487187 | 0.034609943 | 0.055782412 |
| Gm24119 | 26.8554828 | -0.225274921 | 0.28952239 |
| Tmpo | 631.4065621 | 0.624153769 | 0.0866386 |
| Gm26765 | 0.307480012 | -0.054532657 | 1.992054625 |
| Rmst | 270.1421029 | -0.262079981 | 0.120395562 |
| Gm27544 | 10.08854525 | 0.056095554 | 0.300935172 |
| Mir135a-2 | 4.853900475 | -0.193793393 | 0.457398962 |
| Gm27999 | 0 | NA | NA |

| | | | |
|---|---|---|---|
| Gm27608 | 0 | NA | NA |
| Gm27588 | 11.46356049 | -0.026561828 | 0.326919922 |
| Gm27753 | 0 | NA | NA |
| Gm27582 | 0.044191549 | -0.243571806 | 3.724945049 |
| Gm27366 | 0.132101749 | 1.209894437 | 3.72418822 |
| Gm27419 | 0 | NA | NA |
| Mir1251 | 5.125747786 | -0.427204673 | 0.454041905 |
| Gm26700 | 0.516149922 | 1.018684564 | 1.377928678 |
| Gm20757 | 0.625729796 | -0.921153589 | 1.230919965 |
| Gm25884 | 0 | NA | NA |
| Gm22582 | 0 | NA | NA |
| Nedd1 | 87.64473989 | 1.179483985 | 0.170991267 |
| Gm24241 | 0.242925194 | -1.072488203 | 2.100181093 |
| Cfap54 | 546.4457364 | -1.075906656 | 0.180803809 |
| Cdk17 | 2255.31025 | -0.357147513 | 0.069256673 |
| Mir1931 | 9.166638643 | -0.859410856 | 0.371390037 |
| Elk3 | 176.0696239 | 2.477102121 | 0.189367246 |
| Mir7688 | 0.168606834 | -0.623357197 | 2.775232772 |
| Lta4h | 347.4203626 | 0.008343919 | 0.084505968 |
| Hal | 122.7623701 | 6.212625169 | 0.585531296 |
| Amdhd1 | 3.584129096 | 3.400523035 | 0.773587128 |
| Ccdc38 | 1.217260431 | 0.570767368 | 0.840138817 |
| Snrpf | 77.19710455 | 0.350091847 | 0.178004092 |
| 4933408J17Rik | 0.132790949 | -0.724459922 | 2.970952992 |
| Ntn4 | 142.6461492 | -0.43320552 | 0.157163793 |
| Gm8580 | 0.167271369 | 0.718214954 | 2.764694848 |
| Gm15915 | 0.796817125 | 1.878531146 | 1.142916105 |
| Gm15963 | 1.049255278 | -0.716706133 | 0.988061299 |
| Gm3571 | 24.10692972 | 0.206831788 | 0.219843918 |
| Gm26239 | 0 | NA | NA |
| Usp44 | 0.703834089 | 0.469425561 | 1.1252789 |
| Metap2 | 993.8475694 | 0.338670377 | 0.097541864 |
| 4930471D02Rik | 0.044191549 | -0.243571806 | 3.724945049 |
| Gm25962 | 0 | NA | NA |
| Vezt | 1354.242168 | -0.403605804 | 0.121229195 |
| Mir331 | 0 | NA | NA |
| Fgd6 | 264.4343943 | 0.036057793 | 0.115460903 |
| Nr2c1 | 165.6055157 | 0.083758773 | 0.113816567 |
| Gm26122 | 0 | NA | NA |
| Ndufa12 | 388.6068376 | -0.00340448 | 0.093321021 |
| Gm4792 | 0.047948698 | -0.243569582 | 3.724945049 |
| Tmcc3 | 859.2376155 | 0.317367191 | 0.115391849 |
| Gm16155 | 3.245903696 | -0.112430725 | 0.650056297 |
| Tmcc3os | 2.419038749 | 0.709217097 | 0.757789943 |
| Mir7211 | 0 | NA | NA |
| Cep83os | 296.969172 | -0.306550846 | 0.084814364 |
| Cep83 | 388.069701 | -0.081021043 | 0.085014273 |

| | | | |
|---|---|---|---|
| Plxnc1 | 869.6228163 | -0.241800985 | 0.079882041 |
| Gm24186 | 1.18741419 | 0.366319237 | 0.893131754 |
| Gm25266 | 0.715554978 | -0.518774839 | 1.283777992 |
| Cradd | 45.57421372 | 0.513676405 | 0.164722349 |
| 2310039L15Rik | 0 | NA | NA |
| Socs2 | 273.8451504 | 0.017164822 | 0.129409764 |
| 5730420D15Rik | 1.317555436 | 0.323028575 | 0.895972568 |
| Mrpl42 | 211.5063124 | -0.061172284 | 0.11273214 |
| Ube2n | 1238.439884 | -0.170223657 | 0.056482249 |
| Nudt4 | 1713.099017 | -0.180792608 | 0.119308939 |
| Mir3058 | 2.797676517 | -0.390756213 | 0.56946652 |
| Gm23776 | 0 | NA | NA |
| Anapc15-ps | 1.031766027 | 0.714002219 | 1.081337621 |
| 4732465J04Rik | 0.435690537 | 0.602581406 | 1.342869983 |
| Gm24433 | 0 | NA | NA |
| Eea1 | 1239.903993 | 0.211849635 | 0.060463243 |
| Gm5426 | 0 | NA | NA |
| Gm22913 | 0 | NA | NA |
| Gm20091 | 0.131209559 | 1.165289508 | 2.972496041 |
| Gm8601 | 0 | NA | NA |
| Btg1 | 1120.140172 | 0.864033596 | 0.126708604 |
| Gm24587 | 0 | NA | NA |
| mmu-mir-3966 | 0 | NA | NA |
| Dcn | 434.2514662 | 1.860404687 | 0.175606806 |
| Lum | 32.29644331 | 1.003189883 | 0.235837565 |
| Kera | 0.075043205 | 0.237309859 | 3.724945049 |
| Epyc | 1.611010591 | 1.967056222 | 0.908159706 |
| Gm10754 | 7.111199995 | -2.368598213 | 0.497784553 |
| Ccer1 | 0 | NA | NA |
| C030005K15Rik | 6.350231153 | -1.373904196 | 0.54031379 |
| Gm25658 | 1.646274761 | -2.839145245 | 0.920440938 |
| Gm37631 | 0.066050875 | 0.71820487 | 3.724945049 |
| Atp2b1 | 8738.965953 | -0.582892557 | 0.115442128 |
| Poc1b | 99.61505393 | 0.362678708 | 0.117997517 |
| Galnt4 | 92.75288412 | 0.517342628 | 0.150068019 |
| Phxr2 | 0 | NA | NA |
| Gm16239 | 0.066050875 | 0.71820487 | 3.724945049 |
| Dusp6 | 640.3874158 | 0.308016948 | 0.187972409 |
| B530045E10Rik | 1.636950402 | 0.539103745 | 0.896730956 |
| Gad1-ps | 4.900567814 | 0.632327917 | 0.52311327 |
| Csl | 0.893710704 | -0.73701525 | 0.996136838 |
| Kitl | 948.0407053 | 0.171770787 | 0.131179791 |
| Gm22918 | 0 | NA | NA |
| Gm25287 | 0 | NA | NA |
| Gm9476 | 0 | NA | NA |
| Gm8701 | 0 | NA | NA |
| Gm4301 | 0 | NA | NA |

| | | | |
|---|---|---|---|
| Gm4302 | 0 | NA | NA |
| Gm4303 | 0 | NA | NA |
| Gm4305 | 0 | NA | NA |
| Gm8711 | 0 | NA | NA |
| Gm4307 | 0 | NA | NA |
| Gm4308 | 0 | NA | NA |
| Gm8723 | 0 | NA | NA |
| Gm4312 | 0 | NA | NA |
| Gm4781 | 0 | NA | NA |
| Tmtc3 | 401.2372202 | -0.049103312 | 0.075317444 |
| Cep290 | 566.5526866 | -0.073974727 | 0.096573485 |
| 4930430F08Rik | 178.6427657 | -0.251719378 | 0.154328856 |
| 1700017N19Rik | 0.256989293 | 0.620134924 | 2.097725036 |
| Gm38219 | 0 | NA | NA |
| Gm26278 | 0.136194719 | -0.724453471 | 3.724945049 |
| Mgat4c | 312.1695497 | -0.455405279 | 0.093540607 |
| Mir3059 | 1.49611593 | 0.443954645 | 0.880851472 |
| Gm26988 | 0.108767819 | 0.237318756 | 3.724945049 |
| Nts | 88.60440578 | 1.054838106 | 0.237633643 |
| Rassf9 | 16.58629026 | 1.056240522 | 0.29215299 |
| Alx1 | 1.374376221 | -0.854695566 | 0.905566911 |
| Gm17028 | 0 | NA | NA |
| Lrriq1 | 58.42473187 | -0.473422172 | 0.182625627 |
| Gm25376 | 0 | NA | NA |
| Slc6a15 | 616.6613696 | -0.300732693 | 0.085380944 |
| Gm26923 | 0 | NA | NA |
| Gm25143 | 0 | NA | NA |
| Gm6763 | 0 | NA | NA |
| Gm21293 | 0 | NA | NA |
| Gm8764 | 0 | NA | NA |
| Gm21304 | 0 | NA | NA |
| Gm21312 | 0 | NA | NA |
| Gm20765 | 0 | NA | NA |
| Gm4340 | 0 | NA | NA |
| Gm22339 | 0 | NA | NA |
| Mir6411 | 0 | NA | NA |
| Gm22945 | 0 | NA | NA |
| Gm25522 | 0 | NA | NA |
| Gm25587 | 0 | NA | NA |
| Gm25422 | 0.254945911 | -0.243567613 | 2.095323845 |
| Gm36997 | 0.361186056 | 0.250737976 | 1.572115365 |
| Tmtc2 | 369.308983 | -0.252759809 | 0.102619252 |
| Gm15662 | 0 | NA | NA |
| Gm15666 | 12.30954061 | -0.969325662 | 0.403247046 |
| Gm15664 | 0 | NA | NA |
| Gm15663 | 93.64214882 | -0.532444879 | 0.132071064 |
| Mettl25 | 107.6127449 | -0.162696964 | 0.14364835 |

| | | | |
|---|---|---|---|
| n-R5s80 | 1.30631546 | 1.123308904 | 1.180984111 |
| Ccdc59 | 282.5902914 | 0.401491913 | 0.113273821 |
| Gm24237 | 0 | NA | NA |
| Gm25342 | 0 | NA | NA |
| Rps15a-ps1 | 0 | NA | NA |
| C230021G24Rik | 14.38138754 | -0.259643818 | 0.281335065 |
| Ppfia2 | 1751.263262 | -0.424505724 | 0.099450578 |
| Acss3 | 21.10431072 | -0.137014043 | 0.262109675 |
| Gm24745 | 0.047948698 | -0.243569582 | 3.724945049 |
| Lin7a | 1127.089259 | -0.537001399 | 0.105478654 |
| Gm38117 | 0 | NA | NA |
| AC155168.1 | 0.358057361 | -0.700599979 | 3.713097053 |
| Myf5 | 0.154585317 | 1.652042906 | 2.82119606 |
| Myf6 | 0.075043205 | 0.237309859 | 3.724945049 |
| Ptprq | 5.567795988 | -0.156987342 | 0.431650313 |
| Otogl | 7.088242965 | -0.392267676 | 0.408930136 |
| Ppp1r12a | 1620.397248 | 0.237766372 | 0.076216907 |
| Pawr | 30.19438579 | 1.609019312 | 0.264987079 |
| Gm23105 | 0.203286941 | 1.526406207 | 2.603554853 |
| Gm25591 | 0 | NA | NA |
| Syt1 | 12677.56377 | -0.783932507 | 0.114066637 |
| Gm22097 | 0.164132501 | 0.237308425 | 3.397905533 |
| AC125531.1 | 0.108123235 | -0.24357403 | 3.724945049 |
| Gm23041 | 0.2701572 | -0.2435632 | 1.816643769 |
| Nav3 | 1988.147118 | -0.631565941 | 0.116087712 |
| Gm36935 | 26.04041665 | -0.780774115 | 0.412357439 |
| E2f7 | 15.83057167 | 0.872003898 | 0.319404922 |
| 1700020G17Rik | 0 | NA | NA |
| Gm23698 | 0 | NA | NA |
| Gm22186 | 0 | NA | NA |
| Gm22236 | 0 | NA | NA |
| Csrp2 | 26.62848996 | 0.869200442 | 0.225907763 |
| Zdhhc17 | 1450.600249 | -0.284540301 | 0.052524887 |
| Gm5428 | 96.95467252 | 0.534550003 | 0.16595477 |
| Gm22101 | 0 | NA | NA |
| Osbpl8 | 2307.979124 | -0.246588597 | 0.051253236 |
| Bbs10 | 50.75359481 | -0.097737561 | 0.159929996 |
| Nap1l1 | 1669.332245 | -0.154384077 | 0.071545135 |
| Gm25117 | 163.1343521 | 0.004089301 | 0.470657888 |
| Phlda1 | 275.8361962 | 0.878252539 | 0.180610152 |
| Gm28592 | 7.265044534 | 0.397531235 | 0.471708788 |
| 4933440J02Rik | 0 | NA | NA |
| Gm37189 | 0.109727584 | 0.718200421 | 3.724945049 |
| Krr1 | 496.8011956 | 0.015481238 | 0.072547276 |
| Glipr1 | 31.42829398 | 2.92534773 | 0.321571426 |
| Glipr1l1 | 0.138226157 | -0.243563636 | 2.934480157 |
| Glipr1l2 | 0.144013665 | 1.483424291 | 3.538333097 |

| | | | |
|---|---|---|---|
| Gm6808 | 0 | NA | NA |
| Gm15914 | 0 | NA | NA |
| Caps2 | 5.498926737 | 0.39602392 | 0.445358655 |
| Kcnc2 | 1574.614437 | -0.455154871 | 0.085737946 |
| 1700010J16Rik | 0.193455382 | -0.243572074 | 2.632428893 |
| Atxn7l3b | 2117.555783 | -0.198685134 | 0.049608056 |
| Gm26596 | 1.459047308 | -0.250154003 | 0.793240421 |
| Gm22507 | 0.128477747 | 0.718203484 | 3.677131942 |
| Gm38333 | 0 | NA | NA |
| Gm38281 | 0 | NA | NA |
| Trhde | 612.5170607 | -0.880415654 | 0.093132181 |
| 4930473D10Rik | 0 | NA | NA |
| Gm15723 | 23.33627371 | -0.538148782 | 0.25617924 |
| Gm25319 | 0 | NA | NA |
| Gm22070 | 0 | NA | NA |
| Tph2 | 55.63792861 | 0.220817372 | 0.657754445 |
| Tbc1d15 | 653.7365103 | 0.305662229 | 0.097034034 |
| Gm22794 | 0.044544648 | 0.237309859 | 3.724945049 |
| Rab21 | 1138.336811 | 0.792371401 | 0.094676165 |
| Tmem19 | 316.2960843 | -0.053253067 | 0.077538932 |
| Thap2 | 234.4484823 | -0.268938178 | 0.238062719 |
| Zfc3h1 | 1151.166255 | -0.102441002 | 0.101560547 |
| Lgr5 | 129.3954019 | -0.713861549 | 0.27628876 |
| A930009A15Rik | 4.998511455 | -1.142140985 | 0.497575285 |
| Tspan8 | 5.527738138 | 0.708976433 | 0.462555798 |
| Ptprr | 373.9766246 | -0.62648645 | 0.104637431 |
| 4933416C03Rik | 0.062426873 | 0.237312084 | 3.724945049 |
| Ptprb | 234.5662353 | 0.373698096 | 0.197588692 |
| Gm26495 | 0.042716945 | -0.243571806 | 3.724945049 |
| Kcnmb4 | 239.9415874 | -0.617777051 | 0.130732244 |
| Kcnmb4os1 | 0.534709659 | -0.48635464 | 1.555328099 |
| Kcnmb4os2 | 19.7696678 | -0.540480407 | 0.244030572 |
| Cnot2 | 821.4440981 | -0.083633004 | 0.0562371 |
| Gm25190 | 6.442501395 | -0.792892265 | 0.422855567 |
| 5330438D12Rik | 61.66108682 | -0.12090165 | 0.166466876 |
| Gm26579 | 0.044191549 | -0.243571806 | 3.724945049 |
| 1700030O20Rik | 0.066050875 | 0.71820487 | 3.724945049 |
| Myrfl | 6.514367323 | 0.849549409 | 0.523858729 |
| Rab3ip | 355.5556215 | 0.22671978 | 0.082345713 |
| Gm10271 | 0 | NA | NA |
| D630029K05Rik | 3.06831776 | 2.140798639 | 0.731463701 |
| Best3 | 10.8055888 | 0.66621532 | 0.337929314 |
| Lrrc10 | 0.159795471 | 0.733203719 | 2.80534854 |
| Cct2 | 1508.454392 | 0.399891106 | 0.072785156 |
| Frs2 | 1041.72447 | -0.09671203 | 0.063723155 |
| Gm26311 | 0 | NA | NA |
| Yeats4 | 230.0577702 | 0.122852657 | 0.114590889 |

| | | | |
|---|---|---|---|
| 9530003J23Rik | 0.138076337 | 0.718204987 | 3.589802529 |
| Gm10936 | 0 | NA | NA |
| Lyz2 | 917.3809082 | 4.365279179 | 0.450522874 |
| Lyz1 | 2.092478253 | 0.643037956 | 0.7689401 |
| Cpsf6 | 2549.871472 | -0.217223942 | 0.069597101 |
| Gm9002 | 0 | NA | NA |
| Kifc5c-ps | 0 | NA | NA |
| Cpm | 200.0726009 | -0.320742502 | 0.136355041 |
| Gm25709 | 2.906247223 | -0.677053166 | 0.603814152 |
| Mdm2 | 840.5416714 | 1.007114259 | 0.096639399 |
| Slc35e3 | 296.2709981 | -0.097046151 | 0.080673353 |
| Nup107 | 192.1877859 | -0.077849163 | 0.102249442 |
| Rap1b | 1149.527098 | 1.49064382 | 0.125834592 |
| Gm37852 | 0 | NA | NA |
| Mdm1 | 91.75534052 | -0.201930314 | 0.12897022 |
| Il22 | 14.70729829 | 1.44650976 | 0.423156547 |
| Iltifb | 0.488214936 | 1.005730007 | 1.466731296 |
| Gm9040 | 0 | NA | NA |
| Gm9044 | 0 | NA | NA |
| Gm9045 | 0.068097359 | -0.243569582 | 3.724945049 |
| Gm9046 | 0 | NA | NA |
| Gm9048 | 0 | NA | NA |
| Gm9049 | 0.219658751 | -1.513883149 | 3.701869585 |
| Ifng | 22.72435143 | 8.188814229 | 1.02948183 |
| Dyrk2 | 260.899164 | -0.140188255 | 0.110615723 |
| 4932442E05Rik | 0.27373663 | 1.200221346 | 1.795575335 |
| Gm26976 | 0 | NA | NA |
| 1700064J06Rik | 0 | NA | NA |
| Cand1 | 2796.110183 | -0.067811263 | 0.043173131 |
| 1700025F24Rik | 0.942170168 | -0.631327514 | 1.019706287 |
| Grip1os3 | 1.395336821 | -1.813428174 | 0.872294742 |
| Gm16321 | 1.40953533 | -1.88501366 | 0.919076088 |
| Grip1 | 365.1462489 | -0.465261474 | 0.123148641 |
| Grip1os2 | 0 | NA | NA |
| Grip1os1 | 0.072025462 | 0.237312084 | 3.724945049 |
| Helb | 127.7033369 | 1.357571987 | 0.118690494 |
| Irak3 | 30.23155639 | 2.548276058 | 0.279934128 |
| Tmbim4 | 275.0985624 | 0.782839466 | 0.123207947 |
| Llph | 97.51439187 | 0.422614684 | 0.167758495 |
| Gm38141 | 0 | NA | NA |
| Hmga2 | 52.04420527 | 5.283547709 | 0.628289921 |
| 1700006J14Rik | 0.548916986 | 1.175643814 | 1.384501167 |
| 9230105E05Rik | 0 | NA | NA |
| Mir763 | 0.331210058 | 2.146331532 | 1.986775398 |
| Gm24298 | 0 | NA | NA |
| Gm38289 | 0.370157025 | 2.517771483 | 1.614527292 |
| Gm37505 | 0.097139547 | 0.718207094 | 3.724945049 |

| | | | |
|---|---|---|---|
| 4921513I03Rik | 0 | NA | NA |
| Msrb3 | 98.13132724 | 0.207272038 | 0.120616139 |
| Gm15961 | 1.450891998 | -0.507055071 | 0.822159826 |
| Gm15910 | 3.363554134 | 1.135839889 | 0.55504406 |
| Gm4473 | 0.74372259 | -1.170884705 | 1.193258735 |
| Lemd3 | 732.3495551 | -0.175602723 | 0.06732376 |
| Gm16166 | 0.157815785 | 0.718205257 | 3.466717377 |
| Gm23632 | 0.389039412 | -0.953248313 | 1.510088348 |
| Wif1 | 23.54109142 | 0.547448253 | 0.300800022 |
| Tbc1d30 | 832.5656543 | -1.086311898 | 0.099954988 |
| Gns | 1953.99692 | 1.132074645 | 0.105039669 |
| Rassf3 | 242.4931995 | -0.700663423 | 0.123065206 |
| Tbk1 | 841.1038649 | 0.504677363 | 0.091709097 |
| Xpot | 1103.074494 | -0.074277282 | 0.051467143 |
| Gm24330 | 0.750750929 | -0.623015528 | 1.238365177 |
| D930020B18Rik | 4.785986039 | -0.224695562 | 0.499395252 |
| BC048403 | 125.9765068 | -0.017134738 | 0.100889201 |
| C030027H14Rik | 0.046345709 | 0.71820487 | 3.724945049 |
| Srgap1 | 808.6398788 | 0.328159787 | 0.078741721 |
| Tmem5 | 317.9777111 | 0.428688031 | 0.095461195 |
| Gm9081 | 0.90419666 | -0.017007215 | 1.018396255 |
| Avpr1a | 7.942965041 | 1.235007322 | 0.395184989 |
| Mir6412 | 0 | NA | NA |
| Ppm1h | 1340.411078 | 0.183937772 | 0.079304955 |
| Mir8104 | 0 | NA | NA |
| Mirlet7i | 1.909998494 | 1.65032296 | 0.927734629 |
| Mon2 | 1822.582098 | -0.11312141 | 0.045310561 |
| Usp15 | 1257.977968 | 0.129647607 | 0.056842834 |
| Gm27920 | 0 | NA | NA |
| Fam19a2 | 696.2404132 | -0.484931535 | 0.086714799 |
| Gm22958 | 7.051847066 | -1.409368882 | 0.51882105 |
| Gm23777 | 0 | NA | NA |
| Slc16a7 | 304.2321303 | -0.966777981 | 0.152900113 |
| Lrig3 | 67.70650913 | -0.295147939 | 0.144100249 |
| Gm22183 | 0 | NA | NA |
| Mir378d | 0.20330765 | -1.203822874 | 2.208071316 |
| Gm24179 | 0.30098968 | -0.243568757 | 2.036205644 |
| Xrcc6bp1 | 46.49479728 | -0.485701727 | 0.180521009 |
| Ctdsp2 | 1189.150957 | 0.432270567 | 0.089320554 |
| Mir26a-2 | 0.925632266 | -0.626472235 | 1.152112846 |
| Mir546 | 0 | NA | NA |
| Avil | 16.18070083 | 0.648962128 | 0.28250028 |
| Tsfm | 123.6160451 | 0.26358138 | 0.110372286 |
| Mettl21b | 2.262034746 | 0.145664996 | 0.724411745 |
| Mettl1 | 114.9215301 | 0.934059649 | 0.138503701 |
| Cyp27b1 | 5.379193204 | 0.918521707 | 0.476995976 |
| 9-Mar | 204.9397292 | -0.328855597 | 0.111935387 |

| | | | |
|---|---|---|---|
| Cdk4 | 393.057798 | 0.575825438 | 0.129160831 |
| Tspan31 | 608.5951114 | 0.905195309 | 0.107957273 |
| Agap2 | 8823.335821 | -1.124559264 | 0.13950374 |
| Os9 | 1552.713354 | 0.392433156 | 0.079053838 |
| Gm23664 | 0.045882455 | 0.237312084 | 3.724945049 |
| B4galnt1 | 858.1814942 | -0.330509982 | 0.071327437 |
| Slc26a10 | 18.10165712 | -1.118912492 | 0.454395873 |
| Arhgef25 | 709.5156223 | -1.040385557 | 0.10567347 |
| Dtx3 | 1335.531959 | -0.146234381 | 0.051372835 |
| F420014N23Rik | 11.7378327 | -0.102471348 | 0.405228756 |
| Pip4k2c | 1116.600625 | -0.171209084 | 0.061628834 |
| Kif5a | 22627.74193 | -0.314171833 | 0.105136909 |
| Dctn2 | 1693.964189 | -0.045601568 | 0.049109837 |
| Mbd6 | 504.0837198 | -0.144127095 | 0.139249218 |
| Gm20492 | 1.795058199 | -0.557766251 | 0.730821876 |
| Gm4189 | 3.902576099 | -1.370349619 | 0.606815966 |
| Ddit3 | 291.5705915 | 2.171168971 | 0.198348204 |
| Mars | 495.381841 | 0.519188992 | 0.098768013 |
| Gm6283 | 1.594842726 | -1.497706004 | 1.029177932 |
| Arhgap9 | 51.31007067 | 4.263445946 | 0.328924894 |
| Gli1 | 50.51559619 | -0.879924952 | 0.257470801 |
| Inhbe | 0.872322242 | -0.913219132 | 1.166673628 |
| Inhbc | 0.072025462 | 0.237312084 | 3.724945049 |
| R3hdm2 | 2616.368291 | -0.475164176 | 0.073569658 |
| Gm23819 | 0 | NA | NA |
| Stac3 | 24.43958761 | -0.953948259 | 0.405868031 |
| Ndufa4l2 | 7.948266496 | -0.025649974 | 0.365044521 |
| Shmt2 | 276.9083029 | 1.018219091 | 0.094963269 |
| Nxph4 | 143.5244743 | 0.031305926 | 0.226560767 |
| Lrp1 | 13094.28162 | 0.071106302 | 0.084215056 |
| Gm16217 | 0.962459693 | 3.902561862 | 1.157829642 |
| Stat6 | 224.5914028 | 1.994428917 | 0.171522791 |
| Nab2 | 546.0189877 | 0.292447643 | 0.142505151 |
| Gm16229 | 0.705361133 | 1.088396474 | 1.10726489 |
| Tmem194 | 106.2750348 | 0.442534727 | 0.170700499 |
| Gm16230 | 1.3230785 | 1.707769073 | 0.960758898 |
| Myo1a | 8.599191626 | -0.473413793 | 0.340520179 |
| Tac2 | 49.86492223 | 0.352653111 | 0.196276503 |
| Zbtb39 | 235.0509989 | 0.169081366 | 0.130042667 |
| Gpr182 | 35.51224453 | 1.362800356 | 0.237079208 |
| Rdh1 | 7.125358113 | -0.33625889 | 0.522389942 |
| Gm28046 | 0 | NA | NA |
| Rdh9 | 1.759257097 | -0.801774382 | 1.001572806 |
| Rdh16 | 4.45419811 | -0.287639604 | 0.522360248 |
| Rdh18-ps | 0.308514759 | 0.237331356 | 1.777680058 |
| Rdh19 | 0.061920613 | 0.718207094 | 3.724945049 |
| BC089597 | 0.047948698 | -0.243569582 | 3.724945049 |

| | | | |
|---|---|---|---|
| Rdh7 | 0 | NA | NA |
| Sdr9c7 | 0.145038387 | 0.718214452 | 2.888287922 |
| Gm15905 | 0 | NA | NA |
| Gm15900 | 1.613670225 | -1.343843776 | 1.12133307 |
| Gm23996 | 0 | NA | NA |
| Hsd17b6 | 0.048949532 | -0.243569582 | 3.724945049 |
| Prim1 | 70.56909489 | 0.264743783 | 0.13742709 |
| Naca | 1262.918694 | 0.299815555 | 0.064196623 |
| Ptges3 | 1465.432757 | 0.628256583 | 0.094989633 |
| Atp5b | 9103.422248 | -0.061566626 | 0.072618508 |
| Snord59a | 1.560301263 | 0.291565416 | 0.994915719 |
| mmu-mir-677 | 0.075043205 | 0.237309859 | 3.724945049 |
| Mir677 | 0 | NA | NA |
| Baz2a | 1409.129041 | 0.228679792 | 0.085834944 |
| Rbms2 | 301.1430177 | 1.00293423 | 0.141564129 |
| Gls2 | 183.4405106 | -1.006436034 | 0.126735387 |
| Spryd4 | 66.56019931 | 0.419548928 | 0.164774063 |
| Mip | 0 | NA | NA |
| Gm26847 | 5.379666823 | -0.570446762 | 0.565206721 |
| Timeless | 115.0545271 | 1.764312623 | 0.150917874 |
| Apon | 0.725473741 | 2.963959852 | 1.200771481 |
| Apof | 3.028410459 | 2.870755145 | 0.791117042 |
| Stat2 | 2927.543515 | 3.687086895 | 0.106302322 |
| Il23a | 1.801413292 | -0.851369355 | 0.653687823 |
| Gm23241 | 0 | NA | NA |
| Pan2 | 211.7421926 | -0.189032366 | 0.129991894 |
| Cnpy2 | 199.2616733 | 0.513394574 | 0.127149277 |
| Cs | 2382.585503 | -0.004356794 | 0.06624756 |
| Gm23182 | 1.59776671 | -0.049787231 | 0.766635418 |
| Coq10a | 213.594349 | -0.057001168 | 0.083368087 |
| Ankrd52 | 1433.771392 | 0.057505352 | 0.078792206 |
| Gm26308 | 0 | NA | NA |
| Mir6914 | 0.577876106 | 2.023150918 | 1.520776359 |
| mmu-mir-6915 | 0.129647038 | -0.243562299 | 2.992845844 |
| Gm17201 | 0.247536319 | -0.243563245 | 1.84979946 |
| Slc39a5 | 1.555265169 | -2.932659194 | 0.978330585 |
| Nabp2 | 369.4297074 | 0.22598891 | 0.072980792 |
| Gm25361 | 0 | NA | NA |
| Rnf41 | 702.2743574 | -0.121522073 | 0.06386789 |
| Gm26347 | 3.130125749 | -0.99124986 | 0.682296255 |
| Mir8105 | 0.208009692 | 0.811435087 | 2.593857227 |
| Smarcc2 | 3642.728046 | -0.286386002 | 0.071245567 |
| Myl6 | 1464.017352 | 0.468237842 | 0.110974192 |
| Myl6b | 108.9152618 | -0.479922546 | 0.129252646 |
| A430046D13Rik | 90.5350024 | -0.991141388 | 0.164624647 |
| Esyt1 | 580.33528 | 2.135883966 | 0.18665592 |
| Mir6916 | 0 | NA | NA |

| | | | |
|---|---|---|---|
| Gm29585 | 0.520597108 | 0.640882503 | 1.436522982 |
| Zc3h10 | 205.4911794 | 0.271753833 | 0.088246806 |
| Rpl41 | 1706.429966 | 0.737426177 | 0.16048843 |
| Pa2g4 | 643.314172 | 0.478340776 | 0.09068008 |
| AC117232.1 | 0 | NA | NA |
| Erbb3 | 358.7695039 | 0.612859969 | 0.165306445 |
| Mir6917 | 0.044191549 | -0.243571806 | 3.724945049 |
| Rps26 | 1049.123509 | 0.790628379 | 0.102404371 |
| Ikzf4 | 257.6485544 | 0.040819398 | 0.138262763 |
| Gm26876 | 8.215402088 | 0.155282271 | 0.361273215 |
| Suox | 127.1525145 | 0.064708055 | 0.127248098 |
| Rab5b | 1465.571473 | -0.035316702 | 0.062785692 |
| Cdk2 | 38.99389103 | 1.810767572 | 0.237218104 |
| Pmel | 8.731260839 | -0.787965726 | 0.371499979 |
| Dgka | 134.8647648 | 0.690497008 | 0.12756992 |
| Wibg | 245.4724351 | 0.640735364 | 0.128254668 |
| Rpsa-ps2 | 30.10672328 | 0.639611612 | 0.237598813 |
| Mmp19 | 189.6992867 | 5.212725734 | 0.52066833 |
| Tmem198b | 157.6632237 | 0.702025533 | 0.150229444 |
| Dnajc14 | 675.6783784 | 0.258412727 | 0.05502605 |
| Ormdl2 | 72.79417923 | 1.15583103 | 0.213648296 |
| Sarnp | 164.5829855 | 0.163089761 | 0.128051573 |
| Gdf11 | 478.8249761 | 0.531181841 | 0.132668816 |
| Cd63 | 793.3225511 | 2.481008363 | 0.235110829 |
| Rdh5 | 84.87448236 | 1.467308284 | 0.17410686 |
| Bloc1s1 | 41.20713241 | 0.093915292 | 0.264573806 |
| Itga7 | 368.0914646 | 1.314122447 | 0.134738751 |
| Gm25494 | 0.763054649 | -0.171924641 | 1.093052329 |
| Mettl7b | 0.038981395 | 0.718207094 | 3.724945049 |
| 9030616G12Rik | 0 | NA | NA |
| Olfr9 | 0.042716945 | -0.243571806 | 3.724945049 |
| Olfr763 | 0 | NA | NA |
| Olfr765 | 0 | NA | NA |
| Olfr767 | 0 | NA | NA |
| Olfr768 | 0 | NA | NA |
| Olfr769 | 0 | NA | NA |
| Olfr770 | 0 | NA | NA |
| Olfr771 | 0 | NA | NA |
| Gm25807 | 0 | NA | NA |
| Olfr772 | 0 | NA | NA |
| Olfr773 | 0 | NA | NA |
| Olfr774 | 0.045882455 | 0.237312084 | 3.724945049 |
| Olfr775 | 0 | NA | NA |
| Olfr776 | 0 | NA | NA |
| Olfr777 | 0 | NA | NA |
| Gm25764 | 0 | NA | NA |
| Olfr779 | 0 | NA | NA |

| | | | |
|---|---|---|---|
| Olfr780 | 0 | NA | NA |
| Olfr781 | 0 | NA | NA |
| Olfr782 | 0.044544648 | 0.237309859 | 3.724945049 |
| Olfr784 | 0 | NA | NA |
| Olfr786 | 0 | NA | NA |
| Olfr787 | 0 | NA | NA |
| Olfr788 | 0 | NA | NA |
| Olfr790 | 0.08169834 | 0.23732098 | 3.724945049 |
| Olfr791 | 0 | NA | NA |
| Olfr792 | 0 | NA | NA |
| Olfr794 | 0 | NA | NA |
| Gm23252 | 0 | NA | NA |
| Olfr796 | 0 | NA | NA |
| Olfr798 | 0 | NA | NA |
| Olfr799 | 0.046345709 | 0.71820487 | 3.724945049 |
| Olfr800 | 0 | NA | NA |
| Olfr801 | 0 | NA | NA |
| Olfr802 | 0 | NA | NA |
| Olfr803 | 0 | NA | NA |
| Gm24717 | 0 | NA | NA |
| Olfr804 | 0 | NA | NA |
| Olfr805 | 0 | NA | NA |
| Gm24485 | 0 | NA | NA |
| Olfr806 | 0 | NA | NA |
| Olfr807 | 0 | NA | NA |
| Olfr808 | 0 | NA | NA |
| Olfr809 | 0.046345709 | 0.71820487 | 3.724945049 |
| Olfr810 | 0.066050875 | 0.71820487 | 3.724945049 |
| Olfr811 | 0 | NA | NA |
| Olfr812 | 0 | NA | NA |
| Olfr813 | 0 | NA | NA |
| Olfr814 | 0 | NA | NA |
| Olfr815 | 0 | NA | NA |
| Olfr816 | 0.045882455 | 0.237312084 | 3.724945049 |
| Olfr818 | 0 | NA | NA |
| Olfr819 | 0 | NA | NA |
| Olfr247 | 0 | NA | NA |
| Gm10310 | 0 | NA | NA |
| Olfr820 | 0 | NA | NA |
| Olfr821 | 0 | NA | NA |
| Olfr822 | 0 | NA | NA |
| Olfr823 | 0.045882455 | 0.237312084 | 3.724945049 |
| Gm5181 | 0 | NA | NA |
| Olfr824 | 0 | NA | NA |
| Olfr825 | 0.062426873 | 0.237312084 | 3.724945049 |
| Olfr826 | 0 | NA | NA |
| Olfr827 | 0.062426873 | 0.237312084 | 3.724945049 |

| | | | |
|---|---|---|---|
| Neurod4 | 9.261960518 | 0.367589182 | 0.399223883 |
| Tespa1 | 3.163043473 | 4.377121474 | 0.932811482 |
| Vmn2r84 | 31.40616345 | -0.955175663 | 0.28682535 |
| Gm9770 | 48.13481903 | 0.655104955 | 0.160855025 |
| Vmn2r85 | 40.51229935 | -1.08362255 | 0.278420731 |
| Vmn2r86 | 19.28575388 | -1.355174533 | 0.352314526 |
| Vmn2r87 | 47.38212013 | -1.240331904 | 0.297838122 |
| Gm23793 | 0 | NA | NA |
| Gm26445 | 0.178837887 | 0.685271736 | 2.696671491 |
| Gm20746 | 0.242679635 | 1.149639615 | 2.499231879 |
| Pisd-ps1 | 253.7945557 | -0.356132394 | 0.124643924 |
| Sfi1 | 373.2312864 | -0.058466424 | 0.150056176 |
| Gm11399 | 1.590475537 | 2.277401591 | 1.059177406 |
| Gm11400 | 21.45170634 | 0.201220563 | 0.301889976 |
| Drg1 | 586.4306311 | 0.332838407 | 0.074166468 |
| Gm12735 | 2.937817048 | -0.05526247 | 0.514244439 |
| Fau-ps2 | 1.291849242 | -2.068746627 | 0.986623899 |
| Eif4enif1 | 683.4614762 | 0.042138484 | 0.074390181 |
| Patz1 | 528.6289566 | -0.133257919 | 0.082788374 |
| Gm12592 | 46.92358016 | 0.414720626 | 0.230041913 |
| Gm11944 | 38.0840153 | -0.564862929 | 0.22851099 |
| Pik3ip1 | 504.3401145 | 0.574677303 | 0.09251404 |
| Limk2 | 962.7434137 | -0.016652966 | 0.049744312 |
| Gm11948 | 5.24095402 | -1.242560116 | 0.560232915 |
| Rnf185 | 379.3246065 | 0.339157469 | 0.082524084 |
| 8430429K09Rik | 61.46987694 | -0.360313641 | 0.15406802 |
| Gm11945 | 1.421692497 | 0.029999776 | 0.883844671 |
| Gm24439 | 0.228704471 | -1.234915624 | 2.187618024 |
| Pla2g3 | 97.55501564 | 1.446970439 | 0.241348556 |
| Inpp5j | 911.3473185 | -0.51155393 | 0.081570355 |
| Selm | 738.0868402 | 0.3475643 | 0.141372126 |
| Smtn | 69.98786474 | 0.564460349 | 0.184168388 |
| Gm11946 | 0.933952761 | 0.97836053 | 1.152507854 |
| Gm11947 | 0 | NA | NA |
| Gm11949 | 0 | NA | NA |
| Gm11950 | 0 | NA | NA |
| Gm11951 | 0.453565911 | 1.278720653 | 1.530418542 |
| Tug1 | 1777.476593 | 0.25682829 | 0.071326123 |
| Gm27624 | 0 | NA | NA |
| Gm27517 | 0 | NA | NA |
| Gm27937 | 0 | NA | NA |
| Gm27640 | 0 | NA | NA |
| Morc2a | 721.9875827 | 0.218083075 | 0.074304718 |
| Gm26393 | 0.141094079 | 0.718203391 | 3.563850454 |
| Gm16518 | 10.42668312 | -1.024992073 | 0.385800363 |
| Osbp2 | 906.3048646 | -0.838831592 | 0.100726418 |
| Gm24013 | 0.11193333 | 0.718202646 | 3.724945049 |

| | | | |
|---|---|---|---|
| Gm11952 | 0 | NA | NA |
| 4921536K21Rik | 11.5351327 | 1.471874126 | 0.34874872 |
| Dusp18 | 270.8170531 | 0.240845553 | 0.097045102 |
| Slc35e4 | 326.5983649 | 0.69743045 | 0.131767687 |
| Tcn2 | 256.4564119 | 1.093161144 | 0.144079412 |
| 4930556J24Rik | 2.805439532 | -0.527053417 | 0.6342268 |
| Gm11953 | 2.130460637 | -0.881217263 | 0.81408869 |
| Pes1 | 430.3711271 | 0.712686428 | 0.092686839 |
| Gal3st1 | 117.3315211 | 0.01199891 | 0.144912682 |
| Gm11954 | 0.875788929 | -0.648183768 | 1.319217425 |
| Sec14l4 | 4.148966471 | -0.405981117 | 0.587054986 |
| Sec14l3 | 15.8893622 | -1.425574537 | 0.503287107 |
| Mtfp1 | 343.8323402 | -0.676521627 | 0.113951499 |
| Sec14l2 | 346.4809089 | -0.424677808 | 0.111575361 |
| Gm11955 | 0 | NA | NA |
| Gm11957 | 1.061319949 | 0.481291913 | 1.069887919 |
| Rnf215 | 199.4153862 | 0.340785855 | 0.092949688 |
| Mir3060 | 6.092409896 | -0.304672003 | 0.475673511 |
| Ccdc157 | 315.6146368 | 0.001901862 | 0.084739435 |
| Sf3a1 | 976.4741568 | 0.016450054 | 0.059806818 |
| Tbc1d10a | 286.7296832 | 0.811314474 | 0.10981579 |
| Gatsl3 | 28.49626455 | 1.505394485 | 0.293690218 |
| Gm11956 | 0.045882455 | 0.237312084 | 3.724945049 |
| Osm | 17.76436873 | 4.712058026 | 0.542611357 |
| Lif | 163.2509148 | 5.845146482 | 0.514515593 |
| Gm11959 | 0 | NA | NA |
| Gm11958 | 0 | NA | NA |
| Gm24803 | 1.135110426 | -1.120293234 | 0.994014622 |
| Hormad2 | 2.594629735 | 1.594407418 | 0.693611558 |
| Mtmr3 | 1608.347275 | 0.176221326 | 0.06911836 |
| Gm11960 | 2.472144487 | -0.071724156 | 0.670496708 |
| Gm11032 | 0 | NA | NA |
| Gm11961 | 0.364460012 | 1.168397214 | 1.552688198 |
| Ascc2 | 346.5615331 | 0.724259642 | 0.070056708 |
| Uqcr10 | 496.5398793 | 0.125678799 | 0.108885029 |
| Zmat5 | 89.65500592 | 0.367696152 | 0.128538096 |
| Cabp7 | 502.185691 | 0.005618817 | 0.129588425 |
| Nf2 | 1204.431154 | 0.166661027 | 0.054065718 |
| Nipsnap1 | 497.6595648 | -0.401468125 | 0.081704069 |
| Mir6918 | 0.256523886 | -1.18378598 | 1.821889802 |
| Thoc5 | 333.4021351 | 0.084799346 | 0.067343156 |
| AA413626 | 4.943749683 | -1.23002692 | 0.524125906 |
| Nefh | 1154.799543 | -0.259331291 | 0.232447964 |
| Ap1b1 | 2045.497515 | -0.04670648 | 0.045730124 |
| AL645522.1 | 0.147068667 | 0.237310878 | 3.514416608 |
| Gm24958 | 0 | NA | NA |
| Gas2l1 | 586.1682792 | 0.270727094 | 0.067145628 |

| | | | |
|---|---|---|---|
| Rasl10a | 112.452118 | -0.578046438 | 0.263259885 |
| Ewsr1 | 2343.895997 | 0.021035613 | 0.095873468 |
| Rhbdd3 | 147.1104437 | 0.015103449 | 0.123223007 |
| Emid1 | 46.41330166 | -0.646879257 | 0.175111376 |
| Kremen1 | 346.9264262 | 0.897391427 | 0.11579291 |
| Gm25142 | 0 | NA | NA |
| Znrf3 | 372.0960024 | 0.101666552 | 0.093210092 |
| Gm11962 | 0 | NA | NA |
| Gm11963 | 0 | NA | NA |
| Xbp1 | 893.6785691 | 0.612952623 | 0.100662132 |
| Ccdc117 | 216.8499613 | -0.203278319 | 0.101331306 |
| Gm22471 | 0 | NA | NA |
| Ankrd36 | 0.048949532 | -0.243569582 | 3.724945049 |
| Gm11964 | 12.76498303 | 0.147862591 | 0.306868464 |
| Gm11965 | 0.126225412 | 0.718200913 | 3.698754989 |
| Mrps24 | 181.2008852 | 0.376773947 | 0.11234739 |
| Urgcp | 807.8473661 | 0.0754397 | 0.068656438 |
| 2210015D19Rik | 182.3089558 | -0.202740077 | 0.084029892 |
| Dbnl | 1162.267136 | 1.266901404 | 0.12554363 |
| Pgam2 | 10.44641268 | -1.922624843 | 0.375064881 |
| Gm11966 | 36.99000587 | 0.288357048 | 0.204025956 |
| Polm | 78.87651427 | 0.297672037 | 0.13560634 |
| Aebp1 | 394.2398871 | 0.840146399 | 0.177544867 |
| Pold2 | 110.4180631 | 0.367249439 | 0.144186288 |
| Myl7 | 0.376384159 | -0.243564385 | 1.771387359 |
| Gck | 15.14292826 | 0.211749439 | 0.321877642 |
| Gm11967 | 0.249409806 | 0.654476408 | 2.516856909 |
| Ykt6 | 770.8424576 | 0.572321353 | 0.062652837 |
| Camk2b | 7644.2209 | -0.8355952 | 0.132863917 |
| Nudcd3 | 1857.880843 | -0.120842017 | 0.036444067 |
| Rps15a-ps6 | 9.077851534 | -0.312263864 | 0.378632913 |
| Gm11969 | 6.242235417 | -0.436334927 | 0.471773953 |
| A730071L15Rik | 1.959824955 | 1.020232305 | 0.735932025 |
| Npc1l1 | 17.38551341 | 0.650630807 | 0.400843101 |
| Gm11401 | 0.198115332 | -0.261200377 | 3.228165295 |
| Ddx56 | 347.1312207 | 0.467667877 | 0.082202326 |
| Tmed4 | 598.9634342 | 0.286333301 | 0.08607946 |
| n-R5s67 | 0 | NA | NA |
| Ogdh | 3550.241053 | -0.213457162 | 0.063447991 |
| Zmiz2 | 2387.144131 | -0.290100348 | 0.06804436 |
| Mir7651 | 2.015150011 | -0.926229255 | 0.839026351 |
| Ppia | 2699.446977 | -0.052396971 | 0.092781498 |
| H2afv | 161.7851873 | -0.311214346 | 0.139992741 |
| Gm11970 | 0.194008847 | 0.051281511 | 2.632426413 |
| Gm11971 | 0.305478853 | 0.71711122 | 1.768496538 |
| Gm11972 | 0.155921052 | -0.243569951 | 2.831361108 |
| Purb | 8286.574662 | -0.40849831 | 0.084317031 |

| | | | |
|---|---|---|---|
| Gm22123 | 0 | NA | NA |
| Gm11973 | 8.076220047 | 0.559149638 | 0.367085217 |
| Myo1g | 77.4741113 | 5.681393953 | 0.391276854 |
| Gm11974 | 44.13713644 | 1.874517203 | 0.260169259 |
| Gm22457 | 0 | NA | NA |
| Gm24000 | 0.045882455 | 0.237312084 | 3.724945049 |
| Ccm2 | 367.7041191 | 0.186840185 | 0.114489375 |
| Gm11975 | 0 | NA | NA |
| Nacad | 676.8149995 | -0.207475105 | 0.075096191 |
| Tbrg4 | 350.0488541 | 0.341994391 | 0.090335314 |
| Gm24313 | 0.992467499 | -0.783626724 | 0.965617772 |
| Snora5c | 0.582740348 | -1.886164925 | 1.169051081 |
| Wap | 0.066050875 | 0.71820487 | 3.724945049 |
| Gm11978 | 0.38240426 | -1.005636723 | 1.531984065 |
| Ramp3 | 89.90826615 | -0.604525298 | 0.146577809 |
| Gm11980 | 0.066050875 | 0.71820487 | 3.724945049 |
| Gm11981 | 0 | NA | NA |
| Gm11976 | 0 | NA | NA |
| Gm11977 | 0.369792422 | 0.194868827 | 1.533329084 |
| Gm11979 | 0 | NA | NA |
| Gm11982 | 0 | NA | NA |
| Gm11983 | 0.178864798 | 1.594141258 | 3.264221614 |
| Gm11984 | 0 | NA | NA |
| Adcy1 | 9022.462185 | -1.506865972 | 0.152994172 |
| Gm11985 | 0 | NA | NA |
| Igfbp1 | 2.637679718 | 2.17666779 | 0.922725272 |
| Igfbp3 | 602.9842508 | 3.430407821 | 0.317896572 |
| Gm11986 | 0.276998953 | 1.171557012 | 1.812453275 |
| Gm27393 | 0 | NA | NA |
| Gm11987 | 0 | NA | NA |
| Gm24051 | 0 | NA | NA |
| Gm11988 | 0 | NA | NA |
| Gm11989 | 0.796528503 | 0.431021449 | 1.247731017 |
| Tns3 | 1722.397234 | 0.61713205 | 0.131518728 |
| Gm11990 | 0 | NA | NA |
| Pkd1l1 | 3.97064273 | 0.232646389 | 0.523206937 |
| Gm11991 | 0 | NA | NA |
| Hus1 | 255.0186718 | 0.604685049 | 0.118411999 |
| Sun3 | 0.968963158 | 0.266413656 | 0.977329567 |
| Gm11992 | 17.30127652 | 0.504416181 | 0.305078023 |
| Upp1 | 71.12326318 | 3.555139214 | 0.33170073 |
| Gm11993 | 27.56449686 | 0.326574044 | 0.219058086 |
| Abca13 | 10.03132759 | 0.838568945 | 0.398279872 |
| Gm36954 | 0 | NA | NA |
| Eif3s6-ps1 | 4.535504711 | 0.725388111 | 0.526187522 |
| Gm11996 | 0 | NA | NA |
| Gm11997 | 0 | NA | NA |

| | | | |
|---|---|---|---|
| Vwc2 | 143.5725646 | -0.366089189 | 0.103323026 |
| Gm23065 | 0.966893809 | -1.6113851 | 1.0645561 |
| Zpbp | 24.98626369 | -0.679773176 | 0.216147061 |
| Gm12598 | 0.344633608 | -0.243568737 | 1.961162637 |
| Gm26458 | 0.27289602 | -1.604190852 | 1.801859306 |
| Gm12599 | 2.05315699 | 0.230007682 | 0.738740716 |
| 4930415F15Rik | 5.574285272 | 0.984241468 | 0.467683829 |
| Gm11998 | 0.161486377 | 1.199990293 | 2.795640196 |
| 4930512M02Rik | 0.106358623 | -0.24357403 | 3.724945049 |
| Gm11999 | 0.204942615 | 1.292528917 | 2.606106597 |
| Ikzf1 | 243.3512954 | 2.77478657 | 0.16979996 |
| Gm12000 | 0.060174537 | 0.237309859 | 3.724945049 |
| Fignl1 | 18.28682388 | 1.792919811 | 0.33792744 |
| Ddc | 130.1558318 | -0.1025398 | 0.22431498 |
| 1700042O10Rik | 0.263601517 | 1.642893405 | 2.076095547 |
| Gm25058 | 0 | NA | NA |
| Grb10 | 839.2894725 | -0.029835806 | 0.071012678 |
| Gm24036 | 0 | NA | NA |
| Gm12001 | 0.097139547 | 0.718207094 | 3.724945049 |
| Cobl | 1095.669369 | -0.530634083 | 0.128187792 |
| Gm12002 | 1.538964163 | 0.53940193 | 0.705689896 |
| Gm22156 | 0 | NA | NA |
| Gm22789 | 0 | NA | NA |
| Gm12003 | 0.812725683 | 1.421265906 | 1.163435704 |
| Gm12004 | 0 | NA | NA |
| 4930554G24Rik | 0.349921031 | 1.988435067 | 1.927422398 |
| Gm12005 | 0 | NA | NA |
| Gm12006 | 0 | NA | NA |
| 1700046C09Rik | 0 | NA | NA |
| Pom121l12 | 0 | NA | NA |
| Gm12007 | 0 | NA | NA |
| Gm12008 | 0 | NA | NA |
| Gm12009 | 7.023367752 | 0.456793447 | 0.377095033 |
| Gm24707 | 0 | NA | NA |
| Gm25490 | 0 | NA | NA |
| Gm12010 | 0 | NA | NA |
| Gm37319 | 0 | NA | NA |
| Gm12011 | 0 | NA | NA |
| Vstm2a | 820.6459809 | -0.716053214 | 0.085956597 |
| Gm12012 | 0.950406788 | -3.100125769 | 1.264506608 |
| Sec61g | 266.2425794 | 0.906230659 | 0.151431226 |
| Akt2-ps | 59.61674162 | 0.445829047 | 0.140877245 |
| Gm12663 | 89.45054939 | -0.213895921 | 0.112479929 |
| Gm12664 | 0 | NA | NA |
| Gm24714 | 0 | NA | NA |
| Gm25698 | 0 | NA | NA |
| Egfr | 584.8862284 | 0.411664873 | 0.124677262 |

| | | | |
|---|---|---|---|
| Egfros | 2.163206028 | 1.959211265 | 0.725273736 |
| Eldr | 6.358422579 | 2.539588 | 0.520019258 |
| Fbxo48 | 10.03527172 | 0.306990645 | 0.325470395 |
| Plek | 727.3535995 | 2.414704232 | 0.122131004 |
| Gm12014 | 0 | NA | NA |
| Cnrip1 | 500.2213394 | -0.356695272 | 0.076866403 |
| Gm12013 | 1.807586739 | 0.827416695 | 0.765367122 |
| Ppp3r1 | 5062.853256 | -0.601474574 | 0.083390097 |
| Wdr92 | 87.67747582 | 0.065016769 | 0.15238751 |
| Pno1 | 283.54597 | 0.838728073 | 0.122017312 |
| C1d | 398.4995549 | 0.240821804 | 0.12564233 |
| Gm12015 | 0 | NA | NA |
| Gm23959 | 0 | NA | NA |
| Gm12016 | 0 | NA | NA |
| Etaa1 | 152.4498586 | 0.108879201 | 0.105975002 |
| Etaa1os | 0.910758945 | 1.591453072 | 1.124570755 |
| Gm12017 | 1.313957777 | 0.078327544 | 0.891560434 |
| Gm12018 | 0.106112162 | 0.23732098 | 3.724945049 |
| Gm12019 | 0 | NA | NA |
| Gm12020 | 0.546458135 | 0.681856694 | 1.306695628 |
| Gm37869 | 0 | NA | NA |
| Gm12022 | 0.803435494 | 1.214968676 | 1.197497329 |
| Gm12021 | 0.841174947 | -0.256526242 | 1.183334412 |
| Gm12024 | 0.313606012 | -0.243587187 | 3.724945049 |
| Gm12023 | 2.23920711 | 0.544915572 | 1.172948388 |
| Gm23093 | 0 | NA | NA |
| Gm28401 | 1.961032719 | -0.077218265 | 1.03187583 |
| Gm23021 | 0 | NA | NA |
| 8430419K02Rik | 12.33011639 | -0.213643434 | 0.496134871 |
| Meis1 | 395.3527126 | -0.26494472 | 0.104862879 |
| Gm37818 | 22.45192434 | -0.638121412 | 0.305940269 |
| 2900018N21Rik | 18.03296757 | -0.43261403 | 0.360245697 |
| 2610001A08Rik | 15.58260202 | -0.712415021 | 0.347344825 |
| Gm16140 | 0.421709722 | 1.519865342 | 1.433028198 |
| Gm16141 | 1.235942255 | 1.84889555 | 0.911796229 |
| 4933406G16Rik | 0.427771506 | -0.586336407 | 1.534534443 |
| Gm37439 | 0.296404033 | -0.724461431 | 1.797785583 |
| Gm12025 | 0.044191549 | -0.243571806 | 3.724945049 |
| Gm12026 | 0.088534442 | 1.179809211 | 3.723500796 |
| Gm12027 | 0.123841226 | 1.292225235 | 3.719439735 |
| Gm23386 | 0.075043205 | 0.237309859 | 3.724945049 |
| Glns-ps1 | 73.63675014 | -0.785299454 | 0.166528961 |
| Gm12028 | 0 | NA | NA |
| Gm12029 | 0 | NA | NA |
| Spred2 | 1270.960265 | -0.216616077 | 0.081645203 |
| Actr2 | 3727.967054 | 0.260532496 | 0.053964725 |
| Gm12030 | 0 | NA | NA |

| | | | |
|---|---|---|---|
| Gm12031 | 1.129362261 | -1.807071832 | 1.081961525 |
| Gm26253 | 0.30153761 | 0.237330731 | 1.789422496 |
| Rab1 | 1943.110902 | 0.35454201 | 0.080852224 |
| Cep68 | 468.5192915 | -0.110669212 | 0.104043432 |
| Rpl12-ps2 | 0 | NA | NA |
| Slc1a4 | 846.6499244 | 0.561682775 | 0.083628156 |
| Gm12033 | 100.5140019 | -0.055812053 | 0.144195141 |
| Gm12034 | 2.661321104 | -1.018508595 | 0.69587556 |
| Sertad2 | 498.1303247 | 0.237193942 | 0.092413632 |
| Gm12035 | 0 | NA | NA |
| Aftph | 1713.653678 | 0.549516096 | 0.064513392 |
| Gm12036 | 0 | NA | NA |
| Lgalsl | 1035.137919 | 0.084900152 | 0.079329642 |
| Gm12037 | 2.870345996 | 2.00995668 | 0.891069163 |
| Gm12038 | 0.044191549 | -0.243571806 | 3.724945049 |
| Gm22807 | 0 | NA | NA |
| Gm23681 | 0 | NA | NA |
| Peli1 | 911.878915 | 0.887475655 | 0.100255083 |
| 2610024D14Rik | 21.08894881 | 0.142322887 | 0.308509691 |
| Gm22600 | 0 | NA | NA |
| Gm12040 | 0.133633945 | 0.237303512 | 3.724945049 |
| Gm12041 | 0.223001177 | 1.32506149 | 2.54186279 |
| Vps54 | 967.0089129 | 0.768305419 | 0.087510093 |
| Gm23582 | 0 | NA | NA |
| Gm12039 | 0 | NA | NA |
| Gm12042 | 9.247728932 | 0.039635563 | 0.367899847 |
| Ugp2 | 931.9428543 | 0.269599221 | 0.113346831 |
| Mir1933 | 1.993695893 | -1.255603841 | 0.704505321 |
| Gm12043 | 3.417719091 | 0.194405016 | 0.548955522 |
| Gm12044 | 0 | NA | NA |
| Gm12045 | 0 | NA | NA |
| Gm12046 | 0 | NA | NA |
| 4932414J04Rik | 0 | NA | NA |
| Mdh1 | 4531.387594 | -0.359768591 | 0.097314744 |
| Gm22990 | 2.17550669 | -1.412612072 | 0.787217121 |
| Wdpcp | 154.1019773 | -0.065165359 | 0.113881517 |
| Pgam1-ps1 | 0.548058809 | -0.316466196 | 1.143203384 |
| Gm12047 | 0.149223819 | 0.717609421 | 2.865376928 |
| Rpl15-ps1 | 0.048949532 | -0.243569582 | 3.724945049 |
| Otx1 | 56.99576563 | 0.193681516 | 0.156251577 |
| Ehbp1 | 1086.104915 | -0.352499949 | 0.064389383 |
| Rpsa-ps3 | 0 | NA | NA |
| Gm12050 | 0.136201742 | -0.231510441 | 2.944131489 |
| Gm12051 | 0 | NA | NA |
| Tmem17 | 31.45379496 | 0.614590533 | 0.213121224 |
| Gm26829 | 0.106971521 | 0.237307635 | 3.724945049 |
| Gm12052 | 0.878931054 | 0.164261954 | 1.236016352 |

| | | | |
|---|---|---|---|
| Gm12053 | 0.158674157 | -0.724459882 | 2.811010873 |
| Gm12055 | 0.424314013 | -0.77495157 | 1.557633651 |
| Gm12056 | 0 | NA | NA |
| Gm12057 | 7.744039244 | -0.512187405 | 0.355760978 |
| Gm12058 | 1.980218008 | -0.550733233 | 0.807859698 |
| Gm28048 | 0.044544648 | 0.237309859 | 3.724945049 |
| B3gnt2 | 14.36761034 | 0.898766918 | 0.343122962 |
| Gm20456 | 0.099106094 | 1.209894437 | 3.72418822 |
| Gm23772 | 0 | NA | NA |
| 9130230N09Rik | 6.400237851 | 0.734166876 | 0.397672389 |
| Gm24917 | 0 | NA | NA |
| Commd1 | 92.82422243 | 0.238224496 | 0.169314884 |
| Zrsr1 | 521.7180302 | -0.18518182 | 0.101887493 |
| Cct4 | 1159.74844 | 0.605660755 | 0.097150381 |
| Fam161a | 45.44904334 | 0.060200884 | 0.217464964 |
| Gm23740 | 0.228592689 | -0.243563954 | 2.138563777 |
| Gm12059 | 13.03468439 | 0.212194323 | 0.283978603 |
| 1700061J23Rik | 0.046345709 | 0.71820487 | 3.724945049 |
| Gm24398 | 0 | NA | NA |
| Xpo1 | 1829.109104 | 0.561450309 | 0.088125067 |
| Usp34 | 4434.045314 | -0.376420899 | 0.063469123 |
| Gm22753 | 2.309867597 | -1.05991688 | 0.663694012 |
| Ahsa2 | 295.3081191 | -0.309689115 | 0.113132891 |
| AL672049.1 | 0.184646192 | -0.724459678 | 2.707700082 |
| 1700030C12Rik | 0.940659763 | -0.053286313 | 0.995762523 |
| 1700093K21Rik | 3.270068486 | 0.404476923 | 0.690314337 |
| Gm12060 | 7.122622078 | -0.865256321 | 0.462218546 |
| 0610010F05Rik | 731.0037471 | -0.181350631 | 0.066419347 |
| Gm23114 | 5.978366796 | -1.201270103 | 0.673289588 |
| Pex13 | 566.0191946 | 0.359853281 | 0.086662713 |
| Pus10 | 234.206064 | 0.177287171 | 0.102378545 |
| Rel | 254.6607661 | 1.364781977 | 0.169331971 |
| Gm23827 | 0.061920613 | 0.718207094 | 3.724945049 |
| Papolg | 392.4489342 | -0.101547944 | 0.089470784 |
| Gm12061 | 0.546436044 | -1.198266414 | 1.501616417 |
| Gm12062 | 0 | NA | NA |
| AL662810.1 | 0 | NA | NA |
| A830031A19Rik | 4.718729471 | -0.661273535 | 0.486950462 |
| Gm12063 | 6.626652096 | -1.638899267 | 0.460731099 |
| Bcl11a | 1084.849886 | -0.979647914 | 0.154194653 |
| Gm12064 | 0 | NA | NA |
| 4930538E20Rik | 0.409305803 | -0.243564636 | 1.601981006 |
| Gm12065 | 0 | NA | NA |
| Gm12066 | 0.089089296 | 0.237305797 | 3.724945049 |
| Gm12068 | 0.5449234 | 1.413461822 | 1.313471844 |
| 4933430M04Rik | 0.038981395 | 0.718207094 | 3.724945049 |
| Gm12067 | 0 | NA | NA |

| | | | |
|---|---|---|---|
| Gm22787 | 0 | NA | NA |
| Gm25507 | 0 | NA | NA |
| Gm10466 | 0 | NA | NA |
| Gm12665 | 0 | NA | NA |
| 4933427E13Rik | 0.566718995 | 0.23733335 | 1.384811335 |
| Gm24255 | 0 | NA | NA |
| Gm12069 | 0.137762903 | -0.72445853 | 2.937523348 |
| 5730522E02Rik | 70.17051206 | -1.241279741 | 0.258196645 |
| Gm23514 | 0 | NA | NA |
| Mir6372 | 0 | NA | NA |
| Gm22868 | 0 | NA | NA |
| Fancl | 76.3059331 | -0.057580755 | 0.160574771 |
| Gm12713 | 5.450982021 | -0.483104796 | 0.508200789 |
| Vrk2 | 67.55354626 | 1.320919199 | 0.191636259 |
| Gm6899 | 0.12209515 | 0.718202646 | 3.724945049 |
| Gm12071 | 0 | NA | NA |
| Gm12070 | 46.42488409 | -0.193613274 | 0.164347337 |
| Gm12072 | 0.172942268 | -0.724458211 | 3.372002999 |
| Gm12073 | 0 | NA | NA |
| Gm12074 | 0 | NA | NA |
| n-R5s68 | 0 | NA | NA |
| Gm24850 | 0 | NA | NA |
| Gm25387 | 0 | NA | NA |
| Gm12075 | 0 | NA | NA |
| Gm12077 | 0 | NA | NA |
| Gm3810 | 0 | NA | NA |
| Gm12079 | 2.754754232 | 0.065208006 | 0.558389849 |
| Gm12080 | 0 | NA | NA |
| Gm6685 | 30.4764154 | 0.3271613 | 0.191817136 |
| Gm12081 | 0 | NA | NA |
| Ccdc85a | 925.7833576 | -0.713643311 | 0.075656071 |
| 2810471M01Rik | 0.525803784 | -0.718680154 | 1.320412039 |
| Mir216b | 0 | NA | NA |
| mmu-mir-216c | 0 | NA | NA |
| Gm12082 | 0.832920528 | 0.195193039 | 0.99237784 |
| Mir216a | 0.066050875 | 0.71820487 | 3.724945049 |
| Mir217 | 0 | NA | NA |
| Efemp1 | 96.95456267 | 0.678934173 | 0.193214488 |
| Rpsa-ps5 | 0.463552209 | 1.539786658 | 1.467215135 |
| Pnpt1 | 855.1922142 | 1.785642104 | 0.114952307 |
| A630052C17Rik | 26.90946008 | -0.265906406 | 0.253960884 |
| Smek2 | 1087.669138 | 0.428936087 | 0.055558443 |
| Cfap36 | 532.1389956 | -0.188190683 | 0.08400456 |
| Gm12085 | 0 | NA | NA |
| Gm23109 | 0.11197992 | 0.718202646 | 3.724945049 |
| Gm12086 | 0 | NA | NA |
| Gm12087 | 0.124596252 | 0.718200615 | 3.714681068 |

| | | | |
|---|---|---|---|
| Ccdc88a | 2613.854868 | 0.064646942 | 0.058268284 |
| Prorsd1 | 38.98209458 | 1.073631667 | 0.236463079 |
| Gm12089 | 10.3364996 | -0.958333991 | 0.432629767 |
| Mtif2 | 334.856477 | -0.199902711 | 0.075962213 |
| Rps27a | 597.2006918 | 0.585670795 | 0.251769901 |
| Clhc1 | 9.730852973 | 0.471536945 | 0.348866953 |
| Gm12090 | 0 | NA | NA |
| Gm12088 | 0.547473135 | 0.100066967 | 1.334612081 |
| Gm12091 | 2.208204681 | 0.382554742 | 0.697802518 |
| Gm12092 | 0.662335159 | 0.139877897 | 1.103818721 |
| Gm12093 | 0.303276099 | 0.225442916 | 1.620490816 |
| Rtn4 | 7648.216462 | -0.013759315 | 0.071698916 |
| Eml6 | 546.8534648 | -0.667283009 | 0.10533864 |
| 4931440F15Rik | 25.51154845 | -1.256278437 | 0.300281264 |
| Gm12094 | 0.298205325 | 0.063835767 | 1.993872035 |
| Gm22588 | 0 | NA | NA |
| Gm12095 | 0 | NA | NA |
| Sptbn1 | 16443.0819 | -0.355443139 | 0.052790002 |
| Gm29237 | 0.044544648 | 0.237309859 | 3.724945049 |
| Gm8098 | 47.87148983 | -0.515049408 | 0.241563067 |
| 4930505A04Rik | 3.929212317 | -0.399433689 | 0.6341103 |
| Acyp2 | 181.980473 | 0.111318963 | 0.13750384 |
| Gm12096 | 0.263964426 | 0.774521114 | 2.097717396 |
| Gm12098 | 9.953545217 | -0.768420307 | 0.483086417 |
| Gm12097 | 33.54705527 | -0.531879519 | 0.305500564 |
| Gm12099 | 0 | NA | NA |
| Gm12100 | 0.45607993 | -1.291231651 | 1.766126042 |
| Gm12101 | 0.170229941 | 0.743576216 | 2.745538088 |
| Psme4 | 1642.607637 | 0.047505417 | 0.113945006 |
| Gm12103 | 0.045882455 | 0.237312084 | 3.724945049 |
| Gpr75 | 147.2383283 | -0.533507847 | 0.111712234 |
| Gm12104 | 4.359450978 | 0.419154073 | 0.56610527 |
| Erlec1 | 677.4619536 | 0.008794074 | 0.06078681 |
| Asb3 | 233.2081696 | -0.038624553 | 0.099521005 |
| Chac2 | 85.03464214 | 0.13570281 | 0.167446876 |
| Gm12102 | 0 | NA | NA |
| Gm12105 | 4.176201957 | -1.211652301 | 0.546183505 |
| Gm12106 | 0 | NA | NA |
| Gm17332 | 0 | NA | NA |
| Stc2 | 26.77438353 | 1.708096958 | 0.327214846 |
| Bod1 | 387.4332913 | 0.308287706 | 0.081876154 |
| Gm38061 | 0.047948698 | -0.243569582 | 3.724945049 |
| D630024D03Rik | 1.773596836 | 0.405525273 | 0.80739551 |
| Gm12107 | 0 | NA | NA |
| Cpeb4 | 3230.776281 | 0.110102632 | 0.07020625 |
| 4930524B15Rik | 1.33642055 | 2.477488301 | 1.087378178 |
| Nsg2 | 3948.965464 | -0.645345756 | 0.068400399 |

| | | | |
|---|---|---|---|
| Gm12108 | 0.431234269 | 0.746370308 | 1.55334035 |
| Gm12109 | 0.060174537 | 0.237309859 | 3.724945049 |
| Il9r | 0.166808115 | 0.237311459 | 3.385250326 |
| Snrnp25 | 64.99433264 | -0.197178744 | 0.218733506 |
| Rhbdf1 | 175.977258 | 1.473646592 | 0.169027931 |
| Nprl3 | 290.7020933 | -0.214548157 | 0.092366174 |
| Mpg | 101.0618394 | 0.222468543 | 0.125365734 |
| Hba-x | 0.066050875 | 0.71820487 | 3.724945049 |
| Hba-a1 | 2.403392355 | 0.36369956 | 1.265553852 |
| Hbq1b | 0.787662625 | 0.330478376 | 1.282081229 |
| Hba-a2 | 2.268919436 | 0.546643949 | 2.099475704 |
| Hbq1a | 1.119917166 | -2.757264225 | 1.176151675 |
| Sh3pxd2b | 817.112173 | 1.445328988 | 0.135558816 |
| Ubtd2 | 162.9170653 | -0.350157296 | 0.119672505 |
| Efcab9 | 6.264832892 | -0.470517705 | 0.379092793 |
| Stk10 | 237.0838909 | 0.607096204 | 0.132661231 |
| Fbxw11 | 1952.482481 | 0.445887374 | 0.068784074 |
| AL669951.2 | 0 | NA | NA |
| AL669951.1 | 0 | NA | NA |
| Gm8192 | 0.049553047 | 0.71820487 | 3.724945049 |
| 4930555O08Rik | 0 | NA | NA |
| Smim23 | 0 | NA | NA |
| Gm12110 | 0 | NA | NA |
| Gm12111 | 0.20222093 | 0.23732836 | 2.212727188 |
| Gm12112 | 32.64724094 | 0.605214245 | 0.20542217 |
| Gm26052 | 0 | NA | NA |
| Rpsa-ps4 | 1.265922642 | 1.07143801 | 1.083986679 |
| Gm12114 | 0.044191549 | -0.243571806 | 3.724945049 |
| Fgf18 | 14.860062 | -0.700997899 | 0.267393365 |
| Npm1 | 1433.536217 | 0.562307654 | 0.082585508 |
| Gm12115 | 4.703682786 | 0.16186779 | 0.448157988 |
| Tlx3 | 9.956390658 | -0.119555913 | 0.38098622 |
| Gm12116 | 10.20532615 | -1.515185655 | 0.548656584 |
| Ranbp17 | 205.143616 | -0.480684233 | 0.153146135 |
| Gm12117 | 3.560310751 | -0.402583143 | 0.541324054 |
| Anp32-ps | 6.77190338 | -1.059086873 | 0.447104169 |
| Gm12118 | 0.07796279 | 1.072698098 | 3.721829099 |
| Gabrp | 0.357753142 | 0.250810339 | 1.574930754 |
| D130052B06Rik | 0 | NA | NA |
| Kcnip1 | 277.5432354 | -0.433118955 | 0.08181159 |
| Gm12119 | 0.042716945 | -0.243571806 | 3.724945049 |
| Gm12120 | 0.062426873 | 0.237312084 | 3.724945049 |
| Kcnmb1 | 5.253596414 | -0.034952654 | 0.435292031 |
| 4930469K13Rik | 1.763134642 | -2.097995201 | 1.003487733 |
| Lcp2 | 190.2229504 | 3.436020505 | 0.180158669 |
| Gm12121 | 0 | NA | NA |
| Foxi1 | 0.275123771 | 0.785912103 | 1.805080912 |

| | | | |
|---|---|---|---|
| Dock2 | 455.1128129 | 3.706498454 | 0.163977061 |
| 4930403D09Rik | 0.062426873 | 0.237312084 | 3.724945049 |
| Fam196b | 116.3261388 | 0.306049577 | 0.158508559 |
| Spdl1 | 20.02293662 | 2.243876312 | 0.279531476 |
| Gm22022 | 0 | NA | NA |
| Gm12711 | 0 | NA | NA |
| Gm25799 | 0 | NA | NA |
| Slit3 | 590.7003565 | -0.537679907 | 0.145886315 |
| Gm26070 | 1.06710716 | -2.032273854 | 1.333264883 |
| Gm12123 | 0 | NA | NA |
| Gm12122 | 14.36718598 | -1.364611504 | 0.436195714 |
| Mir218-2 | 2.110899703 | -1.227780465 | 0.747246304 |
| Pank3 | 1420.981158 | -0.207310892 | 0.066721203 |
| Mir103-1 | 0.791513252 | 0.279216838 | 1.076988334 |
| Gm12124 | 1.289573466 | 0.049523432 | 0.829609186 |
| Fbll1 | 180.3183456 | -0.197331572 | 0.122078429 |
| Rars | 540.4176717 | 0.848083805 | 0.100996578 |
| Wwc1 | 973.1767011 | -0.209230401 | 0.0687793 |
| Gm12125 | 0 | NA | NA |
| Tenm2 | 1701.648525 | -0.826911996 | 0.109794841 |
| Gm12126 | 0 | NA | NA |
| Gm12127 | 0 | NA | NA |
| Gm22127 | 0 | NA | NA |
| Gm12128 | 0.048949532 | -0.243569582 | 3.724945049 |
| Gm25693 | 0.047948698 | -0.243569582 | 3.724945049 |
| Gm12129 | 0.066050875 | 0.71820487 | 3.724945049 |
| Gm12130 | 1.911894038 | -2.076231 | 0.814627128 |
| Gm23520 | 0.097501745 | 0.23732098 | 3.724945049 |
| Gm12131 | 1.727409906 | -1.444408964 | 0.884464718 |
| Gm24192 | 0 | NA | NA |
| Gm12132 | 0 | NA | NA |
| Gm12133 | 0 | NA | NA |
| Gm12134 | 0 | NA | NA |
| 3110004A20Rik | 0.20299191 | 0.772127408 | 2.607194423 |
| Gm12135 | 0 | NA | NA |
| Gm12136 | 0 | NA | NA |
| Gm12137 | 0 | NA | NA |
| Gm12138 | 2.171605266 | 0.130717527 | 0.734586607 |
| Mat2b | 871.6522152 | -0.494232127 | 0.074432341 |
| Hmmr | 18.26964906 | 1.637835133 | 0.380701396 |
| Nudcd2 | 124.2665417 | 0.312953213 | 0.123988758 |
| Ccng1 | 1174.108608 | 0.509183205 | 0.109716128 |
| Gm24446 | 0 | NA | NA |
| Hmgb1-ps1 | 6.658866634 | -0.191605551 | 0.4008556 |
| Platr13 | 0 | NA | NA |
| Gm12140 | 0 | NA | NA |
| Hspd1-ps3 | 127.1742578 | 0.740318843 | 0.189717603 |

| | | | |
|---|---|---|---|
| Gm12142 | 0 | NA | NA |
| A230004M16Rik | 12.26251621 | 3.290357958 | 0.512032648 |
| Gm12143 | 0 | NA | NA |
| Gabrg2 | 1919.016568 | -0.537966227 | 0.074703764 |
| Gabra1 | 5062.268846 | -0.512940413 | 0.0877553 |
| Gabra6 | 1128.6417 | -0.517965957 | 0.375250491 |
| Gabrb2 | 2861.342192 | -0.634122984 | 0.091066576 |
| Gm25543 | 0 | NA | NA |
| Gm9972 | 0.444159033 | -2.73182252 | 1.509159056 |
| Gm12145 | 0 | NA | NA |
| Atp10b | 25.10547728 | -1.289425416 | 0.211058731 |
| Gm12146 | 1.289012822 | 3.277267936 | 1.081133223 |
| Gm12144 | 0.970234074 | -1.07485754 | 0.970083107 |
| Gm12147 | 0 | NA | NA |
| Mir146 | 0.061920613 | 0.718207094 | 3.724945049 |
| Gm12148 | 0 | NA | NA |
| Pttg1 | 36.67629004 | 3.745213366 | 0.329798321 |
| Slu7 | 647.400309 | 0.297082661 | 0.070221901 |
| Gm12151 | 0 | NA | NA |
| C1qtnf2 | 6.55721845 | -0.320109324 | 0.406661727 |
| Ccnjl | 56.86863016 | -0.543771499 | 0.173882882 |
| Gm12150 | 0 | NA | NA |
| 4933415A04Rik | 9.074042005 | -0.660463964 | 0.347077921 |
| Fabp6 | 0 | NA | NA |
| Gm12149 | 0 | NA | NA |
| Gm12152 | 0 | NA | NA |
| Pwwp2a | 556.3087863 | -0.253899438 | 0.084055194 |
| Ttc1 | 454.7934483 | 0.670644276 | 0.105749416 |
| Adra1b | 167.6708051 | -0.018476425 | 0.127457227 |
| Gm12153 | 0.799763445 | 0.548485725 | 1.049521225 |
| Gm12154 | 0 | NA | NA |
| Gm12155 | 0.038981395 | 0.718207094 | 3.724945049 |
| Il12b | 80.53366365 | 9.020237012 | 1.26970457 |
| Ublcp1 | 282.0163601 | -0.133814888 | 0.100435959 |
| Gm22121 | 0 | NA | NA |
| Gm12156 | 0.547961811 | -0.254330654 | 1.507376359 |
| Rnf145 | 1154.947274 | -0.157327729 | 0.049701884 |
| 4930597A21Rik | 0.434349921 | 1.587441578 | 1.542102508 |
| Gm37695 | 1.050344328 | -1.776255604 | 1.160581439 |
| AL603913.1 | 0 | NA | NA |
| Ebf1 | 264.2278728 | 0.002321207 | 0.12529701 |
| Gm12158 | 0 | NA | NA |
| Gm12159 | 0 | NA | NA |
| Gm12160 | 0 | NA | NA |
| Gm38087 | 0 | NA | NA |
| Gm37312 | 0 | NA | NA |
| Gm12161 | 1.855823375 | -0.603333622 | 0.783017555 |

| | | | |
|---|---|---|---|
| Gm12162 | 0 | NA | NA |
| Gm22284 | 0 | NA | NA |
| F630206G17Rik | 0.072025462 | 0.237312084 | 3.724945049 |
| Gm12163 | 0.042716945 | -0.243571806 | 3.724945049 |
| Gm12164 | 1.520812657 | 2.593669227 | 1.019377867 |
| Gm22751 | 0 | NA | NA |
| Clint1 | 800.9757842 | 0.40095676 | 0.063470453 |
| Lsm11 | 391.468034 | -0.506110298 | 0.110628087 |
| Thg1l | 79.70507417 | 0.315362478 | 0.126345039 |
| Sox30 | 1.344747471 | 1.05890652 | 0.813759787 |
| Gm12165 | 0 | NA | NA |
| Gm12166 | 16.95436423 | 1.096453023 | 0.264771756 |
| Adam19 | 340.8717861 | 0.070151948 | 0.142935449 |
| Mir8100 | 0.635485296 | -0.576770206 | 1.438161594 |
| Gm16033 | 0.647121192 | 0.703374729 | 1.103889658 |
| Gm16034 | 0.200316428 | -0.243565474 | 2.661959317 |
| Nipal4 | 54.85333669 | 0.214657129 | 0.198111639 |
| Mir7239 | 0.223868884 | 0.318316748 | 2.582883142 |
| Cyfip2 | 10488.44967 | -0.698093343 | 0.086490288 |
| Fndc9 | 118.2743247 | 2.732143822 | 0.399391269 |
| Gm12568 | 12.41784929 | -1.377686626 | 0.395872314 |
| Itk | 74.98247743 | 3.692000391 | 0.271315359 |
| Gm12167 | 0.124347486 | 0.719186602 | 3.717245571 |
| Fam71b | 0.55691522 | 1.840897739 | 1.562293894 |
| Med7 | 299.1592794 | 0.945221674 | 0.105983003 |
| 2310031A07Rik | 13.18712137 | 6.998430235 | 0.91617339 |
| Gm12168 | 5.395875324 | 0.249905965 | 0.449867923 |
| Havcr2 | 109.3330086 | 2.754064922 | 0.211804707 |
| Gm12169 | 0.124308499 | 1.48237659 | 3.714822343 |
| Gm12170 | 0 | NA | NA |
| Gm12171 | 0.288794137 | 1.088508301 | 1.78770644 |
| Rpl9-ps2 | 0 | NA | NA |
| BC053393 | 0.265064467 | 0.709590093 | 1.809370946 |
| Dppa1 | 0.143022002 | 0.719176055 | 3.547735711 |
| Gm4926 | 0.476246927 | -1.060133382 | 1.431909383 |
| Gm12173 | 0 | NA | NA |
| Timd2 | 0.095435502 | 0.718202646 | 3.724945049 |
| Gm12174 | 30.95181581 | 0.666774012 | 0.261209041 |
| Havcr1 | 1.532951352 | 4.23735573 | 1.12023586 |
| Timd4 | 7.437687587 | 5.415454197 | 0.889013603 |
| Gm12175 | 0 | NA | NA |
| Sgcd | 134.9314582 | -1.006547914 | 0.132134939 |
| Gm12176 | 0.97553329 | -1.032031997 | 1.08080425 |
| Gm23039 | 4.120196972 | -1.981317299 | 0.60622548 |
| Gm12178 | 2.707360892 | -0.130938557 | 0.603887455 |
| Gm12179 | 0.104637558 | 0.23732098 | 3.724945049 |
| Gm12180 | 0 | NA | NA |

| | | | |
|---|---|---|---|
| Gm12181 | 0.114443069 | 0.23732098 | 3.724945049 |
| Gm24422 | 0 | NA | NA |
| Gm12182 | 0.476797581 | 1.536472372 | 1.315576146 |
| Gm12183 | 218.1818688 | 0.260983137 | 0.113061268 |
| Gnb2l1 | 1496.591963 | 1.048801761 | 0.117894419 |
| Gm25296 | 0 | NA | NA |
| Snord96a | 0 | NA | NA |
| Snord95 | 0 | NA | NA |
| Trim41 | 551.7216766 | 0.230663816 | 0.06455813 |
| Gm12184 | 52.49590673 | 0.47252731 | 0.159657142 |
| Trim7 | 26.45757724 | -0.195008634 | 0.236359122 |
| Gm16170 | 11.86762837 | 1.678203353 | 0.559100322 |
| Irgm1 | 5470.912599 | 7.252663475 | 0.18900998 |
| Gm5431 | 192.2399859 | 6.487794261 | 0.339598424 |
| Gm12185 | 296.7910131 | 10.26514903 | 0.821193163 |
| Psme2b | 176.4926986 | 3.307597233 | 0.191638269 |
| 9930111J21Rik1 | 96.17708593 | 5.03098239 | 0.318535954 |
| Olfr56 | 27.72815539 | 8.39469884 | 1.004241574 |
| Tgtp1 | 654.3535308 | 9.989766081 | 0.575029201 |
| Gm12186 | 0.311786641 | 1.778305851 | 1.926348281 |
| 9930111J21Rik2 | 219.1877081 | 5.189097309 | 0.239429721 |
| Tgtp2 | 962.1752517 | 11.54055165 | 0.71646602 |
| Gm12187 | 0.675604184 | 3.298076605 | 1.324560794 |
| Ifi47 | 1143.870919 | 9.412939663 | 0.438363486 |
| Gm12188 | 0.333018454 | 2.030928584 | 1.700977313 |
| Olfr1396 | 0 | NA | NA |
| Olfr1395 | 0.571466717 | 0.237329657 | 1.725966895 |
| Olfr1394 | 0.066050875 | 0.71820487 | 3.724945049 |
| Btnl9 | 1.653010361 | 4.014673001 | 1.055410732 |
| Zfp62 | 822.4603642 | -0.070612149 | 0.051395649 |
| Mgat1 | 662.2687355 | 0.62968742 | 0.076917528 |
| Olfr1393 | 2.448580025 | -0.644850158 | 0.777340269 |
| Olfr1392 | 1.186358578 | 1.20955927 | 1.089442016 |
| Olfr10 | 0.306560087 | -1.201365536 | 1.605319639 |
| Olfr1391 | 0.124596252 | 0.718200615 | 3.714681068 |
| Olfr1390 | 0.188595572 | 0.237312021 | 3.299076571 |
| Gm12189 | 0 | NA | NA |
| Olfr1389 | 0.061920613 | 0.718207094 | 3.724945049 |
| Olfr1388 | 0 | NA | NA |
| Olfr1387 | 0.046345709 | 0.71820487 | 3.724945049 |
| Olfr1386 | 0 | NA | NA |
| Olfr1385 | 0 | NA | NA |
| Olfr1384 | 0.045882455 | 0.237312084 | 3.724945049 |
| Olfr1383 | 0 | NA | NA |
| Olfr1382 | 0 | NA | NA |
| Olfr1381 | 0 | NA | NA |
| Olfr1380 | 0 | NA | NA |

| | | | |
|---|---|---|---|
| 1700024J04Rik | 0.046345709 | 0.71820487 | 3.724945049 |
| Gm12190 | 0 | NA | NA |
| Flt4 | 66.9739403 | 2.345132183 | 0.220092065 |
| Scgb3a1 | 5.818159514 | 3.820159864 | 0.825833265 |
| Cnot6 | 1091.733663 | 0.104943543 | 0.077598969 |
| Gm12191 | 162.606518 | 0.562498112 | 0.142796562 |
| Gm12192 | 0 | NA | NA |
| Gm23813 | 0 | NA | NA |
| Gm12193 | 0.060174537 | 0.237309859 | 3.724945049 |
| Gfpt2 | 101.9890799 | 0.677924515 | 0.131622593 |
| Mapk9 | 2585.547055 | -0.292590269 | 0.049000175 |
| Gm23492 | 0.042716945 | -0.243571806 | 3.724945049 |
| Rasgef1c | 167.346068 | -0.254309418 | 0.136032495 |
| Rnf130 | 704.3708933 | 0.00098459 | 0.056362683 |
| Mir340 | 5.005408619 | -0.721416067 | 0.507092438 |
| Gm12194 | 0.07796279 | 1.072698098 | 3.721829099 |
| Tbc1d9b | 2240.773314 | -0.237645095 | 0.079534686 |
| Gm12195 | 0 | NA | NA |
| 3010026O09Rik | 75.01584386 | -0.216487039 | 0.145727679 |
| Sqstm1 | 5575.809125 | 0.892811028 | 0.098886055 |
| Mgat4b | 410.2977067 | -0.298748667 | 0.095580865 |
| Mir6919 | 0.119234754 | -0.243576254 | 3.724945049 |
| Ltc4s | 8.403515007 | -2.536580935 | 0.513890047 |
| Gm26542 | 2.8898106 | -0.871866087 | 0.572341815 |
| Maml1 | 333.2785951 | 0.522451481 | 0.127268477 |
| Canx | 5951.612133 | 0.321729871 | 0.081889589 |
| Gm12196 | 0.048949532 | -0.243569582 | 3.724945049 |
| Cby3 | 1.08873821 | 0.712050828 | 1.073861924 |
| Mir804 | 0.587766037 | 0.277475783 | 1.186086228 |
| Hnrnph1 | 2118.885193 | -0.120729826 | 0.085462385 |
| Rufy1 | 315.9868514 | 0.106322474 | 0.084433022 |
| Gm23824 | 0 | NA | NA |
| Gm12198 | 7.905317881 | -0.475838426 | 0.38899884 |
| Adamts2 | 218.5703262 | -0.479844902 | 0.100724728 |
| Gm12199 | 0 | NA | NA |
| Gm12200 | 0.062426873 | 0.237312084 | 3.724945049 |
| Gm23392 | 0 | NA | NA |
| Zfp354c | 737.6052082 | -0.337000343 | 0.077100354 |
| Zfp879 | 31.60590681 | -0.274674154 | 0.218167409 |
| Grm6 | 0 | NA | NA |
| Gm12660 | 0 | NA | NA |
| Zfp454 | 55.6120256 | -0.428752018 | 0.159402636 |
| Zfp2 | 299.1331036 | -0.537883567 | 0.081690814 |
| Zfp354b | 46.21141538 | -0.833890173 | 0.180794586 |
| 4933414I15Rik | 0.060174537 | 0.237309859 | 3.724945049 |
| Prop1 | 0.410687608 | 1.749512452 | 1.712862451 |
| Olfr1379-ps1 | 0.047948698 | -0.243569582 | 3.724945049 |

| | | | |
|---|---|---|---|
| Olfr1378 | 0.11037557 | -0.243571806 | 3.724945049 |
| Gm12661 | 0 | NA | NA |
| Olfr1377 | 0.049553047 | 0.71820487 | 3.724945049 |
| Olfr1376-ps1 | 0 | NA | NA |
| Olfr51 | 0 | NA | NA |
| Olfr54 | 0 | NA | NA |
| Olfr1375 | 0 | NA | NA |
| Zfp354a | 104.6133705 | -0.540131274 | 0.151235501 |
| 9230009I02Rik | 1.062461154 | -1.461052571 | 1.102203934 |
| Gm12202 | 0 | NA | NA |
| Gm12201 | 0.113999573 | 0.23732098 | 3.724945049 |
| Olfr1374-ps1 | 0 | NA | NA |
| Gm12203 | 0 | NA | NA |
| Platr20 | 0.066050875 | 0.71820487 | 3.724945049 |
| Gm12569 | 0.212513282 | 0.237311278 | 3.175662942 |
| BC049762 | 1.440851686 | 3.521705895 | 0.995755185 |
| Clk4 | 881.9365492 | -0.146558239 | 0.089589713 |
| Gm25082 | 22.05723362 | -0.67348179 | 0.286526511 |
| Col23a1 | 236.6432467 | 0.430191966 | 0.145928932 |
| Phykpl | 90.42005935 | -0.545403097 | 0.183179509 |
| Hnrnpab | 1099.124842 | 0.671571807 | 0.12410651 |
| Nhp2 | 191.8689779 | 0.243749857 | 0.109076474 |
| Rmnd5b | 328.920745 | -0.142655733 | 0.072211445 |
| N4bp3 | 118.9340086 | -0.169681625 | 0.105571646 |
| D930048N14Rik | 22.3543129 | -1.016535684 | 0.383782341 |
| 0610009B22Rik | 109.4926551 | -0.054905598 | 0.15305802 |
| Sec24a | 739.2067926 | 0.23091428 | 0.096293601 |
| Gm25291 | 176.1642292 | -0.134008277 | 0.307478515 |
| Sar1b | 418.9258972 | 0.240722873 | 0.089436325 |
| Jade2 | 1658.936098 | 0.617805952 | 0.09591446 |
| Gm26551 | 2.891936054 | -0.777289176 | 0.716878308 |
| Gm26061 | 0 | NA | NA |
| Cdkn2aipnl | 319.6958028 | 0.03685466 | 0.087743724 |
| Ube2b | 1147.001563 | -0.062802859 | 0.061097031 |
| Gm12204 | 0.487092162 | -0.068368522 | 1.420039502 |
| Cdkl3 | 149.4357486 | 0.037754756 | 0.095101656 |
| Gm12205 | 0 | NA | NA |
| Ppp2ca | 3811.689111 | 0.052034311 | 0.08073699 |
| Gm24157 | 3.140872897 | 0.074917185 | 0.562258682 |
| 9430098F02Rik | 0.363523191 | 0.718223035 | 1.945708226 |
| Mir3061 | 0 | NA | NA |
| Olfr1373 | 0 | NA | NA |
| Olfr1372-ps1 | 1.830548353 | 0.426250844 | 0.85074441 |
| Gm12206 | 0.049553047 | 0.71820487 | 3.724945049 |
| Olfr1371 | 0 | NA | NA |
| Skp1a | 2595.731114 | -0.16374295 | 0.07734371 |
| Gm12207 | 10.46687507 | -0.816229672 | 0.407262882 |

| | | | |
|---|---|---|---|
| Tcf7 | 44.21850987 | 0.611282816 | 0.199350658 |
| A630014C17Rik | 2.221200599 | 2.196486525 | 0.813757771 |
| Vdac1 | 3622.033645 | -0.275913409 | 0.054323158 |
| 9530068E07Rik | 1167.821488 | 0.785068868 | 0.094187831 |
| Gm12208 | 2.679228202 | -0.328638782 | 0.675200168 |
| Gm12209 | 0 | NA | NA |
| Gm12210 | 0 | NA | NA |
| Fstl4 | 404.2876281 | -0.307300804 | 0.148961933 |
| Gm11186 | 0.223616349 | 1.102930766 | 1.862306448 |
| Gm11189 | 2.178677355 | 0.752783638 | 0.676134296 |
| Hspa4 | 2862.524315 | 0.152205277 | 0.079874627 |
| Zcchc10 | 74.48748572 | 0.309355143 | 0.130840955 |
| Gm11188 | 0.045882455 | 0.237312084 | 3.724945049 |
| Aff4 | 5179.750789 | -0.022840947 | 0.048792638 |
| Gm25930 | 3.800172548 | -1.623710335 | 0.627139647 |
| mmu-mir-6920 | 1.519428967 | 0.240219192 | 0.815046919 |
| Leap2 | 0.397769051 | -1.088978167 | 1.486678478 |
| Uqcrq | 757.3825943 | 0.195384328 | 0.132734554 |
| Gdf9 | 9.225389834 | 0.021144202 | 0.348821976 |
| Gm10447 | 1.155224236 | -2.09803425 | 1.137382646 |
| Shroom1 | 58.79842685 | -0.445208874 | 0.151724462 |
| Gm12212 | 3.290890966 | -1.517925641 | 0.625894807 |
| Gm9837 | 0.267041986 | -1.033597216 | 1.946458129 |
| Sowaha | 702.0530503 | -1.156959027 | 0.140398267 |
| Gm9945 | 10.46328265 | -1.222658336 | 0.35580666 |
| Rpl9-ps3 | 1.040046165 | -0.843081126 | 0.964670971 |
| 8-Sep | 2499.725176 | -0.491291496 | 0.066046235 |
| A430108G06Rik | 9.283107748 | -0.891053484 | 0.354657258 |
| Kif3a | 1223.698615 | -0.616844564 | 0.077592534 |
| Il4 | 0.170190954 | 1.460732368 | 2.744856262 |
| Il13 | 0 | NA | NA |
| Rps2-ps4 | 0.4561058 | 0.719145478 | 1.524163332 |
| Rad50 | 251.7638459 | 0.285838125 | 0.129300594 |
| Gm22275 | 0 | NA | NA |
| Il5 | 1.025767513 | -1.976372483 | 1.078954114 |
| Gm12214 | 0.972493339 | 1.719617296 | 1.469807503 |
| Gm12213 | 0 | NA | NA |
| Mir7671 | 0.940546779 | 0.515512126 | 1.143250977 |
| Irf1 | 1452.897363 | 6.233939166 | 0.280138881 |
| Gm17334 | 5.090110731 | 2.308126767 | 0.632701811 |
| Gm12216 | 33.70519998 | 2.955148659 | 0.366542657 |
| Slc22a5 | 121.7288286 | 0.017656226 | 0.119031454 |
| Gm12217 | 0 | NA | NA |
| Gm12218 | 0 | NA | NA |
| Gm12220 | 0 | NA | NA |
| Slc22a21 | 25.5234154 | 0.559795517 | 0.305891828 |
| Gm12219 | 2.781362659 | -0.332749047 | 0.782747384 |

| | | | |
|---|---|---|---|
| Slc22a4 | 168.0969617 | 0.106267115 | 0.12011947 |
| Pdlim4 | 206.5108102 | 2.259566267 | 0.163699218 |
| P4ha2 | 113.5551428 | -0.372438769 | 0.154801726 |
| Gm12221 | 0.192293863 | 0.726555744 | 2.249617967 |
| 4933405E24Rik | 0 | NA | NA |
| Gm12222 | 0 | NA | NA |
| Csf2 | 0.705254363 | 3.441780904 | 1.164167581 |
| Gm12223 | 0.986197153 | 0.515485755 | 1.038098911 |
| Il3 | 0.048949532 | -0.243569582 | 3.724945049 |
| Acsl6 | 2735.937845 | -0.391586385 | 0.064184976 |
| Gm12224 | 0.727497348 | -0.351906688 | 1.052969948 |
| Meikin | 5.386405023 | 3.634878753 | 0.824755033 |
| Gm12226 | 3.723226009 | 1.470550964 | 0.596106481 |
| Gm12225 | 0 | NA | NA |
| Fnip1 | 1691.944238 | 0.42810213 | 0.067994509 |
| Gm24198 | 6.506667764 | -0.664544724 | 0.498724323 |
| Rapgef6 | 1160.993809 | -0.080543053 | 0.068045881 |
| Gm26116 | 0 | NA | NA |
| Cdc42se2 | 1261.495251 | -0.204907809 | 0.077758863 |
| Gm12227 | 0.36191905 | 0.211032521 | 1.57007576 |
| Gm12229 | 9.801174999 | -0.456006807 | 0.376869754 |
| Lyrm7os | 0.089089296 | 0.237305797 | 3.724945049 |
| Lyrm7 | 91.39166592 | -0.640269454 | 0.129721033 |
| Gm25426 | 0 | NA | NA |
| Hint1 | 1051.934254 | 0.558948752 | 0.135318938 |
| Gpx3 | 335.6017487 | 1.235574679 | 0.277080589 |
| Tnip1 | 367.6412068 | 0.715521472 | 0.095683311 |
| Gm12230 | 0.092228164 | 0.718202646 | 3.724945049 |
| Anxa6 | 1191.581291 | 0.471989692 | 0.073207396 |
| Ccdc69 | 10.64187061 | 1.246801064 | 0.350510781 |
| Gm2a | 793.8364917 | 0.197315069 | 0.099690507 |
| Gm12231 | 50.21335086 | 0.315414373 | 0.201868026 |
| Mup-ps22 | 0 | NA | NA |
| Slc36a3 | 0.361016523 | 1.641728862 | 3.16310181 |
| Slc36a3os | 0.686935554 | 2.571695437 | 1.247114127 |
| Slc36a2 | 7.128692757 | -2.060548437 | 0.45486892 |
| Slc36a1os | 0.119234754 | -0.243576254 | 3.724945049 |
| Slc36a1 | 1018.826301 | -0.114083981 | 0.05266406 |
| Fat2 | 2711.320312 | -0.436693325 | 0.274438197 |
| Sparc | 5655.720685 | 0.807664146 | 0.164927061 |
| Atox1 | 160.8036094 | 0.296524927 | 0.135060167 |
| Gm12235 | 0 | NA | NA |
| G3bp1 | 943.5798911 | 0.403162855 | 0.074356357 |
| Gm22884 | 0.510616014 | 0.745688327 | 1.444325686 |
| Gm12238 | 0 | NA | NA |
| Mir7652 | 0 | NA | NA |
| Glra1 | 265.7926779 | -0.489273019 | 0.385476509 |

| | | | |
|---|---|---|---|
| Gm12236 | 0.101408268 | 0.717551239 | 3.724898127 |
| Gm12237 | 0.066050875 | 0.71820487 | 3.724945049 |
| Gm12239 | 4.674577041 | -0.711952197 | 0.488580601 |
| Nmur2 | 2.399470382 | 0.267948924 | 0.621709236 |
| Gm12240 | 0.124308499 | 1.48237659 | 3.714822343 |
| Gm24253 | 0 | NA | NA |
| Gm12241 | 0.724742635 | -0.475417417 | 1.217396401 |
| Gm22603 | 0 | NA | NA |
| Gria1 | 3990.825874 | -0.843678984 | 0.110214554 |
| Gm12242 | 0.047948698 | -0.243569582 | 3.724945049 |
| Fam114a2 | 280.8245509 | 0.12648225 | 0.075376151 |
| Mfap3 | 364.5954244 | 0.122714393 | 0.077657287 |
| Gm12243 | 0.840033521 | 1.311855318 | 1.548733298 |
| 4933424L21Rik | 0 | NA | NA |
| Galnt10 | 170.7549866 | 0.172031961 | 0.127059273 |
| 4933426K07Rik | 0 | NA | NA |
| Gm24582 | 0.170988841 | -0.724459866 | 2.745561862 |
| n-R5s69 | 0.143140564 | -0.24357283 | 3.546633223 |
| 2010001A14Rik | 26.6509321 | -0.216688993 | 0.206357998 |
| Sap30l | 239.499411 | 0.103531221 | 0.110251419 |
| Hand1 | 0.104366086 | -0.243576254 | 3.724945049 |
| Gm12244 | 0 | NA | NA |
| Gm12245 | 0.101408268 | 0.717551239 | 3.724898127 |
| Gm26025 | 0 | NA | NA |
| Gm12249 | 0 | NA | NA |
| Larp1 | 2687.565109 | 0.415948226 | 0.066069766 |
| Gm12248 | 0.378899548 | 0.316212855 | 1.737000624 |
| Gm12247 | 0.394465599 | 0.617076858 | 1.737819361 |
| Cnot8 | 374.7702201 | 0.196048612 | 0.061360736 |
| Gemin5 | 376.1235968 | 0.467200643 | 0.070857662 |
| Gm12246 | 2.686282868 | -1.15012243 | 0.786093026 |
| Mrpl22 | 119.8290135 | 0.444702872 | 0.125823784 |
| Gm12250 | 1005.570989 | 9.620287456 | 0.455779561 |
| Igtp | 4765.845661 | 9.413581075 | 0.277002865 |
| Irgm2 | 1836.176375 | 8.531403123 | 0.24820544 |
| 4930438A08Rik | 0.296006519 | 0.671732444 | 2.033812951 |
| Zfp692 | 130.8218891 | -0.733777611 | 0.202281308 |
| Zfp672 | 340.7570754 | -0.160685984 | 0.078979032 |
| 1700047K16Rik | 2.233748514 | 0.737882159 | 0.712283498 |
| Sh3bp5l | 520.8892961 | 0.059100189 | 0.083477935 |
| Lypd8 | 0.598341843 | 2.558781957 | 1.399857638 |
| Gm12251 | 32.14483831 | 0.194115938 | 0.217621737 |
| Gm12252 | 0 | NA | NA |
| 1810065E05Rik | 0.095898756 | 1.192083721 | 3.724434897 |
| Gm12253 | 8.666853529 | 6.836235659 | 0.897998415 |
| 4930504O13Rik | 0 | NA | NA |
| Olfr30 | 0 | NA | NA |

| | | | |
|---|---|---|---|
| Olfr333-ps1 | 0 | NA | NA |
| Olfr332 | 0.045882455 | 0.237312084 | 3.724945049 |
| Olfr331 | 0 | NA | NA |
| Gm12254 | 27.93282506 | 0.744377463 | 0.216309562 |
| Olfr330 | 0.072025462 | 0.237312084 | 3.724945049 |
| Gm12255 | 0 | NA | NA |
| Olfr329-ps | 0.0878635 | -0.72445792 | 3.724945049 |
| Olfr328 | 0.068097359 | -0.243569582 | 3.724945049 |
| Olfr224 | 0 | NA | NA |
| Olfr325 | 0 | NA | NA |
| Olfr324 | 0.389730154 | 1.195668464 | 1.689961696 |
| 2210407C18Rik | 0.08169834 | 0.23732098 | 3.724945049 |
| Olfr323 | 0.122991902 | -0.24357403 | 3.724945049 |
| Trim58 | 1.195637209 | -0.976677624 | 1.12411423 |
| Gm22599 | 0 | NA | NA |
| Olfr322 | 0 | NA | NA |
| Olfr320 | 0 | NA | NA |
| Gm12570 | 0 | NA | NA |
| Olfr319 | 0 | NA | NA |
| Olfr318 | 0 | NA | NA |
| Olfr317 | 0 | NA | NA |
| Olfr316 | 0 | NA | NA |
| Olfr315 | 0.049553047 | 0.71820487 | 3.724945049 |
| Olfr314 | 0 | NA | NA |
| Fam183b | 11.51123108 | 0.117081594 | 0.451212871 |
| Olfr313 | 0 | NA | NA |
| Olfr312 | 0 | NA | NA |
| Olfr311 | 0 | NA | NA |
| Gm12258 | 234.0799246 | -0.372390485 | 0.096252301 |
| Gm12259 | 3.878080432 | 0.256709066 | 0.731765644 |
| 2810021J22Rik | 159.8131489 | -0.178253504 | 0.089245819 |
| Zfp39 | 105.0647483 | 0.432509185 | 0.155362081 |
| Gm12256 | 0.146451277 | -0.243562218 | 2.882358805 |
| Gm12257 | 0.048949532 | -0.243569582 | 3.724945049 |
| Btnl10 | 1.324943677 | -0.363365572 | 1.012514982 |
| Rnf187 | 2464.147731 | -0.090762951 | 0.059198949 |
| Hist3h2ba | 174.4290419 | -0.504768921 | 0.136821371 |
| Hist3h2bb-ps | 13.70298843 | -0.747654796 | 0.297109114 |
| Trim17 | 59.87795607 | 1.253205221 | 0.211108701 |
| Hist3h2a | 106.0988195 | 0.282272306 | 0.154645915 |
| AL662809.1 | 1.743011301 | -0.397925539 | 0.999179223 |
| Gm12260 | 5.27939244 | 0.205229183 | 0.481941221 |
| 2310058D17Rik | 14.57587431 | -0.533575332 | 0.320458947 |
| Trim11 | 219.4000615 | -0.120106683 | 0.118619626 |
| Obscn | 127.4374054 | -1.634762517 | 0.287086654 |
| Gm10435 | 0.809248464 | -0.093808396 | 1.165045917 |
| Iba57 | 77.73534886 | 0.166303173 | 0.142229989 |

| | | | |
|---|---|---|---|
| Gjc2 | 99.21955442 | -0.421161044 | 0.175967571 |
| Guk1 | 493.4726625 | -0.074692214 | 0.115042621 |
| 2610507I01Rik | 3.133265115 | -0.712996904 | 0.590457101 |
| Mrpl55 | 89.18665174 | 0.036675887 | 0.176441901 |
| Gm10434 | 1.747669216 | 1.100049842 | 0.732167084 |
| 2310033P09Rik | 92.90886109 | -0.181112903 | 0.138888293 |
| Arf1 | 2962.793734 | 0.297787575 | 0.069446663 |
| Wnt3a | 0.23148996 | -0.738676785 | 2.542479383 |
| Wnt9a | 76.15545421 | -1.164659925 | 0.164896109 |
| Gm24756 | 0 | NA | NA |
| Prss38 | 1.918719673 | 0.979847315 | 0.940463539 |
| Gm15755 | 0 | NA | NA |
| Snap47 | 2340.344067 | -0.269884166 | 0.052808458 |
| Jmjd4 | 428.2804086 | -0.174980739 | 0.063596989 |
| Zfp867 | 99.85436412 | 0.801583605 | 0.178802719 |
| Gm12263 | 27.6794227 | 0.603970419 | 0.221397594 |
| Gm12262 | 0 | NA | NA |
| Zkscan17 | 266.3947386 | 0.683575392 | 0.125234645 |
| 4933439C10Rik | 96.1816591 | -1.089596853 | 0.184648435 |
| Nlrp3 | 44.3755037 | 2.902670418 | 0.224571794 |
| Olfr222 | 0.26446025 | 0.785512202 | 2.059683277 |
| Gm12261 | 0.418705261 | 1.000633028 | 1.368643271 |
| Olfr223 | 0 | NA | NA |
| Olfr225 | 0 | NA | NA |
| Mprip | 6005.10193 | 0.110202318 | 0.060097024 |
| 1700007J10Rik | 1.636732283 | -1.254602602 | 0.845479039 |
| Gm12264 | 0.290485957 | -1.091709056 | 1.772899275 |
| Pld6 | 8.31364424 | -0.488537534 | 0.384508678 |
| Flcn | 628.2729707 | 0.111201036 | 0.062901239 |
| Gm16062 | 8.661166123 | 0.011102854 | 0.333369944 |
| Gm24939 | 0.352988078 | -1.029648199 | 1.437380652 |
| Cops3 | 618.5280119 | 0.321896653 | 0.082067089 |
| Nt5m | 258.4297601 | -0.071330798 | 0.077631228 |
| Gm12714 | 0.04393175 | -0.243571806 | 3.724945049 |
| 1810063I02Rik | 0.161948271 | -0.243562662 | 3.446645093 |
| Med9os | 2.791248013 | -0.488202015 | 0.563839403 |
| Med9 | 292.2435254 | 0.041840687 | 0.092470772 |
| Rasd1 | 132.7603455 | 0.565974405 | 0.202859968 |
| Pemt | 14.2244874 | 0.118385496 | 0.281959795 |
| Gm27511 | 0 | NA | NA |
| Rai1 | 1366.993626 | -0.406627513 | 0.080004753 |
| 4930412M03Rik | 2.074250591 | 1.621063471 | 0.726737541 |
| Srebf1 | 894.7642396 | 0.468113588 | 0.126273388 |
| Mir6921 | 0 | NA | NA |
| Mir6922 | 0 | NA | NA |
| Tom1l2 | 3217.981596 | -0.304351192 | 0.068853588 |
| Gm12265 | 0.343978253 | 1.749955227 | 1.486689611 |

| | | | |
|---|---|---|---|
| Gm27711 | 2.937723515 | -1.532174729 | 0.719346399 |
| Gm12266 | 0 | NA | NA |
| Lrrc48 | 90.36745919 | -0.039853829 | 0.120750403 |
| Atpaf2 | 218.7768415 | 0.264857055 | 0.074400193 |
| Gid4 | 732.8917463 | -0.451841694 | 0.090215549 |
| Gm12268 | 1.686071008 | -1.298410086 | 0.839926057 |
| Gm12267 | 0.287879534 | 0.639238704 | 1.744251578 |
| Drg2 | 463.374575 | 0.273614049 | 0.064696554 |
| Myo15 | 13.54234393 | 0.625043229 | 0.451750636 |
| Alkbh5 | 1342.8845 | 0.262291345 | 0.068283407 |
| Gm12622 | 0 | NA | NA |
| Gm12628 | 0 | NA | NA |
| Gm26837 | 0 | NA | NA |
| Gm12623 | 0.044191549 | -0.243571806 | 3.724945049 |
| Gm12621 | 0 | NA | NA |
| Gm12627 | 0 | NA | NA |
| Gm12613 | 0 | NA | NA |
| Gm12614 | 0 | NA | NA |
| Gm12615 | 0 | NA | NA |
| Gm12626 | 0 | NA | NA |
| Gm12625 | 0 | NA | NA |
| Gm12619 | 0 | NA | NA |
| Gm12624 | 0 | NA | NA |
| Gm12612 | 0 | NA | NA |
| Gm12616 | 0 | NA | NA |
| Gm12620 | 0 | NA | NA |
| Gm12617 | 0 | NA | NA |
| Llgl1 | 814.2720307 | 0.307839155 | 0.087216317 |
| Flii | 919.6804464 | 0.740420809 | 0.096440919 |
| Mief2 | 152.1320646 | 0.326322157 | 0.110367116 |
| Mir5100 | 0.418037274 | -1.383374235 | 3.284866157 |
| Gm23341 | 0.048949532 | -0.243569582 | 3.724945049 |
| Top3a | 117.1519205 | 0.715858342 | 0.116062313 |
| Smcr8 | 1137.612935 | 0.650102484 | 0.077695275 |
| Shmt1 | 28.85367593 | 0.603308595 | 0.218325156 |
| Gm24945 | 0.047948698 | -0.243569582 | 3.724945049 |
| Gm12611 | 0.097139547 | 0.718207094 | 3.724945049 |
| Gm12618 | 0.132101749 | 1.209894437 | 3.72418822 |
| Dhrs7b | 155.5720981 | 0.037870266 | 0.140440053 |
| Tmem11 | 223.7050137 | 0.511820625 | 0.106438844 |
| Gm26534 | 14.74527245 | -0.408489079 | 0.276971603 |
| Natd1 | 186.5605054 | 0.156655574 | 0.12568765 |
| Map2k3os | 10.95972877 | 1.748276795 | 0.382501016 |
| Map2k3 | 195.4668182 | 1.140331603 | 0.152700207 |
| Kcnj12 | 243.7727264 | -0.545625133 | 0.142396622 |
| Tnfrsf13b | 25.743611 | 2.084884457 | 0.336155731 |
| Gm12269 | 0 | NA | NA |

| | | | |
|---|---|---|---|
| Usp22 | 3131.14818 | -0.089256973 | 0.053299394 |
| Rps13-ps5 | 0.882161971 | 0.182663537 | 1.076410756 |
| Aldh3a1 | 6.417984165 | -0.33821174 | 0.429380271 |
| Aldh3a2 | 629.8388114 | 0.334990708 | 0.097612264 |
| Slc47a2 | 0.919894618 | -0.431538429 | 1.357674661 |
| Gm25929 | 0 | NA | NA |
| Slc47a1 | 39.2732027 | -0.05461024 | 0.246079222 |
| Gm12271 | 0.275857165 | -0.243571112 | 2.464247975 |
| Rnf112 | 1605.26594 | -1.061140373 | 0.13522807 |
| Mfap4 | 9.173601336 | -0.407455757 | 0.352820739 |
| Mapk7 | 210.9337897 | 0.468755099 | 0.10406977 |
| B9d1os | 5.30594644 | -0.770316693 | 0.468236467 |
| B9d1 | 90.68224864 | 0.05573853 | 0.126448202 |
| Epn2 | 1283.27487 | -0.480102845 | 0.062280656 |
| Gm12272 | 1.010954397 | 0.18919954 | 1.267412125 |
| Grap | 28.95643677 | 1.912798547 | 0.25315715 |
| Slc5a10 | 1.756770038 | 0.384537108 | 0.911109647 |
| Fam83g | 11.17234595 | 4.682165407 | 0.632057681 |
| Prpsap2 | 311.9887896 | 0.17756733 | 0.107847569 |
| Ulk2 | 1353.293992 | -0.552287618 | 0.090788958 |
| A530017D24Rik | 232.7074356 | -0.454080946 | 0.079739901 |
| Gm26964 | 0.259490603 | -1.398755635 | 2.447964515 |
| Akap10 | 671.4108366 | -0.120094997 | 0.060126777 |
| Gm23721 | 0 | NA | NA |
| Gm12273 | 0.316938377 | 1.543556756 | 2.03043179 |
| Specc1 | 1898.855375 | -0.01520625 | 0.065022638 |
| Gm12274 | 0.049553047 | 0.71820487 | 3.724945049 |
| Adora2b | 44.79440823 | -0.262702264 | 0.210691236 |
| Zswim7 | 50.11354397 | 0.067666195 | 0.156665456 |
| Ttc19 | 1111.563203 | -0.518613525 | 0.086011496 |
| Gm12275 | 2.211543673 | -1.297319633 | 0.738771518 |
| Ncor1 | 3691.866875 | -0.337352308 | 0.072515878 |
| Gm12276 | 22.41048438 | -0.345617214 | 0.226176492 |
| Pigl | 79.7396089 | -0.11776888 | 0.141321281 |
| Gm12278 | 0.637771319 | -0.716843223 | 1.06530534 |
| Gm12949 | 0 | NA | NA |
| Cenpv | 151.2123072 | -0.112423036 | 0.111585984 |
| Gm12279 | 7.228218235 | -0.664255103 | 0.395362225 |
| Ubb | 6377.498152 | 0.540410658 | 0.104598602 |
| Gm12280 | 0 | NA | NA |
| Trpv2 | 211.9525486 | 0.766331244 | 0.124093903 |
| 2410006H16Rik | 257.1498311 | 1.763896459 | 0.183643086 |
| Snord49b | 2.860297155 | 0.048394742 | 0.599390659 |
| Snord49a | 6.687144773 | 0.032914984 | 0.440845421 |
| Snord65 | 1.151518338 | 0.790657836 | 1.002020624 |
| Lrrc75a | 93.92612434 | -0.161626886 | 0.165174674 |
| Lrrc75aos2 | 8.114142353 | -0.542196326 | 0.430249378 |

| | | | |
|---|---|---|---|
| Lrrc75aos1 | 0 | NA | NA |
| Mmgt2 | 82.28640359 | 0.097182267 | 0.115479795 |
| Gm12281 | 0.339850366 | -0.069830986 | 1.719301308 |
| Zfp287 | 282.6640166 | 0.014615337 | 0.10249338 |
| Gm27194 | 0.484088067 | 0.894635293 | 1.465909823 |
| Wsb2-ps | 33.81592792 | 0.032302917 | 0.196104596 |
| Gm10428 | 0.042716945 | -0.243571806 | 3.724945049 |
| Zfp286 | 136.9541373 | -0.27364857 | 0.112160031 |
| Zfp286it | 0.804776955 | -0.500278353 | 1.201540915 |
| Zfp286os | 0 | NA | NA |
| Gm24151 | 0 | NA | NA |
| Trim16 | 38.08170184 | 1.208621278 | 0.22131066 |
| Fbxw10 | 9.908817 | -0.050691069 | 0.329390219 |
| Tvp23b | 188.0397332 | 0.018404453 | 0.154140056 |
| Tvp23bos | 2.044779979 | 0.531659474 | 0.680764909 |
| Cdrt4os2 | 0 | NA | NA |
| Cdrt4 | 2.232638995 | 1.961272821 | 0.779460984 |
| Cdrt4os1 | 0 | NA | NA |
| Tekt3 | 0.583929304 | -0.02501167 | 1.318029967 |
| Pmp22 | 317.1591218 | 0.344890057 | 0.158879686 |
| B430319H21Rik | 0.78030421 | 1.350059354 | 1.276713038 |
| Gm12285 | 0 | NA | NA |
| Gm12286 | 16.66800335 | -0.394082699 | 0.249967729 |
| B130011K05Rik | 1.258106493 | 0.221443226 | 1.067003666 |
| Gm12287 | 0.044544648 | 0.237309859 | 3.724945049 |
| Gm12288 | 0 | NA | NA |
| Hmgb1-ps3 | 8.21809902 | 0.443874956 | 0.407567993 |
| Hs3st3b1 | 91.21008015 | 0.816218786 | 0.190863861 |
| Cox10 | 247.0159409 | 0.113111254 | 0.084872868 |
| 9630013K17Rik | 0.09689823 | -0.724453471 | 3.724945049 |
| Gm12289 | 0 | NA | NA |
| 2810001G20Rik | 35.83958181 | -0.161338906 | 0.190288561 |
| Gm12290 | 1.421699425 | 1.435291171 | 0.894666291 |
| F930015N05Rik | 5.181464156 | -1.909615835 | 0.577973179 |
| Hs3st3a1 | 37.68402565 | -0.540901218 | 0.247515921 |
| Gm24275 | 0 | NA | NA |
| Gm12291 | 0.332139225 | 0.961316408 | 1.738716932 |
| Gm12292 | 0.254955332 | 1.273369708 | 2.459478168 |
| Elac2 | 339.1600695 | 0.231077811 | 0.086979466 |
| Arhgap44 | 1922.5473 | -0.494732784 | 0.074351195 |
| Gm12294 | 19.04726638 | -1.412304089 | 0.462212061 |
| Gm25991 | 1.580643696 | -1.130627614 | 0.817301186 |
| 1700086D15Rik | 40.20348781 | -1.482207487 | 0.395269185 |
| Gm12293 | 0.046345709 | 0.71820487 | 3.724945049 |
| Myocd | 4.652030534 | 0.787150953 | 0.533945871 |
| Gm12295 | 2.004716341 | 2.675961842 | 0.887162975 |
| Gm26128 | 0 | NA | NA |

| | | | |
|---|---|---|---|
| Map2k4 | 1997.328949 | -0.313711879 | 0.045814314 |
| Mir744 | 16.92311974 | -0.926330512 | 0.292423268 |
| Zkscan6 | 133.418627 | 0.207691017 | 0.109477686 |
| Dnah9 | 161.9890621 | -0.205262186 | 0.128097999 |
| Gm12296 | 7.015874426 | -1.071334791 | 0.501059774 |
| Shisa6 | 795.9631443 | -0.334781588 | 0.132511236 |
| Gm12297 | 1.996095088 | -1.003248557 | 0.722966777 |
| Gm12298 | 0 | NA | NA |
| Pirt | 6.080719795 | -0.261425076 | 0.492857568 |
| 9130409J20Rik | 0 | NA | NA |
| Gm12299 | 0.127580795 | 0.237325162 | 3.007567814 |
| Tmem220 | 8.211970812 | 0.639008351 | 0.341508331 |
| Adprm | 137.339951 | 0.599152684 | 0.122745782 |
| Sco1 | 243.6408562 | 1.338659672 | 0.155202174 |
| Myh3 | 10.50794026 | -0.58071545 | 0.411763539 |
| Mir6923 | 0 | NA | NA |
| 2310065F04Rik | 0.176426445 | 0.237327358 | 2.718482184 |
| Myh2 | 5.858603139 | -0.731190057 | 0.544523927 |
| 9430073C21Rik | 0.072025462 | 0.237312084 | 3.724945049 |
| Myh1 | 7.368640863 | -0.94827559 | 0.451627523 |
| Gm12300 | 1.913586201 | -2.957515205 | 1.017553977 |
| Myh4 | 5.497083594 | -1.3249036 | 0.495876424 |
| Myh8 | 7.157414437 | -1.405737415 | 0.495268822 |
| Myh13 | 1.635466931 | -0.14081876 | 0.817709174 |
| Gm26476 | 0 | NA | NA |
| Gas7 | 7031.634939 | -0.593843152 | 0.106346939 |
| Gm12301 | 0.159545055 | 0.237325453 | 2.806181458 |
| Glp2r | 34.73267854 | 1.33753817 | 0.263747944 |
| Rcvrn | 8.311551091 | -0.806385905 | 0.475979513 |
| Gm12302 | 0.259617586 | 2.294088793 | 2.408644118 |
| Dhrs7c | 0.646105857 | 1.83530477 | 1.368803978 |
| Usp43 | 27.84987753 | -0.587461818 | 0.206217022 |
| 9130017K11Rik | 3.804522088 | -1.193882309 | 0.512826455 |
| Cfap52 | 13.68582642 | 0.539203663 | 0.287846684 |
| Stx8 | 213.3769252 | 0.326260731 | 0.090995711 |
| Gm12303 | 0.044544648 | 0.237309859 | 3.724945049 |
| Gm26157 | 1.267602396 | -0.171179668 | 1.053098589 |
| Gm12304 | 0 | NA | NA |
| C78197 | 2.497471105 | -0.029018714 | 0.636631263 |
| Ntn1 | 541.1663475 | 1.132958484 | 0.154218779 |
| Mir6406 | 0.083526043 | 0.718202646 | 3.724945049 |
| Gm12305 | 1.238804443 | -0.919384118 | 0.844647867 |
| Pik3r5 | 167.1648793 | 3.570341947 | 0.262301391 |
| Pik3r6 | 32.88554754 | 1.16813926 | 0.274514842 |
| Mfsd6l | 0.872192936 | 2.50579725 | 1.231688983 |
| Ccdc42os | 0 | NA | NA |
| Ccdc42 | 2.972495032 | -1.695801548 | 0.644096765 |

| | | | |
|---|---|---|---|
| Myh10 | 3525.532232 | -0.710228991 | 0.091407223 |
| Ndel1 | 867.2338058 | 0.281775873 | 0.070160656 |
| Rnf222 | 1.039519431 | 0.773928598 | 1.128195394 |
| Rpl26 | 246.1278693 | 0.645983829 | 0.160435929 |
| Gm22608 | 0.141071297 | 0.237322348 | 3.564043836 |
| Odf4 | 1.2405255 | -1.445376411 | 1.044637421 |
| Arhgef15 | 134.1597504 | 0.116261623 | 0.156271511 |
| Slc25a35 | 55.2247238 | -0.708511022 | 0.217125833 |
| Rangrf | 38.50784585 | 0.102526159 | 0.203808125 |
| Pfas | 417.0734811 | -0.095362341 | 0.070408541 |
| Mir3062 | 0 | NA | NA |
| Gm25819 | 0 | NA | NA |
| Ctc1 | 356.1546317 | -0.406666972 | 0.098871004 |
| Aurkb | 11.54692778 | 2.645041708 | 0.524728265 |
| 9330160F10Rik | 16.80478976 | -1.433167819 | 0.289699835 |
| 2310047M10Rik | 219.3535084 | -0.098651045 | 0.104258806 |
| Tmem107 | 35.40669845 | -0.270333915 | 0.216699822 |
| Snord118 | 16.03095282 | 0.977323776 | 0.36807904 |
| Gm12306 | 0.293428113 | -0.226022733 | 1.629029963 |
| Gm25371 | 0.515366098 | 0.231507013 | 1.376236547 |
| Vamp2 | 8741.82778 | -0.574481958 | 0.078479913 |
| Per1 | 1912.220295 | -0.267161604 | 0.164347337 |
| 9130213A22Rik | 7.871844516 | -0.729560674 | 0.374232847 |
| Hes7 | 8.964305686 | 0.888419497 | 0.440857017 |
| Aloxe3 | 69.09659255 | 0.123537571 | 0.184280879 |
| Alox12b | 38.03970433 | -1.06544892 | 0.323654977 |
| Alox8 | 33.51618146 | -0.428924791 | 0.205494528 |
| Gucy2e | 40.20224287 | -0.229088121 | 0.234939773 |
| Gm23194 | 0 | NA | NA |
| Cntrobos | 0 | NA | NA |
| Cntrob | 149.0597755 | -0.385292752 | 0.125689555 |
| Trappc1 | 165.2164685 | -0.087384212 | 0.123913573 |
| Kcnab3 | 622.5251017 | -0.561187909 | 0.104929923 |
| Kcnab3os | 3.132524194 | 1.035695624 | 0.639656478 |
| Chd3os | 937.6597823 | -0.141249858 | 0.077137638 |
| Chd3 | 5496.775121 | -1.345997184 | 0.128182594 |
| Gm22442 | 482.6561501 | -0.163850972 | 0.123149341 |
| Gm17305 | 8.479929393 | -0.952959979 | 0.419016497 |
| Cyb5d1 | 166.3102935 | -0.099845049 | 0.091106177 |
| Naa38 | 194.4625342 | -0.119318537 | 0.147735752 |
| Tmem88 | 12.86887999 | 1.383129064 | 0.363242437 |
| Kdm6b | 1048.172561 | 0.607397401 | 0.163252898 |
| Kdm6bos | 0.21690863 | -0.69216663 | 2.210015506 |
| Dnah2 | 63.85404554 | -0.78091768 | 0.149623377 |
| Dnah2os | 1.999759116 | 1.676037763 | 0.805699653 |
| Gm27493 | 0.038981395 | 0.718207094 | 3.724945049 |
| Efnb3 | 1089.578534 | -0.419203323 | 0.080250014 |

| | | | |
|---|---|---|---|
| Wrap53 | 98.47005554 | 0.179536725 | 0.115900102 |
| Trp53 | 281.3127673 | 1.156069592 | 0.127770716 |
| Atp1b2 | 7719.462683 | -0.008759537 | 0.113689702 |
| Shbg | 1.970655031 | 2.917003413 | 0.977595312 |
| Sat2 | 69.39285511 | -0.520692601 | 0.142986484 |
| Gm12308 | 0 | NA | NA |
| Gm12309 | 0 | NA | NA |
| Fxr2 | 1268.108725 | -0.080996227 | 0.05584548 |
| Mir467f | 0 | NA | NA |
| Sox15 | 1.197362358 | -1.555734978 | 1.031122814 |
| Mpdu1 | 282.855519 | 0.346275713 | 0.094298541 |
| Mir1934 | 0.521325481 | -0.620774217 | 1.588700131 |
| Cd68 | 498.24161 | 4.271109947 | 0.339168445 |
| Eif4a1 | 2251.416923 | 0.788880461 | 0.113561953 |
| Gm24029 | 0 | NA | NA |
| Gm25835 | 93.03237133 | 0.429853195 | 0.197968438 |
| Gm22988 | 0 | NA | NA |
| Senp3 | 708.1528605 | 0.180630756 | 0.057284284 |
| Tnfsf13os | 8.601013049 | 1.167793486 | 0.373194381 |
| Tnfsfm13 | 5.612199682 | 1.480603672 | 0.515876879 |
| Tnfsf13 | 5.550421209 | 0.654479354 | 0.429698803 |
| Tnfsf12 | 55.83410333 | 0.896218114 | 0.173922389 |
| Gm12307 | 0.075043205 | 0.237309859 | 3.724945049 |
| Polr2a | 2028.090813 | 0.571544856 | 0.077500125 |
| Slc35g3 | 0.474471242 | -1.001098924 | 1.404118296 |
| Zbtb4 | 2349.635844 | -0.396354931 | 0.055658744 |
| Chrnb1 | 9.482089685 | 2.028287414 | 0.455695706 |
| Fgf11 | 217.0904447 | -0.830162151 | 0.105726696 |
| Tmem102 | 2.228356223 | 0.233296099 | 0.744488898 |
| 4933402P03Rik | 0 | NA | NA |
| Spem1 | 3.349245703 | 0.60634851 | 0.534511404 |
| Nlgn2 | 4433.469855 | -0.020972802 | 0.09290374 |
| Tmem256 | 162.3407789 | 0.184923727 | 0.102997403 |
| Plscr3 | 159.9402606 | 0.8399989 | 0.11401159 |
| Tnk1 | 0.927088552 | -0.28701199 | 1.0594583 |
| Tmem95 | 1.006289673 | 1.425557678 | 0.944491839 |
| Kctd11 | 45.25365547 | 2.12356369 | 0.283062071 |
| Acap1 | 25.63139517 | 7.300009416 | 0.872645947 |
| mmu-mir-6924 | 0 | NA | NA |
| 2810408A11Rik | 1.623343953 | 0.364866287 | 0.826867307 |
| Neurl4 | 1197.492728 | -0.280710387 | 0.05512917 |
| Gps2 | 171.2820456 | 0.409323195 | 0.154933107 |
| Eif5a | 3116.497447 | 0.661587702 | 0.096873871 |
| Ybx2 | 14.27143314 | -1.889947598 | 0.365215877 |
| Slc2a4 | 20.6317304 | -0.732196457 | 0.353693047 |
| Gm12310 | 0 | NA | NA |
| Cldn7 | 0.279349812 | -0.728578385 | 1.791276788 |

| | | | |
|---|---|---|---|
| Elp5 | 407.5792987 | 0.207834707 | 0.100200917 |
| AL596185.1 | 14.58297401 | 0.424546347 | 0.309546854 |
| Ctdnep1 | 632.1150543 | 0.45807107 | 0.070800402 |
| Gabarap | 1356.650847 | 0.945654735 | 0.127063568 |
| Phf23 | 440.621247 | 0.475042718 | 0.090762875 |
| Dvl2 | 217.8185947 | -0.154938062 | 0.133102732 |
| Acadvl | 388.2749648 | 0.413037503 | 0.092019046 |
| Mir324 | 0 | NA | NA |
| Dlg4 | 6142.381317 | -0.810664697 | 0.0996509 |
| Asgr1 | 14.73418541 | 0.195397616 | 0.311337481 |
| Asgr2 | 0.670311827 | 2.697294063 | 1.400733201 |
| Gm12311 | 0 | NA | NA |
| Mgl2 | 3.864736009 | -0.373776102 | 0.443843087 |
| Clec10a | 7.893594261 | 1.24042011 | 0.390324347 |
| Slc16a11 | 154.4850972 | -1.198289063 | 0.177188313 |
| Slc16a13 | 67.2589845 | -0.317205968 | 0.172456425 |
| Bcl6b | 16.16416338 | 1.131841169 | 0.283494258 |
| Mir497b | 0.072025462 | 0.237312084 | 3.724945049 |
| mmu-mir-497a | 0.695279289 | -1.45529342 | 1.249858193 |
| Mir195a | 1.190084389 | -0.112938462 | 0.951981381 |
| 0610010K14Rik | 9.73724523 | -0.74769055 | 0.369739419 |
| Gm21988 | 3.655964835 | 1.321588562 | 0.61761782 |
| Rnasek | 9.847312144 | -0.319078835 | 0.335692145 |
| Alox12 | 4.024321177 | 1.063512294 | 0.596834662 |
| Alox12e | 0.602894331 | -2.370107714 | 1.333468478 |
| Alox15 | 1.707423372 | 2.610124998 | 0.998232075 |
| Gm12312 | 2.105566461 | 0.266439486 | 0.63212569 |
| Pelp1 | 421.1587153 | 0.092124133 | 0.05572219 |
| Gm12313 | 0 | NA | NA |
| Gm12314 | 0.049553047 | 0.71820487 | 3.724945049 |
| Arrb2 | 535.4570107 | 0.747598355 | 0.076934708 |
| Mir7115 | 0 | NA | NA |
| Med11 | 69.16924485 | 0.819504277 | 0.205745941 |
| Cxcl16 | 296.0926 | 4.546847343 | 0.20993759 |
| Zmynd15 | 30.5256007 | 3.74265253 | 0.341200484 |
| Tm4sf5 | 2.723137057 | 0.79439292 | 0.638865102 |
| Vmo1 | 1.167525075 | -0.739469067 | 1.095376727 |
| Gm12317 | 0.120679735 | 0.631141837 | 3.722822053 |
| Gltpd2 | 0.276014283 | 0.730520242 | 2.089029314 |
| Psmb6 | 866.9721086 | 0.677886179 | 0.090160909 |
| Gm10418 | 0.929389619 | -0.384393767 | 0.945101564 |
| Pld2 | 260.2865872 | 0.246091436 | 0.128796671 |
| Mink1 | 3061.414012 | -0.459271346 | 0.091604626 |
| Gm12316 | 0.706900582 | 0.440304053 | 1.270271358 |
| Chrne | 2.081055214 | 0.043087008 | 0.800465423 |
| 4930544D05Rik | 1.506657277 | -1.074914989 | 0.978646775 |
| Gm12315 | 0.418629604 | 0.681702112 | 1.39890186 |

| | | | |
|---|---|---|---|
| Gp1ba | 0.170190954 | 1.460732368 | 2.744856262 |
| Slc25a11 | 758.6093433 | -0.304590544 | 0.072982547 |
| Rnf167 | 591.5226223 | -0.074652496 | 0.061895305 |
| Pfn1 | 1454.63144 | 1.349987622 | 0.130351211 |
| Gm12319 | 3.626546672 | 0.558690845 | 0.596919703 |
| Eno3 | 42.27077307 | -0.780628612 | 0.20249057 |
| Spag7 | 215.7681227 | 0.037260615 | 0.129650304 |
| Camta2 | 2839.805525 | -0.809400532 | 0.094933608 |
| Inca1 | 17.29699935 | -0.361409065 | 0.262500466 |
| Kif1c | 1453.545255 | 0.272694548 | 0.081191834 |
| mmu-mir-6925 | 0 | NA | NA |
| Gm12320 | 0.135956459 | -0.243575439 | 3.642411469 |
| Gm12318 | 0 | NA | NA |
| Gm22760 | 0 | NA | NA |
| Zfp3 | 139.4854402 | -0.128918014 | 0.104896405 |
| Scimp | 44.06491336 | 8.693414578 | 0.879026931 |
| Rabep1 | 1978.442608 | -0.037989325 | 0.062437177 |
| Gm19967 | 38.13002371 | -0.611849412 | 0.233083965 |
| Rpl23a-ps2 | 9.688741338 | -0.836277924 | 0.420824125 |
| Gm12322 | 0.177934686 | -0.243571879 | 2.723282651 |
| Nup88 | 578.9786411 | 0.098537812 | 0.080084821 |
| Rpain | 118.6108747 | 0.052118715 | 0.115585997 |
| C1qbp | 471.9490247 | 0.36774079 | 0.124388473 |
| Dhx33 | 545.9733439 | -0.09810416 | 0.087508218 |
| Gm22297 | 1.938621463 | -0.942671136 | 0.745362617 |
| Derl2 | 390.884449 | 0.449146793 | 0.090011327 |
| Mis12 | 261.7075808 | 0.165824116 | 0.131664988 |
| 6330403K07Rik | 1104.505225 | -0.110799875 | 0.104624459 |
| Gm12321 | 0 | NA | NA |
| Gm15377 | 0 | NA | NA |
| Nlrp1a | 0.174317197 | 1.763489003 | 2.733201371 |
| Nlrp1b | 15.20331131 | 4.429468395 | 0.499774954 |
| Gm23266 | 0 | NA | NA |
| Nlrp1c-ps | 0 | NA | NA |
| Gm23311 | 0 | NA | NA |
| Gm16013 | 0.447195792 | 1.662597811 | 1.509588299 |
| 9230020A06Rik | 0.161949631 | 1.583504458 | 2.7626009 |
| Gm12324 | 1.538502375 | -1.500638632 | 0.732777416 |
| Gm12325 | 0 | NA | NA |
| Gm12326 | 2.668994113 | -1.020689455 | 0.57367088 |
| Wscd1 | 774.0552541 | -0.048170955 | 0.082748311 |
| Gm22927 | 0 | NA | NA |
| 4930401O10Rik | 0.10503227 | 1.179809211 | 3.723500796 |
| Gm23226 | 0 | NA | NA |
| Aipl1 | 1.181230752 | 0.75023264 | 0.911995436 |
| Fam64a | 7.10051421 | 2.154763326 | 0.620218976 |
| Pitpnm3 | 1123.234306 | -0.711130307 | 0.114365553 |

| | | | |
|---|---|---|---|
| 4933427D14Rik | 402.8306143 | -0.356872739 | 0.154067942 |
| Txndc17 | 137.0373374 | 0.630532024 | 0.184682442 |
| Med31 | 91.72905903 | 0.282809141 | 0.149805669 |
| 4930563E22Rik | 22.70421901 | -0.926083616 | 0.259565875 |
| Slc13a5 | 115.5056273 | -0.810257605 | 0.183966805 |
| 1700051A21Rik | 0.48812072 | -0.243565235 | 1.73504208 |
| n-R5s70 | 0 | NA | NA |
| Xaf1 | 848.1965907 | 6.080282727 | 0.169304907 |
| Fbxo39 | 1.50133588 | 3.961866505 | 1.017775129 |
| Gm23224 | 0 | NA | NA |
| Tekt1 | 14.19528596 | 0.092708834 | 0.278791934 |
| Mir6338 | 0 | NA | NA |
| Smtnl2 | 23.5679464 | -0.611497141 | 0.351938624 |
| Ggt6 | 0.550474514 | -1.727173513 | 1.284299283 |
| Mybbp1a | 930.9934562 | 0.898792983 | 0.085569363 |
| Spns2 | 584.9244595 | 0.223620615 | 0.158508115 |
| Spns3 | 6.967261194 | 1.248567608 | 0.485105021 |
| Gm12712 | 1.554484234 | -0.538521761 | 0.862173491 |
| Ube2g1 | 638.5559752 | -0.162518864 | 0.07750612 |
| Gm24143 | 0 | NA | NA |
| Gm25678 | 1.584377093 | -0.892608958 | 0.840530556 |
| Ankfy1 | 1808.623818 | 1.496760736 | 0.078504912 |
| Cyb5d2 | 155.8716509 | -0.158403724 | 0.127072359 |
| Zzef1 | 1733.026892 | 0.107913279 | 0.073840096 |
| Gm22075 | 0 | NA | NA |
| Gm6733 | 29.33660425 | 0.078398956 | 0.267332012 |
| Gm26121 | 4.21018102 | -1.266075903 | 0.580851349 |
| AL663082.1 | 0 | NA | NA |
| Atp2a3 | 446.9867994 | -0.105363493 | 0.264110788 |
| P2rx1 | 3.999946171 | 0.12025831 | 0.563248913 |
| 4732414G09Rik | 8.889423667 | -1.250779814 | 0.377936183 |
| Camkk1 | 1051.478754 | -0.520725634 | 0.132043641 |
| 1200014J11Rik | 480.4026274 | -0.097828717 | 0.086390845 |
| Itgae | 5.448280777 | 1.182483469 | 0.70567051 |
| Gsg2 | 7.452195508 | 2.233711401 | 0.491475103 |
| P2rx5 | 25.59531151 | 0.927151295 | 0.246815126 |
| Emc6 | 323.5210552 | 0.354758347 | 0.107070577 |
| Tax1bp3 | 46.14509597 | 1.81001956 | 0.239597838 |
| Ctns | 210.1024968 | 0.360378992 | 0.114083776 |
| Shpk | 35.71037531 | -0.071223338 | 0.208940258 |
| Gm16021 | 0 | NA | NA |
| Trpv1 | 1.771085664 | 1.450633704 | 0.856184055 |
| Trpv3 | 13.35021764 | -0.03083779 | 0.320525203 |
| Aspa | 279.3669201 | -0.486127534 | 0.18936761 |
| Spata22 | 0.720179912 | -0.925042592 | 1.125387334 |
| Olfr20 | 0 | NA | NA |
| Olfr376 | 0 | NA | NA |

| | | | |
|---|---|---|---|
| Olfr377-ps1 | 0 | NA | NA |
| Olfr1 | 0.108309328 | 0.237310006 | 3.724945049 |
| Olfr75-ps1 | 0 | NA | NA |
| Olfr378 | 0 | NA | NA |
| Olfr379-ps1 | 0 | NA | NA |
| Olfr380 | 0 | NA | NA |
| Olfr381 | 0 | NA | NA |
| Olfr382 | 0 | NA | NA |
| Olfr383-ps1 | 0 | NA | NA |
| Olfr385 | 0 | NA | NA |
| Olfr384 | 0 | NA | NA |
| Gm12327 | 0 | NA | NA |
| Olfr387-ps1 | 0 | NA | NA |
| Zfp735 | 0 | NA | NA |
| Olfr388-ps1 | 0 | NA | NA |
| Gm22381 | 0 | NA | NA |
| Gm12328 | 0.100902008 | 1.15059983 | 3.720931964 |
| Olfr389 | 0 | NA | NA |
| Olfr390 | 0.097139547 | 0.718207094 | 3.724945049 |
| Olfr391-ps | 0 | NA | NA |
| Olfr392 | 0.046345709 | 0.71820487 | 3.724945049 |
| Gm12329 | 0 | NA | NA |
| Olfr393 | 0 | NA | NA |
| Olfr394 | 0 | NA | NA |
| Olfr395 | 0 | NA | NA |
| Olfr396-ps1 | 0 | NA | NA |
| Olfr23 | 0 | NA | NA |
| Olfr22-ps1 | 0 | NA | NA |
| Olfr397 | 0 | NA | NA |
| Olfr398 | 0.042716945 | -0.243571806 | 3.724945049 |
| Olfr139 | 0 | NA | NA |
| Olfr399 | 0 | NA | NA |
| Zfp616 | 0 | NA | NA |
| Gm12331 | 0.304339188 | 1.443904167 | 1.729473692 |
| Olfr400-ps1 | 0 | NA | NA |
| Olfr401 | 0 | NA | NA |
| Olfr402 | 0.044191549 | -0.243571806 | 3.724945049 |
| Olfr403 | 0 | NA | NA |
| Olfr43 | 0 | NA | NA |
| Olfr404-ps1 | 0 | NA | NA |
| Olfr405-ps1 | 0 | NA | NA |
| Olfr406 | 0 | NA | NA |
| Olfr407-ps1 | 0.066050875 | 0.71820487 | 3.724945049 |
| Olfr59 | 0.062426873 | 0.237312084 | 3.724945049 |
| Olfr408-ps1 | 0 | NA | NA |
| Olfr409-ps1 | 0 | NA | NA |
| n-R5s71 | 0 | NA | NA |

| | | | |
|---|---|---|---|
| Gm12332 | 0 | NA | NA |
| Olfr410 | 0.049553047 | 0.71820487 | 3.724945049 |
| Olfr411 | 0.046345709 | 0.71820487 | 3.724945049 |
| Olfr412 | 0.048949532 | -0.243569582 | 3.724945049 |
| Rap1gap2 | 2227.61315 | -0.36464902 | 0.10382564 |
| Gm22993 | 0.552195079 | 1.052446761 | 1.292676114 |
| E130309D14Rik | 443.6514258 | -0.268858923 | 0.069715118 |
| Cluh | 1336.662119 | -0.016522948 | 0.048438673 |
| Pafah1b1 | 8824.289057 | -0.156900598 | 0.057783918 |
| Gm24211 | 0 | NA | NA |
| Gm16032 | 0.075043205 | 0.237309859 | 3.724945049 |
| Mettl16 | 528.4778054 | -0.064445042 | 0.073883928 |
| Mnt | 707.5314908 | -0.071254903 | 0.097638265 |
| Sgsm2 | 1130.427896 | -0.06513675 | 0.081987017 |
| Mir6926 | 0 | NA | NA |
| Tsr1 | 408.3790304 | 0.727668297 | 0.119631255 |
| Snord91a | 2.110678478 | -0.279261077 | 0.747870442 |
| Gm22771 | 1.808802678 | 1.211784632 | 0.868455237 |
| Srr | 740.5480199 | -0.642160784 | 0.113635423 |
| Smg6 | 855.9187194 | -0.107790047 | 0.07611078 |
| Gm22733 | 0 | NA | NA |
| Gm28180 | 0 | NA | NA |
| Gm12333 | 0 | NA | NA |
| Gm12335 | 2.169066299 | -0.654924626 | 0.739045475 |
| Rps12-ps19 | 2.189863395 | 1.603289992 | 0.728638299 |
| Hic1 | 51.99884426 | 0.954174745 | 0.225972417 |
| 1700016P03Rik | 48.88953987 | -0.533341457 | 0.491415628 |
| Mir212 | 22.07890698 | -0.08441777 | 0.703135909 |
| Mir132 | 4.115963635 | 2.374740809 | 0.746507599 |
| Ovca2 | 77.25714499 | 0.407577978 | 0.161465707 |
| Dph1 | 85.00803555 | -0.011874345 | 0.132028533 |
| Rtn4rl1 | 700.6189518 | -0.405380844 | 0.15417344 |
| Rpa1 | 558.9030021 | 0.037923144 | 0.082683858 |
| AL603834.1 | 0.047948698 | -0.243569582 | 3.724945049 |
| Smyd4 | 164.5240954 | 0.007587997 | 0.094303888 |
| Serpinf1 | 126.7245348 | 0.250931035 | 0.198779021 |
| Gm12336 | 0.089062654 | 0.237318756 | 3.724945049 |
| Gm12337 | 0.219222703 | 0.237328821 | 2.177977162 |
| Serpinf2 | 5.648606599 | 2.337296351 | 0.56298844 |
| Wdr81 | 600.0628896 | 0.687928853 | 0.098166638 |
| Mir22hg | 200.1675278 | 0.674535932 | 0.172250558 |
| Tlcd2 | 9.434003272 | 0.54927068 | 0.328287189 |
| Mir22 | 0 | NA | NA |
| Prpf8 | 4246.36712 | 0.229659296 | 0.058532251 |
| Rilp | 13.6021336 | 0.406035232 | 0.283129704 |
| Scarf1 | 30.35591607 | 2.610897263 | 0.281102464 |
| Slc43a2 | 1420.800711 | 0.27399108 | 0.076836499 |

| | | | |
|---|---|---|---|
| Mir3971 | 2.406714335 | -1.156865971 | 0.875034195 |
| Pitpna | 3114.811598 | -0.053954131 | 0.062441125 |
| 4931413K12Rik | 0 | NA | NA |
| Gm12338 | 94.1563446 | 0.047080529 | 0.132395009 |
| Inpp5k | 474.3717951 | -0.093359454 | 0.113790291 |
| Myo1c | 416.9620297 | 1.318553081 | 0.145884178 |
| Crk | 1895.988957 | 0.324353461 | 0.09091898 |
| Ywhae | 6464.299146 | 0.04687519 | 0.062998875 |
| Gm26836 | 0.047948698 | -0.243569582 | 3.724945049 |
| Doc2b | 509.5744778 | -0.702940684 | 0.134885647 |
| Gm12339 | 4.361932539 | -1.601864942 | 0.558557118 |
| Rph3al | 23.25949346 | 0.006635671 | 0.250278347 |
| Gm16075 | 3.522107247 | -1.47283379 | 0.696759303 |
| 1700016K19Rik | 11.13294416 | 0.070109711 | 0.336332008 |
| Fam101b | 120.2091784 | -0.427910586 | 0.15431488 |
| Vps53 | 790.8122356 | -0.045794927 | 0.06609478 |
| Glod4 | 354.7645622 | -0.333878481 | 0.075231637 |
| Gm12340 | 0 | NA | NA |
| Fam57a | 179.7700705 | -0.350294308 | 0.099321709 |
| Gemin4 | 5.229107669 | -0.208371051 | 0.503110648 |
| Dbil5 | 3.336932514 | -0.755432618 | 0.538354017 |
| Rnmtl1 | 118.9831452 | 0.260957696 | 0.110471848 |
| Nxn | 198.759646 | 0.337630108 | 0.128370927 |
| Timm22 | 253.9457515 | 0.567221989 | 0.11811901 |
| Abr | 5618.476176 | -0.312391987 | 0.068678382 |
| Gm12341 | 0.351219581 | 1.153109891 | 1.668396384 |
| Rpl36-ps2 | 4.05888357 | 0.980428785 | 0.619940937 |
| Gm26780 | 0.189892101 | 1.744852741 | 3.278133644 |
| Bhlha9 | 8.304293226 | -0.575618632 | 0.390935274 |
| Tusc5 | 10.29048676 | 1.061001512 | 0.392467383 |
| Gosr1 | 767.1062309 | 0.346638733 | 0.065034862 |
| Cpd | 2040.908043 | 0.067299271 | 0.04723619 |
| Tmigd1 | 0.164253626 | 0.718216032 | 2.777778051 |
| Blmh | 568.2821817 | 0.03987097 | 0.069033942 |
| Gm22611 | 3.32088423 | -0.661443474 | 0.561686067 |
| Slc6a4 | 46.2681185 | 0.611083761 | 0.794009638 |
| Gm12343 | 0 | NA | NA |
| Ccdc55 | 161.7852798 | 0.397353551 | 0.199934757 |
| Gm22772 | 0 | NA | NA |
| Mir423 | 0.516177223 | 2.193513671 | 1.435140837 |
| Efcab5 | 66.97098547 | -0.573644708 | 0.176858882 |
| Gm12344 | 1.005780181 | -1.931959017 | 1.176116855 |
| Gm12345 | 29.19047675 | 0.461678023 | 0.191736384 |
| Ssh2 | 1266.750364 | -0.042886077 | 0.057492174 |
| Gm24989 | 0 | NA | NA |
| Gm12346 | 262.7722694 | 0.128887551 | 0.103940495 |
| Coro6 | 239.6645014 | -1.48986093 | 0.20325269 |

| | | | |
|---|---|---|---|
| Ankrd13b | 701.4661961 | -0.146977076 | 0.093904286 |
| 2210008F06Rik | 16.34395686 | 0.28999939 | 0.293513014 |
| Git1 | 3031.997312 | 0.151066206 | 0.066503948 |
| Trp53i13 | 48.23483047 | 0.607640041 | 0.179092713 |
| Abhd15 | 33.33330842 | 1.473877943 | 0.243241254 |
| Gm10392 | 0.289147236 | 1.538517021 | 2.056177782 |
| Taok1 | 6408.541416 | -0.287638736 | 0.047850936 |
| Gm12734 | 2.565340877 | 1.601852681 | 0.705917369 |
| Gm12732 | 0 | NA | NA |
| Gm12733 | 0 | NA | NA |
| Gm23132 | 0.137483974 | 0.684519582 | 2.929862181 |
| Nufip2 | 1887.858992 | 0.245432392 | 0.060740624 |
| Cryba1 | 0.167183452 | -0.72445987 | 2.765153652 |
| Gm11191 | 0.293994645 | 0.718222164 | 2.058366262 |
| Myo18a | 2948.916186 | 0.300639839 | 0.039630367 |
| Gm10277 | 0.30796268 | 1.144398706 | 1.777277866 |
| Pipox | 16.89892386 | -0.419166741 | 0.25926059 |
| Gm11190 | 4.356108356 | -0.10696689 | 0.453434245 |
| Sez6 | 1855.373666 | -0.872482161 | 0.135983131 |
| Phf12 | 804.2323745 | -0.022573939 | 0.101794689 |
| Dhrs13os | 5.524772491 | -0.353291597 | 0.443752775 |
| Dhrs13 | 73.22834646 | 0.003352443 | 0.137202 |
| Flot2 | 917.7646301 | -0.055875181 | 0.080035038 |
| Mir144 | 0 | NA | NA |
| Mir451a | 0 | NA | NA |
| mmu-mir-451a | 0.164553454 | 0.719098949 | 2.779126835 |
| Eral1 | 174.3755178 | 0.298778994 | 0.083268596 |
| Fam222b | 474.7107881 | 0.05113181 | 0.07039448 |
| Gm12571 | 0.999428373 | 0.747798578 | 1.134308662 |
| Traf4 | 100.814411 | 0.10133698 | 0.147216021 |
| Nek8 | 29.99850749 | 0.393233469 | 0.228723536 |
| Tlcd1 | 129.4641411 | -1.413232371 | 0.163660574 |
| Mir7653 | 0 | NA | NA |
| Rpl23a | 798.9907631 | 0.491734202 | 0.125421064 |
| Snord42a | 0.955875529 | -0.050485294 | 1.058653113 |
| Snord4a | 1.972325891 | -1.157837579 | 0.725451734 |
| Snord42b | 3.467268438 | 0.752983429 | 0.589445574 |
| Rab34 | 131.9099265 | 0.090487341 | 0.148044025 |
| Proca1 | 20.46478121 | -0.963584263 | 0.273921897 |
| Supt6 | 2745.970221 | 0.147851036 | 0.090785838 |
| Sdf2 | 269.515758 | 0.27920699 | 0.095133897 |
| 2610507B11Rik | 4624.687977 | 0.395806287 | 0.056031507 |
| Gm11192 | 0.159819677 | -0.198487585 | 3.46125206 |
| BC030499 | 82.88575948 | -2.191842013 | 0.465354169 |
| Gm11193 | 0.044191549 | -0.243571806 | 3.724945049 |
| Spag5 | 131.8137158 | -0.246164956 | 0.174437586 |
| Aldoc | 6307.64853 | -0.502586918 | 0.148696749 |

| | | | |
|---|---|---|---|
| Pigs | 527.1172141 | -0.079088691 | 0.089137646 |
| Unc119 | 149.7412516 | -0.59852135 | 0.12709502 |
| Foxn1 | 0.172802443 | -0.243571997 | 3.388555944 |
| Slc13a2os | 0 | NA | NA |
| Slc13a2 | 0 | NA | NA |
| Gm11195 | 0.045882455 | 0.237312084 | 3.724945049 |
| Slc46a1 | 58.13216195 | -0.1735602 | 0.162252681 |
| Sarm1 | 378.1139612 | -0.51850938 | 0.081036638 |
| Vtn | 396.8074538 | -0.189369355 | 0.157540544 |
| Sebox | 24.6969001 | -0.295986558 | 0.268007131 |
| Tmem199 | 242.7545441 | 0.137321822 | 0.088858336 |
| Poldip2 | 775.1605727 | -0.10331847 | 0.081827739 |
| Tnfaip1 | 559.6315322 | 0.47385002 | 0.077509041 |
| Ift20 | 265.7227452 | -0.408283824 | 0.07489499 |
| Tmem97 | 90.94084935 | -0.097232662 | 0.146320866 |
| D11Bhm181e | 1.727533833 | -1.298119326 | 1.138117854 |
| Gm22947 | 0.742175153 | -3.44852352 | 1.159392811 |
| Nlk | 1172.726608 | -0.716470738 | 0.09737382 |
| Gm17235 | 0 | NA | NA |
| Gm11196 | 6.623448048 | -0.585548237 | 0.503550748 |
| Gm11197 | 0 | NA | NA |
| Fam58b | 92.53061276 | 0.399189556 | 0.130967013 |
| Gm23840 | 0 | NA | NA |
| Gm23219 | 0 | NA | NA |
| Lyrm9 | 495.4572617 | -0.665046821 | 0.083185255 |
| Nos2 | 95.34328637 | 6.89095856 | 0.772543394 |
| Lgals9 | 1210.742748 | 6.160526312 | 0.261226609 |
| Ksr1 | 731.8760987 | -0.310184657 | 0.088423872 |
| Gm11201 | 0.406995362 | 1.557675301 | 1.671878081 |
| Gm11200 | 0 | NA | NA |
| Wsb1 | 1011.052735 | 0.266610181 | 0.069859305 |
| Gm9964 | 0 | NA | NA |
| Nf1 | 4025.801747 | -0.356688058 | 0.036850857 |
| Gm11198 | 0 | NA | NA |
| Gm11199 | 0 | NA | NA |
| AU040972 | 4.210711181 | -1.367142967 | 0.558236775 |
| Omg | 735.6960469 | -0.674560333 | 0.082460284 |
| Gm21975 | 2.443568159 | 4.040346086 | 0.934455413 |
| Evi2b | 0 | NA | NA |
| Evi2a | 264.6184616 | 0.669015636 | 0.122460655 |
| Rab11fip4 | 3582.291547 | -0.839125656 | 0.095948091 |
| Gm23293 | 1.882325847 | -1.965115403 | 0.950921587 |
| Rab11fip4os1 | 0.112642008 | -0.24357403 | 3.724945049 |
| Gm25867 | 0 | NA | NA |
| Gm24887 | 0 | NA | NA |
| Rab11fip4os2 | 0.045882455 | 0.237312084 | 3.724945049 |
| Gm11205 | 0.304750214 | 0.237329052 | 1.989237162 |

| | | | |
|---|---|---|---|
| Mir193a | 0.044191549 | -0.243571806 | 3.724945049 |
| Mir365-2 | 0.211497531 | 1.671276746 | 2.173517236 |
| 9130204K15Rik | 0.414175607 | 2.435783302 | 1.646706837 |
| Gm24454 | 0.133633945 | 0.237303512 | 3.724945049 |
| Gm11203 | 0 | NA | NA |
| Gm11204 | 0.206494471 | -0.243564123 | 2.202635613 |
| Utp6 | 497.722905 | 0.112999356 | 0.075574023 |
| Suz12 | 822.7645743 | 0.300463911 | 0.070578876 |
| Gm26563 | 1.315499399 | 0.399745793 | 0.986432981 |
| Crlf3 | 219.9408582 | 0.455270532 | 0.097829775 |
| Atad5 | 155.6883303 | 0.043628085 | 0.184276813 |
| Tefm | 62.86592296 | 0.631419108 | 0.153285691 |
| Adap2 | 173.6780421 | 2.5632378 | 0.181335649 |
| Adap2os | 0.240869569 | 1.212829121 | 3.089662278 |
| Rnf135 | 133.3790234 | 2.424520057 | 0.192501674 |
| Rhot1 | 976.0750427 | -0.104782745 | 0.045294168 |
| Rhbdl3 | 250.4651554 | -0.299073979 | 0.142144154 |
| 5730455P16Rik | 801.0455395 | -0.260151257 | 0.070438984 |
| Zfp207 | 1918.849716 | -0.135481955 | 0.077202276 |
| Psmd11 | 1260.672455 | 0.618068419 | 0.087722661 |
| Cdk5r1 | 2328.335231 | -0.72957415 | 0.09842094 |
| Myo1d | 327.8254873 | 0.133152019 | 0.092349975 |
| C030013C21Rik | 2.678984608 | -1.126516107 | 0.71967237 |
| Tmem98 | 68.95410002 | 0.199143025 | 0.214664937 |
| Gm11207 | 0 | NA | NA |
| 4930507D10Rik | 0.549547734 | 0.135272268 | 1.270203395 |
| Spaca3 | 0.045882455 | 0.237312084 | 3.724945049 |
| Asic2 | 625.5974346 | -0.548699885 | 0.071855143 |
| Gm11416 | 0 | NA | NA |
| Gm11417 | 0.04393175 | -0.243571806 | 3.724945049 |
| 4930527B05Rik | 0 | NA | NA |
| 1700071K01Rik | 9.922463048 | -1.740319076 | 0.491135893 |
| Gm11418 | 0 | NA | NA |
| 5530401A14Rik | 0.732425175 | 0.534860504 | 1.115839553 |
| Gm11419 | 2.867374077 | -0.712182352 | 0.683087513 |
| Gm17268 | 0.061920613 | 0.718207094 | 3.724945049 |
| Ccl2 | 1280.074291 | 10.93165935 | 0.818249928 |
| Ccl7 | 524.7321755 | 11.91940972 | 1.030882918 |
| Ccl11 | 7.798742698 | 6.004873222 | 0.914482225 |
| Ccl12 | 226.9053113 | 8.420197594 | 0.651750293 |
| Ccl8 | 52.35493119 | 9.172415823 | 1.020378589 |
| Ccl1 | 0.343821097 | 1.156688638 | 1.999080139 |
| Tmem132e | 318.4462841 | 0.468880871 | 0.130956112 |
| Gm24612 | 0 | NA | NA |
| Gm11426 | 0 | NA | NA |
| Cct6b | 3.620020008 | -0.260931743 | 0.661918676 |
| Gm11421 | 0 | NA | NA |

| | | | |
|---|---|---|---|
| Gm11422 | 0 | NA | NA |
| Gm11420 | 0.621419653 | 1.194189796 | 1.391876344 |
| Zfp830 | 109.1762647 | 0.014873546 | 0.113489642 |
| Gm11423 | 5.437272543 | 0.431533963 | 0.484044602 |
| Lig3 | 405.8341476 | 0.193423683 | 0.066365228 |
| Rffl | 192.7097897 | -0.038578727 | 0.127323036 |
| Rad51d | 202.7225488 | -0.144755329 | 0.081935318 |
| Fndc8 | 0.20460011 | -1.118574121 | 3.233930613 |
| Nle1 | 99.69246992 | 0.298852818 | 0.116584284 |
| Unc45b | 11.03143117 | 1.57567835 | 0.388499628 |
| Unc45bos | 2.092676817 | 1.288333745 | 0.793357613 |
| Slfn5os | 0.252317246 | 1.281389321 | 2.500491872 |
| Slfn5 | 854.2001971 | 6.438341032 | 0.299902888 |
| Gm11424 | 0.243199696 | 0.961162147 | 2.442365941 |
| Slfn9 | 576.8930042 | 7.625102669 | 0.333424954 |
| Mir7679 | 0.98936666 | 3.708466233 | 1.071496621 |
| Slfn8 | 1181.313578 | 8.065561208 | 0.288467038 |
| Gm11425 | 15.12308696 | 4.889051009 | 0.576596743 |
| Slfn10-ps | 120.7759095 | 9.792760814 | 0.912770425 |
| Gm27598 | 0.379788221 | 2.590757739 | 1.84308383 |
| Slfn2 | 559.7972036 | 7.570537943 | 0.379720572 |
| Slfn1 | 299.5776399 | 11.98258739 | 0.921209111 |
| Slfn4 | 2011.837015 | 13.03300378 | 0.839022267 |
| Slfn3 | 17.21960857 | 4.106671844 | 0.502810716 |
| AI662270 | 42.49283015 | 5.639770453 | 0.511487144 |
| Gm11427 | 4.559114128 | 4.326591105 | 0.910415585 |
| Gm23444 | 10.89494943 | 0.558241086 | 0.388743895 |
| Slfn14 | 0.407088326 | 0.265421477 | 1.590775362 |
| AA465934 | 34.77868168 | -0.582597922 | 0.189173847 |
| Snord7 | 2.473802751 | -1.209831077 | 0.632263218 |
| Pex12 | 83.19986102 | 0.65878472 | 0.15896938 |
| Ap2b1 | 4452.733103 | -0.126820619 | 0.053638525 |
| Gm23951 | 0 | NA | NA |
| Rasl10b | 2095.394555 | -0.995140644 | 0.120252747 |
| Gas2l2 | 4.963609251 | -0.14343691 | 0.471386839 |
| 1700020L24Rik | 3.62909452 | 0.831807541 | 0.552086782 |
| Mmp28 | 45.16190099 | 1.446892423 | 0.213753818 |
| Taf15 | 585.2888291 | 0.380511495 | 0.128791821 |
| Heatr9 | 16.53926368 | 7.859222376 | 0.941159368 |
| Gm23247 | 1.077303893 | 1.430028831 | 1.047657756 |
| Ccl5 | 670.2920451 | 11.90199167 | 0.941664581 |
| Ccl9 | 101.7341889 | 6.235858523 | 0.560507555 |
| E230016K23Rik | 2.428642307 | 1.762593337 | 0.675603119 |
| Ccl6 | 71.52556139 | 4.51367988 | 0.384459254 |
| Rpl9-ps1 | 0 | NA | NA |
| Ccl3 | 250.7978955 | 8.185048291 | 0.942900579 |
| Ccl4 | 181.900904 | 8.960043012 | 0.860709357 |

| | | | | |
|---|---|---|---|---|
| Gm11433 | 0.066050875 | | 0.71820487 | 3.724945049 |
| Gm11438 | 0 | NA | | NA |
| Gm11432 | 0.038981395 | | 0.718207094 | 3.724945049 |
| Gm11430 | 0 | NA | | NA |
| Gm11429 | 0 | NA | | NA |
| Wfdc17 | 58.81490581 | | 6.701964951 | 0.679908903 |
| Wfdc18 | 2.91100996 | | 0.218525744 | 0.573946695 |
| Wfdc21 | 0.36705964 | | 0.237305091 | 3.724945049 |
| Heatr6 | 295.8490656 | | 0.138038015 | 0.086072935 |
| Gm22311 | 0 | NA | | NA |
| Gm11434 | 0.042716945 | | -0.243571806 | 3.724945049 |
| Gm12576 | 1.282016488 | | -1.636138025 | 0.945208638 |
| Gm23211 | 0 | NA | | NA |
| Hnf1b | 4.558351918 | | -0.03131939 | 0.514648839 |
| Ddx52 | 333.6973482 | | 0.706979278 | 0.108559056 |
| Synrg | 1329.598341 | | -0.083248877 | 0.056126394 |
| Dusp14 | 294.0225346 | | -0.171368674 | 0.178708129 |
| Tada2a | 253.1517061 | | -0.5022325 | 0.153023313 |
| Acaca | 1627.488358 | | 0.010186774 | 0.059803671 |
| Gm11437 | 0.12092566 | | 0.237307635 | 3.724945049 |
| Gm37391 | 10.91891655 | | -0.53633168 | 0.331676849 |
| Aatf | 235.0359788 | | 0.408198804 | 0.141799631 |
| Lhx1 | 140.9707341 | | -0.339858432 | 0.231157939 |
| Lhx1os | 31.23601852 | | -0.566369839 | 0.225773969 |
| 1700109G15Rik | 0 | NA | | NA |
| Gm11431 | 0 | NA | | NA |
| Mrm1 | 101.0285373 | | 0.169964143 | 0.117829385 |
| Dhrs11 | 57.07916713 | | -0.155165944 | 0.16499406 |
| 4930502E09Rik | 0.377064288 | | 0.272904323 | 3.693238232 |
| Ggnbp2 | 1167.177578 | | 0.096293766 | 0.084284504 |
| Pigw | 46.32962589 | | 0.112127908 | 0.159542043 |
| Myo19 | 116.4020599 | | -0.380972206 | 0.197673394 |
| Gm23564 | 1.319357828 | | 0.249027254 | 0.936313165 |
| Znhit3 | 108.0352272 | | -0.00321459 | 0.120198642 |
| Car4 | 225.7880006 | | -1.253774087 | 0.203208532 |
| Usp32 | 2485.609366 | | -0.345333833 | 0.048991994 |
| Gm11440 | 7.169804879 | | -1.208409301 | 0.398628651 |
| Gm11439 | 9.641612845 | | -0.70975976 | 0.490879005 |
| Rpl13-ps1 | 6.979134821 | | -1.219549596 | 0.526928144 |
| Gm23506 | 0 | NA | | NA |
| 1700125H20Rik | 0.788910745 | | 2.914647655 | 1.157100631 |
| Appbp2 | 1916.277606 | | -0.033109218 | 0.061494354 |
| Akirin1-ps | 0.295845421 | | -1.195086818 | 2.068648647 |
| Appbp2os | 6.2436769 | | -0.422876325 | 0.480074367 |
| Gm22587 | 0 | NA | | NA |
| Ppm1d | 315.9501097 | | -0.16955118 | 0.108697793 |
| Bcas3 | 671.5506855 | | -0.453905841 | 0.070259076 |

| | | | |
|---|---|---|---|
| Gm23765 | 0.716442652 | -0.749385909 | 1.22675903 |
| Bcas3os1 | 0.047948698 | -0.243569582 | 3.724945049 |
| Mir5110 | 0 | NA | NA |
| Bcas3os2 | 0 | NA | NA |
| Gm23473 | 1.868429801 | -1.535581179 | 0.967604245 |
| 2610027K06Rik | 0.597123318 | 1.711764048 | 1.378853591 |
| Tbx2 | 23.31208816 | 0.521283152 | 0.238352183 |
| Gm11444 | 0 | NA | NA |
| Tbx4 | 0.57584922 | -0.399931955 | 1.387077054 |
| Gm11443 | 0 | NA | NA |
| Brip1 | 34.57829007 | 3.310976463 | 0.316487601 |
| Atp5l-ps2 | 2.273527578 | -1.01054738 | 0.867761196 |
| Brip1os | 315.6828442 | 1.482563223 | 0.146233587 |
| Ints2 | 398.3495836 | -0.100329547 | 0.128627006 |
| Med13 | 3263.787859 | 0.215193987 | 0.050326799 |
| Gm25427 | 0 | NA | NA |
| Gm23667 | 0 | NA | NA |
| Gm26439 | 0 | NA | NA |
| Rnft1 | 257.0685552 | 0.069400374 | 0.099754079 |
| Rps6kb1 | 899.7579894 | -0.085733929 | 0.057939666 |
| Tubd1 | 49.92708075 | -0.104942345 | 0.187305286 |
| Vmp1 | 1340.608191 | 0.937221463 | 0.105998905 |
| Mir21a | 0 | NA | NA |
| Gm11478 | 49.42339314 | 0.329070578 | 0.168458041 |
| Ptrh2 | 123.2383261 | 0.221780408 | 0.117563956 |
| Cltc | 12039.16266 | 0.049123214 | 0.056851955 |
| Dhx40 | 393.4698973 | 0.337835047 | 0.084126824 |
| Ypel2 | 824.3745057 | -0.61388844 | 0.110658005 |
| Gm22387 | 0 | NA | NA |
| Gdpd1 | 414.5698482 | -0.216901111 | 0.073412871 |
| Gm11479 | 0.152667883 | -0.243571276 | 2.845214345 |
| Smg8 | 236.0766196 | 0.365168712 | 0.122028268 |
| Gm9378 | 0.044191549 | -0.243571806 | 3.724945049 |
| Prr11 | 19.51012886 | 1.160436224 | 0.288909652 |
| Ska2 | 149.134786 | 0.160650476 | 0.125492798 |
| Mir301 | 1.204075239 | -0.590787178 | 0.855257226 |
| Gm11491 | 0.332371241 | -1.372353956 | 3.691775746 |
| Trim37 | 3977.710216 | -0.79852698 | 0.077404897 |
| Ppm1e | 3157.400212 | -0.417627036 | 0.099344501 |
| Gm11490 | 0 | NA | NA |
| Gm22883 | 12.43333056 | -1.176471329 | 0.492413747 |
| Rad51c | 45.74186524 | -0.17986617 | 0.191326493 |
| Tex14 | 23.09397913 | 1.561045383 | 0.247705417 |
| Rnu1a1 | 44.69699138 | -0.940142385 | 0.500443036 |
| Gm22068 | 772.7219214 | 0.044872328 | 0.166345041 |
| Rnu3b1 | 0.136120942 | 1.15080397 | 3.595683408 |
| Gm23137 | 0 | NA | NA |

| | | | |
|---|---|---|---|
| Rnu3b3 | 0.045882455 | 0.237312084 | 3.724945049 |
| Rnu3b4 | 0 | NA | NA |
| Gm11489 | 0.231904818 | -1.573998253 | 2.137353648 |
| Rnu3b2 | 0 | NA | NA |
| Gm11492 | 0.278403928 | -1.075062943 | 2.018542961 |
| 4-Sep | 1581.659409 | -0.547711904 | 0.088707306 |
| Mtmr4 | 1614.493203 | -0.046386935 | 0.061957647 |
| Hsf5 | 20.47148481 | -0.275297213 | 0.336472821 |
| Rnf43 | 59.17937029 | -0.264696157 | 0.169641292 |
| Gm25968 | 0 | NA | NA |
| 1110028F11Rik | 2.71851656 | -0.329398074 | 0.682258223 |
| Supt4a | 181.1208407 | 0.239621946 | 0.084020679 |
| Mir142hg | 12.62452238 | 4.244398921 | 0.542547196 |
| Mir142b | 0.614441445 | -1.232586532 | 1.194692205 |
| mmu-mir-142a | 0.12092566 | 0.237307635 | 3.724945049 |
| Bzrap1 | 2131.147747 | -1.100235365 | 0.155185079 |
| Mpo | 4.50923424 | 4.628995501 | 0.907698428 |
| Gm11505 | 0.14164095 | 0.719186731 | 3.60310638 |
| Lpo | 20.44324603 | 1.563845063 | 0.270890133 |
| Mks1 | 36.78772966 | 0.593104343 | 0.19285805 |
| Epx | 0.366894703 | 2.441690902 | 1.589294207 |
| Olfr462 | 0.183788242 | -0.243563235 | 2.683430441 |
| Olfr463 | 0.038981395 | 0.718207094 | 3.724945049 |
| Olfr464 | 0.707581183 | -0.345631929 | 1.002269996 |
| Gm11504 | 0 | NA | NA |
| Gm11507 | 0.066050875 | 0.71820487 | 3.724945049 |
| Dynll2 | 5064.124974 | -0.40843325 | 0.076609194 |
| Gm11508 | 7.373820318 | 0.084047694 | 0.395472229 |
| 2010015M23Rik | 13.47151829 | -0.244701702 | 0.300426775 |
| Srsf1 | 2301.674563 | 0.246142376 | 0.04954746 |
| Gm15892 | 24.85441836 | -0.321499961 | 0.291349176 |
| Vezf1 | 819.1262531 | 0.436922218 | 0.087803959 |
| Cuedc1 | 496.2529045 | 0.254918366 | 0.072111619 |
| Mrps23 | 225.4695307 | 0.433322144 | 0.10095859 |
| 1700106J16Rik | 0.47539415 | -1.27201561 | 1.352644305 |
| Gm11509 | 0 | NA | NA |
| Msi2 | 2918.689029 | 0.058175126 | 0.072784822 |
| Mir378b | 0 | NA | NA |
| Gm15893 | 0.758254207 | 0.07464023 | 1.166432196 |
| Gm11510 | 2.984155966 | -0.465147604 | 0.579721683 |
| C030037D09Rik | 74.7814881 | -0.424004604 | 0.154761598 |
| Gm11493 | 0 | NA | NA |
| Akap1 | 422.0526709 | -0.667467762 | 0.098197937 |
| 4930556N13Rik | 0.0878635 | -0.72445792 | 3.724945049 |
| Gm23507 | 0 | NA | NA |
| Scpep1os | 0 | NA | NA |
| Scpep1 | 257.6478549 | 1.755905824 | 0.186880713 |

| | | | |
|---|---|---|---|
| Gm15698 | 0.068097359 | -0.243569582 | 3.724945049 |
| Coil | 170.2213894 | -0.013158726 | 0.091180665 |
| 2210409E12Rik | 0.042716945 | -0.243571806 | 3.724945049 |
| Gm11496 | 0.352842203 | 2.054284545 | 1.965595139 |
| Trim25 | 943.4408156 | 5.280006294 | 0.197442383 |
| Dgke | 1158.574667 | -0.146924758 | 0.070185012 |
| Dgkeos | 4.476524561 | 0.405967054 | 0.491076865 |
| Gm525 | 0 | NA | NA |
| Gm22534 | 0.038981395 | 0.718207094 | 3.724945049 |
| Gm11497 | 0 | NA | NA |
| Gm24734 | 0 | NA | NA |
| Nog | 67.55626535 | -0.755483956 | 0.176793639 |
| Gm2018 | 0.338611762 | -0.676860937 | 2.091678786 |
| 4932411E22Rik | 35.54587762 | -0.994515188 | 0.226569958 |
| Ankfn1 | 45.04732695 | -0.805896141 | 0.176384457 |
| Pctp | 14.03537449 | 0.988112071 | 0.297485743 |
| Tmem100 | 84.40838063 | -0.237787484 | 0.145751888 |
| Gm11494 | 0 | NA | NA |
| n-R5s72 | 0 | NA | NA |
| Mmd | 1332.487735 | -0.656489245 | 0.100253828 |
| Hlf | 2065.341962 | -1.473786648 | 0.132181209 |
| Stxbp4 | 295.2899188 | -0.252394589 | 0.098874024 |
| Gm11511 | 3.871387691 | -0.132829197 | 0.651793885 |
| Gm11512 | 2.280412116 | -0.867757886 | 0.700069787 |
| Cox11 | 152.2450534 | 0.084781227 | 0.122212227 |
| Tom1l1 | 226.4499419 | -0.296882482 | 0.113171977 |
| Gm22762 | 0 | NA | NA |
| 4930405D11Rik | 0.230025872 | -1.163275339 | 2.530754534 |
| Gm11516 | 0 | NA | NA |
| Kif2b | 0 | NA | NA |
| Gm11498 | 0 | NA | NA |
| Gm22702 | 0 | NA | NA |
| Gm11500 | 0 | NA | NA |
| Gm11501 | 0 | NA | NA |
| Gm25113 | 0 | NA | NA |
| Car10 | 1148.248468 | -0.716351593 | 0.088830417 |
| Gm24856 | 0 | NA | NA |
| Gm11502 | 0.044544648 | 0.237309859 | 3.724945049 |
| Gm11506 | 0 | NA | NA |
| Utp18 | 272.3077803 | 0.352415429 | 0.085039309 |
| Mbtd1 | 630.0243432 | -0.393267597 | 0.089354039 |
| Nme2 | 16.60911457 | 0.594149397 | 0.341009029 |
| Gm20390 | 0.049553047 | 0.71820487 | 3.724945049 |
| Nme1 | 441.0854408 | 0.229979606 | 0.078210841 |
| Spag9 | 5268.156952 | -0.606218336 | 0.049683949 |
| Gm11518 | 36.08732431 | 1.118005769 | 0.228100475 |
| B230206L02Rik | 10.54869608 | -1.376397747 | 0.353631586 |

| | | | |
|---|---|---|---|
| Tob1 | 560.157712 | -0.363933577 | 0.083487599 |
| Wfikkn2 | 33.87170838 | -0.109377526 | 0.2174483 |
| Gm21885 | 0.047948698 | -0.243569582 | 3.724945049 |
| Luc7l3 | 1814.337726 | -0.088718337 | 0.097552426 |
| Ankrd40 | 1549.21346 | 0.023260417 | 0.090197939 |
| Abcc3 | 101.8740177 | 2.178171335 | 0.305568259 |
| Cacna1g | 1923.935302 | -0.495676302 | 0.137006388 |
| Spata20 | 1.880027439 | -0.52130335 | 0.717773654 |
| Epn3 | 81.88700119 | -0.847885607 | 0.155492734 |
| Mycbpap | 22.79270077 | -0.149178285 | 0.240704736 |
| Rsad1 | 129.9462632 | -0.376632385 | 0.144492518 |
| Acsf2 | 203.4382502 | -0.360030583 | 0.165544101 |
| Chad | 2.13663281 | -0.083192421 | 0.702159348 |
| Gm11540 | 0 | NA | NA |
| Lrrc59 | 1158.226995 | 0.899092771 | 0.109116252 |
| Eme1 | 6.299531046 | 3.550292047 | 0.72863562 |
| Mrpl27 | 283.0008542 | 0.837339699 | 0.112130913 |
| Xylt2 | 348.7120337 | -0.031364371 | 0.091403553 |
| Gm11542 | 2.690072256 | -0.644587792 | 0.606645632 |
| Gm11541 | 0.211252453 | -0.243563317 | 2.188118052 |
| Cdc34-ps | 2.829894638 | -0.211478324 | 0.595520266 |
| Gm11545 | 0.145451803 | 1.498658476 | 3.529361416 |
| Tmem92 | 0.060174537 | 0.237309859 | 3.724945049 |
| Gm11546 | 0 | NA | NA |
| AA623943 | 0 | NA | NA |
| Gm11543 | 0 | NA | NA |
| Gm11544 | 0 | NA | NA |
| A430060F13Rik | 0 | NA | NA |
| Gm22456 | 0 | NA | NA |
| Col1a1 | 391.7651446 | 0.527056921 | 0.159413883 |
| Gm24877 | 0 | NA | NA |
| Sgca | 2.248524686 | 1.022213751 | 0.832086586 |
| Hils1 | 0 | NA | NA |
| Ppp1r9b | 8227.147287 | -0.585656074 | 0.100808084 |
| Samd14 | 673.600847 | -0.039720107 | 0.117097806 |
| Pdk2 | 946.573687 | -0.587505811 | 0.100200551 |
| Gm11513 | 1.036123199 | -2.121588508 | 1.115852379 |
| Itga3 | 513.8169415 | 0.311835423 | 0.089958983 |
| Dlx3 | 0.948391139 | -0.786926186 | 1.144584802 |
| Dlx4 | 3.24303925 | -0.709060643 | 0.572692171 |
| A730090H04Rik | 3.582880236 | 0.783159435 | 0.670758755 |
| Gm11514 | 1.617110251 | 1.066288415 | 0.821960527 |
| Gm11515 | 0.488879413 | 1.122248634 | 1.462498854 |
| Tac4 | 0.201378205 | -0.724461904 | 2.645200898 |
| Kat7 | 1091.977074 | 0.249721353 | 0.077405652 |
| Gm11520 | 2.683034273 | 0.197884141 | 0.643264577 |
| Gm11521 | 0.240698147 | 0.237299344 | 3.724945049 |

| | | | |
|---|---|---|---|
| Fam117a | 84.85769561 | 0.027079274 | 0.160419266 |
| Gm29477 | 0.19690755 | 0.246781514 | 2.677258802 |
| Slc35b1 | 393.4153695 | 0.593291993 | 0.107341964 |
| Spop | 1831.86833 | 0.114354734 | 0.072646555 |
| Gm11522 | 0 | NA | NA |
| Nxph3 | 195.7791226 | -0.384509368 | 0.125992118 |
| Ngfr | 42.86765523 | 0.701077127 | 0.269727597 |
| Gm11528 | 0 | NA | NA |
| Phb | 513.3109458 | 0.287685419 | 0.11672387 |
| Gm26830 | 2.519878091 | -1.633601295 | 0.721494896 |
| Gm9796 | 6.527712466 | -0.58902566 | 0.402454732 |
| Zfp652os | 6.322034818 | -0.762180974 | 0.526085452 |
| Zfp652 | 1286.698194 | -0.42782967 | 0.05866609 |
| B130006D01Rik | 36.80483133 | -0.799083486 | 0.265218053 |
| Polr2k-ps | 4.087166043 | -0.492818058 | 0.502493464 |
| Phospho1 | 32.25678491 | 0.063137324 | 0.192605554 |
| Abi3 | 109.2635649 | 2.14128139 | 0.18584097 |
| Gngt2 | 37.40057289 | 3.051516106 | 0.302355881 |
| Gm11527 | 0.940895551 | 1.247997812 | 0.977855667 |
| 4833417C18Rik | 16.33142681 | -0.41953612 | 0.306846437 |
| B4galnt2 | 0.576309813 | 2.553330924 | 1.412215775 |
| Gm11534 | 0 | NA | NA |
| Gm24725 | 0 | NA | NA |
| Igf2bp1 | 1.321944539 | 1.265411038 | 1.014555154 |
| Mir3063 | 0 | NA | NA |
| Gm29202 | 0 | NA | NA |
| Gm37179 | 0 | NA | NA |
| Gip | 0 | NA | NA |
| Gm24949 | 0 | NA | NA |
| Snf8 | 318.1547224 | 0.1138072 | 0.10087241 |
| Ube2z | 1093.216647 | 0.281646221 | 0.062706999 |
| Atp5g1 | 503.7452782 | -0.2607991 | 0.088014968 |
| Gm22423 | 0 | NA | NA |
| Calcoco2 | 0.681926656 | 0.67355736 | 1.289888222 |
| Ttll6 | 7.585280821 | -0.502156654 | 0.357570404 |
| Gm11535 | 0 | NA | NA |
| Hoxb13 | 0 | NA | NA |
| Gm27898 | 0 | NA | NA |
| Gm27605 | 0 | NA | NA |
| Gm11539 | 1.7546716 | -0.303923507 | 0.688021693 |
| Gm53 | 0.110242424 | 0.237318756 | 3.724945049 |
| Mir196a-1 | 0 | NA | NA |
| Hoxb9 | 0.072025462 | 0.237312084 | 3.724945049 |
| Hoxb8 | 30.1408604 | 0.947712399 | 1.728818864 |
| Hoxb7 | 3.606077423 | 1.14217455 | 1.427013419 |
| Hoxb5os | 0.345588754 | 1.168281176 | 3.21622731 |
| Hoxb6 | 10.70947983 | 0.61954995 | 1.234177344 |

| | | | |
|---|---|---|---|
| Hoxb5 | 50.63370746 | 0.880465114 | 1.018957142 |
| Mir10a | 0.182096932 | -0.243562086 | 2.691326143 |
| Hoxb4 | 17.2026822 | 1.054963695 | 0.78179062 |
| Hoxb3 | 35.54127179 | 0.64029011 | 0.744536697 |
| Gm11536 | 0.751775467 | 1.482876086 | 1.251113228 |
| Hoxb3os | 4.383958145 | 0.37021207 | 0.526014177 |
| Hoxb2 | 10.86433349 | 1.223310517 | 0.868748415 |
| Hoxb1 | 0.038981395 | 0.718207094 | 3.724945049 |
| Gm11531 | 0.072025462 | 0.237312084 | 3.724945049 |
| Gm11529 | 0.471635551 | -0.634700085 | 1.350722707 |
| Skap1 | 41.01891481 | 1.340212097 | 0.251108445 |
| 2010300F17Rik | 0.313173757 | 0.219049912 | 2.037916367 |
| Gm11537 | 0.487078227 | 2.040831245 | 1.332694146 |
| Snx11 | 313.2983275 | 0.539649359 | 0.080650829 |
| Cbx1 | 382.9098145 | -0.003745007 | 0.095636769 |
| Gm11517 | 5.580449154 | 1.024594536 | 0.47515466 |
| Nfe2l1 | 4454.295115 | 0.168534858 | 0.046028056 |
| Copz2 | 26.60350595 | -0.056108171 | 0.245118695 |
| Mir152 | 0.090665643 | -0.724455696 | 3.724945049 |
| Gm11523 | 0.271332793 | -0.223447886 | 2.072372688 |
| Gm11524 | 0 | NA | NA |
| Cdk5rap3 | 188.9212374 | 0.091519218 | 0.105242057 |
| D030028A08Rik | 76.08496205 | -0.028400749 | 0.142461368 |
| Gm11525 | 3.501911378 | 0.034238712 | 0.594022534 |
| Prr15l | 1.00014491 | 1.909945084 | 1.196841696 |
| Pnpo | 446.5624225 | -0.259081073 | 0.072711491 |
| Sp2 | 265.9732879 | 0.518119743 | 0.133346113 |
| Sp6 | 5.037085609 | -0.391810016 | 0.443968623 |
| Gm11532 | 1.602391015 | 1.458301893 | 0.876843567 |
| Scrn2 | 30.51638522 | -0.129988581 | 0.244753722 |
| Lrrc46 | 5.763699656 | 0.910084085 | 0.442909411 |
| Gm11533 | 0.306021358 | 0.223592098 | 1.799454175 |
| Mrpl10 | 403.0972378 | 0.312180467 | 0.083267886 |
| Gm11574 | 2.609456233 | 0.007785357 | 0.638908739 |
| Osbpl7 | 231.9002821 | 0.122627465 | 0.084832799 |
| Mir8103 | 1.700203044 | -1.574983386 | 0.896688016 |
| Gm11573 | 0 | NA | NA |
| Tbx21 | 34.3699015 | 1.752995934 | 0.425815071 |
| Gm11572 | 0.075043205 | 0.237309859 | 3.724945049 |
| Tbkbp1 | 732.199412 | 0.081055627 | 0.071252314 |
| Mir7235 | 0 | NA | NA |
| Kpnb1 | 2804.956967 | 0.312153538 | 0.061116356 |
| Gm11583 | 2.829785719 | 0.248796934 | 0.657456144 |
| Npepps | 2128.553621 | -0.112994112 | 0.057161161 |
| Gm11592 | 3.097463869 | -0.57482283 | 0.561028782 |
| Mrpl45 | 265.8735872 | 0.245679203 | 0.105162412 |
| Gm11594 | 0.046345709 | 0.71820487 | 3.724945049 |

| | | | |
|---|---|---|---|
| Gpr179 | 53.52322391 | 0.946322493 | 0.269376196 |
| Socs7 | 1787.58141 | -0.234930896 | 0.071665618 |
| Arhgap23 | 1777.390419 | 0.085888558 | 0.063867774 |
| 4933428G20Rik | 1.863773006 | 0.262956229 | 0.90318221 |
| Srcin1 | 3857.360928 | -0.906797783 | 0.090628546 |
| Gm11611 | 105.1460556 | -0.841569942 | 0.2177993 |
| Gm11612 | 0.519932157 | -1.477916754 | 1.420587847 |
| 2410003L11Rik | 0.232084656 | 0.741256983 | 2.132105878 |
| E130012A19Rik | 333.7269305 | -1.414332421 | 0.157866677 |
| Gm11613 | 75.52422125 | -1.11356128 | 0.172268018 |
| Gm24158 | 0 | NA | NA |
| Mllt6 | 2810.779345 | -0.395783146 | 0.061613907 |
| Mir6927 | 0 | NA | NA |
| Cisd3 | 84.26726889 | 0.023098256 | 0.16568341 |
| Pcgf2 | 208.1768514 | -0.473748334 | 0.146500853 |
| Psmb3 | 579.8326605 | 0.858912215 | 0.143650609 |
| Pip4k2b | 1705.574018 | -0.418850455 | 0.064093437 |
| Gm11614 | 0 | NA | NA |
| Mir8102 | 0.581519725 | 0.293097273 | 1.354742568 |
| Cwc25 | 283.5958626 | 0.346457551 | 0.112973839 |
| Atp5l2-ps | 17.66472391 | 0.090758626 | 0.3295988 |
| 1700001P01Rik | 6.295635024 | 0.227173872 | 0.40011497 |
| Rpl23 | 782.0632473 | 0.645939799 | 0.125466571 |
| Gm27326 | 2.560534601 | 0.28453999 | 0.660317338 |
| Snora21 | 15.9140621 | 0.137397989 | 0.282928517 |
| Lasp1 | 1454.162824 | 0.130275081 | 0.081372378 |
| B230217C12Rik | 275.8448458 | -0.956136263 | 0.134576388 |
| Fbxo47 | 12.48405167 | 0.051018466 | 0.293748407 |
| Gm11630 | 0.282072957 | -0.722963945 | 1.785735073 |
| Plxdc1 | 370.3130502 | -0.536433132 | 0.12838709 |
| Gm22461 | 0.138038828 | -0.243577639 | 3.633333205 |
| Gm11633 | 0.1140246 | 0.718202646 | 3.724945049 |
| Arl5c | 23.21351478 | 1.54284045 | 0.310539331 |
| Cacnb1 | 1086.374005 | -0.678816498 | 0.142354707 |
| Gm11629 | 3.893533808 | -0.693315121 | 0.536680718 |
| Rpl19 | 1377.971411 | 0.715329362 | 0.144005152 |
| Gm27597 | 0 | NA | NA |
| Stac2 | 733.8978338 | -0.192813818 | 0.096991327 |
| Gm11631 | 0.045882455 | 0.237312084 | 3.724945049 |
| Fbxl20 | 783.1636922 | -0.464070227 | 0.097691978 |
| Gm11632 | 0 | NA | NA |
| Med1 | 1253.203097 | 0.142387036 | 0.051694743 |
| Gm11604 | 1.649565372 | 0.777889491 | 1.085598721 |
| Cdk12 | 1154.320364 | 0.329573878 | 0.094300212 |
| Gm27399 | 1.487186809 | 0.219826476 | 0.826909281 |
| Mir5119 | 5.066509227 | 0.673771377 | 0.507323253 |
| n-R5s73 | 0 | NA | NA |

| | | | |
|---|---|---|---|
| Gm20644 | 4.541882812 | 0.472171713 | 0.799321106 |
| Neurod2 | 846.9061386 | -0.424957701 | 0.135781421 |
| Ppp1r1b | 2292.267063 | -1.04728532 | 0.258020488 |
| 1700003D09Rik | 3.143667968 | -0.078592448 | 0.779879547 |
| Stard3 | 334.9821919 | 0.904434515 | 0.106378071 |
| Tcap | 15.2625128 | -1.989154232 | 0.313766291 |
| Pnmt | 1.554873959 | 1.095845413 | 0.954089883 |
| Pgap3 | 82.83766852 | -0.063824016 | 0.163756832 |
| Erbb2 | 73.62724615 | 1.584413727 | 0.192725988 |
| Gm25048 | 0 | NA | NA |
| Mien1 | 288.359261 | 0.046363005 | 0.133448503 |
| Gm12352 | 1.915443829 | 1.527747841 | 0.851707946 |
| Grb7 | 32.20120666 | -0.31782801 | 0.202691205 |
| Ikzf3 | 75.08787135 | 5.42707982 | 0.418360051 |
| Gm25106 | 0.044544648 | 0.237309859 | 3.724945049 |
| Zpbp2 | 0.628980615 | 0.225653444 | 1.15358907 |
| Ormdl3 | 555.4362379 | 0.479537328 | 0.07463506 |
| Gm12 | 0 | NA | NA |
| Gm12355 | 8.225470329 | 0.583023945 | 0.345370962 |
| Gsdma3 | 0.182775946 | 0.237309473 | 3.335217586 |
| Gsdma2 | 0 | NA | NA |
| Gsdma | 1.768266679 | 0.539998038 | 0.726886226 |
| Psmd3 | 1360.534359 | 0.554353251 | 0.092014951 |
| Gm12356 | 1.14577023 | 0.793499777 | 0.946958048 |
| Csf3 | 0.907385681 | -0.243566402 | 1.39056566 |
| Med24 | 900.5465924 | -0.233949533 | 0.072289249 |
| Gm22059 | 0.481050399 | 0.239779405 | 1.679239167 |
| Thra | 4319.830123 | -0.64290297 | 0.076556809 |
| Nr1d1 | 1275.640339 | -0.621312763 | 0.090345934 |
| Msl1 | 1734.681355 | -0.367875612 | 0.067573879 |
| Gm12359 | 6.105237519 | 0.171791883 | 0.445619505 |
| Casc3 | 869.1967834 | 0.0632853 | 0.055115517 |
| Rapgefl1 | 1200.68795 | -1.305119765 | 0.144557144 |
| Wipf2 | 1511.917858 | -0.201085552 | 0.048737725 |
| Gm23640 | 0 | NA | NA |
| Cdc6 | 14.16964528 | 1.46433123 | 0.294559565 |
| Rara | 285.8551299 | 1.205515364 | 0.162231553 |
| Gm22061 | 0.570559668 | 1.053195405 | 1.349032633 |
| Gjd3 | 5.273691302 | 0.859719204 | 0.607911122 |
| Top2a | 129.6343859 | 2.916128221 | 0.392373796 |
| Gm26394 | 0 | NA | NA |
| Gm23451 | 0 | NA | NA |
| Igfbp4 | 440.1552601 | 0.675533156 | 0.122650337 |
| Tns4 | 6.644394917 | 0.501551322 | 0.400386597 |
| Ccr7 | 12.21422624 | 4.698354825 | 0.664067274 |
| Smarce1 | 477.9356527 | 0.017977158 | 0.071578275 |
| Krt222 | 403.0542806 | -0.189093735 | 0.069989005 |

| | | | |
|---|---|---|---|
| Krt24 | 0.665888905 | -0.955098813 | 1.588490846 |
| Gm12358 | 0 | NA | NA |
| Krt25 | 4.739584563 | -0.261523224 | 0.473940574 |
| Krt26 | 0.63227042 | 1.56342062 | 1.373005409 |
| Krt27 | 0.305696448 | 0.718222015 | 2.027117056 |
| Krt28 | 1.347390254 | 0.267811201 | 0.960913953 |
| Krt10 | 43.27398083 | -0.747473504 | 0.406377237 |
| Gm11940 | 7.488360699 | 0.230268738 | 0.347721486 |
| Krt12 | 43.87784049 | -1.527462948 | 0.217022648 |
| A830036E02Rik | 268.3212654 | -2.127182283 | 0.359419416 |
| Krt20 | 0.572048112 | -0.912873938 | 1.440480756 |
| Krt23 | 0.336553411 | -0.277494266 | 2.019239084 |
| Krt39 | 0 | NA | NA |
| Krt40 | 0 | NA | NA |
| Krtap3-3 | 0 | NA | NA |
| Krtap3-2 | 0 | NA | NA |
| Gm11939 | 0 | NA | NA |
| Krtap3-1 | 0 | NA | NA |
| Krtap1-5 | 0 | NA | NA |
| Krtap1-4 | 0 | NA | NA |
| Krtap1-3 | 0 | NA | NA |
| Krtap9-3 | 0 | NA | NA |
| Gm11938 | 0 | NA | NA |
| Gm11937 | 0 | NA | NA |
| Krtap2-4 | 0 | NA | NA |
| Gm11562 | 0 | NA | NA |
| Krtap4-1 | 0 | NA | NA |
| Krtap4-2 | 0 | NA | NA |
| Krtap4-7 | 0 | NA | NA |
| Gm11555 | 0 | NA | NA |
| Gm11563 | 0.072025462 | 0.237312084 | 3.724945049 |
| Gm14182 | 0 | NA | NA |
| Krtap4-6 | 0 | NA | NA |
| Gm11557 | 0.202171817 | 1.626940071 | 2.587140394 |
| 4930415H17Rik | 0 | NA | NA |
| Gm14190 | 0 | NA | NA |
| Gm14186 | 0 | NA | NA |
| Gm14193 | 0 | NA | NA |
| Gm14180 | 0 | NA | NA |
| Gm14188 | 0 | NA | NA |
| Gm11595 | 0 | NA | NA |
| Krtap4-8 | 0.046345709 | 0.71820487 | 3.724945049 |
| Krtap4-9 | 0 | NA | NA |
| Gm11596 | 0 | NA | NA |
| Gm11569 | 0 | NA | NA |
| Gm11554 | 0 | NA | NA |
| Krtap4-13 | 0 | NA | NA |

| | | | |
|---|---|---|---|
| Gm11564 | 0 | NA | NA |
| 2300003K06Rik | 0 | NA | NA |
| Gm11556 | 0 | NA | NA |
| Krtap4-16 | 0.075043205 | 0.237309859 | 3.724945049 |
| Gm26324 | 0 | NA | NA |
| Gm11568 | 0 | NA | NA |
| Gm11559 | 0 | NA | NA |
| Krtap9-1 | 0 | NA | NA |
| Gm11567 | 0 | NA | NA |
| Gm11566 | 0 | NA | NA |
| Krtap31-1 | 0.038981395 | 0.718207094 | 3.724945049 |
| Gm11565 | 0 | NA | NA |
| Gm11560 | 17.37014342 | 0.9044288 | 0.356056645 |
| Krtap31-2 | 0 | NA | NA |
| Krtap9-5 | 0 | NA | NA |
| Gm11561 | 0 | NA | NA |
| Gm11553 | 0 | NA | NA |
| Krtap29-1 | 0.072025462 | 0.237312084 | 3.724945049 |
| Krtap16-1 | 0.141781211 | 1.200006512 | 2.911624965 |
| Gm11558 | 0 | NA | NA |
| Krtap17-1 | 0.608261712 | 1.681512611 | 1.33562522 |
| Krt33a | 0 | NA | NA |
| Krt33b | 0 | NA | NA |
| Krt34 | 0 | NA | NA |
| Krt31 | 0 | NA | NA |
| Gm11571 | 0 | NA | NA |
| Krt32 | 0 | NA | NA |
| Krt35 | 0 | NA | NA |
| Krt36 | 0.15956642 | 0.719166641 | 3.418731035 |
| Gm12347 | 0 | NA | NA |
| Krt13 | 0.652975239 | -0.744757502 | 1.71903486 |
| Krt15 | 1.82600251 | -0.802975415 | 0.734525513 |
| Krt19 | 0.608478225 | 0.327522294 | 1.465532462 |
| Krt9 | 26.56616351 | 0.374162572 | 0.290862086 |
| Krt14 | 1.447260382 | 2.307660699 | 1.41445342 |
| Gm11597 | 0 | NA | NA |
| Gm11598 | 0.706088228 | 0.132293287 | 1.269049532 |
| Krt16 | 2.108251394 | 4.086427073 | 1.230746899 |
| Krt17 | 3.233276867 | 2.393627028 | 0.848771474 |
| Krt42 | 0 | NA | NA |
| Gm14206 | 0.67116138 | 1.885078251 | 1.310105695 |
| Gm12349 | 7.416241275 | 6.158348635 | 1.220992166 |
| Eif1 | 1981.137352 | 0.420856403 | 0.095800647 |
| Gast | 0.589408234 | 0.171261672 | 1.21044605 |
| Hap1 | 3690.110612 | 1.120103759 | 0.142416922 |
| Gm20394 | 9.564810652 | 1.455042445 | 0.526231822 |
| Gm10039 | 471.8830643 | -0.12305185 | 0.089127012 |

| | | | |
|---|---|---|---|
| Jup | 876.4788789 | 0.478033986 | 0.098322913 |
| Gm12348 | 0.360464465 | -1.166623093 | 1.365601883 |
| P3h4 | 107.8090644 | 0.20274589 | 0.115078639 |
| Fkbp10 | 103.2641343 | 1.165086825 | 0.16098605 |
| Nt5c3b | 220.2515258 | -0.252239417 | 0.086584873 |
| Klhl10 | 2.707144304 | -0.067120081 | 0.592465772 |
| Gm27374 | 1.132676578 | -1.015502365 | 1.003562138 |
| Klhl11 | 985.9766124 | -0.160543282 | 0.062748778 |
| Acly | 2346.506125 | 0.006022346 | 0.084131214 |
| mmu-mir-7116 | 0.285551684 | -0.693767079 | 2.022601887 |
| Ttc25 | 14.3848452 | 0.177372031 | 0.252206898 |
| Cnp | 4776.985164 | 0.237522572 | 0.115829578 |
| Dnajc7 | 1151.346391 | 0.096900135 | 0.07172014 |
| Nkiras2 | 157.3130789 | 1.058636808 | 0.106715186 |
| Zfp385c | 87.71940538 | -0.052331302 | 0.227918524 |
| Dhx58os | 2.117912561 | 0.081644407 | 0.686618025 |
| Dhx58 | 526.9953047 | 6.16376209 | 0.25098865 |
| Kat2a | 785.4970274 | 0.28891739 | 0.057654944 |
| Hspb9 | 0.468943176 | 2.972163835 | 3.221976663 |
| Rab5c | 1897.413452 | 1.196429718 | 0.128174546 |
| Kcnh4 | 95.02347477 | -1.532220036 | 0.264849302 |
| Hcrt | 38.8486744 | -0.340067915 | 0.204784896 |
| Ghdc | 106.0321 | 1.102414975 | 0.176614123 |
| Gm24358 | 0.048949532 | -0.243569582 | 3.724945049 |
| Stat5b | 602.2188295 | 0.409380492 | 0.105674015 |
| Stat5a | 70.6584695 | 1.816137133 | 0.188437606 |
| Stat3 | 2401.024981 | 2.460183989 | 0.165816233 |
| Ptrf | 400.0738531 | 2.133837662 | 0.180128894 |
| Atp6v0a1 | 5069.409655 | -0.311746574 | 0.063172654 |
| Mir6928 | 2.18332649 | -1.383050974 | 0.791046407 |
| Naglu | 145.2410652 | 1.217013351 | 0.152014983 |
| Hsd17b1 | 3.534966268 | -1.137527631 | 0.602553191 |
| Coasy | 224.3589791 | 0.404156933 | 0.09260389 |
| Mlx | 272.5349141 | 0.490493873 | 0.103364132 |
| Psmc3ip | 15.59208128 | 0.13581832 | 0.282171156 |
| Fam134c | 741.7829735 | 0.257826378 | 0.066098675 |
| Tubg1 | 286.1670221 | -0.101852397 | 0.086525442 |
| Gm27626 | 0 | NA | NA |
| Tubg2 | 274.8310085 | -0.514191735 | 0.085916333 |
| Plekhh3 | 197.5858469 | 0.04254429 | 0.116194216 |
| Cntnap1 | 2524.742535 | -0.960772142 | 0.085109831 |
| Ccr10 | 4.101636231 | 2.17152777 | 0.640151626 |
| Ezh1 | 709.9648608 | -0.154501949 | 0.079788001 |
| Ramp2 | 134.6263258 | 0.684361461 | 0.154386048 |
| Vps25 | 18.38762957 | -0.125896646 | 0.275622109 |
| Wnk4 | 77.9757232 | -0.026386493 | 0.17177964 |
| Gm11615 | 0.280085221 | -0.243565364 | 2.091622978 |

| | | | |
|---|---|---|---|
| Coa3 | 343.5032495 | 0.046325122 | 0.141407094 |
| Cntd1 | 4.658726077 | -0.226243027 | 0.467265725 |
| Becn1 | 989.2074398 | 0.242039754 | 0.068742715 |
| Gm11617 | 0 | NA | NA |
| Psme3 | 1184.471621 | 0.240498102 | 0.065840985 |
| Aoc2 | 39.91492154 | 0.11034402 | 0.295984841 |
| Aoc3 | 5.919817619 | 0.528929078 | 0.377459919 |
| Gm28156 | 0 | NA | NA |
| Gm11618 | 0 | NA | NA |
| Gm11619 | 0 | NA | NA |
| Gm25593 | 0 | NA | NA |
| G6pc | 0.591033593 | -0.527140528 | 1.327431546 |
| Gm25475 | 0 | NA | NA |
| Gm27029 | 8.009665686 | -0.897929153 | 0.440804851 |
| Aarsd1 | 19.81807774 | -0.523026618 | 0.269610307 |
| Ptges3l | 9.728067973 | 0.361292301 | 0.332855771 |
| Mir6929 | 0.072025462 | 0.237312084 | 3.724945049 |
| Rundc1 | 403.5552599 | -0.035482368 | 0.082601536 |
| Rpl27 | 526.9415286 | 0.792400116 | 0.127990485 |
| Ifi35 | 471.7219268 | 6.029762677 | 0.25170241 |
| Gm11626 | 1.005912905 | 0.571097045 | 0.945974823 |
| Vat1 | 770.8644348 | 0.698315815 | 0.162637654 |
| Rnd2 | 698.8704723 | -0.554889357 | 0.14951606 |
| Gm11625 | 0.085327104 | 1.179809211 | 3.723500796 |
| Brca1 | 51.12254382 | 1.033018929 | 0.264382439 |
| Nbr1 | 2218.562145 | 0.090014727 | 0.038590483 |
| Gm27456 | 0 | NA | NA |
| Tmem106a | 300.3975511 | 5.271125338 | 0.327562909 |
| Gm11634 | 0 | NA | NA |
| Gm26316 | 17.08146439 | -0.216774535 | 0.334634389 |
| Gm11635 | 0.049553047 | 0.71820487 | 3.724945049 |
| Rdm1 | 27.87786844 | -0.184659998 | 0.220340765 |
| Gm23849 | 1.056059865 | 0.942141652 | 1.105252375 |
| Gm24950 | 3.757983618 | 0.21309972 | 0.565620244 |
| Gm23971 | 333.7774647 | 0.110711037 | 0.194068311 |
| Rnu2-10 | 3.530281436 | 0.639565875 | 0.596790597 |
| Arl4d | 211.6319964 | 0.604642562 | 0.335735758 |
| Dhx8 | 571.733905 | 0.295491704 | 0.067125491 |
| Etv4 | 83.85799785 | -0.370587669 | 0.142285908 |
| Meox1 | 5.667363994 | 1.099709009 | 0.447446888 |
| Gm11551 | 0.198035107 | -0.243574128 | 3.222442586 |
| Rpl27-ps2 | 0.195392278 | 0.237309854 | 3.242394139 |
| 4930417O22Rik | 0.123841226 | 1.292225235 | 3.719439735 |
| Sost | 2.176193535 | -0.411633555 | 0.64699649 |
| Gm20659 | 0.323889686 | -0.243584415 | 3.724945049 |
| Dusp3 | 1716.840804 | -0.452978471 | 0.081071135 |
| 1700006E09Rik | 0.21733455 | -0.243564037 | 2.170157949 |

| | | | |
|---|---|---|---|
| Mpp3 | 656.0571513 | -0.69468342 | 0.11200333 |
| E130111B04Rik | 2.471955472 | -0.305803212 | 0.648079443 |
| Cd300lg | 0.955408868 | 3.457821958 | 1.096374334 |
| Mpp2 | 1306.923634 | -1.037896032 | 0.101313523 |
| Ppy | 0.061920613 | 0.718207094 | 3.724945049 |
| Pyy | 0.755118542 | 0.624907473 | 1.457651512 |
| Gm11584 | 0.066050875 | 0.71820487 | 3.724945049 |
| Gm11585 | 0.714893038 | 0.176174082 | 1.214054219 |
| Nags | 0.527376817 | 0.452563898 | 1.192714385 |
| Gm11586 | 0 | NA | NA |
| Tmem101 | 154.1546923 | 0.037668597 | 0.108315254 |
| Lsm12 | 620.6014926 | 0.444574299 | 0.090966597 |
| G6pc3 | 331.7313334 | 0.094960508 | 0.100341253 |
| Hdac5 | 1648.304379 | -0.508614881 | 0.10405042 |
| Mir8101 | 0 | NA | NA |
| BC030867 | 13.51292698 | 0.795563244 | 0.366871936 |
| Asb16 | 47.65270219 | -0.727172746 | 0.179498185 |
| Tmub2 | 469.4932909 | -0.037409948 | 0.062918213 |
| Atxn7l3 | 1951.581005 | -0.360690794 | 0.053757056 |
| Ubtf | 1192.509439 | -0.009180688 | 0.069553245 |
| Slc4a1 | 10.15205109 | 0.946666059 | 0.651109888 |
| Bloodlinc | 0.118974955 | -0.243576254 | 3.724945049 |
| Rundc3a | 2248.259499 | -0.199381607 | 0.059502603 |
| Slc25a39 | 589.2097923 | 0.501196131 | 0.100837473 |
| Mir6930 | 0 | NA | NA |
| Grn | 3148.773445 | 3.515101627 | 0.160553996 |
| Fam171a2 | 531.5743851 | -0.615683177 | 0.090662866 |
| Itga2b | 26.87028233 | 1.278066547 | 0.229590096 |
| Gm23563 | 0.090427103 | 0.237307635 | 3.724945049 |
| Gpatch8 | 1536.270915 | -0.251340519 | 0.098353848 |
| Gm11628 | 8.708300683 | -1.44248968 | 0.496567896 |
| Mdk-ps1 | 0 | NA | NA |
| Gm25337 | 4.232730408 | -1.157785108 | 0.544497668 |
| Gm11627 | 10.86367426 | -3.83896529 | 0.573678533 |
| Fzd2 | 124.2975267 | -0.57785255 | 0.268284792 |
| Gm11636 | 0.35856548 | -1.403815463 | 1.758152834 |
| 2810433D01Rik | 2.836445391 | 0.480268838 | 0.537918237 |
| Psmb5-ps | 0 | NA | NA |
| Gm22920 | 0 | NA | NA |
| Gm1564 | 0.341943343 | 0.237330552 | 1.772096005 |
| Ccdc43 | 214.8599409 | 0.250136825 | 0.086389698 |
| Gm11637 | 5.095273339 | -0.005960909 | 0.478336199 |
| Gm16342 | 0.648681289 | -1.981793865 | 1.407351516 |
| Adam11 | 1411.899054 | -0.775004962 | 0.121599428 |
| Gjc1 | 65.81803581 | 0.673053452 | 0.179128139 |
| Gm22683 | 0 | NA | NA |
| Higd1b | 5.836578858 | -0.255782665 | 0.420387632 |

| | | | |
|---|---|---|---|
| Eftud2 | 907.904142 | 0.07768496 | 0.07208716 |
| Gm26668 | 0.400913792 | -0.419109944 | 1.736041663 |
| Ccdc103 | 10.14206194 | 0.050461718 | 0.317741664 |
| Fam187a | 0.4546328 | -1.234488303 | 1.382875985 |
| Gfap | 17257.76253 | 3.234293443 | 0.171368907 |
| Kif18b | 11.6304259 | 2.867485049 | 0.561707657 |
| C1ql1 | 74.68787927 | -0.178739717 | 0.188736828 |
| 2410004I01Rik | 2.828652251 | 0.052387228 | 0.612035835 |
| Dcakd | 326.4044415 | 0.45596197 | 0.102948771 |
| Mir6931 | 0.170867779 | -1.299761979 | 3.70728864 |
| Nmt1 | 1910.096591 | 0.241406989 | 0.046414208 |
| Plcd3 | 130.7093725 | -0.240178951 | 0.101325075 |
| Acbd4 | 248.1541703 | -0.025035957 | 0.097100098 |
| Hexim1 | 470.7862717 | -0.251857707 | 0.103979675 |
| Hexim2 | 107.5740266 | -0.001251375 | 0.149843452 |
| Fmnl1 | 978.2301766 | 0.149641368 | 0.110018431 |
| Gm20511 | 5.853403199 | -0.775526636 | 0.466661428 |
| 1700023F06Rik | 21.3606864 | -0.393077936 | 0.261227186 |
| Spata32 | 11.37911894 | -1.042022513 | 0.406429319 |
| Map3k14 | 131.8227025 | 1.166413537 | 0.18803385 |
| 1700028N14Rik | 3.893030208 | -0.297753688 | 0.522798885 |
| Arhgap27 | 221.5069098 | 0.874739958 | 0.100985755 |
| Arhgap27os3 | 8.082714316 | 2.809641295 | 0.540055419 |
| Arhgap27os2 | 8.179740595 | 0.077073435 | 0.37617677 |
| Arhgap27os1 | 2.613037336 | -0.01843917 | 0.602872594 |
| Plekhm1 | 513.8582558 | 0.753506867 | 0.086223455 |
| Lrrc37a | 0.598753221 | 3.255625801 | 1.332262239 |
| Gm884 | 1.391014422 | 3.762447738 | 1.093239948 |
| Gm11643 | 0.038981395 | 0.718207094 | 3.724945049 |
| Lyzl6 | 0.124596252 | 0.718200615 | 3.714681068 |
| Rprml | 232.2274179 | 0.300859991 | 0.199665707 |
| Gosr2 | 931.0627658 | 0.722113505 | 0.094907375 |
| C130046K22Rik | 12.71673352 | -0.050924949 | 0.271947538 |
| Gm11642 | 9.924921471 | -0.007672672 | 0.341200784 |
| Wnt9b | 31.73347887 | 1.085965361 | 0.2889108 |
| Wnt3 | 86.98556824 | -0.060021646 | 0.217480367 |
| Nsf | 6788.932109 | -0.535556432 | 0.061051826 |
| Gm8738 | 9.128726587 | -0.940540429 | 0.521432515 |
| Arf2 | 510.9708085 | 0.563370983 | 0.106770788 |
| Crhr1 | 208.0710271 | -0.92452555 | 0.111933422 |
| Sppl2c | 0.098502579 | 0.23732098 | 3.724945049 |
| Mapt | 4636.496193 | -0.744764552 | 0.067874683 |
| Gm11659 | 0.104366086 | -0.243576254 | 3.724945049 |
| Gm22743 | 0 | NA | NA |
| AL596383.1 | 0 | NA | NA |
| Kansl1 | 1511.632219 | -0.096563926 | 0.036222756 |
| AL593843.1 | 0.534702693 | 1.919962515 | 1.539977586 |

| | | | |
|---|---|---|---|
| Gm11660 | 0 | NA | NA |
| Cdc27 | 1303.554971 | 0.122008919 | 0.053560125 |
| Myl4 | 65.04662385 | -1.410260783 | 0.223067668 |
| Gm11665 | 0 | NA | NA |
| Gm22000 | 0.267335191 | 1.154036039 | 1.792158276 |
| Gm11661 | 0 | NA | NA |
| Gm11663 | 0.542039732 | 1.257221434 | 1.498651191 |
| Itgb3 | 53.54022327 | 0.497063912 | 0.186038018 |
| Gm11662 | 0.044191549 | -0.243571806 | 3.724945049 |
| Gm11639 | 8.031358769 | -0.343243874 | 0.385393315 |
| Gm11623 | 0.110595523 | 0.718200421 | 3.724945049 |
| Gm11621 | 0.116046057 | -0.724453471 | 3.724945049 |
| Gm11622 | 0.090074004 | -0.24357403 | 3.724945049 |
| Gm11624 | 0.048949532 | -0.243569582 | 3.724945049 |
| Efcab3 | 4.279745736 | -0.258978676 | 0.613186851 |
| Gm25362 | 0 | NA | NA |
| Mettl2 | 133.1323383 | 0.151876506 | 0.126819262 |
| Gm24290 | 0.451210867 | -0.34085325 | 1.449859377 |
| Gm10842 | 0.310932149 | 0.708215857 | 1.7785663 |
| Gm11620 | 9.032629779 | -0.771202012 | 0.334586098 |
| 1700052K11Rik | 167.7122578 | -0.119434449 | 0.095081691 |
| Tlk2 | 1033.874952 | 0.739708773 | 0.101770618 |
| Mrc2 | 195.2873509 | 0.37708401 | 0.152512498 |
| 10-Mar | 11.52093314 | -0.666310473 | 0.340104717 |
| Gm11638 | 3.337525259 | -1.062745351 | 0.704955569 |
| Gm11640 | 0.456339188 | 0.461929289 | 1.321563134 |
| Tanc2 | 3818.814185 | -0.620588509 | 0.105374109 |
| Gm23990 | 3.257021581 | -0.321380422 | 0.586183505 |
| Gm11652 | 25.69936743 | -0.638186619 | 0.285255712 |
| Gm11653 | 35.85399802 | -1.111516261 | 0.293704991 |
| Cyb561 | 385.0963259 | -0.256236266 | 0.068497516 |
| Gm9910 | 2.885346812 | 0.524560314 | 0.585692053 |
| Gm11651 | 0.18135768 | -1.179311526 | 2.732659101 |
| Ace | 389.3701917 | -0.120637762 | 0.111519347 |
| Ace3 | 0.089089296 | 0.237305797 | 3.724945049 |
| Kcnh6 | 35.00512634 | -0.670357161 | 0.185401793 |
| Dcaf7 | 2563.358122 | -0.372072518 | 0.062008618 |
| Gm11646 | 0.25779132 | 1.373353627 | 2.014630796 |
| Taco1os | 4.692172118 | 1.089121585 | 0.465716783 |
| Taco1 | 111.1047001 | 0.142576037 | 0.16121621 |
| Map3k3 | 378.7212756 | 0.224455369 | 0.119214448 |
| Gm11672 | 0.164893722 | 0.23730812 | 3.395459203 |
| Limd2 | 471.2960561 | 0.006517988 | 0.109114973 |
| Gm10840 | 0.825556236 | -0.891701491 | 1.066159689 |
| Strada | 263.1762033 | -0.427614172 | 0.124040245 |
| Gm25556 | 0 | NA | NA |
| Ccdc47 | 1290.145382 | 0.088074248 | 0.066717967 |

| | | | |
|---|---|---|---|
| Ddx42 | 1058.979334 | -0.04278411 | 0.085062521 |
| Ftsj3 | 311.2842433 | 0.708574228 | 0.130174853 |
| Psmc5 | 1264.146288 | 0.4523814 | 0.097822694 |
| Gm23645 | 2.33541923 | -1.148997539 | 0.752314281 |
| Smarcd2 | 283.2968771 | 0.413258342 | 0.115494471 |
| Tcam1 | 4.654232259 | 1.12495515 | 0.499651086 |
| Gh | 110.6277303 | 2.120070627 | 2.421273295 |
| Cd79b | 1.812060818 | 3.622815503 | 0.990222059 |
| Scn4a | 1.34423325 | 3.183232687 | 1.096638908 |
| 2310007L24Rik | 2.639496584 | 0.908745059 | 0.61218452 |
| Icam2 | 17.72490718 | 2.509764474 | 0.322642979 |
| Ern1 | 301.964099 | 0.665011583 | 0.104119588 |
| Snord104 | 16.93328137 | 0.522988706 | 0.268771166 |
| SNORA50 | 0.197850883 | -0.229353099 | 2.220624018 |
| Gm22711 | 0 | NA | NA |
| Tex2 | 1392.018177 | -0.198732925 | 0.050569126 |
| Pecam1 | 139.8937637 | 2.430929296 | 0.153777216 |
| Milr1 | 37.7867777 | 6.524433801 | 0.646891577 |
| Gm25889 | 0.53783135 | 0.69707864 | 1.404419534 |
| Polg2 | 47.1941341 | -0.22141072 | 0.253877148 |
| Ddx5 | 9164.25171 | -0.02867466 | 0.077307976 |
| Mir3064 | 0 | NA | NA |
| Gm25994 | 0 | NA | NA |
| Cep95 | 165.3431975 | -0.264366617 | 0.156560924 |
| Gm11706 | 1.894273405 | 0.030421787 | 0.88798877 |
| Smurf2 | 621.2522263 | 0.266016372 | 0.06079918 |
| Gm27322 | 0.075043205 | 0.237309859 | 3.724945049 |
| Gm11707 | 0.436105163 | 2.213688554 | 1.657043419 |
| Kpna2 | 179.0519059 | 0.359520436 | 0.128759983 |
| Gm11708 | 0 | NA | NA |
| Gm11705 | 2.194360983 | -0.15878558 | 0.720102267 |
| 1810010H24Rik | 17.13476079 | -0.90087637 | 0.273675396 |
| Bptf | 2737.946275 | -0.199834352 | 0.063218824 |
| Gm11715 | 0.139856492 | 0.237322262 | 3.574411025 |
| Nol11 | 402.0846801 | 0.453648583 | 0.089175244 |
| Rps12-ps26 | 3.470784895 | -0.311166522 | 0.642267641 |
| Gm11718 | 0.038981395 | 0.718207094 | 3.724945049 |
| Pitpnc1 | 980.2405448 | -0.663322124 | 0.065457053 |
| Gm11719 | 0 | NA | NA |
| Gm11716 | 0.710007532 | 2.023497323 | 1.185196867 |
| Gm11712 | 0.072025462 | 0.237312084 | 3.724945049 |
| Gm11713 | 0.068097359 | -0.243569582 | 3.724945049 |
| Gm11714 | 1.466365607 | 3.987639418 | 1.045753257 |
| Psmd12 | 782.060693 | 0.551542824 | 0.096227238 |
| Gm11722 | 4.228535416 | -0.215414466 | 0.523174371 |
| C330019F10Rik | 0.3156755 | -0.64125647 | 1.760121728 |
| A830035A12Rik | 0.580962004 | 0.213119369 | 1.288899144 |

| | | | |
|---|---|---|---|
| Gm11649 | 0 | NA | NA |
| Helz | 1652.750466 | -0.509395883 | 0.089856083 |
| Mir6932 | 0.570554915 | 0.626912455 | 1.570706362 |
| Cacng1 | 1.900742604 | -1.05773855 | 0.853009648 |
| Cacng4 | 438.0588544 | -0.647629957 | 0.108681484 |
| Gm11650 | 3.119570997 | -1.630730213 | 0.612902163 |
| Gm23120 | 0 | NA | NA |
| Gm27595 | 0 | NA | NA |
| Cacng5 | 366.3860086 | 0.385759997 | 0.154550809 |
| Prkca | 1907.502606 | -0.618916702 | 0.094540425 |
| Gm11657 | 0 | NA | NA |
| Mir7223 | 0 | NA | NA |
| Gm11655 | 0.048949532 | -0.243569582 | 3.724945049 |
| Apoh | 2.789102959 | 1.369531049 | 0.741573769 |
| Gm11656 | 0.072025462 | 0.237312084 | 3.724945049 |
| Gm11667 | 0 | NA | NA |
| Cep112 | 85.63833006 | -0.691885884 | 0.178116176 |
| Cep112os1 | 0 | NA | NA |
| Olfr1397-ps1 | 0.047948698 | -0.243569582 | 3.724945049 |
| Rpl23a-ps14 | 5.868082076 | -0.608269713 | 0.417076318 |
| Cep112os2 | 13.20886531 | -0.558384414 | 0.462978001 |
| Axin2 | 366.2212925 | -0.605343737 | 0.109579652 |
| Gm11670 | 1.328031477 | 1.944610737 | 1.022430641 |
| Gm11668 | 0.544841187 | 0.214959477 | 1.314229168 |
| Gm24149 | 0.044544648 | 0.237309859 | 3.724945049 |
| E030025P04Rik | 0.237045706 | -0.605220011 | 2.124314614 |
| Gm11658 | 0.044544648 | 0.237309859 | 3.724945049 |
| Rgs9 | 522.9399079 | -1.085475509 | 0.245924407 |
| 1700096J18Rik | 0 | NA | NA |
| Gm11696 | 30.54970298 | 0.481594921 | 0.204622987 |
| Gna13 | 2195.672482 | 1.618603499 | 0.131560477 |
| 9930022D16Rik | 1.783663422 | 2.573163188 | 1.022202113 |
| Amz2 | 356.9352975 | -0.143821355 | 0.117754496 |
| Gm15642 | 0 | NA | NA |
| Slc16a6 | 246.9242258 | 0.649095352 | 0.171588548 |
| Gm25540 | 3.251877087 | -0.431738756 | 0.699094902 |
| Arsg | 200.3840775 | -0.661507288 | 0.130902644 |
| Gm22378 | 0 | NA | NA |
| Wipi1 | 268.8479832 | 0.209405948 | 0.088353781 |
| Gm11685 | 0.075043205 | 0.237309859 | 3.724945049 |
| Prkar1a | 7255.166353 | -0.010255807 | 0.058051535 |
| Fam20a | 97.31167799 | -0.499702648 | 0.13503735 |
| Gm11684 | 0 | NA | NA |
| 1700012B07Rik | 0.10503227 | 1.179809211 | 3.723500796 |
| Gm11683 | 0 | NA | NA |
| 1700023C21Rik | 0 | NA | NA |
| Abca8b | 635.0819168 | -0.262220256 | 0.110250782 |

| | | | |
|---|---|---|---|
| Gm11697 | 0 | NA | NA |
| Abca8a | 98.13962934 | -1.842759892 | 0.175730936 |
| Abca9 | 172.9692926 | -0.960353835 | 0.219255235 |
| Abca6 | 37.25579249 | -0.612620617 | 0.220634304 |
| Abca5 | 1374.997336 | -0.457996778 | 0.057992428 |
| Map2k6 | 151.8243697 | -1.350651166 | 0.1420937 |
| Gm11682 | 0.63216095 | -0.740660522 | 1.182866077 |
| Kcnj16 | 546.3597821 | -0.068189176 | 0.156878363 |
| Kcnj2 | 326.6825132 | -0.494385775 | 0.10150601 |
| Gm11675 | 0.837406104 | 2.245121911 | 1.207084209 |
| Gm11676 | 0 | NA | NA |
| Gm11677 | 0.110870145 | 0.238305784 | 3.724874707 |
| Gm11674 | 0.062426873 | 0.237312084 | 3.724945049 |
| Gm11679 | 0 | NA | NA |
| Gm11680 | 0 | NA | NA |
| BC006965 | 12.81790452 | -1.764126824 | 0.336495258 |
| Sox9 | 1126.317198 | 1.32754238 | 0.133493275 |
| Gm11681 | 4.401306684 | 0.37540242 | 0.502942405 |
| 4933434M16Rik | 0 | NA | NA |
| 2610035D17Rik | 77.49257512 | 0.235306711 | 0.140955233 |
| 4732490B19Rik | 0.42613393 | 2.82930742 | 1.535845159 |
| Slc39a11 | 151.2282737 | 0.46681484 | 0.101760968 |
| Gm16487 | 1.032826873 | -0.458502283 | 0.867175041 |
| Sstr2 | 160.0653886 | 0.373785419 | 0.323712788 |
| Cog1 | 598.811076 | 0.182556733 | 0.067122134 |
| Fam104a | 378.1237319 | 0.315325798 | 0.072188255 |
| Gm11693 | 0.456148335 | 1.195927841 | 1.627281522 |
| D11Wsu47e | 95.6758204 | 0.158454866 | 0.157350384 |
| Cpsf4l | 2.642413763 | 0.707771658 | 0.628321377 |
| Cdc42ep4 | 690.6837532 | 0.637921883 | 0.141622471 |
| Gm11736 | 0.197525616 | 0.701885133 | 2.225104381 |
| Sdk2 | 727.2965037 | -0.479857425 | 0.118900982 |
| 1700025D23Rik | 0.122991902 | -0.24357403 | 3.724945049 |
| 1700001J04Rik | 0.288935699 | 0.860355611 | 2.061810815 |
| 1700092K14Rik | 0.758667428 | -0.420858913 | 1.241099263 |
| Gm11690 | 0.042716945 | -0.243571806 | 3.724945049 |
| Gm11691 | 0.047948698 | -0.243569582 | 3.724945049 |
| Gm11692 | 0 | NA | NA |
| 4932435O22Rik | 2.584340838 | -0.871527017 | 0.695488326 |
| Gm11689 | 0 | NA | NA |
| Rpl38 | 617.2880742 | 0.371679635 | 0.198455301 |
| Ttyh2 | 758.4082506 | -0.244782262 | 0.096909956 |
| Dnaic2 | 33.61650857 | 0.80259039 | 0.225282274 |
| Kif19a | 46.14809591 | -0.311219775 | 0.1775388 |
| Btbd17 | 82.92747472 | -2.350130996 | 0.269792845 |
| Gpr142 | 0.138152447 | 0.237324354 | 2.934963576 |
| Gprc5c | 168.3894624 | -0.214866469 | 0.194228459 |

| | | | |
|---|---|---|---|
| Cd300a | 55.24117626 | 1.300934277 | 0.265837606 |
| Gm11698 | 0.946192865 | 0.479382002 | 1.091333719 |
| Cd300lb | 7.999125163 | 3.576168044 | 0.537010339 |
| Gm11699 | 0 | NA | NA |
| Cd300c | 0.985558492 | 2.855496995 | 2.211908396 |
| Cd300ld | 9.206469351 | 6.606492035 | 0.897107529 |
| AF251705 | 73.50869836 | 2.911025168 | 0.214900694 |
| Gm11709 | 0.932213635 | 3.275655186 | 1.350286968 |
| Gm11710 | 0.854414486 | 2.792604399 | 1.251370123 |
| Gm11711 | 0.991446059 | 3.50549994 | 1.265609447 |
| Cd300lh | 2.74015677 | 3.819846334 | 0.967954471 |
| Cd300e | 5.001744579 | 3.383343938 | 0.649556678 |
| Rab37 | 152.6386923 | -0.368636978 | 0.114670076 |
| Cd300lf | 291.2418054 | 7.320764372 | 0.423793519 |
| Slc9a3r1 | 582.5242563 | 0.912966547 | 0.140364721 |
| Nat9 | 95.23938015 | 0.214272493 | 0.13136249 |
| Tmem104 | 386.5952224 | 0.92704345 | 0.106964142 |
| Grin2c | 638.1417609 | -1.145595274 | 0.163100007 |
| Gm25837 | 0.089089296 | 0.237305797 | 3.724945049 |
| Fdxr | 101.2810521 | -0.390530977 | 0.115684353 |
| Fads6 | 408.9834072 | -0.544475009 | 0.101738362 |
| Otop2 | 12.03184821 | -0.506428368 | 0.490649287 |
| Ush1g | 18.68407142 | 0.365079402 | 0.32781921 |
| Otop3 | 2.248585396 | 2.037394173 | 0.818682651 |
| Hid1 | 846.0541876 | -0.362412379 | 0.078802688 |
| Cdr2l | 554.9718017 | 0.040251985 | 0.082643865 |
| Ict1 | 293.2357144 | 0.037069377 | 0.07312903 |
| Ict1os | 0.802129109 | -1.192331193 | 1.105184702 |
| Atp5h | 982.123738 | -0.113762654 | 0.088111955 |
| Kctd2 | 366.4040735 | -0.1161732 | 0.068930356 |
| Trim80 | 1.020577158 | 0.657849219 | 1.285374457 |
| Mir3968 | 0 | NA | NA |
| Gm11695 | 0.907486357 | -1.160341071 | 1.080341447 |
| Slc16a5 | 0.45393394 | 2.458922711 | 3.337878916 |
| Armc7 | 103.9306984 | 0.75704741 | 0.137999958 |
| Nt5c | 193.1442848 | -0.040067073 | 0.08380446 |
| Gm11694 | 1.069943461 | 1.838743473 | 0.990311501 |
| Hn1 | 386.0221868 | 0.263090173 | 0.111362955 |
| Sumo2 | 322.8270685 | 0.060381604 | 0.112404229 |
| Nup85 | 257.3817318 | -0.222761839 | 0.077227074 |
| Gga3 | 719.1270991 | -0.204057372 | 0.076765207 |
| Gm25364 | 0 | NA | NA |
| Mrps7 | 278.078163 | 0.570768328 | 0.102803391 |
| Mif4gd | 118.7681913 | 0.640962199 | 0.140398405 |
| Slc25a19 | 241.7616655 | -0.416265817 | 0.077663042 |
| Gm26613 | 0.042716945 | -0.243571806 | 3.724945049 |
| Grb2 | 1605.895681 | 0.22623523 | 0.059065637 |

| | | | |
|---|---|---|---|
| Gm11702 | 38.96983336 | -0.434306863 | 0.185489423 |
| Gm11703 | 12.20134384 | 0.936557526 | 0.371064271 |
| Gm23699 | 0 | NA | NA |
| Gm11704 | 0 | NA | NA |
| 2310067B10Rik | 722.2047825 | -0.277679855 | 0.096084301 |
| Mir5621 | 0.180265876 | -1.188459231 | 2.743873342 |
| Caskin2 | 336.692886 | 0.1267772 | 0.111183211 |
| Tsen54 | 70.37095567 | 0.201589991 | 0.125781666 |
| Llgl2 | 24.64085341 | 1.233669221 | 0.241866425 |
| Myo15b | 2.674434539 | 0.227973 | 0.828596019 |
| Recql5 | 210.8391631 | -0.043467161 | 0.153022259 |
| Smim5 | 3.739367707 | 1.405297594 | 0.567805606 |
| Smim6 | 0.320213338 | -1.041058958 | 1.71886891 |
| Recql5os1 | 0.062426873 | 0.237312084 | 3.724945049 |
| Sap30bp | 450.623263 | 0.279803429 | 0.076428511 |
| Sap30bpos | 16.7515477 | 2.2748501 | 0.336985638 |
| Itgb4 | 300.6627537 | 0.038597822 | 0.138457754 |
| B230344G16Rik | 10.95198732 | 0.249739603 | 0.322465775 |
| Galk1 | 66.2841132 | -0.279656509 | 0.171321896 |
| H3f3b | 3094.200091 | 1.445499372 | 0.171182653 |
| Unk | 347.7817187 | 0.259737302 | 0.063638898 |
| Unc13d | 61.4735399 | 2.356510173 | 0.252151927 |
| Wbp2 | 3196.52038 | -0.196155203 | 0.072610408 |
| Trim47 | 116.7135195 | 3.244242254 | 0.227294175 |
| Trim65 | 57.48636768 | -0.194763384 | 0.147428764 |
| Mrpl38 | 328.9494862 | 0.284300429 | 0.072145353 |
| Fbf1 | 655.435408 | -0.332056518 | 0.10057848 |
| Acox1 | 2125.616724 | 0.131804326 | 0.051720633 |
| Ten1 | 127.8018245 | 0.149852911 | 0.112521752 |
| Cdk3-ps | 6.063886428 | -0.413842545 | 0.431255678 |
| Evpl | 18.36159365 | -0.2174986 | 0.226178212 |
| Srp68 | 664.4255585 | 0.455055384 | 0.070127994 |
| Galr2 | 1.093534925 | -1.596239915 | 1.042016179 |
| Exoc7 | 686.1507893 | -0.080541429 | 0.049365832 |
| Gm26730 | 1.17710504 | 0.271493197 | 0.968572868 |
| n-R5s74 | 0 | NA | NA |
| Foxj1 | 234.7792888 | 1.443673935 | 0.163208348 |
| Rnf157 | 2678.61245 | -0.652268242 | 0.081636131 |
| Rps11-ps2 | 3.484535615 | -0.269932259 | 0.537396167 |
| Ubald2 | 148.6109574 | 0.850283088 | 0.134834346 |
| Qrich2 | 0.384783943 | 1.143968091 | 1.56644784 |
| Gm11739 | 0 | NA | NA |
| Prpsap1 | 399.2746756 | 0.095702397 | 0.072412786 |
| Rpl36-ps1 | 0 | NA | NA |
| Gm29292 | 10.23524976 | 3.407280803 | 0.597362902 |
| Sphk1 | 151.140251 | 2.579608603 | 0.363748386 |
| Ube2o | 1590.155959 | -0.369555246 | 0.074213455 |

| | | | |
|---|---|---|---|
| Gm11742 | 0 | NA | NA |
| Aanat | 3.437531617 | -2.10155073 | 0.647943701 |
| Rhbdf2 | 162.7293829 | 5.670202352 | 0.328511038 |
| Cygb | 534.7165052 | -0.644850818 | 0.086487747 |
| Gm11744 | 17.7695872 | -0.974635016 | 0.302559288 |
| 1810032O08Rik | 43.33906545 | 0.559527505 | 0.185305697 |
| Snord1c | 0 | NA | NA |
| Snord1b | 0 | NA | NA |
| Snord1a | 0 | NA | NA |
| St6galnac2 | 21.17771716 | -0.636303761 | 0.238030347 |
| Gm11735 | 0.061920613 | 0.718207094 | 3.724945049 |
| BC018473 | 0 | NA | NA |
| St6galnac1 | 0.21290388 | 1.264634319 | 2.620255742 |
| Mxra7 | 93.40309728 | -0.276580418 | 0.130030966 |
| Jmjd6 | 264.2573602 | 0.185382726 | 0.119409253 |
| Gm27401 | 0 | NA | NA |
| Mettl23 | 60.97083361 | -0.045658353 | 0.153086584 |
| Srsf2 | 1239.606384 | 0.281104186 | 0.072044141 |
| Mfsd11 | 294.976682 | 0.296804412 | 0.078443421 |
| Mgat5b | 569.5109985 | -0.949291638 | 0.115517053 |
| Gm11728 | 0.813341144 | -0.370411156 | 1.110669581 |
| Gm11731 | 0.068097359 | -0.243569582 | 3.724945049 |
| Snhg20 | 102.2173952 | -0.468028103 | 0.152647589 |
| Gm26083 | 0 | NA | NA |
| mmu-mir-6516 | 0 | NA | NA |
| Gm24323 | 0.154433973 | -0.245012343 | 3.500771551 |
| Gm11730 | 1.74719428 | -0.021902579 | 0.68809497 |
| Sec14l1 | 2162.074218 | -0.712821564 | 0.100411173 |
| Gm22667 | 7.017353583 | -0.724182944 | 0.525463488 |
| Gm11745 | 7.216095472 | -1.370433207 | 0.611259492 |
| 9-Sep | 1015.394361 | 1.213578644 | 0.11447421 |
| Gm16045 | 1.18260799 | -1.874016847 | 0.998236724 |
| Gm11729 | 0.295494355 | -0.59400977 | 1.767329605 |
| Gm8624 | 0.24265998 | -0.242738854 | 2.50889915 |
| Gm11732 | 4.722481653 | -0.694674053 | 0.449634481 |
| Gm11733 | 2.291093954 | 1.371736862 | 0.706059033 |
| Gm11734 | 3.524723029 | 0.412332944 | 0.524943262 |
| 2900041M22Rik | 4.279435866 | -0.439581645 | 0.518218582 |
| Tnrc6c | 1645.798208 | -0.625067459 | 0.119728385 |
| Tmc6 | 62.04352302 | 1.202358202 | 0.191629834 |
| Tmc8 | 13.88221168 | 2.525725538 | 0.364457401 |
| Gm11723 | 0.109124069 | -0.24357403 | 3.724945049 |
| Gm11724 | 0.044544648 | 0.237309859 | 3.724945049 |
| 6030468B19Rik | 0.072025462 | 0.237312084 | 3.724945049 |
| Gm20708 | 0.075043205 | 0.237309859 | 3.724945049 |
| Syngr2 | 214.0619962 | 2.348095031 | 0.152685577 |
| Tk1 | 11.63018472 | 2.154141926 | 0.425445692 |

| | | | |
|---|---|---|---|
| Afmid | 13.88472388 | 0.293822732 | 0.317201578 |
| Birc5 | 15.65536197 | 0.984904179 | 0.322739767 |
| Tmem235 | 0.809771811 | -0.044217542 | 1.05319008 |
| Tha1 | 35.97253463 | -0.136518331 | 0.197984311 |
| Gm11725 | 4.41486742 | -0.436530861 | 0.526301883 |
| Gm11726 | 0.061920613 | 0.718207094 | 3.724945049 |
| Socs3 | 775.4504325 | 5.578946633 | 0.414361146 |
| Pgs1 | 896.2403674 | 0.927556897 | 0.089227199 |
| Mir6933 | 0.186399948 | 0.718204068 | 3.299750385 |
| Dnah17 | 7.417164814 | 0.500085712 | 0.355028655 |
| Gm11738 | 6.391237767 | 0.230763755 | 0.58260176 |
| Cyth1 | 1001.744073 | 0.179942084 | 0.059793502 |
| Gm24060 | 17.80235189 | -0.858602569 | 0.289589504 |
| Gm11737 | 8.112751975 | 0.373396683 | 0.48493546 |
| Usp36 | 634.1888172 | 0.2806665 | 0.151981817 |
| Timp2 | 2901.777889 | -0.139032752 | 0.065823972 |
| BC100451 | 15.86145379 | -0.537234019 | 0.338293793 |
| Lgals3bp | 4154.63823 | 5.878890906 | 0.175178632 |
| Cant1 | 623.271791 | 0.442297525 | 0.060396534 |
| C1qtnf1 | 107.3234082 | 1.527360265 | 0.198877501 |
| Gm11747 | 0.291849543 | 0.71779276 | 1.778128226 |
| Gm11748 | 1.545561197 | 0.258800994 | 0.768905779 |
| Engase | 86.50548769 | -0.062996049 | 0.143934055 |
| Rbfox3 | 2325.205722 | -0.799646448 | 0.104455716 |
| Gm11750 | 0.497147205 | 2.836468682 | 1.569869669 |
| Gm25040 | 1.127514739 | -3.006359099 | 1.072815248 |
| Gm11749 | 17.63452542 | -1.137834855 | 0.349098285 |
| Enpp7 | 0.038981395 | 0.718207094 | 3.724945049 |
| Cbx2 | 46.53041809 | -0.941405526 | 0.248190326 |
| Cbx8 | 139.4173976 | -0.448704484 | 0.115820946 |
| Gm26508 | 4.027668366 | 0.711510245 | 0.565667965 |
| Cbx4 | 968.7534903 | 0.196919497 | 0.102832431 |
| Gm11754 | 0.08486385 | 0.717551239 | 3.724898127 |
| Gm11755 | 2.180156761 | -1.781420011 | 0.794715484 |
| Tbc1d16 | 764.1409362 | -0.94148563 | 0.121046013 |
| Gm26888 | 0.416129427 | -0.543861437 | 1.538102891 |
| Mir6934 | 0 | NA | NA |
| Gm11753 | 1.914943955 | -0.386638579 | 0.723441913 |
| Gm11752 | 0.044191549 | -0.243571806 | 3.724945049 |
| Ccdc40 | 32.62461688 | 0.897331353 | 0.219385619 |
| Gaa | 3712.581302 | -0.048862747 | 0.046103386 |
| Eif4a3 | 482.0859025 | 0.454619738 | 0.085753615 |
| Card14 | 10.20988058 | 0.794604884 | 0.364789448 |
| Sgsh | 132.7659391 | 0.154616986 | 0.131320164 |
| Slc26a11 | 53.44579997 | -0.093596038 | 0.158211764 |
| Mir1932 | 5.939787235 | 4.91655228 | 0.913451047 |
| Rnf213 | 13237.76603 | 6.066400634 | 0.164148807 |

| | | | |
|---|---|---|---|
| Endov | 287.30136 | -0.440962422 | 0.127384937 |
| Nptx1 | 4410.742837 | -0.183328149 | 0.106313768 |
| Gm11762 | 16.8669638 | 0.128711381 | 0.272778561 |
| Rptor | 1323.078836 | -0.021660206 | 0.040013272 |
| Gm11761 | 0 | NA | NA |
| Gm23663 | 1.256631142 | 0.894738781 | 0.924573865 |
| Rptoros | 0.455959994 | 1.724955068 | 1.535607409 |
| Chmp6 | 176.3066057 | 0.236686227 | 0.10868234 |
| Gm11767 | 18.81677128 | -1.224429955 | 0.311029339 |
| Gm11766 | 0.898050108 | 1.629818375 | 1.277498796 |
| Baiap2 | 2183.791868 | -0.352226652 | 0.134482287 |
| Aatk | 3564.404415 | -0.412348051 | 0.04617803 |
| Mir338 | 4.047384828 | -0.869451377 | 0.590758701 |
| mmu-mir-3065 | 0.136882968 | 0.663945046 | 3.630339258 |
| Cep131 | 180.3487011 | -0.792088304 | 0.17543049 |
| Gm25948 | 0.585001534 | -0.857093811 | 1.401792383 |
| Enthd2 | 135.2998032 | -0.054783196 | 0.121588958 |
| 1810043H04Rik | 75.99610051 | 0.17942595 | 0.189530946 |
| Slc38a10 | 947.089944 | 0.324611355 | 0.055080087 |
| Gm11769 | 0.670691275 | 0.53806534 | 1.13505878 |
| 2810410L24Rik | 42.09187015 | -1.004924572 | 0.202611107 |
| Gm11770 | 0.567693313 | 0.678236158 | 1.364009791 |
| 2900052L18Rik | 38.54445378 | -1.022986049 | 0.281016736 |
| Bahcc1 | 551.7191568 | -0.365301986 | 0.096638263 |
| Gm11772 | 31.98353754 | 2.459788599 | 0.379823975 |
| AL928567.1 | 0 | NA | NA |
| Actg1 | 4258.481376 | 0.616251102 | 0.116540322 |
| Gm23419 | 0 | NA | NA |
| Mir6935 | 0 | NA | NA |
| 0610009L18Rik | 11.33403993 | -0.601316246 | 0.313436269 |
| Fscn2 | 1.822354792 | -1.452256566 | 0.953657765 |
| 2310003H01Rik | 147.9010322 | -0.199065687 | 0.109188645 |
| Nploc4 | 1106.700852 | 0.72255933 | 0.076497343 |
| Tspan10 | 1.198353109 | 1.516509056 | 1.042491947 |
| Pde6g | 3.767627917 | 0.018376858 | 0.49654541 |
| Oxld1 | 37.49257177 | -0.156948292 | 0.231882888 |
| Ccdc137 | 228.8428678 | 0.759023429 | 0.12556082 |
| Arl16 | 155.9283109 | -0.378356685 | 0.107044233 |
| Hgs | 966.1986603 | 0.446303432 | 0.061779725 |
| Mrpl12 | 411.1473201 | 0.3125863 | 0.121858156 |
| Gm16755 | 0.616632713 | -0.055044717 | 1.308220632 |
| Gm11788 | 0.154696864 | 0.237325224 | 2.833481316 |
| Slc25a10 | 142.1098057 | -0.317827937 | 0.121293559 |
| Gm11789 | 0.038981395 | 0.718207094 | 3.724945049 |
| Gcgr | 1.708960799 | 0.604612835 | 0.76438479 |
| Fam195b | 400.1399544 | 0.291685685 | 0.099727438 |
| Ppp1r27 | 0.150086409 | 0.237305146 | 3.724945049 |

| | | | |
|---|---|---|---|
| P4hb | 1498.417556 | 1.007517 | 0.10350473 |
| Arhgdia | 2780.717337 | 0.817344953 | 0.095287705 |
| Gm3807 | 3.342078322 | 0.016338064 | 0.54365731 |
| Alyref | 326.2916262 | 0.3392604 | 0.142359143 |
| Anapc11 | 300.5463482 | -0.026178249 | 0.110177648 |
| Npb | 4.91649906 | 0.528243375 | 0.486720552 |
| Pcyt2 | 527.1846474 | -0.171368243 | 0.075663288 |
| Sirt7 | 232.8371362 | 0.205776768 | 0.112203526 |
| Mir6936 | 0 | NA | NA |
| Mafg | 947.7400382 | -0.054757038 | 0.086692498 |
| AL663030.1 | 0 | NA | NA |
| Gm17586 | 36.02489538 | 0.120422525 | 0.204412583 |
| Pycr1 | 25.50189758 | 1.156702398 | 0.236522595 |
| Myadml2 | 26.50837914 | -0.432733364 | 0.216089498 |
| Myadml2os | 1.355090691 | -2.088104571 | 1.160793741 |
| Notum | 9.73076196 | -0.513121407 | 0.342770735 |
| Notumos | 2.072861717 | -0.661542317 | 0.712797477 |
| Aspscr1 | 368.7567786 | 0.299804167 | 0.08013969 |
| Gm17178 | 0.221007548 | -1.0869266 | 2.162457626 |
| Stra13 | 57.79830129 | -0.024726838 | 0.174224135 |
| Lrrc45 | 209.0769499 | -0.673456401 | 0.206370447 |
| Rac3 | 88.08912298 | -0.617047924 | 0.137387175 |
| Dcxr | 57.27746725 | 0.342987071 | 0.195281199 |
| Cbr2 | 7.287723491 | -0.83824326 | 0.634985521 |
| Hmga1-rs1 | 524.4408412 | 2.521090927 | 0.261532718 |
| Gm11771 | 0 | NA | NA |
| Rfng | 389.6471229 | -0.248683914 | 0.068768673 |
| Gps1 | 549.026855 | 0.104542973 | 0.085036041 |
| Dus1l | 255.7077501 | 0.230256475 | 0.089102681 |
| Fasn | 4709.556278 | -0.200248285 | 0.0855401 |
| Ccdc57 | 96.87917321 | -0.535068686 | 0.213931644 |
| Gm11773 | 0.216076386 | 0.237307547 | 3.724945049 |
| Slc16a3 | 72.70316503 | 2.177387941 | 0.237238793 |
| Csnk1d | 1494.74923 | 0.229703525 | 0.056960872 |
| Gm28192 | 0 | NA | NA |
| Gm11774 | 0.066050875 | 0.71820487 | 3.724945049 |
| Gm11775 | 0.090277459 | 0.237318756 | 3.724945049 |
| Gm22152 | 0.044191549 | -0.243571806 | 3.724945049 |
| Cd7 | 13.98101131 | 1.844638918 | 0.357520488 |
| Sectm1b | 0.607868506 | 3.287677713 | 1.373104822 |
| Sectm1a | 17.65934462 | 8.0464864 | 0.90594711 |
| Gm11792 | 0 | NA | NA |
| Gm11791 | 0 | NA | NA |
| Gm11793 | 1.517674278 | 1.275306609 | 0.825487804 |
| Tex19.2 | 0 | NA | NA |
| Tex19.1 | 0.038981395 | 0.718207094 | 3.724945049 |
| Uts2r | 2.195192171 | 1.40025468 | 0.755322971 |

| | | | |
|---|---|---|---|
| Gm11790 | 0.802756328 | -0.24356641 | 1.424921788 |
| Ogfod3 | 69.70424457 | 0.290638178 | 0.15833563 |
| Hexdc | 139.714264 | -0.493950848 | 0.110697928 |
| BC017643 | 297.1303026 | 1.253747165 | 0.125832483 |
| Narf | 577.2846721 | -0.830566567 | 0.094511754 |
| Foxk2 | 1306.171416 | -0.061486325 | 0.048218866 |
| Wdr45b | 650.5471699 | 0.402592093 | 0.076849003 |
| Rab40b | 176.4499084 | -0.728931303 | 0.159551781 |
| Mir7236 | 2.550224353 | -0.660887729 | 0.716710512 |
| Gm11866 | 0 | NA | NA |
| Fn3krp | 434.4560457 | -0.362118314 | 0.103561334 |
| Fn3k | 227.4287037 | -0.940308475 | 0.130901776 |
| Tbcd | 616.9440393 | 0.041375322 | 0.062474808 |
| Zfp750 | 17.50032145 | 1.299350386 | 0.31183609 |
| B3gntl1 | 72.02965567 | 0.138413056 | 0.178869447 |
| Metrnl | 107.2840231 | 1.83891145 | 0.206798783 |
| Gm12589 | 0.398050276 | 2.359985058 | 1.597654132 |
| Gm29513 | 0 | NA | NA |
| Gm15643 | 0 | NA | NA |
| Ptchd3 | 0 | NA | NA |
| Gm12586 | 0 | NA | NA |
| Gm12590 | 0 | NA | NA |
| Gm12587 | 0 | NA | NA |
| Gm12591 | 0 | NA | NA |
| Gm9071 | 0 | NA | NA |
| Gm9075 | 0 | NA | NA |
| Rab10os | 387.6915989 | -0.366398934 | 0.099150666 |
| Rab10 | 2113.636369 | 0.409510017 | 0.07835453 |
| Gm26520 | 3.045321195 | -0.703863918 | 0.655324049 |
| Kif3c | 2815.319838 | -0.434512319 | 0.059942916 |
| 1110002L01Rik | 127.452233 | 0.316165861 | 0.11270961 |
| Asxl2 | 1098.710745 | -0.001063024 | 0.076132969 |
| Dtnb | 785.8508435 | -0.608617192 | 0.0860056 |
| Gm26050 | 2.458093412 | -1.092045675 | 0.766870792 |
| Dtnbos | 0.849323088 | -1.222974718 | 1.0704749 |
| Dnmt3a | 1911.391238 | -0.424433083 | 0.091931385 |
| Dnmt3aos | 1.674671121 | 3.874656041 | 1.019066458 |
| Pomc | 33.59521228 | 1.375165394 | 0.910775852 |
| Efr3b | 2749.842645 | -0.077024412 | 0.054653488 |
| Dnajc27 | 1474.757923 | -0.56854201 | 0.064805564 |
| Adcy3 | 597.0799168 | -0.42968464 | 0.079895529 |
| Cenpo | 63.78631368 | 0.258416202 | 0.133952479 |
| Ptrhd1 | 115.3757405 | -0.231035358 | 0.112780502 |
| Ncoa1 | 2595.887549 | -0.061613004 | 0.053143625 |
| Gm23756 | 0 | NA | NA |
| Gm22343 | 0 | NA | NA |
| Gm3625 | 2.933921127 | -0.015511399 | 0.594384715 |

| | | | |
|---|---|---|---|
| Itsn2 | 887.6567088 | 0.023700356 | 0.064368887 |
| Gm17541 | 0.106465261 | 0.718202646 | 3.724945049 |
| Fam228a | 28.71218686 | -1.727049508 | 0.343396622 |
| Fam228b | 74.82727959 | -1.205131167 | 0.188876339 |
| Gm23676 | 0 | NA | NA |
| Gm22003 | 0 | NA | NA |
| Pfn4 | 43.32535029 | -1.143458348 | 0.178032405 |
| Gm6682 | 21.88487653 | 0.482176973 | 0.216806315 |
| Sf3b6 | 380.7093426 | 0.578293875 | 0.146334435 |
| Fkbp1b | 121.655647 | -0.718458175 | 0.125054077 |
| BC068281 | 110.397684 | -0.207822472 | 0.1315178 |
| Mfsd2b | 23.20891546 | 0.044314595 | 0.413765444 |
| Ubxn2a | 760.0083217 | -0.244537551 | 0.078422972 |
| Atad2b | 541.029005 | -0.295410211 | 0.092858326 |
| Gm23167 | 0 | NA | NA |
| Klhl29 | 743.3499042 | -0.395785605 | 0.12060545 |
| 2810032G03Rik | 93.60943587 | -0.744544452 | 0.111886946 |
| Gm5652 | 0.048949532 | -0.243569582 | 3.724945049 |
| Gm23752 | 0.431854921 | -0.243564043 | 1.59886167 |
| 1700101O22Rik | 0.245087052 | 0.237329926 | 2.095767575 |
| Gm9144 | 9.249347624 | 0.826515959 | 0.364081231 |
| Apob | 11.59973813 | 0.558820512 | 0.492885045 |
| Ldah | 329.5450486 | 0.055445121 | 0.110186311 |
| Gdf7 | 3.829800073 | -0.759472827 | 0.604958812 |
| Hs1bp3 | 177.2730511 | 0.981560803 | 0.1127278 |
| Rhob | 3017.567704 | 0.511905678 | 0.115132752 |
| Slc7a15 | 0.061920613 | 0.718207094 | 3.724945049 |
| Pum2 | 3606.55936 | 0.158583956 | 0.066327547 |
| Gm4755 | 0.187019432 | -0.24356566 | 2.718962445 |
| Sdc1 | 95.2175769 | 2.460349831 | 0.232941921 |
| 9930038B18Rik | 0.162099466 | -0.2435696 | 2.789369104 |
| Laptm4a | 2359.716549 | 1.292508917 | 0.119315589 |
| Matn3 | 0.908179219 | -0.628745515 | 0.988133737 |
| Wdr35 | 500.8882983 | -0.319828737 | 0.07037624 |
| Ttc32 | 25.69466073 | 0.181490598 | 0.21737389 |
| Gm7099 | 8.593693136 | 0.916943127 | 0.438630974 |
| 1700022H16Rik | 0 | NA | NA |
| Osr1 | 21.77471978 | -0.304477204 | 0.282057942 |
| Gm23979 | 0 | NA | NA |
| Gm22845 | 0 | NA | NA |
| Nt5c1b | 1.964636517 | 0.423032084 | 0.92100679 |
| Rdh14 | 214.0479375 | -0.004985767 | 0.106199766 |
| Pgk1-rs7 | 21.9025282 | 0.272269274 | 0.235303151 |
| Kcns3 | 56.27639958 | -0.171310103 | 0.172338108 |
| Msgn1 | 0 | NA | NA |
| 1700034J04Rik | 0.087261593 | -0.243576254 | 3.724945049 |
| Gen1 | 22.65896892 | 0.518488563 | 0.28617011 |

| | | | |
|---|---|---|---|
| Smc6 | 783.5087835 | 0.158524823 | 0.065023244 |
| Mir7675 | 1.906635116 | -0.520892984 | 0.710689797 |
| Vsnl1 | 3374.486255 | -0.828167436 | 0.08418109 |
| 4930511A02Rik | 1.096741448 | 0.685945703 | 0.937797377 |
| Rad51ap2 | 31.8005175 | -0.965549794 | 0.270683228 |
| Gm23781 | 0 | NA | NA |
| Gm25610 | 0.211102819 | 0.745356413 | 2.622609005 |
| Gm4294 | 93.43490713 | 0.971000271 | 0.183920684 |
| Gm26171 | 0 | NA | NA |
| Fam49a | 2242.155381 | -0.368165585 | 0.073349279 |
| Gm37414 | 0.066050875 | 0.71820487 | 3.724945049 |
| Gm27952 | 0 | NA | NA |
| Gm25538 | 0 | NA | NA |
| Platr19 | 0 | NA | NA |
| 4930519A11Rik | 0.060174537 | 0.237309859 | 3.724945049 |
| Rpl36-ps3 | 0.314164845 | -0.269387251 | 1.628204234 |
| Mycn | 47.66468018 | -1.424795826 | 0.193393805 |
| Gm24208 | 0.042716945 | -0.243571806 | 3.724945049 |
| Ddx1 | 1202.104352 | 0.095678212 | 0.058785907 |
| Nbas | 1083.615517 | 0.128274488 | 0.051680902 |
| Gm6736 | 3.145663366 | 0.7650569 | 0.57218499 |
| Gm24142 | 0 | NA | NA |
| Gm22170 | 0.191089262 | -0.724458717 | 2.650365316 |
| Gm4929 | 0.355586071 | -0.72218471 | 1.937004552 |
| Fam84a | 632.1102194 | -0.94641098 | 0.117532819 |
| Gm9847 | 0.673365293 | 0.686248403 | 1.134227816 |
| Gm25414 | 0.044191549 | -0.243571806 | 3.724945049 |
| Gm26449 | 0 | NA | NA |
| Gm27682 | 0.049553047 | 0.71820487 | 3.724945049 |
| Gm4804 | 2.483041552 | -1.289836257 | 0.688692723 |
| Trib2 | 460.7235091 | -0.880080402 | 0.148980328 |
| Mir6387 | 0.891236725 | -1.515647135 | 1.129146629 |
| Gm5432 | 2.197748528 | 0.154133538 | 0.632115212 |
| Lpin1 | 478.9431543 | -0.494268575 | 0.109444932 |
| Ntsr2 | 425.5275216 | -1.688462921 | 0.167497449 |
| Greb1 | 74.66542959 | 0.204651237 | 0.147611699 |
| AC122228.1 | 0 | NA | NA |
| E2f6 | 314.0510662 | 0.00785926 | 0.091245503 |
| Rock2 | 3956.78452 | -0.058372782 | 0.07679161 |
| Gm22303 | 1.492607829 | -0.930221477 | 0.795711245 |
| Pqlc3 | 54.75605864 | 2.017679306 | 0.22043752 |
| 2410004P03Rik | 23.25089268 | 0.982532025 | 0.280541053 |
| Kcnf1 | 695.0270978 | -0.934635702 | 0.158691444 |
| Pdia6 | 1198.63775 | 0.957712739 | 0.118577369 |
| Atp6v1c2 | 10.10846665 | -1.983441231 | 0.394581132 |
| Nol10 | 367.5541281 | 0.441483454 | 0.126640055 |
| Mir3066 | 1.524462984 | 2.072940877 | 0.989886782 |

| | | | |
|---|---|---|---|
| Odc1 | 613.484201 | 1.639366097 | 0.169335886 |
| Gm22748 | 7.258283802 | 0.958196133 | 0.436307326 |
| Gm25925 | 0 | NA | NA |
| Hpcal1 | 649.1744098 | -0.154238225 | 0.142518486 |
| B430203G13Rik | 10.4805672 | -0.94541987 | 0.362283827 |
| Gm10480 | 62.44942269 | -0.120498813 | 0.142364147 |
| 5730507C01Rik | 3.246809683 | 0.139545266 | 0.662037853 |
| Gm3944 | 0.500379391 | 0.915885337 | 1.446262159 |
| Gm5784 | 1.293338167 | -0.715267113 | 1.037901814 |
| Gm6802 | 0 | NA | NA |
| Gm9257 | 0.060174537 | 0.237309859 | 3.724945049 |
| Gm22189 | 0 | NA | NA |
| Gm21863 | 0.047948698 | -0.243569582 | 3.724945049 |
| Gm3993 | 0.060174537 | 0.237309859 | 3.724945049 |
| Gm10479 | 0.270569244 | 1.183809591 | 2.073291076 |
| 1700030C10Rik | 18.72189217 | -0.549886125 | 0.267862265 |
| Gm28503 | 0 | NA | NA |
| 3110053B16Rik | 90.02746546 | -0.248732079 | 0.16097604 |
| Gm10478 | 5.792289935 | -0.417420069 | 0.459989371 |
| Zfp125 | 38.28518153 | 0.045992608 | 0.187129375 |
| Gm21989 | 3.535470757 | -1.105757196 | 0.585560779 |
| Gm5977 | 1.165717276 | -3.225963923 | 1.033759751 |
| Asap2 | 769.9183214 | 0.214137031 | 0.086792108 |
| Itgb1bp1 | 249.9318875 | -0.10119987 | 0.092526325 |
| Gm20535 | 1.863766393 | 1.22463696 | 0.959956502 |
| Cpsf3 | 408.1523475 | 0.159822139 | 0.068542117 |
| Gm25821 | 2.089596347 | 0.016143229 | 0.671460457 |
| Iah1 | 73.05919511 | -0.258554256 | 0.151546151 |
| Adam17 | 298.4801547 | 0.871961631 | 0.117577019 |
| Gm22766 | 0 | NA | NA |
| Ywhaq | 1407.12046 | -0.261411312 | 0.075570691 |
| Gm4419 | 4.781372743 | -0.240024452 | 0.483909732 |
| Gm5633 | 5.363150736 | -0.514077102 | 0.44361697 |
| Gm22908 | 0 | NA | NA |
| Gm4425 | 1.228836612 | -0.570389406 | 0.908231567 |
| Gm20486 | 0 | NA | NA |
| Gm6838 | 0.045882455 | 0.237312084 | 3.724945049 |
| Gm10476 | 0.988361678 | -0.63863521 | 1.158627713 |
| Gm17391 | 0.141071297 | 0.237322348 | 3.564043836 |
| 2410018L13Rik | 0.412847196 | 0.581226494 | 1.567599118 |
| Gm26140 | 0 | NA | NA |
| Gm20472 | 0.391014225 | 1.019861856 | 1.433137677 |
| Gm20473 | 0 | NA | NA |
| Gm9285 | 0.258012988 | -0.724461208 | 2.135707359 |
| Gm9286 | 0 | NA | NA |
| Gm20474 | 0.284703419 | -0.94940968 | 1.981522332 |
| Gm10330 | 0.04393175 | -0.243571806 | 3.724945049 |

| | | | |
|---|---|---|---|
| Gm17330 | 0 | NA | NA |
| 9030624G23Rik | 0.15672128 | 0.237326597 | 2.821957917 |
| Gm9312 | 0.046345709 | 0.71820487 | 3.724945049 |
| Gm16372 | 0.480740522 | 0.865231342 | 1.527413031 |
| Taf1b | 209.6766852 | 0.009449006 | 0.085370039 |
| Grhl1 | 110.9007615 | -0.166518002 | 0.172368901 |
| Gm6969 | 32.27026886 | -0.099774941 | 0.299401789 |
| Klf11 | 102.5281956 | 0.483428837 | 0.155911278 |
| Cys1 | 49.24944908 | -1.455972223 | 0.242460101 |
| Rrm2 | 38.30661462 | 2.205805138 | 0.413041371 |
| Gm26211 | 0 | NA | NA |
| Mboat2 | 425.7860528 | -0.628393357 | 0.104367011 |
| Kidins220 | 4110.912429 | -0.424109858 | 0.057122728 |
| Id2 | 1035.883774 | 0.80579221 | 0.148934983 |
| Gm38052 | 0 | NA | NA |
| Gm22527 | 0 | NA | NA |
| Rnf144a | 371.6551165 | -0.454865868 | 0.089725344 |
| Mir6538 | 1.774752459 | -2.200072186 | 0.926352001 |
| Rsad2 | 3584.881863 | 10.18828794 | 0.454638154 |
| Cmpk2 | 2770.91634 | 4.404933926 | 0.266874622 |
| Gm9866 | 90.23738068 | -0.550601769 | 0.14239626 |
| Sox11 | 612.1246314 | 0.467437456 | 0.126183289 |
| Gm24326 | 0 | NA | NA |
| Gm25923 | 0 | NA | NA |
| Gm25564 | 0 | NA | NA |
| Dcdc2c | 0.829077587 | -0.897145994 | 1.351616489 |
| 1700022F17Rik | 0.238210844 | 0.776310309 | 3.037334902 |
| Allc | 2.669291208 | -2.469798285 | 0.792987282 |
| Colec11 | 3.087887419 | 0.940755277 | 0.580466825 |
| Rps7 | 1086.152334 | 0.666802071 | 0.124847003 |
| Rnaseh1 | 69.48037499 | 0.01737985 | 0.159097651 |
| Adi1 | 240.4881119 | -0.13385852 | 0.110495445 |
| Mir6937 | 0 | NA | NA |
| Trappc12 | 622.1510625 | -0.42262326 | 0.08035856 |
| Tssc1 | 236.185517 | -0.123690287 | 0.081181814 |
| Gm23342 | 0 | NA | NA |
| Myt1l | 2705.158939 | -0.816561771 | 0.126997346 |
| Pxdn | 585.0750425 | 0.19408367 | 0.18418547 |
| Tpo | 6.12441587 | -0.571268749 | 0.405059103 |
| Gm15691 | 3.03177422 | -0.006206882 | 0.845942771 |
| Sntg2 | 50.41524398 | -0.428190872 | 0.200095326 |
| 2310016D03Rik | 0.251964224 | 0.740676851 | 2.081917693 |
| Gm25865 | 0 | NA | NA |
| Gm28806 | 0 | NA | NA |
| Gm25243 | 0.038981395 | 0.718207094 | 3.724945049 |
| Tmem18 | 291.1926203 | 0.211476086 | 0.089832139 |
| Fam150b | 13.83564331 | 1.703956799 | 0.315641275 |

| | | | |
|---|---|---|---|
| Acp1 | 179.9360824 | 0.306724938 | 0.103853203 |
| Sh3yl1 | 69.38233294 | -0.31352292 | 0.138834108 |
| Gm25408 | 0 | NA | NA |
| Fam110c | 3.004281714 | 2.362161046 | 0.676758852 |
| 6030469F06Rik | 0.048949532 | -0.243569582 | 3.724945049 |
| Gm24613 | 0.553521388 | -1.476348182 | 1.253843216 |
| Lamb1 | 296.7992853 | -0.233002442 | 0.113073284 |
| Dld | 1183.816382 | 0.140481592 | 0.084444615 |
| Slc26a3 | 2.221929775 | -0.5091518 | 0.774410219 |
| Gm9359 | 0 | NA | NA |
| Cbll1 | 375.0362158 | 0.231891778 | 0.073198755 |
| Gm29542 | 0 | NA | NA |
| Gm29541 | 0 | NA | NA |
| Slc26a4 | 32.11330933 | 0.202208824 | 0.231440629 |
| Bcap29 | 377.0266182 | -0.075619936 | 0.088875004 |
| Dus4l | 47.02044193 | 0.591409605 | 0.174423011 |
| Cog5 | 397.0078528 | -0.223823471 | 0.07684928 |
| Gpr22 | 1006.691692 | 0.147013264 | 0.101544909 |
| Gm25913 | 1.857927479 | -1.140123038 | 0.739485665 |
| Hbp1 | 434.1401043 | 0.092283396 | 0.104136868 |
| Prkar2b | 606.5567347 | -0.88224411 | 0.098547025 |
| Pik3cg | 109.1690308 | 1.584554847 | 0.131251284 |
| Gm23875 | 0 | NA | NA |
| Ccdc71l | 284.7411437 | 0.717508778 | 0.122468965 |
| Nampt | 2161.206624 | 2.122901655 | 0.123729022 |
| Gdap10 | 195.362713 | 1.389645335 | 0.195624434 |
| 4933406C10Rik | 19.31754882 | -0.829401355 | 0.312906767 |
| Sypl | 610.6402324 | 0.281790355 | 0.086981356 |
| Gm24922 | 0 | NA | NA |
| Cdhr3 | 9.236620993 | -0.488348125 | 0.318796461 |
| Gm22904 | 0.044544648 | 0.237309859 | 3.724945049 |
| F730043M19Rik | 202.5005712 | -0.480423985 | 0.138618286 |
| Gm16267 | 0 | NA | NA |
| Atxn7l1 | 715.5692364 | 0.194533447 | 0.087925945 |
| Atxn7l1os1 | 0.109124069 | -0.24357403 | 3.724945049 |
| Atxn7l1os2 | 0 | NA | NA |
| Efcab10 | 8.606127557 | -0.500776967 | 0.365136671 |
| Twistnb | 221.7759397 | 0.489743885 | 0.095342006 |
| Gm23529 | 0 | NA | NA |
| Ferd3l | 0.671239363 | -0.145697074 | 1.250719215 |
| Twist1 | 6.585054892 | 0.767502608 | 0.383905915 |
| Gm18025 | 5.060744911 | -0.117827546 | 0.459190287 |
| Hdac9 | 424.2109994 | -0.986780974 | 0.129356283 |
| 4921508M14Rik | 2.520821878 | -0.916932823 | 0.649245011 |
| Gm24429 | 0 | NA | NA |
| Prps1l1 | 0 | NA | NA |
| AC133459.1 | 0 | NA | NA |

| | | | |
|---|---|---|---|
| Gm24359 | 0.075043205 | 0.237309859 | 3.724945049 |
| Snx13 | 1558.690179 | -0.286868101 | 0.051524005 |
| Mir680-3 | 0 | NA | NA |
| AC132434.1 | 0 | NA | NA |
| Gm26071 | 0 | NA | NA |
| Ahr | 138.7406453 | 0.357348648 | 0.099553434 |
| Agr3 | 0.095295241 | 0.23732098 | 3.724945049 |
| Agr2 | 0.23039375 | 0.23165277 | 2.131410783 |
| Tspan13 | 1131.407557 | -0.264253369 | 0.069685421 |
| Gm5434 | 2.449668862 | 0.870159253 | 0.617721932 |
| Bzw2 | 293.0080855 | -0.32771826 | 0.094612616 |
| Ankmy2 | 458.9611624 | -0.103665835 | 0.064956366 |
| Lrrc72 | 0.323815284 | 0.751908189 | 1.734718826 |
| Sostdc1 | 41.87300568 | -2.373835276 | 0.272195321 |
| Gm25438 | 0.046345709 | 0.71820487 | 3.724945049 |
| D630036H23Rik | 7.277711592 | -0.446186354 | 0.395858366 |
| Ispd | 147.0734498 | 0.319990569 | 0.097199456 |
| Mir5099 | 0 | NA | NA |
| CT010456.1 | 0 | NA | NA |
| Gm29007 | 0 | NA | NA |
| Meox2 | 0.449709139 | -1.928433422 | 1.453760351 |
| Agmo | 60.54347211 | -0.71522596 | 0.188967183 |
| Dgkb | 1454.497695 | -1.36614753 | 0.150110182 |
| Etv1 | 1246.89087 | -0.83722553 | 0.144784406 |
| Gm24069 | 0 | NA | NA |
| Gm24434 | 0 | NA | NA |
| Arl4a | 499.8111158 | 0.291846598 | 0.100006092 |
| Arl4aos | 1.421581723 | -1.601597145 | 1.02616378 |
| Scin | 6.368096109 | 1.274356467 | 0.427266875 |
| Lsmem1 | 17.83132388 | 0.821934827 | 0.287492802 |
| Gm17056 | 0.249945515 | 0.237931501 | 1.847817612 |
| Ifrd1 | 942.5158603 | 1.535624072 | 0.143991475 |
| Gm17024 | 29.97646992 | -0.029676799 | 0.317159079 |
| Mir1938 | 6.231293264 | 0.924980885 | 0.517976593 |
| Gm7008 | 11.68594954 | -0.030682885 | 0.30212537 |
| Zfp277 | 160.924537 | 0.00522483 | 0.132055372 |
| Dock4 | 2536.949238 | -0.483366969 | 0.101282404 |
| Immp2l | 31.50920012 | -0.342977034 | 0.188634372 |
| Gm25497 | 0.147068667 | 0.237310878 | 3.514416608 |
| Lrrn3 | 669.4920803 | -0.369765076 | 0.073982015 |
| Gm25248 | 0 | NA | NA |
| Gm26326 | 0 | NA | NA |
| Dnajb9 | 682.3485262 | 0.66387777 | 0.109560777 |
| Mir5627 | 0.044191549 | -0.243571806 | 3.724945049 |
| Pnpla8 | 1553.715183 | -0.226217887 | 0.065062096 |
| Nrcam | 5819.560715 | -0.020035904 | 0.079088016 |
| Stxbp6 | 733.0217604 | -0.756065092 | 0.106280383 |

| | | | |
|---|---|---|---|
| Gm15901 | 0.132101749 | 1.209894437 | 3.72418822 |
| Gm15902 | 13.4453144 | -1.083677644 | 0.316115708 |
| Gm24731 | 0.429113465 | 1.085838267 | 1.420114679 |
| Gm22076 | 0 | NA | NA |
| Gm9921 | 0.047948698 | -0.243569582 | 3.724945049 |
| Gm24948 | 0 | NA | NA |
| Nova1 | 1267.849016 | -0.383563665 | 0.072293579 |
| Gm25051 | 0 | NA | NA |
| Gm36971 | 0 | NA | NA |
| Gm1818 | 26.89079701 | -0.230553614 | 0.231853673 |
| Gm26454 | 0 | NA | NA |
| Gm7456 | 0 | NA | NA |
| Foxg1 | 626.865741 | -0.368188226 | 0.136510828 |
| 3110039M20Rik | 111.2821837 | -1.141080648 | 0.249271743 |
| RP24-490H13.4 | 48.60029554 | -0.781897325 | 0.242910872 |
| Gm9804 | 13.1091654 | -0.693806926 | 0.421230175 |
| 1810007C17Rik | 0.183032319 | 0.695603381 | 3.319201136 |
| Gm25015 | 0 | NA | NA |
| Gm7481 | 0.044191549 | -0.243571806 | 3.724945049 |
| Prkd1 | 131.3087976 | -0.135938422 | 0.108527949 |
| Gm22088 | 0 | NA | NA |
| Rps11-ps4 | 0.060174537 | 0.237309859 | 3.724945049 |
| G2e3 | 315.7949451 | -0.353217052 | 0.103408281 |
| Scfd1 | 573.8494792 | 0.718034632 | 0.097020599 |
| Coch | 246.9813543 | 0.112718279 | 0.155882119 |
| Strn3 | 1743.354882 | 0.02677473 | 0.054802867 |
| Ap4s1 | 296.6051223 | -0.228897839 | 0.079922695 |
| Gm22663 | 0.270436937 | -0.889034919 | 2.376095253 |
| Hectd1 | 2718.891165 | -0.030173426 | 0.040259233 |
| Heatr5a | 367.9070692 | 0.510595017 | 0.12888576 |
| Gm26517 | 0.276231724 | 1.623936569 | 2.061871023 |
| Dtd2 | 62.41787465 | 0.382089725 | 0.163959668 |
| Gpr33 | 0.154148777 | 1.255550436 | 2.843849521 |
| Nubpl | 61.33971456 | -0.289947536 | 0.149352675 |
| Arhgap5 | 3781.700427 | -0.346650798 | 0.104318028 |
| 1700031P21Rik | 0.115603922 | 1.209894437 | 3.72418822 |
| Gm24859 | 1.957101456 | -0.424870913 | 0.670548394 |
| Gm23296 | 1.92086539 | 0.617946324 | 0.786457362 |
| Akap6 | 5339.749417 | -0.454314432 | 0.063069815 |
| n-R5s58 | 0 | NA | NA |
| Npas3 | 426.0647591 | 0.009573182 | 0.08122694 |
| 1700060O08Rik | 0.07796279 | 1.072698098 | 3.721829099 |
| Egln3 | 112.0375452 | 0.08261065 | 0.120358763 |
| Gm22220 | 0 | NA | NA |
| Gm24692 | 0 | NA | NA |
| Sptssa | 171.7818652 | 0.127561522 | 0.141010737 |
| mmu-mir-1892 | 0 | NA | NA |

| | | | |
|---|---|---|---|
| Gm24971 | 0.26923737 | -0.154540376 | 2.112226549 |
| Eapp | 505.3675571 | 0.063048987 | 0.074224198 |
| Gm22513 | 8.835653036 | -0.104984779 | 0.394789567 |
| Rpl31-ps17 | 1.178692824 | 1.256127607 | 0.979636627 |
| Gm22634 | 17.48739027 | -1.192219978 | 0.522750746 |
| Gm23804 | 1.13792396 | 0.442375328 | 0.965649257 |
| Gm25679 | 43.52464816 | -0.159766716 | 0.251036179 |
| Gm26444 | 126.6439584 | 0.049432106 | 0.152689514 |
| Gm24305 | 91.6640994 | 0.64785591 | 0.204607823 |
| Gm22317 | 529.3228771 | -0.21986734 | 0.239898949 |
| Snx6 | 482.8085486 | 0.823461623 | 0.087665788 |
| Cfl2 | 953.9436151 | 0.432744521 | 0.095390991 |
| Baz1a | 191.7919684 | 2.30769822 | 0.182948148 |
| Gm27014 | 0 | NA | NA |
| Gm20403 | 0 | NA | NA |
| Gm24296 | 0 | NA | NA |
| 2700097O09Rik | 100.6248846 | -0.640617573 | 0.17028877 |
| Srp54a | 0.941593011 | -0.222198616 | 1.001423982 |
| Fam177a | 2.46611504 | -0.707528843 | 0.603293658 |
| Srp54b | 195.0106852 | 0.180738332 | 0.143174475 |
| 1700047I17Rik2 | 3.457287012 | -0.394380788 | 0.591972033 |
| Ppp2r3c | 188.8955074 | 0.357472231 | 0.112216374 |
| 1110008L16Rik | 73.54196958 | 0.288935138 | 0.123138093 |
| Psma6 | 718.3364319 | 0.722246055 | 0.129359034 |
| Rpl18a-ps1 | 0.963303439 | -0.037858565 | 0.923305996 |
| Nfkbia | 657.3079934 | 2.165006485 | 0.28526474 |
| Aldoart2 | 0.61825299 | 0.013614006 | 1.17111078 |
| Insm2 | 3.842056669 | 0.540650712 | 0.520435995 |
| Ralgapa1 | 2973.912485 | -0.413837944 | 0.065341591 |
| Gm9812 | 2.978437241 | -0.038648767 | 0.609134896 |
| Gm23394 | 0 | NA | NA |
| Brms1l | 538.8067484 | -0.254968395 | 0.080991913 |
| Gm22296 | 0 | NA | NA |
| Gm23799 | 0 | NA | NA |
| Gm25917 | 0.075043205 | 0.237309859 | 3.724945049 |
| Gm24685 | 0.291934259 | -0.24596943 | 1.784723596 |
| Mbip | 262.0904343 | -0.16129555 | 0.149328639 |
| Gm6265 | 9.407196128 | -0.182433679 | 0.567195863 |
| Nkx2-1 | 18.9524978 | -0.528427132 | 0.285849999 |
| Gm26973 | 0.859979859 | -1.51359756 | 1.381422431 |
| Gm25760 | 0 | NA | NA |
| C87198 | 0.421608343 | -1.680828493 | 1.599998564 |
| Gm38103 | 1.370820674 | -0.088180531 | 0.845943388 |
| Nkx2-9 | 4.229279226 | -1.130547008 | 0.487585206 |
| Gm15524 | 0 | NA | NA |
| Pax9 | 0.231631338 | 0.713119065 | 2.12291018 |
| Slc25a21 | 5.399330309 | -0.522068404 | 0.431890112 |

| | | | |
|---|---|---|---|
| Prps1l3 | 132.0803981 | 0.305581708 | 0.097596588 |
| Mipol1 | 126.8360654 | -0.140095438 | 0.112017211 |
| Gm11025 | 0 | NA | NA |
| Gm16246 | 0.044544648 | 0.237309859 | 3.724945049 |
| Gm2568 | 0 | NA | NA |
| Gm37291 | 0 | NA | NA |
| Foxa1 | 3.942515582 | -0.610103818 | 0.581075384 |
| 4921506M07Rik | 10.13732351 | -0.961483348 | 0.387185176 |
| 4921518K17Rik | 0.038981395 | 0.718207094 | 3.724945049 |
| Rpl26-ps2 | 0 | NA | NA |
| Sstr1 | 99.2471505 | -0.316367647 | 0.166791663 |
| Clec14a | 49.14502572 | 1.803435065 | 0.207592165 |
| Gm17529 | 0.072025462 | 0.237312084 | 3.724945049 |
| Sec23a | 1353.977473 | -0.2261416 | 0.052253423 |
| Gemin2 | 59.82869919 | 0.405243674 | 0.13904055 |
| Gm22973 | 1.521163403 | 2.499551461 | 1.12920598 |
| Trappc6b | 482.938189 | 0.007754261 | 0.07030352 |
| Pnn | 591.1787483 | -0.15448265 | 0.154643269 |
| Gm5786 | 32.33765274 | 0.221309103 | 0.279364935 |
| Rpl21-ps3 | 1.284833043 | 1.113881226 | 1.043584799 |
| Mia2 | 5.838823621 | -0.463220375 | 0.532966367 |
| Ctage5 | 709.7395958 | 0.125884506 | 0.076302012 |
| Gm24233 | 4.255553331 | -1.453383665 | 0.554173552 |
| Fbxo33 | 714.5346483 | 0.025818151 | 0.117041617 |
| Gm27284 | 0 | NA | NA |
| Gm7719 | 0.04393175 | -0.243571806 | 3.724945049 |
| Gm24377 | 0 | NA | NA |
| Gm22673 | 0 | NA | NA |
| Gm23059 | 0 | NA | NA |
| Gm38172 | 0 | NA | NA |
| Gm7745 | 2.081729379 | 0.496033958 | 0.778288806 |
| Nanog-ps2 | 0 | NA | NA |
| Lrfn5 | 1144.044751 | -0.676285844 | 0.079733277 |
| Gm23213 | 11.15017964 | -0.888629062 | 0.382941651 |
| Gm24750 | 0 | NA | NA |
| Gm24590 | 0 | NA | NA |
| Spanxn4 | 0 | NA | NA |
| Gm7763 | 0 | NA | NA |
| Gm5185 | 0 | NA | NA |
| Gm24921 | 0 | NA | NA |
| Fscb | 0.143022002 | 0.719176055 | 3.547735711 |
| Gm24066 | 0 | NA | NA |
| Gm6395 | 108.7166347 | 0.128539018 | 0.155547286 |
| Gm527 | 45.68336777 | -0.682219479 | 0.153283551 |
| Klhl28 | 276.6149016 | -0.04838618 | 0.084443503 |
| Fam179b | 1164.381407 | -0.092342693 | 0.073016654 |
| Prpf39 | 881.5198811 | -0.042662991 | 0.103321261 |

| | | | |
|---|---|---|---|
| Gm25970 | 3.058502859 | -2.120562967 | 0.693660133 |
| Fkbp3 | 818.3394673 | -0.344194881 | 0.101391029 |
| Fancm | 325.0406225 | -0.455842738 | 0.122522883 |
| Mis18bp1 | 20.94898307 | 1.566784886 | 0.345332111 |
| Wdr20rt | 0.136963818 | 0.718204821 | 3.59954433 |
| Gm26015 | 0 | NA | NA |
| Gm25599 | 0 | NA | NA |
| Gm23948 | 0 | NA | NA |
| Gm22782 | 0 | NA | NA |
| Rpl10l | 0.061920613 | 0.718207094 | 3.724945049 |
| Mdga2 | 1092.640628 | -0.93768252 | 0.106830917 |
| Gm23826 | 1.550082411 | -1.41872294 | 0.849905811 |
| n-R5s60 | 0 | NA | NA |
| Gm26256 | 0 | NA | NA |
| n-R5s61 | 0 | NA | NA |
| Gm15571 | 0.136963818 | 0.718204821 | 3.59954433 |
| Gm24499 | 0.13681464 | 0.237322887 | 2.94378999 |
| Rps29 | 1758.697591 | 0.321868071 | 0.105066679 |
| Rn7s1 | 42650.12024 | 0.682535662 | 0.309748326 |
| AC099934.2 | 0 | NA | NA |
| AC099934.1 | 0 | NA | NA |
| Lrr1 | 2.041322758 | 1.713360853 | 0.90783242 |
| Rpl36al | 266.3361645 | 0.904658815 | 0.159906628 |
| Mgat2 | 450.2752201 | 0.803566538 | 0.111673303 |
| Dnaaf2 | 60.5402635 | 0.505688359 | 0.169189707 |
| 9330151L19Rik | 197.4609258 | -0.43390103 | 0.098303259 |
| Pole2 | 9.423356029 | 1.465919977 | 0.365543157 |
| Klhdc1 | 176.7937862 | -0.620567057 | 0.146924902 |
| Gm15561 | 0 | NA | NA |
| Klhdc2 | 1197.243616 | -0.158437956 | 0.060246483 |
| Nemf | 637.5580402 | 0.066643568 | 0.079006595 |
| AC157822.1 | 0 | NA | NA |
| Rn7s2 | 34914.20044 | 0.290949662 | 0.206979059 |
| AC157822.2 | 0 | NA | NA |
| Gm9887 | 2.974284523 | 0.063645601 | 0.574123764 |
| Arf6 | 1024.329735 | 0.932448681 | 0.093704331 |
| Gm23018 | 0.038981395 | 0.718207094 | 3.724945049 |
| Vcpkmt | 182.6319952 | 0.764649592 | 0.10211397 |
| Sos2 | 821.7158266 | -0.139327673 | 0.05762264 |
| Gm24449 | 1.413161897 | -0.471451229 | 0.787844969 |
| L2hgdh | 176.5242931 | -0.486237535 | 0.119586058 |
| Atp5s | 115.1168851 | -0.623383894 | 0.123530447 |
| Cdkl1 | 209.0876337 | -0.217936871 | 0.108899135 |
| Mir681 | 2.100525097 | -0.754259798 | 0.689243759 |
| 4930512B01Rik | 3.315021913 | -0.456539116 | 0.568453974 |
| Map4k5 | 890.2594534 | -0.311410517 | 0.115062566 |
| 4931403G20Rik | 0.07796279 | 1.072698098 | 3.721829099 |

| | | | |
|---|---|---|---|
| Atl1 | 920.937231 | -0.742442171 | 0.059453395 |
| Gm6316 | 3.637700379 | -0.670414836 | 0.567257114 |
| Gm22058 | 5.202496254 | -0.838885674 | 0.487247359 |
| Gm3086 | 7.899535637 | -0.604967717 | 0.490044546 |
| Sav1 | 344.851752 | 0.942383285 | 0.131645017 |
| Gm23579 | 0 | NA | NA |
| Nin | 902.9475348 | -0.129026517 | 0.138158716 |
| Abhd12b | 8.085658348 | -0.02748698 | 0.412002755 |
| Pygl | 47.59505423 | 0.947603834 | 0.174989721 |
| F730035M05Rik | 0.381387581 | 0.141531205 | 1.541086638 |
| Trim9 | 3338.081355 | -0.432767686 | 0.103994007 |
| Tmx1 | 634.4816424 | 0.504657482 | 0.096404089 |
| Gm24474 | 0.044544648 | 0.237309859 | 3.724945049 |
| Frmd6 | 461.8109092 | -0.193123178 | 0.138903069 |
| Gm24857 | 0 | NA | NA |
| Actr10 | 1289.027537 | -0.100568503 | 0.065569834 |
| Psma3 | 949.5019891 | 0.580618214 | 0.076063842 |
| 3110056K07Rik | 60.20099725 | -0.59472143 | 0.181006525 |
| Arid4a | 748.6510582 | -0.146471715 | 0.107657284 |
| Tomm20l | 0.146692594 | 1.229893553 | 3.521109737 |
| Timm9 | 217.381226 | -0.195220462 | 0.071618706 |
| 2700049A03Rik | 196.0398049 | -0.250702572 | 0.125070634 |
| Dact1 | 166.182804 | 0.148914808 | 0.158309696 |
| 4930474H06Rik | 0.409321926 | -0.29375214 | 1.381306185 |
| 1700083H02Rik | 0 | NA | NA |
| Gm22959 | 0 | NA | NA |
| Daam1 | 1425.073008 | -0.074690287 | 0.048125755 |
| Gm7985 | 3.193668984 | -0.988365226 | 0.668212436 |
| Gpr135 | 77.04711869 | -0.28284182 | 0.13786555 |
| L3hypdh | 19.58973956 | -0.168199868 | 0.258318237 |
| Jkamp | 337.9089615 | -0.21887846 | 0.099975948 |
| Ccdc175 | 2.241016658 | 2.862419454 | 0.892312043 |
| Gm24466 | 0 | NA | NA |
| Rtn1 | 7105.395095 | -0.73322173 | 0.074146297 |
| Lrrc9 | 75.53725959 | 0.11027221 | 0.145636643 |
| AC103353.1 | 0 | NA | NA |
| Pcnxl4 | 483.304744 | -0.232099106 | 0.068291825 |
| Dhrs7 | 149.9058334 | 0.239629258 | 0.116316163 |
| Ppm1a | 1736.427688 | -0.197950798 | 0.061779981 |
| 4930447C04Rik | 136.3889103 | -0.820796292 | 0.193158673 |
| Six6 | 2.974111705 | 0.485276488 | 0.861772983 |
| Gm26709 | 0 | NA | NA |
| Gm38170 | 0 | NA | NA |
| Six1 | 9.772493715 | 1.124008306 | 0.384572663 |
| Six4 | 52.62636027 | -0.600923888 | 0.188668977 |
| Mnat1 | 196.833954 | 0.045483666 | 0.085433679 |
| Gm23910 | 0 | NA | NA |

| | | | |
|---|---|---|---|
| Trmt5 | 87.0705171 | 0.064696422 | 0.184443211 |
| Slc38a6 | 110.6102002 | 0.150197621 | 0.178322028 |
| D830013O20Rik | 0 | NA | NA |
| Tmem30b | 3.209381466 | 0.486290547 | 0.561741818 |
| Prkch | 135.4372229 | 0.821733995 | 0.144322821 |
| Hif1a | 2116.265102 | 0.768665054 | 0.129343321 |
| Gm15283 | 3.904436585 | -1.614953481 | 0.564585394 |
| Gm5068 | 0 | NA | NA |
| Snapc1 | 313.850414 | -0.009012404 | 0.090810732 |
| Syt16 | 456.7941473 | -0.88404448 | 0.1468527 |
| Gm29587 | 0 | NA | NA |
| 1700086L19Rik | 251.9530011 | -0.373991746 | 0.174929744 |
| Dbpht2 | 497.2679655 | -0.401481702 | 0.156457489 |
| Gm11042 | 0.897295135 | -1.405514809 | 1.190609265 |
| Kcnh5 | 625.0782806 | -1.272492611 | 0.146212758 |
| Rhoj | 239.1716956 | 3.561618617 | 0.330547391 |
| Gphb5 | 1.844461247 | 4.181912275 | 1.082801547 |
| Ppp2r5e | 1376.244815 | -0.057555071 | 0.045967705 |
| Gm29236 | 0.285890061 | 2.010053882 | 1.763578364 |
| Wdr89 | 187.0451121 | 0.71111294 | 0.108633832 |
| Sgpp1 | 600.9576085 | 0.248013178 | 0.090045732 |
| Syne2 | 912.2047069 | 0.189527552 | 0.07527702 |
| Mir5101 | 0.042716945 | -0.243571806 | 3.724945049 |
| Esr2 | 7.599516194 | 0.241944413 | 0.356517042 |
| Gm7862 | 4.308722595 | 0.188252879 | 0.498084788 |
| Tex21 | 0.318741927 | 1.629617631 | 1.722515214 |
| Mthfd1 | 536.2563463 | 0.031693624 | 0.069920951 |
| Akap5 | 895.5134347 | -2.13975404 | 0.187040013 |
| Zbtb25 | 191.2371808 | -0.06714616 | 0.090470956 |
| Zbtb1 | 407.9327547 | 0.350201724 | 0.085949393 |
| 4930426I24Rik | 3.588158775 | -1.046897493 | 0.56062022 |
| Hspa2 | 433.2408539 | 0.355472799 | 0.132926181 |
| Ppp1r36 | 10.56443581 | 0.671582032 | 0.300811415 |
| Gm25563 | 0 | NA | NA |
| Gm10451 | 3.439471239 | -0.347844691 | 0.555757392 |
| Mir5135 | 0.047948698 | -0.243569582 | 3.724945049 |
| Plekhg3 | 262.9259336 | -0.31945115 | 0.106503312 |
| Sptb | 3404.075389 | -0.678142289 | 0.093148019 |
| Gm24010 | 0 | NA | NA |
| Churc1 | 76.27497148 | 0.135600696 | 0.146566095 |
| Fntb | 334.4021306 | 0.030064062 | 0.0813863 |
| Gpx2 | 1.498903582 | 0.410113678 | 0.803680708 |
| Rab15 | 1748.719287 | -0.202148449 | 0.124495933 |
| Max | 974.4805696 | 0.980110864 | 0.08982941 |
| Gm22696 | 0 | NA | NA |
| Fut8 | 862.4398163 | -0.15098465 | 0.072810687 |
| Gm24775 | 2.254287688 | -1.883915291 | 0.80203164 |

| | | | |
|---|---|---|---|
| Gm25926 | 5.521549126 | -0.591672572 | 0.487102093 |
| Gm24070 | 0 | NA | NA |
| Gm24994 | 0 | NA | NA |
| Gm6657 | 0 | NA | NA |
| Gphn | 1083.325677 | -0.230449422 | 0.064678512 |
| Fam71d | 7.845798074 | -0.593009028 | 0.438977666 |
| Mpp5 | 811.6373095 | 0.392742668 | 0.097265155 |
| Gm25225 | 0.097139547 | 0.718207094 | 3.724945049 |
| Atp6v1d | 1557.396393 | 0.105523061 | 0.057804146 |
| Eif2s1 | 534.2845462 | 0.936946199 | 0.105392833 |
| Plek2 | 8.61393615 | 2.841841775 | 0.472400459 |
| Tmem229b | 976.9241392 | 0.627559065 | 0.115274586 |
| Plekhh1 | 575.4863948 | -0.459781382 | 0.10286203 |
| Pigh | 108.6090619 | 0.300497012 | 0.130673722 |
| Gm27297 | 0 | NA | NA |
| Arg2 | 76.92116969 | 1.937288003 | 0.28828715 |
| Vti1b | 794.0198813 | -0.090173136 | 0.075206985 |
| Gm17195 | 12.61275266 | -0.777525998 | 0.329759116 |
| Gm17194 | 0 | NA | NA |
| Rdh11 | 314.5614071 | 0.255197922 | 0.071211681 |
| Rdh12 | 16.14501922 | -0.035406494 | 0.272840428 |
| Zfyve26 | 591.7739187 | 0.388256173 | 0.094527618 |
| Rad51b | 1.767988423 | 0.990150749 | 0.816910527 |
| Gm28056 | 0 | NA | NA |
| Zfp36l1 | 660.9038404 | 1.150154594 | 0.119809179 |
| Gm26669 | 0.480640422 | 2.560030851 | 1.326036167 |
| 2310015A10Rik | 112.2249857 | -0.215061863 | 0.103095847 |
| Actn1 | 1100.046826 | 0.853149475 | 0.091707234 |
| Dcaf5 | 1241.49422 | -0.117109621 | 0.06988842 |
| mmu-mir-1843a | 0 | NA | NA |
| Scarna3b | 2.198311352 | -0.316401716 | 0.811787764 |
| Exd2 | 554.3707735 | -0.701995268 | 0.101759657 |
| Gm26777 | 28.6766377 | -1.141291482 | 0.236140687 |
| Galnt16 | 1559.333316 | -0.142557492 | 0.064765315 |
| Gm23817 | 0.07796279 | 1.072698098 | 3.721829099 |
| Erh | 198.2980328 | 0.47866084 | 0.102529691 |
| Slc39a9 | 697.3314638 | 0.258376199 | 0.065690415 |
| Plekhd1os | 1.509627474 | -0.239120678 | 0.877195144 |
| Plekhd1 | 169.5676397 | -0.536684938 | 0.152073832 |
| Ccdc177 | 193.1722448 | -0.917276791 | 0.123476246 |
| Gm26796 | 1.873447992 | 0.334851574 | 0.771563135 |
| Susd6 | 1200.089806 | 0.83875963 | 0.109252535 |
| Gm26545 | 0 | NA | NA |
| Gm23298 | 0.127971488 | 1.23029335 | 3.685533863 |
| 1700052I22Rik | 0 | NA | NA |
| Srsf5 | 1958.310891 | -0.174099366 | 0.085928677 |
| Slc10a1 | 0.066050875 | 0.71820487 | 3.724945049 |

| | | | |
|---|---|---|---|
| Smoc1 | 160.7710477 | 0.643111543 | 0.16714263 |
| Mir3067 | 0.784624355 | -0.10108783 | 1.238302475 |
| Slc8a3 | 615.5791632 | -0.544326719 | 0.101367897 |
| Gm3693 | 13.26815873 | -2.137420074 | 0.468165008 |
| Gm20498 | 37.16701504 | -0.373136097 | 0.188542456 |
| Cox16 | 98.00354183 | -0.206791149 | 0.122128874 |
| Gm3695 | 17.50620631 | -0.709414206 | 0.404306306 |
| Gm4787 | 60.98925875 | -0.825622019 | 0.19400686 |
| Hspe1-ps2 | 0 | NA | NA |
| Adam4 | 66.87698665 | -0.899325319 | 0.253023597 |
| Gm28370 | 1.129831383 | -1.153814298 | 1.008473301 |
| Synj2bp | 590.4251372 | -0.296029205 | 0.105215707 |
| Gm24853 | 0.038981395 | 0.718207094 | 3.724945049 |
| Gm17191 | 9.020757225 | -0.564158231 | 0.371616319 |
| 6530401F13Rik | 4.716167235 | -1.340932432 | 0.564745788 |
| Adam21 | 15.49915397 | -0.432914849 | 0.267281612 |
| Med6 | 165.5733909 | 0.389327205 | 0.121867406 |
| Gm16572 | 0.53222664 | 0.049933216 | 1.584728437 |
| Gm16570 | 4.946459833 | -0.6126635 | 0.570494866 |
| Ttc9 | 455.2096388 | 0.065222981 | 0.086833766 |
| Map3k9 | 1178.392681 | -0.743968614 | 0.100852947 |
| Pcnx | 2449.47711 | -0.353106277 | 0.069717277 |
| n-R5s62 | 0 | NA | NA |
| Sipa1l1 | 4759.581652 | -0.562862981 | 0.127643622 |
| Gm5435 | 1.384865981 | -0.309683696 | 0.861875386 |
| Gm8358 | 0 | NA | NA |
| n-R5s63 | 0 | NA | NA |
| Rgs6 | 645.1774152 | -0.491671806 | 0.105273203 |
| SNORA43 | 0 | NA | NA |
| Gm22149 | 0 | NA | NA |
| 1700085C21Rik | 0.21789665 | 0.237329415 | 2.168530734 |
| 4930423C22Rik | 0.847184061 | -0.227018369 | 1.326220421 |
| Gm29530 | 0.044544648 | 0.237309859 | 3.724945049 |
| 4933436F18Rik | 0 | NA | NA |
| 1700030I03Rik | 2.414365681 | -1.310505885 | 0.670730845 |
| Dpf3 | 155.1908164 | -0.543810918 | 0.142972376 |
| Dcaf4 | 167.3521746 | -0.17041499 | 0.10448842 |
| Zfyve1 | 610.9257158 | -0.089816914 | 0.056871046 |
| Gm26623 | 0.048949532 | -0.243569582 | 3.724945049 |
| Rbm25 | 2301.160026 | -0.169457017 | 0.110498538 |
| Gm29361 | 5.081951061 | -0.180418657 | 0.492396044 |
| Psen1 | 837.0310577 | 0.362082416 | 0.064943451 |
| Papln | 26.40192091 | -1.472851457 | 0.243428212 |
| Numb | 544.7112012 | -0.054465797 | 0.088070352 |
| 2410016O06Rik | 126.434428 | 0.115336328 | 0.131968473 |
| Gm26571 | 3.09834508 | 0.467044851 | 0.572155759 |
| Gm17673 | 1.031598502 | -1.474631609 | 1.268616139 |

| | | | |
|---|---|---|---|
| Acot2 | 62.86110075 | 0.557744834 | 0.157821722 |
| Acot1 | 68.7388213 | 1.30066529 | 0.252159791 |
| Acot4 | 13.42834362 | 0.366159198 | 0.309934833 |
| 4732463B04Rik | 11.2321051 | -1.268856508 | 0.352745731 |
| Acot3 | 12.40614317 | -0.33857243 | 0.367934432 |
| Acot5 | 14.23358867 | -0.064585956 | 0.354336349 |
| Gm8385 | 0 | NA | NA |
| Acot6 | 73.66342787 | -0.194598968 | 0.187224846 |
| Dnal1 | 1112.279329 | -0.815899961 | 0.094961635 |
| Pnma1 | 28.80769789 | -1.127900942 | 0.230426305 |
| Elmsan1 | 799.7188921 | 0.586620585 | 0.171424401 |
| Gm5436 | 39.89268335 | -0.040325528 | 0.209775887 |
| Ptgr2 | 431.9967278 | -0.131543356 | 0.097257359 |
| Zfp410 | 288.7236925 | 0.24392988 | 0.088734977 |
| Fam161b | 141.580014 | -0.341977292 | 0.181797948 |
| Coq6 | 126.6661858 | 0.170335528 | 0.110755313 |
| Entpd5 | 391.9413108 | -0.306651345 | 0.07099335 |
| Ccdc176 | 64.5472327 | -1.110733761 | 0.187988811 |
| AC120402.1 | 0 | NA | NA |
| Rnf113a2 | 0 | NA | NA |
| Aldh6a1 | 576.5073902 | -0.769434262 | 0.128940035 |
| Lin52 | 260.6951333 | -0.041805847 | 0.08873994 |
| Vsx2 | 9.446677945 | 0.087785736 | 0.475513124 |
| Abcd4 | 114.3789344 | -0.179840932 | 0.124923864 |
| Vrtn | 1.178380111 | 1.088785609 | 1.104140905 |
| Syndig1l | 316.8534421 | -0.834894186 | 0.112235231 |
| Npc2 | 1928.697367 | 2.617560211 | 0.159089978 |
| Isca2 | 199.1382904 | 0.144115511 | 0.087964815 |
| Ltbp2 | 65.48633923 | -0.347262586 | 0.213644294 |
| D030025P21Rik | 2.762732937 | -0.59054308 | 0.589901841 |
| Gm17139 | 0.226559374 | 0.2373201 | 2.539258673 |
| Arel1 | 2266.135036 | 0.433813623 | 0.071207057 |
| Gm17193 | 24.019554 | -0.026118156 | 0.319263146 |
| Fcf1 | 140.4858461 | 1.03670808 | 0.136037036 |
| Ylpm1 | 2692.241911 | -0.587200593 | 0.105074067 |
| Prox2os | 42.00672927 | -1.084641144 | 0.29909747 |
| Prox2 | 14.36306314 | 0.284244954 | 0.263201416 |
| Gm38355 | 1.352228905 | -0.251294373 | 0.848144457 |
| Dlst | 1232.377248 | 0.044671246 | 0.05606974 |
| Rps6kl1 | 284.8098805 | -0.280658572 | 0.116948244 |
| Pgf | 33.44763411 | 0.96082189 | 0.258659006 |
| Eif2b2 | 302.5668293 | 0.763619333 | 0.110184321 |
| Mlh3 | 324.2556209 | -0.344054767 | 0.106242851 |
| Mir6938 | 0.348080155 | 0.816099736 | 1.955652706 |
| Acyp1 | 106.7121191 | -0.411239685 | 0.131214596 |
| Zc2hc1c | 44.15230839 | 0.399126358 | 0.166388269 |
| Nek9 | 985.3989254 | 0.593201662 | 0.069527539 |

| | | | |
|---|---|---|---|
| Tmed10 | 1862.85624 | 0.628005684 | 0.097738094 |
| Fos | 757.1228931 | 1.486682236 | 0.462935868 |
| Jdp2 | 344.9332217 | -0.134971093 | 0.09146258 |
| Batf | 16.13625052 | 5.671505214 | 0.772616257 |
| n-R5s64 | 0 | NA | NA |
| AC117240.1 | 0 | NA | NA |
| Mfsd7c | 17.4725598 | -0.14921089 | 0.279308845 |
| 0610007P14Rik | 325.4729982 | -0.079753141 | 0.062642836 |
| Ttll5 | 894.6595886 | -0.427216418 | 0.071082228 |
| Gm26531 | 0.135217741 | 0.237306921 | 3.620320246 |
| Tgfb3 | 281.890318 | 0.901930879 | 0.127722019 |
| Gm37804 | 1.282550361 | 0.193473344 | 0.979597868 |
| Ift43 | 63.1959967 | 0.184624521 | 0.144938717 |
| Gm805 | 0.516561414 | -0.510363916 | 1.324294919 |
| Gpatch2l | 583.809052 | 0.143104889 | 0.067100966 |
| 4732487G21Rik | 0.11657011 | 0.237307635 | 3.724945049 |
| Esrrb | 65.56798319 | -0.0598794 | 0.154890587 |
| Gm27993 | 0 | NA | NA |
| Gm22004 | 0 | NA | NA |
| Vash1 | 371.0212885 | -0.413521543 | 0.071094035 |
| Angel1 | 126.5566759 | -0.038291236 | 0.121567033 |
| Gm17138 | 0 | NA | NA |
| Lrrc74a | 0.176451472 | 0.7182167 | 2.72333919 |
| Gm25888 | 0 | NA | NA |
| Irf2bpl | 1475.011036 | 0.576822085 | 0.111303064 |
| Gm26698 | 1.907481327 | 0.969267782 | 0.743561582 |
| Cipc | 1257.563404 | -0.187212957 | 0.042330921 |
| Zdhhc22 | 154.2953736 | -0.114271371 | 0.114305826 |
| Tmem63c | 504.9714176 | -0.272867301 | 0.11357541 |
| Gm25847 | 6.00824128 | -0.470495246 | 0.543669553 |
| Gm15745 | 1.796415912 | -2.166049575 | 1.006828206 |
| Ngb | 44.68760207 | -0.52176612 | 0.225684252 |
| Pomt2 | 325.0464884 | -0.102770671 | 0.068732377 |
| Gstz1 | 241.2031799 | -0.201508687 | 0.086476355 |
| Tmed8 | 929.2418369 | -0.235808812 | 0.074882441 |
| Samd15 | 21.85261841 | 0.037535208 | 0.261242756 |
| Noxred1 | 12.44760366 | -0.094048163 | 0.340919251 |
| Vipas39 | 515.9231004 | 0.18349824 | 0.055600762 |
| Ahsa1 | 741.40917 | 0.262167713 | 0.088883325 |
| AC151967.1 | 0.038981395 | 0.718207094 | 3.724945049 |
| Ism2 | 2.941330987 | -0.225801054 | 0.571485482 |
| Sptlc2 | 820.5122325 | 0.995544727 | 0.131158073 |
| Gm26764 | 0.618129248 | 0.818731353 | 1.269350431 |
| Alkbh1 | 295.3910853 | 0.34475788 | 0.15280342 |
| mmu-mir-3068 | 0.086930093 | 0.236669296 | 3.724898107 |
| Mir3068 | 0.060174537 | 0.237309859 | 3.724945049 |
| Gm29362 | 0 | NA | NA |

| | | | |
|---|---|---|---|
| Slirp | 288.8250465 | 0.732161828 | 0.126306118 |
| Snw1 | 789.2603872 | 0.606750673 | 0.081431818 |
| Gm8300 | 0 | NA | NA |
| Oog1 | 0 | NA | NA |
| Gm4027 | 0 | NA | NA |
| Gm16381 | 0 | NA | NA |
| Gm21319 | 0 | NA | NA |
| Gm21936 | 0 | NA | NA |
| BB287469 | 0 | NA | NA |
| Gm2001 | 0 | NA | NA |
| Gm2016 | 0 | NA | NA |
| Gm2022 | 0 | NA | NA |
| Gm2035 | 0 | NA | NA |
| Gm2042 | 0 | NA | NA |
| Gm2046 | 0 | NA | NA |
| Gm2075 | 0 | NA | NA |
| Gm6803 | 0 | NA | NA |
| Gm2056 | 0 | NA | NA |
| Gm16368 | 0 | NA | NA |
| RP23-174O6.3 | 0 | NA | NA |
| Gm8587 | 0.789916185 | 0.24167952 | 1.153752204 |
| Gm10436 | 0 | NA | NA |
| Gm8332 | 0 | NA | NA |
| Gm5662 | 0 | NA | NA |
| Gm7104 | 0 | NA | NA |
| Gm5039 | 0.128271896 | -0.243573716 | 3.679089475 |
| Gm10264 | 0 | NA | NA |
| Adck1 | 270.5190383 | 0.097908103 | 0.081057841 |
| Nrxn3 | 5644.729856 | -0.438571866 | 0.105539567 |
| Gm37980 | 0.244108046 | -1.361580503 | 2.485561782 |
| Gm27313 | 0 | NA | NA |
| Gm23989 | 9.194225744 | -0.612249934 | 0.479507974 |
| Dio2 | 725.4291824 | -1.408514026 | 0.187774425 |
| Gm26512 | 30.26736896 | -0.701775935 | 0.2278049 |
| 4930544I03Rik | 0.534296218 | -0.715145155 | 1.497453915 |
| n-R5s65 | 0 | NA | NA |
| Gm23195 | 0 | NA | NA |
| Gm26477 | 0 | NA | NA |
| Cep128 | 104.2602885 | -0.218898634 | 0.124229231 |
| 5430427M07Rik | 0 | NA | NA |
| CR974487.1 | 0 | NA | NA |
| Tshr | 7.627433336 | -0.414982143 | 0.38536514 |
| Gm18827 | 0.108266322 | 1.230376831 | 3.722845817 |
| Gtf2a1 | 1089.805438 | 0.304195002 | 0.062249851 |
| Gm25776 | 271.3150006 | 0.059081992 | 0.223055937 |
| Ston2 | 295.1567159 | 0.297155732 | 0.121101171 |
| Gm8378 | 10.50589885 | -0.810367714 | 0.349748481 |

| | | | |
|---|---|---|---|
| Gm16876 | 0.085327104 | 1.179809211 | 3.723500796 |
| Gm22015 | 0 | NA | NA |
| Sel1l | 2575.426926 | -0.111388506 | 0.038761831 |
| Rpl31-ps1 | 0.104719185 | 0.237305575 | 3.724945049 |
| Gm21614 | 0 | NA | NA |
| Gm23249 | 0 | NA | NA |
| Gm9726 | 14.69313774 | 0.355751548 | 0.281817946 |
| Mir8099-1 | 0.066050875 | 0.71820487 | 3.724945049 |
| Gm26055 | 0.060174537 | 0.237309859 | 3.724945049 |
| Gm22246 | 0.682665822 | 0.252197091 | 1.153118876 |
| Flrt2 | 1031.326902 | -0.583807374 | 0.073776975 |
| 1700019M22Rik | 0 | NA | NA |
| Gm26480 | 0 | NA | NA |
| Gm6863 | 145.6421742 | 0.406188848 | 0.137646368 |
| Gm23558 | 0 | NA | NA |
| Galc | 301.800896 | -0.43743408 | 0.09687198 |
| Gpr65 | 55.39552026 | 5.607514212 | 0.451828301 |
| Kcnk10 | 201.901865 | 0.085030668 | 0.128993284 |
| Gm25486 | 0 | NA | NA |
| Spata7 | 282.5135196 | -0.337417914 | 0.142811775 |
| Ptpn21 | 244.2151489 | 0.675367201 | 0.088808915 |
| Zc3h14 | 1282.947868 | 0.045289074 | 0.058809309 |
| Eml5 | 1567.483707 | -0.793487157 | 0.150763397 |
| Ttc8 | 210.9206023 | -0.592185739 | 0.154047875 |
| Rpl30-ps8 | 10.08383252 | 0.370809119 | 0.320448769 |
| Gm28933 | 0.301731016 | 1.071639968 | 2.028101104 |
| Foxn3 | 762.8771155 | -0.031511393 | 0.053655477 |
| Gm19898 | 0.254453631 | 1.256673259 | 2.971544927 |
| Gm20682 | 16.54715445 | -0.93404186 | 0.386797767 |
| 3300002A11Rik | 4.616394162 | -0.614619711 | 0.548597278 |
| Gm26839 | 2.688780499 | -0.122860946 | 0.671072174 |
| 1700064M15Rik | 1.043907264 | 0.409955729 | 1.083342085 |
| Efcab11 | 4.011499476 | 0.440126777 | 0.543390387 |
| Tdp1 | 173.9666548 | -0.470480817 | 0.159293046 |
| Kcnk13 | 123.3766849 | 1.023122872 | 0.14662194 |
| Gm23291 | 0.044544648 | 0.237309859 | 3.724945049 |
| Psmc1 | 1000.106951 | 0.491663903 | 0.112134091 |
| Nrde2 | 211.3513749 | 0.101189904 | 0.104084395 |
| Gm10433 | 3.50734834 | -0.258979439 | 0.575880456 |
| Calm1 | 27013.61596 | -0.36577293 | 0.082365049 |
| Gm26723 | 0.970285468 | 1.171243909 | 1.120928833 |
| Ttc7b | 1662.607921 | -0.465823881 | 0.069587338 |
| Rps6ka5 | 469.9143002 | -0.647545864 | 0.09102732 |
| Gm25801 | 0.225129614 | 0.237300054 | 3.724945049 |
| 9030617O03Rik | 99.5661288 | 0.264432071 | 0.165976778 |
| Gm8482 | 2.942155206 | 3.23721058 | 1.155187552 |
| Gpr68 | 165.6196338 | 0.03048014 | 0.117652653 |

| | | | |
|---|---|---|---|
| Ccdc88c | 393.2724972 | -0.157350682 | 0.14893546 |
| A630072L19Rik | 0.314467885 | 0.23732882 | 2.033924006 |
| Mir1190 | 0.80558184 | -2.508492695 | 1.193793764 |
| Smek1 | 938.2289803 | 0.320566418 | 0.05259583 |
| D130020L05Rik | 15.63426209 | -0.351736722 | 0.277056799 |
| Kif4-ps | 0 | NA | NA |
| Gm18848 | 0 | NA | NA |
| Gm22749 | 0 | NA | NA |
| Catsperb | 0.068097359 | -0.243569582 | 3.724945049 |
| Gm24542 | 0 | NA | NA |
| Tc2n | 7.321160109 | -0.976255321 | 0.477660588 |
| Gm16285 | 0 | NA | NA |
| Fbln5 | 104.6927819 | 1.280450083 | 0.203975652 |
| Trip11 | 1573.263012 | 0.133745548 | 0.048231843 |
| Gm20701 | 0 | NA | NA |
| Gm20702 | 0 | NA | NA |
| Gm20036 | 4.340709641 | 0.373060115 | 0.534264478 |
| Atxn3 | 451.4130845 | -0.028905409 | 0.072250631 |
| Cpsf2 | 797.9273037 | 0.42988947 | 0.077038953 |
| Gm16339 | 11.18950982 | -0.793823011 | 0.479621 |
| Slc24a4 | 531.3783664 | -0.39377806 | 0.185403409 |
| Gm25203 | 0 | NA | NA |
| Rin3 | 105.1190873 | 2.711724807 | 0.193986068 |
| Lgmn | 2861.87464 | 2.16417913 | 0.132602597 |
| Golga5 | 336.8030054 | 0.164225952 | 0.088577495 |
| Chga | 1338.904208 | 0.319034236 | 0.177087308 |
| Itpk1 | 872.6122047 | 0.64566815 | 0.122681057 |
| Gm37019 | 35.26459873 | -0.516383828 | 0.309378046 |
| Gm20069 | 0.551997393 | 2.088399286 | 1.380248249 |
| Gm28373 | 39.99975009 | -0.252503311 | 0.292987365 |
| Mir1936 | 2.100202168 | 0.328905324 | 0.758367011 |
| Gm28051 | 0.060174537 | 0.237309859 | 3.724945049 |
| Gm21971 | 0 | NA | NA |
| Gm20604 | 131.2845531 | 0.214141585 | 0.115539877 |
| Moap1 | 4.795281827 | 0.401237359 | 0.507601263 |
| Tmem251 | 141.6121926 | 1.123266823 | 0.15583709 |
| AK010878 | 58.12541358 | 0.120077683 | 0.147047759 |
| Ubr7 | 418.4523783 | 0.185351023 | 0.07145921 |
| Btbd7 | 452.2462838 | 0.378223464 | 0.076267159 |
| Cox8c | 0 | NA | NA |
| Unc79 | 1662.520299 | -0.676894538 | 0.115109439 |
| AC122367.1 | 17.42179249 | -1.462154734 | 0.385206965 |
| Gm27976 | 0.999255461 | -0.61200255 | 0.949831466 |
| Prima1 | 64.20214066 | 0.110572186 | 0.164874359 |
| Fam181a | 11.59340835 | -1.030496539 | 0.348416283 |
| Gm29508 | 2.89487079 | -1.913342925 | 0.681841802 |
| Asb2 | 49.37814463 | 0.713409272 | 0.218143753 |

| | | | |
|---|---|---|---|
| Gm15523 | 0.643207634 | -1.013315415 | 1.238962776 |
| Otub2 | 164.6469116 | -0.218133576 | 0.166820186 |
| 9330161L09Rik | 14.42659375 | 0.250337467 | 0.282429615 |
| Ddx24 | 2634.287795 | 0.702420942 | 0.097825033 |
| Ifi27 | 153.7870265 | 0.8111285 | 0.243509853 |
| Ifi27l2a | 1103.207706 | 8.902869177 | 0.367965247 |
| Ifi27l2b | 0.044191549 | -0.243571806 | 3.724945049 |
| Ppp4r4 | 572.2723124 | -0.099474993 | 0.095893516 |
| Serpina10 | 1.913476902 | 4.633750838 | 1.075993297 |
| Gm18501 | 0 | NA | NA |
| Serpina6 | 0.146451277 | -0.243562218 | 2.882358805 |
| Serpina16 | 0 | NA | NA |
| Serpina1f | 0.066050875 | 0.71820487 | 3.724945049 |
| Gm5656 | 0 | NA | NA |
| Gm17142 | 0 | NA | NA |
| Serpina1b | 7.892462027 | 1.28308999 | 0.42246864 |
| Gm17042 | 0 | NA | NA |
| Serpina1d | 0 | NA | NA |
| Gm17043 | 0 | NA | NA |
| Gm8893 | 0.487995331 | -1.480187854 | 1.314872295 |
| Gm6918 | 0 | NA | NA |
| Serpina1a | 0.839309821 | 1.972084202 | 1.188075666 |
| Gm8895 | 0.092691418 | 1.183079489 | 3.724560917 |
| Serpina1c | 0.721092599 | -0.387024332 | 1.202219417 |
| B430119L08Rik | 0.437661561 | 0.795098386 | 1.33669418 |
| Gm17198 | 0 | NA | NA |
| Serpina1e | 0.347991053 | -0.140182396 | 1.710851449 |
| Serpina11 | 2.314302159 | 4.240553322 | 0.966049358 |
| Gm28577 | 0 | NA | NA |
| Serpina9 | 42.26228332 | -1.331496646 | 0.245361394 |
| Gm37496 | 0.085433889 | -0.72445792 | 3.724945049 |
| Serpina12 | 0.359936507 | -0.243577445 | 3.724945049 |
| Serpina4-ps1 | 0.088534442 | 1.179809211 | 3.723500796 |
| Serpina5 | 0.066050875 | 0.71820487 | 3.724945049 |
| Gm23289 | 0 | NA | NA |
| Serpina3a | 0.228000506 | 1.906981631 | 2.476810464 |
| Serpina3b | 0.099106094 | 1.209894437 | 3.72418822 |
| Serpina3c | 2.76240905 | 2.330160389 | 0.915462099 |
| Serpina3d-ps | 0 | NA | NA |
| Serpina3e-ps | 7.226489367 | 3.090041189 | 0.602380729 |
| Serpina3f | 27.21369777 | 6.43901257 | 0.841785422 |
| Gm8918 | 0 | NA | NA |
| Serpina3g | 192.2982361 | 7.339683405 | 0.574436025 |
| Serpina3h | 81.26495241 | 5.673959438 | 0.521913328 |
| Serpina3i | 29.39647476 | 6.890777387 | 0.903060936 |
| Serpina3j | 0 | NA | NA |
| Serpina3m | 31.36521844 | 3.385446489 | 0.409784359 |

| | | | |
|---|---|---|---|
| Serpina3k | 1.059265352 | 0.956686741 | 1.053475882 |
| Serpina3l-ps | 0 | NA | NA |
| Serpina3n | 2233.484408 | 3.458991347 | 0.308463692 |
| Gm24334 | 0 | NA | NA |
| Gsc | 0.456948362 | 0.007430084 | 1.467694231 |
| Gm10000 | 0.122601409 | 0.237307635 | 3.724945049 |
| Dicer1 | 1104.172179 | -0.232770752 | 0.097822615 |
| Clmn | 1628.39178 | -0.953645341 | 0.103773861 |
| 4930408O17Rik | 1.425419836 | -0.144938629 | 0.846156333 |
| Gm28875 | 11.6389744 | 0.625418508 | 0.387429408 |
| Syne3 | 100.3522481 | 0.840122587 | 0.147988631 |
| Scarna13 | 339.6868988 | 0.051724729 | 0.290227758 |
| mmu-mir-3069 | 0 | NA | NA |
| Glrx5 | 236.9685444 | 0.031917687 | 0.102267579 |
| Tcl1b2 | 0.273490436 | 0.237325663 | 2.423326845 |
| Tcl1b1 | 0.12092566 | 0.237307635 | 3.724945049 |
| Tcl1b5 | 0 | NA | NA |
| Tcl1b3 | 0 | NA | NA |
| Tcl1b4 | 0 | NA | NA |
| Tcl1 | 0 | NA | NA |
| Tunar | 293.4716965 | -0.227204448 | 0.101982552 |
| D430019H16Rik | 1275.143419 | -0.188963662 | 0.139666809 |
| Bdkrb2 | 21.21549069 | 2.588861803 | 0.357646133 |
| Bdkrb1 | 6.675953194 | 3.473428329 | 0.760423636 |
| Atg2b | 1471.256828 | -0.27072942 | 0.055728272 |
| Gm22790 | 0 | NA | NA |
| Gskip | 216.9176705 | 0.251896171 | 0.131460687 |
| Ak7 | 31.57741441 | 0.363651139 | 0.204062678 |
| Papola | 1915.550768 | 0.144163626 | 0.081388493 |
| Gm22120 | 0 | NA | NA |
| Vrk1 | 161.0238951 | -0.094319311 | 0.113254088 |
| Gm24986 | 0 | NA | NA |
| Gm17032 | 0.342848402 | 0.818231756 | 1.967429857 |
| 1700121N20Rik | 0 | NA | NA |
| Gm3191 | 0.406963084 | 2.650513801 | 3.680819858 |
| Gm17031 | 0.100902008 | 1.15059983 | 3.720931964 |
| Gm17053 | 0.121578509 | 0.718202646 | 3.724945049 |
| Gm17033 | 0 | NA | NA |
| 4933406K04Rik | 0 | NA | NA |
| 1700013N06Rik | 0 | NA | NA |
| Gm9599 | 0 | NA | NA |
| Gm16087 | 0.365883897 | -2.185737693 | 1.455697607 |
| Gm16086 | 0.197846125 | -0.570038387 | 3.255083671 |
| Gm16085 | 0 | NA | NA |
| Gm16084 | 0 | NA | NA |
| Gm2800 | 0.20562365 | -1.049594134 | 2.592716595 |
| Gm5187 | 0 | NA | NA |

| | | | |
|---|---|---|---|
| Gm17052 | 0.203140137 | 0.261253882 | 2.216123385 |
| 3110018I06Rik | 0 | NA | NA |
| Gm9006 | 0 | NA | NA |
| 4930465M20Rik | 0.148569308 | -0.243563529 | 2.869495678 |
| Gm15208 | 0.15596086 | -1.187811386 | 3.483969583 |
| Bcl11b | 1306.155659 | -1.256400018 | 0.152323898 |
| Setd3 | 1675.172341 | -0.137664984 | 0.053809622 |
| Ccnk | 411.8626785 | 0.421391432 | 0.106997876 |
| Ccdc85c | 333.5260322 | -0.579445866 | 0.082334379 |
| Hhipl1 | 69.09591632 | -0.015958671 | 0.162670695 |
| 4930478K11Rik | 1.119709845 | -0.013013163 | 0.997043859 |
| Cyp46a1 | 1040.049922 | -1.151437632 | 0.153468857 |
| Eml1 | 662.3640817 | -0.096316962 | 0.097937999 |
| Gm15636 | 0 | NA | NA |
| Gm16596 | 7.416129088 | -0.282594173 | 0.374414531 |
| Evl | 1049.201664 | -0.267181185 | 0.055838699 |
| Mir342 | 1.676187767 | -1.122322769 | 1.039616736 |
| Degs2 | 4.353830809 | -0.899805926 | 0.709306863 |
| Gm23872 | 0 | NA | NA |
| Yy1 | 536.7063009 | 0.233277134 | 0.081031647 |
| Slc25a29 | 103.2788333 | -0.462171851 | 0.130748926 |
| Mir345 | 1.700378639 | -0.21584143 | 0.829237558 |
| Slc25a47 | 16.0241689 | -0.157406285 | 0.308835583 |
| Wars | 1271.197342 | 1.038211481 | 0.116063283 |
| Gm22079 | 2.165001819 | -0.514011115 | 0.759760171 |
| Wdr25 | 96.15949304 | -0.045803635 | 0.107694893 |
| Begain | 792.2631617 | -0.804085782 | 0.108812082 |
| Gm28498 | 1.537157928 | -0.999530827 | 0.991843453 |
| Gm22234 | 0 | NA | NA |
| Gm25297 | 0 | NA | NA |
| Dlk1 | 383.0032136 | 0.069647637 | 0.175287719 |
| Gm27528 | 0 | NA | NA |
| Meg3 | 23754.92684 | -1.4918225 | 0.443929357 |
| Gm27596 | 7.046539481 | -1.432507232 | 0.605839966 |
| Gm27300 | 0 | NA | NA |
| Mir1906-1 | 0 | NA | NA |
| Mir770 | 0 | NA | NA |
| Gm26906 | 0.900331428 | -0.67303353 | 1.298875483 |
| Mir673 | 3.122023216 | -1.348633927 | 0.880976752 |
| Mir493 | 16.71408191 | -1.347644836 | 0.489443335 |
| Mir337 | 13.9759078 | -1.523697276 | 0.364205169 |
| mmu-mir-3544 | 0 | NA | NA |
| Mir540 | 13.29036576 | -1.128495338 | 0.441810756 |
| Mir665 | 11.41153131 | -1.379478184 | 0.461818935 |
| Mir3070a | 19.33383777 | -1.345773906 | 0.444290879 |
| Mir3070b | 15.52429422 | -0.89659092 | 0.400405847 |
| Rtl1 | 49.79748458 | 1.735851125 | 0.280164959 |

| | | | |
|---|---|---|---|
| Mir431 | 8.591852434 | -1.206086583 | 0.439367025 |
| Mir433 | 30.4253769 | -1.468391277 | 0.375185577 |
| Mir127 | 8.254821574 | -1.353011377 | 0.444619447 |
| Mir434 | 6.186359681 | -0.955656511 | 0.431498852 |
| Mir432 | 5.368193358 | -1.063065209 | 0.460629976 |
| Mir136 | 0 | NA | NA |
| mmu-mir-136 | 4.439960643 | -1.105229122 | 0.584791127 |
| Gm27350 | 1.495241998 | -2.220972074 | 1.066203834 |
| Rian | 7362.023447 | -0.999829244 | 0.112678785 |
| Mir341 | 46.98560136 | -1.722330002 | 0.310998583 |
| Mir1188 | 40.87071727 | -1.690306108 | 0.396543904 |
| Gm23912 | 5.84972083 | -2.217446972 | 0.654437191 |
| Mir370 | 7.676564023 | -1.180620406 | 0.606434675 |
| Gm24564 | 32.85825865 | -1.044360873 | 0.33513106 |
| Gm23474 | 0 | NA | NA |
| AF357425 | 0 | NA | NA |
| Gm26945 | 332.9786148 | -1.209822197 | 0.206253513 |
| Gm24899 | 0 | NA | NA |
| Gm23600 | 67.41334034 | -0.934747231 | 0.179803742 |
| Gm24895 | 0 | NA | NA |
| DQ267100 | 0 | NA | NA |
| DQ267101 | 11.01360181 | -0.902750243 | 0.327364763 |
| Gm23508 | 0 | NA | NA |
| Gm26922 | 0 | NA | NA |
| DQ267102 | 24.0816795 | -0.970238196 | 0.26741066 |
| Gm25357 | 8.037931983 | -1.631937564 | 0.377015637 |
| Gm27604 | 0 | NA | NA |
| Gm37899 | 685.7979278 | -1.147230264 | 0.293156413 |
| Gm23347 | 0 | NA | NA |
| AF357428 | 5.570026761 | -1.045445103 | 0.574916061 |
| Gm23736 | 2.72795131 | -0.729668079 | 0.602882375 |
| AF357341 | 4.666739356 | -0.891172418 | 0.626492967 |
| Mir882 | 2.263396373 | -0.881529787 | 0.734593858 |
| Gm22981 | 1.907638496 | -1.235011634 | 0.873302899 |
| Gm25856 | 14.93485102 | -1.862631656 | 0.347868886 |
| Gm23787 | 13.95901734 | -1.09832137 | 0.381825948 |
| Gm25854 | 10.14198828 | -0.958752111 | 0.381414615 |
| Gm22205 | 21.52538062 | -0.903315411 | 0.284032826 |
| Gm25224 | 10.64312297 | -0.534689127 | 0.378106858 |
| B830012L14Rik | 195.3433677 | -1.252321224 | 0.356452866 |
| Mir379 | 4.424652029 | -1.205058535 | 0.665426177 |
| Mir411 | 6.496099058 | -1.089070735 | 0.469015969 |
| mmu-mir-299a | 2.459243483 | -1.376474407 | 0.729331915 |
| Mir299b | 0 | NA | NA |
| Gm25750 | 0 | NA | NA |
| mmu-mir-380 | 4.04787115 | -0.964348799 | 0.562187477 |
| Mir1197 | 3.390250948 | -1.282166133 | 0.63873651 |

| | | | |
|---|---|---|---|
| Mir323 | 3.255378368 | -1.566992806 | 0.830349607 |
| Mir758 | 6.502832914 | -1.001032347 | 0.473873513 |
| Mir329 | 5.909624248 | -1.235099239 | 0.555703686 |
| Mir494 | 6.573050005 | -0.915570165 | 0.46627165 |
| Mir679 | 11.72323245 | -1.621533222 | 0.397308636 |
| Mir1193 | 10.23592663 | -1.141379645 | 0.503891705 |
| Mir666 | 14.05147067 | -0.899464824 | 0.400231736 |
| Mir543 | 4.380404366 | -1.65549158 | 0.698741654 |
| Mir495 | 2.202405632 | -0.447608619 | 0.790421772 |
| Mir667 | 9.470118255 | -1.335761705 | 0.384124774 |
| Mir376b | 10.35390795 | -1.165849999 | 0.425863606 |
| Mir654 | 8.477009777 | -0.699720281 | 0.423292941 |
| mmu-mir-376b | 2.209145549 | -0.163166105 | 0.717077355 |
| Gm25252 | 0.047948698 | -0.243569582 | 3.724945049 |
| mmu-mir-376a | 5.585566604 | -0.804841928 | 0.527066 |
| Mir300 | 6.166566501 | -0.96682527 | 0.505460979 |
| Mir381 | 6.601995524 | -1.360818303 | 0.541255395 |
| Mir487b | 4.545755139 | -1.946960829 | 0.699325321 |
| AC121784.1 | 0 | NA | NA |
| Mir539 | 4.506246945 | -1.112762542 | 0.470355715 |
| Gm27379 | 8.227541344 | -1.706229744 | 0.529723022 |
| Mir544 | 5.245180705 | -1.658618192 | 0.55087442 |
| Mirg | 537.7305624 | -0.922056029 | 0.270477551 |
| Mir382 | 2.007426773 | -1.373578178 | 0.707858856 |
| Mir134 | 0 | NA | NA |
| Mir668 | 0 | NA | NA |
| Mir485 | 0 | NA | NA |
| Mir453 | 14.69478301 | -0.984974121 | 0.432413918 |
| Mir154 | 6.301721876 | -0.589057414 | 0.43818594 |
| Mir496a | 7.410274305 | -0.994048059 | 0.425600044 |
| Mir377 | 0 | NA | NA |
| Mir541 | 8.228173234 | -1.446467218 | 0.417753068 |
| mmu-mir-409 | 3.297680665 | -0.627062083 | 0.548138042 |
| Gm24321 | 0 | NA | NA |
| Mir412 | 2.651183465 | -2.052219179 | 0.785382008 |
| Mir369 | 1.213203356 | -1.444160137 | 0.964495801 |
| Mir410 | 0 | NA | NA |
| Mir3072 | 1.399651712 | -1.461996312 | 1.125291924 |
| Mir1247 | 0.141331096 | 0.237322366 | 3.56184083 |
| Dio3 | 10.88659069 | -1.577852334 | 0.371343172 |
| Ppp2r5c | 1467.571291 | -0.242049483 | 0.054370515 |
| Dync1h1 | 16373.17254 | 0.181226206 | 0.045891717 |
| Gm17111 | 0.611020101 | -1.039233263 | 1.415436075 |
| 1700001K19Rik | 0.241252684 | 2.061059537 | 3.028957171 |
| Gm27299 | 0.462939631 | -1.466629527 | 1.388220509 |
| Hsp90aa1 | 8408.664402 | 0.216832986 | 0.11881553 |
| Wdr20 | 372.1537339 | -0.183749369 | 0.097605836 |

| | | | |
|---|---|---|---|
| Mok | 93.83344369 | -0.376924085 | 0.125735127 |
| 4921507G05Rik | 0.309276477 | -1.945858062 | 1.712991017 |
| Gm26912 | 1.084888343 | 1.315306996 | 1.116487797 |
| Zfp839 | 119.0822902 | -0.304062687 | 0.136575968 |
| Mpc1-ps | 240.9835416 | 0.112643268 | 0.113767381 |
| Cinp | 250.286756 | 0.041693498 | 0.087410167 |
| Tecpr2 | 926.178597 | -0.598850529 | 0.079386993 |
| Rps19-ps6 | 18.4757316 | 0.208844219 | 0.254898882 |
| Ankrd9 | 131.61924 | -0.279626992 | 0.098585106 |
| Rcor1 | 394.3624715 | 0.185472933 | 0.088997777 |
| 4930595D18Rik | 0.920001838 | 0.457077892 | 1.079069538 |
| Traf3 | 916.582985 | 0.061401803 | 0.061903226 |
| Amn | 31.33882718 | -1.072172562 | 0.227768291 |
| A230087F16Rik | 11.27729028 | -1.341256274 | 0.368199919 |
| Cdc42bpb | 2200.324733 | -0.09574646 | 0.063930663 |
| A230065H16Rik | 14.31749866 | -0.086450805 | 0.342497705 |
| Exoc3l4 | 26.39184743 | 1.091177415 | 0.264622888 |
| Gm10425 | 4.124014502 | 4.858300172 | 0.996687465 |
| Tnfaip2 | 430.9565934 | 7.212508808 | 0.58545621 |
| Gm266 | 27.84625617 | 0.067310507 | 0.232502559 |
| Eif5 | 3419.805663 | 0.676893558 | 0.107105934 |
| Snora28 | 29.94917886 | -0.633024467 | 0.235361517 |
| 2810029C07Rik | 61.53833547 | -0.526709008 | 0.191792088 |
| Mark3 | 1177.712359 | 0.216355601 | 0.071259342 |
| Ckb | 9080.847841 | -0.25893624 | 0.092992229 |
| Trmt61a | 154.0841474 | 0.503670852 | 0.10387892 |
| Bag5 | 421.2921291 | 0.173102971 | 0.073619898 |
| Apopt1 | 168.4724147 | -0.207338333 | 0.102819627 |
| Gm15995 | 0.042716945 | -0.243571806 | 3.724945049 |
| Gm15996 | 0.04393175 | -0.243571806 | 3.724945049 |
| Klc1 | 4978.643089 | -0.246678008 | 0.075669315 |
| Xrcc3 | 62.34707954 | -0.673777202 | 0.169743901 |
| Zfyve21 | 80.73079552 | -0.161396532 | 0.155913268 |
| Ppp1r13b | 928.8327789 | -0.279091912 | 0.075731978 |
| 2010107E04Rik | 287.253024 | -0.196678164 | 0.16410596 |
| Tdrd9 | 6.653230718 | 0.023287212 | 0.462189134 |
| Rd3l | 8.12228703 | 0.364278226 | 0.444607777 |
| Aspg | 140.2133012 | 3.18857559 | 0.20965846 |
| Mir3073a | 0.372180926 | 1.246631404 | 1.587112566 |
| mmu-mir-3073b | 0 | NA | NA |
| Mir203 | 0 | NA | NA |
| Kif26a | 86.52596024 | -0.293727537 | 0.164687585 |
| Gm38123 | 0.492632247 | 2.2863263 | 1.429851045 |
| A730018C14Rik | 2.250253578 | 0.111828482 | 0.735774877 |
| A530016L24Rik | 0 | NA | NA |
| Tmem179 | 794.5266479 | -0.301659492 | 0.06165727 |
| Inf2 | 1281.719595 | -0.385679876 | 0.085490623 |

| | | | |
|---|---|---|---|
| Adssl1 | 268.3142132 | 0.460274942 | 0.111036292 |
| Siva1 | 44.47611933 | 0.426604963 | 0.200243945 |
| Akt1 | 1539.989558 | 0.110223316 | 0.072757239 |
| Mir6939 | 0.990373052 | 0.482869015 | 1.000474657 |
| Zbtb42 | 29.83589809 | 2.475757632 | 0.291415118 |
| Gm30238 | 24.73544283 | -0.650857282 | 0.351367215 |
| Cep170b | 3968.143558 | -0.446947658 | 0.076411171 |
| Mir6940 | 0.393711421 | -0.968363821 | 1.689503283 |
| Pld4 | 304.2279336 | 3.345102813 | 0.159114787 |
| Ahnak2 | 140.5418072 | 0.919675232 | 0.247485557 |
| BC022687 | 106.0209119 | 0.773137468 | 0.127931023 |
| Cdca4 | 92.32272779 | 0.964563973 | 0.157148221 |
| Gpr132 | 12.16430624 | 6.109957743 | 0.887631732 |
| Jag2 | 271.294481 | -0.482898291 | 0.116834255 |
| Mir6941 | 0.653922745 | -0.839177843 | 1.491040435 |
| Nudt14 | 37.15232537 | -0.095351196 | 0.190013967 |
| Gm26583 | 4.273896503 | 0.082603006 | 0.796432099 |
| Brf1 | 375.6428438 | -0.078035438 | 0.077525107 |
| Btbd6 | 332.183834 | -0.335763327 | 0.069681422 |
| 9230104M06Rik | 1.025204639 | -2.366780349 | 1.274028921 |
| Pacs2 | 2506.266713 | 0.392437187 | 0.092562592 |
| AC073562.1 | 0.303558719 | 1.552525985 | 2.024686499 |
| Tex22 | 3.833641248 | -1.12628704 | 0.554915988 |
| Mta1 | 765.704057 | 0.105794278 | 0.058628134 |
| Crip2 | 531.6473017 | 0.177551393 | 0.101094873 |
| Crip1 | 42.16258074 | 2.075348815 | 0.262178862 |
| 4930427A07Rik | 20.00319577 | 0.498398188 | 0.266949985 |
| RP24-468O24.5 | 0.060174537 | 0.237309859 | 3.724945049 |
| Tmem121 | 107.667371 | -0.073598495 | 0.116752281 |
| Gm25622 | 0 | NA | NA |
| Igha | 7.387870444 | 6.325606672 | 1.804252314 |
| Ighe | 0.136120942 | 1.15080397 | 3.595683408 |
| Ighg2c | 6.374334057 | 3.951721423 | 1.204506133 |
| Ighg2b | 4.144711669 | 5.355392938 | 1.494822244 |
| Gm37944 | 0.048949532 | -0.243569582 | 3.724945049 |
| Ighg1 | 1.048355144 | 2.118452781 | 2.230405001 |
| Ighg3 | 2.594242155 | 4.216504056 | 1.814994872 |
| Gm17421 | 0.038981395 | 0.718207094 | 3.724945049 |
| Gm9134 | 2.929093634 | 0.950751093 | 0.67939438 |
| Gm9136 | 0 | NA | NA |
| Gm5190 | 0.038981395 | 0.718207094 | 3.724945049 |
| Ighd | 2.086187063 | -1.021522324 | 0.798175655 |
| Ighm | 86.67814311 | 0.564221506 | 0.444811465 |
| Gm25747 | 0 | NA | NA |
| Gm24022 | 0 | NA | NA |
| AC073553.1 | 0 | NA | NA |
| Gm22175 | 0 | NA | NA |

| | | | |
|---|---|---|---|
| Ighj4 | 0.495312865 | 0.39792526 | 1.709473648 |
| Ighj3 | 0.07796279 | 1.072698098 | 3.721829099 |
| Ighj2 | 0.566709933 | 1.135635336 | 1.348837573 |
| Ighj1 | 0.937655582 | -0.975351605 | 1.145907584 |
| Ighd4-1 | 0 | NA | NA |
| Ighd3-2 | 0 | NA | NA |
| Ighd5-6 | 0 | NA | NA |
| Ighd2-8 | 0 | NA | NA |
| Ighd5-5 | 0 | NA | NA |
| Ighd2-7 | 0 | NA | NA |
| Ighd5-8 | 0 | NA | NA |
| Ighd5-4 | 0 | NA | NA |
| Ighd2-6 | 0 | NA | NA |
| Ighd5-7 | 0 | NA | NA |
| Ighd5-3 | 0 | NA | NA |
| Ighd2-5 | 0 | NA | NA |
| Gm37327 | 0 | NA | NA |
| Ighd5-2 | 0 | NA | NA |
| Ighd2-4 | 0 | NA | NA |
| Ighd6-2 | 0 | NA | NA |
| Gm37227 | 0 | NA | NA |
| Ighd2-3 | 0 | NA | NA |
| Ighd6-1 | 0 | NA | NA |
| Ighd1-1 | 0 | NA | NA |
| Adam6b | 0.203125505 | -1.113966547 | 2.619597531 |
| Ighd3-1 | 0 | NA | NA |
| Ighd5-1 | 0 | NA | NA |
| Gm16968 | 0 | NA | NA |
| Adam6a | 0 | NA | NA |
| Ighv5-1 | 0 | NA | NA |
| Ighv2-1 | 0 | NA | NA |
| Ighv5-2 | 0 | NA | NA |
| Ighv2-2 | 0 | NA | NA |
| Ighv5-3 | 0 | NA | NA |
| Ighv5-4 | 0.345657073 | 2.555296112 | 1.908296113 |
| Ighv6-1 | 0 | NA | NA |
| Ighv2-3 | 0.072025462 | 0.237312084 | 3.724945049 |
| Ighv5-5 | 0 | NA | NA |
| Ighv5-6 | 0.066050875 | 0.71820487 | 3.724945049 |
| Gm9517 | 109.6899113 | -0.296344757 | 0.134318453 |
| Ighv5-7 | 0 | NA | NA |
| Ighv2-4 | 0 | NA | NA |
| Ighv5-8 | 0 | NA | NA |
| Ighv5-9 | 0.097139547 | 0.718207094 | 3.724945049 |
| Ighv5-10 | 0 | NA | NA |
| Gm37187 | 0 | NA | NA |
| Ighv2-5 | 0 | NA | NA |

| | | | |
|---|---|---|---|
| Ighv5-11 | 0 | NA | NA |
| Ighv5-12 | 0 | NA | NA |
| Ighv2-6 | 0.060174537 | 0.237309859 | 3.724945049 |
| Gm37976 | 0 | NA | NA |
| Gm37418 | 0 | NA | NA |
| Ighv5-9-1 | 0 | NA | NA |
| Gm38203 | 0 | NA | NA |
| Ighv5-12-4 | 0 | NA | NA |
| Gm38205 | 0 | NA | NA |
| Ighv2-9-1 | 0 | NA | NA |
| Gm37722 | 0 | NA | NA |
| Ighv5-13 | 0 | NA | NA |
| Ighv2-6-8 | 0.046345709 | 0.71820487 | 3.724945049 |
| Gm7003 | 0 | NA | NA |
| Ighv2-7 | 0 | NA | NA |
| Rpl26-ps3 | 0 | NA | NA |
| Ighv5-15 | 0 | NA | NA |
| Ighv5-16 | 0 | NA | NA |
| Ighv5-17 | 0.427539584 | 2.305685472 | 3.695649599 |
| Ighv5-18 | 0 | NA | NA |
| Ighv2-8 | 0 | NA | NA |
| Ighv2-9 | 0.631149539 | 3.391443954 | 3.661079862 |
| Ighv5-19 | 0 | NA | NA |
| Ighv7-1 | 0 | NA | NA |
| Ighv7-2 | 0 | NA | NA |
| Ighv14-1 | 0.480014214 | 1.882366523 | 3.703952121 |
| Ighv4-1 | 0.072025462 | 0.237312084 | 3.724945049 |
| Ighv3-1 | 0 | NA | NA |
| Gm37961 | 0 | NA | NA |
| Ighv11-1 | 0 | NA | NA |
| Ighv14-2 | 0.20773507 | 1.199953802 | 2.580740143 |
| Ighv4-2 | 0 | NA | NA |
| Ighv3-2 | 0 | NA | NA |
| Ighv11-2 | 0 | NA | NA |
| Ighv14-3 | 0 | NA | NA |
| Ighv16-1 | 0 | NA | NA |
| Ighv6-2 | 0 | NA | NA |
| Ighv9-1 | 0 | NA | NA |
| Ighv12-1 | 0 | NA | NA |
| Ighv9-2 | 0 | NA | NA |
| Ighv12-2 | 0 | NA | NA |
| Ighv9-3 | 0 | NA | NA |
| Ighv7-3 | 0 | NA | NA |
| Ighv15-1 | 0 | NA | NA |
| Ighv14-4 | 0 | NA | NA |
| Gm37838 | 0 | NA | NA |
| Ighv3-3 | 0 | NA | NA |

| | | | | |
|---|---|---|---|---|
| Ighv7-4 | 0 | NA | NA | |
| Ighv3-4 | 0 | NA | NA | |
| Ighv3-5 | 0 | NA | NA | |
| Gm18551 | 0 | NA | NA | |
| Ighv13-1 | 0 | NA | NA | |
| Ighv3-6 | 0.625300531 | | 2.846423907 | 3.67826431 |
| Ighv9-4 | 0 | NA | NA | |
| Ighv3-7 | 0 | NA | NA | |
| Ighv5-21 | 0 | NA | NA | |
| Ighv3-8 | 0.038981395 | | 0.718207094 | 3.724945049 |
| Ighv8-1 | 0 | NA | NA | |
| Ighv1-1 | 0 | NA | NA | |
| Ighv13-2 | 0 | NA | NA | |
| Ighv12-3 | 0 | NA | NA | |
| Ighv6-3 | 0 | NA | NA | |
| Ighv6-4 | 0 | NA | NA | |
| Ighv6-5 | 0 | NA | NA | |
| Ighv6-6 | 0.211582616 | | 0.771702457 | 2.572998568 |
| Ighv6-7 | 0 | NA | NA | |
| Ighv8-2 | 0 | NA | NA | |
| Ighv1-2 | 0 | NA | NA | |
| mmu-mir-7094-1 | 0 | NA | NA | |
| Ighv10-1 | 0.049553047 | | 0.71820487 | 3.724945049 |
| Ighv1-3 | 0 | NA | NA | |
| Ighv1-4 | 0 | NA | NA | |
| Ighv10-2 | 0 | NA | NA | |
| Ighv1-5 | 0 | NA | NA | |
| Mir7094-2 | 0 | NA | NA | |
| Ighv10-3 | 0 | NA | NA | |
| Ighv1-6 | 0 | NA | NA | |
| Ighv1-7 | 0 | NA | NA | |
| Ighv10-4 | 0 | NA | NA | |
| Ighv1-8 | 0 | NA | NA | |
| Ighv15-2 | 0 | NA | NA | |
| Ighv1-9 | 0.287786674 | | 1.737825039 | 1.999459068 |
| Ighv1-10 | 0 | NA | NA | |
| Ighv1-11 | 0 | NA | NA | |
| Ighv1-12 | 0 | NA | NA | |
| Ighv1-13 | 0 | NA | NA | |
| Ighv1-14 | 0 | NA | NA | |
| Ighv1-15 | 0 | NA | NA | |
| Ighv1-16 | 0 | NA | NA | |
| Ighv1-17-1 | 0 | NA | NA | |
| Ighv1-17 | 0 | NA | NA | |
| Ighv1-18 | 0.194279094 | | 1.292225235 | 3.719439735 |
| Ighv1-19-1 | 0 | NA | NA | |
| Ighv1-19 | 0 | NA | NA | |

| | | | |
|---|---|---|---|
| Ighv1-20 | 0 | NA | NA |
| Ighv1-21-1 | 0 | NA | NA |
| Ighv1-21 | 0 | NA | NA |
| Ighv1-22 | 0 | NA | NA |
| Ighv1-23 | 0.097139547 | 0.718207094 | 3.724945049 |
| Ighv1-24 | 0 | NA | NA |
| Ighv1-25 | 0 | NA | NA |
| Ighv1-26 | 0 | NA | NA |
| Ighv1-27 | 0 | NA | NA |
| Ighv1-28 | 0 | NA | NA |
| Ighv1-29 | 0 | NA | NA |
| Ighv1-30 | 0 | NA | NA |
| Ighv1-31 | 0 | NA | NA |
| Ighv1-32 | 0 | NA | NA |
| Ighv1-33 | 0.097139547 | 0.718207094 | 3.724945049 |
| Ighv1-34 | 0 | NA | NA |
| Ighv1-35 | 0 | NA | NA |
| Ighv1-36 | 0 | NA | NA |
| Ighv1-37 | 0 | NA | NA |
| Ighv1-38 | 0 | NA | NA |
| Ighv1-39 | 0 | NA | NA |
| Ighv1-40 | 0 | NA | NA |
| Ighv1-41 | 0 | NA | NA |
| Ighv1-42 | 0 | NA | NA |
| Ighv1-43 | 0 | NA | NA |
| Ighv1-44 | 0 | NA | NA |
| Ighv1-45 | 0 | NA | NA |
| Ighv1-46 | 0 | NA | NA |
| Ighv1-47 | 0 | NA | NA |
| Ighv8-3 | 0 | NA | NA |
| Ighv8-4 | 0 | NA | NA |
| Ighv1-48 | 0 | NA | NA |
| Ighv1-49 | 0 | NA | NA |
| Ighv8-5 | 0 | NA | NA |
| Ighv1-50 | 0 | NA | NA |
| Ighv1-51 | 0 | NA | NA |
| Ighv1-52 | 0.072025462 | 0.237312084 | 3.724945049 |
| Ighv1-53 | 0 | NA | NA |
| Ighv8-6 | 0 | NA | NA |
| Gm9227 | 0 | NA | NA |
| Ighv1-54 | 0 | NA | NA |
| Ighv1-55 | 0 | NA | NA |
| Gm7142 | 0 | NA | NA |
| Ighv1-56 | 0 | NA | NA |
| Gm7019 | 1.468020652 | 3.008057949 | 1.094989442 |
| Ighv8-7 | 0 | NA | NA |
| Gm30948 | 0 | NA | NA |

| | | | |
|---|---|---|---|
| Ighv1-57 | 0 | NA | NA |
| Ighv8-8 | 0.144050924 | 0.237309606 | 3.724945049 |
| Ighv1-58 | 0 | NA | NA |
| Ighv1-59 | 0 | NA | NA |
| Ighv1-60 | 0 | NA | NA |
| Ighv1-61 | 0 | NA | NA |
| Ighv1-62 | 0 | NA | NA |
| Ighv1-62-1 | 0 | NA | NA |
| Gm19331 | 0 | NA | NA |
| Gm37511 | 0 | NA | NA |
| Gm38184 | 0 | NA | NA |
| Ighv1-62-2 | 0 | NA | NA |
| Ighv1-62-3 | 0 | NA | NA |
| Ighv8-9 | 0 | NA | NA |
| RP24-350F17.8 | 0 | NA | NA |
| Ighv1-63 | 0.046345709 | 0.71820487 | 3.724945049 |
| Gm27722 | 0 | NA | NA |
| Ighv1-64 | 0 | NA | NA |
| Ighv8-10 | 0 | NA | NA |
| RP23-218B2.1 | 0 | NA | NA |
| Ighv1-65 | 0 | NA | NA |
| RP23-218B2.3 | 0.230454321 | 0.238068929 | 2.186165662 |
| Ighv8-11 | 0.066050875 | 0.71820487 | 3.724945049 |
| RP23-218B2.5 | 0 | NA | NA |
| Ighv1-66 | 0 | NA | NA |
| Ighv1-67 | 0.060174537 | 0.237309859 | 3.724945049 |
| Ighv1-68 | 0 | NA | NA |
| RP23-218B2.9 | 0 | NA | NA |
| RP23-218B2.10 | 0 | NA | NA |
| Ighv1-69 | 0 | NA | NA |
| Ighv8-12 | 0 | NA | NA |
| RP23-218B2.13 | 0 | NA | NA |
| Ighv1-70 | 0 | NA | NA |
| Mir6388 | 0 | NA | NA |
| Ighv1-71 | 0 | NA | NA |
| Ighv1-72 | 0 | NA | NA |
| Ighv8-13 | 0 | NA | NA |
| RP23-123N21.1 | 0 | NA | NA |
| Ighv1-73 | 0 | NA | NA |
| Ighv1-74 | 0.136120942 | 1.15080397 | 3.595683408 |
| Ighv8-14 | 0 | NA | NA |
| RP23-123N21.5 | 0 | NA | NA |
| RP23-123N21.6 | 0 | NA | NA |
| Ighv1-75 | 0 | NA | NA |
| Ighv8-15 | 0 | NA | NA |
| Ighv1-76 | 0 | NA | NA |
| Ighv8-16 | 0 | NA | NA |

| | | | |
|---|---|---|---|
| Ighv1-77 | 0.046345709 | 0.71820487 | 3.724945049 |
| Ighv1-78 | 0 | NA | NA |
| RP23-172K2.2 | 0 | NA | NA |
| RP23-172K2.1 | 0 | NA | NA |
| Ighv1-79 | 0 | NA | NA |
| RP23-172K2.4 | 2.956099666 | -0.959812917 | 0.707519764 |
| Ighv1-80 | 0 | NA | NA |
| Ighv1-81 | 0 | NA | NA |
| RP23-172K2.8 | 0 | NA | NA |
| Olfr241-ps1 | 0 | NA | NA |
| Ighv1-82 | 0 | NA | NA |
| Ighv1-83 | 0 | NA | NA |
| Ighv1-84 | 0 | NA | NA |
| RP23-172K2.12 | 5.084652238 | 1.893546421 | 0.545390028 |
| Ighv1-85 | 0.072025462 | 0.237312084 | 3.724945049 |
| Ighv1-86 | 0 | NA | NA |
| Zfp386 | 523.1915969 | 0.115577721 | 0.082537019 |
| Vipr2 | 33.917483 | -0.390880017 | 0.236715352 |
| Gm20658 | 0.517478627 | 2.418294715 | 1.307978932 |
| AC105950.1 | 0 | NA | NA |
| Gm23732 | 0 | NA | NA |
| Wdr60 | 178.5716657 | -0.298950553 | 0.109945204 |
| Gm23190 | 0.086648695 | -0.72445792 | 3.724945049 |
| Gm11027 | 0 | NA | NA |
| Esyt2 | 1140.517075 | -0.18005933 | 0.055283627 |
| Ncapg2 | 136.9875684 | 1.299098068 | 0.172270237 |
| Gm25112 | 0.475792389 | -0.274602773 | 1.467846115 |
| Gm27649 | 0.04393175 | -0.243571806 | 3.724945049 |
| Ptprn2 | 2936.458257 | -0.871532502 | 0.099269507 |
| Gm6750 | 0.877833841 | 0.23070835 | 1.177211182 |
| Gm17807 | 4.466807706 | -2.121602108 | 0.58032151 |
| Gm10421 | 10.74823256 | -0.829750915 | 0.357044103 |
| Gm5441 | 1.251482101 | -0.676285833 | 0.880822704 |
| Mir153 | 1.154465371 | -3.302245431 | 1.107564044 |
| Rapgef5 | 853.7827578 | -0.380131894 | 0.096953868 |
| Gm24741 | 1.59128742 | -0.298537406 | 1.133328549 |
| Cdca7l | 18.95984674 | 0.737065007 | 0.303488365 |
| Gm20613 | 0.046345709 | 0.71820487 | 3.724945049 |
| Dnah11 | 127.3083906 | -0.683502939 | 0.146713927 |
| Sp4 | 671.9126227 | -0.081597144 | 0.096697412 |
| Gm25577 | 0.046345709 | 0.71820487 | 3.724945049 |
| Gm22537 | 0 | NA | NA |
| Sp8 | 103.1593826 | -0.231809473 | 0.573009221 |
| Abcb5 | 0 | NA | NA |
| Itgb8 | 2067.75423 | 0.467341615 | 0.146986448 |
| Gm6768 | 116.6792349 | 0.354773632 | 0.106727643 |
| Macc1 | 2.369395584 | 0.320034689 | 0.653897589 |

| | | | |
|---|---|---|---|
| Gm25675 | 0 | NA | NA |
| Tmem196 | 135.2367115 | -0.530086054 | 0.105049457 |
| Gm27236 | 2.533080183 | -0.448306694 | 0.615219227 |
| Gm17177 | 0 | NA | NA |
| Speer6-ps1 | 0 | NA | NA |
| Gm16505 | 1.0973389 | -2.481270509 | 1.201626155 |
| 2810429I04Rik | 28.4214383 | -0.993490743 | 0.212431586 |
| Gdi2 | 2568.627456 | 0.315031091 | 0.08184989 |
| Fam208b | 1293.848084 | 0.665393253 | 0.092640834 |
| Asb13 | 641.9525158 | 0.414968885 | 0.115259138 |
| Gm23084 | 0.981575226 | 1.58351837 | 1.013329981 |
| Calml3 | 0.554260052 | 0.759641946 | 3.722221816 |
| Calm4 | 0.780001735 | 0.23730057 | 3.724945049 |
| Calm5 | 0.045882455 | 0.237312084 | 3.724945049 |
| AC132239.1 | 0 | NA | NA |
| Net1 | 365.646222 | 0.110567383 | 0.092066125 |
| Tubal3 | 0.038981395 | 0.718207094 | 3.724945049 |
| Ucn3 | 2.74335279 | -0.456396396 | 0.699292583 |
| Gm22460 | 0 | NA | NA |
| Akr1c14 | 23.64608385 | 0.819517799 | 0.368771577 |
| Akr1c18 | 8.892648978 | -0.199755688 | 0.355554597 |
| Akr1c13 | 6.039600961 | 3.080358669 | 0.547697217 |
| Akr1c19 | 0.060174537 | 0.237309859 | 3.724945049 |
| Marcksl1-ps4 | 0 | NA | NA |
| Akr1c12 | 0.642786378 | 1.238296077 | 1.443619308 |
| Akr1c6 | 0.085327104 | 1.179809211 | 3.723500796 |
| Akr1c20 | 0.875592853 | 3.576701821 | 1.469146958 |
| Akr1c21 | 0 | NA | NA |
| Akr1e1 | 194.9019369 | -0.063334314 | 0.116299599 |
| Rpl29-ps2 | 3.103436511 | -0.477885354 | 0.583760181 |
| Gm5444 | 0.134148234 | 0.23732184 | 3.624632408 |
| Gm24790 | 0 | NA | NA |
| Gm26043 | 0 | NA | NA |
| 1700016G22Rik | 0.413724333 | 2.676099005 | 1.586473845 |
| Klf6 | 1264.230787 | 1.844395249 | 0.170478211 |
| Gm26259 | 0 | NA | NA |
| Gm28465 | 0 | NA | NA |
| Pitrm1 | 461.3651909 | 0.248633794 | 0.073544139 |
| Pfkp | 2522.968723 | -0.066645778 | 0.07423186 |
| Gm10029 | 6.775184329 | 1.259228982 | 0.424636748 |
| Gm22689 | 0 | NA | NA |
| Gm9742 | 0.283836568 | 1.381499229 | 2.075593192 |
| Adarb2 | 565.2457729 | -0.759631285 | 0.072747339 |
| Gm10357 | 4.418419051 | 0.627628191 | 0.515834536 |
| Wdr37 | 1368.014662 | -0.224516712 | 0.071810906 |
| Idi1 | 182.2969244 | 0.123291484 | 0.177222777 |
| Gm22750 | 0.171083144 | 1.76201309 | 3.383974148 |

| | | | |
|---|---|---|---|
| Rpl10a-ps2 | 0.785454149 | 2.710635192 | 1.15685818 |
| Gm9745 | 0.104637558 | 0.23732098 | 3.724945049 |
| Gm25274 | 0.066050875 | 0.71820487 | 3.724945049 |
| Gm22115 | 0.115603922 | 1.209894437 | 3.72418822 |
| Idi2 | 0.326649557 | 0.214325087 | 1.474558922 |
| Gm23923 | 0.066050875 | 0.71820487 | 3.724945049 |
| Gtpbp4 | 492.4961443 | 0.557107007 | 0.114333273 |
| Larp4b | 1451.804564 | 0.090063799 | 0.052849098 |
| Gm23553 | 10.68005917 | -0.086879038 | 0.370887548 |
| Gm28155 | 0 NA | | NA |
| Dip2c | 2087.5135 | 0.111121026 | 0.071528243 |
| Gm26601 | 155.8395443 | -0.785105374 | 0.387981297 |
| Gm23496 | 0.048949532 | -0.243569582 | 3.724945049 |
| Zmynd11 | 1907.336184 | -0.351752347 | 0.075575185 |
| Gm24187 | 251.7405501 | 0.90800992 | 0.979984002 |
| Chrm3 | 417.2410317 | -1.01247614 | 0.109173059 |
| Gm26861 | 0.849695755 | -0.60302195 | 1.067468387 |
| Gm23415 | 0 NA | | NA |
| Gm25496 | 0 NA | | NA |
| Ryr2 | 4975.297084 | -1.003238649 | 0.152896188 |
| Gm26956 | 0.116944185 | 1.302646088 | 3.719336636 |
| Mtr | 314.5275728 | 0.084252219 | 0.115405818 |
| Actn2 | 215.0662038 | -1.017818901 | 0.204612379 |
| Gm5445 | 1.336133173 | -0.999616712 | 0.9212743 |
| Heatr1 | 496.059456 | 0.41853687 | 0.089643565 |
| Lgals8 | 902.3953984 | 1.819581744 | 0.141897995 |
| Edaradd | 6.858033054 | -1.060057551 | 0.427972734 |
| Ero1lb | 286.3774935 | -0.134788867 | 0.085843017 |
| Gm25483 | 3.19383198 | -0.292468246 | 0.683442592 |
| Gpr137b-ps | 100.0559785 | 0.394758512 | 0.153797827 |
| Gm24065 | 0 NA | | NA |
| Gm2399 | 0.304316145 | 0.256525124 | 1.59669634 |
| Prl2c3 | 1.422091401 | -1.827159921 | 0.912081756 |
| Prl2c2 | 0.494725653 | -0.828812432 | 1.47086057 |
| Prl2c5 | 0.049553047 | 0.71820487 | 3.724945049 |
| Gm2423 | 35.43526429 | 0.189751235 | 0.249364384 |
| Gm26133 | 0 NA | | NA |
| Gm25804 | 0 NA | | NA |
| Gpr137b | 160.9115362 | 0.100793061 | 0.113626509 |
| Gm23081 | 0 NA | | NA |
| Gm7426 | 0 NA | | NA |
| Nid1 | 422.1371149 | 1.112721461 | 0.145377387 |
| Lyst | 2594.837581 | 0.209384938 | 0.100979972 |
| Gng4 | 505.6373642 | -0.468606705 | 0.253451363 |
| Gm25271 | 3.698276596 | -1.446578604 | 0.678236636 |
| B3galnt2 | 227.2140593 | -0.297203143 | 0.109147887 |
| Gm18856 | 1.228493641 | 2.411854589 | 1.153286141 |

| | | | |
|---|---|---|---|
| Tbce | 525.6659255 | -0.163655267 | 0.103600721 |
| Ggps1 | 552.9079473 | 0.233899735 | 0.080842531 |
| Arid4b | 1466.024357 | 0.01052686 | 0.080840844 |
| Gm26129 | 7.167244977 | -0.559766747 | 0.385265095 |
| n-R5s52 | 0 | NA | NA |
| Hecw1 | 1632.790211 | -0.752304077 | 0.097380854 |
| Mrpl32 | 170.5037384 | 0.64992793 | 0.113716767 |
| Psma2 | 925.2712897 | 0.929496528 | 0.113682189 |
| Gm25444 | 0.046345709 | 0.71820487 | 3.724945049 |
| AW209491 | 188.160879 | -0.202578341 | 0.14553938 |
| Gm6556 | 34.49633554 | -0.644964492 | 0.239467876 |
| Gm26645 | 23.14899875 | -1.103998238 | 0.304720886 |
| Gli3 | 305.6293553 | -0.113134659 | 0.114365139 |
| Inhba | 124.8662631 | 2.240231266 | 0.403866443 |
| Gm22628 | 0.250684468 | -0.766137155 | 2.133838007 |
| Sugct | 24.65360541 | -0.404021995 | 0.234699815 |
| Gm26098 | 0 | NA | NA |
| Gm27043 | 0 | NA | NA |
| Gm25020 | 0 | NA | NA |
| Gm24873 | 1.427842546 | 0.33694632 | 0.873031423 |
| Mplkip | 150.6828418 | 0.2930283 | 0.212872179 |
| Cdk13 | 1371.430534 | 0.017611204 | 0.05933139 |
| Mir466i | 0.78113724 | 2.097415165 | 1.14225298 |
| Gm24018 | 0.047948698 | -0.243569582 | 3.724945049 |
| Gm18859 | 0.219031842 | 1.195339194 | 2.578179985 |
| Rala | 653.2611828 | 0.169116972 | 0.088260376 |
| Yae1d1 | 237.983152 | 0.108884087 | 0.091938169 |
| Gm37888 | 0.089089296 | 0.237305797 | 3.724945049 |
| Vdac3-ps1 | 215.80711 | 0.215962959 | 0.117018091 |
| Pou6f2 | 51.8533924 | -0.311385604 | 0.174972462 |
| Vps41 | 1555.895699 | -0.202760253 | 0.058898367 |
| Gm26172 | 0.049553047 | 0.71820487 | 3.724945049 |
| Amph | 1915.694854 | -0.514799867 | 0.072941369 |
| RP23-167F18.7 | 0 | NA | NA |
| Tcrg-V7 | 0.107803068 | 0.719187033 | 3.724874737 |
| Tcrg-V4 | 0.038981395 | 0.718207094 | 3.724945049 |
| Tcrg-V6 | 0.075043205 | 0.237309859 | 3.724945049 |
| Tcrg-V5 | 0.192276286 | 1.280626648 | 3.260418707 |
| Trgj1 | 0 | NA | NA |
| Tcrg-C1 | 0.641965332 | 2.425886079 | 1.355327488 |
| Tcrg-V3 | 0 | NA | NA |
| Trgj3 | 0 | NA | NA |
| Tcrg-C3 | 0.092228164 | 0.718202646 | 3.724945049 |
| Tcrg-C2 | 0.588064751 | 1.057537607 | 1.470029257 |
| Trgj2 | 0 | NA | NA |
| Trgv2 | 0.097139547 | 0.718207094 | 3.724945049 |
| Tcrg-V1 | 0 | NA | NA |

| | | | |
|---|---|---|---|
| Trgj4 | 0 | NA | NA |
| Tcrg-C4 | 0.312934076 | 1.697531427 | 2.035499258 |
| Stard3nl | 346.4093214 | 0.363035007 | 0.090782093 |
| RP23-306G1.11 | 25.52656834 | -0.449300686 | 0.281450167 |
| RP23-306G1.12 | 0.525666135 | -1.29384395 | 1.362615715 |
| Epdr1 | 994.4722949 | -0.105411796 | 0.072818174 |
| Sfrp4 | 29.48509951 | 1.123290178 | 0.223068855 |
| Nme8 | 0.465692922 | 1.147401019 | 1.372362378 |
| A530099J19Rik | 0.780625029 | 3.013242674 | 1.193393078 |
| Gpr141 | 71.74378744 | 8.12954286 | 0.781667651 |
| Gm5446 | 4.31637155 | 0.952076528 | 0.494916371 |
| Elmo1 | 1541.907887 | -0.09082378 | 0.056193764 |
| Gm7611 | 0 | NA | NA |
| Aoah | 31.82302678 | 4.265701188 | 0.455015332 |
| Gm25605 | 0.583760085 | 2.375020035 | 1.332607234 |
| Olfr1370 | 0 | NA | NA |
| Olfr1369-ps1 | 0 | NA | NA |
| Olfr263 | 0.213549163 | 1.094701859 | 2.569305339 |
| Olfr1368 | 0 | NA | NA |
| Trim27 | 275.0576437 | 0.353660561 | 0.07857696 |
| Gpx5 | 0 | NA | NA |
| Gm11270 | 0 | NA | NA |
| Gpx6 | 2.180372029 | 0.016296331 | 0.657755688 |
| Olfr1367 | 0.046345709 | 0.71820487 | 3.724945049 |
| Zscan12 | 64.42265566 | 0.124487083 | 0.149929333 |
| Gm11271 | 0.280341511 | -0.722977944 | 1.78846901 |
| Zkscan3 | 573.1062161 | -0.39363932 | 0.131989102 |
| Mir6942 | 2.650403497 | -0.82224498 | 0.598467116 |
| Pgbd1 | 43.29911835 | -0.74321237 | 0.175286335 |
| 4930470G03Rik | 1.381930285 | 0.830481439 | 0.907251907 |
| Zscan26 | 980.2046494 | -0.194784502 | 0.057726433 |
| Mir1896 | 0 | NA | NA |
| Nkapl | 5.734391589 | 0.444568796 | 0.468683291 |
| Zkscan4 | 37.6093419 | -0.09970821 | 0.18679741 |
| Gm10065 | 0.298142035 | 0.700197384 | 3.721584259 |
| Gm23779 | 0 | NA | NA |
| AK157302 | 180.0808189 | -0.354016357 | 0.089639163 |
| AL589742.1 | 0 | NA | NA |
| AL589742.2 | 0 | NA | NA |
| Gm11273 | 0 | NA | NA |
| Zfp389 | 4.935521334 | 0.2124991 | 0.462243193 |
| Zkscan8 | 581.7328674 | -0.113611542 | 0.101435692 |
| Olfr1366 | 0.049553047 | 0.71820487 | 3.724945049 |
| Olfr1535 | 0 | NA | NA |
| Olfr1364 | 0.097899064 | -0.724453471 | 3.724945049 |
| Olfr1362 | 0 | NA | NA |
| Olfr11 | 0.091666476 | -0.724455696 | 3.724945049 |

| | | | |
|---|---|---|---|
| Olfr1361 | 0.061920613 | 0.718207094 | 3.724945049 |
| Olfr1360 | 0 | NA | NA |
| Olfr1359 | 0.062426873 | 0.237312084 | 3.724945049 |
| Hist1h2bl | 17.04762427 | 1.575267221 | 0.378537561 |
| Hist1h2ai | 8.571363186 | 1.05400487 | 0.397072685 |
| AL589651.1 | 0.12092566 | 0.237307635 | 3.724945049 |
| Hist1h3h | 40.08083796 | 0.747843211 | 0.230483136 |
| AL589651.2 | 0 | NA | NA |
| Hist1h2aj | 0.635935163 | 1.798508262 | 1.224574289 |
| Hist1h2bm | 22.0061962 | 1.681322702 | 0.330903405 |
| Hist1h4j | 57.33528979 | 0.727483375 | 0.221133817 |
| Hist1h4k | 181.1049781 | 0.291922646 | 0.148772986 |
| Hist1h2ak | 39.92311535 | 1.198519339 | 0.285271736 |
| AL589651.3 | 0.07796279 | 1.072698098 | 3.721829099 |
| Hist1h2bn | 10.00898834 | 1.086624174 | 0.381125377 |
| Gm11274 | 1.966252851 | 1.653003018 | 0.738261919 |
| Hist1h1b | 62.13502671 | 0.987795431 | 0.226002264 |
| Hist1h3i | 77.67695009 | 0.826531068 | 0.239569984 |
| Hist1h2an | 2.07791248 | 1.295766216 | 0.734159744 |
| AL589651.4 | 0 | NA | NA |
| Hist1h2bp | 6.144754156 | 0.819292389 | 0.423054112 |
| Hist1h2bq | 12.73633966 | 1.299841711 | 0.321432583 |
| Hist1h2ao | 3.983646474 | 0.659709297 | 0.522993057 |
| AL589879.1 | 0.072025462 | 0.237312084 | 3.724945049 |
| Hist1h4m | 15.4295438 | 0.870279747 | 0.314819371 |
| Hist1h4n | 166.7008044 | 0.629070369 | 0.165568405 |
| Hist1h2ap | 1.700876845 | 0.897334206 | 0.97471121 |
| AL589879.3 | 0.060174537 | 0.237309859 | 3.724945049 |
| Hist1h2br | 3.568372849 | 1.164350389 | 0.614263455 |
| Gm11278 | 0 | NA | NA |
| Mir1983 | 1.672508104 | 0.630384272 | 0.817233746 |
| Gm11279 | 0.061920613 | 0.718207094 | 3.724945049 |
| AL589879.2 | 0.514605876 | 1.835609355 | 1.424120256 |
| Gm11280 | 0.088534442 | 1.179809211 | 3.723500796 |
| Zfp184 | 35.438407 | -0.57289344 | 0.176738189 |
| Gm11281 | 0 | NA | NA |
| Gm11282 | 0.068097359 | -0.243569582 | 3.724945049 |
| Pom121l2 | 0.476324185 | -0.243564052 | 1.582887117 |
| Prss16 | 7.136397789 | 0.74201706 | 0.535411098 |
| Gm11290 | 0 | NA | NA |
| Gm11291 | 0 | NA | NA |
| Gm11292 | 0.121388914 | 0.718200421 | 3.724945049 |
| AL590614.1 | 0.110870145 | 0.238305784 | 3.724874707 |
| Hist1h2ah | 3.414149285 | 2.277795363 | 0.722901671 |
| Hist1h2bk | 6.592695331 | 1.293339902 | 0.429006817 |
| Hist1h4i | 181.0810821 | 0.984241888 | 0.180150572 |
| Hist1h2ag | 7.85075097 | 0.839563638 | 0.396528371 |

| | | | |
|---|---|---|---|
| AL590614.2 | 0.038981395 | 0.718207094 | 3.724945049 |
| Hist1h2bj | 11.38635759 | 1.456588287 | 0.389531515 |
| Vmn1r-ps92 | 0 | NA | NA |
| Vmn1r-ps93 | 0 | NA | NA |
| Gm11285 | 0 | NA | NA |
| Gm26300 | 0 | NA | NA |
| Vmn1r188 | 0 | NA | NA |
| Vmn1r-ps94 | 0.341912768 | 2.104933275 | 1.976758136 |
| Vmn1r189 | 0 | NA | NA |
| Vmn1r-ps95 | 0 | NA | NA |
| Vmn1r-ps97 | 0 | NA | NA |
| Vmn1r190-ps | 0 | NA | NA |
| Gm11284 | 0.374772292 | 0.704022001 | 1.559683085 |
| Vmn1r-ps99 | 0 | NA | NA |
| Vmn1r191 | 0 | NA | NA |
| Vmn1r192 | 0 | NA | NA |
| Gm25084 | 0 | NA | NA |
| Gm23282 | 0.060174537 | 0.237309859 | 3.724945049 |
| Vmn1r193 | 0 | NA | NA |
| Vmn1r194 | 0.049553047 | 0.71820487 | 3.724945049 |
| Vmn1r195 | 0 | NA | NA |
| Gm11295 | 0.521690028 | -0.162274088 | 1.277621377 |
| Vmn1r196 | 0 | NA | NA |
| Vmn1r-ps100 | 0 | NA | NA |
| Vmn1r197 | 0.066050875 | 0.71820487 | 3.724945049 |
| Vmn1r198 | 0 | NA | NA |
| Vmn1r199 | 0.038981395 | 0.718207094 | 3.724945049 |
| Vmn1r-ps102 | 0 | NA | NA |
| Vmn1r200 | 0.302366008 | 2.249050225 | 1.988401742 |
| Vmn1r-ps103 | 0 | NA | NA |
| Vmn1r-ps104 | 0 | NA | NA |
| Gm11298 | 0.060174537 | 0.237309859 | 3.724945049 |
| Vmn1r201 | 0 | NA | NA |
| Vmn1r-ps106 | 0 | NA | NA |
| Vmn1r-ps107 | 0 | NA | NA |
| Vmn1r-ps108 | 0 | NA | NA |
| Vmn1r202 | 0 | NA | NA |
| Gm11300 | 0.038981395 | 0.718207094 | 3.724945049 |
| Vmn1r203 | 0 | NA | NA |
| Vmn1r204 | 0 | NA | NA |
| Vmn1r205 | 0.044191549 | -0.243571806 | 3.724945049 |
| Vmn1r206 | 1.041252144 | -1.818399111 | 1.085133135 |
| Vmn1r-ps109 | 0.733882059 | -1.53926814 | 1.077537961 |
| Vmn1r-ps110 | 1.137431431 | -0.773282288 | 0.995123314 |
| Gm11307 | 0 | NA | NA |
| Vmn1r-ps111 | 0.097139547 | 0.718207094 | 3.724945049 |
| Gm11309 | 0 | NA | NA |

| | | | |
|---|---|---|---|
| Vmn1r207-ps | 0.447844902 | 1.538300833 | 1.532152092 |
| Vmn1r-ps112 | 0 | NA | NA |
| Vmn1r-ps113 | 1.019683961 | 0.733256058 | 1.094680925 |
| Gm11313 | 0 | NA | NA |
| Vmn1r208 | 0 | NA | NA |
| Gm11314 | 0 | NA | NA |
| Vmn1r209 | 0.344645118 | 0.193588992 | 1.555429846 |
| Vmn1r210 | 0.134880151 | 1.580366464 | 2.92708711 |
| Vmn1r-ps114 | 0 | NA | NA |
| Vmn1r211 | 0 | NA | NA |
| Vmn1r-ps116 | 0 | NA | NA |
| Gm23774 | 0 | NA | NA |
| Vmn1r212 | 0.038981395 | 0.718207094 | 3.724945049 |
| Gm11320 | 0 | NA | NA |
| Gm11321 | 0 | NA | NA |
| Vmn1r-ps120 | 0 | NA | NA |
| Gm11322 | 0 | NA | NA |
| Gm11318 | 0.044544648 | 0.237309859 | 3.724945049 |
| Vmn1r-ps123 | 0 | NA | NA |
| Vmn1r213 | 0.109982625 | 0.237318756 | 3.724945049 |
| Vmn1r-ps125 | 0 | NA | NA |
| Vmn1r214 | 0 | NA | NA |
| Vmn1r-ps126 | 0 | NA | NA |
| Vmn1r215 | 0 | NA | NA |
| Vmn1r216 | 0 | NA | NA |
| Gm11325 | 0 | NA | NA |
| Vmn1r-ps127 | 0 | NA | NA |
| Vmn1r217 | 0 | NA | NA |
| Vmn1r-ps128 | 0 | NA | NA |
| Vmn1r218 | 0 | NA | NA |
| Vmn1r219 | 0 | NA | NA |
| Vmn1r220 | 0.038981395 | 0.718207094 | 3.724945049 |
| Vmn1r-ps132 | 0 | NA | NA |
| Vmn1r221 | 0 | NA | NA |
| Vmn1r222 | 0 | NA | NA |
| Vmn1r223 | 0 | NA | NA |
| Gm11331 | 0 | NA | NA |
| Vmn1r-ps135 | 0 | NA | NA |
| 4930557F10Rik | 0.136120942 | 1.15080397 | 3.595683408 |
| 4933404K08Rik | 0.044544648 | 0.237309859 | 3.724945049 |
| Gm11333 | 0 | NA | NA |
| 4930586N03Rik | 0.124596252 | 0.718200615 | 3.714681068 |
| Zfp322a | 495.523674 | -0.353433225 | 0.112467764 |
| Gm11334 | 0 | NA | NA |
| Abt1 | 128.8824264 | 0.40316888 | 0.129409303 |
| C230035I16Rik | 12.59447301 | 1.694621887 | 0.458689634 |
| Btn1a1 | 18.16397336 | 5.336057976 | 0.940044695 |

| | | | |
|---|---|---|---|
| Btn2a2 | 7.935836238 | -0.589541913 | 0.444701913 |
| AC034285.1 | 0 | NA | NA |
| Gm11335 | 0.44640837 | 0.213853161 | 1.343490659 |
| Hist1h4h | 362.5692348 | -0.19486204 | 0.179245384 |
| Hist1h2af | 13.35131425 | 2.517842604 | 0.517123314 |
| AC034285.3 | 0.10503227 | 1.179809211 | 3.723500796 |
| Hist1h3g | 12.55213016 | 1.46780428 | 0.407357216 |
| Hist1h2bh | 13.53059552 | 1.867863683 | 0.412580244 |
| Hist1h3f | 117.9595612 | 0.90840088 | 0.180806907 |
| Hist1h4f | 133.4200694 | 0.066750925 | 0.185087903 |
| Hist1h1d | 190.8763881 | 0.846738728 | 0.195210967 |
| Hist1h3e | 180.3700906 | 0.452243483 | 0.137978919 |
| Hist1h2ae | 13.93887753 | 0.535221442 | 0.385713863 |
| AC034285.2 | 0.480269614 | -0.235399999 | 1.392178314 |
| Hist1h2bg | 107.7133796 | 0.702945865 | 0.113452212 |
| Hist1h2bf | 17.61300926 | 1.239079902 | 0.363877596 |
| Hist1h2ad | 7.472743362 | 1.114628616 | 0.414116283 |
| AC034285.4 | 0.068097359 | -0.243569582 | 3.724945049 |
| Hist1h3d | 38.7205137 | 0.40869601 | 0.236975491 |
| Hist1h4d | 957.8385019 | 0.585589723 | 0.126462308 |
| Hist1h2be | 153.5786973 | 0.832823566 | 0.149050742 |
| Gm11398 | 0 | NA | NA |
| Hist1h1e | 485.7404665 | 0.42436798 | 0.180644888 |
| Gm22452 | 0 | NA | NA |
| Gm24991 | 0.1140246 | 0.718202646 | 3.724945049 |
| Hist1h2ac | 9.474268641 | 0.456411425 | 0.384243338 |
| AL592149.1 | 0.048949532 | -0.243569582 | 3.724945049 |
| Hist1h2bc | 147.070612 | 1.240407194 | 0.143408804 |
| Hist1h1t | 0.61465116 | 1.729047047 | 1.220888539 |
| Hist1h4c | 207.5103062 | 0.315873241 | 0.167366571 |
| Hfe | 68.88156831 | 1.897916074 | 0.212778026 |
| Gm11338 | 0 | NA | NA |
| Hist1h1c | 665.5210639 | 1.551879984 | 0.166233206 |
| Hist1h3c | 20.18455272 | 2.555757583 | 0.37265821 |
| Gm11336 | 0.090890357 | 0.718200421 | 3.724945049 |
| AL590388.2 | 0.068097359 | -0.243569582 | 3.724945049 |
| Hist1h2bb | 38.92068366 | 0.889535496 | 0.293547397 |
| Hist1h2ab | 19.95108352 | 1.728730882 | 0.268210832 |
| AL590388.1 | 0.038981395 | 0.718207094 | 3.724945049 |
| Hist1h3b | 12.81895394 | 2.124295032 | 0.397675587 |
| 4930558J22Rik | 0.487944759 | -0.913224744 | 1.378816442 |
| Hist1h4b | 52.56315827 | 0.08567112 | 0.235578474 |
| Hist1h4a | 71.08590372 | 0.067262957 | 0.145451209 |
| Hist1h3a | 21.90840992 | 1.080351409 | 0.266124529 |
| Hist1h1a | 77.50845308 | 0.807385844 | 0.199851973 |
| Trim38 | 1.17402693 | 0.310895218 | 0.887293264 |
| Gm11337 | 0 | NA | NA |

| | | | |
|---|---|---|---|
| Slc17a2 | 0.366957926 | 1.84447623 | 1.383360958 |
| Slc17a3 | 0.879126543 | 0.233566363 | 1.347190495 |
| Slc17a1 | 0.066050875 | 0.71820487 | 3.724945049 |
| Slc17a4 | 0 | NA | NA |
| Snrp1c-ps2 | 0 | NA | NA |
| Hist1h2ba | 0.304971633 | 2.165437301 | 1.718719143 |
| Hist1h2aa | 0 | NA | NA |
| AL606464.1 | 0 | NA | NA |
| Scgn | 17.90384641 | 0.878834637 | 0.554164418 |
| Gm11339 | 0 | NA | NA |
| Lrrc16a | 976.9628538 | 0.753674957 | 0.091928841 |
| Gm11343 | 0 | NA | NA |
| Gm11340 | 0.044191549 | -0.243571806 | 3.724945049 |
| Gm23340 | 0 | NA | NA |
| Gm11341 | 0 | NA | NA |
| Cmah | 7.485215113 | 2.034015881 | 0.506751404 |
| Gm11342 | 0.062426873 | 0.237312084 | 3.724945049 |
| Gm11345 | 0.130840244 | -1.076724316 | 2.947200097 |
| Gm11344 | 0 | NA | NA |
| Gm22013 | 0 | NA | NA |
| Fam65b | 791.2498213 | -0.583553315 | 0.115505873 |
| Gm11346 | 3.822611367 | 1.044662963 | 0.546511083 |
| C530050E15Rik | 25.2429927 | -1.002325045 | 0.343389081 |
| 1700016G14Rik | 1.09380521 | 0.251039154 | 0.902257363 |
| Gmnn | 28.95427006 | 0.261598006 | 0.250989039 |
| BC005537 | 3337.74108 | 0.648498329 | 0.051359955 |
| Gm22358 | 0.227031315 | -1.225057273 | 2.14026266 |
| Acot13 | 187.4166705 | -0.392286561 | 0.157431424 |
| Tdp2 | 199.2904976 | -0.113920862 | 0.092182262 |
| D130043K22Rik | 395.2440717 | -0.882935883 | 0.122627417 |
| 4932702P03Rik | 1.792617364 | -1.088721288 | 0.760517659 |
| Aldh5a1 | 1047.224437 | -1.094819547 | 0.065969873 |
| 9330162012Rik | 49.19052179 | -0.528547716 | 0.249073945 |
| Gpld1 | 489.1648644 | -0.641917967 | 0.134639544 |
| Mrs2 | 387.1535434 | -0.262632075 | 0.085690136 |
| Dcdc2a | 119.9340571 | 0.720919239 | 0.163935088 |
| Gm11348 | 0 | NA | NA |
| Nrsn1 | 2860.65622 | -0.34861335 | 0.088697551 |
| Tpi-rs8 | 0.046345709 | 0.71820487 | 3.724945049 |
| Gm11349 | 2.028713949 | 0.079814751 | 0.663843469 |
| Gm11350 | 0 | NA | NA |
| Gm11351 | 0.19500485 | 1.768080507 | 3.216087831 |
| Gm11352 | 0.044544648 | 0.237309859 | 3.724945049 |
| 1700092E19Rik | 0 | NA | NA |
| Gm11353 | 25.4493118 | 0.670872316 | 0.244932426 |
| Hdgfl1 | 0 | NA | NA |
| Gm11354 | 0 | NA | NA |

| | | | | |
|---|---|---|---|---|
| Gm11355 | 0 | NA | NA | |
| Prl | 159.9440114 | 2.2001382 | | 2.871908804 |
| Prl3d1 | 0.046345709 | 0.71820487 | | 3.724945049 |
| Prl3d2 | 0 | NA | NA | |
| Prl3d3 | 0 | NA | NA | |
| Prl3c1 | 0 | NA | NA | |
| Prl3b1 | 0 | NA | NA | |
| Prl3a1 | 0 | NA | NA | |
| 4930511O05Rik | 0 | NA | NA | |
| Gm11356 | 0 | NA | NA | |
| Prl6a1 | 0.046345709 | 0.71820487 | | 3.724945049 |
| Gm11357 | 0 | NA | NA | |
| Gm11358 | 0 | NA | NA | |
| Prl8a2 | 0 | NA | NA | |
| Prl2b1 | 0 | NA | NA | |
| Prl8a6 | 0 | NA | NA | |
| Prl8a8 | 0 | NA | NA | |
| Prl8a9 | 0 | NA | NA | |
| Prl8a1 | 0 | NA | NA | |
| Prl7b1 | 0 | NA | NA | |
| Prl7a1 | 0 | NA | NA | |
| Prl7a2 | 0 | NA | NA | |
| Gm11359 | 0.049553047 | 0.71820487 | | 3.724945049 |
| Prl7d1 | 0.045882455 | 0.237312084 | | 3.724945049 |
| Prl7c1 | 0.062426873 | 0.237312084 | | 3.724945049 |
| Prl2a1 | 0 | NA | NA | |
| Prl2c1 | 0 | NA | NA | |
| Gm22476 | 0 | NA | NA | |
| Gm11360 | 0 | NA | NA | |
| Prl4a1 | 0 | NA | NA | |
| Prl5a1 | 0 | NA | NA | |
| Gm11361 | 310.4479338 | 0.904634743 | | 0.151648705 |
| 2610307P16Rik | 11.882246 | -0.603090863 | | 0.313946963 |
| Gm6081 | 0.66818121 | -0.230921775 | | 1.156428984 |
| Mir6368 | 0.123992736 | -0.243574001 | | 3.720643035 |
| Gm17528 | 0 | NA | NA | |
| Gm11362 | 0.126225412 | 0.718200913 | | 3.698754989 |
| Gm26735 | 7.2496826 | -0.501870098 | | 0.390489619 |
| Sox4 | 422.4467281 | 0.108909207 | | 0.150251845 |
| A330102I10Rik | 83.11002475 | -0.885245042 | | 0.241373854 |
| Gm11363 | 0.134452335 | 0.237310968 | | 3.631824793 |
| Cdkal1 | 296.7585439 | 0.017789304 | | 0.109819518 |
| Gm11364 | 0 | NA | NA | |
| E2f3 | 366.9818336 | -0.011244212 | | 0.071068317 |
| Gm11365 | 0.042716945 | -0.243571806 | | 3.724945049 |
| Gm11368 | 0.105852363 | 0.23732098 | | 3.724945049 |
| Gm11367 | 0 | NA | NA | |

| | | | |
|---|---|---|---|
| Mboat1 | 33.58227926 | 0.525368895 | 0.220658376 |
| Gm23206 | 0 | NA | NA |
| Gm11366 | 0.322434302 | 0.188496645 | 1.699010415 |
| Agtr1a | 5.923613152 | -0.669450211 | 0.429667026 |
| Gm11369 | 0.085327104 | 1.179809211 | 3.723500796 |
| Uqcrfs1 | 840.2137481 | 0.002116037 | 0.112861748 |
| 4930519D14Rik | 0.04393175 | -0.243571806 | 3.724945049 |
| Dusp22 | 130.6997309 | -0.150031754 | 0.116120952 |
| Gm11370 | 0 | NA | NA |
| Irf4 | 27.40671112 | 3.589787254 | 0.358197725 |
| Exoc2 | 757.5578618 | 0.172947958 | 0.045054644 |
| Hus1b | 7.907650439 | -0.731492863 | 0.39610061 |
| Gm5447 | 0.689547818 | 1.281271038 | 1.278141256 |
| Gm11371 | 0.046345709 | 0.71820487 | 3.724945049 |
| Gm11372 | 0 | NA | NA |
| 4930401O12Rik | 0.147083031 | -0.714489443 | 2.875092884 |
| Gm11376 | 0 | NA | NA |
| Gm11373 | 0.313638404 | -0.714956433 | 1.786883446 |
| Gm11374 | 0.506145433 | -1.581972446 | 1.429476306 |
| G630018N14Rik | 1.020023321 | 1.138193294 | 1.400212538 |
| Gm22330 | 0 | NA | NA |
| Gm11375 | 1.368265809 | -2.504136176 | 1.053473597 |
| Gm23351 | 0 | NA | NA |
| A530084C06Rik | 4.406782827 | 0.3795658 | 0.43553895 |
| Foxq1 | 10.28519308 | -0.526138248 | 0.344929273 |
| Gm11377 | 0 | NA | NA |
| 1700018A04Rik | 0.705754361 | 0.831337226 | 1.138517996 |
| Gm11378 | 0 | NA | NA |
| Foxf2 | 22.59941583 | 1.377380013 | 0.329649608 |
| Gm27516 | 0.075043205 | 0.237309859 | 3.724945049 |
| Gm25186 | 0.382576472 | 1.269692523 | 1.751258729 |
| Gm11379 | 0.403183719 | -0.212794485 | 1.439215061 |
| Foxc1 | 125.7748194 | 1.22230022 | 0.196736979 |
| Gmds | 102.4660629 | -0.05000091 | 0.120993244 |
| Gm11380 | 0.044544648 | 0.237309859 | 3.724945049 |
| Gm11381 | 0 | NA | NA |
| Mylk4 | 2.177335128 | -0.779418886 | 0.690505044 |
| Gm27342 | 0 | NA | NA |
| Wrnip1 | 563.9693568 | 0.192242812 | 0.070135922 |
| D930007J09Rik | 3.931524232 | -0.351052727 | 0.479253111 |
| Serpinb1a | 58.83320495 | -1.735971864 | 0.216471667 |
| Serpinb1-ps1 | 0.259499533 | 1.19911226 | 2.506484949 |
| Serpinb1c | 2.097314789 | 3.893201714 | 0.97727298 |
| Gm11382 | 0.61025173 | 2.838462551 | 1.324501827 |
| Serpinb6b | 64.34938334 | 3.772046978 | 0.347265954 |
| Serpinb9 | 320.5775695 | 3.103129929 | 0.227476077 |
| Serpinb9b | 16.14145208 | 6.903793505 | 0.953318829 |

| | | | |
|---|---|---|---|
| Serpinb1b | 57.17042438 | 1.44340689 | 0.338891707 |
| Gm11384 | 0 | NA | NA |
| Gm11383 | 0 | NA | NA |
| Serpinb9c | 0.260452197 | -0.111831034 | 1.813395508 |
| Serpinb9d | 0 | NA | NA |
| Gm11385 | 0 | NA | NA |
| Serpinb9e | 0.061920613 | 0.718207094 | 3.724945049 |
| Serpinb9f | 0 | NA | NA |
| Gm11397 | 0 | NA | NA |
| Serpinb9g | 0 | NA | NA |
| Gm11387 | 0 | NA | NA |
| Gm11386 | 0 | NA | NA |
| Gm11393 | 0 | NA | NA |
| Gm25043 | 0 | NA | NA |
| Gm11388 | 0 | NA | NA |
| Serpinb6d | 0 | NA | NA |
| Gm11389 | 0 | NA | NA |
| Gm26701 | 0 | NA | NA |
| Gm11392 | 0 | NA | NA |
| Gm11394 | 0 | NA | NA |
| Gm11395 | 0 | NA | NA |
| Vmn2r-ps94 | 0 | NA | NA |
| Gm16566 | 0 | NA | NA |
| Serpinb6e | 0.733278584 | -0.455492221 | 1.281015008 |
| Serpinb6c | 3.256068972 | 0.644425237 | 0.578768259 |
| Serpinb6a | 261.2870453 | 0.858897564 | 0.108813054 |
| 1110046J04Rik | 30.45720878 | -0.217435549 | 0.220204853 |
| Nqo2 | 104.6663154 | 0.744918494 | 0.1556362 |
| Ripk1 | 212.9648014 | 1.993053648 | 0.16547806 |
| Gm17083 | 0.214446769 | 0.718215348 | 2.568216429 |
| Gm17084 | 5.185998495 | 0.639797509 | 0.489832086 |
| Bphl | 141.617793 | -0.278946695 | 0.100495231 |
| Tubb2a | 3268.123541 | 0.530485904 | 0.114316253 |
| Tubb2b | 1731.528415 | 1.969778759 | 0.173276741 |
| Psmg4 | 78.61758626 | 1.52199777 | 0.187057524 |
| Slc22a23 | 1092.32252 | 0.08274499 | 0.04948042 |
| Gm24530 | 6.119618873 | -1.146864971 | 0.463081652 |
| Pxdc1 | 58.73051673 | 0.760747984 | 0.205467496 |
| Gm15908 | 0.16390591 | -0.724458634 | 2.782471951 |
| Fam50b | 0.097139547 | 0.718207094 | 3.724945049 |
| Prpf4b | 962.083236 | -0.208597398 | 0.098066914 |
| Fam217a | 2.489500889 | -0.100155236 | 0.656352098 |
| 4933417A18Rik | 0.093141081 | -0.724455696 | 3.724945049 |
| Gm8245 | 0.066050875 | 0.71820487 | 3.724945049 |
| Eci3 | 1.298423698 | 1.02536075 | 0.835978919 |
| Gm16984 | 0.868294842 | -0.167702191 | 1.034873755 |
| Eci2 | 255.3897712 | 0.306996052 | 0.130644386 |

| | | | |
|---|---|---|---|
| 1700011B04Rik | 0.157356115 | 1.255377571 | 2.825615364 |
| Gm22674 | 0 | NA | NA |
| Cdyl | 458.3848708 | 0.161597781 | 0.112870964 |
| Gm22126 | 0 | NA | NA |
| Rpp40 | 47.71410153 | -0.011881386 | 0.199587276 |
| Ppp1r3g | 58.16805527 | -1.581179071 | 0.295249476 |
| Lyrm4 | 177.4291997 | -0.156899863 | 0.110721997 |
| Fars2 | 206.9435174 | -0.048982473 | 0.115371724 |
| Gm26395 | 0.075043205 | 0.237309859 | 3.724945049 |
| Nrn1 | 1029.09811 | -0.485389715 | 0.159628014 |
| F13a1 | 144.5358196 | 3.411180871 | 0.423482419 |
| Ly86 | 116.6552724 | 1.739409664 | 0.277799031 |
| Gm4035 | 0.308888673 | -1.541660199 | 3.685836331 |
| Gm29459 | 0 | NA | NA |
| Gm29458 | 0.240304698 | 0.239875883 | 2.11890435 |
| Gm29590 | 0 | NA | NA |
| Gm37149 | 0 | NA | NA |
| Rreb1 | 710.6869941 | -0.179231252 | 0.17452121 |
| Ssr1 | 2094.214592 | 0.106371071 | 0.04568737 |
| Cage1 | 29.07551637 | -0.506443301 | 0.225974366 |
| Riok1 | 219.0593976 | 0.409509719 | 0.12277487 |
| Gm27387 | 0 | NA | NA |
| Gm10129 | 0.169723681 | 1.276730323 | 3.407267785 |
| Dsp | 96.30191792 | -0.48165119 | 0.170004961 |
| Snrnp48 | 159.0698413 | -0.088798721 | 0.166859203 |
| Bmp6 | 104.867172 | -0.048026051 | 0.196591417 |
| Txndc5 | 655.6323935 | 0.512267371 | 0.08252413 |
| Bloc1s5 | 130.6301522 | -0.006343295 | 0.104222582 |
| Eef1e1 | 223.262343 | 0.629533683 | 0.1244254 |
| Gm22035 | 0.046345709 | 0.71820487 | 3.724945049 |
| Slc35b3 | 116.4993211 | 0.175522423 | 0.121515963 |
| Gm26514 | 39.29548882 | -0.5172532 | 0.227243593 |
| 5033403F01Rik | 3.803373425 | 0.119827785 | 0.556220302 |
| Gm38206 | 0 | NA | NA |
| Gm23972 | 0 | NA | NA |
| Ofcc1 | 0.044191549 | -0.243571806 | 3.724945049 |
| Gm9979 | 0.471418172 | 1.485885759 | 1.472440748 |
| Tfap2a | 38.58151316 | 0.005300736 | 0.236625461 |
| Gm26688 | 1.413787916 | 0.660464305 | 0.920884577 |
| Gm37823 | 0 | NA | NA |
| Gm6245 | 0.59099902 | 0.629335062 | 1.322947073 |
| Gcnt2 | 477.1222298 | 1.287639265 | 0.141774382 |
| Mir5124a | 0.74668639 | 2.443661186 | 1.276046397 |
| A730081D07Rik | 27.84660377 | -0.455922203 | 0.288585156 |
| Pak1ip1 | 411.5182475 | 0.563936053 | 0.116233397 |
| Tmem14c | 229.1318417 | 0.425810549 | 0.101255988 |
| Mak | 31.22020358 | 0.242527746 | 0.203729933 |

| | | | |
|---|---|---|---|
| Gcm2 | 0.462408164 | 1.587515913 | 1.412651479 |
| Sycp2l | 3.696698795 | -0.81553627 | 0.576498152 |
| Elovl2 | 197.6053173 | -0.246277667 | 0.123327167 |
| Smim13 | 2327.736401 | -0.638294708 | 0.085780027 |
| Gm17364 | 0.17692102 | 0.773454306 | 2.723401318 |
| Nedd9 | 334.1506425 | 1.47609239 | 0.18453487 |
| Gm26877 | 0.927545067 | 0.31184019 | 1.014103107 |
| Gm24819 | 0.43165004 | -0.858020664 | 1.318974348 |
| Tmem170b | 1737.683987 | -0.287670458 | 0.048858723 |
| Gm5082 | 0.47229425 | -0.477802513 | 3.672712246 |
| Adtrp | 1.749374567 | 1.877023048 | 1.044844967 |
| 1700061E18Rik | 0.094294407 | 0.23732098 | 3.724945049 |
| Gm28564 | 0.066050875 | 0.71820487 | 3.724945049 |
| Gm37698 | 0 | NA | NA |
| Gm22157 | 0.123841226 | 1.292225235 | 3.719439735 |
| Gm28707 | 0.850789295 | 1.488381692 | 1.246169318 |
| Hivep1 | 1475.754828 | 0.015206642 | 0.061214469 |
| Edn1 | 31.52376167 | 2.153030993 | 0.334752232 |
| Gm23086 | 0 | NA | NA |
| Phactr1 | 3991.264433 | -0.763642853 | 0.117588611 |
| Gm15810 | 0.418886444 | -0.246958233 | 1.408572912 |
| Gm15812 | 0 | NA | NA |
| Gm15809 | 21.27740935 | -0.841422309 | 0.330366433 |
| Gm26086 | 0 | NA | NA |
| Gm15813 | 0.11193333 | 0.718202646 | 3.724945049 |
| Tbc1d7 | 105.3200195 | -0.109993881 | 0.115667909 |
| Gfod1 | 1583.085298 | -0.368243502 | 0.089683739 |
| Sirt5 | 74.72223409 | -0.691198825 | 0.141357171 |
| Nol7 | 275.008259 | 0.175992905 | 0.137929493 |
| Ranbp9 | 1368.855223 | 0.287190994 | 0.076012997 |
| Gm26064 | 7.226595754 | -1.21162486 | 0.502245795 |
| Mcur1 | 233.0888101 | 0.96144503 | 0.109553994 |
| Rnf182 | 236.6714893 | -0.225086163 | 0.106804086 |
| Cd83 | 231.474017 | 0.245237293 | 0.129855543 |
| Gm23117 | 0 | NA | NA |
| Gm5083 | 5.023184258 | -0.12532891 | 0.433261107 |
| Gm27007 | 0 | NA | NA |
| Gm27036 | 0.066050875 | 0.71820487 | 3.724945049 |
| 1700029N11Rik | 0 | NA | NA |
| Jarid2 | 797.878151 | 0.053323687 | 0.087855431 |
| Gm22213 | 0.047948698 | -0.243569582 | 3.724945049 |
| Dtnbp1 | 383.1011531 | 0.257561201 | 0.111245344 |
| Gm23104 | 0 | NA | NA |
| Hsp25-ps1 | 4.708514708 | 2.571147413 | 0.749528717 |
| Gm9817 | 0.309768565 | 1.14074261 | 1.774715175 |
| Gm8513 | 0.148910734 | -0.243572529 | 2.86160923 |
| Gm23755 | 0 | NA | NA |

| | | | |
|---|---|---|---|
| Mylip | 196.9662467 | 0.397824459 | 0.101895115 |
| Gm23387 | 0 | NA | NA |
| Gmpr | 269.5668325 | -0.687979824 | 0.087421409 |
| Atxn1 | 2899.31005 | -0.240134069 | 0.082024569 |
| 5033430I15Rik | 31.65815391 | 0.531259271 | 0.184523561 |
| Gm10113 | 0.097139547 | 0.718207094 | 3.724945049 |
| Stmnd1 | 2.201457459 | 1.672909044 | 0.884874648 |
| Rbm24 | 161.1267455 | -0.706140251 | 0.139887317 |
| Cap2 | 1272.593817 | -0.544629308 | 0.134831457 |
| Gm24915 | 2.015269402 | -2.230508609 | 0.807133409 |
| C78339 | 1105.225121 | -0.000403733 | 0.081048006 |
| Nup153 | 1192.694187 | 0.492558462 | 0.059528291 |
| Kif13a | 663.6183412 | -0.343312247 | 0.085261624 |
| Gm24620 | 4.394585356 | -0.750239596 | 0.49610755 |
| Gm22960 | 0 | NA | NA |
| Gm24769 | 0 | NA | NA |
| 2010001K21Rik | 1.984554007 | 0.65163535 | 0.653997348 |
| Nhlrc1 | 147.1966146 | -0.255462223 | 0.097676118 |
| Tpmt | 47.2986511 | -0.283899246 | 0.173726763 |
| Kdm1b | 181.3782971 | 0.005104645 | 0.118684102 |
| Dek | 884.7800832 | 0.835972283 | 0.121392758 |
| Rnf144b | 226.1613996 | 0.109298135 | 0.114912092 |
| 4931429P17Rik | 0.349470464 | -1.024462413 | 1.772500167 |
| A330033J07Rik | 16.48077188 | -2.35722334 | 0.373042373 |
| Id4 | 466.0183473 | -0.806407576 | 0.161929071 |
| A330048O09Rik | 5.209108024 | -1.488182841 | 0.465980547 |
| Zfp169 | 346.1617771 | -0.427499962 | 0.096674826 |
| A830005F24Rik | 4.046338601 | -0.800616168 | 0.532126864 |
| 6720427I07Rik | 280.6546708 | 0.19332139 | 0.101399239 |
| Mirlet7d | 0 | NA | NA |
| Gm24111 | 0 | NA | NA |
| mmu-let-7f-1 | 12.7179762 | 0.133131013 | 0.297313799 |
| Mirlet7a-1 | 9.148397703 | 0.396587696 | 0.442647808 |
| Gm37238 | 83.05504175 | -0.564013977 | 0.234625679 |
| Gm25232 | 0 | NA | NA |
| Ptpdc1 | 475.0012653 | 0.027695208 | 0.078345206 |
| Barx1 | 0 | NA | NA |
| Phf2 | 871.8668907 | 0.055282525 | 0.06395192 |
| Phf2os1 | 14.57948276 | -1.088084734 | 0.34049434 |
| Fam120a | 2396.718561 | 0.298799576 | 0.058790088 |
| Fam120aos | 129.3350566 | -0.485799247 | 0.148514844 |
| Wnk2 | 2363.287208 | -0.684409111 | 0.112175006 |
| Ninj1 | 228.5158888 | 1.391200357 | 0.172165289 |
| 1110007C09Rik | 111.7269501 | 1.642392233 | 0.184567406 |
| Susd3 | 9.113448631 | 0.705499808 | 0.351052571 |
| Fgd3 | 292.1672564 | 1.301554944 | 0.159367555 |
| Bicd2 | 1037.207963 | -0.017290862 | 0.058044472 |

| | | | |
|---|---|---|---|
| Ippk | 334.3509724 | 0.126120913 | 0.091438437 |
| Cenpp | 10.13615864 | 0.123186887 | 0.411459527 |
| Tes3-ps | 44.94843152 | 0.212848305 | 0.296724248 |
| Ecm2 | 93.28506817 | 1.24160754 | 0.180679845 |
| Aspn | 20.67132123 | -0.718512353 | 0.281757476 |
| Omd | 21.95627783 | 0.165757569 | 0.328515874 |
| Ogn | 102.8893265 | -0.421051626 | 0.219388791 |
| Nol8 | 394.8682349 | 0.182024909 | 0.063829025 |
| Iars | 963.3246765 | 0.512168024 | 0.07368654 |
| Gm25394 | 8.321309178 | -0.022790132 | 0.433198972 |
| Gm23482 | 2.859218534 | -1.638910197 | 0.605431765 |
| Gm7251 | 0.539950272 | 0.706774033 | 1.45781887 |
| Gm8674 | 0 | NA | NA |
| Gm10784 | 0 | NA | NA |
| Gm17611 | 0 | NA | NA |
| Gm24300 | 0 | NA | NA |
| Gm906 | 0 | NA | NA |
| Gm24017 | 0 | NA | NA |
| Gm8739 | 0 | NA | NA |
| Fbxw17 | 179.6250207 | 1.640507613 | 0.162134198 |
| Nutm2 | 0.276311821 | 0.718216777 | 2.43009122 |
| Mir683-1 | 0 | NA | NA |
| Mir683-2 | 0.088123299 | -0.72445792 | 3.724945049 |
| Gm904 | 0.038981395 | 0.718207094 | 3.724945049 |
| Gm23952 | 0 | NA | NA |
| Gm8765 | 0 | NA | NA |
| Gm22199 | 0 | NA | NA |
| Spin1 | 2716.345326 | 0.000509063 | 0.065412132 |
| Nxnl2 | 1.249332652 | 1.072265881 | 1.035260291 |
| Hist1h2al | 110.2101923 | 0.612041619 | 0.164484651 |
| AC160122.1 | 0.260264958 | 1.000223936 | 2.084627883 |
| S1pr3 | 554.2109028 | 3.993713487 | 0.296498781 |
| Shc3 | 425.7683225 | -1.541909125 | 0.149987736 |
| Cks2 | 12.99324003 | 0.586401864 | 0.371545838 |
| Gm22806 | 0.426919779 | 0.529701741 | 1.740535991 |
| Secisbp2 | 291.1100637 | 0.259315364 | 0.087553244 |
| Gm15440 | 205.1826132 | -0.483047783 | 0.117804904 |
| Sema4d | 1108.340496 | 0.645418236 | 0.091070132 |
| Gm23566 | 0 | NA | NA |
| Gadd45g | 275.2348046 | 2.707189324 | 0.324809266 |
| Gm26651 | 0.10503227 | 1.179809211 | 3.723500796 |
| Gm37872 | 0 | NA | NA |
| Diras2 | 2681.114 | -0.709906361 | 0.107271135 |
| Syk | 204.4789825 | 3.039590322 | 0.230101069 |
| Auh | 575.8122667 | -0.265135222 | 0.055938114 |
| n-R5s54 | 0.832946694 | 1.599213728 | 1.300561664 |
| Nfil3 | 206.9757982 | 1.962404762 | 0.345065344 |

| | | | |
|---|---|---|---|
| Ror2 | 25.4305805 | 0.292351287 | 0.225284298 |
| Sptlc1 | 374.6291411 | 0.646074364 | 0.09281458 |
| RP23-340E6.2 | 12.21145091 | -0.745041574 | 0.316776947 |
| RP23-340E6.3 | 0 | NA | NA |
| Msx2 | 19.08535284 | 0.540656837 | 0.330424581 |
| Gm5449 | 0.111006857 | 0.717551239 | 3.724898127 |
| Drd1 | 333.3471759 | -0.822121944 | 0.221540367 |
| Sfxn1 | 561.8273373 | -0.169074229 | 0.066876771 |
| Hrh2 | 63.56667738 | -0.197975259 | 0.173887035 |
| Gm16578 | 0.107803068 | 0.719187033 | 3.724874737 |
| Cplx2 | 9264.934547 | -1.179759791 | 0.121215586 |
| Gm16248 | 0 | NA | NA |
| Gm22668 | 0 | NA | NA |
| Thoc3 | 293.4655856 | 0.208170237 | 0.094866593 |
| Gm2830 | 71.59267597 | 0.279746326 | 0.16184466 |
| Simc1 | 267.6907852 | -0.269180403 | 0.088472545 |
| 4833439L19Rik | 1366.025163 | 0.028600417 | 0.061343094 |
| Arl10 | 135.7640415 | -0.094928408 | 0.128798648 |
| Gm24195 | 1.207241537 | -2.739690012 | 1.122331288 |
| Nop16 | 117.0143181 | 0.379770836 | 0.138448703 |
| Higd2a | 413.1629045 | -0.476563216 | 0.094119156 |
| Cltb | 2228.333823 | 0.762918306 | 0.093590204 |
| Faf2 | 759.1273305 | 0.537076322 | 0.071456383 |
| Rnf44 | 1513.683017 | -0.15442211 | 0.063942418 |
| Cdhr2 | 4.783323946 | -0.741465273 | 0.451124546 |
| Gprin1 | 1231.562756 | -0.510710336 | 0.097123373 |
| Sncb | 3503.722137 | -0.792937292 | 0.101723076 |
| Gm16249 | 0 | NA | NA |
| Eif4e1b | 0 | NA | NA |
| Tspan17 | 212.3703016 | -0.134309974 | 0.130793251 |
| Gm16250 | 0 | NA | NA |
| Gm29431 | 0.504101427 | -0.305826559 | 1.319112213 |
| 5330429C05Rik | 1.674398058 | 1.494836532 | 0.840064936 |
| 4930526F13Rik | 0.237545104 | -0.243569534 | 2.559929356 |
| Unc5a | 875.420532 | -0.616593172 | 0.126296605 |
| Hk3 | 302.6549373 | 6.116307847 | 0.374945595 |
| Uimc1 | 267.0504781 | 0.159664784 | 0.117316538 |
| Gm6344 | 1.345796852 | 1.71596134 | 0.889710445 |
| Zfp346 | 338.7158424 | 0.094570982 | 0.100489593 |
| Gm17617 | 0 | NA | NA |
| Fgfr4 | 13.86132062 | 3.365856771 | 0.471082709 |
| Nsd1 | 4468.214251 | -0.320246232 | 0.069149305 |
| Rab24 | 365.9432809 | -0.112351996 | 0.070405241 |
| Prelid1 | 846.0882492 | 0.391916727 | 0.09129361 |
| Mxd3 | 8.517488434 | -0.406825974 | 0.526391773 |
| Lman2 | 1016.200266 | 0.508861823 | 0.085731065 |
| Rgs14 | 301.6653793 | -0.90582118 | 0.11812722 |

| | | | |
|---|---|---|---|
| Slc34a1 | 0.764795395 | -0.095390439 | 1.136921218 |
| Pfn3 | 0.525579064 | -1.825817958 | 1.429176849 |
| F12 | 5.68547348 | 1.203791995 | 0.61664773 |
| Grk6 | 508.6179697 | 0.060092679 | 0.055881283 |
| Mir6943 | 0 | NA | NA |
| Prr7 | 196.075479 | -0.326537937 | 0.124972268 |
| Dbn1 | 1526.862046 | -0.637751993 | 0.102160631 |
| Mir6944 | 0 | NA | NA |
| Pdlim7 | 426.9912292 | 0.671068975 | 0.087401917 |
| Mir6945 | 0 | NA | NA |
| Dok3 | 92.02924505 | 1.813177443 | 0.228474228 |
| Ddx41 | 338.5120362 | 0.53331994 | 0.108814364 |
| Fam193b | 537.9735023 | -0.711856187 | 0.180655178 |
| Tmed9 | 877.8577826 | 0.773455329 | 0.078428785 |
| B4galt7 | 189.7572043 | -0.073009736 | 0.085271838 |
| Gm15911 | 0.088476398 | -0.243576254 | 3.724945049 |
| Caml | 264.5844916 | 0.580915575 | 0.08045851 |
| Ddx46 | 1095.580132 | -0.04405354 | 0.08007449 |
| B230219D22Rik | 1459.612451 | -0.142816647 | 0.074037337 |
| Txndc15 | 462.6097524 | 0.201358292 | 0.099591587 |
| Pcbd2 | 43.31228221 | -0.098548776 | 0.191164291 |
| 4930451E10Rik | 0 | NA | NA |
| Catsper3 | 9.093479106 | -1.075893696 | 0.364725562 |
| Pitx1 | 1.253999855 | -0.219068105 | 0.911453499 |
| Gm28760 | 0.04393175 | -0.243571806 | 3.724945049 |
| H2afy | 903.7078744 | 0.537566113 | 0.072390142 |
| Gm25148 | 0 | NA | NA |
| Tifab | 116.9424394 | 2.84852628 | 0.296922352 |
| 4930550C17Rik | 0 | NA | NA |
| Neurog1 | 0.267034963 | -2.040308387 | 2.037186825 |
| Cxcl14 | 505.7540529 | -0.580795078 | 0.123304866 |
| Gm24716 | 0 | NA | NA |
| Gm10782 | 0 | NA | NA |
| Slc25a48 | 3.125560887 | 0.447480721 | 0.73677479 |
| Il9 | 0 | NA | NA |
| Fbxl21 | 83.03568542 | -0.321157051 | 0.134473794 |
| Lect2 | 0.045882455 | 0.237312084 | 3.724945049 |
| Tgfbi | 1350.337662 | 6.226710809 | 0.321697627 |
| Gm27698 | 0 | NA | NA |
| Smad5 | 771.3450184 | 0.160033645 | 0.07888342 |
| Gm27301 | 0.049553047 | 0.71820487 | 3.724945049 |
| Gm27834 | 0.26282868 | -0.222241123 | 1.811081714 |
| Gm27271 | 2.902142703 | -1.55738218 | 0.605144445 |
| Trpc7 | 38.7170757 | -0.464811767 | 0.1646873 |
| Gm22777 | 0 | NA | NA |
| Spock1 | 3106.932593 | -0.585152011 | 0.072559416 |
| Klhl3 | 316.2281292 | -0.921996983 | 0.160572063 |

| | | | |
|---|---|---|---|
| Mir874 | 4.532224978 | -1.118483218 | 0.578435762 |
| Hnrnpa0 | 1502.786418 | 0.139434122 | 0.100191961 |
| 1700066J03Rik | 0.225300421 | -0.723175418 | 2.572018836 |
| Idnk | 305.8515994 | 1.752462113 | 0.160028336 |
| Ubqln1 | 2320.123517 | 0.208683237 | 0.06262985 |
| Gkap1 | 124.6432152 | -0.042297309 | 0.140049857 |
| Gm26555 | 4.731241973 | -1.126174501 | 0.580319033 |
| Kif27 | 62.59338873 | -0.311952067 | 0.173982841 |
| Mir6369 | 0.042716945 | -0.243571806 | 3.724945049 |
| Gm24284 | 0 | NA | NA |
| 2210016F16Rik | 128.8048172 | 1.203055164 | 0.16326714 |
| Hnrnpk | 4038.157108 | 0.344698134 | 0.076784579 |
| Mir7-1 | 15.77224037 | 0.075330447 | 0.281439975 |
| Rmi1 | 123.5749391 | 0.370032557 | 0.111318808 |
| Slc28a3 | 2.52226614 | -1.615747503 | 0.751181054 |
| Ntrk2 | 6695.738885 | -0.920464495 | 0.105717959 |
| Agtpbp1 | 2886.668731 | -0.61404366 | 0.068915049 |
| Naa35 | 495.813822 | -0.083310474 | 0.055166945 |
| Golm1 | 357.7061881 | 0.250727605 | 0.117951092 |
| 4921517D22Rik | 0.150248541 | -0.243571377 | 2.859451661 |
| Spata31d1a | 2.564266786 | 0.335594387 | 0.653005016 |
| Spata31d1b | 1.567374066 | 1.101792679 | 0.879370419 |
| Spata31d1d | 3.578253317 | -0.901605904 | 0.639908971 |
| 4930528D03Rik | 0.109124069 | -0.24357403 | 3.724945049 |
| 1700014D04Rik | 13.37826059 | -0.552223688 | 0.309075079 |
| Isca1 | 1058.772678 | -0.174958178 | 0.055673962 |
| Etohd2 | 12.41969927 | -1.192739997 | 0.294975507 |
| Zcchc6 | 1650.158349 | 1.162256981 | 0.098397119 |
| Gas1 | 102.0490903 | 0.073526822 | 0.183469666 |
| Gm5084 | 0.588182734 | -0.748023533 | 1.418672817 |
| Gm24999 | 0 | NA | NA |
| Gm25000 | 0 | NA | NA |
| Dapk1 | 1289.496349 | -0.289376176 | 0.121124239 |
| Gm10312 | 0 | NA | NA |
| BC051665 | 0.048949532 | -0.243569582 | 3.724945049 |
| Ctsll3 | 0 | NA | NA |
| 4930486L24Rik | 7.005733943 | 3.930308199 | 0.695456278 |
| Ctla2b | 52.79599323 | 4.287098074 | 0.36248158 |
| Tpbpb | 0 | NA | NA |
| Gm24594 | 0 | NA | NA |
| Ctla2a | 216.8941751 | 4.10199024 | 0.277105662 |
| Tpbpa | 0 | NA | NA |
| Ctsj | 0 | NA | NA |
| Ctsq | 0 | NA | NA |
| Ctsr | 0 | NA | NA |
| Cts6 | 0 | NA | NA |
| Cts8 | 0 | NA | NA |

| | | | |
|---|---|---|---|
| Gm27704 | 0 | NA | NA |
| Cts7 | 0 | NA | NA |
| Ctsm | 0 | NA | NA |
| Cts3 | 0 | NA | NA |
| Gm19961 | 1.330930655 | 1.32865794 | 0.839551254 |
| Gm23739 | 0 | NA | NA |
| Gm24193 | 0 | NA | NA |
| Gm20563 | 3.002659297 | -0.309450964 | 0.740632177 |
| Gm25649 | 0.044544648 | 0.237309859 | 3.724945049 |
| Gm6478 | 2.109479346 | -0.009194282 | 0.663333063 |
| Zfp808 | 54.99002886 | 0.002255017 | 0.191760403 |
| Gm23713 | 0.185738947 | 0.237327673 | 2.674436203 |
| Gm24539 | 0.901339752 | -0.747587713 | 1.03128407 |
| Gm3604 | 26.94184347 | 0.40781482 | 0.236396385 |
| Gm25933 | 0.061920613 | 0.718207094 | 3.724945049 |
| Gm23855 | 0 | NA | NA |
| Zfp935 | 99.08513468 | 0.082772245 | 0.134321749 |
| Gm22256 | 0.093484797 | 0.237318756 | 3.724945049 |
| Zfp934 | 37.404617 | 0.090090686 | 0.215060022 |
| Gm7664 | 14.88251027 | 0.020779192 | 0.264873999 |
| Gm24095 | 0.244748102 | -0.085981486 | 2.490197097 |
| Gm22615 | 1.125305835 | -0.691844272 | 0.817293017 |
| 6720489N17Rik | 53.55286888 | -0.543945476 | 0.186741767 |
| Gm24020 | 0.042716945 | -0.243571806 | 3.724945049 |
| Platr25 | 30.64885467 | -1.128693392 | 0.307276587 |
| Mir713 | 0 | NA | NA |
| Gm5141 | 99.08605519 | -0.345232725 | 0.163804602 |
| Gm21963 | 0 | NA | NA |
| Gm24588 | 0 | NA | NA |
| Gm25448 | 0 | NA | NA |
| Fbp2 | 0.981774026 | -1.766981642 | 1.111012309 |
| Fbp1 | 2.587088777 | 1.201336538 | 0.671147293 |
| 2010111I01Rik | 498.5900927 | 0.084302684 | 0.090421371 |
| Gm16907 | 5.155791298 | 0.027217394 | 0.456852283 |
| Mir23b | 8.607625547 | 0.262657534 | 0.429090562 |
| Mir27b | 11.22797161 | 0.022045944 | 0.370197387 |
| Mir3074-1 | 0.352569346 | 0.65770984 | 1.585308208 |
| mmu-mir-24-1 | 9.164853348 | -0.022307425 | 0.351351617 |
| Fancc | 74.63926839 | -0.335674948 | 0.135246477 |
| Gm16133 | 0.546112947 | 2.028025765 | 1.383574044 |
| Gm16132 | 1.982187692 | 0.393226499 | 0.788554289 |
| Gm16134 | 0 | NA | NA |
| Ptch1 | 1765.336376 | -0.569982549 | 0.102817942 |
| A930032L01Rik | 2.69933516 | -0.475203605 | 0.564681555 |
| Ercc6l2 | 388.7591555 | 0.23311313 | 0.083506461 |
| Hsd17b3 | 0.044191549 | -0.243571806 | 3.724945049 |
| Slc35d2 | 16.4056275 | 3.243535146 | 0.388244738 |

| | | | |
|---|---|---|---|
| Zfp367 | 307.36151 | 0.168718881 | 0.124981526 |
| Habp4 | 1131.608262 | -0.443509345 | 0.079682937 |
| Cdc14b | 233.8312342 | -0.331889981 | 0.099940406 |
| 1810034E14Rik | 1.766063332 | 2.467370015 | 0.883373204 |
| Aaed1 | 90.98287385 | -0.411142324 | 0.140829008 |
| Gm25654 | 2.492378667 | -0.218349614 | 0.670700656 |
| Ctsl | 1539.723944 | 1.801421061 | 0.179894597 |
| Cdk20 | 89.79521824 | 0.164139456 | 0.119272902 |
| 1190003K10Rik | 0.204782255 | -1.217702835 | 2.205163063 |
| Gm4810 | 1.123243924 | -2.989060037 | 1.068481229 |
| Cntnap3 | 108.0176306 | -0.995083901 | 0.133634471 |
| Spata31 | 0 | NA | NA |
| Spata31d1c | 0 | NA | NA |
| Prss47 | 0.08486385 | 0.717551239 | 3.724898127 |
| Hiatl1 | 800.205226 | 0.220673228 | 0.081734439 |
| Olfr466 | 0 | NA | NA |
| Gm24130 | 0 | NA | NA |
| Nlrp4f | 0.716239967 | 2.333597116 | 1.20578806 |
| Gm10775 | 2.001003253 | 0.97150955 | 0.821766175 |
| Zfp369 | 347.6968427 | -0.326176358 | 0.106612403 |
| Gm5792 | 0 | NA | NA |
| Gm10139 | 0 | NA | NA |
| Gm6901 | 0.223727156 | 0.726215108 | 2.152942531 |
| Gm26639 | 0 | NA | NA |
| Platr2 | 0 | NA | NA |
| Gm6917 | 0 | NA | NA |
| Gm6910 | 5.181120878 | -0.094478187 | 0.464434493 |
| Gm10324 | 0 | NA | NA |
| Gm17514 | 0 | NA | NA |
| Gm10772 | 0 | NA | NA |
| Gm26754 | 0 | NA | NA |
| Gm26715 | 0 | NA | NA |
| Vmn2r-ps104 | 0 | NA | NA |
| 2410141K09Rik | 0 | NA | NA |
| Gm26806 | 0 | NA | NA |
| Gm27501 | 0 | NA | NA |
| Zfp640 | 0 | NA | NA |
| Gm17404 | 0 | NA | NA |
| Gm17449 | 0 | NA | NA |
| Gm10323 | 0 | NA | NA |
| Gm5196 | 0.205773706 | -0.700440041 | 2.189704783 |
| Gm5451 | 26.41010186 | 0.846890838 | 0.243818156 |
| Uqcrb | 583.1177258 | 0.129100821 | 0.100246791 |
| Gm10767 | 12.81661586 | -0.400779857 | 0.279310076 |
| Mterf3 | 229.6188801 | -0.020735644 | 0.080218477 |
| Ptdss1 | 668.8494159 | 0.047114449 | 0.069339722 |
| Zfp712 | 93.19383338 | -0.378662905 | 0.171902722 |

| | | | |
|---|---|---|---|
| Zfp708 | 0.458268618 | -1.284289483 | 1.359973036 |
| Gm28557 | 4.418616242 | -0.479961896 | 0.492222716 |
| Rslcan18 | 4.299918267 | -0.675480972 | 0.461802541 |
| Zfp759 | 131.3437174 | -0.035671982 | 0.157088538 |
| Rsl1 | 30.30590387 | 0.896934173 | 0.279293095 |
| Zfp455 | 45.95318099 | 1.056415755 | 0.27450171 |
| Zfp458 | 240.4665658 | -0.103601697 | 0.133259367 |
| Zfp457 | 21.39111743 | -0.509393916 | 0.278373101 |
| Zfp595 | 65.32548619 | 0.65306748 | 0.199266552 |
| Gm28044 | 0.23851432 | -0.243563887 | 2.112354461 |
| Zfp953 | 36.77585063 | 0.765821162 | 0.207401827 |
| Gm28041 | 0.153723802 | 0.237326469 | 2.839085032 |
| Gm17039 | 2.363953117 | -0.200410364 | 0.685351656 |
| Zfp456 | 19.86136891 | 1.40498883 | 0.266359604 |
| Zfp429 | 33.84957497 | 1.396176413 | 0.218033773 |
| Zfp459 | 66.34388881 | -0.142112394 | 0.156068685 |
| Zfp874a | 90.85288293 | 0.318630468 | 0.153289145 |
| Gm38307 | 0.282611191 | -0.2217227 | 1.640898061 |
| Gm7928 | 0 | NA | NA |
| Zfp874b | 171.5607145 | -0.032352226 | 0.157250132 |
| Zfp58 | 50.28513376 | -0.023209453 | 0.167842934 |
| Gm26965 | 0 | NA | NA |
| Zfp87 | 184.3921584 | 0.020798646 | 0.103124223 |
| Zfp748 | 247.3472852 | 0.246383649 | 0.134345624 |
| 9430065F17Rik | 2.796353279 | -0.534379299 | 0.634856247 |
| Zfp729b | 275.2153557 | 0.473546874 | 0.118230095 |
| Gm37276 | 23.42523615 | -0.176463067 | 0.239611835 |
| Zfp729a | 293.0100012 | 0.232119782 | 0.086718317 |
| Gm26587 | 0 | NA | NA |
| Zfp738 | 303.073871 | -0.022346383 | 0.131846176 |
| Gm26875 | 0 | NA | NA |
| Zfp65 | 240.5209492 | 0.351280639 | 0.11347359 |
| Zfp85os | 44.02786501 | 0.68046665 | 0.198993451 |
| Zfp85 | 74.6551353 | 0.401494236 | 0.161979289 |
| Gm9894 | 0 | NA | NA |
| Zfp493 | 42.58231206 | -0.042608119 | 0.277034096 |
| 4930525G20Rik | 7.15404337 | -0.546842173 | 0.431112531 |
| Gm9625 | 125.9289179 | 1.3687865 | 0.159625584 |
| Zfp273 | 60.68720866 | -0.055569461 | 0.161147724 |
| Gm10037 | 3.305635379 | -0.102092549 | 0.51884233 |
| BC048507 | 0.806474979 | -0.396215127 | 1.185439347 |
| Mtrr | 182.8372363 | 0.200845164 | 0.098649534 |
| Fastkd3 | 102.6350786 | 0.009897697 | 0.120553149 |
| 1700001L19Rik | 42.40692636 | -0.856508164 | 0.174843147 |
| Adcy2 | 1089.553893 | -0.983932962 | 0.090870075 |
| Gm26929 | 0 | NA | NA |
| Gm37122 | 2.65229942 | -1.806878753 | 0.76403192 |

| | | | |
|---|---|---|---|
| Gm26844 | 0 | NA | NA |
| Gm4812 | 0 | NA | NA |
| AC161592.1 | 0 | NA | NA |
| Papd7 | 947.3406657 | 0.869630776 | 0.081564322 |
| Nsun2 | 820.5404197 | 0.303144395 | 0.063373847 |
| A530095I07Rik | 0.197913552 | 0.220998123 | 2.223833422 |
| Srd5a1 | 47.64261977 | -0.278811862 | 0.174164054 |
| Gm17108 | 3.053155272 | 0.051458476 | 0.655644008 |
| Ube2ql1 | 1424.867787 | -0.230275112 | 0.085254201 |
| Med10 | 183.4117316 | 0.499247307 | 0.109520373 |
| Gm26819 | 0.108772582 | 0.718202646 | 3.724945049 |
| 1700084F23Rik | 0.092691418 | 1.183079489 | 3.724560917 |
| AC123920.1 | 0.239188194 | -0.223574406 | 1.837830148 |
| Gm26018 | 0 | NA | NA |
| 1700100L14Rik | 0 | NA | NA |
| Ice1 | 1670.349411 | -0.279830662 | 0.110891187 |
| Adamts16 | 57.86550676 | -1.407596041 | 0.238522784 |
| 8030423J24Rik | 1.260089781 | -0.468282942 | 0.897810673 |
| Gm38191 | 0.247225956 | 0.718211434 | 2.992047273 |
| Mir466f-4 | 0 | NA | NA |
| Irx1 | 50.49064827 | 0.10153118 | 0.209697722 |
| Rpl9-ps4 | 43.5655805 | 0.528410969 | 0.186865818 |
| Gm20554 | 3.453678518 | 0.17607787 | 0.527014038 |
| Irx2 | 81.72096566 | 0.375154885 | 0.211939179 |
| Irx4 | 3.045905106 | 0.644076557 | 0.585221981 |
| Gm10263 | 1.909878088 | -0.385839189 | 0.774704456 |
| Ndufs6 | 294.0100797 | -0.097178312 | 0.077387936 |
| Mrpl36 | 129.8851055 | 0.04589746 | 0.175577615 |
| Lpcat1 | 574.2185807 | 0.555379926 | 0.087989829 |
| Slc6a3 | 74.17620595 | -1.283103911 | 0.295715213 |
| Clptm1l | 878.6178926 | 0.409661005 | 0.063965316 |
| Tert | 17.88891691 | -1.214523231 | 0.295008521 |
| Slc6a18 | 0.178490723 | 0.718216836 | 2.713461433 |
| Slc6a19 | 6.934680317 | 1.197666023 | 0.399244534 |
| Slc6a19os | 0.042716945 | -0.243571806 | 3.724945049 |
| Slc12a7 | 159.1782715 | 1.835992004 | 0.17263589 |
| Nkd2 | 44.99299481 | -0.559130265 | 0.194548878 |
| Gm8062 | 0 | NA | NA |
| Trip13 | 13.34211523 | 1.209800805 | 0.297089394 |
| Gm25197 | 0.235280811 | -1.217126864 | 2.117696254 |
| Brd9 | 453.7906305 | -0.465018301 | 0.092320456 |
| Zdhhc11 | 0.149223819 | 0.717609421 | 2.865376928 |
| Gm10126 | 0.987133562 | -1.582594565 | 1.12503846 |
| Gm15912 | 22.38223563 | -0.764647659 | 0.250297424 |
| Tppp | 4530.77903 | -0.661181312 | 0.080769761 |
| Gm6263 | 0.459143553 | 1.354949814 | 1.383797358 |
| Cep72 | 28.65246828 | -0.120626599 | 0.221683911 |

| | | | |
|---|---|---|---|
| Slc9a3 | 15.35249687 | -1.009895227 | 0.33436944 |
| Exoc3 | 1219.916005 | -0.048475965 | 0.047376965 |
| Ahrr | 8.460366272 | -0.135405852 | 0.348041158 |
| Pdcd6 | 440.5530706 | 0.471421692 | 0.083647568 |
| Sdha | 3363.70453 | -0.187643145 | 0.06120713 |
| Gm16125 | 0 | NA | NA |
| Ccdc127 | 908.4665307 | -0.198471657 | 0.062241862 |
| Lrrc14b | 12.94872141 | 2.564491868 | 0.457266197 |
| Zfp72 | 58.03635434 | -0.45774509 | 0.207483276 |
| Gm10116 | 125.931969 | 2.24446909 | 0.214454 |
| Mir692-3 | 0 | NA | NA |
| Zfp825 | 69.54643972 | 0.213771203 | 0.184249268 |
| Erap1 | 747.9723258 | 3.142570699 | 0.122731652 |
| Cast | 225.7625238 | 1.634890315 | 0.14834886 |
| Hnrnpa1l2-ps | 0.737427016 | 0.217584187 | 1.116835468 |
| Pcsk1 | 401.8836238 | 0.166461466 | 0.164909987 |
| Gm4149 | 23.46034018 | 0.864757324 | 0.653898458 |
| Mir682 | 0 | NA | NA |
| Gm23012 | 0 | NA | NA |
| Ell2 | 509.0942641 | 0.589880594 | 0.099177141 |
| Gm23127 | 1.778293916 | -0.069957435 | 0.778008549 |
| Glrx | 434.8245854 | 1.365750191 | 0.173897268 |
| Rhobtb3 | 441.2117491 | -0.693315012 | 0.128568327 |
| Gm24163 | 0 | NA | NA |
| Gm15528 | 4.932600309 | -0.639856702 | 0.531969218 |
| Gm6311 | 20.93266756 | 0.078271322 | 0.294520079 |
| Gm23319 | 0.214976432 | -0.243563295 | 2.177043894 |
| Spata9 | 2.596037066 | 2.475998606 | 0.843530765 |
| Rfesd | 170.1520903 | 0.067353219 | 0.100890125 |
| Gpr150 | 6.580668916 | -1.207504748 | 0.416826132 |
| Arsk | 95.44909683 | -0.269733402 | 0.137861372 |
| Ttc37 | 526.0109267 | 0.1579811 | 0.067111985 |
| Gm24025 | 0 | NA | NA |
| Mctp1 | 1090.396185 | -0.681196144 | 0.097054508 |
| Gm24519 | 0 | NA | NA |
| Gm21959 | 0 | NA | NA |
| Gm15740 | 8.875000737 | -0.373159634 | 0.39831828 |
| Gm8185 | 13.50337661 | -1.437000242 | 0.381804736 |
| Ankrd32 | 492.724753 | -0.450765613 | 0.082742146 |
| 2210408I21Rik | 307.5143814 | -0.269481214 | 0.08181449 |
| Gm26088 | 0.042716945 | -0.243571806 | 3.724945049 |
| Fam172a | 517.357941 | -0.163443198 | 0.067018543 |
| Pou5f2 | 26.05560058 | -0.39156193 | 0.367452886 |
| Nr2f1 | 507.987299 | -0.719043192 | 0.105921915 |
| A830082K12Rik | 207.7537088 | -1.303111141 | 0.159283937 |
| Gm6317 | 0 | NA | NA |
| Gm25383 | 0 | NA | NA |

| | | | |
|---|---|---|---|
| Gm29318 | 0.046345709 | 0.71820487 | 3.724945049 |
| Gm38349 | 0 | NA | NA |
| Gm24863 | 0.296104695 | -0.243563591 | 1.772080761 |
| Gm28526 | 0.110242424 | 0.237318756 | 3.724945049 |
| Gm24635 | 0 | NA | NA |
| Arrdc3 | 1031.103706 | 0.058069989 | 0.098238669 |
| 1700023H06Rik | 0.473445911 | 1.708120296 | 1.29552742 |
| Gm4211 | 15.12852727 | -0.468650324 | 0.30022172 |
| 9330111N05Rik | 6.889709052 | -1.66385003 | 0.496500362 |
| Gm8399 | 71.44903874 | 0.339459329 | 0.150789604 |
| Gm25241 | 0 | NA | NA |
| Adgrv1 | 98.32611678 | -0.21668538 | 0.155181291 |
| Lysmd3 | 266.3986882 | 0.456096866 | 0.105826839 |
| Polr3g | 69.8321072 | -0.508346258 | 0.130944106 |
| Mblac2 | 357.5721259 | -0.738549928 | 0.102097057 |
| Gm17259 | 17.35136828 | -0.216078789 | 0.273068397 |
| Cetn3 | 274.7140815 | -0.141720134 | 0.138870619 |
| Gm24295 | 0 | NA | NA |
| mmu-mir-3961 | 0 | NA | NA |
| Mef2c | 4656.118685 | -1.265259561 | 0.15867549 |
| C130071C03Rik | 839.3578359 | -0.549204295 | 0.127211277 |
| Gm26803 | 22.51607708 | -0.750652613 | 0.240553742 |
| Gm33366 | 4.40666377 | -0.014786278 | 0.46548579 |
| Mir9-2 | 0 | NA | NA |
| 2810049E08Rik | 34.32347081 | 0.044649787 | 0.263818177 |
| Gm17750 | 9.902264217 | -0.661231938 | 0.343918238 |
| Gm24331 | 0 | NA | NA |
| A230107N01Rik | 12.62433136 | -0.780616135 | 0.372332979 |
| Tmem161b | 374.8315931 | 0.105150399 | 0.098620455 |
| Gm26927 | 0.174715781 | -0.724458669 | 2.726888653 |
| Gm26913 | 0.044191549 | -0.243571806 | 3.724945049 |
| Gm4076 | 2.353414094 | 0.045920368 | 0.683851341 |
| Ccnh | 371.5577082 | -0.211000905 | 0.097650181 |
| Rasa1 | 1059.457239 | -0.133103423 | 0.077209296 |
| Gm37641 | 0 | NA | NA |
| Cox7c | 960.0717164 | -0.01440402 | 0.11129643 |
| Gm22574 | 1.539931973 | 0.223171335 | 0.791616467 |
| Gm27025 | 0 | NA | NA |
| Gm25700 | 0 | NA | NA |
| Gm24935 | 0 | NA | NA |
| Gm21232 | 0 | NA | NA |
| Gm27044 | 0 | NA | NA |
| Gm8526 | 0 | NA | NA |
| Edil3 | 2220.587599 | -0.426253243 | 0.091200512 |
| Hapln1 | 201.6502422 | -1.367239429 | 0.216376155 |
| Gm8546 | 11.32043437 | -0.191113417 | 0.3415013 |
| Vcan | 1385.185601 | 1.199951296 | 0.123459108 |

| | | | |
|---|---|---|---|
| Gm16318 | 0.048949532 | -0.243569582 | 3.724945049 |
| Gm4117 | 32.71244237 | 2.768179031 | 0.293965824 |
| Xrcc4 | 102.7407697 | 0.184043357 | 0.133205347 |
| Gm24498 | 3.65796296 | -1.096994207 | 0.56247977 |
| Tmem167 | 992.695371 | 0.095616445 | 0.056523145 |
| Gm24507 | 5.663741534 | -0.058021881 | 0.419027755 |
| Gm37727 | 1.381073991 | -0.2815065 | 0.900957542 |
| Gm37054 | 0.576775271 | 1.204581581 | 1.434909586 |
| Gm37708 | 0 | NA | NA |
| Gm21726 | 0.255118679 | 1.209391965 | 2.477343878 |
| Gm36966 | 0 | NA | NA |
| AC147633.1 | 0 | NA | NA |
| Atp6ap1l | 69.06133185 | -0.754063748 | 0.201817639 |
| Rps23 | 1420.870961 | 0.569982801 | 0.091481128 |
| Atg10 | 83.32890499 | -0.0638532 | 0.12020529 |
| Gm17450 | 0 | NA | NA |
| A830009L08Rik | 17.31106247 | -2.377316369 | 0.323101541 |
| Gm29540 | 0.823808628 | -2.06251957 | 1.424522383 |
| Ssbp2 | 1111.715029 | -0.391999212 | 0.084404113 |
| 4833422C13Rik | 42.21647888 | 0.237915427 | 0.176163157 |
| Gm22778 | 1.426922496 | -0.985765771 | 0.98141108 |
| Acot12 | 6.478410357 | 0.804590782 | 0.444295419 |
| Gm24597 | 0 | NA | NA |
| Gm27656 | 0 | NA | NA |
| Zcchc9 | 113.9382792 | 0.277633324 | 0.114327526 |
| Ckmt2 | 1.022167862 | 0.170881069 | 0.942195697 |
| 1700119I11Rik | 3.377993944 | -0.626183982 | 0.592767896 |
| Rasgrf2 | 1981.271647 | -0.496975942 | 0.095103085 |
| Msh3 | 384.3197434 | 0.315803387 | 0.103207177 |
| Gm20379 | 0 | NA | NA |
| AC161588.1 | 0 | NA | NA |
| Dhfr | 139.8089927 | 1.967596226 | 0.194281035 |
| C030017D09Rik | 1.139049109 | 0.565814726 | 0.916295026 |
| Ankrd34b | 374.0828976 | -0.542605349 | 0.101924043 |
| Fam151b | 60.4619812 | -0.061474964 | 0.142593481 |
| Zfyve16 | 462.5719769 | -0.163161802 | 0.096413562 |
| Spz1 | 0 | NA | NA |
| Serinc5 | 1004.030055 | -0.25822734 | 0.082208607 |
| Thbs4 | 44.88367789 | -0.665153787 | 0.349345178 |
| Mtx3 | 682.3862295 | -0.558884086 | 0.075719185 |
| Cmya5 | 96.50354882 | -0.72946019 | 0.149810236 |
| Papd4 | 358.1413963 | 0.409161207 | 0.119693878 |
| Gm2076 | 0.653050501 | 1.02990838 | 1.333812202 |
| Homer1 | 3096.738791 | -0.168696885 | 0.242811379 |
| Jmy | 1417.927992 | -0.297561481 | 0.053200125 |
| Bhmt | 2.668832678 | -1.155933961 | 0.739401541 |
| Gm15622 | 0.30042186 | 0.704737745 | 1.6312554 |

| | | | |
|---|---|---|---|
| Bhmt2 | 0.855497761 | -0.688106986 | 1.16359678 |
| Dmgdh | 2.316670105 | -1.599475302 | 0.746029898 |
| AW495222 | 7.0482006 | -1.263590647 | 0.446261273 |
| Gm15620 | 0 | NA | NA |
| Arsb | 966.5384739 | -0.715603215 | 0.128423902 |
| Mir5624 | 0.311503589 | -0.112949063 | 1.76666987 |
| Gm25534 | 0.045882455 | 0.237312084 | 3.724945049 |
| Gm26527 | 0.345934107 | 1.172693949 | 1.728279078 |
| Gm24737 | 0.579409632 | -0.724462214 | 1.438148674 |
| Lhfpl2 | 534.543113 | 1.618276852 | 0.156744397 |
| Gm15907 | 0.284494328 | 1.203225557 | 2.064767643 |
| Scamp1 | 2084.156384 | -0.536151337 | 0.05333448 |
| Gm9776 | 34.06058694 | -0.012785399 | 0.184051221 |
| Ap3b1 | 770.6625569 | 0.276077539 | 0.064411002 |
| Gm25109 | 1.803894262 | -0.745350484 | 0.817122843 |
| Gm36991 | 0.478351411 | 0.237293462 | 3.724945049 |
| Tbca | 287.401201 | 0.710098528 | 0.124056443 |
| Otp | 27.26391552 | 0.286278205 | 0.332362667 |
| Wdr41 | 404.2473427 | 0.020346087 | 0.068871223 |
| Pde8b | 739.5660274 | -0.64881545 | 0.139260249 |
| Gm16243 | 0.075043205 | 0.237309859 | 3.724945049 |
| Zbed3 | 136.959666 | 0.462938131 | 0.156584746 |
| Snora47 | 6.619789565 | -0.479181458 | 0.379616896 |
| Gm22661 | 5.694343269 | -0.254937103 | 0.425198108 |
| Aggf1 | 620.7099307 | -0.004136828 | 0.075223919 |
| Crhbp | 115.8235207 | 0.512187403 | 0.152235317 |
| S100z | 0 | NA | NA |
| F2rl1 | 1.572027216 | -1.104349979 | 0.767421519 |
| F2r | 208.5978153 | 0.398626967 | 0.140133582 |
| Iqgap2 | 782.9040417 | 1.370382022 | 0.106064084 |
| F2rl2 | 16.42119601 | 1.730525814 | 0.327671821 |
| Gm24032 | 0.090427103 | 0.237307635 | 3.724945049 |
| Sv2c | 1728.872108 | 0.564633099 | 0.13626472 |
| Gm17190 | 0.114742407 | -0.24357403 | 3.724945049 |
| Gm23955 | 0.962110563 | 0.439599291 | 1.111184788 |
| Gm29543 | 3.164183009 | 0.732013433 | 0.590551073 |
| Gm25213 | 0 | NA | NA |
| Poc5 | 406.0730432 | -0.100371693 | 0.07848719 |
| Ankdd1b | 3.542632829 | 0.386655503 | 0.602367265 |
| Polk | 176.7680094 | 0.390823529 | 0.120613584 |
| Gm17622 | 0 | NA | NA |
| Col4a3bp | 1491.199334 | -0.018666266 | 0.051065223 |
| Hmgcr | 1361.035643 | -0.111914033 | 0.093659348 |
| Gm27792 | 0 | NA | NA |
| Gcnt4 | 101.3429506 | -0.531436355 | 0.144907357 |
| 1700029F12Rik | 0 | NA | NA |
| Fam169a | 842.6174546 | -0.414563092 | 0.075042879 |

| | | | |
|---|---|---|---|
| Gm6169 | 6.279356964 | 2.491517255 | 0.497741535 |
| Nsa2 | 317.8801227 | 0.59459272 | 0.126946964 |
| Gfm2 | 387.8031722 | -0.195120614 | 0.122008104 |
| Hexb | 1048.089142 | 0.076266828 | 0.171114003 |
| Enc1 | 4686.802237 | -0.71474841 | 0.131717271 |
| C430039J16Rik | 0.150086409 | 0.237305146 | 3.724945049 |
| Lincenc1 | 22.69480111 | 0.585913327 | 0.256123084 |
| Gm8756 | 1.313089905 | -0.149459668 | 0.912402356 |
| 5330416C01Rik | 28.18081706 | 0.344327865 | 0.306900225 |
| Gm26619 | 4.459207448 | 0.559073738 | 0.507561057 |
| Gm10260 | 388.9520094 | 1.117871859 | 0.140206678 |
| Arhgef28 | 438.5791712 | -0.581245845 | 0.091569735 |
| Utp15 | 405.8469762 | 0.08325518 | 0.105330743 |
| Ankra2 | 370.3092848 | -0.13917581 | 0.09189291 |
| Gm21976 | 0.640962652 | 1.459978197 | 1.123908062 |
| Btf3 | 823.2235028 | 0.674298955 | 0.088776864 |
| Gm9828 | 25.61986482 | -0.414276047 | 0.240421916 |
| Foxd1 | 28.76092711 | 0.540866973 | 0.284855673 |
| Gm2445 | 3.175554283 | 0.113419 | 0.546176006 |
| 2310005E17Rik | 0 | NA | NA |
| Gm29501 | 0 | NA | NA |
| Gm10320 | 0.789918879 | 1.714017963 | 1.199081369 |
| Gm37425 | 0 | NA | NA |
| Gm22698 | 0 | NA | NA |
| Tmem174 | 0.124347486 | 0.719186602 | 3.717245571 |
| Tmem171 | 14.34229216 | 4.876465721 | 0.609776021 |
| Fcho2 | 753.4025212 | -0.067791642 | 0.067088321 |
| Tnpo1 | 2400.224482 | 0.652375916 | 0.087347169 |
| Gm5453 | 2.269219555 | 1.452109399 | 0.690890403 |
| 1700024P04Rik | 0 | NA | NA |
| Gm24733 | 0.085433889 | -0.72445792 | 3.724945049 |
| 2310020H05Rik | 0 | NA | NA |
| Gm807 | 7.497632429 | 2.333124952 | 0.52839273 |
| Zfp366 | 15.07922646 | 0.45189945 | 0.291198236 |
| Ptcd2 | 245.1837604 | 0.046206037 | 0.074238247 |
| Mrps27 | 194.2719257 | 0.040922531 | 0.083409388 |
| 6430562O15Rik | 0.528537426 | 1.486927808 | 1.383983551 |
| Map1b | 25160.03304 | -0.322431141 | 0.096027755 |
| Gm26559 | 14.25506631 | -0.974871713 | 0.36595179 |
| 1700012J22Rik | 0.18753903 | -0.243565652 | 2.716639112 |
| Gm24471 | 0 | NA | NA |
| Cartpt | 99.17765322 | -0.063466796 | 0.231922197 |
| Mccc2 | 207.1112753 | -0.546731468 | 0.089828666 |
| Bdp1 | 1242.860492 | 0.066518887 | 0.080829193 |
| BC001981 | 0.782345574 | 1.649340615 | 1.205276798 |
| Serf1 | 47.59636433 | -0.409249916 | 0.178602051 |
| Smn1 | 181.1517505 | 0.38975231 | 0.091911646 |

| | | | |
|---|---|---|---|
| Naip2 | 165.0036826 | 4.507694697 | 0.231228693 |
| Naip5 | 40.97982787 | 3.490735685 | 0.310311785 |
| Naip6 | 36.81044465 | 2.713573099 | 0.284944924 |
| Naip3 | 1.709236637 | 2.785164315 | 0.956259348 |
| Naip1 | 6.360266967 | 0.944189364 | 0.47843012 |
| Rpl31-ps13 | 5.091748836 | 0.919347603 | 0.535442674 |
| Gtf2h2 | 239.663237 | -0.24856047 | 0.126106328 |
| Ocln | 88.83136681 | 0.015555425 | 0.155405737 |
| Marveld2 | 42.42039179 | -0.413652143 | 0.192638913 |
| Gm24261 | 0 | NA | NA |
| Rad17 | 254.2430578 | 0.176223471 | 0.099076362 |
| Ak6 | 71.85688038 | 0.265946582 | 0.138754189 |
| Taf9 | 6.955395866 | -0.834098291 | 0.38504921 |
| Ccdc125 | 6.403646332 | 2.117621195 | 0.447604044 |
| Cdk7 | 820.0282203 | -0.22291619 | 0.055634397 |
| Mrps36 | 129.039131 | 0.111291335 | 0.12802641 |
| Cenph | 6.451376144 | 0.731466232 | 0.58947454 |
| Gm16335 | 0.346789506 | 1.527458421 | 1.435890877 |
| Ccnb1 | 20.0589336 | 1.157583835 | 0.39213106 |
| Slc30a5 | 515.9001532 | -0.062393668 | 0.075384393 |
| Gm29502 | 6.014679714 | 0.388730556 | 0.442151588 |
| Gm37830 | 0.152961209 | 0.219193978 | 2.838062251 |
| Gm10257 | 10.11133075 | 0.294219543 | 0.326245114 |
| Gm6114 | 0 | NA | NA |
| Hmgb1-ps9 | 1.368962209 | 0.473696991 | 1.099110537 |
| 4932411K12Rik | 0.386457151 | 2.101420569 | 1.66073738 |
| Gm36994 | 0 | NA | NA |
| Gm29341 | 0 | NA | NA |
| Pik3r1 | 3272.596283 | 0.248814058 | 0.069095447 |
| Gm36638 | 65.83543421 | -0.418804518 | 0.241937184 |
| Gm38133 | 0 | NA | NA |
| Gm25808 | 0 | NA | NA |
| Cd180 | 333.6593441 | 6.379243715 | 0.279588533 |
| Mast4 | 1713.710742 | -0.320984138 | 0.081494447 |
| 1700099I09Rik | 0.905219458 | 1.401488193 | 1.185690352 |
| Gm17160 | 0.372057435 | 2.295653122 | 1.48134018 |
| Gm6211 | 10.83872605 | -1.904886729 | 0.465070667 |
| Gm5454 | 142.1539982 | -0.84066025 | 0.184589353 |
| n-R5s56 | 0 | NA | NA |
| Gm16416 | 0.30254151 | -1.142388321 | 1.783407285 |
| Gm24441 | 0 | NA | NA |
| Gm24870 | 0 | NA | NA |
| Srek1 | 793.2803317 | -0.29302678 | 0.152252087 |
| Erbb2ip | 3344.991394 | 0.926335396 | 0.132327753 |
| Nln | 399.6247544 | 0.113664599 | 0.079836959 |
| Sgtb | 1145.826265 | -0.725253576 | 0.088970442 |
| Trappc13 | 722.8966951 | 0.061448109 | 0.07137679 |

| | | | |
|---|---|---|---|
| Trim23 | 1039.038422 | -0.149889417 | 0.091216124 |
| Ppwd1 | 182.8657831 | 0.08334881 | 0.098126515 |
| Cenpk | 9.293126097 | 0.89490764 | 0.38755677 |
| Adamts6 | 345.3432011 | 1.510656349 | 0.150776446 |
| Cwc27 | 204.665214 | -0.064994714 | 0.163748895 |
| 2610204G07Rik | 0.047948698 | -0.243569582 | 3.724945049 |
| Srek1ip1 | 192.1915529 | 0.090547277 | 0.125597266 |
| Fam159b | 3.808355539 | 1.212364962 | 0.610369346 |
| Gm4938 | 0.726615386 | -0.253879588 | 1.187866701 |
| Rgs7bp | 3653.625861 | -0.739061513 | 0.099042027 |
| 4933425L06Rik | 0.717517214 | 1.462775532 | 1.108500556 |
| Rnf180 | 161.367705 | -0.019735479 | 0.097200788 |
| Gm25631 | 2.548041448 | -0.384650043 | 0.632166496 |
| Htr1a | 192.2048649 | -0.243660098 | 0.110872544 |
| Dph3b-ps | 0 | NA | NA |
| Gm24527 | 0.388660859 | -1.012792237 | 1.583562953 |
| Gm28989 | 0.486723951 | 0.971396462 | 1.606113869 |
| Ipo11 | 842.5268281 | -0.285446625 | 0.060618015 |
| Gm28988 | 0.166709681 | -0.724459871 | 2.767631179 |
| Dimt1 | 147.6538309 | -0.075316982 | 0.093908071 |
| Kif2a | 1440.730209 | -0.318505559 | 0.065154228 |
| Gm24431 | 2.158765044 | -0.785164672 | 0.841667283 |
| 3830408C21Rik | 12.03259843 | -1.003964116 | 0.340976083 |
| Apoo-ps | 41.16296874 | 0.410224522 | 0.27679139 |
| 1700006H21Rik | 0.426601203 | 2.864632504 | 3.210283662 |
| Zswim6 | 635.8544846 | 0.017906292 | 0.095332108 |
| Gm22118 | 4.848883265 | -0.588459719 | 0.544426584 |
| Smim15 | 306.3096422 | 0.253111482 | 0.105646369 |
| Ndufaf2 | 61.73980177 | -0.197361108 | 0.25481244 |
| Ercc8 | 184.4501039 | -0.265959191 | 0.112135942 |
| Elovl7 | 210.6233394 | -0.215087871 | 0.143697506 |
| Gm2822 | 0.316450815 | -0.670168527 | 1.76190542 |
| Depdc1b | 9.399548456 | 0.312475377 | 0.352308596 |
| Pde4d | 1367.797614 | -0.470922771 | 0.086599706 |
| Gm27572 | 0 | NA | NA |
| Gm28016 | 0 | NA | NA |
| Rps3a3 | 38.33507003 | 0.544966523 | 0.190318714 |
| Mir582 | 2.019407445 | -0.133870691 | 0.733210902 |
| AC161585.1 | 9.306437351 | -1.217558597 | 0.41268976 |
| Mir1904 | 1.272510498 | -0.555859918 | 0.947849636 |
| Rab3c | 4667.9155 | -0.643796509 | 0.092642339 |
| Gm5455 | 0.386702458 | 0.045195316 | 1.677919798 |
| Gapt | 3.214970131 | 4.299984046 | 0.977955065 |
| Plk2 | 1844.679949 | 0.268743089 | 0.140234384 |
| Actbl2 | 1.958193336 | -1.526443059 | 0.700207383 |
| Gm15285 | 0.507331407 | 1.340695981 | 1.352052267 |
| Gm15289 | 0 | NA | NA |

| | | | |
|---|---|---|---|
| Gm6270 | 0 | NA | NA |
| Gpbp1 | 1669.061245 | 0.30531114 | 0.071301546 |
| Gm15290 | 0 | NA | NA |
| Gm15288 | 0 | NA | NA |
| Gm15286 | 0.075043205 | 0.237309859 | 3.724945049 |
| Gm10198 | 0 | NA | NA |
| Gm10736 | 67.77195831 | 0.816459006 | 0.17777449 |
| Mier3 | 595.0131209 | 0.284712963 | 0.091215898 |
| Gm15488 | 0.182466651 | 1.62097259 | 2.67233915 |
| Gm15287 | 0.237498462 | 1.485140168 | 2.099245401 |
| Map3k1 | 549.7994848 | 0.972365554 | 0.156489535 |
| Gm15327 | 2.461737376 | -0.253725177 | 0.699422835 |
| Gm15325 | 0 | NA | NA |
| Gm15324 | 0 | NA | NA |
| Gm15326 | 3.336705128 | 2.851527541 | 0.696730604 |
| Gm9025 | 2.20249356 | 2.947155754 | 0.935163243 |
| Gm15322 | 0.293377407 | -1.282765545 | 3.695996407 |
| Gm15323 | 0.143781519 | -0.710963077 | 3.581286957 |
| Gm37427 | 0 | NA | NA |
| Ankrd55 | 97.25075632 | 0.094076654 | 0.172598497 |
| Gm22768 | 0.696415487 | -1.940546196 | 1.184621705 |
| Il6st | 2388.948021 | 0.523098069 | 0.070771278 |
| Il31ra | 12.44555901 | 0.901572938 | 0.358085075 |
| Ddx4 | 13.81354613 | 1.882193765 | 0.343595646 |
| Slc38a9 | 402.4228987 | -0.423724324 | 0.125684841 |
| Ppap2a | 177.5947723 | 0.092256775 | 0.094777096 |
| Skiv2l2 | 776.3773442 | 0.09424099 | 0.058264236 |
| Gm26943 | 0.235215884 | 1.251453655 | 2.487520524 |
| Dhx29 | 365.0039571 | 0.163897724 | 0.067027441 |
| Ccno | 13.15772638 | -0.120326952 | 0.412560326 |
| Mcidas | 3.060161834 | 0.018509953 | 0.655625758 |
| Cdc20b | 1.498584608 | -0.248141873 | 0.787801021 |
| Mir449c | 0.04393175 | -0.243571806 | 3.724945049 |
| Mir449b | 0.038981395 | 0.718207094 | 3.724945049 |
| Mir449a | 0 | NA | NA |
| Gm10735 | 1.512916718 | 0.41779482 | 0.898947448 |
| Gpx8 | 87.13244342 | 1.753488841 | 0.203755931 |
| Gzma | 199.9537804 | 9.294227136 | 0.793408425 |
| Gm9848 | 0 | NA | NA |
| Gzmk | 39.86666651 | 2.925962596 | 0.357626536 |
| Esm1 | 8.383027344 | 2.112325652 | 0.513504382 |
| BC067074 | 57.61241419 | -0.578827773 | 0.171769397 |
| Gm22385 | 0 | NA | NA |
| Snx18 | 579.4915124 | 0.601570354 | 0.104112468 |
| Hspb3 | 12.0017422 | 0.833248627 | 0.317415611 |
| Arl15 | 416.4383343 | -0.401664542 | 0.093851388 |
| Gm37907 | 0.135812198 | -1.187715541 | 3.646939241 |

| | | | |
|---|---|---|---|
| Ndufs4 | 556.7189796 | 0.165258376 | 0.10171554 |
| Fst | 27.88029129 | 0.272136545 | 0.276231736 |
| Gm25950 | 0 | NA | NA |
| 4930544M13Rik | 0 | NA | NA |
| Mocs2 | 459.8435434 | 0.39255422 | 0.072312359 |
| Itga2 | 10.13867756 | 3.011729083 | 0.48709365 |
| Itga1 | 103.8467935 | 0.835578568 | 0.178402287 |
| Pelo | 139.7216595 | 0.344571796 | 0.110978177 |
| Gm25985 | 0 | NA | NA |
| Gm6404 | 0 | NA | NA |
| Isl1 | 36.11947267 | -0.132696656 | 0.187988535 |
| Gm38193 | 6.0191764 | -0.63982277 | 0.476575509 |
| Gm23198 | 0 | NA | NA |
| Parp8 | 540.5637389 | 0.482340207 | 0.113874015 |
| Gm17509 | 11.37141038 | -0.883612119 | 0.368349586 |
| Emb | 394.0332575 | 0.093667765 | 0.123204566 |
| Gm6421 | 12.74783509 | 0.439238678 | 0.34094543 |
| Hcn1 | 981.9976838 | 0.014645458 | 0.068344119 |
| 4933413L06Rik | 0 | NA | NA |
| Gm26349 | 0.072025462 | 0.237312084 | 3.724945049 |
| Mrps30 | 237.5140506 | 0.349616849 | 0.094292352 |
| B430218F22Rik | 1.974484148 | 0.290157484 | 0.676545512 |
| 1700003P14Rik | 0 | NA | NA |
| Fgf10 | 52.0629943 | -0.802770052 | 0.159703731 |
| Gm16263 | 1.381163524 | -2.190788613 | 1.009045071 |
| AC154241.1 | 0 | NA | NA |
| Nnt | 359.3284767 | -0.267454524 | 0.112111188 |
| Paip1 | 990.7888142 | 0.205766848 | 0.073047546 |
| 4833420G17Rik | 380.1582038 | -0.445518569 | 0.103500896 |
| Gm9752 | 0.269061364 | -0.243585702 | 3.724945049 |
| 3110070M22Rik | 13.70697502 | -0.298614422 | 0.298573747 |
| Gm7120 | 31.73718637 | -0.039521164 | 0.200615198 |
| Gm21967 | 80.16892809 | -0.125079076 | 0.148510972 |
| Ccl28 | 18.99182376 | -0.55395778 | 0.29917335 |
| 1700074H08Rik | 0 | NA | NA |
| Hmgcs1 | 2950.403097 | -0.007024365 | 0.1301002 |
| Nim1k | 238.1724293 | -0.773229408 | 0.133757969 |
| Zfp131 | 574.5633546 | 0.418212415 | 0.087059021 |
| Gm23308 | 0 | NA | NA |
| AF067063 | 0 | NA | NA |
| D13Ertd608e | 1.077945179 | 3.036017143 | 1.395612343 |
| Gm21358 | 0 | NA | NA |
| Gm37619 | 0.171181394 | 0.718216336 | 2.749537022 |
| Tcstv1 | 0 | NA | NA |
| Gm21761 | 0 | NA | NA |
| Gm21181 | 0 | NA | NA |
| B020031M17Rik | 0 | NA | NA |

| | | | |
|---|---|---|---|
| Gm20784 | 1.793435801 | -0.270225181 | 0.713181459 |
| Gm22108 | 0 | NA | NA |
| Gm21370 | 0 | NA | NA |
| Gm21188 | 0.521673782 | 2.887694495 | 1.351027426 |
| Gm24798 | 0 | NA | NA |
| Gm21378 | 13.07532112 | 0.382995401 | 0.306237047 |
| Gm20767 | 0 | NA | NA |
| Gm21818 | 0.095898756 | 1.192083721 | 3.724434897 |
| Gm21762 | 1.958730402 | 4.132632069 | 1.092346171 |
| Gm21731 | 0 | NA | NA |
| AF067061 | 0 | NA | NA |
| Gm20900 | 10.70701964 | 0.281572518 | 0.364380932 |
| BC147527 | 33.1074984 | 8.402665461 | 0.925437714 |
| Tcstv3 | 0 | NA | NA |
| Gm2888 | 0 | NA | NA |
| Gm2897 | 10.21470906 | 0.080570614 | 0.318586693 |
| Gm2904 | 0.731111494 | 1.209138579 | 1.243153522 |
| Gm17051 | 0 | NA | NA |
| Gm10340 | 0.412722675 | -0.412042763 | 1.619108408 |
| Gm17221 | 0 | NA | NA |
| Gm17188 | 0 | NA | NA |
| Gm5795 | 0.0878635 | -0.72445792 | 3.724945049 |
| Gm17222 | 1.483555279 | 2.603491254 | 0.973524401 |
| D830030K20Rik | 12.49083266 | -0.457412612 | 0.306983644 |
| Gm4904 | 0 | NA | NA |
| 1700110I01Rik | 0.924198543 | -0.7841317 | 0.988098027 |
| Gm2956 | 0.84966665 | -1.985852532 | 1.246466966 |
| Gm17223 | 0 | NA | NA |
| Gm10413 | 0 | NA | NA |
| Gm10409 | 0.801287066 | -0.031586351 | 1.41882956 |
| Gm3005 | 2.629406015 | 0.301348048 | 0.84740325 |
| Gm26817 | 0 | NA | NA |
| Gm10408 | 0.044191549 | -0.243571806 | 3.724945049 |
| Gm8323 | 0 | NA | NA |
| Gm8324 | 0 | NA | NA |
| Gm11109 | 0 | NA | NA |
| Gm17107 | 0 | NA | NA |
| Gm3012 | 0.072025462 | 0.237312084 | 3.724945049 |
| Gm3020 | 0.88261066 | 0.494745924 | 1.235676852 |
| Gm2866 | 0 | NA | NA |
| Gm3043 | 0 | NA | NA |
| Gm2871 | 0 | NA | NA |
| Gm2877 | 0 | NA | NA |
| Gm21965 | 0.038981395 | 0.718207094 | 3.724945049 |
| Gm3002 | 2.349117343 | -0.001219793 | 0.606440783 |
| Gm26886 | 0 | NA | NA |
| Gm26570 | 0 | NA | NA |

| | | | |
|---|---|---|---|
| Gm3033 | 0.403254479 | 1.848388269 | 1.646037256 |
| Gm2907 | 0 | NA | NA |
| Gm2916 | 0.044191549 | -0.243571806 | 3.724945049 |
| Gm3029 | 0 | NA | NA |
| Gm3095 | 0.864161856 | -0.061654476 | 1.108273699 |
| Gm17129 | 0 | NA | NA |
| Gm3099 | 0.095898756 | 1.192083721 | 3.724434897 |
| Gm5796 | 5.645051647 | 2.311454271 | 0.554173213 |
| Gm11107 | 0 | NA | NA |
| Gm3115 | 0.045882455 | 0.237312084 | 3.724945049 |
| Gm8108 | 0.608376789 | -1.213522233 | 1.291428235 |
| Gm17152 | 0 | NA | NA |
| Gm3127 | 0.0878635 | -0.72445792 | 3.724945049 |
| Gm2974 | 9.121692315 | 0.816227812 | 0.370426839 |
| Gm11105 | 0 | NA | NA |
| Gm3138 | 0.101408268 | 0.717551239 | 3.724898127 |
| Gm17085 | 0 | NA | NA |
| Gm3149 | 0.060174537 | 0.237309859 | 3.724945049 |
| 2610042L04Rik | 9.926510475 | 1.73358055 | 0.381203125 |
| Gm17086 | 0 | NA | NA |
| Gm3159 | 0.075043205 | 0.237309859 | 3.724945049 |
| Gm3164 | 5.22863904 | 0.71706326 | 0.542950914 |
| Gm3173 | 3.269157551 | 1.524417608 | 0.725455198 |
| Gm17170 | 0 | NA | NA |
| Gm3182 | 0.048949532 | -0.243569582 | 3.724945049 |
| Gm17074 | 0 | NA | NA |
| Gm3030 | 0.11997416 | -0.243571806 | 3.724945049 |
| Gm26655 | 0 | NA | NA |
| Gm3047 | 0.107078754 | 0.236669296 | 3.724898107 |
| Gm8164 | 0 | NA | NA |
| Gm26898 | 0.093141081 | -0.724455696 | 3.724945049 |
| Gm8159 | 0 | NA | NA |
| Gm3239 | 1.60056093 | -0.108211671 | 0.908773823 |
| Gm17073 | 0 | NA | NA |
| Gm7876 | 0.046345709 | 0.71820487 | 3.724945049 |
| Gm3252 | 10.87374526 | 0.582907701 | 0.413034618 |
| Gm3084 | 0 | NA | NA |
| Gm9602 | 0.3514278 | -0.252761832 | 1.744757875 |
| Gm3091 | 0 | NA | NA |
| Gm3269 | 0 | NA | NA |
| Gm3264 | 2.10462685 | 2.107927086 | 0.912808914 |
| Gm26650 | 0 | NA | NA |
| Gm3278 | 0.274437282 | -1.118701382 | 1.607404148 |
| Gm8305 | 0 | NA | NA |
| Gm10410 | 0 | NA | NA |
| Gm8297 | 0 | NA | NA |
| Gm3298 | 0.938975218 | 1.131774172 | 1.024630523 |

| | | | |
|---|---|---|---|
| 4930555G01Rik | 0.088534442 | 1.179809211 | 3.723500796 |
| Gm8281 | 11.41715526 | -0.196528012 | 0.345317579 |
| Gm17069 | 0 | NA | NA |
| Gm8279 | 0.163190422 | 1.22955094 | 3.396300676 |
| Gm26552 | 0 | NA | NA |
| Gm3317 | 1.057189525 | -0.056345866 | 1.216636894 |
| Gm3140 | 0 | NA | NA |
| Gm8271 | 0.068097359 | -0.243569582 | 3.724945049 |
| Gm3326 | 0 | NA | NA |
| Gm3526 | 0.137762903 | -0.72445853 | 2.937523348 |
| Gm3339 | 0 | NA | NA |
| Gm3542 | 0 | NA | NA |
| Gm17126 | 0 | NA | NA |
| Gm3500 | 0.712486263 | 0.14672153 | 1.02657855 |
| Gm3348 | 0 | NA | NA |
| Gm3194 | 0.375475346 | 0.674872203 | 1.718966938 |
| Gm8246 | 0 | NA | NA |
| Gm3187 | 0.044191549 | -0.243571806 | 3.724945049 |
| Gm3488 | 1.283947918 | -0.001974318 | 0.856589559 |
| Gm26893 | 0 | NA | NA |
| Gm26746 | 0 | NA | NA |
| Gm3532 | 0 | NA | NA |
| Gm8266 | 0 | NA | NA |
| Gm26731 | 0 | NA | NA |
| Gm3383 | 0.828062179 | -0.083230874 | 1.200928241 |
| Gm3344 | 0 | NA | NA |
| Gm8265 | 0 | NA | NA |
| Gm9603 | 0.111006857 | 0.717551239 | 3.724898127 |
| Gm3373 | 0.817666292 | 0.758366174 | 1.107507697 |
| Gm3170 | 0 | NA | NA |
| Gm26557 | 0 | NA | NA |
| Gm3424 | 0.342449771 | 0.433004533 | 1.533413365 |
| Gm17158 | 0 | NA | NA |
| Gm8237 | 12.34732519 | 0.97950502 | 0.384337145 |
| Gm17046 | 0.139037128 | 1.464453538 | 3.717743456 |
| Gm3242 | 0 | NA | NA |
| Gm17047 | 0 | NA | NA |
| Gm3248 | 20.30228851 | 0.486926101 | 0.2369883 |
| Gm17048 | 0 | NA | NA |
| Gm3453 | 0.066050875 | 0.71820487 | 3.724945049 |
| Gm3261 | 0 | NA | NA |
| Gm8206 | 2.632186321 | 0.477343997 | 0.594027948 |
| Gm3468 | 14.54750696 | 0.60718252 | 0.364942516 |
| Gm17045 | 0.385383671 | -0.289692325 | 1.601407075 |
| Gm6337 | 0 | NA | NA |
| Gm3275 | 0 | NA | NA |
| Gm3476 | 0.611787247 | -0.001147677 | 1.402618957 |

| | | | |
|---|---|---|---|
| Gm26991 | 0 | NA | NA |
| Gm3208 | 0 | NA | NA |
| Gm21560 | 0 | NA | NA |
| Gm21772 | 0 | NA | NA |
| Gm3411 | 2.644118259 | 0.030857704 | 0.681899973 |
| Gm21103 | 0 | NA | NA |
| Gm3618 | 0 | NA | NA |
| Gm26630 | 0 | NA | NA |
| Gm8340 | 0 | NA | NA |
| Gm16440 | 0.365173462 | -0.658150218 | 1.43778621 |
| Gm3591 | 2.764354947 | 1.657766115 | 0.67721984 |
| Gm3594 | 0 | NA | NA |
| Gm8356 | 0.045882455 | 0.237312084 | 3.724945049 |
| Gm3629 | 1.032577368 | -1.960775264 | 1.095950896 |
| Gm3460 | 0 | NA | NA |
| Gm10251 | 0.095435502 | 0.718202646 | 3.724945049 |
| Gm26680 | 0 | NA | NA |
| Gm3466 | 0 | NA | NA |
| Gm8050 | 0.427185557 | -0.636932964 | 1.529478748 |
| Gm3636 | 17.74053186 | 0.595269657 | 0.253302358 |
| Gm8362 | 0 | NA | NA |
| Gm3642 | 0.049553047 | 0.71820487 | 3.724945049 |
| Gm3667 | 13.76281871 | 0.340352187 | 0.316932086 |
| Gm6356 | 0.072025462 | 0.237312084 | 3.724945049 |
| Gm10406 | 10.70611474 | 1.098101214 | 0.365359763 |
| Gm3685 | 0.098502579 | 0.23732098 | 3.724945049 |
| Gm3696 | 16.41020923 | 0.627958521 | 0.276182815 |
| Gm6676 | 0.092269992 | 0.237318756 | 3.724945049 |
| Gm3512 | 4.130618934 | 1.569258948 | 0.593407143 |
| Gm16434 | 0 | NA | NA |
| Gm3715 | 1.793529573 | 0.856483977 | 0.760189365 |
| Gm5456 | 0.437842326 | 0.206677945 | 1.376783381 |
| Gm3727 | 0.094294407 | 0.23732098 | 3.724945049 |
| Gm3739 | 19.15242329 | 0.810457969 | 0.303494223 |
| Gm5797 | 0.049553047 | 0.71820487 | 3.724945049 |
| Gm8374 | 0.038981395 | 0.718207094 | 3.724945049 |
| Gm10339 | 0 | NA | NA |
| 4933406F09Rik | 0.083526043 | 0.718202646 | 3.724945049 |
| Gm8379 | 0.046345709 | 0.71820487 | 3.724945049 |
| Gm3755 | 0.852403115 | -0.708908015 | 1.045295617 |
| Gm3752 | 3.599181491 | 1.125575103 | 0.732995597 |
| Gm3760 | 0.150455055 | 1.610657435 | 2.838957197 |
| Gm3558 | 7.230339464 | -0.450249326 | 0.356794307 |
| Gm17121 | 0.092140247 | -0.724455696 | 3.724945049 |
| Gm10338 | 0 | NA | NA |
| Gm10128 | 0.356838182 | 1.196023368 | 1.761125647 |
| Gm10044 | 5.427300065 | -0.634620021 | 0.449452764 |

| | | | |
|---|---|---|---|
| Gm16741 | 0.543837322 | -0.66771797 | 1.293515809 |
| Gm26661 | 0.047948698 | -0.243569582 | 3.724945049 |
| Gm8396 | 0.090277459 | 0.237318756 | 3.724945049 |
| Flnb | 1267.997399 | 0.174174264 | 0.150718726 |
| Dnase1l3 | 17.4478056 | 5.198923074 | 0.769560024 |
| Abhd6 | 413.5173793 | -0.325357868 | 0.077963057 |
| Rpp14 | 225.7839547 | -0.009806503 | 0.079881319 |
| Pxk | 710.6699321 | 0.857588189 | 0.086889918 |
| Pdhb | 939.7685917 | -0.245568087 | 0.086369958 |
| Kctd6 | 337.2401318 | -0.136727451 | 0.144340225 |
| Acox2 | 3.533179683 | 0.729173243 | 0.611138459 |
| Fam107a | 4664.005083 | 0.50406192 | 0.182739684 |
| Oit1 | 0 | NA | NA |
| 4930452B06Rik | 90.32014768 | -1.081037637 | 0.148131647 |
| 4930455B14Rik | 0 | NA | NA |
| Gm9800 | 154.1573682 | 0.977383816 | 0.174463123 |
| Fhit | 18.766523 | -0.681126883 | 0.249739874 |
| Hmgb3-ps | 0 | NA | NA |
| Mnd1-ps | 0.293028276 | -1.826555226 | 3.692614927 |
| Gm22053 | 0 | NA | NA |
| Fhitos | 0 | NA | NA |
| Slc25a5-ps | 0.541865536 | -0.225178492 | 1.473709756 |
| Rpl21-ps4 | 0.428504882 | 0.654121507 | 1.72834915 |
| Gm3839 | 0.806310386 | -1.471968658 | 1.148003005 |
| AC158396.1 | 0 | NA | NA |
| Mir5124b | 0 | NA | NA |
| Gm25466 | 0 | NA | NA |
| Ptprg | 1480.140492 | -0.357266894 | 0.060661389 |
| Gm3848 | 0.118974955 | -0.243576254 | 3.724945049 |
| 3830406C13Rik | 205.9760077 | -0.14602468 | 0.108554095 |
| Gm24578 | 0.075043205 | 0.237309859 | 3.724945049 |
| Fezf2 | 140.4500694 | -0.643216691 | 0.129703661 |
| Cadps | 3883.453482 | -0.542267488 | 0.052957079 |
| n-R5s45 | 5.980340237 | -1.981435917 | 0.558807842 |
| Gm5457 | 6.294359381 | -0.530796091 | 0.45772506 |
| Gm5087 | 8.752821209 | -0.125463026 | 0.388101167 |
| Synpr | 975.6483675 | -0.478304414 | 0.167684743 |
| Gm15913 | 0 | NA | NA |
| Sntn | 3.353049333 | 0.426322618 | 0.59324464 |
| Rpl19-ps4 | 1.071806319 | 0.184096135 | 1.091839656 |
| Gm27506 | 0 | NA | NA |
| Gm281 | 0.82274574 | 1.22754841 | 1.030987231 |
| Gm25964 | 0 | NA | NA |
| Gm11100 | 1.662603624 | 3.428321635 | 1.016776594 |
| Thoc7 | 350.5995921 | 0.230706857 | 0.096997137 |
| Atxn7 | 575.3815957 | -0.03308886 | 0.092724033 |
| Psmd6 | 699.5893116 | 0.488737152 | 0.087664188 |

| | | | |
|---|---|---|---|
| Olfr720 | 0 | NA | NA |
| Olfr31 | 0.115957212 | -0.24357403 | 3.724945049 |
| Il3ra | 52.25554781 | 3.106840372 | 0.32104349 |
| Olfr721-ps1 | 0 | NA | NA |
| Slc4a7 | 745.5514613 | 0.192122366 | 0.088139688 |
| Nek10 | 32.25918065 | -1.714504003 | 0.228049281 |
| Lrrc3b | 129.9205039 | -0.501203016 | 0.099925925 |
| B230110C06Rik | 12.80781388 | -1.251299905 | 0.350872628 |
| Oxsm | 84.03786033 | 0.414484881 | 0.154947378 |
| Ngly1 | 371.6785756 | 0.114127767 | 0.070177732 |
| Top2b | 2165.3358 | -0.110475875 | 0.067381404 |
| Rarb | 169.0811627 | -0.717782476 | 0.114598636 |
| Thrb | 548.3553106 | -0.823106767 | 0.098679345 |
| Gm24140 | 0 | NA | NA |
| Gm27851 | 0 | NA | NA |
| Gm6773 | 0.143022002 | 0.719176055 | 3.547735711 |
| CU024892.1 | 0 | NA | NA |
| Nr1d2 | 2201.828554 | -0.420804191 | 0.062771642 |
| Rpl15 | 1078.184859 | 0.552160256 | 0.120753294 |
| Nkiras1 | 524.1011176 | -0.646029932 | 0.082206543 |
| Ube2e1 | 611.3267082 | 0.305608728 | 0.099840261 |
| Gm22350 | 0.766686841 | 0.680922189 | 1.399846961 |
| Gm6781 | 3.825407769 | 0.642842577 | 0.613237306 |
| Ube2e2 | 649.6422344 | -0.358869821 | 0.08387683 |
| Gm20678 | 11.63441953 | -0.39111692 | 0.449162071 |
| Gm20699 | 63.74099188 | -1.040385498 | 0.316771057 |
| Gm22499 | 0 | NA | NA |
| Gm15916 | 0 | NA | NA |
| Gm20677 | 0 | NA | NA |
| Gm8582 | 0.356401535 | -0.237971111 | 1.756649906 |
| Gm8584 | 0.374155823 | -0.240899135 | 1.753231061 |
| Gm9637 | 0 | NA | NA |
| Gm21738 | 1.310113234 | 1.56876572 | 0.898942928 |
| Gm7352 | 0 | NA | NA |
| Gm2244 | 0.202043434 | 0.264367403 | 2.607302291 |
| Gm2237 | 15.97977484 | 0.739635545 | 0.296195222 |
| Gm5458 | 0.372874528 | -1.106867355 | 1.542095856 |
| 4930469B13Rik | 0 | NA | NA |
| Gm17030 | 0.08169834 | 0.23732098 | 3.724945049 |
| Nid2 | 107.7830705 | 1.392530264 | 0.17106994 |
| 2700060E02Rik | 556.5044864 | 0.65243304 | 0.10017701 |
| Gng2 | 1773.216353 | -0.19391428 | 0.065169108 |
| Saysd1 | 59.98590933 | 0.146069233 | 0.147614693 |
| Kcnk5 | 24.84306935 | 1.135238839 | 0.260495756 |
| Gm24912 | 0 | NA | NA |
| Kcnk16 | 0.038981395 | 0.718207094 | 3.724945049 |
| Nudt13 | 124.3202546 | 0.724774147 | 0.128562485 |

| | | | |
|---|---|---|---|
| Ecd | 421.6024906 | 0.280307216 | 0.071916986 |
| Fam149b | 307.5778511 | -0.204266443 | 0.086163707 |
| Dnajc9 | 241.2584147 | 0.267943113 | 0.119368418 |
| Mrps16 | 150.6596876 | 0.529606542 | 0.106915735 |
| Cfap70 | 17.61224759 | 0.086472168 | 0.261737605 |
| 7330404K18Rik | 0 | NA | NA |
| Anxa7 | 995.7442115 | 0.759505206 | 0.084263164 |
| Mss51 | 28.46984622 | -0.313785188 | 0.264153996 |
| Ppp3cb | 3014.123295 | -0.485301861 | 0.060287109 |
| 1810062O18Rik | 11.1715319 | -0.169198013 | 0.330905554 |
| Usp54 | 1368.558277 | -0.069612706 | 0.072149474 |
| 2810402E24Rik | 26.39841974 | -0.321636661 | 0.208873496 |
| Myoz1 | 2.494795275 | 0.349427525 | 0.78454477 |
| Synpo2l | 4.120754503 | 0.332067058 | 0.592039709 |
| Sec24c | 1144.691689 | 0.486510179 | 0.072819503 |
| 6230400D17Rik | 14.90222924 | 0.043768919 | 0.278896562 |
| Mir6946 | 0.27962944 | -0.549028774 | 2.462540246 |
| Fut11 | 188.4790295 | -0.315298949 | 0.093481398 |
| Gm25328 | 3.501233001 | -0.35135491 | 0.581674073 |
| Chchd1 | 206.0508832 | 0.76821358 | 0.112989709 |
| Zswim8 | 1959.999352 | 0.192406654 | 0.072786116 |
| Ndst2 | 264.5758868 | 0.8236337 | 0.111239357 |
| Camk2g | 2341.047828 | -0.756595282 | 0.056359712 |
| Plau | 140.9153563 | 2.51317405 | 0.290400964 |
| Vcl | 823.9348543 | 0.740126948 | 0.113108255 |
| Ap3m1 | 731.0242943 | 0.102885474 | 0.068158776 |
| Adk | 509.0668211 | -0.44201026 | 0.130049872 |
| Gm26372 | 4.248258162 | -0.849615511 | 0.53382917 |
| Gm25864 | 0.26743452 | -0.226601444 | 2.086623697 |
| Kat6b | 1504.25356 | -0.494477445 | 0.079305359 |
| Dupd1 | 0.592353126 | 0.758927555 | 1.341292733 |
| Dusp13 | 5.809763333 | 0.975698861 | 0.46050097 |
| Gm15935 | 0.68377221 | 3.167868916 | 1.389851071 |
| Samd8 | 1631.759024 | 0.118856278 | 0.075253417 |
| Gm23502 | 0.371001759 | -2.036151632 | 3.187383283 |
| Vdac2 | 1632.626953 | 0.153756545 | 0.068071131 |
| Comtd1 | 88.38321911 | 0.115771506 | 0.119418448 |
| A430057M04Rik | 12.38941375 | 0.381241621 | 0.341151186 |
| Gm37590 | 0 | NA | NA |
| Zfp503 | 121.7214339 | 0.008349133 | 0.157671938 |
| C130012C08Rik | 2.040813782 | -0.864386485 | 0.746187219 |
| 1700112E06Rik | 23.04170766 | 0.999114458 | 0.311844822 |
| Gm27532 | 0 | NA | NA |
| Gm5670 | 0 | NA | NA |
| Gm10248 | 1.061926442 | 1.233840834 | 1.127017533 |
| Kcnma1 | 2520.351437 | -0.709215158 | 0.120625604 |
| A830039N20Rik | 615.447544 | -0.746108638 | 0.114262735 |

| | | | |
|---|---|---|---|
| Gm29626 | 0 | NA | NA |
| Gm6158 | 5.934383314 | -0.082631993 | 0.405420738 |
| mmu-mir-7210 | 0 | NA | NA |
| Dlg5 | 927.9578943 | 0.426390689 | 0.083289154 |
| Gm17105 | 11.22543593 | -0.35799556 | 0.399008347 |
| E330034G19Rik | 0.520501943 | -0.338923378 | 1.503710021 |
| Polr3a | 517.7839581 | -0.084041406 | 0.066617295 |
| Rps24 | 2358.715317 | 0.822539661 | 0.106654699 |
| 4930542C16Rik | 0.048949532 | -0.243569582 | 3.724945049 |
| 4931403M11Rik | 0.541236733 | 0.568949568 | 1.56177793 |
| 4930428N03Rik | 0.347568383 | 1.671989832 | 1.725345179 |
| Gm10398 | 0 | NA | NA |
| Gm26660 | 26.92887363 | -0.727670716 | 0.289482751 |
| Zmiz1os1 | 24.20908764 | -0.376057079 | 0.208559411 |
| Zmiz1 | 3086.44194 | 0.350710227 | 0.081494541 |
| Mir3075 | 1.765392608 | -0.930009145 | 0.798578772 |
| Gm26772 | 0.433939115 | 0.656342236 | 1.418596323 |
| Ppif | 256.0875963 | 0.036850453 | 0.079466139 |
| 1700054O19Rik | 0.226439421 | 0.239954661 | 2.155523336 |
| Zcchc24 | 1699.51765 | 0.262506782 | 0.127187289 |
| Gm27185 | 0 | NA | NA |
| Gm23701 | 0 | NA | NA |
| Anxa11 | 152.4094582 | 0.41105692 | 0.09805889 |
| Anxa11os | 1.923336751 | 4.166346839 | 0.980600458 |
| Plac9a | 0 | NA | NA |
| Tmem254a | 0 | NA | NA |
| Cphx1 | 0 | NA | NA |
| Gm19013 | 0 | NA | NA |
| Duxbl1 | 0.094294407 | 0.23732098 | 3.724945049 |
| Gm2260 | 1.263942296 | -0.043707282 | 1.158248094 |
| Gm9780 | 0.038981395 | 0.718207094 | 3.724945049 |
| Tmem254c | 0.63148455 | -1.496258528 | 1.365662241 |
| Gm19014 | 0 | NA | NA |
| Cphx2 | 0 | NA | NA |
| Cphx3 | 0 | NA | NA |
| Duxbl2 | 0 | NA | NA |
| Gm2274 | 0.075043205 | 0.237309859 | 3.724945049 |
| Plac9b | 0 | NA | NA |
| Tmem254b | 0 | NA | NA |
| Gm19015 | 0 | NA | NA |
| Duxbl3 | 0 | NA | NA |
| 4933413J09Rik | 0 | NA | NA |
| Gm16245 | 0.158569248 | 0.718214251 | 2.811594796 |
| Slmapos2 | 0 | NA | NA |
| Slmap | 1697.427193 | -0.350911697 | 0.045384688 |
| Gm24031 | 0 | NA | NA |
| Gm2178 | 0 | NA | NA |

| | | | |
|---|---|---|---|
| Dennd6a | 497.1097104 | -0.174127885 | 0.081679937 |
| Arf4os | 0.854956099 | 0.466676376 | 1.055538656 |
| Arf4 | 1525.733304 | 1.198985041 | 0.129657824 |
| Pde12 | 370.8916435 | 0.323716194 | 0.139214481 |
| Mir7672 | 0 | NA | NA |
| 9930004E17Rik | 4.759985783 | -0.335034435 | 0.49158259 |
| Dnah12 | 39.20255926 | -0.437493942 | 0.218221363 |
| Asb14 | 0.828364824 | -1.224132552 | 1.096887721 |
| Appl1 | 1598.16168 | -0.269750181 | 0.049962655 |
| Hesx1 | 0 | NA | NA |
| Il17rd | 75.9257241 | -0.409213686 | 0.178089205 |
| Arhgef3 | 850.6410935 | -0.230980185 | 0.060752095 |
| Gm23633 | 0 | NA | NA |
| Fam208a | 1486.670546 | -0.08759879 | 0.119952981 |
| Ccdc66 | 229.1061877 | -0.13125755 | 0.125028207 |
| Gm22485 | 2.784173388 | -0.180952937 | 0.729074544 |
| Gm25697 | 0 | NA | NA |
| Erc2 | 1776.688752 | -0.724008325 | 0.112007176 |
| Wnt5a | 214.1936815 | 0.022810387 | 0.089723613 |
| Gm2670 | 2.740285017 | 3.544649317 | 0.927511583 |
| Cacna2d3 | 929.5729355 | -0.848467343 | 0.116530911 |
| Gm23999 | 4.885903765 | -1.900262231 | 0.665982811 |
| Lrtm1 | 0.20890592 | -0.235220952 | 2.608283839 |
| Gm6043 | 0.823931708 | -0.328968306 | 1.0837591 |
| Selk | 760.4563827 | 0.361076281 | 0.108773202 |
| Actr8 | 526.6961103 | -0.11920722 | 0.068411005 |
| mmu-mir-6947 | 1.07422636 | -0.958459459 | 1.137185652 |
| Il17rb | 16.19919013 | 0.085208108 | 0.243373548 |
| Chdh | 62.42317444 | -0.282003435 | 0.237992706 |
| Cacna1d | 1000.382172 | -0.577042402 | 0.148459233 |
| Dcp1a | 375.2765008 | 0.125502332 | 0.067377702 |
| Tkt | 1261.009206 | 0.016136977 | 0.075206182 |
| Mir3076 | 0.281790215 | 0.770862473 | 2.065210462 |
| Prkcd | 783.4000836 | 0.320185946 | 0.099454078 |
| Rft1 | 140.5032926 | 0.248399251 | 0.116317807 |
| Sfmbt1 | 877.7713785 | 0.125281835 | 0.06155847 |
| 1700087M22Rik | 0.048949532 | -0.243569582 | 3.724945049 |
| Gm15667 | 10.02380282 | -0.157666066 | 0.377978273 |
| Tmem110 | 229.3938144 | -0.367094587 | 0.092567226 |
| Mustn1 | 28.82764176 | 2.162687007 | 0.324764571 |
| Itih4 | 5.953922204 | 3.225960991 | 0.698989121 |
| Itih3 | 692.5237228 | -0.153400136 | 0.165545842 |
| Itih1 | 0 | NA | NA |
| Nek4 | 254.004222 | -0.057287911 | 0.08981705 |
| Spcs1 | 557.2353825 | 0.592047906 | 0.146777123 |
| Glt8d1 | 210.2471507 | -0.136247089 | 0.101707585 |
| Gnl3 | 342.5965878 | 0.699191978 | 0.168757886 |

| | | | |
|---|---|---|---|
| Snord69 | 6.499084164 | 0.20874501 | 0.448337216 |
| Gm24916 | 0.901079044 | 1.68401242 | 1.15023352 |
| Snord19 | 6.156095358 | 0.038497069 | 0.526357258 |
| Pbrm1 | 1674.501212 | 0.099517156 | 0.044976569 |
| Gm5459 | 0.740733539 | -0.576130435 | 1.326860213 |
| Smim4 | 39.16029245 | -0.085479617 | 0.173569817 |
| Nt5dc2 | 56.55209549 | 0.465175864 | 0.176295989 |
| Stab1 | 251.2128195 | 0.993053742 | 0.140653928 |
| Nisch | 5669.263495 | -0.156278725 | 0.043236188 |
| Tnnc1 | 26.93316509 | -1.222165278 | 0.250953574 |
| Sema3g | 57.06689575 | 2.258607982 | 0.244393454 |
| Phf7 | 44.60374799 | 0.161618161 | 0.190019931 |
| Bap1 | 1295.484164 | -0.022536798 | 0.066142364 |
| Dnah1 | 99.51919862 | -0.249350237 | 0.124689744 |
| Capn7 | 711.6756629 | -0.174197208 | 0.067829201 |
| Sh3bp5 | 520.8229502 | -0.054506522 | 0.090905189 |
| Mettl6 | 226.22855 | 0.286775046 | 0.104646514 |
| Eaf1 | 382.8875853 | 0.840335149 | 0.121806465 |
| Colq | 11.36104049 | -0.626054169 | 0.360893134 |
| Gm24057 | 0.166923653 | -0.724459871 | 2.766511172 |
| Hacl1 | 56.72216686 | -0.495055284 | 0.226795187 |
| Btd | 112.7781148 | 0.54824357 | 0.140172984 |
| Ankrd28 | 1173.696225 | 0.30082011 | 0.071903126 |
| Gm22876 | 0.047948698 | -0.243569582 | 3.724945049 |
| D830044D21Rik | 0.364803694 | -0.724461113 | 1.744007999 |
| Gm26045 | 0 | NA | NA |
| Galnt15 | 194.1661234 | 2.02030276 | 0.243049105 |
| Dph3 | 331.9002246 | 0.077853099 | 0.096473421 |
| Oxnad1 | 151.3315786 | 0.015931778 | 0.124409348 |
| Msmb | 0 | NA | NA |
| Ncoa4 | 556.7801885 | 0.298990384 | 0.075607707 |
| Timm23 | 256.1091139 | 0.471986195 | 0.118944421 |
| Gm23946 | 1.072725949 | -1.244918652 | 0.947521931 |
| Parg | 736.7172274 | -0.013463028 | 0.053668539 |
| Gm7707 | 0.203659793 | 1.199956923 | 2.597136559 |
| Gm17210 | 0.136194719 | -0.724453471 | 3.724945049 |
| Ogdhl | 1425.208207 | -0.203966549 | 0.069148097 |
| 1700024G13Rik | 0.700875826 | -1.479239838 | 1.157533934 |
| Chat | 45.18468771 | 0.138198568 | 0.202594555 |
| Slc18a3 | 49.59830663 | 0.325307888 | 0.336904845 |
| Gm2990 | 1.893473903 | -1.407403299 | 0.816817837 |
| Ercc6 | 588.9859623 | -0.268058286 | 0.11331923 |
| Prrxl1 | 7.584775115 | 0.074844938 | 0.598495177 |
| 3425401B19Rik | 4.622495198 | -0.36613311 | 0.507993084 |
| Gm28651 | 0.300197857 | 0.10489027 | 1.716990986 |
| 1810011H11Rik | 3.566868347 | 0.472759359 | 0.574987089 |
| Fam170b | 0.045882455 | 0.237312084 | 3.724945049 |

| | | | |
|---|---|---|---|
| Vstm4 | 53.0402276 | 0.633504193 | 0.25854819 |
| Gm7734 | 0.640869663 | -0.461665391 | 1.265188097 |
| Wdfy4 | 177.0804989 | 2.373751896 | 0.184071708 |
| Lrrc18 | 16.544867 | 1.551080753 | 0.331051971 |
| Gm25498 | 0 | NA | NA |
| Arhgap22 | 86.59583876 | 0.67268152 | 0.13991565 |
| Gm15512 | 6.43024678 | -1.124112498 | 0.485881359 |
| Mapk8 | 1298.322725 | -0.458168342 | 0.063608423 |
| Ptpn20 | 4.379735502 | -0.180604449 | 0.462544784 |
| Gm26228 | 0 | NA | NA |
| Gdf10 | 323.3389736 | 0.21616611 | 0.354659907 |
| Gdf2 | 0.41755988 | 0.750895706 | 1.446527949 |
| Rbp3 | 3.927501353 | 1.027594681 | 0.735805634 |
| Zfp488 | 48.54553077 | 0.307811017 | 0.185429253 |
| Gm5460 | 0 | NA | NA |
| A630023A22Rik | 0.092691418 | 1.183079489 | 3.724560917 |
| Antxrl | 0 | NA | NA |
| Anxa8 | 7.103537313 | 1.670892204 | 0.527370623 |
| Gm24998 | 0 | NA | NA |
| Npy4r | 1.230904645 | -0.364590227 | 0.995648136 |
| Rpl23a-ps3 | 207.5274121 | 0.617858723 | 0.13926014 |
| Gprin2 | 18.00709799 | -0.061992689 | 0.297811165 |
| Syt15 | 3.163297818 | 0.070576607 | 0.515990534 |
| Fam35a | 121.8552786 | -0.329438088 | 0.131535717 |
| Glud1 | 2362.144562 | -0.484560477 | 0.104800305 |
| Fam25c | 0.591489534 | 0.821792915 | 1.340404094 |
| Sncg | 85.14162449 | 0.120627343 | 0.327105225 |
| Mmrn2 | 26.86649225 | 2.900237943 | 0.327234137 |
| Bmpr1a | 1071.470205 | 0.042392686 | 0.093709133 |
| Gm17110 | 1.151183522 | -0.43206078 | 1.011169577 |
| 9230112D13Rik | 0.075043205 | 0.237309859 | 3.724945049 |
| Ldb3 | 91.38560832 | -0.765545087 | 0.153309251 |
| Opn4 | 1.339915753 | 2.113076511 | 1.01893836 |
| Wapal | 1670.7941 | 0.293555673 | 0.070491328 |
| A930038B10Rik | 0.504415982 | -1.281965114 | 1.349708473 |
| Grid1 | 726.7856293 | -0.484751101 | 0.066220583 |
| Mir346 | 1.214291556 | -1.091244403 | 1.064789135 |
| Gm27617 | 0.732444829 | -0.049457587 | 1.068829412 |
| 4930596D02Rik | 0 | NA | NA |
| Gm7853 | 0 | NA | NA |
| 4930474N05Rik | 0 | NA | NA |
| Gm24394 | 0 | NA | NA |
| Gm25204 | 0 | NA | NA |
| n-R5s46 | 0.182014725 | 0.23731949 | 2.695472047 |
| Ccser2 | 2137.802388 | -0.101177101 | 0.037907728 |
| Rgr | 0.246474914 | -0.243563134 | 2.09234022 |
| Lrit1 | 0.184428051 | 1.460668374 | 2.661975286 |

| | | | |
|---|---|---|---|
| Lrit2 | 0.338204618 | 1.978922994 | 3.687881963 |
| Cdhr1 | 304.1165299 | -0.004848497 | 0.587948272 |
| 2610528A11Rik | 2.326728918 | 1.159276132 | 0.624076966 |
| Ghitm | 4045.339356 | 0.282367141 | 0.090186063 |
| Gm22469 | 0 | NA | NA |
| Gm23909 | 0 | NA | NA |
| Nrg3 | 1077.56296 | -0.556771385 | 0.135868019 |
| Gm20641 | 0 | NA | NA |
| Nrg3os | 0 | NA | NA |
| Gm27332 | 2.333541239 | -0.538345756 | 0.60950817 |
| Gm20642 | 0.139856492 | 0.237322262 | 3.574411025 |
| 1700109I08Rik | 0 | NA | NA |
| Eif1-ps1 | 3.471245013 | -0.642738911 | 0.5049398 |
| Gm25012 | 0 | NA | NA |
| Gm22191 | 0.04393175 | -0.243571806 | 3.724945049 |
| Gm22726 | 0.094831987 | -0.243571806 | 3.724945049 |
| Gm24388 | 0 | NA | NA |
| Gm5205 | 0.0885994 | -0.24357403 | 3.724945049 |
| Sh2d4b | 40.5303677 | -0.246254243 | 0.252191753 |
| Tspan14 | 417.092454 | 0.44902375 | 0.089461218 |
| Fam213a | 601.8361932 | -0.264760311 | 0.114268251 |
| Dydc2 | 4.886726711 | 0.521221998 | 0.485604212 |
| Dydc1 | 0.704807757 | -0.113676544 | 1.157822216 |
| Mat1a | 4.404352952 | 2.166543348 | 0.682303898 |
| Sftpa1 | 0.045882455 | 0.237312084 | 3.724945049 |
| Mbl1 | 0 | NA | NA |
| Sftpd | 0 | NA | NA |
| Gm7925 | 0 | NA | NA |
| Gm22823 | 0.476740939 | 2.025852771 | 1.652178315 |
| Gm2832 | 0.567915064 | 1.167285775 | 1.2918623 |
| Gm9671 | 0 | NA | NA |
| Gm5798 | 0 | NA | NA |
| Gm2855 | 0 | NA | NA |
| Gm7945 | 0.044544648 | 0.237309859 | 3.724945049 |
| Gm6482 | 0.047948698 | -0.243569582 | 3.724945049 |
| Gm7953 | 0 | NA | NA |
| Gm7954 | 0 | NA | NA |
| Gm2878 | 0 | NA | NA |
| Gm3486 | 0 | NA | NA |
| Gm6490 | 0 | NA | NA |
| Gm17061 | 0 | NA | NA |
| Gm7970 | 0 | NA | NA |
| 1700049E17Rik2 | 0 | NA | NA |
| Gm17159 | 0 | NA | NA |
| Gm3072 | 0 | NA | NA |
| Gm3077 | 0 | NA | NA |
| Gm3676 | 0 | NA | NA |

| | | | |
|---|---|---|---|
| Gm8065 | 0 | NA | NA |
| Gm8068 | 0 | NA | NA |
| Gm7929 | 0 | NA | NA |
| Gm17062 | 0 | NA | NA |
| Gm7980 | 0 | NA | NA |
| 1700049E17Rik1 | 0 | NA | NA |
| 1700024B05Rik | 0 | NA | NA |
| Gm9890 | 0 | NA | NA |
| Gm6401 | 0 | NA | NA |
| Gm3543 | 0 | NA | NA |
| Gm17063 | 0 | NA | NA |
| Gm2930 | 0 | NA | NA |
| Gm7991 | 0 | NA | NA |
| 1700091H14Rik | 0 | NA | NA |
| Gm7951 | 0 | NA | NA |
| Gm17064 | 0 | NA | NA |
| Gm17027 | 0 | NA | NA |
| Gm17166 | 0 | NA | NA |
| Gm3573 | 0.045882455 | 0.237312084 | 3.724945049 |
| Gm17165 | 0 | NA | NA |
| Gm17038 | 0 | NA | NA |
| Gm17026 | 0 | NA | NA |
| Gm25208 | 0 | NA | NA |
| Gm25892 | 0 | NA | NA |
| Gm9611 | 0 | NA | NA |
| Gm7995 | 0 | NA | NA |
| Gm17164 | 0 | NA | NA |
| Gm2951 | 0 | NA | NA |
| Gm2959 | 0 | NA | NA |
| Gm8005 | 0 | NA | NA |
| Gm8011 | 0 | NA | NA |
| Gm2966 | 0 | NA | NA |
| Gm8020 | 0 | NA | NA |
| Gm8024 | 0 | NA | NA |
| Gm8032 | 0.048949532 | -0.243569582 | 3.724945049 |
| Gm17204 | 0 | NA | NA |
| Gm17124 | 0 | NA | NA |
| Gm25228 | 0 | NA | NA |
| Gm23620 | 0.048949532 | -0.243569582 | 3.724945049 |
| Gm3633 | 0 | NA | NA |
| Gm10378 | 0 | NA | NA |
| Gm17162 | 0 | NA | NA |
| Gm5929 | 0 | NA | NA |
| Gm3008 | 0 | NA | NA |
| Gm17161 | 0 | NA | NA |
| Gm3015 | 0 | NA | NA |
| Gm3025 | 0 | NA | NA |

| | | | |
|---|---|---|---|
| Gm10377 | 0 | NA | NA |
| Gm7152 | 0 | NA | NA |
| Gm21977 | 0 | NA | NA |
| Gm3102 | 0 | NA | NA |
| Gm8082 | 0 | NA | NA |
| Gm3111 | 0 | NA | NA |
| Gm10376 | 0 | NA | NA |
| Gm6536 | 0 | NA | NA |
| Gm3123 | 0 | NA | NA |
| Gm8094 | 0.086930093 | 0.236669296 | 3.724898107 |
| Gm8104 | 2.686914903 | -0.671127876 | 0.699328649 |
| Gm7236 | 14.77965937 | 0.550131608 | 0.260159188 |
| Gm8114 | 0 | NA | NA |
| Gm3141 | 0.276783799 | -0.129786273 | 2.481703695 |
| Gm7233 | 2.888901366 | -1.03637799 | 0.708605167 |
| Gm7316 | 0 | NA | NA |
| Gm8122 | 0 | NA | NA |
| Gm8126 | 0.513218277 | 0.737649693 | 1.335145112 |
| Gm17153 | 0 | NA | NA |
| Gm8127 | 0 | NA | NA |
| Gm9732 | 2.036409272 | 1.846536854 | 0.822720074 |
| Gm17155 | 0 | NA | NA |
| Gm17154 | 0 | NA | NA |
| 1700001F09Rik | 0.08486385 | 0.717551239 | 3.724898127 |
| Gm9597 | 0.358995975 | 2.315962152 | 3.694579245 |
| Gm8138 | 0 | NA | NA |
| Gm5799 | 0.049553047 | 0.71820487 | 3.724945049 |
| Gm3216 | 0 | NA | NA |
| Gm17654 | 0 | NA | NA |
| Gm3221 | 0 | NA | NA |
| Gm10375 | 0 | NA | NA |
| Gm8046 | 0.277407222 | 1.608182954 | 1.762283438 |
| Gm8161 | 0 | NA | NA |
| Gm8165 | 0 | NA | NA |
| Gm9672 | 0 | NA | NA |
| Gm17208 | 0.132101749 | 1.209894437 | 3.72418822 |
| Gm7324 | 15.17033637 | 0.664866092 | 0.271010664 |
| Gm16506 | 0 | NA | NA |
| Gm6526 | 0.404621698 | 2.434706208 | 1.6455863 |
| Gm9587 | 0 | NA | NA |
| Gm8180 | 0 | NA | NA |
| Ear1 | 0.104637558 | 0.23732098 | 3.724945049 |
| Ear-ps12 | 0 | NA | NA |
| Ang-ps2 | 0 | NA | NA |
| Ear-ps9 | 0 | NA | NA |
| Ear-ps8 | 0.045882455 | 0.237312084 | 3.724945049 |
| Ang-ps3 | 0 | NA | NA |

| | | | | |
|---|---|---|---|---|
| Ear10 | 0 | NA | NA | |
| Gm8113 | 3.368233017 | -0.189101432 | | 0.64378905 |
| Gm9550 | 0 | NA | NA | |
| Ang5 | 0 | NA | NA | |
| Ear-ps7 | 0 | NA | NA | |
| Ear-ps10 | 0 | NA | NA | |
| Ang6 | 0 | NA | NA | |
| Gm8194 | 0 | NA | NA | |
| Gm3287 | 0 | NA | NA | |
| Ear-ps2 | 0 | NA | NA | |
| Ang-ps1 | 0.046345709 | 0.71820487 | | 3.724945049 |
| Gm16545 | 0 | NA | NA | |
| Ear2 | 0.663448651 | 3.141368109 | | 1.242535301 |
| Gm3327 | 0 | NA | NA | |
| Gm8207 | 0 | NA | NA | |
| 4930503E14Rik | 0.57934571 | 1.872391235 | | 1.298456637 |
| Gm8212 | 0.068097359 | -0.243569582 | | 3.724945049 |
| Gm8220 | 0.072025462 | 0.237312084 | | 3.724945049 |
| Gm8224 | 0 | NA | NA | |
| Gm5930 | 0.049553047 | 0.71820487 | | 3.724945049 |
| Gm8229 | 0 | NA | NA | |
| Gm3370 | 0 | NA | NA | |
| Gm3371 | 0 | NA | NA | |
| Gm8232 | 0.045882455 | 0.237312084 | | 3.724945049 |
| Gm21991 | 0 | NA | NA | |
| Gm17184 | 0.465126201 | 1.597922199 | | 1.750640033 |
| BC061237 | 0.120349074 | 0.237305146 | | 3.724945049 |
| Gm17093 | 0 | NA | NA | |
| Gm17183 | 0 | NA | NA | |
| Gm5624 | 0.172257197 | 0.980031371 | | 2.734354362 |
| Gm8247 | 0.04393175 | -0.243571806 | | 3.724945049 |
| Gm16439 | 6.205147171 | -0.502937218 | | 0.414875701 |
| Gm8256 | 0 | NA | NA | |
| Gm9593 | 0 | NA | NA | |
| Gm8257 | 0 | NA | NA | |
| Gm21754 | 0 | NA | NA | |
| Gm6541 | 0 | NA | NA | |
| Gm8267 | 0.060174537 | 0.237309859 | | 3.724945049 |
| Gm17187 | 0 | NA | NA | |
| Gm17070 | 0 | NA | NA | |
| Ptgdr | 10.70749769 | 0.269299835 | | 0.431706873 |
| Gm16976 | 1.330389483 | 0.509422734 | | 0.899572499 |
| mmu-mir-327 | 0 | NA | NA | |
| 4930431P03Rik | 1.669097723 | 0.942027165 | | 0.831629964 |
| Gm17173 | 1.687842655 | 2.804411541 | | 0.996175417 |
| Ptger2 | 14.92504073 | 2.227479901 | | 0.35466007 |
| Txndc16 | 1225.114205 | -0.275749526 | | 0.068649929 |

| | | | |
|---|---|---|---|
| Gm15765 | 0 | NA | NA |
| Gpr137c | 408.4638035 | -0.578353434 | 0.082322164 |
| Ero1l | 547.1939839 | 0.025080124 | 0.054320718 |
| Psmc6 | 1042.749598 | 0.644405847 | 0.100384501 |
| Styx | 68.64544997 | -0.469879989 | 0.211503026 |
| Gnpnat1 | 150.1275212 | 0.530509952 | 0.151895566 |
| Gm25353 | 0 | NA | NA |
| Fermt2 | 877.773119 | 0.191350306 | 0.073492058 |
| Gm22637 | 3.849995044 | -0.735276098 | 0.578343829 |
| Gm15601 | 0.688336999 | 2.737780858 | 1.216685067 |
| Ddhd1 | 1001.935963 | -0.271650547 | 0.080823866 |
| Mir5131 | 0 | NA | NA |
| A530076I17Rik | 4.579416614 | 1.922536645 | 0.599544108 |
| Gm15516 | 0.119234754 | -0.243576254 | 3.724945049 |
| Gm7206 | 18.53655595 | 0.43193442 | 0.251545672 |
| Gm15218 | 0.089089296 | 0.237305797 | 3.724945049 |
| Gm8317 | 0.802853338 | -0.062506751 | 1.23426241 |
| Gm15220 | 0 | NA | NA |
| Gm6580 | 0.146060149 | 0.555964358 | 3.524366763 |
| Gm37257 | 0 | NA | NA |
| Gm1821 | 160.7341621 | 0.811313541 | 0.165086771 |
| Gm15217 | 0.045882455 | 0.237312084 | 3.724945049 |
| Bmp4 | 27.37682956 | -0.22314593 | 0.238785017 |
| Gm15222 | 0.402965827 | -0.65311795 | 1.532807927 |
| Gm15221 | 0.062426873 | 0.237312084 | 3.724945049 |
| E130120K24Rik | 0.066050875 | 0.71820487 | 3.724945049 |
| Rubie | 0.049553047 | 0.71820487 | 3.724945049 |
| Cdkn3 | 4.411638001 | 1.807409427 | 0.534284649 |
| Cnih1 | 494.7336958 | 0.144792315 | 0.076142709 |
| Gmfb | 1861.18559 | 0.089620364 | 0.091129449 |
| Gm10101 | 2.636333195 | -0.431833824 | 0.621144376 |
| Cgrrf1 | 236.9675109 | 0.460143852 | 0.095823972 |
| Samd4 | 517.3838205 | -0.233530327 | 0.077376457 |
| Mir378c | 0 | NA | NA |
| Gm15562 | 5.089503423 | -0.146865504 | 0.424577427 |
| Gm10371 | 0.119587853 | 0.237305575 | 3.724945049 |
| Gch1 | 83.87125827 | 3.697726204 | 0.315520402 |
| Wdhd1 | 97.88412802 | 0.348541548 | 0.168717294 |
| Gm24378 | 1.312748754 | -0.546938472 | 0.87137326 |
| Socs4 | 478.1944072 | 0.296183369 | 0.072406079 |
| Mapk1ip1l | 616.9594476 | 0.665678252 | 0.073023432 |
| Lgals3 | 777.9121779 | 7.305740187 | 0.458885335 |
| Dlgap5 | 14.65831982 | 3.041691518 | 0.484545774 |
| Fbxo34 | 544.4943191 | -0.223615486 | 0.071875591 |
| Atg14 | 247.0243887 | 0.097634365 | 0.074122521 |
| Ktn1 | 1793.170985 | 0.320857749 | 0.079565379 |
| Gm37874 | 52.01025927 | -0.290970791 | 0.223147763 |

| | | | |
|---|---|---|---|
| Gm6055 | 178.1871439 | 0.43374202 | 0.109402735 |
| Peli2 | 376.3843311 | -0.301470305 | 0.0929529 |
| Gm6498 | 16.31733287 | -0.807588543 | 0.34841043 |
| Gm23881 | 0 | NA | NA |
| Gm3534 | 2.621659813 | 1.526711369 | 0.660916311 |
| Tmem260 | 200.6788583 | -0.344883996 | 0.167694832 |
| Gm23469 | 0 | NA | NA |
| Otx2 | 54.10925475 | 0.1401163 | 0.142753141 |
| Otx2os1 | 2.535154825 | -0.928139519 | 0.616200007 |
| Gm25380 | 0 | NA | NA |
| Gm22699 | 0 | NA | NA |
| Gm23470 | 0.573140788 | -0.226321319 | 1.37631128 |
| Exoc5 | 1321.293313 | 0.180114011 | 0.08149232 |
| Ap5m1 | 326.8043512 | -0.083218223 | 0.076598822 |
| Naa30 | 880.0118064 | 0.133423783 | 0.1041167 |
| 1700011H14Rik | 0 | NA | NA |
| Slc35f4 | 116.8448588 | -0.709069835 | 0.126530577 |
| Gm22116 | 5.008872367 | -2.293292989 | 0.581059482 |
| Gm22949 | 0 | NA | NA |
| 3632451O06Rik | 470.0630739 | -0.75028939 | 0.105172413 |
| Gm16082 | 1.838486464 | -1.575270355 | 0.84339528 |
| Gm22989 | 0 | NA | NA |
| Olfr722 | 0 | NA | NA |
| Olfr723 | 0 | NA | NA |
| Olfr724 | 0 | NA | NA |
| Olfr725 | 0 | NA | NA |
| Gm37194 | 0.41257556 | -0.729765288 | 1.508967265 |
| Olfr726 | 0 | NA | NA |
| Olfr727 | 0 | NA | NA |
| Olfr728 | 0.068097359 | -0.243569582 | 3.724945049 |
| Olfr729 | 0 | NA | NA |
| Olfr730 | 0 | NA | NA |
| Olfr731 | 0.046345709 | 0.71820487 | 3.724945049 |
| Olfr732 | 0 | NA | NA |
| Olfr733 | 0.092881282 | -0.724455696 | 3.724945049 |
| Olfr734 | 0.109869311 | 0.238305784 | 3.724874707 |
| Olfr735 | 0.066050875 | 0.71820487 | 3.724945049 |
| Tlr11 | 0.527131164 | 1.138293169 | 1.652108713 |
| Olfr736 | 0 | NA | NA |
| Olfr738 | 0 | NA | NA |
| Olfr739 | 0 | NA | NA |
| Olfr740 | 0 | NA | NA |
| Olfr741 | 0.048949532 | -0.243569582 | 3.724945049 |
| Olfr742 | 0 | NA | NA |
| Olfr743 | 0 | NA | NA |
| Olfr744 | 0 | NA | NA |
| Olfr745 | 0 | NA | NA |

| | | | |
|---|---|---|---|
| Olfr746 | 0 | NA | NA |
| Olfr747 | 0 | NA | NA |
| Olfr748 | 0 | NA | NA |
| Gm24316 | 0 | NA | NA |
| Olfr749 | 0.172571121 | 1.188801635 | 2.733273164 |
| Ttc5 | 245.1817073 | 0.007639732 | 0.091980415 |
| Gm10916 | 0.925454891 | 1.37764282 | 1.104806091 |
| Ccnb1ip1 | 5.389104856 | -0.654247399 | 0.451943864 |
| Rpph1 | 38912.15563 | 0.369133324 | 0.218597311 |
| Parp2 | 165.0646803 | 0.194080503 | 0.117909175 |
| Tep1 | 233.8406273 | 1.591448297 | 0.1174665 |
| Gm26782 | 38.7123841 | -0.494850548 | 0.22651556 |
| Klhl33 | 73.15949532 | 0.127852749 | 0.18472554 |
| A930018M24Rik | 44.75620921 | -0.013150048 | 0.182797477 |
| Osgep | 153.0925057 | -0.38010414 | 0.099158411 |
| Gm24689 | 0 | NA | NA |
| Apex1 | 280.3220763 | 0.305286361 | 0.099756348 |
| Tmem55b | 636.4352874 | 0.113133786 | 0.057989346 |
| Pnp | 408.8476884 | 2.678242277 | 0.273587485 |
| Gm38316 | 0.038981395 | 0.718207094 | 3.724945049 |
| Pnp2 | 4.854385997 | 2.099839563 | 0.510996231 |
| Rnase10 | 0.957491927 | 0.730785201 | 0.920116605 |
| Rnase9 | 0.066050875 | 0.71820487 | 3.724945049 |
| Rnase11 | 0.038981395 | 0.718207094 | 3.724945049 |
| Rnase12 | 0.062426873 | 0.237312084 | 3.724945049 |
| Olfr750 | 0.092493346 | -0.24357403 | 3.724945049 |
| Rnase4 | 86.11690773 | 0.448801443 | 0.222692723 |
| Ang | 3.984262514 | 1.664168511 | 0.539338178 |
| Eddm3b | 0 | NA | NA |
| Rnase6 | 15.68530113 | 7.572210306 | 0.901065282 |
| Rnase1 | 3.783653538 | -0.383968888 | 0.511574283 |
| Rnase2b | 0.112642008 | -0.24357403 | 3.724945049 |
| Ang2 | 0.049553047 | 0.71820487 | 3.724945049 |
| Ear14 | 0.049553047 | 0.71820487 | 3.724945049 |
| Rnase2a | 0.216167348 | -0.142243255 | 1.865242354 |
| Gm21718 | 0 | NA | NA |
| Gm3719 | 0 | NA | NA |
| Gm16289 | 0 | NA | NA |
| Gm7247 | 0.038981395 | 0.718207094 | 3.724945049 |
| Vmn2r-ps111 | 0 | NA | NA |
| Vmn2r88 | 0 | NA | NA |
| Vmn2r89 | 0 | NA | NA |
| Gm5622 | 0 | NA | NA |
| Gm16290 | 0 | NA | NA |
| Gm17175 | 0.075043205 | 0.237309859 | 3.724945049 |
| Gm17174 | 0 | NA | NA |
| Gm17078 | 0 | NA | NA |

| | | | |
|---|---|---|---|
| Gm4181 | 0 | NA | NA |
| Gm17186 | 0 | NA | NA |
| Gm17079 | 0 | NA | NA |
| Gm5800 | 0.136813032 | -1.18772039 | 3.638145365 |
| Ear-ps3 | 0 | NA | NA |
| Ang4 | 0.095435502 | 0.718202646 | 3.724945049 |
| Ear-ps4 | 0 | NA | NA |
| AY358078 | 0 | NA | NA |
| Gm6047 | 0.182466651 | 1.62097259 | 2.67233915 |
| Gm27375 | 0 | NA | NA |
| Ear-ps5 | 0 | NA | NA |
| Ear6 | 0 | NA | NA |
| Mettl17 | 160.3336998 | -0.435623335 | 0.157544306 |
| Slc39a2 | 25.14959081 | -0.315775494 | 0.265496162 |
| Ndrg2 | 5788.517411 | -0.06753977 | 0.150643881 |
| Tppp2 | 0.205880021 | -0.710941108 | 2.63262206 |
| Rnase13 | 0.16050186 | -0.243570964 | 2.800912406 |
| Arhgef40 | 469.4959225 | 0.531290556 | 0.085580458 |
| Gm16617 | 1.237774572 | -1.109818441 | 1.337480875 |
| Zfp219 | 473.6521586 | 0.119547553 | 0.089533913 |
| Tmem253 | 3.25900046 | 0.579825899 | 0.563021015 |
| Olfr221 | 0.038981395 | 0.718207094 | 3.724945049 |
| Gm22354 | 0.087261593 | -0.243576254 | 3.724945049 |
| Hnrnpc | 1799.363866 | 0.337205664 | 0.073529016 |
| Rpgrip1 | 13.58329738 | 0.065003976 | 0.293229094 |
| Supt16 | 1326.937056 | 0.293685454 | 0.080076575 |
| Gm26590 | 3.018699778 | 1.073544543 | 0.542563616 |
| Chd8 | 2275.929444 | -0.085552563 | 0.076383011 |
| Snord8 | 7.001247873 | -0.070463405 | 0.422378909 |
| RP24-223I19.6 | 3.473627573 | -1.037409976 | 0.551257901 |
| Rab2b | 331.1686235 | -0.071486618 | 0.072824935 |
| Gm23758 | 0 | NA | NA |
| Tox4 | 931.9067073 | 0.636297977 | 0.067592462 |
| Mettl3 | 190.3858702 | 0.033725047 | 0.136958865 |
| Sall2 | 992.2939578 | -0.241360012 | 0.064949362 |
| Gm26328 | 0.151841013 | -0.724459891 | 2.850049718 |
| Olfr1513 | 0 | NA | NA |
| RP24-461L18.3 | 26.78966518 | 0.83278658 | 0.2163864 |
| Olfr1512 | 0 | NA | NA |
| Gm37504 | 0 | NA | NA |
| Olfr1511 | 0 | NA | NA |
| Olfr1510 | 0.062426873 | 0.237312084 | 3.724945049 |
| RP24-461L18.7 | 0.061920613 | 0.718207094 | 3.724945049 |
| Trav1 | 0.280850934 | -0.243568509 | 2.041402103 |
| Olfr1509 | 0.473637868 | -1.835220031 | 1.518815718 |
| Olfr1508 | 0.186324499 | -0.724459786 | 2.716596789 |
| RP23-117I19.7 | 0 | NA | NA |

| | | | |
|---|---|---|---|
| Olfr1507 | 1.123152174 | -0.323620077 | 0.936907482 |
| RP23-117I19.2 | 0.060174537 | 0.237309859 | 3.724945049 |
| RP23-117I19.3 | 0.172571121 | 1.188801635 | 2.733273164 |
| Trav2 | 2.527889093 | 2.412052968 | 1.036345707 |
| Trav3-1 | 0.371273269 | 1.200222299 | 1.73619626 |
| Trav4-1 | 0 | NA | NA |
| Trav5-1 | 0.4844674 | 1.462298334 | 1.371434216 |
| Trav6-1 | 0.59120296 | 2.845550498 | 1.452365067 |
| RP24-447O5.2 | 0 | NA | NA |
| Trav7-1 | 0.214323314 | 1.222688595 | 2.18194044 |
| Trav6-2 | 0.503770797 | 2.394647677 | 1.483098699 |
| Trav7d-2 | 0 | NA | NA |
| Trav4d-2 | 0 | NA | NA |
| Trav6d-3 | 0 | NA | NA |
| Trav7d-3 | 0.231728546 | 1.928525196 | 3.701789631 |
| Trav6d-4 | 0.360686062 | 1.664558047 | 1.561958517 |
| Trav7d-4 | 0.038981395 | 0.718207094 | 3.724945049 |
| RP23-452K12.17 | 0 | NA | NA |
| Trav8d-1 | 0.480868668 | 2.802610321 | 1.563401856 |
| Trav9d-1 | 0.075043205 | 0.237309859 | 3.724945049 |
| Trav6d-5 | 0.045882455 | 0.237312084 | 3.724945049 |
| Trav10d | 0.08486385 | 0.717551239 | 3.724898127 |
| Trav6d-6 | 0.038981395 | 0.718207094 | 3.724945049 |
| Trav11d | 0 | NA | NA |
| Trav7d-5 | 0 | NA | NA |
| Trav12d-1 | 0 | NA | NA |
| Trav13d-1 | 0.061920613 | 0.718207094 | 3.724945049 |
| Trav14d-1 | 0 | NA | NA |
| RP23-452K12.18 | 0.160559904 | 1.165833556 | 2.790729435 |
| Trav15d-1-dv6d-1 | 0 | NA | NA |
| Trav3d-2 | 0 | NA | NA |
| Trav9d-2 | 0 | NA | NA |
| Trav4d-3 | 0 | NA | NA |
| Rps19-ps2 | 0.546944188 | 1.853930455 | 1.471373052 |
| Trav5d-2 | 0 | NA | NA |
| Trav12d-2 | 0.075043205 | 0.237309859 | 3.724945049 |
| Trav9d-3 | 0.216532645 | 2.049607601 | 3.145284541 |
| Trav5d-3 | 0 | NA | NA |
| Trav12d-3 | 0 | NA | NA |
| Trav13d-2 | 0 | NA | NA |
| Trav14d-2 | 0 | NA | NA |
| Trav15d-2-dv6d-2 | 0 | NA | NA |
| RP24-548I6.12 | 0 | NA | NA |
| Trav3d-3 | 0 | NA | NA |
| Trav9d-4 | 0.11147366 | 1.230376831 | 3.722845817 |
| Trav4d-4 | 0 | NA | NA |
| RP24-334B8.4 | 0 | NA | NA |

| | | | |
|---|---|---|---|
| Trav5d-4 | 0 | NA | NA |
| Trav6d-7 | 0.046345709 | 0.71820487 | 3.724945049 |
| Trav7d-6 | 0 | NA | NA |
| RP24-334B8.8 | 0.414202207 | -0.239899092 | 1.540707969 |
| Trav13d-3 | 0 | NA | NA |
| Trav8d-2 | 0.305990011 | 2.52179724 | 1.924155571 |
| Trav16d-dv11 | 0.583486461 | 2.964529631 | 3.371554198 |
| Trav13d-4 | 0.066050875 | 0.71820487 | 3.724945049 |
| Trav14d-3-dv8 | 0.35026241 | 2.37293227 | 1.644907253 |
| Trav15d-3 | 0 | NA | NA |
| Trav7n-4 | 0 | NA | NA |
| Trav9n-1 | 0.141094079 | 0.718203391 | 3.563850454 |
| Trav6n-5 | 0 | NA | NA |
| Trav10n | 0 | NA | NA |
| Trav6n-6 | 0.230315653 | 1.91489812 | 2.106303821 |
| Trav11n | 0 | NA | NA |
| Trav7n-5 | 0 | NA | NA |
| Trav12n-1 | 0 | NA | NA |
| Trav13n-1 | 0.049553047 | 0.71820487 | 3.724945049 |
| Trav14n-1 | 0 | NA | NA |
| RP23-371K2.5 | 0.606297752 | 0.871412304 | 1.251599421 |
| Trav15n-1 | 0 | NA | NA |
| Trav3n-2 | 0 | NA | NA |
| Trav9n-2 | 0 | NA | NA |
| Trav4n-3 | 0 | NA | NA |
| Rps19-ps1 | 0 | NA | NA |
| Trav5n-2 | 0 | NA | NA |
| Trav12n-2 | 0 | NA | NA |
| Trav9n-3 | 0 | NA | NA |
| Trav5n-3 | 0 | NA | NA |
| Trav12n-3 | 0 | NA | NA |
| Trav13n-2 | 0 | NA | NA |
| Trav14n-2 | 0 | NA | NA |
| Trav15n-2 | 0 | NA | NA |
| RP23-125A3.6 | 0 | NA | NA |
| Trav3n-3 | 0 | NA | NA |
| Trav9n-4 | 0.08486385 | 0.717551239 | 3.724898127 |
| Trav4n-4 | 0.12092566 | 0.237307635 | 3.724945049 |
| Trav5n-4 | 0 | NA | NA |
| Trav6n-7 | 0 | NA | NA |
| Trav7n-6 | 0 | NA | NA |
| RP24-334B8.20 | 0 | NA | NA |
| Trav13n-3 | 0 | NA | NA |
| Trav8n-2 | 0.072025462 | 0.237312084 | 3.724945049 |
| Trav16n | 0.219234697 | 1.252219195 | 2.544014054 |
| Trav13n-4 | 0.049553047 | 0.71820487 | 3.724945049 |
| Trav14n-3 | 0.163190422 | 1.22955094 | 3.396300676 |

| | | | |
|---|---|---|---|
| Trav15n-3 | 0 | NA | NA |
| Trav7-2 | 0 | NA | NA |
| Trav4-2 | 0.045882455 | 0.237312084 | 3.724945049 |
| Trav6-3 | 0.15956642 | 0.719166641 | 3.418731035 |
| Trav7-3 | 0.350945957 | 2.013317477 | 1.707095074 |
| Trav6-4 | 0.163577647 | 1.198505612 | 2.784038014 |
| Trav7-4 | 0.394584743 | 1.832033153 | 3.700316041 |
| RP23-230L15.11 | 0.04393175 | -0.243571806 | 3.724945049 |
| Trav8-1 | 0 | NA | NA |
| Trav9-1 | 0.21728767 | 1.453426934 | 2.53000544 |
| Trav6-5 | 0.124347486 | 0.719186602 | 3.717245571 |
| Trav10 | 0.044544648 | 0.237309859 | 3.724945049 |
| Trav6-6 | 0.141781211 | 1.200006512 | 2.911624965 |
| Trav11 | 0 | NA | NA |
| Trav7-5 | 0 | NA | NA |
| Trav12-1 | 0.061920613 | 0.718207094 | 3.724945049 |
| Trav13-1 | 0.353543486 | 1.496253447 | 1.686989079 |
| Trav14-1 | 0 | NA | NA |
| Trav15-1-dv6-1 | 0.099106094 | 1.209894437 | 3.72418822 |
| Trav3-2 | 0 | NA | NA |
| Trav9-2 | 0.219744001 | 1.876864594 | 2.125981944 |
| Trav4-3 | 0 | NA | NA |
| Trav5-2 | 0 | NA | NA |
| Trav12-2 | 0 | NA | NA |
| Trav12-3 | 0.264301748 | 1.209303621 | 2.450606297 |
| Trav13-2 | 0.124853745 | 0.237309606 | 3.724945049 |
| Trav14-2 | 0 | NA | NA |
| Trav15-2-dv6-2 | 0 | NA | NA |
| RP24-100P9.4 | 0 | NA | NA |
| Trav3-3 | 2.07315737 | 0.403792511 | 0.724119833 |
| Trav9-4 | 0.183309527 | 1.199972785 | 2.685766727 |
| Trav4-4-dv10 | 0.315232318 | 0.703156334 | 3.723830706 |
| RP24-100P9.5 | 0 | NA | NA |
| Trav5-4 | 0 | NA | NA |
| Trav6-7-dv9 | 0.09176491 | 0.237310218 | 3.724945049 |
| Trav7-6 | 0 | NA | NA |
| Trav13-3 | 0 | NA | NA |
| Trav8-2 | 0 | NA | NA |
| Trav16 | 0 | NA | NA |
| Trav13-4-dv7 | 0.143485256 | 1.229961212 | 3.547405499 |
| Trav14-3 | 0 | NA | NA |
| Trav15-3 | 0 | NA | NA |
| Trav3-4 | 0.305831651 | -0.724461362 | 2.046051074 |
| Trav12-4 | 0.062426873 | 0.237312084 | 3.724945049 |
| B230359F08Rik | 0.108266322 | 1.230376831 | 3.722845817 |
| Trav17 | 0.511617548 | 2.103805628 | 1.386832996 |
| Trav18 | 0 | NA | NA |

| | | | |
|---|---|---|---|
| Trav19 | 0 | NA | NA |
| Trav20 | 0 | NA | NA |
| Trav21-dv12 | 0.038981395 | 0.718207094 | 3.724945049 |
| Trdv1 | 0.294928389 | 2.241026026 | 3.69705936 |
| RP23-339J4.3 | 0.166286699 | 0.237325754 | 2.769850422 |
| Trav22 | 0 | NA | NA |
| Trdv2-1 | 0 | NA | NA |
| Trdv2-2 | 0.066050875 | 0.71820487 | 3.724945049 |
| Trav23 | 0.0878635 | -0.72445792 | 3.724945049 |
| Trdv3 | 0 | NA | NA |
| RP23-6A14.5 | 0.391491935 | -0.214730127 | 1.399051959 |
| Trdv4 | 0.800063227 | 1.913346311 | 1.155605024 |
| Trdd1 | 0 | NA | NA |
| Gm23704 | 0 | NA | NA |
| Trdd2 | 0.215047464 | 0.742538427 | 2.548074105 |
| Trdj1 | 0.138110619 | 0.718202458 | 3.62359848 |
| RP23-6A14.10 | 1.151756246 | 0.611728478 | 0.908482463 |
| Trdj2 | 0.506381433 | 2.794445094 | 1.676229863 |
| Trdc | 6.909450248 | 5.483984417 | 0.903809377 |
| Trdv5 | 0.261680116 | -0.730832978 | 2.10587865 |
| Traj61 | 0.355005915 | -0.243563508 | 1.585608907 |
| Traj60 | 0 | NA | NA |
| Traj59 | 0.384053089 | 0.249674994 | 1.963098643 |
| Traj58 | 0.060174537 | 0.237309859 | 3.724945049 |
| Traj57 | 0.066050875 | 0.71820487 | 3.724945049 |
| Traj56 | 0.049553047 | 0.71820487 | 3.724945049 |
| Traj55 | 0.038981395 | 0.718207094 | 3.724945049 |
| Traj54 | 0 | NA | NA |
| Traj53 | 0.235191775 | 1.214577565 | 2.111185558 |
| Traj52 | 0.049553047 | 0.71820487 | 3.724945049 |
| Traj51 | 0.143140564 | -0.24357283 | 3.546633223 |
| Traj50 | 0.066050875 | 0.71820487 | 3.724945049 |
| Traj49 | 0.094097695 | 0.718200421 | 3.724945049 |
| Traj48 | 0.268211629 | -0.243563819 | 2.092673895 |
| Traj47 | 0 | NA | NA |
| Traj46 | 0.173853943 | 1.25450333 | 2.738478996 |
| Traj45 | 0.089089296 | 0.237305797 | 3.724945049 |
| Traj44 | 0.378197461 | 1.95253445 | 1.806006581 |
| Traj43 | 0.372057435 | 2.295653122 | 1.48134018 |
| Traj42 | 0.270744093 | 1.605428202 | 2.413549857 |
| Traj41 | 0.062426873 | 0.237312084 | 3.724945049 |
| Traj40 | 0.373373093 | 1.793711551 | 1.550594288 |
| Traj39 | 0.520484338 | 2.97149704 | 1.547201025 |
| Traj38 | 0.304511964 | 2.470755254 | 1.948638501 |
| Traj37 | 0.321009791 | 2.511362702 | 1.669019933 |
| Traj36 | 1.007510561 | 2.811276355 | 1.243328073 |
| Traj35 | 0.11147366 | 1.230376831 | 3.722845817 |

| | | | |
|---|---|---|---|
| Traj34 | 0.174360202 | 0.718216754 | 2.725782262 |
| Traj33 | 0.478372902 | 2.432447492 | 3.477232883 |
| Traj32 | 0.292144397 | 2.355594723 | 3.690600465 |
| Traj31 | 0.534577455 | 1.129804548 | 1.351444649 |
| Traj30 | 0.493327417 | 1.987384147 | 1.370420207 |
| Traj29 | 0.56458922 | 3.236870723 | 1.30551993 |
| Traj28 | 0.229902237 | 1.059823776 | 2.522859681 |
| Traj27 | 0.668611641 | 1.740907856 | 1.260787959 |
| Traj26 | 0.066050875 | 0.71820487 | 3.724945049 |
| Traj25 | 0.632730142 | 2.30585024 | 1.220520598 |
| Traj24 | 0.739637387 | 1.586079827 | 1.394749987 |
| Traj23 | 0.722755445 | 2.270370446 | 1.297020196 |
| Traj22 | 0.569236379 | 1.906195889 | 1.257250278 |
| Traj21 | 0.962101734 | 1.370898275 | 1.178050017 |
| Traj20 | 0.661355215 | 2.426447512 | 1.24273787 |
| Traj19 | 0.223817141 | -0.243568726 | 2.170700296 |
| Traj18 | 0.206905613 | 1.604093267 | 2.172787258 |
| Traj17 | 0.414345469 | 2.346919624 | 1.658731411 |
| Traj16 | 0.11193333 | 0.718202646 | 3.724945049 |
| Traj15 | 0.43445422 | 2.173196039 | 1.690798746 |
| Traj14 | 0.377056126 | 1.193331276 | 3.296216838 |
| Traj13 | 0.208613207 | 1.792485849 | 2.58997165 |
| Traj12 | 0.486308198 | 2.233981486 | 1.488790885 |
| Traj11 | 0.728352697 | 2.713333529 | 1.212608632 |
| Traj9 | 0.154585317 | 1.652042906 | 2.82119606 |
| Traj8 | 0.203604809 | 1.252986864 | 2.604693634 |
| Traj7 | 0.161486377 | 1.199990293 | 2.795640196 |
| Traj6 | 0.490533913 | 2.67373883 | 1.428812904 |
| Traj5 | 0.517714159 | 2.590919352 | 1.392630257 |
| Traj4 | 0.260182639 | 2.003947424 | 2.05447458 |
| Traj3 | 0 | NA | NA |
| Traj2 | 0.364308502 | 2.403266516 | 1.684021165 |
| Traj1 | 0.178447459 | 1.663739594 | 3.327415937 |
| Trac | 42.85427748 | 3.687287813 | 0.36169918 |
| Dad1 | 331.339338 | 0.093677957 | 0.095392997 |
| Abhd4 | 923.6765976 | 0.646592473 | 0.136767888 |
| Olfr49 | 0.04393175 | -0.243571806 | 3.724945049 |
| RP24-378G4.3 | 0.161488833 | 1.352814168 | 3.431296303 |
| RP24-378G4.4 | 0 | NA | NA |
| RP24-378G4.5 | 0 | NA | NA |
| RP24-378G4.6 | 0.093484797 | 0.237318756 | 3.724945049 |
| Oxa1l | 446.5440451 | 0.279406047 | 0.062550513 |
| Slc7a7 | 63.95572232 | 2.35751302 | 0.191910234 |
| Mrpl52 | 179.7423914 | 0.799769719 | 0.116211913 |
| Mmp14 | 516.3861542 | 2.448801631 | 0.190634567 |
| Gm24424 | 0 | NA | NA |
| Lrp10 | 683.6561126 | 1.487983147 | 0.137840362 |

| | | | |
|---|---|---|---|
| Rem2 | 153.9580932 | 1.104085924 | 0.335359697 |
| Prmt5 | 476.4345395 | 0.394039283 | 0.060415514 |
| Haus4 | 37.03329486 | 0.53302062 | 0.201169037 |
| Ajuba | 48.90169317 | 2.051762148 | 0.268990835 |
| Gm24705 | 0 | NA | NA |
| 4931414P19Rik | 61.32594599 | 0.35418636 | 0.140738488 |
| Psmb5 | 352.4206726 | 0.510430596 | 0.100255462 |
| Mir686 | 0 | NA | NA |
| Psmb11 | 6.261784242 | -0.517859357 | 0.521386807 |
| Cdh24 | 130.6564516 | 0.082819368 | 0.13142172 |
| Gm20726 | 0.104106287 | -0.243576254 | 3.724945049 |
| Gm17606 | 1.502658278 | 2.29952197 | 1.112755954 |
| Acin1 | 1506.43145 | -0.016466411 | 0.097833196 |
| mmu-mir-6948 | 0.536873645 | -1.275347673 | 1.245869426 |
| 4930579G18Rik | 0.898085247 | 1.102843956 | 1.045034338 |
| 1700123O20Rik | 137.487668 | 0.689309838 | 0.133319297 |
| Cebpe | 1.473260831 | 4.280132917 | 1.069215493 |
| Slc7a8 | 825.5824292 | 0.181071129 | 0.084791993 |
| Gm8865 | 0 | NA | NA |
| Homez | 148.0548322 | -0.399755699 | 0.153621984 |
| Ppp1r3e | 55.21987148 | -0.38461414 | 0.178519277 |
| Gm37271 | 5.557962796 | -0.991893284 | 0.456793348 |
| Bcl2l2 | 1139.670049 | -0.337407363 | 0.064952172 |
| Gm20521 | 0.375216023 | 0.237295703 | 3.724945049 |
| Pabpn1 | 264.758843 | -0.59034655 | 0.215472905 |
| Slc22a17 | 4291.308412 | -0.577771226 | 0.055053571 |
| Efs | 132.9613014 | 0.084623842 | 0.180148011 |
| Il25 | 0.855745269 | 0.953434912 | 0.970567811 |
| Cmtm5 | 177.6437715 | -0.785064887 | 0.188921396 |
| Myh6 | 15.5941189 | 0.836190753 | 0.377776472 |
| Mir208a | 0 | NA | NA |
| Mhrt | 3.992004508 | -1.391988647 | 0.715844668 |
| Myh7 | 58.29767794 | 1.196109785 | 0.284516669 |
| Gm29015 | 0.163519603 | -0.243571882 | 2.778606197 |
| Mir208b | 0 | NA | NA |
| Gm17428 | 0.038981395 | 0.718207094 | 3.724945049 |
| Ngdn | 176.4773903 | 0.747176 | 0.134779517 |
| Gm10364 | 2.051660529 | -0.051974412 | 0.679923126 |
| Gm20687 | 0.094831987 | -0.243571806 | 3.724945049 |
| Thtpa | 328.5777116 | -0.12737196 | 0.083487774 |
| Zfhx2 | 1084.53136 | -0.778349644 | 0.172723133 |
| Zfhx2os | 47.4886808 | -1.292476718 | 0.385193462 |
| Gm37034 | 35.49877453 | -1.573790237 | 0.353407635 |
| Gm38129 | 19.96999586 | -1.525373578 | 0.363718642 |
| Ap1g2 | 74.95724327 | 0.340919675 | 0.214072685 |
| Jph4 | 3324.850563 | -1.432478417 | 0.129147359 |
| Gm23596 | 0 | NA | NA |

| | | | |
|---|---|---|---|
| Dhrs2 | 1.791652276 | -1.229433169 | 0.756305951 |
| Gm8894 | 12.48756017 | 0.552852058 | 0.320085492 |
| Dhrs4 | 75.51960051 | 0.153383575 | 0.155483662 |
| Lrrc16b | 392.3729988 | -1.083603124 | 0.137624944 |
| Cpne6 | 868.2829338 | -1.096324028 | 0.127712985 |
| Nrl | 2.663415179 | -0.957223083 | 0.611271255 |
| Pck2 | 235.7861946 | 0.207266678 | 0.101938488 |
| A730061H03Rik | 1.918506749 | -0.094103937 | 0.922525188 |
| Dcaf11 | 703.1883374 | 0.07275336 | 0.05257028 |
| Gm15932 | 0.28189265 | -0.243586284 | 3.724945049 |
| Fitm1 | 0.170186936 | 1.71882767 | 3.39900844 |
| Psme1 | 1237.910375 | 3.217896919 | 0.156993101 |
| Emc9 | 130.4834388 | -0.571049117 | 0.137428815 |
| Psme2 | 385.2943989 | 2.891695347 | 0.164503221 |
| Rnf31 | 653.5334504 | 1.448404652 | 0.11027383 |
| Irf9 | 958.7618358 | 4.000048304 | 0.141614233 |
| Rec8 | 6.735313399 | 0.491282387 | 0.429916501 |
| Ipo4 | 859.5313815 | 0.062184769 | 0.047451398 |
| Tm9sf1 | 264.973998 | 0.447942145 | 0.084731741 |
| Tssk4 | 3.653146442 | -0.539118306 | 0.53653438 |
| Mdp1 | 234.6429327 | -0.069350024 | 0.09665681 |
| Nedd8 | 570.9432889 | 0.152414314 | 0.109978426 |
| Gmpr2 | 130.1104825 | -0.088297019 | 0.132268633 |
| Tinf2 | 129.962745 | 0.677420937 | 0.183668074 |
| Gm25887 | 0.119587853 | 0.237305575 | 3.724945049 |
| Tgm1 | 450.7877214 | 8.294879182 | 0.498902939 |
| Rabggta | 331.6413799 | 0.124523001 | 0.065201302 |
| Dhrs1 | 1107.416416 | 1.888776316 | 0.131114127 |
| Nop9 | 182.2591248 | 1.093368488 | 0.143130344 |
| Cideb | 12.60069031 | -1.406950832 | 0.344335808 |
| Ltb4r2 | 1.834185785 | 1.727447482 | 0.813192308 |
| Ltb4r1 | 13.33054419 | 5.768929388 | 0.851334933 |
| Adcy4 | 30.55724621 | 2.574455489 | 0.335754478 |
| Ripk3 | 29.79384862 | 5.076753247 | 0.513483387 |
| Nfatc4 | 25.04774085 | 0.786296144 | 0.227535012 |
| Nynrin | 300.1840239 | -0.092273839 | 0.10578618 |
| Gm26751 | 4.016269843 | -0.956917177 | 0.694314297 |
| Cbln3 | 2048.884425 | -0.127912836 | 0.317386901 |
| Khnyn | 336.7112361 | 0.470059923 | 0.091248959 |
| Sdr39u1 | 401.4242335 | -0.304456803 | 0.094233029 |
| Cma1 | 1.421829526 | 2.374835618 | 0.985105352 |
| Cma2 | 0.212113984 | 0.237311491 | 3.173700036 |
| Gm5801 | 19.00002645 | 0.463777981 | 0.246958347 |
| Mcpt1 | 0.342615221 | 2.378147234 | 1.486465699 |
| Mcpt9 | 0 | NA | NA |
| Mcpt2 | 0.74687904 | 3.126367253 | 3.573231817 |
| Mcpt4 | 0.384964691 | 0.227555075 | 1.437765832 |

| | | | |
|---|---|---|---|
| Mcpt8 | 0 | NA | NA |
| Mir6949 | 0 | NA | NA |
| Ctsg | 0.191334258 | 1.574896507 | 2.660412218 |
| Gzme | 0.045882455 | 0.237312084 | 3.724945049 |
| Gzmd | 0.066050875 | 0.71820487 | 3.724945049 |
| Gzmg | 0 | NA | NA |
| Gzmn | 0 | NA | NA |
| Gzmf | 1.629358998 | 4.644615816 | 1.232229087 |
| Gzmc | 1.857191415 | 3.985649289 | 1.096291997 |
| Gm24118 | 0 | NA | NA |
| Gm4819 | 0 | NA | NA |
| Gzmb | 1148.613576 | 13.3141853 | 1.01104521 |
| Atp12a | 1.447939701 | 0.063137415 | 0.84818221 |
| Rnf17 | 3.730754265 | 0.543242151 | 0.621701127 |
| Gm24392 | 0.066050875 | 0.71820487 | 3.724945049 |
| Cenpj | 130.4607887 | 1.836891335 | 0.204774663 |
| Gm22431 | 0 | NA | NA |
| Parp4 | 406.4978698 | 1.73125318 | 0.110676947 |
| Gm16573 | 0 | NA | NA |
| Mphosph8 | 456.601835 | -0.098317555 | 0.122332242 |
| Gm16973 | 101.7464204 | 0.275722684 | 0.145648153 |
| Pspc1 | 679.2798492 | -0.334881246 | 0.074129587 |
| C030013D06Rik | 71.86212202 | -1.240914502 | 0.302445757 |
| Gm4479 | 0.082913145 | 0.23732098 | 3.724945049 |
| Zmym5 | 534.3723691 | -0.138529246 | 0.080018953 |
| 2410022M11Rik | 46.59279127 | 0.206016184 | 0.165711023 |
| Gm22498 | 0 | NA | NA |
| Gm27635 | 0 | NA | NA |
| Gm22218 | 0 | NA | NA |
| Zmym2 | 1925.381415 | -0.217879131 | 0.072930145 |
| B020004C17Rik | 0.385592413 | 2.118986542 | 3.694977693 |
| Gja3 | 0.839270164 | 1.373061476 | 1.26335611 |
| Gjb2 | 207.1465136 | 1.779444842 | 0.189233074 |
| Gm4491 | 3.32060927 | -2.190437101 | 0.65162387 |
| Gjb6 | 476.1042231 | -0.730744323 | 0.165651114 |
| Cryl1 | 92.72897768 | -0.728418595 | 0.153723885 |
| Gm23593 | 0.852011583 | 0.041302079 | 1.065228471 |
| Ift88 | 288.14491 | -0.218644885 | 0.10057327 |
| Gm15836 | 0.068097359 | -0.243569582 | 3.724945049 |
| Ift88os | 3.791653849 | -1.755302651 | 0.70021429 |
| Il17d | 56.83289221 | -0.869951294 | 0.174002636 |
| N6amt2 | 87.43391111 | -0.121010852 | 0.149045978 |
| 1700039M10Rik | 1.307453931 | -0.655603764 | 1.008965391 |
| Xpo4 | 372.4125732 | 0.080266845 | 0.109917675 |
| Lats2 | 654.4618862 | 0.625693672 | 0.113568068 |
| Gm26440 | 0.104366086 | -0.243576254 | 3.724945049 |
| Gm20430 | 0.779952531 | -1.207982208 | 1.068180254 |

| | | | |
|---|---|---|---|
| Sap18 | 492.5086436 | 0.10031375 | 0.094249655 |
| Mir3077 | 0 | NA | NA |
| Ska3 | 10.9912364 | 1.38421881 | 0.489457205 |
| Mrpl57 | 188.5806294 | -0.03902314 | 0.121115553 |
| Gm22738 | 2.721618137 | -0.885164638 | 0.641080883 |
| Zdhhc20 | 1037.770143 | 0.018354171 | 0.099137098 |
| Micu2 | 298.7097166 | 0.888473459 | 0.113192398 |
| 3110083C13Rik | 6.409078304 | -0.987738354 | 0.510643612 |
| Fgf9 | 261.7121968 | -0.509599714 | 0.082344572 |
| Gm17109 | 0 | NA | NA |
| Gm25614 | 0 | NA | NA |
| Gm9022 | 0.171181394 | 0.718216336 | 2.749537022 |
| Rpl13-ps3 | 2.686924039 | 1.100570162 | 0.707984497 |
| 1700129C05Rik | 0.097139547 | 0.718207094 | 3.724945049 |
| Gm5142 | 0 | NA | NA |
| Rcbtb1 | 723.0799953 | -0.186427877 | 0.083461248 |
| Gm6904 | 9.706043522 | 7.161523265 | 0.91072355 |
| Phf11a | 116.1058508 | 10.10236936 | 0.913133426 |
| Phf11b | 380.8323055 | 8.683401709 | 0.524263446 |
| Phf11d | 768.3922217 | 9.729499329 | 0.546862468 |
| Phf11c | 63.97794148 | 3.010040882 | 0.255851117 |
| Setdb2 | 303.1628553 | 2.19004448 | 0.205603361 |
| Gm16301 | 4.198859835 | 2.577720801 | 0.787992889 |
| Cab39l | 377.7726523 | 0.075010917 | 0.094271089 |
| Cdadc1 | 398.4046235 | -0.189285497 | 0.07608323 |
| Shisa2 | 57.22810011 | 0.4331226 | 0.265026798 |
| Atp8a2 | 620.7074487 | -0.5477022 | 0.094282428 |
| Gm9013 | 6.16443421 | -1.237376995 | 0.492997233 |
| 4930563I02Rik | 0.208097268 | 0.72638144 | 2.198688534 |
| Gm6913 | 0 | NA | NA |
| Gm16261 | 0 | NA | NA |
| Nupl1 | 647.3223968 | 0.085991701 | 0.070272166 |
| Mir719 | 0.1633926 | -0.24356215 | 2.785222606 |
| Mtmr6 | 1302.815769 | -0.08874084 | 0.052087743 |
| Amer2 | 525.7164864 | -0.607308592 | 0.106743785 |
| Gm29266 | 1.126110934 | -2.685282298 | 1.124084272 |
| Spata13 | 889.6645646 | 1.411384188 | 0.131225523 |
| C1qtnf9 | 1.516716675 | 1.494756606 | 0.947166879 |
| Mipepos | 0.63216027 | 0.456315825 | 1.253338844 |
| Mipep | 261.2781114 | -0.104803609 | 0.084231369 |
| Tnfrsf19 | 737.4688333 | 0.767092686 | 0.126875827 |
| Sacs | 3375.54935 | -0.309961227 | 0.069535351 |
| Sgcg | 2.524458954 | 0.100605778 | 0.676219625 |
| 1700109G14Rik | 1.219573449 | -0.093418552 | 1.065569087 |
| Arl11 | 45.06214385 | 4.025360552 | 0.324938067 |
| Ebpl | 74.2502746 | -0.07468457 | 0.151817571 |
| Gm25716 | 0.646183563 | 0.586202022 | 1.213306442 |

| | | | |
|---|---|---|---|
| Kpna3 | 1362.917462 | 0.198090582 | 0.072103646 |
| Spryd7 | 646.9547196 | 0.274013506 | 0.10982359 |
| Trim13 | 93.97118903 | 0.0345994 | 0.144430721 |
| Dleu2 | 509.193176 | 0.084991156 | 0.137709189 |
| Kcnrg | 4.493413694 | -0.678016449 | 0.475751396 |
| Gm27010 | 39.15760298 | -0.242986704 | 0.25955169 |
| Gm27845 | 0.122991902 | -0.24357403 | 3.724945049 |
| Gm26916 | 0.226559374 | 0.2373201 | 2.539258673 |
| Mir16-1 | 0.351804329 | 0.660094489 | 1.475215541 |
| Mir15a | 2.378988063 | -0.120942085 | 0.669437378 |
| Gm27747 | 0.12092566 | 0.237307635 | 3.724945049 |
| Gm27972 | 0 | NA | NA |
| Gm27017 | 1.269792884 | 1.032653245 | 0.818690253 |
| Gm27928 | 0.047948698 | -0.243569582 | 3.724945049 |
| Gm37420 | 92.02312766 | -0.638092778 | 0.228192234 |
| Gm37472 | 129.5225664 | -0.636861396 | 0.240246264 |
| Gm37820 | 1.452044762 | 0.668572492 | 1.017890961 |
| Gm26969 | 0 | NA | NA |
| Dleu7 | 46.2767513 | -0.184812047 | 0.180377788 |
| Rnaseh2b | 84.37813906 | 0.057425133 | 0.132947436 |
| Gucy1b2 | 4.464211597 | 1.669530785 | 0.553147077 |
| Gm4131 | 0.940148893 | 2.862044121 | 1.066644745 |
| Gm24625 | 0 | NA | NA |
| Fam124a | 214.0573691 | -0.228623205 | 0.095303696 |
| Gm23816 | 0 | NA | NA |
| Serpine3 | 0.736171263 | -0.127148989 | 1.152640705 |
| Gm25162 | 0 | NA | NA |
| Ints6 | 590.4517194 | 0.016420316 | 0.118583578 |
| 4931440J10Rik | 4.963742587 | -1.508601968 | 0.537757002 |
| Gm26536 | 1.065495214 | 0.167662386 | 0.929855011 |
| Wdfy2 | 169.1506073 | 0.23956683 | 0.099332456 |
| Mir8098 | 0 | NA | NA |
| Mir6541 | 1.551969969 | -0.102741151 | 0.788801402 |
| n-R5s47 | 0.9747039 | -0.827743943 | 1.13642736 |
| Defb48 | 0 | NA | NA |
| Defb47 | 0.450831908 | 0.332158464 | 1.335031002 |
| Defb43 | 0 | NA | NA |
| Defb30 | 0.101408268 | 0.717551239 | 3.724898127 |
| Defb42 | 4.17522043 | -1.510686284 | 0.742655003 |
| Gm23629 | 0 | NA | NA |
| Ctsb | 9370.229512 | 1.758615722 | 0.145533395 |
| Fdft1 | 819.3075922 | 0.310901995 | 0.117301421 |
| Neil2 | 16.96188054 | 0.11549742 | 0.304059273 |
| Gata4 | 0.136623169 | 0.66393974 | 3.632616033 |
| Blk | 0.439667587 | 1.633835567 | 1.432064927 |
| Gm21430 | 0.110595523 | 0.718200421 | 3.724945049 |
| Fam167a | 87.58736384 | 1.91436729 | 0.20322056 |

| | | | |
|---|---|---|---|
| Tdh | 0.432707378 | -0.874984397 | 1.478443587 |
| Gm17232 | 0.288435958 | 0.25573145 | 2.046791651 |
| Mtmr9 | 857.9831177 | 0.045948509 | 0.052170355 |
| Gm15918 | 7.496844332 | -0.368278531 | 0.3586634 |
| Xkr6 | 411.7970283 | -0.771804618 | 0.158810194 |
| Mir598 | 1.860426698 | -1.710811283 | 0.783647424 |
| Pinx1 | 73.25514319 | -0.065740277 | 0.140761055 |
| Sox7 | 14.05727425 | 1.13633729 | 0.330510467 |
| 4930578I06Rik | 0.15100991 | 0.718202046 | 3.527597384 |
| Rp1l1 | 0.46004984 | 0.72134589 | 1.556588339 |
| Prss55 | 1.808854001 | 2.202923691 | 1.00724922 |
| Prss51 | 0 | NA | NA |
| 4930471C04Rik | 6.910637616 | -0.781529279 | 0.562782361 |
| Prss52 | 0.240888117 | 0.273023763 | 2.521242023 |
| Gm17116 | 16.40399705 | -0.842819387 | 0.389002839 |
| Msra | 295.1015591 | -0.007646396 | 0.090596081 |
| Mir124a-1hg | 1670.155338 | -1.162321857 | 0.360212801 |
| Mir124a-1 | 0 | NA | NA |
| Mir3078 | 0 | NA | NA |
| Kif13b | 733.5243985 | -0.238345368 | 0.106667391 |
| Gm26225 | 1.137833813 | -1.194856626 | 0.986509393 |
| Hmbox1 | 1181.12912 | -0.280843199 | 0.053848466 |
| Gm37183 | 36.08914756 | -0.596682935 | 0.325152744 |
| Gm20111 | 0 | NA | NA |
| Ints9 | 171.8987803 | 0.527441749 | 0.103758544 |
| Extl3 | 2015.132037 | 0.099558759 | 0.062863863 |
| Fzd3 | 1027.303391 | -0.548469529 | 0.076291528 |
| Gm27263 | 1.700332359 | 0.185685661 | 0.818161353 |
| Fbxo16 | 53.77017717 | -0.39628016 | 0.14364247 |
| Zfp395 | 203.6050118 | -0.171727491 | 0.123483171 |
| Pnoc | 87.06028916 | 0.556867572 | 0.143552565 |
| Elp3 | 772.4740043 | 0.114531795 | 0.061112515 |
| Nuggc | 2.07464996 | 4.094336334 | 1.052767518 |
| Scara5 | 27.21228639 | 2.215064209 | 0.373736664 |
| Pbk | 12.44226385 | 1.867410641 | 0.584323474 |
| Esco2 | 8.165872688 | 1.966100088 | 0.531867426 |
| Ccdc25 | 465.8537132 | 0.906518304 | 0.128215281 |
| Scara3 | 275.1443948 | 0.356598002 | 0.100406388 |
| Clu | 7658.013756 | 0.555390671 | 0.108754434 |
| Gulo | 1.585533927 | -0.652454865 | 0.997314861 |
| Adam2 | 0.443182709 | 1.504363253 | 1.741910346 |
| Gm10233 | 0.231561134 | 0.726133901 | 2.131538857 |
| Ephx2 | 103.3385784 | -0.710180162 | 0.141605015 |
| Chrna2 | 9.44470098 | -0.338857027 | 0.346031712 |
| Ptk2b | 3860.082922 | -0.257783561 | 0.15104249 |
| 1700001G11Rik | 4.137181521 | 0.431562926 | 0.500657875 |
| Trim35 | 1878.620081 | 0.373116868 | 0.069128935 |

| | | | |
|---|---|---|---|
| Stmn4 | 1213.698151 | -0.353728745 | 0.098631478 |
| Gm23899 | 0 | NA | NA |
| Adra1a | 340.4045469 | -0.16064931 | 0.113614902 |
| Gm10032 | 0.180493943 | 0.719085619 | 2.699005327 |
| Dpysl2 | 11326.80766 | -0.179737656 | 0.054467923 |
| Gm5464 | 7.318891932 | 0.018375188 | 0.38607016 |
| Pnma2 | 428.5622982 | -0.863434932 | 0.120715253 |
| 4930578I07Rik | 0 | NA | NA |
| Gm24258 | 0 | NA | NA |
| Bnip3l | 1147.218713 | -0.334762671 | 0.098221398 |
| Ppp2r2a | 1306.363493 | 0.393205173 | 0.072052694 |
| Gm27647 | 0 | NA | NA |
| Gm20675 | 0.146692594 | 1.229893553 | 3.521109737 |
| Ebf2 | 68.02274029 | -0.014319053 | 0.178653915 |
| Gm23178 | 0 | NA | NA |
| Gm6878 | 0.454723033 | -1.216658582 | 1.742997423 |
| 1700120O09Rik | 0 | NA | NA |
| 4930438E09Rik | 0 | NA | NA |
| Gm10860 | 0 | NA | NA |
| Gm24981 | 0 | NA | NA |
| Gm20667 | 1.359754244 | 1.157157398 | 1.040469751 |
| Cdca2 | 13.24071265 | 1.223428923 | 0.398397952 |
| Kctd9 | 293.1358824 | -0.164702641 | 0.137793295 |
| Gnrh1 | 6.140945712 | -0.932907974 | 0.478888152 |
| Dock5 | 320.7362463 | 0.317381815 | 0.117456305 |
| Mir6539 | 0.673941775 | -2.765921885 | 1.237110507 |
| Nefm | 2120.125399 | -0.126377768 | 0.22075665 |
| Nefl | 4244.358884 | -0.506052809 | 0.134279088 |
| Gm22319 | 0 | NA | NA |
| Adam7 | 0.28902587 | 0.969425426 | 2.08180942 |
| Adamdec1 | 0.646331757 | 0.782324178 | 1.258362165 |
| Adam28 | 1.311804504 | -0.271859135 | 1.091750114 |
| Stc1 | 48.95481958 | 0.437212785 | 0.199384524 |
| Rps2-ps5 | 0.934890173 | -0.267302175 | 0.969772476 |
| 1700092C10Rik | 0 | NA | NA |
| Nkx2-6 | 0 | NA | NA |
| Nkx3-1 | 12.50312542 | 0.32792587 | 0.333034847 |
| Gm37762 | 0.256447508 | 1.225217374 | 2.469852681 |
| Slc25a37 | 364.4713757 | 0.055456997 | 0.118706117 |
| Gm27222 | 0 | NA | NA |
| Gm20236 | 0.061920613 | 0.718207094 | 3.724945049 |
| Gm31748 | 0 | NA | NA |
| Gm27223 | 0 | NA | NA |
| Synb | 0 | NA | NA |
| Gm16677 | 0 | NA | NA |
| Entpd4 | 0.136880458 | -0.250074019 | 3.635317246 |
| Gm27221 | 0 | NA | NA |

| | | | |
|---|---|---|---|
| Gm28049 | 0.042716945 | -0.243571806 | 3.724945049 |
| Gm21451 | 0 | NA | NA |
| Gm38378 | 0 | NA | NA |
| Gm37094 | 0 | NA | NA |
| Gm37847 | 0.111006857 | 0.717551239 | 3.724898127 |
| Gm27175 | 0.061920613 | 0.718207094 | 3.724945049 |
| Gm27179 | 0 | NA | NA |
| Gm27174 | 0 | NA | NA |
| Gm21685 | 17.74312811 | 0.653222724 | 0.330631626 |
| Gm27176 | 0 | NA | NA |
| Gm27177 | 0.075043205 | 0.237309859 | 3.724945049 |
| Loxl2 | 21.89650722 | 2.421500402 | 0.34115722 |
| Gm37513 | 0 | NA | NA |
| Mir6950 | 0 | NA | NA |
| R3hcc1 | 176.6082625 | -0.006816301 | 0.097402002 |
| Chmp7 | 739.9914465 | -0.026749387 | 0.05876087 |
| 4930480K23Rik | 35.48654686 | -0.852154146 | 0.182801272 |
| Tnfrsf10b | 17.05474737 | 3.103876726 | 0.371140902 |
| Rhobtb2 | 1292.582811 | -0.574454293 | 0.099511481 |
| Pebp4 | 0.585213848 | -0.461143754 | 1.24466285 |
| Egr3 | 501.7132685 | -0.527657026 | 0.320933887 |
| Bin3 | 135.9347836 | 1.069216138 | 0.108740937 |
| Ccar2 | 987.7091471 | -0.184137223 | 0.051833892 |
| 9930012K11Rik | 11.13584856 | 0.025037556 | 0.338627583 |
| Pdlim2 | 142.1719157 | -0.331472403 | 0.136382537 |
| Sorbs3 | 390.0743824 | 0.500733049 | 0.117477451 |
| Gm26908 | 10.16694102 | -0.148049048 | 0.350465519 |
| Ppp3cc | 243.8924496 | -0.093425446 | 0.087829176 |
| AC139294.1 | 1.827339133 | 0.157345768 | 0.743515137 |
| Slc39a14 | 953.2277302 | 1.385452191 | 0.178256183 |
| Gm9174 | 1.024625068 | 1.654091073 | 1.02149244 |
| Piwil2 | 21.10693623 | 1.605430152 | 0.242625493 |
| Gm22725 | 0.106465261 | 0.718202646 | 3.724945049 |
| Gm24890 | 0.088476398 | -0.243576254 | 3.724945049 |
| Polr3d | 217.1158271 | 0.266684417 | 0.114417388 |
| Mir320 | 1.741823475 | -1.744216995 | 0.873695515 |
| Phyhip | 4018.25641 | -0.857197166 | 0.113173951 |
| Bmp1 | 483.1780716 | 0.142294972 | 0.098722628 |
| Sftpc | 3.468929289 | -0.790467756 | 0.580329 |
| Lgi3 | 1208.980655 | -0.249063181 | 0.085757113 |
| Reep4 | 37.19800003 | 0.96066004 | 0.201273607 |
| Hr | 708.6183582 | -0.091316413 | 0.115401709 |
| Nudt18 | 347.76934 | 0.171765806 | 0.076569218 |
| Fam160b2 | 415.787422 | -0.205842138 | 0.093511576 |
| Dmtn | 2327.269897 | -0.680857038 | 0.085195796 |
| Fgf17 | 6.855534712 | 0.117511095 | 0.461660675 |
| Npm2 | 6.446150388 | 0.020772176 | 0.409428207 |

| | | | |
|---|---|---|---|
| Xpo7 | 1222.808954 | -0.239843251 | 0.054094734 |
| Dok2 | 40.29440054 | 2.946831721 | 0.252115304 |
| 2410012E07Rik | 0.066050875 | 0.71820487 | 3.724945049 |
| Gfra2 | 372.4700337 | -0.284552672 | 0.096134466 |
| Gm37455 | 0 | NA | NA |
| Gm9192 | 0 | NA | NA |
| Gm23735 | 0 | NA | NA |
| Fndc3a | 2663.159039 | 1.577057379 | 0.100113073 |
| Gm16549 | 0.514076732 | 1.889183518 | 1.531561676 |
| Cysltr2 | 20.03900042 | 3.571843894 | 0.434715574 |
| Rcbtb2 | 290.6187941 | -0.04690822 | 0.089089092 |
| Gm17233 | 4.303707252 | -1.108998681 | 0.568072681 |
| Rb1 | 791.2567668 | 0.149687466 | 0.069997591 |
| Mir687 | 2.668100091 | -0.421204637 | 0.635614468 |
| Lpar6 | 59.93468125 | 1.150518819 | 0.185415487 |
| Itm2b | 7614.81607 | 0.81324473 | 0.102713018 |
| Med4 | 170.1772693 | 0.008568726 | 0.133074501 |
| Nudt15 | 22.3670934 | -0.240644349 | 0.241147146 |
| Sucla2 | 1247.988688 | -0.174757664 | 0.086900706 |
| Gm21750 | 0.774729477 | 2.081887655 | 1.129184603 |
| Gm24540 | 2.261342538 | 0.708005204 | 0.626147073 |
| Gm16409 | 3.92403765 | 1.201806791 | 0.503533995 |
| Gm22164 | 0 | NA | NA |
| Gm6984 | 34.159564 | -0.223467082 | 0.182798201 |
| Htr2a | 177.8320817 | -0.470258581 | 0.178962948 |
| Esd | 684.0533462 | 1.468995239 | 0.138195279 |
| Lrch1 | 437.4414308 | 0.334847502 | 0.078285979 |
| 5031414D18Rik | 62.98482591 | 2.545198939 | 0.229284905 |
| Lrrc63 | 0.699855268 | 0.581026287 | 1.36089444 |
| Lcp1 | 670.0106953 | 3.404933982 | 0.164709486 |
| Gm15628 | 0 | NA | NA |
| Gm15629 | 0.174360202 | 0.718216754 | 2.725782262 |
| Cpb2 | 0 | NA | NA |
| AC161877.1 | 0 | NA | NA |
| Zc3h13 | 935.6372034 | -0.247490116 | 0.127857339 |
| Siah3 | 24.32705589 | -0.396427772 | 0.224517765 |
| Spert | 0.0885994 | -0.24357403 | 3.724945049 |
| 4930564B18Rik | 0.087261593 | -0.243576254 | 3.724945049 |
| Gm23622 | 0 | NA | NA |
| Cog3 | 540.4936263 | -0.009777729 | 0.073071025 |
| Gm24876 | 5.352175477 | -0.615109409 | 0.50930333 |
| Slc25a30 | 95.05997151 | 0.386743917 | 0.150257352 |
| Gm4285 | 34.11371507 | -0.12846991 | 0.218628114 |
| Tpt1 | 4005.741273 | 1.017158703 | 0.098225226 |
| Snora31 | 5.414963088 | -0.069651244 | 0.425012726 |
| Gtf2f2 | 148.747947 | 0.879512703 | 0.107897162 |
| Kctd4 | 189.0479674 | -1.187386087 | 0.143909919 |

| | | | |
|---|---|---|---|
| Gm25517 | 1.093745531 | -0.846909616 | 0.855989326 |
| Gpalpp1 | 612.613148 | -0.088808805 | 0.057254683 |
| Nufip1 | 200.4613363 | 0.430675325 | 0.108977765 |
| Rps2-ps6 | 4.539665628 | 0.429138189 | 0.464758644 |
| Tsc22d1 | 3622.389764 | -0.231708255 | 0.064968147 |
| 4930444M15Rik | 1.224655951 | -0.072035096 | 0.980521545 |
| Serp2 | 345.0104461 | -0.290243918 | 0.106083467 |
| 1700108F19Rik | 0.956737924 | -1.005216896 | 1.079743742 |
| Lacc1 | 58.05462338 | 1.68867348 | 0.196705273 |
| Ccdc122 | 14.09176844 | 1.359392799 | 0.305715982 |
| Enox1 | 345.9653688 | -0.312752916 | 0.124188971 |
| Gm10132 | 18.18438502 | 0.651427268 | 0.288431028 |
| Gm6994 | 0 | NA | NA |
| n-R5s48 | 4.967990864 | -1.652046188 | 0.613147013 |
| Gm23823 | 0 | NA | NA |
| Gm1587 | 0.104719185 | 0.237305575 | 3.724945049 |
| Dnajc15 | 183.7589994 | 0.392338129 | 0.096611938 |
| Epsti1 | 187.7081649 | 7.104090643 | 0.396132122 |
| Gm19301 | 0 | NA | NA |
| Fam216b | 12.69086337 | 0.413693659 | 0.298725071 |
| Mir1971 | 0 | NA | NA |
| Gm23652 | 0 | NA | NA |
| Tnfsf11 | 1.022937004 | 1.327680828 | 1.015006834 |
| Gm26197 | 0 | NA | NA |
| Akap11 | 5712.428926 | -0.29550213 | 0.059955073 |
| Dgkh | 929.9045385 | -0.298040175 | 0.118252444 |
| Gm26251 | 0.072025462 | 0.237312084 | 3.724945049 |
| Vwa8 | 1020.041174 | -0.479249802 | 0.065078283 |
| Zfp957 | 0 | NA | NA |
| Rgcc | 194.1349241 | 0.388333103 | 0.145236312 |
| Naa16 | 214.9779865 | -0.299876715 | 0.132053067 |
| Mtrf1 | 71.56468441 | 0.213569509 | 0.136110232 |
| Kbtbd7 | 573.16381 | -0.305583 | 0.077325923 |
| Kbtbd6 | 5.414111675 | -0.090234996 | 0.445189978 |
| AC124745.1 | 0 | NA | NA |
| Wbp4 | 689.3854996 | 0.196142067 | 0.088675173 |
| Elf1 | 298.3952992 | 2.441008025 | 0.161141163 |
| Gm27857 | 0 | NA | NA |
| Sugt1 | 598.6834534 | 0.018266326 | 0.083567746 |
| Lect1 | 10.35988659 | 0.509482227 | 0.348355719 |
| Mir759 | 0 | NA | NA |
| Pcdh8 | 429.8832926 | -0.054944504 | 0.158369293 |
| Gm9748 | 3.167342963 | 0.145708871 | 0.506577529 |
| Gm6999 | 2.919730593 | 1.770461384 | 0.665579954 |
| Gm10845 | 0.072025462 | 0.237312084 | 3.724945049 |
| Olfm4 | 11.07339153 | -0.035135495 | 0.325618541 |
| Gm9578 | 0 | NA | NA |

| | | | |
|---|---|---|---|
| Gm22891 | 0 | NA | NA |
| Gm24774 | 0 | NA | NA |
| Pcdh17 | 1873.471252 | -0.011870636 | 0.06462455 |
| Gm23926 | 0 | NA | NA |
| Gm25115 | 0.075043205 | 0.237309859 | 3.724945049 |
| Diap3 | 34.4707947 | 0.811054012 | 0.237532128 |
| Gm23438 | 0 | NA | NA |
| Gm37002 | 0 | NA | NA |
| Gm23278 | 0 | NA | NA |
| Tdrd3 | 499.7196116 | -0.084306344 | 0.056724768 |
| Rps3a2 | 226.2208805 | 0.590799639 | 0.146099429 |
| Gm26405 | 0.047948698 | -0.243569582 | 3.724945049 |
| Gm9274 | 0.048949532 | -0.243569582 | 3.724945049 |
| Pcdh20 | 350.8125221 | -0.835211974 | 0.158380531 |
| Gm36946 | 0 | NA | NA |
| Gm25886 | 0 | NA | NA |
| Gm25415 | 0.048949532 | -0.243569582 | 3.724945049 |
| Gm10110 | 26.30868621 | 0.789615883 | 0.221062821 |
| Gm25495 | 0 | NA | NA |
| Gm23225 | 0 | NA | NA |
| Gm24127 | 0 | NA | NA |
| Gm37341 | 0 | NA | NA |
| Pcdh9 | 3659.38219 | -0.637450447 | 0.073668016 |
| Gm23509 | 13.87461055 | -0.132706897 | 0.415437934 |
| Gm23324 | 0 | NA | NA |
| Gm24680 | 0 | NA | NA |
| 4921530L21Rik | 0.197024265 | -1.191745932 | 2.225999129 |
| Klhl1 | 133.1425135 | 0.526726224 | 0.193816572 |
| Gm15515 | 0.04393175 | -0.243571806 | 3.724945049 |
| Gm24043 | 0 | NA | NA |
| Gm25133 | 0 | NA | NA |
| Gm24770 | 0 | NA | NA |
| Gm27441 | 0 | NA | NA |
| Gm23276 | 0.217428854 | 2.009909443 | 2.547915794 |
| Dach1 | 251.06346 | -0.748562935 | 0.107362672 |
| Gm25831 | 2.466058065 | 0.124477231 | 0.6872856 |
| Gm16331 | 0 | NA | NA |
| Gm26006 | 0 | NA | NA |
| Gm26139 | 0 | NA | NA |
| Gm27034 | 0 | NA | NA |
| Rpl36a-ps1 | 11.21814357 | 0.577604246 | 0.317630837 |
| Mzt1 | 677.8415215 | -0.2044615 | 0.072659727 |
| Bora | 35.47661271 | -0.268705993 | 0.228130399 |
| Dis3 | 309.1793791 | 0.329989123 | 0.131882916 |
| Pibf1 | 250.2980066 | 0.068191507 | 0.068533565 |
| Gm4412 | 0 | NA | NA |
| Klf5 | 148.4473388 | 0.679385334 | 0.168756188 |

| | | | |
|---|---|---|---|
| Gm22984 | 0 | NA | NA |
| Gm25130 | 0 | NA | NA |
| Gm37093 | 0.13998015 | 0.718212532 | 2.923064887 |
| Gm22970 | 0 | NA | NA |
| Gm5958 | 0.163532861 | 0.237326876 | 2.784469947 |
| Klf12 | 318.1322375 | -0.397016108 | 0.090483197 |
| 4930517O19Rik | 0.663801701 | -1.11473989 | 1.578358139 |
| Gm16260 | 0 | NA | NA |
| Gm26367 | 0 | NA | NA |
| Gm24345 | 0 | NA | NA |
| Gm22401 | 0 | NA | NA |
| Gm23538 | 0 | NA | NA |
| Prr30 | 0.448613728 | 0.737912693 | 1.394677123 |
| Gm26173 | 0 | NA | NA |
| Tbc1d4 | 374.3985729 | -0.183900449 | 0.086926215 |
| Commd6 | 158.7771116 | -0.007821056 | 0.113757403 |
| Uchl3 | 201.0806668 | 0.006902683 | 0.09839125 |
| Gm9922 | 0.794139519 | 0.538733386 | 1.022568461 |
| Lmo7 | 977.0007035 | -1.060966063 | 0.184270463 |
| Gm22347 | 0 | NA | NA |
| Gm23764 | 0 | NA | NA |
| Kctd12 | 489.5908408 | -0.429501429 | 0.140133248 |
| AC102815.1 | 0 | NA | NA |
| Gm26778 | 8.453551642 | -0.842870008 | 0.379864894 |
| Mir5130 | 0 | NA | NA |
| Irg1 | 221.3397588 | 10.6058822 | 0.965313505 |
| Cln5 | 245.9919061 | 0.920132281 | 0.13411455 |
| Fbxl3 | 1187.753289 | -0.090953281 | 0.064892908 |
| Mycbp2 | 9169.971231 | -0.333024516 | 0.08381045 |
| Gm4654 | 6.877366695 | -0.458457665 | 0.381042196 |
| Scel | 26.78325797 | 0.156333252 | 0.216845043 |
| Slain1 | 561.6860723 | -0.470972679 | 0.080483322 |
| Gm22877 | 0.140122821 | -0.243570953 | 3.572128793 |
| Slain1os | 0.09349418 | -0.24357403 | 3.724945049 |
| Ednrb | 961.6585733 | -0.301624967 | 0.176307813 |
| Gm37910 | 0.066050875 | 0.71820487 | 3.724945049 |
| Pou4f1 | 42.92036986 | 0.458758434 | 0.336773868 |
| Rnf219 | 139.6039557 | -0.082116273 | 0.095376112 |
| Gm22021 | 0 | NA | NA |
| Gm25724 | 0 | NA | NA |
| Gm24280 | 0 | NA | NA |
| Rbm26 | 1630.5591 | -0.012878993 | 0.095585484 |
| Gm17066 | 187.5789837 | 0.189330257 | 0.125954159 |
| Gm22290 | 0 | NA | NA |
| Ndfip2 | 1106.337769 | -0.194803396 | 0.063170403 |
| Gm24207 | 0 | NA | NA |
| Gm16259 | 0.115603922 | 1.209894437 | 3.72418822 |

| | | | |
|---|---|---|---|
| 4930449E01Rik | 0.297771212 | -0.286843314 | 1.616202106 |
| Mir6390 | 7.744732698 | -2.139678347 | 0.450852027 |
| Gm10076 | 151.4983943 | 0.878716152 | 0.168829355 |
| Gm22406 | 0 | NA | NA |
| Spry2 | 675.0338336 | 0.452271445 | 0.130611585 |
| Gm4775 | 0 | NA | NA |
| Trim52 | 0 | NA | NA |
| Gm23459 | 0 | NA | NA |
| 1700128A07Rik | 0 | NA | NA |
| Gm24634 | 0 | NA | NA |
| Gm17366 | 0 | NA | NA |
| Gm23437 | 0 | NA | NA |
| Gm24818 | 0 | NA | NA |
| Gm24229 | 0 | NA | NA |
| Rpl27a-ps2 | 2.255766892 | -0.329406948 | 0.858924602 |
| Slitrk1 | 1192.574154 | -0.851247565 | 0.094277181 |
| Gm25901 | 0 | NA | NA |
| Gm24369 | 0 | NA | NA |
| n-R5s50 | 0 | NA | NA |
| Gm23242 | 0 | NA | NA |
| Gm25670 | 0 | NA | NA |
| Gm26255 | 0 | NA | NA |
| Slitrk6 | 94.50009073 | -0.250158612 | 0.134702139 |
| Gm22802 | 0 | NA | NA |
| Slitrk5 | 1071.418396 | -0.359782567 | 0.099574554 |
| Gm22764 | 0 | NA | NA |
| Gm4822 | 0 | NA | NA |
| Tpm3-rs7 | 867.6432377 | 1.365769803 | 0.11217833 |
| Gm4487 | 0 | NA | NA |
| Gm37495 | 0 | NA | NA |
| Gm24073 | 0 | NA | NA |
| Mir17hg | 18.52657897 | 1.207969943 | 0.28559728 |
| Mir17 | 0 | NA | NA |
| Mir18 | 0 | NA | NA |
| Mir19a | 0 | NA | NA |
| Mir20a | 0 | NA | NA |
| Mir19b-1 | 0 | NA | NA |
| Mir92-1 | 0 | NA | NA |
| Gpc5 | 280.2541914 | -0.92235693 | 0.135944684 |
| 4930505G20Rik | 0 | NA | NA |
| Gm18367 | 0 | NA | NA |
| Gm20713 | 0.048949532 | -0.243569582 | 3.724945049 |
| Gm38045 | 0 | NA | NA |
| Gpc6 | 374.2491254 | -0.061607321 | 0.083165149 |
| Mir6239 | 0 | NA | NA |
| Dct | 6.057780134 | 0.16049636 | 0.436077919 |
| Gm26773 | 0.075043205 | 0.237309859 | 3.724945049 |

| | | | |
|---|---|---|---|
| Tgds | 62.65483873 | 0.533817616 | 0.196448764 |
| Gpr180 | 179.3319749 | -0.248761707 | 0.086853395 |
| Sox21 | 98.70143058 | -0.969826833 | 0.232088671 |
| Gm4675 | 4.582536418 | -0.753686037 | 0.463321166 |
| Mir6241 | 0.075043205 | 0.237309859 | 3.724945049 |
| Gm9376 | 0 | NA | NA |
| Gm23669 | 0 | NA | NA |
| Gm26791 | 0.061920613 | 0.718207094 | 3.724945049 |
| 1700044C05Rik | 0 | NA | NA |
| Abcc4 | 177.0567019 | 0.437950054 | 0.140922474 |
| Gm23302 | 0 | NA | NA |
| Mir6391 | 0.804474276 | 0.286533716 | 1.149607609 |
| Gm27198 | 0 | NA | NA |
| Gm22379 | 0.042716945 | -0.243571806 | 3.724945049 |
| Cldn10 | 133.9096642 | -1.478183912 | 0.190159656 |
| n-R5s51 | 0 | NA | NA |
| Dzip1 | 1031.570717 | -0.50791322 | 0.075272496 |
| Dnajc3 | 1082.666445 | 0.516800645 | 0.075301252 |
| Uggt2 | 618.685887 | -0.647600321 | 0.089468916 |
| Gm16835 | 39.44922461 | -0.641938749 | 0.274449407 |
| Hs6st3 | 282.2429297 | -0.985770882 | 0.160438481 |
| 1700006F04Rik | 17.63169857 | -0.920974312 | 0.349838688 |
| Oxgr1 | 3.717291957 | 0.105527878 | 0.587547481 |
| Mbnl2 | 3584.706982 | 0.230930646 | 0.068157106 |
| Gm26679 | 5.318848262 | -1.189878987 | 0.60978546 |
| Rap2a | 997.0783507 | -0.350520636 | 0.070192239 |
| Gm17613 | 0.138137327 | 1.112377053 | 3.608654359 |
| Gm25781 | 0 | NA | NA |
| Ipo5 | 1921.943571 | 0.433601828 | 0.062654099 |
| Farp1 | 948.5700461 | -0.247133164 | 0.073443378 |
| B930095G15Rik | 416.2277983 | -1.047349581 | 0.114945543 |
| Stk24 | 646.4051468 | 0.061094296 | 0.08408432 |
| Slc15a1 | 1.066426709 | 1.540109455 | 1.030477785 |
| Dock9 | 2161.334577 | -0.596567921 | 0.091524088 |
| 1810041H14Rik | 1.973916092 | -0.458639138 | 0.764232153 |
| Ubac2 | 353.8238038 | 0.086320636 | 0.092958694 |
| Gpr18 | 9.840232673 | 6.110803233 | 0.933746755 |
| Gpr183 | 14.82188085 | 0.693539397 | 0.310728102 |
| Timm8a2 | 1.438846906 | 1.279201202 | 1.044001814 |
| A330035P11Rik | 10.89891607 | 0.39931359 | 0.302710885 |
| Tm9sf2 | 1465.863616 | 0.202224683 | 0.069480722 |
| Gm6254 | 34.64912996 | 0.345233898 | 0.324598557 |
| Gm17082 | 26.87597815 | -0.287546152 | 0.256691649 |
| Clybl | 80.6719109 | -0.43420531 | 0.14609838 |
| 1700108J01Rik | 0.0878635 | -0.72445792 | 3.724945049 |
| Gm25464 | 0.76305848 | -1.223153382 | 1.084806641 |
| Gm5089 | 8.639524202 | -1.585131705 | 0.483140916 |

| | | | |
|---|---|---|---|
| Zic5 | 171.9911989 | 0.02303337 | 0.144060582 |
| 2610035F20Rik | 11.57667783 | -0.737111621 | 0.320400555 |
| Zic2 | 736.6056247 | -0.03160111 | 0.192415316 |
| Gm10837 | 14.11353788 | -1.088975575 | 0.448091222 |
| Gm9399 | 0.044544648 | 0.237309859 | 3.724945049 |
| Pcca | 484.4393937 | -0.432456083 | 0.11128157 |
| Ggact | 197.5556307 | 0.195843847 | 0.125242821 |
| 4930594M22Rik | 104.9172262 | -0.513243415 | 0.201271544 |
| Tmtc4 | 235.3928922 | -0.147243572 | 0.106355576 |
| Gm15735 | 9.699810692 | -1.209999224 | 0.423825554 |
| Nalcn | 1752.798546 | -0.818517689 | 0.059096808 |
| Itgbl1 | 26.65978906 | -0.847237066 | 0.322144658 |
| Gm9308 | 0 | NA | NA |
| Gm25272 | 0.047948698 | -0.243569582 | 3.724945049 |
| Fgf14 | 940.1745018 | -0.74712792 | 0.091251347 |
| Gm28932 | 1.468722268 | -1.612714103 | 1.008488641 |
| Gm15422 | 16.80940044 | -0.72172201 | 0.366347387 |
| 1700024B18Rik | 0.075043205 | 0.237309859 | 3.724945049 |
| Gm17615 | 0.320086762 | -1.149989943 | 1.600436011 |
| Sepp1 | 2912.542392 | -0.038676923 | 0.136252347 |
| Ccdc152 | 0.495781838 | 0.170677574 | 1.380659584 |
| Ghr | 164.1599138 | -0.236453098 | 0.100151463 |
| Gm22031 | 0 | NA | NA |
| Gm4823 | 0.066050875 | 0.71820487 | 3.724945049 |
| Gm23228 | 0 | NA | NA |
| Fbxo4 | 73.81450778 | 0.605842286 | 0.135672304 |
| AW549877 | 1232.520806 | 0.018818812 | 0.072445051 |
| Gm2093 | 0.919878397 | 1.3695657 | 1.104186152 |
| BC037032 | 34.71017491 | -0.136190285 | 0.268624534 |
| Oxct1 | 2597.175127 | -0.402662999 | 0.087979607 |
| Gm15632 | 0 | NA | NA |
| Plcxd3 | 320.4411957 | -0.633571732 | 0.109355797 |
| C6 | 2.125003216 | 3.968259215 | 1.084383373 |
| Mroh2b | 0.19444009 | -0.724462299 | 2.692294029 |
| C7 | 8.962534464 | 0.332520646 | 0.410339648 |
| Gm25652 | 0 | NA | NA |
| Gm16077 | 0.04393175 | -0.243571806 | 3.724945049 |
| Card6 | 43.77270322 | 1.173916812 | 0.174847764 |
| Rpl37 | 814.5795284 | 0.387668393 | 0.109742515 |
| Snord72 | 5.885017498 | 0.467540744 | 0.448719558 |
| Gm10250 | 114.0624678 | -0.235315988 | 0.110975497 |
| Prkaa1 | 690.6189185 | 0.566709024 | 0.104468506 |
| Gm15938 | 2.619102942 | -0.531780385 | 0.711931206 |
| Ttc33 | 464.303568 | 0.011886252 | 0.070352613 |
| Ptger4 | 41.58960014 | 2.27208059 | 0.2567919 |
| Gm15937 | 14.4175352 | -0.286337655 | 0.332514898 |
| Gm27760 | 0.173816988 | -1.316703932 | 3.321126619 |

| | | | |
|---|---|---|---|
| Gm23139 | 0 | NA | NA |
| Dab2 | 418.7976734 | 1.863959164 | 0.184586965 |
| Gm16311 | 0 | NA | NA |
| C9 | 0.135879625 | -0.243487442 | 2.938858143 |
| Gm23218 | 0 | NA | NA |
| Fyb | 220.4543545 | 3.164037851 | 0.187502968 |
| Gm2245 | 0.436872751 | 1.637169187 | 1.661643543 |
| Gm7666 | 27.5160756 | 0.739727867 | 0.196653202 |
| Rictor | 1871.956267 | 0.106919236 | 0.066026071 |
| Osmr | 628.1045177 | 4.617226923 | 0.294487484 |
| Gm38282 | 0.159330469 | 0.718203216 | 3.465376669 |
| Lifr | 1758.792625 | 0.34532394 | 0.071563294 |
| Egflam | 43.8423358 | -0.046425255 | 0.16392254 |
| Gm16030 | 0 | NA | NA |
| Gm16029 | 0.048949532 | -0.243569582 | 3.724945049 |
| Gm37743 | 0.061920613 | 0.718207094 | 3.724945049 |
| Gdnf | 23.58641988 | 1.857256512 | 0.259862299 |
| Wdr70 | 191.8981038 | 0.072520056 | 0.106045238 |
| Gm27529 | 0.871987541 | 2.553038843 | 1.360147004 |
| Nup155 | 488.0096314 | -0.052290984 | 0.114956557 |
| 2410089E03Rik | 1512.066955 | -0.501881745 | 0.161928538 |
| Gm24144 | 14.79135611 | -1.342709723 | 0.460005669 |
| Nipbl | 1929.6716 | 0.151029474 | 0.063997227 |
| Gm2310 | 0 | NA | NA |
| Gm2348 | 0.104106287 | -0.243576254 | 3.724945049 |
| Slc1a3 | 7685.648105 | 0.340017116 | 0.116785612 |
| Gm37310 | 119.6237137 | 0.177824879 | 0.301424676 |
| RP24-341L14.2 | 0.280881662 | 1.6426832 | 2.072362143 |
| Gm5210 | 0 | NA | NA |
| Ranbp3l | 56.33245746 | -0.988190261 | 0.250731375 |
| Nadk2 | 494.6776195 | -0.258807632 | 0.110292865 |
| Skp2 | 125.2851504 | -0.332383739 | 0.133330578 |
| Lmbrd2 | 1052.783199 | -0.465284675 | 0.053958153 |
| Ugt3a1 | 0.150086409 | 0.237305146 | 3.724945049 |
| Ugt3a2 | 0.171181394 | 0.718216336 | 2.749537022 |
| Capsl | 15.44470158 | 1.087262378 | 0.308240606 |
| Il7r | 194.8832465 | 4.986234434 | 0.445617392 |
| Spef2 | 48.63543957 | 0.035224244 | 0.180032514 |
| Gm26350 | 0 | NA | NA |
| Prlr | 148.8562054 | -1.018468392 | 0.179848613 |
| Gm21973 | 0 | NA | NA |
| Agxt2 | 0.124596252 | 0.718200615 | 3.714681068 |
| Dnajc21 | 254.7197003 | -0.405732223 | 0.107783693 |
| Brix1 | 263.1432847 | 0.57172309 | 0.109464979 |
| Rad1 | 85.02316572 | -0.02837743 | 0.134611343 |
| Ttc23l | 0.267080418 | -0.243563716 | 1.817541942 |
| Gm17044 | 0.038981395 | 0.718207094 | 3.724945049 |

| | | | |
|---|---|---|---|
| Gm8174 | 0.386177252 | 0.16718854 | 1.543169854 |
| Rai14 | 244.3821753 | 1.214932226 | 0.133496028 |
| 4930556M19Rik | 7.969462537 | 0.677292642 | 0.379736472 |
| C1qtnf3 | 3.38967256 | 1.241678681 | 0.612470115 |
| Amacr | 74.0218532 | -0.098103724 | 0.15057028 |
| Slc45a2 | 0.883742299 | -0.244445988 | 1.065754081 |
| Rxfp3 | 34.39250656 | 0.274818772 | 0.198587387 |
| Adamts12 | 17.29016175 | 0.217356288 | 0.274569495 |
| Gm24844 | 0 | NA | NA |
| Tars | 425.9557127 | 1.014900582 | 0.120195029 |
| Npr3 | 146.1473597 | -1.24047526 | 0.139826495 |
| 1700047G03Rik | 0.146692594 | 1.229893553 | 3.521109737 |
| Gm24074 | 0 | NA | NA |
| Sub1 | 1754.400611 | -0.303670438 | 0.079259857 |
| RP24-105K16.2 | 0.060174537 | 0.237309859 | 3.724945049 |
| Zfr | 4001.152899 | 0.080551635 | 0.057345749 |
| Gm25713 | 0 | NA | NA |
| Mir1898 | 18.01255643 | -0.010516209 | 0.254498625 |
| Mtmr12 | 542.6306912 | -0.454928419 | 0.094194147 |
| 1810049J17Rik | 2.812275527 | -0.045375868 | 0.67259005 |
| Golph3 | 1317.912574 | 0.446324006 | 0.091588307 |
| Pdzd2 | 1799.644411 | -0.05595714 | 0.166749968 |
| Gm24135 | 1.998901016 | -1.93944529 | 0.810254281 |
| Gm2606 | 25.26788695 | -0.168947084 | 0.249549378 |
| F830212C03Rik | 14.03902647 | -1.508070589 | 0.425587328 |
| Gm24302 | 0 | NA | NA |
| 6030458C11Rik | 453.2360569 | -0.340345464 | 0.101692795 |
| Drosha | 1331.381267 | -0.228480774 | 0.050114424 |
| AC133189.1 | 0 | NA | NA |
| Gm24664 | 0.315718849 | -0.243566872 | 1.742959166 |
| Cdh6 | 210.1792556 | -0.236586949 | 0.117819272 |
| Gm9630 | 0 | NA | NA |
| Gm24080 | 0 | NA | NA |
| Gm25550 | 0 | NA | NA |
| Cdh9 | 185.9667162 | -1.117374971 | 0.135185093 |
| Gm22032 | 0 | NA | NA |
| Gm22895 | 0 | NA | NA |
| Gm24713 | 0.044544648 | 0.237309859 | 3.724945049 |
| 4921515E04Rik | 0 | NA | NA |
| Gm8318 | 1.912323087 | 0.486099634 | 0.75709006 |
| C030047K22Rik | 5.198307323 | -1.281928476 | 0.529637235 |
| Cdh10 | 687.055273 | -0.873692581 | 0.119833692 |
| Gm25976 | 0 | NA | NA |
| Gm37815 | 0 | NA | NA |
| Acot10 | 1.155605027 | 0.300500164 | 0.965915422 |
| Gm25711 | 4.168298335 | -2.39538322 | 0.665529233 |
| Gm27397 | 3.423615971 | -1.879348055 | 0.626305981 |

| | | | |
|---|---|---|---|
| Cdh12 | 473.520525 | -0.624588252 | 0.110082494 |
| Gm25101 | 4.033556537 | -2.615631692 | 0.694239599 |
| Gm38234 | 6.221313409 | -0.43012693 | 0.402268351 |
| AC166159.1 | 0 | NA | NA |
| Gm27525 | 0 | NA | NA |
| Cdh18 | 540.5763889 | -0.517449546 | 0.103354727 |
| Gm5803 | 0.68701006 | -0.082972565 | 1.064237441 |
| Gm19840 | 13.48907376 | -1.123631579 | 0.302759445 |
| Gm24656 | 2.60150024 | -1.371542222 | 0.670872098 |
| Gm23980 | 0 | NA | NA |
| Gm22810 | 0 | NA | NA |
| Gm2824 | 69.97913716 | -1.759510445 | 0.215774119 |
| 9230109A22Rik | 0.817550754 | 1.655965873 | 1.16854044 |
| Gm38341 | 0.308655657 | 0.718190755 | 3.724945049 |
| Basp1 | 2855.029311 | -0.387447356 | 0.107434989 |
| Gm5468 | 5.212139954 | -0.204356238 | 0.541531945 |
| Myo10 | 988.6576999 | 0.276342116 | 0.119478369 |
| Gm2862 | 0.999928428 | 0.616015059 | 1.151772269 |
| Fam134b | 559.6848899 | -0.004667712 | 0.082398529 |
| Mir7212 | 1.890715168 | -1.041584148 | 0.870871129 |
| Zfp622 | 385.825252 | 0.307577982 | 0.097526707 |
| 11-Mar | 93.58712996 | -0.346554185 | 0.12492852 |
| Fbxl7 | 13.24742036 | -0.337754455 | 0.338266109 |
| Gm6576 | 4.198207553 | 0.778011833 | 0.491144157 |
| Gm23903 | 0 | NA | NA |
| Ank | 1809.490276 | -0.555014555 | 0.050849711 |
| Mir7117 | 0 | NA | NA |
| Otulin | 178.6870836 | 0.440092816 | 0.102863345 |
| Fam105a | 144.2450621 | 0.126719466 | 0.120115497 |
| Trio | 4183.182376 | -0.171608171 | 0.08560867 |
| Dnah5 | 99.5374478 | -0.823933889 | 0.12619989 |
| Mir3964 | 0 | NA | NA |
| Gm23033 | 0 | NA | NA |
| Gm24554 | 0 | NA | NA |
| Gm23327 | 0 | NA | NA |
| Gm26163 | 0 | NA | NA |
| Ctnnd2 | 3344.046455 | -0.095566471 | 0.061784985 |
| Gm23956 | 8.765840549 | -0.991280351 | 0.441396892 |
| Gm24487 | 2.253680491 | -1.015096844 | 0.715526462 |
| Gm26416 | 0.503286714 | 0.718185597 | 3.724945049 |
| Dap | 102.7869284 | 1.395553555 | 0.195672577 |
| Fam136b-ps | 0.047948698 | -0.243569582 | 3.724945049 |
| Ankrd33b | 1536.741971 | -0.039535302 | 0.161717997 |
| Ropn1l | 5.055244084 | 0.705498394 | 0.425140203 |
| Gm6361 | 5.200498561 | -0.10287005 | 0.486631409 |
| 6-Mar | 3828.438788 | -0.460069381 | 0.048032476 |
| Cmbl | 49.64000691 | -1.011672686 | 0.192394708 |

| | | | |
|---|---|---|---|
| Cct5 | 1440.817807 | 0.473380322 | 0.077223773 |
| Fam173b | 140.20352 | -0.538805496 | 0.232630113 |
| Tas2r119 | 0 | NA | NA |
| Snhg18 | 13.92677096 | 0.06918259 | 0.277018756 |
| Snord123 | 0.145446656 | 1.165556206 | 2.878908292 |
| Sema5a | 1219.347837 | -0.360822923 | 0.080471293 |
| Sdc2 | 591.0431138 | 0.162455673 | 0.103858959 |
| Cpq | 169.9525733 | 0.78633687 | 0.17155755 |
| 1700084J12Rik | 0.496918936 | 0.263805659 | 3.575964429 |
| Gm24476 | 0.306418947 | 2.425190707 | 3.688182498 |
| Tspyl5 | 513.6623244 | -0.21148009 | 0.09501841 |
| Mtdh | 1529.028053 | 0.164705211 | 0.061707884 |
| Gm24500 | 0 | NA | NA |
| Laptm4b | 633.1682788 | -0.094118098 | 0.103531628 |
| Gm25809 | 0 | NA | NA |
| Matn2 | 218.3605345 | 0.410374461 | 0.169076762 |
| 9430069I07Rik | 0.203825279 | 1.160606195 | 2.62542815 |
| Rpl30 | 585.149464 | 0.483352538 | 0.081212373 |
| Gm24523 | 0 | NA | NA |
| Erich5 | 14.29563529 | -2.050430163 | 0.397661573 |
| Hrsp12 | 208.6478451 | -0.480220107 | 0.139584805 |
| Pop1 | 119.2294828 | -0.198456268 | 0.158554507 |
| Gm22208 | 2.694090619 | -0.372360473 | 0.589089465 |
| Nipal2 | 159.5133862 | -0.148698686 | 0.095833886 |
| Kcns2 | 254.6488734 | -0.437881889 | 0.098941872 |
| Stk3 | 354.1359327 | 0.211199029 | 0.089944422 |
| Osr2 | 0.114742407 | -0.24357403 | 3.724945049 |
| Vps13b | 2358.901886 | -0.353552407 | 0.070364452 |
| Gm24771 | 4.664523441 | -1.361971679 | 0.534742237 |
| Mir599 | 0 | NA | NA |
| Mir875 | 0 | NA | NA |
| Cox6c | 1187.10824 | -0.243159877 | 0.149442471 |
| Rgs22 | 32.16090155 | -0.19818558 | 0.194552031 |
| Gm25372 | 0 | NA | NA |
| Gm18889 | 2.742579127 | 0.253098007 | 0.563251592 |
| Fbxo43 | 1.809171643 | 0.995344938 | 0.73761289 |
| Gm22979 | 0 | NA | NA |
| Polr2k | 87.53921573 | 0.382203073 | 0.138522963 |
| Spag1 | 177.5961547 | -0.366805043 | 0.132479456 |
| Gm23754 | 0.706791778 | -0.348833618 | 1.085680288 |
| Rnf19a | 663.4722742 | 0.130942299 | 0.100351617 |
| Mir8097 | 0 | NA | NA |
| Rpl7a-ps3 | 1.338063731 | 0.505199471 | 0.882328316 |
| Ankrd46 | 1500.096336 | -0.077729775 | 0.058964348 |
| Snx31 | 4.226100588 | -0.437703223 | 0.478487121 |
| AC137871.1 | 0 | NA | NA |
| Pabpc1 | 2806.56132 | 0.97421566 | 0.076058855 |

| | | | |
|---|---|---|---|
| Ywhaz | 10312.43491 | -0.135738555 | 0.066064934 |
| Gm25303 | 0 | NA | NA |
| Gm10384 | 1.188161708 | -1.577297588 | 0.996105663 |
| Zfp706 | 1437.63576 | -0.158463663 | 0.056456111 |
| Gm26766 | 0.392978793 | 0.303567517 | 1.683149001 |
| Gm16136 | 0.154527273 | 0.237323928 | 2.834454908 |
| Grhl2 | 2.049607807 | 1.22351604 | 0.972427795 |
| Gm16137 | 0 | NA | NA |
| Gm16138 | 0.695937234 | 1.376189725 | 2.417011494 |
| Ncald | 3437.612725 | -0.67486444 | 0.106965834 |
| Gm28221 | 0.522018477 | -0.193809797 | 1.236511343 |
| Gm23654 | 2.696581576 | -0.954288026 | 0.681147965 |
| Gm15941 | 47.58132438 | -1.352404502 | 0.377636985 |
| Gm8668 | 0 | NA | NA |
| 4930447A16Rik | 0.047948698 | -0.243569582 | 3.724945049 |
| Gm3267 | 0 | NA | NA |
| Gm24833 | 0.580994137 | -2.201863492 | 1.289254334 |
| Gm15942 | 0.537603422 | -1.062780396 | 1.358782094 |
| Gm15940 | 9.81108231 | -1.462740414 | 0.416039096 |
| Rps19-ps5 | 1.865250533 | -0.906982872 | 0.925064613 |
| Rrm2b | 427.3554432 | -0.208562009 | 0.068496748 |
| Gm3362 | 13.70112511 | 0.685765205 | 0.30220677 |
| Ubr5 | 4115.687366 | 0.060479818 | 0.060023304 |
| Gm22353 | 0 | NA | NA |
| Odf1 | 0.128271896 | -0.243573716 | 3.679089475 |
| Gm24098 | 0 | NA | NA |
| Gm8705 | 0.498759394 | 1.273914481 | 1.684036444 |
| Klf10 | 430.8187853 | 1.090524427 | 0.251480597 |
| Gm26863 | 0.301278804 | -0.215253814 | 1.605696692 |
| Azin1 | 2466.052833 | 0.301167164 | 0.082107902 |
| Mir6951 | 0.490407978 | -0.709068176 | 1.372257384 |
| 2310043O21Rik | 0.088534442 | 1.179809211 | 3.723500796 |
| Atp6v1c1 | 1712.938684 | -0.055982145 | 0.043985542 |
| Gm26704 | 1.33281409 | 1.467251828 | 0.919831557 |
| Baalc | 276.5131432 | -0.227816463 | 0.090403732 |
| Fzd6 | 96.39112974 | 0.839976351 | 0.133648995 |
| Gm24789 | 0.09176491 | 0.237310218 | 3.724945049 |
| Cthrc1 | 75.42071906 | -0.126994371 | 0.15167418 |
| Slc25a32 | 170.269925 | 0.15795195 | 0.088952366 |
| Dcaf13 | 241.4209333 | 0.590147735 | 0.090258203 |
| n-R5s39 | 0.07796279 | 1.072698098 | 3.721829099 |
| Gm9522 | 11.31287606 | -0.225044284 | 0.332169655 |
| Rims2 | 2518.228734 | -0.668675416 | 0.089015159 |
| Gm25381 | 0 | NA | NA |
| Dcstamp | 2.546063767 | 1.692753013 | 0.682080711 |
| Dpys | 1.614744503 | -0.467905014 | 0.809893688 |
| Gm16291 | 9.213264714 | 3.325850445 | 0.610383349 |

| | | | |
|---|---|---|---|
| Lrp12 | 640.2551409 | -0.219608477 | 0.079496398 |
| 9330182O14Rik | 0 | NA | NA |
| Gm25958 | 0 | NA | NA |
| Gm16294 | 6.25597695 | 0.361509983 | 0.469513521 |
| Zfpm2 | 484.4239322 | -0.248658646 | 0.11497282 |
| 4930555K19Rik | 0 | NA | NA |
| Oxr1 | 3719.638992 | -0.297765276 | 0.051114038 |
| Gm26854 | 0.181780715 | 1.124102266 | 3.311260804 |
| Abra | 0.899802893 | 0.193110045 | 1.139697628 |
| Gm24751 | 0.127971488 | 1.23029335 | 3.685533863 |
| Angpt1 | 263.5197957 | 1.28705435 | 0.16529718 |
| Gm17473 | 35.3169963 | 4.986279264 | 0.524557119 |
| Rspo2 | 100.8944359 | -0.193194255 | 0.15623611 |
| Gm9658 | 1.11165528 | 0.999225949 | 1.031706689 |
| Gm22037 | 0.385685737 | 0.806339126 | 1.709989355 |
| Eif3e | 970.451907 | 0.640250951 | 0.103973286 |
| Gm10373 | 1.964982618 | -0.582682495 | 0.785942458 |
| Emc2 | 503.3907252 | -0.058752694 | 0.098320226 |
| Tmem74 | 82.76716195 | 0.169253414 | 0.143103673 |
| Trhr | 65.62486328 | -0.254706241 | 0.136341301 |
| Gm24903 | 0 | NA | NA |
| Gm25093 | 0 | NA | NA |
| Nudcd1 | 289.802079 | 0.041448633 | 0.127057999 |
| Eny2 | 530.2164271 | -0.094570698 | 0.076256874 |
| Gm22392 | 8.528292636 | -0.242694652 | 0.340367727 |
| Pkhd1l1 | 13.16766861 | -0.603792315 | 0.266655851 |
| Ebag9 | 211.2230902 | -0.031694741 | 0.086348346 |
| Sybu | 684.3366891 | -0.658861034 | 0.065543595 |
| 2310069G16Rik | 44.1689834 | -0.277902813 | 0.17443795 |
| A930017M01Rik | 111.592861 | -0.231043022 | 0.146826746 |
| Gm24768 | 0.351351704 | 0.577309665 | 1.571054222 |
| Gm23343 | 0.299746944 | 0.706629447 | 1.615580847 |
| Gm5471 | 0.105143817 | -0.24357403 | 3.724945049 |
| Kcnv1 | 528.0842402 | -1.723770412 | 0.15230124 |
| Gm5472 | 0.382197862 | -0.721759453 | 1.579808573 |
| Gm26063 | 0 | NA | NA |
| 1700022A22Rik | 0 | NA | NA |
| Gm23562 | 0 | NA | NA |
| 4930548G14Rik | 0.066050875 | 0.71820487 | 3.724945049 |
| Gm23665 | 0 | NA | NA |
| Csmd3 | 1405.512621 | -1.162311675 | 0.115038904 |
| Gm16300 | 8.844286263 | -1.680955694 | 0.386753997 |
| Gm23530 | 0.060174537 | 0.237309859 | 3.724945049 |
| Gm25007 | 0 | NA | NA |
| Gm23185 | 0 | NA | NA |
| Gm23848 | 0 | NA | NA |
| Trps1 | 1156.791181 | -0.114017373 | 0.063099536 |

| | | | |
|---|---|---|---|
| Mir1907 | 3.164287563 | -1.031246243 | 0.631977655 |
| Eif3h | 1020.195418 | 0.399848042 | 0.083767836 |
| Utp23 | 94.48504293 | 0.322240233 | 0.139560533 |
| Rad21 | 1630.277696 | 0.087396474 | 0.089044296 |
| Aard | 24.65755326 | -0.323338399 | 0.240218634 |
| Slc30a8 | 0.5812948 | 0.156470521 | 1.204577193 |
| Gm10020 | 9.983419005 | 0.821661859 | 0.440250346 |
| Gm23545 | 0 | NA | NA |
| Gm23267 | 0 | NA | NA |
| Med30 | 115.6898845 | 0.441189218 | 0.140453504 |
| Ext1 | 627.9625793 | -0.042387089 | 0.06409992 |
| Gm27926 | 0.066050875 | 0.71820487 | 3.724945049 |
| Samd12 | 154.297153 | -0.896607302 | 0.115816523 |
| Gm23200 | 7.643354964 | -1.688856337 | 0.455034641 |
| mmu-mir-28c | 7.291715215 | -1.307616431 | 0.6315718 |
| Gm7489 | 0 | NA | NA |
| Gm26933 | 0 | NA | NA |
| Tnfrsf11b | 55.23446549 | 1.72253232 | 0.244614834 |
| Gm5215 | 0 | NA | NA |
| Colec10 | 0.252184553 | 0.669365403 | 1.817532981 |
| Mal2 | 1117.686592 | -0.556220712 | 0.101985721 |
| Nov | 659.4404828 | -1.031658687 | 0.14079151 |
| Gm7577 | 0 | NA | NA |
| Enpp2 | 3338.021252 | -1.046345034 | 0.137728054 |
| Gm26684 | 4.138921123 | -3.33983636 | 0.902601615 |
| Taf2 | 560.7641421 | -0.254054558 | 0.072081836 |
| Dscc1 | 3.002576494 | 0.707799705 | 0.59773082 |
| Gm9920 | 2.459937312 | -0.784860902 | 0.647891173 |
| Deptor | 493.9836769 | -0.174411704 | 0.084090699 |
| Gm26296 | 0 | NA | NA |
| Col14a1 | 17.16314811 | -0.471643377 | 0.253889442 |
| 1700040F17Rik | 0.075043205 | 0.237309859 | 3.724945049 |
| Mrpl13 | 304.6669654 | 0.426421108 | 0.08235944 |
| Mtbp | 67.98096681 | 1.194388533 | 0.179880201 |
| Sntb1 | 66.74833637 | 0.235045298 | 0.163034196 |
| Has2 | 80.3213104 | 4.695875549 | 0.517519899 |
| Has2os | 3.183229149 | -0.020691164 | 0.595154 |
| Gm26178 | 0 | NA | NA |
| Slc22a22 | 0.334826625 | 1.171776776 | 1.58994219 |
| Zhx2 | 102.0661113 | -0.558980386 | 0.148074521 |
| Gm16006 | 1.944759997 | 1.418335561 | 0.809736931 |
| 9330154K18Rik | 0.788164698 | 1.482174969 | 1.241157932 |
| Derl1 | 846.7041548 | 0.70680386 | 0.090008558 |
| Tbc1d31 | 114.9980324 | -0.403247941 | 0.195796025 |
| Fam83a | 0.697414605 | 1.426202192 | 1.225191352 |
| 9130401M01Rik | 105.150685 | 0.008046628 | 0.114551175 |
| Gm29394 | 4.265622792 | -1.785215532 | 0.62680236 |

| | | | |
|---|---|---|---|
| Gm22299 | 0.817774247 | -1.18016074 | 1.197940557 |
| Zhx1 | 1069.677375 | -0.036148776 | 0.06793646 |
| Atad2 | 218.8401466 | 0.430123284 | 0.16336807 |
| Wdyhv1 | 104.6557226 | 0.203406079 | 0.110429879 |
| Gm23475 | 0 | NA | NA |
| Gm22488 | 1.584432859 | 0.10166359 | 1.00809543 |
| Fbxo32 | 391.5402532 | -0.265320356 | 0.102875381 |
| Gm15943 | 6.572403966 | -0.942250406 | 0.411040852 |
| Klhl38 | 13.89074863 | 0.877602925 | 0.329495516 |
| Anxa13 | 0 | NA | NA |
| D15Ertd621e | 1138.565379 | 0.052904535 | 0.054889525 |
| Fer1l6 | 7.081223484 | 1.250188554 | 0.447702516 |
| Gm23789 | 0 | NA | NA |
| Gm20712 | 48.91645952 | -0.748153884 | 0.183314432 |
| Tmem65 | 1185.359275 | 0.034084174 | 0.061693242 |
| Gm24041 | 11.06577238 | -0.892123859 | 0.330870647 |
| Ube2d4 | 0 | NA | NA |
| Trmt12 | 90.91210585 | 0.034577003 | 0.116038414 |
| Rnf139 | 607.7861192 | 1.008455018 | 0.108400381 |
| Tatdn1 | 116.8025155 | 0.022222293 | 0.129822978 |
| Ndufb9 | 863.2536908 | -0.016170854 | 0.123030456 |
| Mtss1 | 982.1708213 | -0.202284653 | 0.099217244 |
| Gm5045 | 4.498534705 | 0.698144969 | 0.617620749 |
| Zfp572 | 0.238889773 | -1.151582127 | 2.141900865 |
| Sqle | 640.1137635 | 0.090608775 | 0.113419169 |
| E430025E21Rik | 722.4473385 | 0.473625155 | 0.086794515 |
| Nsmce2 | 214.2833592 | 0.105633491 | 0.077924817 |
| Trib1 | 585.2550097 | 1.623639478 | 0.257694294 |
| Gm7083 | 0.061920613 | 0.718207094 | 3.724945049 |
| Gm7713 | 0 | NA | NA |
| 4933412E24Rik | 0 | NA | NA |
| Gm25875 | 0.044544648 | 0.237309859 | 3.724945049 |
| Gm23835 | 0.042716945 | -0.243571806 | 3.724945049 |
| Fam84b | 212.0412458 | 0.297637458 | 0.152879707 |
| 9930014A18Rik | 33.67317411 | -0.336652822 | 0.200697465 |
| A1bg | 0.161114758 | 0.237309167 | 3.43255351 |
| Gm24696 | 0 | NA | NA |
| Gm37947 | 0.038981395 | 0.718207094 | 3.724945049 |
| Myc | 144.727581 | 2.600662102 | 0.276701855 |
| Pvt1 | 146.0044478 | 0.021155905 | 0.142285971 |
| Gm27913 | 0 | NA | NA |
| Gm27782 | 1.314786873 | -0.564819109 | 0.818088651 |
| Gm27718 | 0 | NA | NA |
| Gm27957 | 0.583672784 | -0.829609251 | 1.205259547 |
| H2afy3 | 0.122507438 | 1.112024065 | 3.722822067 |
| Gm24810 | 0 | NA | NA |
| Gm22521 | 0.240278643 | 0.669041827 | 1.840077954 |

| | | | |
|---|---|---|---|
| RP23-55P19.1 | 7.30667397 | 0.424008425 | 0.408894121 |
| Gm25702 | 0.042716945 | -0.243571806 | 3.724945049 |
| Gm37615 | 0 | NA | NA |
| Gm5473 | 0.91428204 | -1.595808651 | 1.131740545 |
| Gsdmc | 0.046345709 | 0.71820487 | 3.724945049 |
| 1700065I16Rik | 0 | NA | NA |
| Gsdmc2 | 0.112396584 | 1.192083721 | 3.724434897 |
| Gsdmcl1 | 0 | NA | NA |
| Gsdmc3 | 0.258721378 | 2.275285631 | 1.99996924 |
| Gsdmcl2 | 0 | NA | NA |
| Gsdmc4 | 1.216796211 | 1.705022084 | 1.03860374 |
| Gsdmcl-ps | 0 | NA | NA |
| Fam49b | 1750.706656 | -0.034788233 | 0.049105863 |
| Gm25628 | 0 | NA | NA |
| Asap1 | 1466.62248 | -0.331181309 | 0.060917092 |
| Gm20717 | 0.527708665 | 2.123783961 | 1.496122669 |
| Adcy8 | 473.5293802 | -0.851173385 | 0.071789631 |
| Gm21798 | 0 | NA | NA |
| Gm21961 | 0 | NA | NA |
| Gm5046 | 0.097139547 | 0.718207094 | 3.724945049 |
| Gm23907 | 0 | NA | NA |
| Efr3a | 2113.727247 | -0.151651169 | 0.065132144 |
| Gm20405 | 22.08023906 | -0.33010054 | 0.403173762 |
| Oc90 | 0.439163765 | 1.913270167 | 1.496381649 |
| Hhla1 | 1.7811158 | -1.184485798 | 1.072400597 |
| Gm2991 | 7.931298069 | 0.665142456 | 0.475233782 |
| Kcnq3 | 2910.472199 | -1.049616086 | 0.092425139 |
| Gm27153 | 0.044544648 | 0.237309859 | 3.724945049 |
| Gm27242 | 2.088118819 | -0.679236074 | 0.902826084 |
| Lrrc6 | 31.55937458 | -0.807947164 | 0.216950953 |
| Tmem71 | 8.010071188 | 4.09572922 | 0.702566972 |
| Phf20l1 | 1418.308876 | -0.074630392 | 0.070880431 |
| Tg | 5.923287473 | 2.647801106 | 0.537727686 |
| Gm17140 | 0 | NA | NA |
| Sla | 260.5870413 | 1.176496096 | 0.113135059 |
| Wisp1 | 38.31189845 | 0.745486401 | 0.314223223 |
| Ndrg1 | 857.5715411 | 0.075304499 | 0.132543689 |
| Gm17035 | 0.30380643 | -0.163139529 | 2.013644718 |
| Gm2895 | 1.340196514 | -0.525414772 | 1.297641954 |
| St3gal1 | 321.0877622 | 0.756410367 | 0.11020266 |
| 1700012I11Rik | 0.522753486 | -1.44758672 | 1.366514577 |
| Zfat | 173.6083377 | -0.274298356 | 0.130042676 |
| Gm10240 | 8.242000484 | 0.739027964 | 0.457676587 |
| Gm37320 | 57.92546311 | -0.378132556 | 0.485932357 |
| Gm26621 | 67.44066586 | -0.275530332 | 0.309381082 |
| Mir30b | 2.936662866 | -0.727235972 | 0.633298175 |
| Gm37296 | 72.21542741 | -0.143527832 | 0.204756831 |

| | | | |
|---|---|---|---|
| Gm20732 | 11.82845525 | 0.628881214 | 0.348292908 |
| Mir30d | 0 | NA | NA |
| Gm25987 | 0 | NA | NA |
| Gm7125 | 2.234104145 | -0.298846645 | 0.631745336 |
| Khdrbs3 | 814.1907786 | -0.209028181 | 0.116553312 |
| Gm24232 | 0 | NA | NA |
| Gm23987 | 0 | NA | NA |
| Fam135b | 709.6683967 | -0.115418507 | 0.117815556 |
| AC157602.1 | 0 | NA | NA |
| Gm22519 | 0.044191549 | -0.243571806 | 3.724945049 |
| Col22a1 | 42.46963243 | 0.98349274 | 0.206936316 |
| Gm16308 | 0.108309328 | 0.237310006 | 3.724945049 |
| Gm23217 | 4.535027026 | -1.036553541 | 0.523623438 |
| Gm24390 | 0 | NA | NA |
| Kcnk9 | 245.3309406 | -0.572262542 | 0.095308568 |
| Trappc9 | 814.4156458 | -0.353166352 | 0.043973155 |
| Gm28020 | 1.075827988 | -1.935626129 | 1.104478897 |
| Peg13 | 3399.486185 | -0.602699401 | 0.052144081 |
| Gm3150 | 0.456099668 | 0.809601016 | 1.41098195 |
| Chrac1 | 76.46489437 | 0.595426467 | 0.165966918 |
| Ago2 | 2822.203106 | 0.052373814 | 0.071901113 |
| Ptk2 | 1504.870551 | -0.215294121 | 0.068610434 |
| Mir151 | 5.981125562 | -1.773393525 | 0.585038293 |
| Gm24787 | 0 | NA | NA |
| Dennd3 | 115.7213391 | 0.953619541 | 0.224416995 |
| Slc45a4 | 787.9133075 | -0.063666059 | 0.072868554 |
| 1700010B13Rik | 0.609775176 | -0.623158645 | 1.275564494 |
| Gpr20 | 1.015418215 | -0.882422699 | 1.112673239 |
| Ptp4a3 | 340.214372 | -0.161488905 | 0.117818449 |
| Gm22106 | 2.287044039 | -1.212655747 | 0.673494047 |
| Mroh5 | 17.84213286 | -0.378496712 | 0.249170437 |
| Gm3244 | 3.864046566 | -0.500322036 | 0.540142002 |
| Gm6569 | 1.915143953 | 0.535653574 | 0.669389848 |
| Gm7935 | 7.955384209 | 0.563461313 | 0.356068622 |
| Gm15387 | 13.22477098 | 0.530651347 | 0.493335312 |
| Adgrb1 | 3078.93178 | -0.568208438 | 0.145851454 |
| Gm26117 | 0 | NA | NA |
| Mroh4 | 0.178490723 | 0.718216836 | 2.713461433 |
| Arc | 2012.660561 | -0.160607614 | 0.498187815 |
| Jrk | 151.5292579 | 0.184628424 | 0.105382804 |
| 4933427E11Rik | 4.797225747 | 0.397262679 | 0.541906161 |
| Psca | 1.592565788 | 3.776975067 | 1.025093745 |
| Gm28976 | 0 | NA | NA |
| Them6 | 413.7288862 | 0.013037245 | 0.079269097 |
| Slurp1 | 1.036256916 | 0.702236707 | 1.209343115 |
| Lypd2 | 2.552977391 | 0.582301328 | 0.649579039 |
| 2300005B03Rik | 0.58378306 | 2.209896632 | 1.634632278 |

| | | | |
|---|---|---|---|
| Lynx1 | 4130.871836 | 0.008859519 | 0.070138678 |
| Ly6d | 10.57966156 | 4.174556607 | 0.711826601 |
| D730001G18Rik | 0.049553047 | 0.71820487 | 3.724945049 |
| Gm17189 | 0.097139547 | 0.718207094 | 3.724945049 |
| Ly6k | 0.732543458 | 1.780773287 | 1.206149603 |
| Gml | 0 | NA | NA |
| Hemt1 | 0 | NA | NA |
| Cyp11b1 | 0.755541319 | 2.152554932 | 1.236133248 |
| Cyp11b2 | 1.063164669 | 1.167703171 | 1.093495901 |
| 2010109I03Rik | 3.337759678 | 2.573153189 | 0.707645739 |
| Gm6610 | 1.171300456 | 4.221602515 | 1.10426189 |
| Gm29126 | 0.728772687 | 3.076676688 | 1.170059304 |
| Ly6e | 7038.794082 | 3.810695634 | 0.168032959 |
| Ly6i | 26.75927167 | 8.017743327 | 1.005929321 |
| Gm8000 | 1.59659966 | 4.340732491 | 1.005418085 |
| Gm37042 | 1.174124634 | 1.424740929 | 0.960503044 |
| Ly6a | 4998.068703 | 9.455739478 | 0.340952344 |
| Gm28068 | 1.632511506 | 2.546914507 | 0.98044104 |
| Gm28069 | 0.045882455 | 0.237312084 | 3.724945049 |
| Ly6c1 | 1373.160618 | 5.595955784 | 0.366204512 |
| Gm5960 | 192.4448412 | 11.53587813 | 1.065777679 |
| Gm28502 | 0.175945213 | 1.166110064 | 2.710746636 |
| Ly6c2 | 840.6845555 | 12.13667846 | 0.92213741 |
| I830127L07Rik | 12.15111785 | 7.514152537 | 0.939589747 |
| Ly6g | 0.512981325 | 2.32236325 | 1.341003638 |
| Gm28117 | 0 | NA | NA |
| BC025446 | 0.044544648 | 0.237309859 | 3.724945049 |
| Gm10238 | 0 | NA | NA |
| Gm28115 | 0 | NA | NA |
| Ly6f | 0 | NA | NA |
| Gm28116 | 0 | NA | NA |
| Gm3454 | 0.07796279 | 1.072698098 | 3.721829099 |
| Gm28113 | 0.038981395 | 0.718207094 | 3.724945049 |
| Gm29517 | 0 | NA | NA |
| Gm28114 | 0.149988252 | 1.037032011 | 3.492945861 |
| Gm3142 | 0 | NA | NA |
| 9030619P08Rik | 0.161532967 | 1.200059132 | 3.449701177 |
| Gm20654 | 0 | NA | NA |
| Ly6h | 944.0802929 | -0.032247428 | 0.096405506 |
| Gpihbp1 | 0.278632317 | 1.087785664 | 1.803623529 |
| Zfp41 | 98.87672504 | 0.298048998 | 0.128142934 |
| 2810039B14Rik | 19.1597757 | -0.049458834 | 0.251102721 |
| Top1mt | 124.7567236 | 0.122987643 | 0.115446332 |
| Gm15945 | 0.585634519 | -0.459789178 | 1.320905131 |
| Rhpn1 | 84.34622614 | -0.138473332 | 0.145881712 |
| Mafa | 7.826514485 | -0.179928431 | 0.403566662 |
| Zc3h3 | 159.3577311 | -0.027205584 | 0.15243558 |

| | | | |
|---|---|---|---|
| Gm23747 | 0 | NA | NA |
| Gm9568 | 0.083526043 | 0.718202646 | 3.724945049 |
| Gsdmd | 138.4714216 | 4.242988597 | 0.230048687 |
| Mroh6 | 1.934735662 | -0.039907371 | 0.706573986 |
| Naprt | 43.80295543 | 0.155396824 | 0.237923817 |
| Eef1d | 688.7842089 | 0.221839322 | 0.065725533 |
| Tigd5 | 16.71972297 | 0.007355536 | 0.259762192 |
| AC116520.1 | 0 | NA | NA |
| Pycrl | 118.6008582 | 0.359551508 | 0.169657113 |
| Tsta3 | 168.0293222 | 0.55807223 | 0.126918367 |
| Zfp623 | 159.2088236 | 0.062744558 | 0.133450588 |
| Zfp707 | 75.27886443 | -0.067357186 | 0.143245278 |
| Ccdc166 | 61.81052859 | 0.106078693 | 0.14927796 |
| Mapk15 | 15.79109506 | -0.085434975 | 0.338076265 |
| Fam83h | 77.85627554 | 0.42435191 | 0.135646674 |
| K230010J24Rik | 12.00267003 | 0.168822608 | 0.334823598 |
| Gm23027 | 0.654384519 | 0.875572582 | 1.381843634 |
| Scrib | 592.0251915 | -0.017170186 | 0.075638308 |
| Mir6952 | 0 | NA | NA |
| Puf60 | 1138.944899 | 0.266767733 | 0.086554421 |
| Nrbp2 | 715.0247032 | -0.730200052 | 0.116741737 |
| BC024139 | 9.779656096 | -0.03618298 | 0.327234489 |
| Plec | 5010.938922 | 0.297786966 | 0.078410674 |
| Gm25720 | 3.45116213 | -0.996552232 | 0.599804266 |
| Mir1942 | 3.221922816 | -1.346242397 | 0.64124203 |
| Parp10 | 866.3362482 | 6.462151231 | 0.213367424 |
| Grina | 4908.811971 | 0.584037529 | 0.085503618 |
| Mir6953 | 0 | NA | NA |
| Gm10872 | 0.972832988 | 3.349214117 | 1.15949029 |
| Spatc1 | 0.136963818 | 0.718204821 | 3.59954433 |
| Mir3079 | 0 | NA | NA |
| Smpd5 | 4.917982605 | 0.41519673 | 0.465171203 |
| Oplah | 309.3643565 | -0.310937442 | 0.117370164 |
| Exosc4 | 178.5313927 | 0.512599106 | 0.113682363 |
| Gpaa1 | 246.0551654 | -0.148351053 | 0.0881496 |
| Rps6-ps1 | 8.203160127 | 0.484103186 | 0.35647663 |
| Cyc1 | 1071.038313 | 0.063064185 | 0.092632372 |
| Sharpin | 316.269203 | 0.353048959 | 0.08362962 |
| Maf1 | 531.081438 | 0.193635512 | 0.081024407 |
| Hgh1 | 82.07062805 | 0.762495808 | 0.188019701 |
| Tssk5 | 6.475749014 | 0.331644358 | 0.478135739 |
| Mroh1 | 1129.595412 | -0.142613674 | 0.086896798 |
| Mir6954 | 0 | NA | NA |
| Bop1 | 410.1160048 | 0.160254065 | 0.06200116 |
| Scx | 29.69111678 | 0.161145036 | 0.220539946 |
| Hsf1 | 313.4887952 | 0.145962984 | 0.09850014 |
| Dgat1 | 157.725041 | -0.142316623 | 0.119274383 |

| | | | |
|---|---|---|---|
| Scrt1 | 1878.856055 | -1.033955795 | 0.078812931 |
| Fbxl6 | 146.2430035 | 0.078424571 | 0.094522113 |
| Slc52a2 | 132.5881942 | 0.225976931 | 0.104991644 |
| Adck5 | 90.29773825 | -0.12422573 | 0.119513818 |
| Cpsf1 | 643.6551908 | 0.180517216 | 0.078050101 |
| Slc39a4 | 6.380351065 | 1.566063588 | 0.488904632 |
| Tonsl | 636.9176432 | 0.064485793 | 0.084730959 |
| Cyhr1 | 972.3595972 | -0.216753985 | 0.056752274 |
| Kifc2 | 2493.40518 | -1.496104284 | 0.139732378 |
| Foxh1 | 0.909626782 | -1.071654808 | 1.097383119 |
| Ppp1r16a | 515.0001235 | -0.267774264 | 0.069693809 |
| Gpt | 84.05780739 | -1.408910856 | 0.191614686 |
| Mfsd3 | 34.06178562 | 0.161586988 | 0.218754661 |
| Recql4 | 21.17447754 | 0.433011964 | 0.299754704 |
| Lrrc14 | 184.1576737 | 0.111824325 | 0.084350733 |
| Lrrc24 | 405.4748953 | -0.315841086 | 0.079146982 |
| Arhgap39 | 936.4403093 | -0.423941054 | 0.093747877 |
| Gm17271 | 0.060174537 | 0.237309859 | 3.724945049 |
| Zfp251 | 456.6486897 | -0.403729507 | 0.066552175 |
| Zfp7 | 92.43301678 | 0.002566445 | 0.118431455 |
| Commd5 | 140.0820505 | 0.375195156 | 0.129466105 |
| Rpl8 | 2132.795094 | 0.628597226 | 0.101771133 |
| Zfp647 | 28.94240596 | -0.094035425 | 0.202372809 |
| 1110038F14Rik | 133.089957 | 0.794885672 | 0.109339935 |
| Mb | 1.797034208 | -0.107555063 | 0.850402088 |
| Apol6 | 78.23562678 | 6.285909241 | 0.496710109 |
| Gm24753 | 2.783881763 | 0.720160684 | 0.728217553 |
| Rbfox2 | 3954.719732 | -0.026855083 | 0.080528591 |
| 1700109K24Rik | 0.04393175 | -0.243571806 | 3.724945049 |
| Gm24056 | 9.581094532 | -1.015055447 | 0.427034442 |
| Apol7a | 18.60379838 | 3.137813574 | 0.55532713 |
| Gm20688 | 0.311586316 | 1.210028468 | 1.97689001 |
| Apol9a | 239.6974814 | 9.12885663 | 0.70466314 |
| Apol7b | 0.961738709 | 1.412052458 | 1.142449435 |
| 4930571N24Rik | 0.157815785 | 0.718205257 | 3.466717377 |
| Gm6506 | 0.409269261 | 1.229307117 | 1.453943663 |
| Apol10a | 0.15100991 | 0.718202046 | 3.527597384 |
| Apol11a | 0 | NA | NA |
| Apol7c | 0.292232717 | 2.171570149 | 1.949434453 |
| 1700025B11Rik | 0.199129965 | 1.667487852 | 2.212011517 |
| Apol10b | 22.02299047 | 7.766622962 | 0.9019808 |
| Gm8221 | 0.475668934 | 0.299443414 | 1.3051666 |
| Apol11b | 3.059439512 | 4.815427351 | 0.96170447 |
| Apol7e | 1.633903828 | 3.455999294 | 1.050611213 |
| Apol9b | 163.6719493 | 9.588893864 | 0.886898283 |
| Apol8 | 54.12186927 | -0.059135647 | 0.157962735 |
| Myh9 | 2978.335968 | 1.652359599 | 0.130478247 |

| | | | |
|---|---|---|---|
| Gm22107 | 5.021758635 | 1.281256424 | 0.460277683 |
| Txn2 | 707.7197206 | 0.123672409 | 0.071965999 |
| Foxred2 | 319.3024116 | -0.181223756 | 0.094707773 |
| Eif3d | 737.7459319 | 0.881726893 | 0.095826172 |
| Gm17638 | 11.65945967 | 0.340953974 | 0.385288613 |
| Cacng2 | 970.2458113 | -0.575987932 | 0.089179344 |
| A730060N03Rik | 73.25342862 | -0.828244554 | 0.171639964 |
| Ift27 | 107.8713168 | -0.215628297 | 0.131964241 |
| Pvalb | 1103.933019 | -0.04969328 | 0.152060079 |
| Ncf4 | 100.2792551 | 4.945794744 | 0.305167812 |
| Csf2rb2 | 172.9194105 | 5.390221991 | 0.287942182 |
| Mir7676-1 | 0 | NA | NA |
| Csf2rb | 284.283327 | 5.363349461 | 0.254620339 |
| Mir7676-2 | 0 | NA | NA |
| Gm26329 | 0 | NA | NA |
| Tex33 | 0.613719218 | 1.906267681 | 1.251655513 |
| Tst | 291.9109073 | 0.989807461 | 0.155781783 |
| Mpst | 172.564482 | 0.06637444 | 0.136817655 |
| Kctd17 | 1094.573162 | -0.213358965 | 0.059571502 |
| Tmprss6 | 5.777471174 | 1.472599241 | 0.515215084 |
| Il2rb | 153.2540098 | 6.709239348 | 0.485953534 |
| C1qtnf6 | 29.05525177 | 0.843037808 | 0.202228406 |
| Sstr3 | 170.7394951 | -0.10207898 | 0.163504424 |
| Gm6723 | 0.648304949 | 0.579858666 | 1.16393623 |
| Rac2 | 192.5000097 | 4.775324994 | 0.245010788 |
| Cyth4 | 257.1621641 | 2.680612729 | 0.146397288 |
| Elfn2 | 1215.602818 | -0.830238323 | 0.121719604 |
| Mfng | 23.1312152 | 1.010666354 | 0.282999719 |
| Card10 | 133.9315844 | 0.13738806 | 0.176863546 |
| Cdc42ep1 | 428.9155534 | 1.165946415 | 0.165882145 |
| Lgals2 | 1.456018128 | 2.192284627 | 0.8549327 |
| Gm26634 | 0.085327104 | 1.179809211 | 3.723500796 |
| Gga1 | 712.034101 | 0.177873447 | 0.068728528 |
| Mir6955 | 1.342239839 | -1.215316356 | 0.902807065 |
| Sh3bp1 | 1228.874401 | -0.465086249 | 0.057475229 |
| 1700027A07Rik | 1.879021917 | -1.231028992 | 0.74690048 |
| Lgals1 | 413.7512805 | 3.494347854 | 0.331930356 |
| Nol12 | 155.5073409 | 0.83743243 | 0.11257831 |
| Triobp | 275.7571062 | 1.84617389 | 0.170283236 |
| Mir6956 | 0 | NA | NA |
| H1f0 | 1010.011615 | 0.182583955 | 0.102999951 |
| Gcat | 70.63897891 | 0.252832896 | 0.149063405 |
| Ankrd54 | 161.076979 | 0.241721791 | 0.14409247 |
| Eif3l | 629.3207194 | 0.51116643 | 0.076639668 |
| Gm25072 | 0 | NA | NA |
| Micall1 | 863.8231058 | 0.191275312 | 0.080716656 |
| 1700088E04Rik | 67.10927695 | 0.633577941 | 0.144903902 |

| | | | |
|---|---|---|---|
| Polr2f | 144.8684693 | 0.52703573 | 0.191678208 |
| Sox10 | 508.9917057 | 0.573926365 | 0.107373102 |
| Gm10863 | 17.85705234 | 0.966158389 | 0.262343872 |
| Pick1 | 359.5554522 | -0.384851173 | 0.090162286 |
| Slc16a8 | 3.295224942 | -4.538748056 | 0.921065237 |
| Lgals1-ps2 | 0 | NA | NA |
| Baiap2l2 | 8.450121399 | -0.664326411 | 0.43613369 |
| Pla2g6 | 246.7582411 | -0.130791695 | 0.10493313 |
| Gm20420 | 0 | NA | NA |
| Maff | 106.5092356 | 3.986219962 | 0.481611389 |
| Tmem184b | 1712.559106 | 1.252079617 | 0.127907878 |
| Mir1943 | 2.514149661 | 0.427159699 | 0.734151317 |
| Csnk1e | 892.3901685 | -0.200478675 | 0.079075217 |
| Gm16059 | 0.387099777 | -0.629435988 | 1.849575414 |
| AC165278.1 | 0 | NA | NA |
| Kcnj4 | 701.1998511 | -0.948255905 | 0.152695438 |
| Kdelr3 | 10.19668411 | 1.954659207 | 0.425903939 |
| Ddx17 | 3680.454381 | -0.124766424 | 0.063133374 |
| Dmc1 | 9.014701878 | -0.65920638 | 0.364313265 |
| Gm27903 | 0.358948382 | -0.684633518 | 1.457837157 |
| Fam227a | 83.10424768 | -1.134299403 | 0.247532243 |
| Cby1 | 177.468213 | -0.523458807 | 0.106576102 |
| Tomm22 | 344.3752573 | 0.328474892 | 0.097482101 |
| Josd1 | 713.1554615 | 0.052482986 | 0.0858801 |
| Gm26884 | 1.046429311 | 1.378821857 | 1.092353308 |
| Gtpbp1 | 638.7878775 | 0.204277987 | 0.062513584 |
| n-R5s40 | 1.378629347 | -0.245196175 | 0.881132649 |
| Sun2 | 786.962264 | 0.247816213 | 0.071563217 |
| Gm16575 | 0.364940181 | 0.371622401 | 1.706349477 |
| Gm16576 | 14.37465387 | 0.232503954 | 0.313346465 |
| Dnal4 | 230.0228909 | -0.262104805 | 0.08315343 |
| Nptxr | 4627.776829 | -1.132279147 | 0.152808962 |
| Npcd | 0.498288516 | -0.731957843 | 1.452616181 |
| Cbx6 | 4151.173169 | -0.648069313 | 0.088600181 |
| Apobec3 | 148.4452752 | 3.58729448 | 0.19365287 |
| Cbx7 | 316.6582599 | -0.437808183 | 0.086596973 |
| Pdgfb | 148.5994786 | 0.443168376 | 0.160102865 |
| Mir7213 | 0 | NA | NA |
| Rpl3 | 1750.858766 | 0.361901945 | 0.09543848 |
| Snord83b | 1.863983961 | -0.057434263 | 0.733347301 |
| Gm22077 | 0 | NA | NA |
| AC161199.1 | 2.341657818 | 0.878212506 | 0.711262268 |
| Gm24204 | 0.553939527 | 0.13251335 | 1.400926531 |
| Syngr1 | 4833.272579 | -0.720168655 | 0.060246915 |
| Tab1 | 241.9134514 | 0.11130424 | 0.085685921 |
| Mgat3 | 1733.281968 | -0.406742351 | 0.071011663 |
| Mief1 | 547.7891883 | 0.360961604 | 0.079952479 |

| | | | |
|---|---|---|---|
| Atf4 | 1610.207181 | 1.438272914 | 0.158158498 |
| Rps19bp1 | 105.055002 | 0.667929814 | 0.149847602 |
| Cacna1i | 1385.507454 | -0.778141319 | 0.120777856 |
| Enthd1 | 1.187168766 | 3.425466423 | 1.073480376 |
| Gm23220 | 0 | NA | NA |
| Grap2 | 30.59568962 | 4.317537132 | 0.415002122 |
| Mir6957 | 0 | NA | NA |
| Fam83f | 18.37852399 | 2.292989956 | 0.390937987 |
| Tnrc6b | 3449.044664 | -0.390646973 | 0.077699136 |
| Mir5113 | 0 | NA | NA |
| Adsl | 284.5366327 | 0.691370272 | 0.077238163 |
| Gm6612 | 1.14679352 | -0.495600212 | 1.070247239 |
| Sgsm3 | 461.0312359 | -0.138893124 | 0.067187136 |
| Mkl1 | 795.5391381 | -0.139475219 | 0.08470852 |
| Gm25131 | 0 | NA | NA |
| 4930483J18Rik | 0.130580445 | -1.07128359 | 3.667847045 |
| Mchr1 | 166.8774582 | 0.19623786 | 0.187347881 |
| Slc25a17 | 303.5824259 | 0.209189111 | 0.097119547 |
| St13 | 1980.343047 | 0.050951098 | 0.064447374 |
| Xpnpep3 | 220.8414576 | 0.205752009 | 0.108568667 |
| Dnajb7 | 0.134148234 | 0.23732184 | 3.624632408 |
| Gm17025 | 0.441788749 | -1.722554535 | 1.237574663 |
| Rbx1 | 792.4993545 | 0.190796958 | 0.081593179 |
| Gm5218 | 0.038981395 | 0.718207094 | 3.724945049 |
| 1110025M09Rik | 12.66128553 | 0.125717908 | 0.298702279 |
| Ep300 | 2415.831866 | 0.038597368 | 0.063616693 |
| Gm23880 | 0 | NA | NA |
| Gm22067 | 12.43328101 | -0.769161504 | 0.275157366 |
| L3mbtl2 | 382.5292509 | -0.101403145 | 0.064205733 |
| Gm18284 | 12.73109718 | -0.042984358 | 0.287843367 |
| Chadl | 117.5180505 | -0.197632711 | 0.132614027 |
| Rangap1 | 1965.883829 | -0.036954422 | 0.065797313 |
| Zc3h7b | 3000.898726 | -0.054465574 | 0.079688795 |
| Gm17597 | 7.602086333 | -1.263834128 | 0.381588512 |
| Tef | 3374.752106 | -0.944772984 | 0.107400102 |
| Gm8444 | 1.234579709 | -0.448442395 | 0.925498532 |
| Tob2 | 667.3737093 | 0.19835151 | 0.111027417 |
| Phf5a | 193.4131225 | 0.77374807 | 0.135169585 |
| Aco2 | 4367.876781 | -0.349995542 | 0.068054619 |
| Polr3h | 303.152543 | -0.085444864 | 0.094112925 |
| Csdc2 | 982.7828558 | -0.457673057 | 0.112123381 |
| Pmm1 | 652.8758253 | -0.226195649 | 0.05797363 |
| Gm5805 | 68.93469726 | 0.904571458 | 0.214635566 |
| 1700029P11Rik | 0.075043205 | 0.237309859 | 3.724945049 |
| Xrcc6 | 153.2673552 | 0.055600459 | 0.116864306 |
| Desi1 | 1377.267412 | -0.064327067 | 0.078212147 |
| Nhp2l1 | 547.7572345 | 0.345330158 | 0.10540979 |

| | | | |
|---|---|---|---|
| 4930407I10Rik | 0.062426873 | 0.237312084 | 3.724945049 |
| Mei1 | 19.77925728 | 0.135015922 | 0.27519436 |
| Ccdc134 | 79.33616336 | 0.230117084 | 0.141406402 |
| Srebf2 | 2183.707136 | -0.346448212 | 0.079071495 |
| Mir33 | 0.155921052 | -0.243569951 | 2.831361108 |
| Shisa8 | 125.8398763 | -0.023082762 | 0.202565814 |
| Tnfrsf13c | 6.674742414 | 1.361401587 | 0.463897397 |
| Cenpm | 13.06399243 | 0.652199718 | 0.324147236 |
| 1500009C09Rik | 1247.839228 | 0.86172693 | 0.137083576 |
| 3-Sep | 4869.520885 | -0.581896114 | 0.069544179 |
| Wbp2nl | 0.428891288 | 1.193237511 | 1.564899853 |
| Naga | 263.3133539 | 0.851924796 | 0.120699248 |
| Fam109b | 4.886605523 | 0.412332099 | 0.471380557 |
| Smdt1 | 561.4413452 | 0.172502714 | 0.10903848 |
| n-R5s41 | 0 | NA | NA |
| Ndufa6 | 517.8924798 | 0.172884596 | 0.10780416 |
| Cyp2d22 | 207.7231852 | 0.253268409 | 0.132817696 |
| Cyp2d11 | 0 | NA | NA |
| Mir7684 | 0 | NA | NA |
| Cyp2d10 | 0 | NA | NA |
| Gm27331 | 0 | NA | NA |
| Gm25990 | 0 | NA | NA |
| Cyp2d9 | 0.049553047 | 0.71820487 | 3.724945049 |
| Gm27622 | 0 | NA | NA |
| Gm27825 | 0 | NA | NA |
| Gm5806 | 3.903672454 | 1.195107817 | 0.596233224 |
| Gm28023 | 0 | NA | NA |
| Cyp2d12 | 0.066050875 | 0.71820487 | 3.724945049 |
| Gm27924 | 0 | NA | NA |
| Gm27742 | 0 | NA | NA |
| Cyp2d34 | 0.18989612 | 1.462508577 | 2.650629334 |
| Gm27463 | 0.194279094 | 1.292225235 | 3.719439735 |
| Cyp2d37-ps | 0.066050875 | 0.71820487 | 3.724945049 |
| Cyp2d40 | 0.493912024 | 2.43791403 | 1.488207873 |
| Cyp2d41-ps | 0.567195248 | 1.179385947 | 1.332672698 |
| Gm27618 | 0 | NA | NA |
| Cyp2d26 | 2.149324495 | 0.984346192 | 0.681284814 |
| Tcf20 | 4578.253049 | -0.701195753 | 0.097138389 |
| Gm20324 | 2.615112804 | 0.629841688 | 0.670027405 |
| Mir3080 | 2.487744732 | -2.531878606 | 0.837039978 |
| Gm29019 | 5.11266864 | -1.262382795 | 0.605126345 |
| AW121686 | 10.56313198 | -0.296237752 | 0.341127138 |
| Nfam1 | 67.01581601 | 2.204695266 | 0.230010851 |
| Serhl | 61.58247766 | 0.755414191 | 0.1801122 |
| Gm17206 | 0.066050875 | 0.71820487 | 3.724945049 |
| Rrp7a | 918.7979819 | 0.39473604 | 0.07516597 |
| Poldip3 | 1119.919797 | 0.44322697 | 0.07712873 |

| | | | |
|---|---|---|---|
| Rnu12 | 255.543187 | -0.028003213 | 0.166057559 |
| Cyb5r3 | 666.3127489 | 0.811305045 | 0.117253389 |
| A4galt | 36.02363458 | 1.705866816 | 0.256592588 |
| Arfgap3 | 242.8767015 | 0.73631071 | 0.077316612 |
| 1700001L05Rik | 36.6738785 | -0.630397079 | 0.204316765 |
| Pacsin2 | 643.9682478 | 0.376477438 | 0.088581601 |
| Gm5417 | 28.18046227 | 0.057200199 | 0.248908505 |
| Ttll1 | 251.422686 | -0.565517578 | 0.110951874 |
| Gm24575 | 0 NA | | NA |
| Bik | 0.708533877 | 3.225678522 | 1.217934215 |
| Mcat | 123.3968876 | 0.566795292 | 0.120450978 |
| Tspo | 459.8898812 | 6.087918745 | 0.343527717 |
| Ttll12 | 215.0437835 | 0.457137431 | 0.110605482 |
| Scube1 | 480.4921202 | -1.424037563 | 0.173339281 |
| Mpped1 | 824.048876 | -1.486621682 | 0.144175843 |
| Efcab6 | 85.58806344 | 0.030530332 | 0.291669949 |
| Rpl31-ps22 | 0 NA | | NA |
| Sult4a1 | 2771.319555 | -0.47264375 | 0.086181905 |
| Pnpla5 | 0 NA | | NA |
| Pnpla3 | 102.6316791 | -1.203949724 | 0.190798981 |
| Mir6392 | 3.244976357 | -1.368205056 | 0.577932126 |
| Samm50 | 528.8198655 | 0.006636463 | 0.070216366 |
| Parvb | 380.2549727 | 0.807755696 | 0.101000254 |
| Gm22890 | 0.046345709 | 0.71820487 | 3.724945049 |
| Parvg | 82.64255846 | 1.473453571 | 0.182063192 |
| 1810041L15Rik | 649.3280647 | -0.352090864 | 0.126342802 |
| Ldoc1l | 614.5341085 | -0.257602104 | 0.083065389 |
| Prr5 | 124.3176796 | 0.310271989 | 0.133819029 |
| Arhgap8 | 19.60055845 | 4.527594551 | 0.57708568 |
| Phf21b | 132.294507 | -0.805701297 | 0.176420002 |
| Gm4217 | 13.23963404 | 0.542910932 | 0.359297056 |
| Gm29666 | 48.12944454 | -0.847413698 | 0.216851965 |
| Nup50 | 602.2802685 | 0.360061627 | 0.081698498 |
| 5031439G07Rik | 1832.693521 | -0.121779091 | 0.056395352 |
| Mir1249 | 0 NA | | NA |
| Upk3a | 0.636985128 | -0.936685327 | 1.321754964 |
| Fam118a | 232.4075255 | 0.276441369 | 0.083657247 |
| Smc1b | 5.078404989 | 1.232665129 | 0.535135592 |
| Gm10923 | 2.764924986 | 1.400413113 | 0.623029413 |
| Ribc2 | 3.994063061 | 0.655705286 | 0.493164241 |
| Fbln1 | 204.5288485 | 1.319282574 | 0.178619406 |
| Atxn10 | 2123.828845 | 0.107529304 | 0.062432326 |
| Gm23517 | 0 NA | | NA |
| 7530416G11Rik | 0 NA | | NA |
| Gm4825 | 0.947696482 | 2.759438794 | 1.123946243 |
| Wnt7b | 268.9427193 | -0.219184496 | 0.132128379 |
| AU022754 | 9.118481155 | -0.161099434 | 0.347703717 |

| | | | |
|---|---|---|---|
| D130051D11Rik | 5.759629403 | -0.291697183 | 0.4563937 |
| Gm37699 | 45.08239792 | -0.635951443 | 0.252988367 |
| Mirlet7c-2 | 5.369208344 | -0.254486522 | 0.494226486 |
| Mirlet7b | 13.61717214 | -0.050925878 | 0.369495602 |
| Ppara | 39.64814777 | -0.650663444 | 0.213728518 |
| Cdpf1 | 48.27012333 | 0.176326496 | 0.185691501 |
| Pkdrej | 11.56756557 | -0.001185141 | 0.33790433 |
| Ttc38 | 59.42181862 | -0.405512224 | 0.158218364 |
| Gtse1 | 17.82322006 | 2.029334674 | 0.343184372 |
| Trmu | 131.3669997 | -0.171194009 | 0.12480325 |
| Celsr1 | 124.8124239 | -0.607093363 | 0.1555602 |
| Gramd4 | 450.6965781 | 0.02555552 | 0.06980285 |
| Gm15722 | 0 | NA | NA |
| Cerk | 1311.25289 | -0.064592078 | 0.083085861 |
| Gm24011 | 0 | NA | NA |
| Gm15569 | 1.568615446 | 0.604149613 | 0.902019098 |
| Gm22818 | 0.772793883 | -2.507881436 | 1.110573481 |
| Tbc1d22a | 375.15338 | 0.136134761 | 0.070673121 |
| Gm23416 | 0.044544648 | 0.237309859 | 3.724945049 |
| n-R5s42 | 0 | NA | NA |
| Fam19a5 | 1108.298912 | -0.312690991 | 0.076217473 |
| Mir6393 | 0 | NA | NA |
| Zdhhc25 | 0 | NA | NA |
| Brd1 | 988.6755636 | -0.100719101 | 0.115091577 |
| Zbed4 | 103.903103 | 0.157932944 | 0.111060286 |
| Alg12 | 119.1075428 | 0.37565678 | 0.111433774 |
| Creld2 | 178.5176635 | 1.356650447 | 0.161221257 |
| Gm23144 | 0 | NA | NA |
| Pim3 | 592.2908598 | 0.83490061 | 0.17172362 |
| Gm20621 | 2.301900616 | -0.623827337 | 0.644292645 |
| Ttll8 | 19.07562425 | -0.806571965 | 0.376958148 |
| Mlc1 | 1260.237927 | -0.189485843 | 0.129611454 |
| Mov10l1 | 0.232306593 | -0.272977932 | 2.17831276 |
| Gm23737 | 0 | NA | NA |
| Panx2 | 872.8564831 | -0.77570491 | 0.099938665 |
| 1810021B22Rik | 9.773541415 | 0.355803455 | 0.349287236 |
| Trabd | 275.7218993 | 0.389348857 | 0.090504811 |
| Selo | 248.0954824 | -0.17914547 | 0.091205192 |
| Tubgcp6 | 650.1368911 | -0.174697443 | 0.083688528 |
| Hdac10 | 82.94525066 | -0.314065875 | 0.15722014 |
| Mapk12 | 96.05029267 | -0.600326634 | 0.170613091 |
| Mapk11 | 187.7470408 | -0.611545351 | 0.153613727 |
| Plxnb2 | 1637.125997 | 0.998391275 | 0.125777008 |
| mmu-mir-7118 | 0 | NA | NA |
| Dennd6b | 661.1196248 | 0.41488716 | 0.18248363 |
| Mir6958 | 0 | NA | NA |
| Gm26798 | 0.47411084 | -1.103168526 | 1.359289921 |

| | | | |
|---|---|---|---|
| Ppp6r2 | 1298.582558 | -0.395909506 | 0.054242029 |
| Sbf1 | 2446.17373 | -0.250130651 | 0.052710768 |
| Mir6959 | 1.302949929 | -1.529392695 | 0.970623525 |
| Adm2 | 0.810290415 | 3.616195999 | 1.207586016 |
| Miox | 0.11193333 | 0.718202646 | 3.724945049 |
| Lmf2 | 236.2729926 | 0.774320967 | 0.115236841 |
| Ncaph2 | 806.3681184 | 0.002778022 | 0.063170379 |
| Sco2 | 38.3484847 | -0.046250993 | 0.177656155 |
| Tymp | 10.59392881 | 0.106445179 | 0.303364426 |
| Odf3b | 13.0856276 | -0.584538638 | 0.29373691 |
| Klhdc7b | 7.376514687 | -1.225443027 | 0.367082181 |
| Syce3 | 0 | NA | NA |
| Chkb | 231.7816191 | 0.022590941 | 0.095950296 |
| Cpt1b | 2.011645178 | -0.052093907 | 0.704993193 |
| C730034F03Rik | 13.21654071 | 0.365806085 | 0.263309383 |
| Mapk8ip2 | 3763.788627 | -0.362499953 | 0.054902171 |
| Arsa | 324.9924385 | -0.080246364 | 0.097767151 |
| C230037L18Rik | 26.54613617 | -1.629086125 | 0.289556694 |
| Shank3 | 2814.903766 | -1.2537654 | 0.133231987 |
| Gm15609 | 24.97572873 | -1.152186658 | 0.35506474 |
| Acr | 23.80281401 | -0.829330237 | 0.447014866 |
| Rabl2 | 143.4438333 | -0.229007148 | 0.134900941 |
| Syt10 | 129.1730868 | -0.732610495 | 0.176974116 |
| Gm24643 | 0 | NA | NA |
| Gm24415 | 0 | NA | NA |
| Alg10b | 588.5402971 | 0.104309228 | 0.081097964 |
| Mir7648 | 0 | NA | NA |
| Cpne8 | 261.9200795 | 0.117615956 | 0.107027694 |
| Gm6746 | 0 | NA | NA |
| Rpl31-ps8 | 113.0537037 | 0.53774729 | 0.242196182 |
| Tcea1-ps1 | 127.5401109 | 0.24582554 | 0.112193042 |
| Kif21a | 3097.142762 | -0.332961959 | 0.06025729 |
| Abcd2 | 509.0775856 | -0.989586111 | 0.098690727 |
| CN725425 | 2.738760097 | -1.13294637 | 0.673955532 |
| Slc2a13 | 1660.411825 | -0.55363455 | 0.100955502 |
| Gm15383 | 16.38285574 | -1.988707571 | 0.392968255 |
| Gm15381 | 25.44622861 | -2.009657857 | 0.313587557 |
| Gm24647 | 4.156244294 | -2.094249819 | 0.666250398 |
| 4933438A12Rik | 0.060174537 | 0.237309859 | 3.724945049 |
| Gm25389 | 0 | NA | NA |
| Lrrk2 | 1067.387671 | -0.335358322 | 0.139261856 |
| Gm15382 | 4.663180715 | -0.429628535 | 0.452297924 |
| Smgc | 0.890372413 | 2.041920676 | 1.181915284 |
| Muc19 | 0.988759962 | 1.343206693 | 1.109789063 |
| Cntn1 | 4874.212289 | -0.953736573 | 0.066866361 |
| Gm26760 | 2.85183316 | -0.997919288 | 0.650611833 |
| Pdzrn4 | 94.36282265 | -0.591684624 | 0.115459186 |

| | | | |
|---|---|---|---|
| Gm37304 | 37.54844209 | -1.492901023 | 0.319611237 |
| Gm4335 | 6.712093293 | 0.065355443 | 0.442489257 |
| Gm24447 | 11246.06972 | 0.14125791 | 0.147220954 |
| Gxylt1 | 308.8367025 | -0.241327762 | 0.098204444 |
| Yaf2 | 648.3196561 | -0.285823562 | 0.076005338 |
| Gm26371 | 0.04393175 | -0.243571806 | 3.724945049 |
| Zcrb1 | 380.1198483 | -0.057283987 | 0.124134865 |
| Pphln1 | 620.7309094 | 0.133674161 | 0.055436851 |
| Prickle1 | 831.5948449 | -0.490698086 | 0.107441891 |
| Adamts20 | 204.9752347 | -0.94778024 | 0.202264739 |
| Pus7l | 33.70876001 | 0.210384491 | 0.186429063 |
| Irak4 | 54.36204711 | 2.139585374 | 0.247857794 |
| Twf1 | 752.74737 | 0.656854891 | 0.082739302 |
| Tmem117 | 131.7499303 | 0.011560583 | 0.102809335 |
| Gm25546 | 0 | NA | NA |
| Gm23129 | 0 | NA | NA |
| 1700129L04Rik | 4.754384054 | 0.294915682 | 0.666700136 |
| Nell2 | 3266.316947 | -0.910491685 | 0.123327833 |
| Gm24668 | 0 | NA | NA |
| Dbx2 | 163.3891668 | 0.542202897 | 0.148676033 |
| Gm8849 | 1.166878852 | 1.903591212 | 1.154526321 |
| A130051J06Rik | 4.097955438 | 2.043221696 | 0.572592837 |
| Ano6 | 622.2228295 | 1.06640188 | 0.100648782 |
| Gm17546 | 0.627185525 | -0.054900198 | 1.128795888 |
| Gm25070 | 0.044544648 | 0.237309859 | 3.724945049 |
| D030018L15Rik | 0.718519848 | 1.586039629 | 1.219911507 |
| 2610037D02Rik | 115.5236633 | -0.571610499 | 0.225661817 |
| 4833422M21Rik | 10.72830272 | 0.290984106 | 0.345656137 |
| Arid2 | 997.3079313 | -0.243627121 | 0.069505599 |
| Gm25397 | 0 | NA | NA |
| Scaf11 | 1520.248971 | 0.309821981 | 0.067434615 |
| Slc38a1 | 3775.966176 | -0.336524597 | 0.048020259 |
| Gm22045 | 0 | NA | NA |
| Slc38a2 | 2617.581037 | 0.117334459 | 0.0766155 |
| Gm38144 | 0.240186774 | 0.741017661 | 2.110992042 |
| Gm8888 | 0 | NA | NA |
| Slc38a4 | 9.499651179 | -0.923843743 | 0.347029625 |
| Amigo2 | 212.2260951 | 0.34371403 | 0.169189202 |
| Pced1b | 100.5813245 | 0.441503247 | 0.134501431 |
| Rpap3 | 291.5755603 | 0.06675272 | 0.078001693 |
| Endou | 46.06882177 | 0.789428438 | 0.210353384 |
| Rapgef3 | 390.02648 | -0.316386986 | 0.113816421 |
| Rapgef3os1 | 0.072025462 | 0.237312084 | 3.724945049 |
| Rapgef3os2 | 3.594151187 | -0.804197934 | 0.605343884 |
| Slc48a1 | 1148.386551 | -0.014496557 | 0.077045175 |
| Hdac7 | 379.135267 | 0.297348574 | 0.164664491 |
| Vdr | 23.71495927 | 2.388696037 | 0.327557843 |

| | | | |
|---|---|---|---|
| Gm22202 | 0 | NA | NA |
| Tmem106c | 169.9404391 | -0.106096717 | 0.142667947 |
| Col2a1 | 22.89251041 | -0.145328519 | 0.284757797 |
| Senp1 | 454.6842185 | 0.566376878 | 0.078853142 |
| Gm25668 | 0.044544648 | 0.237309859 | 3.724945049 |
| Pfkm | 3358.190513 | -0.247842717 | 0.064869836 |
| Gm26513 | 7.485808625 | -0.265907725 | 0.402530108 |
| Asb8 | 417.0210594 | 0.095275667 | 0.095273006 |
| Ccdc184 | 256.2935464 | -0.426983246 | 0.1245943 |
| Olfr288 | 1.13985991 | -0.669101039 | 1.046102799 |
| Olfr287 | 28.52921329 | -0.990770574 | 0.232928892 |
| Olfr286 | 0.16123844 | -1.214248636 | 2.795194053 |
| H1fnt | 0 | NA | NA |
| Zfp641 | 111.9084074 | -0.545280102 | 0.124828879 |
| Olfr285 | 0 | NA | NA |
| Olfr284 | 0 | NA | NA |
| Olfr283 | 0.082913145 | 0.23732098 | 3.724945049 |
| 1700031M16Rik | 0.045882455 | 0.237312084 | 3.724945049 |
| Olfr282 | 0 | NA | NA |
| Olfr281 | 0.061920613 | 0.718207094 | 3.724945049 |
| Lalba | 0.845223607 | 2.381255333 | 1.120790637 |
| Olfr279 | 0.133280846 | -0.243579231 | 3.654655979 |
| Kansl2 | 403.2058431 | 0.236320365 | 0.074376132 |
| Snora34 | 1.248930875 | 1.097228305 | 1.013542961 |
| mmu-mir-1291 | 1.156326783 | -1.004028389 | 0.97642645 |
| Snora2b | 4.632530903 | -1.057741938 | 0.4810425 |
| Ccnt1 | 886.6569998 | -0.02393589 | 0.04568285 |
| 9330020H09Rik | 18.2002576 | -0.331531681 | 0.289072184 |
| 4930415O20Rik | 0.192434788 | 0.772657829 | 2.651844621 |
| Gm29331 | 0.089062654 | 0.237318756 | 3.724945049 |
| Adcy6 | 518.3618143 | -0.309730835 | 0.096864679 |
| Cacnb3 | 1158.604023 | -1.013711397 | 0.097334812 |
| Ddx23 | 691.4598963 | 0.133016484 | 0.093119355 |
| Rnd1 | 252.5003933 | 0.571666579 | 0.218512842 |
| Ccdc65 | 49.36712147 | -0.347003407 | 0.152638467 |
| Fkbp11 | 8.263769739 | 1.358553665 | 0.428185435 |
| Arf3 | 8081.452134 | -0.35585456 | 0.068160776 |
| Wnt10b | 8.994095922 | 0.576046577 | 0.527576184 |
| Wnt1 | 0.424046041 | 0.568140355 | 1.562040175 |
| Ddn | 6153.970663 | -1.50070569 | 0.183543811 |
| B130046B21Rik | 36.69080057 | -0.834100732 | 0.246990685 |
| Prkag1 | 322.4183075 | 0.0968705 | 0.083707459 |
| Kmt2d | 5538.759192 | -0.147559527 | 0.082188092 |
| Rhebl1 | 33.80626636 | -0.377587793 | 0.182932776 |
| Dhh | 5.749732932 | 2.817709766 | 0.561025073 |
| Lmbr1l | 116.8431874 | 0.188113304 | 0.155286846 |
| Tuba1b | 2389.31285 | 0.187950634 | 0.133527011 |

| | | | |
|---|---|---|---|
| Tuba1a | 4753.134624 | -0.067235943 | 0.115932607 |
| 4930578M01Rik | 10.95583967 | -0.969389355 | 0.373800022 |
| Gm8973 | 0.060174537 | 0.237309859 | 3.724945049 |
| Tuba1c | 280.0639814 | 1.340290518 | 0.189250261 |
| Prph | 45.07156546 | 0.624509428 | 0.329903114 |
| Troap | 4.659496152 | 2.183515868 | 0.707605477 |
| Mir6960 | 0 | NA | NA |
| C1ql4 | 6.463082896 | -0.021797316 | 0.396328734 |
| Dnajc22 | 1.159556269 | -1.540353328 | 1.098973881 |
| Spats2 | 344.7749557 | -0.104165856 | 0.063462088 |
| Gm25183 | 0 | NA | NA |
| Gm25057 | 0 | NA | NA |
| Kcnh3 | 619.2177828 | -1.872511117 | 0.166574829 |
| Mcrs1 | 447.4221575 | 0.002802413 | 0.083391275 |
| 1700120C14Rik | 12.4281098 | -0.050894572 | 0.320143988 |
| Gm25999 | 0 | NA | NA |
| Fam186b | 17.25303227 | -0.21806308 | 0.289129681 |
| Prpf40b | 472.5579373 | -0.503260307 | 0.131895696 |
| Fmnl3 | 135.4055067 | 0.802437776 | 0.148544441 |
| Gm22951 | 0.744072464 | 0.600984168 | 1.377544833 |
| Tmbim6 | 4439.906879 | 1.046745923 | 0.122815865 |
| Nckap5l | 263.9144233 | -0.451963862 | 0.10724404 |
| Nckap5los | 0.092140247 | -0.724455696 | 3.724945049 |
| Bcdin3d | 22.52486403 | 0.476681837 | 0.216165787 |
| Faim2 | 4926.385511 | -0.498053238 | 0.06841486 |
| Gm5808 | 0.066050875 | 0.71820487 | 3.724945049 |
| Aqp2 | 0.235388486 | 0.712571808 | 2.118161634 |
| Aqp5 | 2.102526643 | 0.552042013 | 0.767472947 |
| Aqp6 | 109.2123799 | -0.026623967 | 0.224905831 |
| Racgap1 | 122.2464084 | -0.075208386 | 0.124445778 |
| Asic1 | 756.9424049 | -0.358184898 | 0.070659853 |
| Gm17349 | 9.498973506 | -0.357215182 | 0.324832859 |
| Smarcd1 | 1391.542852 | -0.472050864 | 0.081552225 |
| Gm16537 | 2.123900911 | -1.215641414 | 0.72457932 |
| Gpd1 | 1414.234084 | 0.436356385 | 0.103785645 |
| Cox14 | 343.5443005 | -0.051322465 | 0.091992939 |
| Cers5 | 400.0721546 | -0.087989713 | 0.07221912 |
| Gm17058 | 0.41865154 | 1.172989262 | 1.549933302 |
| Lima1 | 217.8937591 | 0.987779013 | 0.111762894 |
| Gm25897 | 0.60093359 | 0.260314803 | 1.332213503 |
| Gm4468 | 0.964311207 | 0.514966473 | 1.087878986 |
| Gm17057 | 0.099106094 | 1.209894437 | 3.72418822 |
| Gm9763 | 0.448689577 | 0.526789956 | 1.406275951 |
| Fam186a | 4.175781228 | -1.030644233 | 0.542407867 |
| Gm21917 | 0.34674037 | -0.727331566 | 1.577287399 |
| Gm23133 | 0 | NA | NA |
| Larp4 | 1360.21083 | 0.396065006 | 0.084961966 |

| | | | |
|---|---|---|---|
| 2310068J16Rik | 5.082496618 | -0.578748651 | 0.50370275 |
| Dip2b | 2171.940491 | -0.295194159 | 0.045002219 |
| Atf1 | 263.8766334 | 0.634309014 | 0.126940134 |
| Tmprss12 | 0.281502737 | -0.210509574 | 2.053121891 |
| Mettl7a1 | 199.3407439 | -0.299894906 | 0.216085767 |
| Mettl7a3 | 1.559777635 | -0.749703496 | 0.962762944 |
| Methig1 | 0 | NA | NA |
| Mettl7a2 | 0.046345709 | 0.71820487 | 3.724945049 |
| Higd1c | 0 | NA | NA |
| Slc11a2 | 350.6928657 | 0.273399588 | 0.096611759 |
| Gm5475 | 0.572323712 | 0.221559756 | 1.448565689 |
| n-R5s43 | 0 | NA | NA |
| 5330439K02Rik | 13.54861615 | -0.380360085 | 0.267481675 |
| Letmd1 | 357.2468629 | 0.125710905 | 0.080628911 |
| Csrnp2 | 483.3671844 | -0.077208022 | 0.129003527 |
| Tfcp2 | 253.237872 | -0.192434924 | 0.109008332 |
| Pou6f1 | 1003.020766 | -0.316741217 | 0.081198804 |
| Gm20680 | 0.523790777 | -0.377293035 | 1.314813861 |
| Gm27209 | 0.085433889 | -0.72445792 | 3.724945049 |
| C330013E15Rik | 16.45784469 | -0.201197523 | 0.27402327 |
| Dazap2 | 1246.964278 | 0.524745249 | 0.107685636 |
| Smagp | 10.67246666 | 1.820259896 | 0.351800844 |
| Bin2 | 77.96195219 | 2.275085542 | 0.222244263 |
| Mir6961 | 0 | NA | NA |
| Cela1 | 27.94372578 | 2.606421569 | 0.282745597 |
| Galnt6 | 328.7283006 | 1.303151112 | 0.189456623 |
| Galnt6os | 0.119234754 | -0.243576254 | 3.724945049 |
| Slc4a8 | 2144.399781 | -0.495242107 | 0.082792074 |
| Scn8a | 4962.265033 | -1.029598897 | 0.113571064 |
| Fignl2 | 63.27470895 | 0.615927896 | 0.166725832 |
| Ankrd33 | 0.864742418 | -0.748650163 | 1.034276234 |
| Acvrl1 | 107.1769413 | 2.277774588 | 0.15364523 |
| Gm16031 | 0.097139547 | 0.718207094 | 3.724945049 |
| Acvr1b | 1299.302991 | -0.436406692 | 0.065926187 |
| mmu-mir-6962 | 0.93249843 | 0.264050515 | 1.016728804 |
| A330009N23Rik | 60.10913543 | -0.192593387 | 0.176951214 |
| Grasp | 286.1441832 | 0.08447341 | 0.216032234 |
| Nr4a1 | 763.9124175 | -0.764079885 | 0.422502516 |
| Atg101 | 182.2922892 | 0.113287431 | 0.105954451 |
| 6030408B16Rik | 2.433334847 | 0.279141608 | 0.6583659 |
| Krt80 | 28.28862397 | 1.21686908 | 0.223242709 |
| Mir1941 | 0.066050875 | 0.71820487 | 3.724945049 |
| Krt7 | 2.609690864 | 0.816879872 | 0.640034692 |
| Krt83 | 0.375360954 | -0.243564871 | 1.616070176 |
| 1700011A15Rik | 0.04393175 | -0.243571806 | 3.724945049 |
| Krt81 | 0 | NA | NA |
| Krt86 | 0.614518174 | 2.10048864 | 1.412262891 |

| | | | |
|---|---|---|---|
| 5430421N21Rik | 0.095897396 | -0.724453471 | 3.724945049 |
| Krt84 | 0 | NA | NA |
| Krt82 | 0 | NA | NA |
| 4732456N10Rik | 0.738905749 | -0.537492104 | 1.446849524 |
| AC103674.1 | 0 | NA | NA |
| Krt75 | 1.991379766 | 4.731811092 | 1.021276952 |
| AC103674.2 | 0 | NA | NA |
| Gm5414 | 0 | NA | NA |
| Gm5478 | 0 | NA | NA |
| AC120787.3 | 0 | NA | NA |
| Krt6b | 0.519670895 | 2.019605008 | 1.729759025 |
| AC120787.2 | 0 | NA | NA |
| Krt6a | 4.257533425 | 5.756230116 | 1.066490888 |
| AC120787.4 | 0 | NA | NA |
| Krt5 | 1.350984256 | 1.969024334 | 1.180771159 |
| AC120787.1 | 0 | NA | NA |
| Krt71 | 0.51940401 | -0.714495451 | 1.4213453 |
| Krt74 | 0 | NA | NA |
| Krt72 | 0.233626327 | 0.718209579 | 3.116229621 |
| Krt73 | 9.672351053 | 0.569810679 | 0.358550319 |
| Krt2 | 16.20338829 | 0.091719612 | 0.87806171 |
| Krt1 | 24.4159562 | -0.303372844 | 0.310429223 |
| Krt77 | 35.13814104 | -1.234563785 | 0.298657138 |
| Krt76 | 0.099106094 | 1.209894437 | 3.72418822 |
| AC139844.1 | 0 | NA | NA |
| Krt4 | 0.254192696 | -0.243571929 | 2.492352056 |
| Krt79 | 0.300336086 | 1.289969094 | 3.719548429 |
| Krt78 | 0.207810967 | -0.723749303 | 2.59344861 |
| Krt8 | 17.9984126 | 2.365413566 | 0.387356053 |
| Krt18 | 21.72902479 | 1.123692189 | 0.345164638 |
| Eif4b | 2876.650182 | 0.056684484 | 0.058466328 |
| Tns2 | 268.4246283 | 0.288163883 | 0.1231947 |
| Spryd3 | 1083.158277 | -0.262698678 | 0.068376416 |
| Igfbp6 | 171.4122214 | 0.14914335 | 0.113601889 |
| Soat2 | 21.31164465 | 5.257706468 | 0.573258312 |
| Csad | 172.5508944 | -0.513209069 | 0.106203906 |
| Zfp740 | 877.8255617 | 0.029365553 | 0.066961888 |
| Itgb7 | 116.7476434 | 5.094509746 | 0.417229535 |
| Rarg | 131.1736488 | 0.17215752 | 0.113153514 |
| Gm9918 | 0.827590884 | -1.113623278 | 1.170416465 |
| Mfsd5 | 351.832301 | 0.556735468 | 0.102134511 |
| Espl1 | 27.92326781 | 1.327926766 | 0.307232873 |
| Pfdn5 | 660.9445207 | 0.136282995 | 0.094306996 |
| Myg1 | 214.9838279 | 0.106111702 | 0.08839825 |
| Aaas | 128.6515988 | 0.104545221 | 0.168925546 |
| Sp7 | 27.79878053 | -0.370107737 | 0.332119203 |
| Gm23590 | 0 | NA | NA |

| | | | |
|---|---|---|---|
| Sp1 | 861.7402076 | 0.551210526 | 0.097836075 |
| Amhr2 | 7.396618114 | 0.381121882 | 0.376453807 |
| Gm26518 | 0.703641397 | -0.814916757 | 1.105157865 |
| Gm24128 | 0.061920613 | 0.718207094 | 3.724945049 |
| Prr13 | 456.4408238 | 0.765283666 | 0.113907046 |
| Pcbp2 | 2462.12792 | 0.3454258 | 0.071018249 |
| Gm27406 | 2.84356551 | 0.696258274 | 0.630238166 |
| Map3k12 | 1206.534751 | -0.672106144 | 0.067458826 |
| Gm10337 | 2.690945547 | 0.320143446 | 0.776805299 |
| Tarbp2 | 121.0001225 | -0.148364732 | 0.115675507 |
| Npff | 8.501785748 | -0.501789594 | 0.576200046 |
| Gm28047 | 0 | NA | NA |
| Atf7 | 596.7832338 | 0.08817741 | 0.068879006 |
| Gm25141 | 0 | NA | NA |
| Rpl39-ps | 40.47241404 | 0.829087329 | 0.240852312 |
| Atp5g2 | 389.3075053 | 0.157855608 | 0.113142427 |
| Mir688 | 0 | NA | NA |
| Calcoco1 | 660.9432622 | -0.266674086 | 0.096392147 |
| Hoxc13 | 0.157352566 | 1.165775181 | 2.808585416 |
| Hoxc12 | 0 | NA | NA |
| Hotair | 0 | NA | NA |
| Gm27337 | 0 | NA | NA |
| Gm27443 | 0 | NA | NA |
| Gm28011 | 0 | NA | NA |
| Gm27535 | 0 | NA | NA |
| Hoxc11 | 0 | NA | NA |
| Hoxc10 | 0 | NA | NA |
| Hoxc5 | 4.717838401 | 0.122259949 | 1.2111556 |
| Mir196a-2 | 0 | NA | NA |
| Hoxc9 | 0.095435502 | 0.718202646 | 3.724945049 |
| Hoxc8 | 0.100902008 | 1.15059983 | 3.720931964 |
| Hoxc6 | 1.521766428 | 0.823204665 | 1.983891129 |
| Mir615 | 0 | NA | NA |
| Hoxc4 | 16.48418288 | 0.218998961 | 1.02276376 |
| 5730585A16Rik | 0.314534502 | -1.345660215 | 3.692614927 |
| Gm28876 | 4.308624721 | 0.041035029 | 0.497310142 |
| Gm10830 | 0.40206694 | 0.167208448 | 1.53083217 |
| Gm28265 | 0.496950623 | -0.680483969 | 1.76451045 |
| D930007P13Rik | 1.440959972 | -1.748203744 | 1.08009382 |
| Smug1 | 140.5464891 | 0.057526667 | 0.117180346 |
| Cbx5 | 3979.323211 | -0.256499031 | 0.065242426 |
| Hnrnpa1 | 1468.269564 | 0.042822102 | 0.093140583 |
| Nfe2 | 3.16484666 | 4.637139915 | 0.953939636 |
| Copz1 | 570.9806916 | 0.567139336 | 0.092429722 |
| Mir148b | 1.892187285 | -0.902422021 | 0.809100172 |
| Gpr84 | 34.27634392 | 3.476758744 | 0.452634973 |
| Zfp385a | 1024.49315 | 0.028022009 | 0.08206242 |

| | | | |
|---|---|---|---|
| Itga5 | 497.2109644 | 4.378531131 | 0.343028905 |
| Mir6963 | 0.095435502 | 0.718202646 | 3.724945049 |
| Gm37322 | 0.248806291 | -0.22996297 | 2.130252479 |
| Gtsf1 | 0.666252615 | 0.291192372 | 1.15336756 |
| BC048502 | 0.169045294 | -0.243573848 | 3.402281771 |
| Nckap1l | 484.3215996 | 2.814328423 | 0.147876161 |
| Pde1b | 1849.40123 | -1.004923741 | 0.124598798 |
| Ppp1r1a | 376.4709638 | -0.691875832 | 0.135930041 |
| Glycam1 | 0.54398204 | -1.417355171 | 1.310943367 |
| Mucl1 | 0.04393175 | -0.243571806 | 3.724945049 |
| Gm22852 | 0 | NA | NA |
| Spt1 | 0.136548098 | -0.724458525 | 2.945560552 |
| Gm23215 | 0 | NA | NA |
| Gm22862 | 0 | NA | NA |
| Gm22598 | 0 | NA | NA |
| Olfr161 | 0 | NA | NA |
| Mefv | 48.54285063 | 9.403564113 | 0.976443716 |
| Tdgf1-ps1 | 0.267098211 | 1.629643063 | 2.459808811 |
| Zfp263 | 304.9461567 | 0.176040073 | 0.104965555 |
| Olfr15 | 0.061920613 | 0.718207094 | 3.724945049 |
| Zfp174 | 93.47116396 | -0.405357862 | 0.173469977 |
| Zfp597 | 424.0273003 | 0.032097741 | 0.105541802 |
| Gm26675 | 8.550105792 | -0.356020928 | 0.409607377 |
| Naa60 | 734.2670764 | -0.274210049 | 0.064488787 |
| Gm20695 | 1.863767 | -1.317912852 | 0.902398904 |
| 1700037C18Rik | 31.49051414 | 0.295327168 | 0.21894047 |
| Cluap1 | 161.5850996 | -0.199069268 | 0.101986557 |
| 1700016D08Rik | 0 | NA | NA |
| Gm15538 | 10.93859038 | -0.817382707 | 0.437405153 |
| Gm23493 | 2.266218546 | -0.027386009 | 0.667961762 |
| Gm15539 | 10.65583297 | -0.671976104 | 0.414853958 |
| Gm15537 | 0.604736192 | 0.712103006 | 1.162664343 |
| Nlrc3 | 40.04733098 | 0.978042045 | 0.207184679 |
| Slx4 | 453.7919637 | -0.124326462 | 0.09177419 |
| Gm15879 | 36.75047168 | -0.42167336 | 0.197646479 |
| Dnase1 | 0.317440118 | -1.165449703 | 1.562758706 |
| Trap1 | 710.530572 | 0.115725824 | 0.054978819 |
| Gm23413 | 3.363014587 | -1.134813334 | 0.571141061 |
| Crebbp | 3534.609602 | -0.046770798 | 0.053027502 |
| Gm37186 | 92.75499785 | -0.77440864 | 0.314266898 |
| Gm5766 | 0.297064053 | 0.718784707 | 1.770719177 |
| Adcy9 | 953.9955112 | -0.845531998 | 0.129605232 |
| Gm6142 | 13.94176219 | -1.223559066 | 0.330358492 |
| Srl | 63.68776743 | -0.679849165 | 0.154876693 |
| Gm15885 | 0 | NA | NA |
| Tfap4 | 61.90644834 | 0.32657992 | 0.176475735 |
| Glis2 | 152.6195355 | 0.24496407 | 0.102228256 |

| | | | |
|---|---|---|---|
| Pam16 | 156.721799 | 0.172742855 | 0.150038809 |
| Coro7 | 608.4531973 | -0.554446816 | 0.10840054 |
| Vasn | 199.7417839 | 1.636930537 | 0.21499972 |
| Dnaja3 | 623.6357018 | 0.184247786 | 0.085269547 |
| Nmral1 | 147.3588111 | 0.229470954 | 0.136805204 |
| Hmox2 | 693.4042527 | 0.477977996 | 0.081443426 |
| Gm15859 | 12.58934561 | -0.132106413 | 0.378052499 |
| Cdip1 | 1931.022938 | -0.314854389 | 0.054151827 |
| Gm15835 | 24.15715004 | -0.878227368 | 0.31370378 |
| 4930562C15Rik | 0.450624642 | 1.995754701 | 1.519778348 |
| Ubald1 | 776.8048839 | -0.257132622 | 0.086729879 |
| Mgrn1 | 1884.371733 | -0.115263658 | 0.050983734 |
| Gm16861 | 17.52762606 | 0.447726053 | 0.272784986 |
| Nudt16l1 | 313.4024066 | 0.062097921 | 0.086114177 |
| Anks3 | 314.2732224 | -0.22264508 | 0.097905962 |
| 4930451G09Rik | 27.70571581 | -0.616870252 | 0.20154757 |
| 12-Sep | 0.042716945 | -0.243571806 | 3.724945049 |
| Smim22 | 0.651416456 | 0.625428878 | 1.309061858 |
| RP23-450O11.3 | 0.373087082 | 1.43516427 | 1.685280486 |
| Rogdi | 663.6085146 | -0.665680656 | 0.078793537 |
| Glyr1 | 1981.897547 | 0.005796382 | 0.054864292 |
| Ubn1 | 976.2780589 | 0.116788362 | 0.080175967 |
| Gm23483 | 0 | NA | NA |
| Ppl | 31.86021987 | 2.090529685 | 0.271876614 |
| Sec14l5 | 164.8073848 | 0.028206226 | 0.130898304 |
| Nagpa | 106.0130434 | 0.740487858 | 0.121096018 |
| AU021092 | 9.404176926 | 1.343882685 | 0.408955282 |
| Mir6364 | 0.049553047 | 0.71820487 | 3.724945049 |
| Gm23689 | 0.15591167 | 0.237325282 | 2.826544723 |
| Gm26095 | 0.09689823 | -0.724453471 | 3.724945049 |
| Alg1 | 199.3302287 | 0.288767421 | 0.108050145 |
| Eef2kmt | 50.93323876 | 0.892960967 | 0.211284598 |
| Gm15983 | 5.210713431 | 0.540064896 | 0.475267608 |
| n-R5s30 | 0 | NA | NA |
| 1700123O21Rik | 37.36951086 | -1.129457627 | 0.386107495 |
| Rbfox1 | 3132.04572 | -0.854790631 | 0.08090708 |
| Gm23476 | 0 | NA | NA |
| Gm26159 | 0 | NA | NA |
| Gm24579 | 0 | NA | NA |
| Tmem114 | 8.994908763 | -1.147658129 | 0.399430279 |
| Gm25782 | 0 | NA | NA |
| Mettl22 | 149.3545907 | 0.036915193 | 0.090606072 |
| Gm23706 | 0 | NA | NA |
| Abat | 3761.014197 | -0.606638484 | 0.085491132 |
| Gm22137 | 0 | NA | NA |
| Gm7541 | 1.437230806 | -0.022108929 | 0.919594993 |
| Tmem186 | 191.5203021 | -0.123645654 | 0.098464442 |

| | | | |
|---|---|---|---|
| Pmm2 | 257.7422973 | 0.287476166 | 0.100593135 |
| Carhsp1 | 736.470531 | 1.239801368 | 0.155507464 |
| RP24-183G19.6 | 0.790581521 | 1.205503888 | 1.565553855 |
| Usp7 | 2331.28947 | -0.03461419 | 0.068645996 |
| Gm25805 | 0 | NA | NA |
| 1810013L24Rik | 1107.486804 | 0.227432571 | 0.066629568 |
| Gm22224 | 0 | NA | NA |
| Gm10247 | 1.54124956 | 1.024867123 | 0.916554367 |
| Grin2a | 4615.108378 | -1.076178318 | 0.161652939 |
| Gm7572 | 0 | NA | NA |
| Gm3890 | 0 | NA | NA |
| Rpl39l | 0.823456907 | 2.120323316 | 1.193486822 |
| Atf7ip2 | 0.994634002 | -0.08697655 | 1.124468238 |
| Gm24667 | 0 | NA | NA |
| Gm22932 | 0 | NA | NA |
| Emp2 | 291.2677059 | 0.961661361 | 0.13291699 |
| Tekt5 | 64.82616325 | -0.708641592 | 0.179031232 |
| Nubp1 | 155.2730509 | 0.506723067 | 0.134704329 |
| Tvp23a | 456.5008628 | 0.851962718 | 0.1072352 |
| Ciita | 53.45720072 | 5.948217071 | 0.518069158 |
| Dexi | 379.7312674 | 0.056619388 | 0.07980978 |
| Clec16a | 1806.749408 | -0.204975359 | 0.067520848 |
| Gm15558 | 0.339464816 | 0.722693558 | 1.763903096 |
| Socs1 | 257.5468929 | 5.304977439 | 0.38176896 |
| Tnp2 | 0.356459053 | 2.089113474 | 1.607544349 |
| Prm3 | 0 | NA | NA |
| Prm2 | 0.134416897 | 1.165350059 | 2.95041533 |
| Prm1 | 0.978261292 | 4.189466907 | 1.163729284 |
| Gm10343 | 5.069691734 | 0.997320736 | 0.531810295 |
| Gm11172 | 0.044544648 | 0.237309859 | 3.724945049 |
| Gm26822 | 0.840740177 | -0.535354444 | 1.331139113 |
| Rmi2 | 18.46361415 | 1.627362886 | 0.285212249 |
| Gm21859 | 0.468571393 | 1.793031708 | 1.400571165 |
| Litaf | 694.90215 | 2.634851358 | 0.257180357 |
| Gm26268 | 0 | NA | NA |
| Gm24961 | 0 | NA | NA |
| Gm4262 | 0.72304365 | 3.0080662 | 1.177187961 |
| Snn | 1735.197761 | -0.465547567 | 0.080181861 |
| Txndc11 | 466.704329 | -0.323029192 | 0.092133266 |
| Zc3h7a | 853.6995849 | 0.621500744 | 0.126709566 |
| Gm23935 | 15251.03865 | 0.5608671 | 0.922177726 |
| Gm24145 | 5.507590669 | 0.048031571 | 0.548496628 |
| 4930509G22Rik | 5.036420442 | -0.149888127 | 0.439982692 |
| Gm7638 | 2.798124148 | -0.423583553 | 0.64847404 |
| Gm6305 | 2.348678771 | 0.442854694 | 0.637647417 |
| Rsl1d1 | 606.0058677 | 0.709453105 | 0.101374702 |
| 2610020C07Rik | 17.10628321 | -0.845775473 | 0.276711961 |

| | | | |
|---|---|---|---|
| Gspt1 | 1482.382663 | 0.176728335 | 0.052142452 |
| Mir1945 | 0.407802109 | 1.359017601 | 1.379609872 |
| Tnfrsf17 | 1.11445967 | 1.340597874 | 1.093348281 |
| Snx29 | 234.388902 | -0.641632124 | 0.120845866 |
| Gm15897 | 0.089089296 | 0.237305797 | 3.724945049 |
| Gm4279 | 0.739397817 | 0.481410495 | 1.148298419 |
| Cpped1 | 428.5321132 | 0.227731697 | 0.081438323 |
| Gm9961 | 0.136963818 | 0.718204821 | 3.59954433 |
| Shisa9 | 635.7404322 | -0.849831399 | 0.101000576 |
| Gm6327 | 0.086930093 | 0.236669296 | 3.724898107 |
| 1700003L19Rik | 0.364102347 | -1.741832977 | 1.637457783 |
| Ercc4 | 327.5978949 | -0.080426734 | 0.116368825 |
| Mkl2 | 3224.420774 | -0.830459076 | 0.122470968 |
| Gm15738 | 0.13861434 | -1.149001357 | 3.614496194 |
| Mir193b | 0.600656357 | -0.716427995 | 1.414405217 |
| Mir365-1 | 1.735414367 | -2.50164886 | 0.953308026 |
| Mir6365 | 0 | NA | NA |
| 2310015D24Rik | 1.269356065 | -0.856215171 | 0.942259092 |
| Gm25276 | 0.11037557 | -0.243571806 | 3.724945049 |
| Parn | 231.9657214 | -0.107649206 | 0.086733098 |
| Bfar | 419.0822541 | 0.373676469 | 0.0964748 |
| 3110001I22Rik | 41.39457374 | 0.12981173 | 0.176381217 |
| Pla2g10 | 0 | NA | NA |
| Pla2g10os | 0.090890357 | 0.718200421 | 3.724945049 |
| Rrn3 | 908.1970389 | 0.255685764 | 0.064604098 |
| Gm27734 | 0.941805692 | -1.809028183 | 1.087959613 |
| Ntan1 | 347.9210283 | -0.037327081 | 0.089097054 |
| Pdxdc1 | 751.5294376 | 0.331948517 | 0.06062957 |
| Mpv17l | 545.4884099 | -0.623925036 | 0.082216026 |
| Gm15950 | 0.843513109 | 0.430385667 | 1.257419695 |
| Gm26213 | 0.252317246 | 1.281389321 | 2.500491872 |
| Ifitm7 | 16.50735806 | 0.854823286 | 0.305522483 |
| 2900011O08Rik | 1845.656536 | -0.401047445 | 0.061136338 |
| Gm15806 | 0.061920613 | 0.718207094 | 3.724945049 |
| Gm15807 | 0 | NA | NA |
| Gm15808 | 7.400018289 | -1.312805051 | 0.601427791 |
| Marf1 | 2957.81909 | -0.459576161 | 0.075159724 |
| Gm27671 | 0 | NA | NA |
| Mir484 | 0.262798536 | 0.237330456 | 1.836165506 |
| Nde1 | 330.4944427 | 1.22393749 | 0.146226887 |
| Myh11 | 25.31456723 | 0.052424248 | 0.293448629 |
| Fopnl | 265.3373334 | -0.007091531 | 0.092512637 |
| Gm15868 | 0 | NA | NA |
| Gm15869 | 19.70131526 | -0.7486292 | 0.277646419 |
| Abcc1 | 333.839765 | -0.186798169 | 0.112382592 |
| Gm23116 | 0.060174537 | 0.237309859 | 3.724945049 |
| Gm1758 | 0.227501076 | 0.712530631 | 2.139256712 |

| | | | |
|---|---|---|---|
| A630010A05Rik | 0 | NA | NA |
| Snai2 | 6.714012416 | 0.098877346 | 0.381963301 |
| Gm25547 | 1.090696383 | 0.890308823 | 0.88940325 |
| Efcab1 | 65.3173956 | -0.645891019 | 0.160305978 |
| Gm7731 | 0 | NA | NA |
| Gm21897 | 0.148910734 | -0.243572529 | 2.86160923 |
| Gm6031 | 0 | NA | NA |
| Ube2v2 | 393.0830429 | -0.382050661 | 0.09802219 |
| Mcm4 | 207.5248428 | 0.839352911 | 0.162188956 |
| Prkdc | 719.6145477 | -0.728478718 | 0.12329942 |
| Mzt2 | 126.2984278 | 0.051031954 | 0.132453463 |
| Cebpd | 477.928814 | 4.410320961 | 0.347619779 |
| Spidr | 147.239883 | 2.948776139 | 0.265752457 |
| n-R5s31 | 0.50758071 | 1.593140026 | 1.270722623 |
| Pkp2 | 124.1564146 | -0.264494352 | 0.13849871 |
| Yars2 | 199.1181058 | 0.273205928 | 0.108175642 |
| Dnm1l | 3082.769836 | -0.12989415 | 0.043832537 |
| Gm7765 | 0 | NA | NA |
| Fgd4 | 1158.129079 | -0.120645194 | 0.084867224 |
| Gm16139 | 0 | NA | NA |
| Olfr19 | 0.447268164 | 0.664944758 | 1.389330541 |
| Spag6 | 66.08212979 | -0.313744912 | 0.159729562 |
| Rpl31-ps12 | 2.913194718 | 0.074386531 | 0.658245283 |
| Gm15714 | 0.095898756 | 1.192083721 | 3.724434897 |
| Igll1 | 0.075043205 | 0.237309859 | 3.724945049 |
| Vpreb1 | 0.913331526 | -0.832385479 | 1.112598545 |
| Top3b | 435.3366088 | 0.006149475 | 0.119880702 |
| Ppm1f | 789.0102064 | -0.232194224 | 0.089515593 |
| Mapk1 | 4968.897064 | -0.495709487 | 0.070311454 |
| Gm22558 | 0 | NA | NA |
| 1700056N10Rik | 40.39149775 | -0.721176646 | 0.319138814 |
| Ypel1 | 118.1725456 | -1.428156974 | 0.211621746 |
| Gm9974 | 6.459321561 | -0.711320583 | 0.432091304 |
| Ppil2 | 680.5363805 | 0.18577359 | 0.053821063 |
| Gm15585 | 2.078701808 | -1.367217165 | 0.814945605 |
| 2610318N02Rik | 4.21806971 | -0.526824044 | 0.485781033 |
| Mir130b | 0.075043205 | 0.237309859 | 3.724945049 |
| Mir301b | 0.118974955 | -0.243576254 | 3.724945049 |
| Sdf2l1 | 144.324589 | 1.120528851 | 0.156766359 |
| Ccdc116 | 27.44048453 | -0.621287966 | 0.239028067 |
| Gm15646 | 0 | NA | NA |
| Ydjc | 88.53278356 | -0.592495998 | 0.13136614 |
| Ube2l3 | 1433.809399 | 0.452711208 | 0.08118744 |
| Gm15648 | 2.296535557 | -0.544166342 | 0.668675878 |
| Gm22129 | 0.093744596 | 0.237318756 | 3.724945049 |
| Rimbp3 | 28.96339831 | -0.773848183 | 0.22739443 |
| Hic2 | 126.3818273 | -0.922314326 | 0.180209026 |

| | | | |
|---|---|---|---|
| Gm26635 | 0.934835317 | -0.235710185 | 1.145657719 |
| Tmem191c | 275.0346229 | -1.335615888 | 0.160179594 |
| Pi4ka | 4405.809757 | -0.13201543 | 0.080706294 |
| Serpind1 | 61.15174061 | 0.880350651 | 0.203204722 |
| Gm24927 | 3.890661782 | -0.010560228 | 0.537906595 |
| Snap29 | 384.8317826 | 0.793867239 | 0.144291638 |
| Crkl | 918.3467102 | 0.565222851 | 0.08284997 |
| Gm22770 | 0 | NA | NA |
| Aifm3 | 673.3284071 | -1.3985325 | 0.175244439 |
| Lztr1 | 1143.994835 | -0.229896571 | 0.052078691 |
| Thap7 | 186.8558046 | 0.647125726 | 0.104480141 |
| Lrrc74b | 11.80310187 | -0.194435454 | 0.29532952 |
| P2rx6 | 28.77111211 | 0.743274673 | 0.234578568 |
| Slc7a4 | 422.6595786 | -0.103732837 | 0.106709207 |
| Smpd4 | 474.152001 | -0.208998777 | 0.143012934 |
| Ccdc74a | 148.3892086 | -0.777390368 | 0.19463121 |
| Med15 | 779.2021573 | 0.481715669 | 0.06681237 |
| Rpl26-ps4 | 0 | NA | NA |
| Klhl22 | 742.0166947 | -0.220939171 | 0.046451344 |
| Scarf2 | 44.76671024 | -0.235785728 | 0.186028128 |
| Car15 | 16.28727946 | -0.184391541 | 0.308618776 |
| Gm20518 | 10.60745708 | -0.755468736 | 0.429336688 |
| Dgcr2 | 954.3027326 | -0.005098153 | 0.070864057 |
| Tssk1 | 0.674642588 | 0.49764467 | 1.139174768 |
| Tssk2 | 0.04393175 | -0.243571806 | 3.724945049 |
| Dgcr14 | 286.5719584 | 0.188566178 | 0.069574926 |
| Gsc2 | 0.267511063 | -1.746225864 | 2.479702835 |
| AA914427 | 7.57215109 | -0.687213248 | 0.371151841 |
| Slc25a1 | 386.8663658 | 0.467775189 | 0.116923936 |
| Vpreb2 | 0.042716945 | -0.243571806 | 3.724945049 |
| Dgcr6 | 244.4688473 | -0.121456651 | 0.108992548 |
| Prodh | 187.7866812 | -1.184308215 | 0.1660347 |
| Gm15954 | 5.737854463 | -1.493642095 | 0.470650271 |
| Rtn4r | 444.6635611 | -0.873789033 | 0.124518823 |
| Mir6366 | 0 | NA | NA |
| Gm7873 | 0.093141081 | -0.724455696 | 3.724945049 |
| Zdhhc8 | 854.0005545 | -0.699372084 | 0.08664945 |
| Ranbp1 | 621.7089953 | 0.89733489 | 0.149916683 |
| Gm25777 | 2.388976987 | -0.817337477 | 0.745992104 |
| Trmt2a | 346.4940552 | 0.381806045 | 0.075821087 |
| Dgcr8 | 527.8985729 | -0.295558681 | 0.105181153 |
| Mir1306 | 0 | NA | NA |
| Gm24572 | 0 | NA | NA |
| Tango2 | 629.321088 | -0.289687028 | 0.094775765 |
| Mir185 | 2.706681505 | -0.12827966 | 0.62401643 |
| Arvcf | 417.1584827 | -0.221757858 | 0.065914304 |
| Gm26170 | 1.640007924 | -0.604722669 | 0.743601042 |

| | | | |
|---|---|---|---|
| Gm26219 | 0 | NA | NA |
| Comt | 659.6866933 | 0.207121953 | 0.057628814 |
| Gm15764 | 1.82265688 | 1.423159348 | 0.732946116 |
| Txnrd2 | 113.8682495 | 0.040260568 | 0.135704199 |
| Gnb1l | 33.08347225 | -0.001057663 | 0.187719715 |
| Gm16314 | 20.96206684 | -0.372790997 | 0.230065768 |
| Tbx1 | 2.659687333 | 0.785695821 | 0.589419276 |
| 4930588K23Rik | 0.132314849 | -1.08153651 | 3.65395166 |
| Gp1bb | 61.84750879 | -0.412446766 | 0.190473904 |
| 5-Sep | 2743.444644 | -0.935409561 | 0.11062046 |
| Cldn5 | 120.886244 | 2.371825972 | 0.185477297 |
| Cdc45 | 22.31723845 | 0.804355591 | 0.276648712 |
| Ufd1l | 707.9940102 | 0.257459218 | 0.055459148 |
| 2510002D24Rik | 76.81866928 | -0.62759235 | 0.122920317 |
| Gm28539 | 0.048949532 | -0.243569582 | 3.724945049 |
| Mrpl40 | 175.5484898 | 0.315362629 | 0.102945495 |
| Hira | 202.5271098 | -0.135926503 | 0.12062058 |
| Gm15799 | 5.251177882 | -0.582647931 | 0.61531663 |
| Gm15798 | 0 | NA | NA |
| RP23-374P12.1 | 0 | NA | NA |
| Gm10088 | 0.415109666 | 0.1935522 | 1.515317547 |
| RP23-247I11.8 | 0.329535318 | 1.626264426 | 1.707163382 |
| Iglc1 | 0.667277359 | 2.790570808 | 2.450564251 |
| Iglj1 | 0 | NA | NA |
| Iglc3 | 1.77022427 | 4.519358488 | 0.999968685 |
| Iglj3p | 0 | NA | NA |
| Iglj3 | 0.072025462 | 0.237312084 | 3.724945049 |
| Iglv1 | 0.611248024 | 2.747443029 | 2.516349021 |
| RP23-247I11.7 | 0 | NA | NA |
| Iglc4 | 0 | NA | NA |
| Iglj4 | 0 | NA | NA |
| Iglc2 | 0.938589659 | 3.797805666 | 1.401800508 |
| Iglj2 | 0 | NA | NA |
| RP23-272P19.8 | 0.046345709 | 0.71820487 | 3.724945049 |
| RP23-272P19.7 | 0.343041433 | 1.656789088 | 1.73610926 |
| Iglv3 | 0.097139547 | 0.718207094 | 3.724945049 |
| RP23-272P19.10 | 1.305037701 | 3.637356867 | 1.295324988 |
| Iglv2 | 0.097139547 | 0.718207094 | 3.724945049 |
| Olfr164 | 0.207953851 | -0.243562013 | 2.579759988 |
| Gm22158 | 0.391389582 | 1.264886637 | 3.360424918 |
| Gm23612 | 0.049553047 | 0.71820487 | 3.724945049 |
| Olfr165 | 0 | NA | NA |
| Gm25762 | 0.144848288 | 0.719115904 | 2.892350495 |
| Olfr166 | 0.296045541 | 0.25558062 | 2.032068974 |
| Olfr167 | 0 | NA | NA |
| Olfr168 | 0 | NA | NA |
| Olfr169 | 0.0878635 | -0.72445792 | 3.724945049 |

| | | | |
|---|---|---|---|
| Olfr170 | 0 | NA | NA |
| Olfr171 | 0 | NA | NA |
| Lamp3 | 0.505840948 | 0.194560696 | 1.341493335 |
| Gm17799 | 0.062426873 | 0.237312084 | 3.724945049 |
| B3gnt5 | 145.4656175 | 1.545417098 | 0.151864146 |
| A930003A15Rik | 2.53052731 | 2.10206792 | 0.734355074 |
| Gm27883 | 0 | NA | NA |
| Klhl6 | 70.61673169 | 3.467333528 | 0.313500571 |
| Klhl24 | 1378.234747 | -0.343653036 | 0.079022175 |
| Yeats2 | 481.9769426 | -0.395623584 | 0.130404498 |
| Gm10241 | 24.31474395 | 0.290442212 | 0.268921089 |
| Map6d1 | 369.8382378 | -1.153201431 | 0.104481663 |
| Parl | 312.4995396 | 0.292912979 | 0.072044214 |
| Cyp2ab1 | 1.187519348 | 3.449795513 | 1.089620422 |
| Abcc5 | 1948.385061 | -0.526045189 | 0.13779432 |
| Gm20614 | 0.26923737 | -0.154540376 | 2.112226549 |
| Gm23332 | 0 | NA | NA |
| Gm20615 | 0.04393175 | -0.243571806 | 3.724945049 |
| Eif2b5 | 489.2421831 | 0.274019472 | 0.063143862 |
| Gm16618 | 0.359481004 | -0.553546107 | 1.759913965 |
| Dvl3 | 617.5796496 | -0.155441334 | 0.073133245 |
| Ap2m1 | 2464.108253 | 0.001403203 | 0.04077435 |
| Gm15760 | 0.195053328 | -0.244160407 | 2.683734912 |
| Abcf3 | 658.1946899 | 0.277398901 | 0.064892449 |
| Vwa5b2 | 365.3797502 | -1.189051964 | 0.203543693 |
| Mir1224 | 0 | NA | NA |
| Alg3 | 92.27766346 | 0.50791256 | 0.126008546 |
| Ece2 | 434.908798 | -0.224956654 | 0.100651175 |
| Camk2n2 | 831.1974061 | -0.373667401 | 0.098727929 |
| Psmd2 | 2330.871484 | 0.49914214 | 0.082136098 |
| Gm4462 | 1.62367863 | 0.030866633 | 0.869130288 |
| Eif4g1 | 4413.780657 | 0.324466299 | 0.045258496 |
| Snord66 | 1.086592418 | 1.689807922 | 1.176022929 |
| Fam131a | 1339.029144 | -1.216424314 | 0.112082474 |
| Clcn2 | 427.9328786 | -0.48061017 | 0.110512527 |
| Polr2h | 104.4427863 | -0.015635946 | 0.116181853 |
| Thpo | 27.14520041 | -1.043465845 | 0.395853606 |
| Chrd | 234.488815 | -0.688709184 | 0.210638736 |
| Gm22741 | 0.075043205 | 0.237309859 | 3.724945049 |
| Rps10-ps2 | 203.4377443 | 0.813684403 | 0.134597303 |
| Gm9697 | 0 | NA | NA |
| Teddm3 | 0.044544648 | 0.237309859 | 3.724945049 |
| AC168061.1 | 0 | NA | NA |
| Ephb3 | 170.6997336 | -0.759676275 | 0.131583616 |
| Gm16863 | 0.106358623 | -0.24357403 | 3.724945049 |
| Mir7680 | 0 | NA | NA |
| Vps8 | 810.4907143 | -0.121750553 | 0.057113592 |

| | | | |
|---|---|---|---|
| 2510009E07Rik | 1017.889882 | 0.092088509 | 0.088666609 |
| Ehhadh | 44.10308811 | 0.04845402 | 0.217527324 |
| Gm24898 | 0 | NA | NA |
| 1300002E11Rik | 152.2564682 | -0.758925832 | 0.18140156 |
| Gm26744 | 2.147345651 | 1.608295184 | 0.84093716 |
| Map3k13 | 489.4826492 | -0.601362475 | 0.08107209 |
| Gm6467 | 24.4487381 | 0.344687012 | 0.214444022 |
| 2610020F03Rik | 0.233260489 | 1.6698275 | 3.039917586 |
| Tmem41a | 97.70389257 | -0.140811921 | 0.131584033 |
| Liph | 10.28461881 | 1.409015165 | 0.341005723 |
| Senp2 | 1234.002812 | -0.050202035 | 0.07276889 |
| Gm4943 | 1.131366322 | -0.338193929 | 0.974148472 |
| Igf2bp2 | 94.65434489 | 3.55275044 | 0.292166525 |
| Gm25810 | 0 | NA | NA |
| Gm5809 | 0.299370719 | 0.541620873 | 1.742633393 |
| Gm26980 | 0 | NA | NA |
| Tra2b | 1042.494832 | 0.390570367 | 0.083018128 |
| Gm15776 | 0.868382071 | 2.996026382 | 1.131216149 |
| Etv5 | 804.3619116 | -0.570165801 | 0.118713326 |
| Dgkg | 917.1223497 | -0.179812888 | 0.101749207 |
| 9230117E06Rik | 0.105852363 | 0.23732098 | 3.724945049 |
| Mir8095 | 6.066797584 | -1.342392778 | 0.484978648 |
| Gm15651 | 2.236386351 | -0.452224846 | 0.641902519 |
| Gm22481 | 0 | NA | NA |
| Mir6362 | 0 | NA | NA |
| Crygs | 0.564059227 | 2.649145925 | 1.35756198 |
| Tbccd1 | 144.3603587 | 0.190429716 | 0.117864478 |
| Dnajb11 | 661.7688257 | 0.839393715 | 0.134535149 |
| Ahsg | 1.513507497 | -0.608684668 | 1.000264953 |
| Fetub | 1.838490909 | 0.712880569 | 0.677363616 |
| Hrg | 0.529344572 | 1.678014388 | 1.421855939 |
| Kng2 | 1.792037434 | 4.131007282 | 1.006364756 |
| Gm6493 | 33.39922536 | 1.039070922 | 0.198228986 |
| Gm23715 | 0 | NA | NA |
| Kng1 | 0.385993897 | 1.834912068 | 1.701169274 |
| Eif4a2 | 5039.606027 | -0.201048286 | 0.063328977 |
| Snord2 | 0 | NA | NA |
| Gm24201 | 0 | NA | NA |
| AC154540.1 | 0 | NA | NA |
| Snora81 | 0 | NA | NA |
| Gm26447 | 0 | NA | NA |
| Gm24616 | 7.869682849 | 0.602806889 | 0.374258314 |
| Rfc4 | 32.64420203 | 0.442259383 | 0.195462401 |
| Adipoq | 0.218718758 | 0.237328752 | 2.162828245 |
| B630019A10Rik | 0.40993559 | -0.64366998 | 1.544717569 |
| Gm6630 | 0.062426873 | 0.237312084 | 3.724945049 |
| Gm15746 | 0 | NA | NA |

| | | | |
|---|---|---|---|
| BC106179 | 11.069689 | -0.44660807 | 0.353413143 |
| St6gal1 | 316.7496297 | 0.291085411 | 0.100242953 |
| Gm23430 | 0 | NA | NA |
| Gm23066 | 0 | NA | NA |
| Rtp1 | 5.214837505 | -0.402011745 | 0.598216305 |
| Masp1 | 117.7213065 | 0.443454928 | 0.111509621 |
| Rtp4 | 910.9235272 | 7.448035697 | 0.23443169 |
| Gm6640 | 0.597217552 | 0.237333239 | 1.392187762 |
| Sst | 654.4591436 | -0.358449453 | 0.150905112 |
| Rtp2 | 3.004889707 | -1.053775956 | 0.706745416 |
| Bcl6 | 818.2508927 | 0.883399392 | 0.134192981 |
| Gm37419 | 0.567687411 | 1.735964275 | 1.403310865 |
| Lppos | 20.64171431 | -0.793327709 | 0.240095292 |
| Lpp | 689.3135003 | 0.676957171 | 0.131827199 |
| Gm24440 | 0 | NA | NA |
| Mir28a | 2.122337162 | -0.693269097 | 0.882715388 |
| Gm22672 | 0.704522364 | -0.511910323 | 1.088068982 |
| A230028O05Rik | 10.09423595 | 3.463090891 | 0.597875898 |
| Tprg | 0.366915913 | 1.083451698 | 1.519958992 |
| Trp63 | 10.83564004 | -0.391082712 | 0.347844698 |
| Gm24928 | 0 | NA | NA |
| Gm27005 | 0.212482393 | -0.242826614 | 2.184520977 |
| Gm4524 | 1.767199681 | -0.72177649 | 0.73153985 |
| P3h2 | 56.59938758 | -0.055261664 | 0.156500105 |
| AU015336 | 6.800499267 | -0.981990827 | 0.418671297 |
| Cldn1 | 56.29871389 | 0.820430207 | 0.181049034 |
| Cldn16 | 0 | NA | NA |
| Tmem207 | 0.473315337 | -1.105399316 | 1.365163361 |
| Il1rap | 1004.716365 | 0.431551651 | 0.06969837 |
| Gm20319 | 0.326503072 | 0.3563481 | 1.540748589 |
| Gmnc | 8.309823615 | -1.033630563 | 0.432286683 |
| Ostn | 1.447793589 | 3.109266002 | 0.982115205 |
| Gm15970 | 0 | NA | NA |
| Uts2b | 1.222794617 | 0.233691711 | 1.149902877 |
| Ccdc50 | 1385.300587 | 0.210164099 | 0.067069053 |
| Fgf12 | 2033.20002 | -0.653285529 | 0.082220926 |
| Mir690 | 11.63860447 | -1.232326297 | 0.400634514 |
| Mb21d2 | 459.0744955 | -0.121559588 | 0.084023855 |
| Hrasls | 202.7297218 | -0.658326408 | 0.087859129 |
| Atp13a5 | 116.1853686 | -1.086829907 | 0.156801105 |
| Gm15752 | 0.667341687 | -1.309538253 | 1.167399917 |
| Atp13a4 | 513.9834346 | 1.431111362 | 0.152490804 |
| Opa1 | 2535.740764 | -0.301654731 | 0.055432395 |
| Gm26834 | 2.849524632 | -0.003450073 | 0.625844443 |
| Gm26569 | 0.312910946 | 2.256522142 | 1.657524181 |
| Gm1968 | 0.303184894 | 2.353374162 | 3.690204768 |
| 4632428C04Rik | 13.7946314 | -0.236119058 | 0.314503766 |

| | | | |
|---|---|---|---|
| Hes1 | 94.25849431 | 0.337870431 | 0.129378984 |
| n-R5s32 | 0 | NA | NA |
| 1700025H01Rik | 0 | NA | NA |
| Gm26562 | 0.090074004 | -0.24357403 | 3.724945049 |
| Cpn2 | 0.207381971 | 0.719063912 | 2.58214438 |
| Lrrc15 | 1.556998443 | 2.14797489 | 0.947779036 |
| Gp5 | 14.94000079 | 0.534600073 | 0.309581259 |
| Atp13a3 | 1577.269787 | 0.023042071 | 0.054544108 |
| Tmem44 | 273.2629563 | -1.55870614 | 0.128441028 |
| Lsg1 | 245.3569069 | 0.607118237 | 0.102309616 |
| Fam43a | 229.047573 | 0.190850093 | 0.093308814 |
| Gm26295 | 0.327620686 | 2.151570642 | 2.012571525 |
| Gm25375 | 0 | NA | NA |
| Xxylt1 | 202.0539679 | -0.128729689 | 0.089592242 |
| Gm15742 | 2.112839908 | -0.849599414 | 0.802663452 |
| Acap2 | 1866.515927 | -0.380085286 | 0.08072718 |
| Ppp1r2 | 1509.036798 | 0.061470616 | 0.079181692 |
| Gm27415 | 0 | NA | NA |
| Apod | 8482.021723 | 3.798033023 | 0.259669334 |
| 1700007E05Rik | 2.68870073 | 0.261651089 | 0.683832966 |
| Bdh1 | 477.7842783 | -0.199751718 | 0.065663327 |
| Gm15743 | 22.43508857 | -0.001491807 | 0.324759393 |
| Dlg1 | 2632.440439 | -0.096557495 | 0.046319796 |
| Gm24879 | 0 | NA | NA |
| Mfi2 | 8.341406001 | -0.413884357 | 0.352370625 |
| Gm15703 | 0.299449808 | 1.607025405 | 2.00867912 |
| Pigz | 89.07597162 | -0.791441173 | 0.155645619 |
| NCBP2-AS2 | 84.09966569 | -0.421098476 | 0.130550914 |
| 0610012G03Rik | 0 | NA | NA |
| Ncbp2 | 349.8016824 | 0.000445002 | 0.125116289 |
| Senp5 | 762.9688528 | 0.149436414 | 0.068905507 |
| Gm15696 | 12.05015047 | -0.850192019 | 0.35950194 |
| Gm26269 | 0.072025462 | 0.237312084 | 3.724945049 |
| Gm15694 | 6.232026559 | 0.518369223 | 0.412419825 |
| Hmgb1-ps6 | 11.46318819 | -0.096020871 | 0.359914018 |
| Pak2 | 1045.01706 | 0.671056605 | 0.082514994 |
| Gm26419 | 0 | NA | NA |
| Pigx | 116.9372743 | 0.248711971 | 0.135849028 |
| Cep19 | 336.7429659 | -0.351083313 | 0.091508179 |
| Gm15729 | 0.971776759 | -0.01488442 | 1.056059917 |
| Nrros | 147.122329 | 1.642092054 | 0.136082499 |
| Bex6 | 0.377709955 | 1.920408501 | 1.513460745 |
| Fbxo45 | 678.3823491 | -0.222252582 | 0.073603648 |
| Gm25848 | 45.86024405 | 0.257939317 | 0.31847193 |
| Wdr53 | 71.48528253 | 0.268952031 | 0.147181598 |
| Mir1946a | 0 | NA | NA |
| Smco1 | 1.487580774 | 0.695542949 | 0.872170525 |

| | | | |
|---|---|---|---|
| Rnf168 | 482.2284669 | 0.06935148 | 0.058861059 |
| Ubxn7 | 1865.438353 | -0.144905209 | 0.075080438 |
| Tm4sf19 | 8.203405253 | 5.535452903 | 0.942750468 |
| Gm20056 | 14.90719541 | 1.403062506 | 0.334976716 |
| Tctex1d2 | 113.226245 | 0.121052215 | 0.104817873 |
| Pcyt1a | 752.3083677 | 0.685496867 | 0.0927444 |
| Slc51a | 0 | NA | NA |
| Zdhhc19 | 0.768268477 | 0.843261488 | 1.102950466 |
| Gm5405 | 0 | NA | NA |
| Tfrc | 1390.85657 | 0.036613866 | 0.113793254 |
| Tnk2 | 2566.202347 | -0.37139755 | 0.048221032 |
| Tnk2os | 0.364228253 | 0.254778247 | 1.571487726 |
| Gm9556 | 0 | NA | NA |
| Muc4 | 0.409426173 | 2.335925149 | 1.622062779 |
| Muc20 | 0.330893424 | 1.213585103 | 1.589865215 |
| Gm933 | 0.097139547 | 0.718207094 | 3.724945049 |
| Gm26769 | 0 | NA | NA |
| 1700021K19Rik | 991.8281154 | 0.095544537 | 0.103683026 |
| Fyttd1 | 1180.673463 | 0.074767776 | 0.094206475 |
| Lrch3 | 977.5154375 | 0.116513997 | 0.064748913 |
| Gm6611 | 33.95866455 | -0.585697373 | 0.34554514 |
| Gm17106 | 0.097139547 | 0.718207094 | 3.724945049 |
| n-R5s33 | 0.130017973 | 0.238301601 | 3.662714184 |
| Iqcg | 37.65510366 | 0.433955299 | 0.192635662 |
| Rpl35a | 394.3677081 | 0.523671201 | 0.093081087 |
| Lmln | 221.22393 | -0.392660777 | 0.089075164 |
| Gm22231 | 0.549303492 | -0.734830023 | 1.278740269 |
| Mir1947 | 0.769571883 | -0.160107497 | 1.210634794 |
| Gm23916 | 0 | NA | NA |
| Osbpl11 | 369.2866785 | 0.305464437 | 0.093001704 |
| Snx4 | 448.4376452 | 0.432988326 | 0.082899854 |
| 1700007L15Rik | 6.048360371 | -0.308006119 | 0.384445565 |
| Zfp148 | 2196.281864 | -0.218320567 | 0.047785074 |
| Slc12a8 | 28.22738804 | 0.016863725 | 0.210432755 |
| Heg1 | 831.6435367 | 0.667013743 | 0.077896752 |
| Gm15658 | 0 | NA | NA |
| Gm15657 | 0.194279094 | 1.292225235 | 3.719439735 |
| Muc13 | 4.915050087 | 4.264321517 | 0.900003625 |
| Itgb5 | 478.5994184 | 0.402473874 | 0.198339094 |
| Umps | 206.1452455 | 0.463119217 | 0.087404661 |
| Gm15829 | 0.247806867 | 0.237328056 | 2.108994839 |
| Kalrn | 4921.446429 | -1.204175608 | 0.183128264 |
| Ropn1 | 0.318826904 | -0.24659801 | 1.714752299 |
| Ccdc14 | 47.35523537 | 0.121959993 | 0.174932166 |
| Mylk | 164.66998 | 0.845107104 | 0.131322756 |
| 1700119H24Rik | 0.164132501 | 0.237308425 | 3.397905533 |
| Hacd2 | 451.6459504 | 0.342512707 | 0.095108547 |

| | | | |
|---|---|---|---|
| Adcy5 | 2609.480313 | -1.267301886 | 0.131799869 |
| Gm25179 | 0 | NA | NA |
| Sec22a | 183.1160017 | -0.017750757 | 0.113517751 |
| Gm25079 | 0 | NA | NA |
| Gm23518 | 0.038981395 | 0.718207094 | 3.724945049 |
| Pdia5 | 30.59877438 | 0.227316581 | 0.220042775 |
| Gm25548 | 0.246683951 | 0.23793457 | 1.853897646 |
| 1600019K03Rik | 0.049553047 | 0.71820487 | 3.724945049 |
| Sema5b | 130.570813 | -0.653560133 | 0.117607858 |
| Gm5963 | 36.49740693 | 0.283345522 | 0.292706935 |
| Dirc2 | 652.6475954 | -0.322741293 | 0.078588331 |
| Gm25967 | 2.125672366 | -0.655536978 | 0.79059667 |
| Hspbap1 | 70.57271391 | -0.086358053 | 0.195673873 |
| Gm26838 | 0 | NA | NA |
| Parp14 | 2858.919116 | 7.191187709 | 0.187499213 |
| Gm10237 | 0.042716945 | -0.243571806 | 3.724945049 |
| Dtx3l | 1910.456235 | 5.696895892 | 0.167145722 |
| Parp9 | 1102.792659 | 5.75337949 | 0.168268606 |
| Gm15564 | 359.9778991 | 0.830783145 | 0.972448086 |
| Kpna1 | 1948.470927 | -0.07331988 | 0.057817523 |
| Wdr5b | 49.21408818 | 0.05210371 | 0.167696105 |
| Fam162a | 149.3833853 | 0.023596946 | 0.118315022 |
| Ccdc58 | 55.1879007 | -0.089739921 | 0.160986685 |
| Csta1 | 0 | NA | NA |
| Gm10092 | 0.045882455 | 0.237312084 | 3.724945049 |
| Gm23080 | 0 | NA | NA |
| Stfa2l1 | 0.423264953 | 2.287710792 | 1.369991725 |
| Gm22885 | 0 | NA | NA |
| Gm5483 | 0.535697641 | 2.438695293 | 1.680381554 |
| Gm25024 | 0 | NA | NA |
| Gm10913 | 0.156524568 | 0.718214077 | 2.823069743 |
| Gm5416 | 0.092228164 | 0.718202646 | 3.724945049 |
| Gm24801 | 0 | NA | NA |
| 2010005H15Rik | 0.699696385 | 2.698923483 | 1.209391976 |
| Gm10813 | 0 | NA | NA |
| Gm24569 | 0 | NA | NA |
| Stfa1 | 0 | NA | NA |
| Gm23306 | 0 | NA | NA |
| Gm4758 | 0 | NA | NA |
| Gm25254 | 0 | NA | NA |
| Gm24437 | 0 | NA | NA |
| BC117090 | 0 | NA | NA |
| Gm25465 | 0.060174537 | 0.237309859 | 3.724945049 |
| BC100530 | 0.115000406 | 0.23732098 | 3.724945049 |
| Gm15845 | 0.075043205 | 0.237309859 | 3.724945049 |
| Gm24218 | 0 | NA | NA |
| Gm22349 | 0 | NA | NA |

| | | | |
|---|---|---|---|
| Stfa2 | 0.143022002 | 0.719176055 | 3.547735711 |
| Gm22263 | 0 | NA | NA |
| Gm25903 | 0 | NA | NA |
| Stfa3 | 0.131209559 | 1.165289508 | 2.972496041 |
| Casr | 17.16410259 | 0.589632738 | 0.283809032 |
| Gm25902 | 0 | NA | NA |
| Cd86 | 81.75402155 | 2.984575331 | 0.20806294 |
| Ildr1 | 14.35142951 | 1.300386022 | 0.366097795 |
| Slc15a2 | 296.0330333 | -1.320012435 | 0.219935911 |
| Gm8425 | 0.186025035 | 0.237327682 | 2.673127101 |
| Gm22617 | 0 | NA | NA |
| Eaf2 | 14.00843118 | 0.911558145 | 0.291430322 |
| Iqcb1 | 366.8902262 | -0.178577953 | 0.084164641 |
| Golgb1 | 1930.593704 | -0.209979752 | 0.107907526 |
| 4930565N06Rik | 0.233785498 | 1.52767907 | 2.489795918 |
| Hcls1 | 216.1493986 | 3.948124309 | 0.197835603 |
| Fbxo40 | 25.85214164 | 0.658489494 | 0.312946395 |
| Polq | 42.09947402 | -0.248966372 | 0.289579285 |
| Stxbp5l | 2108.032249 | -0.491840361 | 0.095197514 |
| Gm15665 | 42.97331767 | -1.023024862 | 0.304522141 |
| Gtf2e1 | 231.0323693 | 0.503334963 | 0.131932307 |
| Rabl3 | 128.7194147 | 0.578803522 | 0.120728898 |
| Hgd | 1.882553937 | 0.266727966 | 0.767042231 |
| Ndufb4 | 405.4328191 | -0.143895153 | 0.136187402 |
| Fstl1 | 650.539503 | 1.074177118 | 0.175010536 |
| n-R5s34 | 0 | NA | NA |
| Lrrc58 | 3637.314835 | 0.041123005 | 0.100268896 |
| Gm15725 | 0 | NA | NA |
| Gpr156 | 12.90207318 | -0.149123916 | 0.472297469 |
| Gm25140 | 0.106056992 | 0.237307635 | 3.724945049 |
| Gsk3b | 4829.972571 | -0.280095066 | 0.045556996 |
| SNORA17 | 0 | NA | NA |
| Gm25617 | 0 | NA | NA |
| Gm27931 | 0 | NA | NA |
| Nr1i2 | 0.200534954 | -0.646301368 | 2.636631645 |
| Maats1 | 18.76607067 | 0.239573551 | 0.240459976 |
| Maats1os | 0.072025462 | 0.237312084 | 3.724945049 |
| Cox17 | 102.4271693 | -0.253932603 | 0.206807944 |
| Gm21987 | 0.2386254 | -0.242851348 | 2.112154837 |
| Popdc2 | 3.459091227 | 0.155073415 | 0.536885926 |
| Pla1a | 56.89416345 | 4.110670822 | 0.359280548 |
| Adprh | 679.8642621 | 0.153520067 | 0.083684595 |
| D930030I03Rik | 2.891043216 | -0.449654218 | 0.590613956 |
| Cd80 | 14.42150522 | 3.151722316 | 0.466319731 |
| Timmdc1 | 207.9193946 | 0.226360234 | 0.131710761 |
| Gm15953 | 0 | NA | NA |
| Poglut1 | 522.8527119 | 0.151081715 | 0.061939904 |

| | | | |
|---|---|---|---|
| Tmem39a | 165.5822775 | 0.408643031 | 0.129932907 |
| Gm17103 | 0.159330469 | 0.718203216 | 3.465376669 |
| Arhgap31 | 855.9405782 | 0.770535047 | 0.094085014 |
| Gm15530 | 5.747139663 | -0.19957855 | 0.427006248 |
| B4galt4 | 168.6399155 | -0.082665755 | 0.098547125 |
| Gm15575 | 0 | NA | NA |
| Upk1b | 9.705317616 | 1.155256485 | 0.440982821 |
| 4930435E12Rik | 0 | NA | NA |
| Gm15803 | 0 | NA | NA |
| Gm15802 | 0 | NA | NA |
| Igsf11 | 559.2207678 | 0.248153847 | 0.137575852 |
| Gm22500 | 7.621091703 | -0.667164798 | 0.506959914 |
| Gm24795 | 0 | NA | NA |
| Gm8609 | 0.167459142 | 1.165958262 | 2.753748546 |
| Lsamp | 5421.197146 | -0.689470219 | 0.063310122 |
| Gm37508 | 3.928946461 | -0.841311971 | 0.600563705 |
| Gm22543 | 16.99085758 | -1.4891499 | 0.364584894 |
| Gm36939 | 48.56639706 | -0.960519623 | 0.241574379 |
| Gm27887 | 0 | NA | NA |
| Gm27717 | 0 | NA | NA |
| Gm26381 | 1.397116197 | -1.914183112 | 0.964121154 |
| AC124193.1 | 0 | NA | NA |
| Gm27587 | 0 | NA | NA |
| Gm28750 | 0 | NA | NA |
| Gap43 | 1479.213936 | -0.402766993 | 0.096033293 |
| Mir6540 | 2.543989632 | -0.945838329 | 0.6790949 |
| 4932412D23Rik | 1.816140692 | -0.305336486 | 0.747906224 |
| Zbtb20 | 6323.085067 | 0.065962434 | 0.083901957 |
| BC002163 | 27.11636185 | -0.349051605 | 0.216134016 |
| Gm25873 | 5.983328485 | 0.029123352 | 0.448649218 |
| Gm15712 | 29.69002061 | -0.706275993 | 0.360364261 |
| Gm37946 | 0 | NA | NA |
| Gm15711 | 21.32018645 | -0.630853452 | 0.318202685 |
| Gm9968 | 1.290099097 | 1.081839669 | 1.012060422 |
| Gm15713 | 0.087930927 | 0.236669296 | 3.724898107 |
| Mir568 | 0 | NA | NA |
| Tigit | 22.00729436 | 8.102933315 | 0.918265253 |
| Drd3 | 8.262393275 | -1.114053812 | 0.444049571 |
| Gm25996 | 0.1580736 | -0.72445813 | 3.474509404 |
| Gm26074 | 0 | NA | NA |
| Qtrtd1 | 200.210793 | -0.04740994 | 0.109062596 |
| Gm25989 | 9.070232471 | -0.071924614 | 0.408073849 |
| 2610015P09Rik | 57.77977805 | -0.592478785 | 0.218718794 |
| Zdhhc23 | 82.33084375 | -0.52461328 | 0.155388358 |
| Gramd1c | 24.07085944 | -0.255656564 | 0.213168524 |
| Mir3470b | 0 | NA | NA |
| Atp6v1a | 5242.393468 | -0.145089293 | 0.059937363 |

| | | | |
|---|---|---|---|
| Naa50 | 1580.032754 | 0.533754791 | 0.090042609 |
| Gm608 | 1399.505967 | -0.379284781 | 0.073093731 |
| Sidt1 | 676.9943462 | -0.590372186 | 0.141921535 |
| Gm26732 | 6.046421774 | -1.671839821 | 0.438195493 |
| Spice1 | 128.0135962 | -0.538800297 | 0.166712784 |
| Cfap44 | 93.24835868 | -0.694402053 | 0.173253516 |
| Boc | 445.6221661 | -0.221564265 | 0.16666777 |
| Mir3081 | 1.772274333 | -0.961190771 | 0.736399362 |
| BC027231 | 116.0115484 | 0.350594973 | 0.164527074 |
| Gtpbp8 | 357.6076025 | 0.063210679 | 0.084335676 |
| Cd200r1 | 51.42872498 | 2.24540313 | 0.300393703 |
| Cd200r4 | 69.15376312 | 4.785671903 | 0.403116843 |
| Cd200r2 | 1.386349714 | 0.670116947 | 0.84610586 |
| Cd200r3 | 0.534659868 | 1.161562023 | 1.346734295 |
| Ccdc80 | 217.6073964 | 2.13093507 | 0.155386706 |
| Slc35a5 | 550.0251731 | 0.03778686 | 0.101155583 |
| Atg3 | 539.1722443 | 0.373396906 | 0.069701903 |
| Gm6030 | 17.33716352 | 0.193159306 | 0.291435719 |
| Btla | 10.07865719 | 2.228286138 | 0.377896557 |
| Cd200 | 1252.555305 | 0.822434317 | 0.104405275 |
| Gm5406 | 0.966602524 | 1.399748954 | 1.084824455 |
| Gm609 | 0.707603298 | -0.479978226 | 1.346042542 |
| Gm16011 | 0.581873452 | 1.082665395 | 1.258195897 |
| Gm6903 | 0.049553047 | 0.71820487 | 3.724945049 |
| Slc9c1 | 0.038981395 | 0.718207094 | 3.724945049 |
| Gcsam | 0.711975465 | -0.047937282 | 1.161067292 |
| BC016579 | 3.578730896 | -0.379675047 | 0.641644837 |
| Tmprss7 | 24.10678365 | -0.445560087 | 0.301898476 |
| Tagln3 | 1506.536885 | -0.34869312 | 0.095709614 |
| Abhd10 | 203.4530973 | -0.084635588 | 0.092617193 |
| Gm15591 | 0.060174537 | 0.237309859 | 3.724945049 |
| Phldb2 | 129.9886621 | 0.154038636 | 0.123070434 |
| Gm15640 | 0.229952095 | 0.663005536 | 2.112891911 |
| Gm15638 | 0.108266322 | 1.230376831 | 3.722845817 |
| Plcxd2 | 1552.118851 | -0.713066947 | 0.078785426 |
| Gm24754 | 0 | NA | NA |
| Cd96 | 3.576396818 | 3.408414511 | 0.886086019 |
| Gm4737 | 112.4227191 | 0.364503815 | 0.164995812 |
| Gm26297 | 0 | NA | NA |
| Pvrl3 | 517.9500622 | -0.317943349 | 0.086503588 |
| Gm6912 | 0 | NA | NA |
| Gm24651 | 0 | NA | NA |
| Gm25318 | 0 | NA | NA |
| Gm25339 | 0 | NA | NA |
| Gm7275 | 0 | NA | NA |
| Gm7204 | 1.57829385 | 3.204492581 | 0.982163957 |
| Dppa4 | 0 | NA | NA |

| | | | |
|---|---|---|---|
| Dppa2 | 0.379773675 | 2.574088852 | 1.586808914 |
| n-R5s36 | 0 | NA | NA |
| Morc1 | 1.278798984 | 1.078556448 | 0.82054802 |
| Trat1 | 6.136963414 | 1.301201531 | 0.462606413 |
| Retnlb | 0 | NA | NA |
| Retnla | 0.060174537 | 0.237309859 | 3.724945049 |
| Retnlg | 1.932622373 | 3.827674801 | 1.057953819 |
| Dzip3 | 1926.33653 | -0.10298666 | 0.094930389 |
| Gm22576 | 7.017922756 | -0.203461735 | 0.420742544 |
| C330027C09Rik | 62.11505808 | 0.540897513 | 0.167167468 |
| Myh15 | 9.416055846 | -0.284189077 | 0.330662335 |
| 1700026J12Rik | 0.047948698 | -0.243569582 | 3.724945049 |
| Gm5407 | 0.075043205 | 0.237309859 | 3.724945049 |
| Gm6931 | 0.04393175 | -0.243571806 | 3.724945049 |
| Ift57 | 340.5122929 | -0.32856998 | 0.086284988 |
| Gm15518 | 0.295614739 | 0.221655062 | 2.085469128 |
| Gm16619 | 0.436453079 | 1.151139832 | 1.439331306 |
| Cd47 | 3446.906187 | 0.668486209 | 0.098742441 |
| AC107830.1 | 0 | NA | NA |
| Gm8824 | 0.047948698 | -0.243569582 | 3.724945049 |
| Bbx | 1297.513501 | 0.758631876 | 0.112202171 |
| G730013B05Rik | 0.417980132 | 0.900174036 | 1.471577252 |
| Ccdc54 | 0.042716945 | -0.243571806 | 3.724945049 |
| Gm9575 | 16.04186629 | 0.124444584 | 0.271370542 |
| 5330426P16Rik | 100.6912749 | -0.264622564 | 0.172936686 |
| Mir6363 | 0 | NA | NA |
| 1700116B05Rik | 0 | NA | NA |
| Gm24925 | 0 | NA | NA |
| Gm6767 | 3.585344364 | 0.528739636 | 0.54421559 |
| Cblb | 789.0152622 | 0.459486671 | 0.092713938 |
| Gm23513 | 0 | NA | NA |
| Alcam | 2206.12951 | -0.479984998 | 0.071762465 |
| Gm25723 | 0 | NA | NA |
| Gm22459 | 0 | NA | NA |
| Gm25041 | 0 | NA | NA |
| Gm25853 | 0 | NA | NA |
| Gm23180 | 0 | NA | NA |
| Gm22977 | 0 | NA | NA |
| Zpld1 | 1.833497833 | -1.330344938 | 0.870527197 |
| Mir5118 | 0 | NA | NA |
| Gm23003 | 0 | NA | NA |
| Gm26369 | 0 | NA | NA |
| Nfkbiz | 257.9280466 | 1.622956846 | 0.164275241 |
| Nxpe3 | 312.8495634 | -0.39401211 | 0.095270138 |
| Cep97 | 386.1470673 | -0.635709036 | 0.147257341 |
| Rpl24 | 16.95551249 | 0.49874149 | 0.307284315 |
| Gm28037 | 0 | NA | NA |

| | | | |
|---|---|---|---|
| Zbtb11os1 | 37.78932254 | 0.184017483 | 0.203847521 |
| Zbtb11 | 1088.837139 | -0.089975577 | 0.069019631 |
| Pcnp | 1051.54738 | 0.205171932 | 0.078217502 |
| Trmt10c | 191.2919896 | 0.554845062 | 0.100129622 |
| Gm22422 | 89.64062021 | -0.082538246 | 0.388438963 |
| Senp7 | 795.4829463 | -0.26246354 | 0.119314391 |
| Gm24047 | 0.105143817 | -0.24357403 | 3.724945049 |
| Impg2 | 63.00063838 | 0.601353839 | 0.143173459 |
| Abi3bp | 266.089939 | -0.164721927 | 0.205808624 |
| Tfg | 756.0033368 | 0.371338349 | 0.056973455 |
| Adgrg7 | 0.562361829 | 1.273340062 | 1.433071037 |
| Gm15693 | 0.044191549 | -0.243571806 | 3.724945049 |
| Tmem45a | 10.44425536 | 2.192755735 | 0.380278837 |
| Gm15839 | 0.181290818 | 0.703161966 | 2.739792952 |
| Gm15840 | 0 | NA | NA |
| Tmem45a2 | 3.654246218 | 1.353828599 | 0.624018493 |
| Tomm70a | 2858.851143 | 0.030029697 | 0.081263461 |
| Nit2 | 67.55208235 | 0.075791733 | 0.14144787 |
| Gm16892 | 2.540529182 | -0.543490786 | 0.613548645 |
| Tbc1d23 | 546.2969052 | 0.204978687 | 0.057244347 |
| Tmem30c | 8.630201669 | -0.42414324 | 0.408867569 |
| Gm26800 | 0.820767868 | 1.491235805 | 1.260452807 |
| Cmss1 | 65.09916204 | -0.026752375 | 0.132018025 |
| Filip1l | 99.78162097 | 0.895501617 | 0.285118447 |
| Col8a1 | 91.62796141 | 0.249456699 | 0.172520604 |
| Gm24039 | 0 | NA | NA |
| Gm37458 | 0.072025462 | 0.237312084 | 3.724945049 |
| Gm37825 | 0 | NA | NA |
| 4930461C15Rik | 1.662681963 | -1.52178725 | 0.878359639 |
| Dcbld2 | 658.0902064 | -0.17015754 | 0.094185509 |
| St3gal6 | 198.8642845 | 0.461131658 | 0.100763874 |
| Gm813 | 1.602843074 | 4.236015232 | 1.050291106 |
| E330017A01Rik | 0 | NA | NA |
| Cpox | 461.6201025 | -0.211316276 | 0.061587633 |
| Gpr15 | 6.100871677 | 6.100548412 | 0.957619518 |
| Cldnd1 | 811.4291424 | 0.294377981 | 0.088129056 |
| Gm27564 | 0 | NA | NA |
| Gm26019 | 0 | NA | NA |
| Olfr172 | 0 | NA | NA |
| Olfr173 | 0 | NA | NA |
| Olfr175-ps1 | 0.049553047 | 0.71820487 | 3.724945049 |
| Olfr177 | 0 | NA | NA |
| Olfr178 | 0 | NA | NA |
| Olfr180 | 0 | NA | NA |
| Olfr181 | 0 | NA | NA |
| Olfr183 | 0 | NA | NA |
| Olfr186 | 0.172571121 | 1.188801635 | 2.733273164 |

| | | | |
|---|---|---|---|
| Olfr187 | 0.106520246 | 0.718200421 | 3.724945049 |
| Olfr190 | 0 | NA | NA |
| Olfr191 | 0 | NA | NA |
| Olfr192 | 0.046345709 | 0.71820487 | 3.724945049 |
| Olfr193 | 0 | NA | NA |
| Olfr194 | 0 | NA | NA |
| Olfr195 | 0.097139547 | 0.718207094 | 3.724945049 |
| Olfr196 | 0 | NA | NA |
| Olfr197 | 0 | NA | NA |
| Olfr198 | 0.194201709 | -0.549026435 | 3.270763994 |
| Olfr199 | 0 | NA | NA |
| Olfr201 | 0 | NA | NA |
| Olfr202 | 0 | NA | NA |
| Olfr203 | 0 | NA | NA |
| Olfr204 | 0.068097359 | -0.243569582 | 3.724945049 |
| Olfr205 | 0.046345709 | 0.71820487 | 3.724945049 |
| Olfr206 | 0.075043205 | 0.237309859 | 3.724945049 |
| Olfr209 | 0.048949532 | -0.243569582 | 3.724945049 |
| Gabrr3 | 1.001170369 | 0.928171745 | 1.122042625 |
| 1700022E09Rik | 0.097139547 | 0.718207094 | 3.724945049 |
| Mina | 105.9311312 | 0.485996687 | 0.134285894 |
| Crybg3 | 125.2170434 | 0.190339507 | 0.110130366 |
| Gm9581 | 73.34347339 | 0.202959078 | 0.124867312 |
| Arl6 | 230.3950377 | 0.088839034 | 0.102894139 |
| Epha6 | 866.3468396 | -1.168343102 | 0.1249305 |
| Gm24755 | 0.366471122 | -0.266398464 | 1.596060386 |
| Gm4660 | 10.78697284 | -1.253043214 | 0.432100948 |
| Gm9017 | 5.516248819 | -0.475579509 | 0.408512289 |
| Gm27001 | 0.095897396 | -0.724453471 | 3.724945049 |
| Gm27784 | 0 | NA | NA |
| Gm17065 | 0 | NA | NA |
| Gm23852 | 0 | NA | NA |
| Gm17054 | 0 | NA | NA |
| n-R5s37 | 0 | NA | NA |
| Gm9027 | 1.878461702 | 0.654911586 | 0.728730456 |
| Gm17192 | 0 | NA | NA |
| Gm24164 | 0 | NA | NA |
| Gm9816 | 144.6420699 | 0.205079563 | 0.110872136 |
| Nsun3 | 164.1841752 | 0.628761522 | 0.097827208 |
| Arl13b | 185.8996067 | 0.175770762 | 0.092645409 |
| Gm25986 | 0 | NA | NA |
| Stx19 | 0.062426873 | 0.237312084 | 3.724945049 |
| Pros1 | 274.6408184 | 2.065161532 | 0.165470542 |
| Epha3 | 257.2072295 | -0.540266909 | 0.156908279 |
| Gm22769 | 4.369109362 | -1.146684109 | 0.510017301 |
| Csnka2ip | 2.872518017 | 0.432553554 | 0.649633388 |
| 4930453N24Rik | 641.6073447 | 0.897757596 | 0.114727883 |

| | | | |
|---|---|---|---|
| Zfp654 | 590.4534688 | 0.127058063 | 0.120642114 |
| Cggbp1 | 1135.996861 | 0.563768677 | 0.094835345 |
| Gm9061 | 0.350048956 | 0.285269514 | 1.51092873 |
| Gm23374 | 2.380614889 | -0.475879173 | 0.710855709 |
| Htr1f | 54.47759132 | -0.69182447 | 0.164696374 |
| Gm22429 | 0 | NA | NA |
| Gm23659 | 0 | NA | NA |
| Pou1f1 | 0.726361874 | 2.808828499 | 3.664952517 |
| Chmp2b | 560.9048391 | 0.074716858 | 0.085511396 |
| Vgll3 | 46.32731703 | -0.350112413 | 0.162638557 |
| Gm22217 | 0.045882455 | 0.237312084 | 3.724945049 |
| Gm22448 | 0 | NA | NA |
| Cadm2 | 4980.463628 | -0.754956944 | 0.091900256 |
| 1700010K23Rik | 0.914990437 | 0.793472193 | 1.218773135 |
| 4933411O13Rik | 0 | NA | NA |
| Gm15828 | 81.5457743 | -0.738436897 | 0.298212867 |
| Gm24681 | 0 | NA | NA |
| Gm23591 | 0 | NA | NA |
| RP24-377L12.1 | 0.046345709 | 0.71820487 | 3.724945049 |
| Speer2 | 0.175920186 | 0.773506405 | 2.728285484 |
| Gbe1 | 229.1812539 | 0.648623307 | 0.093801059 |
| Gm24968 | 0.168853147 | 1.513624709 | 2.756733182 |
| Gm22797 | 0 | NA | NA |
| Robo1 | 968.3837098 | -0.818913503 | 0.092361873 |
| Gm25699 | 0 | NA | NA |
| Robo2 | 1762.408431 | -0.854104481 | 0.108663004 |
| Gm24272 | 0 | NA | NA |
| Gm22163 | 1.874863541 | -2.588325579 | 0.930618633 |
| mmu-mir-691 | 8.12823547 | -0.614673852 | 0.448725284 |
| Gm26307 | 0 | NA | NA |
| Gm26937 | 0.185762564 | -1.566905147 | 3.304313696 |
| Lipi | 0.34836924 | 1.157332615 | 1.55720032 |
| Rbm11 | 76.42304198 | 0.262501555 | 0.190220394 |
| Gm15553 | 0 | NA | NA |
| Hspa13 | 738.3975908 | 0.383997488 | 0.076137586 |
| Gm22923 | 0 | NA | NA |
| Gm15554 | 0.120974994 | -0.243571806 | 3.724945049 |
| Samsn1 | 95.65148686 | 3.326627281 | 0.276378717 |
| Gm15555 | 0 | NA | NA |
| Gm37705 | 0 | NA | NA |
| Gm22268 | 0 | NA | NA |
| Gm26915 | 0 | NA | NA |
| Nrip1 | 1272.517009 | -0.012681199 | 0.090329486 |
| Gm9843 | 41.81605505 | 0.472746732 | 0.198801199 |
| 1700063K16Rik | 0.044544648 | 0.237309859 | 3.724945049 |
| 1700041M19Rik | 0.061920613 | 0.718207094 | 3.724945049 |
| Usp25 | 1277.058717 | 1.545182516 | 0.110974892 |

| | | | |
|---|---|---|---|
| Mir99ahg | 442.0252824 | -0.051243714 | 0.12987065 |
| Gm31258 | 0.728787387 | -1.372094908 | 1.2722172 |
| Gm37466 | 10.46537345 | -0.240064512 | 0.368561815 |
| Gm21816 | 50.06920957 | -0.324453872 | 0.231956253 |
| 9430092D12Rik | 19.231994 | -0.379812929 | 0.276420163 |
| Gm38071 | 38.74659047 | -0.582090609 | 0.240425083 |
| 9430053O09Rik | 25.41324745 | 0.048825412 | 0.222830437 |
| Gm37063 | 47.20854318 | -0.137749672 | 0.206714497 |
| Gm37241 | 6.880611024 | -0.484396678 | 0.510983469 |
| Gm36963 | 34.35472376 | 0.03450586 | 0.225575696 |
| Gm37606 | 95.09549254 | -0.314650977 | 0.218541802 |
| Gm11146 | 0.04393175 | -0.243571806 | 3.724945049 |
| Mir99a | 7.589586068 | 0.234313333 | 0.393642933 |
| Mirlet7c-1 | 8.265456032 | 0.234482889 | 0.379744599 |
| Gm37311 | 115.4089426 | 0.041498447 | 0.205016638 |
| Mir125b-2 | 0.487613095 | 0.423919734 | 1.477041538 |
| C130023A14Rik | 179.7393386 | -0.078557796 | 0.147848873 |
| 9530003O04Rik | 61.28675002 | 0.096993784 | 0.168290703 |
| Gm37694 | 93.40348466 | 0.066545422 | 0.211097268 |
| Gm17333 | 0.423282238 | 1.639558832 | 3.702783046 |
| Gm25025 | 0.066050875 | 0.71820487 | 3.724945049 |
| Gm25916 | 0 | NA | NA |
| Cxadr | 476.9468567 | -0.034183805 | 0.144445197 |
| Btg3 | 111.9560118 | 1.043699923 | 0.238547925 |
| Gm25038 | 0 | NA | NA |
| D16Ertd472e | 110.9694825 | 0.305843002 | 0.12599473 |
| Gm24861 | 0 | NA | NA |
| Chodl | 16.16523681 | 0.027462918 | 0.306447209 |
| Tmprss15 | 0.287833757 | 0.796503725 | 2.035025104 |
| Gm15692 | 0 | NA | NA |
| Gm37709 | 0 | NA | NA |
| Rps19-ps12 | 0 | NA | NA |
| 1700066C05Rik | 0 | NA | NA |
| Gm23083 | 0 | NA | NA |
| Gm24773 | 0 | NA | NA |
| Ncam2 | 1411.136076 | -0.423821301 | 0.077671487 |
| Gm26023 | 0.228677828 | -1.094057945 | 2.143310477 |
| Gm23355 | 0 | NA | NA |
| Gm21833 | 0.117046891 | -0.724453471 | 3.724945049 |
| Rpl21-ps5 | 0 | NA | NA |
| Gm24508 | 0 | NA | NA |
| Gm25292 | 0.193566712 | 1.949803564 | 2.630756597 |
| Gm26667 | 11.04978208 | 6.5224353 | 0.876268302 |
| Mir155 | 0 | NA | NA |
| Mrpl39 | 254.7822395 | 0.32972423 | 0.103244609 |
| Gm26153 | 0.072025462 | 0.237312084 | 3.724945049 |
| Jam2 | 457.3571526 | -0.431597618 | 0.155674489 |

| | | | |
|---|---|---|---|
| Gm25908 | 0.500634304 | -1.440064214 | 1.284582434 |
| Atp5j | 806.9086276 | -0.179849576 | 0.130736614 |
| Gabpa | 471.5751915 | 0.298460552 | 0.059605131 |
| App | 15888.22377 | -0.301716198 | 0.058651529 |
| Gm27295 | 0 | NA | NA |
| Cyyr1 | 66.70143812 | 2.233030436 | 0.182249015 |
| Gm2541 | 17.77359248 | -0.066416722 | 0.273382297 |
| Adamts1 | 613.1894008 | 2.076407766 | 0.303238245 |
| Adamts5 | 105.3116989 | 0.668901089 | 0.199480806 |
| Gm25715 | 0 | NA | NA |
| N6amt1 | 193.0747588 | -0.206920976 | 0.113289335 |
| Ltn1 | 1170.966149 | -0.043890459 | 0.041060448 |
| Gm25896 | 1.990209358 | -0.528280241 | 0.808098225 |
| Rwdd2b | 73.7279705 | -0.035559989 | 0.129340471 |
| Usp16 | 561.8276137 | 0.057715187 | 0.122968618 |
| Cct8 | 1716.316278 | 0.327976955 | 0.087103186 |
| B130034C11Rik | 16.77205567 | -0.82873901 | 0.337590501 |
| Gm24891 | 0 | NA | NA |
| Rpl31-ps4 | 0.092228164 | 0.718202646 | 3.724945049 |
| Map3k7cl | 1.664220238 | 0.433659638 | 0.857895862 |
| Gm22808 | 0 | NA | NA |
| Bach1 | 550.4479081 | 1.324288771 | 0.123975084 |
| Gm23994 | 0 | NA | NA |
| 2810407A14Rik | 5.016221076 | -0.96622307 | 0.826880689 |
| Grik1 | 248.488386 | -0.671815348 | 0.096798815 |
| Cldn17 | 0.086930093 | 0.236669296 | 3.724898107 |
| Cldn8 | 0.133745955 | -0.724460126 | 3.671134981 |
| Krtap24-1 | 0 | NA | NA |
| Gm23587 | 0 | NA | NA |
| 2310079G19Rik | 0 | NA | NA |
| Krtap26-1 | 0 | NA | NA |
| Krtap27-1 | 0 | NA | NA |
| 2310061N02Rik | 0 | NA | NA |
| Krtap13-1 | 0.115603922 | 1.209894437 | 3.72418822 |
| Krtap13 | 0 | NA | NA |
| 2310034C09Rik | 0 | NA | NA |
| AC125199.1 | 0 | NA | NA |
| 2310057N15Rik | 0 | NA | NA |
| Gm5965 | 0 | NA | NA |
| Krtap14 | 0.061920613 | 0.718207094 | 3.724945049 |
| Krtap15 | 0 | NA | NA |
| Krtap19-1 | 0 | NA | NA |
| Krtap19-2 | 0 | NA | NA |
| Krtap19-3 | 0 | NA | NA |
| Krtap19-4 | 0 | NA | NA |
| Krtap19-5 | 0 | NA | NA |
| Krtap19-9a | 0 | NA | NA |

| | | | |
|---|---|---|---|
| Krtap19-9b | 0 | NA | NA |
| Krtap16-3 | 0 | NA | NA |
| AC140194.1 | 0 | NA | NA |
| Krtap22-2 | 0 | NA | NA |
| Gm10229 | 0 | NA | NA |
| Krtap6-1 | 0 | NA | NA |
| Gm10228 | 0 | NA | NA |
| Krtap6-5 | 0 | NA | NA |
| 1110025L11Rik | 0 | NA | NA |
| Krtap6-3 | 0 | NA | NA |
| Gm6358 | 0 | NA | NA |
| Gm10061 | 0 | NA | NA |
| Gm9789 | 0 | NA | NA |
| Gm7735 | 0 | NA | NA |
| 1110057P08Rik | 0 | NA | NA |
| Krtap20-2 | 0 | NA | NA |
| Krtap21-1 | 0 | NA | NA |
| Krtap6-2 | 0 | NA | NA |
| AC121818.1 | 0 | NA | NA |
| Gm23655 | 0.049553047 | 0.71820487 | 3.724945049 |
| Krtap8-1 | 0 | NA | NA |
| Krtap7-1 | 0.106618422 | -0.24357403 | 3.724945049 |
| Krtap11-1 | 0.045882455 | 0.237312084 | 3.724945049 |
| Tiam1 | 2404.733508 | -0.398221476 | 0.130428669 |
| Sod1 | 1216.13554 | 0.101376743 | 0.094947123 |
| Scaf4 | 656.9093113 | -0.000335181 | 0.107482844 |
| Hunk | 320.5353761 | 0.07651751 | 0.068664649 |
| 1110008E08Rik | 0.670543963 | 2.957141979 | 1.310009444 |
| Gm23406 | 0.090074004 | -0.24357403 | 3.724945049 |
| Mis18a | 41.5770124 | 0.26395376 | 0.208562712 |
| Gm17518 | 6.352540116 | -0.677357174 | 0.406607084 |
| Mrap | 10.74916131 | 3.005139342 | 0.618235276 |
| Urb1 | 305.5173951 | 0.004676547 | 0.086694388 |
| Eva1c | 41.11539447 | 0.401146053 | 0.208416037 |
| Gm7831 | 1.072190385 | -1.201969346 | 1.135807649 |
| 1110004E09Rik | 190.7191731 | 0.372164728 | 0.115594986 |
| Synj1 | 5333.078176 | -0.272610805 | 0.061840127 |
| Gm22933 | 0 | NA | NA |
| Gm15965 | 0.045882455 | 0.237312084 | 3.724945049 |
| 4930404I05Rik | 0.517542306 | 1.36498908 | 1.318283872 |
| Paxbp1 | 967.3107701 | -0.110563177 | 0.135615483 |
| 4931406G06Rik | 3.965964068 | 1.629046584 | 0.600477934 |
| 4932438H23Rik | 22.83793654 | -0.078017886 | 0.211404153 |
| H3f3a-ps2 | 2.549826182 | -0.121626552 | 0.682923058 |
| Gm9881 | 5.470790701 | -1.823822386 | 0.532837808 |
| Olig2 | 347.143824 | 0.727286433 | 0.161002733 |
| Gm37016 | 0.089089296 | 0.237305797 | 3.724945049 |

| | | | |
|---|---|---|---|
| Olig1 | 1119.055513 | 0.316755424 | 0.147815453 |
| Gm15966 | 0 | NA | NA |
| Gm24695 | 0 | NA | NA |
| Ifnar2 | 243.5630765 | 1.689936417 | 0.120177576 |
| Gm21970 | 0 | NA | NA |
| Il10rb | 126.5063742 | 2.844160676 | 0.197573863 |
| A930006K02Rik | 5.618572222 | -0.525397027 | 0.458092448 |
| Ifnar1 | 638.5446908 | 0.182675549 | 0.067715581 |
| Gm7856 | 0.270435483 | 0.237298438 | 3.724945049 |
| Mir6367 | 0 | NA | NA |
| Gm7860 | 1.38062972 | 0.900339147 | 0.994820759 |
| Ifngr2 | 476.6547008 | 0.289855521 | 0.065970129 |
| Tmem50b | 796.4973955 | -0.207765332 | 0.059220319 |
| Gm15964 | 4.772112467 | 3.534684813 | 0.740053924 |
| Dnajc28 | 95.3526294 | -0.28397163 | 0.175766985 |
| Gart | 409.6463015 | 0.448420647 | 0.098342526 |
| Son | 6238.367876 | -0.018530503 | 0.06046962 |
| Donson | 181.5744247 | -0.079321476 | 0.097062851 |
| Atp5o | 1115.945799 | -0.05110797 | 0.076211034 |
| Gm10785 | 5.027936496 | -0.128681783 | 0.485936795 |
| Cryzl1 | 541.6613726 | -0.261338292 | 0.079393577 |
| Itsn1 | 2861.680614 | -0.52645169 | 0.05795295 |
| Gm25245 | 0.133144048 | -0.243572169 | 2.968531442 |
| Gm15976 | 0.272598396 | -0.262323621 | 1.814448785 |
| D430001F17Rik | 1.721321887 | 0.178665598 | 0.739260933 |
| Gm27773 | 0 | NA | NA |
| Gm23210 | 0 | NA | NA |
| Mrps6 | 345.9199876 | 2.034255578 | 0.170201727 |
| Slc5a3 | 2337.043536 | 1.053338672 | 0.131542504 |
| Gm16310 | 32.27711759 | -0.44594781 | 0.317201303 |
| Kcne2 | 13.78272213 | -3.44298363 | 0.593153258 |
| Smim11 | 43.31674533 | -0.219400015 | 0.21898316 |
| 1700048M11Rik | 0 | NA | NA |
| 4930563D23Rik | 0.422874972 | 1.133253873 | 1.68148146 |
| Kcne1 | 0.561001133 | 0.196084789 | 1.392869367 |
| Rcan1 | 683.9052406 | 0.949015847 | 0.157059151 |
| 2410124H12Rik | 1.475924581 | -1.090140885 | 0.916516677 |
| Clic6 | 121.7859387 | -1.123598439 | 0.170594847 |
| Runx1 | 156.9186044 | 2.835222862 | 0.181670684 |
| Gm26626 | 3.202063597 | 5.170964067 | 0.912063706 |
| Gm28003 | 0 | NA | NA |
| 1810053B23Rik | 0 | NA | NA |
| Mir802 | 0 | NA | NA |
| 1700029J03Rik | 0.049553047 | 0.71820487 | 3.724945049 |
| Setd4 | 64.56323875 | -0.49922353 | 0.166446211 |
| Cbr1 | 336.8011914 | -0.327080562 | 0.083497606 |
| Gm5678 | 9.467024167 | -0.286690412 | 0.380202346 |

| | | | |
|---|---|---|---|
| Gm22344 | 0 | NA | NA |
| Cbr3 | 89.79952323 | 0.257706831 | 0.162844423 |
| Dopey2 | 1643.428677 | -0.11294638 | 0.064745816 |
| 2310043M15Rik | 4.325146265 | 4.373718078 | 0.846965603 |
| Morc3 | 612.5180941 | 0.990064 | 0.090049402 |
| Chaf1b | 23.6865518 | 2.556537457 | 0.333582221 |
| Cldn14 | 11.98292911 | 0.632334559 | 0.401188754 |
| Sim2 | 1.471461872 | 0.316470866 | 0.797593742 |
| Hlcs | 243.6874178 | -0.451877902 | 0.082067536 |
| Gm23062 | 1.571274675 | -1.356251971 | 0.977214417 |
| Ripply3 | 0.878604591 | 1.142460694 | 1.190141043 |
| Pigp | 188.369765 | 0.243031401 | 0.159839562 |
| Gm15310 | 0.048949532 | -0.243569582 | 3.724945049 |
| Ttc3 | 12670.90535 | -0.693868288 | 0.057737026 |
| Gm23692 | 8.647424846 | -1.451610671 | 0.374756824 |
| Dscr3 | 356.437628 | 0.723021305 | 0.16556736 |
| Dyrk1a | 1886.330775 | 0.220925964 | 0.085742447 |
| Gm15971 | 19.22130654 | -0.728602529 | 0.348401982 |
| Kcnj6 | 616.7000442 | -0.694747714 | 0.118472416 |
| Gm22745 | 0 | NA | NA |
| Gm7976 | 0.168942356 | 0.23732465 | 2.756030556 |
| Kcnj15 | 5.024696343 | 4.455469721 | 0.808014576 |
| Erg | 22.49114717 | 1.291275148 | 0.255673468 |
| Ets2 | 1062.543237 | 0.31250673 | 0.106460812 |
| Gm37259 | 0.066050875 | 0.71820487 | 3.724945049 |
| 2810404F17Rik | 0.08169834 | 0.23732098 | 3.724945049 |
| 1600002D24Rik | 0.782188385 | 3.447809031 | 1.196484254 |
| Gm15340 | 0 | NA | NA |
| Psmg1 | 92.11413496 | 0.23090398 | 0.122485788 |
| Brwd1 | 2327.039944 | -0.463931958 | 0.115219515 |
| Gm15342 | 0 | NA | NA |
| Gm27833 | 5.512382878 | -0.893047284 | 0.513288329 |
| 1700093J21Rik | 4.649104205 | -1.079161071 | 0.541106486 |
| Gm15341 | 0.325655381 | -0.243568467 | 1.95483669 |
| Hmgn1 | 496.6434735 | 0.628713382 | 0.110208552 |
| Wrb | 502.1431527 | -0.426762461 | 0.089181369 |
| Gm15317 | 0.04393175 | -0.243571806 | 3.724945049 |
| Lca5l | 22.78202919 | -0.630853746 | 0.281501757 |
| Sh3bgr | 25.544519 | -0.533714451 | 0.234868399 |
| B3galt5 | 665.4843436 | -0.063484431 | 0.135986764 |
| Igsf5 | 4.211675288 | 0.10994371 | 0.471505726 |
| Itgb2l | 0.181155367 | 0.237326356 | 2.695761639 |
| Pcp4 | 324.4405224 | -1.075336529 | 0.17156293 |
| Dscam | 1116.965166 | -0.662147058 | 0.091021398 |
| Bace2 | 46.32598539 | -0.102721911 | 0.166448265 |
| Mx1 | 1975.925507 | 10.14704559 | 0.469486161 |
| Fam3b | 2.301576675 | 4.780050497 | 0.934894155 |

| | | | |
|---|---|---|---|
| Gm9242 | 1.910909441 | 2.880900675 | 1.01628777 |
| Gm24777 | 0 | NA | NA |
| Mx2 | 1286.777433 | 7.009017721 | 0.263277266 |
| Tmprss2 | 0.798754509 | -1.014353378 | 1.230099108 |
| Ripk4 | 12.75005239 | 0.493792626 | 0.29994491 |
| Prdm15 | 265.211607 | -0.62830003 | 0.148558054 |
| C2cd2 | 212.6215641 | -0.039239386 | 0.095352544 |
| mmu-mir-6964 | 0.047948698 | -0.243569582 | 3.724945049 |
| Zbtb21 | 472.0135521 | 0.346131439 | 0.092067744 |
| C030010L15Rik | 1.895179359 | -0.17186375 | 0.813596987 |
| B230307C23Rik | 102.4124531 | 0.848704709 | 0.135484494 |
| Gm10232 | 17.35894481 | 0.554109628 | 0.303389154 |
| Gm16569 | 0 | NA | NA |
| Gm5479 | 0 | NA | NA |
| Gm25909 | 0.276643538 | -0.562235651 | 2.077652072 |
| Pisd-ps2 | 107.0691569 | -0.227219155 | 0.127249427 |
| Scaf8 | 1244.518299 | 0.016270585 | 0.09538581 |
| Tiam2 | 517.9732827 | -0.398592022 | 0.12016427 |
| Tfb1m | 51.82495722 | -0.146846953 | 0.174011291 |
| Cldn20 | 1.178313167 | 1.450237015 | 1.151704013 |
| Nox3 | 0.272415186 | 0.237330524 | 1.820908668 |
| Gm23186 | 0 | NA | NA |
| Arid1b | 2127.270207 | -0.139323399 | 0.063731041 |
| Gm15599 | 0.896855871 | -0.586669224 | 1.10323569 |
| Gm29050 | 0.223558305 | -0.222437585 | 2.154258091 |
| Tmem242 | 341.4840666 | 0.176525923 | 0.09734267 |
| Ldhal6b | 8.737325657 | -1.309886155 | 0.47248072 |
| Gm22475 | 0.038981395 | 0.718207094 | 3.724945049 |
| Zdhhc14 | 490.2302194 | 0.061378155 | 0.113553891 |
| 3300005D01Rik | 0.549721405 | 1.622090905 | 1.527605805 |
| Gm10231 | 1.299158781 | -0.216509123 | 1.111168569 |
| Gm26595 | 0.095897396 | -0.724453471 | 3.724945049 |
| Gm26622 | 0 | NA | NA |
| Snx9 | 182.1926714 | 0.630548945 | 0.124857288 |
| Synj2 | 857.4048739 | -0.3664716 | 0.091053935 |
| Serac1 | 260.8621082 | -0.556752087 | 0.121447178 |
| Gtf2h5 | 444.1448567 | -0.203052029 | 0.082962704 |
| Tulp4 | 5049.784843 | -0.120849068 | 0.066722172 |
| Gm5812 | 12.25279831 | 0.52850836 | 0.317201541 |
| Gm22774 | 1.94531079 | 2.300565459 | 0.773580917 |
| Gm15590 | 16.93295172 | 2.380957706 | 0.34350041 |
| n-R5s26 | 7.164646703 | -0.291980722 | 0.401818807 |
| Ppp1r2-ps6 | 4.838948672 | 0.821620894 | 0.476029616 |
| Tmem181a | 366.4664287 | -0.702507256 | 0.126958647 |
| Dynlt1a | 11.06668227 | 0.025635455 | 0.338849865 |
| Gm26848 | 0.173022395 | 0.718215388 | 2.735311368 |
| Ppp1r2-ps1 | 3.995967374 | -0.950849775 | 0.551930957 |

| | | | |
|---|---|---|---|
| Dynlt1b | 39.38951723 | 0.282393299 | 0.20469724 |
| Tmem181b-ps | 797.3566581 | -1.274885997 | 0.258653983 |
| Dynlt1c | 50.2770515 | 0.205320517 | 0.210558085 |
| Tmem181c-ps | 423.9445639 | -1.074409687 | 0.269170496 |
| Dynlt1f | 15.90420978 | 1.231743852 | 0.291475167 |
| Sytl3 | 19.32529137 | 3.959536477 | 0.555714245 |
| Ezr | 1074.367285 | 1.642000282 | 0.130042552 |
| Gm2885 | 11.26381867 | -0.604006449 | 0.319497195 |
| Mir692-1 | 0.048949532 | -0.243569582 | 3.724945049 |
| Gm29205 | 0 | NA | NA |
| Rsph3b | 25.73551809 | 0.384632156 | 0.241676526 |
| Tagap1 | 187.261273 | 0.003657914 | 0.085694056 |
| Rnaset2b | 43.79098034 | 1.486688608 | 0.494084176 |
| Gm1604b | 6.225126559 | -1.244785248 | 0.61450111 |
| Gm25119 | 0 | NA | NA |
| Rps6ka2 | 480.6686781 | -0.561595998 | 0.09258996 |
| Tcp10a | 0.109727584 | 0.718200421 | 3.724945049 |
| Ttll2 | 0.198340327 | 0.742299072 | 2.231534896 |
| Gm9992 | 2.241760946 | 3.589458762 | 0.987964792 |
| Gm26057 | 0 | NA | NA |
| Gm25872 | 0 | NA | NA |
| Fndc1 | 124.9694906 | -0.461294562 | 0.172237841 |
| E430024P14Rik | 37.33337678 | -0.877327142 | 0.226653752 |
| Tagap | 96.63924738 | 2.107871014 | 0.164203848 |
| Rsph3a | 115.5642771 | 0.332630658 | 0.100093414 |
| Gm1604a | 4.428521402 | -1.478468436 | 0.605735454 |
| Gm22416 | 0 | NA | NA |
| Rnaset2a | 127.2262442 | 1.2511678 | 0.440470946 |
| Fgfr1op | 286.1447221 | -0.012880817 | 0.101367287 |
| Ccr6 | 7.182727363 | -3.803896468 | 0.69046434 |
| Mpc1 | 409.8670687 | -0.219114087 | 0.067259631 |
| 4930506C21Rik | 16.13295338 | 0.016341082 | 0.246242808 |
| Sft2d1 | 202.5780256 | 0.264735848 | 0.135043251 |
| Prr18 | 521.3498503 | -0.712117621 | 0.161858207 |
| T2 | 5.357486873 | -0.805716528 | 0.447445208 |
| Gm16702 | 11.86087499 | -0.497867382 | 0.296359054 |
| T | 0.764042974 | 2.062169608 | 1.113712331 |
| Pde10a | 2524.291417 | -1.376588899 | 0.183564975 |
| Gm15425 | 0 | NA | NA |
| Gm17087 | 0.0878635 | -0.72445792 | 3.724945049 |
| Gm15426 | 0.048949532 | -0.243569582 | 3.724945049 |
| 1700010I14Rik | 16.46611213 | 0.936375467 | 0.265266355 |
| 6530411M01Rik | 2.907305987 | -1.619634247 | 0.622510174 |
| Gm8492 | 0.83813809 | -0.071035106 | 1.120354328 |
| Gm17748 | 3.088466909 | 0.127212089 | 0.584332229 |
| Gm17728 | 0 | NA | NA |
| Pabpc6 | 0.088534442 | 1.179809211 | 3.723500796 |

| | | | |
|---|---|---|---|
| Qk | 8468.123674 | -0.178819548 | 0.12083361 |
| 1700110C19Rik | 10.49805674 | -1.015262783 | 0.444490661 |
| Pacrg | 86.69560762 | -0.159710646 | 0.13193328 |
| A230009B12Rik | 11.05179156 | -0.141735657 | 0.351882612 |
| Gm16168 | 29.62534574 | -0.733622551 | 0.320735473 |
| Gm16169 | 3.038906809 | 1.470129218 | 0.680385272 |
| Park2 | 207.7195687 | -1.182586602 | 0.232152513 |
| Gm3145 | 3.839486989 | -0.739176879 | 0.563869012 |
| Gm28505 | 0.112396584 | 1.192083721 | 3.724434897 |
| Gm37402 | 18.41567043 | -1.086645581 | 0.314713116 |
| Gm37332 | 0.321071396 | 2.241026026 | 3.69705936 |
| Gm28118 | 0 | NA | NA |
| Gm10513 | 0.405560335 | 1.382536539 | 1.490489213 |
| Agpat4 | 470.2599501 | 0.26226808 | 0.109777279 |
| Tdgf1-ps2 | 2.738925537 | -0.723969685 | 0.638289382 |
| Map3k4 | 824.0897494 | -0.259964553 | 0.059808799 |
| Gm24794 | 0 | NA | NA |
| 4732491K20Rik | 48.6473752 | -0.353260764 | 0.176772163 |
| Plg | 1.917939655 | 0.882998036 | 0.753022021 |
| Slc22a3 | 26.84679186 | -1.633484075 | 0.2274525 |
| Slc22a2 | 15.39931173 | 0.483499682 | 0.32718748 |
| Slc22a1 | 2.772496349 | 0.546033528 | 0.646561567 |
| Igf2r | 1261.515538 | 0.39817877 | 0.065527131 |
| Airn | 54.16339736 | -1.018100401 | 0.237143583 |
| Gm23833 | 6.456263911 | -1.436133206 | 0.507466276 |
| Gm16574 | 6.618925518 | -1.467067748 | 0.494194446 |
| Gm7162 | 0.709135383 | -2.417751272 | 1.330499215 |
| Mas1 | 77.7093898 | -0.929316678 | 0.210118371 |
| Gm24085 | 0 | NA | NA |
| Mrgprh | 1.689186202 | -0.427233888 | 0.671171162 |
| Pnldc1 | 15.6955681 | -0.768744261 | 0.364089571 |
| Mrpl18 | 276.4221612 | 0.143035904 | 0.091850039 |
| Tcp1 | 1522.478764 | 0.324657037 | 0.06553954 |
| Gm26130 | 0 | NA | NA |
| Snora20 | 1.9528199 | -1.827509428 | 0.801913898 |
| Acat3 | 48.71270492 | -0.076954802 | 0.191819102 |
| Acat2 | 268.88389 | -0.309799403 | 0.085963611 |
| Gm15946 | 1.917568322 | -0.281033775 | 0.664457684 |
| Wtap | 592.5866412 | 0.200298191 | 0.074007474 |
| Sod2 | 2289.929684 | -0.233765328 | 0.066970559 |
| Gpr31b | 8.443869178 | 6.569402513 | 0.901140011 |
| Tcp10b | 7.010379163 | 0.147882537 | 0.620840605 |
| Unc93a | 6.630768804 | 4.241335632 | 0.703509087 |
| Gm23050 | 0 | NA | NA |
| Smok2a | 0.046345709 | 0.71820487 | 3.724945049 |
| Smok2b | 0 | NA | NA |
| Gm26500 | 0.250452015 | 0.725943726 | 2.083611474 |

| | | | |
|---|---|---|---|
| Tcp10c | 0.224267024 | 0.726209466 | 2.151436481 |
| Gm8597 | 4.592470227 | 0.887306297 | 0.550935685 |
| Gm25730 | 0.075043205 | 0.237309859 | 3.724945049 |
| Rnu6 | 0.503427435 | -0.668299511 | 1.50312522 |
| Tcte2 | 120.1207726 | -0.785064247 | 0.209498613 |
| Gm16050 | 0.048949532 | -0.243569582 | 3.724945049 |
| Gm8603 | 0 | NA | NA |
| Smok4a | 0 | NA | NA |
| Gm26375 | 1.918309794 | -0.931467176 | 0.747988161 |
| Gm24867 | 0 | NA | NA |
| Gm16046 | 10.12097898 | -0.795752343 | 0.493406216 |
| Gm16052 | 12.43848109 | -0.890165678 | 0.298313881 |
| Gm16049 | 10.2923578 | -0.716433954 | 0.340442354 |
| Mllt4 | 2109.491735 | -0.191821806 | 0.082126933 |
| Gm3355 | 0.441163179 | -0.174639713 | 1.415947888 |
| Gm7168 | 0.355971994 | 2.58972749 | 1.858095125 |
| Gm7356 | 0 | NA | NA |
| Gm7358 | 0 | NA | NA |
| 4930488N24Rik | 0.072025462 | 0.237312084 | 3.724945049 |
| Dact2 | 350.5464997 | -0.580905663 | 0.139589727 |
| Smoc2 | 139.6069329 | 0.915899024 | 0.211975841 |
| Thbs2 | 224.1080342 | 1.165853986 | 0.181412637 |
| Gm3222 | 8.619157282 | -0.342977058 | 0.34791205 |
| Gm23352 | 0 | NA | NA |
| Wdr27 | 15.49212616 | -0.717722992 | 0.325174207 |
| Gm7393 | 0 | NA | NA |
| 1600012H06Rik | 204.8958293 | 0.363315551 | 0.094842861 |
| Phf10 | 695.9833845 | -0.004426518 | 0.076977456 |
| Gm3448 | 0.554247143 | 1.715262461 | 1.352237167 |
| Ermard | 167.2744434 | 1.375283989 | 0.151221196 |
| Tcte3 | 0 | NA | NA |
| Gm28869 | 9.40787293 | 2.401094179 | 0.430126561 |
| Gm28873 | 3.819986632 | 0.586261796 | 0.526712486 |
| Gm5091 | 0 | NA | NA |
| Dll1 | 136.4076956 | 0.627373013 | 0.146883896 |
| A930024N18Rik | 109.5960546 | 0.628460585 | 0.158876372 |
| Fam120b | 782.860338 | -0.233604298 | 0.090945473 |
| Psmb1 | 1128.534319 | 0.600774298 | 0.119140361 |
| Tbp | 458.9414412 | -0.246449941 | 0.131074435 |
| Pdcd2 | 107.0321996 | 0.534461844 | 0.137266497 |
| Prdm9 | 37.41141876 | -0.339673238 | 0.17996908 |
| Gm6686 | 0 | NA | NA |
| Chd1 | 779.4457597 | 0.219394792 | 0.091138614 |
| Rgmb | 695.6204429 | -0.129245616 | 0.076411545 |
| Gm25598 | 0 | NA | NA |
| BC002059 | 100.3680048 | 0.117195408 | 0.147910498 |
| Zfp960 | 12.66666622 | -0.509817862 | 0.443531116 |

| | | | |
|---|---|---|---|
| Zfp97 | 21.19205506 | 0.365523705 | 0.259682368 |
| Gm6712 | 39.55212394 | 0.099062235 | 0.191381802 |
| Oaz1-ps | 23.5739542 | 0.059650594 | 0.243384589 |
| Riok2 | 310.94823 | 0.434257627 | 0.076170771 |
| Gm26873 | 0.801515891 | -0.243198837 | 1.24956599 |
| Lix1 | 332.7540407 | -1.353175484 | 0.163500989 |
| Lnpep | 2780.673188 | 0.202554682 | 0.056160098 |
| Vmn2r90 | 0.038981395 | 0.718207094 | 3.724945049 |
| Gm23837 | 0 NA | NA | |
| Mir99b | 6.303641128 | -1.601765509 | 0.454737286 |
| Gm25927 | 0.066050875 | 0.71820487 | 3.724945049 |
| mmu-let-7e | 6.339667104 | -1.058331488 | 0.426794642 |
| Spaca6 | 577.2041235 | -0.585651458 | 0.291429839 |
| Mir125a | 0 NA | NA | |
| Has1 | 11.28386909 | 0.542174777 | 0.331777277 |
| Fpr1 | 8.843961322 | 7.002460397 | 0.915621013 |
| Fpr2 | 37.5235979 | 3.80379173 | 0.457579156 |
| Fpr3 | 0.406865304 | 2.966020135 | 1.500725672 |
| Gm7535 | 0.668988238 | 0.555071109 | 1.151057877 |
| Fpr-rs4 | 0.097139547 | 0.718207094 | 3.724945049 |
| Vmn2r124 | 0.527106987 | 2.43743869 | 1.392641206 |
| Vmn2r91 | 0.129871752 | 1.198487306 | 2.991078732 |
| Vmn2r92 | 0.150961315 | 1.165658083 | 2.845512047 |
| Vmn2r94 | 0 NA | NA | |
| Vmn2r93 | 0 NA | NA | |
| Vmn2r95 | 0.066050875 | 0.71820487 | 3.724945049 |
| Vmn2r-ps114 | 0 NA | NA | |
| Vmn2r96 | 0 NA | NA | |
| Vmn2r-ps115 | 0 NA | NA | |
| Vmn2r-ps116 | 0 NA | NA | |
| Vmn2r-ps117 | 0 NA | NA | |
| Vmn2r97 | 0 NA | NA | |
| Vmn2r-ps118 | 0 NA | NA | |
| Vmn2r98 | 0 NA | NA | |
| Vmn2r-ps119 | 0 NA | NA | |
| Vmn2r-ps120 | 0 NA | NA | |
| Vmn2r99 | 0.062426873 | 0.237312084 | 3.724945049 |
| Vmn2r-ps121 | 0 NA | NA | |
| Vmn2r100 | 0 NA | NA | |
| Vmn2r101 | 0 NA | NA | |
| Vmn2r102 | 0 NA | NA | |
| Vmn2r-ps122 | 0 NA | NA | |
| Vmn2r103 | 0 NA | NA | |
| Vmn2r-ps123 | 0 NA | NA | |
| Vmn2r-ps124 | 0 NA | NA | |
| Vmn2r-ps125 | 0 NA | NA | |
| Vmn2r104 | 0 NA | NA | |

| | | | |
|---|---|---|---|
| Vmn2r-ps126 | 0 | NA | NA |
| Fpr-rs7 | 0 | NA | NA |
| Fpr-rs6 | 0 | NA | NA |
| Vmn2r105 | 0 | NA | NA |
| Vmn2r106 | 0 | NA | NA |
| Vmn2r-ps127 | 0 | NA | NA |
| Vmn2r107 | 0 | NA | NA |
| Vmn1r-ps141 | 0 | NA | NA |
| Vmn1r224 | 0 | NA | NA |
| Vmn1r-ps142 | 0 | NA | NA |
| Vmn1r-ps143 | 0 | NA | NA |
| Vmn2r108 | 0 | NA | NA |
| Vmn1r225 | 0 | NA | NA |
| Vmn2r109 | 0.225617294 | 1.251911154 | 2.520825197 |
| Gm5145 | 2.222558862 | 0.532201719 | 0.658702385 |
| Vmn2r110 | 0.133946075 | 0.7191812 | 3.626570343 |
| Fpr-rs3 | 0 | NA | NA |
| Vmn1r226 | 0 | NA | NA |
| Vmn1r227 | 0 | NA | NA |
| Vmn1r228 | 0.088534442 | 1.179809211 | 3.723500796 |
| Vmn1r229 | 0 | NA | NA |
| Vmn1r230 | 0 | NA | NA |
| Vmn1r231 | 0.108767819 | 0.237318756 | 3.724945049 |
| Vmn1r232 | 0.192963196 | 0.981890925 | 2.637065715 |
| Vmn1r-ps144 | 0 | NA | NA |
| Ppp2r1a | 4099.317594 | -0.070519335 | 0.060599063 |
| Vmn1r-ps145 | 0 | NA | NA |
| Vmn1r233 | 0 | NA | NA |
| Zfp160 | 354.561668 | 0.296255367 | 0.083716729 |
| Gm5681 | 0 | NA | NA |
| Vmn1r-ps146 | 0 | NA | NA |
| Gm5092 | 0.092140247 | -0.724455696 | 3.724945049 |
| Gm6811 | 0.490429956 | 1.295821939 | 1.312554123 |
| Gm7732 | 0.049553047 | 0.71820487 | 3.724945049 |
| Gm7736 | 0 | NA | NA |
| Gm7740 | 0 | NA | NA |
| Gm7742 | 0 | NA | NA |
| Vmn1r-ps147 | 0 | NA | NA |
| Vmn1r234 | 0 | NA | NA |
| Vmn1r-ps148 | 0 | NA | NA |
| Vmn1r235 | 0.11193333 | 0.718202646 | 3.724945049 |
| Vmn1r-ps149 | 0 | NA | NA |
| Gm26384 | 1.87298797 | 1.339099012 | 0.662595661 |
| Vmn1r236 | 0 | NA | NA |
| Vmn1r237 | 0.097139547 | 0.718207094 | 3.724945049 |
| Vmn1r-ps150 | 0 | NA | NA |
| Zfp677 | 55.30162814 | 0.826920445 | 0.165805046 |

| | | | |
|---|---|---|---|
| Zfp54 | 27.41881594 | 0.795781877 | 0.257613828 |
| Zfp51 | 105.8080167 | 0.855152593 | 0.131372971 |
| Gm26753 | 1.899580325 | 0.522381102 | 0.743311015 |
| Zfp53 | 100.822271 | 0.369055502 | 0.139010426 |
| 9330136K24Rik | 11.4954026 | -1.019811111 | 0.43312134 |
| Zfp52 | 67.11583658 | 0.21241726 | 0.171846651 |
| Zfp948 | 235.9906384 | 0.122245293 | 0.121405012 |
| 3110052M02Rik | 172.7372577 | -0.013074861 | 0.104744891 |
| Gm10509 | 18.52972673 | 0.380451039 | 0.247629099 |
| Gm10226 | 26.0094674 | 0.378532002 | 0.219473537 |
| Zfp760 | 279.5658373 | -0.1263807 | 0.148372178 |
| Zfp229 | 105.1619278 | 0.243178212 | 0.178629373 |
| 4930515G13Rik | 0 | NA | NA |
| Gm9818 | 0.049553047 | 0.71820487 | 3.724945049 |
| Zfp820 | 16.99717544 | 0.655280094 | 0.274559411 |
| 2210404O09Rik | 62.10937004 | 0.287316704 | 0.145099206 |
| Zfp942 | 313.3301051 | -0.081709149 | 0.097076006 |
| Zfp943 | 306.7466701 | -0.017910436 | 0.100734839 |
| Gm7805 | 0 | NA | NA |
| Gm7809 | 5.578788543 | 1.466034318 | 0.456392202 |
| Gm9772 | 0 | NA | NA |
| Gm7052 | 0.038981395 | 0.718207094 | 3.724945049 |
| Zfp947 | 19.48152888 | 0.52359799 | 0.249140065 |
| Gm4944 | 183.4875709 | 0.479513928 | 0.11602793 |
| Zfp944 | 179.0922888 | 0.112518272 | 0.102198528 |
| Zfp758 | 147.1029375 | 0.485149149 | 0.104215098 |
| Zfp946 | 267.7372296 | 0.11627712 | 0.093983211 |
| Vmn2r111 | 0.571153248 | 3.311501009 | 1.49037385 |
| Vmn2r112 | 0 | NA | NA |
| Gm9805 | 0.259702268 | 0.740457362 | 2.063779476 |
| Gm5224 | 0 | NA | NA |
| Gm5493 | 1.253500652 | 0.010894712 | 0.8269982 |
| Gm16386 | 36.63223039 | -0.327811885 | 0.209801855 |
| Zfp945 | 660.5015324 | 1.27391491 | 0.120951109 |
| Vmn2r-ps128 | 0 | NA | NA |
| Vmn2r113 | 0.044544648 | 0.237309859 | 3.724945049 |
| Vmn2r-ps129 | 0 | NA | NA |
| Gm18415 | 0 | NA | NA |
| Gm18396 | 0 | NA | NA |
| Vmn2r-ps130 | 0 | NA | NA |
| Gm18397 | 0 | NA | NA |
| Gm18416 | 0 | NA | NA |
| Vmn2r-ps131 | 0 | NA | NA |
| Vmn2r-ps132 | 0.415262948 | 0.199882788 | 1.363218369 |
| Zfp40 | 220.4414794 | -0.15242776 | 0.130079818 |
| Vmn2r-ps133 | 0 | NA | NA |
| Vmn2r114 | 0 | NA | NA |

| | | | |
|---|---|---|---|
| Vmn2r115 | 0 | NA | NA |
| Vmn2r116 | 0 | NA | NA |
| Vmn2r-ps134 | 0 | NA | NA |
| Vmn2r117 | 0.156477978 | 0.718214073 | 2.823333319 |
| Casp16 | 0.468340293 | -0.162921828 | 1.387819537 |
| Gm26695 | 3.060190583 | 0.078019901 | 0.708512657 |
| Zfp213 | 78.04727618 | 0.484727206 | 0.138077608 |
| Zfp13 | 172.7206419 | -0.141760705 | 0.105105692 |
| Zscan10 | 0.304370145 | 0.237299194 | 3.724945049 |
| Mmp25 | 15.55909309 | 4.86865216 | 0.622568995 |
| Ccdc64b | 0.263919366 | 0.718222839 | 2.067926716 |
| Thoc6 | 57.82215382 | 0.96176192 | 0.176528542 |
| Hcfc1r1 | 508.7659468 | -0.13026971 | 0.097071645 |
| Tnfrsf12a | 409.2070675 | 3.618699618 | 0.393688325 |
| Cldn6 | 0.270371171 | -0.243563199 | 1.81628852 |
| Cldn9 | 1.372057613 | 0.589270471 | 0.809127167 |
| 1520401A03Rik | 0.641047896 | -1.222220736 | 1.352755459 |
| Pkmyt1 | 17.10398538 | 0.32149979 | 0.403990253 |
| Paqr4 | 569.8115097 | -0.236505217 | 0.099846588 |
| Kremen2 | 6.737691786 | -0.014122519 | 0.420389678 |
| 9530082P21Rik | 164.7586404 | 4.233539264 | 0.248485134 |
| Flywch1 | 2434.68631 | -0.054523897 | 0.059201362 |
| Gm16275 | 0.151377979 | 1.628752834 | 2.835614948 |
| Flywch2 | 102.393296 | -0.610694704 | 0.140802064 |
| D930048G16Rik | 12.89624846 | -1.531771908 | 0.345166064 |
| Srrm2 | 8917.247412 | -0.122774304 | 0.097965282 |
| Mir5125 | 32.86136734 | -1.356968241 | 0.354646143 |
| Tceb2 | 859.0037201 | 0.096046906 | 0.098333059 |
| Prss33 | 1.317762752 | -2.086207623 | 1.184447018 |
| Prss41 | 25.3916789 | -0.288182769 | 0.210384646 |
| Prss32 | 1.380959146 | -0.683974236 | 0.874739107 |
| Gm15947 | 0.088534442 | 1.179809211 | 3.723500796 |
| Prss21 | 0 | NA | NA |
| Dcpp1 | 0 | NA | NA |
| Dcpp2 | 0 | NA | NA |
| Dcpp3 | 0 | NA | NA |
| Sbp | 0 | NA | NA |
| Sbpl | 0 | NA | NA |
| Prss30 | 3.148853658 | 0.797041676 | 0.488270769 |
| Prss22 | 4.912676513 | 1.117292742 | 0.429875328 |
| Gm27700 | 0 | NA | NA |
| Gm25092 | 0 | NA | NA |
| Gm5225 | 0 | NA | NA |
| Prss27 | 0.22024148 | 1.26320404 | 2.1616408 |
| Kctd5 | 261.9043685 | 0.670128671 | 0.110617375 |
| Pdpk1 | 2152.703641 | -0.236024875 | 0.072885496 |
| Amdhd2 | 151.6535398 | 0.805024754 | 0.157924674 |

| | | | |
|---|---|---|---|
| RP23-27H19.7 | 0 | NA | NA |
| Atp6v0c | 323.8959897 | -0.132212988 | 0.128419356 |
| Tbc1d24 | 880.7733529 | -0.463299003 | 0.110673017 |
| BC028777 | 3.029716819 | -0.39039299 | 0.560361496 |
| 1600002H07Rik | 75.97695628 | 0.30975734 | 0.163403196 |
| Ccnf | 89.28741565 | 1.00612285 | 0.177916696 |
| Mir5134 | 0.102891481 | -0.243576254 | 3.724945049 |
| Mir6965 | 0.049553047 | 0.71820487 | 3.724945049 |
| Abca17 | 4.624081096 | 0.016154741 | 0.478559629 |
| Gm24427 | 0 | NA | NA |
| Abca3 | 1345.901901 | -0.205063075 | 0.06608133 |
| Gm25618 | 0 | NA | NA |
| D330041H03Rik | 54.50038671 | -1.193870567 | 0.212400455 |
| Rnps1 | 386.2518119 | 0.110764901 | 0.064755297 |
| Eci1 | 99.86578218 | 0.064974439 | 0.1714173 |
| Dnase1l2 | 15.71919709 | -1.192340738 | 0.555009808 |
| E4f1 | 179.8701777 | -0.071014535 | 0.102386039 |
| Pgp | 366.4948428 | -0.059094748 | 0.120188863 |
| Mlst8 | 286.6817983 | 0.043027086 | 0.069088152 |
| Bricd5 | 7.409630445 | -1.115805202 | 0.387182532 |
| Caskin1 | 1583.967537 | -0.905955173 | 0.070751772 |
| Traf7 | 402.3587103 | 0.199783936 | 0.066783856 |
| Rab26 | 113.4819396 | -1.264327752 | 0.168409026 |
| Rab26os | 18.9116278 | -0.054187829 | 0.273037563 |
| Snord60 | 0 | NA | NA |
| Pkd1 | 2223.361591 | -0.509035897 | 0.139652253 |
| Gm24589 | 0 | NA | NA |
| Tsc2 | 1269.544961 | -0.036720983 | 0.048605088 |
| Nthl1 | 19.09256242 | 0.603443258 | 0.285172586 |
| Slc9a3r2 | 379.4006015 | 0.305071201 | 0.079276423 |
| Npw | 2.00579695 | 2.815633809 | 0.872723786 |
| Zfp598 | 484.0735371 | 0.259010346 | 0.112629372 |
| Syngr3 | 1091.758285 | -0.474395854 | 0.063259692 |
| Gfer | 118.1883123 | 0.712807647 | 0.132090915 |
| Noxo1 | 16.22553485 | -0.702933689 | 0.271800474 |
| Tbl3 | 217.5084557 | 0.414838933 | 0.104540424 |
| Rnf151 | 7.038343901 | -0.025271436 | 0.362428416 |
| Rps2 | 374.06437 | 0.493288245 | 0.134973311 |
| Snhg9 | 0.044191549 | -0.243571806 | 3.724945049 |
| Snora78 | 10.19340988 | 0.335566325 | 0.410834006 |
| Snora64 | 0 | NA | NA |
| Ndufb10 | 578.8159123 | -0.163875442 | 0.100823994 |
| Rpl3l | 1.739503778 | -0.347637274 | 0.808647324 |
| Msrb1 | 303.5260477 | 2.222210965 | 0.197450782 |
| Hs3st6 | 1.483788266 | -1.210344848 | 0.996388177 |
| Meiob | 0.700302977 | 2.382982285 | 1.196324634 |
| Hagh | 454.8144511 | -0.050967759 | 0.08081781 |

| | | | |
|---|---|---|---|
| Fahd1 | 88.20798843 | -0.194061193 | 0.139659161 |
| Igfals | 0.769314682 | -0.460358565 | 3.657516129 |
| Nubp2 | 229.2468458 | 0.89929103 | 0.13159204 |
| Spsb3 | 316.2208862 | 0.445114459 | 0.112167043 |
| Eme2 | 125.0526026 | 0.224318446 | 0.132456493 |
| Mapk8ip3 | 5120.798928 | -0.148353473 | 0.049945111 |
| Mrps34 | 292.3664307 | 0.424480586 | 0.103216789 |
| Nme3 | 0.049553047 | 0.71820487 | 3.724945049 |
| Hn1l | 155.596658 | 1.98797207 | 0.197594535 |
| Cramp1l | 709.8548994 | -0.237386158 | 0.100047227 |
| Ift140 | 319.2473791 | -0.603830554 | 0.128008439 |
| Tmem204 | 25.86076114 | -0.438786353 | 0.222045228 |
| Telo2 | 173.173594 | -0.27416871 | 0.120495306 |
| Ptx4 | 2.281795086 | -0.642077138 | 0.793986993 |
| Clcn7 | 760.3513654 | 0.794384582 | 0.100376769 |
| Ccdc154 | 1.912657768 | 1.233151918 | 0.707388109 |
| BC003965 | 211.743186 | -0.151459926 | 0.107860148 |
| Unkl | 277.8387526 | -0.29080246 | 0.126669884 |
| Gnptg | 438.2214292 | -0.04716913 | 0.072452902 |
| Tsr3 | 147.8211637 | 0.513509323 | 0.131906259 |
| Baiap3 | 567.9275415 | 0.13452841 | 0.160102354 |
| Mir3547 | 0 | NA | NA |
| Ube2i | 715.8321189 | 0.119980255 | 0.106841411 |
| Prss34 | 0.061920613 | 0.718207094 | 3.724945049 |
| Gm25921 | 0 | NA | NA |
| Prss28 | 0 | NA | NA |
| Prss29 | 0 | NA | NA |
| Gm5430 | 0.892004414 | -0.330941467 | 1.075625002 |
| Tpsab1 | 3.207720559 | 0.965346948 | 0.554317613 |
| Tpsb2 | 1.275377428 | -0.635519394 | 0.857679754 |
| Tpsg1 | 0.042716945 | -0.243571806 | 3.724945049 |
| Cacna1h | 727.8784213 | -0.472682199 | 0.129354475 |
| Tekt4 | 20.49047743 | 0.346323421 | 0.38283129 |
| Sstr5 | 11.97499174 | -0.581421163 | 0.319387008 |
| Sox8 | 497.4054169 | -0.910265226 | 0.116770255 |
| 2810468N07Rik | 100.8375544 | -1.184102431 | 0.193087121 |
| Lmf1 | 167.6280357 | -0.32032852 | 0.136979955 |
| Gm22645 | 0 | NA | NA |
| Gng13 | 254.443737 | -0.409828854 | 0.202718001 |
| Chtf18 | 22.59824114 | 0.642740546 | 0.331369348 |
| Rpusd1 | 275.4402358 | -0.473775439 | 0.087741194 |
| Mslnl | 3.752562866 | 0.224848478 | 0.504673528 |
| Gm25083 | 0 | NA | NA |
| Msln | 26.85201184 | 2.434399497 | 0.348434608 |
| Narfl | 208.0663513 | 0.213416037 | 0.076808822 |
| Haghl | 395.0534574 | -0.895393488 | 0.099370583 |
| Mir6966 | 0 | NA | NA |

| | | | |
|---|---|---|---|
| Ccdc78 | 1.842483344 | -0.656594036 | 0.698226427 |
| Fam173a | 248.2616466 | -0.27291777 | 0.121316666 |
| Metrn | 475.4136771 | 0.022196667 | 0.259757647 |
| Gm26694 | 2.65610448 | 2.017310596 | 0.727942827 |
| Fbxl16 | 5262.479825 | -1.327886647 | 0.127914641 |
| Wdr24 | 242.1513502 | 0.206522634 | 0.079208091 |
| Jmjd8 | 285.0629585 | -0.156293271 | 0.088959764 |
| Stub1 | 735.6538842 | -0.025841347 | 0.089799016 |
| Mir3082 | 1.578690045 | -0.82409368 | 0.816748422 |
| Rhbdl1 | 30.55610411 | -1.0955783 | 0.208047862 |
| Gm20683 | 0 | NA | NA |
| Rhot2 | 307.1541338 | -0.280840138 | 0.087790608 |
| Wdr90 | 125.7964112 | -0.759813428 | 0.195618863 |
| Fam195a | 55.95550801 | 0.041834224 | 0.155634364 |
| 0610011F06Rik | 119.2220752 | -0.558866358 | 0.120747514 |
| Wfikkn1 | 12.97584509 | -0.628845699 | 0.624038018 |
| Rab40c | 356.9302055 | -0.250897285 | 0.107788378 |
| Gm10012 | 55.0896093 | -0.317067385 | 0.202389514 |
| Gm16278 | 14.17840066 | 0.07127782 | 0.321960299 |
| Pigq | 882.6485576 | -0.036719706 | 0.100869473 |
| Nhlrc4 | 4.440385187 | -0.805510329 | 0.480502785 |
| A930017K11Rik | 39.98571024 | -0.369551854 | 0.223225704 |
| D630044L22Rik | 2.092098959 | -1.226962528 | 0.80861814 |
| Capn15 | 519.3883034 | -0.321355188 | 0.100637123 |
| 1700022N22Rik | 1.65831888 | 1.5326204 | 0.791012654 |
| Rab11fip3 | 1324.716031 | -0.557113283 | 0.082591887 |
| Decr2 | 128.9572076 | -0.551014854 | 0.118699295 |
| Nme4 | 11.32066076 | -0.312169249 | 0.319531552 |
| Gm8186 | 9.968334961 | 0.381408598 | 0.35915372 |
| Tmem8 | 155.1022499 | 0.046422821 | 0.105620356 |
| Mrpl28 | 258.4561109 | 0.177429743 | 0.099855515 |
| Axin1 | 402.0340246 | 0.423583041 | 0.089268647 |
| Pdia2 | 0.090277459 | 0.237318756 | 3.724945049 |
| Arhgdig | 330.2245481 | -0.544564775 | 0.098325946 |
| Rgs11 | 126.1618791 | -1.318226052 | 0.290444512 |
| Itfg3 | 218.9037408 | 0.307275265 | 0.138326965 |
| Gm23123 | 4.679423458 | 1.14350126 | 0.605758452 |
| Luc7l | 1216.152592 | -0.068027861 | 0.070926069 |
| Gm16580 | 14.51804168 | 0.099760136 | 0.262318183 |
| Hba-ps4 | 0.121388914 | 0.718200421 | 3.724945049 |
| Neurl1b | 522.8334351 | -1.88414847 | 0.176358446 |
| Dusp1 | 584.3483468 | 0.493086819 | 0.313880107 |
| Gm8225 | 0.51166646 | 0.414798795 | 1.261061387 |
| 1700049J03Rik | 1.147468153 | -2.313277627 | 1.050983923 |
| Ergic1 | 1341.076534 | 0.028240785 | 0.054611748 |
| Gm22420 | 0 | NA | NA |
| Gm17218 | 0 | NA | NA |

| | | | |
|---|---|---|---|
| Atp6v0e | 321.5954019 | 0.812926706 | 0.128822577 |
| Crebrf | 2489.558306 | 0.157567949 | 0.091399578 |
| Bnip1 | 161.7726201 | -0.006552196 | 0.157707044 |
| Nkx2-5 | 0 | NA | NA |
| Gm17382 | 7.241949452 | 0.139419112 | 0.414729125 |
| Gm20468 | 0.607877193 | -0.15170055 | 1.268461368 |
| Kifc5b | 20.39272261 | 0.07735873 | 0.267255855 |
| Gm27775 | 0 | NA | NA |
| Phf1 | 420.9294313 | -0.066246936 | 0.084274801 |
| Mir6968 | 0.061920613 | 0.718207094 | 3.724945049 |
| Cuta | 316.9454618 | 0.265431303 | 0.107031305 |
| Syngap1 | 4379.193043 | -1.016434595 | 0.135715881 |
| Mir3083 | 0 | NA | NA |
| Zbtb9 | 0 | NA | NA |
| Ggnbp1 | 24.84235746 | -1.209530524 | 0.26112941 |
| Bak1 | 376.0418335 | 2.388064773 | 0.1639567 |
| Itpr3 | 158.6747885 | 1.39281969 | 0.166023277 |
| Mir7677 | 0 | NA | NA |
| Itpr3os | 0 | NA | NA |
| Uqcc2 | 404.6628392 | 0.089557847 | 0.10597893 |
| Ip6k3 | 19.80013244 | -0.515891163 | 0.2972318 |
| Lemd2 | 337.854379 | 0.48212261 | 0.085204982 |
| Gm26724 | 0.046345709 | 0.71820487 | 3.724945049 |
| mmu-mir-7214 | 0.072025462 | 0.237312084 | 3.724945049 |
| Mir7216 | 0 | NA | NA |
| Gm10505 | 1.106753324 | 2.075564093 | 1.039901294 |
| mmu-mir-7217 | 0 | NA | NA |
| mmu-mir-7218 | 0 | NA | NA |
| Grm4 | 1362.166969 | -0.459779127 | 0.169244156 |
| Gm16005 | 0 | NA | NA |
| Gm9703 | 3.905880329 | 0.968805481 | 0.529415402 |
| Hmga1 | 483.005091 | 2.307957643 | 0.263360159 |
| AI413582 | 165.7897589 | -0.149936006 | 0.125902828 |
| Nudt3 | 2683.122826 | -0.091155867 | 0.064921576 |
| Rps10 | 731.2072688 | 0.671276432 | 0.115115434 |
| Gm15420 | 1.006594715 | 0.347266938 | 1.069503098 |
| Pacsin1 | 4703.755317 | -0.491457852 | 0.075036543 |
| Spdef | 9.032787089 | 0.233670179 | 0.599469346 |
| Gm15458 | 0.243671637 | 0.162562113 | 2.123491562 |
| D17Wsu92e | 3051.297663 | 0.286675296 | 0.07808022 |
| Rps2-ps9 | 0.733389001 | 0.513160906 | 1.499447432 |
| Snrpc | 219.5654359 | 0.762303468 | 0.088908972 |
| Uhrf1bp1 | 565.7822905 | 0.03787345 | 0.099195677 |
| Taf11 | 229.7296066 | -0.067044084 | 0.084330866 |
| Anks1 | 569.0372512 | 0.165828714 | 0.064758304 |
| Gm15597 | 0 | NA | NA |
| Gm15598 | 0.962678808 | -0.516803801 | 1.11000521 |

| | | | |
|---|---|---|---|
| Tcp11 | 4.048470876 | 0.20773442 | 0.530500092 |
| 4930526A20Rik | 2.494837672 | -0.046643724 | 0.712990486 |
| Scube3 | 58.8908513 | -1.012934436 | 0.240436551 |
| Zfp523 | 600.1388987 | -0.536093768 | 0.106003697 |
| Def6 | 48.95250913 | 2.214240741 | 0.22285154 |
| Ppard | 357.9241621 | 0.46989712 | 0.070731436 |
| Fance | 139.6977314 | 0.140539411 | 0.110988318 |
| Rpl10a | 628.5301939 | 0.613870736 | 0.092053983 |
| Tead3 | 55.35341921 | 1.412960852 | 0.256560708 |
| Tulp1 | 1.121467505 | 2.331204331 | 1.407798224 |
| Fkbp5 | 903.5710338 | 0.845851785 | 0.172181947 |
| Gm22146 | 0 | NA | NA |
| Gm20109 | 2.702534539 | 0.085133951 | 0.65359491 |
| E230001N04Rik | 14.53897155 | 0.630210897 | 0.292339245 |
| Gm25195 | 0 | NA | NA |
| Armc12 | 0.464894546 | -0.085312082 | 1.502345566 |
| Clpsl2 | 0 | NA | NA |
| Clps | 0.10503227 | 1.179809211 | 3.723500796 |
| Mir6969 | 0 | NA | NA |
| Lhfpl5 | 32.4049498 | -0.141340298 | 0.24339631 |
| Srpk1 | 762.4953308 | -0.243871176 | 0.082831124 |
| Slc26a8 | 26.07395248 | -1.363464672 | 0.241148159 |
| Mapk14 | 951.4501698 | -0.009727034 | 0.062588292 |
| Mapk13 | 6.079458248 | -0.048260198 | 0.446731166 |
| Brpf3 | 576.6212084 | 0.042204193 | 0.080668367 |
| Pnpla1os | 0.18989612 | 1.462508577 | 2.650629334 |
| Pnpla1 | 2.646101737 | 0.328275493 | 0.638087496 |
| 4930539E08Rik | 83.46186464 | 0.265226033 | 0.166696896 |
| 1700030A11Rik | 0.307834097 | 0.752739745 | 1.756072748 |
| Gm16191 | 0.066050875 | 0.71820487 | 3.724945049 |
| Pxt1 | 0.042716945 | -0.243571806 | 3.724945049 |
| Kctd20 | 733.6283113 | 0.000465961 | 0.055654591 |
| Stk38 | 479.5171983 | -0.055581678 | 0.11567052 |
| Gm23887 | 7.551778246 | -1.083879981 | 0.519675888 |
| Gm16196 | 0.628440879 | 0.614256307 | 1.337928366 |
| Gm16195 | 0.038981395 | 0.718207094 | 3.724945049 |
| Srsf3 | 1460.401065 | 0.366455388 | 0.086580922 |
| Trp53cor1 | 8.461310733 | 1.687963151 | 0.437251202 |
| Gm27599 | 0.04393175 | -0.243571806 | 3.724945049 |
| Cdkn1a | 962.23884 | 3.138518076 | 0.358233731 |
| Gm16194 | 0.143847454 | 0.719116779 | 2.898586705 |
| Rab44 | 2.105186647 | 2.807396852 | 0.856670128 |
| Rpl35a-ps3 | 1.427311429 | 1.129608468 | 0.98834656 |
| Cpne5 | 630.1572259 | -0.984259683 | 0.247092099 |
| Ppil1 | 83.28828306 | 0.595443584 | 0.151869619 |
| BC004004 | 500.5361452 | 0.288806653 | 0.06191553 |
| Gm10045 | 2.961333848 | -0.543676139 | 0.650360071 |

| | | | |
|---|---|---|---|
| Pi16 | 15.76972868 | 0.995313831 | 0.271481467 |
| Mtch1 | 2170.561962 | -0.315939734 | 0.065683354 |
| Fgd2 | 41.20869392 | -0.07265902 | 0.312688699 |
| Gm26885 | 6.791476013 | 1.352259182 | 0.464399945 |
| Pim1 | 294.2735208 | 3.293995789 | 0.307187482 |
| Gm17657 | 0 | NA | NA |
| Tmem217 | 0.817974253 | -1.463247136 | 1.450224431 |
| Tbc1d22b | 597.8773199 | -0.183336465 | 0.06378293 |
| Gm28052 | 0 | NA | NA |
| Tbc1d22bos | 0 | NA | NA |
| Mir7667 | 0.28873988 | -1.606105062 | 1.782469188 |
| Rnf8 | 73.9322441 | -0.129477317 | 0.165852225 |
| Gm28043 | 40.56409608 | 0.37276033 | 0.342644027 |
| Cmtr1 | 326.8605501 | 0.234402365 | 0.11915503 |
| Ccdc167 | 100.0491434 | -0.755125848 | 0.131322813 |
| Mdga1 | 427.208838 | -0.753592031 | 0.142925689 |
| Gm16758 | 7.249476259 | -1.305493381 | 0.393677362 |
| Gm22327 | 0 | NA | NA |
| Zfand3 | 884.6551979 | 0.655164842 | 0.083101942 |
| Btbd9 | 1023.020761 | 0.160162627 | 0.059553517 |
| Gm25049 | 0 | NA | NA |
| Gm9874 | 0.097139547 | 0.718207094 | 3.724945049 |
| Glo1 | 537.9219683 | -0.14340056 | 0.092290967 |
| 1700097N02Rik | 0.406736653 | 1.24775055 | 1.548651137 |
| Dnah8 | 41.46003794 | 1.402731683 | 0.25605649 |
| Gm24661 | 0 | NA | NA |
| Glp1r | 10.69878445 | 0.225111028 | 0.292052272 |
| Umodl1 | 10.985597 | -0.375647708 | 0.594420573 |
| Gm24026 | 0 | NA | NA |
| Abcg1 | 1340.501293 | 0.641577349 | 0.076059846 |
| Gm25447 | 2.050495351 | 0.375937217 | 0.78055929 |
| Tff3 | 1.029781442 | -1.516377161 | 1.053206982 |
| Tff2 | 0 | NA | NA |
| Gm15318 | 0.237699072 | 1.742403233 | 2.113982826 |
| Tff1 | 0.320114016 | 1.657179491 | 1.989075608 |
| Tmprss3 | 1.885701583 | -1.692320528 | 0.747868013 |
| Ubash3a | 6.422161135 | 3.132205037 | 0.606599676 |
| Rsph1 | 24.15190014 | 0.04821914 | 0.252382301 |
| Slc37a1 | 77.30229299 | 0.50526606 | 0.135536647 |
| Pde9a | 243.102937 | -0.039391063 | 0.139688652 |
| Wdr4 | 235.3192353 | 0.339161706 | 0.102544421 |
| Ndufv3 | 462.2896515 | -0.107753194 | 0.088729278 |
| Gm38099 | 1.2404959 | -1.070354027 | 0.916017832 |
| 4833413E03Rik | 0.117046891 | -0.724453471 | 3.724945049 |
| Pknox1 | 214.0257755 | 0.206140137 | 0.089237082 |
| Gm24970 | 0.09349418 | -0.24357403 | 3.724945049 |
| Cbs | 185.6562434 | -0.514304776 | 0.155063516 |

| | | | |
|---|---|---|---|
| U2af1 | 321.058565 | 0.65595441 | 0.101555572 |
| Cryaa | 0.372981075 | 0.297660169 | 1.547827606 |
| Sik1 | 432.2003957 | 0.182289223 | 0.188037513 |
| Hsf2bp | 3.787275169 | 0.80441438 | 0.58283108 |
| Gm17572 | 0 | NA | NA |
| Gm25883 | 0 | NA | NA |
| Rrp1b | 198.546731 | -0.131223001 | 0.144997832 |
| Pdxk-ps | 1.082653786 | -1.320426912 | 0.90350668 |
| Gm17276 | 31.48193691 | -0.492775655 | 0.256999548 |
| Notch3 | 246.8071195 | -0.898316889 | 0.199481947 |
| Ephx3 | 2.938467936 | 0.639780973 | 0.600544278 |
| Brd4 | 1978.347052 | 0.083252432 | 0.074537269 |
| Gm26549 | 15.20514642 | -0.773029973 | 0.387166805 |
| Akap8 | 845.3496397 | -0.201720531 | 0.132690979 |
| Akap8l | 594.3929537 | -0.505818698 | 0.138757372 |
| Wiz | 755.6427441 | -0.137150004 | 0.062784901 |
| Gm26858 | 0.274350536 | -0.247235395 | 2.087708211 |
| Gm22201 | 0.044191549 | -0.243571806 | 3.724945049 |
| Rasal3 | 63.09606561 | 2.064307053 | 0.262826965 |
| Pglyrp2 | 9.032459669 | 6.789213631 | 0.905688624 |
| Cyp4f39 | 0 | NA | NA |
| Cyp4f17 | 39.89199084 | 0.144994881 | 0.193650467 |
| Cyp4f16 | 69.22286846 | 1.161675259 | 0.183565623 |
| Cyp4f37 | 0.44264929 | 2.826124122 | 1.367974698 |
| Cyp4f40 | 0.108772582 | 0.718202646 | 3.724945049 |
| Cyp4f15 | 57.06117773 | -2.307745105 | 0.238249241 |
| Zfp871 | 1423.833684 | -0.247308587 | 0.060450898 |
| Gm17115 | 5.126403706 | -0.864792308 | 0.472955807 |
| Zfp811 | 109.0107342 | -0.40474847 | 0.166725306 |
| Gm23294 | 1.960852514 | -0.834599593 | 0.718673226 |
| Gm25555 | 1.60505521 | -2.749709078 | 1.041228138 |
| Gm26693 | 0.210884633 | 1.190411796 | 2.175374416 |
| Zfp799 | 476.0807173 | -0.462561907 | 0.094801811 |
| Zfp870 | 141.8172216 | -0.189295765 | 0.092937834 |
| Cyp4f14 | 50.57166942 | -0.770102726 | 0.199556124 |
| Cyp4f13 | 93.77020389 | 0.108892788 | 0.150603372 |
| Cyp4f41-ps | 1.046059986 | 0.454191475 | 1.060116917 |
| Gm25667 | 0.310541583 | 0.237309481 | 3.724945049 |
| Zfp472 | 71.14659937 | 0.265962352 | 0.14761868 |
| Zfp952 | 219.1287848 | 0.000435389 | 0.088567722 |
| Zfp763 | 281.1564641 | -0.299328073 | 0.098423562 |
| 4921501E09Rik | 0.687686063 | 0.157252372 | 1.173446267 |
| Zfp563 | 131.0379676 | -0.386634767 | 0.12343172 |
| Morc2b | 49.27469275 | -0.948389259 | 0.361705344 |
| Olfr55 | 0.072025462 | 0.237312084 | 3.724945049 |
| Olfr239 | 0.04393175 | -0.243571806 | 3.724945049 |
| Gm4461 | 9.889894451 | -2.42853178 | 0.417162072 |

| | | | |
|---|---|---|---|
| Zfp955a | 185.3926737 | -0.295631608 | 0.130746567 |
| Olfr63 | 0.330364319 | 1.531636342 | 1.706787558 |
| Zfp955b | 228.1232927 | -0.261697944 | 0.086812756 |
| Zfp422-rs1 | 1.757111958 | -0.453277283 | 0.797309153 |
| Zfp81 | 498.1008011 | -0.205250354 | 0.08495 |
| Zfp101 | 191.0430263 | -0.579216758 | 0.10113116 |
| CT030734.1 | 0 | NA | NA |
| Actl9 | 0 | NA | NA |
| Adamts10 | 190.1840441 | -0.541357444 | 0.146686205 |
| Myo1f | 224.7996339 | 3.738928188 | 0.224511505 |
| Zfp414 | 163.0997787 | 0.133729407 | 0.10417914 |
| Gm20507 | 3.129648303 | -0.983358833 | 0.685939676 |
| Pram1 | 0.821047175 | 0.295294443 | 1.059980888 |
| Hnrnpm | 845.7194937 | 0.265293315 | 0.145225029 |
| 2-Mar | 559.2133353 | -0.212170012 | 0.062841822 |
| Gm22943 | 0 | NA | NA |
| Rab11b | 1217.267332 | -0.149798299 | 0.076446313 |
| Gm17251 | 14.95472732 | 0.352938914 | 0.270819863 |
| Angptl4 | 147.9950521 | 3.744746832 | 0.440642903 |
| 4931413I07Rik | 0.260232477 | 1.495587004 | 2.451991008 |
| Kank3 | 84.23068527 | -0.291506402 | 0.194866427 |
| Rps28 | 433.7597313 | 0.469183148 | 0.169175959 |
| Ndufa7 | 437.6705464 | 0.058547679 | 0.096104512 |
| Cd320 | 102.4821458 | -0.215871161 | 0.128783978 |
| Gm4134 | 0.061920613 | 0.718207094 | 3.724945049 |
| Gm7029 | 1.064012846 | 1.059918251 | 1.05448116 |
| Kifc1 | 18.16257989 | 0.114317125 | 0.256546402 |
| Gm27740 | 0 | NA | NA |
| Platr17 | 2.945475634 | 3.31275879 | 0.929975431 |
| BC051226 | 10.35737011 | -0.178023007 | 0.366153322 |
| Daxx | 959.7967857 | 2.744630039 | 0.175383586 |
| Tapbp | 4153.144377 | 4.356211681 | 0.161563576 |
| Zbtb22 | 294.4051857 | -0.096939187 | 0.078737083 |
| Gm19412 | 30.80291434 | 1.192840576 | 0.238858415 |
| Rgl2 | 188.6076046 | -0.197768096 | 0.131373042 |
| Pfdn6 | 222.2854681 | -0.001618635 | 0.14049652 |
| Wdr46 | 114.1398418 | 1.072173287 | 0.146333281 |
| B3galt4 | 35.09101575 | 0.175739288 | 0.176578096 |
| Rps18 | 253.3875219 | 0.805307144 | 0.201569612 |
| Vps52 | 738.2973603 | -0.175283405 | 0.064972049 |
| Gm23111 | 0.99397625 | -0.553731697 | 1.198776244 |
| H2-K2 | 47.37552283 | 6.98419642 | 0.688931498 |
| Gm4152 | 0 | NA | NA |
| H2-K1 | 2006.445401 | 7.034623469 | 0.184935826 |
| Ring1 | 223.6236612 | 0.202567016 | 0.082869491 |
| Gm26940 | 0.048949532 | -0.243569582 | 3.724945049 |
| Mir219a-1 | 0.047948698 | -0.243569582 | 3.724945049 |

| | | | |
|---|---|---|---|
| mmu-mir-219c | 1.228891946 | -0.575018564 | 0.801271204 |
| H2-Ke6 | 1.732078677 | -0.494082165 | 0.710455599 |
| Gm20427 | 1.185090727 | 2.005076687 | 0.9799451 |
| Slc39a7 | 92.16732074 | 0.666674919 | 0.154458007 |
| Rxrb | 612.8230062 | 0.009639569 | 0.056518688 |
| Col11a2 | 1625.713743 | 2.530756817 | 0.245469374 |
| H2-Pb | 0.204910391 | -1.671415104 | 2.623494331 |
| BC051537 | 0.820567722 | -1.589529984 | 1.299291307 |
| H2-Pa | 0 | NA | NA |
| H2-Oa | 2.48498024 | 2.24074215 | 1.014624654 |
| Brd2 | 3845.362694 | 0.716508279 | 0.080111253 |
| Gm20493 | 0 | NA | NA |
| H2-DMa | 136.095749 | 1.15597762 | 0.161290578 |
| H2-DMb2 | 12.8997608 | 3.163843401 | 0.507841082 |
| H2-DMb1 | 15.23037961 | 5.077931084 | 0.657771417 |
| Psmb9 | 510.8582445 | 7.768310382 | 0.323220541 |
| Tap1 | 2068.383747 | 7.259124783 | 0.196364227 |
| Psmb8 | 1292.910119 | 8.037423827 | 0.290215318 |
| Gm20496 | 0.676590931 | 2.740430178 | 1.437695312 |
| Tap2 | 1370.634535 | 5.397899954 | 0.174290061 |
| Gm15821 | 0.89598866 | 3.197961877 | 1.357386465 |
| H2-Ob | 4.820269621 | 1.464625049 | 0.523302299 |
| Gm20506 | 0.159280631 | 1.185012522 | 3.461052012 |
| H2-Ab1 | 139.2632816 | 4.255612939 | 0.288933575 |
| H2-Aa | 90.90688896 | 4.533416921 | 0.348249168 |
| Gm20513 | 0.289806465 | 1.549959474 | 1.968503316 |
| H2-Eb1 | 122.6882474 | 5.614898798 | 0.523301809 |
| H2-Eb2 | 1.375804068 | 1.649969231 | 0.963735857 |
| H2-Ea-ps | 95.63606511 | 4.924414764 | 0.386026142 |
| Btnl2 | 0.44006443 | 1.838134555 | 1.473712121 |
| Btnl1 | 1.318555613 | 3.704042308 | 1.123021641 |
| Gm15320 | 0 | NA | NA |
| Gm15321 | 0 | NA | NA |
| BC051142 | 7.899042157 | -0.505343962 | 0.375480903 |
| Btnl4 | 0.635806793 | 3.113856345 | 1.33858642 |
| Gm15332 | 0 | NA | NA |
| Btnl5-ps | 0 | NA | NA |
| Gm15331 | 0 | NA | NA |
| Btnl6 | 0 | NA | NA |
| Gm15329 | 0 | NA | NA |
| Btnl7-ps | 0.643414179 | 2.848729793 | 1.188451 |
| Notch4 | 57.84156358 | 1.402796057 | 0.184308147 |
| Gm20462 | 0 | NA | NA |
| Gpsm3 | 60.20938841 | 3.293191758 | 0.281576713 |
| Pbx2 | 551.6966075 | -0.268702178 | 0.068970111 |
| Ager | 5.903964042 | 0.065469261 | 0.569680779 |
| Rnf5 | 318.1261628 | -0.401029423 | 0.074685656 |

| | | | |
|---|---|---|---|
| Gm20463 | 0.104106287 | -0.243576254 | 3.724945049 |
| Agpat1 | 1285.566767 | -0.077415194 | 0.061334254 |
| Egfl8 | 5.399883896 | -1.477963365 | 0.489260062 |
| Gm20460 | 0 | NA | NA |
| Ppt2 | 144.3949312 | -0.199083421 | 0.094950607 |
| Prrt1 | 1088.589987 | -0.541305485 | 0.119107701 |
| Gm20461 | 0.042716945 | -0.243571806 | 3.724945049 |
| Fkbpl | 63.97067545 | 0.362310424 | 0.142581768 |
| Atf6b | 618.9161125 | 0.307095818 | 0.079755951 |
| Tnxb | 123.7697874 | -0.247166281 | 0.196196403 |
| Gm24389 | 0.068097359 | -0.243569582 | 3.724945049 |
| Cyp21a2-ps | 0.84670602 | 1.462839439 | 1.288471421 |
| C4b | 4516.244341 | 4.245358729 | 0.142931776 |
| Stk19-ps1 | 0.301345325 | 1.298531221 | 2.002123968 |
| Skiv2l-ps1 | 0.092269992 | 0.237318756 | 3.724945049 |
| Gm27471 | 0 | NA | NA |
| Tnxa | 0.049553047 | 0.71820487 | 3.724945049 |
| Cyp21a1 | 0.116944185 | 1.302646088 | 3.719336636 |
| C4a | 141.065081 | 2.93055193 | 0.214654545 |
| Stk19 | 195.7826537 | 0.369438122 | 0.166656843 |
| Dxo | 129.6180202 | 0.394114658 | 0.127082711 |
| Skiv2l | 714.6748977 | 0.077395581 | 0.055706836 |
| Mir6971 | 0 | NA | NA |
| Mir6970 | 0 | NA | NA |
| Nelfe | 167.1258369 | 0.427641082 | 0.088319229 |
| Cfb | 85.69436929 | 10.10361412 | 0.883381686 |
| Gm20547 | 0 | NA | NA |
| Mir6972 | 0 | NA | NA |
| C2 | 78.60544563 | 3.557229939 | 0.255005285 |
| Zbtb12 | 59.55277683 | -0.156056159 | 0.236975449 |
| Ehmt2 | 1573.434019 | -0.362921982 | 0.072702733 |
| Slc44a4 | 2.383600578 | -0.7094388 | 0.690351266 |
| Neu1 | 530.836773 | 0.254041111 | 0.079229717 |
| 1110038B12Rik | 128.359516 | 1.486778087 | 0.170695169 |
| Snord52 | 0 | NA | NA |
| Gm25744 | 0 | NA | NA |
| Gm8696 | 0.535596869 | 2.315049807 | 1.449203259 |
| Hspa1b | 205.5992703 | 2.270184896 | 0.618708276 |
| Hspa1a | 168.1854328 | 2.222648568 | 0.571972848 |
| Gm20481 | 0.568897959 | 1.978301803 | 1.480405312 |
| Hspa1l | 8.258336928 | 2.087022178 | 0.421093542 |
| Gm10501 | 57.48590907 | -0.765322709 | 0.174031903 |
| Lsm2 | 72.82678909 | 0.25316188 | 0.127224898 |
| Vars | 655.1030319 | 0.392179655 | 0.067084891 |
| Vwa7 | 91.21471042 | -0.635238438 | 0.1948978 |
| Sapcd1 | 2.507914302 | -0.406553434 | 0.74225172 |
| Msh5 | 15.17733262 | 0.074731674 | 0.382605423 |

| | | | |
|---|---|---|---|
| Gm23421 | 0 | NA | NA |
| Clic1 | 397.1204603 | 3.507802638 | 0.210357535 |
| Ddah2 | 89.30682263 | -0.018064493 | 0.157607514 |
| G6b | 1.225313711 | -1.46074366 | 0.975539267 |
| Ly6g6c | 0.500444145 | 2.327073288 | 1.449198194 |
| Ly6g6d | 2.936279196 | -1.376364301 | 0.617958166 |
| Ly6g6e | 18.02936065 | 1.238305009 | 0.448222632 |
| Ly6g6f | 3.98262746 | -0.464572949 | 0.51928903 |
| Abhd16a | 1009.474395 | 0.335629114 | 0.103733454 |
| Ly6g5c | 0.083526043 | 0.718202646 | 3.724945049 |
| Ly6g5b | 3.278244484 | 0.07620742 | 0.534619525 |
| Csnk2b | 658.9798502 | 0.435052431 | 0.067980719 |
| Gpank1 | 122.9329401 | 0.214099083 | 0.127332901 |
| Gm20522 | 3.138335889 | 0.199730868 | 0.547570964 |
| D17H6S53E | 194.6956857 | 0.23299243 | 0.092692784 |
| Apom | 0.669558094 | 2.062837052 | 1.233562434 |
| Bag6 | 2358.238324 | -0.071686999 | 0.046459193 |
| Prrc2a | 6282.854268 | -0.082312047 | 0.076330316 |
| Gm23442 | 7.065932701 | -0.177466778 | 0.444432466 |
| Gm17705 | 6.475434284 | -0.829959359 | 0.431058818 |
| Aif1 | 114.5567538 | 2.542458709 | 0.196856312 |
| Ncr3-ps | 0.383232264 | 0.237329819 | 1.737333312 |
| Lst1 | 34.76546749 | 2.842251784 | 0.283923377 |
| Ltb | 23.43129296 | 4.167978289 | 0.443712692 |
| Mir6973a | 0.196800764 | 1.969753326 | 2.170503945 |
| Tnf | 84.48636142 | 7.399232098 | 0.730352584 |
| Lta | 7.270911819 | 0.316639988 | 0.565891374 |
| Mir6974 | 0 | NA | NA |
| Nfkbil1 | 124.1921778 | 0.760603007 | 0.156159014 |
| Gm16181 | 0.333516142 | 0.272427187 | 1.725989429 |
| Atp6v1g2 | 2556.244658 | -0.876345741 | 0.094508358 |
| Ddx39b | 1299.932362 | 0.142264753 | 0.070970907 |
| Gm22589 | 0 | NA | NA |
| Mir6975 | 0.574701389 | 0.249354763 | 1.289070948 |
| Gm25128 | 2.512166706 | -0.79784383 | 0.684456657 |
| Mir8094 | 0.904930538 | -1.566194317 | 1.178573139 |
| H2-D1 | 7783.618613 | 5.760078224 | 0.188087358 |
| Gm18733 | 1.69076615 | 0.937673403 | 0.704401511 |
| Gm18734 | 0.938757343 | 0.409132421 | 1.094832968 |
| H2-Q1 | 113.5987717 | 4.774255195 | 0.29656096 |
| Gm8741 | 9.465940354 | -0.307821803 | 0.356781259 |
| H2-Q2 | 32.27579733 | 5.41304134 | 0.499978854 |
| H2-Q3 | 0.044544648 | 0.237309859 | 3.724945049 |
| Gm11131 | 30.17597747 | 8.175963443 | 0.860632674 |
| H2-Q4 | 2741.587666 | 8.778868906 | 0.27728287 |
| H2-Q5 | 203.0857095 | 9.431989873 | 0.737080169 |
| Gm8750 | 0.56888582 | 1.009556795 | 1.238032159 |

| | | | |
|---|---|---|---|
| H2-Q6 | 1065.781777 | 10.36618656 | 0.506120169 |
| H2-Q7 | 469.1606159 | 10.11004812 | 0.62548177 |
| Gm8752 | 0.72059279 | 1.736605777 | 1.25303555 |
| H2-Q10 | 15.54474821 | 3.602959033 | 0.453000422 |
| Pou5f1 | 0 | NA | NA |
| Gm19553 | 0 | NA | NA |
| Tcf19 | 44.18076177 | 0.690049016 | 0.321995172 |
| Cchcr1 | 54.80302612 | 0.974384167 | 0.192814176 |
| Psors1c2 | 0.771204239 | -0.548616812 | 1.245868517 |
| Cdsn | 7.227748083 | 2.312672778 | 0.440383542 |
| Gm9613 | 0.0885994 | -0.24357403 | 3.724945049 |
| 2300002M23Rik | 0 | NA | NA |
| Sfta2 | 0.116217011 | -0.24357403 | 3.724945049 |
| Gm9573 | 0.122991902 | -0.24357403 | 3.724945049 |
| Dpcr1 | 0.220032676 | 1.195301988 | 2.574515385 |
| Gm20483 | 0.193987248 | -0.745348799 | 2.240590368 |
| Vars2 | 218.6187569 | -0.594272844 | 0.093285333 |
| Gtf2h4 | 76.61225677 | 0.191314836 | 0.140692414 |
| Ddr1 | 1349.901542 | 0.926009884 | 0.113280743 |
| Gm23864 | 3.332665857 | -0.231916079 | 0.542144248 |
| Gm4577 | 0.622983114 | -0.243564102 | 1.358677777 |
| Gm20443 | 0.385495486 | 0.713775374 | 1.530017028 |
| Gm20442 | 0.201858732 | 0.718217268 | 2.604406118 |
| 4833427F10Rik | 0.08486385 | 0.717551239 | 3.724898127 |
| Ier3 | 124.9554162 | 1.359562844 | 0.251622163 |
| Flot1 | 628.8586158 | 0.023650204 | 0.089499615 |
| Tubb5 | 3619.132628 | -0.089441853 | 0.093200419 |
| Mdc1 | 303.644182 | 0.195791186 | 0.11347632 |
| Ppp1r18os | 5.57648041 | 1.279999536 | 0.468464028 |
| Nrm | 18.32685686 | 0.142787414 | 0.245117851 |
| Ppp1r18 | 331.6778556 | 2.760170972 | 0.207580002 |
| Gm28033 | 0.098502579 | 0.23732098 | 3.724945049 |
| Dhx16 | 338.5688034 | 0.513586661 | 0.098777936 |
| 2310061I04Rik | 406.7426439 | -0.212268794 | 0.082526307 |
| Gm16279 | 3.489032565 | -0.543131147 | 0.655095347 |
| Atat1 | 349.1959485 | -0.52624781 | 0.07890024 |
| Mrps18b | 109.9134655 | 0.608286628 | 0.121092084 |
| Ppp1r10 | 838.888173 | 0.417154443 | 0.081498289 |
| Mir1894 | 0 | NA | NA |
| Gm9840 | 3.211017017 | 0.480010454 | 0.56172265 |
| Gm8801 | 0.654072159 | -0.972117362 | 1.175813739 |
| Abcf1 | 700.9733728 | 0.254661521 | 0.085196271 |
| Mir877 | 0 | NA | NA |
| Prr3 | 210.8648339 | 0.23941511 | 0.126757704 |
| Gnl1 | 1616.148979 | -0.016235445 | 0.068341682 |
| Gm20508 | 2.081650308 | -0.256691978 | 0.665005486 |
| A930015D03Rik | 113.1905163 | 1.718082124 | 0.167570859 |

| | | | |
|---|---|---|---|
| H2-T24 | 905.0134114 | 6.795704828 | 0.264033826 |
| H2-T23 | 3145.49009 | 7.780953377 | 0.211323407 |
| H2-T22 | 1287.107806 | 5.110433872 | 0.16736451 |
| Gm6034 | 25.06496248 | 4.257868408 | 0.433402512 |
| Gm11127 | 246.9603189 | 6.357107166 | 0.290682073 |
| Gm8810 | 46.0581403 | 6.388570382 | 0.591305833 |
| H2-Bl | 5.276086801 | 5.891938923 | 0.926522827 |
| Gm20530 | 0 | NA | NA |
| Gm8815 | 1.211501154 | 3.930691599 | 1.066846762 |
| C920025E04Rik | 19.33663716 | 6.394776067 | 0.848958646 |
| Gm7030 | 5.681141433 | 5.998315665 | 0.948595034 |
| BC023719 | 0.374687972 | 1.442394305 | 1.530935682 |
| H2-T10 | 170.5753034 | 4.307687003 | 0.223483298 |
| Gm19684 | 10.16906943 | 6.155730466 | 0.895308173 |
| Gm9574 | 0.115603922 | 1.209894437 | 3.72418822 |
| Gm10499 | 19.27648584 | 3.958374356 | 0.504254 |
| 2410017I17Rik | 0.142780685 | -0.70560352 | 2.899062601 |
| Gm17782 | 0.038981395 | 0.718207094 | 3.724945049 |
| Gm6659 | 0.124347486 | 0.719186602 | 3.717245571 |
| Gm8909 | 5.891852665 | 4.477764642 | 0.8421903 |
| Gm20478 | 1.20453066 | 3.595346689 | 1.372267105 |
| Gm6623 | 0.15621502 | 0.774550521 | 2.832050821 |
| Gm20495 | 0 | NA | NA |
| H2-T3 | 0.239012627 | 1.577311796 | 2.097643552 |
| Gm4246 | 0.068097359 | -0.243569582 | 3.724945049 |
| Gm20447 | 0 | NA | NA |
| Gm8835 | 0.696575882 | 3.612315218 | 1.220383687 |
| Gm5682 | 0.219744001 | 1.876864594 | 2.125981944 |
| Gm20392 | 0.238833577 | 0.237322268 | 2.490474962 |
| Gm10074 | 1.752125518 | 1.306487786 | 0.837228407 |
| Gm20545 | 0.204225461 | 0.23731275 | 3.221243153 |
| Gm29402 | 0 | NA | NA |
| Gm18604 | 0 | NA | NA |
| Gm20391 | 0 | NA | NA |
| Gm22453 | 0 | NA | NA |
| Gm4252 | 0 | NA | NA |
| Gm20546 | 0.114742407 | -0.24357403 | 3.724945049 |
| Gm6633 | 0.13733263 | -1.191829832 | 3.63430361 |
| Rpp21 | 97.94756067 | 0.058113591 | 0.122734528 |
| Trim39 | 333.3226075 | -0.254419456 | 0.130872718 |
| H2-M10.2 | 0.048949532 | -0.243569582 | 3.724945049 |
| H2-M10.1 | 0 | NA | NA |
| H2-M10.3 | 0 | NA | NA |
| Stk-ps1 | 0 | NA | NA |
| H2-M10.5-ps1 | 0 | NA | NA |
| Lgals1-ps1 | 0 | NA | NA |
| Rpl9-ps9 | 0 | NA | NA |

| | | | |
|---|---|---|---|
| Obox3-ps8 | 0.062426873 | 0.237312084 | 3.724945049 |
| H2-M10.4 | 0 | NA | NA |
| Gm20452 | 0.139037128 | 1.464453538 | 3.717743456 |
| Gm20451 | 0.072025462 | 0.237312084 | 3.724945049 |
| Gm20454 | 0 | NA | NA |
| Gm20453 | 0 | NA | NA |
| H2-M11 | 0.066050875 | 0.71820487 | 3.724945049 |
| Gm8868 | 0 | NA | NA |
| Gm20450 | 0 | NA | NA |
| H2-M9 | 0.541282201 | 3.277560769 | 1.294475368 |
| Rpl9-ps5 | 0.163190422 | 1.22955094 | 3.396300676 |
| H2-M1 | 0.163289894 | 1.708362142 | 3.406546227 |
| Gm8877 | 0 | NA | NA |
| Gm20549 | 0 | NA | NA |
| Gm20523 | 0 | NA | NA |
| Gm20525 | 0 | NA | NA |
| Gm20526 | 0 | NA | NA |
| Gm4271 | 0 | NA | NA |
| Gm20401 | 0 | NA | NA |
| Gm8878 | 0.22323753 | 0.237325468 | 2.561679775 |
| H2-M10.5 | 0.066050875 | 0.71820487 | 3.724945049 |
| H2-M10.6 | 0 | NA | NA |
| Trim26 | 648.1818355 | 1.574778452 | 0.101391268 |
| Gm25973 | 0 | NA | NA |
| Trim15 | 1.517773654 | 4.449824371 | 1.0697393 |
| Trim10 | 2.069903214 | 3.335693465 | 1.068771219 |
| Trim40 | 1.721525315 | -1.179265397 | 0.854413194 |
| Trim31 | 0.590394638 | 1.860930817 | 1.255563069 |
| 1700031A10Rik | 0.066050875 | 0.71820487 | 3.724945049 |
| Rnf39 | 43.61237388 | 0.085399398 | 0.247406132 |
| Ppp1r11 | 632.5865216 | 0.473481598 | 0.0968677 |
| Znrd1 | 116.024339 | 0.474175655 | 0.136046082 |
| Znrd1as | 20.09275304 | 0.467007492 | 0.249707791 |
| Gm27735 | 0 | NA | NA |
| H2-M6-ps | 4.48790112 | -1.245804819 | 0.598034397 |
| 2410137M14Rik | 2.456754703 | -1.908500197 | 0.809404162 |
| H2-M5 | 10.19594007 | -1.716219916 | 0.43088789 |
| Zfp57 | 63.14501023 | -0.862847318 | 0.223766184 |
| Mog | 603.8852982 | -0.265116026 | 0.156154949 |
| Gm22388 | 0.655055538 | -1.744907657 | 1.201374766 |
| Gabbr1 | 8020.403664 | -0.472684276 | 0.070777274 |
| Olfr90 | 0.046345709 | 0.71820487 | 3.724945049 |
| Olfr91 | 0.106520246 | 0.718200421 | 3.724945049 |
| Olfr92 | 0.119587853 | 0.237305575 | 3.724945049 |
| Gm4831 | 0.046345709 | 0.71820487 | 3.724945049 |
| Gm19807 | 2.281184893 | 1.810978121 | 0.73667775 |
| Olfr93 | 0.291730981 | 1.629907502 | 1.754787612 |

| | | | |
|---|---|---|---|
| Vhl-ps1 | 3.944458201 | 5.446322236 | 0.945321969 |
| Olfr753-ps1 | 11.84344878 | 7.303593692 | 0.931833452 |
| Gm20497 | 17.12714754 | 7.652012649 | 0.937442206 |
| Gm7031 | 0.038981395 | 0.718207094 | 3.724945049 |
| Ubd | 60.38255723 | 9.782358652 | 1.039861448 |
| Olfr94 | 0.247779669 | 1.172881041 | 1.834514665 |
| Olfr95 | 0.082913145 | 0.23732098 | 3.724945049 |
| Olfr96 | 0.046345709 | 0.71820487 | 3.724945049 |
| Olfr97 | 0 | NA | NA |
| Gm20429 | 1.971464702 | 4.365547357 | 1.033860167 |
| Olfr754-ps1 | 0.99187381 | 3.237988683 | 1.157883115 |
| Olfr98 | 0 | NA | NA |
| Gm25104 | 0.045882455 | 0.237312084 | 3.724945049 |
| H2-M3 | 192.7482628 | 4.72457897 | 0.233075892 |
| Olfr755-ps1 | 6.424303794 | 3.343958269 | 0.690224188 |
| Olfr99 | 0 | NA | NA |
| Olfr756-ps1 | 0 | NA | NA |
| Olfr101 | 0.110870145 | 0.238305784 | 3.724874707 |
| Olfr102 | 0.068097359 | -0.243569582 | 3.724945049 |
| Olfr103 | 0 | NA | NA |
| Olfr104-ps | 0 | NA | NA |
| Olfr757-ps1 | 0 | NA | NA |
| Olfr105-ps | 0 | NA | NA |
| Olfr758-ps1 | 0 | NA | NA |
| Olfr106-ps | 0 | NA | NA |
| Olfr107 | 0 | NA | NA |
| Olfr759-ps1 | 0 | NA | NA |
| Gm18808 | 0 | NA | NA |
| Olfr108 | 0 | NA | NA |
| Olfr760-ps1 | 0 | NA | NA |
| Olfr109 | 0.108772582 | 0.718202646 | 3.724945049 |
| H2-M2 | 1.126195538 | 3.156880814 | 1.285338982 |
| Olfr110 | 0.784115673 | -0.497595645 | 1.230330645 |
| Olfr111 | 1.850398219 | 1.072879691 | 0.833766342 |
| Olfr112 | 0.062426873 | 0.237312084 | 3.724945049 |
| Olfr113 | 0 | NA | NA |
| CR974439.2 | 0 | NA | NA |
| Olfr114 | 0 | NA | NA |
| Olfr115 | 0.046345709 | 0.71820487 | 3.724945049 |
| CR974439.1 | 0 | NA | NA |
| Olfr116 | 0.046345709 | 0.71820487 | 3.724945049 |
| Olfr117 | 0 | NA | NA |
| Olfr118 | 0.061920613 | 0.718207094 | 3.724945049 |
| Olfr119 | 0 | NA | NA |
| Gm7335 | 43.44352658 | -0.369290564 | 0.19927985 |
| Olfr120 | 0.155118291 | 1.200062356 | 3.496459569 |
| Olfr121 | 0 | NA | NA |

| | | | |
|---|---|---|---|
| Olfr122 | 0.08486385 | 0.717551239 | 3.724898127 |
| Olfr123 | 0 | NA | NA |
| Gm20524 | 0.042716945 | -0.243571806 | 3.724945049 |
| Olfr124 | 0 | NA | NA |
| Olfr125 | 0 | NA | NA |
| Gm9577 | 0.148659141 | 1.513781604 | 3.716315176 |
| Olfr126 | 0 | NA | NA |
| Gm18737 | 0.194588561 | -0.243564226 | 2.240900713 |
| Gm18520 | 0 | NA | NA |
| Olfr127 | 0 | NA | NA |
| Olfr128 | 0 | NA | NA |
| Gm20543 | 0 | NA | NA |
| Gm20542 | 0 | NA | NA |
| Gm5967 | 0 | NA | NA |
| Olfr761 | 0 | NA | NA |
| Gm18522 | 0 | NA | NA |
| Gm20541 | 1.213329596 | 0.216878762 | 1.036247028 |
| Olfr129 | 0 | NA | NA |
| Olfr130 | 0.068097359 | -0.243569582 | 3.724945049 |
| Gm18833 | 0 | NA | NA |
| Olfr131 | 0.048949532 | -0.243569582 | 3.724945049 |
| Olfr132 | 0.042716945 | -0.243571806 | 3.724945049 |
| Olfr133 | 0.061920613 | 0.718207094 | 3.724945049 |
| Olfr134 | 0 | NA | NA |
| Olfr135 | 0 | NA | NA |
| Olfr138 | 0 | NA | NA |
| Olfr137 | 0 | NA | NA |
| Olfr762-ps1 | 0 | NA | NA |
| Olfr136 | 0.089062654 | 0.237318756 | 3.724945049 |
| Esp36 | 0 | NA | NA |
| Gm20409 | 0 | NA | NA |
| Gm6726 | 0 | NA | NA |
| Esp34 | 0 | NA | NA |
| Esp31 | 0.04393175 | -0.243571806 | 3.724945049 |
| Esp24 | 0 | NA | NA |
| Gm21903 | 0 | NA | NA |
| Esp23 | 0 | NA | NA |
| Esp18 | 0 | NA | NA |
| Gm26429 | 0 | NA | NA |
| Esp16 | 0 | NA | NA |
| Esp15 | 0 | NA | NA |
| Gm26917 | 335746.0181 | 0.75474988 | 0.10482809 |
| Rn18s-rs5 | 102.7885986 | 0.352242805 | 0.819911173 |
| Yam1 | 128743.8113 | 0.715924531 | 0.635052632 |
| Esp38 | 0 | NA | NA |
| Gm26376 | 0 | NA | NA |
| Tgif2-ps2 | 0.524319023 | 0.278242951 | 1.357611241 |

| | | | |
|---|---|---|---|
| Pgk2 | 0 | NA | NA |
| Crisp3 | 0 | NA | NA |
| Crisp1 | 0.331267484 | 1.09564947 | 1.728839436 |
| Esp8 | 0 | NA | NA |
| Esp6Esp5 | 0 | NA | NA |
| Esp6 | 0 | NA | NA |
| Esp4 | 0 | NA | NA |
| Esp3 | 0 | NA | NA |
| Esp1 | 0 | NA | NA |
| Crisp2 | 0.293737151 | 1.133789136 | 3.717753581 |
| Rhag | 0.351655166 | 1.168348944 | 3.720612353 |
| Gm20574 | 0 | NA | NA |
| 9130008F23Rik | 2.649375936 | 1.569800982 | 0.611046304 |
| Glyatl3 | 0.199876958 | -0.680910717 | 2.647758279 |
| Gm23078 | 0.852119224 | -1.007579931 | 1.104067614 |
| Cenpq | 63.31617002 | 1.152682267 | 0.167720359 |
| Mut | 485.5759982 | -0.295165954 | 0.100457587 |
| Gm6771 | 0.062426873 | 0.237312084 | 3.724945049 |
| Ptchd4 | 384.0058745 | -0.215611817 | 0.079365394 |
| Opn5 | 0.714665336 | 0.520465681 | 1.161220429 |
| Adgrf4 | 19.83835518 | 0.665531258 | 0.251788478 |
| Adgrf2 | 18.64465502 | -0.396257617 | 0.300097755 |
| Cd2ap | 541.0115779 | 0.316087981 | 0.141626771 |
| Tnfrsf21 | 965.9785103 | -0.320573537 | 0.06724196 |
| Adgrf1 | 1.021366362 | 0.517671896 | 1.120683112 |
| Adgrf5 | 194.5698432 | 1.253427506 | 0.182232647 |
| Gm25135 | 0.066050875 | 0.71820487 | 3.724945049 |
| Mep1a | 1.39389272 | 2.854434206 | 1.110903661 |
| Ankrd66 | 10.82149774 | 2.918808343 | 0.464877438 |
| Pla2g7 | 1523.964145 | 0.424175167 | 0.129657607 |
| 1700071M16Rik | 0.275776889 | -0.622106591 | 2.051565649 |
| Tdrd6 | 25.59334993 | -0.937010397 | 0.288805876 |
| Slc25a27 | 391.4686422 | -0.734933391 | 0.134656914 |
| Cyp39a1 | 17.12697954 | -0.352475696 | 0.284696141 |
| Gm25783 | 0 | NA | NA |
| Rcan2 | 1857.55946 | -0.235760366 | 0.101508841 |
| Gm24532 | 0.062426873 | 0.237312084 | 3.724945049 |
| Enpp5 | 2499.132028 | -0.460914476 | 0.050884261 |
| Enpp4 | 696.8122402 | 1.594471996 | 0.166642025 |
| Clic5 | 79.94705142 | 1.950161169 | 0.205284961 |
| Runx2 | 205.0751855 | 0.882115874 | 0.168691398 |
| Runx2os3 | 0 | NA | NA |
| n-R5s27 | 0.766229633 | 1.249280046 | 1.190999434 |
| Runx2os2 | 0.092269992 | 0.237318756 | 3.724945049 |
| Runx2os1 | 7.863194607 | 1.138873598 | 0.452983626 |
| Supt3 | 134.9145071 | 0.109238037 | 0.122019096 |
| Gm24979 | 0 | NA | NA |

| | | | |
|---|---|---|---|
| Cdc5l | 544.4221431 | 0.149653283 | 0.06524478 |
| B230354K17Rik | 184.8456956 | -0.259794768 | 0.127885637 |
| Spats1 | 16.79904519 | -0.11768801 | 0.277136358 |
| Gm9104 | 1.30365031 | -0.616680297 | 0.92368407 |
| Aars2 | 207.7781551 | -0.250551959 | 0.090085514 |
| Tcte1 | 13.23124629 | -0.262684034 | 0.31784672 |
| Tmem151b | 1924.460039 | -0.557161744 | 0.086668221 |
| Nfkbie | 174.6593782 | 2.656086631 | 0.271282748 |
| Slc35b2 | 231.0133132 | 0.547140487 | 0.147395083 |
| Hsp90ab1 | 12795.80252 | 0.259351436 | 0.08346035 |
| Slc29a1 | 163.1308914 | -0.157149794 | 0.10130288 |
| Gm17080 | 1.296318896 | -0.301135156 | 0.854136687 |
| Gm7325 | 0 | NA | NA |
| Gm25008 | 0 | NA | NA |
| Capn11 | 4.929061397 | -0.196826683 | 0.527077033 |
| Tmem63b | 1144.513863 | -0.658075955 | 0.073516477 |
| Gm16172 | 0.25133133 | 0.756075996 | 2.431336391 |
| Mrpl14 | 161.5517627 | 0.346341091 | 0.133963457 |
| 1600014C23Rik | 0.431841237 | -0.711688933 | 1.395536285 |
| F630040K05Rik | 2.147486072 | -3.882820439 | 0.943072707 |
| Vegfa | 569.830768 | -0.511756183 | 0.100509897 |
| Mrps18a | 383.939429 | 0.331124722 | 0.079275959 |
| Rsph9 | 56.95556391 | -0.245498678 | 0.141141813 |
| Mad2l1bp | 60.28485607 | 0.523443725 | 0.157562861 |
| Gtpbp2 | 597.9990834 | 0.322020226 | 0.082835844 |
| Polh | 84.99680479 | 0.386474974 | 0.150554038 |
| Xpo5 | 708.0881202 | 0.241493174 | 0.058842722 |
| Mir693 | 0.626433637 | -1.008842042 | 1.227907215 |
| Gm22474 | 0 | NA | NA |
| Polr1c | 186.1516209 | 0.453752424 | 0.110686472 |
| Yipf3 | 497.8450709 | 0.331787365 | 0.07198553 |
| Lrrc73 | 196.1398283 | -0.896926653 | 0.118324119 |
| Tjap1 | 262.6446001 | 0.404493013 | 0.107016552 |
| Gm26785 | 0.440626437 | 1.596229982 | 1.700308928 |
| Dlk2 | 119.3259875 | -1.17137258 | 0.18101967 |
| Abcc10 | 137.5955981 | -0.475788226 | 0.168884692 |
| Zfp318 | 978.2373152 | -0.327652506 | 0.094333333 |
| Gm23037 | 4.835503307 | -0.412403644 | 0.589295411 |
| Crip3 | 13.66803183 | -1.706862777 | 0.370324073 |
| Slc22a7 | 3.396474255 | 0.938125732 | 0.551603941 |
| Gm5093 | 4.877282298 | 0.076825741 | 0.550043817 |
| Ttbk1 | 1698.649847 | -0.388626545 | 0.08945013 |
| Gm26904 | 0.300318634 | -0.163263496 | 1.775528842 |
| Dnph1 | 20.31270367 | 0.327776205 | 0.250464664 |
| Cul9 | 1048.399803 | -0.625040772 | 0.076319746 |
| Srf | 574.4040805 | -0.020861365 | 0.135511241 |
| mmu-mir-6976 | 0.656458977 | -0.231571461 | 1.307154503 |

| | | | |
|---|---|---|---|
| Ptk7 | 194.7380974 | 0.18123962 | 0.08869394 |
| Klc4 | 177.2125805 | -0.204521669 | 0.095242312 |
| Mrpl2 | 228.4860238 | 0.257813546 | 0.115867427 |
| Cul7 | 465.2672236 | -0.142197334 | 0.094244504 |
| Rrp36 | 135.9417813 | 0.775680655 | 0.099727874 |
| Klhdc3 | 1295.864183 | -0.254074031 | 0.05006441 |
| Mea1 | 401.4088604 | 0.211284713 | 0.1232162 |
| Ppp2r5d | 1040.331458 | -0.116683966 | 0.066433915 |
| Pex6 | 410.5596741 | -0.123189747 | 0.131327708 |
| Gnmt | 15.57555665 | 0.281435609 | 0.32494657 |
| Cnpy3 | 552.2456984 | 0.366873777 | 0.079463322 |
| Ptcra | 0.771865818 | 0.228463336 | 1.099404767 |
| 2310039H08Rik | 29.97597163 | 0.687063321 | 0.220712592 |
| Rpl7l1 | 408.9885597 | 0.711331297 | 0.103725805 |
| Gltscr1l | 1089.761893 | -0.305979877 | 0.082356135 |
| A330017A19Rik | 35.0263174 | -1.252806176 | 0.267904953 |
| Tbcc | 165.35841 | 0.457059507 | 0.129692757 |
| Prph2 | 0.359258722 | 0.642662993 | 1.458915818 |
| Ubr2 | 1461.655788 | -0.073197266 | 0.055170484 |
| Gm23657 | 4.719513391 | -0.762415167 | 0.496452695 |
| Gm16494 | 2.052894931 | 1.065057313 | 0.904153609 |
| Gm4945 | 0 | NA | NA |
| Trerf1 | 592.2320734 | -1.274458314 | 0.155254693 |
| Gm26216 | 1.145909063 | -2.019849795 | 0.976815887 |
| Mrps10 | 197.4832179 | 0.07056769 | 0.09971521 |
| Guca1b | 0.605175792 | 1.098883239 | 1.249924832 |
| Guca1a | 2.149392939 | 3.252907779 | 1.117663721 |
| Gm5814 | 2.49590342 | 1.796436583 | 0.723603898 |
| 1700001C19Rik | 0.29699235 | -0.243563108 | 1.774962751 |
| AI661453 | 6.189948269 | 1.013106863 | 0.435603405 |
| Taf8 | 186.7542863 | 0.324454347 | 0.091818533 |
| Ccnd3 | 416.7992574 | 1.17614603 | 0.128486843 |
| Bysl | 253.0965139 | 0.377922547 | 0.089703697 |
| Med20 | 240.4788844 | 0.106720459 | 0.119929774 |
| Gm20517 | 21.19396069 | -0.793310958 | 0.34274968 |
| Kpna2-ps | 1.554960773 | 0.68081767 | 0.826781696 |
| Usp49 | 129.8291219 | -0.267041152 | 0.12634646 |
| Tomm6 | 2.522522481 | -1.302357195 | 0.836582759 |
| Gm21981 | 0.57942775 | -0.243564171 | 1.372884017 |
| Tomm6os | 39.84955966 | -0.618949077 | 0.174409411 |
| Prickle4 | 1.072312625 | -1.664414151 | 1.141570604 |
| Frs3 | 230.2853897 | -0.188665897 | 0.110617283 |
| Frs3os | 0.363326102 | -0.054953738 | 1.688397364 |
| Pgc | 0.329750641 | 1.150547912 | 1.787994845 |
| Tfeb | 165.6362251 | 0.018231188 | 0.128919828 |
| Mdfi | 11.42660599 | 0.153044777 | 0.310420412 |
| Gm25201 | 0.048949532 | -0.243569582 | 3.724945049 |

| | | | |
|---|---|---|---|
| n-R5s28 | 0 | NA | NA |
| Foxp4 | 579.0805347 | -0.464780375 | 0.105303073 |
| Gm15556 | 0.196245641 | 1.710950971 | 3.230301063 |
| Gm5228 | 0 | NA | NA |
| Gm14871 | 0 | NA | NA |
| 1700122O11Rik | 0 | NA | NA |
| 1700067P10Rik | 0 | NA | NA |
| 9830107B12Rik | 0 | NA | NA |
| A530064D06Rik | 30.08457509 | 7.373711993 | 0.914357925 |
| Trem1 | 13.84162536 | 7.615528603 | 0.977502914 |
| Trem3 | 3.961576269 | 5.931049911 | 0.929763217 |
| Treml4 | 2.160387598 | 3.620168347 | 1.1380968 |
| Treml2 | 120.9667195 | 6.503459234 | 0.393679387 |
| B430306N03Rik | 110.3507723 | 9.219514124 | 0.898866635 |
| Trem2 | 83.40596837 | 0.372492036 | 0.487519268 |
| Treml1 | 1.368647145 | -0.482423822 | 0.845183938 |
| Nfya | 266.8544357 | 0.478366859 | 0.126920402 |
| Oard1 | 77.30484208 | 0.245774839 | 0.144958202 |
| Apobec2 | 1.336996185 | 0.218338691 | 0.881818878 |
| Tspo2 | 0 | NA | NA |
| Unc5cl | 6.179264061 | 0.109698805 | 0.411861896 |
| Gm24071 | 0 | NA | NA |
| Gm23642 | 0 | NA | NA |
| Lrfn2 | 299.7160112 | -0.317289864 | 0.137112771 |
| AC154247.1 | 0.135598227 | -1.1790103 | 2.942835517 |
| 1700008K24Rik | 0 | NA | NA |
| Gm22978 | 0.046345709 | 0.71820487 | 3.724945049 |
| Gm20540 | 0.410699371 | 0.15534364 | 1.450495098 |
| Mocs1 | 175.5053655 | 0.631041829 | 0.116523587 |
| Daam2 | 1383.444506 | -0.371312657 | 0.099785644 |
| Kif6 | 28.85323937 | -0.222519312 | 0.240676964 |
| Gm16555 | 0 | NA | NA |
| Gm16554 | 0.075043205 | 0.237309859 | 3.724945049 |
| Rftn1 | 212.415448 | 0.97009724 | 0.138433512 |
| Gm37173 | 0 | NA | NA |
| AY702103 | 0 | NA | NA |
| Dazl | 26.34726813 | -0.862204628 | 0.276813883 |
| Plcl2 | 1538.397032 | -0.425366726 | 0.082832848 |
| Gm7334 | 7.373441669 | 1.461634243 | 0.443629935 |
| Tbc1d5 | 596.1325065 | -0.139938318 | 0.061965286 |
| Gm25177 | 2.526566463 | -0.172864684 | 0.671815765 |
| Satb1 | 1280.976831 | -0.595145483 | 0.109379548 |
| E430014B02Rik | 30.56272651 | -0.210858832 | 0.326936997 |
| C230085N15Rik | 40.73930876 | -0.68051147 | 0.337500498 |
| Gm36931 | 44.82250833 | -0.597336771 | 0.235480572 |
| 5830444F18Rik | 19.30851287 | -0.73907454 | 0.390011122 |
| Gm37248 | 39.69177475 | -0.70541751 | 0.337399684 |

| | | | |
|---|---|---|---|
| Gm6934 | 12.03475686 | -1.20788473 | 0.452789621 |
| Gm37593 | 18.0879042 | -1.330620686 | 0.428448901 |
| Gm37266 | 10.80255973 | -1.497548031 | 0.41894827 |
| Gm37176 | 9.616469211 | -1.476857139 | 0.529399749 |
| Gm19585 | 5.697516218 | 4.076998568 | 0.823597538 |
| Gm27217 | 0.396216368 | 0.272302979 | 1.764869129 |
| Kcnh8 | 23.78866619 | -0.28173444 | 0.254366363 |
| Efhb | 28.27811259 | 2.339407672 | 0.281977422 |
| Rab5a | 1148.748334 | 0.343472853 | 0.085641088 |
| Pp2d1 | 0.305447682 | 0.718226064 | 1.771549079 |
| Kat2b | 803.7724207 | 0.641710548 | 0.10000687 |
| Gm6919 | 0.108772582 | 0.718202646 | 3.724945049 |
| Sgol1 | 6.962193728 | 2.173985793 | 0.671559088 |
| Gm19518 | 0.186767869 | -0.243573031 | 3.290881976 |
| 4932415M13Rik | 0 | NA | NA |
| Ck-ps1 | 0.277663158 | 0.718193929 | 3.724945049 |
| RP24-173D2.6 | 0 | NA | NA |
| RP24-173D2.8 | 0 | NA | NA |
| RP24-173D2.9 | 0 | NA | NA |
| Sult1c2 | 0.362371572 | -1.175421894 | 1.732696631 |
| Mrps36-ps1 | 2.335139537 | 1.234870501 | 0.801075501 |
| RP24-173D2.10 | 0 | NA | NA |
| RP24-173D2.11 | 0 | NA | NA |
| RP23-416K11.2 | 0 | NA | NA |
| RP23-416K11.3 | 0 | NA | NA |
| RP23-416K11.4 | 0 | NA | NA |
| Sult1c1 | 1.161929334 | -0.318205089 | 1.171188959 |
| RP23-416K11.5 | 0 | NA | NA |
| Gm37966 | 0 | NA | NA |
| Slc5a7 | 146.1741277 | -0.027217305 | 0.246986265 |
| 1700025K24Rik | 0.046345709 | 0.71820487 | 3.724945049 |
| Trp53-ps | 0 | NA | NA |
| Gm26291 | 0 | NA | NA |
| Gm17377 | 2.328145098 | -0.211617419 | 0.712395056 |
| St6gal2 | 374.6218881 | -1.259099947 | 0.161778314 |
| Gm26547 | 0.778497894 | 0.527316223 | 1.330147288 |
| Vmn2r118 | 1.673678531 | 1.152459405 | 0.776554771 |
| Pot1b | 63.94887526 | 0.098644914 | 0.154295269 |
| Gm22654 | 0 | NA | NA |
| Adgre4 | 3.084562812 | 3.783989873 | 0.946657634 |
| Gm18417 | 1.232924928 | 1.299543646 | 1.140326899 |
| Zfp119a | 72.0705982 | 0.370243986 | 0.136892564 |
| Zfp959 | 70.36865212 | 0.265713306 | 0.136420521 |
| Rpl21-ps6 | 2.92891158 | 0.824481862 | 0.683085843 |
| Zfp119b | 52.43127221 | 0.961121124 | 0.194381559 |
| Ebi3 | 12.63508742 | 1.518063386 | 0.341489277 |
| Gm16712 | 2.434106523 | -0.208115648 | 0.614492141 |

| | | | |
|---|---|---|---|
| Ccdc94 | 63.31143036 | 0.491610731 | 0.161962706 |
| Shd | 180.526558 | -0.729288517 | 0.0884885 |
| Fsd1 | 267.0610619 | -0.430771421 | 0.084924446 |
| Stap2 | 23.62948936 | -0.571242483 | 0.213530028 |
| Mpnd | 574.8071085 | -0.147808827 | 0.059442909 |
| Gm16276 | 0.401552702 | -1.41142361 | 1.502003895 |
| Sh3gl1 | 328.504642 | 0.23701648 | 0.084835602 |
| Chaf1a | 59.75683498 | 0.499330913 | 0.216454458 |
| Ubxn6 | 593.1748788 | -0.127912284 | 0.052104249 |
| Hdgfrp2 | 630.2550697 | -0.10304753 | 0.077002029 |
| Plin4 | 580.4887041 | 1.57387273 | 0.281427639 |
| Plin5 | 6.445794394 | -0.238863292 | 0.408131852 |
| Lrg1 | 47.22509705 | 4.28756675 | 0.404890054 |
| Sema6b | 1185.955188 | -0.84587022 | 0.094871384 |
| Tnfaip8l1 | 28.12050717 | -0.814431774 | 0.224323481 |
| AC026385.1 | 0.119587853 | 0.237305575 | 3.724945049 |
| Mydgf | 321.3040017 | 0.815605168 | 0.121624951 |
| Dpp9 | 687.0448644 | -0.024031203 | 0.055589325 |
| Mir7b | 2.668141924 | -0.645381006 | 0.635069792 |
| Fem1a | 839.6826125 | 0.078790195 | 0.077744956 |
| Ticam1 | 133.3080346 | 1.304160686 | 0.144556804 |
| Plin3 | 209.6287355 | 0.876240932 | 0.196124266 |
| Arrdc5 | 0 | NA | NA |
| Uhrf1 | 83.37304998 | 3.489347193 | 0.33411665 |
| Kdm4b | 809.6209241 | -0.362047596 | 0.081057655 |
| Ptprs | 5796.573917 | -0.740415342 | 0.087589877 |
| Mir6977 | 0 | NA | NA |
| Znrf4 | 0.092691418 | 1.183079489 | 3.724560917 |
| Safb2 | 645.9551208 | -0.222482062 | 0.178856083 |
| Safb | 970.0585309 | -0.156845534 | 0.159935135 |
| 2410015M20Rik | 237.0366201 | -0.111565177 | 0.141605435 |
| Rpl36 | 557.6166036 | 0.600899581 | 0.151122303 |
| Lonp1 | 634.6361425 | 0.282724781 | 0.074733191 |
| Catsperd | 6.596655121 | -0.681962241 | 0.418095714 |
| Ranbp3 | 724.4876957 | 0.186944243 | 0.058434725 |
| Vmac | 91.17240515 | -0.448177171 | 0.123112005 |
| Ndufa11 | 437.8651622 | -0.08650357 | 0.106936551 |
| Nrtn | 25.07553935 | -0.056634935 | 0.23210116 |
| Dus3l | 477.2858719 | -0.33003706 | 0.091220692 |
| Prr22 | 6.373319375 | -0.290305611 | 0.404888781 |
| Rfx2 | 53.12569894 | 1.131267376 | 0.172729894 |
| Acsbg2 | 0.382360723 | 1.851666359 | 1.651868503 |
| 1700061G19Rik | 1.170313971 | 0.851743456 | 1.129710746 |
| Mllt1 | 783.4489317 | 0.006398195 | 0.069812705 |
| Acer1 | 0.046345709 | 0.71820487 | 3.724945049 |
| Clpp | 295.2646874 | 0.298054657 | 0.087369956 |
| Alkbh7 | 82.81251037 | -0.244703444 | 0.181625709 |

| | | | |
|---|---|---|---|
| Pspn | 0.342416647 | -0.647263458 | 1.984346592 |
| Gtf2f1 | 546.3864564 | 0.791864576 | 0.131493989 |
| Khsrp | 1672.094708 | 0.205302775 | 0.040844885 |
| Slc25a41 | 1.408439352 | -0.564762627 | 0.779937915 |
| Gm17168 | 0.17329739 | -1.314315834 | 3.323993654 |
| Slc25a23 | 2717.008405 | -0.930021277 | 0.08981116 |
| Crb3 | 6.90059268 | 0.474868314 | 0.443168695 |
| Dennd1c | 54.0186804 | 3.525705202 | 0.28146015 |
| Tubb4a | 6699.351768 | -0.737488398 | 0.095294354 |
| Gm11110 | 11.24476628 | -0.260060186 | 0.336390635 |
| Tnfsf9 | 20.88192311 | 1.472611246 | 0.340161675 |
| Cd70 | 3.132506103 | 0.611136505 | 0.538735676 |
| Tnfsf14 | 6.752466611 | 5.824515023 | 1.005765285 |
| C3 | 1371.649863 | 6.652767394 | 0.4732168 |
| Mir6978 | 0 | NA | NA |
| Gpr108 | 284.67282 | 0.587072343 | 0.097460884 |
| Trip10 | 80.25548548 | 1.199166044 | 0.173856804 |
| Vav1 | 298.6647182 | 4.104696094 | 0.191240358 |
| Adgre1 | 147.8605273 | 2.418761592 | 0.244616932 |
| Vmn2r120 | 0 | NA | NA |
| Gm21834 | 0.060174537 | 0.237309859 | 3.724945049 |
| Gm25102 | 0 | NA | NA |
| Cntnap5c | 204.371656 | -0.117147659 | 0.149397226 |
| Rpl7a-ps5 | 12.52156343 | 0.872134504 | 0.457910086 |
| Gm5815 | 49.22368419 | -0.369733179 | 0.189602761 |
| Pdzph1 | 37.42674745 | 0.111466806 | 0.240074044 |
| Nudt12os | 0.099106094 | 1.209894437 | 3.72418822 |
| Nudt12 | 83.70631604 | 0.245782466 | 0.171584509 |
| Gm23769 | 0 | NA | NA |
| Gm23203 | 0 | NA | NA |
| Gm29051 | 0 | NA | NA |
| Gm23447 | 0 | NA | NA |
| Gm23773 | 0 | NA | NA |
| Obox6-ps1 | 0 | NA | NA |
| Gm4518 | 4.468190482 | -0.257698681 | 0.514824537 |
| Gm27023 | 0.131193343 | -0.71236593 | 2.978303909 |
| Gm25800 | 0 | NA | NA |
| Efna5 | 380.5177091 | -0.297767276 | 0.087255124 |
| Fbxl17 | 855.8478773 | -0.387372267 | 0.061068727 |
| Gm25348 | 2.594574602 | -0.890183685 | 0.630618665 |
| Gm24730 | 0 | NA | NA |
| 4930405O22Rik | 44.37612019 | -0.924961125 | 0.279802325 |
| Gm17228 | 21.45301415 | -0.871171556 | 0.302643412 |
| Gm24813 | 0 | NA | NA |
| A930002H24Rik | 1.337378336 | 0.281038429 | 1.005648102 |
| Fer | 419.2667044 | -0.030968068 | 0.062823856 |
| Pja2 | 6703.040193 | -0.431464387 | 0.059321406 |

| | | | |
|---|---|---|---|
| Gm24194 | 3.223858552 | -1.159162776 | 0.705570379 |
| Mir6420 | 0 | NA | NA |
| AU016765 | 0 | NA | NA |
| Man2a1 | 771.0017978 | 0.806422057 | 0.109801708 |
| Gm17133 | 0 | NA | NA |
| 4930583I09Rik | 0 | NA | NA |
| Gm25344 | 0 | NA | NA |
| Tmem232 | 14.97100193 | -0.0116351 | 0.263123604 |
| Gm25731 | 0 | NA | NA |
| Vapa | 1462.977574 | 0.451333327 | 0.090568309 |
| Txndc2 | 4.780619236 | 0.588799355 | 0.501713229 |
| Rab31 | 1383.126595 | 0.86285484 | 0.128803841 |
| Ppp4r1 | 548.0723031 | 0.497606312 | 0.086201432 |
| Ralbp1 | 849.2774884 | -0.006701668 | 0.083765121 |
| Twsg1 | 633.6219113 | 0.898996465 | 0.106008798 |
| Ankrd12 | 2379.847845 | -0.329444091 | 0.07240007 |
| Ndufv2 | 1210.518907 | 0.077424112 | 0.095973276 |
| Gm23264 | 0.127717592 | 0.237323721 | 3.006584758 |
| Wash1 | 277.8590539 | 0.375204155 | 0.077007101 |
| Ddx11 | 26.93519898 | 0.033635834 | 0.21788511 |
| Mtcl1 | 1748.393845 | -0.987245486 | 0.141220769 |
| Gm4705 | 22.01963504 | -0.829916428 | 0.272275528 |
| Rab12 | 884.2363581 | 0.046468807 | 0.085757265 |
| Themis3 | 0.652038697 | 1.116761791 | 1.306392448 |
| Ptprm | 817.855665 | -0.295453837 | 0.067059053 |
| Mir5709 | 2.098507799 | -0.138060736 | 0.74663796 |
| Gm22277 | 0 | NA | NA |
| Lrrc30 | 0.047948698 | -0.243569582 | 3.724945049 |
| Lama1 | 126.6409752 | 0.088433758 | 0.178767515 |
| Arhgap28 | 21.82346039 | 1.715066171 | 0.260000632 |
| Gm17097 | 0.284087786 | 1.19912272 | 1.80160308 |
| L3mbtl4 | 19.07932733 | -0.613965626 | 0.248262334 |
| Gm15974 | 0.045882455 | 0.237312084 | 3.724945049 |
| Tmem200c | 195.2513106 | 0.171423509 | 0.095546587 |
| Epb4.1l3 | 2609.206492 | -0.609935207 | 0.070715637 |
| 2410021H03Rik | 0.229803803 | 0.161911872 | 2.159244993 |
| Zbtb14 | 266.5595265 | -0.012000234 | 0.085534155 |
| A930029G22Rik | 17.70809633 | -0.518798589 | 0.265106748 |
| C030034I22Rik | 43.35133415 | 0.294010821 | 0.189684585 |
| A330050F15Rik | 100.5892871 | 0.185130019 | 0.128528875 |
| Gm22967 | 0 | NA | NA |
| Dlgap1 | 5541.017591 | -0.597455899 | 0.104256015 |
| 5031415H12Rik | 2.165235617 | 2.114174219 | 0.729254128 |
| Gm20703 | 2.008244415 | 0.636060586 | 1.123263769 |
| Tgif1 | 173.2492349 | 4.502704355 | 0.286946192 |
| Gm28727 | 2.554427972 | 1.873258181 | 0.71956521 |
| Mir1195 | 0 | NA | NA |

| | | | |
|---|---|---|---|
| Gm26510 | 1.053857694 | 2.51513489 | 1.121989169 |
| Gm9320 | 7.10377163 | 0.7469493 | 0.428268532 |
| Gm26561 | 0.274682661 | 0.718190974 | 3.724945049 |
| Gm16519 | 0 | NA | NA |
| Myl12b | 1231.826391 | -0.136660552 | 0.100986712 |
| Myl12a | 318.06527 | 1.990722874 | 0.187172779 |
| Myom1 | 11.02360487 | -0.156419723 | 0.323276979 |
| Lpin2 | 1181.411907 | 0.167209357 | 0.060102234 |
| Emilin2 | 338.4431336 | 3.356993716 | 0.292729538 |
| Smchd1 | 1270.998503 | 0.619981893 | 0.071765288 |
| Gm37639 | 37.94753101 | 0.006719169 | 0.258985625 |
| Gm38220 | 18.1195503 | 0.202769405 | 0.417103135 |
| Gm4707 | 30.94223545 | 0.06735574 | 0.329441938 |
| Ndc80 | 7.673135682 | 2.36778632 | 0.557731882 |
| Gm23115 | 0.060174537 | 0.237309859 | 3.724945049 |
| Mettl4-ps1 | 0.604171475 | 0.401682202 | 1.079068688 |
| Spdya | 9.137166137 | -1.0449722 | 0.368826738 |
| Trmt61b | 64.18131499 | 0.37867929 | 0.152616273 |
| Wdr43 | 540.8743764 | 0.861080149 | 0.106050928 |
| Snord92 | 1.968787839 | -0.322150236 | 0.906136869 |
| Snord53 | 3.604104618 | -0.236619252 | 0.715765613 |
| Gm22858 | 0.576946659 | -2.062442511 | 3.565006659 |
| Fam179a | 77.43913187 | 0.974186988 | 0.179615487 |
| Gm15641 | 0 | NA | NA |
| BC027072 | 1.833725576 | 0.547249833 | 0.727829425 |
| Clip4 | 625.3662385 | 0.072921085 | 0.055088617 |
| Alk | 126.1837875 | 0.118515987 | 0.111607592 |
| Gm24736 | 0.908322817 | -1.719742694 | 1.107451984 |
| Gm26963 | 0 | NA | NA |
| Ypel5 | 905.935313 | 0.515557254 | 0.07974793 |
| Lbh | 1015.797448 | 1.25295079 | 0.135493774 |
| Gm23649 | 0.075043205 | 0.237309859 | 3.724945049 |
| Lclat1 | 432.1555874 | 0.057714193 | 0.08075651 |
| Gm9311 | 0.089089296 | 0.237305797 | 3.724945049 |
| Gm25406 | 3.791534426 | -0.97748313 | 0.505952151 |
| Capn13 | 0.17782045 | 0.70160365 | 2.294087018 |
| Galnt14 | 193.3014098 | -0.15666969 | 0.151130003 |
| D630014O11Rik | 13.27744998 | -0.874777274 | 0.433951254 |
| Ehd3 | 1261.372702 | 0.541820611 | 0.085421046 |
| Xdh | 746.6583068 | 5.850902826 | 0.267902797 |
| Gm24475 | 0 | NA | NA |
| Srd5a2 | 1.0712781 | -0.363819572 | 0.967334814 |
| Memo1 | 287.5100735 | 0.125381198 | 0.084354111 |
| Dpy30 | 145.3608656 | 0.060509031 | 0.120600358 |
| Spast | 787.6234006 | -0.092081813 | 0.05461144 |
| Slc30a6 | 112.2934621 | 0.214630632 | 0.131149906 |
| Nlrc4 | 11.8016311 | 2.350370031 | 0.371237881 |

| | | | |
|---|---|---|---|
| Yipf4 | 489.5110183 | 0.129092769 | 0.097274184 |
| Birc6 | 5732.824248 | 0.115070708 | 0.064939198 |
| Gm26749 | 0.249552943 | 0.717769878 | 1.84829206 |
| Ttc27 | 207.7543264 | 0.293721003 | 0.079706873 |
| Ltbp1 | 508.8403771 | 2.430640125 | 0.182307459 |
| Gm6276 | 0.110870145 | 0.238305784 | 3.724874707 |
| Rasgrp3 | 150.64243 | -0.152176588 | 0.187538312 |
| Fam98a | 451.495364 | 0.288545771 | 0.088020125 |
| Gm24126 | 0 | NA | NA |
| Gm26124 | 0 | NA | NA |
| Gm16391 | 0 | NA | NA |
| Gm17538 | 12.6024622 | -0.489767899 | 0.30311983 |
| Gm5229 | 0.072025462 | 0.237312084 | 3.724945049 |
| Crim1 | 1289.41108 | -0.087958131 | 0.07765943 |
| Gm25069 | 0.582565216 | -0.642380244 | 1.367310588 |
| Fez2 | 418.1889862 | 0.504438824 | 0.093064388 |
| Gm10093 | 94.11779407 | 1.144791141 | 0.161736342 |
| Vit | 19.4303546 | -0.410500969 | 0.238966995 |
| Gm22562 | 0.092493346 | -0.24357403 | 3.724945049 |
| Strn | 1780.921145 | -0.571563768 | 0.060305921 |
| Heatr5b | 1002.913214 | -0.402363381 | 0.060037154 |
| Gpatch11 | 308.9091273 | -0.441412834 | 0.124170849 |
| Gm6548 | 198.422454 | 4.181322431 | 0.202884482 |
| Eif2ak2 | 2234.053102 | 4.98900515 | 0.16607383 |
| n-R5s29 | 0.136963818 | 0.718204821 | 3.59954433 |
| Gm26637 | 5.907155527 | 4.28352375 | 0.883880436 |
| Sult6b1 | 1.803778753 | 1.012794284 | 0.881003123 |
| Cebpzos | 71.15640056 | -0.156686833 | 0.154085632 |
| Cebpz | 422.2918863 | 0.178222318 | 0.088540208 |
| Ndufaf7 | 198.2430765 | -0.402693541 | 0.08143548 |
| Prkd3 | 324.5446214 | 0.42546584 | 0.116508021 |
| Gm26957 | 0 | NA | NA |
| Qpct | 101.3108535 | -0.197819464 | 0.159035048 |
| Cdc42ep3 | 151.9232374 | -0.295516503 | 0.119978498 |
| Gm17315 | 8.227418531 | -0.410758535 | 0.369657357 |
| Rmdn2 | 100.9181395 | -0.090584918 | 0.112579564 |
| 4921513D11Rik | 17.43713135 | -0.041683981 | 0.272378244 |
| Cyp1b1 | 282.3238744 | 3.183391886 | 0.22987413 |
| Gm37555 | 0.136963818 | 0.718204821 | 3.59954433 |
| Atl2 | 973.0030504 | -0.370497281 | 0.069132503 |
| Hnrnpll | 974.8453895 | -0.010467768 | 0.085485564 |
| Gm22215 | 0 | NA | NA |
| Gm23125 | 0 | NA | NA |
| Galm | 27.71090595 | 0.368093017 | 0.300628133 |
| Gm25706 | 3.582960898 | -0.427728935 | 0.546954758 |
| Srsf7 | 905.8057965 | 0.196459676 | 0.072103049 |
| Ttc39d | 1.912265302 | 0.611511304 | 0.709177723 |

| | | | |
|---|---|---|---|
| Gemin6 | 32.80368467 | 0.235498285 | 0.202855127 |
| Dhx57 | 544.389297 | -0.422067173 | 0.104481423 |
| Morn2 | 84.79043694 | 0.057316964 | 0.165106538 |
| Arhgef33 | 272.4084982 | -0.170483893 | 0.302953504 |
| Gm10190 | 0.322675615 | -0.688343563 | 1.496939308 |
| Sos1 | 1511.796088 | -0.143529083 | 0.052544916 |
| Cdkl4 | 159.4655411 | -0.82862079 | 0.165202259 |
| Map4k3 | 1163.46542 | 0.015024307 | 0.050793992 |
| Gm9959 | 0.271824581 | -0.201163116 | 1.781450756 |
| Gm9386 | 0.459282249 | 0.711067407 | 1.508609389 |
| Tmem178 | 593.6338004 | -0.479565566 | 0.090851766 |
| Thumpd2 | 64.74403033 | 0.490579675 | 0.151653685 |
| Slc8a1 | 2319.537813 | -0.289825501 | 0.087632977 |
| Gm11096 | 0.461313532 | -0.692378097 | 1.370210424 |
| Rpl31-ps25 | 0 | NA | NA |
| Gm27359 | 0 | NA | NA |
| Gm22386 | 0 | NA | NA |
| Gm23400 | 0 | NA | NA |
| Gm6594 | 14.12482822 | 0.960125848 | 0.318819126 |
| Gm29052 | 0 | NA | NA |
| Pkdcc | 80.91185254 | -0.03115473 | 0.145232834 |
| Gm24240 | 0 | NA | NA |
| Rpl21-ps7 | 2.037653348 | 0.522245919 | 0.753158813 |
| Eml4 | 575.2161808 | -0.16937674 | 0.079617703 |
| Cox7a2l | 853.8877004 | -0.054674751 | 0.078880423 |
| Kcng3 | 81.41433254 | -1.139112532 | 0.188818309 |
| Mta3 | 891.3237956 | -0.215518582 | 0.058204 |
| Haao | 4.556506885 | 3.508713334 | 0.954465968 |
| C430042M11Rik | 1.362936493 | 0.782932276 | 0.828400909 |
| Gm23905 | 0.061920613 | 0.718207094 | 3.724945049 |
| Gm37549 | 0.452867449 | 1.20363687 | 1.848481458 |
| Zfp36l2 | 928.29596 | 2.038101353 | 0.226059811 |
| Thada | 445.9157641 | -0.172045772 | 0.09138923 |
| Gm24492 | 0.916014489 | -0.045782245 | 0.949626724 |
| Gm24722 | 0.550570057 | 0.531872415 | 1.33824865 |
| Plekhh2 | 185.6708911 | -0.392737966 | 0.110237045 |
| Gm26066 | 0.13205505 | -1.080010628 | 3.656016608 |
| Dync2li1 | 99.93135035 | -0.239596932 | 0.142700191 |
| Abcg5 | 0.732414602 | 1.64042298 | 1.422940901 |
| Abcg8 | 0.118974955 | -0.243576254 | 3.724945049 |
| Lrpprc | 1213.010164 | -0.209681905 | 0.072851649 |
| 1110020A21Rik | 4.802739388 | -0.425501856 | 0.432295337 |
| Ppm1b | 1207.845512 | -0.367423253 | 0.062794656 |
| Slc3a1 | 11.21614055 | 0.069239179 | 0.392824675 |
| Prepl | 4734.893635 | -0.518086855 | 0.053981099 |
| Camkmt | 72.64996259 | -0.174698608 | 0.14098955 |
| Rpl31-ps16 | 2.862600439 | 0.075326487 | 0.663625735 |

| | | | |
|---|---|---|---|
| Gm27178 | 0 | NA | NA |
| Six3os1 | 52.19109397 | -0.510009108 | 0.225331176 |
| Gm27435 | 0.089062654 | 0.237318756 | 3.724945049 |
| Gm27801 | 0.250814404 | -1.548900481 | 1.784277208 |
| Six3 | 109.2161158 | -0.544990577 | 0.150571158 |
| Gm27610 | 0.226876767 | -1.574924574 | 2.148259335 |
| Gm27283 | 0.388809915 | -1.579644358 | 1.718958247 |
| Gm27275 | 0.04393175 | -0.243571806 | 3.724945049 |
| Gm27502 | 0.090665643 | -0.724455696 | 3.724945049 |
| Gm27427 | 0.359940881 | -0.882091968 | 1.711769764 |
| Gm28528 | 1.48446612 | -0.331651859 | 0.920712679 |
| Six2 | 7.803849726 | 0.696601366 | 0.355139409 |
| Gm29418 | 0.434220697 | -0.727593856 | 1.401509288 |
| Srbd1 | 196.7237653 | -0.062676431 | 0.115524981 |
| Prkce | 4692.067451 | -0.520703111 | 0.093379511 |
| 2010106C02Rik | 0 | NA | NA |
| Gm10309 | 10.12416963 | 4.009685852 | 0.78370353 |
| Gm22235 | 0 | NA | NA |
| Epas1 | 1364.101885 | 0.1618695 | 0.114040708 |
| Gm29168 | 0.150086409 | 0.237305146 | 3.724945049 |
| Tmem247 | 0 | NA | NA |
| Atp6v1e2 | 0.434133818 | 0.343715658 | 1.544507561 |
| Rhoq | 718.01171 | 0.492369689 | 0.10537561 |
| Pigf | 100.958299 | 0.111865341 | 0.150589791 |
| Cript | 263.4740334 | 0.278418035 | 0.140895366 |
| Gm24648 | 0 | NA | NA |
| Gm5499 | 18.0195093 | 0.230918076 | 0.26113868 |
| Socs5 | 1211.318638 | -0.456178673 | 0.092796329 |
| Gm22346 | 0.044191549 | -0.243571806 | 3.724945049 |
| Mcfd2 | 543.0470171 | 0.677128814 | 0.085685529 |
| 4833418N02Rik | 13.1105544 | -0.135017983 | 0.291888859 |
| Ttc7 | 196.4810743 | 0.917564431 | 0.146455887 |
| Gm15978 | 0.6240965 | -1.199808209 | 1.22945889 |
| 0610012D04Rik | 0.332525094 | 1.054173008 | 1.700876742 |
| 1700011E24Rik | 3.15672782 | 0.87897097 | 0.576180228 |
| Calm2 | 8582.912101 | -0.456674708 | 0.104391749 |
| Gm26005 | 0.04393175 | -0.243571806 | 3.724945049 |
| Gm28676 | 0.161026707 | 1.711863256 | 3.451443504 |
| Epcam | 12.12751263 | 3.080843153 | 0.482665263 |
| Msh2 | 333.7442204 | -0.362069236 | 0.097773007 |
| Kcnk12 | 130.455902 | -0.129597375 | 0.136140906 |
| Rpl36-ps4 | 0.224455361 | -0.243577557 | 3.146467167 |
| Msh6 | 331.2527847 | 0.051471798 | 0.07660589 |
| Fbxo11 | 1810.721455 | -0.119633757 | 0.066424647 |
| Gm4832 | 0.217012189 | 1.638157874 | 2.152597871 |
| Gm26612 | 0.40569442 | -1.493434057 | 1.411856804 |
| Gm22542 | 0.253603277 | -0.216623787 | 2.120224239 |

| | | | |
|---|---|---|---|
| Gm23947 | 0 | NA | NA |
| Foxn2 | 537.5500919 | 0.14364227 | 0.092992562 |
| Ppp1r21 | 678.815562 | -0.35868355 | 0.057455412 |
| Gm38109 | 8.730948186 | -0.979471259 | 0.407535641 |
| Ston1 | 93.35916316 | 0.476978392 | 0.176025251 |
| Gtf2a1l | 2.669826932 | -0.230008725 | 0.586662698 |
| Lhcgr | 3.114871368 | -1.624049769 | 0.678548047 |
| Fshr | 0.045882455 | 0.237312084 | 3.724945049 |
| Gm10184 | 3.494817459 | 0.667719409 | 0.547371292 |
| Nrxn1 | 7760.329357 | -0.758074734 | 0.098224285 |
| Gm24510 | 0 | NA | NA |
| Gm10493 | 3.600385196 | -0.504064886 | 0.534014307 |
| Gm19815 | 7.90218533 | -0.738127212 | 0.543684014 |
| Gm20491 | 88.08114114 | -0.95627521 | 0.314409405 |
| 4930480K15Rik | 0.097899064 | -0.724453471 | 3.724945049 |
| AC151286.1 | 0.277228681 | -2.065548249 | 3.683251455 |
| Gm10308 | 1.537590905 | -1.081941316 | 0.852937785 |
| Gm27475 | 0 | NA | NA |
| Gm6741 | 17.12533191 | -0.437783973 | 0.355925238 |
| Gm15404 | 0.089089296 | 0.237305797 | 3.724945049 |
| Gm15403 | 0 | NA | NA |
| Gm24244 | 0 | NA | NA |
| AC131325.1 | 0 | NA | NA |
| Gm27039 | 0 | NA | NA |
| Adcyap1 | 181.5271029 | 0.221245034 | 0.182093217 |
| Gm25935 | 0 | NA | NA |
| Gm22593 | 0.196742721 | 0.712364957 | 2.231534681 |
| Mettl4 | 150.9550503 | -0.118781604 | 0.161654854 |
| 2700099C18Rik | 6.495084589 | 1.141622164 | 0.451616228 |
| Gm27038 | 3.867668655 | -1.762224599 | 0.636063711 |
| Gm1976 | 94.15986379 | -0.844512952 | 0.121802734 |
| Gm23964 | 0.139856492 | 0.237322262 | 3.574411025 |
| Gm26734 | 6.887014156 | -0.421394765 | 0.419818134 |
| Gm23102 | 0 | NA | NA |
| Gm20939 | 23.37788906 | 0.703390498 | 0.353722155 |
| Vmn1r-ps151 | 0 | NA | NA |
| Vmn1r-ps152 | 0.092691418 | 1.183079489 | 3.724560917 |
| Vmn1r238 | 0.090890357 | 0.718200421 | 3.724945049 |
| Crem | 361.0165932 | 1.540563653 | 0.163409634 |
| Gm6225 | 15.32231194 | 0.630519971 | 0.371408189 |
| Cul2 | 847.8838813 | 0.29267419 | 0.080720447 |
| G430049J08Rik | 0.148708979 | 1.198497773 | 2.868467391 |
| Bambi | 60.79279935 | 0.677528848 | 0.177248521 |
| Gm23643 | 0 | NA | NA |
| Gm24632 | 0 | NA | NA |
| Lyzl1 | 0 | NA | NA |
| Gm10557 | 201.5185117 | 0.413152181 | 0.146758244 |

| | | | |
|---|---|---|---|
| Map3k8 | 99.88469274 | 2.232669352 | 0.183886863 |
| Gm26865 | 0.332096027 | -0.374840273 | 1.686898156 |
| Mtpap | 376.5909295 | 0.202950806 | 0.095696783 |
| 9430020K01Rik | 1287.14579 | -0.411564065 | 0.111469721 |
| Gm10556 | 0.381271471 | 1.527733027 | 1.649781887 |
| Svil | 295.6710269 | 0.43393612 | 0.103886231 |
| Gm26682 | 7.326093239 | 0.737201415 | 0.466946703 |
| Zfp438 | 70.78775138 | -0.065956421 | 0.148792164 |
| Gm10125 | 71.18097263 | -0.506592337 | 0.188481024 |
| Gm26575 | 19.11664666 | -0.648073168 | 0.33126114 |
| Gm2238 | 59.71255148 | -0.922386294 | 0.194993791 |
| Zeb1 | 1461.053986 | -0.061570033 | 0.051780916 |
| Gm16147 | 0 | NA | NA |
| Gm22709 | 4.178956302 | -0.304483542 | 0.516841986 |
| Arhgap12 | 782.7285597 | -0.036708641 | 0.075502314 |
| Kif5b | 4777.179878 | 0.265773276 | 0.115622223 |
| Gm17036 | 1.215361072 | 1.479263389 | 1.036025737 |
| Rpl27-ps3 | 9.972102879 | 0.93507357 | 0.368923272 |
| Gm6291 | 0 | NA | NA |
| Epc1 | 763.0696826 | 0.035504621 | 0.074534501 |
| Gm28529 | 13.22339979 | 0.009419257 | 0.35135496 |
| Mir1893 | 0 | NA | NA |
| 4921524L21Rik | 0.072025462 | 0.237312084 | 3.724945049 |
| Rab18 | 1241.658341 | 0.142663324 | 0.08046939 |
| Mkx | 204.0449348 | -0.411408973 | 0.123512807 |
| Armc4 | 20.81306997 | 0.087740261 | 0.245902017 |
| Mpp7 | 246.8020213 | -0.471194481 | 0.102734751 |
| Fabp5l2 | 0.240849226 | 0.236870678 | 1.864799325 |
| Gm24292 | 0 | NA | NA |
| Wac | 1913.852717 | 0.239157732 | 0.063230523 |
| Gm7502 | 0.768676318 | -2.245996347 | 1.185245137 |
| Gm5819 | 9.13781552 | -0.53006765 | 0.330823934 |
| Gm26119 | 0 | NA | NA |
| Gm37148 | 0 | NA | NA |
| Fzd8 | 66.68569282 | 0.188948636 | 0.152164953 |
| Gjd4 | 0.815409053 | 0.906392039 | 1.053672175 |
| Ccny | 1305.979739 | -0.094850296 | 0.05154165 |
| Gm22765 | 0 | NA | NA |
| Cetn1 | 0.038981395 | 0.718207094 | 3.724945049 |
| Colec12 | 279.2173052 | 1.081143379 | 0.143996987 |
| Gm17430 | 0 | NA | NA |
| Gm7497 | 4.858631947 | 0.512287376 | 0.591359578 |
| Gm23350 | 0 | NA | NA |
| Gm23637 | 0 | NA | NA |
| Gm5501 | 0 | NA | NA |
| Thoc1 | 432.2063309 | -0.037453705 | 0.079741934 |
| Usp14 | 1395.726703 | -0.131154388 | 0.050411569 |

| | | | |
|---|---|---|---|
| Rock1 | 1035.477942 | 0.359834481 | 0.056724775 |
| Gm2531 | 25.92851566 | -0.503662942 | 0.315787012 |
| Gm26244 | 55.4930898 | 0.358999757 | 0.191398994 |
| Greb1l | 215.658039 | -0.646582003 | 0.145962117 |
| AC124740.1 | 0 | NA | NA |
| Gm10349 | 1.210176943 | -0.389373227 | 0.942096198 |
| Gm24894 | 2.156355716 | -0.617471565 | 0.659076552 |
| Esco1 | 664.6422751 | 0.150930665 | 0.081085068 |
| Snrpd1 | 236.8850658 | 0.408366917 | 0.131344416 |
| Abhd3 | 409.5174636 | -0.741893276 | 0.145182579 |
| 4930563E18Rik | 1.299310135 | -1.293497967 | 0.855632944 |
| Mib1 | 3835.229447 | -0.330057775 | 0.060370022 |
| Gm25116 | 4.118368001 | -1.228806176 | 0.709446449 |
| Gm21747 | 0.049553047 | 0.71820487 | 3.724945049 |
| Mir133a-1 | 0 | NA | NA |
| Mir1b | 7.73100226 | -0.881930818 | 0.381485867 |
| mmu-mir-1a-2 | 0 | NA | NA |
| 1010001N08Rik | 0.099155932 | 0.718202646 | 3.724945049 |
| Gata6 | 1.608823659 | 0.730603014 | 0.848214808 |
| Rbbp8 | 116.5073145 | 0.910690185 | 0.11838316 |
| Gm6277 | 35.32926637 | -0.274883207 | 0.243986502 |
| Cables1 | 273.0530706 | -0.736357084 | 0.120303299 |
| Mir1901 | 6.131714793 | 0.495435557 | 0.429701717 |
| Tmem241 | 88.72054137 | -0.532692263 | 0.159372673 |
| Riok3 | 978.9154087 | 0.590940341 | 0.084180744 |
| Gm25289 | 0.089062654 | 0.237318756 | 3.724945049 |
| 3110002H16Rik | 281.027959 | 0.086742198 | 0.091157651 |
| Npc1 | 1148.204684 | 0.170595558 | 0.08683151 |
| Gm15956 | 8.077703934 | 0.13355774 | 0.379958699 |
| Ankrd29 | 171.8859524 | -0.55886363 | 0.089006134 |
| Gm16072 | 0.87006192 | -1.119348403 | 1.100667724 |
| Lama3 | 110.321609 | -0.408898838 | 0.166794013 |
| Gm29201 | 0.436251183 | -1.264025283 | 1.726684964 |
| Gm29199 | 0 | NA | NA |
| Gm29200 | 0.582012116 | 1.810625573 | 1.251940963 |
| Gm22340 | 0.038981395 | 0.718207094 | 3.724945049 |
| Ttc39c | 360.5212832 | 0.241519159 | 0.104900632 |
| Mir1948 | 2.898584899 | -1.116204821 | 0.654255321 |
| Cabyr | 18.12948923 | 0.830806245 | 0.444117989 |
| Gm7599 | 1.737466599 | -1.349753014 | 0.92705015 |
| Osbpl1a | 2161.262522 | -0.80570738 | 0.069947881 |
| Gm5687 | 0.269295688 | 0.525511902 | 2.060259505 |
| Gm26407 | 27.03744251 | -1.250251669 | 0.287875449 |
| Impact | 3627.068778 | -0.175784901 | 0.083382428 |
| Hrh4 | 0.228099782 | 1.1839131 | 3.115458725 |
| Gm22251 | 0 | NA | NA |
| Zfp521 | 803.5141072 | 0.289443932 | 0.103549637 |

| | | | |
|---|---|---|---|
| Gm5160 | 3.237902636 | 0.28079644 | 0.598077388 |
| Gm6457 | 44.70599757 | 0.506829539 | 0.157498365 |
| Ss18 | 416.8945723 | 0.330319963 | 0.0966998 |
| Gm24924 | 13.08384193 | -0.006408588 | 0.291246388 |
| Psma8 | 5.559946366 | 0.999794444 | 0.476203352 |
| Taf4b | 120.7625759 | 0.559763064 | 0.138404006 |
| Kctd1 | 603.8212261 | -0.855622487 | 0.099767701 |
| Gm6304 | 6.355730337 | -0.813945139 | 0.443646139 |
| Aqp4 | 4262.513783 | 1.03838373 | 0.140896418 |
| Chst9 | 8.958957809 | -1.094757794 | 0.375693956 |
| Gm10036 | 42.03144848 | 0.86560514 | 0.260372103 |
| Gm24228 | 0.072025462 | 0.237312084 | 3.724945049 |
| Gm4835 | 1.43143147 | 0.048458778 | 0.838194565 |
| Gm22389 | 1.98025533 | -0.161811238 | 0.817922864 |
| Gm24385 | 0 | NA | NA |
| Gm7665 | 11.35479647 | 4.725033395 | 0.712561203 |
| Cdh2 | 2013.677982 | 0.140442967 | 0.062216555 |
| Gm15485 | 0 | NA | NA |
| Gm15328 | 30.8296316 | -1.409248836 | 0.287540812 |
| Gm7670 | 0 | NA | NA |
| 1700001G01Rik | 0.046345709 | 0.71820487 | 3.724945049 |
| Gm22470 | 0 | NA | NA |
| Gm24015 | 0 | NA | NA |
| Gm26235 | 0.048949532 | -0.243569582 | 3.724945049 |
| Gm25392 | 0 | NA | NA |
| Gm24238 | 0 | NA | NA |
| Dsc3 | 13.88762389 | 0.407023874 | 0.303187822 |
| Dsc2 | 11.39730211 | -0.148248281 | 0.368536224 |
| Dsc1 | 2.506558332 | 3.977426271 | 1.250978728 |
| Dsg1c | 4.094471938 | -2.302207196 | 0.780360342 |
| Dsg1a | 1.903639553 | -1.986619749 | 3.021840169 |
| Dsg1b | 0.835693444 | -0.104411181 | 1.340454303 |
| Dsg4 | 0 | NA | NA |
| Dsg3 | 1.159328825 | 1.865101413 | 1.099390603 |
| Dsg2 | 36.65644163 | 0.143895775 | 0.193843975 |
| Gm16090 | 0.646850989 | -0.044089943 | 1.355344115 |
| Ttr | 2265.392118 | -2.815892283 | 0.335448551 |
| Gm10269 | 7.797321979 | -0.02536714 | 0.407819833 |
| B4galt6 | 2045.964961 | -0.429480992 | 0.062708195 |
| Trappc8 | 1142.654653 | 0.245099843 | 0.05329236 |
| Gm24959 | 4.935114692 | -0.401175283 | 0.452487834 |
| Rnf125 | 36.46586033 | 3.497366089 | 0.402089039 |
| Rnf138 | 267.4053105 | 0.242554388 | 0.087425014 |
| Mep1b | 0 | NA | NA |
| Gm6378 | 0.155469778 | 0.237325261 | 2.829060383 |
| Garem | 494.2667245 | -0.397029894 | 0.099761478 |
| Gm24372 | 0.044191549 | -0.243571806 | 3.724945049 |

| | | | |
|---|---|---|---|
| Gm22886 | 0 | NA | NA |
| Klhl14 | 23.8882674 | -0.713840858 | 0.244612375 |
| 4930426D05Rik | 6.412194933 | -0.966836424 | 0.42982514 |
| Ccdc178 | 1.26832265 | 1.440872632 | 1.064791609 |
| Asxl3 | 371.9529472 | -0.340532186 | 0.117398375 |
| Nol4 | 882.8164278 | -0.982914373 | 0.095323043 |
| Gm22495 | 0 | NA | NA |
| Dtna | 1930.500953 | -0.151224185 | 0.084539991 |
| Gm15972 | 1.928713869 | -1.086042433 | 0.825379616 |
| Mapre2 | 3079.479868 | -0.487045515 | 0.060441207 |
| Mir6360 | 0 | NA | NA |
| Gm23207 | 0 | NA | NA |
| Zfp397 | 704.3356492 | -0.039835765 | 0.08304926 |
| Zscan30 | 8.824857606 | -0.210838262 | 0.327910205 |
| Zfp35 | 243.5789265 | 0.332084269 | 0.093872967 |
| Zfp191 | 825.6271123 | 0.434247543 | 0.097138409 |
| Ino80c | 294.4170576 | 0.559254875 | 0.109225891 |
| Galnt1 | 801.5304355 | 0.339000435 | 0.089014363 |
| Gm7701 | 14.47583842 | -0.823030208 | 0.28392994 |
| Mir187 | 0.214710016 | 1.850837512 | 3.129502904 |
| 2700062C07Rik | 106.228182 | 0.190836244 | 0.114567988 |
| Rprd1a | 820.2179591 | -0.158132325 | 0.069046895 |
| Gm23606 | 0.061920613 | 0.718207094 | 3.724945049 |
| Slc39a6 | 544.4852359 | 0.213476161 | 0.093415113 |
| Elp2 | 877.5310991 | -0.208299069 | 0.049762343 |
| Mocos | 101.3743352 | 3.425284765 | 0.210621936 |
| n-R5s23 | 0.045882455 | 0.237312084 | 3.724945049 |
| Gm9955 | 0.957795399 | -0.75261903 | 1.00484807 |
| Fhod3 | 695.6586359 | -0.284172929 | 0.130749191 |
| Gm24294 | 1.176033076 | -0.818600077 | 0.969789469 |
| Tpgs2 | 457.1009357 | -0.601120661 | 0.075294215 |
| AW554918 | 270.5192402 | -0.32357678 | 0.078839171 |
| Gm16558 | 5.405763959 | -1.429725656 | 0.505006281 |
| Celf4 | 4385.110905 | -0.409004761 | 0.090894392 |
| Gm25845 | 9.017131998 | -1.782790315 | 0.385040983 |
| Gm7847 | 0 | NA | NA |
| Gm23674 | 0 | NA | NA |
| Gm26106 | 0 | NA | NA |
| Gm7729 | 7.595035811 | -0.442425994 | 0.397805508 |
| Gm24246 | 0 | NA | NA |
| Pik3c3 | 522.6692759 | -0.258970219 | 0.057914795 |
| Rpl29-ps5 | 0.839167439 | -0.379877321 | 1.00265086 |
| Gm22870 | 0 | NA | NA |
| Gm7936 | 0.203976984 | -0.635344147 | 2.157428343 |
| Rit2 | 621.7940845 | -0.464077676 | 0.092352383 |
| Gm23283 | 0 | NA | NA |
| Syt4 | 3383.452968 | 0.078589009 | 0.077638889 |

| | | | |
|---|---|---|---|
| Gm26658 | 4.064855943 | 0.240975834 | 0.502343183 |
| Gm25396 | 0 | NA | NA |
| Slc25a46 | 1147.053562 | 0.077611149 | 0.080877027 |
| B930094E09Rik | 4.156831401 | 0.663981877 | 0.516617755 |
| Sap130 | 827.4019423 | -0.074809886 | 0.055299106 |
| Ammecr1l | 554.7126231 | 0.287073017 | 0.077699284 |
| Gm26533 | 2.98239301 | -0.004459894 | 0.567246031 |
| Polr2d | 122.288999 | 0.496313437 | 0.117940832 |
| Wdr33 | 970.9585418 | 0.050903604 | 0.053584001 |
| Gm6665 | 1.827239475 | 1.362212733 | 0.775683504 |
| Gm26823 | 0.062426873 | 0.237312084 | 3.724945049 |
| Sft2d3 | 54.7958328 | -0.034885012 | 0.1499573 |
| Lims2 | 61.79027259 | 0.118512918 | 0.16104958 |
| Gpr17 | 643.1742382 | -0.47128156 | 0.284605495 |
| Myo7b | 0.405683672 | -0.789985188 | 1.595166951 |
| Gm16344 | 0.401548357 | -0.177910996 | 1.534592212 |
| Gm16345 | 0.075043205 | 0.237309859 | 3.724945049 |
| Iws1 | 844.5236566 | -0.044149204 | 0.077474971 |
| Proc | 0.048949532 | -0.243569582 | 3.724945049 |
| Gm26717 | 0.046345709 | 0.71820487 | 3.724945049 |
| Map3k2 | 1196.837555 | -0.023609335 | 0.088652113 |
| Ercc3 | 374.8626789 | 0.332010059 | 0.066363674 |
| Gm22325 | 0 | NA | NA |
| A830052D11Rik | 8.708922263 | -1.798586921 | 0.409111885 |
| Bin1 | 1973.364051 | -0.374192576 | 0.041141029 |
| Gypc | 57.79268309 | 2.868338436 | 0.301384027 |
| Gm25826 | 0 | NA | NA |
| 4930455D15Rik | 0.382692223 | 1.64145076 | 1.546741997 |
| Tslp | 1.429800265 | 2.019261398 | 0.953597436 |
| Gm26378 | 0.072025462 | 0.237312084 | 3.724945049 |
| Wdr36 | 397.0988872 | 0.257561275 | 0.062828514 |
| Mir6361 | 0 | NA | NA |
| 2310026I22Rik | 3.939202308 | -0.507904665 | 0.523151168 |
| Camk4 | 6160.809613 | -1.02296676 | 0.157867433 |
| Gm22814 | 0 | NA | NA |
| Stard4 | 406.2855849 | -0.347608163 | 0.09950134 |
| Nrep | 1304.95571 | -0.817574906 | 0.156709111 |
| Gm10549 | 1.094754251 | -1.092887314 | 0.959059712 |
| Epb4.1l4aos | 32.18986031 | 0.664908881 | 0.241987136 |
| Gm23639 | 3.703464063 | 1.095365806 | 0.529664616 |
| Epb4.1l4a | 142.1446161 | 0.26760098 | 0.129290141 |
| Gm22788 | 0 | NA | NA |
| Gm24432 | 0 | NA | NA |
| Gm23411 | 0 | NA | NA |
| Gm10548 | 0.669147434 | 1.821233072 | 1.253786541 |
| Apc | 7569.545602 | -0.526492615 | 0.074992762 |
| Srp19 | 313.7947171 | 0.242749201 | 0.090839004 |

| | | | |
|---|---|---|---|
| Reep5 | 3174.602423 | -0.17138981 | 0.064476472 |
| Pkd2l2 | 2.995780043 | 0.200122895 | 0.52334038 |
| Fam13b | 1185.027783 | 0.113749965 | 0.064578296 |
| n-R5s24 | 0.131453142 | -0.714386264 | 2.976894461 |
| Wnt8a | 0.202010919 | 1.156346953 | 2.194321124 |
| Nme5 | 50.66745939 | -0.489307254 | 0.16266472 |
| 4933408B17Rik | 16.70085405 | -0.644133486 | 0.321549268 |
| Brd8 | 925.5748569 | -0.113242209 | 0.074599654 |
| Kif20a | 22.72681451 | 2.305478724 | 0.415995422 |
| Cdc23 | 555.3324902 | -0.182854198 | 0.072802128 |
| n-R5s25 | 0 | NA | NA |
| Gfra3 | 0 | NA | NA |
| Cdc25c | 5.025978301 | 1.085614066 | 0.477037096 |
| Gm3550 | 0.207873913 | 1.07512317 | 2.592691985 |
| 2010110K18Rik | 0.294784161 | 0.718223719 | 2.01714337 |
| Fam53c | 1037.28919 | 0.041996581 | 0.054946427 |
| Kdm3b | 1100.794087 | 0.039211945 | 0.050977644 |
| Gm26538 | 0.390497984 | 0.674513476 | 1.548740505 |
| Reep2 | 1197.302419 | -0.273029576 | 0.068205811 |
| Egr1 | 3179.097212 | 0.03997565 | 0.353898174 |
| Etf1 | 1390.357339 | 0.875914844 | 0.099604845 |
| Hspa9 | 2869.344892 | 0.689674643 | 0.10679982 |
| Gm22200 | 4.566833944 | -0.132132802 | 0.505766332 |
| Gm26109 | 3.869151513 | -0.65810562 | 0.590389085 |
| Ctnna1 | 941.2579512 | 0.94729212 | 0.102193549 |
| Lrrtm2 | 1096.201602 | -0.276878534 | 0.078864162 |
| Sil1 | 202.6507413 | 0.196661081 | 0.096499004 |
| Gm5239 | 0.164165709 | -0.724458634 | 2.781083781 |
| Snhg4 | 73.54980766 | -0.527352214 | 0.145712125 |
| Gm28285 | 6.985251871 | 0.36788969 | 0.363567404 |
| Mir1949 | 0 | NA | NA |
| mmu-mir-1949 | 0 | NA | NA |
| Snora74a | 0 | NA | NA |
| AC121821.1 | 0.048949532 | -0.243569582 | 3.724945049 |
| Matr3 | 4954.108798 | -0.155340294 | 0.068682782 |
| Gm36989 | 219.4274089 | -0.658198286 | 0.135693448 |
| Paip2 | 1058.0764 | -0.082142434 | 0.095685475 |
| Slc23a1 | 0.196018198 | -0.24356877 | 2.634006312 |
| Mzb1 | 4.18320868 | 1.436501552 | 0.623385542 |
| Prob1 | 99.4461201 | -0.617425051 | 0.150007215 |
| Spata24 | 30.05154039 | -0.183015447 | 0.206344028 |
| Dnajc18 | 1508.902544 | -0.106988013 | 0.045143904 |
| 1700066B19Rik | 1.640932998 | 0.514198469 | 0.782917868 |
| Ecscr | 35.18133015 | 2.538785948 | 0.266804152 |
| Tmem173 | 105.9553805 | 2.890354787 | 0.208080959 |
| Gm16490 | 0 | NA | NA |
| Ube2d2a | 1153.052153 | 0.233332053 | 0.07663964 |

| | | | |
|---|---|---|---|
| Cxxc5 | 830.8617964 | -0.112723998 | 0.062430828 |
| Gm29417 | 3.083648494 | -0.644233231 | 0.531414092 |
| Gm23882 | 0 | NA | NA |
| Psd2 | 681.0294117 | -0.945187661 | 0.143074627 |
| Nrg2 | 144.6574193 | -0.632890408 | 0.124744835 |
| Pura | 8433.573605 | -0.55484148 | 0.077347781 |
| Cystm1 | 401.2118129 | 0.069510502 | 0.099418306 |
| Pfdn1 | 311.4709643 | 0.642820252 | 0.134239188 |
| Hbegf | 582.7227112 | 3.21383611 | 0.315936766 |
| Slc4a9 | 2.54483004 | -1.447854243 | 0.751314583 |
| Ankhd1 | 1581.43401 | 0.011648629 | 0.084568674 |
| Gm6542 | 84.43238 | 0.479724578 | 0.140149187 |
| Eif4ebp3 | 7.191909191 | -0.681119478 | 0.396202442 |
| Sra1 | 541.6420646 | -0.051531094 | 0.068235941 |
| Slc35a4 | 487.0228943 | 0.444841243 | 0.07533742 |
| E230025N22Rik | 1.365445969 | 0.303056446 | 0.838478019 |
| Cd14 | 75.49450281 | 2.974145536 | 0.328784995 |
| Tmco6 | 58.12891111 | -0.059895188 | 0.171298721 |
| Ndufa2 | 320.0471109 | 0.229565423 | 0.116815662 |
| Ik | 860.0388227 | 0.197968004 | 0.117755132 |
| Wdr55 | 90.11495044 | 0.431948853 | 0.138780215 |
| Dnd1 | 14.61022796 | 0.531125865 | 0.29988499 |
| Hars | 700.2311977 | 0.295948497 | 0.124999046 |
| Hars2 | 250.6053811 | -0.15180031 | 0.107324095 |
| Zmat2 | 1268.32983 | -0.051131726 | 0.062569143 |
| Vaultrc5 | 18.77737098 | 1.564868598 | 0.409384347 |
| Gm23424 | 0.049553047 | 0.71820487 | 3.724945049 |
| Gm6756 | 1.105122771 | 0.520646321 | 0.993239518 |
| Pcdha1 | 15.4569917 | -1.463523617 | 0.277232419 |
| Gm37751 | 3.355297506 | -0.851357507 | 0.56165905 |
| Pcdha2 | 32.39827103 | -0.824993744 | 0.199647759 |
| Pcdha3 | 47.11146249 | -1.133712938 | 0.184147301 |
| Pcdha4 | 76.22283242 | -0.429955333 | 0.161551022 |
| Gm38225 | 2.632135046 | -0.376472379 | 0.75404414 |
| Pcdha5 | 55.26370248 | -1.046555466 | 0.19029991 |
| Gm37753 | 3.33123091 | -1.632873051 | 0.601311645 |
| Pcdha6 | 52.26043541 | -1.109636909 | 0.168930049 |
| Gm37013 | 0 | NA | NA |
| Pcdha7 | 45.39741467 | -1.231977256 | 0.205665384 |
| Gm37388 | 0 | NA | NA |
| Gm36858 | 26.49116316 | -1.388485246 | 0.288414613 |
| Pcdha8 | 34.08858227 | -1.166116838 | 0.31514079 |
| Pcdha9 | 61.46070133 | -1.022311424 | 0.172159776 |
| Pcdha11 | 154.8993467 | -1.184808724 | 0.174321695 |
| Gm10545 | 8.703390065 | -1.212401844 | 0.35271734 |
| Gm18150 | 13.89624275 | -1.17489251 | 0.42927605 |
| Pcdha12 | 117.7268123 | -0.908214168 | 0.139465642 |

| | | | |
|---|---|---|---|
| Gm19035 | 45.21062936 | -1.264794382 | 0.287702693 |
| Pcdhac1 | 92.62921019 | -0.696662136 | 0.126736618 |
| Pcdhac2 | 545.5711865 | -0.459074343 | 0.097666365 |
| Gm10544 | 2.182918281 | 0.178089501 | 0.679384889 |
| Gm38097 | 0.07796279 | 1.072698098 | 3.721829099 |
| Pcdhb1 | 5.985971091 | -0.910430992 | 0.436854107 |
| Pcdhb2 | 60.01309998 | -0.937951624 | 0.179757824 |
| Pcdhb3 | 83.43922558 | -0.798647027 | 0.132048346 |
| Gm18529 | 8.138222198 | -0.636184875 | 0.334716197 |
| Pcdhb4 | 60.73868439 | -0.709356835 | 0.166759052 |
| Pcdhb5 | 51.69657475 | -0.891637201 | 0.178782422 |
| Pcdhb6 | 68.6643995 | -0.954791865 | 0.164268299 |
| Pcdhb7 | 114.8261978 | -0.69663601 | 0.147776808 |
| Pcdhb8 | 89.35877465 | -0.590991301 | 0.162522114 |
| Pcdhb9 | 170.4389826 | -0.48508558 | 0.106976067 |
| Pcdhb10 | 105.3601154 | -0.454328475 | 0.12900178 |
| Pcdhb11 | 148.8037279 | -0.754629137 | 0.122638625 |
| Gm37446 | 0.821410149 | 0.915878304 | 1.405468223 |
| Gm20162 | 21.72530242 | -0.906429067 | 0.3233428 |
| Pcdhb12 | 117.9366013 | -0.740756444 | 0.160594799 |
| Pcdhb13 | 90.40139946 | -0.857918537 | 0.173377038 |
| Pcdhb14 | 293.0642983 | -0.664162681 | 0.118759952 |
| Pcdhb15 | 108.4402084 | -0.798292881 | 0.157230196 |
| Pcdhb16 | 347.3551605 | -0.523939632 | 0.080384611 |
| Pcdhb17 | 319.6024478 | -0.659971019 | 0.110979374 |
| Pcdhb18 | 229.3400325 | -0.548073115 | 0.091685128 |
| Pcdhb19 | 338.4815422 | -0.76222256 | 0.140471986 |
| Pcdhb20 | 206.2975924 | -0.612387952 | 0.1254264 |
| Pcdhb21 | 135.2304553 | -0.492912943 | 0.130728825 |
| Pcdhb22 | 238.2686593 | -0.536290788 | 0.081730732 |
| 4930517L18Rik | 0.044544648 | 0.237309859 | 3.724945049 |
| Slc25a2 | 10.03606488 | -0.023373444 | 0.356571637 |
| Taf7 | 183.7126313 | 0.705092888 | 0.116082699 |
| Gm8242 | 6.947208237 | 0.314086893 | 0.443516981 |
| 3222401L13Rik | 32.8963234 | 0.263582821 | 0.195816525 |
| Pcdhga1 | 89.5175488 | -0.088661417 | 0.128926202 |
| Gm37118 | 0.066050875 | 0.71820487 | 3.724945049 |
| Pcdhga2 | 78.94781358 | -0.662616322 | 0.147626612 |
| Pcdhga3 | 73.82529401 | -0.431168855 | 0.15160498 |
| Pcdhgb1 | 82.71764577 | -0.402188412 | 0.153811619 |
| Pcdhga4 | 165.6903328 | -0.378076633 | 0.096526343 |
| Pcdhgb2 | 147.8807373 | -0.090307344 | 0.137857024 |
| Pcdhga5 | 172.6956286 | -0.32817419 | 0.114016307 |
| Gm24401 | 7.516116141 | -1.197311344 | 0.560736294 |
| Pcdhga6 | 188.1446477 | -0.386971353 | 0.112347531 |
| Gm37165 | 0.949659951 | 0.889522955 | 1.084497261 |
| Pcdhga7 | 242.4294219 | -0.505258505 | 0.104605966 |

| | | | |
|---|---|---|---|
| Pcdhgb4 | 169.0788982 | -0.671013647 | 0.157125681 |
| Pcdhga8 | 424.9442205 | -0.518760266 | 0.093725608 |
| Pcdhgb5 | 111.8894427 | -0.299119938 | 0.133317281 |
| Pcdhga9 | 152.843188 | -0.516866107 | 0.103713844 |
| Pcdhgb6 | 97.63560197 | -0.641923437 | 0.193192683 |
| Gm26672 | 13.40694889 | -0.259219552 | 0.297227496 |
| Pcdhga10 | 152.3565402 | -0.164995794 | 0.139071621 |
| Pcdhgb7 | 116.7201272 | -0.511674579 | 0.140482083 |
| Pcdhga11 | 97.79522649 | -0.529579634 | 0.183803751 |
| Pcdhgb8 | 105.1432484 | -0.557313258 | 0.168823626 |
| Pcdhga12 | 164.3031905 | -0.876058483 | 0.164669375 |
| BC037039 | 16.91683977 | 0.684915143 | 0.272278166 |
| Pcdhgc3 | 1154.35759 | -0.120020435 | 0.107212442 |
| Pcdhgc5 | 972.9292026 | -0.603184377 | 0.104389829 |
| Gm38182 | 0 | NA | NA |
| Gm29994 | 1.935845755 | 0.826064697 | 0.95559692 |
| Diap1 | 993.6614968 | 0.720068522 | 0.095033275 |
| Mir6979 | 0.451714327 | 1.854363462 | 1.467423798 |
| Hdac3 | 567.0461211 | -0.227242501 | 0.070742023 |
| Rell2 | 1076.046952 | -0.326346675 | 0.086790643 |
| Fchsd1 | 97.53749871 | 0.774519624 | 0.119022582 |
| Arap3 | 55.62706146 | 0.841470204 | 0.30725836 |
| Mir6981 | 0 | NA | NA |
| Mir6980 | 0 | NA | NA |
| Pcdh1 | 2873.63663 | -0.500390646 | 0.110945535 |
| 2010320O07Rik | 1.189163173 | -0.473656027 | 1.017718129 |
| 1700086O06Rik | 44.53713014 | -0.116003606 | 0.183537813 |
| Gm10268 | 1.309431784 | 0.395728786 | 0.875926884 |
| 0610009O20Rik | 346.6488783 | -0.071914054 | 0.07784947 |
| Pcdh12 | 4.608338074 | 1.483631999 | 0.510445744 |
| Rnf14 | 3142.218856 | 0.150106587 | 0.071919389 |
| Gm23205 | 10.03026786 | -1.391864319 | 0.508651009 |
| Gnpda1 | 269.096148 | 1.060995855 | 0.109524997 |
| Gm24690 | 0 | NA | NA |
| Ndfip1 | 2572.872965 | 0.072216162 | 0.070663922 |
| Spry4 | 260.3722342 | 0.008990774 | 0.152455524 |
| Gm27369 | 0 | NA | NA |
| Gm27804 | 0 | NA | NA |
| Arhgap26 | 2536.520982 | -0.0540487 | 0.095879954 |
| Gm19774 | 0.900419082 | -1.278903758 | 1.035787986 |
| Gm15335 | 0 | NA | NA |
| Gm8302 | 0.044544648 | 0.237309859 | 3.724945049 |
| Fgf1 | 1013.624655 | -0.410848964 | 0.122867526 |
| Gm15334 | 1.714908477 | -1.202819437 | 1.106819806 |
| Gm15336 | 0.09349418 | -0.24357403 | 3.724945049 |
| Gm15333 | 0 | NA | NA |
| Gm15337 | 14.68375219 | 0.220993535 | 0.346289004 |

| | | | |
|---|---|---|---|
| Nr3c1 | 1778.771107 | -0.131906781 | 0.053208937 |
| Pabpc2 | 0.066050875 | 0.71820487 | 3.724945049 |
| Yipf5 | 514.5136962 | 0.511296091 | 0.084662273 |
| 2900055J20Rik | 110.5064999 | -1.463913568 | 0.202747435 |
| Kctd16 | 294.0495108 | -1.193836279 | 0.191528426 |
| Rps19-ps13 | 0.270136144 | -0.243301623 | 2.100416214 |
| Gm22515 | 0 | NA | NA |
| Gm22483 | 0.046345709 | 0.71820487 | 3.724945049 |
| AC149064.1 | 0 | NA | NA |
| Prelid2 | 7.690273632 | 1.585186186 | 0.408524441 |
| Grxcr2 | 0.311588305 | -1.015194801 | 1.987600934 |
| Sh3rf2 | 92.9302299 | -0.654700544 | 0.278638109 |
| Gm5689 | 0.669351884 | 2.368894162 | 1.27650581 |
| Plac8l1 | 0.74423012 | 2.392725674 | 1.456148785 |
| Lars | 747.0783727 | 0.665211243 | 0.113444285 |
| Rps19-ps4 | 0.931580594 | 1.090623857 | 1.137323835 |
| Rbm27 | 1071.37103 | 0.086149557 | 0.063068729 |
| Gm22697 | 0 | NA | NA |
| Pou4f3 | 1.201894167 | 1.693914594 | 0.998886953 |
| Gm16415 | 0.834345451 | 1.520372039 | 1.261593075 |
| Tcerg1 | 1111.337889 | 0.043482246 | 0.087228002 |
| Gm27457 | 2.628954575 | -0.879448597 | 0.661907577 |
| Gpr151 | 33.37780463 | 2.512744559 | 0.290373772 |
| Ppp2r2b | 978.0231163 | -0.520754751 | 0.066918642 |
| 4933407I08Rik | 0.075043205 | 0.237309859 | 3.724945049 |
| Gm22264 | 0 | NA | NA |
| Gm8181 | 9.04075607 | 0.008396167 | 0.46258983 |
| Stk32a | 162.2987704 | 0.338965176 | 0.14140465 |
| Dpysl3 | 999.9963044 | 0.091004753 | 0.068545814 |
| Gm3650 | 17.99992091 | 0.312491123 | 0.355375847 |
| Eif3j2 | 165.1918275 | 0.583261393 | 0.118531445 |
| Gm27325 | 0 | NA | NA |
| Jakmip2 | 846.4346633 | -0.289978652 | 0.077717235 |
| Spink1 | 0 | NA | NA |
| Gm23894 | 0 | NA | NA |
| Scgb3a2 | 0.047948698 | -0.243569582 | 3.724945049 |
| Gm94 | 0.18352982 | 0.237308059 | 3.724945049 |
| Spink5 | 0.975055063 | -0.193493246 | 3.566193858 |
| Spink14 | 0 | NA | NA |
| Spink6 | 0 | NA | NA |
| Spink12 | 0 | NA | NA |
| Gm37797 | 0 | NA | NA |
| Gm10267 | 0 | NA | NA |
| Spinkl | 0 | NA | NA |
| Spink11 | 0.047948698 | -0.243569582 | 3.724945049 |
| Gm10542 | 0 | NA | NA |
| Npy6r | 0.261451683 | -0.129265576 | 2.08654729 |

| | | | |
|---|---|---|---|
| Myot | 0.836501452 | 0.243395887 | 1.116798492 |
| Dcp2 | 1781.048542 | 0.779450144 | 0.094885232 |
| Mcc | 632.7057759 | -0.014441051 | 0.092200015 |
| A930012L18Rik | 13.61962428 | -1.108579862 | 0.295269441 |
| RP23-342A15.2 | 0 | NA | NA |
| Ythdc2 | 676.2351253 | -0.316228251 | 0.084048661 |
| Gm26928 | 0 | NA | NA |
| Kcnn2 | 557.7293241 | -1.040727499 | 0.109846692 |
| Gm10540 | 0.30732292 | 0.237331854 | 1.605764792 |
| A330093E20Rik | 2.019082719 | -0.994243121 | 0.919327337 |
| Gm26262 | 0 | NA | NA |
| Gm38337 | 0.335887269 | 0.237331373 | 1.576837564 |
| Trim36 | 95.57263784 | -0.568907183 | 0.172002515 |
| 1700018A14Rik | 0.75323169 | 2.432634884 | 1.340447322 |
| Pggt1b | 266.2078042 | 0.411318712 | 0.089113264 |
| Ccdc112 | 108.5123134 | -0.241946602 | 0.156332704 |
| Gm24617 | 0.282138413 | 1.625534534 | 3.717603603 |
| Mospd4 | 3.756273683 | -0.151672962 | 0.535054071 |
| Gm24076 | 0.776630353 | 0.717758585 | 1.370014436 |
| Fem1c | 864.4434414 | 0.346796365 | 0.083709938 |
| Ticam2 | 12.34676471 | 1.692924376 | 0.402965988 |
| Tmed7 | 1507.489644 | 0.44493281 | 0.107319634 |
| Eif1a | 755.0916563 | 1.193253251 | 0.149445382 |
| Cdo1 | 339.5541898 | 0.593984466 | 0.148567008 |
| Atg12 | 650.7230075 | 0.403468774 | 0.086575115 |
| Ap3s1 | 461.3279815 | 0.231982209 | 0.086629176 |
| Lvrn | 10.74167226 | -0.386994371 | 0.362614904 |
| Gm23610 | 0 | NA | NA |
| Arl14epl | 1.575196263 | 1.491521919 | 0.894021635 |
| Commd10 | 179.0565603 | -0.002472703 | 0.089464511 |
| Gm22791 | 0 | NA | NA |
| Gm25036 | 0 | NA | NA |
| Sema6a | 831.0817601 | -0.210318669 | 0.059305243 |
| Eno1b | 521.4844324 | 0.265924236 | 0.122872079 |
| Gm5839 | 0 | NA | NA |
| 1700044K03Rik | 0 | NA | NA |
| Gm25440 | 0 | NA | NA |
| Dtwd2 | 52.5847649 | -0.193757377 | 0.151924014 |
| Gm16283 | 2.090780166 | -0.176752367 | 0.670267162 |
| 1700065O20Rik | 0 | NA | NA |
| Dmxl1 | 3104.946172 | -0.038423186 | 0.049781069 |
| Tnfaip8 | 137.5382232 | 1.367550773 | 0.146255137 |
| C030005K06Rik | 2.787674368 | -3.760211663 | 0.887224266 |
| Hsd17b4 | 542.6081984 | -0.256588803 | 0.085814289 |
| Cd63-ps | 108.0511914 | 2.401994917 | 0.208675372 |
| Gm8529 | 0 | NA | NA |
| Fam170a | 0 | NA | NA |

| | | | |
|---|---|---|---|
| Hdhd1a | 0.113999573 | 0.23732098 | 3.724945049 |
| Prr16 | 138.718701 | -0.882443288 | 0.139116121 |
| Gm25739 | 0 | NA | NA |
| Gm25785 | 0 | NA | NA |
| Gm4950 | 20.25727402 | 0.581546455 | 0.251106366 |
| Gm22259 | 0 | NA | NA |
| Ftmt | 0 | NA | NA |
| Srfbp1 | 140.6912451 | 0.297091064 | 0.113223314 |
| Gm27026 | 0 | NA | NA |
| Lox | 47.85031176 | 2.538484767 | 0.342717551 |
| Zfp474 | 3.585501729 | 0.589699369 | 0.511748794 |
| 1700034E13Rik | 0.108266322 | 1.230376831 | 3.722845817 |
| Gykl1 | 0 | NA | NA |
| Sncaip | 113.2002606 | -0.087892538 | 0.110159379 |
| Snx2 | 781.8559442 | 0.923967716 | 0.084628438 |
| Snx24 | 137.1094418 | 0.088862726 | 0.100476589 |
| Ppic | 20.57672658 | 0.901272985 | 0.231145622 |
| Prdm6 | 12.21164467 | 0.243827313 | 0.314884751 |
| Cep120 | 755.1587019 | -0.152024591 | 0.060065607 |
| Csnk1g3 | 932.5335226 | -0.142126942 | 0.0604667 |
| Gm5507 | 5.787175505 | -0.714106573 | 0.420083324 |
| Redrum | 0.747472414 | 2.567322621 | 1.147792474 |
| Zfp608 | 685.0116038 | -0.13141561 | 0.072633228 |
| Gm26742 | 0.583001974 | 0.382184994 | 1.133015955 |
| Gm4221 | 12.78354649 | -1.450932061 | 0.372379608 |
| Gm24183 | 0 | NA | NA |
| Gm34073 | 0.428599116 | -1.430222795 | 3.327984098 |
| Gm22597 | 0 | NA | NA |
| Gm37828 | 0 | NA | NA |
| Gm37337 | 0.587544172 | 0.835094904 | 1.268367232 |
| Gm24514 | 0.493679557 | -1.975776735 | 1.435516245 |
| Gm26959 | 0 | NA | NA |
| Gm8614 | 0.29391207 | -1.211271379 | 2.017889165 |
| Gm23335 | 0 | NA | NA |
| Gm6944 | 0 | NA | NA |
| Gm22484 | 0.068097359 | -0.243569582 | 3.724945049 |
| Gm23555 | 0 | NA | NA |
| Gm25476 | 0 | NA | NA |
| Gramd3 | 170.2177708 | 0.55331464 | 0.110497297 |
| Aldh7a1 | 399.3017201 | -0.132607579 | 0.076457757 |
| Mir1258 | 0 | NA | NA |
| Phax | 381.2748711 | 0.280166291 | 0.10208033 |
| Tex43 | 2.674966445 | -0.344732778 | 0.56124724 |
| Lmnb1 | 214.1019998 | 0.94487293 | 0.145643569 |
| 3-Mar | 67.75458388 | 0.582233283 | 0.177671456 |
| Gm15345 | 0.999978248 | 1.841712929 | 1.11762733 |
| C330018D20Rik | 87.14994236 | -0.240526432 | 0.137936904 |

| | | | |
|---|---|---|---|
| Megf10 | 704.5314188 | 0.229000391 | 0.125975254 |
| Prrc1 | 265.116747 | 0.315057501 | 0.081671326 |
| 4930511M06Rik | 2.334218344 | 1.572495015 | 0.68917714 |
| Ctxn3 | 52.35061762 | 0.233621158 | 0.190824396 |
| 1700066O22Rik | 0.436302845 | 1.753976696 | 1.378169843 |
| 1700011I03Rik | 2.706585767 | 0.662111666 | 0.66176167 |
| Gm26038 | 0 | NA | NA |
| AC102232.1 | 0.04393175 | -0.243571806 | 3.724945049 |
| Slc12a2 | 1824.003985 | -0.518887169 | 0.088399234 |
| Fbn2 | 140.4520598 | -0.744171279 | 0.169268163 |
| Gm26507 | 13.7398474 | -1.257815809 | 0.310428533 |
| Gm25660 | 0 | NA | NA |
| Slc27a6 | 3.148605881 | 1.16314312 | 0.601131808 |
| Gm23270 | 0 | NA | NA |
| Isoc1 | 459.7670982 | 0.717782776 | 0.124915675 |
| Adamts19 | 19.03947336 | -0.751453733 | 0.343830816 |
| A730017C20Rik | 501.0808718 | 0.41790418 | 0.109639509 |
| Chsy3 | 230.6243843 | -0.667808417 | 0.156425204 |
| Mir6355 | 0 | NA | NA |
| Gm26028 | 0 | NA | NA |
| Gm23576 | 0 | NA | NA |
| Gm4951 | 925.2780923 | 9.812201975 | 0.448273597 |
| Gm5970 | 122.1217523 | 8.907892253 | 0.811675511 |
| Gm4841 | 260.7014249 | 11.70959322 | 0.920515771 |
| F830016B08Rik | 944.1534938 | 10.49761114 | 0.530301075 |
| Iigp1 | 5189.009943 | 10.27040214 | 0.274833713 |
| Gm6960 | 0.926812825 | 2.928930459 | 1.171580252 |
| BC023105 | 164.7922582 | 10.68556801 | 0.900900148 |
| Smim3 | 173.6181185 | 2.279086677 | 0.231260227 |
| Dctn4 | 1434.033031 | 0.452659924 | 0.059198429 |
| Rbm22 | 390.7352864 | 0.388751577 | 0.142878728 |
| Myoz3 | 0.060174537 | 0.237309859 | 3.724945049 |
| Synpo | 1700.826908 | -0.280245525 | 0.12929633 |
| Ndst1 | 2081.012663 | 0.754934062 | 0.090858889 |
| Rps14 | 1385.960924 | 0.760404863 | 0.137454017 |
| Gm8731 | 0.109869311 | 0.238305784 | 3.724874707 |
| Cd74 | 447.6522266 | 4.278454661 | 0.329470657 |
| Mir5107 | 0.060174537 | 0.237309859 | 3.724945049 |
| Tcof1 | 457.500707 | 0.68967285 | 0.094476713 |
| Arsi | 22.1297461 | 1.641594644 | 0.291082774 |
| Camk2a | 16056.95831 | -1.416441283 | 0.175329861 |
| Mir7220 | 4.383661021 | -2.274828349 | 0.619361824 |
| Mir6982 | 3.753094389 | -2.326997455 | 0.710060923 |
| Slc6a7 | 440.8262577 | -0.949579775 | 0.143470028 |
| Cdx1 | 0.329049156 | -1.389183999 | 1.701879364 |
| Pdgfrb | 307.8251967 | 0.439118382 | 0.090567175 |
| Csf1r | 863.5867635 | -0.709383114 | 0.301297086 |

| | | | |
|---|---|---|---|
| Mir6983 | 0 | NA | NA |
| Hmgxb3 | 688.8088613 | -0.14065414 | 0.059189762 |
| Gm26403 | 2.513420025 | -0.790808678 | 0.73727885 |
| Slc26a2 | 293.7967227 | 0.524206244 | 0.101661861 |
| Pde6a | 3.821803709 | -0.054953162 | 0.490601639 |
| Rps2-ps10 | 1236.213371 | 0.398690865 | 0.129625799 |
| Ppargc1b | 407.0618076 | -0.882887335 | 0.127848087 |
| Gm25301 | 0 | NA | NA |
| mmu-mir-378a | 3.818950962 | -1.481792703 | 0.526300038 |
| Gm25834 | 0.038981395 | 0.718207094 | 3.724945049 |
| Arhgef37 | 140.4268379 | 0.550103139 | 0.140373161 |
| Csnk1a1 | 4557.884024 | 0.256831103 | 0.06690307 |
| Bvht | 5.444855971 | 2.036046701 | 0.583541782 |
| Mir145a | 0.194631057 | 0.237309881 | 3.243494927 |
| Mir143 | 0.147068667 | 0.237310878 | 3.514416608 |
| Mir143hg | 8.471314973 | -0.934964036 | 0.466057036 |
| Il17b | 2.212656516 | 1.194286202 | 0.85204315 |
| Gm38165 | 0.662591016 | 0.609755598 | 1.316173126 |
| Pcyox1l | 207.9956365 | -0.398854768 | 0.136529825 |
| Grpel2 | 247.9051696 | -0.057857872 | 0.096224901 |
| 1500015A07Rik | 35.15775905 | -0.445873013 | 0.189871134 |
| Afap1l1 | 138.1820362 | -0.24273456 | 0.095853574 |
| Ablim3 | 445.2700515 | -0.982638424 | 0.116221783 |
| Gm37774 | 0 | NA | NA |
| Sh3tc2 | 44.0296305 | 0.157527224 | 0.218309954 |
| Adrb2 | 16.87312987 | -0.620549698 | 0.263747832 |
| Gm9949 | 1.519386227 | 0.912390838 | 0.930904765 |
| Htr4 | 92.71409538 | -0.36067135 | 0.144322069 |
| Fbxo38 | 754.8506995 | 0.232940005 | 0.057077733 |
| Spink10 | 11.52016113 | 0.133605541 | 0.318798909 |
| Spink7 | 0.099106094 | 1.209894437 | 3.72418822 |
| Spink13 | 0.546984718 | -0.043578931 | 1.216345121 |
| Gm17732 | 5.889985807 | -0.575988401 | 0.443799137 |
| 2700046A07Rik | 9.548929167 | -0.232301907 | 0.458241658 |
| Apcdd1 | 206.8351242 | -0.442352156 | 0.165080954 |
| Napg | 1597.555114 | -0.283737766 | 0.056183782 |
| Piezo2 | 47.03296206 | 0.546071129 | 0.198269101 |
| Gm26972 | 0 | NA | NA |
| Gm23581 | 0 | NA | NA |
| Gm22560 | 0 | NA | NA |
| Gm22210 | 0 | NA | NA |
| Gm24848 | 0 | NA | NA |
| Txnl1 | 825.0032602 | 0.468670102 | 0.096180748 |
| Wdr7 | 3498.348225 | -0.251846427 | 0.045134592 |
| Gm24384 | 0 | NA | NA |
| St8sia3os | 1.428069919 | 1.504732824 | 0.970650927 |
| St8sia3 | 1740.780446 | -0.377139338 | 0.081225626 |

| | | | |
|---|---|---|---|
| Onecut2 | 194.3572374 | -0.225783161 | 0.155519495 |
| Amd-ps3 | 30.2089109 | 0.151292194 | 0.188231807 |
| Fech | 621.1316025 | -0.080346485 | 0.067604589 |
| Nars | 1964.183135 | 0.511788717 | 0.088235047 |
| Atp8b1 | 51.70323717 | 1.029836028 | 0.203698139 |
| Gm23016 | 0 | NA | NA |
| Gm24504 | 0 | NA | NA |
| Nedd4l | 2403.215903 | -0.170711993 | 0.105137375 |
| Mir122 | 0.049553047 | 0.71820487 | 3.724945049 |
| Alpk2 | 11.61777028 | -0.303025906 | 0.353123787 |
| Gm22567 | 0.046345709 | 0.71820487 | 3.724945049 |
| Malt1 | 178.3264262 | 1.020852846 | 0.180150244 |
| Zfp532 | 1029.175328 | -0.11603484 | 0.060567071 |
| Gm37643 | 15.88324691 | -0.957682333 | 0.354439338 |
| Oacyl | 19.6943626 | -0.667978481 | 0.387068796 |
| Sec11c | 480.2888366 | 0.083668791 | 0.086814693 |
| Grp | 19.80146899 | -0.833851715 | 0.254835527 |
| Gm25150 | 0 | NA | NA |
| Rax | 0.423125362 | 0.051605724 | 1.490560692 |
| Raxos1 | 0 | NA | NA |
| Cplx4 | 0.598788582 | 0.97533791 | 1.283780424 |
| Lman1 | 670.4253144 | 0.466907338 | 0.063404988 |
| Ccbe1 | 53.63741432 | -1.080867232 | 0.221187882 |
| Mir694 | 0.217748738 | -1.530389524 | 2.460727054 |
| Gm15958 | 0.904555577 | -2.232247576 | 1.108010404 |
| Pmaip1 | 46.65352456 | 2.637647558 | 0.208929665 |
| Gm24679 | 0 | NA | NA |
| Gm22219 | 0 | NA | NA |
| Mc4r | 36.60476633 | 0.135990412 | 0.226929639 |
| Gnal | 2244.602178 | -0.582631524 | 0.095233248 |
| Gm23512 | 0.291502873 | -1.596673129 | 2.009766309 |
| Mppe1 | 130.6858995 | 0.421175308 | 0.108892967 |
| Impa2 | 34.6996934 | -0.505792111 | 0.224084416 |
| B430212C06Rik | 2.198202818 | -0.687120217 | 0.945695888 |
| Cidea | 13.25459769 | -0.633675551 | 0.322427098 |
| Tubb6 | 218.1172078 | 4.570435977 | 0.332048723 |
| Afg3l2 | 820.858718 | -0.076154757 | 0.077717723 |
| Gm24900 | 0 | NA | NA |
| Slmo1 | 256.90097 | -0.554470943 | 0.102574587 |
| Spire1 | 2405.877136 | -0.222373561 | 0.071943068 |
| Gm17669 | 0.12092566 | 0.237307635 | 3.724945049 |
| Cep76 | 301.4880058 | 0.197671559 | 0.160456073 |
| Psmg2 | 96.05648597 | 0.103909645 | 0.157927644 |
| Ptpn2 | 330.6143394 | 0.368002342 | 0.095661776 |
| Gm26910 | 20.45356711 | -0.591224438 | 0.312163879 |
| Seh1l | 391.9333236 | 0.512492635 | 0.064900676 |
| Cep192 | 279.5064016 | 0.326002539 | 0.127140107 |

| | | | |
|---|---|---|---|
| Ldlrad4 | 528.0018019 | -0.100243002 | 0.09765626 |
| Fam210a | 679.3179832 | -0.751909507 | 0.071609369 |
| Mir7219 | 0 | NA | NA |
| Rnmt | 878.6655392 | -0.172850969 | 0.050671938 |
| Mc5r | 18.63850153 | 0.412518857 | 0.328016941 |
| Mc2r | 1.528578462 | 2.789445812 | 1.099262746 |
| Mir6356 | 0 | NA | NA |
| 4930546C10Rik | 0.160559904 | 1.165833556 | 2.790729435 |
| Mir145b | 0 | NA | NA |
| Gm23536 | 0.335787014 | -0.222591191 | 1.730419906 |
| Gm4842 | 0 | NA | NA |
| Tcf4 | 6112.811251 | -0.227713865 | 0.08799673 |
| Gm24845 | 5.630675305 | -1.340008701 | 0.497497232 |
| Gm22508 | 0 | NA | NA |
| Ccdc68 | 4.327574142 | 0.27648177 | 0.499957466 |
| 1700061H18Rik | 0.072025462 | 0.237312084 | 3.724945049 |
| 4930448D08Rik | 0.605479419 | 0.237197489 | 1.20080369 |
| Rab27b | 340.9708684 | -0.350249369 | 0.101764544 |
| Dynap | 0.072025462 | 0.237312084 | 3.724945049 |
| 4930503L19Rik | 92.03394658 | 0.503213117 | 0.130503086 |
| Stard6 | 8.232924272 | -0.058946905 | 0.340108743 |
| Poli | 92.01233665 | 0.287612818 | 0.122433498 |
| Mir6357 | 0.407891471 | -0.681179459 | 1.447662419 |
| Mbd2 | 568.7990903 | 1.227469894 | 0.12518435 |
| Dcc | 395.337002 | -1.001262459 | 0.123446622 |
| Gm25509 | 0 | NA | NA |
| n-R5-8s1 | 0 | NA | NA |
| Mex3c | 685.3858209 | 0.562108641 | 0.09427644 |
| Gm26438 | 0 | NA | NA |
| Gm22065 | 0.042716945 | -0.243571806 | 3.724945049 |
| Smad4 | 842.5159027 | 0.216611677 | 0.063021716 |
| Elac1 | 466.4413671 | -0.277360166 | 0.085866576 |
| Me2 | 344.6597954 | 0.148797648 | 0.071311939 |
| Mro | 101.9368696 | -0.685159997 | 0.121052698 |
| Mapk4 | 1333.921859 | 0.088234502 | 0.097864775 |
| D730045A05Rik | 2.135082683 | 3.168542301 | 0.890403075 |
| Gm9925 | 7.46780898 | -1.238781593 | 0.45435226 |
| Ska1 | 2.02584455 | 2.946015326 | 0.989623258 |
| Cxxc1 | 352.5600067 | 0.121177879 | 0.089149229 |
| Mbd1 | 382.0312851 | 0.257648726 | 0.138844737 |
| Cfap53 | 11.98097365 | 0.78870461 | 0.490857835 |
| Gm14328 | 4.493268881 | 0.81461019 | 0.517412241 |
| Myo5b | 428.355463 | -0.247252447 | 0.109544635 |
| 1700120E14Rik | 0 | NA | NA |
| Scarna17 | 10.66151498 | -0.281381887 | 0.387913249 |
| Gm23119 | 43.96335493 | 0.155658025 | 0.172428376 |
| Acaa2 | 104.6941341 | -0.647723992 | 0.17518874 |

| | | | |
|---|---|---|---|
| Gm24559 | 0 | NA | NA |
| Gm25594 | 0 | NA | NA |
| Lipg | 453.85713 | 4.57199464 | 0.427941099 |
| Rpl17 | 766.1645459 | 0.542958205 | 0.076902208 |
| Snord58b | 1.376506768 | -0.213430803 | 0.927027435 |
| Gm23301 | 2.714581473 | -0.140758028 | 0.646711032 |
| Gm26202 | 2.364995654 | -0.173439876 | 0.71884319 |
| BC031181 | 600.9122168 | 0.06030438 | 0.094176026 |
| Dym | 677.9260573 | -0.375902333 | 0.058828209 |
| Gm27781 | 0.317560373 | -0.319194067 | 3.702639598 |
| Gm8807 | 0 | NA | NA |
| 2010010A06Rik | 0 | NA | NA |
| Gm20544 | 9.279688315 | 0.164100883 | 0.33562066 |
| Smad7 | 616.9288546 | 0.340192841 | 0.080494659 |
| Ctif | 1554.441727 | -0.278549625 | 0.066615868 |
| Gm10532 | 5.803090283 | 1.042091813 | 0.516860473 |
| 1700034B16Rik | 0 | NA | NA |
| Zbtb7c | 113.4256275 | -0.15089646 | 0.137164208 |
| Mir6358 | 0 | NA | NA |
| Smad2 | 378.1534108 | 0.142532619 | 0.079858778 |
| Skor2 | 31.77545133 | -0.321968197 | 0.30110102 |
| Ier3ip1 | 232.6003113 | 0.292996972 | 0.092917006 |
| Hdhd2 | 753.876256 | -0.148602674 | 0.059657656 |
| Katnal2 | 40.6261297 | -0.684262758 | 0.204706462 |
| Pias2 | 1238.28436 | -0.144129606 | 0.053042039 |
| Gm7276 | 15.00176566 | -1.036983924 | 0.361466876 |
| St8sia5 | 623.4113925 | -0.540170066 | 0.107596974 |
| Gm15957 | 0 | NA | NA |
| Loxhd1 | 23.89937881 | 2.061513039 | 0.637102481 |
| Rnf165 | 750.5636098 | -1.055402689 | 0.160805393 |
| 8030462N17Rik | 630.218415 | -0.06670109 | 0.067304003 |
| Gm26014 | 0 | NA | NA |
| 4930465K10Rik | 2.048990233 | -1.010540718 | 0.666059726 |
| A330094K24Rik | 6.372749315 | 0.825127841 | 0.456775926 |
| Haus1 | 42.89733926 | -0.153363465 | 0.164505412 |
| Atp5a1 | 6848.041245 | -0.101186294 | 0.06909834 |
| F830208F22Rik | 1.770308639 | 0.783200993 | 0.7627764 |
| Pstpip2 | 100.7389169 | -0.351931904 | 0.134213595 |
| Gm23954 | 0.113999573 | 0.23732098 | 3.724945049 |
| Epg5 | 1099.858229 | -0.140165928 | 0.088623714 |
| Siglec15 | 0.39439978 | 0.930398638 | 1.602238663 |
| Slc14a1 | 350.1029007 | 0.826927236 | 0.124030007 |
| Slc14a2 | 4.084512964 | -0.0755982 | 0.537960845 |
| Gm6133 | 2.464583914 | -1.515110989 | 0.692189021 |
| Gm26430 | 0 | NA | NA |
| Gm23895 | 0 | NA | NA |
| Setbp1 | 1103.372977 | -0.356811914 | 0.088035938 |

| | | | |
|---|---|---|---|
| Gm25824 | 0 | NA | NA |
| Gm26977 | 0.099106094 | 1.209894437 | 3.72418822 |
| Pard6g | 127.4897943 | 0.069056743 | 0.144618021 |
| Gm21886 | 0.269331125 | -0.21319046 | 2.073908625 |
| Adnp2 | 487.4784465 | 0.344028699 | 0.096758976 |
| Rbfa | 151.797274 | -0.124953305 | 0.091451586 |
| Gm16286 | 543.6190761 | 0.47780135 | 0.098182541 |
| Txnl4a | 164.8076413 | 0.229242055 | 0.111038103 |
| Hsbp1l1 | 8.924943958 | -0.612051925 | 0.352326866 |
| Pqlc1 | 221.7266294 | 0.34594174 | 0.101220677 |
| Kcng2 | 77.54725198 | -0.289096461 | 0.195954683 |
| Gm37015 | 4.004740963 | -1.443040555 | 0.652200775 |
| Gm26676 | 5.505563926 | 1.521572326 | 0.522718601 |
| D330025C20Rik | 2.748297996 | 0.570650882 | 0.68756538 |
| Ctdp1 | 381.7265928 | 0.195274911 | 0.098760067 |
| Nfatc1 | 215.3281497 | 1.793296915 | 0.135442058 |
| Atp9b | 951.6403139 | 0.144235451 | 0.073366708 |
| Gm25071 | 1.130492263 | 0.055810312 | 0.867685452 |
| Sall3 | 188.0210551 | -0.318362172 | 0.175406084 |
| Gm27238 | 1.264061948 | -1.633828971 | 0.92123663 |
| Gm27239 | 2.200041218 | -1.864957942 | 0.86351025 |
| Gm25718 | 0 | NA | NA |
| 4930594M17Rik | 0.045882455 | 0.237312084 | 3.724945049 |
| Gm26012 | 0 | NA | NA |
| Mir5127 | 0 | NA | NA |
| Gm17383 | 86.5616375 | 0.158369812 | 0.172201936 |
| Gm25060 | 0 | NA | NA |
| Galr1 | 70.21229872 | 0.307854553 | 0.257283474 |
| Mbp | 30062.83706 | -0.423183967 | 0.176851299 |
| Rpl21-ps8 | 1.385867572 | 1.34390668 | 0.825239348 |
| Zfp236 | 860.9371065 | -0.456588587 | 0.109311146 |
| Gm10524 | 101.1240389 | -0.51096508 | 0.125597982 |
| Mir5112 | 0.320492979 | -0.55289372 | 1.539493636 |
| 2210420H20Rik | 0.04393175 | -0.243571806 | 3.724945049 |
| 4930445N18Rik | 0.552122516 | -0.322802948 | 1.42664383 |
| Zfp516 | 555.826877 | 0.406048306 | 0.09166289 |
| 1700095A13Rik | 0 | NA | NA |
| Gm24720 | 0 | NA | NA |
| Tshz1 | 687.2218015 | -0.148822596 | 0.140511507 |
| Zadh2 | 324.0628898 | -0.270323955 | 0.109094865 |
| Zfp407 | 728.9401691 | -0.147455294 | 0.098898349 |
| Gm37216 | 25.25091243 | -0.252595711 | 0.299739343 |
| Cndp1 | 9.704862909 | 1.220727589 | 0.423218498 |
| Gm7289 | 1.584876195 | 1.401628039 | 0.869442821 |
| Cndp2 | 787.0114979 | 1.705473374 | 0.161709182 |
| D030046N08Rik | 4.544185544 | -0.442661923 | 0.489556304 |
| Fam69c | 47.9173264 | -0.737229526 | 0.186528393 |

| | | | |
|---|---|---|---|
| Gm22014 | 0 | NA | NA |
| Gm25005 | 0 | NA | NA |
| Gm22819 | 0 | NA | NA |
| Cyb5a | 343.7727805 | -0.027501183 | 0.076507642 |
| Gm16146 | 0.241190471 | 0.754107443 | 3.036905984 |
| Fbxo15 | 2.684576295 | 0.74407405 | 0.609478163 |
| Timm21 | 78.28501918 | -0.23337893 | 0.201910213 |
| Gm25453 | 0 | NA | NA |
| Neto1 | 920.988223 | -0.352579092 | 0.11316496 |
| Cbln2 | 282.7539475 | -0.660371041 | 0.110875539 |
| Gm5096 | 0.150520013 | 0.237326327 | 2.857840489 |
| Gm24987 | 0 | NA | NA |
| Socs6 | 448.8133279 | 0.241725855 | 0.096935101 |
| Rttn | 192.02834 | -0.009998962 | 0.097610448 |
| Mir6359 | 0.075043205 | 0.237309859 | 3.724945049 |
| Cd226 | 7.086717468 | 4.377307267 | 0.842013278 |
| 9330132A10Rik | 749.6793612 | -0.441098846 | 0.098183257 |
| Dok6 | 388.457898 | -0.568577137 | 0.105552633 |
| Gm16296 | 5.46580188 | -0.388697387 | 0.439242129 |
| Gm16297 | 0 | NA | NA |
| Gm22694 | 0 | NA | NA |
| Gm27037 | 0 | NA | NA |
| Gm23859 | 0 | NA | NA |
| Eif3s6-ps2 | 6.153886045 | 0.564238205 | 0.497472645 |
| Gm25418 | 0 | NA | NA |
| Tmx3 | 1329.967682 | 0.211708306 | 0.089585531 |
| 1700030N03Rik | 0.163190422 | 1.22955094 | 3.396300676 |
| Gm25431 | 0 | NA | NA |
| Ighmbp2 | 158.9101558 | 0.18525724 | 0.114633554 |
| Mrpl21 | 165.3835846 | 0.385541208 | 0.114524666 |
| Mir6984 | 0.950361064 | -2.517466717 | 1.119633132 |
| Cpt1a | 573.796439 | 1.131406824 | 0.143119014 |
| Mtl5 | 3.167783942 | -0.53568079 | 0.625200841 |
| Gal | 59.25410188 | 1.701482752 | 0.363227057 |
| Ppp6r3 | 1233.355573 | 0.127724708 | 0.061278653 |
| Gm23940 | 0 | NA | NA |
| Lrp5 | 186.6160727 | 0.760090414 | 0.150792253 |
| 1810055G02Rik | 195.6912383 | 0.151939923 | 0.097041761 |
| Suv420h1 | 939.8295677 | -0.198702957 | 0.073382971 |
| Gm16066 | 1.927026077 | -0.828641568 | 0.775856051 |
| Chka | 1064.49316 | -0.369240618 | 0.168868049 |
| Tcirg1 | 611.8661545 | 3.460381287 | 0.171815253 |
| Ndufs8 | 433.8570073 | -0.188120419 | 0.111375639 |
| Aldh3b1 | 78.94818307 | 1.281550549 | 0.227948616 |
| Unc93b1 | 424.5399235 | 2.887316832 | 0.158176496 |
| Aldh3b3 | 1.060607173 | 0.845676356 | 0.965602222 |
| Aldh3b2 | 8.359127856 | -0.813607396 | 0.36984052 |

| | | | |
|---|---|---|---|
| Acy3 | 20.90646341 | 0.302041051 | 0.245052973 |
| Tbx10 | 0.479449896 | 0.504573601 | 1.414467474 |
| Nudt8 | 43.46713457 | -0.489167034 | 0.175334899 |
| Gm16312 | 0.283672814 | 0.664548546 | 1.622485146 |
| Doc2g | 55.67602762 | -0.385554022 | 0.614734645 |
| Ndufv1 | 1083.51582 | -0.163554714 | 0.065090362 |
| Gstp1 | 541.1525819 | -0.168382694 | 0.139031951 |
| Gstp2 | 1.939495377 | -0.384236766 | 0.75283898 |
| BC021614 | 1.429393014 | 3.244054722 | 1.118324355 |
| Gm23927 | 0 | NA | NA |
| Cabp2 | 3.994848174 | -0.206228446 | 0.685810943 |
| Cdk2ap2 | 150.3217526 | 0.444626012 | 0.131946187 |
| Pitpnm1 | 2202.207616 | -0.494547807 | 0.07005453 |
| Aip | 570.1830399 | -0.005059234 | 0.08228415 |
| Tmem134 | 248.6189701 | 0.21927075 | 0.097089717 |
| Cabp4 | 3.477720489 | 3.609500644 | 0.922624158 |
| Gpr152 | 2.789673889 | 1.304312579 | 0.686430946 |
| Coro1b | 812.1320772 | 1.045973316 | 0.092338203 |
| Ptprcap | 24.66290501 | 4.653794189 | 0.523955073 |
| Rps6kb2 | 211.501265 | -0.412386767 | 0.14882251 |
| Carns1 | 117.9629228 | -0.791968731 | 0.123148075 |
| Gm17552 | 4.682108124 | 0.173732681 | 0.479421101 |
| Tbc1d10c | 30.14122667 | 2.112439038 | 0.261724884 |
| Ppp1ca | 1432.944828 | 0.156959087 | 0.083876194 |
| Rad9a | 127.978386 | 0.959980193 | 0.120113512 |
| Clcf1 | 180.1899159 | 5.317959846 | 0.383842167 |
| Gm26115 | 0.127364494 | -0.243565133 | 3.724945049 |
| Pold4 | 108.3674615 | 0.888376571 | 0.137490799 |
| Ssh3 | 269.6560211 | -0.103283647 | 0.151273813 |
| Mir6985 | 0 | NA | NA |
| Ankrd13d | 332.9753029 | -0.975334023 | 0.111790773 |
| Adrbk1 | 2117.90304 | -0.254827922 | 0.059511998 |
| Kdm2a | 1799.114924 | 0.051482785 | 0.062007587 |
| Rhod | 32.92487569 | 2.628481182 | 0.316436158 |
| A930001C03Rik | 2.701377055 | 1.24473201 | 0.651532734 |
| Syt12 | 590.78774 | 0.238213689 | 0.11650838 |
| 2010003K11Rik | 0.302393035 | 2.005931201 | 1.86912769 |
| Pcx | 931.2379665 | -0.513696104 | 0.104823488 |
| Lrfn4 | 397.610308 | 0.292627621 | 0.093269218 |
| Rce1 | 129.1447744 | 0.518320931 | 0.146068057 |
| Mir6986 | 0.044191549 | -0.243571806 | 3.724945049 |
| Gm960 | 25.50013179 | -0.767160451 | 0.226850641 |
| Sptbn2 | 9930.18535 | -0.581124465 | 0.103389837 |
| Gm21992 | 41.25034136 | -0.613476477 | 0.184163727 |
| Rbm4b | 188.1038022 | -0.164259128 | 0.086496421 |
| Gm37206 | 190.853055 | -0.505368714 | 0.201158116 |
| Rbm4 | 178.0962253 | -0.49721402 | 0.173967684 |

| | | | |
|---|---|---|---|
| Rbm14 | 411.7655184 | -0.228566999 | 0.084927855 |
| Gm21844 | 0.416305032 | 2.246707107 | 1.432208788 |
| Ccs | 117.7739688 | 0.226473508 | 0.136773676 |
| Ccdc87 | 18.82604018 | -0.244779249 | 0.271151082 |
| Ctsf | 447.0332469 | -0.670255068 | 0.085553088 |
| Actn3 | 10.89038947 | -0.359394063 | 0.323309384 |
| Zdhhc24 | 191.9093747 | 0.085645386 | 0.091179987 |
| Bbs1 | 425.8384116 | -0.443635238 | 0.077623674 |
| Gm25334 | 0 | NA | NA |
| Dpp3 | 394.3142871 | 0.25011299 | 0.080634927 |
| Peli3 | 250.3912601 | -0.485847828 | 0.091119039 |
| Mrpl11 | 282.4075369 | 0.054163575 | 0.087561555 |
| Npas4 | 211.216604 | -0.309890955 | 0.599457252 |
| Slc29a2 | 100.667391 | -0.353094479 | 0.118286185 |
| B4gat1 | 647.79448 | -0.666920144 | 0.062964775 |
| Brms1 | 180.8653628 | 0.809767596 | 0.10577288 |
| Rin1 | 659.0041585 | 0.030653113 | 0.180808358 |
| Cd248 | 53.27825101 | 0.825367081 | 0.198471648 |
| Tmem151a | 1868.678076 | -0.235275564 | 0.069341863 |
| Yif1a | 156.5502458 | 0.781854717 | 0.147666754 |
| Cnih2 | 897.9476727 | -0.420646684 | 0.099131009 |
| Rab1b | 1115.051491 | 0.221222852 | 0.097538717 |
| Klc2 | 2105.216524 | -0.410982067 | 0.070019191 |
| Pacs1 | 821.0770873 | -0.51816794 | 0.071186986 |
| Gm27468 | 0.909851095 | -2.184250589 | 1.067852701 |
| Sf3b2 | 2394.762425 | 0.088000238 | 0.061465971 |
| Gal3st3 | 390.9376513 | -0.919093086 | 0.111430536 |
| Catsper1 | 0 | NA | NA |
| Cst6 | 23.01852267 | -0.822226668 | 0.253414535 |
| Banf1 | 258.7929532 | 0.276528872 | 0.094156331 |
| Eif1ad | 254.8369414 | 0.557115717 | 0.107837388 |
| Sart1 | 490.457391 | 0.227502737 | 0.147940925 |
| Tsga10ip | 0.362047852 | -0.242093081 | 1.586077597 |
| 4930481A15Rik | 2.199134214 | -0.76292593 | 0.948799893 |
| Gm6293 | 0.045882455 | 0.237312084 | 3.724945049 |
| Drap1 | 882.2497186 | 0.215485107 | 0.09093829 |
| AI837181 | 578.5093112 | 0.613257444 | 0.115411939 |
| Gm25432 | 0 | NA | NA |
| Fosl1 | 112.2972541 | 8.279959736 | 0.828971213 |
| Ccdc85b | 470.1620099 | -0.119970275 | 0.105629758 |
| Fibp | 216.1359764 | -0.009409654 | 0.07589575 |
| Ctsw | 32.65133826 | 4.412864094 | 0.42019935 |
| Efemp2 | 186.5996474 | 1.547581402 | 0.155277987 |
| Mus81 | 195.7583366 | -0.180939391 | 0.115195224 |
| Cfl1 | 6985.640524 | 0.299308866 | 0.069324523 |
| Snx32 | 463.4435586 | -0.743391779 | 0.08357897 |
| 1700020D05Rik | 15.41874029 | -1.111520039 | 0.416718748 |

| | | | |
|---|---|---|---|
| Ovol1 | 1.251933375 | 3.350336503 | 1.335106517 |
| Ap5b1 | 86.41281867 | 0.843063457 | 0.136598667 |
| Rnaseh2c | 82.70313133 | -0.578730706 | 0.174141783 |
| Kat5 | 511.7440207 | 0.106589123 | 0.06440936 |
| Rela | 372.9586236 | 1.154936151 | 0.162921129 |
| Sipa1 | 217.0465711 | 1.427778417 | 0.151362357 |
| Pcnxl3 | 996.3058461 | -0.082214986 | 0.0720543 |
| Mir6987 | 0 | NA | NA |
| Map3k11 | 329.8028705 | 0.632821893 | 0.109120647 |
| Kcnk7 | 0.811426433 | 2.355659523 | 1.227721679 |
| Ehbp1l1 | 284.648246 | 0.47732744 | 0.10827393 |
| Gm16538 | 0.233003758 | 2.148481989 | 2.52304222 |
| Fam89b | 0.325597809 | -1.521518584 | 1.71557966 |
| Sssca1 | 172.9191299 | -0.023406319 | 0.101170921 |
| Ltbp3 | 514.8847917 | -0.298787512 | 0.074290844 |
| Scyl1 | 711.1667025 | 0.123984611 | 0.072473533 |
| Malat1 | 158003.9038 | -0.26114019 | 0.226882376 |
| Gm20417 | 3.344837808 | 0.216164554 | 0.928049009 |
| Gm27694 | 0 | NA | NA |
| Gm37376 | 17.03796044 | -0.150730471 | 0.969267384 |
| Gm27702 | 0 | NA | NA |
| Neat1 | 2624.857955 | 1.380145072 | 0.289207193 |
| Gm27533 | 0 | NA | NA |
| Gm27505 | 0 | NA | NA |
| Gm28006 | 0 | NA | NA |
| Frmd8os | 2.120407615 | 0.203911421 | 0.714401802 |
| Frmd8 | 356.4512706 | 1.618110589 | 0.186562484 |
| Slc25a45 | 48.19350137 | 1.238318231 | 0.228295029 |
| Tigd3 | 29.86989943 | -0.375061102 | 0.208627181 |
| Dpf2 | 712.9918355 | 0.371470497 | 0.066841152 |
| Cdc42ep2 | 113.290494 | -0.298683426 | 0.159257346 |
| Pola2 | 84.75758802 | 0.913495507 | 0.148494538 |
| Slc22a20 | 0.141071297 | 0.237322348 | 3.564043836 |
| Capn1 | 318.3485814 | 0.811083911 | 0.100586216 |
| Gm10814 | 14.06531991 | -0.202254688 | 0.305771408 |
| Gm8034 | 7.330356508 | -0.283250247 | 0.39075024 |
| Syvn1 | 680.7779014 | 0.516596426 | 0.082679856 |
| Mir6988 | 0 | NA | NA |
| Mrpl49 | 285.2862106 | 0.055663426 | 0.090852497 |
| Fau | 1011.116782 | 0.608999726 | 0.100176777 |
| Znhit2 | 165.2069863 | 0.238865264 | 0.09841216 |
| Tm7sf2 | 103.8839401 | -1.232994706 | 0.14699919 |
| Vps51 | 374.0248763 | -0.044259925 | 0.07079735 |
| Gm28374 | 0.132725991 | 0.717601035 | 2.971216202 |
| BC048609 | 0.046345709 | 0.71820487 | 3.724945049 |
| Zfpl1 | 181.431277 | 0.250943059 | 0.111824088 |
| Cdca5 | 6.165523022 | 2.526383948 | 0.585703428 |

| | | | |
|---|---|---|---|
| Naaladl1 | 0.666154845 | -1.696189621 | 1.203482544 |
| Sac3d1 | 83.16707754 | 0.133297548 | 0.14017505 |
| Snx15 | 211.7518732 | -0.381130409 | 0.080662572 |
| Arl2 | 383.7572647 | -0.045143407 | 0.098004852 |
| Batf2 | 180.1949883 | 7.562442764 | 0.535364301 |
| n-R5s19 | 0 | NA | NA |
| Gm5510 | 2.351247315 | 0.817717867 | 0.643363269 |
| 1700123I01Rik | 7.996023092 | 2.827563671 | 0.490321112 |
| Gpha2 | 0.377319533 | -0.243563846 | 1.582959984 |
| Ppp2r5b | 1407.304023 | -0.266496001 | 0.053257681 |
| Atg2a | 650.2802182 | -0.160099118 | 0.060639758 |
| Mir194-2 | 0.099106094 | 1.209894437 | 3.72418822 |
| Mir192 | 0.254293348 | 0.237327089 | 2.472801316 |
| Gm14963 | 2.026491629 | -0.302244146 | 0.71452278 |
| Ehd1 | 703.3125589 | 0.677329594 | 0.118310077 |
| Cdc42bpg | 176.7180393 | -0.287772221 | 0.153126188 |
| Men1 | 585.1800558 | -0.109687944 | 0.052905842 |
| Map4k2 | 638.3859077 | -0.700505222 | 0.098425173 |
| Mir6989 | 0 | NA | NA |
| Gm14966 | 12.5179983 | -0.621324437 | 0.312157359 |
| Sf1 | 2104.915733 | 0.247577232 | 0.082484731 |
| Pygm | 267.9090404 | -0.958715598 | 0.17072338 |
| Rasgrp2 | 263.4585251 | -1.498245305 | 0.198274598 |
| Gm14965 | 2.017609801 | 0.456286484 | 0.596763659 |
| Nrxn2 | 6243.867174 | -0.499920298 | 0.0763781 |
| Gm14964 | 23.8586541 | -0.452003446 | 0.269856496 |
| Slc22a12 | 5.133294896 | 0.78431097 | 0.562951099 |
| Gm14967 | 0 | NA | NA |
| Gm26460 | 0.404356784 | 1.708512988 | 1.523818162 |
| Hmgb1-ps4 | 0.663061751 | 0.44403559 | 1.175190471 |
| 2410152P15Rik | 0 | NA | NA |
| Gm14968 | 0.11037557 | -0.243571806 | 3.724945049 |
| Rps6ka4 | 374.0853682 | -0.036123503 | 0.079537278 |
| mmu-mir-5046 | 0 | NA | NA |
| Ccdc88b | 121.8708714 | 3.261574918 | 0.190345182 |
| Prdx5 | 1538.036727 | 0.328705334 | 0.085979132 |
| Trmt112 | 108.60505 | 0.612956246 | 0.162883457 |
| Esrra | 457.3522009 | -0.318417695 | 0.083793773 |
| Mir6990 | 0 | NA | NA |
| Tex40 | 21.48008408 | -1.219289029 | 0.315824868 |
| Kcnk4 | 51.05303909 | -1.567056381 | 0.218941394 |
| Gpr137 | 725.8468719 | -0.191927216 | 0.059855345 |
| Bad | 198.4137653 | 0.063860939 | 0.119826164 |
| Plcb3 | 323.134626 | 0.568026095 | 0.120490131 |
| Ppp1r14b | 149.5878577 | 1.071128221 | 0.166382443 |
| Fkbp2 | 300.3276835 | -0.149020075 | 0.283476741 |
| Vegfb | 398.1291403 | -0.358863409 | 0.106322568 |

| | | | |
|---|---|---|---|
| Dnajc4 | 95.01009773 | -0.129498819 | 0.132987275 |
| Nudt22 | 67.37727446 | 0.516516462 | 0.145498401 |
| Trpt1 | 56.60693119 | -0.447564335 | 0.147608877 |
| Fermt3 | 175.2517023 | 3.231385806 | 0.16765068 |
| Stip1 | 1604.279865 | 0.24007902 | 0.096263777 |
| Macrod1 | 69.5949187 | -0.413660786 | 0.144196239 |
| Flrt1 | 608.6636325 | -0.905431409 | 0.1264131 |
| Otub1 | 1632.950069 | -0.1762963 | 0.065778422 |
| Cox8a | 2092.571017 | -0.316145997 | 0.074665142 |
| Naa40 | 276.9429562 | -0.093381581 | 0.121733449 |
| Gm22113 | 0 | NA | NA |
| Gm23997 | 0 | NA | NA |
| Rcor2 | 69.31615873 | -1.412634695 | 0.189566483 |
| Mark2 | 1468.643762 | -0.232952336 | 0.078510471 |
| Gm17227 | 0.066050875 | 0.71820487 | 3.724945049 |
| AI846148 | 273.2784663 | -0.013030199 | 0.100553655 |
| 1700105P06Rik | 6.754696947 | -0.32415968 | 0.412897019 |
| Rab11b-ps2 | 191.3767199 | -0.030366951 | 0.125385695 |
| 2700081O15Rik | 415.5788108 | -0.438758831 | 0.103796399 |
| Mir6991 | 0.662460401 | -0.954747411 | 1.461671565 |
| Rtn3 | 9090.002618 | -0.567006488 | 0.051424145 |
| Atl3 | 817.3787042 | 0.238400276 | 0.06221005 |
| Pla2g16 | 615.7304892 | 0.938720011 | 0.131320445 |
| Lgals12 | 1.476747611 | -2.137159344 | 0.939254885 |
| Hrasls5 | 1.687958028 | 1.271647865 | 1.129026926 |
| Gm16437 | 26.12384685 | 0.278015467 | 0.261646451 |
| Slc22a19 | 0 | NA | NA |
| Gm15967 | 0 | NA | NA |
| Gm15968 | 0 | NA | NA |
| Slc22a26 | 0.04393175 | -0.243571806 | 3.724945049 |
| Slc22a27 | 0 | NA | NA |
| Gm24951 | 0 | NA | NA |
| Slc22a28 | 0 | NA | NA |
| Slc22a29 | 1.523828694 | 0.427596493 | 0.751035065 |
| Slc22a30 | 0.083526043 | 0.718202646 | 3.724945049 |
| Gm6425 | 1.341472408 | 0.410182058 | 0.805662163 |
| Slc22a8 | 69.67438683 | -1.808484143 | 0.289131061 |
| Slc22a6 | 62.43237234 | -1.157612207 | 0.269664408 |
| Chrm1 | 1319.762724 | -1.374739779 | 0.177111015 |
| Gm22820 | 0.118974955 | -0.243576254 | 3.724945049 |
| Slc3a2 | 1681.539151 | 1.575919486 | 0.142404373 |
| Gm25855 | 7.213846394 | 0.337294974 | 0.40246032 |
| Gm25894 | 7.96667369 | 0.668362717 | 0.441386594 |
| Gm24452 | 7.467891793 | 0.249439961 | 0.428509874 |
| Gm24453 | 6.941141185 | 1.225945503 | 0.39977532 |
| Gm22744 | 7.168190812 | 0.705152194 | 0.40897164 |
| Gm22680 | 5.248670073 | 1.023220692 | 0.494763866 |

| | | | |
|---|---|---|---|
| Gm23246 | 6.637239202 | -0.042285613 | 0.409227777 |
| Snord22 | 166.9869455 | 0.588257267 | 0.242207877 |
| Wdr74 | 196.0682104 | 0.428398681 | 0.113786063 |
| 1700092M07Rik | 1.245725704 | -1.4777562 | 1.034568868 |
| Stx5a | 534.5973207 | -0.041532393 | 0.080767597 |
| Mir6992 | 0 | NA | NA |
| Nxf1 | 945.8212657 | -0.066516781 | 0.067999764 |
| Taf6l | 170.5846256 | 0.653784324 | 0.110779299 |
| Polr2g | 319.7014694 | 0.306355888 | 0.078256175 |
| Zbtb3 | 40.55560979 | 0.289617398 | 0.186449035 |
| Ttc9c | 517.0751682 | 0.566614753 | 0.07846961 |
| Hnrnpul2 | 2234.773551 | -0.20426532 | 0.111083622 |
| Gng3 | 1184.138002 | -0.520349472 | 0.071247116 |
| Bscl2 | 767.6870937 | 0.063778847 | 0.068669039 |
| Lrrn4cl | 17.94000658 | 0.802726988 | 0.245909622 |
| Ubxn1 | 553.7697328 | 0.474358043 | 0.091896525 |
| Uqcc3 | 121.6763236 | 0.281625529 | 0.140532339 |
| Lbhd1 | 49.53080462 | -0.099323727 | 0.178098788 |
| Snora57 | 16.08879982 | 1.290181084 | 0.333307606 |
| Mir5136 | 0.568319789 | -1.75512309 | 1.299218027 |
| 1810009A15Rik | 2.938865308 | -0.380331529 | 0.633726808 |
| Ints5 | 306.8553324 | 0.5387867 | 0.101958609 |
| Ganab | 1308.618353 | 0.244659802 | 0.061817041 |
| B3gat3 | 910.6532437 | -0.119861555 | 0.091396225 |
| Rom1 | 80.3855847 | 0.477664454 | 0.162272317 |
| Eml3 | 209.4625949 | 0.379529026 | 0.108204126 |
| Mta2 | 687.4573254 | 0.577892014 | 0.080112323 |
| Tut1 | 264.2884532 | 0.320841349 | 0.094187506 |
| Eef1g | 1677.123862 | 0.537631338 | 0.083554239 |
| Ahnak | 4057.702084 | 2.697875917 | 0.173306726 |
| Scgb1a1 | 0 | NA | NA |
| Asrgl1 | 1213.125555 | -0.674983578 | 0.119402011 |
| Gm9750 | 3.69099669 | 0.343380051 | 0.53178285 |
| Gm24543 | 0 | NA | NA |
| Pcna-ps2 | 9.410251089 | 1.033579531 | 0.348942756 |
| Gm2617 | 0.087261593 | -0.243576254 | 3.724945049 |
| Stxbp3-ps | 23.11005125 | 1.078447057 | 0.238092919 |
| Gm29231 | 0.2218918 | -1.084728494 | 3.137147691 |
| Gm6445 | 1.943633249 | -0.443112048 | 0.732757592 |
| Scgb2a2 | 0.152314784 | -0.724459891 | 2.847275348 |
| Gm22369 | 0 | NA | NA |
| Incenp | 144.1045022 | -0.120891637 | 0.126891588 |
| Gm24484 | 0.061920613 | 0.718207094 | 3.724945049 |
| Fth1 | 13460.26008 | 1.089425183 | 0.167225675 |
| Best1 | 1.289165993 | 2.244031198 | 1.060169321 |
| Rab3il1 | 131.0561169 | 0.870492244 | 0.171329149 |
| Fads3 | 258.367544 | 0.162573026 | 0.08757719 |

| | | | |
|---|---|---|---|
| Fads2 | 1025.070797 | -0.157679574 | 0.11921997 |
| Fads1 | 1910.999182 | -0.313299143 | 0.098292977 |
| Mir6993 | 1.831497397 | -0.717098278 | 0.676913174 |
| Gm10143 | 2.039768525 | -0.737449042 | 0.794323468 |
| Fen1 | 177.2425644 | 0.838040473 | 0.146781321 |
| Tmem258 | 152.1698681 | 0.44752618 | 0.164977817 |
| Myrf | 1402.777588 | 0.939430661 | 0.136805141 |
| Dagla | 1427.846811 | -0.640322024 | 0.113413993 |
| Syt7 | 3169.726564 | -0.989284771 | 0.103923767 |
| Gm26859 | 2.155457834 | -0.832824394 | 0.675845795 |
| Lrrc10b | 214.7262662 | -1.265446333 | 0.216475357 |
| Ppp1r32 | 18.15485456 | 0.67546775 | 0.228696027 |
| Sdhaf2 | 369.9013947 | 0.463196195 | 0.072094505 |
| Cpsf7 | 982.7429653 | 0.039850558 | 0.079759979 |
| Tmem216 | 43.25074293 | 0.531483437 | 0.180561518 |
| Tmem138 | 84.89562639 | 0.258265178 | 0.119550992 |
| Cyb561a3 | 70.26552882 | 0.911077803 | 0.192670721 |
| Tkfc | 98.20098442 | -0.215122937 | 0.131484631 |
| Ddb1 | 2897.624334 | 0.181463141 | 0.055155763 |
| Vwce | 6.549817523 | 0.805828793 | 0.461253424 |
| Pga5 | 0.066050875 | 0.71820487 | 3.724945049 |
| Vps37c | 286.9571528 | 0.424103049 | 0.099409508 |
| Cd5 | 21.9583943 | 3.937715596 | 0.425617701 |
| A430093F15Rik | 5.018136537 | 3.935986466 | 0.824967573 |
| Cd6 | 55.03509925 | 1.945329559 | 0.224230814 |
| Slc15a3 | 354.5772757 | 5.176670457 | 0.303287129 |
| Tmem132a | 2470.423893 | -0.386519565 | 0.093405483 |
| Tmem109 | 292.8556633 | -0.13681213 | 0.120547981 |
| Prpf19 | 1647.79328 | 0.180969882 | 0.06564846 |
| Zp1 | 0.369197529 | 0.261462252 | 1.453018841 |
| Ptgdr2 | 2.273986841 | 0.613150994 | 0.659960491 |
| Ccdc86 | 181.1409562 | 1.393204925 | 0.138071485 |
| Ms4a10 | 0.044544648 | 0.237309859 | 3.724945049 |
| Ms4a15 | 28.11515156 | -0.021318755 | 1.129009275 |
| Ms4a18 | 0.263774563 | 2.227043398 | 2.376523115 |
| Gm28347 | 2.460283742 | 5.08462951 | 0.955248706 |
| AW112010 | 268.3498837 | 7.08492316 | 0.45575982 |
| Ms4a8a | 62.44774592 | 9.428268331 | 1.06077495 |
| Gm336 | 28.43566127 | 0.736590801 | 0.207067799 |
| 1700017D01Rik | 0.314055832 | 1.60254248 | 1.730742929 |
| Gm6501 | 23.72387051 | 0.800900381 | 0.269654401 |
| 1700025F22Rik | 0.715113835 | 3.070083763 | 1.396251822 |
| Ms4a13 | 0 | NA | NA |
| 4930526L06Rik | 0.358247177 | 0.242378919 | 1.578371256 |
| Ms4a12 | 0.470940457 | 2.940134128 | 1.531913554 |
| Ms4a1 | 1.60259925 | 3.785092825 | 0.986807211 |
| Ms4a5 | 0 | NA | NA |

| | | | |
|---|---|---|---|
| Ms4a14 | 14.20065461 | 5.78966623 | 0.806315272 |
| Ms4a7 | 116.0378029 | 5.981372835 | 0.388434937 |
| Gm28935 | 0.610760527 | 2.576923879 | 1.451001053 |
| Ms4a4a | 77.63966184 | 4.985029665 | 0.416446693 |
| Gm5692 | 0.269752699 | 2.100162329 | 1.769126485 |
| Ms4a4c | 622.4266349 | 10.47372131 | 0.671555568 |
| Gm5511 | 2.864211081 | 1.273979077 | 0.677896289 |
| Ms4a4b | 229.0039815 | 9.589353362 | 0.796541185 |
| Gm37387 | 7.980888526 | 6.193026286 | 0.945716343 |
| Ms4a6c | 638.4433901 | 7.167498381 | 0.332962928 |
| BE692007 | 14.17246462 | 5.397346623 | 0.789103437 |
| Gm8369 | 26.70344983 | 5.269365266 | 0.575114937 |
| Gm19261 | 0.679959982 | 3.158537799 | 1.178573902 |
| Ms4a6b | 404.189003 | 5.907461261 | 0.305720305 |
| Ms4a4d | 12.73072494 | 7.440820803 | 0.900892423 |
| Gm10212 | 1.028441291 | 3.263914252 | 2.112425317 |
| Gm38275 | 1.398451413 | 3.464625582 | 1.036022944 |
| Ms4a6d | 780.7598453 | 7.196970962 | 0.381285533 |
| Gm24599 | 0 | NA | NA |
| Ms4a2 | 1.450910466 | 2.371841405 | 0.985729216 |
| Ms4a3 | 0.922047842 | 2.147406047 | 1.144867736 |
| Oosp2 | 0.454235466 | 2.002611612 | 1.36858175 |
| Oosp1 | 0.420049603 | 2.72593407 | 1.596106554 |
| Gm26183 | 0 | NA | NA |
| Gm25443 | 0 | NA | NA |
| Oosp3 | 0 | NA | NA |
| Gm19224 | 0.308730509 | 1.632004418 | 3.716508703 |
| Gif | 0.046345709 | 0.71820487 | 3.724945049 |
| Mrpl16 | 136.9670042 | 0.092803836 | 0.117425271 |
| Stx3 | 268.7199862 | 0.318551937 | 0.114191587 |
| Olfr1417 | 0.045882455 | 0.237312084 | 3.724945049 |
| Olfr1418 | 0 | NA | NA |
| Olfr1419 | 0.385885129 | -0.404191203 | 1.460723925 |
| Gm6365 | 6.926587766 | 0.933762958 | 0.398364281 |
| Olfr1420 | 0.092228164 | 0.718202646 | 3.724945049 |
| Patl1 | 512.1726233 | 0.604838304 | 0.08313914 |
| mmu-mir-6994 | 0.26492599 | 0.717778447 | 1.820983113 |
| Gm22272 | 0 | NA | NA |
| Osbp | 1002.556198 | -0.012336823 | 0.047050229 |
| Olfr1423 | 0 | NA | NA |
| Olfr1424 | 0.049553047 | 0.71820487 | 3.724945049 |
| Olfr1425 | 0.045882455 | 0.237312084 | 3.724945049 |
| Olfr1426 | 0 | NA | NA |
| Olfr1427 | 0 | NA | NA |
| Olfr1428 | 0.090277459 | 0.237318756 | 3.724945049 |
| Olfr76 | 0.045882455 | 0.237312084 | 3.724945049 |
| Olfr1431 | 0 | NA | NA |

| | | | |
|---|---|---|---|
| Olfr1432 | 0 | NA | NA |
| Olfr262 | 0 | NA | NA |
| Gm26864 | 0 | NA | NA |
| Olfr235 | 0.119234754 | -0.243576254 | 3.724945049 |
| Olfr1434 | 0 | NA | NA |
| Olfr1436 | 0 | NA | NA |
| Olfr1437 | 0.435432306 | 2.734264239 | 3.679082036 |
| Olfr1440 | 0.060174537 | 0.237309859 | 3.724945049 |
| Olfr1441 | 0.105852363 | 0.23732098 | 3.724945049 |
| Pfpl | 0 | NA | NA |
| Mpeg1 | 2850.264192 | 4.747983763 | 0.186473226 |
| Dtx4 | 733.6416179 | 0.526543267 | 0.094943959 |
| Fam111a | 54.9166158 | 3.569380976 | 0.361057496 |
| Gm16274 | 2.331142904 | -0.366089576 | 0.700184104 |
| A330040F15Rik | 6.392297317 | 0.336854525 | 0.426215668 |
| Gm4952 | 2.655904937 | 4.713087964 | 0.937627896 |
| Glyat | 2.115680473 | 0.707860882 | 0.774728278 |
| Gm24521 | 0.066050875 | 0.71820487 | 3.724945049 |
| Olfr1442 | 0.259089178 | 2.114906188 | 2.017823773 |
| Olfr1443 | 0.092691418 | 1.183079489 | 3.724560917 |
| Keg1 | 0.403815225 | 2.788641999 | 3.67649238 |
| Gm15962 | 0 | NA | NA |
| Cntf | 4.709263031 | -0.297561582 | 0.476239538 |
| Zfp91 | 2396.758906 | 0.136559818 | 0.061437453 |
| Lpxn | 113.3029764 | 5.28354562 | 0.328325794 |
| Gm5244 | 0.635603962 | 1.979674481 | 1.309597724 |
| Olfr1444 | 0.544199285 | 2.365723614 | 1.242716319 |
| Olfr1445 | 0.066050875 | 0.71820487 | 3.724945049 |
| Olfr1446 | 0 | NA | NA |
| Olfr1447 | 0 | NA | NA |
| Olfr1448 | 0.092228164 | 0.718202646 | 3.724945049 |
| Olfr1449 | 0.066050875 | 0.71820487 | 3.724945049 |
| Olfr1450 | 0 | NA | NA |
| Olfr1451 | 0 | NA | NA |
| Olfr1452-ps1 | 0 | NA | NA |
| Olfr1453 | 0 | NA | NA |
| Olfr1454 | 0 | NA | NA |
| Olfr1456-ps1 | 0 | NA | NA |
| Olfr1457 | 0 | NA | NA |
| Olfr1458 | 0.049553047 | 0.71820487 | 3.724945049 |
| Olfr1459 | 0 | NA | NA |
| Olfr1461 | 0 | NA | NA |
| Olfr1462 | 0 | NA | NA |
| Olfr1463 | 0 | NA | NA |
| Gm22443 | 0 | NA | NA |
| Olfr1465 | 0 | NA | NA |
| Olfr1466 | 0.062426873 | 0.237312084 | 3.724945049 |

| | | | |
|---|---|---|---|
| Olfr1467 | 0 | NA | NA |
| Olfr1469 | 0 | NA | NA |
| Olfr1471 | 0 | NA | NA |
| Olfr1472 | 0 | NA | NA |
| Olfr1474 | 0 | NA | NA |
| Olfr1475 | 0 | NA | NA |
| Olfr1477 | 0 | NA | NA |
| Olfr1480 | 0 | NA | NA |
| Olfr1484 | 0.068097359 | -0.243569582 | 3.724945049 |
| Gm22323 | 0 | NA | NA |
| Gm23931 | 0 | NA | NA |
| Olfr1487 | 0 | NA | NA |
| Olfr1489 | 0 | NA | NA |
| Gm24124 | 0 | NA | NA |
| Olfr1490 | 0 | NA | NA |
| Olfr1491 | 0 | NA | NA |
| Olfr1494 | 0.061920613 | 0.718207094 | 3.724945049 |
| Olfr1495 | 0 | NA | NA |
| Olfr1496 | 0 | NA | NA |
| Olfr1497 | 0.04393175 | -0.243571806 | 3.724945049 |
| Olfr1499 | 0 | NA | NA |
| Olfr1500 | 0 | NA | NA |
| Olfr1501 | 0.044191549 | -0.243571806 | 3.724945049 |
| Olfr1502 | 0 | NA | NA |
| Olfr1503-ps1 | 0 | NA | NA |
| Olfr1504 | 0 | NA | NA |
| Olfr1505 | 0 | NA | NA |
| Gm26993 | 0 | NA | NA |
| Tle4 | 1024.20378 | 0.145067898 | 0.069327253 |
| Gm37997 | 0 | NA | NA |
| Gm26026 | 0 | NA | NA |
| Gm5513 | 1.595119001 | 1.087402909 | 0.820009409 |
| Gm24319 | 0 | NA | NA |
| Psat1 | 926.2091933 | 0.161926722 | 0.125989362 |
| Gm3329 | 0 | NA | NA |
| Cep78 | 106.2535858 | -0.535873732 | 0.124880795 |
| C130060C02Rik | 7.941562867 | -1.104739271 | 0.47772008 |
| Gnaq | 3032.47275 | -0.250749115 | 0.038029113 |
| E030024N20Rik | 20.01620722 | -0.361358663 | 0.22448972 |
| Gm10819 | 108.4716668 | -0.067719982 | 0.128390566 |
| Gm17068 | 25.77906067 | -0.324540386 | 0.376145198 |
| Mir496b | 0 | NA | NA |
| Gna14 | 16.24861454 | 1.915850778 | 0.463039247 |
| Vps13a | 1858.188761 | -0.447157385 | 0.073451088 |
| Gm9806 | 0.46212231 | 1.209816184 | 1.542466877 |
| Foxb2 | 1.423250569 | 0.273045962 | 0.929793334 |
| Prune2 | 1550.090265 | 0.283815669 | 0.1140846 |

| | | | |
|---|---|---|---|
| Gm27970 | 0 | NA | NA |
| Gm27796 | 0 | NA | NA |
| Gcnt1 | 110.511412 | 0.175248886 | 0.189241755 |
| Rfk | 726.378061 | 0.244653112 | 0.108708051 |
| Pcsk5 | 144.4692259 | -0.193510789 | 0.189806191 |
| Gm22880 | 0 | NA | NA |
| Gm23063 | 0 | NA | NA |
| Gm23061 | 0 | NA | NA |
| Ostf1 | 237.4482545 | 1.752189744 | 0.14509636 |
| Gm8250 | 0.075043205 | 0.237309859 | 3.724945049 |
| Nmrk1 | 21.71250768 | -0.175461502 | 0.236910727 |
| 2410127L17Rik | 57.66373482 | -0.015029477 | 0.154916739 |
| D030056L22Rik | 161.6691014 | 0.029359926 | 0.102931832 |
| Gm23238 | 1902.741476 | 0.430072132 | 0.398663822 |
| Trpm6 | 41.27246436 | 1.946739299 | 0.226628978 |
| Gm24190 | 0 | NA | NA |
| Rorb | 1457.656088 | 0.204800118 | 0.083676806 |
| Gm25437 | 0 | NA | NA |
| Gm22684 | 0 | NA | NA |
| Anxa1 | 128.5436956 | 3.458744227 | 0.30456126 |
| 1500015L24Rik | 2.063333161 | -0.881598083 | 0.749426902 |
| Gm6684 | 0 | NA | NA |
| Aldh1a1 | 547.0244569 | 0.780860449 | 0.187509459 |
| Aldh1a7 | 11.46744072 | -0.864794694 | 0.49417649 |
| Tmc1 | 1.743158711 | -0.049725798 | 0.86561744 |
| Zfand5 | 1899.320038 | -0.073959929 | 0.074088643 |
| Gda | 1770.327303 | -0.513167045 | 0.104779621 |
| Gm3443 | 0.062426873 | 0.237312084 | 3.724945049 |
| 1110059E24Rik | 203.1103973 | 0.278171284 | 0.133388371 |
| Abhd17b | 456.3664533 | 0.093715922 | 0.076637223 |
| Tmem2 | 509.6691455 | 0.668088816 | 0.10569402 |
| Gm5514 | 10.26669978 | -0.508418377 | 0.364604023 |
| Trpm3 | 1663.018852 | -0.685337455 | 0.081649694 |
| Gm22506 | 0 | NA | NA |
| Gm27151 | 4.77538895 | -0.830494485 | 0.49279899 |
| Mir204 | 3.450472164 | -1.10364984 | 0.714488084 |
| C330002G04Rik | 1.481948713 | 2.298163882 | 0.927740568 |
| 2410080I02Rik | 2.836846521 | -0.337664168 | 0.668011156 |
| Klf9 | 2856.577684 | 0.078310306 | 0.121906147 |
| Mir1192 | 10.09841093 | -0.596493838 | 0.405271203 |
| Smc5 | 682.7510276 | -0.123901035 | 0.08883027 |
| Gm24263 | 0.146692594 | 1.229893553 | 3.521109737 |
| Mamdc2 | 41.82610674 | -0.322547269 | 0.222679185 |
| 1700028P14Rik | 7.326566073 | 0.89636022 | 0.393105049 |
| Gm9493 | 43.04589625 | 0.642892455 | 0.197686427 |
| Gm22144 | 0 | NA | NA |
| Gm21542 | 0 | NA | NA |

| | | | |
|---|---|---|---|
| Gm6563 | 117.5227222 | 0.455678572 | 0.145401274 |
| Ptar1 | 312.4373579 | -0.167030864 | 0.083839453 |
| Gm9938 | 199.0723875 | -0.189298607 | 0.093816942 |
| Apba1 | 2979.940385 | -0.373132514 | 0.068230297 |
| Fam189a2 | 83.27374125 | -0.088483555 | 0.147826074 |
| 1700021P04Rik | 0.123841226 | 1.292225235 | 3.719439735 |
| Tjp2 | 802.6063976 | 1.120823902 | 0.109280183 |
| Fxn | 57.39510889 | 0.20247131 | 0.155577562 |
| Gm16216 | 0.110870145 | 0.238305784 | 3.724874707 |
| Pip5k1b | 239.2249923 | -0.48408894 | 0.110811723 |
| Pip5k1bos | 0.196777805 | -0.712620893 | 2.227175217 |
| Fam122a | 94.19792559 | 0.108589087 | 0.159913546 |
| Tmem252 | 56.70084035 | 5.166780765 | 0.432565007 |
| Pgm5 | 91.2960709 | 2.050771285 | 0.217094473 |
| Gm17800 | 0.452161616 | 1.003655877 | 1.381776019 |
| Gm10053 | 118.3751616 | 0.315462876 | 0.158741414 |
| Foxd4 | 0.431881386 | 1.190745401 | 1.714314669 |
| Cbwd1 | 142.8925874 | 0.378160526 | 0.104130494 |
| mmu-mir-3084-1 | 0 | NA | NA |
| Gm23658 | 0.086648695 | -0.72445792 | 3.724945049 |
| Dock8 | 246.4602774 | 2.535643472 | 0.17318518 |
| Gm24252 | 0 | NA | NA |
| Kank1 | 435.2904436 | 0.630187689 | 0.132670074 |
| Gm26207 | 0 | NA | NA |
| Dmrt1 | 0 | NA | NA |
| Dmrt3 | 3.281054849 | 1.306084116 | 0.589892281 |
| 2610016A17Rik | 0.172734917 | -0.243562117 | 2.736751389 |
| Dmrt2 | 7.322241276 | -1.086836195 | 0.409592153 |
| Gm8825 | 0 | NA | NA |
| Gm25176 | 0.061920613 | 0.718207094 | 3.724945049 |
| Smarca2 | 2885.991631 | -0.680428926 | 0.069791099 |
| Gm20616 | 0.038981395 | 0.718207094 | 3.724945049 |
| 4931403E22Rik | 1.142959485 | 0.389147973 | 0.925873949 |
| Gm815 | 0.098502579 | 0.23732098 | 3.724945049 |
| Vldlr | 1529.815537 | -0.277457865 | 0.062483345 |
| Kcnv2 | 1.799299276 | 0.376604149 | 0.86282309 |
| D19Bwg1357e | 318.8741132 | 0.325414808 | 0.097215133 |
| Gm24782 | 0 | NA | NA |
| Rfx3 | 1251.313148 | -0.671889621 | 0.11726546 |
| Gm8412 | 0.124853745 | 0.237309606 | 3.724945049 |
| Gm20407 | 0 | NA | NA |
| Gm28228 | 0.047948698 | -0.243569582 | 3.724945049 |
| Glis3 | 334.0083546 | 1.862389669 | 0.119528325 |
| 4933413C19Rik | 10.33626457 | 0.512581316 | 0.372675957 |
| D930032P07Rik | 0.131806241 | -0.243579363 | 3.668090746 |
| Rps15a-ps2 | 0.107078754 | 0.236669296 | 3.724898107 |
| Gm6788 | 2.966347917 | 0.373315993 | 0.55805544 |

| | | | |
|---|---|---|---|
| Gm26942 | 0.038981395 | 0.718207094 | 3.724945049 |
| Slc1a1 | 1257.835077 | -0.435290999 | 0.080453506 |
| Gm5822 | 3.84996234 | 0.231548076 | 0.536263113 |
| 4430402I18Rik | 33.49194021 | -1.058049998 | 0.209282713 |
| Ppapdc2 | 422.5883542 | -0.345469126 | 0.083820648 |
| Cdc37l1 | 944.3468794 | -0.217662215 | 0.059617888 |
| Gm10136 | 20.53681307 | 0.56600975 | 0.236206492 |
| Ak3 | 872.3625232 | 0.207741578 | 0.117750167 |
| 1700018L02Rik | 10.4047421 | 0.006384857 | 0.393997966 |
| Rcl1 | 158.5573071 | 0.199607549 | 0.089261998 |
| Mir101b | 0.39937024 | -1.601324029 | 1.452789668 |
| Gm5518 | 30.36473408 | 0.244763698 | 0.275372718 |
| Jak2 | 1349.483123 | 1.148762967 | 0.097377917 |
| Insl6 | 6.983240532 | 2.152758027 | 0.619225934 |
| Rln1 | 4.858786034 | 1.331153952 | 0.570015804 |
| Plgrkt | 208.1764301 | 0.735440255 | 0.105149254 |
| Cd274 | 1796.888883 | 7.294057158 | 0.274202516 |
| Pdcd1lg2 | 5.725591314 | 5.248275379 | 0.904076042 |
| A930007I19Rik | 28.01513506 | 1.222119479 | 0.217964605 |
| Ric1 | 1103.079595 | 0.032400818 | 0.084740162 |
| Ermp1 | 862.3818722 | -0.378613696 | 0.081287758 |
| Mlana | 1.722373549 | 3.447561635 | 1.040369962 |
| 9930021J03Rik | 1199.788204 | -0.210995169 | 0.049340972 |
| Ranbp6 | 784.8181587 | -0.086143273 | 0.091879569 |
| Il33 | 391.409633 | 2.059101621 | 0.289053707 |
| Gm26046 | 0 | NA | NA |
| Trpd52l3 | 0 | NA | NA |
| Uhrf2 | 805.07859 | 0.179072251 | 0.067167218 |
| Gldc | 269.8156168 | -0.199171234 | 0.170145655 |
| Rpl31-ps20 | 0 | NA | NA |
| Mbl2 | 0.229843471 | -0.136361492 | 2.545197784 |
| Gm24108 | 0 | NA | NA |
| Gm24883 | 0 | NA | NA |
| Ppp1r2-ps3 | 1.44902218 | 1.183569245 | 0.902033434 |
| Dkk1 | 0.239409371 | -1.154275891 | 2.141382825 |
| Prkg1 | 254.0169383 | -0.74082247 | 0.126834307 |
| Gm23426 | 0 | NA | NA |
| Cstf2t | 1157.615467 | -0.004336717 | 0.054492373 |
| A1cf | 0.581447236 | 2.553201207 | 1.326266039 |
| Asah2 | 367.16758 | 0.375265923 | 0.145054876 |
| Sgms1 | 487.606945 | -0.100791065 | 0.093805789 |
| 2700046G09Rik | 25.94331831 | -0.395357043 | 0.233719573 |
| Rpl9-ps6 | 47.62432018 | 0.479128758 | 0.177424273 |
| Minpp1 | 627.6333895 | 0.166785892 | 0.081204572 |
| Papss2 | 133.0147862 | 0.224397675 | 0.178129865 |
| Atad1 | 1547.98136 | 0.353199875 | 0.101581836 |
| Pten | 3804.391786 | -0.332692251 | 0.069965786 |

| | | | |
|---|---|---|---|
| Gm22482 | 0.072025462 | 0.237312084 | 3.724945049 |
| Rnls | 22.89813653 | 0.5645736 | 0.232724934 |
| mmu-mir-3970 | 0 | NA | NA |
| Gm7237 | 0.127971488 | 1.23029335 | 3.685533863 |
| Gm8975 | 0 | NA | NA |
| Lipo4 | 0.195256443 | 0.237327512 | 2.230306796 |
| Lipo1 | 94.81852167 | 0.916259877 | 0.120639599 |
| Gm27440 | 0 | NA | NA |
| Gm8978 | 0 | NA | NA |
| Gm8980 | 0.107078754 | 0.236669296 | 3.724898107 |
| Lipo2 | 1.691184024 | 0.606580573 | 0.925746769 |
| Gm5097 | 0.090537259 | 0.237318756 | 3.724945049 |
| Gm5519 | 0.296824366 | 1.231287729 | 2.372115562 |
| Gm8988 | 0 | NA | NA |
| Gm23095 | 0.106971521 | 0.237307635 | 3.724945049 |
| Lipf | 0.066050875 | 0.71820487 | 3.724945049 |
| Lipk | 0.091880448 | -0.724455696 | 3.724945049 |
| Lipn | 0.251288325 | 1.772852883 | 2.091451752 |
| Gm23334 | 0 | NA | NA |
| Lipm | 1.074639982 | -0.107055827 | 1.144212839 |
| Ankrd22 | 0.527224786 | -1.779768013 | 1.523054295 |
| Stambpl1 | 217.0575449 | 0.036417756 | 0.087460976 |
| Acta2 | 30.64504911 | 1.24270496 | 0.288866215 |
| Fas | 57.72959901 | 3.693963213 | 0.338917501 |
| Gm27818 | 0.914190395 | 2.036940182 | 1.355597824 |
| Gm23060 | 0 | NA | NA |
| Ch25h | 163.957365 | 7.099261237 | 0.497962488 |
| Gm26902 | 14.57275529 | 0.377564472 | 0.381649299 |
| Lipa | 315.4234948 | 1.020746297 | 0.152515415 |
| Ifit2 | 10029.24711 | 7.040518123 | 0.281707869 |
| Ifit3 | 3311.123546 | 8.58214004 | 0.263608007 |
| Gm14446 | 387.1233819 | 5.36424839 | 0.46394922 |
| Ifit3b | 2018.601927 | 7.029622299 | 0.258205742 |
| 2010002M12Rik | 153.7706214 | 4.014763973 | 0.25062224 |
| Ifit1 | 4083.025541 | 9.198264423 | 0.286222931 |
| Slc16a12 | 39.40020055 | -1.040825978 | 0.219374525 |
| Gm27476 | 0 | NA | NA |
| Gm22082 | 0 | NA | NA |
| Pank1 | 222.2087571 | -0.549086485 | 0.105437531 |
| Mir107 | 0.848910726 | -2.358856714 | 1.359492734 |
| Kif20b | 26.85031262 | 1.1814194 | 0.259794595 |
| Mir1950 | 0 | NA | NA |
| Gm5248 | 0 | NA | NA |
| Gm24006 | 0 | NA | NA |
| Gm23075 | 0 | NA | NA |
| Gm22853 | 0 | NA | NA |
| Gm25611 | 0.204668643 | -1.044336943 | 2.595258787 |

| | | | |
|---|---|---|---|
| A830019P07Rik | 16.97608067 | 0.096846957 | 0.302581271 |
| Htr7 | 177.6287066 | 0.385490007 | 0.110381654 |
| Rpp30 | 118.3865456 | 0.435630932 | 0.118358465 |
| Ankrd1 | 0.384174469 | 1.561841333 | 1.707460405 |
| Pcgf5 | 290.0103938 | 1.524965026 | 0.110709499 |
| Gm23170 | 0 | NA | NA |
| Hectd2 | 578.0733075 | -0.032319124 | 0.104783568 |
| 1500017E21Rik | 5.192905494 | -0.966040423 | 0.463523474 |
| Ppp1r3c | 367.6091572 | -0.430183889 | 0.178592369 |
| Tnks2 | 2533.779749 | 0.059264943 | 0.059012971 |
| Fgfbp3 | 45.89989236 | -0.210970468 | 0.196281455 |
| Btaf1 | 981.4939569 | 0.134778791 | 0.08987462 |
| Cpeb3 | 1630.611145 | -0.308961066 | 0.05072851 |
| Gm22714 | 7.127945666 | -1.029435922 | 0.453941719 |
| A330032B11Rik | 4.137844924 | -0.834382345 | 0.644189388 |
| Gm23918 | 1.204149766 | 0.28741496 | 1.022651555 |
| 5-Mar | 1036.41543 | 1.258931812 | 0.120973914 |
| Gm25268 | 0 | NA | NA |
| Ide | 826.1150129 | 0.170693353 | 0.067227518 |
| Rpl10-ps6 | 9.176167568 | -0.726175178 | 0.34802237 |
| Kif11 | 42.94126884 | 1.231044913 | 0.273202924 |
| Gm38345 | 1.850005102 | 2.177984321 | 0.852577031 |
| Hhex | 20.51235511 | 1.548283075 | 0.332679435 |
| Gm23026 | 0 | NA | NA |
| Exoc6 | 443.4655257 | -0.134530062 | 0.077343617 |
| Cyp26c1 | 3.181018617 | -0.10689723 | 0.67461404 |
| Cyp26a1 | 1.890572446 | 0.351354226 | 0.740193003 |
| Myof | 324.4012162 | 2.58653601 | 0.274867658 |
| I830134H01Rik | 0.540537234 | -2.094435572 | 1.269207492 |
| Cep55 | 11.79181242 | 2.497587012 | 0.416822447 |
| Gm23300 | 0 | NA | NA |
| Ffar4 | 0.430805093 | -0.17205632 | 1.611830765 |
| Rbp4 | 74.47465664 | -0.031530055 | 0.184497895 |
| Pde6c | 1.452976049 | 0.537657017 | 0.82143287 |
| Fra10ac1 | 83.34982365 | -0.23319549 | 0.173277291 |
| Lgi1 | 1139.244008 | -0.587166609 | 0.093091797 |
| RP24-417P12.1 | 0.567844938 | 0.031490229 | 1.288951995 |
| Slc35g1 | 45.27465032 | 0.056847902 | 0.18290863 |
| Gm24930 | 0 | NA | NA |
| Plce1 | 890.1324761 | 0.969692043 | 0.135342062 |
| Gm28991 | 0 | NA | NA |
| Noc3l | 300.5445623 | 0.31840305 | 0.078343149 |
| Tbc1d12 | 321.3981898 | -0.059300342 | 0.08960677 |
| Gm25838 | 2.422369998 | -0.956188327 | 0.668519293 |
| Hells | 136.3095865 | 0.388158969 | 0.142912431 |
| Cyp2c55 | 1.660109277 | 0.784227431 | 0.734215859 |
| Gm23800 | 0 | NA | NA |

| | | | |
|---|---|---|---|
| Cyp2c65 | 0.792025016 | 1.900156708 | 1.194368398 |
| Cyp2c66 | 0.061920613 | 0.718207094 | 3.724945049 |
| Cyp2c53-ps | 0.038981395 | 0.718207094 | 3.724945049 |
| Cyp2c29 | 0.220032676 | 1.195301988 | 2.574515385 |
| Cyp2c38 | 0.045882455 | 0.237312084 | 3.724945049 |
| Cyp2c39 | 0.206165386 | 0.319485876 | 2.649120405 |
| Cyp2c67 | 0 | NA | NA |
| Cyp2c68 | 0.045882455 | 0.237312084 | 3.724945049 |
| Cyp2c40 | 0.138763737 | -0.724458535 | 2.930963593 |
| Cyp2c69 | 0.075043205 | 0.237309859 | 3.724945049 |
| Cyp2c37 | 0.045882455 | 0.237312084 | 3.724945049 |
| Cyp2c54 | 0.045882455 | 0.237312084 | 3.724945049 |
| Cyp2c50 | 0.153424464 | 0.684814535 | 2.831062225 |
| Cyp2c70 | 0.309994989 | -0.62972627 | 1.720159622 |
| Gm16470 | 3.026502306 | 0.264548116 | 0.614201447 |
| Pdlim1 | 63.17543641 | 0.752087633 | 0.18036985 |
| Sorbs1 | 1628.853159 | 0.472580168 | 0.093076986 |
| Aldh18a1 | 344.8097266 | 0.774978215 | 0.115639213 |
| Gm15801 | 18.90733818 | 0.927444962 | 0.251924455 |
| Gm27042 | 1.18341529 | 0.90070116 | 0.833330865 |
| Tctn3 | 216.4391839 | -0.48366065 | 0.102369658 |
| Entpd1 | 233.1810674 | 1.266553788 | 0.183158771 |
| Gm16027 | 0.848235736 | -1.42118008 | 1.23679135 |
| Gm8783 | 30.04226012 | 0.871251037 | 0.190435252 |
| Ccnj | 128.7530246 | 0.004115307 | 0.127982691 |
| E030044B06Rik | 1.397451337 | 0.68357885 | 0.938807405 |
| Mir8092 | 0 | NA | NA |
| Zfp518a | 366.4925213 | 0.508108582 | 0.114103819 |
| Blnk | 124.657933 | 1.417619423 | 0.193599953 |
| Dntt | 1.408550033 | 4.242697253 | 1.158280761 |
| Opalin | 167.6099803 | -0.036194456 | 0.136948125 |
| Tll2 | 11.98457804 | 0.155654972 | 0.300429966 |
| Tm9sf3 | 2559.714795 | -0.036269043 | 0.058419826 |
| Pik3ap1 | 444.6212228 | 4.94443174 | 0.194561689 |
| Gm9788 | 0.072025462 | 0.237312084 | 3.724945049 |
| Lcor | 579.5255258 | -0.394291648 | 0.113912839 |
| Gm340 | 323.3061025 | -0.003984595 | 0.113811933 |
| Mir8091 | 15.23441784 | 0.382441329 | 0.297737538 |
| AI606181 | 120.0613128 | -0.076757709 | 0.126291546 |
| Slit1 | 868.1752054 | -0.773427744 | 0.099843615 |
| Gm22601 | 0.086648695 | -0.72445792 | 3.724945049 |
| Arhgap19 | 33.0013039 | -0.30153648 | 0.183003492 |
| Frat1 | 29.17957322 | 0.441799455 | 0.209195082 |
| Frat2 | 52.5338819 | -0.581316026 | 0.15687147 |
| Rrp12 | 552.962244 | 0.529408919 | 0.085103606 |
| Pgam1 | 1666.384237 | 0.045553526 | 0.061153152 |
| Exosc1 | 140.0755093 | 0.26758139 | 0.09701554 |

| | | | |
|---|---|---|---|
| Zdhhc16 | 217.4843809 | 0.290883605 | 0.086945859 |
| Mms19 | 424.2345625 | 0.169511758 | 0.116099738 |
| Gm23113 | 0 | NA | NA |
| Ubtd1 | 125.2316124 | 1.336297291 | 0.152562066 |
| Ankrd2 | 4.166437747 | 0.479134476 | 0.539995107 |
| Hoga1 | 9.277047816 | 0.80064916 | 0.402206681 |
| 4933411K16Rik | 1.522479514 | -2.607798283 | 1.003122569 |
| Morn4 | 316.6990762 | -0.430618713 | 0.101433566 |
| Pi4k2a | 1103.365836 | 0.933776363 | 0.106631687 |
| Avpi1 | 112.560822 | 0.028452474 | 0.128545701 |
| Marveld1 | 84.73861597 | 0.048654689 | 0.124394507 |
| Zfyve27 | 786.8275757 | -0.0874731 | 0.074384452 |
| Sfrp5 | 34.79157235 | -1.210601718 | 0.355867183 |
| Golga7b | 962.3374335 | -0.258817894 | 0.06974205 |
| Gm26339 | 1.445692662 | 1.272793992 | 0.978286633 |
| Mir3085 | 0.135858026 | -1.180985464 | 2.941468349 |
| Crtac1 | 458.2694328 | 0.172303014 | 0.102698146 |
| R3hcc1l | 325.8532248 | 0.082266877 | 0.092543374 |
| Gm16541 | 0.261817563 | 0.719232662 | 2.461197138 |
| Loxl4 | 7.914160501 | 2.708654113 | 0.471594988 |
| Gm25216 | 0 | NA | NA |
| Pyroxd2 | 33.56925608 | -1.024004855 | 0.245933509 |
| Hps1 | 90.94188656 | 0.240808074 | 0.149187355 |
| Gm16244 | 3.388720562 | -1.510397378 | 0.705497734 |
| Hpse2 | 1.819823472 | -0.189217283 | 0.83426107 |
| Gm22135 | 0 | NA | NA |
| Gm6776 | 3.379576263 | -0.319489089 | 0.494535208 |
| Cnnm1 | 1191.472548 | -0.56306349 | 0.081076618 |
| Got1 | 2463.067521 | -0.192628466 | 0.063092934 |
| Gm20467 | 0.07796279 | 1.072698098 | 3.721829099 |
| Nkx2-3 | 0.04393175 | -0.243571806 | 3.724945049 |
| Gm22646 | 0 | NA | NA |
| Slc25a28 | 398.2739226 | 0.89921025 | 0.13534349 |
| Entpd7 | 424.4745962 | 0.108059622 | 0.094941812 |
| Cox15 | 306.6258307 | 0.699489837 | 0.117171472 |
| Cutc | 59.7964683 | 0.905004104 | 0.189359811 |
| Abcc2 | 1.945998912 | 0.373893031 | 0.725142298 |
| Dnmbp | 248.0196757 | -0.03764998 | 0.108457541 |
| Gm24400 | 2.333191932 | -2.272652889 | 0.779649383 |
| n-R5s21 | 0 | NA | NA |
| Cpn1 | 2.984315308 | 0.411211965 | 0.538680468 |
| Cyp2c44 | 1.49985572 | 0.033637668 | 0.696827825 |
| Erlin1 | 312.6811813 | 0.215397093 | 0.07739245 |
| Chuk | 597.7287007 | 0.2090644 | 0.085382392 |
| Cwf19l1 | 148.8001304 | -0.024529618 | 0.100104894 |
| Gm24336 | 70.39117647 | 0.259454923 | 0.143394356 |
| Bloc1s2 | 164.8635678 | 0.480152556 | 0.119522247 |

| | | | |
|---|---|---|---|
| Pkd2l1 | 8.957326903 | 2.590395293 | 0.444545698 |
| Scd3 | 92.32162226 | -0.432550099 | 0.151207473 |
| Scd2 | 13180.9979 | -1.095694061 | 0.125890758 |
| Mir5114 | 0 | NA | NA |
| Scd4 | 17.24753904 | 0.313648138 | 0.26213914 |
| Scd1 | 2469.814438 | -0.549837502 | 0.142023786 |
| Wnt8b | 3.609855578 | -1.158028868 | 0.569827325 |
| Sec31b | 4.76996336 | 0.388241925 | 0.484607324 |
| Gm20538 | 0.061920613 | 0.718207094 | 3.724945049 |
| Ndufb8 | 791.3026735 | -0.068726218 | 0.119519004 |
| Hif1an | 511.8430662 | -0.047423381 | 0.07128012 |
| Gm26644 | 1.498020843 | -0.532989212 | 0.835516494 |
| Pax2 | 56.35874315 | 0.233510076 | 0.302550755 |
| Gm20395 | 0 | NA | NA |
| Mir6405 | 0.042716945 | -0.243571806 | 3.724945049 |
| 1700039E22Rik | 0.097899064 | -0.724453471 | 3.724945049 |
| Fam178a | 1155.011791 | -0.02449342 | 0.091431506 |
| Gm25482 | 3.723338729 | -0.002590379 | 0.669669924 |
| Sema4g | 448.4563521 | -0.785093194 | 0.107276869 |
| Mrpl43 | 278.1537674 | 0.341928961 | 0.095261559 |
| Peo1 | 143.5811533 | -0.145014156 | 0.110753736 |
| Lzts2 | 639.0048677 | 0.510380112 | 0.090554376 |
| Pdzd7 | 38.25076259 | -0.808275814 | 0.323304684 |
| Sfxn3 | 897.777185 | -0.174054648 | 0.043216205 |
| Kazald1 | 26.8175025 | 0.150508156 | 0.210851617 |
| Tlx1 | 0.675592865 | -0.147584616 | 1.171312748 |
| Tlx1os | 0 | NA | NA |
| Lbx1 | 26.55161603 | 0.206209445 | 0.329336264 |
| Gm29595 | 9.837264003 | 0.034472518 | 0.321246213 |
| Gm29593 | 4.239070863 | 0.476044116 | 0.504980972 |
| Gm29594 | 0.57177649 | -1.007306648 | 1.271085784 |
| Gm28578 | 19.28251988 | 0.217207005 | 0.364349111 |
| Btrc | 1150.587465 | -0.280338539 | 0.065207123 |
| Gm6807 | 12.07198287 | 0.628604759 | 0.3105901 |
| Poll | 98.22017834 | 0.115819549 | 0.136931301 |
| Dpcd | 221.260172 | 0.290526997 | 0.141859121 |
| Fbxw4 | 248.8152368 | 0.113481718 | 0.10073722 |
| Fgf8 | 0.275408967 | -0.71221843 | 2.070152497 |
| Npm3 | 55.30170471 | 0.878267263 | 0.192265899 |
| Gm15491 | 0.865681968 | 0.926247405 | 0.988790685 |
| Mgea5 | 3029.551855 | 0.265129392 | 0.106202872 |
| Kcnip2 | 682.9003887 | -1.591847712 | 0.160791548 |
| 9130011E15Rik | 279.4855959 | 0.07358061 | 0.077708047 |
| 4930505N22Rik | 0.892093866 | 0.346414124 | 0.899189412 |
| Hps6 | 95.93381243 | 0.570166611 | 0.154641233 |
| Ldb1 | 957.9423255 | -0.303200993 | 0.077368131 |
| Pprc1 | 454.9862607 | 0.507732193 | 0.088859623 |

| | | | |
|---|---|---|---|
| Nolc1 | 518.4413524 | 0.184448576 | 0.110537846 |
| Elovl3 | 0.899697409 | 3.333479613 | 1.177342054 |
| Pitx3 | 0.694683682 | 0.224915671 | 1.250046183 |
| Gbf1 | 1560.460353 | 0.070336749 | 0.050709201 |
| Gm8974 | 0.178004931 | -0.244115601 | 2.299166908 |
| Gm16726 | 0.21650593 | 2.065999099 | 2.130658308 |
| Nfkb2 | 234.2002576 | 3.229047539 | 0.241033501 |
| Psd | 2382.095847 | -1.315179937 | 0.120333903 |
| Fbxl15 | 156.8858204 | -0.048932016 | 0.113607862 |
| Cuedc2 | 723.9696748 | -0.05401684 | 0.082367627 |
| Gm26792 | 1.450243399 | -0.176486468 | 0.791717771 |
| Tmem180 | 197.5786801 | -0.386977732 | 0.10740919 |
| Mir146b | 0.495786544 | 1.012098611 | 1.315920594 |
| 2310034G01Rik | 6.1931368 | -0.826359564 | 0.487883395 |
| Gm24610 | 1.876595299 | -2.063476449 | 0.794911362 |
| Actr1a | 1762.40465 | 0.126185288 | 0.051004546 |
| Sufu | 283.8186178 | 0.100149653 | 0.093709234 |
| Trim8 | 1307.724394 | 0.098116995 | 0.071328681 |
| Arl3 | 347.6175142 | -0.331859906 | 0.118189998 |
| Sfxn2 | 104.3476033 | 0.339572002 | 0.153587162 |
| Wbp1l | 711.1586369 | 0.873692573 | 0.097617615 |
| Cyp17a1 | 0.280011045 | 0.718798921 | 1.796497879 |
| Gm6967 | 0.062426873 | 0.237312084 | 3.724945049 |
| 2010012O05Rik | 189.8383209 | -0.286437186 | 0.093159259 |
| As3mt | 170.4653533 | 0.438014841 | 0.133767454 |
| Cnnm2 | 423.4727026 | -0.386530566 | 0.069169631 |
| Nt5c2 | 724.1452932 | -0.135988512 | 0.078805876 |
| Ina | 2102.910469 | -0.175412995 | 0.113217106 |
| Pcgf6 | 132.4800137 | -0.507103013 | 0.149070735 |
| Gm23140 | 0.042716945 | -0.243571806 | 3.724945049 |
| Taf5 | 105.5145098 | 0.241510777 | 0.164787132 |
| Usmg5 | 854.0068898 | 0.106843361 | 0.140848408 |
| Pdcd11 | 426.8069572 | 0.414049547 | 0.087773887 |
| Calhm2 | 55.28273003 | 0.058290818 | 0.249728508 |
| Calhm1 | 0 | NA | NA |
| Calhm3 | 0.222639008 | 1.13028641 | 2.551709879 |
| Gm6970 | 2.783210482 | -0.727409002 | 0.672493916 |
| Neurl1a | 1174.052599 | -0.897067345 | 0.104017307 |
| Sh3pxd2a | 901.8376151 | -0.477475569 | 0.080041027 |
| Mir6995 | 1.143944469 | -1.649400483 | 1.090654341 |
| Obfc1 | 85.46752745 | 0.030403436 | 0.130295205 |
| Gm19557 | 1.777092953 | 1.459115264 | 0.835314676 |
| Slk | 1543.68933 | 0.036735185 | 0.074036669 |
| Gm16068 | 0.225284295 | -1.184793932 | 1.883450619 |
| Col17a1 | 13.5201549 | 0.057950275 | 0.312139235 |
| Gm26085 | 0 | NA | NA |
| Gm24805 | 0 | NA | NA |

| | | | |
|---|---|---|---|
| Sfr1 | 586.6289326 | 0.246873015 | 0.099459617 |
| Cfap43 | 93.17411037 | 1.84184223 | 0.188542741 |
| Mir8090 | 0.046345709 | 0.71820487 | 3.724945049 |
| Gsto1 | 187.9833352 | -0.003591447 | 0.110704379 |
| Gsto2 | 5.405473811 | -0.230775026 | 0.408201862 |
| Gm22240 | 0 | NA | NA |
| Cfap58 | 6.680071795 | 1.071628089 | 0.438066831 |
| Sorcs3 | 648.2072618 | 0.29646343 | 0.142097008 |
| Gm23857 | 0 | NA | NA |
| Gm27935 | 0 | NA | NA |
| Rpl13a-ps1 | 0.376112881 | 0.716738138 | 3.307048878 |
| Sorcs1 | 756.9127055 | -0.002975648 | 0.155045622 |
| Gm27749 | 0 | NA | NA |
| Gm26629 | 0.136194719 | -0.724453471 | 3.724945049 |
| Gm25659 | 0 | NA | NA |
| Gm25644 | 0 | NA | NA |
| Rpl31-ps18 | 0 | NA | NA |
| Ins1 | 0 | NA | NA |
| Xpnpep1 | 575.104864 | 0.547705605 | 0.084188564 |
| Mir6407 | 1.567823505 | -0.912987672 | 0.922176941 |
| 1700054A03Rik | 0.092691418 | 1.183079489 | 3.724560917 |
| Add3 | 1815.758105 | 0.40910007 | 0.094567228 |
| Gm24724 | 0 | NA | NA |
| Mxi1 | 685.6681876 | -0.544646993 | 0.070735349 |
| Gm10197 | 0.330485341 | 0.770905196 | 1.777504449 |
| Smndc1 | 390.3324799 | 0.57091425 | 0.122378773 |
| 5830416P10Rik | 35.48233613 | 0.264919578 | 0.242810347 |
| 4833407H14Rik | 1.343421229 | 1.930388336 | 1.087889371 |
| Dusp5 | 436.4752899 | 1.007003714 | 0.264395823 |
| Nutf2-ps1 | 25.95248433 | 0.620660915 | 0.263499262 |
| Smc3 | 928.9007826 | -0.136999325 | 0.087990742 |
| Rbm20 | 75.31453989 | -0.405157653 | 0.131797015 |
| Gm16298 | 0.11037557 | -0.243571806 | 3.724945049 |
| Gm16299 | 1.831283359 | -0.283647386 | 0.85051284 |
| Pdcd4 | 830.0440107 | -0.576552656 | 0.077286691 |
| Bbip1 | 292.1551775 | -0.031609152 | 0.103715046 |
| Shoc2 | 875.8097201 | 0.094560479 | 0.056334382 |
| Adra2a | 223.1903651 | -0.748302547 | 0.157828867 |
| Gpam | 844.7644142 | 0.15126421 | 0.087342762 |
| Tectb | 0.98697613 | 0.1858959 | 1.04507381 |
| Mir6715 | 0 | NA | NA |
| Gucy2g | 23.72729764 | -1.538869257 | 0.269683312 |
| Acsl5 | 698.4865369 | 0.946014355 | 0.069469166 |
| Zdhhc6 | 305.077262 | 0.382476893 | 0.083363536 |
| Vti1a | 524.7365872 | -0.124432511 | 0.063361839 |
| 4930552P12Rik | 3.0018596 | 2.949148398 | 0.766074473 |
| Tcf7l2 | 729.2267912 | 0.015780517 | 0.084158932 |

| | | | |
|---|---|---|---|
| Ppnr | 24.84417134 | -0.294901012 | 0.394375099 |
| Gm22271 | 0 | NA | NA |
| Habp2 | 0.044544648 | 0.237309859 | 3.724945049 |
| Nrap | 37.22294834 | 1.920762789 | 0.280640181 |
| Gm17197 | 0.54074031 | 0.086708702 | 1.243707989 |
| Casp7 | 137.0862569 | 3.091537761 | 0.179133913 |
| Plekhs1 | 3.487526041 | 0.827028524 | 0.564859954 |
| Dclre1a | 154.6814328 | 0.61198656 | 0.104664837 |
| Nhlrc2 | 312.5790796 | 0.479454843 | 0.129675251 |
| Adrb1 | 210.0560459 | -0.569626789 | 0.104106015 |
| Gm6990 | 0.88041345 | -1.095888449 | 1.02006141 |
| Ccdc186 | 1033.921743 | -0.089699692 | 0.060611844 |
| Tdrd1 | 1.956606073 | 0.097618782 | 0.762611195 |
| Vwa2 | 0.389464624 | 1.50334758 | 1.539268844 |
| Afap1l2 | 288.9123761 | 0.45247579 | 0.138247852 |
| Ablim1 | 1533.129369 | -0.29817894 | 0.064092352 |
| B230217O12Rik | 56.18862831 | -0.675978537 | 0.154785959 |
| Fam160b1 | 577.2204722 | -0.134867736 | 0.073715303 |
| Trub1 | 145.5468936 | -0.354985564 | 0.149029084 |
| Gm26874 | 0 | NA | NA |
| Atrnl1 | 1851.542762 | -0.479724466 | 0.085629865 |
| Mir5623 | 3.511196682 | 0.249590952 | 0.778738419 |
| Gm16277 | 0.097501745 | 0.23732098 | 3.724945049 |
| Gfra1 | 699.0851246 | 0.869821668 | 0.093259192 |
| Ccdc172 | 1.84083352 | 0.718764659 | 0.729480015 |
| 1810007D17Rik | 0.526835603 | 0.569643649 | 1.534435552 |
| Pnlip | 3.471747287 | 0.021735537 | 0.541201334 |
| 1810018F18Rik | 0 | NA | NA |
| Pnliprp1 | 0.093484797 | 0.237318756 | 3.724945049 |
| Pnliprp2 | 0.200944149 | -0.242815215 | 2.220206551 |
| 1700019N19Rik | 1.010445843 | 1.863343635 | 1.061440043 |
| Gm22561 | 0 | NA | NA |
| Gm17150 | 0 | NA | NA |
| Hspa12a | 2209.460681 | -0.76216331 | 0.080446629 |
| Mir3086 | 1.922482983 | -2.549610912 | 0.948349766 |
| Eno4 | 32.39598207 | 0.179781351 | 0.184871058 |
| Shtn1 | 427.0917043 | -0.523105535 | 0.106741611 |
| Vax1 | 9.031124418 | 0.417917855 | 0.403710725 |
| Kcnk18 | 0.044191549 | -0.243571806 | 3.724945049 |
| Gm29261 | 1.407425821 | -0.293379036 | 0.972585042 |
| Slc18a2 | 109.7658993 | -0.176134966 | 0.301397872 |
| Pdzd8 | 1685.906693 | -0.037459369 | 0.063462054 |
| Rps12-ps3 | 13.1319493 | 0.898646269 | 0.320242265 |
| Emx2os | 17.8875887 | -1.389264925 | 0.332752054 |
| Emx2 | 54.25553675 | -0.413138204 | 0.17214854 |
| 2700089I24Rik | 0.185374628 | -1.198364947 | 2.266154646 |
| Gm17203 | 36.89967017 | -0.502826751 | 0.2402119 |

| | | | |
|---|---|---|---|
| Rab11fip2 | 973.2053457 | -0.285486082 | 0.069618189 |
| Gm4219 | 3.097868212 | 0.597120285 | 0.554308856 |
| E330013P04Rik | 2.49875752 | -0.107368726 | 0.56920451 |
| Fam204a | 247.5678764 | 0.149344335 | 0.1058603 |
| 4933412A08Rik | 0.53212218 | 0.701265659 | 1.337499108 |
| Gm28351 | 0 | NA | NA |
| Prlhr | 4.925174813 | -0.347565038 | 0.434228073 |
| Cacul1 | 885.5461787 | 0.443103488 | 0.07886701 |
| Gm25238 | 3.512009492 | -0.358785429 | 0.593871323 |
| Nanos1 | 296.0425935 | -0.28350392 | 0.106159905 |
| Eif3a | 2000.929003 | 0.46970591 | 0.118218584 |
| Snora19 | 2.074201168 | 0.203796875 | 0.690650883 |
| Gm26158 | 3.094791288 | 1.565441491 | 0.648432889 |
| Fam45a | 537.0189958 | 0.26860272 | 0.106757966 |
| Gm15959 | 0 | NA | NA |
| Sfxn4 | 143.5137547 | -0.529381628 | 0.144603929 |
| Prdx3 | 403.3254737 | 0.232477549 | 0.123461816 |
| Grk5 | 237.0149773 | -0.249761714 | 0.091124437 |
| Gm9529 | 0.17796667 | -0.21293993 | 2.712315561 |
| Gm22520 | 0 | NA | NA |
| 5830428H23Rik | 346.0755543 | -0.498704858 | 0.223101078 |
| Gm22365 | 0 | NA | NA |
| Gm7102 | 0.828685883 | -0.527181425 | 1.101283323 |
| Gm6020 | 0 | NA | NA |
| Csf2ra | 189.2145181 | 1.312858685 | 0.151363326 |
| Gm21060 | 0 | NA | NA |
| Gm37392 | 0 | NA | NA |
| Gm27306 | 0 | NA | NA |
| Fam171a1 | 789.1479404 | -0.122876092 | 0.076706318 |
| Nmt2 | 1499.866317 | -0.041700553 | 0.070663479 |
| Gm22005 | 0.075043205 | 0.237309859 | 3.724945049 |
| Rpp38 | 43.57699851 | 0.324715932 | 0.204175714 |
| Acbd7 | 3.006698544 | 0.15239158 | 0.736428865 |
| Olah | 0.742270129 | 1.558873516 | 1.225688406 |
| Gm37525 | 0.038981395 | 0.718207094 | 3.724945049 |
| Meig1 | 9.115918498 | 0.37716572 | 0.320541394 |
| Dclre1c | 259.4107841 | 0.635296916 | 0.152743039 |
| Suv39h2 | 54.79723866 | -0.384216585 | 0.158362115 |
| Gm13184 | 2.921456852 | 1.304841698 | 0.765175748 |
| Hspa14 | 484.0326706 | 0.127613233 | 0.11422046 |
| Cdnf | 15.49217047 | 0.069546298 | 0.266615171 |
| Gm13186 | 0.075043205 | 0.237309859 | 3.724945049 |
| Gm13182 | 0.275763028 | 0.661977244 | 1.956963858 |
| Fam107b | 448.5992859 | 1.032646331 | 0.175647121 |
| Gm13183 | 0 | NA | NA |
| Gm13181 | 0.648853987 | 0.338444683 | 1.249083293 |
| Gm13185 | 0.04393175 | -0.243571806 | 3.724945049 |

| | | | |
|---|---|---|---|
| Gm13180 | 0.088736197 | -0.243576254 | 3.724945049 |
| Gm13191 | 0.045882455 | 0.237312084 | 3.724945049 |
| Frmd4a | 1510.085395 | 0.53810253 | 0.062559106 |
| Gm2639 | 0.385855529 | -0.951618201 | 3.262688888 |
| Gm13188 | 0 | NA | NA |
| Gm10862 | 12.42465393 | -0.938316359 | 0.363184766 |
| Gm38085 | 10.53103825 | -0.792987514 | 0.469776944 |
| 1700080N15Rik | 0 | NA | NA |
| Gm13189 | 2.288528955 | -2.010630939 | 0.92017421 |
| Gm37814 | 13.6296294 | -1.140041425 | 0.474247501 |
| Gm37107 | 25.13242011 | -1.623925798 | 0.372461367 |
| Gm38348 | 18.53952862 | -1.557472234 | 0.434088021 |
| Gm13187 | 1.991338835 | 0.857502929 | 0.660476358 |
| Gm36988 | 22.65632828 | -1.217910681 | 0.336224596 |
| Gm23691 | 0 | NA | NA |
| Gm13175 | 0 | NA | NA |
| Gm13179 | 0.804355203 | 2.34115509 | 1.16939368 |
| Prpf18 | 625.4248675 | 0.157616076 | 0.064804448 |
| Gm13196 | 0.049553047 | 0.71820487 | 3.724945049 |
| Bend7 | 58.21736025 | 0.06548155 | 0.140224878 |
| Gm13176 | 0 | NA | NA |
| Sephs1 | 350.7542985 | -0.065556078 | 0.080488809 |
| Phyh | 939.6459375 | 0.603583779 | 0.083866141 |
| Gm13194 | 2.056906148 | 0.496272803 | 0.938417943 |
| Gm13193 | 0 | NA | NA |
| Gm13192 | 14.54002108 | -0.046149432 | 0.269028684 |
| Ucma | 8.601202928 | 0.858039116 | 0.383842766 |
| Mcm10 | 26.1128149 | 1.899982283 | 0.402384893 |
| Optn | 314.9999699 | 0.232405745 | 0.091194717 |
| 4930551O13Rik | 0.150455055 | 1.610657435 | 2.838957197 |
| Gm26776 | 1.257166043 | 0.50849845 | 1.15252479 |
| Ccdc3 | 115.3722469 | 0.624697318 | 0.112176605 |
| Gm13198 | 1.380091768 | -0.583428807 | 0.852823215 |
| Gm23118 | 0 | NA | NA |
| Gm13197 | 0.068097359 | -0.243569582 | 3.724945049 |
| Camk1d | 1938.800003 | -0.200677983 | 0.091806823 |
| Gm13216 | 0.097501745 | 0.23732098 | 3.724945049 |
| Cdc123 | 504.6193172 | 0.036282926 | 0.070588541 |
| Nudt5 | 187.3923638 | -0.106961797 | 0.127692434 |
| Gm13199 | 8.870008164 | 0.020073613 | 0.375681322 |
| Sec61a2 | 789.7387573 | -0.313406146 | 0.112817458 |
| Gm13267 | 0.3753498 | 1.678905232 | 1.607187487 |
| Dhtkd1 | 85.77264844 | -0.537617069 | 0.153088386 |
| Upf2 | 873.5434251 | 0.257564384 | 0.079462494 |
| Proser2 | 48.48109559 | -0.320625216 | 0.162392994 |
| Gm13384 | 0 | NA | NA |
| Gm10857 | 0.297167372 | 1.528008401 | 1.729450466 |

| | | | |
|---|---|---|---|
| Echdc3 | 8.401856911 | 1.044629254 | 0.371817337 |
| A230108P19Rik | 0.266247585 | -1.154830068 | 2.501021303 |
| Gm13383 | 0.729279448 | 1.419629392 | 1.219331304 |
| Usp6nl | 655.1216898 | 0.037779978 | 0.071324828 |
| Gm13388 | 0.223035371 | 0.718216941 | 2.568721411 |
| Gm13391 | 5.144468292 | 1.255625852 | 0.507441223 |
| Gm38386 | 2.604362067 | 1.630803804 | 0.670109868 |
| Celf2 | 6175.024994 | -0.64435261 | 0.120537612 |
| Gm13389 | 0.654952557 | 0.45610898 | 1.396302688 |
| Gm10855 | 169.6995702 | -1.546979911 | 0.498646769 |
| Gm37340 | 0.066050875 | 0.71820487 | 3.724945049 |
| Gm10115 | 101.9444587 | -1.790712967 | 0.471978407 |
| Gm26315 | 191.8007175 | -0.175371798 | 0.251872606 |
| Gm26565 | 0.531771505 | -1.456867968 | 1.380909145 |
| Gm24340 | 0 | NA | NA |
| Gm28641 | 0.190780345 | -1.461403718 | 3.283197437 |
| Gm13210 | 0.681349771 | 0.819814079 | 1.423769239 |
| Gm13211 | 0.138642343 | 0.718199153 | 3.607279928 |
| Gm13254 | 0.062426873 | 0.237312084 | 3.724945049 |
| Gm24534 | 0 | NA | NA |
| Gm13255 | 0 | NA | NA |
| Gm13217 | 27.36624143 | -0.287248296 | 0.192948722 |
| 1700061F12Rik | 0 | NA | NA |
| Gm23963 | 0 | NA | NA |
| Gm13221 | 0 | NA | NA |
| Gm13219 | 0 | NA | NA |
| Gm13220 | 0 | NA | NA |
| Gm13218 | 0 | NA | NA |
| Gm37866 | 0 | NA | NA |
| Gata3 | 51.8974266 | 0.536069566 | 0.16176053 |
| Gm13256 | 0.066050875 | 0.71820487 | 3.724945049 |
| 4930412O13Rik | 0.725537781 | 1.398987958 | 1.567777763 |
| 9230102O04Rik | 0.685158741 | -1.819545316 | 1.126317639 |
| Gm13262 | 0 | NA | NA |
| Taf3 | 355.5580828 | 0.004571745 | 0.064946495 |
| C630004M23Rik | 3.262821873 | -0.021404067 | 0.679874017 |
| Atp5c1 | 1526.152544 | -0.053506934 | 0.079331073 |
| Gm23608 | 6.323166952 | -1.307335704 | 0.427206419 |
| Kin | 91.12058367 | 0.124207205 | 0.12823999 |
| Itih2 | 90.97624014 | 1.846429813 | 0.231053553 |
| Itih5 | 570.7931782 | -0.081239695 | 0.154726001 |
| Gm13261 | 0.038981395 | 0.718207094 | 3.724945049 |
| Sfmbt2 | 174.706509 | 0.918617845 | 0.125208453 |
| Gm22072 | 0.909783392 | -0.390533786 | 1.14767941 |
| Gm22677 | 0.580675992 | 2.697116908 | 1.661844763 |
| Mir466m | 0.940874839 | -1.238141117 | 1.082158404 |
| Mir466f-1 | 0.953625643 | 0.098850111 | 1.159299312 |

| | | | |
|---|---|---|---|
| Mir669f | 1.177175292 | 1.517638567 | 0.903147123 |
| Mir669e | 0.868322942 | 0.922796796 | 1.138220675 |
| Mir669b | 1.131281954 | 0.734081871 | 1.087286973 |
| Gm23578 | 1.362203202 | 2.191817977 | 1.045391508 |
| Mir669d | 0.802169344 | -0.076379131 | 1.128199079 |
| Mir466f-2 | 0.667839854 | 1.614583005 | 1.350815158 |
| Mir669l | 0.610120006 | -2.107734251 | 1.187398815 |
| Mir669d-2 | 0.709755772 | -0.171495558 | 1.19569726 |
| Mir466f-3 | 0.856346278 | -0.09377191 | 1.223138556 |
| Mir297a-2 | 0.78950392 | 0.702244461 | 1.097190458 |
| Mir466o | 1.891729189 | 0.33924734 | 0.746997321 |
| Gm26073 | 1.013961535 | 0.506415137 | 1.011211654 |
| Mir467c | 0.194094924 | 1.169565035 | 2.224736307 |
| mmu-mir-466b-1 | 0.068097359 | -0.243569582 | 3.724945049 |
| Mir669a-3 | 0.890972926 | 0.269157942 | 1.415585259 |
| Mir669k | 1.409213138 | -0.982322017 | 0.852149215 |
| Gm26491 | 1.682866984 | 0.255673101 | 0.91508958 |
| Mir467a-1 | 0.921542032 | 0.298464536 | 1.116771209 |
| Mir466b-8 | 0.330565531 | 1.090455616 | 1.715153751 |
| Mir669a-1 | 1.495866474 | 0.537267585 | 0.858389073 |
| Mir669g | 0.495109938 | 0.274100643 | 1.368417564 |
| Gm25701 | 0.561285994 | 0.898369589 | 1.220552141 |
| Mir669j | 1.067120961 | 1.892221872 | 1.284104461 |
| Mir467a-2 | 0.612405649 | 0.906650357 | 1.269094838 |
| mmu-mir-466e | 0.75911489 | 2.600027829 | 1.384360652 |
| mmu-mir-669a-4 | 2.460216864 | 0.071375737 | 0.645786831 |
| Gm24315 | 0.616248996 | 0.260755953 | 1.24952781 |
| Gm25451 | 0.626970746 | -2.121947801 | 1.34223351 |
| Mir467b | 1.221212496 | 0.00050716 | 0.851954189 |
| mmu-mir-466c-1 | 1.101702777 | 0.828473396 | 1.008592867 |
| Gm24761 | 0.825397108 | 2.216590418 | 1.08971406 |
| Gm23651 | 1.10380611 | -0.749202354 | 1.049580462 |
| Mir467a-3 | 0 | NA | NA |
| Mir466c-2 | 0 | NA | NA |
| Gm23108 | 0.643541246 | 1.693467297 | 1.58008862 |
| Gm25680 | 0 | NA | NA |
| Gm22775 | 0.32589515 | 1.211913735 | 1.768951046 |
| Mir467a-4 | 0.173861546 | 1.847947637 | 2.700164252 |
| Mir466b-4 | 0 | NA | NA |
| Gm26060 | 0 | NA | NA |
| Gm26431 | 0.062426873 | 0.237312084 | 3.724945049 |
| Gm26229 | 0.045882455 | 0.237312084 | 3.724945049 |
| Mir467a-5 | 0 | NA | NA |
| Mir466b-5 | 0 | NA | NA |
| Mir669p-1 | 0 | NA | NA |
| Gm24167 | 0.038981395 | 0.718207094 | 3.724945049 |
| Gm24640 | 0 | NA | NA |

| | | | |
|---|---|---|---|
| Gm23288 | 0.060174537 | 0.237309859 | 3.724945049 |
| Mir467a-6 | 0 | NA | NA |
| Gm25285 | 0.094831987 | -0.243571806 | 3.724945049 |
| Gm24673 | 0.458955183 | 0.227613535 | 1.493121012 |
| Gm22635 | 0.309435895 | -1.667716719 | 3.702082823 |
| Gm23569 | 0.231961655 | 0.712553981 | 2.127211619 |
| Mir467a-7 | 0 | NA | NA |
| Mir466b-6 | 0 | NA | NA |
| Gm26426 | 0 | NA | NA |
| Gm24488 | 0.108266322 | 1.230376831 | 3.722845817 |
| Gm25158 | 0.038981395 | 0.718207094 | 3.724945049 |
| Mir467a-8 | 0 | NA | NA |
| Mir466b-7 | 0.158320866 | 0.733224281 | 2.813533472 |
| Mir669p-2 | 0 | NA | NA |
| Gm24028 | 0.085433889 | -0.72445792 | 3.724945049 |
| Gm22626 | 0.305016863 | 0.366463585 | 2.023236213 |
| Gm22166 | 0.060174537 | 0.237309859 | 3.724945049 |
| Mir467a-9 | 0.048949532 | -0.243569582 | 3.724945049 |
| Mir466b-2 | 0.150451996 | 0.237323705 | 2.858243839 |
| Gm26156 | 1.082398122 | -0.686274633 | 0.855220725 |
| Gm23571 | 0.95850512 | -0.467392892 | 1.057631267 |
| Gm25316 | 0.639171633 | -1.100184964 | 1.155122739 |
| Gm25022 | 0.159191956 | -0.24356343 | 2.808135645 |
| Gm25943 | 0.342780899 | 0.643573279 | 1.582772374 |
| Gm23988 | 0.869956765 | 1.630811619 | 1.124815975 |
| Gm25673 | 0.68128338 | 2.23421096 | 1.295130373 |
| Gm25753 | 0.533146218 | 0.197154595 | 1.29666663 |
| Mir467a-10 | 0 | NA | NA |
| Mir466b-3 | 0.655224956 | -1.058399845 | 1.30348322 |
| Gm26398 | 0.832395869 | 0.946783923 | 1.078910803 |
| Mir467e | 0.572335319 | 0.64522372 | 1.341263836 |
| Mir466p | 1.677881506 | 0.109063902 | 0.76040381 |
| Gm22051 | 0.979291814 | 1.252812143 | 1.200876078 |
| Gm22678 | 0.741352231 | -0.835312993 | 1.159623176 |
| Mir467d | 0.189382193 | -0.710859249 | 2.701715785 |
| Mir466 | 0.816872591 | 0.041547318 | 1.055806294 |
| Gm22091 | 0.829130969 | -1.527186806 | 1.076258459 |
| Mir297c | 0.615674597 | 0.439101124 | 1.262873087 |
| Mir669c | 0.542133415 | -0.243565895 | 1.749908809 |
| Gm24342 | 0.492872509 | -1.161942741 | 1.272061937 |
| Gm25263 | 0.34438493 | 0.217315484 | 1.493973311 |
| Mir669a-2 | 0.768366922 | 0.564055109 | 1.148865539 |
| Gm22239 | 0.500671887 | 1.223348754 | 1.572335322 |
| Gm22925 | 0.21460851 | -0.230645797 | 2.598321842 |
| Gm23542 | 0.931192987 | 0.548591993 | 1.029068933 |
| Mir297b | 0.635701853 | -0.664255071 | 1.154445175 |
| Mir466d | 0.422902843 | 0.277774564 | 1.581086594 |

| | | | |
|---|---|---|---|
| Gm26092 | 0.669827454 | -0.674158609 | 1.450687857 |
| Mir669m-1 | 0.636524932 | 1.283183837 | 1.150669001 |
| Mir669m-2 | 0.70834526 | -0.048799275 | 1.197403657 |
| Mir466n | 0.60136891 | 0.762825026 | 1.186910834 |
| Mir669o | 1.139979212 | 0.901958647 | 1.017564104 |
| Mir466g | 0.512204714 | -0.474581663 | 1.233797304 |
| Mir466h | 0.440637317 | 0.237318447 | 3.40099462 |
| Mir297a-3 | 0.589191525 | -0.738950113 | 1.402508061 |
| Mir466l | 0.747968401 | 0.995791346 | 1.320803852 |
| Mir297a-4 | 0.343551435 | 1.703662542 | 1.736201265 |
| Gm24806 | 0.654401101 | 0.63336064 | 1.235578498 |
| Mir669i | 0.597540007 | -0.379833064 | 1.157193649 |
| Mir669h | 0.264963494 | -0.621491402 | 2.075074973 |
| Gm18547 | 0 | NA | NA |
| Gm13265 | 0.236919184 | 0.237320365 | 2.511382982 |
| Gm13263 | 0.565796537 | 0.237330663 | 1.551323019 |
| Gm37343 | 0 | NA | NA |
| Gm23877 | 0 | NA | NA |
| Gm13264 | 0.423046101 | 1.151195706 | 1.660088068 |
| Gm26478 | 0 | NA | NA |
| Gm13297 | 0 | NA | NA |
| Gm13294 | 1.544067245 | -0.074365293 | 0.876099554 |
| Prkcq | 193.2675698 | 0.685381237 | 0.122225248 |
| Gm36932 | 3.868039749 | 0.811306794 | 0.528760607 |
| Gm37520 | 0.177524535 | 1.789751687 | 2.721783096 |
| Gm37766 | 3.664845353 | 1.265705848 | 0.708515787 |
| Gm38171 | 1.860369092 | 4.11568617 | 1.043195842 |
| Gm37851 | 0 | NA | NA |
| 8030442B05Rik | 2.076510791 | 0.792755815 | 0.739014856 |
| Gm13293 | 46.58195302 | 1.127922796 | 0.184214695 |
| Gm37730 | 0.669761723 | -2.243022839 | 1.260721329 |
| 4933403L11Rik | 0.631690809 | -1.81212292 | 1.130650223 |
| Gm13291 | 0.045882455 | 0.237312084 | 3.724945049 |
| Gm13292 | 11.7837924 | -0.015505545 | 0.299974763 |
| Gm13296 | 0 | NA | NA |
| Pfkfb3 | 910.8026235 | 1.053431896 | 0.130404784 |
| Gm10851 | 1.168248907 | 4.136756116 | 1.072410412 |
| Gm37975 | 7.09582318 | -1.854774936 | 0.517433723 |
| Rbm17 | 378.7991564 | 0.420854396 | 0.130306633 |
| Gm13260 | 0.138087489 | 1.595384126 | 3.611428798 |
| Gm17490 | 0 | NA | NA |
| Il2ra | 71.84653869 | 5.724481738 | 0.445400906 |
| Gm37881 | 0.909342915 | 3.705133369 | 2.527594565 |
| Il15ra | 192.4286151 | 2.224389092 | 0.219988612 |
| Fbxo18 | 817.0147377 | -0.258970131 | 0.135690196 |
| Ankrd16 | 279.6460949 | -0.490358685 | 0.176814552 |
| Gm34768 | 0.194631057 | 0.237309881 | 3.243494927 |

| | | | |
|---|---|---|---|
| Gm13310 | 0.075043205 | 0.237309859 | 3.724945049 |
| Itga8 | 97.06132719 | -1.63767965 | 0.22474409 |
| Gm37701 | 1.141257353 | -3.013786479 | 1.02539227 |
| E030013I19Rik | 6.292898879 | -1.005384947 | 0.509188687 |
| Fam188a | 576.4656474 | 0.403516768 | 0.0886823 |
| Gm37289 | 14.48869372 | 0.253365332 | 0.356166596 |
| Gm37255 | 40.61645279 | 0.129854149 | 0.314427597 |
| Gm38014 | 40.65078467 | -0.323126672 | 0.327902072 |
| Gm13321 | 0 | NA | NA |
| Gm37565 | 0.527338585 | 3.28370678 | 3.145285826 |
| Pter | 24.4696158 | 0.132256416 | 0.217230278 |
| C1ql3 | 535.3698455 | -0.92366883 | 0.082514127 |
| Gm37824 | 3.481665934 | -0.620352759 | 0.576958048 |
| Gm37811 | 9.289570475 | -1.100931325 | 0.352343224 |
| Gm37356 | 2.692269002 | -1.579093469 | 0.713066292 |
| Gm37742 | 0.85289683 | -0.88937349 | 1.264269445 |
| Rsu1 | 237.1157615 | 0.161459622 | 0.115775035 |
| Gm38156 | 11.10230631 | -0.439949393 | 0.397655179 |
| Gm37780 | 17.42481316 | -0.405075297 | 0.271567141 |
| Gm27540 | 0 | NA | NA |
| Gm37160 | 10.15377686 | -0.703850477 | 0.310810488 |
| Gm13270 | 8.36952636 | 0.160261618 | 0.377658833 |
| Cubn | 24.44027687 | 0.083026097 | 0.248095129 |
| Trdmt1 | 114.6086781 | 0.698729809 | 0.126456271 |
| Vim | 4040.904579 | 4.330067984 | 0.283115514 |
| St8sia6 | 68.51491012 | -0.156604139 | 0.160834322 |
| Gm37126 | 0 | NA | NA |
| Hacd1 | 81.82130967 | -0.262579003 | 0.137555105 |
| D930036K23Rik | 55.51995555 | -0.563433031 | 0.209932909 |
| Gm37894 | 18.6674481 | -1.173347275 | 0.309845274 |
| Stamos | 3.400319483 | 0.414264058 | 0.528961502 |
| Stam | 1143.237138 | 0.153673317 | 0.058543159 |
| Gm37499 | 111.4820659 | -0.367975618 | 0.129316311 |
| Tmem236 | 7.691974964 | 2.228365195 | 0.448962237 |
| Gm13320 | 0.278513079 | 0.237328245 | 2.101938298 |
| Mrc1 | 62.46851479 | -0.835492108 | 0.278843593 |
| Mir511 | 0.109727584 | 0.718200421 | 3.724945049 |
| Gm37697 | 0.16123844 | -1.214248636 | 2.795194053 |
| Slc39a12 | 432.812937 | -1.274549496 | 0.175970904 |
| Gm13266 | 1.979745462 | -0.347680011 | 0.781222664 |
| Cacnb2 | 1007.083002 | -0.322565285 | 0.151576876 |
| Gm10849 | 2.270217088 | -0.816589627 | 0.791889154 |
| Gm38105 | 15.2328292 | -0.791294241 | 0.322502383 |
| Gm13268 | 1.66036403 | 1.849101219 | 0.930547792 |
| Gm13312 | 24.04166377 | -0.625324266 | 0.448760626 |
| 2900092N22Rik | 49.08839729 | -0.732744403 | 0.321303115 |
| Gm13314 | 0.346172528 | -0.835277977 | 1.688081421 |

| | | | |
|---|---|---|---|
| Gm13311 | 0.601079571 | -0.257372272 | 1.409275538 |
| Gm10848 | 3.555860714 | 0.323070657 | 0.661974196 |
| Gm37006 | 66.77607684 | -0.674611853 | 0.351831721 |
| Gm13315 | 24.30272673 | -0.869523182 | 0.43904993 |
| Gm38273 | 9.78169311 | 0.21964455 | 0.376863721 |
| Gm13269 | 0.198547815 | 1.143005721 | 2.601213951 |
| Gm13316 | 0.175141309 | 0.701564674 | 2.304192594 |
| Gm37437 | 53.34605896 | -0.818777831 | 0.380133513 |
| Gm37891 | 58.0044897 | -0.255806076 | 0.264938024 |
| Nsun6 | 110.3532385 | -0.095095427 | 0.1558779 |
| Gm36930 | 6.501557468 | -0.670903869 | 0.415932665 |
| Arl5b | 609.464288 | -0.098177943 | 0.101616649 |
| Gm38257 | 38.93688588 | -0.705315982 | 0.294759251 |
| Gm13313 | 0.521845371 | 0.619021405 | 1.318939028 |
| Gm13367 | 0 | NA | NA |
| Gm13366 | 0 | NA | NA |
| Gm13365 | 0.11997416 | -0.243571806 | 3.724945049 |
| Gm13368 | 0 | NA | NA |
| Malrd1 | 1.059665568 | 2.429848298 | 1.152384599 |
| Gm37595 | 0 | NA | NA |
| Gm6587 | 0 | NA | NA |
| Gm37416 | 2.356264561 | 0.064374637 | 0.689971738 |
| Plxdc2 | 861.0341044 | -0.486609378 | 0.074384391 |
| Gm24933 | 0.072025462 | 0.237312084 | 3.724945049 |
| Gm13326 | 1.471932881 | -1.698475877 | 0.998012672 |
| Gm13325 | 0 | NA | NA |
| Gm13322 | 3.121057985 | -2.209670066 | 0.687377889 |
| Nebl | 1622.729757 | 0.004020823 | 0.06341617 |
| Gm13324 | 0.045882455 | 0.237312084 | 3.724945049 |
| Mir6419 | 0 | NA | NA |
| Gm13323 | 0 | NA | NA |
| H2afb1 | 0.165156969 | 1.598327176 | 3.401540795 |
| Gm13329 | 0.121578509 | 0.718202646 | 3.724945049 |
| A930004D18Rik | 38.49094849 | 0.564659424 | 0.177021552 |
| Gm13330 | 0.172363982 | -0.700047083 | 2.780806472 |
| Skida1 | 111.6125948 | -0.174534789 | 0.11918313 |
| 1810059C17Rik | 0.960935127 | 0.901319388 | 0.932714118 |
| Mllt10 | 761.3719482 | -0.279237599 | 0.095032176 |
| Mir7655 | 0.321829426 | -2.149999884 | 1.66780851 |
| Gm23638 | 0 | NA | NA |
| Gm25147 | 6.022279029 | -0.845256475 | 0.471129803 |
| Gm13351 | 0.9210764 | 2.437579335 | 2.150329205 |
| Dnajc1 | 546.8891029 | 0.048904288 | 0.068723578 |
| Gm13350 | 4.932017945 | -2.389070355 | 0.583446052 |
| Gm13349 | 8.865425504 | -0.803147153 | 0.39703129 |
| Gm13353 | 0 | NA | NA |
| Gm13355 | 0.097139547 | 0.718207094 | 3.724945049 |

| | | | |
|---|---|---|---|
| Gm13352 | 0.195196415 | 1.110259803 | 2.208870759 |
| Commd3 | 408.9733762 | 0.245193448 | 0.105723313 |
| Bmi1 | 684.4399484 | 0.374773053 | 0.095852816 |
| Spag6l | 18.02665997 | -1.124529945 | 0.300952944 |
| Gm20539 | 2.239606267 | 0.130377093 | 0.636657417 |
| Gm13333 | 0.068097359 | -0.243569582 | 3.724945049 |
| Carlr | 12.90806688 | 0.692290326 | 0.295057639 |
| Pip4k2a | 1173.833449 | 0.114142882 | 0.111969158 |
| 4930426L09Rik | 9.715138516 | 0.232507179 | 0.317496635 |
| Gm13332 | 0 | NA | NA |
| Armc3 | 8.367447745 | -0.502726864 | 0.358023758 |
| 4930447M23Rik | 25.2336097 | -0.998285835 | 0.233263137 |
| Msrb2 | 108.7981829 | 0.193777514 | 0.127929625 |
| Gm13344 | 0 | NA | NA |
| Ptf1aos | 0 | NA | NA |
| Ptf1a | 0.167536551 | -0.243570703 | 2.763312762 |
| 1810010K12Rik | 2.426680169 | -1.937882707 | 0.704113142 |
| 4921504E06Rik | 0.234904022 | 1.950034461 | 2.102426917 |
| Gm13345 | 0 | NA | NA |
| Gm13348 | 0.146089079 | 0.238291249 | 3.522464378 |
| Gm13343 | 0.231405101 | -0.24356407 | 2.545874405 |
| Otud1 | 292.6909858 | 1.033447533 | 0.16306254 |
| Gm13346 | 0.525489152 | 2.775019797 | 1.561144519 |
| Gm13347 | 0.240250011 | 1.207935541 | 2.114649114 |
| Gm24670 | 0 | NA | NA |
| BX571892.1 | 0.123992736 | -0.243574001 | 3.720643035 |
| Etl4 | 1406.655128 | -0.822997948 | 0.09614808 |
| Gm25859 | 5.580134033 | -1.583713458 | 0.527845627 |
| Gm13361 | 0.132101749 | 1.209894437 | 3.72418822 |
| Gm13363 | 9.874902262 | -0.867071529 | 0.383552246 |
| Gm13360 | 0.049553047 | 0.71820487 | 3.724945049 |
| Gm13328 | 2.887214733 | -0.077657945 | 0.615080134 |
| Gm23970 | 0.864953071 | -1.451072459 | 1.090778385 |
| Gm17171 | 1.518325143 | 0.831535038 | 0.898733503 |
| Gm13335 | 0.728721824 | -1.696881315 | 1.305294657 |
| Gm27446 | 0.851828701 | -1.607682568 | 1.084516213 |
| Gm13362 | 0.31548088 | 0.169799016 | 1.709105884 |
| Arhgap21 | 2959.631058 | -0.450624477 | 0.083516824 |
| Gm13380 | 18.50560142 | -0.996216021 | 0.380061446 |
| Gm13375 | 61.63895035 | -0.868031418 | 0.165925353 |
| Gm13377 | 0.2958732 | -1.434463539 | 1.724710447 |
| Gm13376 | 0.075043205 | 0.237309859 | 3.724945049 |
| Gm25368 | 0 | NA | NA |
| Enkur | 19.33843805 | 0.348009261 | 0.259358336 |
| Gm23007 | 0.130861843 | -0.243562293 | 2.984314958 |
| Thnsl1 | 325.2700769 | -0.309471355 | 0.115438866 |
| Gm13378 | 1.494280535 | 0.073389315 | 0.816532335 |

| | | | |
|---|---|---|---|
| Gm13327 | 3.973417911 | 0.015529596 | 0.514227718 |
| Gpr158 | 2965.115641 | -0.418870386 | 0.096903073 |
| Gm13379 | 0 | NA | NA |
| Gm24886 | 0 | NA | NA |
| Gm13337 | 5.556964523 | 0.222999095 | 0.482028122 |
| Myo3a | 0.925384949 | -0.475301865 | 0.947210437 |
| Gm13342 | 2.124844546 | 1.231012197 | 0.71918704 |
| Gm13341 | 22.54937325 | -0.323777157 | 0.337535735 |
| Gm13340 | 244.019484 | -0.176199047 | 0.143065962 |
| Gm13339 | 4.846303826 | -0.205358851 | 0.877751935 |
| Gad2 | 3413.524295 | 0.098167817 | 0.088169452 |
| mmu-mir-3967 | 2.839739068 | 0.062921187 | 0.639727462 |
| Gm23636 | 1.903429752 | 0.502687455 | 0.751089576 |
| Gm13338 | 0.254763098 | -0.243651485 | 1.831079368 |
| 1700092C17Rik | 0.045882455 | 0.237312084 | 3.724945049 |
| Gm13356 | 0.169165009 | 0.719161184 | 3.351346564 |
| Apbb1ip | 159.2402503 | 2.076464588 | 0.124647356 |
| A130006I12Rik | 3.281667885 | -0.608723806 | 0.533128053 |
| Pdss1 | 223.7617087 | -0.086750554 | 0.131341513 |
| Gm25544 | 3.664064784 | -0.652795081 | 0.649966398 |
| Gm14441 | 0 | NA | NA |
| Abi1 | 1539.731744 | 0.143374197 | 0.049784095 |
| Gm13357 | 12.25840608 | -0.456879206 | 0.391490099 |
| Acbd5 | 986.3592934 | -0.287083143 | 0.075401059 |
| Mastl | 28.61317683 | 0.960806133 | 0.308149354 |
| Yme1l1 | 1605.641626 | 0.22683421 | 0.064666214 |
| Gm13423 | 14.64943441 | -0.315217171 | 0.292178356 |
| 4931423N10Rik | 14.857103 | 0.469975745 | 0.320809574 |
| Gm13422 | 10.18410332 | 0.259577946 | 0.327238577 |
| Nxph2 | 18.63269461 | -0.621226634 | 0.243647391 |
| Gm38309 | 0.149576918 | 1.19849824 | 2.863264735 |
| Spopl | 699.9338885 | -0.162043329 | 0.093470245 |
| Gm13331 | 22.66616647 | 0.050696478 | 0.325524137 |
| Sh3d2c-ps1 | 0.219384069 | -0.703807056 | 1.896297501 |
| Gm26323 | 0 | NA | NA |
| Gm13392 | 0.616639959 | -0.749710916 | 1.273009635 |
| Hnmt | 147.4068612 | -0.599923531 | 0.124832997 |
| Tbpl2 | 0 | NA | NA |
| Il1f8 | 0 | NA | NA |
| Gm13410 | 44.62557519 | 0.228899987 | 0.162770049 |
| Il1f9 | 2.482099328 | 3.984512242 | 1.006134319 |
| Il1f6 | 0 | NA | NA |
| Gm37703 | 0 | NA | NA |
| Gm13409 | 0.143485256 | 1.229961212 | 3.547405499 |
| Il1f5 | 0.291332888 | 0.783778086 | 3.720845901 |
| Il1f10 | 0 | NA | NA |
| Gm13411 | 0.061920613 | 0.718207094 | 3.724945049 |

| | | | |
|---|---|---|---|
| Il1rn | 704.4699326 | 12.66831101 | 1.063225429 |
| Psd4 | 122.4570999 | 4.183286826 | 0.26527035 |
| Pax8 | 14.32991156 | 0.857443519 | 0.498660273 |
| Gm13415 | 0.371620636 | 2.115779348 | 1.70329581 |
| Gm27193 | 0 | NA | NA |
| Gm13418 | 0 | NA | NA |
| Gm13417 | 0.292450921 | -0.233136408 | 2.077152858 |
| Gm38383 | 0 | NA | NA |
| Gm13416 | 20.11579464 | -0.28122941 | 0.232451883 |
| Gm25455 | 0 | NA | NA |
| Cacna1b | 2031.063943 | -0.763341601 | 0.13064099 |
| Gm13459 | 0 | NA | NA |
| Ehmt1 | 682.9022921 | -0.313332376 | 0.105002873 |
| Gm38147 | 36.95672247 | -0.868598783 | 0.308109375 |
| Gm37139 | 53.01011326 | -0.938798236 | 0.301964765 |
| Arrdc1 | 89.73200459 | 0.316545221 | 0.124369106 |
| Gm22875 | 0.044544648 | 0.237309859 | 3.724945049 |
| Zmynd19 | 269.41162 | -0.019396029 | 0.121074781 |
| Dph7 | 184.0627944 | 0.186819136 | 0.088651076 |
| Mrpl41 | 329.4748938 | 0.089725061 | 0.119708544 |
| Pnpla7 | 332.8704237 | 0.975715658 | 0.111201163 |
| Gm13370 | 5.147326582 | 0.200028733 | 0.461196356 |
| Gm13369 | 0.218114541 | 0.263957289 | 2.544355846 |
| Gm25518 | 0 | NA | NA |
| Gm24875 | 0 | NA | NA |
| Nsmf | 2802.591237 | -0.578936203 | 0.092108539 |
| Gm24328 | 0 | NA | NA |
| Mir7664 | 1.286742338 | -2.560120912 | 1.087280129 |
| Entpd8 | 0.593920364 | 2.955697986 | 1.293306285 |
| Noxa1 | 0 | NA | NA |
| Gm13387 | 1.538924929 | -1.770641202 | 0.830609945 |
| Nrarp | 119.8106717 | -1.877027515 | 0.262889191 |
| Gm38287 | 0.509714849 | 0.767579888 | 1.609131796 |
| Tor4a | 84.46660317 | 2.547202844 | 0.21958016 |
| Nelfb | 451.1973938 | 0.21558806 | 0.065496053 |
| Gm22572 | 0 | NA | NA |
| 4933433C11Rik | 0.117650406 | 0.23732098 | 3.724945049 |
| Fam166a | 0.395870588 | -0.243563819 | 1.478958502 |
| Tubb4b | 1827.495376 | 0.169059184 | 0.098763524 |
| Slc34a3 | 1.160340464 | 0.432870315 | 0.915841413 |
| Rnf224 | 0.060174537 | 0.237309859 | 3.724945049 |
| Cysrt1 | 0.419553954 | -0.791013624 | 1.56711349 |
| Rnf208 | 937.0051333 | -0.717968394 | 0.066492246 |
| Ndor1 | 276.8690761 | 0.196480788 | 0.146696086 |
| Tmem203 | 61.6019059 | -0.399022583 | 0.167755281 |
| Tprn | 478.8306303 | 0.32716908 | 0.145019213 |
| Ssna1 | 171.2042094 | 0.181099569 | 0.120607866 |

| | | | |
|---|---|---|---|
| Anapc2 | 780.9388462 | -0.030601479 | 0.073356907 |
| Tmem210 | 0.270178086 | 1.823786162 | 3.707629481 |
| Lrrc26 | 0.469296467 | 1.623126293 | 3.264037629 |
| Grin1 | 5090.601218 | -0.693408917 | 0.120205039 |
| Grin1os | 5.022424966 | -0.799021339 | 0.500290018 |
| AA543186 | 4.244450911 | -0.740753939 | 0.52949308 |
| Man1b1 | 806.4183462 | 0.152399318 | 0.060983323 |
| Gm25220 | 2.58962508 | -1.000163961 | 0.679049353 |
| Dpp7 | 187.4441202 | 0.463254262 | 0.10643399 |
| Uap1l1 | 215.8935137 | 1.13791029 | 0.141721551 |
| Sapcd2 | 9.831372002 | -0.765151872 | 0.395401885 |
| Entpd2 | 57.81792083 | -0.323364577 | 0.199396834 |
| Npdc1 | 1413.945948 | -0.072059336 | 0.049955198 |
| Fut7 | 5.453509397 | 1.007623692 | 0.639675295 |
| Abca2 | 5084.226177 | -0.303961337 | 0.070094428 |
| mmu-mir-3087 | 0 | NA | NA |
| BC029214 | 267.2170635 | 0.073938806 | 0.086497171 |
| Clic3 | 1.915678769 | 0.582866974 | 0.808602951 |
| Ptgds | 4437.782382 | -0.733948285 | 0.159981425 |
| Lcn12 | 0.07796279 | 1.072698098 | 3.721829099 |
| C8g | 3.739276094 | -0.598110063 | 0.5316723 |
| Fbxw5 | 594.4106511 | -0.122254899 | 0.059131686 |
| Traf2 | 170.0261398 | 1.115561187 | 0.131877996 |
| Gm13433 | 0 | NA | NA |
| Gm13432 | 0.086930093 | 0.236669296 | 3.724898107 |
| Edf1 | 525.3660091 | 0.22402079 | 0.20071111 |
| Mamdc4 | 22.73292264 | -0.845300717 | 0.300239167 |
| Phpt1 | 63.01003723 | -1.023935529 | 0.16084527 |
| Gm996 | 1271.384592 | -1.447026354 | 0.129436301 |
| Gm26702 | 5.898309265 | -1.169924192 | 0.599125517 |
| Rabl6 | 1439.377497 | -0.291193407 | 0.056479298 |
| Fcnaos | 8.104967773 | -0.541270143 | 0.361563535 |
| Ccdc183 | 0.262568881 | -0.220732528 | 1.811193303 |
| Tmem141 | 79.20590439 | -0.419562577 | 0.175349728 |
| Fcna | 2.11389961 | -2.037325342 | 0.740271371 |
| Lcn8 | 0.091666476 | -0.724455696 | 3.724945049 |
| Lcn5 | 0.13733263 | -1.191829832 | 3.63430361 |
| Lcn6 | 0.208472373 | -0.708698314 | 2.637394356 |
| Lcn10 | 0.112396584 | 1.192083721 | 3.724434897 |
| Obp2a | 0.229116515 | 0.179807596 | 2.173657491 |
| A230005M16Rik | 4.640067633 | -0.822013556 | 0.471514497 |
| Bmyc | 286.5842141 | -0.299611009 | 0.097856395 |
| Obp2b | 1.10611924 | 1.403339692 | 1.128169266 |
| Lcn3 | 0.068097359 | -0.243569582 | 3.724945049 |
| Lcn11 | 0 | NA | NA |
| Gm13538 | 0.049553047 | 0.71820487 | 3.724945049 |
| Gm13539 | 0.066050875 | 0.71820487 | 3.724945049 |

| | | | |
|---|---|---|---|
| Glt6d1 | 0 | NA | NA |
| Lcn9 | 0.042716945 | -0.243571806 | 3.724945049 |
| Sohlh1 | 0.142434329 | -0.243563591 | 2.907370952 |
| Kcnt1 | 1559.190496 | -0.709463049 | 0.18232827 |
| Camsap1 | 1832.116154 | -0.054442731 | 0.069177569 |
| Gm13542 | 4.306872891 | -1.053767046 | 0.612448471 |
| Ubac1 | 350.8632011 | -0.051624527 | 0.06550667 |
| Nacc2 | 2614.976181 | -0.094538292 | 0.101646341 |
| Gm13554 | 0.124596252 | 0.718200615 | 3.714681068 |
| C330006A16Rik | 755.4666773 | -0.012849645 | 0.075291134 |
| 1810012K08Rik | 1.703298555 | -1.189408155 | 0.918255997 |
| Gm13553 | 0.097139547 | 0.718207094 | 3.724945049 |
| Lhx3 | 3.930402932 | 0.64302063 | 0.525923706 |
| Gm27196 | 0.679199126 | 0.160082995 | 1.181425407 |
| Qsox2 | 316.2840522 | 0.256279586 | 0.080533768 |
| C030048H21Rik | 9.452175138 | -1.346349562 | 0.377552945 |
| 4932418E24Rik | 35.63244048 | -1.800375246 | 0.25111167 |
| Gpsm1 | 740.3008245 | -0.428979615 | 0.090923371 |
| Dnlz | 309.3938586 | 0.16663633 | 0.108686698 |
| Card9 | 28.78353682 | 1.77302493 | 0.264994929 |
| Gm13562 | 6.216571396 | -1.206972454 | 0.437991617 |
| Snapc4 | 357.5941975 | -0.125152998 | 0.173134232 |
| Gm13563 | 26.0969972 | -0.813052807 | 0.27159343 |
| Sdccag3 | 585.8077495 | -0.083178246 | 0.072911853 |
| Pmpca | 864.0585262 | 0.091720258 | 0.065839302 |
| Inpp5e | 354.6391807 | -0.028686316 | 0.090256852 |
| Sec16a | 1481.474974 | 0.296942052 | 0.045210362 |
| 0610009E02Rik | 13.44346893 | -0.702507401 | 0.330465022 |
| Gm13568 | 2.19974275 | 0.272178606 | 0.666715928 |
| Notch1 | 1250.567747 | 1.369105976 | 0.118298184 |
| Mir6996 | 0.216707685 | 0.726289085 | 2.17296241 |
| Gm13358 | 0 | NA | NA |
| Egfl7 | 168.1436807 | 1.234425321 | 0.124005323 |
| Gm20532 | 0 | NA | NA |
| Mir126a | 0 | NA | NA |
| mmu-mir-126b | 0 | NA | NA |
| Agpat2 | 19.91019668 | 2.422822114 | 0.321054413 |
| Fam69b | 271.5122921 | -0.340141526 | 0.072696969 |
| Snhg7 | 0.151994617 | 0.237326393 | 2.849149087 |
| Snora43 | 0 | NA | NA |
| Snora17 | 8.121164419 | 0.278073855 | 0.365509641 |
| Snhg7os | 0.566020539 | -0.5017541 | 1.294046366 |
| Lcn4 | 0.485200684 | 2.513078955 | 1.626154655 |
| Gm13359 | 0 | NA | NA |
| Abo | 0.44885175 | 0.920952361 | 1.354481551 |
| Surf6 | 237.2764312 | 0.390864934 | 0.090990867 |
| Med22 | 318.481323 | 0.74708843 | 0.105671953 |

| | | | |
|---|---|---|---|
| Rpl7a | 773.724984 | 0.716705758 | 0.132152903 |
| Gm23969 | 2.942191425 | 0.584716508 | 0.670847742 |
| Gm22879 | 0 | NA | NA |
| Gm24134 | 0 | NA | NA |
| Surf1 | 388.1526091 | 0.360008147 | 0.083494471 |
| Surf2 | 198.1272666 | 0.084831423 | 0.08426359 |
| Surf4 | 907.7278515 | 0.960577959 | 0.104912418 |
| Stkld1 | 2.796069476 | 0.372554813 | 0.686730018 |
| Rexo4 | 357.7792058 | 0.018412352 | 0.10632774 |
| Adamts13 | 7.574428338 | -0.784393376 | 0.380828943 |
| Cacfd1 | 416.4720264 | 0.195929067 | 0.071991151 |
| Slc2a6 | 213.3264522 | -0.311072282 | 0.1023141 |
| Gm13397 | 0.26409243 | 2.102070284 | 2.011061824 |
| Gm13398 | 0.795847279 | 1.870319462 | 1.238182979 |
| Tmem8c | 3.290439168 | 0.346191285 | 0.653851372 |
| Adamtsl2 | 40.03111391 | -0.339072668 | 0.210235319 |
| Fam163b | 723.0454923 | -1.34241095 | 0.148644959 |
| Dbhos | 1.810426623 | -0.166741471 | 0.720841107 |
| Dbh | 21.92637974 | 0.003059439 | 0.346066 |
| Sardh | 177.0436753 | 0.585853739 | 0.164492409 |
| Sardhos | 0 | NA | NA |
| Vav2 | 234.323641 | 0.10127545 | 0.094341111 |
| AA645442 | 0 | NA | NA |
| Gm24049 | 0 | NA | NA |
| Brd3 | 790.2310845 | -0.271059697 | 0.067868902 |
| Mir7578 | 0.414979285 | -0.257500459 | 1.485146209 |
| Wdr5 | 327.3142271 | 0.309260629 | 0.100421038 |
| Gm25541 | 8.286915501 | -0.239148157 | 0.387263984 |
| Gm13421 | 4.157031413 | -0.544293373 | 0.51256458 |
| Rxra | 493.0791904 | 0.299381535 | 0.080461533 |
| 2810430I11Rik | 3.721301919 | 0.132680943 | 0.578584377 |
| Col5a1 | 208.2168104 | 0.12022538 | 0.14475419 |
| Gm13372 | 0 | NA | NA |
| Fcnb | 0.737107465 | 1.496936626 | 1.10520643 |
| Gm13371 | 0.299853729 | -1.167055122 | 1.787187062 |
| F730016J06Rik | 3.372121501 | -0.695121506 | 0.581400871 |
| Olfm1 | 4803.122293 | -0.524812354 | 0.095152517 |
| Gm13373 | 2.464667239 | 0.128604588 | 0.676582358 |
| Ppp1r26 | 345.0378359 | -0.62582554 | 0.067583083 |
| 1700007K13Rik | 15.71640074 | -0.12887449 | 0.299645133 |
| Mrps2 | 321.9683543 | 0.425551195 | 0.092156356 |
| Rpsa-ps9 | 16.7483217 | 0.541092746 | 0.293737436 |
| Gbgt1 | 30.60507794 | 0.739060311 | 0.227772623 |
| Gm10134 | 5.669552113 | 0.51572465 | 0.475016542 |
| Ralgds | 1434.688426 | 0.467390741 | 0.103749347 |
| Cel | 0.371662472 | -1.61077481 | 1.612385808 |
| Gm13381 | 0.066050875 | 0.71820487 | 3.724945049 |

| | | | |
|---|---|---|---|
| Gtf3c5 | 232.8597514 | 0.435360174 | 0.08577584 |
| Gm22675 | 0 | NA | NA |
| Gm22824 | 0.072025462 | 0.237312084 | 3.724945049 |
| Gfi1b | 0.585889909 | 1.358874475 | 1.534487404 |
| Tsc1 | 1195.112856 | -0.29618952 | 0.085864593 |
| 1700026L06Rik | 2.561385301 | 0.289050301 | 0.66635284 |
| Ak8 | 10.0543632 | -0.269731324 | 0.343014165 |
| Mir3088 | 0 | NA | NA |
| Gm13385 | 0 | NA | NA |
| Gm13386 | 0.066050875 | 0.71820487 | 3.724945049 |
| Gtf3c4 | 803.8265441 | 0.238333225 | 0.056397899 |
| Ddx31 | 110.419689 | 0.153747394 | 0.110541224 |
| Barhl1 | 48.18788692 | -0.355297618 | 0.249736486 |
| 1700101E01Rik | 8.146087367 | 0.682501199 | 0.338518907 |
| Gm13394 | 2432.869465 | -0.092057349 | 0.075004002 |
| Ttf1 | 162.3829832 | 0.068993658 | 0.137872482 |
| Gm13393 | 0.138076337 | 0.718204987 | 3.589802529 |
| Gm13396 | 0.120349074 | 0.237305146 | 3.724945049 |
| Setx | 1922.416912 | -0.249014466 | 0.067420302 |
| Ntng2 | 269.5695528 | -0.559901319 | 0.125880443 |
| 6530402F18Rik | 198.2560696 | -0.335745963 | 0.10114628 |
| Med27 | 188.5821441 | 0.049194391 | 0.102031503 |
| Gm13402 | 0.13742593 | -0.724459914 | 2.939744481 |
| Gm24976 | 1.183229494 | -1.435167953 | 1.102672521 |
| Mir133c | 0 | NA | NA |
| Gm13401 | 1.918047756 | -1.444442089 | 0.772418275 |
| Rapgef1 | 1745.697021 | -0.110194962 | 0.059217995 |
| Gm9823 | 0.449135467 | 2.558909396 | 1.318483196 |
| Gm13420 | 1.634298146 | -0.760934374 | 0.833422293 |
| Gm13547 | 0.049553047 | 0.71820487 | 3.724945049 |
| Trub2 | 328.3662552 | 0.268978661 | 0.083084431 |
| Coq4 | 113.9900504 | 0.083257748 | 0.103908292 |
| Slc27a4 | 871.144694 | -0.104317111 | 0.052427534 |
| Urm1 | 399.0492176 | 0.475266994 | 0.087390457 |
| Mir219a-2 | 4.701385445 | -0.151456942 | 0.638849882 |
| mmu-mir-219b | 2.33080191 | 0.152369981 | 0.763871782 |
| 2600006K01Rik | 2.48771953 | 0.014047684 | 0.623951546 |
| Cercam | 101.0078076 | 0.252255124 | 0.145659226 |
| Odf2 | 917.3723799 | -0.52942261 | 0.148287859 |
| Gm23865 | 0 | NA | NA |
| Gle1 | 489.083346 | -0.230662099 | 0.069105516 |
| Sptan1 | 16083.58909 | -0.347661552 | 0.070490524 |
| Mir6997 | 1.080983412 | -2.306224241 | 1.169556052 |
| Wdr34 | 186.8631446 | -0.114994993 | 0.12153842 |
| Set | 1857.674102 | 0.285441909 | 0.079551455 |
| Pkn3 | 10.32177187 | 3.072576896 | 0.441543127 |
| Zdhhc12 | 49.78225683 | 0.761654261 | 0.151686974 |

| | | | |
|---|---|---|---|
| Zer1 | 1722.618374 | -0.367356268 | 0.055017331 |
| Tbc1d13 | 421.3439125 | 0.346275614 | 0.087664555 |
| Slc39a1-ps | 5.938480901 | 1.459815247 | 0.473704116 |
| Endog | 111.3222115 | 0.131353618 | 0.136193996 |
| D2Wsu81e | 251.2755507 | 0.026334826 | 0.09139346 |
| Ccbl1 | 92.26571604 | -0.845506503 | 0.147338445 |
| 1700084E18Rik | 9.307150498 | 0.067241321 | 0.345101044 |
| Lrrc8a | 523.7153376 | 0.554237189 | 0.102824938 |
| Gm28035 | 4.247830997 | -0.839720919 | 0.514531655 |
| Phyhd1 | 42.27143342 | 0.566215351 | 0.255404448 |
| Gm28038 | 0 | NA | NA |
| Dolk | 191.5944489 | 0.123700927 | 0.106570795 |
| Nup188 | 388.5180733 | 0.425040178 | 0.090231945 |
| Gm14471 | 0 | NA | NA |
| Sh3glb2 | 1245.303918 | -0.505599699 | 0.067344885 |
| Fam73b | 803.1893246 | -0.158863989 | 0.06372719 |
| AL954388.1 | 0 | NA | NA |
| Dolpp1 | 175.3732175 | 0.020635993 | 0.088626548 |
| Crat | 430.9212061 | -0.16768745 | 0.080551135 |
| Gm16323 | 10.6181011 | -0.464866313 | 0.407034165 |
| Ppp2r4 | 1830.296797 | -0.026591782 | 0.073821725 |
| Ier5l | 217.6479524 | 0.443188311 | 0.167664725 |
| Gm14487 | 1.256366316 | 1.071012669 | 1.488949747 |
| Cstad | 12.63152331 | -0.550416981 | 0.329840149 |
| 9330198N18Rik | 2.48591285 | 0.165613393 | 0.792201243 |
| Gm14486 | 1.55717633 | 1.851827429 | 1.011652911 |
| Gm14488 | 4.29828934 | 2.399643408 | 0.570573018 |
| Mir3089 | 0.367473112 | -1.470143495 | 1.508726601 |
| 1700001O22Rik | 34.98746724 | -0.125638947 | 0.20757956 |
| 4930527E20Rik | 0.124415284 | -0.243565133 | 3.724945049 |
| Ntmt1 | 195.850967 | 0.343015428 | 0.120886101 |
| Asb6 | 292.993291 | 0.419906462 | 0.082982945 |
| Prrx2 | 13.36258994 | 1.311680602 | 0.31219725 |
| Ptges | 99.72996419 | 1.926193572 | 0.285345464 |
| Gm13400 | 1.094212761 | 1.422873186 | 1.064571056 |
| Tor1b | 368.7015473 | 0.144091489 | 0.075671491 |
| Tor1a | 257.103606 | 0.236951904 | 0.084839163 |
| BC005624 | 537.4115427 | -0.072069038 | 0.099577334 |
| Usp20 | 792.5786421 | -0.454379364 | 0.070150625 |
| Fnbp1 | 2552.528974 | -0.322156741 | 0.071547403 |
| D330023K18Rik | 31.26496909 | -0.292012081 | 0.214362189 |
| Gpr107 | 862.5963975 | 0.305522506 | 0.048260163 |
| Gm22516 | 4.36673596 | -0.138587838 | 0.525955898 |
| Gm13405 | 0.116046057 | -0.724453471 | 3.724945049 |
| Ncs1 | 3353.155418 | -0.170926755 | 0.10564957 |
| Hmcn2 | 32.42021316 | 1.376871012 | 0.276963995 |
| Ass1 | 313.9716636 | 0.795058432 | 0.132686034 |

| | | | |
|---|---|---|---|
| Gm13426 | 0 | NA | NA |
| Fubp3 | 562.5375217 | 0.225542427 | 0.080514119 |
| Mir6998 | 1.273153199 | -0.932661473 | 1.059547964 |
| Gm13425 | 3.449187758 | -0.322546939 | 0.584627804 |
| Prdm12 | 5.973420824 | -0.223683043 | 0.447004789 |
| Gm13424 | 0 | NA | NA |
| Exosc2 | 146.3618478 | 0.484752769 | 0.101793397 |
| Gm13427 | 1.245568681 | -0.054720046 | 0.874173258 |
| Abl1 | 1015.182759 | -0.023300991 | 0.052823939 |
| Qrfp | 1.096023364 | 1.539787082 | 0.988844159 |
| Fibcd1 | 171.8501216 | -1.256958485 | 0.144085596 |
| Gm13441 | 0.122601409 | 0.237307635 | 3.724945049 |
| Lamc3 | 85.86094398 | -0.236839805 | 0.183028833 |
| Aif1l | 75.90635778 | 0.215770836 | 0.215674663 |
| Gm13442 | 0.192796986 | 0.261585525 | 2.249927941 |
| Nup214 | 817.8687593 | 0.148326661 | 0.073562838 |
| Gm13608 | 10.74285294 | -0.476506441 | 0.485678087 |
| Gm13609 | 0.288698636 | 0.761870787 | 2.056046409 |
| Fam78a | 86.39529847 | -0.346554547 | 0.142774213 |
| Gm16534 | 0.731853201 | -1.445702286 | 1.254487591 |
| Mir7674 | 0.20460011 | -1.118574121 | 3.233930613 |
| Ppapdc3 | 209.3998458 | -0.103722485 | 0.093450027 |
| Prrc2b | 8454.239442 | -0.381470255 | 0.069808036 |
| Gm27805 | 3.918894907 | -1.336328568 | 0.585850327 |
| Gm22192 | 0.800855801 | -1.228379355 | 1.160080114 |
| Gm25632 | 3.196964654 | -0.793170815 | 0.747099102 |
| Gm13610 | 1.740926027 | -0.334311707 | 0.690742208 |
| Pomt1 | 154.4461912 | -0.132081857 | 0.167601038 |
| Uck1 | 264.0364267 | 0.163746766 | 0.095440775 |
| Gm13611 | 4.282612097 | 1.454645675 | 0.583987189 |
| Swi5 | 679.3934042 | 0.225360073 | 0.075362947 |
| Golga2 | 1066.72792 | -0.02657148 | 0.082746502 |
| Dnm1 | 10321.66057 | -0.694576124 | 0.084466453 |
| Gm23363 | 0.044544648 | 0.237309859 | 3.724945049 |
| Mir199b | 0.042716945 | -0.243571806 | 3.724945049 |
| Ciz1 | 515.2909118 | 0.349120869 | 0.076866743 |
| 1110008P14Rik | 542.4132862 | -0.715077019 | 0.097489311 |
| Lcn2 | 343.4844549 | 6.903685139 | 0.805823582 |
| Ptges2 | 252.589667 | 0.098660977 | 0.088919634 |
| Gm13413 | 0.473410537 | 0.892754924 | 1.391167123 |
| Slc25a25 | 955.9872403 | 0.300546994 | 0.13152866 |
| Naif1 | 76.9182076 | 0.40074158 | 0.147950188 |
| Gm37169 | 0 | NA | NA |
| Gm13412 | 0 | NA | NA |
| Fam102a | 1133.257023 | 0.104975479 | 0.088541136 |
| Dpm2 | 258.7392704 | 0.298301253 | 0.082862823 |
| 9430097D07Rik | 4.085120842 | -0.144418181 | 0.519255486 |

| | | | |
|---|---|---|---|
| Pip5kl1 | 31.38216028 | -0.94387452 | 0.288514249 |
| St6galnac4 | 520.5645504 | 0.828440445 | 0.089116107 |
| Gm23369 | 0.044544648 | 0.237309859 | 3.724945049 |
| St6galnac6 | 748.2103617 | -0.867810326 | 0.089265694 |
| Ak1 | 1088.578713 | -0.172390638 | 0.071382906 |
| Eng | 245.6138286 | 2.365232273 | 0.172280234 |
| Mir1954 | 0.415365334 | 2.301309367 | 3.225002104 |
| AL772271.1 | 0 | NA | NA |
| Gm26236 | 0.112642008 | -0.24357403 | 3.724945049 |
| Fpgs | 67.89073458 | 0.561430273 | 0.176011467 |
| Mir8093 | 0 | NA | NA |
| Cdk9 | 551.2940997 | 0.166823437 | 0.070944982 |
| Mir2861 | 0 | NA | NA |
| mmu-mir-3960 | 0 | NA | NA |
| Sh2d3c | 241.4715273 | 0.136902773 | 0.116175855 |
| 6330409D20Rik | 20.79074414 | -0.753837831 | 0.230636732 |
| Ttc16 | 12.6100887 | 0.247273246 | 0.333764872 |
| Tor2a | 191.8598868 | 0.393035486 | 0.097775949 |
| Ptrh1 | 37.42700904 | 1.219600905 | 0.211679331 |
| 1700019L03Rik | 4.625372364 | -0.043755735 | 0.44997241 |
| Stxbp1 | 9943.087999 | -0.337650684 | 0.062087736 |
| Gm24165 | 0 | NA | NA |
| Gm13524 | 5.575737892 | 0.244192005 | 0.461152468 |
| Gm13523 | 5.508606278 | 0.318390217 | 0.474858006 |
| Fam129b | 816.2277192 | 1.980762506 | 0.172581156 |
| Lrsam1 | 401.4005097 | -0.312820515 | 0.074743457 |
| Rpl12 | 1113.211754 | 0.695721974 | 0.098895137 |
| Snora65 | 0 | NA | NA |
| Slc2a8 | 138.0519622 | -0.825045646 | 0.12754155 |
| Garnl3 | 1194.21391 | -0.518129837 | 0.128710785 |
| Gm38299 | 48.15575876 | -0.777884383 | 0.417322726 |
| Ralgps1 | 1094.760979 | -0.405833687 | 0.098238092 |
| Gm13528 | 0.134209209 | -0.240821775 | 3.665582479 |
| Gm23546 | 4.82094354 | -0.780262677 | 0.506374467 |
| Angptl2 | 62.62522016 | 0.253513582 | 0.161243544 |
| Gm13537 | 0.097139547 | 0.718207094 | 3.724945049 |
| Gm25998 | 3.056188022 | -0.865123494 | 0.654796182 |
| Zbtb34 | 281.4807306 | -0.308318912 | 0.098733513 |
| Gm13536 | 1.068960359 | -0.285306813 | 0.982746284 |
| Zbtb43 | 400.5586369 | -0.011888291 | 0.104310171 |
| Gm13530 | 1.493521525 | -1.767537333 | 0.928760261 |
| Lmx1b | 29.823212 | 0.461815078 | 0.376704604 |
| Gm38011 | 0 | NA | NA |
| C130021I20Rik | 27.33416534 | 0.225226711 | 0.378740717 |
| Gm13529 | 0.108772582 | 0.718202646 | 3.724945049 |
| 9430024E24Rik | 0.066050875 | 0.71820487 | 3.724945049 |
| Mvb12b | 1603.937987 | -0.0268115 | 0.058620821 |

| | | | |
|---|---|---|---|
| Nron | 0.158932157 | -0.243563432 | 2.809576835 |
| Gm13403 | 3.213919713 | 0.611684028 | 0.621757545 |
| Gm13404 | 2.447136968 | 0.15674162 | 0.618870755 |
| C230014O12Rik | 11.75502483 | -0.25043439 | 0.285196351 |
| Gm38389 | 0.847399432 | -2.926102802 | 1.155618969 |
| Pbx3 | 410.290469 | -0.284635897 | 0.158328747 |
| C79798 | 0.252257838 | 1.731297673 | 2.47551742 |
| Mapkap1 | 523.1598314 | -0.024516561 | 0.093580256 |
| Gm13407 | 0 | NA | NA |
| Gm13406 | 0 | NA | NA |
| Gm13408 | 58.61577562 | 0.522479959 | 0.195236582 |
| Gm13414 | 23.60150067 | 0.297604203 | 0.274534017 |
| Gapvd1 | 1564.103693 | 0.011876743 | 0.042745153 |
| Hspa5 | 4684.153117 | 1.051989241 | 0.123290383 |
| Rabepk | 186.7520506 | 0.059453199 | 0.09812124 |
| Fbxw2 | 731.5136731 | 0.028551131 | 0.077257099 |
| Gm13419 | 0.060174537 | 0.237309859 | 3.724945049 |
| RP23-446N16.6 | 0.320849018 | -0.256473937 | 1.744244045 |
| Psmd5 | 541.5249382 | 0.239909082 | 0.066988567 |
| Cutal | 4.992381895 | 0.578895832 | 0.444343743 |
| Phf19 | 22.86602922 | 0.901676472 | 0.382718088 |
| Gm13448 | 0 | NA | NA |
| Gm24905 | 0 | NA | NA |
| Traf1 | 68.5058879 | 2.141383003 | 0.230219067 |
| Gm13450 | 0.044544648 | 0.237309859 | 3.724945049 |
| Hc | 5.496526028 | 1.18081795 | 0.44370544 |
| Gm13449 | 0.579463306 | 0.781078223 | 1.493754912 |
| AI182371 | 3.278154351 | 1.000879913 | 0.601957082 |
| Cntrl | 338.5002484 | 0.625335324 | 0.153580673 |
| Rab14 | 2869.966926 | -0.012040035 | 0.03831763 |
| Gm13605 | 0 | NA | NA |
| Gsn | 990.1649303 | 0.335015458 | 0.134845602 |
| RP23-186B18.8 | 6.833187758 | 0.276834143 | 0.44259719 |
| Stom | 265.4412871 | 1.357795434 | 0.144231326 |
| Gm13443 | 73.68671835 | 0.067941653 | 0.138785188 |
| 4930568D16Rik | 0 | NA | NA |
| 4930402F06Rik | 0.092691418 | 1.183079489 | 3.724560917 |
| Ggta1 | 165.7903113 | 3.051372559 | 0.194205238 |
| RP23-458A10.4 | 2.488423977 | 2.181832882 | 0.836788246 |
| Gm13444 | 0.047948698 | -0.243569582 | 3.724945049 |
| Gm13445 | 0 | NA | NA |
| Gm13447 | 0.202171817 | 1.626940071 | 2.587140394 |
| Gm13446 | 33.55988347 | -1.581118702 | 0.219978906 |
| Dab2ip | 1883.686994 | -0.457293792 | 0.077121721 |
| Ttll11 | 316.1783459 | -0.18107926 | 0.114132944 |
| RP23-366N13.3 | 20.18499025 | -1.239159328 | 0.413944209 |
| 9030204H09Rik | 0.047948698 | -0.243569582 | 3.724945049 |

| | | | |
|---|---|---|---|
| RP23-366N13.2 | 0.655197851 | 1.537007836 | 1.290043889 |
| Ndufa8 | 520.9048915 | 0.058315132 | 0.104220568 |
| Morn5 | 2.567728017 | -0.371208856 | 0.544722245 |
| RP23-169L4.6 | 0 | NA | NA |
| Lhx6 | 117.6126763 | -0.488081365 | 0.130380725 |
| Gm27953 | 0 | NA | NA |
| Rbm18 | 518.3893294 | 0.201205702 | 0.076447937 |
| Mrrf | 178.907885 | 0.245112663 | 0.099601578 |
| Gm13428 | 0 | NA | NA |
| Gm13429 | 0.948980887 | 0.405470161 | 1.087088623 |
| Gm13431 | 0.366933406 | 0.709657068 | 1.731498455 |
| Ptgs1 | 131.6150916 | 0.234723906 | 0.222411847 |
| RP23-454A14.10 | 0.048949532 | -0.243569582 | 3.724945049 |
| Olfr334-ps1 | 0.427301431 | 1.825028395 | 3.361678582 |
| Olfr335-ps | 0.092140247 | -0.724455696 | 3.724945049 |
| Olfr336-ps1 | 0.11657011 | 0.237307635 | 3.724945049 |
| Olfr337-ps1 | 0.194279094 | 1.292225235 | 3.719439735 |
| Olfr338 | 0.195866688 | 1.276112028 | 3.248877671 |
| Gm13430 | 9.735155595 | -0.356119024 | 0.38555331 |
| Olfr339 | 0 | NA | NA |
| Olfr340 | 0 | NA | NA |
| Olfr341 | 0 | NA | NA |
| Olfr342 | 0.194279094 | 1.292225235 | 3.719439735 |
| Olfr343-ps1 | 0 | NA | NA |
| Olfr344 | 0.046345709 | 0.71820487 | 3.724945049 |
| Olfr345 | 0 | NA | NA |
| Olfr346 | 0 | NA | NA |
| Olfr347 | 0 | NA | NA |
| Gm13438 | 0 | NA | NA |
| Olfr348 | 0 | NA | NA |
| Olfr50 | 0 | NA | NA |
| Gm13436 | 42.25227907 | 1.1328613 | 0.216188041 |
| Olfr3 | 0 | NA | NA |
| Olfr349-ps1 | 0 | NA | NA |
| Olfr350 | 0 | NA | NA |
| Olfr351 | 0.097139547 | 0.718207094 | 3.724945049 |
| Olfr352 | 0 | NA | NA |
| Olfr353 | 0 | NA | NA |
| Olfr354 | 0 | NA | NA |
| Gm13440 | 0 | NA | NA |
| Olfr355 | 0 | NA | NA |
| Olfr356 | 0 | NA | NA |
| Olfr357 | 0 | NA | NA |
| Olfr358 | 0 | NA | NA |
| Olfr359-ps1 | 0 | NA | NA |
| Gm13439 | 0 | NA | NA |
| Olfr360 | 0 | NA | NA |

| | | | |
|---|---|---|---|
| Gm13437 | 0 | NA | NA |
| Olfr361 | 0 | NA | NA |
| Gm13434 | 0 | NA | NA |
| Olfr362 | 0 | NA | NA |
| Olfr363-ps | 0 | NA | NA |
| Olfr364-ps1 | 0 | NA | NA |
| Olfr365 | 0 | NA | NA |
| Gm13435 | 0 | NA | NA |
| Olfr366 | 0.068097359 | -0.243569582 | 3.724945049 |
| Olfr367-ps | 0 | NA | NA |
| Gm22306 | 0 | NA | NA |
| Olfr368 | 0 | NA | NA |
| Pdcl | 328.2885349 | 0.21142613 | 0.122219747 |
| Rc3h2 | 3083.559831 | 0.025595785 | 0.037452478 |
| Snord90 | 1.013447778 | -0.426900464 | 1.066132973 |
| Zbtb6 | 434.5484348 | 0.422163529 | 0.075488369 |
| Zbtb26 | 221.5652181 | 0.002862229 | 0.128171875 |
| Rabgap1 | 1677.895686 | -0.16503659 | 0.043084486 |
| Strbp | 4912.750144 | -1.39225109 | 0.131891071 |
| Gpr21 | 129.8264781 | -0.363302529 | 0.15341908 |
| Gm13543 | 0 | NA | NA |
| Mir5128 | 8.038112018 | -1.844481752 | 0.443411272 |
| Crb2 | 102.1842241 | 0.147228716 | 0.160682556 |
| Dennd1a | 974.3781944 | 0.266663771 | 0.060881246 |
| AL929186.1 | 4.630423038 | -0.464275646 | 0.51911507 |
| Gm13556 | 0.04393175 | -0.243571806 | 3.724945049 |
| Lhx2 | 320.8917483 | -0.511652757 | 0.125323119 |
| Gm27197 | 1.480046296 | -0.687646644 | 1.044804229 |
| Gm13584 | 0 | NA | NA |
| Gm13589 | 3.470889967 | 0.260403768 | 0.531182203 |
| Gm25081 | 0.13733263 | -1.191829832 | 3.63430361 |
| Gm13588 | 0.045882455 | 0.237312084 | 3.724945049 |
| Nek6 | 512.9966335 | 1.7793537 | 0.138214986 |
| Psmb7 | 966.8015143 | 0.486539006 | 0.085519447 |
| Gm13586 | 0.536876103 | 0.989195579 | 1.427332337 |
| Gm27634 | 1.100154067 | -0.044069997 | 1.149796001 |
| Adgrd2-ps | 0 | NA | NA |
| Gm13587 | 0 | NA | NA |
| Nr5a1 | 5.753592802 | 0.020165769 | 0.509614516 |
| Nr5a1os | 0.104719185 | 0.237305575 | 3.724945049 |
| Nr6a1 | 153.9253641 | -0.512088857 | 0.14261241 |
| Rpl15-ps4 | 0 | NA | NA |
| Nr6a1os | 2.499784555 | -1.793559918 | 0.776611866 |
| Mir181a-2 | 0.139226991 | -0.243563626 | 2.927945967 |
| Mir181b-2 | 0.377368388 | 0.254138341 | 1.720871248 |
| Olfml2a | 59.91228772 | -0.25127075 | 0.208593859 |
| Gm13473 | 0 | NA | NA |

| | | | |
|---|---|---|---|
| Gm13474 | 0.110595523 | 0.718200421 | 3.724945049 |
| Gm13496 | 0.873180464 | -0.281660708 | 0.947588437 |
| Wdr38 | 12.8811223 | 0.061463722 | 0.29356188 |
| Rpl35 | 950.799867 | 0.758706264 | 0.103117311 |
| Arpc5l | 615.8797353 | 0.406891751 | 0.098853719 |
| Golga1 | 507.3639855 | -0.184674343 | 0.085644261 |
| Gm26127 | 0 | NA | NA |
| Scai | 1810.878198 | -0.427692463 | 0.074104146 |
| Gm16523 | 10.92727314 | -0.502231337 | 0.42029765 |
| Ppp6c | 892.2623954 | 0.397774675 | 0.082788182 |
| Gm13511 | 6.921421994 | -1.252519067 | 0.469083216 |
| Gm13504 | 0 | NA | NA |
| Vmn2r-ps2 | 0.066050875 | 0.71820487 | 3.724945049 |
| Gm13506 | 0 | NA | NA |
| Gm13454 | 0 | NA | NA |
| Gm13451 | 0 | NA | NA |
| Gm13452 | 0.066050875 | 0.71820487 | 3.724945049 |
| Gm13453 | 7.860772947 | 0.351271384 | 0.351500696 |
| Gm13457 | 0 | NA | NA |
| Gm13458 | 0.061920613 | 0.718207094 | 3.724945049 |
| Gm13455 | 0 | NA | NA |
| Gm24467 | 0 | NA | NA |
| Gm13456 | 339.2160491 | 0.722523345 | 0.114530781 |
| Lrp1b | 2840.147083 | -0.892644511 | 0.122906996 |
| Gm27348 | 0.290738398 | -0.678605226 | 1.769415157 |
| Gm13462 | 0 | NA | NA |
| Gm13460 | 0 | NA | NA |
| Gm13461 | 25.32005373 | 0.345467861 | 0.214967633 |
| Gm13463 | 0 | NA | NA |
| Mir6336 | 9.1407384 | -0.867037334 | 0.386430685 |
| Gm24350 | 0 | NA | NA |
| Gm13464 | 14.15448784 | 0.016171862 | 0.305962127 |
| Kynu | 6.170416941 | 2.779349639 | 0.527687261 |
| Arhgap15 | 136.0233683 | 1.034900209 | 0.162783332 |
| Arhgap15os | 0.169875191 | -0.243569355 | 2.748385943 |
| Gm22867 | 0.042716945 | -0.243571806 | 3.724945049 |
| Gtdc1 | 447.6665794 | -0.560465823 | 0.097789188 |
| Gm13477 | 0.271375346 | -0.243568522 | 2.084009298 |
| Zeb2 | 3739.770028 | 0.113116064 | 0.097034138 |
| Mir5129 | 1.526907099 | -0.69130008 | 0.737389284 |
| Zeb2os | 56.92489478 | 1.765694133 | 0.205808352 |
| Gm27761 | 0 | NA | NA |
| Gm27676 | 0.743251506 | 1.668286446 | 1.173120599 |
| Gm27404 | 0 | NA | NA |
| Gm27645 | 0 | NA | NA |
| Gm28643 | 0.172790758 | 0.795772189 | 3.387088085 |
| Vis1 | 2.225936112 | 2.303216046 | 0.741576352 |

| | | | |
|---|---|---|---|
| Gm13475 | 0.044191549 | -0.243571806 | 3.724945049 |
| Gm13479 | 1.673299741 | -0.209848844 | 0.760174852 |
| Gm13478 | 0 | NA | NA |
| 1700019E08Rik | 0 | NA | NA |
| Gm13465 | 0.044191549 | -0.243571806 | 3.724945049 |
| Gm13467 | 0 | NA | NA |
| Gm13466 | 0 | NA | NA |
| Gm23072 | 0.060174537 | 0.237309859 | 3.724945049 |
| Gm13469 | 0 | NA | NA |
| Gm13470 | 0.459694823 | 1.619841743 | 1.512299854 |
| Gm25264 | 0 | NA | NA |
| Gm25959 | 0.869297867 | 2.836052921 | 1.084188245 |
| Gm13481 | 2.425955829 | 0.942993879 | 0.717388306 |
| Gm13471 | 0 | NA | NA |
| Gm13482 | 0 | NA | NA |
| Gm25338 | 0.121578509 | 0.718202646 | 3.724945049 |
| Gm13472 | 0.062426873 | 0.237312084 | 3.724945049 |
| Gm13489 | 0.524554913 | -1.888285276 | 1.371588677 |
| Acvr2a | 849.6809722 | -0.376591102 | 0.063867859 |
| Orc4 | 1017.373273 | 0.207416769 | 0.089720048 |
| Mbd5 | 1674.173319 | -0.377494734 | 0.096731207 |
| Gm13510 | 17.67851426 | -0.584259554 | 0.301593096 |
| Gm13509 | 582.147492 | 0.06677687 | 0.097636994 |
| Gm13508 | 0 | NA | NA |
| Gm25215 | 0.07796279 | 1.072698098 | 3.721829099 |
| Epc2 | 915.7051038 | 0.171751623 | 0.098760459 |
| Kif5c | 11473.79799 | -0.381217743 | 0.086535381 |
| Gm13525 | 21.45150064 | -1.027939282 | 0.446018285 |
| Lypd6b | 138.4529101 | 0.212324427 | 0.162411375 |
| Gm13480 | 0.55522199 | 1.180831499 | 1.36001553 |
| Gm25223 | 0 | NA | NA |
| Lypd6 | 201.0019211 | -0.369625207 | 0.114500142 |
| Mmadhc | 325.9114616 | 0.409754708 | 0.094168209 |
| Gm13483 | 20.39339472 | -0.60406816 | 0.265283148 |
| Gm13486 | 0.761951221 | -1.330114068 | 1.139376494 |
| Gm13488 | 0.134148234 | 0.23732184 | 3.624632408 |
| Gm13487 | 0.658756006 | 0.474192476 | 1.260627994 |
| Gm13485 | 1.138498193 | -0.531370808 | 1.024448153 |
| Gm13484 | 0 | NA | NA |
| Gm13498 | 2.899184114 | 0.682662793 | 0.627416688 |
| Rnd3 | 459.0834406 | 1.101211178 | 0.165560002 |
| Gm28639 | 0 | NA | NA |
| Gm13500 | 0 | NA | NA |
| Gm13497 | 0.338153501 | 0.690752345 | 1.485248987 |
| Gm13499 | 0.047948698 | -0.243569582 | 3.724945049 |
| 1700057H21Rik | 0.253674139 | 0.252551829 | 2.108404788 |
| Gm23505 | 0.049553047 | 0.71820487 | 3.724945049 |

| | | | |
|---|---|---|---|
| Gm13494 | 0 | NA | NA |
| Gm13491 | 0.045882455 | 0.237312084 | 3.724945049 |
| Tas2r134 | 0 | NA | NA |
| Gm13490 | 0.088383099 | -0.72445792 | 3.724945049 |
| Gm13492 | 0 | NA | NA |
| Gm13493 | 88.56531709 | -0.060403311 | 0.130795272 |
| Rbm43 | 483.6855949 | 2.291835116 | 0.155979714 |
| Gm13495 | 1.701798027 | 3.52849277 | 1.19946653 |
| Nmi | 215.7031363 | 4.315946667 | 0.263918698 |
| Tnfaip6 | 134.9876113 | 0.158224773 | 0.176890924 |
| Rif1 | 1558.481454 | -0.127332857 | 0.070101425 |
| Neb | 246.6161145 | -0.209644495 | 0.161531253 |
| Gm22095 | 0 | NA | NA |
| Gm13521 | 0 | NA | NA |
| Gm13522 | 0.21580698 | -0.685787784 | 2.583860003 |
| Arl5a | 1365.664992 | -0.055749629 | 0.064774095 |
| A430018G15Rik | 10.01027238 | -0.349148987 | 0.348176601 |
| Cacnb4 | 3963.510753 | -0.637314403 | 0.0679164 |
| Gm13541 | 0 | NA | NA |
| Bloc1s2-ps | 5.732035973 | 0.300073623 | 0.446042079 |
| Stam2 | 454.7228158 | 0.52252154 | 0.076212922 |
| Gm13548 | 1.65657422 | -0.281949751 | 0.85762536 |
| Fmnl2 | 1786.717828 | 0.929123288 | 0.11609506 |
| Gm26603 | 0.266304556 | 1.36146117 | 3.715728684 |
| Prpf40a | 1253.218375 | 0.412234072 | 0.094976279 |
| Arl6ip6 | 155.8469731 | 0.304665251 | 0.102216143 |
| Gm25898 | 0 | NA | NA |
| Gm13501 | 0 | NA | NA |
| Gm13503 | 0 | NA | NA |
| Gm13502 | 9.659148329 | 1.170431548 | 0.351063465 |
| Gm23315 | 0 | NA | NA |
| Rprm | 258.4970102 | -0.58395757 | 0.095061484 |
| Gm22298 | 0.222122807 | -0.243564144 | 2.578296719 |
| Gm13505 | 0 | NA | NA |
| Gm14035 | 0 | NA | NA |
| Gm14034 | 0.195609542 | 0.718214769 | 2.623112067 |
| Galnt13 | 988.7175182 | -0.58695937 | 0.104263907 |
| Gm24549 | 0 | NA | NA |
| Gm25489 | 4.670143882 | -1.556439543 | 0.558737575 |
| Gm14032 | 0.804512061 | -0.065010122 | 1.163232818 |
| Gm14033 | 39.47582969 | -0.281140708 | 0.238709877 |
| Gm26794 | 31.41234798 | -0.993959809 | 0.206921879 |
| Kcnj3 | 1639.12147 | -0.892219223 | 0.094431024 |
| Gm13517 | 0.255816663 | 0.740567145 | 2.072795991 |
| Gm13615 | 0 | NA | NA |
| Gm13518 | 0.061920613 | 0.718207094 | 3.724945049 |
| Mir195b | 0 | NA | NA |

| | | | |
|---|---|---|---|
| Gm13527 | 0 | NA | NA |
| Nr4a2 | 294.6002765 | -1.016478379 | 0.194232141 |
| BB557941 | 2.213086149 | -1.661969056 | 0.837967767 |
| 4930555B11Rik | 0.042716945 | -0.243571806 | 3.724945049 |
| Gm13526 | 0 | NA | NA |
| A930012O16Rik | 7.994888912 | 0.358050396 | 0.379438298 |
| Gpd2 | 1202.471657 | -0.471217844 | 0.08787975 |
| Gm13535 | 0 | NA | NA |
| Gm13531 | 0 | NA | NA |
| Gm13532 | 2.612576842 | 1.830919514 | 0.663145418 |
| Gm13533 | 0 | NA | NA |
| Gm25388 | 0 | NA | NA |
| Galnt5 | 0.79124893 | 0.376750615 | 1.286200316 |
| Ermn | 1375.435044 | 0.898257243 | 0.176553906 |
| Gm13534 | 0.908099955 | -0.16488007 | 0.912942239 |
| Cytip | 92.50638567 | 5.66416761 | 0.399525237 |
| Gm13546 | 1.506085986 | 2.384166119 | 0.99224259 |
| Acvr1c | 451.2463121 | -0.519707287 | 0.170095191 |
| Gm13544 | 2.162167769 | 0.448930561 | 0.742918517 |
| Gm13545 | 0.038981395 | 0.718207094 | 3.724945049 |
| Acvr1 | 465.7691716 | 0.274699104 | 0.093916065 |
| Gm17409 | 0 | NA | NA |
| Upp2 | 286.6269545 | -0.883423278 | 0.102542963 |
| Gm13558 | 10.32835312 | -1.055515615 | 0.350429166 |
| Gm13559 | 1.310073478 | -0.943043513 | 0.969841547 |
| Ccdc148 | 215.227496 | -0.818868981 | 0.111121929 |
| Gm11099 | 0.095897396 | -0.724453471 | 3.724945049 |
| Gm13560 | 0.045882455 | 0.237312084 | 3.724945049 |
| Gm13557 | 1.02920101 | -1.204880108 | 1.504381751 |
| Gm13550 | 2.191863138 | -0.721358499 | 0.765301153 |
| Pkp4 | 2887.099339 | -0.433267416 | 0.058555283 |
| 5330411J11Rik | 0.17957102 | 0.773316785 | 2.710633433 |
| Gm13551 | 0 | NA | NA |
| Gm13549 | 0 | NA | NA |
| Dapl1 | 3.189629895 | -1.543934071 | 0.661080269 |
| Gm13552 | 6.37558536 | 0.332551563 | 0.390905811 |
| Tanc1 | 1294.003133 | 0.240207766 | 0.086982805 |
| Wdsub1 | 122.8229018 | 0.229216204 | 0.125830039 |
| Baz2b | 1280.473641 | -0.275692596 | 0.099857033 |
| Gm13574 | 8.429661362 | -0.984568875 | 0.375307204 |
| Gm13572 | 1.149049355 | 1.875824241 | 1.110211401 |
| Gm13570 | 9.56573459 | -0.27465945 | 0.357526719 |
| Gm13620 | 2.37993816 | -0.096326949 | 0.59087406 |
| Ldha-ps | 0.208487022 | 0.531317491 | 2.578957643 |
| 7-Mar | 1192.427979 | 0.153477826 | 0.095806864 |
| Cd302 | 130.1829803 | 0.599176417 | 0.162736694 |
| Gm13573 | 2.148767599 | 0.005380466 | 0.671580181 |

| | | | |
|---|---|---|---|
| Ly75 | 45.62619954 | 1.790630226 | 0.227105544 |
| Gm13571 | 1.148968944 | 1.877492952 | 1.05934986 |
| Gm13580 | 0.165156969 | 1.598327176 | 3.401540795 |
| Pla2r1 | 18.48570985 | 1.701485479 | 0.302671867 |
| Gm13583 | 0 | NA | NA |
| Gm13566 | 0 | NA | NA |
| Gm13567 | 0 | NA | NA |
| Itgb6 | 5.350769152 | 2.335092123 | 0.559710652 |
| Rbms1 | 693.4012186 | 1.250110753 | 0.123910634 |
| Gm13582 | 0.21728767 | 1.453426934 | 2.53000544 |
| Gm13579 | 0 | NA | NA |
| Gm13581 | 0 | NA | NA |
| Gm26422 | 0 | NA | NA |
| Gm22338 | 0 | NA | NA |
| Tank | 472.8096782 | 0.677571205 | 0.084706075 |
| Gm13626 | 0.359719716 | 0.278583139 | 1.580067463 |
| Psmd14 | 942.4841296 | 0.47210549 | 0.074370069 |
| Gm22403 | 0 | NA | NA |
| Gm25966 | 3.561922538 | -0.160605141 | 0.550642814 |
| Tbr1 | 776.6407633 | -1.417437035 | 0.181700907 |
| Gm13555 | 0 | NA | NA |
| Slc4a10 | 3386.482564 | -0.381497607 | 0.083098664 |
| Dpp4 | 80.82233867 | 1.432816797 | 0.168925278 |
| Gm13561 | 3.915321939 | 0.826589088 | 0.611334977 |
| Gm13565 | 0 | NA | NA |
| Gm13564 | 0.340218975 | 0.231098164 | 2.026787287 |
| Gcg | 1.314123019 | 1.352950745 | 1.009644394 |
| Fap | 70.46597968 | 1.134419958 | 0.319117913 |
| Ifih1 | 2560.45954 | 6.231957976 | 0.240526236 |
| Gca | 214.646939 | 0.675631081 | 0.121738124 |
| Gm13590 | 0.316156321 | 0.238002069 | 2.002529216 |
| Kcnh7 | 1930.150615 | -1.168737713 | 0.156536295 |
| Gm22020 | 4.453022388 | -1.399191347 | 0.680110232 |
| Gm13569 | 0.048949532 | -0.243569582 | 3.724945049 |
| Gm23503 | 0 | NA | NA |
| Fign | 177.2882869 | -0.505916872 | 0.092075741 |
| Gm13575 | 0 | NA | NA |
| Gm13577 | 10.38401885 | 0.235499103 | 0.324015573 |
| Gm13576 | 0 | NA | NA |
| Gm13578 | 0.248954575 | 0.201607492 | 1.836620317 |
| Grb14 | 287.0919978 | -0.417224129 | 0.095545968 |
| Cobll1 | 254.6797965 | 0.627678869 | 0.131180698 |
| Gm13594 | 1.366457952 | -0.587078252 | 0.841826577 |
| Gm13592 | 0 | NA | NA |
| Gm13593 | 0 | NA | NA |
| Slc38a11 | 11.09653691 | -1.242480724 | 0.383699689 |
| Mir6337 | 0 | NA | NA |

| | | | |
|---|---|---|---|
| Gm24138 | 0.150086409 | 0.237305146 | 3.724945049 |
| Scn3a | 1030.005389 | -0.435582035 | 0.074798868 |
| Gm13595 | 0.072025462 | 0.237312084 | 3.724945049 |
| Scn2a1 | 6263.540254 | -0.888042558 | 0.11070899 |
| Csrnp3 | 2276.632116 | -0.269509603 | 0.080433947 |
| Gm13619 | 2.502007469 | -0.668328045 | 0.643890305 |
| Gm13617 | 8.016718732 | 3.997244159 | 0.711315203 |
| Galnt3 | 26.28681886 | 0.14901113 | 0.243811455 |
| Gm28638 | 0.099106094 | 1.209894437 | 3.72418822 |
| Gm13616 | 0 | NA | NA |
| Ttc21b | 309.885117 | -0.579790996 | 0.169173678 |
| Gm13618 | 0.367269401 | -0.980960591 | 1.538258858 |
| Scn1a | 3205.479907 | -0.826811363 | 0.06614045 |
| Gm13630 | 13.73615189 | -0.133142904 | 0.318069422 |
| Gm13629 | 78.07883946 | -0.636205995 | 0.162352424 |
| Rpl35-ps1 | 0.152586256 | -0.243562192 | 2.845690286 |
| Scn9a | 113.7734046 | -0.048405475 | 0.173078414 |
| Gm13627 | 0.809136868 | -2.143781852 | 1.183661684 |
| Gm23046 | 0.7960769 | 1.168353277 | 1.295123085 |
| Scn7a | 68.38610524 | -0.120365357 | 0.181359952 |
| Gm13600 | 0.068097359 | -0.243569582 | 3.724945049 |
| Gm13602 | 0.090277459 | 0.237318756 | 3.724945049 |
| Gm13598 | 0 | NA | NA |
| Gm1322 | 0 | NA | NA |
| Gm13599 | 0 | NA | NA |
| Gm21830 | 0 | NA | NA |
| Gm23328 | 0 | NA | NA |
| Gm26727 | 0 | NA | NA |
| Xirp2 | 1.65068616 | 0.011521597 | 0.744982614 |
| Gm13601 | 10.77684248 | -0.955948227 | 0.406491336 |
| B3galt1 | 1121.981432 | 0.366921375 | 0.085806763 |
| Mir7224 | 5.441005428 | -0.327520771 | 0.408351579 |
| Gm26743 | 0.083526043 | 0.718202646 | 3.724945049 |
| Gm13604 | 0.106618422 | -0.24357403 | 3.724945049 |
| Gm13603 | 9.345943703 | 0.222748945 | 0.433555874 |
| Mir6335 | 8.512614594 | -0.398787501 | 0.442868437 |
| Gm37964 | 115.6859084 | -0.249647628 | 0.233328202 |
| Gm13607 | 9.156164614 | -0.670089586 | 0.468198968 |
| Stk39 | 1177.70026 | -0.053075802 | 0.058965508 |
| Gm13596 | 0.577259679 | -1.405025712 | 1.281643137 |
| Gm26321 | 0.044191549 | -0.243571806 | 3.724945049 |
| Gm13597 | 10.00332192 | 1.396767305 | 0.356231283 |
| 4933409G03Rik | 0.150086409 | 0.237305146 | 3.724945049 |
| 4932414N04Rik | 0.339353417 | 1.471086221 | 1.547698945 |
| Gm13612 | 0.421611163 | -0.179826467 | 1.466996783 |
| Cers6 | 1364.374441 | 0.14864362 | 0.048505225 |
| Gm37159 | 76.06150798 | -0.19463248 | 0.234940016 |

| | | | |
|---|---|---|---|
| Gm38236 | 46.00811634 | -0.544481946 | 0.308259089 |
| Gm37651 | 29.9890416 | -0.428218706 | 0.336756321 |
| Gm37747 | 50.62951173 | -0.107127352 | 0.250895631 |
| Gm37666 | 22.07432209 | -0.341056819 | 0.316936958 |
| Gm38377 | 75.00233876 | -0.181244674 | 0.196619405 |
| Nostrin | 27.43524618 | 1.083776878 | 0.254367556 |
| Spc25 | 14.006815 | 2.373388518 | 0.424338552 |
| G6pc2 | 0.378093966 | 16.68573949 | 3.683449809 |
| Abcb11 | 1.943768998 | 2.416236912 | 0.907002655 |
| Gm13613 | 0 | NA | NA |
| Gm13614 | 0 | NA | NA |
| Dhrs9 | 8.333878881 | 4.325425596 | 0.695082555 |
| Lrp2 | 28.81441089 | 0.405690175 | 0.289208318 |
| Bbs5 | 149.8333186 | -0.23293715 | 0.118347972 |
| Klhl41 | 8.40861694 | -0.176885239 | 0.386276258 |
| Fastkd1 | 167.3175697 | -0.451163374 | 0.091103709 |
| Ppig | 1069.962896 | -0.127856278 | 0.080512389 |
| Gm13622 | 1.277078827 | 0.509819639 | 1.05886815 |
| Ccdc173 | 33.00873947 | -0.303229322 | 0.201261 |
| Gm13623 | 0.146451277 | -0.243562218 | 2.882358805 |
| Phospho2 | 316.3809453 | 0.272897465 | 0.122845548 |
| Gm13624 | 0.441815039 | -0.24357922 | 3.406154375 |
| Klhl23 | 607.9612626 | -0.550536044 | 0.093411454 |
| Ssb | 1044.348789 | 0.257061176 | 0.092251845 |
| Mettl5 | 79.04134513 | 0.238676605 | 0.128768899 |
| Gm13625 | 2.609485436 | -0.57248144 | 0.618516524 |
| Mettl5os | 3.090821495 | -1.011071815 | 0.607266766 |
| Ubr3 | 4154.724246 | -0.351066914 | 0.068965431 |
| Gm13631 | 0.295881152 | 0.813238491 | 2.049360465 |
| Myo3b | 31.90249792 | 0.276003599 | 0.261649075 |
| Rpl9-ps7 | 51.73231683 | 0.526671242 | 0.202190648 |
| Sp5 | 11.17023444 | 1.044535305 | 0.319724747 |
| Gad1os | 2.192682401 | 1.396095496 | 0.650000451 |
| Erich2os | 3.966806659 | 2.876875686 | 0.623183483 |
| Erich2 | 9.894614024 | 1.690104828 | 0.362094367 |
| Gm37026 | 0.230844305 | -0.152631994 | 2.125624719 |
| Gad1 | 4252.286034 | -0.200308217 | 0.092210301 |
| Gm16510 | 0.089814205 | -0.24357403 | 3.724945049 |
| Gm26558 | 1.60906251 | 0.803959756 | 0.86851862 |
| Gorasp2 | 1047.261539 | 0.354300825 | 0.068609539 |
| Tlk1 | 1317.27295 | -0.13981325 | 0.056991887 |
| Gm13633 | 0.219273179 | 1.591795885 | 2.133770018 |
| Gm13632 | 0.086908494 | -0.72445792 | 3.724945049 |
| Mettl8 | 114.1898919 | -0.523811698 | 0.159058334 |
| Dcaf17 | 286.7511595 | -0.335748831 | 0.130808058 |
| Cybrd1 | 58.06687379 | 0.395338558 | 0.281258436 |
| Dync1i2 | 1658.410193 | -0.014400307 | 0.043032456 |

| | | | |
|---|---|---|---|
| Slc25a12 | 1679.553867 | -0.108112081 | 0.0462057 |
| Gm13641 | 0.124853745 | 0.237309606 | 3.724945049 |
| Hat1 | 224.6790855 | 0.843585845 | 0.130830364 |
| Gm13642 | 0.843921636 | 1.616463905 | 1.212414372 |
| Gm23166 | 0 | NA | NA |
| Metap1d | 226.2128202 | -0.506623677 | 0.093988768 |
| Gm23253 | 0.412462835 | 0.186558279 | 1.49993992 |
| Dlx1as | 32.54297881 | -1.755408034 | 0.295767769 |
| Dlx1 | 196.3712922 | -0.523013316 | 0.299231505 |
| Dlx2 | 66.09861157 | -0.461712909 | 0.266140612 |
| Platr26 | 0.08486385 | 0.717551239 | 3.724898127 |
| Gm13647 | 0.342086031 | -0.243584783 | 3.724945049 |
| Itga6 | 388.4782796 | 0.524369076 | 0.139637868 |
| Gm13663 | 0.038981395 | 0.718207094 | 3.724945049 |
| Gm17250 | 1.158557209 | 1.001081921 | 1.055105918 |
| Gm13662 | 0 | NA | NA |
| Gm13739 | 0 | NA | NA |
| Pdk1 | 468.9053974 | -0.347231406 | 0.152657057 |
| Gm13746 | 0 | NA | NA |
| Rapgef4 | 3912.521623 | -0.578791192 | 0.112502572 |
| Gm26408 | 0 | NA | NA |
| Rapgef4os1 | 0.258731086 | -0.242869406 | 2.063229988 |
| Rapgef4os2 | 0.479549184 | -0.626371886 | 1.419752363 |
| Rapgef4os3 | 1.315840983 | -0.041920013 | 1.082291927 |
| Zak | 235.4261635 | 0.254187915 | 0.087262978 |
| Zakit | 0.427144302 | -0.483546397 | 1.382109505 |
| Rpl36a-ps4 | 0.431106946 | 0.198382954 | 1.379730368 |
| Cdca7 | 32.93658422 | 0.027311856 | 0.192089032 |
| Rps13-ps8 | 0 | NA | NA |
| Eif1-ps3 | 0 | NA | NA |
| Pex13-ps | 0.061920613 | 0.718207094 | 3.724945049 |
| Ak3l2-ps | 18.80802623 | -0.252102925 | 0.245090928 |
| 8430437L04Rik | 0.222986776 | 1.155651747 | 2.149646621 |
| 6430710C18Rik | 1.790214168 | -0.026672589 | 0.685487451 |
| Sp3 | 1150.979175 | 0.382768161 | 0.087018738 |
| Gm25960 | 2.212404147 | -0.646203614 | 0.71261978 |
| Hspe1-ps5 | 0 | NA | NA |
| Gm11084 | 3.542390089 | 0.101079764 | 0.500120272 |
| Sp3os | 52.06250527 | -0.602427295 | 0.176512755 |
| Ola1 | 736.2602298 | -0.212280703 | 0.069536431 |
| Gm13665 | 0 | NA | NA |
| Gm13666 | 0.293512495 | 1.282243191 | 2.021458828 |
| Gm13664 | 0 | NA | NA |
| Sp9 | 90.07522838 | -0.867995623 | 0.245617813 |
| Cir1 | 188.1924414 | -0.195158813 | 0.109791869 |
| Scrn3 | 297.9077961 | -0.196426188 | 0.109909979 |
| Gpr155 | 760.0828086 | -0.683682796 | 0.070867266 |

| | | | |
|---|---|---|---|
| Gm13703 | 0.773921687 | 1.635876455 | 1.275023707 |
| Gm13709 | 0.142443712 | -0.724458029 | 3.596298704 |
| D230022J07Rik | 3.614518035 | -0.423579805 | 0.513648854 |
| Gm13707 | 1.060348673 | -0.064680827 | 0.910609184 |
| Wipf1 | 548.8928039 | 1.559733335 | 0.143166995 |
| Gm13708 | 0 | NA | NA |
| Chrna1 | 3.856870735 | 1.68154769 | 0.67217289 |
| Chrna1os | 4.004686001 | 0.054822654 | 0.512800375 |
| Chn1os3 | 3.866965461 | 0.48866455 | 0.505052871 |
| Chn1 | 4895.40377 | -0.886218636 | 0.117640744 |
| Chn1os1 | 6.863703603 | -0.772998255 | 0.412850495 |
| Gm13640 | 0 | NA | NA |
| Atf2 | 1720.92591 | -0.194845508 | 0.057054154 |
| Gm27421 | 0 | NA | NA |
| Gm10822 | 3.167618981 | -0.485122538 | 0.528630631 |
| Atp5g3 | 1417.808151 | -0.082569042 | 0.094524348 |
| Gm13668 | 0 | NA | NA |
| A630050E04Rik | 0.102891481 | -0.243576254 | 3.724945049 |
| Gm28640 | 0 | NA | NA |
| Gm13667 | 0 | NA | NA |
| 4930441J16Rik | 0.04393175 | -0.243571806 | 3.724945049 |
| Lnp | 1061.691288 | -0.269892946 | 0.072400226 |
| 4930445N08Rik | 0 | NA | NA |
| Mrpl23-ps1 | 10.74443234 | 0.372772274 | 0.445214801 |
| Gm22975 | 0 | NA | NA |
| Gm13672 | 0 | NA | NA |
| Evx2 | 6.495853999 | 0.023456009 | 0.37882242 |
| Hoxd13 | 0.292684482 | 0.714701903 | 1.632696722 |
| Hoxd12 | 0 | NA | NA |
| Hoxd11 | 0 | NA | NA |
| Gm28309 | 0 | NA | NA |
| Hoxd10 | 0 | NA | NA |
| Gm14396 | 0 | NA | NA |
| Gm28793 | 0 | NA | NA |
| Hoxd9 | 0 | NA | NA |
| AL928644.1 | 0 | NA | NA |
| Hoxd3os1 | 0.132812548 | 0.218958578 | 2.965535997 |
| Hoxd8 | 0.583508756 | 1.216262137 | 3.393076145 |
| Hoxd3 | 4.727112919 | -0.070720758 | 0.652849382 |
| Hoxd4 | 1.678255395 | -0.519151845 | 0.934669579 |
| Mir10b | 0.042716945 | -0.243571806 | 3.724945049 |
| Gm28230 | 0.047948698 | -0.243569582 | 3.724945049 |
| Haglr | 0.983322447 | -0.896844737 | 1.14779807 |
| Gm14424 | 0.086930093 | 0.236669296 | 3.724898107 |
| Gm14426 | 0 | NA | NA |
| Hoxd1 | 1.509597968 | 1.551414222 | 0.998788858 |
| Mtx2 | 366.8248721 | -0.025282247 | 0.080387634 |

| | | | |
|---|---|---|---|
| Gm14425 | 13.47600813 | -0.657615938 | 0.331421842 |
| Gm14427 | 0 | NA | NA |
| n-R5s198 | 0 | NA | NA |
| Gm13652 | 1.412897596 | -0.022879452 | 0.774657229 |
| 2600014E21Rik | 28.30992819 | -0.909347703 | 0.2266282 |
| A330043C09Rik | 1.006727404 | 1.176743972 | 1.084352917 |
| Rps6-ps4 | 71.2506606 | 0.730212785 | 0.143777384 |
| Gm13653 | 0.886214715 | 0.528427818 | 1.067569532 |
| Gm37243 | 0 | NA | NA |
| Gm13658 | 0.044544648 | 0.237309859 | 3.724945049 |
| Gm13660 | 0 | NA | NA |
| Gm13655 | 0.042716945 | -0.243571806 | 3.724945049 |
| Gm24574 | 0.04393175 | -0.243571806 | 3.724945049 |
| Hnrnpa3 | 910.2237922 | -0.032920595 | 0.077048629 |
| Nfe2l2 | 396.3091985 | 1.740051943 | 0.156097167 |
| Gm24996 | 0.882521852 | -0.191987023 | 0.963257375 |
| E030042O20Rik | 0.923975021 | 1.640305631 | 1.011145029 |
| Gm13656 | 0 | NA | NA |
| Gm13657 | 0.629646617 | 2.53940742 | 1.439629272 |
| Gm13669 | 1.634157898 | 2.990006739 | 1.035524771 |
| Gm13670 | 1.659798175 | -1.185720172 | 0.785973201 |
| Agps | 976.4418691 | 0.068766922 | 0.058390213 |
| Ttc30b | 125.5045713 | 0.18135517 | 0.119935532 |
| Ttc30a2 | 2.514664855 | 0.106713595 | 0.652608997 |
| Ttc30a1 | 46.83376903 | 0.220924525 | 0.186956104 |
| Pde11a | 85.82156519 | -0.429101172 | 0.124118629 |
| AL845373.1 | 0 | NA | NA |
| Cyct | 0 | NA | NA |
| Rbm45 | 248.2191173 | 0.222288955 | 0.07930752 |
| Osbpl6 | 1518.253489 | -0.487467197 | 0.089310971 |
| Gm23164 | 1.915350769 | -1.222473124 | 0.841164398 |
| Prkra | 162.1113476 | 0.155843955 | 0.093798075 |
| Dfnb59 | 0.622664233 | -1.500733524 | 1.359477554 |
| Fkbp7 | 23.5112496 | 0.247726898 | 0.21665665 |
| Plekha3 | 466.6346452 | 0.248823425 | 0.077045631 |
| Ttn | 182.6205874 | -0.303286688 | 0.222622585 |
| Gm13938 | 1.375616079 | -0.435184654 | 1.068068448 |
| Ccdc141 | 278.830304 | -0.060377237 | 0.116085821 |
| AL928681.1 | 1.008651653 | -1.851133391 | 1.117304653 |
| Sestd1 | 1270.998036 | -0.436771256 | 0.075646752 |
| Gm13650 | 19.02386222 | -0.794547575 | 0.390137392 |
| Gm13651 | 3.760012592 | 0.3430658 | 0.531869996 |
| Gm13943 | 0 | NA | NA |
| Gm13649 | 0.072025462 | 0.237312084 | 3.724945049 |
| Zfp385b | 633.2440097 | -0.655800717 | 0.100453178 |
| Gm13944 | 5.255555274 | 0.286854689 | 0.602337393 |
| Gm13661 | 1.194926524 | -0.930881294 | 1.045291524 |

| | | | |
|---|---|---|---|
| Gm13727 | 0.386897144 | -1.291981643 | 1.385494013 |
| Cwc22 | 371.0030043 | 0.080233261 | 0.106170532 |
| Gm13726 | 0 | NA | NA |
| 4930440I19Rik | 0.335959249 | 1.421841708 | 2.017601087 |
| Gm14461 | 0.674729951 | 2.070820237 | 1.218813152 |
| Gm14460 | 0 | NA | NA |
| Gm23149 | 0 | NA | NA |
| Gm14464 | 0.757266105 | 1.913155822 | 1.218848923 |
| Gm14463 | 0.136194719 | -0.724453471 | 3.724945049 |
| Gm17014 | 0 | NA | NA |
| Ube2e3 | 827.2373707 | -0.081137146 | 0.083865396 |
| Gm37004 | 1.348672834 | 2.342194286 | 0.91530764 |
| Gm14466 | 0.233003758 | 2.148481989 | 2.52304222 |
| Gm14469 | 0.066050875 | 0.71820487 | 3.724945049 |
| Gm14465 | 0.075043205 | 0.237309859 | 3.724945049 |
| Itga4 | 452.6472453 | 1.002422437 | 0.139425144 |
| Cerkl | 55.52887644 | -0.654363888 | 0.270755657 |
| Gm22606 | 0 | NA | NA |
| Gm13673 | 0.097139547 | 0.718207094 | 3.724945049 |
| Neurod1 | 741.6134161 | -0.64275829 | 0.28470513 |
| Gm26755 | 3.778983902 | 1.26666298 | 0.59310804 |
| Ssfa2 | 1354.956421 | 1.254716496 | 0.138149639 |
| Ppp1r1c | 7.321389695 | 1.316868198 | 0.433647791 |
| Pde1a | 1335.258322 | -0.48471739 | 0.139588777 |
| Gm13674 | 0 | NA | NA |
| Prdx6b | 0 | NA | NA |
| Dnajc10 | 1132.405827 | 0.013013051 | 0.050211682 |
| Gm13752 | 0 | NA | NA |
| Frzb | 132.7433967 | -1.024250856 | 0.177282914 |
| Gm13687 | 0 | NA | NA |
| Nckap1 | 5774.673254 | -0.322122191 | 0.046718751 |
| Gm13689 | 37.9418997 | -0.814162976 | 0.305207109 |
| Gm24461 | 0 | NA | NA |
| Gm13688 | 12.81351646 | 0.054396245 | 0.303296321 |
| Dusp19 | 119.9719043 | -0.34350569 | 0.114187565 |
| Nup35 | 181.7556722 | 0.361913642 | 0.127239909 |
| mmu-mir-684-1 | 0.866431109 | 1.037496659 | 1.122631827 |
| Gm13690 | 0.627304132 | -0.176370741 | 1.32305132 |
| Gm25420 | 0.262453343 | 0.762050116 | 2.103818725 |
| Gm4735 | 399.2680756 | 0.182995577 | 0.114283474 |
| Gm22652 | 0 | NA | NA |
| Gm13700 | 0 | NA | NA |
| Gm13755 | 0.332110845 | 0.683307314 | 1.598753998 |
| Gm13756 | 0 | NA | NA |
| Gm23452 | 0 | NA | NA |
| Gm23900 | 0 | NA | NA |
| Zfp804a | 500.2371726 | 0.243616399 | 0.126781175 |

| | | | |
|---|---|---|---|
| Gm13677 | 0 | NA | NA |
| A130030D18Rik | 0.048949532 | -0.243569582 | 3.724945049 |
| Gm13675 | 0.141830814 | -1.161222926 | 3.589786304 |
| Gm13676 | 0 | NA | NA |
| Gm13678 | 0 | NA | NA |
| Fsip2 | 0.617250809 | -0.702635174 | 1.335249532 |
| Gm28675 | 0.121388914 | 0.718200421 | 3.724945049 |
| Gm13680 | 10.62432231 | 0.756562495 | 0.322918113 |
| Gm13679 | 0 | NA | NA |
| Gm13682 | 0 | NA | NA |
| Gm13681 | 0 | NA | NA |
| Gm13683 | 0 | NA | NA |
| Gm13685 | 0 | NA | NA |
| Zc3h15 | 953.3117506 | 0.151604299 | 0.104542753 |
| Gm13686 | 0 | NA | NA |
| Itgav | 2001.084212 | 0.262407075 | 0.077462437 |
| Gm13684 | 0.113999573 | 0.23732098 | 3.724945049 |
| Dnmt3l-ps1 | 0.097139547 | 0.718207094 | 3.724945049 |
| Fam171b | 2724.632699 | -0.504423374 | 0.086018943 |
| Zswim2 | 0.405375818 | 0.237332723 | 1.606645898 |
| Gm13698 | 0 | NA | NA |
| Gm13693 | 0 | NA | NA |
| Gm13695 | 0 | NA | NA |
| Gm13696 | 0 | NA | NA |
| Gm13694 | 0.047948698 | -0.243569582 | 3.724945049 |
| Gm13697 | 0.061920613 | 0.718207094 | 3.724945049 |
| Gm13691 | 0 | NA | NA |
| Gm13692 | 0 | NA | NA |
| Gm13702 | 0 | NA | NA |
| Gm25972 | 0.044191549 | -0.243571806 | 3.724945049 |
| Gm13711 | 0.180501107 | -0.243571141 | 2.711307924 |
| Calcrl | 175.101438 | 0.665423303 | 0.123214359 |
| Tfpi | 63.47998308 | 0.916592973 | 0.145690723 |
| Gm13710 | 7.944981639 | -0.706111029 | 0.506741707 |
| Ctnnd1 | 284.3298198 | 0.023915688 | 0.092326089 |
| Gm28635 | 0 | NA | NA |
| Btbd18 | 0.061920613 | 0.718207094 | 3.724945049 |
| 2700094K13Rik | 93.3688872 | -0.013229892 | 0.110268895 |
| Tmx2 | 1256.227034 | 0.123457458 | 0.057633419 |
| Med19 | 347.2590899 | 0.631194548 | 0.089862308 |
| Zdhhc5 | 1049.31998 | 0.421282786 | 0.063467732 |
| n-R5s200 | 2.639998601 | -0.599263361 | 0.688744226 |
| Gm27942 | 0.072025462 | 0.237312084 | 3.724945049 |
| Clp1 | 87.12695073 | 0.429815114 | 0.160031599 |
| Ypel4 | 577.9281882 | -0.291561357 | 0.115870283 |
| Mir130a | 0.283042562 | 0.075011513 | 3.698929216 |
| Gm19426 | 6.293410423 | 0.078126014 | 0.451441085 |

| | | | |
|---|---|---|---|
| Serping1 | 1376.102164 | 5.445250096 | 0.281059494 |
| Ube2l6 | 958.9461895 | 6.043080347 | 0.28625847 |
| Smtnl1 | 0.1140246 | 0.718202646 | 3.724945049 |
| Timm10 | 121.5002785 | 0.339601893 | 0.121362343 |
| Slc43a1 | 4.475148185 | 1.840284551 | 0.572313837 |
| Rtn4rl2 | 239.9515241 | -0.118712074 | 0.189643993 |
| Gm24332 | 0.173129629 | 0.685170389 | 2.724725116 |
| Slc43a3 | 68.69465978 | 3.088228458 | 0.33206463 |
| Prg2 | 0.484678492 | 1.998376259 | 3.689457286 |
| Prg3 | 0.205912129 | 1.252872323 | 2.595364951 |
| P2rx3 | 14.94560384 | 0.001113649 | 0.29486743 |
| Ssrp1 | 920.1403945 | 0.45679381 | 0.068127981 |
| Tnks1bp1 | 1348.991537 | 0.550341798 | 0.083670085 |
| 4930443O20Rik | 0.247670153 | -1.091040403 | 2.092819555 |
| Aplnr | 18.49192457 | 1.13372651 | 0.261398002 |
| Gm13712 | 0.447666499 | 1.474808332 | 1.721471971 |
| Gm13713 | 6.805315018 | 2.570673091 | 0.497631241 |
| Lrrc55 | 258.0702094 | 0.030463661 | 0.126649838 |
| Gm13716 | 0.932468966 | 0.389361832 | 1.305219379 |
| Gm13715 | 0.832268342 | 2.661071139 | 1.127489729 |
| Gm13714 | 0.047948698 | -0.243569582 | 3.724945049 |
| Olfr987 | 0.046345709 | 0.71820487 | 3.724945049 |
| Olfr988 | 0.042716945 | -0.243571806 | 3.724945049 |
| Olfr990-ps1 | 0 | NA | NA |
| Olfr992 | 0 | NA | NA |
| Olfr993 | 0 | NA | NA |
| Gm23716 | 0 | NA | NA |
| Olfr994 | 0.042716945 | -0.243571806 | 3.724945049 |
| Olfr995 | 0 | NA | NA |
| 4833423E24Rik | 3.225753529 | -0.262928296 | 0.528803249 |
| Olfr996 | 0 | NA | NA |
| Olfr997-ps1 | 0 | NA | NA |
| Olfr998 | 0.038981395 | 0.718207094 | 3.724945049 |
| Olfr999-ps1 | 0 | NA | NA |
| Olfr1000 | 0 | NA | NA |
| Olfr1001-ps1 | 0.046345709 | 0.71820487 | 3.724945049 |
| Olfr1002 | 0 | NA | NA |
| Olfr1003-ps1 | 0 | NA | NA |
| Olfr1004-ps1 | 0 | NA | NA |
| Olfr154 | 0 | NA | NA |
| Olfr1006 | 0.048949532 | -0.243569582 | 3.724945049 |
| Olfr1008 | 0 | NA | NA |
| Olfr1009 | 0 | NA | NA |
| Olfr1010 | 0.038981395 | 0.718207094 | 3.724945049 |
| Olfr1012 | 0.089062654 | 0.237318756 | 3.724945049 |
| Olfr1013 | 0 | NA | NA |
| Olfr1014 | 0 | NA | NA |

| | | | |
|---|---|---|---|
| Olfr1015 | 0 | NA | NA |
| Olfr1016 | 0.04393175 | -0.243571806 | 3.724945049 |
| Olfr1017-ps1 | 0 | NA | NA |
| Gm13720 | 0 | NA | NA |
| Olfr1018 | 0 | NA | NA |
| Olfr1019 | 0.062426873 | 0.237312084 | 3.724945049 |
| Olfr1020 | 0 | NA | NA |
| Olfr1021-ps1 | 0 | NA | NA |
| Olfr1022 | 0 | NA | NA |
| Gm24037 | 0 | NA | NA |
| Olfr1023 | 0 | NA | NA |
| Olfr1024 | 0 | NA | NA |
| Olfr1025-ps1 | 0 | NA | NA |
| Olfr1026 | 0 | NA | NA |
| Olfr1027-ps1 | 0 | NA | NA |
| Olfr1028 | 0 | NA | NA |
| Olfr1029 | 0.108772582 | 0.718202646 | 3.724945049 |
| Olfr1030 | 0 | NA | NA |
| Olfr1031 | 0 | NA | NA |
| Olfr1032 | 0 | NA | NA |
| Olfr1033 | 0 | NA | NA |
| Olfr1034 | 0 | NA | NA |
| Olfr1035-ps1 | 0 | NA | NA |
| Olfr1036 | 0 | NA | NA |
| Olfr1037 | 0 | NA | NA |
| Gm13722 | 0.13742593 | -0.724459914 | 2.939744481 |
| Olfr1038-ps | 0 | NA | NA |
| Olfr1039 | 0 | NA | NA |
| Olfr1040 | 0 | NA | NA |
| Olfr1042 | 0 | NA | NA |
| Olfr1043 | 0 | NA | NA |
| Olfr1044 | 0 | NA | NA |
| Olfr52 | 0.108772582 | 0.718202646 | 3.724945049 |
| Olfr1045 | 0 | NA | NA |
| Olfr1046 | 0 | NA | NA |
| Olfr1047 | 0 | NA | NA |
| Olfr1048 | 0 | NA | NA |
| Olfr1049 | 0 | NA | NA |
| Olfr1050-ps1 | 0 | NA | NA |
| Olfr1051 | 0 | NA | NA |
| Olfr1052 | 0.044544648 | 0.237309859 | 3.724945049 |
| Olfr1053 | 0 | NA | NA |
| Gm13719 | 2.60552406 | 0.435889082 | 0.678595439 |
| Olfr1054 | 0 | NA | NA |
| Olfr1055 | 0 | NA | NA |
| Olfr1056 | 0 | NA | NA |
| Olfr1057 | 0 | NA | NA |

| | | | |
|---|---|---|---|
| Olfr1058 | 0 | NA | NA |
| Olfr1060 | 0 | NA | NA |
| Olfr1061 | 0 | NA | NA |
| Olfr1062 | 0 | NA | NA |
| Olfr1063-ps1 | 0 | NA | NA |
| Olfr1065 | 0 | NA | NA |
| Olfr1066 | 0 | NA | NA |
| Olfr1069-ps1 | 0 | NA | NA |
| Olfr228 | 0 | NA | NA |
| Olfr1073-ps1 | 0 | NA | NA |
| Olfr1076 | 0 | NA | NA |
| Olfr1077-ps1 | 0 | NA | NA |
| Olfr1078-ps1 | 0 | NA | NA |
| Olfr1079 | 0 | NA | NA |
| Olfr1080 | 0 | NA | NA |
| Olfr1081-ps1 | 0 | NA | NA |
| Olfr1082 | 0 | NA | NA |
| Olfr1083-ps | 0 | NA | NA |
| Olfr1084 | 0 | NA | NA |
| Olfr1085 | 0 | NA | NA |
| Olfr1086 | 0 | NA | NA |
| Olfr1087 | 0 | NA | NA |
| Olfr1089 | 0 | NA | NA |
| Olfr1090 | 0 | NA | NA |
| Olfr1093 | 0 | NA | NA |
| BX284685.1 | 0 | NA | NA |
| Olfr141 | 0 | NA | NA |
| Olfr1094 | 0 | NA | NA |
| Olfr1095 | 0 | NA | NA |
| Olfr1096-ps1 | 0 | NA | NA |
| Gm13723 | 0 | NA | NA |
| Olfr1097 | 0 | NA | NA |
| Olfr1098 | 0 | NA | NA |
| Olfr1099 | 0.048949532 | -0.243569582 | 3.724945049 |
| Olfr1100 | 0 | NA | NA |
| Olfr1101 | 0 | NA | NA |
| Olfr1102 | 0 | NA | NA |
| Olfr1103-ps1 | 0 | NA | NA |
| Olfr1104 | 0 | NA | NA |
| Olfr1105 | 0 | NA | NA |
| Olfr1106 | 0 | NA | NA |
| Olfr1107 | 0 | NA | NA |
| Olfr1109 | 0.046345709 | 0.71820487 | 3.724945049 |
| Olfr259 | 0 | NA | NA |
| Gm13733 | 0 | NA | NA |
| Olfr1110 | 0 | NA | NA |
| Gm13734 | 0 | NA | NA |

| | | | |
|---|---|---|---|
| Olfr1111 | 0 | NA | NA |
| Nlk-ps1 | 0.21574249 | -0.7244605 | 2.174796322 |
| Olfr1112 | 0 | NA | NA |
| Gm13730 | 0 | NA | NA |
| Olfr1113 | 0 | NA | NA |
| Gm13735 | 0 | NA | NA |
| Olfr1115 | 0 | NA | NA |
| Gm27438 | 0 | NA | NA |
| Gm13732 | 0 | NA | NA |
| Olfr1116 | 0 | NA | NA |
| Gm13728 | 0.042716945 | -0.243571806 | 3.724945049 |
| Olfr1117-ps1 | 0 | NA | NA |
| Gm13729 | 0 | NA | NA |
| Olfr1118 | 0 | NA | NA |
| Gm13737 | 0 | NA | NA |
| Gm13731 | 0 | NA | NA |
| Olfr1120 | 0 | NA | NA |
| Olfr1121 | 0 | NA | NA |
| Olfr1122 | 0 | NA | NA |
| Gm1826 | 8.12848589 | 0.253685299 | 0.346099286 |
| Olfr1123 | 0 | NA | NA |
| Olfr1124 | 0 | NA | NA |
| Olfr1126 | 0 | NA | NA |
| Pramel7 | 0 | NA | NA |
| Pramel6 | 0 | NA | NA |
| Gm13738 | 0 | NA | NA |
| Olfr1127-ps1 | 0 | NA | NA |
| Gm13736 | 22.02889899 | 0.443286519 | 0.274181233 |
| Olfr153 | 0 | NA | NA |
| Olfr1128 | 0 | NA | NA |
| Gm13740 | 0.18866314 | -0.685656973 | 2.661691149 |
| Gm13744 | 0 | NA | NA |
| Gm13743 | 3.080495725 | 0.289524173 | 0.592438126 |
| Olfr1129 | 0 | NA | NA |
| Olfr1130 | 0 | NA | NA |
| Olfr1131 | 0 | NA | NA |
| Olfr1132 | 0 | NA | NA |
| Gm13742 | 0 | NA | NA |
| Olfr1133 | 0 | NA | NA |
| Gm13741 | 0 | NA | NA |
| Olfr1134 | 0 | NA | NA |
| Olfr1135 | 0 | NA | NA |
| Olfr1136 | 0 | NA | NA |
| Olfr1137 | 0 | NA | NA |
| Olfr1138 | 0 | NA | NA |
| Olfr1140 | 0 | NA | NA |
| Olfr1141 | 0 | NA | NA |

| | | | |
|---|---|---|---|
| Olfr152 | 0 | NA | NA |
| Olfr1143 | 0 | NA | NA |
| Olfr1144-ps1 | 0 | NA | NA |
| Olfr1145 | 0 | NA | NA |
| Olfr1146-ps1 | 0 | NA | NA |
| Olfr1147-ps1 | 0 | NA | NA |
| Olfr1148 | 0 | NA | NA |
| Olfr1149-ps1 | 0 | NA | NA |
| Olfr1150-ps1 | 0 | NA | NA |
| Olfr1151 | 0 | NA | NA |
| Olfr1152 | 0 | NA | NA |
| Gm13745 | 0 | NA | NA |
| Olfr1153 | 0 | NA | NA |
| Olfr1154 | 0 | NA | NA |
| Olfr1155 | 0 | NA | NA |
| Olfr1156 | 0 | NA | NA |
| Olfr1157 | 0 | NA | NA |
| Olfr74 | 0 | NA | NA |
| Olfr1158 | 0 | NA | NA |
| Olfr1159-ps1 | 0 | NA | NA |
| Olfr1160 | 0 | NA | NA |
| Olfr1161 | 0 | NA | NA |
| Olfr73 | 0 | NA | NA |
| Gm13747 | 0 | NA | NA |
| Olfr1162 | 0 | NA | NA |
| Olfr1163 | 0 | NA | NA |
| Olfr1164 | 0 | NA | NA |
| Olfr1165-ps | 0.046345709 | 0.71820487 | 3.724945049 |
| Olfr1166 | 0 | NA | NA |
| Olfr1167 | 0 | NA | NA |
| Olfr1168 | 0.046345709 | 0.71820487 | 3.724945049 |
| Olfr1169-ps1 | 0 | NA | NA |
| Olfr1170 | 0 | NA | NA |
| Olfr1171-ps1 | 0 | NA | NA |
| Gm13753 | 0 | NA | NA |
| Olfr1172-ps1 | 0 | NA | NA |
| Olfr1173 | 0 | NA | NA |
| Olfr1174-ps | 0 | NA | NA |
| Olfr1175-ps | 0 | NA | NA |
| Olfr1176 | 0.061920613 | 0.718207094 | 3.724945049 |
| Olfr1177-ps | 0 | NA | NA |
| Olfr1178 | 0 | NA | NA |
| Olfr1179 | 0 | NA | NA |
| Olfr1180 | 0 | NA | NA |
| Olfr1181 | 0 | NA | NA |
| Olfr1182 | 0 | NA | NA |
| Gm13757 | 0 | NA | NA |

| | | | |
|---|---|---|---|
| Olfr1183 | 0 | NA | NA |
| Olfr1184 | 0 | NA | NA |
| Olfr1185-ps1 | 0 | NA | NA |
| Olfr1186 | 0 | NA | NA |
| Olfr1187-ps1 | 0 | NA | NA |
| Olfr1188 | 0 | NA | NA |
| Gm13758 | 0 | NA | NA |
| Olfr1189 | 0 | NA | NA |
| Olfr475-ps1 | 0 | NA | NA |
| Olfr1191-ps1 | 0.062426873 | 0.237312084 | 3.724945049 |
| Gm13759 | 0 | NA | NA |
| Olfr1192-ps1 | 0 | NA | NA |
| Olfr1193 | 0.042716945 | -0.243571806 | 3.724945049 |
| Olfr1195 | 0.131862677 | -0.243562287 | 2.97735462 |
| Olfr1196 | 0 | NA | NA |
| Olfr1197 | 0 | NA | NA |
| Olfr1198 | 0 | NA | NA |
| Olfr1199 | 0 | NA | NA |
| Gm13760 | 0 | NA | NA |
| Olfr1200 | 0 | NA | NA |
| Olfr1201 | 0 | NA | NA |
| Olfr1202 | 0 | NA | NA |
| Olfr1205 | 0 | NA | NA |
| Olfr1204 | 0 | NA | NA |
| Olfr1206 | 0 | NA | NA |
| Olfr1208 | 0 | NA | NA |
| Olfr1209 | 0 | NA | NA |
| Olfr1211 | 0.068097359 | -0.243569582 | 3.724945049 |
| Gm13763 | 0 | NA | NA |
| Olfr1212 | 0 | NA | NA |
| Olfr1213 | 0 | NA | NA |
| Gm13762 | 0 | NA | NA |
| Olfr1214 | 0 | NA | NA |
| Olfr1215 | 0 | NA | NA |
| Olfr1216 | 0 | NA | NA |
| Olfr1217 | 0 | NA | NA |
| Olfr1218 | 0 | NA | NA |
| Olfr1219 | 0.045882455 | 0.237312084 | 3.724945049 |
| Olfr1220 | 0.095435502 | 0.718202646 | 3.724945049 |
| Olfr1221 | 0 | NA | NA |
| Gm13764 | 0.048949532 | -0.243569582 | 3.724945049 |
| Olfr1222 | 0.048949532 | -0.243569582 | 3.724945049 |
| Olfr1223 | 0 | NA | NA |
| Olfr1224-ps1 | 0 | NA | NA |
| Olfr1225 | 0 | NA | NA |
| Gm13761 | 0 | NA | NA |
| Olfr1226 | 0 | NA | NA |

| | | | |
|---|---|---|---|
| Olfr1228 | 0.062426873 | 0.237312084 | 3.724945049 |
| Olfr1229 | 0 | NA | NA |
| Olfr1230 | 0 | NA | NA |
| Olfr1231 | 0.046345709 | 0.71820487 | 3.724945049 |
| Olfr1232 | 0 | NA | NA |
| Olfr1233 | 0.048949532 | -0.243569582 | 3.724945049 |
| Olfr1234 | 0.048949532 | -0.243569582 | 3.724945049 |
| Olfr1235-ps1 | 0 | NA | NA |
| Olfr1238 | 0 | NA | NA |
| Olfr1239 | 0 | NA | NA |
| Olfr1240 | 0 | NA | NA |
| Olfr1241 | 0 | NA | NA |
| Olfr1242 | 0 | NA | NA |
| Olfr1243 | 0.106618422 | -0.24357403 | 3.724945049 |
| Olfr1245 | 0 | NA | NA |
| Olfr1246 | 0.045882455 | 0.237312084 | 3.724945049 |
| Olfr1247 | 0 | NA | NA |
| Olfr1248 | 0 | NA | NA |
| Olfr1249 | 0 | NA | NA |
| Olfr1250 | 0.042716945 | -0.243571806 | 3.724945049 |
| Olfr1251 | 0 | NA | NA |
| Olfr1252 | 0 | NA | NA |
| Olfr1253 | 0 | NA | NA |
| Olfr1254 | 0.04393175 | -0.243571806 | 3.724945049 |
| Olfr1255 | 0 | NA | NA |
| Olfr1256 | 0 | NA | NA |
| Olfr48 | 0.156220659 | -1.190320593 | 2.817542717 |
| Olfr1257 | 0 | NA | NA |
| Olfr1258 | 0.254495662 | 1.772500168 | 2.083803576 |
| Olfr1259 | 0 | NA | NA |
| Olfr1260 | 0.042716945 | -0.243571806 | 3.724945049 |
| Olfr1261 | 0.198041555 | 1.570941826 | 2.587932611 |
| Olfr1262 | 0.31782415 | 2.175537539 | 3.698236668 |
| Olfr1263 | 0 | NA | NA |
| Olfr1264 | 0.068097359 | -0.243569582 | 3.724945049 |
| Olfr1265 | 0.097139547 | 0.718207094 | 3.724945049 |
| Olfr140 | 0.229124067 | 1.267476167 | 2.560995054 |
| Olfr1267-ps1 | 0.049553047 | 0.71820487 | 3.724945049 |
| Olfr1268-ps1 | 0.202674043 | 1.045489697 | 2.606063585 |
| Olfr1269 | 0.045882455 | 0.237312084 | 3.724945049 |
| Olfr32 | 0.631605561 | -0.376533484 | 1.338247294 |
| Gm4222 | 0 | NA | NA |
| Olfr1270 | 0.066050875 | 0.71820487 | 3.724945049 |
| Gm13768 | 4.397907185 | -0.305988847 | 0.573203113 |
| Olfr1506 | 0 | NA | NA |
| Gm18953 | 1.627769934 | 1.361763081 | 0.815176849 |
| Olfr142 | 0.123841226 | 1.292225235 | 3.719439735 |

| | | | |
|---|---|---|---|
| Olfr1271 | 0 | NA | NA |
| Olfr1272 | 0 | NA | NA |
| Olfr1273-ps | 0.044544648 | 0.237309859 | 3.724945049 |
| Gm13769 | 0 | NA | NA |
| Gm13767 | 0.326846603 | 1.16080624 | 1.579319892 |
| Olfr1274-ps | 0.163996116 | 0.719099421 | 2.782099506 |
| Ptprj | 2106.140895 | 0.117524089 | 0.077312166 |
| 4933423P22Rik | 0.57303442 | 1.808476451 | 1.394171915 |
| Nup160 | 358.4498607 | 0.172854976 | 0.107538503 |
| Fnbp4 | 883.9475744 | 0.335435313 | 0.155221029 |
| Rpl30-ps3 | 4.003505758 | 0.884196205 | 0.585003997 |
| Agbl2 | 38.0546618 | 0.022148206 | 0.198257326 |
| Gm13772 | 2.379780159 | -0.924096246 | 0.770459204 |
| Mtch2 | 1065.574097 | -0.089000686 | 0.054117897 |
| C1qtnf4 | 729.131802 | -0.882701098 | 0.096062815 |
| Ndufs3 | 556.9779417 | -0.064761454 | 0.053829218 |
| Kbtbd4 | 295.8165819 | 0.131430079 | 0.078482936 |
| Ptpmt1 | 185.7476871 | 0.137157524 | 0.090858568 |
| Gm17661 | 14.75789179 | -0.745994847 | 0.342246512 |
| Celf1 | 3282.919782 | -0.228672447 | 0.069347699 |
| Gm13777 | 0.066050875 | 0.71820487 | 3.724945049 |
| Rapsn | 7.482500097 | -0.06544848 | 0.41576715 |
| Gm13778 | 1.349943917 | -0.582783559 | 0.882422034 |
| Psmc3 | 1077.312818 | 0.382228846 | 0.091817629 |
| Slc39a13 | 259.1457734 | 0.013533074 | 0.08060052 |
| Spi1 | 202.8166777 | 3.944174409 | 0.226617915 |
| Mybpc3 | 140.415035 | -0.504930495 | 0.245582567 |
| Madd | 2828.091406 | -0.402418732 | 0.077495474 |
| Nr1h3 | 80.38358335 | 2.14926794 | 0.220560795 |
| Gm13783 | 1.952547309 | -1.290306524 | 0.727462926 |
| Gm23390 | 0 | NA | NA |
| Acp2 | 985.571155 | 0.319190616 | 0.044841219 |
| Ddb2 | 90.31751625 | -0.616155795 | 0.211996001 |
| A330069E16Rik | 10.08996547 | 0.456105565 | 0.319055817 |
| Gm17281 | 10.85400332 | -1.268618938 | 0.371537171 |
| Pacsin3 | 255.622615 | -0.509223469 | 0.122581744 |
| Arfgap2 | 730.4068622 | 0.451188172 | 0.074523629 |
| 1110051M20Rik | 549.9156413 | -0.334976267 | 0.057617899 |
| Gm13787 | 4.652905194 | -0.782839421 | 0.527068331 |
| Gm22071 | 0.442430462 | 0.491297358 | 3.326896204 |
| Lrp4 | 768.9458671 | 0.018555204 | 0.142322587 |
| Gm13770 | 1.121024803 | -0.380112779 | 1.012204288 |
| Ckap5 | 3392.946938 | 0.162300971 | 0.056581333 |
| Snord67 | 0.174512195 | -1.319653964 | 3.342629336 |
| F2 | 4.096713608 | 1.502809021 | 0.575835314 |
| Gm13771 | 3.006370356 | 0.008984127 | 0.630191 |
| Zfp408 | 162.7132082 | 0.610487195 | 0.171953716 |

| | | | |
|---|---|---|---|
| Arhgap1 | 635.3928673 | 0.300312169 | 0.062756378 |
| Atg13 | 1068.978477 | -0.03882538 | 0.051258342 |
| Harbi1 | 67.29079701 | -0.10770419 | 0.156538944 |
| Gm13776 | 0 | NA | NA |
| Ambra1 | 1182.397324 | -0.080034923 | 0.069139469 |
| Gm10805 | 2.718864041 | -0.411901384 | 0.731955509 |
| Chrm4 | 179.7457586 | -0.880291432 | 0.165038893 |
| Mdk | 159.9674338 | 0.798527986 | 0.183212577 |
| Dgkz | 3070.276697 | -0.585553629 | 0.091848002 |
| Mir6999 | 2.840504449 | -1.298554369 | 0.72827467 |
| Gm9821 | 0.506210706 | 1.258887809 | 1.658375308 |
| Creb3l1 | 136.2000294 | 2.599424958 | 0.230732198 |
| Gm13780 | 0 | NA | NA |
| 2900072N19Rik | 2.504782524 | -1.638005406 | 0.783584665 |
| Phf21a | 888.8724875 | -0.141096176 | 0.082476875 |
| Mir1955 | 9.919958007 | -0.116089735 | 0.345119317 |
| Gm13786 | 17.47248729 | -0.404247113 | 0.302153086 |
| Gm13785 | 11.39462224 | -0.954403435 | 0.344401791 |
| Gyltl1b | 4.391866736 | -0.594463069 | 0.488930247 |
| Pex16 | 172.030486 | 0.314740631 | 0.100435965 |
| 1700029I15Rik | 16.98714461 | -0.537037042 | 0.280408115 |
| Mapk8ip1 | 3383.018323 | -0.221852865 | 0.08377689 |
| Mir7000 | 0.999544483 | -0.955286184 | 1.121968022 |
| Cry2 | 1477.051092 | -0.557645642 | 0.124158351 |
| D930015M05Rik | 1.112373992 | -0.166845348 | 1.006725573 |
| Slc35c1 | 209.4076668 | 0.26142637 | 0.097733255 |
| Gm13814 | 0.592656722 | 0.005656789 | 1.211783114 |
| Gm13817 | 0 | NA | NA |
| Gm13815 | 0.046345709 | 0.71820487 | 3.724945049 |
| Mir7221 | 0 | NA | NA |
| Mir7222 | 0 | NA | NA |
| Chst1 | 1513.4659 | -0.615746964 | 0.104480815 |
| Gm13816 | 0.371244445 | 1.313557381 | 1.626148026 |
| Gm13791 | 2.292177245 | 0.930896104 | 0.760863257 |
| Gm13792 | 0.47588132 | -1.578755371 | 1.735843654 |
| Syt13 | 2064.734027 | -0.262661732 | 0.077704765 |
| Prdm11 | 417.3168277 | -0.645966747 | 0.113263423 |
| 4631405J19Rik | 0.851482706 | -2.939913329 | 1.105482211 |
| Gm13802 | 0 | NA | NA |
| Trp53i11 | 736.2819127 | 0.358930477 | 0.114045634 |
| Tspan18 | 97.99565433 | -0.209317349 | 0.146225466 |
| Gm23380 | 0 | NA | NA |
| Gm13807 | 0 | NA | NA |
| Cd82 | 303.0056944 | 0.896410825 | 0.141626163 |
| Mir7001 | 0.756442295 | -0.21625294 | 1.348497265 |
| Gm10804 | 0.060174537 | 0.237309859 | 3.724945049 |
| Gm10803 | 0 | NA | NA |

| | | | |
|---|---|---|---|
| Alx4 | 48.87403417 | 0.675660696 | 0.356929804 |
| Ext2 | 641.1248268 | 0.209014628 | 0.07604967 |
| Gm13818 | 0 | NA | NA |
| Accs | 46.96754752 | 0.016236373 | 0.195537662 |
| Accsl | 3.620414628 | 0.409602606 | 0.634169457 |
| Gm1335 | 0.929738683 | 0.910942355 | 1.181767243 |
| Gm13889 | 181.9860749 | 0.998280094 | 0.342636635 |
| AL772372.1 | 0 | NA | NA |
| Gm27027 | 1.373972637 | -1.157940351 | 0.884884119 |
| 4921507L20Rik | 0.891453265 | 3.651398542 | 1.146687023 |
| Alkbh3 | 168.1665116 | 0.138864479 | 0.139448891 |
| Hsd17b12 | 914.0604724 | 0.718918356 | 0.126712447 |
| Gm23630 | 0.694735342 | 1.056813065 | 1.223435745 |
| Mir129-2 | 0.549517402 | 0.114944929 | 1.275687777 |
| mmu-mir-129b | 0 | NA | NA |
| E530001K10Rik | 26.89108908 | -0.641790993 | 0.285247941 |
| Mir670 | 0.304542382 | -1.337608332 | 2.009338117 |
| Ttc17 | 942.8773192 | -0.190374022 | 0.095515764 |
| Itpa-ps1 | 137.8875215 | 0.415521052 | 0.121195674 |
| 2810002D19Rik | 26.44289622 | -0.134907246 | 0.220836922 |
| Api5 | 1200.933468 | 0.28165465 | 0.064499885 |
| AL672251.1 | 0 | NA | NA |
| Gm13788 | 0 | NA | NA |
| Gm24318 | 0 | NA | NA |
| Gm13793 | 0 | NA | NA |
| Gm26396 | 0 | NA | NA |
| Gm13794 | 0 | NA | NA |
| Gm13798 | 0 | NA | NA |
| Gm13797 | 0 | NA | NA |
| Gm13799 | 0 | NA | NA |
| Gm37175 | 0 | NA | NA |
| Gm13801 | 0 | NA | NA |
| Lrrc4c | 1313.773726 | -0.555832773 | 0.084384417 |
| 4930445B16Rik | 0 | NA | NA |
| Gm17075 | 11.93369344 | -0.418704633 | 0.314114479 |
| Gm17076 | 16.91552809 | -1.176444315 | 0.364595333 |
| Gm22957 | 5.852693499 | -2.086863093 | 0.553411053 |
| Gm22313 | 8.145373654 | -1.830718135 | 0.617156191 |
| Gm23317 | 0 | NA | NA |
| BX813336.1 | 0 | NA | NA |
| Gm13803 | 11.5628337 | -0.4023029 | 0.350662458 |
| Gm13804 | 0 | NA | NA |
| Gm13806 | 0 | NA | NA |
| Gm10801 | 0.412640929 | -1.119629606 | 1.555320814 |
| Gm10800 | 8.185264119 | -1.140016137 | 0.471242862 |
| Gm29235 | 0 | NA | NA |
| Gm13809 | 0 | NA | NA |

| | | | |
|---|---|---|---|
| Gm13808 | 0 | NA | NA |
| Gm13811 | 0.109727584 | 0.718200421 | 3.724945049 |
| Gm13813 | 0 | NA | NA |
| Gm13812 | 0.047948698 | -0.243569582 | 3.724945049 |
| Gm13810 | 0 | NA | NA |
| B230118H07Rik | 178.272976 | -0.117137641 | 0.149223757 |
| Gm20693 | 0 | NA | NA |
| Rag2 | 0.139883403 | 1.411723814 | 3.551352091 |
| Rag1 | 5.95628265 | 0.414870249 | 0.425133286 |
| Traf6 | 437.7135397 | 0.388979252 | 0.107329104 |
| Prr5l | 253.5858244 | 0.680400892 | 0.137521272 |
| Gm13905 | 0.083526043 | 0.718202646 | 3.724945049 |
| Commd9 | 191.447391 | 0.424792118 | 0.116275191 |
| Ldlrad3 | 167.5338321 | 0.130285616 | 0.125527551 |
| Gm13919 | 2.169032929 | 1.573378616 | 0.870873172 |
| Gm13920 | 4.969440295 | 3.009093063 | 0.674183426 |
| Gm13921 | 0.311841693 | 0.237330958 | 1.755529323 |
| Gm13866 | 1.228132247 | -0.671498002 | 1.003662311 |
| Trim44 | 3706.544855 | -0.194866496 | 0.046809886 |
| Gm13868 | 13.87276866 | -0.462712376 | 0.408024364 |
| Fjx1 | 429.7281807 | -0.590508879 | 0.09289113 |
| Pamr1 | 172.126028 | -0.634500312 | 0.151511042 |
| Slc1a2 | 36923.0076 | -1.40630938 | 0.160622854 |
| Gm13872 | 1.821369377 | 0.136552304 | 0.882823639 |
| Cd44 | 1546.027959 | 5.170697722 | 0.333429099 |
| Gm23731 | 0 | NA | NA |
| Pdhx | 517.8134996 | 0.025002018 | 0.068718256 |
| Apip | 121.8280582 | 0.756690331 | 0.111666898 |
| Gm10038 | 4.822154624 | 0.494285699 | 0.51409572 |
| Gm13874 | 0.503556157 | 2.458905797 | 1.689782704 |
| Gm13869 | 0.217279565 | -0.243563282 | 2.170317414 |
| Gm13873 | 0 | NA | NA |
| Ehf | 2.553683234 | 1.118675016 | 0.614875638 |
| A930006I01Rik | 0 | NA | NA |
| Gm37220 | 0 | NA | NA |
| BC016548 | 0.652016589 | -0.344131001 | 1.154901562 |
| Elf5 | 2.42823633 | -2.866769541 | 0.835433056 |
| Cat | 444.9583296 | -0.187681769 | 0.114912999 |
| Gm13996 | 0 | NA | NA |
| Abtb2 | 322.9374325 | 0.983953224 | 0.141156723 |
| Nat10 | 403.567961 | -0.028763256 | 0.170481007 |
| Caprin1 | 3734.751293 | 0.502391287 | 0.052336433 |
| 4930547E08Rik | 0.171199454 | 0.718206393 | 3.387969756 |
| Gm13878 | 0 | NA | NA |
| Gm13880 | 2.206276467 | 0.396490519 | 0.621777195 |
| Gm13877 | 0 | NA | NA |
| Gm13875 | 0 | NA | NA |

| | | | |
|---|---|---|---|
| Gm22757 | 0 | NA | NA |
| Gm13879 | 0 | NA | NA |
| Gm22761 | 0 | NA | NA |
| Gm13876 | 0 | NA | NA |
| Gm13881 | 0.097899064 | -0.724453471 | 3.724945049 |
| Lmo2 | 263.3668701 | 0.911647768 | 0.11139705 |
| 4931422A03Rik | 0.485755142 | 0.237333428 | 1.415645104 |
| Fbxo3 | 1097.464659 | -0.224474615 | 0.054464838 |
| Gm24644 | 12.13107012 | -1.027933308 | 0.379645564 |
| Gm10912 | 3.939700132 | -0.258255448 | 0.436603386 |
| Gm10799 | 0.09689823 | -0.724453471 | 3.724945049 |
| Cd59b | 12.1589774 | -0.073335778 | 0.297833591 |
| Cd59a | 104.2603775 | 0.669627023 | 0.215015918 |
| A930018P22Rik | 1.374894827 | 1.052947575 | 0.859547976 |
| D430041D05Rik | 2766.772851 | -0.965676122 | 0.150653714 |
| Gm22501 | 3.530580087 | -1.638377033 | 0.68421178 |
| Gm13883 | 6.270791699 | 0.669745243 | 0.546672357 |
| Gm25554 | 0 | NA | NA |
| Gm13882 | 5.232096263 | 0.206797134 | 0.441870166 |
| Hipk3 | 2248.948924 | -0.133602786 | 0.062094149 |
| Mir1902 | 0 | NA | NA |
| Gm13870 | 0 | NA | NA |
| Gm13884 | 0.326587101 | 1.112697679 | 1.98566164 |
| Platr14 | 2.284861264 | -0.75131104 | 0.727075479 |
| Gm22439 | 0 | NA | NA |
| Gm13885 | 0.266822755 | -1.248666532 | 2.399007414 |
| Cstf3 | 425.9326561 | -0.045212273 | 0.074429467 |
| Tcp11l1 | 978.2370623 | -0.328628194 | 0.071448895 |
| Pin1rt1 | 0.178918687 | -0.560686608 | 2.708509187 |
| Depdc7 | 17.9411153 | -0.11722587 | 0.280656277 |
| Qser1 | 622.2252455 | 0.004297067 | 0.065354754 |
| Gm13886 | 11.02501732 | -0.108518601 | 0.326447395 |
| Prrg4 | 12.12341216 | 1.415268522 | 0.333823106 |
| Ccdc73 | 1.808963643 | -0.138305775 | 0.787304643 |
| Gm13887 | 0.209356743 | 0.237328535 | 2.19052869 |
| Eif3m | 484.4513724 | 0.32319414 | 0.081217271 |
| Gm13888 | 0 | NA | NA |
| Wt1os | 0.172571121 | 1.188801635 | 2.733273164 |
| Gm11060 | 0 | NA | NA |
| Gm27715 | 0 | NA | NA |
| Gm27538 | 0 | NA | NA |
| Gm27636 | 0 | NA | NA |
| Gm27308 | 0 | NA | NA |
| Wt1 | 11.92317521 | 3.671591106 | 0.524369412 |
| Them7 | 0.493638219 | 0.230659886 | 1.401445733 |
| Rcn1 | 399.3853322 | 0.847622262 | 0.127724981 |
| 4930527A07Rik | 0 | NA | NA |

| | | | |
|---|---|---|---|
| Pax6os1 | 4.443791399 | -1.170846579 | 0.555595251 |
| Gm13890 | 0.122991902 | -0.24357403 | 3.724945049 |
| Rpl10-ps1 | 5.581713243 | 0.823583294 | 0.407183954 |
| Paupar | 4.745398909 | -0.461719069 | 0.455962647 |
| Pax6 | 311.7227136 | -0.353681209 | 0.161515971 |
| Elp4 | 237.3869113 | -0.118740064 | 0.084486118 |
| Gm13954 | 0.303403639 | 0.718222014 | 2.031766505 |
| Immp1l | 117.1741609 | 0.708275553 | 0.193907683 |
| Dnajc24 | 115.5254544 | 0.115787739 | 0.099863319 |
| Gm22002 | 0.104637558 | 0.23732098 | 3.724945049 |
| n-R5s201 | 0 | NA | NA |
| Dcdc5 | 20.38028449 | -0.872141823 | 0.269333431 |
| Gm13992 | 0.047948698 | -0.243569582 | 3.724945049 |
| Gm29053 | 0 | NA | NA |
| Gm14013 | 0.459558249 | -1.586081209 | 1.327170979 |
| Gm14015 | 1.812101644 | -1.864796548 | 0.86849464 |
| Gm14014 | 0.072025462 | 0.237312084 | 3.724945049 |
| BB218582 | 7.281320362 | -1.542294655 | 0.455028672 |
| Mpped2 | 264.5495323 | -0.752720346 | 0.100834812 |
| Gm13901 | 0.066050875 | 0.71820487 | 3.724945049 |
| Gm13902 | 0.292336004 | -0.695053139 | 1.612916721 |
| Arl14ep | 637.6380912 | 0.348824776 | 0.079132266 |
| Gm13900 | 0.111006857 | 0.717551239 | 3.724898127 |
| Gm13904 | 0.527697161 | -1.388500625 | 1.385836323 |
| Fshb | 0.248616998 | 2.225260438 | 3.00105206 |
| Gm24885 | 0 | NA | NA |
| Gm13903 | 0 | NA | NA |
| Kcna4 | 692.1728847 | -0.53059913 | 0.115655881 |
| Gm9864 | 1.048421568 | 0.251771322 | 0.916893328 |
| Gm23439 | 0 | NA | NA |
| Gm23749 | 0.105852363 | 0.23732098 | 3.724945049 |
| Gm13912 | 1.945135077 | 3.739903043 | 0.973155335 |
| Gm13916 | 0.123845245 | 1.035807111 | 3.720871489 |
| Gm13915 | 0.038981395 | 0.718207094 | 3.724945049 |
| Gm13913 | 0 | NA | NA |
| Gm13914 | 0 | NA | NA |
| Rpl35a-ps4 | 1.664460293 | 1.137919424 | 0.83223079 |
| Gm13910 | 140.7366961 | -0.012171797 | 0.153962032 |
| Gm13923 | 0 | NA | NA |
| Mettl15 | 54.80204305 | -0.120950291 | 0.167089766 |
| Gm24115 | 0.713863726 | 0.444502754 | 1.05266899 |
| Kif18a | 32.64308567 | 0.326318785 | 0.21158669 |
| Gm13924 | 0 | NA | NA |
| Gm13925 | 0 | NA | NA |
| Gm20638 | 0.165580463 | 0.23732449 | 2.773571271 |
| Bdnf | 331.5846942 | 0.425344134 | 0.216606602 |
| Gm13932 | 0.182620985 | 0.718216066 | 2.688869866 |

| | | | |
|---|---|---|---|
| Lin7c | 1365.827837 | -0.06811073 | 0.063206712 |
| Platr8 | 0 | NA | NA |
| Lgr4 | 521.05221 | -0.270388914 | 0.099060226 |
| Ccdc34 | 111.7137007 | -0.3994741 | 0.147812637 |
| Ccdc34os | 0 | NA | NA |
| Gm13936 | 1.575920251 | -0.132399313 | 0.973110835 |
| Gm13937 | 0.157911564 | -0.724458612 | 2.815266154 |
| Gm22997 | 0.40088837 | -0.243563459 | 1.475392019 |
| Bbox1 | 28.0529557 | -1.238591075 | 0.271137733 |
| Fibin | 60.99905906 | 0.351841613 | 0.215722551 |
| Gm13935 | 0.04393175 | -0.243571806 | 3.724945049 |
| Gm10794 | 0.046345709 | 0.71820487 | 3.724945049 |
| Slc5a12 | 0.290078826 | -0.279544126 | 1.809020424 |
| Ano3 | 1378.79017 | -1.413244682 | 0.166528357 |
| Muc15 | 15.98576495 | 2.066233637 | 0.30574638 |
| Gm13942 | 0 | NA | NA |
| Gm13941 | 0 | NA | NA |
| Gm15130 | 0 | NA | NA |
| 4930430A15Rik | 0.123841226 | 1.292225235 | 3.719439735 |
| Gm13961 | 21.20792997 | 0.078387311 | 0.219993182 |
| Olfr1275 | 0 | NA | NA |
| Gm13962 | 0 | NA | NA |
| Olfr1276 | 0.075043205 | 0.237309859 | 3.724945049 |
| Olfr1277 | 0 | NA | NA |
| Olfr1278 | 0 | NA | NA |
| Olfr1279 | 0 | NA | NA |
| Olfr1280 | 0 | NA | NA |
| Olfr1281 | 0 | NA | NA |
| Olfr1282 | 0 | NA | NA |
| Olfr1283 | 0.038981395 | 0.718207094 | 3.724945049 |
| Olfr1284 | 0 | NA | NA |
| Olfr1285 | 0 | NA | NA |
| Olfr1286 | 0 | NA | NA |
| Olfr1287 | 0 | NA | NA |
| Olfr1288 | 0.11197992 | 0.718202646 | 3.724945049 |
| Olfr1289 | 0 | NA | NA |
| Olfr1290 | 0.062426873 | 0.237312084 | 3.724945049 |
| Olfr1291-ps1 | 0 | NA | NA |
| Olfr1292-ps1 | 0 | NA | NA |
| Olfr1293-ps | 0 | NA | NA |
| Olfr1294 | 0.134986936 | -0.220861911 | 3.655468937 |
| Olfr1295 | 0.08486385 | 0.717551239 | 3.724898127 |
| Olfr1296-ps1 | 0 | NA | NA |
| Olfr1297 | 0 | NA | NA |
| Olfr1298 | 0 | NA | NA |
| Olfr1299 | 0 | NA | NA |
| Olfr1300-ps1 | 0 | NA | NA |

| | | | |
|---|---|---|---|
| Gm13922 | 0 | NA | NA |
| Gm13930 | 0 | NA | NA |
| Olfr1301 | 0.046345709 | 0.71820487 | 3.724945049 |
| Olfr1302 | 0 | NA | NA |
| Olfr1303 | 0.046345709 | 0.71820487 | 3.724945049 |
| Gm26421 | 0 | NA | NA |
| Olfr268-ps1 | 0.068097359 | -0.243569582 | 3.724945049 |
| Olfr1305 | 0 | NA | NA |
| Olfr1306 | 0 | NA | NA |
| Gm13928 | 0 | NA | NA |
| Olfr1307 | 0 | NA | NA |
| Olfr1308 | 0 | NA | NA |
| Olfr1309 | 0 | NA | NA |
| Gm13929 | 0 | NA | NA |
| Olfr1310 | 0.048949532 | -0.243569582 | 3.724945049 |
| Olfr1311 | 0 | NA | NA |
| Olfr1312 | 0 | NA | NA |
| Olfr1313 | 0.066050875 | 0.71820487 | 3.724945049 |
| Olfr1314 | 0 | NA | NA |
| Olfr1315-ps1 | 0 | NA | NA |
| Olfr1316 | 0.049553047 | 0.71820487 | 3.724945049 |
| Olfr1317 | 0 | NA | NA |
| Olfr1318 | 0 | NA | NA |
| Lpcat4 | 780.3235412 | -0.752215401 | 0.095037705 |
| Nutm1 | 0.463200932 | 1.015679229 | 1.499770166 |
| Nop10 | 173.0751968 | -0.211935333 | 0.117673667 |
| Slc12a6 | 925.7716682 | -0.712742899 | 0.083876835 |
| Gm21985 | 0.288125237 | -0.243584957 | 3.724945049 |
| Emc4 | 460.0520838 | 0.056396332 | 0.074196488 |
| Katnbl1 | 227.6757466 | 0.288969102 | 0.105022005 |
| Gm13940 | 0 | NA | NA |
| Emc7 | 1191.037724 | 0.138687096 | 0.065851494 |
| n-R5s202 | 0 | NA | NA |
| Chrm5 | 22.04456264 | -0.68687223 | 0.239844699 |
| Aven | 92.20401975 | 0.184452347 | 0.110167829 |
| Gm22676 | 0.183828322 | -0.542678823 | 2.663972021 |
| AL691423.1 | 0.566015516 | -2.375179303 | 1.457441368 |
| Ryr3 | 1324.13524 | -0.871101739 | 0.270570128 |
| Tmco5b | 1.585639959 | 3.310905398 | 1.06243756 |
| Gm26505 | 0.194120612 | 0.718217805 | 2.642167127 |
| Fmn1 | 837.5690741 | -0.476142602 | 0.075694709 |
| Gm13967 | 4.347705094 | -0.21456908 | 0.591127445 |
| Gm13963 | 0.697976008 | 3.474534727 | 1.302533348 |
| Gm13965 | 7.79987822 | -0.909930271 | 0.409824735 |
| Gm13966 | 0.044191549 | -0.243571806 | 3.724945049 |
| AL691437.1 | 0 | NA | NA |
| Gm13964 | 0.060174537 | 0.237309859 | 3.724945049 |

| | | | |
|---|---|---|---|
| 4930533B01Rik | 0.045882455 | 0.237312084 | 3.724945049 |
| Grem1 | 36.57688901 | 0.083588212 | 0.225127848 |
| Scg5 | 2876.760524 | -0.37385439 | 0.095638007 |
| Arhgap11a | 55.95224419 | 0.644048769 | 0.242556026 |
| Gm21596 | 8.155034204 | 0.06293845 | 0.391341589 |
| Gjd2 | 128.1457705 | -0.396019767 | 0.132207868 |
| A530058N18Rik | 102.4745166 | -1.056599891 | 0.238721451 |
| Actc1 | 0.709411339 | 3.400369751 | 1.209555018 |
| C130080G10Rik | 3.984159091 | 0.033086101 | 0.543321098 |
| Aqr | 833.2625079 | 0.164035479 | 0.04587685 |
| Mir7002 | 0.230049058 | -1.173529547 | 2.117076221 |
| Zfp770 | 474.7123103 | -0.086857956 | 0.087796752 |
| Gm25103 | 0.044544648 | 0.237309859 | 3.724945049 |
| Gm29234 | 0.986742042 | -0.908322179 | 1.04316883 |
| Dph6 | 238.7373716 | -0.262672858 | 0.116052297 |
| Gm13972 | 0.263371022 | -0.229200414 | 1.816772981 |
| Gm13975 | 0.044191549 | -0.243571806 | 3.724945049 |
| Gm13974 | 2.444458607 | -0.918001093 | 0.685644268 |
| Gm28494 | 0 | NA | NA |
| Gm28492 | 0.508786284 | -1.2045366 | 1.272041676 |
| Gm13973 | 0 | NA | NA |
| Gm28493 | 0.257385133 | 1.917654578 | 2.027399582 |
| 4930528P14Rik | 0.479723347 | 0.620962108 | 1.343005172 |
| Gm13976 | 0.060174537 | 0.237309859 | 3.724945049 |
| 3110099E03Rik | 2.086914652 | 0.741981393 | 0.657277014 |
| Gm13977 | 0.219276728 | 1.700730588 | 2.590888607 |
| BC052040 | 104.7146314 | -0.165027271 | 0.140344301 |
| Mir1951 | 0.245748936 | -0.10280023 | 2.04460657 |
| Meis2 | 1352.514286 | -0.599623162 | 0.141557584 |
| 2700033N17Rik | 6.560169035 | 0.212668156 | 0.390427976 |
| G630016G05Rik | 19.18698179 | -1.465025073 | 0.32779058 |
| 2810405F15Rik | 1.233431567 | -1.103772195 | 1.033859767 |
| Gm13991 | 44.848571 | 0.106596221 | 0.163523201 |
| Gm13990 | 0.730890257 | -0.10182669 | 1.249292811 |
| Tmco5 | 9.517001854 | 0.550004789 | 0.366346882 |
| D330050G23Rik | 15.28741942 | -0.311321066 | 0.25579886 |
| Gm24914 | 0 | NA | NA |
| Gm25514 | 0.127971488 | 1.23029335 | 3.685533863 |
| Gm29340 | 5.225310162 | 1.813188358 | 0.516495475 |
| Gm13981 | 0.117907917 | 0.237310006 | 3.724945049 |
| Spred1 | 3571.650489 | 0.277455684 | 0.088266911 |
| Mir674 | 5.596050554 | -0.544933598 | 0.418318924 |
| Rap1a-ps2 | 0.847725382 | 0.844701983 | 1.023650531 |
| Gm25189 | 0.284781514 | 0.238025105 | 2.060972187 |
| Fam98b | 385.3078931 | 0.065392889 | 0.086339337 |
| Rasgrp1 | 4351.652765 | -0.714931535 | 0.139241779 |
| Gm13982 | 0.169723681 | 1.276730323 | 3.407267785 |

| | | | |
|---|---|---|---|
| Gm28183 | 0.166952883 | 1.610951781 | 2.74653892 |
| Gm13985 | 0.090890357 | 0.718200421 | 3.724945049 |
| 4930412B13Rik | 0 | NA | NA |
| Gm13986 | 9.239123008 | 1.640175986 | 0.415586094 |
| Gm36979 | 0 | NA | NA |
| Gm25308 | 0.048949532 | -0.243569582 | 3.724945049 |
| Thbs1 | 1597.215543 | 5.562240206 | 0.457712431 |
| Gm29233 | 0.136201742 | -0.231510441 | 2.944131489 |
| Fsip1 | 6.681599266 | 0.188397805 | 0.39049275 |
| Gm22800 | 0 | NA | NA |
| Gm29232 | 0.049553047 | 0.71820487 | 3.724945049 |
| A930104D05Rik | 0.722394163 | -0.435086974 | 1.25291186 |
| Gpr176 | 129.1472356 | -0.309907649 | 0.119085692 |
| 1700054M17Rik | 0.459216712 | -1.821299427 | 1.477438974 |
| Gm22651 | 0 | NA | NA |
| Eif2ak4 | 429.1232767 | -0.368211503 | 0.138407441 |
| Srp14 | 522.9120673 | 0.279741126 | 0.104768235 |
| Gm13983 | 0 | NA | NA |
| Bmf | 64.79741328 | 0.130535133 | 0.152817728 |
| Gm14016 | 0 | NA | NA |
| Bub1b | 14.01725363 | 2.684397286 | 0.442319387 |
| Pak6 | 596.5307493 | -0.46706887 | 0.150371055 |
| Ankrd63 | 286.9989274 | -1.66292609 | 0.333724597 |
| Plcb2 | 54.89159527 | 1.269130634 | 0.252518079 |
| Inafm2 | 508.6639642 | 0.05137783 | 0.087944749 |
| A430105I19Rik | 127.3481097 | 0.651611356 | 0.140329842 |
| Phgr1 | 0.264111009 | 0.718192361 | 3.724945049 |
| Disp2 | 4917.700072 | -0.269586545 | 0.059087633 |
| Knstrn | 21.59877154 | 1.150199943 | 0.388916968 |
| Gm14091 | 1.226992653 | -1.407269344 | 1.063973367 |
| Ivd | 475.053727 | -0.432757237 | 0.082608073 |
| Bahd1 | 560.8711464 | -0.007525129 | 0.089489638 |
| Chst14 | 36.12922769 | 1.086400552 | 0.205621882 |
| Gm14089 | 0.273591019 | 1.15691822 | 2.025264683 |
| Gm14088 | 0.141331096 | 0.237322366 | 3.56184083 |
| Ccdc32 | 448.3274695 | 0.719044677 | 0.118551472 |
| Rpusd2 | 91.45199017 | 0.099870919 | 0.144465866 |
| Casc5 | 31.52527001 | 2.379816029 | 0.362941096 |
| Rad51 | 19.29236894 | 0.918233718 | 0.337093495 |
| Rmdn3 | 789.494809 | 0.603415032 | 0.102411337 |
| Gchfr | 7.98091292 | -0.900248857 | 0.408283964 |
| Dnajc17 | 78.13003113 | 0.154358897 | 0.141512917 |
| Gm14137 | 2.178995086 | 4.608531854 | 0.974045524 |
| Gm14138 | 1.456796327 | -0.395833804 | 0.818501979 |
| Zfyve19 | 182.5094088 | 0.167665577 | 0.087679669 |
| Ppp1r14d | 0.089814205 | -0.24357403 | 3.724945049 |
| Spint1 | 12.20675764 | 1.02143917 | 0.3047022 |

| | | | |
|---|---|---|---|
| Rhov | 153.8616767 | 1.840639536 | 0.169140311 |
| Gm14175 | 0 | NA | NA |
| Vps18 | 675.7131133 | 0.265134869 | 0.054498877 |
| Gm14207 | 0.327809541 | 0.718222761 | 1.951230145 |
| Dll4 | 18.4801647 | 0.216441046 | 0.401121629 |
| Chac1 | 175.096179 | 2.275365476 | 0.260896744 |
| Ino80 | 813.323168 | 0.295936736 | 0.063643262 |
| Gm14208 | 0.083526043 | 0.718202646 | 3.724945049 |
| Exd1 | 16.27157243 | -0.091917381 | 0.260652776 |
| Chp1 | 1163.742042 | -0.185275684 | 0.055218014 |
| 1700020I14Rik | 6679.70898 | -0.961718815 | 0.088358098 |
| Oip5 | 7.956782189 | -0.364430659 | 0.408127901 |
| Nusap1 | 22.39643515 | 2.020227435 | 0.411958464 |
| n-R5s204 | 0 | NA | NA |
| Ndufaf1 | 192.2611336 | 0.249242424 | 0.076748828 |
| H3f3c | 22.18271704 | 0.236268487 | 0.248990818 |
| Gm14383 | 0.267580615 | -0.243563209 | 1.820952685 |
| Rtf1 | 860.2757225 | -0.209544987 | 0.080317766 |
| Gm25857 | 1.36269325 | -2.198830557 | 1.085156385 |
| Itpka | 639.0705542 | -1.089672501 | 0.134645928 |
| Ltk | 62.0682925 | -1.24399166 | 0.15807205 |
| Rpap1 | 285.1701272 | -0.152688299 | 0.094292404 |
| Gm23814 | 0 | NA | NA |
| Tyro3 | 1444.38765 | -0.976844742 | 0.086665394 |
| Gm13998 | 0 | NA | NA |
| Mga | 2323.037544 | -0.041531834 | 0.065861518 |
| Mapkbp1 | 844.8660816 | -0.461941736 | 0.11240039 |
| Mir7665 | 0.088383099 | -0.72445792 | 3.724945049 |
| Gm13999 | 0 | NA | NA |
| Jmjd7 | 2.162814427 | 0.394391811 | 0.707143204 |
| Gm28042 | 0.10503227 | 1.179809211 | 3.723500796 |
| Pla2g4b | 5.643927168 | -0.415841577 | 0.401616692 |
| Sptbn5 | 13.1047588 | -0.484881331 | 0.354348512 |
| Ehd4 | 458.2061878 | 2.188484528 | 0.157065324 |
| Pla2g4e | 246.4736302 | -0.612508902 | 0.141218781 |
| Gm13997 | 1.380678606 | -0.292425606 | 0.859727457 |
| Pla2g4d | 1.305860104 | 1.806509068 | 1.105293062 |
| Pla2g4f | 1.649860538 | 0.893219449 | 0.80510001 |
| Vps39 | 1015.694296 | -0.017054215 | 0.05560259 |
| Tmem87a | 799.4634507 | 0.33562705 | 0.065253522 |
| Ganc | 158.6786428 | 0.243924172 | 0.109798009 |
| Capn3 | 66.72612438 | -0.819352481 | 0.246682063 |
| Zfp106 | 4291.41446 | -0.239569693 | 0.048224741 |
| Gm26899 | 5.004844821 | 0.096651587 | 0.494511035 |
| Snap23 | 220.6488251 | 0.401000069 | 0.140419938 |
| Lrrc57 | 243.5709634 | 0.070458541 | 0.088479432 |
| Haus2 | 232.8508983 | 0.560272102 | 0.118793725 |

| | | | |
|---|---|---|---|
| Stard9 | 314.5610356 | -0.52075023 | 0.173663889 |
| n-R5s205 | 0 | NA | NA |
| Cdan1 | 293.1784353 | -0.399560837 | 0.156985765 |
| Ttbk2 | 1910.365418 | -0.36929755 | 0.065240227 |
| AV039307 | 6.926568459 | -0.758844562 | 0.400414369 |
| Ubr1 | 1554.190815 | -0.220780257 | 0.046660306 |
| Tmem62 | 383.5196607 | 0.013032606 | 0.072271933 |
| AL844548.1 | 1.862757 | -0.547893105 | 0.763629111 |
| Gm22865 | 3.966952568 | 0.085354365 | 0.679138055 |
| Ccndbp1 | 483.2632691 | -0.152234298 | 0.073854672 |
| Epb4.2 | 1.132377036 | 1.507146789 | 1.102061917 |
| Tgm5 | 0.516597926 | 1.314087043 | 1.448638169 |
| Gm14213 | 0 | NA | NA |
| Tgm7 | 0.64993548 | -0.283743169 | 1.040852021 |
| Lcmt2 | 75.2947259 | 0.626489764 | 0.160174713 |
| Gm23858 | 0.195161473 | -0.20882949 | 2.671768033 |
| Adal | 331.7277646 | -0.068897443 | 0.063438969 |
| Zscan29 | 154.5557525 | 1.040413497 | 0.129454985 |
| Tubgcp4 | 399.137152 | -0.187689874 | 0.088067913 |
| Trp53bp1 | 1866.10298 | -0.339092726 | 0.075589896 |
| Map1a | 23447.45946 | -0.62683792 | 0.068379296 |
| Gm14978 | 2.354129177 | -0.889940669 | 0.736995675 |
| Ppip5k1 | 1660.739469 | -0.194543174 | 0.084565905 |
| Ckmt1 | 1627.233016 | -0.660349503 | 0.066937078 |
| Strc | 2.337199581 | 0.393170053 | 0.705283914 |
| Catsper2 | 52.60120326 | -0.471187107 | 0.195705022 |
| Gm14001 | 3.014229995 | -1.498116771 | 0.56155478 |
| Gm14000 | 0 | NA | NA |
| Pdia3 | 2693.142216 | 0.949362921 | 0.1029795 |
| Ell3 | 0.594313001 | -0.678136482 | 1.300828392 |
| Serinc4 | 3.508801613 | 0.556803479 | 0.552413255 |
| Serf2 | 753.224674 | 0.209461398 | 0.069852349 |
| Gm22953 | 0.938485331 | 1.317733544 | 1.109863038 |
| Hypk | 31.77497691 | -0.747646424 | 0.302448179 |
| Mfap1b | 106.0742756 | 0.443008022 | 0.131807915 |
| Gm14017 | 2.307785566 | -0.321748026 | 0.717715858 |
| Gm14018 | 0.899881821 | -0.662233843 | 0.938793206 |
| Mfap1a | 281.266986 | 0.028864768 | 0.063902394 |
| Wdr76 | 70.42691052 | -0.03160555 | 0.13311742 |
| Frmd5 | 605.5814555 | 0.010600014 | 0.073793329 |
| Gm14021 | 44.18792118 | -1.042043827 | 0.263870905 |
| Gm14020 | 0.04393175 | -0.243571806 | 3.724945049 |
| Gm14019 | 0.838491377 | 1.519700348 | 1.111141581 |
| Casc4 | 1001.124057 | -0.185502455 | 0.064807999 |
| Gm23983 | 0 | NA | NA |
| Mageb3 | 1.086684801 | -0.197579866 | 0.867534523 |
| Ctdspl2 | 624.7685435 | 0.112680792 | 0.086283834 |

| | | | |
|---|---|---|---|
| Eif3j1 | 235.1963187 | 0.170609754 | 0.130143115 |
| AL845457.1 | 0 | NA | NA |
| Spg11 | 412.0809888 | 0.125530652 | 0.093620376 |
| Patl2 | 2.452851275 | -0.226248341 | 0.675167826 |
| B2m | 9604.418411 | 5.775493261 | 0.161424175 |
| Trim69 | 8.404189488 | 5.628995588 | 0.874667026 |
| 4933406J08Rik | 0.139037128 | 1.464453538 | 3.717743456 |
| Gm14050 | 0 | NA | NA |
| Gm14058 | 0.04393175 | -0.243571806 | 3.724945049 |
| Rps12-ps10 | 0.709725355 | 1.23910537 | 1.386259068 |
| Sord | 185.688388 | -0.266138611 | 0.086928774 |
| Duox2 | 2.808703517 | 0.946539678 | 0.668604523 |
| Duoxa2 | 0.276302848 | 0.237329286 | 2.039329957 |
| Duoxa1 | 3.074242624 | 1.688428257 | 0.975893295 |
| Duox1 | 2.917555992 | -0.845870486 | 0.671830471 |
| Shf | 467.0710296 | -0.118158169 | 0.112853374 |
| Gm24409 | 0 | NA | NA |
| Slc28a2 | 188.3325738 | 11.10235108 | 1.057298981 |
| Gm14086 | 0 | NA | NA |
| Bambi-ps1 | 12.14750959 | 0.615464767 | 0.369438488 |
| Gm14085 | 41.28799092 | 9.32933362 | 0.914192377 |
| Gatm | 1118.00039 | 0.365421066 | 0.154627199 |
| Spata5l1 | 53.6862279 | 0.920903909 | 0.194703207 |
| AA467197 | 88.13508126 | 7.257031752 | 0.644665145 |
| Mir147 | 0 | NA | NA |
| Slc30a4 | 1022.722383 | -0.00272693 | 0.083257671 |
| 4930417H01Rik | 4.957477421 | -0.629227718 | 0.549142473 |
| Bloc1s6os | 0.578202636 | -0.517877153 | 1.200948698 |
| Bloc1s6 | 473.7227365 | 0.880098533 | 0.103301449 |
| Sqrdl | 62.88662421 | 1.347191384 | 0.214434313 |
| Gm13988 | 0.046345709 | 0.71820487 | 3.724945049 |
| Sema6d | 1571.33555 | -0.249854548 | 0.077346695 |
| 4930517E11Rik | 0.136309558 | 0.23730798 | 3.614873719 |
| 4930583P06Rik | 0.286786342 | 1.658093888 | 1.989386936 |
| Gm13994 | 0 | NA | NA |
| Slc24a5 | 5.913197406 | -1.324691338 | 0.47114596 |
| Myef2 | 1104.874051 | -0.358215196 | 0.08819554 |
| Gm22352 | 0 | NA | NA |
| Gm14004 | 11.63604071 | -0.746457296 | 0.324846988 |
| Ctxn2 | 55.00778503 | -0.421152454 | 0.180249209 |
| Slc12a1 | 4.418703675 | -0.46245092 | 0.464621463 |
| Gm14002 | 3.728554256 | -0.398662455 | 0.564934436 |
| Dut | 174.499205 | -0.051926546 | 0.103293589 |
| A530010F05Rik | 0 | NA | NA |
| Gm14003 | 0.116944185 | 1.302646088 | 3.719336636 |
| Fbn1 | 344.1427708 | 1.250575818 | 0.131448553 |
| Gm9913 | 2.575840985 | 1.243432862 | 0.656441063 |

| | | | |
|---|---|---|---|
| Cep152 | 85.54108715 | -0.04841768 | 0.152423897 |
| Shc4 | 125.3363377 | 0.394805536 | 0.141944718 |
| Eid1 | 1534.982027 | -0.445072173 | 0.071920287 |
| Secisbp2l | 3169.337731 | -0.090942186 | 0.043204803 |
| Cops2 | 1452.800378 | 0.352046279 | 0.108350768 |
| Galk2 | 221.865611 | -0.004621778 | 0.08917025 |
| Fam227b | 26.91957889 | -1.962729629 | 0.318444323 |
| Fgf7 | 42.37332393 | -0.422028186 | 0.236641632 |
| Gm26496 | 0.514662899 | 0.504534036 | 1.257423487 |
| Dtwd1 | 64.96816098 | 0.323613858 | 0.137439067 |
| Gm17555 | 0 | NA | NA |
| Gm24739 | 0 | NA | NA |
| Atp8b4 | 80.28755012 | 7.584290601 | 0.637521926 |
| Gm26697 | 1.020000067 | 0.135520503 | 0.931725624 |
| Slc27a2 | 66.76116308 | -0.821835846 | 0.146942938 |
| Hdc | 76.84591766 | 2.368267192 | 0.269663258 |
| Gabpb1 | 280.3852881 | 0.273591162 | 0.116662983 |
| Gm23753 | 1.578568981 | -2.189691709 | 0.973749475 |
| Gm27003 | 13.76199491 | -0.804517058 | 0.354993566 |
| Usp8 | 1065.714832 | 0.205970508 | 0.053992401 |
| Usp50 | 5.50178608 | 0.283214881 | 0.466793554 |
| Gm10774 | 0 | NA | NA |
| Trpm7 | 1310.402208 | -0.140709665 | 0.100468411 |
| Gm14212 | 0 | NA | NA |
| Sppl2a | 1346.162496 | 1.188706831 | 0.101127221 |
| Mir3473g | 5.710082525 | 0.41705886 | 0.617889311 |
| Ap4e1 | 265.198369 | -0.012850841 | 0.12739067 |
| Gm24081 | 0 | NA | NA |
| Blvra | 139.0631896 | 0.670274135 | 0.133863782 |
| Ncaph | 24.30005576 | 1.397985266 | 0.282144444 |
| Itpripl1 | 51.57497973 | 1.614441192 | 0.196833156 |
| 1810024B03Rik | 6.848270579 | -0.767897566 | 0.425616453 |
| Snrnp200 | 2169.627635 | 0.246891935 | 0.054866116 |
| Ciao1 | 447.7927703 | 0.245897564 | 0.097841447 |
| Tmem127 | 1631.67063 | 0.233131749 | 0.06199274 |
| Gm10766 | 3.235620746 | -1.286808563 | 0.566339508 |
| Stard7 | 1005.190084 | -0.201201487 | 0.053092048 |
| Dusp2 | 53.83102264 | 4.303311937 | 0.412616745 |
| Astl | 1.38321721 | 2.657217688 | 1.075190098 |
| Adra2b | 12.20916545 | -2.152041871 | 0.402268093 |
| Gpat2 | 10.37835042 | -1.142153127 | 0.357941459 |
| Fahd2a | 142.0280488 | -0.702572958 | 0.109148538 |
| Kcnip3 | 821.4776332 | 0.12092488 | 0.07848804 |
| Gm22613 | 5.411512881 | -1.143406947 | 0.539202423 |
| Prom2 | 9.157968272 | -0.096599946 | 0.360759081 |
| Gm14229 | 0 | NA | NA |
| Zfp661 | 72.87159097 | -0.14386641 | 0.13271321 |

| | | | |
|---|---|---|---|
| Mrps5 | 469.1048793 | -0.063793097 | 0.066965383 |
| Mal | 1616.595918 | -0.42283079 | 0.095506337 |
| Gm22889 | 0 | NA | NA |
| Gm14245 | 0 | NA | NA |
| Gm14244 | 0.04393175 | -0.243571806 | 3.724945049 |
| Mall | 5.752169691 | 0.885960934 | 0.4628773 |
| Nphp1 | 102.8958306 | 0.017121684 | 0.110342364 |
| 1500011K16Rik | 147.7564642 | 0.184096724 | 0.179567669 |
| Bub1 | 15.64735813 | 2.392271281 | 0.444013209 |
| Acoxl | 0 | NA | NA |
| Gm22411 | 0.044191549 | -0.243571806 | 3.724945049 |
| Gm23101 | 0 | NA | NA |
| Bcl2l11 | 158.1317849 | 1.584551715 | 0.180057809 |
| Gm14009 | 0.120974994 | -0.243571806 | 3.724945049 |
| Gm14005 | 19.18807827 | 4.616986889 | 0.558730995 |
| Gm14012 | 0.247225956 | 0.718211434 | 2.992047273 |
| Gm14010 | 0.594047536 | 2.174461524 | 1.349673292 |
| Gm29010 | 0 | NA | NA |
| Gm14006 | 0.075043205 | 0.237309859 | 3.724945049 |
| Gm14008 | 0.04393175 | -0.243571806 | 3.724945049 |
| Gm355 | 0 | NA | NA |
| Gm14007 | 0 | NA | NA |
| Anapc1 | 1911.795479 | 0.094188002 | 0.05871569 |
| Mertk | 498.8880327 | 0.746209834 | 0.134306686 |
| Gm14011 | 0 | NA | NA |
| Gm23172 | 3.964087094 | -0.929623707 | 0.755857278 |
| Tmem87b | 666.3521244 | 0.050883441 | 0.075012985 |
| Fbln7 | 48.08116317 | -0.055346203 | 0.199623005 |
| Gm22859 | 0 | NA | NA |
| Zc3h8 | 36.68443589 | -0.345151553 | 0.245684993 |
| Gm14028 | 0 | NA | NA |
| 4933427J07Rik | 0.358075244 | -1.812208238 | 1.691566301 |
| Gm10762 | 30.64394088 | 0.138150383 | 0.208469212 |
| Zc3h6 | 404.6004255 | -0.120438934 | 0.110489814 |
| Gm14027 | 11.52035073 | -0.660592121 | 0.321005558 |
| Gm14025 | 1.764317597 | 1.314252836 | 0.81134904 |
| Gm14022 | 5.545450415 | -1.2007876 | 0.445826474 |
| Ttl | 570.9256818 | -0.508240292 | 0.095869962 |
| Polr1b | 151.5190305 | 0.177747725 | 0.150056256 |
| Chchd5 | 87.19880141 | 0.517191062 | 0.155669524 |
| AI847159 | 3.26828567 | 1.324770193 | 0.547920019 |
| Gm14026 | 27.00074528 | -0.203159798 | 0.23453059 |
| Gm14029 | 5.503856369 | 1.388772629 | 0.436133548 |
| Gm4430 | 83.58514794 | -0.133378343 | 0.127962018 |
| Slc20a1 | 1626.484812 | 0.165112376 | 0.079729472 |
| 9830144P21Rik | 0.631002787 | 1.606393215 | 1.278333243 |
| A730036I17Rik | 9.097055571 | -0.659550764 | 0.39879714 |

| | | | |
|---|---|---|---|
| Gm14024 | 0.180493943 | 0.719085619 | 2.699005327 |
| Ckap2l | 17.79315673 | 1.488004521 | 0.474534436 |
| Gm14023 | 2.610220779 | -0.131361643 | 0.641505465 |
| Il1a | 25.73372477 | 0.827896825 | 0.335042773 |
| Gm25703 | 0 | NA | NA |
| Gm14039 | 0 | NA | NA |
| Il1b | 17.42109151 | 4.760826893 | 0.533585958 |
| Il1bos | 0.58991373 | 0.201276354 | 1.297758367 |
| Spcs2-ps | 12.01699771 | 0.860716104 | 0.291102279 |
| F830045P16Rik | 7.997923877 | -1.828015862 | 0.543766548 |
| Gm14040 | 0.25468147 | -0.188782661 | 2.103565842 |
| Gm14041 | 2.468877162 | -1.259766939 | 0.711477857 |
| Sirpa | 3103.836334 | -0.420792888 | 0.08404698 |
| Pdyn | 260.8278238 | -0.327832646 | 0.151898733 |
| Gm14046 | 0.049553047 | 0.71820487 | 3.724945049 |
| Stk35 | 654.0656122 | -0.080596065 | 0.064315812 |
| Gm28196 | 0.664320874 | -0.44500455 | 1.562404454 |
| Tgm3 | 28.71390692 | -0.503235563 | 0.273519523 |
| Mir6339 | 0 | NA | NA |
| Gm22424 | 0.046345709 | 0.71820487 | 3.724945049 |
| Gm24665 | 0.038981395 | 0.718207094 | 3.724945049 |
| AU015228 | 0.066050875 | 0.71820487 | 3.724945049 |
| Tgm6 | 0.648101363 | -1.497127986 | 1.145535428 |
| Snrpb | 546.91693 | 0.518714637 | 0.106378095 |
| Tmc2 | 0.617199072 | -2.231839252 | 1.312878688 |
| Nop56 | 546.9667433 | 0.215316714 | 0.102019944 |
| Snord110 | 0.459163063 | -1.708664489 | 1.276257475 |
| Gm24451 | 0 | NA | NA |
| Gm23187 | 0 | NA | NA |
| Gm23650 | 0 | NA | NA |
| Snord57 | 0 | NA | NA |
| Idh3b | 1254.967299 | -0.223444624 | 0.057266669 |
| Ebf4 | 86.18724845 | -0.941399368 | 0.129371244 |
| Gm14044 | 0.90691343 | 0.901982601 | 1.009387284 |
| Cpxm1 | 57.79122351 | -0.662811831 | 0.274852299 |
| Gm14042 | 0.049553047 | 0.71820487 | 3.724945049 |
| Gm28372 | 0 | NA | NA |
| 1700020A23Rik | 0.230076657 | 0.718205339 | 3.121436614 |
| Tmem239 | 0.066050875 | 0.71820487 | 3.724945049 |
| Pced1a | 228.252696 | -0.246534465 | 0.09896962 |
| Vps16 | 502.9920582 | -0.106068585 | 0.063288291 |
| Ptpra | 1842.435841 | 0.063977453 | 0.06183583 |
| Gm23876 | 0 | NA | NA |
| 4930473A02Rik | 0.931693893 | 0.993390489 | 1.061899645 |
| Mrps26 | 372.6054324 | 0.10363773 | 0.126271724 |
| Oxt | 55.53220185 | -0.199384399 | 0.529405895 |
| Avp | 105.9017596 | -0.224865999 | 0.310903236 |

| | | | |
|---|---|---|---|
| Ubox5 | 144.7706665 | 0.02086951 | 0.108881091 |
| Fastkd5 | 53.00231664 | 0.046013685 | 0.164745994 |
| Lzts3 | 1886.840697 | -1.205470075 | 0.136509603 |
| Ddrgk1 | 422.8448348 | 0.256671476 | 0.088445645 |
| Itpa | 134.0024633 | 0.134688719 | 0.117275686 |
| Slc4a11 | 82.95192658 | 2.251892591 | 0.234704434 |
| Gm24893 | 1.856422655 | -0.806569721 | 0.760145181 |
| 4930402H24Rik | 1406.23195 | -0.587230752 | 0.111694631 |
| Gm14057 | 0.232395771 | 1.183255052 | 2.122081047 |
| A730017L22Rik | 0.834338752 | 0.561132931 | 1.142312966 |
| Gm14048 | 32.5064405 | 1.064024374 | 0.210453413 |
| Atrn | 2962.760329 | -0.643014199 | 0.071894743 |
| Gm14049 | 10.85113572 | -0.953221785 | 0.359925397 |
| Gfra4 | 152.1835605 | -0.595262226 | 0.13613583 |
| Mir6973b | 0 | NA | NA |
| Adam33 | 45.18950367 | -0.530992293 | 0.21406518 |
| Siglec1 | 680.3250049 | 6.58328675 | 0.217249096 |
| Hspa12b | 36.83173324 | 1.077061459 | 0.205428087 |
| Gm11037 | 0.127971488 | 1.23029335 | 3.685533863 |
| 1700037H04Rik | 296.061265 | -0.527096788 | 0.087957968 |
| Gm14232 | 0.292476601 | -0.243566209 | 1.790231676 |
| Spef1 | 54.9579452 | -0.361557626 | 0.152262942 |
| Cenpb | 1691.510661 | 0.26540915 | 0.087271987 |
| Cdc25b | 96.01317364 | -0.799258855 | 0.157067864 |
| Ap5s1 | 93.61776372 | -0.043155362 | 0.115107659 |
| Mavs | 109.308156 | 0.645569854 | 0.189863445 |
| Pank2 | 601.5473895 | -0.131408205 | 0.066494381 |
| Mir103-2 | 2.477700388 | -0.614068873 | 0.608576744 |
| Rnf24 | 607.3774196 | -0.208003809 | 0.064065117 |
| Gm14233 | 1.848697502 | -1.604613699 | 0.762356743 |
| Gm14284 | 25.84299431 | 0.88416111 | 0.207985822 |
| Gm22902 | 0 | NA | NA |
| Gm14283 | 0 | NA | NA |
| Gm14304 | 0.048949532 | -0.243569582 | 3.724945049 |
| Smox | 612.36022 | 0.888449006 | 0.115422995 |
| Adra1d | 160.5552724 | -0.805302421 | 0.124865647 |
| Gm14285 | 9.998759481 | -0.734394974 | 0.416929117 |
| Gm14280 | 0.350616066 | 0.237330443 | 1.702125356 |
| 5330413P13Rik | 17.0164514 | -0.531850944 | 0.303616579 |
| Gm14282 | 0 | NA | NA |
| Gm14281 | 0.072025462 | 0.237312084 | 3.724945049 |
| Erv3 | 13.96015691 | 3.307987867 | 0.471994332 |
| Lamr1-ps1 | 0 | NA | NA |
| Prnp | 7268.086947 | 0.435951284 | 0.091815331 |
| Gm28045 | 0.311063295 | 2.031058787 | 1.738808305 |
| Prnd | 0.302678705 | 0.718225365 | 1.786544143 |
| Rassf2 | 1100.292024 | 0.35605808 | 0.113511118 |

| | | | |
|---|---|---|---|
| Slc23a2 | 2945.124527 | 0.090634501 | 0.071346451 |
| Gm14052 | 0.072025462 | 0.237312084 | 3.724945049 |
| Gm14051 | 0.790290772 | -0.077384068 | 1.322719928 |
| Tmem230 | 195.6462545 | 0.210810441 | 0.099904011 |
| Pcna | 440.6784249 | 0.998756979 | 0.156482514 |
| AV099323 | 3.007737845 | -0.835009943 | 0.622419086 |
| Cds2 | 6744.842328 | -0.517866737 | 0.060816453 |
| Prokr2 | 44.48347877 | -0.114093469 | 0.202818673 |
| 4921508D12Rik | 3.197768018 | 3.507464567 | 0.925271532 |
| Gm14095 | 0 NA | NA | |
| 1700026D11Rik | 0.269634638 | 1.199122435 | 1.824400259 |
| Gpcpd1 | 1339.978017 | -0.252688786 | 0.09331505 |
| AU019990 | 0.221462189 | 0.249407702 | 2.207548918 |
| Rpl23a-ps4 | 0 NA | NA | |
| Gm14099 | 0 NA | NA | |
| Gm14098 | 0.075043205 | 0.237309859 | 3.724945049 |
| 1110034G24Rik | 59.26919426 | -0.223953051 | 0.185158482 |
| Gm14097 | 0.654170826 | 1.730456188 | 1.324147621 |
| Gm22245 | 1.105592398 | -1.679163074 | 0.992636358 |
| Chgb | 2627.906697 | -0.164815107 | 0.25294823 |
| Trmt6 | 322.2692483 | 0.179365912 | 0.081720344 |
| Mcm8 | 58.14385559 | -0.132402455 | 0.177622785 |
| Gm14104 | 1.507066517 | -0.343837043 | 0.788543063 |
| Crls1 | 253.7650353 | 0.321956243 | 0.106760186 |
| Lrrn4 | 5.522564292 | -0.6093855 | 0.576216315 |
| Fermt1 | 3.449481863 | -0.890995704 | 0.690143236 |
| Gm14102 | 0.088736197 | -0.243576254 | 3.724945049 |
| Gm14101 | 0 NA | NA | |
| Gm25704 | 0 NA | NA | |
| Gm14103 | 0 NA | NA | |
| Gm14094 | 0.049553047 | 0.71820487 | 3.724945049 |
| Gm27449 | 0 NA | NA | |
| A430048G15Rik | 0.11037557 | -0.243571806 | 3.724945049 |
| Gm14100 | 0 NA | NA | |
| Bmp2 | 72.0830543 | 0.544010786 | 0.175558406 |
| Mir3090 | 0 NA | NA | |
| Gm25258 | 0 NA | NA | |
| Hao1 | 8.182416264 | 3.044314736 | 0.873600869 |
| Tmx4 | 4150.07463 | -0.125499928 | 0.06662246 |
| Gm14037 | 0.044191549 | -0.243571806 | 3.724945049 |
| Gm14038 | 8.079079625 | -0.489948792 | 0.376696975 |
| Plcb1 | 2558.946918 | -0.918967241 | 0.097547917 |
| Gm14036 | 21.28777894 | -0.1484303 | 0.241672965 |
| 4930545L23Rik | 0.686786262 | 0.866019895 | 1.231432371 |
| 9630028H03Rik | 3.015362371 | -0.616868685 | 0.665523974 |
| Platr3 | 0.132101749 | 1.209894437 | 3.72418822 |
| Plcb4 | 1658.213635 | -0.381478193 | 0.059947181 |

| | | | |
|---|---|---|---|
| Gm14209 | 16.51388486 | -1.095980826 | 0.347556503 |
| Gm14211 | 0.046345709 | 0.71820487 | 3.724945049 |
| Gm14210 | 0 | NA | NA |
| Gm26227 | 0.166709681 | -0.724459871 | 2.767631179 |
| Lamp5 | 675.5566326 | -1.07497253 | 0.154308398 |
| Pak7 | 618.9095195 | -0.719626069 | 0.120051752 |
| Gm14218 | 0 | NA | NA |
| Gm14056 | 0.369423471 | 1.901930455 | 1.503553419 |
| Gm14054 | 0.515606452 | 1.645834415 | 1.760782956 |
| Ankef1 | 28.51623535 | -0.502017108 | 0.203039949 |
| Gm36967 | 0.044544648 | 0.237309859 | 3.724945049 |
| Snap25 | 16400.01604 | -0.752235355 | 0.076324792 |
| Gm14053 | 0.15906016 | 1.290286593 | 3.433492162 |
| Mkks | 278.0206276 | -0.18745417 | 0.148134398 |
| Slx4ip | 166.0135647 | 0.157026574 | 0.118188093 |
| Jag1 | 129.0559854 | 0.498594166 | 0.130869862 |
| Gm28214 | 0.110595523 | 0.718200421 | 3.724945049 |
| Gm14055 | 0.284116082 | 1.220680246 | 2.011004084 |
| Gm14064 | 0 | NA | NA |
| Gm14062 | 10.87638177 | 3.898158296 | 0.897306444 |
| Btbd3 | 2584.985912 | -0.614389388 | 0.108029446 |
| Gm14061 | 4.211247599 | -0.69797614 | 0.531275858 |
| Mir1952 | 0 | NA | NA |
| Gm14065 | 0.143485256 | 1.229961212 | 3.547405499 |
| Gm14067 | 0 | NA | NA |
| Gm14069 | 0 | NA | NA |
| Gm14066 | 25.91994233 | -1.05587245 | 0.232818972 |
| Gm14071 | 0 | NA | NA |
| Sptlc3 | 1.944739822 | 0.3650263 | 0.863609759 |
| Gm27385 | 0 | NA | NA |
| Ism1 | 46.24466388 | -0.68138447 | 0.177965729 |
| Tasp1 | 470.1320558 | -0.166912382 | 0.089884493 |
| Gm14070 | 0 | NA | NA |
| Rps19-ps7 | 1.19142324 | 1.164496225 | 0.974782055 |
| Gm14073 | 0.066050875 | 0.71820487 | 3.724945049 |
| Esf1 | 372.9243209 | 0.241020399 | 0.089476724 |
| Gm17374 | 0 | NA | NA |
| Ndufaf5 | 256.6486657 | -0.008171126 | 0.080186852 |
| Sel1l2 | 0 | NA | NA |
| Macrod2 | 727.1380872 | -0.728893731 | 0.090996817 |
| Gm23846 | 0 | NA | NA |
| Flrt3 | 365.9243577 | -0.627869531 | 0.078896353 |
| Calr-ps | 0.108772582 | 0.718202646 | 3.724945049 |
| Zcchc9-ps | 30.12202849 | -1.348714735 | 0.384666783 |
| Macrod2os2 | 0 | NA | NA |
| Macrod2os1 | 0 | NA | NA |
| Gm25583 | 2.242176843 | -2.321104943 | 0.846858949 |

| | | | |
|---|---|---|---|
| Fau-ps1 | 8.271865471 | -1.68017915 | 0.532627502 |
| Gm23179 | 0.743559953 | -2.246914042 | 1.292712092 |
| Rpl26-ps5 | 0 | NA | NA |
| Rbmx2-ps | 0.144013665 | 1.483424291 | 3.538333097 |
| Kif16b | 346.7941092 | -0.286775531 | 0.102674051 |
| Gm22310 | 0.089089296 | 0.237305797 | 3.724945049 |
| Kif16bos | 0.612922179 | 0.65992478 | 1.266809124 |
| Snrpb2 | 243.6495243 | 0.49312052 | 0.13421647 |
| Otor | 0.348607679 | 2.650348489 | 1.892361484 |
| Gm14083 | 0 | NA | NA |
| Gm14082 | 0 | NA | NA |
| Gm11644 | 0 | NA | NA |
| Gm11645 | 3.276776717 | -1.498826691 | 0.747378515 |
| Pcsk2os1 | 65.67909268 | -0.802424322 | 0.176009278 |
| Pcsk2 | 3501.594654 | -0.983226823 | 0.105260827 |
| Pcsk2os2 | 0.594248913 | 0.310444603 | 1.466149787 |
| Bfsp1 | 5.877618629 | 0.147274879 | 0.41062284 |
| Rps13-ps6 | 0.132101749 | 1.209894437 | 3.72418822 |
| Gm11673 | 0 | NA | NA |
| Gm11686 | 0.0878635 | -0.72445792 | 3.724945049 |
| Dstn | 1131.896076 | 0.180918914 | 0.090937147 |
| Rrbp1 | 1060.324141 | 1.663786567 | 0.17124867 |
| Banf2 | 0 | NA | NA |
| Banf2os | 0.710948324 | -0.754641265 | 1.169861325 |
| Gm11687 | 1.624027788 | 1.233009756 | 0.811933777 |
| Gm5535 | 4.621059661 | 0.235952531 | 0.431552981 |
| 4930444E06Rik | 0.044191549 | -0.243571806 | 3.724945049 |
| Snx5 | 1040.74114 | 1.042208855 | 0.097119086 |
| Gm11688 | 3.796974204 | 0.609714508 | 0.521426291 |
| Gm26165 | 17.87795363 | 0.613965077 | 0.290322368 |
| Snord17 | 426.2071553 | 0.298667882 | 0.144534929 |
| Mgme1 | 58.73843915 | 0.355681657 | 0.151645241 |
| Ovol2 | 23.1269052 | 0.090931903 | 0.302647069 |
| 1700108N11Rik | 1.600434186 | 0.934711164 | 0.840055288 |
| Gm11700 | 0 | NA | NA |
| Gm11701 | 1.140052013 | -0.205231339 | 0.987021189 |
| Csrp2bp | 346.2940874 | -0.024829947 | 0.065418461 |
| Pet117 | 0 | NA | NA |
| Zfp133-ps | 17.17923113 | -0.503280622 | 0.253896316 |
| Dzank1 | 3640.794771 | -0.779570113 | 0.06224274 |
| Polr3f | 438.0816624 | 0.233825677 | 0.116484955 |
| Rbbp9 | 270.2515903 | -0.199023245 | 0.135001751 |
| Sec23b | 663.5913093 | 0.776536035 | 0.09040571 |
| Gm561 | 84.95980064 | 0.103691189 | 0.15075849 |
| Dtd1 | 350.4218442 | -0.26819816 | 0.073318894 |
| Gm24823 | 0 | NA | NA |
| Gm11743 | 12.81133691 | -0.740114541 | 0.415369029 |

| | | | |
|---|---|---|---|
| Scp2d1 | 0.431157666 | 1.652440871 | 1.561539495 |
| Gm11746 | 0 | NA | NA |
| Slc24a3 | 1355.879502 | -0.594145728 | 0.077798396 |
| Gm14093 | 14.45402812 | -0.903937823 | 0.332508521 |
| Gm22731 | 3.697861772 | -1.723158086 | 0.637257142 |
| Gm22390 | 0 | NA | NA |
| Gm11759 | 0.254720735 | 0.717772792 | 1.838858596 |
| Gm14092 | 0.048949532 | -0.243569582 | 3.724945049 |
| BC039771 | 5.098733496 | 0.43671568 | 0.499760106 |
| Gm11763 | 0.066050875 | 0.71820487 | 3.724945049 |
| Rin2 | 761.9091662 | 1.429440447 | 0.123767873 |
| Gm25026 | 1.82714634 | -0.493559199 | 0.706013455 |
| Gm11764 | 0 | NA | NA |
| Naa20 | 524.4015188 | 0.829306055 | 0.099379036 |
| Crnkl1 | 313.9592317 | 0.221833667 | 0.073310934 |
| Cfap61 | 40.89814708 | -0.94620482 | 0.187836483 |
| Insm1 | 43.73815878 | -0.615530769 | 0.192143024 |
| Ralgapa2 | 804.7755793 | -0.451457772 | 0.151633164 |
| A930019D19Rik | 0.11147366 | 1.230376831 | 3.722845817 |
| Gm14117 | 89.00856018 | -0.175077123 | 0.133309471 |
| 4933406D12Rik | 0 | NA | NA |
| Gm14111 | 0.398068774 | 1.15831352 | 3.723962117 |
| Gm14114 | 1.234305601 | -0.454208628 | 0.881007593 |
| Kiz | 194.5668815 | -0.01532807 | 0.100400199 |
| Xrn2 | 1167.834356 | 0.527737471 | 0.08755375 |
| Nkx2-4 | 1.324777524 | 1.646428953 | 1.231277665 |
| Gm14116 | 0.119587853 | 0.237305575 | 3.724945049 |
| Nkx2-2 | 88.16107763 | 0.591773582 | 0.141142568 |
| Nkx2-2os | 6.403733378 | -0.041021538 | 0.40204717 |
| 6430503K07Rik | 6.531065735 | -0.409241326 | 0.453080976 |
| Gm22261 | 0 | NA | NA |
| Gm14115 | 0.048949532 | -0.243569582 | 3.724945049 |
| Gm14113 | 3.843754305 | 0.05931067 | 0.468310805 |
| Gm14110 | 0 | NA | NA |
| AI646519 | 0 | NA | NA |
| Pax1 | 0.59047672 | 0.553992629 | 1.281466589 |
| Gm14109 | 0 | NA | NA |
| Gm14107 | 0 | NA | NA |
| Gm14106 | 0.066050875 | 0.71820487 | 3.724945049 |
| A530006G24Rik | 0 | NA | NA |
| Gm25516 | 0 | NA | NA |
| 9030622O22Rik | 0.80144685 | -1.41255907 | 1.245014873 |
| Foxa2 | 2.785346695 | 0.321607094 | 0.833787065 |
| Gm27420 | 0 | NA | NA |
| Gm14108 | 0 | NA | NA |
| Gm24221 | 0 | NA | NA |
| Sstr4 | 86.59783816 | -1.164617201 | 0.155027864 |

| | | | |
|---|---|---|---|
| Thbd | 190.3799444 | 1.234134891 | 0.173501035 |
| Gm14119 | 0.109727584 | 0.718200421 | 3.724945049 |
| Cd93 | 290.705625 | 4.106359526 | 0.210534018 |
| Gm14120 | 2.718509683 | -1.824330336 | 0.968522896 |
| Gm25866 | 0 | NA | NA |
| Nxt1 | 62.52770177 | 0.636513171 | 0.189955704 |
| Gzf1 | 464.5606654 | 0.017165817 | 0.062187668 |
| Napb | 4683.537772 | -0.353425131 | 0.049660445 |
| Cstl1 | 7.743612027 | 0.766502642 | 0.387677635 |
| Cst11 | 0.068097359 | -0.243569582 | 3.724945049 |
| 8030411F24Rik | 0.116944185 | 1.302646088 | 3.719336636 |
| Cst12 | 0.585343961 | 1.588861409 | 1.569621431 |
| Cst8 | 0.132101749 | 1.209894437 | 3.72418822 |
| Cst13 | 0.138076337 | 0.718204987 | 3.589802529 |
| Cst9 | 0.038981395 | 0.718207094 | 3.724945049 |
| 9230104L09Rik | 0.693909001 | 1.295677402 | 1.188566556 |
| Cst3 | 6207.776889 | -0.377911773 | 0.125102795 |
| Gm38221 | 0.060174537 | 0.237309859 | 3.724945049 |
| Gm14129 | 0 | NA | NA |
| Gm1330 | 0 | NA | NA |
| Gm10750 | 0 | NA | NA |
| Gm14135 | 0 | NA | NA |
| Gm14132 | 0.875345121 | -1.211440935 | 1.140022797 |
| Gm14131 | 0.1140246 | 0.718202646 | 3.724945049 |
| Cst10 | 0 | NA | NA |
| Gm24551 | 0.086648695 | -0.72445792 | 3.724945049 |
| Gm14130 | 2.711335961 | 0.311285701 | 0.748381634 |
| Gm14133 | 2.358317775 | -0.221244766 | 0.735440611 |
| Syndig1 | 375.012134 | -0.32190454 | 0.086883247 |
| C530025M09Rik | 3.362260024 | -0.641060231 | 0.550109805 |
| Gm14140 | 1.613620997 | 0.447011329 | 0.719829752 |
| Zfp120 | 248.9582029 | 0.164631284 | 0.132104922 |
| Gm10770 | 0 | NA | NA |
| Gm14139 | 0.236020217 | -1.078077901 | 2.493450191 |
| Zfp937 | 71.56539432 | -0.107422288 | 0.158918843 |
| Gm21994 | 0.580893185 | 0.980064406 | 1.546906504 |
| Gm14124 | 2.427470934 | -0.776737535 | 0.802666632 |
| Gm14125 | 0 | NA | NA |
| 3300002I08Rik | 53.22250888 | -0.065688823 | 0.199073618 |
| Gm10130 | 6.196226755 | 0.680990769 | 0.415950836 |
| Gm14126 | 0.606327527 | -1.164975684 | 1.33734221 |
| Gm14122 | 12.30982907 | -0.674178419 | 0.302716907 |
| Zfp442 | 70.96873342 | -0.00704698 | 0.173512204 |
| Gm14127 | 0.190396633 | 0.718208389 | 3.269807176 |
| Zfp345 | 0.580755766 | 1.129566445 | 1.289830001 |
| Gm14121 | 87.55734381 | -0.257809951 | 0.169390431 |
| Gm14128 | 0 | NA | NA |

| | | | |
|---|---|---|---|
| Cst7 | 47.58322733 | 6.086185164 | 0.621007655 |
| Apmap | 559.647641 | 0.197655163 | 0.091261844 |
| Gm14123 | 1.105496479 | 2.349327001 | 1.075634469 |
| Acss1 | 208.0155599 | -0.791745775 | 0.119933712 |
| E130215H24Rik | 2.738071508 | -1.135371371 | 0.675761536 |
| Vsx1 | 2.657225507 | -1.145687421 | 0.658096881 |
| Platr15 | 0.232351398 | -0.680203956 | 2.548595388 |
| Gm28450 | 0 | NA | NA |
| Gm26681 | 0.300525961 | 2.314851091 | 3.691932167 |
| Entpd6 | 438.5268923 | -0.315056522 | 0.073208334 |
| Pygb | 1906.36878 | -0.251691445 | 0.105708439 |
| Abhd12 | 2275.162201 | -0.011490187 | 0.048630725 |
| Gm23598 | 0 | NA | NA |
| Gins1 | 16.63953435 | 1.610671847 | 0.330084016 |
| Gm14143 | 0.083526043 | 0.718202646 | 3.724945049 |
| Ninl | 67.5515232 | -0.171136721 | 0.16598422 |
| Gm14142 | 0.423021739 | -0.240769554 | 1.71531797 |
| Gm14141 | 0 | NA | NA |
| Nanp | 8.071585247 | -0.231005347 | 0.482184926 |
| Gm22187 | 0 | NA | NA |
| Gm24199 | 0 | NA | NA |
| Gm14151 | 0 | NA | NA |
| Gm14147 | 0 | NA | NA |
| Gm14145 | 0 | NA | NA |
| Gm22517 | 0 | NA | NA |
| Gm14149 | 0.579212071 | -1.10842932 | 1.332684175 |
| Gm14144 | 0 | NA | NA |
| Gm23005 | 0 | NA | NA |
| Gm24077 | 0 | NA | NA |
| 4930442J19Rik | 0 | NA | NA |
| Gm23261 | 0 | NA | NA |
| Gm14148 | 0.134276636 | 0.684459443 | 2.951368184 |
| Gm14152 | 0 | NA | NA |
| Gm24868 | 0 | NA | NA |
| Gm14150 | 75.86947207 | 0.24397653 | 0.216538468 |
| Gm14146 | 0 | NA | NA |
| 4921509C19Rik | 0.322748876 | 1.906772647 | 3.700711097 |
| Nsfl1c | 781.3049052 | 0.330600899 | 0.059514587 |
| Gm14167 | 3.663554111 | -0.292652125 | 0.541529399 |
| Fkbp1a | 3786.330142 | -0.042402442 | 0.067483636 |
| Sdcbp2 | 21.61712664 | 3.491489154 | 0.434121072 |
| Snph | 3369.871354 | -0.446631447 | 0.09232487 |
| Rad21l | 5.277545605 | -0.396060527 | 0.459022064 |
| Tmem74bos | 12.83654197 | 0.087818495 | 0.338142806 |
| Gm14159 | 1.598224206 | -1.353340871 | 0.905606677 |
| Tmem74b | 29.74572494 | -0.902340917 | 0.224599724 |
| Psmf1 | 502.2961145 | 0.514938413 | 0.099266464 |

| | | | |
|---|---|---|---|
| 4930556L07Rik | 0.192951122 | 0.237321161 | 2.643859218 |
| Gm14153 | 0 | NA | NA |
| Gm14155 | 0 | NA | NA |
| Rspo4 | 1.536911596 | -0.588400779 | 0.808629239 |
| Rspo4os | 0.417636614 | 0.237333904 | 1.472330291 |
| Angpt4 | 4.455728532 | 3.25771563 | 0.701150916 |
| Gm14154 | 0.934869149 | 0.28162358 | 1.257660629 |
| Mir1953 | 0.11193333 | 0.718202646 | 3.724945049 |
| Fam110a | 57.87769406 | 0.228449197 | 0.150631078 |
| Slc52a3 | 22.60185597 | 2.584561136 | 0.283118234 |
| AA387200 | 2.676545693 | 0.236391752 | 0.589510302 |
| Scrt2 | 297.4913887 | 0.283865919 | 0.142927839 |
| Srxn1 | 1496.550893 | 1.470196944 | 0.11133562 |
| Tcf15 | 3.917975245 | 0.264152793 | 0.509017698 |
| Rps15a-ps7 | 16.65743553 | 0.91654493 | 0.265492816 |
| Csnk2a1 | 1499.289359 | -0.052915508 | 0.038260688 |
| Tbc1d20 | 711.5116405 | 0.298458427 | 0.054974027 |
| Rbck1 | 775.7363636 | 0.037369077 | 0.065149349 |
| Trib3 | 48.90459884 | 4.23739891 | 0.398288017 |
| Gm14164 | 0.458177261 | -0.441848985 | 1.496176528 |
| Gm14165 | 3.039451885 | 1.10421287 | 0.608942509 |
| Nrsn2 | 894.8007339 | -0.205555267 | 0.08694646 |
| Sox12 | 175.956736 | -0.487650551 | 0.142995134 |
| Zcchc3 | 147.4086379 | 0.120538432 | 0.118883027 |
| 6820408C15Rik | 27.31515061 | -0.073743364 | 0.191576616 |
| Gm27343 | 3.046712631 | -0.88686581 | 0.548054902 |
| Gm14158 | 0.634981691 | -0.741663315 | 1.206066523 |
| Defb23 | 0.097139547 | 0.718207094 | 3.724945049 |
| Gm14156 | 0 | NA | NA |
| Defb20 | 0.545971167 | -1.230288498 | 1.32061789 |
| Defb22 | 0.061920613 | 0.718207094 | 3.724945049 |
| Defb26 | 0 | NA | NA |
| Defb28 | 0 | NA | NA |
| Defb29 | 0 | NA | NA |
| Gm14161 | 0.118371171 | 0.718202646 | 3.724945049 |
| Gm17416 | 0 | NA | NA |
| Defb21 | 0.04393175 | -0.243571806 | 3.724945049 |
| Defb19 | 0 | NA | NA |
| Defb45 | 0.07796279 | 1.072698098 | 3.721829099 |
| Defb36 | 0 | NA | NA |
| Gm14163 | 0 | NA | NA |
| Defb25 | 0.234671876 | -0.243564023 | 2.172263027 |
| Rem1 | 3.041196478 | -0.862111271 | 0.609886487 |
| 1700030C14Rik | 0.290525896 | -0.724461378 | 1.805594899 |
| H13 | 832.2285858 | 0.405122206 | 0.072434557 |
| Mcts2 | 80.0283442 | 0.053187533 | 0.121951181 |
| Gm26841 | 0.794703889 | -0.4682824 | 1.108713764 |

| | | | |
|---|---|---|---|
| Id1 | 158.7587211 | 1.959209547 | 0.205927175 |
| Gm14162 | 0.044191549 | -0.243571806 | 3.724945049 |
| Cox4i2 | 12.07102929 | 0.04427328 | 0.328735438 |
| Gm23802 | 0.93732221 | -1.918943399 | 1.092367144 |
| Bcl2l1 | 1656.202641 | 0.814153961 | 0.112173934 |
| Tpx2 | 97.61010764 | 2.373247056 | 0.236582847 |
| Mir5622 | 0 | NA | NA |
| Mylk2 | 13.81209412 | 7.737747762 | 0.964446904 |
| Foxs1 | 7.663944057 | 2.464253336 | 0.48108984 |
| Dusp15 | 161.0227663 | -0.918106783 | 0.121088153 |
| Ttll9 | 15.9748206 | -0.063622649 | 0.326784327 |
| Gm14199 | 0.260717194 | 1.638578725 | 2.039123502 |
| Pdrg1 | 314.6393574 | 0.973044694 | 0.090366358 |
| Xkr7 | 77.04392738 | -0.734535364 | 0.159832924 |
| Ccm2l | 14.10545208 | 0.449934738 | 0.275070558 |
| Hck | 217.2183995 | 4.715708136 | 0.218941634 |
| Gm25643 | 0 | NA | NA |
| Tm9sf4 | 988.450612 | 0.078813441 | 0.04241609 |
| Tspyl3 | 233.7002 | -0.519256531 | 0.094714229 |
| Plagl2 | 263.3156821 | 1.066079291 | 0.113365225 |
| Pofut1 | 393.8680983 | -0.050114926 | 0.084094623 |
| Kif3b | 984.8683119 | 0.039350386 | 0.045433796 |
| 2500004C02Rik | 39.85926624 | -0.824380167 | 0.223442398 |
| Asxl1 | 918.9589658 | -0.085794575 | 0.083041663 |
| Nol4l | 699.5239319 | -0.563761841 | 0.125660707 |
| Gm14472 | 0.227745227 | -0.243570935 | 2.58737708 |
| 4930404H24Rik | 0.825833058 | 0.042290595 | 1.052483277 |
| Commd7 | 204.4885226 | 0.421006039 | 0.092579788 |
| Dnmt3bos | 0 | NA | NA |
| Dnmt3b | 18.90429361 | -0.046414972 | 0.285738471 |
| Gm14490 | 1.779788623 | 1.623495457 | 0.759742705 |
| Mapre1 | 1211.391518 | 0.234583661 | 0.070953002 |
| Efcab8 | 3.424510338 | 2.001775461 | 0.895107651 |
| Gm14494 | 80.10444224 | 0.130066986 | 0.131658532 |
| Sun5 | 0.124596252 | 0.718200615 | 3.714681068 |
| Bpifb2 | 0.955311604 | 0.362757725 | 1.031705568 |
| Bpifb6 | 1.501918467 | 0.82685685 | 0.816811057 |
| Bpifb3 | 1.988936712 | -0.15791372 | 1.472858256 |
| Bpifb4 | 3.358904097 | 0.443584645 | 1.203102342 |
| Bpifa6 | 0.496470236 | -0.278783185 | 3.678519652 |
| Bpifa2 | 0 | NA | NA |
| Gm21850 | 0 | NA | NA |
| Gm14497 | 0.15596086 | -1.187811386 | 3.483969583 |
| Bpifa3 | 0 | NA | NA |
| Bpifa1 | 1.270782233 | -0.362997474 | 1.078520859 |
| Bpifa5 | 0.243228626 | 0.867111339 | 3.022466272 |
| Bpifb1 | 0.700162942 | 0.995007119 | 1.097242634 |

| | | | |
|---|---|---|---|
| Bpifb5 | 0.765485319 | -0.682062252 | 1.160454361 |
| Bpifb9a | 1.362283058 | -0.68894176 | 0.925660962 |
| Gm14197 | 1.834732021 | -0.024027956 | 0.695429317 |
| Bpifb9b | 3.934499224 | -1.142874247 | 1.489660133 |
| Cdk5rap1 | 105.428526 | 0.014966934 | 0.135859799 |
| Snta1 | 269.5638173 | -0.161663481 | 0.112945997 |
| Cbfa2t2 | 875.8606125 | -0.120406652 | 0.05059502 |
| Necab3 | 257.0570188 | -1.66146345 | 0.173837901 |
| 1700003F12Rik | 3.150202034 | -0.657769338 | 0.702796799 |
| Actl10 | 0.266983968 | 0.290665739 | 1.813687922 |
| E2f1 | 73.04566475 | -0.636048193 | 0.228084266 |
| Pxmp4 | 135.3858191 | -0.531625839 | 0.115281102 |
| 4930519P11Rik | 1.196740622 | -0.575648534 | 0.947672218 |
| Zfp341 | 104.6788915 | -0.516229697 | 0.130489514 |
| Gm14198 | 0.162949104 | -0.243562152 | 2.787607662 |
| Chmp4b | 2004.387491 | 1.011048457 | 0.136539481 |
| Rpl5-ps2 | 5.302412506 | 0.947736333 | 0.425685566 |
| Gm14215 | 0 | NA | NA |
| Gm14216 | 0.099155932 | 0.718202646 | 3.724945049 |
| Gm14214 | 0 | NA | NA |
| Raly | 850.5641207 | 0.629613444 | 0.086685101 |
| a | 0.140640116 | 0.719187302 | 3.611523994 |
| Eif2s2 | 723.9443759 | 0.76580985 | 0.115058636 |
| Gm14226 | 0.71216146 | 1.364522364 | 1.298637265 |
| Ahcy | 112.8096952 | 0.512053421 | 0.147448301 |
| 2310005A03Rik | 0.04393175 | -0.243571806 | 3.724945049 |
| Itch | 1259.365133 | 0.137706952 | 0.062206412 |
| Gm14227 | 0.141071297 | 0.237322348 | 3.564043836 |
| Dynlrb1 | 713.8695448 | -0.047864354 | 0.088378759 |
| Map1lc3a | 781.5540749 | -0.185465046 | 0.121666362 |
| Pigu | 261.7083534 | 0.150062569 | 0.109388408 |
| Mir695 | 0 | NA | NA |
| Trp53inp2 | 2971.079894 | 0.042661236 | 0.106867417 |
| Ncoa6 | 1310.679018 | -0.149772426 | 0.068907603 |
| Gm14239 | 28.68396613 | -0.879514878 | 0.304498023 |
| Gm14238 | 0.886238366 | -0.845369774 | 1.103505454 |
| Ggt7 | 845.8338809 | -0.432203527 | 0.073215666 |
| Acss2 | 635.0895078 | -0.344492829 | 0.06644025 |
| Mipep-ps | 1.837594399 | 0.297421575 | 0.809041385 |
| Gm20637 | 2.163212633 | -0.897004535 | 0.719644583 |
| Acss2os | 0.277114854 | -0.298530038 | 2.054940167 |
| Gss | 143.0865772 | 0.097803129 | 0.123562974 |
| Gssos2 | 1.802868249 | -0.266994968 | 0.913025876 |
| Gssos1 | 1.992642508 | -1.911498053 | 1.000630048 |
| Gm14257 | 0.038981395 | 0.718207094 | 3.724945049 |
| Myh7b | 88.10465837 | -0.32557978 | 0.272615069 |
| Mir499 | 0 | NA | NA |

| | | | |
|---|---|---|---|
| Trpc4ap | 1703.710218 | 0.121776073 | 0.054743375 |
| Gm22334 | 1.228515858 | 0.015034539 | 1.054555544 |
| Edem2 | 274.0937995 | 0.3702649 | 0.089302934 |
| Gm17581 | 0 | NA | NA |
| Procr | 21.11771009 | 4.131664832 | 0.446854131 |
| Mmp24 | 793.763571 | -0.310633285 | 0.133039956 |
| BC029722 | 543.0851477 | -0.1833406 | 0.092345815 |
| Eif6 | 475.8961878 | 1.066151888 | 0.118136963 |
| Gm16098 | 1.335672075 | 1.930820348 | 0.960849876 |
| Fam83c | 0.240161549 | 1.34798213 | 3.035217049 |
| Uqcc1 | 374.2692931 | 0.249895489 | 0.083624702 |
| Gm15557 | 0.53525542 | 0.25627681 | 1.432519954 |
| Gdf5 | 4.286069218 | 0.752705914 | 0.605223137 |
| Cep250 | 337.5416032 | -0.338793389 | 0.158602909 |
| 6430550D23Rik | 7.007522312 | -0.712376099 | 0.401968372 |
| Ergic3 | 697.3852742 | 0.183998061 | 0.089271653 |
| Fer1l4 | 0.699011511 | 3.099509098 | 2.561566208 |
| Gm14240 | 0 | NA | NA |
| Spag4 | 20.11396154 | -0.97122078 | 0.308782313 |
| Cpne1 | 7.154347536 | -1.115912969 | 0.454606203 |
| Gm28036 | 0.689591474 | -0.016764327 | 1.051493541 |
| Gm37274 | 40.70937969 | -0.821770524 | 0.326055549 |
| Rbm12 | 333.4415097 | -0.112984111 | 0.083242739 |
| Nfs1 | 286.4122944 | -0.092006993 | 0.089499726 |
| Romo1 | 128.8625568 | -0.226756138 | 0.178991458 |
| Rbm39 | 3275.570639 | 0.13484301 | 0.088178373 |
| Phf20 | 1427.32006 | -0.287098891 | 0.051668845 |
| Scand1 | 367.1199098 | 0.707912571 | 0.175965278 |
| Cnbd2 | 68.49456786 | 0.326630059 | 0.134672035 |
| Gm14251 | 0.489024281 | 2.474713512 | 1.621449742 |
| Gm22759 | 0 | NA | NA |
| Gm14252 | 0.106056992 | 0.237307635 | 3.724945049 |
| Gm14253 | 61.94935748 | 0.750609247 | 0.198362068 |
| Gm14170 | 0.179599912 | 0.18250098 | 2.721541985 |
| Gm22316 | 0.234323835 | 1.155422934 | 2.538723173 |
| Rpl21-ps9 | 0.666172099 | -1.501017126 | 1.201738726 |
| 2900097C17Rik | 2086.303339 | -0.516584572 | 0.092373521 |
| Gm14173 | 0.760931553 | 0.349746548 | 0.977852569 |
| Gm22758 | 0 | NA | NA |
| Epb4.1l1 | 5881.508901 | -0.46998924 | 0.078144552 |
| Gm14225 | 0 | NA | NA |
| Gm14224 | 0.038981395 | 0.718207094 | 3.724945049 |
| Aar2 | 389.6193864 | 0.303752047 | 0.071278741 |
| Gm14168 | 20.52960382 | 0.341364351 | 0.303682098 |
| Gm14169 | 37.69901258 | -0.791335549 | 0.206953849 |
| Dlgap4 | 2838.730653 | -0.326175302 | 0.101056644 |
| Gm14174 | 0.748568137 | -2.18584661 | 1.15330589 |

| | | | |
|---|---|---|---|
| Gm14172 | 0 | NA | NA |
| 4930405A21Rik | 7.408513515 | 0.049795323 | 0.361445869 |
| Myl9 | 37.34594635 | -0.215456337 | 0.263962525 |
| Gm14231 | 0.108266322 | 1.230376831 | 3.722845817 |
| Gm14230 | 5.976257995 | 1.948517868 | 0.489276142 |
| Tgif2 | 38.14849271 | 2.283063819 | 0.318611535 |
| 5430405H02Rik | 85.38075778 | 0.125744455 | 0.172423337 |
| Gm37790 | 6.47799121 | 0.000379409 | 0.438892391 |
| 4930518I15Rik | 26.40534196 | -0.215830287 | 0.252082662 |
| Gm27651 | 0 | NA | NA |
| 1110008F13Rik | 336.8881828 | 0.527981689 | 0.110633632 |
| Sla2 | 15.40209609 | 2.324638851 | 0.390119894 |
| Gm14278 | 0.888132063 | -0.728109013 | 1.105683733 |
| Gm14248 | 0 | NA | NA |
| Gm14247 | 0.534422126 | -2.447959824 | 1.651600044 |
| Ndrg3 | 3682.906368 | -0.801933375 | 0.078136903 |
| Gm27653 | 9.457828482 | -1.140629069 | 0.459033679 |
| Gm14276 | 8.337030673 | -0.383019361 | 0.353451168 |
| Dsn1 | 45.4075692 | 0.312638686 | 0.203662364 |
| Soga1 | 1303.081876 | -0.294425278 | 0.120128815 |
| Tldc2 | 4.0710617 | -0.9830167 | 0.497261046 |
| Samhd1 | 1799.230623 | 4.128769808 | 0.185260165 |
| Rbl1 | 152.3466098 | 2.484534327 | 0.164641708 |
| Gm14277 | 3.35648137 | 0.5417721 | 0.502825613 |
| Mroh8 | 16.34303778 | -0.627559383 | 0.284645776 |
| Rpn2 | 1339.45037 | 0.580345709 | 0.076809858 |
| Ghrh | 3.638464397 | 0.946508034 | 0.609685653 |
| Manbal | 220.2483533 | 0.343248311 | 0.128761775 |
| Src | 1335.75724 | -0.011613863 | 0.06177751 |
| Gm14279 | 2.044822302 | 1.583692594 | 0.770928914 |
| Gm14287 | 40.93809312 | -0.15497726 | 0.255470533 |
| Gm14286 | 22.25133751 | -1.256093482 | 0.355053208 |
| Blcap | 862.3755201 | 0.055866624 | 0.062681287 |
| Nnat | 1274.354139 | 0.013952408 | 0.167601567 |
| Gm23134 | 19.82322296 | -0.498978078 | 0.450553724 |
| Platr27 | 0.11147366 | 1.230376831 | 3.722845817 |
| Gm25407 | 0 | NA | NA |
| Gm23463 | 0 | NA | NA |
| Ctnnbl1 | 279.0876754 | -0.035044923 | 0.079823424 |
| Gm14176 | 7.679034499 | 0.283632909 | 0.353794769 |
| Vstm2l | 466.6791405 | -0.598642144 | 0.09828874 |
| Tti1 | 247.1897018 | 0.155846579 | 0.119215756 |
| Rprd1b | 501.6807887 | 0.056801468 | 0.065357157 |
| 2010009K17Rik | 0.075043205 | 0.237309859 | 3.724945049 |
| Tgm2 | 2012.035572 | 5.944506622 | 0.332753379 |
| Gm20412 | 1.717954688 | 3.59900263 | 1.367638433 |
| D630003M21Rik | 59.49929145 | -0.700577503 | 0.176955674 |

| | | | |
|---|---|---|---|
| 1700060C20Rik | 0.29895891 | 1.381014202 | 2.041101338 |
| Mir7685 | 0 | NA | NA |
| Bpi | 0.044544648 | 0.237309859 | 3.724945049 |
| Lbp | 183.630573 | 2.162801131 | 0.249685089 |
| Snhg17 | 148.0843631 | 0.432747094 | 0.18666818 |
| Gm24411 | 0 | NA | NA |
| Gm26003 | 0 | NA | NA |
| Gm23925 | 0 | NA | NA |
| Snhg11 | 9158.007078 | -1.263936222 | 0.334602707 |
| Gm25187 | 0 | NA | NA |
| Gm25129 | 29.82494218 | -0.911941117 | 0.299148737 |
| Gm14201 | 0 | NA | NA |
| Gm14202 | 0 | NA | NA |
| Ralgapb | 2672.82956 | -0.244164261 | 0.052592318 |
| Gm14200 | 0.166639798 | 0.718214905 | 2.767997355 |
| Gm24379 | 4.859614805 | -0.520368376 | 0.454752656 |
| Adig | 1.688935393 | -0.479084521 | 0.664372329 |
| Arhgap40 | 1.707670524 | 0.231918602 | 0.787802862 |
| Gm14204 | 129.1982071 | -0.603408742 | 0.144305606 |
| Slc32a1 | 1229.442361 | -0.332565035 | 0.062224118 |
| Gm14205 | 1.579080267 | 0.975958688 | 0.812096364 |
| Actr5 | 113.0741844 | 0.200677832 | 0.17005743 |
| Mir3474 | 0.201505633 | 0.237327064 | 2.605863055 |
| Ppp1r16b | 2425.103286 | -0.296949771 | 0.077397614 |
| Gm26141 | 1.537427175 | -0.279498769 | 0.775774293 |
| Fam83d | 51.99511303 | 0.257207182 | 0.217932325 |
| Dhx35 | 216.8272346 | 0.153622478 | 0.11030376 |
| AL669913.1 | 0.255320825 | 0.237320997 | 2.417314357 |
| Gm25090 | 0 | NA | NA |
| Gm14219 | 0.112288909 | -0.724455696 | 3.724945049 |
| Gm11445 | 0 | NA | NA |
| Gm11446 | 0 | NA | NA |
| Gm14223 | 0 | NA | NA |
| 4933416E03Rik | 0 | NA | NA |
| Gm14220 | 0 | NA | NA |
| 9130015L21Rik | 0 | NA | NA |
| Gm826 | 0.8433133 | 1.076312262 | 1.251532215 |
| Mafb | 493.6009209 | 2.445716293 | 0.171966141 |
| Gm14221 | 0.304607851 | 2.099788164 | 1.981031123 |
| Gm14222 | 0 | NA | NA |
| Gm23968 | 0.062426873 | 0.237312084 | 3.724945049 |
| Top1 | 1394.007593 | -0.159157545 | 0.06285326 |
| Plcg1 | 938.5103693 | -0.253685416 | 0.068295781 |
| Gm27206 | 3.426664286 | 0.304055262 | 0.576940257 |
| Zhx3 | 1591.941502 | -0.230421684 | 0.052893386 |
| Gm16751 | 10.07322116 | -0.006041114 | 0.309549587 |
| Gm11447 | 0.089814205 | -0.24357403 | 3.724945049 |

| | | | |
|---|---|---|---|
| Lpin3 | 58.0580359 | 3.738036909 | 0.257131554 |
| Emilin3 | 11.35137183 | -0.106157624 | 0.332638093 |
| Gm14228 | 0.449436652 | 0.237294043 | 3.724945049 |
| Chd6 | 2195.755601 | -0.416259502 | 0.068986561 |
| Gm25561 | 0 | NA | NA |
| Gm14243 | 0.04393175 | -0.243571806 | 3.724945049 |
| Gm14242 | 0 | NA | NA |
| Gm5270 | 0.106618422 | -0.24357403 | 3.724945049 |
| Gm11448 | 0.467761363 | -0.089514686 | 1.373070742 |
| Gm14241 | 0 | NA | NA |
| Gm11449 | 0.644287204 | 0.28909765 | 1.284420514 |
| Gm29229 | 0.291544834 | 0.210157196 | 1.581536393 |
| Ptprt | 2587.731476 | -0.765648877 | 0.076741682 |
| Ptprtos | 0.352863859 | 2.187946718 | 1.87241383 |
| 9430021M05Rik | 23.91964892 | -1.385488678 | 0.262873323 |
| Gm14246 | 8.592019774 | -0.901002502 | 0.431419024 |
| Gm22936 | 0 | NA | NA |
| Gm11450 | 0 | NA | NA |
| Gm24478 | 0 | NA | NA |
| Gm14255 | 0.044191549 | -0.243571806 | 3.724945049 |
| Gm11451 | 16.98984007 | 0.396325645 | 0.30936149 |
| Gm11452 | 0.150711795 | 0.237323719 | 2.856704362 |
| Srsf6 | 1346.747418 | 0.268717926 | 0.082030311 |
| L3mbtl1 | 55.09161921 | -1.988820241 | 0.370130119 |
| Gm11453 | 0.038981395 | 0.718207094 | 3.724945049 |
| Sgk2 | 21.73993536 | -0.993913613 | 0.297432312 |
| Ift52 | 322.3681409 | 0.145485233 | 0.075543891 |
| Mybl2 | 17.83251159 | 0.92015027 | 0.292809552 |
| Mir7003 | 0.049553047 | 0.71820487 | 3.724945049 |
| Gtsf1l | 0.489956056 | 0.646805813 | 1.363939873 |
| Gm22174 | 0 | NA | NA |
| Gm11454 | 2.386488514 | -0.205753198 | 0.663928493 |
| Gm14254 | 0 | NA | NA |
| Tox2 | 401.2220022 | -0.209331405 | 0.104263903 |
| Jph2 | 9.352170419 | -0.079287169 | 0.339716604 |
| Oser1 | 111.6600522 | 0.586519288 | 0.132778541 |
| 2900093K20Rik | 14.33683216 | -0.129869259 | 0.264223841 |
| Gdap1l1 | 668.5251384 | -0.013170323 | 0.090218692 |
| Fitm2 | 424.7360197 | -0.030931064 | 0.06510397 |
| 2310001K24Rik | 0.198365723 | -0.577757461 | 2.624989092 |
| R3hdml | 0.349069061 | -0.147592965 | 1.75577197 |
| Hnf4aos | 0 | NA | NA |
| Hnf4a | 0.497304619 | 0.879467093 | 1.48687912 |
| Gm25881 | 0 | NA | NA |
| Ttpal | 655.7681734 | -0.162654507 | 0.088841949 |
| Serinc3 | 3221.802251 | 0.636970982 | 0.072005833 |
| 0610039K10Rik | 0.721809532 | 1.114855944 | 1.156092953 |

| | | | |
|---|---|---|---|
| Pkig | 413.2230987 | 0.113101297 | 0.076918654 |
| Gm16316 | 3.804176175 | 4.600099684 | 0.918185226 |
| Ada | 48.39031174 | 3.260953699 | 0.283993792 |
| Wisp2 | 23.40200403 | 3.858014818 | 0.490801969 |
| Kcnk15 | 8.160985404 | -0.154378764 | 0.481225245 |
| Rims4 | 1059.645651 | -0.401454971 | 0.082702624 |
| Ywhab | 4440.302886 | -0.181606521 | 0.058996892 |
| Pabpc1l | 10.92999753 | 1.964611898 | 0.489967501 |
| Tomm34 | 543.6160348 | -0.245999044 | 0.069915578 |
| Stk4 | 685.9737304 | 0.338551336 | 0.076305375 |
| Gm11455 | 4.303662858 | -1.107437444 | 0.592694007 |
| n-R5s207 | 0 | NA | NA |
| Kcns1 | 75.78496726 | -0.726067137 | 0.201163776 |
| Wfdc5 | 0.042716945 | -0.243571806 | 3.724945049 |
| A730032A03Rik | 0.162822621 | 1.512421763 | 3.396659703 |
| Wfdc12 | 0.231631338 | 0.713119065 | 2.12291018 |
| Wfdc15a | 0.066050875 | 0.71820487 | 3.724945049 |
| Wfdc15b | 0 | NA | NA |
| Svs2 | 0 | NA | NA |
| Svs3b | 0.141331096 | 0.237322366 | 3.56184083 |
| Svs4 | 0 | NA | NA |
| Svs3a | 0 | NA | NA |
| Svs6 | 0.038981395 | 0.718207094 | 3.724945049 |
| Svs5 | 0.282943117 | 0.895835863 | 2.020629928 |
| Slpi | 106.5214453 | 4.017850327 | 0.52012526 |
| Mir7678 | 0.061920613 | 0.718207094 | 3.724945049 |
| Matn4 | 90.53204317 | -0.558026257 | 0.172761198 |
| Rbpjl | 13.41464962 | -0.873065282 | 0.299375553 |
| Sdc4 | 2785.760239 | 2.356161167 | 0.251929018 |
| Sys1 | 225.5397579 | 0.325185297 | 0.080915056 |
| Gm14302 | 1.474490841 | -0.698683358 | 0.886860306 |
| Gm20458 | 0.478777135 | -0.210100535 | 1.325288521 |
| Trp53tg5 | 0.109982625 | 0.237318756 | 3.724945049 |
| Dbndd2 | 527.4245514 | 0.049640243 | 0.093136143 |
| Pigt | 917.1375041 | 0.228464999 | 0.083104433 |
| Gm14317 | 0.567797178 | -0.572322308 | 1.299458837 |
| Wfdc2 | 6.468348977 | 0.740864625 | 0.438871968 |
| Spint3 | 0 | NA | NA |
| Wfdc6a | 0.062426873 | 0.237312084 | 3.724945049 |
| Eppin | 0.044544648 | 0.237309859 | 3.724945049 |
| Wfdc8 | 0.123845245 | 1.035807111 | 3.720871489 |
| Gm25512 | 0 | NA | NA |
| Wfdc6b | 0 | NA | NA |
| Gm24473 | 0 | NA | NA |
| Wfdc16 | 0.038981395 | 0.718207094 | 3.724945049 |
| Gm11456 | 0 | NA | NA |
| Wfdc9 | 0.046345709 | 0.71820487 | 3.724945049 |

| | | | |
|---|---|---|---|
| Wfdc10 | 0.171131556 | 1.200054296 | 3.383981483 |
| Wfdc11 | 0 | NA | NA |
| Wfdc13 | 0 | NA | NA |
| Spint4 | 0 | NA | NA |
| Spint5 | 0 | NA | NA |
| Wfdc3 | 9.036228213 | 0.256416838 | 0.331105696 |
| Dnttip1 | 245.2841238 | 0.508594823 | 0.107224886 |
| Gm11457 | 0.366597908 | -0.109927724 | 1.563674928 |
| Gm22596 | 5.545541213 | -0.182803092 | 0.487557069 |
| Ube2c | 12.17535567 | 2.192914064 | 0.474415785 |
| Tnnc2 | 0.984947528 | 1.273365181 | 1.198887973 |
| Snx21 | 316.0422822 | -0.450379912 | 0.07037165 |
| Acot8 | 121.1509639 | 0.481696535 | 0.108605405 |
| Zswim3 | 75.20523749 | -0.236839931 | 0.174404943 |
| Zswim1 | 246.2299447 | -0.031466871 | 0.074025222 |
| Spata25 | 2.304449883 | 1.041394032 | 0.853602428 |
| Neurl2 | 13.0448324 | -0.203003153 | 0.299021721 |
| Ctsa | 1093.099412 | 0.803284333 | 0.092337661 |
| Pltp | 377.8729552 | 0.440772678 | 0.17951047 |
| Pcif1 | 806.4495592 | 0.136876627 | 0.092559676 |
| Zfp335 | 457.5970439 | 0.071042477 | 0.101722122 |
| Zfp335os | 35.33703191 | 0.130633352 | 0.199134248 |
| Mmp9 | 18.18569822 | 0.10753918 | 0.256208827 |
| Slc12a5 | 6336.594774 | -0.90611302 | 0.07352494 |
| Ncoa5 | 554.329722 | 0.150826234 | 0.101026302 |
| Cd40 | 64.99215734 | 6.46011799 | 0.525367178 |
| 1700025C18Rik | 0 | NA | NA |
| Cdh22 | 506.5456601 | -0.337298532 | 0.105545082 |
| Gm14397 | 0 | NA | NA |
| Gm25569 | 0 | NA | NA |
| Slc35c2 | 388.3991378 | 0.101999218 | 0.062809335 |
| Elmo2 | 1659.397843 | -0.242662563 | 0.07153689 |
| Zfp663 | 0.749070597 | -0.450386956 | 1.254296493 |
| Zfp334 | 573.5096397 | -0.193799506 | 0.060487328 |
| Ocstamp | 4.715999993 | 3.721002666 | 0.926028662 |
| Slc13a3 | 348.6971527 | 0.07421793 | 0.129859761 |
| Gm14437 | 0 | NA | NA |
| Trp53rka | 121.9068042 | 0.211364138 | 0.139656635 |
| Slc2a10 | 29.04143457 | 1.060823212 | 0.271620439 |
| Eya2 | 64.95457155 | 0.282296027 | 0.169154402 |
| Gm28163 | 0 | NA | NA |
| Gm11460 | 0.325497449 | -1.56771057 | 1.552329276 |
| Zmynd8 | 1768.666945 | -0.091218947 | 0.094557369 |
| BC046401 | 0.509709817 | 0.074436848 | 1.249954383 |
| Gm11465 | 2.089642419 | -0.29279091 | 0.704980858 |
| Gm22692 | 0.117907917 | 0.237310006 | 3.724945049 |
| Gm11461 | 0 | NA | NA |

| | | | |
|---|---|---|---|
| Gm11462 | 0 | NA | NA |
| Platr29 | 0 | NA | NA |
| Gm11464 | 0.663210203 | 0.545400935 | 1.176269617 |
| Ncoa3 | 801.9318103 | 0.310778802 | 0.092381759 |
| Sulf2 | 1316.311321 | 0.620048263 | 0.063443155 |
| Gm22687 | 0 | NA | NA |
| Gm11466 | 0 | NA | NA |
| Gm11467 | 0 | NA | NA |
| Gm11468 | 0.423089777 | 0.237132781 | 1.475878136 |
| Gm14268 | 0.409889595 | 1.980048773 | 1.603371388 |
| 5031425F14Rik | 0.951684498 | 1.360804951 | 1.028023199 |
| Gm11469 | 0.922271222 | 1.241012113 | 1.014078769 |
| Gm14267 | 0.809026523 | -0.868739693 | 1.189428302 |
| Prex1 | 2399.621753 | 0.633575896 | 0.071218007 |
| Gm23152 | 1.245355593 | 0.394880216 | 1.020405544 |
| Gm25265 | 0 | NA | NA |
| Gm24477 | 0 | NA | NA |
| Trp53rkb | 54.71816161 | -0.469709075 | 0.144343635 |
| Arfgef2 | 1429.215814 | 0.053805219 | 0.043597119 |
| Cse1l | 859.0610123 | 0.042084133 | 0.06925446 |
| Gm17096 | 7.764297659 | -0.485714023 | 0.537867872 |
| Stau1 | 911.3828237 | 0.222470982 | 0.06951597 |
| Gm11470 | 0.342301797 | 0.718193163 | 3.724945049 |
| Ddx27 | 291.9835414 | 0.244292562 | 0.12115673 |
| Znfx1 | 5280.077714 | 3.114872186 | 0.166999171 |
| 1500012F01Rik | 269.2484598 | 1.499442848 | 0.146936847 |
| Gm27494 | 0 | NA | NA |
| Gm25878 | 0 | NA | NA |
| Gm23201 | 0 | NA | NA |
| Snord12 | 0 | NA | NA |
| Gm11472 | 0.906216104 | 0.213393142 | 1.281933531 |
| Gm11471 | 0.062426873 | 0.237312084 | 3.724945049 |
| Gm14290 | 607.9128014 | -0.987942454 | 0.176164029 |
| Kcnb1 | 1138.507184 | -0.85624425 | 0.148759628 |
| 1110018N20Rik | 50.53414942 | -0.63045901 | 0.21253473 |
| Ptgis | 12.61769256 | 0.117801647 | 0.273626292 |
| Gm14291 | 10.76029111 | -0.414670689 | 0.386464656 |
| Gm11473 | 0.150086409 | 0.237305146 | 3.724945049 |
| B4galt5 | 1227.37508 | 1.182610726 | 0.106816687 |
| Slc9a8 | 501.9560524 | 0.219848721 | 0.075412784 |
| Spata2 | 691.9053702 | -0.196367757 | 0.088641895 |
| Rnf114 | 1225.538309 | 1.843422812 | 0.122606039 |
| Gm11474 | 0.049553047 | 0.71820487 | 3.724945049 |
| Snai1 | 13.66662012 | 1.813888651 | 0.345392113 |
| Gm11476 | 0.445249545 | -0.722770434 | 1.368809821 |
| Gm11475 | 0 | NA | NA |
| Ube2v1 | 21.15999022 | 0.130552533 | 0.236935171 |

| | | | |
|---|---|---|---|
| Gm20431 | 0 | NA | NA |
| Tmem189 | 432.2688695 | 0.542064406 | 0.093170093 |
| Gm22355 | 0.959597608 | -0.115539306 | 1.069866097 |
| Gm14320 | 1.589793752 | -0.838360833 | 0.744074279 |
| Cebpb | 250.3011784 | 3.423023845 | 0.287140535 |
| A530013C23Rik | 6.464581192 | 0.807486257 | 0.492240975 |
| Gm14321 | 4.54434231 | -0.627764381 | 0.523904164 |
| 9230111E07Rik | 2.522272708 | -0.674018414 | 0.661414692 |
| 1200007C13Rik | 1.134624916 | 2.573458576 | 1.342363633 |
| Gm14319 | 0.048949532 | -0.243569582 | 3.724945049 |
| Ptpn1 | 822.4208813 | 0.832690289 | 0.105932593 |
| Fam65c | 5.824423608 | -0.292389247 | 0.536884279 |
| Gm14236 | 0.314827273 | 0.237332193 | 1.609835843 |
| Gm24327 | 0 | NA | NA |
| Pard6b | 76.59336974 | 0.125921046 | 0.203731312 |
| Gm14235 | 0.229025006 | 0.702317088 | 2.13114309 |
| E130018N17Rik | 1.228602664 | 0.348125012 | 1.156471518 |
| Gm24561 | 2.66823802 | -1.375008961 | 0.654065643 |
| Adnp | 84.6346395 | -0.005678987 | 0.17977147 |
| Gm20716 | 2.018522609 | -0.111156855 | 0.692803837 |
| Dpm1 | 240.6952631 | -0.543508171 | 0.191681148 |
| Mocs3 | 110.2099467 | 0.334314929 | 0.149899528 |
| Gm14237 | 0.075043205 | 0.237309859 | 3.724945049 |
| Kcng1 | 163.4779742 | -0.882608379 | 0.200203203 |
| Gm14262 | 0 | NA | NA |
| Gm22704 | 0.118974955 | -0.243576254 | 3.724945049 |
| Gm14234 | 0.075043205 | 0.237309859 | 3.724945049 |
| Nfatc2 | 337.8978929 | 0.222310881 | 0.086116674 |
| Atp9a | 3839.633382 | -0.285020209 | 0.038849417 |
| mmu-mir-7004 | 0.834414618 | -0.141703191 | 1.193706297 |
| Sall4 | 2.654068103 | -0.948736343 | 0.68937983 |
| Gm14261 | 0.095897396 | -0.724453471 | 3.724945049 |
| Gm25214 | 0.150086409 | 0.237305146 | 3.724945049 |
| Zfp64 | 184.749999 | 0.16537566 | 0.110261298 |
| 1700017J07Rik | 0.04393175 | -0.243571806 | 3.724945049 |
| Gm14260 | 0 | NA | NA |
| Gm9873 | 2.521204246 | -0.303017777 | 0.641952925 |
| Gm26469 | 0.075043205 | 0.237309859 | 3.724945049 |
| Gm14258 | 0.182681975 | 1.156412857 | 2.723875067 |
| 9430093N23Rik | 0 | NA | NA |
| Gm14250 | 0 | NA | NA |
| 4930529I22Rik | 0 | NA | NA |
| Gm14249 | 0.152853976 | 0.237319074 | 2.829491075 |
| Gm11011 | 0 | NA | NA |
| Gm34294 | 5.209546091 | -0.555975698 | 0.509232549 |
| Tshz2 | 720.7148521 | -0.486244161 | 0.10702111 |
| Gm26883 | 2.358426798 | 0.4872537 | 0.638916567 |

| | | | |
|---|---|---|---|
| AY702102 | 0 | NA | NA |
| A630075F10Rik | 0 | NA | NA |
| Zfp217 | 142.5419381 | 1.82345069 | 0.158136861 |
| Gm17619 | 0.100902008 | 1.15059983 | 3.720931964 |
| Gm14270 | 11.75654133 | 0.793465102 | 0.294531259 |
| Bcas1 | 1958.212372 | -0.054136257 | 0.141961663 |
| Bcas1os2 | 1.937199938 | -0.658587915 | 0.751145529 |
| Bcas1os1 | 3.671955489 | 0.541918732 | 0.543515592 |
| Gm14269 | 52.69193393 | 0.299824191 | 0.168473146 |
| Cyp24a1 | 1.170018213 | 0.627972299 | 1.057545604 |
| Pfdn4 | 118.4952192 | 0.477720991 | 0.194007516 |
| Gm16796 | 10.39617679 | 0.523498188 | 0.359304463 |
| Gm14637 | 2.883279791 | 1.172493868 | 0.701175907 |
| 4930470P17Rik | 0.106465261 | 0.718202646 | 3.724945049 |
| Gm14264 | 0.597452826 | 1.602501356 | 1.391826693 |
| Dok5 | 201.9773433 | 0.509153895 | 0.146482968 |
| Gm14263 | 0 | NA | NA |
| Gm14266 | 0.20369824 | 0.718208928 | 3.19658972 |
| Gm14640 | 0 | NA | NA |
| 1700007M16Rik | 0.088534442 | 1.179809211 | 3.723500796 |
| Gm26489 | 0 | NA | NA |
| 1700028P15Rik | 0 | NA | NA |
| Gm14272 | 0 | NA | NA |
| Gm14641 | 0 | NA | NA |
| Cbln4 | 171.4002324 | -0.540933415 | 0.137093367 |
| Gm14271 | 0.25568908 | 1.239968177 | 1.82592321 |
| Gm14274 | 0 | NA | NA |
| AL844846.1 | 0 | NA | NA |
| Mc3r | 7.192941036 | 0.672570425 | 0.441381488 |
| Gm14275 | 0 | NA | NA |
| Gm14273 | 0.454387946 | 1.6212663 | 1.479019735 |
| Fam210b | 297.4704023 | -0.145987655 | 0.097821227 |
| Aurka | 15.8043844 | 1.368280509 | 0.322455356 |
| Cstf1 | 247.1233043 | 0.250596469 | 0.079906711 |
| Cass4 | 5.704398591 | 1.839032396 | 0.533673569 |
| Gm14455 | 6.260709098 | -0.285717573 | 0.4295664 |
| Rtfdc1 | 1259.401244 | 0.517103558 | 0.064203305 |
| Gcnt7 | 0.721712508 | 2.138333217 | 1.250937884 |
| Fam209 | 5.869341775 | -0.216052118 | 0.544818631 |
| Gm14303 | 72.23225202 | 0.472106291 | 0.19219541 |
| Tfap2c | 3.754962934 | -1.071676843 | 0.652970488 |
| Gm22773 | 0 | NA | NA |
| Bmp7 | 187.5782872 | 0.423599916 | 0.195008773 |
| Spo11 | 3.986507545 | -0.137057049 | 0.531391581 |
| Rae1 | 377.0248253 | 0.168982288 | 0.08176984 |
| Rbm38 | 50.55568089 | 0.759113901 | 0.188967315 |
| Gm20490 | 0 | NA | NA |

| | | | |
|---|---|---|---|
| Gm14453 | 0 | NA | NA |
| Ctcfl | 0.389334344 | 0.983951735 | 1.432187837 |
| Ctcflos | 0.803203798 | 1.710396809 | 1.281114044 |
| Pck1 | 0.852421781 | 1.612314151 | 1.371964232 |
| Zbp1 | 1311.11427 | 10.64101437 | 0.533315477 |
| Pmepa1 | 867.4379167 | 1.072270891 | 0.158371729 |
| Pmepa1os | 1.650675276 | 2.21773222 | 0.899343005 |
| Gm14642 | 0 | NA | NA |
| 1700021F07Rik | 0 | NA | NA |
| Ankrd60 | 0.203084296 | -0.680948678 | 2.634616364 |
| Ppp4r1l-ps | 195.0349838 | -0.352273816 | 0.155492019 |
| Rab22a | 871.4199496 | 0.230481295 | 0.073829317 |
| Mir6340 | 0 | NA | NA |
| 1700010B08Rik | 0.15795635 | 0.237325379 | 2.815015565 |
| Vapb | 1573.7862 | -0.254062658 | 0.061674249 |
| Gm14371 | 0 | NA | NA |
| Gm14336 | 0 | NA | NA |
| Atp5k-ps2 | 1.191345417 | 0.540490662 | 0.933786857 |
| Vamp7-ps | 9.408148121 | 0.092415079 | 0.409470117 |
| Stx16 | 905.3860616 | -0.25827546 | 0.071102315 |
| Npepl1 | 158.3376538 | 0.511826991 | 0.134559225 |
| Gm10714 | 1.585474913 | -2.718093811 | 0.933351698 |
| Gm27591 | 0 | NA | NA |
| Mir296 | 0.229181457 | -0.243566376 | 2.603735228 |
| Mir298 | 0.18819539 | 1.162950383 | 3.281668061 |
| Nespas | 11.25411321 | -0.154900275 | 0.370100447 |
| Gnas | 10595.54034 | -0.02071807 | 0.079658171 |
| Gm27899 | 0.248714005 | -1.650463295 | 1.8126423 |
| Gm27869 | 0.247908139 | -0.243564003 | 2.146724066 |
| Gm27849 | 0.638233968 | -2.159266713 | 2.443621008 |
| Gm20721 | 37.27970981 | 0.081077668 | 0.265640204 |
| Nelfcd | 290.7180538 | -0.021734325 | 0.065484655 |
| Ctsz | 1102.778525 | 2.924248352 | 0.149405579 |
| Rpl30-ps5 | 0 | NA | NA |
| Tubb1 | 0.687228199 | 2.165178143 | 1.200663722 |
| Atp5e | 358.6752708 | -0.030479979 | 0.136477366 |
| Slmo2 | 386.0940015 | 0.337920684 | 0.099078338 |
| Gm14385 | 0 | NA | NA |
| Vmn1r-ps1 | 0.042716945 | -0.243571806 | 3.724945049 |
| Zfp831 | 392.3466882 | -0.798501542 | 0.190355707 |
| Edn3 | 25.09549224 | -2.702841975 | 0.429003778 |
| Gm14617 | 0.43206172 | 1.517643699 | 3.290707453 |
| Gm14618 | 0.490955376 | 1.513564324 | 3.694534144 |
| Gm14616 | 0 | NA | NA |
| Gm14444 | 1.336470638 | -0.235343807 | 0.961277358 |
| Gm16354 | 2.08236629 | -1.466169207 | 0.964947592 |
| Gm14445 | 0.131100043 | -1.074360889 | 3.663654906 |

| | | | |
|---|---|---|---|
| Gm14393 | 21.09049984 | -0.603945524 | 0.323299568 |
| Gm14439 | 0.167183452 | -0.72445987 | 2.765153652 |
| Gm14387 | 3.01954408 | -0.578552887 | 0.56184017 |
| Gm14392 | 2.964599863 | -0.643217013 | 0.591093897 |
| Gm14399 | 171.4815682 | -0.621090319 | 0.141959856 |
| Gm16355 | 0 | NA | NA |
| Gm14443 | 181.6545351 | -0.750290666 | 0.199586049 |
| Gm14391 | 24.59774181 | -0.466204934 | 0.259515301 |
| Gm14442 | 0.132475575 | 0.237325435 | 2.973122108 |
| Gm14438 | 2.078804666 | -0.002143477 | 0.661037862 |
| Gm14440 | 1.419637689 | 0.219430454 | 0.746375605 |
| Gm14298 | 0 | NA | NA |
| Gm4631 | 1.996170162 | -0.340059295 | 0.678948456 |
| Gm4723 | 1.755952801 | -2.36574062 | 0.988564447 |
| Gm14297 | 0 | NA | NA |
| Gm4724 | 1.988766949 | -0.228241458 | 0.724229905 |
| Gm8923 | 0.572800032 | 0.780972187 | 1.435846468 |
| Gm11007 | 1.812554613 | -0.534174505 | 0.74608558 |
| Gm16359 | 0 | NA | NA |
| Gm4245 | 2.26497711 | 0.34881456 | 0.725678462 |
| Gm2007 | 0.319719969 | 0.683315584 | 2.065928481 |
| Gm16358 | 0 | NA | NA |
| Gm14288 | 19.90464905 | -0.572556825 | 0.313546765 |
| Rps8-ps2 | 2.50324314 | 0.549377255 | 0.741836025 |
| Gm2020 | 0.047948698 | -0.243569582 | 3.724945049 |
| Gm6710 | 28.49017795 | 0.452274357 | 0.246429237 |
| Gm16356 | 0 | NA | NA |
| Gm8898 | 2.984855315 | -0.582296292 | 0.579966155 |
| Gm2026 | 2.773167693 | -0.377406122 | 0.710191488 |
| Gm11008 | 0.226297694 | 0.718221951 | 2.13634781 |
| Gm16357 | 0.13861434 | -1.149001357 | 3.614496194 |
| Gm2004 | 6.050452602 | -0.688599322 | 0.478373654 |
| Gm6182 | 0.19897497 | -0.724459706 | 2.220827427 |
| 2210418O10Rik | 24.2680911 | -0.671330212 | 0.320292893 |
| Gm17258 | 4.979667379 | -0.2094907 | 0.639087265 |
| Gm1980 | 0.288549458 | 0.242200592 | 1.640418519 |
| Gm11009 | 0.13733263 | -1.191829832 | 3.63430361 |
| Gm14435 | 0.646335998 | -0.394501049 | 1.18083164 |
| Gm16362 | 0 | NA | NA |
| Gm14434 | 1.597696658 | 0.709621258 | 0.972396926 |
| Gm14431 | 1.363863261 | -1.633109526 | 1.020437887 |
| Gm14307 | 0 | NA | NA |
| Gm14308 | 2.466484677 | 0.391955131 | 0.810214688 |
| Gm14306 | 4.680219553 | -0.398095707 | 0.497640929 |
| Gm14305 | 173.9135921 | -0.39344937 | 0.126915014 |
| Gm14301 | 0.675521499 | 0.410020883 | 1.460594601 |
| Gm14390 | 0.066050875 | 0.71820487 | 3.724945049 |

| | | | |
|---|---|---|---|
| Gm14295 | 67.51412393 | -0.191293158 | 0.25685192 |
| Gm14408 | 0.284066144 | 0.084371705 | 2.070056406 |
| Gm14405 | 1.448471955 | -1.325561017 | 0.883283531 |
| Gm14296 | 30.11651545 | -0.472475239 | 0.305263554 |
| Mup-ps25 | 0 | NA | NA |
| Gm14417 | 0.204225461 | 0.23731275 | 3.221243153 |
| Gm14419 | 4.87970569 | -0.330011418 | 0.466472167 |
| Gm14402 | 1.138899903 | -0.356042972 | 0.872633246 |
| Gm14421 | 5.323853793 | -0.625838472 | 0.443832006 |
| Gm14401 | 0.702992681 | -0.202857677 | 1.176932415 |
| Gm14407 | 3.984100659 | -0.996119078 | 0.540620569 |
| Gm14415 | 18.81040368 | -0.912111059 | 0.320288834 |
| Gm14411 | 16.94595019 | -0.487899533 | 0.322139384 |
| Gm14416 | 3.389752464 | -0.652672159 | 0.554710544 |
| Gm14413 | 14.03696361 | -1.13842916 | 0.301171007 |
| Gm14410 | 143.7066193 | -0.5671062 | 0.150661362 |
| Gm14400 | 0.439370512 | 0.588753799 | 1.291207718 |
| Gm14409 | 0.902861002 | -1.92212541 | 1.118024432 |
| Gm14412 | 64.87765343 | -0.573694483 | 0.240797263 |
| Gm14418 | 19.13716893 | -0.002262673 | 0.261343378 |
| Gm14404 | 0.1140246 | 0.718202646 | 3.724945049 |
| Gm14414 | 1.844525372 | 0.8702632 | 0.700759739 |
| Gm14420 | 72.73691108 | -0.405466422 | 0.189809481 |
| Gm14403 | 33.59942204 | -0.645722217 | 0.239160752 |
| Gm16363 | 0.136194719 | -0.724453471 | 3.724945049 |
| Gm14406 | 1.816762255 | -1.521883017 | 0.82964126 |
| Gm14324 | 0.130017973 | 0.238301601 | 3.662714184 |
| Gm14322 | 38.40290494 | -0.320922096 | 0.202592885 |
| Gm14325 | 156.7524774 | 0.082000618 | 0.170906878 |
| Gm14429 | 5.50850624 | -0.257817286 | 0.467739121 |
| Gm14323 | 1.346967727 | -1.321229853 | 1.058747952 |
| Gm14327 | 9.478232069 | -0.255833313 | 0.353266244 |
| C330013J21Rik | 6.088359734 | 0.110970223 | 0.57017466 |
| Gm14326 | 207.3808432 | -0.212232017 | 0.114280937 |
| Gm14428 | 0.751182044 | 0.922595592 | 1.22711007 |
| Etohi1 | 168.5832989 | -0.404392883 | 0.123182149 |
| Gm26869 | 1.499946283 | -0.383949557 | 0.701681449 |
| Zfp931 | 152.8081414 | -0.089002806 | 0.161724058 |
| Phactr3 | 1341.248435 | -0.803152867 | 0.100640354 |
| Sycp2 | 51.90291932 | -0.870838919 | 0.186403194 |
| Ppp1r3d | 90.63451093 | -0.315147588 | 0.169152365 |
| Fam217b | 760.5922785 | -0.633644014 | 0.08658569 |
| Cdh26 | 2.606367607 | -0.100299047 | 0.554583506 |
| Gm14314 | 0 | NA | NA |
| Gm14292 | 70.00844022 | 0.345095389 | 0.128102353 |
| Gm14293 | 0 | NA | NA |
| Gm14294 | 0 | NA | NA |

| | | | |
|---|---|---|---|
| 4930591A17Rik | 0 | NA | NA |
| Cdh4 | 374.0015759 | -0.4823571 | 0.090020623 |
| Gm14300 | 0.042716945 | -0.243571806 | 3.724945049 |
| Taf4a | 305.5847678 | -0.195651609 | 0.098116018 |
| 4921531C22Rik | 29.89290028 | -0.265109918 | 0.274230709 |
| Gm14299 | 0.10503227 | 1.179809211 | 3.723500796 |
| Lsm14b | 804.4044537 | -0.592111772 | 0.072740488 |
| Psma7 | 1015.88698 | 0.937443212 | 0.11644894 |
| Ss18l1 | 608.8536376 | -0.500301166 | 0.066647563 |
| Mtg2 | 112.4777664 | 0.089479949 | 0.111814259 |
| Hrh3 | 541.8584486 | -1.493288825 | 0.147213135 |
| Gm17180 | 0.40273248 | -0.897731486 | 3.380031657 |
| Osbpl2 | 658.7052841 | -0.129501006 | 0.052791288 |
| Adrm1 | 690.902229 | 0.600968668 | 0.083576335 |
| Lama5 | 192.3097236 | 0.759326749 | 0.195141472 |
| Mir7005 | 0 | NA | NA |
| Rps21 | 796.5226521 | 0.292803069 | 0.147250881 |
| Mir3091 | 3.756198497 | -0.52741625 | 0.597862771 |
| Cables2 | 296.6979083 | -0.103570844 | 0.084619337 |
| Rbbp8nl | 0.042716945 | -0.243571806 | 3.724945049 |
| Gata5 | 0.107803068 | 0.719187033 | 3.724874737 |
| Gata5os | 0 | NA | NA |
| B230312C02Rik | 9.425892428 | -1.467335642 | 0.379641922 |
| Mir1a-1 | 0.044544648 | 0.237309859 | 3.724945049 |
| Mir133a-2 | 0.230754588 | -1.849515512 | 2.536465953 |
| Slco4a1 | 65.52497335 | 0.301310697 | 0.15557622 |
| Ntsr1 | 81.28326963 | -1.192298788 | 0.14091704 |
| Mrgbp | 164.3286957 | 0.292800459 | 0.088108846 |
| Ogfr | 1008.536141 | 2.219837454 | 0.132242623 |
| Col9a3 | 196.0069601 | -0.311030147 | 0.117840403 |
| Tcfl5 | 5.628819471 | -0.137990295 | 0.428131932 |
| Dido1 | 1582.103518 | -0.211584468 | 0.089678143 |
| Gm22502 | 0 | NA | NA |
| Gid8 | 550.3352372 | 0.340776431 | 0.058849657 |
| Slc17a9 | 7.930149013 | 0.60756812 | 0.569224594 |
| Gm14343 | 0 | NA | NA |
| Bhlhe23 | 1.454032994 | -0.609744078 | 0.919889551 |
| Gm14340 | 2.077246534 | 0.272040187 | 0.62297474 |
| Gm14342 | 4.321485189 | -0.462849346 | 0.517741119 |
| Gm14344 | 0.429573641 | 0.280358992 | 1.450018324 |
| Gm22842 | 0.112288909 | -0.724455696 | 3.724945049 |
| Gm14339 | 0.202674043 | 1.045489697 | 2.606063585 |
| Mir124a-3 | 0.458109599 | 1.052945242 | 1.762288146 |
| Gm27032 | 102.6108423 | -0.779236998 | 0.192956834 |
| Ythdf1 | 817.9793704 | 0.379594333 | 0.072503502 |
| Birc7 | 1.578816077 | -2.913619132 | 0.919842502 |
| Nkain4 | 83.76982434 | -2.11304484 | 0.254216628 |

| | | | |
|---|---|---|---|
| Gm14341 | 1.152403953 | -1.303430498 | 0.986139397 |
| Arfgap1 | 898.465573 | -0.073220083 | 0.095242072 |
| 9230112E08Rik | 145.4154449 | -0.514826487 | 0.195988066 |
| Col20a1 | 78.79971225 | 0.695902076 | 0.225862977 |
| Chrna4 | 653.8085059 | -0.310725368 | 0.124777342 |
| Kcnq2 | 4581.893317 | -0.527441431 | 0.127912841 |
| Eef1a2 | 7678.32573 | -0.309836973 | 0.074220208 |
| Gm24892 | 0 | NA | NA |
| Ppdpf | 209.3303033 | 0.014106498 | 0.094772366 |
| Ptk6 | 1.572767014 | 2.673796914 | 0.891227516 |
| Srms | 1.881316667 | 0.783638371 | 0.765345037 |
| Gm14398 | 0.631396267 | 1.502197067 | 1.328445204 |
| BC051628 | 0.900425091 | 1.54403827 | 1.144953919 |
| Helz2 | 1488.152321 | 6.092097694 | 0.278643067 |
| Gmeb2 | 338.4791566 | 0.363171357 | 0.117824814 |
| Gm7645 | 4.486974988 | -0.383263828 | 0.54087748 |
| Stmn3 | 2848.78216 | -0.370304378 | 0.106347198 |
| Rtel1 | 189.1322798 | -0.128029122 | 0.152358463 |
| Arfrp1 | 263.798925 | 0.093109822 | 0.098396958 |
| Zgpat | 398.0436084 | -0.268897303 | 0.122084785 |
| Lime1 | 3.160497059 | -1.194854231 | 0.70661985 |
| Slc2a4rg-ps | 70.84153722 | -0.640258429 | 0.352053995 |
| Zbtb46 | 247.6912476 | -0.463452144 | 0.135648911 |
| Gm22708 | 0.139376635 | -0.243570432 | 2.924218065 |
| Abhd16b | 0 | NA | NA |
| Tpd52l2 | 603.1097156 | 0.196369038 | 0.062054489 |
| Gm25184 | 2.339668925 | -1.565468445 | 0.831494642 |
| Dnajc5 | 4139.797779 | -0.155283865 | 0.042676899 |
| Uckl1 | 187.8382027 | 0.14541778 | 0.127478592 |
| Uckl1os | 3.735935155 | -0.423925925 | 0.538484942 |
| Znf512b | 722.5500756 | -0.294653845 | 0.11467173 |
| Mir7006 | 0 | NA | NA |
| Prpf6 | 758.9138183 | 0.313855143 | 0.081798619 |
| Samd10 | 162.7504342 | -0.2916419 | 0.100102698 |
| Gm25879 | 0 | NA | NA |
| Sox18 | 27.0834614 | 1.1493537 | 0.226386243 |
| Tcea2 | 231.4654535 | -0.884722185 | 0.14911019 |
| Rgs19 | 143.057256 | 0.539892519 | 0.114498699 |
| Lkaaear1 | 0 | NA | NA |
| Oprl1 | 406.903062 | -0.441930947 | 0.086466956 |
| Gm14489 | 0 | NA | NA |
| Myt1 | 485.0253362 | -0.534934318 | 0.135748514 |
| Pcmtd2 | 953.1475173 | -0.44835393 | 0.088622134 |
| Polr3k | 406.1158035 | -0.302877809 | 0.060681258 |
| Gm5466 | 17.7294434 | -0.320153048 | 0.267566797 |
| Gm14496 | 0.046345709 | 0.71820487 | 3.724945049 |
| Gm28002 | 0 | NA | NA |

| | | | |
|---|---|---|---|
| Gm27460 | 0 | NA | NA |
| Gm37338 | 0 | NA | NA |
| Gm8747 | 0 | NA | NA |
| Hnf4g | 1.172273398 | 2.473803132 | 1.128141151 |
| Gm37000 | 0.047948698 | -0.243569582 | 3.724945049 |
| Gm23237 | 0 | NA | NA |
| Rps24-ps2 | 0.229688899 | -1.221261407 | 2.548865832 |
| Gm2071 | 0 | NA | NA |
| Gm2077 | 0.293065915 | 0.273161822 | 1.779570523 |
| Gm8775 | 6.987551743 | 0.686062043 | 0.388446882 |
| Gm23976 | 0.841305426 | -0.71641718 | 1.208846252 |
| n-R5s194 | 0 | NA | NA |
| Gm34023 | 0 | NA | NA |
| Gm38131 | 0 | NA | NA |
| Gm37029 | 0 | NA | NA |
| Gm22944 | 0 | NA | NA |
| Gm37763 | 0 | NA | NA |
| Gm37801 | 0 | NA | NA |
| 1110015O18Rik | 13.63005333 | -0.46923497 | 0.316755937 |
| Gm6140 | 0 | NA | NA |
| Gm17308 | 1.32255022 | 0.154329581 | 0.988757047 |
| 2700069I18Rik | 10.07409941 | -0.504701365 | 0.352317419 |
| Gm10748 | 0.529004033 | 0.717328356 | 1.413932263 |
| Zfhx4 | 931.4280682 | -0.090945271 | 0.073282547 |
| 4930555M17Rik | 0 | NA | NA |
| Pex2 | 255.6278662 | 0.329853665 | 0.104665789 |
| Gm8797 | 20.70001869 | 0.835305865 | 0.33111288 |
| Gm37350 | 0.062426873 | 0.237312084 | 3.724945049 |
| Gm25352 | 0 | NA | NA |
| Gm25144 | 0 | NA | NA |
| Gm17934 | 0.363515313 | -0.724462012 | 1.985488265 |
| Gm6162 | 0.35319577 | 0.131662914 | 1.569196267 |
| Gm38298 | 0 | NA | NA |
| Gm23147 | 0.044544648 | 0.237309859 | 3.724945049 |
| Gm18658 | 0 | NA | NA |
| Gm5272 | 0 | NA | NA |
| Gm38291 | 0 | NA | NA |
| 1700008P02Rik | 0.048949532 | -0.243569582 | 3.724945049 |
| Olfr289-ps1 | 0.152964131 | 0.237325139 | 2.843489581 |
| Gm22074 | 0 | NA | NA |
| Gm8823 | 0 | NA | NA |
| Pkia | 2228.089358 | 0.060982655 | 0.083162499 |
| Gm37918 | 0 | NA | NA |
| Gm8826 | 0.880107612 | 0.660678332 | 1.105543289 |
| Gm23592 | 0 | NA | NA |
| Zc2hc1a | 623.5639214 | -0.241809003 | 0.071416566 |
| Il7 | 17.75567427 | 3.945799372 | 0.453198197 |

| | | | |
|---|---|---|---|
| Gm16685 | 14.13537915 | 5.180749175 | 0.802879667 |
| Gm29747 | 0.364508673 | 2.462782321 | 1.571389843 |
| 1700010I02Rik | 0.136120942 | 1.15080397 | 3.595683408 |
| Gm7103 | 0.4787828 | 1.562551428 | 1.497916257 |
| Gm5841 | 5.450194582 | 1.346611999 | 0.467679489 |
| Gm24614 | 0 | NA | NA |
| 1700003N22Rik | 1.26814241 | -1.160437391 | 0.974023203 |
| Stmn2 | 2333.139224 | -0.435475537 | 0.068365085 |
| Gm18822 | 4.048703541 | 0.368682815 | 0.539998216 |
| Gm32496 | 1.115586983 | 0.237330523 | 1.43279214 |
| Hey1 | 408.0829732 | -0.130733183 | 0.069238983 |
| Gm38007 | 0.33971137 | -1.831866842 | 1.646392649 |
| Gm23670 | 0 | NA | NA |
| C230057A21Rik | 0.383304402 | 1.116787926 | 1.665922681 |
| Gm37647 | 0.199458489 | -0.243565486 | 2.665489557 |
| Mrps28 | 80.2242516 | 0.66747738 | 0.157156353 |
| Gm15467 | 0 | NA | NA |
| Gm15466 | 6.37553239 | -0.835941167 | 0.457225316 |
| Tpd52 | 690.3315485 | 0.424645508 | 0.071124867 |
| Gm37616 | 0 | NA | NA |
| 4930539M17Rik | 0.310536081 | -1.152850963 | 1.601437158 |
| Gm8860 | 0.282898957 | 0.23732945 | 2.066165417 |
| Zbtb10 | 267.8821127 | 0.001065605 | 0.100249065 |
| Gm22604 | 0.061920613 | 0.718207094 | 3.724945049 |
| C030034L19Rik | 1.859233056 | 0.100191679 | 0.785934426 |
| Zfp704 | 1430.007501 | -0.797611797 | 0.091814613 |
| Gm38001 | 0 | NA | NA |
| Gm17806 | 0 | NA | NA |
| Pag1 | 670.6925854 | -0.380990037 | 0.113088119 |
| Gm16337 | 0 | NA | NA |
| Gm24786 | 3.953076191 | -1.274713462 | 0.560747871 |
| Gm6236 | 0 | NA | NA |
| Gm17877 | 0.154612032 | 1.66937168 | 3.490354585 |
| Fabp5 | 368.7894691 | 0.421676095 | 0.129322543 |
| Gm38335 | 0.498917054 | 1.461874774 | 1.383964038 |
| Gm9833 | 92.22302499 | -0.062546908 | 0.165503072 |
| Gm37308 | 0 | NA | NA |
| Pmp2 | 0 | NA | NA |
| Gm37389 | 0 | NA | NA |
| Fabp9 | 0 | NA | NA |
| Fabp4 | 3.144700407 | 4.386638868 | 0.945992456 |
| Gm38310 | 0.547572158 | 3.311297008 | 1.490784983 |
| Gm37421 | 0 | NA | NA |
| Gm37384 | 0 | NA | NA |
| Fabp12 | 1.277907819 | 2.458203646 | 1.039462148 |
| Gm37316 | 0.044544648 | 0.237309859 | 3.724945049 |
| Gm38364 | 0 | NA | NA |

| | | | |
|---|---|---|---|
| Impa1 | 639.4989438 | 0.034063722 | 0.056061557 |
| Gm38303 | 0 | NA | NA |
| Slc10a5 | 0.840861136 | 0.527421176 | 1.033533541 |
| Zfand1 | 152.3432722 | -0.554073508 | 0.137280259 |
| Chmp4c | 3.570328421 | 0.483023655 | 0.550321291 |
| Snx16 | 391.7704708 | 0.116181102 | 0.106879408 |
| Gm37831 | 0.049553047 | 0.71820487 | 3.724945049 |
| Gm18692 | 0 | NA | NA |
| Gm37244 | 0.092140247 | -0.724455696 | 3.724945049 |
| Gm22795 | 0 | NA | NA |
| Gm21631 | 0 | NA | NA |
| Gm37832 | 0 | NA | NA |
| Gm22547 | 0.047948698 | -0.243569582 | 3.724945049 |
| Gm8919 | 0 | NA | NA |
| Gm10745 | 0 | NA | NA |
| Gm37996 | 0 | NA | NA |
| Gm2429 | 0 | NA | NA |
| Gm8935 | 0.243796007 | 0.210251092 | 1.847899763 |
| Gm8939 | 0 | NA | NA |
| Gm38317 | 9.626198242 | 0.013606252 | 0.340788079 |
| Gm37813 | 0.0885994 | -0.24357403 | 3.724945049 |
| Gm37197 | 0.044544648 | 0.237309859 | 3.724945049 |
| Gm2464 | 10.69073118 | -0.332527325 | 0.31345331 |
| Gm38112 | 362.4382461 | -0.576387499 | 0.143580096 |
| Gm37164 | 1.862656823 | 0.243151786 | 0.734433066 |
| Gm2474 | 11.24932943 | -1.410609653 | 0.366296936 |
| Gm38400 | 64.08375772 | -0.564029472 | 0.176108738 |
| Gm19114 | 0.044544648 | 0.237309859 | 3.724945049 |
| Gm25981 | 24.11327504 | -0.56570891 | 0.296609651 |
| Ralyl | 450.4153493 | -0.480157155 | 0.077402788 |
| Gm22026 | 0.132101749 | 1.209894437 | 3.72418822 |
| Gm23112 | 0 | NA | NA |
| Gm38311 | 0.286827043 | 0.221913361 | 2.064176379 |
| Gm6300 | 1.130984576 | -0.032602902 | 0.977994617 |
| Gm37358 | 0 | NA | NA |
| Gm8955 | 0.038981395 | 0.718207094 | 3.724945049 |
| Slc7a12 | 0 | NA | NA |
| Gm8957 | 0.226376398 | 0.741428046 | 2.147568127 |
| Lrrcc1 | 419.00625 | -0.337289344 | 0.091814634 |
| Gm17119 | 0 | NA | NA |
| E2f5 | 109.6456114 | 0.242429279 | 0.120315007 |
| 1810022K09Rik | 264.8506355 | 0.336838744 | 0.117237109 |
| Gm38151 | 0.061920613 | 0.718207094 | 3.724945049 |
| Car13 | 144.2580462 | 2.769834557 | 0.242630399 |
| Gm5843 | 0 | NA | NA |
| Car1 | 0.04393175 | -0.243571806 | 3.724945049 |
| Gm16399 | 0.953016895 | 0.113436822 | 1.229435086 |

| | | | |
|---|---|---|---|
| Car3 | 3.829701666 | -1.256443122 | 0.539931937 |
| Car2 | 1336.217829 | -0.256076448 | 0.182157056 |
| A930001A20Rik | 0.572612294 | 1.223962125 | 1.311765439 |
| Gm7132 | 0 | NA | NA |
| Gm33819 | 0.263469446 | 0.718934585 | 2.105945703 |
| Gm9733 | 1.473887978 | 4.097182278 | 1.116064838 |
| Sirpb1a | 36.66926605 | 8.627755461 | 0.907701165 |
| Sirpb1b | 7.832067487 | 6.705794503 | 0.947953338 |
| Gm37468 | 0 | NA | NA |
| Sirpb1c | 25.99497885 | 8.07273953 | 1.021429426 |
| Gm6321 | 0.066050875 | 0.71820487 | 3.724945049 |
| Gm7343 | 0.430172323 | 2.904178799 | 1.546368585 |
| Gm5150 | 10.40898798 | 7.319901667 | 0.921879 |
| Gm24694 | 0 | NA | NA |
| Gm37836 | 0 | NA | NA |
| Gm25863 | 0 | NA | NA |
| Ythdf3 | 1623.468186 | 0.133799187 | 0.048792975 |
| Gm26485 | 0 | NA | NA |
| Gm5283 | 0.46900631 | 0.997597721 | 1.310826933 |
| Gm24963 | 0.253979914 | -1.204035645 | 1.83096817 |
| Gm6350 | 0.222549579 | -0.243563358 | 1.900001963 |
| Gm30340 | 13.55485612 | -0.334253861 | 0.312138328 |
| Gm19694 | 0 | NA | NA |
| Gm38154 | 0 | NA | NA |
| Mir124-2hg | 730.7247633 | -0.490513524 | 0.092962884 |
| Mir124a-2 | 2.558881719 | -0.441809252 | 0.623658381 |
| Gm23441 | 0 | NA | NA |
| RP24-84E6.5 | 0.192405858 | 0.981841522 | 2.639510113 |
| Cypt12 | 0.109982625 | 0.237318756 | 3.724945049 |
| Gm5844 | 33.81437118 | 0.295613781 | 0.194016066 |
| Bhlhe22 | 439.9718104 | -0.531151783 | 0.159383393 |
| Gm23726 | 0 | NA | NA |
| Cyp7b1 | 183.0659675 | 0.326806436 | 0.134663436 |
| Gm23686 | 3.150048424 | -0.768118217 | 0.618738815 |
| RP23-77D22.2 | 2.298046648 | 0.591550083 | 0.698975405 |
| Gm30667 | 0 | NA | NA |
| 4930433B08Rik | 0 | NA | NA |
| Gm6369 | 0 | NA | NA |
| RP23-427B4.3 | 0 | NA | NA |
| Gm37317 | 2.158753835 | -1.203475622 | 0.864929465 |
| RP23-427B4.2 | 0.137548931 | -0.724458529 | 2.938932991 |
| Gm30341 | 0 | NA | NA |
| Armc1 | 876.5532839 | -0.087094764 | 0.07952889 |
| Gm37979 | 14.15384062 | -0.809861637 | 0.314565296 |
| Mtfr1 | 148.0536149 | 0.444281313 | 0.132561825 |
| Pde7a | 422.8626837 | -0.502693721 | 0.107277503 |
| Gm16093 | 2.00382768 | -1.158968367 | 0.791740225 |

| | | | |
|---|---|---|---|
| Gm22282 | 0.538570099 | -1.483574481 | 1.31773399 |
| Dnajc5b | 1.101603561 | 3.665713242 | 1.024359177 |
| Gm23330 | 0.225375977 | -0.242838997 | 2.147468582 |
| Gm6141 | 0.044191549 | -0.243571806 | 3.724945049 |
| 1700064H15Rik | 1.062875418 | -2.58871065 | 1.09959249 |
| Trim55 | 5.56456899 | 4.14394048 | 1.007401041 |
| BC002189 | 0 | NA | NA |
| Crh | 42.16080477 | 0.521185843 | 0.283120394 |
| Gm18864 | 0.120349074 | 0.237305146 | 3.724945049 |
| RP23-105K2.2 | 0 | NA | NA |
| 4632415L05Rik | 336.8514353 | 0.02509733 | 0.072493772 |
| Gm37523 | 0.176667124 | -0.243563549 | 2.304379988 |
| Gm17771 | 0.368202395 | 2.222641892 | 1.469249079 |
| Cp | 857.3323739 | 2.639632261 | 0.251849454 |
| Hps3 | 339.45544 | 0.076280934 | 0.084961071 |
| Hltf | 534.1407693 | 0.011723484 | 0.071821101 |
| Gyg | 294.265161 | 0.566456779 | 0.127070516 |
| Cpa3 | 1.364377633 | 0.522262664 | 0.802998213 |
| Mir7007 | 0 | NA | NA |
| Cpb1 | 1.12517466 | 1.794317701 | 0.980271733 |
| RP24-190F20.3 | 0.04393175 | -0.243571806 | 3.724945049 |
| Agtr1b | 1.030384379 | 0.229793277 | 1.059110795 |
| Gm18426 | 0.046345709 | 0.71820487 | 3.724945049 |
| RP24-190F20.4 | 0 | NA | NA |
| Gm31466 | 0 | NA | NA |
| Gm18491 | 0 | NA | NA |
| Gm38323 | 0 | NA | NA |
| Gm18493 | 0 | NA | NA |
| Gm37480 | 0.122601409 | 0.237307635 | 3.724945049 |
| Gm7488 | 13.25965715 | -0.156551522 | 0.304143168 |
| RP23-105J10.1 | 0.180386268 | -1.335976324 | 3.333134146 |
| Gm29137 | 0.309067702 | 0.517977895 | 3.709899797 |
| RP23-458I4.1 | 0 | NA | NA |
| RP23-458I4.2 | 0 | NA | NA |
| RP23-458I4.3 | 0 | NA | NA |
| 7530428D23Rik | 1.976191175 | -0.778527003 | 0.734278615 |
| RP23-105M18.2 | 4.557716346 | -0.962324344 | 0.537273352 |
| RP23-105M18.1 | 0 | NA | NA |
| RP24-487F9.4 | 40.33606232 | -0.626845391 | 0.167234087 |
| Tbl1xr1 | 1839.527346 | -0.368900387 | 0.071770592 |
| Gm5842 | 2.255157929 | 0.41889812 | 0.634874667 |
| Rprl2 | 0.403845505 | 0.272507101 | 1.554384773 |
| Gm38324 | 0 | NA | NA |
| Gm37636 | 0 | NA | NA |
| Gm26354 | 0 | NA | NA |
| Spin3-ps | 2.893792528 | 2.016793449 | 0.835222978 |
| Gm37889 | 0.097139547 | 0.718207094 | 3.724945049 |

| | | | |
|---|---|---|---|
| Gm36983 | 1.90140223 | 3.280946992 | 1.043190206 |
| Gm37021 | 1.244068253 | 3.903678124 | 1.067271913 |
| Gm37492 | 0 | NA | NA |
| 4930412E21Rik | 0.700626182 | 2.30790019 | 1.232491934 |
| Naaladl2 | 18.35136831 | -0.119765604 | 0.283444375 |
| Gm7536 | 51.1862964 | 0.814890592 | 0.195814347 |
| Gm24704 | 0 | NA | NA |
| Gm24062 | 0 | NA | NA |
| RP23-152K14.2 | 3.653026081 | -0.104740364 | 0.689518553 |
| Gm37136 | 0.341775432 | -0.810247837 | 2.023376244 |
| Gm38002 | 0 | NA | NA |
| Gm10259 | 3.08375672 | 0.567940414 | 0.600325613 |
| Nlgn1 | 2154.133087 | -0.269262301 | 0.09449221 |
| Gm24250 | 0 | NA | NA |
| Gm37573 | 125.0600198 | -0.476601453 | 0.193717481 |
| Gm6197 | 0.660619988 | 0.594881448 | 1.344633043 |
| A830092H15Rik | 35.52709241 | -0.476906916 | 0.267280391 |
| Gm4855 | 1.087037337 | -0.833223973 | 1.171575832 |
| Gm25773 | 0.04393175 | -0.243571806 | 3.724945049 |
| Spata16 | 0.312179078 | -0.698404384 | 1.753199291 |
| Gm37659 | 0 | NA | NA |
| AC108829.1 | 0 | NA | NA |
| Gm7558 | 0 | NA | NA |
| Ect2 | 26.23418059 | 1.968043674 | 0.410953566 |
| Gm25152 | 0 | NA | NA |
| 1700125G22Rik | 0.384795782 | -1.627507084 | 1.433229927 |
| Nceh1 | 1766.692954 | 0.113621002 | 0.075019028 |
| Gm37191 | 0.357797561 | -0.697106009 | 1.961653179 |
| RP23-131B7.3 | 0.060174537 | 0.237309859 | 3.724945049 |
| Tnfsf10 | 460.2181555 | 6.837995017 | 0.367959526 |
| Ghsr | 15.57801817 | 0.476761913 | 0.32385863 |
| Fndc3b | 605.684463 | 0.817222599 | 0.07885585 |
| RP24-62O16.3 | 0 | NA | NA |
| Mir3092 | 1.750169107 | 0.129988046 | 0.706014859 |
| Tmem212 | 11.59725055 | -0.521616176 | 0.331299741 |
| Gm26040 | 0 | NA | NA |
| Pld1 | 212.9629629 | -0.003511746 | 0.118666121 |
| Gm37098 | 0 | NA | NA |
| Tnik | 3513.493683 | -0.749329991 | 0.130130827 |
| Gm37537 | 40.89549279 | -1.262392666 | 0.312843585 |
| Mir466q | 6.638222057 | -1.261376761 | 0.446492979 |
| Slc2a2 | 4.994744966 | 2.701934818 | 0.558457178 |
| 1700112D23Rik | 5.161670123 | -0.157749276 | 0.46630028 |
| Gm6505 | 17.67248122 | 0.205275969 | 0.253981318 |
| Eif5a2 | 1024.993172 | -0.495271796 | 0.122263695 |
| Gm32950 | 1.23849881 | -0.496798245 | 0.969886135 |
| Rpl22l1 | 357.7709375 | 0.312518078 | 0.145046603 |

| | | | |
|---|---|---|---|
| Gm33051 | 0.1140246 | 0.718202646 | 3.724945049 |
| Gm37867 | 0 | NA | NA |
| Gm37924 | 0 | NA | NA |
| Gm1527 | 0.240368072 | 2.150082938 | 2.497391186 |
| Egfem1 | 145.805512 | -1.087984347 | 0.148170735 |
| Gm38029 | 61.7718118 | -1.97429209 | 0.376545017 |
| Gm37965 | 30.75954022 | -1.535376631 | 0.269422201 |
| Mir551b | 1.726387458 | -2.705994546 | 1.02310807 |
| Mir21b | 0.755943838 | -1.936875835 | 1.13903497 |
| Gm37557 | 0.247197661 | 0.71822182 | 2.116562565 |
| Gm33206 | 0.045882455 | 0.237312084 | 3.724945049 |
| Mannr | 0.312486075 | 0.237330473 | 1.796057737 |
| Gm37281 | 0.306751849 | -0.693371102 | 1.60214119 |
| Mecom | 32.25235124 | 1.245228491 | 0.265401392 |
| Mecomos | 0.404625785 | 0.865304639 | 1.77652635 |
| Gm38066 | 1.468378907 | 1.189854715 | 1.087365795 |
| Gm38197 | 1.573498709 | 0.860878469 | 0.874974637 |
| Gm10258 | 0.786098488 | -0.416722862 | 1.251381817 |
| Gm38362 | 0.449676329 | 0.225384934 | 1.375335561 |
| Gm37024 | 0 | NA | NA |
| Gm37652 | 2.839259381 | 1.291956271 | 0.796291499 |
| Gm38258 | 0.044191549 | -0.243571806 | 3.724945049 |
| Gm38153 | 0.060174537 | 0.237309859 | 3.724945049 |
| 1600017P15Rik | 2.396959378 | -0.234980053 | 0.662366764 |
| Actrt3 | 12.26637337 | 0.050529916 | 0.323612739 |
| Mynn | 359.7065454 | 0.171222285 | 0.070583366 |
| Lrrc34 | 2.945986073 | -1.232419441 | 0.635224975 |
| Lrriq4 | 0.139037128 | 1.464453538 | 3.717743456 |
| Lrrc31 | 0.163642604 | -0.243570845 | 2.783881602 |
| Gm15462 | 0.843690133 | -1.368838003 | 1.076642888 |
| Samd7 | 0.989999884 | 2.442738565 | 1.094765038 |
| 4933429H19Rik | 2.000591198 | 0.328061638 | 0.764618513 |
| Sec62 | 1867.564255 | -0.046554901 | 0.058736707 |
| Gm38080 | 0.150086409 | 0.237305146 | 3.724945049 |
| Gpr160 | 12.95102122 | 1.213995436 | 0.363404927 |
| Phc3 | 2283.871593 | 0.13777384 | 0.072795738 |
| 9530022L04Rik | 9.381402519 | -0.740724994 | 0.351391554 |
| Gm2979 | 0 | NA | NA |
| Prkci | 784.3797283 | -0.089611335 | 0.056044104 |
| Mir7008 | 0.429612315 | -0.537726879 | 1.377089181 |
| Skil | 1820.158226 | 0.676888193 | 0.09505592 |
| Cldn11 | 3469.821532 | 1.057983481 | 0.142294702 |
| Slc7a14 | 2555.251066 | 0.078769121 | 0.061848109 |
| Gm15496 | 15.67699621 | 0.208897983 | 0.270813634 |
| Gm38025 | 1.47797824 | -0.663539381 | 0.906277508 |
| Gm6558 | 0 | NA | NA |
| Kcnmb2 | 45.73165487 | -0.499921967 | 0.178366299 |

| | | | |
|---|---|---|---|
| Gm37834 | 14.36667755 | -1.235200617 | 0.316792623 |
| Gm37770 | 0 | NA | NA |
| Zmat3 | 1732.179252 | -0.653631223 | 0.061651619 |
| Gm36990 | 0.044544648 | 0.237309859 | 3.724945049 |
| Pik3ca | 1418.733806 | -0.024478004 | 0.041424 |
| Gm37123 | 26.09608814 | -0.774795838 | 0.330307164 |
| Kcnmb3 | 0.414197093 | -1.341333973 | 1.663505696 |
| Zfp639 | 349.0890839 | 0.09496939 | 0.090026678 |
| Gm37592 | 0 | NA | NA |
| Mfn1 | 1164.365532 | 0.305994848 | 0.069155746 |
| Gnb4 | 421.880452 | 0.123892455 | 0.107138238 |
| Gm37640 | 0.609267175 | -0.15851787 | 1.260434309 |
| Actl6a | 159.1766196 | 0.150803169 | 0.151214765 |
| Gm37353 | 0.579969052 | -1.66122237 | 1.240482983 |
| Mrpl47 | 199.0986341 | 0.467514059 | 0.094853126 |
| Ndufb5 | 763.5212508 | -0.13022008 | 0.076703731 |
| Usp13 | 553.4368463 | -0.053350584 | 0.089618718 |
| Gm15574 | 0 | NA | NA |
| Gm37459 | 3.866617621 | -0.1454615 | 0.584598622 |
| Pex5l | 2341.167297 | -0.316833916 | 0.081013331 |
| Gm23214 | 0 | NA | NA |
| 4930419G24Rik | 1.068758569 | 0.908067925 | 0.987418036 |
| Gm37443 | 87.11784037 | -0.985229256 | 0.31498754 |
| Gm19445 | 130.8739924 | -0.97138141 | 0.295156377 |
| Gm37128 | 0.066050875 | 0.71820487 | 3.724945049 |
| Gm27974 | 0 | NA | NA |
| Gm37031 | 0.110870145 | 0.238305784 | 3.724874707 |
| Gm24780 | 0 | NA | NA |
| Gm37398 | 0.221866436 | -0.69933098 | 2.162516774 |
| Gm5845 | 1.420437351 | 1.407896074 | 0.884358826 |
| RP23-373G12.3 | 0.234927675 | -0.756690633 | 1.872171324 |
| RP23-373G12.1 | 1.69071356 | -0.974786079 | 0.881995529 |
| RP23-373G12.4 | 0.110242424 | 0.237318756 | 3.724945049 |
| Ttc14 | 2035.534113 | -0.637160896 | 0.131771176 |
| Ccdc39 | 244.5153087 | -0.818819753 | 0.186040989 |
| RP23-373G12.5 | 10.01486383 | 0.453912126 | 0.41486895 |
| RP23-373G12.2 | 0 | NA | NA |
| RP24-457B22.2 | 0 | NA | NA |
| Gm23175 | 0 | NA | NA |
| Gm9791 | 24.4382261 | 1.445402748 | 0.269432737 |
| RP24-457B22.3 | 1.01049493 | -2.10903737 | 1.116016412 |
| Fxr1 | 1205.298799 | 0.133493253 | 0.05343584 |
| RP23-418O21.3 | 27.80606419 | -0.451262978 | 0.226681495 |
| Dnajc19 | 148.9300614 | -0.413285161 | 0.138587845 |
| RP23-418O21.5 | 200.8974819 | -0.403336599 | 0.165917165 |
| Sox2ot | 356.2706365 | -0.335969276 | 0.102316567 |
| RP23-418O21.1 | 0.658880446 | -0.501589325 | 1.23713586 |

| | | | |
|---|---|---|---|
| Gm20446 | 0 | NA | NA |
| RP24-396D16.3 | 2.524695083 | -2.525629584 | 0.850174828 |
| Gm20515 | 12.14791881 | -0.251151056 | 0.362634111 |
| Gm29135 | 0 | NA | NA |
| Gm27403 | 0 | NA | NA |
| RP24-277H8.2 | 0.177597096 | 1.095593133 | 2.263528593 |
| RP24-277H8.3 | 0.133275802 | 0.684440622 | 2.958200925 |
| Mir1897 | 0 | NA | NA |
| mmu-mir-1897 | 0 | NA | NA |
| RP24-277H8.1 | 54.03438621 | -0.964950802 | 0.228270073 |
| Sox2 | 458.0693419 | 0.45486161 | 0.126401514 |
| Gm27611 | 0 | NA | NA |
| RP23-423J10.6 | 3.912046527 | 1.325224017 | 0.616021291 |
| RP23-423J10.7 | 0.80598721 | -1.137093295 | 1.030844726 |
| RP23-423J10.4 | 0 | NA | NA |
| RP23-423J10.2 | 0.090665643 | -0.724455696 | 3.724945049 |
| Mir6378 | 0 | NA | NA |
| RP23-477L20.1 | 1.570552818 | 1.498228943 | 0.834785605 |
| Gm25442 | 0.241676147 | -1.165713399 | 1.82625298 |
| RP24-141D20.1 | 2.674854522 | -0.39325472 | 0.659355305 |
| RP23-112A11.3 | 2.926346155 | -0.124232138 | 0.595827714 |
| Ube2l3-ps1 | 0 | NA | NA |
| RP23-112A11.1 | 0.159330469 | 0.718203216 | 3.465376669 |
| Gm25696 | 0 | NA | NA |
| Atp11b | 1725.242478 | -0.038278454 | 0.106902892 |
| RP23-331L5.1 | 12.60277648 | 0.569644437 | 0.433360248 |
| Gm7741 | 2.653371563 | -0.825658224 | 0.751835627 |
| Dcun1d1 | 831.5482228 | -0.03495011 | 0.063672274 |
| Gm15952 | 9.495619415 | -0.206650292 | 0.411619751 |
| RP23-111M12.3 | 21.60654816 | -0.83992279 | 0.402766768 |
| Mccc1 | 363.2980275 | -0.509451227 | 0.147781061 |
| RP23-111M12.2 | 53.110291 | -0.857157796 | 0.289448677 |
| AC116677.4 | 0.320207964 | -0.527851034 | 1.716633908 |
| Mccc1os | 32.77909455 | -0.255075443 | 0.196722699 |
| Ccdc144b | 0.1140246 | 0.718202646 | 3.724945049 |
| Acad9 | 366.3759439 | -0.177379056 | 0.065215794 |
| RP23-218L17.3 | 0 | NA | NA |
| RP23-218L17.2 | 12.80704659 | -0.292902406 | 0.272131698 |
| D3Ertd254e | 555.8188512 | -0.164629792 | 0.093442048 |
| Qrfpr | 20.7155208 | -0.21366116 | 0.237645172 |
| RP24-389H4.1 | 1.921125148 | -0.441501467 | 0.801140094 |
| RP24-389H4.2 | 1.29860622 | -2.92947243 | 1.053875852 |
| RP24-363L5.1 | 0.700674929 | 1.98144828 | 1.205343603 |
| RP23-165D1.3 | 0.038981395 | 0.718207094 | 3.724945049 |
| RP23-165D1.2 | 0 | NA | NA |
| Anxa5 | 799.2156265 | 1.175404293 | 0.144549653 |
| Mir7009 | 0 | NA | NA |

| | | | |
|---|---|---|---|
| 1810062G17Rik | 1.725215873 | -0.604739133 | 0.773100201 |
| Gm11548 | 0.199639456 | 0.718205476 | 3.225149067 |
| Gm11549 | 318.3834355 | 0.632216601 | 0.284172646 |
| RP23-151D16.2 | 14.17505885 | 2.428637642 | 0.434174979 |
| Gm11550 | 0 | NA | NA |
| Exosc9 | 304.3832438 | 0.364143387 | 0.10187104 |
| Ccna2 | 45.411053 | 1.490235615 | 0.292449519 |
| Bbs7 | 206.6797326 | -0.562600873 | 0.134789321 |
| Trpc3 | 351.384491 | -0.220631275 | 0.113803891 |
| Gm12377 | 0 | NA | NA |
| Gm12376 | 0 | NA | NA |
| 4932438A13Rik | 5270.27695 | -0.142561232 | 0.067598441 |
| Gm12531 | 0.264018088 | 1.195737513 | 2.09756407 |
| Adad1 | 0.098502579 | 0.23732098 | 3.724945049 |
| Gm12582 | 0 | NA | NA |
| Gm12583 | 0.046345709 | 0.71820487 | 3.724945049 |
| Gm12584 | 0 | NA | NA |
| Il2 | 0.52475134 | 1.109656733 | 1.296119678 |
| Gm12532 | 1.446123796 | -0.720762343 | 0.874273999 |
| Il21 | 4.872996577 | 4.844287654 | 0.903110406 |
| Gm12534 | 11.60045621 | 6.516953746 | 0.97142696 |
| RP23-243E17.13 | 4.833786608 | 5.345070946 | 1.027202751 |
| Gm24169 | 0.075043205 | 0.237309859 | 3.724945049 |
| Cetn4 | 69.0187925 | 0.268411144 | 0.155155551 |
| RP23-328P19.3 | 0 | NA | NA |
| Bbs12 | 67.61927239 | 0.436989071 | 0.202187405 |
| Gm12540 | 6.941958254 | -0.178499081 | 0.446097026 |
| Fgf2os | 0.170190954 | 1.460732368 | 2.744856262 |
| RP23-137F6.5 | 13.65974216 | -0.058080468 | 0.34817051 |
| Fgf2 | 254.8186985 | 1.725049069 | 0.224265143 |
| RP23-328P19.4 | 0.559063306 | -0.194307439 | 1.248227935 |
| Nudt6 | 28.00862341 | -0.32887697 | 0.220332395 |
| Spata5 | 377.9700528 | 0.488966186 | 0.090662247 |
| RP23-465A17.7 | 107.3750371 | -0.069925807 | 0.253203835 |
| Gm12564 | 0 | NA | NA |
| Gm12565 | 0.230175392 | 1.41897741 | 2.102972987 |
| 4930594O21Rik | 0.042716945 | -0.243571806 | 3.724945049 |
| RP23-382L1.6 | 4.839488669 | -0.802573537 | 0.567500202 |
| Spry1 | 135.2416231 | 0.733169075 | 0.197975228 |
| RP23-382L1.4 | 8.157939626 | -0.691544968 | 0.408133581 |
| RP23-382L1.3 | 1.313092963 | -1.434233588 | 0.978657377 |
| Gm5148 | 59.44844248 | -0.8278433 | 0.220434781 |
| Rps23-ps1 | 35.46711007 | 0.734783291 | 0.189630879 |
| Rpl31-ps15 | 0.453368568 | 0.213866059 | 1.552103362 |
| RP23-382L1.7 | 0.0878635 | -0.72445792 | 3.724945049 |
| Gm26404 | 0 | NA | NA |
| RP23-175A22.1 | 0.138375801 | -0.243570473 | 2.930747511 |

| | | | |
|---|---|---|---|
| RP23-175A22.2 | 0.167536551 | -0.243570703 | 2.763312762 |
| RP23-112C14.1 | 0.320788412 | 0.668852211 | 1.732694419 |
| Gm7799 | 0 | NA | NA |
| RP23-112C14.2 | 0 | NA | NA |
| Gm22899 | 0 | NA | NA |
| RP24-537K9.3 | 0.194322771 | 0.237326828 | 2.636226981 |
| RP24-537K9.1 | 6.837594151 | 0.401651682 | 0.393899722 |
| RP24-537K9.2 | 0 | NA | NA |
| RP24-392E13.1 | 0 | NA | NA |
| Gm25574 | 0 | NA | NA |
| RP23-334N1.5 | 0 | NA | NA |
| RP23-334N1.3 | 0.614372065 | -1.2097267 | 1.437791613 |
| RP23-334N1.1 | 2.007620227 | -4.374030356 | 1.032624684 |
| RP23-334N1.2 | 0.353731249 | -2.085449472 | 1.646120903 |
| Ankrd50 | 1197.125447 | 0.150179993 | 0.066274109 |
| Gm24462 | 0 | NA | NA |
| RP23-33I24.2 | 0 | NA | NA |
| RP23-33I24.1 | 0 | NA | NA |
| C230034O21Rik | 45.21372464 | -0.906826232 | 0.180469351 |
| Fat4 | 722.5535298 | -0.676197684 | 0.101369892 |
| RP23-194E5.1 | 0 | NA | NA |
| RP23-382L1.5 | 0.075043205 | 0.237309859 | 3.724945049 |
| RP23-49O22.2 | 0.068097359 | -0.243569582 | 3.724945049 |
| Gm9845 | 9.009532912 | 2.125322568 | 0.401576417 |
| n-R5s195 | 0 | NA | NA |
| RP24-85O24.1 | 0.162691105 | -0.724458629 | 2.788998794 |
| Gm23796 | 0 | NA | NA |
| Gm23831 | 0 | NA | NA |
| RP23-256D7.1 | 0 | NA | NA |
| RP23-256D7.2 | 0.608405602 | -1.540538176 | 1.568234337 |
| 1700017G19Rik | 0.863538605 | 0.16436808 | 0.93168226 |
| Gm23810 | 0 | NA | NA |
| Intu | 226.3559068 | 0.055656632 | 0.088896433 |
| Gm16114 | 0 | NA | NA |
| Slc25a31 | 2.917089469 | 0.200217358 | 0.783107818 |
| Hspa4l | 1914.656164 | -0.31040412 | 0.072789528 |
| Plk4 | 7.648973505 | 0.440589216 | 0.363814697 |
| AC146980.1 | 0 | NA | NA |
| Mfsd8 | 158.5183234 | -0.325412215 | 0.103679322 |
| 3110057O12Rik | 161.8450865 | -0.646848043 | 0.193574252 |
| Gm10731 | 0.47056225 | -0.243564994 | 1.598839944 |
| 6430590A07Rik | 35.09747937 | -0.879583884 | 0.322800989 |
| 1700034I23Rik | 0.704088048 | -0.543160712 | 1.216432825 |
| Larp1b | 264.3748089 | -0.192229477 | 0.161461377 |
| Gm22043 | 4.896223194 | -0.700952887 | 0.487252733 |
| RP23-235G13.3 | 37.44091296 | -0.384736631 | 0.345535931 |
| RP23-235G13.4 | 1.041424966 | 1.49744751 | 1.1405238 |

| | | | |
|---|---|---|---|
| Pgrmc2 | 739.9745203 | 0.014391206 | 0.07982823 |
| RP23-144N19.1 | 0.207481443 | 1.276479684 | 2.601747941 |
| RP23-149I5.1 | 0 | NA | NA |
| RP23-325M18.3 | 0 | NA | NA |
| Gm25487 | 0 | NA | NA |
| RP23-325M18.5 | 0.106618422 | -0.24357403 | 3.724945049 |
| Platr4 | 0.110814304 | -0.724455696 | 3.724945049 |
| RP23-325M18.2 | 0.374922842 | 1.199122684 | 1.738986582 |
| RP23-325M18.1 | 0.284631292 | -2.057801814 | 2.351506599 |
| Jade1 | 693.3829749 | -0.242366802 | 0.067889282 |
| Gm31266 | 2.246458349 | -0.669983563 | 0.763644546 |
| Sclt1 | 316.1906754 | -0.738037085 | 0.123530386 |
| Gm38047 | 3.02071253 | -0.690005719 | 0.548338167 |
| D3Ertd751e | 153.9838688 | -0.897202213 | 0.155791047 |
| Gm37268 | 18.97111997 | -1.005247935 | 0.326566146 |
| Gm7899 | 12.35707777 | 0.443376582 | 0.327938897 |
| Gm37348 | 0.897478403 | 0.118514235 | 1.213688228 |
| Gm37846 | 1.825474869 | -1.75402843 | 0.764612572 |
| Gm38274 | 0 | NA | NA |
| Gm37533 | 0.066050875 | 0.71820487 | 3.724945049 |
| Gm38044 | 0 | NA | NA |
| Gm9723 | 0 | NA | NA |
| Gm18865 | 0 | NA | NA |
| Gm36976 | 0.179149492 | -0.243571833 | 2.717452308 |
| Gm29739 | 0 | NA | NA |
| Gm25714 | 0 | NA | NA |
| Gm32175 | 0 | NA | NA |
| Gm37746 | 0 | NA | NA |
| Gm37365 | 0 | NA | NA |
| Gm38128 | 0 | NA | NA |
| Gm37597 | 0 | NA | NA |
| Gm24997 | 0 | NA | NA |
| Gm27989 | 0 | NA | NA |
| Gm38124 | 15.15835603 | -1.060301209 | 0.296842125 |
| 5930409G06Rik | 29.06761066 | -0.98814419 | 0.232260298 |
| 2610316D01Rik | 48.48921872 | -0.805710281 | 0.190647996 |
| Gm36996 | 0.744898608 | -1.401760111 | 1.106245279 |
| Pcdh10 | 3146.497889 | -0.681623855 | 0.08058735 |
| 1700027H10Rik | 0.202816535 | 1.003054481 | 3.205965041 |
| Gm2136 | 79.20150034 | -0.745959497 | 0.133346 |
| Gm37345 | 48.63821402 | -0.410638283 | 0.21830879 |
| Gm37835 | 0 | NA | NA |
| Gm38166 | 0.402076344 | -0.245250374 | 1.766412339 |
| Gm32340 | 0.086930093 | 0.236669296 | 3.724898107 |
| Gm25796 | 0 | NA | NA |
| Gm23337 | 0 | NA | NA |
| Gm26081 | 0 | NA | NA |

| | | | |
|---|---|---|---|
| Gm18843 | 0 | NA | NA |
| Pabpc4l | 7.77852774 | 0.383444092 | 0.446739856 |
| Gm37445 | 0 | NA | NA |
| Mir6379 | 0 | NA | NA |
| Gm10356 | 11.08110355 | 0.160007297 | 0.315084619 |
| Gm5274 | 0 | NA | NA |
| Gm23891 | 0 | NA | NA |
| Rpl23a-ps5 | 0.044544648 | 0.237309859 | 3.724945049 |
| Gm10730 | 0 | NA | NA |
| Gm2229 | 0 | NA | NA |
| Gm7977 | 1.207653455 | 0.345194189 | 0.925506283 |
| Gm37295 | 0 | NA | NA |
| 1700018B24Rik | 0 | NA | NA |
| Gm37190 | 0.042716945 | -0.243571806 | 3.724945049 |
| Gm23511 | 0 | NA | NA |
| Gm31415 | 0 | NA | NA |
| Gm37819 | 0.126225412 | 0.718200913 | 3.698754989 |
| Gm37684 | 0 | NA | NA |
| Gm20557 | 0.243142889 | 1.621374458 | 2.12417188 |
| Gm5846 | 20.89105308 | 0.572403439 | 0.250458483 |
| Gm22480 | 0 | NA | NA |
| Gm37424 | 0 | NA | NA |
| Gm38261 | 0.543563254 | 1.659017045 | 1.401121852 |
| Gm38088 | 0 | NA | NA |
| Gm2328 | 0 | NA | NA |
| Gm29741 | 0 | NA | NA |
| Gm23096 | 0 | NA | NA |
| Pcdh18 | 119.7553779 | -0.358966926 | 0.163551832 |
| Gm37550 | 0.066050875 | 0.71820487 | 3.724945049 |
| Slc7a11 | 1346.770682 | 1.077699458 | 0.147366948 |
| Gm37854 | 0 | NA | NA |
| Gm37826 | 0 | NA | NA |
| Gm37692 | 0 | NA | NA |
| Gm2345 | 0 | NA | NA |
| Gm37498 | 31.49812525 | 0.687417169 | 0.298591565 |
| Gm6209 | 0.762494802 | 0.508730162 | 1.134813674 |
| 5830415G21Rik | 0.198884431 | 1.258521403 | 3.225284387 |
| Gm38237 | 0.062426873 | 0.237312084 | 3.724945049 |
| Gm37199 | 2.566445083 | 3.235472358 | 0.834443543 |
| Gm37461 | 0.509388139 | 1.818193163 | 1.489704304 |
| Gm24503 | 0.088534442 | 1.179809211 | 3.723500796 |
| Gm37548 | 1.788766466 | 1.743123172 | 0.918924775 |
| Gm37843 | 13.03457715 | 4.705271925 | 0.648734214 |
| Gm37209 | 0.2407585 | 1.111458469 | 2.075178798 |
| Gm29230 | 0.044191549 | -0.243571806 | 3.724945049 |
| Gm38246 | 0 | NA | NA |
| Ccrn4l | 783.7619654 | 1.105344429 | 0.127150012 |

| | | | |
|---|---|---|---|
| Gm38357 | 107.4530042 | 0.84486498 | 0.231376973 |
| Elf2 | 758.9211253 | 0.342293809 | 0.078109728 |
| 4930577N17Rik | 6.225444594 | -0.636862064 | 0.452037891 |
| Gm24282 | 0 | NA | NA |
| Gm37399 | 7.483535934 | 0.154980753 | 0.3978482 |
| Gm37203 | 0.04393175 | -0.243571806 | 3.724945049 |
| Gm5103 | 0.703642298 | 0.068619319 | 1.358386591 |
| Mgarp | 3.540390614 | 0.626277897 | 0.598022231 |
| Gm9442 | 4.775535441 | -0.509594302 | 0.545931167 |
| Ndufc1 | 324.928662 | -0.114405982 | 0.09936417 |
| Naa15 | 1426.402046 | 0.248614367 | 0.070132577 |
| Gm37646 | 5.007836009 | -0.28723319 | 0.554933401 |
| Gm38160 | 2.044538346 | 0.583330352 | 0.745131546 |
| Rab33b | 363.4884665 | 0.049075288 | 0.094754402 |
| Gm37968 | 16.21162866 | -0.708231392 | 0.337899433 |
| Setd7 | 3529.863564 | -0.464641693 | 0.086030235 |
| Gm38200 | 38.03688965 | -0.448213248 | 0.301014939 |
| 5031434O11Rik | 65.52831371 | -1.013915842 | 0.166522029 |
| Gm38334 | 0 | NA | NA |
| 5430433H01Rik | 0.38231627 | -0.243587317 | 3.724945049 |
| Gm38247 | 0 | NA | NA |
| Mgst2 | 4.6105502 | 2.921508763 | 0.68006771 |
| Gm37261 | 0.485780372 | 1.866369466 | 1.259724833 |
| Maml3 | 325.2409999 | -0.190616114 | 0.094028408 |
| Gm10729 | 0.120349074 | 0.237305146 | 3.724945049 |
| Gm37342 | 22.56178576 | -0.953064763 | 0.254610693 |
| Gm37229 | 0.104366086 | -0.243576254 | 3.724945049 |
| n-R5s196 | 0.213343419 | 0.718218105 | 2.570977616 |
| 3110080O07Rik | 13.48253032 | -1.379993136 | 0.43366998 |
| Gm38000 | 2.827666343 | -0.653251025 | 0.662056612 |
| Gm37675 | 6.636235977 | -1.17032495 | 0.478704468 |
| Gm38252 | 9.035061384 | -1.257709093 | 0.45430371 |
| Gm10728 | 9.423739219 | -2.188362742 | 0.484391598 |
| Gm20089 | 0.225870551 | 0.3181867 | 2.575825623 |
| Gm38055 | 26.67308699 | -1.194168161 | 0.317042734 |
| Gm37526 | 8.63944635 | -0.906080156 | 0.38644685 |
| Gm37465 | 35.32081551 | -1.261971913 | 0.321786619 |
| C130089K02Rik | 104.1019996 | -1.042821092 | 0.313884931 |
| 5430420F09Rik | 30.46786594 | -0.931180035 | 0.29048132 |
| Gm37202 | 0 | NA | NA |
| Gm30074 | 0.368444795 | 1.064614741 | 1.425263719 |
| Gm38132 | 0 | NA | NA |
| Gm37170 | 1.831335546 | 0.044524541 | 0.758274233 |
| Olfr1398-ps1 | 0 | NA | NA |
| Foxo1 | 708.0426262 | 0.182189255 | 0.106373876 |
| Gm20402 | 0.491186624 | -0.724470769 | 3.355377818 |
| Gm38034 | 38.7415979 | -0.384602609 | 0.321297258 |

| | | | |
|---|---|---|---|
| Gm38098 | 1.419478831 | -0.01187986 | 0.839686645 |
| Gm30173 | 1.249461252 | -0.481064584 | 0.870004778 |
| Gm10293 | 11.59886104 | 0.13184358 | 0.30995678 |
| Gm22798 | 7.790214668 | -0.258230919 | 0.441061858 |
| Gm2447 | 1.231911519 | -0.466608901 | 1.265281418 |
| Gm30292 | 0.646295659 | 1.903441388 | 1.329733199 |
| Gm20750 | 0.229084632 | 0.216946891 | 2.564343408 |
| Gm37007 | 0 | NA | NA |
| Cog6 | 435.3182281 | -0.198909649 | 0.068520306 |
| RP23-256O22.3 | 10.07518621 | 0.055234773 | 0.337145236 |
| Lhfp | 547.2260578 | 0.420058735 | 0.101282873 |
| RP23-372C7.5 | 0.218578798 | -0.308098384 | 2.567633543 |
| RP23-372C7.4 | 86.73692889 | -0.754704108 | 0.291836443 |
| RP23-372C7.2 | 134.5349933 | -0.863090005 | 0.361950998 |
| RP23-372C7.7 | 21.66084825 | -0.614318627 | 0.330492271 |
| RP23-372C7.6 | 0 | NA | NA |
| RP23-372C7.8 | 30.07124974 | -0.640203626 | 0.376177885 |
| RP23-335E20.1 | 0 | NA | NA |
| RP23-335E20.2 | 0.902201473 | 0.286772522 | 1.147914406 |
| Nhlrc3 | 292.4384838 | 1.0782211 | 0.108088549 |
| Gm16206 | 0.175945213 | 1.166110064 | 2.710746636 |
| Proser1 | 377.6447841 | -0.115793958 | 0.115269799 |
| Stoml3 | 10.96884363 | -1.32877347 | 0.796084324 |
| RP23-335E20.3 | 0.440404983 | -0.567440519 | 1.546646291 |
| Gm23675 | 0.061920613 | 0.718207094 | 3.724945049 |
| Frem2 | 82.40792598 | -0.711464035 | 0.138384832 |
| RP24-291E8.1 | 0 | NA | NA |
| RP24-291E8.2 | 16.32404086 | 0.831433215 | 0.285767928 |
| RP24-291E8.4 | 0.325009107 | -0.243564832 | 1.802373943 |
| RP24-291E8.3 | 0.344979474 | 0.727361586 | 1.785292223 |
| RP24-211P15.1 | 0.713939428 | -0.714904691 | 1.183457088 |
| Gm24851 | 0 | NA | NA |
| Gm10985 | 0 | NA | NA |
| Ufm1 | 702.309027 | 0.196462292 | 0.061166711 |
| RP23-391P1.1 | 0 | NA | NA |
| Gm8109 | 0.115603922 | 1.209894437 | 3.72418822 |
| RP23-234G15.2 | 0 | NA | NA |
| Trpc4 | 278.9395309 | -0.389785716 | 0.101490502 |
| Postn | 86.52036441 | 0.823516559 | 0.274445174 |
| Gm5641 | 12.39303311 | -0.14148919 | 0.447954523 |
| Gm26678 | 0 | NA | NA |
| RP23-382K4.1 | 0 | NA | NA |
| Supt20 | 511.0808974 | 0.087855241 | 0.100183225 |
| Gm21958 | 0.229069447 | 0.718215456 | 2.551127553 |
| Exosc8 | 123.3892681 | -0.023442794 | 0.116006537 |
| Alg5 | 141.422727 | 0.393522136 | 0.128729608 |
| Smad9 | 371.890028 | -0.447723808 | 0.102050652 |

| | | | |
|---|---|---|---|
| Rfxap | 213.8530842 | -0.075763286 | 0.114477861 |
| Gm10254 | 54.85742217 | 0.613269441 | 0.185263733 |
| Sertm1 | 270.0278678 | 0.396061844 | 0.167012826 |
| Ccna1 | 5.128124108 | 0.370464005 | 0.581274771 |
| RP23-137P16.2 | 0.532719758 | -0.241608179 | 1.579461791 |
| RP23-137P16.1 | 8.16035312 | -0.127609193 | 0.388531885 |
| Spg20 | 505.3869147 | -0.128931699 | 0.07919232 |
| Ccdc169 | 8.621555654 | 0.185129361 | 0.384529685 |
| Gm22029 | 0.119587853 | 0.237305575 | 3.724945049 |
| Sohlh2 | 0.439952846 | 1.282973331 | 3.220679284 |
| Dclk1 | 10037.27044 | -0.418180405 | 0.119387541 |
| Gm25132 | 4.787591675 | -2.089304085 | 0.577082809 |
| RP24-372J8.2 | 44.9034577 | -1.195927618 | 0.318467414 |
| RP24-372J8.3 | 0.907267504 | 1.189527601 | 0.953062525 |
| Gm21954 | 0.194279094 | 1.292225235 | 3.719439735 |
| RP24-555P13.2 | 122.1621789 | -1.196749379 | 0.242374756 |
| RP24-555P13.4 | 139.4117742 | -1.215768435 | 0.262321848 |
| Gm25404 | 0.125889889 | -0.243563781 | 3.019818789 |
| WI1-49P9.1 | 0.657052355 | 2.369910575 | 1.19428231 |
| WI1-49P9.2 | 2.567514962 | 3.120479228 | 0.896180165 |
| Nbea | 6573.173349 | -0.693233071 | 0.087720191 |
| Mab21l1 | 155.2590693 | -0.094417403 | 0.139696114 |
| RP24-549I13.1 | 0.075043205 | 0.237309859 | 3.724945049 |
| RP24-549I13.2 | 0.126225412 | 0.718200913 | 3.698754989 |
| Gm22784 | 14.91209993 | -1.013757262 | 0.291660329 |
| RP24-293C5.5 | 80.47609759 | -1.086165845 | 0.275610424 |
| RP24-80L24.3 | 126.7577974 | -0.942139132 | 0.390317756 |
| RP24-80L24.2 | 91.10912563 | -0.939293099 | 0.421593917 |
| RP24-80L24.1 | 244.6441849 | -1.003525231 | 0.297613323 |
| Gm25727 | 0 | NA | NA |
| RP24-111L5.2 | 0.108266322 | 1.230376831 | 3.722845817 |
| Gm22269 | 0.061920613 | 0.718207094 | 3.724945049 |
| Gm8177 | 1.764273044 | 1.006252267 | 0.808184867 |
| RP24-218M19.4 | 0.066050875 | 0.71820487 | 3.724945049 |
| RP24-218M19.5 | 0 | NA | NA |
| Tm4sf1 | 103.6301723 | 2.356517709 | 0.237379817 |
| RP23-234F9.2 | 0.925167547 | 0.151856817 | 1.033770375 |
| Tm4sf4 | 0.133813382 | 0.218970433 | 2.958683677 |
| Wwtr1 | 465.1289583 | 1.962832703 | 0.14628818 |
| Gm16016 | 0 | NA | NA |
| Gm26671 | 2.020290962 | 0.16415112 | 0.69946669 |
| Mir6377 | 0 | NA | NA |
| RP23-132D7.2 | 73.00516792 | -0.261388712 | 0.172519593 |
| Commd2 | 168.5798793 | 0.277921784 | 0.107465311 |
| RP23-132D7.4 | 6.836736274 | -1.253130275 | 0.466642099 |
| Ankub1 | 37.54081804 | -0.081754172 | 0.262251894 |
| RP23-132D7.5 | 0 | NA | NA |

| | | | |
|---|---|---|---|
| RP24-352B14.1 | 0.588070563 | 0.780584151 | 1.447494347 |
| Rnf13 | 1345.661625 | -0.023041662 | 0.108388562 |
| RP24-352B14.2 | 0.154696864 | 0.237325224 | 2.833481316 |
| RP24-352B14.3 | 0.210855403 | -1.179371778 | 2.60734478 |
| Pfn2 | 3248.658378 | -0.334996408 | 0.079954128 |
| Gm24531 | 0.132790949 | -0.724459922 | 2.970952992 |
| RP23-1B20.1 | 0.115957212 | -0.24357403 | 3.724945049 |
| RP23-1B20.2 | 6.805132405 | -1.481956103 | 0.486596987 |
| RP23-1B20.3 | 0.497476243 | 1.400481938 | 1.519305044 |
| RP23-1B20.4 | 0.383276703 | 0.80268803 | 1.738319072 |
| Gm5537 | 1.183520696 | -1.286196963 | 0.982291312 |
| RP23-375H15.2 | 20.27877935 | -1.921872378 | 0.326788021 |
| 1700007F19Rik | 6.275202345 | -0.957233567 | 0.479591729 |
| Gm26166 | 0 | NA | NA |
| RP24-376N4.1 | 0 | NA | NA |
| RP23-413C3.1 | 0 | NA | NA |
| Tsc22d2 | 1435.051753 | 0.183583352 | 0.069917833 |
| Serp1 | 878.7409205 | 1.132612784 | 0.13099842 |
| Eif2a | 501.9842971 | 0.190054709 | 0.071594779 |
| RP23-413C3.4 | 13.02020907 | -0.24645654 | 0.309368554 |
| Selt | 2800.757624 | 0.510944099 | 0.095031567 |
| Erich6 | 23.0190923 | -0.181246552 | 0.24379731 |
| RP24-275F18.1 | 4.381802891 | -0.424616707 | 0.571984994 |
| RP24-275F18.2 | 3.555894033 | -1.315129466 | 0.547773495 |
| Siah2 | 196.4969752 | -0.047739799 | 0.135366187 |
| RP24-275F18.4 | 3.656147857 | -0.776380217 | 0.51206536 |
| 4930593A02Rik | 1.120771396 | -0.997287268 | 1.004481194 |
| Fam188b2 | 11.97822246 | -0.003443609 | 0.363071941 |
| Clrn1 | 6.849902671 | -0.019728616 | 0.393628146 |
| Gm22491 | 0 | NA | NA |
| Gm25572 | 0.089089296 | 0.237305797 | 3.724945049 |
| Gm10071 | 7.465476509 | 0.584769741 | 0.392335626 |
| Gm27793 | 0.08169834 | 0.23732098 | 3.724945049 |
| RP24-493L18.4 | 21.6695156 | -1.007121664 | 0.26373925 |
| Med12l | 1187.74355 | -0.850356188 | 0.094250547 |
| Gpr171 | 49.42350281 | 8.295031879 | 0.905539429 |
| P2ry14 | 82.16983007 | 3.28758196 | 0.261208224 |
| Gpr87 | 0.049553047 | 0.71820487 | 3.724945049 |
| P2ry13 | 94.23721791 | 0.628214757 | 0.188984504 |
| P2ry12 | 174.5045895 | -2.196573635 | 0.23099481 |
| RP24-427G24.5 | 16.25700884 | -1.355183923 | 0.415554607 |
| Igsf10 | 258.1667321 | 1.459362965 | 0.184784425 |
| Gm38186 | 1.889693139 | 1.074575213 | 0.842927891 |
| Gm37715 | 0.038981395 | 0.718207094 | 3.724945049 |
| Gm8276 | 1.405995652 | 0.719908404 | 1.048294215 |
| Gm38288 | 0 | NA | NA |
| Gm37771 | 0 | NA | NA |

| | | | |
|---|---|---|---|
| Gm4856 | 0.075043205 | 0.237309859 | 3.724945049 |
| Gm5709 | 0 | NA | NA |
| Gm37926 | 0 | NA | NA |
| Gm5538 | 0 | NA | NA |
| Gm38208 | 0 | NA | NA |
| Gm37579 | 0 | NA | NA |
| Gm5539 | 2.248417857 | 0.467799335 | 0.69797914 |
| 4930449A18Rik | 0.066050875 | 0.71820487 | 3.724945049 |
| Gm2756 | 0 | NA | NA |
| Gm8298 | 0.066050875 | 0.71820487 | 3.724945049 |
| C130079G13Rik | 0.049553047 | 0.71820487 | 3.724945049 |
| Gm9696 | 0.049553047 | 0.71820487 | 3.724945049 |
| Aadacl2 | 0 | NA | NA |
| Gm18427 | 0 | NA | NA |
| Aadac | 0.048949532 | -0.243569582 | 3.724945049 |
| Sucnr1 | 0.697362497 | 3.010456114 | 1.456161981 |
| Gm24382 | 0 | NA | NA |
| RP23-254F12.1 | 1.05607488 | -0.649006975 | 1.187935724 |
| Mbnl1 | 2752.745771 | 0.265651354 | 0.049906854 |
| Gm37589 | 184.5056286 | -0.027693021 | 0.199398936 |
| Gm37488 | 59.18839802 | -0.161320353 | 0.20917455 |
| Gm38280 | 0.072025462 | 0.237312084 | 3.724945049 |
| Gm33707 | 0.263895486 | -1.5203379 | 3.704486237 |
| Gm38326 | 0 | NA | NA |
| Gm36985 | 0.325677113 | 0.707943309 | 1.60134381 |
| Gm8325 | 112.612681 | 0.030905468 | 0.290163803 |
| Gm37035 | 1.517989225 | -2.049672214 | 0.98413277 |
| P2ry1 | 94.52974778 | -0.729765126 | 0.135192194 |
| Gm37719 | 0.066050875 | 0.71820487 | 3.724945049 |
| Gm19901 | 0.038981395 | 0.718207094 | 3.724945049 |
| Gm37696 | 26.28877038 | -0.304092872 | 0.230196761 |
| Rap2b | 869.3150327 | 0.0852769 | 0.0815252 |
| B430305J03Rik | 35.36276701 | 0.100730869 | 0.195131139 |
| Gm8349 | 2.822290259 | -0.65551112 | 0.572822342 |
| Gm9700 | 0.470561891 | -1.644921253 | 1.542804782 |
| Gm37404 | 0.15100991 | 0.718202046 | 3.527597384 |
| 9330121J05Rik | 22.77484088 | 0.532359538 | 0.263006433 |
| Arhgef26 | 647.6477621 | 0.317037417 | 0.109779103 |
| Dhx36 | 1564.510516 | 0.111241811 | 0.063341683 |
| Gpr149 | 48.93600836 | -0.199367832 | 0.185530986 |
| Gm18382 | 0.246177344 | -0.788169888 | 2.49861285 |
| Gm9701 | 0.309747539 | 0.980254627 | 3.720736542 |
| Gm22433 | 0 | NA | NA |
| Gm8388 | 2.905967886 | 2.041943921 | 0.660998552 |
| Mme | 252.6871563 | -0.650169767 | 0.092449751 |
| Gm34240 | 0 | NA | NA |
| Gm34302 | 0.608629602 | -1.132125665 | 1.280048998 |

| | | | |
|---|---|---|---|
| Gm37279 | 0 | NA | NA |
| Gm24946 | 0 | NA | NA |
| Plch1 | 346.9457733 | 0.283623395 | 0.092982291 |
| Gm23500 | 0 | NA | NA |
| Gm16162 | 0 | NA | NA |
| Gm16129 | 0 | NA | NA |
| Gm34379 | 0.085327104 | 1.179809211 | 3.723500796 |
| E130311K13Rik | 62.02924702 | -0.230505063 | 0.191982359 |
| Slc33a1 | 460.1328625 | 0.325119147 | 0.084234388 |
| Gm26850 | 5.346930916 | 0.122724072 | 0.450431443 |
| Gmps | 2065.915186 | -0.07449027 | 0.05440935 |
| Vmn2r-ps3 | 0.09176491 | 0.237310218 | 3.724945049 |
| Vmn2r1 | 7.920741964 | 0.84564746 | 0.39946402 |
| Gm22165 | 0.115603922 | 1.209894437 | 3.72418822 |
| Vmn2r2 | 2.02639444 | 0.198644219 | 0.710529664 |
| Gm5139 | 0.217472118 | 1.155507983 | 2.165423537 |
| Gm18294 | 0.049553047 | 0.71820487 | 3.724945049 |
| Vmn2r3 | 0.204914957 | 1.193798894 | 2.202071459 |
| Gm23897 | 0.049553047 | 0.71820487 | 3.724945049 |
| Vmn2r-ps4 | 0.124853745 | 0.237309606 | 3.724945049 |
| Vmn2r-ps5 | 0.174149299 | 1.185086784 | 3.359371254 |
| Vmn2r4 | 0.145501641 | 1.198496036 | 2.88801119 |
| Vmn2r-ps6 | 0 | NA | NA |
| Vmn2r-ps7 | 0 | NA | NA |
| Gm20656 | 0.439965521 | -0.721445256 | 1.578390959 |
| Vmn2r-ps8 | 0 | NA | NA |
| Vmn2r5 | 0.344244685 | 0.801692526 | 1.834817094 |
| Vmn2r6 | 0.549943695 | 0.841846925 | 1.466971081 |
| Vmn2r-ps9 | 0.898522788 | 0.340526407 | 1.072837758 |
| Gm20657 | 0.48442959 | -0.626878162 | 1.339284473 |
| Vmn2r-ps10 | 0 | NA | NA |
| Vmn2r-ps11 | 0.791552024 | 0.230663087 | 1.087050098 |
| Gm26939 | 0 | NA | NA |
| Vmn2r7 | 3.934381616 | -0.515583618 | 0.573578654 |
| Vmn2r-ps12 | 0.044544648 | 0.237309859 | 3.724945049 |
| Gm20698 | 0 | NA | NA |
| Gm25222 | 0.066050875 | 0.71820487 | 3.724945049 |
| Vmn2r-ps13 | 0 | NA | NA |
| Gm38048 | 50.77510117 | -0.483537799 | 0.2221502 |
| Kcnab1 | 1328.585016 | -0.515266636 | 0.09373488 |
| A330015K06Rik | 13.82607597 | -1.048558575 | 0.361588294 |
| A730090N16Rik | 2.24970321 | 0.908798604 | 0.778354057 |
| Ssr3 | 1935.252712 | 0.695563328 | 0.092500522 |
| Gm5540 | 6.757178092 | 0.37749496 | 0.376425147 |
| Gm34780 | 0.183309527 | 1.199972785 | 2.685766727 |
| Gm37009 | 0.786503428 | -1.867410602 | 1.206634233 |
| 4931440P22Rik | 18.83552398 | 1.175686933 | 0.290290435 |

| | | | |
|---|---|---|---|
| Gm22279 | 0 | NA | NA |
| Tiparp | 657.7234625 | 1.311637913 | 0.233736977 |
| Gm5847 | 0.982102764 | 1.326785992 | 1.184726666 |
| Gm38233 | 0 | NA | NA |
| Mir8120 | 0.138110619 | 0.718202458 | 3.62359848 |
| Lekr1 | 106.5364298 | -0.425525397 | 0.193412458 |
| Gm37776 | 35.53555177 | -1.123400977 | 0.317150965 |
| Gm25755 | 0 | NA | NA |
| Gm37359 | 0.166952883 | 1.610951781 | 2.74653892 |
| Gm37131 | 0.161076545 | 1.165842935 | 2.787893623 |
| Gm37037 | 0.659599365 | 0.876578501 | 1.227217122 |
| Ccnl1 | 947.3859406 | 0.530875323 | 0.136000502 |
| Gm37305 | 112.0234715 | 0.056236089 | 0.105031065 |
| Gm35106 | 0.709932405 | 1.61128606 | 1.257522933 |
| Gm37078 | 4.562323805 | -0.917892449 | 0.442256444 |
| Gm19024 | 0 | NA | NA |
| Gm6546 | 0 | NA | NA |
| Veph1 | 5.036914537 | -0.692215567 | 0.517154042 |
| Gm37822 | 0 | NA | NA |
| Gm26442 | 0 | NA | NA |
| Gm36973 | 0.066050875 | 0.71820487 | 3.724945049 |
| Tpi-rs2 | 0 | NA | NA |
| Gm37330 | 0 | NA | NA |
| Ptx3 | 208.6062493 | 7.086302148 | 0.652148578 |
| Gm17952 | 0.276493613 | -0.724460678 | 1.796298024 |
| Gm6555 | 0 | NA | NA |
| Gm38356 | 0.174174109 | 0.237326083 | 2.729571827 |
| RP23-103D17.4 | 0 | NA | NA |
| Shox2 | 134.0978986 | -0.362664907 | 0.145996828 |
| Rsrc1 | 525.7525617 | -0.338247614 | 0.063233987 |
| Gm22295 | 1.86515265 | -0.32346345 | 0.749252173 |
| Gm17402 | 0 | NA | NA |
| Mlf1 | 15.4417281 | 0.936359583 | 0.311748474 |
| Gfm1 | 616.4629507 | 0.026043211 | 0.049768168 |
| Lxn | 276.0893367 | -0.353549679 | 0.158545844 |
| Gm37453 | 18.12819328 | -0.57204034 | 0.278723494 |
| Rarres1 | 12.46569881 | -0.62184959 | 0.319909841 |
| Gm35299 | 0.137416547 | -0.240825018 | 3.63730774 |
| 4930402C01Rik | 0 | NA | NA |
| Gm25919 | 0 | NA | NA |
| Mfsd1 | 388.2294903 | 0.973196076 | 0.112354271 |
| Gm8515 | 6.537093261 | 0.825322902 | 0.37865895 |
| Gm412 | 0 | NA | NA |
| Gm37601 | 0.128477747 | 0.718203484 | 3.677131942 |
| Gm18717 | 0.045882455 | 0.237312084 | 3.724945049 |
| Iqcj | 18.38164937 | -1.953693353 | 0.369533464 |
| Schip1 | 1612.457602 | -0.206220898 | 0.088053203 |

| | | | |
|---|---|---|---|
| Gm10292 | 11.84136725 | -1.134473862 | 0.485831223 |
| Gm10723 | 64.31874068 | -1.147989401 | 0.35196644 |
| Gm22073 | 0 | NA | NA |
| Il12a | 10.96521039 | 0.367787914 | 0.355964279 |
| Gm10040 | 0.437154627 | -0.625090793 | 1.330949025 |
| 4930535E02Rik | 0 | NA | NA |
| Gm35584 | 0.044191549 | -0.243571806 | 3.724945049 |
| Gm7270 | 0 | NA | NA |
| Gm17641 | 6.199809403 | -1.305237309 | 0.557856693 |
| 1110032F04Rik | 89.81938694 | -0.415936673 | 0.186445559 |
| Ift80 | 422.7464304 | -0.360713077 | 0.12498155 |
| Gm20689 | 16.19484603 | 1.024463548 | 0.291746161 |
| Gm20690 | 0.088383099 | -0.72445792 | 3.724945049 |
| Gm18588 | 78.97133165 | 0.544052389 | 0.164671216 |
| Gm20691 | 11.60463875 | -0.024759265 | 0.274040951 |
| Smc4 | 201.8629177 | 1.070359976 | 0.139366579 |
| Mir15b | 0.132199999 | 0.23730893 | 3.642365845 |
| Mir16-2 | 0.358973896 | 0.205355929 | 1.423708122 |
| Trim59 | 149.282414 | -0.794897122 | 0.142985083 |
| Kpna4 | 1744.547512 | 0.508437062 | 0.057740218 |
| Gm22009 | 385.5335551 | -0.428615984 | 0.27800026 |
| Gm37558 | 126.3325027 | -0.021792599 | 0.216488381 |
| Gm1647 | 0.255219096 | -0.265378238 | 1.832121749 |
| Arl14 | 0.146692594 | 1.229893553 | 3.521109737 |
| Ppm1l | 3121.106828 | -0.434403762 | 0.07670725 |
| Gm17214 | 0 | NA | NA |
| Gm17212 | 0 | NA | NA |
| Gm17213 | 0 | NA | NA |
| Gm8565 | 1.479948724 | -1.719899379 | 0.986773366 |
| B3galnt1 | 333.6927738 | -0.307963792 | 0.089899724 |
| Gm37380 | 0.038981395 | 0.718207094 | 3.724945049 |
| Gm25621 | 0 | NA | NA |
| Rpl32-ps | 0.038981395 | 0.718207094 | 3.724945049 |
| Nmd3 | 566.0817292 | 0.374891767 | 0.101430506 |
| Sptssb | 37.86867121 | -0.446857319 | 0.186174094 |
| Gm17895 | 0 | NA | NA |
| Gm23484 | 0 | NA | NA |
| Gm37784 | 14.73966199 | 0.394843044 | 0.294018613 |
| Otol1 | 0.139883403 | 1.411723814 | 3.551352091 |
| Gm37681 | 0.143485256 | 1.229961212 | 3.547405499 |
| Gm37585 | 0 | NA | NA |
| Gm23477 | 0 | NA | NA |
| Gm6631 | 0 | NA | NA |
| Gm37213 | 0 | NA | NA |
| Gm10780 | 0 | NA | NA |
| Gm6634 | 0.207012398 | -0.243565388 | 2.635059287 |
| Gm8599 | 0 | NA | NA |

| | | | |
|---|---|---|---|
| Gm38305 | 0 | NA | NA |
| Gm19194 | 0 | NA | NA |
| Gm8604 | 0 | NA | NA |
| Gm37617 | 0 | NA | NA |
| Gm3351 | 0 | NA | NA |
| Gm37603 | 0 | NA | NA |
| Gm27465 | 0 | NA | NA |
| Gm37901 | 0 | NA | NA |
| Gm22451 | 0 | NA | NA |
| Sis | 3.661596079 | -1.089871574 | 0.72879907 |
| Gm37056 | 46.70151825 | -0.85391998 | 0.163240416 |
| Slitrk3 | 736.758703 | -0.838308394 | 0.084953318 |
| Gm20754 | 35.42307591 | -1.051052971 | 0.223115981 |
| Gm24544 | 0.953051642 | -2.618034281 | 1.204803149 |
| Gm37725 | 9.175969031 | -1.030945876 | 0.380273559 |
| Gm22538 | 0 | NA | NA |
| Gm38353 | 21.39725853 | -0.906483167 | 0.424948315 |
| 4930509J09Rik | 0 | NA | NA |
| Bche | 53.21720014 | -0.466532949 | 0.168925689 |
| Gm8625 | 0 | NA | NA |
| Gm20356 | 0 | NA | NA |
| Gm6098 | 0 | NA | NA |
| Gm37050 | 0 | NA | NA |
| Gm8643 | 0 | NA | NA |
| Gm37464 | 5.385716294 | -0.474474324 | 0.400908052 |
| Zbbx | 20.8812694 | -0.657518055 | 0.261426369 |
| Serpini2 | 1.184533449 | 2.966860299 | 1.064385033 |
| Wdr49 | 44.34530745 | 2.249510034 | 0.301422758 |
| Gm17836 | 0 | NA | NA |
| Pdcd10 | 402.9200995 | 0.591786014 | 0.129479619 |
| Serpini1 | 2225.756587 | -0.725564612 | 0.095386417 |
| Gm37672 | 32.21841053 | -1.265400328 | 0.267041466 |
| Platr10 | 0.223571151 | -0.243571348 | 2.553503579 |
| Gm37193 | 0 | NA | NA |
| Gm25846 | 0.062426873 | 0.237312084 | 3.724945049 |
| Gm29133 | 0 | NA | NA |
| Golim4 | 319.1997903 | -0.101080187 | 0.093489088 |
| Gm37685 | 0 | NA | NA |
| Gm37256 | 0.048949532 | -0.243569582 | 3.724945049 |
| Gm18428 | 0 | NA | NA |
| Fstl5 | 496.3711575 | -0.460349364 | 0.100583873 |
| Gm37897 | 17.02018816 | -1.010068331 | 0.322716054 |
| Gm37290 | 0.044544648 | 0.237309859 | 3.724945049 |
| Gm19231 | 0.236516182 | 0.237330568 | 2.111948259 |
| Gm23644 | 0 | NA | NA |
| Dnmt3a-ps1 | 0.044544648 | 0.237309859 | 3.724945049 |
| Gm37806 | 0.072025462 | 0.237312084 | 3.724945049 |

| | | | |
|---|---|---|---|
| Gm38195 | 0 | NA | NA |
| Gm37576 | 0 | NA | NA |
| RP23-257I21.2 | 0 | NA | NA |
| Gm6677 | 0 | NA | NA |
| Gm8684 | 0.072025462 | 0.237312084 | 3.724945049 |
| Gm6680 | 0 | NA | NA |
| RP23-399J17.1 | 4.954402907 | 0.665785204 | 0.524518405 |
| Gm15442 | 3.380166461 | 0.119545151 | 0.533224303 |
| Gm18952 | 0 | NA | NA |
| Gm24749 | 0 | NA | NA |
| Gm17953 | 0.338292759 | 2.441795379 | 3.689344679 |
| Gm5277 | 0.066050875 | 0.71820487 | 3.724945049 |
| Gm10291 | 12.97480729 | 0.017872607 | 0.339831969 |
| Gm9762 | 88.84803059 | 0.421215504 | 0.179929206 |
| Gm23440 | 0 | NA | NA |
| Rapgef2 | 2966.610995 | -0.163226271 | 0.07702599 |
| Gm24150 | 0 | NA | NA |
| RP24-253I2.4 | 0.238666214 | 1.232561363 | 2.534197894 |
| 6430573P05Rik | 8.415433566 | 0.183391641 | 0.361997538 |
| Gm17359 | 0.372745468 | 0.718229673 | 1.486677229 |
| Fnip2 | 1106.231662 | 1.305866761 | 0.147011814 |
| Gm3513 | 13.83725982 | 0.274923386 | 0.326167402 |
| 4930589L23Rik | 0.532739632 | 0.70490142 | 1.194985988 |
| Gm35067 | 0.044544648 | 0.237309859 | 3.724945049 |
| Ppid | 668.4459743 | -0.074669111 | 0.074398936 |
| Etfdh | 350.8915493 | 0.023304398 | 0.103371138 |
| Gm37973 | 5.608718336 | -0.218400127 | 0.428227221 |
| 4930579G24Rik | 15.15182425 | 0.064523134 | 0.267949767 |
| Rxfp1 | 93.03486081 | -1.928873414 | 0.253875658 |
| Gm26984 | 4.172577245 | -1.811704344 | 0.616253808 |
| Tmem144 | 92.24781446 | -1.04376553 | 0.179391847 |
| Gm26420 | 0 | NA | NA |
| Gm36569 | 0.140714442 | -0.243572565 | 3.582146371 |
| Fam198b | 79.79527804 | -1.079660299 | 0.173699964 |
| A330069K06Rik | 1.631297931 | -0.529471532 | 1.022889997 |
| Gm19066 | 0.045882455 | 0.237312084 | 3.724945049 |
| RP23-121N19.4 | 0.091880448 | -0.724455696 | 3.724945049 |
| Gm38295 | 0.117760149 | -0.243576254 | 3.724945049 |
| Gm37892 | 0.469035539 | -1.161921191 | 1.563546141 |
| Gm37971 | 2.078197752 | 0.649071499 | 0.6484561 |
| Gm6706 | 0.213298592 | 2.049674178 | 2.13630919 |
| Gria2 | 8463.13547 | -0.78239338 | 0.11423397 |
| Gm25749 | 0 | NA | NA |
| Glrb | 2044.481699 | -0.311476653 | 0.066810401 |
| RP24-445D5.2 | 0 | NA | NA |
| Gm29808 | 0.687463866 | 0.380474003 | 1.316250912 |
| RP23-187I18.1 | 0.148659141 | 1.513781604 | 3.716315176 |

| | | | |
|---|---|---|---|
| Pdgfc | 119.0878968 | 0.287796127 | 0.117272026 |
| Gm16000 | 0.137483974 | 0.684519582 | 2.929862181 |
| Gm37300 | 0 | NA | NA |
| Gm4857 | 0 | NA | NA |
| RP23-144D4.1 | 0 | NA | NA |
| A830029E22Rik | 4.685490163 | 0.449385996 | 0.504935115 |
| Gm22531 | 0 | NA | NA |
| Gm9989 | 1.473237501 | -1.037153223 | 0.865873405 |
| Ctso | 423.7109783 | 0.774289971 | 0.114132932 |
| Tdo2 | 9.666145708 | -0.193062487 | 0.360417842 |
| Asic5 | 0.518949687 | -0.243563595 | 1.446218512 |
| Gucy1b3 | 1608.78633 | -0.579957107 | 0.091025968 |
| Gucy1a3 | 860.8261742 | -1.17973511 | 0.1097621 |
| Mir7010 | 0.40751274 | -0.562655666 | 1.421426981 |
| Map9 | 1296.963444 | -0.577380279 | 0.070740065 |
| Npy2r | 123.6004455 | 0.698340816 | 0.206135086 |
| Gm35323 | 0 | NA | NA |
| Gm29999 | 0.156071933 | -0.724458118 | 3.489245271 |
| RP23-469J3.5 | 0.062426873 | 0.237312084 | 3.724945049 |
| RP23-223I2.3 | 0 | NA | NA |
| Gm22038 | 0 | NA | NA |
| RP23-223I2.2 | 0 | NA | NA |
| Rbm46 | 23.88977253 | 0.347390765 | 0.242244316 |
| Rbm46os | 0.56520525 | -0.671192875 | 1.373801598 |
| Lrat | 10.45023781 | -0.284458979 | 0.481319801 |
| Gm30097 | 0.374039128 | -0.232778163 | 1.602952323 |
| Fgg | 0.874079576 | 1.578055812 | 1.20312762 |
| Fga | 10.18503337 | 5.975173659 | 0.987212053 |
| Fgb | 1.174237097 | 2.342292926 | 1.423910901 |
| Plrg1 | 372.6327839 | 0.423118419 | 0.095874992 |
| Gm10710 | 19.45997954 | -1.290363618 | 0.287205599 |
| Dchs2 | 240.9884021 | -0.828593377 | 0.115284864 |
| Gm38096 | 1.042898341 | 1.173851538 | 1.291072643 |
| 1700028M03Rik | 0.205907367 | 0.719065078 | 2.58804216 |
| Gm22342 | 0.083172944 | 0.23732098 | 3.724945049 |
| Rpl28-ps2 | 0 | NA | NA |
| Gm24584 | 0 | NA | NA |
| Sfrp2 | 19.78329196 | -0.668599833 | 0.268573573 |
| Gm26771 | 5.06408741 | -0.075026016 | 0.469703021 |
| Gm7115 | 0 | NA | NA |
| Tlr2 | 267.1921433 | 4.453657261 | 0.246093523 |
| D930015E06Rik | 251.151783 | -0.617951611 | 0.150220825 |
| Gm36981 | 2.682934602 | 2.327575769 | 0.807343629 |
| Mnd1 | 6.972706414 | 0.349729653 | 0.578050443 |
| Trim2 | 5153.10411 | -0.311308653 | 0.06134805 |
| Gm6525 | 0.265820157 | -2.04581302 | 2.43893295 |
| 4930565D16Rik | 0.075043205 | 0.237309859 | 3.724945049 |

| | | | |
|---|---|---|---|
| Fhdc1 | 188.3832 | 0.465836878 | 0.102966707 |
| Arfip1 | 76.98979055 | 0.203732538 | 0.123735524 |
| Gm37240 | 3.253759892 | -1.103718651 | 0.58966719 |
| Gm8813 | 4.188906716 | -0.466218812 | 0.513283698 |
| Gm38241 | 5.062387905 | -0.858897517 | 0.481829257 |
| Tigd4 | 1.127018203 | -0.095564884 | 1.011159024 |
| Tmem154 | 14.23682184 | 3.400367387 | 0.504245264 |
| Fbxw7 | 2169.618475 | -0.337847652 | 0.067301579 |
| Gm37726 | 37.63354527 | -0.110509837 | 0.282415524 |
| AC124496.1 | 0 | NA | NA |
| Gm38041 | 0 | NA | NA |
| Gm38313 | 0 | NA | NA |
| 1700036G14Rik | 0.202694752 | -1.652845375 | 2.186041579 |
| RP23-94M23.3 | 2.853198453 | -1.251397011 | 0.599358087 |
| Mir466k | 0 | NA | NA |
| RP24-389G6.1 | 1.736188672 | -0.50620755 | 0.699856767 |
| Gm15535 | 15.003366 | -0.994185651 | 0.500584953 |
| Gatb | 159.5508096 | -0.009335735 | 0.108035138 |
| Fam160a1 | 54.14219974 | -0.473422238 | 0.150097886 |
| Glt28d2 | 6.204872336 | -0.700114821 | 0.43830724 |
| RP23-329L9.3 | 0.488911873 | -0.666699981 | 1.433360492 |
| Gm26204 | 0 | NA | NA |
| Gm38030 | 0 | NA | NA |
| Gm9790 | 12.80588666 | 0.046678296 | 0.336103738 |
| Gm25876 | 0 | NA | NA |
| Gm24501 | 0 | NA | NA |
| Gm27811 | 0 | NA | NA |
| Sh3d19 | 475.5047945 | -0.033656643 | 0.112614603 |
| Prss48 | 0.087261593 | -0.243576254 | 3.724945049 |
| Gm25188 | 1.524225208 | -0.918277153 | 0.962978551 |
| Rps3a1 | 1477.132456 | 0.510058021 | 0.095879116 |
| Snord73a | 1.006316758 | 0.267939179 | 0.924198506 |
| Rnu73b | 4.371881307 | -0.178926131 | 0.519742155 |
| Gm37933 | 3.2042328 | 0.565852937 | 0.646681319 |
| Gm37493 | 0 | NA | NA |
| Gm23025 | 0 | NA | NA |
| Lrba | 1713.427863 | -0.60544296 | 0.072112286 |
| Gm37288 | 0 | NA | NA |
| Gm26343 | 0 | NA | NA |
| Gm23850 | 3.443253566 | -1.111225224 | 0.685930073 |
| Gm25039 | 5.355852112 | -0.323306278 | 0.430613152 |
| Mab21l2 | 76.13297546 | 0.849806531 | 0.433381942 |
| Gm3788 | 95.68068835 | 1.512570177 | 0.285862826 |
| Gm37876 | 2.813292308 | -0.520332667 | 0.741880753 |
| Dclk2 | 1006.44227 | -0.232074541 | 0.07641059 |
| Gm37025 | 0.066050875 | 0.71820487 | 3.724945049 |
| Gm23376 | 0 | NA | NA |

| | | | |
|---|---|---|---|
| Gm37207 | 0.045882455 | 0.237312084 | 3.724945049 |
| Cd1d2 | 0.436249931 | 2.947350276 | 1.358514297 |
| Cd1d1 | 24.21988694 | 2.054674726 | 0.246635207 |
| Gm8869 | 0 | NA | NA |
| Gm18403 | 0 | NA | NA |
| Gm37575 | 0 | NA | NA |
| Gm24472 | 0 | NA | NA |
| Kirrel | 258.042677 | 1.191266261 | 0.148934969 |
| Gm37855 | 0.445528452 | 1.575980164 | 1.400925618 |
| Gm10705 | 72.15716491 | 0.584989948 | 0.153917267 |
| Fcrls | 15.92947883 | -4.307700978 | 0.514096934 |
| Cd5l | 3.379428938 | 5.587549838 | 1.027925562 |
| Fcrl1 | 1.476596612 | 4.396267621 | 1.003934116 |
| Fcrl5 | 1.398885825 | 1.833218516 | 0.973306151 |
| Etv3 | 674.3690235 | 0.745531037 | 0.114487393 |
| Etv3l | 0.263217861 | 0.786452883 | 2.111187172 |
| Gm25167 | 0 | NA | NA |
| Gm24109 | 0 | NA | NA |
| Arhgef11 | 1362.693994 | -0.193562019 | 0.040144896 |
| Lrrc71 | 7.798021523 | 0.325371497 | 0.439843153 |
| Pear1 | 78.16134709 | 1.101002697 | 0.14545856 |
| Ntrk1 | 25.65422587 | 0.340285404 | 0.268125665 |
| Insrr | 23.94499803 | 1.700112569 | 0.249420043 |
| Sh2d2a | 48.27836269 | 7.999924152 | 0.890009556 |
| Prcc | 536.6201964 | 0.148999095 | 0.061902899 |
| Hdgf | 1352.852318 | 0.353720341 | 0.058347367 |
| Mrpl24 | 277.5276893 | 0.336404699 | 0.09070048 |
| Rrnad1 | 232.3898889 | -0.346178986 | 0.123974708 |
| Isg20l2 | 217.9624788 | 0.555287017 | 0.0918893 |
| Crabp2 | 45.98201362 | 2.12358697 | 0.273841057 |
| Gm3745 | 0 | NA | NA |
| Nes | 844.5112463 | 4.708500296 | 0.407716754 |
| Bcan | 1604.188113 | -1.643775442 | 0.148597233 |
| Hapln2 | 312.1547389 | 0.354341027 | 0.169092742 |
| RP23-240E15.3 | 115.0424175 | 0.32072479 | 0.198338594 |
| Gpatch4 | 115.8497267 | 0.586503597 | 0.184528611 |
| Apoa1bp | 265.3851978 | 0.111701051 | 0.102401502 |
| Ttc24 | 0.042716945 | -0.243571806 | 3.724945049 |
| Iqgap3 | 18.40657348 | 2.354633249 | 0.478600909 |
| Mef2d | 2901.836846 | -0.393620538 | 0.077305222 |
| 1700113A16Rik | 8.541957373 | -0.481965906 | 0.360548758 |
| AI849053 | 5.915945368 | -1.468466824 | 0.453580223 |
| AW047730 | 113.9278857 | -1.338419137 | 0.198928695 |
| Gm3764 | 811.9285128 | -0.341579512 | 0.180703917 |
| Mir3093 | 13.97251529 | -0.10972295 | 0.291623989 |
| Gm25641 | 0.574683629 | -0.987518883 | 1.230827665 |
| mmu-mir-9-1 | 11.97633615 | -0.521871686 | 0.325544887 |

| | | | |
|---|---|---|---|
| Rhbg | 2.720079777 | 4.61813984 | 1.007173661 |
| Gm38392 | 2.080370332 | -0.755210125 | 0.760795879 |
| Tsacc | 7.84413125 | 0.141098889 | 0.375674853 |
| Gm37116 | 2.788796254 | 0.281784936 | 0.701051487 |
| Cct3 | 1136.589819 | 0.676831775 | 0.08342073 |
| Glmp | 457.6296344 | 0.850853356 | 0.125808469 |
| Tmem79 | 9.024460536 | 0.518495779 | 0.328395182 |
| Smg5 | 1241.608425 | 0.367378133 | 0.069819553 |
| Paqr6 | 85.91657556 | -0.932503886 | 0.170740126 |
| Bglap3 | 2.927464552 | 0.331625082 | 0.569564987 |
| Gm21956 | 0.097139547 | 0.718207094 | 3.724945049 |
| Bglap2 | 0.555158567 | 1.189395665 | 1.187667368 |
| Gm6821 | 0.068097359 | -0.243569582 | 3.724945049 |
| Bglap | 0.801792645 | 2.049533364 | 1.228248441 |
| Pmf1 | 65.8586388 | 0.815422269 | 0.18367271 |
| Slc25a44 | 1273.147583 | 0.279473935 | 0.064425027 |
| RP24-298F19.6 | 2.520287764 | 1.848029273 | 0.675369541 |
| Sema4a | 442.3120329 | 0.241076711 | 0.099804216 |
| Mir7011 | 0 | NA | NA |
| Gm5979 | 0 | NA | NA |
| Lmna | 660.5220702 | 1.189295595 | 0.168578961 |
| Gm37584 | 6.065917723 | -1.215398668 | 0.42406601 |
| Mex3a | 75.1280634 | -0.026909255 | 0.158457067 |
| Mir1905 | 0.318342637 | -1.636370806 | 3.682092106 |
| Rab25 | 1.64905476 | 0.317031647 | 0.767707273 |
| Lamtor2 | 189.4900279 | 0.014150212 | 0.113521322 |
| Ubqln4 | 1064.387744 | -0.264571178 | 0.062964975 |
| Ssr2 | 493.374432 | 0.550972015 | 0.08625797 |
| Gm10704 | 39.754764 | 0.72252964 | 0.234057051 |
| Arhgef2 | 1402.732759 | -0.049597124 | 0.064383279 |
| Gm20652 | 1.311025493 | -0.484584148 | 0.941586362 |
| Rxfp4 | 0 | NA | NA |
| RP23-199B2.8 | 2.30646632 | -0.015036169 | 0.715154121 |
| 2810403A07Rik | 753.6655769 | 0.067628602 | 0.096945006 |
| Gm25945 | 6.725632374 | -0.11829001 | 0.418640577 |
| Rit1 | 282.0700927 | 0.420359727 | 0.107715506 |
| Gm17146 | 5.547323716 | -1.433098846 | 0.477728255 |
| Gm10253 | 5.431051021 | -0.29406135 | 0.622131771 |
| Syt11 | 7534.20102 | -0.420849268 | 0.051345219 |
| 5830417I10Rik | 63.92099027 | -0.17202274 | 0.147329318 |
| 1500004A13Rik | 4.959186419 | -0.595527279 | 0.490974252 |
| Gon4l | 808.0428841 | -0.109212071 | 0.073150508 |
| RP23-51N3.3 | 0.176220594 | -0.724458674 | 2.719488399 |
| Gm23054 | 2.17116889 | -0.566227787 | 0.696461233 |
| Msto1 | 187.4319594 | 0.268492035 | 0.085255987 |
| n-R5s197 | 0.163066018 | -0.243578385 | 3.449621945 |
| Dap3 | 627.5926466 | -0.168109454 | 0.055065072 |

| | | | |
|---|---|---|---|
| Ash1l | 4532.149695 | -0.300259108 | 0.061521494 |
| Gm26465 | 0 | NA | NA |
| Gm24403 | 0 | NA | NA |
| RP24-394C15.7 | 0 | NA | NA |
| Rusc1 | 959.1680519 | -0.496499611 | 0.064682272 |
| Fdps | 34.50620784 | 0.076176068 | 0.201118041 |
| Gm22935 | 1.101484168 | -0.742923626 | 1.267173307 |
| RP24-394C15.4 | 0 | NA | NA |
| Pklr | 0.541363579 | -1.903431528 | 1.222069555 |
| Hcn3 | 139.9590133 | -0.070808049 | 0.128124618 |
| Clk2 | 311.1271416 | -0.25534964 | 0.145001297 |
| Scamp3 | 397.6184147 | 0.423590516 | 0.093349654 |
| Gm16069 | 1.160335885 | -1.342190136 | 1.026716301 |
| Fam189b | 352.3484391 | -0.326949773 | 0.069378432 |
| Gm23269 | 0 | NA | NA |
| Gba | 494.4063209 | 1.262673899 | 0.116599762 |
| Mtx1 | 203.9215083 | 0.030163842 | 0.085256783 |
| RP23-350K3.2 | 104.7257883 | 0.428681234 | 0.110499119 |
| Thbs3 | 91.591488 | 0.012490115 | 0.193475215 |
| Mir92b | 0.280267168 | 0.613265721 | 1.796298609 |
| Muc1 | 17.55319324 | 2.877610903 | 0.398305356 |
| Trim46 | 576.4645691 | -0.470914388 | 0.127355602 |
| Krtcap2 | 231.132102 | 0.761859281 | 0.105888969 |
| Dpm3 | 102.4945791 | 0.548608325 | 0.132747936 |
| Slc50a1 | 178.718743 | 0.183141292 | 0.101424222 |
| Efna1 | 37.54655563 | 0.542336928 | 0.201702288 |
| Gm22642 | 0 | NA | NA |
| Efna3 | 366.6839132 | -0.534424177 | 0.147251853 |
| Gm15998 | 1.818279517 | 1.474874187 | 1.017065767 |
| Efna4 | 3.626077548 | 1.099571758 | 0.530667086 |
| 4731419I09Rik | 0.772156528 | 0.425740655 | 1.083958489 |
| Adam15 | 928.1765902 | 0.337452051 | 0.075812999 |
| Dcst1 | 1.444845127 | 0.796612437 | 0.860148262 |
| Gm10702 | 17.15096642 | -1.086875202 | 0.303922581 |
| Zbtb7b | 489.1333668 | 1.096667496 | 0.133980264 |
| Gm15417 | 31.01182027 | 0.012873276 | 0.189285562 |
| Lenep | 0.241023442 | -0.653889742 | 1.826648313 |
| Flad1 | 214.0339016 | 0.061853882 | 0.086711746 |
| Cks1b | 21.556343 | 1.065017609 | 0.364642247 |
| Shc1 | 291.0704285 | 1.18918461 | 0.122576635 |
| Pygo2 | 447.2134043 | 0.047726535 | 0.096348027 |
| Pbxip1 | 1167.4111 | 1.301781697 | 0.128313831 |
| Pmvk | 242.2372548 | 0.018499425 | 0.120841524 |
| Kcnn3 | 548.2658225 | -0.451055129 | 0.095752945 |
| Adar | 3619.123941 | 1.007085204 | 0.090967838 |
| Chrnb2 | 737.8398015 | -0.566845792 | 0.070276026 |
| 4632404H12Rik | 215.8736812 | -0.19263346 | 0.134345881 |

| | | | |
|---|---|---|---|
| Ube2q1 | 886.7801256 | 0.058294337 | 0.054521043 |
| RP24-122I22.4 | 0 | NA | NA |
| She | 25.99300754 | 1.637499887 | 0.292324649 |
| Il6ra | 238.4815568 | 2.327247098 | 0.131178812 |
| RP24-122I22.6 | 1.56556854 | 1.362042746 | 0.943360294 |
| Atp8b2 | 1163.874269 | -0.402504088 | 0.09786916 |
| Aqp10-ps | 0.173853943 | 1.25450333 | 2.738478996 |
| Gm24046 | 0.066050875 | 0.71820487 | 3.724945049 |
| Hax1 | 320.365027 | 0.808638499 | 0.115517997 |
| RP23-114P22.5 | 1.940516017 | -0.60796673 | 0.723010635 |
| Ubap2l | 1956.985286 | 0.07564532 | 0.068381421 |
| Gm24608 | 5.024933871 | -1.702185855 | 0.606191133 |
| Mir7669 | 4.045153519 | -1.280883978 | 0.61345569 |
| 4933434E20Rik | 645.1823872 | 0.296267993 | 0.088997262 |
| Gm16540 | 6.216095886 | -0.969822036 | 0.509782975 |
| Mir190b | 0.194279094 | 1.292225235 | 3.719439735 |
| Tpm3 | 824.8814748 | 0.895219509 | 0.106072529 |
| Nup210l | 14.08624278 | -0.10161646 | 0.342159744 |
| Gm23723 | 0.35561486 | 0.638707989 | 1.575476688 |
| RP24-257C17.2 | 1.986149933 | 0.61836831 | 0.702569399 |
| Gm24185 | 0 | NA | NA |
| Rps27 | 1242.923509 | 0.659373227 | 0.143238248 |
| Rab13 | 74.74244885 | 1.775653936 | 0.205186406 |
| Jtb | 232.6321241 | 0.384503639 | 0.09743135 |
| Creb3l4 | 5.167911745 | 0.704457991 | 0.489839228 |
| Slc39a1 | 919.2059152 | 1.659340873 | 0.134285173 |
| Crtc2 | 364.8494084 | 0.469858425 | 0.115281293 |
| Dennd4b | 682.1714222 | -0.228804102 | 0.064589226 |
| Mir7012 | 0 | NA | NA |
| RP23-358O22.1 | 2.468095164 | 0.215819362 | 0.622905693 |
| Gatad2b | 2548.376701 | -0.00504659 | 0.063308085 |
| Slc27a3 | 56.99060069 | -0.037088266 | 0.167093967 |
| Ints3 | 947.6918643 | 0.174403876 | 0.083208703 |
| Npr1 | 184.6399887 | 1.085293841 | 0.208865379 |
| Gm16048 | 1.106761898 | 0.048228817 | 0.884471324 |
| RP24-333B11.3 | 0.066050875 | 0.71820487 | 3.724945049 |
| Ilf2 | 670.5056944 | -0.058764623 | 0.083554144 |
| Snapin | 374.92744 | -0.054860322 | 0.077241166 |
| Chtop | 1458.724535 | 0.237795682 | 0.075687149 |
| S100a1 | 401.8878366 | 0.230493286 | 0.151679881 |
| S100a13 | 63.775755 | 0.982995577 | 0.165362204 |
| S100a14 | 1.045271382 | -0.288699012 | 1.300281881 |
| S100a16 | 533.8051986 | 1.631488052 | 0.169511777 |
| Gm24593 | 0 | NA | NA |
| Gm23464 | 0 | NA | NA |
| S100a3 | 9.213372739 | 2.108892426 | 0.431000152 |
| S100a2 | 0.475453923 | -1.441742408 | 1.41284587 |

| | | | |
|---|---|---|---|
| RP23-326D6.5 | 0 | NA | NA |
| S100a4 | 14.92865454 | 3.67192857 | 0.497145776 |
| S100a5 | 23.65689645 | -0.349490712 | 1.455973823 |
| S100a6 | 362.9019992 | 3.972104657 | 0.273682318 |
| RP24-174D10.3 | 0 | NA | NA |
| S100a7a | 5.924278019 | 1.843616322 | 0.458686672 |
| S100a8 | 13.16848317 | 5.50263819 | 0.809729721 |
| S100a9 | 10.76590429 | 3.88263346 | 0.58032335 |
| RP24-174D10.4 | 0 | NA | NA |
| RP24-174D10.5 | 0 | NA | NA |
| Pglyrp4 | 0.060174537 | 0.237309859 | 3.724945049 |
| RP24-174D10.6 | 2.240082864 | -0.234671786 | 0.67040187 |
| Gm5849 | 0 | NA | NA |
| RP23-111A11.1 | 0 | NA | NA |
| RP23-309N14.1 | 0 | NA | NA |
| RP24-296O9.1 | 0 | NA | NA |
| 4930529C04Rik | 0.183030605 | -0.243564335 | 2.280962878 |
| 9130204L05Rik | 0 | NA | NA |
| Gm23697 | 0 | NA | NA |
| Pglyrp3 | 2.336438979 | 4.650058267 | 1.043272625 |
| Mif-ps3 | 0.176744298 | -0.244116297 | 2.303885067 |
| Gm25145 | 0 | NA | NA |
| Lor | 15.81042869 | -0.401785488 | 0.311384802 |
| RP23-381I16.5 | 0 | NA | NA |
| RP23-381I16.3 | 0 | NA | NA |
| RP23-381I16.4 | 0 | NA | NA |
| Prr9 | 0 | NA | NA |
| Lelp1 | 0 | NA | NA |
| RP23-381I16.6 | 0.291617713 | -0.724460872 | 1.768942416 |
| Sprr2a1 | 0 | NA | NA |
| Sprr2a2 | 0 | NA | NA |
| Gm5850 | 0.099106094 | 1.209894437 | 3.72418822 |
| Gm37087 | 0.047948698 | -0.243569582 | 3.724945049 |
| Gm5851 | 0.36081128 | 1.997940866 | 1.665878557 |
| Gm37752 | 0 | NA | NA |
| Sprr2a3 | 0 | NA | NA |
| Sprr2b | 0.207368832 | 1.226334974 | 2.637089982 |
| Gm37428 | 0 | NA | NA |
| Sprr2c-ps | 0 | NA | NA |
| Sprr2d | 0 | NA | NA |
| Sprr2e | 0.186228844 | 0.718206409 | 3.289587508 |
| Sprr2f | 0 | NA | NA |
| Sprr2g | 0.097139547 | 0.718207094 | 3.724945049 |
| Sprr2h | 0 | NA | NA |
| Sprr2i | 0 | NA | NA |
| Sprr2j-ps | 0 | NA | NA |
| Gm9774 | 28.33359312 | 0.687363574 | 0.225238494 |

| | | | |
|---|---|---|---|
| Sprr2k | 0 | NA | NA |
| Sprr1b | 0.278361807 | -0.243576532 | 3.724945049 |
| Sprr3 | 0 | NA | NA |
| Gm37377 | 0 | NA | NA |
| Sprr1a | 21.48993395 | 5.057037148 | 0.823268864 |
| Sprr4 | 0 | NA | NA |
| 2310046K23Rik | 0.227721369 | -0.243573166 | 3.11953315 |
| Ivl | 0 | NA | NA |
| Smcp | 0 | NA | NA |
| 2210017I01Rik | 0 | NA | NA |
| Lce6a | 0.09176491 | 0.237310218 | 3.724945049 |
| Lce1a1 | 0.475368968 | 0.237303829 | 3.724945049 |
| Lce1b | 0.18352982 | 0.237308059 | 3.724945049 |
| Rpl13-ps4 | 0 | NA | NA |
| Lce1a2 | 0 | NA | NA |
| Lce1c | 0.27529473 | 0.237306482 | 3.724945049 |
| Lce1d | 0.45492314 | -0.724468639 | 3.724945049 |
| Gm4202 | 299.4188111 | 0.305986218 | 0.172786986 |
| Lce1e | 0 | NA | NA |
| Lce1f | 0.373862045 | 2.635230816 | 1.612558125 |
| Gm38119 | 0 | NA | NA |
| Lce1g | 0.655362467 | 3.230599131 | 2.414384756 |
| Lce1h | 0.27529473 | 0.237306482 | 3.724945049 |
| Lce1i | 0.17962841 | -0.724461859 | 3.374012434 |
| Lce1j | 0.09176491 | 0.237310218 | 3.724945049 |
| Lce1k | 0 | NA | NA |
| Kprp | 0.596471915 | 0.237302484 | 3.724945049 |
| Gm38201 | 0 | NA | NA |
| Lce1l | 0.09176491 | 0.237310218 | 3.724945049 |
| 2310050C09Rik | 0.31922648 | -0.243580115 | 3.724945049 |
| Lce3a | 0 | NA | NA |
| Lce3b | 0 | NA | NA |
| Gm38078 | 0 | NA | NA |
| Lce3c | 0.342841614 | 0.739473624 | 2.030805745 |
| Gm24481 | 0 | NA | NA |
| Lce3d | 0 | NA | NA |
| Gm22577 | 0 | NA | NA |
| Lce3e | 0 | NA | NA |
| Lce3f | 0.038981395 | 0.718207094 | 3.724945049 |
| Crct1 | 1.039606943 | 3.474601977 | 1.126948416 |
| Lce1m | 0.045882455 | 0.237312084 | 3.724945049 |
| Gm38040 | 0.139839889 | 0.237324449 | 2.92397156 |
| Gm4858 | 0 | NA | NA |
| Gm37361 | 0 | NA | NA |
| Crnn | 0 | NA | NA |
| Gm37501 | 0 | NA | NA |
| Flg2 | 0.73411928 | 0.237301116 | 3.724945049 |

| | | | |
|---|---|---|---|
| Gm18432 | 0.092691418 | 1.183079489 | 3.724560917 |
| Flg | 1.695700131 | -0.243594542 | 3.724945049 |
| Rplp0-ps1 | 2.415407705 | 1.701551626 | 0.696974669 |
| Gm37922 | 0 | NA | NA |
| Gm37245 | 0.045882455 | 0.237312084 | 3.724945049 |
| Gm37913 | 0.038981395 | 0.718207094 | 3.724945049 |
| Hrnr | 2.939544198 | -0.200328595 | 3.716988201 |
| AC123859.1 | 0 | NA | NA |
| Gm37857 | 0 | NA | NA |
| Rptn | 0.124596252 | 0.718200615 | 3.714681068 |
| Gm18296 | 0.119587853 | 0.237305575 | 3.724945049 |
| Gm22752 | 0.140442044 | -0.724458015 | 3.613090186 |
| Gm5278 | 0.680250709 | -0.609682805 | 1.553652699 |
| Tchh | 32.61220021 | 0.159583711 | 0.24567641 |
| Tchhl1 | 0.282572244 | -0.224239229 | 2.070043404 |
| Gm36953 | 0 | NA | NA |
| S100a11 | 216.4239304 | 4.682416996 | 0.391159448 |
| Gm29950 | 0 | NA | NA |
| S100a10 | 413.4047504 | 2.684479048 | 0.195406193 |
| Gm5541 | 0 | NA | NA |
| Gm38207 | 0 | NA | NA |
| Tdpoz2 | 0 | NA | NA |
| Tdpoz1 | 0 | NA | NA |
| Gm18433 | 0 | NA | NA |
| Gm37596 | 0 | NA | NA |
| Gm38086 | 0 | NA | NA |
| Gm5852 | 0 | NA | NA |
| Gm5773 | 0 | NA | NA |
| Tdpoz4 | 0 | NA | NA |
| Tdpoz3 | 0 | NA | NA |
| Gm37430 | 0 | NA | NA |
| Gm9113 | 0.044544648 | 0.237309859 | 3.724945049 |
| Gm9116 | 0 | NA | NA |
| Gm37185 | 0 | NA | NA |
| Gm9117 | 0 | NA | NA |
| Gm37104 | 0 | NA | NA |
| Tdpoz5 | 0 | NA | NA |
| Gm37807 | 0 | NA | NA |
| Gm37481 | 0 | NA | NA |
| Gm9121 | 0.045882455 | 0.237312084 | 3.724945049 |
| Gm9124 | 0 | NA | NA |
| Gm37085 | 0 | NA | NA |
| Gm9125 | 0 | NA | NA |
| Gm37081 | 0 | NA | NA |
| Gm10697 | 0 | NA | NA |
| Gm37778 | 0 | NA | NA |
| Gm37155 | 0 | NA | NA |

| | | | |
|---|---|---|---|
| Gm5542 | 0 | NA | NA |
| Gm9131 | 0 | NA | NA |
| Gm37882 | 0 | NA | NA |
| Gm5543 | 0 | NA | NA |
| Gm37001 | 0 | NA | NA |
| Gm16560 | 0 | NA | NA |
| Gm10696 | 0.045882455 | 0.237312084 | 3.724945049 |
| Gm5286 | 0 | NA | NA |
| Gm16584 | 0 | NA | NA |
| Gm4982 | 0.038981395 | 0.718207094 | 3.724945049 |
| Gm9141 | 0 | NA | NA |
| Gm4778 | 0 | NA | NA |
| Gm16561 | 0 | NA | NA |
| Them4 | 202.6068946 | -0.405392998 | 0.089064443 |
| RP23-148G12.2 | 0 | NA | NA |
| Them5 | 0.249936274 | -0.726505664 | 2.498159312 |
| RP23-148G12.3 | 4.416068108 | -0.603031866 | 0.506700329 |
| C2cd4d | 4.518953556 | 0.195077236 | 0.457951844 |
| RP23-148G12.5 | 0.049553047 | 0.71820487 | 3.724945049 |
| Rorc | 43.04669428 | -0.503408623 | 0.242377827 |
| RP23-298M23.3 | 0.086648695 | -0.72445792 | 3.724945049 |
| Lingo4 | 68.20621523 | -0.475753714 | 0.245436267 |
| 1700040D17Rik | 7.265577632 | -0.793931736 | 0.364171433 |
| Tdrkh | 464.6804853 | -0.438451419 | 0.087006674 |
| RP23-298M23.7 | 44.92890297 | -1.110396255 | 0.258846811 |
| RP23-298M23.8 | 0.072025462 | 0.237312084 | 3.724945049 |
| Oaz3 | 2.190110068 | 1.868007545 | 1.198937269 |
| Mrpl9 | 334.7621041 | 0.639969487 | 0.087381709 |
| RP23-298M23.9 | 0.961922688 | -0.028576294 | 0.932010192 |
| Riiad1 | 12.40794134 | -0.297085125 | 0.3318833 |
| Celf3 | 927.0754436 | -0.99412039 | 0.142591029 |
| Snx27 | 1992.373795 | -0.040031922 | 0.057278117 |
| RP23-298M23.11 | 19.53939973 | -0.406506277 | 0.353781263 |
| RP23-298M23.12 | 7.754593511 | -0.50490127 | 0.495703205 |
| RP24-276A6.1 | 23.82234211 | -0.747938843 | 0.331672571 |
| AC127252.1 | 0.108123235 | -0.24357403 | 3.724945049 |
| Rpl31-ps11 | 1.676330775 | -0.310415647 | 0.873949246 |
| Tuft1 | 38.81743166 | 0.900549511 | 0.221827871 |
| Gm10972 | 0 | NA | NA |
| Gm15234 | 0.403742549 | 0.224694974 | 1.487932675 |
| BC021767 | 1.492922606 | 0.137104172 | 0.907678732 |
| Selenbp2 | 2.718408089 | 0.245931653 | 0.695845101 |
| Gm15264 | 3.338245402 | -0.573428585 | 0.658002624 |
| RP23-449N19.1 | 0 | NA | NA |
| Cgn | 50.96569556 | -1.224479543 | 0.182502343 |
| Gm15263 | 0.664861716 | -2.415077065 | 1.385456711 |
| Pogz | 1033.996817 | -0.215839323 | 0.062179163 |

| | | | |
|---|---|---|---|
| Psmb4 | 959.4736378 | 0.836495287 | 0.11487269 |
| Gm26279 | 0.212513282 | 0.237311278 | 3.175662942 |
| Selenbp1 | 39.74177677 | -0.070700473 | 0.258618899 |
| Rfx5 | 386.701994 | 0.143369872 | 0.075110007 |
| B230398E01Rik | 27.22757332 | 2.916262036 | 0.49433178 |
| Pi4kb | 609.4035235 | -0.055347574 | 0.063652415 |
| A730011C13Rik | 177.9491982 | 3.642368525 | 0.213363587 |
| Gm15265 | 34.26348499 | 0.021785317 | 0.276048929 |
| Mir7013 | 0.881607526 | -1.747836854 | 1.14485815 |
| Zfp687 | 719.9016467 | 0.327345501 | 0.089124528 |
| 4930481B07Rik | 9.171066605 | 0.180690263 | 0.339023446 |
| Psmd4 | 779.7769061 | 0.571420997 | 0.097179865 |
| Pip5k1a | 758.4226424 | 0.101018872 | 0.086325088 |
| Gm24920 | 5.105149463 | -1.462169212 | 0.580522294 |
| Gm22301 | 0 | NA | NA |
| Vps72 | 306.4355309 | 0.358617242 | 0.066290634 |
| Tmod4 | 6.657288501 | -1.170434419 | 0.441983176 |
| Scnm1 | 114.2649021 | 0.496080802 | 0.114000174 |
| Lysmd1 | 47.85101064 | -0.230852471 | 0.155318375 |
| Mir8099-2 | 0.049553047 | 0.71820487 | 3.724945049 |
| Tnfaip8l2 | 38.08952461 | 3.317902931 | 0.276505183 |
| Gm9169 | 2.545959807 | 0.832174262 | 0.632867567 |
| Sema6c | 68.04043718 | -0.925499645 | 0.193754427 |
| AC140190.1 | 0 | NA | NA |
| Gabpb2 | 547.3167726 | -0.108086455 | 0.110212988 |
| Gm16740 | 8.225987562 | -0.085045864 | 0.360826894 |
| Mllt11 | 2083.533929 | -0.317110038 | 0.072520583 |
| Cdc42se1 | 925.5629768 | 1.118906156 | 0.11067726 |
| Gm128 | 1.475208495 | -1.512596808 | 0.809582553 |
| Bnipl | 3.731006451 | -1.787255861 | 0.584674514 |
| Prune | 303.723384 | -0.01645018 | 0.075951347 |
| Fam63a | 346.9866243 | 0.365277601 | 0.100102319 |
| Anxa9 | 8.904347822 | -0.038087394 | 0.362578127 |
| 6330562C20Rik | 7.360490999 | 0.692913798 | 0.382395331 |
| Cers2 | 766.0749113 | 0.809091678 | 0.106942238 |
| Setdb1 | 649.1102505 | -0.133044659 | 0.092200218 |
| RP23-407B16.2 | 0 | NA | NA |
| Gm37500 | 1.278257442 | -0.961221475 | 1.060090202 |
| RP23-407B16.3 | 0 | NA | NA |
| Gm5070 | 0.368600158 | -0.867381309 | 1.486765048 |
| RP23-407B16.4 | 0.048949532 | -0.243569582 | 3.724945049 |
| Gm4349 | 3.988126414 | -1.397372623 | 0.508631722 |
| Arnt | 674.0580543 | -0.155397226 | 0.110206519 |
| RP23-422N18.1 | 29.38944538 | -1.015173592 | 0.272618365 |
| Ctsk | 101.6290436 | 2.007765173 | 0.19682354 |
| Ctss | 2705.512219 | 3.477284805 | 0.182585897 |
| Hmgb1-ps5 | 4.681085585 | 4.88965913 | 0.938114217 |

| | | | |
|---|---|---|---|
| Hormad1 | 0.108266322 | 1.230376831 | 3.722845817 |
| Golph3l | 291.0130091 | 0.746133742 | 0.109999052 |
| RP23-422N18.3 | 2.848305637 | -0.326773854 | 0.713833779 |
| Rps10-ps1 | 80.90969664 | 0.757543767 | 0.186457342 |
| Ensa | 2238.110929 | 0.113421502 | 0.07336622 |
| RP23-231L15.4 | 47.75963671 | -0.596187113 | 0.263279506 |
| Mcl1 | 1783.072468 | 1.764123127 | 0.142169604 |
| Adamtsl4 | 317.4111096 | 1.298494203 | 0.142270792 |
| Ecm1 | 388.3138544 | 4.218713633 | 0.342740488 |
| Mir7014 | 0 | NA | NA |
| Tars2 | 297.5330298 | 0.017274857 | 0.087839226 |
| Rprd2 | 1419.095113 | -0.222713526 | 0.055131978 |
| RP23-218K6.1 | 0.165206806 | 1.198506451 | 2.775355574 |
| Gm26361 | 3.436253382 | -1.550235564 | 0.638387035 |
| RP23-218K6.4 | 0.306564387 | 0.718222451 | 2.005374229 |
| Prpf3 | 300.0577214 | 0.231784495 | 0.114488688 |
| Gm26594 | 0.250472921 | 1.720117518 | 2.484005275 |
| Mrps21 | 159.8335192 | -0.06067209 | 0.120249736 |
| C920021L13Rik | 8.391545128 | 0.048365099 | 0.416395791 |
| Ciart | 209.0128952 | -1.341089485 | 0.198086722 |
| BC028528 | 36.09890605 | 1.768529506 | 0.194832231 |
| Aph1a | 300.5160635 | 0.572017928 | 0.102316827 |
| Car14 | 134.9063813 | -0.520747425 | 0.13762251 |
| Anp32e | 879.4154652 | -0.073183354 | 0.06194535 |
| RP23-49O17.4 | 9.354061232 | -1.408023741 | 0.416053972 |
| RP23-49O17.5 | 0 | NA | NA |
| Plekho1 | 424.3660083 | 0.948361085 | 0.12235049 |
| Vps45 | 380.4768023 | -0.06519008 | 0.095046615 |
| RP23-261M19.2 | 0.092691418 | 1.183079489 | 3.724560917 |
| Otud7b | 1265.281876 | -0.052250708 | 0.061478865 |
| Mtmr11 | 145.3552089 | 1.31091943 | 0.157159615 |
| Gm17690 | 21.29907499 | -0.01589489 | 0.259629689 |
| Sf3b4 | 445.2184839 | 0.763734989 | 0.113036167 |
| Gm22027 | 1.636315489 | -0.5753025 | 0.768634881 |
| Sv2a | 5873.135983 | -0.462655139 | 0.041133563 |
| Bola1 | 50.41452498 | -0.028620507 | 0.18144429 |
| Gm15444 | 1.046383212 | 0.361645413 | 0.980394073 |
| Hist2h2ab | 34.08697731 | -0.186328876 | 0.301508857 |
| Hist2h2ac | 86.71087739 | 0.535497923 | 0.171575488 |
| AC125099.2 | 1.307878832 | 0.588979313 | 0.943617508 |
| Hist2h2be | 54.11456325 | -0.729410173 | 0.210461569 |
| Hist2h3c2 | 19.58774007 | 0.755918671 | 0.23798428 |
| Gm20632 | 14.19037399 | -0.052238589 | 0.310977528 |
| Hist2h2aa2 | 13.3101306 | 0.603550555 | 0.284847991 |
| AC125099.3 | 0 | NA | NA |
| Gm20633 | 28.70123197 | -0.0151797 | 0.223470279 |
| Hist2h2aa1 | 0.135217741 | 0.237306921 | 3.620320246 |

| | | | |
|---|---|---|---|
| Gm20634 | 1.134122097 | 2.891758416 | 1.066699406 |
| AC125099.1 | 0 | NA | NA |
| Hist2h3c1 | 0 | NA | NA |
| Gm20627 | 9.369992915 | 0.229818091 | 0.405314895 |
| Hist2h4 | 368.633472 | 0.368517757 | 0.154357466 |
| Gm38227 | 4.279540226 | 0.249885087 | 0.48035796 |
| Gm20628 | 12.09656833 | 0.118246213 | 0.330441157 |
| Hist2h3b | 50.50959981 | 0.771715475 | 0.289525012 |
| Hist2h2bb | 201.6384358 | 0.812606456 | 0.156175095 |
| Fcgr1 | 583.6067119 | 5.742410694 | 0.285610567 |
| Gm24830 | 0 | NA | NA |
| Platr30 | 0 | NA | NA |
| Gm26553 | 0 | NA | NA |
| Rnu1b1 | 6.056136525 | 0.694554109 | 0.53652316 |
| Rnu1b2 | 0.421929236 | 0.240014793 | 1.485875834 |
| Gm17575 | 0 | NA | NA |
| Gm26689 | 0 | NA | NA |
| 4930477E14Rik | 0 | NA | NA |
| Gm26654 | 2.969550979 | 2.139667977 | 0.592913264 |
| Pafah1b1-ps1 | 0.705160482 | 0.01250669 | 1.197999765 |
| Gm21960 | 0.551586351 | 0.614118673 | 1.237592967 |
| Terc | 7.890901767 | 1.702670064 | 0.465752617 |
| Gm24136 | 0 | NA | NA |
| BC107364 | 0.047948698 | -0.243569582 | 3.724945049 |
| Gm26232 | 0 | NA | NA |
| RP23-404J7.8 | 2.243072942 | 0.270392769 | 0.772131051 |
| RP23-404J7.10 | 0 | NA | NA |
| Rnu1b6 | 0.046345709 | 0.71820487 | 3.724945049 |
| RP23-404J7.9 | 0 | NA | NA |
| Gm28001 | 0 | NA | NA |
| Gm25890 | 0 | NA | NA |
| Gm22614 | 1.309217219 | 2.384395499 | 1.090088977 |
| Hfe2 | 3.500549162 | -1.064287796 | 0.514114259 |
| Gm15441 | 0.831325084 | 0.910808374 | 1.172599167 |
| Txnip | 698.8955186 | 2.51602117 | 0.186583763 |
| Gm16253 | 0.327232518 | 1.881189855 | 2.226562402 |
| Polr3gl | 127.7419174 | -0.32325135 | 0.124622445 |
| Ankrd34a | 497.5908006 | -0.718929567 | 0.108020649 |
| Lix1l | 406.2282066 | 0.796788769 | 0.150684864 |
| 6330549D23Rik | 6.560446392 | -0.240557347 | 0.457262513 |
| Rbm8a | 404.2810953 | 0.377751092 | 0.121304924 |
| Pex11b | 270.344613 | -0.122945265 | 0.0771165 |
| Itga10 | 23.71082201 | -0.249867116 | 0.273376682 |
| RP23-204O17.2 | 0.060174537 | 0.237309859 | 3.724945049 |
| Ankrd35 | 72.85459675 | -0.447472323 | 0.174452219 |
| Pias3 | 236.294852 | 0.043948302 | 0.100112429 |
| Nudt17 | 21.37927619 | -1.149732835 | 0.241962852 |

| | | | |
|---|---|---|---|
| Polr3c | 240.883843 | 0.313304612 | 0.094195866 |
| Gm22581 | 3.657530093 | -0.710103681 | 0.620576388 |
| RP23-204O17.4 | 9.840503469 | -0.36878972 | 0.378505132 |
| Rnf115 | 868.8846253 | 0.596413324 | 0.096976718 |
| RP23-395B13.2 | 10.83277785 | -0.454816853 | 0.43197238 |
| Cd160 | 2.898004674 | 4.683155736 | 0.9451731 |
| Pdzk1 | 2.857891658 | 1.327277341 | 0.632543851 |
| 4930442L01Rik | 0.150086409 | 0.237305146 | 3.724945049 |
| Gpr89 | 501.075764 | -0.152990437 | 0.060638008 |
| Gja5 | 5.28099507 | 1.094012111 | 0.431023208 |
| Gja8 | 0 | NA | NA |
| RP24-86K23.1 | 0 | NA | NA |
| RP23-97A19.1 | 0.188929682 | 0.237328747 | 2.260134566 |
| RP23-97A19.2 | 0.09349418 | -0.24357403 | 3.724945049 |
| Gm24435 | 0 | NA | NA |
| Acp6 | 115.9459266 | -0.639884079 | 0.181587321 |
| Bcl9 | 929.9899206 | 0.092644756 | 0.060687922 |
| RP23-394K8.5 | 0.094831987 | -0.243571806 | 3.724945049 |
| RP23-394K8.3 | 12.87106279 | -2.097346631 | 0.401966602 |
| Olfr1402 | 0 | NA | NA |
| RP23-290C4.1 | 0 | NA | NA |
| Chd1l | 117.2581915 | 0.189819938 | 0.133609475 |
| RP24-316D12.2 | 2.928805847 | -0.659469345 | 0.603748466 |
| Fmo5 | 26.93201772 | -0.681174314 | 0.275206956 |
| Prkab2 | 567.66263 | 0.301318968 | 0.12618132 |
| Gm17651 | 0 | NA | NA |
| Pde4dip | 3395.191701 | -0.302574747 | 0.093524516 |
| Mir7225 | 0 | NA | NA |
| RP24-124P9.3 | 0.671139028 | -0.742959872 | 1.173038007 |
| Gm15999 | 0.08486385 | 0.717551239 | 3.724898127 |
| Sec22b | 837.8239496 | 0.259440259 | 0.062864583 |
| Gm5544 | 1.267397472 | 2.904528677 | 1.014568787 |
| Notch2 | 1217.583416 | 1.240839599 | 0.133494642 |
| RP23-97M20.1 | 25.75372241 | -0.000150232 | 0.269465315 |
| RP23-3C4.2 | 11.28696547 | -0.433040743 | 0.346998229 |
| Adam30 | 0.456191127 | 1.220406776 | 3.271474321 |
| RP23-3C4.5 | 0.046345709 | 0.71820487 | 3.724945049 |
| Reg4 | 0.197005158 | 0.267412562 | 3.277479704 |
| RP23-348H21.1 | 0 | NA | NA |
| Hmgcs2 | 85.88327543 | -0.244595247 | 0.287817494 |
| RP23-348H21.2 | 2.607862967 | -0.545391844 | 0.812317482 |
| Phgdh | 330.7849704 | 0.246344562 | 0.181837645 |
| Zfp697 | 358.3939029 | -0.797805045 | 0.135873453 |
| Gm4450 | 0 | NA | NA |
| RP23-381G13.3 | 0.131100043 | -1.074360889 | 3.663654906 |
| RP23-381G13.4 | 0 | NA | NA |
| Hsd3b4 | 0 | NA | NA |

| | | | |
|---|---|---|---|
| RP23-381G13.6 | 0 | NA | NA |
| Gm10681 | 0 | NA | NA |
| RP24-285G1.2 | 0.109727584 | 0.718200421 | 3.724945049 |
| RP24-285G1.1 | 0 | NA | NA |
| Hsd3b5 | 0 | NA | NA |
| Gm12399 | 0 | NA | NA |
| Gm12400 | 0 | NA | NA |
| Hsd3b2 | 0.319559307 | -0.663827566 | 1.561071868 |
| Hsd3b3 | 0.048949532 | -0.243569582 | 3.724945049 |
| Hsd3b6 | 0 | NA | NA |
| Gm12427 | 0 | NA | NA |
| Gm12428 | 10.43905456 | -0.551887003 | 0.36547309 |
| Gm12431 | 0 | NA | NA |
| Gm12432 | 0.046345709 | 0.71820487 | 3.724945049 |
| Gm12433 | 0.907786306 | -1.720147558 | 1.225711404 |
| Hsd3b1 | 0.244145947 | 0.625011252 | 1.815580732 |
| Hao2 | 0.110595523 | 0.718200421 | 3.724945049 |
| 5730437C11Rik | 0 | NA | NA |
| Gm12440 | 0.266043621 | -0.2435714 | 2.474534405 |
| Gm12441 | 0 | NA | NA |
| Gm22341 | 0 | NA | NA |
| Gm24216 | 0 | NA | NA |
| Wars2 | 117.2102137 | -0.232825846 | 0.153309644 |
| RP23-200E11.1 | 19.63971965 | -0.816402437 | 0.303398677 |
| RP23-200E11.3 | 0 | NA | NA |
| Mir6481 | 0 | NA | NA |
| RP23-200E11.2 | 15.98219274 | -0.826121907 | 0.334376882 |
| Tbx15 | 21.3154785 | 0.410776316 | 0.293401141 |
| RP23-200E11.4 | 0 | NA | NA |
| Gm12449 | 0 | NA | NA |
| M6pr-ps | 7.110781671 | 1.157467048 | 0.489705932 |
| Gm12448 | 0 | NA | NA |
| Gm22505 | 0 | NA | NA |
| Spag17 | 14.25668977 | 0.06102057 | 0.29900427 |
| Rpl36-ps7 | 0 | NA | NA |
| Tra2b-ps | 0 | NA | NA |
| Spag17os | 4.312461576 | -0.85958091 | 0.572833706 |
| Wdr3 | 327.3352847 | 0.139391121 | 0.084053583 |
| Gdap2 | 404.7187721 | -0.082253069 | 0.058324797 |
| Gm25802 | 0 | NA | NA |
| Gm12464 | 0 | NA | NA |
| 4930406D18Rik | 0 | NA | NA |
| RP23-335A19.3 | 0 | NA | NA |
| Fam46c | 258.8118225 | 3.316363288 | 0.180337272 |
| Gm12474 | 1.5464145 | 0.563089423 | 0.850719077 |
| RP23-356K2.5 | 3.883316289 | 2.127384488 | 0.70550061 |
| Man1a2 | 2251.936784 | -0.258643362 | 0.066976832 |

| | | | |
|---|---|---|---|
| RP23-356K2.6 | 73.44238526 | -0.770812664 | 0.21464158 |
| RP23-267O21.4 | 93.44924604 | -0.578804067 | 0.213675193 |
| RP23-267O21.3 | 152.3385727 | -0.574242598 | 0.244354531 |
| RP23-267O21.2 | 27.15683575 | -0.645540336 | 0.270859643 |
| Gm12485 | 0 | NA | NA |
| RP23-267O21.5 | 2.75295346 | 4.767436427 | 0.940429149 |
| Gm23465 | 0 | NA | NA |
| Vtcn1 | 9.162540417 | 3.298354678 | 0.652404731 |
| Trim45 | 89.1891917 | -0.15109345 | 0.126972747 |
| Ttf2 | 41.18065889 | 1.590537847 | 0.232414989 |
| Gm12488 | 0 | NA | NA |
| Cd101 | 11.43243995 | 2.343065614 | 0.429044992 |
| Ptgfrn | 610.398054 | 0.423025521 | 0.090526354 |
| RP23-434L7.5 | 6.750937273 | -1.186296198 | 0.488090897 |
| Gm12486 | 0 | NA | NA |
| Tpt1-ps1 | 0.068097359 | -0.243569582 | 3.724945049 |
| Gm12490 | 0.045882455 | 0.237312084 | 3.724945049 |
| Cd2 | 30.67976892 | 7.378748928 | 0.86884962 |
| Gm10355 | 2.030855555 | -0.200154413 | 0.802666466 |
| RP23-347G20.3 | 0 | NA | NA |
| Igsf3 | 593.9020174 | 0.097573831 | 0.102638549 |
| RP23-347G20.4 | 4.3771345 | -0.58426791 | 0.512439164 |
| RP23-347G20.5 | 2.646385505 | -1.574177245 | 0.688256365 |
| RP23-347G20.6 | 0 | NA | NA |
| Atp1a1 | 7569.239803 | -0.297114766 | 0.076586098 |
| RP23-103P11.2 | 13.65833158 | -1.100667502 | 0.347861106 |
| RP23-103P11.1 | 32.33019698 | -1.29380106 | 0.328937699 |
| RP24-115G23.1 | 0.049553047 | 0.71820487 | 3.724945049 |
| RP24-115G23.2 | 0.117046891 | -0.724453471 | 3.724945049 |
| Mab21l3 | 2.984303933 | 2.202872477 | 0.684761808 |
| Slc22a15 | 163.9002955 | -0.140958842 | 0.117135177 |
| Nhlh2 | 53.60793837 | -0.121106471 | 0.179440986 |
| RP23-226J22.2 | 0 | NA | NA |
| RP23-226J22.3 | 0 | NA | NA |
| Casq2 | 54.88658604 | 0.148956592 | 0.215644491 |
| RP24-89C11.1 | 0.931900496 | -0.760483473 | 0.944933195 |
| RP24-89C11.2 | 3.004992861 | -1.105549415 | 0.645321604 |
| Vangl1 | 127.7913766 | 0.280227612 | 0.113817148 |
| Gm16160 | 0.666323006 | 0.249665473 | 1.070407889 |
| RP23-191N23.1 | 0.144846928 | -1.027946598 | 3.528682026 |
| RP23-63A19.1 | 24.83821061 | -0.870406861 | 0.221341795 |
| RP24-409P22.1 | 3.653697379 | -1.271451656 | 0.581489397 |
| RP24-409P22.2 | 0 | NA | NA |
| Ngf | 23.55789567 | 1.588711253 | 0.223064965 |
| RP23-63A19.3 | 5.480294069 | 1.37665364 | 0.449171487 |
| RP24-89D17.1 | 0.141094079 | 0.718203391 | 3.563850454 |
| RP24-89D17.3 | 0.352483205 | -0.243109739 | 1.757305561 |

| | | | |
|---|---|---|---|
| RP24-89D17.2 | 0 | NA | NA |
| RP23-183L19.1 | 0.184015763 | 1.259045766 | 3.310013623 |
| Tspan2os | 4.207560025 | -0.234261886 | 0.504516643 |
| Tspan2 | 1566.869316 | -1.037099199 | 0.097756645 |
| Tshb | 0.532135551 | 1.03952108 | 1.329175194 |
| RP23-183L19.3 | 0.133946075 | 0.7191812 | 3.626570343 |
| Sycp1 | 43.46231598 | 0.620739534 | 0.320452277 |
| RP23-183L19.5 | 0 | NA | NA |
| RP23-183L19.6 | 0 | NA | NA |
| RP23-119F18.1 | 0 | NA | NA |
| Nr1h5 | 0.37220402 | 1.194469422 | 1.561504783 |
| Gm22826 | 0.042716945 | -0.243571806 | 3.724945049 |
| Sike1 | 543.965291 | -0.238861665 | 0.074078973 |
| Csde1 | 5133.717668 | 0.389089111 | 0.076565065 |
| Nras | 746.5356171 | 0.65387379 | 0.090982712 |
| RP23-106G24.2 | 13.15425346 | -0.696602772 | 0.319589967 |
| Ampd1 | 2.035201819 | 3.082212029 | 1.031337283 |
| Gm23820 | 0 | NA | NA |
| RP24-409D4.3 | 0.140443404 | 1.16653613 | 2.909819564 |
| Dennd2c | 11.7142143 | 1.410540489 | 0.326406241 |
| Bcas2 | 429.1205886 | -0.083835323 | 0.064354348 |
| Gm27024 | 0 | NA | NA |
| 2410024N13Rik | 0 | NA | NA |
| RP23-70G12.2 | 0 | NA | NA |
| Trim33 | 1476.330058 | -0.096069817 | 0.062445501 |
| RP23-70G12.3 | 47.1675988 | -0.88011611 | 0.348607135 |
| Gm25009 | 0 | NA | NA |
| Gm25869 | 2.797214178 | 0.773037894 | 0.568136811 |
| Gm25199 | 0 | NA | NA |
| Syt6 | 477.8102627 | 0.272303419 | 0.173873993 |
| RP23-184N6.1 | 10.82643969 | -0.632625082 | 0.42836001 |
| Atg4a-ps | 7.430379725 | 0.664265836 | 0.365583498 |
| Gm25325 | 0 | NA | NA |
| Gm15986 | 0 | NA | NA |
| Olfml3 | 289.0413337 | 1.602137067 | 0.161856867 |
| Gm10964 | 0 | NA | NA |
| Hipk1 | 3690.970568 | 0.225586389 | 0.064036741 |
| RP24-559G13.2 | 18.01004 | -0.621585625 | 0.253450632 |
| Gm15886 | 0.872687401 | 1.019491405 | 1.123655129 |
| Gm16520 | 20.13453085 | -0.746053631 | 0.31611499 |
| Dclre1b | 139.879516 | 0.174249899 | 0.1022746 |
| Gm15471 | 0.37827039 | 0.658340672 | 1.496336178 |
| Ap4b1 | 32.06009994 | 0.136953573 | 0.204283511 |
| RP24-180F7.1 | 1.345434513 | -2.315798516 | 1.048917195 |
| Bcl2l15 | 59.73746658 | -0.282575818 | 0.210987017 |
| Ptpn22 | 140.1073571 | 2.959767724 | 0.352729094 |
| Gm15472 | 7.402758977 | 1.373461919 | 0.628506736 |

| | | | |
|---|---|---|---|
| RP24-180F7.2 | 12.50366051 | 1.22643768 | 0.39980266 |
| Gm15602 | 0.044544648 | 0.237309859 | 3.724945049 |
| Rsbn1 | 982.5522449 | 0.052827512 | 0.055475851 |
| Phtf1os | 24.52540072 | -0.202694914 | 0.208291217 |
| Phtf1 | 538.6159203 | -0.350792796 | 0.097704826 |
| Magi3 | 898.9838019 | -0.38778843 | 0.087137061 |
| RP24-535M14.2 | 5.341278601 | -1.986334366 | 0.521737728 |
| RP24-535M14.1 | 11.88755228 | -0.954707536 | 0.325289111 |
| Gm23830 | 0 | NA | NA |
| RP23-314M6.1 | 7.461541511 | -1.254552243 | 0.417251546 |
| RP23-333I5.9 | 6.272582582 | -0.780666213 | 0.579346227 |
| RP23-333I5.8 | 0.141094079 | 0.718203391 | 3.563850454 |
| RP23-333I5.7 | 34.16158652 | -1.086459142 | 0.320138035 |
| RP23-333I5.6 | 17.9477799 | -0.791815938 | 0.27578665 |
| RP23-333I5.5 | 22.39591524 | -1.196412832 | 0.363723035 |
| RP23-333I5.4 | 10.78090146 | -1.610446374 | 0.367021358 |
| RP23-333I5.3 | 57.87337 | -0.928155831 | 0.257775979 |
| RP23-333I5.2 | 44.8221039 | -0.730071012 | 0.294342959 |
| RP23-333I5.1 | 53.3220374 | -0.915437737 | 0.26697783 |
| Gm23077 | 0 | NA | NA |
| Gm5546 | 0.706390846 | -1.906423811 | 1.210122549 |
| Lrig2 | 847.580948 | -0.189472453 | 0.06460393 |
| RP23-357D17.5 | 0.044191549 | -0.243571806 | 3.724945049 |
| RP23-357D17.6 | 0.315700266 | -0.243564193 | 1.802497714 |
| RP23-357D17.4 | 84.69226509 | -0.727228914 | 0.269696787 |
| RP23-357D17.3 | 51.88363519 | -1.129872372 | 0.260499813 |
| RP23-38L11.4 | 32.89618772 | -0.665064907 | 0.212897073 |
| Gm26091 | 0.643663771 | -0.092751743 | 1.217931206 |
| RP23-38L11.3 | 7.967952249 | -0.224188268 | 0.425003774 |
| Slc16a1 | 692.2362281 | 0.676215581 | 0.113102457 |
| RP23-38L11.1 | 26.81542658 | -0.417238815 | 0.283720125 |
| RP23-38L11.2 | 8.709900708 | -0.725829543 | 0.44741592 |
| Gm29561 | 0 | NA | NA |
| Gm29560 | 0.214789954 | 0.771545301 | 2.560687178 |
| Gm24223 | 0 | NA | NA |
| Gm29572 | 0 | NA | NA |
| Gm6485 | 49.19972077 | 1.408679348 | 0.215739458 |
| RP24-503L4.2 | 0.134880151 | 1.580366464 | 2.92708711 |
| Fam19a3 | 6.79823111 | 0.146184583 | 0.466897065 |
| Ppm1j | 22.89084466 | 0.978330578 | 0.358646876 |
| Rhoc | 431.5423868 | 3.095663544 | 0.190646108 |
| Mov10 | 458.979696 | 2.640572668 | 0.148704981 |
| Capza1 | 545.1140594 | 0.906529572 | 0.091975315 |
| RP24-503L4.4 | 28.08267805 | -0.423138708 | 0.3350157 |
| RP24-503L4.3 | 42.80149299 | -0.420802982 | 0.34311652 |
| St7l | 574.3995767 | 0.001664393 | 0.066558488 |
| RP24-274H19.1 | 0.085327104 | 1.179809211 | 3.723500796 |

| | | | |
|---|---|---|---|
| Wnt2b | 46.92012226 | 1.045465244 | 0.247415094 |
| RP24-274H19.3 | 0.479138389 | -1.590952973 | 1.407428003 |
| RP24-274H19.5 | 3.753741655 | -0.289705977 | 0.550261858 |
| Cttnbp2nl | 873.2904401 | 1.349279929 | 0.081376362 |
| 4930564D02Rik | 0.045882455 | 0.237312084 | 3.724945049 |
| Kcnd3os | 14.28524435 | -1.453387345 | 0.377224078 |
| Kcnd3 | 1457.767906 | -0.265235153 | 0.068749847 |
| RP23-443O1.1 | 42.61810582 | -1.456787976 | 0.257415758 |
| Ddx20 | 175.0365538 | 0.179452524 | 0.108059703 |
| Fam212b | 966.0315667 | -0.014137181 | 0.173921601 |
| RP24-500I24.6 | 43.0223173 | -0.214082131 | 0.207100109 |
| Rap1a | 552.0763281 | 0.294117179 | 0.107104781 |
| RP24-500I24.4 | 37.31668493 | -0.616348792 | 0.275725716 |
| RP24-500I24.3 | 48.64596038 | -0.384992709 | 0.253574579 |
| RP24-500I24.2 | 32.60373694 | -0.575605814 | 0.258866168 |
| RP24-500I24.5 | 33.83971767 | -0.188162218 | 0.265156685 |
| RP24-500I24.7 | 2.098595915 | 0.309138817 | 0.665885854 |
| Adora3 | 9.886871762 | 1.48429341 | 0.411618125 |
| Tmigd3 | 0.650450437 | 2.070574672 | 1.373271627 |
| I830077J02Rik | 45.37358814 | 2.490703642 | 0.293898181 |
| Atp5f1 | 2177.75384 | 0.067045587 | 0.084373234 |
| Wdr77 | 315.9051542 | 0.053115867 | 0.074718583 |
| RP23-169H9.3 | 8.021508243 | -0.27910445 | 0.446698995 |
| Ovgp1 | 35.01689641 | -0.447255627 | 0.302899722 |
| Pifo | 4.561976979 | -0.626031944 | 0.471526896 |
| RP23-169H9.6 | 8.449170836 | -2.03287182 | 0.570050752 |
| RP23-169H9.5 | 0 | NA | NA |
| Chil5 | 6.28406316 | 1.674833131 | 0.43534765 |
| Gm4540 | 10.64549809 | 0.047417008 | 0.298308627 |
| RP23-169H9.8 | 0 | NA | NA |
| RP23-62K10.1 | 0.969048608 | -2.004352299 | 1.160791135 |
| Chia1 | 8.605265352 | -1.679852497 | 0.462293273 |
| Chil3 | 390.2692537 | 12.70340912 | 1.320531357 |
| RP23-169H9.9 | 0 | NA | NA |
| Chil4 | 0.092691418 | 1.183079489 | 3.724560917 |
| RP23-191E1.1 | 0.325695176 | 2.581414813 | 1.869788021 |
| RP23-191E1.2 | 0 | NA | NA |
| Chil6 | 1.21043771 | 1.51157427 | 1.089847887 |
| Gm38244 | 0 | NA | NA |
| 2010016I18Rik | 10.3751689 | 3.909444563 | 0.626886112 |
| Dennd2d | 26.9763545 | 2.938623762 | 0.312759514 |
| Cept1 | 592.7431419 | 0.818831186 | 0.115949243 |
| Dram2 | 391.8607469 | 0.802515384 | 0.112356235 |
| RP23-164N7.2 | 0 | NA | NA |
| RP23-164N7.3 | 46.01383439 | 0.204766101 | 0.291881817 |
| Gm38271 | 1.062454588 | 1.401036872 | 1.099755734 |
| Lrif1 | 343.0611023 | 0.179122245 | 0.099363078 |

| | | | |
|---|---|---|---|
| Gm38253 | 70.59424867 | -0.042067458 | 0.229820988 |
| Cd53 | 595.4541458 | 4.399183686 | 0.291436639 |
| Olfr266 | 0.317967701 | -0.243564166 | 1.799472047 |
| RP24-481E4.4 | 1.491118976 | 0.779055672 | 0.995210938 |
| Gm27008 | 4.962650854 | -0.888132184 | 0.495796227 |
| Kcna3 | 229.4345153 | 0.216611843 | 0.096310852 |
| AI504432 | 766.1594696 | 0.198867749 | 0.093356832 |
| A930002I21Rik | 0.066050875 | 0.71820487 | 3.724945049 |
| Kcna2 | 5020.598395 | -0.622153106 | 0.074023857 |
| Kcna10 | 0.10503227 | 1.179809211 | 3.723500796 |
| Cym | 1.145893073 | 2.624628087 | 1.095971708 |
| A630076J17Rik | 0 | NA | NA |
| Prok1 | 4.595448637 | 0.675317865 | 0.649441258 |
| Lamtor5 | 258.6284024 | 0.401902879 | 0.136385121 |
| Slc16a4 | 15.80845376 | -0.328002259 | 0.327168278 |
| Rbm15 | 884.0175483 | 0.266218569 | 0.066650743 |
| RP23-332I1.7 | 0.048949532 | -0.243569582 | 3.724945049 |
| Gm24383 | 0.089089296 | 0.237305797 | 3.724945049 |
| Kcnc4 | 618.4655694 | -0.463892136 | 0.081539557 |
| Slc6a17 | 6021.526622 | -0.139224055 | 0.110189207 |
| Ubl4b | 0.42233865 | 1.072294012 | 1.513650744 |
| Alx3 | 13.73531081 | 1.827774638 | 0.471485328 |
| Strip1 | 672.0896654 | -0.050797176 | 0.072946639 |
| Gm10961 | 0.150451996 | 0.237323705 | 2.858243839 |
| Ahcyl1 | 4251.746979 | -0.386017778 | 0.076873947 |
| Csf1 | 1357.448556 | 2.982696705 | 0.229251399 |
| RP23-455E11.2 | 0.180126469 | -1.336007643 | 3.33473513 |
| RP23-455E11.1 | 0.049553047 | 0.71820487 | 3.724945049 |
| RP23-313F7.2 | 1.373329198 | -0.15109984 | 0.862831927 |
| RP23-313F7.8 | 0 | NA | NA |
| Eps8l3 | 0.290796633 | -1.110156872 | 1.58261049 |
| RP23-313F7.4 | 0 | NA | NA |
| 4933431E20Rik | 826.6688943 | -1.405464359 | 0.119689894 |
| Gstm5 | 640.7472393 | -0.697104964 | 0.131212566 |
| RP23-313F7.6 | 0.038981395 | 0.718207094 | 3.724945049 |
| Gm12489 | 0 | NA | NA |
| Gstm7 | 114.4264593 | -0.433049417 | 0.186104406 |
| Gstm6 | 23.14962032 | -0.16787018 | 0.25882416 |
| Gm12494 | 1.001132476 | -0.000985558 | 0.983827069 |
| Gstm3 | 0.425236713 | 0.874128406 | 1.683492002 |
| Gm12497 | 0 | NA | NA |
| Gstm2 | 20.7015664 | 0.110807302 | 0.316043134 |
| RP23-313F7.7 | 0.044544648 | 0.237309859 | 3.724945049 |
| Gstm1 | 1044.038813 | -0.744872306 | 0.181229428 |
| Gm12498 | 0.194528622 | 1.200042521 | 3.242027088 |
| Gm12502 | 2.704774846 | 0.680803849 | 0.665807099 |
| Gm12499 | 0 | NA | NA |

| | | | |
|---|---|---|---|
| Gm25592 | 0 | NA | NA |
| Gstm4 | 56.39273402 | 0.058779011 | 0.177289353 |
| Ampd2 | 616.2955637 | -0.425358525 | 0.071387567 |
| Gm12500 | 6.670440749 | -0.263489531 | 0.365804694 |
| Gnat2 | 1.625657436 | 1.302054961 | 0.822183237 |
| Gnai3 | 600.7292191 | 0.890514386 | 0.132210924 |
| Gpr61 | 149.8257791 | -0.77236241 | 0.12849965 |
| Gm12524 | 0.213105667 | 1.175348221 | 2.596470818 |
| Amigo1 | 1089.407811 | -0.394218908 | 0.068281977 |
| Cyb561d1 | 232.812123 | 0.086925987 | 0.079935384 |
| Atxn7l2 | 171.8889948 | -0.810277207 | 0.166978125 |
| Sypl2 | 13.67443026 | -2.532889601 | 0.352864136 |
| RP23-89M15.15 | 1.384376546 | -3.80122844 | 1.086568003 |
| Gm12501 | 0.378789762 | -1.249226634 | 1.639481725 |
| 1700010K24Rik | 1.755101273 | -0.07108805 | 0.732628164 |
| Psma5 | 407.5269465 | 0.788877067 | 0.120240113 |
| Gm12523 | 1.090677807 | 0.095567041 | 0.915313675 |
| Sort1 | 4425.028415 | -1.181991587 | 0.104094935 |
| RP23-27B23.9 | 51.99739385 | -1.436904079 | 0.385131945 |
| Gm12525 | 7.569909558 | -0.892221033 | 0.4834722 |
| Mybphl | 0.67227865 | -2.0307303 | 1.188129857 |
| Gm12522 | 10.72785558 | -1.6816562 | 0.413975547 |
| Psrc1 | 41.89133445 | -0.104450721 | 0.229490772 |
| Celsr2 | 4197.124515 | -1.151853062 | 0.121838772 |
| Sars | 1527.302422 | 0.446571986 | 0.082704483 |
| Gm22942 | 0 | NA | NA |
| RP23-37G19.3 | 0.129625439 | -1.065581744 | 3.675612039 |
| 5330417C22Rik | 475.2428445 | -0.580730953 | 0.130242897 |
| Gm23336 | 0 | NA | NA |
| Gm37859 | 19.39367035 | -1.150096308 | 0.476322277 |
| 1700013F07Rik | 7.727506971 | -0.389947532 | 0.371321737 |
| Scarna2 | 566.7431568 | 0.218969458 | 0.235659171 |
| RP23-128G10.7 | 0 | NA | NA |
| Tmem167b | 537.5116706 | 0.321988486 | 0.066259332 |
| Gm22860 | 0.047948698 | -0.243569582 | 3.724945049 |
| Taf13 | 217.8213017 | 0.310033108 | 0.117850839 |
| Wdr47 | 1240.073746 | -0.395946352 | 0.073675573 |
| Gm12517 | 0 | NA | NA |
| Clcc1 | 305.3641183 | 0.402211472 | 0.081544872 |
| Gpsm2 | 151.0149954 | 0.332586314 | 0.161278117 |
| Gm27244 | 0 | NA | NA |
| Aknad1 | 0.537146556 | 1.232403254 | 1.291965454 |
| Stxbp3 | 425.8909719 | 1.112190235 | 0.109751119 |
| Fndc7 | 3.245667884 | 3.149165323 | 0.761448857 |
| RP23-316M11.8 | 0.10503227 | 1.179809211 | 3.723500796 |
| Prpf38b | 771.447708 | 0.122473698 | 0.115372589 |
| Henmt1 | 3.246767699 | -0.248685545 | 0.609411328 |

| | | | |
|---|---|---|---|
| Gm9857 | 0.597180582 | 0.19793274 | 1.360364441 |
| Fam102b | 647.6814876 | -0.343951721 | 0.111638684 |
| RP23-316M11.10 | 0 | NA | NA |
| AL671921.1 | 0 | NA | NA |
| RP23-288B16.2 | 0 | NA | NA |
| Slc25a54 | 0 | NA | NA |
| RP23-288B16.4 | 8.294058351 | -0.711229161 | 0.401775883 |
| Slc25a24 | 72.40842471 | 2.309521793 | 0.228209528 |
| Gm13865 | 0 | NA | NA |
| Vav3 | 309.6866449 | 0.587837034 | 0.077146039 |
| Ntng1 | 634.8071448 | -0.891896001 | 0.094336247 |
| Gm25681 | 0 | NA | NA |
| Gm12535 | 0 | NA | NA |
| RP23-327M1.2 | 2.031725428 | 1.895671735 | 0.843487952 |
| Prmt6 | 217.8360219 | 0.468938124 | 0.103582148 |
| RP23-327M1.1 | 30.13062775 | 0.664294762 | 0.216092064 |
| RP23-440G4.1 | 0 | NA | NA |
| RP23-440G4.2 | 0.155610039 | 0.718213998 | 2.828260953 |
| RP23-43A4.1 | 0 | NA | NA |
| Gm26076 | 0 | NA | NA |
| RP24-366N1.1 | 0.538110248 | 1.266180311 | 1.575426078 |
| RP24-223F21.1 | 0.075043205 | 0.237309859 | 3.724945049 |
| RP24-230D18.1 | 0 | NA | NA |
| RP23-14C8.1 | 0 | NA | NA |
| RP23-14C8.2 | 0 | NA | NA |
| RP24-93G19.1 | 0.12092566 | 0.237307635 | 3.724945049 |
| Gm25519 | 0 | NA | NA |
| RP24-325G15.1 | 0 | NA | NA |
| RP24-325G15.2 | 0.39879887 | 0.974545941 | 1.735214585 |
| RP23-117K10.1 | 0 | NA | NA |
| RP24-221P14.1 | 0 | NA | NA |
| RP23-430D14.1 | 5.464530712 | -0.052517974 | 0.481094918 |
| RP24-141M5.1 | 0 | NA | NA |
| Gm5548 | 0.097139547 | 0.718207094 | 3.724945049 |
| RP23-470O13.1 | 0 | NA | NA |
| Amy2b | 0.038981395 | 0.718207094 | 3.724945049 |
| Amy2-ps1 | 0 | NA | NA |
| Amy2a5 | 0.143022002 | 0.719176055 | 3.547735711 |
| Amy2a4 | 0.089089296 | 0.237305797 | 3.724945049 |
| Amy2a3 | 0.060174537 | 0.237309859 | 3.724945049 |
| Amy2a2 | 0 | NA | NA |
| Amy2a1 | 0.175778459 | 1.213873066 | 2.722156148 |
| Amy1 | 192.4654876 | -0.43306778 | 0.187614928 |
| Rnpc3 | 532.6768979 | -0.30124517 | 0.083215361 |
| RP24-237D17.3 | 0 | NA | NA |
| RP24-182K13.1 | 0.088383099 | -0.72445792 | 3.724945049 |
| RP23-256J12.1 | 0.075043205 | 0.237309859 | 3.724945049 |

| | | | |
|---|---|---|---|
| Col11a1 | 184.655142 | 0.342961742 | 0.148202502 |
| RP23-281B17.2 | 0 | NA | NA |
| Gm23750 | 0 | NA | NA |
| RP23-292P14.1 | 0 | NA | NA |
| Olfm3 | 941.997379 | -0.602444745 | 0.147235663 |
| RP23-231J6.1 | 0.046345709 | 0.71820487 | 3.724945049 |
| Gm22337 | 0 | NA | NA |
| Gm25347 | 0 | NA | NA |
| RP23-238A12.1 | 0.12092566 | 0.237307635 | 3.724945049 |
| S1pr1 | 1328.879257 | 0.545860719 | 0.098786821 |
| Gm9889 | 3.625155807 | 0.650659877 | 0.595335805 |
| A930005H10Rik | 72.61233888 | -0.355873751 | 0.133170583 |
| Dph5 | 126.6011858 | 0.491170861 | 0.121275138 |
| RP24-155C13.8 | 0.199959441 | 0.237325476 | 2.66342588 |
| RP24-155C13.6 | 0.510666263 | -0.866539955 | 1.305749431 |
| RP24-155C13.7 | 0.595761752 | 0.742152447 | 1.37172703 |
| Slc30a7 | 561.2209546 | 0.291981507 | 0.088351563 |
| RP24-245I16.1 | 1.76977757 | -1.174410821 | 1.009758426 |
| RP24-155C13.3 | 4.122231334 | 0.471635225 | 0.53422611 |
| Mir669n | 0 | NA | NA |
| Extl2 | 975.1969821 | -0.113521734 | 0.07959883 |
| Gm26344 | 0 | NA | NA |
| Vcam1 | 1522.902751 | 3.213582866 | 0.215617464 |
| RP24-245I16.3 | 1.059433203 | -0.098733626 | 0.99092346 |
| RP23-93F11.2 | 1.275102148 | 2.438587009 | 1.09833825 |
| Gm26544 | 0.239854152 | 0.240623648 | 2.528885923 |
| RP23-93F11.4 | 0.265934997 | -1.11372022 | 2.10415641 |
| Gpr88 | 1159.493482 | -1.418276811 | 0.155103269 |
| Cdc14a | 108.7934714 | 0.753699147 | 0.127813711 |
| Gm29151 | 0 | NA | NA |
| Gm22361 | 0 | NA | NA |
| Gm26358 | 0 | NA | NA |
| Rtca | 414.5859796 | 0.2737248 | 0.110921924 |
| Gm26530 | 0.207691825 | 1.136717191 | 2.608990766 |
| Dbt | 216.6118277 | 0.191557957 | 0.102520319 |
| Gm4596 | 16.93547575 | 0.554765099 | 0.271277863 |
| RP23-453M11.3 | 3.08271666 | -3.073857954 | 0.890787437 |
| Gm9761 | 55.52035785 | 0.429009426 | 0.156881846 |
| Lrrc39 | 5.281818806 | 0.075888984 | 0.502798201 |
| RP23-453M11.4 | 10.63896767 | 0.520542395 | 0.444492 |
| Trmt13 | 134.1213703 | -0.055614655 | 0.133068253 |
| Sass6 | 170.9486267 | 0.915611302 | 0.112210486 |
| RP24-565B3.4 | 8.173995324 | -0.392818858 | 0.362867857 |
| Hiat1 | 617.2974484 | -0.076960079 | 0.065978273 |
| RP24-565B3.2 | 6.147691891 | 0.237021363 | 0.446354169 |
| RP24-565B3.5 | 0 | NA | NA |
| RP24-565B3.3 | 0.186953752 | 0.237327712 | 2.668894796 |

| | | | |
|---|---|---|---|
| Slc35a3 | 453.1222322 | 0.170333336 | 0.083822584 |
| Agl | 702.330067 | -0.346843158 | 0.075179359 |
| RP24-423B3.1 | 0.424532791 | 1.151927074 | 1.463611043 |
| RP24-423B3.2 | 0.346647033 | -0.25841231 | 1.612164452 |
| RP24-539I8.1 | 0 | NA | NA |
| CH25-626J19.1 | 0.097139547 | 0.718207094 | 3.724945049 |
| Frrs1 | 66.10065566 | 2.586323702 | 0.224162697 |
| Palmd | 159.5277226 | 0.014134462 | 0.162593192 |
| RP24-370B22.3 | 26.73605718 | -1.432118053 | 0.372903621 |
| 4930455H04Rik | 0.11657011 | 0.237307635 | 3.724945049 |
| RP24-341P14.2 | 0 | NA | NA |
| 1700061I17Rik | 0 | NA | NA |
| RP24-83G17.1 | 0.038981395 | 0.718207094 | 3.724945049 |
| RP24-83G17.2 | 0 | NA | NA |
| RP23-173L24.3 | 0.766169159 | 0.671142798 | 1.440189741 |
| RP23-173L24.2 | 1.114955615 | 2.676430314 | 1.167843956 |
| Lppr4 | 2571.916126 | -0.405155106 | 0.136432667 |
| Gm22408 | 0.596872081 | 0.68621607 | 1.449220504 |
| Lppr5 | 515.7492512 | -0.32257276 | 0.11289473 |
| Snx7 | 152.0630282 | 0.128335949 | 0.111811824 |
| Gm24811 | 0.346782673 | -0.685910217 | 1.55577586 |
| CH25-626J19.2 | 0 | NA | NA |
| RP24-294N3.1 | 0 | NA | NA |
| Gm26871 | 573.8481393 | -0.56743156 | 0.258084762 |
| Mir137 | 0 | NA | NA |
| Gm24155 | 3.962088627 | -1.279521172 | 0.666157356 |
| Gm9916 | 100.2258943 | -0.963557968 | 0.391180446 |
| Gm37773 | 129.487562 | -0.421775435 | 0.173564948 |
| Gm37928 | 218.6973413 | -0.57293299 | 0.246871888 |
| Gm32444 | 100.5808754 | -0.275022085 | 0.192832522 |
| Dpyd | 164.1135791 | 0.248003051 | 0.120258625 |
| RP24-173A7.2 | 9.88657072 | -0.162117983 | 0.359566869 |
| Gm23432 | 0 | NA | NA |
| RP23-235E15.1 | 152.0784296 | 0.789431731 | 0.146249722 |
| Ptbp2 | 2043.522665 | -0.181237344 | 0.117916533 |
| RP23-235E15.5 | 196.3079525 | -0.320703774 | 0.260877563 |
| RP23-235E15.6 | 474.5268742 | -0.187370952 | 0.211569671 |
| RP23-235E15.7 | 0 | NA | NA |
| Gm24744 | 0 | NA | NA |
| Gm26137 | 0 | NA | NA |
| RP23-427K12.1 | 0 | NA | NA |
| Gm23733 | 0 | NA | NA |
| 6530403H02Rik | 55.37544644 | -1.165976193 | 0.20478154 |
| Gm25013 | 0.075043205 | 0.237309859 | 3.724945049 |
| RP23-324P1.1 | 0.575937436 | -1.110196163 | 1.296142963 |
| RP24-295P14.2 | 0.374268822 | -0.715208349 | 3.68104514 |
| RP24-295P14.1 | 0.066050875 | 0.71820487 | 3.724945049 |

| | | | |
|---|---|---|---|
| Rwdd3 | 95.46200367 | -0.413518936 | 0.194400079 |
| RP24-210J6.4 | 0.060174537 | 0.237309859 | 3.724945049 |
| Gm5710 | 0 | NA | NA |
| Tmem56 | 921.2771509 | -0.72222656 | 0.072738517 |
| RP24-291L17.1 | 8.847422939 | -0.975828885 | 0.46646466 |
| RP24-291L17.2 | 0.233612452 | 0.718216207 | 2.540177828 |
| Alg14 | 122.1469618 | 0.251604369 | 0.125283316 |
| Gm26027 | 1.314748965 | -0.350757377 | 0.975145858 |
| RP24-277J22.6 | 0 | NA | NA |
| RP24-277J22.4 | 33.98944694 | 4.057012084 | 0.496518701 |
| Cnn3 | 2221.060378 | 2.393290966 | 0.170110722 |
| RP24-277J22.3 | 26.71658222 | 1.009450471 | 0.333425026 |
| Slc44a3 | 22.72182752 | 4.300160404 | 0.535957893 |
| A530020G20Rik | 0.979355264 | 2.053326245 | 1.056795892 |
| A730020M07Rik | 12.26765879 | 0.861663737 | 0.323338839 |
| 4930432M17Rik | 0 | NA | NA |
| RP24-404P10.3 | 0.136120942 | 1.15080397 | 3.595683408 |
| RP24-404P10.4 | 0 | NA | NA |
| RP24-404P10.2 | 5.861481922 | 0.13558573 | 0.416173949 |
| F3 | 702.2491587 | 1.059608766 | 0.150619098 |
| RP24-404P10.8 | 9.75452295 | -1.114544289 | 0.362320652 |
| Abcd3 | 888.7244778 | -0.102345764 | 0.084994539 |
| RP24-404P10.7 | 19.74428529 | -0.926657231 | 0.250575785 |
| RP24-404P10.6 | 21.98336171 | -0.504145406 | 0.307935769 |
| RP24-404P10.9 | 0.234491734 | 1.723339577 | 2.118440779 |
| RP23-54L16.1 | 0 | NA | NA |
| Arhgap29 | 378.6264762 | 0.71613823 | 0.081635178 |
| RP23-54L16.2 | 5.712404952 | -1.499331131 | 0.443689732 |
| Abca4 | 76.95354951 | -0.684537669 | 0.18829821 |
| RP24-471B15.3 | 0.334549462 | 0.246393399 | 1.688475854 |
| RP24-471B15.2 | 0 | NA | NA |
| RP24-471B15.1 | 0.04393175 | -0.243571806 | 3.724945049 |
| Gm4609 | 10.08219113 | 0.003558574 | 0.347628189 |
| Gm22903 | 0.085433889 | -0.72445792 | 3.724945049 |
| Gclm | 1174.677346 | 0.633177334 | 0.105220658 |
| Gm17494 | 67.68281203 | -1.12917584 | 0.280927239 |
| Gm6061 | 18.81787436 | 0.121547865 | 0.257542226 |
| Dnttip2 | 377.7972389 | 0.769205485 | 0.118999855 |
| RP23-71G16.7 | 1.463879734 | -0.905064764 | 0.866126677 |
| Mir760 | 0 | NA | NA |
| Bcar3 | 197.0241314 | 0.480362183 | 0.12227822 |
| RP23-71G16.6 | 1.51534681 | 3.523923614 | 1.016575425 |
| RP23-192O12.4 | 0.11657011 | 0.237307635 | 3.724945049 |
| Gm25153 | 0.892441177 | -2.074899738 | 1.127796161 |
| Fnbp1l | 1523.630982 | -0.599011708 | 0.101500356 |
| mmu-mir-7657 | 0 | NA | NA |
| RP23-1A12.4 | 0.048949532 | -0.243569582 | 3.724945049 |

| | | | |
|---|---|---|---|
| RP23-1A12.2 | 29.85261811 | -1.131603823 | 0.284099905 |
| RP23-1A12.5 | 0.187986659 | 1.057580841 | 3.306965487 |
| RP23-1A12.1 | 0.044544648 | 0.237309859 | 3.724945049 |
| Pde5a | 353.2225523 | -0.324974852 | 0.165381049 |
| RP24-182A8.1 | 4.735351793 | -0.564006126 | 0.464681468 |
| RP24-182A8.2 | 0.332478248 | -1.715066883 | 1.947363378 |
| RP24-70L3.3 | 0.663651821 | 0.961886562 | 1.204382204 |
| 4933405D12Rik | 1.378174623 | 0.357847103 | 0.837014234 |
| Fabp2 | 0 | NA | NA |
| Gm10959 | 0.205306308 | -0.243571185 | 2.610168303 |
| 1810037I17Rik | 393.3493054 | 0.698519171 | 0.134500531 |
| Usp53 | 810.7985606 | 0.436042106 | 0.114006486 |
| RP23-48L9.1 | 1.574470888 | -0.154268575 | 0.756932651 |
| Myoz2 | 0.617569467 | 1.021367717 | 1.36396444 |
| Gm22066 | 0.060174537 | 0.237309859 | 3.724945049 |
| RP23-48L9.2 | 0 | NA | NA |
| Synpo2 | 278.5160077 | -0.388451343 | 0.133023699 |
| RP23-48L9.4 | 0.075043205 | 0.237309859 | 3.724945049 |
| RP23-48L9.3 | 2.800041622 | -1.026814897 | 0.607734257 |
| RP24-488I7.1 | 9.542608144 | -1.083203334 | 0.419974604 |
| RP24-488I7.2 | 12.44003729 | -0.547697964 | 0.367794693 |
| Sec24d | 210.6853567 | 1.452162835 | 0.130668333 |
| Mettl14 | 164.8105688 | -0.007318797 | 0.094955678 |
| Gm22713 | 0.066050875 | 0.71820487 | 3.724945049 |
| RP23-113O9.4 | 1.285708346 | 0.267045833 | 1.026510662 |
| Prss12 | 146.351687 | -0.414236612 | 0.161328237 |
| RP23-113O9.3 | 0.319644482 | -0.232352042 | 1.600091901 |
| Snhg8 | 70.39141616 | 0.304044577 | 0.340389839 |
| Snora24 | 0 | NA | NA |
| RP24-149B7.2 | 0.066050875 | 0.71820487 | 3.724945049 |
| Ndst3 | 624.8561942 | -0.582603425 | 0.166457372 |
| RP24-374F22.1 | 20.67476562 | -1.047035387 | 0.293361335 |
| Gm24456 | 0 | NA | NA |
| Gm15400 | 0.119587853 | 0.237305575 | 3.724945049 |
| Gm15399 | 0.060174537 | 0.237309859 | 3.724945049 |
| RP24-107N14.2 | 0 | NA | NA |
| Gm9372 | 0.194279094 | 1.292225235 | 3.719439735 |
| RP24-107N14.4 | 0 | NA | NA |
| Tram1l1 | 242.7416911 | -0.366255192 | 0.089593806 |
| Gm4617 | 19.97291991 | 1.421970134 | 0.353651917 |
| RP23-412L13.3 | 0 | NA | NA |
| RP23-412L13.5 | 0 | NA | NA |
| 1700006A11Rik | 0.255320825 | 0.237320997 | 2.417314357 |
| Gm22575 | 0.066050875 | 0.71820487 | 3.724945049 |
| RP23-412L13.4 | 0 | NA | NA |
| Gm22900 | 0 | NA | NA |
| 1700003H04Rik | 0 | NA | NA |

| | | | |
|---|---|---|---|
| RP24-332L11.2 | 0 | NA | NA |
| RP23-24O21.1 | 0.192951104 | -0.243571233 | 3.266798291 |
| Gm23673 | 0.219094128 | 0.237313857 | 3.147696964 |
| Gm9381 | 0.088383099 | -0.72445792 | 3.724945049 |
| RP23-446O11.1 | 0 | NA | NA |
| Ndst4 | 306.8135137 | -0.528794627 | 0.138169422 |
| Ugt8a | 1013.869147 | -0.625649405 | 0.224626966 |
| Gm26275 | 0.551648795 | 0.313012409 | 1.697689541 |
| RP24-297G3.1 | 0 | NA | NA |
| RP23-301O12.1 | 0.159882028 | 0.237326729 | 2.804321407 |
| Arsj | 51.42050542 | 1.107301733 | 0.20972181 |
| Gm17225 | 0.384085174 | 1.094500408 | 3.28526054 |
| RP23-60D3.4 | 6.256257677 | 0.500574477 | 0.563780418 |
| RP23-60D3.3 | 21.83940584 | 0.205439722 | 0.339302754 |
| Camk2d | 2946.282952 | 0.267024227 | 0.049355007 |
| RP23-293K21.1 | 122.6972469 | 0.000991408 | 0.236338963 |
| RP23-293K21.2 | 86.00755271 | 0.243061825 | 0.212810479 |
| RP23-293K21.3 | 29.22029167 | -0.06205708 | 0.264809185 |
| RP23-293K21.4 | 16.36398547 | 0.335529314 | 0.393267466 |
| RP23-293K21.5 | 4.108293731 | -0.054871065 | 0.56003634 |
| Gm15551 | 0 | NA | NA |
| RP23-293K21.7 | 52.63659607 | 0.172275508 | 0.27123587 |
| RP23-293K21.8 | 19.23585701 | 0.187459917 | 0.293102882 |
| Ank2 | 12996.91815 | -0.389532257 | 0.054690311 |
| RP24-423L4.5 | 54.25629816 | -0.857941046 | 0.292966379 |
| RP24-423L4.4 | 84.96617208 | -0.863868332 | 0.246731187 |
| RP24-423L4.3 | 17.18518708 | -0.931438772 | 0.312890938 |
| Gm25501 | 0 | NA | NA |
| RP24-423L4.2 | 66.13758907 | -1.075815669 | 0.333378096 |
| RP24-423L4.1 | 97.87426476 | -0.956591969 | 0.278793178 |
| RP23-50E10.7 | 69.57910019 | -0.590954053 | 0.227819007 |
| Gm16958 | 3.547171809 | 0.785849873 | 0.531356216 |
| RP23-50E10.5 | 105.9269057 | -0.482375411 | 0.259204264 |
| RP23-50E10.4 | 141.5973348 | -0.512173229 | 0.211576679 |
| RP23-50E10.3 | 170.8359331 | -0.97159599 | 0.372021789 |
| RP23-50E10.2 | 42.82266199 | -1.112035549 | 0.342974376 |
| RP23-50E10.1 | 29.57400322 | -1.26383241 | 0.41635454 |
| RP24-163F11.1 | 44.61355707 | -0.721497658 | 0.318315513 |
| RP24-260H22.2 | 2.710738714 | -0.785044622 | 0.607474555 |
| RP24-260H22.1 | 2.472653692 | -1.796652321 | 0.80483565 |
| AC157817.1 | 0 | NA | NA |
| RP24-166J20.2 | 0 | NA | NA |
| Larp7 | 118.9071558 | 0.175594724 | 0.148394169 |
| Mir302b | 0 | NA | NA |
| Mir302c | 0 | NA | NA |
| Mir302a | 0 | NA | NA |
| Mir302d | 0 | NA | NA |

| | | | |
|---|---|---|---|
| Mir367 | 0 | NA | NA |
| Zgrf1 | 113.3311546 | 0.309632863 | 0.149633094 |
| RP23-182M12.2 | 0 | NA | NA |
| Neurog2 | 4.696444918 | -0.252905331 | 0.481805305 |
| Alpk1 | 333.345234 | 1.174577519 | 0.162919802 |
| Gm16238 | 2.994207206 | 1.018772094 | 0.621722201 |
| RP23-182M12.4 | 9.869931754 | 1.142331187 | 0.462260517 |
| RP23-182M12.3 | 4.2109437 | 0.931775132 | 0.659766757 |
| Gm23279 | 0 | NA | NA |
| Tifa | 352.3487698 | 1.836594108 | 0.164223795 |
| Ap1ar | 641.0884169 | -0.261852629 | 0.080557683 |
| RP23-173N16.4 | 70.08836606 | -0.596759949 | 0.353913717 |
| RP23-173N16.3 | 2.767049531 | -0.446325276 | 0.568258439 |
| RP23-173N16.5 | 0.803713496 | -0.730969176 | 1.391475544 |
| RP23-173N16.8 | 0.269164771 | 1.237208448 | 2.060544012 |
| 5730508B09Rik | 54.45665569 | 2.534000214 | 0.238965119 |
| RP24-285A14.1 | 0.588650639 | 0.729466026 | 1.474996493 |
| RP24-285A14.2 | 0.048949532 | -0.243569582 | 3.724945049 |
| Gm10650 | 0 | NA | NA |
| RP24-380G1.1 | 1.298106709 | 0.537999696 | 0.817515559 |
| RP24-380G1.2 | 0 | NA | NA |
| Gm22293 | 0 | NA | NA |
| RP24-196G4.1 | 0 | NA | NA |
| RP23-61G16.1 | 0.062426873 | 0.237312084 | 3.724945049 |
| RP24-261K17.1 | 0 | NA | NA |
| Gm9387 | 8.054043807 | -0.179593904 | 0.353647788 |
| Gm28543 | 0 | NA | NA |
| Pitx2 | 11.30362075 | -0.548005365 | 0.329842298 |
| Gm27195 | 0.265852463 | 0.728349929 | 1.827080215 |
| RP23-446B17.3 | 0.066050875 | 0.71820487 | 3.724945049 |
| Enpep | 24.04966124 | -0.030779779 | 0.274884677 |
| RP24-140K12.2 | 0 | NA | NA |
| RP24-140K12.4 | 0 | NA | NA |
| RP23-257P11.1 | 0 | NA | NA |
| RP23-257P11.2 | 0.095897396 | -0.724453471 | 3.724945049 |
| Rpl7a-ps7 | 2.979957284 | 0.116332811 | 0.626499231 |
| RP23-257P11.4 | 0.174406792 | 0.718206751 | 3.367085639 |
| Elovl6 | 655.176359 | -0.487799889 | 0.076607307 |
| RP23-3L20.3 | 30.90056436 | -1.60915128 | 0.253322436 |
| Egf | 35.62468515 | -0.141960549 | 0.238347462 |
| Gm24515 | 0.098502579 | 0.23732098 | 3.724945049 |
| RP23-3L20.6 | 0.263299413 | -1.27192617 | 2.097131309 |
| RP23-3L20.8 | 0 | NA | NA |
| RP23-3L20.5 | 1.405849262 | -1.033315137 | 0.879541008 |
| Lrit3 | 3.350603553 | 1.771457148 | 0.87226979 |
| Rrh | 25.69789624 | 0.188418089 | 0.475403652 |
| Gar1 | 114.8672286 | 0.203270339 | 0.203820188 |

| | | | |
|---|---|---|---|
| Cfi | 0.976809389 | 1.773726548 | 1.313345569 |
| Pla2g12a | 138.6047462 | -0.180202898 | 0.111802239 |
| Casp6 | 39.32840354 | 0.332412468 | 0.190232396 |
| Ccdc109b | 33.80637658 | 1.639495342 | 0.255257616 |
| Sec24b | 993.3623414 | 0.056962296 | 0.048805085 |
| Gm9396 | 8.082634732 | 0.928634044 | 0.429508466 |
| RP23-123C15.1 | 0 | NA | NA |
| Col25a1 | 767.761363 | -0.6691099 | 0.106709439 |
| RP23-123C15.2 | 0.147068667 | 0.237310878 | 3.514416608 |
| Gm22682 | 2.066582291 | -0.559216613 | 0.717689418 |
| RP23-105B8.2 | 34.01273389 | -0.837487873 | 0.320937102 |
| RP23-105B8.3 | 9.17532523 | -1.25935903 | 0.392904144 |
| RP23-105B8.4 | 6.910068968 | -1.146811814 | 0.446890739 |
| Etnppl | 214.5413129 | -1.224296312 | 0.189174677 |
| Ostc | 293.8716669 | 0.774120114 | 0.147810747 |
| RP23-246L14.2 | 8.0379673 | -0.541658508 | 0.443241798 |
| Rpl34 | 837.1533585 | 0.822393052 | 0.131357747 |
| RP23-246L14.3 | 2.763783036 | -0.996926921 | 0.60480635 |
| RP23-246L14.4 | 0.231616962 | -0.72446114 | 2.127076405 |
| RP23-246L14.5 | 0.678557026 | -0.613514711 | 1.258069107 |
| RP23-349J3.1 | 0 | NA | NA |
| RP23-349J3.2 | 0.048949532 | -0.243569582 | 3.724945049 |
| RP23-349J3.3 | 0.086908494 | -0.72445792 | 3.724945049 |
| Gm26090 | 0 | NA | NA |
| RP23-349J3.4 | 6.268689147 | 6.507584612 | 0.917841675 |
| RP23-349J3.5 | 0.049553047 | 0.71820487 | 3.724945049 |
| RP23-349J3.6 | 0 | NA | NA |
| RP23-349J3.7 | 0.075043205 | 0.237309859 | 3.724945049 |
| RP23-349J3.8 | 18.61381697 | 0.185240841 | 0.236937494 |
| RP24-176P19.1 | 0.066050875 | 0.71820487 | 3.724945049 |
| RP24-176P19.3 | 3.678638758 | -0.061092265 | 0.467347967 |
| Lef1 | 120.0536891 | -0.355322445 | 0.148101399 |
| RP24-176P19.6 | 5.471034591 | -1.390293104 | 0.456527767 |
| Hadh | 222.3007882 | -0.065421164 | 0.145196091 |
| RP24-434E24.5 | 0.259720851 | 0.553060111 | 2.015970751 |
| Cyp2u1 | 190.2662416 | -0.7404631 | 0.097615638 |
| 4930534D22Rik | 5.431134866 | -0.503428417 | 0.415886868 |
| Sgms2 | 102.4694722 | 1.077554304 | 0.195380864 |
| RP23-86I19.3 | 0 | NA | NA |
| Papss1 | 709.3146802 | -0.291297924 | 0.08370145 |
| RP24-188G23.3 | 0.068097359 | -0.243569582 | 3.724945049 |
| RP23-292C19.1 | 0.108772582 | 0.718202646 | 3.724945049 |
| Olfr375-ps1 | 0 | NA | NA |
| Gm26103 | 0 | NA | NA |
| Gm22421 | 0.047948698 | -0.243569582 | 3.724945049 |
| Dkk2 | 4.638032695 | -0.690208935 | 0.425137 |
| RP23-435E17.1 | 0 | NA | NA |

| | | | |
|---|---|---|---|
| Gm25307 | 0 | NA | NA |
| Gm5549 | 0 | NA | NA |
| RP24-450N10.1 | 0.627611986 | 0.277899631 | 1.244471378 |
| Aimp1 | 305.8677058 | 0.213381058 | 0.118323055 |
| Tbck | 868.7732137 | -0.295223999 | 0.06381324 |
| RP24-450N10.3 | 11.22849639 | -0.989037425 | 0.337697969 |
| Gm22322 | 0 | NA | NA |
| RP23-447N11.5 | 18.07808909 | -0.61811938 | 0.285871202 |
| RP23-447N11.2 | 0.788050755 | -1.660423128 | 1.102197324 |
| RP23-447N11.3 | 31.71702897 | -0.393080777 | 0.248434624 |
| Npnt | 229.9057745 | -0.497766554 | 0.136931637 |
| Gm23196 | 0 | NA | NA |
| Gstcd | 129.9010644 | 0.225180212 | 0.134027281 |
| RP23-10D15.4 | 0 | NA | NA |
| RP23-10D15.3 | 6.257617145 | -0.484208988 | 0.559236555 |
| RP23-10D15.2 | 9.313879906 | -0.358433742 | 0.417944041 |
| Ints12 | 209.4893335 | 0.61531668 | 0.101367687 |
| Arhgef38 | 3.684370567 | 0.314427735 | 0.48810727 |
| RP23-335C9.1 | 28.10162786 | -0.233171604 | 0.214474766 |
| Ppa2 | 314.4988618 | -0.461894538 | 0.075677788 |
| Gm25042 | 0.060174537 | 0.237309859 | 3.724945049 |
| Tet2 | 1445.087248 | -0.066648195 | 0.150101003 |
| RP24-333H9.2 | 2.368693662 | -0.091879688 | 0.628476871 |
| RP24-333H9.1 | 0 | NA | NA |
| Gm23011 | 0 | NA | NA |
| Gm37228 | 0 | NA | NA |
| RP24-289J19.2 | 0 | NA | NA |
| RP23-437C24.1 | 0.044544648 | 0.237309859 | 3.724945049 |
| RP23-437C24.2 | 0.116046057 | -0.724453471 | 3.724945049 |
| Gm26691 | 49.78358579 | -1.130855719 | 0.236495605 |
| AC123608.1 | 1.334504357 | -2.707874472 | 1.044595618 |
| RP23-304A10.3 | 0 | NA | NA |
| RP23-304A10.1 | 0 | NA | NA |
| RP23-304A10.2 | 4.293971641 | -1.068836603 | 0.504959158 |
| Cxxc4 | 699.6168967 | -0.490438289 | 0.080933925 |
| Mir1895 | 0 | NA | NA |
| RP23-384A11.6 | 0.213180531 | 1.157027057 | 2.596008878 |
| RP23-384A11.5 | 0.844415878 | -1.745710465 | 1.082004712 |
| RP23-384A11.4 | 0.116217011 | -0.24357403 | 3.724945049 |
| RP23-424L10.1 | 0 | NA | NA |
| Gm26820 | 0.046345709 | 0.71820487 | 3.724945049 |
| Tacr3 | 61.25689593 | 0.074752643 | 0.175550439 |
| RP23-465M13.2 | 0.165933562 | 0.718213875 | 2.765757588 |
| RP23-465M13.3 | 0.044544648 | 0.237309859 | 3.724945049 |
| RP23-465M13.4 | 0 | NA | NA |
| RP23-93G15.1 | 0 | NA | NA |
| Gm24048 | 0 | NA | NA |

| | | | |
|---|---|---|---|
| RP23-93G15.2 | 0 | NA | NA |
| RP23-453I14.1 | 0 | NA | NA |
| Cenpe | 78.90502966 | 1.752970806 | 0.199731599 |
| Bdh2 | 27.37528258 | 1.347872031 | 0.335433369 |
| Slc9b2 | 28.12143682 | -0.394788714 | 0.224859556 |
| Slc9b1 | 1.329624306 | -1.485081739 | 0.843498298 |
| Cisd2 | 673.1771875 | -0.165085153 | 0.058446676 |
| RP24-372J15.5 | 30.69028919 | -0.722313559 | 0.256956747 |
| RP24-372J15.4 | 100.7051577 | -0.689214111 | 0.197854961 |
| 4930539J05Rik | 36.18476307 | -0.520733206 | 0.263580565 |
| Ube2d3 | 3995.493788 | 0.43124577 | 0.079820409 |
| RP23-167C5.2 | 0 | NA | NA |
| Manba | 173.3995577 | -0.111606851 | 0.101442434 |
| Gm16100 | 2.944243352 | -0.092639231 | 0.642091004 |
| Oaz2-ps | 15.79173318 | -0.02388717 | 0.250439832 |
| Nfkb1 | 567.2400552 | 0.937265419 | 0.124190294 |
| RP23-167C5.1 | 62.27141224 | -0.150213511 | 0.36367355 |
| Slc39a8 | 39.47806482 | 1.057499968 | 0.206515062 |
| RP24-349N15.3 | 0.185615945 | 0.237326522 | 2.674999822 |
| RP24-349N15.4 | 1.794573941 | 0.731797329 | 0.751982232 |
| RP24-349N15.5 | 1.687301136 | -0.934042271 | 0.773280527 |
| RP24-349N15.6 | 0 | NA | NA |
| RP24-349N15.2 | 0.623481245 | -0.942395793 | 1.263380549 |
| RP24-349N15.1 | 0.362626786 | 0.739836875 | 1.741825213 |
| RP24-326F21.2 | 0.246385664 | 0.71822404 | 2.07558433 |
| Bank1 | 11.35624835 | 0.568645655 | 0.34358658 |
| RP24-335I21.1 | 0.047948698 | -0.243569582 | 3.724945049 |
| RP24-335I21.2 | 0.075043205 | 0.237309859 | 3.724945049 |
| 1700030L20Rik | 0.443470786 | 0.780576026 | 1.504618463 |
| Ppp3ca | 9465.004104 | -0.73623959 | 0.112371877 |
| RP24-164P24.2 | 107.1091577 | -0.792602145 | 0.382038894 |
| RP24-164P24.5 | 0.044191549 | -0.243571806 | 3.724945049 |
| AC124177.1 | 0 | NA | NA |
| RP23-174N20.1 | 0.068097359 | -0.243569582 | 3.724945049 |
| RP23-174N20.3 | 0.099155932 | 0.718202646 | 3.724945049 |
| RP23-174N20.2 | 0 | NA | NA |
| Gm26107 | 0 | NA | NA |
| RP23-77E5.1 | 0.132101749 | 1.209894437 | 3.72418822 |
| Emcn | 39.45898547 | 1.998245662 | 0.271540196 |
| Gm4861 | 0.22850592 | -1.242835959 | 2.556398893 |
| Gm22468 | 0.075043205 | 0.237309859 | 3.724945049 |
| Ddit4l | 50.60904352 | -1.397169373 | 0.191079177 |
| RP24-134B9.2 | 0.132574648 | -1.082845688 | 3.713921268 |
| Gm21962 | 0.087261593 | -0.243576254 | 3.724945049 |
| RP24-134B9.3 | 0 | NA | NA |
| Gm25080 | 0 | NA | NA |
| Mir6380 | 0 | NA | NA |

| | | | |
|---|---|---|---|
| RP24-380C2.1 | 0 | NA | NA |
| RP24-380C2.2 | 1.550909849 | 1.010894992 | 1.0347525 |
| H2afz | 499.3132004 | 0.118123421 | 0.096736562 |
| Dnajb14 | 1867.871122 | -0.468999389 | 0.049341193 |
| RP23-209N19.2 | 0 | NA | NA |
| RP23-209N19.1 | 20.11453365 | -0.824424382 | 0.347469199 |
| Gm15460 | 2.001670347 | -0.514713093 | 0.723286131 |
| Gm25920 | 0 | NA | NA |
| RP23-209N19.4 | 0.841319663 | 0.443263472 | 1.123438329 |
| Lamtor3 | 343.7921853 | 0.117210276 | 0.082417456 |
| Dapp1 | 62.05077585 | 1.83391565 | 0.217799592 |
| RP23-209N19.6 | 0.453752633 | 0.221158621 | 1.406047299 |
| Gm5105 | 1.256408573 | 0.213016945 | 0.860990215 |
| 1110002E22Rik | 61.32006928 | 0.565763771 | 0.18094709 |
| Mttp | 190.0120384 | -0.140186928 | 0.107787594 |
| RP24-553B2.3 | 0.388301457 | 2.779538615 | 3.676627848 |
| Trmt10a | 78.50456046 | -0.074369464 | 0.147874179 |
| 4930579F01Rik | 0.038981395 | 0.718207094 | 3.724945049 |
| 0610031O16Rik | 0 | NA | NA |
| Adh7 | 0.650173001 | -0.766234316 | 1.363074808 |
| Adh1 | 2.324828864 | 1.160303959 | 0.780867172 |
| Gm16559 | 0.066050875 | 0.71820487 | 3.724945049 |
| Adh6a | 0.234786912 | 0.755649615 | 3.056246368 |
| Adh6b | 0.808183924 | 0.105350686 | 1.033638506 |
| Gm23467 | 0 | NA | NA |
| Adh6-ps1 | 0 | NA | NA |
| Adh4 | 0.195798127 | -1.266543985 | 3.71166609 |
| Adh5 | 412.48678 | 0.176379099 | 0.096999455 |
| Metap1 | 515.9695598 | 0.073188568 | 0.051206559 |
| RP24-129H9.2 | 0.047948698 | -0.243569582 | 3.724945049 |
| RP24-129H9.1 | 0.20323857 | 1.27652582 | 2.63933021 |
| Eif4e | 1261.786848 | 0.134849161 | 0.064939429 |
| Mir1956 | 0 | NA | NA |
| RP24-129H9.7 | 0 | NA | NA |
| RP24-129H9.6 | 0.597553932 | -2.356827388 | 1.347053396 |
| RP24-129H9.8 | 0 | NA | NA |
| RP24-93M7.1 | 0 | NA | NA |
| RP24-93M7.2 | 0.061920613 | 0.718207094 | 3.724945049 |
| Tspan5 | 1750.479894 | -0.128919416 | 0.079967725 |
| Gm16060 | 0.740177848 | 0.851353551 | 1.196701875 |
| Rap1gds1 | 3054.008212 | -0.281922211 | 0.049464118 |
| RP24-428N8.5 | 39.16231688 | -0.699166616 | 0.291356143 |
| Gm4862 | 0 | NA | NA |
| Stpg2 | 4.136924382 | -0.189158396 | 0.518138767 |
| RP23-276I11.1 | 0.091666476 | -0.724455696 | 3.724945049 |
| Gm24260 | 1.280600893 | 0.995978836 | 1.037343752 |
| RP23-4F17.1 | 21.90285516 | -0.479977777 | 0.260713661 |

| | | | |
|---|---|---|---|
| RP23-434M21.1 | 0.112396584 | 1.192083721 | 3.724434897 |
| RP23-434M21.2 | 0.115957212 | -0.24357403 | 3.724945049 |
| Mir7j | 0.266304556 | 1.36146117 | 3.715728684 |
| RP24-221D2.1 | 0 | NA | NA |
| RP24-63C3.1 | 0.099155932 | 0.718202646 | 3.724945049 |
| RP23-201A9.1 | 0 | NA | NA |
| RP23-460F15.1 | 0 | NA | NA |
| RP23-205G9.1 | 0 | NA | NA |
| Gm6214 | 0 | NA | NA |
| RP23-148J20.2 | 0 | NA | NA |
| RP23-148J20.3 | 0.226297694 | 0.718221951 | 2.13634781 |
| RP23-148J20.4 | 0.157356115 | 1.255377571 | 2.825615364 |
| Pdha2 | 0 | NA | NA |
| Gm15688 | 15.75159018 | -0.506844221 | 0.301989865 |
| Unc5c | 1697.764115 | -0.322305957 | 0.089498523 |
| Gm15689 | 1.903171754 | 2.376767377 | 0.998455878 |
| RP23-97D22.3 | 43.75796597 | -0.511477661 | 0.283405021 |
| Bmpr1b | 318.2842301 | -0.255972903 | 0.155487941 |
| RP23-100B7.1 | 0.094097695 | 0.718200421 | 3.724945049 |
| Pdlim5 | 665.5298056 | 1.090549323 | 0.107637385 |
| Gbp5 | 1952.644448 | 8.402595184 | 0.427167582 |
| Gbp7 | 3441.7084 | 7.121136456 | 0.243012166 |
| RP23-100J23.5 | 0 | NA | NA |
| Gbp3 | 3129.531274 | 7.621493386 | 0.250069595 |
| Gbp2b | 4052.460821 | 9.104312528 | 0.345085745 |
| Gbp2 | 2406.349028 | 7.746365741 | 0.287366256 |
| Mir6381 | 0.56675494 | 2.500901075 | 1.251974685 |
| RP24-314I8.2 | 0 | NA | NA |
| RP24-314I8.3 | 0.091666476 | -0.724455696 | 3.724945049 |
| Ccbl2 | 63.67129629 | -0.148133106 | 0.14951147 |
| Gm15540 | 0.047948698 | -0.243569582 | 3.724945049 |
| Gtf2b | 311.3568098 | 0.450419181 | 0.0771748 |
| Gm24457 | 0 | NA | NA |
| Pkn2 | 955.7411507 | 0.55406679 | 0.080518643 |
| Gm25965 | 0 | NA | NA |
| RP24-445M20.2 | 0.255858694 | 1.291643236 | 2.132292767 |
| 9530052C20Rik | 8.046682377 | -0.636500573 | 0.375895711 |
| RP24-445M20.3 | 0 | NA | NA |
| Gm2574 | 15.8626213 | -0.09497683 | 0.281680942 |
| RP23-49B23.1 | 0.306809375 | 1.649855054 | 1.716691972 |
| RP23-49B23.2 | 0.297888126 | -0.215203329 | 1.751678899 |
| RP24-377E5.1 | 10.93206221 | 0.763794019 | 0.362569257 |
| RP24-337A2.1 | 0.793655395 | 1.926543335 | 1.430411965 |
| RP24-337A2.2 | 1.407833646 | 0.36172583 | 0.795067557 |
| RP24-337A2.3 | 0.046345709 | 0.71820487 | 3.724945049 |
| RP23-402L6.2 | 0.617903777 | -1.697415621 | 1.192866864 |
| RP23-402L6.1 | 1.629102473 | -1.459066356 | 0.850475247 |

| | | | |
|---|---|---|---|
| RP24-273H20.2 | 0.222190708 | -1.217671171 | 2.15133339 |
| Gm38006 | 0 | NA | NA |
| RP24-273H20.3 | 0.068097359 | -0.243569582 | 3.724945049 |
| RP24-273H20.4 | 3.820552698 | -2.38148812 | 0.705430774 |
| Gm24728 | 0 | NA | NA |
| RP23-229C5.1 | 0.29112196 | -0.693810761 | 1.614058653 |
| RP23-229C5.2 | 0.131251387 | 0.717600267 | 2.981415988 |
| RP24-224P21.1 | 0.177934686 | -0.243571879 | 2.723282651 |
| RP23-327I19.1 | 2.34560619 | -1.170767493 | 0.664854737 |
| RP24-224P21.2 | 0.044544648 | 0.237309859 | 3.724945049 |
| RP23-327I19.2 | 0.038981395 | 0.718207094 | 3.724945049 |
| Lmo4 | 1293.515492 | 0.114611838 | 0.075356227 |
| RP23-461P7.1 | 43.89163664 | -0.938821878 | 0.246935661 |
| RP23-461P7.2 | 0 | NA | NA |
| RP24-232N19.1 | 0.421951652 | 0.237327346 | 1.771276567 |
| RP24-224M8.1 | 22.99821006 | -1.146587765 | 0.289556888 |
| Hs2st1 | 927.2177343 | -0.185185413 | 0.054202823 |
| RP23-225M12.2 | 46.28994543 | -1.008302085 | 0.25240993 |
| RP23-225M12.1 | 14.69975535 | -0.97244291 | 0.37267277 |
| 15-Sep | 1591.561157 | 0.621802821 | 0.124022691 |
| Gm24406 | 0 | NA | NA |
| Rpl9-ps8 | 0.045882455 | 0.237312084 | 3.724945049 |
| Sh3glb1 | 1627.178665 | 0.711340155 | 0.083925195 |
| RP24-321M14.4 | 0.899213277 | -0.614477092 | 1.041128264 |
| Clca3a1 | 41.15834729 | 1.442130668 | 0.551213962 |
| RP24-321M14.5 | 0.600994565 | 1.033873002 | 1.296397779 |
| RP24-271G16.1 | 1.394083107 | -1.998809373 | 0.992822868 |
| Clca3a2 | 1.360869654 | 0.88826346 | 0.950992073 |
| RP24-271G16.2 | 0.93582689 | -0.348893969 | 0.951443584 |
| Clca3b | 0.093141081 | -0.724455696 | 3.724945049 |
| Gm16215 | 0.201808035 | 1.466409448 | 3.188838746 |
| Clca4c-ps | 0.221289295 | 0.237320238 | 2.545524212 |
| Clca4b | 0 | NA | NA |
| Gm16233 | 0.233058144 | -0.685249192 | 1.863179717 |
| Clca4a | 0 | NA | NA |
| RP24-277E17.1 | 0.140122821 | -0.243570953 | 3.572128793 |
| Clca1 | 0.274296751 | 1.865203426 | 3.710574064 |
| Clca2 | 0.311801716 | -0.243580258 | 3.724945049 |
| Odf2l | 262.3245237 | -0.022240612 | 0.1073478 |
| Col24a1 | 47.56753531 | -1.216025042 | 0.24538504 |
| RP23-307M9.1 | 0 | NA | NA |
| RP23-307M9.2 | 0.046345709 | 0.71820487 | 3.724945049 |
| Znhit6 | 139.6637941 | -0.075297625 | 0.137664773 |
| Cyr61 | 545.855505 | 3.545635811 | 0.586936327 |
| Gm17501 | 9.319672861 | 0.641388779 | 0.337162116 |
| RP24-267M22.1 | 0 | NA | NA |
| Ddah1 | 1181.961696 | 0.474488268 | 0.105591253 |

| | | | |
|---|---|---|---|
| Rpl36a-ps2 | 93.94039235 | 0.453769682 | 0.164454222 |
| Bcl10 | 200.6606098 | 0.960293001 | 0.113919712 |
| 2410004B18Rik | 209.7691853 | 0.345885651 | 0.121995522 |
| Syde2 | 65.37702174 | -0.553350872 | 0.148978344 |
| RP24-267M11.2 | 0 | NA | NA |
| Wdr63 | 17.44671922 | -0.307316459 | 0.293704841 |
| Gm25627 | 0 | NA | NA |
| Mcoln3 | 1.139337436 | 1.614294848 | 0.970183423 |
| Mcoln2 | 13.29155609 | 6.526705162 | 0.883816407 |
| Lpar3 | 6.729774449 | 0.567887971 | 0.388307487 |
| RP24-455A5.3 | 0 | NA | NA |
| RP24-364E17.1 | 0.399538249 | 2.265581726 | 1.633034642 |
| Gm22078 | 0 | NA | NA |
| Gm22901 | 0 | NA | NA |
| Ssx2ip | 795.7799406 | -0.414044178 | 0.080067199 |
| Ctbs | 130.2632904 | 0.511433239 | 0.14470639 |
| Spata1 | 56.80384134 | -0.096491195 | 0.180548832 |
| Gng5 | 243.1666755 | 1.53546457 | 0.155538973 |
| Rpf1 | 164.2090017 | 0.26757198 | 0.114271684 |
| RP24-100G24.2 | 0 | NA | NA |
| Uox | 7.769118165 | 1.566491676 | 0.433901315 |
| Dnase2b | 0.061920613 | 0.718207094 | 3.724945049 |
| Gm16325 | 1.444443621 | 1.332107153 | 0.811039122 |
| 4930503B20Rik | 2.396082498 | 4.074364506 | 0.989087611 |
| Samd13 | 1.10620882 | 0.775319964 | 0.890255146 |
| RP24-86C15.1 | 0.038981395 | 0.718207094 | 3.724945049 |
| Prkacb | 5477.753168 | -0.374204242 | 0.047380792 |
| RP24-409N2.3 | 0.044544648 | 0.237309859 | 3.724945049 |
| Gm10288 | 30.25828668 | 0.708956708 | 0.24797279 |
| Ttll7 | 4265.777113 | -0.698142778 | 0.07195989 |
| Gm29771 | 2.774619275 | -1.660340229 | 0.689394377 |
| Gm6074 | 0 | NA | NA |
| RP24-293H7.2 | 0 | NA | NA |
| Gm37803 | 0 | NA | NA |
| Gm37041 | 0.332175987 | 1.261837681 | 3.720999938 |
| Gm36888 | 0 | NA | NA |
| RP24-184C18.1 | 0.044191549 | -0.243571806 | 3.724945049 |
| Gm23131 | 0 | NA | NA |
| RP23-290J2.1 | 0 | NA | NA |
| Gm38068 | 0 | NA | NA |
| RP24-196A21.2 | 0 | NA | NA |
| Adgrl2 | 1106.274129 | -0.728136638 | 0.127427589 |
| RP24-224A19.1 | 3.670148074 | -1.366507594 | 0.551898411 |
| RP24-224A19.2 | 2.247096996 | -1.706139479 | 0.714478988 |
| Gm25127 | 1.189027437 | 0.438329387 | 0.842204851 |
| RP23-7K18.3 | 0.083172944 | 0.23732098 | 3.724945049 |
| Gm10287 | 19.5831566 | -1.41771619 | 0.377538086 |

| | | | |
|---|---|---|---|
| RP23-473G18.1 | 38.52654686 | -1.596561031 | 0.241497822 |
| RP24-512E12.1 | 0.471094982 | 0.732309172 | 1.452325476 |
| Gm26468 | 0.506521265 | -0.221050052 | 1.321830722 |
| RP24-248L23.1 | 0 | NA | NA |
| Rpsa-ps10 | 128.4715849 | 0.540304941 | 0.129271908 |
| RP24-134A5.1 | 4.381038091 | 1.083705063 | 0.492816507 |
| RP24-257N6.1 | 0 | NA | NA |
| Mir3963 | 0 | NA | NA |
| RP24-232H5.1 | 0 | NA | NA |
| RP24-232H5.2 | 0.216927137 | 0.237330228 | 2.165803523 |
| RP24-232H5.3 | 0.510381651 | -0.761158605 | 1.35115346 |
| RP23-239A13.1 | 0.433950198 | 1.667517635 | 1.336026866 |
| RP23-239A13.2 | 0.049553047 | 0.71820487 | 3.724945049 |
| Adgrl4 | 62.50733234 | 0.134182483 | 0.226640385 |
| Gm24458 | 0.493466105 | -0.371052539 | 1.431382226 |
| Ifi44 | 3535.414574 | 10.3051289 | 0.357777386 |
| Ifi44l | 509.0915789 | 9.843201082 | 0.612726766 |
| Ptgfr | 13.2398592 | -0.34775542 | 0.365859668 |
| RP24-267D17.1 | 0.162302921 | 0.237325578 | 2.79109695 |
| RP24-267D17.2 | 0 | NA | NA |
| Gm25056 | 0 | NA | NA |
| RP23-245O13.3 | 0.112396584 | 1.192083721 | 3.724434897 |
| RP23-245O13.1 | 0.13307909 | 1.198489135 | 2.968793202 |
| RP23-245O13.2 | 0.060174537 | 0.237309859 | 3.724945049 |
| RP24-499N1.1 | 0.129258854 | 0.717599224 | 2.995409112 |
| Gipc2 | 25.12877053 | -0.049854291 | 0.204071955 |
| RP24-499N1.2 | 2.888459128 | 0.04186167 | 0.681296824 |
| Gm16198 | 0.109869311 | 0.238305784 | 3.724874707 |
| D630002J18Rik | 0.719682426 | 0.417934713 | 1.161542678 |
| Dnajb4 | 820.1226806 | -0.263931598 | 0.078576258 |
| Fubp1 | 2019.104171 | -0.160779241 | 0.092178246 |
| Nexn | 60.15238185 | -0.677068575 | 0.187622392 |
| RP23-181C16.4 | 5.705262924 | -0.727688562 | 0.473053263 |
| Fam73a | 1902.314384 | -0.33054903 | 0.055562792 |
| Usp33 | 1820.049344 | -0.288203947 | 0.065754033 |
| RP24-511F2.1 | 7.968776044 | -0.546287147 | 0.360826209 |
| RP24-511F2.2 | 7.38870447 | -0.000793273 | 0.419579796 |
| RP24-511F2.3 | 58.00030583 | -1.336124999 | 0.378253579 |
| RP24-511F2.4 | 45.38790308 | -0.204600966 | 0.220184914 |
| Zzz3 | 889.1435612 | -0.224600992 | 0.068055123 |
| Ak5 | 942.1680984 | -0.742114231 | 0.129773936 |
| AI115009 | 13.76623626 | -0.78692223 | 0.27817117 |
| Pigk | 806.3801702 | -0.136168922 | 0.045909207 |
| Rpl29-ps1 | 0.333447913 | 0.718225951 | 1.73228833 |
| St6galnac5 | 598.912246 | -0.770830519 | 0.125698901 |
| Gm16232 | 3.327150153 | -2.115260632 | 0.70910374 |
| Gm16213 | 1.998664546 | 4.13158741 | 1.018450906 |

| | | | |
|---|---|---|---|
| Gm16231 | 0.342796299 | 1.318403919 | 1.767701169 |
| RP23-394P9.1 | 0.038981395 | 0.718207094 | 3.724945049 |
| RP24-117O8.1 | 6.160904613 | -0.850530793 | 0.570363849 |
| St6galnac3 | 220.6741086 | -0.364379534 | 0.112958233 |
| RP24-303G10.2 | 9.238644217 | -0.836496635 | 0.547048468 |
| RP24-303G10.1 | 48.22872844 | -0.820917899 | 0.311244285 |
| Gm27405 | 0.046345709 | 0.71820487 | 3.724945049 |
| RP24-104D12.1 | 4.330890979 | -0.806554581 | 0.598446786 |
| RP24-104D12.2 | 7.42840581 | -0.399024829 | 0.497949779 |
| Gm22136 | 0 | NA | NA |
| Gm22206 | 0 | NA | NA |
| RP24-186A10.2 | 0 | NA | NA |
| Gm10042 | 0 | NA | NA |
| 5730460C07Rik | 0 | NA | NA |
| Gm25757 | 0 | NA | NA |
| Asb17os | 1.419315775 | 2.981803748 | 1.217048116 |
| Asb17 | 0.04393175 | -0.243571806 | 3.724945049 |
| Msh4 | 11.18809076 | -0.301820442 | 0.320476354 |
| Gm24395 | 0 | NA | NA |
| Rabggtb | 426.99162 | 0.541394233 | 0.09964593 |
| Snord45b | 0 | NA | NA |
| Gm24494 | 6.926426429 | 0.133029005 | 0.429736334 |
| Snord45c | 0 | NA | NA |
| RP23-403C5.3 | 0.748230394 | -0.212036086 | 1.120794094 |
| Acadm | 231.7967069 | -0.349267379 | 0.127807964 |
| Slc44a5 | 191.2716275 | -0.68103867 | 0.154200171 |
| Gm3076 | 3.513132756 | -0.599932284 | 0.655246373 |
| RP24-334O8.1 | 5.951295928 | -1.83685978 | 0.624039132 |
| RP24-527P22.4 | 0 | NA | NA |
| Lhx8 | 17.86896063 | -0.048575539 | 0.266090855 |
| AI606473 | 4.337481582 | -0.743171268 | 0.503064741 |
| Tyw3 | 99.42373595 | -0.439602978 | 0.131615596 |
| Cryz | 179.0070392 | -0.333087005 | 0.089992853 |
| Gm38139 | 1.950077631 | -1.561488415 | 0.750117316 |
| Erich3 | 339.8935089 | -0.30431427 | 0.152054118 |
| Gm20389 | 0.04393175 | -0.243571806 | 3.724945049 |
| Tnni3k | 1.221396067 | 1.321563504 | 1.148642317 |
| Gm37890 | 0 | NA | NA |
| Fpgt | 339.8878321 | 0.451725532 | 0.089972319 |
| Gm26456 | 0 | NA | NA |
| Lrriq3 | 12.459711 | -0.124685181 | 0.279796369 |
| Gm37482 | 1.621103067 | -0.087011685 | 0.716449087 |
| Gm5285 | 0 | NA | NA |
| Scp2-ps1 | 0.068097359 | -0.243569582 | 3.724945049 |
| Gm37449 | 0.554117557 | -0.676060782 | 1.361457071 |
| Gm37324 | 0.156124879 | 0.237325292 | 2.825334012 |
| Gm23038 | 0 | NA | NA |

| | | | |
|---|---|---|---|
| Gm37102 | 0.042716945 | -0.243571806 | 3.724945049 |
| Gm37515 | 0.408926061 | -0.575840531 | 1.430707372 |
| RP23-322I5.1 | 0.044191549 | -0.243571806 | 3.724945049 |
| RP23-322I5.2 | 0 | NA | NA |
| RP23-322I5.3 | 0 | NA | NA |
| RP23-187E3.1 | 0 | NA | NA |
| 4930570G19Rik | 328.3713293 | -0.127264582 | 0.170435562 |
| Negr1 | 3447.761138 | -0.477188412 | 0.091626931 |
| RP23-354G14.1 | 240.3499704 | -1.005622278 | 0.314549907 |
| RP23-354G14.2 | 93.63872821 | -1.016825025 | 0.260224742 |
| RP23-354G14.3 | 137.6961956 | -1.014918795 | 0.270334578 |
| RP23-354G14.4 | 101.0414782 | -1.064068425 | 0.284439738 |
| Gm15578 | 21.23400752 | -1.364247877 | 0.297356761 |
| Gm15577 | 12.27858454 | 0.933374244 | 0.400432745 |
| RP24-176M3.1 | 45.69791764 | -0.97543523 | 0.339057348 |
| RP23-103L12.1 | 51.84091573 | -1.30234811 | 0.266217782 |
| 4930566N20Rik | 0.044544648 | 0.237309859 | 3.724945049 |
| Gm24373 | 15.34325161 | -1.284349162 | 0.484169322 |
| Gm23725 | 0 | NA | NA |
| Gm22458 | 0.088383099 | -0.72445792 | 3.724945049 |
| Gm23379 | 0 | NA | NA |
| RP23-64P1.1 | 0.244387265 | 2.019757066 | 2.046368005 |
| RP23-210I6.1 | 1.128024331 | 1.285070938 | 1.108026918 |
| Zranb2 | 1982.94177 | -0.243477797 | 0.081525808 |
| Mir186 | 29.35376171 | -0.600905738 | 0.242100081 |
| Ptger3 | 53.06310274 | -0.101459489 | 0.182219541 |
| RP24-411A22.2 | 0.062426873 | 0.237312084 | 3.724945049 |
| RP23-287M3.1 | 0.332723394 | 0.545019666 | 1.534908184 |
| RP23-287M3.3 | 0 | NA | NA |
| Cth | 54.84234108 | 1.420159825 | 0.191184406 |
| RP23-287M3.2 | 3.057191251 | -0.558435087 | 0.580279084 |
| 1810013D15Rik | 0.328075509 | 0.237330623 | 1.77573068 |
| Ankrd13c | 1568.866954 | 0.224042795 | 0.06491168 |
| RP24-68G17.2 | 78.17925785 | -0.780805931 | 0.301058887 |
| Srsf11 | 1647.601996 | -0.116428673 | 0.121045317 |
| RP24-68G17.3 | 48.08364435 | -0.129322395 | 0.26935844 |
| Lrrc40 | 326.5754133 | 0.003361647 | 0.07356293 |
| RP24-68G17.4 | 1897.008325 | -1.131368455 | 0.229488287 |
| Lrrc7 | 2770.36321 | -1.021302293 | 0.219662824 |
| RP24-213M16.2 | 0.56553622 | 1.937043139 | 3.472769516 |
| RP24-213M16.1 | 0.354487963 | 0.250879875 | 1.577648743 |
| RP24-481I21.3 | 0.26582355 | -1.530681685 | 2.467868246 |
| RP24-481I21.2 | 0 | NA | NA |
| Gm9423 | 0 | NA | NA |
| RP23-324D2.1 | 2.321564718 | -4.037187231 | 1.06540261 |
| RP24-209L5.1 | 0.316660866 | 2.077527337 | 1.935250374 |
| Depdc1a | 5.945976878 | 2.268595977 | 0.651914192 |

| | | | |
|---|---|---|---|
| Rpe65 | 5.700803298 | -3.97537493 | 0.738366178 |
| Wls | 574.2412302 | 0.232866041 | 0.103007014 |
| Wl1-1749A12.1 | 12.79191494 | -0.852167629 | 0.397985937 |
| Gm23941 | 0 | NA | NA |
| Gm27878 | 0 | NA | NA |
| Vmn1r-ps2 | 0 | NA | NA |
| Gm11812 | 0 | NA | NA |
| Vmn1r2 | 0 | NA | NA |
| Vmn1r3 | 0 | NA | NA |
| AI838599 | 1.421121277 | -1.302770197 | 0.8156751 |
| Gm11783 | 0.04393175 | -0.243571806 | 3.724945049 |
| Gm11786 | 0 | NA | NA |
| Gm11784 | 0 | NA | NA |
| Gm11785 | 0.044544648 | 0.237309859 | 3.724945049 |
| Gm11787 | 2.521183062 | 2.188763418 | 0.753829346 |
| Mir3471-2 | 0 | NA | NA |
| Tmem68 | 334.4267812 | -0.173395839 | 0.069366595 |
| Tgs1 | 548.3289277 | 0.4025372 | 0.090582426 |
| Gm22541 | 0.302788994 | -0.220691268 | 1.802614547 |
| 2210414B05Rik | 0.54041206 | 2.006611789 | 1.516378958 |
| Lyn | 538.8172436 | 2.531631502 | 0.137006175 |
| Gm26857 | 5.106476976 | 1.268858189 | 0.445908686 |
| Gm11805 | 0.72046455 | -1.207590789 | 1.338244355 |
| Gm22781 | 2.17130535 | -0.085350179 | 0.750994868 |
| 9430025C20Rik | 38.03548056 | -0.000751592 | 0.289454426 |
| Rps20 | 938.4544041 | 0.818748406 | 0.09800748 |
| Gm24016 | 0 | NA | NA |
| Gm24125 | 0 | NA | NA |
| Mos | 0.721375344 | -0.465286375 | 1.250743471 |
| Plag1 | 101.48303 | -0.568850849 | 0.157681012 |
| Chchd7 | 221.4013722 | -0.44820618 | 0.106940936 |
| Gm11808 | 144.4071686 | 0.316659326 | 0.211478593 |
| Rps10-ps3 | 0 | NA | NA |
| Sdr16c5 | 0.092228164 | 0.718202646 | 3.724945049 |
| Sdr16c6 | 0.09176491 | 0.237310218 | 3.724945049 |
| Penk | 1553.820092 | -0.133146409 | 0.221795601 |
| A830012C17Rik | 4.915491126 | -0.515904361 | 0.474965997 |
| Gm11780 | 0.085433889 | -0.72445792 | 3.724945049 |
| Gm26436 | 0 | NA | NA |
| Impad1 | 2842.589061 | -0.049484431 | 0.084927219 |
| Gm11779 | 0 | NA | NA |
| Gm11782 | 0 | NA | NA |
| Gm11781 | 0 | NA | NA |
| 4930423M02Rik | 0 | NA | NA |
| Fam110b | 526.7316921 | -0.232416242 | 0.079003822 |
| Gm24337 | 0 | NA | NA |
| Gm11796 | 0 | NA | NA |

| | | | |
|---|---|---|---|
| Gm11797 | 0.624599851 | -1.273751187 | 3.61217587 |
| Ubxn2b | 545.8183538 | -0.428481181 | 0.124383288 |
| Gm11798 | 0 | NA | NA |
| Cyp7a1 | 0.264608745 | -1.520485425 | 2.469844052 |
| 4930430E12Rik | 36.79472905 | 8.051124028 | 0.924499852 |
| Sdcbp | 2354.562233 | 1.187766706 | 0.126633933 |
| Gm37689 | 8.169802865 | 0.472154124 | 0.453840497 |
| Nsmaf | 844.8221621 | 0.32558782 | 0.112145922 |
| BX510318.1 | 42.81714402 | -0.15748453 | 0.280558374 |
| Gm11803 | 0 | NA | NA |
| Gm11801 | 0 | NA | NA |
| Gm22473 | 0 | NA | NA |
| Tox | 323.9376703 | -0.388557356 | 0.124220612 |
| Gm11802 | 0.913351404 | 2.5409098 | 1.113211089 |
| Gm11804 | 0 | NA | NA |
| 8430436N08Rik | 0.23059471 | 0.718222388 | 2.144949844 |
| Gm11795 | 0.166709681 | -0.724459871 | 2.767631179 |
| Gm11794 | 0.481267046 | 1.749312275 | 1.361476041 |
| Gm23860 | 0 | NA | NA |
| Gm11800 | 0 | NA | NA |
| Car8 | 1324.707954 | -0.16371323 | 0.304716386 |
| Gm11799 | 5.921281838 | -1.712391483 | 0.448894203 |
| Gm25355 | 0 | NA | NA |
| Gm37386 | 0.066050875 | 0.71820487 | 3.724945049 |
| Gm11810 | 0.196054161 | -0.244105808 | 2.235824846 |
| Rab2a | 2207.111245 | -0.166257938 | 0.067574416 |
| Gm11809 | 0 | NA | NA |
| Chd7 | 1522.223342 | 0.023581656 | 0.136008154 |
| Rps18-ps2 | 0.644285774 | 0.736317589 | 1.39893529 |
| Gm23423 | 0.075043205 | 0.237309859 | 3.724945049 |
| Gm26548 | 1.161740484 | 0.928854495 | 0.916573629 |
| Clvs1 | 941.1106066 | -0.308649355 | 0.079029299 |
| Gm11816 | 3.30561101 | 1.488239489 | 0.634908978 |
| Gm11817 | 0.206997213 | 0.237320372 | 2.599503519 |
| Asph | 1754.934329 | -0.42054542 | 0.104425544 |
| Gm25677 | 0.044191549 | -0.243571806 | 3.724945049 |
| Gm24152 | 0 | NA | NA |
| 4930412C18Rik | 9.306510091 | -0.701216379 | 0.34218464 |
| Gdf6 | 2.455729561 | -1.863972917 | 0.988170528 |
| 4930448K20Rik | 0 | NA | NA |
| Gm11834 | 0 | NA | NA |
| Gm11835 | 0 | NA | NA |
| Gm11814 | 0.850521408 | 0.658270754 | 0.997992364 |
| 1700123O12Rik | 0 | NA | NA |
| Gm11815 | 0.853137171 | -2.519556011 | 1.103455507 |
| Gm11813 | 0 | NA | NA |
| Gm12920 | 0.044191549 | -0.243571806 | 3.724945049 |

| | | | |
|---|---|---|---|
| Mir3471-1 | 0 | NA | NA |
| Gm12918 | 39.37875012 | 0.493452055 | 0.207048326 |
| Gm12919 | 1.164100891 | 0.409021323 | 0.929777204 |
| AL672160.1 | 0.087930927 | 0.236669296 | 3.724898107 |
| 2610301B20Rik | 374.8592291 | 0.3092653 | 0.107898261 |
| Rps11-ps3 | 0.170186936 | 1.71882767 | 3.39900844 |
| Plekhf2 | 240.1964193 | 1.972506375 | 0.144798307 |
| Ndufaf6 | 73.7669564 | 0.011339924 | 0.166674698 |
| Gm11827 | 1.479390131 | -0.592526401 | 1.006522857 |
| Gm11829 | 0 | NA | NA |
| mmu-mir-684-2 | 0.974719181 | -0.005558147 | 1.2041748 |
| Gm11830 | 0.15906016 | 1.290286593 | 3.433492162 |
| Trp53inp1 | 416.2178088 | 0.235852868 | 0.125677068 |
| Ccne2 | 114.245643 | 0.241412603 | 0.150190708 |
| Ints8 | 627.572806 | 0.087699198 | 0.089962668 |
| Dpy19l4 | 659.7126594 | -0.30182653 | 0.070549629 |
| Gm22940 | 3.154500895 | 0.581386834 | 0.827015678 |
| Gm26663 | 0.326609485 | 0.251503048 | 1.602572799 |
| Esrp1 | 1.320956406 | -0.0201151 | 1.023019128 |
| Gm11821 | 0.370194986 | 0.712748946 | 1.601111071 |
| Gm11822 | 0 | NA | NA |
| 1110037F02Rik | 957.3336285 | 0.163757736 | 0.053193464 |
| Gm11831 | 0 | NA | NA |
| Rad54b | 15.9562118 | 0.809022185 | 0.32009759 |
| Fsbp | 18.80141561 | -0.141356331 | 0.256491611 |
| Gm11832 | 0 | NA | NA |
| Gm11833 | 0 | NA | NA |
| Gem | 164.20295 | 3.586534314 | 0.364496229 |
| Cdh17 | 2.087534489 | 3.011708179 | 0.878783124 |
| Gm22477 | 0.061920613 | 0.718207094 | 3.724945049 |
| Gm25002 | 0 | NA | NA |
| Pdp1 | 1783.616816 | -0.369842779 | 0.126375074 |
| 1700123M08Rik | 21.10087427 | -0.600152097 | 0.240544277 |
| Gm10604 | 0.068097359 | -0.243569582 | 3.724945049 |
| Gm26895 | 2.384116323 | -1.061428038 | 0.696135118 |
| Gm11840 | 0.048949532 | -0.243569582 | 3.724945049 |
| Gm11842 | 0.297840172 | -0.728314311 | 1.763109119 |
| Tmem67 | 313.6983145 | 0.5855039 | 0.104816938 |
| Rbm12b2 | 274.5941422 | -0.124489994 | 0.083111857 |
| Gm11839 | 15.15531981 | -0.955529643 | 0.373275987 |
| Gm11841 | 7.772074058 | -0.599231145 | 0.450390788 |
| Gm11843 | 1.645711418 | 1.046443153 | 0.706768849 |
| Rbm12b1 | 103.5381567 | -0.189741499 | 0.17113956 |
| Fam92a | 431.8026252 | -0.104195549 | 0.072601449 |
| Gm11847 | 135.4062284 | 0.133541391 | 0.122889016 |
| Gm11846 | 0.044191549 | -0.243571806 | 3.724945049 |
| Gm37985 | 1.26802963 | 1.101264475 | 0.860602224 |

| | | | |
|---|---|---|---|
| n-R5s182 | 0 | NA | NA |
| Triqk | 145.0715167 | -0.659734952 | 0.13762627 |
| Gm11820 | 0.208450558 | -0.72446052 | 2.196616084 |
| Gm26250 | 0 | NA | NA |
| Gm11819 | 0 | NA | NA |
| Gm11826 | 3.374863659 | 0.399572815 | 0.563570319 |
| Gm11825 | 0.044544648 | 0.237309859 | 3.724945049 |
| Gm23162 | 0 | NA | NA |
| Runx1t1 | 591.5908782 | -0.767200892 | 0.097875372 |
| Gm11823 | 0.253691411 | 2.191682051 | 2.43059992 |
| Gm24068 | 0 | NA | NA |
| Gm11824 | 3.085586268 | 2.352882908 | 0.86088 |
| 1700120G11Rik | 0 | NA | NA |
| Gm24908 | 0 | NA | NA |
| Gm11836 | 2.257326888 | 0.028359549 | 0.731780542 |
| Slc26a7 | 38.39735321 | -0.558080249 | 0.222282856 |
| Lrrc69 | 6.895850206 | 0.244248297 | 0.47507342 |
| Gm11838 | 0 | NA | NA |
| Otud6b | 619.2074641 | -0.091388537 | 0.062848018 |
| Tmem55a | 534.2645095 | 0.155647549 | 0.091212213 |
| Gm11837 | 8.354147893 | 0.21070436 | 0.356344924 |
| Necab1 | 1188.342868 | -0.956644119 | 0.112678808 |
| Gm11844 | 0 | NA | NA |
| Gm11845 | 0 | NA | NA |
| Tmem64 | 564.4500706 | -0.469816818 | 0.085546802 |
| Gm24317 | 0 | NA | NA |
| Gm11856 | 0 | NA | NA |
| Gm11859 | 0.314457495 | 0.226391261 | 1.786471123 |
| Gm11857 | 0 | NA | NA |
| Gm11858 | 0.914915251 | -1.218224444 | 1.141096662 |
| Gm11855 | 11.73213907 | -0.581346341 | 0.34657062 |
| Gm24567 | 0 | NA | NA |
| Calb1 | 2166.896649 | -0.51526387 | 0.156278958 |
| Gm26148 | 0.061920613 | 0.718207094 | 3.724945049 |
| Decr1 | 158.7523432 | 0.250582792 | 0.131972242 |
| Mir6400 | 0.947067764 | 1.15948577 | 0.881925208 |
| Nbn | 245.1479976 | 0.430573125 | 0.080453053 |
| Gm11869 | 0.066050875 | 0.71820487 | 3.724945049 |
| Osgin2 | 252.2581438 | 0.225188992 | 0.097858982 |
| Gm11870 | 26.42157704 | -0.398477241 | 0.268522576 |
| Gm11874 | 4.610084374 | 0.134138279 | 0.512912194 |
| Ripk2 | 184.8884919 | 1.376373947 | 0.180509271 |
| Gm22690 | 1.641288957 | -0.252858398 | 0.792685529 |
| A530072M11Rik | 6.322029942 | 0.312150721 | 0.371332592 |
| Gm11865 | 0.577397264 | 0.827436937 | 1.333657534 |
| Gm37355 | 0.136963818 | 0.718204821 | 3.59954433 |
| Gm11864 | 0.066050875 | 0.71820487 | 3.724945049 |

| | | | | |
|---|---|---|---|---|
| Gm11861 | 0.117650406 | | 0.23732098 | 3.724945049 |
| Gm24978 | 0.911472295 | | -0.496599521 | 1.136569859 |
| Gm11862 | 0 | NA | | NA |
| Gm23975 | 0 | NA | | NA |
| Gm11848 | 0.044191549 | | -0.243571806 | 3.724945049 |
| Gm11850 | 0 | NA | | NA |
| Gm11849 | 0.314045685 | | 0.219169048 | 1.728138005 |
| Gm23384 | 1.375344072 | | 1.650758156 | 1.077551297 |
| Gm11851 | 0.112396584 | | 1.192083721 | 3.724434897 |
| Mmp16 | 803.4952648 | | -0.432760721 | 0.094182731 |
| Gm11860 | 0 | NA | | NA |
| Gm11852 | 0.075043205 | | 0.237309859 | 3.724945049 |
| Gm11867 | 0.108266322 | | 1.230376831 | 3.722845817 |
| Gm11854 | 0 | NA | | NA |
| Gm11853 | 0 | NA | | NA |
| Gm25971 | 0 | NA | | NA |
| Gm24419 | 0 | NA | | NA |
| Gm11868 | 68.16477173 | | 0.243613946 | 0.163404085 |
| Cnbd1 | 0 | NA | | NA |
| Gm12392 | 0 | NA | | NA |
| Gm12393 | 0 | NA | | NA |
| Cngb3 | 8.051515703 | | 5.401717161 | 0.961114145 |
| Gm11877 | 0 | NA | | NA |
| Gm25655 | 0.263406493 | | 1.696453761 | 2.070862568 |
| Cpne3 | 505.9295095 | | 1.093088715 | 0.11475664 |
| Rmdn1 | 159.5744036 | | -0.475782861 | 0.099676459 |
| Gm12353 | 18.78038428 | | -0.383059775 | 0.274203255 |
| Wwp1 | 1103.425899 | | -0.249009462 | 0.0632498 |
| Gm12354 | 0 | NA | | NA |
| Slc7a13 | 0.367515839 | | 2.415465226 | 1.683660775 |
| Atp6v0d2 | 1.127083422 | | 2.441382064 | 1.072024175 |
| 4930480G23Rik | 0.075043205 | | 0.237309859 | 3.724945049 |
| Ttpa | 39.70977185 | | -0.69717947 | 0.253729321 |
| 4930598A11Rik | 0.729378188 | | 1.448914819 | 1.265623904 |
| Ggh | 127.1892972 | | 0.818849078 | 0.133907491 |
| Nkain3 | 142.3883735 | | -0.656811266 | 0.116484631 |
| Gm11872 | 3.989504448 | | 0.415089848 | 0.522465961 |
| Gm11873 | 0.255186079 | | -1.165183556 | 1.807034923 |
| Gm11871 | 0 | NA | | NA |
| Gm22817 | 0 | NA | | NA |
| Gm25977 | 0 | NA | | NA |
| Gm11875 | 0 | NA | | NA |
| Gm11876 | 0 | NA | | NA |
| Prdm13 | 1.264085207 | | -0.796033843 | 0.967905404 |
| Ccnc | 534.707253 | | 0.217772789 | 0.08308245 |
| Tstd3 | 155.9862153 | | -0.003184397 | 0.098955934 |
| Usp45 | 1038.908871 | | 0.110595461 | 0.055047438 |

| | | | |
|---|---|---|---|
| Pnisr | 1930.889929 | -0.750026418 | 0.141517655 |
| Coq3 | 168.8413342 | 0.023983275 | 0.113183591 |
| Faxc | 940.4861022 | -0.487730677 | 0.090447911 |
| Gm11885 | 0.049553047 | 0.71820487 | 3.724945049 |
| Gm11880 | 0 | NA | NA |
| Gm11882 | 0 | NA | NA |
| Fbxl4 | 229.5017738 | -0.140429112 | 0.096912355 |
| Gm24607 | 2.356080661 | 0.012556649 | 0.837042867 |
| Gm37900 | 25.72877726 | -0.122533697 | 0.36368136 |
| Pou3f2 | 455.224007 | -0.994499478 | 0.129247606 |
| Gm11878 | 0.318934723 | 0.655907957 | 1.581236262 |
| Gm11881 | 0 | NA | NA |
| Gm11879 | 0 | NA | NA |
| Gm24078 | 0 | NA | NA |
| 1700025O08Rik | 0 | NA | NA |
| Gm11884 | 0.044544648 | 0.237309859 | 3.724945049 |
| Gm11888 | 0 | NA | NA |
| Gm11889 | 2.706150538 | -0.975035843 | 0.648111917 |
| Gm25978 | 0.086648695 | -0.72445792 | 3.724945049 |
| Gm11890 | 1.145011917 | 0.629460972 | 1.301314244 |
| Gm28448 | 1.698407129 | -2.375811483 | 0.984301521 |
| Gm11892 | 0.26738533 | 1.219471665 | 2.089224018 |
| Gm11891 | 0 | NA | NA |
| Gm27243 | 8.985200294 | -1.989554627 | 0.438609003 |
| Mms22l | 56.96292425 | 0.818686837 | 0.267385031 |
| Klhl32 | 321.673524 | -0.087004946 | 0.092281914 |
| Gm11899 | 0.512549389 | -0.153219615 | 1.41614614 |
| Ndufaf4 | 345.0672365 | -0.077664983 | 0.102105031 |
| C230012O17Rik | 12.79852409 | -0.048978528 | 0.310555438 |
| Gpr63 | 190.3210986 | -0.523804116 | 0.153865243 |
| Gm11906 | 2.251356483 | -0.898544887 | 0.631506346 |
| Fhl5 | 2.616435038 | 0.401077683 | 0.679120265 |
| Ufl1 | 454.4895708 | -0.086956355 | 0.074330851 |
| Prdx6-ps2 | 0.044191549 | -0.243571806 | 3.724945049 |
| Gm11897 | 0 | NA | NA |
| Gm11894 | 0.044544648 | 0.237309859 | 3.724945049 |
| Gm25975 | 0.109124069 | -0.24357403 | 3.724945049 |
| Gm11896 | 0 | NA | NA |
| Gm11893 | 3.313083698 | 0.917076205 | 0.543414897 |
| Gm24977 | 0 | NA | NA |
| Fut9 | 4215.601891 | -0.711914611 | 0.102193526 |
| Gm24042 | 0 | NA | NA |
| Gm11898 | 0 | NA | NA |
| Gm26067 | 0 | NA | NA |
| Gm11895 | 0 | NA | NA |
| Gm24965 | 0 | NA | NA |
| Manea | 627.7666335 | -0.017289907 | 0.11183741 |

| | | | |
|---|---|---|---|
| Bhmt-ps1 | 0 | NA | NA |
| Gm11903 | 0 | NA | NA |
| Gng2-ps1 | 0.3346585 | -0.775465785 | 1.757484089 |
| Gm11905 | 0 | NA | NA |
| 4930548K13Rik | 0 | NA | NA |
| Gm11904 | 0 | NA | NA |
| Gm11902 | 0 | NA | NA |
| Gm11901 | 0.045882455 | 0.237312084 | 3.724945049 |
| Gm25173 | 0 | NA | NA |
| Gm11914 | 3.923996896 | 0.173812749 | 0.560639297 |
| Tpm3-rs2 | 0 | NA | NA |
| Gm11911 | 0 | NA | NA |
| Gm11907 | 0.192582685 | -0.708793343 | 2.67885232 |
| Gm25378 | 0 | NA | NA |
| Gm11908 | 0 | NA | NA |
| Gm11909 | 0 | NA | NA |
| Gm11910 | 0 | NA | NA |
| Gm11912 | 0 | NA | NA |
| Gm11913 | 0 | NA | NA |
| Epha7 | 823.6749401 | -0.60756216 | 0.141489679 |
| Gm11915 | 4.082662799 | -0.381649324 | 0.498794762 |
| Gm11917 | 0.126225412 | 0.718200913 | 3.698754989 |
| Gm11916 | 0 | NA | NA |
| Gm11918 | 0 | NA | NA |
| Gm11919 | 0 | NA | NA |
| Gm11923 | 0 | NA | NA |
| Gm11925 | 0 | NA | NA |
| Gm11924 | 0.272022412 | 1.280116177 | 2.443962013 |
| 4930556G01Rik | 0.046345709 | 0.71820487 | 3.724945049 |
| Gm22831 | 0 | NA | NA |
| Gm11920 | 0.174512195 | -1.319653964 | 3.342629336 |
| Gm25705 | 0.044544648 | 0.237309859 | 3.724945049 |
| Gm11922 | 0.132199999 | 0.23730893 | 3.642365845 |
| Map3k7 | 821.1190469 | 0.152444457 | 0.054662721 |
| Gm11926 | 0 | NA | NA |
| Gm26254 | 0 | NA | NA |
| Gm11927 | 0.564158409 | 1.014042998 | 1.545561704 |
| Gm11928 | 0 | NA | NA |
| Gm11929 | 0 | NA | NA |
| Bach2 | 252.756904 | -0.182466208 | 0.081478491 |
| Bach2it1 | 12.10687002 | -1.248428561 | 0.40092583 |
| Gm24371 | 4.271398782 | -1.474380175 | 0.582878369 |
| Bach2os | 18.90185043 | 4.444989973 | 0.461142798 |
| Gja10 | 0.072025462 | 0.237312084 | 3.724945049 |
| Gm11933 | 0.233612452 | 0.718216207 | 2.540177828 |
| Casp8ap2 | 404.3248676 | -0.015188181 | 0.095377887 |
| Gm11936 | 0 | NA | NA |

| | | | |
|---|---|---|---|
| Mdn1 | 1768.721666 | -0.184623428 | 0.11800598 |
| Lyrm2 | 76.45699897 | 0.140327894 | 0.181177803 |
| Ankrd6 | 449.0953625 | -0.294399852 | 0.094567691 |
| Gm11941 | 0 | NA | NA |
| Gm11942 | 24.81150096 | 0.624199647 | 0.267708206 |
| Rragd | 1716.34318 | -0.523923893 | 0.098242541 |
| Gm11943 | 0.062426873 | 0.237312084 | 3.724945049 |
| 4933421O10Rik | 104.7690149 | -0.455527345 | 0.126802518 |
| Ube2j1 | 1400.842074 | 0.124562279 | 0.065228769 |
| Gm11931 | 0 | NA | NA |
| Gabrr2 | 12.63021313 | -0.032211405 | 0.315338422 |
| Gabrr1 | 1.88379093 | 1.612625281 | 0.822470416 |
| Pm20d2 | 35.06690641 | -0.788583989 | 0.197920521 |
| Gm11930 | 0 | NA | NA |
| Srsf12 | 183.1024447 | -0.610272017 | 0.102743944 |
| Pnrc1 | 961.8622753 | -0.345372046 | 0.097544595 |
| Gm11934 | 0.906232762 | 0.191165562 | 1.07291934 |
| Rngtt | 426.6224941 | 0.007715829 | 0.061634615 |
| Mir8118 | 1.705950494 | 0.893782476 | 0.824198148 |
| Gm11935 | 0 | NA | NA |
| Gm23304 | 0.144050924 | 0.237309606 | 3.724945049 |
| Gm24341 | 0 | NA | NA |
| Cnr1 | 3191.885636 | -0.325899934 | 0.103259815 |
| Gm22010 | 0 | NA | NA |
| Spaca1 | 0.461160426 | 0.235262188 | 1.33267203 |
| Gm25623 | 0 | NA | NA |
| Gm12753 | 0.179340113 | 0.184531913 | 2.722406815 |
| Gm12751 | 94.08612612 | 0.434115841 | 0.128247477 |
| Gm12752 | 0.088534442 | 1.179809211 | 3.723500796 |
| Gm12754 | 0.046345709 | 0.71820487 | 3.724945049 |
| Gm23466 | 0 | NA | NA |
| Gm12351 | 0 | NA | NA |
| Gm12350 | 35.03170047 | 0.110585796 | 0.201274217 |
| Akirin2 | 531.2879269 | 0.261032247 | 0.071049661 |
| Orc3 | 556.144257 | 0.027058621 | 0.098554359 |
| Rars2 | 246.197839 | -0.19237414 | 0.098129578 |
| Slc35a1 | 169.2461004 | -0.210108404 | 0.10646052 |
| Gm22849 | 0.748256776 | 0.578779196 | 1.223343462 |
| 1700003M02Rik | 17.88018956 | -0.200703562 | 0.284133538 |
| Gm136 | 1.409557441 | -2.230275481 | 1.153154447 |
| Smim8 | 84.47012772 | 0.465650171 | 0.130664286 |
| Zfp292 | 1650.746143 | -0.066487743 | 0.094668307 |
| Gm12363 | 0 | NA | NA |
| Cga | 2.158593695 | 1.796442034 | 1.494223442 |
| Platr9 | 0 | NA | NA |
| Gm12361 | 0.042716945 | -0.243571806 | 3.724945049 |
| Gm12362 | 0.090537259 | 0.237318756 | 3.724945049 |

| | | | |
|---|---|---|---|
| Mob3b | 188.1401066 | -0.295200418 | 0.159305565 |
| Gm12364 | 0 | NA | NA |
| Ifnk | 0.038981395 | 0.718207094 | 3.724945049 |
| 3110043O21Rik | 349.8244095 | 0.179128866 | 0.075621474 |
| Gm12366 | 24.68870437 | 0.385623766 | 0.232195133 |
| Gm12367 | 4.278993874 | -0.667060634 | 0.519451982 |
| Gm12368 | 0 | NA | NA |
| Gm12365 | 0 | NA | NA |
| Lingo2 | 668.3004972 | -0.766179511 | 0.123549367 |
| Gm12369 | 25.01436732 | -1.235066413 | 0.275915263 |
| Gm12370 | 0 | NA | NA |
| Mir876 | 9.32634141 | -0.822890725 | 0.332840762 |
| Mir873a | 8.772029568 | -0.705549231 | 0.368062941 |
| mmu-mir-873a | 0 | NA | NA |
| Gm23314 | 0 | NA | NA |
| AL844160.1 | 0 | NA | NA |
| Gm12371 | 46.23436975 | -0.256648396 | 0.196437415 |
| Gm12372 | 1.336268985 | 1.037884797 | 0.843147454 |
| Rpl23a-ps6 | 0 | NA | NA |
| Gm12374 | 2.339895364 | -0.241335264 | 0.685276069 |
| Gm22732 | 0.04393175 | -0.243571806 | 3.724945049 |
| Gm12375 | 0 | NA | NA |
| Gm12378 | 0 | NA | NA |
| Gm12379 | 0 | NA | NA |
| Gm12380 | 0.504525611 | 0.103499215 | 1.357038112 |
| Gm12381 | 0 | NA | NA |
| Gm12382 | 0 | NA | NA |
| Gm25581 | 0 | NA | NA |
| Gm12383 | 0 | NA | NA |
| Gm23607 | 0 | NA | NA |
| Gm17094 | 0.175174943 | 0.237326123 | 2.724622258 |
| 1700009N14Rik | 0 | NA | NA |
| Gm12385 | 0.045882455 | 0.237312084 | 3.724945049 |
| Gm12384 | 0.119234754 | -0.243576254 | 3.724945049 |
| Gm12386 | 0 | NA | NA |
| Gm22850 | 0 | NA | NA |
| Gm12387 | 0.097139547 | 0.718207094 | 3.724945049 |
| Gm26087 | 0 | NA | NA |
| Gm12388 | 0.062426873 | 0.237312084 | 3.724945049 |
| Gm12389 | 0 | NA | NA |
| Aco1 | 373.8664707 | -0.114560464 | 0.095679682 |
| Ddx58 | 2204.242023 | 6.222782708 | 0.227831546 |
| Topors | 450.6972402 | 0.228383033 | 0.071976233 |
| Toporsos | 15.19562275 | 0.410515929 | 0.313593525 |
| Ndufb6 | 365.4765144 | -0.098670147 | 0.10530997 |
| 4930509K18Rik | 0 | NA | NA |
| Gm23338 | 0.239936926 | 0.237320696 | 2.454644493 |

| | | | |
|---|---|---|---|
| Tmem215 | 126.9200701 | -0.5767636 | 0.207409322 |
| 4933428C19Rik | 0.268438559 | 0.691329555 | 1.807842089 |
| Gm12391 | 0 | NA | NA |
| Aptx | 267.7610802 | -0.246970874 | 0.091472807 |
| Gm12390 | 0.717871974 | -1.034831588 | 1.279615647 |
| Gm6297 | 15.05970898 | -0.617278292 | 0.280773337 |
| Dnaja1 | 2433.261954 | 0.336724255 | 0.102459229 |
| Mir207 | 0.128150834 | -1.056605085 | 3.714811905 |
| Smu1 | 643.1185477 | 0.244886899 | 0.078225501 |
| Gm12396 | 8.729157278 | 0.686617444 | 0.335649089 |
| B4galt1 | 382.463159 | 2.847946458 | 0.195355042 |
| Mir5123 | 0.217530652 | 1.222589782 | 2.172555094 |
| Gm24112 | 0.24398126 | -1.585252391 | 1.806934748 |
| Gm25931 | 0.653402772 | 2.659358251 | 1.41078445 |
| Gm12397 | 7.076756442 | 0.022875456 | 0.383467083 |
| Spink4 | 0.044544648 | 0.237309859 | 3.724945049 |
| Bag1 | 715.7527563 | 0.429803021 | 0.090753801 |
| Chmp5 | 658.9006136 | 0.450725555 | 0.114702038 |
| Nfx1 | 1354.303035 | -0.136287793 | 0.091620918 |
| Mir3094 | 0.798730366 | 0.583811853 | 1.111099452 |
| Aqp7 | 0.174839421 | -0.71915759 | 2.306333002 |
| Gm25637 | 0 | NA | NA |
| Aqp3 | 1.154822103 | 2.061460165 | 1.095999328 |
| Nol6 | 1475.373276 | -0.050914754 | 0.04637608 |
| Gm12402 | 0.230106605 | -1.101438274 | 2.560976671 |
| Ube2r2 | 789.6006133 | 0.289486239 | 0.064385963 |
| Gm12403 | 11.82053707 | -0.722884022 | 0.409575108 |
| Gm24650 | 0 | NA | NA |
| Ubap2 | 693.4431244 | 0.170098212 | 0.084495166 |
| Gm22888 | 2.630926952 | -0.925151664 | 0.698314165 |
| Gm24837 | 2.20124 | -0.650568587 | 0.759115783 |
| Gm23090 | 0.341439828 | 0.279935831 | 1.596078972 |
| Gm12401 | 0.164132501 | 0.237308425 | 3.397905533 |
| Dcaf12 | 631.7207173 | 0.341865037 | 0.085445588 |
| n-R5s183 | 0 | NA | NA |
| Gm26049 | 4.518367824 | -0.566405656 | 0.616492146 |
| Gm26084 | 0.449030389 | -2.815084022 | 1.450219674 |
| Ubap1 | 662.4155343 | 0.523956258 | 0.070751003 |
| Kif24 | 28.6589064 | 0.74544256 | 0.199189065 |
| Nudt2 | 96.39035252 | 0.104838095 | 0.116265398 |
| AI464131 | 299.7303309 | -1.036395408 | 0.200362886 |
| 1110017D15Rik | 25.98366852 | 0.133873491 | 0.198844548 |
| Fam219aos | 0.808497328 | -0.050039671 | 1.162047521 |
| Fam219a | 1771.430822 | -0.214049554 | 0.051627979 |
| Gm12404 | 1.315597923 | 0.032794512 | 0.82491269 |
| Dnaic1 | 23.7871792 | -0.485422699 | 0.275605143 |
| Enho | 315.6348979 | -0.513790883 | 0.13030383 |

| | | | |
|---|---|---|---|
| Gm12406 | 1.108658207 | -1.16440331 | 0.98588686 |
| Gm12405 | 0.625648376 | -3.086164499 | 1.521776276 |
| Cntfr | 660.2301978 | -0.440584285 | 0.105152708 |
| Il11ra1 | 183.2536711 | 0.068638451 | 0.103980218 |
| Rpp25l | 90.6293161 | 0.553667779 | 0.125169236 |
| Dctn3 | 483.0857412 | 0.041976727 | 0.069514833 |
| Arid3c | 1.591794818 | 0.086226616 | 0.988832824 |
| Sigmar1 | 271.0497585 | 0.112497448 | 0.084940799 |
| Galt | 238.390805 | -0.532932283 | 0.086023934 |
| Ccl27a | 57.35970255 | -1.73890538 | 0.164453508 |
| Gm12407 | 0.693596057 | 2.124832367 | 1.252753739 |
| Gm10601 | 0.106056992 | 0.237307635 | 3.724945049 |
| Gm13307 | 0 | NA | NA |
| Gm13302 | 0 | NA | NA |
| Gm21541 | 0 | NA | NA |
| Ccl19-ps1 | 0 | NA | NA |
| Gm20878 | 0 | NA | NA |
| Gm20938 | 0 | NA | NA |
| Gm13303 | 0 | NA | NA |
| Gm21093 | 0.097139547 | 0.718207094 | 3.724945049 |
| Gm37530 | 0.382550573 | -1.291739185 | 1.3879946 |
| Gm3892 | 0 | NA | NA |
| Gm3893 | 0 | NA | NA |
| Gm13304 | 0.049553047 | 0.71820487 | 3.724945049 |
| Gm13299 | 0.097501745 | 0.23732098 | 3.724945049 |
| Ccl19-ps3 | 0.089089296 | 0.237305797 | 3.724945049 |
| Gm13306 | 1.067961214 | 0.665190312 | 1.053565684 |
| Gm13305 | 0.062426873 | 0.237312084 | 3.724945049 |
| 1700045I11Rik | 0 | NA | NA |
| Gm17167 | 0.21074972 | 0.264474914 | 2.623964103 |
| Gm13301 | 0 | NA | NA |
| Gm5859 | 0 | NA | NA |
| Gm17081 | 0 | NA | NA |
| Gm10600 | 0 | NA | NA |
| Ccl21b | 0 | NA | NA |
| Gm21980 | 0 | NA | NA |
| Gm21953 | 8.239491219 | -1.374568965 | 0.382760385 |
| Gm21598 | 0 | NA | NA |
| Gm21586 | 0.193603971 | 0.71820869 | 3.251616718 |
| Gm3883 | 0 | NA | NA |
| Gm2163 | 0 | NA | NA |
| Gm13298 | 0.089814205 | -0.24357403 | 3.724945049 |
| Gm10593 | 0.444623086 | 0.718200753 | 3.258894588 |
| Gm10591 | 0 | NA | NA |
| Gm2564 | 0 | NA | NA |
| Ccl27b | 1.785252223 | 0.219371587 | 0.75604342 |
| Il11ra2 | 0.424152111 | -0.083894472 | 1.664452084 |

| | | | |
|---|---|---|---|
| Gm21955 | 83.35882431 | -1.262699825 | 0.214304099 |
| 4930578G10Rik | 26.07152981 | -1.870189668 | 0.3920822 |
| Gm12395 | 6.47297608 | -1.825545857 | 0.652519075 |
| Ccl19 | 26.89383427 | 6.541236334 | 0.866959752 |
| Ccl21a | 1.739286695 | 2.080656314 | 0.821437056 |
| Gm12394 | 0.61514822 | -1.205804284 | 1.227580805 |
| Gm12429 | 2.41835347 | -2.062615739 | 0.822045896 |
| BC049635 | 0.963622687 | 1.386275076 | 1.072721785 |
| N28178 | 2446.431538 | -1.240839991 | 0.150817254 |
| Dnajb5 | 1123.231011 | -0.286725241 | 0.165439961 |
| 1700022I11Rik | 5.07266936 | 4.493197152 | 0.8098172 |
| Vcp | 2874.0526 | 0.408008591 | 0.065395434 |
| Fancg | 91.45811176 | -0.26098129 | 0.168892767 |
| Pigo | 262.034385 | -0.121882068 | 0.082309869 |
| Stoml2 | 348.4296267 | 0.256739592 | 0.095030396 |
| Gm26643 | 0.357756446 | -1.725280656 | 1.352728745 |
| Fam214b | 535.3240349 | 0.750936732 | 0.081055297 |
| Gm26881 | 11.80662906 | -1.166268645 | 0.329946742 |
| Unc13b | 466.7629952 | -0.551805307 | 0.185589211 |
| Gm23709 | 0 | NA | NA |
| Gm25010 | 0 | NA | NA |
| Atp8b5 | 7.649160145 | 0.534940253 | 0.344122533 |
| Rusc2 | 1981.730262 | -0.209776221 | 0.070841456 |
| Fam166b | 3.554918289 | -1.240045952 | 0.627801621 |
| Tesk1 | 834.3782588 | -0.065764032 | 0.064219518 |
| Cd72 | 162.2518301 | 6.341679824 | 0.377672093 |
| Sit1 | 2.004396688 | 3.467101169 | 1.013164109 |
| Rmrp | 17396.1871 | 0.417569452 | 0.169896527 |
| Ccdc107 | 211.9356722 | 0.188600638 | 0.150789016 |
| Arhgef39 | 4.559760072 | 0.155270572 | 0.508016826 |
| Gm12454 | 3.511636557 | -0.15745868 | 0.580912817 |
| Car9 | 11.98985167 | -0.311995972 | 0.351597236 |
| Tpm2 | 58.51175456 | -0.304970394 | 0.199239768 |
| Tln1 | 1972.671443 | 1.870791229 | 0.130468848 |
| Creb3 | 390.8704704 | 0.898034662 | 0.126984075 |
| Gba2 | 788.9852365 | -0.313541781 | 0.053786487 |
| Rgp1 | 609.9554589 | 0.253164383 | 0.055419632 |
| Msmp | 0.198131738 | -0.243563397 | 2.22921086 |
| Gm12472 | 0.580871419 | 1.647871659 | 1.406213909 |
| Gm25262 | 9.682546495 | -0.48017321 | 0.41436815 |
| Gm12473 | 1.944300039 | 0.725211628 | 0.768555271 |
| Npr2 | 447.7307936 | -0.334028658 | 0.096603707 |
| Spag8 | 14.00717955 | 0.267099687 | 0.31628021 |
| Hint2 | 86.66528165 | 0.209410176 | 0.153538454 |
| Fam221b | 17.22801957 | -0.662427163 | 0.269939859 |
| Gm12481 | 74.19163858 | 0.410844638 | 0.175500312 |
| Tmem8b | 757.5108737 | -0.74262254 | 0.081308591 |

| | | | |
|---|---|---|---|
| Gm23257 | 4.230344492 | -0.729279515 | 0.496815041 |
| Olfr70 | 0 | NA | NA |
| Olfr71 | 0 | NA | NA |
| Hrct1 | 4.20546811 | 1.722371743 | 0.591075281 |
| 5430416O09Rik | 3.015746376 | 0.447899641 | 0.56910102 |
| A630077J23Rik | 0.321964759 | 0.769031624 | 1.998609257 |
| Olfr269-ps1 | 0 | NA | NA |
| Olfr159 | 0 | NA | NA |
| Olfr29-ps1 | 0 | NA | NA |
| Gm12492 | 0 | NA | NA |
| Olfr156 | 0.048949532 | -0.243569582 | 3.724945049 |
| Olfr157 | 0 | NA | NA |
| Olfr155 | 0.08169834 | 0.23732098 | 3.724945049 |
| Reck | 121.3301511 | -0.113075601 | 0.187601476 |
| Gm27265 | 2.022028428 | -0.513711621 | 0.787928068 |
| Glipr2 | 555.7186848 | 5.440647266 | 0.354490919 |
| Ccin | 0.952506814 | 2.118487071 | 1.179324087 |
| Clta | 1302.037201 | 0.215045083 | 0.080604261 |
| Gne | 417.5634539 | -0.005295687 | 0.077834357 |
| Gm22706 | 0.185638728 | 0.718215363 | 2.668820032 |
| Gm12503 | 0.506185493 | 1.618993942 | 1.345027416 |
| Gm12504 | 0.323274704 | -0.243564124 | 1.623916085 |
| Rnf38 | 803.2801551 | -0.158466668 | 0.068960286 |
| Mir5106 | 2.834065591 | 0.22401878 | 0.626358938 |
| Melk | 12.45686811 | 2.847272951 | 0.430233874 |
| Gm12677 | 0 | NA | NA |
| Gm12455 | 0 | NA | NA |
| Gm24376 | 0 | NA | NA |
| Pax5 | 62.51624943 | 1.535617418 | 0.194278407 |
| Mir5120 | 0.090665643 | -0.724455696 | 3.724945049 |
| Gm12462 | 5.015296868 | 0.921081699 | 0.498654179 |
| 4930517J16Rik | 1.109399417 | 1.336966724 | 1.201564964 |
| 5730488B01Rik | 1.238426347 | 0.184043884 | 0.946046866 |
| Gm12463 | 0.758386275 | -0.057880918 | 1.021771962 |
| Zcchc7 | 970.4223863 | -0.48583926 | 0.147501808 |
| Gm22639 | 3.577120001 | -0.463536674 | 0.500184098 |
| Gm12678 | 0 | NA | NA |
| Gm12493 | 0 | NA | NA |
| Gm12679 | 2.474621302 | 1.165447769 | 0.767980655 |
| Grhpr | 156.0778864 | -0.724129589 | 0.126246423 |
| Zbtb5 | 270.4928444 | 0.8085324 | 0.111005351 |
| 1700055D18Rik | 14.55774235 | 0.547652536 | 0.281365789 |
| Polr1e | 68.48577099 | 0.513125761 | 0.148035083 |
| Fbxo10 | 581.7003137 | -0.24544962 | 0.071310742 |
| Tomm5 | 134.9204341 | 0.389382324 | 0.146767494 |
| Gm12566 | 0 | NA | NA |
| Frmpd1 | 273.0835449 | 0.383835453 | 0.139313299 |

| | | | |
|---|---|---|---|
| Frmpd1os | 0.15456712 | -0.7244586 | 2.83422604 |
| Trmt10b | 109.9286799 | -0.060822653 | 0.10405888 |
| Exosc3 | 78.17585261 | 0.699991128 | 0.163435964 |
| Gm12585 | 0 | NA | NA |
| Dcaf10 | 447.5914563 | 0.291000552 | 0.106804095 |
| Slc25a51 | 825.9827367 | 0.024686389 | 0.046355938 |
| Gm25692 | 0 | NA | NA |
| Shb | 131.2113877 | 0.409422373 | 0.120802012 |
| Gm22518 | 0.171044682 | 0.237327163 | 2.745278379 |
| Gm22055 | 0 | NA | NA |
| Gm12411 | 5.818705562 | -0.359738058 | 0.474781662 |
| Gm12408 | 2.792582105 | -0.799800897 | 0.586521328 |
| Gm829 | 0.154612032 | 1.66937168 | 3.490354585 |
| Gm12410 | 0.044191549 | -0.243571806 | 3.724945049 |
| Gm12409 | 0.36880298 | 1.100698747 | 1.574424242 |
| Gm12412 | 0.044544648 | 0.237309859 | 3.724945049 |
| Aldh1b1 | 125.3782894 | 1.395716999 | 0.180799856 |
| Igfbpl1 | 40.81830737 | 1.596876774 | 0.266955927 |
| Stra6l | 52.92012026 | 6.601188315 | 0.564177155 |
| E230008N13Rik | 12.72585994 | 0.562920731 | 0.345877119 |
| Tdrd7 | 1191.587 | 1.889291865 | 0.12533677 |
| Tmod1 | 407.2062115 | -0.675686248 | 0.129261485 |
| Tstd2 | 221.2458736 | 0.376300946 | 0.105186676 |
| Gm23250 | 0.397571503 | -0.137186088 | 3.259168755 |
| Ncbp1 | 647.6804023 | 0.115488069 | 0.07120735 |
| Xpa | 75.64072436 | 0.078850653 | 0.178540038 |
| Gm12444 | 2.312647122 | -0.584940912 | 0.682159168 |
| Gm12445 | 0 | NA | NA |
| Gm12446 | 0.091666476 | -0.724455696 | 3.724945049 |
| Gm12447 | 2.955086367 | -1.079648198 | 0.644666706 |
| Foxe1 | 0.36004552 | 1.203083784 | 1.758955206 |
| 5830415F09Rik | 17.35131692 | -0.068317729 | 0.250284146 |
| Hemgn | 1.129829066 | 1.766144701 | 1.339923281 |
| Gm12443 | 0.104366086 | -0.243576254 | 3.724945049 |
| Anp32b | 407.7977659 | 0.701452373 | 0.121232914 |
| Nans | 132.4373756 | 0.442665633 | 0.144428029 |
| Trim14 | 69.63745902 | 5.178728196 | 0.343339555 |
| Gm22247 | 0.26079097 | -0.243563865 | 2.453645846 |
| Gm16731 | 0.468050163 | 2.348693656 | 1.368279679 |
| Coro2a | 311.7321339 | -0.142113659 | 0.080926599 |
| Tbc1d2 | 54.86226986 | 0.955245844 | 0.202015342 |
| Gabbr2 | 3104.900051 | -0.719984753 | 0.117990064 |
| Gm568 | 0.609243813 | -0.538150095 | 1.427531832 |
| Anks6 | 149.640635 | -0.625150376 | 0.196708373 |
| Gm12425 | 0 | NA | NA |
| Gm12422 | 0.506756615 | -0.243565509 | 1.492015699 |
| Galnt12 | 14.31060596 | -0.243527418 | 0.333450344 |

| | | | |
|---|---|---|---|
| Gm12424 | 0.961043033 | -1.381932948 | 1.095323693 |
| Gm12423 | 0 | NA | NA |
| Gm12426 | 1.368259282 | -0.323912649 | 0.862238911 |
| Col15a1 | 61.55752535 | -0.556758619 | 0.179631474 |
| Tgfbr1 | 481.364332 | -0.038487887 | 0.145091521 |
| Gm12430 | 5.138410126 | 0.225940205 | 0.417704951 |
| Alg2 | 1382.70197 | -0.110521139 | 0.063892494 |
| Sec61b | 158.1254426 | 1.037404631 | 0.156649055 |
| Gm27601 | 0 | NA | NA |
| Gm27668 | 0 | NA | NA |
| Gm38343 | 0.066050875 | 0.71820487 | 3.724945049 |
| Gm22670 | 0 | NA | NA |
| Gm12434 | 0 | NA | NA |
| Nr4a3 | 542.7975049 | -0.605952054 | 0.309416004 |
| Stx17 | 355.1670537 | -0.001843943 | 0.074281472 |
| Erp44 | 524.436621 | 0.522971616 | 0.092373571 |
| Mir1958 | 0.694929783 | -2.595832067 | 1.234773237 |
| Invs | 182.2944049 | 0.022843207 | 0.112762372 |
| Gm12435 | 13.93294907 | 1.197372015 | 0.280568111 |
| Tex10 | 345.3321695 | 0.111431207 | 0.076576222 |
| Msantd3 | 126.8937499 | -0.02670843 | 0.113163069 |
| Tmeff1 | 650.557545 | -0.839329985 | 0.113757647 |
| Murc | 22.05245246 | -0.645370357 | 0.246959856 |
| Gm12439 | 2.996720251 | -0.877496585 | 0.625577627 |
| Gm12438 | 0 | NA | NA |
| Gm24573 | 0 | NA | NA |
| Gm23746 | 0 | NA | NA |
| Gm12436 | 0.044544648 | 0.237309859 | 3.724945049 |
| Gm12442 | 0 | NA | NA |
| Lppr1 | 119.1937067 | -1.058789831 | 0.152516495 |
| Gm22685 | 0.467201629 | -1.861831948 | 1.471147055 |
| Gm12453 | 0.624449418 | -1.362925592 | 1.294178472 |
| Rbpsuh-ps1 | 0.090074004 | -0.24357403 | 3.724945049 |
| Acnat2 | 0.218471152 | 0.717751605 | 1.911199781 |
| Gm12451 | 0.314054052 | -0.243576782 | 3.724945049 |
| Gm12452 | 0.075043205 | 0.237309859 | 3.724945049 |
| Acnat1 | 2.07728567 | 0.91420836 | 0.814286171 |
| Gm26424 | 0 | NA | NA |
| Baat | 6.000090479 | 0.292947616 | 0.468686021 |
| Mrpl50 | 361.7674449 | 0.083971739 | 0.090095852 |
| Zfp189 | 205.1948466 | -0.040130617 | 0.100673688 |
| Aldob | 16.11872095 | -0.944377417 | 0.335706989 |
| Tmem246 | 593.6062876 | -0.25115472 | 0.074543906 |
| Rnf20 | 1042.085228 | -0.066913858 | 0.07106794 |
| Grin3a | 866.8310369 | -0.440230052 | 0.076018967 |
| Ppp3r2 | 15.47209461 | -0.752663218 | 0.344790149 |
| Gm12459 | 3.099509663 | -2.391983963 | 0.694549577 |

| | | | |
|---|---|---|---|
| Gm12458 | 0.048949532 | -0.243569582 | 3.724945049 |
| Gm25485 | 0 | NA | NA |
| Gm12460 | 0.083526043 | 0.718202646 | 3.724945049 |
| Cylc2 | 0 | NA | NA |
| Amd-ps4 | 32.8894957 | -0.146607963 | 0.217881078 |
| Gm22563 | 0 | NA | NA |
| Gm23043 | 0 | NA | NA |
| Gm24943 | 0 | NA | NA |
| Gm22042 | 0.482432794 | 1.666534541 | 1.413893611 |
| Gm12461 | 0 | NA | NA |
| C630028M04Rik | 0.154174174 | -0.242934529 | 2.834967975 |
| Gm12465 | 0 | NA | NA |
| Gm12466 | 0.144050924 | 0.237309606 | 3.724945049 |
| Gm12467 | 0 | NA | NA |
| Gm12475 | 1.233559488 | 2.826225354 | 1.158478448 |
| Smc2os | 3.507361272 | -0.735371929 | 0.610165289 |
| Smc2 | 184.4916281 | 0.066583443 | 0.17070952 |
| Tmc4-ps | 0 | NA | NA |
| Vma21-ps | 0.038981395 | 0.718207094 | 3.724945049 |
| Toporsl | 0 | NA | NA |
| 4930552N02Rik | 0 | NA | NA |
| Gm12482 | 0.399060758 | 1.209724135 | 1.671356624 |
| Olfr275 | 0 | NA | NA |
| Olfr273 | 0.062426873 | 0.237312084 | 3.724945049 |
| Cct3-ps1 | 0.33824293 | 1.623203087 | 1.885291768 |
| Olfr272 | 0.046345709 | 0.71820487 | 3.724945049 |
| Olfr271-ps1 | 0.118974955 | -0.243576254 | 3.724945049 |
| Olfr270 | 0 | NA | NA |
| Nipsnap3a | 0 | NA | NA |
| Nipsnap3b | 127.01825 | 1.013510638 | 0.186875814 |
| Abca1 | 2154.257843 | 1.023404042 | 0.145612616 |
| 4930412L05Rik | 1.32543013 | 0.966543495 | 1.035866982 |
| 4930522O17Rik | 0.038981395 | 0.718207094 | 3.724945049 |
| AI427809 | 3.06421064 | 1.964358501 | 0.728101341 |
| Gm12495 | 0 | NA | NA |
| Gm12496 | 0.503001268 | -0.446843914 | 1.453860242 |
| Slc44a1 | 1434.826822 | 0.173965138 | 0.086215392 |
| 1700060J05Rik | 0.251076704 | 0.838298884 | 2.493560933 |
| Fsd1l | 701.9750228 | -0.437547774 | 0.061495308 |
| Gm12470 | 15.34197034 | -0.981141759 | 0.36882852 |
| Fktn | 705.1094383 | -0.41834949 | 0.062382638 |
| Gm25445 | 0 | NA | NA |
| Tal2 | 3.478949006 | 1.784510401 | 0.629185657 |
| Rps15a-ps8 | 0 | NA | NA |
| Tmem38b | 167.1596389 | 0.414711235 | 0.111268941 |
| n-R5s185 | 0 | NA | NA |
| Gm12468 | 0.068097359 | -0.243569582 | 3.724945049 |

| | | | |
|---|---|---|---|
| Gm23745 | 0.047948698 | -0.243569582 | 3.724945049 |
| Gm24177 | 0 | NA | NA |
| Gm12484 | 0 | NA | NA |
| Gm12469 | 0.094831987 | -0.243571806 | 3.724945049 |
| Gm25687 | 0 | NA | NA |
| Tpt1-ps2 | 0 | NA | NA |
| Gm29067 | 0 | NA | NA |
| Gm12478 | 0 | NA | NA |
| Gm12480 | 0.086930093 | 0.236669296 | 3.724898107 |
| Gm12477 | 0 | NA | NA |
| Gm24628 | 0 | NA | NA |
| Gm12479 | 0 | NA | NA |
| Zfp462 | 1321.864971 | -0.490401001 | 0.076469596 |
| Gm12515 | 0.097139547 | 0.718207094 | 3.724945049 |
| Gm12514 | 0.145451803 | 1.498658476 | 3.529361416 |
| Gm12509 | 0 | NA | NA |
| Gm12508 | 0.167320703 | 0.237327023 | 2.764437523 |
| Gm12512 | 0 | NA | NA |
| Gm25419 | 0 | NA | NA |
| Rad23b | 2280.331224 | 0.352217422 | 0.073702538 |
| Gm12516 | 0 | NA | NA |
| Gm12513 | 0 | NA | NA |
| Gm12505 | 0.048949532 | -0.243569582 | 3.724945049 |
| Gm12510 | 0 | NA | NA |
| Gm22150 | 0 | NA | NA |
| Gm12507 | 0 | NA | NA |
| Klf4 | 85.90030529 | 1.969293541 | 0.271016098 |
| Gm12511 | 0.772413307 | 0.571373892 | 1.230288022 |
| Gm12506 | 1.081462396 | 0.126500987 | 0.867513989 |
| Gm12519 | 0.384295504 | 0.304118739 | 1.564783657 |
| Gm12520 | 0 | NA | NA |
| 2310081O03Rik | 1.122999083 | 4.050869849 | 1.098963368 |
| Gm12518 | 0 | NA | NA |
| Gm26144 | 0.110595523 | 0.718200421 | 3.724945049 |
| Actl7b | 1.554701746 | -0.40285845 | 0.868758379 |
| Gm26657 | 0 | NA | NA |
| Actl7a | 0.04393175 | -0.243571806 | 3.724945049 |
| Ikbkap | 924.710914 | -0.379195468 | 0.088712649 |
| Gm22952 | 4.197905476 | -1.253694794 | 0.553567255 |
| Fam206a | 278.4213948 | -0.038746806 | 0.090141413 |
| Ctnnal1 | 338.824844 | -0.091803475 | 0.131419289 |
| Tmem245 | 2131.110561 | -0.537837436 | 0.091911376 |
| Mir32 | 0.646815127 | -2.164534106 | 1.412959653 |
| Gm25053 | 0.048949532 | -0.243569582 | 3.724945049 |
| Gm22300 | 0.061920613 | 0.718207094 | 3.724945049 |
| Frrs1l | 3263.550938 | -0.977531492 | 0.103729884 |
| Epb4.1l4b | 724.2344794 | -0.858219717 | 0.084855808 |

| | | | |
|---|---|---|---|
| Gm12530 | 6.592824524 | 3.594193042 | 0.68507584 |
| Ptpn3 | 752.3310739 | -0.682247441 | 0.11733895 |
| Gm12536 | 3.13007401 | -0.347158177 | 0.608745527 |
| 1700042G15Rik | 0 | NA | NA |
| Gm12537 | 23.16944423 | -0.168519401 | 0.227543991 |
| Gm12538 | 1.143639422 | 0.360276408 | 1.208947215 |
| Palm2 | 230.289783 | -0.613558417 | 0.11668743 |
| Gm12539 | 0 | NA | NA |
| Pakap | 9.557701205 | -1.119055245 | 0.359511677 |
| Akap2 | 22.84245767 | 0.173005769 | 0.251580924 |
| Gm12526 | 0.201955748 | 1.192557315 | 2.220435447 |
| Gm24277 | 1.91044629 | -0.666931045 | 0.784344747 |
| Gm26138 | 0 | NA | NA |
| D630039A03Rik | 3.846205949 | 0.17538354 | 0.563934222 |
| Txn1 | 650.0859016 | 1.543043435 | 0.16410663 |
| Txndc8 | 0 | NA | NA |
| Svep1 | 270.025923 | -0.831832362 | 0.217901611 |
| Gm26889 | 1.802555565 | -0.653636949 | 0.70490408 |
| Musk | 19.66503011 | -0.126085043 | 0.228974112 |
| Lpar1 | 579.1101207 | -0.178489062 | 0.13732852 |
| Mir3095 | 0.494865952 | -0.929774577 | 1.402591883 |
| Gm12577 | 4.644629704 | 0.063315294 | 0.566007095 |
| Gm12578 | 0 | NA | NA |
| Gm12579 | 0.044191549 | -0.243571806 | 3.724945049 |
| Gm12580 | 0.526179982 | 0.577911547 | 1.188716167 |
| Olfr267 | 2.961215905 | 1.354903253 | 0.540140479 |
| AI314180 | 2945.683157 | 0.101523317 | 0.074350182 |
| Gm26400 | 0 | NA | NA |
| Zkscan16 | 430.9535622 | -1.039067333 | 0.098836673 |
| Ptgr1 | 21.48653641 | 1.763180407 | 0.378391519 |
| Dnajc25 | 87.02722299 | 0.133054356 | 0.162475409 |
| Gm20503 | 0 | NA | NA |
| Gng10 | 14.26907646 | -0.172639469 | 0.282751092 |
| AI481877 | 0.167924218 | 1.199985076 | 2.761407179 |
| Gm12593 | 0.308655657 | 0.718190755 | 3.724945049 |
| Gm12594 | 0 | NA | NA |
| Rpsa-ps11 | 1.26617739 | 0.777851915 | 0.842650229 |
| Ugcg | 1300.167576 | 0.106807944 | 0.057285857 |
| Gm12596 | 0.346136167 | 2.071697297 | 1.648880817 |
| Susd1 | 107.3121581 | -0.570071273 | 0.165201409 |
| Gm12527 | 0.551374303 | -1.67907683 | 1.267685396 |
| Ptbp3 | 1030.313491 | 1.571955745 | 0.117041058 |
| Gm12529 | 0 | NA | NA |
| 1700018M17Rik | 1.139841548 | 1.224430834 | 1.131220626 |
| Hsdl2 | 568.5576942 | 0.64213966 | 0.117755822 |
| E130308A19Rik | 348.4769703 | -0.272818005 | 0.068456891 |
| Inip | 231.4780127 | 0.127916532 | 0.0853813 |

| | | | |
|---|---|---|---|
| Snx30 | 979.4902318 | -0.139246025 | 0.066946469 |
| Gm12543 | 7.392584082 | -0.316480112 | 0.454398711 |
| Gm12542 | 0.038981395 | 0.718207094 | 3.724945049 |
| Slc46a2 | 0 | NA | NA |
| Mup4 | 0.187777844 | -1.444274885 | 3.298122832 |
| Mup6 | 0.095435502 | 0.718202646 | 3.724945049 |
| Mup7 | 0.045882455 | 0.237312084 | 3.724945049 |
| Mup-ps1 | 0 | NA | NA |
| Mup2 | 0 | NA | NA |
| Mup-ps2 | 0 | NA | NA |
| Mup8 | 0 | NA | NA |
| Mup-ps3 | 0 | NA | NA |
| Mup-ps29 | 0 | NA | NA |
| Mup-ps28 | 0 | NA | NA |
| Mup9 | 0 | NA | NA |
| Mup-ps4 | 0 | NA | NA |
| Mup-ps26 | 0 | NA | NA |
| Mup1 | 0 | NA | NA |
| Gm13773 | 0 | NA | NA |
| Mup-ps6 | 0 | NA | NA |
| Mup-ps24 | 0 | NA | NA |
| Mup10 | 0 | NA | NA |
| Mup-ps5 | 0 | NA | NA |
| Mup11 | 0 | NA | NA |
| Mup-ps7 | 0 | NA | NA |
| Mup12 | 0 | NA | NA |
| Mup-ps8 | 0 | NA | NA |
| Gm21286 | 0 | NA | NA |
| Mup-ps9 | 0 | NA | NA |
| Mup13 | 0 | NA | NA |
| Mup-ps10 | 0 | NA | NA |
| Mup14 | 0 | NA | NA |
| Mup-ps11 | 0 | NA | NA |
| Mup15 | 0 | NA | NA |
| Mup-ps12 | 0 | NA | NA |
| Mup16 | 0 | NA | NA |
| Mup-ps13 | 0 | NA | NA |
| Mup17 | 0 | NA | NA |
| Mup-ps14 | 0 | NA | NA |
| Mup18 | 0 | NA | NA |
| Mup-ps15 | 0 | NA | NA |
| Gm12573 | 0 | NA | NA |
| Mup19 | 0 | NA | NA |
| Gm13775 | 0 | NA | NA |
| Mup-ps16 | 0 | NA | NA |
| Mup5 | 0.135858026 | -1.180985464 | 2.941468349 |
| Mup-ps23 | 0 | NA | NA |

| | | | |
|---|---|---|---|
| Mup-ps17 | 0 | NA | NA |
| Mup-ps18 | 0 | NA | NA |
| Mup-ps19 | 0 | NA | NA |
| Mup-ps20 | 0 | NA | NA |
| Mup-ps21 | 0 | NA | NA |
| Mup20 | 0 | NA | NA |
| Mup3 | 1.273710888 | -0.086584281 | 0.960015286 |
| Gm12909 | 0 | NA | NA |
| Gm12910 | 0 | NA | NA |
| Mup21 | 0.088476398 | -0.243576254 | 3.724945049 |
| Zfp37 | 301.4106421 | 0.176210739 | 0.07807157 |
| Slc31a2 | 164.8023296 | 1.932438802 | 0.212213783 |
| Gm23148 | 0 | NA | NA |
| Fkbp15 | 744.3892095 | 0.325398951 | 0.074291977 |
| Slc31a1 | 348.3933879 | 0.724486479 | 0.113124976 |
| Cdc26 | 169.15404 | 0.164586546 | 0.10568838 |
| Prpf4 | 294.0494031 | 0.241936839 | 0.074833102 |
| Rnf183 | 1.151952751 | 2.816100104 | 1.20540607 |
| Wdr31 | 53.77822627 | -0.513169544 | 0.156336411 |
| Bspry | 7.199256456 | 1.25438916 | 0.406552544 |
| Hdhd3 | 119.7174087 | -0.235981798 | 0.141698381 |
| Alad | 452.599456 | -0.076732543 | 0.084104535 |
| Pole3 | 241.0396122 | 0.481430595 | 0.098184514 |
| 4933430I17Rik | 8.456414667 | 1.129521693 | 0.372493375 |
| Rgs3 | 280.2490358 | 0.153036461 | 0.079433159 |
| Gm11209 | 0.195038611 | -0.199365204 | 3.238350286 |
| Gm11210 | 0.707441185 | 1.793946282 | 1.23622732 |
| Gm11211 | 0.066050875 | 0.71820487 | 3.724945049 |
| Gm24117 | 0 | NA | NA |
| Gm25391 | 0 | NA | NA |
| Gm11481 | 0 | NA | NA |
| Gm11480 | 0 | NA | NA |
| Zfp618 | 184.4983496 | -0.173543275 | 0.107764344 |
| Ambp | 0.720892307 | 0.912470404 | 1.485132957 |
| Kif12 | 0.229693479 | 0.237328807 | 2.141518339 |
| Col27a1 | 179.4777456 | -0.978875654 | 0.198041644 |
| Mir455 | 0.26766946 | -0.729181198 | 2.079953896 |
| Orm1 | 0.137647365 | 0.237309079 | 3.724945049 |
| Gm11212 | 0 | NA | NA |
| Orm3 | 0.157314084 | 0.718207559 | 3.435173412 |
| Orm2 | 0.751984476 | 2.873141615 | 1.183581565 |
| Akna | 219.0544516 | 1.98249173 | 0.117609889 |
| Aknaos | 0.425011668 | 0.680945007 | 1.557369634 |
| Whrn | 563.7172746 | -0.627574033 | 0.083271678 |
| Atp6v1g1 | 617.2868754 | 0.507683647 | 0.113229224 |
| 6330416G13Rik | 190.6195337 | 0.301509895 | 0.116835623 |
| Gm11483 | 0 | NA | NA |

| | | | |
|---|---|---|---|
| Gm24448 | 0 | NA | NA |
| Gm11213 | 0 | NA | NA |
| 1700018C11Rik | 0 | NA | NA |
| Gm24234 | 0 | NA | NA |
| Gm11482 | 1.060662479 | 0.56902497 | 1.030167759 |
| Gm11214 | 0 | NA | NA |
| Tnfsf15 | 4.678869644 | 3.233807932 | 0.685084272 |
| Tnfsf8 | 49.08655836 | 6.253231305 | 0.549014679 |
| Rpl17-ps4 | 3.619380876 | 0.679079104 | 0.478813508 |
| Tnc | 854.476176 | 2.8778134 | 0.319859842 |
| 8030451A03Rik | 1.099630792 | 1.304825844 | 1.026366764 |
| Gm11216 | 0.038981395 | 0.718207094 | 3.724945049 |
| Gm11217 | 0 | NA | NA |
| Gm23950 | 0 | NA | NA |
| Gm11218 | 0.060174537 | 0.237309859 | 3.724945049 |
| Gm11219 | 0 | NA | NA |
| Pappa | 141.7302013 | 1.242488142 | 0.215455232 |
| Astn2 | 1043.79549 | -0.541690177 | 0.086742649 |
| Trim32 | 1762.038928 | -0.303435616 | 0.059885102 |
| Gm11484 | 0 | NA | NA |
| Gm25480 | 0 | NA | NA |
| Gm11220 | 0 | NA | NA |
| Tlr4 | 178.7502272 | 2.701187784 | 0.239358841 |
| Gm11403 | 0 | NA | NA |
| Hmgb1-rs18 | 0 | NA | NA |
| AL589845.1 | 0 | NA | NA |
| Gm11249 | 12.40046496 | 0.340627737 | 0.36052131 |
| Gm11751 | 0.126225412 | 0.718200913 | 3.698754989 |
| Gm23738 | 0 | NA | NA |
| Tcp1-ps1 | 0.143022002 | 0.719176055 | 3.547735711 |
| Gm12911 | 0 | NA | NA |
| Brinp1 | 2163.011251 | -0.095362104 | 0.116873829 |
| Gm24697 | 0 | NA | NA |
| Gm11404 | 1.959659191 | 0.071968194 | 1.045637121 |
| Gm11221 | 11.46002724 | 0.888653358 | 0.348738648 |
| Gm11222 | 0.623048784 | 0.495068709 | 1.25906175 |
| Gm11223 | 93.55249628 | -0.812398589 | 0.159342416 |
| Gm11224 | 5.294292594 | 0.578596539 | 0.447235834 |
| Gm11225 | 0.044191549 | -0.243571806 | 3.724945049 |
| Cdk5rap2 | 668.7817866 | 0.649674043 | 0.098381535 |
| Gm11226 | 21.32553314 | -0.225784851 | 0.261820443 |
| Gm26320 | 0.207279166 | 0.237325733 | 2.634010914 |
| Megf9 | 2380.976153 | -0.652437159 | 0.082973504 |
| Gm11227 | 0 | NA | NA |
| Gm11228 | 0 | NA | NA |
| Gm11229 | 0 | NA | NA |
| Rps18-ps1 | 0.478648982 | 1.980763254 | 1.39706757 |

| | | | |
|---|---|---|---|
| Gm11231 | 0.345321076 | -0.7227525 | 1.748533964 |
| Gm11232 | 0.115000406 | 0.23732098 | 3.724945049 |
| Gm11233 | 0 | NA | NA |
| Gm11234 | 0 | NA | NA |
| Tle1 | 623.9206616 | -0.5361144 | 0.097107607 |
| Gm11250 | 0 | NA | NA |
| C630043F03Rik | 39.86748014 | -0.700017968 | 0.200173743 |
| Gm11235 | 0.263040051 | 0.730531029 | 1.805398239 |
| Aldoart1 | 2.66134452 | -0.587542463 | 0.631038619 |
| Gm5287 | 0 | NA | NA |
| Gm25653 | 0 | NA | NA |
| Gm11251 | 0 | NA | NA |
| Gm25769 | 0 | NA | NA |
| Gm11487 | 0 | NA | NA |
| Gm11486 | 0 | NA | NA |
| Gm11485 | 0.097139547 | 0.718207094 | 3.724945049 |
| Gm11488 | 14.4078922 | -0.3134014 | 0.336488091 |
| Gm24170 | 0 | NA | NA |
| Gm11238 | 0 | NA | NA |
| Gm11237 | 0 | NA | NA |
| Gm11236 | 0 | NA | NA |
| Gm11239 | 0 | NA | NA |
| Gm428 | 0 | NA | NA |
| Rasef | 2.980195284 | -0.138869691 | 0.634259435 |
| Gm11240 | 0.068097359 | -0.243569582 | 3.724945049 |
| Gm13867 | 0.251494677 | 0.718217178 | 2.510715589 |
| Gm11758 | 0 | NA | NA |
| Gm11757 | 0 | NA | NA |
| Gm13871 | 0 | NA | NA |
| Gm11756 | 0 | NA | NA |
| 2310002L09Rik | 0.092493346 | -0.24357403 | 3.724945049 |
| Frmd3 | 144.1270821 | -0.381176342 | 0.125236206 |
| Gm11760 | 4.795815343 | -0.965941501 | 0.537289589 |
| Gm11405 | 10.60243355 | 0.323983109 | 0.359409436 |
| Kdm4c | 648.3243287 | -0.144504119 | 0.085991228 |
| A230083N12Rik | 2.600875038 | -0.442890229 | 0.745378997 |
| Gm26379 | 0 | NA | NA |
| Gm11254 | 0.066050875 | 0.71820487 | 3.724945049 |
| Gm11258 | 0 | NA | NA |
| Gm11253 | 0 | NA | NA |
| Gm11257 | 0 | NA | NA |
| Gm12912 | 0.235437891 | -0.228806217 | 2.562082607 |
| Tmem261 | 148.8567893 | -0.01334577 | 0.113777583 |
| Gm22864 | 0 | NA | NA |
| Gm11255 | 0 | NA | NA |
| Gm11259 | 0 | NA | NA |
| Gm11256 | 0 | NA | NA |

| | | | |
|---|---|---|---|
| Gm11241 | 0.044544648 | 0.237309859 | 3.724945049 |
| Ptprd | 4811.876378 | -0.991785432 | 0.099801641 |
| Gm11252 | 0 | NA | NA |
| Gm11242 | 0.10503227 | 1.179809211 | 3.723500796 |
| Gm11243 | 0 | NA | NA |
| Gm11244 | 7.743757494 | -1.406465062 | 0.413858849 |
| Gm11245 | 0 | NA | NA |
| Gm11246 | 0.158325629 | 1.199992869 | 2.813062165 |
| Gm23159 | 10.5795326 | -1.347204836 | 0.432850283 |
| B230208B08Rik | 26.21483297 | -1.036882446 | 0.274461048 |
| Gm11260 | 0.431719086 | -0.225072208 | 1.479239473 |
| Gm11261 | 0.068097359 | -0.243569582 | 3.724945049 |
| Gm22262 | 0 | NA | NA |
| Gm11262 | 0.136963818 | 0.718204821 | 3.59954433 |
| Gm25836 | 0 | NA | NA |
| Gm11263 | 32.27155379 | 0.661709784 | 0.235710827 |
| Gm11409 | 0 | NA | NA |
| Gm11407 | 1.154096829 | -3.183029393 | 1.012948073 |
| Gm11408 | 0.245813264 | 0.71822219 | 2.073589709 |
| Gm11410 | 0.40441216 | 0.669931977 | 1.674209972 |
| Gm11406 | 0 | NA | NA |
| Tyrp1 | 7.568395809 | 1.019389399 | 0.443015175 |
| Lurap1l | 131.167045 | -0.475055117 | 0.115941624 |
| Gm27452 | 1.818514136 | -0.854136833 | 0.916187048 |
| Mpdz | 900.4834607 | -0.156759728 | 0.073026845 |
| Gm25803 | 3.471866242 | -0.757142117 | 0.537119588 |
| Gm11765 | 0 | NA | NA |
| Gm11411 | 0.049553047 | 0.71820487 | 3.724945049 |
| Gm11412 | 0.477185738 | -1.340585445 | 1.294774511 |
| n-R5s187 | 0 | NA | NA |
| Gm12914 | 0 | NA | NA |
| Gm11264 | 0.922800452 | -0.070078897 | 1.143269097 |
| Gm22402 | 0 | NA | NA |
| Gm5860 | 3.25720724 | -0.492658619 | 0.575104066 |
| Nfib | 2628.924338 | -0.418930417 | 0.097971327 |
| n-R5s188 | 0 | NA | NA |
| Gm11266 | 18.26086893 | -0.951851322 | 0.276762729 |
| Gm11267 | 0.708875568 | 0.138039521 | 1.093690802 |
| Gm11268 | 0.189675868 | 0.237327798 | 2.656640733 |
| Gm11269 | 0.044544648 | 0.237309859 | 3.724945049 |
| Zdhhc21 | 1208.957435 | -0.256667592 | 0.052657976 |
| Cer1 | 0.481690985 | -1.93648002 | 1.390588077 |
| Frem1 | 91.57009446 | -0.902532105 | 0.176674323 |
| Gm24884 | 0 | NA | NA |
| Gm11248 | 0 | NA | NA |
| Gm11185 | 2.941272167 | -0.274982014 | 0.65761929 |
| Ttc39b | 1497.137622 | 0.485736183 | 0.099501631 |

| | | | |
|---|---|---|---|
| Gm23412 | 0.392867355 | 2.050304495 | 1.922916048 |
| Gm11413 | 1.891350131 | -0.304007177 | 0.864192681 |
| Gm11184 | 0 | NA | NA |
| Snapc3 | 534.9796752 | -0.167089695 | 0.098043073 |
| Psip1 | 2060.877521 | -0.137274948 | 0.076586697 |
| Gm11414 | 0.092493346 | -0.24357403 | 3.724945049 |
| Gm10154 | 11.65614726 | 0.845585547 | 0.498372565 |
| Ccdc171 | 44.96192503 | -0.040055552 | 0.174713342 |
| Gm25899 | 0 | NA | NA |
| Gm27046 | 0.323557773 | 0.782382288 | 1.730895835 |
| Cbx3-ps3 | 0.066050875 | 0.71820487 | 3.724945049 |
| Gm26968 | 0.264809544 | -0.712178301 | 2.093082436 |
| Gm11415 | 0 | NA | NA |
| 6030471H07Rik | 0 | NA | NA |
| Gm12416 | 6.021123113 | 0.922097602 | 0.41877442 |
| Gm12414 | 0.092269992 | 0.237318756 | 3.724945049 |
| Bnc2 | 94.00517019 | -0.089108039 | 0.160982406 |
| Gm12421 | 0.473868817 | -0.721725804 | 1.463228218 |
| Gm12420 | 0 | NA | NA |
| Gm12415 | 0.047948698 | -0.243569582 | 3.724945049 |
| Cntln | 269.0535505 | -0.548362187 | 0.091632288 |
| Gm12418 | 9.82225973 | -0.211146647 | 0.306041079 |
| Gm12419 | 0.575593679 | 0.737138111 | 1.179801871 |
| Gm12417 | 0 | NA | NA |
| Sh3gl2 | 2324.421738 | -0.49116062 | 0.059780261 |
| Gm12413 | 0 | NA | NA |
| Gm12680 | 8.074609009 | -0.083731888 | 0.442399554 |
| Gm26069 | 1.027412341 | -0.67680768 | 0.956496577 |
| Gm25811 | 0 | NA | NA |
| Adamtsl1 | 125.2776745 | 0.580775785 | 0.136385781 |
| Gm12550 | 0.075043205 | 0.237309859 | 3.724945049 |
| Saxo1 | 0.416337256 | -1.323024001 | 1.714813016 |
| Gm26017 | 0 | NA | NA |
| Saxo1os | 0 | NA | NA |
| Rraga | 797.0060883 | 0.140720374 | 0.086240385 |
| Haus6 | 193.2480707 | 0.054517314 | 0.087909595 |
| Scarna8 | 0.537445765 | -1.882430918 | 1.241188883 |
| Gm12551 | 0.925310778 | 3.826465969 | 1.258352321 |
| Plin2 | 668.7166373 | 4.655041892 | 0.354541717 |
| Dennd4c | 657.3655427 | 0.586977866 | 0.072446115 |
| Gm12630 | 2.006602685 | 1.321262235 | 1.141456802 |
| Rps6 | 1228.330895 | 0.715002111 | 0.069191276 |
| Acer2 | 61.2692064 | 0.610571927 | 0.19757813 |
| Slc24a2 | 8155.610505 | -0.921090796 | 0.103684427 |
| Gm12600 | 0.561663198 | 0.327936807 | 1.203647633 |
| Gm23154 | 0 | NA | NA |
| Gm12604 | 0.075043205 | 0.237309859 | 3.724945049 |

| | | | |
|---|---|---|---|
| Mllt3 | 907.8146563 | 0.066000729 | 0.07159747 |
| Gm12631 | 3.345683444 | 0.055849829 | 0.54545911 |
| Gm12646 | 0.591515762 | 1.195706381 | 1.394099368 |
| Focad | 1053.950382 | -0.538938531 | 0.108970104 |
| Mir491 | 10.7430229 | -1.061278785 | 0.336044631 |
| Gm12645 | 3.761156174 | -0.473158675 | 0.547976514 |
| Hacd4 | 63.03965297 | 1.596169556 | 0.225220673 |
| Ifnb1 | 44.66037375 | 9.007097746 | 1.213913409 |
| Ifna15 | 0.599468885 | 2.019964945 | 1.546875122 |
| Ifna14 | 0.481692262 | 2.230878659 | 1.645810735 |
| Ifna-ps1 | 0 | NA | NA |
| Ifna9 | 0.316561663 | 2.484428409 | 1.8862611 |
| Mrpl48-ps | 38.64384611 | 0.12945886 | 0.244655349 |
| Ifna12 | 0.176391548 | -0.243570398 | 2.718652661 |
| Gm12601 | 0.215725693 | 0.096368704 | 2.175168995 |
| C87499 | 0.779996687 | 1.882630844 | 1.192041738 |
| Ifna13 | 0.075043205 | 0.237309859 | 3.724945049 |
| Gm13281 | 0.046345709 | 0.71820487 | 3.724945049 |
| Ifna16 | 0.145446656 | 1.165556206 | 2.878908292 |
| Ifna2 | 8.251847137 | 5.514882829 | 1.067698984 |
| Ifnab | 1.397532748 | 3.769977394 | 1.057674173 |
| Gm13273 | 0.313664056 | 1.216494719 | 3.720482438 |
| Gm13282 | 2.459004477 | -0.296719843 | 0.782116984 |
| Klhl9 | 1871.695579 | -0.051441475 | 0.087355876 |
| Gm26566 | 4.232091902 | 0.053113296 | 0.491649681 |
| Gm13274 | 0.044191549 | -0.243571806 | 3.724945049 |
| Gm13271 | 0.08486385 | 0.717551239 | 3.724898127 |
| Gm16686 | 0 | NA | NA |
| Gm13286 | 0 | NA | NA |
| Gm10583 | 0.144244773 | -0.243562779 | 3.581436307 |
| Gm13283 | 0 | NA | NA |
| Gm10582 | 0 | NA | NA |
| Gm13284 | 0 | NA | NA |
| Gm10581 | 0 | NA | NA |
| Gm13290 | 0 | NA | NA |
| Gm10580 | 0 | NA | NA |
| Gm13289 | 0 | NA | NA |
| Gm13272 | 0 | NA | NA |
| Ifnz | 0 | NA | NA |
| Gm26525 | 0.092269992 | 0.237318756 | 3.724945049 |
| Gm13276 | 0 | NA | NA |
| Gm13277 | 0 | NA | NA |
| Gm13278 | 0 | NA | NA |
| Gm13275 | 0 | NA | NA |
| Gm26867 | 0.045882455 | 0.237312084 | 3.724945049 |
| Gm13279 | 0 | NA | NA |
| Gm13285 | 0 | NA | NA |

| | | | |
|---|---|---|---|
| Gm13287 | 0 | NA | NA |
| Gm13288 | 0 | NA | NA |
| Ifna7 | 0.305381988 | 1.201083703 | 2.442844655 |
| Ifna11 | 0.275757984 | 0.726495856 | 3.724360356 |
| Ifna6 | 0.418480079 | 0.718190415 | 3.724945049 |
| Ifna5 | 11.73825278 | 6.928563929 | 1.277956129 |
| Ifna4 | 14.65900129 | 6.924793756 | 1.402434863 |
| Ifna1 | 2.156732396 | 4.247693153 | 1.314725415 |
| 4930553M12Rik | 0.266535354 | 0.768954872 | 2.105555535 |
| Ifne | 0.083526043 | 0.718202646 | 3.724945049 |
| Mir31 | 1.252737907 | 0.148191576 | 1.037492826 |
| Gm12603 | 0.048949532 | -0.243569582 | 3.724945049 |
| Gm12602 | 0 | NA | NA |
| Mtap | 151.2844411 | 0.347016032 | 0.148815308 |
| Gm12607 | 0 | NA | NA |
| Gm26490 | 0 | NA | NA |
| Gm12606 | 0.046345709 | 0.71820487 | 3.724945049 |
| Cdkn2a | 2.014914513 | 3.610328749 | 1.012882193 |
| Cdkn2b | 6.964193097 | 2.083974887 | 0.536465524 |
| Gm12609 | 0.942397274 | 0.712895907 | 1.146808853 |
| Gm12610 | 0.811625864 | 1.627408014 | 1.325798315 |
| Gm12608 | 0 | NA | NA |
| Gm27580 | 0.090427103 | 0.237307635 | 3.724945049 |
| Dmrta1 | 4.432480129 | -0.836586364 | 0.49136025 |
| Gm12629 | 0 | NA | NA |
| Zfp352 | 0 | NA | NA |
| Gm12633 | 0.176993581 | 0.220699569 | 2.297185952 |
| Gm12634 | 0 | NA | NA |
| Gm12635 | 0 | NA | NA |
| Gm12636 | 0 | NA | NA |
| Gm12632 | 0 | NA | NA |
| Gm25244 | 0.236157963 | 0.237320405 | 2.511414526 |
| Gm12643 | 0 | NA | NA |
| Gm12644 | 0.116217011 | -0.24357403 | 3.724945049 |
| Elavl2 | 1479.215199 | -0.024747347 | 0.122441157 |
| Gm12653 | 0 | NA | NA |
| Mir6402 | 6.56355345 | -0.582305944 | 0.441261496 |
| Gm12668 | 0.048949532 | -0.243569582 | 3.724945049 |
| Gm12670 | 0.219222703 | 0.237328821 | 2.177977162 |
| Gm12667 | 0.171083144 | 1.76201309 | 3.383974148 |
| Gm12669 | 20.03569431 | 0.459532193 | 0.229480034 |
| Gm12671 | 13.22935111 | 0.370686951 | 0.284032999 |
| 4930577H14Rik | 0 | NA | NA |
| Izumo3 | 0 | NA | NA |
| Gm12666 | 1.628724105 | -0.739219075 | 0.93819077 |
| Gm12638 | 0 | NA | NA |
| Gm12637 | 0 | NA | NA |

| | | | |
|---|---|---|---|
| Gm22605 | 0 | NA | NA |
| Gm12639 | 0 | NA | NA |
| Gm12641 | 0 | NA | NA |
| Gm12642 | 0 | NA | NA |
| Tusc1 | 50.66388116 | 0.153807653 | 0.169164316 |
| Gm12649 | 0.203196539 | 0.718218497 | 2.595971176 |
| Gm23443 | 0 | NA | NA |
| Gm12640 | 0 | NA | NA |
| Gm24796 | 0 | NA | NA |
| Gm22619 | 0 | NA | NA |
| Gm12650 | 0 | NA | NA |
| Gm12647 | 0 | NA | NA |
| Gm12648 | 0.255640523 | 0.703139007 | 2.106863157 |
| Gm23836 | 0 | NA | NA |
| Gm12654 | 0 | NA | NA |
| Gm26410 | 0.038981395 | 0.718207094 | 3.724945049 |
| Gm12655 | 23.30350443 | 0.03810068 | 0.222916266 |
| Gm22804 | 0 | NA | NA |
| Gm12656 | 0 | NA | NA |
| Caap1 | 152.6588818 | -0.05575091 | 0.114869015 |
| Gm10306 | 0.49643671 | -1.856755059 | 1.431072529 |
| Plaa | 900.5907788 | 0.539571205 | 0.08435123 |
| Gm12657 | 14.22801902 | 0.181182156 | 0.287575791 |
| Ift74 | 232.3827732 | 0.173987083 | 0.089954881 |
| Lrrc19 | 0.551940858 | 2.537798693 | 1.462049155 |
| Mir872 | 1.137124893 | -0.481185809 | 0.968956762 |
| Tek | 47.11216226 | 0.810774389 | 0.202730772 |
| Eqtn | 11.24146221 | 0.625466798 | 0.377760342 |
| Gm12693 | 29.98890503 | 0.104111708 | 0.192623422 |
| Mysm1 | 851.8413644 | 0.249074448 | 0.085080581 |
| Gm12694 | 0.203088869 | 0.237324761 | 2.638249046 |
| Gm12703 | 1.47600346 | 1.205720051 | 0.875766291 |
| Gm12704 | 0.126912322 | 0.555057727 | 3.69382869 |
| Jun | 1515.464005 | 2.048003501 | 0.231585366 |
| Junos | 29.43953063 | 0.36270008 | 0.228182858 |
| n-R5s189 | 0.792393139 | -1.155998875 | 1.161387695 |
| 4930551L18Rik | 0.046345709 | 0.71820487 | 3.724945049 |
| Gm28096 | 1.499901305 | -1.226609705 | 0.922162593 |
| Gm12708 | 1.926663799 | 2.412969884 | 0.839981864 |
| Gm29064 | 0 | NA | NA |
| Gm830 | 0.19500349 | 0.267447742 | 3.288425011 |
| Gm12707 | 0 | NA | NA |
| Gm25004 | 0 | NA | NA |
| Fggy | 21.91263025 | -0.482885403 | 0.27455671 |
| Gm20651 | 0 | NA | NA |
| 9530080O11Rik | 18.68130564 | -0.041209561 | 0.244466931 |
| Hook1 | 1088.16238 | -0.358579712 | 0.111343001 |

| | | | |
|---|---|---|---|
| Cyp2j13 | 7.229154167 | -0.36253671 | 0.420765272 |
| Hspe1-ps6 | 0.042716945 | -0.243571806 | 3.724945049 |
| Cyp2j12 | 25.97268065 | -0.155113982 | 0.211316392 |
| Cyp2j7-ps3 | 0 | NA | NA |
| Cyp2j7-ps2 | 0 | NA | NA |
| Cyp2j7-ps1 | 0 | NA | NA |
| Cyp2j7 | 0.068097359 | -0.243569582 | 3.724945049 |
| Cyp2j15-ps | 0 | NA | NA |
| Cyp2j11 | 0.27158854 | 0.68358377 | 2.070242552 |
| Cyp2j14-ps | 0 | NA | NA |
| Cyp2j8 | 2.219411473 | -0.228399401 | 0.717324172 |
| Cyp2j6 | 245.6472783 | 0.105016321 | 0.130795596 |
| Cyp2j9 | 418.0043846 | 0.126785323 | 0.149764235 |
| Cyp2j5 | 0.490888186 | 1.091377452 | 1.656509381 |
| Gm12695 | 0.642811059 | -1.434297339 | 1.167246761 |
| Gm27579 | 0 | NA | NA |
| Gm27521 | 0 | NA | NA |
| Gm10192 | 0 | NA | NA |
| Gm12696 | 28.72011168 | 0.157856442 | 0.200206317 |
| Gm12697 | 3.534004104 | -1.737770245 | 0.756875167 |
| E130114P18Rik | 46.0958998 | -0.199300823 | 0.172587317 |
| Gm12676 | 1.650824257 | -0.402774932 | 1.176680848 |
| Nfia | 1906.233348 | -0.015309877 | 0.082201646 |
| Gm12691 | 1.556603733 | -0.485069521 | 0.815614411 |
| Gm12692 | 0.373707575 | -1.982531008 | 1.50959638 |
| Gm12787 | 0 | NA | NA |
| Gm24203 | 0 | NA | NA |
| 0610025J13Rik | 0.422542347 | -0.219575826 | 1.448759136 |
| Gm12788 | 0 | NA | NA |
| Tm2d1 | 205.9564824 | 0.005798361 | 0.10557866 |
| Gm24960 | 0 | NA | NA |
| Inadl | 702.0612203 | -0.322872798 | 0.128363157 |
| Gm22625 | 0.042716945 | -0.243571806 | 3.724945049 |
| Gm26154 | 4.026424441 | -0.212591121 | 0.607608832 |
| Gm12846 | 13.02039719 | -1.335262184 | 0.382677931 |
| L1td1 | 1.260641324 | -0.494338163 | 0.855748061 |
| Gm17376 | 0.049553047 | 0.71820487 | 3.724945049 |
| Kank4 | 103.4907107 | -0.336691449 | 0.162120004 |
| Kank4os | 0.479560471 | 0.606926417 | 1.708549056 |
| I0C0044D17Rik | 0.072025462 | 0.237312084 | 3.724945049 |
| Gm12851 | 0 | NA | NA |
| Usp1 | 392.4990393 | 0.644580768 | 0.103618283 |
| Dock7 | 964.8693113 | -0.160061768 | 0.082594496 |
| Angptl3 | 2.293039322 | 1.81226829 | 0.780972919 |
| Gm12852 | 6.635996975 | -1.294155169 | 0.550409102 |
| Gm12705 | 0.112642008 | -0.24357403 | 3.724945049 |
| Gm12681 | 0.364815146 | 1.54993673 | 1.583868392 |

| | | | |
|---|---|---|---|
| Atg4c | 539.9595983 | -0.52388771 | 0.088555703 |
| Gm10305 | 27.16950442 | -0.86533091 | 0.284187061 |
| Gm12689 | 9.012307495 | -0.48834781 | 0.433289072 |
| Gm37556 | 0 | NA | NA |
| Gm12685 | 0 | NA | NA |
| Gm12686 | 0 | NA | NA |
| Gm12684 | 0 | NA | NA |
| Gm12683 | 0 | NA | NA |
| Gm37117 | 0 | NA | NA |
| Gm12690 | 0 | NA | NA |
| Gm12687 | 0 | NA | NA |
| Gm12688 | 2.558789337 | -0.289454582 | 0.805351027 |
| Foxd3 | 16.0637295 | 1.784178379 | 1.061954342 |
| Gm23366 | 0 | NA | NA |
| Gm12682 | 0 | NA | NA |
| Alg6 | 154.9068683 | -0.431626255 | 0.159739397 |
| Itgb3bp | 53.12261853 | 0.571327895 | 0.164204751 |
| Efcab7 | 42.69675528 | 0.929095841 | 0.187096564 |
| Gm26425 | 0.198326984 | 0.237328063 | 2.234447446 |
| Pgm2 | 636.1507098 | -0.122523114 | 0.082438012 |
| Gm12702 | 0 | NA | NA |
| Ror1 | 118.724752 | -0.13503101 | 0.153774499 |
| Gm12701 | 11.03703998 | -1.619890086 | 0.435219219 |
| Gm12706 | 0.200296414 | 0.237329899 | 2.216693753 |
| Ube2u | 0.310183902 | 0.688928987 | 1.77269257 |
| Ube2uos | 0 | NA | NA |
| Ccdc50-ps | 0.574153598 | -1.709614329 | 1.287722216 |
| Gm12700 | 0 | NA | NA |
| Cachd1 | 422.3160165 | 0.112353322 | 0.074897871 |
| Gm10577 | 0 | NA | NA |
| Gm10576 | 0 | NA | NA |
| Raver2 | 272.6356933 | -0.964871073 | 0.188684599 |
| Gm22100 | 0.061920613 | 0.718207094 | 3.724945049 |
| Jak1 | 2519.077659 | 0.409084087 | 0.062908398 |
| Gm24468 | 0.165470308 | 0.237318778 | 2.761559324 |
| Gm12785 | 2.331474832 | -0.337752038 | 0.703966862 |
| Gm25124 | 0 | NA | NA |
| Gm12801 | 0 | NA | NA |
| Gm12796 | 0.250633908 | 0.691603379 | 1.832681367 |
| Gm12793 | 0.066050875 | 0.71820487 | 3.724945049 |
| E130102H24Rik | 106.6384263 | 0.372385929 | 0.212748728 |
| Mir101a | 0 | NA | NA |
| 0610043K17Rik | 25.29227341 | -0.080868605 | 0.26392824 |
| Gm12795 | 0 | NA | NA |
| Gm12798 | 2.114628594 | -0.567832836 | 0.792615669 |
| Ak4 | 432.3296171 | -0.74822673 | 0.147471412 |
| Dnajc6 | 4085.768198 | -0.509121914 | 0.057777828 |

| | | | |
|---|---|---|---|
| Gm22533 | 11.8805187 | -1.597577157 | 0.390173347 |
| Leprot | 268.8974141 | 0.648713454 | 0.1161944 |
| Lepr | 95.56977508 | 0.714767507 | 0.144136119 |
| B020004J07Rik | 0.102891481 | -0.243576254 | 3.724945049 |
| Gm12791 | 3.064345978 | 0.307823047 | 0.550845961 |
| Gm12797 | 0.048949532 | -0.243569582 | 3.724945049 |
| C130073F10Rik | 0 | NA | NA |
| Gm12792 | 0 | NA | NA |
| Gm12800 | 0.092691418 | 1.183079489 | 3.724560917 |
| Gm12794 | 0.223364959 | 1.199942 | 2.521503358 |
| Gm12790 | 0.047948698 | -0.243569582 | 3.724945049 |
| Gm12789 | 0.194279094 | 1.292225235 | 3.719439735 |
| Gm12799 | 4.949076095 | -0.106616742 | 0.430190663 |
| Pde4b | 1759.695653 | 0.367633005 | 0.094323932 |
| Gm24869 | 0 | NA | NA |
| Sgip1 | 2873.427705 | -0.583772733 | 0.075978305 |
| Gm12709 | 0.447471404 | -1.401491033 | 1.555409019 |
| Tctex1d1 | 5.63377873 | 0.45236351 | 0.489343628 |
| BB031773 | 3.502581246 | -3.369575408 | 0.909226217 |
| Insl5 | 5.684128039 | -0.537408374 | 0.444934701 |
| Wdr78 | 212.7570655 | -0.261311568 | 0.159779191 |
| Gm12710 | 0.045882455 | 0.237312084 | 3.724945049 |
| Mier1 | 957.8111048 | 0.244829049 | 0.059704551 |
| Slc35d1 | 196.800066 | -0.227270125 | 0.103461517 |
| 4921539E11Rik | 0.165933562 | 0.718213875 | 2.765757588 |
| 4930456L15Rik | 0.038981395 | 0.718207094 | 3.724945049 |
| Oma1 | 95.40654656 | 0.4146551 | 0.171088616 |
| Gm12718 | 0.250798881 | 0.725940307 | 2.082776451 |
| Gm23064 | 0 | NA | NA |
| Gm12720 | 0.060174537 | 0.237309859 | 3.724945049 |
| Gm12716 | 0 | NA | NA |
| Gm12715 | 393.1600641 | 0.664387044 | 0.107493227 |
| Gm22801 | 0 | NA | NA |
| Gm25877 | 0 | NA | NA |
| Dab1 | 1235.718576 | -0.576370553 | 0.078069881 |
| Gm12719 | 0 | NA | NA |
| Gm12717 | 3.781002861 | -2.601737074 | 0.763368007 |
| C8b | 6.711931076 | 0.330454335 | 1.036560222 |
| C8a | 0.626920446 | 1.174862345 | 1.174310346 |
| Gm17662 | 0.225129614 | 0.237300054 | 3.724945049 |
| 1700024P16Rik | 18.7794585 | -0.893434828 | 0.350797342 |
| Gm12721 | 0.110242424 | 0.237318756 | 3.724945049 |
| Prkaa2 | 803.30275 | -0.578373176 | 0.051311902 |
| Ppap2b | 1987.101825 | -0.628156944 | 0.131593087 |
| Gm12722 | 2.944400151 | 0.306699131 | 0.600966221 |
| Gm12723 | 0 | NA | NA |
| Gm12726 | 0.106112162 | 0.23732098 | 3.724945049 |

| | | | |
|---|---|---|---|
| Gm12728 | 0.174579622 | -0.717341564 | 2.306965692 |
| Gm24363 | 0 | NA | NA |
| Gm12729 | 0.086930093 | 0.236669296 | 3.724898107 |
| Gm12727 | 0.640223289 | 1.240410844 | 1.35654814 |
| Gm26047 | 0 | NA | NA |
| Hspd1-ps4 | 0.60453213 | 2.0007945 | 1.384170158 |
| Gm12724 | 0.146823232 | -0.712492187 | 2.876272087 |
| Gm12731 | 0 | NA | NA |
| Gm12730 | 0.072025462 | 0.237312084 | 3.724945049 |
| Gm37253 | 0 | NA | NA |
| Usp24 | 2364.206895 | 0.270925716 | 0.075401537 |
| Pcsk9 | 22.35786243 | -1.554612495 | 0.279495284 |
| Bsnd | 0.044191549 | -0.243571806 | 3.724945049 |
| Tmem61 | 0 | NA | NA |
| 2700068H02Rik | 1.22619721 | -0.8457048 | 1.123674875 |
| Dhcr24 | 1131.153985 | -0.260040987 | 0.113449308 |
| Gm12744 | 0.280446515 | -0.651375049 | 1.797077456 |
| BC055111 | 2.227745575 | 0.320909855 | 0.67074648 |
| Ttc22 | 5.229776616 | -0.621284598 | 0.427719912 |
| Pars2 | 65.0940951 | 0.031289059 | 0.14764394 |
| Ttc4 | 363.5749445 | -0.072169468 | 0.060659745 |
| Mroh7 | 34.09442817 | -1.662682069 | 0.312121625 |
| Fam151a | 2.097884303 | 1.408149462 | 0.729566479 |
| Acot11 | 491.6282229 | -0.779142603 | 0.146489875 |
| Gm12745 | 0 | NA | NA |
| Gm12746 | 0.048949532 | -0.243569582 | 3.724945049 |
| Ssbp3 | 1314.52194 | -0.337080646 | 0.099555859 |
| Gm12786 | 0.08486385 | 0.717551239 | 3.724898127 |
| Mrpl37 | 409.2304581 | 0.042938969 | 0.078999151 |
| Cyb5rl | 41.42110935 | -0.304999939 | 0.182118671 |
| Cdcp2 | 0.295634611 | 1.284610721 | 2.042973107 |
| Gm12802 | 0.337683083 | 1.138815782 | 1.723125619 |
| Tceanc2 | 245.3997247 | 0.267144959 | 0.095598381 |
| Tmem59 | 1671.351481 | 0.287042773 | 0.073515423 |
| Ldlrad1 | 1.155427534 | 0.606538223 | 1.064278653 |
| Lrrc42 | 182.0305453 | 0.154553464 | 0.108821947 |
| Hspb11 | 48.05584592 | 0.162615267 | 0.147760696 |
| Gm12850 | 2.64217102 | -1.466120105 | 0.683021813 |
| Dio1 | 0.200957741 | 1.97380807 | 3.229647234 |
| Yipf1 | 215.0974917 | 0.048417098 | 0.081814675 |
| Gm12870 | 0.418005171 | -0.623676268 | 1.428023086 |
| Gm12869 | 0 | NA | NA |
| Ndc1 | 286.6300775 | -0.119624214 | 0.083734043 |
| Gm24804 | 0 | NA | NA |
| Glis1 | 17.53901492 | 0.649624595 | 0.288267798 |
| Dmrtb1 | 26.37534565 | 1.091410857 | 0.236291977 |
| Gm12899 | 0 | NA | NA |

| | | | |
|---|---|---|---|
| 1700047F07Rik | 0.364322736 | -1.792322603 | 1.652447643 |
| Gm25123 | 0 | NA | NA |
| Gm12906 | 0.718440912 | 2.024418636 | 1.343427271 |
| Gm12907 | 0 | NA | NA |
| 4933424M12Rik | 0.383698231 | 1.718198569 | 1.694097783 |
| Lrp8os3 | 15.16761459 | -0.637390243 | 0.319018662 |
| Lrp8 | 1123.019012 | -0.100790688 | 0.097924105 |
| Lrp8os2 | 23.3868331 | -0.855366937 | 0.2470455 |
| Lrp8os1 | 0.049553047 | 0.71820487 | 3.724945049 |
| Magoh | 160.5650247 | 0.825808932 | 0.143984184 |
| 0610037L13Rik | 286.4495153 | -0.209943089 | 0.086648045 |
| Cpt2 | 89.15985385 | 0.649600036 | 0.191699563 |
| 4930407G08Rik | 0.217432726 | -1.22539114 | 2.167380711 |
| Slc1a7 | 1.139422845 | 0.869606648 | 0.985224757 |
| Podn | 28.84068978 | -0.777822083 | 0.286666772 |
| Gm23354 | 0.487093236 | -0.885325576 | 1.452756241 |
| Scp2 | 720.4369207 | -0.086664887 | 0.083810499 |
| Mir6397 | 0 | NA | NA |
| Echdc2 | 50.39875141 | -1.035252149 | 0.201305553 |
| Zyg11a | 0.650364221 | 1.206955372 | 1.34735783 |
| Zyg11b | 3195.301796 | -0.518273752 | 0.047231532 |
| Gm12742 | 0 | NA | NA |
| Gm12740 | 0 | NA | NA |
| Coa7 | 154.9104418 | -0.246535718 | 0.11467131 |
| Fam159a | 1.112155548 | -0.4308671 | 1.082913523 |
| Gpx7 | 31.9655395 | 0.84587958 | 0.208223367 |
| Gm12738 | 0.049553047 | 0.71820487 | 3.724945049 |
| Zcchc11 | 1174.50692 | 0.062426277 | 0.094968619 |
| Gm24967 | 6.345655539 | 0.094319914 | 0.438465781 |
| Gm12739 | 0.090277459 | 0.237318756 | 3.724945049 |
| Gm23138 | 4.286477608 | -0.902213569 | 0.61895548 |
| Gm12737 | 23.64047355 | -0.806905665 | 0.363936409 |
| Gm25827 | 0.189729909 | 0.742577392 | 2.26039112 |
| Gm22345 | 0.183160349 | 0.662407564 | 2.259866273 |
| Prpf38a | 397.5696076 | 0.863430788 | 0.095321134 |
| Orc1 | 6.089138634 | 2.638262529 | 0.549145037 |
| Gm12743 | 18.25140362 | -0.283629972 | 0.294359181 |
| Cc2d1b | 307.3753207 | 0.317708731 | 0.105277587 |
| Gm20731 | 6.967362313 | -0.374808007 | 0.435125821 |
| Gm17354 | 78.27363649 | -0.70608643 | 0.191813463 |
| Zfyve9 | 2056.889909 | -0.164630501 | 0.068593893 |
| 3110021N24Rik | 19.00627441 | -0.965341165 | 0.266995557 |
| Gm12741 | 0 | NA | NA |
| Btf3l4 | 356.4128181 | -0.199561148 | 0.086609246 |
| Txndc12 | 312.2245455 | 0.638378854 | 0.120251181 |
| A730015C16Rik | 2.046195117 | 0.304994635 | 0.713528051 |
| Kti12 | 133.8561621 | 0.167468163 | 0.102102234 |

| | | | |
|---|---|---|---|
| Rab3b | 288.1114516 | -0.784721196 | 0.142759384 |
| Gm12736 | 0 | NA | NA |
| 8030443G20Rik | 0.272094973 | 0.68830177 | 2.080499469 |
| Gm23589 | 0.08486385 | 0.717551239 | 3.724898127 |
| Nrd1 | 1613.530245 | -0.079632528 | 0.05286389 |
| Mir761 | 0.40746677 | -2.043979392 | 1.3973592 |
| Osbpl9 | 801.6924287 | 0.079521795 | 0.068229172 |
| Gm23743 | 1.374639076 | -1.569489761 | 0.844368857 |
| Calr4 | 3.300543358 | -0.854105617 | 0.512294729 |
| Eps15 | 2866.84323 | -0.458598098 | 0.092062555 |
| Gm12749 | 6.563129853 | 0.625533249 | 0.441521998 |
| Ttc39aos1 | 29.58635856 | -0.516273949 | 0.243349166 |
| Ttc39a | 61.40979687 | -0.690465424 | 0.185314011 |
| Rnf11 | 2222.627844 | -0.146567648 | 0.08425488 |
| 4930522H14Rik | 0 | NA | NA |
| Gm22497 | 0 | NA | NA |
| Gm12811 | 0.075043205 | 0.237309859 | 3.724945049 |
| Gm12803 | 0 | NA | NA |
| 9630013D21Rik | 4.08627299 | -0.72491748 | 0.601724046 |
| Cdkn2c | 32.18597743 | 0.807792693 | 0.284173785 |
| Faf1 | 667.4892566 | 0.162603346 | 0.070388923 |
| Gm12808 | 0 | NA | NA |
| Dmrta2os | 0.681178858 | -1.267455458 | 1.23011319 |
| Dmrta2 | 37.20715882 | 0.755530404 | 0.202995607 |
| Elavl4 | 1395.565762 | -0.144967354 | 0.079316258 |
| Agbl4 | 173.4577221 | -1.071180032 | 0.170486406 |
| Gm12806 | 4.671720804 | -0.781809497 | 0.528256245 |
| Gm12807 | 1.537250626 | 0.010391983 | 0.805890259 |
| Gm12804 | 7.93214406 | -0.651382257 | 0.408655388 |
| Gm12805 | 2.823428998 | 2.987818098 | 0.963802758 |
| Bend5 | 101.6608683 | -0.06541332 | 0.172961549 |
| n-R5s191 | 0 | NA | NA |
| Spata6 | 105.6804718 | 1.364108567 | 0.140406213 |
| Gm12961 | 0.049553047 | 0.71820487 | 3.724945049 |
| Gm12960 | 11.3450459 | 0.156356942 | 0.386921602 |
| Slc5a9 | 0.45131048 | 2.802789326 | 1.626100148 |
| Skint8 | 0.165933562 | 0.718213875 | 2.765757588 |
| Skint7 | 0.891009806 | 0.521517005 | 1.15008639 |
| Gm25844 | 0 | NA | NA |
| Skint1 | 0.404673549 | 1.966604447 | 1.522426561 |
| Skint4 | 0 | NA | NA |
| Gm12820 | 0 | NA | NA |
| Gm12815 | 0 | NA | NA |
| Skint3 | 0 | NA | NA |
| Skint9 | 0 | NA | NA |
| Gm12819 | 0 | NA | NA |
| Gm12821 | 0.143485256 | 1.229961212 | 3.547405499 |

| | | | |
|---|---|---|---|
| Skint2 | 0.106112162 | 0.23732098 | 3.724945049 |
| Gm12814 | 0 | NA | NA |
| Gm24099 | 0 | NA | NA |
| Gm12817 | 0 | NA | NA |
| Skint10 | 0 | NA | NA |
| Skint6 | 0 | NA | NA |
| Gm12823 | 0.15269957 | -0.231642517 | 2.841177074 |
| Gm12816 | 17.15975745 | 0.54547415 | 0.270544892 |
| Gm12818 | 0.762181831 | -0.036287263 | 1.151616666 |
| Gm12813 | 0 | NA | NA |
| Skint5 | 0 | NA | NA |
| Gm12825 | 0 | NA | NA |
| Skint11 | 0 | NA | NA |
| Gm15741 | 0 | NA | NA |
| Trabd2b | 136.8658561 | -0.007315518 | 0.13715182 |
| Gm28864 | 0.044544648 | 0.237309859 | 3.724945049 |
| Gm12829 | 0.792430516 | -0.151721884 | 1.297100552 |
| 9130410C08Rik | 0.223616349 | 1.102930766 | 1.862306448 |
| Gm12830 | 0.072025462 | 0.237312084 | 3.724945049 |
| Gm23230 | 0 | NA | NA |
| Foxd2 | 5.895850479 | 0.316770604 | 0.415837968 |
| Foxd2os | 3.321918732 | 1.854229184 | 0.624482139 |
| Foxe3 | 0.279606198 | 2.155515886 | 3.700692188 |
| Cmpk1 | 803.5483932 | 0.265056413 | 0.099467468 |
| Gm24045 | 1.058755838 | -0.698442375 | 1.215282207 |
| Stil | 19.48329208 | 0.325916741 | 0.305633785 |
| Tal1 | 51.3179016 | 0.296049684 | 0.165535124 |
| Pdzk1ip1 | 3.920075909 | 0.758990482 | 0.518297191 |
| Cyp4x1 | 182.705986 | -0.68202988 | 0.102001126 |
| Cyp4x1os | 7.5998123 | -0.610748625 | 0.402761082 |
| Cyp4a28-ps | 0 | NA | NA |
| Cyp4a29-ps1 | 0 | NA | NA |
| Gm12834 | 0.117650406 | 0.23732098 | 3.724945049 |
| Cyp4a29 | 0.128150834 | -1.056605085 | 3.714811905 |
| Cyp4a12a | 0.066050875 | 0.71820487 | 3.724945049 |
| Cyp4a30-ps | 0 | NA | NA |
| Cyp4a12b | 2.349144555 | 4.5414167 | 0.990374723 |
| Gm12837 | 0 | NA | NA |
| Cyp4a30b | 0 | NA | NA |
| Gm12838 | 0 | NA | NA |
| Gm12833 | 0 | NA | NA |
| Cyp4a14 | 0.301400513 | 2.063605848 | 2.360996073 |
| Cyp4a10 | 0.11147366 | 1.230376831 | 3.722845817 |
| Cyp4b1-ps1 | 0 | NA | NA |
| Cyp4a31 | 0 | NA | NA |
| Cyp4b1-ps2 | 0 | NA | NA |
| Cyp4a32 | 0 | NA | NA |

| | | | |
|---|---|---|---|
| Cyp4b1 | 3.175114547 | -0.742819646 | 0.572537059 |
| Gm12848 | 0 | NA | NA |
| Gm12849 | 0 | NA | NA |
| Efcab14 | 1217.01994 | 0.434120453 | 0.095375238 |
| Tex38 | 1.456205946 | 0.214473302 | 0.87681663 |
| Atpaf1 | 375.0164648 | -0.354243918 | 0.07288465 |
| Gm12847 | 4.394170468 | -0.644238789 | 0.639293388 |
| Mob3c | 80.03911941 | 1.368377266 | 0.171470148 |
| Mknk1 | 268.8412447 | 0.053534972 | 0.076312949 |
| Gm29434 | 0.203831703 | -0.24357123 | 2.616105226 |
| Kncn | 2.503085161 | 0.862785999 | 0.683300224 |
| 6430628N08Rik | 3.312938502 | 1.03629159 | 0.590235919 |
| Dmbx1 | 11.05645536 | 0.06647484 | 0.326674365 |
| Faah | 605.5629304 | -1.010909039 | 0.102675977 |
| Nsun4 | 130.3320412 | 0.468140858 | 0.130230908 |
| Uqcrh | 768.3040327 | 0.007896174 | 0.149797171 |
| Gm12854 | 9.36533067 | 4.017681475 | 0.621930378 |
| Lrrc41 | 643.5531935 | 0.604266521 | 0.090265777 |
| Rad54l | 35.66809566 | 0.89910004 | 0.301566354 |
| Pomgnt1 | 564.1232995 | -0.109904555 | 0.061158921 |
| Lurap1 | 633.8593911 | -0.535733669 | 0.154626457 |
| Tspan1 | 1.564275958 | 0.054263502 | 0.676108358 |
| 1700042G07Rik | 0.107078754 | 0.236669296 | 3.724898107 |
| Gm12951 | 1.402628926 | -0.243814746 | 0.859275639 |
| Llph-ps1 | 0.397229082 | -1.45745384 | 1.67100973 |
| Pik3r3 | 1226.032443 | -0.323576626 | 0.070451454 |
| Mast2 | 1537.82531 | -0.061138182 | 0.054352618 |
| Gm12950 | 0 | NA | NA |
| Ipp | 171.4951214 | -0.28492162 | 0.097258398 |
| Tmem69 | 92.62743766 | 0.377761418 | 0.149404355 |
| Gm12953 | 2.041941669 | 0.508000005 | 0.793869275 |
| Gpbp1l1 | 655.1959422 | -0.012942981 | 0.068177062 |
| C530005A16Rik | 44.26883829 | -0.563028483 | 0.194775838 |
| Ccdc17 | 25.14108292 | -0.460094891 | 0.231275465 |
| Nasp | 221.6735445 | 0.189928349 | 0.128638707 |
| Akr1a1 | 2348.648558 | 0.691424492 | 0.097846643 |
| Gm25137 | 0 | NA | NA |
| Prdx1 | 1494.062489 | 1.10824506 | 0.172482544 |
| Mmachc | 150.583502 | -0.035013339 | 0.105282282 |
| Ccdc163 | 28.56972285 | 0.001096314 | 0.249209181 |
| Tesk2 | 80.19330176 | 0.077460746 | 0.118600367 |
| Gm27688 | 0.669412454 | -0.769536118 | 1.176057452 |
| Toe1 | 130.6486763 | 0.525126738 | 0.103568354 |
| Mutyh | 16.21670338 | -0.073634637 | 0.270380713 |
| Hpdl | 10.50466602 | 0.200518857 | 0.345488762 |
| Gm12996 | 2.848165926 | -1.133039058 | 0.620721351 |
| Rpl36-ps8 | 0 | NA | NA |

| | | | |
|---|---|---|---|
| Gm12994 | 0.047948698 | -0.243569582 | 3.724945049 |
| Gm12993 | 8.790287878 | -0.958122159 | 0.357097102 |
| Zswim5 | 280.6732534 | -0.25481251 | 0.10361753 |
| Gm26355 | 2.145613104 | -1.460298277 | 0.754858523 |
| Gm25435 | 4.493034422 | -0.007117226 | 0.502015955 |
| Gm22092 | 0.088123299 | -0.72445792 | 3.724945049 |
| Gm12998 | 0.333118728 | 0.718191893 | 3.724945049 |
| Urod | 271.8023691 | 0.050890006 | 0.091727883 |
| Hectd3 | 678.4462804 | -0.148382509 | 0.055117302 |
| 1700021J08Rik | 0.083172944 | 0.23732098 | 3.724945049 |
| Gm12997 | 0 | NA | NA |
| Eif2b3 | 158.1507724 | -0.002606099 | 0.084619549 |
| Gm22335 | 0.400019583 | -0.724396828 | 1.499392096 |
| Ptch2 | 34.03195546 | -0.559307893 | 0.239469705 |
| Btbd19 | 68.25176527 | 0.542471445 | 0.132249332 |
| Tctex1d4 | 5.254348095 | -0.391742387 | 0.453116263 |
| Plk3 | 168.7442851 | 0.741085633 | 0.132955298 |
| Best4-ps | 0.256651301 | -0.557733292 | 2.067892374 |
| Gm13015 | 0.494552988 | -0.365570232 | 1.278969298 |
| Rps8 | 1469.97404 | 0.560447552 | 0.112095438 |
| Gm22980 | 0.6528699 | 0.60707177 | 1.325496122 |
| Snord38a | 0 | NA | NA |
| Gm26330 | 0 | NA | NA |
| Snord55 | 5.259902018 | 1.153173539 | 0.47902349 |
| Gm22398 | 0 | NA | NA |
| Kif2c | 10.61551923 | 1.810850915 | 0.452922529 |
| Gm25099 | 10.85160868 | 0.898557563 | 0.390853241 |
| Gm23143 | 1.628265577 | 0.501166566 | 0.829341403 |
| Gm23287 | 0.580091356 | 1.95670043 | 1.541204224 |
| Gm1661 | 2.47324152 | -0.31702864 | 0.658411178 |
| 1700012C08Rik | 0 | NA | NA |
| Tmem53 | 15.28288558 | 0.747024461 | 0.269667558 |
| Rnf220 | 1260.662527 | -0.118906829 | 0.060951845 |
| Gm12843 | 2.662738786 | 1.079283556 | 0.617408292 |
| Gm12828 | 0.106520246 | 0.718200421 | 3.724945049 |
| Gm12827 | 1.000155967 | -0.753328968 | 1.149624756 |
| Gm12826 | 0.267394713 | 0.756313675 | 1.824423707 |
| Eri3 | 969.1178252 | -0.281729734 | 0.053020348 |
| Gm25559 | 3.182023478 | -1.102127649 | 0.716476521 |
| Dmap1 | 212.1899085 | 0.248860793 | 0.096046494 |
| Gm12840 | 0.644938119 | 1.74160577 | 1.293746665 |
| Gm12845 | 0.176646398 | 1.281666154 | 3.35399331 |
| Gm12844 | 4.659505038 | 0.514310441 | 0.469363795 |
| Klf17 | 0.667883529 | -0.075572719 | 1.392040654 |
| Gm12842 | 0.226488584 | -0.684194756 | 1.868523929 |
| Slc6a9 | 973.6643676 | -0.234641959 | 0.139021717 |
| Gm17114 | 0.046345709 | 0.71820487 | 3.724945049 |

| | | | |
|---|---|---|---|
| Ccdc24 | 3.944649576 | -0.421979723 | 0.478332325 |
| B4galt2 | 391.215399 | -0.940602391 | 0.086447887 |
| Gm12841 | 2.986054906 | -0.970498394 | 0.590610965 |
| Atp6v0b | 1543.475499 | 0.177374568 | 0.066619769 |
| Dph2 | 90.95128011 | 0.532359697 | 0.112869687 |
| Ipo13 | 1183.676242 | -0.007907394 | 0.055788289 |
| Artn | 7.776272826 | 0.28834945 | 0.479252227 |
| St3gal3 | 444.5428053 | -0.033955018 | 0.06176834 |
| 9530034E10Rik | 0.241524581 | 0.787445787 | 2.114341428 |
| Kdm4a | 879.391759 | 0.393311936 | 0.067176792 |
| Gm23382 | 0 | NA | NA |
| Ptprf | 1597.948165 | 0.051203138 | 0.046236567 |
| Mir7226 | 0 | NA | NA |
| Hyi | 25.72014861 | 0.10342786 | 0.213435524 |
| Szt2 | 768.797515 | -0.320498965 | 0.123336825 |
| Med8 | 175.0860242 | 0.793970497 | 0.13900001 |
| Elovl1 | 262.3958902 | 1.320846321 | 0.13777692 |
| Cdc20 | 20.13486822 | 1.356519239 | 0.317121703 |
| Gm12858 | 0.124415284 | -0.243565133 | 3.724945049 |
| Mpl | 4.268644651 | -0.166669267 | 0.628586652 |
| Tie1 | 38.04113095 | 1.444099609 | 0.20852624 |
| Gm12859 | 0 | NA | NA |
| Gm12857 | 0 | NA | NA |
| 2610528J11Rik | 0.045882455 | 0.237312084 | 3.724945049 |
| Tmem125 | 85.91567833 | -0.327118133 | 0.190459183 |
| Cfap57 | 12.09672597 | -0.469798438 | 0.284749032 |
| Ebna1bp2 | 374.5410988 | 0.129502318 | 0.088866866 |
| Gm12853 | 0 | NA | NA |
| Olfr62 | 0.260073237 | 2.24256775 | 3.694036856 |
| Olfr1343-ps1 | 0 | NA | NA |
| Olfr1342 | 0.045882455 | 0.237312084 | 3.724945049 |
| Olfr1341 | 0.138076337 | 0.718204987 | 3.589802529 |
| Olfr1340 | 0.163066018 | -0.243578385 | 3.449621945 |
| Olfr1339 | 0.060174537 | 0.237309859 | 3.724945049 |
| Olfr1338 | 0 | NA | NA |
| Olfr1337 | 0 | NA | NA |
| Gm12855 | 0 | NA | NA |
| Olfr1335 | 0 | NA | NA |
| Olfr1333 | 0 | NA | NA |
| Olfr1332-ps1 | 0 | NA | NA |
| Olfr1331 | 0 | NA | NA |
| Olfr1330 | 0 | NA | NA |
| Olfr1329 | 0 | NA | NA |
| Olfr1328 | 0.212007022 | 0.718208831 | 3.149720606 |
| Gm12856 | 0.270533821 | 2.134499984 | 2.409115729 |
| Gm12863 | 0.232395771 | 1.183255052 | 2.122081047 |
| Lao1 | 0.148650935 | -0.243572541 | 2.863172849 |

| | | | |
|---|---|---|---|
| Gm12861 | 0 | NA | NA |
| Gm24352 | 0 | NA | NA |
| Gm12865 | 0.488817361 | 1.060683161 | 1.498603806 |
| Gm12864 | 0 | NA | NA |
| Gm12862 | 0.177984204 | 1.200050841 | 3.339898728 |
| Gm12866 | 0.218085611 | 0.531996412 | 2.542083288 |
| Slc2a1 | 824.8283364 | 0.928695542 | 0.109927616 |
| Mir1957a | 1.374216805 | 0.566728717 | 0.987408296 |
| Gm12867 | 0.088534442 | 1.179809211 | 3.723500796 |
| Gm12868 | 0.130524232 | -0.243571427 | 3.657871338 |
| Zfp691 | 52.30840471 | 0.375556234 | 0.144569754 |
| Ermap | 7.770417513 | 0.934930122 | 0.384275197 |
| Ccdc23 | 99.45512681 | 1.352064089 | 0.182439276 |
| AU022252 | 374.3096476 | -0.431318607 | 0.107731796 |
| 4930538K18Rik | 9.22375745 | -0.860359768 | 0.334843793 |
| Gm12898 | 0.061920613 | 0.718207094 | 3.724945049 |
| P3h1 | 112.7927286 | 0.349462044 | 0.135221159 |
| Gm12927 | 0.131862677 | -0.243562287 | 2.97735462 |
| Cldn19 | 10.33246628 | 1.157652335 | 0.408846765 |
| Ybx1 | 865.8775829 | 0.79319706 | 0.127914909 |
| Gm37817 | 0.046345709 | 0.71820487 | 3.724945049 |
| Ppih | 73.62859532 | -0.245825295 | 0.168915798 |
| Ccdc30 | 150.4461269 | -0.414646965 | 0.200519084 |
| Ldha-ps2 | 11.09854899 | -0.420893091 | 0.309591316 |
| Ppcs | 48.56475404 | 0.418049808 | 0.15668843 |
| Zmynd12 | 4.300457922 | 0.107883386 | 0.472144964 |
| Rimkla | 301.1007889 | -0.354653532 | 0.128034833 |
| Gm12954 | 0.144050924 | 0.237309606 | 3.724945049 |
| Gm12956 | 0.301349932 | -0.243563199 | 1.630883096 |
| Gm12955 | 1.414308396 | -0.71282031 | 0.901362191 |
| Gm12957 | 0.066050875 | 0.71820487 | 3.724945049 |
| Rnu6-ps2 | 0 | NA | NA |
| AA415398 | 75.1633106 | -0.530793811 | 0.144145324 |
| Foxj3 | 1047.552468 | 0.005853097 | 0.076167353 |
| Gm12959 | 0 | NA | NA |
| Guca2a | 0.577807171 | -0.23667394 | 1.352226815 |
| Guca2b | 0.198152624 | 1.757323469 | 3.707396009 |
| Hivep3 | 1525.201888 | 0.082048457 | 0.108812591 |
| Edn2 | 0.478833344 | -0.186460906 | 1.326263035 |
| Foxo6 | 102.9223404 | -0.818653137 | 0.141149073 |
| Foxo6os | 0.889815719 | 1.807741533 | 1.135857886 |
| Scmh1 | 665.7514625 | -0.345098182 | 0.093265068 |
| Slfnl1 | 5.927427543 | -1.358144341 | 0.511277923 |
| Ctps | 504.9997313 | 0.441812195 | 0.108620061 |
| Gm8439 | 0.147068667 | 0.237310878 | 3.514416608 |
| Gm12860 | 1.253221006 | 0.187410276 | 0.857140375 |
| Cited4 | 44.06063388 | 0.075823874 | 0.197398065 |

| | | | |
|---|---|---|---|
| Kcnq4 | 121.7819068 | 0.156802159 | 0.160292561 |
| Nfyc | 366.6066488 | 0.038047119 | 0.088351617 |
| Mir30c-1 | 1.678046897 | -1.333889379 | 0.803142591 |
| mmu-mir-30e | 0 | NA | NA |
| Mir30e | 2.389660156 | 0.354521665 | 0.681108563 |
| Gm23936 | 0.479339789 | -0.629866102 | 1.355832594 |
| Rims3 | 2593.580875 | -0.792027845 | 0.086319875 |
| Gm12871 | 0.804633503 | 0.837196436 | 1.210105998 |
| Gm24678 | 0.394656821 | 0.237310862 | 3.34824039 |
| Exo5 | 86.65934631 | 0.402511148 | 0.146287693 |
| Zfp69 | 17.8840471 | 0.249535017 | 0.265549743 |
| Smap2 | 1633.072005 | -0.168587614 | 0.077193629 |
| mmu-mir-7015 | 1.055430728 | -1.274140868 | 0.957828787 |
| Col9a2 | 128.0701482 | -0.778042339 | 0.136314483 |
| Zmpste24 | 545.8456237 | 0.429236306 | 0.077330989 |
| Gm26186 | 0 | NA | NA |
| Gm23540 | 0.845517282 | 0.611927328 | 1.138186637 |
| Tmco2 | 0.224241997 | 0.246553956 | 2.571903766 |
| Gm12882 | 0 | NA | NA |
| Gm12881 | 0.075043205 | 0.237309859 | 3.724945049 |
| Gm12879 | 0 | NA | NA |
| Gm22982 | 0 | NA | NA |
| Rlf | 853.6771386 | 0.007451193 | 0.11607649 |
| Gm12890 | 0 | NA | NA |
| Gm12888 | 0 | NA | NA |
| Gm12889 | 0 | NA | NA |
| Gm12885 | 0 | NA | NA |
| Gm12884 | 0 | NA | NA |
| Gm12883 | 0.060174537 | 0.237309859 | 3.724945049 |
| Gm12886 | 0 | NA | NA |
| Gm12880 | 0 | NA | NA |
| Gm26162 | 0 | NA | NA |
| Gm12887 | 0 | NA | NA |
| Gm12878 | 0 | NA | NA |
| Gm12892 | 61.92315877 | -0.194803864 | 0.171295698 |
| Gm12896 | 0 | NA | NA |
| Gm12897 | 0 | NA | NA |
| Gm12893 | 0 | NA | NA |
| Gm12895 | 0 | NA | NA |
| Gm12877 | 0.066050875 | 0.71820487 | 3.724945049 |
| Gm12874 | 0 | NA | NA |
| Gm12875 | 0 | NA | NA |
| Gm12876 | 0.038981395 | 0.718207094 | 3.724945049 |
| 9530002B09Rik | 0 | NA | NA |
| Gm12872 | 0 | NA | NA |
| Gm12873 | 0.136623169 | 0.66393974 | 3.632616033 |
| Ppt1 | 1536.560467 | 0.29271259 | 0.054268069 |

| | | | |
|---|---|---|---|
| Cap1 | 2420.151979 | 0.676774888 | 0.06559085 |
| Gm12891 | 0 | NA | NA |
| Mfsd2a | 324.2124731 | 1.094905731 | 0.152983998 |
| 4933421A08Rik | 2.139806348 | 1.267743339 | 0.714601308 |
| Mycl | 128.6415578 | -0.930069615 | 0.190717808 |
| Trit1 | 181.2275785 | 0.169186377 | 0.170233468 |
| Gm12923 | 0.164553454 | 0.719098949 | 2.779126835 |
| Bmp8b | 1.325872221 | 0.702930258 | 0.820897653 |
| Gm17244 | 11.29059836 | -0.504182952 | 0.314213033 |
| Oxct2b | 0.96273357 | -1.649982556 | 1.151319088 |
| Ppie | 115.1280594 | 0.901273305 | 0.13530994 |
| Hpcal4 | 5044.886741 | -0.565115485 | 0.163746209 |
| Nt5c1a | 53.31890087 | -0.255846383 | 0.160872433 |
| Heyl | 92.72271586 | 0.001197357 | 0.189368646 |
| Gm12901 | 0.083526043 | 0.718202646 | 3.724945049 |
| Pabpc4 | 351.2920336 | 0.436977069 | 0.103451098 |
| Gm12900 | 0 | NA | NA |
| Gm25788 | 10.10737276 | 0.454927238 | 0.340031595 |
| Gm22154 | 9.014346252 | -0.009568263 | 0.335240458 |
| Bmp8a | 2.710969876 | 1.259311855 | 0.696192118 |
| Oxct2a | 0.793872245 | -1.482669064 | 1.199575635 |
| Macf1 | 11900.70528 | -0.199047111 | 0.06141224 |
| Gm12924 | 0.060174537 | 0.237309859 | 3.724945049 |
| Gm23548 | 0 | NA | NA |
| Mir6398 | 0 | NA | NA |
| Gm12926 | 0 | NA | NA |
| Gm12925 | 2.717467516 | 0.421886388 | 0.568901938 |
| Gm12922 | 4.003874815 | -0.184512048 | 0.541346012 |
| Ndufs5 | 423.3196819 | -0.109462108 | 0.096464675 |
| Akirin1 | 729.4963505 | 0.551202431 | 0.097647552 |
| Rhbdl2 | 5.725445938 | 0.420088118 | 0.42211054 |
| 1700121C08Rik | 6.402408373 | -0.868588842 | 0.516915019 |
| Gm12904 | 0.192693611 | 0.237326772 | 2.64331282 |
| 4933427I04Rik | 0.332356096 | 0.718190717 | 3.724945049 |
| Gm26606 | 53.85320551 | -0.46788254 | 0.239789098 |
| D130007C19Rik | 26.17483582 | -0.722437152 | 0.29759392 |
| Gm12903 | 0 | NA | NA |
| Mycbp | 64.1662213 | 0.065695783 | 0.13635019 |
| Gm12905 | 20.80298438 | -0.023820998 | 0.29149233 |
| Rragc | 863.0862859 | 0.492711228 | 0.087936621 |
| 4930535I16Rik | 1.505311134 | 0.361444095 | 0.904310087 |
| Gm12902 | 7.717556106 | -1.027377186 | 0.403696199 |
| Gm37667 | 0.274851136 | 1.700199074 | 3.715468711 |
| Gm12916 | 0.134481855 | -0.243575541 | 3.655499445 |
| 1700021L23Rik | 0.106971521 | 0.237307635 | 3.724945049 |
| 4933435F18Rik | 0.562550613 | -0.078205569 | 1.431319597 |
| Gm22983 | 0 | NA | NA |

| | | | |
|---|---|---|---|
| 1700057H15Rik | 1.747775439 | -0.432729111 | 0.831411885 |
| Gm27247 | 15.84162281 | -0.62224859 | 0.327091931 |
| 4933407E24Rik | 0 | NA | NA |
| 3100002H09Rik | 4.323581818 | -0.798804078 | 0.653114539 |
| Pou3f1 | 370.5457929 | 0.538349392 | 0.129848273 |
| Utp11l | 181.9343619 | 0.417120239 | 0.08280855 |
| Gm24480 | 0 | NA | NA |
| Fhl3 | 24.63805473 | 1.611867161 | 0.248993177 |
| Sf3a3 | 314.4333351 | 0.465631484 | 0.092297941 |
| Gm12917 | 6.099625106 | -0.58479529 | 0.599315718 |
| Mir697 | 1.448093177 | -0.725469588 | 0.874465174 |
| Gm12915 | 2.377012089 | -0.118848103 | 0.674705576 |
| Gm24721 | 0.117760149 | -0.243576254 | 3.724945049 |
| Inpp5b | 535.9436542 | 0.224583223 | 0.073286694 |
| Mir698 | 0.286901049 | 0.241069013 | 2.051807113 |
| Mtf1 | 507.9230992 | -0.061587109 | 0.092041552 |
| n-R5s192 | 1.205568873 | -1.868815098 | 1.039247192 |
| 1110065P20Rik | 188.1940773 | -0.277017218 | 0.110376557 |
| Yrdc | 152.5953946 | 0.608548331 | 0.19458619 |
| Maneal | 554.488513 | -0.357521163 | 0.082549274 |
| Epha10 | 353.1436211 | -1.08554914 | 0.172441011 |
| 1700125G02Rik | 0.77578221 | -0.466304203 | 1.085879305 |
| Cdca8 | 18.02129889 | 0.784798481 | 0.330568319 |
| 9930104L06Rik | 211.7189511 | 0.154489891 | 0.080181574 |
| Gm12930 | 0.650770932 | -1.169622942 | 1.183138429 |
| Rspo1 | 34.80671183 | -1.092887317 | 0.263250772 |
| Gnl2 | 487.0670719 | 0.366532753 | 0.103161578 |
| Gm12931 | 0 | NA | NA |
| Dnali1 | 15.85509031 | 0.570858712 | 0.280791811 |
| Snip1 | 158.6932189 | 0.432387772 | 0.103346306 |
| Gm12932 | 3.658838888 | 0.307453314 | 0.536692126 |
| Meaf6 | 803.9924443 | -0.597701682 | 0.103634576 |
| Zc3h12a | 71.74715942 | 3.14490678 | 0.23998847 |
| 1700041M05Rik | 0 | NA | NA |
| Grik3 | 1453.049993 | -0.517204493 | 0.081425805 |
| Ftl2 | 1.736617851 | 3.510569658 | 1.091700143 |
| Mir692-2 | 0.541671891 | 1.372898848 | 1.449303211 |
| 2610028E06Rik | 11.39828975 | 1.271430712 | 0.376843865 |
| Csf3r | 194.1912691 | 2.764156534 | 0.192416446 |
| Mrps15 | 201.7560872 | 0.098182093 | 0.105995658 |
| Oscp1 | 268.9501632 | -0.683212291 | 0.095252253 |
| Rpl28-ps3 | 19.78160788 | 0.765246537 | 0.268582965 |
| Lsm10 | 59.05863203 | 0.816968493 | 0.171205576 |
| Stk40 | 956.9610872 | 1.834127573 | 0.18956635 |
| Gm12946 | 0.363304151 | 1.65821487 | 1.659847299 |
| Eva1b | 26.65916579 | 1.606219823 | 0.252093598 |
| Sh3d21 | 14.96427342 | 0.704800613 | 0.295417616 |

| | | | |
|---|---|---|---|
| Thrap3 | 1545.929057 | 0.135634357 | 0.05932558 |
| Gm25323 | 1.633037017 | -1.081943534 | 0.945897827 |
| Map7d1 | 3375.866482 | 0.304877797 | 0.076113952 |
| Trappc3 | 541.5073919 | 0.168693876 | 0.073087712 |
| Col8a2 | 71.84796312 | -0.588273143 | 0.220962443 |
| Adprhl2 | 208.9240687 | 0.389114229 | 0.105932596 |
| Tekt2 | 9.162665626 | -0.18954525 | 0.379013988 |
| Gm12945 | 0.40938346 | 0.68702353 | 1.430387242 |
| Ago3 | 1555.337107 | -0.653103489 | 0.104230303 |
| Ago1 | 1116.199882 | -0.31839849 | 0.073037792 |
| Gm12929 | 1.033962832 | 1.993789496 | 1.06603415 |
| Gm12928 | 17.68241393 | 0.184702177 | 0.270010444 |
| Ago4 | 190.9073592 | -0.54425492 | 0.125446386 |
| Gm12935 | 0.038981395 | 0.718207094 | 3.724945049 |
| Gm12934 | 4.064846038 | -1.010627708 | 0.639504287 |
| Clspn | 29.12900793 | 1.149586735 | 0.404972157 |
| Mir7119 | 0 | NA | NA |
| Gm12936 | 0.066050875 | 0.71820487 | 3.724945049 |
| 5730409E04Rik | 892.9733379 | -0.477046638 | 0.06410066 |
| Gm12933 | 1.508740896 | -0.292215617 | 0.809896016 |
| Psmb2 | 628.3734742 | 0.921837505 | 0.109088216 |
| Tfap2e | 0.181508466 | 0.71821599 | 2.694094898 |
| Ncdn | 11330.12844 | -0.511308851 | 0.078538272 |
| AU040320 | 846.8326345 | 0.133293138 | 0.049882061 |
| Gm12937 | 0 | NA | NA |
| Gm24161 | 0 | NA | NA |
| Gm25604 | 3.572392357 | -0.238268976 | 0.559784016 |
| Zmym4 | 1413.511916 | -0.006128538 | 0.045151413 |
| Gm12939 | 0 | NA | NA |
| Sfpq | 2483.364451 | -0.338421411 | 0.09380179 |
| Gm12940 | 393.2352481 | -0.823389011 | 0.254467701 |
| Gm12941 | 12.76876565 | -1.269289512 | 0.522751111 |
| Zmym1 | 160.873064 | -0.316215436 | 0.13866573 |
| Gm12944 | 1.541842 | 0.407772823 | 0.800819608 |
| Zmym6 | 362.4092715 | -0.301575953 | 0.117783344 |
| Gm25600 | 0 | NA | NA |
| Gm12942 | 27.99737021 | -0.07577912 | 0.247497206 |
| Gm12943 | 0.338503588 | -1.397714321 | 1.518886239 |
| Dlgap3 | 2650.38369 | -1.067046624 | 0.123325269 |
| Smim12 | 221.697071 | 0.047082255 | 0.088376277 |
| Gm22221 | 0 | NA | NA |
| Gm12948 | 0 | NA | NA |
| Gja4 | 31.84417143 | 1.505092215 | 0.255995304 |
| Gjb3 | 2.20869273 | 2.594352422 | 0.829042462 |
| Gjb4 | 1.851903907 | 4.336517262 | 1.032275665 |
| Gjb5 | 0.85820172 | 3.197736629 | 1.31468538 |
| Gm12947 | 0 | NA | NA |

| | | | |
|---|---|---|---|
| 4930471C06Rik | 0.072025462 | 0.237312084 | 3.724945049 |
| 1700080G11Rik | 0.674986302 | 0.541002841 | 1.209118713 |
| A630031M04Rik | 0.919688069 | 1.646292844 | 1.079008383 |
| 1700112K13Rik | 0.099155932 | 0.718202646 | 3.724945049 |
| CK137956 | 8.83914943 | -0.263306865 | 0.339564846 |
| Csmd2 | 1037.327866 | -0.453400764 | 0.118919831 |
| Csmd2os | 0.11657011 | 0.237307635 | 3.724945049 |
| Hmgb4 | 0.309883016 | 2.467318587 | 3.687546276 |
| Hmgb4os | 3.44465028 | -1.246371141 | 0.664571926 |
| Zscan20 | 123.2244835 | -0.102511232 | 0.157438141 |
| Tlr12 | 8.189155563 | -0.281283149 | 0.543354064 |
| Gm12958 | 0 | NA | NA |
| Phc2 | 1259.457039 | 0.749148759 | 0.093875515 |
| A3galt2 | 16.37679573 | 0.940648126 | 0.318268002 |
| Zfp362 | 336.5354883 | -0.252823263 | 0.09149509 |
| Gm12968 | 0.072025462 | 0.237312084 | 3.724945049 |
| Gm12969 | 0.192749185 | -1.055188118 | 3.283777405 |
| Trim62 | 671.0414608 | -0.170629352 | 0.155967952 |
| Azin2 | 295.6170531 | -0.054494229 | 0.087669 |
| Gm15904 | 0.046345709 | 0.71820487 | 3.724945049 |
| Ak2 | 283.4084168 | 1.232043364 | 0.107843661 |
| 1700086P04Rik | 1.876204568 | 0.617866635 | 0.742106655 |
| Rnf19b | 1117.961967 | 1.700401256 | 0.162095042 |
| Tmem54 | 5.394435422 | -0.932381944 | 0.482165163 |
| Hpca | 2746.890777 | -1.462718566 | 0.156920584 |
| Gm15906 | 1.230936373 | -0.201478243 | 0.903346511 |
| Fndc5 | 973.7975932 | -0.524734449 | 0.095811906 |
| S100pbp | 678.3393157 | -0.379564863 | 0.068965379 |
| Yars | 969.2323716 | 0.489131165 | 0.083897873 |
| C77080 | 318.9680605 | -0.424497757 | 0.111331635 |
| Gm12976 | 2.076702053 | -1.536267961 | 0.702357186 |
| Gm22994 | 0 | NA | NA |
| Sync | 16.23627725 | -0.147099668 | 0.252459056 |
| Rbbp4 | 1422.281671 | 0.195103313 | 0.071633272 |
| Gm26722 | 3.033581745 | 0.929920002 | 0.555824489 |
| Zbtb8os | 112.0160394 | -0.194868906 | 0.112353444 |
| Zbtb8a | 48.71610654 | -0.086866739 | 0.175876265 |
| Gm12980 | 0.075043205 | 0.237309859 | 3.724945049 |
| Zbtb8b | 38.62991341 | -1.020255653 | 0.241849375 |
| Gm24621 | 0 | NA | NA |
| Bsdc1 | 654.7636716 | 0.125450213 | 0.065968742 |
| Tssk3 | 1.78535976 | -1.175456854 | 0.973286084 |
| Fam229a | 1.511769237 | -0.252387965 | 0.810176559 |
| Gm12979 | 0 | NA | NA |
| Marcksl1 | 1179.090896 | 0.685611932 | 0.128381238 |
| Hdac1 | 211.8204638 | 1.044548529 | 0.115484998 |
| Lck | 123.7661852 | 4.075928464 | 0.268017836 |

| | | | |
|---|---|---|---|
| Mir8119 | 0.156477978 | 0.718214073 | 2.823333319 |
| Fam167b | 2.785268692 | 1.018076206 | 0.668118401 |
| Eif3i | 596.7175796 | 0.357624503 | 0.085843658 |
| Tmem234 | 475.3810905 | 0.160984474 | 0.055875722 |
| Dcdc2b | 6.370853605 | -1.554400493 | 0.438420645 |
| Iqcc | 93.52464096 | -0.397995437 | 0.149060345 |
| Ccdc28b | 159.6210152 | -0.332327798 | 0.128291975 |
| Txlna | 633.3282999 | 0.401127296 | 0.067181805 |
| Kpna6 | 1687.26505 | 0.05551873 | 0.050673255 |
| Tmem39b | 67.54451293 | -0.066656802 | 0.168917488 |
| Mir7016 | 0.152586256 | -0.243562192 | 2.845690286 |
| Khdrbs1 | 1365.931483 | 0.016094211 | 0.062046219 |
| Gm12967 | 0 | NA | NA |
| Gm28017 | 0 | NA | NA |
| Ptp4a2 | 1712.805092 | 0.405020489 | 0.085231669 |
| Gm10571 | 11.18052223 | -0.83579456 | 0.354919882 |
| Gm23845 | 0.075043205 | 0.237309859 | 3.724945049 |
| Gm12966 | 92.3427904 | 0.518704992 | 0.194076821 |
| E330017L17Rik | 0.177020223 | 0.237324565 | 2.765325947 |
| Spocd1 | 2.098579804 | -0.812186758 | 0.814539434 |
| 1700003M07Rik | 24.71805353 | -0.755243188 | 0.279050929 |
| Adgrb2 | 3754.218321 | -0.678260427 | 0.129088719 |
| Gm12963 | 2.392410068 | -1.433084877 | 0.768211447 |
| Col16a1 | 419.3408002 | 1.990639904 | 0.152812665 |
| Pef1 | 447.0267585 | 0.305876847 | 0.094991751 |
| Hcrtr1 | 16.529312 | -0.040347903 | 0.294921403 |
| Tinagl1 | 47.82351247 | 2.105891279 | 0.249424455 |
| Gm853 | 1.50545653 | -1.787301984 | 0.91069662 |
| Serinc2 | 80.11973323 | 0.266991144 | 0.163601289 |
| Gm10570 | 4.998772346 | 0.145701954 | 0.43007706 |
| Fabp3 | 388.7351392 | -0.560551153 | 0.105013714 |
| Zcchc17 | 445.3923863 | 0.053395304 | 0.075388685 |
| Snrnp40 | 199.2041572 | 0.246706543 | 0.080912306 |
| Nkain1 | 233.0882025 | 0.070023898 | 0.123938145 |
| Pum1 | 1545.221209 | -0.20605579 | 0.056385409 |
| Snord85 | 3.381283184 | -0.20329202 | 0.578308751 |
| Gm12971 | 8.510907694 | -1.094515453 | 0.401535146 |
| Gm24091 | 2.986937453 | -0.55883942 | 0.536598572 |
| Gm26516 | 0.225317292 | 0.532376345 | 2.559353476 |
| Gm23711 | 1.081483968 | -0.485692015 | 0.883061615 |
| Gm12972 | 0.415245445 | -0.136607071 | 1.433256191 |
| Sdc3 | 6090.612101 | 1.407581736 | 0.122367195 |
| Gm12970 | 0.131862677 | -0.243562287 | 2.97735462 |
| Gm26716 | 0.436796958 | 2.492318836 | 1.590551441 |
| Gm22660 | 0 | NA | NA |
| Laptm5 | 869.0779485 | 2.386914177 | 0.165135595 |
| Matn1 | 10.36744075 | 3.104088102 | 0.501261616 |

| | | | |
|---|---|---|---|
| Gm12973 | 0.042716945 | -0.243571806 | 3.724945049 |
| A930031H19Rik | 0.159888962 | -0.724460061 | 3.461347753 |
| Gm25261 | 0 | NA | NA |
| Gm831 | 0.963234904 | 1.834478335 | 1.069710663 |
| Gm12962 | 0.066050875 | 0.71820487 | 3.724945049 |
| Gm16080 | 0.066050875 | 0.71820487 | 3.724945049 |
| Ptpru | 445.2308485 | -0.71872345 | 0.090529602 |
| Mecr | 135.1425178 | -0.06367785 | 0.122043176 |
| Srsf4 | 621.5934249 | 0.05850695 | 0.075049363 |
| A930004J17Rik | 191.8201737 | -0.509810855 | 0.239902933 |
| Gm12992 | 46.74477129 | -0.390085622 | 0.201020479 |
| Gm12978 | 0 | NA | NA |
| Tmem200b | 45.76488705 | -0.145396897 | 0.230195391 |
| Epb4.1 | 1119.732651 | -0.366945201 | 0.137173952 |
| Gm27445 | 1.407831039 | -1.267327612 | 0.852577757 |
| Gm13063 | 0.62973509 | 1.204841109 | 1.176975449 |
| Gm10300 | 1.165032807 | 0.398727812 | 1.021624925 |
| Gm13214 | 0 | NA | NA |
| Gm37711 | 81.49722938 | -0.312079317 | 0.218242599 |
| Oprd1 | 249.9746901 | -0.197202087 | 0.15461747 |
| Gm13215 | 7.792582945 | -0.176641182 | 0.390580213 |
| Ythdf2 | 645.7985944 | 0.499043747 | 0.104814747 |
| Gm22033 | 0.373327567 | 1.552389869 | 3.698703546 |
| Rps15a-ps4 | 8.387413765 | 0.786145625 | 0.373459304 |
| Gmeb1 | 299.6588852 | 0.120781936 | 0.11055264 |
| Gm13252 | 3.851277915 | -0.685901534 | 0.515429751 |
| Gm28874 | 1.719071184 | -1.300311766 | 0.881825788 |
| Gm28872 | 0.044544648 | 0.237309859 | 3.724945049 |
| Rnu11 | 24.6768285 | 0.394364193 | 0.258506228 |
| Taf12 | 193.1512937 | 0.737174894 | 0.087484724 |
| Rab42 | 6.121133301 | -0.413892352 | 0.398142642 |
| Snhg12 | 177.2717645 | 1.147522751 | 0.16144463 |
| snopsi28S-3327 | 0 | NA | NA |
| Snora16a | 0 | NA | NA |
| Snora44 | 0 | NA | NA |
| Snora61 | 0 | NA | NA |
| Snord99 | 0 | NA | NA |
| Trnau1ap | 131.2572649 | 0.242064015 | 0.106876799 |
| Rcc1 | 103.1344041 | 0.249922949 | 0.127705536 |
| Snhg3 | 63.93668591 | 0.589147377 | 0.145570654 |
| Gm17300 | 0.711576293 | -0.68527603 | 1.280776817 |
| Snora73b | 425.1513821 | 0.320502303 | 0.211311207 |
| Snora73a | 0.690105543 | 0.026955575 | 1.267482263 |
| Phactr4 | 351.1978573 | 0.991805287 | 0.142456956 |
| Gm25707 | 0 | NA | NA |
| Gm23055 | 0 | NA | NA |
| Med18 | 61.32883924 | 0.243622763 | 0.142045526 |

| | | | |
|---|---|---|---|
| Sesn2 | 307.2550336 | 1.627780084 | 0.174879722 |
| Gm12981 | 4.426758316 | 0.39048615 | 0.46694229 |
| Atpif1 | 785.2790718 | -0.27587466 | 0.132198711 |
| Gm12999 | 2.573520391 | -0.57101675 | 0.6668069 |
| Dnajc8 | 648.8436146 | 0.153873279 | 0.088267821 |
| Gm24762 | 0 | NA | NA |
| Ptafr | 215.886901 | 4.932980235 | 0.375179238 |
| Eya3 | 516.9045441 | 0.252672076 | 0.071606166 |
| Xkr8 | 74.57135612 | 0.031597583 | 0.136733923 |
| Smpdl3b | 33.7182008 | 0.304604519 | 0.219669828 |
| Rpa2 | 85.55301726 | 0.181337369 | 0.156658374 |
| Themis2 | 329.3702901 | 6.36579619 | 0.354909894 |
| Gm13022 | 0 | NA | NA |
| Ppp1r8 | 368.0675459 | 0.328680034 | 0.080090685 |
| Gm22767 | 11.91181268 | -0.070793114 | 0.341009029 |
| Stx12 | 1297.871004 | 0.061829187 | 0.062680935 |
| Gm13033 | 2.338964691 | 0.534891042 | 0.745329461 |
| Fam76a | 504.2948798 | 0.013285007 | 0.074765657 |
| Gm24913 | 0 | NA | NA |
| Fgr | 92.45807742 | 6.211867918 | 0.453221365 |
| Ahdc1 | 1531.214203 | -0.292933783 | 0.097110249 |
| Gm26615 | 5.273376998 | -1.102517242 | 0.576702252 |
| Wasf2 | 693.9247449 | 1.42248508 | 0.119885537 |
| Gm24636 | 0 | NA | NA |
| Mir7017 | 0.269321163 | -0.243585702 | 3.724945049 |
| Gpr3 | 98.5239047 | 0.157837271 | 0.253802376 |
| Cd164l2 | 13.0383265 | 0.272817325 | 0.295042392 |
| Map3k6 | 67.01031991 | 2.006822904 | 0.380496622 |
| Sytl1 | 19.69897721 | -0.568155217 | 0.279656395 |
| Tmem222 | 442.6011969 | 0.252045037 | 0.088954205 |
| Gm13259 | 0.441858849 | -1.398096045 | 1.374143519 |
| Wdtc1 | 1171.200647 | 0.817862118 | 0.089902496 |
| Slc9a1 | 963.2912325 | 0.376369493 | 0.070137067 |
| Mir5122 | 0 | NA | NA |
| Gm13257 | 0.719481786 | 2.283114812 | 1.292471519 |
| Fam46b | 4.767060263 | 1.816997433 | 0.617402508 |
| Trnp1 | 753.4664742 | -0.184237246 | 0.116024282 |
| Kdf1 | 1.792464502 | 0.314322392 | 0.810219873 |
| Nudc | 578.9021612 | 0.255993509 | 0.169262437 |
| Nr0b2 | 0.783927492 | -2.275341794 | 1.102407565 |
| Gm23158 | 0.049553047 | 0.71820487 | 3.724945049 |
| Gpatch3 | 56.4655971 | 0.713888083 | 0.17313656 |
| Gpn2 | 58.10242144 | 0.69361032 | 0.191398232 |
| Sfn | 14.04179457 | 0.699885825 | 0.389520965 |
| Zdhhc18 | 588.2128307 | 0.90065469 | 0.089206143 |
| Gm13213 | 0.097899064 | -0.724453471 | 3.724945049 |
| Gm22662 | 0 | NA | NA |

| | | | |
|---|---|---|---|
| Pigv | 97.35604423 | 0.840543448 | 0.173279545 |
| Arid1a | 2860.653318 | -0.059801526 | 0.064408626 |
| mmu-mir-7227 | 2.041220482 | -1.535167126 | 0.746492082 |
| Gm25270 | 0 | NA | NA |
| Gm12974 | 0.15596086 | -1.187811386 | 3.483969583 |
| Rps6ka1 | 359.3082904 | 0.058849211 | 0.114481182 |
| Gm12977 | 0 | NA | NA |
| Gm12985 | 2.189170689 | -0.361761527 | 0.705327945 |
| Gm22523 | 2.911568235 | -1.819076681 | 0.772040005 |
| Hmgn2 | 174.4388056 | 0.305444602 | 0.169028669 |
| Dhdds | 602.5039709 | -0.134960716 | 0.060599013 |
| Lin28a | 0.305648149 | 1.16019121 | 1.734082216 |
| Gm13061 | 0 | NA | NA |
| Zfp683 | 0.393365814 | 2.175855783 | 1.43073075 |
| Aim1l | 11.08135394 | 0.589245697 | 0.293465673 |
| Cd52 | 180.6648992 | 5.819693432 | 0.379037573 |
| Ubxn11 | 97.44407537 | -0.50757255 | 0.204066461 |
| Sh3bgrl3 | 569.2295487 | 0.296480321 | 0.088187055 |
| Cep85 | 164.0264212 | 0.195386379 | 0.09503036 |
| Gm27619 | 0.047948698 | -0.243569582 | 3.724945049 |
| Gm7534 | 0 | NA | NA |
| Gm13131 | 1.316625395 | -0.244982272 | 0.789113814 |
| Catsper4 | 0.735041978 | 1.035508907 | 1.263908383 |
| Cnksr1 | 18.73766402 | 2.933392637 | 0.425883836 |
| Zfp593 | 102.2427118 | 1.00213061 | 0.170080021 |
| E130218I03Rik | 0.926475344 | 2.427705939 | 1.227803467 |
| Grrp1 | 25.39142091 | 2.676729586 | 0.282130695 |
| Pdik1l | 224.1146395 | 0.145885768 | 0.086581969 |
| Trim63 | 3.249610149 | 1.740175919 | 0.706589012 |
| Gm13195 | 0 | NA | NA |
| Slc30a2 | 1.413987658 | 1.157563287 | 0.859288565 |
| Extl1 | 547.7075812 | -0.886586336 | 0.187493595 |
| Pafah2 | 65.33746165 | 0.229557984 | 0.203569719 |
| 1700021N21Rik | 2.462361163 | -0.010922275 | 0.649350942 |
| Stmn1 | 1361.893308 | -1.024504444 | 0.092892942 |
| Gm23634 | 0.446933609 | -0.761421085 | 1.710754355 |
| Gm13250 | 0 | NA | NA |
| Paqr7 | 388.1743221 | -0.92203512 | 0.101677418 |
| Aunip | 1.169101948 | 0.664285985 | 1.083804386 |
| Mtfr1l | 639.4674001 | 0.396370015 | 0.06914521 |
| Sepn1 | 472.582935 | 0.307270759 | 0.080753636 |
| Man1c1 | 640.4685701 | 0.17057332 | 0.114373335 |
| Mir6403 | 1.273845189 | -1.982761145 | 0.93566681 |
| Ldlrap1 | 252.7861691 | 1.247408958 | 0.163418429 |
| Tmem57 | 886.6149259 | -0.285311672 | 0.075734625 |
| Rhd | 1.579581474 | -0.827179934 | 0.76304621 |
| Tmem50a | 553.5327818 | 0.430644987 | 0.105475874 |

| | | | |
|---|---|---|---|
| Rsrp1 | 2711.383038 | 0.27669888 | 0.091839399 |
| Syf2 | 379.4524818 | 0.572576841 | 0.134738191 |
| Gm16225 | 0 | NA | NA |
| Runx3 | 97.20798678 | 7.957265218 | 0.714107371 |
| Gm16224 | 0 | NA | NA |
| Clic4 | 3776.355182 | 2.403507138 | 0.232979186 |
| Gm12983 | 0.072025462 | 0.237312084 | 3.724945049 |
| Gm12984 | 0 | NA | NA |
| Gm12982 | 0 | NA | NA |
| Srrm1 | 1767.897914 | -0.06321851 | 0.09516549 |
| Ncmap | 11.39922601 | -0.750046162 | 0.34453052 |
| Gm12991 | 0 | NA | NA |
| Gm12990 | 3.613492829 | -0.439383321 | 0.584579363 |
| Rcan3 | 591.8608699 | -0.15280289 | 0.094805278 |
| Mir700 | 0.366383937 | -1.506129194 | 1.7191649 |
| Gm15979 | 4.479156939 | -0.868833752 | 0.560541567 |
| Nipal3 | 1375.805614 | -0.178741551 | 0.065167711 |
| Gm37270 | 0.048949532 | -0.243569582 | 3.724945049 |
| Stpg1 | 13.32983798 | -0.977841865 | 0.305826558 |
| Gm25317 | 0 | NA | NA |
| Grhl3 | 12.09498146 | 0.254347427 | 0.307832671 |
| Gm23106 | 0.133280846 | -0.243579231 | 3.654655979 |
| 1700029M20Rik | 0.369889 | 1.084471068 | 1.573017357 |
| Ifnlr1 | 9.250335689 | 1.56308612 | 0.366071913 |
| Gm12989 | 39.77014755 | 0.66760724 | 0.23701307 |
| Il22ra1 | 3.4240698 | 1.844283235 | 0.691473677 |
| Gm12988 | 0.337511725 | -0.149257963 | 1.566363218 |
| Myom3 | 10.58892132 | -2.121849533 | 0.399473718 |
| Gm13000 | 0 | NA | NA |
| Srsf10 | 1164.791877 | 0.2264166 | 0.077913689 |
| Pnrc2 | 595.9273528 | 0.593426577 | 0.102571527 |
| Gm13006 | 0 | NA | NA |
| Cnr2 | 15.77841223 | 3.482784239 | 0.509378433 |
| Fuca1 | 282.912463 | 0.531966317 | 0.093667549 |
| Hmgcl | 258.9557325 | 0.42829692 | 0.09368686 |
| Gale | 89.88904888 | 0.279552978 | 0.123883087 |
| Lypla2 | 893.641742 | 1.22013913 | 0.130338889 |
| Pithd1 | 571.8837247 | 0.425796273 | 0.080511374 |
| Gm13007 | 0.235668049 | -0.724459919 | 2.542840095 |
| Tceb3 | 686.4047522 | 0.642987225 | 0.078060252 |
| Gm13008 | 0.394976806 | -0.243563855 | 1.599456159 |
| Gm24362 | 2.378997451 | -0.619077217 | 0.827749476 |
| Rpl11 | 1068.809872 | 0.593122129 | 0.137430539 |
| Gm13009 | 0 | NA | NA |
| Gm26001 | 0 | NA | NA |
| Id3 | 323.3694017 | 1.162654469 | 0.161636845 |
| E2f2 | 28.44497551 | 1.201660182 | 0.282114765 |

| | | | |
|---|---|---|---|
| Asap3 | 274.1575355 | 2.629614404 | 0.191939352 |
| Tcea3 | 4.626838346 | -0.185586257 | 0.475736599 |
| Zfp46 | 472.2796557 | -0.224631431 | 0.12627171 |
| Gm13013 | 0 | NA | NA |
| Hnrnpr | 1542.097552 | -0.365788581 | 0.078444561 |
| Gm27022 | 0 | NA | NA |
| Htr1d | 41.88636825 | 0.292961409 | 0.201035791 |
| 6030445D17Rik | 4.632584323 | 0.918224209 | 0.433400764 |
| Luzp1 | 1615.824644 | -0.007128322 | 0.061621874 |
| Kdm1a | 902.2554757 | 0.414534681 | 0.071668429 |
| Gm12987 | 8.499369334 | -0.484485242 | 0.510564322 |
| Gm12986 | 0 | NA | NA |
| Gm22180 | 0 | NA | NA |
| 4930549C01Rik | 0 | NA | NA |
| Lactbl1 | 0.276289708 | 0.718217053 | 2.444991654 |
| Gm37583 | 0.251203948 | -0.601197334 | 2.087522995 |
| Ephb2 | 367.915314 | -0.351109427 | 0.101380806 |
| C1qb | 684.4735841 | 1.844360179 | 0.122873961 |
| C1qc | 767.5830084 | 1.828691515 | 0.154052825 |
| C1qa | 854.8487578 | 1.498728721 | 0.170301819 |
| Epha8 | 193.1729168 | -0.787561158 | 0.105952398 |
| Zbtb40 | 195.6296531 | -0.313718216 | 0.170057168 |
| Gm23834 | 0.094097695 | 0.718200421 | 3.724945049 |
| 1700037C06Rik | 0 | NA | NA |
| Gm13001 | 0.262972053 | 0.237330792 | 1.813525 |
| Gm13003 | 0.049553047 | 0.71820487 | 3.724945049 |
| Gm25399 | 0 | NA | NA |
| Gm23263 | 0 | NA | NA |
| Gm13005 | 0.48771898 | -2.037099534 | 1.420468066 |
| Gm13002 | 0.904140992 | -0.513706491 | 0.980595598 |
| Wnt4 | 161.8183686 | 0.145716487 | 0.139253085 |
| Cdc42 | 2845.278538 | 0.490662008 | 0.088166752 |
| Gm25772 | 0 | NA | NA |
| 2810405F17Rik | 18.73872419 | -0.250741421 | 0.351485222 |
| Rpl31-ps10 | 1.205425868 | -0.054908493 | 0.975847256 |
| Cela3a | 0 | NA | NA |
| Cela3b | 0 | NA | NA |
| Gm13010 | 0.090890357 | 0.718200421 | 3.724945049 |
| 1700013G24Rik | 0 | NA | NA |
| Hspg2 | 519.9526523 | 1.469881301 | 0.141887103 |
| Mir7018 | 0 | NA | NA |
| Ldlrad2 | 0.178810541 | -0.645565375 | 2.733025976 |
| Usp48 | 1402.55254 | -0.473102439 | 0.124831523 |
| Rap1gap | 2225.616142 | -0.678793733 | 0.094546915 |
| Rap1gapos | 0.214365142 | 0.726314033 | 2.179830127 |
| Alpl | 62.50808463 | 0.476636803 | 0.170961515 |
| Ece1 | 1063.140229 | 0.629285257 | 0.08911356 |

| | | | |
|---|---|---|---|
| Gm13012 | 0.199351704 | 1.51582692 | 2.615059144 |
| Eif4g3 | 3297.877854 | 0.022163099 | 0.072105263 |
| 1700095J12Rik | 0 | NA | NA |
| Mir6399 | 2.27686558 | 0.085665355 | 0.800634483 |
| 2310026L22Rik | 1.951947418 | 0.808618211 | 0.747661995 |
| Hp1bp3 | 1903.997062 | 0.006215265 | 0.067456849 |
| Sh2d5 | 591.0024521 | 0.211528165 | 0.134464738 |
| Kif17 | 150.6944098 | -0.069020339 | 0.206786713 |
| Ddost | 796.5474421 | 0.758829997 | 0.086663867 |
| Pink1 | 2455.55194 | -0.291744442 | 0.095032482 |
| Mir7019 | 0 | NA | NA |
| Gm27646 | 0 | NA | NA |
| Cda | 12.16163641 | 0.098867796 | 0.336784839 |
| Fam43b | 205.4390261 | -0.421917589 | 0.142158802 |
| AB041806 | 16.25226339 | -0.996709458 | 0.302071687 |
| Rpl38-ps1 | 1.247502082 | 0.693474249 | 0.959330967 |
| Mul1 | 449.0477894 | 0.379171725 | 0.11536398 |
| Camk2n1 | 9074.818375 | -0.94509385 | 0.123163757 |
| Vwa5b1 | 72.56742202 | -0.995915174 | 0.187818697 |
| Gm22961 | 0.044191549 | -0.243571806 | 3.724945049 |
| Ubxn10 | 30.26165497 | 0.583593796 | 0.198845515 |
| Pla2g2c | 2.097114995 | 0.360036014 | 0.630472615 |
| Pla2g2f | 0.721011721 | -1.109392101 | 1.350326585 |
| Pla2g2d | 2.315173619 | 0.870116627 | 0.70788812 |
| Pla2g5 | 22.3998674 | -0.138624248 | 0.266092983 |
| Gm25280 | 0.126225412 | 0.718200913 | 3.698754989 |
| Pla2g2a | 0 | NA | NA |
| Gm13030 | 0.123845245 | 1.035807111 | 3.720871489 |
| Pla2g2e | 6.503439099 | 6.4202113 | 0.924019199 |
| Otud3 | 118.1387416 | -0.029316735 | 0.134396932 |
| Rnf186 | 0.191999622 | -0.243571491 | 3.27205423 |
| Tmco4 | 88.13299696 | 2.393384776 | 0.215811498 |
| Htr6 | 39.32244043 | -0.751746316 | 0.201278742 |
| Nbl1 | 578.5377914 | 0.437461589 | 0.128857554 |
| Minos1 | 533.822241 | 0.12915566 | 0.101481247 |
| Hspe1-ps4 | 0 | NA | NA |
| Gm16287 | 5.315121335 | 0.339544619 | 0.458815758 |
| Capzb | 1556.163037 | 0.428061584 | 0.064618194 |
| Pqlc2 | 134.0855149 | 0.201854684 | 0.119321394 |
| Akr7a5 | 131.3216444 | -0.202720976 | 0.125218211 |
| Gm25732 | 0 | NA | NA |
| Gm24245 | 1.005478281 | -0.450566294 | 2.262465375 |
| Gm13019 | 0 | NA | NA |
| Ccnd3-ps | 1.016942265 | 0.962645719 | 0.990085524 |
| Mrto4 | 197.6742609 | 0.471373846 | 0.106946277 |
| Emc1 | 1429.8396 | 0.089937489 | 0.071332632 |
| Ubr4 | 6917.676742 | 0.460894712 | 0.050714113 |

| | | | |
|---|---|---|---|
| 2700016F22Rik | 3.508143679 | 0.064788176 | 0.54480458 |
| Iffo2 | 505.8284133 | 0.750765636 | 0.156225891 |
| Gm25571 | 0 | NA | NA |
| Gm21969 | 1.469723789 | -0.885507868 | 0.920026982 |
| Aldh4a1 | 91.67888502 | 0.395969526 | 0.130495886 |
| Mir7020 | 0.163190422 | 1.22955094 | 3.396300676 |
| Tas1r2 | 0.187591875 | 0.237324135 | 2.704135342 |
| Pax7 | 25.98741359 | -0.311343575 | 0.236065488 |
| Gm1667 | 0.227050272 | 0.673471025 | 2.173296379 |
| Gm13028 | 0.115000406 | 0.23732098 | 3.724945049 |
| 4933427I22Rik | 0.61103801 | 0.088337447 | 1.35815846 |
| Klhdc7a | 224.24311 | -0.329774238 | 0.193682157 |
| Mir2139 | 0 | NA | NA |
| Igsf21 | 330.4037649 | -0.806840705 | 0.084914861 |
| Gm13027 | 0.336994846 | 0.718753894 | 1.717995599 |
| Gm13029 | 0.895881185 | -0.051489296 | 1.172449263 |
| Gm9867 | 0 | NA | NA |
| Gm13016 | 1.130346611 | 0.51278331 | 1.125159593 |
| Gm13017 | 0.248149725 | 2.088277992 | 2.390389312 |
| Gm13021 | 0 | NA | NA |
| Gm13026 | 0.049553047 | 0.71820487 | 3.724945049 |
| Arhgef10l | 696.9675906 | -0.071564718 | 0.081291036 |
| Gm13025 | 0.470217137 | -0.709122586 | 1.28975348 |
| Gm23045 | 0 | NA | NA |
| Rcc2 | 1056.188668 | 0.272992766 | 0.059840155 |
| Gm25951 | 0 | NA | NA |
| Padi6 | 0.601311955 | -0.867616258 | 1.240676405 |
| Padi4 | 0.880251114 | 2.662809384 | 1.208877134 |
| Padi3 | 0.34999831 | 0.548188333 | 1.509320281 |
| Gm13032 | 0.111006857 | 0.717551239 | 3.724898127 |
| Padi1 | 3.269897503 | -0.058304373 | 0.586026757 |
| 4930515B02Rik | 0.062426873 | 0.237312084 | 3.724945049 |
| Padi2 | 199.5704136 | -1.458610711 | 0.232956774 |
| Gm26226 | 0.275809893 | -1.873223112 | 1.971958652 |
| Gm13031 | 7.163063276 | -1.3467466 | 0.424079331 |
| Sdhb | 803.5181823 | 0.061807195 | 0.072264711 |
| Atp13a2 | 1396.203774 | -0.345496461 | 0.066016488 |
| Mfap2 | 17.61408699 | -0.572627076 | 0.254259027 |
| Crocc | 384.0657555 | -1.282994431 | 0.178277844 |
| Necap2 | 429.1644651 | 1.161267223 | 0.121848008 |
| Gm13047 | 0 | NA | NA |
| Spata21 | 3.445363304 | -1.047600043 | 0.652056297 |
| Szrd1 | 732.8062499 | 0.92843345 | 0.116654433 |
| 4921514A10Rik | 0.424480717 | 1.65095541 | 1.515502857 |
| Fbxo42 | 663.8042497 | 0.104930732 | 0.079273514 |
| Rsg1 | 15.89625249 | -0.881623077 | 0.315775541 |
| C630004L07Rik | 0.566706395 | -1.137356123 | 1.306435263 |

| | | | |
|---|---|---|---|
| Arhgef19 | 124.7706795 | -0.683429136 | 0.199383506 |
| Gm13055 | 0.062426873 | 0.237312084 | 3.724945049 |
| Gm13056 | 0.112396584 | 1.192083721 | 3.724434897 |
| Epha2 | 56.26488158 | 2.508581197 | 0.270989166 |
| Gm25690 | 0 | NA | NA |
| Gm13074 | 0 | NA | NA |
| Gm13076 | 0 | NA | NA |
| Fam131c | 33.32296123 | -0.464839111 | 0.211470263 |
| Clcnka | 0.814794334 | 1.401551341 | 1.235764047 |
| Gm13075 | 0.049553047 | 0.71820487 | 3.724945049 |
| Clcnkb | 0.117046891 | -0.724453471 | 3.724945049 |
| Hspb7 | 5.320519917 | 3.145545625 | 0.747123901 |
| Gm694 | 13.84823866 | 2.129958434 | 0.387080724 |
| Zbtb17 | 276.1721208 | 0.287779363 | 0.107111647 |
| Spen | 2269.995177 | -0.208232727 | 0.070385048 |
| B330016D10Rik | 28.20952649 | -0.192518943 | 0.247115208 |
| Fblim1 | 31.34586486 | 2.12158443 | 0.281135781 |
| Tmem82 | 7.023389443 | 0.81785661 | 0.395814297 |
| AI507597 | 0.761152721 | 1.699018579 | 1.384179342 |
| Slc25a34 | 67.64740712 | -0.515936768 | 0.288783048 |
| Plekhm2 | 1488.934527 | 0.24029362 | 0.097247394 |
| Ddi2 | 701.2059972 | -0.078782308 | 0.075190942 |
| Rsc1a1 | 0 | NA | NA |
| Agmat | 14.27242461 | 0.437387297 | 0.345607365 |
| Dnajc16 | 741.3636435 | -0.135711246 | 0.059764135 |
| Casp9 | 369.2534085 | -0.373207549 | 0.069527967 |
| Cela2a | 0.706300188 | -0.254039121 | 1.523898415 |
| Ctrc | 0 | NA | NA |
| Ctrcos | 0 | NA | NA |
| Efhd2 | 2399.369846 | 1.192811496 | 0.095854469 |
| Gm29367 | 15.45718579 | -0.328302479 | 0.273501394 |
| Fhad1os2 | 0.207873529 | 1.1552549 | 2.194146315 |
| Fhad1 | 63.64227438 | 1.443656308 | 0.225385454 |
| Gm13059 | 0 | NA | NA |
| Fhad1os1 | 0 | NA | NA |
| 4930455G09Rik | 2.445960431 | 3.23911111 | 0.881183183 |
| Tmem51 | 80.10815602 | 1.195748879 | 0.196962775 |
| Gm13053 | 1.791442492 | 0.500539661 | 0.744726778 |
| Tmem51os1 | 3.504632306 | 0.521373417 | 0.56140527 |
| Kazn | 1222.293754 | -0.083348742 | 0.067360817 |
| Gm13062 | 0.107803068 | 0.719187033 | 3.724874737 |
| Gm13052 | 20.91063599 | -1.255160999 | 0.479733986 |
| Gm37624 | 0 | NA | NA |
| Prdm2 | 1439.09533 | -0.082475488 | 0.051951917 |
| mmu-mir-7021 | 0.613445446 | 1.155540082 | 1.377405895 |
| Gm13038 | 0 | NA | NA |
| Gm13039 | 0.205676178 | 1.19183487 | 2.643985805 |

| | | | |
|---|---|---|---|
| Gm22039 | 0 | NA | NA |
| Gm13050 | 32.91936983 | 0.221695203 | 0.190716816 |
| Pdpn | 594.7265377 | 3.402292658 | 0.256361255 |
| Gm26313 | 0 | NA | NA |
| Lrrc38 | 24.86342066 | 0.206135555 | 0.207523122 |
| Anp32b-ps1 | 18.32970893 | 0.530690106 | 0.323949568 |
| Pramel1 | 0.631666884 | 1.416642991 | 1.264067544 |
| Gm13045 | 0.09176491 | 0.237310218 | 3.724945049 |
| Pramef8 | 229.9222564 | 0.660215322 | 0.102712162 |
| Oog4 | 0 | NA | NA |
| Gm13042 | 1.564646527 | 0.929979153 | 0.739397069 |
| Gm13046 | 0 | NA | NA |
| Gm13041 | 0 | NA | NA |
| Gm13043 | 0 | NA | NA |
| Gm13040 | 0 | NA | NA |
| BC080695 | 0 | NA | NA |
| Gm13057 | 0 | NA | NA |
| Gm13058 | 0 | NA | NA |
| Gm13080 | 5.317189742 | 0.128991194 | 0.465585963 |
| Gm13083 | 0 | NA | NA |
| Gm13088 | 0 | NA | NA |
| Gm13085 | 0 | NA | NA |
| Gm13082 | 2.119392185 | -0.492294139 | 0.731490878 |
| Gm13089 | 0 | NA | NA |
| Gm13078 | 0 | NA | NA |
| Gm13087 | 0 | NA | NA |
| Gm13081 | 0 | NA | NA |
| Gm13023 | 0 | NA | NA |
| Gm13084 | 0 | NA | NA |
| Gm13103 | 0 | NA | NA |
| Pramef6 | 0 | NA | NA |
| Gm13100 | 0 | NA | NA |
| Gm13106 | 0 | NA | NA |
| Pramef25 | 0 | NA | NA |
| Gm13101 | 0 | NA | NA |
| Pramef17 | 0 | NA | NA |
| Gm13108 | 0 | NA | NA |
| Gm13104 | 0 | NA | NA |
| Pramel4 | 0 | NA | NA |
| Gm13102 | 0 | NA | NA |
| Oog3 | 0 | NA | NA |
| Oog2 | 0 | NA | NA |
| C87977 | 0 | NA | NA |
| Gm13123 | 21.87672395 | 0.048618293 | 0.230439347 |
| Pramel5 | 0 | NA | NA |
| Gm13120 | 0 | NA | NA |
| Gm13130 | 0 | NA | NA |

| | | | |
|---|---|---|---|
| Gm13128 | 0 | NA | NA |
| Gm13122 | 0 | NA | NA |
| Gm13119 | 0 | NA | NA |
| Gm13125 | 0 | NA | NA |
| Pramef12 | 0 | NA | NA |
| Pramef12os | 0 | NA | NA |
| 1700012P22Rik | 0.273864066 | -0.767436272 | 2.397202529 |
| Gm9944 | 0.048949532 | -0.243569582 | 3.724945049 |
| Aadacl3 | 0 | NA | NA |
| Gm13118 | 0 | NA | NA |
| 9430007A20Rik | 0.068097359 | -0.243569582 | 3.724945049 |
| Gm13121 | 0.042716945 | -0.243571806 | 3.724945049 |
| Gm13126 | 0.046345709 | 0.71820487 | 3.724945049 |
| Gm13124 | 0 | NA | NA |
| Gm13127 | 28.94697736 | -0.090386972 | 0.213888066 |
| Gm13177 | 0 | NA | NA |
| Gm436 | 0 | NA | NA |
| Gm13178 | 0.072025462 | 0.237312084 | 3.724945049 |
| Gm25938 | 0 | NA | NA |
| Gm438 | 0.169902121 | 0.718202065 | 3.402973925 |
| Prdx2-rs3 | 0.044191549 | -0.243571806 | 3.724945049 |
| Dhrs3 | 254.6119046 | 1.454223947 | 0.126374764 |
| Gm38074 | 1.747383901 | -0.394258962 | 0.818883407 |
| Vps13d | 3201.77473 | -0.331727876 | 0.071831724 |
| Tnfrsf1b | 457.2834194 | 4.656742688 | 0.295997974 |
| Tnfrsf8 | 21.18745374 | 0.824987254 | 0.232261292 |
| Gm13024 | 0.288426457 | 0.264931877 | 2.06139142 |
| Gm13224 | 0 | NA | NA |
| Platr16 | 0 | NA | NA |
| Gm13230 | 0.097899064 | -0.724453471 | 3.724945049 |
| Gm13223 | 0.093141081 | -0.724455696 | 3.724945049 |
| Smarca5-ps | 7.147813683 | 0.077417396 | 0.387740463 |
| Gm13229 | 0 | NA | NA |
| Gm13225 | 0.120679735 | 0.631141837 | 3.722822053 |
| Gm21978 | 0.060174537 | 0.237309859 | 3.724945049 |
| Gm13228 | 0 | NA | NA |
| Gm13226 | 9.748555663 | 0.816720108 | 0.365926858 |
| Gm13222 | 0 | NA | NA |
| Gm13212 | 8.306006014 | 3.820475335 | 0.60199765 |
| Gm26763 | 0 | NA | NA |
| Gm13234 | 0.712482738 | -0.047958947 | 1.052233437 |
| Gm13231 | 1.074501331 | 0.335922595 | 1.184043389 |
| Gm13249 | 0 | NA | NA |
| Gm13242 | 0.150086409 | 0.237305146 | 3.724945049 |
| Gm13239 | 0 | NA | NA |
| Gm13232 | 0 | NA | NA |
| Gm13244 | 0.205478627 | 1.922714726 | 3.178544584 |

| | | | |
|---|---|---|---|
| Gm13236 | 0 | NA | NA |
| Gm13243 | 1.675672734 | 0.42875987 | 0.772581113 |
| Vmn2r-ps15 | 0.062426873 | 0.237312084 | 3.724945049 |
| Znf41-ps | 66.78666031 | 0.866915995 | 0.211841761 |
| Gm13238 | 8.42982011 | -0.052463823 | 0.382158255 |
| Gm13246 | 0 | NA | NA |
| Gm13235 | 2.295964185 | 1.884721356 | 0.830420809 |
| Gm13036 | 0 | NA | NA |
| Gm13034 | 8.179144896 | 0.34101845 | 0.354017596 |
| Gm13035 | 0 | NA | NA |
| Gm13051 | 0 | NA | NA |
| Gm26573 | 0 | NA | NA |
| Zfp600 | 0.463671963 | -1.932624043 | 1.311115087 |
| Gm13165 | 0 | NA | NA |
| Gm13170 | 0 | NA | NA |
| Gm13167 | 0 | NA | NA |
| Gm13169 | 0.687288638 | -1.64570781 | 1.191508572 |
| Gm13166 | 0.061920613 | 0.718207094 | 3.724945049 |
| Gm13168 | 0.090665643 | -0.724455696 | 3.724945049 |
| Vmn2r-ps17 | 0 | NA | NA |
| Gm13251 | 42.78458666 | 0.737172372 | 0.183924027 |
| Gm13241 | 5.328295998 | 0.610284962 | 0.433807265 |
| Gm13245 | 1.259358607 | 0.244688327 | 0.93519664 |
| Gm13247 | 3.086345102 | 0.319845424 | 0.844269833 |
| Gm26880 | 0 | NA | NA |
| 1700095A21Rik | 0 | NA | NA |
| Gm13237 | 0.2312517 | -0.243563998 | 2.173638425 |
| Gm13233 | 0.066050875 | 0.71820487 | 3.724945049 |
| Gm13240 | 0.060174537 | 0.237309859 | 3.724945049 |
| Vmn2r-ps16 | 0.194279094 | 1.292225235 | 3.719439735 |
| Gm13248 | 6.591240675 | 1.015772503 | 0.5220622 |
| Gm21411 | 0.046345709 | 0.71820487 | 3.724945049 |
| Gm20760 | 0.333659237 | 1.061701942 | 1.730009445 |
| Gm13140 | 10.11168674 | 0.157199674 | 0.340580824 |
| Gm13149 | 0 | NA | NA |
| Gm13150 | 0 | NA | NA |
| C230088H06Rik | 0 | NA | NA |
| Rex2 | 0.066050875 | 0.71820487 | 3.724945049 |
| Gm13153 | 0 | NA | NA |
| 4933438K21Rik | 0 | NA | NA |
| Gm13160 | 0.210629592 | 1.638094857 | 2.171652041 |
| Gm13142 | 2.102244023 | 1.073665293 | 0.666054123 |
| Gm13162 | 1.043892331 | 1.502871818 | 1.0024099 |
| Vmn2r-ps18 | 0.642632734 | -0.157731964 | 1.240694629 |
| Gm13139 | 85.76215223 | 0.687852995 | 0.134986498 |
| Gm26624 | 9.940525084 | 0.146061025 | 0.338818648 |
| Gm13146 | 6.463804552 | -0.286117088 | 0.390124132 |

| | | | |
|---|---|---|---|
| Gm13163 | 0 | NA | NA |
| Gm13147 | 1.544988629 | 0.189422086 | 0.713251147 |
| Gm13158 | 0.256692543 | -0.702891098 | 1.82158712 |
| Gm13135 | 13.09013645 | 0.404399481 | 0.362164237 |
| Gm13144 | 0 | NA | NA |
| Gm13151 | 2.603243443 | -0.018792558 | 0.657730466 |
| Gm13145 | 0.352653913 | 0.860769418 | 1.954023473 |
| Gm8178 | 0 | NA | NA |
| Gm13161 | 3.441294106 | -0.567248824 | 0.599894975 |
| Vmn2r-ps19 | 2.482748844 | -0.176218725 | 0.606918857 |
| Gm13152 | 33.96153438 | 0.149033433 | 0.235518063 |
| Gm15951 | 0 | NA | NA |
| Vmn2r-ps20 | 4.516610571 | -0.455820654 | 0.474706861 |
| Gm13141 | 0 | NA | NA |
| Gm16503 | 25.68570846 | -0.524678957 | 0.339929171 |
| Gm13154 | 0.973397315 | 0.720358814 | 0.998794655 |
| 2610305D13Rik | 21.91887679 | 0.975746271 | 0.267030446 |
| Gm13155 | 0.092140247 | -0.724455696 | 3.724945049 |
| Gm16211 | 0.547750696 | 1.016927425 | 1.566399231 |
| Zfp534 | 0.518969261 | 1.038343625 | 1.322423306 |
| Gm13137 | 0 | NA | NA |
| Gm13136 | 14.06422963 | 0.507746077 | 0.259937937 |
| Gm13148 | 0.33580603 | 2.492824674 | 1.645955527 |
| Gm13157 | 104.6671989 | 1.539435547 | 0.143548957 |
| Gm20707 | 14.6133478 | 1.328833376 | 0.369425431 |
| Rps19-ps3 | 30.20379719 | -0.071390871 | 0.211102337 |
| Zfp933 | 375.230941 | -0.192342434 | 0.082234202 |
| Miip | 134.4730676 | -0.231554306 | 0.166225909 |
| Fv1 | 52.36557907 | -0.655140955 | 0.163906343 |
| Mfn2 | 2014.021368 | -0.173337987 | 0.037580061 |
| Gm13156 | 0.836555448 | -0.378036866 | 1.015952697 |
| Plod1 | 436.6411955 | 0.929595496 | 0.117759609 |
| 2510039O18Rik | 318.750318 | 0.770828726 | 0.101792432 |
| Nppb | 0.91071486 | 1.787761123 | 1.12557338 |
| Nppa | 2.986404308 | 0.133877415 | 0.739455212 |
| Gm13054 | 12.47324529 | 0.335340186 | 0.370748661 |
| Clcn6 | 1341.943323 | -0.024654325 | 0.047117629 |
| Gm13207 | 0.045882455 | 0.237312084 | 3.724945049 |
| Mthfr | 731.1208707 | 1.788009085 | 0.115573877 |
| Gm13202 | 5.417309824 | 0.313309078 | 0.453725484 |
| Gm13201 | 0.498362359 | -0.983390362 | 1.564850332 |
| Agtrap | 214.8520272 | 1.056482368 | 0.144324926 |
| Draxin | 11.61009598 | 0.104811783 | 0.332806915 |
| Mad2l2 | 105.7912503 | 1.33332324 | 0.160612632 |
| Fbxo6 | 181.4057765 | 1.024230279 | 0.1139461 |
| Mir7022 | 0.141872876 | 0.237324562 | 2.910933148 |
| Fbxo44 | 836.5623358 | -0.224297854 | 0.054201369 |

| | | | |
|---|---|---|---|
| Fbxo2 | 551.7280945 | -0.437193429 | 0.115313948 |
| Ptchd2 | 379.3452522 | -0.71513084 | 0.12676598 |
| Gm13206 | 0.047948698 | -0.243569582 | 3.724945049 |
| Gm13200 | 2.555034347 | -1.361772905 | 0.765211512 |
| Ubiad1 | 175.5010009 | 0.637548978 | 0.119999142 |
| Gm23303 | 1.89929672 | 0.478303008 | 0.778307536 |
| Mtor | 2020.048309 | -0.24685802 | 0.035975212 |
| Angptl7 | 17.72480738 | 1.334945428 | 0.327776089 |
| Gm24002 | 0.589151384 | -0.979685764 | 1.220307909 |
| Gm23318 | 1.238646735 | -0.18251239 | 0.835480403 |
| Exosc10 | 484.0039437 | 0.367009119 | 0.080540262 |
| Gm13204 | 0 | NA | NA |
| Gm13209 | 0.331673312 | 2.431474473 | 1.684761418 |
| Srm | 303.9037999 | 0.112344921 | 0.088690543 |
| Masp2 | 31.81998135 | -0.472931569 | 0.280606348 |
| Tardbp | 2054.279046 | -0.051724075 | 0.074351483 |
| Gm572 | 0.786597823 | 0.465588119 | 1.112830994 |
| Gm13203 | 0 | NA | NA |
| Casz1 | 60.05755845 | 0.417563758 | 0.177194503 |
| Gm15969 | 0 | NA | NA |
| Gm13205 | 27.32691937 | -1.045251706 | 0.228943005 |
| Gm17029 | 1.365296087 | 1.217346078 | 0.902357735 |
| Pex14 | 521.3504447 | 0.337955844 | 0.088295622 |
| Gm9506 | 0 | NA | NA |
| Dffa | 268.8330888 | -0.351733862 | 0.072720638 |
| Cort | 13.59908539 | -1.292689999 | 0.323192852 |
| Apitd1 | 26.0334167 | -0.282809177 | 0.214009075 |
| Rpsa-ps12 | 20.84511033 | 0.196683542 | 0.335031979 |
| Pgd | 591.4610587 | 1.227116135 | 0.098992696 |
| Kif1b | 15537.60889 | -0.386130247 | 0.050632383 |
| Ube4b | 1942.055481 | -0.086171017 | 0.04567072 |
| Ube4bos3 | 0.106056992 | 0.237307635 | 3.724945049 |
| Ube4bos2 | 0.038981395 | 0.718207094 | 3.724945049 |
| Ube4bos1 | 0.415914162 | 0.478133072 | 3.33403506 |
| Rbp7 | 1.09427696 | 1.683019048 | 1.132986957 |
| Trmt112-ps2 | 31.55425196 | 0.411668716 | 0.226333392 |
| Nmnat1 | 53.5119218 | -0.413583107 | 0.16951734 |
| Gm13097 | 0.04393175 | -0.243571806 | 3.724945049 |
| Lzic | 141.3885089 | 0.394472071 | 0.112533887 |
| Gm13071 | 9.973492174 | -0.523628705 | 0.318944786 |
| Gm24381 | 0 | NA | NA |
| Ctnnbip1 | 373.0545822 | -0.260109665 | 0.099230189 |
| Mir5616 | 0.908927132 | 1.853502045 | 1.207627012 |
| Gm13066 | 0.3242121 | 0.613240568 | 1.783489725 |
| Clstn1 | 10198.28505 | -0.817169103 | 0.08192107 |
| Pik3cd | 275.4671803 | 1.135000093 | 0.117891283 |
| Mir7023 | 0 | NA | NA |

| | | | |
|---|---|---|---|
| Tmem201 | 968.9539549 | -0.376154114 | 0.089901852 |
| Slc25a33 | 270.7812472 | -0.049227097 | 0.119619966 |
| Gm13068 | 0.185281328 | -1.567553191 | 2.686846338 |
| Gm16188 | 0.247276735 | 2.182825213 | 1.996544723 |
| Gm13073 | 0.234101041 | 0.717760958 | 1.878164381 |
| Gm13065 | 0.044544648 | 0.237309859 | 3.724945049 |
| Gm13064 | 0.150086409 | 0.237305146 | 3.724945049 |
| n-R5s193 | 0 | NA | NA |
| AL626808.1 | 0 | NA | NA |
| Spsb1 | 673.0771038 | 2.059359896 | 0.186374903 |
| Gm13070 | 0.062426873 | 0.237312084 | 3.724945049 |
| H6pd | 247.2321755 | 0.531795608 | 0.136666778 |
| Gm13067 | 8.956120355 | -0.673838096 | 0.376702958 |
| Mir34a | 0.164553454 | 0.719098949 | 2.779126835 |
| Gm37795 | 24.19206811 | -0.569349139 | 0.331219047 |
| Gpr157 | 52.57716929 | 0.929949254 | 0.192292511 |
| Gm13093 | 6.364908752 | -1.538062259 | 0.4726802 |
| Slc2a5 | 12.42258672 | -0.34100327 | 0.294268825 |
| Slc2a7 | 0 | NA | NA |
| Car6 | 1.391048348 | 4.498317708 | 1.184192372 |
| Gm13094 | 0.173900533 | 1.254500896 | 2.738247499 |
| Eno1 | 2763.278328 | 0.102180859 | 0.0664374 |
| Rere | 3154.94565 | -0.0889784 | 0.061099514 |
| Gm13092 | 5.881560967 | 0.798089167 | 0.638665431 |
| Gm23209 | 6.850924464 | -1.551849994 | 0.547158147 |
| Gm13091 | 33.11693011 | 0.780879998 | 0.26607215 |
| Slc45a1 | 569.9271968 | -0.197946913 | 0.063551145 |
| Gm16079 | 0 | NA | NA |
| Gm13049 | 0 | NA | NA |
| 1700045H11Rik | 0.109982625 | 0.237318756 | 3.724945049 |
| Rps15a-ps3 | 6.113901122 | 0.515341447 | 0.395212615 |
| Errfi1 | 790.4060845 | 1.085735017 | 0.236306287 |
| Park7 | 548.339284 | 0.021997979 | 0.102164909 |
| Tnfrsf9 | 42.91515327 | 4.762453264 | 0.593050343 |
| Camta1 | 4644.702699 | -0.515805538 | 0.05366957 |
| Gm23405 | 0 | NA | NA |
| Uts2 | 1.009670171 | 2.989283314 | 1.180403596 |
| Per3 | 705.3281863 | -1.511097221 | 0.23828173 |
| Vamp3 | 420.1066725 | 0.832756424 | 0.119349078 |
| Gm13090 | 0.329009797 | 0.881797274 | 1.509784535 |
| 4930589P08Rik | 0.062426873 | 0.237312084 | 3.724945049 |
| Gm37141 | 0.038981395 | 0.718207094 | 3.724945049 |
| Dnajc11 | 693.2058939 | 0.29802501 | 0.062752476 |
| Thap3 | 106.1287797 | 0.043026452 | 0.126955012 |
| Phf13 | 200.4894639 | 0.349285169 | 0.163953567 |
| Klhl21 | 246.8014045 | 0.230771074 | 0.110340751 |
| Zbtb48 | 85.35332904 | 0.254108282 | 0.159986453 |

| | | | |
|---|---|---|---|
| Tas1r1 | 4.276683327 | 0.981925809 | 0.603842043 |
| Nol9 | 273.9583489 | 0.335448315 | 0.081362179 |
| Plekhg5 | 606.8905356 | -0.504547229 | 0.166132339 |
| Tnfrsf25 | 29.08316862 | -1.437817937 | 0.600668368 |
| Espn | 20.79395242 | -0.484430139 | 0.229778267 |
| Gm13077 | 2.185424001 | 1.006150934 | 0.682937139 |
| Hes2 | 0.293168959 | 0.309873513 | 2.040869581 |
| Acot7 | 1614.713023 | -0.21111351 | 0.072910806 |
| Gm27500 | 0 | NA | NA |
| Gm13096 | 2.546102757 | -1.834177756 | 0.722417553 |
| Gpr153 | 338.048994 | -0.718828737 | 0.091827138 |
| Hes3 | 57.0044881 | 0.133695001 | 0.263914783 |
| Icmt | 1103.865666 | -0.402089444 | 0.118276185 |
| Rnf207 | 32.84951904 | -1.357607485 | 0.268253087 |
| Rpl22 | 661.306753 | 0.507907232 | 0.117863659 |
| Chd5 | 2858.157954 | -0.884736706 | 0.095613626 |
| Gm13645 | 1.241205481 | -2.620628873 | 1.123046034 |
| Kcnab2 | 3430.815311 | -0.68740711 | 0.08007702 |
| Gm16333 | 0.211164147 | 1.166722342 | 2.556588043 |
| Gm16334 | 14.13596111 | -1.0546537 | 0.355063324 |
| Nphp4 | 205.6488101 | -0.75133493 | 0.192715368 |
| Gm13172 | 0 | NA | NA |
| Gm833 | 0.678447637 | 1.80306331 | 1.351656537 |
| Gm25779 | 0 | NA | NA |
| Gm13173 | 0.184770788 | 0.718214619 | 2.691102959 |
| Gm13174 | 0.2360389 | 1.079679865 | 2.132223921 |
| Ajap1 | 418.755463 | -0.40517397 | 0.115948101 |
| Gm25880 | 0 | NA | NA |
| Gm13115 | 0.044191549 | -0.243571806 | 3.724945049 |
| BC039966 | 5.183547717 | -1.19422748 | 0.497167127 |
| A430005L14Rik | 129.8028141 | 0.392084102 | 0.11975762 |
| Dffb | 64.42929542 | -0.29890839 | 0.204832943 |
| Cep104 | 594.1324898 | 0.14163979 | 0.069901374 |
| Lrrc47 | 577.7083269 | 0.20766483 | 0.09598504 |
| Smim1 | 121.5585543 | -0.152589353 | 0.144591363 |
| Ccdc27 | 0.478716937 | -0.820888695 | 1.479631375 |
| Gm13117 | 0 | NA | NA |
| Gm13114 | 0 | NA | NA |
| Trp73 | 20.9206457 | -0.890920202 | 0.305772898 |
| Trp73os | 0.141071297 | 0.237322348 | 3.564043836 |
| Gm26036 | 0 | NA | NA |
| Wrap73 | 158.2566248 | -0.114860488 | 0.097940466 |
| Tprgl | 931.6745304 | -0.242994722 | 0.090104244 |
| Megf6 | 28.54359773 | 0.031666801 | 0.203639091 |
| Gm13132 | 0 | NA | NA |
| Arhgef16 | 13.58180626 | 0.081226964 | 0.297338539 |
| Prdm16 | 164.5019658 | 0.018669471 | 0.124591713 |

| | | | |
|---|---|---|---|
| B230104I21Rik | 0.989733885 | -2.523149946 | 1.222184612 |
| Gm13133 | 0.742042093 | 2.400629521 | 1.271847401 |
| Gm13134 | 0.593269415 | 1.144457578 | 1.188029387 |
| Gm13111 | 52.71521894 | -1.683831654 | 0.275938528 |
| Gm27202 | 0 | NA | NA |
| Gm13110 | 1.937464243 | -0.197914444 | 0.762068435 |
| Actrt2 | 0 | NA | NA |
| Gm13113 | 1.711017467 | 0.54762266 | 0.795753992 |
| Ttc34 | 32.00110063 | -0.267797103 | 0.189374005 |
| Gm13112 | 3.984283232 | -1.016563694 | 0.545112588 |
| Mmel1 | 12.70547386 | -0.140020745 | 0.316967575 |
| Fam213b | 258.5981968 | -0.537841797 | 0.135694623 |
| Tnfrsf14 | 53.00596033 | 3.786054108 | 0.283492358 |
| Gm20421 | 6.460871859 | -0.905095727 | 0.442519206 |
| Hes5 | 56.4435291 | -2.641513516 | 0.343608828 |
| Pank4 | 318.1852538 | -0.103358061 | 0.073013921 |
| Plch2 | 799.0502262 | -1.274061046 | 0.148662828 |
| Gm10564 | 2.918309484 | -1.627540866 | 0.851675016 |
| Pex10 | 129.4440069 | -0.240515736 | 0.118241457 |
| Rer1 | 559.9984354 | 0.684220033 | 0.089567044 |
| Morn1 | 92.30872075 | -0.790699691 | 0.232812044 |
| Ski | 2357.896731 | 0.019127921 | 0.090298245 |
| 2610002J02Rik | 142.9826346 | 0.614229773 | 0.116723283 |
| Prkcz | 1482.735323 | -0.701766795 | 0.107056179 |
| Gm27200 | 1.590510877 | 0.064185143 | 0.913877532 |
| Gabrd | 409.0842723 | -1.058683932 | 0.163098129 |
| 5830444B04Rik | 0.822034208 | 0.559870037 | 1.052977336 |
| Cfap74 | 161.3175935 | -0.752501244 | 0.185711194 |
| Tmem52 | 14.34551441 | 1.565953226 | 0.360369951 |
| Gnb1 | 6791.840275 | -0.146664699 | 0.056712249 |
| Gm13171 | 0.045882455 | 0.237312084 | 3.724945049 |
| Nadk | 615.6859532 | 0.441948986 | 0.092628428 |
| Slc35e2 | 942.3227332 | -0.203336177 | 0.060134872 |
| Gm16023 | 7.870775316 | 0.382162187 | 0.437274561 |
| Gm10563 | 0.110242424 | 0.237318756 | 3.724945049 |
| Gm16024 | 3.554695965 | -0.84390227 | 0.671092283 |
| Cdk11b | 500.6534106 | 0.500699436 | 0.131480206 |
| Mmp23 | 18.56308676 | 0.001261633 | 0.268161024 |
| Mib2 | 880.1110826 | -0.26313612 | 0.066924342 |
| AL670236.1 | 0 | NA | NA |
| B930041F14Rik | 0 | NA | NA |
| Ssu72 | 463.6570633 | 0.335325301 | 0.079058738 |
| Gm22573 | 0.048949532 | -0.243569582 | 3.724945049 |
| 1500002C15Rik | 18.92769097 | -0.542750527 | 0.250283809 |
| Tmem240 | 359.2263579 | -0.752134018 | 0.108561646 |
| Atad3a | 323.5868663 | 0.209203806 | 0.103570905 |
| Atad3aos | 2.955322402 | -0.720436996 | 0.577109026 |

| | | | |
|---|---|---|---|
| Vwa1 | 254.228511 | 0.835775518 | 0.110153734 |
| Gm26840 | 23.01521584 | 0.517959568 | 0.225145823 |
| Tmem88b | 1074.722227 | 0.172063293 | 0.137654708 |
| Gm27743 | 0 | NA | NA |
| Ankrd65 | 0.81397646 | -1.120330603 | 1.095133031 |
| Mrpl20 | 295.0369231 | 0.391766502 | 0.13809926 |
| Ccnl2 | 1143.506995 | 0.136947163 | 0.132800958 |
| Aurkaip1 | 435.2140168 | 0.347500146 | 0.112267438 |
| Mxra8os | 0.102891481 | -0.243576254 | 3.724945049 |
| Mxra8 | 155.5730168 | 1.098624232 | 0.181722277 |
| Dvl1 | 770.5694415 | -0.059270687 | 0.075513956 |
| Tas1r3 | 8.894659495 | -0.086556744 | 0.354156566 |
| Cptp | 215.1678806 | 0.223771578 | 0.100145349 |
| Cpsf3l | 232.6024494 | 0.669490157 | 0.074859814 |
| Pusl1 | 121.0895384 | -0.40248457 | 0.134635017 |
| Acap3 | 1132.056167 | -0.511896054 | 0.107558397 |
| Gm22991 | 0 | NA | NA |
| Ube2j2 | 313.5792266 | 0.398044057 | 0.096836146 |
| Fam132a | 40.80633962 | -1.238499228 | 0.249385604 |
| B3galt6 | 79.13773638 | 0.201594599 | 0.1669965 |
| Sdf4 | 2638.964062 | -0.023232967 | 0.055197537 |
| Tnfrsf4 | 12.95165872 | 1.248301387 | 0.465222653 |
| Gm37090 | 21.13329923 | -1.231119584 | 0.297591734 |
| Gm16008 | 35.8777959 | -0.348769651 | 0.218033116 |
| Gm25982 | 1.203677616 | 0.197882783 | 1.017694276 |
| Tnfrsf18 | 8.73841548 | 6.69095497 | 0.963589223 |
| Ttll10 | 1.729677859 | -0.267604389 | 0.767209308 |
| Gm13648 | 0.840567365 | -1.112972244 | 1.138544079 |
| Mir429 | 0 | NA | NA |
| Mir200a | 0.075043205 | 0.237309859 | 3.724945049 |
| Mir200b | 0.044191549 | -0.243571806 | 3.724945049 |
| 9430015G10Rik | 166.5177331 | -0.662117674 | 0.145904697 |
| Agrn | 2086.461589 | 0.240226375 | 0.094233451 |
| Isg15 | 2298.739811 | 10.05779218 | 0.440094005 |
| AW011738 | 181.0853581 | 2.782778573 | 0.217144111 |
| Perm1 | 7.861063657 | 1.330015764 | 0.405541044 |
| Plekhn1 | 36.96362493 | 0.938120324 | 0.193038342 |
| Klhl17 | 254.2446718 | -0.535944598 | 0.120713712 |
| Mir7658 | 0 | NA | NA |
| Noc2l | 659.9224954 | 0.350973732 | 0.073854091 |
| Samd11 | 37.37872384 | -0.097849135 | 0.214928404 |
| Vmn2r-ps159 | 0.208220181 | -0.769418513 | 3.172055944 |
| Gm20782 | 0 | NA | NA |
| Vmn1r-ps5 | 0 | NA | NA |
| Gm3027 | 0 | NA | NA |
| RP24-481N17.5 | 0 | NA | NA |
| RP24-481N17.7 | 0 | NA | NA |

| | | | |
|---|---|---|---|
| RP24-481N17.8 | 0 | NA | NA |
| Gm6486 | 0 | NA | NA |
| Gm3053 | 0 | NA | NA |
| RP23-128P4.1 | 0 | NA | NA |
| Gm8715 | 0 | NA | NA |
| RP23-135F23.1 | 0 | NA | NA |
| Gm15772 | 876.5883103 | 0.680201086 | 0.165290538 |
| RP23-135F23.2 | 0 | NA | NA |
| Cdk6 | 185.3456827 | 2.02523827 | 0.175047833 |
| RP23-454P8.1 | 0.082913145 | 0.23732098 | 3.724945049 |
| RP23-454P8.3 | 0.295949093 | 1.731823648 | 2.042708398 |
| Fam133b | 414.4885661 | -0.152060703 | 0.090406543 |
| RP23-434H17.4 | 7.609880518 | -0.744383405 | 0.407509189 |
| n-R5s169 | 0 | NA | NA |
| 1700109H08Rik | 123.3221518 | -0.6439902 | 0.378611562 |
| Rbm48 | 131.5051051 | 0.220679342 | 0.137076027 |
| RP23-434H17.3 | 18.64070939 | -0.840574279 | 0.313329033 |
| Pex1 | 339.3891102 | -0.350322155 | 0.131863337 |
| RP23-434H17.5 | 1.254627161 | 0.133836822 | 0.879319676 |
| RP23-434H17.6 | 46.52130311 | -0.799733829 | 0.154017144 |
| Gatad1 | 720.1271276 | -0.043940953 | 0.085392775 |
| Tmbim7 | 0.149576918 | 1.19849824 | 2.863264735 |
| Gm26238 | 0.120349074 | 0.237305146 | 3.724945049 |
| Ankib1 | 1655.656652 | 0.139982945 | 0.059620088 |
| Gm22926 | 0 | NA | NA |
| RP23-425I8.2 | 106.4878435 | -0.69024484 | 0.246377249 |
| Krit1 | 1289.563298 | -0.32426546 | 0.106975209 |
| Lrrd1 | 7.878715828 | 0.39019753 | 0.33766144 |
| Rps4x-ps | 7.918913376 | 0.831075527 | 0.364017395 |
| Mterf1a | 31.30236157 | 0.020060872 | 0.240448905 |
| Akap9 | 2881.578872 | -0.191713191 | 0.100000056 |
| Wdr46-ps | 30.12730886 | -0.782099944 | 0.279426804 |
| RP23-369O21.1 | 30.72591236 | -0.577092249 | 0.263737791 |
| Cyp51 | 835.8806558 | 0.257254937 | 0.137656398 |
| Mterf1b | 11.84616759 | -0.319490437 | 0.347373973 |
| Fzd1 | 479.4036875 | 0.334288287 | 0.14002572 |
| Gm15750 | 1.582330525 | -0.655050085 | 0.870770288 |
| RP23-114F4.1 | 8.620768254 | -0.783368495 | 0.472254977 |
| AC026231.1 | 0 | NA | NA |
| Cdk14 | 2684.227082 | -0.899056303 | 0.082320024 |
| RP23-114F4.2 | 25.16791745 | -1.566687755 | 0.360158619 |
| RP23-114F4.3 | 17.31984568 | -1.034913292 | 0.309377335 |
| Gm25037 | 1.520499152 | 1.490157011 | 0.971626151 |
| RP23-114F4.4 | 36.06826517 | -1.156492159 | 0.413956641 |
| RP23-282L6.2 | 14.6407167 | -1.098054816 | 0.340559716 |
| RP23-282L6.1 | 20.90691181 | -1.10760925 | 0.314259101 |
| RP23-407F12.1 | 44.77564562 | -0.907349344 | 0.245149821 |

| | | | |
|---|---|---|---|
| RP23-396K2.1 | 12.0291979 | 0.179601639 | 0.331910434 |
| RP23-396K2.2 | 12.35285954 | 0.347466495 | 0.356925493 |
| 1700015F17Rik | 1.819296732 | 0.619781188 | 0.752239106 |
| Cldn12 | 558.7465676 | 0.029439114 | 0.076510637 |
| Gtpbp10 | 131.1217465 | 0.164327505 | 0.160818734 |
| RP23-119M19.2 | 0 | NA | NA |
| Gm8773 | 3.281282651 | 1.024060918 | 0.550149405 |
| Cfap69 | 276.8459991 | -0.511003532 | 0.116145441 |
| Steap2 | 544.5451271 | -0.143026455 | 0.11178529 |
| RP23-261D18.1 | 0.350305046 | -1.205778921 | 1.764454447 |
| Steap1 | 8.981601178 | 0.30918508 | 0.358876113 |
| Gm26825 | 3.449447184 | 0.181464314 | 0.582116744 |
| Gm15459 | 4780.466862 | 0.402790631 | 0.104141966 |
| RP23-261D18.2 | 0 | NA | NA |
| Gm23838 | 0 | NA | NA |
| RP24-323L24.1 | 0.269679299 | -0.999128353 | 1.590586128 |
| RP24-323L24.2 | 0 | NA | NA |
| Gm24686 | 0 | NA | NA |
| RP24-338C23.1 | 0.161949631 | 1.583504458 | 2.7626009 |
| RP24-338C23.2 | 0.322473324 | 0.73162861 | 1.767917875 |
| RP23-189N11.1 | 0.187072314 | -0.724459847 | 2.668356397 |
| RP23-74J16.1 | 1.533852754 | -1.910284369 | 1.025592684 |
| Zfp804b | 64.00592618 | -1.023850851 | 0.142623289 |
| Gm24525 | 5.266732454 | -1.254362823 | 0.491834007 |
| Tubb4b-ps1 | 53.69871365 | 0.34637329 | 0.184278985 |
| RP24-72E12.3 | 27.34862605 | -1.683827813 | 0.33781543 |
| RP24-72E12.2 | 24.54249589 | -1.129200393 | 0.29015918 |
| RP24-72E12.4 | 0 | NA | NA |
| 4921511H03Rik | 0 | NA | NA |
| Gm4960 | 0.287827876 | 0.221924985 | 2.062203635 |
| RP23-465J5.1 | 0.130017973 | 0.238301601 | 3.662714184 |
| Gm8802 | 0.060174537 | 0.237309859 | 3.724945049 |
| Gm22196 | 0 | NA | NA |
| RP23-212F6.1 | 0 | NA | NA |
| Steap4 | 41.56055496 | 4.221025173 | 0.694027915 |
| Gm15731 | 0.761917678 | 2.080581888 | 1.219745286 |
| Sri | 562.266319 | 0.312407321 | 0.088950943 |
| Gm15730 | 0.062426873 | 0.237312084 | 3.724945049 |
| Adam22 | 4541.002357 | -0.902516564 | 0.092234901 |
| Gm21759 | 0.083172944 | 0.23732098 | 3.724945049 |
| Gm24196 | 4.833961456 | -1.60931692 | 0.544540516 |
| Gm23993 | 0 | NA | NA |
| RP23-201E13.1 | 54.61168029 | -1.360352883 | 0.283577143 |
| Dbf4 | 47.80200343 | 1.148962114 | 0.259316663 |
| Slc25a40 | 206.1233218 | -0.20885168 | 0.098081412 |
| Rundc3b | 435.3356137 | -0.416800671 | 0.095746806 |
| RP24-159A20.4 | 29.56068002 | -1.172284176 | 0.290192991 |

| | | | |
|---|---|---|---|
| Abcb1a | 172.8364366 | 0.781963409 | 0.160805532 |
| RP24-223N5.1 | 5.526269997 | 0.026966659 | 0.509760535 |
| RP23-162P5.1 | 1.143206505 | -0.824995924 | 0.927875041 |
| Gm5106 | 1.599390384 | 0.649080689 | 0.874615269 |
| Abcb1b | 71.1266548 | 1.371880224 | 0.169358988 |
| RP23-338P12.2 | 0.602373977 | 1.649285301 | 1.377942321 |
| Abcb4 | 48.46738024 | -1.496013189 | 0.187116894 |
| RP23-338P12.3 | 11.71418499 | 2.08395197 | 0.3666602 |
| Crot | 573.9541497 | 0.730411908 | 0.122041665 |
| Gm15610 | 11.65246704 | 0.652691574 | 0.453668425 |
| Gm37153 | 0.09689823 | -0.724453471 | 3.724945049 |
| Gm27941 | 0 NA | NA | |
| Gm15611 | 0.120349074 | 0.237305146 | 3.724945049 |
| Tmem243 | 183.1811946 | 1.095384536 | 0.127951246 |
| Dmtf1 | 881.7559076 | -0.163929637 | 0.099459609 |
| Gm24666 | 1.171812973 | -0.131322754 | 0.85586375 |
| Gm7332 | 10.24110484 | 1.014089373 | 0.366903625 |
| Gm15734 | 0.410151891 | -0.276858381 | 1.563841803 |
| 9330182L06Rik | 795.1077193 | -0.186662324 | 0.057727347 |
| Gm15733 | 3.241155436 | -1.276820659 | 0.60735302 |
| Mir879 | 0.599062483 | -0.525754911 | 1.317589903 |
| Grm3 | 803.8407714 | -1.760080203 | 0.129447714 |
| RP24-118C7.3 | 12.97903413 | -2.569569816 | 0.366526321 |
| Gm23312 | 0 NA | NA | |
| RP23-193N17.1 | 0.49396662 | 0.265132965 | 1.465755527 |
| RP23-193C5.2 | 0.302344664 | 2.089901222 | 3.702587768 |
| Gm6586 | 0.313880683 | 1.72868081 | 1.781582291 |
| RP23-193C5.3 | 0.217113707 | 0.263982495 | 2.548098786 |
| Gm10482 | 0.386285301 | 1.173453022 | 1.568691179 |
| Gm28685 | 0.419836731 | 1.281230695 | 1.366927889 |
| Gm5152 | 0.268081508 | 1.637406458 | 2.017854505 |
| RP23-412L2.1 | 0.048949532 | -0.243569582 | 3.724945049 |
| Gm17091 | 0.90048758 | -2.264615267 | 1.088906837 |
| RP24-383C4.1 | 0.834920584 | -2.87713555 | 1.24725653 |
| RP23-281K10.3 | 0 NA | NA | |
| RP23-281K10.1 | 0 NA | NA | |
| RP23-281K10.2 | 0 NA | NA | |
| Gm6455 | 0 NA | NA | |
| RP24-113C21.2 | 0 NA | NA | |
| RP23-307D8.3 | 0 NA | NA | |
| Gm8871 | 0 NA | NA | |
| Gm8857 | 0 NA | NA | |
| RP24-79B1.2 | 0 NA | NA | |
| RP23-307D8.4 | 0 NA | NA | |
| RP24-79B1.3 | 0 NA | NA | |
| RP23-307D8.5 | 0 NA | NA | |
| Gm8879 | 0 NA | NA | |

| | | | |
|---|---|---|---|
| RP23-92D22.3 | 0 | NA | NA |
| RP23-307D8.7 | 0 | NA | NA |
| Gm5861 | 0 | NA | NA |
| RP23-307D8.9 | 0 | NA | NA |
| Gm8890 | 0 | NA | NA |
| RP24-79B1.4 | 0 | NA | NA |
| RP23-307D8.8 | 0 | NA | NA |
| Speer1 | 0 | NA | NA |
| RP24-390F5.2 | 0 | NA | NA |
| RP23-307D8.10 | 0 | NA | NA |
| Gm8897 | 0 | NA | NA |
| RP23-299O17.3 | 0 | NA | NA |
| RP23-299O17.4 | 0.739845808 | 0.621345486 | 1.389968935 |
| RP23-307D8.11 | 0 | NA | NA |
| Gm8906 | 0 | NA | NA |
| RP23-299O17.5 | 0 | NA | NA |
| RP23-307D8.12 | 0 | NA | NA |
| Gm6460 | 0 | NA | NA |
| RP23-395E24.4 | 0 | NA | NA |
| RP23-307D8.6 | 0 | NA | NA |
| Gm8922 | 0 | NA | NA |
| 1700003C15Rik | 0 | NA | NA |
| RP23-395E24.3 | 0 | NA | NA |
| Gm8926 | 0 | NA | NA |
| RP23-408K8.3 | 0 | NA | NA |
| RP23-307D8.13 | 0 | NA | NA |
| Gm6465 | 0 | NA | NA |
| RP24-341O14.4 | 0 | NA | NA |
| RP23-307D8.14 | 0 | NA | NA |
| 4933402N22Rik | 0 | NA | NA |
| RP24-341O14.3 | 0.201351563 | -0.692409618 | 2.201968076 |
| Gm29066 | 0 | NA | NA |
| Gm38242 | 0 | NA | NA |
| RP24-235L14.3 | 0.152964131 | 0.237325139 | 2.843489581 |
| RP23-364E18.2 | 0.046345709 | 0.71820487 | 3.724945049 |
| Sema3d | 247.8894151 | 0.464352074 | 0.152266947 |
| RP23-237H9.3 | 0 | NA | NA |
| RP23-237H9.2 | 4.98564735 | -0.251377784 | 0.438374835 |
| RP24-346C14.1 | 0 | NA | NA |
| Gm23243 | 0 | NA | NA |
| Gm10481 | 11.05583305 | 0.049772254 | 0.310972289 |
| RP23-30B3.2 | 0 | NA | NA |
| Gm10108 | 1.450945253 | 1.720215258 | 0.845808448 |
| Sema3a | 401.2948366 | -0.433343181 | 0.09168272 |
| RP23-30B3.4 | 0 | NA | NA |
| RP23-10F17.2 | 15.6478416 | -1.092113021 | 0.363874454 |
| RP23-10F17.3 | 7.940230631 | -0.442021692 | 0.371326938 |

| | | | |
|---|---|---|---|
| RP23-10F17.4 | 8.41488845 | -1.221415488 | 0.453745029 |
| RP23-192D21.1 | 16.18937152 | -0.510626355 | 0.251291336 |
| RP23-192D21.2 | 0 | NA | NA |
| RP24-405C17.3 | 0 | NA | NA |
| RP24-405C17.4 | 0.101408268 | 0.717551239 | 3.724898127 |
| Gm6647 | 0.217530652 | 1.222589782 | 2.172555094 |
| Speer3 | 0 | NA | NA |
| Gm6650 | 0 | NA | NA |
| RP24-405C17.5 | 0 | NA | NA |
| RP24-405C17.6 | 0 | NA | NA |
| RP24-405C17.7 | 0 | NA | NA |
| Gm24054 | 0 | NA | NA |
| Sema3e | 550.5002642 | 0.27516142 | 0.115608123 |
| RP24-194J1.1 | 17.56400965 | -0.380015789 | 0.291755972 |
| Gm4128 | 0 | NA | NA |
| RP24-481P22.1 | 12.20307049 | 0.307210367 | 0.321630888 |
| Pclo | 8266.158298 | -0.877866181 | 0.108135335 |
| Gm26918 | 1.104625153 | -1.304812314 | 1.068739184 |
| RP23-342M4.2 | 153.5905569 | -1.071965004 | 0.236494795 |
| RP23-342M4.3 | 100.8300785 | -1.054264434 | 0.234479109 |
| RP23-342M4.4 | 73.09923858 | -1.040753222 | 0.195326617 |
| RP23-342M4.5 | 112.0484156 | -1.227753549 | 0.258884753 |
| RP23-342M4.6 | 56.2336579 | -1.398162315 | 0.283370886 |
| RP23-342M4.7 | 8.648288359 | -1.000114268 | 0.374709101 |
| Gm26954 | 0.454340154 | 0.217613459 | 1.371704231 |
| RP23-451C1.2 | 88.75190297 | -0.669391857 | 0.216770841 |
| RP23-451C1.1 | 51.00169416 | -0.807443833 | 0.267187021 |
| Gm9758 | 0.061920613 | 0.718207094 | 3.724945049 |
| Speer4e | 0.072025462 | 0.237312084 | 3.724945049 |
| RP23-239L21.4 | 0.368800866 | -1.637688986 | 1.553252988 |
| Gm10354 | 0.066050875 | 0.71820487 | 3.724945049 |
| Gm17019 | 0.062426873 | 0.237312084 | 3.724945049 |
| RP24-152F23.2 | 0.163328881 | 1.142321958 | 2.769442353 |
| Gm8579 | 0.295956696 | 2.238693612 | 1.677163455 |
| Gm21847 | 0.231556444 | 1.637815325 | 2.111331104 |
| RP24-152F23.3 | 0.112396584 | 1.192083721 | 3.724434897 |
| RP23-465P18.1 | 0.409112724 | 0.803298303 | 1.687251256 |
| Gm21655 | 0 | NA | NA |
| Speer4d | 0 | NA | NA |
| 4930572O03Rik | 0 | NA | NA |
| Speer4cos | 0.174149299 | 1.185086784 | 3.359371254 |
| Speer4c | 0.220272213 | 1.091888084 | 2.175258066 |
| RP23-77E18.1 | 0 | NA | NA |
| RP23-77E18.2 | 0 | NA | NA |
| RP23-77E18.3 | 0 | NA | NA |
| RP23-77E18.5 | 0.109124069 | -0.24357403 | 3.724945049 |
| RP23-77E18.6 | 0 | NA | NA |

| | | | |
|---|---|---|---|
| Gm3510 | 30.33973269 | -1.098298223 | 0.225333206 |
| Cacna2d1 | 2942.12003 | -0.864998329 | 0.098566565 |
| RP23-123P10.5 | 0.04393175 | -0.243571806 | 3.724945049 |
| RP24-220P21.4 | 77.60211899 | -1.030592889 | 0.255987895 |
| RP24-372A2.2 | 0 | NA | NA |
| Gm8984 | 0 | NA | NA |
| Hgf | 116.1833121 | 0.536911538 | 0.162367302 |
| RP24-295A5.2 | 4.076536194 | 0.16752973 | 0.552856959 |
| RP24-295A5.3 | 3.112827446 | 0.463274297 | 0.671827998 |
| Gm8991 | 9.192051872 | 1.009657703 | 0.370118548 |
| Gm28710 | 0.299775621 | 1.089946151 | 1.762152014 |
| RP24-363D4.1 | 1.258311994 | 0.534818541 | 0.89228832 |
| Gm6673 | 0 | NA | NA |
| Gm3495 | 0 | NA | NA |
| Gm21999 | 0 | NA | NA |
| Speer4f | 0 | NA | NA |
| RP23-76L18.2 | 0 | NA | NA |
| Sema3c | 248.8209663 | -0.592650531 | 0.126781073 |
| Cd36 | 46.54322315 | 2.511965285 | 0.549978724 |
| Gnat3 | 0.097139547 | 0.718207094 | 3.724945049 |
| Gnai1 | 1646.477261 | -0.627328206 | 0.075199396 |
| RP24-302O2.1 | 0 | NA | NA |
| RP24-459D3.1 | 0.038981395 | 0.718207094 | 3.724945049 |
| RP24-332E15.1 | 0 | NA | NA |
| Gm23822 | 0.060174537 | 0.237309859 | 3.724945049 |
| RP23-350E1.2 | 51.82168262 | -0.341494429 | 0.155555674 |
| 4921504A21Rik | 48.57949567 | -0.421994993 | 0.193319092 |
| RP23-350E1.3 | 0 | NA | NA |
| Magi2 | 1985.599496 | -0.65054308 | 0.106831322 |
| RP24-388K6.6 | 0.090665643 | -0.724455696 | 3.724945049 |
| Gm25335 | 0 | NA | NA |
| Gm23570 | 4.591749267 | -1.931910674 | 0.573095078 |
| Gm25761 | 0 | NA | NA |
| RP24-531D1.4 | 0.048949532 | -0.243569582 | 3.724945049 |
| Gm22755 | 8.57913668 | -0.383288862 | 0.375525242 |
| RP23-281M20.1 | 6.048354155 | -1.691672884 | 0.526440914 |
| Gm29254 | 8.172606336 | -1.098165889 | 0.433383029 |
| RP23-343F16.2 | 0.689035189 | 0.211288192 | 1.178154134 |
| Phtf2 | 298.2117683 | -0.426303795 | 0.105584138 |
| Rpl31-ps5 | 0.299266739 | 0.718782919 | 1.767544496 |
| Tmem60 | 129.5434812 | -0.199983554 | 0.137188099 |
| Rsbn1l | 755.8552143 | 0.194560682 | 0.07014528 |
| A630072M18Rik | 116.1077269 | 0.452265529 | 0.136091727 |
| Ptpn12 | 569.2757863 | 0.519427937 | 0.090514723 |
| RP23-234P16.4 | 0.061920613 | 0.718207094 | 3.724945049 |
| Gm22490 | 0 | NA | NA |
| Rpl31-ps21 | 0 | NA | NA |

| | | | |
|---|---|---|---|
| RP23-234P16.2 | 0.792202851 | -0.522915971 | 1.18349725 |
| RP23-234P16.3 | 0.272741579 | 0.237330755 | 1.811420715 |
| Gsap | 232.3497055 | 1.128799932 | 0.145429955 |
| RP23-250D22.2 | 3.212737209 | 2.461641952 | 0.740701503 |
| RP23-250D22.3 | 0 | NA | NA |
| Ccdc146 | 10.59368012 | -0.3044533 | 0.332783306 |
| Fgl2 | 737.5148379 | 5.448263717 | 0.274336563 |
| Fam185a | 169.3008059 | -0.415675904 | 0.129954641 |
| Fbxl13 | 5.258800805 | -1.165616415 | 0.47624899 |
| Lrrc17 | 1.924082696 | -0.351100114 | 0.766836996 |
| RP23-6C10.3 | 0.285998089 | 0.199025115 | 2.095067819 |
| RP23-6C10.4 | 1.597009952 | 0.52657307 | 0.796131046 |
| Armc10 | 162.4313593 | 0.037062797 | 0.096345935 |
| Napepld | 190.2960083 | -0.543083999 | 0.112891245 |
| Gm15719 | 0.801310233 | -1.004043195 | 1.079250838 |
| Pmpcb | 597.1185698 | 0.458651818 | 0.076323828 |
| Gm15715 | 0.120974994 | -0.243571806 | 3.724945049 |
| Dnajc2 | 320.2722277 | 0.370745421 | 0.123980403 |
| Psmc2 | 881.0786388 | 0.523745508 | 0.084378516 |
| Slc26a5 | 5.142851575 | -1.367949918 | 0.478837851 |
| Reln | 2935.282946 | -0.668551198 | 0.128055208 |
| Gm16111 | 0 | NA | NA |
| Gm10475 | 1.275826341 | 0.663495616 | 1.001159412 |
| Dnaja1-ps | 7.533972655 | -0.741695459 | 0.49774736 |
| RP23-205D3.1 | 11.79089672 | -1.513421358 | 0.406081568 |
| Gm16113 | 6.656687351 | -0.990166686 | 0.432672043 |
| Gm16112 | 0 | NA | NA |
| Gm16110 | 0 | NA | NA |
| Gm25459 | 0 | NA | NA |
| RP23-143M9.2 | 0.121578509 | 0.718202646 | 3.724945049 |
| n-R5s170 | 0 | NA | NA |
| RP24-86E4.1 | 1.834562507 | 1.107125957 | 0.889403237 |
| Orc5 | 252.3162393 | 0.012594349 | 0.098392093 |
| Gm15421 | 12.14808016 | 0.338896805 | 0.304733771 |
| Gm22887 | 3.183676125 | -0.237132396 | 0.63745776 |
| 6030443J06Rik | 71.11363277 | -0.668273589 | 0.226486944 |
| RP23-258C13.1 | 3.322405445 | -0.46709959 | 0.558472795 |
| Rpl17-ps5 | 55.4808862 | 0.626061518 | 0.168452928 |
| Gm24512 | 0 | NA | NA |
| RP24-324L19.2 | 13.10249877 | 0.356751269 | 0.373532356 |
| RP24-324L19.3 | 161.9541949 | 0.236190569 | 0.11532006 |
| RP24-324L19.4 | 19.8545456 | 0.766986934 | 0.228775125 |
| AC164538.1 | 0.163066018 | -0.243578385 | 3.449621945 |
| Lhfpl3 | 375.6236598 | -0.17320538 | 0.098584346 |
| Gm21846 | 71.52268729 | -0.712297671 | 0.318564998 |
| RP23-211E12.6 | 0.049553047 | 0.71820487 | 3.724945049 |
| Gm6564 | 0 | NA | NA |

| | | | |
|---|---|---|---|
| RP23-240A11.2 | 0.172588697 | 0.712228794 | 2.317759978 |
| Gm28022 | 0 | NA | NA |
| RP23-443N7.4 | 0.181654796 | 1.679598083 | 3.311020364 |
| RP23-443N7.5 | 0 | NA | NA |
| 5031425E22Rik | 328.5648936 | -0.299237538 | 0.164775171 |
| RP23-443N7.6 | 0 | NA | NA |
| RP23-443N7.10 | 6.257156534 | -0.131774871 | 0.548237207 |
| RP23-443N7.7 | 0.066050875 | 0.71820487 | 3.724945049 |
| Kmt2e | 2575.672994 | -0.299427669 | 0.078436798 |
| Gm25219 | 2.780058051 | -1.150492892 | 0.722358448 |
| Srpk2 | 2903.573116 | -0.465892234 | 0.046999729 |
| Gm24009 | 0.699061148 | -0.941001856 | 1.35784194 |
| RP23-110C17.1 | 13.01527202 | -0.336325135 | 0.285330765 |
| RP23-110C17.2 | 20.4004732 | 0.666400244 | 0.266750733 |
| Mir3096 | 0.548185132 | 0.706222461 | 1.407825128 |
| Pus7 | 222.5424342 | -0.362254746 | 0.139711448 |
| Rint1 | 243.1730144 | -0.014298278 | 0.097622613 |
| RP23-354E8.1 | 9.469473219 | -0.082536475 | 0.367615105 |
| 4933427G23Rik | 0.868720842 | -0.603458985 | 1.332247946 |
| Gm22054 | 0.593565611 | -0.517089728 | 1.274340713 |
| Tomm7 | 416.4948252 | 0.059569952 | 0.108754123 |
| 2700038G22Rik | 16.39405462 | -0.036392063 | 0.304457775 |
| Snord93 | 2.473235781 | -0.469844711 | 0.614337818 |
| Fam126a | 374.7933423 | 0.067399255 | 0.093104163 |
| RP23-170E15.2 | 3.709199715 | -0.489460957 | 0.539540827 |
| RP23-451E8.3 | 0 | NA | NA |
| Klhl7 | 1038.582517 | -0.186859615 | 0.063030415 |
| RP23-451E8.2 | 13.50599551 | 0.270974045 | 0.276763362 |
| RP23-451E8.7 | 63.10412136 | -1.0005977 | 0.266247386 |
| Nupl2 | 116.6198412 | -0.096136669 | 0.133546494 |
| Gm5575 | 2.4388784 | 0.55826501 | 0.69279995 |
| RP23-451E8.8 | 0 | NA | NA |
| RP23-451E8.9 | 0.083526043 | 0.718202646 | 3.724945049 |
| Gm9836 | 12.95280587 | -0.089639942 | 0.272641723 |
| Kcnh2 | 620.4579014 | -0.352948852 | 0.108945172 |
| Gm15589 | 0 | NA | NA |
| Nos3 | 66.34634559 | 1.773325948 | 0.190545887 |
| Gm15587 | 5.162842253 | -2.49895678 | 0.639963366 |
| Atg9b | 48.1694312 | -0.462216851 | 0.179328502 |
| Abcb8 | 361.5327256 | -0.428758029 | 0.085704224 |
| Asic3 | 3.0483297 | -0.308275274 | 0.849064253 |
| Cdk5 | 656.490936 | -0.344475919 | 0.054799308 |
| Slc4a2 | 638.8880438 | 0.683999921 | 0.11697676 |
| Fastk | 605.3488211 | -0.105225499 | 0.061517626 |
| Gm22333 | 0 | NA | NA |
| Tmub1 | 167.7441116 | 0.088147183 | 0.111149992 |
| RP24-239E2.2 | 2.781790008 | -0.55029013 | 0.545291299 |

| | | | |
|---|---|---|---|
| Gm10472 | 2.635270342 | 0.263131001 | 0.679525231 |
| Agap3 | 2182.413688 | -0.357659562 | 0.065017738 |
| Gbx1 | 12.22572069 | -0.312886288 | 0.29446498 |
| Asb10 | 0.545953117 | -1.50787227 | 1.257579586 |
| 4931409K22Rik | 0.666655512 | -0.660048878 | 1.170644093 |
| Abcf2 | 862.9471066 | 0.206454391 | 0.069241974 |
| Gm10221 | 170.6058012 | 0.128006374 | 0.150346262 |
| Chpf2 | 822.9892482 | 0.374465362 | 0.081472098 |
| Smarcd3 | 501.6330187 | -0.51318768 | 0.083657568 |
| Mir671 | 0 | NA | NA |
| 1700022A21Rik | 0 | NA | NA |
| Nub1 | 1661.721015 | 0.847629006 | 0.082290691 |
| Gm26648 | 1.867746029 | 0.131493588 | 0.70964849 |
| Wdr86 | 10.62884738 | -0.986623591 | 0.444748989 |
| Crygn | 1.666884929 | -0.400430952 | 0.835594856 |
| Rheb | 967.3691609 | 0.376445914 | 0.103417036 |
| Gm25363 | 7.580824313 | -0.912876081 | 0.385304205 |
| Prkag2 | 719.0261586 | -0.472204461 | 0.112395461 |
| Prkag2os2 | 1.088852924 | 0.444893892 | 1.037775344 |
| Prkag2os1 | 0 | NA | NA |
| 1500035N22Rik | 3.98331357 | -1.203923266 | 0.527384818 |
| 2900005J15Rik | 14.08399665 | -0.854748778 | 0.334547724 |
| RP24-84G9.2 | 1.333500337 | -1.797068559 | 0.946828982 |
| Gm26051 | 0 | NA | NA |
| Galntl5 | 0.110595523 | 0.718200421 | 3.724945049 |
| Gm26356 | 0 | NA | NA |
| Galnt11 | 598.673327 | -0.393297615 | 0.071814064 |
| E130116L18Rik | 0 | NA | NA |
| Kmt2c | 3673.91328 | -0.210865202 | 0.079009792 |
| RP23-171F16.5 | 0.112029109 | -0.724455696 | 3.724945049 |
| Hspe1-ps3 | 0.038981395 | 0.718207094 | 3.724945049 |
| Gm20432 | 5.780307253 | -0.337963329 | 0.533084161 |
| Gm18343 | 0 | NA | NA |
| 4831440E17Rik | 12.8027681 | -1.706390117 | 0.343813758 |
| Gm22656 | 0.372618932 | 0.688971948 | 1.53886579 |
| Cct8l1 | 0.82877443 | 0.003869884 | 1.097653249 |
| 1700096K18Rik | 15.86316258 | -0.107595303 | 0.277338777 |
| RP23-405O11.3 | 0.04393175 | -0.243571806 | 3.724945049 |
| Gm10062 | 3.869150979 | -1.922229746 | 0.618621577 |
| RP24-121I11.1 | 0 | NA | NA |
| RP24-121I11.4 | 17.85167491 | -1.514281915 | 0.296079315 |
| Xrcc2 | 97.35748713 | -0.529641354 | 0.146105957 |
| RP24-121I11.5 | 0.245367999 | 0.236873241 | 1.856129601 |
| Actr3b | 678.1794346 | -0.624258451 | 0.075130332 |
| RP23-56M18.4 | 0 | NA | NA |
| RP23-56M18.5 | 0.232282403 | 0.718215903 | 2.551194947 |
| Gm21671 | 0.275686258 | 0.139921644 | 1.808825036 |

| | | | |
|---|---|---|---|
| RP23-56M18.8 | 24.28531059 | -1.0111458 | 0.337065829 |
| RP23-56M18.6 | 0.144833758 | 0.684656385 | 2.882712487 |
| Gm21698 | 0 | NA | NA |
| Gm1979 | 0.35614329 | 0.754380958 | 1.740729038 |
| Gm5067 | 39.82603458 | -0.906652053 | 0.242344115 |
| Gm5862 | 0 | NA | NA |
| Speer4a | 0.739238851 | 1.529273594 | 1.286278428 |
| Gm7347 | 0.182828849 | 1.122216213 | 2.253829118 |
| RP24-162F23.9 | 8.999765542 | -1.30189421 | 0.410004868 |
| Gm10471 | 0.138573873 | 1.200070582 | 3.628829444 |
| 5031410I06Rik | 0 | NA | NA |
| Gm10220 | 0.108266322 | 1.230376831 | 3.722845817 |
| Gm7361 | 6.292791096 | -0.526886989 | 0.395103863 |
| RP24-362M8.2 | 0.097139547 | 0.718207094 | 3.724945049 |
| 4930584F24Rik | 0 | NA | NA |
| RP24-165O21.1 | 0 | NA | NA |
| Dpp6 | 3300.38036 | -0.398711988 | 0.063398575 |
| RP23-474A22.1 | 0 | NA | NA |
| Gm16056 | 0 | NA | NA |
| Gm16057 | 0 | NA | NA |
| RP23-478E6.1 | 84.06891195 | -1.064414123 | 0.311082341 |
| RP23-478E6.3 | 79.08630011 | -1.312260193 | 0.311099501 |
| Speer4b | 0.378318939 | -0.243563821 | 1.7144404 |
| Gm16058 | 0.092228164 | 0.718202646 | 3.724945049 |
| Paxip1 | 503.8218817 | -0.252466471 | 0.080061556 |
| RP23-397C5.2 | 0.110870145 | 0.238305784 | 3.724874707 |
| RP23-397C5.1 | 0 | NA | NA |
| Htr5a | 258.8451686 | -0.087306652 | 0.123437512 |
| RP23-397C5.5 | 4.181737927 | 0.110979502 | 0.511790306 |
| RP23-397C5.4 | 1.516892051 | 0.148847725 | 0.840008747 |
| RP23-256A5.2 | 3.457092485 | -0.52472972 | 0.595874495 |
| Gm26608 | 2.045375739 | -0.15645399 | 0.672210697 |
| RP23-256A5.4 | 1.504629277 | -1.644200379 | 0.912874548 |
| Insig1 | 653.7555455 | 0.108883202 | 0.138446914 |
| En2 | 252.7745475 | -0.307670827 | 0.281848172 |
| RP23-358F24.3 | 0.075043205 | 0.237309859 | 3.724945049 |
| Cnpy1 | 429.1835849 | -1.128843012 | 0.296279021 |
| Rbm33 | 1626.312609 | -0.401944424 | 0.125380822 |
| Shh | 57.43406997 | 0.411934447 | 0.157225424 |
| 9530036O11Rik | 6.185525992 | -1.359509564 | 0.480778509 |
| Gm26894 | 16.72981752 | -1.37411625 | 0.285567564 |
| RP23-94F17.1 | 1.563650339 | 0.290632431 | 0.736400366 |
| RP23-94F17.2 | 0.559314221 | -1.180920478 | 1.487016172 |
| Rnf32 | 99.31368546 | -0.840444035 | 0.15564146 |
| Gm7420 | 6.011270768 | -2.162916851 | 0.676533812 |
| Lmbr1 | 450.4254583 | -0.691673274 | 0.079138354 |
| RP23-396H16.2 | 45.39463601 | -1.128665529 | 0.258153098 |

| | | | |
|---|---|---|---|
| Gm23801 | 1.101786575 | -0.671968012 | 0.862443565 |
| 4632411P08Rik | 8.069396325 | -0.472000229 | 0.383063744 |
| Nom1 | 466.9739484 | 0.359840217 | 0.073641788 |
| Mnx1 | 0.180050447 | 0.719165083 | 3.327036943 |
| Gm24538 | 0 | NA | NA |
| Gm6083 | 0 | NA | NA |
| Ube3c | 1443.177322 | -0.06832249 | 0.046974542 |
| RP24-224N3.5 | 0.401137426 | -1.532042485 | 1.518162756 |
| Gm26346 | 0.288530209 | 0.238794541 | 1.764048745 |
| RP24-224N3.4 | 0 | NA | NA |
| Gm5129 | 0.619575296 | -1.264261822 | 1.154429227 |
| Dnajb6 | 825.5467343 | 0.161665327 | 0.057574987 |
| RP24-224N3.8 | 0.381425259 | 1.554745497 | 1.537218212 |
| RP24-224N3.9 | 0 | NA | NA |
| RP24-176O2.1 | 0 | NA | NA |
| RP24-176O2.2 | 0.121388914 | 0.718200421 | 3.724945049 |
| RP24-176O2.4 | 0.21891683 | 1.527064226 | 2.542683672 |
| RP24-418J2.4 | 0.04393175 | -0.243571806 | 3.724945049 |
| Il6 | 43.66316143 | 9.0068779 | 1.156263541 |
| Tyms | 45.1820026 | 0.24990721 | 0.202807793 |
| RP24-418J2.5 | 0.550684032 | -0.214725462 | 1.338733539 |
| 3110082J24Rik | 7.185718025 | -1.71110467 | 0.461699232 |
| Gareml | 196.41537 | -1.220554047 | 0.157670743 |
| Hadha | 925.8447827 | 0.5069447 | 0.087529962 |
| Hadhb | 496.8674058 | -0.200114314 | 0.084302156 |
| Mir1960 | 2.414716579 | -0.496305803 | 0.776365699 |
| Adgrf3 | 4.779796961 | 1.017463002 | 0.574244113 |
| Ept1 | 1152.189136 | -0.340255065 | 0.050237185 |
| Gm16288 | 0.621982117 | 1.043442648 | 1.314253634 |
| Drc1 | 103.3566621 | 0.175424199 | 0.184696702 |
| WI1-2080J1.2 | 1.385170032 | -3.021761294 | 1.028700871 |
| Otof | 183.3860885 | -2.389823901 | 0.216616922 |
| 1700001C02Rik | 2.869726659 | -0.918565594 | 0.627068703 |
| Cib4 | 0.038981395 | 0.718207094 | 3.724945049 |
| Gm25666 | 0 | NA | NA |
| RP23-446J15.1 | 0.136419714 | 0.237324253 | 2.946414804 |
| Gm9899 | 195.7601427 | -0.420211968 | 0.176002955 |
| Kcnk3 | 462.3649798 | -0.203774362 | 0.119633603 |
| Slc35f6 | 587.4667661 | 1.185021621 | 0.096030186 |
| Mir5625 | 0 | NA | NA |
| Cenpa | 31.10520093 | 0.723698579 | 0.26645655 |
| Dpysl5 | 464.621931 | -0.310965148 | 0.090780498 |
| RP23-342N8.1 | 3.407592442 | -2.70213354 | 0.691197883 |
| Mapre3 | 1596.309188 | -0.323665376 | 0.064994734 |
| RP23-342N8.2 | 49.78286253 | -0.918971441 | 0.327946153 |
| Gm15461 | 0 | NA | NA |
| Tmem214 | 593.0310336 | 0.267403896 | 0.06099255 |

| | | | |
|---|---|---|---|
| RP23-113M20.5 | 2.578624624 | -0.196001485 | 0.853509282 |
| Agbl5 | 130.4690619 | -0.40570944 | 0.142623359 |
| RP23-113M20.4 | 16.2135406 | -0.63264569 | 0.260912595 |
| Ost4 | 231.3485647 | 0.376830998 | 0.147835608 |
| Emilin1 | 69.27241042 | 2.060813817 | 0.188795509 |
| Khk | 59.85891304 | -0.318769222 | 0.166541361 |
| Cgref1 | 143.7238615 | 0.37280003 | 0.192583739 |
| Abhd1 | 97.11033058 | -0.537918449 | 0.134846799 |
| Preb | 614.623067 | 0.036565179 | 0.065528218 |
| Tcf23 | 3.89979361 | 1.242605159 | 0.567577699 |
| Slc5a6 | 215.3588507 | -0.46677847 | 0.090267566 |
| Atraid | 300.310359 | 0.036433028 | 0.108556178 |
| Cad | 344.3964736 | 0.282910242 | 0.076891907 |
| Slc30a3 | 365.9223408 | -1.245361105 | 0.150120328 |
| Gm9924 | 4.140852306 | -0.467401265 | 0.482483145 |
| Dnajc5g | 0.288733551 | 1.585265222 | 2.009257491 |
| Trim54 | 0.834511584 | -1.942333933 | 1.122037595 |
| Gm15469 | 0 | NA | NA |
| Ucn | 1.659798835 | -1.46313022 | 1.24140734 |
| Mpv17 | 189.2554659 | -0.130592784 | 0.102799088 |
| Gm29609 | 14.60460443 | -1.026828974 | 0.312289419 |
| Gtf3c2 | 322.248693 | -0.306604749 | 0.099277231 |
| Eif2b4 | 322.6252131 | 0.392790013 | 0.078183544 |
| Snx17 | 719.475255 | 0.390146246 | 0.071926012 |
| Zfp513 | 310.9180716 | 0.510698394 | 0.098781796 |
| Ppm1g | 655.6864786 | 0.315486548 | 0.077334455 |
| RP24-86I23.4 | 34.84001459 | -0.519367521 | 0.369396948 |
| Gm9970 | 1.14535829 | 0.47325225 | 0.935302779 |
| Nrbp1 | 774.3138472 | 0.273389147 | 0.062143975 |
| Krtcap3 | 6.967827923 | 0.271019846 | 0.380916811 |
| Ift172 | 621.7871443 | -0.409046679 | 0.082219748 |
| Fndc4 | 773.732385 | 0.036074063 | 0.087997132 |
| Gckr | 6.742714648 | -0.612223195 | 0.399149028 |
| RP23-383E21.3 | 0 | NA | NA |
| RP23-383E21.5 | 0 | NA | NA |
| RP23-383E21.4 | 0.38059087 | 1.208147893 | 1.718114936 |
| Zfp512 | 587.0033922 | -0.079715327 | 0.064546866 |
| RP23-383E21.1 | 18.84380616 | -0.916883086 | 0.319944841 |
| 4930548H24Rik | 1.368337452 | 0.615646748 | 0.964584649 |
| 4930566F21Rik | 0.183097766 | -0.243562083 | 2.686642813 |
| Gpn1 | 250.7486316 | 0.105289895 | 0.077323128 |
| Supt7l | 184.2270805 | 0.123038927 | 0.098505969 |
| Slc4a1ap | 405.404987 | -0.051767105 | 0.069503913 |
| RP24-80F7.3 | 2.627266017 | 3.448278902 | 0.822657673 |
| RP24-80F7.5 | 6.412813486 | 2.297406489 | 0.449284382 |
| RP24-80F7.6 | 3.22349573 | -0.192328785 | 0.726273098 |
| Mrpl33 | 188.1679828 | 0.42812711 | 0.116655926 |

| | | | |
|---|---|---|---|
| RP24-80F7.7 | 0 | NA | NA |
| Rbks | 30.53545577 | 0.852203193 | 0.213391929 |
| Mir3473e | 1.131450813 | -0.026177786 | 0.93793709 |
| Bre | 293.3718634 | 0.1212891 | 0.070018359 |
| RP24-80F7.8 | 3.022895243 | 0.102823296 | 0.65252424 |
| Gm24436 | 2.112795873 | -0.136655993 | 0.819752342 |
| RP23-181I1.4 | 6.442610278 | 0.098889097 | 0.378229368 |
| Gm17130 | 0.230862969 | 0.726141081 | 2.133407852 |
| Gm10463 | 2.064454091 | 1.017147681 | 0.710744367 |
| Fosl2 | 1087.482308 | 1.13059607 | 0.309951056 |
| Gm26335 | 0 | NA | NA |
| RP23-415E17.3 | 0 | NA | NA |
| Plb1 | 16.55472255 | 0.595215537 | 0.288990372 |
| Gm15615 | 0 | NA | NA |
| RP23-262J12.5 | 0.060174537 | 0.237309859 | 3.724945049 |
| RP23-262J12.4 | 0 | NA | NA |
| Gm5864 | 1.456934103 | 0.526088865 | 0.829115779 |
| Ppp1cb | 4028.577039 | 0.015019631 | 0.087460495 |
| RP23-262J12.3 | 12.26687181 | -0.801019136 | 0.327355055 |
| RP23-262J12.2 | 50.45321847 | -0.794803145 | 0.285954969 |
| Gm15613 | 0 | NA | NA |
| Gm15614 | 0.985784616 | -1.239909674 | 1.344227616 |
| RP23-149A1.7 | 0.075043205 | 0.237309859 | 3.724945049 |
| RP23-149A1.5 | 0.156588133 | 0.707833931 | 2.818547658 |
| RP23-149A1.6 | 21.7617041 | -0.604777854 | 0.25499396 |
| Yes1 | 334.4835077 | -0.078313811 | 0.093829279 |
| RP23-149A1.2 | 1.316494506 | 0.682804405 | 0.883182572 |
| RP23-353M5.2 | 7.746880428 | -0.964068215 | 0.43249101 |
| Gm2420 | 4.384803307 | -1.746033655 | 0.560036161 |
| Pisd | 289.4875556 | 0.314203415 | 0.076673627 |
| Gm20671 | 0 | NA | NA |
| RP23-353M5.1 | 0 | NA | NA |
| Prr14l | 1555.512628 | -0.252224841 | 0.054823576 |
| RP23-118L1.4 | 11.68546513 | -1.833416661 | 0.424730512 |
| RP23-118L1.5 | 0 | NA | NA |
| Depdc5 | 821.0906113 | -0.367128668 | 0.075147394 |
| Gm24003 | 1.689516722 | -2.78717902 | 1.0117781 |
| RP23-69J7.1 | 15.99339509 | -1.247578148 | 0.405124286 |
| Ywhah | 4942.475359 | -0.293022373 | 0.074752175 |
| RP23-69J7.4 | 0 | NA | NA |
| Rpl35a-ps5 | 4.14321256 | 0.400747319 | 0.535293824 |
| Slc5a1 | 30.58515113 | 0.95932641 | 0.306954701 |
| Gm15936 | 0.038981395 | 0.718207094 | 3.724945049 |
| 5033423K11Rik | 0.310320078 | 1.511235446 | 1.958736152 |
| Spon2 | 4.327007151 | 2.920441012 | 0.837192343 |
| Ctbp1 | 1818.873768 | 0.035623435 | 0.07415769 |
| RP24-406N3.3 | 0 | NA | NA |

| | | | |
|---|---|---|---|
| Gm20465 | 0 | NA | NA |
| Maea | 757.2802516 | 0.36225657 | 0.076749154 |
| Uvssa | 220.4651291 | -0.116451615 | 0.132094716 |
| Nkx1-1 | 0.498903533 | 0.758683947 | 1.46122053 |
| RP24-94L5.3 | 0.038981395 | 0.718207094 | 3.724945049 |
| Fam53a | 291.3243653 | 0.301321484 | 0.078050297 |
| Gm9903 | 6.764630232 | -0.002224419 | 0.396906174 |
| Slbp | 223.0429402 | 0.426520603 | 0.094620917 |
| RP23-391P21.2 | 0.046345709 | 0.71820487 | 3.724945049 |
| Tmem129 | 242.2710187 | 0.179345965 | 0.096384056 |
| Tacc3 | 66.98387769 | 0.708766221 | 0.211493817 |
| RP23-391P21.5 | 0 | NA | NA |
| RP23-391P21.4 | 17.89361774 | -0.928116406 | 0.281736242 |
| Fgfr3 | 1284.08441 | -0.179596487 | 0.134509641 |
| Letm1 | 945.73928 | -0.059757103 | 0.048990445 |
| Gm10059 | 2.485764843 | 0.02751 | 0.709174055 |
| RP23-391P21.3 | 0 | NA | NA |
| Whsc1 | 2442.311656 | -0.140194237 | 0.045275598 |
| Nelfa | 291.9774525 | 0.201631734 | 0.0899036 |
| Mir7024 | 0 | NA | NA |
| RP24-294K15.5 | 2.887331315 | 0.589239555 | 0.597332126 |
| RP24-294K15.4 | 13.77895346 | 1.096240803 | 0.364602994 |
| Gm1673 | 134.0090932 | 0.210404183 | 0.162715765 |
| RP24-294K15.7 | 56.16396208 | -0.566271634 | 0.162892764 |
| Nat8l | 3474.475194 | -0.893628276 | 0.07028068 |
| Poln | 11.42523575 | -0.313365661 | 0.283938859 |
| RP23-60E7.2 | 0.517722191 | 1.147696266 | 1.342145684 |
| RP23-60E7.3 | 0.042716945 | -0.243571806 | 3.724945049 |
| RP23-60E7.1 | 0.122601409 | 0.237307635 | 3.724945049 |
| Haus3 | 129.8840804 | 0.212946938 | 0.107499797 |
| Gm25918 | 0 | NA | NA |
| Mxd4 | 934.5782134 | -0.68994426 | 0.116922601 |
| RP24-230P10.6 | 0.165470308 | 0.237318778 | 2.761559324 |
| Zfyve28 | 544.0876334 | -0.905951185 | 0.129482715 |
| Gm15513 | 73.95229317 | -1.285706788 | 0.308841841 |
| RP23-201D22.6 | 0 | NA | NA |
| RP23-201D22.4 | 0.535802326 | 1.838860926 | 1.376045732 |
| RP23-201D22.5 | 0.419184927 | -0.243574657 | 3.363637513 |
| Rnf4 | 1197.186287 | 0.616079056 | 0.083174846 |
| Fam193a | 1067.198729 | -0.036325338 | 0.069638399 |
| RP23-429E19.4 | 44.14393845 | -0.677489668 | 0.283255378 |
| Gm26931 | 0.110870145 | 0.238305784 | 3.724874707 |
| RP23-429E19.5 | 37.84079439 | -0.550619572 | 0.242386941 |
| Tnip2 | 144.594261 | 0.791678985 | 0.16456857 |
| Sh3bp2 | 240.9989662 | 3.203370204 | 0.176029731 |
| Gm22847 | 1.623325927 | 2.349898038 | 0.960796609 |
| Add1 | 3389.415062 | -0.35183413 | 0.051331422 |

| | | | |
|---|---|---|---|
| Mir7036b | 0 | NA | NA |
| RP24-229D18.2 | 13.74406399 | -0.442146383 | 0.454677972 |
| Mfsd10 | 128.2944897 | 0.135545167 | 0.156298157 |
| Nop14 | 276.7616417 | 0.81968088 | 0.140433396 |
| Gm15522 | 0.123992736 | -0.243574001 | 3.720643035 |
| Grk4 | 52.31774275 | -0.358606858 | 0.190647422 |
| RP23-44L13.1 | 0.042716945 | -0.243571806 | 3.724945049 |
| Mir467g | 0 | NA | NA |
| Htt | 3899.683035 | 0.024931011 | 0.065656966 |
| Msantd1 | 17.05411229 | 0.207216521 | 0.268506684 |
| Rgs12 | 414.5610989 | -0.363597683 | 0.093439439 |
| Mir3097 | 0.476896902 | -0.243393085 | 1.483785158 |
| RP23-352G19.2 | 0.30060319 | 0.706640579 | 1.614670333 |
| Hgfac | 2.316699699 | -0.091856784 | 0.645152932 |
| Dok7 | 162.3460107 | -0.371186942 | 0.13997449 |
| Lrpap1 | 1373.379439 | 0.2956832 | 0.101747956 |
| Adra2c | 169.5646234 | -1.010636024 | 0.122179457 |
| RP24-276N7.5 | 0.097139547 | 0.718207094 | 3.724945049 |
| E130018O15Rik | 0 | NA | NA |
| Hmx1 | 0.038981395 | 0.718207094 | 3.724945049 |
| 4930442P19Rik | 0 | NA | NA |
| RP24-276N7.7 | 0 | NA | NA |
| Cpz | 3.532213938 | 0.832265741 | 0.554981648 |
| RP23-26P10.1 | 28.70925171 | 0.250165476 | 0.310419214 |
| Plk-ps1 | 0 | NA | NA |
| RP23-26P10.2 | 15.37615298 | -0.223454387 | 0.313710636 |
| Trmt44 | 83.81309991 | 0.025881387 | 0.163627019 |
| 4931431C16Rik | 1.224874833 | 1.349107568 | 1.018248643 |
| Acox3 | 452.6412082 | -0.12814831 | 0.083857254 |
| Htra3 | 58.84703746 | 0.939182666 | 0.204777341 |
| Sh3tc1 | 32.64252681 | 1.761236722 | 0.250248607 |
| Gm15650 | 0.369202626 | 0.652126819 | 1.742868043 |
| Ablim2 | 1153.537665 | -0.845752904 | 0.130658321 |
| Afap1 | 482.3908662 | 0.035602634 | 0.068569757 |
| H2af-ps | 0.473803596 | -0.59545663 | 1.437687055 |
| Sorcs2 | 714.7892678 | -0.072852445 | 0.077130523 |
| Psapl1 | 0.670061835 | 1.002052192 | 1.220109895 |
| Mir7689 | 0.235888945 | 0.69030369 | 2.154631984 |
| 2210406O10Rik | 10.33515927 | -0.006816562 | 0.306817874 |
| Grpel1 | 361.9003875 | 0.57959727 | 0.112965044 |
| Tada2b | 496.4167252 | 0.16333505 | 0.107541077 |
| Ccdc96 | 19.96485888 | 0.774919842 | 0.250700646 |
| Tbc1d14 | 875.8556229 | 0.048802537 | 0.052996999 |
| RP23-463H10.2 | 0 | NA | NA |
| D5Ertd579e | 1861.527523 | -0.270265608 | 0.055643892 |
| RP23-463H10.1 | 0 | NA | NA |
| Gm24878 | 6.521421862 | -0.010034165 | 0.467719076 |

| | | | |
|---|---|---|---|
| Gm25767 | 0 | NA | NA |
| Bloc1s4 | 95.95579344 | 0.613012373 | 0.157138896 |
| RP23-382O15.2 | 0.348252234 | 0.206601073 | 1.463271833 |
| Mrfap1 | 2250.019853 | 0.181646656 | 0.077193958 |
| Man2b2 | 514.8886766 | 0.633322647 | 0.108690087 |
| Ppp2r2c | 4810.614684 | -0.422473542 | 0.099457946 |
| Ppp2r2cos | 0.150455055 | 1.610657435 | 2.838957197 |
| RP23-16N23.2 | 5.207155389 | -1.258929393 | 0.52209449 |
| Wfs1 | 1615.855355 | 0.661584786 | 0.134373881 |
| Jakmip1 | 619.5416328 | -0.920050316 | 0.187301505 |
| Gm26647 | 0.176363682 | 0.773483308 | 2.72611693 |
| Gm25551 | 0 | NA | NA |
| RP23-106D12.1 | 30.55419668 | -1.03196042 | 0.289828326 |
| Gm1043 | 54.30076238 | -0.498334424 | 0.226957886 |
| Crmp1 | 1273.345192 | -0.720525635 | 0.073938723 |
| Evc | 77.51767331 | -0.098036609 | 0.201536164 |
| RP23-339H11.3 | 0 | NA | NA |
| Evc2 | 92.04870115 | -0.137162968 | 0.134013973 |
| Gm15732 | 0 | NA | NA |
| Stk32b | 51.08340483 | -0.020567453 | 0.17570692 |
| Cytl1 | 1.87157638 | -1.12707668 | 0.722637837 |
| RP24-199D21.1 | 0 | NA | NA |
| Msx1 | 35.29830261 | -0.208681413 | 0.257761554 |
| RP24-292O13.1 | 0.159287055 | -0.243571011 | 2.807608812 |
| RP23-453O15.1 | 0.961470315 | -0.721465901 | 0.978412932 |
| RP23-453O15.4 | 0.066050875 | 0.71820487 | 3.724945049 |
| Stx18 | 243.7354772 | 0.266528703 | 0.08470901 |
| RP24-535K22.2 | 24.02896761 | -0.30325102 | 0.223991857 |
| Nsg1 | 2866.052638 | -0.339047546 | 0.085421584 |
| Zbtb49 | 139.0035971 | -0.430881776 | 0.138019706 |
| Lyar | 86.46303146 | 0.525433187 | 0.21698894 |
| Tmem128 | 186.1267044 | 0.66463537 | 0.126822448 |
| Otop1 | 1.049274302 | 0.589735647 | 0.923900578 |
| Gm26560 | 1.795106397 | 2.021624314 | 0.787104696 |
| Drd5 | 35.97084852 | 0.43586615 | 0.18916681 |
| RP23-426E16.4 | 0.045882455 | 0.237312084 | 3.724945049 |
| Slc2a9 | 46.88057285 | 2.916112077 | 0.250253215 |
| Gm15796 | 0.265425963 | 1.163913365 | 2.075800997 |
| Wdr1 | 2375.805282 | 0.995949988 | 0.103844293 |
| Gm22249 | 0 | NA | NA |
| 4930421P07Rik | 0 | NA | NA |
| Zfp518b | 296.1034572 | 0.040872148 | 0.076046014 |
| Gm22779 | 0 | NA | NA |
| Clnk | 4.40624827 | 2.847838603 | 0.650015733 |
| Clnkos | 0.326535978 | -0.633738051 | 1.744633913 |
| RP24-258B22.1 | 0.156480458 | -1.191887415 | 3.480826907 |
| RP24-263A3.3 | 0 | NA | NA |

| | | | |
|---|---|---|---|
| RP24-263A3.2 | 0.133021046 | -0.243579254 | 3.657009941 |
| RP24-263A3.1 | 0 | NA | NA |
| Gm25505 | 0.048949532 | -0.243569582 | 3.724945049 |
| RP23-477D20.1 | 0.049553047 | 0.71820487 | 3.724945049 |
| RP23-477D20.4 | 0.771325962 | 0.252932735 | 1.158240862 |
| RP23-477D20.3 | 0.202816535 | 1.003054481 | 3.205965041 |
| RP23-477D20.2 | 2.015324298 | -0.686432394 | 0.781654189 |
| Gm25377 | 0 | NA | NA |
| RP23-2C16.2 | 158.46767 | -0.343358506 | 0.119927282 |
| Hs3st1 | 277.9990456 | -0.398589505 | 0.103664096 |
| Gm7816 | 17.85736311 | 0.415286021 | 0.26045114 |
| RP23-318O24.1 | 0.038981395 | 0.718207094 | 3.724945049 |
| Gm2810 | 23.07760637 | -0.330607166 | 0.259220798 |
| Gm23022 | 0 | NA | NA |
| RP23-477O15.2 | 0 | NA | NA |
| RP23-477O15.1 | 3.151818414 | -0.27661354 | 0.886886181 |
| RP24-377B20.1 | 0 | NA | NA |
| Rab28 | 662.797046 | 0.116045766 | 0.056634722 |
| Nkx3-2 | 0.153659334 | 0.237323881 | 2.839457797 |
| Bod1l | 1615.891892 | -0.170744769 | 0.11290608 |
| RP24-188B2.2 | 1.139214769 | 0.323262774 | 1.050783889 |
| Mir6414 | 0 | NA | NA |
| RP24-188B2.1 | 0.100902008 | 1.15059983 | 3.720931964 |
| Gm16223 | 2.960191199 | -0.052039291 | 0.572074716 |
| RP23-146B9.1 | 0.099155932 | 0.718202646 | 3.724945049 |
| RP24-167K2.2 | 0.184373794 | -1.191730714 | 2.724223369 |
| RP24-560A18.4 | 0.659798846 | -1.643818277 | 1.191102968 |
| RP24-560A18.1 | 1.041030589 | -1.891842939 | 0.985239565 |
| RP24-560A18.3 | 0.172203646 | -0.724459864 | 2.739420422 |
| RP24-560A18.2 | 14.93512307 | -0.909781482 | 0.409369566 |
| Gm7854 | 28.25369218 | -0.599458939 | 0.268324306 |
| Cpeb2 | 1599.741892 | -0.268287396 | 0.064618802 |
| RP24-230L5.2 | 66.96472975 | -0.533503049 | 0.217984533 |
| RP24-230L5.3 | 79.78188433 | -0.963868588 | 0.212541547 |
| RP24-230L5.4 | 0.343955217 | 0.224740337 | 1.756696329 |
| RP24-230L5.5 | 0.150451996 | 0.237323705 | 2.858243839 |
| RP23-15G1.1 | 0.062426873 | 0.237312084 | 3.724945049 |
| C1qtnf7 | 8.52901898 | -0.394422233 | 0.336667776 |
| RP23-15G1.2 | 2.619111478 | -1.243752645 | 0.643996545 |
| Gm15866 | 0.088534442 | 1.179809211 | 3.723500796 |
| RP23-15G1.3 | 0.396976673 | -0.747345912 | 1.571170484 |
| Cc2d2a | 374.1500514 | -0.32542343 | 0.121027699 |
| Fbxl5 | 1545.557561 | -0.042676028 | 0.071793541 |
| RP23-336O18.1 | 33.11233713 | 0.063564784 | 0.318451573 |
| RP23-336O18.2 | 3.500141698 | -0.477261835 | 0.540350501 |
| RP23-336O18.3 | 11.15943119 | 7.171343663 | 0.9828493 |
| RP23-336O18.4 | 0.062426873 | 0.237312084 | 3.724945049 |

| | | | |
|---|---|---|---|
| Bst1 | 59.92677726 | 5.904257272 | 0.497892366 |
| Cd38 | 164.7434097 | 1.818769295 | 0.20940906 |
| Gm16014 | 2.851290958 | 2.241121899 | 0.705676297 |
| Gm16015 | 0 | NA | NA |
| Gm7879 | 31.12801235 | 0.042434873 | 0.215269476 |
| Gm7882 | 0 | NA | NA |
| Gm5292 | 0 | NA | NA |
| RP23-58C20.3 | 0 | NA | NA |
| RP23-58C20.4 | 0 | NA | NA |
| Fgfbp1 | 20.48618477 | -0.11626431 | 0.306956368 |
| Prom1 | 116.7320341 | 0.092626656 | 0.1971811 |
| RP24-230J14.2 | 3.662822882 | -0.524384817 | 0.636396777 |
| Gm16401 | 1.371463738 | -1.232372104 | 1.057844408 |
| RP24-230J14.1 | 0.385783664 | 0.766239835 | 1.794451709 |
| RP24-230J14.3 | 0.273584676 | 0.718804454 | 1.806800059 |
| RP24-230J14.4 | 0.127971488 | 1.23029335 | 3.685533863 |
| RP24-230J14.5 | 0.770607605 | 0.856747649 | 1.122566325 |
| Tapt1 | 762.3494051 | 0.238216762 | 0.07487325 |
| RP24-230J14.8 | 46.64492421 | -0.607707699 | 0.285973743 |
| RP24-230J14.7 | 0 | NA | NA |
| RP24-230J14.9 | 0 | NA | NA |
| RP24-230J14.10 | 12.44662364 | -0.568533095 | 0.292838112 |
| RP24-137F10.3 | 0 | NA | NA |
| RP24-137F10.4 | 0.038981395 | 0.718207094 | 3.724945049 |
| RP24-137F10.1 | 0.921219545 | -0.581213298 | 1.093480717 |
| RP24-137F10.2 | 0.094097695 | 0.718200421 | 3.724945049 |
| Ldb2 | 272.9695285 | -0.807542579 | 0.165545218 |
| RP24-188E19.2 | 17.17285275 | -1.699259062 | 0.504987585 |
| RP24-188E19.3 | 7.245259879 | -2.230535374 | 0.552070679 |
| RP24-188E19.4 | 5.313268904 | -0.739871851 | 0.704048573 |
| RP24-188E19.5 | 0 | NA | NA |
| RP24-247A21.1 | 40.44753983 | -1.466650048 | 0.380322816 |
| RP24-496C22.1 | 0.159422358 | 0.774378149 | 2.814104785 |
| RP24-496C22.2 | 19.70483131 | -1.657808004 | 0.365379516 |
| RP24-496C22.3 | 8.581378291 | -1.508210501 | 0.465895598 |
| RP24-496C22.4 | 7.00608203 | -1.532305538 | 0.4562458 |
| RP24-496C22.5 | 14.99026705 | -2.101519811 | 0.435080073 |
| RP24-496C22.6 | 21.60948068 | -1.903041383 | 0.339163072 |
| RP23-133N3.1 | 0 | NA | NA |
| 4930431F12Rik | 0.1567526 | 0.265588935 | 2.825354309 |
| Gm10205 | 0 | NA | NA |
| 4930435H24Rik | 4.37057135 | 2.305642703 | 0.635013821 |
| RP23-194K16.1 | 0.718177591 | 1.200223358 | 1.242797459 |
| RP23-33C9.1 | 1.272457209 | 0.850929643 | 0.893455686 |
| Qdpr | 1058.229875 | -0.721496834 | 0.118732317 |
| AC092711.4 | 2.958230677 | -0.904337027 | 0.559501101 |
| Clrn2 | 1.123782183 | -1.940146008 | 1.032012812 |

| | | | |
|---|---|---|---|
| Lap3 | 1007.429224 | 1.881060122 | 0.149060886 |
| Med28 | 563.4186552 | 0.394875062 | 0.086344459 |
| Fam184b | 121.4810835 | -1.222342982 | 0.209259575 |
| RP23-7D3.3 | 16.87252161 | -0.106308906 | 0.420983233 |
| RP23-118K12.1 | 0.502057302 | 1.589375289 | 1.318831841 |
| 9630001P10Rik | 17.90177705 | -0.88583956 | 0.282445662 |
| RP23-7E4.8 | 0.664660227 | -0.28996345 | 1.23126497 |
| RP23-7E4.7 | 0.154585317 | 1.652042906 | 2.82119606 |
| RP23-7E4.9 | 3.048203817 | -1.226237177 | 0.708043821 |
| Ncapg | 19.54545852 | 2.663675408 | 0.381316833 |
| Lcorl | 806.219256 | 0.00592539 | 0.067797719 |
| Gm36401 | 0.944820375 | 0.732767846 | 1.058217656 |
| RP23-7E4.3 | 66.02500977 | -0.066333641 | 0.17497099 |
| Gm29036 | 24.05605521 | -0.06494465 | 0.273226977 |
| RP23-7E4.4 | 46.18654444 | -0.070057411 | 0.214202251 |
| AC158786.1 | 0.746606991 | 0.987502004 | 1.256168538 |
| RP23-228M24.1 | 0 | NA | NA |
| Gm7931 | 9.771161355 | 0.806281631 | 0.360192662 |
| RP24-400K16.1 | 0 | NA | NA |
| RP23-277H1.1 | 0 | NA | NA |
| RP23-287A8.1 | 0.858277593 | -2.47560044 | 1.229134408 |
| RP23-319E24.2 | 0 | NA | NA |
| RP23-319E24.1 | 0 | NA | NA |
| RP23-195A18.4 | 0 | NA | NA |
| Slit2 | 695.3750179 | -0.645184676 | 0.089393782 |
| 4930518C09Rik | 0 | NA | NA |
| Gm20475 | 5.876575912 | -1.08366876 | 0.412240752 |
| RP23-195A18.3 | 20.06045718 | -0.752657582 | 0.332194238 |
| RP23-219K16.2 | 8.550972595 | -1.230243285 | 0.385214538 |
| RP23-219K16.1 | 10.624249 | -1.175177375 | 0.419080049 |
| Mir218-1 | 2.27602632 | -2.328080397 | 0.768224692 |
| RP23-79E5.4 | 4.747594266 | -0.959775168 | 0.517981022 |
| RP23-79E5.5 | 0 | NA | NA |
| Pacrgl | 220.5840451 | -0.218582979 | 0.143821026 |
| Kcnip4 | 719.2995326 | -0.953193606 | 0.085932335 |
| RP23-14C20.2 | 34.79027562 | -1.561452336 | 0.322892 |
| Gm20700 | 0 | NA | NA |
| RP24-174D16.1 | 87.81495278 | -1.391783075 | 0.235208112 |
| Gm19031 | 0 | NA | NA |
| RP24-170C14.2 | 202.1229481 | -1.378602865 | 0.239829775 |
| RP23-309G12.4 | 37.2300334 | -1.659736121 | 0.35536254 |
| RP23-309G12.3 | 161.4099445 | -1.230664725 | 0.279488328 |
| RP23-309G12.2 | 70.38172967 | -1.354390403 | 0.272754037 |
| RP23-79E5.7 | 138.4994086 | -1.243609696 | 0.287261163 |
| RP23-309G12.1 | 115.2856778 | -1.026872602 | 0.262949809 |
| Gm10025 | 0 | NA | NA |
| Gm10048 | 309.3515181 | -1.53566016 | 0.334910749 |

| | | | |
|---|---|---|---|
| RP23-187D22.9 | 187.5255152 | -1.614269785 | 0.374136398 |
| RP23-187D22.8 | 194.8703213 | -1.249310916 | 0.221338484 |
| RP23-187D22.7 | 88.63220688 | -1.312033743 | 0.308220344 |
| RP23-187D22.6 | 375.4133243 | -1.414876855 | 0.293313023 |
| RP23-187D22.5 | 181.2108272 | -1.433441679 | 0.245002085 |
| RP23-187D22.4 | 188.0951161 | -1.631930856 | 0.359262933 |
| RP23-187D22.10 | 319.4630086 | -1.294385857 | 0.295374111 |
| RP23-187D22.11 | 0.476944705 | -2.455199005 | 1.444169572 |
| RP23-187D22.3 | 106.6585838 | -1.571088028 | 0.430696491 |
| RP23-187D22.1 | 388.4487423 | -1.472272108 | 0.29835873 |
| RP23-140B23.2 | 209.5621235 | -1.5022494 | 0.34561817 |
| RP23-140B23.1 | 603.2557397 | -1.28861876 | 0.253971614 |
| RP23-394G5.2 | 44.14745288 | -1.521277036 | 0.378743488 |
| RP23-394G5.1 | 396.9657275 | -1.335741143 | 0.309861826 |
| RP24-158C2.3 | 251.5456547 | -1.504790097 | 0.296883998 |
| Gm19932 | 0 | NA | NA |
| RP24-158C2.4 | 0.083526043 | 0.718202646 | 3.724945049 |
| RP24-253A20.1 | 476.3543519 | -1.536140181 | 0.307603693 |
| Gm5555 | 28.60310479 | 0.366262087 | 0.208253913 |
| RP24-79O7.3 | 0.322897824 | 0.237332241 | 1.601978208 |
| RP24-79O7.1 | 1.170209419 | -1.256550889 | 0.88294153 |
| Adgra3 | 255.2158286 | -0.288961121 | 0.106655434 |
| RP24-315J23.1 | 0.596915583 | -0.31977714 | 1.37266258 |
| RP24-315J23.2 | 0 | NA | NA |
| RP24-478I4.1 | 0.047948698 | -0.243569582 | 3.724945049 |
| Mir8117 | 0 | NA | NA |
| Gm22618 | 0.389682156 | -0.634309577 | 1.531256637 |
| RP23-21O1.1 | 0.042716945 | -0.243571806 | 3.724945049 |
| RP23-21O1.3 | 0.04393175 | -0.243571806 | 3.724945049 |
| RP23-21O1.2 | 0.038981395 | 0.718207094 | 3.724945049 |
| RP23-125B15.1 | 0 | NA | NA |
| RP24-363A3.1 | 0.195174816 | 0.231463797 | 2.236723735 |
| RP24-312A3.1 | 0 | NA | NA |
| RP24-441G10.1 | 0 | NA | NA |
| AC119939.1 | 0 | NA | NA |
| Ppargc1a | 1065.622016 | -0.196137383 | 0.062005577 |
| RP23-385G21.4 | 13.87657411 | -0.720293444 | 0.339069111 |
| RP23-385G21.3 | 24.14107729 | -0.636525608 | 0.259628842 |
| RP23-385G21.2 | 0 | NA | NA |
| RP23-385G21.1 | 28.94227565 | -0.43526365 | 0.250182808 |
| RP24-296L6.1 | 199.7747531 | -0.813864813 | 0.366936317 |
| RP24-296L6.2 | 266.7259097 | -0.779924544 | 0.23397421 |
| RP24-296L6.3 | 71.01041919 | -1.220432438 | 0.345789183 |
| RP24-296L6.4 | 241.104923 | -0.896715032 | 0.306889011 |
| RP24-299M10.1 | 94.15749016 | -1.618676151 | 0.462776159 |
| RP24-299M10.2 | 0.645477656 | -2.409003438 | 1.317579242 |
| Mir6417 | 1.584881738 | -0.555046576 | 0.827656222 |

| | | | |
|---|---|---|---|
| RP24-309H3.1 | 30.5102578 | -0.858348681 | 0.287021506 |
| Dhx15 | 1662.063983 | 0.267371953 | 0.098507686 |
| RP24-309H3.3 | 27.29790927 | -0.45820297 | 0.240591564 |
| 9230114K14Rik | 31.71347572 | -0.389532917 | 0.202489766 |
| RP24-309H3.5 | 0.319554274 | -0.243564207 | 1.629736656 |
| RP24-309H3.6 | 0.802626749 | 0.592705963 | 1.112842773 |
| RP24-237I4.4 | 0 | NA | NA |
| Gm25601 | 0.045882455 | 0.237312084 | 3.724945049 |
| Sod3 | 146.4300929 | 0.906138597 | 0.148596622 |
| Ccdc149 | 659.1518796 | -0.358407903 | 0.075804002 |
| RP24-237I4.3 | 0.09689823 | -0.724453471 | 3.724945049 |
| Lgi2 | 1350.453356 | -0.749017715 | 0.07556218 |
| RP23-181F22.4 | 0 | NA | NA |
| RP23-181F22.3 | 4.780205594 | -1.76607165 | 0.601326207 |
| Gm5866 | 0.405467889 | -1.02086589 | 1.564295806 |
| 8030423F21Rik | 2.177346141 | 1.038418784 | 0.668154889 |
| Sepsecs | 236.6757126 | 0.150837027 | 0.099737375 |
| Pi4k2b | 58.07000918 | 1.710602428 | 0.210366324 |
| Zcchc4 | 173.3575901 | -0.217542325 | 0.166680132 |
| Anapc4 | 711.191152 | 0.15556916 | 0.052117232 |
| RP24-373A18.2 | 0 | NA | NA |
| Gm23532 | 0 | NA | NA |
| RP24-373A18.1 | 0 | NA | NA |
| RP24-373A18.3 | 0.457291341 | 2.990740256 | 1.681639794 |
| Slc34a2 | 1.437183928 | 3.433989071 | 1.06864843 |
| Gm17182 | 0 | NA | NA |
| Sel1l3 | 908.2236204 | -0.641348431 | 0.126832861 |
| n-R5s171 | 0 | NA | NA |
| Smim20 | 129.4033044 | 0.418434787 | 0.140843072 |
| RP23-425N12.2 | 79.50346624 | -0.379760334 | 0.154069564 |
| n-R5s172 | 0.379461129 | -1.087388011 | 1.717138877 |
| RP23-425N12.1 | 0.485522898 | 0.176884845 | 1.483370519 |
| Gm26486 | 0 | NA | NA |
| Rbpj | 346.6721826 | 0.436705043 | 0.100440439 |
| RP24-540G23.3 | 0.236145685 | -0.545868074 | 2.11430247 |
| RP23-137N6.2 | 0.29199547 | 0.727910435 | 1.785213019 |
| Cckar | 6.464590576 | 0.747087682 | 0.479571065 |
| Tbc1d19 | 456.9575057 | -0.287421441 | 0.104067967 |
| RP24-196M24.2 | 141.7804511 | -0.472767025 | 0.215588745 |
| RP24-196M24.3 | 10.28263209 | -0.222155807 | 0.388897047 |
| Stim2 | 905.9735999 | -0.224713452 | 0.061686481 |
| Gm15820 | 0 | NA | NA |
| RP24-482F14.1 | 1.962603442 | -0.029998753 | 0.747961366 |
| RP23-148F6.1 | 0 | NA | NA |
| RP23-187K3.1 | 6.300906923 | 0.342672209 | 0.403618353 |
| RP24-201P24.1 | 0.044544648 | 0.237309859 | 3.724945049 |
| Gm24235 | 0 | NA | NA |

| | | | |
|---|---|---|---|
| RP23-267P6.1 | 0 | NA | NA |
| Gm25513 | 0 | NA | NA |
| RP24-133H21.1 | 0 | NA | NA |
| RP23-317I8.3 | 0 | NA | NA |
| RP23-317I8.2 | 0 | NA | NA |
| Gm15975 | 34.36835703 | 0.648685503 | 0.239138027 |
| RP23-317I8.1 | 0.181740481 | -1.200076803 | 2.280596453 |
| RP23-162F3.1 | 0.072025462 | 0.237312084 | 3.724945049 |
| RP23-162F3.2 | 1.035655884 | -0.99324696 | 1.081149632 |
| RP24-301M6.1 | 0.124853745 | 0.237309606 | 3.724945049 |
| RP24-394H13.1 | 0.707701802 | -0.143440103 | 1.47224611 |
| Gm23896 | 0 | NA | NA |
| Gm24902 | 0 | NA | NA |
| Gm23015 | 0 | NA | NA |
| RP23-17A11.1 | 0 | NA | NA |
| RP24-499N22.1 | 0 | NA | NA |
| RP24-499N22.2 | 0 | NA | NA |
| RP23-297D16.1 | 0 | NA | NA |
| RP23-297D16.2 | 0.038981395 | 0.718207094 | 3.724945049 |
| 4932441J04Rik | 57.67652023 | -0.446824682 | 0.151863488 |
| Gm8121 | 18.9482281 | 0.533179188 | 0.242809986 |
| RP24-189M4.2 | 0 | NA | NA |
| RP24-189M4.1 | 0.956833923 | -0.065115054 | 0.939949539 |
| Pcdh7 | 3274.629757 | -0.518663377 | 0.092700547 |
| RP23-458G12.1 | 132.1876841 | -0.698363105 | 0.132113827 |
| RP23-458G12.4 | 95.53520872 | -0.443102538 | 0.208442064 |
| RP23-458G12.3 | 115.8575128 | -0.769315515 | 0.17839494 |
| RP23-458G12.2 | 78.02550846 | -0.780702288 | 0.222315317 |
| RP23-358D12.4 | 15.63086198 | -0.488315576 | 0.403490499 |
| RP23-358D12.1 | 30.61318993 | -0.274299609 | 0.276319187 |
| RP23-358D12.3 | 67.73431069 | -0.668306084 | 0.175677082 |
| RP23-358D12.2 | 74.52970784 | -0.699023874 | 0.230057562 |
| RP24-231P10.2 | 87.86336388 | -0.679339658 | 0.202266185 |
| RP24-231P10.3 | 68.50219268 | -1.053724239 | 0.262023264 |
| RP24-231P10.4 | 91.67305281 | -0.678179561 | 0.234912921 |
| Gm37720 | 43.33085034 | -0.63624527 | 0.188925265 |
| RP24-231P10.5 | 54.31474321 | -0.626081955 | 0.206357061 |
| RP24-340K5.2 | 33.48457548 | -0.297453001 | 0.229131503 |
| RP24-340K5.4 | 0 | NA | NA |
| RP24-340K5.3 | 0 | NA | NA |
| RP24-502C10.1 | 0 | NA | NA |
| Gm22536 | 0 | NA | NA |
| RP24-375D24.2 | 0 | NA | NA |
| 1700029E06Rik | 0 | NA | NA |
| RP24-353N4.4 | 0 | NA | NA |
| Gm6632 | 0 | NA | NA |
| RP24-353N4.3 | 0 | NA | NA |

| | | | |
|---|---|---|---|
| Gm22308 | 0 | NA | NA |
| RP23-324H16.1 | 0 | NA | NA |
| RP23-324H16.2 | 3.023846459 | -0.397739624 | 0.615006832 |
| RP23-340N4.1 | 0 | NA | NA |
| RP24-267O6.1 | 1.051957033 | -2.803858915 | 1.15812854 |
| RP24-543J12.1 | 0 | NA | NA |
| RP24-543J12.2 | 0.062426873 | 0.237312084 | 3.724945049 |
| RP24-543J12.3 | 0 | NA | NA |
| RP24-543J12.4 | 0.075043205 | 0.237309859 | 3.724945049 |
| RP23-183O3.1 | 0.136194719 | -0.724453471 | 3.724945049 |
| Cbfa2t2-ps1 | 30.26364068 | -0.418637823 | 0.19738191 |
| Gm23331 | 0 | NA | NA |
| RP23-419P12.1 | 0 | NA | NA |
| RP23-84N24.1 | 0 | NA | NA |
| Gm23708 | 0 | NA | NA |
| RP24-390N7.1 | 2.35460177 | -0.147719453 | 0.599921295 |
| RP24-390N7.2 | 0 | NA | NA |
| RP24-390N7.3 | 0 | NA | NA |
| RP24-300K9.1 | 0 | NA | NA |
| RP23-210A16.1 | 0 | NA | NA |
| RP23-86N20.1 | 0 | NA | NA |
| G6pd2 | 0.044191549 | -0.243571806 | 3.724945049 |
| RP23-480E8.1 | 0 | NA | NA |
| Gm22273 | 0 | NA | NA |
| RP24-447I18.1 | 0.236971717 | -0.724459691 | 2.110625472 |
| Gm24079 | 0 | NA | NA |
| Arap2 | 2567.739427 | 0.440661916 | 0.072610753 |
| RP23-387F14.1 | 16.56268708 | 0.020970378 | 0.283229084 |
| RP23-387F14.2 | 57.3555258 | 0.092848782 | 0.188544845 |
| Dthd1 | 4.062000091 | 3.265599839 | 0.730189195 |
| RP23-37I24.1 | 105.4712107 | -0.889751955 | 0.168653764 |
| Nwd2 | 487.7321807 | -0.857174163 | 0.160945207 |
| Gm3687 | 4.714620213 | -1.284810734 | 0.522161507 |
| 0610040J01Rik | 36.24267069 | 0.054745147 | 0.190227646 |
| Rell1 | 246.5447793 | 0.87204267 | 0.120561033 |
| Gm15819 | 0.356802427 | 1.563231376 | 3.710948037 |
| RP23-192C21.2 | 0.04393175 | -0.243571806 | 3.724945049 |
| Gm25608 | 0 | NA | NA |
| RP23-192C21.3 | 1.013052095 | 2.214424447 | 1.070867681 |
| Pgm1 | 176.2902017 | 1.20565581 | 0.138933771 |
| Gm23967 | 0 | NA | NA |
| Gm22395 | 0 | NA | NA |
| Gm22559 | 0 | NA | NA |
| Tbc1d1 | 746.5571268 | 0.876427641 | 0.099409166 |
| Gm15824 | 0.060174537 | 0.237309859 | 3.724945049 |
| RP23-339I6.1 | 13.2226886 | 0.705722817 | 0.31477055 |
| Gm15823 | 0 | NA | NA |

| | | | |
|---|---|---|---|
| Gm6044 | 4.011017284 | -0.434338502 | 0.578468275 |
| RP23-339I6.2 | 0.245047857 | 0.718222864 | 2.107737545 |
| RP23-339I6.3 | 0.541557618 | 1.132712261 | 1.513961029 |
| RP23-195B3.4 | 0.062426873 | 0.237312084 | 3.724945049 |
| 1700027F09Rik | 0 | NA | NA |
| RP23-195B3.3 | 0 | NA | NA |
| RP23-195B3.5 | 10.29678862 | -4.942927748 | 0.702675015 |
| RP24-448C16.4 | 0.540262476 | -0.504287131 | 1.338667117 |
| RP24-448C16.6 | 0.192501058 | -0.629765963 | 2.663543715 |
| RP24-448C16.5 | 1.291588001 | -0.426478954 | 0.964374774 |
| RP24-448C16.7 | 3.668071862 | -0.180306665 | 0.514865189 |
| RP24-448C16.9 | 0.116217011 | -0.24357403 | 3.724945049 |
| RP24-448C16.8 | 0 | NA | NA |
| Gm25306 | 0.049553047 | 0.71820487 | 3.724945049 |
| Gm26761 | 0.503421469 | -1.187903903 | 1.378795365 |
| Gm20033 | 20.30300836 | -0.652922314 | 0.242517106 |
| Klf3 | 642.6073925 | 0.771907855 | 0.110561932 |
| RP23-326O2.2 | 26.4782624 | -0.288286275 | 0.226933101 |
| Tlr1 | 85.37836272 | 2.027513161 | 0.210214892 |
| Tlr6 | 42.58363877 | 2.694397181 | 0.270298752 |
| Fam114a1 | 72.52929912 | 1.35362533 | 0.177990652 |
| Mir574 | 0.038981395 | 0.718207094 | 3.724945049 |
| RP24-87P2.3 | 1.276825249 | 0.550953986 | 0.953628986 |
| Tmem156 | 6.703461548 | 2.974435518 | 0.567478713 |
| Klhl5 | 474.7177188 | -0.133422825 | 0.108100035 |
| Gm25154 | 2.287715794 | 0.731299171 | 0.867033492 |
| Wdr19 | 272.9715081 | -0.426062976 | 0.117171985 |
| Rfc1 | 611.6308179 | 0.526580197 | 0.07683444 |
| RP23-344O11.3 | 13.50952572 | -0.708187941 | 0.396490631 |
| Klb | 1.57634325 | -0.669293858 | 0.78750577 |
| Rpl9 | 1617.17554 | 0.508132683 | 0.087769663 |
| Lias | 326.7940019 | 0.078880834 | 0.085605248 |
| RP23-477N9.1 | 7.915089792 | -0.937664609 | 0.520280739 |
| RP23-477N9.5 | 5.012299285 | -0.772002379 | 0.472471487 |
| Ugdh | 393.579355 | 0.524254746 | 0.091166432 |
| AC136513.6 | 0.044544648 | 0.237309859 | 3.724945049 |
| RP23-477N9.2 | 5.168412192 | -0.611528133 | 0.496520532 |
| Smim14 | 1287.523276 | -0.330617641 | 0.054654595 |
| RP23-477N9.4 | 25.33295597 | -1.180286276 | 0.345074142 |
| RP23-477N9.6 | 5.250217998 | -1.031879169 | 0.449688406 |
| RP23-477N9.3 | 33.76352782 | -1.441925504 | 0.422122578 |
| Ube2k | 1596.82342 | -0.203057163 | 0.062124683 |
| RP23-403P24.1 | 88.56705528 | -0.882337326 | 0.291362302 |
| RP23-403P24.3 | 0 | NA | NA |
| Pds5a | 1522.438719 | -0.030824305 | 0.065221272 |
| RP23-403P24.5 | 11.79713159 | -0.641972851 | 0.333845671 |
| RP23-403P24.6 | 4.819094623 | 0.145604473 | 0.472129571 |

| | | | |
|---|---|---|---|
| Gm24399 | 0.0878635 | -0.72445792 | 3.724945049 |
| RP23-403P24.7 | 16.35270107 | -0.498004127 | 0.362449428 |
| RP23-403P24.8 | 29.36603598 | -0.731982286 | 0.274900374 |
| N4bp2os | 32.02296958 | -0.053310895 | 0.201760959 |
| N4bp2 | 387.2265267 | -0.123784309 | 0.11658478 |
| Gm24346 | 4.621381795 | -1.252461802 | 0.549992629 |
| RP23-42O8.7 | 30.0274551 | -0.609227208 | 0.339093466 |
| RP23-42O8.6 | 0.112396584 | 1.192083721 | 3.724434897 |
| RP23-42O8.5 | 0.165156969 | 1.598327176 | 3.401540795 |
| RP23-42O8.4 | 0 | NA | NA |
| Rhoh | 32.07602456 | 2.32573523 | 0.241774268 |
| RP23-42O8.9 | 0.263966226 | 0.236883522 | 1.822623217 |
| Chrna9 | 0.40746857 | 2.29049474 | 1.467684812 |
| RP23-92N2.1 | 0 | NA | NA |
| RP23-92N2.2 | 0 | NA | NA |
| 9130230L23Rik | 1.235688375 | 4.169487597 | 1.039692592 |
| Gm25952 | 0 | NA | NA |
| Rbm47 | 123.9754224 | 2.718985103 | 0.208243596 |
| RP24-351O18.5 | 7.439759207 | 2.595178296 | 0.563715543 |
| RP24-351O18.4 | 9.722078793 | 2.572171207 | 0.46131815 |
| Gm26725 | 0.204218301 | 0.66222277 | 2.57764078 |
| Gm22929 | 0 | NA | NA |
| 1700126H18Rik | 0 | NA | NA |
| RP24-351O18.2 | 0 | NA | NA |
| Gm15794 | 0 | NA | NA |
| RP24-351O18.7 | 0.049553047 | 0.71820487 | 3.724945049 |
| Nsun7 | 120.513026 | 0.720221048 | 0.174930804 |
| Apbb2 | 1193.808036 | -0.069574698 | 0.07236316 |
| RP23-21D6.2 | 18.43062459 | -0.864983092 | 0.30449659 |
| Gm23183 | 8.321784305 | -0.453023719 | 0.462781105 |
| RP23-21D6.1 | 24.68721265 | -0.67012055 | 0.340767782 |
| Gm16273 | 0.062426873 | 0.237312084 | 3.724945049 |
| RP23-157G2.2 | 24.13982844 | -0.750359117 | 0.279227293 |
| RP23-157G2.1 | 9.142804473 | -1.072172317 | 0.405370932 |
| RP24-360B3.2 | 21.21818474 | -0.68432901 | 0.367933873 |
| RP24-360B3.3 | 23.13608884 | -0.820332103 | 0.327263592 |
| RP24-360B3.1 | 33.09598643 | -0.498029481 | 0.366861298 |
| Uchl1os | 3.562613421 | -0.545015662 | 0.575490647 |
| Uchl1 | 2844.203176 | -0.215394354 | 0.077144383 |
| RP23-472J13.3 | 20.85386141 | -0.395957061 | 0.529841649 |
| RP23-352L15.1 | 0.061920613 | 0.718207094 | 3.724945049 |
| Limch1 | 1533.433457 | -0.329748444 | 0.055500714 |
| Gm6517 | 9.113272248 | -1.325677485 | 0.390023579 |
| RP24-79P8.1 | 29.56015194 | -0.769045096 | 0.28278201 |
| RP24-383P24.2 | 11.74058244 | -0.956293675 | 0.355667765 |
| Gm15949 | 0 | NA | NA |
| Gm15948 | 0.038981395 | 0.718207094 | 3.724945049 |

| | | | |
|---|---|---|---|
| RP24-383P24.3 | 0.808719243 | 0.020632067 | 1.028187749 |
| Gm23841 | 0 | NA | NA |
| Phox2b | 13.13116622 | 0.64267098 | 0.552780607 |
| RP24-95M11.1 | 1.073023658 | 0.440939734 | 1.096211864 |
| RP24-353A8.1 | 1.943439641 | 1.91673617 | 0.767485069 |
| Tmem33 | 1537.582746 | 0.061707636 | 0.064934596 |
| Slc30a9 | 1760.179162 | -0.497510555 | 0.056586138 |
| RP24-353A8.2 | 0.708786783 | -0.47671826 | 1.204968322 |
| Bend4 | 204.8953353 | -0.436993003 | 0.11200936 |
| Gm26756 | 4.883969858 | -1.379607824 | 0.507644635 |
| RP24-103L16.3 | 7.272572362 | -1.476003281 | 0.549797202 |
| RP24-103L16.4 | 0 | NA | NA |
| RP24-103L16.5 | 15.55852251 | -0.810309539 | 0.267874245 |
| RP23-213N8.1 | 1.075240245 | -0.416421977 | 0.952899161 |
| Shisa3 | 72.51937258 | 0.364532888 | 0.535915273 |
| Atp8a1 | 4334.315932 | -0.204875381 | 0.060746048 |
| Gm23617 | 2.388208885 | -0.436855175 | 0.65534915 |
| RP23-213N8.2 | 52.3886502 | -0.668311271 | 0.281870416 |
| Gm15478 | 2.676788521 | -0.738868968 | 0.674448992 |
| RP24-274I18.1 | 56.96681602 | -0.762083481 | 0.304902958 |
| RP23-130C18.1 | 59.36905083 | -0.571778575 | 0.277285366 |
| RP23-130C18.2 | 29.63687175 | -0.593438357 | 0.302833541 |
| Gm15477 | 9.54473322 | 0.242673266 | 0.357263236 |
| RP23-130C18.3 | 0.310726904 | 0.307691265 | 2.067812291 |
| Gm21905 | 0.178178593 | 0.237301575 | 3.724945049 |
| RP23-130C18.4 | 1.374506326 | 1.723635807 | 1.377135366 |
| Gm5108 | 0.225572436 | 0.71768769 | 2.199023238 |
| RP23-387L12.2 | 0.395257705 | 0.100552781 | 1.714076798 |
| RP23-387L12.3 | 1.482312138 | -0.304165711 | 0.832064044 |
| Grxcr1 | 0.175796035 | 0.726744679 | 2.308680797 |
| RP23-387L12.4 | 0.038981395 | 0.718207094 | 3.724945049 |
| RP24-224K15.1 | 0 | NA | NA |
| Gm23605 | 0 | NA | NA |
| RP24-441E22.1 | 0.086648695 | -0.72445792 | 3.724945049 |
| RP23-72G21.1 | 2.775994898 | -0.461486564 | 0.648364474 |
| Kctd8 | 186.6518599 | -0.478683452 | 0.097352709 |
| RP23-319P12.6 | 23.8423184 | -0.883571191 | 0.247826428 |
| RP23-319P12.5 | 17.40396653 | -1.146267588 | 0.28740506 |
| RP23-319P12.4 | 57.79128105 | -0.988625525 | 0.257393943 |
| RP23-319P12.3 | 56.51122244 | -0.790493226 | 0.262499231 |
| RP23-319P12.2 | 47.23969796 | -1.212388338 | 0.271763538 |
| RP23-319P12.1 | 45.02692642 | -1.256686598 | 0.306149549 |
| RP23-114M24.1 | 151.2873273 | -0.875831491 | 0.226682304 |
| Gm24368 | 0.334883495 | 0.197332555 | 1.578985564 |
| Gm23932 | 0 | NA | NA |
| Gm5867 | 0.309731683 | -0.243563031 | 1.750531252 |
| RP24-351E22.1 | 0.956841293 | 0.241308392 | 1.137508874 |

| | | | |
|---|---|---|---|
| Yipf7 | 2.071883479 | 1.397449488 | 0.721181192 |
| RP24-351E22.2 | 0.331853251 | -0.215240547 | 1.764682824 |
| Guf1 | 544.3541319 | -0.349678179 | 0.10942498 |
| Gnpda2 | 249.4753375 | 0.070737118 | 0.128741918 |
| Gm26044 | 0 | NA | NA |
| Gm23661 | 0 | NA | NA |
| RP23-236G19.1 | 0 | NA | NA |
| Gm23067 | 0 | NA | NA |
| Gm26072 | 0 | NA | NA |
| RP23-332O18.1 | 0 | NA | NA |
| RP23-332O18.2 | 0.480717083 | -1.144129232 | 1.526529995 |
| Gabrg1 | 384.6081965 | -0.310788427 | 0.135975675 |
| Gabra2 | 1383.339494 | -0.767903478 | 0.083008537 |
| Cox7b2 | 1.615076682 | -0.049025379 | 0.706056847 |
| Gabra4 | 1311.463824 | -0.845914665 | 0.110132293 |
| Gabrb1 | 2188.449478 | -0.495757101 | 0.080994288 |
| RP24-156O12.1 | 9.874941751 | -0.890452088 | 0.328664051 |
| Gm15617 | 0.150086409 | 0.237305146 | 3.724945049 |
| Gm15616 | 12.70121426 | -0.925442018 | 0.292005085 |
| Commd8 | 452.1455523 | 0.212914422 | 0.072941096 |
| RP23-27G7.2 | 0 | NA | NA |
| Atp10d | 139.6844161 | 0.809710035 | 0.142151456 |
| Corin | 18.43000873 | 0.052510736 | 0.303188428 |
| Gm25768 | 0.088383099 | -0.72445792 | 3.724945049 |
| Gm20647 | 0 | NA | NA |
| Gm19560 | 0 | NA | NA |
| Gm23712 | 0.116046057 | -0.724453471 | 3.724945049 |
| AU023070 | 0.14021869 | -0.243563615 | 2.921525367 |
| Nfxl1 | 313.9107044 | 0.080478247 | 0.07388139 |
| RP24-316L21.3 | 23.03118454 | -2.72E-05 | 0.22439061 |
| RP24-506B15.1 | 0.413335106 | 0.668516257 | 1.459606399 |
| Zar1 | 3.771757384 | 0.169837285 | 0.540136953 |
| Gm5868 | 0.655263372 | 1.274116435 | 1.45293673 |
| Cnga1 | 0.310549208 | 2.19189667 | 3.694476578 |
| RP24-506B15.2 | 5.852374332 | 1.22897418 | 0.438982518 |
| RP24-506B15.3 | 0 | NA | NA |
| Nipal1 | 7.542068972 | 4.389641265 | 0.856902266 |
| Txk | 24.20729952 | 3.77208906 | 0.508427499 |
| RP23-415O9.1 | 0 | NA | NA |
| Tec | 81.72636874 | 2.747664916 | 0.291599383 |
| RP23-204A8.1 | 6.319016411 | 1.059339188 | 0.437788391 |
| Gm9870 | 6.265701809 | -0.04009514 | 0.410489185 |
| Slain2 | 498.0036851 | 0.137760112 | 0.07508105 |
| Gm10135 | 0.045882455 | 0.237312084 | 3.724945049 |
| Gm24776 | 0 | NA | NA |
| Slc10a4 | 54.26526042 | -0.046898163 | 0.186467421 |
| Gm15847 | 1.549834172 | -2.807613725 | 1.369078659 |

| | | | |
|---|---|---|---|
| AC108848.6 | 1.313571363 | -0.212002493 | 0.866921963 |
| Fryl | 1628.389783 | -0.039764084 | 0.061285409 |
| RP23-300K5.1 | 35.04837016 | -0.655886999 | 0.260847672 |
| 1700071G01Rik | 0 | NA | NA |
| RP23-318I19.2 | 17.43330629 | -0.562296573 | 0.32583115 |
| Gm17207 | 5.90520374 | -0.767274995 | 0.41293647 |
| RP23-153H17.2 | 0.097139547 | 0.718207094 | 3.724945049 |
| 1700025M24Rik | 0.068097359 | -0.243569582 | 3.724945049 |
| RP23-153H17.3 | 0 | NA | NA |
| Ociad1 | 3089.545592 | -0.209938848 | 0.046691632 |
| RP23-153H17.6 | 0.3552573 | 1.751624623 | 2.011465178 |
| RP23-153H17.5 | 470.3152913 | -0.762333962 | 0.087112157 |
| Ociad2 | 527.2792456 | 0.098723727 | 0.149870923 |
| RP23-153H17.7 | 29.63938725 | 0.212679444 | 0.268580664 |
| RP23-153H17.9 | 0.315779812 | 1.298078541 | 3.707347035 |
| Cwh43 | 14.92583306 | -0.323039822 | 0.32016012 |
| RP24-274B10.1 | 0.149988742 | -0.243571388 | 2.860996817 |
| Gm15653 | 0 | NA | NA |
| Dcun1d4 | 1162.429234 | -0.462089937 | 0.067386837 |
| Lrrc66 | 2.285793377 | -0.425353857 | 0.83857605 |
| Sgcb | 773.7364474 | 0.449799811 | 0.092762177 |
| RP23-105O12.2 | 2.772617781 | -0.23551194 | 0.589840979 |
| Spata18 | 12.87588817 | -0.227217227 | 0.349277982 |
| RP23-276J7.1 | 0 | NA | NA |
| RP23-276J7.2 | 0.066050875 | 0.71820487 | 3.724945049 |
| Usp46 | 1284.997462 | -0.476968501 | 0.0734476 |
| Usp46os1 | 0.374638785 | -0.674466457 | 1.584197982 |
| Usp46os2 | 0.045882455 | 0.237312084 | 3.724945049 |
| CH25-194I22.2 | 46.11457756 | -1.428109338 | 0.312118383 |
| CH25-194I22.3 | 0 | NA | NA |
| CH25-194I22.4 | 0.285862058 | 0.217107517 | 1.776712065 |
| Snora26 | 6.265788682 | -0.058784513 | 0.405953769 |
| Dancr | 0 | NA | NA |
| RP23-422P4.2 | 0.766635478 | 0.798151402 | 1.034434524 |
| Rasl11b | 390.2960293 | -0.556901491 | 0.138199586 |
| Scfd2 | 235.5014393 | -0.276408396 | 0.07976437 |
| RP23-422P4.3 | 0.472351266 | 0.460051599 | 1.556602553 |
| Gm15981 | 4.746611391 | -0.671972814 | 0.512340939 |
| Gm15982 | 6.659240228 | -0.563320494 | 0.446200664 |
| RP24-465E5.2 | 15.61428043 | -0.728890031 | 0.401899602 |
| RP24-465E5.1 | 9.836186905 | -0.793713299 | 0.461483491 |
| Fip1l1 | 1364.710559 | -0.138300026 | 0.069900511 |
| RP24-245O20.1 | 42.86441423 | -0.509143088 | 0.207388308 |
| Lnx1 | 208.7467131 | -0.750913089 | 0.092074285 |
| Gm15984 | 0.100902008 | 1.15059983 | 3.720931964 |
| Gm15985 | 0 | NA | NA |
| RP23-88L6.1 | 0 | NA | NA |

| | | | |
|---|---|---|---|
| RP24-245O20.2 | 0 | NA | NA |
| Gm6116 | 0.042716945 | -0.243571806 | 3.724945049 |
| RP23-382I10.2 | 10.57589096 | 0.450041959 | 0.370868262 |
| Chic2 | 373.5242777 | 0.138195885 | 0.104955809 |
| RP23-382I10.1 | 56.62750935 | -0.655199888 | 0.275254048 |
| RP23-382I10.3 | 0.095897396 | -0.724453471 | 3.724945049 |
| Gsx2 | 1.492637219 | -0.484219221 | 0.912221732 |
| RP23-55P22.2 | 0 | NA | NA |
| RP23-55P22.3 | 0.097139547 | 0.718207094 | 3.724945049 |
| Olfr1401-ps1 | 0 | NA | NA |
| Pdgfra | 784.1705611 | 0.11657274 | 0.108776408 |
| RP23-55P22.6 | 0.141219523 | -0.243563604 | 2.915099341 |
| RP23-55P22.5 | 0 | NA | NA |
| Mir7025 | 0.116046057 | -0.724453471 | 3.724945049 |
| RP23-55P22.7 | 0 | NA | NA |
| RP23-149H20.2 | 0 | NA | NA |
| Gm22084 | 0 | NA | NA |
| RP23-149H20.3 | 0 | NA | NA |
| Gm24502 | 0 | NA | NA |
| Kit | 814.734425 | -0.265726788 | 0.11407743 |
| RP24-273B9.2 | 0.092228164 | 0.718202646 | 3.724945049 |
| RP24-273B9.1 | 0.222682676 | 1.157320706 | 2.136262342 |
| RP24-273B9.3 | 0 | NA | NA |
| RP23-329A21.3 | 0.435363789 | -1.035082042 | 1.381908612 |
| Gm38180 | 0.072025462 | 0.237312084 | 3.724945049 |
| Gm28865 | 0.066050875 | 0.71820487 | 3.724945049 |
| Gm25374 | 0 | NA | NA |
| Kdr | 411.3836948 | 4.001994169 | 0.295106425 |
| Gm24738 | 0 | NA | NA |
| RP24-445F15.2 | 0 | NA | NA |
| RP24-445F15.1 | 2.139707088 | -0.454422975 | 0.725091361 |
| Srd5a3 | 137.1421468 | 0.488854375 | 0.112549885 |
| Tmem165 | 351.682121 | 0.302367557 | 0.095668814 |
| Apc-ps1 | 7.578865732 | -0.391339359 | 0.516391365 |
| Clock | 2177.681423 | -0.059555943 | 0.060795921 |
| RP23-386C10.8 | 47.58777522 | 0.196851585 | 0.204845768 |
| RP23-386C10.5 | 89.03088876 | -0.551824533 | 0.287380788 |
| Gm7467 | 7.824312757 | -0.89466652 | 0.395026202 |
| Pdcl2 | 0.114443069 | 0.23732098 | 3.724945049 |
| 4930432L08Rik | 0.99774422 | 0.357723071 | 1.201424663 |
| Nmu | 1.605616479 | 0.251638449 | 0.881870572 |
| RP24-186A17.1 | 0 | NA | NA |
| RP24-186A17.2 | 0.068097359 | -0.243569582 | 3.724945049 |
| Gm7271 | 1.031658297 | 0.464221994 | 1.114026413 |
| Exoc1 | 794.8721072 | 0.036382374 | 0.097994738 |
| Cep135 | 200.4137152 | -0.030779657 | 0.113600054 |
| A730089K16Rik | 11.31091771 | -0.2317953 | 0.329342623 |

| | | | |
|---|---|---|---|
| C530008M17Rik | 974.9986135 | -0.412800946 | 0.072367462 |
| Gm7494 | 0.044191549 | -0.243571806 | 3.724945049 |
| Aasdh | 129.6955173 | -0.571632781 | 0.170131524 |
| Ppat | 241.3568791 | -0.268678847 | 0.085240129 |
| 2310040G07Rik | 1.39740727 | 0.586845153 | 0.997716011 |
| Paics | 931.5296296 | 0.147681753 | 0.091771329 |
| Srp72 | 1194.911166 | 0.196358564 | 0.063407836 |
| Arl9 | 3.563487532 | 1.36091113 | 0.567502009 |
| 1700112J05Rik | 0 | NA | NA |
| 1700023E05Rik | 5.201511052 | -0.02182972 | 0.452558699 |
| Hopx | 352.4399641 | 0.639028768 | 0.140164356 |
| Hopxos | 6.015739958 | 0.902836683 | 0.469608807 |
| Gm15831 | 0.13681464 | 0.237322887 | 2.94378999 |
| RP23-3L23.1 | 0 | NA | NA |
| Spink2 | 0.769070381 | 2.519957559 | 1.225700371 |
| RP24-439P5.1 | 0 | NA | NA |
| Rest | 302.1062253 | 1.424189203 | 0.135646626 |
| Mir5098 | 0.366116714 | 0.541318844 | 1.540387108 |
| Noa1 | 216.3475351 | 0.237413468 | 0.084158432 |
| Polr2b | 926.3349238 | 0.053216052 | 0.056511655 |
| RP24-439P5.3 | 1.692103223 | -1.500756529 | 0.881696692 |
| Igfbp7 | 165.7570609 | 1.935583094 | 0.156349974 |
| RP24-89O12.2 | 0 | NA | NA |
| RP24-89O12.4 | 0.126225412 | 0.718200913 | 3.698754989 |
| RP24-89O12.3 | 0 | NA | NA |
| RP24-89O12.5 | 0 | NA | NA |
| Pea15b | 0.138038828 | -0.243577639 | 3.633333205 |
| Gm22011 | 0 | NA | NA |
| RP24-358F13.1 | 0.17692102 | 0.773454306 | 2.723401318 |
| RP23-465P13.1 | 0 | NA | NA |
| Mthfr-ps1 | 0.438166815 | -0.312718574 | 1.444892056 |
| RP23-135B1.1 | 0 | NA | NA |
| Gm25756 | 0 | NA | NA |
| RP24-318J14.1 | 0 | NA | NA |
| RP24-187F3.3 | 0 | NA | NA |
| Gm22553 | 0 | NA | NA |
| Gm23429 | 0 | NA | NA |
| Gm24304 | 0 | NA | NA |
| Gm22974 | 0 | NA | NA |
| Gm2199 | 0.805896106 | 1.179660185 | 1.097407411 |
| Adgrl3 | 3423.719416 | -0.211919167 | 0.083100522 |
| RP24-248M2.5 | 137.1470277 | -0.266042118 | 0.142317634 |
| RP24-74E24.5 | 20.74210032 | -0.417153891 | 0.321705871 |
| AC104196.1 | 2.384345449 | -0.302350725 | 0.692355716 |
| Gm16054 | 1.328319734 | -0.060977515 | 1.130277388 |
| RP24-82B9.1 | 52.62095602 | -0.428149977 | 0.167902063 |
| RP23-393A24.1 | 0.106465261 | 0.718202646 | 3.724945049 |

| | | | |
|---|---|---|---|
| 1700031L13Rik | 0.049553047 | 0.71820487 | 3.724945049 |
| RP23-151G22.1 | 9.983061651 | 1.230514087 | 0.34217631 |
| Mir1187 | 0 | NA | NA |
| Gm24129 | 0 | NA | NA |
| RP24-558F5.1 | 0 | NA | NA |
| Tecrl | 1.615357459 | -0.541467315 | 0.784055617 |
| Gm15626 | 0.088383099 | -0.72445792 | 3.724945049 |
| AC120867.1 | 0 | NA | NA |
| Gm25765 | 0 | NA | NA |
| Gm25806 | 0 | NA | NA |
| Gm24792 | 0 | NA | NA |
| RP23-189P23.1 | 0 | NA | NA |
| Epha5 | 1720.415688 | -0.556066729 | 0.094820152 |
| Hmgn2-ps1 | 6.552657376 | 0.579510112 | 0.409766891 |
| RP24-166L23.1 | 2.846346598 | -0.616074763 | 0.795785729 |
| RP24-138J15.1 | 0 | NA | NA |
| RP24-273G7.1 | 0 | NA | NA |
| RP23-227I15.1 | 0.342219405 | -1.196749819 | 1.718645153 |
| Gm24626 | 0 | NA | NA |
| RP23-258M22.2 | 1.514822285 | -0.721346506 | 0.728797806 |
| Cenpc1 | 463.6409797 | -0.301814562 | 0.062283335 |
| RP23-19G21.1 | 0.150451996 | 0.237323705 | 2.858243839 |
| Stap1 | 5.819746149 | 4.318750602 | 0.791118914 |
| Gm24524 | 0.085327104 | 1.179809211 | 3.723500796 |
| Uba6 | 668.4127751 | 0.016771451 | 0.064303368 |
| Gnrhr | 0.310823584 | 0.66434229 | 1.728134864 |
| Tmprss11c | 0.04393175 | -0.243571806 | 3.724945049 |
| Gm25473 | 0 | NA | NA |
| Tmprss11d | 1.632697194 | -0.404045202 | 0.828869558 |
| Tmprss11a | 3.616981247 | -4.833444856 | 0.919511524 |
| Tmprss11g | 0.130746305 | 0.703205782 | 3.666340321 |
| Tmprss11f | 0.18352982 | 0.237308059 | 3.724945049 |
| Tmprss11b | 2.079765874 | 3.478701877 | 1.011844185 |
| RP23-314N3.1 | 0.090074004 | -0.24357403 | 3.724945049 |
| Tmprss11e | 0.166952883 | 1.610951781 | 2.74653892 |
| Gm16579 | 0.090277459 | 0.237318756 | 3.724945049 |
| RP24-567O8.4 | 0 | NA | NA |
| RP24-567O8.3 | 0.115000406 | 0.23732098 | 3.724945049 |
| Ythdc1 | 1204.211079 | 0.127253137 | 0.107903258 |
| RP24-158N23.1 | 0 | NA | NA |
| Ugt2b34 | 1.309248412 | 3.830835019 | 1.129563064 |
| Gm25211 | 0 | NA | NA |
| Ugt2b1 | 0 | NA | NA |
| Gm24121 | 0 | NA | NA |
| RP23-59B8.1 | 0 | NA | NA |
| RP23-227D7.1 | 0.092228164 | 0.718202646 | 3.724945049 |
| RP23-59B8.2 | 33.07073811 | 0.610325615 | 0.23827837 |

| | | | |
|---|---|---|---|
| Ugt2b35 | 0.394550875 | 1.160148643 | 1.460243282 |
| RP23-59B8.3 | 0 | NA | NA |
| RP23-227D7.2 | 0 | NA | NA |
| Ugt2b36 | 0.256447508 | 1.225217374 | 2.469852681 |
| RP23-227D7.3 | 0 | NA | NA |
| RP23-67B9.4 | 0 | NA | NA |
| Ugt2b5 | 0.491974008 | 1.905076031 | 1.307778619 |
| RP23-67B9.2 | 0.045882455 | 0.237312084 | 3.724945049 |
| RP23-227D7.4 | 0 | NA | NA |
| RP23-67B9.1 | 0.072025462 | 0.237312084 | 3.724945049 |
| Ugt2b37 | 0.046345709 | 0.71820487 | 3.724945049 |
| RP23-67B9.3 | 0.075043205 | 0.237309859 | 3.724945049 |
| Ugt2a3 | 0.072025462 | 0.237312084 | 3.724945049 |
| RP23-227D7.7 | 0 | NA | NA |
| RP23-67B9.5 | 0 | NA | NA |
| RP23-227D7.8 | 0.093141081 | -0.724455696 | 3.724945049 |
| RP23-227D7.6 | 0 | NA | NA |
| RP23-227D7.5 | 0 | NA | NA |
| Ugt2b38 | 0.123841226 | 1.292225235 | 3.719439735 |
| Ugt2a2 | 0.062426873 | 0.237312084 | 3.724945049 |
| Ugt2a1 | 0 | NA | NA |
| RP23-227D7.10 | 0 | NA | NA |
| Sult1b1 | 0.042716945 | -0.243571806 | 3.724945049 |
| Sult1d1 | 1.935656465 | -0.234290461 | 1.090578892 |
| Sult1e1 | 0.061920613 | 0.718207094 | 3.724945049 |
| mmu-mir-3969 | 0 | NA | NA |
| RP24-571N6.2 | 0 | NA | NA |
| RP24-571N6.3 | 0 | NA | NA |
| Csn1s1 | 0.044544648 | 0.237309859 | 3.724945049 |
| Csn2 | 0 | NA | NA |
| Gm25066 | 0 | NA | NA |
| RP23-440C1.3 | 0.266205562 | 1.208749481 | 1.803586166 |
| Gm24314 | 0 | NA | NA |
| Csn1s2a | 0.075043205 | 0.237309859 | 3.724945049 |
| Csn1s2b | 0 | NA | NA |
| Prr27 | 0 | NA | NA |
| RP23-440C1.5 | 0 | NA | NA |
| Gm7337 | 0 | NA | NA |
| Odam | 0 | NA | NA |
| 1700066N21Rik | 0 | NA | NA |
| RP23-423C8.4 | 0 | NA | NA |
| Csn3 | 0.488468934 | 2.93071371 | 1.284260768 |
| BC051076 | 0.204292078 | -1.281279935 | 3.71120435 |
| 2310003L06Rik | 0.63240349 | 1.333589412 | 1.237275253 |
| Gm28434 | 0 | NA | NA |
| Cabs1 | 0.109869311 | 0.238305784 | 3.724874707 |
| Smr3a | 0 | NA | NA |

| | | | |
|---|---|---|---|
| RP24-324D14.5 | 0 | NA | NA |
| RP24-324D14.6 | 0 | NA | NA |
| Smr2 | 0 | NA | NA |
| Gm11116 | 0 | NA | NA |
| RP23-50G20.2 | 0 | NA | NA |
| Gm11115 | 0 | NA | NA |
| RP23-50G20.3 | 0 | NA | NA |
| RP23-50G20.5 | 0 | NA | NA |
| RP23-50G20.4 | 0 | NA | NA |
| Gm7714 | 0 | NA | NA |
| Prol1 | 0 | NA | NA |
| Amtn | 0 | NA | NA |
| RP23-265M4.3 | 1.532002984 | -1.491010815 | 1.013853857 |
| RP23-265M4.4 | 0 | NA | NA |
| Ambn | 0 | NA | NA |
| Enam | 0 | NA | NA |
| Jchain | 15.78987055 | 1.540465508 | 0.582295887 |
| RP24-132B11.1 | 0.854816446 | 0.332675074 | 0.983959043 |
| Utp3 | 218.2673712 | 0.585913876 | 0.1837293 |
| Rufy3 | 2816.589347 | -0.07427207 | 0.050865407 |
| Grsf1 | 1026.519618 | 0.087652316 | 0.061259961 |
| Mob1b | 844.5626815 | 0.156185742 | 0.120131944 |
| Gm15682 | 3.374206779 | 1.209210584 | 0.617373859 |
| Dck | 405.0431763 | 1.100357767 | 0.110854804 |
| RP24-242L17.1 | 22.06035809 | -1.044890691 | 0.377740716 |
| RP24-242L17.2 | 0 | NA | NA |
| Mkrn1-ps1 | 1.85792581 | 0.365830797 | 0.679334764 |
| Slc4a4 | 4189.981023 | -0.671106168 | 0.132912084 |
| Gc | 1.073168736 | 0.964403522 | 1.147212496 |
| RP24-365I5.1 | 0.045882455 | 0.237312084 | 3.724945049 |
| RP24-365I5.2 | 0 | NA | NA |
| Npffr2 | 7.733873509 | -0.842582516 | 0.460037469 |
| Adamts3 | 373.7060012 | -0.724656192 | 0.211253785 |
| Gm25320 | 0.186976534 | 0.718216355 | 2.668791307 |
| Gm25758 | 0 | NA | NA |
| Eif5al3-ps | 2.402331814 | 1.555950809 | 0.778773211 |
| Cox18 | 236.2894485 | 0.752682665 | 0.109766159 |
| Ankrd17 | 4578.10987 | 0.181376705 | 0.062587741 |
| Gm9958 | 14.24586082 | 0.309889084 | 0.249991015 |
| RP23-279P6.5 | 0 | NA | NA |
| Alb | 7.938965767 | 0.216522229 | 0.592897028 |
| Afp | 3.204684021 | 1.792054516 | 0.650609808 |
| RP23-279P6.3 | 0.060174537 | 0.237309859 | 3.724945049 |
| Afm | 0.397459543 | 1.917753916 | 1.684571988 |
| RP24-63O24.1 | 0.290146865 | -0.672292599 | 2.027255783 |
| RP24-63O24.4 | 0 | NA | NA |
| 5830473C10Rik | 0.606733601 | 0.574417776 | 1.287142397 |

| | | | |
|---|---|---|---|
| Rassf6 | 13.93087578 | -0.7377581 | 0.300247052 |
| RP24-168A7.1 | 0.237346126 | 0.237328964 | 2.121312375 |
| Cxcl5 | 7.088183414 | 1.31853841 | 0.48243652 |
| Ppbp | 1.785008614 | 1.131395102 | 0.822789283 |
| Pf4 | 3.421604734 | -0.3134518 | 0.572172288 |
| Cxcl3 | 3.225786153 | 5.73806684 | 1.006622426 |
| Cxcl15 | 0.068097359 | -0.243569582 | 3.724945049 |
| Gm24322 | 0 | NA | NA |
| RP24-168A7.7 | 0 | NA | NA |
| RP23-307P18.1 | 0 | NA | NA |
| Cxcl1 | 39.73533034 | 6.034344289 | 1.043240916 |
| Gm22816 | 0.235449268 | 0.718217112 | 2.543787894 |
| Cxcl2 | 19.17032838 | 7.243100232 | 1.166480634 |
| RP23-307P18.3 | 0 | NA | NA |
| Mthfd2l | 111.8726162 | -0.259624624 | 0.119525741 |
| Epgn | 0.633611781 | -1.054045352 | 1.32632263 |
| RP23-42K10.4 | 0.186036187 | 1.136436328 | 2.703027039 |
| Ereg | 4.701452776 | 6.237596968 | 1.039853344 |
| Areg | 0.714414789 | 3.151948573 | 1.254050252 |
| Gm23092 | 5.083391071 | 0.053089058 | 0.519865276 |
| RP23-258C6.2 | 6.099374268 | 0.539772946 | 0.433968615 |
| Btc | 22.83547764 | 2.381120502 | 0.306927034 |
| Parm1 | 754.069321 | -0.225519796 | 0.058436448 |
| RP23-118H9.3 | 3.970608911 | 0.446176986 | 0.544257042 |
| RP23-17D18.3 | 4.213956554 | 0.084801065 | 0.459045541 |
| Gm28271 | 0.521165882 | 0.857536796 | 1.4933322 |
| 1700063O14Rik | 0.323348989 | -0.195771692 | 1.719978378 |
| RP24-266G11.2 | 0.541747793 | 1.807159583 | 1.525842701 |
| RP24-266G11.4 | 0 | NA | NA |
| Gm5558 | 5.850622695 | -0.203339819 | 0.46405595 |
| RP23-442L7.2 | 0.329047954 | 1.203599098 | 1.766593391 |
| Rchy1 | 427.1403147 | 0.566986782 | 0.08713464 |
| RP23-442L7.3 | 26.15808167 | -0.285556088 | 0.314862806 |
| Thap6 | 30.14759305 | 0.888994704 | 0.187703813 |
| Gm24113 | 0.102891481 | -0.243576254 | 3.724945049 |
| Gm1045 | 3.240137224 | 1.088558064 | 0.661232717 |
| Cdkl2 | 400.6574654 | -0.61640376 | 0.093006995 |
| Gm26582 | 7.944179279 | -0.340522829 | 0.352669454 |
| G3bp2 | 4286.859093 | 0.530682697 | 0.102616401 |
| Gm22728 | 6.520177888 | 0.61341596 | 0.503554568 |
| Gm24931 | 1.274348854 | -1.208521392 | 0.937869044 |
| Uso1 | 1247.998467 | 0.180971747 | 0.055168279 |
| Gm15710 | 147.9457653 | 0.619047269 | 0.155923673 |
| RP23-321C24.2 | 0 | NA | NA |
| Gm25290 | 1.111053339 | -1.766619501 | 1.031479489 |
| U90926 | 0.097139547 | 0.718207094 | 3.724945049 |
| Ppef2 | 8.480238973 | -0.45096892 | 0.341482384 |

| | | | |
|---|---|---|---|
| Naaa | 229.011174 | 0.699268767 | 0.144409576 |
| Sdad1 | 341.6351534 | 0.502119245 | 0.108124925 |
| RP23-447A7.2 | 0.049553047 | 0.71820487 | 3.724945049 |
| Gm23031 | 0.045882455 | 0.237312084 | 3.724945049 |
| Cxcl9 | 597.6617564 | 11.87039528 | 0.936315244 |
| Art3 | 40.01616527 | 0.992194786 | 0.198280383 |
| Cxcl10 | 3859.437308 | 12.34550471 | 0.71309469 |
| Cxcl11 | 885.873838 | 9.799566456 | 0.606219781 |
| Gm2673 | 0.689365755 | 3.386342918 | 1.146832218 |
| Nup54 | 228.7748306 | 0.471729508 | 0.122090283 |
| Gm26039 | 0 | NA | NA |
| Scarb2 | 2989.989026 | 0.653324977 | 0.085520868 |
| Mir6415 | 0 | NA | NA |
| Fam47e | 1.965355678 | -0.633899544 | 0.692701467 |
| Stbd1 | 13.78635077 | 0.896803864 | 0.365160571 |
| Ccdc158 | 6.083259113 | -0.099240291 | 0.386468012 |
| Shroom3 | 115.9678624 | 1.05998509 | 0.18335781 |
| Gm20500 | 0.399554685 | -1.227005032 | 1.546698086 |
| Mir1961 | 0 | NA | NA |
| 4932430I15Rik | 0 | NA | NA |
| RP23-9D24.2 | 1.173754143 | -0.806633517 | 0.942847695 |
| Gm25521 | 0.066050875 | 0.71820487 | 3.724945049 |
| Sowahb | 261.7368774 | -0.163456768 | 0.131782378 |
| Gm22915 | 1.677596521 | -1.388817287 | 0.857008538 |
| RP24-470E5.2 | 19.84950728 | -0.913817687 | 0.386297226 |
| RP24-470E5.3 | 3.144773293 | -1.464869452 | 0.660292999 |
| 11-Sep | 1923.005761 | -0.045593133 | 0.046631301 |
| RP24-470E5.4 | 138.2382822 | -1.013741078 | 0.156165887 |
| Ccni | 2707.469122 | -0.143747873 | 0.087404663 |
| RP24-470E5.9 | 57.10845697 | -0.900571969 | 0.23923722 |
| Gm2986 | 3.888599492 | 0.229222444 | 0.573652976 |
| 2010109A12Rik | 6.375499996 | -0.828816837 | 0.544855067 |
| Ccng2 | 404.8129049 | -0.590651221 | 0.104937986 |
| Kat2b-ps | 1.175757053 | 2.391754692 | 0.974920076 |
| RP23-388L12.2 | 0 | NA | NA |
| RP23-173B14.3 | 0 | NA | NA |
| Gm16427 | 0 | NA | NA |
| C87414 | 0 | NA | NA |
| RP23-388L12.3 | 0 | NA | NA |
| RP23-388L12.1 | 0 | NA | NA |
| RP23-57N5.1 | 0 | NA | NA |
| RP23-173B14.4 | 0 | NA | NA |
| RP23-57N5.3 | 0 | NA | NA |
| Gm16429 | 0 | NA | NA |
| RP23-173B14.2 | 0 | NA | NA |
| RP23-173B14.1 | 0 | NA | NA |
| BC061212 | 0 | NA | NA |

| | | | |
|---|---|---|---|
| Gm6351 | 0 | NA | NA |
| Gm3106 | 0 | NA | NA |
| Gm16522 | 0 | NA | NA |
| Gm6502 | 0 | NA | NA |
| AA792892 | 0 | NA | NA |
| RP23-432M17.3 | 0 | NA | NA |
| Gm7682 | 0 | NA | NA |
| Gm3139 | 0 | NA | NA |
| Gm3147 | 0 | NA | NA |
| Gm6509 | 0 | NA | NA |
| Gm6205 | 0 | NA | NA |
| RP23-133M13.3 | 0 | NA | NA |
| RP23-432M17.5 | 0 | NA | NA |
| Gm3183 | 0 | NA | NA |
| RP24-85B14.2 | 0 | NA | NA |
| Gm7647 | 0 | NA | NA |
| Gm6367 | 0 | NA | NA |
| RP23-365E3.3 | 0 | NA | NA |
| RP24-85B14.3 | 0 | NA | NA |
| Gm7942 | 0 | NA | NA |
| Gm16513 | 0 | NA | NA |
| Gm10424 | 0 | NA | NA |
| RP23-320H13.5 | 0 | NA | NA |
| Gm3259 | 0 | NA | NA |
| RP23-365E3.2 | 0 | NA | NA |
| RP23-320H13.4 | 0 | NA | NA |
| RP23-320H13.3 | 0 | NA | NA |
| D5Ertd577e | 0 | NA | NA |
| Gm3286 | 0 | NA | NA |
| Gm3176 | 0 | NA | NA |
| RP23-23P3.2 | 0 | NA | NA |
| RP23-23P3.1 | 0 | NA | NA |
| Gm7978 | 0 | NA | NA |
| Gm7982 | 0 | NA | NA |
| E330014E10Rik | 0 | NA | NA |
| RP24-322E6.3 | 0 | NA | NA |
| Gm5559 | 314.382447 | 0.020025803 | 0.129828464 |
| Gm25937 | 0 | NA | NA |
| Cxcl13 | 78.29238741 | 6.749804705 | 0.657736064 |
| RP23-57A17.2 | 106.8115065 | 1.750179182 | 0.208645453 |
| Cnot6l | 1535.89927 | 0.317438522 | 0.072493873 |
| Gm2840 | 2.621041726 | 0.316826419 | 0.581557824 |
| Mrpl1 | 300.8320522 | 0.194645045 | 0.077307574 |
| Fras1 | 497.6484057 | -1.277719174 | 0.173259509 |
| RP23-381C14.2 | 0.206671645 | -0.242712299 | 2.636505309 |
| RP23-381C14.5 | 0.680615379 | 0.611909316 | 1.182002305 |
| RP23-480P21.2 | 0.07796279 | 1.072698098 | 3.721829099 |

| | | | |
|---|---|---|---|
| Anxa3 | 185.5288032 | 3.7648284 | 0.235742995 |
| RP23-480P21.3 | 7.010785502 | 1.257436429 | 0.563510484 |
| RP24-142B15.2 | 0.147290723 | 0.701519566 | 2.864649326 |
| RP24-142B15.3 | 0.270258959 | 1.678086604 | 1.796003949 |
| RP24-142B15.4 | 0.176646398 | 1.281666154 | 3.35399331 |
| Gm24623 | 0 | NA | NA |
| RP24-142B15.5 | 2.750814746 | 4.076997266 | 0.95808001 |
| RP24-142B15.6 | 0.520715546 | 2.860724403 | 1.460743662 |
| RP24-142B15.7 | 1.115544105 | 0.551747386 | 0.849245714 |
| RP24-142B15.8 | 0.895791331 | -0.085272195 | 0.971291332 |
| RP24-142B15.9 | 0.252544957 | 1.293440875 | 2.086455436 |
| RP24-142B15.11 | 5.79381536 | 0.837850973 | 0.459044714 |
| Bmp2k | 887.8164416 | 1.382932483 | 0.10681407 |
| RP24-142B15.10 | 11.84488042 | 0.391224152 | 0.441300534 |
| Paqr3 | 238.9609811 | -0.429284248 | 0.149991356 |
| RP23-45P1.1 | 1.824918576 | -1.874546618 | 0.866315077 |
| RP23-28F1.3 | 1.733684669 | 0.65756639 | 0.919155732 |
| Gm5560 | 26.0495091 | 0.151806716 | 0.257137637 |
| RP23-28F1.4 | 0 | NA | NA |
| Naa11 | 0.23191841 | 0.510901202 | 2.569307743 |
| 4930467D21Rik | 11.74611447 | -2.083518531 | 0.411941128 |
| Gk2 | 0.04393175 | -0.243571806 | 3.724945049 |
| Antxr2 | 68.57005387 | 1.836334407 | 0.198641699 |
| RP23-178N8.7 | 1.361532721 | 1.769152191 | 0.968026375 |
| RP23-178N8.3 | 0.042716945 | -0.243571806 | 3.724945049 |
| RP23-178N8.4 | 0 | NA | NA |
| RP23-178N8.5 | 0.202865515 | 0.735120215 | 2.211509827 |
| Gm8048 | 0.123841226 | 1.292225235 | 3.719439735 |
| Prdm8 | 199.5149206 | -0.688045676 | 0.137767876 |
| A730035I17Rik | 9.47161779 | -0.208004471 | 0.387753207 |
| Fgf5 | 19.39898948 | -0.67175183 | 0.281857179 |
| 1700007G11Rik | 3.95741196 | 0.092441573 | 0.504739973 |
| RP23-425F8.1 | 1.376239274 | -0.377384921 | 0.980567593 |
| RP24-240P8.1 | 0.208122295 | 1.206543051 | 2.195383563 |
| mmu-mir-703 | 13.41816306 | 0.295173941 | 0.282279426 |
| Gm24279 | 0.141779851 | -0.698448454 | 3.595043549 |
| Gm11111 | 0.419266243 | 1.130481541 | 1.378472124 |
| 1700010H22Rik | 0 | NA | NA |
| Bmp3 | 131.1088595 | -1.090725937 | 0.160051933 |
| Prkg2 | 141.6564417 | -0.502025898 | 0.135323207 |
| Gm3470 | 1.546431532 | -0.141340253 | 0.807058687 |
| RP24-548B7.1 | 0.424196832 | 2.459223761 | 1.601312933 |
| Rasgef1b | 664.9745574 | 0.524396511 | 0.072601643 |
| RP23-253I14.3 | 18.40400765 | -0.351768991 | 0.287486166 |
| A930011G23Rik | 101.3913818 | -0.625734837 | 0.292833813 |
| RP23-380D12.1 | 0.13861434 | -1.149001357 | 3.614496194 |
| Gm16227 | 6.434094977 | -0.593851783 | 0.445959895 |

| | | | |
|---|---|---|---|
| Gm16228 | 3.733916757 | -1.197032645 | 0.585030796 |
| 4930405H06Rik | 0 | NA | NA |
| Gm25969 | 0 | NA | NA |
| Gm16226 | 0 | NA | NA |
| 4930522N08Rik | 0 | NA | NA |
| RP23-438O22.1 | 33.98607118 | -1.365913641 | 0.366356689 |
| RP23-57P23.3 | 0.038981395 | 0.718207094 | 3.724945049 |
| Gm26252 | 0 | NA | NA |
| RP23-57P23.4 | 0.879409403 | 0.20510171 | 1.268781331 |
| Gm22838 | 0.115603922 | 1.209894437 | 3.72418822 |
| RP23-57P23.5 | 0.082913145 | 0.23732098 | 3.724945049 |
| Hnrnpd | 2150.330006 | -0.043038977 | 0.079677258 |
| Gm17092 | 3.730625592 | 0.707922569 | 0.545727502 |
| 4930524J08Rik | 4.763368001 | 0.942104544 | 0.482888483 |
| Hnrnpdl | 2079.483625 | 0.057926893 | 0.084254918 |
| Enoph1 | 302.3695605 | -0.246075723 | 0.094484472 |
| Gm26363 | 0 | NA | NA |
| Tmem150c | 388.2756464 | -0.417493702 | 0.072495739 |
| Tmem150cos | 0.555352798 | 2.351877316 | 1.348908919 |
| Gm9932 | 0.452014659 | 1.034428518 | 1.48581123 |
| 2310034O05Rik | 0.531171913 | 0.743882953 | 1.495421957 |
| Gm8091 | 0.417821482 | -0.952783015 | 1.334688394 |
| Sec31a | 1485.020543 | 0.153678076 | 0.039978996 |
| 5430416N02Rik | 76.01050414 | 0.303213245 | 0.175711083 |
| Gm27553 | 0.170081438 | -0.631816917 | 2.769839891 |
| Gm27997 | 0.038981395 | 0.718207094 | 3.724945049 |
| Gm28030 | 0 | NA | NA |
| Gm27286 | 0.243228626 | 0.867111339 | 3.022466272 |
| RP23-325M7.4 | 4.625370955 | 0.136543195 | 0.538152536 |
| Lin54 | 315.1002167 | 0.212882782 | 0.097018042 |
| Gm23222 | 0.640027454 | -0.64005722 | 1.436741508 |
| RP23-313F5.1 | 0 | NA | NA |
| Gm23173 | 0 | NA | NA |
| Cops4 | 680.1477187 | 0.286950991 | 0.063246785 |
| Plac8 | 627.7400569 | 11.68774863 | 0.920169342 |
| RP23-313F5.2 | 0 | NA | NA |
| Coq2 | 258.0378681 | 0.212824746 | 0.082593754 |
| Gm15777 | 0.230076657 | 0.718205339 | 3.121436614 |
| Hpse | 70.70159234 | 3.441857318 | 0.243594984 |
| Helq | 242.9045317 | -0.226628407 | 0.152067423 |
| Mrps18c | 124.8778561 | 0.779541694 | 0.147169882 |
| RP23-7M16.1 | 22.40968198 | -0.344580736 | 0.278683856 |
| Fam175a | 72.7916956 | -0.314990768 | 0.146003962 |
| RP23-7M16.2 | 5.191824483 | -1.414967928 | 0.575445141 |
| RP23-7M16.3 | 1.089839937 | 1.410229961 | 1.079773403 |
| Gm26286 | 0.158374819 | -0.243563437 | 2.812678226 |
| 4930458D05Rik | 1.568767244 | 0.929460785 | 0.78688513 |

| | | | |
|---|---|---|---|
| Agpat9 | 94.53062151 | 1.648977709 | 0.212604336 |
| RP23-99A9.1 | 2.620592534 | 0.485894001 | 0.61569099 |
| RP23-99A9.2 | 0.075043205 | 0.237309859 | 3.724945049 |
| RP23-302F24.1 | 0.045882455 | 0.237312084 | 3.724945049 |
| RP23-336N16.1 | 0 | NA | NA |
| Gm20484 | 0 | NA | NA |
| Nkx6-1 | 10.48168021 | 0.812331779 | 0.522640438 |
| RP24-292F8.2 | 0 | NA | NA |
| RP24-292F8.1 | 1.593328872 | 0.62516422 | 0.784640439 |
| Cds1 | 1169.603167 | -0.121873062 | 0.073154871 |
| RP24-292F8.3 | 17.79350931 | -1.051695401 | 0.340151229 |
| WI1-1010L24.1 | 36.41421054 | -0.9937801 | 0.291708679 |
| Wdfy3 | 5687.595008 | -0.351861711 | 0.060466483 |
| Gm20548 | 3.946887117 | 0.653599793 | 0.535314605 |
| RP24-491C20.4 | 0.127971488 | 1.23029335 | 3.685533863 |
| RP23-409I5.1 | 28.85031586 | -0.699017443 | 0.338627357 |
| RP23-409I5.2 | 1.952620281 | -0.756207475 | 0.705260909 |
| RP23-409I5.3 | 0.494178821 | 0.252514952 | 1.459818051 |
| RP23-409I5.4 | 2.040397007 | 0.225352988 | 0.661356535 |
| RP23-260N17.2 | 0.650505631 | -1.221951879 | 1.206071016 |
| RP23-260N17.1 | 0.199035941 | -0.243574084 | 3.217008196 |
| Arhgap24 | 215.5874552 | 0.81000557 | 0.108446721 |
| Mapk10 | 3784.726759 | -0.502459404 | 0.07617504 |
| RP23-286M22.2 | 10.7259886 | -1.165869072 | 0.37569808 |
| RP23-286M22.1 | 125.4642517 | -0.849826497 | 0.262296243 |
| RP24-212G3.4 | 0 | NA | NA |
| RP24-212G3.3 | 0 | NA | NA |
| RP24-270K7.2 | 0 | NA | NA |
| RP24-270K7.1 | 0 | NA | NA |
| RP24-270K7.3 | 2.638949413 | 1.930766549 | 0.735006735 |
| Ptpn13 | 508.0720606 | 0.516346635 | 0.094337163 |
| RP24-102B13.3 | 0 | NA | NA |
| Gm23085 | 0.072025462 | 0.237312084 | 3.724945049 |
| Slc10a6 | 36.57358822 | 7.810645902 | 1.169286191 |
| Gm15844 | 0 | NA | NA |
| RP24-102B13.5 | 0.358939267 | -1.721912061 | 1.493595023 |
| 1700016H13Rik | 1.060754565 | 2.297168033 | 1.092868462 |
| Gm25721 | 0 | NA | NA |
| Aff1 | 1183.051492 | 1.75117029 | 0.154117221 |
| Klhl8 | 320.9854992 | -0.543195392 | 0.084744441 |
| RP23-337H1.2 | 4.609961798 | -1.227343198 | 0.602421417 |
| Gm24897 | 0 | NA | NA |
| Hsd17b13 | 1.339240761 | -0.154788101 | 0.953687704 |
| Gm8200 | 0 | NA | NA |
| Hsd17b11 | 306.1707364 | -0.08650466 | 0.096714472 |
| RP23-102H7.7 | 6.166014486 | -0.753585611 | 0.416453097 |
| Nudt9 | 452.8860333 | 0.555541094 | 0.103322946 |

| | | | |
|---|---|---|---|
| Mir5619 | 0.706746895 | -0.592592838 | 1.189127951 |
| Gm17660 | 2.469786297 | -1.125761058 | 0.797861815 |
| RP23-102H7.9 | 2.342408071 | -4.23016218 | 1.010509063 |
| Sparcl1 | 15907.62479 | -0.854509883 | 0.098520744 |
| Gm26703 | 13.10610984 | -0.938733436 | 0.36335829 |
| Dspp | 1.924149456 | -0.259363037 | 0.791890874 |
| Dmp1 | 39.86314599 | -0.132835878 | 0.231063959 |
| RP24-191K3.1 | 0.046345709 | 0.71820487 | 3.724945049 |
| Gm23616 | 0 | NA | NA |
| Ibsp | 1.789549342 | 0.935083527 | 0.777915893 |
| Mepe | 0 | NA | NA |
| Spp1 | 1206.760665 | 3.629139868 | 0.321197938 |
| RP24-299A7.2 | 2.453753307 | 2.161756982 | 0.774545048 |
| Pkd2 | 529.8119712 | -0.181856586 | 0.113108675 |
| BC005561 | 622.8318008 | -0.321355578 | 0.1164345 |
| D930016D06Rik | 256.987905 | -0.435324516 | 0.153389601 |
| Gm8258 | 4.524171958 | 0.502981902 | 0.496980652 |
| Zfp951 | 41.98758719 | -0.547570916 | 0.209767279 |
| Abcg3 | 1.390608906 | 2.466181371 | 1.035165971 |
| Gm24097 | 0 | NA | NA |
| RP23-370D2.2 | 0 | NA | NA |
| RP23-370D2.5 | 0 | NA | NA |
| RP23-370D2.4 | 0.124853745 | 0.237309606 | 3.724945049 |
| Gbp8 | 237.2535119 | 8.629725189 | 0.600565133 |
| Gbp9 | 1809.026709 | 7.559042039 | 0.252326162 |
| Gbp4 | 4374.309881 | 10.47736036 | 0.383272492 |
| Gbp6 | 1754.483529 | 8.212302205 | 0.280820554 |
| Gbp10 | 886.6277886 | 10.25929787 | 0.557751937 |
| Gbp11 | 1003.541595 | 6.384959272 | 0.310402264 |
| Gm26519 | 1.456550475 | 1.882412979 | 0.948656137 |
| RP24-63G23.2 | 1.81533978 | 2.457392031 | 0.969442305 |
| Lrrc8b | 1659.203879 | -0.496500051 | 0.067124016 |
| RP23-41J13.2 | 3.634356604 | 1.100796978 | 0.619238606 |
| Lrrc8c | 697.388411 | 0.621692751 | 0.107646964 |
| C230066G23Rik | 8.141609263 | -0.495073508 | 0.449265058 |
| Lrrc8d | 580.2514977 | 0.126414971 | 0.079450271 |
| Rps15a-ps5 | 17.05466322 | 1.057379995 | 0.276662246 |
| Lrrc8dos | 0.047948698 | -0.243569582 | 3.724945049 |
| Gm10359 | 23.82378434 | -0.202314293 | 0.256618849 |
| RP24-354P12.2 | 0.324285546 | 0.718190289 | 3.724945049 |
| Zfp326 | 429.4665804 | -0.478859961 | 0.097190565 |
| RP24-354P12.3 | 23.94047403 | -0.575785437 | 0.311429608 |
| RP24-400H4.1 | 0 | NA | NA |
| Gm25672 | 0 | NA | NA |
| RP24-476K21.1 | 0.34546946 | 1.919200287 | 1.660697777 |
| RP23-424O22.1 | 0.204914957 | 1.193798894 | 2.202071459 |
| RP23-424O22.2 | 0.075043205 | 0.237309859 | 3.724945049 |

| | | | |
|---|---|---|---|
| RP23-424O22.3 | 0 | NA | NA |
| RP24-289G19.1 | 0.04393175 | -0.243571806 | 3.724945049 |
| RP24-289G19.3 | 0.147068667 | 0.237310878 | 3.514416608 |
| RP24-289G19.2 | 0.089089296 | 0.237305797 | 3.724945049 |
| RP24-289G19.4 | 0 | NA | NA |
| Barhl2 | 85.11172079 | -0.108838388 | 0.239248729 |
| Gm26872 | 0.927992339 | 0.617045367 | 1.102172923 |
| Gm28050 | 8.739896192 | -0.135566327 | 0.376321581 |
| Gm28039 | 0.075043205 | 0.237309859 | 3.724945049 |
| Gm29464 | 0.340386612 | 1.810885516 | 1.668718359 |
| Gm17304 | 0 | NA | NA |
| RP23-167H23.3 | 0.042716945 | -0.243571806 | 3.724945049 |
| Zfp644 | 2759.468483 | -0.431532408 | 0.075481227 |
| RP23-167H23.2 | 69.78784834 | -0.844754875 | 0.320097891 |
| Gm24191 | 0 | NA | NA |
| RP24-512F21.1 | 0 | NA | NA |
| Hfm1 | 28.3914873 | -0.206743285 | 0.208641673 |
| Gm27527 | 0.338550938 | 0.237332292 | 1.585738527 |
| Olfr719-ps | 0 | NA | NA |
| Cdc7 | 158.8450571 | -0.661732097 | 0.192869354 |
| RP23-24K16.1 | 0.066050875 | 0.71820487 | 3.724945049 |
| Tgfbr3 | 375.510489 | -0.004614666 | 0.119371015 |
| RP23-24K16.3 | 17.70335292 | -1.197757827 | 0.472103546 |
| RP23-24K16.2 | 5.270225733 | -1.524525455 | 0.624853541 |
| Gm8145 | 0 | NA | NA |
| RP24-357G2.2 | 11.97838418 | -1.865749474 | 0.439474152 |
| Gm22557 | 0 | NA | NA |
| Brdt | 172.1838994 | -0.039557152 | 0.108579499 |
| Gm8152 | 0.483474337 | -0.065442505 | 1.446027943 |
| RP24-140D11.1 | 19.5481605 | -1.203279009 | 0.255346569 |
| n-R5s173 | 0.044191549 | -0.243571806 | 3.724945049 |
| Ephx4 | 172.3772472 | -1.139855402 | 0.127363601 |
| Gm17202 | 1.45035625 | -2.994422429 | 1.088594892 |
| Lpcat2b | 2.065408989 | -1.446040872 | 0.707901741 |
| Btbd8 | 41.15341184 | -0.59419886 | 0.160189751 |
| Gm9727 | 10.68833693 | 0.535264832 | 0.344365251 |
| A830010M20Rik | 1635.382598 | -0.493875668 | 0.090179726 |
| RP23-32E20.1 | 2.258596541 | 1.084966383 | 0.805851089 |
| 1700028K03Rik | 21.85631515 | -0.749267937 | 0.332826236 |
| Glmn | 205.8721023 | -0.398380121 | 0.090994125 |
| Rpap2 | 173.555107 | 0.006153575 | 0.094127008 |
| A930041C12Rik | 0.201858732 | 0.718217268 | 2.604406118 |
| Gm26692 | 1.12480443 | -1.456194427 | 1.063630528 |
| RP23-32E20.3 | 0.156258025 | -0.243568418 | 2.817094184 |
| RP24-140D11.4 | 0.388935867 | 2.175034127 | 1.621690912 |
| Gfi1 | 6.525819315 | 5.020793191 | 0.933849235 |
| A430072P03Rik | 0.11147366 | 1.230376831 | 3.722845817 |

| | | | |
|---|---|---|---|
| Evi5 | 1027.982555 | -0.428626638 | 0.061609906 |
| RP24-140D11.6 | 51.7496199 | -1.10160068 | 0.209457094 |
| Ube2d2b | 0 | NA | NA |
| Gm16007 | 0 | NA | NA |
| RP24-140D11.5 | 1.387064387 | 0.86871037 | 0.859876105 |
| Rpl5 | 1390.886114 | 0.465343773 | 0.095532129 |
| Snord21 | 0 | NA | NA |
| Gm22270 | 2.785641484 | -0.455918115 | 0.567936654 |
| Gm26387 | 2.437479595 | -0.000630133 | 0.659577477 |
| Fam69a | 302.41774 | -0.268067151 | 0.088370599 |
| Gm9850 | 0.572539054 | -1.264828082 | 1.292254893 |
| Rps19-ps8 | 0 | NA | NA |
| Mtf2 | 757.8828192 | -0.374786846 | 0.08746099 |
| Tmed5 | 501.0379422 | 0.906761439 | 0.113945293 |
| Ccdc18 | 25.54476345 | 0.107808957 | 0.238675616 |
| RP23-46J11.4 | 0 | NA | NA |
| RP23-46J11.5 | 0 | NA | NA |
| RP23-46J11.6 | 0 | NA | NA |
| Dr1 | 585.9554827 | 0.458452552 | 0.096899908 |
| RP24-317M4.1 | 6.093084322 | -0.843717755 | 0.514540382 |
| Pigg | 279.9406767 | -0.106164721 | 0.087987308 |
| RP24-317M4.2 | 2.428646146 | -1.108339633 | 0.624855687 |
| RP24-317M4.3 | 9.503775188 | -0.990686075 | 0.364589004 |
| RP24-317M4.6 | 137.5888963 | -0.103681373 | 0.16887089 |
| Gm10419 | 867.0367941 | -0.067861194 | 0.121190532 |
| Pde6b | 10.02136167 | 4.359805102 | 0.536663936 |
| Atp5k | 305.7650685 | -0.33750346 | 0.182209879 |
| Mfsd7a | 18.35894452 | 1.934820869 | 0.289525915 |
| Pcgf3 | 699.3355955 | -0.044936146 | 0.069331754 |
| Gm10086 | 0.202977511 | 1.123041874 | 2.185177258 |
| Cplx1 | 5792.979381 | -0.489118469 | 0.079886622 |
| RP24-529E15.1 | 10.79973995 | 0.380133403 | 0.575404745 |
| Gak | 1581.201267 | 0.168503991 | 0.057215024 |
| RP24-529E15.3 | 21.02397872 | -0.532204819 | 0.292891795 |
| Tmem175 | 304.8091594 | -0.490408163 | 0.101650913 |
| Dgkq | 523.7346149 | -0.469594292 | 0.100869021 |
| Idua | 131.045456 | 0.629587737 | 0.128189822 |
| Slc26a1 | 12.6609391 | -0.326618492 | 0.324479376 |
| Fgfrl1 | 247.8383437 | 1.308316021 | 0.150462882 |
| Rnf212 | 0.048949532 | -0.243569582 | 3.724945049 |
| 1700047L14Rik | 0.220636191 | 2.023934468 | 2.539537598 |
| Tmed11 | 0 | NA | NA |
| Vmn2r8 | 0 | NA | NA |
| Gm18594 | 0 | NA | NA |
| Vmn2r9 | 0 | NA | NA |
| Gm20629 | 0.046345709 | 0.71820487 | 3.724945049 |
| Gm10417 | 1.121165163 | -0.498488763 | 0.974914706 |

| | | | |
|---|---|---|---|
| Vmn2r-ps21 | 0 | NA | NA |
| RP23-37G13.2 | 0.042716945 | -0.243571806 | 3.724945049 |
| RP23-37G13.3 | 0 | NA | NA |
| Gm18595 | 0 | NA | NA |
| Vmn2r10 | 0 | NA | NA |
| Gm5871 | 0.397993176 | 1.69751829 | 1.706524431 |
| Vmn2r11 | 0 | NA | NA |
| Vmn2r12 | 0 | NA | NA |
| Vmn2r13 | 0 | NA | NA |
| Vmn2r14 | 0 | NA | NA |
| Gm8473 | 0 | NA | NA |
| Vmn2r15 | 0 | NA | NA |
| Vmn2r16 | 0 | NA | NA |
| Gm7171 | 0 | NA | NA |
| Vmn2r17 | 0.044191549 | -0.243571806 | 3.724945049 |
| RP23-410M21.3 | 0 | NA | NA |
| Gm8488 | 0.787704992 | 1.505101543 | 1.085257589 |
| RP23-203O7.3 | 2.482507492 | -0.240863987 | 0.566478478 |
| Crlf2 | 173.2274311 | 1.121512565 | 0.122068571 |
| Vmn2r-ps23 | 0.148650935 | -0.243572541 | 2.863172849 |
| Gm28000 | 0 | NA | NA |
| 5430403G16Rik | 36.45249734 | -0.247534151 | 0.258493325 |
| 4930522L14Rik | 39.79845386 | -0.053452583 | 0.189913039 |
| Gm8508 | 4.624103064 | -0.061120106 | 0.464515335 |
| RP23-396K22.3 | 1.154557212 | -1.653365709 | 1.027853626 |
| RP23-396K22.4 | 4.399392508 | -0.874082495 | 0.630721493 |
| Gm26779 | 37.79669302 | -0.694664829 | 0.193404355 |
| Gm26808 | 0.439601053 | 0.976878097 | 1.51120383 |
| Gm15446 | 190.6866757 | -0.449915247 | 0.132091057 |
| 1010001B22Rik | 0.952639941 | -0.8895088 | 1.021935136 |
| Zfp932 | 186.9135822 | -0.318075939 | 0.126949554 |
| Gm17655 | 4.402479275 | -0.286724677 | 0.492236089 |
| Plcxd1 | 122.7290074 | -0.096039387 | 0.144889864 |
| Gtpbp6 | 242.7828586 | 0.211730292 | 0.07278173 |
| Zfp605 | 230.5230773 | 0.053161228 | 0.082136024 |
| Gm15484 | 2.365693163 | 0.715729549 | 0.616683375 |
| Chfr | 523.8452094 | -0.341839026 | 0.11282973 |
| RP24-374B14.4 | 1.731816556 | 0.850470155 | 0.767696709 |
| Gm15787 | 10.57349902 | 2.024824164 | 0.37535762 |
| Golga3 | 1104.05704 | 0.220345153 | 0.065756281 |
| RP24-374B14.6 | 9.781179111 | -0.845490475 | 0.388525531 |
| RP24-374B14.8 | 3.53955661 | -0.013007737 | 0.627324038 |
| Ankle2 | 833.7400944 | 0.575581425 | 0.065090513 |
| Pgam5 | 509.9939935 | 0.164017056 | 0.067437349 |
| Gm26718 | 13.31072805 | -0.512677572 | 0.308569598 |
| Gm15788 | 2.377542658 | -0.230147481 | 0.660912922 |
| Pxmp2 | 26.9648925 | -0.12972507 | 0.266258157 |

| | | | |
|---|---|---|---|
| Pole | 51.3992184 | 1.914434114 | 0.324452736 |
| Gm26277 | 0 | NA | NA |
| Gm15791 | 0.092493346 | -0.24357403 | 3.724945049 |
| Gm15792 | 0.061920613 | 0.718207094 | 3.724945049 |
| P2rx2 | 1.822501977 | -0.206054411 | 0.788888622 |
| Lrcol1 | 0.046345709 | 0.71820487 | 3.724945049 |
| Fbrsl1 | 654.9798202 | -0.321400031 | 0.081889376 |
| Gm26711 | 1.656227335 | -0.088082081 | 0.88780305 |
| RP23-231C16.1 | 2.276862309 | -0.205794098 | 0.720955415 |
| RP23-231C16.2 | 0 | NA | NA |
| A630023P12Rik | 2.678264972 | -0.172073875 | 0.698800212 |
| Galnt9 | 582.7762908 | -0.474837444 | 0.095370471 |
| Noc4l | 266.4007122 | 1.162359615 | 0.154532421 |
| Ddx51 | 235.8120389 | 0.357380734 | 0.100111418 |
| RP23-393E10.2 | 29.91262942 | 0.358400575 | 0.22314915 |
| Ep400 | 2474.898703 | 0.094930657 | 0.039894535 |
| Mir7026 | 1.550196443 | -0.368829731 | 0.848732458 |
| Gm22716 | 3.382263423 | -0.749648101 | 0.559018241 |
| Gm26079 | 0.044544648 | 0.237309859 | 3.724945049 |
| Gm22583 | 0 | NA | NA |
| Gm15559 | 13.41128988 | 0.272497188 | 0.274798913 |
| Pus1 | 142.3904629 | 0.330717731 | 0.100965587 |
| Gm26515 | 2.087315227 | -0.316179339 | 0.681824783 |
| Ulk1 | 1100.014041 | -0.235436285 | 0.054873788 |
| Hscb | 47.36481145 | 0.441333743 | 0.192103048 |
| Chek2 | 15.72251656 | 0.441541274 | 0.273559496 |
| RP24-207A18.2 | 0.348192234 | 1.115625355 | 1.584132297 |
| Ttc28 | 598.1288249 | 0.069816293 | 0.056850118 |
| Gm15989 | 3.684955975 | -0.812740909 | 0.568255678 |
| RP23-326H18.1 | 27.43115472 | -0.124849115 | 0.216736838 |
| Mir701 | 2.293891663 | -1.416901039 | 0.732579267 |
| Gm15988 | 0 | NA | NA |
| RP24-230N16.2 | 6.359331415 | -1.061003164 | 0.472678139 |
| RP23-72O23.3 | 4.724619101 | -0.87855135 | 0.524954511 |
| RP23-72O23.4 | 1.873630482 | -1.30186599 | 0.927298918 |
| Gm27602 | 0 | NA | NA |
| RP23-131F7.2 | 0.716273392 | -3.077333298 | 1.166238713 |
| Gm27680 | 0 | NA | NA |
| Pitpnb | 878.8123659 | 0.044943268 | 0.065151673 |
| RP23-131F7.5 | 0.255202296 | 0.691710223 | 1.824364618 |
| Mn1 | 649.1153886 | -1.007639284 | 0.144241317 |
| RP23-131F7.4 | 14.92077947 | -1.598531465 | 0.329643106 |
| C130026L21Rik | 34.50140241 | -0.601872014 | 0.266729666 |
| RP24-572L22.2 | 3.151250669 | -1.270125841 | 0.621382971 |
| Gm26897 | 0.18833836 | 0.529869577 | 2.663963407 |
| E130006D01Rik | 3.354483374 | -0.193784351 | 0.519689108 |
| Gm16019 | 0 | NA | NA |

| | | | |
|---|---|---|---|
| RP24-340B11.1 | 0 | NA | NA |
| RP23-427D2.1 | 0.576279854 | -0.552983655 | 1.241241927 |
| RP24-317K11.4 | 0 | NA | NA |
| 1700028D13Rik | 0 | NA | NA |
| Miat | 1661.236509 | -2.047769724 | 0.491091554 |
| Gm27998 | 0 | NA | NA |
| Gm27882 | 0 | NA | NA |
| Gm26953 | 0.344925633 | 1.319169243 | 1.758077751 |
| Gm27627 | 0 | NA | NA |
| Gm27357 | 0 | NA | NA |
| Cryba4 | 4.194020461 | 1.001751709 | 0.620063756 |
| Crybb1 | 14.29264834 | -2.575256139 | 0.583077481 |
| Tpst2 | 280.8629906 | 0.752436456 | 0.095041183 |
| Tfip11 | 581.9359828 | 0.272126993 | 0.06397861 |
| Srrd | 100.6125371 | -0.004961901 | 0.118447976 |
| Hps4 | 178.5209335 | -0.199742253 | 0.133331716 |
| Gm6583 | 0.632098713 | 2.078152519 | 1.399231021 |
| RP23-459I1.3 | 20.2973942 | -1.308043852 | 0.437050679 |
| RP23-459I1.5 | 0 | NA | NA |
| Gm20636 | 0.231217418 | -0.243563208 | 2.131487463 |
| Asphd2 | 618.5894786 | -0.532089152 | 0.082545858 |
| Sez6l | 3346.420789 | -0.178331684 | 0.089489756 |
| Gm6588 | 0 | NA | NA |
| Gm27004 | 3.168227339 | 0.425073158 | 0.612584874 |
| 1700034G24Rik | 0.106465261 | 0.718202646 | 3.724945049 |
| RP23-156K23.3 | 0.158818843 | -0.196148845 | 2.807459993 |
| RP23-156K23.4 | 7.736953527 | -1.668684285 | 0.475291733 |
| RP24-304N8.2 | 2.801170497 | -1.583056254 | 0.759364643 |
| Myo18b | 28.0640786 | 0.833311113 | 0.22638154 |
| 1700095B10Rik | 0 | NA | NA |
| Adrbk2 | 1355.458738 | -0.851727597 | 0.082424521 |
| Gm22740 | 0 | NA | NA |
| Crybb2 | 0.278057897 | 1.209174331 | 2.412826006 |
| RP23-4P1.1 | 0.92508897 | 0.984000393 | 1.015258435 |
| Crybb3 | 5.690583048 | 0.288194402 | 0.399379683 |
| 2900026A02Rik | 622.1930781 | -0.554284817 | 0.093946236 |
| RP23-4P1.6 | 0.061920613 | 0.718207094 | 3.724945049 |
| RP23-4P1.5 | 5.731549506 | -0.599022052 | 0.449607155 |
| RP23-4P1.3 | 0.48780114 | -1.011182557 | 1.287763012 |
| RP23-4P1.7 | 0.066050875 | 0.71820487 | 3.724945049 |
| RP23-4P1.2 | 1.950711236 | -2.145092627 | 1.043255705 |
| Tmem211 | 0.590173444 | 0.028883058 | 1.201126091 |
| Sgsm1 | 1205.098046 | -0.323518511 | 0.081571403 |
| RP24-456F12.1 | 1.004172246 | 2.167029511 | 1.132799594 |
| RP24-456F12.2 | 0.236337014 | 1.631657247 | 2.149159569 |
| mmu-mir-7229 | 0 | NA | NA |
| mmu-mir-7230 | 0 | NA | NA |

| | | | |
|---|---|---|---|
| Aym1 | 0.330879691 | 0.237328172 | 2.020593963 |
| Wscd2 | 877.7252593 | -0.772912719 | 0.130354917 |
| Cmklr1 | 152.364205 | 3.403819695 | 0.191710576 |
| Gm15736 | 0.075043205 | 0.237309859 | 3.724945049 |
| 1700069L16Rik | 1.680905595 | 0.56644716 | 0.962495188 |
| Ficd | 177.6330377 | 0.299733361 | 0.104904237 |
| Sart3 | 480.3375866 | 0.100636111 | 0.096068506 |
| Gm3511 | 0.172107867 | 0.718215321 | 2.739902689 |
| Iscu | 446.2315743 | 0.078410772 | 0.090788495 |
| Tmem119 | 184.4126201 | -0.802816225 | 0.222248237 |
| RP24-555M20.7 | 0 | NA | NA |
| Selplg | 268.6005576 | 1.231514026 | 0.160438209 |
| Coro1c | 1185.306564 | 0.53797288 | 0.096778509 |
| Gm17122 | 0.291024678 | 0.28155918 | 2.054139395 |
| RP24-555M20.4 | 0.261813223 | -1.171744231 | 2.497637454 |
| RP24-555M20.3 | 33.32994184 | -0.122858807 | 0.357611311 |
| RP24-555M20.5 | 0.345648411 | 1.028366416 | 1.551699962 |
| Gm22056 | 0 | NA | NA |
| Ssh1 | 648.036513 | 0.25152605 | 0.061592129 |
| Dao | 29.66175715 | -0.878161878 | 0.325977255 |
| Svop | 1130.54196 | -0.557588977 | 0.078912815 |
| Usp30 | 451.0626031 | 0.037223238 | 0.106670922 |
| RP24-226L20.2 | 8.375339354 | -0.810888948 | 0.417017124 |
| RP24-226L20.1 | 11.87863406 | -1.18136653 | 0.575056349 |
| Alkbh2 | 30.67206188 | 0.602809356 | 0.186368368 |
| Ung | 32.04545787 | 0.701559205 | 0.238192624 |
| Acacb | 113.2622031 | 0.611848493 | 0.132266538 |
| Gm13784 | 0 | NA | NA |
| Foxn4 | 0.07796279 | 1.072698098 | 3.721829099 |
| Myo1h | 18.55275259 | 0.001001742 | 0.331285975 |
| Kctd10 | 678.4952583 | 0.946574488 | 0.119821094 |
| Ube3b | 1199.562226 | -0.105201786 | 0.075493887 |
| Mir7027 | 0.775147523 | -0.651151206 | 1.13417863 |
| Mmab | 164.6481927 | -0.571033984 | 0.091995997 |
| Mvk | 173.1109817 | 0.043379818 | 0.12966952 |
| Gm13790 | 0.110814304 | -0.724455696 | 3.724945049 |
| Fam222a | 196.5946661 | -0.070128287 | 0.126217721 |
| Trpv4 | 24.56884945 | 0.418732553 | 0.256452491 |
| Gltp | 531.1234222 | 0.796365467 | 0.122959532 |
| Tchp | 125.3804559 | -0.103671088 | 0.180857066 |
| RP23-3M10.11 | 0.104637558 | 0.23732098 | 3.724945049 |
| Mir7028 | 0.496352305 | -1.449490399 | 1.386230189 |
| Git2 | 916.5459927 | 0.050456443 | 0.099225313 |
| 4930515G01Rik | 7.627541774 | -0.676079028 | 0.401678604 |
| Ankrd13a | 914.9811008 | 0.778454607 | 0.103563749 |
| 1500011B03Rik | 398.2607319 | -0.042397886 | 0.086222809 |
| Gm20499 | 1.052193156 | 0.619302155 | 1.150292474 |

| | | | |
|---|---|---|---|
| 2610524H06Rik | 87.22289935 | 0.258220725 | 0.132541829 |
| Ywhaq-ps2 | 0.12209515 | 0.718202646 | 3.724945049 |
| RP24-297N9.13 | 1.537880876 | 0.692363204 | 0.823677072 |
| Mir6240 | 0 | NA | NA |
| 4930519G04Rik | 0.461107361 | 0.153263994 | 1.417563378 |
| Gm9936 | 3.314927513 | 1.245156945 | 0.620297563 |
| Gm13823 | 0.097139547 | 0.718207094 | 3.724945049 |
| Oasl2 | 5567.265836 | 8.812686857 | 0.170967487 |
| Gm13822 | 5.72139037 | 6.456853088 | 0.959250515 |
| Oasl1 | 994.9398756 | 10.74104198 | 0.61092907 |
| 2210016L21Rik | 838.2899106 | -0.203737415 | 0.070586903 |
| Hnf1a | 0.04393175 | -0.243571806 | 3.724945049 |
| Hnf1aos1 | 0.088476398 | -0.243576254 | 3.724945049 |
| Hnf1aos2 | 0.044544648 | 0.237309859 | 3.724945049 |
| RP24-556H22.2 | 0.075043205 | 0.237309859 | 3.724945049 |
| Sppl3 | 907.0167297 | -0.208818427 | 0.063712362 |
| RP24-556H22.3 | 18.34444885 | -0.299764075 | 0.325672163 |
| Gm13827 | 0.049553047 | 0.71820487 | 3.724945049 |
| Gm10401 | 0.066050875 | 0.71820487 | 3.724945049 |
| Rpl37rt | 126.6981566 | 0.522885985 | 0.162488798 |
| Acads | 60.73886381 | 0.510514641 | 0.189690117 |
| Unc119b | 384.4903987 | 0.619785139 | 0.122695002 |
| Mlec | 1723.019123 | 0.374517608 | 0.112292231 |
| Cabp1 | 800.6070211 | -0.908256349 | 0.125226703 |
| Gm13828 | 0.405693472 | -2.434372363 | 1.609780954 |
| Pop5 | 169.6778782 | -0.319533406 | 0.109958225 |
| Rnf10 | 2262.437902 | 0.246347133 | 0.066172671 |
| Gm13831 | 0.044544648 | 0.237309859 | 3.724945049 |
| Coq5 | 405.2808921 | 0.265360543 | 0.077917482 |
| Gm13829 | 0 | NA | NA |
| Dynll1 | 1360.661757 | 0.071917705 | 0.114697979 |
| Gm13830 | 22.05592487 | -0.856823586 | 0.288220486 |
| Srsf9 | 360.4762658 | 0.063052885 | 0.083384053 |
| Gatc | 179.8265356 | -0.243399006 | 0.102937433 |
| Triap1 | 96.42262351 | 0.104492795 | 0.127172932 |
| Cox6a1 | 2038.657447 | 0.041639876 | 0.111794877 |
| RP24-132L16.11 | 0.287284012 | 0.691764028 | 1.777664448 |
| 4930401G09Rik | 0.097899064 | -0.724453471 | 3.724945049 |
| Gm13832 | 4.797855873 | 0.71302245 | 0.472131162 |
| RP24-132L16.12 | 3.340097371 | -2.081691136 | 0.61790653 |
| RP24-132L16.13 | 6.416277231 | -1.847511787 | 0.525975108 |
| Msi1 | 624.614844 | 0.067959195 | 0.096519473 |
| RP24-132L16.14 | 0.383028346 | -0.619318237 | 1.448720818 |
| Pla2g1b | 0.110595523 | 0.718200421 | 3.724945049 |
| Sirt4 | 90.54918992 | -0.5561084 | 0.19674202 |
| Gm24265 | 261.4064373 | 0.224594049 | 0.214331413 |
| Gm24407 | 234.2932087 | -0.092590439 | 0.179500854 |

| | | | |
|---|---|---|---|
| Gm13836 | 0.227885852 | 1.138658693 | 2.505164943 |
| Pxn | 603.300534 | 0.814864466 | 0.124224968 |
| Gm13840 | 1.630127001 | 0.621696583 | 0.744022394 |
| Rplp0 | 924.8935804 | 1.322863746 | 0.130670537 |
| Gcn1l1 | 1454.726854 | -0.02192915 | 0.072928321 |
| Mir7029 | 0.140122821 | -0.243570953 | 3.572128793 |
| RP23-228F6.9 | 3.230897769 | -0.305051552 | 0.612384167 |
| Rab35 | 708.4090342 | 0.169343205 | 0.067442436 |
| Mir7030 | 0.697123821 | -0.286709694 | 1.187292213 |
| Ccdc64 | 496.8343264 | -0.572819658 | 0.102666004 |
| Gm13841 | 108.5809223 | 0.456110006 | 0.165810895 |
| Cit | 2100.160842 | -0.869183494 | 0.134860618 |
| RP23-265I19.3 | 0.244575677 | 0.23732928 | 2.093685082 |
| RP23-265I19.2 | 0.484822013 | -1.159673102 | 1.569119149 |
| Gm14508 | 2.937257927 | -1.136219975 | 0.613401597 |
| Prkab1 | 201.4925811 | 0.64651514 | 0.128570382 |
| Gm13842 | 0 | NA | NA |
| Gm17420 | 0 | NA | NA |
| Tmem233 | 0.047948698 | -0.243569582 | 3.724945049 |
| C330018A13Rik | 4.719781364 | -1.411552601 | 0.575467806 |
| Ccdc60 | 12.74564928 | 0.658921547 | 0.3295571 |
| Gm14507 | 0.354637581 | -1.612239618 | 1.63157277 |
| Gm13837 | 0.143618953 | 0.68463387 | 2.890309179 |
| Gm13839 | 0.393020699 | 1.260710985 | 1.689812692 |
| Gm13838 | 0.066050875 | 0.71820487 | 3.724945049 |
| B230112J18Rik | 0.528863779 | 0.254631974 | 1.751018068 |
| RP24-79A6.9 | 0.038981395 | 0.718207094 | 3.724945049 |
| 4930562A09Rik | 0.146089079 | 0.238291249 | 3.522464378 |
| Hspb8 | 433.011455 | 2.717538235 | 0.208607549 |
| Srrm4os | 0.683645101 | -1.675213993 | 1.133775598 |
| Srrm4 | 850.8466253 | -0.53133561 | 0.091727688 |
| RP24-79A6.8 | 32.11370528 | -1.269985457 | 0.324468294 |
| RP23-312N12.2 | 10.26602459 | -0.855364265 | 0.439138146 |
| RP23-312N12.1 | 15.57120856 | -2.119265545 | 0.428357615 |
| Gstm2-ps1 | 32.88268672 | -0.623007304 | 0.252508832 |
| RP23-418H8.3 | 0.072025462 | 0.237312084 | 3.724945049 |
| RP23-280O16.1 | 0.060174537 | 0.237309859 | 3.724945049 |
| 4930569F06Rik | 0 | NA | NA |
| RP23-73P16.2 | 0.598047226 | 2.955045596 | 1.205822183 |
| Suds3 | 810.9948176 | -0.134732545 | 0.058301829 |
| 9530046B11Rik | 1.655761812 | -0.789832611 | 0.801833528 |
| Taok3 | 787.4816357 | 0.129740529 | 0.068377767 |
| RP23-376H8.2 | 15.63613524 | -0.313437151 | 0.354320886 |
| n-R5s174 | 1.199099511 | -0.476073094 | 1.001635381 |
| Pebp1 | 1757.043143 | -0.042666841 | 0.072631663 |
| RP23-76J15.3 | 0 | NA | NA |
| RP23-76J15.4 | 0.215166026 | -0.243568332 | 2.556929146 |

| | | | |
|---|---|---|---|
| BC051077 | 1.218454842 | -0.494914143 | 1.046021617 |
| Gm10399 | 0.929291888 | 1.340994254 | 1.073750278 |
| Vsig10 | 55.95498491 | 0.991280671 | 0.231235726 |
| Gm15727 | 0 | NA | NA |
| Wsb2 | 1799.690181 | 0.109412826 | 0.066481628 |
| Rfc5 | 71.308591 | 0.048012927 | 0.16865382 |
| Gm15728 | 37.43701224 | 0.564568233 | 0.206563341 |
| Ksr2 | 1532.422263 | -0.924376415 | 0.078401679 |
| RP23-429P17.1 | 4.731857296 | -1.336478101 | 0.510314838 |
| Nos1 | 1164.626249 | -0.744446561 | 0.082644412 |
| Gm37028 | 0.69250253 | -1.259534905 | 1.393359524 |
| Gm26995 | 0.399888529 | -0.689605564 | 1.443348872 |
| Gm25343 | 0.046345709 | 0.71820487 | 3.724945049 |
| Fbxo21 | 1283.047223 | -0.525155025 | 0.063153169 |
| RP23-42D10.2 | 0.853574741 | 1.338979428 | 1.11270918 |
| Tesc | 196.9276744 | -0.876368422 | 0.167770301 |
| Fbxw8 | 652.4874552 | 0.523300902 | 0.097129056 |
| Gm9754 | 24.46512519 | -1.307173653 | 0.440705754 |
| Gm26411 | 1.298260455 | -0.341897745 | 0.99140516 |
| Hrk | 446.6006905 | -0.844869692 | 0.193936584 |
| Rnft2 | 775.8299915 | -0.652447998 | 0.082229522 |
| Gm24050 | 1.923134663 | -0.364048973 | 0.683765379 |
| 2410131K14Rik | 480.8992711 | 0.505604006 | 0.099413312 |
| Gm25076 | 4.628642042 | -0.099946076 | 0.445939135 |
| Gm28563 | 0.587040768 | -1.877299491 | 1.158535406 |
| Gm26455 | 0.133946075 | 0.7191812 | 3.626570343 |
| Gm15754 | 0.066050875 | 0.71820487 | 3.724945049 |
| Med13l | 2276.940129 | -0.041652404 | 0.066478376 |
| RP24-122N24.1 | 0 | NA | NA |
| RP24-122N24.2 | 76.64565332 | -0.495784837 | 0.299620705 |
| RP24-122N24.3 | 27.41095622 | -0.679888489 | 0.302410405 |
| RP24-122N24.4 | 64.02437309 | -0.53580866 | 0.319060776 |
| RP23-155F13.2 | 26.31192174 | -0.860106852 | 0.340808438 |
| RP23-155F13.3 | 28.57824406 | -0.642748505 | 0.3582608 |
| RP24-372H3.1 | 0 | NA | NA |
| RP24-372H3.2 | 0 | NA | NA |
| RP24-372H3.4 | 0 | NA | NA |
| RP24-372H3.3 | 0 | NA | NA |
| 1700081H04Rik | 0.049553047 | 0.71820487 | 3.724945049 |
| RP23-316P20.1 | 0.048949532 | -0.243569582 | 3.724945049 |
| n-R5s175 | 0 | NA | NA |
| RP23-73O10.1 | 0 | NA | NA |
| Tbx3os1 | 2.722982093 | -0.609454394 | 0.724610035 |
| RP23-44J10.1 | 0.387852398 | -1.869722392 | 1.49615485 |
| Gm16063 | 0 | NA | NA |
| Gm16064 | 0 | NA | NA |
| Tbx3 | 28.477325 | 0.091649446 | 0.23544709 |

| | | | |
|---|---|---|---|
| Tbx3os2 | 0.046345709 | 0.71820487 | 3.724945049 |
| RP24-150D8.1 | 0 | NA | NA |
| RP24-150D8.2 | 0 | NA | NA |
| Tbx5 | 0.360567166 | 1.85974319 | 1.974746816 |
| RP24-150D8.3 | 0 | NA | NA |
| RP24-69A21.1 | 0 | NA | NA |
| n-R5s176 | 0 | NA | NA |
| Rbm19 | 227.3684793 | 0.698226632 | 0.076975067 |
| Gm10390 | 1.46581892 | 1.572623369 | 0.881762566 |
| RP23-213H20.1 | 1.459375539 | 2.765608913 | 1.049271727 |
| Gm26474 | 0 | NA | NA |
| RP23-213H20.2 | 0.303117673 | 0.237299767 | 3.724945049 |
| Gm27199 | 3.757776822 | -1.00259895 | 0.558341673 |
| Lhx5 | 49.17951797 | -0.18414591 | 0.206726618 |
| Sdsl | 34.31511951 | 0.208083051 | 0.196837672 |
| Sds | 0.913087876 | -0.929767423 | 1.35182343 |
| Gm15690 | 5.787489139 | -0.912868834 | 0.494209821 |
| Plbd2 | 910.6907376 | 0.206688547 | 0.080242984 |
| Slc8b1 | 149.2518986 | 2.18248962 | 0.170458649 |
| Tpcn1 | 461.0027236 | -1.090967622 | 0.151095392 |
| RP23-297M4.7 | 22.51234608 | -0.761117936 | 0.269698171 |
| Iqcd | 5.452284118 | -1.439746562 | 0.517876426 |
| Rita1 | 142.064277 | -0.468536513 | 0.134160412 |
| Ddx54 | 525.9668418 | 0.60811372 | 0.090607293 |
| RP23-297M4.4 | 3.794821076 | 0.149760332 | 0.564731914 |
| Ccdc42b | 3.366237703 | -0.19767692 | 0.543338045 |
| RP23-297M4.1 | 0 | NA | NA |
| Gm15749 | 0.639699895 | 0.805223264 | 1.487868987 |
| Rasal1 | 506.8043477 | -1.285171066 | 0.133216014 |
| Dtx1 | 1090.783309 | 0.036322794 | 0.074141939 |
| Oas2 | 477.0806149 | 9.003266242 | 0.46171123 |
| RP24-547J21.3 | 0.066050875 | 0.71820487 | 3.724945049 |
| Oas3 | 836.3377262 | 12.10159005 | 0.874641582 |
| RP24-547J21.4 | 1.429586369 | 3.99503793 | 1.071440746 |
| Oas1e | 0.736605981 | 1.875709613 | 1.327478716 |
| Oas1c | 32.60080593 | 2.574628642 | 0.263136803 |
| Oas1b | 318.3948597 | 7.550093912 | 0.436779757 |
| Oas1f | 0.049553047 | 0.71820487 | 3.724945049 |
| Gm22907 | 0 | NA | NA |
| Oas1h | 0.100902008 | 1.15059983 | 3.720931964 |
| Oas1g | 272.4663882 | 9.916496098 | 0.770902169 |
| Oas1a | 611.2994765 | 10.13911568 | 0.583929587 |
| Oas1d | 0.674169638 | 3.43937509 | 1.195689425 |
| Rph3a | 3838.839031 | -0.81817568 | 0.089095932 |
| RP23-39M18.2 | 36.68051303 | -2.251963715 | 0.340988585 |
| Ptpn11 | 2663.867196 | 0.143990733 | 0.074520622 |
| RP23-257E17.2 | 45.93026392 | -1.17901837 | 0.340237428 |

| | | | |
|---|---|---|---|
| Rpl6 | 1338.143399 | 0.640191185 | 0.084740677 |
| Gm26205 | 0 | NA | NA |
| Gm15800 | 6892.751409 | -0.862071059 | 0.090237012 |
| Gm24671 | 2.5200291 | -0.83556442 | 0.683632842 |
| Trafd1 | 1684.87789 | 2.567273615 | 0.15271398 |
| RP23-50I16.1 | 5.493864583 | 0.962454147 | 0.481548448 |
| RP23-50I16.2 | 19.99381265 | 0.259381288 | 0.254918328 |
| Naa25 | 948.0018726 | 0.662508003 | 0.087220706 |
| RP23-50I16.3 | 14.51075867 | -0.077835528 | 0.362665689 |
| Erp29 | 574.7376244 | 0.180982585 | 0.100558355 |
| Tmem116 | 26.92021731 | -0.203828168 | 0.212670072 |
| Gm15546 | 0 | NA | NA |
| Adam1b | 0.227186609 | 1.364281994 | 2.549472411 |
| RP24-435H23.1 | 2.953870943 | -0.397128897 | 0.687828619 |
| Adam1a | 36.9795004 | -0.444100635 | 0.281138302 |
| Mapkapk5 | 369.1571913 | 0.171615295 | 0.093065957 |
| Gm15547 | 4.040079771 | -1.127091272 | 0.727537955 |
| Aldh2 | 693.3685642 | -0.318318244 | 0.117451848 |
| RP23-181P9.3 | 1.724244418 | 0.307633343 | 0.766989438 |
| Acad12 | 28.57697775 | -0.458670206 | 0.232384079 |
| Acad10 | 41.29551714 | 0.021965949 | 0.191961458 |
| Brap | 823.207007 | 0.168630283 | 0.079492301 |
| Atxn2 | 1833.502759 | -0.269005709 | 0.069680428 |
| RP23-18K20.2 | 63.62259654 | -0.57809715 | 0.361657413 |
| Gm16552 | 0 | NA | NA |
| RP23-18K20.3 | 37.1706614 | -0.907478466 | 0.329933461 |
| Sh2b3 | 323.628142 | 0.234537704 | 0.105499022 |
| Mir7031 | 0.877654538 | -0.134203716 | 1.061722212 |
| RP23-431M7.2 | 0.523632775 | 2.255418859 | 1.426308753 |
| RP23-431M7.4 | 0.25563639 | -0.71214286 | 2.113943058 |
| Fam109a | 123.4761975 | -0.022052772 | 0.109088989 |
| Cux2 | 1234.512826 | -0.968951493 | 0.129558644 |
| AC113302.1 | 0 | NA | NA |
| RP23-254C8.1 | 0 | NA | NA |
| Gm15637 | 0.291530733 | -0.226232216 | 1.786380053 |
| Myl2 | 1.403419482 | -1.742352954 | 1.025390002 |
| Ccdc63 | 13.49297597 | 0.244763395 | 0.354597208 |
| Ppp1cc | 926.7286209 | 0.031933289 | 0.069278782 |
| RP24-347B12.1 | 50.60541787 | -0.842932476 | 0.325264443 |
| Hvcn1 | 77.26580927 | 1.577009427 | 0.236338364 |
| Tctn1 | 251.1741939 | -0.135671249 | 0.100261758 |
| Gm15481 | 0 | NA | NA |
| Pptc7 | 900.5757088 | -0.146992181 | 0.061574423 |
| Gm15842 | 8.407355864 | 0.181977343 | 0.454573207 |
| Rad9b | 39.0160455 | -0.409056968 | 0.200883066 |
| Vps29 | 780.350795 | -0.258115533 | 0.086968655 |
| Fam216a | 407.2222889 | 0.112007519 | 0.067212086 |

| | | | |
|---|---|---|---|
| Gpn3 | 109.4263101 | 0.035787328 | 0.131378177 |
| RP23-407I6.1 | 0.395669761 | -0.567893577 | 1.649184744 |
| Arpc3 | 620.2922037 | 0.332292982 | 0.092427536 |
| RP23-407I6.2 | 78.71060985 | -0.653078681 | 0.155767981 |
| Anapc7 | 355.9121168 | 0.242335336 | 0.082922194 |
| Gm15846 | 0 | NA | NA |
| Atp2a2 | 11857.11526 | -0.241682223 | 0.060985445 |
| RP23-407I6.6 | 93.54004702 | -1.259175408 | 0.410396671 |
| RP23-407I6.5 | 69.66663032 | -1.325380751 | 0.309537512 |
| RP23-407I6.4 | 0.863835993 | 1.627204353 | 1.325404978 |
| Gm22965 | 1.295159943 | 1.402659285 | 1.058255399 |
| Ift81 | 401.1137358 | -0.303359714 | 0.08306163 |
| RP23-407I6.7 | 1.063558902 | -2.907420002 | 1.077407666 |
| RP24-114E20.2 | 0.51473784 | -0.247388942 | 1.259478685 |
| P2rx7 | 217.2536391 | 1.994981248 | 0.164681602 |
| Mir8115 | 0 | NA | NA |
| Gm10064 | 0 | NA | NA |
| P2rx4 | 293.0467081 | 1.525508686 | 0.146339863 |
| Camkk2 | 1870.397667 | -1.751808357 | 0.14174851 |
| RP23-31J24.2 | 7.791943915 | -1.206841291 | 0.427674958 |
| Anapc5 | 1309.787107 | -0.280469107 | 0.054744374 |
| Rnf34 | 645.0189091 | 0.234008108 | 0.066805502 |
| Kdm2b | 295.4828908 | -0.463635168 | 0.093676054 |
| RP23-214I6.2 | 6.36596584 | -1.791522026 | 0.613325473 |
| RP23-214I6.5 | 0 | NA | NA |
| A930024E05Rik | 27.86517634 | 0.007162523 | 0.20007038 |
| RP23-214I6.3 | 0.145501641 | 1.198496036 | 2.88801119 |
| Orai1 | 101.2846242 | 1.02411153 | 0.152816178 |
| Morn3 | 1.434593181 | 0.532920108 | 0.783754839 |
| RP23-214I6.4 | 5.158449836 | -0.210176278 | 0.646568389 |
| Gm26745 | 0.042716945 | -0.243571806 | 3.724945049 |
| Gm6444 | 0.573335652 | -0.64330823 | 1.259542818 |
| Tmem120b | 108.3890549 | -0.405792774 | 0.144849672 |
| Rhof | 259.5705259 | -0.351049028 | 0.143468776 |
| RP23-371A16.4 | 14.62755803 | -2.526859024 | 0.43364757 |
| AI480526 | 273.446315 | -0.850436541 | 0.221833352 |
| Gm38102 | 1.104086423 | 3.06855105 | 1.094903404 |
| RP23-371A16.3 | 8.618846306 | 0.448511903 | 0.37105382 |
| Setd1b | 1298.15226 | -0.154564751 | 0.103045415 |
| Psmd9 | 209.0408427 | 0.265742027 | 0.089751747 |
| Hpd | 0.5402512 | 1.503666606 | 1.448529183 |
| RP23-297I15.1 | 0 | NA | NA |
| Wdr66 | 48.81208703 | -1.052655543 | 0.168116719 |
| Rpl35a-ps6 | 0.817924277 | 0.525981338 | 1.142036102 |
| Gm15857 | 0.981461204 | -0.991422868 | 1.100142198 |
| Gm15860 | 13.45882266 | -0.718806302 | 0.27709433 |
| Bcl7a | 835.7958974 | -0.977856932 | 0.108016274 |

| | | | |
|---|---|---|---|
| Gm23458 | 4.576786753 | -1.528310343 | 0.534297374 |
| RP23-370G17.1 | 16.44557487 | -1.325600502 | 0.362797326 |
| Gm15747 | 1.272170519 | 2.042774665 | 1.0782548 |
| Mlxip | 530.2039296 | 0.116074222 | 0.1022757 |
| RP23-370G17.2 | 23.88538577 | -0.437197711 | 0.299076971 |
| RP23-370G17.3 | 10.05076668 | 0.17511161 | 0.40303317 |
| RP23-370G17.4 | 55.71562266 | -0.299595322 | 0.252904928 |
| Rpl31-ps6 | 0.955200391 | -2.034317337 | 1.060189531 |
| Il31 | 0.27099349 | 1.770707551 | 2.046397532 |
| Lrrc43 | 5.510592113 | -0.025187199 | 0.52860536 |
| Gm15751 | 2.413806088 | -1.683470675 | 0.667480559 |
| Diablo | 222.7606092 | 0.267544536 | 0.09700525 |
| B3gnt4 | 5.682107971 | -0.258432914 | 0.393807428 |
| Gm24682 | 0.386251008 | -1.9293626 | 1.721162037 |
| Mir7647 | 1.111319392 | -1.223903478 | 1.010360669 |
| Vps33a | 1223.976526 | 0.111587496 | 0.052861161 |
| RP23-121M13.2 | 3.773545612 | -0.739081182 | 0.579198067 |
| Clip1 | 1856.591368 | -0.272035333 | 0.088753362 |
| Gm24840 | 0 | NA | NA |
| RP23-121M13.3 | 21.46809472 | -1.038858878 | 0.321777752 |
| RP23-121M13.4 | 2.51877741 | 0.339486803 | 0.695061917 |
| Zcchc8 | 358.9731995 | -0.068787379 | 0.100766842 |
| Rsrc2 | 977.6377703 | -0.110999385 | 0.079093575 |
| Kntc1 | 15.85629698 | 1.76200194 | 0.397749939 |
| Hcar2 | 55.90105025 | 8.765439535 | 0.887263865 |
| Hcar1 | 26.46370296 | 0.634463051 | 0.264770269 |
| Denr | 363.9918925 | 0.1663381 | 0.107584982 |
| RP24-492O4.5 | 0.261825437 | -1.080113759 | 2.073931698 |
| Ccdc62 | 24.55556878 | -0.668428473 | 0.247745402 |
| Hip1r | 946.4062031 | -0.213418609 | 0.068429993 |
| Mir7032 | 0 | NA | NA |
| Vps37b | 406.419246 | 0.325812142 | 0.104902456 |
| Gm37939 | 0.497466187 | -0.175586639 | 1.510159029 |
| RP24-492O4.9 | 0.226827357 | -0.242840368 | 2.143469669 |
| RP24-492O4.8 | 0.200931026 | 1.950293203 | 2.148509737 |
| Abcb9 | 408.729264 | -0.754958271 | 0.076419342 |
| Gm16001 | 32.41362582 | -1.006397544 | 0.19923421 |
| Ogfod2 | 183.0496373 | 0.092580426 | 0.083977035 |
| Arl6ip4 | 210.5407291 | 0.12306868 | 0.138289589 |
| Pitpnm2 | 2668.846213 | -0.995666304 | 0.14204516 |
| Pitpnm2os2 | 1.19134753 | -0.483217165 | 0.873751767 |
| Pitpnm2os1 | 0.048949532 | -0.243569582 | 3.724945049 |
| RP24-257M14.2 | 0 | NA | NA |
| Mphosph9 | 491.0917071 | -0.540119038 | 0.169580692 |
| Gm23171 | 0 | NA | NA |
| RP24-257M14.3 | 50.23293639 | -0.984693831 | 0.349169239 |
| RP23-180G16.2 | 6.931404989 | -0.857946632 | 0.45392093 |

| | | | |
|---|---|---|---|
| 2810006K23Rik | 85.50115535 | -0.205057196 | 0.209674551 |
| Cdk2ap1 | 265.3510822 | 0.162304457 | 0.097305799 |
| Sbno1 | 3030.840915 | -0.295090589 | 0.052901389 |
| Gm37415 | 6.225233596 | -0.593687879 | 0.462373903 |
| Gm27486 | 0 | NA | NA |
| Setd8 | 525.0620955 | 0.614909563 | 0.101158516 |
| Rilpl2 | 121.8049509 | 0.56085498 | 0.116971912 |
| Gm15621 | 0.143485256 | 1.229961212 | 3.547405499 |
| Snrnp35 | 68.78650209 | 0.421276081 | 0.212175849 |
| Rilpl1 | 373.8418609 | -0.454561852 | 0.094316726 |
| RP23-311A21.2 | 18.59305175 | -1.169813841 | 0.29770567 |
| Tmed2 | 550.4698307 | 0.721536234 | 0.089539848 |
| Ddx55 | 213.2561564 | -0.415460954 | 0.137421437 |
| Eif2b1 | 210.495533 | 0.253868031 | 0.098241792 |
| Gtf2h3 | 153.3355758 | 0.034323033 | 0.134678266 |
| Tctn2 | 238.196889 | -0.102882247 | 0.125265309 |
| Atp6v0a2 | 1102.977081 | 0.666417196 | 0.101667996 |
| Gm24854 | 0 | NA | NA |
| WI1-2065I1.2 | 0.075043205 | 0.237309859 | 3.724945049 |
| Dnah10 | 37.63473061 | 0.163329039 | 0.210847103 |
| RP23-437D10.1 | 3.569206524 | 0.411961768 | 0.689541444 |
| Ccdc92 | 896.6833926 | -0.295833108 | 0.078999364 |
| Zfp664 | 1184.705232 | -0.122473276 | 0.062479158 |
| Fam101a | 19.01723143 | 4.583957865 | 0.540875824 |
| Ncor2 | 4028.896925 | 0.126841294 | 0.074277762 |
| RP24-291G12.3 | 0.185374628 | -1.198364947 | 2.266154646 |
| RP24-291G12.2 | 0.590920558 | 0.661503897 | 1.324783849 |
| RP23-363M4.1 | 0.852297073 | 0.260497484 | 1.246040984 |
| Scarb1 | 198.4544971 | 0.147466417 | 0.095032904 |
| RP23-363M4.2 | 6.758329482 | -0.58963911 | 0.441420545 |
| RP24-105J12.1 | 0 | NA | NA |
| Ubc | 623.813315 | 1.146720968 | 0.1472441 |
| Gm10382 | 3.55196556 | -0.247057824 | 0.541297503 |
| Dhx37 | 317.3772995 | 0.637230838 | 0.083008216 |
| RP24-105J12.3 | 0.046345709 | 0.71820487 | 3.724945049 |
| Bri3bp | 1674.728707 | -0.824133339 | 0.066994212 |
| RP24-324M10.2 | 0.427937583 | -1.244446069 | 1.346067333 |
| Aacs | 461.1003154 | 0.187614394 | 0.091476689 |
| Gm27551 | 0.589769752 | -0.38779419 | 1.224706229 |
| Tmem132b | 1925.698646 | -0.899405152 | 0.120361011 |
| Gm4868 | 0 | NA | NA |
| RP24-332C17.1 | 1.392497718 | 1.285911958 | 1.010828613 |
| RP24-332C17.2 | 0 | NA | NA |
| RP24-332C17.3 | 0.274664082 | 2.268192365 | 2.025446229 |
| RP24-332C17.4 | 5.259892424 | 0.624275275 | 0.4706079 |
| Gm23151 | 0 | NA | NA |
| Gm37700 | 0 | NA | NA |

| | | | |
|---|---|---|---|
| RP23-414E14.1 | 0.0878635 | -0.72445792 | 3.724945049 |
| Gm24839 | 0 | NA | NA |
| RP23-74L20.1 | 1.146741161 | -0.019132822 | 1.004306502 |
| RP23-74L20.2 | 0.044191549 | -0.243571806 | 3.724945049 |
| RP23-145H8.1 | 0 | NA | NA |
| RP23-145H8.2 | 0.512350924 | 1.513194577 | 1.505705966 |
| RP23-282A16.1 | 0.230183376 | 0.718216341 | 2.508489011 |
| RP23-37N15.1 | 0.075043205 | 0.237309859 | 3.724945049 |
| RP23-37N15.3 | 0 | NA | NA |
| RP23-37N15.4 | 0.066050875 | 0.71820487 | 3.724945049 |
| RP23-37N15.2 | 0 | NA | NA |
| Tmem132c | 179.9200518 | -0.079200148 | 0.113438261 |
| Tmem132cos | 0 | NA | NA |
| RP23-71M14.1 | 1.788302827 | -1.234116869 | 0.970114316 |
| WI1-2285D16.1 | 0.302970058 | 0.718222726 | 1.998383114 |
| Slc15a4 | 300.9485309 | 0.109497091 | 0.109379294 |
| WI1-1024F3.1 | 0.545152564 | 0.632115031 | 1.314685345 |
| Glt1d1 | 207.4632902 | -0.653581308 | 0.125023009 |
| Tmem132d | 473.83412 | -0.747347847 | 0.145307999 |
| Gm17132 | 0.461821276 | -1.960761442 | 1.529232043 |
| Gm17131 | 0.522940275 | 0.837099539 | 1.473528398 |
| RP23-378C15.2 | 8.098685304 | -1.160176791 | 0.52847509 |
| RP23-378C15.1 | 6.802413532 | -2.0674169 | 0.514836937 |
| WI1-2255B14.1 | 0.141177696 | 0.237325886 | 2.91536683 |
| CH36-12I15.2 | 13.22303069 | 2.11512355 | 0.35940845 |
| CH36-12I15.3 | 7.173227082 | 2.822632325 | 0.60441149 |
| CH36-12I15.1 | 0 | NA | NA |
| CH36-12I15.4 | 0.075043205 | 0.237309859 | 3.724945049 |
| 5930412G12Rik | 23.11388567 | -0.855268452 | 0.223121133 |
| RP24-258A20.1 | 1.894888719 | -0.499859895 | 0.707617724 |
| Fzd10 | 39.99764841 | -0.444313003 | 0.173618542 |
| Piwil1 | 0.809884792 | -0.528491054 | 1.095830127 |
| RP23-156F20.2 | 0 | NA | NA |
| Rimbp2 | 1419.490251 | -0.555898877 | 0.22328174 |
| RP23-156F20.4 | 0.134481855 | -0.243575541 | 3.655499445 |
| RP23-156F20.5 | 1.664368785 | -1.06813699 | 0.748620175 |
| Stx2 | 147.9774137 | 0.407799313 | 0.120052201 |
| Ran | 1531.557904 | 0.442927841 | 0.089805546 |
| Adgrd1 | 48.98301064 | 0.711692837 | 0.173216402 |
| Rps16-ps2 | 75.19513727 | 0.622187468 | 0.160974081 |
| RP24-158G19.1 | 0 | NA | NA |
| RP24-81J14.1 | 2.60471737 | 0.403297847 | 0.799170424 |
| Sfswap | 880.7266452 | -0.270774695 | 0.137140939 |
| RP24-161C20.4 | 61.43406924 | -0.725009684 | 0.355460748 |
| Mmp17 | 1263.227049 | -0.545814258 | 0.134631828 |
| Gm6139 | 8.767336692 | 1.037826151 | 0.39215211 |
| Zfp11 | 110.3957661 | -0.094869098 | 0.134803507 |

| | | | |
|---|---|---|---|
| 14-Sep | 0.250988417 | 1.181866011 | 2.47690198 |
| Mrps17 | 416.7636034 | -0.222356489 | 0.062615123 |
| Gbas | 690.6986552 | -0.119314678 | 0.068788552 |
| Gm15903 | 27.46139881 | -0.322727873 | 0.324480118 |
| Psph | 139.9382442 | 0.519304068 | 0.146774023 |
| Cct6a | 1504.12042 | 0.466163731 | 0.087833208 |
| Gm23245 | 2.474741284 | 0.029445904 | 0.610095931 |
| Snora15 | 0 | NA | NA |
| Sumf2 | 86.25597758 | 0.418029151 | 0.120863592 |
| Phkg1 | 43.76477553 | -1.704865385 | 0.267304025 |
| WI1-535J9.1 | 0.523518094 | -2.738358451 | 1.282366332 |
| Chchd2 | 843.0146962 | 0.291990905 | 0.117402051 |
| Zbed5 | 99.19524039 | 0.270589838 | 0.155922935 |
| Nupr1l | 8.851478164 | -1.007731617 | 0.350509215 |
| Vkorc1l1 | 758.1542901 | -0.166964858 | 0.062773945 |
| RP23-189N6.2 | 51.40024564 | -0.95428101 | 0.318375529 |
| RP23-189N6.3 | 26.74839225 | -0.56214179 | 0.368709345 |
| Gusb | 191.859854 | 1.453007657 | 0.137627526 |
| Asl | 102.1205115 | 0.352451088 | 0.134133946 |
| Crcp | 187.2081582 | 0.716834736 | 0.111377486 |
| Tpst1 | 399.6196954 | 0.496238404 | 0.090938041 |
| RP23-353L10.2 | 7.510072057 | -0.076157235 | 0.492303739 |
| Kctd7 | 165.5730604 | -0.310000656 | 0.101244572 |
| Rabgef1 | 458.137125 | 0.101197091 | 0.070721259 |
| Gm15920 | 252.9729714 | 0.154934085 | 0.106125443 |
| Gm15921 | 0.048949532 | -0.243569582 | 3.724945049 |
| Tmem248 | 679.0584884 | 0.486875349 | 0.103983886 |
| AC113029.6 | 6.876401235 | -0.086149818 | 0.422804494 |
| RP23-206J17.2 | 14.12832924 | -0.096867703 | 0.434045039 |
| Sbds | 553.098637 | 0.539502265 | 0.119407549 |
| Tyw1 | 213.1214519 | -0.0269251 | 0.131586483 |
| Caln1 | 4353.134745 | -0.267136375 | 0.095370601 |
| A330070K13Rik | 2.26626686 | -1.778493722 | 0.864743638 |
| RP24-571K14.1 | 0 | NA | NA |
| Gm23761 | 4.136581771 | -0.520209387 | 0.543551281 |
| RP24-559P3.1 | 12.49495635 | 0.175090094 | 0.344066201 |
| Wbscr17 | 856.0949554 | -0.37517921 | 0.083046638 |
| RP23-323D23.1 | 6.092562558 | -1.502714054 | 0.498750454 |
| RP23-323D23.2 | 13.43160181 | -1.125560773 | 0.415900298 |
| RP23-323D23.3 | 19.70298692 | -1.200845977 | 0.427925069 |
| RP23-445H19.1 | 18.79099024 | -1.362284409 | 0.290002222 |
| RP23-114L5.4 | 5.83927098 | -0.494514339 | 0.465102839 |
| Auts2 | 2085.675582 | -0.773013866 | 0.126594588 |
| Gm27266 | 0 | NA | NA |
| RP23-120C8.7 | 31.89874726 | -1.435100883 | 0.403303435 |
| RP23-120C8.6 | 19.56297173 | -1.536175368 | 0.363357654 |
| RP23-120C8.9 | 0.108767819 | 0.237318756 | 3.724945049 |

| | | | |
|---|---|---|---|
| RP23-120C8.5 | 11.30893109 | -1.326715651 | 0.482226784 |
| RP23-120C8.4 | 44.54237424 | -1.258633787 | 0.356264786 |
| RP23-120C8.3 | 22.2103361 | -1.179224106 | 0.364345188 |
| RP23-120C8.2 | 22.94359978 | -1.627246441 | 0.316070492 |
| RP24-95O4.7 | 0 | NA | NA |
| RP24-95O4.6 | 73.13605096 | -1.323529266 | 0.300103488 |
| RP24-95O4.5 | 20.70201435 | -1.660744342 | 0.514399698 |
| RP24-95O4.4 | 58.37291526 | -1.504688324 | 0.345221116 |
| RP24-95O4.9 | 0 | NA | NA |
| RP24-95O4.3 | 42.64488512 | -0.909354821 | 0.255639648 |
| RP24-95O4.2 | 132.5110994 | -1.471744969 | 0.370878734 |
| RP24-95O4.1 | 237.0691924 | -1.485290521 | 0.322001094 |
| RP23-378O9.2 | 72.35690436 | -1.905148005 | 0.32987693 |
| RP23-378O9.1 | 291.0443093 | -1.40547639 | 0.290732497 |
| RP24-147H20.3 | 37.48355553 | -1.510815586 | 0.323335402 |
| RP24-147H20.2 | 50.07029278 | -1.61521043 | 0.252911464 |
| RP23-276C23.3 | 53.66935453 | -1.58370737 | 0.332198989 |
| RP23-276C23.2 | 105.0397251 | -1.396955067 | 0.232978989 |
| RP23-276C23.1 | 58.80184518 | -1.225134661 | 0.338842038 |
| RP24-399A15.2 | 87.32223591 | -1.586630578 | 0.309868654 |
| RP24-399A15.3 | 100.1496488 | -1.463578709 | 0.286996488 |
| RP24-399A15.5 | 46.61621938 | -1.439903927 | 0.384850221 |
| Gm28616 | 0.118974955 | -0.243576254 | 3.724945049 |
| RP24-399A15.6 | 0.098502579 | 0.23732098 | 3.724945049 |
| Gm38373 | 0 | NA | NA |
| Gm25722 | 0 | NA | NA |
| RP23-455A10.3 | 0 | NA | NA |
| Gm10051 | 4.533521042 | 0.800149416 | 0.602172889 |
| RP23-455A10.2 | 0 | NA | NA |
| RP23-455A10.1 | 0.061920613 | 0.718207094 | 3.724945049 |
| RP23-409D19.1 | 5.435068931 | -0.753613788 | 0.429482282 |
| RP23-409D19.2 | 0 | NA | NA |
| RP24-199E1.1 | 0.733529363 | 0.182891436 | 1.123871188 |
| Gm15627 | 4.57824934 | -1.03985089 | 0.515453798 |
| Gatsl2 | 1933.057981 | -0.434239965 | 0.076609506 |
| Wbscr16 | 129.6578973 | 0.657221341 | 0.118247682 |
| B230377A18Rik | 10.36497119 | 0.942349859 | 0.374082277 |
| Gtf2ird2 | 207.0338271 | -0.112437984 | 0.100233873 |
| Gm24139 | 2.685875012 | -0.685709192 | 0.675968685 |
| Ncf1 | 204.7455163 | 1.879925388 | 0.199392602 |
| Gtf2i | 2503.932034 | -0.354059288 | 0.103360925 |
| RP23-15K13.2 | 110.8580539 | -1.194823408 | 0.371317588 |
| RP23-15K13.3 | 5.451269312 | -1.034854493 | 0.566367122 |
| Mir3965 | 0 | NA | NA |
| Gtf2ird1 | 384.0610061 | -0.097903386 | 0.13850275 |
| n-R5s178 | 0 | NA | NA |
| Gm26340 | 0 | NA | NA |

| | | | |
|---|---|---|---|
| RP24-359H20.2 | 0.585862523 | -0.676580468 | 1.245046834 |
| Gm16020 | 0 | NA | NA |
| RP24-359H20.3 | 3.016053333 | -0.433184939 | 0.504501323 |
| Clip2 | 1985.813075 | 0.29289248 | 0.050969534 |
| RP23-85H24.1 | 29.77219559 | -0.595771966 | 0.317594139 |
| RP23-85H24.2 | 8.735753554 | -0.999912068 | 0.36490179 |
| Syna | 7.492197562 | 0.128798195 | 0.417198753 |
| mmu-mir-7228 | 0.285083685 | -0.398868857 | 1.763266355 |
| Rfc2 | 247.4000499 | 0.254400984 | 0.084725336 |
| Lat2 | 113.3855065 | 3.596209802 | 0.246274064 |
| Eif4h | 3070.987643 | 0.092067287 | 0.049088823 |
| Limk1 | 829.834934 | -0.33120617 | 0.121340414 |
| Gm10369 | 1.019349322 | 0.183359786 | 1.416392693 |
| Eln | 72.96738645 | 0.080109956 | 0.187778134 |
| RP23-315E2.1 | 8.223192194 | -1.556145915 | 0.421944541 |
| Wbscr28 | 0.045882455 | 0.237312084 | 3.724945049 |
| Cldn13 | 0 | NA | NA |
| RP23-201C9.3 | 0 | NA | NA |
| RP23-201C9.2 | 0.038981395 | 0.718207094 | 3.724945049 |
| Wbscr27 | 62.20310215 | -0.785364698 | 0.167932427 |
| Cldn4 | 0.565252138 | 2.2465443 | 3.077357899 |
| Cldn3 | 2.938076242 | 0.381581141 | 0.63421124 |
| Wbscr25 | 2.760772158 | -1.077997585 | 0.672564801 |
| Rps29-ps | 0 | NA | NA |
| RP23-38B15.1 | 4.804143474 | -0.616905042 | 0.540601872 |
| Abhd11 | 157.6817239 | 0.050343364 | 0.117014031 |
| Abhd11os | 7.140605062 | -0.513096684 | 0.515053982 |
| Stx1a | 708.4200384 | -1.929541823 | 0.161748849 |
| Wbscr22 | 194.0156334 | 0.119478578 | 0.07952905 |
| Gm25492 | 29.10926416 | -0.127241285 | 0.204305793 |
| Dnajc30 | 252.3546012 | -0.150115215 | 0.087017436 |
| Vps37d | 164.5301101 | -0.137005735 | 0.12268966 |
| Mlxipl | 13.0557667 | -1.323449751 | 0.382353497 |
| RP23-240H6.1 | 0 | NA | NA |
| Tbl2 | 380.9054715 | 0.026518433 | 0.065367445 |
| Bcl7b | 442.8768574 | 0.306016027 | 0.057037161 |
| Baz1b | 2277.625964 | -0.110649684 | 0.08720799 |
| Gm25315 | 0 | NA | NA |
| Fzd9 | 41.32503692 | -0.765531166 | 0.189644296 |
| RP23-240H6.2 | 0.095295241 | 0.23732098 | 3.724945049 |
| Fkbp6 | 0.371024948 | 0.904203521 | 1.692447917 |
| Trim50 | 0.160978641 | -1.214251972 | 2.79661493 |
| Nsun5 | 116.6234585 | 0.109421233 | 0.175658007 |
| Pom121 | 1177.537542 | 0.08768045 | 0.072983444 |
| Mir7033 | 0 | NA | NA |
| Hip1 | 1047.747718 | 0.665098782 | 0.114575007 |
| RP23-333I24.3 | 8.728400981 | -0.360535099 | 0.358624066 |

| | | | |
|---|---|---|---|
| Ccl26 | 0 | NA | NA |
| Ccl24 | 1.573001602 | -0.780828203 | 0.712587023 |
| RP23-423A22.3 | 0.045882455 | 0.237312084 | 3.724945049 |
| Rhbdd2 | 635.3255966 | -0.290957109 | 0.055670333 |
| Gm15607 | 0.101408268 | 0.717551239 | 3.724898127 |
| Gm21839 | 0.540306067 | -0.159304689 | 1.299279871 |
| RP23-67P7.1 | 0.422515209 | 0.672054688 | 1.396402663 |
| Por | 973.1641497 | 0.668733782 | 0.101055964 |
| Gm16061 | 0.898502897 | 0.552663396 | 1.078302229 |
| Tmem120a | 119.540289 | 0.739352208 | 0.118667818 |
| Mir7034 | 0 | NA | NA |
| Styxl1 | 3.717676903 | -0.291294559 | 0.536222787 |
| Mdh2 | 1871.37566 | -0.076737256 | 0.05915125 |
| Srrm3 | 617.0965085 | -0.706359055 | 0.144949562 |
| Gm23268 | 0 | NA | NA |
| Srrm3os | 0.79881978 | 1.577814554 | 3.645582969 |
| Gm22450 | 0.295673938 | -0.245378918 | 1.619543012 |
| Hspb1 | 217.0812965 | 3.908890923 | 0.440271594 |
| RP23-11P12.2 | 1.422147357 | 0.758269173 | 1.416772974 |
| Ywhag | 12717.10653 | 0.056628641 | 0.068637197 |
| Ssc4d | 19.88563498 | 0.572692431 | 0.259224772 |
| Zp3 | 0.564341384 | -0.932726954 | 1.382114288 |
| Dtx2 | 165.5707377 | 0.903784554 | 0.133489228 |
| Upk3b | 0.665037802 | 1.427939278 | 1.358494439 |
| Upk3bl | 6.950616643 | 2.915383193 | 0.530625659 |
| Rasa4 | 403.5442416 | 2.478396764 | 0.136554391 |
| Gm22214 | 0 | NA | NA |
| Gm15701 | 0.196892959 | -0.24410536 | 2.233059386 |
| Mir7035 | 0 | NA | NA |
| Polr2j | 169.450244 | 0.046725286 | 0.102027136 |
| RP23-314O1.1 | 1.911784405 | 0.286086051 | 0.671527301 |
| Lrwd1 | 120.9503839 | -0.135624639 | 0.111245639 |
| Alkbh4 | 76.72992867 | 0.47560315 | 0.123817233 |
| Orai2 | 675.4374974 | -0.477854347 | 0.086694259 |
| Prkrip1 | 213.2039114 | 0.435224176 | 0.130714516 |
| RP23-284P20.2 | 0 | NA | NA |
| Sh2b2 | 93.36965878 | 0.551536754 | 0.160727305 |
| Cux1 | 1881.15298 | -0.266513173 | 0.07084103 |
| Gm16599 | 1.050839697 | -1.03180689 | 0.98793237 |
| A430110C17Rik | 51.6013035 | -0.847877866 | 0.306121837 |
| mmu-mir-721 | 2.23838086 | -1.416645878 | 0.77107097 |
| Gm38082 | 75.94798503 | -0.61627407 | 0.281592304 |
| Myl10 | 0.304278192 | 0.237298954 | 3.724945049 |
| Col26a1 | 25.30413262 | -1.150374359 | 0.224877449 |
| Gm20485 | 0.218750444 | 0.389044063 | 2.589750937 |
| RP24-456B4.2 | 0 | NA | NA |
| Ift22 | 390.4508874 | -0.057593594 | 0.085899342 |

| | | | |
|---|---|---|---|
| 4933404O12Rik | 126.181175 | -0.438606445 | 0.109011379 |
| Fis1 | 968.7424033 | 0.264341498 | 0.080194244 |
| Cldn15 | 7.096013574 | 1.262976409 | 0.502898637 |
| Znhit1 | 217.8120474 | 0.412364479 | 0.099489319 |
| Plod3 | 660.8862868 | 1.537981874 | 0.11117369 |
| Mir702 | 0 | NA | NA |
| Gm7284 | 0.113979814 | -0.243571806 | 3.724945049 |
| Gm26989 | 0 | NA | NA |
| Vgf | 2915.612919 | 0.937961478 | 0.163482338 |
| Ap1s1 | 1188.779745 | 0.241963073 | 0.070282087 |
| Serpine1 | 1051.265589 | 7.079140017 | 0.510540146 |
| Trim56 | 1329.516906 | 3.884545877 | 0.166795954 |
| Muc3 | 0.044544648 | 0.237309859 | 3.724945049 |
| Gm3054 | 0.061920613 | 0.718207094 | 3.724945049 |
| CAAA01077340.1 | 0.185747766 | -0.243562532 | 3.292273734 |
| CAAA01154573.1 | 3.791680059 | 1.347857756 | 0.570347806 |
| Muc3a | 21.091838 | -1.126750049 | 0.476185788 |
| WI1-166E14.1 | 15.34798663 | -1.321313003 | 0.320198946 |
| Ache | 1111.230768 | 0.229440149 | 0.105250983 |
| Mir8116 | 1.629432974 | -0.061623748 | 0.892771902 |
| Ufsp1 | 91.40840618 | 0.27791607 | 0.160961855 |
| Srrt | 944.0836702 | 0.170317238 | 0.074559443 |
| Mir7036 | 0 | NA | NA |
| RP24-210N1.1 | 0.752770407 | 1.732259435 | 1.232739481 |
| Trip6 | 103.7948532 | 1.610290189 | 0.213804694 |
| Slc12a9 | 407.3671883 | 0.468779217 | 0.085125914 |
| RP24-210N1.3 | 0.151841013 | -0.724459891 | 2.850049718 |
| Ephb4 | 94.20008279 | 1.065964567 | 0.149608947 |
| Zan | 8.749737854 | -0.385442262 | 0.400839504 |
| RP24-210N1.4 | 0 | NA | NA |
| Epo | 0.229076001 | -1.650624631 | 2.528990758 |
| Pop7 | 101.0875345 | 0.227313745 | 0.134374081 |
| RP23-45G4.2 | 6.33503719 | 0.205556626 | 0.403315661 |
| RP23-45G4.3 | 32.98247416 | -0.701569788 | 0.219329356 |
| Gigyf1 | 841.5272401 | -0.297659908 | 0.121993589 |
| Gnb2 | 2014.783637 | 0.325884526 | 0.062329384 |
| Mir6418 | 0 | NA | NA |
| Gm17112 | 0.842992085 | 0.227630884 | 0.970771965 |
| Actl6b | 409.7728831 | -0.823987655 | 0.110315977 |
| Tfr2 | 33.22771086 | -0.51093951 | 0.231708462 |
| Mospd3 | 114.8382337 | -0.080653497 | 0.110125208 |
| RP23-286M3.4 | 4.875430065 | 0.178839463 | 0.480866053 |
| Pcolce | 87.87249262 | 0.519317575 | 0.190221663 |
| Fbxo24 | 2.595989607 | -1.205583242 | 0.697643463 |
| RP23-286M3.5 | 4.924586763 | -0.33996093 | 0.451247814 |
| Gm16089 | 9.122415521 | 0.173498175 | 0.350203984 |
| Lrch4 | 4.483355963 | -0.954228854 | 0.61025194 |

| | | | |
|---|---|---|---|
| Gm20605 | 6.061787305 | -0.203225443 | 0.471884511 |
| Sap25 | 0.075043205 | 0.237309859 | 3.724945049 |
| Irs3 | 1.938977309 | -0.217934447 | 0.654695933 |
| Agfg2 | 460.5243141 | -0.098591085 | 0.078775049 |
| Gm15498 | 0.332718546 | 1.604354814 | 1.994073304 |
| Gm15753 | 9.170347486 | 2.76692412 | 0.45411184 |
| Nyap1 | 848.2957187 | -0.411270378 | 0.077508884 |
| Tsc22d4 | 1384.328407 | 0.966925034 | 0.145687258 |
| Ppp1r35 | 80.27309234 | 0.388965543 | 0.151510884 |
| Mepce | 500.0316796 | 0.007338809 | 0.071736001 |
| Zcwpw1 | 45.4661166 | -0.568868435 | 0.191644319 |
| Pilra | 8.121710499 | 2.329461958 | 0.482052223 |
| Pilrb1 | 1.522009338 | 4.005438147 | 1.087805585 |
| RP23-425N17.3 | 0.609942678 | 2.801473395 | 1.365333886 |
| Pilrb2 | 1.815499725 | 2.6244671 | 0.932793477 |
| RP23-425N17.5 | 0.143022002 | 0.719176055 | 3.547735711 |
| RP23-425N17.6 | 0 | NA | NA |
| RP23-131D6.3 | 0 | NA | NA |
| Cyp3a13 | 18.8351161 | -0.644159824 | 0.348595279 |
| Gm15502 | 0.499480636 | -0.345073865 | 1.433839641 |
| RP23-131D6.4 | 4.460458762 | 0.061737992 | 0.530622079 |
| Gjc3 | 574.2169864 | -0.317230015 | 0.130279062 |
| Azgp1 | 0.218824723 | -0.724459763 | 2.584608499 |
| Gm4963 | 0.586951709 | -0.223212645 | 1.285731445 |
| Gm29127 | 0 | NA | NA |
| Smok3a | 0.044544648 | 0.237309859 | 3.724945049 |
| Smok3b | 0 | NA | NA |
| Smok3c | 0 | NA | NA |
| Zkscan1 | 1601.280236 | -0.007585159 | 0.047847899 |
| RP23-147H4.1 | 0 | NA | NA |
| Zscan21 | 197.9428209 | 0.107833853 | 0.09233302 |
| Zfp113 | 154.778503 | 0.058633494 | 0.166618025 |
| Cops6 | 678.0499843 | 0.242751835 | 0.088784763 |
| Mcm7 | 172.4442146 | 0.947203456 | 0.143714956 |
| Mir25 | 0 | NA | NA |
| Mir93 | 0 | NA | NA |
| Mir106b | 0.072025462 | 0.237312084 | 3.724945049 |
| Ap4m1 | 196.8680612 | -0.068109306 | 0.112608232 |
| Taf6 | 295.6377165 | 0.192245942 | 0.104792772 |
| Cnpy4 | 86.00518766 | 0.077461041 | 0.140495599 |
| Mblac1 | 28.62003678 | 0.093775418 | 0.214310925 |
| Gm454 | 0 | NA | NA |
| RP24-86H20.4 | 0 | NA | NA |
| Gm22956 | 0 | NA | NA |
| Nxpe5 | 0.64031546 | 1.530048213 | 1.239014771 |
| Lamtor4 | 172.5096594 | 0.006878082 | 0.116492444 |
| BC037034 | 543.4878643 | -0.018507049 | 0.05790614 |

| | | | |
|---|---|---|---|
| 6330418K02Rik | 28.7450452 | 0.081389277 | 0.223556271 |
| Gal3st4 | 0 | NA | NA |
| Gpc2 | 31.36245639 | -0.484950389 | 0.214602692 |
| Stag3 | 27.52049443 | 0.149907016 | 0.272276402 |
| RP24-95J4.1 | 0.060174537 | 0.237309859 | 3.724945049 |
| Gm10874 | 0.888510269 | 1.303687082 | 1.144990584 |
| Got2-ps1 | 146.5220016 | 0.022584523 | 0.102654137 |
| Gm25525 | 0.061920613 | 0.718207094 | 3.724945049 |
| Zfp157 | 295.429195 | -0.169631188 | 0.09640213 |
| Gm5564 | 8.712984693 | 0.395430019 | 0.367039723 |
| RP24-255F19.2 | 1.435448476 | 0.429004391 | 0.821782623 |
| 1700123K08Rik | 2.221046269 | -0.212875016 | 0.899639365 |
| Zfp68 | 529.1369275 | -0.114395991 | 0.06736189 |
| Gm24082 | 0 | NA | NA |
| Gm15497 | 0.148708979 | 1.198497773 | 2.868467391 |
| RP24-255F19.5 | 0.796574019 | 3.320212197 | 1.332533424 |
| A430033K04Rik | 179.5683617 | -0.403748886 | 0.115491885 |
| Gm38232 | 0.132101749 | 1.209894437 | 3.72418822 |
| Fam20c | 570.0411155 | 0.185333127 | 0.096241759 |
| Gm5294 | 0.042716945 | -0.243571806 | 3.724945049 |
| Pdgfa | 337.774443 | 0.054425442 | 0.0835475 |
| 6330403L08Rik | 206.3643988 | -0.920563555 | 0.136084358 |
| Prkar1b | 4853.393054 | -0.330717126 | 0.083462289 |
| Gm15672 | 16.59824677 | -0.339740022 | 0.307351321 |
| Dnaaf5 | 128.0720811 | 0.9191538 | 0.118648684 |
| Sun1 | 919.0838976 | -0.426737503 | 0.108388122 |
| RP23-430G6.3 | 0.066050875 | 0.71820487 | 3.724945049 |
| Get4 | 323.1872879 | 0.535305073 | 0.114052457 |
| Adap1 | 1303.85996 | -0.413363775 | 0.069135916 |
| Gm27316 | 0 | NA | NA |
| Cox19 | 240.9171869 | -0.176056708 | 0.076782237 |
| Cyp2w1 | 0 | NA | NA |
| 3110082I17Rik | 68.15138838 | 0.268182636 | 0.125216711 |
| Mir339 | 1.20342686 | -0.35133556 | 1.030124996 |
| Gpr146 | 293.4623801 | 0.437810131 | 0.097529887 |
| C130050O18Rik | 9.082211208 | 2.140464463 | 0.42532903 |
| Gper1 | 14.62948997 | -0.10916103 | 0.319206174 |
| Gm26285 | 0.142244466 | 1.482178313 | 3.552423338 |
| Zfand2a | 664.2777979 | 0.69110531 | 0.099621381 |
| RP23-400D9.3 | 0.170157162 | 0.237324706 | 2.749797554 |
| Uncx | 53.72924227 | -0.339703132 | 0.194429878 |
| RP23-92L4.4 | 0 | NA | NA |
| Micall2 | 117.2123074 | 2.401786243 | 0.291299682 |
| Ints1 | 1337.189342 | -0.173918761 | 0.055927415 |
| Mir7037 | 0 | NA | NA |
| Mafk | 342.8849901 | 2.397437206 | 0.205759209 |
| Tmem184a | 2.905166453 | 1.536300285 | 0.652019383 |

| | | | |
|---|---|---|---|
| Gm26938 | 0.061920613 | 0.718207094 | 3.724945049 |
| Psmg3 | 54.0547785 | 0.474598424 | 0.159025197 |
| Gm24146 | 2.128603593 | 1.136231881 | 0.770693495 |
| RP23-168E11.1 | 0.493577707 | -0.97703476 | 1.409664446 |
| Elfn1 | 266.0295 | -0.4648157 | 0.132948056 |
| RP23-31L10.2 | 0 | NA | NA |
| Mad1l1 | 200.5910728 | 0.008991899 | 0.099041173 |
| RP23-31L10.4 | 9.073372328 | -1.060370584 | 0.346535013 |
| Gm16122 | 0.099106094 | 1.209894437 | 3.72418822 |
| Gm16121 | 0.641775973 | 0.275983974 | 1.480687144 |
| RP24-97G6.1 | 10.02733677 | -1.318795829 | 0.373581918 |
| Gm16120 | 0.194043089 | 0.194157179 | 2.227684541 |
| RP23-307I19.1 | 0.437638936 | 0.670950678 | 1.884513209 |
| Nudt1 | 23.04760792 | 0.543771385 | 0.252390735 |
| Ftsj2 | 69.98359942 | -0.002406808 | 0.14972993 |
| Snx8 | 233.8617382 | 0.974100034 | 0.126703725 |
| Eif3b | 1148.304761 | 0.804672873 | 0.083525285 |
| Mir7038 | 0.519085648 | -1.8925155 | 1.428934234 |
| RP23-307I19.6 | 0 | NA | NA |
| RP23-307I19.5 | 0.947788321 | 1.16330757 | 1.050071431 |
| Chst12 | 156.329386 | -0.116496351 | 0.105746172 |
| RP23-462C22.2 | 1.646628032 | 2.899578242 | 1.012368422 |
| RP23-462C22.1 | 0.174406792 | 0.718206751 | 3.367085639 |
| Grifin | 1.541974883 | 0.867405075 | 0.762246432 |
| RP23-462C22.5 | 1.34756889 | -0.962590657 | 1.066402504 |
| Lfng | 314.5144813 | 0.639451931 | 0.174514378 |
| Ttyh3 | 2824.785642 | -0.242311203 | 0.092795787 |
| Iqce | 386.6329369 | -0.513029621 | 0.141961639 |
| Brat1 | 216.0259619 | 0.03857594 | 0.109813856 |
| Amz1 | 137.7511797 | -0.651818828 | 0.15811633 |
| Gna12 | 984.1548664 | -0.065243571 | 0.118154673 |
| Card11 | 40.90469028 | 5.213576662 | 0.443307779 |
| AC161570.1 | 0 | NA | NA |
| Gm26113 | 0 | NA | NA |
| Sdk1 | 231.3078279 | -0.260368244 | 0.09871356 |
| Gm6433 | 1.643128883 | 0.269780129 | 0.876034006 |
| RP23-49G7.2 | 0 | NA | NA |
| RP23-286C5.1 | 0 | NA | NA |
| RP23-286C5.2 | 0.061920613 | 0.718207094 | 3.724945049 |
| Gm16035 | 0 | NA | NA |
| Gm24131 | 0 | NA | NA |
| Gm16036 | 0 | NA | NA |
| Gm26970 | 0 | NA | NA |
| Foxk1 | 1176.476401 | -0.037355098 | 0.075973157 |
| Ap5z1 | 229.1563663 | 0.172710046 | 0.114187505 |
| Radil | 157.9503125 | -0.688572078 | 0.129680514 |
| Papolb | 13.6158541 | -1.111235217 | 0.330042631 |

| | | | |
|---|---|---|---|
| RP23-106N8.2 | 0 | NA | NA |
| Mmd2 | 1010.130574 | -0.704225099 | 0.156871305 |
| RP23-106N8.4 | 0 | NA | NA |
| Wipi2 | 1453.522254 | 0.010327104 | 0.058180656 |
| Slc29a4 | 200.9815486 | -0.372564139 | 0.117819209 |
| Tnrc18 | 2698.838952 | 0.027129677 | 0.066827616 |
| Gm15770 | 0.972120366 | 2.933631562 | 1.266243838 |
| Fbxl18 | 720.5681582 | -0.470401335 | 0.071573072 |
| D430018E03Rik | 19.83389332 | -1.535543005 | 0.329805706 |
| Actb | 25647.77243 | 1.001316147 | 0.113812605 |
| Fscn1 | 1883.804362 | -0.115905079 | 0.08730472 |
| Rnf216 | 737.7813341 | 0.00334663 | 0.046367899 |
| RP23-393G8.5 | 4.643209367 | -0.603386658 | 0.467263613 |
| RP23-393G8.10 | 0.927313021 | -0.011904561 | 1.073578627 |
| Olfr718-ps1 | 0 | NA | NA |
| RP23-301L9.8 | 0.114443069 | 0.23732098 | 3.724945049 |
| AC123686.1 | 0 | NA | NA |
| Rbakdn | 0.061920613 | 0.718207094 | 3.724945049 |
| Rbak | 178.0893661 | -0.140214326 | 0.142314438 |
| 4933411G11Rik | 0.556792772 | 0.012518188 | 1.305305275 |
| Spdyb | 0.075043205 | 0.237309859 | 3.724945049 |
| 4930448H16Rik | 0 | NA | NA |
| Zfp12 | 444.4098317 | -0.038783332 | 0.09350452 |
| Zfp316 | 378.1393857 | -0.243318472 | 0.099961863 |
| RP23-301L9.3 | 37.38971463 | -0.740885328 | 0.288129118 |
| Zfp853 | 32.64981703 | -1.40565107 | 0.223959402 |
| E130309D02Rik | 214.9676355 | 0.210577892 | 0.106283616 |
| RP23-301L9.6 | 1.497168834 | -0.475842201 | 0.952335993 |
| Zdhhc4 | 177.8279622 | 0.07290405 | 0.121525507 |
| 0610040B10Rik | 20.77576342 | 0.004312208 | 0.308910131 |
| RP23-301L9.4 | 0.904055358 | -1.35718286 | 1.328796578 |
| Grid2ip | 247.2010809 | -0.816100223 | 0.22274434 |
| Gm20635 | 4.77646696 | -0.610514732 | 0.50002558 |
| RP23-372N19.4 | 0.76879839 | -1.850790264 | 1.177354929 |
| RP23-372N19.2 | 152.8195759 | -0.853471106 | 0.217578272 |
| 9530056K15Rik | 3.981355527 | -1.424237888 | 0.571847527 |
| Kdelr2 | 375.7129238 | 0.800769705 | 0.128108894 |
| RP23-158A20.2 | 0 | NA | NA |
| RP23-158A20.4 | 0 | NA | NA |
| Daglb | 512.0781547 | -0.111532829 | 0.078820421 |
| Rac1 | 2571.473266 | 0.299374648 | 0.076853951 |
| RP23-158A20.7 | 0.049553047 | 0.71820487 | 3.724945049 |
| Fam220a | 378.2842679 | -0.011281691 | 0.081659809 |
| Cyth3 | 478.6545393 | -0.091147239 | 0.073357704 |
| Usp42 | 463.5942125 | 0.326344994 | 0.069951601 |
| D130017N08Rik | 140.3508044 | -0.745916435 | 0.153378519 |
| RP23-151C5.2 | 0 | NA | NA |

| | | | |
|---|---|---|---|
| RP23-151C5.3 | 4.718276104 | -1.437945321 | 0.560704096 |
| Eif2ak1 | 746.7429788 | 0.185974665 | 0.061498609 |
| Ankrd61 | 4.124592769 | -0.4417297 | 0.470894134 |
| Aimp2 | 134.5489307 | 0.318354492 | 0.121722898 |
| Pms2 | 217.1478713 | -0.607065542 | 0.104253933 |
| Gm17135 | 0.441552168 | -1.196895079 | 1.470247765 |
| PMS2 | 0.78876463 | 0.235391087 | 1.224649232 |
| Rsph10b | 55.58878597 | -0.692244677 | 0.19189462 |
| Ccz1 | 450.6103287 | 0.50791801 | 0.067106546 |
| Gm24311 | 0 | NA | NA |
| Ocm | 1.753517396 | 0.904139118 | 0.794791598 |
| RP24-194F23.5 | 0.296409788 | -0.751325329 | 3.719653481 |
| Lmtk2 | 3277.074586 | -0.328543858 | 0.067737481 |
| Gm26925 | 0 | NA | NA |
| Bhlha15 | 4.240558544 | 1.181364287 | 0.627999906 |
| Tecpr1 | 1209.265373 | -0.548978654 | 0.067214219 |
| RP24-82B23.5 | 8.247112194 | -0.979984553 | 0.353807614 |
| 2900089D17Rik | 4.905083075 | -1.366265166 | 0.461365465 |
| Bri3 | 199.49671 | 0.782780618 | 0.126186736 |
| Baiap2l1 | 221.2967602 | -0.739023454 | 0.125554212 |
| Gm15708 | 7.40733758 | 0.56342266 | 0.425753802 |
| RP24-182D2.1 | 0.265735694 | 1.329573413 | 3.714216621 |
| Gm15709 | 0 | NA | NA |
| Nptx2 | 612.3205461 | 1.940431803 | 0.316889972 |
| Gm6272 | 0.315729261 | 0.233534293 | 1.609531157 |
| Tmem130 | 1967.351054 | -0.270694097 | 0.08816136 |
| Trrap | 3060.283518 | -0.317102554 | 0.076647952 |
| Gm23995 | 1.477838792 | -0.900642346 | 0.829153928 |
| Smurf1 | 700.542789 | 0.435559193 | 0.068051055 |
| Mir7039 | 0.22969359 | 1.166625474 | 3.129888948 |
| Kpna7 | 0.7084488 | 1.066162179 | 1.332868569 |
| RP23-424M12.7 | 0 | NA | NA |
| Gm4871 | 0 | NA | NA |
| RP23-424M12.4 | 0 | NA | NA |
| RP23-424M12.5 | 0.075043205 | 0.237309859 | 3.724945049 |
| 1700018F24Rik | 0.068097359 | -0.243569582 | 3.724945049 |
| Arpc1a | 1136.742611 | 0.136764217 | 0.068550239 |
| Arpc1b | 585.2155934 | 3.299545198 | 0.16618163 |
| Pdap1 | 676.5616892 | 0.322865573 | 0.131552957 |
| Bud31 | 300.4083312 | 0.63610447 | 0.089101112 |
| Ptcd1 | 337.0823285 | 0.504261971 | 0.079768475 |
| Cpsf4 | 256.7055373 | 0.043977382 | 0.091304619 |
| Atp5j2 | 504.0996134 | -0.089801169 | 0.242973291 |
| Zkscan14 | 94.39392942 | 0.130301063 | 0.121263332 |
| Zkscan5 | 178.3410506 | 0.155644121 | 0.123575801 |
| Zfp655 | 776.8726219 | 0.562111123 | 0.099128146 |
| RP23-60E24.5 | 0 | NA | NA |

| | | | |
|---|---|---|---|
| Zscan25 | 198.2716498 | 0.382775077 | 0.163734239 |
| RP23-60E24.3 | 23.00508971 | 0.513733006 | 0.431217527 |
| Cyp3a57 | 13.25184546 | 4.226697638 | 0.595724434 |
| RP24-443E20.1 | 0.066050875 | 0.71820487 | 3.724945049 |
| RP24-443E20.2 | 0.066050875 | 0.71820487 | 3.724945049 |
| Cyp3a16 | 0 | NA | NA |
| RP24-127J3.1 | 0.120974994 | -0.243571806 | 3.724945049 |
| Cyp3a41b | 0 | NA | NA |
| RP23-379D24.1 | 0 | NA | NA |
| RP23-379D24.2 | 0 | NA | NA |
| RP24-443E20.4 | 0 | NA | NA |
| Cyp3a41a | 0 | NA | NA |
| RP23-102C14.1 | 0 | NA | NA |
| Cyp3a44 | 0 | NA | NA |
| RP23-102C14.2 | 0.060174537 | 0.237309859 | 3.724945049 |
| Cyp3a11 | 0.492793603 | -1.97052752 | 1.507541056 |
| RP23-161K8.1 | 0 | NA | NA |
| RP23-161K8.2 | 0 | NA | NA |
| RP23-161K8.3 | 0 | NA | NA |
| Cyp3a25 | 0.261272317 | 1.879600272 | 2.011158969 |
| RP23-53G23.1 | 0 | NA | NA |
| RP23-53G23.2 | 0 | NA | NA |
| RP23-53G23.3 | 0 | NA | NA |
| Cyp3a59 | 0 | NA | NA |
| RP23-441I24.9 | 0.282778275 | 1.205901682 | 2.008574658 |
| Gm6309 | 0.038981395 | 0.718207094 | 3.724945049 |
| 1700001J03Rik | 1.161484436 | 0.842431311 | 1.072990273 |
| RP23-441I24.5 | 22.34345423 | -0.12545006 | 0.50626639 |
| RP23-441I24.4 | 2.948678684 | 0.452363956 | 0.697153358 |
| RP23-441I24.3 | 9.915980733 | -0.285240867 | 0.438797774 |
| Rnf6 | 700.4219294 | 0.071504059 | 0.068707894 |
| Cdk8 | 479.7397194 | -0.375671764 | 0.105081605 |
| Gm15739 | 20.56479697 | -0.773233357 | 0.279939771 |
| Wasf3 | 1567.401421 | -0.499308256 | 0.074210186 |
| Gm6408 | 0.779579221 | 0.972078678 | 1.249419401 |
| Gm6370 | 0.038981395 | 0.718207094 | 3.724945049 |
| 4930449I24Rik | 0 | NA | NA |
| Gm3402 | 0 | NA | NA |
| Gm3404 | 0 | NA | NA |
| Gm3409 | 0 | NA | NA |
| Gm3415 | 0 | NA | NA |
| Gm8494 | 20.48103304 | 0.728969383 | 0.228917079 |
| Gpr12 | 122.0110685 | -0.8764972 | 0.161562909 |
| RP23-134K6.2 | 0.089089296 | 0.237305797 | 3.724945049 |
| RP23-134K6.3 | 0 | NA | NA |
| Usp12 | 869.2011401 | 0.445855165 | 0.070764329 |
| RP24-126A19.2 | 0.044191549 | -0.243571806 | 3.724945049 |

| | | | |
|---|---|---|---|
| Rpl21 | 82.06727034 | 0.23857557 | 0.141593511 |
| Gm25091 | 0 | NA | NA |
| Gm23202 | 0.066050875 | 0.71820487 | 3.724945049 |
| Rasl11a | 102.228073 | 2.192219118 | 0.307199282 |
| Gtf3a | 110.5136172 | 0.074275789 | 0.125325078 |
| Mtif3 | 139.0778404 | 0.30026625 | 0.094953343 |
| Lnx2 | 154.2635579 | 0.26448588 | 0.127030733 |
| Polr1d | 179.4983994 | 0.277737816 | 0.122663769 |
| Rps10-ps4 | 0.048949532 | -0.243569582 | 3.724945049 |
| Gsx1 | 3.152922583 | -0.017373564 | 0.543622072 |
| Gm24556 | 0 | NA | NA |
| 2210019I11Rik | 0 | NA | NA |
| Pdx1 | 0.046345709 | 0.71820487 | 3.724945049 |
| Gm27033 | 0 | NA | NA |
| Mir7k | 0 | NA | NA |
| Cdx2 | 0 | NA | NA |
| Urad | 0 | NA | NA |
| RP24-510G5.6 | 0 | NA | NA |
| Flt3 | 155.671954 | -0.721604035 | 0.1531579 |
| Gm26597 | 1.472170236 | -0.950830576 | 0.878154465 |
| Gm6054 | 0.773509025 | -0.224318596 | 1.360343237 |
| RP24-544J6.3 | 0.075043205 | 0.237309859 | 3.724945049 |
| RP24-544J6.4 | 1.088238703 | 0.232358968 | 0.849529959 |
| RP24-544J6.5 | 2.007455813 | -2.640133857 | 0.957952001 |
| RP24-544J6.6 | 0.135435006 | 0.718198586 | 3.635348467 |
| Pan3 | 1329.249898 | -0.316824632 | 0.1004311 |
| RP24-544J6.7 | 8.762375666 | -1.594714643 | 0.485693186 |
| Flt1 | 324.459667 | 1.530410897 | 0.198164762 |
| RP23-270L23.1 | 0.088534442 | 1.179809211 | 3.723500796 |
| RP23-270L23.2 | 0 | NA | NA |
| Pomp | 809.6499027 | 0.871251549 | 0.094718139 |
| RP23-270L23.4 | 0.636753058 | -1.271726555 | 1.253523779 |
| Slc46a3 | 67.21071847 | 0.195709066 | 0.144585669 |
| Gm15721 | 25.84402214 | -0.768221 | 0.228302769 |
| Mtus2 | 777.696009 | -0.812419703 | 0.108752556 |
| Gm10167 | 12.31383881 | 0.244613754 | 0.28870245 |
| RP24-71D19.1 | 4.780781177 | -1.607427121 | 0.622449527 |
| Slc7a1 | 875.5929286 | 0.989766913 | 0.16224731 |
| RP24-307I24.1 | 0.318558491 | 0.467825204 | 1.712942493 |
| Ubl3 | 1852.018664 | 0.008608162 | 0.059309513 |
| RP24-61O18.3 | 0 | NA | NA |
| RP24-61O18.1 | 0.068097359 | -0.243569582 | 3.724945049 |
| RP24-61O18.2 | 0 | NA | NA |
| 2210417A02Rik | 0 | NA | NA |
| n-R5s179 | 0 | NA | NA |
| 4930505K14Rik | 0 | NA | NA |
| Gm15407 | 0.046345709 | 0.71820487 | 3.724945049 |

| | | | |
|---|---|---|---|
| Katnal1 | 1032.752 | -0.113401857 | 0.067669711 |
| RP23-188F20.1 | 0 | NA | NA |
| Gm15410 | 0.390771115 | 0.689409592 | 1.52630587 |
| Gm15406 | 5.088172819 | 0.67640045 | 0.476786536 |
| 8430423G03Rik | 0.143485256 | 1.229961212 | 3.547405499 |
| Gm15409 | 0.354937333 | 0.237330648 | 1.69222271 |
| RP23-134H19.3 | 52.59786348 | 0.483102134 | 0.194855986 |
| Gm15411 | 7.066285288 | -0.466283631 | 0.457932672 |
| Gm15408 | 2.719793829 | 1.795021506 | 0.731347303 |
| Hmgb1 | 340.808951 | 0.119324785 | 0.101726958 |
| 5730422E09Rik | 12.70416494 | -0.520750409 | 0.363741887 |
| RP23-408A1.6 | 1.653178163 | -0.851411154 | 0.865488104 |
| RP23-408A1.4 | 3.467320147 | 3.165383489 | 0.86385166 |
| Gm23362 | 0 | NA | NA |
| Gm8615 | 17.8882258 | 0.002402944 | 0.239770976 |
| RP23-408A1.8 | 0.278466489 | 0.237330646 | 1.810174079 |
| Uspl1 | 565.005295 | 0.075335556 | 0.094072853 |
| 2310047D07Rik | 2.953893651 | -0.913349617 | 0.610242568 |
| Gm29264 | 0.433555451 | -0.272983531 | 1.469001176 |
| AC122546.1 | 0 | NA | NA |
| Gm24105 | 0 | NA | NA |
| Alox5ap | 91.57756619 | 3.452548952 | 0.242671081 |
| n-R5s180 | 0 | NA | NA |
| WI1-1993I8.1 | 5.494432944 | 0.811769628 | 0.459632716 |
| BC028471 | 1.239154484 | -0.223132702 | 0.937415913 |
| Gm20488 | 0.478946765 | -0.178738747 | 1.532705023 |
| RP23-375E13.2 | 0.173926504 | 0.701546944 | 2.308837118 |
| Medag | 75.60539664 | 0.889569865 | 0.210619886 |
| RP23-375E13.4 | 0 | NA | NA |
| RP23-375E13.3 | 0.045882455 | 0.237312084 | 3.724945049 |
| Tex26 | 5.173350472 | 0.833309037 | 0.52473705 |
| RP23-375E13.6 | 1.443591694 | 1.773482997 | 0.923476994 |
| Gm15997 | 2.38686755 | -1.498149105 | 0.686003392 |
| Gm2566 | 0.852004391 | -0.933792975 | 1.030939453 |
| Wdr95 | 6.751900483 | 0.212952222 | 0.417062258 |
| Hsph1 | 3279.88901 | 0.506365344 | 0.228690574 |
| RP23-256B18.1 | 1.058337859 | 2.270222141 | 1.157111014 |
| Gm5566 | 1.331862033 | 2.467589418 | 0.997981416 |
| B3glct | 236.8677375 | -0.248856261 | 0.110137025 |
| RP23-346P15.1 | 0 | NA | NA |
| RP23-346P15.3 | 0.483795822 | 1.488501023 | 1.633959963 |
| RP23-346P15.4 | 0 | NA | NA |
| RP23-337K7.2 | 0 | NA | NA |
| RP23-337K7.1 | 0.29963536 | 0.580441052 | 1.572043422 |
| RP23-337K7.3 | 0 | NA | NA |
| Rxfp2 | 8.210362686 | 0.130810167 | 0.338894052 |
| RP23-301H20.1 | 0 | NA | NA |

| | | | |
|---|---|---|---|
| RP23-301H20.2 | 0 | NA | NA |
| Fry | 3743.235337 | -0.821977131 | 0.079231483 |
| RP23-327P24.3 | 2.277130459 | -2.465898393 | 0.857887424 |
| RP23-327P24.2 | 0 | NA | NA |
| RP24-333I1.2 | 4.567880012 | -1.486941025 | 0.529918879 |
| RP24-333I1.3 | 1.611190043 | -0.585879824 | 0.745695071 |
| Zar1l | 2.866525085 | -0.870770358 | 0.581866772 |
| Gm15933 | 0.686705509 | -0.820995383 | 1.134937857 |
| Brca2 | 165.9286918 | -0.220828445 | 0.169913584 |
| Gm25284 | 0.075043205 | 0.237309859 | 3.724945049 |
| Gm23641 | 0.146089079 | 0.238291249 | 3.522464378 |
| RP24-333I1.6 | 1.253964362 | -1.014699775 | 0.869237567 |
| RP24-333I1.5 | 4.525610808 | -1.320058064 | 0.494754216 |
| RP24-333I1.7 | 13.83704783 | -0.195948966 | 0.290387933 |
| N4bp2l1 | 393.7981301 | -0.36681766 | 0.09645744 |
| RP23-34P11.1 | 29.55790751 | -0.661789917 | 0.388604576 |
| RP23-34P11.2 | 251.9245356 | -0.267591521 | 0.170784852 |
| N4bp2l2 | 1453.675956 | -0.086039248 | 0.081087281 |
| Pds5b | 2134.751181 | -0.307491592 | 0.07033915 |
| Gm24630 | 4.821230019 | -1.602097175 | 0.546434704 |
| Gm26013 | 0.049553047 | 0.71820487 | 3.724945049 |
| RP23-336F4.1 | 29.09809687 | -0.612538272 | 0.269316762 |
| RP23-336F4.4 | 4.80270818 | 0.094415865 | 0.481987846 |
| RP23-336F4.5 | 0 | NA | NA |
| Kl | 209.3353533 | -1.229422483 | 0.217766574 |
| RP23-412C8.2 | 0.322046281 | 2.042580287 | 1.727691993 |
| Stard13 | 272.414666 | 0.408984482 | 0.082584277 |
| RP24-155K17.1 | 0.139883403 | 1.411723814 | 3.551352091 |
| RP24-343A12.4 | 0 | NA | NA |
| RP24-343A12.3 | 0 | NA | NA |
| D730045B01Rik | 4.778829002 | -0.597926776 | 0.522917648 |
| 1700028E10Rik | 23.33760472 | -0.362152606 | 0.221145403 |
| Vmn2r-ps25 | 0.141684195 | 0.718205511 | 3.558854591 |
| Vmn2r-ps26 | 0.233449105 | 0.718219013 | 2.502791986 |
| Vmn2r18 | 0.476067163 | 2.7075336 | 1.583393202 |
| RP24-211L21.1 | 0.060174537 | 0.237309859 | 3.724945049 |
| Rfc3 | 87.13724546 | 1.501458238 | 0.197877557 |
| RP24-211L21.2 | 0 | NA | NA |
| RP24-211L21.3 | 0.154433973 | -0.245012343 | 3.500771551 |
| Gm24242 | 0.09689823 | -0.724453471 | 3.724945049 |
| Gm26463 | 0 | NA | NA |
| Gm23551 | 0 | NA | NA |
| Vmn1r-ps6 | 0 | NA | NA |
| Vmn1r-ps7 | 0.046345709 | 0.71820487 | 3.724945049 |
| RP24-328P2.7 | 0.256207099 | -1.167285757 | 2.111180689 |
| RP24-328P2.8 | 0.219080626 | 0.260757134 | 2.168083021 |
| RP23-368D12.2 | 0 | NA | NA |

| | | | |
|---|---|---|---|
| RP23-368D12.3 | 0 | NA | NA |
| RP23-258F1.1 | 0.066050875 | 0.71820487 | 3.724945049 |
| Gm18222 | 0 | NA | NA |
| RP23-253G12.8 | 136.7192892 | 5.525702955 | 0.33999303 |
| RP24-328P2.5 | 839.9270636 | 6.030526198 | 0.200153463 |
| RP23-253G12.10 | 0.128477747 | 0.718203484 | 3.677131942 |
| RP23-253G12.9 | 311.8525769 | 5.671271575 | 0.252410977 |
| Samd9l | 3774.434653 | 6.962041593 | 0.206671421 |
| Hepacam2 | 3.625877892 | 0.046499838 | 0.488126197 |
| Ccdc132 | 896.7331518 | -0.07135406 | 0.075305031 |
| RP23-318M11.2 | 26.8210992 | -0.357985274 | 0.351391011 |
| Calcr | 25.19477301 | 0.066027611 | 0.255690545 |
| Mir653 | 0 | NA | NA |
| Mir489 | 0.044191549 | -0.243571806 | 3.724945049 |
| Tfpi2 | 9.186610503 | 6.64715006 | 0.906507093 |
| Gngt1 | 0 | NA | NA |
| Gng11 | 162.5242613 | 1.390468526 | 0.178383647 |
| Bet1 | 205.6506794 | 0.382781977 | 0.130051013 |
| Gm8602 | 0 | NA | NA |
| Gm24442 | 0 | NA | NA |
| Gm37883 | 0 | NA | NA |
| Col1a2 | 642.3682694 | 1.158183372 | 0.135087061 |
| Gm26696 | 3.592646241 | -0.68501221 | 0.547702314 |
| Casd1 | 867.3601419 | -0.349695094 | 0.086231021 |
| Sgce | 237.5338859 | -0.137318201 | 0.115251676 |
| Peg10 | 1587.476363 | 1.173944076 | 0.220671161 |
| Gm20714 | 0 | NA | NA |
| Gm26828 | 0 | NA | NA |
| Gm9835 | 0.856801049 | 2.868293372 | 1.143769282 |
| Ppp1r9a | 4137.715075 | -0.745681627 | 0.104433764 |
| 2210017G18Rik | 30.57888805 | -1.260768291 | 0.345699437 |
| Gm26814 | 84.80524745 | -0.855105754 | 0.275737724 |
| Gm26736 | 22.89958731 | -1.401642953 | 0.350091043 |
| Gm26556 | 14.27493484 | -0.988410292 | 0.307238335 |
| Pon1 | 0.759396657 | -1.434742624 | 1.110433247 |
| Pon3 | 56.36395702 | 0.538156664 | 0.228224477 |
| Pon2 | 309.9460515 | 0.115299796 | 0.110117889 |
| Asb4 | 38.68988062 | 1.083870786 | 0.284391186 |
| Pdk4 | 314.1644295 | 0.713133715 | 0.189890667 |
| Dync1i1 | 1162.518359 | -0.563313621 | 0.061630221 |
| 1700019G24Rik | 1.274692649 | 1.541812609 | 0.925823064 |
| Slc25a13 | 101.7915475 | 0.013181963 | 0.176375452 |
| Gm20619 | 0.313425824 | 1.206230349 | 2.046332542 |
| Gm8649 | 0 | NA | NA |
| Gm27018 | 0 | NA | NA |
| Gm8652 | 0.294205253 | 0.669168049 | 1.610483038 |
| Gm20617 | 0.379053103 | 1.191543876 | 1.708025176 |

| | | | | |
|---|---|---|---|---|
| Gm20685 | 0 | NA | | NA |
| Shfm1 | 204.0907547 | | 0.594649199 | 0.248971475 |
| Gm20618 | 0.47603017 | | -0.948691116 | 1.305208181 |
| Gm18006 | 0 | NA | | NA |
| Gm5873 | 4.742073203 | | 0.20707932 | 0.455085836 |
| Dlx6os1 | 337.2803174 | | -1.180140808 | 0.262310952 |
| Gm27699 | 0.095295241 | | 0.23732098 | 3.724945049 |
| Gm27905 | 0 | NA | | NA |
| Dlx6os3 | 6.490635638 | | -1.559910124 | 0.474313162 |
| Dlx6 | 64.91335236 | | -0.566526393 | 0.202831923 |
| Dlx6os2 | 8.021264403 | | -1.369194522 | 0.519488572 |
| Dlx5 | 52.02918498 | | -0.534526335 | 0.21156676 |
| Gm22155 | 0 | NA | | NA |
| Gm26774 | 0.048949532 | | -0.243569582 | 3.724945049 |
| Sdhaf3 | 139.5460954 | | -0.490279315 | 0.112862041 |
| Gm27190 | 0 | NA | | NA |
| Gm8669 | 0 | NA | | NA |
| Gm19272 | 0.25372768 | | 1.240871483 | 2.070088427 |
| Tac1 | 258.728839 | | -0.653344045 | 0.151846976 |
| Gm8676 | 0 | NA | | NA |
| Asns | 800.622713 | | 0.405082581 | 0.089711524 |
| C1galt1 | 215.3562405 | | 0.494838738 | 0.121090948 |
| Gm22276 | 0.301804762 | | -0.153660576 | 2.014459629 |
| Gm9825 | 372.9565206 | | 0.238065649 | 0.085173079 |
| Col28a1 | 8.368185802 | | -0.770798409 | 0.388464908 |
| Mios | 475.6118824 | | -0.106705472 | 0.06921817 |
| Rpa3 | 69.13831145 | | 0.004156874 | 0.143118582 |
| Umad1 | 363.7124902 | | -0.329079331 | 0.079547841 |
| Gm16055 | 0.775287707 | | -1.422087006 | 1.42137716 |
| Gm8428 | 0 | NA | | NA |
| Gm16043 | 0.136120942 | | 1.15080397 | 3.595683408 |
| Gm24059 | 4.143222737 | | -0.918653359 | 0.541839567 |
| A430035B10Rik | 32.4009088 | | -0.429266543 | 0.250244352 |
| Gm26075 | 0.250670796 | | -0.734330346 | 2.480260474 |
| Glcci1 | 81.25189137 | | -0.697727962 | 0.131306671 |
| Gm16042 | 0.476187739 | | 2.108310498 | 1.666193974 |
| Ica1 | 575.9737311 | | -0.45103995 | 0.123049124 |
| Nxph1 | 529.823522 | | -0.403608684 | 0.076308057 |
| Gm26058 | 0.921802213 | | -1.046601265 | 0.935451695 |
| Gm27894 | 0 | NA | | NA |
| Ndufa4 | 1081.345019 | | -0.377114507 | 0.140893311 |
| Phf14 | 900.5267977 | | -0.237836765 | 0.063946839 |
| Gm6578 | 0.045882455 | | 0.237312084 | 3.724945049 |
| Thsd7a | 1278.572731 | | -0.436884326 | 0.057443583 |
| Gm16123 | 0 | NA | | NA |
| n-R5s160 | 0 | NA | | NA |
| Gm15529 | 0 | NA | | NA |

| | | | |
|---|---|---|---|
| Gm5300 | 0.044544648 | 0.237309859 | 3.724945049 |
| Tmem106b | 2433.505759 | -0.400593845 | 0.076726304 |
| Gm24985 | 0 | NA | NA |
| Vwde | 1.568186543 | 2.331585813 | 1.075555118 |
| 1700016P04Rik | 0.132574648 | -1.082845688 | 3.713921268 |
| Gm25737 | 0.154696864 | 0.237325224 | 2.833481316 |
| Tmem168 | 343.227461 | 0.525663368 | 0.095568887 |
| B630005N14Rik | 527.4785247 | 0.107660195 | 0.09063781 |
| Gpr85 | 431.2033423 | 0.069630631 | 0.08941514 |
| 2610001J05Rik | 197.506035 | 0.407769711 | 0.105246957 |
| 1110019D14Rik | 33.23543665 | -0.287609161 | 0.208056344 |
| Gm23760 | 0.097899064 | -0.724453471 | 3.724945049 |
| Gm23013 | 0 | NA | NA |
| Ppp1r3a | 6.253343815 | -1.036098112 | 0.44783034 |
| Foxp2 | 446.2894329 | -0.662624763 | 0.09096696 |
| Gm25470 | 1.281282525 | -0.468351131 | 1.1473963 |
| Mdfic | 320.1052487 | 3.557167129 | 0.232005264 |
| Gm29589 | 0 | NA | NA |
| RP23-244M15.1 | 0.212513282 | 0.237311278 | 3.175662942 |
| Tfec | 15.16942446 | 5.065412716 | 0.720505647 |
| Gm5721 | 0 | NA | NA |
| Gm15473 | 0.646370819 | 2.216926232 | 1.467051365 |
| Gm4876 | 26.61523984 | 0.349764384 | 0.223124876 |
| Tes | 60.86189865 | 2.744167226 | 0.253406401 |
| D830026I12Rik | 0 | NA | NA |
| Gm15581 | 0.736762124 | -2.323364194 | 1.452811265 |
| Cav2 | 214.4350625 | 0.7150065 | 0.139926348 |
| Cav1 | 153.6111711 | 1.726742547 | 0.16530715 |
| Gm29591 | 0 | NA | NA |
| Gm25021 | 0 | NA | NA |
| Met | 239.4772955 | 1.853722335 | 0.17098432 |
| Capza2 | 2658.372187 | 0.502968546 | 0.079694531 |
| St7 | 410.6774712 | -0.366482403 | 0.111928128 |
| Gm27755 | 0 | NA | NA |
| Gm27762 | 0 | NA | NA |
| Gm27960 | 2.550499828 | 0.299333708 | 0.605146794 |
| Gm27365 | 1.826179369 | -0.599643794 | 0.706102127 |
| Gm27542 | 1.845381307 | -0.588090804 | 0.742169252 |
| Gm28008 | 1.623521978 | 0.06212117 | 0.730345739 |
| Gm13898 | 0.097139547 | 0.718207094 | 3.724945049 |
| Gm13899 | 0 | NA | NA |
| RP23-479D21.3 | 13.97715435 | -0.147248949 | 0.336252745 |
| RP23-479D21.4 | 0.045882455 | 0.237312084 | 3.724945049 |
| Gm26738 | 0.238158741 | 1.209559574 | 2.530286746 |
| Gm27444 | 0 | NA | NA |
| Gm27402 | 3.526582245 | -0.949885697 | 0.618102867 |
| RP23-86H7.3 | 81.37030569 | -0.997962691 | 0.291802568 |

| | | | |
|---|---|---|---|
| Wnt2 | 29.40825172 | -0.192084939 | 0.313773091 |
| Asz1 | 0.284445263 | -0.243568748 | 2.047426847 |
| RP23-275N13.1 | 0 | NA | NA |
| Gm15593 | 0.075043205 | 0.237309859 | 3.724945049 |
| Gm15596 | 0.396988867 | 0.60818064 | 1.470508579 |
| Cftr | 19.0807837 | -0.65085202 | 0.30969957 |
| Gm15592 | 0 | NA | NA |
| Gm15595 | 0.243349506 | -0.834651148 | 2.446611182 |
| Cttnbp2 | 1527.7495 | -0.917483025 | 0.139731777 |
| Gm15594 | 1.482657567 | 2.982124508 | 0.997018859 |
| RP23-16J8.6 | 154.7156367 | -1.64877227 | 0.351269792 |
| Gm26233 | 3.766699628 | -0.57194837 | 0.575016563 |
| Gm23648 | 0 | NA | NA |
| RP23-148L8.1 | 0 | NA | NA |
| Gm5874 | 1.63591125 | 0.659724746 | 0.852569608 |
| Gm26809 | 2.591679042 | 0.721849734 | 0.636100474 |
| RP23-148L8.3 | 35.09407007 | 0.048778627 | 0.189157239 |
| Lsm8 | 195.2002428 | -0.092181502 | 0.090918499 |
| Ankrd7 | 2.04319782 | 0.986331925 | 0.751966472 |
| RP24-230L6.3 | 0.39809594 | 1.266875908 | 1.765106758 |
| Gm22718 | 0 | NA | NA |
| RP23-18G14.1 | 0.045882455 | 0.237312084 | 3.724945049 |
| RP23-179A12.2 | 0 | NA | NA |
| RP23-179A12.1 | 0 | NA | NA |
| Gm24213 | 0 | NA | NA |
| Mir6370 | 0 | NA | NA |
| Gm22828 | 0 | NA | NA |
| RP23-302D18.3 | 0 | NA | NA |
| RP23-302D18.1 | 0 | NA | NA |
| RP23-302D18.2 | 0.044191549 | -0.243571806 | 3.724945049 |
| RP23-441A23.1 | 0 | NA | NA |
| Gm23960 | 0 | NA | NA |
| Gm26373 | 0 | NA | NA |
| Kcnd2 | 2260.738386 | -0.764939504 | 0.104978942 |
| RP23-307L24.2 | 1439.777736 | -1.227630895 | 0.330596045 |
| RP23-94H23.1 | 58.13546571 | -1.445685576 | 0.308261145 |
| RP23-301C7.2 | 101.8565513 | -0.812419406 | 0.2434899 |
| Tspan12 | 200.3279207 | -0.245044691 | 0.132879506 |
| RP24-169J24.2 | 0.152959369 | -0.233631232 | 2.840073126 |
| RP24-169J24.1 | 0.164540623 | -0.243578283 | 3.439350116 |
| Ing3 | 280.4244847 | -0.037582562 | 0.073867391 |
| Gm26719 | 0 | NA | NA |
| Cped1 | 74.68292297 | 0.102059038 | 0.147048295 |
| RP24-279C2.3 | 0 | NA | NA |
| Wnt16 | 10.34745373 | 0.738536519 | 0.318343795 |
| Fam3c | 626.0288257 | -0.51171131 | 0.093808898 |
| RP23-369P2.2 | 0 | NA | NA |

| | | | |
|---|---|---|---|
| Gm8927 | 0.687801143 | 2.695098821 | 1.327847471 |
| RP24-241H23.2 | 1.331472478 | -0.227102399 | 0.923437584 |
| RP24-241H23.1 | 0.086648695 | -0.72445792 | 3.724945049 |
| Gm23961 | 0 | NA | NA |
| Gm25942 | 0 | NA | NA |
| RP23-369P2.3 | 0.047948698 | -0.243569582 | 3.724945049 |
| Ptprz1 | 4928.564296 | -0.425526457 | 0.134742573 |
| Aass | 39.94585083 | -0.804357988 | 0.233765579 |
| RP24-301O17.2 | 0 | NA | NA |
| RP24-301O17.3 | 0 | NA | NA |
| RP24-301O17.1 | 0 | NA | NA |
| Fezf1 | 7.883156094 | 0.637798852 | 0.449348404 |
| RP23-124H2.4 | 3.80210492 | -0.411122227 | 0.486226113 |
| Cadps2 | 1566.7278 | -0.632492815 | 0.128413799 |
| Gm27959 | 0 | NA | NA |
| Gm24217 | 2.108983506 | -1.175566937 | 0.793214607 |
| Rnf133 | 16.93650597 | -1.091491311 | 0.430142698 |
| Rnf148 | 13.88597797 | -0.683609786 | 0.402749639 |
| Tas2r118 | 0 | NA | NA |
| Slc13a1 | 10.97794222 | 0.462729137 | 0.370287567 |
| Gm20706 | 0 | NA | NA |
| Gm26386 | 0.072025462 | 0.237312084 | 3.724945049 |
| Iqub | 42.66293401 | -0.119394525 | 0.171873128 |
| Ndufa5 | 270.339159 | -0.161478418 | 0.218960363 |
| Asb15 | 1.031776346 | 0.378031718 | 0.994669863 |
| RP23-392I2.4 | 0 | NA | NA |
| Lmod2 | 0.965679557 | -1.981156431 | 1.109318881 |
| Wasl | 1854.805742 | -0.19417725 | 0.055145573 |
| RP23-392I2.3 | 73.97553726 | -0.299186702 | 0.183082323 |
| Hyal6 | 0.086908494 | -0.72445792 | 3.724945049 |
| Hyal4 | 0 | NA | NA |
| Spam1 | 0 | NA | NA |
| RP24-255M7.1 | 0 | NA | NA |
| Hyal5 | 0 | NA | NA |
| Gm22267 | 0 | NA | NA |
| Tmem229a | 986.0139867 | -1.194716017 | 0.161716135 |
| RP23-339M5.2 | 0.278190828 | 1.470944277 | 2.414659645 |
| Gm24443 | 0 | NA | NA |
| Gm22529 | 0 | NA | NA |
| Gpr37 | 655.1826324 | -0.678774786 | 0.151350767 |
| Pot1a | 253.1759545 | -0.383575917 | 0.154317162 |
| RP23-302O24.1 | 0.342793059 | -0.541814677 | 1.686740721 |
| RP23-302O24.2 | 0 | NA | NA |
| Gm24691 | 0 | NA | NA |
| RP23-308F24.1 | 0 | NA | NA |
| Grm8 | 291.4031537 | -0.614093313 | 0.128977306 |
| RP23-414I9.1 | 5.120409944 | -0.60641536 | 0.492291614 |

| | | | |
|---|---|---|---|
| Gm26310 | 0 | NA | NA |
| Mir592 | 0 | NA | NA |
| RP23-23C4.6 | 185.154323 | 0.031275802 | 0.136162682 |
| 6530409C15Rik | 8.058619829 | -0.74103309 | 0.401341744 |
| RP23-23C4.5 | 118.802725 | -0.406942568 | 0.253067236 |
| RP23-23C4.4 | 75.79663319 | -0.511260126 | 0.226952086 |
| Zfp800 | 452.6169934 | 0.263124802 | 0.068185065 |
| Gm5303 | 1.49686415 | -0.915910101 | 0.831311486 |
| Gcc1 | 343.3632238 | 0.346994002 | 0.089847219 |
| Arf5 | 1016.757531 | 0.06601877 | 0.075223918 |
| Fscn3 | 1.623741216 | -0.671826732 | 0.808208854 |
| Pax4 | 0.249106506 | 0.709481059 | 1.837811495 |
| RP23-434O14.5 | 0 | NA | NA |
| Snd1 | 1136.070214 | 0.318580075 | 0.074383901 |
| Gm37321 | 29.66800731 | -0.280718297 | 0.300347917 |
| Gm37978 | 39.62155469 | -0.10272488 | 0.303470472 |
| Lrrc4 | 977.950627 | -0.678882909 | 0.10454856 |
| Gm37940 | 70.33009473 | -0.350746103 | 0.274573311 |
| Gm3294 | 93.79479471 | -1.209475438 | 0.278286736 |
| Mir129-1 | 0 | NA | NA |
| Lep | 0.088534442 | 1.179809211 | 3.723500796 |
| Gm25589 | 0 | NA | NA |
| Rbm28 | 422.2309319 | 0.018338477 | 0.114896377 |
| Gm24303 | 1.400035301 | -1.701024411 | 1.118867798 |
| Prrt4 | 24.29690244 | -1.240813316 | 0.24685666 |
| Impdh1 | 348.4065021 | -0.664618507 | 0.077676556 |
| Gm26021 | 0 | NA | NA |
| Hilpda | 32.36285512 | 0.830981827 | 0.208735498 |
| Fam71f2 | 3.490009053 | 0.668531903 | 0.592656735 |
| Fam71f1 | 0.2407585 | 1.111458469 | 2.075178798 |
| Gm17172 | 0.045882455 | 0.237312084 | 3.724945049 |
| Calu | 1240.173328 | 0.96638776 | 0.116116976 |
| Opn1sw | 0.679991162 | -0.901847221 | 1.280736935 |
| Ccdc136 | 845.6284117 | -0.647556364 | 0.100898316 |
| Gm26220 | 0.160408561 | -0.72446006 | 3.457615423 |
| AC044807.1 | 0.581411716 | -1.724973972 | 1.253308418 |
| Flnc | 1393.252646 | 4.974589093 | 0.376809389 |
| Atp6v1f | 586.7322244 | 0.116156977 | 0.092581453 |
| Gm9047 | 2.295347566 | -0.341007365 | 0.617184758 |
| Kcp | 10.30552492 | -0.499577364 | 0.318742865 |
| Gm26484 | 0 | NA | NA |
| Gm26627 | 0.072025462 | 0.237312084 | 3.724945049 |
| Irf5 | 262.9012068 | 4.258552654 | 0.21819604 |
| Tnpo3 | 1004.219965 | 0.01144919 | 0.075563983 |
| RP23-446B13.7 | 31.58542221 | -0.531458939 | 0.271554182 |
| Tspan33 | 165.1105126 | -0.706716954 | 0.128237811 |
| Smo | 219.4811244 | 0.112050488 | 0.142418881 |

| | | | |
|---|---|---|---|
| Gm13717 | 1.21856855 | -1.949922794 | 0.954049957 |
| Ahcyl2 | 2068.520444 | -0.377079303 | 0.045140549 |
| RP23-353E8.7 | 0.12092566 | 0.237307635 | 3.724945049 |
| Strip2 | 859.1687429 | -0.495885482 | 0.076370623 |
| Gm13724 | 2.453329902 | -0.426778787 | 0.646743727 |
| Gm24536 | 0.309201626 | -0.696633147 | 1.568196628 |
| Gm13725 | 0.309323177 | 0.238006957 | 2.014555284 |
| Smkr-ps | 0.763164319 | 0.142780715 | 1.175257345 |
| Gm23049 | 0 | NA | NA |
| Nrf1 | 255.4170055 | 0.056408808 | 0.093823772 |
| Gm25580 | 0 | NA | NA |
| Mir182 | 0 | NA | NA |
| Mir96 | 0 | NA | NA |
| Mir183 | 0 | NA | NA |
| Gm37046 | 0.075043205 | 0.237309859 | 3.724945049 |
| Ube2h | 1345.992283 | -0.173835996 | 0.058704515 |
| 1700095J07Rik | 0 | NA | NA |
| Gm25625 | 0.044191549 | -0.243571806 | 3.724945049 |
| Zc3hc1 | 183.2490365 | 0.465926437 | 0.096344004 |
| n-R5s161 | 0 | NA | NA |
| Klhdc10 | 1743.124813 | -0.091894911 | 0.042273934 |
| 1700012C14Rik | 0.900590653 | -0.523060678 | 0.983401046 |
| Tmem209 | 379.778664 | 0.532692631 | 0.091807983 |
| Ssmem1 | 1.951291546 | -0.922118723 | 0.698180422 |
| Cpa2 | 12.99980896 | -0.154382271 | 0.32243736 |
| Cpa4 | 11.88556507 | 3.359322818 | 0.550284179 |
| Gm13781 | 0.062426873 | 0.237312084 | 3.724945049 |
| Cpa5 | 0.592694123 | 0.359638251 | 1.253614112 |
| Cpa1 | 0.176646398 | 1.281666154 | 3.35399331 |
| Cep41 | 83.12317599 | 0.825654918 | 0.132583825 |
| Gm13782 | 0.049553047 | 0.71820487 | 3.724945049 |
| Gm14528 | 0 | NA | NA |
| Mest | 610.9026533 | -1.260340588 | 0.119201969 |
| Gm27522 | 0 | NA | NA |
| Gm27568 | 0 | NA | NA |
| Mir335 | 5.501266716 | -0.83619779 | 0.539117455 |
| Copg2 | 853.6229713 | -0.119348247 | 0.059323003 |
| 4930412F09Rik | 0.271012069 | 0.237298623 | 3.724945049 |
| Tsga13 | 1.195665945 | 3.230282881 | 1.063397895 |
| Klf14 | 3.978034112 | 0.470426213 | 0.533018505 |
| Mir29a | 6.497893077 | 0.001874239 | 0.520156359 |
| Mir29b-1 | 0.108266322 | 1.230376831 | 3.722845817 |
| Gm27019 | 20.56482463 | 0.153105825 | 0.376158983 |
| Gm13834 | 35.42125562 | -0.196419029 | 0.338667816 |
| Lincpint | 345.1173544 | -0.791599265 | 0.217778019 |
| Gm13833 | 0.042716945 | -0.243571806 | 3.724945049 |
| 5330406M23Rik | 159.3696281 | -1.039174098 | 0.380212133 |

| | | | |
|---|---|---|---|
| Gm13835 | 176.8303318 | 0.713598346 | 0.131514434 |
| Gm37728 | 114.4100306 | -1.20217179 | 0.422379566 |
| 2210408F21Rik | 141.5258413 | -0.436442521 | 0.162635869 |
| RP23-400L4.7 | 44.53988991 | -0.907128789 | 0.419451782 |
| Gm14532 | 12.88187382 | -0.730700458 | 0.32938387 |
| Mkln1os | 23.92253865 | -0.800227799 | 0.233433494 |
| Gm13844 | 13.1985606 | -0.950034161 | 0.481044056 |
| Mkln1 | 2442.640936 | -0.144358127 | 0.086976116 |
| Podxl | 278.0855086 | 1.752716409 | 0.12832518 |
| Gm6117 | 0.646697505 | -1.265914094 | 1.266371522 |
| RP24-414G20.1 | 3.468425366 | -0.464890188 | 0.598001501 |
| Gm13846 | 0.441802452 | -0.724462587 | 1.728964224 |
| Gm13848 | 0.595326517 | 1.405113987 | 1.56633625 |
| RP24-414G20.4 | 0.125911488 | 0.555009803 | 3.703517916 |
| Gm13849 | 0.554297541 | -1.660217581 | 1.341938859 |
| 1700012A03Rik | 0.048949532 | -0.243569582 | 3.724945049 |
| Gm13850 | 0 | NA | NA |
| Plxna4 | 2374.903231 | -0.536407646 | 0.101213992 |
| Plxna4os3 | 0 | NA | NA |
| Plxna4os2 | 0.093141081 | -0.724455696 | 3.724945049 |
| Plxna4os1 | 3.620095065 | 2.578069927 | 0.742036747 |
| Gm13852 | 0.049553047 | 0.71820487 | 3.724945049 |
| RP23-225G5.2 | 0.573159608 | -1.162000219 | 1.511079103 |
| Chchd3 | 512.9631055 | 0.441283928 | 0.096447232 |
| Gm14540 | 0 | NA | NA |
| RP24-246G10.3 | 30.7240258 | -0.81022324 | 0.363913031 |
| Gm22321 | 0 | NA | NA |
| Gm1401 | 0 | NA | NA |
| Exoc4 | 1197.152527 | -0.103416776 | 0.067713322 |
| Gm13854 | 0 | NA | NA |
| RP23-4H14.1 | 15.19819234 | -0.808776753 | 0.426266297 |
| RP24-365C10.2 | 0 | NA | NA |
| Gm13853 | 0.045882455 | 0.237312084 | 3.724945049 |
| Lrguk | 29.47287266 | -0.401477754 | 0.21783513 |
| Gm13856 | 0.085327104 | 1.179809211 | 3.723500796 |
| Slc35b4 | 730.9802596 | -0.283473561 | 0.054715975 |
| Gm13855 | 0 | NA | NA |
| Gm14546 | 0.956739943 | -0.085443317 | 1.047887202 |
| Akr1b3 | 294.7538566 | 0.593543716 | 0.088880998 |
| Akr1b8 | 26.07990468 | 3.561315019 | 0.368137815 |
| Gm13859 | 0.259720851 | 0.553060111 | 2.015970751 |
| Gm13858 | 0.289603658 | 1.598935388 | 2.038145796 |
| Akr1b10 | 145.9333712 | 0.169961961 | 0.100570676 |
| Akr1b7 | 1.910658871 | 0.478334321 | 0.755171593 |
| Gm13857 | 0.038981395 | 0.718207094 | 3.724945049 |
| Bpgm | 500.8527338 | 0.208415 | 0.080167676 |
| Gm13860 | 0 | NA | NA |

| | | | |
|---|---|---|---|
| Cald1 | 388.6951145 | 0.811355702 | 0.120107238 |
| Gm13861 | 2.144383846 | -0.835695321 | 0.841954793 |
| Npn2 | 0 | NA | NA |
| Gm14547 | 0 | NA | NA |
| Agbl3 | 107.7106235 | 0.342552908 | 0.153051969 |
| Gm13864 | 2.929070206 | 0.957358256 | 0.571604897 |
| Gm13862 | 0.066050875 | 0.71820487 | 3.724945049 |
| Gm13863 | 0 | NA | NA |
| RP23-386A17.7 | 1.526274188 | 0.84574118 | 0.941449727 |
| Tmem140 | 313.938631 | 5.894462383 | 0.270255529 |
| 3110062M04Rik | 36.44138757 | 0.009429452 | 0.22777343 |
| Wdr91 | 325.7975239 | 0.105556578 | 0.079673811 |
| Stra8 | 0.124347486 | 0.719186602 | 3.717245571 |
| 2010107G12Rik | 6.64236274 | 1.082837059 | 0.468045917 |
| Cnot4 | 879.3333849 | -0.008990008 | 0.068972859 |
| Nup205 | 597.9465125 | 0.142564851 | 0.086239547 |
| Gm23435 | 0.60633683 | -1.652310513 | 1.192387954 |
| 1810058I24Rik | 158.214639 | 0.497048495 | 0.131259412 |
| Slc13a4 | 187.4589476 | -0.619256966 | 0.2065804 |
| Fam180a | 13.25868466 | -1.464826033 | 0.345438431 |
| RP23-338E5.5 | 0.361225529 | 0.718865292 | 1.931443906 |
| Mtpn | 3362.617321 | -0.008002735 | 0.059292973 |
| Gm23053 | 1.826002588 | -0.031906762 | 0.743746433 |
| RP23-20C5.1 | 0 | NA | NA |
| RP23-13B24.1 | 13.51087949 | -0.232594095 | 0.296649991 |
| Gm23273 | 0 | NA | NA |
| RP23-352D5.1 | 6.269730523 | -0.158186572 | 0.45778259 |
| 9330158H04Rik | 22.23221858 | 0.121272683 | 0.226487007 |
| Chrm2 | 536.4163306 | 0.089776655 | 0.121392355 |
| Mir490 | 3.003392332 | -1.113650013 | 0.694511198 |
| RP23-81D20.2 | 0 | NA | NA |
| RP23-5K15.1 | 0 | NA | NA |
| Gm25111 | 0.298614356 | -0.243568581 | 2.01181534 |
| Ptn | 1821.983259 | -0.593886215 | 0.134753499 |
| Dgki | 1466.320099 | -0.465002182 | 0.114799577 |
| 1700111E14Rik | 0.088736197 | -0.243576254 | 3.724945049 |
| RP23-151F20.5 | 2.105924665 | 1.464023711 | 0.659467498 |
| Creb3l2 | 335.1776498 | 0.472908433 | 0.09945771 |
| Akr1d1 | 0.049553047 | 0.71820487 | 3.724945049 |
| RP24-549J13.2 | 0 | NA | NA |
| RP23-245J24.1 | 9.865094166 | 0.813475259 | 0.342773521 |
| Gm15487 | 1555.447171 | 0.087878593 | 0.070844271 |
| RP23-459F14.2 | 0.11197992 | 0.718202646 | 3.724945049 |
| Trim24 | 495.6674005 | -0.1542836 | 0.10961157 |
| Svopl | 1.88997817 | 2.197324601 | 0.827058628 |
| Atp6v0a4 | 3.783295768 | 1.176334426 | 0.552129787 |
| Tmem213 | 0 | NA | NA |

| | | | |
|---|---|---|---|
| Gm23231 | 0 | NA | NA |
| D630045J12Rik | 1061.614314 | -0.924075764 | 0.133884317 |
| RP23-387J8.3 | 0 | NA | NA |
| Zc3hav1l | 80.97671505 | -0.818294418 | 0.186769459 |
| Zc3hav1 | 1446.894863 | 5.146643745 | 0.173859749 |
| RP24-142K18.4 | 0.189025308 | 0.22087251 | 2.253764058 |
| Rpl30-ps1 | 9.381830078 | 0.443919743 | 0.348931727 |
| Ttc26 | 133.6908354 | -0.260205322 | 0.117770712 |
| Gm16563 | 1.549622433 | -0.959633257 | 0.836893881 |
| Ubn2 | 2325.889383 | -0.378546715 | 0.101507206 |
| Mir7670 | 0 | NA | NA |
| 1110001J03Rik | 118.1303116 | -0.127888754 | 0.14729699 |
| Gm15697 | 2.902282796 | -0.026284367 | 0.690142765 |
| Luc7l2 | 4182.925321 | -0.306764974 | 0.112110609 |
| Gm26699 | 1.935704857 | -1.20119362 | 0.878207528 |
| Klrg2 | 32.43324731 | 2.450889733 | 0.314058723 |
| Clec2l | 411.0156647 | -0.412321853 | 0.079329877 |
| Hipk2 | 3020.872203 | 0.287520447 | 0.099744252 |
| Tbxas1 | 29.76369512 | 1.331633573 | 0.280904447 |
| RP23-346B17.4 | 0.320565116 | 0.718189471 | 3.724945049 |
| Gm23010 | 0.282660793 | -2.093698044 | 3.682238457 |
| 1700025N23Rik | 9.679120195 | 5.864297573 | 0.932459089 |
| Parp12 | 1518.309522 | 4.996207397 | 0.156133989 |
| 4930599N23Rik | 26.63738886 | 4.630446964 | 0.428605515 |
| RP23-345F4.3 | 0.255942976 | 2.259817959 | 3.017298019 |
| Kdm7a | 1749.004419 | -0.057591548 | 0.073959486 |
| RP24-173B18.1 | 0.413211764 | 1.899954636 | 1.506043511 |
| Gm26008 | 0.222111871 | 0.237311956 | 3.129813646 |
| Slc37a3 | 628.5897833 | -0.169144828 | 0.071038133 |
| Rab19 | 12.00148128 | 6.210011786 | 0.882550214 |
| Mkrn1 | 1161.354 | 0.170327756 | 0.049406256 |
| Gm10244 | 4.940723394 | 0.271863366 | 0.467168844 |
| Gm25402 | 0.04393175 | -0.243571806 | 3.724945049 |
| Dennd2a | 408.9845452 | 0.493875166 | 0.093172686 |
| 8030453O22Rik | 22.33756815 | -0.30707941 | 0.30053067 |
| Adck2 | 176.9877061 | -0.210839762 | 0.095773962 |
| Gm16272 | 1.598725798 | 0.846506761 | 0.783828848 |
| RP23-366P9.8 | 0.091666476 | -0.724455696 | 3.724945049 |
| Ndufb2 | 352.2685807 | -0.227883381 | 0.092886787 |
| Braf | 3301.479618 | -0.381365454 | 0.077567177 |
| Fth-ps2 | 26.73179484 | -1.024047318 | 0.402517137 |
| Mrps33 | 137.537649 | -0.014178494 | 0.135028674 |
| RP23-380O16.6 | 0 | NA | NA |
| RP23-380O16.7 | 0.107078754 | 0.236669296 | 3.724898107 |
| Gm26833 | 7.290008566 | 0.686909597 | 0.42214373 |
| Tmem178b | 2878.275615 | -0.651599884 | 0.08443173 |
| Gm37995 | 95.94723699 | -1.130426977 | 0.312502444 |

| | | | |
|---|---|---|---|
| Gm37673 | 72.41137234 | -1.341790639 | 0.287207311 |
| Gm26638 | 42.96957209 | -1.694888955 | 0.410723948 |
| Gm26807 | 1.057496005 | -1.339802611 | 1.371462959 |
| Gm37003 | 29.74312774 | -1.14363565 | 0.377747678 |
| Agk | 341.3868807 | -0.415785066 | 0.08382015 |
| E330009J07Rik | 1236.299548 | -0.924053718 | 0.140589607 |
| Wee2 | 3.520209586 | 0.933916251 | 0.534639871 |
| Ssbp1 | 232.2371204 | 0.082038404 | 0.136848486 |
| Tas2r137 | 4.276107861 | -0.085314641 | 0.513792364 |
| Tas2r108 | 4.194868342 | 0.878893544 | 0.570152228 |
| Prss37 | 0 | NA | NA |
| RP24-503P15.7 | 0 | NA | NA |
| Olfr461 | 0 | NA | NA |
| Olfr460 | 0.474871048 | 1.164849407 | 1.413770553 |
| Clec5a | 38.15951351 | 2.548906183 | 0.270091578 |
| Tas2r138 | 0 | NA | NA |
| Mgam | 2.716524655 | 1.459526076 | 0.732522656 |
| Mgam2-ps | 0.962702212 | 3.317032473 | 1.102566988 |
| Moxd2 | 0 | NA | NA |
| Gm38276 | 0 | NA | NA |
| Trbv1 | 1.116438897 | 2.833776175 | 1.143241069 |
| Prss58 | 0 | NA | NA |
| 1700074P13Rik | 0 | NA | NA |
| Gm4744 | 0 | NA | NA |
| Gm29588 | 0 | NA | NA |
| 1810009J06Rik | 0 | NA | NA |
| Gm2663 | 0 | NA | NA |
| Gm38015 | 0 | NA | NA |
| Gm37878 | 0 | NA | NA |
| Gm38294 | 0 | NA | NA |
| 2210010C04Rik | 0.16656219 | 0.597911793 | 2.779636975 |
| Trbv2 | 1.408278968 | 3.910946604 | 1.101025619 |
| Trbv3 | 0.461584485 | 1.629948099 | 1.68059839 |
| 5830405F06Rik | 0.95870128 | 2.18071719 | 1.143878884 |
| Trbv5 | 0.150914725 | 1.165657226 | 2.845788074 |
| Trbv6 | 0 | NA | NA |
| Trbv7 | 0 | NA | NA |
| Gm37610 | 0 | NA | NA |
| Trbv8 | 0 | NA | NA |
| Trbv9 | 0 | NA | NA |
| Trbv10 | 0 | NA | NA |
| Trbv11 | 0 | NA | NA |
| Trbv12-1 | 0 | NA | NA |
| Trbv13-1 | 2.762763099 | 5.3585167 | 0.989759149 |
| Trbv12-2 | 0.921345027 | 3.34849012 | 1.082931396 |
| Trbv13-2 | 3.038727124 | 4.873788081 | 0.985695877 |
| Trbv12-3 | 0 | NA | NA |

| | | | |
|---|---|---|---|
| Trbv13-3 | 1.194558986 | 3.953768129 | 1.107723631 |
| Trbv14 | 1.5023977 | 3.719292136 | 1.79770907 |
| Trbv15 | 0.168389893 | 1.080949625 | 2.764823542 |
| Trbv16 | 0.74889667 | 1.917681487 | 1.47403684 |
| Trbv17 | 1.696056422 | 4.190508489 | 1.080838542 |
| Trbv18 | 0 | NA | NA |
| Trbv19 | 0.811168495 | 3.325118833 | 1.163971646 |
| Trbv20 | 0.334925437 | 1.972611946 | 1.980464618 |
| Trbv21 | 0 | NA | NA |
| Trbv22 | 0.097139547 | 0.718207094 | 3.724945049 |
| Trbv23 | 0 | NA | NA |
| Trbv24 | 0.038981395 | 0.718207094 | 3.724945049 |
| Trbv25 | 0 | NA | NA |
| Trbv26 | 0.038981395 | 0.718207094 | 3.724945049 |
| Gm2719 | 0.3093278 | -0.238993779 | 1.785103355 |
| Trbv27 | 0 | NA | NA |
| Trbv28 | 0 | NA | NA |
| Trbv29 | 1.593432551 | 3.777867014 | 1.163304329 |
| Trbv30 | 0.049553047 | 0.71820487 | 3.724945049 |
| Try4 | 0.044544648 | 0.237309859 | 3.724945049 |
| Gm18964 | 0 | NA | NA |
| Try5 | 0 | NA | NA |
| Try10 | 0 | NA | NA |
| Prss3 | 0 | NA | NA |
| Gm5771 | 0 | NA | NA |
| Gm38306 | 0 | NA | NA |
| Gm5409 | 0 | NA | NA |
| Gm10334 | 0.980132196 | -1.094200802 | 1.077864621 |
| Prss1 | 0 | NA | NA |
| Gm37301 | 0 | NA | NA |
| Gm29813 | 0.146823232 | -0.712492187 | 2.876272087 |
| Gm37841 | 0 | NA | NA |
| Gm38018 | 0 | NA | NA |
| Prss2 | 0.046345709 | 0.71820487 | 3.724945049 |
| Trbd1 | 0.960611118 | 3.314009111 | 1.096269082 |
| Trbj1-1 | 0.739650571 | 2.739917251 | 1.158080061 |
| Trbj1-2 | 0.363286374 | 2.688950746 | 3.679631232 |
| Trbj1-3 | 0.347715489 | 2.561502899 | 1.447102126 |
| Trbj1-4 | 0.471163529 | 2.780138779 | 1.408656983 |
| Trbj1-5 | 0.450331108 | 1.614315125 | 1.426152519 |
| Trbj1-6 | 0.583857846 | 3.159918036 | 1.28924178 |
| Trbj1-7 | 0.161076545 | 1.165842935 | 2.787893623 |
| Trbc1 | 24.37840666 | 7.59179972 | 0.89525639 |
| Trbd2 | 0.440524946 | 1.765580671 | 1.686409622 |
| Trbj2-1 | 0.112396584 | 1.192083721 | 3.724434897 |
| Trbj2-2 | 0.218053095 | 1.392588641 | 2.131916016 |
| Trbj2-3 | 0.696513874 | -1.315216412 | 1.148930741 |

| | | | |
|---|---|---|---|
| Trbj2-4 | 0.088736197 | -0.243576254 | 3.724945049 |
| Trbj2-5 | 0.202417642 | -0.243564318 | 2.653394741 |
| Trbj2-6 | 0.354669972 | 2.480703459 | 1.842032297 |
| Trbj2-7 | 1.180533128 | -1.371921748 | 0.999570481 |
| Trbc2 | 69.31285373 | 1.352466633 | 0.210915429 |
| Trbv31 | 1.290124216 | 2.717356789 | 1.174973818 |
| Gm27549 | 0 | NA | NA |
| Ephb6 | 693.8526302 | -0.716075213 | 0.133837192 |
| Trpv6 | 9.753117248 | -0.148052905 | 0.391524255 |
| Trpv5 | 0 | NA | NA |
| 1700034O15Rik | 0.14896402 | 0.717609292 | 2.866934839 |
| Kel | 0.227427828 | 0.23732637 | 2.559553268 |
| Gm37527 | 0 | NA | NA |
| Olfr459 | 0 | NA | NA |
| Pip | 0.038981395 | 0.718207094 | 3.724945049 |
| Sval2 | 0.163190422 | 1.22955094 | 3.396300676 |
| Sval1 | 0.173394273 | 1.740859167 | 3.379052784 |
| Sval3 | 0.066050875 | 0.71820487 | 3.724945049 |
| Sva | 0.066050875 | 0.71820487 | 3.724945049 |
| Tas2r139 | 0 | NA | NA |
| Tas2r144 | 0.049553047 | 0.71820487 | 3.724945049 |
| Gstk1 | 46.48690246 | -0.287346795 | 0.190182827 |
| Tmem139 | 0.247571123 | -1.165553575 | 2.480837065 |
| Casp2 | 320.4543346 | 0.438680883 | 0.127370461 |
| Gm25108 | 3.821797685 | 0.446982716 | 0.554851001 |
| Clcn1 | 102.7118201 | -0.374701788 | 0.286299732 |
| Fam131b | 1482.538637 | -0.634609683 | 0.088494379 |
| Gm24611 | 0 | NA | NA |
| Zyx | 868.7508899 | 1.240669519 | 0.169016007 |
| Epha1 | 4.796176745 | 0.568695985 | 0.46674329 |
| 2010310C07Rik | 14.52291164 | -0.313340472 | 0.410688878 |
| Tas2r143 | 4.118545241 | -0.317171822 | 0.557327964 |
| Tas2r135 | 5.600410292 | 0.366248112 | 0.553503813 |
| Tas2r126 | 2.194003788 | -0.107512246 | 0.71930367 |
| Olfr458 | 0.062426873 | 0.237312084 | 3.724945049 |
| Olfr457 | 0 | NA | NA |
| Olfr456 | 0 | NA | NA |
| Gm4877 | 0.146848595 | -1.036216968 | 3.718157501 |
| Olfr455 | 0 | NA | NA |
| Tcaf3 | 0.038981395 | 0.718207094 | 3.724945049 |
| Tcaf2 | 17.75907395 | 1.048082003 | 0.267896978 |
| Tcaf1 | 3402.596378 | -0.505035072 | 0.08157117 |
| Olfr453 | 0.410820671 | 1.661370342 | 1.537140536 |
| Olfr38 | 0.225129614 | 0.237300054 | 3.724945049 |
| Gm22363 | 0.272364125 | 1.701176406 | 1.99524632 |
| Olfr452 | 0.180050447 | 0.719165083 | 3.327036943 |
| Olfr450 | 0.211502678 | 1.844922517 | 2.522003129 |

| | | | |
|---|---|---|---|
| Gm23982 | 0.194279094 | 1.292225235 | 3.719439735 |
| Olfr449 | 0.134148234 | 0.23732184 | 3.624632408 |
| Olfr448 | 0.07796279 | 1.072698098 | 3.721829099 |
| Olfr447 | 0 | NA | NA |
| Olfr446 | 0.048949532 | -0.243569582 | 3.724945049 |
| Olfr444 | 0 | NA | NA |
| Gm17472 | 0 | NA | NA |
| Gm20730 | 0 | NA | NA |
| Olfr443-ps1 | 0.336338362 | 1.97650831 | 1.696312305 |
| Olfr441 | 0 | NA | NA |
| Olfr237-ps1 | 0 | NA | NA |
| Olfr437 | 0 | NA | NA |
| Olfr13 | 0 | NA | NA |
| Olfr435 | 0.038981395 | 0.718207094 | 3.724945049 |
| Gm21827 | 0 | NA | NA |
| Olfr434 | 0.061920613 | 0.718207094 | 3.724945049 |
| Olfr47 | 0 | NA | NA |
| Arhgef5 | 73.22831166 | 1.465514549 | 0.176468652 |
| Nobox | 0.405536581 | -0.750309289 | 1.654250943 |
| Tpk1 | 61.31573283 | 0.055609769 | 0.174050971 |
| Gm15550 | 0.332944463 | 0.291362643 | 1.522917421 |
| Gm22939 | 0 | NA | NA |
| Cntnap2 | 2123.978884 | -0.559876289 | 0.062834221 |
| Gm22839 | 8.429271064 | -0.46859086 | 0.378203502 |
| Gm25161 | 0 | NA | NA |
| Rbpsuh-rs3 | 97.81620039 | 0.850406247 | 0.132393366 |
| Cul1 | 1508.243995 | 0.09776051 | 0.059633509 |
| A930035D04Rik | 12.16101713 | 0.230545517 | 0.292150334 |
| Ezh2 | 210.0637519 | -0.001434874 | 0.106213404 |
| Gm26625 | 0.182019239 | 1.106583154 | 2.712073064 |
| Gm25283 | 0 | NA | NA |
| Rny3 | 804.362109 | -0.710734912 | 0.320673039 |
| Rpl31-ps7 | 0 | NA | NA |
| Rny1 | 909.9952825 | -0.573794432 | 0.417976302 |
| Pdia4 | 818.8143374 | 1.067542941 | 0.125268473 |
| Mir704 | 0.123992736 | -0.243574001 | 3.720643035 |
| Rpl35a-ps7 | 0.314214724 | 0.237328524 | 1.752182381 |
| Zfp786 | 50.22326823 | -0.070397133 | 0.170893821 |
| Zfp398 | 324.873339 | -0.424560526 | 0.145541663 |
| Zfp282 | 307.4764343 | -0.324620599 | 0.109941323 |
| Zfp212 | 200.032767 | 0.025667036 | 0.134529052 |
| Zfp783 | 130.5839755 | -0.177354339 | 0.160739582 |
| Zfp956 | 173.5071749 | -0.290981336 | 0.143934964 |
| Zfp777 | 460.5844228 | 0.326694189 | 0.100975714 |
| Gm24563 | 1.500420242 | 0.43196516 | 0.790125601 |
| Zfp746 | 384.3199641 | 0.033865018 | 0.087677615 |
| Gm16630 | 7.347387118 | -0.15469048 | 0.406275919 |

| | | | |
|---|---|---|---|
| Gm22122 | 0 | NA | NA |
| Krba1 | 549.5583407 | -0.719000419 | 0.135291631 |
| Gm24325 | 1.916854426 | -1.686366188 | 0.782564545 |
| Zfp467 | 330.9730189 | -0.57110466 | 0.089081304 |
| Sspo | 38.70866699 | -0.685545986 | 0.34943136 |
| Gm10243 | 43.3226042 | 0.992268176 | 0.202804186 |
| Gm7887 | 75.72156547 | 0.077740804 | 0.158044529 |
| 1700026J14Rik | 0.552687697 | -0.843362954 | 1.348565082 |
| Atp6v0e2 | 1764.72768 | -0.368980545 | 0.067663266 |
| Lrrc61 | 226.76103 | -0.083653801 | 0.110617905 |
| Rarres2 | 195.195546 | 3.080567301 | 0.321610526 |
| Gm5111 | 0.049553047 | 0.71820487 | 3.724945049 |
| Repin1 | 624.3197349 | 0.169419391 | 0.089792324 |
| Zfp775 | 56.68412911 | -0.27681467 | 0.15798663 |
| AI854703 | 220.6699541 | -0.513421317 | 0.116861394 |
| Gimap8 | 51.49836854 | 2.920478455 | 0.27963441 |
| Gimap9 | 17.15776395 | 3.442896522 | 0.391061054 |
| Gimap4 | 77.7956503 | 7.020115518 | 0.573600785 |
| Gimap6 | 85.26288905 | 2.986590748 | 0.26000138 |
| Gm28053 | 0.321006207 | 1.947249551 | 2.354797057 |
| Gimap7 | 21.5292461 | 7.198834756 | 0.911261092 |
| Gimap1os | 2.672780519 | 3.467942481 | 0.947558361 |
| Gimap1 | 14.27405217 | 3.954852679 | 0.516849894 |
| Gimap5 | 14.86559326 | 3.806854599 | 0.519586873 |
| Gimap3 | 25.51805368 | 7.29880887 | 0.894191619 |
| Tmem176b | 421.451915 | 1.571746083 | 0.185589053 |
| Tmem176a | 202.5557947 | 1.706678722 | 0.189085669 |
| Gm7932 | 3.063139515 | 2.573892642 | 0.838294252 |
| Aoc1 | 3.736083012 | 3.562781805 | 0.86996592 |
| 1600015I10Rik | 0 | NA | NA |
| Gm24348 | 0 | NA | NA |
| Gm10242 | 0.061920613 | 0.718207094 | 3.724945049 |
| Doxl2 | 0.106112162 | 0.23732098 | 3.724945049 |
| Svs1 | 0.044191549 | -0.243571806 | 3.724945049 |
| Gpnmb | 840.8388242 | 6.617213402 | 0.437938652 |
| Malsu1 | 255.5915214 | -0.006813948 | 0.096215415 |
| Igf2bp3 | 6.6227068 | 3.637933046 | 0.859901762 |
| Tra2a | 624.172519 | 0.261557246 | 0.088447643 |
| Ccdc126 | 154.9854233 | 0.265285035 | 0.113911753 |
| Fam221a | 18.46478838 | 0.226695653 | 0.269172648 |
| Stk31 | 5.37787014 | 0.102051429 | 0.442078375 |
| Gm17060 | 0.066050875 | 0.71820487 | 3.724945049 |
| Npy | 399.7067098 | 0.440316329 | 0.18533291 |
| Gm21786 | 0.284351963 | -0.720685959 | 3.721442299 |
| Mpp6 | 900.1739491 | 0.064862064 | 0.055590852 |
| Dfna5 | 75.89232883 | -0.424463791 | 0.142378793 |
| Osbpl3 | 392.5077654 | -0.547585364 | 0.111933835 |

| | | | |
|---|---|---|---|
| Gm26289 | 0 | NA | NA |
| Cycs | 696.5473769 | 0.394492248 | 0.146654278 |
| 5430402O13Rik | 7.75721647 | -1.139505204 | 0.388705977 |
| Mir6371 | 0.990815542 | 0.476298754 | 1.221648549 |
| 4921507P07Rik | 15.81567947 | -0.169084833 | 0.2597167 |
| Gm4782 | 0.19378329 | 0.194150365 | 2.228559436 |
| Gm8038 | 0.04393175 | -0.243571806 | 3.724945049 |
| Npvf | 3.377670371 | 0.294905777 | 0.565554351 |
| G930045G22Rik | 0 | NA | NA |
| Gm29004 | 0.141094079 | 0.718203391 | 3.563850454 |
| Mir148a | 0.208883516 | 1.209863461 | 2.634551626 |
| Gm26677 | 0.297812842 | 0.236901192 | 1.769393438 |
| Nfe2l3 | 68.43159579 | 0.667957074 | 0.161836199 |
| Hnrnpa2b1 | 3001.421392 | -0.021180221 | 0.091506305 |
| Cbx3 | 448.3183731 | -0.193686045 | 0.105552092 |
| Snx10 | 1492.510951 | 0.722905393 | 0.104399065 |
| Gm22914 | 0.310734401 | 0.627278641 | 1.739830171 |
| Skap2 | 251.8896519 | 1.664860233 | 0.146463344 |
| Halr1 | 0.212468866 | 1.193833687 | 2.178974378 |
| Rps8-ps3 | 0 | NA | NA |
| Rps19-ps14 | 0 | NA | NA |
| Hoxa1 | 1.953182484 | 0.174840996 | 0.843289504 |
| Hotairm1 | 2.721387227 | -0.594384587 | 0.610925571 |
| Gm27867 | 0 | NA | NA |
| Gm27861 | 0.467025382 | -0.342962776 | 1.451985284 |
| Gm27868 | 0 | NA | NA |
| Gm27477 | 0.460648582 | -1.035786348 | 1.275739171 |
| Hoxa2 | 5.976910094 | 0.799019649 | 0.65735157 |
| Gm28308 | 0 | NA | NA |
| Hoxaas2 | 3.211323041 | 0.051907868 | 0.700543509 |
| Hoxa3 | 12.50440182 | 0.379395086 | 0.776962173 |
| Gm15050 | 0.11193333 | 0.718202646 | 3.724945049 |
| Hoxaas3 | 0.798143448 | 1.631051149 | 1.447069977 |
| 5730596B20Rik | 0.683443395 | -1.433205843 | 1.311815668 |
| Gm29430 | 0.397176565 | -0.243566537 | 1.901396828 |
| Hoxa4 | 7.87519976 | -0.033057078 | 0.661744572 |
| Hoxa5 | 22.4157456 | -0.022675217 | 0.920155756 |
| Hoxa6 | 2.164192679 | 0.164118633 | 1.64608027 |
| Hoxa7 | 1.260199239 | -0.567494873 | 1.096969715 |
| Hoxa9 | 0.137647365 | 0.237309079 | 3.724945049 |
| AC015583.1 | 0 | NA | NA |
| Mir196b | 0 | NA | NA |
| Hoxa10 | 0 | NA | NA |
| Hoxa11 | 0 | NA | NA |
| Hoxa11os | 0 | NA | NA |
| Gm28032 | 0 | NA | NA |
| Gm27492 | 0 | NA | NA |

| | | | |
|---|---|---|---|
| Gm27695 | 0 | NA | NA |
| Gm27817 | 0 | NA | NA |
| Gm27454 | 0 | NA | NA |
| Gm27543 | 0 | NA | NA |
| 9530018H14Rik | 0 | NA | NA |
| Hoxa13 | 0 | NA | NA |
| Gm27280 | 0 | NA | NA |
| Gm27985 | 0 | NA | NA |
| Hottip | 0 | NA | NA |
| Gm27383 | 0 | NA | NA |
| Gm27866 | 0 | NA | NA |
| 5730457N03Rik | 3.810690039 | 0.149198029 | 0.529646948 |
| Evx1 | 6.587797916 | 0.753375874 | 0.525270016 |
| Gm8129 | 27.45075818 | 0.248984008 | 0.352447795 |
| Gm15054 | 0 | NA | NA |
| Gm37351 | 0.106358623 | -0.24357403 | 3.724945049 |
| Hibadh | 577.34931 | 0.291530892 | 0.101505537 |
| Gm24355 | 3.634340466 | -0.393317351 | 0.595443227 |
| Tax1bp1 | 2381.399223 | 0.664964204 | 0.070273442 |
| Jazf1 | 226.345128 | -0.074674724 | 0.104807198 |
| Gm15572 | 3.883676966 | -1.376791782 | 0.601717713 |
| Gm26215 | 6.191745285 | -0.448332405 | 0.442682607 |
| Gm15573 | 0.191192637 | -0.598851582 | 2.65601198 |
| Gm25818 | 0.106618422 | -0.24357403 | 3.724945049 |
| Creb5 | 153.7701789 | 0.995450583 | 0.204610805 |
| Tril | 1025.612528 | 0.303419561 | 0.167110666 |
| Gm16499 | 0.523988269 | 1.306087119 | 1.402327326 |
| Gm22910 | 0 | NA | NA |
| Cpvl | 0.864647981 | 1.785540704 | 1.078287964 |
| Chn2 | 1528.702195 | -0.393515018 | 0.220102031 |
| 9530036M11Rik | 0.430808853 | 1.234924349 | 1.550304175 |
| Gm15526 | 0.194037777 | -0.197508093 | 2.634480038 |
| Gm15527 | 0.824096613 | 1.039380012 | 1.161335576 |
| 9130019P16Rik | 5.722762008 | -0.429289511 | 0.459438338 |
| Prr15 | 15.96290937 | 1.085215336 | 0.326134262 |
| Wipf3 | 1686.195845 | -0.839989416 | 0.151424084 |
| Scrn1 | 2778.202593 | -0.347693015 | 0.052834431 |
| Fkbp14 | 126.0321638 | 0.385244015 | 0.157772204 |
| Plekha8 | 312.5409318 | -0.563854849 | 0.07581041 |
| Gm15863 | 0.061920613 | 0.718207094 | 3.724945049 |
| Gm28402 | 0.435961187 | -0.227223811 | 1.44029365 |
| Mturn | 1138.29535 | -1.036954258 | 0.102052792 |
| Znrf2 | 429.2306728 | 0.139040054 | 0.07200468 |
| Nod1 | 393.3170085 | 3.075628862 | 0.180587848 |
| Gm24230 | 0 | NA | NA |
| Ggct | 96.47110576 | 0.401101345 | 0.213581921 |
| Gars | 1477.498831 | 0.614538704 | 0.093235111 |

| | | | |
|---|---|---|---|
| Crhr2 | 14.43709135 | -0.714992141 | 0.298732275 |
| Inmt | 2.148441887 | 0.562061062 | 0.701839536 |
| Fam188b | 89.47598892 | -0.242936717 | 0.150222779 |
| Gm25458 | 0 | NA | NA |
| Aqp1 | 74.91150286 | 2.157063545 | 0.555249871 |
| Ghrhr | 0.426601203 | 2.864632504 | 3.210283662 |
| Adcyap1r1 | 2280.821534 | -0.267864117 | 0.049694028 |
| AC069309.1 | 0.075043205 | 0.237309859 | 3.724945049 |
| Neurod6 | 272.3533864 | -1.581367751 | 0.189869181 |
| Ccdc129 | 12.62732475 | 0.25897892 | 0.289240962 |
| Ppp1r17 | 219.4288715 | -0.137670935 | 0.298155052 |
| Pde1c | 512.5279336 | -0.617624146 | 0.109579926 |
| AC079956.1 | 1.707215572 | -0.89222393 | 0.737002924 |
| Gm24709 | 0 | NA | NA |
| Gm22493 | 0.217296566 | -1.057518501 | 2.166387933 |
| Lsm5 | 26.01436472 | 0.20705114 | 0.212154039 |
| Avl9 | 1503.371839 | 0.411666724 | 0.083941827 |
| Kbtbd2 | 937.4819014 | 0.183239603 | 0.063783958 |
| Fkbp9 | 377.8815204 | 0.225206709 | 0.121117508 |
| Nt5c3 | 889.5108846 | 1.691141047 | 0.139776458 |
| Vmn1r4 | 1.941638245 | 0.043866516 | 0.779871626 |
| Vmn1r5 | 0.356992291 | -0.243574256 | 3.321892865 |
| Vmn1r6 | 0.118371171 | 0.718202646 | 3.724945049 |
| Vmn1r7 | 0 | NA | NA |
| Vmn1r8 | 0 | NA | NA |
| Vmn1r9 | 0 | NA | NA |
| Vmn1r10 | 0 | NA | NA |
| Vmn1r11 | 0 | NA | NA |
| Vmn1r12 | 0 | NA | NA |
| Vmn1r-ps8 | 0 | NA | NA |
| Vmn1r13 | 0 | NA | NA |
| Vmn1r14 | 0 | NA | NA |
| Vmn1r15 | 0 | NA | NA |
| Vmn1r-ps9 | 0 | NA | NA |
| Vmn1r-ps10 | 0 | NA | NA |
| Vmn1r16 | 0 | NA | NA |
| Vmn1r17 | 0 | NA | NA |
| Vmn1r18 | 0 | NA | NA |
| Vmn1r19 | 0 | NA | NA |
| Vmn1r20 | 0 | NA | NA |
| Vmn1r-ps11 | 0 | NA | NA |
| 4930533I22Rik | 0.062426873 | 0.237312084 | 3.724945049 |
| Ppm1k | 1833.8061 | 0.202420645 | 0.106821712 |
| Gm27896 | 0 | NA | NA |
| Herc6 | 2713.472928 | 4.910284819 | 0.168539667 |
| Gm37397 | 18.0739852 | -0.366528002 | 0.24769637 |
| Pyurf | 116.2739024 | -0.296188066 | 0.111368245 |

| | | | |
|---|---|---|---|
| Gm26712 | 0 | NA | NA |
| Lancl2 | 881.9561078 | -0.311209519 | 0.063059524 |
| Vopp1 | 778.6664814 | -0.216860054 | 0.081897186 |
| Vmn1r21 | 0.060174537 | 0.237309859 | 3.724945049 |
| Gm25726 | 0 | NA | NA |
| Vmn1r-ps12 | 0 | NA | NA |
| Vmn1r22 | 0 | NA | NA |
| Vmn1r23 | 0 | NA | NA |
| Gm20660 | 0 | NA | NA |
| Vmn1r24 | 0 | NA | NA |
| Vmn1r25 | 0 | NA | NA |
| Vmn1r26 | 0 | NA | NA |
| Rps15-ps2 | 2.860642787 | -0.585329112 | 0.775115062 |
| Vmn1r-ps14 | 0 | NA | NA |
| Vmn1r-ps15 | 0.09176491 | 0.237310218 | 3.724945049 |
| Gm6695 | 0.119587853 | 0.237305575 | 3.724945049 |
| Vmn1r-ps17 | 0 | NA | NA |
| Gm20718 | 0.110595523 | 0.718200421 | 3.724945049 |
| Vmn1r-ps18 | 0 | NA | NA |
| Vmn1r27 | 0 | NA | NA |
| Vmn1r-ps19 | 0 | NA | NA |
| Vmn1r28 | 0.068097359 | -0.243569582 | 3.724945049 |
| Vmn1r29 | 0.139856492 | 0.237322262 | 3.574411025 |
| Gm20674 | 0 | NA | NA |
| Vmn1r-ps20 | 0 | NA | NA |
| Vmn1r30 | 0.075043205 | 0.237309859 | 3.724945049 |
| Vmn1r-ps21 | 0 | NA | NA |
| Gm20673 | 0 | NA | NA |
| Vmn1r31 | 0 | NA | NA |
| Vmn1r-ps22 | 0.047948698 | -0.243569582 | 3.724945049 |
| Vmn1r-ps23 | 0.066050875 | 0.71820487 | 3.724945049 |
| Gm20672 | 0 | NA | NA |
| Abcg2 | 150.9866917 | 0.436576877 | 0.114753066 |
| Herc3 | 2335.927333 | -0.078367302 | 0.08676086 |
| Nap1l5 | 4015.777592 | -0.017988047 | 0.128740225 |
| Fam13a | 397.9236056 | -0.130457168 | 0.069850189 |
| BB365896 | 2.654833662 | 1.053325782 | 0.735113482 |
| Gm22367 | 0 | NA | NA |
| Tigd2 | 330.9779313 | 0.682093701 | 0.110413246 |
| Gprin3 | 236.7557592 | -0.811242982 | 0.125150861 |
| Gm22622 | 0 | NA | NA |
| Gm5570 | 0.128070691 | 0.718198228 | 3.723419581 |
| Gm22723 | 0 | NA | NA |
| n-R5s162 | 0 | NA | NA |
| A530053G22Rik | 0.106618422 | -0.24357403 | 3.724945049 |
| Snca | 1013.675009 | -0.655844464 | 0.126423688 |
| Mmrn1 | 0.717105192 | 2.967675074 | 1.210417871 |

| | | | |
|---|---|---|---|
| A730020E08Rik | 23.88982311 | -0.383909922 | 0.278325088 |
| Ccser1 | 527.4917736 | -0.547273406 | 0.10580991 |
| Gm22212 | 0 | NA | NA |
| Grid2 | 1329.763725 | -0.503612445 | 0.118163708 |
| Gm25205 | 1.741084885 | -0.486291333 | 0.957060171 |
| Atoh1 | 0.260468428 | 1.558713868 | 2.020704085 |
| Gm15534 | 0.038981395 | 0.718207094 | 3.724945049 |
| Smarcad1 | 613.7764423 | 0.049698984 | 0.066522804 |
| Hpgds | 46.4750562 | -0.421915488 | 0.270828494 |
| Gm17147 | 2.881706403 | 0.362704275 | 0.70393128 |
| C130060K24Rik | 9.901210978 | 0.434845464 | 0.366131258 |
| Tnip3 | 4.68769279 | 4.855114388 | 1.00498116 |
| Ndnf | 237.9647372 | -0.435982184 | 0.163315946 |
| Gm15490 | 0.045882455 | 0.237312084 | 3.724945049 |
| Gm16838 | 1.376958878 | -0.395907922 | 0.879672445 |
| Prdm5 | 99.54165155 | 0.137101975 | 0.113477281 |
| Gm23543 | 0 | NA | NA |
| Gm20445 | 3.495574212 | -1.518720824 | 0.599217319 |
| 4930544G11Rik | 0 | NA | NA |
| Gm23625 | 0 | NA | NA |
| Mad2l1 | 71.55826495 | -0.061134949 | 0.174519292 |
| Vmn1r32 | 0 | NA | NA |
| Vmn1r33 | 0.066050875 | 0.71820487 | 3.724945049 |
| Vmn1r34 | 0.225129614 | 0.237300054 | 3.724945049 |
| Vmn1r35 | 0.038981395 | 0.718207094 | 3.724945049 |
| Vmn1r36 | 0 | NA | NA |
| Vmn1r37 | 0 | NA | NA |
| Vmn1r-ps24 | 0.066050875 | 0.71820487 | 3.724945049 |
| Vmn1r38 | 0 | NA | NA |
| Vmn1r39 | 0.047948698 | -0.243569582 | 3.724945049 |
| RP23-263B18.4 | 379.7491701 | 0.706472727 | 0.155882574 |
| 4930597O21Rik | 2.946845261 | 0.79176565 | 0.59466496 |
| Gng12 | 1315.427679 | 0.988243191 | 0.118668429 |
| Gm15644 | 4.54732579 | -0.145863011 | 0.530514763 |
| Gm25260 | 0.532344628 | 0.531776954 | 1.539859007 |
| Gadd45a | 98.08999083 | 1.863758219 | 0.272312186 |
| E230016M11Rik | 92.89630384 | -0.710765966 | 0.190551809 |
| A430010J10Rik | 11.54166846 | -0.60662451 | 0.356913404 |
| Gm27015 | 0.824269312 | 0.255037787 | 1.385901974 |
| Gm8566 | 29.8603066 | 0.252297327 | 0.219822484 |
| Serbp1 | 2115.70978 | -0.151198682 | 0.111371278 |
| Il12rb2 | 27.958596 | 4.581611328 | 0.589862762 |
| Gm4761 | 0 | NA | NA |
| Il23r | 2.075698228 | 1.454091972 | 0.923680847 |
| RP23-123I10.3 | 0.062426873 | 0.237312084 | 3.724945049 |
| Tacstd2 | 4.381073283 | 1.763763948 | 0.598993938 |
| Igkv2-137 | 0.061920613 | 0.718207094 | 3.724945049 |

| | | | |
|---|---|---|---|
| RP23-123I10.4 | 0 | NA | NA |
| Igkv1-136 | 0 | NA | NA |
| Igkv1-135 | 0.427013396 | 2.641813523 | 1.539463762 |
| RP24-322E13.1 | 0 | NA | NA |
| RP24-322E13.2 | 0 | NA | NA |
| Igkv14-134-1 | 0 | NA | NA |
| Igkv17-134 | 0 | NA | NA |
| Igkv1-133 | 0.216076386 | 0.237307547 | 3.724945049 |
| RP23-255N3.3 | 0 | NA | NA |
| RP23-255N3.4 | 0 | NA | NA |
| RP23-255N3.6 | 0 | NA | NA |
| Igkv1-132 | 0.132101749 | 1.209894437 | 3.72418822 |
| Igkv1-131 | 0 | NA | NA |
| RP23-255N3.8 | 0 | NA | NA |
| Igkv14-130 | 0 | NA | NA |
| Gm22425 | 0 | NA | NA |
| Gm24911 | 0 | NA | NA |
| Igkv9-129 | 0 | NA | NA |
| Igkv9-128 | 0 | NA | NA |
| Igkv17-127 | 0.143022002 | 0.719176055 | 3.547735711 |
| Igkv14-126-1 | 0 | NA | NA |
| RP23-234A12.5 | 0 | NA | NA |
| Igkv14-126 | 0 | NA | NA |
| Gm25910 | 0 | NA | NA |
| Igkv11-125 | 0.062426873 | 0.237312084 | 3.724945049 |
| Igkv9-124 | 0 | NA | NA |
| Igkv9-123 | 0 | NA | NA |
| Igkv1-122 | 0 | NA | NA |
| Igkv17-121 | 0.097139547 | 0.718207094 | 3.724945049 |
| Igkv9-120 | 0.291418641 | 1.927454675 | 3.70657633 |
| Igkv9-119 | 0 | NA | NA |
| Igkv14-118-2 | 0 | NA | NA |
| Igkv14-118-1 | 0 | NA | NA |
| RP23-185M15.10 | 0 | NA | NA |
| RP23-185M15.7 | 0 | NA | NA |
| Igkv11-118 | 0 | NA | NA |
| Igkv1-117 | 0.45022207 | 2.963647719 | 3.671267695 |
| RP24-76K4.8 | 0 | NA | NA |
| RP24-76K4.7 | 0.141094079 | 0.718203391 | 3.563850454 |
| Igkv2-116 | 0.060174537 | 0.237309859 | 3.724945049 |
| Igkv1-115 | 0 | NA | NA |
| Igkv11-114 | 0 | NA | NA |
| Igkv2-113 | 0 | NA | NA |
| RP24-76K4.1 | 0 | NA | NA |
| Igkv2-112 | 0.157314084 | 0.718207559 | 3.435173412 |
| Igkv14-111 | 0.94930219 | 3.402695791 | 1.116702358 |
| Gm25436 | 0 | NA | NA |

| | | | |
|---|---|---|---|
| Igkv1-110 | 0.099106094 | 1.209894437 | 3.72418822 |
| RP24-76K4.12 | 0 | NA | NA |
| RP24-76K4.11 | 0 | NA | NA |
| Igkv2-109 | 0.106971521 | 0.237307635 | 3.724945049 |
| Igkv1-108 | 0 | NA | NA |
| Igkv2-107 | 0 | NA | NA |
| Igkv19-93 | 0 | NA | NA |
| Igkv2-105 | 0 | NA | NA |
| RP23-29F15.2 | 0 | NA | NA |
| Igkv16-104 | 0.862499085 | 1.82550048 | 3.711599481 |
| Igkv15-103 | 0.066050875 | 0.71820487 | 3.724945049 |
| Igkv15-102 | 0 | NA | NA |
| Igkv20-101-2 | 0 | NA | NA |
| Igkv15-101-1 | 0 | NA | NA |
| Igkv15-101 | 0 | NA | NA |
| Igkv14-100 | 0 | NA | NA |
| Gm25833 | 0 | NA | NA |
| Igkv1-99 | 0 | NA | NA |
| Igkv12-98 | 0 | NA | NA |
| Igkv15-97 | 0 | NA | NA |
| Igkv10-96 | 0.163190422 | 1.22955094 | 3.396300676 |
| Igkv2-95-2 | 0 | NA | NA |
| Igkv2-95-1 | 0 | NA | NA |
| Igkv10-95 | 0 | NA | NA |
| Igkv10-94 | 0.066050875 | 0.71820487 | 3.724945049 |
| Igkv2-93-1 | 0 | NA | NA |
| RP24-405O23.8 | 0 | NA | NA |
| AC120404.1 | 0 | NA | NA |
| Igkv4-92 | 0 | NA | NA |
| RP23-211L5.2 | 0 | NA | NA |
| Igkv4-91 | 0 | NA | NA |
| Igkv4-90 | 0 | NA | NA |
| Igkv13-89-1 | 0 | NA | NA |
| Igkv12-89 | 0.175102337 | 1.440451655 | 3.306427423 |
| Igkv1-88 | 0.108772582 | 0.718202646 | 3.724945049 |
| RP23-211L5.7 | 0 | NA | NA |
| IGKV1D-42 | 0 | NA | NA |
| RP23-211L5.9 | 0.045882455 | 0.237312084 | 3.724945049 |
| Igkv13-87 | 0.049553047 | 0.71820487 | 3.724945049 |
| RP23-211L5.12 | 0.088534442 | 1.179809211 | 3.723500796 |
| Igkv4-86 | 0 | NA | NA |
| Igkv12-47 | 0.097139547 | 0.718207094 | 3.724945049 |
| Igkv13-84 | 0.060174537 | 0.237309859 | 3.724945049 |
| Igkv4-56 | 0 | NA | NA |
| Igkv13-82 | 0 | NA | NA |
| Igkv4-81 | 0 | NA | NA |
| Igkv13-80-1 | 0 | NA | NA |

| | | | |
|---|---|---|---|
| Igkv4-80 | 0.097139547 | 0.718207094 | 3.724945049 |
| Igkv4-79 | 0 | NA | NA |
| Igkv13-78-1 | 0 | NA | NA |
| Igkv4-78 | 0 | NA | NA |
| RP23-344N8.12 | 0 | NA | NA |
| Igkv4-77 | 0 | NA | NA |
| Igkv13-76 | 0 | NA | NA |
| Igkv4-75 | 0 | NA | NA |
| Igkv13-74-1 | 0 | NA | NA |
| Igkv4-74 | 0.062426873 | 0.237312084 | 3.724945049 |
| Igkv13-73-1 | 0 | NA | NA |
| Igkv4-73 | 0.046345709 | 0.71820487 | 3.724945049 |
| Igkv4-72 | 0.182573437 | -0.243564306 | 3.311478753 |
| Igkv13-71-1 | 0 | NA | NA |
| Igkv4-71 | 0 | NA | NA |
| Igkv4-70 | 0.046345709 | 0.71820487 | 3.724945049 |
| Igkv4-69 | 0 | NA | NA |
| Igkv4-68 | 0.060174537 | 0.237309859 | 3.724945049 |
| Igkv12-67 | 0 | NA | NA |
| Rprl1 | 0 | NA | NA |
| Igkv12-66 | 0 | NA | NA |
| Igkv4-65 | 0 | NA | NA |
| Igkv13-64 | 0 | NA | NA |
| Igkv4-63 | 0 | NA | NA |
| Igkv13-62-1 | 0 | NA | NA |
| Igkv4-62 | 0 | NA | NA |
| Igkv13-61-1 | 0 | NA | NA |
| Igkv4-61 | 0.266304556 | 1.36146117 | 3.715728684 |
| Igkv4-59 | 0 | NA | NA |
| Igkv4-60 | 0 | NA | NA |
| Igkv4-58 | 0 | NA | NA |
| Rn7s6 | 476.0174306 | 0.289892388 | 0.334346588 |
| AC156953.1 | 0 | NA | NA |
| Igkv13-57-2 | 0 | NA | NA |
| Igkv4-57-1 | 0.097139547 | 0.718207094 | 3.724945049 |
| Igkv13-57-1 | 0 | NA | NA |
| Igkv4-57 | 0.102891481 | -0.243576254 | 3.724945049 |
| Igkv13-56-1 | 0 | NA | NA |
| Igkv13-55-1 | 0 | NA | NA |
| Igkv4-55 | 0.097139547 | 0.718207094 | 3.724945049 |
| Igkv13-54-1 | 0 | NA | NA |
| Igkv4-54 | 0 | NA | NA |
| Igkv4-53 | 0 | NA | NA |
| Gm11143 | 0 | NA | NA |
| RP23-24I14.7 | 0 | NA | NA |
| Igkv4-50 | 0 | NA | NA |
| Igkv12-49 | 0 | NA | NA |

| | | | |
|---|---|---|---|
| Igkv5-48 | 0.183032319 | 0.695603381 | 3.319201136 |
| Igkv12-46 | 0.771993125 | 3.298765275 | 3.513920301 |
| Igkv5-45 | 0.049553047 | 0.71820487 | 3.724945049 |
| Igkv12-44 | 0.240624804 | 1.893902921 | 3.03496855 |
| Igkv5-43 | 0.240624804 | 1.893902921 | 3.03496855 |
| Igkv12-42 | 0 | NA | NA |
| Igkv12-41 | 0.194279094 | 1.292225235 | 3.719439735 |
| Igkv5-40-1 | 0 | NA | NA |
| Igkv12-40 | 0 | NA | NA |
| Igkv5-39 | 0.975696798 | 2.75435191 | 2.648219456 |
| RP24-475M8.10 | 0 | NA | NA |
| Igkv12-38 | 0.099106094 | 1.209894437 | 3.72418822 |
| Igkv5-37 | 0 | NA | NA |
| Igkv18-36 | 0 | NA | NA |
| Igkv1-35 | 0 | NA | NA |
| Gm23694 | 0 | NA | NA |
| RP24-314D8.6 | 0 | NA | NA |
| Igkv8-34 | 0.066050875 | 0.71820487 | 3.724945049 |
| Igkv7-33 | 0.093831153 | -0.243571806 | 3.724945049 |
| Igkv6-32 | 0.998588786 | 1.527620321 | 3.675501912 |
| Igkv8-31 | 0 | NA | NA |
| Igkv8-30 | 0.203286941 | 1.526406207 | 2.603554853 |
| Amd-ps2 | 0.04393175 | -0.243571806 | 3.724945049 |
| Igkv6-29 | 0 | NA | NA |
| Igkv8-28 | 0 | NA | NA |
| Gm24096 | 0 | NA | NA |
| Igkv8-27 | 0.112396584 | 1.192083721 | 3.724434897 |
| Igkv8-26 | 0 | NA | NA |
| RP23-64J23.3 | 0.049553047 | 0.71820487 | 3.724945049 |
| Igkv6-25 | 0.097139547 | 0.718207094 | 3.724945049 |
| Igkv8-24 | 0 | NA | NA |
| Igkv8-23-1 | 0 | NA | NA |
| Igkv6-23 | 1.206331356 | 4.280336937 | 3.647219182 |
| RP23-64J23.7 | 0 | NA | NA |
| Gm11065 | 0 | NA | NA |
| Igkv8-22 | 0 | NA | NA |
| Igkv8-21 | 0.277388482 | 0.237330532 | 1.787163803 |
| Amd-ps5 | 0.048949532 | -0.243569582 | 3.724945049 |
| Igkv6-20 | 0.038981395 | 0.718207094 | 3.724945049 |
| Igkv8-19 | 0.269481931 | -0.883869421 | 2.410497298 |
| Igkv8-18 | 0 | NA | NA |
| RP23-64J23.14 | 0.089814205 | -0.24357403 | 3.724945049 |
| Igkv6-17 | 0.659092009 | 3.152227755 | 3.667902572 |
| Igkv8-16 | 0.097139547 | 0.718207094 | 3.724945049 |
| RP23-175B24.3 | 0 | NA | NA |
| Igkv6-15 | 0.421418085 | 1.843448336 | 1.664973325 |
| RP23-175B24.4 | 0.066050875 | 0.71820487 | 3.724945049 |

| | | | |
|---|---|---|---|
| Gm10360 | 0 | NA | NA |
| RP23-175B24.5 | 5.844746325 | -0.416280167 | 0.379860197 |
| Igkv6-14 | 0 | NA | NA |
| RP23-175B24.8 | 0 | NA | NA |
| Igkv6-13 | 0.044544648 | 0.237309859 | 3.724945049 |
| RP23-175B24.10 | 0 | NA | NA |
| RP23-175B24.13 | 0.042716945 | -0.243571806 | 3.724945049 |
| RP23-175B24.14 | 0 | NA | NA |
| Igkv3-12-1 | 0 | NA | NA |
| Igkv3-12 | 0.232585366 | 1.152901533 | 2.538730861 |
| Igkv3-11 | 0 | NA | NA |
| Igkv3-10 | 0.300744356 | 1.802836861 | 3.711870465 |
| Igkv3-9 | 0 | NA | NA |
| Igkv3-8 | 0 | NA | NA |
| Igkv3-7 | 4.685552657 | 5.795968833 | 1.560888412 |
| Igkv3-6 | 0 | NA | NA |
| Igkv3-5 | 0.169165009 | 0.719161184 | 3.351346564 |
| Igkv3-4 | 0.266304556 | 1.36146117 | 3.715728684 |
| Igkv3-3 | 0 | NA | NA |
| Igkv3-2 | 0.309883016 | 2.467318587 | 3.687546276 |
| Igkv3-1 | 0 | NA | NA |
| RP23-435I4.11 | 0.318875346 | 1.869770846 | 3.709404448 |
| Igkj1 | 0.169165009 | 0.719161184 | 3.351346564 |
| Igkj2 | 0 | NA | NA |
| Igkj3 | 0.062426873 | 0.237312084 | 3.724945049 |
| Igkj4 | 0.254453631 | 1.256673259 | 2.971544927 |
| Igkj5 | 0.189830966 | 1.668992102 | 3.258054319 |
| Igkc | 42.12345985 | 5.907522364 | 1.051625606 |
| Rpia | 130.920887 | -0.141335292 | 0.103802021 |
| Eif2ak3 | 463.4739881 | 0.034856866 | 0.07110906 |
| Gm23346 | 0.975898478 | 0.037839043 | 0.980364594 |
| RP24-122N23.3 | 0.09349418 | -0.24357403 | 3.724945049 |
| Tex37 | 0 | NA | NA |
| Foxi3 | 0.042716945 | -0.243571806 | 3.724945049 |
| Gm23485 | 0 | NA | NA |
| Rpl34-ps1 | 54.38793798 | 0.815487407 | 0.25116767 |
| Thnsl2 | 33.95158729 | 0.135412066 | 0.212888996 |
| Fabp1 | 0.161948271 | -0.243562662 | 3.446645093 |
| Smyd1 | 16.55316792 | -1.476197623 | 0.316947837 |
| Mir8112 | 5.618825651 | 0.977676982 | 0.496250706 |
| Krcc1 | 217.6208105 | 1.185059272 | 0.173571909 |
| Gm1070 | 1.970056041 | 1.438412632 | 0.813739767 |
| Cd8b1 | 33.23125073 | 8.532941678 | 0.892659816 |
| Cd8a | 74.76319009 | 8.605621681 | 0.88978395 |
| Rmnd5a | 2145.404065 | -0.400427207 | 0.062086781 |
| Rnf103 | 778.7289262 | -0.168424627 | 0.062803105 |
| Chmp3 | 861.0926457 | -0.080096199 | 0.073462605 |

| | | | |
|---|---|---|---|
| Kdm3a | 752.2610037 | 0.175541551 | 0.076369227 |
| Reep1 | 1109.512451 | -0.2152494 | 0.073435727 |
| Mrpl35 | 321.4157534 | 0.130986258 | 0.073308917 |
| Immt | 1466.512924 | 0.185052846 | 0.059265464 |
| Ptcd3 | 534.5440444 | -0.015993543 | 0.075171763 |
| Gm22486 | 336.2636012 | -0.140882245 | 0.32949797 |
| Polr1a | 650.2299052 | 0.437332493 | 0.05491636 |
| Gm26628 | 0.134276636 | 0.684459443 | 2.951368184 |
| St3gal5 | 1041.7803 | -0.328933495 | 0.055259784 |
| Gm29438 | 74.24082104 | -1.295920503 | 0.310765784 |
| Atoh8 | 37.89726758 | 1.160178368 | 0.246085329 |
| Sftpb | 0.285479113 | 0.718794291 | 1.787992454 |
| Usp39 | 316.7202377 | 0.528237389 | 0.074852504 |
| 0610030E20Rik | 238.8730958 | -0.041405799 | 0.086721132 |
| Tmem150a | 75.99670554 | 0.321271373 | 0.154578554 |
| Rnf181 | 551.102173 | 0.490503567 | 0.09348532 |
| Vamp5 | 21.09757441 | 1.599145404 | 0.30971948 |
| Vamp8 | 157.0775941 | 2.131986185 | 0.176235422 |
| Ggcx | 228.9491815 | 0.01021266 | 0.092136223 |
| Mat2a | 2291.964425 | 0.579540922 | 0.123340803 |
| Sh2d6 | 8.896846792 | 0.819304477 | 0.381268641 |
| Capg | 296.8370465 | 6.17624027 | 0.429833924 |
| Elmod3 | 206.9098534 | -0.198306596 | 0.091882853 |
| Retsat | 77.16422045 | -0.384834972 | 0.178330148 |
| Tgoln1 | 2850.702626 | 0.514840816 | 0.076008015 |
| Gm20536 | 0.787340167 | -1.591118921 | 1.198509791 |
| Tcf7l1 | 89.55502137 | 0.532351432 | 0.139699508 |
| Gm15401 | 1.324332992 | 1.202125999 | 0.984086506 |
| Gm37969 | 4.217504979 | -0.868453277 | 0.496397871 |
| Gm38114 | 1.798572027 | -0.736541917 | 0.772562323 |
| Gm38320 | 0.230331698 | 0.237324042 | 2.566107632 |
| Gm15402 | 0.090074004 | -0.24357403 | 3.724945049 |
| Gm37736 | 8.749811602 | -0.766478954 | 0.395487095 |
| Kcmf1 | 755.6815892 | 0.446997274 | 0.079023859 |
| Gm26640 | 0.344365504 | 1.230369784 | 3.722845822 |
| Tmsb10 | 1562.811079 | 1.080975526 | 0.218379827 |
| Dnah6 | 180.1945128 | -0.786697098 | 0.173412103 |
| Gm24533 | 0.239208914 | -0.724460444 | 2.110573275 |
| Suclg1 | 851.7330975 | -0.005416346 | 0.057004633 |
| Gm24370 | 0 | NA | NA |
| 4931417E11Rik | 0 | NA | NA |
| Gm26179 | 0 | NA | NA |
| Gm23924 | 0 | NA | NA |
| Gm22254 | 0 | NA | NA |
| Gm8993 | 0 | NA | NA |
| Gm9001 | 4.797213872 | 0.228508589 | 0.521937102 |
| AC161363.1 | 0 | NA | NA |

| | | | |
|---|---|---|---|
| Gm22283 | 0 | NA | NA |
| Gm9008 | 0.868283866 | 0.040199994 | 1.084988058 |
| Ctnna2 | 1519.198385 | -0.56835473 | 0.05976445 |
| Lrrtm1 | 446.8085958 | -0.576762799 | 0.110432394 |
| Gm21045 | 0 | NA | NA |
| Gm5576 | 0.929768241 | -0.094386676 | 0.980197921 |
| Reg3b | 0.091880448 | -0.724455696 | 3.724945049 |
| Reg3d | 0 | NA | NA |
| Reg3a | 0 | NA | NA |
| Reg2 | 0 | NA | NA |
| Reg1 | 0 | NA | NA |
| Reg3g | 0.501377686 | 2.43043679 | 1.397109704 |
| Gm21392 | 0 | NA | NA |
| Gm23727 | 0 | NA | NA |
| Gm29005 | 0 | NA | NA |
| Gm20594 | 0 | NA | NA |
| Gm4409 | 0.396083486 | -1.344253654 | 1.526726686 |
| Lrrtm4 | 669.3441651 | -0.963280562 | 0.121976515 |
| Gm26264 | 0 | NA | NA |
| Gm22530 | 0 | NA | NA |
| mmu-mir-7232 | 0.163166504 | -0.723831518 | 2.784882282 |
| mmu-mir-468 | 0 | NA | NA |
| 1700009C05Rik | 0.362568279 | -1.606280096 | 1.550894516 |
| Gcfc2 | 109.5450915 | -0.148046228 | 0.182775234 |
| Mrpl19 | 289.1779516 | 0.055938418 | 0.068280381 |
| Eva1a | 89.53357422 | -0.453236092 | 0.169518152 |
| Gm15864 | 0 | NA | NA |
| Tacr1 | 94.67186864 | 0.124467412 | 0.123163881 |
| Pole4 | 302.9621574 | 0.309389733 | 0.114305119 |
| Hk2 | 352.4423473 | 2.055254119 | 0.198904253 |
| Gm17034 | 3.741498389 | 5.834459913 | 0.950768362 |
| 2310069B03Rik | 0.29717078 | 1.776591104 | 2.016152136 |
| Sema4f | 443.9451415 | -0.308349678 | 0.09753583 |
| M1ap | 2.42047183 | 1.544702316 | 0.885242951 |
| Dok1 | 94.45580167 | 3.771203821 | 0.293249421 |
| Loxl3 | 99.54709547 | 2.025684708 | 0.19910991 |
| Mir7040 | 0 | NA | NA |
| Htra2 | 198.8763925 | 0.450679264 | 0.08305155 |
| Aup1 | 620.1822418 | 0.797341252 | 0.05551638 |
| Dqx1 | 10.03751516 | -0.045910696 | 0.404835949 |
| Tlx2 | 0.090665643 | -0.724455696 | 3.724945049 |
| Gm37092 | 0.240333628 | 0.691358771 | 1.852219835 |
| Pcgf1 | 96.34584019 | 0.336690332 | 0.10909448 |
| Lbx2 | 0.810336573 | 0.916641465 | 1.107280575 |
| Mir3470a | 0 | NA | NA |
| Ccdc142 | 74.38601428 | -0.377287725 | 0.179694837 |
| Ccdc142os | 12.6558528 | -0.029409684 | 0.306694191 |

| | | | |
|---|---|---|---|
| Mrpl53 | 0 | NA | NA |
| Mogs | 256.0958102 | 0.026655697 | 0.103915648 |
| Wbp1 | 156.9620067 | -0.20712328 | 0.125280301 |
| Ino80b | 119.6851225 | 0.272113579 | 0.165455007 |
| Rtkn | 581.9726703 | 0.350503248 | 0.081258465 |
| Wdr54 | 115.1061269 | -0.577582471 | 0.11715923 |
| 1700003E16Rik | 151.7522411 | -0.455396968 | 0.107458534 |
| Dctn1 | 3696.021488 | -0.500462729 | 0.05886959 |
| Gm15624 | 21.19334217 | -1.645978754 | 0.401365328 |
| Slc4a5 | 42.25336671 | -1.678379512 | 0.264613389 |
| Mthfd2 | 159.228204 | 1.740410603 | 0.1710621 |
| Mob1a | 606.7587231 | 1.355441287 | 0.129968779 |
| Bola3 | 181.7266574 | 0.22464388 | 0.093675604 |
| Tet3 | 1814.49476 | -0.144337552 | 0.105717613 |
| Mir6374 | 7.73301019 | -1.489153944 | 0.529195473 |
| B230319C09Rik | 9.836627767 | 0.333889429 | 0.33083657 |
| Dguok | 113.566417 | -0.18834809 | 0.167131152 |
| Actg2 | 1.148058676 | 1.53555005 | 0.969383906 |
| Stambp | 312.4465512 | 0.060263372 | 0.083432877 |
| Gm24990 | 0 | NA | NA |
| Clec4f | 0.297646291 | 1.155870254 | 1.795877463 |
| Cd207 | 0.357781856 | 2.107253821 | 1.526650388 |
| Vax2os | 1.069186981 | -1.09893517 | 1.033422714 |
| Gm27772 | 0 | NA | NA |
| Gm27631 | 0 | NA | NA |
| Gm27893 | 0 | NA | NA |
| Vax2 | 0.095898756 | 1.192083721 | 3.724434897 |
| Gm25054 | 0 | NA | NA |
| Atp6v1b1 | 0.540009155 | -1.510964685 | 1.572763479 |
| 1700124L16Rik | 10.03157174 | -1.099931651 | 0.399809062 |
| Ankrd53 | 0 | NA | NA |
| Tex261 | 481.6030204 | 0.537315484 | 0.106628513 |
| Nagk | 216.7968119 | 0.101116925 | 0.100801311 |
| Paip2b | 685.0019845 | 0.110012824 | 0.07516905 |
| Gm23014 | 0 | NA | NA |
| Gm5138 | 12.43742249 | 0.189451682 | 0.28879694 |
| Zfp638 | 2492.303382 | -0.098086106 | 0.079338139 |
| Dysf | 208.5045205 | 2.139163781 | 0.238018842 |
| Gm15475 | 0.044544648 | 0.237309859 | 3.724945049 |
| Cyp26b1 | 126.4459904 | -0.651651129 | 0.173063481 |
| Exoc6b | 1761.121593 | -0.113110165 | 0.047351858 |
| Gm7507 | 0.706207703 | -0.208169845 | 1.225951935 |
| Gm26353 | 0.038981395 | 0.718207094 | 3.724945049 |
| Npm3-ps1 | 4.293674138 | 0.460369708 | 0.591400716 |
| Gm5878 | 0.534626289 | 0.386673665 | 1.284232744 |
| Spr | 209.6465649 | 0.185603116 | 0.09228805 |
| Gm10444 | 0.134148234 | 0.23732184 | 3.624632408 |

| | | | |
|---|---|---|---|
| Gm22893 | 0 | NA | NA |
| Spr-ps1 | 0.674932609 | -1.013278043 | 1.429913286 |
| Emx1 | 100.6339851 | 0.023423857 | 0.17105228 |
| Sfxn5 | 2038.190871 | -0.730265971 | 0.114793314 |
| Rab11fip5 | 1169.630522 | 0.180290069 | 0.081012908 |
| Mir705 | 0 | NA | NA |
| Noto | 0 | NA | NA |
| Smyd5 | 326.4950938 | 0.245971819 | 0.074927412 |
| Pradc1 | 68.96069805 | -0.777397166 | 0.150081102 |
| Cct7 | 1480.602166 | 0.450640024 | 0.075930154 |
| Fbxo41 | 1325.807268 | -0.924477671 | 0.119477167 |
| Egr4 | 333.4082471 | -0.130866675 | 0.534015533 |
| Alms1 | 761.6584805 | -0.522621952 | 0.130539566 |
| RP23-27D8.2 | 34.95142274 | -1.22908531 | 0.251489511 |
| RP24-478I22.8 | 43.9950079 | -0.593711514 | 0.243140209 |
| Gm4477 | 0.754021959 | -0.019383738 | 1.016394506 |
| Alms1-ps1 | 10.38534064 | -0.524829542 | 0.299414202 |
| Cml3 | 8.402099551 | -0.91055582 | 0.385185279 |
| Gm27317 | 0.061920613 | 0.718207094 | 3.724945049 |
| Alms1-ps2 | 11.88847144 | -0.251309479 | 0.320767248 |
| mmu-mir-6375 | 0.167183452 | -0.72445987 | 2.765153652 |
| Gm11128 | 2.945191291 | -1.904718845 | 0.591026505 |
| Cml5 | 13.56449452 | -0.216769908 | 0.353236121 |
| Nat8 | 14.92973809 | -0.038972465 | 0.291085978 |
| Gm9769 | 437.2277849 | 0.622488777 | 0.084229989 |
| Cml2 | 0.711965957 | -1.421352891 | 1.230268733 |
| RP23-448G18.6 | 0 | NA | NA |
| 1700019G17Rik | 48.23925206 | 0.538495615 | 0.196069117 |
| Cml1 | 59.08754969 | 0.021904564 | 0.181782092 |
| Tprkb | 600.9053318 | 0.044754037 | 0.138592323 |
| RP23-345O17.4 | 0 | NA | NA |
| Nat8b | 0.082913145 | 0.23732098 | 3.724945049 |
| RP23-345O17.6 | 0.298750603 | 0.237330868 | 1.779430853 |
| Dusp11 | 1203.32624 | -0.102253277 | 0.12216542 |
| RP23-345O17.5 | 10.75741223 | -0.488229181 | 0.341905922 |
| RP23-345O17.7 | 10.85736135 | 0.177369771 | 0.325471046 |
| Figla | 0.579661142 | -0.744266858 | 1.485218109 |
| Add2 | 1996.526052 | -0.343129848 | 0.115336226 |
| Tgfa | 554.9848714 | 0.443268491 | 0.092077191 |
| Gm7545 | 0.358406704 | 0.729020509 | 1.733766709 |
| Gm10443 | 50.80569197 | 0.248456051 | 0.234214851 |
| Fam136a | 140.0252418 | 0.093270617 | 0.154751231 |
| Snrpg | 133.4774109 | 0.530009733 | 0.103150236 |
| Pcyox1 | 1206.150805 | -0.296572176 | 0.057956447 |
| Tia1 | 1452.959773 | -0.567882987 | 0.126499704 |
| C87436 | 200.9232409 | -0.170118334 | 0.116134949 |
| A430078I02Rik | 3.729996922 | -0.576956771 | 0.546047728 |

| | | | |
|---|---|---|---|
| 2310040G24Rik | 5.093180958 | -0.004872655 | 0.427452538 |
| Gm28720 | 0.502304703 | 0.674321621 | 1.395747216 |
| Gm28719 | 0.796860523 | 0.590077922 | 1.354571267 |
| Pcbp1 | 1630.30914 | 0.510611094 | 0.082780579 |
| 1600020E01Rik | 80.68127648 | 0.163490096 | 0.154255543 |
| Gm22184 | 0 | NA | NA |
| Asprv1 | 17.06775386 | 0.874912229 | 0.291066146 |
| Mxd1 | 914.3853629 | 1.927207933 | 0.189358428 |
| Snrnp27 | 334.3660018 | 0.371419483 | 0.110551036 |
| Gmcl1 | 364.6549906 | -0.413808391 | 0.092922782 |
| Anxa4 | 269.2860603 | 3.607510268 | 0.233371263 |
| 2610306M01Rik | 36.58485778 | 0.105408152 | 0.180345626 |
| Aak1 | 5511.101378 | -0.42424755 | 0.077436402 |
| Nfu1 | 197.0210827 | 0.122983985 | 0.090704342 |
| Gfpt1 | 1366.619225 | 0.091328394 | 0.054168333 |
| D6Ertd527e | 0.429069343 | 0.718228328 | 1.582414148 |
| Antxr1 | 334.4251876 | 0.158692745 | 0.121546627 |
| n-R5s164 | 0 | NA | NA |
| Gkn1 | 0.539210579 | 2.528817749 | 1.366771052 |
| Gkn2 | 0 | NA | NA |
| Gkn3 | 2.877452801 | -3.402174687 | 0.830786095 |
| Bmp10 | 1.435473298 | 4.291210702 | 1.002791959 |
| Arhgap25 | 146.1837619 | 1.56577164 | 0.156425138 |
| Prokr1 | 11.83858572 | 3.783664582 | 0.524364366 |
| Aplf | 74.79263325 | -0.06872114 | 0.146213753 |
| E230015B07Rik | 2.634345682 | -0.620551153 | 0.604944554 |
| 1810020O05Rik | 22.22824979 | -1.167566096 | 0.258733458 |
| Efcc1 | 88.67690409 | -0.861856791 | 0.187820954 |
| Gm5879 | 12.16014049 | 0.54108812 | 0.31910892 |
| Gp9 | 0.777280298 | -0.427075096 | 1.191655217 |
| Rab43 | 434.7988563 | 0.915682345 | 0.084621243 |
| Isy1 | 190.4889355 | 0.041428815 | 0.094155694 |
| Cnbp | 2353.702497 | 0.606489748 | 0.106122273 |
| Copg1 | 2888.377927 | -0.199515247 | 0.031982538 |
| Gm26636 | 0.2969049 | 0.237327654 | 1.964299551 |
| Hmces | 147.1776525 | 0.15959782 | 0.135711605 |
| Rps19-ps9 | 0 | NA | NA |
| H1fx | 161.3903661 | -0.634383939 | 0.122173189 |
| Gm5577 | 38.22855657 | -1.099047805 | 0.280609417 |
| AC153872.1 | 0.355821277 | 0.625055056 | 1.454358679 |
| Rab7 | 2929.399181 | 0.435363141 | 0.07697738 |
| Rpn1 | 1428.832432 | 0.52815007 | 0.069839617 |
| Mir6376 | 0 | NA | NA |
| Gata2 | 41.94893332 | 1.776186989 | 0.244071661 |
| Dnajb8 | 0 | NA | NA |
| 1700031F10Rik | 0.520495925 | 1.191609935 | 1.609376855 |
| Eefsec | 187.7376768 | -0.022189145 | 0.102774517 |

| | | | |
|---|---|---|---|
| Gm25178 | 0 | NA | NA |
| Ruvbl1 | 250.3254379 | 0.267956284 | 0.09700238 |
| Sec61a1 | 1211.826051 | 1.064122281 | 0.101503846 |
| Kbtbd12 | 4.967738157 | 0.999407183 | 0.553646694 |
| Gm26588 | 16.08933923 | -0.97925719 | 0.378562463 |
| Mgll | 1645.494617 | 0.027609321 | 0.090186791 |
| Abtb1 | 225.610541 | -0.30991752 | 0.090083387 |
| Podxl2 | 1494.724341 | -0.235187364 | 0.082912954 |
| Gm15612 | 6.463977267 | -0.363810111 | 0.473015271 |
| Mcm2 | 112.1005328 | 1.330237227 | 0.22380235 |
| Tpra1 | 240.1461408 | -0.18043841 | 0.117857934 |
| 4933427D06Rik | 0.766530834 | 1.789004823 | 1.292597396 |
| Gm1965 | 0.219739983 | 2.071179841 | 2.555741374 |
| Gm6507 | 0 | NA | NA |
| Gm839 | 0.209172349 | 1.705675153 | 2.585387636 |
| Gm25961 | 0.088534442 | 1.179809211 | 3.723500796 |
| Gm26811 | 0.627488786 | -0.416753218 | 1.317082941 |
| Plxna1 | 1720.837352 | -0.198300574 | 0.070541331 |
| Chchd6 | 271.9148547 | -0.246978518 | 0.101175094 |
| Txnrd3 | 60.18168024 | -0.075409238 | 0.185137744 |
| Vmn1r-ps25 | 0 | NA | NA |
| Vmn1r40 | 0.046345709 | 0.71820487 | 3.724945049 |
| Vmn1r41 | 0 | NA | NA |
| Vmn1r-ps26 | 0 | NA | NA |
| Vmn1r-ps27 | 0 | NA | NA |
| Vmn1r42 | 0 | NA | NA |
| Vmn1r43 | 0.049553047 | 0.71820487 | 3.724945049 |
| Vmn1r44 | 0 | NA | NA |
| Vmn1r-ps28 | 0 | NA | NA |
| Vmn1r45 | 0 | NA | NA |
| Vmn1r46 | 0 | NA | NA |
| Vmn1r-ps29 | 0 | NA | NA |
| Vmn1r47 | 0 | NA | NA |
| Vmn1r48 | 0 | NA | NA |
| Vmn1r-ps31 | 0 | NA | NA |
| Gm20534 | 0 | NA | NA |
| Vmn1r49 | 0 | NA | NA |
| Vmn1r50 | 0 | NA | NA |
| Vmn1r51 | 0.060174537 | 0.237309859 | 3.724945049 |
| Vmn1r-ps32 | 0 | NA | NA |
| Vmn1r52 | 0 | NA | NA |
| V1ra8 | 0 | NA | NA |
| Vmn1r53 | 0.265601376 | -0.685290612 | 1.793603868 |
| Gm20426 | 1.074962736 | 0.768356462 | 1.050693361 |
| Gm4287 | 0.044544648 | 0.237309859 | 3.724945049 |
| Vmn1r-ps34 | 0 | NA | NA |
| Vmn1r54 | 0 | NA | NA |

| | | | |
|---|---|---|---|
| BC048671 | 0 | NA | NA |
| Chst13 | 0.497746636 | 1.144861273 | 1.35252206 |
| Uroc1 | 0.105143817 | -0.24357403 | 3.724945049 |
| Zxdc | 516.6253474 | -0.09779043 | 0.086839942 |
| Ccdc37 | 25.13015435 | -1.178910889 | 0.232683338 |
| Klf15 | 358.4959565 | 0.139141668 | 0.136308314 |
| Aldh1l1 | 1458.014872 | 1.387745191 | 0.11079773 |
| Gm15756 | 0.143022002 | 0.719176055 | 3.547735711 |
| Slc41a3 | 95.87477886 | 0.037521988 | 0.142207097 |
| Iqsec1 | 4998.243591 | -0.21085119 | 0.064316839 |
| Gm24816 | 4.887144399 | -0.874409508 | 0.660287356 |
| Rpl21-ps12 | 7.418190298 | -0.752272714 | 0.478678346 |
| Gm25185 | 0 | NA | NA |
| Ppp1r2-ps2 | 19.04760772 | 0.396456777 | 0.295019418 |
| Nup210 | 552.2987281 | 0.039954015 | 0.087176394 |
| Hdac11 | 1873.279407 | -0.520206832 | 0.067667699 |
| Gm14573 | 0.08169834 | 0.23732098 | 3.724945049 |
| Fbln2 | 117.7950445 | 1.223271224 | 0.242849198 |
| Wnt7a | 199.3147353 | 0.024047096 | 0.184900368 |
| Gm4575 | 4.420079801 | -0.517457717 | 0.477785528 |
| 4930471M09Rik | 1.669552124 | 0.514823744 | 0.744321402 |
| 1810044D09Rik | 5.670729865 | -0.096702527 | 0.41791252 |
| Chchd4 | 184.8478106 | 0.10701786 | 0.096820079 |
| Tmem43 | 388.1633489 | 1.038573853 | 0.08919213 |
| Xpc | 207.6063262 | -0.018490936 | 0.107908249 |
| Lsm3 | 147.4136989 | 0.205317685 | 0.108840228 |
| Slc6a6 | 1683.442733 | -0.034344003 | 0.07420208 |
| Grip2 | 311.1559795 | -0.658471359 | 0.109901318 |
| Ccdc174 | 303.9255113 | -0.134897587 | 0.110990776 |
| 4930590J08Rik | 9.858077027 | 0.599726376 | 0.398843237 |
| Fgd5 | 79.82046982 | 0.51689081 | 0.171867067 |
| 4930466I24Rik | 0.042716945 | -0.243571806 | 3.724945049 |
| Nr2c2 | 2086.98739 | -0.334031357 | 0.057322004 |
| Mrps25 | 334.8028407 | -0.286558395 | 0.072644589 |
| Rbsn | 899.6634951 | 0.300955532 | 0.091165773 |
| Trh | 83.76368859 | 1.203448639 | 0.408082304 |
| Prickle2 | 2847.184781 | -0.627555499 | 0.106941943 |
| Gm17797 | 0.118974955 | -0.243576254 | 3.724945049 |
| Adamts9 | 272.0878833 | 1.509974389 | 0.199000575 |
| A730049H05Rik | 9.582230866 | 3.73491833 | 0.580269715 |
| Gm15737 | 0 | NA | NA |
| 9530026P05Rik | 37.55723895 | -0.363368042 | 0.214800021 |
| Gm5313 | 0 | NA | NA |
| Gm25094 | 0 | NA | NA |
| Gm22312 | 0 | NA | NA |
| Magi1 | 1204.640072 | -0.610451271 | 0.070668156 |
| Gm23035 | 0 | NA | NA |

| | | | |
|---|---|---|---|
| Gm22840 | 0 | NA | NA |
| Slc25a26 | 101.4816024 | -0.286449528 | 0.122202598 |
| Lrig1 | 1206.313659 | 0.488911069 | 0.124621521 |
| Mir7041 | 0.248387756 | -0.685788557 | 1.83325957 |
| Gm7833 | 0.121388914 | 0.718200421 | 3.724945049 |
| Gm23364 | 0 | NA | NA |
| Gm24601 | 0 | NA | NA |
| Gm26501 | 0 | NA | NA |
| Kbtbd8os | 0.511579953 | -1.405186738 | 1.235870528 |
| Kbtbd8 | 179.6004661 | -0.132488561 | 0.123277408 |
| AY512915 | 0.311830279 | 0.718223437 | 2.001638361 |
| Gm10234 | 0 | NA | NA |
| Suclg2 | 141.9514173 | -0.287781134 | 0.149833783 |
| Fam19a1 | 272.9938384 | -0.482362369 | 0.142835123 |
| 1700123L14Rik | 0.257669488 | -0.384724312 | 2.084788683 |
| Gm22971 | 0 | NA | NA |
| Gm26011 | 0 | NA | NA |
| Fam19a4 | 11.35413066 | -0.035362793 | 0.311174177 |
| Eogt | 164.467491 | 0.042308083 | 0.131426368 |
| Tmf1 | 950.0693584 | 0.131800053 | 0.068299454 |
| Gm26175 | 2.7824929 | 0.213319273 | 0.622629744 |
| Uba3 | 755.4137509 | 0.09464866 | 0.087672288 |
| Arl6ip5 | 1177.848838 | 0.429794026 | 0.063476243 |
| Lmod3 | 0.289573199 | -0.260101323 | 2.06205345 |
| Frmd4b | 362.2741639 | 0.426624021 | 0.088229212 |
| Gm25852 | 1.201575708 | -0.167708469 | 0.824139992 |
| Gm23510 | 0.586937805 | -2.070430011 | 1.62127154 |
| Gm24565 | 0 | NA | NA |
| Gm23703 | 0 | NA | NA |
| Mitf | 247.2340933 | 0.564837462 | 0.113152145 |
| Gm15531 | 0.049553047 | 0.71820487 | 3.724945049 |
| Gm765 | 8.927308765 | -1.54714564 | 0.377589937 |
| Foxp1 | 1512.202388 | -0.560582345 | 0.123762511 |
| AC155839.1 | 0.110870145 | 0.238305784 | 3.724874707 |
| Gm20696 | 52.11740566 | -0.854490291 | 0.268787311 |
| Gm20705 | 26.36886051 | -0.972462997 | 0.38434823 |
| Gm22328 | 2.778696684 | -0.292894853 | 0.628253176 |
| Gm24936 | 0 | NA | NA |
| Eif4e3 | 223.9200148 | 0.273793531 | 0.088605875 |
| Gpr27 | 69.76717276 | -0.740786529 | 0.194945245 |
| Prok2 | 9.10031652 | 0.7632325 | 0.393399899 |
| Gm26748 | 2.205804803 | -1.637501501 | 0.718097074 |
| Tpt1-ps3 | 354.4684848 | 1.156398944 | 0.166910164 |
| RP23-191I2.1 | 0 | NA | NA |
| Gm24248 | 0 | NA | NA |
| Rybp | 419.1411725 | 0.181907573 | 0.081549658 |
| Gm23234 | 0.066050875 | 0.71820487 | 3.724945049 |

| | | | |
|---|---|---|---|
| 1700049E22Rik | 0 | NA | NA |
| Shq1 | 52.12462995 | 0.080869945 | 0.140287188 |
| Gm6565 | 46.92634586 | 0.010985281 | 0.183166016 |
| Gxylt2 | 11.76095793 | -0.638054288 | 0.330645648 |
| Gm15576 | 0.092881282 | -0.724455696 | 3.724945049 |
| Ppp4r2 | 799.647895 | 0.29163812 | 0.07960907 |
| Pdzrn3 | 253.9254666 | -0.322118471 | 0.130655138 |
| Gm26911 | 0.672570739 | -2.583160394 | 1.299531507 |
| Gm9871 | 0.521928645 | -0.240669206 | 1.5738359 |
| Cntn3 | 401.4213994 | -0.39986734 | 0.141999656 |
| Mir6373 | 0 | NA | NA |
| Gm26022 | 0 | NA | NA |
| Chl1 | 3209.185577 | -0.150410613 | 0.075867893 |
| Gm24784 | 4.963107029 | -0.398998243 | 0.568232272 |
| Gm25656 | 0 | NA | NA |
| RP24-151A23.1 | 0 | NA | NA |
| Cntn6 | 392.3293275 | -0.594206654 | 0.111223718 |
| Gm23339 | 0 | NA | NA |
| Gm25570 | 0 | NA | NA |
| Gm15631 | 17.2682059 | -0.80409187 | 0.352364051 |
| Cntn4 | 819.1636369 | -0.475545453 | 0.073251548 |
| Il5ra | 0.538620109 | 2.850511004 | 1.289128031 |
| Trnt1 | 529.4811257 | 0.386582347 | 0.076608681 |
| Crbn | 1050.434189 | 0.18561942 | 0.089850533 |
| Gm22418 | 0 | NA | NA |
| Gm27690 | 0 | NA | NA |
| Lrrn1 | 1083.285535 | -0.354787885 | 0.075843875 |
| Setmar | 60.36555304 | 0.20559201 | 0.140825486 |
| Sumf1 | 349.8000208 | 0.797385195 | 0.088958102 |
| Gm4604 | 20.23814071 | 0.370480848 | 0.246597425 |
| Itpr1 | 13527.53245 | -0.691163199 | 0.165176355 |
| Gm27813 | 4.625054459 | -1.586478577 | 0.493543138 |
| mmu-mir-7661 | 0.67439107 | -0.851215518 | 1.113116015 |
| Gm16433 | 10.96864868 | 0.592373723 | 0.330286111 |
| 0610040F04Rik | 4.191962974 | -1.195716434 | 0.513082239 |
| Gm38167 | 1.527506968 | -0.81226837 | 0.890506501 |
| Gm17055 | 0.461373441 | -1.087490763 | 1.656777033 |
| Bhlhe40 | 1442.70058 | 0.185402152 | 0.129998718 |
| Arl8b | 2174.485167 | 0.058246738 | 0.055162194 |
| Edem1 | 637.9057285 | 1.506403801 | 0.136060523 |
| Gm20387 | 22.36903969 | -0.49010986 | 0.261980259 |
| Grm7 | 1035.879049 | -0.504986708 | 0.14415896 |
| Gm20386 | 13.98619218 | -0.943872516 | 0.323685557 |
| Gm22093 | 0.0878635 | -0.72445792 | 3.724945049 |
| Gm24005 | 0 | NA | NA |
| Gm26362 | 0 | NA | NA |
| Gm25454 | 0 | NA | NA |

| | | | |
|---|---|---|---|
| 1700054K19Rik | 0 | NA | NA |
| n-R5s165 | 0 | NA | NA |
| Lmcd1 | 50.45369243 | 0.753543942 | 0.217065386 |
| 5031434C07Rik | 0.668421634 | -1.688800755 | 1.181584324 |
| Ssu2 | 1.179082066 | 0.517751699 | 1.064035733 |
| Cav3 | 2.338446252 | 2.132424306 | 0.822868591 |
| Oxtr | 56.02683163 | -0.497302036 | 0.147920575 |
| Gm27944 | 0 | NA | NA |
| Gm15519 | 37.45729361 | 0.650818668 | 0.179309994 |
| Gm23002 | 0.075043205 | 0.237309859 | 3.724945049 |
| Gm5578 | 12.94208794 | 0.564282072 | 0.330925286 |
| Rad18 | 111.1939575 | -0.391682417 | 0.17533414 |
| Gm26799 | 2.930040241 | 0.592396085 | 0.677574644 |
| Srgap3 | 4773.252643 | -0.529263085 | 0.071958366 |
| Gm6134 | 0.068097359 | -0.243569582 | 3.724945049 |
| Gm23244 | 0 | NA | NA |
| Thumpd3 | 249.1100717 | 0.544007073 | 0.08523128 |
| Gm22591 | 0.068097359 | -0.243569582 | 3.724945049 |
| Gt(ROSA)26Sor | 121.8807241 | 0.410578897 | 0.139619335 |
| Setd5 | 2417.574525 | -0.185543366 | 0.072811952 |
| Lhfpl4 | 1265.24945 | -0.531176043 | 0.079823555 |
| Mtmr14 | 178.5916463 | 0.274669227 | 0.085751643 |
| Gm16161 | 0.217535639 | 0.408281962 | 3.169413733 |
| Cpne9 | 367.3936013 | -1.204958248 | 0.13206383 |
| Brpf1 | 657.1446642 | 0.231694983 | 0.116060623 |
| Gm15492 | 1.140702544 | -1.469013096 | 1.148925726 |
| Ogg1 | 60.44940537 | 0.497278164 | 0.17614949 |
| Camk1 | 512.5663802 | -0.221084237 | 0.09110169 |
| Tada3 | 319.0164571 | -0.091480635 | 0.072664296 |
| Arpc4 | 1219.326062 | 0.25898378 | 0.082491686 |
| Ttll3 | 119.1716164 | -0.328002409 | 0.197796726 |
| Rpusd3 | 80.45315769 | -0.487152803 | 0.150580614 |
| Gm24387 | 0.061920613 | 0.718207094 | 3.724945049 |
| Cidec | 0.392922208 | 0.724805278 | 1.6710349 |
| Jagn1 | 177.5032672 | 0.491853064 | 0.14098093 |
| Il17re | 7.25222539 | -0.217282761 | 0.450761608 |
| Il17rc | 69.78172622 | 2.235255664 | 0.228611404 |
| Creld1 | 601.1655518 | -0.198493934 | 0.06927692 |
| Prrt3 | 474.5330712 | -0.638692819 | 0.117327046 |
| Emc3 | 603.6486607 | 0.471826798 | 0.080839578 |
| Gm23365 | 0.654623538 | 0.67258082 | 1.275163011 |
| Fancd2 | 38.49030515 | 0.374395916 | 0.179278419 |
| Fancd2os | 0.484250087 | 0.862391528 | 1.360926957 |
| Brk1 | 853.6369896 | -0.03147582 | 0.063044625 |
| Vhl | 414.2383988 | 0.207206171 | 0.086281048 |
| 1700015O11Rik | 0.140088945 | -1.149030578 | 3.601861891 |
| Irak2 | 312.5390426 | 0.93118042 | 0.147783552 |

| | | | |
|---|---|---|---|
| Tatdn2 | 686.0036907 | 0.546966952 | 0.10173626 |
| mmu-mir-7042 | 1.150390536 | 0.596256579 | 0.889542609 |
| Gm26982 | 73.25146826 | -0.766064043 | 0.22206361 |
| Ghrl | 1.379464875 | 0.246480593 | 0.808277092 |
| Sec13 | 453.3719348 | 0.764922354 | 0.08359954 |
| Atp2b2 | 10812.54371 | -1.129265392 | 0.114742075 |
| Gm15082 | 0 | NA | NA |
| Gm15083 | 4.119074061 | -0.7146628 | 0.505918397 |
| Slc6a11 | 2799.545472 | -1.399806159 | 0.160371352 |
| Slc6a1 | 4290.118131 | -0.806945565 | 0.085808438 |
| Hrh1 | 148.4312325 | 0.337310495 | 0.12588643 |
| Atg7 | 275.6563131 | 0.260210881 | 0.095803029 |
| Vgll4 | 362.9008528 | 0.278509145 | 0.093978303 |
| Tamm41 | 94.56214301 | -0.373887335 | 0.152105378 |
| Gm24767 | 0 | NA | NA |
| Gm17733 | 5.222249821 | -0.043180997 | 0.410270785 |
| Syn2 | 4179.711114 | -0.248645224 | 0.087876499 |
| Gm17482 | 8.270587696 | -0.860101515 | 0.361400704 |
| Timp4 | 415.701942 | 0.504644766 | 0.201563436 |
| Pparg | 54.34159874 | 0.61630866 | 0.210441691 |
| Tsen2 | 155.5268573 | -0.411260925 | 0.111504213 |
| Mkrn2os | 0.394336369 | 0.684173546 | 1.580663311 |
| Mkrn2 | 337.3089779 | 0.157786733 | 0.10387936 |
| Raf1 | 1017.633135 | 0.173170927 | 0.090781638 |
| Gm14335 | 18.11564472 | -0.119813637 | 0.365556143 |
| Gm24008 | 0 | NA | NA |
| Tmem40 | 16.46128129 | 1.232247269 | 0.417385083 |
| Gm27853 | 0 | NA | NA |
| Cand2 | 86.32620824 | 0.253506224 | 0.162649573 |
| Mir7660 | 0.118974955 | -0.243576254 | 3.724945049 |
| Rpl32 | 874.900214 | 0.72507677 | 0.142708572 |
| Snora7a | 4.44476562 | 0.345586533 | 0.590211494 |
| Efcab12 | 50.83190883 | -0.402219465 | 0.237741475 |
| Mbd4 | 111.5763811 | -0.421487574 | 0.177078335 |
| Ift122 | 349.0430765 | -0.074175549 | 0.112283153 |
| Rho | 4.082856454 | 0.415623975 | 0.483433032 |
| H1foo | 0.271840384 | 1.173843315 | 2.045681495 |
| Plxnd1 | 1058.405309 | -0.37345903 | 0.093868068 |
| Tmcc1 | 1022.727567 | -0.438578078 | 0.055109242 |
| Gm20404 | 1.065143116 | 1.25997335 | 1.066984461 |
| Gm8203 | 110.3300294 | 0.259026652 | 0.136117015 |
| Gm8213 | 4.570295035 | 0.489823333 | 0.486661395 |
| Fam21 | 1941.589181 | 0.130921141 | 0.082928781 |
| Zfand4 | 120.8501776 | -0.069142235 | 0.123476084 |
| 8-Mar | 943.7508335 | 0.296690465 | 0.079846461 |
| Gm23527 | 0 | NA | NA |
| Alox5 | 9.062762248 | -0.854561306 | 0.38631242 |

| | | | |
|---|---|---|---|
| Olfr211 | 0.287786674 | 1.737825039 | 1.999459068 |
| Olfr212 | 0.265357793 | 1.144092151 | 1.797908657 |
| Olfr213 | 0.046345709 | 0.71820487 | 3.724945049 |
| Gm5580 | 0.092493346 | -0.24357403 | 3.724945049 |
| Olfr214 | 0 | NA | NA |
| Olfr215 | 0 | NA | NA |
| Zfp422 | 293.8653895 | 0.264231248 | 0.068995727 |
| Rassf4 | 281.0708732 | 0.252273567 | 0.151947837 |
| Mir7043 | 0 | NA | NA |
| 8430408G22Rik | 5.938959585 | 1.163766321 | 0.5970675 |
| Tmem72 | 34.03285026 | -0.77688681 | 0.240731154 |
| Gm22882 | 0 | NA | NA |
| Cxcl12 | 287.0988006 | 0.348568228 | 0.123286571 |
| Rpl28-ps4 | 0 | NA | NA |
| Gm9946 | 0.160978641 | -1.214251972 | 2.79661493 |
| 1700030F04Rik | 1.580617114 | -1.312897988 | 0.830646868 |
| Gm7292 | 1492.722712 | -0.584326468 | 0.075687107 |
| Gm6253 | 0.669418093 | -0.924595644 | 1.377404722 |
| Zfp637 | 220.273934 | 0.008062921 | 0.085259049 |
| Zfp239 | 252.8739187 | -0.766423292 | 0.112602537 |
| Gm29509 | 0.066050875 | 0.71820487 | 3.724945049 |
| Hnrnpf | 625.6970897 | 1.109621559 | 0.116348845 |
| 4933440N22Rik | 4.749668983 | 1.946138183 | 0.553502561 |
| Fxyd4 | 0.432778942 | 0.582962078 | 1.364410894 |
| Gm25535 | 0.060174537 | 0.237309859 | 3.724945049 |
| Gm4875 | 25.05432516 | 0.02033186 | 0.200367735 |
| Rasgef1a | 1668.685123 | -0.774166875 | 0.107838782 |
| Mir7044 | 0.497274054 | 0.002459976 | 1.3479309 |
| Csgalnact2 | 271.8412438 | 0.006013364 | 0.074454834 |
| Gm23908 | 2.493933935 | -0.903138235 | 0.638714499 |
| Ret | 333.8009791 | -0.030155522 | 0.148080644 |
| Bms1 | 452.1647011 | 0.429442555 | 0.118983072 |
| Zfp248 | 186.5369703 | -0.305665283 | 0.103114688 |
| Gm25953 | 0 | NA | NA |
| Zfp9 | 323.1489127 | -0.070977468 | 0.080003945 |
| Ankrd26 | 319.7692467 | -0.549299626 | 0.174724351 |
| Cacna1c | 2127.985368 | -0.786903557 | 0.124344009 |
| Gm18981 | 18.01703932 | -1.452352543 | 0.407762364 |
| Gm25905 | 0 | NA | NA |
| Dcp1b | 230.6682507 | -0.420857598 | 0.118772934 |
| Cacna2d4 | 45.15394803 | -0.647085695 | 0.159248652 |
| Lrtm2 | 701.9863131 | -0.313811972 | 0.072199936 |
| Adipor2 | 1326.364367 | 0.619779148 | 0.110438551 |
| Wnt5b | 23.55176808 | 1.068468468 | 0.239639593 |
| Fbxl14 | 457.0418174 | 0.473643746 | 0.09145129 |
| Erc1 | 1392.520042 | -0.344982975 | 0.083737776 |
| 3110021A11Rik | 3.049669005 | -0.884346917 | 0.608848557 |

| | | | |
|---|---|---|---|
| Gm15532 | 0 | NA | NA |
| 4930540M05Rik | 1.752268337 | 1.086979864 | 0.779688263 |
| Rad52 | 228.0315541 | -0.030055066 | 0.093924489 |
| Wnk1 | 6814.715123 | 0.474062602 | 0.062904148 |
| Gm16199 | 3.97190282 | 0.093373988 | 0.563069283 |
| Mir706 | 1.782601153 | -0.060911702 | 0.70719343 |
| Ninj2 | 7.372786058 | -0.268453646 | 0.363296789 |
| Gm25327 | 0 | NA | NA |
| B4galnt3 | 7.654890081 | -1.344446886 | 0.374608233 |
| D330020A13Rik | 0 | NA | NA |
| Ccdc77 | 82.48745712 | -0.190104602 | 0.174640048 |
| Kdm5a | 1440.026259 | 0.204580841 | 0.084401318 |
| Gm24592 | 0 | NA | NA |
| Il17ra | 446.1211285 | 2.017744572 | 0.187481433 |
| Cecr6 | 417.0501185 | -1.158229964 | 0.168311193 |
| Cecr5 | 216.0841903 | -0.488020515 | 0.145526071 |
| Gm20692 | 2.176719344 | -2.302179402 | 0.858309425 |
| Gm24391 | 0.075043205 | 0.237309859 | 3.724945049 |
| Cecr2 | 227.0044623 | -0.049864988 | 0.155457707 |
| Slc25a18 | 412.4554696 | -1.000516793 | 0.143874844 |
| Atp6v1e1 | 2143.722746 | -0.208847453 | 0.066626971 |
| Gm25193 | 1.884175004 | -0.691136017 | 0.834529451 |
| Bcl2l13 | 678.3631799 | -0.019313524 | 0.071404315 |
| Bid | 179.7781668 | 1.958010899 | 0.172242653 |
| Mical3 | 1830.192808 | -0.54948352 | 0.123184778 |
| Gm4651 | 0.182574094 | 0.237323313 | 2.706657689 |
| Pex26 | 225.9169415 | -0.096658846 | 0.111082966 |
| Tuba8 | 200.697007 | 0.649354303 | 0.163944113 |
| Gm15856 | 6.178163136 | 6.010811413 | 0.969792542 |
| Usp18 | 2014.636858 | 9.833639205 | 0.403102696 |
| Slc6a13 | 90.14607231 | -0.18654169 | 0.252549415 |
| Gm24855 | 0 | NA | NA |
| Slc6a12 | 25.52923883 | 0.679887861 | 0.296274163 |
| Iqsec3 | 2651.332656 | -0.152759004 | 0.097212412 |
| A2m | 460.9649802 | 2.524458136 | 0.261452508 |
| Mug1 | 1.274577395 | 0.979039326 | 1.182430196 |
| Mug2 | 0.087930927 | 0.236669296 | 3.724898107 |
| Gm10319 | 0 | NA | NA |
| Mug-ps1 | 0.15956642 | 0.719166641 | 3.418731035 |
| Gm27703 | 0 | NA | NA |
| Klrg1 | 4.976324902 | 2.387350928 | 0.627961551 |
| Gm27826 | 0.044544648 | 0.237309859 | 3.724945049 |
| M6pr | 1445.217025 | 0.955160162 | 0.106472721 |
| Phc1 | 481.579932 | -0.164264226 | 0.083640908 |
| 1700063H04Rik | 0 | NA | NA |
| Rimklb | 372.2860085 | -0.183022536 | 0.061412452 |
| Gm8430 | 34.0404713 | 0.655930651 | 0.285683546 |

| | | | |
|---|---|---|---|
| Mfap5 | 3.28087796 | 1.841514127 | 0.735997189 |
| Aicda | 0 | NA | NA |
| Gm16556 | 1.225054178 | 4.146401181 | 1.065181208 |
| Gm16557 | 0.258216092 | 1.592790931 | 2.032402786 |
| Gm8459 | 0.839467773 | 3.695958746 | 2.281167729 |
| Apobec1 | 484.8295906 | 5.664859732 | 0.236305606 |
| Gdf3 | 0.810901572 | 1.482863426 | 1.230929836 |
| Gm26168 | 0 | NA | NA |
| Dppa3 | 0.038981395 | 0.718207094 | 3.724945049 |
| Gm10420 | 14.33523795 | 0.315060213 | 0.288403366 |
| Gm10224 | 1.550258836 | 0.921757056 | 0.797209234 |
| Nanog | 0.725797938 | 1.511359201 | 1.544115538 |
| Slc2a3 | 1520.555383 | -0.282344465 | 0.08260612 |
| AC163108.1 | 10.5610363 | 0.270868695 | 0.328535442 |
| Gm26826 | 2.946188958 | -0.357242438 | 0.600891577 |
| Foxj2 | 640.279476 | -0.05107283 | 0.064744253 |
| mmu-mir-7231 | 1.544805485 | -1.771828698 | 0.990585488 |
| C3ar1 | 498.6296836 | 5.093404325 | 0.299467706 |
| Necap1 | 1570.429647 | -0.067809172 | 0.06385447 |
| Gm5316 | 0 | NA | NA |
| Clec4a1 | 94.94432573 | 5.718216275 | 0.419088625 |
| Clec4a3 | 123.1859983 | 5.33291603 | 0.466343209 |
| Clec4a4 | 1.253602477 | 4.250369349 | 1.041059779 |
| Clec4b1 | 0.237601816 | 0.237329794 | 2.114704841 |
| Clec4a2 | 17.55388936 | 2.239167004 | 0.399990019 |
| Clec4b2 | 0.180312491 | 0.685297793 | 2.689604957 |
| Gm22946 | 0 | NA | NA |
| Gm5883 | 0.359325923 | -0.732411682 | 1.989870213 |
| Clec4n | 6.544164894 | 2.393931904 | 0.486965545 |
| Clec4d | 34.33899703 | 9.047361331 | 1.16417362 |
| Clec4e | 72.26271789 | 9.708874153 | 1.10633311 |
| Vmn2r19 | 0 | NA | NA |
| Vmn2r20 | 0 | NA | NA |
| Gm22833 | 0 | NA | NA |
| Vmn2r-ps31 | 0 | NA | NA |
| Vmn2r21 | 0 | NA | NA |
| Gm25891 | 0 | NA | NA |
| Gm22722 | 0 | NA | NA |
| Vmn2r-ps32 | 0 | NA | NA |
| Vmn2r22 | 0 | NA | NA |
| Gm25065 | 0 | NA | NA |
| Vmn2r23 | 0.060174537 | 0.237309859 | 3.724945049 |
| Gm22724 | 0 | NA | NA |
| Vmn2r24 | 0 | NA | NA |
| Gm23079 | 0 | NA | NA |
| Vmn2r25 | 0 | NA | NA |
| A330044P14Rik | 0.221456285 | 1.263084442 | 2.158168584 |

| | | | |
|---|---|---|---|
| Gm22763 | 0 | NA | NA |
| Vmn2r26 | 0 | NA | NA |
| Gm24843 | 0.072025462 | 0.237312084 | 3.724945049 |
| Vmn2r-ps33 | 0 | NA | NA |
| Gm22916 | 0 | NA | NA |
| Vmn2r-ps34 | 0 | NA | NA |
| Gm23554 | 0 | NA | NA |
| Vmn2r27 | 0.851816096 | 1.603886428 | 1.041787729 |
| Gm25311 | 0.355991514 | -1.285033125 | 1.631500949 |
| Cd163 | 12.17190331 | -1.832169308 | 0.370140673 |
| 1700013D24Rik | 0.329404011 | 1.787115388 | 1.952032897 |
| Pex5 | 534.6967276 | 0.109299232 | 0.07242903 |
| Clstn3 | 4659.131254 | -0.493778443 | 0.060843669 |
| C1rl | 16.99094215 | 2.709397176 | 0.344470746 |
| C1ra | 434.4635122 | 6.401877574 | 0.326146622 |
| C1s1 | 241.4617169 | 6.208707139 | 0.434074729 |
| Gm16567 | 0 | NA | NA |
| C1rb | 42.44241018 | 5.520451134 | 0.533155411 |
| C1s2 | 22.96225106 | 3.733598577 | 0.434092486 |
| Lpcat3 | 223.387876 | 0.67339354 | 0.099065773 |
| Emg1 | 187.4943886 | 0.387386911 | 0.095599794 |
| Phb2 | 583.7592251 | 0.402790996 | 0.075462703 |
| Gm23547 | 0.819923432 | -0.770362412 | 1.054905776 |
| Gm15884 | 0.260697633 | 0.238043878 | 2.112155273 |
| Gm29008 | 0.068097359 | -0.243569582 | 3.724945049 |
| Mir141 | 0 | NA | NA |
| Mir200c | 0 | NA | NA |
| Ptpn6 | 244.4682648 | 4.151569095 | 0.225726104 |
| Gm20531 | 0.660431168 | 1.495279253 | 1.216159913 |
| Grcc10 | 122.9669071 | -0.176100708 | 0.117659253 |
| Atn1 | 2476.146781 | -0.222653384 | 0.07505695 |
| Rnu7 | 0 | NA | NA |
| Eno2 | 8072.573921 | -0.426029148 | 0.057440174 |
| Lrrc23 | 22.31127345 | -0.135012832 | 0.249104951 |
| Spsb2 | 56.84758583 | 1.557304502 | 0.210385006 |
| Tpi1 | 3615.771022 | -0.201491981 | 0.059379714 |
| Usp5 | 1925.798522 | -0.027386229 | 0.04578681 |
| Cdca3 | 13.51901529 | 1.5541859 | 0.43948096 |
| Gnb3 | 5.506721047 | 1.351541771 | 0.622452161 |
| P3h3 | 139.641216 | -0.503261177 | 0.181925912 |
| Gpr162 | 1473.024775 | -0.344498499 | 0.072881419 |
| Cd4 | 50.03247416 | -0.684752598 | 0.222125989 |
| Lag3 | 118.9770841 | 2.547095995 | 0.226880562 |
| Ptms | 3060.359574 | -0.37390639 | 0.079284927 |
| A230083G16Rik | 12.40694247 | -0.567605307 | 0.307447714 |
| Mlf2 | 3556.102244 | -0.08197765 | 0.06142817 |
| Cops7a | 930.3160726 | -0.18298179 | 0.067978901 |

| | | | |
|---|---|---|---|
| Gm23851 | 0 | NA | NA |
| Pianp | 1373.810968 | -0.304823299 | 0.063893188 |
| Zfp384 | 433.8857635 | 0.111163338 | 0.072663475 |
| Ing4 | 294.8789085 | -0.15658296 | 0.079868836 |
| Acrbp | 30.74231455 | -0.593323095 | 0.278365581 |
| Lpar5 | 6.566982277 | 1.911050584 | 0.437356333 |
| Chd4 | 3499.470299 | 0.106481625 | 0.057872325 |
| Mir7045 | 0 | NA | NA |
| Gm23751 | 1.854332816 | 0.039869077 | 0.692892722 |
| Nop2 | 279.2923541 | 0.667550687 | 0.136870872 |
| Iffo1 | 569.9560514 | -0.260891262 | 0.107149435 |
| Gapdh | 740.2095212 | -0.062621733 | 0.076531502 |
| Ncapd2 | 168.4681777 | -0.408739456 | 0.156267753 |
| Scarna10 | 0.038981395 | 0.718207094 | 3.724945049 |
| Gm28967 | 0 | NA | NA |
| Mrpl51 | 347.6131604 | 0.344831296 | 0.082054167 |
| Mir3098 | 14.8885832 | -0.640794381 | 0.347191326 |
| Vamp1 | 2358.40077 | -0.370298405 | 0.164230207 |
| Tapbpl | 330.3516113 | 3.446540951 | 0.174848425 |
| Cd27 | 20.08776085 | 7.383216874 | 0.891055887 |
| Mir8113 | 0 | NA | NA |
| Gm24175 | 0 | NA | NA |
| Tuba3a | 0.099106094 | 1.209894437 | 3.72418822 |
| Ltbr | 255.90804 | 2.5058794 | 0.198781038 |
| Scnn1a | 38.07952501 | 0.155150081 | 0.188151314 |
| Tnfrsf1a | 469.1495194 | 3.317251898 | 0.187846667 |
| Plekhg6 | 3.530240572 | -0.170787448 | 0.526925836 |
| Cd9 | 682.5462011 | 1.227403834 | 0.132240663 |
| Gm28809 | 1.776315563 | 2.727142628 | 0.895656088 |
| Gm26728 | 0.409864138 | 0.802272856 | 1.6747275 |
| Vwf | 172.1922304 | 2.149580858 | 0.172207292 |
| Ano2 | 81.25229014 | -1.524860896 | 0.154805897 |
| Gm16303 | 0.512522738 | 1.263921218 | 1.411223506 |
| Gm24947 | 0.271265366 | -1.211780194 | 2.089321436 |
| Ntf3 | 14.00482386 | -1.181899717 | 0.313534864 |
| Gm29009 | 0 | NA | NA |
| Kcna5 | 77.10056723 | -0.392219996 | 0.165929425 |
| Gm33320 | 17.12675634 | -0.297995355 | 0.306924132 |
| Kcna1 | 5474.951842 | -0.319565918 | 0.094861832 |
| Kcna6 | 1135.188091 | -0.371645688 | 0.094100611 |
| Gm26673 | 76.86612292 | 0.091006463 | 0.22861027 |
| Ndufa9 | 671.0675705 | -0.132218151 | 0.070441095 |
| RP23-222G22.5 | 0.062426873 | 0.237312084 | 3.724945049 |
| Akap3 | 1.281806 | -0.470307144 | 0.852671099 |
| Dyrk4 | 3.739156486 | 1.083571447 | 0.640882823 |
| Rad51ap1 | 20.13187148 | 0.28091383 | 0.332143679 |
| RP23-222G22.9 | 0 | NA | NA |

| | | | |
|---|---|---|---|
| D6Wsu163e | 358.0575039 | 0.082493865 | 0.079062829 |
| RP23-8L20.9 | 0 | NA | NA |
| RP23-8L20.8 | 0.141823039 | 0.704039222 | 3.596944413 |
| Fgf6 | 0.246418769 | 1.197997911 | 1.838700562 |
| Fgf23 | 4.436253295 | -0.129723611 | 0.528980111 |
| Tigar | 254.7556436 | -0.18994148 | 0.092180105 |
| Platr31 | 1.918110815 | -0.211332568 | 0.725136044 |
| RP23-8L20.6 | 0.061920613 | 0.718207094 | 3.724945049 |
| Ccnd2 | 1274.440512 | 0.253622549 | 0.134909717 |
| 9330179D12Rik | 23.50546626 | -0.060493105 | 0.221519315 |
| RP23-387P23.5 | 2.073082813 | -1.033542662 | 0.818887999 |
| RP23-403P6.1 | 26.03534595 | -0.161469274 | 0.208516341 |
| Gm4968 | 0.813324618 | 0.000963844 | 1.0676125 |
| RP23-387P23.7 | 0 | NA | NA |
| Gm7308 | 60.57587847 | -0.539792984 | 0.181254805 |
| RP23-387P23.3 | 0.11147366 | 1.230376831 | 3.722845817 |
| RP23-387P23.4 | 2.402022068 | -0.812572231 | 0.678058735 |
| RP23-387P23.9 | 0.145088245 | 0.23829189 | 3.530638546 |
| RP23-387P23.8 | 0 | NA | NA |
| RP23-223J11.1 | 0.049553047 | 0.71820487 | 3.724945049 |
| Parp11 | 870.7509703 | 2.522758398 | 0.182615297 |
| RP23-422D12.3 | 30.48963593 | 1.347980605 | 0.326221195 |
| Cracr2a | 10.36781213 | -0.633294292 | 0.32453196 |
| RP23-42M8.2 | 6.20278278 | -0.381206734 | 0.464165807 |
| Prmt8 | 579.4218534 | -0.886576559 | 0.149596189 |
| RP23-42M8.9 | 0 | NA | NA |
| RP23-42M8.10 | 14.99285838 | -0.975793876 | 0.320047601 |
| RP23-42M8.5 | 0.045882455 | 0.237312084 | 3.724945049 |
| RP23-42M8.7 | 0.12092566 | 0.237307635 | 3.724945049 |
| mmu-mir-7233 | 0 | NA | NA |
| RP24-473H20.4 | 0.306891002 | 1.714894839 | 1.744070442 |
| Gm23098 | 0.762875166 | -0.183306916 | 1.10127057 |
| RP24-473H20.3 | 0.711610403 | 1.908367611 | 1.199697034 |
| Tspan11 | 99.16132127 | -0.190347798 | 0.161718521 |
| Gm15870 | 0.825566729 | 2.072909848 | 1.150990572 |
| Tspan9 | 1288.655582 | 0.295495357 | 0.134397903 |
| Gm26338 | 0.326202429 | 0.232274551 | 1.816484719 |
| Gm10010 | 121.3586711 | -1.413972706 | 0.185582445 |
| Gm4691 | 17.793641 | -0.452535574 | 0.288682755 |
| Tead4 | 29.15459458 | 0.553502372 | 0.25753277 |
| Gm21893 | 0 | NA | NA |
| Gm26951 | 0.061920613 | 0.718207094 | 3.724945049 |
| Rpl18-ps2 | 5.774417445 | 1.602322168 | 0.428708161 |
| Tulp3 | 551.9556277 | 1.224919325 | 0.10910429 |
| Gm26770 | 0.55622196 | 0.127510744 | 1.225457805 |
| Rhno1 | 92.43520098 | 0.193819247 | 0.11783882 |
| Foxm1 | 45.1673141 | 1.125587527 | 0.369456004 |

| | | | |
|---|---|---|---|
| 4933413G19Rik | 17.62518862 | -0.621546011 | 0.348565469 |
| Gm15862 | 1.729221609 | 0.12299526 | 0.780958169 |
| Nrip2 | 216.9234121 | -0.336146678 | 0.09313517 |
| Itfg2 | 280.4983282 | 0.165733976 | 0.103277191 |
| Fkbp4 | 1131.070024 | 0.020095141 | 0.082006897 |
| Gm10069 | 14.56716468 | -0.452364515 | 0.323877823 |
| Pzp | 1.32803898 | 0.467689409 | 1.03393068 |
| Gm8724 | 0.141071297 | 0.237322348 | 3.564043836 |
| BC048546 | 30.70108197 | -0.34431248 | 0.199257843 |
| Klrb1a | 2.241070021 | 4.264276666 | 1.01024422 |
| Gm5884 | 150.0674224 | 0.429222593 | 0.120616275 |
| Clec2j | 0 | NA | NA |
| Clec2h | 0.169629266 | 0.117186178 | 2.763788639 |
| Klrb1 | 0 | NA | NA |
| Klrb1c | 5.966816195 | 6.326104456 | 0.976476604 |
| Klrb1b | 4.841237096 | 3.21955014 | 0.900417842 |
| Gm26656 | 0.322709507 | 0.718222766 | 1.971788094 |
| BC035044 | 6.979976624 | 0.388753323 | 0.480921062 |
| Clec2i | 12.44756223 | 5.334845165 | 0.854381854 |
| Gm27336 | 0.319483352 | 1.620019128 | 1.970253129 |
| Clec2g | 0.594455495 | -0.324345195 | 1.478768616 |
| Gm15987 | 6.297883917 | 4.799006887 | 0.872952392 |
| Mir7649 | 0 | NA | NA |
| Gm8778 | 0.092691418 | 1.183079489 | 3.724560917 |
| BC064078 | 21.92509358 | 1.440075316 | 0.307091677 |
| Gm16187 | 0.11197992 | 0.718202646 | 3.724945049 |
| Clec2f | 1.610080848 | 1.224955139 | 0.916048766 |
| Gm16186 | 0 | NA | NA |
| Klrb1f | 4.97974666 | 4.902679807 | 0.916910025 |
| Clec2e | 0.752785761 | 2.161306617 | 1.311828747 |
| Klrb1-ps1 | 0 | NA | NA |
| Gm26160 | 0.088534442 | 1.179809211 | 3.723500796 |
| Clec2d | 395.2909725 | 4.191776588 | 0.262518111 |
| Gm27514 | 0.12209515 | 0.718202646 | 3.724945049 |
| 2310001H17Rik | 41.45082234 | 4.401958262 | 0.362332254 |
| Cd69 | 49.45403992 | 9.244529832 | 0.890925539 |
| 4922502D21Rik | 0.201858732 | 0.718217268 | 2.604406118 |
| Clec12a | 44.34900859 | 3.997001409 | 0.379017829 |
| Clec12b | 2.028096924 | 4.115581077 | 1.099947324 |
| Clec1b | 1.612175364 | 0.972478037 | 0.891408408 |
| Clec9a | 4.424159844 | 2.413171689 | 0.770927477 |
| Clec1a | 25.40223335 | 1.615355706 | 0.295353393 |
| Clec7a | 77.62764879 | 3.882789657 | 0.309062548 |
| Olr1 | 7.563519298 | 6.344566548 | 0.905382532 |
| Tmem52b | 0.046345709 | 0.71820487 | 3.724945049 |
| 1700101I11Rik | 14.17165844 | 0.900038996 | 0.295375501 |
| Gabarapl1 | 4493.649597 | -0.137967226 | 0.059417402 |

| | | | |
|---|---|---|---|
| Klre1 | 4.71579917 | 5.112047497 | 0.920491773 |
| Klrd1 | 23.39137982 | 7.31125929 | 0.873988121 |
| Klrk1 | 186.6532185 | 9.261426173 | 0.760014667 |
| Klrc3 | 5.985488163 | 6.279168479 | 0.926257131 |
| Klrc2 | 17.16294782 | 7.373828683 | 0.907360115 |
| Klrc1 | 33.05721631 | 8.567249671 | 0.877405774 |
| Gm7309 | 0 | NA | NA |
| Mir680-1 | 0.066050875 | 0.71820487 | 3.724945049 |
| Klri1 | 2.598038951 | 5.308180354 | 0.97975729 |
| Klri2 | 8.227621471 | 5.979692385 | 0.989344295 |
| Gm156 | 0.097139547 | 0.718207094 | 3.724945049 |
| Gm16242 | 0.044544648 | 0.237309859 | 3.724945049 |
| Klra17 | 6.058344776 | 6.088823125 | 0.992103989 |
| Gm17631 | 0 | NA | NA |
| Gm23977 | 0 | NA | NA |
| Klra5 | 0.677147933 | 3.162944166 | 1.402810346 |
| Gm15854 | 0 | NA | NA |
| Gm22555 | 0 | NA | NA |
| Klra6 | 0 | NA | NA |
| Klra4 | 0.378042144 | 2.34546093 | 1.65531192 |
| Gm24676 | 0 | NA | NA |
| Klra11-ps | 0 | NA | NA |
| Gm22621 | 0 | NA | NA |
| Klra8 | 0.110595523 | 0.718200421 | 3.724945049 |
| Gm24712 | 0 | NA | NA |
| Klra14-ps | 0 | NA | NA |
| Gm23552 | 0 | NA | NA |
| Klra9 | 0.604622052 | 3.05991298 | 1.216808561 |
| Klra7 | 3.947549917 | 5.850425366 | 0.933878309 |
| Klra10 | 0.285107744 | 2.038594153 | 2.393662212 |
| Klra13-ps | 0.52061208 | 3.10310241 | 1.319403497 |
| Gm24072 | 0 | NA | NA |
| Klra3 | 12.4313916 | 6.445419724 | 0.906767081 |
| Klra1 | 0.328294527 | 1.901087936 | 1.683669081 |
| Gm6584 | 0 | NA | NA |
| Gm15861 | 0 | NA | NA |
| Gm20997 | 0 | NA | NA |
| Gm27013 | 0.044544648 | 0.237309859 | 3.724945049 |
| Gm8816 | 0 | NA | NA |
| Gm26996 | 0.113979814 | -0.243571806 | 3.724945049 |
| Gm26914 | 0 | NA | NA |
| Gm27047 | 0 | NA | NA |
| Gm26919 | 0.161486377 | 1.199990293 | 2.795640196 |
| Gm6600 | 0 | NA | NA |
| Gm8837 | 0 | NA | NA |
| Gm5581 | 0.915657815 | 3.84053447 | 1.36087087 |
| Klra2 | 81.98899445 | 6.989281811 | 0.606431891 |

| | | | |
|---|---|---|---|
| Gm5582 | 1.579793576 | -0.456331803 | 0.795834878 |
| Magohb | 84.69062322 | 0.02004777 | 0.159883594 |
| Styk1 | 21.45282092 | 1.922687397 | 0.300634009 |
| Ybx3 | 331.3395029 | 0.716755248 | 0.104254304 |
| Gm22362 | 7.679914887 | -0.06499374 | 0.416381585 |
| 5430401F13Rik | 0.172107867 | 0.718215321 | 2.739902689 |
| Gm6619 | 0 | NA | NA |
| Gm22190 | 0 | NA | NA |
| Gm16571 | 0 | NA | NA |
| Hmgb1l | 0 | NA | NA |
| Tas2r130 | 0 | NA | NA |
| Tas2r107 | 0 | NA | NA |
| Tas2r106 | 0 | NA | NA |
| Tas2r104 | 0 | NA | NA |
| Tas2r105 | 0 | NA | NA |
| Tas2r114 | 0 | NA | NA |
| Gm4736 | 0 | NA | NA |
| Prpmp5 | 0 | NA | NA |
| Gm8882 | 0.046345709 | 0.71820487 | 3.724945049 |
| n-R5s166 | 0 | NA | NA |
| Prh1 | 0 | NA | NA |
| n-R5s167 | 0 | NA | NA |
| Prp2 | 0 | NA | NA |
| A630073D07Rik | 0 | NA | NA |
| Tas2r120 | 0 | NA | NA |
| Tas2r121 | 0 | NA | NA |
| Tas2r122 | 0 | NA | NA |
| Tas2r115 | 0 | NA | NA |
| Tas2r124 | 0 | NA | NA |
| Tas2r102 | 0 | NA | NA |
| Tas2r136 | 0.061920613 | 0.718207094 | 3.724945049 |
| Tas2r117 | 0 | NA | NA |
| Tas2r123 | 0 | NA | NA |
| Tas2r116 | 0.061920613 | 0.718207094 | 3.724945049 |
| Tas2r110 | 0 | NA | NA |
| Tas2r113 | 0 | NA | NA |
| Tas2r125 | 0 | NA | NA |
| Tas2r129 | 0 | NA | NA |
| Tas2r131 | 0 | NA | NA |
| Tas2r109 | 0 | NA | NA |
| Tas2r103 | 0 | NA | NA |
| Tas2r140 | 0 | NA | NA |
| Smim10l1 | 1843.017742 | -0.079628633 | 0.060255599 |
| Gm23403 | 0 | NA | NA |
| Gm25019 | 0.075043205 | 0.237309859 | 3.724945049 |
| 5530400C23Rik | 0 | NA | NA |
| Gm5885 | 0 | NA | NA |

| | | | |
|---|---|---|---|
| Gm23375 | 0.502960085 | 1.148508911 | 1.520822661 |
| Kap | 0.042716945 | -0.243571806 | 3.724945049 |
| Etv6 | 690.3323098 | 1.347008246 | 0.128574641 |
| Gm17088 | 0.202171817 | 1.626940071 | 2.587140394 |
| Gm17089 | 0.375570109 | 2.540840786 | 1.622213777 |
| Bcl2l14 | 0.379111656 | 1.153099391 | 1.45961221 |
| Lrp6 | 1132.17589 | -0.463861519 | 0.064173851 |
| Mansc1 | 117.9240851 | -0.127192658 | 0.128676525 |
| Loh12cr1 | 180.7292652 | -0.121159734 | 0.093752207 |
| Dusp16 | 411.5721016 | 0.628998807 | 0.125305706 |
| Crebl2 | 520.8784886 | -0.156829906 | 0.111963632 |
| Gpr19 | 174.2410689 | 0.182783132 | 0.130293408 |
| 2810454H06Rik | 8.20098561 | -0.572881493 | 0.384998186 |
| Cdkn1b | 860.406846 | 0.118695953 | 0.100638617 |
| 1190002F15Rik | 1.018292001 | 2.063613418 | 1.168022315 |
| Apold1 | 80.86165162 | 0.468675265 | 0.260249493 |
| Ddx47 | 401.4348954 | 0.172210789 | 0.069111231 |
| Gprc5a | 6.993744504 | 2.878218779 | 0.604414682 |
| Gprc5d | 0.543589123 | 0.82525739 | 1.422595353 |
| Hebp1 | 102.192585 | 0.093193062 | 0.169239057 |
| Gm22881 | 0 | NA | NA |
| 8430419L09Rik | 1464.187594 | -0.454362658 | 0.055394254 |
| Gsg1 | 0.811303554 | 1.103634009 | 1.239172749 |
| Pbp2 | 0 | NA | NA |
| Dynlt1-ps1 | 35.59685521 | 0.789083754 | 0.266110419 |
| Emp1 | 465.3937255 | 5.130245761 | 0.417907066 |
| Gm25136 | 0.672405421 | 1.086099982 | 1.220301681 |
| Grin2b | 9373.268553 | -1.336367799 | 0.189604339 |
| Gm14329 | 0.341881936 | -1.062983581 | 1.562612399 |
| Gm22892 | 0.112396584 | 1.192083721 | 3.724434897 |
| Gm38202 | 40.04288314 | -1.523062486 | 0.347371036 |
| Gm37182 | 48.96267409 | -1.779825186 | 0.411701782 |
| Gm14330 | 0.038981395 | 0.718207094 | 3.724945049 |
| Gm26653 | 7.145952665 | -1.050061414 | 0.449584433 |
| Gm26105 | 0.110595523 | 0.718200421 | 3.724945049 |
| Gm8994 | 0.107803068 | 0.719187033 | 3.724874737 |
| Gm15476 | 0.04393175 | -0.243571806 | 3.724945049 |
| E330021D16Rik | 0 | NA | NA |
| Gm25882 | 0 | NA | NA |
| Gm6728 | 11.37417031 | 0.180375509 | 0.310348178 |
| Rpl36a-ps3 | 8.907631286 | 0.682574881 | 0.351691824 |
| Atf7ip | 2145.155086 | 0.280959821 | 0.075809521 |
| 5830415B17Rik | 15.21012998 | 0.208492559 | 0.310267118 |
| Plbd1 | 20.58495972 | 3.872720374 | 0.443902682 |
| Gucy2c | 13.73199816 | 0.515976921 | 0.319732744 |
| Hist4h4 | 361.153536 | -0.427012442 | 0.158876946 |
| H2afj | 164.4299334 | 0.186033749 | 0.128667936 |

| | | | |
|---|---|---|---|
| AC122804.1 | 2.891209199 | 0.91771039 | 0.597423718 |
| Wbp11 | 918.8633835 | 0.067396944 | 0.086161695 |
| BC049715 | 12.24740255 | 0.186927069 | 0.319107578 |
| Smco3 | 28.46706308 | -0.616549195 | 0.200662056 |
| Art4 | 1.936338333 | -0.102786258 | 0.73824204 |
| Mgp | 118.7015475 | 1.30886561 | 0.180921099 |
| Erp27 | 1.418067754 | 1.637439449 | 1.077533735 |
| Arhgdib | 195.7130349 | 2.598547409 | 0.172443995 |
| Pde6h | 2.208047999 | 0.180318893 | 1.014757089 |
| Rerg | 82.27360041 | -0.745881888 | 0.141819583 |
| Ptpro | 585.5542697 | -0.054363991 | 0.161809818 |
| Eps8 | 778.7034068 | 0.526578914 | 0.122156381 |
| Strap | 1329.576707 | 0.307005401 | 0.083364522 |
| Dera | 50.46973726 | 1.342347452 | 0.230010807 |
| Gm24308 | 0 | NA | NA |
| Slc15a5 | 0.807800175 | -1.013696352 | 1.191836218 |
| Mgst1 | 196.4190543 | 2.06815831 | 0.221295709 |
| Lmo3 | 872.8499117 | -1.371317159 | 0.124447541 |
| A830011K09Rik | 12.61465114 | -0.694070686 | 0.367138602 |
| B230110G15Rik | 1.657582323 | -3.143886596 | 1.032227949 |
| Igbp1b | 1.236116855 | -3.205498702 | 1.067410395 |
| Gm25434 | 0 | NA | NA |
| Rergl | 0.244560259 | -1.635384651 | 2.108054029 |
| Pik3c2g | 12.46784403 | 2.402751786 | 0.386051356 |
| Gm37269 | 0.211089261 | 1.18296644 | 2.182449302 |
| Plcz1 | 6.588952628 | 0.134108516 | 0.384690644 |
| Capza3 | 0.060174537 | 0.237309859 | 3.724945049 |
| Gm24174 | 0 | NA | NA |
| n-R5s168 | 0 | NA | NA |
| Plekha5 | 1362.818818 | -0.462708943 | 0.108081917 |
| Gm25780 | 21.43207649 | -1.049122361 | 0.330673499 |
| Aebp2 | 1104.796083 | 0.108191345 | 0.062872727 |
| Gm11077 | 0.333293459 | 1.012413838 | 1.700776948 |
| Rpl38-ps2 | 307.4974746 | 0.404415107 | 0.090398851 |
| Gm28524 | 1.176308512 | -0.721614454 | 1.00571749 |
| Gm28523 | 12.22868002 | -0.519243112 | 0.315436585 |
| Pde3a | 315.7616379 | 0.893544389 | 0.103660909 |
| Slco1c1 | 209.1116899 | -0.708735291 | 0.22913924 |
| Slco1b2 | 3.268651982 | -0.783480353 | 0.577042423 |
| Gm5724 | 0.34217064 | -1.114523573 | 1.567286496 |
| Gm23069 | 0 | NA | NA |
| Slco1a4 | 155.850488 | -1.025809643 | 0.345445478 |
| Slco1a1 | 0.113999573 | 0.23732098 | 3.724945049 |
| Gm22036 | 0 | NA | NA |
| Gm6614 | 0.602494748 | 2.079943827 | 1.393848103 |
| Gm23775 | 0 | NA | NA |
| Slco1a6 | 0.857960599 | 1.767713286 | 1.194755174 |

| | | | |
|---|---|---|---|
| Gm18159 | 0.046345709 | 0.71820487 | 3.724945049 |
| Gm20400 | 0.439621597 | 1.915271256 | 1.525205511 |
| Slco1a5 | 10.05483302 | 2.122479762 | 0.430341482 |
| Gm25430 | 0 | NA | NA |
| Iapp | 0.092228164 | 0.718202646 | 3.724945049 |
| Pyroxd1 | 173.2075479 | 0.094545042 | 0.084859206 |
| Recql | 183.7198923 | -0.222001731 | 0.145510808 |
| Golt1b | 467.436909 | 0.370667211 | 0.063360063 |
| B230216G23Rik | 18.91388396 | -0.642626203 | 0.264469275 |
| Gys2 | 1.458192719 | 2.122692376 | 1.15952597 |
| Ldhb | 2909.672582 | -0.469521517 | 0.072428356 |
| Kcnj8 | 29.13393712 | 1.075528619 | 0.213806299 |
| Abcc9 | 80.16356092 | -0.089467151 | 0.16670077 |
| 5330439B14Rik | 0 | NA | NA |
| Cmas | 639.4281021 | -0.225530721 | 0.06578809 |
| Gm766 | 1.893318208 | -1.359007151 | 0.883678218 |
| Gm7457 | 1.3052917 | 0.201207824 | 0.858733325 |
| St8sia1 | 1060.037098 | -0.909344196 | 0.063589651 |
| C2cd5 | 774.2859654 | -0.671741806 | 0.121660709 |
| Etnk1 | 6574.447317 | 0.457294659 | 0.090925148 |
| D6Ertd474e | 2.829004364 | -0.301866767 | 0.609947387 |
| Gm23272 | 0 | NA | NA |
| 4930579D09Rik | 0 | NA | NA |
| 1700060C16Rik | 0.044191549 | -0.243571806 | 3.724945049 |
| Sox5 | 636.5935809 | -0.364825518 | 0.096510644 |
| Sox5os1 | 0.085433889 | -0.72445792 | 3.724945049 |
| Sox5os2 | 0 | NA | NA |
| Sox5it | 4.538093024 | -0.466034915 | 0.531493049 |
| Sox5os3 | 0.137647365 | 0.237309079 | 3.724945049 |
| Sox5os4 | 0 | NA | NA |
| Sox5os5 | 0 | NA | NA |
| Gm22792 | 0 | NA | NA |
| Bcat1 | 1548.257675 | 0.261007736 | 0.103294491 |
| Gm23071 | 1.299875216 | -1.820517168 | 0.998194258 |
| Gm26666 | 7.30636117 | -0.858880663 | 0.41163897 |
| Gm15687 | 0.219710209 | 0.734628013 | 2.160637886 |
| Lrmp | 42.64411814 | 1.845679311 | 0.218366495 |
| Casc1 | 19.07366442 | -0.210687799 | 0.260056133 |
| Gm15543 | 0.049553047 | 0.71820487 | 3.724945049 |
| Lyrm5 | 128.9520887 | -0.073690795 | 0.123547008 |
| Kras | 1419.910694 | 0.061149722 | 0.071897251 |
| Gm15706 | 24.72915098 | -0.80289629 | 0.233334724 |
| Rps25-ps1 | 12.76755632 | 0.790252113 | 0.32832829 |
| Gm22793 | 0 | NA | NA |
| 2010013B24Rik | 0.275148976 | 1.192062318 | 2.06642393 |
| Gm15707 | 0.08486385 | 0.717551239 | 3.724898127 |
| Lmntd1 | 13.42183498 | -0.335159963 | 0.333953363 |

| | | | |
|---|---|---|---|
| Gm23498 | 0 | NA | NA |
| 1700073E17Rik | 0.253253743 | 1.300974525 | 2.470000538 |
| Gm15499 | 0 | NA | NA |
| Gm25373 | 0 | NA | NA |
| Tuba3b | 0.095898756 | 1.192083721 | 3.724434897 |
| Gm15704 | 2.261169167 | -0.654125001 | 0.634451537 |
| Rassf8 | 399.0644075 | 0.043123115 | 0.109057413 |
| Bhlhe41 | 914.9264729 | -0.173407402 | 0.06231237 |
| Sspn | 140.0205592 | -0.405808833 | 0.189383365 |
| Sspnos | 0.044191549 | -0.243571806 | 3.724945049 |
| Itpr2 | 644.2115075 | 0.132422417 | 0.098273345 |
| Gm15720 | 1.860207608 | -0.794999507 | 0.807263869 |
| Asun | 359.4234607 | 0.001981069 | 0.105197874 |
| Fgfr1op2 | 883.1771317 | 0.100072287 | 0.052570087 |
| Tm7sf3 | 427.7210704 | -0.04140312 | 0.079379239 |
| Med21 | 211.1955998 | 0.807208881 | 0.156745736 |
| Gm6654 | 0.926048393 | 2.089447953 | 1.065775049 |
| 4930479D17Rik | 2.935438062 | -0.245260828 | 0.559130491 |
| Stk38l | 426.5635259 | 0.304980907 | 0.082453678 |
| Arntl2 | 48.93835431 | -0.611233141 | 0.22101487 |
| Smco2 | 1.092136027 | 2.70770279 | 1.125703502 |
| Ppfibp1 | 368.3784692 | 0.382940893 | 0.083707052 |
| 1700034J05Rik | 0.062426873 | 0.237312084 | 3.724945049 |
| Rep15 | 3.085232335 | -0.465576434 | 0.530637497 |
| Mrps35 | 175.5694965 | -0.133262209 | 0.102965837 |
| Mansc4 | 2.80060554 | 0.987900602 | 0.598281576 |
| Klhl42 | 434.0929622 | -0.358508657 | 0.072095063 |
| Gm7571 | 0.356315666 | 0.733666583 | 1.58319482 |
| E330012B07Rik | 0.576204374 | 1.868441238 | 1.49902659 |
| Gm15767 | 0.045882455 | 0.237312084 | 3.724945049 |
| Pthlh | 19.04363101 | -0.143098056 | 0.293817081 |
| Gm15766 | 2.701807514 | 0.186037861 | 0.611616838 |
| Gm6288 | 5.185232385 | 0.472834631 | 0.462703285 |
| Ccdc91 | 322.5136679 | 0.021784658 | 0.101025692 |
| 1700049E15Rik | 0.04393175 | -0.243571806 | 3.724945049 |
| Gm15762 | 0.295243939 | -1.010346107 | 1.749139951 |
| Far2os2 | 3.668828368 | 1.46913849 | 0.684836141 |
| Far2 | 187.1712709 | -0.923518696 | 0.12234429 |
| Far2os1 | 2.690587419 | 0.671345022 | 0.572518446 |
| Ergic2 | 794.7085187 | 0.040915309 | 0.07622925 |
| Gm17216 | 1.982138501 | -1.530838672 | 0.913772244 |
| Tmtc1 | 1255.055725 | -0.457630067 | 0.0854075 |
| Gm15763 | 0.045882455 | 0.237312084 | 3.724945049 |
| Rps4l | 60.78732413 | 0.37380919 | 0.262782084 |
| Gm6313 | 0.072025462 | 0.237312084 | 3.724945049 |
| Gm23456 | 0.106358623 | -0.24357403 | 3.724945049 |
| 4930528G23Rik | 0.514450355 | -2.155472882 | 1.486346682 |

| | | | |
|---|---|---|---|
| Gm15778 | 0 | NA | NA |
| Ipo8 | 948.9724006 | -0.119354049 | 0.052741633 |
| Caprin2 | 102.1573326 | -0.887476394 | 0.159315739 |
| Gm7618 | 0.320079734 | 1.639296172 | 1.708290133 |
| Fam60a | 49.2104772 | 1.087608655 | 0.176238125 |
| 3010003L21Rik | 2.014334011 | 2.265836875 | 0.89954368 |
| Gm25539 | 0 | NA | NA |
| Dennd5b | 2078.167419 | -0.40329417 | 0.052004246 |
| Gm26540 | 0.259164831 | 0.29271656 | 1.826965837 |
| Gm15779 | 0 | NA | NA |
| Gm15781 | 0 | NA | NA |
| Gm22578 | 4.906737206 | -0.993215241 | 0.486464916 |
| Gm15780 | 0 | NA | NA |
| Gm23462 | 2.97460163 | -0.956441006 | 0.670684526 |
| Gm10203 | 0.141071297 | 0.237322348 | 3.564043836 |
| Mettl20 | 22.19794556 | 0.624276719 | 0.324219413 |
| Gm10011 | 0.371635155 | -0.243564261 | 1.601546068 |
| Amn1 | 149.0049197 | 0.002934422 | 0.111946137 |
| 2810474O19Rik | 949.1809365 | 1.81529054 | 0.146271524 |
| Gm15782 | 0.129871752 | 1.198487306 | 2.991078732 |
| Gm15784 | 6.345417981 | 0.050532424 | 0.373516343 |
| Gm15786 | 0.200503209 | 0.718218166 | 2.61511129 |
| Gm15783 | 0.047948698 | -0.243569582 | 3.724945049 |
| Bicd1 | 1658.556008 | -0.381533465 | 0.050395586 |
| Gm15785 | 0.570614426 | -1.724982247 | 1.263330722 |
| Gm24307 | 1.150914894 | 0.420421404 | 0.905504751 |
| Gm21814 | 6.725415989 | -0.338963936 | 0.389559014 |
| Gm3104 | 0 | NA | NA |
| Gm7353 | 46.36268977 | 0.13489533 | 0.163674819 |
| AU018091 | 0.640879238 | 3.297127647 | 1.200921464 |
| Mir290a | 0 | NA | NA |
| mmu-mir-290b | 0 | NA | NA |
| Mir291a | 0 | NA | NA |
| Mir292 | 0 | NA | NA |
| mmu-mir-292b | 0 | NA | NA |
| Mir291b | 0 | NA | NA |
| Mir293 | 0 | NA | NA |
| Mir294 | 0 | NA | NA |
| Mir295 | 0 | NA | NA |
| Nlrp12 | 0.605827841 | -0.194664691 | 1.240583199 |
| Myadm | 554.9737066 | -0.021595417 | 0.089542688 |
| Prkcg | 1983.083602 | -1.209341068 | 0.159281765 |
| Cacng7 | 1812.960324 | -0.187882032 | 0.078390058 |
| Cacng8 | 593.3089663 | -0.638370027 | 0.102640253 |
| Gm23450 | 0 | NA | NA |
| Cacng6 | 9.245691091 | 1.167536598 | 0.385380413 |
| Tarm1 | 15.12490142 | 6.398721943 | 1.189383578 |

| | | | |
|---|---|---|---|
| Gm25100 | 0 | NA | NA |
| Oscar | 1.834747314 | 1.931912628 | 0.929137277 |
| Ndufa3 | 436.48222 | 0.008914179 | 0.11830782 |
| Tfpt | 142.0055102 | 0.209615059 | 0.129728955 |
| Prpf31 | 410.4576804 | 0.563708016 | 0.072931608 |
| Gm15927 | 8.516043887 | -1.019634198 | 0.381727216 |
| Cnot3 | 493.1599569 | -0.011748665 | 0.076673463 |
| Mir3572 | 0 | NA | NA |
| Leng1 | 69.24448306 | 0.349800185 | 0.177736109 |
| Tmc4 | 26.68951419 | 0.083381685 | 0.281004889 |
| Gm15929 | 0.377779179 | 1.326675169 | 3.150272027 |
| Mboat7 | 1337.109882 | 0.023759502 | 0.051666879 |
| Tsen34 | 403.0156256 | 0.107638476 | 0.121698369 |
| Rps9 | 1343.617836 | 0.752867357 | 0.141109903 |
| Pirb | 100.508228 | 5.216408765 | 0.345948335 |
| Gm15922 | 4.272688704 | 5.811532861 | 0.904850044 |
| Gm15930 | 0 | NA | NA |
| Gm15925 | 2.457332247 | 5.03448621 | 1.110799867 |
| Gm15448 | 3.00180885 | 5.481341974 | 1.041782141 |
| Pira2 | 10.48735136 | 7.161391311 | 0.893160033 |
| Gm10693 | 0.291730981 | 1.629907502 | 1.754787612 |
| Gm14548 | 29.20745553 | 6.848123647 | 0.849705344 |
| Lilra6 | 22.5183229 | 6.466219426 | 0.777227483 |
| Gm7363 | 4.253190973 | 0.893053484 | 0.478474996 |
| Gm15928 | 0.849955171 | -2.116640685 | 1.169849355 |
| Lair1 | 165.5340521 | 1.272912056 | 0.212774885 |
| 9430041J12Rik | 62.38036442 | -0.219083104 | 0.154400669 |
| Gm23741 | 0.042716945 | -0.243571806 | 3.724945049 |
| Ttyh1 | 4104.837726 | -0.711308026 | 0.116221875 |
| D030047H15Rik | 2.31889725 | -1.487252258 | 0.705174404 |
| Leng8 | 1590.8448 | -0.85252511 | 0.285752531 |
| Leng9 | 55.71701833 | -0.052955392 | 0.14651465 |
| Cdc42ep5 | 10.88113561 | 0.580713498 | 0.350322122 |
| Gm5319 | 0 | NA | NA |
| Gm15926 | 0 | NA | NA |
| Lilra5 | 0.951293484 | -0.109622805 | 1.101142322 |
| Gm15923 | 0 | NA | NA |
| Gm15931 | 19.55735248 | 5.426342994 | 0.640434169 |
| Ncr1 | 7.234741561 | 6.225325552 | 0.90697936 |
| Gp6 | 0.386555381 | 2.571445334 | 3.683869617 |
| Gm25349 | 0.061920613 | 0.718207094 | 3.724945049 |
| Gm24752 | 0.04393175 | -0.243571806 | 3.724945049 |
| Rdh13 | 296.6943263 | -0.484040173 | 0.11308047 |
| Gm24605 | 0 | NA | NA |
| Gm15494 | 14.28857811 | -2.360348699 | 0.35573143 |
| Eps8l1 | 3.37593979 | -1.1486848 | 0.610337016 |
| Ppp1r12c | 945.1906909 | 0.033674591 | 0.079083645 |

| | | | |
|---|---|---|---|
| Tnnt1 | 15.65154636 | -1.038349832 | 0.274028412 |
| Tnni3 | 10.79028321 | -0.180979464 | 0.454087024 |
| Dnaaf3 | 15.48723177 | 1.254296663 | 0.316055135 |
| Gm15873 | 0.239904818 | 1.87075327 | 2.531082556 |
| Syt5 | 799.4033589 | -0.454594892 | 0.116905322 |
| Ptprh | 6.302761948 | 1.185226241 | 0.464487433 |
| Gm23577 | 0 | NA | NA |
| Tmem86b | 45.98200731 | -0.228309888 | 0.255921507 |
| Ppp6r1 | 1889.707859 | 0.375628558 | 0.062683266 |
| Hspbp1 | 559.9998375 | 0.155953637 | 0.081645513 |
| Brsk1 | 2659.133262 | -0.819460966 | 0.084099215 |
| Tmem150b | 4.110811422 | -0.282347535 | 0.49333292 |
| Gm24537 | 0 | NA | NA |
| Suv420h2 | 136.5439044 | -0.533110869 | 0.169077259 |
| Cox6b2 | 14.20000006 | -1.218082817 | 0.344431613 |
| Fam71e2 | 0.747386132 | 2.798987957 | 1.199662611 |
| Il11 | 12.48613724 | 2.91547972 | 0.540134269 |
| Tmem190 | 0.273106308 | 0.932841234 | 1.760687163 |
| Tmem238 | 19.96996773 | 0.166803719 | 0.244546096 |
| Rpl28 | 949.3495926 | 0.617376178 | 0.099677599 |
| Ube2s | 303.8563974 | 0.381214003 | 0.114758933 |
| Shisa7 | 1255.260455 | -1.172440779 | 0.237359287 |
| Gm15525 | 0 | NA | NA |
| Isoc2b | 8.838166061 | -0.526774966 | 0.383193254 |
| Isoc2a | 117.0217193 | -0.249323196 | 0.121885752 |
| Zfp628 | 234.1479967 | 0.477246196 | 0.141054931 |
| Nat14 | 284.6791148 | -0.311011746 | 0.114674966 |
| Ssc5d | 87.37008735 | 1.815001794 | 0.176544669 |
| Sbk2 | 2.02355919 | 0.003379187 | 0.741785448 |
| Sbk3 | 14.88936924 | -1.16712355 | 0.377719938 |
| Zfp579 | 184.6131844 | -0.305346004 | 0.105715496 |
| Fiz1 | 274.5806166 | 0.464421785 | 0.081795821 |
| Zfp524 | 43.21558028 | -0.002259596 | 0.201442374 |
| Zfp865 | 368.9050981 | 0.138637646 | 0.075977428 |
| 4632433K11Rik | 17.57441645 | -0.228182119 | 0.286821482 |
| Zfp784 | 56.52918824 | 0.268139569 | 0.160948154 |
| Gm15510 | 3.40753209 | -0.427177492 | 0.518735563 |
| Zfp580 | 125.3343534 | -0.02825019 | 0.136127053 |
| Ccdc106 | 206.5384023 | -0.35736782 | 0.111572036 |
| U2af2 | 1303.244093 | 0.346506925 | 0.054987992 |
| Epn1 | 2656.292776 | -0.384743958 | 0.072957378 |
| Rfpl4 | 0 | NA | NA |
| Rasl2-9 | 0.99583754 | -0.046931228 | 1.033271756 |
| Gm26217 | 0 | NA | NA |
| Vmn1r55 | 0.048949532 | -0.243569582 | 3.724945049 |
| Vmn1r56 | 0 | NA | NA |
| Vmn1r57 | 0.085327104 | 1.179809211 | 3.723500796 |

| | | | |
|---|---|---|---|
| Gm23942 | 0 | NA | NA |
| Nlrp2 | 0 | NA | NA |
| Gm5065 | 0.07796279 | 1.072698098 | 3.721829099 |
| Vmn1r58 | 0 | NA | NA |
| Vmn1r59 | 0 | NA | NA |
| Vmn2r28 | 0 | NA | NA |
| Vmn1r60 | 0 | NA | NA |
| Vmn1r-ps40 | 0 | NA | NA |
| Vmn1r61 | 0 | NA | NA |
| AC161762.1 | 0 | NA | NA |
| Vmn1r62 | 0 | NA | NA |
| Vmn1r63 | 0 | NA | NA |
| Vmn1r64 | 0 | NA | NA |
| Gm20399 | 0 | NA | NA |
| Vmn1r-ps45 | 0 | NA | NA |
| Vmn1r-ps46 | 0 | NA | NA |
| Vmn1r65 | 0.115603922 | 1.209894437 | 3.72418822 |
| Gm5321 | 0.234623521 | 1.156840198 | 2.103128903 |
| Nlrp4c | 0 | NA | NA |
| Zfp787 | 156.8006384 | 0.367372305 | 0.099800664 |
| Zfp444 | 319.2682102 | -0.051565745 | 0.061297129 |
| Galp | 0.278580731 | 1.920033141 | 1.753038086 |
| Zscan5b | 0.398992304 | 2.192579091 | 1.618105269 |
| Gm6792 | 0.123841226 | 1.292225235 | 3.719439735 |
| Zfp667 | 214.0871925 | -0.182017973 | 0.10647246 |
| Zfp583 | 81.78065755 | -0.377637117 | 0.136263308 |
| Zfp78 | 26.1290204 | -0.384944632 | 0.224277543 |
| Zfp28 | 117.9628295 | -0.189397284 | 0.099625992 |
| Smim17 | 178.4907971 | -0.920876953 | 0.171379021 |
| Gm16306 | 2.355736216 | -1.154608993 | 0.766688132 |
| Gm16307 | 0 | NA | NA |
| Gm20715 | 6.848240459 | -1.333848679 | 0.442813509 |
| Olfr1344 | 0 | NA | NA |
| Olfr1336 | 0 | NA | NA |
| Olfr1346 | 0.047948698 | -0.243569582 | 3.724945049 |
| Olfr5 | 0.046345709 | 0.71820487 | 3.724945049 |
| Olfr1347 | 0.072025462 | 0.237312084 | 3.724945049 |
| Olfr1348 | 0 | NA | NA |
| n-R5s150 | 0 | NA | NA |
| Olfr1349 | 0.115957212 | -0.24357403 | 3.724945049 |
| Olfr1350 | 0 | NA | NA |
| Zim1 | 102.3587915 | -0.007199928 | 0.18787447 |
| Peg3 | 8284.965525 | -0.490980877 | 0.07740369 |
| Usp29 | 1064.211711 | -0.254219548 | 0.099889911 |
| Gm27675 | 0 | NA | NA |
| Mir3099 | 1.920827858 | -1.021570934 | 0.814259535 |
| Gm22649 | 0 | NA | NA |

| | | | |
|---|---|---|---|
| Aurkc | 1.107598099 | -1.11050652 | 0.997306041 |
| Zfp954 | 110.343462 | 0.628197967 | 0.141789682 |
| Zfp773 | 25.68791501 | -0.228015904 | 0.234553527 |
| Zfp418 | 91.99907945 | -0.189027047 | 0.155034243 |
| Zfp772 | 42.36616452 | -0.419693033 | 0.163702002 |
| Vmn2r29 | 3.485300844 | -0.938527873 | 0.546921637 |
| Gm18190 | 0.639457996 | -0.418548743 | 1.269202856 |
| Clcn4-2 | 1846.754651 | -0.274310029 | 0.056940935 |
| Mir5620 | 0.831589853 | -0.314315878 | 1.1444964 |
| Vmn2r30 | 0.25221175 | -0.243564665 | 2.131649039 |
| Gm3912 | 1.415212796 | -0.504747616 | 0.871934628 |
| Vmn2r31 | 0.094294407 | 0.23732098 | 3.724945049 |
| Vmn2r-ps36 | 0 | NA | NA |
| Vmn2r32 | 0 | NA | NA |
| Gm3563 | 0 | NA | NA |
| Vmn2r33 | 0 | NA | NA |
| Gm7648 | 0 | NA | NA |
| Gm7651 | 0 | NA | NA |
| Vmn2r-ps37 | 0.072025462 | 0.237312084 | 3.724945049 |
| Vmn2r34 | 0 | NA | NA |
| Vmn2r-ps38 | 0 | NA | NA |
| Vmn2r35 | 0.093141081 | -0.724455696 | 3.724945049 |
| Vmn2r-ps40 | 0.038981395 | 0.718207094 | 3.724945049 |
| Vmn2r36 | 0 | NA | NA |
| Gm7230 | 0 | NA | NA |
| Gm3978 | 0.160408561 | -0.72446006 | 3.457615423 |
| Gm20438 | 0.038981395 | 0.718207094 | 3.724945049 |
| Gm18200 | 0.398333997 | -0.627398217 | 1.526094049 |
| Gm4073 | 0 | NA | NA |
| Gm3675 | 0 | NA | NA |
| 2810047C21Rik1 | 0.042716945 | -0.243571806 | 3.724945049 |
| Gm3665 | 0 | NA | NA |
| Vmn2r41 | 0.042716945 | -0.243571806 | 3.724945049 |
| Vmn2r42 | 0 | NA | NA |
| Vmn2r43 | 0 | NA | NA |
| Gm3654 | 0 | NA | NA |
| Vmn2r-ps45 | 0.124853745 | 0.237309606 | 3.724945049 |
| Gm7696 | 0.676743848 | -0.078095484 | 1.141362717 |
| Vmn2r44 | 0.263039019 | -0.243564584 | 2.106825953 |
| Vmn2r-ps46 | 0 | NA | NA |
| Vmn2r45 | 0.068097359 | -0.243569582 | 3.724945049 |
| Vmn2r-ps47 | 0 | NA | NA |
| Vmn2r-ps49 | 0 | NA | NA |
| Gm3584 | 0.047948698 | -0.243569582 | 3.724945049 |
| Gm3956 | 0 | NA | NA |
| Gm3585 | 0 | NA | NA |
| Gm3962 | 0.333123609 | -0.98624477 | 1.68046324 |

| | | | |
|---|---|---|---|
| Gm18191 | 0.264294608 | 0.612581613 | 1.782244687 |
| Vmn2r-ps44 | 0 | NA | NA |
| Gm3644 | 0 | NA | NA |
| Gm20482 | 0 | NA | NA |
| Vmn2r-ps43 | 0 | NA | NA |
| Vmn2r40 | 0 | NA | NA |
| Vmn2r-ps42 | 0 | NA | NA |
| Vmn2r39 | 0 | NA | NA |
| Zfp419 | 0.038981395 | 0.718207094 | 3.724945049 |
| Vmn2r38 | 0.068097359 | -0.243569582 | 3.724945049 |
| Gm5583 | 0 | NA | NA |
| Gm18201 | 0 | NA | NA |
| Vmn2r-ps41 | 0 | NA | NA |
| Vmn2r37 | 0 | NA | NA |
| Gm4011 | 0 | NA | NA |
| Gm3623 | 0 | NA | NA |
| Gm3616 | 0 | NA | NA |
| Gm20287 | 0 | NA | NA |
| Gm3611 | 0 | NA | NA |
| Gm3972 | 0.099106094 | 1.209894437 | 3.72418822 |
| Gm18199 | 0.40494807 | 0.735190669 | 1.717889494 |
| Gm10302 | 0 | NA | NA |
| Gm3683 | 0 | NA | NA |
| Gm20345 | 0 | NA | NA |
| Vmn2r46 | 0.048949532 | -0.243569582 | 3.724945049 |
| Vmn2r47 | 0 | NA | NA |
| Vmn2r-ps51 | 0 | NA | NA |
| Vmn2r-ps52 | 0 | NA | NA |
| Vmn2r48 | 0 | NA | NA |
| Vmn2r49 | 0 | NA | NA |
| Vmn2r50 | 0.038981395 | 0.718207094 | 3.724945049 |
| Vmn2r51 | 0 | NA | NA |
| Vmn2r52 | 0 | NA | NA |
| Gm20679 | 0 | NA | NA |
| Vmn1r66 | 0 | NA | NA |
| Vmn1r-ps47 | 0 | NA | NA |
| Vmn1r-ps48 | 0 | NA | NA |
| Nlrp4d | 0.062426873 | 0.237312084 | 3.724945049 |
| Vmn1r67 | 0 | NA | NA |
| Zik1 | 251.415385 | 0.013075519 | 0.091649237 |
| Vmn1r68 | 0 | NA | NA |
| Vmn1r-ps49 | 0 | NA | NA |
| Vmn1r69 | 0 | NA | NA |
| Vmn1r-ps50 | 0 | NA | NA |
| Vmn1r70 | 0 | NA | NA |
| Vmn1r-ps51 | 0 | NA | NA |
| Gm6900 | 2.818881342 | 0.720674 | 0.619326927 |

| | | | |
|---|---|---|---|
| Vmn1r-ps52 | 0 | NA | NA |
| Nlrp4b | 0 | NA | NA |
| Vmn1r71 | 0 | NA | NA |
| Zscan4b | 0 | NA | NA |
| Mir297-1 | 0 | NA | NA |
| Zscan4c | 0 | NA | NA |
| Zscan4d | 0 | NA | NA |
| Gm26248 | 0 | NA | NA |
| Zscan4e | 0 | NA | NA |
| Gm23087 | 0 | NA | NA |
| Zscan4f | 0 | NA | NA |
| Vmn1r-ps53 | 0 | NA | NA |
| Vmn1r72 | 0 | NA | NA |
| Vmn1r73 | 0 | NA | NA |
| Vmn1r-ps54 | 0 | NA | NA |
| Vmn1r74 | 0 | NA | NA |
| Vmn1r-ps55 | 0 | NA | NA |
| Vmn1r75 | 0 | NA | NA |
| Vmn1r-ps56 | 0 | NA | NA |
| Vmn1r76 | 0 | NA | NA |
| Vmn1r-ps57 | 0 | NA | NA |
| Gm24414 | 1.280135606 | 0.272952758 | 0.847807637 |
| Vmn1r-ps58 | 0 | NA | NA |
| Vmn1r-ps59 | 0 | NA | NA |
| Vmn1r-ps60 | 0 | NA | NA |
| Vmn1r77 | 0 | NA | NA |
| Gm10179 | 23.18691661 | -0.437954552 | 0.245954315 |
| Vmn1r-ps61 | 0 | NA | NA |
| Vmn1r-ps62 | 0 | NA | NA |
| Vmn1r-ps63 | 0 | NA | NA |
| Vmn1r78 | 0 | NA | NA |
| Vmn1r79 | 0 | NA | NA |
| Vmn1r80 | 0 | NA | NA |
| Vmn1r-ps64 | 0 | NA | NA |
| Vmn1r-ps65 | 0 | NA | NA |
| Vmn1r81 | 0 | NA | NA |
| Vmn1r82 | 0 | NA | NA |
| Vmn1r83 | 0 | NA | NA |
| Gm23521 | 0 | NA | NA |
| Vmn1r84 | 0.047948698 | -0.243569582 | 3.724945049 |
| Vmn1r-ps66 | 0 | NA | NA |
| Vmn1r-ps67 | 0 | NA | NA |
| Vmn1r-ps68 | 0 | NA | NA |
| Zfp551 | 75.69228729 | -0.301843118 | 0.12222987 |
| Vmn2r-ps53 | 0 | NA | NA |
| Zfp606 | 312.4159778 | -0.157390963 | 0.071883894 |
| 2900092C05Rik | 0.326351934 | 1.332166066 | 1.671010596 |

| | | | |
|---|---|---|---|
| Gm15877 | 0.593305459 | 0.778339182 | 1.282359613 |
| Vmn2r53 | 2.096357237 | -0.485256615 | 0.71736707 |
| Gm15773 | 0.596424136 | 0.002176907 | 1.290640466 |
| Vmn2r54 | 1.895285101 | 0.824808273 | 0.845037463 |
| Vmn2r55 | 1.524911348 | -0.682170474 | 0.954621209 |
| Vmn2r56 | 2.422070769 | 0.022495357 | 1.031648536 |
| Gm6946 | 0 | NA | NA |
| Zscan18 | 180.8411845 | -0.334086839 | 0.147736795 |
| Zfp329 | 519.818062 | -0.360298157 | 0.124573649 |
| Zfp110 | 270.5868238 | 1.136629758 | 0.114852803 |
| Zfp128 | 118.8763104 | -0.169603221 | 0.134394595 |
| Zscan22 | 156.5051589 | 0.081460556 | 0.158115991 |
| Rps5 | 1620.953915 | 1.262406427 | 0.129567568 |
| Rnf225 | 104.6098484 | 3.52029885 | 0.22255196 |
| Gm26325 | 0 | NA | NA |
| 1810019N24Rik | 6.340267528 | 1.343032293 | 0.479431104 |
| Gm29638 | 0.046345709 | 0.71820487 | 3.724945049 |
| Zfp324 | 270.4925259 | 1.07215539 | 0.109048036 |
| Zfp446 | 119.2751431 | 0.314487879 | 0.145397837 |
| Slc27a5 | 10.7500977 | -0.439031178 | 0.342423507 |
| Zbtb45 | 249.6021384 | 0.112940829 | 0.100273829 |
| Trim28 | 1232.544845 | 0.329418895 | 0.072839577 |
| Chmp2a | 510.643161 | 0.582002501 | 0.108309576 |
| Ube2m | 1024.417635 | 0.268336407 | 0.082533344 |
| Mzf1 | 76.8160005 | -1.210559337 | 0.195637386 |
| Vmn1r85 | 0 | NA | NA |
| Vmn1r86 | 0 | NA | NA |
| Vmn1r87 | 0 | NA | NA |
| Vmn1r88 | 0.289463044 | -0.243568266 | 2.051711593 |
| Gm4879 | 6.771121092 | 0.727371023 | 0.46217311 |
| Vmn1r89 | 0.099106094 | 1.209894437 | 3.72418822 |
| Vmn1r-ps69 | 0 | NA | NA |
| 6330408A02Rik | 197.7921584 | -0.333068182 | 0.181957809 |
| Lig1 | 167.3687248 | 0.358325654 | 0.125808353 |
| Pla2g4c | 22.31652208 | 1.600273285 | 0.303155283 |
| Rps8-ps4 | 2.214157104 | 0.671164407 | 0.669392564 |
| Gm24909 | 0.201760502 | 0.74218997 | 2.220505766 |
| Gm23499 | 0 | NA | NA |
| Cabp5 | 0.609661125 | -0.646411452 | 1.265840032 |
| Bsph1 | 0.222333927 | 0.71822328 | 2.143130283 |
| Bsph2-ps | 0 | NA | NA |
| Bsph2 | 0 | NA | NA |
| Sult2a5 | 0 | NA | NA |
| Zbed4-ps1 | 0 | NA | NA |
| Kat6b-ps1 | 0 | NA | NA |
| Tmem167-ps1 | 0 | NA | NA |
| Sult2a-ps1 | 0 | NA | NA |

| | | | |
|---|---|---|---|
| Sult2a2 | 0 | NA | NA |
| Zbed4-ps2 | 0.047948698 | -0.243569582 | 3.724945049 |
| Sult2a1 | 0.169165009 | 0.719161184 | 3.351346564 |
| Kat6b-ps2 | 0 | NA | NA |
| Sult2a-ps2 | 0 | NA | NA |
| Tmem167-ps2 | 0 | NA | NA |
| Sult2a4 | 0 | NA | NA |
| Zbed4-ps3 | 0.112642008 | -0.24357403 | 3.724945049 |
| Tmem167-ps3 | 0 | NA | NA |
| Sult2a3 | 0 | NA | NA |
| Sult2a-ps3 | 0 | NA | NA |
| Sult2a-ps4 | 0 | NA | NA |
| Rpap3-ps1 | 0 | NA | NA |
| Sult2a6 | 0 | NA | NA |
| Rpap3-ps2 | 0 | NA | NA |
| Phf20-ps | 17.46056627 | 0.237637384 | 0.246770482 |
| Obox8 | 0 | NA | NA |
| Mrps36-ps3 | 0 | NA | NA |
| Crs-ps | 0 | NA | NA |
| Sult2a8 | 0.047948698 | -0.243569582 | 3.724945049 |
| Sult2a7 | 0.119587853 | 0.237305575 | 3.724945049 |
| Obox4-ps1 | 0.155414793 | 0.237326542 | 2.82937401 |
| Obox4-ps2 | 0.183158741 | -0.698418772 | 2.72428501 |
| Nlrp5-ps | 183.9697913 | -0.236533626 | 0.175227095 |
| Obox4-ps3 | 0 | NA | NA |
| Vmn1r90 | 3.490537078 | -0.087369382 | 0.573926421 |
| Zfp-ps | 0 | NA | NA |
| Ranbp2-ps1 | 0 | NA | NA |
| Obox3-ps7 | 4.106584469 | -0.845525252 | 0.534219853 |
| Obox7 | 0.129647038 | -0.243562299 | 2.992845844 |
| Obox3-ps1 | 0 | NA | NA |
| RP24-194K24.10 | 0 | NA | NA |
| Gtpbp4-ps1 | 0 | NA | NA |
| Ranbp2-ps2 | 0 | NA | NA |
| Obox4-ps36 | 0 | NA | NA |
| Obox4-ps37 | 0 | NA | NA |
| Gm26056 | 0 | NA | NA |
| Obox3-ps2 | 0 | NA | NA |
| RP23-349I13.7 | 0 | NA | NA |
| Gtpbp4-ps2 | 0 | NA | NA |
| Ranbp2-ps3 | 0 | NA | NA |
| Obox4-ps38 | 0 | NA | NA |
| Obox4-ps39 | 0 | NA | NA |
| Ranbp2-ps4 | 0 | NA | NA |
| Obox4-ps6 | 0 | NA | NA |
| Gm24701 | 0 | NA | NA |
| Obox3-ps3 | 0 | NA | NA |

| | | | |
|---|---|---|---|
| RP23-263E7.8 | 0 | NA | NA |
| Gtpbp4-ps3 | 0 | NA | NA |
| Ranbp2-ps5 | 0 | NA | NA |
| Obox4-ps8 | 0 | NA | NA |
| Obox4-ps9 | 0 | NA | NA |
| Obox3-ps4 | 0 | NA | NA |
| RP23-263E7.7 | 0 | NA | NA |
| Gtpbp4-ps4 | 0 | NA | NA |
| Ranbp2-ps6 | 0 | NA | NA |
| Obox4-ps11 | 0 | NA | NA |
| Obox4-ps12 | 0 | NA | NA |
| Obox4-ps13 | 0 | NA | NA |
| Obox4-ps14 | 0 | NA | NA |
| Obox4-ps15 | 0 | NA | NA |
| Obox4-ps16 | 0 | NA | NA |
| Obox4-ps17 | 0 | NA | NA |
| Obox4-ps18 | 0 | NA | NA |
| Obox4-ps19 | 0 | NA | NA |
| Obox4-ps20 | 0 | NA | NA |
| Obox4-ps21 | 0 | NA | NA |
| Obox4-ps22 | 0 | NA | NA |
| Obox4-ps23 | 0 | NA | NA |
| Mrip-ps | 35.07510409 | 0.473605868 | 0.356227807 |
| Obox3-ps5 | 0 | NA | NA |
| RP24-84K14.12 | 0 | NA | NA |
| Gtpbp4-ps5 | 0 | NA | NA |
| RP24-105I10.4 | 0 | NA | NA |
| Obox4-ps26 | 0 | NA | NA |
| Ranbp2-ps7 | 0 | NA | NA |
| Obox2 | 0 | NA | NA |
| Obox1 | 0 | NA | NA |
| Ranbp2-ps8 | 0 | NA | NA |
| Obox4-ps27 | 0 | NA | NA |
| Obox4-ps28 | 0 | NA | NA |
| Obox4-ps29 | 0 | NA | NA |
| Obox3 | 0 | NA | NA |
| Obox3-ps6 | 0 | NA | NA |
| Obox4-ps30 | 0 | NA | NA |
| Ranbp2-ps9 | 0 | NA | NA |
| Ranbp2-ps10 | 0 | NA | NA |
| Obox4-ps31 | 0 | NA | NA |
| Gm25354 | 0 | NA | NA |
| RP23-140B24.1 | 0 | NA | NA |
| RP23-140B24.2 | 0 | NA | NA |
| Obox4-ps33 | 0 | NA | NA |
| Gm23714 | 0 | NA | NA |
| Obox4-ps34 | 0 | NA | NA |

| | | | |
|---|---|---|---|
| RP24-370G23.12 | 0 | NA | NA |
| Obox5 | 0 | NA | NA |
| RP24-370G23.13 | 0 | NA | NA |
| RP24-370G23.14 | 0 | NA | NA |
| Obox4-ps35 | 0.220980774 | 1.68041995 | 3.10409395 |
| Obox6 | 0.123845245 | 1.035807111 | 3.720871489 |
| Crx | 0.603215896 | 0.22632817 | 2.628214393 |
| Crxos | 0.501748666 | -2.168678045 | 1.283847841 |
| RP24-466N11.6 | 0 | NA | NA |
| Sepw1 | 5104.264334 | 0.899142844 | 0.096666049 |
| Gltscr2 | 737.8364871 | 0.554782058 | 0.093392279 |
| Snord23 | 0.926190085 | 1.693135497 | 1.141327781 |
| Gm24576 | 0.169505627 | 0.219382274 | 2.747555816 |
| Ehd2 | 172.4468587 | 2.703100373 | 0.190937356 |
| Gltscr1 | 438.3260183 | 0.155040338 | 0.077075439 |
| Zfp541 | 3.78555653 | -0.636489843 | 0.51470418 |
| Napa | 1603.381598 | -0.20427108 | 0.056383397 |
| Kptn | 155.1087845 | -0.725474481 | 0.10546508 |
| Slc8a2 | 2081.402497 | -1.526899892 | 0.140435637 |
| Meis3 | 261.2098945 | 0.015712395 | 0.103185292 |
| Dhx34 | 160.4271404 | 0.471558033 | 0.087826254 |
| C5ar2 | 2.109974762 | 2.384683373 | 0.898170143 |
| Gm26473 | 0 | NA | NA |
| C5ar1 | 78.4729806 | 5.526084444 | 0.399122826 |
| Inafm1 | 160.8892379 | -0.016617511 | 0.109160775 |
| Ccdc9 | 366.0675831 | 0.015711484 | 0.102057859 |
| Bbc3 | 74.98142202 | 0.316177768 | 0.147016691 |
| Sae1 | 546.3005839 | 0.022479578 | 0.061735985 |
| Zc3h4 | 1014.843218 | 0.090314706 | 0.076470072 |
| Tmem160 | 153.6281965 | 0.155275027 | 0.148449969 |
| Npas1 | 76.47707439 | -0.12759971 | 0.14522955 |
| n-R5s151 | 0 | NA | NA |
| Arhgap35 | 4302.533817 | -0.040281132 | 0.051559094 |
| Gm17826 | 0.07796279 | 1.072698098 | 3.721829099 |
| Gm17981 | 12.95166994 | -0.468116161 | 0.384009294 |
| Gm29443 | 4.384705177 | -1.329161203 | 0.524277192 |
| Gm29442 | 0.132101749 | 1.209894437 | 3.72418822 |
| Gm24708 | 0 | NA | NA |
| Ceacam15 | 0.205464717 | 0.727598501 | 2.642335502 |
| Ceacam9 | 1.215943724 | -1.09684229 | 0.900026934 |
| Ap2s1 | 780.4435061 | 0.000955009 | 0.097484667 |
| Slc1a5 | 49.69459424 | 1.546653802 | 0.237381895 |
| Fkrp | 486.4658568 | -0.182696969 | 0.077167065 |
| Strn4 | 1924.374574 | -0.281991519 | 0.078878322 |
| Prkd2 | 82.10841754 | 1.013150343 | 0.17922917 |
| 9330104G04Rik | 30.06220494 | -0.73609088 | 0.197661871 |
| Dact3 | 1250.275623 | -0.973729042 | 0.099702331 |

| | | | |
|---|---|---|---|
| Gng8 | 8.57217951 | -0.709065843 | 0.354035874 |
| Ptgir | 4.875732241 | 4.580446443 | 0.890838373 |
| Calm3 | 11988.4177 | -0.520364628 | 0.069419598 |
| Pnmal2 | 4095.934161 | -0.530266826 | 0.074507933 |
| Gm25815 | 0 | NA | NA |
| Pnmal1 | 642.282333 | -0.552710002 | 0.104940853 |
| Ccdc8 | 31.97129462 | -0.439097109 | 0.247944968 |
| Ppp5c | 936.7635144 | -0.136201527 | 0.053146729 |
| Hif3a | 169.2274098 | -0.596400841 | 0.2015423 |
| Psg16 | 99.26004857 | -0.02527106 | 0.189406605 |
| Ceacam3 | 2.261397335 | -0.85124181 | 0.74831475 |
| Psg29 | 2.218822718 | 0.335439875 | 0.744720049 |
| Gm24520 | 0 | NA | NA |
| Gm25068 | 0 | NA | NA |
| Ceacam5 | 0 | NA | NA |
| Gm6065 | 0.197744397 | -0.731744894 | 2.224405099 |
| Ceacam14 | 0 | NA | NA |
| Gm5155 | 0 | NA | NA |
| Ceacam11 | 0 | NA | NA |
| Ceacam13 | 0 | NA | NA |
| Gm22414 | 0 | NA | NA |
| Ceacam12 | 0 | NA | NA |
| Igfl3 | 0 | NA | NA |
| Mill1 | 0 | NA | NA |
| Psg18 | 0 | NA | NA |
| Gm24715 | 0 | NA | NA |
| Psg28 | 0.047948698 | -0.243569582 | 3.724945049 |
| Psg26 | 0 | NA | NA |
| Psg25 | 0.049553047 | 0.71820487 | 3.724945049 |
| Gm26062 | 0 | NA | NA |
| Psg27 | 0 | NA | NA |
| Psg23 | 0.066050875 | 0.71820487 | 3.724945049 |
| Psg21 | 0 | NA | NA |
| Psg20 | 0.124347486 | 0.719186602 | 3.717245571 |
| Psg22 | 0 | NA | NA |
| Gm24464 | 0 | NA | NA |
| Psg19 | 0 | NA | NA |
| Psg17 | 1.789359587 | 2.16970397 | 0.977574276 |
| Mill2 | 14.66817284 | 3.100763482 | 0.488374022 |
| Pglyrp1 | 28.41791276 | 1.759433248 | 0.281102846 |
| Ccdc61 | 68.71592428 | 0.056822228 | 0.172433005 |
| Nova2 | 1854.132093 | -0.835617134 | 0.106300565 |
| Gm26821 | 0.975505809 | 1.384055584 | 1.006792325 |
| Nanos2 | 10.4316941 | -0.992397767 | 0.443682237 |
| Gm10676 | 6.752775585 | -0.487000286 | 0.384617919 |
| Mypop | 377.759842 | -0.561840679 | 0.087541401 |
| Mypopos | 3.140367151 | 1.19566823 | 0.63201329 |

| | | | |
|---|---|---|---|
| Irf2bp1 | 595.038875 | 0.035440431 | 0.076736334 |
| Foxa3 | 0.267510113 | 1.353903804 | 2.390275537 |
| Sympk | 1388.597211 | -0.058553678 | 0.059160716 |
| Rsph6a | 1.359012152 | -0.25567125 | 0.805070505 |
| Dmwd | 973.7957556 | -0.065883814 | 0.078770808 |
| Dmpk | 178.6755614 | -0.63301673 | 0.205138363 |
| Mir3100 | 1.552752546 | -0.31293052 | 0.827639754 |
| Six5 | 17.71455458 | -0.56971242 | 0.274875735 |
| Gm4969 | 2.587362481 | -0.693690524 | 0.843301036 |
| Fbxo46 | 157.8854747 | 0.119312991 | 0.146619462 |
| Qpctl | 177.5083971 | 0.581505686 | 0.126318036 |
| Snrpd2 | 249.1707862 | 0.400397731 | 0.122992068 |
| Gipr | 9.076618891 | 0.091140934 | 0.42315653 |
| Eml2 | 1469.733199 | -0.451348907 | 0.071758963 |
| Mir330 | 0 | NA | NA |
| Gpr4 | 26.0798917 | 0.19360221 | 0.222417785 |
| Opa3 | 585.1491548 | 0.127000888 | 0.063484292 |
| Gm25134 | 1.373839161 | -1.413304264 | 0.913977185 |
| Vasp | 344.2888316 | 2.000913437 | 0.143538558 |
| 1700058P15Rik | 0 | NA | NA |
| Ppm1n | 0.85008157 | 1.874694614 | 1.263637141 |
| Rtn2 | 574.5812076 | -0.188363826 | 0.101216836 |
| Gm26802 | 0.561122788 | 2.705886014 | 1.587356969 |
| Fosb | 461.8672414 | 1.815679755 | 0.612604392 |
| Ercc1 | 157.0107263 | 0.82652593 | 0.120209113 |
| Cd3eap | 115.4671044 | 0.386292392 | 0.119411266 |
| Ppp1r13l | 62.86165383 | 1.061118702 | 0.163468689 |
| Ercc2 | 283.3150772 | -0.002298842 | 0.082827809 |
| Gm26852 | 0.604815377 | 1.092827493 | 1.189048467 |
| Mir343 | 1.738846634 | -2.031024965 | 0.902158453 |
| Klc3 | 30.34969473 | 0.930307812 | 0.215575868 |
| Ckm | 1.573181334 | 0.863156217 | 0.822853996 |
| A930016O22Rik | 2.771915612 | -1.616414067 | 0.654268384 |
| Mark4 | 736.3036542 | 0.025783888 | 0.06867192 |
| Gm23099 | 0 | NA | NA |
| Exoc3l2 | 2.258282039 | 0.061811782 | 0.620443186 |
| Bloc1s3 | 84.9281982 | 0.048402906 | 0.120045118 |
| Trappc6a | 46.60324668 | -0.086096934 | 0.178248639 |
| Nkpd1 | 9.973239763 | -0.209379696 | 0.323197328 |
| Ppp1r37 | 1313.266029 | 0.022271809 | 0.073170729 |
| Gemin7 | 188.0515057 | 0.617181147 | 0.103397176 |
| Zfp296 | 23.1978082 | 1.571850149 | 0.279458081 |
| Clasrp | 352.6239218 | -0.336020403 | 0.176605506 |
| Gm26890 | 51.60349904 | -0.813455646 | 0.351547831 |
| Relb | 145.3464235 | 1.944440876 | 0.238735352 |
| Clptm1 | 1928.605293 | 0.202645651 | 0.064478707 |
| Apoc2 | 34.46812154 | 6.253678321 | 0.625215684 |

| | | | |
|---|---|---|---|
| Apoc4 | 0.250169628 | -0.243564681 | 2.136490353 |
| Apoc1 | 11.42902807 | 1.001851117 | 0.368217319 |
| Apoe | 8648.195503 | -0.144302805 | 0.177180045 |
| Tomm40 | 595.2610474 | 0.043191624 | 0.065248311 |
| Gm20512 | 1.197749286 | -2.286929984 | 1.11069438 |
| Pvrl2 | 109.3428072 | 1.009579563 | 0.163492856 |
| Bcam | 112.1762543 | 0.037725617 | 0.147867537 |
| Gm26600 | 0 | NA | NA |
| Cblc | 0.325981489 | -0.226491801 | 1.503645015 |
| Bcl3 | 136.661948 | 6.075187709 | 0.437715408 |
| Gm16175 | 0.542127486 | -0.661817576 | 1.160501569 |
| Gm16174 | 1.534203801 | 3.55046036 | 1.04896823 |
| Gm19345 | 14.35855545 | -0.595193048 | 0.291030787 |
| Ceacam16 | 1.967062318 | 0.758568392 | 0.811031592 |
| Ceacam19 | 1.492916405 | 4.425433031 | 1.027192159 |
| Pvr | 193.8546322 | 1.580261362 | 0.179931904 |
| Gm16184 | 0 | NA | NA |
| Gm16251 | 0.990102919 | -1.415952581 | 1.049869714 |
| Igsf23 | 0.400433843 | 2.638949811 | 1.587647965 |
| Rpl7a-ps8 | 0.1140246 | 0.718202646 | 3.724945049 |
| Ceacam20 | 0.152347716 | 0.718215089 | 2.84428374 |
| Nlrp9b | 0 | NA | NA |
| n-R5s152 | 0 | NA | NA |
| Vmn1r91 | 0 | NA | NA |
| Gm16442 | 0 | NA | NA |
| Vmn1r93 | 0 | NA | NA |
| Vmn1r94 | 0 | NA | NA |
| Vmn1r95 | 0 | NA | NA |
| Gm6164 | 0 | NA | NA |
| Gm4498 | 0 | NA | NA |
| Gm4133 | 0 | NA | NA |
| Vmn1r-ps79 | 0 | NA | NA |
| Gm5728 | 0 | NA | NA |
| Vmn1r100 | 0 | NA | NA |
| Vmn1r101 | 0 | NA | NA |
| Gm10670 | 0 | NA | NA |
| Vmn1r103 | 0 | NA | NA |
| Vmn1r104 | 0 | NA | NA |
| Gm5726 | 0 | NA | NA |
| Gm4141 | 0 | NA | NA |
| Vmn1r107 | 0 | NA | NA |
| Vmn1r-ps71 | 0 | NA | NA |
| Vmn1r111 | 0 | NA | NA |
| Gm10668 | 0 | NA | NA |
| Vmn1r112 | 0 | NA | NA |
| Vmn1r113 | 0 | NA | NA |
| Vmn1r114 | 0 | NA | NA |

| | | | |
|---|---|---|---|
| Vmn1r115 | 0 | NA | NA |
| Vmn1r116 | 0 | NA | NA |
| Vmn1r117 | 0 | NA | NA |
| Vmn1r118 | 0 | NA | NA |
| Gm4526 | 0 | NA | NA |
| Vmn1r119 | 0 | NA | NA |
| Vmn1r-ps73 | 0 | NA | NA |
| Vmn1r120 | 0 | NA | NA |
| Vmn1r-ps74 | 0 | NA | NA |
| Vmn1r121 | 0 | NA | NA |
| Vmn1r122 | 0 | NA | NA |
| Vmn1r123 | 0 | NA | NA |
| Gm5157 | 0 | NA | NA |
| Vmn1r124 | 0 | NA | NA |
| Vmn1r125 | 0 | NA | NA |
| Vmn1r126 | 0 | NA | NA |
| Vmn1r127 | 0 | NA | NA |
| Vmn1r128 | 0 | NA | NA |
| Vmn1r129 | 0 | NA | NA |
| Vmn1r-ps75 | 0 | NA | NA |
| Gm6882 | 0 | NA | NA |
| Vmn1r130 | 0 | NA | NA |
| Vmn1r-ps76 | 0 | NA | NA |
| Vmn1r131 | 0 | NA | NA |
| Gm4545 | 0 | NA | NA |
| Vmn1r132 | 0 | NA | NA |
| Gm8600 | 0 | NA | NA |
| Gm4175 | 0 | NA | NA |
| Gm4177 | 0 | NA | NA |
| Vmn1r135 | 0 | NA | NA |
| Gm5725 | 0 | NA | NA |
| Vmn1r-ps77 | 0 | NA | NA |
| Gm5891 | 0 | NA | NA |
| Vmn1r137 | 0 | NA | NA |
| Vmn1r-ps78 | 0 | NA | NA |
| Vmn1r138 | 0 | NA | NA |
| Gm6176 | 0 | NA | NA |
| Vmn1r139 | 0 | NA | NA |
| AC127227.1 | 0 | NA | NA |
| Gm16451 | 0 | NA | NA |
| Gm10666 | 0 | NA | NA |
| Vmn1r142 | 0 | NA | NA |
| Vmn1r143 | 0 | NA | NA |
| Gm8653 | 0 | NA | NA |
| Gm8453 | 0 | NA | NA |
| Gm4557 | 0 | NA | NA |
| Gm4558 | 0 | NA | NA |

| | | | |
|---|---|---|---|
| Gm4187 | 0 | NA | NA |
| Vmn1r-ps70 | 0 | NA | NA |
| Gm8660 | 0 | NA | NA |
| Vmn1r148 | 0 | NA | NA |
| Vmn1r149 | 0 | NA | NA |
| Gm10665 | 0 | NA | NA |
| Vmn1r151 | 0 | NA | NA |
| Vmn1r152 | 0 | NA | NA |
| Gm8677 | 0 | NA | NA |
| Gm4201 | 0 | NA | NA |
| Gm4565 | 0 | NA | NA |
| Vmn1r155 | 0 | NA | NA |
| Vmn1r-ps80 | 0 | NA | NA |
| Gm8693 | 0 | NA | NA |
| Gm4567 | 0 | NA | NA |
| Vmn1r157 | 0 | NA | NA |
| Vmn1r158 | 0 | NA | NA |
| Gm4207 | 0 | NA | NA |
| Vmn1r159 | 0 | NA | NA |
| Vmn1r160 | 0 | NA | NA |
| Gm8708 | 0 | NA | NA |
| Gm4214 | 0 | NA | NA |
| Gm4216 | 0 | NA | NA |
| Vmn1r163 | 0 | NA | NA |
| Gm8720 | 0 | NA | NA |
| Gm16456 | 0 | NA | NA |
| Gm10662 | 0 | NA | NA |
| Vmn1r165 | 0 | NA | NA |
| Vmn1r-ps83 | 0 | NA | NA |
| Vmn1r166 | 0 | NA | NA |
| Gm6902 | 0 | NA | NA |
| Nlrp4e | 0 | NA | NA |
| Nlrp5 | 1.484051332 | -0.422064015 | 0.909066914 |
| Gm25196 | 0.066050875 | 0.71820487 | 3.724945049 |
| Vmn1r-ps84 | 0 | NA | NA |
| Vmn1r167 | 0 | NA | NA |
| Vmn1r168 | 0.047948698 | -0.243569582 | 3.724945049 |
| Vmn1r169 | 0 | NA | NA |
| Vmn1r170 | 0 | NA | NA |
| Vmn1r171 | 0 | NA | NA |
| Vmn1r172 | 0 | NA | NA |
| Vmn1r173 | 0 | NA | NA |
| Vmn1r174 | 0 | NA | NA |
| Vmn1r-ps85 | 0 | NA | NA |
| Vmn1r175 | 0 | NA | NA |
| Vmn1r176 | 0 | NA | NA |
| Vmn1r177 | 0 | NA | NA |

| | | | |
|---|---|---|---|
| Vmn1r178 | 0 | NA | NA |
| Vmn1r179 | 0 | NA | NA |
| Gm10175 | 13.75409139 | 0.475028515 | 0.327542505 |
| Vmn1r180 | 0 | NA | NA |
| Vmn1r181 | 0 | NA | NA |
| V1rd19 | 0 | NA | NA |
| Vmn1r183 | 0 | NA | NA |
| Ptma-ps1 | 0 | NA | NA |
| Zfp180 | 620.834939 | -0.033454748 | 0.091890951 |
| Zfp112 | 130.5214199 | -0.215477251 | 0.164886818 |
| Zfp235 | 143.0420723 | -0.307032446 | 0.122923094 |
| Zfp114 | 11.47062778 | -0.408144853 | 0.354447608 |
| Zfp111 | 197.4067323 | -0.063525902 | 0.159152533 |
| Zfp109 | 90.1039288 | -0.474133914 | 0.149550772 |
| Zfp108 | 66.9010371 | -0.501462642 | 0.211653681 |
| Zfp93 | 102.2549038 | -0.102663054 | 0.136365493 |
| Zfp61 | 139.4922126 | -0.140066757 | 0.130680984 |
| Zfp94 | 74.342771 | -0.006493855 | 0.147116398 |
| Gm15465 | 0 | NA | NA |
| Tescl | 0 | NA | NA |
| Gm27592 | 0.061920613 | 0.718207094 | 3.724945049 |
| Lypd5 | 0.324847777 | 0.734922895 | 3.72378749 |
| Kcnn4 | 8.685732437 | 3.005481022 | 0.508961482 |
| Smg9 | 288.8311134 | 0.10966125 | 0.077892759 |
| Irgc1 | 0.04393175 | -0.243571806 | 3.724945049 |
| Plaur | 88.94456294 | 5.292092809 | 0.486089602 |
| Cadm4 | 2550.068439 | 0.067970033 | 0.088748297 |
| Zfp428 | 224.1959431 | -0.060366514 | 0.099922757 |
| Irgq | 1232.318714 | -0.437995195 | 0.04984012 |
| Pinlyp | 0.104366086 | -0.243576254 | 3.724945049 |
| Xrcc1 | 220.1632784 | 8.75E-05 | 0.097087061 |
| Zfp575 | 72.63402446 | -0.919455836 | 0.129293986 |
| Ethe1 | 83.5924299 | 0.248098793 | 0.156332019 |
| Phldb3 | 6.837825038 | 3.856763327 | 0.650541759 |
| Gm26550 | 0 | NA | NA |
| Lypd3 | 0.412745228 | -1.615016044 | 3.704286444 |
| Tex101 | 0 | NA | NA |
| BC049730 | 0 | NA | NA |
| Gm4763 | 0 | NA | NA |
| Cd177 | 15.47993706 | 7.998633199 | 0.961866585 |
| Ceacam10 | 1.306976768 | 0.464973502 | 1.127805597 |
| Gm5893 | 0 | NA | NA |
| Gm9844 | 66.59610774 | 1.291747073 | 0.197530865 |
| Lypd4 | 0 | NA | NA |
| Dmrtc2 | 0.042716945 | -0.243571806 | 3.724945049 |
| Rps19 | 1020.693303 | 0.659470445 | 0.087499965 |
| Cd79a | 3.200457292 | 0.902563452 | 0.54879399 |

| | | | |
|---|---|---|---|
| Arhgef1 | 583.8558966 | 0.347661814 | 0.119772715 |
| Rabac1 | 736.5228606 | 0.110391754 | 0.120830971 |
| Mir7046 | 0 | NA | NA |
| Atp1a3 | 26298.87918 | -0.431504617 | 0.062712401 |
| Mir7047 | 0.284641305 | -1.334865199 | 1.71003639 |
| Grik5 | 1931.46398 | -0.795793045 | 0.108420526 |
| Zfp574 | 425.7976918 | 0.160263324 | 0.084054171 |
| Pou2f2 | 170.0044398 | 0.257400408 | 0.159021074 |
| Mir6537 | 0 | NA | NA |
| D930028M14Rik | 1.834100151 | 0.333227174 | 0.74810035 |
| Dedd2 | 172.9180783 | 0.116923835 | 0.131421232 |
| Mir7048 | 0.228760906 | -0.968563628 | 2.10987967 |
| Zfp526 | 147.8424307 | -0.153373366 | 0.115398527 |
| Gsk3a | 1893.258904 | -0.069364668 | 0.047824141 |
| 9130221H12Rik | 75.48946259 | -0.245344294 | 0.1302972 |
| Erf | 724.7532469 | 0.411460756 | 0.126215342 |
| Cic | 3271.904299 | 0.027295433 | 0.058401077 |
| Pafah1b3 | 85.99027182 | 0.332028532 | 0.138154449 |
| Prr19 | 0.750844096 | 1.20883431 | 1.124144224 |
| Tmem145 | 351.2431402 | -1.044756574 | 0.147403679 |
| Megf8 | 2644.487032 | -0.795453525 | 0.083089955 |
| Cnfn | 0.044191549 | -0.243571806 | 3.724945049 |
| Lipe | 172.463792 | 1.005135357 | 0.133857187 |
| 4732471J01Rik | 9.165417789 | -0.868403752 | 0.407781116 |
| Cxcl17 | 0.234440768 | 1.505856806 | 2.117020646 |
| Gm15495 | 0.148650935 | -0.243572541 | 2.863172849 |
| Ceacam1 | 24.75964592 | 1.817728208 | 0.269295091 |
| Ceacam2 | 1.575759893 | 0.167408463 | 1.022342344 |
| Erich4 | 0.961427984 | -1.850270154 | 1.202844061 |
| Atp5sl | 334.5795391 | -0.260024289 | 0.067883944 |
| B3gnt8 | 13.86552076 | 2.939122246 | 0.438067827 |
| Bckdha | 305.8692003 | -0.111481536 | 0.123046285 |
| Exosc5 | 96.23120801 | 0.216287817 | 0.129483887 |
| Tmem91 | 111.1486534 | -1.020324052 | 0.148572151 |
| B9d2 | 96.55601536 | -0.007622992 | 0.14023225 |
| Tgfb1 | 180.5550002 | 2.593834373 | 0.193312946 |
| Ccdc97 | 599.3683796 | 0.403065143 | 0.071475748 |
| Hnrnpul1 | 1763.970679 | 0.32979796 | 0.063054091 |
| Axl | 1705.534697 | 2.154472025 | 0.131234121 |
| Cyp2s1 | 45.56318474 | -0.931432655 | 0.209631557 |
| Gm26707 | 1.31640046 | -2.831278301 | 1.036319207 |
| Gm4613 | 2.523943987 | 1.14459359 | 0.658990423 |
| Gm24865 | 0 | NA | NA |
| Gm23921 | 0 | NA | NA |
| Gm6434 | 0 | NA | NA |
| Cyp2b10 | 0.605952604 | 2.335254743 | 1.252497665 |
| Cyp2b13 | 0.046345709 | 0.71820487 | 3.724945049 |

| | | | |
|---|---|---|---|
| Rnf170-ps | 17.15945834 | -0.464375336 | 0.245077787 |
| Cyp2b9 | 0 | NA | NA |
| Gm8902 | 0.060174537 | 0.237309859 | 3.724945049 |
| Vmn1r-ps86 | 0 | NA | NA |
| Vmn1r184 | 0 | NA | NA |
| Cyp2a4 | 0.118974955 | -0.243576254 | 3.724945049 |
| Nlrp9c | 0.044544648 | 0.237309859 | 3.724945049 |
| Nlrp4a | 0 | NA | NA |
| Gm25947 | 0 | NA | NA |
| Nlrp9a | 0 | NA | NA |
| Vmn1r185 | 0 | NA | NA |
| Cyp2b23 | 0 | NA | NA |
| Cyp2b19 | 0.186596897 | -0.243571703 | 3.291160745 |
| Cyp2g1 | 8.028728541 | -0.657836262 | 1.623718735 |
| Cyp2a5 | 11.27708144 | -0.882978518 | 1.162342579 |
| Cyp2a22 | 0.038981395 | 0.718207094 | 3.724945049 |
| Cyp2a12 | 0.191971534 | 0.26463041 | 2.650118462 |
| Cyp2f2 | 13.0302823 | 0.811488936 | 0.436736052 |
| Gm15883 | 0.044544648 | 0.237309859 | 3.724945049 |
| Cyp2t4 | 0.092269992 | 0.237318756 | 3.724945049 |
| Egln2 | 888.7193016 | 0.261246267 | 0.076253483 |
| Rab4b | 72.36944043 | -0.048743322 | 0.167121294 |
| Gm21983 | 0 | NA | NA |
| Mir3101 | 0 | NA | NA |
| Mia | 1.133325952 | -1.401653661 | 0.978455523 |
| Snrpa | 438.9269378 | 0.302295245 | 0.083131935 |
| BC024978 | 169.2292599 | 0.055581618 | 0.096796516 |
| Mir1191 | 8.542722226 | -0.37932003 | 0.366656923 |
| Itpkc | 197.4348915 | 1.284398723 | 0.215736695 |
| Adck4 | 262.1854901 | 0.838490806 | 0.078768669 |
| Gm15567 | 1.049380316 | -0.632743914 | 0.948942564 |
| Numbl | 1053.281901 | -0.058723202 | 0.095508004 |
| Gm15570 | 0 | NA | NA |
| Ltbp4 | 207.6280549 | -1.060306988 | 0.150368137 |
| Shkbp1 | 92.29759634 | 0.369353879 | 0.173245901 |
| Gm20479 | 0 | NA | NA |
| Sptbn4 | 1935.821563 | -0.643421923 | 0.132920901 |
| Blvrb | 91.00347656 | 0.235840608 | 0.121710427 |
| Pgam1-ps2 | 38.75767613 | -0.030112804 | 0.17387565 |
| Sertad3 | 55.81009826 | 1.435693952 | 0.197491093 |
| Sertad1 | 143.2528654 | 1.60825237 | 0.23196255 |
| Prx | 13.03989321 | 1.727688034 | 0.342540048 |
| Gm15541 | 1.986167605 | 2.358163848 | 0.982490583 |
| Hipk4 | 1.250864467 | 0.775472242 | 0.830746706 |
| Pld3 | 3221.190747 | -0.655599247 | 0.074853179 |
| 2310022A10Rik | 301.4687265 | 0.765681361 | 0.10283809 |
| Akt2 | 915.7419638 | 0.34964139 | 0.075869716 |

| | | | |
|---|---|---|---|
| Ttc9b | 367.5397197 | -0.971186368 | 0.09465238 |
| Map3k10 | 1510.75745 | -0.465348248 | 0.078580288 |
| C030039L03Rik | 80.29137703 | -0.287091837 | 0.163151142 |
| Zfp60 | 578.4836708 | -0.021697969 | 0.174623721 |
| Zfp626 | 57.40632837 | 0.214477765 | 0.151206617 |
| Zfp59 | 63.21053847 | 0.325937937 | 0.16724262 |
| Zfp607 | 29.83338546 | 0.529645393 | 0.237448265 |
| 1700049G17Rik | 131.7655322 | 0.016548512 | 0.099673055 |
| Gm26891 | 2.658590328 | 0.718086202 | 0.825631258 |
| Gm10651 | 26.89522568 | -0.062610871 | 0.230502507 |
| Zfp780b | 147.3540878 | 0.040803632 | 0.138824763 |
| Zfp850 | 137.3735265 | 0.032924037 | 0.107631004 |
| Psmc4 | 769.5473873 | 0.663544363 | 0.086202998 |
| Fcgbp | 5.334611605 | -0.66035519 | 0.446305729 |
| Gm10935 | 0.257434735 | -0.282591297 | 2.117967881 |
| Gm24356 | 0 | NA | NA |
| 9530053A07Rik | 2.222860927 | 1.023666469 | 0.678351567 |
| Fbl | 235.6745308 | 0.787653546 | 0.156332857 |
| Dyrk1b | 419.6037486 | -0.786615278 | 0.104003333 |
| Eid2 | 336.2126705 | 0.002855023 | 0.094447552 |
| Eid2b | 286.8257227 | -0.17612368 | 0.099820154 |
| BC089491 | 5.091535971 | -1.529758193 | 0.500983343 |
| Dll3 | 12.77550445 | -0.09620884 | 0.320277498 |
| Timm50 | 228.1838604 | 0.585212853 | 0.102023592 |
| Supt5 | 1576.058628 | 0.304947861 | 0.057641421 |
| Rps16 | 917.1611222 | 0.762787837 | 0.097170612 |
| AF357399 | 3.515687709 | 0.922388995 | 0.563392226 |
| Plekhg2 | 230.709035 | 2.696052199 | 0.199517019 |
| Zfp36 | 478.3772212 | 4.379757436 | 0.332239575 |
| Med29 | 93.91206282 | 0.537412597 | 0.125534844 |
| Paf1 | 477.8677575 | 0.155855891 | 0.079145525 |
| Pak4 | 1507.799238 | 0.648629276 | 0.095512525 |
| Gmfg | 19.76928172 | 3.135601377 | 0.388810607 |
| Lrfn1 | 754.1318081 | -0.505416477 | 0.100700768 |
| 1700028B04Rik | 0.752450377 | 0.662045149 | 1.299311837 |
| Ifnl2 | 0.046345709 | 0.71820487 | 3.724945049 |
| Ifnl3 | 0.046345709 | 0.71820487 | 3.724945049 |
| Sycn | 0 | NA | NA |
| Nccrp1 | 4.109619456 | -0.381374242 | 0.484979041 |
| Mir7049 | 0.300003454 | -0.226860965 | 2.020595256 |
| Papl | 10.97390351 | -0.112554184 | 0.30299179 |
| Fbxo27 | 104.6688318 | -0.17906689 | 0.118721561 |
| Fbxo17 | 17.81551388 | 0.547848172 | 0.244751444 |
| Mrps12 | 129.792039 | 0.068320122 | 0.123648099 |
| Sars2 | 103.3753545 | 0.04118213 | 0.110650146 |
| Gm6537 | 10.517279 | -0.812725619 | 0.389254909 |
| Nfkbib | 264.1176441 | 0.900246379 | 0.159206046 |

| | | | |
|---|---|---|---|
| Sirt2 | 1431.100394 | 0.12972209 | 0.064642862 |
| Rinl | 62.25372522 | 2.340489092 | 0.251750321 |
| Hnrnpl | 2413.616906 | -0.060881871 | 0.059673133 |
| Ech1 | 280.1649813 | 0.054429846 | 0.100984753 |
| Lgals4 | 13.26484918 | -0.00786342 | 0.312005123 |
| Lgals7 | 1.097858384 | -0.595824097 | 1.384860243 |
| Capn12 | 3.372768181 | -1.510941986 | 0.639260957 |
| Actn4 | 1770.860706 | 0.835339329 | 0.089871332 |
| Eif3k | 526.8853122 | 0.42133868 | 0.113120747 |
| Map4k1 | 43.16772284 | 2.442511061 | 0.240037767 |
| Ryr1 | 194.7438043 | -1.635551829 | 0.224842914 |
| AC165142.1 | 0.214156338 | 0.702115857 | 2.173286021 |
| Gm26604 | 0.400939804 | -1.649725048 | 1.463263636 |
| Mir1963 | 0 | NA | NA |
| Rasgrp4 | 17.50432713 | 1.912458262 | 0.294039066 |
| Gm16282 | 0 | NA | NA |
| Fam98c | 181.7263762 | -0.611686225 | 0.104863462 |
| 1110035H17Rik | 2.173831266 | -0.646896337 | 0.685954769 |
| Spred3 | 426.8502981 | -0.0770453 | 0.111695274 |
| Ggn | 9.283367035 | 0.844968916 | 0.312669784 |
| Psmd8 | 837.7336857 | 0.595131905 | 0.104597889 |
| Catsperg1 | 14.9802426 | -0.687278796 | 0.335947068 |
| Gm17113 | 0 | NA | NA |
| Kcnk6 | 62.8823065 | 0.722874869 | 0.194235254 |
| Yif1b | 285.5672007 | 0.128802916 | 0.10658485 |
| 2200002D01Rik | 2.362804395 | 1.214192558 | 0.59329457 |
| Spint2 | 203.5653366 | 0.024063691 | 0.106764078 |
| Ppp1r14a | 171.4189243 | 0.054749661 | 0.144112589 |
| Dpf1 | 418.7210972 | -0.799887252 | 0.116149482 |
| Sipa1l3 | 1249.288852 | -0.237725282 | 0.075033012 |
| 4932431P20Rik | 0.682519493 | -1.206449498 | 1.308564289 |
| 4930432E11Rik | 0.998548739 | -3.302566957 | 1.123589286 |
| Gm29326 | 0.495715849 | -0.996127659 | 1.443183622 |
| A330087D11Rik | 0.362932359 | -2.459726483 | 1.580125934 |
| n-R5s153 | 0 | NA | NA |
| Catsperg2 | 7.493041037 | -0.917961416 | 0.403483801 |
| Gm23103 | 0 | NA | NA |
| Zfp84 | 275.1985705 | 0.050675987 | 0.08564331 |
| Mir1964 | 1.817730626 | -0.821783071 | 0.712505184 |
| Zfp30 | 158.5141845 | -0.475431744 | 0.101435086 |
| Zfp790 | 215.9787552 | -0.060909284 | 0.115685697 |
| Zfp940 | 138.395624 | -0.429025953 | 0.108221632 |
| Gm26554 | 0 | NA | NA |
| Gm17199 | 0 | NA | NA |
| Zfp420 | 147.2350297 | -0.084786046 | 0.114527382 |
| Zfp27 | 203.8758977 | -0.262733664 | 0.080384112 |
| Mir7668 | 0.042716945 | -0.243571806 | 3.724945049 |

| | | | |
|---|---|---|---|
| Zfp383 | 132.1721319 | -0.338254891 | 0.101957541 |
| Zfp74 | 213.3163599 | -0.274451249 | 0.092891504 |
| C230062I16Rik | 7.143933271 | 0.411308934 | 0.425528681 |
| Gm10169 | 7.433721481 | 0.599722899 | 0.409470374 |
| Zfp568 | 112.9697705 | 0.104404961 | 0.129526376 |
| Zfp14 | 53.58896386 | -0.157437314 | 0.170746257 |
| Zfp82 | 6.950672932 | -1.146884478 | 0.392093108 |
| Gm26920 | 0.228152142 | 0.718216712 | 2.561309204 |
| Zfp566 | 31.53288666 | -0.224158461 | 0.188028067 |
| Zfp260 | 833.8589085 | 0.161220782 | 0.068597427 |
| Gm23368 | 0 | NA | NA |
| Zfp382 | 241.7030479 | -0.056169725 | 0.093387908 |
| Zfp146 | 303.8588219 | 0.176887059 | 0.073369812 |
| Gm5113 | 98.06848606 | -0.739549153 | 0.141528893 |
| Cox7a1 | 8.620735387 | 0.296034338 | 0.345667571 |
| Capns1 | 1143.806175 | 0.736362579 | 0.113415207 |
| Gm26810 | 0.29029136 | 0.683529244 | 1.781240272 |
| Tbcb | 367.5355379 | 0.206088644 | 0.0859514 |
| Polr2i | 115.9016702 | 0.370844464 | 0.133953205 |
| Ovol3 | 0.088736197 | -0.243576254 | 3.724945049 |
| Wdr62 | 63.98026938 | 2.01513644 | 0.249704522 |
| Rps12-ps5 | 0.252860697 | -0.746384882 | 1.836154608 |
| Thap8 | 1.268210335 | 0.409313923 | 0.866876509 |
| Clip3 | 6878.216135 | -0.210387666 | 0.079651655 |
| Alkbh6 | 168.5814911 | -0.367563409 | 0.095801197 |
| Gm24757 | 0 | NA | NA |
| Syne4 | 17.72551575 | -0.107058438 | 0.254408496 |
| Sdhaf1 | 75.1581409 | 0.315591373 | 0.126808646 |
| E130208F15Rik | 4.251243736 | 0.620087049 | 0.544771492 |
| Lrfn3 | 312.5931955 | 0.001373277 | 0.089002157 |
| Gm25259 | 0.679915734 | 0.189407683 | 1.154722094 |
| Tyrobp | 264.1135694 | 2.848407672 | 0.19902191 |
| Hcst | 5.467031529 | 5.13632565 | 0.88549229 |
| Nfkbid | 33.79332328 | 0.936503173 | 0.256344226 |
| Aplp1 | 8728.071125 | -0.475613977 | 0.065989324 |
| Kirrel2 | 45.1116197 | -2.20558248 | 0.282104667 |
| Nphs1 | 8.109329026 | -1.77006444 | 0.445513034 |
| Nphs1os | 0.917433238 | -3.28612243 | 1.159717206 |
| Prodh2 | 0.146848595 | -1.036216968 | 3.718157501 |
| Arhgap33os | 1.185039339 | -1.814747412 | 1.023629775 |
| Arhgap33 | 1004.610349 | -1.449263558 | 0.159912234 |
| Proser3 | 27.82645344 | -0.118670212 | 0.255832968 |
| Hspb6 | 57.72339347 | 0.998329165 | 0.17412977 |
| mmu-mir-9769 | 0 | NA | NA |
| Lin37 | 148.1215379 | 0.153751216 | 0.125199773 |
| Psenen | 209.1857129 | 0.6062359 | 0.097205916 |
| U2af1l4 | 195.8717764 | 0.166935251 | 0.078131474 |

| | | | |
|---|---|---|---|
| Gm17104 | 0.139856492 | 0.237322262 | 3.574411025 |
| Igflr1 | 0 | NA | NA |
| Kmt2b | 1103.022709 | -0.122270672 | 0.137137681 |
| Mir3569 | 0.55981669 | -0.604524915 | 1.123398385 |
| Zbtb32 | 35.12199024 | 8.811902566 | 0.89827558 |
| Upk1a | 5.454212958 | -0.117900254 | 0.503811708 |
| Cox6b1 | 1385.504029 | -0.138343678 | 0.070384048 |
| Etv2 | 0.288531392 | 2.053111372 | 1.732500644 |
| Rbm42 | 1.341866388 | -0.155876301 | 0.858395808 |
| Gm21982 | 0.195772763 | -1.171546878 | 2.219885219 |
| Gm26610 | 2.644816022 | -0.895864185 | 0.633109035 |
| Haus5 | 8.540474638 | -0.177636348 | 0.367950301 |
| Pmis2 | 0 | NA | NA |
| 2200002J24Rik | 0.066050875 | 0.71820487 | 3.724945049 |
| Gm4883 | 0 | NA | NA |
| Atp4a | 2.461184311 | -1.159173224 | 0.689532212 |
| Tmem147 | 361.1132233 | 0.179174652 | 0.081887058 |
| Tmem147os | 88.86610623 | -0.815063782 | 0.186837011 |
| Gapdhs | 0.221390349 | -1.141846804 | 2.608989397 |
| Sbsn | 30.54518145 | -0.318507595 | 0.254065196 |
| Gm26935 | 15.53492519 | -1.099732741 | 0.354776363 |
| Dmkn | 16.49677372 | 1.393547719 | 0.497622738 |
| Krtdap | 1.379211929 | 1.180658261 | 3.708194212 |
| Ffar2 | 3.090575541 | 4.488178591 | 1.078258853 |
| Ffar3 | 0.43344934 | -2.759774044 | 1.534030007 |
| Ffar1 | 0.265062405 | -0.138792144 | 2.434989369 |
| Cd22 | 18.32792618 | 3.277900952 | 0.467061921 |
| Gm29439 | 0.095898756 | 1.192083721 | 3.724434897 |
| Gm29440 | 0.110242424 | 0.237318756 | 3.724945049 |
| Mag | 2336.670031 | 0.292974118 | 0.128910224 |
| Hamp2 | 0.159795471 | 0.733203719 | 2.80534854 |
| Hamp | 0.755877001 | -1.084963198 | 1.337415507 |
| Usf2 | 555.4235933 | -0.063618347 | 0.060081271 |
| Gm17077 | 7.799693549 | -1.030205268 | 0.490365726 |
| Gm4673 | 75.79538166 | -0.526533516 | 0.234832725 |
| Lsr | 56.83401911 | 2.354937255 | 0.23548262 |
| Fam187b | 1.876867459 | -0.2852868 | 0.790641202 |
| Fxyd5 | 213.0889357 | 2.622094453 | 0.222283496 |
| Mir7050 | 0 | NA | NA |
| Fxyd7 | 269.8762191 | -1.657192178 | 0.142362564 |
| Fxyd1 | 309.5145642 | 0.73491242 | 0.174293865 |
| Lgi4 | 451.055952 | 0.828883283 | 0.125450577 |
| Fxyd3 | 1.806277867 | 1.995119041 | 0.789238087 |
| Hpn | 35.43406477 | -0.455663984 | 0.280788789 |
| Scn1b | 2784.178309 | -0.412928228 | 0.076530112 |
| Gramd1a | 871.9470282 | -0.149241513 | 0.096719771 |
| G630030J09Rik | 3.656513056 | -0.256559507 | 0.572807936 |

| | | | |
|---|---|---|---|
| Gm10640 | 1.251626804 | 0.048262853 | 0.845134316 |
| Gm9569 | 0 | NA | NA |
| Gm37292 | 9.829030887 | -0.47396602 | 0.351041159 |
| Scgb1b1-ps | 0 | NA | NA |
| Scgb2b1 | 0 | NA | NA |
| Scgb1b2 | 0 | NA | NA |
| Scgb2b2 | 0 | NA | NA |
| Scgb2b3 | 0 | NA | NA |
| Scgb1b3 | 0 | NA | NA |
| Scgb2b4-ps | 0 | NA | NA |
| Scgb1b4-ps | 0 | NA | NA |
| Scgb1b28-ps | 0 | NA | NA |
| Gm29492 | 0 | NA | NA |
| Scgb2b5-ps | 0 | NA | NA |
| Scgb1b5-ps | 0 | NA | NA |
| Scgb2b28-ps | 0 | NA | NA |
| Rpl23a-ps8 | 0 | NA | NA |
| Scgb2b6 | 0 | NA | NA |
| Scgb1b6-ps | 0 | NA | NA |
| Gm4373 | 0 | NA | NA |
| Rpl23a-ps9 | 0 | NA | NA |
| Scgb2b7 | 0 | NA | NA |
| Scgb1b7 | 0 | NA | NA |
| Gm9114 | 0 | NA | NA |
| Gm29627 | 0 | NA | NA |
| Scgb2b8-ps | 0 | NA | NA |
| Scgb1b8-ps | 0 | NA | NA |
| Gm6669 | 0 | NA | NA |
| Gm9120 | 0 | NA | NA |
| Gm21673 | 0 | NA | NA |
| Scgb2b9-ps | 0 | NA | NA |
| Scgb1b9-ps | 0 | NA | NA |
| Gm5329 | 0 | NA | NA |
| Scgb2b29-ps | 0 | NA | NA |
| Gm9127 | 0 | NA | NA |
| Gm29262 | 0 | NA | NA |
| Gm29263 | 0 | NA | NA |
| Scgb2b10 | 0 | NA | NA |
| Scgb1b10 | 0 | NA | NA |
| Rpl23a-ps10 | 0 | NA | NA |
| Scgb2b30 | 0 | NA | NA |
| Gm9135 | 0 | NA | NA |
| Gm28256 | 0 | NA | NA |
| Gm28257 | 0 | NA | NA |
| Scgb2b11 | 0 | NA | NA |
| Scgb1b11 | 0 | NA | NA |
| Scgb2b12 | 0 | NA | NA |

| | | | |
|---|---|---|---|
| Scgb1b12 | 0 | NA | NA |
| Gm9142 | 0 | NA | NA |
| Gm28862 | 0 | NA | NA |
| Rpl23a-ps11 | 0 | NA | NA |
| Scgb1b29 | 0 | NA | NA |
| Gm9146 | 0 | NA | NA |
| Gm5116 | 0 | NA | NA |
| Scgb2b13-ps | 0 | NA | NA |
| Scgb1b13-ps | 0 | NA | NA |
| Gm5327 | 0 | NA | NA |
| Gm9149 | 0 | NA | NA |
| Gm21135 | 0 | NA | NA |
| Scgb2b14-ps | 0 | NA | NA |
| Scgb1b14-ps | 0 | NA | NA |
| Gm24766 | 0 | NA | NA |
| Gm5058 | 0 | NA | NA |
| Gm29429 | 0 | NA | NA |
| Scgb2b31-ps | 0 | NA | NA |
| Gm4398 | 0 | NA | NA |
| Gm29432 | 0 | NA | NA |
| Gm28408 | 0 | NA | NA |
| Scgb2b15 | 0 | NA | NA |
| Scgb1b15 | 0 | NA | NA |
| Scgb2b16-ps | 0 | NA | NA |
| Scgb1b16-ps | 0 | NA | NA |
| Gm25817 | 0 | NA | NA |
| Gm9156 | 0 | NA | NA |
| Gm28409 | 0 | NA | NA |
| Scgb2b32-ps | 0 | NA | NA |
| Gm4405 | 0 | NA | NA |
| Gm28383 | 0 | NA | NA |
| Gm28384 | 0 | NA | NA |
| Scgb2b17 | 0 | NA | NA |
| Scgb1b17 | 0 | NA | NA |
| Gm28992 | 0 | NA | NA |
| Rpl23a-ps7 | 0 | NA | NA |
| Scgb2b33 | 0 | NA | NA |
| Gm9162 | 0 | NA | NA |
| Gm28559 | 0 | NA | NA |
| Gm6003 | 0 | NA | NA |
| Scgb2b18 | 0 | NA | NA |
| Scgb1b18 | 0 | NA | NA |
| Scgb2b19 | 0 | NA | NA |
| Scgb1b19 | 0 | NA | NA |
| Gm9167 | 0 | NA | NA |
| Scgb2b20 | 0 | NA | NA |
| Scgb1b20 | 0 | NA | NA |

| | | | |
|---|---|---|---|
| Gm5732 | 0 | NA | NA |
| Gm28794 | 0 | NA | NA |
| Gm5731 | 0 | NA | NA |
| Scgb1b34-ps | 0 | NA | NA |
| Scgb2b21 | 0 | NA | NA |
| Scgb1b21 | 0 | NA | NA |
| Gm12770 | 0 | NA | NA |
| Gm12771 | 0 | NA | NA |
| Scgb2b22-ps | 0 | NA | NA |
| Scgb1b22-ps | 0 | NA | NA |
| Scgb2b23-ps | 0 | NA | NA |
| Scgb1b23-ps | 0 | NA | NA |
| Gm12763 | 0.048949532 | -0.243569582 | 3.724945049 |
| Rpl23a-ps12 | 0 | NA | NA |
| Scgb2b24 | 0 | NA | NA |
| Scgb1b24 | 0 | NA | NA |
| Gm12782 | 0 | NA | NA |
| Gm28614 | 0 | NA | NA |
| Gm26443 | 0 | NA | NA |
| Gm12768 | 0 | NA | NA |
| Scgb2b25-ps | 0 | NA | NA |
| Scgb1b25-ps | 0 | NA | NA |
| Gm12757 | 0 | NA | NA |
| Gm28615 | 0 | NA | NA |
| Gm12765 | 0 | NA | NA |
| Scgb2b26 | 0 | NA | NA |
| Scgb1b26-ps | 0 | NA | NA |
| Gm28184 | 0 | NA | NA |
| Gm12773 | 0 | NA | NA |
| Gm12755 | 0 | NA | NA |
| Scgb2b27 | 0.046345709 | 0.71820487 | 3.724945049 |
| Scgb1b27 | 0 | NA | NA |
| Gm12760 | 0 | NA | NA |
| Gm12779 | 0.044544648 | 0.237309859 | 3.724945049 |
| Scgb1b30 | 0 | NA | NA |
| Wtip | 46.54900976 | 0.928317938 | 0.170744566 |
| Uba2 | 890.9651073 | 0.112393286 | 0.059130111 |
| Gm12778 | 0.649479285 | 1.434773769 | 1.326891114 |
| Gm12783 | 0 | NA | NA |
| Pdcd2l | 117.7724749 | 0.532027987 | 0.139295647 |
| Gpi1 | 4331.119929 | -0.273502926 | 0.049453057 |
| Gm26762 | 2.694167648 | 0.275715804 | 0.678826292 |
| Gm12762 | 6.006325422 | -0.534826063 | 0.410450383 |
| Gm12758 | 0.954952387 | 0.142654802 | 0.902874582 |
| 4931406P16Rik | 686.0018774 | 0.233263859 | 0.059894208 |
| Gm6096 | 0.066050875 | 0.71820487 | 3.724945049 |
| Lsm14a | 923.1856796 | 0.450060672 | 0.057692149 |

| | | | |
|---|---|---|---|
| Gm12774 | 0 | NA | NA |
| Gm12764 | 10.54452331 | 3.016331669 | 0.56423917 |
| Gm12766 | 0.212513282 | 0.237311278 | 3.175662942 |
| Gm12780 | 0 | NA | NA |
| Gm12784 | 0.530195705 | -1.280521222 | 1.403449874 |
| Kctd15 | 124.850857 | 0.232927209 | 0.112266313 |
| Gm12781 | 0.50824631 | 0.292758572 | 1.238780062 |
| Gm12756 | 2.131963076 | -1.776954691 | 0.736176901 |
| Chst8 | 122.9302308 | -0.386585603 | 0.125862342 |
| Pepd | 269.5084426 | 0.302336064 | 0.094482104 |
| Gm25922 | 0.089089296 | 0.237305797 | 3.724945049 |
| Cebpg | 578.773901 | 0.77567992 | 0.093869253 |
| Gm25033 | 0 | NA | NA |
| Gm22535 | 0 | NA | NA |
| Cebpa | 259.903502 | 1.64472675 | 0.198644503 |
| Slc7a10 | 228.5227835 | -1.553898884 | 0.181478578 |
| Lrp3 | 882.6663157 | -0.593664943 | 0.077231711 |
| Gm24505 | 0.219162254 | 0.237328761 | 2.161553971 |
| Gpatch1 | 413.9756821 | 0.167033347 | 0.095669828 |
| Rhpn2 | 165.8571245 | 0.588609423 | 0.16198837 |
| C230052I12Rik | 49.69667937 | 1.077572054 | 0.192514164 |
| Cep89 | 116.2122036 | 0.14006311 | 0.136750557 |
| Gm18254 | 0 | NA | NA |
| Slc7a9 | 0.108767819 | 0.237318756 | 3.724945049 |
| Tdrd12 | 4.458748399 | 0.149351554 | 0.493686246 |
| Gm18558 | 0.451259361 | -1.10712139 | 1.424984508 |
| Nudt19 | 464.3454165 | 0.14863992 | 0.105832241 |
| B230322F03Rik | 17.22929159 | 0.33744993 | 0.302661443 |
| Rgs9bp | 10.39343307 | -0.68134863 | 0.41699331 |
| Ankrd27 | 324.283844 | -0.072941502 | 0.147261388 |
| Pdcd5 | 124.9127266 | 0.562152341 | 0.219995876 |
| Dpy19l3 | 892.6871314 | -0.052796208 | 0.097151825 |
| Zfp507 | 543.6392318 | 0.005389554 | 0.071271933 |
| Gm26790 | 0.186516865 | 1.199970255 | 2.670993307 |
| E130304I02Rik | 2.861485906 | -0.664656934 | 0.573077561 |
| Gm28514 | 1.65996821 | 0.246861676 | 0.712769541 |
| Rpl17-ps9 | 11.99013794 | 0.470829884 | 0.302027264 |
| Gm9211 | 0.12092566 | 0.237307635 | 3.724945049 |
| Gm23292 | 2.206361005 | -1.444750329 | 0.719197459 |
| Gm25247 | 0 | NA | NA |
| Gm29129 | 0 | NA | NA |
| Gm38285 | 5.376801637 | -0.293276488 | 0.491831247 |
| Tshz3 | 524.5764167 | -0.380926643 | 0.134925513 |
| Gm37452 | 12.71943701 | -1.249830394 | 0.418293518 |
| Gm38340 | 32.55178755 | -1.127804623 | 0.374359441 |
| E230020A03Rik | 24.88402293 | -0.953137182 | 0.327879332 |
| Gm37827 | 13.69016405 | -1.160171514 | 0.455343323 |

| | | | |
|---|---|---|---|
| Gm24550 | 0 | NA | NA |
| Gm28078 | 0 | NA | NA |
| Gm28076 | 0.144050924 | 0.237309606 | 3.724945049 |
| Gm28075 | 0 | NA | NA |
| Gm28077 | 0.313601826 | -0.243566835 | 1.792795329 |
| Zfp536 | 718.5400894 | -0.200174565 | 0.076923302 |
| 4930505M18Rik | 0 | NA | NA |
| Gm22302 | 0 | NA | NA |
| Uri1 | 620.2418666 | -0.125493632 | 0.075215989 |
| Ccne1 | 105.1069268 | -0.120232143 | 0.122931914 |
| Gm22203 | 0 | NA | NA |
| 1600014C10Rik | 402.5956352 | 1.235961924 | 0.133206942 |
| Plekhf1 | 106.4439667 | 0.534900635 | 0.155702006 |
| Pop4 | 57.53508151 | 0.263442231 | 0.192878481 |
| Gm28581 | 0 | NA | NA |
| Gm9220 | 0 | NA | NA |
| Gm4451 | 0 | NA | NA |
| Gm28453 | 0.075043205 | 0.237309859 | 3.724945049 |
| Gm22638 | 0 | NA | NA |
| Gm6818 | 0 | NA | NA |
| Gm28583 | 0 | NA | NA |
| Gm6605 | 0 | NA | NA |
| Gm6819 | 0 | NA | NA |
| Gm29087 | 0.10503227 | 1.179809211 | 3.723500796 |
| Gm5591 | 0 | NA | NA |
| Gm4454 | 0 | NA | NA |
| Gm20749 | 0 | NA | NA |
| Gm29086 | 0 | NA | NA |
| Gm25286 | 0 | NA | NA |
| Gm21115 | 0 | NA | NA |
| Gm21276 | 0 | NA | NA |
| Gm21122 | 0 | NA | NA |
| Gm29085 | 0 | NA | NA |
| 1700010N08Rik | 0.146060149 | 0.555964358 | 3.524366763 |
| Gm24797 | 0 | NA | NA |
| Gm21129 | 0 | NA | NA |
| Gm21136 | 0 | NA | NA |
| Gm21142 | 0 | NA | NA |
| Gm21152 | 0 | NA | NA |
| Gm25671 | 0 | NA | NA |
| Gm28451 | 0 | NA | NA |
| Gm1982 | 0 | NA | NA |
| Gm1988 | 0 | NA | NA |
| Gm6124 | 0 | NA | NA |
| Gm16387 | 0 | NA | NA |
| Gm9226 | 0 | NA | NA |
| Gm23307 | 0 | NA | NA |

| | | | |
|---|---|---|---|
| Gm29258 | 0 | NA | NA |
| Gm9271 | 0 | NA | NA |
| Gm5114 | 0 | NA | NA |
| Gm27437 | 0 | NA | NA |
| Gm5590 | 0.783781752 | -0.724657877 | 1.239001948 |
| Zfp939 | 78.05214499 | 0.179456641 | 0.157930156 |
| Gm28455 | 4.111696988 | 0.570706767 | 0.50337001 |
| Zfp619 | 83.85373399 | 0.008944986 | 0.141153666 |
| Gm33989 | 0.779068608 | -0.44030253 | 1.072841574 |
| Gm37494 | 103.4889581 | -0.609354456 | 0.150171212 |
| Gm4790 | 216.6323139 | -0.348097662 | 0.150065167 |
| 6230415J03Rik | 105.8274719 | -0.641664538 | 0.191017271 |
| Gm28582 | 30.69961378 | -0.393333883 | 0.222990472 |
| Gm17983 | 0 | NA | NA |
| Gm26526 | 0.0878635 | -0.72445792 | 3.724945049 |
| Gm2058 | 91.57356217 | 0.168157896 | 0.242991586 |
| Gm26327 | 0.206849446 | -0.243571977 | 2.595103581 |
| Gm2021 | 0.044544648 | 0.237309859 | 3.724945049 |
| Gm9230 | 0 | NA | NA |
| Gm37302 | 0.167459142 | 1.165958262 | 2.753748546 |
| 4930558N11Rik | 0 | NA | NA |
| 4930435C17Rik | 0 | NA | NA |
| Gm38199 | 0 | NA | NA |
| Gm6851 | 4.478289484 | 0.435938571 | 0.514242225 |
| 4933404I11Rik | 0 | NA | NA |
| Gm24910 | 0 | NA | NA |
| Gm28807 | 0.144050924 | 0.237309606 | 3.724945049 |
| 4921507H08Rik | 0 | NA | NA |
| 4933402C06Rik | 0.04393175 | -0.243571806 | 3.724945049 |
| A230077H06Rik | 28.4887614 | -1.263743756 | 0.29326198 |
| Gm28307 | 0.072025462 | 0.237312084 | 3.724945049 |
| Gm25784 | 0 | NA | NA |
| Vstm2b | 175.071028 | -0.287040861 | 0.135833543 |
| Gm29293 | 0.161114758 | 0.237309167 | 3.43255351 |
| 4930433I11Rik | 0.134148234 | 0.23732184 | 3.624632408 |
| Gm4884 | 0 | NA | NA |
| 1700110I07Rik | 0.044191549 | -0.243571806 | 3.724945049 |
| Gm28061 | 0 | NA | NA |
| Gm9239 | 0 | NA | NA |
| Gm6820 | 0 | NA | NA |
| Gm2128 | 0 | NA | NA |
| Gm28060 | 0 | NA | NA |
| Gm28667 | 0 | NA | NA |
| Gm5896 | 0.091880448 | -0.724455696 | 3.724945049 |
| Gm6866 | 0 | NA | NA |
| Gm9244 | 0 | NA | NA |
| Gm5592 | 0 | NA | NA |

| | | | |
|---|---|---|---|
| Gm28059 | 0 | NA | NA |
| Gm2108 | 0 | NA | NA |
| Gm17791 | 0.292366188 | -0.809344554 | 2.05189631 |
| Gm9246 | 0 | NA | NA |
| AI987944 | 114.8081952 | 0.519159531 | 0.147341801 |
| Gm17102 | 8.180380778 | -0.840090514 | 0.391087139 |
| Gm20449 | 4.437436714 | -1.035612177 | 0.558139113 |
| Vmn2r57 | 0.538693099 | -1.686177215 | 1.387497526 |
| AW146154 | 60.96357997 | -0.080105417 | 0.182066149 |
| Gm15470 | 0.332094496 | 0.23732819 | 2.018636342 |
| Gm6871 | 0 | NA | NA |
| Gm17217 | 0 | NA | NA |
| 2610021A01Rik | 362.0305132 | -0.013427274 | 0.071728107 |
| Zfp788 | 649.6253934 | -0.141144549 | 0.073370756 |
| Gm9252 | 0 | NA | NA |
| Vmn2r-ps54 | 6.472406225 | 1.934279404 | 0.490522815 |
| Gm6872 | 0.060174537 | 0.237309859 | 3.724945049 |
| Cd9-ps | 0.178447459 | 1.663739594 | 3.327415937 |
| Vmn2r-ps55 | 0.046345709 | 0.71820487 | 3.724945049 |
| Vmn2r-ps56 | 0.066050875 | 0.71820487 | 3.724945049 |
| Vmn2r58 | 0 | NA | NA |
| Vmn2r-ps57 | 0 | NA | NA |
| Vmn2r59 | 0 | NA | NA |
| Hmgb1-ps7 | 2.510100352 | 0.02656278 | 0.630754769 |
| Gm7224 | 23.28790071 | -0.116539211 | 0.219903282 |
| Gm5593 | 0.933648904 | 1.295381337 | 1.253654736 |
| Vmn2r60 | 0 | NA | NA |
| Gm2308 | 2.058388305 | 0.391955177 | 0.686196513 |
| Vmn2r61 | 0 | NA | NA |
| Vmn2r-ps58 | 0 | NA | NA |
| Gm5594 | 0 | NA | NA |
| Vmn2r-ps59 | 0 | NA | NA |
| Vmn2r-ps60 | 0 | NA | NA |
| 4933421I07Rik | 0 | NA | NA |
| Gm6887 | 0.072025462 | 0.237312084 | 3.724945049 |
| Zfp141 | 173.0700645 | 0.065829904 | 0.095221239 |
| Gm19246 | 0.428879182 | -0.55924351 | 1.357754566 |
| Gm21028 | 1.10003083 | 0.704553088 | 1.029584544 |
| Gm7221 | 16.48036244 | -0.368364057 | 0.253428166 |
| Gm24938 | 1.200098617 | 0.018860877 | 1.03986637 |
| 9830147E19Rik | 96.87745312 | -0.061408319 | 0.181496968 |
| Gm5595 | 96.44755503 | -0.233888697 | 0.125975453 |
| Gm17067 | 0 | NA | NA |
| Gm18128 | 0 | NA | NA |
| Gm6004 | 0 | NA | NA |
| Vmn2r62 | 0.088383099 | -0.72445792 | 3.724945049 |
| Gm2381 | 0.372704184 | -0.243563794 | 1.590117799 |

| | | | |
|---|---|---|---|
| Gm7211 | 0 | NA | NA |
| Vmn2r63 | 0 | NA | NA |
| Gm9266 | 0 | NA | NA |
| Gm9294 | 4.49822579 | 1.239337668 | 0.590307768 |
| Gm23256 | 0 | NA | NA |
| Gm9268 | 0 | NA | NA |
| RP23-367H8.3 | 0 | NA | NA |
| Gm10351 | 0 | NA | NA |
| Zfp936 | 0.092140247 | -0.724455696 | 3.724945049 |
| Gm22707 | 0 | NA | NA |
| EU599041 | 0.135598227 | -1.1790103 | 2.942835517 |
| Zfp715 | 456.2127529 | 0.275064509 | 0.080712192 |
| Siglecf | 5.415502606 | -0.035036086 | 0.595393021 |
| Gm10298 | 0 | NA | NA |
| Siglecg | 1.312064872 | 3.69501747 | 1.103181682 |
| Lim2 | 0.046345709 | 0.71820487 | 3.724945049 |
| Nkg7 | 96.73496261 | 10.19261176 | 0.945356209 |
| Cldnd2 | 1.387261314 | 3.714341473 | 1.192271642 |
| Etfb | 181.9463623 | 0.293799861 | 0.130237998 |
| Mir7051 | 0.307454254 | 0.718776366 | 1.756035224 |
| Vsig10l | 33.09847067 | 0.740797291 | 0.18590978 |
| Iglon5 | 396.8950678 | -0.328855025 | 0.100498652 |
| 4931406B18Rik | 0.154787072 | 0.237323942 | 2.832963972 |
| Cd33 | 51.25040222 | -0.365738932 | 0.334765959 |
| Zfp658 | 98.3557729 | -0.258314169 | 0.172852253 |
| Zfp719 | 304.4942731 | 0.32104675 | 0.157050701 |
| Zfp819 | 3.834671984 | 0.817496768 | 0.672623517 |
| Ceacam18 | 0.539507586 | 2.66943708 | 1.271702672 |
| Siglece | 5.430287587 | 0.266110932 | 0.557612179 |
| Ctu1 | 87.06700889 | 0.171873452 | 0.148984373 |
| Klk14 | 0 | NA | NA |
| 2310002F09Rik | 9.427952603 | -0.818460654 | 0.356083332 |
| Klk13 | 0.134945109 | 0.237324167 | 2.95629408 |
| Klk12 | 0 | NA | NA |
| Klk11 | 0.225356219 | -0.730495427 | 2.1322701 |
| Klk10 | 1.594085209 | 0.964640945 | 0.70814136 |
| Klk9 | 11.12317148 | 6.904590076 | 1.001949734 |
| Klk8 | 23.43242316 | 2.090480464 | 0.322572692 |
| Klk7 | 6.053054786 | 5.304211444 | 1.077484117 |
| Klk6 | 383.1545088 | 2.622031496 | 0.331267908 |
| Klk5 | 0.178695003 | 0.208646342 | 2.712882698 |
| Klk4 | 0.195922627 | 1.650451042 | 2.218601548 |
| Klk15 | 0 | NA | NA |
| Klk1b7-ps | 0.097139547 | 0.718207094 | 3.724945049 |
| Klk1b8 | 0.138087489 | 1.595384126 | 3.611428798 |
| Klk1b1 | 0.049553047 | 0.71820487 | 3.724945049 |
| Klk1b9 | 0.07796279 | 1.072698098 | 3.721829099 |

| | | | |
|---|---|---|---|
| Klk1b11 | 0.276660002 | 0.672650728 | 1.827318484 |
| Klk1b26 | 0.046345709 | 0.71820487 | 3.724945049 |
| Klk1b27 | 1.612931274 | 3.488550788 | 1.070505785 |
| Klk1b21 | 0 | NA | NA |
| Klk1b22 | 0 | NA | NA |
| Klk1b16 | 0.166231529 | 0.237308199 | 3.426626645 |
| Klk1b24 | 0.038981395 | 0.718207094 | 3.724945049 |
| Klk1b3 | 0 | NA | NA |
| Klk1b4 | 0 | NA | NA |
| Klk1b5 | 0 | NA | NA |
| Gm10109 | 2.250427361 | 0.377446676 | 0.687812237 |
| Klk1 | 11.42971091 | 6.180032929 | 0.912032561 |
| 1700028J19Rik | 6.192777873 | 0.553600436 | 0.448371411 |
| Gm25386 | 0 | NA | NA |
| 2410002F23Rik | 502.3878778 | -0.072290707 | 0.065626443 |
| Gm28496 | 4.360017817 | 0.386967925 | 0.655199911 |
| Gm26247 | 0 | NA | NA |
| Snord88c | 1.895772021 | -1.096688909 | 0.907800933 |
| Snord88a | 0 | NA | NA |
| Gm23991 | 0 | NA | NA |
| Acpt | 0 | NA | NA |
| Gm15517 | 0 | NA | NA |
| Clec11a | 34.26572508 | -0.765177662 | 0.191533643 |
| Shank1 | 9409.0605 | -1.398730458 | 0.132584831 |
| 1700008O03Rik | 12.39141503 | 0.289963962 | 0.315852695 |
| Syt3 | 466.7523524 | -0.944636147 | 0.101179111 |
| Lrrc4b | 2121.923621 | -0.80066571 | 0.080166415 |
| Aspdh | 27.56526144 | -0.962605324 | 0.244682353 |
| Josd2 | 428.3256145 | 0.001062098 | 0.106219772 |
| Mir7052 | 5.276354033 | -0.998639947 | 0.432264923 |
| Emc10 | 1263.340695 | 0.031100953 | 0.075797974 |
| Fam71e1 | 39.06557393 | -0.695728438 | 0.194959471 |
| Mybpc2 | 3.531971173 | 0.511862163 | 0.559271847 |
| Spib | 15.69420495 | 4.749720879 | 0.612035376 |
| Pold1 | 95.92363125 | 0.648893788 | 0.189904892 |
| Mir7053 | 0.189266655 | 0.237326601 | 2.687034879 |
| Nr1h2 | 489.1957701 | 0.644824863 | 0.10155545 |
| Napsa | 53.67902564 | 7.098253888 | 0.666154663 |
| Gm15396 | 3.09078215 | 0.123847644 | 0.631826087 |
| Kcnc3 | 3773.256925 | -0.529784775 | 0.117958729 |
| Myh14 | 523.8678887 | -0.467865555 | 0.158165228 |
| 2310016G11Rik | 0.744214146 | 0.434682258 | 1.030825987 |
| Izumo2 | 0 | NA | NA |
| Zfp473 | 10.07782813 | -0.222663927 | 0.308691945 |
| Vrk3 | 217.5211013 | 0.505608169 | 0.126792936 |
| Atf5 | 717.0131176 | 2.362825224 | 0.175579547 |
| Nup62 | 303.2451805 | 0.619796763 | 0.115456909 |

| | | | |
|---|---|---|---|
| Il4i1 | 16.51100415 | -0.026113275 | 0.286981316 |
| Tbc1d17 | 424.275241 | 0.057038649 | 0.081042883 |
| Akt1s1 | 507.5705319 | 0.590130704 | 0.065518395 |
| Mir707 | 0 | NA | NA |
| Pnkp | 197.282501 | 0.228585963 | 0.08687602 |
| Ptov1 | 973.5278823 | -0.244908407 | 0.055240909 |
| Med25 | 617.720589 | 0.265706252 | 0.073295436 |
| Fuz | 99.72731422 | -0.480954407 | 0.138951024 |
| Ap2a1 | 2601.373962 | -0.187343584 | 0.050722504 |
| Tsks | 3.95301002 | -1.431528792 | 0.612931443 |
| Cpt1c | 718.0796799 | -0.412887719 | 0.073617187 |
| Prmt1 | 509.9239553 | 0.361533426 | 0.095120483 |
| Gm15545 | 9.287909809 | -0.275181504 | 0.353251037 |
| Bcl2l12 | 25.31009217 | 0.635705523 | 0.28478899 |
| Irf3 | 149.8132699 | 0.365214665 | 0.168239261 |
| Scaf1 | 2253.505891 | -0.166888293 | 0.056533145 |
| Mir7054 | 0.567171362 | -2.099541788 | 1.23776187 |
| Rras | 111.4151899 | 1.856183153 | 0.189738755 |
| Prr12 | 1649.371309 | -0.38443486 | 0.065397777 |
| Prrg2 | 52.66422563 | 0.637730324 | 0.152654387 |
| Nosip | 442.5713574 | 0.54400053 | 0.089221151 |
| Rcn3 | 57.62462159 | 1.442931673 | 0.204382264 |
| Fcgrt | 111.4072777 | -0.058419023 | 0.190634013 |
| Mir150 | 0 | NA | NA |
| Rps11 | 1103.617122 | 0.564250711 | 0.098551288 |
| Snord35b | 0.134597393 | -1.178997129 | 2.949594298 |
| Rpl13a | 2064.540611 | 0.73736975 | 0.144533517 |
| Snord35a | 0.656197145 | -1.565103037 | 1.250302205 |
| Snord34 | 0 | NA | NA |
| Snord33 | 0 | NA | NA |
| mmu-mir-5121 | 0 | NA | NA |
| Snord32a | 0 | NA | NA |
| Flt3l | 50.76228596 | 1.865754364 | 0.230936819 |
| Aldh16a1 | 113.740617 | 0.828010731 | 0.129739656 |
| Pih1d1 | 223.2642417 | 0.264192723 | 0.089968564 |
| Slc17a7 | 6082.783442 | -0.800879925 | 0.121728921 |
| mmu-mir-7055 | 0.339810401 | -1.779788195 | 1.708695433 |
| Gfy | 10.55721659 | 0.18464573 | 0.641286118 |
| Pth2 | 1.157525169 | -0.032846747 | 0.791766094 |
| Ccdc155 | 18.99181208 | 0.008332489 | 0.271323498 |
| Dkkl1 | 52.36991393 | -1.213334463 | 0.215523427 |
| Tead2 | 38.54700613 | 0.424797066 | 0.228498333 |
| Cd37 | 67.79623972 | 2.359582193 | 0.237638504 |
| Slc6a16 | 3.556493767 | 0.104546195 | 0.509800478 |
| Gm23585 | 0 | NA | NA |
| 1700039E15Rik | 0.62192417 | -0.255555399 | 1.389436325 |
| Trpm4 | 109.1332828 | -0.166847601 | 0.174568021 |

| | | | |
|---|---|---|---|
| Rpl14-ps1 | 281.4107973 | 0.693950954 | 0.1337096 |
| Hrc | 0.548344614 | 1.115971687 | 1.226748029 |
| Ppfia3 | 1660.177301 | -0.624923594 | 0.104164232 |
| Mtag2 | 16.72535298 | -0.26455806 | 0.331575327 |
| Lin7b | 211.863161 | -1.660314866 | 0.192873877 |
| Snrnp70 | 1046.054046 | -0.354246754 | 0.178875655 |
| Gm22478 | 0.046345709 | 0.71820487 | 3.724945049 |
| Kcna7 | 0.228985044 | 0.7988754 | 2.565006469 |
| Ntf5 | 1.628131443 | 1.599141855 | 0.876932342 |
| Lhb | 9.980967548 | 1.145867479 | 0.570884414 |
| Ruvbl2 | 254.9844317 | 0.336281191 | 0.107700836 |
| Gys1 | 207.1643521 | 0.151916915 | 0.122151837 |
| Ftl1 | 3794.633164 | 2.310049108 | 0.193441589 |
| Gm22133 | 0 | NA | NA |
| Bax | 270.9969915 | 0.804837178 | 0.098256966 |
| Dhdh | 156.4792992 | -0.114681089 | 0.148361471 |
| Tulp2 | 0.658348016 | -0.457955425 | 1.284712707 |
| Nucb1 | 959.7799318 | 0.51342498 | 0.067974416 |
| Ppp1r15a | 417.9443 | 2.199640117 | 0.354341755 |
| Plekha4 | 287.7791967 | 3.98258556 | 0.318770045 |
| Gm16022 | 0.072025462 | 0.237312084 | 3.724945049 |
| Hsd17b14 | 2.84382331 | 0.010485562 | 0.644211919 |
| 0610005C13Rik | 1.694978974 | 0.019073487 | 0.696073204 |
| Bcat2 | 107.8855267 | -0.207658061 | 0.153806326 |
| Fgf21 | 1.959085883 | 5.010223983 | 0.975954525 |
| Fut1 | 0.956979021 | 0.707082935 | 1.063943063 |
| Izumo1 | 0.702218406 | 1.087596602 | 1.347917733 |
| Rasip1 | 78.58288542 | 1.64817929 | 0.221574527 |
| Gm16047 | 2.391421094 | 0.270014851 | 0.739985072 |
| Mamstr | 31.70904901 | -0.087124005 | 0.263284164 |
| Fut2 | 5.566764691 | 1.571820567 | 0.435565721 |
| Sec1 | 5.465091639 | -0.373384931 | 0.494080771 |
| Ntn5 | 29.63111439 | -1.258913461 | 0.330875646 |
| Car11 | 810.7850525 | -1.167960018 | 0.138937061 |
| Dbp | 869.3346952 | -1.815379105 | 0.153929485 |
| Sphk2 | 380.1522572 | 0.82727738 | 0.109746114 |
| Rpl18 | 1024.378528 | 0.95650593 | 0.125370097 |
| Fam83e | 2.171234967 | -0.586790911 | 0.638274533 |
| Spaca4 | 0 | NA | NA |
| Sult2b1 | 46.17871227 | 0.927650842 | 0.258036517 |
| Lmtk3 | 2315.447368 | -0.970257824 | 0.110216368 |
| Cyth2 | 643.2901348 | -0.073085645 | 0.07765225 |
| Kcnj14 | 17.48480922 | 0.7152119 | 0.398497951 |
| Grwd1 | 158.3457012 | 0.704920521 | 0.153944217 |
| Grin2d | 305.5974088 | -0.294167926 | 0.093385101 |
| Kdelr1 | 538.7074619 | 0.321949893 | 0.07349803 |
| Gm22098 | 0 | NA | NA |

| | | | |
|---|---|---|---|
| Syngr4 | 0.321511422 | 0.680226962 | 1.779309434 |
| Tmem143 | 157.483315 | -0.352285981 | 0.1167947 |
| Emp3 | 69.37531286 | 3.148146924 | 0.285804262 |
| Ccdc114 | 21.307866 | 0.420365912 | 0.238603148 |
| Abcc6 | 6.348614184 | 0.396640346 | 0.387534754 |
| Nomo1 | 1371.14784 | -0.43022706 | 0.065805953 |
| Kcnj11 | 116.36677 | -0.723690729 | 0.12884862 |
| Abcc8 | 310.3174424 | -1.224835385 | 0.123170974 |
| Ush1c | 7.85454541 | -0.198290724 | 0.381167682 |
| Otog | 2.40295838 | -1.461642706 | 0.74863774 |
| Gm38176 | 0 | NA | NA |
| Myod1 | 0.509027769 | -1.120552494 | 1.444247398 |
| Kcnc1 | 3456.691312 | -0.567578423 | 0.075595249 |
| Sergef | 366.9267911 | -0.152688608 | 0.104708352 |
| Gm22969 | 1.792854059 | -1.65125969 | 0.906720895 |
| Gm15700 | 0 | NA | NA |
| Tph1 | 2.962511132 | 0.481614531 | 0.920411847 |
| Saal1 | 161.7785274 | 0.176064705 | 0.093239646 |
| Saa3 | 441.1512112 | 11.66578272 | 1.116718283 |
| Saa4 | 0.385868569 | 2.270964391 | 1.62062496 |
| Saa1 | 4.201556626 | 2.582994479 | 0.80012419 |
| Saa2 | 1.450828853 | 3.189579762 | 1.126103193 |
| Hps5 | 293.9049292 | 0.341627791 | 0.115331078 |
| Gtf2h1 | 540.0561687 | 0.385184997 | 0.086358806 |
| Gm9392 | 24.65709606 | 0.543732932 | 0.227526048 |
| Ldha | 1608.193775 | 0.448972538 | 0.081139975 |
| Ldhc | 0.061920613 | 0.718207094 | 3.724945049 |
| Tsg101 | 606.9962445 | 0.175644672 | 0.073598518 |
| Uevld | 206.960226 | -0.527868376 | 0.115433443 |
| Gm9999 | 0.062426873 | 0.237312084 | 3.724945049 |
| Spty2d1 | 614.2646612 | 0.964772737 | 0.12005436 |
| Tmem86a | 239.3545487 | 1.520293101 | 0.139526837 |
| Ptpn5 | 1663.361268 | -0.252634763 | 0.132988354 |
| Mir7056 | 0.65404267 | -0.477092738 | 1.19696596 |
| Gm14377 | 1.857966922 | 3.390145935 | 1.029265935 |
| Mrgpra6 | 0 | NA | NA |
| Mrgpra9 | 0.097139547 | 0.718207094 | 3.724945049 |
| Mrgprx3-ps | 0 | NA | NA |
| Mrgpra1 | 0 | NA | NA |
| Gm29479 | 0 | NA | NA |
| Gm25326 | 0.128477747 | 0.718203484 | 3.677131942 |
| Mrgpra2a | 0.092691418 | 1.183079489 | 3.724560917 |
| Mrgpra2b | 0.175945213 | 1.166110064 | 2.710746636 |
| Gm7255 | 0 | NA | NA |
| Mrgprc4-ps | 0 | NA | NA |
| Mrgpra14 | 0 | NA | NA |
| Gm9322 | 0 | NA | NA |

| | | | |
|---|---|---|---|
| Mrgpra3 | 0 | NA | NA |
| Mrgprc1-ps | 0 | NA | NA |
| Gm2707 | 0 | NA | NA |
| Gm28288 | 0 | NA | NA |
| Mrgprc3-ps | 0 | NA | NA |
| Gm5733 | 0 | NA | NA |
| Gm22427 | 0 | NA | NA |
| Gm18348 | 0 | NA | NA |
| Gm29563 | 0 | NA | NA |
| Mrgprc7-ps | 0 | NA | NA |
| Gm18349 | 0 | NA | NA |
| Mrgpra18-ps | 0 | NA | NA |
| Gm18350 | 0 | NA | NA |
| Gm5734 | 0 | NA | NA |
| Mrgprc2-ps | 0 | NA | NA |
| Mrgpra4 | 0 | NA | NA |
| Mrgprx1 | 0 | NA | NA |
| Mrgprb7-ps | 0 | NA | NA |
| Gm28666 | 0 | NA | NA |
| Mrgprb12-ps | 0 | NA | NA |
| Gm18351 | 0 | NA | NA |
| Gm28665 | 0 | NA | NA |
| Mrgprb5 | 0 | NA | NA |
| Gm18352 | 0 | NA | NA |
| Mrgprb4 | 0 | NA | NA |
| Gm26950 | 0 | NA | NA |
| Gm18356 | 0 | NA | NA |
| Mrgprb13 | 0 | NA | NA |
| Gm18357 | 0 | NA | NA |
| Mrgprb8 | 0.066050875 | 0.71820487 | 3.724945049 |
| Mrgprb1 | 0.19039241 | -1.084179639 | 2.660990838 |
| Mrgprx2 | 0.10503227 | 1.179809211 | 3.723500796 |
| Mrgprb2 | 0.044544648 | 0.237309859 | 3.724945049 |
| Mrgprb9-ps | 0 | NA | NA |
| Gm18353 | 0 | NA | NA |
| Mrgprb3 | 0 | NA | NA |
| Mrgprb11 | 0.045882455 | 0.237312084 | 3.724945049 |
| Zdhhc13 | 253.9183394 | 0.077445981 | 0.092440434 |
| Csrp3 | 0.531680062 | 3.207113779 | 1.478342513 |
| E2f8 | 18.28425174 | 2.047325863 | 0.324090953 |
| Gm2788 | 0.049553047 | 0.71820487 | 3.724945049 |
| Nav2 | 2595.787216 | 0.352084466 | 0.102438341 |
| Gm37613 | 24.19551003 | -1.371933479 | 0.402497317 |
| Gm38059 | 24.62141553 | -0.752764615 | 0.437788946 |
| Dbx1 | 0.097139547 | 0.718207094 | 3.724945049 |
| Htatip2 | 67.03302061 | 0.964610124 | 0.199262961 |
| Prmt3 | 302.2229905 | 0.05819789 | 0.087914237 |

| | | | |
|---|---|---|---|
| Gm26856 | 0 | NA | NA |
| Slc6a5 | 485.8678254 | -0.189806338 | 0.561948156 |
| Nell1 | 633.6668325 | -0.372865299 | 0.102110236 |
| Nell1os | 0.048949532 | -0.243569582 | 3.724945049 |
| 4933405O20Rik | 9.680996758 | -1.323448598 | 0.417426048 |
| Ano5 | 52.81759069 | -0.270923726 | 0.194277725 |
| Slc17a6 | 1077.091468 | -0.30521193 | 0.1544356 |
| Gm7336 | 39.57013241 | -0.014379432 | 0.181425072 |
| Gm29296 | 0.142694128 | -0.243563589 | 2.905728216 |
| Fancf | 37.44894291 | 0.177309649 | 0.235845963 |
| Gas2 | 177.6414515 | -0.138953642 | 0.13027722 |
| Svip | 477.7009006 | -0.493003405 | 0.095569314 |
| 1700015G11Rik | 0.161582804 | 0.718215627 | 2.799906529 |
| Gm22211 | 0 | NA | NA |
| Gm6181 | 1.4136546 | 0.228844369 | 0.828937926 |
| Gm22444 | 0 | NA | NA |
| Mir6238 | 0 | NA | NA |
| Gm27921 | 0 | NA | NA |
| Gm25956 | 0 | NA | NA |
| Gm5776 | 44.67075326 | -0.095724666 | 0.190677973 |
| Luzp2 | 603.6111141 | -0.815438232 | 0.108539056 |
| Gm24116 | 0 | NA | NA |
| Gm26482 | 0 | NA | NA |
| Siglech | 79.78094538 | -0.781695228 | 0.304147423 |
| Tubgcp5 | 553.9336148 | 0.029750913 | 0.068097865 |
| Cyfip1 | 1268.126837 | 0.454441994 | 0.083897581 |
| Gm17907 | 0.149576918 | 1.19849824 | 2.863264735 |
| Nipa2 | 555.4063321 | 0.353059996 | 0.078827384 |
| A230056P14Rik | 25.86046502 | -0.043459478 | 0.223076101 |
| Nipa1 | 678.816936 | -0.935077809 | 0.107884627 |
| Gm15888 | 0 | NA | NA |
| Rps12l1 | 1.168735097 | 1.861932376 | 1.230817618 |
| Herc2 | 5466.518983 | -0.59836397 | 0.083790078 |
| Oca2 | 2.497417949 | 0.097345031 | 0.631323926 |
| Gabrg3 | 564.6589 | -0.43982115 | 0.098482744 |
| Gm9962 | 23.2644548 | -0.425033475 | 0.224606795 |
| Gabra5 | 375.6717185 | -0.899624006 | 0.109077329 |
| Gabrb3 | 2705.952044 | -0.629356672 | 0.07745138 |
| Mir6389 | 0 | NA | NA |
| Gm23911 | 0.328919656 | 0.520952223 | 1.732945401 |
| Gm26288 | 0 | NA | NA |
| Atp10a | 221.527901 | 2.371479216 | 0.202769349 |
| Gm6226 | 2.39103255 | 0.913031059 | 0.721380911 |
| Ube3a | 3913.118757 | 0.064481951 | 0.073998442 |
| Snhg14 | 6561.535641 | -0.892828949 | 0.125069869 |
| Gm25499 | 5.73832025 | -0.756116321 | 0.704060605 |
| Gm24966 | 65.59095034 | -1.140476882 | 0.241056079 |

| | | | |
|---|---|---|---|
| Gm22496 | 12.77173823 | -1.637410185 | 0.435250318 |
| Gm25648 | 15.8722383 | -1.036142486 | 0.477402855 |
| Gm24306 | 4.90255571 | -0.152329242 | 0.485693966 |
| Gm25934 | 2.147180949 | -0.679006676 | 0.73491029 |
| Gm25156 | 8.000524403 | -1.069561515 | 0.467097931 |
| Gm26488 | 4.065985139 | -1.289253311 | 0.599788467 |
| Gm24220 | 8.515480277 | -1.994641655 | 0.451616231 |
| Gm24653 | 7.944433476 | -1.338910027 | 0.413049264 |
| Gm24969 | 3.545771104 | -1.120141069 | 0.624622951 |
| Gm24417 | 0.567759171 | -2.162421777 | 1.308528657 |
| Gm25121 | 85.00767218 | -1.171283141 | 0.273929627 |
| Gm24657 | 0.475242527 | -2.52014461 | 1.657691792 |
| Gm26230 | 0.075043205 | 0.237309859 | 3.724945049 |
| Gm23922 | 0.156071933 | -0.724458118 | 3.489245271 |
| Gm25314 | 2.261769448 | -0.65820498 | 0.817888551 |
| Gm26096 | 0.262111692 | 0.718216628 | 2.46860757 |
| Gm22996 | 6.119234212 | -1.044574744 | 0.595894315 |
| Gm26499 | 0.650021907 | -1.099629615 | 1.392981545 |
| Gm25646 | 2.157091062 | -0.366879152 | 0.707613377 |
| Gm26498 | 21.92470555 | -0.874223441 | 0.346813613 |
| Gm23660 | 2.095785588 | -1.273394494 | 0.735526999 |
| Gm23933 | 2.60823589 | -1.324772364 | 0.674857146 |
| Gm24862 | 0.637223532 | -1.10258587 | 1.212059174 |
| Gm24866 | 6.740698188 | -1.434114982 | 0.464864028 |
| Gm22348 | 0 | NA | NA |
| Gm26466 | 4.89789051 | -0.627162277 | 0.591469233 |
| Gm24799 | 3.843037622 | -1.63319645 | 0.667203112 |
| Gm24088 | 1.480005993 | -2.439953979 | 1.020164097 |
| Gm25463 | 3.198438298 | -1.208894812 | 0.747495047 |
| Gm25229 | 2.607201442 | -2.084603783 | 0.729976292 |
| Gm24528 | 0.50461959 | -0.099103449 | 1.265311008 |
| Gm23356 | 0.770457139 | 0.08479661 | 1.345344097 |
| Gm25449 | 0.090074004 | -0.24357403 | 3.724945049 |
| Gm24872 | 0.860548126 | -2.461482311 | 1.062646536 |
| Gm26337 | 0.156716517 | -0.243563452 | 2.821984814 |
| Gm23145 | 0.371627904 | 0.240635127 | 1.585780192 |
| Gm22130 | 0.047948698 | -0.243569582 | 3.724945049 |
| Gm25988 | 0 | NA | NA |
| Gm25221 | 0 | NA | NA |
| Gm26389 | 0.044191549 | -0.243571806 | 3.724945049 |
| Gm23471 | 2.996745481 | -0.44178112 | 0.612384477 |
| Gm25120 | 0.150248541 | -0.243571377 | 2.859451661 |
| Gm24418 | 0 | NA | NA |
| Gm23687 | 0 | NA | NA |
| Gm22393 | 0 | NA | NA |
| Gm24639 | 0.168184286 | -0.724459869 | 2.759948137 |
| Gm24566 | 0 | NA | NA |

| | | | |
|---|---|---|---|
| Gm22111 | 0 | NA | NA |
| Gm23076 | 0.042716945 | -0.243571806 | 3.724945049 |
| Gm25874 | 0 | NA | NA |
| Gm25088 | 0.11657011 | 0.237307635 | 3.724945049 |
| Gm22510 | 5.686408051 | -0.835966753 | 0.541924632 |
| Gm24983 | 1.441787318 | -0.631866776 | 0.876512808 |
| Gm24027 | 3.476611664 | -0.611841392 | 0.618791297 |
| Gm22524 | 0.172163037 | 0.719092542 | 2.73973509 |
| Gm22630 | 0.258404708 | 0.71822288 | 2.078647503 |
| Gm25741 | 0.657611471 | -0.342780181 | 1.121114796 |
| Gm26059 | 0.559779376 | -1.895760348 | 1.36196231 |
| Gm25175 | 0.444383067 | -1.068613009 | 1.571309206 |
| Gm22511 | 1.321027671 | 0.445538989 | 0.927094261 |
| Gm25077 | 0.739785724 | 1.763865889 | 1.329239767 |
| Gm22834 | 0.106971521 | 0.237307635 | 3.724945049 |
| Gm24926 | 1.408313197 | 0.184945121 | 0.944075401 |
| Gm26200 | 0.042716945 | -0.243571806 | 3.724945049 |
| Gm22632 | 0.060174537 | 0.237309859 | 3.724945049 |
| Gm23254 | 4.913973528 | -0.627247408 | 0.509000906 |
| Gm26390 | 0.52711509 | -0.255400271 | 1.427688824 |
| Gm26366 | 1.250668422 | -1.582197026 | 1.093570931 |
| Gm26336 | 0.636553247 | -1.747901265 | 1.223513591 |
| Gm26283 | 0.137685729 | -0.724459914 | 2.938031474 |
| Gm25098 | 0.174523186 | -0.690816038 | 3.368236204 |
| Gm24702 | 0.239175706 | 0.237312413 | 3.01480951 |
| Gm26432 | 0.437372945 | -0.730059089 | 1.502248943 |
| Gm25462 | 6.852308181 | -0.747563153 | 0.531914985 |
| Gm24495 | 12.03897181 | -1.425638076 | 0.379429924 |
| Gm25523 | 2.19067646 | -1.448223474 | 0.873852871 |
| Gm22255 | 1.065694062 | -0.202704044 | 1.058632405 |
| Gm23449 | 8.027250868 | -0.999122346 | 0.420725956 |
| Gm23310 | 0.295362062 | -2.209125501 | 1.951579377 |
| Gm25461 | 0.060174537 | 0.237309859 | 3.724945049 |
| Gm24310 | 0.210314706 | 0.237328558 | 2.187621999 |
| Gm22909 | 0.331212083 | -1.188284642 | 1.981708256 |
| Gm25823 | 1.425569217 | -1.715943088 | 0.878998135 |
| Gm25647 | 0.32719657 | 0.332794537 | 1.760762732 |
| Gm22252 | 2.966653922 | -0.882313266 | 0.673455112 |
| Gm26392 | 0.252618675 | -0.53461931 | 2.505827185 |
| Gm24219 | 2.893504144 | -1.128637222 | 0.621916832 |
| Gm25230 | 1.071417476 | 1.232472597 | 1.109844915 |
| Gm22627 | 0.53540973 | -1.675583812 | 1.271907377 |
| Gm24585 | 0.104637558 | 0.23732098 | 3.724945049 |
| Gm23305 | 0.469128522 | 0.143033387 | 1.579917078 |
| Gm24206 | 0.186417525 | 0.237327935 | 2.265609519 |
| Gm25085 | 2.147377591 | -1.215916684 | 0.764743627 |
| Gm25209 | 0.361503534 | -1.126066943 | 1.557038143 |

| | | | |
|---|---|---|---|
| Gm23682 | 0.137685729 | -0.724459914 | 2.938031474 |
| Gm24654 | 0.322713358 | -1.07411572 | 1.733024434 |
| Gm26334 | 0.362177786 | -1.169718189 | 1.56294878 |
| Gm25742 | 0.531759696 | -0.723184906 | 1.435167749 |
| Gm25585 | 0 | NA | NA |
| Gm25840 | 0 | NA | NA |
| Gm25089 | 0.347269729 | -0.724466952 | 3.29115655 |
| Gm24040 | 0.042716945 | -0.243571806 | 3.724945049 |
| Gm22050 | 0.315224781 | -0.980766354 | 1.549575357 |
| Gm23286 | 0 | NA | NA |
| Gm24021 | 0 | NA | NA |
| Gm22274 | 0 | NA | NA |
| Gm24652 | 0.045882455 | 0.237312084 | 3.724945049 |
| Gm23618 | 0 | NA | NA |
| Gm23688 | 0 | NA | NA |
| Gm24570 | 0 | NA | NA |
| Gm26467 | 0 | NA | NA |
| Gm25194 | 0.147068667 | 0.237310878 | 3.514416608 |
| Gm22640 | 0 | NA | NA |
| Gm24952 | 0 | NA | NA |
| Gm25087 | 0 | NA | NA |
| Gm23944 | 0 | NA | NA |
| Gm22629 | 0.292758571 | 0.237328231 | 2.052738961 |
| Gm26374 | 0.198152624 | 1.757323469 | 3.707396009 |
| Gm26284 | 0 | NA | NA |
| Gm25452 | 0 | NA | NA |
| Gm25984 | 0 | NA | NA |
| Gm25146 | 0.176667124 | -0.243563549 | 2.304379988 |
| Gm24759 | 1.076799075 | -0.314643428 | 0.993152644 |
| Gm24100 | 1.19629966 | -1.982993131 | 1.0055696 |
| Gm22253 | 0.244208214 | 0.718220219 | 2.455587708 |
| Gm25017 | 0.060174537 | 0.237309859 | 3.724945049 |
| Gm23575 | 0.210511418 | -0.243563321 | 2.190351522 |
| Gm22391 | 0.323669746 | -1.921081557 | 1.503900664 |
| Gm22449 | 1.252922778 | -0.150072344 | 0.94070774 |
| Gm24137 | 0.11657011 | 0.237307635 | 3.724945049 |
| Gm22912 | 0.442601433 | -0.67812266 | 1.538770173 |
| Gm25710 | 0.427556563 | -1.321009621 | 1.50726979 |
| Gm26434 | 7.062431358 | -1.008383199 | 0.424073032 |
| Gm23560 | 8.541564187 | -0.38964013 | 0.421758745 |
| Gm25870 | 11.09216655 | -0.755957466 | 0.348004926 |
| Gm23862 | 2.015428526 | -1.790052875 | 0.773156072 |
| Gm26504 | 0.522929342 | -0.25831832 | 1.388414532 |
| Gm22285 | 0 | NA | NA |
| Gm22131 | 0.271448477 | -0.3676098 | 2.416077242 |
| Gm24153 | 0.725964322 | -0.141326564 | 1.109913764 |
| Gm26365 | 0 | NA | NA |

| | | | |
|---|---|---|---|
| Gm24609 | 0.334054859 | -1.163360302 | 3.708526686 |
| Gm24264 | 0.41844861 | -0.384116439 | 3.289031767 |
| Gm25155 | 0.246032045 | -0.724461202 | 2.099389713 |
| Gm25816 | 0.401944266 | -1.057259897 | 1.509215742 |
| Gm23141 | 0.141693294 | -0.243563599 | 2.912076028 |
| Gm23767 | 0.37340313 | -1.574925659 | 1.737470897 |
| Gm22417 | 0.221310845 | -1.296684975 | 3.162314925 |
| Gm22584 | 1.21151113 | -0.81482807 | 0.927743198 |
| Gm26270 | 0.134148234 | 0.23732184 | 3.624632408 |
| Gm23089 | 0 | NA | NA |
| Gm24711 | 0.104637558 | 0.23732098 | 3.724945049 |
| Gm26188 | 0 | NA | NA |
| Gm23265 | 0.938116572 | -0.215586115 | 1.246692019 |
| Gm24518 | 0.272816007 | -1.167309158 | 2.074360538 |
| Gm25944 | 0.061920613 | 0.718207094 | 3.724945049 |
| Gm23047 | 0.652779004 | -0.77897041 | 1.228392001 |
| Gm23953 | 0 | NA | NA |
| Gm26332 | 0 | NA | NA |
| Gm26032 | 0 | NA | NA |
| Gm22863 | 0 | NA | NA |
| Gm24618 | 0 | NA | NA |
| Gm22046 | 0 | NA | NA |
| Gm24760 | 0.092140247 | -0.724455696 | 3.724945049 |
| Gm25350 | 0.472357773 | -1.08436895 | 1.447031224 |
| Gm26502 | 0 | NA | NA |
| Gm23313 | 0 | NA | NA |
| Gm25471 | 0.129625439 | -1.065581744 | 3.675612039 |
| Gm22776 | 0 | NA | NA |
| Gm23446 | 0.198993444 | -0.727577853 | 2.222930108 |
| Gm25210 | 0 | NA | NA |
| Gm22128 | 0 | NA | NA |
| Gm22851 | 0.048949532 | -0.243569582 | 3.724945049 |
| Gm25074 | 0.045882455 | 0.237312084 | 3.724945049 |
| Gm25157 | 0.086648695 | -0.72445792 | 3.724945049 |
| Gm22631 | 0.573189294 | -1.371081234 | 1.191169961 |
| Gm25474 | 0.37594025 | -1.129517035 | 1.431687331 |
| Gm22941 | 0.150983267 | 0.719110569 | 2.855208279 |
| Gm26246 | 0.123992736 | -0.243574001 | 3.720643035 |
| Gm22047 | 0 | NA | NA |
| Gm25597 | 0.783837108 | -2.417910861 | 1.136743282 |
| Gm26097 | 0.21090123 | -1.17338211 | 2.172089159 |
| Gm22258 | 0.307376478 | -1.097821221 | 1.74928669 |
| Gm26201 | 0 | NA | NA |
| Gm23619 | 0 | NA | NA |
| Gm22812 | 0.249740077 | -1.520008321 | 2.513367833 |
| Gm26433 | 0.126225412 | 0.718200913 | 3.698754989 |
| Gm23357 | 0.67569888 | -0.246093227 | 1.327069209 |

| | | | |
|---|---|---|---|
| Gm25615 | 0.379570916 | 0.652195939 | 1.569274929 |
| Gm22110 | 0.872838561 | -0.742340778 | 1.083471576 |
| Gm23724 | 0 | NA | NA |
| Gm22188 | 0.093141081 | -0.724455696 | 3.724945049 |
| Gm23549 | 0 | NA | NA |
| Snord116l2 | 0.075043205 | 0.237309859 | 3.724945049 |
| Gm26223 | 0 | NA | NA |
| Gm23524 | 0.112288909 | -0.724455696 | 3.724945049 |
| Gm26136 | 0 | NA | NA |
| Gm24658 | 0.115000406 | 0.23732098 | 3.724945049 |
| Gm23359 | 0.04393175 | -0.243571806 | 3.724945049 |
| Gm24742 | 0.049553047 | 0.71820487 | 3.724945049 |
| Snord116l1 | 0.083526043 | 0.718202646 | 3.724945049 |
| Gm22173 | 0.200209858 | 0.726467642 | 2.223473249 |
| Gm23696 | 0.675101473 | -1.265729877 | 1.407931385 |
| Gm26094 | 1.506060395 | -0.068482324 | 0.848077947 |
| Gm22289 | 0.897799824 | -1.093261405 | 1.051240967 |
| Gm22373 | 29.41355203 | -1.292993018 | 0.302832486 |
| Gm38393 | 1891.788183 | -0.89323314 | 0.150775008 |
| Snord64 | 0 | NA | NA |
| Snord107 | 0 | NA | NA |
| Snrpn | 3523.734492 | -0.472379183 | 0.054891359 |
| Snurf | 0 | NA | NA |
| Gm7367 | 30.85968858 | 0.958032661 | 0.247479551 |
| A230006K03Rik | 161.3268707 | -0.818685358 | 0.173452532 |
| Ube2nl | 15.82980231 | -0.269065051 | 0.413200627 |
| Gm25533 | 0.514202509 | 1.760516247 | 1.563297525 |
| Gm23962 | 0.605864455 | 0.581331572 | 1.215899309 |
| Mir344d-1 | 0.868248373 | 0.167745326 | 1.019372016 |
| Mir344d-2 | 0.131862677 | -0.243562287 | 2.97735462 |
| Gm23490 | 9.098233414 | -0.632041239 | 0.363482964 |
| Gm22466 | 0.385656404 | 0.215635879 | 1.483828365 |
| Mir344d-3 | 1.914600676 | -2.65263724 | 1.01406071 |
| Gm23917 | 0.903269115 | -0.699688844 | 1.142898135 |
| Mir344e | 0.296439348 | -1.449491353 | 1.538312073 |
| Mir344h-1 | 0.876086486 | -0.370348991 | 1.183577895 |
| Mir344h-2 | 2.928954034 | -0.533020361 | 0.662790832 |
| Gm23873 | 4.451125399 | -0.440708294 | 0.513314875 |
| Mir344b | 5.840177946 | -2.119261284 | 0.554851595 |
| Mir344c | 9.878672328 | -1.200311123 | 0.337373651 |
| Mir344 | 15.06924853 | -0.865918319 | 0.345470566 |
| Mir344-2 | 22.73106971 | -1.022287109 | 0.270731541 |
| Mir344g | 49.18631136 | -0.367369571 | 0.17706611 |
| Gm23431 | 7.186751093 | -0.98457016 | 0.445282428 |
| Mir344f | 6.212148508 | -0.667551326 | 0.418066096 |
| Gm24923 | 13.62120263 | -0.900587898 | 0.345801328 |
| Mir344i | 13.24250092 | -1.360979954 | 0.425160229 |

| | | | |
|---|---|---|---|
| Gm9801 | 35.25565351 | -0.9716318 | 0.300173368 |
| Ndn | 2072.502296 | -0.048808999 | 0.104838826 |
| Magel2 | 65.47304857 | -0.356823249 | 0.185603646 |
| Mkrn3 | 19.82500287 | -0.292874421 | 0.246752433 |
| Peg12 | 5.656883354 | 0.610614762 | 0.457249717 |
| Atp5l-ps1 | 1.396333172 | -0.438974891 | 0.813999153 |
| Gm22494 | 0 | NA | NA |
| Chrna7 | 175.1769349 | -0.162660202 | 0.08651768 |
| 4930554H23Rik | 0.967147681 | 1.364189057 | 0.98152077 |
| Otud7a | 560.9803126 | -0.299699096 | 0.12742996 |
| Gm5898 | 84.2128684 | 0.21857644 | 0.130999131 |
| Gm22228 | 3.911342548 | -0.851051657 | 0.505808786 |
| Gm3307 | 7.681223622 | -1.746799699 | 0.485742366 |
| Klf13 | 2862.082524 | -0.083182364 | 0.074893083 |
| Gm27252 | 12.51525409 | 1.686392165 | 0.357328099 |
| E030018B13Rik | 0.724732643 | 3.418571691 | 1.269592832 |
| Gm20670 | 0.072025462 | 0.237312084 | 3.724945049 |
| Trpm1 | 3.26130212 | 1.616825661 | 0.601181252 |
| Mir211 | 0 | NA | NA |
| Mir6416 | 0 | NA | NA |
| Mtmr10 | 533.9198899 | 0.381496015 | 0.092221756 |
| Gm20457 | 18.61572913 | -0.236990713 | 0.257212923 |
| Fan1 | 158.3085833 | -0.019024934 | 0.11153706 |
| Mphosph10 | 195.4413562 | 0.39722392 | 0.10495059 |
| Mcee | 82.01444615 | -0.331424409 | 0.149700873 |
| Apba2 | 1821.774854 | -0.553083809 | 0.083649352 |
| Fam189a1 | 495.1526885 | -0.84830483 | 0.112651943 |
| Ndnl2 | 270.3594914 | -0.236185791 | 0.12294826 |
| Gm25249 | 1.017734822 | -2.590552342 | 1.178930092 |
| Tjp1 | 2187.437935 | 0.493861014 | 0.050133542 |
| Gm7546 | 0 | NA | NA |
| BC046251 | 8.000213675 | -0.611716039 | 0.397951776 |
| Tarsl2 | 377.8388073 | -0.379802568 | 0.075654971 |
| Tm2d3 | 199.3078344 | -0.005209144 | 0.090847636 |
| Gm7551 | 1.00142042 | 0.827449423 | 0.991795199 |
| 1810008I18Rik | 0.117907917 | 0.237310006 | 3.724945049 |
| Gm20684 | 1.680189541 | -1.442841931 | 0.956920158 |
| Pcsk6 | 300.8982086 | -0.43882316 | 0.135554417 |
| Snrpa1 | 232.9850694 | 0.640363466 | 0.134336844 |
| Vimp | 381.3181123 | 0.834252713 | 0.102576363 |
| Gm26827 | 0.677567072 | 1.61882295 | 1.12881122 |
| Chsy1 | 358.5432631 | 0.806608916 | 0.094334102 |
| Gm23042 | 0 | NA | NA |
| Lrrk1 | 209.5656891 | 1.863223529 | 0.146168649 |
| Gm10974 | 0 | NA | NA |
| Gm23957 | 0.352640022 | 1.656667146 | 1.724759357 |
| Mir7057 | 0.172107867 | 0.718215321 | 2.739902689 |

| | | | |
|---|---|---|---|
| Aldh1a3 | 20.34501332 | 0.99745795 | 0.328249873 |
| Asb7 | 386.1973764 | 0.023206196 | 0.077795162 |
| Lins | 133.4462556 | 0.34025919 | 0.095634528 |
| Cers3 | 5.123955725 | 0.781238712 | 0.530643642 |
| Adamts17 | 97.65468606 | -1.261574161 | 0.201521888 |
| 1700112J16Rik | 8.55332328 | -0.979423385 | 0.401096762 |
| Lysmd4 | 213.3922501 | -0.336137739 | 0.090966287 |
| Mef2a | 2662.223246 | -0.059142867 | 0.061642243 |
| Lrrc28 | 238.448027 | -0.318887844 | 0.098549128 |
| Gm23233 | 1.460122852 | -0.351311686 | 0.765428263 |
| Ttc23 | 41.9836744 | 0.428108122 | 0.197224464 |
| Synm | 824.232063 | -0.062215519 | 0.126597085 |
| Gm23844 | 0 | NA | NA |
| Gm7583 | 0.042716945 | -0.243571806 | 3.724945049 |
| 4833412C05Rik | 2.535954356 | -1.425502807 | 0.676786468 |
| Igf1r | 783.90039 | -0.219999434 | 0.078523875 |
| Pgpep1l | 5.087731363 | 0.325923785 | 0.453499639 |
| Gm16157 | 0.108309328 | 0.237310006 | 3.724945049 |
| Fam169b | 17.53390412 | 2.859632323 | 0.374189616 |
| Gm16158 | 0 | NA | NA |
| Gm5334 | 0 | NA | NA |
| Arrdc4 | 222.7753849 | 2.116462654 | 0.239390587 |
| Gm23795 | 0 | NA | NA |
| Gm5342 | 0 | NA | NA |
| Gm23815 | 0.0885994 | -0.24357403 | 3.724945049 |
| Gm7627 | 0 | NA | NA |
| Gm24120 | 0 | NA | NA |
| Gm29327 | 0 | NA | NA |
| Nr2f2 | 456.5175412 | 0.059921554 | 0.113321301 |
| B130024G19Rik | 28.43045277 | -0.736147282 | 0.291898636 |
| Gm28258 | 0.072025462 | 0.237312084 | 3.724945049 |
| Gm7656 | 3.484008338 | -1.512504753 | 0.628178379 |
| Gm24880 | 0.045882455 | 0.237312084 | 3.724945049 |
| 6030442E23Rik | 0.042716945 | -0.243571806 | 3.724945049 |
| Gm10172 | 0 | NA | NA |
| Gm10295 | 0.372348714 | -0.800498752 | 3.289136785 |
| Gm29295 | 0.243433062 | -0.236589482 | 1.85055791 |
| Gm29328 | 0.046345709 | 0.71820487 | 3.724945049 |
| Mctp2 | 119.8191935 | 1.547818358 | 0.157506135 |
| Gm37620 | 0 | NA | NA |
| Gm28745 | 0 | NA | NA |
| Rpl17-ps10 | 146.566573 | 0.533679817 | 0.113079261 |
| Gm28744 | 0 | NA | NA |
| Kansl2-ps | 68.64857961 | 0.631217528 | 0.144153475 |
| Gm7693 | 0 | NA | NA |
| Gm5335 | 0.044544648 | 0.237309859 | 3.724945049 |
| Gm28746 | 0.114742407 | -0.24357403 | 3.724945049 |

| | | | |
|---|---|---|---|
| A730056A06Rik | 25.12540099 | -1.490769842 | 0.26568677 |
| Rgma | 807.0540894 | -0.33723319 | 0.111968012 |
| Chd2 | 1727.684492 | -0.074653484 | 0.105964105 |
| 1810026B05Rik | 562.1011433 | -0.119570058 | 0.109781489 |
| Gm26176 | 0 | NA | NA |
| Gm23020 | 0.068097359 | -0.243569582 | 3.724945049 |
| A830073O21Rik | 30.67817576 | -0.979522836 | 0.300268693 |
| Fam174b | 232.8015944 | -0.224373806 | 0.124156783 |
| Gm10619 | 0 | NA | NA |
| mmu-mir-7234 | 0 | NA | NA |
| St8sia2 | 68.7914267 | 0.46062514 | 0.210889332 |
| Gm38143 | 0.044191549 | -0.243571806 | 3.724945049 |
| Gm27459 | 0 | NA | NA |
| Slco3a1 | 783.7089768 | 0.440688696 | 0.082694286 |
| Gm7580 | 0 | NA | NA |
| Gm22326 | 0 | NA | NA |
| Sv2b | 4509.045788 | -0.486663958 | 0.114773809 |
| Gm24858 | 0.075043205 | 0.237309859 | 3.724945049 |
| 1500012K07Rik | 0.790435591 | -1.508523151 | 1.168000053 |
| Gm10161 | 0 | NA | NA |
| Akap13 | 1197.678966 | 1.065178024 | 0.082197547 |
| Gm25282 | 0.642535093 | 0.704599855 | 1.295876528 |
| Gm23251 | 0.141684195 | 0.718205511 | 3.558854591 |
| AU020206 | 84.50291907 | 3.657293638 | 0.249046751 |
| Gm26729 | 0 | NA | NA |
| Klhl25 | 139.2928838 | 0.764636214 | 0.151530973 |
| Gm5597 | 1.352145875 | 0.452385657 | 0.890079207 |
| Agbl1 | 4.71615247 | -0.921637424 | 0.510123015 |
| Gm23239 | 0.182574094 | 0.237323313 | 2.706657689 |
| AC134536.1 | 0 | NA | NA |
| Ntrk3 | 1530.084009 | -0.387403516 | 0.112608174 |
| Gm9885 | 28.00701669 | 0.345743066 | 0.244587255 |
| Mrpl46 | 163.3248249 | 0.329230998 | 0.122387355 |
| Mrps11 | 173.3279118 | 0.231128529 | 0.095374758 |
| Det1 | 52.30670483 | 0.33727305 | 0.177059753 |
| Mir7-2 | 0.175313191 | -0.722740566 | 2.770426814 |
| Aen | 320.6797765 | 1.151267157 | 0.123521836 |
| Isg20 | 337.9177596 | 8.459693773 | 0.548993342 |
| Gm26633 | 15.61936475 | -0.601968359 | 0.257397715 |
| Acan | 162.5228664 | 1.165391173 | 0.222936926 |
| Hapln3 | 8.674975699 | -0.615510732 | 0.349347464 |
| Mfge8 | 724.7290691 | -1.648887279 | 0.1962495 |
| Abhd2 | 3069.461115 | 0.509568048 | 0.072769916 |
| Gm24224 | 0 | NA | NA |
| Rlbp1 | 85.04677146 | -0.625883379 | 0.164436824 |
| Fanci | 15.02295619 | 1.376963951 | 0.334922489 |
| Gm16017 | 0 | NA | NA |

| | | | |
|---|---|---|---|
| Polg | 722.9823603 | -0.216653792 | 0.096288698 |
| AI854517 | 340.1156097 | -0.311979572 | 0.131606411 |
| Mir9-3 | 6.978027697 | -1.80370702 | 0.494173706 |
| A330074H02Rik | 52.62118314 | -0.79304712 | 0.30839945 |
| 2900037B21Rik | 15.38072168 | -1.441669349 | 0.384342043 |
| Gm37608 | 67.71388935 | -1.048250425 | 0.327390295 |
| Rhcg | 17.15633722 | 2.194423512 | 0.393947613 |
| Ticrr | 11.11653513 | 2.034408701 | 0.434211736 |
| Kif7 | 33.31114625 | 0.132478806 | 0.176843616 |
| 9330171B17Rik | 15.94460044 | -0.722089171 | 0.30779129 |
| Plin1 | 0.616538614 | 1.053992845 | 1.299567906 |
| Pex11a | 89.1075553 | 0.308905805 | 0.139950083 |
| Wdr93 | 5.435520937 | -0.197148659 | 0.437119098 |
| Mesp1 | 0.291110193 | -0.268019941 | 1.627414325 |
| Mesp2 | 2.119758866 | -0.281008718 | 0.792504945 |
| Anpep | 165.5971786 | 2.541217235 | 0.214701763 |
| Ap3s2 | 947.0018652 | -0.23332558 | 0.058617934 |
| Gm26646 | 0.14985154 | 0.661163679 | 2.846050679 |
| 5430400D12Rik | 13.24663712 | -0.686105523 | 0.330479859 |
| 2610034B18Rik | 234.4523726 | 0.247017893 | 0.112157141 |
| Gm24541 | 0 | NA | NA |
| Zfp710 | 414.4888839 | 0.727057579 | 0.093037569 |
| Idh2 | 278.3167953 | -0.522021142 | 0.137642952 |
| Sema4b | 443.5342128 | 0.799275982 | 0.112840351 |
| Cib1 | 109.0301844 | 0.978167761 | 0.153628854 |
| Gdpgp1 | 63.61861299 | -0.311943305 | 0.167931413 |
| Gm15504 | 0 | NA | NA |
| Ttll13 | 6.876810528 | -0.60391781 | 0.514006893 |
| Ngrn | 276.3196493 | 0.308423195 | 0.098697563 |
| Vps33b | 168.9997345 | -0.137342141 | 0.110448772 |
| Prc1 | 39.35090745 | 2.417197374 | 0.436091441 |
| 6330403N20Rik | 1.079531651 | 0.474978913 | 0.987424001 |
| Rccd1 | 59.77890577 | -0.37760712 | 0.157673204 |
| Unc45a | 445.147613 | 0.347409844 | 0.111930918 |
| Hddc3 | 174.8477751 | -0.215694702 | 0.149026547 |
| Man2a2 | 2375.928341 | -0.664443033 | 0.072576473 |
| Fes | 119.6055901 | 3.067247785 | 0.194264043 |
| Furin | 1024.487266 | 1.398868924 | 0.137195244 |
| Gm15544 | 0.049553047 | 0.71820487 | 3.724945049 |
| Blm | 98.86775975 | -0.01840255 | 0.133831446 |
| Gm17257 | 13.53931913 | 0.850561419 | 0.283462727 |
| Crtc3 | 380.3991371 | 0.109507762 | 0.111286089 |
| Gm15880 | 1.230449842 | 2.220314956 | 1.032261812 |
| Iqgap1 | 1218.575044 | 2.436816142 | 0.180424152 |
| Zscan2 | 88.28572028 | -0.298901002 | 0.134475223 |
| Wdr73 | 209.5163078 | -0.198623581 | 0.098031636 |
| Nmb | 36.76065315 | 0.034017573 | 0.315877866 |

| | | | |
|---|---|---|---|
| Sec11a | 259.8445855 | 0.24807511 | 0.094946685 |
| Gm16171 | 0 | NA | NA |
| Zfp592 | 595.8256381 | 0.090778778 | 0.072639122 |
| Alpk3 | 21.69556852 | -0.036362186 | 0.500475241 |
| Platr32 | 1.295334173 | -1.042563085 | 1.078327515 |
| Slc28a1 | 1.668031982 | 0.658693781 | 0.751720925 |
| Gm6415 | 6.092582732 | -0.111699566 | 0.370219994 |
| Pde8a | 237.4167335 | -0.477623652 | 0.121785723 |
| Gm7180 | 6.599784284 | 0.179631792 | 0.448704659 |
| Rps17 | 912.2411316 | 0.498459717 | 0.114835715 |
| Cpeb1 | 643.0184324 | -0.538903133 | 0.117946855 |
| Rpl7a-ps9 | 0 | NA | NA |
| Ap3b2 | 1542.537639 | -0.268719738 | 0.056060746 |
| BC048679 | 0.097899064 | -0.724453471 | 3.724945049 |
| 2900076A07Rik | 53.12144243 | -0.496739347 | 0.181948933 |
| Mir1839 | 0 | NA | NA |
| mmu-mir-1839 | 0 | NA | NA |
| Gm37829 | 0.980535863 | -1.260208565 | 1.127343908 |
| Fsd2 | 3.637652488 | -1.451142668 | 0.544369315 |
| Whamm | 307.5779694 | 0.898345376 | 0.116300436 |
| Homer2 | 622.3176929 | -1.547449116 | 0.161559048 |
| Gm26149 | 0 | NA | NA |
| Fam103a1 | 318.4598396 | -0.152179908 | 0.083707337 |
| 3110040N11Rik | 45.08078921 | -0.079086657 | 0.172324258 |
| Btbd1 | 1030.704126 | -0.31631423 | 0.10183368 |
| Gm10160 | 0.442186748 | -0.031525936 | 1.51072891 |
| Tm6sf1 | 282.4071646 | 0.303754168 | 0.098542032 |
| Hdgfrp3 | 1850.998309 | -0.276981823 | 0.065065947 |
| Gm25907 | 0 | NA | NA |
| Bnc1 | 0.212279271 | 1.124586116 | 2.583711293 |
| 4833418N17Rik | 0 | NA | NA |
| Sh3gl3 | 330.4796501 | -0.432277611 | 0.080127142 |
| Adamtsl3 | 118.3270371 | 1.094217982 | 0.13445289 |
| Saxo2 | 36.73756128 | -0.268899896 | 0.224562391 |
| Eftud1 | 227.2937603 | 0.262273146 | 0.080573924 |
| 4933406J10Rik | 0.25679625 | 0.718222116 | 2.093734043 |
| n-R5s154 | 0 | NA | NA |
| 1700010L04Rik | 0.351906323 | 1.000071045 | 1.551421626 |
| Mex3b | 220.6262723 | -0.598770017 | 0.132056598 |
| Gm22934 | 0 | NA | NA |
| A530021J07Rik | 0.162691105 | -0.724458629 | 2.788998794 |
| Gm10610 | 1.084114685 | 2.041209796 | 1.219873211 |
| Tmc3 | 14.78000229 | 1.493400846 | 0.334064798 |
| Gm16638 | 33.35336936 | -0.73004776 | 0.26186435 |
| Stard5 | 261.421678 | 1.41710028 | 0.137026458 |
| Il16 | 509.4594508 | -0.365016211 | 0.193097893 |
| 2310044K18Rik | 1.104418013 | -1.023414709 | 1.257065052 |

| | | | |
|---|---|---|---|
| Gm7964 | 331.670112 | 0.409267083 | 0.093412224 |
| 1700026D08Rik | 9.589069244 | 0.371570672 | 0.386681179 |
| Mesdc1 | 387.3285222 | 0.053935979 | 0.097708282 |
| Mesdc2 | 527.7692794 | 0.777623274 | 0.098152954 |
| Cemip | 295.0602538 | -0.098736936 | 0.081103356 |
| Abhd17c | 249.4441419 | 0.018021592 | 0.082259143 |
| Gm26708 | 1.414184224 | -0.801308799 | 1.031027824 |
| Gm22177 | 0 | NA | NA |
| Arnt2 | 3090.055291 | -0.607020271 | 0.066347917 |
| Gm25790 | 1.94490296 | -0.547228028 | 0.826767524 |
| Gm2115 | 123.9262083 | -0.312216498 | 0.150144866 |
| Fah | 224.6003258 | -0.153977765 | 0.139727939 |
| Zfand6 | 399.9024983 | 0.199783324 | 0.118377174 |
| 2610206C17Rik | 0.927099338 | 0.998365037 | 1.028281887 |
| Gm18958 | 0.18866986 | 0.23732874 | 2.26103487 |
| Vmn2r-ps62 | 0 | NA | NA |
| Gm6155 | 2.358756262 | 1.225551275 | 0.676377293 |
| Olfr291 | 0.046345709 | 0.71820487 | 3.724945049 |
| Gm8018 | 0.038981395 | 0.718207094 | 3.724945049 |
| Olfr290 | 0.092881282 | -0.724455696 | 3.724945049 |
| Gm18560 | 0 | NA | NA |
| Vmn2r65 | 0 | NA | NA |
| Vmn2r66 | 0 | NA | NA |
| Vmn2r-ps64 | 0 | NA | NA |
| Vmn2r-ps65 | 0 | NA | NA |
| Vmn2r-ps66 | 0 | NA | NA |
| Vmn2r-ps67 | 0 | NA | NA |
| Gm18358 | 0 | NA | NA |
| Vmn2r67 | 0 | NA | NA |
| Vmn2r-ps68 | 0 | NA | NA |
| Vmn2r68 | 0 | NA | NA |
| Gm20414 | 0 | NA | NA |
| Gm20415 | 0 | NA | NA |
| Vmn2r-ps69 | 0 | NA | NA |
| Vmn2r69 | 0 | NA | NA |
| Gm18359 | 0 | NA | NA |
| Vmn2r-ps70 | 0 | NA | NA |
| Gm18360 | 0 | NA | NA |
| Vmn2r70 | 0 | NA | NA |
| Vmn2r-ps71 | 0 | NA | NA |
| Vmn2r-ps72 | 0 | NA | NA |
| Vmn2r71 | 0 | NA | NA |
| Vmn2r-ps73 | 0 | NA | NA |
| Vmn2r-ps74 | 0 | NA | NA |
| Vmn2r72 | 0 | NA | NA |
| Gm18779 | 0 | NA | NA |
| Gm18354 | 0 | NA | NA |

| | | | |
|---|---|---|---|
| Vmn2r-ps75 | 0 | NA | NA |
| Vmn2r-ps76 | 0 | NA | NA |
| Vmn2r-ps77 | 0 | NA | NA |
| Vmn2r73 | 0.08486385 | 0.717551239 | 3.724898127 |
| Vmn2r-ps78 | 0 | NA | NA |
| Vmn2r74 | 0 | NA | NA |
| Vmn2r-ps79 | 0 | NA | NA |
| Vmn2r-ps80 | 0 | NA | NA |
| Vmn2r-ps81 | 0 | NA | NA |
| Vmn2r-ps82 | 0 | NA | NA |
| Vmn2r-ps83 | 0 | NA | NA |
| Vmn2r-ps84 | 0 | NA | NA |
| Vmn2r75 | 0 | NA | NA |
| Vmn2r-ps85 | 0 | NA | NA |
| Vmn2r-ps86 | 0 | NA | NA |
| Vmn2r76 | 0 | NA | NA |
| Olfr310 | 0 | NA | NA |
| Gm20480 | 0 | NA | NA |
| Gm23192 | 0 | NA | NA |
| Olfr309 | 0.297212607 | 1.277416442 | 1.773462536 |
| Olfr308 | 0 | NA | NA |
| Olfr307 | 0 | NA | NA |
| Olfr306-ps1 | 0 | NA | NA |
| Olfr305 | 0 | NA | NA |
| Olfr304 | 0 | NA | NA |
| Olfr303 | 0 | NA | NA |
| Olfr301 | 0 | NA | NA |
| Olfr300-ps1 | 0 | NA | NA |
| Olfr299 | 0 | NA | NA |
| Olfr298 | 0.066050875 | 0.71820487 | 3.724945049 |
| Olfr297 | 0 | NA | NA |
| Gm17919 | 0 | NA | NA |
| Olfr296-ps1 | 0 | NA | NA |
| Olfr295 | 0 | NA | NA |
| Olfr294 | 0 | NA | NA |
| Olfr293 | 0 | NA | NA |
| Olfr292 | 0 | NA | NA |
| Folh1 | 228.0677052 | 0.377788645 | 0.177339727 |
| Vmn2r77 | 0.046345709 | 0.71820487 | 3.724945049 |
| Vmn2r-ps87 | 0.08486385 | 0.717551239 | 3.724898127 |
| Gm19203 | 0 | NA | NA |
| Vmn2r78 | 0.360847685 | -1.581162243 | 1.992523351 |
| Vmn2r79 | 0.061920613 | 0.718207094 | 3.724945049 |
| Nox4 | 23.51386303 | 2.438253116 | 0.319741363 |
| Tyr | 0.679694067 | -0.924620851 | 1.245817343 |
| Grm5 | 3152.369327 | -0.892735152 | 0.105490253 |
| Gm24249 | 0.097139547 | 0.718207094 | 3.724945049 |

| | | | |
|---|---|---|---|
| Ctsc | 748.4761849 | 3.849512536 | 0.181783492 |
| Gm26522 | 0.075043205 | 0.237309859 | 3.724945049 |
| Gm25345 | 0.066050875 | 0.71820487 | 3.724945049 |
| Rab38 | 4.431782484 | -0.734966408 | 0.587970977 |
| Gm15661 | 0 | NA | NA |
| Rps13-ps2 | 13.83618781 | 0.483219532 | 0.359684688 |
| Gm8183 | 0.155358952 | -0.724459886 | 2.829692658 |
| Tmem135 | 828.7159243 | -0.134220099 | 0.057657562 |
| Gm24659 | 0.092269992 | 0.237318756 | 3.724945049 |
| Gm22171 | 0.104637558 | 0.23732098 | 3.724945049 |
| Gm28748 | 6.9950682 | -0.600137606 | 0.42144372 |
| Fzd4 | 116.5445208 | -0.006059097 | 0.185999966 |
| Prss23 | 214.4898237 | 1.358094988 | 0.189932239 |
| Prss23os | 0.31866137 | 0.662973275 | 1.978005995 |
| Gm2546 | 6.356979297 | -0.49224269 | 0.418864304 |
| A230065N10Rik | 4.255075643 | 0.123046979 | 0.501832413 |
| Me3 | 830.2853921 | -0.341193174 | 0.122905964 |
| Gm15744 | 0.161532967 | 1.200059132 | 3.449701177 |
| Ccdc81 | 14.73968261 | -0.139791764 | 0.249026465 |
| l7Rn6 | 302.6286143 | -0.113141801 | 0.114858965 |
| Eed | 245.5434659 | 0.445752509 | 0.088753411 |
| Gm26529 | 7.028223412 | -0.452817807 | 0.370061632 |
| Gm5341 | 0.068097359 | -0.243569582 | 3.724945049 |
| 2310010J17Rik | 13.38140942 | 0.203597116 | 0.279862689 |
| Picalm | 2942.135727 | 0.566938837 | 0.064217829 |
| Ccdc83 | 1.068251668 | 1.93499629 | 1.103721013 |
| n-R5s155 | 0 | NA | NA |
| Sytl2 | 471.4003508 | -0.823476771 | 0.099547523 |
| Ccdc89 | 10.74032172 | 0.102227886 | 0.314068531 |
| Crebzf | 1021.356941 | 0.063800543 | 0.055702328 |
| Tmem126a | 179.1913439 | 0.609511858 | 0.143542726 |
| Tmem126b | 260.3332693 | 0.092637904 | 0.126279627 |
| Dlg2 | 4221.663311 | -0.539404524 | 0.10171605 |
| Gm27470 | 0.068097359 | -0.243569582 | 3.724945049 |
| Gm27660 | 0 | NA | NA |
| Gm24552 | 6.758948741 | -0.998318218 | 0.391738343 |
| Gm22080 | 0 | NA | NA |
| Gm23928 | 0 | NA | NA |
| Ccdc90b | 140.9402912 | 0.347933394 | 0.148466977 |
| Ankrd42 | 228.9912909 | -0.309783065 | 0.124979496 |
| Gm37008 | 21.96798092 | -0.0935523 | 0.249480886 |
| Gm26944 | 56.56169118 | -0.015680752 | 0.26951062 |
| 6430511E19Rik | 167.4764364 | -0.105270691 | 0.227456806 |
| Pcf11 | 1375.530546 | -0.13213747 | 0.079448704 |
| Gm26981 | 0 | NA | NA |
| 4632427E13Rik | 515.8795252 | -0.245290974 | 0.179293687 |
| Rab30 | 322.4076359 | -0.311891496 | 0.075976787 |

| | | | |
|---|---|---|---|
| Ddias | 20.7592005 | 1.522581072 | 0.347801624 |
| Prcp | 124.058315 | 0.057877372 | 0.11815383 |
| Gm26862 | 2.576837399 | 0.580134204 | 0.637318976 |
| Gm9934 | 11.67004998 | 0.076803735 | 0.305092378 |
| Fam181b | 200.5250975 | -0.276603706 | 0.168054245 |
| Gm25860 | 0 | NA | NA |
| Gm15501 | 19.75181214 | 0.41779458 | 0.261725927 |
| Gm5899 | 1.074124899 | -0.095332974 | 0.997601915 |
| Gm23884 | 0.681297893 | -0.545592705 | 1.235060218 |
| Gm25684 | 0 | NA | NA |
| Gm9966 | 3.042324184 | -2.125725166 | 0.670024692 |
| Tenm4 | 3151.27742 | -0.290158478 | 0.125859788 |
| Mir708 | 5.493531071 | -2.231898694 | 0.553526378 |
| Mir6394 | 3.129376264 | -1.554689765 | 0.627276221 |
| Gm15412 | 0 | NA | NA |
| Rps11-ps5 | 0 | NA | NA |
| Gm15414 | 0.304874617 | 2.042114001 | 3.703499636 |
| Gm15413 | 2.589705772 | 4.986064326 | 1.054727715 |
| Gm15416 | 0.568378176 | 1.665203615 | 1.227178134 |
| Gm25712 | 0 | NA | NA |
| Gm15415 | 0.44530367 | 1.678109675 | 1.500926528 |
| Gm22226 | 0 | NA | NA |
| C230038L03Rik | 4.597572509 | 0.042802352 | 0.586654732 |
| Nars2 | 182.0977735 | -0.061523026 | 0.091080705 |
| Gab2 | 640.0412359 | 1.103928767 | 0.123212169 |
| Usp35 | 246.8318203 | -0.323660923 | 0.08811278 |
| Kctd21 | 137.2874877 | -0.070909433 | 0.223853672 |
| Alg8 | 69.82562007 | 0.471329844 | 0.152876034 |
| Ndufc2 | 551.0014102 | 0.287956155 | 0.139347975 |
| Gm16053 | 0.824460844 | -1.849416657 | 1.055589495 |
| Thrsp | 209.116264 | 0.663924349 | 0.137749469 |
| Kctd14 | 10.62226215 | -1.078410211 | 0.328579557 |
| Ints4 | 375.7085532 | 0.257620671 | 0.096574308 |
| Gm24412 | 1.785420374 | -2.110755816 | 0.964884514 |
| Aamdc | 120.3980682 | -0.212963408 | 0.10218692 |
| Rsf1 | 1617.979498 | -0.127398455 | 0.058026 |
| Rsf1os1 | 0 | NA | NA |
| Rsf1os2 | 1.842452978 | -0.467867252 | 0.695198146 |
| Clns1a | 571.5874786 | 0.022200909 | 0.072012366 |
| Gm9990 | 8.809819601 | -0.911116095 | 0.452326079 |
| Aqp11 | 58.08426219 | -1.28323809 | 0.176723665 |
| Pak1 | 1544.109605 | -0.443332009 | 0.076327851 |
| Gdpd4 | 2.28172379 | -1.023694888 | 0.849937762 |
| Myo7a | 84.81893064 | 0.508252307 | 0.123071926 |
| Capn5 | 668.0892524 | 0.420767899 | 0.099386936 |
| Omp | 79.7450718 | -0.221777442 | 0.736920024 |
| Gm16938 | 29.29590882 | 0.14100701 | 0.192554697 |

| | | | |
|---|---|---|---|
| B3gnt6 | 6.185203872 | 0.193709506 | 0.426903802 |
| Acer3 | 413.2037177 | 0.501449938 | 0.094830566 |
| Tsku | 55.8336496 | 0.633179251 | 0.224274937 |
| Gucy2d | 0.893137065 | 0.88632616 | 1.074314648 |
| Lrrc32 | 74.30774312 | 1.681089395 | 0.26857188 |
| Gm23040 | 0.075043205 | 0.237309859 | 3.724945049 |
| A630091E08Rik | 0 | NA | NA |
| 2210018M11Rik | 690.3005469 | -0.410853221 | 0.103921481 |
| Gm15506 | 8.670677481 | -1.111339641 | 0.380881262 |
| Prkrir | 815.4013897 | -0.035775439 | 0.073090884 |
| Wnt11 | 8.862266067 | 0.056320509 | 0.378358247 |
| Gm8398 | 0.117760149 | -0.243576254 | 3.724945049 |
| Uvrag | 562.8884481 | 0.489770055 | 0.088075111 |
| Gm23479 | 0 | NA | NA |
| Dgat2 | 363.7510314 | -0.537522746 | 0.112748447 |
| Mogat2 | 0.144988549 | 1.20006742 | 3.575474141 |
| Gm26705 | 2.196663442 | 0.891955022 | 0.67830981 |
| Map6 | 1290.520615 | -0.393864246 | 0.07450757 |
| Serpinh1 | 377.8077043 | 2.258019247 | 0.20660724 |
| Gdpd5 | 476.3039452 | 0.019098774 | 0.120824361 |
| Klhl35 | 5.303271472 | -0.362258675 | 0.463751142 |
| Rps3 | 2214.045192 | 0.77478237 | 0.110272194 |
| Snord15b | 0 | NA | NA |
| Snord15a | 45.47276232 | 0.893223079 | 0.20691056 |
| n-R5s156 | 0 | NA | NA |
| Arrb1 | 3455.425929 | -0.229511477 | 0.073397931 |
| Mir326 | 2.043157693 | -0.843275865 | 0.723151406 |
| Gm4980 | 50.35484282 | -0.46277778 | 0.19758814 |
| Gm10605 | 21.76352257 | -0.236833095 | 0.233825898 |
| Gm25409 | 0 | NA | NA |
| Slco2b1 | 87.4821162 | -0.928765978 | 0.323183624 |
| Gm15635 | 0.464401238 | 0.713111587 | 1.565628935 |
| Olfr520 | 0.044544648 | 0.237309859 | 3.724945049 |
| Olfr521 | 0 | NA | NA |
| F730035P03Rik | 0.645028293 | 0.735982414 | 1.296625461 |
| Neu3 | 60.27959619 | 0.659020188 | 0.138057099 |
| Spcs2 | 666.5559383 | 0.665814407 | 0.107795547 |
| Xrra1 | 7.264420559 | -0.932813216 | 0.402497506 |
| Rnf169 | 651.1755701 | -0.415771082 | 0.079672153 |
| Chrdl2 | 5.288126676 | -1.032596083 | 0.48670297 |
| Pold3 | 335.8617301 | 0.633871926 | 0.072888561 |
| Lipt2 | 20.2504011 | -0.110685571 | 0.235979783 |
| Kcne3 | 1.196981447 | 0.569991309 | 0.870515177 |
| Pgm2l1 | 3618.346017 | -1.024520418 | 0.110617792 |
| Gpx2-ps1 | 5.150345274 | -1.213740721 | 0.585589298 |
| P4ha3 | 123.5044264 | 1.645135126 | 0.188008674 |
| Ppme1 | 1499.055743 | 0.065451683 | 0.058932362 |

| | | | |
|---|---|---|---|
| C2cd3 | 640.0813776 | 0.088352243 | 0.137692642 |
| Gm10603 | 1.405100988 | -0.972275638 | 0.853622476 |
| Gm37716 | 3.584490054 | -1.683598482 | 0.657418898 |
| Ucp3 | 15.93899765 | -1.806099404 | 0.352891507 |
| Ucp2 | 1246.309197 | 2.661123371 | 0.159218009 |
| Dnajb13 | 7.766840755 | 1.012337623 | 0.392085747 |
| Coa4 | 46.56779514 | 0.724589777 | 0.205205224 |
| Mrpl48 | 372.182467 | -0.184436582 | 0.073387289 |
| Cox20-ps | 0.105143817 | -0.24357403 | 3.724945049 |
| Rab6a | 4454.768753 | -0.214472688 | 0.051816482 |
| Gm14382 | 10.15270512 | -1.063842441 | 0.4290807 |
| Plekhb1 | 2340.080905 | -0.107170526 | 0.243501582 |
| Fam168a | 4198.767291 | -0.264895781 | 0.046648072 |
| Gm3200 | 5.569043595 | 0.134222085 | 0.457159902 |
| Relt | 198.9215136 | 0.35507031 | 0.122619995 |
| Arhgef17 | 2322.379047 | -0.43619957 | 0.063327071 |
| Mir3102 | 0 | NA | NA |
| P2ry6 | 83.22755977 | 2.88402977 | 0.24287238 |
| P2ry2 | 41.59914068 | 4.475384175 | 0.432671157 |
| Fchsd2 | 883.8523491 | -0.049367927 | 0.064634835 |
| Gm6341 | 2.08700676 | -1.403933953 | 0.725857163 |
| Gm15673 | 11.22830619 | -1.137849281 | 0.367433236 |
| Atg16l2 | 153.6349008 | -0.289046144 | 0.160120176 |
| Stard10 | 401.7502125 | -0.186406647 | 0.090895075 |
| Gm20476 | 0.089814205 | -0.24357403 | 3.724945049 |
| Arap1 | 603.6362813 | 1.318555576 | 0.096805955 |
| Pde2a | 2506.890893 | -0.642776712 | 0.12435098 |
| Mir139 | 0.438507067 | -1.172379132 | 1.545448565 |
| Art2a-ps | 2.02887955 | 4.383946461 | 1.076507719 |
| Gm7027 | 32.67737545 | 0.510544447 | 0.236015692 |
| Art2b | 0.341803366 | 2.083994575 | 3.69593078 |
| Clpb | 506.8075599 | -0.227678704 | 0.072089508 |
| Phox2a | 2.085398638 | 0.135252092 | 0.752021181 |
| Inppl1 | 617.7038875 | 0.562853036 | 0.11164058 |
| Gm10602 | 1.695248615 | 0.043928982 | 0.813590709 |
| Folr2 | 8.630795641 | 2.432130212 | 0.448112394 |
| Folr1 | 27.39965009 | -1.214518187 | 0.261975623 |
| Gm22655 | 0.092269992 | 0.237318756 | 3.724945049 |
| Anapc15 | 157.008238 | 0.059059986 | 0.093687516 |
| Tomt | 1.048945322 | 0.485666856 | 0.873969368 |
| Lamtor1 | 591.6362557 | 0.427412947 | 0.098708913 |
| Lrrc51 | 20.02074397 | 0.681610993 | 0.242828318 |
| Numa1 | 1753.65198 | 0.17344726 | 0.083175849 |
| Il18bp | 580.6338207 | 3.863550483 | 0.196826786 |
| Rnf121 | 426.15743 | 0.416655946 | 0.093642361 |
| Xndc1 | 64.93653681 | -0.380072891 | 0.210980682 |
| Xntrpc | 2.817151494 | 0.772350142 | 0.687216304 |

| | | | |
|---|---|---|---|
| Trpc2 | 0 | NA | NA |
| Art5 | 0.775564919 | 0.88363731 | 1.00473161 |
| Art1 | 0.377906432 | -0.157090272 | 1.558035192 |
| Chrna10 | 6.942902888 | 0.050597568 | 0.452972282 |
| Nup98 | 911.2468387 | 0.554068775 | 0.094747697 |
| Pgap2 | 160.2875351 | 0.76888115 | 0.148679935 |
| Rhog | 475.1594366 | 1.38773888 | 0.149359515 |
| Stim1 | 903.1380357 | -0.354213822 | 0.064741121 |
| Rrm1 | 306.720304 | 0.411280805 | 0.122860039 |
| Olfr543 | 0 | NA | NA |
| Olfr544 | 0 | NA | NA |
| Olfr545 | 0 | NA | NA |
| Olfr547 | 0 | NA | NA |
| Olfr548-ps1 | 0 | NA | NA |
| Olfr549 | 0 | NA | NA |
| Trim21 | 602.064124 | 5.712141389 | 0.187169233 |
| Olfr550 | 0.446685102 | 0.730131813 | 1.66955423 |
| Olfr551 | 0 | NA | NA |
| Olfr552 | 0.044544648 | 0.237309859 | 3.724945049 |
| Olfr553 | 0 | NA | NA |
| Olfr554 | 0.152980968 | 0.684806416 | 2.833643346 |
| Olfr555 | 0 | NA | NA |
| Olfr556 | 0.166709681 | -0.724459871 | 2.767631179 |
| Trim68 | 38.7507144 | 0.253235167 | 0.181719148 |
| Olfr557 | 0.253342923 | 0.718222183 | 2.092743516 |
| Olfr558 | 0.693780966 | -0.75806675 | 1.11980514 |
| Olfr33 | 0 | NA | NA |
| Olfr559 | 0 | NA | NA |
| Gm24210 | 0 | NA | NA |
| Olfr78 | 4.264554404 | -3.616449557 | 0.873519952 |
| Olfr561 | 0 | NA | NA |
| Olfr564 | 0 | NA | NA |
| Olfr566 | 0.049553047 | 0.71820487 | 3.724945049 |
| Olfr568 | 0.044544648 | 0.237309859 | 3.724945049 |
| Olfr569 | 0 | NA | NA |
| Olfr570 | 0 | NA | NA |
| Olfr571 | 0 | NA | NA |
| Olfr572 | 0 | NA | NA |
| Olfr573-ps1 | 0 | NA | NA |
| Olfr574 | 0.224720342 | 0.675836698 | 2.133697034 |
| Olfr575 | 0 | NA | NA |
| Olfr576 | 0 | NA | NA |
| Olfr577 | 0 | NA | NA |
| Olfr578 | 0 | NA | NA |
| Olfr582 | 0 | NA | NA |
| Olfr583 | 0 | NA | NA |
| Olfr584 | 0.046345709 | 0.71820487 | 3.724945049 |

| | | | |
|---|---|---|---|
| Olfr585 | 0.062426873 | 0.237312084 | 3.724945049 |
| Olfr586 | 0 | NA | NA |
| Olfr587-ps1 | 0.046345709 | 0.71820487 | 3.724945049 |
| Gm15120 | 0.07796279 | 1.072698098 | 3.721829099 |
| Olfr588-ps1 | 0 | NA | NA |
| Gm15116 | 0 | NA | NA |
| Olfr589 | 0 | NA | NA |
| Olfr590-ps1 | 0 | NA | NA |
| Olfr591 | 0 | NA | NA |
| Olfr592 | 0 | NA | NA |
| Olfr593 | 0.044191549 | -0.243571806 | 3.724945049 |
| Olfr594 | 0 | NA | NA |
| Olfr595-ps1 | 0 | NA | NA |
| Usp17ld | 0 | NA | NA |
| Gm25171 | 0 | NA | NA |
| Gm23550 | 0 | NA | NA |
| Gm23973 | 0 | NA | NA |
| Olfr596 | 0 | NA | NA |
| Olfr597 | 0 | NA | NA |
| Olfr598 | 0 | NA | NA |
| Olfr599 | 0.097139547 | 0.718207094 | 3.724945049 |
| Olfr600 | 0 | NA | NA |
| Olfr601 | 0 | NA | NA |
| Olfr602-ps1 | 0 | NA | NA |
| Olfr603 | 0 | NA | NA |
| Usp17lc | 0 | NA | NA |
| Gm25774 | 0.116217011 | -0.24357403 | 3.724945049 |
| Olfr605 | 0 | NA | NA |
| Olfr606 | 0 | NA | NA |
| Gm15121 | 0 | NA | NA |
| Olfr607 | 0 | NA | NA |
| Olfr608 | 0 | NA | NA |
| Olfr609 | 0 | NA | NA |
| Olfr610 | 0 | NA | NA |
| Olfr611 | 0 | NA | NA |
| Olfr612 | 0 | NA | NA |
| Olfr613 | 0 | NA | NA |
| Olfr615 | 0 | NA | NA |
| Olfr616 | 0.04393175 | -0.243571806 | 3.724945049 |
| Gm26506 | 0.044544648 | 0.237309859 | 3.724945049 |
| Olfr617 | 0 | NA | NA |
| Olfr618 | 0 | NA | NA |
| Olfr619 | 0 | NA | NA |
| Olfr620 | 0 | NA | NA |
| Olfr621-ps1 | 0 | NA | NA |
| Olfr622 | 0 | NA | NA |
| Olfr623 | 0 | NA | NA |

| | | | |
|---|---|---|---|
| Olfr624 | 0 | NA | NA |
| Olfr625-ps1 | 0 | NA | NA |
| Olfr626-ps1 | 0 | NA | NA |
| Olfr260-ps1 | 0 | NA | NA |
| Olfr243 | 0 | NA | NA |
| Olfr628 | 0.061920613 | 0.718207094 | 3.724945049 |
| Olfr629 | 0 | NA | NA |
| Olfr630 | 0 | NA | NA |
| Olfr69 | 0 | NA | NA |
| Olfr68 | 0 | NA | NA |
| Olfr67 | 0.045882455 | 0.237312084 | 3.724945049 |
| Hbb-bt | 10.74344224 | 0.777446612 | 0.693767267 |
| Hbb-bs | 223.1020574 | 1.162771969 | 0.69370834 |
| Gm15115 | 0 | NA | NA |
| Hbb-bh3 | 0 | NA | NA |
| Hbb-bh2 | 0 | NA | NA |
| Hbb-bh1 | 0 | NA | NA |
| Hbb-bh0 | 0 | NA | NA |
| Hbb-y | 0 | NA | NA |
| Olfr66 | 0 | NA | NA |
| Dnajc19-ps | 11.1957499 | 0.506478492 | 0.331445891 |
| Olfr64 | 0 | NA | NA |
| Olfr65 | 0.042716945 | -0.243571806 | 3.724945049 |
| Olfr631 | 0 | NA | NA |
| Olfr632 | 0.042716945 | -0.243571806 | 3.724945049 |
| Olfr633 | 0 | NA | NA |
| Olfr634-ps1 | 0 | NA | NA |
| Gm4886 | 0 | NA | NA |
| Olfr635 | 0.092691418 | 1.183079489 | 3.724560917 |
| Olfr636-ps1 | 0.038981395 | 0.718207094 | 3.724945049 |
| Olfr637-ps1 | 0 | NA | NA |
| 4930516K23Rik | 0 | NA | NA |
| Olfr638 | 0 | NA | NA |
| Olfr639 | 0 | NA | NA |
| Olfr640 | 0 | NA | NA |
| Olfr641 | 0 | NA | NA |
| Olfr642 | 0 | NA | NA |
| Olfr643 | 0.062426873 | 0.237312084 | 3.724945049 |
| Olfr644 | 0 | NA | NA |
| Olfr645 | 0 | NA | NA |
| Olfr646 | 0 | NA | NA |
| 4931431F19Rik | 0.042716945 | -0.243571806 | 3.724945049 |
| Ubqln3 | 0 | NA | NA |
| Ubqlnl | 0 | NA | NA |
| E030002O03Rik | 0.160237607 | -1.206626386 | 2.799265637 |
| Olfr647-ps1 | 0 | NA | NA |
| Olfr648 | 0 | NA | NA |

| | | | |
|---|---|---|---|
| Olfr649 | 0.060174537 | 0.237309859 | 3.724945049 |
| Olfr650-ps1 | 0.524522254 | 2.928189047 | 1.507631992 |
| Trim6 | 12.76726646 | 3.325446766 | 0.440236499 |
| Trim34a | 130.1339841 | 5.32630733 | 0.258067404 |
| Gm15135 | 0 | NA | NA |
| Trim5 | 3.073902706 | 3.527013312 | 0.84801357 |
| Gm24846 | 0 | NA | NA |
| Trim12a | 0.470008961 | 2.046092047 | 1.635362152 |
| Trim34b | 43.17410611 | 5.826095375 | 0.508021097 |
| Gm15133 | 0.044544648 | 0.237309859 | 3.724945049 |
| Trim12c | 411.4065939 | 4.349905935 | 0.274605384 |
| Gm27995 | 0 | NA | NA |
| Trim30b | 23.44767613 | 8.420247411 | 0.942990705 |
| Trim30c | 106.4359115 | 7.429079269 | 0.784467222 |
| Trim30a | 1972.4724 | 8.317916154 | 0.293743215 |
| Gm25405 | 0.545822006 | -1.252523743 | 1.190758524 |
| Trim30d | 381.5932247 | 8.556616313 | 0.494833389 |
| Gm16464 | 7.703381671 | 0.359958966 | 0.421571634 |
| Gm6574 | 0 | NA | NA |
| Trim30e-ps1 | 0 | NA | NA |
| Olfr651 | 0.060174537 | 0.237309859 | 3.724945049 |
| Olfr652 | 0.038981395 | 0.718207094 | 3.724945049 |
| Olfr653 | 0 | NA | NA |
| Olfr654 | 0.212522253 | 0.718222226 | 2.170709238 |
| Olfr655 | 0.733075257 | 0.935737811 | 1.238251116 |
| Olfr656 | 0 | NA | NA |
| Olfr657 | 0 | NA | NA |
| Olfr658 | 7.112035581 | 1.929697872 | 0.517276481 |
| Usp-ps | 0 | NA | NA |
| Olfr659 | 0 | NA | NA |
| Olfr660-ps1 | 0 | NA | NA |
| Olfr661 | 0 | NA | NA |
| Olfr663 | 0 | NA | NA |
| Olfr664 | 0 | NA | NA |
| Usp17le | 0 | NA | NA |
| Gm4887 | 2.328839425 | -0.161716227 | 0.769984095 |
| Usp17lb | 0 | NA | NA |
| Usp17la | 0 | NA | NA |
| Olfr665 | 0 | NA | NA |
| Olfr666 | 0.045882455 | 0.237312084 | 3.724945049 |
| Olfr667 | 0.038981395 | 0.718207094 | 3.724945049 |
| Olfr668 | 0.044544648 | 0.237309859 | 3.724945049 |
| Olfr669 | 0 | NA | NA |
| Gm5900 | 87.04897631 | 0.341248393 | 0.134658797 |
| Olfr670 | 0 | NA | NA |
| Olfr671 | 0.038981395 | 0.718207094 | 3.724945049 |
| Olfr672 | 0.101408268 | 0.717551239 | 3.724898127 |

| | | | |
|---|---|---|---|
| Olfr675 | 0 | NA | NA |
| Olfr676 | 0 | NA | NA |
| Olfr677 | 0 | NA | NA |
| Olfr678 | 0 | NA | NA |
| Olfr679 | 0 | NA | NA |
| Olfr680-ps1 | 0.149335367 | -0.724459895 | 2.864896933 |
| Olfr681 | 0 | NA | NA |
| Olfr682-ps1 | 0.201268616 | 0.718215723 | 2.619264982 |
| Olfr683 | 0.362463993 | 2.467552355 | 1.603190423 |
| Olfr684 | 0.256199707 | 1.811336593 | 2.973144861 |
| Olfr685 | 0.357595806 | 0.741839776 | 3.31635367 |
| Olfr686 | 0.169924525 | -0.711094185 | 3.387745846 |
| Olfr687 | 0 | NA | NA |
| Olfr688 | 0 | NA | NA |
| Olfr689 | 0 | NA | NA |
| Olfr690 | 5.292052314 | -1.418712545 | 0.554201112 |
| Olfr691 | 5.347759076 | -1.00759501 | 0.636089436 |
| Olfr692 | 0.049553047 | 0.71820487 | 3.724945049 |
| Fam160a2 | 1185.683241 | -0.935006368 | 0.101181465 |
| Gm5901 | 0.044544648 | 0.237309859 | 3.724945049 |
| Cnga4 | 8.850792061 | -0.948144096 | 0.384873664 |
| Cckbr | 263.3216545 | -0.614973551 | 0.168022549 |
| Prkcdbp | 205.5345052 | 1.623048558 | 0.199979472 |
| Smpd1 | 1173.492348 | 0.148603411 | 0.069181241 |
| Apbb1 | 2731.609977 | -0.446573372 | 0.062635187 |
| Hpx | 16.70994554 | 3.579040224 | 0.502765243 |
| Trim3 | 1334.924561 | -0.312158493 | 0.084193628 |
| Arfip2 | 801.3768636 | 0.115657489 | 0.069878727 |
| Timm10b | 130.0419406 | -0.042004392 | 0.103092418 |
| Dnhd1 | 163.944155 | 0.123978402 | 0.197549839 |
| Gm22504 | 0.072025462 | 0.237312084 | 3.724945049 |
| Mrps36-ps2 | 0.9406005 | 2.204282818 | 1.131431693 |
| Rrp8 | 163.2337327 | 0.650517692 | 0.125449361 |
| Ilk | 832.9278343 | 0.790058599 | 0.105281188 |
| Taf10 | 390.2000004 | 0.462523833 | 0.112044284 |
| Tpp1 | 810.0873032 | 0.528095355 | 0.117684978 |
| Dchs1 | 413.6250274 | -0.172921961 | 0.104012247 |
| Gm15645 | 4.914941411 | -0.10120809 | 0.463373376 |
| Mrpl17 | 334.4169704 | 0.616664055 | 0.114192052 |
| 9130208D14Rik | 37.17479923 | 7.929362875 | 0.852334165 |
| Gm4070 | 48.02494048 | 6.443966779 | 0.565096755 |
| Gm20663 | 0.150903271 | -0.243571349 | 2.855571776 |
| Gm18852 | 4.812241163 | 5.518377904 | 0.908441289 |
| Gm27228 | 0 | NA | NA |
| Gm8979 | 1.384150549 | 4.175490312 | 1.013981859 |
| Gm27149 | 0.066050875 | 0.71820487 | 3.724945049 |
| Gm8982 | 0 | NA | NA |

| | | | |
|---|---|---|---|
| Gvin1 | 6.195143366 | 6.398011638 | 0.95920527 |
| Gm18853 | 1.133034309 | 4.040969754 | 1.101804968 |
| Gm27150 | 0 | NA | NA |
| Gm8989 | 0.700564593 | 3.188628559 | 1.258667915 |
| Gm4759 | 67.3121842 | 7.747691276 | 0.772457578 |
| Gm22372 | 0 | NA | NA |
| Gm8995 | 675.8586595 | 6.256756963 | 0.219538657 |
| Gm1966 | 19.76971248 | 7.254106796 | 0.891502778 |
| Olfr693 | 0.090277459 | 0.237318756 | 3.724945049 |
| Olfr694 | 0 | NA | NA |
| Olfr695 | 0 | NA | NA |
| Olfr697 | 0 | NA | NA |
| Olfr698 | 0 | NA | NA |
| Olfr699 | 0.049553047 | 0.71820487 | 3.724945049 |
| Olfr700 | 0 | NA | NA |
| Olfr701 | 0.038981395 | 0.718207094 | 3.724945049 |
| Olfr702 | 0 | NA | NA |
| Olfr703 | 0.085433889 | -0.72445792 | 3.724945049 |
| Olfr704 | 0 | NA | NA |
| Olfr705 | 0.136194719 | -0.724453471 | 3.724945049 |
| Olfr706 | 0 | NA | NA |
| Olfr707 | 0.068097359 | -0.243569582 | 3.724945049 |
| Olfr1532-ps1 | 0 | NA | NA |
| Olfr709-ps1 | 0 | NA | NA |
| Olfr710 | 0.04393175 | -0.243571806 | 3.724945049 |
| Olfr6 | 0 | NA | NA |
| Olfr711 | 0 | NA | NA |
| Olfr2 | 0 | NA | NA |
| Olfr713 | 0 | NA | NA |
| Olfr714 | 0 | NA | NA |
| Olfr17 | 0.048949532 | -0.243569582 | 3.724945049 |
| Gm10081 | 0 | NA | NA |
| Olfr715 | 0.097139547 | 0.718207094 | 3.724945049 |
| Olfr716 | 0.092691418 | 1.183079489 | 3.724560917 |
| Nlrp14 | 0 | NA | NA |
| Gm3765 | 0.678124035 | 2.924652805 | 1.368597963 |
| Rbmxl2 | 0 | NA | NA |
| Platr28 | 0.049553047 | 0.71820487 | 3.724945049 |
| Gm24932 | 0.060174537 | 0.237309859 | 3.724945049 |
| Syt9 | 336.6996381 | -0.254427213 | 0.08649159 |
| Olfml1 | 105.9701042 | -0.006736054 | 0.175133106 |
| Ppfibp2 | 154.4178595 | 0.564305176 | 0.124876177 |
| 5330417H12Rik | 0 | NA | NA |
| Cyb5r2 | 9.463141173 | 2.320189656 | 0.425924045 |
| Ovch2 | 0 | NA | NA |
| Olfr467 | 0 | NA | NA |
| Olfr469 | 0 | NA | NA |

| | | | |
|---|---|---|---|
| Olfr470 | 0 | NA | NA |
| Gm10156 | 22.50808326 | 1.404749507 | 0.347946105 |
| Olfr472 | 0.092881282 | -0.724455696 | 3.724945049 |
| Olfr473 | 0.038981395 | 0.718207094 | 3.724945049 |
| Olfr474 | 0 | NA | NA |
| Olfr476 | 0 | NA | NA |
| Olfr477 | 0.165236907 | 0.238279123 | 3.378321226 |
| Olfr478 | 0 | NA | NA |
| Olfr479 | 0 | NA | NA |
| Olfr480 | 0 | NA | NA |
| Olfr481 | 0 | NA | NA |
| Olfr482 | 0 | NA | NA |
| Olfr483 | 0.062426873 | 0.237312084 | 3.724945049 |
| Gm25424 | 0 | NA | NA |
| Olfr484 | 0 | NA | NA |
| Olfr485 | 0 | NA | NA |
| Olfr486 | 0 | NA | NA |
| Olfr487 | 0.123841226 | 1.292225235 | 3.719439735 |
| Olfr488 | 0 | NA | NA |
| Olfr490 | 0 | NA | NA |
| Olfr491 | 0 | NA | NA |
| Olfr492 | 0 | NA | NA |
| Olfr493 | 0 | NA | NA |
| Olfr494 | 0 | NA | NA |
| Olfr495 | 0 | NA | NA |
| Olfr497 | 0.049553047 | 0.71820487 | 3.724945049 |
| Olfr498 | 0 | NA | NA |
| Olfr502 | 0 | NA | NA |
| Olfr503 | 0 | NA | NA |
| Olfr504 | 0 | NA | NA |
| Olfr506 | 0 | NA | NA |
| Olfr507 | 0 | NA | NA |
| Olfr508 | 0 | NA | NA |
| Olfr509 | 0 | NA | NA |
| Olfr510 | 0.046345709 | 0.71820487 | 3.724945049 |
| Olfr512 | 0 | NA | NA |
| Olfr513 | 0 | NA | NA |
| Olfr514 | 0 | NA | NA |
| Olfr516 | 0 | NA | NA |
| Olfr517 | 0 | NA | NA |
| Olfr518 | 0 | NA | NA |
| Olfr519 | 0 | NA | NA |
| Nlrp10 | 10.06809784 | 0.510500004 | 0.337588403 |
| Eif3f | 1059.444892 | 0.447976254 | 0.076266745 |
| Gm26599 | 4.712628857 | 2.092451102 | 0.798493842 |
| Tub | 1801.451592 | -0.793964747 | 0.111654139 |
| Ric3 | 1280.403782 | 0.255108209 | 0.072182922 |

| | | | |
|---|---|---|---|
| Gm25450 | 3.817416342 | -0.714242573 | 0.580424741 |
| Lmo1 | 71.3151019 | 0.378314771 | 0.136840163 |
| Stk33 | 14.06609556 | -0.234743122 | 0.285749355 |
| Gm9105 | 5.082316306 | 0.456066997 | 0.453553682 |
| 1700095J03Rik | 3.157210742 | 0.280888926 | 0.640956776 |
| Trim66 | 632.3889432 | 0.033143704 | 0.105468834 |
| Rpl27a | 1290.635761 | 0.601440618 | 0.102199964 |
| Snora3 | 0 | NA | NA |
| AC124457.1 | 0.875090066 | -0.020430261 | 1.104179008 |
| Gm24888 | 1.185922749 | -0.071437629 | 0.790288782 |
| St5 | 362.0690004 | 0.235053764 | 0.085102572 |
| Akip1 | 31.41517011 | 1.215501159 | 0.225907295 |
| BC051019 | 3.117097637 | 1.96268781 | 0.646951368 |
| Ascl3 | 1.052193792 | 1.00304969 | 0.927286943 |
| Tmem9b | 679.663783 | 0.048419924 | 0.07383378 |
| Nrip3 | 1443.710783 | -0.619682824 | 0.112022372 |
| Gm22822 | 1.435595059 | -0.886025694 | 0.925910482 |
| Scube2 | 22.45075123 | -0.559265832 | 0.314642006 |
| Dennd5a | 3399.692085 | -0.351717235 | 0.068741068 |
| Tmem41b | 489.61422 | 0.239557124 | 0.099803141 |
| Gm24842 | 0.126912322 | 0.555057727 | 3.69382869 |
| 1600010M07Rik | 2.455553585 | 2.998892585 | 0.845398693 |
| Gm23100 | 0 | NA | NA |
| Ipo7 | 1735.407786 | 0.331114624 | 0.075747531 |
| Gm25636 | 6.97985599 | 0.193113631 | 0.390876182 |
| Snora23 | 31.16499087 | -0.040016707 | 0.228146976 |
| AA474408 | 22.01653641 | -0.654290826 | 0.257350682 |
| Zfp143 | 153.6133789 | 0.476847533 | 0.093961704 |
| Wee1 | 194.9507975 | -0.274473784 | 0.119740704 |
| Swap70 | 161.8502858 | 0.915502591 | 0.107708958 |
| Sbf2 | 1399.071276 | 0.026638162 | 0.07277212 |
| Gm9064 | 0 | NA | NA |
| Gm17219 | 0.046345709 | 0.71820487 | 3.724945049 |
| Gm10087 | 0 | NA | NA |
| Gm28863 | 0.430307537 | 0.670850727 | 1.436953515 |
| Adm | 37.32539147 | 1.633558192 | 0.236763654 |
| Ampd3 | 786.3935543 | 1.868621036 | 0.162286258 |
| Rnf141 | 819.8137337 | -0.195353954 | 0.108684805 |
| AC122844.1 | 0.956902809 | 0.210581772 | 1.206802713 |
| Lyve1 | 8.973244537 | -0.721622458 | 0.410077458 |
| Mrvi1 | 43.82669296 | -0.38329352 | 0.182319907 |
| Gm16336 | 0.075043205 | 0.237309859 | 3.724945049 |
| Ctr9 | 691.8584163 | 0.036766641 | 0.059270892 |
| Eif4g2 | 8499.079975 | 0.441220955 | 0.082328523 |
| Gm23262 | 0 | NA | NA |
| Galnt18 | 280.6267264 | -0.726598333 | 0.130252678 |
| Gm25787 | 0 | NA | NA |

| | | | |
|---|---|---|---|
| Gm23345 | 0.066050875 | 0.71820487 | 3.724945049 |
| Usp47 | 1697.490691 | 0.330761382 | 0.062757246 |
| Dkk3 | 1672.01163 | -0.283564533 | 0.122421142 |
| Mical2 | 2673.306519 | -0.801825327 | 0.14147415 |
| Micalcl | 33.92286921 | -1.533661501 | 0.271462008 |
| Parva | 887.2698577 | 0.087448872 | 0.086464264 |
| Parvaos | 2.062454624 | 1.191712612 | 0.721318972 |
| 2310014F06Rik | 4.100390628 | 0.50339975 | 0.491109195 |
| Gm24154 | 0.181894773 | 0.264898351 | 2.695899273 |
| Tead1 | 787.7264761 | 1.267082345 | 0.121815595 |
| AC126052.1 | 1.51359592 | -0.920223359 | 0.830612787 |
| Rassf10 | 10.84181486 | -1.027470997 | 0.334025543 |
| AC104930.1 | 0 | NA | NA |
| Gm23662 | 0 | NA | NA |
| Arntl | 352.8719286 | 0.23170205 | 0.127633613 |
| Btbd10 | 732.5892807 | -0.237299608 | 0.078100352 |
| Gm15994 | 0.277092755 | 2.213303804 | 2.021992352 |
| Pth | 0 | NA | NA |
| Far1 | 1848.391941 | -0.226843113 | 0.06408883 |
| Far1os | 0.286281128 | -0.724462368 | 3.724945049 |
| Gm23700 | 0 | NA | NA |
| Gm5600 | 0.213133906 | -0.221384427 | 2.170898092 |
| Gm29507 | 0.046345709 | 0.71820487 | 3.724945049 |
| Spon1 | 1904.191878 | 0.651331053 | 0.107880743 |
| Rras2 | 177.862388 | 0.321342874 | 0.118922007 |
| Copb1 | 1041.818964 | 0.564764705 | 0.086315554 |
| Psma1 | 653.5961011 | 0.67606674 | 0.106751135 |
| 4933406I18Rik | 91.43073966 | -0.512378617 | 0.12709385 |
| Pde3b | 366.815521 | -0.013053273 | 0.096604232 |
| Cyp2r1 | 3.701896043 | 1.450284798 | 0.561048745 |
| Calca | 61.65923974 | 2.650160697 | 0.247937409 |
| Gm15500 | 180.1198848 | 0.754470819 | 0.164357407 |
| Calcb | 5.761881877 | 2.015701534 | 0.585967191 |
| Insc | 59.233201 | 0.110508258 | 0.251878539 |
| Gm24982 | 0 | NA | NA |
| Rps4l-ps | 0.061920613 | 0.718207094 | 3.724945049 |
| Gm22669 | 0 | NA | NA |
| Gm27684 | 0.263733274 | 0.237328279 | 2.116539665 |
| Sox6 | 489.52169 | 0.27514613 | 0.086128662 |
| Sox6os | 0.611953183 | -1.479672897 | 1.236521551 |
| 1700003G18Rik | 0.77082754 | -0.779525298 | 1.043310349 |
| Gm4353 | 19.37501129 | 0.297792558 | 0.290791261 |
| 1110004F10Rik | 588.1908301 | 0.232268554 | 0.092624355 |
| Plekha7 | 339.2168702 | -0.11753099 | 0.10036247 |
| Rps13 | 763.6410431 | 0.76108674 | 0.101943751 |
| Snord14a | 3.97447326 | 0.215781195 | 0.573909319 |
| Pik3c2a | 1083.338475 | 0.131865982 | 0.066522474 |

| | | | |
|---|---|---|---|
| Nucb2 | 157.5399997 | 0.465468251 | 0.170934239 |
| n-R5s157 | 0 | NA | NA |
| Gm25423 | 0 | NA | NA |
| RP24-246N21.1 | 393.097115 | 0.69355683 | 0.12102277 |
| Gm26054 | 0 | NA | NA |
| Xylt1 | 443.4184742 | 0.092943199 | 0.063920688 |
| Gm24063 | 0 | NA | NA |
| Rps15a | 1296.320441 | 0.480385277 | 0.074847746 |
| Arl6ip1 | 2631.753197 | 0.636455554 | 0.118408035 |
| Smg1 | 4666.36924 | 0.045663231 | 0.103780084 |
| Gm25683 | 0 | NA | NA |
| 4930583K01Rik | 0.241683003 | 0.71822246 | 2.095750154 |
| Gm23229 | 0.044544648 | 0.237309859 | 3.724945049 |
| Syt17 | 179.5473353 | -0.72360027 | 0.156178107 |
| Itpripl2 | 154.921355 | 2.412819876 | 0.16816359 |
| Coq7 | 126.5384451 | 0.121535166 | 0.150226795 |
| Tmc7 | 194.611435 | -0.932600136 | 0.119929007 |
| Tmc5 | 2.207900702 | -0.507004656 | 0.802475029 |
| Gde1 | 1198.036269 | -0.019117919 | 0.058784438 |
| Ccp110 | 802.7760504 | -0.726875211 | 0.081069649 |
| 9030624J02Rik | 685.4574909 | 0.166171656 | 0.069418244 |
| Gm26147 | 1.083234348 | -1.581131728 | 0.972696823 |
| Knop1 | 359.1550831 | 0.34518486 | 0.069282941 |
| Iqck | 21.26371758 | 0.757100759 | 0.239854274 |
| Gprc5b | 1912.548548 | -0.639127001 | 0.094379657 |
| Gpr139 | 17.82217913 | -0.536423177 | 0.293718901 |
| Gm23321 | 0 | NA | NA |
| Gp2 | 0.244487964 | -0.243563258 | 1.85554965 |
| Umod | 0 | NA | NA |
| Pdilt | 0.092881282 | -0.724455696 | 3.724945049 |
| Gm37518 | 0.130524232 | -0.243571427 | 3.657871338 |
| Gm26967 | 0.180739367 | -0.72445825 | 3.322505087 |
| Acsm5 | 5.692917749 | 0.025269201 | 0.481942359 |
| Acsm2 | 0.062426873 | 0.237312084 | 3.724945049 |
| Acsm1 | 0.038981395 | 0.718207094 | 3.724945049 |
| Acsm4 | 1.49381288 | -0.80897374 | 0.785641599 |
| Thumpd1 | 343.5482368 | 0.319263424 | 0.106113428 |
| Gm5601 | 215.6914571 | 0.083433338 | 0.094165684 |
| Acsm3 | 0.623781035 | 0.897632204 | 1.334840631 |
| Eri2 | 94.72177517 | 0.159785636 | 0.149758361 |
| 2610020H08Rik | 22.09157058 | 0.530700201 | 0.274116715 |
| Dcun1d3 | 429.5927485 | 0.203933415 | 0.091328242 |
| Lyrm1 | 64.24767704 | -0.025473754 | 0.14658762 |
| Dnah3 | 28.90499188 | -0.1703135 | 0.213684974 |
| Gm25217 | 0.280224789 | 0.60700547 | 1.801075827 |
| Tmem159 | 54.2618588 | 0.925778145 | 0.162086451 |
| Zp2 | 1.125736089 | 1.351168426 | 1.144966442 |

| | | | |
|---|---|---|---|
| Anks4b | 4.77461678 | 1.881634208 | 0.606866312 |
| Crym | 493.688269 | 1.184323905 | 0.124716599 |
| Abca14 | 0.666404725 | 1.840284367 | 1.171448293 |
| Abca15 | 0.612941558 | 2.035335147 | 1.294839256 |
| Abca16 | 0.254958917 | 2.166073837 | 2.048074188 |
| E130201H02Rik | 1.649955311 | 0.756040268 | 0.836181103 |
| Uqcrc2 | 1539.021604 | 0.027511294 | 0.078570147 |
| Pdzd9 | 1.255624142 | 1.95505819 | 1.003195119 |
| 9030407P20Rik | 49.63574211 | -0.482302471 | 0.178510493 |
| BC030336 | 1048.447558 | 0.011531863 | 0.065188538 |
| Vwa3a | 56.37564545 | -0.612419044 | 0.171120495 |
| Eef2k | 574.705251 | -0.285179557 | 0.10594005 |
| Gm15774 | 0.046345709 | 0.71820487 | 3.724945049 |
| Polr3e | 380.9951259 | -0.190833499 | 0.107747839 |
| Cdr2 | 189.8166021 | 0.277955462 | 0.175233025 |
| 4933427G17Rik | 3.26880693 | -0.398034939 | 0.626128789 |
| Gm9234 | 7.527412146 | 0.528738917 | 0.364487359 |
| Gm25924 | 0 | NA | NA |
| Gm9905 | 21.38643968 | 0.93838887 | 0.255553788 |
| Mettl9 | 576.9325256 | 0.264895509 | 0.111848817 |
| Gm27991 | 0 | NA | NA |
| Igsf6 | 72.32097277 | 4.864102422 | 0.407508629 |
| Otoa | 4.95839639 | 0.125139463 | 0.491861878 |
| Hs3st2 | 198.3223821 | -0.100398491 | 0.311597069 |
| Usp31 | 2536.972672 | -0.260495251 | 0.07877116 |
| 1700069B07Rik | 1.411393777 | 0.813117928 | 0.830724712 |
| Scnn1g | 16.54571043 | 1.617599194 | 0.321657316 |
| Scnn1b | 3.3334194 | 3.025491217 | 0.753265483 |
| Cog7 | 494.7223238 | -0.183142412 | 0.066148197 |
| Gga2 | 393.4606856 | 0.141786935 | 0.096480428 |
| Ears2 | 110.4529885 | 0.360509615 | 0.119958814 |
| Ubfd1 | 1136.037908 | 0.09016529 | 0.062785508 |
| Ndufab1 | 625.6134519 | -0.038349598 | 0.079849151 |
| Palb2 | 22.09319308 | 0.44899996 | 0.256644464 |
| Gm16326 | 0.136194719 | -0.724453471 | 3.724945049 |
| Dctn5 | 636.704053 | 0.055307295 | 0.055173257 |
| Gm15489 | 0.496441948 | -0.288984951 | 1.538886242 |
| Plk1 | 22.12491667 | 1.119592305 | 0.304051818 |
| Ern2 | 3.478004289 | -0.92592819 | 0.575137172 |
| Chp2 | 6.555594633 | -0.738571659 | 0.399703757 |
| Gm23614 | 0 | NA | NA |
| Prkcb | 4097.896829 | -1.158621518 | 0.120092695 |
| Gm14388 | 23.59541605 | -0.140191599 | 0.252853613 |
| Gm26414 | 4.07614265 | -1.707873037 | 0.641631533 |
| Gm14389 | 0 | NA | NA |
| Cacng3 | 479.8047837 | -1.39588741 | 0.166861005 |
| Rbbp6 | 1656.438353 | 0.039258765 | 0.073004562 |

| | | | |
|---|---|---|---|
| Tnrc6a | 1721.479868 | -0.157620429 | 0.09432277 |
| Slc5a11 | 15.52195762 | 0.047513711 | 0.27070494 |
| Arhgap17 | 309.4984358 | 0.971934746 | 0.139726718 |
| Lcmt1 | 343.5567309 | -0.270693457 | 0.069167217 |
| Gm23559 | 0 | NA | NA |
| Aqp8 | 0.525182132 | -0.243564 | 1.549215714 |
| Zkscan2 | 230.1962317 | -0.565346189 | 0.161163344 |
| Gm23788 | 0.167772976 | 0.778749666 | 3.41346025 |
| Gm26974 | 9.009306514 | 0.362419862 | 0.328874185 |
| Gm27040 | 8.361238006 | 0.131037609 | 0.402499778 |
| Hs3st4 | 521.3318301 | -0.162679034 | 0.087835512 |
| Gm25759 | 6.10632452 | -0.928844935 | 0.545053506 |
| Gm15338 | 14.24273356 | -1.424327541 | 0.438224376 |
| Gm15339 | 0 | NA | NA |
| Gm23168 | 0.097139547 | 0.718207094 | 3.724945049 |
| 3100003L05Rik | 1.759517327 | 2.204549906 | 0.891154904 |
| Gm21957 | 0 | NA | NA |
| 4930533L02Rik | 0 | NA | NA |
| 4933440M02Rik | 0.422952957 | -0.796853038 | 3.703775843 |
| 4930571K23Rik | 0 | NA | NA |
| Kdm8 | 38.64691656 | 0.086368346 | 0.249721768 |
| Nsmce1 | 119.9047304 | 1.078443301 | 0.168638404 |
| Il4ra | 356.9425988 | 3.452681812 | 0.194342473 |
| Il21r | 78.43361135 | 4.036462761 | 0.261292655 |
| Gtf3c1 | 2212.631377 | -0.217926451 | 0.062839831 |
| D430042O09Rik | 320.9984056 | -0.036152855 | 0.074830553 |
| Gsg1l | 413.6031104 | -0.992943682 | 0.160957788 |
| Xpo6 | 1490.157562 | 0.241629918 | 0.066872107 |
| Gm17137 | 0.048949532 | -0.243569582 | 3.724945049 |
| Gm17136 | 0.044544648 | 0.237309859 | 3.724945049 |
| Sbk1 | 1551.156001 | -0.359111509 | 0.14366911 |
| Gm10155 | 8.990861825 | 0.50874324 | 0.356787066 |
| Lat | 33.16436472 | 3.723672695 | 0.356616777 |
| Mir7058 | 0 | NA | NA |
| Spns1 | 414.4773662 | -0.019128439 | 0.080218641 |
| Nfatc2ip | 98.35474023 | 0.622125483 | 0.138734511 |
| Cd19 | 0.620398479 | 3.277578814 | 1.469374456 |
| Rabep2 | 121.2584971 | 0.04236435 | 0.106857434 |
| Atp2a1 | 15.93822915 | -0.161204311 | 0.273539706 |
| Sh2b1 | 913.839647 | 0.188910006 | 0.057811235 |
| Tufm | 383.5821207 | 0.05994599 | 0.105415814 |
| Atxn2l | 1896.105242 | 0.195399752 | 0.084274719 |
| Eif3c | 2058.465012 | 0.636669891 | 0.084535291 |
| Gm24596 | 0.884909101 | -0.49998993 | 1.057219056 |
| G730046D07Rik | 2.910496466 | 4.479144203 | 0.939253992 |
| Gm25579 | 0 | NA | NA |
| Cln3 | 171.9378215 | 1.063334814 | 0.10351752 |

| | | | |
|---|---|---|---|
| Mir7059 | 0.228481116 | -0.677352646 | 2.168157963 |
| Apobr | 49.34344806 | 4.410277542 | 0.395648216 |
| Il27 | 13.85132785 | 6.342143741 | 0.898947243 |
| Nupr1 | 177.6816824 | 4.144264061 | 0.278912784 |
| 2510046G10Rik | 4.55499157 | -0.727500429 | 0.459713646 |
| Ccdc101 | 124.2951638 | -0.178629877 | 0.099830707 |
| Sult1a1 | 88.10259738 | -0.268766578 | 0.259164981 |
| Gm6939 | 1.140676537 | 0.717513706 | 1.041681485 |
| Slx1b | 109.1217166 | -0.180902437 | 0.123029317 |
| Bola2 | 141.4468845 | 0.436654589 | 0.145862011 |
| Coro1a | 641.2179666 | 0.334865826 | 0.088011907 |
| Mapk3 | 1099.213407 | -0.372365224 | 0.093347774 |
| Gdpd3 | 17.57179256 | -0.657110636 | 0.281744225 |
| Gm9967 | 7.556454724 | -0.829591697 | 0.457029229 |
| Ypel3 | 1100.394795 | -0.237398369 | 0.071864021 |
| Tbx6 | 9.328334444 | -0.66213878 | 0.412184021 |
| Ppp4c | 390.6480823 | 0.958538807 | 0.122527304 |
| Aldoa | 12436.05844 | 0.013714418 | 0.049784327 |
| Gm15676 | 14.57694585 | 0.058426519 | 0.287162917 |
| Fam57b | 780.1775602 | -0.559907121 | 0.068330185 |
| 4930451I11Rik | 0.843446786 | -1.838710464 | 1.026216589 |
| Doc2a | 204.7161822 | -1.284959327 | 0.165550511 |
| Ino80e | 331.245082 | 0.340521996 | 0.072417067 |
| Hirip3 | 134.4899788 | -0.360870977 | 0.141098514 |
| Taok2 | 2099.502443 | -0.366857737 | 0.048099736 |
| Tmem219 | 154.4928956 | 1.08325305 | 0.157014504 |
| Gm20650 | 0.099106094 | 1.209894437 | 3.72418822 |
| Gm23445 | 0.097899064 | -0.724453471 | 3.724945049 |
| Kctd13 | 647.9350141 | -0.688029045 | 0.087493205 |
| Gm21984 | 106.5590713 | 0.230000521 | 0.198295135 |
| Asphd1 | 163.222168 | -0.410052021 | 0.111870468 |
| Sez6l2 | 4188.453118 | -0.302181114 | 0.072482809 |
| D830044I16Rik | 2.339863245 | 1.168529254 | 0.662227504 |
| Cdipt | 964.396228 | 0.094068329 | 0.04474883 |
| Mvp | 599.1504578 | 3.808562124 | 0.266496693 |
| Pagr1b | 0.194631057 | 0.237309881 | 3.243494927 |
| Pagr1a | 269.1231763 | -0.308822182 | 0.145942535 |
| Prrt2 | 12.36009762 | -0.998126644 | 0.40956498 |
| Maz | 1509.464906 | -0.241588876 | 0.073277742 |
| Kif22 | 26.64433543 | 0.914368792 | 0.258881236 |
| Gm25333 | 0 | NA | NA |
| AC122863.1 | 0 | NA | NA |
| Zg16 | 0.046345709 | 0.71820487 | 3.724945049 |
| Gm22747 | 0 | NA | NA |
| AI467606 | 37.52684094 | 2.081663787 | 0.259848521 |
| Qprt | 3.756627285 | 0.698453288 | 0.550685468 |
| Spn | 82.89016059 | 6.006618351 | 0.424679207 |

| | | | |
|---|---|---|---|
| Gm25735 | 0 | NA | NA |
| Cd2bp2 | 785.0302282 | 0.197898829 | 0.059801349 |
| Tbc1d10b | 882.4067378 | 0.319011271 | 0.067224371 |
| Mylpf | 4.837556069 | 0.709593047 | 0.487405708 |
| 1-Sep | 35.87249771 | 3.485717479 | 0.328837687 |
| Zfp553 | 281.7915856 | -0.080059368 | 0.082200042 |
| Zfp771 | 217.3863754 | 0.181692532 | 0.133390484 |
| Dctpp1 | 50.21803819 | 0.089861029 | 0.163390745 |
| Sephs2 | 242.8260877 | 0.205341817 | 0.104402029 |
| Gm17511 | 0.08486385 | 0.717551239 | 3.724898127 |
| Itgal | 262.0288419 | 4.241717073 | 0.218880539 |
| Zfp768 | 163.5362928 | 0.653400307 | 0.098558642 |
| Zfp747 | 52.03124678 | 0.353242573 | 0.154574735 |
| 9130019O22Rik | 50.28528677 | 0.17501381 | 0.170881655 |
| E430018J23Rik | 67.77893546 | 0.067389068 | 0.146806116 |
| Smt3h2-ps2 | 0 | NA | NA |
| Zfp764 | 37.42744171 | 0.330934456 | 0.184531969 |
| Zfp688 | 66.55993122 | 0.437586796 | 0.141806331 |
| Zfp689 | 64.00074658 | 0.231729561 | 0.139958895 |
| B130055M24Rik | 80.27483843 | -0.515536766 | 0.185468536 |
| Prr14 | 312.8086992 | 0.385456421 | 0.071916092 |
| Fbrs | 544.6972203 | 0.285914336 | 0.097787334 |
| Mir7060 | 0.754347455 | 0.472816923 | 1.032005551 |
| AC136146.1 | 0 | NA | NA |
| 1700008J07Rik | 27.76791627 | 0.273440656 | 0.207330255 |
| Srcap | 3720.548078 | -0.159622094 | 0.059267206 |
| Snora30 | 11.72211257 | -0.297552116 | 0.308634259 |
| Gm28198 | 0.198870357 | -0.243564353 | 2.667920877 |
| Tmem265 | 0 | NA | NA |
| Phkg2 | 467.0988482 | -0.099502072 | 0.072228599 |
| Gm166 | 30.45493143 | -0.32367279 | 0.198166557 |
| Rnf40 | 789.9745989 | 0.349500661 | 0.065773974 |
| 1700120K04Rik | 1.43115425 | -1.117061183 | 0.96700316 |
| Zfp629 | 457.8583134 | -0.125113408 | 0.064146476 |
| Bcl7c | 293.7258915 | 0.222134068 | 0.078948468 |
| Mir762 | 1.435335593 | -0.109531421 | 0.892901028 |
| Ctf1 | 11.31378371 | -0.007941424 | 0.30141867 |
| Ctf2 | 0 | NA | NA |
| Fbxl19 | 610.6258965 | -0.638212646 | 0.098095069 |
| Orai3 | 179.2087718 | 0.63396294 | 0.146962038 |
| Setd1a | 391.9559766 | -0.045102125 | 0.126497877 |
| Hsd3b7 | 112.3828733 | 1.085558557 | 0.139168823 |
| Stx1b | 6333.347927 | -0.359684389 | 0.074169935 |
| Stx4a | 301.4532526 | 0.536650998 | 0.143623402 |
| Zfp668 | 218.1486246 | -0.207988819 | 0.106720233 |
| Zfp646 | 586.462955 | 0.328176825 | 0.072728987 |
| Prss53 | 5.985399039 | -0.727610146 | 0.388684739 |

| | | | |
|---|---|---|---|
| Gm21974 | 0.049553047 | 0.71820487 | 3.724945049 |
| Vkorc1 | 34.89639478 | 0.133701905 | 0.251175371 |
| Bckdk | 313.1884233 | 0.081173696 | 0.086125521 |
| Kat8 | 146.4017955 | -0.239691711 | 0.098465789 |
| Prss8 | 2.192463596 | -1.520730657 | 0.740895033 |
| Prss36 | 34.47699352 | -0.341430427 | 0.183249761 |
| Fus | 2383.412294 | -0.111241367 | 0.118648679 |
| Gm26690 | 0.089089296 | 0.237305797 | 3.724945049 |
| B230325K18Rik | 116.590507 | -0.05548773 | 0.277503074 |
| Pycard | 48.5114067 | 5.144260037 | 0.425274067 |
| Gm15533 | 0 | NA | NA |
| Trim72 | 39.26137666 | 1.83870656 | 0.390640797 |
| Gm22809 | 0 | NA | NA |
| Itgam | 625.9394924 | 2.504762977 | 0.152537224 |
| Itgax | 181.6100497 | 4.75448293 | 0.372737544 |
| Itgad | 28.13752969 | 1.594742268 | 0.307637854 |
| Cox6a2 | 20.40538094 | 0.192960808 | 0.244877401 |
| 9130023H24Rik | 70.01739671 | 0.719210862 | 0.148581936 |
| Armc5 | 244.6664953 | 0.751710113 | 0.09765172 |
| Tgfb1i1 | 119.9274198 | 0.867812531 | 0.169655204 |
| Slc5a2 | 1.211082793 | 1.386185594 | 0.974916277 |
| BC017158 | 206.5939799 | -0.157224975 | 0.083469812 |
| Mir3103 | 0 | NA | NA |
| Rgs10 | 178.444261 | -0.387288035 | 0.127672535 |
| Gm15503 | 2.556904923 | -1.330096603 | 0.739485378 |
| Tial1 | 1499.062975 | -0.191500749 | 0.070574094 |
| Gm24365 | 0 | NA | NA |
| Bag3 | 352.4742549 | 1.706571152 | 0.157452703 |
| Gm7258 | 0 | NA | NA |
| Inpp5f | 1743.240792 | 0.016241573 | 0.073497917 |
| RP24-143L4.2 | 0 | NA | NA |
| Gm16044 | 15.72144794 | -1.042430794 | 0.388376594 |
| Mcmbp | 481.2577654 | 0.485029346 | 0.067743291 |
| Gm23783 | 0 | NA | NA |
| Sec23ip | 696.8616121 | 0.363948034 | 0.073489408 |
| n-R5s158 | 0 | NA | NA |
| Gm25778 | 0.04393175 | -0.243571806 | 3.724945049 |
| Ppapdc1a | 43.3421754 | -0.166384838 | 0.24294622 |
| Wdr11 | 633.8080906 | -0.117373868 | 0.06516229 |
| Gm23847 | 0 | NA | NA |
| Fgfr2 | 819.5441903 | 0.017976334 | 0.096335395 |
| Gm5903 | 24.16642656 | 0.412227772 | 0.220283187 |
| Ate1 | 1165.905196 | -0.051948005 | 0.057070677 |
| Gm4275 | 0.372589196 | 1.671517586 | 1.563096663 |
| Gm27884 | 0.4630892 | -0.650372594 | 1.565804053 |
| Nsmce4a | 299.1185009 | 0.486084948 | 0.116667483 |
| Tacc2 | 858.9102761 | -0.59355293 | 0.075563443 |

| | | | |
|---|---|---|---|
| Etos1 | 0 | NA | NA |
| Btbd16 | 23.45110209 | 1.962811042 | 0.272233355 |
| Gm5602 | 13.63298114 | -0.058965023 | 0.303300205 |
| Plekha1 | 1609.865415 | -0.044101159 | 0.071412826 |
| mmu-mir-7061 | 2.359287033 | 0.007479637 | 0.807969037 |
| Htra1 | 1343.622786 | 0.543355874 | 0.14795643 |
| Dmbt1 | 0.462783582 | 2.354853249 | 3.377870757 |
| 4933402N03Rik | 0.061920613 | 0.718207094 | 3.724945049 |
| 1700029B22Rik | 0 | NA | NA |
| 5430419D17Rik | 1.767964381 | -1.047317633 | 0.735693162 |
| Cuzd1 | 2.321753028 | 0.833736339 | 0.646128946 |
| 1700007K09Rik | 3.665048522 | 0.147548985 | 0.691676616 |
| Fam24a | 0 | NA | NA |
| 2310057M21Rik | 207.8650457 | 0.08147464 | 0.129403644 |
| Pstk | 158.2756739 | -0.550781993 | 0.11167169 |
| Ikzf5 | 441.795929 | 0.348157862 | 0.082236366 |
| Acadsb | 717.8726284 | -0.164620866 | 0.085566942 |
| Hmx3 | 4.229700681 | 1.684191679 | 0.661413226 |
| Hmx2 | 1.376538491 | -0.999164362 | 0.902983987 |
| Bub3 | 556.6642844 | 0.383904872 | 0.081501535 |
| Gm16477 | 27.01530151 | 0.541365752 | 0.221992421 |
| Gpr26 | 2155.295409 | -1.190407128 | 0.178032502 |
| Cpxm2 | 34.2720618 | 0.530976219 | 0.199371413 |
| n-R5s159 | 0.155551505 | 0.718215147 | 2.833462086 |
| Gm15677 | 0.494008362 | 0.657084482 | 1.513204397 |
| Chst15 | 526.7601878 | -0.257429678 | 0.089812094 |
| Oat | 670.5788141 | -0.053808407 | 0.097792392 |
| Nkx1-2 | 0.886311283 | -0.908222991 | 1.058729687 |
| Gm16764 | 0 | NA | NA |
| Lhpp | 114.0965916 | -0.3149882 | 0.123063475 |
| Gm15582 | 0.853586773 | 2.354025801 | 1.091397918 |
| Fam53b | 689.2067909 | -0.344628083 | 0.083927458 |
| Mettl10 | 199.6988259 | -0.129746458 | 0.095910785 |
| Fam175b | 351.9015338 | 0.355759922 | 0.072763267 |
| 1500002F19Rik | 8.219991273 | -0.310364024 | 0.359513266 |
| Zranb1 | 1066.949998 | -0.025667057 | 0.076847934 |
| Gm15718 | 1.397387052 | -0.101046562 | 0.968361606 |
| Ctbp2 | 328.6048763 | 0.004309259 | 0.089887938 |
| Tex36 | 0.64913551 | 0.40246491 | 1.181088923 |
| Gm28747 | 0.061920613 | 0.718207094 | 3.724945049 |
| Edrf1 | 475.154792 | -0.065116448 | 0.123555263 |
| Mmp21 | 0.96439918 | 0.029672919 | 1.103962167 |
| Uros | 150.8894574 | -0.240690286 | 0.113835837 |
| Bccip | 345.0185436 | 0.438697202 | 0.081266775 |
| Dhx32 | 391.5252559 | 0.004014057 | 0.074496883 |
| Gm15483 | 5.172158321 | -1.067222704 | 0.458244895 |
| Fank1 | 22.86474296 | -1.100239287 | 0.237990512 |

| | | | |
|---|---|---|---|
| Adam12 | 174.7221536 | 1.919325034 | 0.187878009 |
| Gm23631 | 0.152314784 | -0.724459891 | 2.847275348 |
| A130023I24Rik | 1.325949706 | 2.74985025 | 0.95466754 |
| D7Ertd443e | 78.15056778 | -0.335207401 | 0.193190403 |
| Dock1 | 717.8821439 | 0.372561012 | 0.078978218 |
| Fam196a | 171.3787744 | -0.136074672 | 0.123997356 |
| Gm24581 | 0 | NA | NA |
| 1700120G07Rik | 0.35282422 | 2.586627861 | 3.683635885 |
| Nps | 2.222986028 | 0.821022621 | 0.746265331 |
| 4930544L04Rik | 0.117650406 | 0.23732098 | 3.724945049 |
| Foxi2 | 0.120349074 | 0.237305146 | 3.724945049 |
| Clrn3 | 0 | NA | NA |
| Ptpre | 396.1280751 | -0.1865861 | 0.096108604 |
| Mir1962 | 1.121758522 | -0.63413116 | 0.970471175 |
| 5830432E09Rik | 5.577809048 | 3.858443494 | 0.721821438 |
| Mki67 | 177.6667962 | 2.672761982 | 0.256875918 |
| Gm9341 | 0 | NA | NA |
| Gm24444 | 0 | NA | NA |
| Mgmt | 8.081835135 | 0.482228289 | 0.343341972 |
| Ebf3 | 140.1280894 | -0.142842527 | 0.164666852 |
| 9430038I01Rik | 31.06055986 | 0.054921078 | 0.241596794 |
| Glrx3 | 666.4075028 | 0.338801976 | 0.105485211 |
| Gm25798 | 0 | NA | NA |
| Tcerg1l | 114.589252 | -0.169880776 | 0.133943986 |
| Mapk1ip1 | 278.2027142 | -0.483584142 | 0.110333678 |
| Ppp2r2d | 650.2346655 | 0.286513574 | 0.116239041 |
| Bnip3 | 859.8363811 | -0.128856291 | 0.086727166 |
| Gm9358 | 0.765055553 | -1.853607264 | 1.18195417 |
| Jakmip3 | 408.0351068 | -0.576153105 | 0.139983501 |
| Dpysl4 | 522.9033322 | -0.78721402 | 0.108294138 |
| Stk32c | 865.5787329 | -0.1819776 | 0.075752172 |
| Gm27975 | 0 | NA | NA |
| Lrrc27 | 79.21060963 | -0.393625541 | 0.142499909 |
| Pwwp2b | 187.2201883 | 0.316703066 | 0.103124529 |
| Inpp5a | 914.9916766 | -0.057080276 | 0.104796011 |
| Nkx6-2 | 248.995156 | 1.34501368 | 0.170922632 |
| Cfap46 | 121.7543508 | -0.864719826 | 0.16821777 |
| Gm4459 | 68.82999537 | 0.170019626 | 0.158016267 |
| Rpl21-ps13 | 0 | NA | NA |
| Mir6401 | 0 | NA | NA |
| Adgra1 | 707.4917962 | -0.515540346 | 0.071790134 |
| Kndc1 | 2147.099969 | -0.392517931 | 0.071533409 |
| Utf1 | 1.176348267 | 1.163015107 | 0.987417778 |
| Gm16201 | 8.156929821 | 0.572117889 | 0.388493108 |
| Gm2044 | 0.046345709 | 0.71820487 | 3.724945049 |
| mmu-mir-7686 | 0 | NA | NA |
| Mir202 | 0 | NA | NA |

| | | | |
|---|---|---|---|
| 6430531B16Rik | 3.025363873 | -0.837723623 | 0.656566603 |
| Adam8 | 325.8439531 | 4.528152252 | 0.490008071 |
| Mir7062 | 0 | NA | NA |
| Tubgcp2 | 416.699423 | -0.021868434 | 0.074092667 |
| Zfp511 | 95.06049784 | 0.301660537 | 0.121183198 |
| Msx3 | 0.762104533 | -1.27817872 | 1.156473896 |
| Caly | 767.6017605 | -0.844134766 | 0.074814428 |
| Prap1 | 0 | NA | NA |
| Fuom | 141.6570441 | -0.113077539 | 0.112504181 |
| Echs1 | 365.0393526 | 0.02686479 | 0.099570715 |
| Paox | 52.9831637 | 0.456833686 | 0.156479223 |
| Mtg1 | 103.7910862 | -0.259488432 | 0.125060849 |
| Sprn | 1168.69905 | -0.66170241 | 0.143719039 |
| Olfr522 | 0 | NA | NA |
| Olfr523 | 0.038981395 | 0.718207094 | 3.724945049 |
| Olfr524 | 0.042716945 | -0.243571806 | 3.724945049 |
| Cd163l1 | 0.514333851 | -0.998539639 | 3.230881657 |
| 5830411N06Rik | 0.560014729 | 0.596387923 | 1.284053797 |
| Olfr525 | 0.068097359 | -0.243569582 | 3.724945049 |
| Olfr527 | 0.1567526 | 0.265588935 | 2.825354309 |
| Olfr60 | 0.072025462 | 0.237312084 | 3.724945049 |
| Olfr530 | 0.095435502 | 0.718202646 | 3.724945049 |
| Olfr531 | 0.095898756 | 1.192083721 | 3.724434897 |
| Olfr532 | 0.042716945 | -0.243571806 | 3.724945049 |
| Olfr533 | 0.120974994 | -0.243571806 | 3.724945049 |
| Olfr535 | 0 | NA | NA |
| Olfr536 | 0.638507312 | 0.295077422 | 1.398538289 |
| Olfr538 | 0 | NA | NA |
| Olfr46 | 0.111376404 | -0.243571806 | 3.724945049 |
| Olfr61 | 0.224318018 | -1.772280945 | 2.570571478 |
| Olfr53 | 0.178643884 | -0.243569231 | 2.71272524 |
| Olfr539 | 0.207883488 | -1.200158497 | 2.193075849 |
| Olfr45 | 0.066050875 | 0.71820487 | 3.724945049 |
| Olfr541 | 0 | NA | NA |
| Cyp2e1 | 1.071126076 | 0.187064878 | 1.219797351 |
| Syce1 | 4.365775959 | 0.695153995 | 0.590925107 |
| Zfp941 | 465.5985334 | 0.240517716 | 0.103424817 |
| Urah | 8.499830305 | 0.981027963 | 0.391220911 |
| C330022C24Rik | 1.303324972 | 3.427804318 | 1.13566803 |
| Scgb1c1 | 1.058089029 | -0.689032823 | 0.981931574 |
| Odf3 | 0.418471408 | -1.564325709 | 1.624090349 |
| Bet1l | 215.3921604 | 0.467574611 | 0.097935165 |
| Ric8 | 517.4466835 | 0.393730887 | 0.07408708 |
| Sirt3 | 296.3129455 | -0.290414109 | 0.111696497 |
| Psmd13 | 560.5188265 | 0.586902811 | 0.097237656 |
| Gm15542 | 83.64766115 | 0.616451576 | 0.172301785 |
| Cox8b | 4.291398722 | -2.316182839 | 0.613069634 |

| | | | |
|---|---|---|---|
| Nlrp6 | 8.710158728 | -0.444915141 | 0.430441331 |
| Athl1 | 114.0507651 | 1.613054985 | 0.160474231 |
| Ifitm5 | 0.496272132 | 2.304524884 | 1.305766643 |
| Ifitm2 | 214.7202724 | 2.875901091 | 0.233123955 |
| Ifitm1 | 100.7816241 | 3.386660485 | 0.272570052 |
| Ifitm3 | 2374.688831 | 6.763345423 | 0.270573454 |
| Ifitm6 | 40.88379849 | 5.958944571 | 0.68560842 |
| Gm17387 | 0.276680467 | 2.073613567 | 1.752697307 |
| B4galnt4 | 713.1671909 | -0.174470675 | 0.108315387 |
| Pkp3 | 31.38673572 | 1.037357528 | 0.257464225 |
| Sigirr | 8.814058262 | 2.99391566 | 0.499348534 |
| Ano9 | 0.759896141 | 2.181403699 | 1.169979019 |
| Gm24962 | 0 | NA | NA |
| Ptdss2 | 397.5875048 | -0.157279683 | 0.081486632 |
| Rnh1 | 785.7956867 | 2.548907093 | 0.192688046 |
| Gm22019 | 0.061920613 | 0.718207094 | 3.724945049 |
| Hras | 652.7768577 | -0.23678036 | 0.081265416 |
| Lrrc56 | 44.78247708 | -0.189741229 | 0.214286629 |
| Lmntd2 | 1.504148252 | 0.498638235 | 0.804599154 |
| Rassf7 | 15.32468764 | 0.592386414 | 0.278428587 |
| Mir210 | 0.042716945 | -0.243571806 | 3.724945049 |
| Phrf1 | 827.2765508 | 0.215916422 | 0.055889899 |
| Irf7 | 4351.088714 | 10.39195662 | 0.317539063 |
| Cdhr5 | 0.766219994 | 2.77044641 | 1.109878495 |
| Sct | 31.79202062 | 6.329789491 | 0.818106522 |
| Drd4 | 5.233666748 | 3.72372992 | 0.761369315 |
| Deaf1 | 397.411066 | -0.466705454 | 0.082454713 |
| Tmem80 | 167.3778283 | -0.297918917 | 0.107476451 |
| Eps8l2 | 83.71994531 | 0.051013803 | 0.149783874 |
| B230206H07Rik | 17.64932275 | -1.093050375 | 0.376910545 |
| Gm25725 | 0 | NA | NA |
| Taldo1 | 760.0746265 | 0.573903882 | 0.114591035 |
| Gm4535 | 4.201928798 | 0.292975355 | 0.493668902 |
| Pddc1 | 201.5225707 | -0.309030723 | 0.110051957 |
| Cend1 | 1948.915175 | -0.388690712 | 0.105627343 |
| Slc25a22 | 2538.937697 | 0.559033677 | 0.125681017 |
| Pidd1 | 23.70766093 | 0.796915618 | 0.258710406 |
| Rplp2 | 1141.878145 | 0.723961648 | 0.093566954 |
| Snora52 | 0 | NA | NA |
| Pnpla2 | 482.1622727 | 1.591809143 | 0.195427546 |
| Cracr2b | 10.14241748 | -0.118451833 | 0.324387081 |
| Gm10575 | 13.15716283 | 0.777263821 | 0.294523336 |
| Cd151 | 360.834311 | 1.900186891 | 0.196062832 |
| Polr2l | 159.7689273 | 0.64723381 | 0.158075427 |
| Tspan4 | 266.6277742 | 1.464739059 | 0.220259764 |
| Chid1 | 475.4024328 | -0.289969095 | 0.119168272 |
| Ap2a2 | 2741.755715 | -0.07013383 | 0.057328429 |

| | | | |
|---|---|---|---|
| Gm16982 | 0.330744509 | 0.738518292 | 1.783519541 |
| Mir7063 | 2.38820333 | -0.36430119 | 0.674782065 |
| Muc6 | 0.236992836 | 1.54848761 | 2.06726318 |
| Muc2 | 8.969982396 | 2.146987165 | 0.5263699 |
| Muc5ac | 0.957274486 | 3.388319023 | 1.214776741 |
| Muc5b | 2.416483532 | -0.42698508 | 1.066048819 |
| Tollip | 1722.402167 | 0.11167406 | 0.063291846 |
| Gm20501 | 7.216052579 | -0.855816227 | 0.400294959 |
| Brsk2 | 1833.13049 | -0.66260734 | 0.086441911 |
| Mir3104 | 0.562632007 | -1.704970946 | 1.202484128 |
| Mob2 | 221.1798062 | 0.056619837 | 0.080340487 |
| Gm25416 | 0.290864751 | -0.270438753 | 2.045403122 |
| Dusp8 | 1691.653108 | -0.639932553 | 0.079435779 |
| Gm25370 | 0.286235171 | 0.23621144 | 1.594459476 |
| Gm4553 | 0 | NA | NA |
| Krtap5-2 | 0 | NA | NA |
| Gm10153 | 0 | NA | NA |
| Krtap5-3 | 0.115000406 | 0.23732098 | 3.724945049 |
| Gm7579 | 0 | NA | NA |
| Krtap5-5 | 0 | NA | NA |
| Gm2431 | 0.097139547 | 0.718207094 | 3.724945049 |
| Gm4559 | 0 | NA | NA |
| Gm10013 | 0 | NA | NA |
| Krtap5-1 | 0.044544648 | 0.237309859 | 3.724945049 |
| Krtap5-4 | 0 | NA | NA |
| Gm5054 | 0.042716945 | -0.243571806 | 3.724945049 |
| Ifitm10 | 182.7874866 | 0.521445826 | 0.116311889 |
| Ctsd | 5752.611281 | 1.332254583 | 0.170285696 |
| Gm26143 | 0.044544648 | 0.237309859 | 3.724945049 |
| Syt8 | 0.543884849 | 1.166560539 | 1.190786441 |
| Tnni2 | 2.708583151 | 3.061443742 | 0.852887684 |
| Lsp1 | 166.316609 | 3.287879426 | 0.191285076 |
| Prr33 | 0.642141737 | -1.672671873 | 1.171185982 |
| Tnnt3 | 1.402656864 | 0.721503383 | 0.908061052 |
| Mrpl23 | 104.8815584 | -0.054561402 | 0.121255165 |
| Nctc1 | 0 | NA | NA |
| H19 | 64.97075824 | 3.813064444 | 0.350066981 |
| Gm27483 | 0.184422046 | 1.199971906 | 2.680605796 |
| Mir675 | 0 | NA | NA |
| Gm27786 | 0.110595523 | 0.718200421 | 3.724945049 |
| Igf2 | 848.6181956 | -0.46079109 | 0.180219663 |
| Mir483 | 0 | NA | NA |
| Igf2os | 4.725164502 | 0.345364171 | 0.537672933 |
| Ins2 | 0.386429078 | 0.576385407 | 1.445930409 |
| Gm7290 | 0.08486385 | 0.717551239 | 3.724898127 |
| Gm6471 | 0 | NA | NA |
| Th | 127.7817552 | -0.093586995 | 0.260613658 |

| | | | |
|---|---|---|---|
| Ascl2 | 2.369287799 | 0.926056446 | 0.926755959 |
| Tspan32os | 6.17140379 | -0.318158605 | 0.411887292 |
| Tspan32 | 1.376815522 | 2.249900772 | 0.997973356 |
| R74862 | 40.83695903 | -0.81315058 | 0.203279526 |
| Cd81 | 3734.509123 | 0.179251864 | 0.106476601 |
| Trpm5 | 0.502574849 | 1.595765034 | 3.385761615 |
| Tssc4 | 336.1484664 | 0.582045799 | 0.08611929 |
| Kcnq1 | 3.259092159 | 0.841448948 | 0.557042959 |
| Gm38321 | 0 | NA | NA |
| Gm37235 | 0.046345709 | 0.71820487 | 3.724945049 |
| Gm28821 | 0 | NA | NA |
| Kcnq1ot1 | 4980.818381 | -0.716482183 | 0.250627929 |
| Gm27395 | 0.113999573 | 0.23732098 | 3.724945049 |
| Gm27389 | 0.179789776 | -1.515970277 | 2.234383842 |
| Gm27803 | 0 | NA | NA |
| Gm27539 | 0.072025462 | 0.237312084 | 3.724945049 |
| Gm37364 | 1.783218599 | 0.972541697 | 0.913297561 |
| 4933417O13Rik | 0.382744327 | -2.590557315 | 1.361539479 |
| Gm38095 | 0.038981395 | 0.718207094 | 3.724945049 |
| Gm38065 | 0 | NA | NA |
| Gm27901 | 0.136963818 | 0.718204821 | 3.59954433 |
| Cdkn1c | 70.29508147 | -0.414631699 | 0.150052956 |
| Slc22a18 | 7.798117052 | 1.52921912 | 0.495139325 |
| Phlda2 | 0.124308499 | 1.48237659 | 3.714822343 |
| Nap1l4 | 877.3075028 | -0.081435335 | 0.08337274 |
| Gm23297 | 5.403433764 | -1.000122043 | 0.522250173 |
| Cars | 570.4979151 | 0.944484373 | 0.113724876 |
| Mir7064 | 0.434125078 | -1.915944709 | 1.254300128 |
| Gm15580 | 0 | NA | NA |
| Gm15579 | 0.072025462 | 0.237312084 | 3.724945049 |
| Tnfrsf26 | 16.36698174 | 5.318592683 | 0.766122444 |
| Tnfrsf22 | 17.56781374 | 1.060036529 | 0.25817875 |
| Tnfrsf23 | 21.91112602 | 1.808446805 | 0.292442125 |
| Osbpl5 | 343.6236498 | 0.18944242 | 0.073548384 |
| Mrgprg | 0.045882455 | 0.237312084 | 3.724945049 |
| Mrgpre | 90.26427259 | 0.134255658 | 0.138866042 |
| Gm22064 | 0.123841226 | 1.292225235 | 3.719439735 |
| Nadsyn1 | 56.75566904 | 0.04799575 | 0.186155223 |
| Dhcr7 | 321.2309411 | -0.111264502 | 0.104064029 |
| Gm498 | 0.049553047 | 0.71820487 | 3.724945049 |
| Mir6404 | 0 | NA | NA |
| Shank2 | 2746.162572 | -0.677717791 | 0.11720204 |
| Mir3105 | 0.102891481 | -0.243576254 | 3.724945049 |
| Gm14372 | 0.097899064 | -0.724453471 | 3.724945049 |
| Cttn | 1215.2151 | 0.500275131 | 0.075303267 |
| Gm14376 | 6.097514919 | -0.815508748 | 0.603091558 |
| Ppfia1 | 744.1652023 | -0.053579263 | 0.082013923 |

| | | | |
|---|---|---|---|
| Faddos | 0.049553047 | 0.71820487 | 3.724945049 |
| Fadd | 59.54792491 | 1.236760323 | 0.179290772 |
| Ano1 | 32.22433977 | -0.845093199 | 0.23259625 |
| Gm10152 | 0 | NA | NA |
| Fgf3 | 4.236074083 | -0.544685742 | 0.540867701 |
| Fgf4 | 0 | NA | NA |
| Gm26793 | 3.941161089 | -0.666753402 | 0.557884975 |
| Fgf15 | 0.844302062 | 0.880989414 | 1.281713939 |
| Oraov1 | 196.1845682 | 0.097941567 | 0.079283033 |
| Ccnd1 | 896.4298962 | 2.394458392 | 0.243404199 |
| Tpcn2 | 87.43800306 | 0.793668221 | 0.19736106 |
| Mrgprf | 52.34489822 | -0.052555567 | 0.252887764 |
| Mrgprd | 0.203212381 | 0.677280635 | 2.635685699 |
| Gm7346 | 0 | NA | NA |
| Gm25775 | 0 | NA | NA |
| Insr | 1190.786105 | -0.141710118 | 0.056356205 |
| Gm16180 | 0 | NA | NA |
| A430078G23Rik | 4.533747209 | 3.139208891 | 0.795370325 |
| Arhgef18 | 750.8432807 | -0.057720245 | 0.072417088 |
| Pex11g | 11.94680139 | 0.772972764 | 0.380158293 |
| 1700019B03Rik | 3.816282625 | 1.086878849 | 0.637046842 |
| Zfp358 | 245.1757097 | 0.275074632 | 0.112021778 |
| Mcoln1 | 510.2888465 | -0.19242888 | 0.058382928 |
| Pnpla6 | 618.1933405 | -0.293639065 | 0.091080942 |
| C330021F23Rik | 0.687005404 | -2.463217544 | 1.163298932 |
| Rps23-ps2 | 34.30525878 | 0.743885749 | 0.191526115 |
| Camsap3 | 321.1724317 | -0.477111505 | 0.096654121 |
| Xab2 | 460.0137195 | 0.319284924 | 0.070420524 |
| Gm22204 | 0.066050875 | 0.71820487 | 3.724945049 |
| Pet100 | 155.4280518 | 0.038782256 | 0.113981748 |
| Pcp2 | 337.4770539 | -0.053320532 | 0.413316133 |
| Stxbp2 | 289.3398245 | 0.146841987 | 0.141345956 |
| Retn | 3.230460487 | 1.433106165 | 0.630294774 |
| Mcemp1 | 19.01625202 | 4.425920507 | 0.626775636 |
| Fcor | 19.48329625 | -0.436611898 | 0.310646896 |
| Trappc5 | 363.4451168 | 0.352108255 | 0.087512499 |
| Fcer2a | 1.327801032 | -0.811916441 | 0.959326092 |
| Clec4g | 0.624219342 | 1.02262958 | 1.295815448 |
| Gm16553 | 0.238000031 | -0.243571794 | 2.485699892 |
| Gm23886 | 0 | NA | NA |
| Cd209a | 1.799118211 | 1.426690248 | 0.7432651 |
| Gm24934 | 0 | NA | NA |
| Rprl3 | 2178.394221 | 0.448531684 | 0.183328763 |
| Gm7381 | 0.061920613 | 0.718207094 | 3.724945049 |
| Cd209e | 0 | NA | NA |
| Cd209d | 0.062426873 | 0.237312084 | 3.724945049 |
| Cd209b | 0.371584567 | -2.531955487 | 1.596794342 |

| | | | |
|---|---|---|---|
| Cd209c | 0.190986542 | 0.246832495 | 2.702582407 |
| Gm7389 | 8.537660985 | 0.723757851 | 0.40150309 |
| Gm16589 | 85.35553407 | 1.866387046 | 0.243712391 |
| Gm26750 | 0 | NA | NA |
| Cd209f | 0.631752858 | -1.739210279 | 1.173786562 |
| Gm23888 | 0 | NA | NA |
| Cd209g | 0.34920187 | -0.745629295 | 1.663145215 |
| 4932443L11Rik | 1.299081245 | 0.404284519 | 0.937402738 |
| Evi5l | 1072.773425 | -0.527869776 | 0.075139212 |
| Prr36 | 298.1056952 | -1.258002397 | 0.123280885 |
| Lrrc8e | 1.436219828 | 2.394761387 | 0.970368406 |
| Map2k7 | 1090.223813 | -0.173392428 | 0.058949113 |
| Tgfbr3l | 14.69813157 | -1.313334931 | 0.4184727 |
| Snapc2 | 202.0178337 | 0.610966601 | 0.118429415 |
| Ctxn1 | 1372.549575 | -1.441113019 | 0.124313654 |
| Timm44 | 376.9137615 | -0.240187437 | 0.103528831 |
| Gm26102 | 0 | NA | NA |
| Elavl1 | 1614.161581 | 0.045701822 | 0.053110214 |
| Ccl25 | 51.47610217 | -0.674284412 | 0.196669382 |
| Cers4 | 639.9032004 | -0.50000216 | 0.124379131 |
| Gm36956 | 0.066050875 | 0.71820487 | 3.724945049 |
| Zfp958 | 156.442319 | 0.086843542 | 0.113468062 |
| Gm7461 | 2.204253537 | -0.033123313 | 0.684724497 |
| Gm6410 | 0.983694817 | -0.768174885 | 1.142594294 |
| Shcbp1 | 11.59037295 | 1.039940754 | 0.455353093 |
| Gm9457 | 0.283539495 | 1.209123794 | 2.398473448 |
| Rpl21-ps14 | 0.417309641 | -0.373290269 | 1.741672399 |
| n-R5s93 | 0 | NA | NA |
| Slc10a2 | 0 | NA | NA |
| n-R5s94 | 0 | NA | NA |
| Gm1840 | 36.14079938 | 0.20352669 | 0.174007828 |
| Gm22565 | 0 | NA | NA |
| Gm24288 | 0 | NA | NA |
| Gm17215 | 0.061920613 | 0.718207094 | 3.724945049 |
| Gm26427 | 0.673557475 | -0.217851052 | 1.245604616 |
| Gm27585 | 0 | NA | NA |
| Gm27545 | 0 | NA | NA |
| Gm25169 | 0 | NA | NA |
| Gm28262 | 0 | NA | NA |
| Efnb2 | 602.2040592 | -0.602241324 | 0.073621652 |
| 4921522P10Rik | 0.133946075 | 0.7191812 | 3.626570343 |
| Arglu1 | 500.6182152 | -0.294954883 | 0.070404158 |
| Mir7654 | 0 | NA | NA |
| Fam155a | 942.9312957 | -0.637609507 | 0.068269382 |
| n-R5s95 | 14.75670795 | -1.142918156 | 0.537819375 |
| AC099760.1 | 0 | NA | NA |
| Gm10217 | 0.045882455 | 0.237312084 | 3.724945049 |

| | | | |
|---|---|---|---|
| Gm10067 | 0 | NA | NA |
| Lig4 | 157.1064013 | -0.165205899 | 0.134592358 |
| Abhd13 | 722.6949606 | 0.146578252 | 0.084257348 |
| Tnfsf13b | 20.67774298 | 1.051894944 | 0.254540565 |
| Myo16 | 882.108668 | -0.021518723 | 0.112539718 |
| Rps16-ps3 | 0 | NA | NA |
| 3930402G23Rik | 10.57984639 | 3.740579719 | 0.541272312 |
| Irs2 | 1258.437026 | 0.289990043 | 0.130586466 |
| 9530052E02Rik | 11.90754286 | -0.72522328 | 0.35821641 |
| Gm15418 | 2.963241567 | -1.897034484 | 0.764455916 |
| Col4a1 | 1092.321152 | 2.080917413 | 0.16381171 |
| Col4a2 | 772.5542793 | 1.119312049 | 0.159601639 |
| Gm15419 | 0.932741312 | 0.095711872 | 0.958537042 |
| Rab20 | 71.74611719 | 5.559898365 | 0.443831046 |
| E230013L22Rik | 6.9128239 | 2.345358981 | 0.493383709 |
| Carkd | 279.2439548 | 0.010815297 | 0.071750743 |
| Cars2 | 87.5486135 | -0.098891638 | 0.126665228 |
| Ing1 | 228.1766961 | -0.323406516 | 0.102282912 |
| Ankrd10 | 320.9466642 | -0.281711489 | 0.113990476 |
| mmu-mir-7238 | 0 | NA | NA |
| 1700016D06Rik | 0.194626871 | 0.237320667 | 2.660960796 |
| Gm27854 | 0 | NA | NA |
| 1700128E19Rik | 0.068097359 | -0.243569582 | 3.724945049 |
| Arhgef7 | 1845.245703 | -0.22270276 | 0.067263458 |
| Gm15875 | 0 | NA | NA |
| Tex29 | 5.082148535 | -1.080961807 | 0.483464012 |
| A230072I06Rik | 0.719797104 | -2.181245958 | 1.263884822 |
| Gm29175 | 0.569306118 | -0.356748877 | 1.508953055 |
| Gm22207 | 0.197462999 | -1.54767071 | 2.627104102 |
| Gm5607 | 118.2838747 | -1.106419477 | 0.137945965 |
| Sox1 | 235.5645892 | -0.599535511 | 0.156931655 |
| Gm25239 | 0 | NA | NA |
| D630011A20Rik | 0 | NA | NA |
| Spaca7 | 0 | NA | NA |
| Tubgcp3 | 535.0497003 | -0.15688056 | 0.08330828 |
| Gm26483 | 0 | NA | NA |
| 4931415C17Rik | 18.23396489 | -0.241135367 | 0.284039672 |
| Gm15348 | 0.194797293 | -1.268452491 | 3.207102014 |
| Atp11a | 1406.61443 | 0.144636316 | 0.071391946 |
| Gm15350 | 0.251121349 | 1.156407898 | 2.061297004 |
| Gm15347 | 0.159763836 | -1.214267593 | 2.803296028 |
| Mcf2l | 2158.382309 | -0.548690126 | 0.071692539 |
| Gm15353 | 0.04393175 | -0.243571806 | 3.724945049 |
| Gm15351 | 6.732411598 | -1.253870345 | 0.461613873 |
| mmu-mir-9768 | 0 | NA | NA |
| Gm15349 | 0 | NA | NA |
| Gm15352 | 1.728865685 | -0.248297943 | 0.97174054 |

| | | | |
|---|---|---|---|
| F7 | 20.47438508 | 5.265602374 | 0.583666883 |
| F10 | 60.6296221 | 7.839875113 | 0.898712886 |
| Proz | 0.405129103 | 1.348871667 | 1.527819745 |
| Gm17023 | 0.048949532 | -0.243569582 | 3.724945049 |
| Pcid2 | 338.1239794 | 0.002074696 | 0.093218406 |
| Gm17022 | 0 | NA | NA |
| Cul4a | 779.7913125 | 0.076454421 | 0.053448935 |
| Mir7065 | 0.26402779 | -1.475253656 | 3.700112569 |
| Lamp1 | 3957.92154 | 0.890237572 | 0.12442009 |
| Grtp1 | 32.55675301 | 0.192948841 | 0.218802804 |
| Mir1968 | 0.779555773 | -1.112054524 | 1.124158656 |
| 2810030D12Rik | 0.867078918 | -2.171537189 | 1.127672182 |
| Adprhl1 | 5.011785424 | -0.1520212 | 0.509557576 |
| Dcun1d2 | 379.2848758 | -0.415979679 | 0.096080611 |
| Gm25014 | 4.915973584 | -0.824978218 | 0.445371743 |
| Tmco3 | 370.903991 | -0.321283099 | 0.091564939 |
| Tfdp1 | 412.9010695 | 0.14338666 | 0.084905879 |
| Atp4b | 0 | NA | NA |
| Grk1 | 1.091106207 | 1.774209176 | 1.298954394 |
| Tmem255b | 12.33555406 | -0.098840152 | 0.336499455 |
| Gas6 | 898.867002 | -0.451431634 | 0.078530532 |
| 1700029H14Rik | 5.973066547 | -0.237569954 | 0.622230655 |
| Rasa3 | 1272.496422 | 0.306014906 | 0.075236445 |
| 4932443I19Rik | 8.590972356 | 0.246746906 | 0.351737858 |
| Cdc16 | 437.6639685 | 0.323579067 | 0.081841242 |
| Upf3a | 175.6020734 | 0.283088617 | 0.167247425 |
| AF366264 | 0.110595523 | 0.718200421 | 3.724945049 |
| Champ1 | 447.8929496 | 0.107590911 | 0.068563785 |
| AC139186.1 | 0 | NA | NA |
| Gm24698 | 0.093141081 | -0.724455696 | 3.724945049 |
| Coprs | 191.5374814 | 0.066440045 | 0.120405796 |
| Gm7676 | 0.12936564 | -1.064030098 | 3.677739829 |
| Fbxo25 | 406.2625769 | -0.361474356 | 0.071403239 |
| Tdrp | 156.0986126 | 0.006186043 | 0.110590737 |
| Erich1 | 48.89166254 | 0.49054585 | 0.157057209 |
| Gm10699 | 61.82645713 | -0.712423641 | 0.176793623 |
| Dlgap2 | 896.9164479 | -0.613231181 | 0.183232298 |
| Gm3160 | 8.920971897 | -2.274322473 | 0.576695145 |
| Gm26184 | 0 | NA | NA |
| AC130213.1 | 3.509323303 | -2.825182656 | 0.69077179 |
| C030037F17Rik | 0 | NA | NA |
| Cln8 | 468.4529787 | 0.495085137 | 0.097635886 |
| Arhgef10 | 845.0217701 | 0.374299162 | 0.111335576 |
| Gm16350 | 0.676711058 | 1.963372754 | 1.353579255 |
| Kbtbd11 | 1707.888606 | -0.194327841 | 0.093378623 |
| 5830468F06Rik | 0.645565975 | 3.319878634 | 1.323274198 |
| Gm37844 | 22.02823964 | -0.452031203 | 0.23124043 |

| | | | |
|---|---|---|---|
| BB014433 | 0 | NA | NA |
| Myom2 | 7.622785529 | 0.738805853 | 0.415762637 |
| Gm15678 | 0 | NA | NA |
| Gm24231 | 0 | NA | NA |
| Gm24222 | 0 | NA | NA |
| Gm25871 | 1.401691129 | -1.094946114 | 0.987712475 |
| Gm25255 | 0 | NA | NA |
| Csmd1 | 2110.968194 | -0.854865851 | 0.239613656 |
| Mir3106 | 1.89102523 | -1.458626182 | 0.933713255 |
| Gm16346 | 0 | NA | NA |
| Gm16347 | 0 | NA | NA |
| Gm25278 | 0 | NA | NA |
| Gm25665 | 0 | NA | NA |
| Gm24469 | 0 | NA | NA |
| Mcph1 | 236.5966961 | -0.385884798 | 0.100722854 |
| Gm25763 | 0 | NA | NA |
| Gm16725 | 0 | NA | NA |
| Gm17149 | 0.323327586 | -0.724461993 | 2.019867436 |
| Angpt2 | 48.84955761 | 2.431074523 | 0.310316689 |
| Agpat5 | 782.015753 | -0.491686763 | 0.069093242 |
| Xkr5 | 3.737801717 | 1.648983143 | 0.802777483 |
| Gm22551 | 0.097139547 | 0.718207094 | 3.724945049 |
| Defb40 | 0 | NA | NA |
| Defb37 | 0 | NA | NA |
| Defb38 | 0 | NA | NA |
| Gm15366 | 0.11193333 | 0.718202646 | 3.724945049 |
| Defb39 | 0 | NA | NA |
| Defb12 | 0 | NA | NA |
| Defb34 | 0 | NA | NA |
| Spag11b | 0.25135723 | -0.712126095 | 2.124018352 |
| Spag11a | 0.093744596 | 0.237318756 | 3.724945049 |
| Defb14 | 0 | NA | NA |
| Defb4 | 0 | NA | NA |
| Defa-ps12 | 0 | NA | NA |
| Defb6 | 0.199888145 | -0.243569408 | 2.628735217 |
| Defb46 | 0 | NA | NA |
| Defb5 | 0 | NA | NA |
| Gm15357 | 0 | NA | NA |
| Defb3 | 0 | NA | NA |
| Defa-ps13 | 0 | NA | NA |
| Defa-ps14 | 0 | NA | NA |
| Defb8 | 0 | NA | NA |
| Defa-ps15 | 0 | NA | NA |
| Rpl19-ps11 | 69.27011738 | 0.234708686 | 0.219579557 |
| Gm27288 | 0 | NA | NA |
| Defb7 | 0 | NA | NA |
| Gm15358 | 0 | NA | NA |

| | | | |
|---|---|---|---|
| Gm27852 | 0 | NA | NA |
| Gm16425 | 10.9074589 | 0.291247211 | 0.334419895 |
| Gm15359 | 1.904847302 | 1.241680527 | 0.737592122 |
| Gm6483 | 149.5723312 | -0.119499233 | 0.125367401 |
| 4930467E23Rik | 2.550781979 | 0.823332679 | 0.656441924 |
| Gm26853 | 52.27869009 | -0.097478616 | 0.228832144 |
| Gm21769 | 152.8336227 | -0.296378392 | 0.167650197 |
| Gm21811 | 150.012751 | -0.033520413 | 0.215750609 |
| 6820431F20Rik | 191.9164182 | -0.042902669 | 0.119553401 |
| Gm26804 | 0 | NA | NA |
| Gm15319 | 17.21563751 | 0.874231796 | 0.360433168 |
| 2610005L07Rik | 198.0277139 | -0.181751073 | 0.213277639 |
| Gm21092 | 212.4801284 | -0.086760532 | 0.099514046 |
| Gm21182 | 0.374378724 | -0.161653154 | 1.69641367 |
| Gm21119 | 1.839005393 | -0.39762893 | 0.819934225 |
| Gm20946 | 1.715564851 | -0.242585232 | 0.779054808 |
| Gm20797 | 0.04393175 | -0.243571806 | 3.724945049 |
| Gm20945 | 0.044191549 | -0.243571806 | 3.724945049 |
| Gm20796 | 3.51668492 | 1.539647745 | 0.677883474 |
| Defb33 | 0.682587184 | 3.35709861 | 1.388978833 |
| Gm15056 | 287.1653067 | 11.85469852 | 1.090526932 |
| Gm6040 | 1.098746849 | 3.034924373 | 1.132416824 |
| AY761185 | 1.275844484 | 3.756485838 | 1.072484761 |
| Defa21 | 0 | NA | NA |
| Gm23142 | 0 | NA | NA |
| Defa23 | 0 | NA | NA |
| Gm10104 | 0 | NA | NA |
| Defa-ps3 | 0 | NA | NA |
| Defa25 | 0 | NA | NA |
| Gm6689 | 0 | NA | NA |
| Gm14851 | 0 | NA | NA |
| Gm24568 | 0 | NA | NA |
| Gm2869 | 0.109727584 | 0.718200421 | 3.724945049 |
| Defa-ps4 | 0 | NA | NA |
| Gm15284 | 0.075043205 | 0.237309859 | 3.724945049 |
| Defa22 | 0 | NA | NA |
| Gm26240 | 0 | NA | NA |
| Defa-rs7 | 0 | NA | NA |
| Gm15293 | 0 | NA | NA |
| Defa-ps5 | 0 | NA | NA |
| Defa-ps6 | 0 | NA | NA |
| Defa-ps7 | 0 | NA | NA |
| Gm15303 | 0 | NA | NA |
| Gm15292 | 0 | NA | NA |
| Defa-ps8 | 0 | NA | NA |
| Defa3 | 0 | NA | NA |
| Defa5 | 0 | NA | NA |

| | | | |
|---|---|---|---|
| Gm15300 | 0 | NA | NA |
| Gm15299 | 0 | NA | NA |
| Defa-rs1 | 0 | NA | NA |
| Gm23561 | 0 | NA | NA |
| Gm15302 | 0 | NA | NA |
| Defa-ps16 | 0 | NA | NA |
| Defa2 | 0 | NA | NA |
| Gm21002 | 0 | NA | NA |
| Gm21498 | 0 | NA | NA |
| Gm15305 | 0 | NA | NA |
| Defa-ps18 | 0 | NA | NA |
| Gm15309 | 0 | NA | NA |
| Gm7849 | 0 | NA | NA |
| Gm23568 | 0 | NA | NA |
| Gm15304 | 0 | NA | NA |
| Defa-ps17 | 0 | NA | NA |
| Defa20 | 0 | NA | NA |
| Gm15308 | 0 | NA | NA |
| Gm24423 | 0 | NA | NA |
| Gm14850 | 0 | NA | NA |
| Gm15311 | 0 | NA | NA |
| Gm6696 | 0 | NA | NA |
| Gm15312 | 0 | NA | NA |
| Gm7861 | 0 | NA | NA |
| Gm24829 | 0 | NA | NA |
| Gm15314 | 0 | NA | NA |
| Defa26 | 0 | NA | NA |
| Gm15313 | 0 | NA | NA |
| Defa17 | 0 | NA | NA |
| Gm15315 | 0 | NA | NA |
| Defa-ps9 | 0 | NA | NA |
| Defa-ps1 | 0 | NA | NA |
| AY761184 | 0.101408268 | 0.717551239 | 3.724898127 |
| Gm22447 | 0 | NA | NA |
| Gm7869 | 0.109869311 | 0.238305784 | 3.724874707 |
| Defa-ps10 | 0 | NA | NA |
| Defa24 | 0.046345709 | 0.71820487 | 3.724945049 |
| Defa-ps11 | 0 | NA | NA |
| Defb1 | 2.378549416 | 0.349546257 | 0.695691851 |
| Gm15316 | 0.049553047 | 0.71820487 | 3.724945049 |
| Defb50 | 0 | NA | NA |
| Defb2 | 0.062426873 | 0.237312084 | 3.724945049 |
| Defb10 | 0.392953962 | -0.663215728 | 1.513404692 |
| Defb9 | 0.27226464 | -0.607186614 | 2.040937322 |
| Defb11 | 0.092881282 | -0.724455696 | 3.724945049 |
| Defb15 | 0 | NA | NA |
| Defb35 | 0 | NA | NA |

| | | | |
|---|---|---|---|
| Defb13 | 0 | NA | NA |
| C86695 | 0 | NA | NA |
| Ccdc70 | 0 | NA | NA |
| Atp7b | 61.90632721 | -0.115126312 | 0.221086892 |
| Alg11 | 609.1938378 | 0.254606869 | 0.076669594 |
| Nek5 | 10.77738955 | 0.786072073 | 0.333623333 |
| Nek3 | 31.0393675 | -0.019200594 | 0.262302865 |
| Ckap2 | 17.47088672 | 1.801501912 | 0.521359472 |
| Vps36 | 310.0223612 | 0.139887011 | 0.088876927 |
| Thsd1 | 30.08606779 | 0.447987022 | 0.262086121 |
| Rpl30-ps2 | 0 | NA | NA |
| Tpte | 0.493569458 | -1.316623647 | 1.370745271 |
| Slc25a15 | 143.2256164 | -0.098787443 | 0.116519154 |
| 1810012K16Rik | 2.758229591 | 0.171491766 | 0.586269907 |
| Mrps31 | 170.4153543 | 0.228687258 | 0.088743446 |
| Smim19 | 208.6627446 | -0.025583195 | 0.104443642 |
| Slc20a2 | 881.8761026 | 0.624110873 | 0.10919921 |
| Gm17491 | 71.03296188 | -0.484820445 | 0.25565894 |
| Vdac3 | 786.9911644 | 0.180681339 | 0.0869809 |
| Dkk4 | 0.121388914 | 0.718200421 | 3.724945049 |
| Polb | 409.3894603 | -0.029053878 | 0.072258118 |
| Gm26909 | 0.178821284 | 0.701618166 | 2.290359596 |
| Ikbkb | 547.644006 | 0.067145281 | 0.096896544 |
| Gm15346 | 0 | NA | NA |
| Plat | 1098.82342 | 2.335441298 | 0.18488249 |
| Ap3m2 | 1163.088985 | -0.174250793 | 0.077411764 |
| 1700041G16Rik | 3.442441086 | 2.110785482 | 0.606538491 |
| Gm24735 | 0.372097677 | 0.916192619 | 1.498075668 |
| Kat6a | 2202.941112 | 0.409225969 | 0.07068254 |
| Ank1 | 2360.60079 | -0.474204827 | 0.091416833 |
| AC126445.1 | 0 | NA | NA |
| Gm15815 | 3.535656746 | -2.08904113 | 0.690343098 |
| Gm15816 | 4.564354496 | -0.237730815 | 0.49754254 |
| Mir486 | 1.840863337 | -1.511215317 | 0.782751976 |
| mmu-mir-486b | 0 | NA | NA |
| Nkx6-3 | 1.853806923 | 0.545450693 | 0.814013654 |
| Agpat6 | 875.7847179 | 0.080755816 | 0.064384071 |
| B230112G18Rik | 0.914342362 | -0.964454464 | 1.037382006 |
| Gins4 | 158.5957338 | -0.013987839 | 0.093657799 |
| Golga7 | 765.1713379 | 0.297747201 | 0.104220501 |
| Gm24335 | 0 | NA | NA |
| Sfrp1 | 332.0038061 | 1.299278889 | 0.189337444 |
| Zmat4 | 554.6012764 | -0.427823598 | 0.099109129 |
| A730045E13Rik | 1.027651671 | -0.802780973 | 1.213588145 |
| Gm26714 | 0.777682211 | 0.210903012 | 1.40465448 |
| 1810011O10Rik | 28.96321082 | 1.131659366 | 0.30483199 |
| Ido2 | 21.23577249 | 0.512376701 | 0.244566847 |

| | | | |
|---|---|---|---|
| Ido1 | 30.3065116 | 0.598695135 | 0.251595435 |
| Adam18 | 1.533690469 | 2.30533126 | 0.958339154 |
| Adam3 | 2.60343246 | 1.259481232 | 0.644171884 |
| Gm26208 | 0 | NA | NA |
| Adam5 | 3.408984752 | 0.821553336 | 0.576207803 |
| Adam32 | 15.27834067 | 0.264536091 | 0.283403289 |
| Gm23898 | 0 | NA | NA |
| Adam9 | 978.4277452 | 0.865038625 | 0.088401013 |
| Tm2d2 | 316.9476182 | -0.001499196 | 0.079359557 |
| Htra4 | 16.56292703 | 2.403618817 | 0.490807643 |
| Gm16933 | 1.72427518 | 0.776747548 | 0.774172416 |
| Plekha2 | 504.2642388 | 2.004861413 | 0.110686486 |
| Tacc1 | 2873.630983 | -0.433798755 | 0.08954131 |
| Mir8108 | 0 | NA | NA |
| Gm10689 | 1.990043323 | 0.268403581 | 0.777489281 |
| Fgfr1 | 1712.090942 | 0.862011665 | 0.111612157 |
| Gm16159 | 0.092228164 | 0.718202646 | 3.724945049 |
| Letm2 | 106.132572 | -0.427075818 | 0.142342419 |
| D830025C05Rik | 3.021300136 | -0.298202081 | 0.550583524 |
| Whsc1l1 | 2360.967649 | 0.24974006 | 0.080250446 |
| Gm23184 | 0 | NA | NA |
| Gm24992 | 3.122516278 | 0.148584804 | 0.554992497 |
| Gm23094 | 7.166830998 | -0.417614889 | 0.594074354 |
| Ppapdc1b | 162.0488265 | -0.313492979 | 0.131795367 |
| Ddhd2 | 814.1538713 | -0.592682718 | 0.079269567 |
| Gm17484 | 32.71518712 | 0.133631292 | 0.286474332 |
| Bag4 | 936.8776659 | -0.473075937 | 0.064503176 |
| Lsm1 | 143.0297487 | 0.179295054 | 0.117846035 |
| Star | 22.62371583 | 0.244435685 | 0.286269565 |
| Ash2l | 496.3350927 | 0.0051912 | 0.060221363 |
| Kcnu1 | 15.88801623 | -0.600368819 | 0.275245595 |
| Hgsnat | 490.0921398 | -0.390838777 | 0.0718347 |
| Pomk | 265.4581434 | -0.347461045 | 0.080507401 |
| Fnta | 475.5451638 | 0.035082978 | 0.069897948 |
| Hook3 | 2198.855902 | -0.332877392 | 0.041995245 |
| 1700047A11Rik | 0.066050875 | 0.71820487 | 3.724945049 |
| Rnf170 | 300.8605897 | -0.552811044 | 0.102544665 |
| Thap1 | 73.51708713 | 0.306678272 | 0.127910238 |
| AC154102.1 | 0 | NA | NA |
| Gm22227 | 0 | NA | NA |
| 2310008N11Rik | 0.04393175 | -0.243571806 | 3.724945049 |
| Zfp703 | 588.8755344 | 0.688470093 | 0.09049168 |
| Erlin2 | 723.3406109 | 0.315022827 | 0.081241832 |
| Proscos | 5.115524656 | 0.392215562 | 0.465366708 |
| Prosc | 626.0658266 | 0.13337715 | 0.065296098 |
| Adgra2 | 197.3133489 | 0.601382639 | 0.123207431 |
| Brf2 | 100.3354801 | 0.228018131 | 0.136835046 |

| | | | |
|---|---|---|---|
| Rab11fip1 | 53.92462846 | 1.123862665 | 0.213950761 |
| 4933416M07Rik | 0.124596252 | 0.718200615 | 3.714681068 |
| Got1l1 | 2.891827676 | -0.228072499 | 0.620849311 |
| Adrb3 | 6.872911924 | -1.013399853 | 0.433949214 |
| Eif4ebp1 | 90.4572564 | 4.020587634 | 0.275574783 |
| Tex24 | 0.451016279 | 1.795798069 | 1.491789578 |
| Chrnb3 | 25.45046697 | -0.974659741 | 0.238484075 |
| Chrna6 | 16.6337043 | -1.138647758 | 0.337977611 |
| Gm24410 | 0 | NA | NA |
| Poteg | 0 | NA | NA |
| Gm1698 | 3.756750159 | 0.252851998 | 0.519441839 |
| AC111034.1 | 0 | NA | NA |
| Gm24202 | 0 | NA | NA |
| Gm8100 | 1.189584463 | -0.589058466 | 0.924382888 |
| Gm26795 | 1.493046057 | 2.06022592 | 1.013528473 |
| Unc5d | 798.2940023 | -0.104683415 | 0.077750188 |
| 4933433F19Rik | 0 | NA | NA |
| Dusp26 | 500.6484086 | 0.558654006 | 0.109003672 |
| Rnf122 | 133.8921804 | 0.333333005 | 0.134197068 |
| Snord13 | 7.690137191 | 0.885393099 | 0.493414679 |
| Tti2 | 237.0865655 | 0.162022153 | 0.099550584 |
| Mak16 | 240.9495093 | 0.462016706 | 0.095941827 |
| Fut10 | 112.6731054 | 1.05366385 | 0.135436128 |
| Gm8051 | 0 | NA | NA |
| Gm23539 | 0.319751656 | 0.800037581 | 1.799538669 |
| Gm24184 | 0.141071297 | 0.237322348 | 3.564043836 |
| Gm5117 | 3.857935733 | 0.99791781 | 0.515609913 |
| Nrg1 | 264.3991212 | 0.71812641 | 0.14440629 |
| Gm26578 | 5.176536363 | 0.385680259 | 0.450963642 |
| Gm22140 | 0 | NA | NA |
| Gm5908 | 9.444744683 | 0.402377474 | 0.354897966 |
| Gm3985 | 0.861087095 | -0.961019023 | 1.217100659 |
| Gm6877 | 13.36504497 | 0.610313826 | 0.316645598 |
| Wrn | 282.207455 | 0.176011976 | 0.108567462 |
| Purg | 493.6010051 | -0.836348368 | 0.089750741 |
| 5930422O12Rik | 8.011746571 | -1.03958003 | 0.389413352 |
| Tex15 | 53.01275634 | 0.29588224 | 0.245172329 |
| Ppp2cb | 866.8510396 | 0.07496172 | 0.083197301 |
| Ubxn8 | 210.3585288 | 0.223503328 | 0.085054027 |
| Gsr | 438.4005393 | 0.938830212 | 0.088296059 |
| Gm24727 | 28.80332305 | 0.405830421 | 0.330036683 |
| 1700104B16Rik | 0.424137699 | 0.064764228 | 1.566833426 |
| Gtf2e2 | 144.048718 | 0.053264314 | 0.099202309 |
| Smim18 | 19.2016788 | -0.702967562 | 0.272745565 |
| Rbpms | 30.315901 | 1.301855416 | 0.260041136 |
| Gm26632 | 0 | NA | NA |
| Gm26978 | 0.712265815 | 1.038787637 | 1.221723074 |

| | | | |
|---|---|---|---|
| Gm9951 | 0.166952883 | 1.610951781 | 2.74653892 |
| Dctn6 | 437.7144974 | -0.285062048 | 0.074374695 |
| Gm26768 | 0.185862605 | 0.718215322 | 2.686445724 |
| Gm10131 | 44.98302574 | 0.755183173 | 0.201051425 |
| Mboat4 | 2.274616701 | 0.71516909 | 0.676618024 |
| Leprotl1 | 852.1230496 | 0.586060254 | 0.088051841 |
| Saraf | 1822.362344 | 0.38507848 | 0.095808048 |
| Mir6395 | 0 | NA | NA |
| Gm29243 | 0.741879394 | 2.444170234 | 1.163828795 |
| Gm23177 | 0.047948698 | -0.243569582 | 3.724945049 |
| Gm6100 | 0 | NA | NA |
| Gm8254 | 0 | NA | NA |
| Gm4889 | 0 | NA | NA |
| Gm29174 | 0.095435502 | 0.718202646 | 3.724945049 |
| Dusp4 | 233.7081911 | 1.233405717 | 0.204058397 |
| Tnks | 1939.237959 | 0.066489783 | 0.05548785 |
| Gm9939 | 2.193369661 | 0.119735505 | 0.615121909 |
| Rpl31-ps23 | 0 | NA | NA |
| Ppp1r3b | 56.58455372 | 1.573760998 | 0.253453844 |
| Eri1 | 199.30024 | 0.333031985 | 0.126663105 |
| Gm16793 | 8.177702489 | 0.324121956 | 0.370877878 |
| Mfhas1 | 794.683006 | -0.105374268 | 0.084890175 |
| Gm22216 | 0.304980231 | -0.724460655 | 1.760607303 |
| AC129211.1 | 0 | NA | NA |
| Cldn23 | 10.064079 | 0.964722929 | 0.44336601 |
| Gm22030 | 0 | NA | NA |
| Rps12l2 | 3.933964785 | 1.906818126 | 0.886385191 |
| D8Ertd82e | 662.5180749 | -0.335713344 | 0.094073582 |
| Lonrf1 | 282.0614182 | -0.459425859 | 0.119936392 |
| Gm5787 | 0.203967006 | 0.742120014 | 2.21351486 |
| Gm26150 | 0.099155932 | 0.718202646 | 3.724945049 |
| 6430573F11Rik | 301.8566133 | -0.133042825 | 0.183923517 |
| Dlc1 | 1005.579271 | -0.324101569 | 0.066665136 |
| AI429214 | 46.9821826 | 0.272941482 | 0.160554377 |
| Gm25126 | 0 | NA | NA |
| Sgcz | 92.85252959 | -0.908194343 | 0.163803428 |
| Gm16205 | 2.502004683 | -2.057324545 | 0.884894836 |
| Gm16204 | 0 | NA | NA |
| Mir383 | 2.557294913 | -1.097618541 | 0.689074496 |
| Gm23128 | 0 | NA | NA |
| Tusc3 | 992.2313355 | -0.301189447 | 0.077270684 |
| Msr1 | 385.467979 | 8.026177434 | 0.467955282 |
| Fgf20 | 6.705059015 | -0.48280758 | 0.386679848 |
| Micu3 | 1360.461354 | -0.401348976 | 0.065935605 |
| Mir7666 | 6.340121748 | -1.023503371 | 0.471225496 |
| Zdhhc2 | 665.2779763 | -0.299837779 | 0.099350937 |
| Cnot7 | 1005.736534 | -0.108651273 | 0.057265089 |

| | | | |
|---|---|---|---|
| Vps37a | 864.8152951 | 0.140669282 | 0.071221621 |
| Mtmr7 | 611.4343167 | -0.385326709 | 0.064654776 |
| Adam24 | 0.11193333 | 0.718202646 | 3.724945049 |
| Adam25 | 0 | NA | NA |
| Adam20 | 0 | NA | NA |
| Adam39 | 0 | NA | NA |
| Gm5345 | 0.200503209 | 0.718218166 | 2.61511129 |
| Gm26584 | 0.132101749 | 1.209894437 | 3.72418822 |
| Slc7a2 | 630.8632614 | 0.65287494 | 0.13544395 |
| Pdgfrl | 37.56964625 | 1.938630686 | 0.261631669 |
| Mtus1 | 1041.21851 | -0.001762231 | 0.079955341 |
| B430010I23Rik | 0.577003193 | 1.361839137 | 1.320061643 |
| Gm16192 | 0.177745132 | -0.244115745 | 2.300135904 |
| Gm16193 | 1.147365236 | 0.467440444 | 1.006133291 |
| Fgl1 | 0.121388914 | 0.718200421 | 3.724945049 |
| Gm16348 | 0.652886586 | 3.332004777 | 3.663057269 |
| Pcm1 | 2899.225793 | -0.069831541 | 0.060742581 |
| Asah1 | 694.5391705 | 0.566929924 | 0.091648817 |
| Frg1 | 216.5532014 | 0.236672256 | 0.106941089 |
| Gapdh-ps14 | 0.166952883 | 1.610951781 | 2.74653892 |
| Smarce1-ps1 | 16.22242323 | 0.328193626 | 0.263356388 |
| 2810404M03Rik | 0.402944817 | 0.098526798 | 1.685812034 |
| Zfp353-ps | 0 | NA | NA |
| Gm6180 | 51.37583421 | 0.334919279 | 0.147550299 |
| Gm23531 | 0 | NA | NA |
| Gm26181 | 0 | NA | NA |
| Triml1 | 0.169875191 | -0.243569355 | 2.748385943 |
| Triml2 | 0.045882455 | 0.237312084 | 3.724945049 |
| Zfp42 | 0.483255338 | 2.324368598 | 1.307649714 |
| Gm22986 | 0 | NA | NA |
| Adam26b | 0 | NA | NA |
| Adam26a | 0 | NA | NA |
| Gm5346 | 0.068097359 | -0.243569582 | 3.724945049 |
| Adam34 | 0.137685729 | -0.724459914 | 2.938031474 |
| Gm24591 | 0 | NA | NA |
| Gm37972 | 0 | NA | NA |
| Gm26241 | 0 | NA | NA |
| Gm26089 | 0 | NA | NA |
| Fat1 | 3170.327197 | -0.227428084 | 0.080052016 |
| Gm2366 | 0.106971521 | 0.237307635 | 3.724945049 |
| AY512931 | 0.183958792 | 0.718217382 | 2.679745607 |
| Mtnr1a | 1.009506168 | -0.669471837 | 1.035358606 |
| F11 | 1.357931586 | 2.048309678 | 1.077887013 |
| Klkb1 | 0.521565614 | 2.438712827 | 1.360899864 |
| Cyp4v3 | 176.7598727 | 1.435255461 | 0.15048934 |
| Fam149a | 309.9705733 | -0.124855845 | 0.1014732 |
| Tlr3 | 1099.900133 | 3.707075936 | 0.192872509 |

| | | | |
|---|---|---|---|
| Sorbs2 | 2803.02876 | -0.633825146 | 0.131177674 |
| Gm23604 | 0 | NA | NA |
| Gm16351 | 0.27308476 | 1.593156749 | 1.998903325 |
| Sorbs2os | 1.437005151 | -0.632226153 | 0.814081742 |
| Gm16352 | 7.186401275 | -1.020242556 | 0.396511263 |
| Gm25309 | 0 | NA | NA |
| Pdlim3 | 13.56838278 | -0.425128093 | 0.362919664 |
| Ccdc110 | 5.316494593 | -0.030314524 | 0.429069916 |
| Gm15878 | 0.075043205 | 0.237309859 | 3.724945049 |
| 1700029J07Rik | 30.24045965 | -0.525976125 | 0.272018115 |
| Ufsp2 | 315.566229 | -0.11024683 | 0.11883229 |
| Ankrd37 | 31.55324993 | -0.964449561 | 0.20664907 |
| Lrp2bp | 13.15399262 | 0.39727967 | 0.327205049 |
| Snx25 | 564.8783107 | -0.372204494 | 0.055084034 |
| Gm23030 | 0 | NA | NA |
| Cfap97 | 360.4521644 | -0.13436142 | 0.074512505 |
| Gm15634 | 3.396395574 | -0.300752217 | 0.588267832 |
| Slc25a4 | 4403.300027 | -0.217784088 | 0.084239901 |
| Actg-ps1 | 1.057130107 | 1.381678739 | 1.047906811 |
| Gm23812 | 0 | NA | NA |
| Gm10313 | 0.79470004 | 1.388541132 | 1.015588082 |
| Gm24669 | 0 | NA | NA |
| Helt | 1.906358267 | 2.889913714 | 0.990842036 |
| Acsl1 | 635.4438796 | -0.030958742 | 0.090058371 |
| Cenpu | 23.99853257 | -0.71224966 | 0.229248138 |
| Primpol | 71.59947947 | 0.329258985 | 0.150452053 |
| Casp3 | 141.296778 | 1.061819422 | 0.148459693 |
| Gm16675 | 21.12817838 | 4.847888886 | 0.523209178 |
| Irf2 | 579.6276244 | 1.75334812 | 0.150682613 |
| Enpp6 | 117.2534975 | -0.270099643 | 0.126483487 |
| 4930579M01Rik | 0.29175089 | 0.75908417 | 1.775015858 |
| Stox2 | 2896.450262 | -0.304235721 | 0.063361865 |
| 4930448N21Rik | 21.17633594 | -0.916879726 | 0.328761732 |
| Gm10083 | 4.406530871 | -3.351355032 | 0.761643797 |
| Gm22857 | 0 | NA | NA |
| Trappc11 | 834.2127417 | -0.017087401 | 0.057337329 |
| Rwdd4a | 348.2097546 | 0.134310092 | 0.083031992 |
| Gm27409 | 0 | NA | NA |
| Gm8623 | 3.204879861 | -0.065902669 | 0.551632118 |
| Ing2 | 227.0032211 | -0.530122421 | 0.106335777 |
| AA386476 | 7.480509175 | -0.967800558 | 0.428368674 |
| Cdkn2aip | 219.8046481 | 0.204677341 | 0.104697302 |
| E030037K01Rik | 3.293407775 | -0.491986286 | 0.552904338 |
| n-R5s96 | 0.212081199 | -0.243570985 | 2.583581921 |
| Cldn24 | 0.048949532 | -0.243569582 | 3.724945049 |
| Cldn22 | 1.967947156 | -0.648813595 | 0.836470973 |
| Wwc2 | 675.3607201 | -0.179543728 | 0.064928956 |

| | | | |
|---|---|---|---|
| n-R5s97 | 0 | NA | NA |
| Dctd | 53.07648255 | -0.424255722 | 0.186538139 |
| Tenm3 | 1960.745608 | -0.84828505 | 0.113205946 |
| Gm19744 | 64.58441548 | 0.62194373 | 0.263703207 |
| Gm25830 | 0.249706104 | -0.243563802 | 1.848218408 |
| 4930555F03Rik | 2.112369234 | -1.327948745 | 0.707910516 |
| Gm23986 | 0.048949532 | -0.243569582 | 3.724945049 |
| AC140437.1 | 0.038981395 | 0.718207094 | 3.724945049 |
| Gm24929 | 0 | NA | NA |
| Gm9892 | 89.20159694 | 0.974612237 | 0.164935637 |
| Gm6463 | 0 | NA | NA |
| Gm25813 | 1.893990199 | 0.476319591 | 0.832127333 |
| Gm23666 | 0 | NA | NA |
| Gm27964 | 0 | NA | NA |
| Gm22841 | 0 | NA | NA |
| Aga | 79.49866868 | 1.13692464 | 0.151007322 |
| Neil3 | 7.05498645 | 1.863098613 | 0.699616847 |
| Vegfc | 39.65073296 | -0.044280148 | 0.168372469 |
| Spcs3 | 857.5164723 | 0.035450119 | 0.077926322 |
| Asb5 | 7.025391348 | 0.502580787 | 0.408850495 |
| 5330439A09Rik | 2.898308479 | -0.774755254 | 0.748506958 |
| Spata4 | 0.398944595 | -0.262787821 | 1.688762139 |
| Wdr17 | 405.2573486 | -1.253078223 | 0.134179689 |
| Gm15758 | 0.38984224 | -0.630811548 | 1.456295622 |
| n-R5s98 | 6.806331161 | -0.878109538 | 0.403077588 |
| Gpm6a | 9377.570103 | -0.82638012 | 0.084371856 |
| Adam29 | 0.15906016 | 1.290286593 | 3.433492162 |
| Glra3 | 117.655716 | 0.065099831 | 0.130767462 |
| Hpgd | 129.4286017 | 2.597696929 | 0.250849501 |
| Gm24375 | 0.075043205 | 0.237309859 | 3.724945049 |
| Cep44 | 122.059907 | -0.236078036 | 0.151093094 |
| Fbxo8 | 283.448663 | 0.00108317 | 0.118319423 |
| Gm25992 | 6.166697826 | -0.505488795 | 0.434237699 |
| Gm29173 | 0.088736197 | -0.243576254 | 3.724945049 |
| AV026068 | 0.061920613 | 0.718207094 | 3.724945049 |
| Hand2 | 0.062426873 | 0.237312084 | 3.724945049 |
| Gm28063 | 0 | NA | NA |
| 5033428I22Rik | 0 | NA | NA |
| Scrg1 | 178.3231663 | -0.40363351 | 0.213097101 |
| Sap30 | 157.6272062 | 1.549340982 | 0.152167261 |
| 2500002B13Rik | 14.59964785 | -0.099797028 | 0.305394144 |
| Hmgb2 | 86.7028358 | 2.104072771 | 0.291969672 |
| Galnt7 | 420.2620518 | 0.349837961 | 0.071035828 |
| Gm25379 | 0 | NA | NA |
| Gm10284 | 11.17616707 | -0.597321212 | 0.300451574 |
| Galntl6 | 211.6445591 | -0.936681912 | 0.109214721 |
| Gm26905 | 1.736582332 | -1.654876247 | 0.851097217 |

| | | | |
|---|---|---|---|
| Gm15882 | 0 | NA | NA |
| Gm15881 | 19.66782202 | -1.650082099 | 0.311140833 |
| BC030500 | 65.02402097 | -0.76470358 | 0.166824863 |
| Gm24847 | 0 | NA | NA |
| Gm26164 | 0 | NA | NA |
| Gm10283 | 0 | NA | NA |
| Aadat | 2.615189709 | -0.105137971 | 0.613058443 |
| Mfap3l | 976.8064727 | -0.193639617 | 0.082132778 |
| Gm23329 | 0.066050875 | 0.71820487 | 3.724945049 |
| 2700029M09Rik | 270.2446424 | 0.037409792 | 0.118503777 |
| Clcn3 | 3034.91923 | -0.342759401 | 0.043269803 |
| B230317F23Rik | 9.640489829 | -1.072473597 | 0.402405881 |
| Nek1 | 677.7359105 | -0.564524215 | 0.151239623 |
| Gm16179 | 0 | NA | NA |
| Gm16178 | 0.095898756 | 1.192083721 | 3.724434897 |
| Sh3rf1 | 531.7020943 | -0.4944647 | 0.101308815 |
| 1700001D01Rik | 0 | NA | NA |
| Gm10015 | 0.445602023 | 0.627081567 | 1.465505326 |
| Cbr4 | 83.07961058 | -0.06215817 | 0.146679624 |
| Palld | 202.5425458 | 2.041409872 | 0.203316418 |
| 4930512H18Rik | 0.155551505 | 0.718215147 | 2.833462086 |
| Ddx60 | 1763.240931 | 7.769466293 | 0.223613025 |
| Anxa10 | 0.200514362 | 1.617596658 | 2.640387911 |
| Gm22324 | 0 | NA | NA |
| Spock3 | 1216.4572 | -0.733127354 | 0.098882055 |
| Gm24236 | 0 | NA | NA |
| Gm5350 | 0 | NA | NA |
| Gm24580 | 0 | NA | NA |
| Sgol2b | 2.142608394 | -0.934220607 | 0.66810939 |
| Tll1 | 166.2460114 | -0.018245126 | 0.190221721 |
| Gm22586 | 0 | NA | NA |
| Mir710 | 0.327710148 | 1.211186266 | 1.970831531 |
| Cpe | 12511.93844 | -0.265877725 | 0.093929406 |
| Msmo1 | 801.7074196 | 0.569401941 | 0.141827026 |
| Klhl2 | 1600.801606 | 0.003592408 | 0.09143703 |
| Gm26210 | 0 | NA | NA |
| Tmem192 | 214.6384359 | 1.071402454 | 0.138358099 |
| Trim75 | 0 | NA | NA |
| Trim60 | 0 | NA | NA |
| Apela | 0.083526043 | 0.718202646 | 3.724945049 |
| Gm10663 | 0 | NA | NA |
| BC030870 | 0 | NA | NA |
| Gm24333 | 0 | NA | NA |
| Gm25403 | 0 | NA | NA |
| 1-Mar | 492.5184607 | 0.163057734 | 0.067876595 |
| Gm23515 | 0 | NA | NA |
| Gm15355 | 0.044191549 | -0.243571806 | 3.724945049 |

| | | | |
|---|---|---|---|
| Gm26123 | 1.969797064 | -0.582554652 | 0.729132474 |
| Gm24205 | 4.278889304 | 0.018130322 | 0.526934285 |
| Gm15354 | 0.20829534 | 1.831782594 | 2.531443297 |
| Gm15356 | 0.306626524 | 1.517386078 | 1.73968374 |
| Tma16 | 219.0930399 | 0.79417098 | 0.184672714 |
| Tktl2 | 2.437974107 | -1.512073399 | 0.822205243 |
| Gm16330 | 0 | NA | NA |
| Npy5r | 36.55808565 | -1.000106702 | 0.198247316 |
| Npy1r | 157.9670103 | -1.102788396 | 0.147646833 |
| Naf1 | 250.5788205 | 0.218312205 | 0.114291967 |
| Gm25275 | 0 | NA | NA |
| Gm24274 | 0 | NA | NA |
| Gm24677 | 0 | NA | NA |
| Gm10660 | 0 | NA | NA |
| Nat1 | 7.474725557 | -1.29449445 | 0.422555106 |
| Nat2 | 24.83237878 | 2.332378458 | 0.267020416 |
| Gm9755 | 12.27831489 | 0.390370348 | 0.298789779 |
| Nat3 | 0.254414262 | 1.319626129 | 2.462869205 |
| AC141868.1 | 0 | NA | NA |
| Psd3 | 7016.150575 | -0.62460591 | 0.104908934 |
| Gm15992 | 24.57258576 | -1.483617134 | 0.320541632 |
| Gm15991 | 12.26391693 | -1.218091139 | 0.315106793 |
| Gm22018 | 0.176406687 | -0.243567884 | 2.71092078 |
| Gm21807 | 5.900355672 | 0.175537635 | 0.380010146 |
| Gm26659 | 0.106520246 | 0.718200421 | 3.724945049 |
| Sh2d4a | 4.03121477 | 2.019471216 | 0.657422046 |
| Csgalnact1 | 201.9711671 | -0.243955634 | 0.115421332 |
| 1700125H03Rik | 0.107078754 | 0.236669296 | 3.724898107 |
| Gm15656 | 0 | NA | NA |
| Gm15654 | 0 | NA | NA |
| Ints10 | 560.1978323 | -0.591443441 | 0.119723525 |
| Gm15549 | 5.491253898 | -1.035318993 | 0.601632595 |
| Lpl | 272.7124532 | -0.740974581 | 0.131746072 |
| Slc18a1 | 6.521337389 | 0.858169624 | 0.395135978 |
| Gm15716 | 0 | NA | NA |
| Atp6v1b2 | 5572.862093 | 0.155848595 | 0.048778817 |
| Gm15717 | 14.06246178 | -0.609293202 | 0.403453395 |
| Lzts1 | 982.0473404 | -1.175920066 | 0.163349523 |
| Zfp930 | 119.9920175 | 0.349786907 | 0.134719786 |
| D130040H23Rik | 96.03445661 | -0.78842095 | 0.143801527 |
| Gm10033 | 37.55209856 | -0.079632648 | 0.232176751 |
| Gm10311 | 21.9424117 | 0.231737768 | 0.236325962 |
| Gm25661 | 0.130524232 | -0.243571427 | 3.657871338 |
| RP23-468H22.1 | 1.045653151 | 2.866277117 | 1.241371319 |
| RP23-468H22.3 | 0.584793824 | 1.000427893 | 1.25613002 |
| RP23-468H22.2 | 0.794752679 | 2.788751568 | 1.288437039 |
| Zfp868 | 254.8017484 | -0.015864408 | 0.085918234 |

| | | | |
|---|---|---|---|
| Zfp964 | 24.06935087 | -0.559777773 | 0.221834241 |
| Zfp869 | 250.7494571 | 0.352031479 | 0.087876172 |
| Zfp963 | 46.44832303 | -0.230940376 | 0.160312306 |
| Gm20422 | 0 | NA | NA |
| Zfp866 | 322.7321497 | -0.330826453 | 0.113493957 |
| Atp13a1 | 689.8934044 | 0.55973724 | 0.086787451 |
| Gmip | 150.8857527 | 0.867700586 | 0.141349218 |
| Lpar2 | 13.32151877 | 0.286041346 | 0.292882541 |
| Pbx4 | 59.57232151 | -0.840681149 | 0.150644813 |
| Cilp2 | 10.39023466 | 0.143212519 | 0.337736105 |
| Yjefn3 | 43.79218575 | -0.645185926 | 0.194082819 |
| Ndufa13 | 488.3269015 | -0.174805133 | 0.343453433 |
| Tssk6 | 2.109268871 | 0.818612096 | 0.673468961 |
| Gatad2a | 755.2849878 | 1.126912849 | 0.10199188 |
| Mau2 | 1031.702982 | 0.033111503 | 0.065638311 |
| Sugp1 | 554.3624155 | 0.1730273 | 0.060494024 |
| Tm6sf2 | 17.28368832 | -0.832197085 | 0.462592871 |
| Hapln4 | 1233.584627 | -0.421881869 | 0.075002854 |
| Ncan | 3935.767652 | 0.622461591 | 0.169688904 |
| Mir7066 | 0.192965486 | -0.724461459 | 2.252205086 |
| Gm25906 | 0 | NA | NA |
| Rfxank | 136.35992 | 0.677016885 | 0.10640288 |
| Nr2c2ap | 176.3814956 | 1.503223724 | 0.151555676 |
| 2310045N01Rik | 134.2125858 | 0.364380334 | 0.101738531 |
| Mef2b | 0.295756179 | -0.780801048 | 2.076351649 |
| Tmem161a | 299.8727589 | 0.135059622 | 0.09443524 |
| Slc25a42 | 283.822492 | -0.624574751 | 0.16254281 |
| Armc6 | 187.6904792 | 0.056443736 | 0.100550581 |
| Sugp2 | 1115.657998 | -0.496524128 | 0.095065107 |
| Homer3 | 488.6342012 | -0.034247215 | 0.201359788 |
| Ddx49 | 235.0236882 | 0.535837553 | 0.087841542 |
| Cope | 659.244897 | 0.224637847 | 0.074814992 |
| Cers1 | 345.2784473 | -0.23415193 | 0.091721071 |
| Gdf1 | 0 | NA | NA |
| Upf1 | 1210.834756 | 0.513304842 | 0.060109708 |
| Comp | 10.12622545 | -1.495115528 | 0.445634728 |
| Crtc1 | 1987.206553 | -0.602430462 | 0.100465794 |
| Klhl26 | 487.3158746 | 0.020589546 | 0.074535352 |
| Tmem59l | 1446.494139 | -0.261231601 | 0.055932634 |
| Crlf1 | 37.87736275 | 1.496636725 | 0.217219373 |
| 2810428I15Rik | 262.175928 | 0.234020103 | 0.106168949 |
| Uba52 | 19.53769797 | 0.29604336 | 0.26361221 |
| Kxd1 | 196.5815617 | 0.202942458 | 0.083832476 |
| Fkbp8 | 2500.961044 | 0.06591383 | 0.076503678 |
| Ell | 288.2178684 | 0.261678188 | 0.120869235 |
| Isyna1 | 251.2867603 | -0.036435212 | 0.143020127 |
| Ssbp4 | 876.1567567 | -0.397501786 | 0.064402857 |

| | | | |
|---|---|---|---|
| Lrrc25 | 65.01020252 | 5.092600501 | 0.437886828 |
| Gdf15 | 12.20491556 | 6.099912029 | 0.918535715 |
| Pgpep1 | 166.5791501 | 0.552008485 | 0.167186202 |
| Lsm4 | 218.808746 | 0.463824503 | 0.095956835 |
| Jund | 1476.72973 | 0.443012504 | 0.137886225 |
| Gm11175 | 17.16041749 | -0.32935122 | 0.285416961 |
| Gm3336 | 0.36285418 | 0.244496723 | 1.709178269 |
| Pde4c | 6.152153976 | -0.949531815 | 0.449592975 |
| Rab3a | 3789.848654 | -0.696888769 | 0.055883862 |
| Mpv17l2 | 124.5250829 | 0.380482571 | 0.14440141 |
| Ifi30 | 138.4088315 | 2.212948625 | 0.214750088 |
| Pik3r2 | 1054.806414 | -0.505271642 | 0.076101988 |
| 2010320M18Rik | 39.80095612 | 0.392979315 | 0.189349762 |
| Mast3 | 1971.583413 | -1.276198581 | 0.118160924 |
| Mir7240 | 0.07796279 | 1.072698098 | 3.721829099 |
| Il12rb1 | 265.2067133 | 4.349768412 | 0.240306775 |
| Arrdc2 | 123.0827081 | 0.42349856 | 0.184656037 |
| Kcnn1 | 121.7377044 | -0.471113485 | 0.129774506 |
| A230052G05Rik | 2.786635864 | -0.904057609 | 0.655817413 |
| Ccdc124 | 301.1606489 | -0.03958007 | 0.113453323 |
| Slc5a5 | 42.07648303 | -1.050608613 | 0.193396695 |
| Rpl18a | 1403.238223 | 0.804395348 | 0.13363378 |
| Mir7067 | 0 | NA | NA |
| Snora68 | 19.80479055 | 0.308088803 | 0.282442181 |
| Map1s | 619.2559287 | 0.077089287 | 0.068988265 |
| Mir1969 | 0.386072962 | 0.634797266 | 3.258544709 |
| Gm10654 | 0.749724794 | 1.572778463 | 1.11510696 |
| 1700026F02Rik | 0.167183452 | -0.72445987 | 2.765153652 |
| Gm17576 | 0.045882455 | 0.237312084 | 3.724945049 |
| Gm27628 | 0.044544648 | 0.237309859 | 3.724945049 |
| Haus8 | 58.62861715 | 0.648861071 | 0.134724498 |
| Myo9b | 1047.53723 | 0.884409834 | 0.093740419 |
| Use1 | 300.4227806 | 0.12808415 | 0.097319704 |
| Ocel1 | 128.1701314 | 0.21716635 | 0.138007875 |
| Nr2f6 | 250.0485222 | 0.271607523 | 0.119131017 |
| Ushbp1 | 11.95745292 | 1.012883271 | 0.360238721 |
| Babam1 | 547.0408888 | 0.04689231 | 0.075209024 |
| Ankle1 | 4.029650605 | 0.013939834 | 0.57215708 |
| Abhd8 | 1731.379142 | -0.379965911 | 0.083349086 |
| Mrpl34 | 112.3878888 | -0.303818876 | 0.117236378 |
| Dda1 | 548.7093606 | 0.205386829 | 0.06821526 |
| Ano8 | 471.5065914 | -0.60676348 | 0.098731527 |
| Gtpbp3 | 220.5550811 | -0.147071997 | 0.081881175 |
| Plvap | 140.3657839 | -0.162315916 | 0.12639197 |
| Bst2 | 1856.786934 | 8.427849553 | 0.311906746 |
| Mvb12a | 194.6062646 | 0.793927033 | 0.099547371 |
| Tmem221 | 4.789566606 | 3.810259885 | 0.881610836 |

| | | | |
|---|---|---|---|
| Nxnl1 | 1.098494234 | 0.298314039 | 1.01942481 |
| Slc27a1 | 744.5497192 | -0.056735301 | 0.11120006 |
| Pgls | 165.2612445 | 0.364238087 | 0.109264157 |
| Fam129c | 18.63060221 | -0.077611946 | 0.272763149 |
| Colgalt1 | 425.2055677 | -0.016825843 | 0.106905199 |
| Unc13a | 2849.774368 | -0.980300646 | 0.134653981 |
| Mir6769b | 0.250814404 | -1.548900481 | 1.784277208 |
| Jak3 | 179.3964878 | 2.527692842 | 0.125893906 |
| Insl3 | 0.723819272 | 2.711528344 | 1.233373198 |
| B3gnt3 | 7.857962099 | 2.476415561 | 0.428036212 |
| Fcho1 | 729.7161296 | -0.810420062 | 0.101196075 |
| Zfp709 | 191.7633267 | 0.227727233 | 0.142226324 |
| Zfp882 | 121.9521505 | -0.362297981 | 0.150402297 |
| Zfp617 | 199.055627 | 0.284306919 | 0.124086532 |
| Ccnb2-ps | 2.364122105 | 0.148774742 | 0.778565894 |
| Zfp961 | 151.8196963 | -0.073545609 | 0.142689848 |
| Cyp4f18 | 43.06846795 | 7.982393089 | 0.882738385 |
| Gm25052 | 0 | NA | NA |
| Olfr372 | 0 | NA | NA |
| Olfr373 | 0 | NA | NA |
| Olfr374 | 0 | NA | NA |
| Gm24747 | 0 | NA | NA |
| Tpm4 | 485.8967539 | 2.002703539 | 0.17583111 |
| Gm26586 | 0.363579626 | -0.351636738 | 1.684244108 |
| Gm16091 | 1.601615687 | -1.065956397 | 0.809296151 |
| Rab8a | 470.4426857 | 0.839614196 | 0.08427446 |
| Hsh2d | 15.58486664 | 6.633636553 | 0.874292726 |
| Cib3 | 0.321217965 | 1.176677003 | 3.721251982 |
| Gm11034 | 0 | NA | NA |
| Fam32a | 524.8981796 | 0.375619511 | 0.119480572 |
| Ap1m1 | 552.8521804 | 0.085193224 | 0.058081324 |
| Gm10282 | 57.72901061 | 0.622634773 | 0.15818617 |
| Klf2 | 122.111689 | 2.309734259 | 0.213254047 |
| Eps15l1 | 756.2958603 | -0.095607081 | 0.07993846 |
| Calr3 | 22.05166792 | 1.085172291 | 0.243599309 |
| Gm3835 | 0.038981395 | 0.718207094 | 3.724945049 |
| 1700030K09Rik | 65.49681279 | 0.189793679 | 0.163486383 |
| Cherp | 754.5952941 | 0.297466468 | 0.070688224 |
| Mir7068 | 0 | NA | NA |
| Slc35e1 | 803.4949084 | 0.156517523 | 0.064904653 |
| Gm17435 | 24.35316493 | 1.371509792 | 0.27844308 |
| Med26 | 154.280037 | 0.321797058 | 0.123569277 |
| Gm28019 | 1.785542434 | -1.022405329 | 0.855338987 |
| Smim7 | 545.215953 | 0.006983332 | 0.106247421 |
| Tmem38a | 713.1160059 | -0.634733228 | 0.083907924 |
| Nwd1 | 810.0404047 | -0.970933821 | 0.192539469 |
| Gm26237 | 0.846376972 | -1.547184265 | 1.137432246 |

| | | | |
|---|---|---|---|
| Sin3b | 1202.221482 | 0.291423397 | 0.088962982 |
| F2rl3 | 3.0074759 | 1.398201126 | 0.562234003 |
| Large | 1767.595343 | -0.307012895 | 0.073772676 |
| Mir28b | 0 | NA | NA |
| Gm22532 | 8.167798302 | -0.164394787 | 0.453144335 |
| Gm23255 | 0 | NA | NA |
| Crry-ps | 1.276549371 | 1.093243679 | 0.931783212 |
| Gm27864 | 0 | NA | NA |
| Gm11033 | 0 | NA | NA |
| Isx | 0.612261413 | 0.38797829 | 1.516705098 |
| Hmgxb4 | 272.0156998 | 0.042569671 | 0.132092929 |
| Tom1 | 187.7576606 | 0.324741432 | 0.096460659 |
| Hmox1 | 1627.230529 | 6.191416088 | 0.45658661 |
| Mcm5 | 102.0398356 | 1.559887365 | 0.21289595 |
| Rasd2 | 1068.740531 | -0.96037655 | 0.146003619 |
| Gm7984 | 22.42806742 | 0.677016661 | 0.259750758 |
| Gm10358 | 5.720807359 | 0.131527069 | 0.423161696 |
| 1700007B14Rik | 4.206508402 | -0.326860695 | 0.571584014 |
| Gm38270 | 0 | NA | NA |
| Gm7901 | 51.21551474 | 0.382757792 | 0.254648659 |
| Gm22544 | 0 | NA | NA |
| Gm22509 | 0 | NA | NA |
| Gm7069 | 0.097139547 | 0.718207094 | 3.724945049 |
| Gm10649 | 41.3073234 | -0.766564605 | 0.214725255 |
| Gm27355 | 0 | NA | NA |
| Gm5353 | 7.21028118 | -0.918865011 | 0.482070227 |
| Nr3c2 | 828.7814193 | -0.341828062 | 0.06406456 |
| Gm24838 | 0 | NA | NA |
| Gm23260 | 3.058609062 | -0.846337679 | 0.590003956 |
| Arhgap10 | 375.9267285 | -0.104206206 | 0.0752638 |
| 0610038B21Rik | 3.891239497 | 0.444952142 | 0.518774431 |
| Prmt10 | 268.6822526 | -0.088881274 | 0.078778483 |
| Gm23468 | 0 | NA | NA |
| Tmem184c | 547.9950958 | -0.694502336 | 0.059481817 |
| 4933431K23Rik | 0.309777098 | -0.718453973 | 1.61058947 |
| 1700092C02Rik | 0 | NA | NA |
| Ednra | 46.53120417 | 0.080086805 | 0.162285947 |
| Gm23981 | 0 | NA | NA |
| Ttc29 | 3.962620848 | -0.569537326 | 0.598838102 |
| Pou4f2 | 9.952480325 | 0.089662544 | 0.343991938 |
| Rbmxl1 | 229.4117921 | 0.276654422 | 0.090235456 |
| Slc10a7 | 127.4608844 | 0.83083459 | 0.121227917 |
| Lsm6 | 350.1356414 | -0.044760744 | 0.094665014 |
| Gm8054 | 0.763271679 | 1.166648945 | 1.262263031 |
| Gm22044 | 0 | NA | NA |
| Zfp827 | 1018.675046 | -0.343285017 | 0.068262301 |
| 1700011L22Rik | 0.586370721 | -0.136498906 | 1.220529092 |

| | | | |
|---|---|---|---|
| Mmaa | 175.5439761 | -0.491012128 | 0.112625529 |
| Gm4890 | 7.177265976 | 0.157069085 | 0.438916169 |
| Smad1 | 552.5127933 | 0.218941053 | 0.099405797 |
| Gm8079 | 0.101408268 | 0.717551239 | 3.724898127 |
| Otud4 | 1259.849304 | 0.159100873 | 0.084877013 |
| Tpd52-ps | 12.01690052 | 0.372427028 | 0.31633402 |
| Abce1 | 1227.066757 | 0.347937081 | 0.078199596 |
| Gm27006 | 0.04393175 | -0.243571806 | 3.724945049 |
| Anapc10 | 137.3995736 | 0.11950089 | 0.108315636 |
| Hhip | 182.8551312 | 0.051664175 | 0.110388389 |
| Gm27820 | 0 | NA | NA |
| Gm5909 | 4.376335103 | 0.30338887 | 0.454826182 |
| Gypa | 2.705694399 | 1.202324014 | 0.839722522 |
| 4930505O20Rik | 0.533083002 | 1.020327584 | 1.286320387 |
| Frem3 | 59.61918439 | 0.994673214 | 0.265660281 |
| Smarca5 | 1563.50047 | 0.405129845 | 0.074508682 |
| Gab1 | 641.4208176 | 0.315584497 | 0.100330257 |
| Gm27048 | 0 | NA | NA |
| Gm4891 | 0.083526043 | 0.718202646 | 3.724945049 |
| Usp38 | 593.7032144 | 0.438173471 | 0.094015534 |
| Gm9725 | 0.99585445 | 1.837158559 | 1.043725627 |
| Gm4899 | 0.173022395 | 0.718215388 | 2.735311368 |
| Gm5354 | 0 | NA | NA |
| Inpp4b | 528.8835549 | 0.553652268 | 0.084692731 |
| 4930579O11Rik | 0 | NA | NA |
| Gm17169 | 0 | NA | NA |
| Gm17072 | 10.76767044 | 0.034478435 | 0.372869692 |
| Gm25267 | 0 | NA | NA |
| Il15 | 30.29114704 | 3.887368271 | 0.332507147 |
| Gm25138 | 0 | NA | NA |
| Gm8167 | 0 | NA | NA |
| Zfp330 | 179.4385362 | 0.356390745 | 0.117352081 |
| Rnf150 | 857.5833068 | -0.867394172 | 0.122382255 |
| Gm10645 | 5.503045566 | -0.23057253 | 0.411250958 |
| Tbc1d9 | 1881.448684 | 0.208111843 | 0.073107233 |
| Ucp1 | 2.960476708 | 1.001366328 | 0.802881912 |
| Elmod2 | 319.9756207 | -0.012406089 | 0.073655311 |
| Mgat4d | 0.674779921 | 2.035022049 | 1.341762256 |
| Gm15660 | 0 | NA | NA |
| Clgn | 175.9442896 | -0.591299721 | 0.119835205 |
| Scoc | 643.4231515 | -0.369250493 | 0.100695059 |
| Olfr370 | 0.049553047 | 0.71820487 | 3.724945049 |
| Ndufb7 | 290.6417338 | 0.068815928 | 0.207575336 |
| Tecr | 2387.710836 | -0.382889219 | 0.067201909 |
| Dnajb1 | 549.0730292 | 0.079555846 | 0.172825389 |
| Gipc1 | 557.0990177 | 0.292461943 | 0.064689214 |
| Ptger1 | 17.78461194 | 0.248567827 | 0.330832118 |

| | | | |
|---|---|---|---|
| Pkn1 | 501.3936003 | 0.565273908 | 0.096937658 |
| Ddx39 | 165.8939195 | 1.444997718 | 0.143831439 |
| Adgre5 | 155.9202038 | 2.416143459 | 0.136695618 |
| Mir1668 | 0 | NA | NA |
| Gm22024 | 0 | NA | NA |
| Gm26721 | 9.148635249 | -2.696342649 | 0.54728468 |
| Adgrl1 | 7570.249026 | -1.024731308 | 0.102629656 |
| AC156028.1 | 0.257982025 | -0.243563247 | 1.837527087 |
| AC156028.2 | 0.523560496 | -1.417497433 | 1.312711335 |
| Gm10644 | 10.70655375 | 0.307379491 | 0.432637543 |
| Asf1b | 15.09267688 | 2.172905797 | 0.397834001 |
| Prkaca | 1912.644388 | -0.377673132 | 0.051718418 |
| Samd1 | 227.0946354 | -0.160470884 | 0.103456607 |
| 1700067K01Rik | 1.769403265 | -1.248061343 | 0.743548726 |
| Mir8111 | 0 | NA | NA |
| 2210011C24Rik | 4.04060031 | -0.856996579 | 0.496706562 |
| Mir1199 | 0 | NA | NA |
| Palm3 | 9.849211078 | -0.673897434 | 0.352064059 |
| Il27ra | 17.46056049 | 3.07014997 | 0.373923158 |
| Rln3 | 9.200686208 | 0.564218642 | 0.854005767 |
| Gm10643 | 25.86145265 | -0.887777448 | 0.245672808 |
| C330011M18Rik | 25.69760837 | -0.837703749 | 0.216516745 |
| Rfx1 | 271.1011036 | -0.161880395 | 0.119364387 |
| Mir709 | 0.78198097 | -0.736448762 | 1.31426676 |
| Dcaf15 | 175.8113705 | 0.132621133 | 0.119716995 |
| Podnl1 | 7.966600571 | 3.219793354 | 0.554162097 |
| Cc2d1a | 577.8296229 | 0.05960494 | 0.049028516 |
| 4930432K21Rik | 28.89408785 | -0.225297221 | 0.208050539 |
| Gm25110 | 0 | NA | NA |
| Nanos3 | 8.912972197 | -1.013517067 | 0.39535933 |
| Mir181d | 2.185676227 | -1.813758499 | 0.733564076 |
| Mir181c | 0.986137052 | -0.269113576 | 0.929183411 |
| Gm26887 | 0.282313603 | 0.245514186 | 1.786539438 |
| Gm37352 | 38.80286291 | 0.845697336 | 0.21285335 |
| Mir23a | 1.248589509 | 2.447302154 | 1.10385036 |
| Mir27a | 0.444870878 | 2.451433038 | 1.35594002 |
| Mir24-2 | 0 | NA | NA |
| mmu-mir-3074-2 | 0.551097043 | 0.201121316 | 1.51905776 |
| Gm26532 | 9.112776141 | 0.819423789 | 0.386652777 |
| Zswim4 | 345.6336222 | 0.629601539 | 0.153526521 |
| D8Ertd738e | 305.0049009 | 0.428628734 | 0.122410344 |
| Mri1 | 76.67094331 | 0.154107489 | 0.145816852 |
| Ccdc130 | 95.20490353 | 0.582204086 | 0.120836684 |
| Gm23626 | 0 | NA | NA |
| Cacna1a | 3274.65666 | -0.73474919 | 0.1263819 |
| Gm16183 | 0.17506888 | -0.243569334 | 2.730117347 |
| Ier2 | 148.7573661 | 1.188154509 | 0.274905306 |

| | | | |
|---|---|---|---|
| Gm26664 | 2.727852729 | -1.786591288 | 0.648917123 |
| Nacc1 | 1589.037921 | 0.226035891 | 0.062079008 |
| Trmt1 | 439.8140685 | -0.023625466 | 0.094373548 |
| Nfix | 4316.174095 | -0.007834106 | 0.089986348 |
| Lyl1 | 28.85883109 | 1.483910148 | 0.315973015 |
| G430095P16Rik | 2.042153182 | 1.062668451 | 0.895801345 |
| Dand5 | 53.8269615 | -0.207277609 | 0.141889791 |
| Gadd45gip1 | 231.7276792 | 0.168946177 | 0.106996089 |
| Rad23a | 572.6689497 | 0.066945831 | 0.060972026 |
| Calr | 4607.859221 | 1.025271984 | 0.107976451 |
| 1700122E12Rik | 0.913679305 | 0.997300693 | 1.14715279 |
| Farsa | 355.8983591 | 0.585592486 | 0.107279525 |
| Mir7069 | 0 | NA | NA |
| Syce2 | 15.66997929 | 0.415443894 | 0.301903348 |
| Gcdh | 224.1995156 | -0.323096427 | 0.082074036 |
| Klf1 | 0.981329651 | 0.35969203 | 1.085237064 |
| Dnase2a | 91.10452649 | 1.228583059 | 0.178720683 |
| Mast1 | 1321.949102 | -0.910459427 | 0.081583581 |
| Gm24197 | 0 | NA | NA |
| Rtbdn | 11.67166441 | -2.4407582 | 0.4209633 |
| Rnaseh2a | 132.5774289 | -0.262091849 | 0.094227954 |
| Prdx2 | 1409.970294 | -0.046144805 | 0.07349235 |
| Junb | 1137.476461 | 1.281845139 | 0.331594345 |
| Hook2 | 101.5563122 | 0.203865855 | 0.14822911 |
| Best2 | 0.602809315 | 1.355426392 | 1.264097551 |
| Asna1 | 647.8422268 | 0.25129611 | 0.081492305 |
| 2310036O22Rik | 524.574075 | 0.343489806 | 0.106344529 |
| Tnpo2 | 1895.024094 | -0.273884023 | 0.053838192 |
| Gm25506 | 0.455189762 | -2.162827672 | 1.351079521 |
| A230103J11Rik | 33.92778836 | -0.144198952 | 0.192233754 |
| Fbxw9 | 120.3613756 | 0.141868885 | 0.109429028 |
| Mir7070 | 0 | NA | NA |
| Gm5741 | 3.225606799 | -0.573889017 | 0.634896316 |
| Dhps | 207.0919154 | 0.570661973 | 0.093063268 |
| Wdr83 | 132.9258923 | -0.124712886 | 0.109111881 |
| Wdr83os | 100.509984 | 0.230845689 | 0.140354852 |
| Man2b1 | 495.4554757 | 0.428479446 | 0.143085165 |
| Zfp791 | 32.04972151 | -0.162911897 | 0.203740938 |
| Gm23388 | 0 | NA | NA |
| Cks1brt | 0 | NA | NA |
| Olfr371 | 0 | NA | NA |
| Vps35 | 2002.604974 | 0.389454916 | 0.08782249 |
| Orc6 | 141.8270405 | 0.047539869 | 0.124540767 |
| Gm22336 | 0.044544648 | 0.237309859 | 3.724945049 |
| Mylk3 | 3.257485277 | -0.691627974 | 0.567897676 |
| AA672651 | 0 | NA | NA |
| 4921524J17Rik | 80.96275031 | -0.138487372 | 0.144118071 |

| | | | |
|---|---|---|---|
| 1700051O22Rik | 0.111006857 | 0.717551239 | 3.724898127 |
| Gpt2 | 699.6609475 | -0.278093438 | 0.15032756 |
| Dnaja2 | 1599.493447 | 0.188319285 | 0.078654843 |
| Neto2 | 1594.195169 | 0.125027922 | 0.092065585 |
| Mir8109 | 0.42925394 | 2.012231887 | 1.686922092 |
| Itfg1 | 2429.683998 | -0.332435893 | 0.081172861 |
| Phkb | 740.2327041 | 0.057269782 | 0.102477847 |
| Gm16167 | 0 | NA | NA |
| Hm629797 | 14.11744488 | -0.767459343 | 0.360482233 |
| Gm23904 | 0 | NA | NA |
| Gm24490 | 0.060174537 | 0.237309859 | 3.724945049 |
| Abcc12 | 27.6985328 | 0.754797261 | 0.244263501 |
| AC132147.1 | 0 | NA | NA |
| Lonp2 | 859.6725007 | 0.181233245 | 0.080168683 |
| Gm16177 | 0.509424666 | -0.204906761 | 1.287935085 |
| Gm22305 | 0.045882455 | 0.237312084 | 3.724945049 |
| Siah1a | 351.8844398 | 0.120431442 | 0.113672645 |
| Gm10638 | 6.090970552 | 0.236713982 | 0.399189788 |
| N4bp1 | 1537.795251 | 0.962547539 | 0.093585137 |
| 2010110E17Rik | 5.305919521 | 2.635307527 | 0.602443132 |
| Gm27167 | 3.969393191 | 1.616394591 | 0.560313222 |
| Gm27168 | 0.090665643 | -0.724455696 | 3.724945049 |
| Gm24781 | 0 | NA | NA |
| Rps13-ps1 | 58.84124857 | 0.819848037 | 0.184936575 |
| Gm5118 | 1.461301642 | -0.201483792 | 0.920556994 |
| Gm27246 | 0.76513295 | 0.571975591 | 1.070675399 |
| Gm10637 | 0.117650406 | 0.23732098 | 3.724945049 |
| Gm27169 | 0.24523014 | 1.744427389 | 2.50859869 |
| Gm27207 | 0.273479446 | -0.605951804 | 2.461054195 |
| Gm27208 | 0 | NA | NA |
| 4930535O05Rik | 0 | NA | NA |
| Gm19872 | 0 | NA | NA |
| Cbln1 | 1077.308355 | -0.351685941 | 0.238425094 |
| Gm2694 | 162.596044 | -0.985094979 | 0.253026402 |
| Gm27016 | 0.046345709 | 0.71820487 | 3.724945049 |
| 1700120C18Rik | 0 | NA | NA |
| 4933402J07Rik | 1.368974137 | -0.193162781 | 1.257606106 |
| Gm24841 | 0.060174537 | 0.237309859 | 3.724945049 |
| Zfp423 | 479.0759597 | -0.385477189 | 0.086603826 |
| Gm2716 | 0.511053402 | 2.245671671 | 1.322182405 |
| Cnep1r1 | 300.1426235 | 0.27085781 | 0.083521668 |
| Heatr3 | 504.4058116 | 0.022365021 | 0.060724638 |
| Gm9988 | 0.95664278 | -2.478672459 | 1.109428345 |
| Mir7071 | 0 | NA | NA |
| 9430002A10Rik | 3.207016025 | -0.814275207 | 0.592863269 |
| Papd5 | 693.6320357 | -0.040992702 | 0.07172569 |
| Adcy7 | 236.1657784 | 0.74611269 | 0.124851359 |

| | | | |
|---|---|---|---|
| Brd7 | 454.0573306 | 0.013998835 | 0.092700711 |
| Nkd1 | 108.148034 | -0.806756603 | 0.142103186 |
| Snx20 | 40.8375519 | 3.263207401 | 0.280220985 |
| Nod2 | 50.86176278 | 3.082519013 | 0.252033178 |
| Cyld | 2133.072429 | -0.485046601 | 0.063596564 |
| Rps6-ps2 | 0.516489969 | -0.355864729 | 1.391548904 |
| Mir8110 | 0 NA | | NA |
| Sall1 | 352.191393 | 0.10996252 | 0.153237589 |
| Gm6625 | 0 NA | | NA |
| Gm5356 | 0.530154252 | 1.63985367 | 1.263869649 |
| Gm26331 | 1.106943244 | 0.680095487 | 0.937130448 |
| Gm24212 | 0 NA | | NA |
| Tox3 | 156.4269144 | -0.572659385 | 0.114670698 |
| AC129321.1 | 0 NA | | NA |
| Chd9 | 2318.475297 | -0.192759523 | 0.071522192 |
| Gm6658 | 1.730564267 | 0.865224726 | 0.861488285 |
| Gm22428 | 0.075043205 | 0.237309859 | 3.724945049 |
| Rbl2 | 619.2188564 | -0.241870478 | 0.074396596 |
| Aktip | 891.7499773 | -0.3371373 | 0.062867854 |
| Gm23408 | 0 NA | | NA |
| Rpgrip1l | 501.0190372 | -0.505662368 | 0.148706803 |
| Fto | 1709.590573 | -0.372880604 | 0.047965322 |
| Gm3235 | 0.204751453 | 0.701986129 | 2.201934228 |
| Irx3 | 54.79817873 | 0.528916952 | 0.224390981 |
| Irx3os | 5.187223665 | -1.072815106 | 0.443061393 |
| Gm28511 | 0.044544648 | 0.237309859 | 3.724945049 |
| Gm28515 | 0.165933562 | 0.718213875 | 2.765757588 |
| Gm28516 | 0.048949532 | -0.243569582 | 3.724945049 |
| Gm24254 | 0 NA | | NA |
| Crnde | 9.401908757 | 0.932741068 | 0.380753256 |
| Gm21817 | 2.549468396 | 0.292510938 | 0.696422201 |
| Irx5 | 30.17646328 | 0.556768526 | 0.216972458 |
| Gm29177 | 0 NA | | NA |
| Irx6 | 3.738898168 | 0.20626885 | 0.526179039 |
| Gm3272 | 14.76791741 | 0.277055683 | 0.306410169 |
| Mmp2 | 39.1239749 | 0.331861417 | 0.179514544 |
| Lpcat2 | 68.00775929 | -0.936172444 | 0.218272858 |
| Capns2 | 4.884769305 | -1.420077217 | 0.649349874 |
| Gm22151 | 0 NA | | NA |
| Slc6a2 | 13.59669148 | -1.31150002 | 0.300594157 |
| Ces1a | 0.156716517 | -0.243563452 | 2.821984814 |
| Ces1b | 0 NA | | NA |
| Ces1c | 0.230645793 | -0.243563908 | 1.885418553 |
| Ces1d | 1.445763283 | -1.994642543 | 1.050291588 |
| Ces1e | 0 NA | | NA |
| Ces1f | 0.742894727 | -2.756923613 | 1.200765732 |
| Ces1g | 0.379165397 | 1.395557927 | 1.488805069 |

| | | | |
|---|---|---|---|
| Ces1h | 0 | NA | NA |
| Ces5a | 9.541882804 | -0.062531422 | 0.387910157 |
| Gm26843 | 19.49403685 | -0.870043818 | 0.297092161 |
| Gnao1 | 5592.419096 | -0.571255932 | 0.063317269 |
| Gm24159 | 0.183166439 | -0.24357018 | 2.686322637 |
| Amfr | 1992.652925 | 0.370612762 | 0.070183669 |
| Nudt21 | 505.4581536 | -0.108622742 | 0.079241119 |
| Ogfod1 | 1369.192929 | -0.288104879 | 0.054634762 |
| Bbs2 | 229.083479 | -0.580836229 | 0.104440118 |
| Mt4 | 0.564343484 | 2.087197229 | 1.528743719 |
| Mt3 | 1914.641927 | 0.677283979 | 0.124419684 |
| Mt2 | 2941.273043 | 2.262611926 | 0.319401316 |
| Mt1 | 3127.581112 | 1.428277312 | 0.231860834 |
| Nup93 | 580.310508 | 0.214910202 | 0.128761469 |
| Mir138-2 | 5.437333865 | 0.290305212 | 0.58443421 |
| Slc12a3 | 11.0278551 | 1.287749925 | 0.400832716 |
| Gm15889 | 0.042716945 | -0.243571806 | 3.724945049 |
| Gm15890 | 0.82545695 | 2.094430845 | 1.16872659 |
| Herpud1 | 583.7203636 | 0.246907241 | 0.09493039 |
| Ap3s1-ps2 | 8.423014622 | 0.285332657 | 0.360383521 |
| 9330175E14Rik | 54.28032715 | 7.366135546 | 0.744910818 |
| Nlrc5 | 1448.823358 | 8.647755835 | 0.252146988 |
| Mir7072 | 0.934615587 | 3.014985855 | 1.371005829 |
| Cpne2 | 360.9580537 | 1.224584199 | 0.145887502 |
| Fam192a | 433.1676574 | 0.552655984 | 0.077896712 |
| Rspry1 | 440.5534813 | -0.033845875 | 0.09083206 |
| Arl2bp | 939.3975919 | -0.029961793 | 0.074061819 |
| Pllp | 348.7112223 | -0.618188087 | 0.173211041 |
| Gm22846 | 0 | NA | NA |
| Ccl22 | 3.029817251 | -0.798237675 | 0.628824732 |
| Cx3cl1 | 4526.862607 | -0.60562924 | 0.09436346 |
| Ccl17 | 8.845676815 | -0.425914319 | 0.471227034 |
| Ciapin1 | 565.2127423 | 0.106826153 | 0.083869329 |
| Gm7418 | 0.075043205 | 0.237309859 | 3.724945049 |
| Coq9 | 367.4866646 | -0.276682282 | 0.074153372 |
| Polr2c | 334.1966378 | 0.567960085 | 0.106274692 |
| Dok4 | 119.6154433 | -0.310026299 | 0.125443819 |
| Ccdc102a | 31.06208967 | 1.713898872 | 0.295038033 |
| Adgrg5 | 33.49054987 | 4.429412451 | 0.416541194 |
| Gm10286 | 0.154272232 | 0.718212858 | 2.830050023 |
| Adgrg1 | 915.8714062 | 0.135282652 | 0.137674928 |
| Adgrg3 | 4.584657955 | 2.623632639 | 0.674252921 |
| Drc7 | 36.74700655 | 0.296035489 | 0.226040917 |
| Katnb1 | 374.5637383 | -0.079396799 | 0.085163915 |
| Kifc3 | 519.7752177 | -0.022867059 | 0.092823707 |
| Cngb1 | 16.48700249 | -0.16861131 | 0.288369365 |
| Tepp | 2.391530947 | -1.287559399 | 0.802138409 |

| | | | |
|---|---|---|---|
| Zfp319 | 384.2111316 | 0.230887642 | 0.083194913 |
| Usb1 | 184.5297308 | 1.835676456 | 0.147842738 |
| Mmp15 | 359.2212613 | -0.126411481 | 0.109670604 |
| Cfap20 | 234.0850033 | -0.213174373 | 0.087487464 |
| Csnk2a2 | 377.4618607 | 0.324048667 | 0.101131503 |
| Ccdc113 | 14.24627225 | 0.352596026 | 0.287347848 |
| Prss54 | 0.189486273 | 0.23732666 | 2.657487022 |
| Gm5912 | 6.84788527 | 0.321417983 | 0.390730628 |
| Gins3 | 30.02162402 | 0.403815839 | 0.227454578 |
| Ndrg4 | 14045.15733 | -0.794849644 | 0.062269976 |
| Setd6 | 77.58215835 | -0.02981901 | 0.162080622 |
| Cnot1 | 2334.865287 | 0.063584858 | 0.045317164 |
| Gm26493 | 9.543263168 | -0.844857511 | 0.390551736 |
| mmu-mir-7073 | 0.68581662 | -1.091097716 | 1.093291754 |
| SNORA76 | 0 | NA | NA |
| Gm26265 | 0 | NA | NA |
| 4930513N10Rik | 11.81172818 | 0.395664616 | 0.31441892 |
| Gm10094 | 48.65651831 | 0.719927373 | 0.239447002 |
| Slc38a7 | 365.4018507 | 0.414011523 | 0.07340894 |
| Got2 | 2359.910276 | 0.044634565 | 0.052865772 |
| Gm22780 | 0 | NA | NA |
| Gm24132 | 0 | NA | NA |
| Gm25160 | 0 | NA | NA |
| Gm26399 | 0.072025462 | 0.237312084 | 3.724945049 |
| Gm5913 | 8.128038844 | 0.672630751 | 0.363489865 |
| Gm7191 | 0 | NA | NA |
| Gm23494 | 0 | NA | NA |
| Gm15679 | 0.097899064 | -0.724453471 | 3.724945049 |
| Cdh8 | 694.165117 | -0.878068517 | 0.096603922 |
| Gm24693 | 0.415711858 | 1.182503169 | 1.535760186 |
| Gm15681 | 35.39573601 | -1.44240518 | 0.329703492 |
| Gm15680 | 7.35783165 | -0.935598229 | 0.414350079 |
| A330008L17Rik | 88.53438246 | -1.004402642 | 0.220386784 |
| Gm8688 | 2.373068043 | -0.750069259 | 0.825154114 |
| Gm36942 | 0 | NA | NA |
| Gm15210 | 118.5129343 | 0.629752809 | 0.1611551 |
| Gm5742 | 0.038981395 | 0.718207094 | 3.724945049 |
| Gm22223 | 0 | NA | NA |
| Gm24602 | 0 | NA | NA |
| Cdh11 | 712.29886 | -0.569948224 | 0.086024488 |
| Gm26301 | 4.260983332 | -1.276432079 | 0.603195701 |
| Gm8730 | 715.9366467 | 1.324542494 | 0.128657434 |
| Cdh5 | 215.0038206 | 2.579210136 | 0.160567293 |
| Bean1 | 97.56300219 | -0.464570726 | 0.111124662 |
| Gm10631 | 0.136963818 | 0.718204821 | 3.59954433 |
| Tk2 | 352.362705 | 0.253299693 | 0.08185655 |
| Cklf | 22.85197058 | 0.818822362 | 0.233660084 |

| | | | |
|---|---|---|---|
| Cmtm2a | 0.047948698 | -0.243569582 | 3.724945049 |
| Cmtm1 | 0.461873605 | 0.930701897 | 1.490799979 |
| Gm11020 | 0 | NA | NA |
| Cmtm2b | 0.088534442 | 1.179809211 | 3.723500796 |
| Cmtm3 | 162.243504 | 1.721332063 | 0.182158236 |
| Cmtm4 | 1562.719887 | -0.637477907 | 0.077039539 |
| Dync1li2 | 3022.19108 | -0.397512819 | 0.050904681 |
| Ccdc79 | 18.23009573 | -0.495727732 | 0.266548245 |
| Gm22148 | 0 | NA | NA |
| Nae1 | 544.9477949 | -0.025231933 | 0.076482598 |
| Car7 | 137.94085 | -0.406045872 | 0.122930806 |
| Pdp2 | 131.2365206 | -0.23148515 | 0.114226081 |
| Cdh16 | 0 | NA | NA |
| Rrad | 32.51512905 | 1.748589399 | 0.294379952 |
| Fam96b | 134.4805842 | 0.624620789 | 0.138187793 |
| 1700082M22Rik | 0.890885745 | -1.696625369 | 1.159824808 |
| AC156564.1 | 0 | NA | NA |
| Ces2a | 0.086648695 | -0.72445792 | 3.724945049 |
| 4932416K20Rik | 0 | NA | NA |
| Ces2b | 0.087930927 | 0.236669296 | 3.724898107 |
| Matr3-ps2 | 0 | NA | NA |
| Ces2c | 0 | NA | NA |
| Ces2d-ps | 0 | NA | NA |
| Ces2e | 0.208319007 | 0.702035566 | 2.190873233 |
| Ces2f | 0 | NA | NA |
| Ces2g | 3.472075296 | 0.116913522 | 0.553076336 |
| Ces2h | 0 | NA | NA |
| Ces3a | 0.090890357 | 0.718200421 | 3.724945049 |
| Ces3b | 0.093831153 | -0.243571806 | 3.724945049 |
| Ces4a | 0.088534442 | 1.179809211 | 3.723500796 |
| Gm3830 | 0 | NA | NA |
| Cbfb | 387.4469817 | 0.931546809 | 0.1322454 |
| Gm22063 | 3.344407031 | -0.375374173 | 0.559223767 |
| Gm24629 | 1.944273076 | 0.428189361 | 0.766263576 |
| D230025D16Rik | 379.8537354 | -0.268881823 | 0.085569466 |
| B3gnt9 | 40.27416652 | -0.23699049 | 0.188471201 |
| Tradd | 49.19104427 | 0.769085138 | 0.210732199 |
| Fbxl8 | 12.45441639 | 0.884293047 | 0.332936813 |
| Hsf4 | 105.9523221 | -0.370384493 | 0.135995754 |
| Nol3 | 174.8745832 | -0.339816312 | 0.11995902 |
| 4931428F04Rik | 216.122387 | -0.644254768 | 0.101511178 |
| Exoc3l | 19.59593397 | 0.184603227 | 0.258705058 |
| E2f4 | 317.7432688 | 0.605097157 | 0.072784714 |
| Elmo3 | 30.0446705 | 0.474653217 | 0.201839452 |
| Gm27320 | 4.749146687 | -0.367926047 | 0.611292743 |
| Mir328 | 3.399306995 | -0.464488373 | 0.665901735 |
| Lrrc29 | 48.99782354 | -0.534297014 | 0.238203804 |

| | | | |
|---|---|---|---|
| Tmem208 | 199.915556 | 0.676155102 | 0.108979369 |
| Fhod1 | 83.53890257 | 1.594541897 | 0.217477558 |
| Slc9a5 | 310.0134594 | -0.961043148 | 0.155140865 |
| Plekhg4 | 37.98167613 | -0.144470789 | 0.360828867 |
| Kctd19 | 6.042255962 | 5.549857279 | 0.9130859 |
| Lrrc36 | 11.57992328 | -0.173933333 | 0.299398987 |
| Tppp3 | 487.5931413 | 0.705608639 | 0.157260889 |
| Zdhhc1 | 228.3370938 | -0.460431234 | 0.096321749 |
| Hsd11b2 | 0.385063269 | -0.910309446 | 1.402017953 |
| Atp6v0d1 | 1618.375995 | -0.004432198 | 0.048883006 |
| Gm38250 | 12.77996492 | -0.273684584 | 0.357408403 |
| Agrp | 1.967041363 | 0.406897784 | 0.821827339 |
| Fam65a | 1008.594529 | -0.156323888 | 0.094875291 |
| Mir1966 | 0 | NA | NA |
| Gm5914 | 15.31483393 | -0.584329756 | 0.319608391 |
| Gm38384 | 0 | NA | NA |
| Ctcf | 822.3735919 | -0.088036912 | 0.066916768 |
| Gm5915 | 1.873087375 | 0.113805631 | 0.70243345 |
| Rltpr | 568.0764324 | -0.467624089 | 0.085526347 |
| Acd | 255.1203144 | 0.207326397 | 0.083500062 |
| Pard6a | 178.3545304 | -0.225812901 | 0.133662656 |
| Enkd1 | 47.5336699 | -0.01990839 | 0.172798602 |
| 4933405L10Rik | 1.137565759 | -0.786144409 | 0.905100948 |
| Gfod2 | 210.1736066 | 0.090717206 | 0.098454163 |
| Ranbp10 | 548.0493242 | 0.114281362 | 0.062533617 |
| Tsnaxip1 | 3.0628478 | -0.76732465 | 0.534093879 |
| Cenpt | 114.9710339 | 0.399341085 | 0.114847553 |
| Thap11 | 254.668566 | 0.068637188 | 0.082384456 |
| Nutf2 | 250.6042831 | 0.166820724 | 0.109055374 |
| Edc4 | 794.3036961 | -0.129845737 | 0.106260009 |
| Nrn1l | 0.575698224 | -0.495158591 | 1.308553362 |
| Pskh1 | 139.9944564 | 0.272998728 | 0.107646317 |
| Ctrl | 5.684503014 | 0.863022478 | 0.448621308 |
| Gm16156 | 0.885244197 | -0.526938803 | 1.122988774 |
| Psmb10 | 1108.11009 | 3.445356252 | 0.185087096 |
| Lcat | 338.7334527 | 0.915319522 | 0.180905084 |
| Slc12a4 | 501.7697974 | 1.033165832 | 0.118244295 |
| Mir7074 | 0.042716945 | -0.243571806 | 3.724945049 |
| Dpep3 | 0.127971488 | 1.23029335 | 3.685533863 |
| Dpep2 | 27.40815167 | 7.235695477 | 0.915787682 |
| Gm22972 | 0 | NA | NA |
| Dus2 | 119.8929111 | -0.086258888 | 0.12899723 |
| Ddx28 | 135.9796744 | 0.681019986 | 0.122263821 |
| Nfatc3 | 441.1062357 | 0.65993633 | 0.074843614 |
| Gm27956 | 0 | NA | NA |
| Esrp2 | 1.734636384 | 0.191799203 | 0.951798992 |
| 1810019D21Rik | 1.035937776 | 1.856272724 | 1.238981724 |

| | | | |
|---|---|---|---|
| Pla2g15 | 252.7616184 | 0.368360164 | 0.11333841 |
| Slc7a6 | 332.464804 | -0.216417738 | 0.131182668 |
| Slc7a6os | 176.7031225 | 0.422480797 | 0.129712673 |
| Prmt7 | 349.6739077 | 0.07109064 | 0.072619457 |
| Gm26786 | 0.213709657 | 0.718218858 | 2.562476354 |
| Smpd3 | 843.1766415 | -1.115272157 | 0.147755894 |
| Gm10629 | 2.127429481 | -2.02846128 | 0.804011175 |
| Zfp90 | 174.4767697 | 0.41509092 | 0.094659564 |
| 6030452D12Rik | 0.119587853 | 0.237305575 | 3.724945049 |
| Cdh3 | 29.60771109 | 1.140005632 | 0.265849283 |
| Gm10073 | 77.24332829 | 0.656855716 | 0.201172919 |
| 1110028F18Rik | 0.166497232 | 1.729737333 | 3.388675328 |
| Cdh1 | 87.82624501 | 0.868080211 | 0.174758045 |
| Tango6 | 76.05254567 | -0.137852811 | 0.175277588 |
| Has3 | 26.28930354 | -1.230099396 | 0.20826481 |
| Chtf8 | 670.8970226 | 0.253814219 | 0.107869637 |
| Cirh1a | 288.9954583 | 0.378795849 | 0.082955036 |
| Rpl10-ps2 | 1.60060964 | 2.193843793 | 0.893504955 |
| Sntb2 | 331.1315353 | -0.179751658 | 0.105067564 |
| Gm22085 | 0 | NA | NA |
| Gm16208 | 0.439742618 | -1.1786829 | 1.743977545 |
| Vps4a | 870.3055482 | -0.093135112 | 0.053942109 |
| Gm16209 | 1.711796903 | -0.112911334 | 0.779266516 |
| Pdf | 11.11514947 | -0.097148692 | 0.325291509 |
| Cog8 | 288.9214874 | 0.333503534 | 0.096511232 |
| Nip7 | 304.7822817 | 0.41346115 | 0.095402299 |
| Tmed6 | 0.075043205 | 0.237309859 | 3.724945049 |
| Terf2 | 412.9827996 | 0.014451786 | 0.067976211 |
| Mir7075 | 0 | NA | NA |
| C630050I24Rik | 0 | NA | NA |
| Cyb5b | 1556.047896 | -0.011721011 | 0.051083107 |
| Rps18-ps3 | 31.42787796 | 0.752716986 | 0.207737029 |
| Nfat5 | 2785.630128 | -0.002835059 | 0.081789297 |
| Nqo1 | 48.27701717 | -0.116433131 | 0.219334187 |
| Nob1 | 223.6934552 | 0.128385552 | 0.097277258 |
| Wwp2 | 543.3421393 | 0.542247192 | 0.081196356 |
| Rps26-ps1 | 30.55227829 | 0.533073252 | 0.205675309 |
| Mir140 | 0.882355932 | -0.546666244 | 1.183888619 |
| Psmd7 | 734.7468457 | 0.561996502 | 0.094847769 |
| Gm23646 | 0.066050875 | 0.71820487 | 3.724945049 |
| Gm22825 | 0.0885994 | -0.24357403 | 3.724945049 |
| Zfhx3 | 1136.418799 | -0.212705248 | 0.091077825 |
| Gm38042 | 32.44911505 | -0.898294199 | 0.351408125 |
| Gm38318 | 5.263687666 | -0.402500412 | 0.482858867 |
| Gm20686 | 0 | NA | NA |
| Mir3108 | 0.47188914 | 0.314729397 | 1.344760371 |
| D030068K23Rik | 6.705867478 | -0.044731846 | 0.381219265 |

| | | | |
|---|---|---|---|
| Gm23163 | 0 | NA | NA |
| Pmfbp1 | 9.435034997 | -0.427015749 | 0.465378644 |
| Dhx38 | 483.563221 | 0.476079643 | 0.097807121 |
| Txnl4b | 126.0385784 | -0.282734134 | 0.108908311 |
| Hp | 55.13014879 | 8.115524793 | 1.043169675 |
| Dhodh | 150.7597164 | -0.142847992 | 0.115231346 |
| Pkd1l3 | 15.56061734 | -0.23460269 | 0.321575747 |
| Ist1 | 750.1340784 | 0.460559606 | 0.07745317 |
| Zfp821 | 241.0196306 | 0.058758874 | 0.100973701 |
| Atxn1l | 914.3276386 | 0.144524379 | 0.088620761 |
| Ap1g1 | 1809.349129 | 0.132185538 | 0.047093132 |
| Snord71 | 7.711840907 | -0.550577221 | 0.388587783 |
| Gm25321 | 0 | NA | NA |
| Gm17344 | 3.933162639 | -0.258608937 | 0.486166936 |
| Phlpp2 | 846.0254909 | -0.447698218 | 0.07367585 |
| Gm16349 | 3.039183673 | -1.574553027 | 0.759767699 |
| Marveld3 | 4.627756683 | 0.898035577 | 0.482373085 |
| Tat | 1.816527333 | -0.32656379 | 0.728812133 |
| Chst4 | 0.843638514 | 1.316774037 | 1.01901183 |
| Zfp612 | 420.3106796 | -0.271398163 | 0.091522864 |
| Gm17720 | 0 | NA | NA |
| Gm21964 | 0.038981395 | 0.718207094 | 3.724945049 |
| Calb2 | 675.5295577 | -0.421791787 | 0.134926863 |
| Cmtr2 | 86.97394878 | 0.161927361 | 0.123278759 |
| Gm26832 | 0 | NA | NA |
| Hydin | 49.41383929 | -0.803587816 | 0.205118039 |
| Gm23627 | 0 | NA | NA |
| Gm22701 | 0 | NA | NA |
| AC069308.1 | 0 | NA | NA |
| AC069308.2 | 0 | NA | NA |
| Vac14 | 425.0913073 | 0.432865307 | 0.071437386 |
| Mtss1l | 3340.251647 | -0.300713688 | 0.106302368 |
| Il34 | 286.7182791 | -0.468532878 | 0.111850989 |
| Gm15894 | 3.21488869 | 0.905588635 | 0.635195756 |
| Gm15895 | 1.066917243 | 0.51103727 | 1.100834969 |
| Sf3b3 | 1283.257394 | 0.672655436 | 0.069230774 |
| Snord111 | 4.735066032 | 0.270643548 | 0.548046227 |
| Gm22193 | 2.867962316 | -0.27306425 | 0.642058457 |
| Cog4 | 549.572635 | 0.438663864 | 0.074858538 |
| Gm26816 | 0.327007429 | 0.718887983 | 1.984158798 |
| Fuk | 171.2700549 | -0.559962135 | 0.131504694 |
| St3gal2 | 1026.922896 | -0.459538848 | 0.065386821 |
| Gm26132 | 2.688917129 | -0.705834318 | 0.674139543 |
| Ddx19a | 222.5239366 | 0.342040338 | 0.096114381 |
| Ddx19b | 400.244091 | -0.02209803 | 0.093818192 |
| Mir3473d | 0.67306248 | -0.45367356 | 1.205442699 |
| Aars | 2524.454962 | 0.850965277 | 0.100765004 |

| | | | |
|---|---|---|---|
| Gm24103 | 0 | NA | NA |
| Clec18a | 9.750147804 | -0.470580064 | 0.335668397 |
| Pdpr | 569.8637507 | -0.160333769 | 0.062490627 |
| 9430091E24Rik | 22.80186376 | -0.769918177 | 0.245485281 |
| Glg1 | 3651.900435 | -0.164470641 | 0.060188727 |
| Gm23193 | 0 | NA | NA |
| Rfwd3 | 350.9863053 | 0.213410379 | 0.089402322 |
| Mlkl | 209.3845513 | 8.45982279 | 0.616683464 |
| Fa2h | 504.9529093 | -0.014026696 | 0.168036098 |
| Gm24793 | 0 | NA | NA |
| Wdr59 | 307.9043311 | -0.152469997 | 0.071030434 |
| Mir1957b | 0.047948698 | -0.243569582 | 3.724945049 |
| Znrf1 | 1233.211656 | -0.261272772 | 0.079891946 |
| Ldhd | 179.3875016 | -0.313047197 | 0.097540233 |
| Zfp1 | 213.1490368 | 0.343482439 | 0.112765533 |
| Ctrb1 | 0.649033156 | 2.736744938 | 1.318607136 |
| Mir7076 | 0 | NA | NA |
| Bcar1 | 823.9151968 | 0.107901638 | 0.106106136 |
| Cfdp1 | 449.2752954 | 0.186465238 | 0.130197157 |
| Tmem170 | 50.89811066 | 0.008370642 | 0.15464978 |
| Gm23853 | 0 | NA | NA |
| Chst5 | 6.122444543 | 0.662994116 | 0.446106526 |
| Tmem231 | 128.7512586 | -0.462870028 | 0.107609868 |
| Gabarapl2 | 905.821491 | -0.03986482 | 0.081196712 |
| Adat1 | 132.6219706 | -0.681151119 | 0.211832427 |
| Kars | 1033.948333 | 1.171844359 | 0.120220803 |
| Terf2ip | 508.2144975 | 0.19182553 | 0.081380259 |
| Gm6793 | 13.31818528 | 0.587648573 | 0.425879818 |
| Gm22827 | 0 | NA | NA |
| Gm26994 | 1.945104103 | 0.214300612 | 0.68206211 |
| Cntnap4 | 605.8576847 | -0.44312543 | 0.07791732 |
| Gm22664 | 1.761579563 | -0.726392982 | 0.826029056 |
| Gm25766 | 0 | NA | NA |
| Gm24291 | 0 | NA | NA |
| Mon1b | 489.7840867 | 0.031821673 | 0.073433906 |
| Syce1l | 2.596219419 | 0.264007581 | 0.740428447 |
| Adamts18 | 110.6812341 | -0.370379709 | 0.205756883 |
| Nudt7 | 34.66451103 | 0.688758821 | 0.227112921 |
| Vat1l | 1139.282187 | -0.057317465 | 0.10887063 |
| 4933408N05Rik | 0.707695957 | 0.761204529 | 1.463770057 |
| Clec3a | 0.137470077 | 0.237309658 | 3.595099416 |
| Wwox | 259.2877687 | -0.296402338 | 0.075142445 |
| Gm16116 | 0 | NA | NA |
| Gm16117 | 0.066050875 | 0.71820487 | 3.724945049 |
| Gm16118 | 0.567921522 | 2.859230537 | 1.227192628 |
| Gm22556 | 0 | NA | NA |
| Gm23677 | 1.922491501 | -1.002152764 | 0.706930708 |

| | | | |
|---|---|---|---|
| Maf | 623.4348012 | 0.513494103 | 0.104891027 |
| Gm15655 | 1.118566861 | 0.840012527 | 0.877729216 |
| Dynlrb2 | 11.99619067 | 0.250778782 | 0.287966557 |
| Cdyl2 | 447.4685583 | -0.013874315 | 0.089523077 |
| Gm10620 | 0 | NA | NA |
| Cmc2 | 85.51804082 | -0.454603311 | 0.122915921 |
| Cenpn | 11.50529104 | 0.130299666 | 0.386356629 |
| Atmin | 1056.148415 | -0.09696333 | 0.07640353 |
| 1700030J22Rik | 85.33078503 | -0.543782883 | 0.199755902 |
| Gcsh | 322.5962855 | -0.069435347 | 0.112958486 |
| Pkd1l2 | 5.568506049 | 2.157971798 | 0.48003493 |
| Gm15395 | 0 | NA | NA |
| Bco1 | 1.165159028 | 1.86549102 | 1.035806089 |
| Gm20694 | 0 | NA | NA |
| Gan | 644.4361812 | -0.255320763 | 0.08379992 |
| Cmip | 3352.599209 | 0.140691089 | 0.074353748 |
| Mir7077 | 0.151026272 | -0.243569982 | 2.852050277 |
| Mir7078 | 0.769887329 | 1.238763046 | 1.213003017 |
| Plcg2 | 185.0113621 | 2.18696009 | 0.175110623 |
| Gm27361 | 0 | NA | NA |
| Sdr42e1 | 58.48922315 | -0.156985842 | 0.179124214 |
| Hsd17b2 | 4.082108608 | -0.277603361 | 0.513413727 |
| Mphosph6 | 123.4766612 | 0.215730564 | 0.120102017 |
| Gm10617 | 0.340027382 | -2.419886016 | 1.424142586 |
| Gm25200 | 0 | NA | NA |
| Cdh13 | 744.3496955 | -0.629678991 | 0.065883841 |
| Gm24035 | 0 | NA | NA |
| Gm22999 | 0.35553296 | -1.525372937 | 1.712411637 |
| Hsbp1 | 1254.7106 | 0.151934705 | 0.053767399 |
| Mlycd | 201.1556644 | 0.114922879 | 0.098213201 |
| Osgin1 | 24.0618504 | 2.802056203 | 0.377065949 |
| Necab2 | 468.9419142 | -0.780880853 | 0.149701722 |
| Slc38a8 | 4.05865859 | -0.959084626 | 0.615058208 |
| Mbtps1 | 1032.591658 | -0.099792544 | 0.068890271 |
| Hsdl1 | 556.8575977 | 0.058636119 | 0.080863038 |
| Dnaaf1 | 4.031485904 | -0.03173558 | 0.543356686 |
| Taf1c | 112.5897251 | 0.395606493 | 0.108502064 |
| Rps13-ps4 | 4.918628032 | 0.129508802 | 0.485857585 |
| Adad2 | 0.577193238 | -0.069365099 | 1.285634787 |
| Kcng4 | 392.3468948 | 0.005869123 | 0.174516002 |
| Wfdc1 | 23.0470312 | 0.409976083 | 0.293798244 |
| Atp2c2 | 14.74911433 | 0.919988383 | 0.344787599 |
| Tldc1 | 91.32362537 | 0.05846546 | 0.112550055 |
| Cotl1 | 860.7212699 | 1.108536399 | 0.112110896 |
| Klhl36 | 75.98809112 | 0.026764926 | 0.131745414 |
| Usp10 | 645.8447964 | 0.014510963 | 0.056707607 |
| Gm20388 | 0 | NA | NA |

| | | | |
|---|---|---|---|
| Crispld2 | 132.7086816 | 1.998506734 | 0.29390014 |
| Gm15684 | 0.208379788 | 0.717452451 | 2.639584027 |
| Zdhhc7 | 282.094203 | 0.263973231 | 0.102163332 |
| Gm15898 | 0 | NA | NA |
| A130014A01Rik | 33.99075599 | -0.252697918 | 0.168569558 |
| Gm38379 | 3.088015912 | -1.838191812 | 0.590792256 |
| 6430548M08Rik | 3271.759844 | -0.14597664 | 0.095376162 |
| Fam92b | 3.336165937 | -0.207107143 | 0.554627673 |
| A330074K22Rik | 139.4201663 | -0.274837513 | 0.151362313 |
| Gse1 | 1501.615146 | 0.108440586 | 0.093709102 |
| Gm22715 | 0 | NA | NA |
| Gm26971 | 9.453064617 | -0.718464823 | 0.511381613 |
| 1700016A09Rik | 5.586702209 | -0.171796658 | 0.461986383 |
| Mir7687 | 1.495561828 | -2.764514479 | 1.098110255 |
| Gins2 | 20.25107559 | 1.52186975 | 0.366596781 |
| Gm10614 | 0.913620126 | -0.309022102 | 0.996716597 |
| Gm26739 | 0.232719486 | 0.23732887 | 2.133424764 |
| 1190005I06Rik | 8.331890292 | -0.828549916 | 0.368016423 |
| Gm27021 | 0.176744647 | 0.237319179 | 2.719535928 |
| Gm26537 | 0 | NA | NA |
| Emc8 | 634.0511647 | -0.126348774 | 0.072477506 |
| Cox4i1 | 2787.69123 | -0.243741513 | 0.091990731 |
| Gm33142 | 0.833731302 | 1.542460923 | 1.221859116 |
| Gm24239 | 0 | NA | NA |
| Gm18709 | 0.119587853 | 0.237305575 | 3.724945049 |
| Irf8 | 490.9940863 | 4.299269442 | 0.190627424 |
| Gm26878 | 0.075043205 | 0.237309859 | 3.724945049 |
| Gm28324 | 0 | NA | NA |
| Fendrr | 2.814059987 | 1.25383557 | 0.687675959 |
| Foxf1 | 17.00115138 | 2.513796103 | 0.385318985 |
| Gm27530 | 0.136963818 | 0.718204821 | 3.59954433 |
| Mthfsd | 357.8554343 | -0.284764469 | 0.113515967 |
| Foxc2 | 30.92999064 | 0.380679396 | 0.255177086 |
| AC127554.1 | 0 | NA | NA |
| Foxl1 | 1.257454774 | 0.718406514 | 1.045835559 |
| Gm37609 | 0.320352182 | -0.77878809 | 1.943260801 |
| 5033426O07Rik | 1.128616957 | -0.256512302 | 1.094404165 |
| Gm26815 | 0 | NA | NA |
| Gm26747 | 6.573009254 | -0.56730689 | 0.379377882 |
| Gm26784 | 0.109869311 | 0.238305784 | 3.724874707 |
| 1700018B08Rik | 0.229212118 | -0.243570082 | 2.547047007 |
| Gm27045 | 0.361384634 | -0.63709112 | 1.773524649 |
| 1700030M09Rik | 0.55198573 | 1.999369095 | 1.272156676 |
| Fbxo31 | 1046.295583 | -0.501473888 | 0.07487092 |
| Gm17786 | 1.168706595 | 0.464147853 | 0.904848102 |
| Map1lc3b | 1461.563508 | 0.168270178 | 0.077285922 |
| Zcchc14 | 1352.76978 | -0.367069417 | 0.097075334 |

| | | | |
|---|---|---|---|
| Gm27030 | 11.36160826 | -1.317419421 | 0.340243768 |
| Gm23299 | 0.157818009 | -0.186853641 | 3.472940599 |
| Jph3 | 2592.754649 | -0.786484651 | 0.080776383 |
| Klhdc4 | 215.8889794 | 0.560074553 | 0.141988197 |
| Gm20406 | 0 | NA | NA |
| Gm24283 | 0.323040168 | 1.049337984 | 1.743213369 |
| Gm26812 | 0.227390921 | 0.71822274 | 2.128167085 |
| 9530085P06Rik | 0.295432452 | 1.758271432 | 1.773903645 |
| Slc7a5 | 545.5188095 | 1.426743829 | 0.136824402 |
| BC048644 | 2.632535955 | 0.406271131 | 0.577147552 |
| Car5a | 0.305395298 | -0.929456877 | 1.783827178 |
| Gm26999 | 0.075043205 | 0.237309859 | 3.724945049 |
| Gm37497 | 0 | NA | NA |
| Gm27011 | 0.490536136 | 1.157968843 | 1.573580021 |
| Banp | 309.1391948 | -0.557688081 | 0.172424162 |
| Gm17709 | 4.900990937 | 1.281125647 | 0.576263773 |
| Mir7237 | 0.780673905 | -1.386351047 | 1.103293038 |
| Gm22 | 32.71914304 | 0.817380529 | 0.202602678 |
| Zfpm1 | 722.8421094 | 0.812786021 | 0.174570706 |
| Trhr2 | 33.68978649 | -0.070087471 | 0.261661007 |
| Gm20681 | 0.36185533 | 1.377395289 | 1.525396811 |
| Zc3h18 | 664.5062256 | 0.265590722 | 0.086188379 |
| Il17c | 0.855647594 | 1.955915342 | 1.434169995 |
| Cyba | 233.8961265 | 4.056348666 | 0.232426675 |
| Mvd | 137.2979585 | -0.199747419 | 0.117567682 |
| 9330133O14Rik | 42.62923582 | -0.211947635 | 0.174897741 |
| Snai3 | 6.952624755 | 1.505095018 | 0.462218009 |
| Rnf166 | 241.1004881 | 0.126151132 | 0.090134633 |
| Gm24366 | 2.393340753 | -1.684872703 | 0.706659355 |
| Ctu2 | 170.7851 | 0.206217433 | 0.097615973 |
| Piezo1 | 341.3561795 | 2.222912428 | 0.160426341 |
| Gm26497 | 0.280709254 | -0.745060788 | 1.788608539 |
| Cdt1 | 76.18453122 | 2.72899769 | 0.288244562 |
| Aprt | 172.1584049 | 1.056346197 | 0.138097052 |
| Galns | 112.6437773 | 1.322501425 | 0.154662122 |
| Trappc2l | 257.9503243 | 0.009321758 | 0.078189644 |
| Pabpn1l | 0.259153158 | -0.218268419 | 2.089255896 |
| Mir6396 | 0 | NA | NA |
| Cbfa2t3 | 329.1625399 | -0.470465759 | 0.164367011 |
| Gm15899 | 22.41855246 | 0.655997689 | 0.230542225 |
| Gm10612 | 0.141684195 | 0.718205511 | 3.558854591 |
| Acsf3 | 163.1806245 | -0.320558496 | 0.114492901 |
| Gm16378 | 9.858353745 | -0.07641425 | 0.306621091 |
| Cdh15 | 97.85341112 | 0.035871583 | 0.237378861 |
| Ankrd11 | 2871.409642 | -0.127065876 | 0.07973419 |
| Gm24445 | 2.473368846 | -0.390368495 | 0.72673885 |
| 2810013P06Rik | 123.5382447 | -0.226298317 | 0.110837089 |

| | | | |
|---|---|---|---|
| Spg7 | 394.1502427 | -0.179804824 | 0.08666135 |
| Rpl13 | 1506.813565 | 0.559513666 | 0.113217745 |
| Snord68 | 0 | NA | NA |
| Mir7079 | 0 | NA | NA |
| Cpne7 | 380.383942 | -0.961744401 | 0.173141853 |
| Mir7080 | 0 | NA | NA |
| Sult5a1 | 0.948467964 | 0.146838743 | 1.039926401 |
| Dpep1 | 2.460632423 | 2.01968771 | 0.836826338 |
| Chmp1a | 416.7948105 | 0.518816865 | 0.089643546 |
| Rps12-ps9 | 5.02878481 | 0.753270636 | 0.565364043 |
| Spata33 | 6.260054198 | -0.323690291 | 0.418259516 |
| Cdk10 | 231.2422657 | -0.500741362 | 0.093654287 |
| Spata2l | 264.1685552 | -0.850404355 | 0.159799533 |
| 4933417D19Rik | 2.606279457 | -0.714626234 | 0.635838351 |
| Vps9d1 | 418.2806154 | -0.152770885 | 0.099296453 |
| Zfp276 | 246.8177323 | 0.059806246 | 0.114853024 |
| Fanca | 30.7735149 | 2.492375347 | 0.289011989 |
| Spire2 | 197.4572324 | -0.010793503 | 0.105879862 |
| Tcf25 | 4948.09245 | -0.153958029 | 0.0564195 |
| Mc1r | 2.754460718 | 2.348125463 | 0.723935707 |
| Gm38092 | 0.099155932 | 0.718202646 | 3.724945049 |
| Tubb3 | 2462.821606 | -0.341077609 | 0.097896602 |
| Def8 | 517.338147 | -0.237462621 | 0.060629299 |
| Gm25428 | 0 | NA | NA |
| Afg3l1 | 423.4960045 | 0.426807394 | 0.063087949 |
| Dbndd1 | 119.4149113 | -0.6574686 | 0.109031961 |
| Gas8 | 123.9867013 | 0.33750321 | 0.180367907 |
| Gm22314 | 2.308872772 | -0.158554669 | 0.71972299 |
| n-R5s100 | 0 | NA | NA |
| n-R5s101 | 0 | NA | NA |
| n-R5s102 | 0 | NA | NA |
| n-R5s103 | 0 | NA | NA |
| n-R5s104 | 0 | NA | NA |
| n-R5s105 | 0 | NA | NA |
| n-R5s106 | 0 | NA | NA |
| n-R5s107 | 0 | NA | NA |
| n-R5s108 | 0 | NA | NA |
| n-R5s109 | 0 | NA | NA |
| n-R5s110 | 0 | NA | NA |
| n-R5s111 | 0 | NA | NA |
| n-R5s112 | 0 | NA | NA |
| n-R5s113 | 0 | NA | NA |
| n-R5s114 | 0 | NA | NA |
| n-R5s115 | 0 | NA | NA |
| Gm25018 | 0 | NA | NA |
| n-R5s117 | 0 | NA | NA |
| n-R5s118 | 0 | NA | NA |

| | | | |
|---|---|---|---|
| n-R5s119 | 0 | NA | NA |
| n-R5s120 | 0 | NA | NA |
| n-R5s121 | 0 | NA | NA |
| n-R5s122 | 0 | NA | NA |
| n-R5s123 | 0 | NA | NA |
| n-R5s124 | 0 | NA | NA |
| n-R5s125 | 0 | NA | NA |
| Gm24312 | 0 | NA | NA |
| n-R5s127 | 0 | NA | NA |
| n-R5s128 | 0 | NA | NA |
| n-R5s129 | 0 | NA | NA |
| n-R5s130 | 0 | NA | NA |
| n-R5s131 | 0 | NA | NA |
| Gm26391 | 0 | NA | NA |
| n-R5s133 | 0 | NA | NA |
| n-R5s134 | 0 | NA | NA |
| Rn5s | 0 | NA | NA |
| n-R5s136 | 0 | NA | NA |
| Gm24089 | 0 | NA | NA |
| n-R5s138 | 0 | NA | NA |
| n-R5s139 | 0 | NA | NA |
| Gm22109 | 0 | NA | NA |
| n-R5s141 | 0 | NA | NA |
| n-R5s142 | 0 | NA | NA |
| n-R5s143 | 0 | NA | NA |
| n-R5s144 | 0 | NA | NA |
| Gm25212 | 0 | NA | NA |
| n-R5s146 | 0 | NA | NA |
| Gm22291 | 0 | NA | NA |
| Gm23284 | 0 | NA | NA |
| n-R5s149 | 0 | NA | NA |
| Gm23647 | 0 | NA | NA |
| Rhou | 659.2759545 | 0.58046097 | 0.125248299 |
| 4732419C18Rik | 5.763193032 | 1.247310022 | 0.504677971 |
| Rab4a | 299.8235972 | -0.464046945 | 0.078633382 |
| 2810455O05Rik | 58.83641319 | -0.85421726 | 0.261464687 |
| Ccsap | 351.2028742 | -0.360148862 | 0.148815027 |
| Gm4342 | 0.738388108 | 1.469470715 | 1.219208241 |
| AC147266.1 | 3.758891091 | 0.069867378 | 0.535906939 |
| Acta1 | 38.57868714 | 0.022185828 | 0.174797215 |
| Nup133 | 281.5212268 | -0.178155461 | 0.07080565 |
| Abcb10 | 231.5059475 | -0.049854682 | 0.110446372 |
| Taf5l | 237.4112607 | 0.378752322 | 0.111810566 |
| Urb2 | 291.7584138 | 0.147793531 | 0.08720934 |
| Mir1967 | 0.881693066 | 0.545895569 | 1.09401791 |
| Galnt2 | 11.54263694 | -0.028603801 | 0.303285575 |
| 1810008B01Rik | 0.299839612 | -0.720056397 | 1.575758354 |

| | | | |
|---|---|---|---|
| Gm15775 | 0.11147366 | 1.230376831 | 3.722845817 |
| Gm24459 | 0 | NA | NA |
| Pgbd5 | 1529.511435 | -0.282615072 | 0.126381549 |
| Cog2 | 399.9630909 | 0.082593961 | 0.081168916 |
| Agt | 480.964286 | -0.638101494 | 0.259575641 |
| Capn9 | 3.44454021 | 1.504384729 | 0.610504262 |
| 2310022B05Rik | 690.1050064 | -1.08996811 | 0.186184313 |
| Ttc13 | 501.1809921 | -0.275317973 | 0.132702545 |
| Arv1 | 50.9902722 | -0.100785428 | 0.171568281 |
| Fam89a | 4.128209337 | 0.996166833 | 0.539803711 |
| Trim67 | 432.3924619 | 0.075733058 | 0.133551318 |
| Gm16163 | 3.256114358 | -0.920870565 | 0.684985275 |
| 2810004N23Rik | 135.240311 | 0.388776054 | 0.113466115 |
| Gnpat | 451.9781084 | -0.245334309 | 0.075707423 |
| Exoc8 | 423.6774759 | -0.124679529 | 0.06965665 |
| Sprtn | 158.0541678 | 0.26631693 | 0.11816536 |
| Egln1 | 684.0866431 | -0.195181827 | 0.085076905 |
| Tsnax | 775.1924979 | -0.164750475 | 0.080066481 |
| Disc1 | 23.96939849 | -0.252412922 | 0.215948921 |
| Gm22197 | 0.208494586 | -0.243564262 | 2.629256435 |
| Gm16237 | 0.184167273 | -0.243570149 | 2.681673463 |
| Sipa1l2 | 1159.737547 | 0.210910982 | 0.084556585 |
| Map10 | 80.06219714 | 0.827043061 | 0.129660533 |
| Ntpcr | 128.5931328 | 0.020115046 | 0.16355782 |
| Pcnxl2 | 1292.263944 | -0.560043724 | 0.121943142 |
| A730098A19Rik | 4.29871648 | -0.299339093 | 0.494942703 |
| BC021891 | 32.34379101 | -0.587443645 | 0.297355022 |
| Kcnk1 | 670.0428367 | -0.029372054 | 0.09289751 |
| Slc35f3 | 269.0447631 | -0.747148429 | 0.170516199 |
| Coa6 | 108.6067078 | 0.261148857 | 0.151836047 |
| Gm17296 | 160.9155127 | -0.731892534 | 0.185173484 |
| Gm6091 | 0 | NA | NA |
| Gm26759 | 0.554140925 | -1.393875658 | 1.548951075 |
| Irf2bp2 | 1675.480812 | 0.553503567 | 0.116497068 |
| Tomm20 | 2682.24209 | -0.508123793 | 0.046946495 |
| Gm26397 | 173.2653863 | 0.016124979 | 0.174707726 |
| Rbm34 | 225.3370361 | 0.193838582 | 0.116558515 |
| Gm22730 | 0 | NA | NA |
| Pard3 | 532.7837827 | -0.102624476 | 0.06903011 |
| Gm6921 | 2.797109942 | -0.51995627 | 0.577954015 |
| Gm22968 | 0.822547242 | -0.927516886 | 1.1276958 |
| Mir21c | 0.121578509 | 0.718202646 | 3.724945049 |
| Nrp1 | 639.8097654 | 0.687355042 | 0.103165502 |
| Mir1903 | 0 | NA | NA |
| Gm26002 | 0 | NA | NA |
| Itgb1 | 1279.915761 | 0.986128998 | 0.100866326 |
| Gm3952 | 0.143618953 | 0.68463387 | 2.890309179 |

| | | | |
|---|---|---|---|
| Gm26100 | 0.368403345 | 1.669814594 | 1.713670053 |
| Ccdc7 | 0.294450523 | 0.462448713 | 1.728428269 |
| Gm22545 | 0 | NA | NA |
| 1700008F21Rik | 0.249972158 | 0.237328829 | 2.09832297 |
| Gm22374 | 0.159888962 | -0.724460061 | 3.461347753 |
| Gm10999 | 0.583900451 | 1.846202342 | 1.365028086 |
| 2610044O15Rik8 | 322.1898291 | -0.42273746 | 0.131288564 |
| Gm21399 | 59.65666709 | 1.055350684 | 0.164531201 |
| Gm26870 | 1.94439448 | 1.844774428 | 1.014133061 |
| Gm10722 | 0.242739364 | 0.676283093 | 2.085877574 |
| Gm11168 | 0.318248637 | 0.304523493 | 2.036557635 |
| Gm10721 | 0.187213551 | 0.23732772 | 2.667715587 |
| Gm10720 | 0 | NA | NA |
| Gm10719 | 0.044544648 | 0.237309859 | 3.724945049 |
| Gm10718 | 0.690752925 | -0.993429655 | 1.267736865 |
| Gm10717 | 0.158171364 | -0.724458613 | 2.813813679 |
| Gm17535 | 0 | NA | NA |
| Gm10715 | 0 | NA | NA |
| mmu-mir-101c | 0 | NA | NA |
| 4930433N12Rik | 0 | NA | NA |
| Gm27679 | 0 | NA | NA |
| Alkbh8 | 460.6917347 | -0.250528485 | 0.058674071 |
| Gm24918 | 2.468298956 | 0.947638193 | 0.706252779 |
| Gm23275 | 1.098824885 | -0.816220981 | 1.267916257 |
| Cwf19l2 | 426.6056693 | 0.497846006 | 0.115362976 |
| Gucy1a2 | 1402.939796 | -0.644112112 | 0.096212279 |
| Gm22195 | 0.192501058 | -0.629765963 | 2.663543715 |
| Aasdhppt | 467.0250061 | -0.093135801 | 0.088488543 |
| Kbtbd3 | 165.3891345 | -0.065149259 | 0.133948763 |
| Gm23992 | 0.044191549 | -0.243571806 | 3.724945049 |
| Msantd4 | 720.9495236 | 0.280366681 | 0.069609927 |
| Gria4 | 2416.785679 | -0.724201001 | 0.085455494 |
| Gm23811 | 8.724062623 | -1.722446636 | 0.407162665 |
| Gm24896 | 0 | NA | NA |
| Casp1 | 190.837744 | 4.153085097 | 0.245452194 |
| Casp4 | 185.281841 | 6.934595786 | 0.468612402 |
| Casp12 | 173.1181621 | 3.760085421 | 0.231332171 |
| Pdgfd | 56.87235133 | -0.465224038 | 0.236277889 |
| Ddi1 | 0 | NA | NA |
| Gm23097 | 0 | NA | NA |
| Dync2h1 | 1480.682915 | -0.529148179 | 0.10100116 |
| Dcun1d5 | 410.911556 | 0.146482759 | 0.090723814 |
| Mmp13 | 18.19977551 | 4.78745352 | 0.617021479 |
| Mmp12 | 7.267951943 | 3.080239832 | 0.586317875 |
| Mmp1b | 0 | NA | NA |
| Mmp3 | 11.53096597 | 6.704228895 | 0.952935732 |
| Mmp1a | 0.181654796 | 1.679598083 | 3.311020364 |

| | | | |
|---|---|---|---|
| Mmp10 | 0.099106094 | 1.209894437 | 3.72418822 |
| Mmp8 | 40.56306267 | 8.22462483 | 1.19135512 |
| Mmp27 | 4.159708068 | 6.058288082 | 0.998149793 |
| Mmp20 | 0.336931769 | 1.205424134 | 1.762671907 |
| AV064505 | 0 | NA | NA |
| Mmp7 | 0.107803068 | 0.719187033 | 3.724874737 |
| Gm10709 | 0.713745664 | 0.524162015 | 1.042619696 |
| Tmem123 | 239.3847869 | 1.19871238 | 0.133957308 |
| Mir5618 | 0.424774107 | -0.533829452 | 1.38201678 |
| Birc2 | 772.896626 | 0.534070627 | 0.107231742 |
| Birc3 | 154.0540683 | 3.557677708 | 0.275503968 |
| Yap1 | 390.0541373 | 0.72223485 | 0.101649221 |
| Gm20416 | 0.109982625 | 0.237318756 | 3.724945049 |
| 9230110C19Rik | 76.10268791 | -0.080116129 | 0.136551897 |
| Cep126 | 298.1066087 | -0.207293504 | 0.085758553 |
| 1700128F08Rik | 0.650208918 | -0.125702518 | 1.203131828 |
| Mir1899 | 0.060174537 | 0.237309859 | 3.724945049 |
| Trpc6 | 122.6815528 | -0.837178065 | 0.155892888 |
| Pgr | 476.6556311 | -0.895008633 | 0.090168016 |
| Gm16485 | 192.3635005 | -0.982188132 | 0.138745725 |
| Arhgap42 | 263.2227951 | -0.445161417 | 0.081751991 |
| Gm16833 | 0 | NA | NA |
| Gm22815 | 0 | NA | NA |
| Cntn5 | 249.9584538 | -1.140055804 | 0.128232434 |
| Gm26195 | 0.87738752 | -1.459111552 | 1.200934488 |
| Mir6237 | 0 | NA | NA |
| Gm24496 | 0.646963711 | -1.144341034 | 1.213387316 |
| Gm25086 | 0.47668666 | -0.196294317 | 1.699137801 |
| Gm25365 | 0 | NA | NA |
| Gm21070 | 0 | NA | NA |
| Gm29176 | 0 | NA | NA |
| Jrkl | 160.5671298 | 0.083500291 | 0.137801861 |
| Ccdc82 | 240.4966967 | -0.224246339 | 0.095382028 |
| Gm26189 | 1.402298525 | 1.761775471 | 0.899817808 |
| Phxr4 | 43.51084174 | 0.068598654 | 0.329191638 |
| Gm17571 | 0 | NA | NA |
| Maml2 | 397.5331643 | 0.660503068 | 0.086942635 |
| Gm22643 | 0 | NA | NA |
| Mtmr2 | 890.4922701 | -0.13434632 | 0.103515668 |
| Cep57 | 215.6600884 | 0.235360983 | 0.120564819 |
| Fam76b | 356.5380025 | 0.010955841 | 0.143921554 |
| Sesn3 | 970.3136834 | -0.056686389 | 0.083234848 |
| Endod1 | 2711.618671 | 1.158940929 | 0.17972634 |
| Kdm4dl | 4.695898157 | -0.046843674 | 0.56551781 |
| Kdm4d | 25.20782128 | -0.287470322 | 0.227451848 |
| Cwc15 | 455.3458575 | 0.511399097 | 0.13097798 |
| Amotl1 | 1488.185642 | 1.149268628 | 0.112422287 |

| | | | |
|---|---|---|---|
| Piwil4 | 14.13799992 | 5.49572583 | 0.747824896 |
| Gm10706 | 3.101177844 | 0.169590976 | 0.523819001 |
| Gm15586 | 0 | NA | NA |
| Fut4 | 18.43119656 | 1.822241278 | 0.295778108 |
| 1700012B09Rik | 5.943573362 | 0.185402935 | 0.428488532 |
| Ankrd49 | 112.3845942 | 0.660662011 | 0.126863265 |
| Mre11a | 230.8957433 | 0.184633195 | 0.090050758 |
| Gm16379 | 0.870448589 | 1.078757639 | 1.077071867 |
| Gm16302 | 2.36738784 | 0.716080883 | 0.642512387 |
| Gpr83 | 316.7124723 | 0.220085962 | 0.125147648 |
| Izumo1r | 3.609677315 | 5.475488059 | 0.922962358 |
| Panx1 | 176.4686585 | 0.643538796 | 0.113239458 |
| Hephl1 | 1.803729066 | -0.16443595 | 0.882757206 |
| Gm22658 | 0.042716945 | -0.243571806 | 3.724945049 |
| Vstm5 | 75.69295347 | -0.274440196 | 0.175826282 |
| Med17 | 334.3072139 | 0.415183859 | 0.075253542 |
| 4931406C07Rik | 230.1835539 | 0.299024564 | 0.117900755 |
| Taf1d | 188.0883961 | 0.426662096 | 0.151723234 |
| Gm25500 | 1.755293934 | -0.558597995 | 0.822719758 |
| mmu-mir-7650 | 0 | NA | NA |
| Gm24455 | 11.03006032 | 0.193527398 | 0.327983009 |
| Gm24299 | 7.628192187 | -0.612914157 | 0.391523751 |
| AC154296.1 | 0.044191549 | -0.243571806 | 3.724945049 |
| Gm25791 | 5.041681076 | 0.282755129 | 0.515369659 |
| Gm22620 | 6.972143724 | -0.017052743 | 0.411910835 |
| Gm24357 | 17.62472295 | 1.345003158 | 0.282776992 |
| Gm23455 | 26.8443768 | 1.603342718 | 0.263571733 |
| Gm22579 | 6.383805412 | -0.078148945 | 0.457045226 |
| Cep295 | 648.5597578 | -0.483864218 | 0.113568615 |
| Scarna9 | 8.758301848 | -1.144033841 | 0.372030806 |
| Gm24339 | 46.69042643 | -0.655314179 | 0.277618171 |
| Gm16568 | 0.41713571 | 0.171965893 | 1.291451138 |
| Smco4 | 17.55463808 | 0.351646041 | 0.2868739 |
| Ccdc67 | 5.175431821 | 0.461624602 | 0.459321904 |
| Slc36a4 | 909.8448179 | -0.302295261 | 0.073959972 |
| Mtnr1b | 0.066050875 | 0.71820487 | 3.724945049 |
| Fat3 | 3920.904863 | -0.964453852 | 0.132528003 |
| Gm5611 | 1.991326778 | -0.314565099 | 0.760496749 |
| CT943659.1 | 0 | NA | NA |
| Chordc1 | 659.9625695 | 0.130348585 | 0.1099622 |
| Naalad2 | 9.142264245 | -0.964322716 | 0.464906237 |
| Ubtfl1 | 0.110595523 | 0.718200421 | 3.724945049 |
| Gm5612 | 0.507208347 | 0.611374396 | 1.400164931 |
| Mbd3l2 | 0.241190471 | 0.754107443 | 3.036905984 |
| Zfp558 | 31.78569487 | -0.396800678 | 0.210321832 |
| Gm22854 | 0 | NA | NA |
| Mbd3l1 | 0.181654796 | 1.679598083 | 3.311020364 |

| | | | |
|---|---|---|---|
| Muc16 | 0.130524232 | -0.243571427 | 3.657871338 |
| Gm25295 | 0 | NA | NA |
| Gm24546 | 0 | NA | NA |
| Olfr24 | 0 | NA | NA |
| Olfr828 | 0 | NA | NA |
| Olfr829 | 0.068097359 | -0.243569582 | 3.724945049 |
| Olfr830 | 0 | NA | NA |
| Olfr831-ps1 | 0 | NA | NA |
| Olfr832 | 0 | NA | NA |
| Olfr833-ps1 | 0 | NA | NA |
| Olfr834 | 0 | NA | NA |
| Olfr835 | 0.046345709 | 0.71820487 | 3.724945049 |
| Olfr836 | 0 | NA | NA |
| Olfr837 | 0 | NA | NA |
| Olfr843 | 0 | NA | NA |
| Olfr844 | 0 | NA | NA |
| Olfr845 | 0 | NA | NA |
| Olfr846 | 0.049553047 | 0.71820487 | 3.724945049 |
| Olfr847 | 0 | NA | NA |
| Gm23048 | 0 | NA | NA |
| Olfr849 | 0 | NA | NA |
| Olfr850 | 0 | NA | NA |
| Olfr851 | 0 | NA | NA |
| Olfr853 | 0 | NA | NA |
| Olfr854 | 0 | NA | NA |
| Olfr855 | 0 | NA | NA |
| Zfp317 | 272.3857193 | 0.291774755 | 0.084306667 |
| Olfr857 | 1.183877471 | 0.512375406 | 1.002408169 |
| Olfr58 | 0.474412771 | 0.730191501 | 1.606693391 |
| Olfr859 | 0.318756133 | 0.260412138 | 1.776492402 |
| Olfr860 | 0 | NA | NA |
| Olfr862 | 0 | NA | NA |
| Olfr77 | 0 | NA | NA |
| Gm7808 | 2.442408819 | 0.143289913 | 0.594614172 |
| Olfr866 | 0 | NA | NA |
| AC121953.1 | 0 | NA | NA |
| Gm26231 | 0 | NA | NA |
| Olfr867 | 0.092691418 | 1.183079489 | 3.724560917 |
| Olfr868 | 0.089062654 | 0.237318756 | 3.724945049 |
| Olfr869 | 0 | NA | NA |
| Olfr870 | 0 | NA | NA |
| Gm25914 | 0 | NA | NA |
| Gm26290 | 0 | NA | NA |
| Olfr871 | 0 | NA | NA |
| Olfr872 | 0 | NA | NA |
| Gm25425 | 0.049553047 | 0.71820487 | 3.724945049 |
| Olfr39 | 0 | NA | NA |

| | | | |
|---|---|---|---|
| Olfr873 | 0 | NA | NA |
| Olfr18 | 0.954195627 | 1.233226261 | 1.274306515 |
| Zfp560 | 108.4598925 | -0.264144772 | 0.142575644 |
| Gm26733 | 0 | NA | NA |
| Zfp26 | 822.179251 | 0.079190614 | 0.089706811 |
| Zfp426 | 384.7268472 | -0.068817774 | 0.117068596 |
| Gm23316 | 0 | NA | NA |
| Zfp266 | 1050.28284 | -0.031518371 | 0.069497055 |
| Zfp846 | 184.0291169 | -0.465317693 | 0.120952318 |
| Fbxl12os | 22.46068426 | -0.730397236 | 0.261285281 |
| Gm26521 | 12.01662456 | -0.022558283 | 0.385136361 |
| Fbxl12 | 213.0329372 | -0.274528965 | 0.16239605 |
| Ubl5 | 521.6630355 | 0.156740694 | 0.111648016 |
| Pin1 | 657.0847573 | 0.075374134 | 0.068018305 |
| Olfm2 | 557.2935379 | -0.442659929 | 0.086710123 |
| Col5a3 | 212.1757937 | 3.172020919 | 0.236214505 |
| Gm26274 | 0 | NA | NA |
| Rdh8 | 0.705140734 | 2.901665101 | 1.442405063 |
| A230050P20Rik | 616.357269 | 0.360028993 | 0.12580682 |
| Angptl6 | 38.80084079 | -0.591123643 | 0.177564966 |
| Ppan | 173.1887125 | 0.509329854 | 0.148021159 |
| Gm24067 | 0 | NA | NA |
| Gm23008 | 1.246341358 | 0.166764872 | 0.951882731 |
| Eif3g | 495.9541041 | 0.492894482 | 0.105668113 |
| Dnmt1 | 868.320093 | 0.24729328 | 0.096813785 |
| Mir7081 | 0.867462339 | -1.098924591 | 1.121720742 |
| S1pr2 | 119.737217 | 2.873304199 | 0.231968841 |
| Mrpl4 | 356.3497304 | 0.263467325 | 0.066781503 |
| Icam1 | 316.3122786 | 4.966553808 | 0.376664782 |
| Icam4 | 2.361925876 | -1.336330481 | 0.730172874 |
| Icam5 | 1112.549133 | -1.5928352 | 0.231366445 |
| Mir1900 | 0.298647883 | 1.012836524 | 2.074434771 |
| Gm26592 | 23.65836997 | -0.038589413 | 0.220284038 |
| Zglp1 | 9.660883422 | 0.075807875 | 0.354103121 |
| Fdx1l | 166.5463135 | 0.210289999 | 0.114486785 |
| Raver1 | 630.7586862 | 0.429736803 | 0.073813363 |
| Mir7082 | 0 | NA | NA |
| 1700084C06Rik | 2.554977676 | -0.24289766 | 0.598098976 |
| Tyk2 | 239.0143666 | 0.65017047 | 0.155153983 |
| Cdc37 | 916.0704561 | 0.162432113 | 0.092773798 |
| Pde4a | 1129.44041 | -0.656346368 | 0.100996324 |
| Gm16754 | 3.260813172 | 0.453511788 | 0.561599659 |
| Keap1 | 857.7412683 | 0.418386448 | 0.066268577 |
| S1pr5 | 222.4130938 | 0.094960077 | 0.142604744 |
| Atg4d | 243.847421 | 0.035068729 | 0.082857516 |
| Kri1 | 243.0174812 | 0.223416433 | 0.110698602 |
| Gm37379 | 20.17791531 | -0.575566039 | 0.289973329 |

| | | | |
|---|---|---|---|
| Cdkn2d | 273.9573276 | 0.143806127 | 0.086749608 |
| Ap1m2 | 9.647799368 | -0.124087994 | 0.378684271 |
| Slc44a2 | 583.7110967 | 0.936117808 | 0.119923589 |
| Ilf3 | 1100.802251 | 0.116903299 | 0.053547963 |
| Gm16853 | 1.441198087 | -0.273386782 | 0.83980065 |
| Qtrt1 | 87.06778592 | 0.211344988 | 0.15154478 |
| Dnm2 | 1057.753246 | 0.71230742 | 0.10394525 |
| Mir199a-1 | 0.092691418 | 1.183079489 | 3.724560917 |
| Tmed1 | 82.01844689 | 0.529397152 | 0.147285021 |
| AB124611 | 105.0329745 | 6.714577804 | 0.437011333 |
| Gm20519 | 0.907036939 | -1.007008918 | 1.111105625 |
| Carm1 | 707.8332732 | -0.071327542 | 0.061374528 |
| Yipf2 | 186.1268908 | -0.265839034 | 0.095963762 |
| 1810026J23Rik | 718.2922187 | 0.369924453 | 0.113753882 |
| Mir1946b | 0.611133411 | -2.045757956 | 1.350050244 |
| Smarca4 | 2424.773633 | 0.097394024 | 0.051541444 |
| Gm27373 | 1.398170896 | 0.39149546 | 0.846103327 |
| Gm26511 | 0.192512644 | -0.735351246 | 2.691637281 |
| Ldlr | 610.6285346 | 0.84153685 | 0.181377836 |
| Spc24 | 9.830363938 | 0.513461616 | 0.38529927 |
| Kank2 | 390.6615536 | 0.477460959 | 0.144151744 |
| Dock6 | 147.3513096 | -0.283458835 | 0.13841337 |
| Mir7083 | 0 | NA | NA |
| Gm6484 | 0.140389662 | 1.036581329 | 3.571608646 |
| Rab3d | 110.9505493 | 0.176291941 | 0.162592224 |
| Tmem205 | 101.5694773 | 0.406902161 | 0.18899215 |
| Ccdc159 | 62.58867534 | -0.704300606 | 0.198753925 |
| Lppr2 | 1319.490486 | -0.350878682 | 0.095897066 |
| Swsap1 | 101.2171663 | -0.161806747 | 0.126982584 |
| Epor | 34.63035605 | -0.341006289 | 0.241581482 |
| Rgl3 | 100.740861 | -0.5965623 | 0.215349036 |
| Ccdc151 | 32.36726646 | -0.48124802 | 0.233219177 |
| Prkcsh | 759.8445769 | 0.55707386 | 0.076282777 |
| Elavl3 | 2565.303786 | -0.655634265 | 0.07090419 |
| Zfp653 | 159.2068104 | -0.220060698 | 0.105885053 |
| Gm16845 | 10.87916454 | -0.891339651 | 0.352308793 |
| Ecsit | 303.4651564 | -0.182970503 | 0.069354735 |
| Cnn1 | 3.417651458 | -0.105337127 | 0.500477091 |
| Elof1 | 264.1837638 | 0.357817029 | 0.112488439 |
| Mir7084 | 0 | NA | NA |
| Acp5 | 37.28268211 | 4.64484 | 0.399273426 |
| Pigyl | 121.7874717 | 0.105455081 | 0.146169066 |
| Rpl15-ps3 | 5.54092659 | 1.102852951 | 0.424849663 |
| Zfp872 | 0.324168737 | 1.860060639 | 1.67484964 |
| 1810064F22Rik | 0.089062654 | 0.237318756 | 3.724945049 |
| Zfp809 | 131.6943064 | 0.153726339 | 0.114175987 |
| Zfp599 | 85.70332244 | -0.107494341 | 0.171137343 |

| | | | |
|---|---|---|---|
| Zfp810 | 217.69277 | -0.126919741 | 0.115118278 |
| Gm10182 | 10.2804556 | 0.463660011 | 0.322852768 |
| Anln | 682.8292644 | 0.246419399 | 0.182227566 |
| 9530077C05Rik | 75.06799202 | -0.065175868 | 0.170454914 |
| Rp9 | 229.1720738 | 0.127594782 | 0.075272177 |
| Gm17545 | 0 | NA | NA |
| Bbs9 | 315.6451564 | -0.394419428 | 0.089464958 |
| Gm2976 | 0 | NA | NA |
| Gm16165 | 1.13143437 | -0.354549285 | 0.818438676 |
| Gm16164 | 0 | NA | NA |
| Gm27639 | 0 | NA | NA |
| Bmper | 91.07485983 | -0.264895235 | 0.115491712 |
| Npsr1 | 33.2953994 | -0.454747325 | 0.200052378 |
| Gm15600 | 0.84374619 | 3.317644469 | 1.09719765 |
| Gm10701 | 1.895990101 | 2.862458382 | 0.993272401 |
| Gm22380 | 0.229628521 | 1.6670716 | 2.121064275 |
| Dpy19l1 | 977.1122373 | -0.42054893 | 0.059786978 |
| Dpy19l2 | 3.565532606 | -0.823568337 | 0.62803168 |
| Cypt4 | 0 | NA | NA |
| Gm38179 | 0.167183452 | -0.72445987 | 2.765153652 |
| Tbx20 | 7.101997061 | 1.040397135 | 0.524994769 |
| Gm23893 | 0 | NA | NA |
| Gm10181 | 0 | NA | NA |
| Gm10180 | 2.742928344 | 0.014767205 | 0.681849073 |
| Herpud2 | 587.9118386 | 0.154764416 | 0.084727017 |
| Gm29642 | 3.768175155 | -0.375219341 | 0.503488243 |
| Gm29375 | 1.122258044 | 0.142923555 | 0.908167714 |
| 7-Sep | 3678.656398 | -0.211194377 | 0.07626498 |
| Eepd1 | 188.7348402 | 0.312210469 | 0.09202018 |
| Gm25861 | 0 | NA | NA |
| Gm25346 | 0 | NA | NA |
| Gm26190 | 0 | NA | NA |
| B3gat1 | 1268.762376 | -1.119926822 | 0.113802697 |
| Glb1l2 | 10.70062985 | 0.031679994 | 0.374129517 |
| Glb1l3 | 0.449952257 | 0.271895264 | 1.29498094 |
| Gm1110 | 2.025517295 | -0.34898201 | 0.867990857 |
| Acad8 | 225.253486 | -0.160922964 | 0.102232523 |
| Thyn1 | 253.8863181 | 0.151672257 | 0.124720255 |
| Vps26b | 1272.753477 | -0.168315971 | 0.05990622 |
| Ncapd3 | 405.8601585 | -0.491625802 | 0.167946077 |
| Jam3 | 390.9541926 | 0.737405018 | 0.131115559 |
| Gm29172 | 0.038981395 | 0.718207094 | 3.724945049 |
| Igsf9b | 818.461646 | -0.302080071 | 0.239054684 |
| Spata19 | 4.643680173 | 0.647087255 | 0.580127918 |
| Opcml | 3000.095646 | -0.733281306 | 0.083053329 |
| Gm15606 | 43.86298618 | -1.955387428 | 0.527532961 |
| AC157511.1 | 4.057004922 | -1.172043685 | 0.554697813 |

| | | | |
|---|---|---|---|
| Ntm | 1996.252357 | -0.80075232 | 0.080036402 |
| Gm15520 | 1.48836363 | 1.559646622 | 1.018978239 |
| Gm15521 | 42.12449345 | -1.574732427 | 0.319182158 |
| Gm37605 | 0 | NA | NA |
| Gm26435 | 0 | NA | NA |
| Snx19 | 1050.451712 | 0.102907946 | 0.048655793 |
| Adamts15 | 127.7747721 | -0.254902203 | 0.118432972 |
| Adamts8 | 29.93733091 | 0.179911798 | 0.230625396 |
| Zbtb44 | 1100.679016 | -0.391681771 | 0.075230438 |
| Gm17508 | 0 | NA | NA |
| St14 | 17.50112768 | 1.119809034 | 0.275875786 |
| Gm16071 | 0.620902317 | 2.629689637 | 1.289393002 |
| Aplp2 | 5929.023008 | -0.435592492 | 0.06394968 |
| Gm7244 | 0.710933469 | -2.089944295 | 1.184057601 |
| Prdm10 | 216.453307 | -0.760130079 | 0.13548433 |
| Nfrkb | 530.6418161 | -0.003085725 | 0.141554207 |
| Tmem45b | 0.191334258 | 1.574896507 | 2.660412218 |
| Barx2 | 29.27701895 | 0.930590892 | 0.307677712 |
| Gm3756 | 26.83663404 | 0.083481519 | 0.254656878 |
| AC160635.1 | 2.239755095 | -0.927273887 | 0.670759248 |
| Arhgap32 | 5811.857867 | -1.06271947 | 0.098259972 |
| Kcnj5 | 21.9094618 | 0.386986003 | 0.273962309 |
| Kcnj1 | 0.551016524 | 2.64915421 | 1.461651809 |
| Fli1 | 127.9880946 | 2.195792435 | 0.201828625 |
| Gm22060 | 0.733464214 | 1.389617929 | 1.20112185 |
| Gm38292 | 5.098635657 | 0.516447093 | 0.456443076 |
| Gm38346 | 11.05146628 | 1.023475794 | 0.444173479 |
| Gm27201 | 0.169165009 | 0.719161184 | 3.351346564 |
| Gm37474 | 11.13391284 | 1.320106755 | 0.51244869 |
| Gm37082 | 13.07759707 | 0.996273759 | 0.462752233 |
| Gm27163 | 0 | NA | NA |
| Ets1 | 209.3380529 | 2.485457484 | 0.20832378 |
| Gm27240 | 0 | NA | NA |
| Gm3331 | 0 | NA | NA |
| Gm37167 | 0.620529807 | -1.418656081 | 1.285517738 |
| Gm27162 | 0 | NA | NA |
| Gm37192 | 0.061920613 | 0.718207094 | 3.724945049 |
| Gm27166 | 0 | NA | NA |
| Gm37758 | 2.309104371 | 0.78246068 | 0.676539589 |
| Gm10698 | 69.88234991 | 0.735399988 | 0.166321043 |
| Gm25439 | 0 | NA | NA |
| Gm27165 | 0 | NA | NA |
| Gm27164 | 0 | NA | NA |
| 7630403G23Rik | 0.458133289 | -0.733016216 | 1.540311426 |
| Gm27161 | 0 | NA | NA |
| Gm38354 | 0 | NA | NA |
| Kirrel3 | 816.5410815 | -0.393072793 | 0.068756932 |

| | | | |
|---|---|---|---|
| Gm38217 | 71.70747919 | -1.43925564 | 0.299799505 |
| Gm37038 | 54.63741103 | -1.27779399 | 0.278680278 |
| Gm26662 | 0.042716945 | -0.243571806 | 3.724945049 |
| Kirrel3os | 3.153914063 | -0.217911683 | 0.673746753 |
| Gm23702 | 2.111071361 | -1.348070283 | 0.770511503 |
| St3gal4 | 323.0715017 | 0.61949882 | 0.09583744 |
| 4930581F22Rik | 31.67819825 | -0.160073898 | 0.210783905 |
| Dcps | 130.3926271 | 0.411541099 | 0.128711609 |
| Gm24262 | 0 | NA | NA |
| Tirap | 88.4573936 | 1.685033995 | 0.192582198 |
| Srpr | 878.7068951 | 0.702668144 | 0.071648409 |
| Foxred1 | 265.905042 | 0.221036392 | 0.096356142 |
| Fam118b | 211.7892874 | -0.083768051 | 0.102620719 |
| Rpusd4 | 78.94005876 | 0.449743712 | 0.153091791 |
| Gm5614 | 0 | NA | NA |
| 2610105M22Rik | 0.044191549 | -0.243571806 | 3.724945049 |
| Cdon | 259.9464149 | -0.587014119 | 0.146751027 |
| Gm25630 | 1.242022914 | -1.738009797 | 0.902888071 |
| Gm24188 | 0 | NA | NA |
| Gm1113 | 0 | NA | NA |
| Ddx25 | 512.2202113 | -0.278499173 | 0.05479882 |
| Pus3 | 127.6005767 | 0.232940214 | 0.106330652 |
| Hyls1 | 86.77207718 | -0.277925051 | 0.147938795 |
| Pate2 | 15.21999782 | -0.478619838 | 0.327948906 |
| Gm6762 | 0 | NA | NA |
| Gm10105 | 0 | NA | NA |
| Pate4 | 0 | NA | NA |
| Gm17365 | 0 | NA | NA |
| Gm17252 | 0 | NA | NA |
| Gm17677 | 0.297465678 | 0.237331021 | 1.773341359 |
| Gm27235 | 0 | NA | NA |
| Gm17727 | 0 | NA | NA |
| D730048I06Rik | 0 | NA | NA |
| Gm36974 | 1.090715414 | -0.909960837 | 0.944177534 |
| 9230110F15Rik | 0 | NA | NA |
| Gm3428 | 0.042716945 | -0.243571806 | 3.724945049 |
| 9230113P08Rik | 1.958122126 | -0.349452597 | 0.701131407 |
| Gm27187 | 0 | NA | NA |
| Gm5916 | 0 | NA | NA |
| Gm3434 | 0 | NA | NA |
| Gm3867 | 0.061920613 | 0.718207094 | 3.724945049 |
| Gm7257 | 0 | NA | NA |
| Gm9513 | 0 | NA | NA |
| Gm27160 | 0 | NA | NA |
| Gm5615 | 0 | NA | NA |
| Gm27218 | 0 | NA | NA |
| Gm17689 | 0.194279094 | 1.292225235 | 3.719439735 |

| | | | |
|---|---|---|---|
| A630095E13Rik | 0.217428854 | 2.009909443 | 2.547915794 |
| 1700027I24Rik | 0.357600009 | 2.389139793 | 1.647075383 |
| Acrv1 | 0.095898756 | 1.192083721 | 3.724434897 |
| Chek1 | 39.39245048 | 0.454700646 | 0.197262956 |
| Stt3a | 958.0649057 | 0.473545456 | 0.066958655 |
| Gm26787 | 0.495189146 | -0.7117464 | 1.348299891 |
| Ei24 | 869.5859806 | 0.004486701 | 0.069857496 |
| Gm27219 | 6.744631513 | 0.858982229 | 0.414325125 |
| Fez1 | 1689.638317 | -0.299949315 | 0.060907365 |
| Pknox2 | 477.999823 | -0.834228378 | 0.107672421 |
| Gm3896 | 0.132101749 | 1.209894437 | 3.72418822 |
| n-R5s82 | 0 | NA | NA |
| Gm25273 | 0 | NA | NA |
| Tmem218 | 64.43814503 | -0.652377935 | 0.13676604 |
| Slc37a2 | 140.0947805 | 2.669916567 | 0.260598549 |
| Gm10177 | 68.34373515 | 0.786904847 | 0.200329106 |
| Ccdc15 | 56.68857134 | -0.467486505 | 0.189692777 |
| Hepacam | 1455.687833 | 0.289884704 | 0.127706532 |
| Robo4 | 52.91085951 | 2.183251512 | 0.25269494 |
| Robo3 | 261.3456688 | 0.46752987 | 0.425221225 |
| Msantd2 | 153.0368494 | -0.076150876 | 0.102026725 |
| Esam | 83.18252856 | 1.377953475 | 0.164068213 |
| Vsig2 | 19.39362468 | 1.734272874 | 0.345104598 |
| Nrgn | 5315.665183 | -1.269310282 | 0.205654581 |
| Spa17 | 8.072628227 | 0.037481743 | 0.395966793 |
| Siae | 198.8133421 | 0.101978175 | 0.132539343 |
| Tbrg1 | 650.3765193 | 1.407279626 | 0.123668 |
| Panx3 | 0.252375374 | -0.24356391 | 2.128498805 |
| Gm8343 | 0 | NA | NA |
| Tpt1-ps5 | 7.622765194 | 1.024037597 | 0.420819897 |
| Olfr160 | 0 | NA | NA |
| Olfr874 | 0 | NA | NA |
| Olfr875 | 0.072025462 | 0.237312084 | 3.724945049 |
| Olfr876 | 0.066050875 | 0.71820487 | 3.724945049 |
| Gm24764 | 0 | NA | NA |
| Olfr877 | 0.162452565 | 0.237319607 | 2.783670453 |
| Olfr145 | 0.104637558 | 0.23732098 | 3.724945049 |
| Olfr878 | 0.170434537 | -0.243564467 | 2.328336882 |
| Olfr881 | 0 | NA | NA |
| Olfr883 | 0 | NA | NA |
| Olfr884 | 0.092140247 | -0.724455696 | 3.724945049 |
| Olfr885 | 0 | NA | NA |
| Olfr887 | 0.072025462 | 0.237312084 | 3.724945049 |
| Olfr888 | 0 | NA | NA |
| Olfr889 | 0 | NA | NA |
| Olfr890 | 0 | NA | NA |
| Olfr891 | 0 | NA | NA |

| | | | |
|---|---|---|---|
| Gm17798 | 0 | NA | NA |
| Olfr892-ps1 | 0 | NA | NA |
| Olfr893 | 0 | NA | NA |
| Olfr894 | 0 | NA | NA |
| Olfr143 | 0 | NA | NA |
| Olfr895 | 0 | NA | NA |
| Olfr25 | 0.111376404 | -0.243571806 | 3.724945049 |
| Olfr898 | 0 | NA | NA |
| Olfr250 | 0.072025462 | 0.237312084 | 3.724945049 |
| Olfr251 | 0 | NA | NA |
| Olfr147 | 0.068097359 | -0.243569582 | 3.724945049 |
| Olfr901 | 0.114443069 | 0.23732098 | 3.724945049 |
| Olfr902 | 0 | NA | NA |
| Olfr904 | 0 | NA | NA |
| Olfr905 | 0 | NA | NA |
| Olfr906 | 0 | NA | NA |
| Olfr907 | 0 | NA | NA |
| Gm23271 | 0 | NA | NA |
| Olfr908 | 0 | NA | NA |
| Olfr911-ps1 | 0 | NA | NA |
| Olfr910 | 0 | NA | NA |
| Olfr912 | 0 | NA | NA |
| Olfr913 | 0 | NA | NA |
| Olfr914 | 0 | NA | NA |
| Olfr916 | 0 | NA | NA |
| Olfr917 | 0 | NA | NA |
| Olfr918 | 0 | NA | NA |
| Olfr919 | 0 | NA | NA |
| Vwa5a | 1119.696084 | 3.0185467 | 0.165595179 |
| Olfr920 | 22.44487698 | 2.634930056 | 0.502629272 |
| Olfr921 | 0.704586541 | 2.7181242 | 1.495038958 |
| Olfr922 | 0.673070397 | 0.798211739 | 1.035562719 |
| Olfr923 | 0.176917541 | 0.24695619 | 2.76696777 |
| Olfr924 | 0.326455024 | -1.674765783 | 1.663865731 |
| Olfr26 | 0.232584034 | 0.237328867 | 2.133784112 |
| Olfr926 | 0.106056992 | 0.237307635 | 3.724945049 |
| Olfr930 | 0 | NA | NA |
| Olfr933 | 0 | NA | NA |
| Olfr934 | 0 | NA | NA |
| Olfr935 | 0 | NA | NA |
| Olfr146 | 0 | NA | NA |
| Olfr936 | 0 | NA | NA |
| Gm25529 | 0 | NA | NA |
| Olfr937 | 0 | NA | NA |
| Gm25032 | 0 | NA | NA |
| Olfr938 | 0 | NA | NA |
| Olfr27 | 0 | NA | NA |

| | | | |
|---|---|---|---|
| Olfr943 | 0 | NA | NA |
| Olfr944 | 0 | NA | NA |
| Olfr1537 | 0 | NA | NA |
| Olfr945 | 0 | NA | NA |
| Olfr948 | 0 | NA | NA |
| Olfr951 | 0 | NA | NA |
| Olfr952 | 0 | NA | NA |
| Olfr954 | 0 | NA | NA |
| Olfr955 | 0 | NA | NA |
| Olfr44 | 0.046345709 | 0.71820487 | 3.724945049 |
| Olfr957 | 0 | NA | NA |
| Gm24278 | 0 | NA | NA |
| Olfr958 | 0 | NA | NA |
| Olfr959 | 0.090427103 | 0.237307635 | 3.724945049 |
| AW551984 | 246.7130081 | 0.183379133 | 0.154410038 |
| Olfr148 | 0 | NA | NA |
| Olfr960 | 0 | NA | NA |
| Olfr961 | 0 | NA | NA |
| Olfr963 | 0 | NA | NA |
| Olfr964-ps1 | 0.048949532 | -0.243569582 | 3.724945049 |
| Olfr149 | 0 | NA | NA |
| Olfr965 | 0 | NA | NA |
| Olfr150 | 0 | NA | NA |
| Olfr967 | 0 | NA | NA |
| Olfr968 | 0.046345709 | 0.71820487 | 3.724945049 |
| Olfr969 | 0 | NA | NA |
| Olfr970 | 0 | NA | NA |
| Olfr971 | 0 | NA | NA |
| Olfr972 | 0 | NA | NA |
| Olfr229 | 0 | NA | NA |
| Olfr974 | 0.049553047 | 0.71820487 | 3.724945049 |
| Olfr975 | 0 | NA | NA |
| Olfr976 | 0.111006857 | 0.717551239 | 3.724898127 |
| Olfr978 | 0 | NA | NA |
| Olfr979 | 0.046345709 | 0.71820487 | 3.724945049 |
| Olfr980 | 0.09176491 | 0.237310218 | 3.724945049 |
| Olfr981 | 0 | NA | NA |
| 1700001J11Rik | 0.049553047 | 0.71820487 | 3.724945049 |
| Olfr982 | 0 | NA | NA |
| Olfr983 | 0 | NA | NA |
| Olfr984 | 0.072025462 | 0.237312084 | 3.724945049 |
| Olfr985 | 0.09176491 | 0.237310218 | 3.724945049 |
| Tmem225 | 0 | NA | NA |
| Olfr986 | 0 | NA | NA |
| Zfp202 | 72.13492 | 0.429821747 | 0.142942918 |
| Scn3b | 970.5865091 | -1.183453926 | 0.106092282 |
| Gramd1b | 3247.828285 | -0.440603744 | 0.080864194 |

| | | | |
|---|---|---|---|
| 1700110K17Rik | 1.343476617 | 0.016129624 | 0.905272518 |
| Gm17540 | 0 | NA | NA |
| Gm21915 | 0 | NA | NA |
| Clmp | 308.4342746 | -0.041111094 | 0.126740363 |
| Gm16095 | 0.148850706 | 0.661143511 | 2.852067089 |
| Gm16096 | 0.692884898 | 2.539253404 | 1.269977517 |
| Hspa8 | 7490.665187 | 0.395468583 | 0.079975195 |
| Snord14c | 0 | NA | NA |
| Snord14d | 0 | NA | NA |
| Snord14e | 0 | NA | NA |
| Bsx | 2.789926683 | 0.453457554 | 0.597635538 |
| 4931429I11Rik | 11.72828212 | 0.48487697 | 0.324805901 |
| Crtam | 123.3542397 | 0.533210022 | 0.247837194 |
| Ubash3b | 620.5872832 | -0.021511985 | 0.093040739 |
| 1700063D05Rik | 6.117005653 | -0.763448148 | 0.402922891 |
| Gm25401 | 0 | NA | NA |
| 2610203C20Rik | 984.4296156 | -0.108496247 | 0.145461843 |
| Mir100 | 2.750782513 | -1.010012812 | 0.735113161 |
| Mirlet7a-2 | 2.112107905 | 0.050381602 | 0.739248569 |
| Gm28119 | 36.18489172 | 0.041675865 | 0.24017645 |
| Mir125b-1 | 0 | NA | NA |
| Gm8707 | 28.79399415 | -0.275987824 | 0.21704719 |
| Sorl1 | 7353.79371 | -0.374741585 | 0.075681481 |
| Gm16214 | 0 | NA | NA |
| Sc5d | 491.2699012 | -0.0149108 | 0.104628165 |
| Gm26272 | 0 | NA | NA |
| Tecta | 9.746868474 | 1.612221456 | 0.441962019 |
| Tbcel | 691.6961961 | -0.268421242 | 0.05914221 |
| AC136903.1 | 0 | NA | NA |
| Gm16322 | 0.291180489 | 0.718190974 | 3.724945049 |
| Grik4 | 287.3585913 | -0.515448841 | 0.098603753 |
| Gm24420 | 0 | NA | NA |
| Gm27509 | 1.432503634 | -2.727547123 | 1.029268877 |
| Arhgef12 | 4563.170901 | -0.060972863 | 0.038397822 |
| Tmem136 | 181.8275258 | -0.437824992 | 0.106260646 |
| Pou2f3 | 2.121136815 | 1.586591412 | 0.779503349 |
| Oaf | 220.6241672 | 1.802773649 | 0.164757149 |
| D630033O11Rik | 2.615308024 | 0.534878615 | 0.593744384 |
| Trim29 | 4.261450001 | 1.306907863 | 0.577787116 |
| Gm28215 | 0 | NA | NA |
| Pvrl1 | 1033.013847 | -0.400169929 | 0.079446481 |
| Gm3898 | 0.044544648 | 0.237309859 | 3.724945049 |
| Gm23326 | 0.048949532 | -0.243569582 | 3.724945049 |
| Thy1 | 4661.495848 | -0.30132022 | 0.077945929 |
| Rnf26 | 340.15711 | -0.046251895 | 0.077673414 |
| Usp2 | 947.8979254 | -0.276315854 | 0.062835443 |
| Mfrp | 54.45536832 | -0.955490961 | 0.26286087 |

| | | | |
|---|---|---|---|
| C1qtnf5 | 0 | NA | NA |
| Gm20444 | 3.364322678 | -0.832840781 | 0.558100235 |
| Gm26737 | 7.342448606 | 0.423685675 | 0.402607882 |
| Gm10687 | 8.115134709 | -0.524602743 | 0.367451498 |
| Mcam | 257.1599992 | 0.384083066 | 0.153713052 |
| Cbl | 967.2924709 | 0.061787401 | 0.072866238 |
| Ccdc153 | 11.25998896 | -1.475711248 | 0.38297704 |
| Pdzd3 | 3.822296293 | -0.455468614 | 0.604892254 |
| Nlrx1 | 97.08268411 | 0.470246364 | 0.144679123 |
| Abcg4 | 564.6710316 | -0.731874035 | 0.085514519 |
| Hinfp | 148.4591195 | 0.074104874 | 0.133983601 |
| C2cd2l | 1532.300989 | -0.564678697 | 0.109523423 |
| Dpagt1 | 146.9363667 | 0.328782555 | 0.113194485 |
| H2afx | 329.2993814 | 0.233994023 | 0.09905459 |
| AC122428.1 | 1.088551472 | -1.906906396 | 1.199864553 |
| Hmbs | 132.3982402 | 0.213047612 | 0.127692475 |
| Vps11 | 770.3516232 | 0.237331317 | 0.066295905 |
| Gm22141 | 0.240440413 | -1.173536932 | 2.14647057 |
| Gm10080 | 110.1720267 | 0.33876211 | 0.166115275 |
| Hyou1 | 2036.023468 | 0.557564446 | 0.080969966 |
| Gm26306 | 0 | NA | NA |
| Slc37a4 | 109.665464 | -0.455146683 | 0.133489886 |
| Mir7085 | 0.487060322 | -1.066283036 | 1.433422395 |
| Trappc4 | 251.412174 | 0.373210088 | 0.126065867 |
| Rps25 | 1102.353914 | 0.45041335 | 0.131793939 |
| Ccdc84 | 64.93599903 | -0.305706663 | 0.162236775 |
| Foxr1 | 0.310849204 | 1.296189755 | 3.719250335 |
| Upk2 | 0.044191549 | -0.243571806 | 3.724945049 |
| Gm9830 | 5.961436335 | 0.295783197 | 0.378948521 |
| Gm22540 | 5.014127157 | -2.054780197 | 0.540972702 |
| C030014I23Rik | 47.83992227 | -0.761443555 | 0.168521363 |
| Bcl9l | 785.1067627 | -0.203109399 | 0.081987327 |
| Cxcr5 | 2.247268683 | 4.402611345 | 1.007663603 |
| Ddx6 | 4111.857025 | 0.108162695 | 0.05350328 |
| Treh | 0.042716945 | -0.243571806 | 3.724945049 |
| Phldb1 | 1542.489038 | -0.095534059 | 0.071009704 |
| Gm24166 | 0 | NA | NA |
| Arcn1 | 1826.393337 | 0.4651732 | 0.081200819 |
| Ift46 | 313.3153656 | -0.232162372 | 0.075188021 |
| Tmem25 | 356.7404894 | -0.908153279 | 0.096713338 |
| Ttc36 | 3.625487916 | 0.156525812 | 0.585715109 |
| Kmt2a | 3944.583612 | -0.34532597 | 0.093762463 |
| Gm26249 | 0 | NA | NA |
| Atp5l | 692.9294976 | -0.049911683 | 0.138282349 |
| Ube4a | 1166.455959 | 0.148376533 | 0.057550719 |
| Cd3g | 29.68427331 | 7.39811294 | 0.905537772 |
| Cd3d | 21.88467821 | 7.849624949 | 0.875223762 |

| | | | |
|---|---|---|---|
| Cd3e | 72.02395476 | 9.153351972 | 0.880855307 |
| Mpzl2 | 55.99719323 | 1.213758524 | 0.235029165 |
| Mpzl3 | 77.05801845 | 0.435777077 | 0.148036798 |
| Amica1 | 164.8869608 | 4.568525614 | 0.345687761 |
| Gm10684 | 5.682893444 | -1.03360461 | 0.505873479 |
| Scn2b | 3837.721104 | -0.696908781 | 0.097977739 |
| Scn4b | 2000.069508 | -1.28165176 | 0.126467709 |
| Tmprss4 | 3.058856891 | 0.500288691 | 0.568837806 |
| Gm17099 | 0.153914375 | -0.24084143 | 3.503921201 |
| BC049352 | 7.258084432 | -0.861345435 | 0.497102865 |
| Il10ra | 224.5975288 | 2.659598419 | 0.151122918 |
| Gm20612 | 0.047948698 | -0.243569582 | 3.724945049 |
| Mir7086 | 0.58926593 | 0.748893562 | 1.381736844 |
| 1700003G13Rik | 0 | NA | NA |
| Tmprss13 | 0.459044381 | 2.308506757 | 1.495152632 |
| Fxyd6 | 526.6237023 | -0.256225039 | 0.110362347 |
| Fxyd2 | 9.473000343 | -0.182867795 | 0.361953644 |
| 4833428L15Rik | 0.565850957 | -1.82712728 | 1.140713821 |
| Dscaml1 | 663.8123879 | -0.769601716 | 0.115005822 |
| Gm22069 | 0 | NA | NA |
| Cep164 | 262.2380998 | -0.310317449 | 0.134008495 |
| Gm7286 | 0.119587853 | 0.237305575 | 3.724945049 |
| Bace1 | 1452.453558 | -0.136618626 | 0.044276972 |
| Gm16536 | 127.1699055 | -0.206540122 | 0.106580005 |
| Rnf214 | 859.5718938 | -0.266033234 | 0.047934898 |
| Gm24646 | 2.996628699 | -0.10944429 | 0.925485782 |
| Pcsk7 | 359.8709397 | 0.421144441 | 0.085653821 |
| Tagln | 130.3382176 | 3.999393068 | 0.438325573 |
| Sidt2 | 826.3607848 | 0.02785919 | 0.085030986 |
| Mir7087 | 1.036669434 | -0.344818494 | 1.166297257 |
| Pafah1b2 | 2591.382288 | 0.016887609 | 0.066904538 |
| Sik3 | 2827.253407 | 0.449436812 | 0.081584793 |
| Apoa1 | 4.148602219 | 0.245888504 | 0.556170389 |
| Apoc3 | 5.512392282 | 1.246981909 | 0.554455947 |
| Apoa4 | 0.183366429 | 0.674767185 | 2.264892113 |
| Gm10680 | 0 | NA | NA |
| Apoa5 | 0.33367842 | 1.904994053 | 1.94560346 |
| Zpr1 | 322.0139224 | 0.404804274 | 0.073169738 |
| Bud13 | 129.6764734 | 0.309113475 | 0.101464132 |
| 4931429L15Rik | 0.137762903 | -0.72445853 | 2.937523348 |
| 4930448E22Rik | 1.471695147 | 1.522365144 | 0.922127241 |
| Gm22659 | 0.246177344 | -0.788169888 | 2.49861285 |
| Gm22805 | 0 | NA | NA |
| 2900052N01Rik | 275.6261956 | -0.112091217 | 0.161632223 |
| Gm4791 | 0.045882455 | 0.237312084 | 3.724945049 |
| Gm22286 | 0.075043205 | 0.237309859 | 3.724945049 |
| Cadm1 | 2092.339405 | -0.143929192 | 0.066662644 |

| | | | |
|---|---|---|---|
| Nxpe2 | 2.03192395 | 2.601021679 | 0.988523922 |
| Nxpe4 | 71.31690674 | 0.02867388 | 0.139445065 |
| Gm5616 | 4.81606905 | 0.397473166 | 0.446576402 |
| Rexo2 | 393.6205041 | 0.690905101 | 0.101720566 |
| Rbm7 | 414.2773601 | 1.062092164 | 0.111841585 |
| Gm5617 | 29.90601096 | 0.271589987 | 0.233968134 |
| Gm23653 | 0 | NA | NA |
| Nnmt | 5.720995077 | 3.374254317 | 0.634130342 |
| Zbtb16 | 1098.476404 | -0.293231006 | 0.176369258 |
| Htr3a | 33.96213922 | 0.003867885 | 0.224724412 |
| Htr3b | 0.066050875 | 0.71820487 | 3.724945049 |
| Usp28 | 333.1237609 | -0.655298004 | 0.119249855 |
| Cldn21 | 0 | NA | NA |
| RP23-136A1.2 | 0 | NA | NA |
| Zw10 | 188.7291143 | 0.159189964 | 0.091370484 |
| Tmprss5 | 18.34493739 | -0.192064593 | 0.246851628 |
| Gm4894 | 0.044544648 | 0.237309859 | 3.724945049 |
| Drd2 | 144.2458334 | -0.994917414 | 0.225602307 |
| Ankk1 | 0.421296087 | 0.613827315 | 1.412914664 |
| Ttc12 | 60.41204789 | 0.125000473 | 0.171418345 |
| Ncam1 | 4558.689867 | -0.369548255 | 0.05666363 |
| Gm11149 | 1.147320409 | -0.916236552 | 0.981535124 |
| Gm25633 | 0 | NA | NA |
| Gm8907 | 0.242574275 | 0.6568138 | 2.526122525 |
| Rpl10-ps3 | 4.878838789 | 0.863257296 | 0.436873516 |
| Plet1os | 0.85456564 | -1.532468492 | 1.10432852 |
| Plet1 | 4.418087894 | 0.315027785 | 0.533439227 |
| Pts | 145.2821715 | -0.092841533 | 0.13266327 |
| Bco2 | 15.34273135 | -0.037281927 | 0.267333949 |
| Tex12 | 0.048949532 | -0.243569582 | 3.724945049 |
| Il18 | 136.7701011 | 0.65391157 | 0.137247617 |
| Sdhd | 647.1493877 | 0.192900713 | 0.072838729 |
| Timm8b | 499.1712965 | -0.007159627 | 0.148916966 |
| AU019823 | 218.1162646 | 0.296114162 | 0.117313113 |
| Pih1d2 | 29.01466556 | -0.020444434 | 0.191011527 |
| Dlat | 1112.83225 | -0.2447147 | 0.059313107 |
| Dixdc1 | 1202.736387 | -0.732446562 | 0.075138627 |
| 2310030G06Rik | 16.94180656 | 0.171055195 | 0.253564455 |
| Hspb2 | 5.620585044 | 1.706701567 | 0.523301559 |
| Cryab | 871.6678754 | 1.169061747 | 0.199613478 |
| 1110032A03Rik | 144.5010807 | -0.023640939 | 0.112509911 |
| Fdxacb1 | 52.09799359 | -0.104779553 | 0.152924586 |
| Alg9 | 194.0505543 | -0.000242446 | 0.085503679 |
| Ppp2r1b | 243.7699036 | 0.840093902 | 0.102092771 |
| Gm25558 | 0.177937505 | -1.118419011 | 2.732390334 |
| Sik2 | 790.3337553 | -0.13111427 | 0.104797874 |
| Layn | 62.49321233 | 1.895050547 | 0.24697139 |

| | | | |
|---|---|---|---|
| 4833427G06Rik | 5.495872342 | 1.11041712 | 0.468237353 |
| Mir34c | 0 | NA | NA |
| Mir34b | 0 | NA | NA |
| Btg4 | 0.260468428 | 1.558713868 | 2.020704085 |
| Pou2af1 | 4.531271601 | 3.188960295 | 0.797468987 |
| Gm684 | 7.066067338 | 0.071693004 | 0.374587495 |
| 2010007H06Rik | 9.722359338 | -0.830933802 | 0.360585883 |
| 1810046K07Rik | 0.216202823 | -0.244095225 | 2.173239048 |
| Gm7293 | 4.644993497 | -0.571885416 | 0.540133552 |
| Arhgap20 | 1285.799934 | -0.742888672 | 0.08856665 |
| Arhgap20os | 1.805117362 | -0.644544882 | 0.983113346 |
| Fdx1 | 101.2393919 | 0.467574486 | 0.149528074 |
| Gm6981 | 16.53353035 | 0.294939938 | 0.286206551 |
| Rdx | 1573.201998 | 0.509353025 | 0.091201866 |
| Gm27686 | 0 | NA | NA |
| Zc3h12c | 704.8556827 | 0.617509306 | 0.121282649 |
| Mir6244 | 0.506103766 | 2.549130456 | 3.28515521 |
| Gm25562 | 0 | NA | NA |
| AI593442 | 2277.537907 | -1.080721613 | 0.101109336 |
| Gm22242 | 0 | NA | NA |
| Ddx10 | 340.9961152 | 0.040718401 | 0.086840752 |
| 4930510E17Rik | 0 | NA | NA |
| Exph5 | 843.2123267 | -0.582982398 | 0.185364812 |
| Kdelc2 | 102.2218059 | 0.017867758 | 0.122800358 |
| Mir130c | 0.049553047 | 0.71820487 | 3.724945049 |
| 4930550C14Rik | 3.372172578 | 0.820321292 | 0.540895373 |
| Atm | 883.8572749 | 0.765740079 | 0.097442157 |
| Npat | 649.966786 | -0.113679449 | 0.095227115 |
| Acat1 | 830.5020792 | -0.171388122 | 0.065130814 |
| Cul5 | 1233.233245 | 0.094072481 | 0.081145034 |
| Gm22304 | 0 | NA | NA |
| Gm16124 | 9.053711666 | -0.559047524 | 0.387302802 |
| Rab39 | 22.16151509 | 0.823765942 | 0.224056811 |
| Rps6-ps3 | 1.738215179 | 0.501456588 | 0.778891863 |
| Slc35f2 | 23.18056578 | 0.171959936 | 0.226780126 |
| Sln | 3.22564905 | 0.420269295 | 0.606702703 |
| Gm16380 | 0.31602048 | -0.724461607 | 1.61289492 |
| Elmod1 | 1587.467376 | -0.711476931 | 0.063548842 |
| Tnfaip8l3 | 95.49101603 | 0.427506095 | 0.142902911 |
| Cyp19a1 | 1.568920191 | -0.364553995 | 0.816211455 |
| Gldnos | 0.038981395 | 0.718207094 | 3.724945049 |
| Gldn | 61.66008331 | 2.147914345 | 0.261343158 |
| Dmxl2 | 4593.55088 | -0.534778157 | 0.067877864 |
| Gm25980 | 0 | NA | NA |
| Sh2d7 | 0.788119377 | 1.603496264 | 1.466698802 |
| Cib2 | 155.6054035 | -0.511055641 | 0.135873635 |
| Idh3a | 1642.060698 | -0.257717418 | 0.066128071 |

| | | | |
|---|---|---|---|
| Acsbg1 | 1565.688373 | -0.061142527 | 0.147298222 |
| Dnaja4 | 456.041311 | 0.171825323 | 0.118178446 |
| Mir5710 | 0.417007765 | -1.005964058 | 1.42005813 |
| Wdr61 | 268.6245129 | 0.292977685 | 0.091914415 |
| Crabp1 | 9.084845767 | -0.855770061 | 0.386918264 |
| Ireb2 | 1512.434034 | -0.128075533 | 0.069066887 |
| Hykk | 65.99315051 | -0.442910118 | 0.174013257 |
| AY074887 | 0.108309328 | 0.237310006 | 3.724945049 |
| Psma4 | 828.6826824 | 0.842147383 | 0.0959672 |
| Chrna5 | 21.98151757 | -0.635053014 | 0.246074728 |
| Chrna3 | 28.52374879 | -0.391366751 | 0.244757102 |
| Chrnb4 | 16.87880904 | 0.332498677 | 0.333881386 |
| Ube2q2 | 800.5171951 | -0.15515469 | 0.06988368 |
| Fbxo22 | 362.6766692 | 0.076720854 | 0.06852985 |
| Nrg4 | 36.91795082 | 1.892153694 | 0.280682386 |
| Gm15625 | 0 | NA | NA |
| AI118078 | 183.6671056 | -0.494852047 | 0.140852043 |
| Etfa | 264.841253 | 0.343242217 | 0.137114857 |
| Isl2 | 0.471858061 | 0.489559772 | 1.386056706 |
| Gm17226 | 0 | NA | NA |
| Scaper | 701.5265708 | -0.559836321 | 0.081307213 |
| Gm23407 | 0.627732484 | -1.918748371 | 1.246829411 |
| 4930563M21Rik | 0.038981395 | 0.718207094 | 3.724945049 |
| Rcn2 | 1128.177669 | -0.310417234 | 0.088979214 |
| Pstpip1 | 94.70433045 | 3.928205194 | 0.261628404 |
| Tspan3 | 2389.846793 | -0.069329312 | 0.089330713 |
| Peak1 | 1406.181793 | 0.130383357 | 0.071998497 |
| Gm24270 | 405.7586766 | 0.973477787 | 0.860059213 |
| Gm29322 | 6.151976361 | -1.071819792 | 0.43977101 |
| Peak1os | 8.730896443 | 3.496059501 | 0.488744484 |
| Gm37842 | 2.222650432 | 4.653845036 | 1.046058961 |
| Gm36940 | 30.43733313 | -0.738544408 | 0.292617866 |
| Hmg20a | 798.0476359 | -0.18415654 | 0.061016246 |
| Gm26868 | 0.201261165 | -0.243564167 | 2.21910279 |
| Lingo1 | 2194.009453 | -0.746168425 | 0.187752375 |
| Odf3l1 | 0.839566529 | -0.151189543 | 1.367455356 |
| Cspg4 | 408.2241713 | 0.504524551 | 0.091930029 |
| Snx33 | 323.7784099 | 1.141679281 | 0.144454996 |
| Imp3 | 174.2311039 | 0.273804709 | 0.143173876 |
| Snupn | 97.8417329 | 0.203569895 | 0.124914504 |
| Ptpn9 | 554.0054497 | 0.466535645 | 0.090834155 |
| Gm10658 | 5.404131218 | -0.324599538 | 0.517824908 |
| Sin3a | 642.6567267 | 0.067522287 | 0.060116949 |
| 2700012I20Rik | 1.975685927 | 0.223149299 | 0.766937964 |
| Man2c1os | 25.71371313 | -0.228026784 | 0.263883194 |
| Man2c1 | 469.3784322 | -0.480330924 | 0.101860751 |
| Neil1 | 55.964215 | -0.32305262 | 0.195803847 |

| | | | |
|---|---|---|---|
| Commd4 | 212.8717519 | 0.016463985 | 0.086807377 |
| 1700041C23Rik | 0.044544648 | 0.237309859 | 3.724945049 |
| Trcg1 | 0.119587853 | 0.237305575 | 3.724945049 |
| 1700017B05Rik | 450.6317575 | 1.509668398 | 0.15480919 |
| Gm5121 | 0.778482992 | 1.824275982 | 1.141718798 |
| Ppcdc | 133.777483 | 0.130173655 | 0.136936726 |
| Scamp5 | 3375.801194 | -0.387101566 | 0.050359571 |
| Gm6166 | 226.0029141 | 0.517146217 | 0.122241033 |
| Rpp25 | 85.87965751 | -0.479398123 | 0.163423239 |
| Cox5a | 985.6266964 | 0.081199223 | 0.101406683 |
| Gm26631 | 4.769631826 | -0.482881077 | 0.690181779 |
| Fam219b | 230.0016874 | 0.082478704 | 0.085452231 |
| Mpi | 358.5215663 | -0.144804127 | 0.070987399 |
| Scamp2 | 323.5294745 | 1.144761484 | 0.12015382 |
| Ulk3 | 153.3484004 | -0.354137968 | 0.148403434 |
| Cplx3 | 72.27239396 | -1.14798773 | 0.231957116 |
| Lman1l | 1.399662294 | 3.916749063 | 1.077769023 |
| Csk | 512.8013575 | 0.795458216 | 0.114924224 |
| Cyp1a2 | 0.530050215 | -0.469549152 | 1.475554973 |
| Cyp1a1 | 0.692555501 | -0.038279761 | 1.146696184 |
| Edc3 | 308.5919229 | 0.076011537 | 0.101741959 |
| Clk3 | 437.4151506 | 0.026637833 | 0.102371413 |
| Gm17231 | 0.756247029 | 0.79020492 | 3.204875992 |
| Arid3b | 117.0600692 | -0.598248993 | 0.215918451 |
| Ubl7 | 525.5396743 | 0.022276904 | 0.078220378 |
| Sema7a | 1597.276098 | -0.163540701 | 0.058157499 |
| Gm17322 | 37.01146921 | -1.363957517 | 0.250037129 |
| Cyp11a1 | 16.97152848 | -1.617399345 | 0.308915464 |
| Ccdc33 | 7.670793784 | 0.552548518 | 0.38999042 |
| Gm16130 | 0.257325455 | -0.553324669 | 2.065025894 |
| Stra6 | 207.0557773 | 3.020728249 | 0.261924715 |
| Gm16131 | 0 | NA | NA |
| Islr | 283.8085694 | 0.851157124 | 0.176236046 |
| Islr2 | 662.2018967 | -0.026138063 | 0.179762503 |
| 1600029O15Rik | 9.217140603 | 0.840974227 | 0.35957412 |
| Pml | 1126.656421 | 3.517411969 | 0.17594452 |
| Mir6385 | 0.090427103 | 0.237307635 | 3.724945049 |
| Stoml1 | 580.7474093 | 0.730966729 | 0.111684786 |
| Loxl1 | 78.19031371 | 1.778308772 | 0.213581749 |
| Tbc1d21 | 0 | NA | NA |
| 1700072B07Rik | 0.112288909 | -0.724455696 | 3.724945049 |
| 4930461G14Rik | 0.112396584 | 1.192083721 | 3.724434897 |
| 6030419C18Rik | 1394.926258 | -0.496444096 | 0.084384148 |
| Cd276 | 125.0400392 | 0.461846334 | 0.107564565 |
| Gm10657 | 8.005046079 | -0.40330736 | 0.390926143 |
| Nptn | 4736.976647 | -0.433596259 | 0.046680382 |
| Rec114 | 2.938016276 | 1.118716073 | 0.613620436 |

| | | | |
|---|---|---|---|
| Gm20620 | 0 | NA | NA |
| Hcn4 | 140.1825926 | -0.127914457 | 0.118420641 |
| Neo1 | 1678.641993 | -0.072024932 | 0.044199033 |
| Gm7589 | 1.467463761 | -0.598339743 | 0.845699134 |
| Adpgk | 228.1330696 | 0.699880149 | 0.078830366 |
| Bbs4 | 465.2733797 | 0.054845862 | 0.062712464 |
| Arih1 | 1854.328923 | 0.26010619 | 0.047178653 |
| Rps11-ps1 | 39.22874844 | -0.232238732 | 0.234414944 |
| Tmem202 | 11.87611272 | 0.798489981 | 0.321468055 |
| Hexa | 567.217733 | 0.818474117 | 0.148945327 |
| Celf6 | 249.5287256 | -0.107318628 | 0.09863637 |
| Parp6 | 1004.151157 | -0.321379313 | 0.078267681 |
| Pkm | 10594.62968 | 0.252810403 | 0.062102344 |
| Gramd2 | 45.41966732 | -1.656408164 | 0.259011125 |
| Senp8 | 100.069545 | 0.018298691 | 0.193844224 |
| Myo9a | 3366.723381 | -0.451349782 | 0.071936834 |
| Gm7613 | 0 | NA | NA |
| Gm2735 | 0.060174537 | 0.237309859 | 3.724945049 |
| Nr2e3 | 0.25324161 | -1.47777566 | 2.98720403 |
| Gm7616 | 0.135217741 | 0.237306921 | 3.620320246 |
| Gm15507 | 0.044544648 | 0.237309859 | 3.724945049 |
| Thsd4 | 209.2641097 | 0.200757552 | 0.155928648 |
| Gm23668 | 0.21890733 | -1.243213754 | 2.590555603 |
| Gm22099 | 0 | NA | NA |
| Lrrc49 | 843.7713801 | -0.263750642 | 0.066754584 |
| B930082K07Rik | 9.514292225 | -0.133741533 | 0.337436335 |
| Larp6 | 390.5730594 | -0.258748323 | 0.09343596 |
| 1700036A12Rik | 5.346398063 | -0.191084351 | 0.50091371 |
| Uaca | 260.6470978 | 2.587913317 | 0.238482225 |
| Gm9869 | 0.554399409 | 2.332147565 | 1.581529628 |
| Gm10655 | 0.545210175 | -0.630629172 | 1.420012425 |
| Tle3 | 1433.600019 | 0.90556998 | 0.13029837 |
| Rplp1 | 2533.538003 | 0.601968369 | 0.127445035 |
| Kif23 | 18.40282969 | 1.209554062 | 0.37988027 |
| Paqr5 | 21.19486437 | 0.471814737 | 0.292424133 |
| Gm23221 | 0 | NA | NA |
| Glce | 558.6811353 | -0.093086414 | 0.06579122 |
| Mir5133 | 0 | NA | NA |
| Gm25603 | 0.463251208 | 2.154033376 | 1.420365102 |
| Spesp1 | 1.257510835 | 4.35155958 | 1.073146488 |
| Anp32a | 1220.827071 | 0.060147057 | 0.091558716 |
| Coro2b | 2114.073909 | -0.351265975 | 0.084125581 |
| n-R5s84 | 0 | NA | NA |
| Itga11 | 34.17670704 | 0.594394148 | 0.243876197 |
| Fem1b | 2918.112869 | 0.365958629 | 0.071984943 |
| Gm26609 | 0.048949532 | -0.243569582 | 3.724945049 |
| Cln6 | 139.097147 | 0.404296192 | 0.098406875 |

| | | | |
|---|---|---|---|
| Gm10653 | 0.777406778 | 0.563811243 | 1.164077446 |
| Calml4 | 9.674253487 | -0.803445301 | 0.386121368 |
| Pias1 | 838.3302677 | -0.014065897 | 0.069578743 |
| Gm24526 | 0 | NA | NA |
| Gm25064 | 0.490308497 | 2.187347345 | 1.318916751 |
| Skor1 | 43.41229914 | 0.639777076 | 0.231508376 |
| Map2k5 | 366.7989628 | -0.32815015 | 0.074172019 |
| 2300009A05Rik | 131.9294766 | -0.301146809 | 0.099541077 |
| Gm16759 | 25.37684091 | -1.255508091 | 0.374257295 |
| Iqch | 1.370360641 | -0.335846927 | 0.858750997 |
| Gm25995 | 0.525324978 | -0.139383221 | 1.662127136 |
| Gm6257 | 8.599489497 | 0.414554687 | 0.414522745 |
| Aagab | 585.5754181 | 0.090792857 | 0.065532504 |
| Smad3 | 973.5426315 | 0.326260399 | 0.09394465 |
| Smad6 | 31.70011363 | 0.839783197 | 0.238021946 |
| 1700055C04Rik | 0 | NA | NA |
| Gm25606 | 0 | NA | NA |
| 1110036E04Rik | 0.062426873 | 0.237312084 | 3.724945049 |
| Scarletltr | 0 | NA | NA |
| Lctl | 0.650092409 | -1.600259128 | 1.264862621 |
| Zwilch | 32.2726367 | -0.107891074 | 0.253132088 |
| Rpl4 | 3612.185091 | 0.528010148 | 0.084948202 |
| Gm23344 | 0.928033173 | -0.275380015 | 0.958986453 |
| Snord16a | 3.328548066 | 0.565232494 | 0.609382064 |
| Gm22571 | 2.155635849 | -0.699853409 | 0.678831806 |
| Gm23136 | 2.475450049 | -0.636092217 | 0.745185568 |
| Gm22455 | 1.074104802 | 0.391150414 | 0.941557479 |
| Snapc5 | 90.39216346 | -0.25673882 | 0.132768864 |
| Map2k1 | 2292.231272 | 0.246961075 | 0.089480704 |
| Gm24106 | 0 | NA | NA |
| Uchl4 | 10.29624034 | 0.106727064 | 0.332369439 |
| Tipin | 120.2139083 | 0.312042345 | 0.131780176 |
| Gm24289 | 1.279592076 | -0.44425106 | 0.933246325 |
| Dis3l | 359.2069684 | -0.182472097 | 0.088278893 |
| Megf11 | 632.3694475 | -0.16430426 | 0.117436645 |
| Rab11a | 1359.033138 | -0.014067326 | 0.087160337 |
| AC147560.1 | 0 | NA | NA |
| Dennd4a | 1982.107441 | -0.042270848 | 0.072107383 |
| Gm24347 | 0 | NA | NA |
| Slc24a1 | 2.426791053 | 0.948794876 | 0.647984676 |
| Vwa9 | 254.7088933 | 0.337522781 | 0.08252422 |
| Gm25640 | 1.475874304 | 0.137543871 | 0.849593825 |
| Hacd3 | 1635.485628 | -0.139537995 | 0.067272793 |
| Dpp8 | 2380.590348 | -0.418125104 | 0.055020896 |
| Igdcc4 | 562.3809832 | -1.119714606 | 0.149709091 |
| Igdcc3 | 23.91639695 | -1.345898267 | 0.246234223 |
| Gm25313 | 3.277280862 | 1.315398931 | 0.657539485 |

| | | | |
|---|---|---|---|
| Rnu5g | 4.132676428 | 1.344909889 | 0.527229728 |
| Parp16 | 35.8728912 | -0.045114503 | 0.18364063 |
| Cilp | 2.578207035 | 0.966957431 | 0.749748109 |
| Clpx | 458.2865452 | 0.306547482 | 0.084031672 |
| Gm16218 | 0 | NA | NA |
| Pdcd7 | 276.9287299 | -0.174022022 | 0.079979125 |
| Ubap1l | 57.59547211 | -0.621882993 | 0.334854871 |
| Kbtbd13 | 1.752461279 | -0.512241433 | 0.93408777 |
| Rasl12 | 22.15572106 | -0.299175575 | 0.227643335 |
| Slc51b | 0.090890357 | 0.718200421 | 3.724945049 |
| Mtfmt | 124.7687732 | -0.267067047 | 0.127677056 |
| Spg21 | 367.5189745 | 1.083335887 | 0.137312357 |
| Ankdd1a | 14.47040722 | -1.906946686 | 0.361012807 |
| Plekho2 | 544.0730231 | 1.431028644 | 0.148885113 |
| Pif1 | 5.794504373 | -0.304210962 | 0.482858427 |
| Rbpms2 | 39.33216484 | 1.352550067 | 0.213063692 |
| Oaz2 | 881.559079 | -0.173219185 | 0.080947817 |
| Zfp609 | 1323.683813 | -0.180296235 | 0.057655654 |
| Trip4 | 581.8480211 | 0.303019514 | 0.056269078 |
| n-R5s85 | 0 | NA | NA |
| Gm16073 | 0.045882455 | 0.237312084 | 3.724945049 |
| 2810417H13Rik | 14.58011455 | 3.602264376 | 0.553672601 |
| Gm23248 | 0 | NA | NA |
| Csnk1g1 | 818.8766249 | -0.268977787 | 0.093726709 |
| Gm22657 | 0 | NA | NA |
| Ppib | 745.1195212 | 0.943611239 | 0.110486416 |
| Snx22 | 48.15551827 | -0.039906578 | 0.18720194 |
| Snx1 | 927.1821691 | 0.974642523 | 0.083096932 |
| Fam96a | 202.4742243 | 0.582094599 | 0.108411457 |
| Dapk2 | 23.66058682 | 0.378272553 | 0.24046856 |
| Herc1 | 7489.340024 | -0.437075828 | 0.095518754 |
| Gm23428 | 0 | NA | NA |
| Gm28379 | 0.527778125 | 0.955067547 | 1.397439199 |
| Fbxl22 | 15.26684527 | -0.950918286 | 0.311715804 |
| Usp3 | 603.065053 | -0.378796604 | 0.077865977 |
| Gm15563 | 0 | NA | NA |
| Car12 | 301.1637606 | 0.375893537 | 0.151564532 |
| Aph1b | 309.2262609 | -0.508076791 | 0.085028435 |
| Gm10647 | 6.118905949 | -1.035800307 | 0.527139792 |
| BC050972 | 1.031670571 | -0.699138181 | 0.950131238 |
| Ppp1r2-ps4 | 7.934796858 | 0.020054895 | 0.396991443 |
| Aph1c | 99.73584743 | -0.122682226 | 0.126072484 |
| Gm17098 | 1.018413619 | -0.88210421 | 1.09896922 |
| Rab8b | 840.6987422 | 0.933054777 | 0.11438034 |
| Rps27l | 178.3460037 | 1.569193908 | 0.175473965 |
| Lactb | 129.5314519 | -0.186199811 | 0.109457853 |
| Gm24225 | 0 | NA | NA |

| | | | |
|---|---|---|---|
| Tpm1 | 1521.522177 | 0.043424937 | 0.117942442 |
| Gm22145 | 0 | NA | NA |
| Tln2 | 1967.481299 | -0.804450885 | 0.092112791 |
| Mir190a | 0.401901456 | 0.218530128 | 1.543916655 |
| Mir7241 | 0.17692102 | 0.773454306 | 2.723401318 |
| Mir7242 | 0.162691105 | -0.724458629 | 2.788998794 |
| M5C1000I18Rik | 30.04157137 | -1.436546803 | 0.232344586 |
| C2cd4b | 11.79896043 | 1.285729218 | 0.40949444 |
| C2cd4a | 1.024692053 | 2.098665531 | 1.166239253 |
| Vps13c | 2530.722944 | -0.421723256 | 0.096649461 |
| Gm28195 | 0 | NA | NA |
| Gm22382 | 0 | NA | NA |
| Gm22169 | 0 | NA | NA |
| Gm22962 | 0.044191549 | -0.243571806 | 3.724945049 |
| Rora | 3475.748826 | -0.248646763 | 0.071543765 |
| AC127262.1 | 19.3378465 | -2.461599148 | 0.405941109 |
| Gm15511 | 0.358080405 | 0.578747246 | 1.564402896 |
| Ice2 | 215.338252 | -0.214242065 | 0.119593672 |
| Gm4978 | 0.090277459 | 0.237318756 | 3.724945049 |
| Anxa2 | 640.8248345 | 3.979551624 | 0.344616073 |
| Mir3109 | 0.728690676 | 1.576475482 | 1.144997653 |
| Gm16144 | 0 | NA | NA |
| Foxb1 | 12.10284569 | -0.752988919 | 0.309050415 |
| B230323A14Rik | 7.674232682 | -1.731766463 | 0.487782432 |
| Gm28731 | 0.659754175 | 0.949923959 | 1.229951725 |
| Bnip2 | 412.8066768 | 0.766466299 | 0.115364295 |
| Gtf2a2 | 313.4260604 | -0.134317798 | 0.100191305 |
| Gcnt3 | 0.134276636 | 0.684459443 | 2.951368184 |
| Fam81a | 750.583868 | -1.446039613 | 0.146020063 |
| Gm24257 | 0 | NA | NA |
| Gm26849 | 0.04393175 | -0.243571806 | 3.724945049 |
| Myo1e | 393.0843238 | 1.637676848 | 0.13783254 |
| Mir5626 | 0.091666476 | -0.724455696 | 3.724945049 |
| Ccnb2 | 19.87963883 | 0.876370885 | 0.324492381 |
| Rnf111 | 823.3856292 | 0.273464758 | 0.065222008 |
| Gm24615 | 0 | NA | NA |
| Gm23588 | 0 | NA | NA |
| Gm23730 | 0.04393175 | -0.243571806 | 3.724945049 |
| Gm22229 | 0 | NA | NA |
| Sltm | 725.1236 | -0.025965291 | 0.115747285 |
| Fam63b | 1775.64688 | -0.206571949 | 0.09241241 |
| Gm10642 | 11.00079997 | 0.266447609 | 0.328304371 |
| Adam10 | 1465.376635 | 0.245924307 | 0.092636233 |
| Lipc | 1.303781876 | 0.330444069 | 1.065717841 |
| Gm25149 | 0 | NA | NA |
| Gm3436 | 6.640611746 | 1.139419028 | 0.389641047 |
| Aqp9 | 16.64346168 | 1.381003098 | 0.32075858 |

| | | | |
|---|---|---|---|
| Aldh1a2 | 316.9562088 | 2.198049868 | 0.201279646 |
| Polr2m | 1314.180415 | 0.273360604 | 0.081447597 |
| Gcom1 | 0 | NA | NA |
| Myzap | 9.380444241 | 0.154732934 | 0.350281513 |
| Cgnl1 | 362.6537632 | -0.078572832 | 0.124825645 |
| Gm27220 | 0 | NA | NA |
| Gm37660 | 16.37804138 | -0.547775292 | 0.349226711 |
| Tcf12 | 1303.134971 | 0.362142508 | 0.068014445 |
| Gm38057 | 46.80930091 | -1.014853835 | 0.303245251 |
| Gm38111 | 32.02976615 | -0.971117466 | 0.320080924 |
| Gm18821 | 0 | NA | NA |
| Gm37879 | 56.51000279 | -0.2694098 | 0.250747982 |
| Gm27152 | 0 | NA | NA |
| Gm37663 | 62.49855959 | -0.428421701 | 0.179995206 |
| Gm37023 | 32.44644427 | -0.53332869 | 0.336289133 |
| A430027C01Rik | 29.11976123 | -0.610920209 | 0.286471132 |
| Gm27450 | 0 | NA | NA |
| Gm37137 | 44.47354403 | -0.625210444 | 0.281575929 |
| Rpl15-ps2 | 4.494967837 | 0.854756462 | 0.600986825 |
| Gm25045 | 0.207243203 | 0.237320053 | 2.611868359 |
| Gm7863 | 0 | NA | NA |
| Gm7866 | 2.795202431 | 0.534543705 | 0.650398621 |
| Gm27159 | 0.158742293 | 1.563747326 | 3.440567392 |
| Gm27188 | 5.84902935 | -0.179805601 | 0.510147285 |
| Gm7870 | 0.124347486 | 0.719186602 | 3.717245571 |
| Gm25163 | 0.730530645 | -1.846388655 | 1.249383892 |
| Gm27253 | 3.522083342 | -1.496325317 | 0.618018024 |
| Zfp280d | 1238.243116 | -0.32487328 | 0.107321508 |
| Gm19353 | 21.12517525 | -1.095171526 | 0.315721272 |
| Gm17828 | 1.657074577 | 0.200108108 | 0.761323429 |
| Gm37983 | 73.73482861 | -0.709682866 | 0.240456356 |
| Gm37303 | 69.68360701 | -0.787422324 | 0.240041872 |
| Gm37581 | 38.67009967 | -0.840065314 | 0.304707565 |
| BC065403 | 0.430924493 | -0.698808187 | 1.464756975 |
| Gm37150 | 0.370677815 | 1.231541656 | 1.722514586 |
| Gm27230 | 0.066050875 | 0.71820487 | 3.724945049 |
| Gm27255 | 0.075043205 | 0.237309859 | 3.724945049 |
| Gm27254 | 0.194277958 | -0.243576487 | 3.254509263 |
| Mns1 | 77.00500091 | 0.470489092 | 0.158907642 |
| Gm36948 | 4.487847835 | 0.879879391 | 0.626019597 |
| Gm27231 | 0.460076145 | -1.254959695 | 1.617497861 |
| Tex9 | 177.6432418 | -0.188732732 | 0.128035446 |
| Gm28010 | 0.172182752 | 0.718209228 | 3.331044598 |
| 4930509E16Rik | 0.362648911 | 0.789252486 | 1.727061378 |
| Rfx7 | 1180.076257 | -0.496083626 | 0.077016554 |
| Gm19427 | 12.05712943 | -0.983022693 | 0.353694834 |
| Gm37105 | 45.12253491 | -0.977871163 | 0.340833103 |

| | | | |
|---|---|---|---|
| Gm18014 | 0 | NA | NA |
| Gm27241 | 0.220148986 | 0.240721873 | 2.596317291 |
| Gm27232 | 0.138573873 | 1.200070582 | 3.628829444 |
| Gm23057 | 0.072025462 | 0.237312084 | 3.724945049 |
| Nedd4 | 6315.797856 | -0.05572709 | 0.048768379 |
| A530064N14Rik | 50.37861278 | -0.815261666 | 0.33235472 |
| Gm7265 | 195.1401439 | 0.446086991 | 0.097515474 |
| Gm37955 | 4.498464866 | -1.277764204 | 0.545835146 |
| Gm36936 | 103.529168 | -0.964428796 | 0.287168501 |
| Gm27204 | 0 | NA | NA |
| Prtg | 50.92406102 | -1.27106348 | 0.214953372 |
| Gm27203 | 0 | NA | NA |
| A730062M13Rik | 2.892325926 | -2.688033466 | 0.758364581 |
| Gm37234 | 1.821448544 | -1.223023988 | 0.896938181 |
| Pygo1 | 611.8774595 | -1.03734843 | 0.100235762 |
| Dyx1c1 | 5.660739534 | -0.114778752 | 0.428550349 |
| Dyx1c1Ccpg1 | 0 | NA | NA |
| Gm23567 | 0.220111735 | 0.260725583 | 2.165130518 |
| Gm37710 | 0.046345709 | 0.71820487 | 3.724945049 |
| Ccpg1os | 38.04226344 | -0.118994024 | 0.210909675 |
| Ccpg1 | 158.4319805 | -0.493560979 | 0.112437319 |
| Gm27353 | 16.47684853 | -1.566839173 | 0.348629116 |
| Pigb | 210.9882647 | -0.056732469 | 0.089660497 |
| Gm38192 | 17.09336393 | -1.192088429 | 0.287125285 |
| 2310009A05Rik | 48.02464468 | -0.05010259 | 0.22106479 |
| Rab27a | 175.4812393 | 0.556455027 | 0.119406129 |
| Gm27186 | 0.89763465 | 0.283090067 | 1.25901378 |
| Khdc3 | 3.338837625 | -1.125742205 | 0.606399644 |
| Gm20509 | 0 | NA | NA |
| Rsl24d1 | 296.3744691 | 0.465294306 | 0.080360144 |
| Gm38215 | 2.049787317 | -0.473440543 | 0.749486696 |
| Gm5745 | 0.038981395 | 0.718207094 | 3.724945049 |
| Gm27211 | 0.328170597 | 1.799839574 | 3.693610719 |
| Unc13c | 1536.822218 | -1.137450592 | 0.098212439 |
| Gm37611 | 8.958307541 | -1.660357572 | 0.461721655 |
| Gm27148 | 1.654347319 | -2.945974672 | 1.03190089 |
| Gm25105 | 4.734499335 | -1.337227761 | 0.519150118 |
| Gm37326 | 31.32772203 | -1.625882786 | 0.329519622 |
| n-R5s86 | 0 | NA | NA |
| Wdr72 | 4.387779186 | 0.887813845 | 0.561225457 |
| Gm24141 | 0 | NA | NA |
| Nr1h2-ps | 0 | NA | NA |
| Gm28622 | 0.044544648 | 0.237309859 | 3.724945049 |
| Gm27189 | 0 | NA | NA |
| Gm27233 | 0 | NA | NA |
| Gm22315 | 0 | NA | NA |
| Gm16551 | 2.498792093 | 0.261240884 | 0.713665159 |

| | | | |
|---|---|---|---|
| Gm20649 | 0 | NA | NA |
| Gm28182 | 0.458414965 | -0.451690286 | 1.346811356 |
| Onecut1 | 8.345816899 | -0.441024414 | 0.380665191 |
| Fam214a | 701.9545256 | -0.435477127 | 0.086031178 |
| Arpp19 | 2304.068667 | -0.701884371 | 0.123089237 |
| Myo5a | 9406.867657 | -0.369120851 | 0.065419349 |
| Gm9719 | 0.038981395 | 0.718207094 | 3.724945049 |
| Gm16353 | 0 | NA | NA |
| Myo5c | 38.56207624 | -0.772248657 | 0.349628055 |
| Gnb5 | 588.9032945 | -0.575524991 | 0.077188425 |
| Bcl2l10 | 2.578686406 | -1.073952002 | 0.669152119 |
| Mapk6 | 1348.202811 | 0.385517227 | 0.095691817 |
| 4933433G15Rik | 7.660142694 | 0.037396146 | 0.427184731 |
| A130057D12Rik | 0.39127578 | 0.723706966 | 1.567518952 |
| Leo1 | 251.7819994 | 0.35695475 | 0.180376528 |
| Tmod3 | 491.2591432 | 1.080313643 | 0.094287754 |
| Tmod2 | 9543.757063 | -0.495592979 | 0.061223888 |
| Lysmd2 | 376.8025865 | 0.301373698 | 0.10691384 |
| Scg3 | 2648.530245 | -0.019421501 | 0.079337226 |
| Gm37871 | 0.090665643 | -0.724455696 | 3.724945049 |
| Gm3671 | 0.843494912 | 0.551110539 | 1.13548107 |
| Bmp5 | 56.01127082 | 0.638039192 | 0.218244134 |
| Gm26903 | 0.668743806 | -0.975852051 | 1.482082344 |
| AC079245.1 | 0 | NA | NA |
| Gm25300 | 0 | NA | NA |
| Hmgcll1 | 224.7683895 | -0.556204363 | 0.118236592 |
| Gm38083 | 27.55755119 | -0.972504744 | 0.265972524 |
| Gfral | 1.244634631 | -0.641152847 | 0.920511494 |
| Hcrtr2 | 58.13001265 | -0.273937061 | 0.232021537 |
| Gm27234 | 0.038981395 | 0.718207094 | 3.724945049 |
| Fam83b | 2.476850474 | 0.039077854 | 0.685944628 |
| Gm27156 | 0 | NA | NA |
| Gm27245 | 0 | NA | NA |
| Gm22938 | 0 | NA | NA |
| Gm37408 | 0 | NA | NA |
| Gm27155 | 0 | NA | NA |
| Gm23791 | 0 | NA | NA |
| Gm18486 | 0 | NA | NA |
| Tinag | 0.568818845 | 0.662860249 | 1.171631687 |
| Gm27157 | 0 | NA | NA |
| Gm27158 | 0 | NA | NA |
| Gm19541 | 0.139037128 | 1.464453538 | 3.717743456 |
| Gm5746 | 0.044544648 | 0.237309859 | 3.724945049 |
| Mlip | 470.9327951 | -0.843653352 | 0.108193346 |
| Gm27210 | 1.42879052 | 0.383166903 | 0.849802593 |
| Gm38254 | 2.89654368 | -0.593721239 | 0.649005834 |
| 4930590H14Rik | 0 | NA | NA |

| | | | |
|---|---|---|---|
| 5730403I07Rik | 4.072737608 | -0.138182849 | 0.571501909 |
| Lrrc1 | 172.5600245 | -0.770094461 | 0.125557757 |
| Gm19569 | 3.396866737 | 0.579169523 | 0.602255701 |
| Gm21013 | 0.284055481 | -1.745302754 | 2.435667199 |
| Gm25250 | 0 | NA | NA |
| Klhl31 | 1.14819629 | -0.395936692 | 1.009183945 |
| Gclc | 631.4586054 | 0.050339763 | 0.054545706 |
| Elovl5 | 1099.939201 | 0.17515227 | 0.101599627 |
| Gm23609 | 0 | NA | NA |
| Gcm1 | 0.062426873 | 0.237312084 | 3.724945049 |
| Fbxo9 | 758.0697948 | -0.156969785 | 0.090909553 |
| Ick | 452.7014818 | 0.040241617 | 0.086861682 |
| Rn7sk | 139206.5501 | 0.120114867 | 0.172334527 |
| C920006O11Rik | 1.914692995 | 0.535612946 | 0.776589397 |
| Gsta4 | 217.5686189 | -0.419896897 | 0.156932211 |
| Gsta1 | 0.047948698 | -0.243569582 | 3.724945049 |
| Gm10639 | 0.087930927 | 0.236669296 | 3.724898107 |
| Omt2a | 0 | NA | NA |
| Gm8074 | 0 | NA | NA |
| Omt2b | 0 | NA | NA |
| Gsta2 | 0.44449992 | 2.149781203 | 1.664512788 |
| Dppa5a | 0.045882455 | 0.237312084 | 3.724945049 |
| Ooep | 0.251515219 | 0.23733065 | 1.833800462 |
| Gm8087 | 0.090074004 | -0.24357403 | 3.724945049 |
| 4930542C12Rik | 0 | NA | NA |
| Ddx43 | 4.317706999 | 0.016855891 | 0.517134784 |
| Gm8093 | 1.913696796 | 2.655349825 | 0.939392924 |
| Mb21d1 | 40.10846614 | 2.74318679 | 0.266001586 |
| Gm17324 | 2.766932339 | 0.442421296 | 0.661180157 |
| Mto1 | 264.8336796 | 0.00098062 | 0.076260898 |
| Eef1a1 | 14473.51984 | 0.667610391 | 0.094531211 |
| Gm26377 | 0 | NA | NA |
| Slc17a5 | 273.2142388 | -0.289781461 | 0.103617956 |
| Cd109 | 135.8248537 | 3.182366402 | 0.263713784 |
| 4921509A06Rik | 0.07796279 | 1.072698098 | 3.721829099 |
| Gm8116 | 61.44867496 | 0.366148761 | 0.13778506 |
| Gm28620 | 0 | NA | NA |
| Col12a1 | 350.8884372 | 1.253055546 | 0.157565845 |
| 4930562D21Rik | 0 | NA | NA |
| Cox7a2 | 993.8036579 | -0.130553568 | 0.132484824 |
| Tmem30a | 5117.156797 | -0.016119108 | 0.090120396 |
| 4930429F24Rik | 2.804294664 | 0.991413661 | 0.560362692 |
| Filip1 | 118.2003987 | -0.901633372 | 0.131352224 |
| Gm20477 | 0.402709178 | -1.484987492 | 1.329874223 |
| Gm3211 | 0 | NA | NA |
| Gm25433 | 0 | NA | NA |
| Senp6 | 1335.192463 | 0.037399165 | 0.055586589 |

| | | | |
|---|---|---|---|
| Gm20648 | 0 | NA | NA |
| Gm17477 | 0 | NA | NA |
| Myo6 | 1685.660187 | -0.359808543 | 0.080128082 |
| Gm27227 | 0 | NA | NA |
| Impg1 | 1.92279121 | -0.992588667 | 0.766369317 |
| Gm27224 | 0.046345709 | 0.71820487 | 3.724945049 |
| Gm27225 | 0.281850154 | -0.593901688 | 1.758885992 |
| Gm8151 | 0.792246753 | 0.915507532 | 1.117055537 |
| Gm26713 | 0.150086409 | 0.237305146 | 3.724945049 |
| Gm27226 | 0 | NA | NA |
| Htr1b | 292.4427632 | -0.572020209 | 0.132553558 |
| D430036J16Rik | 57.68366793 | -1.324429593 | 0.186306103 |
| Gm9531 | 81.22717314 | 0.351386785 | 0.128890912 |
| Gm26907 | 0.36944391 | 0.659636548 | 3.271147628 |
| Mei4 | 18.14348477 | -0.618574268 | 0.249670666 |
| Gm37837 | 2.253692409 | -1.230657874 | 0.827036086 |
| Gm15896 | 1.074470558 | -1.429799401 | 1.240672247 |
| Gm23748 | 0.232637982 | -0.724462006 | 2.547631377 |
| 1700063H06Rik | 1.312371618 | -1.246143016 | 1.089362022 |
| Gm22836 | 0 | NA | NA |
| Gm24393 | 0 | NA | NA |
| Gm27249 | 0.180969744 | -0.724462065 | 2.734435593 |
| Irak1bp1 | 163.4822182 | -0.140880852 | 0.102698411 |
| Phip | 3482.802041 | 0.141485901 | 0.086584634 |
| Gm27248 | 0.875572801 | 0.303523932 | 1.340791312 |
| Hmgn3 | 287.8474579 | -0.242188747 | 0.077033752 |
| Gm2065 | 1.354044546 | 1.926492205 | 1.05022116 |
| Gm29054 | 6.179723687 | -1.825314122 | 0.460809487 |
| Gm38279 | 0.200264842 | 0.663060731 | 2.20130791 |
| Gm27216 | 0 | NA | NA |
| Lca5 | 153.3277019 | -0.351841164 | 0.113090438 |
| Rps27a-ps2 | 369.3263099 | 0.61067095 | 0.141397461 |
| Gm2087 | 1.000715493 | 0.23988527 | 1.345498932 |
| Gm37307 | 1.831656741 | -0.819017466 | 0.927354629 |
| Gm28552 | 0.11657011 | 0.237307635 | 3.724945049 |
| Gm28477 | 0.334445592 | -0.243563573 | 1.976837606 |
| Sh3bgrl2 | 558.0448667 | 0.067812786 | 0.085463033 |
| Gm2109 | 2.113095361 | -1.534201245 | 0.846338976 |
| Elovl4 | 771.8863038 | 0.092212966 | 0.101131205 |
| Ttk | 15.5422173 | 1.533593788 | 0.402929288 |
| Bckdhb | 244.9808363 | -0.06517702 | 0.074254809 |
| Gm29163 | 0 | NA | NA |
| Gm29164 | 0 | NA | NA |
| Gm26146 | 0.204782255 | -1.217702835 | 2.205163063 |
| Gm8226 | 4.794882048 | 0.73653218 | 0.486571542 |
| Gm26126 | 0 | NA | NA |
| 4930554C24Rik | 0.203235021 | 1.156859971 | 2.192141084 |

| | | | |
|---|---|---|---|
| Gm28625 | 0 | NA | NA |
| Gm8228 | 8.163884718 | -0.070223551 | 0.38524846 |
| Gm28070 | 0 | NA | NA |
| Gm11114 | 0 | NA | NA |
| Fam46a | 425.1094954 | 2.53378105 | 0.183486111 |
| Gm36927 | 0 | NA | NA |
| Gm25125 | 0 | NA | NA |
| Ibtk | 540.332865 | 0.045462075 | 0.054355894 |
| 4933431K14Rik | 24.95481328 | 0.365576751 | 0.207340196 |
| Gm22830 | 0.143846094 | -1.022239892 | 3.718482058 |
| Gm5619 | 19.25933968 | 0.959090731 | 0.383972484 |
| Gm18766 | 0.416300838 | -1.163802423 | 1.54346192 |
| 9330154J02Rik | 12.67333215 | -0.034464642 | 0.28378783 |
| Tpbg | 242.3331958 | 0.62158646 | 0.116304315 |
| Ube2cbp | 42.39662063 | 0.122782413 | 0.205234645 |
| Gm16202 | 0.329441611 | 0.251360451 | 1.720256333 |
| Dopey1 | 1570.3345 | -0.347801589 | 0.109865763 |
| Gm17530 | 6.903802255 | -0.053881694 | 0.362822503 |
| Pgm3 | 195.6218196 | 0.078363592 | 0.090370716 |
| Rwdd2a | 134.250853 | -0.714913414 | 0.125692895 |
| Me1 | 717.0322805 | 0.454847685 | 0.103006915 |
| Gm38230 | 5.852527318 | -1.583440443 | 0.490095084 |
| A330041J22Rik | 15.09013069 | -0.188197899 | 0.249043496 |
| Gm25949 | 0 | NA | NA |
| Gm22138 | 0.271189253 | -0.722085259 | 3.723649597 |
| Gm37484 | 1.311065665 | -2.019720877 | 0.93672192 |
| Prss35 | 55.61444896 | -0.079322185 | 0.178245673 |
| Snap91 | 4128.306371 | -0.480094565 | 0.063540239 |
| Gm37941 | 94.46575361 | -0.769977308 | 0.272405199 |
| Ripply2 | 33.98125597 | -0.531818643 | 0.209480329 |
| Cyb5r4 | 351.6163489 | -0.062077514 | 0.078680209 |
| Gm8282 | 3.540589861 | -0.241059083 | 0.516455501 |
| Mrap2 | 94.81455933 | 0.520008335 | 0.206823831 |
| Gm37012 | 4.127795301 | 0.343108888 | 0.485742636 |
| Cep162 | 341.6360393 | -0.127546103 | 0.106073651 |
| Gm24649 | 0.157258859 | -0.243568389 | 2.811434299 |
| Gm9540 | 0 | NA | NA |
| Gm37375 | 0.154920219 | -0.243569986 | 2.837065107 |
| Tbx18 | 38.37151821 | 0.209180458 | 0.180100536 |
| D030062O11Rik | 0.22856443 | 1.125686174 | 2.156383772 |
| Gm37374 | 0.748906342 | 0.412402683 | 1.048794935 |
| Gm9541 | 0 | NA | NA |
| Gm25528 | 0 | NA | NA |
| Gm23176 | 0 | NA | NA |
| Gm28877 | 0.115603922 | 1.209894437 | 3.72418822 |
| Gm2324 | 0 | NA | NA |
| Gm5368 | 0 | NA | NA |

| | | | |
|---|---|---|---|
| Gm28231 | 0 | NA | NA |
| Gm5066 | 2.101151607 | -0.029142452 | 0.842562661 |
| Nt5e | 129.2471619 | 0.653994182 | 0.155545692 |
| Snx14 | 532.7999784 | -0.429160163 | 0.069935685 |
| Gm10163 | 3.877733872 | -0.260795624 | 0.504221634 |
| Gm20537 | 1.59229893 | -0.194566666 | 0.759686003 |
| Gm28229 | 64.56740978 | -0.751265263 | 0.177297988 |
| Syncrip | 1823.979902 | 0.065930292 | 0.066009808 |
| Snhg5 | 101.6600101 | 0.597922316 | 0.156809998 |
| Snord50a | 0 | NA | NA |
| Snord50b | 0 | NA | NA |
| Gm22985 | 0 | NA | NA |
| Zfp949 | 311.469506 | -0.075627996 | 0.094982407 |
| Gm26461 | 289.3437332 | 0.077901203 | 0.198552605 |
| Trim43a | 11.4592027 | 1.294655855 | 0.343256315 |
| 9430037G07Rik | 40.90789348 | -0.189396797 | 0.217120465 |
| Gm29603 | 0.044544648 | 0.237309859 | 3.724945049 |
| Gm23017 | 0 | NA | NA |
| Gm26567 | 0 | NA | NA |
| Gm29415 | 1.336334664 | 0.027096149 | 0.973235679 |
| Gm38214 | 23.29150933 | -1.049802003 | 0.31779085 |
| Gm37226 | 40.31519145 | -1.078422207 | 0.342763272 |
| Mthfsl | 134.439119 | -0.179888985 | 0.157295603 |
| Gm10634 | 2.499553143 | -0.618589193 | 0.602281142 |
| Bcl2a1d | 28.6456985 | 5.844369991 | 0.713369484 |
| Gm37560 | 0.141331096 | 0.237322366 | 3.56184083 |
| Gm26082 | 0.238313115 | 0.768146604 | 2.147196579 |
| Gm37614 | 0.29572682 | -0.722855647 | 1.764980694 |
| Gm29562 | 2.519137971 | 0.24151391 | 0.68855441 |
| Trim43c | 0 | NA | NA |
| 9330159M07Rik | 56.07294295 | -0.110592239 | 0.199103011 |
| Gm9775 | 0 | NA | NA |
| Gm26135 | 0 | NA | NA |
| Gm26611 | 0 | NA | NA |
| Gm2396 | 3.793477816 | 2.137449498 | 0.721411295 |
| Gm23707 | 0 | NA | NA |
| Bcl2a1a | 13.58752781 | 6.904507345 | 0.89267533 |
| Gm37833 | 3.739270958 | -0.89337333 | 0.602492914 |
| Gm37993 | 0.55378357 | 1.563851146 | 1.400476226 |
| Gm37471 | 0.168528002 | 0.23732463 | 2.758169185 |
| Gm24463 | 0.710913347 | 0.956021639 | 1.203361292 |
| Gm19498 | 0 | NA | NA |
| Trim43b | 0.095295241 | 0.23732098 | 3.724945049 |
| 4930579C12Rik | 0.595794717 | 2.177430961 | 1.339736512 |
| Gm26423 | 0 | NA | NA |
| Bcl2a1b | 8.616394077 | 4.758222121 | 0.891348725 |
| Gm29094 | 0.444881751 | 2.84199794 | 1.564376805 |

| | | | |
|---|---|---|---|
| Mthfs | 70.61009405 | -0.16654213 | 0.247612719 |
| Gm38091 | 0.04393175 | -0.243571806 | 3.724945049 |
| AF529169 | 248.5876439 | 0.012699156 | 0.175697545 |
| Tmed3 | 223.4678697 | 0.641699856 | 0.118465411 |
| Ankrd34c | 157.1958196 | -0.791269791 | 0.127076438 |
| Gm29608 | 0.210489921 | 0.264481081 | 2.624963033 |
| Mir184 | 0 | NA | NA |
| 1700017I07Rik | 0 | NA | NA |
| 4930524O08Rik | 0.357611442 | 0.237329923 | 1.970779538 |
| Rasgrf1 | 5020.678698 | -0.37662975 | 0.076448331 |
| Gm23759 | 0.203875117 | 0.71821508 | 2.62924432 |
| Gm17944 | 0.117650406 | 0.23732098 | 3.724945049 |
| Ctsh | 415.3048284 | 3.564791715 | 0.207325299 |
| Morf4l1 | 2238.38623 | 0.050277918 | 0.065652935 |
| Gm26882 | 0.372996712 | 0.237329245 | 1.966514151 |
| Gm6223 | 7.567416792 | 0.566565184 | 0.343538439 |
| 9430062P05Rik | 1.427283908 | 0.532820023 | 0.920304607 |
| Adamts7 | 9.932952153 | -0.117148834 | 0.343515086 |
| Tbc1d2b | 421.9794416 | 0.335177827 | 0.087650864 |
| Gm16200 | 6.214365972 | -1.050643233 | 0.415774776 |
| Gm22717 | 0 | NA | NA |
| Gm22866 | 0.060174537 | 0.237309859 | 3.724945049 |
| Gm2497 | 0 | NA | NA |
| Gm29605 | 0 | NA | NA |
| Gm29604 | 0.044544648 | 0.237309859 | 3.724945049 |
| Gm28703 | 3.32610108 | -1.618354361 | 0.703251172 |
| Gm37665 | 0.317495958 | -0.090202785 | 1.7518824 |
| Gm20561 | 0.047948698 | -0.243569582 | 3.724945049 |
| Gm29408 | 0.136071997 | -1.190208711 | 2.941811982 |
| Gm5620 | 30.93367375 | 0.00330699 | 0.225376308 |
| Gm36960 | 3.033497949 | 0.390425915 | 0.671446242 |
| Gm29602 | 0 | NA | NA |
| Gm37812 | 6.137587838 | -0.215552723 | 0.387155527 |
| Gm37457 | 5.572788009 | -1.003648043 | 0.473030942 |
| Gm37040 | 5.798168318 | 0.214926062 | 0.506515659 |
| Gm28652 | 0 | NA | NA |
| A730094K22Rik | 3.673889323 | -0.086503216 | 0.69665454 |
| Zic1 | 2091.234844 | -0.516759416 | 0.184273222 |
| Zic4 | 124.4589485 | -0.436414629 | 0.14137393 |
| Gm37477 | 5.296492507 | -0.209658188 | 0.467322822 |
| Gm29478 | 1.174654913 | -1.149762074 | 0.930287621 |
| Gm37987 | 0.060174537 | 0.237309859 | 3.724945049 |
| Gm26101 | 0.062426873 | 0.237312084 | 3.724945049 |
| Plscr5 | 1.323501177 | -1.115251609 | 0.880631796 |
| Plscr1 | 60.43889054 | 1.978220898 | 0.282853407 |
| Mir6386 | 0 | NA | NA |
| Plscr2 | 121.3806642 | 2.794771994 | 0.240458715 |

| | | | |
|---|---|---|---|
| Gm29139 | 0 | NA | NA |
| 1700057G04Rik | 0.707934375 | 1.257583179 | 1.174452634 |
| Gm15493 | 0 | NA | NA |
| Plscr4 | 44.4848455 | 0.005018394 | 0.182812732 |
| Gm25417 | 0.083526043 | 0.718202646 | 3.724945049 |
| Gm38391 | 19.9576536 | -0.076378445 | 0.245716605 |
| Gm37545 | 0 | NA | NA |
| Plod2 | 158.0764622 | 0.610096929 | 0.107165471 |
| Gm2495 | 0 | NA | NA |
| Gm29396 | 0 | NA | NA |
| 4933400C23Rik | 0.454057983 | -1.179847089 | 1.496541601 |
| Gm28054 | 0.110242424 | 0.237318756 | 3.724945049 |
| Gm29397 | 3.384086637 | 0.896791526 | 0.535206143 |
| 1700034K08Rik | 0 | NA | NA |
| Gm9621 | 0 | NA | NA |
| Gm24200 | 0 | NA | NA |
| Gm5370 | 0 | NA | NA |
| Gm28934 | 0 | NA | NA |
| 1190002N15Rik | 756.6875991 | -0.191478987 | 0.139395819 |
| Gm24178 | 0.049553047 | 0.71820487 | 3.724945049 |
| Gm29395 | 0.832501709 | -1.604650243 | 1.15844795 |
| Slc9a9 | 216.5651415 | -0.307699297 | 0.122116258 |
| Gm16262 | 0.060174537 | 0.237309859 | 3.724945049 |
| Mir7656 | 0.276333946 | -1.209613235 | 1.800088756 |
| Chst2 | 1637.467997 | 0.360993664 | 0.107351513 |
| Gm37805 | 7.969835414 | 0.121739071 | 0.368876368 |
| Gm8449 | 2.212778765 | 0.940522085 | 0.895720399 |
| Gm19193 | 0 | NA | NA |
| U2surp | 1949.775422 | -0.057592872 | 0.085123855 |
| Gm28424 | 12.26829614 | -0.695365173 | 0.27396724 |
| Paqr9 | 964.539708 | -0.398726171 | 0.098875108 |
| Gm16126 | 0.766553141 | -0.123538137 | 1.408830062 |
| Gm28164 | 0.74026972 | -1.110856081 | 1.335950893 |
| Gm25582 | 0.857373094 | -1.629559974 | 1.109816963 |
| Gm16127 | 0.660063185 | 0.213357835 | 1.450837339 |
| Pcolce2 | 60.65372678 | -0.482738962 | 0.170466232 |
| Itpa-ps2 | 0.066050875 | 0.71820487 | 3.724945049 |
| Trpc1 | 585.9086551 | -0.286313839 | 0.065174338 |
| Pls1 | 183.634994 | -1.008415329 | 0.121077674 |
| 1700065D16Rik | 3.065206492 | -0.065953507 | 0.560587797 |
| Atr | 564.2015883 | -0.147583291 | 0.097102918 |
| Gm24004 | 0 | NA | NA |
| Gm16794 | 29.63475761 | 0.01597493 | 0.201626061 |
| Xrn1 | 1619.277144 | 0.41181944 | 0.083446263 |
| C78334 | 0.042716945 | -0.243571806 | 3.724945049 |
| Gm28085 | 0 | NA | NA |
| Gm38375 | 61.21253001 | 0.268252241 | 0.263158414 |

| | | | |
|---|---|---|---|
| Gk5 | 59.46593611 | -0.010335378 | 0.154639671 |
| Tfdp2 | 637.9968549 | -0.625923857 | 0.086183278 |
| Gm37195 | 34.22903916 | -1.250903076 | 0.311048464 |
| Gm28924 | 0.049553047 | 0.71820487 | 3.724945049 |
| Atp1b3 | 1332.613662 | 0.481463582 | 0.080261777 |
| BC043934 | 5.01349092 | -0.049161556 | 0.504860767 |
| Rnf7 | 496.7963806 | 0.200185292 | 0.096344992 |
| Gm27289 | 0 | NA | NA |
| Gm28729 | 2.066931119 | 0.739615437 | 0.687361251 |
| Gm8495 | 0.930891567 | -2.005960755 | 1.00314801 |
| Rasa2 | 837.5261912 | 0.111688152 | 0.08376713 |
| n-R5s87 | 2.429268922 | -0.504691121 | 0.632891528 |
| Gm19280 | 0 | NA | NA |
| Zbtb38 | 1623.281997 | 0.046269353 | 0.039932693 |
| A930006L05Rik | 0.095435502 | 0.718202646 | 3.724945049 |
| C430002N11Rik | 6.432897433 | 5.743197276 | 0.95917345 |
| Gm28688 | 0.279349467 | 2.385553838 | 1.983067444 |
| Gm8520 | 7.409839226 | -0.092604291 | 0.44027442 |
| Pxylp1 | 198.9385142 | -0.376070242 | 0.120903249 |
| Gm10123 | 609.9706294 | -0.072565952 | 0.094680624 |
| Gm8524 | 2.21064614 | 0.295433538 | 0.65425826 |
| Spsb4 | 74.46297335 | 0.047227994 | 0.129696403 |
| Gm16010 | 1.553703116 | -0.158410186 | 0.682257529 |
| Gm18291 | 0.371128506 | 2.154591324 | 1.944151191 |
| Gm19325 | 18.24486811 | -0.522572451 | 0.330892908 |
| Gm26767 | 0.08169834 | 0.23732098 | 3.724945049 |
| Slc25a36 | 1229.888798 | -0.185984823 | 0.060967738 |
| Gm16185 | 0.038981395 | 0.718207094 | 3.724945049 |
| Rpl7a-ps10 | 2.493931419 | 1.466180492 | 0.828933326 |
| Gm5371 | 0 | NA | NA |
| Gm29241 | 0 | NA | NA |
| Gm29240 | 0.044544648 | 0.237309859 | 3.724945049 |
| A030012G06Rik | 0.136880458 | -0.250074019 | 3.635317246 |
| Trim42 | 0.23239932 | 1.222140377 | 2.131267972 |
| Gm15891 | 0.046345709 | 0.71820487 | 3.724945049 |
| Gm25607 | 0.495675163 | 1.093725313 | 1.642217822 |
| Clstn2 | 1155.089609 | -0.251156758 | 0.085480499 |
| Gm37661 | 32.39089962 | -0.659412013 | 0.286961914 |
| 4921534H16Rik | 0.120974994 | -0.243571806 | 3.724945049 |
| 2610303G11Rik | 0 | NA | NA |
| Nmnat3 | 20.17326344 | 0.423024576 | 0.219741133 |
| Gm28530 | 3.500655289 | 1.555229806 | 0.574928526 |
| Rbp1 | 140.0093858 | 1.84312357 | 0.185858156 |
| Rbp2 | 0.656451805 | 1.122301022 | 1.181606626 |
| Gm37113 | 0 | NA | NA |
| 4930579K19Rik | 5.920659574 | -0.410518486 | 0.479445211 |
| Copb2 | 1299.911645 | 0.619591525 | 0.077068005 |

| | | | |
|---|---|---|---|
| Mrps22 | 205.9148466 | 0.424920759 | 0.088427849 |
| Gm37637 | 13.96030438 | -0.078085265 | 0.272280036 |
| Gm27919 | 0 | NA | NA |
| Pisrt1 | 0 | NA | NA |
| E330023G01Rik | 0.110595523 | 0.718200421 | 3.724945049 |
| Gm37282 | 0 | NA | NA |
| Gm37917 | 0.58604645 | -0.052324603 | 1.433769193 |
| Gm38226 | 0.474407165 | -0.647309142 | 1.346372868 |
| Prr23a | 0.228844266 | -0.24356322 | 2.137880943 |
| Gm6406 | 0 | NA | NA |
| 2410012M07Rik | 0 | NA | NA |
| Gm5920 | 2.082893185 | 0.706445469 | 0.688566051 |
| 7420426K07Rik | 0 | NA | NA |
| Gm28266 | 0 | NA | NA |
| Foxl2os | 2.17543179 | 0.684140849 | 0.730594057 |
| Foxl2 | 2.235327434 | -0.031341705 | 0.630553578 |
| Faim | 196.4395141 | -0.083594581 | 0.106498088 |
| Gm1123 | 1.879715592 | 0.791871752 | 0.835227108 |
| Pik3cb | 544.1368007 | 0.121515402 | 0.065529141 |
| Gm2950 | 0.061920613 | 0.718207094 | 3.724945049 |
| Gm6432 | 0.11197992 | 0.718202646 | 3.724945049 |
| Gm38327 | 0.266212691 | -0.695556941 | 3.704249029 |
| Cep70 | 260.9692196 | -0.682785164 | 0.158984245 |
| 4930422M22Rik | 0.158279039 | 1.199992907 | 2.813322116 |
| Gm37249 | 36.58656883 | -1.351861868 | 0.310484004 |
| Esyt3 | 26.59934933 | -0.432681534 | 0.230633205 |
| 1700008A23Rik | 0.963690934 | -0.456563706 | 1.08393411 |
| Gm28087 | 0 | NA | NA |
| Mras | 2060.728902 | 0.11079281 | 0.063807661 |
| Nme9 | 2.199217757 | -0.385247796 | 0.681304712 |
| Armc8 | 1137.205985 | -0.048400903 | 0.079136237 |
| Dbr1 | 168.1373829 | 0.597756867 | 0.141421206 |
| A4gnt | 0.979608328 | -1.089784172 | 1.018877048 |
| Dzip1l | 286.0060264 | -0.744362513 | 0.154930273 |
| Cldn18 | 0.262323822 | 0.237303174 | 3.724945049 |
| Gm16004 | 0 | NA | NA |
| Gm23949 | 0 | NA | NA |
| Sox14 | 8.749592595 | -0.483748625 | 0.375950642 |
| Gm23594 | 0 | NA | NA |
| Gm8641 | 0.504639468 | 1.042155649 | 1.347477561 |
| 4930519F24Rik | 0.061920613 | 0.718207094 | 3.724945049 |
| 4930540E01Rik | 0 | NA | NA |
| Gm28167 | 0 | NA | NA |
| Gm28267 | 0 | NA | NA |
| Gm28166 | 0.08486385 | 0.717551239 | 3.724898127 |
| Il20rb | 65.86335234 | 0.294363996 | 0.26371586 |
| Nck1 | 358.9705822 | 0.114603321 | 0.078250644 |

| | | | |
|---|---|---|---|
| Gm28586 | 8.399474289 | -0.64609189 | 0.49703734 |
| Slc35g2 | 192.5839218 | 0.010360105 | 0.152736206 |
| Stag1 | 1284.878169 | -0.015712095 | 0.069194812 |
| Gm8661 | 0.119234754 | -0.243576254 | 3.724945049 |
| Gm37204 | 117.4552302 | -0.962252854 | 0.269469609 |
| Gm38297 | 15.65520555 | -1.167301996 | 0.406542449 |
| Gm3081 | 62.39629019 | -0.373429779 | 0.238387545 |
| Pccb | 410.7052558 | -0.501064821 | 0.075936666 |
| Gm3096 | 0 | NA | NA |
| Msl2 | 1119.25172 | 0.024663376 | 0.067262694 |
| Gm37553 | 42.26828888 | -0.499279524 | 0.256904253 |
| Ppp2r3a | 1068.136217 | 0.133311392 | 0.070374792 |
| Gm37962 | 38.30761735 | -0.540897789 | 0.268985175 |
| Gm37621 | 78.87355947 | -0.462732413 | 0.221742732 |
| Gm37953 | 20.22364659 | -0.451610165 | 0.291845361 |
| Gm38344 | 18.31202231 | -0.628581699 | 0.332813326 |
| Gm37314 | 38.77458494 | -0.551287679 | 0.243363573 |
| Gm24338 | 0.290020514 | 0.233418599 | 1.636755953 |
| Gm3121 | 0 | NA | NA |
| Gm28660 | 0 | NA | NA |
| Gm28979 | 0.404019895 | 1.760146503 | 1.466077277 |
| Gm24638 | 0 | NA | NA |
| Gm29385 | 0 | NA | NA |
| Gm5161 | 0.106056992 | 0.237307635 | 3.724945049 |
| Gm29521 | 6.846922448 | -0.791121213 | 0.463183044 |
| Gm37460 | 0.652148592 | 1.029363338 | 1.366708072 |
| Ephb1 | 568.496847 | -0.717272761 | 0.072715748 |
| 9630041A04Rik | 1.37032714 | 0.789892729 | 0.943541614 |
| Gm37683 | 0.138485956 | -0.243563634 | 2.932778697 |
| Gm37562 | 15.3341162 | -1.023564334 | 0.3805246 |
| n-R5s88 | 0 | NA | NA |
| Gm29388 | 6.105350049 | -1.716994594 | 0.599681931 |
| Gm37945 | 0.049553047 | 0.71820487 | 3.724945049 |
| Gm37260 | 0.11197992 | 0.718202646 | 3.724945049 |
| Mir7243 | 1.350235793 | -1.652355535 | 0.845955114 |
| Gm26846 | 1.767979395 | -1.362248396 | 0.886157324 |
| Gm22894 | 0.570356529 | 0.404165387 | 1.240835063 |
| Ky | 45.80996683 | -0.999707903 | 0.22987357 |
| Gm25669 | 0.261093266 | 0.718194452 | 3.724945049 |
| Gm7364 | 0.312630927 | 0.237297968 | 3.724945049 |
| Gm29143 | 0 | NA | NA |
| Gm29387 | 0.088534442 | 1.179809211 | 3.723500796 |
| Cep63 | 192.8660548 | -0.309542862 | 0.098488084 |
| Anapc13 | 138.6441374 | -0.114613931 | 0.155358511 |
| 4930533D04Rik | 0.164893722 | 0.23730812 | 3.395459203 |
| Gm36957 | 0.137470077 | 0.237309658 | 3.595099416 |
| Amotl2 | 701.5681414 | 1.277943966 | 0.181922772 |

| | | | |
|---|---|---|---|
| Gm5627 | 0.038981395 | 0.718207094 | 3.724945049 |
| Gm36958 | 0.813964436 | -1.17349543 | 1.15706956 |
| Gm28621 | 0 | NA | NA |
| Ryk | 428.6544998 | 0.035131672 | 0.073589087 |
| Slco2a1 | 62.94329372 | 0.983691326 | 0.177863208 |
| Gm25842 | 0.189042777 | 0.237326645 | 2.65947082 |
| n-R5s89 | 0 | NA | NA |
| Gm22595 | 0 | NA | NA |
| Rab6b | 7282.57726 | -0.718338335 | 0.087173807 |
| Gm37166 | 45.91226253 | -1.795198614 | 0.403088571 |
| Srprb | 192.082496 | 0.349859897 | 0.101430674 |
| Gm20425 | 0 | NA | NA |
| Trf | 281.060478 | 1.856107865 | 0.208855883 |
| 1300017J02Rik | 2.855215039 | 0.721688718 | 0.59541377 |
| Topbp1 | 318.7285085 | 0.100602331 | 0.092088594 |
| Cdv3 | 1213.725262 | 0.777425938 | 0.094528776 |
| Gm29600 | 0.038981395 | 0.718207094 | 3.724945049 |
| 5830418P13Rik | 5.81979985 | -0.857394286 | 0.459001145 |
| Bfsp2 | 26.48732468 | 0.093523472 | 0.252656488 |
| Gm37938 | 2.10811511 | -0.306922271 | 0.943374477 |
| Gm37546 | 0.280931382 | -0.218399353 | 3.714291231 |
| Gm16252 | 0.115957212 | -0.24357403 | 3.724945049 |
| Tmem108 | 366.5574461 | -0.35640293 | 0.068567222 |
| Gm26360 | 2.674678528 | -0.219962388 | 0.664480067 |
| Gm37649 | 24.6588879 | -1.502269801 | 0.417804083 |
| 4932413F04Rik | 0.450128165 | -0.111723949 | 1.398864599 |
| Gm38032 | 30.93777638 | -1.454834962 | 0.323436991 |
| Gm5372 | 0 | NA | NA |
| Gm29154 | 0.341479653 | -1.170692686 | 3.722689234 |
| Nphp3 | 143.1369062 | -0.241915466 | 0.140975055 |
| Gm28305 | 0 | NA | NA |
| Gm37200 | 6.440447106 | -0.17317183 | 0.40023298 |
| Uba5 | 589.0417371 | 0.12591258 | 0.064701411 |
| Gm37750 | 0.049553047 | 0.71820487 | 3.724945049 |
| Acad11 | 231.7177883 | -0.343897084 | 0.120820853 |
| Ackr4 | 2.634801621 | 0.096821735 | 0.607647483 |
| Dnajc13 | 1701.736079 | 0.403178814 | 0.068977218 |
| Hmgb1-rs16 | 11.38856345 | -0.504707559 | 0.45056557 |
| Gm37563 | 1.931745907 | 3.21496872 | 0.989832256 |
| Acpp | 6.437954073 | 0.594296974 | 0.38810067 |
| Gm37188 | 0.048949532 | -0.243569582 | 3.724945049 |
| Gm28996 | 0 | NA | NA |
| Gm28548 | 0 | NA | NA |
| Mir2136 | 0 | NA | NA |
| Gm25904 | 0 | NA | NA |
| Cpne4 | 536.3195774 | -1.060698646 | 0.156707836 |
| Gm23391 | 1.103801871 | -2.919119666 | 1.081995058 |

| | | | |
|---|---|---|---|
| Mrpl3 | 481.1686626 | -0.021356576 | 0.059334989 |
| Gm38077 | 99.57854452 | -0.847496541 | 0.311951637 |
| Nudt16 | 104.3868709 | -0.172934878 | 0.125526468 |
| 1700080E11Rik | 0.3422147 | 2.308926681 | 1.89526122 |
| Nek11 | 22.44919175 | 0.513059544 | 0.270231298 |
| Gm37582 | 0.092140247 | -0.724455696 | 3.724945049 |
| Gm20643 | 0 | NA | NA |
| Gm15619 | 0 | NA | NA |
| Aste1 | 73.3563129 | 0.304754868 | 0.138957023 |
| Atp2c1 | 2093.602576 | -0.228277779 | 0.058808966 |
| Gm19667 | 0.638749378 | 1.199122655 | 1.648874669 |
| Gm7436 | 0.368385343 | 0.237621136 | 1.799701206 |
| Gm5621 | 1.423724791 | 0.060529003 | 0.88097493 |
| Gm28555 | 0 | NA | NA |
| Pik3r4 | 491.4021009 | -0.297339962 | 0.073352338 |
| Gm22720 | 0 | NA | NA |
| Col6a6 | 14.261268 | 1.048440666 | 0.323822434 |
| Gm38314 | 0 | NA | NA |
| Col6a5 | 28.81713568 | -0.41464334 | 0.268651006 |
| Col6a4 | 6.820926848 | 0.268995524 | 0.364567466 |
| Gm24631 | 0 | NA | NA |
| Gm36993 | 0 | NA | NA |
| Gm9451 | 0.136331796 | -1.191826756 | 3.64305592 |
| Gm29208 | 0 | NA | NA |
| D030055H07Rik | 7.771284628 | -3.439232975 | 0.555222762 |
| Glyctk | 39.68295307 | -0.185754509 | 0.220054425 |
| Mir135a-1 | 0.193036943 | -0.187869281 | 3.246599635 |
| Wdr82 | 1508.867638 | -0.080519224 | 0.057870614 |
| Mirlet7g | 2.956734474 | -0.185053039 | 0.740949017 |
| Ppm1m | 99.17848393 | 0.269099628 | 0.198956811 |
| Twf2 | 271.214745 | -0.145363107 | 0.099806573 |
| Tlr9 | 362.3896947 | 6.399857271 | 0.380817844 |
| Alas1 | 600.2556519 | 0.209764027 | 0.064215241 |
| 4930500F10Rik | 7.519553513 | -0.835234762 | 0.417838317 |
| Poc1a | 35.43149234 | -0.176793932 | 0.224880969 |
| Gm29123 | 0.049553047 | 0.71820487 | 3.724945049 |
| Dusp7 | 640.9294056 | -0.106420327 | 0.084350269 |
| Rpl29 | 8.095074919 | 0.187440774 | 0.384291178 |
| Acy1 | 102.5946641 | -0.289888205 | 0.139262557 |
| Abhd14a | 66.26385201 | -0.50105678 | 0.16563246 |
| Abhd14b | 60.06866735 | -1.048601627 | 0.221363135 |
| Gm28959 | 0.650039631 | -1.115073841 | 1.178203855 |
| Pcbp4 | 1192.577122 | -0.325637367 | 0.064159441 |
| Gpr62 | 124.4874142 | 0.466371329 | 0.145763987 |
| Parp3 | 396.144577 | 4.063497636 | 0.151204247 |
| Rrp9 | 173.3168163 | 0.715500465 | 0.096276582 |
| Iqcf1 | 0.403414764 | 1.286887589 | 1.480419134 |

| | | | |
|---|---|---|---|
| Iqcf3 | 0.731923588 | -2.197475057 | 1.182684729 |
| Iqcf5 | 0.458372996 | -2.188300168 | 1.612005388 |
| Iqcf4 | 0 | NA | NA |
| Iqcf6 | 0 | NA | NA |
| Gm26185 | 0 | NA | NA |
| Gm28111 | 0 | NA | NA |
| Grm2 | 419.4914956 | -0.944942602 | 0.133108328 |
| Tex264 | 547.2087676 | 0.334389562 | 0.079619625 |
| Rad54l2 | 692.1787233 | 0.095729415 | 0.094818448 |
| Gm22384 | 0.206359549 | -0.724462198 | 2.64344885 |
| Vprbp | 629.97184 | 0.053381929 | 0.08513239 |
| Manf | 616.8659586 | 0.819720372 | 0.108528409 |
| Gm37612 | 1.882200288 | -1.061317879 | 0.769043122 |
| Rbm15b | 823.0495333 | -0.096095727 | 0.068086153 |
| 4930524O07Rik | 4.402341252 | 0.119518654 | 0.605645144 |
| Dock3 | 4718.325882 | -0.787182604 | 0.1127685 |
| Gm17205 | 0.109124069 | -0.24357403 | 3.724945049 |
| Gm22454 | 0 | NA | NA |
| Gm17117 | 0 | NA | NA |
| Gm23595 | 0 | NA | NA |
| AA543401 | 0.072025462 | 0.237312084 | 3.724945049 |
| Gm22238 | 0 | NA | NA |
| 1700039M15Rik | 0.049553047 | 0.71820487 | 3.724945049 |
| Mapkapk3 | 63.6118118 | 1.974411552 | 0.219741069 |
| Gm17040 | 0.094097695 | 0.718200421 | 3.724945049 |
| Cish | 89.05641637 | 0.570936485 | 0.171592781 |
| Gm17041 | 0.049553047 | 0.71820487 | 3.724945049 |
| Hemk1 | 115.6953326 | -0.007243914 | 0.113054747 |
| 6430571L13Rik | 122.123982 | -0.307858423 | 0.1675927 |
| 4930429P21Rik | 0.791934515 | -1.826512869 | 1.231905729 |
| Cacna2d2 | 1015.043882 | -0.19041594 | 0.073349715 |
| Gm17141 | 0.093831153 | -0.243571806 | 3.724945049 |
| Gm17118 | 0 | NA | NA |
| Tmem115 | 377.5379067 | 0.53611039 | 0.094523653 |
| Cyb561d2 | 55.18192089 | 0.41661613 | 0.203312939 |
| Nprl2 | 184.5566623 | -0.278569911 | 0.086052469 |
| Zmynd10 | 22.83363798 | -0.643964026 | 0.26587869 |
| Gm34106 | 0.182655267 | 0.7182151 | 2.701208497 |
| Rassf1 | 176.3853711 | 1.120269606 | 0.133037723 |
| Gm9917 | 9.204737304 | -0.855408132 | 0.48122462 |
| Tusc2 | 617.3101563 | -0.165950429 | 0.07122924 |
| Hyal2 | 95.99414798 | 0.890030195 | 0.125909542 |
| Gm16343 | 0.186399948 | 0.718204068 | 3.299750385 |
| Hyal1 | 60.01938957 | 1.000440185 | 0.191266095 |
| Nat6 | 87.56719576 | 0.780570519 | 0.159587822 |
| Hyal3 | 5.312334351 | 0.646922187 | 0.535824234 |
| Ifrd2 | 67.19084161 | -0.051756357 | 0.215584156 |

| | | | |
|---|---|---|---|
| Lsmem2 | 2.036564983 | 1.704128944 | 0.850124944 |
| Gm38150 | 3.547776317 | -1.549112396 | 0.585989206 |
| Sema3b | 58.08865837 | 0.145160138 | 0.196429852 |
| Gnai2 | 2404.56764 | 1.021022486 | 0.114734667 |
| Gm19721 | 19.767715 | -0.049724015 | 0.410778417 |
| Slc38a3 | 598.0789621 | -0.427841464 | 0.180594891 |
| Gnat1 | 1.440190137 | -0.940141565 | 0.90849786 |
| A930036K24Rik | 0.18066559 | 1.166193762 | 2.687923556 |
| Sema3f | 139.5279565 | -0.422980551 | 0.110507968 |
| Rbm5 | 1611.009065 | -0.440271105 | 0.158464776 |
| Gm37850 | 0 | NA | NA |
| Rbm6 | 1030.666851 | -0.377578807 | 0.1235148 |
| Gm37436 | 36.96383849 | -0.768723563 | 0.252407204 |
| Mon1a | 192.9155728 | 0.127800764 | 0.097715793 |
| Gm23856 | 0.487068913 | -0.462680672 | 1.513361617 |
| Mst1r | 16.57438961 | 0.3267328 | 0.256850611 |
| Actl11 | 0.123841226 | 1.292225235 | 3.719439735 |
| Gm37974 | 0.217247303 | -0.724460911 | 2.596047673 |
| Camkv | 2202.703855 | -1.381758612 | 0.156680296 |
| Traip | 114.0875043 | -0.754269414 | 0.246788314 |
| Uba7 | 688.1155965 | 6.553820578 | 0.197506673 |
| Gm20661 | 4.55440648 | 1.664497712 | 0.571496517 |
| Fam212a | 16.26337642 | 0.508720738 | 0.254835715 |
| Gm24944 | 1.177026929 | 2.926488895 | 1.146037451 |
| 4921523L03Rik | 0.198550338 | -0.243567932 | 2.214317373 |
| Cdhr4 | 13.4482997 | 0.947301968 | 0.365791795 |
| Gm20662 | 0.066050875 | 0.71820487 | 3.724945049 |
| 4930535L15Rik | 1.219811481 | 0.19336271 | 0.853283218 |
| Ip6k1 | 2129.145135 | 0.081871083 | 0.053133699 |
| Gm38134 | 0.127971488 | 1.23029335 | 3.685533863 |
| Gmppb | 158.7886363 | 1.165213019 | 0.146803941 |
| Rnf123 | 811.9292402 | -0.535049242 | 0.071848992 |
| Gm20529 | 0.04393175 | -0.243571806 | 3.724945049 |
| Amigo3 | 37.0261778 | 1.812648891 | 0.303610108 |
| Mst1 | 0.439905485 | 1.80674003 | 1.380665547 |
| Mir7088 | 0 | NA | NA |
| Apeh | 282.2906194 | 0.196823377 | 0.082040795 |
| Bsn | 11059.17198 | -0.778990316 | 0.121449793 |
| AC137678.1 | 2.386774017 | -0.878633391 | 0.9335399 |
| 4930447F24Rik | 14.97539824 | 2.54141974 | 0.39534495 |
| Gm37201 | 2.168403122 | 0.362332873 | 0.865852671 |
| Gm37247 | 41.41364108 | -1.183296758 | 0.362039564 |
| Gm37230 | 0.567387455 | -0.892427541 | 1.293051597 |
| Dag1 | 1797.643256 | 0.304427175 | 0.082628267 |
| Nicn1 | 612.9850749 | -0.506376712 | 0.072450063 |
| Amt | 190.2692267 | -0.510959026 | 0.138193582 |
| Tcta | 207.060682 | 0.389859855 | 0.109471582 |

| | | | |
|---|---|---|---|
| Rhoa | 1859.627478 | 0.669735796 | 0.1066189 |
| Gm37568 | 3.725366168 | -0.236290346 | 0.545416316 |
| Gm37401 | 0.049553047 | 0.71820487 | 3.724945049 |
| Gpx1 | 610.2857368 | 1.286664577 | 0.145461516 |
| Usp4 | 1105.767034 | 0.182047154 | 0.05336991 |
| Rps2-ps11 | 0.578661668 | 1.540819215 | 1.243998929 |
| Gm37963 | 32.08673899 | -0.879428374 | 0.341067553 |
| 1700102P08Rik | 15.20131833 | -0.152129454 | 0.267814075 |
| Ccdc36 | 5.280767892 | -0.566115073 | 0.463923719 |
| Gm25456 | 0.042716945 | -0.243571806 | 3.724945049 |
| BC048562 | 0 | NA | NA |
| Klhdc8b | 159.0275307 | -0.842150385 | 0.125797386 |
| Ccdc71 | 275.4617965 | 0.195236122 | 0.075337028 |
| Gm37475 | 3.601477103 | -0.227168664 | 0.539965954 |
| Lamb2 | 362.809454 | 0.179564981 | 0.148389872 |
| Usp19 | 1729.388652 | 0.358794318 | 0.050060998 |
| Qars | 489.2628777 | 0.351753892 | 0.063074936 |
| Qrich1 | 1514.838001 | 0.092629439 | 0.069203966 |
| Gm38366 | 45.47164434 | -0.634149475 | 0.34017042 |
| Gm37678 | 6.993644233 | -1.036389803 | 0.476898767 |
| Impdh2 | 294.2527551 | 0.393585743 | 0.097258621 |
| Ndufaf3 | 182.2571026 | -0.090951481 | 0.08132568 |
| 4833445I07Rik | 30.38428203 | -0.659971366 | 0.208633186 |
| Gm24083 | 0 | NA | NA |
| mmu-mir-191 | 0 | NA | NA |
| Mir425 | 0 | NA | NA |
| Dalrd3 | 336.6293249 | -0.075640191 | 0.077020583 |
| Wdr6 | 1862.952255 | -0.56472977 | 0.09014274 |
| P4htm | 622.680632 | -0.178518864 | 0.078534598 |
| Gm38163 | 3.058745156 | -1.354811469 | 0.640908797 |
| Arih2 | 618.9684878 | 0.376841723 | 0.082449062 |
| Gm26298 | 0 | NA | NA |
| Slc25a20 | 138.8267023 | 0.865514723 | 0.154110338 |
| Prkar2a | 906.3458615 | 0.087354808 | 0.068008064 |
| Gm24259 | 0.258327161 | 1.693545259 | 3.021726448 |
| Ip6k2 | 843.8946416 | 0.025221794 | 0.086469348 |
| Gm23034 | 0.062426873 | 0.237312084 | 3.724945049 |
| Nckipsd | 578.0213764 | -0.231899944 | 0.080472766 |
| Gm35025 | 6.065554424 | -0.502908446 | 0.459663374 |
| Gm37714 | 0.267221877 | 1.096605092 | 1.778545358 |
| Celsr3 | 1375.868327 | -0.690142134 | 0.162012186 |
| Gm23156 | 0.616331352 | -1.149267746 | 1.660667587 |
| Slc26a6 | 70.00938289 | -0.223691154 | 0.159623916 |
| Tmem89 | 0.562660809 | -0.701496929 | 1.377235106 |
| Uqcrc1 | 1892.13114 | 0.021233636 | 0.067336964 |
| Gm37284 | 0 | NA | NA |
| Col7a1 | 41.89192254 | 0.362834862 | 0.255466085 |

| | | | |
|---|---|---|---|
| Mir711 | 0 | NA | NA |
| Ucn2 | 10.62182747 | 6.825987481 | 1.132016559 |
| Gm38004 | 0.533662682 | 1.141843433 | 1.329710995 |
| Pfkfb4 | 187.9455917 | 0.498459674 | 0.10650953 |
| Shisa5 | 1519.355689 | 3.00426073 | 0.147409225 |
| Trex1 | 520.4087808 | 3.166743752 | 0.194970908 |
| Atrip | 28.91987549 | -0.492005507 | 0.394065034 |
| Tma7 | 371.0569712 | 0.187589468 | 0.089443241 |
| Ccdc51 | 52.13264183 | 0.166142961 | 0.263307361 |
| Gm7628 | 6.615983682 | -0.116072131 | 0.463376629 |
| Plxnb1 | 1318.803774 | -0.801971765 | 0.124340791 |
| Fbxw21 | 0 | NA | NA |
| Fbxw13 | 0.127364494 | -0.243565133 | 3.724945049 |
| Fbxw20 | 0.066050875 | 0.71820487 | 3.724945049 |
| Fbxw14 | 0.088383099 | -0.72445792 | 3.724945049 |
| Gm22821 | 0 | NA | NA |
| Gm25484 | 0 | NA | NA |
| Gm24641 | 0 | NA | NA |
| Fbxw22 | 0.038981395 | 0.718207094 | 3.724945049 |
| Fbxw16 | 0.066050875 | 0.71820487 | 3.724945049 |
| RP23-62L19.6 | 0 | NA | NA |
| RP23-62L19.5 | 0 | NA | NA |
| Fbxw19 | 0.038981395 | 0.718207094 | 3.724945049 |
| RP24-187P11.4 | 0.450885768 | 1.372171841 | 1.619779443 |
| RP24-187P11.2 | 9.209079983 | 2.839151062 | 0.553496141 |
| RP24-187P11.5 | 0.238621995 | 1.611275356 | 2.496608988 |
| Fbxw15 | 0.204053819 | -0.724459671 | 2.62442978 |
| RP24-187P11.3 | 0.046345709 | 0.71820487 | 3.724945049 |
| Fbxw24 | 0.124596252 | 0.718200615 | 3.714681068 |
| Fbxw25 | 0.109982625 | 0.237318756 | 3.724945049 |
| Fbxw18 | 0.15906016 | 1.290286593 | 3.433492162 |
| RP24-344B20.5 | 0 | NA | NA |
| Fbxw26 | 0.166952883 | 1.610951781 | 2.74653892 |
| RP23-357H5.5 | 0.394548002 | -0.223572796 | 1.757656987 |
| Spink8 | 16.32657011 | 0.002226613 | 0.310416347 |
| 3000002C10Rik | 56.32280498 | -0.549496453 | 0.184286904 |
| Nme6 | 38.63028386 | 0.617262102 | 0.204944181 |
| Camp | 2.243591458 | 4.66810345 | 1.020991057 |
| Cdc25a | 165.9711309 | 0.380491992 | 0.108577246 |
| Mir7089 | 0.220081591 | 0.71822207 | 2.16660067 |
| RP24-421E18.9 | 0 | NA | NA |
| Map4 | 7553.036783 | 0.240090948 | 0.080654501 |
| RP24-421E18.8 | 10.03429169 | -0.353241895 | 0.328302827 |
| RP24-421E18.10 | 0 | NA | NA |
| RP24-421E18.5 | 78.64983153 | -0.503517356 | 0.127155341 |
| AC161246.1 | 0 | NA | NA |
| RP24-421E18.7 | 15.20321581 | 0.531437565 | 0.353163907 |

| | | | |
|---|---|---|---|
| Dhx30 | 1202.629578 | -0.048620745 | 0.050500391 |
| Gm27855 | 0.367828087 | -1.619082867 | 1.530278457 |
| Smarcc1 | 1075.459101 | 0.058619328 | 0.050337934 |
| Gm23377 | 0 | NA | NA |
| Cspg5 | 2416.656234 | -0.278545778 | 0.078580033 |
| RP24-285I4.2 | 0.569412722 | -0.801914389 | 1.205015132 |
| Mir6236 | 2.610552123 | 0.453736516 | 1.071646699 |
| RP24-330A11.9 | 0.916112882 | -2.301154619 | 1.143164979 |
| Elp6 | 89.70397602 | 0.469328058 | 0.119765917 |
| RP24-330A11.12 | 0.042716945 | -0.243571806 | 3.724945049 |
| Scap | 1121.694434 | -0.096694587 | 0.048288881 |
| 5830462I19Rik | 0.695310992 | 2.30499581 | 1.421356974 |
| Ptpn23 | 992.3070621 | 0.472752904 | 0.092447028 |
| Ngp | 7.348605377 | 4.167568684 | 0.906672333 |
| Klhl18 | 377.384243 | -0.093607427 | 0.076023748 |
| Kif9 | 65.23232709 | 0.38192104 | 0.155288214 |
| Setd2 | 2113.532458 | -0.069145509 | 0.070779659 |
| RP24-363K20.7 | 41.47908056 | -0.324906567 | 0.231627451 |
| Nradd | 21.00591379 | 1.799409443 | 0.316792172 |
| Nbeal2 | 229.3785941 | 0.530519207 | 0.149149653 |
| Mir8107 | 0 | NA | NA |
| RP24-179L19.4 | 0.417968807 | -0.243563898 | 1.540943312 |
| Ccdc12 | 189.7858103 | 0.378222113 | 0.120754672 |
| RP24-179L19.2 | 0.211345377 | -0.325108818 | 2.637775574 |
| Pth1r | 56.35433901 | 0.058269122 | 0.155938401 |
| Myl3 | 1.171543132 | -0.040878289 | 1.095321773 |
| Prss42 | 0.145501641 | 1.198496036 | 2.88801119 |
| Prss44 | 0.225957929 | 1.183168856 | 2.139540998 |
| Prss43 | 1.895398054 | 1.009619146 | 0.89028715 |
| Prss45 | 3.856439033 | 1.653333995 | 0.704284086 |
| Prss46 | 14.47717373 | 7.917910165 | 0.941606288 |
| Prss50 | 6.552872895 | 0.968286738 | 0.445487053 |
| Tmie | 70.89016146 | 0.254979109 | 0.17206835 |
| Als2cl | 108.0650459 | -0.503987377 | 0.142553109 |
| RP23-265F20.14 | 0.087930927 | 0.236669296 | 3.724898107 |
| Gm590 | 0.446134451 | 2.798756339 | 1.512506319 |
| Gm24269 | 0 | NA | NA |
| Tdgf1 | 2.307514072 | -0.488330216 | 0.712454293 |
| Lrrc2 | 11.32380672 | -0.469576095 | 0.369100085 |
| RP24-166N8.8 | 0.109982625 | 0.237318756 | 3.724945049 |
| Rtp3 | 1.735450405 | 3.779991291 | 0.984962793 |
| 1700061E17Rik | 0.467110691 | -0.501794966 | 1.639249822 |
| Gm10030 | 2.141449619 | 0.004976608 | 0.648271056 |
| Ltf | 6.616994115 | 3.515863369 | 0.658309489 |
| RP24-166N8.9 | 4.949110328 | 5.557309462 | 0.912364 |
| Ccrl2 | 163.4421883 | 6.430145552 | 0.456042632 |
| Rp31-ps19 | 0 | NA | NA |

| | | | |
|---|---|---|---|
| Gm26409 | 0.044544648 | 0.237309859 | 3.724945049 |
| Lrrfip2 | 487.0826255 | 0.135610114 | 0.091587192 |
| Mlh1 | 123.8432082 | -0.058894904 | 0.130867064 |
| Gm24860 | 1.095639389 | -0.102772401 | 0.995638506 |
| Epm2aip1 | 2946.703591 | -0.062613485 | 0.067039058 |
| Trank1 | 3806.887753 | -0.656504042 | 0.277307937 |
| RP23-361O11.4 | 10.70607882 | -1.183055301 | 0.408678513 |
| Dclk3 | 344.3535978 | -1.140389689 | 0.149594696 |
| RP24-362N9.3 | 0.122601409 | 0.237307635 | 3.724945049 |
| RP24-362N9.4 | 0 | NA | NA |
| Stac | 39.84967809 | -0.381743498 | 0.189770505 |
| n-R5s90 | 0.892725456 | 0.798367508 | 1.071368172 |
| Arpp21 | 4354.712088 | -0.992488174 | 0.150240582 |
| Mir128-2 | 2.542567827 | -1.008068644 | 0.684147016 |
| 2310075C17Rik | 2.414579911 | 1.19915956 | 0.724797623 |
| 2900079G21Rik | 54.14502989 | -0.040442071 | 0.167134902 |
| Gm27847 | 0 | NA | NA |
| Gm24957 | 0 | NA | NA |
| Gm22492 | 0 | NA | NA |
| Pdcd6ip | 1431.871971 | 0.428661446 | 0.060577626 |
| Gm25059 | 0 | NA | NA |
| Clasp2 | 4322.484509 | -0.644337032 | 0.054806473 |
| Ubp1 | 888.9733183 | -0.143653713 | 0.11626579 |
| Fbxl2 | 321.4966191 | -0.725888437 | 0.099102734 |
| Susd5 | 65.24506384 | -0.829090748 | 0.177136184 |
| 4921528I07Rik | 0.068097359 | -0.243569582 | 3.724945049 |
| Bcl2a1c | 0.061920613 | 0.718207094 | 3.724945049 |
| 4930520O04Rik | 10.94558613 | -0.526168215 | 0.335521567 |
| Crtap | 52.46044785 | 1.287186136 | 0.23592678 |
| Glb1 | 239.9047864 | 0.073529012 | 0.088793004 |
| Tmppe | 289.5667612 | 0.086240664 | 0.074291569 |
| Ccr4 | 0.577241561 | 1.770362296 | 1.487177242 |
| Trim71 | 15.97479315 | -0.641648349 | 0.385388215 |
| C130032M10Rik | 0.045882455 | 0.237312084 | 3.724945049 |
| Cnot10 | 446.9158187 | -0.112923421 | 0.072560768 |
| Gm23232 | 2.256171834 | -1.091095574 | 0.878037287 |
| Dync1li1 | 1024.902468 | 0.067767887 | 0.052973112 |
| Cmtm6 | 352.6464989 | 0.981072604 | 0.096135304 |
| Cmtm7 | 42.27447401 | 2.314609165 | 0.28237291 |
| Gm9888 | 1.350285312 | 1.350111462 | 0.987419827 |
| Cmtm8 | 7.699545734 | 0.614217155 | 0.422915346 |
| Gm23889 | 0 | NA | NA |
| Gpd1l | 938.6073185 | -0.74124978 | 0.079971993 |
| Osbpl10 | 236.991532 | -0.735987164 | 0.139768194 |
| Rps27rt | 78.90118146 | 0.773222156 | 0.225085633 |
| Gm4665 | 0.066050875 | 0.71820487 | 3.724945049 |
| Gm26962 | 0.061920613 | 0.718207094 | 3.724945049 |

| | | | |
|---|---|---|---|
| Gm27002 | 10.03766149 | -1.134857467 | 0.347253332 |
| Stt3b | 1311.019778 | 0.177841419 | 0.06524103 |
| Mir467h | 0 | NA | NA |
| Gm26303 | 0 | NA | NA |
| Gm25510 | 0 | NA | NA |
| Gm22243 | 0 | NA | NA |
| Gm5921 | 6.413797967 | 0.563213274 | 0.436232145 |
| n-R5s92 | 0 | NA | NA |
| Gadl1 | 3.709958051 | 0.099463226 | 0.488099227 |
| Gm16142 | 0.404993149 | 0.188202107 | 1.329095041 |
| Gm9385 | 43.14016071 | 0.709786895 | 0.249689182 |
| Tgfbr2 | 420.264414 | 1.980804263 | 0.110444962 |
| Rbms3 | 651.4525007 | 0.124739087 | 0.096677606 |
| Gm18489 | 0.042716945 | -0.243571806 | 3.724945049 |
| Gm20397 | 0 | NA | NA |
| Gm17396 | 26.24111455 | 0.629651982 | 0.249492183 |
| Gm20396 | 0 | NA | NA |
| Gm26614 | 3.2216295 | -0.106530476 | 0.652614288 |
| Zcwpw2 | 38.02155974 | -0.16165407 | 0.197615598 |
| Gm22008 | 0 | NA | NA |
| Azi2 | 1280.08819 | 0.712900413 | 0.096712039 |
| Cmc1 | 102.3559624 | -0.009913842 | 0.114663374 |
| Gm17399 | 4.930845136 | -0.675561712 | 0.498832717 |
| Gm22198 | 0 | NA | NA |
| 2610509F24Rik | 0.046345709 | 0.71820487 | 3.724945049 |
| Platr11 | 0.154746054 | -1.179258444 | 2.823763514 |
| Gm22479 | 0.124853745 | 0.237309606 | 3.724945049 |
| Eomes | 133.6882393 | -0.102580964 | 0.417526345 |
| Golga4 | 1496.655227 | 0.012263798 | 0.129606275 |
| Itga9 | 160.656506 | 0.015332567 | 0.100581324 |
| Gm10157 | 51.89187838 | 0.194098573 | 0.151032253 |
| Gm2415 | 124.2848123 | -1.363578945 | 0.159217437 |
| Gm16295 | 0.648502457 | -0.855159486 | 1.178265052 |
| Ctdspl | 341.7777662 | -0.736544432 | 0.134490215 |
| Mir26a-1 | 1.512289855 | 0.073052575 | 1.082011739 |
| Vill | 6.901825543 | 1.467442416 | 0.405918685 |
| Plcd1 | 118.7634359 | 0.459772055 | 0.130533275 |
| Dlec1 | 29.71330161 | -0.768496989 | 0.222759068 |
| Acaa1b | 17.68007261 | 0.284266093 | 0.256484951 |
| Gm10608 | 0.106465261 | 0.718202646 | 3.724945049 |
| Slc22a14 | 0.191792366 | 1.616133477 | 2.227787421 |
| Slc22a13 | 1.365498172 | -0.902830185 | 0.887613378 |
| Slc22a13b-ps | 14.19279457 | -1.0130594 | 0.352989325 |
| Oxsr1 | 850.5060215 | 0.030306338 | 0.06863813 |
| Gm22729 | 1.308834292 | -0.528177743 | 0.802819255 |
| Myd88 | 293.492393 | 4.076461837 | 0.232022522 |
| Acaa1a | 291.2994478 | 0.450243811 | 0.103086016 |

| | | | |
|---|---|---|---|
| Xylb | 75.68413251 | 0.078626992 | 0.149225224 |
| Acvr2b | 67.3176329 | -0.763639179 | 0.139945923 |
| 4930516B21Rik | 0 | NA | NA |
| Exog | 250.3105798 | 0.129192611 | 0.086677133 |
| Scn5a | 99.30242439 | -0.567760279 | 0.134298339 |
| Scn10a | 7.212801384 | -0.875426721 | 0.69064824 |
| Scn11a | 3.528093251 | 1.194885905 | 0.53256532 |
| Gm2449 | 4.853795414 | 0.0262935 | 0.479016323 |
| Wdr48 | 797.9104278 | -0.055334352 | 0.049849678 |
| Cx3cr1 | 232.9636481 | -2.374380087 | 0.485596373 |
| Gorasp1 | 319.6341586 | 0.091245876 | 0.130982166 |
| Ttc21a | 29.13279244 | -0.432475148 | 0.277323245 |
| Csrnp1 | 561.5797647 | 2.858372194 | 0.347954025 |
| Xirp1 | 5.455029321 | 4.33923498 | 0.902863042 |
| Ccr8 | 0.637648044 | 1.825451192 | 1.291119975 |
| Slc25a38 | 245.895353 | 0.495125625 | 0.102115282 |
| Rpsa | 2894.845799 | 0.6373981 | 0.097012907 |
| Gm24044 | 12.71611818 | -0.308361635 | 0.360814663 |
| Gm26448 | 99.87717075 | 0.377529842 | 0.182810184 |
| Snora62 | 1.810539642 | 1.133905122 | 0.750777938 |
| Mobp | 12397.77805 | 0.610207021 | 0.158712687 |
| Gm37111 | 46.80149452 | -0.532720899 | 0.193294304 |
| Myrip | 677.5660742 | -0.659732458 | 0.146434488 |
| Gm15566 | 4.5585197 | -2.268609727 | 0.709071998 |
| Gm15565 | 3.283342549 | -1.632941207 | 0.729803969 |
| Eif1b | 736.4044258 | 0.32394786 | 0.094417617 |
| Entpd3 | 153.1227425 | 0.241771313 | 0.11106202 |
| Rpl14 | 1081.536881 | 0.520448778 | 0.11259866 |
| 5830454E08Rik | 6.315262053 | 0.064937034 | 0.369473873 |
| Gm23624 | 0 | NA | NA |
| 1700020M21Rik | 0.275587006 | 2.213841587 | 2.391099463 |
| 4930593C16Rik | 1.144931742 | 0.265202433 | 0.977897798 |
| Ctnnb1 | 5088.011416 | 0.550668511 | 0.090553067 |
| mmu-mir-7090 | 0.464416953 | -0.969856995 | 1.266808085 |
| Ulk4 | 49.64989384 | -0.768569326 | 0.194496874 |
| Gm22987 | 0 | NA | NA |
| Gm17157 | 0 | NA | NA |
| Trak1 | 1974.275711 | -0.423300235 | 0.072222836 |
| Cck | 1108.958497 | -1.057464752 | 0.122458958 |
| Lyzl4 | 13.61068275 | 0.228378308 | 0.356010643 |
| Lyzl4os | 0.219234697 | 1.252219195 | 2.544014054 |
| Vipr1 | 229.8541341 | 0.073009017 | 0.218750546 |
| Sec22c | 369.7565343 | -0.449480006 | 0.072688813 |
| Deb1 | 212.2253457 | -0.179515965 | 0.085350868 |
| Nktr | 2807.808864 | -0.321288571 | 0.170972773 |
| E530011L22Rik | 71.35472267 | -0.87659256 | 0.153400579 |
| Zfp651 | 995.5133322 | -0.290605148 | 0.083553128 |

| | | | |
|---|---|---|---|
| Klhl40 | 81.7128723 | 1.527097398 | 0.226961103 |
| Hhatl | 134.4694746 | -0.647630559 | 0.147405669 |
| Ccdc13 | 49.28911322 | -0.420640799 | 0.185859088 |
| Gm17163 | 0.831375036 | -0.921407458 | 1.178533467 |
| Higd1a | 526.1847965 | -0.074620538 | 0.079395789 |
| Ackr2 | 29.87282185 | -0.25471196 | 0.313900033 |
| Cyp8b1 | 0.044191549 | -0.243571806 | 3.724945049 |
| 1700048O20Rik | 89.81562415 | 0.013376949 | 0.175230599 |
| Fam198a | 45.49996011 | -1.071731232 | 0.257530376 |
| Pomgnt2 | 324.6867476 | -0.343508282 | 0.088693475 |
| Snrk | 1696.951144 | -0.25904298 | 0.107021062 |
| Ano10 | 333.863595 | 0.731489057 | 0.078597411 |
| Mir8106 | 0.421183792 | 0.237330214 | 1.667257599 |
| Gm26797 | 96.16516166 | 3.967246855 | 0.308270038 |
| Abhd5 | 237.0914825 | 0.445211909 | 0.120431875 |
| A730085K08Rik | 3.727958404 | -0.465472281 | 0.532883057 |
| 9530059O14Rik | 357.5582499 | -1.027640992 | 0.342688753 |
| Mir138-1 | 4.764516023 | -1.19164833 | 0.50499048 |
| Topaz1 | 1.867918734 | 2.049810135 | 0.744405819 |
| Tcaim | 210.4247602 | -0.125174914 | 0.102301515 |
| Gm25341 | 1.808478971 | -0.449733868 | 0.74424623 |
| Zfp445 | 1117.035592 | -0.426778889 | 0.107749666 |
| A530083I20Rik | 14.49954602 | -0.289829123 | 0.328212245 |
| Zkscan7 | 42.10827637 | -0.033893918 | 0.193795421 |
| Zfp105 | 88.67340489 | 0.565605589 | 0.123588228 |
| 1110059G10Rik | 126.7733861 | 0.186019426 | 0.11248793 |
| Kif15 | 30.3785014 | 1.655989509 | 0.336927174 |
| Tmem42 | 63.48058778 | -0.335194961 | 0.132370847 |
| Tgm4 | 6.449499428 | -0.406130422 | 0.399231239 |
| Zdhhc3 | 963.4259801 | 0.094507166 | 0.051808608 |
| Exosc7 | 112.7568782 | -0.324580719 | 0.098733842 |
| Clec3b | 48.60893226 | -0.986151419 | 0.208808751 |
| Gm23323 | 0.148038037 | -0.72173093 | 3.549417237 |
| Cdcp1 | 2.585897959 | -0.176626853 | 0.60588371 |
| Tmem158 | 284.2320044 | -0.180664096 | 0.106033371 |
| Scp2-ps2 | 91.28446377 | 0.293348922 | 0.160482969 |
| Lars2 | 5438.700325 | 0.693014354 | 0.817350254 |
| Limd1 | 249.5361358 | 1.315154072 | 0.122888744 |
| Sacm1l | 1325.845621 | 0.099770758 | 0.067114982 |
| Slc6a20b | 4.677638664 | 0.551115686 | 0.493082559 |
| Gm17120 | 4.624289095 | 0.766752839 | 0.912456495 |
| Slc6a20a | 203.7172933 | 0.237725257 | 0.201061901 |
| Gm17200 | 1.771524358 | 0.723492259 | 0.909249079 |
| Ccr9 | 18.10541206 | 3.588791064 | 0.477004189 |
| Gm10052 | 130.6631812 | 0.249180313 | 0.12856144 |
| Lztfl1 | 833.9950152 | -0.2964968 | 0.063351639 |
| Gm17020 | 0.049553047 | 0.71820487 | 3.724945049 |

| | | | |
|---|---|---|---|
| Gm17021 | 0.832525973 | 0.61559013 | 1.028041296 |
| Fyco1 | 809.1842188 | 0.200851707 | 0.084988877 |
| Cxcr6 | 20.53480371 | 8.035557918 | 0.89792 |
| Xcr1 | 0.667406772 | 1.481615805 | 2.607875391 |
| Ccr1 | 181.3036799 | 6.812604811 | 0.654100948 |
| Ccr1l1 | 0.461281393 | 2.214902889 | 1.484557983 |
| Ccr3 | 1.785934789 | 3.532389298 | 1.108757606 |
| Ccr2 | 315.4917145 | 7.165783589 | 0.447290837 |
| Ccr5 | 468.5098695 | 4.268186799 | 0.303485928 |
| 2010315B03Rik | 245.0633303 | -0.1649371 | 0.121668038 |
| Nlrp4g | 0.042716945 | -0.243571806 | 3.724945049 |
| Ppp2r3d | 158.3671903 | 0.022421094 | 0.116536502 |
| Gm21836 | 0 | NA | NA |
| SNORA28 | 0 | NA | NA |
| RP24-571H12.4 | 0 | NA | NA |
| RP24-571H12.6 | 0 | NA | NA |
| AC168054.1 | 0 | NA | NA |
| WI1-2477B13.3 | 0 | NA | NA |
| CAAA01180111.1 | 0 | NA | NA |
| CAAA01202134.1 | 0 | NA | NA |
| CAAA01125382.1 | 0 | NA | NA |
| AC109138.1 | 0 | NA | NA |
| AC109138.3 | 0 | NA | NA |
| AC109138.2 | 0 | NA | NA |
| AC116115.1 | 0 | NA | NA |
| mmu-mir-505 | 0 | NA | NA |
| CAAA01000182.1 | 0 | NA | NA |
| RP23-6C3.5 | 0 | NA | NA |
| RP23-6C3.4 | 0 | NA | NA |
| Hspa9-ps1 | 7.785073125 | 0.99818268 | 0.376665634 |
| RP23-299A23.1 | 0 | NA | NA |
| RP23-6C3.2 | 0 | NA | NA |
| RP23-6C3.3 | 0 | NA | NA |
| RP23-445P15.4 | 0 | NA | NA |
| RP23-445P15.5 | 0 | NA | NA |
| RP23-445P15.2 | 0 | NA | NA |
| RP23-163F11.2 | 0 | NA | NA |
| RP23-346N8.3 | 0 | NA | NA |
| CU041226.1 | 0 | NA | NA |
| RP23-402O17.1 | 0 | NA | NA |
| RP23-16C4.1 | 0 | NA | NA |
| AL593853.1 | 0 | NA | NA |
| RP23-199A2.7 | 0 | NA | NA |
| SNORA77 | 0 | NA | NA |
| mmu-mir-1306 | 0 | NA | NA |
| mmu-mir-3618 | 0 | NA | NA |
| mmu-mir-185 | 0 | NA | NA |

| | | | |
|---|---|---|---|
| AC084822.2 | 0 | NA | NA |
| U1 | 0 | NA | NA |
| AC084822.1 | 0 | NA | NA |
| RP23-285E22.4 | 0 | NA | NA |
| RP23-348F1.1 | 0 | NA | NA |
| 7SK | 0 | NA | NA |
| RP23-58H5.2 | 0 | NA | NA |
| RP23-58H5.5 | 0 | NA | NA |
| RP23-58H5.6 | 0 | NA | NA |
| RP23-58H5.8 | 0 | NA | NA |
| OTTMUSG0000001013 | 0 | NA | NA |
| RP24-364D22.1 | 0 | NA | NA |
| AC132586.1 | 0 | NA | NA |
| AC245272.1 | 0 | NA | NA |
| AC245272.3 | 0 | NA | NA |
| mmu-mir-290a | 0 | NA | NA |
| mmu-mir-291a | 0 | NA | NA |
| mmu-mir-292a | 0 | NA | NA |
| mmu-mir-291b | 0 | NA | NA |
| mmu-mir-293 | 0 | NA | NA |
| mmu-mir-294 | 0 | NA | NA |
| mmu-mir-295 | 0 | NA | NA |
| AC245272.2 | 0 | NA | NA |
| mmu-mir-935 | 0 | NA | NA |
| Mico1 | 0 | NA | NA |
| MEG3_1 | 0 | NA | NA |
| MEG3_2 | 0 | NA | NA |
| mmu-mir-1906-1 | 0 | NA | NA |
| mmu-mir-770 | 0 | NA | NA |
| AC152063.1 | 0 | NA | NA |
| mmu-mir-673 | 0 | NA | NA |
| mmu-mir-493 | 0 | NA | NA |
| mmu-mir-337 | 0 | NA | NA |
| mmu-mir-540 | 0 | NA | NA |
| mmu-mir-665 | 0 | NA | NA |
| mmu-mir-3070-1 | 0 | NA | NA |
| mmu-mir-3070-2 | 0 | NA | NA |
| mmu-mir-431 | 0 | NA | NA |
| mmu-mir-433 | 0 | NA | NA |
| mmu-mir-127 | 0 | NA | NA |
| mmu-mir-434 | 0 | NA | NA |
| mmu-mir-432 | 0 | NA | NA |
| mmu-mir-3071 | 0 | NA | NA |
| MEG8_1 | 0 | NA | NA |
| mmu-mir-341 | 0 | NA | NA |
| mmu-mir-1188 | 0 | NA | NA |
| AC152063.2 | 0 | NA | NA |

| | | | | |
|---|---|---|---|---|
| mmu-mir-370 | 0 | NA | NA | |
| SNORD113 | 0 | NA | NA | |
| RP23-204I16.2 | 0 | NA | NA | |
| RP23-204I16.3 | 0 | NA | NA | |
| mmu-mir-882 | 0 | NA | NA | |
| CH36-472P4.1 | 0 | NA | NA | |
| AC233742.1 | 0 | NA | NA | |
| CH36-472P4.2 | 0 | NA | NA | |
| CH36-472P4.3 | 0 | NA | NA | |
| CH36-169F23.1 | 0 | NA | NA | |
| CH36-169F23.2 | 0 | NA | NA | |
| AC240971.1 | 0 | NA | NA | |
| CH36-169F23.4 | 0 | NA | NA | |
| CH36-169F23.3 | 0 | NA | NA | |
| CH36-169F23.5 | 0 | NA | NA | |
| RP24-152F23.1 | 0 | NA | NA | |
| RP24-152F23.4 | 0 | NA | NA | |
| RP24-152F23.5 | 0 | NA | NA | |
| AC244694.1 | 0 | NA | NA | |
| RP24-505E20.1 | 0 | NA | NA | |
| RP24-410B18.1 | 0 | NA | NA | |
| RP24-410B18.2 | 0 | NA | NA | |
| RP24-410B18.3 | 0 | NA | NA | |
| RP24-410B18.5 | 0 | NA | NA | |
| AC202135.1 | 0 | NA | NA | |
| RP24-410B18.4 | 0 | NA | NA | |
| Rhox5 | 0.049553047 | | 0.71820487 | 3.724945049 |
| Rhox6 | 0 | NA | NA | |
| Rhox7a | 0 | NA | NA | |
| Rhox8 | 0.068097359 | | -0.243569582 | 3.724945049 |
| Rhox7b | 0 | NA | NA | |
| Rhox9 | 0.095435502 | | 0.718202646 | 3.724945049 |
| RP23-202F23.7 | 0 | NA | NA | |
| Rhox11-ps | 0 | NA | NA | |
| AC154679.1 | 0 | NA | NA | |
| AC182748.2 | 0 | NA | NA | |
| AC182748.1 | 0 | NA | NA | |
| U6 | 0 | NA | NA | |
| RP24-176A1.1 | 0 | NA | NA | |
| RP24-176A1.2 | 0 | NA | NA | |
| AC127556.1 | 0 | NA | NA | |
| mmu-mir-194-2 | 0 | NA | NA | |
| mmu-mir-192 | 0 | NA | NA | |
| AC127556.5 | 0 | NA | NA | |
| mmu-mir-6989 | 0 | NA | NA | |
| AC127556.10 | 0 | NA | NA | |
| AC167245.6 | 0 | NA | NA | |

| | | | |
|---|---|---|---|
| AC167245.5 | 0 | NA | NA |
| AC124394.1 | 0 | NA | NA |
| Gm14423 | 0 | NA | NA |
| AC124394.3 | 0 | NA | NA |
| AC124394.4 | 0 | NA | NA |
| RP23-188E3.2 | 0 | NA | NA |
| RP23-188E3.1 | 0 | NA | NA |
| BX890553.1 | 0 | NA | NA |
| RP23-324J18.1 | 0 | NA | NA |
| RP23-324J18.2 | 0 | NA | NA |
| AC164883.2 | 0 | NA | NA |
| AC164883.1 | 0 | NA | NA |
| CH25-484K21.2 | 0 | NA | NA |
| CH25-484K21.3 | 0 | NA | NA |
| CH25-484K21.5 | 0 | NA | NA |
| CH25-484K21.6 | 0 | NA | NA |
| Gm9919 | 0 | NA | NA |
| FO907130.1 | 0 | NA | NA |
| SNORD103 | 0 | NA | NA |
| RP23-157N11.1 | 0 | NA | NA |
| RP23-157N11.4 | 0 | NA | NA |
| RP23-157N11.7 | 0 | NA | NA |
| RP23-157N11.6 | 0 | NA | NA |
| RP23-146J17.3 | 0 | NA | NA |
| AC125149.4 | 0 | NA | NA |
| AC125149.1 | 0 | NA | NA |
| AC125149.2 | 0 | NA | NA |
| AC125149.3 | 0 | NA | NA |
| AC133103.4 | 0 | NA | NA |
| AC133103.6 | 0 | NA | NA |
| AC133103.7 | 0 | NA | NA |
| AC133103.2 | 0 | NA | NA |
| AC133103.5 | 0 | NA | NA |
| AC133103.1 | 0 | NA | NA |
| AC133103.3 | 0 | NA | NA |
| AC168977.2 | 0 | NA | NA |
| AC168977.1 | 0 | NA | NA |
| DHRSX | 0 | NA | NA |
| U2 | 0 | NA | NA |
| Vmn1r186 | 0 | NA | NA |
| AC102264.1 | 0 | NA | NA |
| AC132444.1 | 0 | NA | NA |
| AC132444.3 | 0 | NA | NA |
| AC132444.5 | 0 | NA | NA |
| AC132444.4 | 0 | NA | NA |
| AC132444.2 | 0 | NA | NA |
| Csprs | 0 | NA | NA |

| | | | |
|---|---|---|---|
| AC132444.6 | 0 | NA | NA |
| Vamp7 | 0 | NA | NA |
| CAAA01165726.1 | 0 | NA | NA |
| Spry3 | 0 | NA | NA |
| Tmlhe | 0 | NA | NA |
| CR524822.1 | 0 | NA | NA |
| AC087559.4 | 0 | NA | NA |
| AC087559.5 | 0 | NA | NA |
| AC087559.1 | 0 | NA | NA |
| AC087559.3 | 0 | NA | NA |
| AC087559.2 | 0 | NA | NA |
| Ccl21c | 0 | NA | NA |
| AC140325.2 | 0 | NA | NA |
| AC140325.1 | 0 | NA | NA |
| AC140325.3 | 0 | NA | NA |
| AC140325.4 | 0 | NA | NA |
| CAAA01098150.1 | 0 | NA | NA |
| CAAA01064564.1 | 0 | NA | NA |
| CAAA01205117.2 | 0 | NA | NA |
| CAAA01205117.1 | 0 | NA | NA |
| Vmn2r122 | 0 | NA | NA |
| Gm2002 | 0 | NA | NA |
| CR974586.2 | 0 | NA | NA |
| CR974586.4 | 0 | NA | NA |
| CR974586.8 | 0 | NA | NA |
| CR974586.5 | 0 | NA | NA |
| CR974586.7 | 0 | NA | NA |
| CR974586.6 | 0 | NA | NA |
| CR974586.3 | 0 | NA | NA |
| CR974586.1 | 0 | NA | NA |
| Scya19 | 0 | NA | NA |
| CT868723.1 | 0 | NA | NA |
| Gm10931 | 0 | NA | NA |
| CT868723.2 | 0 | NA | NA |
| CAAA01147332.1 | 0 | NA | NA |
| AC164084.2 | 0 | NA | NA |
| AC164084.3 | 0 | NA | NA |
| AC164084.1 | 0 | NA | NA |
| AC165294.1 | 0 | NA | NA |
| AC165294.2 | 0 | NA | NA |
| AC165294.3 | 0 | NA | NA |
| AC125178.1 | 0 | NA | NA |
| AC125178.3 | 0 | NA | NA |
| AC125178.2 | 0 | NA | NA |
| AC123873.3 | 0 | NA | NA |
| AC123873.1 | 0 | NA | NA |
| AC123873.2 | 0 | NA | NA |

| | | | |
|---|---|---|---|
| AC126035.1 | 0 | NA | NA |
| Gm16367 | 0 | NA | NA |
| AC163611.1 | 0 | NA | NA |
| AC140365.1 | 0 | NA | NA |
| AC124606.2 | 0 | NA | NA |
| AC124606.1 | 0 | NA | NA |
| AC133095.2 | 0 | NA | NA |
| AC133095.1 | 0 | NA | NA |
| AC234645.1 | 0 | NA | NA |
| PISD | 0 | NA | NA |
| mt-Tf | 0 | NA | NA |
| mt-Rnr1 | 119593.2604 | 0.276700092 | 0.923923878 |
| mt-Tv | 0 | NA | NA |
| mt-Rnr2 | 65951.93075 | 0.549500659 | 0.794778659 |
| mt-Tl1 | 208.0973662 | 0.334691734 | 0.219914296 |
| mt-Nd1 | 34373.33605 | -0.400348876 | 0.157397455 |
| mt-Ti | 143.1190915 | -0.588590433 | 0.336428977 |
| mt-Tq | 55.05421077 | -0.208042458 | 0.192705424 |
| mt-Tm | 149.670499 | -0.556404291 | 0.244456945 |
| mt-Nd2 | 20368.9686 | -0.331412281 | 0.224881873 |
| mt-Tw | 186.9799014 | 0.041083148 | 0.187244929 |
| mt-Ta | 11.02334367 | 0.207434388 | 0.345347108 |
| mt-Tn | 0.72655531 | 0.194112794 | 1.286450347 |
| mt-Tc | 1.449755302 | 1.284603293 | 0.978542784 |
| mt-Ty | 0.807376506 | -0.211001706 | 1.324940186 |
| mt-Co1 | 112876.7948 | -0.060865799 | 0.120041532 |
| mt-Ts1 | 1.308396679 | 1.513086774 | 1.035164758 |
| mt-Td | 0.616049689 | 0.623999074 | 1.313475995 |
| mt-Co2 | 1688.977157 | -0.215236469 | 0.128255757 |
| mt-Tk | 0.097139547 | 0.718207094 | 3.724945049 |
| mt-Atp8 | 110.3540989 | -0.298204883 | 0.42925608 |
| mt-Atp6 | 624.2525603 | -0.154040513 | 0.197237198 |
| mt-Co3 | 1101.033194 | -0.154405446 | 0.147082414 |
| mt-Tg | 0 | NA | NA |
| mt-Nd3 | 119.756487 | -0.226835839 | 0.173366154 |
| mt-Tr | 0 | NA | NA |
| mt-Nd4l | 129.7203189 | -0.389119826 | 0.21643647 |
| mt-Nd4 | 21602.70211 | -0.350596277 | 0.143062135 |
| mt-Th | 0.134005754 | -0.72445992 | 2.962652837 |
| mt-Ts2 | 0.119587853 | 0.237305575 | 3.724945049 |
| mt-Tl2 | 0.731462286 | -0.228997562 | 1.306676543 |
| mt-Nd5 | 23114.44776 | -0.439590207 | 0.210333045 |
| mt-Nd6 | 1598.689532 | 0.469596512 | 0.154969727 |
| mt-Te | 5.226567033 | -0.04417262 | 0.481148541 |
| mt-Cytb | 37159.01855 | -0.452125754 | 0.140694911 |
| mt-Tt | 18.73040197 | 0.26434659 | 0.358842797 |
| mt-Tp | 56.15899216 | 0.044532635 | 0.21692565 |

| | | | |
|---|---|---|---|
| Gm21950 | 0 | NA | NA |
| Gm21364 | 0 | NA | NA |
| Gm14331 | 0 | NA | NA |
| Gm14333 | 0 | NA | NA |
| Gm14332 | 0 | NA | NA |
| Gm14334 | 0 | NA | NA |
| Gm14352 | 0 | NA | NA |
| Gm14346 | 0 | NA | NA |
| Gm14353 | 0 | NA | NA |
| Gm14360 | 0 | NA | NA |
| Gm14361 | 0 | NA | NA |
| Gm14345 | 0 | NA | NA |
| Gm14351 | 0 | NA | NA |
| Gm14357 | 0 | NA | NA |
| Gm14359 | 0 | NA | NA |
| Gm14355 | 0 | NA | NA |
| Spin2-ps1 | 0 | NA | NA |
| Gm3701 | 0 | NA | NA |
| Gm15251 | 0 | NA | NA |
| Gm15257 | 0 | NA | NA |
| Gm3706 | 0 | NA | NA |
| Gm15258 | 0 | NA | NA |
| Gm14350 | 0 | NA | NA |
| Gm14349 | 0 | NA | NA |
| Gm14358 | 0 | NA | NA |
| Gm14362 | 0 | NA | NA |
| Gm14356 | 0 | NA | NA |
| Gm14347 | 0 | NA | NA |
| Gm10921 | 0 | NA | NA |
| Gm14348 | 0 | NA | NA |
| Gm14364 | 0 | NA | NA |
| Gm14368 | 0 | NA | NA |
| Gm15260 | 0 | NA | NA |
| Gm10922 | 0 | NA | NA |
| Gm6019 | 0 | NA | NA |
| Gm9427 | 0 | NA | NA |
| Gm15256 | 0 | NA | NA |
| Gm5925 | 0 | NA | NA |
| Gm4725 | 0 | NA | NA |
| Gm9498 | 0 | NA | NA |
| Gm15252 | 0 | NA | NA |
| Gm3750 | 0 | NA | NA |
| Gm4726 | 0 | NA | NA |
| Gm3757 | 0 | NA | NA |
| Gm15259 | 0 | NA | NA |
| Gm15254 | 0 | NA | NA |
| Gm3763 | 0 | NA | NA |

| | | | |
|---|---|---|---|
| Gm14370 | 0 | NA | NA |
| Gm14367 | 0 | NA | NA |
| Gm14365 | 0 | NA | NA |
| Gm9428 | 0 | NA | NA |
| Gm5924 | 0 | NA | NA |
| Gm14369 | 0 | NA | NA |
| Gm14363 | 0 | NA | NA |
| Mycs | 0 | NA | NA |
| Gm14366 | 0 | NA | NA |
| Gm14375 | 0 | NA | NA |
| Gm14374 | 0 | NA | NA |
| Gm14373 | 4.562379813 | 0.71236944 | 0.516009744 |
| Gm14452 | 0.11997416 | -0.243571806 | 3.724945049 |
| Gm14451 | 0.112642008 | -0.24357403 | 3.724945049 |
| Gm14448 | 2.617244783 | -1.38464295 | 0.593874936 |
| Gm14450 | 0 | NA | NA |
| Gm26618 | 11.11537949 | -1.007064522 | 0.329285532 |
| Nudt11 | 271.955026 | 0.068695078 | 0.102619996 |
| Rpl19-ps12 | 2.084829622 | 0.006699792 | 0.870543631 |
| Gm27360 | 0 | NA | NA |
| AU022751 | 17.60759048 | -1.513580732 | 0.392295897 |
| n-R5s2 | 0 | NA | NA |
| Nudt10 | 171.3843856 | -0.154914725 | 0.099103714 |
| Bmp15 | 3.216309252 | 0.411775899 | 0.684566665 |
| Shroom4 | 90.11915289 | 0.269196322 | 0.153325527 |
| Dgkk | 132.1695717 | 0.312301344 | 0.213865428 |
| Ccnb3 | 0.629652017 | 3.467557868 | 1.361048818 |
| Akap4 | 0 | NA | NA |
| Gm14380 | 0.089089296 | 0.237305797 | 3.724945049 |
| Clcn5 | 513.7578379 | 0.012947604 | 0.0947649 |
| Gm26401 | 0 | NA | NA |
| Mir500 | 0.181074517 | 0.16772247 | 3.340104567 |
| Mir501 | 0.726672012 | -0.281113587 | 1.030916882 |
| Mir362 | 0.060174537 | 0.237309859 | 3.724945049 |
| Mir188 | 0.206496347 | -0.70323865 | 2.184500223 |
| Mir532 | 0.516362146 | -2.737326012 | 1.255532869 |
| Usp27x | 183.7584339 | 0.144391581 | 0.129465795 |
| Gm14379 | 5.282615923 | 0.6581587 | 0.453786176 |
| 2010204K13Rik | 54.17774226 | -0.311692974 | 0.1566554 |
| Gm14381 | 0.045882455 | 0.237312084 | 3.724945049 |
| Gm14395 | 0 | NA | NA |
| Gm14394 | 0.287498921 | 2.37074578 | 1.959901931 |
| Gm23082 | 0.046345709 | 0.71820487 | 3.724945049 |
| Ppp1r3f | 110.0662308 | -0.364065999 | 0.112156514 |
| Gm36995 | 5.062940487 | -0.658839408 | 0.499460753 |
| Ppp1r3fos | 7.493797218 | -0.078252705 | 0.373623409 |
| Foxp3 | 2.783566837 | 2.133126838 | 0.833730541 |

| | | | |
|---|---|---|---|
| Ccdc22 | 22.02494333 | 0.097575224 | 0.22019829 |
| Cacna1f | 2.348133422 | 1.652362446 | 0.750025649 |
| Syp | 9951.334057 | -0.527210143 | 0.068775783 |
| Gm14703 | 9.786129742 | -1.038848623 | 0.558730859 |
| Prickle3 | 15.66947633 | 0.225118994 | 0.254364523 |
| Plp2 | 46.01570745 | 2.311544769 | 0.223398018 |
| Magix | 1.802146662 | -0.240759392 | 0.737933453 |
| Gm6079 | 0 | NA | NA |
| Gpkow | 510.2906317 | 0.29694445 | 0.079298444 |
| Gm14720 | 0 | NA | NA |
| Wdr45 | 250.0876241 | -0.165098496 | 0.071020008 |
| Praf2 | 530.2469731 | -0.202808084 | 0.064192363 |
| Ccdc120 | 144.7907492 | -0.466581154 | 0.14036553 |
| Smt3h2-ps | 1.463749607 | -1.177134646 | 1.09206126 |
| Tfe3 | 940.6091959 | 0.535005555 | 0.060487642 |
| Gripap1 | 833.0662841 | 6.39E-05 | 0.116090878 |
| Mir1198 | 4.250207662 | -1.375851229 | 0.546537489 |
| Kcnd1 | 104.6643679 | -1.009517893 | 0.142655817 |
| Otud5 | 1030.482185 | 0.300116383 | 0.04738796 |
| Pim2 | 309.3189051 | -0.377305943 | 0.08985362 |
| Slc35a2 | 203.1775785 | 0.219310016 | 0.102997108 |
| Pqbp1 | 363.0226281 | -0.241345404 | 0.073433823 |
| Timm17b | 118.9161447 | -0.126917793 | 0.150723315 |
| Gm10491 | 0.046345709 | 0.71820487 | 3.724945049 |
| Gm10490 | 0.314722824 | 0.263085455 | 1.710947864 |
| Pcsk1n | 5132.141306 | -0.122375297 | 0.101753593 |
| Eras | 0.25849572 | 0.786007684 | 2.073274158 |
| Hdac6 | 414.7095717 | -0.366221145 | 0.083857255 |
| Psmb6-ps | 0 | NA | NA |
| Gata1 | 0.556661178 | 1.875081089 | 1.561296022 |
| Glod5 | 2.482897828 | -0.405673087 | 0.659858987 |
| Gm9429 | 0.112029109 | -0.724455696 | 3.724945049 |
| Rpl22-ps1 | 1.865863811 | 0.705859869 | 0.752929703 |
| Gm14799 | 1.762166695 | -0.354151712 | 0.848476653 |
| Gm14820 | 0.044544648 | 0.237309859 | 3.724945049 |
| Gm7079 | 50.37483144 | 0.139133673 | 0.297082594 |
| Suv39h1 | 269.5752616 | -0.241791775 | 0.095643317 |
| Was | 67.79222777 | 3.749472949 | 0.269848722 |
| Gm22083 | 0 | NA | NA |
| Gm6787 | 1.653410964 | -0.994356528 | 0.810234399 |
| Psmb7-ps2 | 0.216874018 | -1.204562229 | 2.593170119 |
| Wdr13 | 1616.98844 | -0.690759529 | 0.07853969 |
| Rbm3 | 813.7615764 | 0.597939616 | 0.132526855 |
| Rbm3os | 4.093244608 | 0.290108443 | 0.535442598 |
| Tbc1d25 | 154.8391368 | -0.010326683 | 0.105642028 |
| Ebp | 209.4144051 | 0.636749898 | 0.095049444 |
| Porcn | 473.2143967 | -0.720028844 | 0.114458823 |

| | | | |
|---|---|---|---|
| Ftsj1 | 170.553967 | -0.098833089 | 0.085281064 |
| Slc38a5 | 13.85939591 | 0.864480074 | 0.329487856 |
| Ssxb10 | 0 | NA | NA |
| Ssxb9 | 0 | NA | NA |
| Ssxb1 | 0 | NA | NA |
| Ssxb2 | 0 | NA | NA |
| Gm14459 | 0 | NA | NA |
| Ssxb6 | 0 | NA | NA |
| Ssxb3 | 0.117907917 | 0.237310006 | 3.724945049 |
| Gm14500 | 0 | NA | NA |
| Gm6797 | 0 | NA | NA |
| Gm6798 | 0 | NA | NA |
| Gm9430 | 0.048949532 | -0.243569582 | 3.724945049 |
| Ssxb8 | 0.176646398 | 1.281666154 | 3.35399331 |
| Ssx9 | 0 | NA | NA |
| Ssxb5 | 0 | NA | NA |
| Gm14502 | 0.848591665 | -1.417188213 | 1.148211004 |
| Gm6592 | 0 | NA | NA |
| Gm5751 | 0.339985237 | -0.208074561 | 1.740125057 |
| B630019K06Rik | 470.2877301 | 0.11586303 | 0.134001249 |
| Gm14504 | 0.188390742 | -1.069709803 | 3.306242977 |
| Gm5752 | 0 | NA | NA |
| Gm16479 | 0 | NA | NA |
| Gm14510 | 0 | NA | NA |
| Gm5634 | 0 | NA | NA |
| Gm14509 | 0 | NA | NA |
| Gm14511 | 0 | NA | NA |
| Gm14457 | 0 | NA | NA |
| Gm14458 | 0 | NA | NA |
| Gm9431 | 0.150711795 | 0.237323719 | 2.856704362 |
| Gm14865 | 0 | NA | NA |
| Fthl17 | 0 | NA | NA |
| Gm14503 | 0 | NA | NA |
| Gm14499 | 0 | NA | NA |
| Gm14864 | 0.066050875 | 0.71820487 | 3.724945049 |
| Gm5379 | 0 | NA | NA |
| Gm5635 | 0 | NA | NA |
| Gm16480 | 0 | NA | NA |
| Gm6826 | 0 | NA | NA |
| 4930402K13Rik | 0 | NA | NA |
| Gm5753 | 0 | NA | NA |
| Lancl3 | 181.9738499 | -0.176401526 | 0.097053759 |
| Gm14862 | 0.045882455 | 0.237312084 | 3.724945049 |
| Xk | 302.6762856 | -0.478986907 | 0.092707111 |
| 1700012L04Rik | 1.468848822 | -1.296364106 | 0.795181509 |
| Gm25481 | 0.305717649 | -0.998132011 | 1.732081029 |
| Gm6829 | 5.233925965 | 0.843448872 | 0.462902626 |

| | | | |
|---|---|---|---|
| Gm14863 | 0.303434138 | 1.17134256 | 1.739739513 |
| Gm14501 | 0.106056992 | 0.237307635 | 3.724945049 |
| Cybb | 408.2087086 | 6.394140844 | 0.247813313 |
| Gm25728 | 0.044191549 | -0.243571806 | 3.724945049 |
| Gm5132 | 0.273271296 | 0.237330367 | 2.030826423 |
| Dynlt3 | 1172.505651 | -0.062889229 | 0.098707329 |
| Hypm | 1.020378237 | -1.441740626 | 1.096412251 |
| 4930557A04Rik | 0.934106423 | -0.727624599 | 1.048371688 |
| Sytl5 | 63.53451429 | -0.566440303 | 0.164973647 |
| Gm5754 | 0.681090533 | 0.806889849 | 1.350001159 |
| Srpx | 4.924570278 | 0.339127865 | 0.473814491 |
| Rpgr | 331.1611277 | -0.63611198 | 0.144612797 |
| Otc | 0.279231709 | -0.65089106 | 2.081982346 |
| 4933416E14Rik | 0.989536815 | -1.344536741 | 1.217707725 |
| Tspan7 | 5607.471727 | -0.858240849 | 0.086487136 |
| Gm25885 | 9.492997069 | -1.740516058 | 0.385119145 |
| Gm5931 | 6.023005853 | 0.323837919 | 0.43490589 |
| Gm14470 | 0.083526043 | 0.718202646 | 3.724945049 |
| Gm14480 | 0.946116905 | -1.308604289 | 1.063141417 |
| Gm10489 | 4.50082467 | -0.533806798 | 0.485977644 |
| Mid1ip1 | 964.1006633 | 0.230784113 | 0.107961668 |
| Gm14493 | 2.463509541 | -0.985315049 | 0.681515168 |
| Gm14473 | 0 | NA | NA |
| Gm14485 | 0.048949532 | -0.243569582 | 3.724945049 |
| Gm14481 | 0.306362211 | 0.718223197 | 2.031452835 |
| Gm14483 | 0.04393175 | -0.243571806 | 3.724945049 |
| Gm14474 | 0 | NA | NA |
| Gm14477 | 0 | NA | NA |
| Gm14476 | 0 | NA | NA |
| Gm14484 | 0 | NA | NA |
| Gm14479 | 0 | NA | NA |
| Gm14482 | 0 | NA | NA |
| Gm14478 | 0 | NA | NA |
| Gm14475 | 0 | NA | NA |
| Gm4906 | 0 | NA | NA |
| Gm14515 | 0 | NA | NA |
| Gm14513 | 0 | NA | NA |
| Gm26314 | 0 | NA | NA |
| Rpl23a-ps13 | 0 | NA | NA |
| Gm14491 | 1.107727442 | -1.483592375 | 1.020019433 |
| Gm14512 | 0 | NA | NA |
| Bcor | 487.1376583 | -0.005318267 | 0.123385295 |
| Gm14521 | 0 | NA | NA |
| Gm14635 | 0.042716945 | -0.243571806 | 3.724945049 |
| Rpl30-ps11 | 0.439488226 | 1.997685593 | 1.481577663 |
| Atp6ap2 | 2034.077482 | 0.056770933 | 0.064369787 |
| Gm9432 | 2.504601602 | 0.621786484 | 0.743377896 |

| | | | |
|---|---|---|---|
| Gm4984 | 0.075043205 | 0.237309859 | 3.724945049 |
| 1810030O07Rik | 260.8927199 | 0.079746905 | 0.073656773 |
| Gm16265 | 0.351984013 | 1.533548101 | 1.76217258 |
| Med14 | 1262.541672 | -0.330961686 | 0.042652056 |
| Gm14634 | 22.48773761 | -1.039571603 | 0.260189747 |
| Gm25063 | 0.044544648 | 0.237309859 | 3.724945049 |
| Gm14524 | 0 | NA | NA |
| AA414768 | 46.27663079 | 0.571203366 | 0.193651144 |
| Gm14534 | 0.163190422 | 1.22955094 | 3.396300676 |
| 5730405O15Rik | 64.71272437 | -0.704917489 | 0.176681751 |
| Usp9x | 6600.196581 | -0.097427667 | 0.045972498 |
| Gm15027 | 0.158569248 | 0.718214251 | 2.811594796 |
| Gm14536 | 0 | NA | NA |
| Rpl3-ps1 | 956.7057751 | 0.266522231 | 0.106580088 |
| n-R5s3 | 0 | NA | NA |
| Llph-ps2 | 8.542226199 | 0.772630031 | 0.411663231 |
| 2010308F09Rik | 1.66644364 | -0.309051834 | 0.797668136 |
| Gm16481 | 0 | NA | NA |
| Ddx3x | 4505.87406 | 0.35117428 | 0.100382152 |
| Gm5380 | 0.854198329 | -0.427166929 | 1.015122911 |
| Gm7129 | 0.097899064 | -0.724453471 | 3.724945049 |
| Gm9434 | 0.22969359 | 1.166625474 | 3.129888948 |
| Nyx | 9.174112265 | -0.621955649 | 0.431992987 |
| Gm14506 | 0 | NA | NA |
| Cask | 1849.363507 | 0.100239114 | 0.068191992 |
| Gpr34 | 42.59505678 | -4.306979826 | 0.694056391 |
| Gm25246 | 0 | NA | NA |
| Gpr82 | 1.132182296 | 1.730351589 | 1.167004519 |
| Gm5381 | 0 | NA | NA |
| Gm25552 | 0 | NA | NA |
| Gm5382 | 0 | NA | NA |
| Gm14867 | 0 | NA | NA |
| Gm25202 | 0 | NA | NA |
| Gm14505 | 0.222175754 | 0.727460723 | 2.579270603 |
| Gm9435 | 0 | NA | NA |
| Ppp1r2-ps9 | 0 | NA | NA |
| Gm14519 | 0.126242988 | 0.237323625 | 3.017245153 |
| Gm14518 | 0 | NA | NA |
| Gm14517 | 0 | NA | NA |
| Gm25207 | 0 | NA | NA |
| Gm22464 | 0 | NA | NA |
| Gm23516 | 0 | NA | NA |
| Drr1 | 0.510920245 | 0.320586352 | 1.310147699 |
| Gm14520 | 0 | NA | NA |
| Gm14516 | 0.07796279 | 1.072698098 | 3.721829099 |
| Cypt1 | 0.038981395 | 0.718207094 | 3.724945049 |
| Csnk2a1-ps | 0.061920613 | 0.718207094 | 3.724945049 |

| | | | |
|---|---|---|---|
| Maoa | 471.3466681 | 0.117369345 | 0.098405724 |
| Maob | 199.4044433 | 0.291645764 | 0.141003525 |
| Ndp | 53.73532709 | -0.78302062 | 0.286547581 |
| Efhc2 | 28.39597078 | -0.170551304 | 0.244337047 |
| Gm14514 | 0 | NA | NA |
| Gm22294 | 0.044191549 | -0.243571806 | 3.724945049 |
| Fundc1 | 517.0185679 | 0.190008312 | 0.136855583 |
| Gm5073 | 0.100902008 | 1.15059983 | 3.720931964 |
| Gm9436 | 0 | NA | NA |
| Gm5755 | 0 | NA | NA |
| Gm5123 | 0 | NA | NA |
| Dusp21 | 0 | NA | NA |
| Kdm6a | 813.7693481 | -0.079957305 | 0.08349661 |
| n-R5s5 | 0 | NA | NA |
| Gm14883 | 0 | NA | NA |
| 4930578C19Rik | 7.057036865 | 1.149968998 | 0.509543037 |
| Gm6923 | 0 | NA | NA |
| Gm5383 | 0 | NA | NA |
| Mir221 | 8.635094361 | 0.519276009 | 0.408453933 |
| Mir222 | 2.111772912 | 0.169465918 | 0.754801188 |
| Gm14636 | 161.0641105 | 0.007939889 | 0.184281424 |
| Gm26652 | 38.71122557 | -0.765391015 | 0.279957607 |
| Gm22783 | 0.140090306 | 0.685641014 | 2.912101186 |
| Gm14885 | 0 | NA | NA |
| BC049702 | 0 | NA | NA |
| Gm14531 | 0 | NA | NA |
| Gm23678 | 0 | NA | NA |
| Gm14530 | 0.578381701 | 0.968175963 | 1.528418134 |
| Rpl36-ps10 | 0.038981395 | 0.718207094 | 3.724945049 |
| Gm5384 | 0.309182497 | -1.125015904 | 1.782419036 |
| Chst7 | 15.98671876 | -0.106437615 | 0.293633406 |
| Slc9a7 | 557.1947993 | -0.406517283 | 0.068851508 |
| Gm9083 | 2.845117016 | 0.418220754 | 0.538837362 |
| Gm14529 | 0 | NA | NA |
| Rp2h | 271.8265236 | 0.20710997 | 0.134108688 |
| Gm9085 | 0.090427103 | 0.237307635 | 3.724945049 |
| Gm14537 | 3.667719452 | -0.803797902 | 0.598447169 |
| Jade3 | 162.3751509 | 0.09709857 | 0.121694542 |
| Gm14884 | 0.062426873 | 0.237312084 | 3.724945049 |
| Gm14538 | 0 | NA | NA |
| Rgn | 3.195312812 | 0.904561275 | 0.624770655 |
| Gm14539 | 108.3056856 | 0.07047636 | 0.165813157 |
| Gm23628 | 1.296764287 | 0.042629956 | 0.96569536 |
| Ndufb11 | 499.4740233 | -0.00344963 | 0.137792786 |
| Rbm10 | 598.4934086 | -0.159180699 | 0.072340728 |
| Uba1 | 4405.966393 | 0.485860099 | 0.072847666 |
| Cdk16 | 2633.067354 | -0.260144509 | 0.058288842 |

| | | | |
|---|---|---|---|
| Usp11 | 1238.553824 | -0.700347404 | 0.060545685 |
| Araf | 1405.997007 | -0.220620714 | 0.051745217 |
| Gm26065 | 0 | NA | NA |
| Gm24824 | 0 | NA | NA |
| Gm15028 | 0.262683741 | 1.878630148 | 2.44852992 |
| Syn1 | 5420.391119 | -0.597599958 | 0.09389254 |
| Mir5617 | 0 | NA | NA |
| Timp1 | 225.2635421 | 6.820461433 | 0.52536711 |
| Gm15029 | 0 | NA | NA |
| Cfp | 118.3901945 | 0.535324122 | 0.208368293 |
| Elk1 | 876.8854966 | 0.190118373 | 0.132281058 |
| Uxt | 53.85454816 | 0.470354541 | 0.160456059 |
| A230072C01Rik | 65.61270515 | -0.274836635 | 0.137894817 |
| Zfp182 | 201.9227118 | -0.203424825 | 0.116592718 |
| Spaca5 | 0.708400758 | 2.35066652 | 1.233267287 |
| Zfp300 | 69.84368967 | 0.176632519 | 0.158069767 |
| Ssxa1 | 0 | NA | NA |
| Gm1848 | 2.174616668 | 0.034946611 | 0.72826061 |
| Gm21876 | 0 | NA | NA |
| 4930453H23Rik | 0.062426873 | 0.237312084 | 3.724945049 |
| Gm6938 | 0.624139293 | -1.199110559 | 1.404562753 |
| Gm5124 | 229.7670475 | -0.163762939 | 0.116851124 |
| Gm26593 | 1.567651568 | 1.48876247 | 0.930687068 |
| Agtr2 | 19.19641764 | -0.034315205 | 0.300242582 |
| Gm22592 | 0 | NA | NA |
| Gm9439 | 0.758270043 | 0.16306707 | 1.175599689 |
| Gm14558 | 0 | NA | NA |
| Slc6a14 | 1.053279316 | 0.539187126 | 1.077446718 |
| Gm25553 | 0.068097359 | -0.243569582 | 3.724945049 |
| Gm28269 | 0 | NA | NA |
| Gm28268 | 0.530320922 | 0.206704146 | 1.356794837 |
| Gm24535 | 0 | NA | NA |
| Gm14563 | 0 | NA | NA |
| Gm14561 | 0 | NA | NA |
| Gm6071 | 0 | NA | NA |
| Gm25812 | 0 | NA | NA |
| Gm14562 | 0 | NA | NA |
| Gm15030 | 0 | NA | NA |
| Ppp1r2-ps7 | 0 | NA | NA |
| Gm26131 | 0 | NA | NA |
| Klhl13 | 551.8407492 | -0.499813162 | 0.097283477 |
| Gm14523 | 0.533370488 | -1.882580565 | 1.204862488 |
| Gm7164 | 0.047948698 | -0.243569582 | 3.724945049 |
| Wdr44 | 445.6653234 | 0.014345117 | 0.068857576 |
| Gm4789 | 0 | NA | NA |
| Gm4907 | 0.266401621 | 0.875488724 | 2.475088273 |
| Gm4985 | 0 | NA | NA |

| | | | | |
|---|---|---|---|---|
| Gm25299 | 0 | NA | NA | |
| Gm4732 | 0 | NA | NA | |
| Gm9440 | 0 | NA | NA | |
| Gm6956 | 0 | NA | NA | |
| Gm27192 | 0 | NA | NA | |
| Gm5934 | 0.046345709 | | 0.71820487 | 3.724945049 |
| Gm4297 | 0 | NA | NA | |
| Gm27910 | 0 | NA | NA | |
| Gm27839 | 0 | NA | NA | |
| Gm5935 | 0 | NA | NA | |
| Gm2759 | 0 | NA | NA | |
| CR938728.1 | 0 | NA | NA | |
| Gm2768 | 0 | NA | NA | |
| Gm5169 | 0 | NA | NA | |
| Gm1989 | 0 | NA | NA | |
| Gm1993 | 0 | NA | NA | |
| Gm2003 | 0 | NA | NA | |
| Gm27892 | 0 | NA | NA | |
| Gm7350 | 0 | NA | NA | |
| E330010L02Rik | 0 | NA | NA | |
| Gm14533 | 0 | NA | NA | |
| Gm5168 | 0 | NA | NA | |
| Gm2012 | 0 | NA | NA | |
| Gm26302 | 0 | NA | NA | |
| Gm2030 | 0 | NA | NA | |
| Gm2005 | 0 | NA | NA | |
| Slx | 0 | NA | NA | |
| Gm27341 | 0 | NA | NA | |
| Gm14525 | 0 | NA | NA | |
| Gm6113 | 0 | NA | NA | |
| Gm6121 | 0 | NA | NA | |
| Gm7365 | 0 | NA | NA | |
| Gm7375 | 0 | NA | NA | |
| Gm10230 | 0 | NA | NA | |
| BX813306.1 | 0 | NA | NA | |
| Gm2092 | 0 | NA | NA | |
| CR589930.1 | 0 | NA | NA | |
| Gm2101 | 0 | NA | NA | |
| Gm7386 | 0 | NA | NA | |
| Gm10058 | 0 | NA | NA | |
| Gm2117 | 0 | NA | NA | |
| Gm7391 | 0 | NA | NA | |
| Gm4836 | 0 | NA | NA | |
| Gm7398 | 0 | NA | NA | |
| Gm10147 | 0 | NA | NA | |
| BX813332.1 | 0 | NA | NA | |
| Gm2165 | 0 | NA | NA | |

| | | | | |
|---|---|---|---|---|
| Gm7408 | 0 | NA | NA | |
| Gm10096 | 0 | NA | NA | |
| Gm2200 | 0 | NA | NA | |
| Gm7415 | 0 | NA | NA | |
| Gm26818 | 0 | NA | NA | |
| Gm2066 | 0 | NA | NA | |
| Gm7421 | 0 | NA | NA | |
| Gm15432 | 1.975319427 | | 1.203824352 | 0.802489171 |
| Gm14527 | 0 | NA | NA | |
| BX294196.1 | 0 | NA | NA | |
| Gm3657 | 0 | NA | NA | |
| Gm3669 | 0 | NA | NA | |
| Gm10488 | 0 | NA | NA | |
| BX294192.1 | 0 | NA | NA | |
| E330016L19Rik | 0 | NA | NA | |
| Gm3681 | 0 | NA | NA | |
| Gm14632 | 0 | NA | NA | |
| Gm7437 | 0 | NA | NA | |
| Gm14974 | 0 | NA | NA | |
| Gm7438 | 0 | NA | NA | |
| Gm10487 | 0 | NA | NA | |
| Gm21447 | 0 | NA | NA | |
| Spin2f | 0 | NA | NA | |
| Gm2784 | 0 | NA | NA | |
| Gm15280 | 0 | NA | NA | |
| Gm15276 | 0 | NA | NA | |
| Gm2777 | 0 | NA | NA | |
| Gm21883 | 0 | NA | NA | |
| Spin2e | 0 | NA | NA | |
| Gm21608 | 0 | NA | NA | |
| Gm21637 | 0 | NA | NA | |
| Gm21645 | 0 | NA | NA | |
| Gm2799 | 0 | NA | NA | |
| Gmcl1l | 0 | NA | NA | |
| Gm5926 | 0 | NA | NA | |
| Gm21951 | 0 | NA | NA | |
| Gm21657 | 0 | NA | NA | |
| Gm21789 | 0 | NA | NA | |
| Gm2825 | 0 | NA | NA | |
| Spin2-ps6 | 0 | NA | NA | |
| Gm15275 | 0 | NA | NA | |
| Gm15274 | 0 | NA | NA | |
| Gm2863 | 0 | NA | NA | |
| Gm2182 | 0 | NA | NA | |
| Gm2854 | 0 | NA | NA | |
| Gm21979 | 0 | NA | NA | |
| Gm21990 | 0 | NA | NA | |

| | | | |
|---|---|---|---|
| Gm2892 | 0 | NA | NA |
| Gm2223 | 89.01610747 | 0.304004095 | 0.149256641 |
| Gm2231 | 0 | NA | NA |
| Gm2913 | 0 | NA | NA |
| Gm15273 | 0 | NA | NA |
| Gm15281 | 0 | NA | NA |
| Gm15279 | 0 | NA | NA |
| Gm7023 | 0 | NA | NA |
| Gm2927 | 0 | NA | NA |
| Gm2933 | 0 | NA | NA |
| Gm2964 | 0 | NA | NA |
| Gm21870 | 0 | NA | NA |
| Gm21681 | 0 | NA | NA |
| Spin2g | 0 | NA | NA |
| Gm21699 | 0 | NA | NA |
| Gm2288 | 0 | NA | NA |
| Gm14552 | 0 | NA | NA |
| Gm10486 | 0 | NA | NA |
| Gm2309 | 0 | NA | NA |
| Gm2318 | 0 | NA | NA |
| Gm14553 | 0 | NA | NA |
| Gm14819 | 0 | NA | NA |
| Gm4764 | 0 | NA | NA |
| Gm14554 | 0.066050875 | 0.71820487 | 3.724945049 |
| Gm25624 | 0 | NA | NA |
| Dock11 | 371.5272742 | 0.533741979 | 0.083396677 |
| Il13ra1 | 318.8934833 | 2.832716187 | 0.191387425 |
| Zcchc12 | 687.811863 | -0.46339442 | 0.132098337 |
| Gm14560 | 0 | NA | NA |
| Gm6023 | 0.589180756 | 2.674006852 | 1.497290865 |
| Gm14559 | 0 | NA | NA |
| Lonrf3 | 281.4852912 | 0.42596506 | 0.133160963 |
| Gm25047 | 24.14149978 | 0.498539114 | 0.245848221 |
| Gm14568 | 0.274682661 | 0.718190974 | 3.724945049 |
| AL450395.1 | 0 | NA | NA |
| Gm6268 | 0.075043205 | 0.237309859 | 3.724945049 |
| Gm22364 | 0 | NA | NA |
| Gm14569 | 4.657189074 | 1.125288185 | 0.508927588 |
| Pgrmc1 | 2005.442762 | -0.042452584 | 0.078815786 |
| Akap17b | 352.8596161 | -0.401925706 | 0.07182883 |
| Gm6274 | 10.81761153 | 0.584744079 | 0.380552559 |
| Slc25a43 | 7.740311858 | 2.107174225 | 0.52334421 |
| Gm23613 | 0.350819806 | 2.278737161 | 1.934226257 |
| Slc25a5 | 1945.709052 | 0.177717995 | 0.074249279 |
| Gm14549 | 0 | NA | NA |
| 2310010G23Rik | 9.327370753 | 0.123217901 | 0.404354581 |
| C330007P06Rik | 639.7924073 | 0.180865509 | 0.075340032 |

| | | | | |
|---|---|---|---|---|
| Gm14541 | | 2.652685225 | 0.316641306 | 0.780352197 |
| Ube2a | | 329.5173645 | 0.184054003 | 0.098713887 |
| Nkrf | | 449.5931371 | -0.249035162 | 0.078988517 |
| Gm15008 | | 0.250168298 | -1.034794603 | 2.073054895 |
| | 6-Sep | 721.875096 | -0.384908631 | 0.065157325 |
| Gm14542 | | 0 | NA | NA |
| Sowahd | | 0.087930927 | 0.236669296 | 3.724898107 |
| Gm26099 | | 0 | NA | NA |
| Gm24809 | | 0 | NA | NA |
| Rpl39 | | 722.6800404 | 0.720578565 | 0.127250136 |
| Snora69 | | 1.244281449 | -0.302330597 | 0.882816906 |
| Upf3b | | 273.0236683 | -0.246441206 | 0.119319812 |
| Nkap | | 229.4254487 | 0.205202688 | 0.083903929 |
| Akap14 | | 1.404017228 | 0.018980369 | 1.19627982 |
| Ndufa1 | | 279.6891087 | -0.22984654 | 0.13368423 |
| Rnf113a1 | | 34.43701755 | 0.208433853 | 0.19635222 |
| Gm9 | | 0 | NA | NA |
| Rhox1 | | 0 | NA | NA |
| Rhox2a | | 0 | NA | NA |
| Rhox3a | | 0 | NA | NA |
| Rhox4a | | 0 | NA | NA |
| Rhox3a2 | | 0 | NA | NA |
| Rhox4a2 | | 0 | NA | NA |
| Rhox2b | | 0 | NA | NA |
| Rhox3b-ps | | 0 | NA | NA |
| Rhox4b | | 0 | NA | NA |
| Rhox2c | | 0 | NA | NA |
| Rhox3c | | 0 | NA | NA |
| Rhox4c | | 0 | NA | NA |
| Rhox2d | | 0 | NA | NA |
| Gm26494 | | 0 | NA | NA |
| Rhox3d-ps | | 0 | NA | NA |
| Rhox4d | | 0 | NA | NA |
| Rhox2e | | 0 | NA | NA |
| Rhox3e | | 0 | NA | NA |
| Rhox4e | | 0 | NA | NA |
| Rhox2f | | 0 | NA | NA |
| Rhox3f | | 0 | NA | NA |
| Rhox4f | | 0 | NA | NA |
| Rhox3g | | 0 | NA | NA |
| Dppa3-ps | | 0 | NA | NA |
| Gm24493 | | 0 | NA | NA |
| Rhox2g | | 0 | NA | NA |
| Rhox4g | | 0 | NA | NA |
| Rhox3h | | 0 | NA | NA |
| Gm24147 | | 0 | NA | NA |
| Rhox2h | | 0 | NA | NA |

| | | | |
|---|---|---|---|
| RP23-457N24.12 | 0 | NA | NA |
| Rhox2-ps | 0 | NA | NA |
| Rhox11-ps1 | 0 | NA | NA |
| Rhox11-ps2 | 0 | NA | NA |
| Btg1-ps1 | 0.062426873 | 0.237312084 | 3.724945049 |
| Btg1-ps2 | 0 | NA | NA |
| Rhox10 | 0 | NA | NA |
| Rhox11 | 0 | NA | NA |
| Rhox11-ps3 | 0 | NA | NA |
| Rhox12 | 0.066050875 | 0.71820487 | 3.724945049 |
| Rhox13 | 0 | NA | NA |
| Zfp36l1-ps | 0.671662539 | 2.892940135 | 1.199776526 |
| Zbtb33 | 539.0021361 | 0.248784639 | 0.10129916 |
| Tmem255a | 487.0062224 | -0.037953483 | 0.111606157 |
| Mir3110 | 0 | NA | NA |
| Rhox7-ps1 | 0.075043205 | 0.237309859 | 3.724945049 |
| Atp1b4 | 0.134880151 | 1.580366464 | 2.92708711 |
| Gm40 | 0 | NA | NA |
| Gm4853 | 2.301314035 | 0.572923777 | 0.653118516 |
| Lamp2 | 2710.25613 | 1.661791484 | 0.126720349 |
| Gm7598 | 7.758147147 | -0.386950677 | 0.396022127 |
| Cul4b | 778.5921849 | 0.194042603 | 0.059608183 |
| Mcts1 | 310.2787056 | -0.11167966 | 0.095015005 |
| C1galt1c1 | 195.6813856 | 0.479667484 | 0.189815357 |
| Ap3s1-ps1 | 43.23259583 | 0.456012032 | 0.162499523 |
| Gm14565 | 0.060174537 | 0.237309859 | 3.724945049 |
| Gm25642 | 0 | NA | NA |
| Gm14564 | 0 | NA | NA |
| 6030498E09Rik | 0.374199036 | -0.764729284 | 1.490311566 |
| Gm6024 | 0 | NA | NA |
| Gm14567 | 9.440869076 | 0.183361175 | 0.356335564 |
| Gm16423 | 0 | NA | NA |
| Gm14551 | 0 | NA | NA |
| Gm14566 | 0 | NA | NA |
| Gm26437 | 0 | NA | NA |
| Cypt15 | 0.048949532 | -0.243569582 | 3.724945049 |
| Gm14550 | 0 | NA | NA |
| Gm14570 | 0 | NA | NA |
| Gm14571 | 0.04393175 | -0.243571806 | 3.724945049 |
| Cypt14 | 0 | NA | NA |
| Gm14643 | 0.048949532 | -0.243569582 | 3.724945049 |
| Gm26194 | 0 | NA | NA |
| Gm7189 | 0 | NA | NA |
| Gm16383 | 0 | NA | NA |
| Gm7190 | 0 | NA | NA |
| Gm4986 | 0.708087976 | 1.594252508 | 1.286782745 |
| Gm14575 | 0 | NA | NA |

| | | | |
|---|---|---|---|
| Gm14574 | 0 | NA | NA |
| Gm14576 | 0 | NA | NA |
| Gm5385 | 0 | NA | NA |
| Gm14577 | 0 | NA | NA |
| Gria3 | 2075.810427 | -1.152654699 | 0.100693524 |
| Gm22413 | 0 | NA | NA |
| Gm14614 | 0 | NA | NA |
| Gm14613 | 0 | NA | NA |
| Thoc2 | 1453.981551 | 0.062614701 | 0.062477886 |
| Gm7657 | 0 | NA | NA |
| Gm5135 | 0 | NA | NA |
| AL954355.1 | 0 | NA | NA |
| Xiap | 1927.944407 | 0.032691107 | 0.077529949 |
| Gm14615 | 0.068097359 | -0.243569582 | 3.724945049 |
| Gm37564 | 0.632215448 | -1.244631698 | 1.141298368 |
| Stag2 | 1734.40788 | 0.163573888 | 0.081396711 |
| Gm14619 | 0.123841226 | 1.292225235 | 3.719439735 |
| RP23-280E8.2 | 0.589937099 | 1.9730679 | 1.315942133 |
| Sh2d1a | 7.357494345 | 4.352192595 | 0.781392416 |
| Tenm1 | 2192.810765 | -0.685770847 | 0.102355071 |
| Gm23705 | 2.363489277 | -1.187858272 | 0.702601786 |
| Gm14578 | 7.50102364 | -0.935885429 | 0.54207136 |
| Gm24763 | 0 | NA | NA |
| Gm14587 | 8.400665238 | -1.596517444 | 0.52473175 |
| Gm14588 | 0 | NA | NA |
| Gm14579 | 0 | NA | NA |
| Gm362 | 0 | NA | NA |
| Gm14580 | 0 | NA | NA |
| Gm14628 | 0 | NA | NA |
| Gm14603 | 0 | NA | NA |
| Dcaf12l2 | 11.20309874 | -0.459990297 | 0.304746538 |
| Itpa-ps3 | 0 | NA | NA |
| Gm14602 | 0 | NA | NA |
| Dcaf12l1 | 439.0135136 | -0.611124822 | 0.075893794 |
| Gm7700 | 0 | NA | NA |
| Prr32 | 4.737155157 | -3.701951073 | 0.832410067 |
| Gm14606 | 0 | NA | NA |
| Gm14605 | 0 | NA | NA |
| Gm5386 | 0 | NA | NA |
| Gm14652 | 0 | NA | NA |
| Gm25078 | 0 | NA | NA |
| Gm14653 | 0 | NA | NA |
| 4930515L19Rik | 0 | NA | NA |
| Actrt1 | 0 | NA | NA |
| Gm14654 | 0.049553047 | 0.71820487 | 3.724945049 |
| Gm4987 | 0 | NA | NA |
| Gm29242 | 0 | NA | NA |

| | | | |
|---|---|---|---|
| Gm7722 | 0.112396584 | 1.192083721 | 3.724434897 |
| Gm14609 | 0.13648314 | 0.684500845 | 2.936510035 |
| Gm14610 | 0.187229393 | 0.210760517 | 2.696756358 |
| Gm4908 | 0 | NA | NA |
| Gm14608 | 0.401923326 | -0.640870142 | 1.42605383 |
| Gm6480 | 0.088476398 | -0.243576254 | 3.724945049 |
| Smarca1 | 656.7356124 | -0.665610506 | 0.086862541 |
| Gm14648 | 4.891543041 | 0.574526654 | 0.517377034 |
| Ocrl | 1059.362513 | -0.379241587 | 0.049046893 |
| Apln | 136.796465 | -1.583061244 | 0.182807364 |
| Xpnpep2 | 2.216825317 | -3.405266945 | 0.909180689 |
| Sash3 | 86.39783744 | 3.574398402 | 0.243217378 |
| Zdhhc9 | 642.249467 | -0.225508587 | 0.07174334 |
| Gm24038 | 0 | NA | NA |
| Utp14a | 243.5098185 | 0.440778376 | 0.097052897 |
| 9530027J09Rik | 0.791221454 | -1.592326234 | 1.101746831 |
| Gm22528 | 0 | NA | NA |
| Gm7212 | 0 | NA | NA |
| Gm22612 | 0 | NA | NA |
| Bcorl1 | 414.200989 | -0.615776557 | 0.108558445 |
| Elf4 | 210.3602389 | 4.267903731 | 0.201284295 |
| Aifm1 | 270.3649458 | -0.183664085 | 0.09048861 |
| Rab33a | 124.0995529 | -0.260233334 | 0.111551803 |
| Zfp280c | 445.3017861 | -0.285650004 | 0.100723571 |
| Slc25a14 | 322.987207 | -0.451184536 | 0.078513687 |
| Gm27828 | 1.991846454 | -2.428737182 | 0.915063865 |
| Gpr119 | 0.257827269 | 0.275224014 | 1.836990111 |
| Rbmx2 | 38.78282912 | 0.347442358 | 0.230900844 |
| Gm595 | 0.555008095 | 1.810260802 | 1.553152623 |
| Gm7219 | 0 | NA | NA |
| Enox2 | 126.1449739 | 0.021037189 | 0.132077628 |
| Gm15034 | 0 | NA | NA |
| Gm15036 | 0 | NA | NA |
| Gm14696 | 0.319846959 | 0.702385891 | 1.790294815 |
| Gm14697 | 0.131474741 | 0.237325381 | 2.980045311 |
| Arhgap36 | 111.713745 | 0.331078187 | 0.230702651 |
| Gm15037 | 0 | NA | NA |
| Gm14719 | 6.72125269 | -0.434947212 | 0.387007463 |
| Olfr1320 | 0 | NA | NA |
| Olfr1321 | 0 | NA | NA |
| Igsf1 | 120.6751455 | -1.633259606 | 0.164259921 |
| n-R5s6 | 0 | NA | NA |
| Olfr1322 | 0.062426873 | 0.237312084 | 3.724945049 |
| Gm15033 | 0 | NA | NA |
| Olfr1323 | 0.233510017 | 1.675700291 | 2.527199776 |
| Gm23385 | 0 | NA | NA |
| Gm7803 | 2.841226851 | 1.675161319 | 0.610475269 |

| | | | |
|---|---|---|---|
| Gm14612 | 0.038981395 | 0.718207094 | 3.724945049 |
| Gm22288 | 0.45637343 | 0.600450368 | 1.351430205 |
| Olfr1324 | 0.046345709 | 0.71820487 | 3.724945049 |
| Gm14620 | 0 | NA | NA |
| Firre | 150.6157184 | -0.973988224 | 0.215180772 |
| Stk26 | 85.23139706 | -0.442097197 | 0.143353424 |
| Frmd7 | 56.57679548 | 0.008603556 | 0.652394264 |
| Rap2c | 496.5610328 | 0.712516501 | 0.111067005 |
| Gm5387 | 0 | NA | NA |
| Mbnl3 | 8.934417775 | 1.342988048 | 0.38805075 |
| Gm7834 | 0 | NA | NA |
| Gm5388 | 0.810280881 | 0.593066251 | 1.151599254 |
| Gm23417 | 0 | NA | NA |
| Gm14621 | 0 | NA | NA |
| Hs6st2 | 613.6901847 | -0.478363652 | 0.085519292 |
| Gm14581 | 4.970320586 | -0.501632493 | 0.5221448 |
| Usp26 | 0 | NA | NA |
| Gm7840 | 0 | NA | NA |
| Gm22539 | 0.072025462 | 0.237312084 | 3.724945049 |
| 1700080O16Rik | 0 | NA | NA |
| Gpc4 | 223.3113565 | 0.603421395 | 0.092017072 |
| Gpc3 | 54.37477559 | -0.517742558 | 0.162972421 |
| Gm14582 | 0 | NA | NA |
| mmu-mir-6384 | 0.085433889 | -0.72445792 | 3.724945049 |
| Mir717 | 0 | NA | NA |
| A630012P03Rik | 15.99725564 | 7.803794757 | 0.91679815 |
| Mir363 | 0 | NA | NA |
| Mir92-2 | 0 | NA | NA |
| Mir19b-2 | 0 | NA | NA |
| Mir20b | 0 | NA | NA |
| Mir18b | 0 | NA | NA |
| Mir106a | 0 | NA | NA |
| Kis2 | 0.066050875 | 0.71820487 | 3.724945049 |
| Gm14586 | 42.19938222 | 0.604507376 | 0.222121669 |
| Gm7851 | 0 | NA | NA |
| Ccdc160 | 20.72630267 | -0.111317331 | 0.254005251 |
| Gm14607 | 0.060174537 | 0.237309859 | 3.724945049 |
| Gm6539 | 0 | NA | NA |
| Rps2-ps13 | 156.9088428 | 0.414575334 | 0.153283794 |
| Phf6 | 556.50205 | 0.361681889 | 0.082273847 |
| Hprt | 957.8882762 | -0.094543144 | 0.076803765 |
| Mir450b | 0 | NA | NA |
| Mir450-1 | 0 | NA | NA |
| Mir450-2 | 0 | NA | NA |
| Mir542 | 0 | NA | NA |
| Gm28730 | 0.097899064 | -0.724453471 | 3.724945049 |
| Mir351 | 0 | NA | NA |

| | | | |
|---|---|---|---|
| Mir503 | 0.047948698 | -0.243569582 | 3.724945049 |
| Mir322 | 0 | NA | NA |
| C430049B03Rik | 3.09373086 | -0.798481753 | 0.570706157 |
| Plac1 | 0 | NA | NA |
| Fam122b | 69.8401722 | -0.407490545 | 0.214681561 |
| Fam122c | 0 | NA | NA |
| Gm23718 | 0 | NA | NA |
| Gm14585 | 0 | NA | NA |
| Mospd1 | 150.5973759 | 0.299649502 | 0.11062239 |
| Gm14584 | 111.7814324 | -0.004040922 | 0.118371119 |
| Etd | 0 | NA | NA |
| Gm14597 | 0.48133134 | 1.642822511 | 1.481089536 |
| Cxx1c | 578.0914899 | -0.528100755 | 0.10775345 |
| Gm14592 | 0.289161111 | -0.322702417 | 1.687854192 |
| Gm14600 | 0 | NA | NA |
| Cxx1a | 186.6998316 | -0.312447918 | 0.094109264 |
| Cxx1b | 332.64126 | -0.347419192 | 0.078740935 |
| 4933416I08Rik | 0.129625439 | -1.065581744 | 3.675612039 |
| AW822252 | 37.35059821 | -0.64981447 | 0.248089806 |
| 4930502E18Rik | 2.580172256 | -0.952531281 | 0.6579035 |
| 1700013H16Rik | 0.066050875 | 0.71820487 | 3.724945049 |
| Gm6025 | 0 | NA | NA |
| Gm14593 | 0 | NA | NA |
| Zfp36l3 | 0.434206664 | -2.357736664 | 1.43981746 |
| Xlr | 8.912034157 | 1.419849978 | 0.414841221 |
| Gm14599 | 0.044544648 | 0.237309859 | 3.724945049 |
| Gm14589 | 0 | NA | NA |
| Gm1862 | 2.065160235 | -0.031342657 | 0.648002834 |
| Gm14591 | 0.119587853 | 0.237305575 | 3.724945049 |
| Gm15482 | 0.887798776 | -1.946343909 | 1.229484051 |
| Gm14601 | 0 | NA | NA |
| Gm14644 | 0 | NA | NA |
| Gm22550 | 0 | NA | NA |
| Gm16428 | 0 | NA | NA |
| Gm7916 | 1.902020358 | -0.165945642 | 0.790132254 |
| Gm14598 | 0.066050875 | 0.71820487 | 3.724945049 |
| Gm6591 | 0 | NA | NA |
| Gm14596 | 0 | NA | NA |
| Gm14594 | 0 | NA | NA |
| Gm16405 | 0 | NA | NA |
| Gm16430 | 0 | NA | NA |
| Gm14595 | 0 | NA | NA |
| Gm14590 | 0 | NA | NA |
| Gm16431 | 0 | NA | NA |
| Gm7950 | 0 | NA | NA |
| Gm7958 | 0 | NA | NA |
| Gm7974 | 0 | NA | NA |

| | | | |
|---|---|---|---|
| Slxl1 | 0 | NA | NA |
| Gm6660 | 0 | NA | NA |
| Gm14625 | 0 | NA | NA |
| Gm16404 | 0 | NA | NA |
| Gm14622 | 0 | NA | NA |
| Gm6664 | 0 | NA | NA |
| Gm14626 | 0 | NA | NA |
| Gm7927 | 0 | NA | NA |
| Gm14623 | 0 | NA | NA |
| Gm14624 | 0 | NA | NA |
| Gm22103 | 0.038981395 | 0.718207094 | 3.724945049 |
| Gm14638 | 0 | NA | NA |
| 3830403N18Rik | 1.591056397 | 2.363175973 | 1.038109798 |
| Gm14639 | 1.176771261 | 1.015963804 | 1.125835254 |
| Gm773 | 0 | NA | NA |
| Gm14627 | 0 | NA | NA |
| 1600025M17Rik | 0 | NA | NA |
| Zfp449 | 65.59676692 | -0.070032334 | 0.133047687 |
| Gm2155 | 3.901369078 | -0.92057846 | 0.56268226 |
| Smim10l2a | 178.3604988 | -0.723351926 | 0.112170275 |
| Gm14633 | 2.099444623 | -0.356983209 | 0.741637717 |
| Gm2174 | 0.09349418 | -0.24357403 | 3.724945049 |
| Ddx26b | 661.1565071 | -0.707633599 | 0.132503526 |
| Gm10477 | 0 | NA | NA |
| Gm648 | 0.240593192 | -1.051187001 | 2.543642901 |
| Mmgt1 | 503.4834972 | -0.130580406 | 0.063587226 |
| Mir5116 | 0 | NA | NA |
| Slc9a6 | 1377.918565 | -0.625449906 | 0.057851145 |
| Gm26312 | 0 | NA | NA |
| Fhl1 | 841.3587556 | -0.186397859 | 0.059385544 |
| Gm23258 | 1.567065625 | -1.108574054 | 0.80264771 |
| Mtap7d3 | 3.192641839 | 1.141917464 | 0.625279056 |
| Adgrg4 | 0.589632284 | 1.424196557 | 1.388164606 |
| Brs3 | 2.988176321 | -0.426669218 | 0.74092679 |
| Htatsf1 | 484.4912738 | -0.115931102 | 0.128892238 |
| Vgll1 | 0 | NA | NA |
| Gm14718 | 0.17796667 | -0.21293993 | 2.712315561 |
| Cd40lg | 0.63650561 | 2.120470484 | 1.350164343 |
| Arhgef6 | 229.7557389 | 0.044371471 | 0.085286016 |
| Rbmx | 637.3674608 | -0.442506833 | 0.095078525 |
| Snord61 | 0.989940444 | -0.138812282 | 1.069894478 |
| Gm364 | 0.352737941 | 2.515092076 | 3.685508193 |
| Gpr101 | 120.5546104 | -0.704363154 | 0.152342683 |
| Gm14630 | 0.097139547 | 0.718207094 | 3.724945049 |
| Gm5389 | 0 | NA | NA |
| Gm14629 | 0 | NA | NA |
| Gm8107 | 0.038981395 | 0.718207094 | 3.724945049 |

| | | | |
|---|---|---|---|
| Gm8111 | 0 | NA | NA |
| Gm14631 | 0 | NA | NA |
| Gm25023 | 0 | NA | NA |
| Zic3 | 63.76716955 | -0.368006144 | 0.156629216 |
| Gm14649 | 0 | NA | NA |
| Gm14647 | 0 | NA | NA |
| Gm25851 | 0 | NA | NA |
| Gm14645 | 0 | NA | NA |
| Gm16436 | 0.046345709 | 0.71820487 | 3.724945049 |
| Gm14650 | 0.045882455 | 0.237312084 | 3.724945049 |
| Gm7071 | 0 | NA | NA |
| AL713977.1 | 0 | NA | NA |
| 4930550L24Rik | 0.32299001 | 1.328405168 | 1.925400248 |
| Fgf13 | 698.6388779 | -1.25051333 | 0.104663885 |
| Gm26351 | 1.389516516 | -1.686508952 | 1.085452594 |
| Mir504 | 1.452681833 | -0.62076027 | 0.820222635 |
| Gm26487 | 0 | NA | NA |
| Gm14890 | 0.104106287 | -0.243576254 | 3.724945049 |
| AL670091.1 | 0 | NA | NA |
| AL672036.1 | 0 | NA | NA |
| Gm14887 | 0 | NA | NA |
| F9 | 3.199446176 | 2.436367911 | 0.921348404 |
| Gm5637 | 10.86361093 | 3.077114606 | 0.449591864 |
| Gm14891 | 0.066050875 | 0.71820487 | 3.724945049 |
| Mcf2 | 115.7996196 | -0.621920897 | 0.186178027 |
| Atp11c | 543.5317974 | 0.156695465 | 0.082533031 |
| Gm22287 | 0 | NA | NA |
| Mir505 | 2.930256519 | -0.419039238 | 0.610851066 |
| Gm7073 | 0 | NA | NA |
| Gm715 | 17.31431649 | -0.648715178 | 0.241374938 |
| Gm14661 | 0.27928303 | -0.716348936 | 2.075376776 |
| Gm14651 | 0 | NA | NA |
| Gm4909 | 0 | NA | NA |
| Gm8163 | 0.296485438 | 0.259905725 | 2.092361166 |
| Gm14660 | 0 | NA | NA |
| Sox3 | 13.97423569 | 0.274832009 | 0.287447807 |
| Gm14662 | 1.313401919 | -1.186569024 | 0.968068162 |
| Gm7076 | 0 | NA | NA |
| Rpl7l1-ps1 | 0 | NA | NA |
| Gm14664 | 0.089814205 | -0.24357403 | 3.724945049 |
| Gm14663 | 0.061920613 | 0.718207094 | 3.724945049 |
| C230004F18Rik | 1115.002913 | -0.693271386 | 0.161767007 |
| Gm24396 | 21.15961452 | -0.530838965 | 0.464518794 |
| Cdr1 | 62.47634879 | 0.271655979 | 0.484257869 |
| Gm14665 | 0.120974994 | -0.243571806 | 3.724945049 |
| Gm14666 | 0.43984171 | 1.158058185 | 1.455835877 |
| Gm14821 | 0.106112162 | 0.23732098 | 3.724945049 |

| | | | |
|---|---|---|---|
| Gm14655 | 0.060174537 | 0.237309859 | 3.724945049 |
| Ldoc1 | 14.5236453 | -0.89744279 | 0.287556814 |
| Gm7327 | 29.50965119 | 0.064916725 | 0.303460425 |
| Gm6743 | 0.441426087 | 0.411931458 | 1.481182915 |
| Gm25412 | 0 | NA | NA |
| Gm8215 | 0 | NA | NA |
| Gm5390 | 0 | NA | NA |
| Gm14656 | 0 | NA | NA |
| Gm16411 | 0.044191549 | -0.243571806 | 3.724945049 |
| 4933402E13Rik | 0 | NA | NA |
| Gm14679 | 0 | NA | NA |
| 4931400O07Rik | 0 | NA | NA |
| 1700019B21Rik | 0 | NA | NA |
| Gm14678 | 0 | NA | NA |
| Gm14677 | 0 | NA | NA |
| Gm14680 | 169.5303447 | -0.320668179 | 0.094135701 |
| Gm8244 | 0 | NA | NA |
| Gm6754 | 0 | NA | NA |
| Gm14659 | 0 | NA | NA |
| Gm14657 | 0 | NA | NA |
| Gm8260 | 0 | NA | NA |
| Gm6760 | 0 | NA | NA |
| 3830417A13Rik | 0 | NA | NA |
| Slitrk4 | 1198.015236 | -0.109361083 | 0.098470805 |
| Gm24598 | 1.413287166 | -1.4317448 | 0.869418173 |
| Gm14676 | 0 | NA | NA |
| Gm26111 | 0 | NA | NA |
| Gm14667 | 0 | NA | NA |
| Gm14673 | 0 | NA | NA |
| Ctag2 | 0 | NA | NA |
| Gm14669 | 0 | NA | NA |
| Gm14670 | 0 | NA | NA |
| Gm14668 | 0 | NA | NA |
| Gm14671 | 0 | NA | NA |
| Gm14674 | 0 | NA | NA |
| 4930447F04Rik | 0.792081435 | -0.006974503 | 1.190293476 |
| Gm23028 | 0 | NA | NA |
| Slitrk2 | 714.9978112 | -0.155581041 | 0.096640097 |
| Gm23000 | 0.301026439 | 1.214841724 | 2.043748766 |
| Gm8310 | 0.044544648 | 0.237309859 | 3.724945049 |
| Mir743 | 0 | NA | NA |
| Mir743b | 0 | NA | NA |
| Gm14681 | 55.50849165 | 0.708578475 | 0.144875069 |
| Mir742 | 0 | NA | NA |
| Mir883a | 0 | NA | NA |
| Mir883b | 0 | NA | NA |
| Mir471 | 0 | NA | NA |

| | | | |
|---|---|---|---|
| Mir741 | 0 | NA | NA |
| Mir463 | 0 | NA | NA |
| Mir880 | 0 | NA | NA |
| Mir878 | 0 | NA | NA |
| Mir881 | 0 | NA | NA |
| Mir871 | 0 | NA | NA |
| Mir470 | 0 | NA | NA |
| mmu-mir-465d | 0 | NA | NA |
| Mir465c-1 | 0 | NA | NA |
| Mir465b-1 | 0 | NA | NA |
| Mir465c-2 | 0 | NA | NA |
| Mir465b-2 | 0 | NA | NA |
| Mir465 | 0 | NA | NA |
| Gm6783 | 0 | NA | NA |
| Gm14682 | 0.089089296 | 0.237305797 | 3.724945049 |
| Gm8312 | 0 | NA | NA |
| 1700036O09Rik | 0 | NA | NA |
| Gm26276 | 0 | NA | NA |
| Gm14687 | 0 | NA | NA |
| Gm1140 | 0 | NA | NA |
| Gm14692 | 0 | NA | NA |
| Rpl7a-ps13 | 0 | NA | NA |
| 4933436I01Rik | 0 | NA | NA |
| Mir201 | 0 | NA | NA |
| Mir547 | 0 | NA | NA |
| Gm14688 | 0 | NA | NA |
| mmu-mir-509 | 0 | NA | NA |
| Gm4910 | 0 | NA | NA |
| Gm5391 | 0 | NA | NA |
| Gm6806 | 0.624564003 | 2.690435742 | 1.349446583 |
| Gm22332 | 0 | NA | NA |
| Gm14701 | 0 | NA | NA |
| Gm5638 | 0.108772582 | 0.718202646 | 3.724945049 |
| Fmr1os | 1.106201151 | 0.291156421 | 1.155450093 |
| Gm27979 | 0 | NA | NA |
| Gm27838 | 0 | NA | NA |
| Fmr1 | 1356.393382 | 0.394855821 | 0.076301326 |
| Gm22435 | 0 | NA | NA |
| Fmr1nb | 0.234997993 | 1.192781842 | 2.125031746 |
| Gm14699 | 0 | NA | NA |
| AC055766.1 | 0 | NA | NA |
| Gm14698 | 5.615076243 | -0.050702595 | 0.399849289 |
| Gm6812 | 0 | NA | NA |
| Gm8344 | 0 | NA | NA |
| n-R5s7 | 0 | NA | NA |
| Gm14705 | 0.713101696 | -1.805914052 | 1.260918914 |
| Aff2 | 361.4846116 | -0.360827615 | 0.088164661 |

| | | | |
|---|---|---|---|
| Gm8172 | 2.258517424 | 0.483181597 | 0.727643085 |
| 1700111N16Rik | 0.153668717 | -0.243571235 | 2.839403532 |
| 1700020N15Rik | 0.116217011 | -0.24357403 | 3.724945049 |
| Ids | 5255.302793 | -0.599408935 | 0.070711963 |
| 1110012L19Rik | 65.12692351 | 0.386108807 | 0.185278437 |
| 4930567H17Rik | 0 | NA | NA |
| Gm16407 | 0 | NA | NA |
| Gm14704 | 0 | NA | NA |
| Gm6823 | 0.187945415 | 1.067735365 | 2.675122609 |
| Gm16441 | 0 | NA | NA |
| Gm14722 | 0 | NA | NA |
| Gm14721 | 0 | NA | NA |
| BC023829 | 354.2798382 | -0.046131751 | 0.094864744 |
| Gm14727 | 0 | NA | NA |
| Gm14729 | 0 | NA | NA |
| Gm6038 | 0 | NA | NA |
| Gm8401 | 0.88643884 | 1.402605659 | 1.06174805 |
| Gm14726 | 0 | NA | NA |
| Gm14724 | 0 | NA | NA |
| Gm14725 | 0 | NA | NA |
| Gm14728 | 0.144050924 | 0.237309606 | 3.724945049 |
| Gm14732 | 0 | NA | NA |
| Gm14731 | 0 | NA | NA |
| Rplp1-ps1 | 0.046345709 | 0.71820487 | 3.724945049 |
| Gm22785 | 0 | NA | NA |
| Gm8410 | 0.503546025 | -1.661130632 | 1.518951132 |
| Gm6062 | 0.075043205 | 0.237309859 | 3.724945049 |
| Mamld1 | 657.3894617 | -0.095024279 | 0.0948845 |
| n-R5s8 | 8.139267235 | -0.838467575 | 0.399159903 |
| Gm14730 | 1.043158539 | -0.381759392 | 1.11759586 |
| Mtm1 | 330.8783525 | -0.072851605 | 0.135524825 |
| Mtmr1 | 659.908289 | -0.020068234 | 0.074868838 |
| Cd99l2 | 1057.193581 | -0.073621884 | 0.055532877 |
| Gm16189 | 0 | NA | NA |
| Gm8421 | 0 | NA | NA |
| Gm23945 | 0 | NA | NA |
| Hmgb3 | 158.3977876 | -0.432995244 | 0.172893437 |
| Gpr50 | 5.432542524 | -0.341909303 | 0.398579445 |
| Gm14723 | 0 | NA | NA |
| Rpl30-ps10 | 27.35008078 | 0.464501369 | 0.217047201 |
| Vma21 | 809.8468664 | 0.069737504 | 0.10151667 |
| Gm1141 | 6.002430894 | 0.126593588 | 0.418031341 |
| Prrg3 | 424.0651492 | -0.677879497 | 0.1083423 |
| Fate1 | 0.616736355 | -0.130935337 | 1.189648256 |
| Cnga2 | 3.751482543 | -0.185861116 | 0.743136024 |
| Magea4 | 0 | NA | NA |
| Gabre | 35.49517603 | 0.320777108 | 0.249150806 |

| | | | |
|---|---|---|---|
| Mir224 | 0.097139547 | 0.718207094 | 3.724945049 |
| Mir452 | 0.132574648 | -1.082845688 | 3.713921268 |
| Magea10 | 0 | NA | NA |
| Gabra3 | 880.1056003 | -0.933165382 | 0.108232393 |
| Gm14735 | 0 | NA | NA |
| Mir767 | 0.267172041 | -1.96396915 | 1.977856494 |
| Mir105 | 0.580314088 | -1.670077099 | 1.508072884 |
| Gm23461 | 0 | NA | NA |
| Gm14736 | 0 | NA | NA |
| Gabrq | 110.4557477 | -0.084297893 | 0.176473864 |
| Magea9-ps | 0 | NA | NA |
| Cetn2 | 224.1480122 | -0.12790137 | 0.087371865 |
| Nsdhl | 258.6958972 | 0.175294802 | 0.0932636 |
| Gm14684 | 0.411174732 | -2.238028173 | 1.632728895 |
| Zfp185 | 38.8421771 | 1.008793809 | 0.201880707 |
| Mir2137 | 0.131193343 | -0.71236593 | 2.978303909 |
| Pnma5 | 2.351169846 | -0.977254118 | 0.724815143 |
| Pnma3 | 162.7565536 | -1.121672987 | 0.108847028 |
| Xlr4a | 1.613791847 | 2.3083355 | 1.114407219 |
| Xlr3a | 10.70218797 | 0.43952211 | 0.339671952 |
| Gm20455 | 0 | NA | NA |
| Xlr5a | 0 | NA | NA |
| Gm14685 | 1.153352726 | 0.747005021 | 0.936853733 |
| DXBay18 | 1.088618379 | 1.429147838 | 0.984755143 |
| Xlr5b | 0.044544648 | 0.237309859 | 3.724945049 |
| Spin2d | 0 | NA | NA |
| Xlr3b | 10.24179207 | 1.00672252 | 0.854605017 |
| Xlr4b | 4.01574399 | 4.489809473 | 0.961415104 |
| F8a | 148.8077908 | 0.136756061 | 0.146903332 |
| Xlr4c | 1.800570423 | 4.123931068 | 1.059444202 |
| Xlr3c | 1.249216686 | 3.310409367 | 1.134586233 |
| Xlr5c | 0 | NA | NA |
| Gm5639 | 0.465319851 | -1.201159105 | 1.442065969 |
| Gm8501 | 0 | NA | NA |
| Zfp275 | 599.4099204 | 0.153435642 | 0.072166681 |
| Gm18336 | 1.728761539 | 1.591292391 | 0.945382615 |
| Gm26726 | 0.597994088 | 0.72566574 | 1.605583713 |
| Gm37741 | 0.240513512 | 0.237320228 | 2.504347404 |
| Zfp92 | 61.40182776 | 0.541241004 | 0.163324435 |
| Trex2 | 0.567095806 | 1.016082706 | 1.292067175 |
| Haus7 | 77.87449152 | 0.231004705 | 0.117637139 |
| Bgn | 275.5973356 | 1.581732683 | 0.182166471 |
| Atp2b3 | 1878.820538 | -0.80802213 | 0.092961672 |
| Dusp9 | 7.63352917 | 0.759304474 | 0.401931355 |
| Pnck | 377.2175805 | -0.912982572 | 0.076790645 |
| Slc6a8 | 1645.774035 | 0.712233703 | 0.063960991 |
| Bcap31 | 456.597487 | 0.369644385 | 0.105974368 |

| | | | |
|---|---|---|---|
| Abcd1 | 224.9704374 | 0.887271102 | 0.117818208 |
| Gm25226 | 0 | NA | NA |
| Plxnb3 | 370.4438468 | 0.003703595 | 0.119042994 |
| Srpk3 | 30.04135515 | -0.634220149 | 0.235875535 |
| Idh3g | 816.0853577 | -0.274435519 | 0.049895043 |
| Ssr4 | 382.7763301 | 0.473868251 | 0.100719144 |
| Pdzd4 | 1049.261182 | -0.534612411 | 0.084339417 |
| Gm14817 | 2.398130883 | -0.874072333 | 0.835593217 |
| Gm23615 | 0.046345709 | 0.71820487 | 3.724945049 |
| L1cam | 1414.37122 | -0.9242762 | 0.066546032 |
| Arhgap4 | 118.1046268 | 0.772580619 | 0.173681774 |
| Avpr2 | 0.170941961 | 1.167085536 | 2.735055532 |
| Naa10 | 112.0862865 | -0.522804681 | 0.116238704 |
| Renbp | 87.59544553 | 1.917659556 | 0.182257179 |
| Hcfc1 | 2331.747261 | 0.011718105 | 0.05432422 |
| Gm8545 | 0.428564514 | 0.779253696 | 1.474449054 |
| Irak1 | 580.922968 | 0.342793025 | 0.067348889 |
| Mir5132 | 0 | NA | NA |
| Mir718 | 0.042716945 | -0.243571806 | 3.724945049 |
| Mecp2 | 3093.872843 | -0.165249059 | 0.050794323 |
| Opn1mw | 3.912128538 | -0.149271101 | 0.552711065 |
| Gm6758 | 1.061458158 | 0.392298185 | 0.913935815 |
| Tex28 | 0.045882455 | 0.237312084 | 3.724945049 |
| Rpl3-ps2 | 16.62506886 | 0.721677584 | 0.271085242 |
| Tktl1 | 1.027298454 | 1.788288781 | 1.258512205 |
| Flna | 2140.441872 | 2.025475648 | 0.140522451 |
| Emd | 326.2260813 | 0.679674698 | 0.116091153 |
| Gm14686 | 0 | NA | NA |
| Rpl10 | 1410.009698 | 0.727843928 | 0.096252383 |
| Snora70 | 0 | NA | NA |
| Dnase1l1 | 38.9238987 | 2.673357057 | 0.263451587 |
| Mir7091 | 0 | NA | NA |
| Gm14846 | 0 | NA | NA |
| Taz | 323.2193373 | -0.381714875 | 0.128120608 |
| Atp6ap1 | 2828.059804 | -0.010551881 | 0.055740598 |
| Gdi1 | 4514.466658 | -0.420804734 | 0.044952853 |
| Fam50a | 152.1449703 | 0.2921927 | 0.103924034 |
| mmu-mir-7092 | 0.044191549 | -0.243571806 | 3.724945049 |
| Plxna3 | 360.6800962 | -0.473168375 | 0.091756764 |
| Lage3 | 80.66907545 | 0.286282196 | 0.147796821 |
| Ubl4 | 481.5895494 | 0.029860072 | 0.078518489 |
| Slc10a3 | 39.38722088 | 1.398762212 | 0.226404069 |
| Fam3a | 178.954657 | -0.068967398 | 0.129984399 |
| Ikbkg | 603.2886534 | 0.336603978 | 0.079289993 |
| G6pdx | 686.6621862 | 1.564348668 | 0.149683077 |
| Gm6880 | 0.066050875 | 0.71820487 | 3.724945049 |
| Gm6881 | 0 | NA | NA |

| | | | |
|---|---|---|---|
| Gm15361 | 20.55718455 | -0.171090874 | 0.2559407 |
| Gm15362 | 0 | NA | NA |
| Gm15364 | 0 | NA | NA |
| Gm26125 | 0 | NA | NA |
| Gm4989 | 0 | NA | NA |
| Olfr1326-ps1 | 0 | NA | NA |
| Olfr1325 | 0 | NA | NA |
| Gm26417 | 0 | NA | NA |
| Gm5640 | 1.316030826 | -0.217380232 | 0.93728305 |
| Xlr5d-ps | 0.066050875 | 0.71820487 | 3.724945049 |
| Spin2-ps2 | 0 | NA | NA |
| Xlr3d-ps | 0.16282664 | 1.267090096 | 3.411118614 |
| Gm15365 | 0.152853976 | 0.237319074 | 2.829491075 |
| Gm22351 | 0 | NA | NA |
| Xlr4d-ps | 0 | NA | NA |
| Gm6890 | 0 | NA | NA |
| Gm14755 | 0 | NA | NA |
| Olfr1327-ps1 | 0 | NA | NA |
| Gm14758 | 0 | NA | NA |
| Gm14695 | 0 | NA | NA |
| Gm8617 | 0 | NA | NA |
| Spin2-ps3 | 0 | NA | NA |
| Xlr5e-ps | 0 | NA | NA |
| Gm5936 | 1.015210553 | 1.32434804 | 1.02516521 |
| Gm14694 | 0 | NA | NA |
| Gm6897 | 0 | NA | NA |
| Gm14756 | 0 | NA | NA |
| Spin2-ps4 | 0 | NA | NA |
| Xlr3e-ps | 0.045882455 | 0.237312084 | 3.724945049 |
| Xlr4e-ps | 0.114443069 | 0.23732098 | 3.724945049 |
| Gm15363 | 0 | NA | NA |
| Gab3 | 34.85498717 | 0.215008765 | 0.200144451 |
| Gm25629 | 0.361882898 | 0.249845146 | 1.901100364 |
| Gm15367 | 0 | NA | NA |
| Dkc1 | 249.3852457 | 0.141292117 | 0.087602078 |
| Gm25520 | 1.152914642 | -1.098459978 | 0.978411252 |
| Mpp1 | 513.8380435 | 0.829872053 | 0.096744932 |
| Gm15378 | 0.117046891 | -0.724453471 | 3.724945049 |
| Fam220-ps | 0.313388536 | 0.237329427 | 2.041201642 |
| Smim9 | 2.883149428 | 1.878155144 | 0.707108642 |
| Gm8503 | 0.097139547 | 0.718207094 | 3.724945049 |
| F8 | 100.3533705 | -0.220412832 | 0.152910633 |
| Gm6039 | 5.682746135 | 0.338516332 | 0.414943379 |
| Gm8522 | 1.810659408 | -1.606222204 | 0.83282149 |
| Fundc2 | 533.8816821 | -0.096466602 | 0.105513816 |
| Mtcp1 | 186.8893314 | -0.312474566 | 0.130727156 |
| Brcc3 | 516.7646188 | -0.051668059 | 0.074783695 |

| | | | |
|---|---|---|---|
| Gm15379 | 11.17773374 | 0.135632032 | 0.313899606 |
| Gm15385 | 2.75264805 | 0.773921802 | 0.756593555 |
| Gm8666 | 2.699454663 | 1.10776612 | 0.578899789 |
| Vbp1 | 703.5174424 | 0.094681187 | 0.094251545 |
| Gm15384 | 1.213939271 | -0.558052044 | 0.968081206 |
| Rab39b | 757.2087334 | 0.204624117 | 0.088770167 |
| Gm15063 | 0.98952171 | 3.235831379 | 1.208611206 |
| Gm25421 | 0 | NA | NA |
| Gm15065 | 1.438276361 | 0.703145695 | 0.725560622 |
| 4933407K13Rik | 98.06458889 | -0.04760859 | 0.257746093 |
| Gm7153 | 0.141331096 | 0.237322366 | 3.56184083 |
| Pls3 | 1303.00074 | -0.116471248 | 0.061966994 |
| Gm23320 | 0 | NA | NA |
| Oat-rs1 | 0 | NA | NA |
| Gm14715 | 0 | NA | NA |
| Gm7155 | 0 | NA | NA |
| Gm8692 | 1.34351349 | 0.441153142 | 0.868729967 |
| Gm14710 | 0 | NA | NA |
| Gm5397 | 0 | NA | NA |
| Gm14711 | 0.062426873 | 0.237312084 | 3.724945049 |
| Gm14707 | 0 | NA | NA |
| Gm14717 | 0 | NA | NA |
| Cldn34b3 | 0.209261529 | 1.185614193 | 2.617572548 |
| Gm14712 | 0 | NA | NA |
| Magea7-ps | 0.046345709 | 0.71820487 | 3.724945049 |
| Gm14709 | 0 | NA | NA |
| Cldn34b4 | 0 | NA | NA |
| Gm5793 | 0.038981395 | 0.718207094 | 3.724945049 |
| Cldn34d | 0.044191549 | -0.243571806 | 3.724945049 |
| Gm14716 | 0 | NA | NA |
| Gm14713 | 0 | NA | NA |
| Gm26041 | 0.046345709 | 0.71820487 | 3.724945049 |
| Gm23280 | 0 | NA | NA |
| Gm14706 | 0.190396633 | 0.718208389 | 3.269807176 |
| Gm14714 | 0 | NA | NA |
| Gm5398 | 2.350384682 | -1.071706668 | 0.786188172 |
| Gm5757 | 0 | NA | NA |
| Gm5937 | 0.810948363 | 0.25155045 | 1.151098002 |
| Tbl1x | 1255.905752 | 0.29151545 | 0.053187283 |
| Gm23121 | 6.016926621 | -0.144239171 | 0.408807805 |
| Cldn34-ps | 0 | NA | NA |
| Gm14746 | 0.224947322 | -1.168296805 | 2.157985137 |
| Gm14748 | 0 | NA | NA |
| Gm8722 | 2.37024928 | 0.183449805 | 0.600065003 |
| Prkx | 384.5278709 | 1.203642928 | 0.138747709 |
| Gm14742 | 0.093141081 | -0.724455696 | 3.724945049 |
| Pbsn | 0 | NA | NA |

| | | | |
|---|---|---|---|
| Gm14744 | 0.130840244 | -1.076724316 | 2.947200097 |
| 5430402E10Rik | 0.090665643 | -0.724455696 | 3.724945049 |
| Gm14750 | 0 | NA | NA |
| Gm14753 | 0 | NA | NA |
| Gm16458 | 0 | NA | NA |
| Obp1a | 0.143846094 | -1.022239892 | 3.718482058 |
| Gm5938 | 0 | NA | NA |
| Obp1b | 0.091880448 | -0.724455696 | 3.724945049 |
| Gm14743 | 0.116046057 | -0.724453471 | 3.724945049 |
| Gm8749 | 0 | NA | NA |
| 4930480E11Rik | 0.066050875 | 0.71820487 | 3.724945049 |
| Gm14749 | 0 | NA | NA |
| Gm8754 | 0 | NA | NA |
| Gm14752 | 0 | NA | NA |
| Gm14745 | 0 | NA | NA |
| Prrg1 | 108.3946446 | -1.125630496 | 0.155470774 |
| Gm22843 | 1.022987396 | -2.417459688 | 1.116347589 |
| Gm8757 | 0 | NA | NA |
| Gm8761 | 0 | NA | NA |
| Gm14747 | 0 | NA | NA |
| Fam47c | 0 | NA | NA |
| Gm8770 | 0.097139547 | 0.718207094 | 3.724945049 |
| Gm5939 | 0.62385056 | 0.987375897 | 1.298725037 |
| Gm23556 | 0 | NA | NA |
| Gm14751 | 0 | NA | NA |
| Rps24-ps3 | 11.66239001 | 0.441585885 | 0.303682636 |
| Gm7173 | 57.33673308 | -0.035271484 | 0.159580482 |
| Mageb16 | 1.972874096 | 0.766063807 | 0.784566787 |
| Gm4772 | 0.201649101 | -0.690705636 | 2.200631607 |
| Gm4773 | 0.396881693 | -0.243564447 | 1.604822613 |
| Gm14760 | 16.61937256 | 0.398547998 | 0.260622156 |
| Gm6952 | 0 | NA | NA |
| Gm14768 | 0 | NA | NA |
| Gm5940 | 3.69570757 | 0.649081824 | 0.550550558 |
| Gm8806 | 5.085933841 | 1.141203531 | 0.48278881 |
| Gm6954 | 0 | NA | NA |
| Gm14733 | 0 | NA | NA |
| Gm26775 | 0.462969528 | -1.453603743 | 1.489201447 |
| Tmem47 | 1344.966627 | -0.060708698 | 0.128210241 |
| Gm14754 | 0 | NA | NA |
| 4930595M18Rik | 0.072025462 | 0.237312084 | 3.724945049 |
| Gm8822 | 164.0860792 | 0.682903519 | 0.130949579 |
| Gm26174 | 0 | NA | NA |
| Gm22686 | 0 | NA | NA |
| Rpl10-ps4 | 0 | NA | NA |
| Gm24706 | 0 | NA | NA |
| Gm14740 | 12.83618259 | -2.498996302 | 0.819005316 |

| | | | |
|---|---|---|---|
| Gm14739 | 0 | NA | NA |
| Gm24470 | 0 | NA | NA |
| Gm14741 | 0 | NA | NA |
| Gm23671 | 0 | NA | NA |
| Dmd | 2142.472415 | -0.426213611 | 0.100711179 |
| n-R5s9 | 0.908782567 | 0.642105538 | 1.1189952 |
| Tsga8 | 0 | NA | NA |
| Gm22580 | 0.283050572 | 0.225170817 | 1.64303825 |
| Gm14769 | 8.178611007 | -2.023449888 | 0.536183666 |
| Gm14770 | 2.258554499 | -1.110247847 | 0.652071213 |
| Gm14771 | 0 | NA | NA |
| Gm16463 | 0.470240119 | 1.943669278 | 1.510998111 |
| 1600014K23Rik | 0 | NA | NA |
| Gm14765 | 0 | NA | NA |
| Tab3 | 521.751977 | 0.244978743 | 0.090736132 |
| AL645586.1 | 2.461055123 | -0.870454032 | 0.805615347 |
| Gyk | 504.0204661 | 0.146341905 | 0.056165007 |
| Gm14767 | 0.044544648 | 0.237309859 | 3.724945049 |
| Gm14764 | 0 | NA | NA |
| Gm14766 | 0.576946983 | -0.382766275 | 1.161854373 |
| Gm6964 | 0 | NA | NA |
| Gm14761 | 0.106520246 | 0.718200421 | 3.724945049 |
| Gm14762 | 0.702946434 | -0.779104695 | 1.312743602 |
| 5430427O19Rik | 37.85834025 | 4.038000074 | 0.329359867 |
| Gm14772 | 0 | NA | NA |
| Samt3 | 0.047948698 | -0.243569582 | 3.724945049 |
| Nr0b1 | 0.363180662 | 1.005093828 | 1.530392108 |
| Mageb4 | 0.097139547 | 0.718207094 | 3.724945049 |
| Gm8839 | 0 | NA | NA |
| Mageb10-ps | 0 | NA | NA |
| Gm8844 | 0.25781401 | -0.252790842 | 1.827323946 |
| Mageb8-ps | 0 | NA | NA |
| Gm41 | 0 | NA | NA |
| Mageb6-ps | 0 | NA | NA |
| Mageb7-ps | 0 | NA | NA |
| n-R5s10 | 0 | NA | NA |
| Il1rapl1 | 848.3376803 | -0.764297461 | 0.111725216 |
| Gm14775 | 0 | NA | NA |
| Gm8855 | 54.40271974 | -0.937211391 | 0.254893574 |
| Gm14776 | 77.87756624 | -1.380744625 | 0.300522087 |
| Gm24812 | 2.806226544 | -1.531839763 | 0.738004145 |
| Gm14774 | 28.40504812 | -1.301394906 | 0.296418966 |
| Gm25006 | 0 | NA | NA |
| Gm8858 | 0 | NA | NA |
| Gm14793 | 0 | NA | NA |
| Gm8864 | 0.154920219 | -0.243569986 | 2.837065107 |
| Gm14795 | 0 | NA | NA |

| | | | |
|---|---|---|---|
| Gm14792 | 0 | NA | NA |
| Smek3-ps | 0 | NA | NA |
| Gm14794 | 0.838406277 | 0.038086555 | 1.044467241 |
| Gm8876 | 0 | NA | NA |
| Gm27000 | 21.60956527 | -1.503288366 | 0.402260866 |
| Mir1906-2 | 0 | NA | NA |
| Gm27335 | 0 | NA | NA |
| Gm5759 | 0 | NA | NA |
| Gm8880 | 0.146451277 | -0.243562218 | 2.882358805 |
| Gm16412 | 0.045882455 | 0.237312084 | 3.724945049 |
| Gm14791 | 0 | NA | NA |
| Gm14796 | 0 | NA | NA |
| Gm14797 | 0 | NA | NA |
| Gm649 | 0 | NA | NA |
| Gm14805 | 0 | NA | NA |
| Pet2 | 0 | NA | NA |
| Gm14804 | 0 | NA | NA |
| Gm6973 | 0 | NA | NA |
| 4932429P05Rik | 0 | NA | NA |
| Gm7108 | 0.816803644 | -1.103898523 | 1.321670928 |
| Gm24954 | 0 | NA | NA |
| Nanog-ps1 | 0 | NA | NA |
| 4930415L06Rik | 0 | NA | NA |
| Gm6027 | 0 | NA | NA |
| Gm4746 | 0 | NA | NA |
| Gm6977 | 10.13847199 | 0.89760567 | 0.410246699 |
| Gm44 | 0 | NA | NA |
| Gm14773 | 0 | NA | NA |
| Gm24835 | 0 | NA | NA |
| Gm14786 | 0.042716945 | -0.243571806 | 3.724945049 |
| Gm8908 | 0 | NA | NA |
| Gm14788 | 0 | NA | NA |
| Mageb2 | 0.045882455 | 0.237312084 | 3.724945049 |
| Gm14783 | 0.291614092 | 0.718789178 | 1.778722295 |
| Gm5072 | 0 | NA | NA |
| Gm8914 | 0 | NA | NA |
| Gm14787 | 0 | NA | NA |
| Gm8916 | 0 | NA | NA |
| Gm14785 | 0 | NA | NA |
| 1700084M14Rik | 0 | NA | NA |
| Mageb17-ps | 0 | NA | NA |
| Gm14781 | 0 | NA | NA |
| Mageb5 | 0 | NA | NA |
| Gm14782 | 0 | NA | NA |
| Gm7113 | 0 | NA | NA |
| Gm16465 | 0.297592784 | -0.243563564 | 1.623738006 |
| Gm5071 | 0 | NA | NA |

| | | | |
|---|---|---|---|
| Gm14784 | 0 | NA | NA |
| Mageb1 | 0.062426873 | 0.237312084 | 3.724945049 |
| Gm4911 | 0 | NA | NA |
| Gm4912 | 0.306841164 | 2.059308053 | 1.969528009 |
| Mageb18 | 0.274198742 | -0.243564503 | 1.81942087 |
| Gm14777 | 0 | NA | NA |
| Gm5941 | 0 | NA | NA |
| Gm6982 | 0 | NA | NA |
| Gm25332 | 0 | NA | NA |
| Gm24973 | 0.33162218 | -1.447128189 | 1.720545792 |
| 1700003E24Rik | 0 | NA | NA |
| Gm24123 | 0 | NA | NA |
| BC061195 | 0 | NA | NA |
| Gm23454 | 0 | NA | NA |
| Arx | 155.2305208 | -0.487046574 | 0.171512286 |
| Pola1 | 222.1831865 | 0.124925963 | 0.125240273 |
| Gm14780 | 0 | NA | NA |
| Gm23557 | 1.675207292 | -0.886778426 | 0.718229145 |
| Gm6985 | 0 | NA | NA |
| Pcyt1b | 616.1736765 | -0.035641537 | 0.067691173 |
| Pdk3 | 429.9123672 | 0.034465281 | 0.065664898 |
| Supt20-ps | 0 | NA | NA |
| Gm15164 | 0 | NA | NA |
| AU015836 | 0.093831153 | -0.243571806 | 3.724945049 |
| Gm14790 | 0 | NA | NA |
| Gm15165 | 0.044191549 | -0.243571806 | 3.724945049 |
| Gm6987 | 0 | NA | NA |
| Gm14798 | 0.269031123 | 0.237331212 | 1.815493784 |
| Zfx | 737.8114658 | 0.221720948 | 0.093147565 |
| Eif2s3x | 977.9631196 | 0.387303349 | 0.073751966 |
| Klhl15 | 561.3122414 | -0.148202087 | 0.064196252 |
| Mir7673 | 0 | NA | NA |
| Gm5400 | 0.277195307 | 1.668343358 | 2.051685278 |
| Gm14826 | 0 | NA | NA |
| Gm14825 | 0.060174537 | 0.237309859 | 3.724945049 |
| 4932442L08Rik | 2.026572498 | 0.986079629 | 0.759393288 |
| Apoo | 208.2198411 | -0.554948915 | 0.111526093 |
| AL627302.1 | 0 | NA | NA |
| Gm9003 | 0 | NA | NA |
| Gm14827 | 54.27178075 | -0.96381938 | 0.39536144 |
| Gm24595 | 0 | NA | NA |
| Maged1 | 4444.520797 | -0.226095153 | 0.055206064 |
| Gm9005 | 0 | NA | NA |
| Gm14831 | 0.038981395 | 0.718207094 | 3.724945049 |
| Gm5401 | 0.15966113 | 0.871795722 | 3.444418384 |
| Gm9009 | 0.046345709 | 0.71820487 | 3.724945049 |
| Gspt2 | 293.6766047 | -0.254142025 | 0.08442495 |

| | | | |
|---|---|---|---|
| Gm5402 | 0.046345709 | 0.71820487 | 3.724945049 |
| Zxdb | 233.650166 | 0.040148857 | 0.098550944 |
| Gm7061 | 8.775414701 | -0.876855074 | 0.461937904 |
| Gm14830 | 3.497870509 | -2.110100154 | 0.675178745 |
| Rpl13-ps5 | 0.088383099 | -0.72445792 | 3.724945049 |
| Gm26617 | 0.179489757 | 0.237327731 | 2.290582425 |
| Zxda | 5.296744549 | 0.176760116 | 0.430622 |
| Gm371 | 0.049553047 | 0.71820487 | 3.724945049 |
| Spin4 | 40.24504316 | 1.361869793 | 0.238975338 |
| Arhgef9 | 4283.077243 | -0.815248856 | 0.101908567 |
| Gm6092 | 0.047948698 | -0.243569582 | 3.724945049 |
| Gm14807 | 0.075043205 | 0.237309859 | 3.724945049 |
| Gm3712 | 0.119587853 | 0.237305575 | 3.724945049 |
| Gm14806 | 0.045882455 | 0.237312084 | 3.724945049 |
| Amer1 | 313.2545163 | 0.042025006 | 0.100651408 |
| Asb12 | 0.245805814 | -0.241733758 | 2.122612749 |
| Gm14818 | 0 | NA | NA |
| Gm9043 | 0 | NA | NA |
| Zc4h2 | 203.86252 | -0.321521863 | 0.101035245 |
| Zc3h12b | 477.9337351 | -0.622916963 | 0.117626831 |
| 1700010D01Rik | 0 | NA | NA |
| Las1l | 521.4809968 | 0.252725921 | 0.11727644 |
| Msn | 2032.338517 | 3.798915214 | 0.188482627 |
| Gm24718 | 0 | NA | NA |
| Gm14925 | 0.293430937 | -1.189930328 | 2.033018266 |
| Gm9051 | 0 | NA | NA |
| F630028O10Rik | 26.3940196 | 3.25783976 | 0.445469388 |
| Mir223 | 0 | NA | NA |
| Vsig4 | 0.106465261 | 0.718202646 | 3.724945049 |
| Hsf3 | 1.054375976 | -0.072309223 | 1.04501065 |
| Gm14933 | 0.042716945 | -0.243571806 | 3.724945049 |
| Heph | 82.76888646 | 0.521500183 | 0.188275804 |
| Rpl17-ps8 | 13.95711591 | 0.600108304 | 0.27663373 |
| Gm14935 | 0.695213882 | -0.7164976 | 1.160787882 |
| Gpr165 | 370.7971054 | 0.360900252 | 0.130462036 |
| Gm16394 | 0 | NA | NA |
| Gm14803 | 0 | NA | NA |
| Gm9059 | 0 | NA | NA |
| Gm9060 | 0 | NA | NA |
| Pgr15l | 28.47859877 | 0.7428007 | 0.283061734 |
| Gm14800 | 0 | NA | NA |
| Eda2r | 27.60206714 | 3.537166827 | 0.397451379 |
| Gm9062 | 0 | NA | NA |
| Gm14802 | 0.124347486 | 0.719186602 | 3.717245571 |
| Ar | 593.9010294 | -0.073300529 | 0.123298228 |
| Gm14811 | 0 | NA | NA |
| Gm14815 | 0.75340593 | 0.339267473 | 1.292193272 |

| | | | |
|---|---|---|---|
| Gm14810 | 2.604223114 | 1.234106057 | 0.847980922 |
| Ophn1 | 626.2746632 | -0.591909474 | 0.090300676 |
| Gm27554 | 3.06361838 | -1.083163559 | 0.690906633 |
| Gm26000 | 0.038981395 | 0.718207094 | 3.724945049 |
| Yipf6 | 767.4798748 | 0.280682686 | 0.07033312 |
| Gm14813 | 0.038981395 | 0.718207094 | 3.724945049 |
| Stard8 | 229.0060904 | -0.272297406 | 0.159729558 |
| Gm5760 | 0 | NA | NA |
| Efnb1 | 173.9851955 | 0.292890898 | 0.125974156 |
| Gm14812 | 1.182411106 | -0.245322014 | 3.722873619 |
| Gm14809 | 0.233612452 | 0.718216207 | 2.540177828 |
| Gm14808 | 0.179754939 | -0.724462075 | 2.740145766 |
| Pja1 | 1595.105195 | -0.191507743 | 0.082843948 |
| Gm14816 | 0.266066471 | 0.207772176 | 2.088875421 |
| Pcna-ps1 | 0.075043205 | 0.237309859 | 3.724945049 |
| Tmem28 | 262.17291 | -0.272817436 | 0.104918827 |
| Eda | 61.73840336 | -0.797087099 | 0.170791955 |
| Gm14832 | 1.244840331 | -1.746557138 | 1.087701568 |
| Gm14829 | 1.581404248 | -2.210024571 | 0.939767163 |
| Gm14828 | 1.203205628 | -0.27518939 | 0.876103475 |
| Mir676 | 0 | NA | NA |
| Awat2 | 2.157021785 | 1.095274295 | 0.735233934 |
| Otud6a | 0 | NA | NA |
| Igbp1 | 392.0809371 | 0.201154321 | 0.075108885 |
| Dgat2l6 | 5.685626011 | 0.038025309 | 0.506647698 |
| Awat1 | 0.271018715 | 1.269594465 | 1.808828413 |
| Gm6175 | 0 | NA | NA |
| P2ry4 | 1.586926032 | 2.624927635 | 1.078054109 |
| Arr3 | 2.257394224 | -1.898664262 | 0.718237965 |
| Pdzd11 | 261.6832867 | 0.534066814 | 0.08406099 |
| Kif4 | 26.59334305 | 1.410847235 | 0.352231631 |
| Gdpd2 | 50.75686142 | -1.353340156 | 0.279554281 |
| Gm14866 | 1.936928019 | 0.960306343 | 0.784361387 |
| Gm14902 | 0.875624081 | 2.297252349 | 1.175498179 |
| Gm22160 | 0.092691418 | 1.183079489 | 3.724560917 |
| Dlg3 | 1696.534007 | -0.660868959 | 0.089301898 |
| Tex11 | 4.291613721 | -0.044241778 | 0.565778591 |
| Gm24577 | 0 | NA | NA |
| Gm27510 | 0 | NA | NA |
| Gm14901 | 0 | NA | NA |
| Gm9078 | 0 | NA | NA |
| Slc7a3 | 112.9406466 | 0.715843027 | 0.165743195 |
| Snx12 | 585.2370213 | -0.056052148 | 0.067471582 |
| Gm24061 | 0.522088832 | 0.484260176 | 1.282824714 |
| Gm14842 | 0 | NA | NA |
| Gm22472 | 0.04393175 | -0.243571806 | 3.724945049 |
| Foxo4 | 217.5431392 | -0.044423918 | 0.149380576 |

| | | | |
|---|---|---|---|
| Gm614 | 0.87838318 | 2.668184172 | 1.201885357 |
| Gm20489 | 0 | NA | NA |
| Il2rg | 109.8692796 | 6.370216616 | 0.395535355 |
| Med12 | 591.9911033 | 0.302499899 | 0.119949729 |
| Gm21986 | 6.701345333 | -0.474187911 | 0.460563828 |
| Nlgn3 | 1054.489164 | -0.775041351 | 0.093087147 |
| Gjb1 | 235.1220583 | 0.668243643 | 0.28353075 |
| Zmym3 | 1188.784781 | -0.564766107 | 0.060444237 |
| Nono | 1296.303821 | 0.441381159 | 0.056207817 |
| Itgb1bp2 | 18.97263099 | -0.307255368 | 0.432815438 |
| Gm14843 | 1.315890906 | -1.118010102 | 1.098533158 |
| Gm6184 | 0.809862658 | -1.18731146 | 1.309416155 |
| Taf1 | 1383.236683 | -0.103610851 | 0.045713117 |
| Ogt | 3519.482034 | -0.613553946 | 0.125397836 |
| Gm14845 | 0.98816996 | -0.540119848 | 1.118497197 |
| Gm9089 | 0.045882455 | 0.237312084 | 3.724945049 |
| Cxcr3 | 9.848512992 | 6.218246907 | 0.894933274 |
| Tpt1-ps6 | 6.439785805 | 1.011463137 | 0.406035397 |
| Gm9095 | 0 | NA | NA |
| Gm4779 | 0.781258755 | -1.566464317 | 1.068567773 |
| 8030474K03Rik | 0 | NA | NA |
| Gm9097 | 0 | NA | NA |
| H3f3a-ps1 | 4.580768557 | -0.501175218 | 0.612180212 |
| Nhsl2 | 1474.969298 | -0.804384632 | 0.085488405 |
| Gm25828 | 0 | NA | NA |
| Gm27843 | 0 | NA | NA |
| Rgag4 | 516.2445288 | 0.638757756 | 0.13472907 |
| Gm5125 | 0 | NA | NA |
| Pin4 | 43.34765536 | 0.500236857 | 0.297500794 |
| Gm16471 | 0.089089296 | 0.237305797 | 3.724945049 |
| Ercc6l | 15.19884806 | 3.09008693 | 0.461933999 |
| Gm9103 | 0.062426873 | 0.237312084 | 3.724945049 |
| Rps4x | 1364.482336 | 0.753265091 | 0.115194053 |
| Gm14859 | 0 | NA | NA |
| Cited1 | 20.65218563 | 0.599323783 | 0.275828264 |
| Gm14858 | 2.504509112 | -3.301046448 | 0.932916342 |
| Gm26318 | 0.128271896 | -0.243573716 | 3.679089475 |
| Hdac8 | 196.3053901 | -0.453146287 | 0.116151116 |
| Phka1 | 482.7901274 | -0.401830166 | 0.082364871 |
| Gm9109 | 0 | NA | NA |
| Gm14840 | 0 | NA | NA |
| Gm3880 | 0 | NA | NA |
| Gm9112 | 0 | NA | NA |
| Dmrtc1b | 0 | NA | NA |
| Gm9115 | 4.289267201 | 1.943570709 | 0.534204279 |
| Gm14839 | 0 | NA | NA |
| Gm4234 | 0 | NA | NA |

| | | | |
|---|---|---|---|
| Gm9119 | 0 | NA | NA |
| Gm5166 | 0 | NA | NA |
| Gm25358 | 0 | NA | NA |
| Dmrtc1c1 | 0 | NA | NA |
| Gm9126 | 0 | NA | NA |
| Gm16511 | 0 | NA | NA |
| Gm24509 | 0 | NA | NA |
| Dmrtc1c2 | 0 | NA | NA |
| Gm26152 | 0 | NA | NA |
| 1700031F05Rik | 0 | NA | NA |
| Gm9133 | 0 | NA | NA |
| Dmrtc1a | 21.57087845 | -0.842472775 | 0.244117373 |
| 1700011M02Rik | 0.046345709 | 0.71820487 | 3.724945049 |
| Gm14841 | 0 | NA | NA |
| Gm14837 | 0.193036943 | -0.187869281 | 3.246599635 |
| Gm21998 | 0.296810931 | -0.243563109 | 1.775226324 |
| Pabpc1l2b-ps | 6.125206409 | 0.27235578 | 0.404692559 |
| Gm9143 | 0.044191549 | -0.243571806 | 3.724945049 |
| Pabpc1l2a-ps | 3.182304534 | -0.374304662 | 0.543909144 |
| Gm3928 | 0.473680182 | 0.219325689 | 1.333322418 |
| Gm14838 | 0.200176296 | -1.092374469 | 3.251120522 |
| Gm6206 | 202.4849468 | 0.362462512 | 0.121396456 |
| 4930519F16Rik | 7.945419585 | -0.806642112 | 0.382516188 |
| Nap1l2 | 1056.694576 | -0.289293781 | 0.067400223 |
| Gm9050 | 0 | NA | NA |
| Gm14847 | 0 | NA | NA |
| Gm19200 | 0.512704799 | -2.816981962 | 1.492776705 |
| Gm14848 | 4.393179665 | 0.143366667 | 1.384153803 |
| Gm3931 | 0.286297534 | -0.724734198 | 2.057902363 |
| Cdx4 | 2.430558831 | -0.622571958 | 1.17423849 |
| Gm9516 | 0 | NA | NA |
| Chic1 | 1013.690449 | -0.010006669 | 0.065032856 |
| Gm26952 | 0 | NA | NA |
| Tsx | 2.523036181 | 0.69768577 | 0.658042975 |
| Gm26992 | 0 | NA | NA |
| Tsix | 4.036643334 | 0.126334017 | 0.628236271 |
| Xist | 7370.143953 | -0.296994731 | 0.215637538 |
| Gm27733 | 0 | NA | NA |
| Gm27927 | 0 | NA | NA |
| Gm27520 | 0.255832186 | -1.568230554 | 1.783000394 |
| Gm14861 | 0.384616848 | -1.181068229 | 1.47631189 |
| Rps12-ps1 | 0.281032441 | -0.805900377 | 2.034103399 |
| Jpx | 90.00391533 | -0.627906099 | 0.2176068 |
| Gm27875 | 0 | NA | NA |
| Gm9670 | 11.86610392 | -1.305568042 | 0.379149564 |
| Gm14860 | 4.132034165 | -1.051375689 | 0.510555909 |
| Gm9785 | 0.044544648 | 0.237309859 | 3.724945049 |

| | | | |
|---|---|---|---|
| Gm9157 | 5.137711797 | -1.153782493 | 0.547984889 |
| Ftx | 498.1238306 | -0.828109399 | 0.206195735 |
| Mir421 | 14.61544255 | -0.960677195 | 0.379850601 |
| Mir374b | 9.615598131 | -0.845397736 | 0.460811397 |
| mmu-mir-374c | 0 | NA | NA |
| Gm38020 | 450.5061 | -0.974587066 | 0.27701284 |
| Gm27798 | 0.19090462 | 0.718208435 | 3.267044772 |
| Gm9159 | 0.296935004 | -1.128371656 | 3.721675807 |
| Gm27987 | 0.094097695 | 0.718200421 | 3.724945049 |
| Gm27285 | 0 | NA | NA |
| Gm27321 | 0.436808164 | -0.100375094 | 1.555107393 |
| Zcchc13 | 0.559888142 | 1.117293835 | 1.257858182 |
| Gm14874 | 0.823502631 | -2.439067195 | 1.103002168 |
| Gm14875 | 0 | NA | NA |
| Gm3943 | 0.062426873 | 0.237312084 | 3.724945049 |
| Slc16a2 | 413.5769034 | -0.074084309 | 0.089090188 |
| Gm6222 | 51.51232969 | 0.183937284 | 0.197304785 |
| Gm37956 | 12.17933112 | -1.450609525 | 0.347761668 |
| Gm14893 | 0 | NA | NA |
| Gm9164 | 0.119587853 | 0.237305575 | 3.724945049 |
| Gm9166 | 0.719693239 | 1.234427799 | 1.604689225 |
| Rlim | 2303.194065 | 0.424568189 | 0.09817807 |
| Gm14892 | 0 | NA | NA |
| Gm23378 | 0 | NA | NA |
| Gm22266 | 0.086648695 | -0.72445792 | 3.724945049 |
| Gm24993 | 0 | NA | NA |
| C77370 | 997.2398713 | -0.710651133 | 0.083559541 |
| Mir672 | 0.913602969 | -1.431679094 | 1.012652965 |
| Abcb7 | 785.2393539 | -0.105314523 | 0.081113675 |
| Vcp-rs | 650.7603801 | 0.462710925 | 0.079213859 |
| Gm6238 | 4.907369357 | -1.295304355 | 0.52875599 |
| Uprt | 137.0040339 | 0.182561911 | 0.130502784 |
| Zdhhc15 | 117.8220007 | -0.219814816 | 0.127520458 |
| Gm22430 | 0 | NA | NA |
| Gm14822 | 2.347436586 | -0.180616478 | 0.700434849 |
| 1700121L16Rik | 0.182267886 | 0.237326398 | 2.690523896 |
| Magee2 | 43.93040372 | -0.284278304 | 0.163541803 |
| Gm6285 | 2.055059636 | 1.173778958 | 0.745246522 |
| Gm22090 | 0.110870145 | 0.238305784 | 3.724874707 |
| Gm9673 | 0 | NA | NA |
| 5530601H04Rik | 228.1153272 | -0.398299541 | 0.105783193 |
| Pbdc1 | 243.3133995 | 0.094460008 | 0.103015764 |
| Magee1 | 1870.496208 | -0.475121505 | 0.059384815 |
| Gm6292 | 2.268236772 | -0.548634903 | 0.699827107 |
| E530001F21Rik | 15.1906836 | -0.262564194 | 0.325512458 |
| Rpl30-ps9 | 19.84181199 | -0.596941253 | 0.283805456 |
| Gm14824 | 57.64436276 | -0.903295293 | 0.227771225 |

| | | | |
|---|---|---|---|
| Mir384 | 25.25469777 | -0.412773106 | 0.222909635 |
| 5330434G04Rik | 854.7814095 | -0.594656996 | 0.162257278 |
| Mir325 | 9.585432933 | -0.101868254 | 0.486712783 |
| Rab9b-ps1 | 0.047948698 | -0.243569582 | 3.724945049 |
| Cypt2 | 0 | NA | NA |
| Gm14823 | 0 | NA | NA |
| Gm14855 | 0 | NA | NA |
| Gm14854 | 0 | NA | NA |
| Gm7117 | 0.290452866 | 0.237331432 | 1.781614801 |
| Fgf16 | 4.769070217 | -0.975557878 | 0.48267674 |
| Atrx | 4307.043563 | -0.440803687 | 0.085407436 |
| Gm26020 | 0 | NA | NA |
| Gm14853 | 16.35826218 | -0.886342234 | 0.389114688 |
| Gm23656 | 0 | NA | NA |
| Magt1 | 450.0307998 | 0.532307005 | 0.119292504 |
| Cox7b | 1087.530571 | -0.135975685 | 0.118271327 |
| Atp7a | 328.8971482 | 0.920712477 | 0.102931758 |
| Tlr13 | 410.2003387 | 4.903372309 | 0.301001317 |
| Gm14856 | 0 | NA | NA |
| Pgk1 | 2368.453568 | 0.215842823 | 0.06526337 |
| Taf9b | 306.5549042 | 0.539062028 | 0.11855575 |
| Fnd3c2 | 0.045882455 | 0.237312084 | 3.724945049 |
| Gm14852 | 0 | NA | NA |
| Gm14857 | 0.631966691 | 1.287672342 | 1.212427786 |
| Gm5133 | 0 | NA | NA |
| Mir6382 | 0 | NA | NA |
| Gm14876 | 0 | NA | NA |
| Gm6325 | 0.676409176 | 1.979778601 | 1.241028518 |
| Fndc3c1 | 1.066871938 | 0.691924086 | 0.886211984 |
| Gm27428 | 0.038981395 | 0.718207094 | 3.724945049 |
| Cysltr1 | 35.52481597 | 2.266691962 | 0.328306085 |
| Gm5127 | 6.92021651 | 3.40942774 | 0.563008029 |
| Gm24622 | 0.231961655 | 0.712553981 | 2.127211619 |
| Gm14877 | 0.75733687 | 0.59172651 | 1.445565157 |
| Gm14878 | 0.046345709 | 0.71820487 | 3.724945049 |
| Gm14879 | 0 | NA | NA |
| Gm9200 | 1.650574175 | -0.073331906 | 0.783652264 |
| Gm24491 | 0 | NA | NA |
| Gm22102 | 0 | NA | NA |
| Gm14882 | 0 | NA | NA |
| Gm14881 | 0 | NA | NA |
| Gm14880 | 0 | NA | NA |
| Zcchc5 | 3.290042265 | -0.88224681 | 0.592601795 |
| Lpar4 | 34.6581693 | 0.059035189 | 0.231910437 |
| P2ry10 | 28.76235556 | 7.837782002 | 0.870885488 |
| A630033H20Rik | 13.48921018 | 4.264275107 | 0.571127589 |
| Gm14834 | 0 | NA | NA |

| | | | |
|---|---|---|---|
| Gm14836 | 0 | NA | NA |
| Gpr174 | 17.71929983 | 4.866861176 | 0.573486624 |
| Gm9208 | 0.060174537 | 0.237309859 | 3.724945049 |
| Gm14833 | 0 | NA | NA |
| Gm5392 | 0 | NA | NA |
| Itm2a | 152.8149106 | -0.756424101 | 0.247286583 |
| Gm14835 | 0 | NA | NA |
| 4933401B06Rik | 0 | NA | NA |
| Tbx22 | 1.078738829 | -2.46636913 | 1.045118146 |
| 2610002M06Rik | 1022.333748 | -0.065841108 | 0.092762782 |
| Fam46d | 2.160810062 | 1.564714069 | 0.890167224 |
| Gm732 | 0 | NA | NA |
| Gm14849 | 0 | NA | NA |
| Gm378 | 0 | NA | NA |
| Gm9218 | 0 | NA | NA |
| Gm379 | 0.046345709 | 0.71820487 | 3.724945049 |
| Brwd3 | 697.4322742 | -0.189183975 | 0.080042702 |
| 2810403D21Rik | 30.21590391 | 0.230646988 | 0.189312066 |
| Gm14870 | 0.48610956 | 0.721932319 | 1.435418764 |
| 5730416F02Rik | 0.253264426 | 0.740639729 | 2.078818472 |
| Hmgn5 | 36.85669999 | 0.053528214 | 0.253246723 |
| Gm23322 | 0.212513282 | 0.237311278 | 3.175662942 |
| Gm25797 | 0 | NA | NA |
| Sh3bgrl | 620.2824578 | 0.307757012 | 0.13280481 |
| Gm26415 | 0 | NA | NA |
| Gm14869 | 0 | NA | NA |
| Gm6377 | 31.64666742 | 6.458504812 | 0.67838187 |
| Gm14868 | 0 | NA | NA |
| Gm4784 | 0.759591101 | 2.546733858 | 1.159882436 |
| Gm4991 | 0 | NA | NA |
| Gm7134 | 0 | NA | NA |
| Gm9225 | 0 | NA | NA |
| Gm14886 | 0 | NA | NA |
| Gm26108 | 0.089814205 | -0.24357403 | 3.724945049 |
| Pou3f4 | 86.02711921 | -0.310224738 | 0.141550807 |
| Cylc1 | 0 | NA | NA |
| Gm10112 | 0 | NA | NA |
| Rps6ka6 | 115.5535552 | 0.085302534 | 0.136559065 |
| Hdx | 271.1100276 | -0.568175311 | 0.107084325 |
| Gm7340 | 0 | NA | NA |
| Gm14898 | 0 | NA | NA |
| Gm14899 | 0 | NA | NA |
| Tex16 | 4.590268127 | 0.579235215 | 0.465849572 |
| Gm14900 | 0 | NA | NA |
| Gm6335 | 3.683185528 | 0.566930999 | 0.617869304 |
| 4933403O08Rik | 4.019211381 | 0.520308602 | 0.499943103 |
| Gm5393 | 0 | NA | NA |

| | | | |
|---|---|---|---|
| Apool | 63.23518737 | 1.123110102 | 0.178084207 |
| Satl1 | 0 | NA | NA |
| Gm6418 | 0.152980968 | 0.684806416 | 2.833643346 |
| Gm14908 | 0.684617987 | 1.92614287 | 1.382129742 |
| 2010106E10Rik | 0.828534694 | 0.201027415 | 1.14006165 |
| Zfp711 | 267.1925573 | -0.106682923 | 0.097182754 |
| Pof1b | 1.108407561 | 0.578673733 | 1.443042504 |
| Gm14921 | 0 | NA | NA |
| Gm14922 | 0 | NA | NA |
| Gm14923 | 0 | NA | NA |
| Gm6427 | 0 | NA | NA |
| Gm6429 | 0 | NA | NA |
| Gm14937 | 0 | NA | NA |
| Gm14938 | 0 | NA | NA |
| Gm14936 | 0.124596252 | 0.718200615 | 3.714681068 |
| Gm14939 | 1.913926814 | -0.436219941 | 0.887029586 |
| Chm | 978.6370897 | -0.00816316 | 0.066631092 |
| Mir361 | 1.21006551 | 0.053803204 | 0.972873511 |
| Gm7405 | 0 | NA | NA |
| Dach2 | 22.20089386 | 0.751956151 | 0.230529063 |
| Gm24831 | 0 | NA | NA |
| Gm23985 | 0 | NA | NA |
| Gm4915 | 0 | NA | NA |
| Gm14896 | 0 | NA | NA |
| Gm7416 | 0 | NA | NA |
| Gm7146 | 0.194279094 | 1.292225235 | 3.719439735 |
| Klhl4 | 212.9303465 | -0.423897025 | 0.13651587 |
| Gm14895 | 3.529479033 | 0.060298827 | 0.526439646 |
| Gm1866 | 7.003298379 | 0.017714439 | 0.389565277 |
| Gm14894 | 0.088476398 | -0.243576254 | 3.724945049 |
| Gm5942 | 0 | NA | NA |
| Gm14897 | 0.045882455 | 0.237312084 | 3.724945049 |
| Ube2dnl1 | 0 | NA | NA |
| Ube2dnl2 | 0 | NA | NA |
| 4930555B12Rik | 0 | NA | NA |
| Gm4914 | 0 | NA | NA |
| Gm7429 | 0.963943105 | 0.876346231 | 0.92647625 |
| Gm5943 | 0.089062654 | 0.237318756 | 3.724945049 |
| Gm14905 | 0 | NA | NA |
| Gm25366 | 0 | NA | NA |
| Gm27414 | 0 | NA | NA |
| Gm6461 | 0 | NA | NA |
| Gm5944 | 0.066050875 | 0.71820487 | 3.724945049 |
| Gm14903 | 0 | NA | NA |
| Gm14906 | 0 | NA | NA |
| Gm14907 | 0 | NA | NA |
| Gm24683 | 0 | NA | NA |

| | | | |
|---|---|---|---|
| Gm14904 | 0 | NA | NA |
| Gm24624 | 0 | NA | NA |
| Cpxcr1 | 0 | NA | NA |
| H2afb2 | 0 | NA | NA |
| Gm24438 | 0 | NA | NA |
| Gm16373 | 0.048949532 | -0.243569582 | 3.724945049 |
| Gm14919 | 0 | NA | NA |
| Gm14920 | 0 | NA | NA |
| Gm14918 | 0 | NA | NA |
| Gm14913 | 0 | NA | NA |
| Gm6491 | 0 | NA | NA |
| Gm28579 | 0 | NA | NA |
| Gm14911 | 0 | NA | NA |
| Gm14914 | 0.265235714 | 0.237330249 | 2.048392787 |
| Gm14917 | 0 | NA | NA |
| Gm6500 | 0 | NA | NA |
| Gm14916 | 0 | NA | NA |
| Gm14912 | 0 | NA | NA |
| Gm14915 | 0 | NA | NA |
| Gm14909 | 0 | NA | NA |
| Gm14910 | 0 | NA | NA |
| Tgif2lx2 | 0 | NA | NA |
| Tgif2lx1 | 0 | NA | NA |
| Gm7485 | 0 | NA | NA |
| Gm14926 | 0 | NA | NA |
| Gm26182 | 0 | NA | NA |
| Gm14929 | 0 | NA | NA |
| Pabpc5 | 33.05411456 | -0.105815847 | 0.201102281 |
| Gm4913 | 0.123841226 | 1.292225235 | 3.719439735 |
| Gm16420 | 0 | NA | NA |
| Gm5863 | 0.479236126 | -0.180484899 | 1.333093663 |
| Gm14928 | 0.622864522 | -1.000874944 | 1.320216627 |
| Pcdh11x | 761.4248623 | -0.504563309 | 0.098702835 |
| H2afb3 | 0 | NA | NA |
| Gm14931 | 0 | NA | NA |
| Gm14927 | 1.075015726 | -0.696824354 | 0.932842986 |
| Gm14930 | 0 | NA | NA |
| Gm4993 | 0.279479172 | 0.583591589 | 2.485743045 |
| Gm22590 | 0.042716945 | -0.243571806 | 3.724945049 |
| Gm5394 | 0 | NA | NA |
| Gm25825 | 0 | NA | NA |
| Gm5648 | 1.049721047 | 2.972772406 | 1.133632331 |
| Gm14932 | 0.047948698 | -0.243569582 | 3.724945049 |
| Gm23277 | 0 | NA | NA |
| Gm14943 | 0 | NA | NA |
| Gm14945 | 0 | NA | NA |
| Gm4992 | 0.108772582 | 0.718202646 | 3.724945049 |

| | | | |
|---|---|---|---|
| Gm24783 | 0 | NA | NA |
| Gm16421 | 0.491147404 | 0.222974268 | 1.396683456 |
| Gm14946 | 0.141684195 | 0.718205511 | 3.558854591 |
| Gm14942 | 0.160761659 | -0.243570954 | 2.799488298 |
| Nap1l3 | 532.8242286 | -0.26909158 | 0.122890062 |
| Gm14947 | 0 | NA | NA |
| Gm14940 | 0 | NA | NA |
| Gm17521 | 0 | NA | NA |
| Cldn34c1 | 0.062426873 | 0.237312084 | 3.724945049 |
| Astx6 | 0 | NA | NA |
| Srsx | 0 | NA | NA |
| Gm17577 | 0 | NA | NA |
| Gm14951 | 0 | NA | NA |
| Rps12-ps21 | 0 | NA | NA |
| Astx2 | 0 | NA | NA |
| Gm17412 | 0 | NA | NA |
| Rps12-ps22 | 0.049553047 | 0.71820487 | 3.724945049 |
| Cldn34c2 | 0 | NA | NA |
| Gm14950 | 0 | NA | NA |
| Gm17467 | 0 | NA | NA |
| Rps12-ps23 | 0 | NA | NA |
| Cldn34c3 | 0 | NA | NA |
| Astx5 | 0 | NA | NA |
| Rps12-ps20 | 0 | NA | NA |
| Vmn2r121 | 0 | NA | NA |
| Rps12-ps6 | 0.072025462 | 0.237312084 | 3.724945049 |
| Astx1a | 0 | NA | NA |
| Rps12-ps11 | 0 | NA | NA |
| Gm17584 | 0 | NA | NA |
| Astx4a | 0 | NA | NA |
| Rps12-ps12 | 0 | NA | NA |
| Gm17469 | 0 | NA | NA |
| Astx4b | 0 | NA | NA |
| Rps12-ps18 | 0 | NA | NA |
| Astx1b | 0 | NA | NA |
| Rps12-ps15 | 0 | NA | NA |
| Gm17361 | 0 | NA | NA |
| Gm21616 | 0 | NA | NA |
| Rps12-ps17 | 0.094831987 | -0.243571806 | 3.724945049 |
| Astx4c | 0 | NA | NA |
| Rps12-ps13 | 0 | NA | NA |
| Gm17693 | 0 | NA | NA |
| Astx1c | 0 | NA | NA |
| Rps12-ps14 | 0 | NA | NA |
| Gm17522 | 0 | NA | NA |
| Astx4d | 0 | NA | NA |
| Rps12-ps16 | 0 | NA | NA |

| | | | |
|---|---|---|---|
| Gm17267 | 0 | NA | NA |
| Astx3 | 0 | NA | NA |
| Gm14957 | 0.04393175 | -0.243571806 | 3.724945049 |
| Gm14971 | 0 | NA | NA |
| Gm7746 | 0 | NA | NA |
| 4932411N23Rik | 0 | NA | NA |
| Gm23586 | 0 | NA | NA |
| Gm381 | 0 | NA | NA |
| Gm14973 | 0 | NA | NA |
| Gm14972 | 0 | NA | NA |
| Gm382 | 0 | NA | NA |
| Gm22139 | 0 | NA | NA |
| 4921511C20Rik | 0 | NA | NA |
| Gm22636 | 0 | NA | NA |
| Gm14970 | 0.307002753 | 1.243495601 | 1.752068715 |
| Cldn34c4 | 0.191938586 | 0.79531826 | 3.270846535 |
| Gm25061 | 0 | NA | NA |
| Gm5403 | 0.038981395 | 0.718207094 | 3.724945049 |
| Mdm4-ps | 2.28277985 | 0.211290591 | 0.641674674 |
| Gm14975 | 0 | NA | NA |
| Gm4916 | 0 | NA | NA |
| Gm38390 | 0 | NA | NA |
| Gm26475 | 0 | NA | NA |
| Gm24522 | 0 | NA | NA |
| 4930558G05Rik | 0 | NA | NA |
| Gm26029 | 0 | NA | NA |
| Gm22225 | 0 | NA | NA |
| Gm14986 | 0 | NA | NA |
| Gm14985 | 0 | NA | NA |
| Gm14983 | 0 | NA | NA |
| Diap2 | 874.9384135 | 0.124116411 | 0.074704845 |
| Gm14984 | 0 | NA | NA |
| Gm14987 | 0 | NA | NA |
| Gm7781 | 0.389466168 | 0.324474554 | 1.405634807 |
| Gm14982 | 0.044191549 | -0.243571806 | 3.724945049 |
| Gm26209 | 0 | NA | NA |
| Gm25170 | 0 | NA | NA |
| Gm14976 | 0 | NA | NA |
| Gm7790 | 0 | NA | NA |
| Gm14977 | 0 | NA | NA |
| Gm4994 | 0 | NA | NA |
| Gm7058 | 0 | NA | NA |
| Gm14981 | 0 | NA | NA |
| Gm26406 | 0 | NA | NA |
| Gm23768 | 0 | NA | NA |
| Gm22143 | 0 | NA | NA |
| Gm7820 | 0 | NA | NA |

| | | | |
|---|---|---|---|
| Gm7821 | 0 | NA | NA |
| Gm7823 | 0 | NA | NA |
| Gm14979 | 0 | NA | NA |
| Gm22552 | 0 | NA | NA |
| Gm14980 | 0.189785611 | -0.24356998 | 2.656151365 |
| Pcdh19 | 966.848733 | -0.627518626 | 0.062121041 |
| Gm26851 | 2.30478967 | -0.990179851 | 0.652020191 |
| Tnmd | 3.21833566 | 0.214017394 | 0.546350955 |
| Tspan6 | 133.5839573 | -0.133833356 | 0.140357627 |
| 4930570D08Rik | 0 | NA | NA |
| Srpx2 | 13.98697368 | 0.129522344 | 0.272650518 |
| Sytl4 | 25.13300853 | 0.529788492 | 0.2708789 |
| Gm7841 | 0.106465261 | 0.718202646 | 3.724945049 |
| Cstf2 | 1299.774295 | -0.196531871 | 0.075520903 |
| Mir3112 | 0 | NA | NA |
| Nox1 | 3.798596485 | -0.333051095 | 0.537068011 |
| Xkrx | 14.27994078 | -0.438603529 | 0.264469684 |
| Arl13a | 0.299311364 | 2.291397375 | 3.695464448 |
| Trmt2b | 186.6964886 | -0.192865333 | 0.119031826 |
| Tmem35 | 346.804519 | -0.363756916 | 0.103444522 |
| Cenpi | 15.04443857 | 1.080306774 | 0.346504783 |
| Drp2 | 836.9736946 | -0.873364783 | 0.110050609 |
| Taf7l | 0.599737706 | 2.094631778 | 1.453164845 |
| Gm16410 | 0.92854447 | 0.943570371 | 1.18498272 |
| Gm23124 | 0.094831987 | -0.243571806 | 3.724945049 |
| Timm8a1 | 33.17250175 | 0.645402513 | 0.235586666 |
| Btk | 35.98129176 | 2.812642805 | 0.239635398 |
| Gm15154 | 0 | NA | NA |
| Rpl36a | 325.2517739 | 0.340978979 | 0.204326821 |
| Gla | 403.723937 | 2.078484975 | 0.165824016 |
| Hnrnph2 | 1054.6625 | 0.244490562 | 0.092225416 |
| Gm7855 | 0.152402701 | 0.718213715 | 2.846761649 |
| Gm7857 | 0.072025462 | 0.237312084 | 3.724945049 |
| B230119M05Rik | 0.783798686 | 0.620310841 | 1.21902015 |
| Armcx4 | 981.8027628 | -0.508472688 | 0.100943731 |
| Armcx1 | 541.5903477 | -0.291628809 | 0.081079935 |
| Gm16397 | 0.442616626 | -1.181802727 | 1.538305517 |
| Gm10344 | 1.729071564 | 0.970568035 | 0.803561606 |
| Armcx6 | 82.02743705 | 0.939734171 | 0.210904327 |
| Armcx3 | 414.6905456 | -0.20844876 | 0.077923798 |
| Mir7093 | 0.745433557 | -1.486477044 | 1.374361615 |
| Armcx2 | 966.547524 | -0.11300502 | 0.07547098 |
| Gm7222 | 0 | NA | NA |
| Gm15018 | 0 | NA | NA |
| Nxf2 | 1.760497888 | 1.32526752 | 0.808389729 |
| Zmat1 | 230.6846648 | -0.404842509 | 0.15489876 |
| 3632454L22Rik | 43.94593524 | -0.862594004 | 0.189949639 |

| | | | |
|---|---|---|---|
| Gm6207 | 0 | NA | NA |
| Gm15020 | 0 | NA | NA |
| Gm25868 | 0 | NA | NA |
| Gm15023 | 0.072025462 | 0.237312084 | 3.724945049 |
| Tceal6 | 310.5221001 | -0.563843779 | 0.127923203 |
| Gm15022 | 0 | NA | NA |
| Pramel | 0 | NA | NA |
| Pramel3 | 0.045882455 | 0.237312084 | 3.724945049 |
| Gm25936 | 0 | NA | NA |
| Gm15017 | 0 | NA | NA |
| Gm6215 | 0 | NA | NA |
| Gm5128 | 0 | NA | NA |
| Gm22006 | 0 | NA | NA |
| Gm6221 | 0 | NA | NA |
| Gm7903 | 0 | NA | NA |
| Gm24416 | 0 | NA | NA |
| Gm7905 | 0 | NA | NA |
| AV320801 | 0 | NA | NA |
| Gm22233 | 0 | NA | NA |
| Gm15015 | 0 | NA | NA |
| Gm6228 | 0 | NA | NA |
| Nxf7 | 11.01788405 | 1.281293073 | 0.345191962 |
| Prame | 0 | NA | NA |
| Gm15021 | 0.141317957 | 0.718202955 | 3.59624077 |
| Tcp11x2 | 0.28975322 | 1.799916207 | 2.357436394 |
| Tmsb15a | 0.072025462 | 0.237312084 | 3.724945049 |
| Gm15019 | 0 | NA | NA |
| Gprasp1 | 6518.488455 | -0.512714766 | 0.064032694 |
| Armcx5 | 396.6884651 | -0.166283084 | 0.172276334 |
| mmu-mir-7646 | 0 | NA | NA |
| Mir1970 | 0 | NA | NA |
| Gprasp2 | 1597.674091 | -0.188287578 | 0.111353387 |
| Bhlhb9 | 800.6765123 | 0.071003826 | 0.090943218 |
| Gm15016 | 2.223121331 | -0.099269203 | 0.772062891 |
| Arxes2 | 329.4047032 | -0.305466887 | 0.09877407 |
| Arxes1 | 156.3845472 | 0.064806808 | 0.141015878 |
| Bex2 | 883.5005139 | -0.068064322 | 0.167212063 |
| Nxf3 | 2.723995911 | -0.16039652 | 0.64577774 |
| 5730412P04Rik | 0.591118678 | 1.574033035 | 1.301061117 |
| Bex4 | 82.98433973 | -0.327159255 | 0.154952004 |
| Tceal8 | 272.1310813 | 0.362631346 | 0.142054139 |
| Mir6383 | 0 | NA | NA |
| Tceal5 | 256.9790592 | -0.539546386 | 0.233130595 |
| Bex1 | 254.7050612 | -0.231259091 | 0.143930729 |
| Tceal7 | 6.33875668 | -0.678553793 | 0.46799891 |
| Wbp5 | 588.6657451 | 0.472184971 | 0.100780438 |
| Ngfrap1 | 809.4390861 | -0.11884361 | 0.155682945 |

| | | | |
|---|---|---|---|
| n-R5s12 | 0.061920613 | 0.718207094 | 3.724945049 |
| Nsa2-ps2 | 0 | NA | NA |
| Gm14997 | 0 | NA | NA |
| Gm4917 | 0.046345709 | 0.71820487 | 3.724945049 |
| Gm7973 | 0.061920613 | 0.718207094 | 3.724945049 |
| Gm14994 | 0.090427103 | 0.237307635 | 3.724945049 |
| Gm14999 | 0 | NA | NA |
| Gm14998 | 0 | NA | NA |
| Gm14993 | 0 | NA | NA |
| Gm14992 | 0 | NA | NA |
| Gm14996 | 0 | NA | NA |
| Gm14990 | 0 | NA | NA |
| Gm14995 | 0.106112162 | 0.23732098 | 3.724945049 |
| Kir3dl2 | 0.620158706 | -0.24238849 | 1.284250353 |
| Gm14991 | 0.047948698 | -0.243569582 | 3.724945049 |
| Kir3dl1 | 0 | NA | NA |
| Gm15006 | 0 | NA | NA |
| Gm15003 | 0 | NA | NA |
| Gm15005 | 0 | NA | NA |
| Gm15004 | 0 | NA | NA |
| Tceal3 | 620.6626587 | -0.20813878 | 0.120362513 |
| Gm15010 | 0.784324088 | -1.054335554 | 1.290347844 |
| Gm23068 | 0 | NA | NA |
| Tceal1 | 253.1864694 | -0.095305929 | 0.123797519 |
| Morf4l2 | 2139.902612 | 0.370630535 | 0.084201651 |
| BC065397 | 34.23957864 | -0.801656128 | 0.245436715 |
| Glra4 | 6.335166068 | -0.918668874 | 0.495961494 |
| Gm15009 | 0 | NA | NA |
| Plp1 | 16662.56891 | -0.387788209 | 0.142936922 |
| Rab9b | 322.6084878 | -0.331948435 | 0.109622018 |
| Gm15026 | 0 | NA | NA |
| Gm15025 | 0 | NA | NA |
| H2bfm | 0 | NA | NA |
| Gm6275 | 0.048949532 | -0.243569582 | 3.724945049 |
| Tmsb15l | 3.858449061 | -0.112023789 | 0.604690443 |
| Tmsb15b2 | 0.857879072 | -2.292799694 | 1.148753154 |
| Tmsb15b1 | 3.641358966 | -1.743607831 | 0.603750561 |
| Slc25a53 | 178.3064943 | -0.416178216 | 0.131342387 |
| Zcchc18 | 1940.436389 | -0.586394929 | 0.069269785 |
| Gm15038 | 0 | NA | NA |
| Fam199x | 561.8101836 | -0.133644702 | 0.089476632 |
| Gm25770 | 0 | NA | NA |
| Gm8019 | 0 | NA | NA |
| Gm5761 | 0.368947036 | -0.24356434 | 1.608971141 |
| Gm15039 | 0 | NA | NA |
| Gm15042 | 0 | NA | NA |
| Gm15045 | 0 | NA | NA |

| | | | |
|---|---|---|---|
| Gm15043 | 0 | NA | NA |
| Gm15044 | 0 | NA | NA |
| Esx1 | 0 | NA | NA |
| Gm15040 | 0.045882455 | 0.237312084 | 3.724945049 |
| Gm15041 | 0 | NA | NA |
| Gm23679 | 0 | NA | NA |
| Gm8036 | 0.810529001 | -0.202904502 | 1.006864204 |
| Il1rapl2 | 78.53886856 | -0.948583806 | 0.157833668 |
| Gm14988 | 0 | NA | NA |
| Gm25393 | 0 | NA | NA |
| Gm14989 | 0 | NA | NA |
| Gm24675 | 0 | NA | NA |
| Tex13a | 0 | NA | NA |
| Gm22057 | 0 | NA | NA |
| Gm23958 | 0 | NA | NA |
| Gm15000 | 0 | NA | NA |
| Gm15001 | 0 | NA | NA |
| Nrk | 62.1717717 | -0.15672914 | 0.259699498 |
| Serpina7 | 0.108772582 | 0.718202646 | 3.724945049 |
| 4930513O06Rik | 0 | NA | NA |
| 4933428M09Rik | 0 | NA | NA |
| Gm15007 | 0 | NA | NA |
| Mum1l1 | 62.92430637 | -0.384451463 | 0.142636366 |
| Platr21 | 0.194631057 | 0.237309881 | 3.243494927 |
| Gm15002 | 0 | NA | NA |
| Trap1a | 0 | NA | NA |
| Gm8061 | 0 | NA | NA |
| D330045A20Rik | 6.283406809 | 0.115598959 | 0.406968413 |
| Rnf128 | 220.8248196 | 0.257492129 | 0.108231595 |
| Gm24380 | 0 | NA | NA |
| Gm6322 | 0 | NA | NA |
| Tbc1d8b | 118.5380032 | 0.208819798 | 0.132486131 |
| Gm15013 | 2.788702176 | 0.199765243 | 0.614527401 |
| Ripply1 | 0.677889948 | 1.503933789 | 1.336008296 |
| Gm15012 | 0.066050875 | 0.71820487 | 3.724945049 |
| Cldn2 | 23.20106824 | -0.767121315 | 0.279174185 |
| Morc4 | 90.84373692 | 0.307031943 | 0.164975542 |
| Rbm41 | 210.9730729 | 0.096318117 | 0.10091485 |
| Gm15011 | 0 | NA | NA |
| Nup62cl | 3.861479097 | 0.127794604 | 0.579859067 |
| Pih1h3b | 0.733953745 | -1.083103285 | 1.109107724 |
| Gm6298 | 0.042716945 | -0.243571806 | 3.724945049 |
| Gm15046 | 0.104637558 | 0.23732098 | 3.724945049 |
| Mir3475 | 0.844812178 | -0.436330184 | 1.317865881 |
| Frmpd3 | 67.41637008 | -0.556205501 | 0.154430988 |
| Gm8097 | 0.313170507 | -0.243564451 | 2.088068835 |
| Prps1 | 548.0358506 | -0.133203996 | 0.063075674 |

| | | | |
|---|---|---|---|
| Tsc22d3 | 1422.675399 | 0.718955944 | 0.154626641 |
| Gm15047 | 0 | NA | NA |
| Gm22025 | 0 | NA | NA |
| Mid2 | 552.0542967 | -0.074724849 | 0.075554906 |
| Gm23199 | 2.503876714 | 0.654731082 | 0.671191525 |
| Eif2c5 | 0 | NA | NA |
| Gm6088 | 0.360750703 | 1.406024352 | 1.505853078 |
| Tex13 | 0 | NA | NA |
| Gm7091 | 0 | NA | NA |
| Vsig1 | 0.30513904 | 0.237311308 | 3.724945049 |
| Psmd10 | 105.8155227 | 0.63719525 | 0.150507145 |
| Atg4a | 98.38075259 | 0.323398466 | 0.172368861 |
| Col4a6 | 3.411580873 | 0.860926957 | 0.602171518 |
| Gm15077 | 0 | NA | NA |
| Gm15078 | 0 | NA | NA |
| Gm15079 | 0 | NA | NA |
| Col4a5 | 123.2002736 | 1.156073159 | 0.203520934 |
| Gm15268 | 0.88858162 | 0.914471257 | 1.154324828 |
| Gm15282 | 0 | NA | NA |
| Irs4 | 86.40347737 | -0.499053952 | 0.239926697 |
| Gm15295 | 0.642988239 | 0.126492486 | 1.345374417 |
| Gm15297 | 0 | NA | NA |
| Hmgn2l6 | 0 | NA | NA |
| Gm15294 | 0.045882455 | 0.237312084 | 3.724945049 |
| Gm15298 | 0.198041555 | 1.570941826 | 2.587932611 |
| Gm6368 | 8.824265348 | 0.357809773 | 0.361770143 |
| Gm5762 | 0 | NA | NA |
| Gm6373 | 0 | NA | NA |
| Gucy2f | 26.45992321 | 0.251848505 | 0.218433152 |
| Gm6382 | 25.01211358 | 0.571337938 | 0.226306019 |
| Gm15014 | 0 | NA | NA |
| Nxt2 | 433.560056 | -0.060567347 | 0.090720046 |
| Gm5643 | 2.51713653 | -0.145059344 | 0.604239614 |
| Kcne1l | 125.6411286 | 1.694179034 | 0.243759924 |
| Acsl4 | 1433.382608 | 0.093069282 | 0.072220929 |
| Gm6067 | 0 | NA | NA |
| Gm7123 | 0 | NA | NA |
| Gm15031 | 0 | NA | NA |
| Gm25915 | 0.060174537 | 0.237309859 | 3.724945049 |
| Gm4995 | 0.090537259 | 0.237318756 | 3.724945049 |
| Gm15032 | 0 | NA | NA |
| Gm5763 | 8.930037149 | 0.257140322 | 0.345798429 |
| Tmem164 | 493.7209216 | -0.17953059 | 0.08015371 |
| Mir652 | 2.416261644 | -0.657461831 | 0.745827056 |
| Gm15049 | 0.046345709 | 0.71820487 | 3.724945049 |
| Ammecr1 | 116.0557192 | 0.316556884 | 0.116254688 |
| AL731548.1 | 1.863015902 | -0.227709346 | 0.693942074 |

| | | | |
|---|---|---|---|
| Rgag1 | 33.41144712 | -0.376233257 | 0.228966077 |
| Gm4760 | 0.046345709 | 0.71820487 | 3.724945049 |
| Chrdl1 | 139.3962317 | 0.448575759 | 0.177037516 |
| Gm23325 | 0 | NA | NA |
| Pak3 | 2356.012352 | -0.150524246 | 0.102141941 |
| Capn6 | 31.3985991 | -0.029999155 | 0.227295287 |
| BX530055.1 | 0 | NA | NA |
| Dcx | 707.4354374 | -0.536393811 | 0.160789152 |
| Gm15048 | 0.045882455 | 0.237312084 | 3.724945049 |
| A730046J19Rik | 10.6199982 | 0.166504232 | 0.345436223 |
| Gm8199 | 0.207515118 | 0.264226477 | 2.585168276 |
| Gm8202 | 0 | NA | NA |
| Gm15067 | 0 | NA | NA |
| Gm15066 | 0.044544648 | 0.237309859 | 3.724945049 |
| Mir680-2 | 0 | NA | NA |
| Alg13 | 168.7701936 | -0.22396951 | 0.110393193 |
| Gm2214 | 0.090665643 | -0.724455696 | 3.724945049 |
| Trpc5 | 161.8271037 | -0.862343807 | 0.097310752 |
| Trpc5os | 0.750003179 | 2.219871798 | 2.540357748 |
| Gm15071 | 0 | NA | NA |
| Mageb16-ps1 | 3.715911654 | -1.523205852 | 0.587770193 |
| Zcchc16 | 32.00016923 | -0.99278618 | 0.210428299 |
| Gm26024 | 2.396750964 | -2.828667501 | 0.92564401 |
| Gm8216 | 4.713227208 | -1.978572954 | 0.582892616 |
| Mir3472 | 0.066050875 | 0.71820487 | 3.724945049 |
| Gm25097 | 0 | NA | NA |
| Gm15073 | 0.066050875 | 0.71820487 | 3.724945049 |
| Gm6441 | 0.044191549 | -0.243571806 | 3.724945049 |
| Lhfpl1 | 2.080192563 | 0.602854354 | 0.734270243 |
| Amot | 561.2978999 | -0.266036386 | 0.104897237 |
| Gm5644 | 2.782059835 | -0.898009419 | 0.724000001 |
| Gm15060 | 0 | NA | NA |
| Gm27659 | 0.042716945 | -0.243571806 | 3.724945049 |
| Gm15058 | 0.106618422 | -0.24357403 | 3.724945049 |
| Gm6446 | 0.038981395 | 0.718207094 | 3.724945049 |
| Gm15057 | 0.867677903 | 0.026836358 | 0.988163225 |
| Gm6447 | 0 | NA | NA |
| Gm15062 | 0 | NA | NA |
| Gm4918 | 0 | NA | NA |
| Hmgb1-ps2 | 15.03640688 | 0.056383922 | 0.264408253 |
| Gm15061 | 0 | NA | NA |
| Gm26151 | 0.325583935 | -0.902256417 | 1.987255747 |
| Gm15076 | 0 | NA | NA |
| Gm8261 | 2.625663271 | 0.192499282 | 0.723729832 |
| Gm15075 | 1.268895825 | 0.034698493 | 0.880678205 |
| Gm15074 | 0 | NA | NA |
| Gm4996 | 0 | NA | NA |

| | | | |
|---|---|---|---|
| Htr2c | 718.8775726 | -1.070697054 | 0.112141199 |
| Snora35 | 4.50470804 | -0.440186573 | 0.533143619 |
| Gm22650 | 4.136795173 | -1.135035805 | 0.603883583 |
| Mir764 | 3.164642609 | -1.019113782 | 0.583663283 |
| Mir1912 | 2.763607753 | -0.417112318 | 0.587333168 |
| Mir1264 | 5.146865135 | -1.251198664 | 0.546819588 |
| Mir1298 | 1.82841141 | -0.612890033 | 0.965611686 |
| Gm24627 | 1.916860434 | -0.429608138 | 0.808424363 |
| Mir448 | 0.601491876 | -2.939008963 | 1.381601374 |
| Il13ra2 | 7.235467095 | 0.227710128 | 0.388271839 |
| Vmn1r239-ps | 0 | NA | NA |
| Lrch2 | 586.4938171 | -0.290912522 | 0.088642448 |
| Gm25107 | 111.0918953 | 0.118156236 | 0.208066575 |
| Gm16400 | 0 | NA | NA |
| Gm15167 | 0 | NA | NA |
| Gm8289 | 0 | NA | NA |
| Gm15128 | 0 | NA | NA |
| Gm15081 | 0 | NA | NA |
| Gm15080 | 0 | NA | NA |
| Gm15108 | 0 | NA | NA |
| Gm15110 | 0 | NA | NA |
| Gm8303 | 0 | NA | NA |
| Gm15107 | 0 | NA | NA |
| Gm15109 | 0 | NA | NA |
| Gm15111 | 0 | NA | NA |
| Gm15113 | 0 | NA | NA |
| Gm8320 | 0 | NA | NA |
| Gm15114 | 0 | NA | NA |
| Gm15106 | 0 | NA | NA |
| Gm15112 | 0 | NA | NA |
| Gm8331 | 0 | NA | NA |
| Gm8334 | 0 | NA | NA |
| Gm15127 | 0 | NA | NA |
| Gm15125 | 0 | NA | NA |
| Gm15126 | 0 | NA | NA |
| Kars-ps1 | 0 | NA | NA |
| Luzp4 | 0.048949532 | -0.243569582 | 3.724945049 |
| Gm15099 | 0 | NA | NA |
| Gm15084 | 0 | NA | NA |
| Gm15094 | 0 | NA | NA |
| Gm15096 | 0 | NA | NA |
| Ott | 0 | NA | NA |
| Gm15092 | 0 | NA | NA |
| Gm15088 | 0 | NA | NA |
| Gm15090 | 0 | NA | NA |
| Gm8346 | 0 | NA | NA |
| Gm15093 | 0 | NA | NA |

| | | | |
|---|---|---|---|
| Gm15098 | 0 | NA | NA |
| Gm15089 | 0 | NA | NA |
| Gm8366 | 0 | NA | NA |
| Gm15100 | 0 | NA | NA |
| Gm15085 | 0 | NA | NA |
| Gm15086 | 0 | NA | NA |
| Gm10439 | 0 | NA | NA |
| Gm15101 | 0 | NA | NA |
| Gm8383 | 0 | NA | NA |
| Gm15097 | 0 | NA | NA |
| Gm15095 | 0 | NA | NA |
| Gm15087 | 0 | NA | NA |
| Gm15091 | 0 | NA | NA |
| Gm15102 | 0 | NA | NA |
| Gm15103 | 0 | NA | NA |
| Gm8402 | 0 | NA | NA |
| Gm5946 | 2.587997239 | 1.251934798 | 0.699574394 |
| Gm15104 | 1.895634264 | -0.669116656 | 0.74345795 |
| Gm15105 | 0.604613898 | -0.655329107 | 1.586334572 |
| Tmem29 | 269.2719752 | -0.479528485 | 0.109738075 |
| Gm15129 | 0 | NA | NA |
| 3110067C02Rik | 4.62715315 | -0.449434607 | 0.480297559 |
| Apex2 | 300.6728437 | -0.414759605 | 0.183444173 |
| Alas2 | 56.13115503 | 2.001089215 | 0.803652209 |
| Mir3620 | 3.543203576 | -1.309006465 | 0.601154792 |
| Pfkfb1 | 11.63733137 | 0.312473374 | 0.344099156 |
| Tro | 2279.729976 | -0.108225542 | 0.092813005 |
| Gm15132 | 0 | NA | NA |
| Gm15131 | 0 | NA | NA |
| Maged2 | 434.343474 | 0.030828607 | 0.115719587 |
| Gm24907 | 0.178529143 | -1.502132721 | 2.705292286 |
| Itih5l-ps | 0.139037128 | 1.464453538 | 3.717743456 |
| Gm27191 | 0 | NA | NA |
| Gm8424 | 0.517778811 | 1.748295803 | 1.363277115 |
| Gnl3l | 3189.542655 | -0.373424802 | 0.055206594 |
| Gm15152 | 0 | NA | NA |
| Fgd1 | 278.5392714 | -0.186168598 | 0.080764158 |
| Tsr2 | 790.8463341 | -0.037263528 | 0.063043844 |
| Gm15149 | 0 | NA | NA |
| Rps12-ps7 | 0 | NA | NA |
| Gm15138 | 7.688685255 | -0.42356375 | 0.459150526 |
| Wnk3 | 948.2720462 | -0.365570825 | 0.067175131 |
| Rpl21-ps15 | 49.38234276 | 0.379065546 | 0.210321767 |
| Gm26441 | 0.12209515 | 0.718202646 | 3.724945049 |
| A230072E10Rik | 43.55964103 | 0.568863368 | 0.182481592 |
| Fam120c | 1332.354665 | 0.164707282 | 0.071318656 |
| Gm22248 | 2.415551224 | -1.677001196 | 0.781061056 |

| | | | |
|---|---|---|---|
| Gm6451 | 2.057373881 | 0.203665731 | 0.695394729 |
| Gm15148 | 0.123992736 | -0.243574001 | 3.720643035 |
| Phf8 | 763.8243954 | 0.095371372 | 0.092283546 |
| Gm24460 | 0 | NA | NA |
| Gm5395 | 1.549805528 | 1.492782251 | 1.026455205 |
| Gm15151 | 0.220031754 | 1.209745473 | 2.592709249 |
| Gm4919 | 0 | NA | NA |
| n-R5s13 | 0 | NA | NA |
| Gm15161 | 0 | NA | NA |
| Huwe1 | 8349.182081 | -0.364470682 | 0.05212621 |
| Mir3113 | 0 | NA | NA |
| Mirlet7f-2 | 0.623206311 | 0.017438941 | 1.268401641 |
| Mir98 | 2.834035009 | 0.086646632 | 0.59234887 |
| Gm15191 | 3.302051428 | 0.272372056 | 0.507982274 |
| n-R5s14 | 0 | NA | NA |
| Hsd17b10 | 170.8953565 | 0.437303489 | 0.122593031 |
| Ribc1 | 4.147795559 | 0.842418577 | 0.474570188 |
| Smc1a | 891.0140031 | 0.206086346 | 0.103937365 |
| Gm15267 | 3.127372856 | 0.479974224 | 0.544091897 |
| Gm15266 | 0 | NA | NA |
| Iqsec2 | 2091.297293 | -0.809589924 | 0.099731408 |
| Kdm5c | 1433.211927 | 0.290047857 | 0.056545724 |
| Kantr | 325.3470963 | -0.477699636 | 0.087655171 |
| n-R5s15 | 0 | NA | NA |
| Tspyl2 | 1239.902915 | 0.154298354 | 0.108007447 |
| Gpr173 | 222.5226893 | -0.268401617 | 0.087206799 |
| 3010001F23Rik | 25.65934517 | -1.069231806 | 0.366422554 |
| Gm4750 | 1.060705812 | 1.731822624 | 1.108753265 |
| Gm8464 | 0.320494736 | -0.217672008 | 1.742948413 |
| Gm7158 | 0 | NA | NA |
| Cldn34a | 0.10503227 | 1.179809211 | 3.723500796 |
| Gm8475 | 0.097139547 | 0.718207094 | 3.724945049 |
| Shroom2 | 979.6660779 | -0.79469521 | 0.091068533 |
| Gpr143 | 0.64528552 | 2.268068587 | 1.43021343 |
| Gm24687 | 0 | NA | NA |
| Rpl7a-ps12 | 0.049553047 | 0.71820487 | 3.724945049 |
| Gm6472 | 317.0452057 | 0.760589339 | 0.163439634 |
| Gm26035 | 140.1127517 | 0.499898325 | 0.24790541 |
| Usp51 | 28.56090422 | 0.29054375 | 0.211559009 |
| Mageh1 | 200.6419644 | -0.1268207 | 0.137505414 |
| 9530051G07Rik | 22.2217983 | -0.521783037 | 0.268587865 |
| Gm5396 | 0.044544648 | 0.237309859 | 3.724945049 |
| Gm26960 | 1.043034268 | -1.619139548 | 1.122917202 |
| Foxr2 | 13.36216574 | -0.458477133 | 0.314791408 |
| Rragb | 646.2289516 | -0.379876182 | 0.072442755 |
| Gm15153 | 0.202306569 | -0.698458055 | 2.641239443 |
| Klf8 | 93.85799211 | 0.150675479 | 0.129574175 |

| | | | |
|---|---|---|---|
| Gm7150 | 0 | NA | NA |
| Ubqln2 | 2353.389334 | -0.293921129 | 0.05353823 |
| Gm15166 | 0 | NA | NA |
| Cypt3 | 0 | NA | NA |
| Gm25793 | 0 | NA | NA |
| Gm8534 | 0 | NA | NA |
| Kctd12b | 83.33022565 | 0.264660025 | 0.152283686 |
| 2210013O21Rik | 216.6134074 | 0.174488441 | 0.098416896 |
| Smt3h1-ps | 0.284401297 | -1.193149127 | 2.051614158 |
| Spin2c | 37.12736702 | -0.451864771 | 0.214993678 |
| Gm25895 | 0 | NA | NA |
| Samt1 | 0 | NA | NA |
| 4921511M17Rik | 0 | NA | NA |
| Gm10057 | 0 | NA | NA |
| Gm15143 | 0 | NA | NA |
| Gm5947 | 0 | NA | NA |
| Gm15140 | 0 | NA | NA |
| Gm5645 | 0 | NA | NA |
| Gm16445 | 0 | NA | NA |
| Gm6620 | 0 | NA | NA |
| Gm15147 | 0 | NA | NA |
| Gm5060 | 0 | NA | NA |
| Gm15141 | 0 | NA | NA |
| 4930524N10Rik | 0 | NA | NA |
| Gm5646 | 0 | NA | NA |
| Gm8569 | 0 | NA | NA |
| Samt4 | 0 | NA | NA |
| Samt2 | 0 | NA | NA |
| Gm15142 | 0 | NA | NA |
| Gm22359 | 0 | NA | NA |
| Gm8577 | 0.253240508 | -1.093085126 | 2.097824694 |
| Cldn34b1 | 0 | NA | NA |
| Magea6 | 0 | NA | NA |
| Magea3 | 0 | NA | NA |
| Gm6645 | 1.418926095 | -0.869349874 | 0.869511696 |
| Gm25257 | 0 | NA | NA |
| Magea8 | 0 | NA | NA |
| Magea2 | 0 | NA | NA |
| Gm24268 | 0 | NA | NA |
| Magea5 | 0 | NA | NA |
| Magea1 | 0 | NA | NA |
| Cldn34b2 | 0 | NA | NA |
| Gm25429 | 0.066050875 | 0.71820487 | 3.724945049 |
| Gm4997 | 0 | NA | NA |
| Amd-ps1 | 0.844289582 | -1.525586513 | 1.238948664 |
| Gm8595 | 0.04393175 | -0.243571806 | 3.724945049 |
| Gm15157 | 0.250163086 | 0.240574235 | 2.496469752 |

| | | | |
|---|---|---|---|
| Sat1 | 845.2085452 | 2.138724113 | 0.157819359 |
| Gm15159 | 0 | NA | NA |
| Acot9 | 212.0243755 | 0.66740284 | 0.121856868 |
| Gm650 | 0 | NA | NA |
| Prdx4 | 114.043658 | 0.904104767 | 0.215483185 |
| Gm15158 | 0 | NA | NA |
| Gm8606 | 0.172182752 | 0.718209228 | 3.331044598 |
| n-R5s16 | 0 | NA | NA |
| Ptchd1 | 278.1680322 | -0.271488914 | 0.091888518 |
| Gm15156 | 5.651205916 | 1.036318757 | 0.434783268 |
| Gm15155 | 12.34800111 | -1.557300921 | 0.331288311 |
| Gm15160 | 1.808882573 | -0.86772185 | 0.806187934 |
| Gm23404 | 0 | NA | NA |
| 4930503H13Rik | 1.186325201 | -2.837314291 | 1.02039488 |
| Gm7312 | 0 | NA | NA |
| Gm15162 | 0.389425067 | 0.237310554 | 3.297689459 |
| Gm15163 | 28.67019377 | -0.314254803 | 0.209819151 |
| Gm22666 | 0 | NA | NA |
| Phex | 65.77163163 | 0.407315869 | 0.194192211 |
| Gm23798 | 0 | NA | NA |
| Gm25795 | 0.325008436 | 1.076295602 | 2.051917921 |
| Gm15172 | 0.412251148 | -1.879233977 | 1.672423068 |
| Gm1947 | 0.704217346 | 2.559674185 | 1.306172775 |
| Gm24281 | 0 | NA | NA |
| Sms | 958.8198068 | -0.454493964 | 0.057566102 |
| Mbtps2 | 467.2768864 | -0.117013769 | 0.064226626 |
| Gm6568 | 0.666747433 | 2.018003787 | 1.263354124 |
| Yy2 | 0 | NA | NA |
| Gm15176 | 0 | NA | NA |
| Gm15175 | 0.320922609 | 1.23176036 | 1.742001407 |
| Gm15173 | 0.066050875 | 0.71820487 | 3.724945049 |
| Gm23381 | 0 | NA | NA |
| Gm8644 | 0.329848065 | 1.85737421 | 1.872398093 |
| Gm15171 | 0 | NA | NA |
| Gm8648 | 0 | NA | NA |
| Smpx | 10.49823735 | -0.693460312 | 0.328350437 |
| Gm15169 | 2.861039839 | 2.455285063 | 0.811941191 |
| Klhl34 | 1.755040647 | 0.044942131 | 0.772261742 |
| Cnksr2 | 3647.587889 | -0.817199192 | 0.126439756 |
| Gm8659 | 0.100902008 | 1.15059983 | 3.720931964 |
| Gm15170 | 0.046345709 | 0.71820487 | 3.724945049 |
| Gm8662 | 0 | NA | NA |
| Gm15177 | 0 | NA | NA |
| Gm22522 | 0 | NA | NA |
| Gm15168 | 0 | NA | NA |
| Gm5764 | 0.044544648 | 0.237309859 | 3.724945049 |
| Gm15185 | 0 | NA | NA |

| | | | |
|---|---|---|---|
| Gm15188 | 0 | NA | NA |
| Gm15189 | 0 | NA | NA |
| Rps6ka3 | 1265.248659 | 0.29599427 | 0.072455714 |
| Eif1ax | 1820.91249 | -0.480408059 | 0.069688203 |
| Gm15187 | 0 | NA | NA |
| Map7d2 | 1767.738311 | -0.250490689 | 0.090129285 |
| Gm23806 | 1.294438978 | -0.573957045 | 0.920695971 |
| A830080D01Rik | 225.115431 | 0.259834009 | 0.088796837 |
| Gm15190 | 9.37392129 | 0.513558482 | 0.338556647 |
| Gm15186 | 6.883933299 | 0.028127939 | 0.4500607 |
| Sh3kbp1 | 688.8231907 | 0.028933101 | 0.059199808 |
| Map3k15 | 25.20550808 | 0.625768581 | 0.251105422 |
| Gm15194 | 0 | NA | NA |
| Gm8682 | 0 | NA | NA |
| Pdha1 | 1985.030852 | -0.375480528 | 0.052553338 |
| Gm15195 | 0 | NA | NA |
| Gm15192 | 0 | NA | NA |
| Gm15196 | 0.075043205 | 0.237309859 | 3.724945049 |
| Gm15193 | 0 | NA | NA |
| Adgrg2 | 34.96479159 | 0.937215643 | 0.245058533 |
| Gm15241 | 0.873806335 | 1.518214868 | 1.178676784 |
| Phka2 | 591.6410219 | -0.109343296 | 0.131410441 |
| Gm15243 | 0.261996957 | 0.2373302 | 2.055677216 |
| Ppef1 | 10.0328463 | 1.339028185 | 0.377726217 |
| Gm15242 | 0.08486385 | 0.717551239 | 3.724898127 |
| Gm15244 | 0.061920613 | 0.718207094 | 3.724945049 |
| Rs1 | 1.55354348 | -0.916976607 | 0.891953243 |
| Cdkl5 | 3363.729212 | -0.640239642 | 0.103911574 |
| Gja6 | 33.35128538 | -0.908592794 | 0.291460054 |
| Scml2 | 17.41690607 | -0.224051674 | 0.256445617 |
| Gm5419 | 0 | NA | NA |
| Gm15262 | 0 | NA | NA |
| Prkaca-ps1 | 0.150961315 | 1.165658083 | 2.845512047 |
| Rai2 | 127.3185232 | -0.062604248 | 0.098650165 |
| Scml1 | 0.926668879 | -0.340216762 | 1.077110382 |
| Gm15205 | 1.789402658 | -0.833776231 | 0.714485337 |
| Nhs | 212.8785834 | -0.172971749 | 0.092657556 |
| Gm15202 | 0.888501202 | -0.399340786 | 1.127738751 |
| Gm26317 | 0.531781488 | 1.146859638 | 1.43463889 |
| Gm23794 | 0.04393175 | -0.243571806 | 3.724945049 |
| Gm27490 | 0 | NA | NA |
| Gm23867 | 0.167772976 | 0.778749666 | 3.41346025 |
| Gm15204 | 0.395626157 | 0.663927829 | 1.494294117 |
| Reps2 | 2262.978211 | -0.68058691 | 0.078557093 |
| Gm7331 | 62.14126764 | 0.414625813 | 0.173618818 |
| Gm15206 | 2.290494867 | -1.735509432 | 0.75102726 |
| Gm15203 | 0 | NA | NA |

| | | | |
|---|---|---|---|
| Gm22257 | 0 | NA | NA |
| Gm25557 | 0 | NA | NA |
| Gm8737 | 0 | NA | NA |
| Gm15201 | 8.230899355 | -0.753428006 | 0.397567384 |
| Gm23786 | 0 | NA | NA |
| Rbbp7 | 1339.028074 | 0.540733734 | 0.106902318 |
| Txlng | 376.7131286 | -0.23087067 | 0.098281038 |
| Gm16459 | 0 | NA | NA |
| Gm15207 | 0 | NA | NA |
| Gm23901 | 0.060174537 | 0.237309859 | 3.724945049 |
| Syap1 | 318.0290325 | 0.335183578 | 0.085144816 |
| Mir3473a | 0.568149222 | -1.696348034 | 1.410386779 |
| Gm15209 | 0 | NA | NA |
| Ctps2 | 433.4122222 | -0.085906916 | 0.062845722 |
| S100g | 0.048949532 | -0.243569582 | 3.724945049 |
| Gm23501 | 0 | NA | NA |
| Gm22746 | 0 | NA | NA |
| Gm15215 | 0 | NA | NA |
| Gm15212 | 0 | NA | NA |
| Gm15214 | 0 | NA | NA |
| Gm26007 | 0 | NA | NA |
| Gm7199 | 0 | NA | NA |
| Grpr | 12.17739777 | 0.161757642 | 0.298977838 |
| Rnf138rt1 | 7.321455992 | 0.265282293 | 0.377218119 |
| Gm15213 | 0 | NA | NA |
| Ap1s2 | 658.6125027 | 0.184870272 | 0.106189819 |
| Zrsr2 | 430.8776927 | -0.008504845 | 0.085498999 |
| Car5b | 61.00096164 | 0.181716525 | 0.188181206 |
| Siah1b | 50.98963704 | 0.121390197 | 0.158653283 |
| Tmem27 | 1.459315314 | 1.085594697 | 0.905627737 |
| n-R5s18 | 0.046345709 | 0.71820487 | 3.724945049 |
| Ace2 | 16.8793861 | 0.337630447 | 0.355041185 |
| Bmx | 4.886164753 | -0.01184673 | 0.481042784 |
| Gm25367 | 0 | NA | NA |
| Pir | 44.18337336 | -0.285164927 | 0.177374831 |
| Gm22023 | 0.437539382 | -0.877149 | 1.653910799 |
| Figf | 22.19425402 | -0.069417501 | 0.298094342 |
| Piga | 134.018528 | -0.265698221 | 0.114578121 |
| Asb11 | 36.5181848 | 0.160892841 | 0.241314327 |
| Gm15211 | 0.30015424 | 1.829125579 | 1.987472111 |
| Asb9 | 0.209460102 | 1.182943972 | 2.187386132 |
| Gm15216 | 0 | NA | NA |
| Mospd2 | 329.2346432 | 0.174550189 | 0.095539081 |
| Fancb | 25.83413697 | -0.343920614 | 0.233314706 |
| Gm17604 | 0 | NA | NA |
| Glra2 | 135.3189216 | -0.089786676 | 0.143759688 |
| Gm15236 | 0.167734623 | 1.145446775 | 3.392587305 |

| | | | |
|---|---|---|---|
| Gm22062 | 0 | NA | NA |
| Gm15224 | 0 | NA | NA |
| Gm15225 | 0 | NA | NA |
| Gemin8 | 60.76505143 | -0.425953909 | 0.17033534 |
| Gm15227 | 0 | NA | NA |
| Gpm6b | 8313.7276 | -0.507220969 | 0.111765712 |
| Ofd1 | 183.0566898 | 0.247811426 | 0.109411235 |
| Trappc2 | 241.3367123 | -0.39623689 | 0.103226245 |
| Rab9 | 264.7892433 | 0.52678137 | 0.094852189 |
| Gm15229 | 2.091380692 | 5.10666915 | 1.049392191 |
| Gm15223 | 0.211816017 | 0.71822132 | 2.18128993 |
| Tceanc | 44.56431763 | 0.878807744 | 0.204552244 |
| Egfl6 | 12.75317809 | -0.774254696 | 0.339649699 |
| Gm15226 | 0.151064894 | 0.718212558 | 2.848768331 |
| Gm1720 | 0.332179105 | -0.608199234 | 1.572648959 |
| Rpl7a-ps11 | 9.849636779 | 0.664666627 | 0.349142297 |
| Gm15230 | 0.199910846 | 0.67520288 | 2.208306885 |
| Gm15231 | 0 | NA | NA |
| AL671888.1 | 0.061920613 | 0.718207094 | 3.724945049 |
| Gm22368 | 0 | NA | NA |
| Gm8814 | 0.110595523 | 0.718200421 | 3.724945049 |
| Gm8817 | 0.926280627 | 2.424251304 | 1.155235034 |
| Gm15232 | 1.683504739 | 0.272867373 | 0.903670043 |
| Gm15228 | 2.037836299 | 3.649417367 | 0.980957047 |
| Tmsb4x | 4681.12954 | 0.679222571 | 0.106178407 |
| Tlr8 | 77.00154128 | 5.964587762 | 0.584931508 |
| Tlr7 | 273.9150659 | 3.400609216 | 0.207294802 |
| Prps2 | 362.3824202 | -0.104472484 | 0.098759108 |
| Gm15239 | 0.195968864 | 0.237310703 | 3.246343675 |
| Gm26368 | 0.238567011 | 1.850643534 | 2.543006132 |
| Gm6744 | 0.523620968 | -0.243583646 | 3.148671252 |
| Frmpd4 | 1476.981339 | -0.58659358 | 0.071080162 |
| Gm15238 | 0.110870145 | 0.238305784 | 3.724874707 |
| Gm24772 | 0 | NA | NA |
| Gm15240 | 0 | NA | NA |
| Gm15237 | 0 | NA | NA |
| Gm8832 | 0.092228164 | 0.718202646 | 3.724945049 |
| Gm7209 | 0 | NA | NA |
| Gm23197 | 0 | NA | NA |
| Msl3 | 245.0695052 | 0.079778394 | 0.074217134 |
| Gm15233 | 0 | NA | NA |
| Arhgap6 | 136.4378488 | -0.671715446 | 0.131439383 |
| Gm15261 | 0.130746305 | 0.703205782 | 3.666340321 |
| Amelx | 0.092691418 | 1.183079489 | 3.724560917 |
| Hccs | 283.4725314 | 0.201444875 | 0.101044998 |
| Gm27980 | 0.850639233 | -0.101397326 | 1.238374964 |
| Gm15246 | 24.67077973 | -0.889939739 | 0.303921008 |

| | | | |
|---|---|---|---|
| BC022960 | 47.81857404 | -0.83216475 | 0.174616714 |
| Gm15245 | 0.062426873 | 0.237312084 | 3.724945049 |
| Mid1 | 46.97430459 | 1.044419261 | 0.195559858 |
| Gm25651 | 0 | NA | NA |
| 4933400A11Rik | 0 | NA | NA |
| G530011O06Rik | 0.809500859 | 2.916405868 | 1.372553779 |
| Gm15726 | 1.372866643 | -0.062204817 | 0.821443297 |
| Gm15247 | 0.097139547 | 0.718207094 | 3.724945049 |
| Gm21887 | 20.27296318 | -0.313315336 | 0.375092415 |
| Asmt | 1.358880654 | 0.185628199 | 0.92846701 |
| Gm15069 | 0 | NA | NA |
| Gm15068 | 0 | NA | NA |
| Gm29277 | 0 | NA | NA |
| Gm29089 | 0 | NA | NA |
| Gm2098 | 0.060174537 | 0.237309859 | 3.724945049 |
| Zfy1 | 0.044191549 | -0.243571806 | 3.724945049 |
| Uba1y | 4.871321793 | 0.826795906 | 0.527671441 |
| Gm28588 | 0 | NA | NA |
| Gm28587 | 0.444799355 | -0.992197077 | 3.105741306 |
| Rhoay-ps3 | 0.088383099 | -0.72445792 | 3.724945049 |
| Kdm5d | 320.9905738 | -0.335804133 | 0.186814149 |
| n-R5s1 | 0 | NA | NA |
| AC145392.1 | 0 | NA | NA |
| Eif2s3y | 295.8023547 | 0.088773811 | 0.1601928 |
| Gm29650 | 9.742440598 | -0.727261863 | 0.57753803 |
| Tspy-ps | 2.062938699 | -0.551835679 | 0.991085141 |
| Uty | 395.1440632 | 0.051301913 | 0.125757277 |
| Ddx3y | 555.3045698 | 0.425636108 | 0.145938094 |
| Gm4017 | 4.097680658 | -0.360364605 | 0.638073319 |
| Gm18665 | 3.611403245 | 0.041822493 | 0.6021817 |
| Usp9y | 1.14946738 | -1.697613942 | 3.650689424 |
| Gm25565 | 0 | NA | NA |
| Gm17790 | 0 | NA | NA |
| Gm29167 | 0 | NA | NA |
| Gm8446 | 0.334381441 | -1.25273902 | 1.743550105 |
| Gm28534 | 0 | NA | NA |
| Gm28975 | 0 | NA | NA |
| Gm28974 | 0 | NA | NA |
| Gm18797 | 0 | NA | NA |
| Rhoay-ps1 | 0 | NA | NA |
| Zfy2 | 0 | NA | NA |
| Uba1y-ps2 | 0 | NA | NA |
| Gm18796 | 0 | NA | NA |
| Gm29552 | 0 | NA | NA |
| Gm29550 | 0 | NA | NA |
| Gm29551 | 0 | NA | NA |
| Gm8480 | 0 | NA | NA |

| | | | | |
|---|---|---|---|---|
| Gm29553 | 0 | NA | NA | |
| Gm22119 | 0 | NA | NA | |
| Gm28630 | 0 | NA | NA | |
| Gm28629 | 0 | NA | NA | |
| Gm6026 | 0 | NA | NA | |
| Uba1y-ps1 | 0 | NA | NA | |
| Sry | 0 | NA | NA | |
| Gm18798 | 0 | NA | NA | |
| Gm16501 | 0 | NA | NA | |
| Gm28628 | 0 | NA | NA | |
| Gm4064 | 0 | NA | NA | |
| Gm6137 | 0 | NA | NA | |
| Gm8498 | 0 | NA | NA | |
| Rbmy | 0 | NA | NA | |
| Gm10256 | 0 | NA | NA | |
| Gm8502 | 0 | NA | NA | |
| Gm8506 | 0 | NA | NA | |
| Gm10352 | 0 | NA | NA | |
| Gm8510 | 0 | NA | NA | |
| Gm29289 | 0 | NA | NA | |
| Gm21677 | 0 | NA | NA | |
| Gm29288 | 0 | NA | NA | |
| Gm21693 | 0 | NA | NA | |
| Gm21139 | 0 | NA | NA | |
| Gm21704 | 0 | NA | NA | |
| Gm21147 | 0 | NA | NA | |
| Gm21708 | 0 | NA | NA | |
| Gm21156 | 0 | NA | NA | |
| Rhoay-ps2 | 0 | NA | NA | |
| Gm2316 | 0 | NA | NA | |
| Gm3376 | 0 | NA | NA | |
| Gm2325 | 0 | NA | NA | |
| Gm28649 | 0 | NA | NA | |
| Gm8521 | 0 | NA | NA | |
| Gm18177 | 0 | NA | NA | |
| Gm28429 | 0 | NA | NA | |
| Gm28413 | 0 | NA | NA | |
| Gm28412 | 0 | NA | NA | |
| Gm8525 | 0 | NA | NA | |
| Gm28414 | 0 | NA | NA | |
| Gm28648 | 0 | NA | NA | |
| Gm28647 | 0 | NA | NA | |
| Gm29435 | 0 | NA | NA | |
| Vmn2r-ps139 | 0.046345709 | 0.71820487 | | 3.724945049 |
| Gm29038 | 0 | NA | NA | |
| Gm28191 | 0 | NA | NA | |
| Gm28186 | 0 | NA | NA | |

| | | | |
|---|---|---|---|
| Gm21064 | 0 | NA | NA |
| Gm28241 | 0 | NA | NA |
| Gm28242 | 0 | NA | NA |
| Gm37222 | 0 | NA | NA |
| Gm28356 | 0 | NA | NA |
| Gm28357 | 0 | NA | NA |
| Gm29352 | 0 | NA | NA |
| Gm29351 | 0 | NA | NA |
| Gm29350 | 0 | NA | NA |
| Gm29349 | 0 | NA | NA |
| Gm29355 | 0 | NA | NA |
| Gm20918 | 0 | NA | NA |
| Gm29354 | 0 | NA | NA |
| Gm29353 | 0 | NA | NA |
| Gm21820 | 0 | NA | NA |
| Gm29357 | 0 | NA | NA |
| Gm29356 | 0 | NA | NA |
| Gm29161 | 0 | NA | NA |
| Gm28767 | 0 | NA | NA |
| Gm29160 | 0 | NA | NA |
| Gm20915 | 0 | NA | NA |
| Gm28766 | 0 | NA | NA |
| Gm29159 | 0 | NA | NA |
| Gm27663 | 0 | NA | NA |
| Gm29158 | 0 | NA | NA |
| Gm21854 | 0 | NA | NA |
| Gm28768 | 0 | NA | NA |
| Gm28769 | 0 | NA | NA |
| Gm20914 | 0 | NA | NA |
| Gm28171 | 0 | NA | NA |
| Gm28173 | 0 | NA | NA |
| Gm37236 | 0 | NA | NA |
| Gm29194 | 0.066050875 | 0.71820487 | 3.724945049 |
| Gm21778 | 0 | NA | NA |
| Gm29195 | 0 | NA | NA |
| Gm29196 | 0 | NA | NA |
| Gm28435 | 0 | NA | NA |
| Gm28436 | 0 | NA | NA |
| Gm29197 | 0 | NA | NA |
| Gm29198 | 0 | NA | NA |
| Gm29193 | 0 | NA | NA |
| Gm28430 | 0 | NA | NA |
| Gm21746 | 0 | NA | NA |
| Gm28593 | 0 | NA | NA |
| Gm20873 | 0 | NA | NA |
| Gm28916 | 0 | NA | NA |
| Gm28915 | 0 | NA | NA |

| | | | |
|---|---|---|---|
| Gm28918 | 0.060174537 | 0.237309859 | 3.724945049 |
| Gm28917 | 0 | NA | NA |
| Gm28919 | 0 | NA | NA |
| Gm21719 | 0 | NA | NA |
| Gm28921 | 0 | NA | NA |
| Gm28920 | 0 | NA | NA |
| Gm28444 | 0 | NA | NA |
| Gm28443 | 0 | NA | NA |
| Gm20830 | 0 | NA | NA |
| Gm28442 | 0 | NA | NA |
| Gm28445 | 0 | NA | NA |
| Gm20788 | 0 | NA | NA |
| Gm28999 | 0 | NA | NA |
| Gm21244 | 0.139376635 | -0.243570432 | 2.924218065 |
| Gm28997 | 0 | NA | NA |
| Gm28998 | 0 | NA | NA |
| Gm28398 | 0 | NA | NA |
| Gm28397 | 0 | NA | NA |
| Gm28400 | 0 | NA | NA |
| Gm28399 | 0 | NA | NA |
| Gm28395 | 0 | NA | NA |
| Gm21788 | 0 | NA | NA |
| Gm28394 | 0 | NA | NA |
| Gm28396 | 0 | NA | NA |
| Gm28393 | 0 | NA | NA |
| Gm20826 | 0.204257686 | -1.138338946 | 2.624259596 |
| Vmn2r-ps149 | 0 | NA | NA |
| Gm28576 | 0 | NA | NA |
| Gm28575 | 0 | NA | NA |
| Gm28574 | 0 | NA | NA |
| Gm28573 | 0 | NA | NA |
| Gm28572 | 0 | NA | NA |
| Gm28571 | 0 | NA | NA |
| Gm28570 | 0 | NA | NA |
| Gm29049 | 0 | NA | NA |
| Gm20825 | 0 | NA | NA |
| Gm29047 | 0 | NA | NA |
| Gm29048 | 0 | NA | NA |
| Gm29045 | 0 | NA | NA |
| Gm29046 | 0 | NA | NA |
| Gm29043 | 0 | NA | NA |
| Gm29044 | 0 | NA | NA |
| Gm37951 | 0 | NA | NA |
| Gm20824 | 0 | NA | NA |
| Gm28146 | 0 | NA | NA |
| Gm28145 | 0 | NA | NA |
| Gm28148 | 0 | NA | NA |

| | | | |
|---|---|---|---|
| Gm28147 | 0 | NA | NA |
| Gm28142 | 0 | NA | NA |
| Gm28141 | 0 | NA | NA |
| Gm28144 | 0 | NA | NA |
| Gm28143 | 0 | NA | NA |
| Gm28150 | 0 | NA | NA |
| Gm28149 | 0 | NA | NA |
| Gm28098 | 0 | NA | NA |
| Gm28330 | 0 | NA | NA |
| Gm28331 | 0 | NA | NA |
| Gm28099 | 0 | NA | NA |
| Gm20815 | 0 | NA | NA |
| Gm28328 | 0 | NA | NA |
| Gm28329 | 0 | NA | NA |
| Gm28270 | 0 | NA | NA |
| Gm28327 | 0 | NA | NA |
| Gm28097 | 0 | NA | NA |
| Gm29365 | 0 | NA | NA |
| Gm29364 | 0 | NA | NA |
| Gm29363 | 0 | NA | NA |
| Gm21292 | 0 | NA | NA |
| Gm21721 | 0 | NA | NA |
| Gm29366 | 0 | NA | NA |
| Gm29281 | 0 | NA | NA |
| Gm29330 | 0 | NA | NA |
| Gm29329 | 0 | NA | NA |
| Gm21812 | 0 | NA | NA |
| Gm29535 | 0 | NA | NA |
| Gm29533 | 0 | NA | NA |
| Gm29534 | 0 | NA | NA |
| Gm21874 | 0 | NA | NA |
| Gm28849 | 0 | NA | NA |
| Gm28847 | 0 | NA | NA |
| Gm28848 | 0 | NA | NA |
| Gm28846 | 0 | NA | NA |
| Gm29471 | 0 | NA | NA |
| Gm29474 | 0 | NA | NA |
| Gm29475 | 0 | NA | NA |
| Gm20821 | 0 | NA | NA |
| Gm28506 | 0 | NA | NA |
| Gm28507 | 0 | NA | NA |
| Gm21301 | 0 | NA | NA |
| Gm29142 | 0 | NA | NA |
| Gm21310 | 0 | NA | NA |
| Gm28509 | 0 | NA | NA |
| Gm28508 | 0 | NA | NA |
| Gm20834 | 0 | NA | NA |

| | | | |
|---|---|---|---|
| Gm28510 | 0 | NA | NA |
| Gm29525 | 0 | NA | NA |
| Gm29526 | 0 | NA | NA |
| Gm20737 | 0.044544648 | 0.237309859 | 3.724945049 |
| Gm21775 | 0 | NA | NA |
| Gm21900 | 0 | NA | NA |
| Gm29527 | 0 | NA | NA |
| Gm20775 | 1.978445198 | 1.44314189 | 0.752903783 |
| Gm29522 | 0 | NA | NA |
| Gm29523 | 0 | NA | NA |
| Gm29524 | 0 | NA | NA |
| Gm29528 | 0 | NA | NA |
| Gm20777 | 0 | NA | NA |
| Gm28955 | 0 | NA | NA |
| Gm28954 | 0 | NA | NA |
| Gm28953 | 0 | NA | NA |
| Gm20776 | 0 | NA | NA |
| Gm28958 | 0 | NA | NA |
| Gm20828 | 0 | NA | NA |
| Gm28957 | 0 | NA | NA |
| Gm28956 | 0 | NA | NA |
| Gm28952 | 0 | NA | NA |
| Gm28951 | 0 | NA | NA |
| Gm28390 | 0 | NA | NA |
| Gm28391 | 0 | NA | NA |
| Gm28389 | 0 | NA | NA |
| Gm28655 | 0 | NA | NA |
| Gm28654 | 0 | NA | NA |
| Gm28657 | 0 | NA | NA |
| Gm28656 | 0 | NA | NA |
| Gm29554 | 0 | NA | NA |
| Gm29556 | 0 | NA | NA |
| Gm20812 | 0 | NA | NA |
| Gm35108 | 0 | NA | NA |
| Gm37739 | 0 | NA | NA |
| Gm21736 | 0 | NA | NA |
| Gm21379 | 0 | NA | NA |
| Gm36928 | 0 | NA | NA |
| Gm20807 | 0 | NA | NA |
| Gm38084 | 0 | NA | NA |
| Gm37231 | 0 | NA | NA |
| Gm21891 | 0 | NA | NA |
| Gm21425 | 0 | NA | NA |
| Gm37252 | 0 | NA | NA |
| Gm21828 | 0 | NA | NA |
| Gm21440 | 0 | NA | NA |
| Gm21454 | 0 | NA | NA |

| | | | | |
|---|---|---|---|---|
| Gm37147 | 0 | NA | NA | |
| Gm38159 | 0 | NA | NA | |
| Gm21725 | 0 | NA | NA | |
| Gm37870 | 0 | NA | NA | |
| Gm37378 | 0 | NA | NA | |
| Gm21866 | 0 | NA | NA | |
| Gm20773 | 0 | NA | NA | |
| Gm36398 | 0 | NA | NA | |
| Gm37441 | 0 | NA | NA | |
| Gm37561 | 0 | NA | NA | |
| Gm36728 | 0 | NA | NA | |
| Gm32295 | 0 | NA | NA | |
| Gm20836 | 0 | NA | NA | |
| Gm37263 | 0 | NA | NA | |
| Gm21767 | 0 | NA | NA | |
| Gm37130 | 0 | NA | NA | |
| Gm37467 | 0 | NA | NA | |
| Gm20928 | 0 | NA | NA | |
| Gm38363 | 0 | NA | NA | |
| Gm30174 | 0 | NA | NA | |
| Gm30353 | 0 | NA | NA | |
| Gm30686 | 0 | NA | NA | |
| Gm30737 | 0 | NA | NA | |
| Gm36987 | 0 | NA | NA | |
| Gm31186 | 0 | NA | NA | |
| Gm37654 | 0 | NA | NA | |
| Gm37044 | 0 | NA | NA | |
| Gm31511 | 0 | NA | NA | |
| Gm37572 | 0 | NA | NA | |
| Gm37344 | 0 | NA | NA | |
| Gm37059 | 0 | NA | NA | |
| Gm31942 | 0 | NA | NA | |
| Gm37440 | 0 | NA | NA | |
| Gm38185 | 0 | NA | NA | |
| Gm37395 | 0 | NA | NA | |
| Gm37577 | 0 | NA | NA | |
| Gm20822 | 0 | NA | NA | |
| Gm38183 | 0 | NA | NA | |
| Gm38149 | 0 | NA | NA | |
| Gm38145 | 0 | NA | NA | |
| Gm32033 | 0 | NA | NA | |
| Gm32447 | 0 | NA | NA | |
| Gm20770 | 0 | NA | NA | |
| Gm38016 | 0 | NA | NA | |
| Gm37158 | 0.04393175 | | -0.243571806 | 3.724945049 |
| Gm36998 | 0 | NA | NA | |
| Gm33112 | 0 | NA | NA | |

| | | | |
|---|---|---|---|
| Gm21809 | 0 | NA | NA |
| Gm34015 | 0 | NA | NA |
| Gm20877 | 0 | NA | NA |
| Gm33191 | 0 | NA | NA |
| Gm38003 | 0 | NA | NA |
| Gm20781 | 0 | NA | NA |
| Rbm31y | 0 | NA | NA |
| Gm37325 | 0 | NA | NA |
| Gm20833 | 0 | NA | NA |
| Gm37865 | 0 | NA | NA |
| Gm36941 | 0 | NA | NA |
| Gm21904 | 0 | NA | NA |
| Gm37937 | 0 | NA | NA |
| Gm20772 | 0 | NA | NA |
| Gm20831 | 0 | NA | NA |
| Gm37286 | 0 | NA | NA |
| Srsy | 0 | NA | NA |
| Gm38028 | 0 | NA | NA |
| Gm21764 | 0 | NA | NA |
| Ssty1 | 0 | NA | NA |
| Gm38072 | 0 | NA | NA |
| Gm20851 | 0 | NA | NA |
| Gm30312 | 0 | NA | NA |
| Gm37433 | 0 | NA | NA |
| Gm37473 | 0 | NA | NA |
| Gm37656 | 0 | NA | NA |
| Gm38127 | 0 | NA | NA |
| Gm30638 | 0 | NA | NA |
| Gm38361 | 0 | NA | NA |
| Gm31260 | 0 | NA | NA |
| Gm21660 | 0 | NA | NA |
| Gm38209 | 0 | NA | NA |
| Gm21822 | 0 | NA | NA |
| Gm37734 | 0 | NA | NA |
| Gm38013 | 0 | NA | NA |
| Gm21678 | 0 | NA | NA |
| Gm37948 | 0 | NA | NA |
| Gm37442 | 0 | NA | NA |
| Gm34550 | 0 | NA | NA |
| Gm37544 | 0 | NA | NA |
| Gm34716 | 0 | NA | NA |
| Gm37635 | 0 | NA | NA |
| Gm35070 | 0 | NA | NA |
| Gm35134 | 0 | NA | NA |
| Gm37998 | 0 | NA | NA |
| Gm38094 | 0 | NA | NA |
| Gm21763 | 0 | NA | NA |

| | | | | |
|---|---|---|---|---|
| Gm38369 | 0 | NA | NA | |
| Gm35843 | 0 | NA | NA | |
| Gm38168 | 0 | NA | NA | |
| Gm37740 | 0 | NA | NA | |
| Gm27701 | 0 | NA | NA | |
| Gm37723 | 0 | NA | NA | |
| Gm36261 | 0 | NA | NA | |
| Gm37898 | 0 | NA | NA | |
| Gm36467 | 0 | NA | NA | |
| Gm37927 | 0 | NA | NA | |
| Gm36891 | 0 | NA | NA | |
| Gm36929 | 0 | NA | NA | |
| Gm38174 | 0 | NA | NA | |
| Gm29866 | 0 | NA | NA | |
| Gm36950 | 0 | NA | NA | |
| Gm20910 | 0 | NA | NA | |
| Gm30598 | 0 | NA | NA | |
| Gm20909 | 0 | NA | NA | |
| Gm20865 | 0 | NA | NA | |
| Gm37462 | 0 | NA | NA | |
| Gm37658 | 0 | NA | NA | |
| Gm32347 | 0 | NA | NA | |
| Gm32419 | 0 | NA | NA | |
| Gm38370 | 0 | NA | NA | |
| Gm38054 | 0 | NA | NA | |
| Gm37476 | 0 | NA | NA | |
| Gm37369 | 0 | NA | NA | |
| Gm38181 | 0 | NA | NA | |
| Gm31335 | 0 | NA | NA | |
| Gm37690 | 0 | NA | NA | |
| Gm38107 | 0 | NA | NA | |
| Gm31571 | 0 | NA | NA | |
| Gm32114 | 0 | NA | NA | |
| Gm37075 | 0 | NA | NA | |
| Gm37921 | 0 | NA | NA | |
| Gm32180 | 0 | NA | NA | |
| Gm38371 | 0 | NA | NA | |
| Gm37657 | 0 | NA | NA | |
| Gm37448 | 0 | NA | NA | |
| Gm33163 | 0 | NA | NA | |
| Gm21065 | 0.044191549 | -0.243571806 | | 3.724945049 |
| Gm36978 | 0 | NA | NA | |
| Gm37721 | 0 | NA | NA | |
| Gm37129 | 0 | NA | NA | |
| Gm37990 | 0 | NA | NA | |
| Gm21366 | 0 | NA | NA | |
| Gm33954 | 0 | NA | NA | |

| | | | |
|---|---|---|---|
| Gm37538 | 0 | NA | NA |
| Gm21871 | 0 | NA | NA |
| Gm34274 | 0 | NA | NA |
| Gm34217 | 0 | NA | NA |
| Gm21469 | 0 | NA | NA |
| Gm21661 | 0 | NA | NA |
| Gm37952 | 0 | NA | NA |
| Gm21907 | 0 | NA | NA |
| Gm37627 | 0 | NA | NA |
| Gm20925 | 0 | NA | NA |
| Gm20926 | 0 | NA | NA |
| Gm37454 | 0 | NA | NA |
| Gm38138 | 0 | NA | NA |
| Gm29740 | 0 | NA | NA |
| Gm37840 | 0 | NA | NA |
| Gm37574 | 0 | NA | NA |
| Gm21768 | 0 | NA | NA |
| Gm37172 | 0 | NA | NA |
| Gm35670 | 0 | NA | NA |
| Gm20840 | 0 | NA | NA |
| Gm35955 | 0 | NA | NA |
| Gm37687 | 0 | NA | NA |
| Gm38296 | 0 | NA | NA |
| Gm38290 | 0 | NA | NA |
| Gm38121 | 0 | NA | NA |
| Gm36047 | 0 | NA | NA |
| Gm36345 | 0 | NA | NA |
| Gm37157 | 0 | NA | NA |
| Gm38017 | 0 | NA | NA |
| Gm36468 | 0 | NA | NA |
| Gm21845 | 0 | NA | NA |
| Gm20809 | 0 | NA | NA |
| Gm37434 | 0 | NA | NA |
| Gm37765 | 0 | NA | NA |
| Gm38229 | 0 | NA | NA |
| Gm36866 | 0 | NA | NA |
| Gm30045 | 0 | NA | NA |
| Gm21921 | 0 | NA | NA |
| Gm37112 | 0 | NA | NA |
| Gm37287 | 0 | NA | NA |
| Gm30175 | 0 | NA | NA |
| Gm21350 | 0 | NA | NA |
| Gm21412 | 0 | NA | NA |
| Gm37071 | 0 | NA | NA |
| Gm37547 | 0 | NA | NA |
| Gm36968 | 0 | NA | NA |
| Gm38035 | 0 | NA | NA |

| | | | |
|---|---|---|---|
| Gm20857 | 0 | NA | NA |
| Gm30705 | 0 | NA | NA |
| Gm21773 | 0 | NA | NA |
| Gm37451 | 0 | NA | NA |
| Gm37793 | 0 | NA | NA |
| Gm30858 | 0 | NA | NA |
| Gm21791 | 0 | NA | NA |
| Gm21427 | 0 | NA | NA |
| Gm37875 | 0 | NA | NA |
| Gm37237 | 0 | NA | NA |
| Gm38350 | 0 | NA | NA |
| Gm20858 | 0 | NA | NA |
| Gm31422 | 0 | NA | NA |
| Gm37798 | 0 | NA | NA |
| Gm37318 | 0 | NA | NA |
| Gm31572 | 0 | NA | NA |
| Gm21890 | 0 | NA | NA |
| Gm21443 | 0 | NA | NA |
| Gm37346 | 0 | NA | NA |
| Gm38136 | 0 | NA | NA |
| Gm37991 | 0 | NA | NA |
| Gm37808 | 0 | NA | NA |
| Gm33571 | 0 | NA | NA |
| Gm32181 | 0 | NA | NA |
| Gm28662 | 0 | NA | NA |
| Gm28780 | 0 | NA | NA |
| Gm21755 | 0 | NA | NA |
| Gm20895 | 0 | NA | NA |
| Gm28853 | 0 | NA | NA |
| Gm28852 | 0 | NA | NA |
| Gm28855 | 0 | NA | NA |
| Gm28854 | 0 | NA | NA |
| Gm28858 | 0 | NA | NA |
| Gm20890 | 0 | NA | NA |
| Gm28245 | 0 | NA | NA |
| Gm28246 | 0 | NA | NA |
| Gm28247 | 0 | NA | NA |
| Gm28248 | 0 | NA | NA |
| Gm20892 | 0 | NA | NA |
| Gm28249 | 0 | NA | NA |
| Gm28250 | 0 | NA | NA |
| Gm28251 | 0 | NA | NA |
| Gm28243 | 0 | NA | NA |
| Gm28244 | 0 | NA | NA |
| Gm21771 | 0 | NA | NA |
| Gm21476 | 0 | NA | NA |
| Gm29021 | 0 | NA | NA |

| | | | |
|---|---|---|---|
| Gm28880 | 0.044191549 | -0.243571806 | 3.724945049 |
| Gm28452 | 0 | NA | NA |
| Gm21488 | 0 | NA | NA |
| Gm28454 | 0 | NA | NA |
| Gm28878 | 0 | NA | NA |
| Gm28879 | 0 | NA | NA |
| Gm29369 | 0 | NA | NA |
| Gm21506 | 0 | NA | NA |
| Gm29399 | 0 | NA | NA |
| Gm29400 | 0 | NA | NA |
| Gm29401 | 0 | NA | NA |
| Gm20891 | 0 | NA | NA |
| Gm20893 | 0 | NA | NA |
| Gm28799 | 0 | NA | NA |
| Gm28798 | 0 | NA | NA |
| Gm28797 | 0 | NA | NA |
| Gm28796 | 0 | NA | NA |
| Gm20894 | 0 | NA | NA |
| Gm28938 | 0 | NA | NA |
| Gm28937 | 0 | NA | NA |
| Gm28936 | 0 | NA | NA |
| Gm21642 | 0 | NA | NA |
| Gm21804 | 0 | NA | NA |
| Gm29122 | 0 | NA | NA |
| Gm28939 | 0 | NA | NA |
| Gm28983 | 0 | NA | NA |
| Gm28985 | 0 | NA | NA |
| Gm20810 | 0 | NA | NA |
| Gm20811 | 0 | NA | NA |
| Gm28725 | 0 | NA | NA |
| Gm28984 | 0 | NA | NA |
| Gm28987 | 0 | NA | NA |
| Gm28986 | 0 | NA | NA |
| Gm29171 | 0 | NA | NA |
| Gm28129 | 0 | NA | NA |
| Gm28127 | 0 | NA | NA |
| Gm28128 | 0 | NA | NA |
| Gm21853 | 0 | NA | NA |
| Gm28132 | 0 | NA | NA |
| Gm28130 | 0 | NA | NA |
| Gm28131 | 0 | NA | NA |
| Gm21679 | 0 | NA | NA |
| Gm29317 | 0 | NA | NA |
| Gm29316 | 0 | NA | NA |
| Gm29315 | 0 | NA | NA |
| Gm28216 | 0 | NA | NA |
| Gm28217 | 0 | NA | NA |

| | | | |
|---|---|---|---|
| Gm28093 | 0 | NA | NA |
| Gm28092 | 0 | NA | NA |
| Gm28091 | 0 | NA | NA |
| Gm28090 | 0 | NA | NA |
| Gm29580 | 0 | NA | NA |
| Gm28089 | 0 | NA | NA |
| Gm28088 | 0 | NA | NA |
| Gm29579 | 0 | NA | NA |
| Gm29582 | 0 | NA | NA |
| Gm21529 | 0 | NA | NA |
| Gm29581 | 0 | NA | NA |
| Gm21539 | 0 | NA | NA |
| Gm29578 | 0 | NA | NA |
| Gm21908 | 0 | NA | NA |
| Gm21780 | 0 | NA | NA |
| Gm28486 | 0 | NA | NA |
| Gm21737 | 0 | NA | NA |
| Gm28487 | 0 | NA | NA |
| Gm28489 | 0 | NA | NA |
| Gm28488 | 0 | NA | NA |
| Gm28491 | 0 | NA | NA |
| Gm28490 | 0 | NA | NA |
| Gm21562 | 0 | NA | NA |
| Gm28298 | 0 | NA | NA |
| Gm21572 | 0 | NA | NA |
| Gm28297 | 0 | NA | NA |
| Gm21753 | 0 | NA | NA |
| Gm21723 | 0 | NA | NA |
| Gm28296 | 0 | NA | NA |
| Gm21821 | 0 | NA | NA |
| Gm28295 | 0 | NA | NA |
| Gm21842 | 0 | NA | NA |
| Gm28170 | 0 | NA | NA |
| Gm21744 | 0 | NA | NA |
| Gm28764 | 0 | NA | NA |
| Gm28762 | 0 | NA | NA |
| Gm28763 | 0 | NA | NA |
| Gm21094 | 0 | NA | NA |
| Gm21588 | 0 | NA | NA |
| Gm28761 | 0 | NA | NA |
| Gm21599 | 0 | NA | NA |
| Gm28765 | 0 | NA | NA |
| Gm21810 | 0 | NA | NA |
| Gm21916 | 0 | NA | NA |
| Gm29191 | 0 | NA | NA |
| Gm21756 | 0 | NA | NA |
| Gm28210 | 0 | NA | NA |

| | | | |
|---|---|---|---|
| Gm29074 | 0 | NA | NA |
| Gm28211 | 0 | NA | NA |
| Gm28518 | 0 | NA | NA |
| Gm28519 | 0 | NA | NA |
| Gm28520 | 0 | NA | NA |
| Gm20838 | 0 | NA | NA |
| Gm21626 | 0 | NA | NA |
| Gm28521 | 0 | NA | NA |
| Gm21633 | 0 | NA | NA |
| Gm28522 | 0 | NA | NA |
| Gm21745 | 0 | NA | NA |
| Gm28517 | 0 | NA | NA |
| Gm29532 | 0 | NA | NA |
| Gm29531 | 0 | NA | NA |
| Gm28464 | 0 | NA | NA |
| Gm29656 | 0 | NA | NA |
| Gm29655 | 0 | NA | NA |
| Gm21647 | 0 | NA | NA |
| Gm29653 | 0 | NA | NA |
| Gm29654 | 0 | NA | NA |
| Gm29651 | 0 | NA | NA |
| Gm29652 | 0 | NA | NA |
| Gm29080 | 0 | NA | NA |
| Gm21184 | 0 | NA | NA |
| Gm21201 | 0 | NA | NA |
| Gm29081 | 0 | NA | NA |
| Gm29077 | 0 | NA | NA |
| Gm29076 | 0 | NA | NA |
| Gm29079 | 0 | NA | NA |
| Gm29078 | 0 | NA | NA |
| Gm20853 | 0 | NA | NA |
| Gm29075 | 0 | NA | NA |
| Gm29225 | 0 | NA | NA |
| Gm29226 | 0 | NA | NA |
| Gm29227 | 0 | NA | NA |
| Gm28339 | 0 | NA | NA |
| Gm20855 | 0 | NA | NA |
| Gm29386 | 0 | NA | NA |
| Gm29384 | 0 | NA | NA |
| Gm21896 | 0 | NA | NA |
| Gm29280 | 0 | NA | NA |
| Gm20799 | 0 | NA | NA |
| Gm29279 | 0 | NA | NA |
| Gm28260 | 0 | NA | NA |
| Gm21795 | 0 | NA | NA |
| Gm28261 | 0 | NA | NA |
| Gm28264 | 0 | NA | NA |

| | | | |
|---|---|---|---|
| Gm21872 | 0 | NA | NA |
| Gm20798 | 0 | NA | NA |
| Gm28463 | 0 | NA | NA |
| Gm28584 | 0 | NA | NA |
| Gm28585 | 0 | NA | NA |
| Gm20896 | 0 | NA | NA |
| Gm28470 | 0 | NA | NA |
| Gm28469 | 0 | NA | NA |
| Gm21898 | 0 | NA | NA |
| Gm28468 | 0 | NA | NA |
| Gm20898 | 0 | NA | NA |
| Gm29333 | 0 | NA | NA |
| Gm28332 | 0 | NA | NA |
| Gm28333 | 0 | NA | NA |
| Gm28334 | 0 | NA | NA |
| Gm28335 | 0 | NA | NA |
| Gm28336 | 0 | NA | NA |
| Gm28337 | 0 | NA | NA |
| Gm28338 | 0 | NA | NA |
| Gm28812 | 0 | NA | NA |
| Gm28811 | 0 | NA | NA |
| Gm28810 | 0 | NA | NA |
| Gm20835 | 0 | NA | NA |
| Gm29027 | 0 | NA | NA |
| Gm29028 | 0 | NA | NA |
| Gm29029 | 0 | NA | NA |
| Gm29030 | 0 | NA | NA |
| Gm29023 | 0 | NA | NA |
| Gm29024 | 0 | NA | NA |
| Gm29025 | 0 | NA | NA |
| Gm29026 | 0 | NA | NA |
| Gm28213 | 0 | NA | NA |
| Gm28212 | 0 | NA | NA |
| Gm21799 | 0 | NA | NA |
| Gm28227 | 0 | NA | NA |
| Gm20905 | 0 | NA | NA |
| Gm29411 | 0 | NA | NA |
| Gm28222 | 0 | NA | NA |
| Gm21835 | 0 | NA | NA |
| Gm28820 | 0 | NA | NA |
| Gm28223 | 0 | NA | NA |
| Gm29412 | 0 | NA | NA |
| Gm29413 | 0 | NA | NA |
| Gm29671 | 0 | NA | NA |
| Gm21878 | 0 | NA | NA |
| Gm28162 | 0 | NA | NA |
| Gm20922 | 0 | NA | NA |

| | | | |
|---|---|---|---|
| Gm20738 | 0 | NA | NA |
| Gm28545 | 0 | NA | NA |
| Gm28547 | 0 | NA | NA |
| Gm28546 | 0 | NA | NA |
| Gm21924 | 0 | NA | NA |
| Gm21735 | 0 | NA | NA |
| Gm28771 | 0 | NA | NA |
| Gm28772 | 0 | NA | NA |
| Gm28773 | 0 | NA | NA |
| Gm20897 | 0 | NA | NA |
| Gm28774 | 0 | NA | NA |
| Gm21732 | 0 | NA | NA |
| Gm28775 | 0 | NA | NA |
| Gm21765 | 0 | NA | NA |
| Gm20899 | 12.70112357 | -0.084990573 | 0.330869435 |
| Gm28776 | 0 | NA | NA |
| Gm21787 | 0 | NA | NA |
| Gm28770 | 0 | NA | NA |
| Gm20901 | 0 | NA | NA |
| Gm28176 | 0 | NA | NA |
| Gm28786 | 0 | NA | NA |
| Gm29360 | 0 | NA | NA |
| Gm29496 | 0 | NA | NA |
| Gm29497 | 0 | NA | NA |
| Gm29499 | 0 | NA | NA |
| Gm29498 | 0 | NA | NA |
| Gm36782 | 0 | NA | NA |
| Gm29839 | 0 | NA | NA |
| Gm29495 | 0 | NA | NA |
| Gm28371 | 0.044191549 | -0.243571806 | 3.724945049 |
| Gm28595 | 0 | NA | NA |
| Gm37032 | 0 | NA | NA |
| Gm21865 | 0 | NA | NA |
| Gm29433 | 0 | NA | NA |
| Gm29215 | 0 | NA | NA |
| Gm21882 | 0 | NA | NA |
| Gm21801 | 0 | NA | NA |
| Gm28850 | 0 | NA | NA |
| Gm21914 | 0 | NA | NA |
| Gm28617 | 0 | NA | NA |
| Gm28618 | 0 | NA | NA |
| Gm28619 | 0 | NA | NA |
| Gm21852 | 0 | NA | NA |
| Gm20793 | 0 | NA | NA |
| Gm27466 | 0 | NA | NA |
| Gm28018 | 0 | NA | NA |
| Gm28993 | 0 | NA | NA |

| | | | | |
|---|---|---|---|---|
| Gm28994 | 0 | NA | NA | |
| Gm21797 | 0 | NA | NA | |
| Gm20795 | 0 | NA | NA | |
| Gm28596 | 0 | NA | NA | |
| Gm20832 | 0 | NA | NA | |
| Gm28597 | 0 | NA | NA | |
| Gm28428 | 0 | NA | NA | |
| Gm20827 | 0 | NA | NA | |
| Gm28427 | 0 | NA | NA | |
| Gm21800 | 0 | NA | NA | |
| Gm28426 | 0 | NA | NA | |
| Gm28425 | 0 | NA | NA | |
| Gm21729 | 0 | NA | NA | |
| Gm28834 | 0 | NA | NA | |
| Gm28833 | 0 | NA | NA | |
| Gm21774 | 0 | NA | NA | |
| Gm29537 | 0 | NA | NA | |
| Gm29274 | 0 | NA | NA | |
| Gm29275 | 0.04393175 | -0.243571806 | | 3.724945049 |
| Gm29276 | 0 | NA | NA | |
| Gm21805 | 0 | NA | NA | |
| Gm29645 | 0 | NA | NA | |
| Gm28312 | 0 | NA | NA | |
| Gm29165 | 0 | NA | NA | |
| Gm29042 | 0 | NA | NA | |
| Gm28311 | 0 | NA | NA | |
| Gm28310 | 0 | NA | NA | |
| Gm28504 | 0 | NA | NA | |
| Gm28567 | 0 | NA | NA | |
| Gm28568 | 0 | NA | NA | |
| Gm29286 | 0 | NA | NA | |
| Gm29285 | 0 | NA | NA | |
| Gm28280 | 0 | NA | NA | |
| Gm21241 | 0 | NA | NA | |
| Gm28278 | 0 | NA | NA | |
| Gm28279 | 0 | NA | NA | |
| Gm21739 | 0 | NA | NA | |
| Gm28886 | 0 | NA | NA | |
| Gm29426 | 0 | NA | NA | |
| Gm28885 | 0 | NA | NA | |
| Gm21856 | 0 | NA | NA | |
| Gm28887 | 0 | NA | NA | |
| Gm29444 | 0 | NA | NA | |
| Gm21783 | 0 | NA | NA | |
| Gm29445 | 0 | NA | NA | |
| Gm28457 | 0 | NA | NA | |
| Gm21779 | 0 | NA | NA | |

| | | | |
|---|---|---|---|
| Gm28458 | 0 | NA | NA |
| Gm28459 | 0 | NA | NA |
| Gm28461 | 0 | NA | NA |
| Gm28460 | 0 | NA | NA |
| Gm28851 | 0 | NA | NA |
| Gm28456 | 0 | NA | NA |
| Gm29073 | 0 | NA | NA |
| Gm29648 | 0 | NA | NA |
| Gm29647 | 0 | NA | NA |
| Gm29646 | 0 | NA | NA |
| Gm28549 | 0 | NA | NA |
| Gm29211 | 0 | NA | NA |
| Gm29212 | 0 | NA | NA |
| Gm21247 | 0 | NA | NA |
| Gm29210 | 0 | NA | NA |
| Gm29213 | 0 | NA | NA |
| Gm28550 | 0 | NA | NA |
| Gm28604 | 0 | NA | NA |
| Gm28603 | 0 | NA | NA |
| Gm28565 | 0 | NA | NA |
| Gm28566 | 0 | NA | NA |
| Gm28697 | 0 | NA | NA |
| Gm28554 | 0 | NA | NA |
| Gm21256 | 0 | NA | NA |
| Gm29056 | 0 | NA | NA |
| Gm29098 | 0 | NA | NA |
| Gm21170 | 0 | NA | NA |
| Gm29373 | 0 | NA | NA |
| Gm29370 | 0 | NA | NA |
| Gm21751 | 0 | NA | NA |
| Gm29166 | 0 | NA | NA |
| Gm28194 | 0 | NA | NA |
| Gm20792 | 0 | NA | NA |
| Gm28754 | 0 | NA | NA |
| Gm28753 | 0 | NA | NA |
| Gm21191 | 0 | NA | NA |
| Gm21271 | 0 | NA | NA |
| Gm21281 | 0 | NA | NA |
| Gm28193 | 0 | NA | NA |
| Gm20802 | 0 | NA | NA |
| Gm29070 | 0 | NA | NA |
| Gm28553 | 0 | NA | NA |
| Gm28751 | 0 | NA | NA |
| Gm28752 | 0 | NA | NA |
| Gm28726 | 0 | NA | NA |
| Gm21308 | 0 | NA | NA |
| Gm29267 | 0 | NA | NA |

| | | | |
|---|---|---|---|
| Gm29268 | 0 | NA | NA |
| Gm29269 | 0 | NA | NA |
| Gm29270 | 0 | NA | NA |
| Gm29271 | 0 | NA | NA |
| Gm21209 | 0 | NA | NA |
| Gm28682 | 0 | NA | NA |
| Gm28569 | 0 | NA | NA |
| Gm28681 | 0 | NA | NA |
| Gm28684 | 0 | NA | NA |
| Gm21316 | 0 | NA | NA |
| Gm21923 | 0 | NA | NA |
| Gm28683 | 0 | NA | NA |
| Gm28680 | 0 | NA | NA |
| Gm28679 | 0 | NA | NA |
| Gm28870 | 0 | NA | NA |
| Gm28871 | 0 | NA | NA |
| Gm29512 | 0 | NA | NA |
| Gm29511 | 0 | NA | NA |
| Gm20880 | 0 | NA | NA |
| Gm20881 | 0 | NA | NA |
| Gm29121 | 0 | NA | NA |
| Gm29120 | 0 | NA | NA |
| Gm29117 | 0 | NA | NA |
| Gm20883 | 0 | NA | NA |
| Gm29119 | 0 | NA | NA |
| Gm29118 | 0 | NA | NA |
| Gm29116 | 0 | NA | NA |
| Gm20889 | 0 | NA | NA |
| Gm20887 | 0 | NA | NA |
| Gm29320 | 0 | NA | NA |
| Gm21849 | 0 | NA | NA |
| Gm29321 | 0 | NA | NA |
| Gm21117 | 0 | NA | NA |
| Gm29319 | 0 | NA | NA |
| Gm28317 | 0 | NA | NA |
| Gm28318 | 0 | NA | NA |
| Gm20884 | 0 | NA | NA |
| Gm28313 | 0 | NA | NA |
| Gm28314 | 0 | NA | NA |
| Gm28315 | 0 | NA | NA |
| Gm28316 | 0 | NA | NA |
| Gm21387 | 0 | NA | NA |
| Gm21396 | 0 | NA | NA |
| Gm28944 | 0 | NA | NA |
| Gm28943 | 0 | NA | NA |
| Gm28946 | 0 | NA | NA |
| Gm28945 | 0 | NA | NA |

| | | | |
|---|---|---|---|
| Gm28947 | 0 | NA | NA |
| Gm20920 | 0 | NA | NA |
| Gm28949 | 0 | NA | NA |
| Gm28948 | 0 | NA | NA |
| Gm28950 | 0 | NA | NA |
| Gm21245 | 0 | NA | NA |
| Gm29169 | 0 | NA | NA |
| Gm21899 | 0 | NA | NA |
| Gm29063 | 0 | NA | NA |
| Gm28737 | 0 | NA | NA |
| Gm28738 | 0 | NA | NA |
| Gm20923 | 0 | NA | NA |
| Gm28197 | 0 | NA | NA |
| Gm21249 | 0 | NA | NA |
| Gm28736 | 0 | NA | NA |
| Gm21840 | 0 | NA | NA |
| Gm28735 | 0 | NA | NA |
| Gm21258 | 0 | NA | NA |
| Gm28734 | 0 | NA | NA |
| Gm28232 | 0 | NA | NA |
| Gm28732 | 0 | NA | NA |
| Gm28733 | 0 | NA | NA |
| Gm21268 | 0 | NA | NA |
| Gm21275 | 0 | NA | NA |
| Gm28233 | 0 | NA | NA |
| Gm28235 | 0 | NA | NA |
| Gm28234 | 0 | NA | NA |
| Gm21902 | 0 | NA | NA |
| Gm28236 | 0 | NA | NA |
| Gm21285 | 0 | NA | NA |
| Gm20747 | 0 | NA | NA |
| Gm29250 | 0 | NA | NA |
| Gm29251 | 0 | NA | NA |
| Gm21906 | 0 | NA | NA |
| Gm29252 | 0 | NA | NA |
| Gm29248 | 0 | NA | NA |
| Gm29249 | 0 | NA | NA |
| Gm29484 | 0 | NA | NA |
| Gm21920 | 0 | NA | NA |
| Gm28908 | 0 | NA | NA |
| Gm28907 | 0 | NA | NA |
| Gm21749 | 0 | NA | NA |
| Gm20929 | 0 | NA | NA |
| Gm28910 | 0 | NA | NA |
| Gm28909 | 0 | NA | NA |
| Gm28276 | 0 | NA | NA |
| Gm28274 | 0 | NA | NA |

| | | | |
|---|---|---|---|
| Gm28275 | 0 | NA | NA |
| Gm21302 | 0 | NA | NA |
| Gm28290 | 0 | NA | NA |
| Gm28292 | 0 | NA | NA |
| Gm28291 | 0 | NA | NA |
| Gm28293 | 0 | NA | NA |
| Gm21740 | 0 | NA | NA |
| Gm21912 | 0 | NA | NA |
| Gm28698 | 0 | NA | NA |
| Gm29304 | 0 | NA | NA |
| Gm29305 | 0 | NA | NA |
| Gm29303 | 0 | NA | NA |
| Gm21894 | 0 | NA | NA |
| Gm29307 | 0 | NA | NA |
| Gm29557 | 0 | NA | NA |
| Gm23052 | 0 | NA | NA |
| Gm20927 | 0 | NA | NA |
| Gm20930 | 0 | NA | NA |
| Gm29306 | 0 | NA | NA |
| Gm29300 | 0 | NA | NA |
| Gm29301 | 0 | NA | NA |
| Gm20931 | 0 | NA | NA |
| Gm29559 | 0 | NA | NA |
| Gm29558 | 0 | NA | NA |
| Gm29555 | 0 | NA | NA |
| Gm21733 | 0 | NA | NA |
| Gm21330 | 0 | NA | NA |
| Gm28355 | 0 | NA | NA |
| Gm28354 | 0 | NA | NA |
| Gm28353 | 0 | NA | NA |
| Gm28352 | 0 | NA | NA |
| Gm21858 | 0 | NA | NA |
| Gm29244 | 0 | NA | NA |
| Gm28200 | 0 | NA | NA |
| Gm28201 | 0 | NA | NA |
| Gm28202 | 0 | NA | NA |
| Gm28203 | 0 | NA | NA |
| Gm21340 | 0 | NA | NA |
| Gm28204 | 0 | NA | NA |
| Gm28205 | 0 | NA | NA |
| Gm28206 | 0 | NA | NA |
| Gm28207 | 0 | NA | NA |
| Gm28208 | 0 | NA | NA |
| Gm21851 | 0 | NA | NA |
| Gm21819 | 0 | NA | NA |
| Gm29224 | 0 | NA | NA |
| Gm29223 | 0 | NA | NA |

| | | | |
|---|---|---|---|
| Gm29222 | 0 | NA | NA |
| Gm29221 | 0 | NA | NA |
| Gm29220 | 0 | NA | NA |
| Gm21815 | 0 | NA | NA |
| Gm29407 | 0 | NA | NA |
| Gm29405 | 0 | NA | NA |
| Gm29406 | 0 | NA | NA |
| Gm20913 | 0 | NA | NA |
| Gm29404 | 0 | NA | NA |
| Gm29403 | 0 | NA | NA |
| Gm28350 | 0 | NA | NA |
| Gm28964 | 0 | NA | NA |
| Sly | 0 | NA | NA |
| Gm28165 | 0 | NA | NA |
| Gm28709 | 0 | NA | NA |
| Gm28965 | 0 | NA | NA |
| Gm21868 | 0 | NA | NA |
| Gm21943 | 0 | NA | NA |
| Gm28532 | 0 | NA | NA |
| Gm28533 | 0 | NA | NA |
| Gm28966 | 0 | NA | NA |
| Gm28961 | 0 | NA | NA |
| Gm28962 | 0 | NA | NA |
| Gm28963 | 0 | NA | NA |
| Gm29466 | 0 | NA | NA |
| Gm29467 | 0 | NA | NA |
| Gm21724 | 0 | NA | NA |
| Gm20963 | 0 | NA | NA |
| Gm28326 | 0 | NA | NA |
| Gm28325 | 0 | NA | NA |
| Gm21405 | 0 | NA | NA |
| Gm28284 | 0 | NA | NA |
| Gm29368 | 0 | NA | NA |
| Gm21861 | 0 | NA | NA |
| Gm21413 | 0 | NA | NA |
| Gm28073 | 0 | NA | NA |
| Gm21419 | 0 | NA | NA |
| Gm28074 | 0 | NA | NA |
| Gm21428 | 0 | NA | NA |
| Gm28072 | 0 | NA | NA |
| Gm21758 | 0 | NA | NA |
| Gm29672 | 0 | NA | NA |
| Gm29131 | 0 | NA | NA |
| Gm20871 | 0 | NA | NA |
| Gm29130 | 0 | NA | NA |
| Gm20978 | 0 | NA | NA |
| Gm29132 | 0 | NA | NA |

| | | | |
|---|---|---|---|
| Gm21450 | 0 | NA | NA |
| Gm21843 | 0 | NA | NA |
| Gm28889 | 0 | NA | NA |
| Gm28890 | 0 | NA | NA |
| Gm28891 | 0 | NA | NA |
| Gm28691 | 0 | NA | NA |
| Gm28692 | 0 | NA | NA |
| Gm20794 | 0 | NA | NA |
| Gm28690 | 0 | NA | NA |
| Gm28689 | 0 | NA | NA |
| Gm28687 | 0 | NA | NA |
| Gm20987 | 0 | NA | NA |
| Gm21918 | 0 | NA | NA |
| 1700040F15Rik | 0 | NA | NA |
| Gm29207 | 0 | NA | NA |
| Gm29204 | 0 | NA | NA |
| Gm21489 | 0 | NA | NA |
| Gm21832 | 0 | NA | NA |
| Gm21573 | 0 | NA | NA |
| Gm28179 | 0 | NA | NA |
| Gm21577 | 0 | NA | NA |
| Gm29206 | 0 | NA | NA |
| Gm29203 | 0 | NA | NA |
| Gm21497 | 0 | NA | NA |
| Gm29614 | 0 | NA | NA |
| Gm29613 | 0 | NA | NA |
| Gm29612 | 0 | NA | NA |
| Gm29343 | 0 | NA | NA |
| Gm29342 | 0 | NA | NA |
| Gm20803 | 0 | NA | NA |
| Gm20804 | 0 | NA | NA |
| Gm28598 | 0 | NA | NA |
| Gm28599 | 0 | NA | NA |
| Gm28600 | 0 | NA | NA |
| Gm21518 | 0 | NA | NA |
| Gm28185 | 0 | NA | NA |
| Gm29037 | 0 | NA | NA |
| Gm20823 | 0 | NA | NA |
| Gm21877 | 0 | NA | NA |
| Gm28835 | 0 | NA | NA |
| Gm29548 | 0 | NA | NA |
| Gm29547 | 0 | NA | NA |
| Gm29546 | 0 | NA | NA |
| Gm29545 | 0 | NA | NA |
| Gm29544 | 0 | NA | NA |
| Gm29209 | 0 | NA | NA |
| Gm29549 | 0 | NA | NA |

| | | | |
|---|---|---|---|
| Gm21530 | 0 | NA | NA |
| Gm21617 | 0 | NA | NA |
| Gm28589 | 0 | NA | NA |
| Gm29420 | 0 | NA | NA |
| Gm29421 | 0 | NA | NA |
| Gm21627 | 0 | NA | NA |
| Gm29422 | 0 | NA | NA |
| Gm28808 | 0 | NA | NA |
| Gm21634 | 0 | NA | NA |
| Gm28446 | 0 | NA | NA |
| Gm29060 | 0 | NA | NA |
| Gm29061 | 0 | NA | NA |
| Gm20885 | 0 | NA | NA |
| Gm29062 | 0 | NA | NA |
| Gm29057 | 0 | NA | NA |
| Gm29625 | 0 | NA | NA |
| Gm21654 | 0 | NA | NA |
| Gm29058 | 0 | NA | NA |
| Gm29059 | 0 | NA | NA |
| Gm29628 | 0 | NA | NA |
| Gm20882 | 0 | NA | NA |
| Gm20907 | 0 | NA | NA |
| Gm29624 | 0 | NA | NA |
| Gm29623 | 0 | NA | NA |
| Gm29622 | 0 | NA | NA |
| Gm29265 | 0 | NA | NA |
| Gm20908 | 0 | NA | NA |
| Gm29259 | 0 | NA | NA |
| Gm28860 | 0 | NA | NA |
| Gm28252 | 0 | NA | NA |
| Gm28859 | 0 | NA | NA |
| Gm20921 | 0 | NA | NA |
| Gm28253 | 0 | NA | NA |
| Gm21826 | 0 | NA | NA |
| Gm28254 | 0 | NA | NA |
| Gm28861 | 0 | NA | NA |
| Gm28255 | 0 | NA | NA |
| Gm21171 | 0 | NA | NA |
| Gm20924 | 0 | NA | NA |
| Gm28990 | 0 | NA | NA |
| Gm21848 | 0 | NA | NA |
| Gm29457 | 0 | NA | NA |
| Gm20736 | 0 | NA | NA |
| Gm28558 | 0 | NA | NA |
| Gm28560 | 0 | NA | NA |
| Gm28561 | 0 | NA | NA |
| Gm29584 | 0 | NA | NA |

| | | | |
|---|---|---|---|
| Gm20936 | 0 | NA | NA |
| Gm20934 | 0 | NA | NA |
| Gm29586 | 0 | NA | NA |
| Gm21895 | 0 | NA | NA |
| Gm28134 | 0 | NA | NA |
| Gm28135 | 0 | NA | NA |
| Gm20933 | 0 | NA | NA |
| Gm28133 | 0 | NA | NA |
| Gm28137 | 0 | NA | NA |
| Gm28138 | 0 | NA | NA |
| Gm21837 | 0 | NA | NA |
| Gm28136 | 0 | NA | NA |
| Gm21910 | 0 | NA | NA |
| Gm29377 | 0 | NA | NA |
| Gm21889 | 0 | NA | NA |
| Gm29378 | 0 | NA | NA |
| Gm29379 | 0 | NA | NA |
| Gm29380 | 0 | NA | NA |
| Gm29381 | 0 | NA | NA |
| Gm28795 | 0 | NA | NA |
| Gm20852 | 0 | NA | NA |
| Gm29003 | 0 | NA | NA |
| Gm28749 | 0 | NA | NA |
| Gm29001 | 0 | NA | NA |
| Gm29002 | 0 | NA | NA |
| Ssty2 | 0 | NA | NA |
| Gm29000 | 0 | NA | NA |
| Gm28646 | 0 | NA | NA |
| Gm28612 | 0 | NA | NA |
| Gm28613 | 0 | NA | NA |
| Gm28611 | 0 | NA | NA |
| Gm21114 | 0 | NA | NA |
| Gm29633 | 0 | NA | NA |
| Gm21794 | 0 | NA | NA |
| Gm29219 | 0 | NA | NA |
| Gm29632 | 0 | NA | NA |
| Gm29631 | 0 | NA | NA |
| Gm29637 | 0 | NA | NA |
| Gm20874 | 0 | NA | NA |
| Gm29082 | 0 | NA | NA |
| Gm29217 | 0 | NA | NA |
| Gm29636 | 0 | NA | NA |
| Gm29635 | 0 | NA | NA |
| Gm29634 | 0 | NA | NA |
| Gm29639 | 0 | NA | NA |
| Gm29218 | 0 | NA | NA |
| Gm21806 | 0 | NA | NA |

| | | | | |
|---|---|---|---|---|
| Gm20935 | 0 | NA | NA | |
| Gm29662 | 0 | NA | NA | |
| Gm29660 | 0 | NA | NA | |
| Gm29661 | 0 | NA | NA | |
| Gm29659 | 0 | NA | NA | |
| Gm20937 | 0 | NA | NA | |
| Gm29450 | 0 | NA | NA | |
| Gm29449 | 0 | NA | NA | |
| Gm21867 | 0 | NA | NA | |
| Gm29448 | 0 | NA | NA | |
| Gm21127 | 0 | NA | NA | |
| Gm20816 | 0 | NA | NA | |
| Gm29447 | 0 | NA | NA | |
| Gm29446 | 0 | NA | NA | |
| Gm29452 | 0 | NA | NA | |
| Gm29451 | 0.044191549 | -0.243571806 | | 3.724945049 |
| Gm20817 | 0 | NA | NA | |
| Gm28633 | 0 | NA | NA | |
| Gm29410 | 0 | NA | NA | |
| Gm28819 | 0 | NA | NA | |
| Gm28462 | 0 | NA | NA | |
| Gm20818 | 0 | NA | NA | |
| Gm21151 | 0 | NA | NA | |
| Gm29409 | 0 | NA | NA | |
| Gm28632 | 0 | NA | NA | |
| Gm28101 | 0 | NA | NA | |
| Gm33815 | 0 | NA | NA | |
| Gm21160 | 0 | NA | NA | |
| Gm28637 | 0 | NA | NA | |
| Gm28636 | 0 | NA | NA | |
| Gm20932 | 0 | NA | NA | |
| Gm29134 | 0 | NA | NA | |
| Gm28303 | 0 | NA | NA | |
| Gm28304 | 0 | NA | NA | |
| Gm28824 | 0 | NA | NA | |
| Gm28081 | 0 | NA | NA | |
| Gm28080 | 0 | NA | NA | |
| Gm21524 | 0 | NA | NA | |
| Gm28079 | 0 | NA | NA | |
| Gm28302 | 0 | NA | NA | |
| Gm28082 | 0 | NA | NA | |
| Gm21672 | 0 | NA | NA | |
| Gm29644 | 0 | NA | NA | |
| Gm21118 | 0 | NA | NA | |
| Gm29423 | 0 | NA | NA | |
| Gm29424 | 0 | NA | NA | |
| 4932431L22Rik | 0 | NA | NA | |

| | | | |
|---|---|---|---|
| Gm29425 | 0 | NA | NA |
| Gm20888 | 0 | NA | NA |
| Gm28238 | 0 | NA | NA |
| Gm28433 | 0 | NA | NA |
| Gm28431 | 0 | NA | NA |
| Gm28432 | 0 | NA | NA |
| Gm20886 | 0 | NA | NA |
| Gm21808 | 0 | NA | NA |
| Gm28840 | 0 | NA | NA |
| Gm28841 | 0 | NA | NA |
| Gm28842 | 0 | NA | NA |
| Gm28843 | 0 | NA | NA |
| Gm28837 | 0 | NA | NA |
| Gm28838 | 0 | NA | NA |
| Gm28839 | 0 | NA | NA |
| Gm20866 | 0 | NA | NA |
| Gm20868 | 0 | NA | NA |
| Gm28844 | 0 | NA | NA |
| Gm21841 | 0 | NA | NA |
| Gm28668 | 0 | NA | NA |
| Gm20869 | 0 | NA | NA |
| Gm28481 | 0 | NA | NA |
| Gm28482 | 0 | NA | NA |
| Gm28485 | 0 | NA | NA |
| Gm20870 | 0 | NA | NA |
| Gm28483 | 0 | NA | NA |
| Gm28484 | 0 | NA | NA |
| Gm28480 | 0 | NA | NA |
| Gm20842 | 0 | NA | NA |
| Gm29179 | 0 | NA | NA |
| Gm21880 | 0 | NA | NA |
| Gm29180 | 0 | NA | NA |
| Gm29181 | 0 | NA | NA |
| Gm20843 | 0 | NA | NA |
| Gm29182 | 0 | NA | NA |
| Gm20844 | 0 | NA | NA |
| Gm29178 | 0 | NA | NA |
| Gm21757 | 0 | NA | NA |
| Gm28968 | 0 | NA | NA |
| Gm28495 | 0 | NA | NA |
| Gm20846 | 0 | NA | NA |
| Gm20847 | 0 | NA | NA |
| Gm20848 | 0 | NA | NA |
| Gm28969 | 0 | NA | NA |
| Gm21766 | 0 | NA | NA |
| Gm20800 | 0 | NA | NA |
| Gm21683 | 0 | NA | NA |

| | | | |
|---|---|---|---|
| Gm28677 | 0 | NA | NA |
| Gm21855 | 0 | NA | NA |
| Gm29255 | 0 | NA | NA |
| Gm20902 | 0 | NA | NA |
| Gm28678 | 0 | NA | NA |
| Gm20903 | 0 | NA | NA |
| Gm28272 | 0 | NA | NA |
| Gm29256 | 0 | NA | NA |
| Gm28259 | 0 | NA | NA |
| Gm28273 | 0 | NA | NA |
| Gm21715 | 0 | NA | NA |
| Gm29617 | 0 | NA | NA |
| Gm29615 | 0 | NA | NA |
| Gm29616 | 0 | NA | NA |
| Gm21784 | 0 | NA | NA |
| Gm20805 | 0 | NA | NA |
| Gm29618 | 0 | NA | NA |
| Gm29109 | 0 | NA | NA |
| Gm29108 | 0 | NA | NA |
| Gm29110 | 0 | NA | NA |
| Gm28781 | 0 | NA | NA |
| Gm21913 | 0 | NA | NA |
| Gm28174 | 0 | NA | NA |
| Gm22832 | 0 | NA | NA |
| Gm21076 | 0 | NA | NA |
| Gm29416 | 0 | NA | NA |
| Gm25359 | 0 | NA | NA |
| Gm21869 | 0 | NA | NA |
| Gm20813 | 0 | NA | NA |
| Gm28790 | 0 | NA | NA |
| Gm29272 | 0 | NA | NA |
| Gm28789 | 0 | NA | NA |
| Gm20814 | 0 | NA | NA |
| Gm28787 | 0 | NA | NA |
| Gm28788 | 0 | NA | NA |
| Gm29278 | 0 | NA | NA |
| Gm21922 | 0 | NA | NA |
| Gm21111 | 0 | NA | NA |
| Gm28608 | 0 | NA | NA |
| Gm28607 | 0 | NA | NA |
| Gm28606 | 0 | NA | NA |
| Gm20850 | 0 | NA | NA |
| Gm29574 | 0 | NA | NA |
| Gm29575 | 0 | NA | NA |
| Gm21875 | 0 | NA | NA |
| Gm20904 | 0 | NA | NA |
| Gm29576 | 0 | NA | NA |

| | | | |
|---|---|---|---|
| Gm29577 | 0 | NA | NA |
| Gm29573 | 0 | NA | NA |
| Gm28377 | 0 | NA | NA |
| Gm21776 | 0 | NA | NA |
| Gm21155 | 0 | NA | NA |
| Gm21163 | 0 | NA | NA |
| Gm28122 | 0 | NA | NA |
| Gm28121 | 0 | NA | NA |
| Gm21173 | 0 | NA | NA |
| Gm28123 | 0 | NA | NA |
| Gm28120 | 0 | NA | NA |
| Gm29313 | 0 | NA | NA |
| Gm21180 | 0 | NA | NA |
| Gm20859 | 0 | NA | NA |
| Gm29314 | 0 | NA | NA |
| Gm29310 | 0 | NA | NA |
| Gm29311 | 0 | NA | NA |
| Gm21638 | 0 | NA | NA |
| Gm29312 | 0 | NA | NA |
| Gm29308 | 0 | NA | NA |
| Gm29309 | 0 | NA | NA |
| Gm28702 | 0 | NA | NA |
| Gm28701 | 0 | NA | NA |
| Gm28700 | 0 | NA | NA |
| Gm28699 | 0 | NA | NA |
| Gm28705 | 0 | NA | NA |
| Gm28704 | 0 | NA | NA |
| Gm28540 | 0 | NA | NA |
| Gm21650 | 0 | NA | NA |
| Gm28541 | 0 | NA | NA |
| Gm28542 | 0 | NA | NA |
| Gm28537 | 0 | NA | NA |
| Gm28538 | 0 | NA | NA |
| Gm29569 | 0 | NA | NA |
| Gm20867 | 0 | NA | NA |
| Gm21658 | 0 | NA | NA |
| Gm29568 | 0 | NA | NA |
| Gm29567 | 0 | NA | NA |
| Gm21198 | 0 | NA | NA |
| Gm29566 | 0 | NA | NA |
| Gm29565 | 0 | NA | NA |
| Gm29564 | 0 | NA | NA |
| Gm29147 | 0 | NA | NA |
| Gm29146 | 0 | NA | NA |
| Gm28664 | 0 | NA | NA |
| Gm28663 | 0 | NA | NA |
| Gm29149 | 0 | NA | NA |

| | | | |
|---|---|---|---|
| Gm29148 | 0 | NA | NA |
| Gm29145 | 0 | NA | NA |
| Gm29144 | 0 | NA | NA |
| Gm28817 | 0 | NA | NA |
| Gm28816 | 0 | NA | NA |
| Gm20808 | 0 | NA | NA |
| Gm20806 | 0 | NA | NA |
| Gm28815 | 0 | NA | NA |
| Gm28814 | 0 | NA | NA |
| Gm28813 | 0 | NA | NA |
| Gm20919 | 0 | NA | NA |
| Gm20917 | 0 | NA | NA |
| Gm28421 | 0 | NA | NA |
| Gm28420 | 0 | NA | NA |
| Gm28423 | 0 | NA | NA |
| Gm28422 | 0 | NA | NA |
| Gm20916 | 0 | NA | NA |
| Gm29033 | 0 | NA | NA |
| Gm29034 | 0 | NA | NA |
| Gm21823 | 0 | NA | NA |
| Gm21242 | 0 | NA | NA |
| Gm29031 | 0 | NA | NA |
| Gm29032 | 0 | NA | NA |
| Gm29606 | 0 | NA | NA |
| Gm29607 | 0 | NA | NA |
| Gm21248 | 0 | NA | NA |
| Gm29189 | 0 | NA | NA |
| Gm21873 | 0 | NA | NA |
| Gm29190 | 0 | NA | NA |
| Gm21760 | 0 | NA | NA |
| Gm28159 | 0 | NA | NA |
| Gm28160 | 0 | NA | NA |
| Gm28157 | 0 | NA | NA |
| Gm28158 | 0 | NA | NA |
| Gm21257 | 0 | NA | NA |
| Gm28161 | 0 | NA | NA |
| Gm28695 | 0 | NA | NA |
| Gm28696 | 0 | NA | NA |
| Gm20791 | 0 | NA | NA |
| Gm20841 | 0 | NA | NA |
| Gm28899 | 0 | NA | NA |
| Gm21722 | 0 | NA | NA |
| Gm28898 | 0 | NA | NA |
| Gm28897 | 0 | NA | NA |
| Gm28896 | 0 | NA | NA |
| Gm21282 | 0 | NA | NA |
| Gm28895 | 0 | NA | NA |

| | | | |
|---|---|---|---|
| Gm28894 | 0 | NA | NA |
| Gm21829 | 0 | NA | NA |
| Gm29324 | 0 | NA | NA |
| Gm20911 | 0 | NA | NA |
| Gm28739 | 0 | NA | NA |
| Gm28741 | 0 | NA | NA |
| Gm20912 | 0 | NA | NA |
| Gm20801 | 0 | NA | NA |
| Gm28742 | 0 | NA | NA |
| Gm21919 | 0 | NA | NA |
| Gm28743 | 0 | NA | NA |
| Gm28124 | 0 | NA | NA |
| Gm21790 | 0 | NA | NA |
| Gm21728 | 0 | NA | NA |
| Gm28126 | 0 | NA | NA |
| Gm28125 | 0 | NA | NA |
| Gm21825 | 0 | NA | NA |
| Gm21317 | 0 | NA | NA |
| Gm29468 | 0 | NA | NA |
| Gm29469 | 0 | NA | NA |
| Gm29472 | 0 | NA | NA |
| Gm29473 | 0 | NA | NA |
| Gm29470 | 0 | NA | NA |
| Gm21333 | 0 | NA | NA |
| Gm29382 | 0 | NA | NA |
| Gm28224 | 0 | NA | NA |
| Gm28226 | 0 | NA | NA |
| Gm28225 | 0 | NA | NA |
| Gm21344 | 0 | NA | NA |
| Gm20849 | 0 | NA | NA |
| Gm28829 | 0 | NA | NA |
| Gm21901 | 0 | NA | NA |
| Gm28827 | 0 | NA | NA |
| Gm28828 | 0 | NA | NA |
| Gm28831 | 0 | NA | NA |
| Gm28832 | 0 | NA | NA |
| Gm21864 | 0 | NA | NA |
| Gm21802 | 0 | NA | NA |
| Gm28658 | 0 | NA | NA |
| Gm29238 | 0 | NA | NA |
| Gm21088 | 0 | NA | NA |
| Gm29239 | 0 | NA | NA |
| Gm25729 | 0 | NA | NA |
| Gm21741 | 0 | NA | NA |
| Gm20862 | 0 | NA | NA |
| Gm22624 | 0 | NA | NA |
| Gm20863 | 0 | NA | NA |

| | | | |
|---|---|---|---|
| Gm20864 | 0 | NA | NA |
| Gm29476 | 0 | NA | NA |
| Gm21717 | 0 | NA | NA |
| Gm28472 | 0 | NA | NA |
| Gm21095 | 0 | NA | NA |
| Gm28473 | 0 | NA | NA |
| Gm28474 | 0 | NA | NA |
| Gm28475 | 0 | NA | NA |
| Gm28476 | 0 | NA | NA |
| Gm21380 | 0 | NA | NA |
| Gm21782 | 0 | NA | NA |
| Gm21394 | 0 | NA | NA |
| Gm28471 | 0 | NA | NA |
| Gm21824 | 0 | NA | NA |
| Gm29338 | 0 | NA | NA |
| Gm21409 | 0 | NA | NA |
| Gm29339 | 0 | NA | NA |
| Gm28759 | 0 | NA | NA |
| Gm28758 | 0 | NA | NA |
| Gm29071 | 0 | NA | NA |
| Gm21879 | 0 | NA | NA |
| Gm28757 | 0 | NA | NA |
| Gm20856 | 0 | NA | NA |
| Gm28348 | 0 | NA | NA |
| Gm28973 | 0 | NA | NA |
| Gm21734 | 0 | NA | NA |
| Gm28972 | 0 | NA | NA |
| Gm28970 | 0 | NA | NA |
| Gm28971 | 0 | NA | NA |
| Gm24828 | 0 | NA | NA |
| Gm21785 | 0 | NA | NA |
| Gm20854 | 0 | NA | NA |
| Gm28349 | 0 | NA | NA |
| Gm21892 | 0 | NA | NA |
| Gm28346 | 0 | NA | NA |
| Gm28345 | 0 | NA | NA |
| Gm29090 | 0 | NA | NA |
| Gm28152 | 0 | NA | NA |
| Gm23770 | 0 | NA | NA |
| Gm21881 | 0 | NA | NA |
| Gm21435 | 0 | NA | NA |
| Gm29505 | 0 | NA | NA |
| Gm21862 | 0 | NA | NA |
| Gm28823 | 0 | NA | NA |
| Gm28220 | 0 | NA | NA |
| Gm20820 | 0 | NA | NA |
| Gm28219 | 0 | NA | NA |

| | | | | |
|---|---|---|---|---|
| Gm28218 | 0 | NA | NA | |
| Gm23863 | 0 | NA | NA | |
| Gm20861 | 0 | NA | NA | |
| Gm20860 | 0 | NA | NA | |
| Gm28057 | 0 | NA | NA | |
| Gm21796 | 0 | NA | NA | |
| Gm29302 | 0 | NA | NA | |
| Gm28440 | 0 | NA | NA | |
| Gm29299 | 0 | NA | NA | |
| Gm29298 | 0 | NA | NA | |
| Gm29092 | 0 | NA | NA | |
| Gm29091 | 0 | NA | NA | |
| Gm29297 | 0 | NA | NA | |
| Gm21462 | 0 | NA | NA | |
| Gm21470 | 0 | NA | NA | |
| Gm28366 | 0 | NA | NA | |
| Gm21752 | 0 | NA | NA | |
| Gm28365 | 0 | NA | NA | |
| Gm21477 | 0 | NA | NA | |
| Gm28368 | 0 | NA | NA | |
| Gm28369 | 0 | NA | NA | |
| Gm28367 | 0 | NA | NA | |
| Gm29115 | 0 | NA | NA | |
| Gm21813 | 0 | NA | NA | |
| Gm20906 | 0 | NA | NA | |
| Gm29111 | 0 | NA | NA | |
| Gm28110 | 0 | NA | NA | |
| Gm28109 | 0 | NA | NA | |
| Gm28108 | 0 | NA | NA | |
| Gm28107 | 0 | NA | NA | |
| Gm28106 | 0 | NA | NA | |
| Gm28105 | 0 | NA | NA | |
| Gm28104 | 0 | NA | NA | |
| Gm28103 | 0 | NA | NA | |
| Gm28102 | 0 | NA | NA | |
| Gm28716 | 0 | NA | NA | |
| Gm28717 | 0 | NA | NA | |
| Gm28718 | 0 | NA | NA | |
| Gm29162 | 0.046345709 | 0.71820487 | | 3.724945049 |
| Gm21777 | 0 | NA | NA | |
| Gm28866 | 0 | NA | NA | |
| Gm28405 | 0 | NA | NA | |
| Gm21792 | 0 | NA | NA | |
| Gm28406 | 0 | NA | NA | |
| Gm28978 | 0 | NA | NA | |
| Gm29436 | 0 | NA | NA | |
| Gm20819 | 0 | NA | NA | |

| | | | |
|---|---|---|---|
| Gm29437 | 0 | NA | NA |
| Gm28977 | 0 | NA | NA |
| Gm28407 | 0 | NA | NA |
| Gm21287 | 0 | NA | NA |
| Gm29390 | 0 | NA | NA |
| Gm29391 | 0 | NA | NA |
| Gm21720 | 0 | NA | NA |
| Gm29392 | 0 | NA | NA |
| Gm29393 | 0 | NA | NA |
| Gm21294 | 0 | NA | NA |
| Gm28672 | 0 | NA | NA |
| Gm21838 | 0 | NA | NA |
| Gm28670 | 0 | NA | NA |
| Gm28671 | 0 | NA | NA |
| Gm28673 | 0 | NA | NA |
| Gm28674 | 0 | NA | NA |
| Gm28531 | 0 | NA | NA |
| Gm21996 | 0 | NA | NA |
| Gm21803 | 0 | NA | NA |
| Gm28930 | 0 | NA | NA |
| Gm20879 | 0 | NA | NA |
| Gm29504 | 0 | NA | NA |
| Gm20837 | 0 | NA | NA |
| Gm29503 | 0 | NA | NA |
| Gm28300 | 0 | NA | NA |
| Gm28301 | 0 | NA | NA |
| Gm21857 | 2.299789819 | 1.00812413 | 0.689743561 |
| Gm21860 | 0.344895107 | 1.595654113 | 1.549418196 |
| Erdr1 | 44.24775491 | 0.014385544 | 0.216812008 |
| Gm21742 | 0 | NA | NA |
| Gm21748 | 0 | NA | NA |

| stat | pvalue | padj |
|---|---|---|
| NA | NA | NA |
| NA | NA | NA |
| -4.810765573 | 1.50E-06 | 6.88E-06 |
| -0.287752335 | 0.773536326 | NA |
| -3.189187032 | 0.001426735 | 0.003970998 |
| -3.775108708 | 0.000159938 | 0.000533482 |
| -2.769235167 | 0.005618806 | 0.013647003 |
| NA | NA | NA |
| -2.675678889 | 0.007457806 | 0.017527051 |
| 0.896254689 | 0.370116753 | NA |
| -3.247112438 | 0.001165823 | 0.00331296 |
| -3.39307705 | 0.000691122 | 0.002053999 |
| 3.788902685 | 0.000151314 | 0.00050681 |
| NA | NA | NA |
| NA | NA | NA |
| -0.065389369 | 0.947863988 | NA |
| 0.192810118 | 0.847107679 | NA |
| -0.533881092 | 0.593423804 | 0.692466099 |
| NA | NA | NA |
| -2.105338792 | 0.035261828 | 0.068344707 |
| 0.806300917 | 0.420069362 | 0.531779088 |
| 0.348332884 | 0.7275902 | NA |
| 0.414461455 | 0.67853619 | NA |
| NA | NA | NA |
| -1.307174664 | 0.191153398 | NA |
| 0.063708882 | 0.949202033 | NA |
| 0.446269038 | 0.655402918 | 0.74320369 |
| -0.300350002 | 0.763910197 | 0.829530248 |
| -0.065389369 | 0.947863988 | NA |
| -1.105983861 | 0.268733505 | NA |
| 1.075739687 | 0.282043693 | 0.389063303 |
| 1.518609545 | 0.128860807 | 0.205950506 |
| -1.942233725 | 0.052108812 | 0.095945532 |
| 0.733984897 | 0.462957943 | 0.572408482 |
| 0.324874675 | 0.745275925 | NA |
| 4.127107588 | 3.67E-05 | 0.00013689 |
| -0.875381644 | 0.381366284 | 0.493925183 |
| 3.181197948 | 0.001466674 | 0.004071744 |
| 4.200515726 | 2.66E-05 | 0.000101565 |
| -0.462321258 | 0.643850958 | 0.733527058 |
| -1.503091475 | 0.132815457 | 0.211010638 |
| 0.639749403 | 0.522335532 | NA |
| -0.069891852 | 0.944279738 | 0.962232702 |
| 3.217435572 | 0.001293421 | 0.003632259 |
| NA | NA | NA |
| -1.979270942 | 0.047785509 | 0.089033147 |

| | | |
|---|---|---|
| -0.853357289 | 0.3934612 | 0.505398471 |
| -6.335318118 | 2.37E-10 | 1.72E-09 |
| NA | NA | NA |
| NA | NA | NA |
| NA | NA | NA |
| 0.268631646 | 0.788213157 | 0.849288649 |
| 0.192809521 | 0.847108147 | NA |
| -3.153381212 | 0.001613909 | 0.00444402 |
| -2.029773518 | 0.042379566 | 0.080190546 |
| 0.155456793 | 0.876461214 | NA |
| 0.431593367 | 0.66603698 | NA |
| NA | NA | NA |
| -2.358717868 | 0.018338192 | 0.038783687 |
| -3.932134044 | 8.42E-05 | 0.000295237 |
| NA | NA | NA |
| -0.065389966 | 0.947863513 | NA |
| 1.852006948 | 0.064024825 | 0.114124866 |
| NA | NA | NA |
| NA | NA | NA |
| NA | NA | NA |
| NA | NA | NA |
| 0.11217595 | 0.910683898 | 0.938505351 |
| NA | NA | NA |
| -2.575871715 | 0.009998774 | 0.022786971 |
| 0.136392141 | 0.891511286 | 0.924073525 |
| NA | NA | NA |
| NA | NA | NA |
| 0.31685483 | 0.751353758 | NA |
| -3.708198756 | 0.000208739 | 0.000683686 |
| NA | NA | NA |
| NA | NA | NA |
| NA | NA | NA |
| 0.192809521 | 0.847108147 | NA |
| NA | NA | NA |
| NA | NA | NA |
| -5.91910271 | 3.24E-09 | 2.09E-08 |
| -1.780348159 | 0.075019 | 0.130688027 |
| -2.016993607 | 0.04369618 | 0.08240773 |
| -4.426265718 | 9.59E-06 | 3.92E-05 |
| -0.114374582 | 0.908940862 | 0.937172095 |
| -1.133350128 | 0.257067249 | 0.361624978 |
| -1.037790894 | 0.299367417 | 0.408280197 |
| 0.320071032 | 0.748914485 | NA |
| -0.072137591 | 0.942492411 | 0.960959104 |
| -0.090026302 | 0.928266314 | NA |
| 0.223746869 | 0.822954274 | NA |
| -1.6478084 | 0.099391995 | 0.165763701 |

| | | |
|---|---|---|
| -4.078679465 | 4.53E-05 | 0.000165837 |
| -1.854889769 | 0.063611969 | 0.113534483 |
| NA | NA | NA |
| -0.214011832 | 0.830537831 | NA |
| NA | NA | NA |
| NA | NA | NA |
| NA | NA | NA |
| -0.845779001 | 0.397676042 | 0.509499773 |
| 5.072236608 | 3.93E-07 | 1.96E-06 |
| 1.288268125 | 0.197652645 | 0.292771079 |
| -1.626342772 | 0.103876758 | 0.171891792 |
| 0.329899765 | 0.741475701 | 0.81178973 |
| -6.524309493 | 6.83E-11 | 5.29E-10 |
| NA | NA | NA |
| -0.023178083 | 0.981508221 | 0.987227277 |
| 5.882410576 | 4.04E-09 | 2.58E-08 |
| -1.341917229 | 0.179622835 | 0.270694175 |
| 5.822139292 | 5.81E-09 | 3.64E-08 |
| -0.125565959 | 0.90007551 | 0.930292358 |
| NA | NA | NA |
| 1.142703293 | 0.253161802 | 0.357203415 |
| 1.497646363 | 0.134225154 | 0.212865463 |
| -0.388385392 | 0.697730853 | 0.777534537 |
| 2.007499944 | 0.044696457 | 0.084028264 |
| 0.111866421 | 0.910929321 | NA |
| -1.475622815 | 0.140045177 | 0.22050266 |
| NA | NA | NA |
| NA | NA | NA |
| 3.592874808 | 0.00032705 | 0.00103281 |
| -4.747787426 | 2.06E-06 | 9.24E-06 |
| NA | NA | NA |
| 1.881236441 | 0.059939763 | 0.107996851 |
| NA | NA | NA |
| -1.695488041 | 0.089982877 | 0.152593045 |
| -1.316376521 | 0.188047699 | 0.281055347 |
| -2.128952653 | 0.033258178 | 0.064984558 |
| -0.828450996 | 0.407415136 | 0.519264254 |
| -0.234825393 | 0.814344258 | NA |
| -0.065389369 | 0.947863988 | NA |
| NA | NA | NA |
| 0.192809521 | 0.847108147 | NA |
| NA | NA | NA |
| NA | NA | NA |
| -6.843808458 | 7.71E-12 | 6.50E-11 |
| NA | NA | NA |
| NA | NA | NA |
| 0.063708285 | 0.949202508 | NA |

| | | |
|---|---|---|
| -10.99335666 | 4.11E-28 | 1.29E-26 |
| -3.612216978 | 0.00030359 | 0.000963956 |
| NA | NA | NA |
| -2.050371088 | 0.040328232 | 0.076749974 |
| 0.137333993 | 0.890766802 | NA |
| NA | NA | NA |
| NA | NA | NA |
| -0.885351482 | 0.375967092 | NA |
| NA | NA | NA |
| 0.192808924 | 0.847108615 | NA |
| NA | NA | NA |
| 6.728102146 | 1.72E-11 | 1.41E-10 |
| 1.855280718 | 0.06355615 | 0.113447406 |
| NA | NA | NA |
| NA | NA | NA |
| NA | NA | NA |
| 0.254117618 | 0.799404693 | NA |
| -1.94104029 | 0.052253391 | 0.096176363 |
| -2.37674016 | 0.017466387 | 0.037207121 |
| -2.621898724 | 0.008744141 | 0.02019244 |
| NA | NA | NA |
| -3.196384608 | 0.001391615 | 0.003881448 |
| -4.348828127 | 1.37E-05 | 5.45E-05 |
| -1.652301278 | 0.098473149 | 0.164472369 |
| -3.299189099 | 0.000969646 | 0.00280192 |
| -2.009201242 | 0.044515796 | 0.083738943 |
| -0.666629026 | 0.505009124 | 0.611526121 |
| 0.192810118 | 0.847107679 | NA |
| 7.679366713 | 1.60E-14 | 1.78E-13 |
| 0.130775086 | 0.895953232 | 0.927678887 |
| -0.533982936 | 0.59335334 | 0.692416155 |
| 1.509217734 | 0.13124315 | 0.20890404 |
| NA | NA | NA |
| 0.951433933 | 0.341384142 | 0.452898981 |
| -0.194487022 | 0.845794556 | NA |
| NA | NA | NA |
| NA | NA | NA |
| -4.467165816 | 7.93E-06 | 3.28E-05 |
| NA | NA | NA |
| -2.751654702 | 0.005929501 | 0.014306478 |
| -3.033896414 | 0.002414173 | 0.006374802 |
| 0.525237434 | 0.599418145 | 0.697282901 |
| -2.685887612 | 0.007233742 | 0.017047411 |
| -0.349025383 | 0.727070253 | 0.800711389 |
| 0.023082742 | 0.981584272 | 0.987264098 |
| -1.34941294 | 0.177204367 | 0.267710363 |
| NA | NA | NA |

| | | | |
|---|---|---|---|
| | 0.835601059 | 0.403379373 | 0.515035814 |
| | -7.018324079 | 2.25E-12 | 2.00E-11 |
| | -4.221852519 | 2.42E-05 | 9.28E-05 |
| | -3.673161913 | 0.000239568 | 0.000774926 |
| NA | NA | NA | |
| | -0.41847347 | 0.67560098 | NA |
| | 0.192809521 | 0.847108147 | NA |
| | 0.192808924 | 0.847108615 | NA |
| | -1.738570998 | 0.082110252 | 0.141213384 |
| | -1.076955508 | 0.281500143 | 0.388506208 |
| | -1.703860681 | 0.088407118 | 0.150344329 |
| | 0.192810118 | 0.847107679 | NA |
| NA | NA | NA | |
| | -0.150638759 | 0.880260685 | 0.915863364 |
| | 6.635425394 | 3.24E-11 | 2.58E-10 |
| | 5.900106884 | 3.63E-09 | 2.33E-08 |
| | 0.31571572 | 0.752218293 | 0.82031191 |
| NA | NA | NA | |
| | -6.324490666 | 2.54E-10 | 1.84E-09 |
| | 0.282999869 | 0.777176938 | NA |
| | -1.711322614 | 0.087021574 | 0.148334922 |
| | 2.300718535 | 0.021407546 | 0.044323837 |
| | -1.603668519 | 0.108787139 | 0.178471657 |
| | 0.095515416 | 0.923905446 | NA |
| | -0.23036245 | 0.817810137 | NA |
| | 1.734647339 | 0.082803289 | 0.142219607 |
| | 3.7740944 | 0.00016059 | 0.000535514 |
| NA | NA | NA | |
| | 2.923685836 | 0.003459137 | 0.008808851 |
| | -0.232595903 | 0.816075209 | NA |
| NA | NA | NA | |
| | -10.75477168 | 5.63E-27 | 1.63E-25 |
| | 0.473754222 | 0.635675182 | NA |
| | -6.07808306 | 1.22E-09 | 8.21E-09 |
| | -0.86623014 | 0.386363968 | 0.498643401 |
| | 0.192809521 | 0.847108147 | NA |
| | 0.210360696 | 0.833386166 | NA |
| | 0.341077186 | 0.733045475 | 0.80538127 |
| | 0.192810118 | 0.847107679 | NA |
| NA | NA | NA | |
| | 1.424975695 | 0.154164227 | 0.23881239 |
| | -3.596343157 | 0.000322722 | 0.00102056 |
| | -5.692168009 | 1.25E-08 | 7.57E-08 |
| | -9.061270739 | 1.29E-19 | 2.23E-18 |
| | -0.857469882 | 0.391185253 | 0.503153488 |
| NA | NA | NA | |
| | -3.166620337 | 0.001542215 | 0.00426302 |

| | | |
|---|---|---|
| NA | NA | NA |
| -0.355466115 | 0.722240426 | 0.796970324 |
| -3.504268407 | 0.000457864 | 0.001405485 |
| NA | NA | NA |
| -0.065388772 | 0.947864464 | NA |
| NA | NA | NA |
| NA | NA | NA |
| 0.102220301 | 0.918581814 | NA |
| NA | NA | NA |
| NA | NA | NA |
| NA | NA | NA |
| NA | NA | NA |
| NA | NA | NA |
| -0.039588875 | 0.968420897 | 0.978407366 |
| -1.749075283 | 0.080278007 | 0.13849049 |
| 0.405082664 | 0.685416749 | 0.76757108 |
| -1.765428164 | 0.077491834 | 0.134387902 |
| 0.192808327 | 0.847109082 | NA |
| NA | NA | NA |
| 2.69681761 | 0.00700056 | 0.016575917 |
| NA | NA | NA |
| -0.194488217 | 0.845793621 | NA |
| -0.147420768 | 0.882799914 | 0.917894814 |
| NA | NA | NA |
| -1.289789105 | 0.197123892 | NA |
| 0.111708329 | 0.911054675 | NA |
| NA | NA | NA |
| NA | NA | NA |
| -1.738208705 | 0.082174046 | 0.14130368 |
| NA | NA | NA |
| NA | NA | NA |
| -0.07451834 | 0.940597949 | NA |
| 8.735074104 | 2.43E-18 | 3.79E-17 |
| NA | NA | NA |
| -0.065388772 | 0.947864464 | NA |
| NA | NA | NA |
| -1.712855117 | 0.086739197 | 0.147924106 |
| -3.050659302 | 0.002283395 | 0.006069171 |
| NA | NA | NA |
| -1.572506085 | 0.115833229 | 0.188168855 |
| 0.700784865 | 0.483437285 | 0.591853032 |
| 12.86329263 | 7.24E-38 | 3.63E-36 |
| -0.112406574 | 0.910501043 | NA |
| -1.323850476 | 0.185552708 | 0.278040247 |
| -6.667043725 | 2.61E-11 | 2.10E-10 |
| -0.726138334 | 0.467753971 | 0.576970328 |
| -3.210011615 | 0.001327296 | 0.003717796 |

| | | |
|---|---|---|
| -1.49668706 | 0.134474705 | 0.213126187 |
| 0.087905337 | 0.929951915 | 0.952440625 |
| 0.634701 | 0.525623442 | NA |
| 0.192809521 | 0.847108147 | NA |
| 0.063708285 | 0.949202508 | NA |
| NA | NA | NA |
| NA | NA | NA |
| 0.192810118 | 0.847107679 | NA |
| NA | NA | NA |
| NA | NA | NA |
| NA | NA | NA |
| NA | NA | NA |
| -7.498347539 | 6.46E-14 | 6.76E-13 |
| -4.113665418 | 3.89E-05 | 0.000144448 |
| 2.454192796 | 0.014120125 | 0.030896647 |
| -4.455581854 | 8.37E-06 | 3.45E-05 |
| -0.263926973 | 0.791836191 | 0.85190887 |
| NA | NA | NA |
| NA | NA | NA |
| -9.557932313 | 1.20E-21 | 2.43E-20 |
| -0.065388772 | 0.947864464 | NA |
| -0.689537294 | 0.490485213 | 0.59796624 |
| -0.463364847 | 0.643102871 | NA |
| 0.110009718 | 0.912401668 | NA |
| NA | NA | NA |
| -0.065389369 | 0.947863988 | NA |
| 0.609554017 | 0.54215728 | 0.646230617 |
| -1.906683152 | 0.05656163 | 0.102906645 |
| NA | NA | NA |
| -0.509188592 | 0.610620038 | 0.706700863 |
| 0.246772625 | 0.805084198 | NA |
| -5.902025839 | 3.59E-09 | 2.31E-08 |
| 0.192809521 | 0.847108147 | NA |
| 0.663450952 | 0.507041782 | 0.613489429 |
| -7.807959163 | 5.81E-15 | 6.77E-14 |
| -3.176357711 | 0.001491369 | 0.004133698 |
| -2.171581303 | 0.029887259 | 0.05916467 |
| -0.065390563 | 0.947863038 | NA |
| -1.666546134 | 0.095604687 | 0.160538396 |
| -0.905783442 | 0.365050487 | 0.4775481 |
| NA | NA | NA |
| 3.033290294 | 0.002419028 | 0.006384254 |
| -8.218028915 | 2.07E-16 | 2.74E-15 |
| -1.985378574 | 0.047102368 | 0.087912644 |
| 0.510816551 | 0.609479518 | 0.705935786 |
| -2.756432432 | 0.005843568 | 0.014126378 |
| -1.560260624 | 0.118698304 | 0.192112419 |

| | | | |
|---|---|---|---|
| | 0.116311233 | 0.907405883 | 0.935974996 |
| | -0.692851245 | 0.488402909 | 0.596298952 |
| | -0.737313223 | 0.460931893 | 0.570326031 |
| | -1.779803217 | 0.075108171 | 0.130834263 |
| | 1.017948246 | 0.308702552 | 0.418593904 |
| | -7.075836875 | 1.49E-12 | 1.35E-11 |
| NA | NA | NA | |
| | 0.507275069 | 0.611961829 | NA |
| | 0.192809521 | 0.847108147 | NA |
| | 0.260395747 | 0.794558523 | NA |
| | -7.01422231 | 2.31E-12 | 2.05E-11 |
| | 0.2972438 | 0.766280385 | NA |
| | -3.964372907 | 7.36E-05 | 0.00026075 |
| | -3.053780048 | 0.002259777 | 0.00601278 |
| | -3.494213061 | 0.000475461 | 0.001455399 |
| | -5.045778991 | 4.52E-07 | 2.23E-06 |
| | -4.486945728 | 7.23E-06 | 3.00E-05 |
| | -4.062590533 | 4.85E-05 | 0.000177204 |
| | -5.1924868 | 2.08E-07 | 1.07E-06 |
| | -3.789459728 | 0.000150975 | 0.000505878 |
| NA | NA | NA | |
| | -5.988496651 | 2.12E-09 | 1.39E-08 |
| | -4.63540912 | 3.56E-06 | 1.55E-05 |
| | -5.175697796 | 2.27E-07 | 1.17E-06 |
| | -6.350668411 | 2.14E-10 | 1.57E-09 |
| | -4.859083864 | 1.18E-06 | 5.47E-06 |
| | -4.683497344 | 2.82E-06 | 1.24E-05 |
| | -4.04507417 | 5.23E-05 | 0.000190157 |
| | -6.276562395 | 3.46E-10 | 2.47E-09 |
| | -6.082649124 | 1.18E-09 | 7.99E-09 |
| | -5.541992499 | 2.99E-08 | 1.73E-07 |
| | -6.428284344 | 1.29E-10 | 9.69E-10 |
| | -5.009757497 | 5.45E-07 | 2.65E-06 |
| | -5.437246161 | 5.41E-08 | 3.02E-07 |
| | -3.169174048 | 0.001528728 | 0.004229476 |
| NA | NA | NA | |
| NA | NA | NA | |
| NA | NA | NA | |
| NA | NA | NA | |
| NA | NA | NA | |
| NA | NA | NA | |
| | -0.065389369 | 0.947863988 | NA |
| NA | NA | NA | |
| | 0.063708217 | 0.949202562 | NA |
| NA | NA | NA | |
| | 1.271008971 | 0.203725457 | 0.300081472 |
| | -0.065388772 | 0.947864464 | NA |

| | | |
|---|---|---|
| -0.085312598 | 0.932012876 | NA |
| NA | NA | NA |
| NA | NA | NA |
| NA | NA | NA |
| NA | NA | NA |
| NA | NA | NA |
| -0.065389369 | 0.947863988 | NA |
| 2.869654129 | 0.00410921 | 0.010301836 |
| NA | NA | NA |
| NA | NA | NA |
| -0.194487022 | 0.845794556 | NA |
| -1.052298981 | 0.292662399 | NA |
| -1.287103645 | 0.198058165 | 0.293284972 |
| 0.675435901 | 0.499398863 | NA |
| -2.165389634 | 0.030357859 | 0.059961487 |
| -4.456025519 | 8.35E-06 | 3.45E-05 |
| -3.691226883 | 0.000223175 | 0.000726596 |
| 6.07684363 | 1.23E-09 | 8.27E-09 |
| NA | NA | NA |
| NA | NA | NA |
| 0.192810118 | 0.847107679 | NA |
| NA | NA | NA |
| NA | NA | NA |
| NA | NA | NA |
| -0.065388772 | 0.947864464 | NA |
| NA | NA | NA |
| NA | NA | NA |
| 0.629519727 | 0.529008858 | NA |
| NA | NA | NA |
| NA | NA | NA |
| NA | NA | NA |
| 0.192809521 | 0.847108147 | NA |
| NA | NA | NA |
| NA | NA | NA |
| 2.404838656 | 0.016179607 | 0.034848599 |
| NA | NA | NA |
| NA | NA | NA |
| -3.659452521 | 0.000252755 | 0.000814005 |
| -0.065388772 | 0.947864464 | NA |
| -2.496175101 | 0.012554061 | 0.027864846 |
| -1.60707546 | 0.108037816 | 0.177430676 |
| -1.944636615 | 0.051818729 | 0.095495714 |
| 0.199526594 | 0.841850842 | NA |
| -4.396107217 | 1.10E-05 | 4.46E-05 |
| -0.189010623 | 0.850084491 | NA |
| NA | NA | NA |
| -1.652412326 | 0.098450524 | 0.164445554 |

| | | |
|---|---|---|
| -0.090308947 | 0.92804171 | NA |
| -0.328131109 | 0.742812534 | 0.812897767 |
| -0.738943534 | 0.459941286 | NA |
| 0.391055383 | 0.695756299 | 0.775914155 |
| -0.18697769 | 0.851678125 | 0.896866688 |
| -0.144837628 | 0.884839073 | NA |
| NA | NA | NA |
| 0.480482063 | 0.630884655 | 0.722933796 |
| NA | NA | NA |
| NA | NA | NA |
| -0.404950762 | 0.685513704 | 0.767645312 |
| -0.194488217 | 0.845793621 | NA |
| NA | NA | NA |
| -0.433074869 | 0.66496038 | 0.75101337 |
| NA | NA | NA |
| -3.714367304 | 0.000203713 | 0.000668712 |
| NA | NA | NA |
| -0.755557335 | 0.449914644 | 0.559849046 |
| 0.399535292 | 0.689498824 | 0.770831907 |
| -2.269356008 | 0.023246685 | 0.047639146 |
| -1.33261928 | 0.182656775 | 0.274506099 |
| -0.920486551 | 0.357318558 | NA |
| -2.625592809 | 0.008649822 | 0.020004163 |
| -0.98892118 | 0.322701707 | 0.433583323 |
| -4.704969764 | 2.54E-06 | 1.13E-05 |
| -3.901591478 | 9.56E-05 | 0.000331751 |
| -2.716133604 | 0.006604926 | 0.015752348 |
| -3.859806832 | 0.000113477 | 0.0003885 |
| NA | NA | NA |
| NA | NA | NA |
| -2.261635211 | 0.023719951 | 0.048466152 |
| -1.665670132 | 0.095779134 | 0.1607235 |
| 0.352684266 | 0.724325156 | 0.79871943 |
| 0.222947504 | 0.823576364 | 0.87632768 |
| -0.813864819 | 0.415722397 | 0.527342975 |
| NA | NA | NA |
| 0.276206047 | 0.782389813 | NA |
| NA | NA | NA |
| -3.150464553 | 0.00163011 | 0.004483207 |
| -2.061566836 | 0.039248994 | 0.074935341 |
| NA | NA | NA |
| 1.659548031 | 0.09700541 | 0.162421929 |
| NA | NA | NA |
| 5.296337836 | 1.18E-07 | 6.31E-07 |
| 8.379496383 | 5.32E-17 | 7.39E-16 |
| 2.601934426 | 0.009269958 | 0.021281027 |
| 1.092174308 | 0.274756493 | 0.38144961 |

| | | | |
|---|---|---|---|
| | 0.913517788 | 0.360970283 | NA |
| NA | NA | NA | |
| | -0.267825953 | 0.788833293 | NA |
| NA | NA | NA | |
| NA | NA | NA | |
| | -0.387626123 | 0.698292734 | 0.77795287 |
| | -4.43226562 | 9.32E-06 | 3.82E-05 |
| | 0.063708882 | 0.949202033 | NA |
| | 0.240059543 | 0.810284097 | NA |
| | 0.874450475 | 0.38187298 | 0.494376781 |
| | 3.526639015 | 0.00042087 | 0.001300701 |
| | -9.995383182 | 1.60E-23 | 3.70E-22 |
| NA | NA | NA | |
| | -1.357382682 | 0.174659648 | 0.264600593 |
| | -0.981791992 | 0.326202334 | 0.437277444 |
| | -1.365825705 | 0.171993676 | 0.261162443 |
| | -2.094766621 | 0.036191744 | 0.069876531 |
| | -0.475226149 | 0.634625791 | 0.726090577 |
| | -0.893572312 | 0.371550766 | 0.484320158 |
| | 1.230553245 | 0.218490001 | NA |
| | -0.39252633 | 0.69466936 | NA |
| | 0.063708882 | 0.949202033 | NA |
| NA | NA | NA | |
| NA | NA | NA | |
| | 0.400954005 | 0.688453988 | 0.770145399 |
| NA | NA | NA | |
| NA | NA | NA | |
| | -0.577871317 | 0.563351002 | 0.665534317 |
| NA | NA | NA | |
| NA | NA | NA | |
| | -0.423848218 | 0.67167651 | NA |
| NA | NA | NA | |
| | -0.065388772 | 0.947864464 | NA |
| | -0.195150768 | 0.845274919 | 0.891844675 |
| | -0.697622945 | 0.485413028 | NA |
| | -0.129201837 | 0.897197943 | NA |
| | 0.162635435 | 0.870805486 | 0.90999755 |
| | -0.301950037 | 0.762690151 | 0.828600828 |
| | 13.59020102 | 4.58E-42 | 2.68E-40 |
| | 8.615126982 | 6.99E-18 | 1.04E-16 |
| | 0.342031543 | 0.732327155 | 0.804871726 |
| | 3.410917957 | 0.000647446 | 0.001933377 |
| | -0.065389369 | 0.947863988 | NA |
| | 2.173462711 | 0.029745509 | 0.058940002 |
| | -1.669371812 | 0.095043716 | 0.159789375 |
| | -0.938705872 | 0.347881778 | 0.459473426 |
| | -2.876461853 | 0.004021608 | 0.010104482 |

| | | | |
|---|---|---|---|
| NA | NA | NA | |
| | -3.887006441 | 0.000101488 | 0.000350427 |
| | 0.262055901 | 0.793278345 | NA |
| | -0.232929539 | 0.81581612 | 0.870373642 |
| | -0.612751332 | 0.540040775 | 0.644321493 |
| | 5.500913821 | 3.78E-08 | 2.16E-07 |
| | 0.526733202 | 0.598378876 | 0.696559541 |
| | 0.713928352 | 0.475271485 | 0.58419905 |
| | -0.253761763 | 0.799679616 | 0.857722749 |
| NA | NA | NA | |
| | -0.858569649 | 0.390577989 | 0.502719692 |
| | -4.306722048 | 1.66E-05 | 6.52E-05 |
| | 0.943340644 | 0.345506693 | 0.457011706 |
| | -3.944732835 | 7.99E-05 | 0.00028132 |
| | -1.461612767 | 0.143847354 | 0.22550493 |
| | 8.288432398 | 1.15E-16 | 1.55E-15 |
| NA | NA | NA | |
| | -0.446614342 | 0.655153539 | 0.743021888 |
| | -2.853936841 | 0.004318112 | 0.010769463 |
| | -1.553624467 | 0.12027402 | 0.194323439 |
| | -1.273756447 | 0.202749745 | 0.299064536 |
| | 0.120039149 | 0.904452135 | NA |
| NA | NA | NA | |
| | 0.714166805 | 0.47512404 | 0.584046495 |
| | -2.522887707 | 0.011639556 | 0.026070523 |
| | 1.887247245 | 0.059127078 | 0.106740049 |
| NA | NA | NA | |
| | -11.04910858 | 2.21E-28 | 7.02E-27 |
| | -0.065389369 | 0.947863988 | NA |
| NA | NA | NA | |
| | 2.222814495 | 0.026228311 | 0.052853605 |
| | 2.101235433 | 0.035620304 | 0.068911441 |
| | -5.225814062 | 1.73E-07 | 9.04E-07 |
| | -3.186532248 | 0.001439894 | 0.004005396 |
| | -5.667667611 | 1.45E-08 | 8.68E-08 |
| | 0.192809521 | 0.847108147 | NA |
| | 0.39931459 | 0.689661418 | NA |
| | -4.827263711 | 1.38E-06 | 6.36E-06 |
| | -3.795227866 | 0.000147508 | 0.000494989 |
| | -6.073660569 | 1.25E-09 | 8.42E-09 |
| | -1.593036741 | 0.111151949 | 0.181731876 |
| | 13.69284513 | 1.12E-42 | 6.77E-41 |
| | 7.110108647 | 1.16E-12 | 1.07E-11 |
| | -2.446334003 | 0.014431724 | 0.031465683 |
| | 0.969652943 | 0.332219514 | NA |
| | 0.320051528 | 0.748929269 | NA |
| NA | NA | NA | |

| | | |
|---|---|---|
| 7.40934412 | 1.27E-13 | 1.29E-12 |
| 2.892656849 | 0.003819984 | 0.009641444 |
| -3.366689943 | 0.000760762 | 0.002243122 |
| -3.286918368 | 0.001012902 | 0.002917148 |
| 0.201524559 | 0.840288428 | NA |
| -7.432968744 | 1.06E-13 | 1.09E-12 |
| 0.529576976 | 0.596405261 | 0.694779098 |
| NA | NA | NA |
| 1.803912428 | 0.071245036 | 0.125158848 |
| NA | NA | NA |
| NA | NA | NA |
| NA | NA | NA |
| -6.314146181 | 2.72E-10 | 1.96E-09 |
| -10.22942071 | 1.46E-24 | 3.62E-23 |
| NA | NA | NA |
| 0.069729964 | 0.944408592 | NA |
| 2.187659991 | 0.028694378 | 0.057137653 |
| 3.554676821 | 0.000378444 | 0.001180074 |
| -2.698837903 | 0.006958206 | 0.01648967 |
| 1.036830243 | 0.299814979 | 0.408712429 |
| -0.093881882 | 0.925202986 | NA |
| NA | NA | NA |
| -5.243439836 | 1.58E-07 | 8.28E-07 |
| -0.148901947 | 0.881631005 | NA |
| -2.172204096 | 0.029840272 | 0.059090349 |
| 5.238171578 | 1.62E-07 | 8.50E-07 |
| -6.337600995 | 2.33E-10 | 1.70E-09 |
| -2.941402547 | 0.003267296 | 0.008369147 |
| NA | NA | NA |
| 1.246967532 | 0.212409403 | 0.310320892 |
| 2.32054398 | 0.020311469 | 0.042357728 |
| 0.820410484 | 0.411982141 | 0.523633422 |
| 0.777301178 | 0.4369811 | 0.547301546 |
| 7.425429317 | 1.12E-13 | 1.15E-12 |
| 2.769400054 | 0.005615963 | 0.013643362 |
| 0.561589829 | 0.574395512 | 0.675616526 |
| -2.274390157 | 0.022942539 | 0.047115458 |
| -1.876168501 | 0.060632146 | 0.109008951 |
| -0.614299882 | 0.53901718 | 0.643314897 |
| -0.065389369 | 0.947863988 | NA |
| -0.138546859 | 0.889808239 | NA |
| 2.814812291 | 0.004880573 | 0.012049788 |
| -1.833950747 | 0.066661306 | 0.118209882 |
| 6.509444524 | 7.54E-11 | 5.81E-10 |
| -4.024114636 | 5.72E-05 | 0.000206382 |
| 3.476115798 | 0.000508733 | 0.001549851 |
| 8.4473023 | 2.98E-17 | 4.23E-16 |

| | | | |
|---|---|---|---|
| | 1.278617267 | 0.201031867 | 0.297013665 |
| | 2.939823515 | 0.003283992 | 0.008407316 |
| | -0.302141151 | 0.762544463 | NA |
| | 1.778987009 | 0.075241892 | 0.131003377 |
| | 7.116274878 | 1.11E-12 | 1.02E-11 |
| | 9.702861891 | 2.93E-22 | 6.16E-21 |
| | 0.063708882 | 0.949202033 | NA |
| | -1.231806732 | 0.218021288 | 0.316847985 |
| | 0.324874675 | 0.745275925 | NA |
| | -1.79921613 | 0.071984499 | 0.126183347 |
| NA | NA | NA | |
| | -0.122730807 | 0.902320268 | 0.932150576 |
| | 1.499223484 | 0.133815665 | 0.212323682 |
| | -0.733491866 | 0.463258487 | 0.572723498 |
| NA | NA | NA | |
| | 0.063708285 | 0.949202508 | NA |
| | 0.202714101 | 0.839358501 | NA |
| | 0.989868403 | 0.322238445 | 0.433286505 |
| NA | NA | NA | |
| NA | NA | NA | |
| | 0.338599155 | 0.73491172 | NA |
| NA | NA | NA | |
| | -1.53887941 | 0.123833739 | 0.199277259 |
| | 0.30922356 | 0.757151473 | NA |
| | -1.219660053 | 0.222593771 | NA |
| | -5.50917513 | 3.61E-08 | 2.06E-07 |
| NA | NA | NA | |
| | -6.752444169 | 1.45E-11 | 1.20E-10 |
| | -3.796078361 | 0.000147003 | 0.000493427 |
| | -2.833434397 | 0.004605076 | 0.011427437 |
| | -4.745515144 | 2.08E-06 | 9.35E-06 |
| | -2.380061784 | 0.017309736 | 0.03693629 |
| NA | NA | NA | |
| | -1.770299954 | 0.076677186 | 0.13314117 |
| | -0.40232487 | 0.687444954 | NA |
| | -0.211679286 | 0.832357249 | 0.882642855 |
| | -0.898130785 | 0.369115828 | 0.481672967 |
| | -1.872543959 | 0.061131386 | 0.109764605 |
| | -3.321156265 | 0.000896453 | 0.002605763 |
| | 4.18483601 | 2.85E-05 | 0.00010834 |
| | 3.119004097 | 0.001814634 | 0.004933771 |
| | -0.065388772 | 0.947864464 | NA |
| NA | NA | NA | |
| NA | NA | NA | |
| | -2.067945833 | 0.038645117 | 0.073968592 |
| | -2.964157345 | 0.003035131 | 0.007837544 |
| | -3.125034922 | 0.00177784 | 0.004844782 |

| | | |
|---|---|---|
| -1.915963107 | 0.055369792 | 0.101043007 |
| -1.245854959 | 0.212817648 | 0.310728912 |
| -1.414528055 | 0.157206916 | NA |
| -1.149579176 | 0.250317239 | 0.353939696 |
| -1.695998671 | 0.089886132 | 0.152470173 |
| NA | NA | NA |
| -0.340756924 | 0.733286582 | 0.80538127 |
| NA | NA | NA |
| NA | NA | NA |
| -0.11230281 | 0.910583314 | NA |
| 1.038654742 | 0.298965337 | 0.407865176 |
| 0.081455619 | 0.935079619 | 0.955854074 |
| NA | NA | NA |
| 1.267355274 | 0.205028285 | 0.301577962 |
| 1.382240474 | 0.166897877 | 0.254722796 |
| NA | NA | NA |
| -1.75800174 | 0.078747213 | 0.136262893 |
| -3.133436154 | 0.001727725 | 0.004722612 |
| -4.625728905 | 3.73E-06 | 1.62E-05 |
| -0.899947557 | 0.36814816 | 0.480635742 |
| -0.681982031 | 0.495250313 | 0.602337966 |
| NA | NA | NA |
| NA | NA | NA |
| 2.491993487 | 0.012702837 | 0.028164762 |
| 0.419958351 | 0.67451588 | 0.758042945 |
| -2.100992758 | 0.035641602 | 0.068947314 |
| -2.513905787 | 0.011940233 | 0.026653351 |
| 0.507275069 | 0.611961829 | NA |
| -1.341148495 | 0.179872245 | 0.271004773 |
| 0.063707688 | 0.949202983 | NA |
| NA | NA | NA |
| NA | NA | NA |
| NA | NA | NA |
| 10.313098 | 6.15E-25 | 1.56E-23 |
| NA | NA | NA |
| 8.715728124 | 2.89E-18 | 4.46E-17 |
| 2.79693453 | 0.005158999 | 0.012652335 |
| -0.065388772 | 0.947864464 | NA |
| 0.372603537 | 0.70944354 | NA |
| NA | NA | NA |
| NA | NA | NA |
| NA | NA | NA |
| 0.417818162 | 0.676080071 | NA |
| NA | NA | NA |
| 5.263119797 | 1.42E-07 | 7.48E-07 |
| NA | NA | NA |
| 1.385283938 | 0.165965663 | 0.25351645 |

| | | | |
|---|---|---|---|
| | 0.083798905 | 0.933216319 | NA |
| | -0.078634545 | 0.937323309 | NA |
| | -4.619997049 | 3.84E-06 | 1.66E-05 |
| NA | NA | NA | |
| NA | NA | NA | |
| | 0.362780088 | 0.716769167 | NA |
| NA | NA | NA | |
| NA | NA | NA | |
| | -5.258179609 | 1.45E-07 | 7.67E-07 |
| | -0.065389369 | 0.947863988 | NA |
| | -0.982248123 | 0.325977625 | 0.437069849 |
| | -4.009597359 | 6.08E-05 | 0.000218232 |
| | -2.70972147 | 0.006733973 | 0.016028363 |
| | -0.936359609 | 0.349088067 | 0.460750596 |
| | -2.932154283 | 0.003366195 | 0.008595782 |
| | -1.064349329 | 0.287170479 | 0.394510108 |
| | 0.279538226 | 0.779831807 | NA |
| | 2.226788112 | 0.025961438 | 0.052379075 |
| | 0.228443079 | 0.819301802 | 0.873015075 |
| NA | NA | NA | |
| NA | NA | NA | |
| | 1.994689095 | 0.0460768 | 0.086247554 |
| | 1.688518752 | 0.091311692 | 0.15443825 |
| | 1.077657106 | 0.281186807 | 0.388202593 |
| NA | NA | NA | |
| NA | NA | NA | |
| NA | NA | NA | |
| NA | NA | NA | |
| NA | NA | NA | |
| NA | NA | NA | |
| NA | NA | NA | |
| NA | NA | NA | |
| NA | NA | NA | |
| NA | NA | NA | |
| NA | NA | NA | |
| NA | NA | NA | |
| NA | NA | NA | |
| NA | NA | NA | |
| NA | NA | NA | |
| NA | NA | NA | |
| NA | NA | NA | |
| | -0.065388772 | 0.947864464 | NA |
| NA | NA | NA | |
| NA | NA | NA | |
| NA | NA | NA | |
| NA | NA | NA | |
| NA | NA | NA | |

| | | |
|---|---|---|
| 0.317642674 | 0.750756 | NA |
| -2.159657129 | 0.030799222 | 0.060720269 |
| -1.293123755 | 0.19596826 | 0.290740653 |
| -2.530438876 | 0.011391993 | 0.025603088 |
| 0.094087818 | 0.925039397 | NA |
| -3.833007393 | 0.000126586 | 0.000429854 |
| -2.728991354 | 0.006352837 | 0.01521059 |
| 0.650855921 | 0.515139498 | 0.620467814 |
| 4.449166881 | 8.62E-06 | 3.55E-05 |
| -0.432410491 | 0.665443095 | 0.751395926 |
| 0.352770413 | 0.724260567 | NA |
| 4.954762933 | 7.24E-07 | 3.47E-06 |
| 2.527781287 | 0.011478583 | 0.025769929 |
| -0.319010171 | 0.7497188 | NA |
| 0.214912324 | 0.829835676 | 0.880976671 |
| -2.780268715 | 0.005431393 | 0.013248979 |
| -0.048598266 | 0.961239452 | 0.973530124 |
| -2.271617632 | 0.023109615 | 0.047400939 |
| NA | NA | NA |
| NA | NA | NA |
| 0.611152837 | 0.541098403 | NA |
| 6.977640666 | 3.00E-12 | 2.64E-11 |
| 4.089156196 | 4.33E-05 | 0.000159035 |
| 0.143776291 | 0.885677126 | 0.919892878 |
| 43.910005 | 0 | 0 |
| 0.192808924 | 0.847108615 | NA |
| -13.28214836 | 2.94E-40 | 1.62E-38 |
| -0.065389369 | 0.947863988 | NA |
| NA | NA | NA |
| 4.086809212 | 4.37E-05 | 0.000160557 |
| 0.192810118 | 0.847107679 | NA |
| -0.976691266 | 0.32872202 | 0.439666141 |
| -1.451280778 | 0.146701692 | 0.229156426 |
| 1.804475672 | 0.07115677 | NA |
| 0.951510318 | 0.341345383 | 0.452894609 |
| -0.881073693 | 0.378277935 | 0.490915991 |
| -2.181033592 | 0.029180932 | 0.057968153 |
| -2.149333472 | 0.031607975 | 0.062128759 |
| -3.860957848 | 0.000112943 | 0.000386992 |
| -0.362858557 | 0.716710546 | 0.792720503 |
| 6.371909564 | 1.87E-10 | 1.37E-09 |
| -0.556434738 | 0.577913697 | 0.678765888 |
| 0.903298684 | 0.366367395 | 0.47887534 |
| -1.091694645 | 0.27496734 | 0.381636493 |
| 0.877887117 | 0.380004981 | 0.492620918 |
| NA | NA | NA |
| -1.419998048 | 0.155608249 | 0.240676913 |

| | | |
|---|---|---|
| -3.047897209 | 0.002304487 | 0.006116789 |
| NA | NA | NA |
| 3.68080423 | 0.000232499 | 0.000753914 |
| -0.065389369 | 0.947863988 | NA |
| -1.910819253 | 0.056027813 | 0.102072651 |
| -0.533387036 | 0.593765689 | 0.692768154 |
| -1.503075714 | 0.132819521 | 0.211010638 |
| 1.118423205 | 0.263386288 | 0.368876387 |
| 0.192810118 | 0.847107679 | NA |
| NA | NA | NA |
| NA | NA | NA |
| -1.912175358 | 0.055853706 | 0.101785093 |
| -0.065389966 | 0.947863513 | NA |
| 10.62752168 | 2.22E-26 | 6.19E-25 |
| NA | NA | NA |
| -7.57423631 | 3.61E-14 | 3.88E-13 |
| -0.065388772 | 0.947864464 | NA |
| -0.959676618 | 0.337217995 | 0.44847996 |
| -4.354641361 | 1.33E-05 | 5.32E-05 |
| 0.083955171 | 0.933092074 | NA |
| -3.268044606 | 0.001082933 | 0.003098161 |
| 0.068991756 | 0.944996182 | 0.962531718 |
| -6.7776689 | 1.22E-11 | 1.02E-10 |
| -4.695637509 | 2.66E-06 | 1.18E-05 |
| 0.192808924 | 0.847108615 | NA |
| 2.753522044 | 0.00589578 | 0.01423335 |
| 0.732733268 | 0.463721129 | NA |
| -0.697775083 | 0.485317864 | 0.593661721 |
| NA | NA | NA |
| -0.403417362 | 0.686641217 | 0.768667188 |
| 2.134557152 | 0.032797201 | 0.064209146 |
| 1.273486628 | 0.202845414 | 0.299099895 |
| 0.339908183 | 0.733925674 | 0.805814784 |
| -2.873478185 | 0.004059791 | 0.010191212 |
| 4.802450872 | 1.57E-06 | 7.15E-06 |
| 3.799857961 | 0.000144779 | 0.000486288 |
| -0.476178973 | 0.633946878 | NA |
| 0.184624604 | 0.853523486 | 0.898134596 |
| 1.593174245 | 0.111121107 | 0.181705188 |
| 0.511581024 | 0.608944269 | NA |
| -2.959514595 | 0.003081241 | 0.007941049 |
| -1.951367765 | 0.051013313 | 0.094205639 |
| -3.398503594 | 0.000677556 | 0.002016075 |
| 1.520472117 | 0.128392361 | 0.20542499 |
| 3.332721297 | 0.00086001 | 0.002511206 |
| NA | NA | NA |
| -0.065388772 | 0.947864464 | NA |

| | | |
|---|---|---|
| 0.388660194 | 0.697527532 | 0.77734257 |
| -1.814834414 | 0.06954936 | 0.122558133 |
| 2.28651177 | 0.022224336 | 0.045821675 |
| -2.102987049 | 0.035466902 | 0.068683679 |
| -0.95442232 | 0.339869913 | 0.451344647 |
| NA | NA | NA |
| NA | NA | NA |
| 1.337117794 | 0.18118419 | NA |
| NA | NA | NA |
| NA | NA | NA |
| -4.019317024 | 5.84E-05 | 0.000210206 |
| -6.596829562 | 4.20E-11 | 3.32E-10 |
| -4.589005017 | 4.45E-06 | 1.91E-05 |
| NA | NA | NA |
| 0.063708285 | 0.949202508 | NA |
| 1.651166546 | 0.098704572 | 0.164814906 |
| NA | NA | NA |
| 0.178047637 | 0.858685564 | 0.90185044 |
| -2.755297247 | 0.005863884 | 0.014168646 |
| 5.405024599 | 6.48E-08 | 3.58E-07 |
| 2.968340397 | 0.002994125 | 0.007744442 |
| NA | NA | NA |
| NA | NA | NA |
| NA | NA | NA |
| -0.646805567 | 0.517757789 | NA |
| 0.063708285 | 0.949202508 | NA |
| 15.22797692 | 2.31E-52 | 1.87E-50 |
| 0.192636473 | 0.847243678 | NA |
| 0.533131522 | 0.59394254 | 0.692939002 |
| 2.106211802 | 0.035185958 | 0.068229357 |
| 7.924001477 | 2.30E-15 | 2.78E-14 |
| 1.510419761 | 0.130936349 | 0.20856159 |
| NA | NA | NA |
| -0.136828103 | 0.891166669 | 0.923833166 |
| 0.203595243 | 0.838669811 | NA |
| -1.111675455 | 0.266277717 | 0.372010005 |
| -1.23374675 | 0.217297287 | 0.31596091 |
| 3.884600104 | 0.000102498 | 0.000353526 |
| -4.233625964 | 2.30E-05 | 8.85E-05 |
| 1.340227207 | 0.180171489 | 0.271373954 |
| 2.686860601 | 0.007212705 | 0.017001036 |
| 1.820304926 | 0.068712582 | 0.121340195 |
| -0.8533309 | 0.393475829 | 0.505398471 |
| 0.429952402 | 0.667230265 | 0.752524379 |
| -0.321135838 | 0.748107453 | 0.816856769 |
| -1.704613803 | 0.088266475 | 0.150150189 |
| NA | NA | NA |

| | | |
|---|---|---|
| NA | NA | NA |
| 0.303799102 | 0.761280951 | 0.827357136 |
| 2.523330731 | 0.011624901 | 0.02604469 |
| 5.122125186 | 3.02E-07 | 1.53E-06 |
| NA | NA | NA |
| 17.65242183 | 9.75E-70 | 1.23E-67 |
| NA | NA | NA |
| 18.19286632 | 5.88E-74 | 8.27E-72 |
| 7.18123109 | 6.91E-13 | 6.53E-12 |
| 1.540638393 | 0.123404818 | 0.19863811 |
| 0.272389173 | 0.785322797 | 0.846794021 |
| -3.48591325 | 0.00049046 | 0.001498378 |
| 0.192808924 | 0.847108615 | NA |
| 0.769723836 | 0.441463728 | NA |
| 0.08765515 | 0.930150767 | NA |
| NA | NA | NA |
| -0.18243076 | 0.855244688 | 0.899406518 |
| 0.141851683 | 0.88719716 | 0.920898437 |
| -3.089937732 | 0.002001985 | 0.005396042 |
| 0.063708285 | 0.949202508 | NA |
| NA | NA | NA |
| -2.218757412 | 0.026503233 | 0.053304614 |
| -0.929524025 | 0.352617581 | 0.464502818 |
| NA | NA | NA |
| -0.07562122 | 0.939720454 | NA |
| 2.43685174 | 0.014815752 | 0.032215992 |
| -0.065389369 | 0.947863988 | NA |
| 0.398161514 | 0.690511133 | 0.771550094 |
| 0.192808924 | 0.847108615 | NA |
| 0.320051528 | 0.748929269 | NA |
| 3.406080408 | 0.000659028 | 0.001964681 |
| 0.110705143 | 0.911850167 | NA |
| -2.24416984 | 0.024821476 | 0.050404149 |
| -1.144671643 | 0.252345202 | 0.356384383 |
| 2.948882003 | 0.003189257 | 0.008186556 |
| -1.057525418 | 0.290271861 | 0.397848246 |
| NA | NA | NA |
| -1.601792635 | 0.109201472 | 0.179034051 |
| 1.042995444 | 0.296950403 | 0.405558388 |
| -2.894640474 | 0.00379593 | 0.009582666 |
| -3.369535831 | 0.000752949 | 0.002222181 |
| -0.364808922 | 0.715254045 | NA |
| -2.270905325 | 0.02315271 | 0.047472984 |
| -2.951408047 | 0.003163287 | 0.008128443 |
| NA | NA | NA |
| -8.317593393 | 8.98E-17 | 1.22E-15 |
| -0.010907849 | 0.991296968 | 0.993917766 |

| | | |
|---|---|---|
| -6.910069319 | 4.84E-12 | 4.19E-11 |
| -0.535556359 | 0.592265197 | 0.691468719 |
| -2.028541852 | 0.042504976 | 0.080385309 |
| NA | NA | NA |
| 5.841779321 | 5.16E-09 | 3.26E-08 |
| -0.783689902 | 0.433222089 | 0.543945267 |
| -8.280339526 | 1.23E-16 | 1.66E-15 |
| -0.020065788 | 0.983990892 | 0.988810503 |
| -2.622417771 | 0.008730833 | 0.020169152 |
| 1.414396397 | 0.157245547 | NA |
| -3.571773317 | 0.000354572 | 0.001113125 |
| 0.115484181 | 0.908061359 | NA |
| NA | NA | NA |
| NA | NA | NA |
| 7.747522606 | 9.37E-15 | 1.07E-13 |
| 13.03475065 | 7.76E-39 | 4.08E-37 |
| 0.31685483 | 0.751353758 | NA |
| 0.675738624 | 0.499206609 | NA |
| 2.210135528 | 0.027095758 | 0.054326083 |
| 0.192809521 | 0.847108147 | NA |
| -0.084501006 | 0.932658103 | NA |
| -1.658808853 | 0.097154316 | 0.162636482 |
| 7.674884924 | 1.66E-14 | 1.84E-13 |
| 1.71535938 | 0.086279358 | 0.147192692 |
| -0.065388772 | 0.947864464 | NA |
| 10.28812193 | 7.97E-25 | 2.00E-23 |
| 4.747795931 | 2.06E-06 | 9.24E-06 |
| 1.652765918 | 0.098378513 | 0.164347205 |
| NA | NA | NA |
| NA | NA | NA |
| NA | NA | NA |
| 0.333816862 | 0.738517758 | NA |
| NA | NA | NA |
| NA | NA | NA |
| NA | NA | NA |
| 0.330493631 | 0.741027004 | NA |
| NA | NA | NA |
| 0.192809521 | 0.847108147 | NA |
| NA | NA | NA |
| 4.787806162 | 1.69E-06 | 7.67E-06 |
| 2.24066865 | 0.025047546 | 0.0508014 |
| -0.670702579 | 0.502410019 | 0.6092516 |
| 0.523576119 | 0.600573395 | 0.698302229 |
| NA | NA | NA |
| 1.256843907 | 0.208810163 | 0.305930601 |
| NA | NA | NA |
| -0.065389369 | 0.947863988 | NA |

| | | | |
|---|---|---|---|
| | 0.138584883 | 0.88977819 | NA |
| | 0.382660899 | 0.70197121 | 0.781246689 |
| | 0.450180844 | 0.652580047 | NA |
| | -0.014917394 | 0.988098083 | 0.991703557 |
| NA | NA | NA | |
| | 0.265141849 | 0.79090019 | 0.851164385 |
| | 6.408297728 | 1.47E-10 | 1.09E-09 |
| | 1.015974988 | 0.309641297 | 0.419594049 |
| | -3.403050522 | 0.00066638 | 0.001985416 |
| | 0.469385103 | 0.638794395 | NA |
| | -0.558602029 | 0.576433355 | 0.677440458 |
| NA | NA | NA | |
| NA | NA | NA | |
| NA | NA | NA | |
| NA | NA | NA | |
| | -3.376245958 | 0.000734822 | 0.002174065 |
| | -3.765990354 | 0.00016589 | 0.000552232 |
| | -5.323990805 | 1.02E-07 | 5.47E-07 |
| | 0.075823031 | 0.939559893 | NA |
| | 3.783294641 | 0.000154766 | 0.00051747 |
| NA | NA | NA | |
| | 0.209074615 | 0.834389991 | NA |
| NA | NA | NA | |
| | 1.619277521 | 0.105387565 | 0.173874547 |
| | -1.98486235 | 0.047159788 | 0.088004883 |
| | 0.192809521 | 0.847108147 | NA |
| | 0.192809521 | 0.847108147 | NA |
| | -0.194487022 | 0.845794556 | NA |
| | -0.19448762 | 0.845794089 | NA |
| | -0.065389966 | 0.947863513 | NA |
| | -3.866697156 | 0.000110319 | 0.000378742 |
| | -0.090199405 | 0.928128757 | NA |
| NA | NA | NA | |
| | 0.904323871 | 0.365823693 | 0.478399531 |
| | -12.77430677 | 2.28E-37 | 1.11E-35 |
| NA | NA | NA | |
| | -0.471825095 | 0.637051637 | 0.728100569 |
| NA | NA | NA | |
| | -1.879616384 | 0.060160379 | 0.108291122 |
| | 1.180950887 | 0.237622234 | 0.339634795 |
| | 0.556606987 | 0.577795979 | 0.678659472 |
| | 1.394896395 | 0.163047084 | NA |
| | -1.616348853 | 0.10601891 | 0.174693483 |
| | -3.06477143 | 0.002178364 | 0.005819045 |
| | -0.429148782 | 0.667814954 | 0.752912405 |
| | 1.806229247 | 0.070882538 | NA |
| | -0.036196681 | 0.971125532 | 0.980052902 |

| | | |
|---|---|---|
| 0.288947695 | 0.772621404 | 0.836449557 |
| 0.892022167 | 0.372381051 | 0.485199029 |
| -0.130162487 | 0.896437873 | 0.927873623 |
| NA | NA | NA |
| NA | NA | NA |
| -1.955882757 | 0.050478967 | 0.093377322 |
| -0.525512242 | 0.599227146 | 0.697157918 |
| 1.128070838 | 0.259290004 | 0.364308076 |
| NA | NA | NA |
| 2.873161539 | 0.004063863 | 0.010199027 |
| -0.007018097 | 0.994400415 | 0.996031949 |
| 4.169387091 | 3.05E-05 | 0.000115391 |
| -0.612600463 | 0.540140552 | 0.644348394 |
| 1.117965269 | 0.263581827 | 0.369088287 |
| NA | NA | NA |
| 3.274935728 | 0.00105686 | 0.003030497 |
| NA | NA | NA |
| -0.09562842 | 0.923815693 | NA |
| NA | NA | NA |
| NA | NA | NA |
| NA | NA | NA |
| NA | NA | NA |
| 0.063707019 | 0.949203516 | NA |
| 0.537899024 | 0.590646762 | 0.690062008 |
| -1.276843195 | 0.201657612 | 0.297776588 |
| -0.095025761 | 0.924294365 | NA |
| 0.262922253 | 0.792610504 | 0.852485429 |
| -0.67543553 | 0.499399099 | 0.60649973 |
| 4.4404245 | 8.98E-06 | 3.69E-05 |
| -0.169240834 | 0.865607213 | 0.906246133 |
| NA | NA | NA |
| NA | NA | NA |
| NA | NA | NA |
| NA | NA | NA |
| NA | NA | NA |
| NA | NA | NA |
| NA | NA | NA |
| NA | NA | NA |
| NA | NA | NA |
| NA | NA | NA |
| NA | NA | NA |
| NA | NA | NA |
| -2.442656656 | 0.0145796 | 0.0317632 |
| -2.016313893 | 0.043767163 | 0.0825043 |
| 0.836906288 | 0.402645245 | 0.514378119 |
| -1.26028012 | 0.207568329 | 0.304592558 |
| -12.50822152 | 6.73E-36 | 3.06E-34 |

| | | |
|---|---|---|
| -1.6272037 | 0.103693842 | 0.17162668 |
| 5.937071919 | 2.90E-09 | 1.88E-08 |
| 1.159602618 | 0.246210635 | 0.349257704 |
| -2.819851845 | 0.004804583 | 0.011875057 |
| -3.563014574 | 0.00036662 | 0.001146562 |
| 1.647905157 | 0.099372135 | 0.165751091 |
| 1.831522176 | 0.067022641 | 0.118808548 |
| -1.965111098 | 0.04940138 | 0.091661716 |
| -0.266184652 | 0.790097003 | 0.850695917 |
| 3.331794926 | 0.000862878 | 0.002518699 |
| 1.370658333 | 0.170481489 | 0.259196482 |
| 7.173246593 | 7.32E-13 | 6.90E-12 |
| 4.125845453 | 3.69E-05 | 0.00013754 |
| 0.063708285 | 0.949202508 | NA |
| 0.063707688 | 0.949202983 | NA |
| 0.251471875 | 0.801449304 | 0.859075675 |
| 0.793620098 | 0.427416629 | 0.538736943 |
| 0.668750982 | 0.50365434 | 0.610187228 |
| -12.22426733 | 2.31E-34 | 9.79E-33 |
| -0.966751735 | 0.333668158 | 0.444862856 |
| 0.31685483 | 0.751353758 | NA |
| NA | NA | NA |
| NA | NA | NA |
| NA | NA | NA |
| NA | NA | NA |
| NA | NA | NA |
| NA | NA | NA |
| NA | NA | NA |
| -13.92717601 | 4.33E-44 | 2.82E-42 |
| -2.813092599 | 0.004906752 | 0.012104869 |
| -1.820441541 | 0.068691791 | 0.121315585 |
| -0.579619379 | 0.562171321 | 0.664297431 |
| -1.574136756 | 0.115455837 | 0.187682135 |
| -3.259439176 | 0.001116327 | 0.003183873 |
| -3.030538651 | 0.002441179 | 0.006436608 |
| -2.091916386 | 0.036445996 | 0.070329449 |
| -4.083569262 | 4.43E-05 | 0.00016267 |
| NA | NA | NA |
| NA | NA | NA |
| -0.065388772 | 0.947864464 | NA |
| -0.922359766 | 0.356340949 | 0.468224786 |
| NA | NA | NA |
| 5.015335043 | 5.29E-07 | 2.58E-06 |
| 1.565121485 | 0.117554427 | 0.190507433 |
| -0.363328061 | 0.716359834 | 0.792437647 |
| -0.065389369 | 0.947863988 | NA |
| 0.276024982 | 0.782528878 | 0.844795893 |

| | | | |
|---|---|---|---|
| NA | NA | NA | |
| NA | NA | NA | |
| NA | NA | NA | |
| | -0.065389369 | 0.947863988 | NA |
| | 0.096545474 | 0.92308736 | NA |
| NA | NA | NA | |
| | -3.082897296 | 0.002049959 | 0.005510145 |
| | 7.865247307 | 3.68E-15 | 4.37E-14 |
| | 0.438862535 | 0.660761145 | 0.747586288 |
| NA | NA | NA | |
| | 0.377087183 | 0.706108816 | 0.784426064 |
| NA | NA | NA | |
| | 3.182255566 | 0.001461328 | 0.004059605 |
| | 4.694905794 | 2.67E-06 | 1.18E-05 |
| | 1.351551928 | 0.176518697 | 0.266867782 |
| | 1.281185817 | 0.200128407 | 0.295877969 |
| | -0.383716286 | 0.701188741 | 0.780449759 |
| NA | NA | NA | |
| NA | NA | NA | |
| | 0.109127625 | 0.913101264 | NA |
| | -0.219498892 | 0.826261443 | NA |
| NA | NA | NA | |
| NA | NA | NA | |
| | 0.392023906 | 0.69504055 | NA |
| | 0.204183227 | 0.838210318 | NA |
| | 0.828064868 | 0.407633764 | 0.519451844 |
| | 4.504912317 | 6.64E-06 | 2.78E-05 |
| | -4.901170024 | 9.53E-07 | 4.49E-06 |
| | -0.757502033 | 0.448749149 | 0.558732089 |
| | -1.534598052 | 0.124882591 | 0.200681257 |
| | 1.908144307 | 0.056372567 | 0.102603648 |
| | -2.491479282 | 0.012721239 | 0.028195573 |
| | -2.97641303 | 0.002916418 | 0.007559853 |
| | 3.605830368 | 0.000311156 | 0.000986228 |
| | -3.837404216 | 0.000124342 | 0.000422748 |
| | 0.452944158 | 0.650588953 | NA |
| | -3.391242159 | 0.000695766 | 0.002066084 |
| | 0.146247849 | 0.883725734 | NA |
| | 2.699041225 | 0.006953956 | 0.016481159 |
| | 5.155290958 | 2.53E-07 | 1.29E-06 |
| | 1.792653989 | 0.073028268 | 0.127655391 |
| | 1.066942967 | 0.285997591 | 0.393109264 |
| NA | NA | NA | |
| NA | NA | NA | |
| | 0.063708285 | 0.949202508 | NA |
| NA | NA | NA | |
| | -0.095223702 | 0.924137144 | NA |

| | | | |
|---|---|---|---|
| NA | NA | NA | |
| NA | NA | NA | |
| | 0.192809521 | 0.847108147 | NA |
| | 2.350115031 | 0.01876761 | 0.039541522 |
| NA | NA | NA | |
| | -3.061395945 | 0.002203076 | 0.005876903 |
| | -2.00317686 | 0.045158308 | 0.084769196 |
| NA | NA | NA | |
| | -0.180065661 | 0.85710102 | NA |
| | 3.077910343 | 0.002084576 | 0.005591185 |
| | 3.239154057 | 0.001198848 | 0.003395244 |
| | 3.513922568 | 0.000441542 | 0.001360554 |
| | 0.497915394 | 0.618543677 | NA |
| NA | NA | NA | |
| NA | NA | NA | |
| | 7.885791231 | 3.13E-15 | 3.73E-14 |
| | 0.192808924 | 0.847108615 | NA |
| | 1.993665828 | 0.046188586 | 0.086424483 |
| | 0.543583719 | 0.586727966 | NA |
| | -7.277467146 | 3.40E-13 | 3.31E-12 |
| | 0.254628458 | 0.799010077 | 0.85719509 |
| | 9.449609488 | 3.40E-21 | 6.63E-20 |
| | -4.333291591 | 1.47E-05 | 5.83E-05 |
| | -3.450998149 | 0.000558517 | 0.001688776 |
| | -0.088402443 | 0.92955682 | 0.952113893 |
| | 17.07510974 | 2.27E-65 | 2.60E-63 |
| | 7.367708718 | 1.74E-13 | 1.74E-12 |
| | -0.343678669 | 0.731087955 | NA |
| | 1.885521839 | 0.059359419 | 0.107089971 |
| | 0.731532105 | 0.464454202 | 0.573889948 |
| | -1.792309159 | 0.073083457 | 0.127734022 |
| | 4.430826054 | 9.39E-06 | 3.84E-05 |
| | 6.893980157 | 5.43E-12 | 4.67E-11 |
| | 2.229826934 | 0.025758935 | 0.051999849 |
| | -1.134172184 | 0.25672233 | 0.361290435 |
| | -3.137926587 | 0.001701475 | 0.00465756 |
| | 2.937428294 | 0.003309467 | 0.008464741 |
| | 1.304881934 | 0.191933054 | 0.285821296 |
| | -2.176378719 | 0.029526954 | 0.058562571 |
| | -1.131581775 | 0.257810306 | 0.362507272 |
| | 0.261092137 | 0.794021455 | 0.853526177 |
| NA | NA | NA | |
| | -2.463261333 | 0.013767952 | 0.030234414 |
| | 0.06371127 | 0.949200131 | NA |
| | 1.415627848 | 0.156884493 | 0.24233639 |
| | -0.530764445 | 0.595582021 | 0.694013873 |
| | -7.077590628 | 1.47E-12 | 1.34E-11 |

| | | |
|---|---|---|
| 1.223487942 | 0.221145456 | 0.320410183 |
| 2.507862957 | 0.012146377 | 0.027053195 |
| NA | NA | NA |
| -5.746047495 | 9.14E-09 | 5.60E-08 |
| -0.065388772 | 0.947864464 | NA |
| -0.324348095 | 0.745674511 | 0.814960834 |
| 0.063708285 | 0.949202508 | NA |
| 0.7793916 | 0.435749071 | 0.54629598 |
| 2.706206415 | 0.006805673 | 0.016171064 |
| 6.784778503 | 1.16E-11 | 9.69E-11 |
| -2.829461497 | 0.004662641 | 0.011550466 |
| 1.893490306 | 0.058292696 | 0.105507715 |
| -2.202259207 | 0.027647004 | 0.055316108 |
| -0.052977631 | 0.95774973 | NA |
| -7.202338184 | 5.92E-13 | 5.64E-12 |
| -3.370190766 | 0.000751162 | 0.002217952 |
| 0.827010931 | 0.408230866 | 0.520080347 |
| NA | NA | NA |
| 4.613868124 | 3.95E-06 | 1.71E-05 |
| -11.6701884 | 1.81E-31 | 6.76E-30 |
| 1.259295735 | 0.207923533 | 0.304962645 |
| -0.101253237 | 0.919349437 | 0.9446741 |
| 0.191964722 | 0.84776984 | 0.893723585 |
| -1.112460997 | 0.265939991 | 0.371621147 |
| 3.295494266 | 0.000982487 | 0.002836735 |
| -0.580071533 | 0.56186638 | 0.664031139 |
| 4.353987186 | 1.34E-05 | 5.33E-05 |
| 5.911670313 | 3.39E-09 | 2.18E-08 |
| -0.946579399 | 0.343853145 | 0.455209405 |
| -5.745445632 | 9.17E-09 | 5.62E-08 |
| NA | NA | NA |
| 1.621277693 | 0.104958097 | 0.173348807 |
| -8.825398498 | 1.09E-18 | 1.75E-17 |
| 1.412153094 | 0.157904888 | 0.24346183 |
| 2.028154762 | 0.042544455 | 0.080453892 |
| -1.16791072 | 0.24284278 | 0.345535972 |
| -0.389750628 | 0.696720955 | 0.776618161 |
| -6.091398721 | 1.12E-09 | 7.59E-09 |
| NA | NA | NA |
| -1.068110605 | 0.285470623 | 0.39261983 |
| 0.06371127 | 0.949200131 | NA |
| -11.62080736 | 3.23E-31 | 1.19E-29 |
| 0.454979602 | 0.649123914 | NA |
| 0.324874675 | 0.745275925 | NA |
| 0.06371127 | 0.949200131 | NA |
| NA | NA | NA |
| NA | NA | NA |

| | | |
|---|---|---|
| -0.401495214 | 0.688055561 | NA |
| 1.414760329 | 0.157138779 | 0.242624369 |
| NA | NA | NA |
| -1.25511186 | 0.209438156 | NA |
| NA | NA | NA |
| NA | NA | NA |
| -7.258224435 | 3.92E-13 | 3.79E-12 |
| -2.808202162 | 0.004981894 | 0.012270893 |
| -1.693117302 | 0.090433142 | 0.153221521 |
| -0.485475065 | 0.627339396 | NA |
| NA | NA | NA |
| 0.063708285 | 0.949202508 | NA |
| -0.502030651 | 0.615645961 | 0.710805304 |
| -3.26054641 | 0.001111978 | 0.003172552 |
| -0.157230409 | 0.875063264 | NA |
| -1.991067251 | 0.046473494 | 0.086905608 |
| 3.506005958 | 0.000454885 | 0.00139737 |
| NA | NA | NA |
| -2.923281706 | 0.00346363 | 0.008818498 |
| -2.266238137 | 0.023436807 | 0.047985544 |
| -0.438957902 | 0.66069204 | 0.747541893 |
| 0.552312704 | 0.580734128 | NA |
| NA | NA | NA |
| -2.657160081 | 0.007880201 | 0.018383954 |
| 0.313465947 | 0.753926688 | 0.821504724 |
| 1.154520047 | 0.248287024 | 0.351525778 |
| 1.486513108 | 0.137143452 | NA |
| 2.042405465 | 0.041111328 | 0.078032649 |
| 0.109178353 | 0.91306103 | NA |
| -0.077373275 | 0.938326601 | 0.958058216 |
| 3.632602181 | 0.000280577 | 0.000896573 |
| 0.192810118 | 0.847107679 | NA |
| NA | NA | NA |
| 7.555351504 | 4.18E-14 | 4.46E-13 |
| -0.186971533 | 0.851682953 | 0.896866688 |
| 9.708789676 | 2.77E-22 | 5.82E-21 |
| 1.62396917 | 0.104382388 | NA |
| 3.0799348 | 0.002070459 | 0.005558086 |
| 1.411052329 | 0.158229185 | 0.243841676 |
| -0.412630698 | 0.679877206 | 0.76269839 |
| 1.616425595 | 0.106002328 | 0.174681564 |
| NA | NA | NA |
| -0.176367936 | 0.860004893 | NA |
| 0.835811791 | 0.403260792 | NA |
| NA | NA | NA |
| 5.45653581 | 4.86E-08 | 2.73E-07 |
| NA | NA | NA |

| | | | |
|---|---|---|---|
| | 0.293532392 | 0.769115244 | 0.83366337 |
| | 1.484196162 | 0.137756888 | 0.217515449 |
| | -0.514692702 | 0.606767774 | 0.703509155 |
| | 0.095722367 | 0.923741076 | 0.947630605 |
| | 1.119560711 | 0.262901006 | 0.368279168 |
| | 2.87427108 | 0.004049612 | 0.010169739 |
| | 0.06371127 | 0.949200131 | NA |
| | 5.426533995 | 5.75E-08 | 3.20E-07 |
| | 1.616827232 | 0.105915579 | 0.174583875 |
| | 0.975634283 | 0.329245729 | 0.440107909 |
| | 1.189553167 | 0.234222062 | 0.335541336 |
| NA | NA | NA | |
| NA | NA | NA | |
| | -7.513893913 | 5.74E-14 | 6.04E-13 |
| NA | NA | NA | |
| NA | NA | NA | |
| | 4.164937998 | 3.11E-05 | 0.000117505 |
| NA | NA | NA | |
| NA | NA | NA | |
| NA | NA | NA | |
| | -3.468223361 | 0.000523912 | 0.001592798 |
| | 0.425562139 | 0.670426934 | NA |
| | 7.307485465 | 2.72E-13 | 2.68E-12 |
| | 1.979513027 | 0.047758274 | 0.088995653 |
| | -0.394504716 | 0.693208446 | NA |
| | 0.193076837 | 0.846898793 | NA |
| | -0.411265037 | 0.680878204 | 0.763581766 |
| | -2.651246693 | 0.008019524 | 0.0186707 |
| NA | NA | NA | |
| | 1.281546071 | 0.200001928 | 0.295743385 |
| | -2.471576009 | 0.013451895 | 0.029615704 |
| | -2.149110578 | 0.031625635 | 0.06215371 |
| | 0.063710673 | 0.949200606 | NA |
| NA | NA | NA | |
| | 0.30922356 | 0.757151473 | NA |
| | 0.89065545 | 0.373114042 | NA |
| NA | NA | NA | |
| | 0.106035613 | 0.915554096 | NA |
| | 3.073972603 | 0.002112288 | 0.005658844 |
| | 0.038510891 | 0.969280348 | 0.978742716 |
| | -1.625653261 | 0.104023438 | 0.172100381 |
| NA | NA | NA | |
| NA | NA | NA | |
| NA | NA | NA | |
| | -0.065389369 | 0.947863988 | NA |
| | 0.192809521 | 0.847108147 | NA |
| | 0.510446462 | 0.609738713 | NA |

| | | |
|---|---|---|
| NA | NA | NA |
| -0.777288685 | 0.436988469 | 0.547301546 |
| NA | NA | NA |
| NA | NA | NA |
| -5.112654245 | 3.18E-07 | 1.60E-06 |
| -0.065388772 | 0.947864464 | NA |
| -4.409531031 | 1.04E-05 | 4.22E-05 |
| NA | NA | NA |
| 1.041433838 | 0.297674245 | 0.40639175 |
| 0.063707019 | 0.949203516 | NA |
| NA | NA | NA |
| 3.030218116 | 0.002443772 | 0.00644073 |
| -0.959046979 | 0.337535079 | 0.448720152 |
| NA | NA | NA |
| -0.20144474 | 0.840350834 | 0.88803434 |
| -2.610245002 | 0.00904774 | 0.020826276 |
| -3.217155386 | 0.001294685 | 0.0036354 |
| -4.307377668 | 1.65E-05 | 6.50E-05 |
| -0.522813248 | 0.601104219 | NA |
| 8.010450826 | 1.14E-15 | 1.42E-14 |
| -0.736228881 | 0.461591419 | 0.571000949 |
| -1.67408496 | 0.094113903 | 0.158385732 |
| -0.982090663 | 0.326055185 | 0.437135619 |
| -2.001018024 | 0.045390448 | 0.085147486 |
| 6.262291748 | 3.79E-10 | 2.69E-09 |
| 0.153462498 | 0.878033562 | NA |
| 9.133062973 | 6.66E-20 | 1.18E-18 |
| -0.044640706 | 0.964393697 | 0.975777113 |
| NA | NA | NA |
| 6.022434083 | 1.72E-09 | 1.14E-08 |
| NA | NA | NA |
| 7.261593411 | 3.83E-13 | 3.70E-12 |
| 8.753333098 | 2.07E-18 | 3.24E-17 |
| 0.324874675 | 0.745275925 | NA |
| 8.662653638 | 4.61E-18 | 6.98E-17 |
| 0.192810118 | 0.847107679 | NA |
| 0.31685483 | 0.751353758 | NA |
| 1.66795642 | 0.095324378 | 0.160099925 |
| 8.060785838 | 7.58E-16 | 9.56E-15 |
| 0.324874675 | 0.745275925 | NA |
| 6.938931201 | 3.95E-12 | 3.44E-11 |
| 6.148084068 | 7.84E-10 | 5.39E-09 |
| 1.407234384 | 0.159357899 | 0.245203677 |
| 1.120869477 | 0.262343425 | NA |
| 2.97662635 | 0.00291439 | 0.007555378 |
| 4.14921537 | 3.34E-05 | 0.000125156 |
| -1.71538835 | 0.08627405 | 0.147192692 |

| | | |
|---|---|---|
| 4.973997395 | 6.56E-07 | 3.16E-06 |
| NA | NA | NA |
| 22.91009304 | 3.69E-116 | 1.15E-113 |
| 6.227175142 | 4.75E-10 | 3.34E-09 |
| 22.13514408 | 1.45E-108 | 3.99E-106 |
| 4.620473938 | 3.83E-06 | 1.66E-05 |
| 1.459425414 | 0.144448055 | 0.226267903 |
| 23.26130054 | 1.09E-119 | 3.46E-117 |
| NA | NA | NA |
| 8.80354658 | 1.33E-18 | 2.12E-17 |
| NA | NA | NA |
| NA | NA | NA |
| 1.25215755 | 0.210512461 | NA |
| NA | NA | NA |
| 0.978089126 | 0.328030246 | 0.439045883 |
| -2.35543105 | 0.01850123 | 0.039075672 |
| -4.939111141 | 7.85E-07 | 3.74E-06 |
| -1.126989027 | 0.25974712 | 0.364759337 |
| 5.51250487 | 3.54E-08 | 2.03E-07 |
| 0.09635447 | 0.923239051 | NA |
| 1.053767989 | 0.291989152 | 0.399825205 |
| -0.665120771 | 0.505973251 | 0.612403794 |
| 7.537095097 | 4.81E-14 | 5.10E-13 |
| 5.259123269 | 1.45E-07 | 7.64E-07 |
| -0.468803726 | 0.639209937 | 0.729947381 |
| 8.721728828 | 2.74E-18 | 4.24E-17 |
| -0.630440966 | 0.528406115 | 0.633324216 |
| -0.978008081 | 0.328070327 | NA |
| -0.83024019 | 0.406402996 | 0.518226451 |
| 1.920841416 | 0.054751703 | 0.100090208 |
| 6.311103149 | 2.77E-10 | 2.00E-09 |
| 1.83005788 | 0.067241284 | 0.119115438 |
| NA | NA | NA |
| NA | NA | NA |
| 4.378585487 | 1.19E-05 | 4.80E-05 |
| NA | NA | NA |
| 0.156713226 | 0.875470863 | NA |
| NA | NA | NA |
| -0.554917596 | 0.578951026 | 0.679729076 |
| -1.586838757 | 0.112549169 | 0.183632468 |
| 0.78708332 | 0.431233093 | 0.542197669 |
| -0.065389369 | 0.947863988 | NA |
| NA | NA | NA |
| 5.587976705 | 2.30E-08 | 1.35E-07 |
| -0.297090195 | 0.76639765 | 0.831506598 |
| -0.087659735 | 0.930147124 | NA |
| 0.156284267 | 0.875808957 | 0.913357702 |

| | | |
|---|---|---|
| NA | NA | NA |
| 0.673782828 | 0.500449394 | 0.607509982 |
| 1.803300917 | 0.071340969 | 0.125274621 |
| -3.154158988 | 0.001609614 | 0.00443512 |
| -0.065389369 | 0.947863988 | NA |
| -0.654363798 | 0.512877452 | 0.618509307 |
| -3.18161252 | 0.001464576 | 0.004066993 |
| -2.810932495 | 0.004939815 | 0.012173237 |
| -1.973909018 | 0.048392089 | 0.089962418 |
| -1.552863004 | 0.120455867 | 0.194554452 |
| -3.736457961 | 0.000186631 | 0.000616031 |
| 3.492737609 | 0.000478096 | 0.001463106 |
| -3.189895649 | 0.001423242 | 0.003962596 |
| 0.063708882 | 0.949202033 | NA |
| -1.274984114 | 0.202314865 | 0.298511027 |
| -3.8820797 | 0.000103567 | 0.000356965 |
| -0.215013851 | 0.82975652 | 0.88093 |
| NA | NA | NA |
| NA | NA | NA |
| NA | NA | NA |
| NA | NA | NA |
| 1.230553245 | 0.218490001 | NA |
| 3.52093533 | 0.000430027 | 0.001326545 |
| 6.899695643 | 5.21E-12 | 4.50E-11 |
| 3.28582856 | 0.001016829 | 0.002926774 |
| NA | NA | NA |
| 0.354627914 | 0.722868361 | NA |
| 1.816445289 | 0.069302093 | 0.122191561 |
| 0.865297337 | 0.386875612 | 0.499097854 |
| -6.377846812 | 1.80E-10 | 1.32E-09 |
| -0.678911602 | 0.497193874 | NA |
| -2.875896646 | 0.004028816 | 0.010121576 |
| NA | NA | NA |
| -4.577214342 | 4.71E-06 | 2.02E-05 |
| NA | NA | NA |
| -0.711412203 | 0.476828839 | NA |
| -7.068785625 | 1.56E-12 | 1.42E-11 |
| -2.248090874 | 0.024570396 | 0.049930747 |
| -0.443234805 | 0.657595907 | NA |
| 0.912257851 | 0.361632997 | 0.47386001 |
| 16.46109261 | 6.98E-61 | 7.19E-59 |
| -2.955048391 | 0.0031262 | 0.008045321 |
| -0.71167137 | 0.476668301 | 0.585561225 |
| 0.820000696 | 0.412215711 | 0.523877087 |
| 2.501583276 | 0.012363936 | 0.027493721 |
| -2.473244804 | 0.013389239 | 0.029501114 |
| 1.385623925 | 0.165861769 | 0.253388676 |

| | | | |
|---|---|---|---|
| | -1.807223557 | 0.070727429 | 0.124406715 |
| | -2.611904315 | 0.009003945 | 0.020735003 |
| | 0.625921037 | 0.531366736 | 0.635964848 |
| NA | NA | NA | |
| | 0.06371127 | 0.949200131 | NA |
| | 0.192809521 | 0.847108147 | NA |
| | 0.573193348 | 0.566513788 | NA |
| | 0.192810118 | 0.847107679 | NA |
| | 0.192810118 | 0.847107679 | NA |
| NA | NA | NA | |
| | 6.190689627 | 5.99E-10 | 4.16E-09 |
| | -0.065389369 | 0.947863988 | NA |
| | 2.342659822 | 0.019146832 | 0.040228758 |
| | 2.352956482 | 0.018624814 | 0.039293584 |
| | -0.091738921 | 0.926905474 | NA |
| | 2.088962809 | 0.036711071 | 0.070775484 |
| | 1.824093392 | 0.068137954 | 0.120427411 |
| | 0.324459073 | 0.745590502 | 0.814940187 |
| | 1.193838803 | 0.23254103 | NA |
| | -0.038193522 | 0.969533386 | 0.978801301 |
| | 1.020461421 | 0.307509677 | 0.41727026 |
| | 1.123941463 | 0.26103786 | NA |
| | 0.748031863 | 0.454440939 | 0.564275062 |
| | 2.271243373 | 0.023132249 | 0.047435703 |
| | -0.13520322 | 0.892451201 | NA |
| | -0.114845519 | 0.908567569 | 0.936941589 |
| | 0.370415585 | 0.711072863 | 0.788296296 |
| | -0.099432242 | 0.920795084 | 0.945538501 |
| | 0.192808327 | 0.847109082 | NA |
| | 2.141474955 | 0.032235756 | 0.063238575 |
| | -0.555720496 | 0.578401943 | NA |
| NA | NA | NA | |
| | 5.009202879 | 5.47E-07 | 2.66E-06 |
| | 0.192810118 | 0.847107679 | NA |
| | 0.192810118 | 0.847107679 | NA |
| | -0.194488217 | 0.845793621 | NA |
| | 2.141571357 | 0.032227991 | 0.063228297 |
| NA | NA | NA | |
| | -0.065388772 | 0.947864464 | NA |
| NA | NA | NA | |
| NA | NA | NA | |
| NA | NA | NA | |
| NA | NA | NA | |
| | -0.019972361 | 0.984065421 | 0.988845698 |
| | -2.138530378 | 0.032473721 | 0.063665485 |
| | 0.063708882 | 0.949202033 | NA |
| | 0.192809521 | 0.847108147 | NA |

| | | | |
|---|---|---|---|
| | 0.181680723 | 0.855833295 | NA |
| | -2.463033129 | 0.013776719 | 0.030243051 |
| | -2.454218915 | 0.0141191 | 0.030896647 |
| | -1.348192116 | 0.1775966 | 0.268173437 |
| | -1.747739597 | 0.080509129 | 0.138850927 |
| | 0.437089463 | 0.662046469 | NA |
| | -2.795806992 | 0.005177031 | 0.012690342 |
| | -2.851150481 | 0.004356135 | 0.010856722 |
| | 1.148437558 | 0.250787976 | 0.354465134 |
| | -0.698853214 | 0.484643769 | 0.59306901 |
| | -3.523513223 | 0.000425866 | 0.001315208 |
| | -1.819118633 | 0.068893333 | 0.121625061 |
| | 0.192808924 | 0.847108615 | NA |
| | 4.483088206 | 7.36E-06 | 3.05E-05 |
| | -0.065389369 | 0.947863988 | NA |
| | -0.065388772 | 0.947864464 | NA |
| | 1.098579073 | 0.27195171 | 0.378416238 |
| NA | NA | NA | |
| | 0.740324274 | 0.459103255 | NA |
| | 1.132676243 | 0.257350238 | 0.36198238 |
| | 2.990789696 | 0.002782571 | 0.00724667 |
| | 0.192810118 | 0.847107679 | NA |
| | -2.124635822 | 0.033617012 | 0.065580608 |
| NA | NA | NA | |
| NA | NA | NA | |
| | -3.688005748 | 0.000226019 | 0.000735184 |
| | 0.924450665 | 0.355251727 | 0.467176162 |
| | 0.382329229 | 0.702217177 | NA |
| NA | NA | NA | |
| | 2.466130533 | 0.013658155 | 0.030019637 |
| | 2.025914858 | 0.04277351 | 0.080813781 |
| | 0.85698741 | 0.391451842 | NA |
| | -4.659055079 | 3.18E-06 | 1.39E-05 |
| | 0.97185934 | 0.33112053 | 0.442023833 |
| | 2.037173104 | 0.041632698 | 0.07895639 |
| | -0.065389966 | 0.947863513 | NA |
| | 5.843313729 | 5.12E-09 | 3.23E-08 |
| | 0.06353712 | 0.949338802 | NA |
| | 1.655935384 | 0.097734905 | 0.163490165 |
| | 0.798630193 | 0.424504875 | 0.536038595 |
| | 4.579735894 | 4.66E-06 | 1.99E-05 |
| | -0.313953449 | 0.753556395 | 0.821208488 |
| | 0.249979153 | 0.80260347 | 0.860091829 |
| | -1.17765161 | 0.238935526 | 0.341032989 |
| | -6.33267148 | 2.41E-10 | 1.75E-09 |
| NA | NA | NA | |
| NA | NA | NA | |

| | | | |
|---|---|---|---|
| | 0.316754367 | 0.751429992 | NA |
| | -2.78718734 | 0.005316773 | 0.012992174 |
| | -6.166474531 | 6.98E-10 | 4.81E-09 |
| | -0.088545518 | 0.929443108 | NA |
| | -5.457476339 | 4.83E-08 | 2.72E-07 |
| | -1.001219766 | 0.316720572 | 0.427267216 |
| | -1.194252236 | 0.232379315 | 0.333321161 |
| NA | NA | NA | |
| | -2.248531498 | 0.024542319 | 0.049889892 |
| | -0.135115751 | 0.892520357 | NA |
| NA | NA | NA | |
| | -2.735221107 | 0.006233839 | 0.014962887 |
| NA | NA | NA | |
| | 0.330823454 | 0.740777844 | NA |
| | -3.685466735 | 0.000228284 | 0.000742071 |
| | -1.037962026 | 0.299287735 | 0.408221572 |
| NA | NA | NA | |
| NA | NA | NA | |
| NA | NA | NA | |
| NA | NA | NA | |
| NA | NA | NA | |
| NA | NA | NA | |
| NA | NA | NA | |
| NA | NA | NA | |
| NA | NA | NA | |
| NA | NA | NA | |
| NA | NA | NA | |
| | 1.110935007 | 0.266596326 | 0.372392769 |
| | -0.847035245 | 0.396975477 | 0.508823032 |
| | -0.817623954 | 0.413571946 | 0.525174055 |
| NA | NA | NA | |
| | -0.085782536 | 0.93163929 | NA |
| | 3.592500347 | 0.00032752 | 0.001034166 |
| | 0.299446355 | 0.764599498 | 0.830206725 |
| | -1.277769934 | 0.201330559 | 0.297409341 |
| | 3.178196331 | 0.001481944 | 0.004110033 |
| | 6.573380552 | 4.92E-11 | 3.87E-10 |
| | -2.006677117 | 0.044784054 | 0.084155021 |
| | 7.859122512 | 3.87E-15 | 4.57E-14 |
| | 0.344242911 | 0.730663614 | 0.803396163 |
| | -0.171177426 | 0.864084257 | 0.905433929 |
| | 13.95475453 | 2.94E-44 | 1.93E-42 |
| | 1.868201161 | 0.061734037 | NA |
| | 0.43565198 | 0.663089251 | NA |
| | -5.81889323 | 5.92E-09 | 3.71E-08 |
| | 0.172804329 | 0.862805234 | 0.904507246 |
| | 0.192808924 | 0.847108615 | NA |

| | | | |
|---|---|---|---|
| | 0.949596438 | 0.342317349 | 0.453652031 |
| | -0.07397841 | 0.941027564 | 0.959886935 |
| | -7.780973812 | 7.20E-15 | 8.32E-14 |
| NA | NA | NA | |
| | -0.003315953 | 0.997354257 | 0.998111913 |
| | -0.987614481 | 0.323341492 | 0.434256454 |
| | -0.215233129 | 0.829585563 | 0.880823222 |
| | -0.502634077 | 0.615221566 | 0.710515194 |
| | -4.681903955 | 2.84E-06 | 1.25E-05 |
| | -0.257324468 | 0.796928306 | NA |
| | -1.389565379 | 0.164660895 | 0.251922174 |
| | -2.462315776 | 0.013804307 | 0.030292986 |
| | -0.095290854 | 0.924083807 | 0.947850385 |
| | 5.43718062 | 5.41E-08 | 3.02E-07 |
| | 1.589100451 | 0.112037716 | 0.18292916 |
| | 8.615933937 | 6.94E-18 | 1.03E-16 |
| | -2.549226973 | 0.010796201 | 0.024415333 |
| | -4.074375393 | 4.61E-05 | 0.00016886 |
| | -4.914422674 | 8.90E-07 | 4.21E-06 |
| | -1.554805379 | 0.119992428 | 0.193968643 |
| | -4.299281599 | 1.71E-05 | 6.73E-05 |
| | -1.247633107 | 0.212165448 | 0.310047943 |
| | -0.106131009 | 0.915478408 | NA |
| | -1.951155547 | 0.051038545 | 0.094231378 |
| | -1.039393119 | 0.298621943 | 0.407467555 |
| | 4.040960716 | 5.32E-05 | 0.000193384 |
| NA | NA | NA | |
| | -1.741329094 | 0.081625909 | 0.140505915 |
| | 0.129456818 | 0.896996192 | NA |
| | -0.551288949 | 0.581435613 | 0.681942429 |
| | -1.114805689 | 0.264933698 | 0.370545958 |
| | -0.041033067 | 0.967269535 | 0.977647039 |
| | -0.229988164 | 0.818100967 | 0.872181072 |
| | -3.040869291 | 0.002358962 | 0.006242497 |
| | 7.739602039 | 9.97E-15 | 1.14E-13 |
| | 0.514345524 | 0.607010439 | NA |
| | -0.047727531 | 0.961933393 | 0.973823966 |
| NA | NA | NA | |
| | 0.541356021 | 0.588262214 | NA |
| NA | NA | NA | |
| | -0.194488217 | 0.845793621 | NA |
| | -0.968195257 | 0.332946863 | 0.444130249 |
| | -1.694446967 | 0.090180381 | 0.152876054 |
| | 5.629897756 | 1.80E-08 | 1.07E-07 |
| | 0.25204008 | 0.801010085 | 0.858751977 |
| | 0.33176097 | 0.740069758 | 0.810817913 |
| NA | NA | NA | |

| | | |
|---|---|---|
| NA | NA | NA |
| -0.069958086 | 0.944227021 | 0.962218155 |
| NA | NA | NA |
| NA | NA | NA |
| NA | NA | NA |
| NA | NA | NA |
| 0.063708285 | 0.949202508 | NA |
| NA | NA | NA |
| NA | NA | NA |
| 0.573193348 | 0.566513788 | NA |
| NA | NA | NA |
| -2.000647883 | 0.045430349 | 0.085215951 |
| -0.712470033 | 0.476173762 | 0.585078261 |
| 0.495984725 | 0.619905188 | NA |
| NA | NA | NA |
| NA | NA | NA |
| 4.284933307 | 1.83E-05 | 7.15E-05 |
| -1.617431439 | 0.105785184 | 0.174427085 |
| 0.09122233 | 0.927315932 | NA |
| 0.201524559 | 0.840288428 | NA |
| -0.192857988 | 0.847070188 | 0.893174101 |
| -1.125555754 | 0.260353605 | 0.36530359 |
| -2.649801475 | 0.008053908 | 0.01873855 |
| -0.3518252 | 0.724969359 | 0.79917204 |
| -5.484079723 | 4.16E-08 | 2.36E-07 |
| -2.732457015 | 0.006286387 | 0.015068787 |
| NA | NA | NA |
| 3.578163067 | 0.000346018 | 0.001088179 |
| -5.074498996 | 3.89E-07 | 1.94E-06 |
| 0.192809521 | 0.847108147 | NA |
| 1.137546238 | 0.255310007 | NA |
| -0.109710303 | 0.912639129 | NA |
| NA | NA | NA |
| -0.666682924 | 0.504974688 | NA |
| 0.167180473 | 0.867228048 | NA |
| 2.703045316 | 0.006870738 | 0.016314825 |
| NA | NA | NA |
| 0.080644518 | 0.935724661 | 0.956279123 |
| 0.640701726 | 0.521716492 | 0.627001671 |
| 0.063708285 | 0.949202508 | NA |
| 0.063708285 | 0.949202508 | NA |
| -7.298254004 | 2.92E-13 | 2.86E-12 |
| 1.363318926 | 0.17278202 | 0.262184538 |
| -2.924232685 | 0.003453065 | 0.008795179 |
| -1.408486336 | 0.15898711 | 0.244738461 |
| NA | NA | NA |
| -1.681648592 | 0.092637002 | 0.156394453 |

| | | |
|---|---|---|
| -2.141736005 | 0.032214732 | 0.063207238 |
| NA | NA | NA |
| NA | NA | NA |
| NA | NA | NA |
| NA | NA | NA |
| NA | NA | NA |
| NA | NA | NA |
| 0.064259733 | 0.948763415 | 0.965010012 |
| NA | NA | NA |
| 0.063708882 | 0.949202033 | NA |
| -0.052328221 | 0.958267167 | 0.971148562 |
| NA | NA | NA |
| NA | NA | NA |
| NA | NA | NA |
| NA | NA | NA |
| NA | NA | NA |
| NA | NA | NA |
| NA | NA | NA |
| NA | NA | NA |
| 4.320636094 | 1.56E-05 | 6.15E-05 |
| 0.418577516 | 0.675524924 | NA |
| NA | NA | NA |
| -0.065389369 | 0.947863988 | NA |
| NA | NA | NA |
| NA | NA | NA |
| NA | NA | NA |
| 0.541356021 | 0.588262214 | NA |
| 0.98930624 | 0.322513332 | 0.433471614 |
| -0.524044634 | 0.600247496 | 0.698053005 |
| NA | NA | NA |
| -1.975376275 | 0.048225463 | 0.08969929 |
| NA | NA | NA |
| 0.063707194 | 0.949203376 | NA |
| 0.192809521 | 0.847108147 | NA |
| -3.402825913 | 0.000666928 | 0.001986782 |
| NA | NA | NA |
| -0.265134095 | 0.790906163 | 0.851164385 |
| -0.517658029 | 0.604696887 | 0.701742573 |
| 4.203182585 | 2.63E-05 | 0.000100513 |
| 1.83865815 | 0.065965483 | 0.117158591 |
| -0.648868019 | 0.516423688 | 0.621681731 |
| 1.093122622 | 0.274339965 | 0.381040415 |
| NA | NA | NA |
| NA | NA | NA |
| -1.52062835 | 0.128353128 | 0.205387981 |
| -1.908367797 | 0.056343696 | 0.102558557 |
| 0.875493761 | 0.381305303 | 0.493871759 |

| | | | |
|---|---|---|---|
| | 9.545555386 | 1.35E-21 | 2.73E-20 |
| | 0.263726495 | 0.791990678 | NA |
| NA | NA | NA | |
| | 5.20441337 | 1.95E-07 | 1.01E-06 |
| | 7.249458446 | 4.18E-13 | 4.04E-12 |
| | 9.261381736 | 2.02E-20 | 3.68E-19 |
| | 0.320071032 | 0.748914485 | NA |
| | -0.235186148 | 0.814064257 | 0.869247402 |
| | -2.320828412 | 0.020296106 | 0.042337824 |
| | -4.09428714 | 4.23E-05 | 0.000155828 |
| | -3.584261916 | 0.000338033 | 0.001064941 |
| | 0.669669065 | 0.503068776 | NA |
| | 2.218984686 | 0.026487766 | 0.05328207 |
| | 1.912302953 | 0.055837347 | 0.101770101 |
| | 1.095974989 | 0.27308972 | 0.379788626 |
| | 0.49917203 | 0.617658197 | NA |
| | -0.920902911 | 0.357101119 | 0.468928231 |
| | -0.479341889 | 0.631695427 | 0.723570672 |
| | 3.180010692 | 0.001472696 | 0.004086197 |
| | 3.529379426 | 0.000416535 | 0.001288419 |
| NA | NA | NA | |
| | 0.401171491 | 0.688293869 | NA |
| NA | NA | NA | |
| | -0.065388772 | 0.947864464 | NA |
| | -0.065389966 | 0.947863513 | NA |
| | 0.063708285 | 0.949202508 | NA |
| NA | NA | NA | |
| | -0.065393756 | 0.947860495 | NA |
| | 0.063708882 | 0.949202033 | NA |
| NA | NA | NA | |
| | 0.49606901 | 0.619845723 | NA |
| | 0.31685483 | 0.751353758 | NA |
| | -0.415932138 | 0.677459662 | NA |
| NA | NA | NA | |
| | 1.012718233 | 0.311194766 | NA |
| NA | NA | NA | |
| | 5.699154366 | 1.20E-08 | 7.29E-08 |
| | 2.222605528 | 0.026242411 | 0.052867647 |
| | 3.784725909 | 0.000153878 | 0.000514916 |
| NA | NA | NA | |
| NA | NA | NA | |
| | 0.063708285 | 0.949202508 | NA |
| NA | NA | NA | |
| | 0.330493631 | 0.741027004 | NA |
| NA | NA | NA | |
| NA | NA | NA | |
| | -0.065389369 | 0.947863988 | NA |

| | | | |
|---|---|---|---|
| NA | NA | NA | |
| NA | NA | NA | |
| NA | NA | NA | |
| NA | NA | NA | |
| | -0.065388772 | 0.947864464 | NA |
| NA | NA | NA | |
| | -0.44278398 | 0.657921992 | NA |
| | -4.194188072 | 2.74E-05 | 0.00010433 |
| NA | NA | NA | |
| NA | NA | NA | |
| NA | NA | NA | |
| | 2.03392405 | 0.041959254 | 0.079485364 |
| | -1.515316014 | 0.129692395 | 0.2070149 |
| | -2.503865545 | 0.012284471 | 0.027331583 |
| | 1.004651186 | 0.315064841 | 0.425630018 |
| | -0.065388772 | 0.947864464 | NA |
| | -1.662670778 | 0.096378355 | 0.161566554 |
| NA | NA | NA | |
| | -3.060741405 | 0.002207897 | 0.005888509 |
| | 0.083043937 | 0.933816603 | NA |
| NA | NA | NA | |
| NA | NA | NA | |
| NA | NA | NA | |
| | -0.740700273 | 0.458875194 | 0.568371244 |
| NA | NA | NA | |
| NA | NA | NA | |
| NA | NA | NA | |
| NA | NA | NA | |
| NA | NA | NA | |
| NA | NA | NA | |
| NA | NA | NA | |
| | 1.219874848 | 0.222512322 | 0.321977754 |
| NA | NA | NA | |
| | -3.482446938 | 0.000496854 | 0.001516801 |
| | -9.007969333 | 2.10E-19 | 3.57E-18 |
| NA | NA | NA | |
| | -0.771655919 | 0.440318246 | NA |
| | 0.506380903 | 0.612589278 | 0.708259487 |
| NA | NA | NA | |
| NA | NA | NA | |
| NA | NA | NA | |
| NA | NA | NA | |
| NA | NA | NA | |
| NA | NA | NA | |
| NA | NA | NA | |
| NA | NA | NA | |
| NA | NA | NA | |

| | | |
|---|---|---|
| NA | NA | NA |
| NA | NA | NA |
| 0.549531246 | 0.582640928 | NA |
| NA | NA | NA |
| NA | NA | NA |
| NA | NA | NA |
| NA | NA | NA |
| NA | NA | NA |
| -0.065389369 | 0.947863988 | NA |
| NA | NA | NA |
| 0.192809521 | 0.847108147 | NA |
| NA | NA | NA |
| -0.62693057 | 0.530704748 | 0.635454451 |
| 4.709025032 | 2.49E-06 | 1.10E-05 |
| 0.216157829 | 0.828864724 | NA |
| -6.10362118 | 1.04E-09 | 7.06E-09 |
| NA | NA | NA |
| -0.195029726 | 0.845369677 | 0.891903027 |
| NA | NA | NA |
| 2.790518068 | 0.005262376 | 0.012874331 |
| NA | NA | NA |
| NA | NA | NA |
| NA | NA | NA |
| NA | NA | NA |
| 0.358973738 | 0.719614735 | 0.794984192 |
| NA | NA | NA |
| -1.737223949 | 0.082347649 | 0.141553579 |
| NA | NA | NA |
| NA | NA | NA |
| NA | NA | NA |
| -1.393044261 | 0.163606407 | 0.250539514 |
| 0.401135131 | 0.688320637 | 0.770144563 |
| 11.00800113 | 3.50E-28 | 1.10E-26 |
| NA | NA | NA |
| -0.210973165 | 0.832908212 | 0.882927856 |
| 0.953885974 | 0.340141364 | 0.451657206 |
| NA | NA | NA |
| 0.997318343 | 0.318610013 | 0.429240998 |
| -0.34093621 | 0.733151604 | NA |
| -0.116065114 | 0.907600937 | 0.936099038 |
| -1.99836148 | 0.045677485 | 0.085608943 |
| -4.295990838 | 1.74E-05 | 6.83E-05 |
| -0.646497727 | 0.517957069 | 0.623261643 |
| -3.316674862 | 0.000910956 | 0.002644539 |
| NA | NA | NA |
| NA | NA | NA |
| 0.063708285 | 0.949202508 | NA |

| | | |
|---|---|---|
| 0.089388503 | 0.92877316 | NA |
| NA | NA | NA |
| -2.224026238 | 0.026146679 | 0.052714052 |
| -0.45820353 | 0.646806224 | 0.735889081 |
| NA | NA | NA |
| -2.093436265 | 0.036310228 | 0.070094478 |
| 5.578622899 | 2.42E-08 | 1.42E-07 |
| 2.888049717 | 0.003876386 | 0.009775913 |
| -1.347806099 | 0.177720756 | 0.268344725 |
| 1.743368402 | 0.081269284 | 0.140004274 |
| 0.486620152 | 0.62652754 | 0.719874993 |
| 0.063708285 | 0.949202508 | NA |
| 0.485676506 | 0.627196543 | NA |
| 0.832717931 | 0.40500383 | 0.516784556 |
| 0.149865936 | 0.880870387 | NA |
| NA | NA | NA |
| 0.192808924 | 0.847108615 | NA |
| 0.063708285 | 0.949202508 | NA |
| NA | NA | NA |
| 0.652778213 | 0.513899271 | NA |
| 0.063708882 | 0.949202033 | NA |
| NA | NA | NA |
| -4.290357683 | 1.78E-05 | 6.99E-05 |
| NA | NA | NA |
| -1.004369084 | 0.315200746 | 0.425696519 |
| 0.485754458 | 0.627141268 | 0.720259977 |
| -1.173236547 | 0.24070096 | 0.343055747 |
| 0.324874675 | 0.745275925 | NA |
| -1.175308444 | 0.239871337 | 0.342048629 |
| NA | NA | NA |
| 0.192810118 | 0.847107679 | NA |
| -1.682557758 | 0.092460735 | 0.156129006 |
| NA | NA | NA |
| 8.17794873 | 2.89E-16 | 3.77E-15 |
| NA | NA | NA |
| NA | NA | NA |
| NA | NA | NA |
| NA | NA | NA |
| NA | NA | NA |
| NA | NA | NA |
| NA | NA | NA |
| NA | NA | NA |
| NA | NA | NA |
| NA | NA | NA |
| -0.085070848 | 0.932205066 | NA |
| 0.31685483 | 0.751353758 | NA |
| -0.065390563 | 0.947863038 | NA |

| | | |
|---|---|---|
| 0.192808327 | 0.847109082 | NA |
| NA | NA | NA |
| -8.914594723 | 4.90E-19 | 8.11E-18 |
| -3.354032142 | 0.000796431 | 0.002338104 |
| -3.492685574 | 0.000478189 | 0.001463212 |
| -3.265959123 | 0.00109094 | 0.003117861 |
| -4.725512618 | 2.30E-06 | 1.03E-05 |
| -3.341354926 | 0.000833706 | 0.002438665 |
| -2.734821287 | 0.006241415 | 0.014979636 |
| -4.565757511 | 4.98E-06 | 2.12E-05 |
| NA | NA | NA |
| -4.161382967 | 3.16E-05 | 0.000119044 |
| -4.812037423 | 1.49E-06 | 6.84E-06 |
| -3.041264772 | 0.002355866 | 0.006236628 |
| NA | NA | NA |
| -2.546412745 | 0.010883642 | 0.024577549 |
| -2.672529089 | 0.007528185 | 0.017668117 |
| -3.23929039 | 0.001198275 | 0.003394005 |
| 0.192810118 | 0.847107679 | NA |
| 7.42375274 | 1.14E-13 | 1.16E-12 |
| NA | NA | NA |
| 6.544333669 | 5.98E-11 | 4.65E-10 |
| NA | NA | NA |
| 1.544810216 | 0.122392176 | 0.197236425 |
| -12.64319998 | 1.22E-36 | 5.75E-35 |
| 1.575396919 | 0.115164855 | 0.187289736 |
| -0.341149938 | 0.732990708 | 0.80538127 |
| 0.083606531 | 0.933369274 | 0.954768603 |
| -0.178297977 | 0.858488968 | 0.901804189 |
| NA | NA | NA |
| NA | NA | NA |
| -3.716316533 | 0.000202148 | 0.000664186 |
| -1.348377699 | 0.177536933 | 0.268115689 |
| -2.122756989 | 0.033774221 | 0.065864335 |
| 2.089817126 | 0.03663423 | 0.070638224 |
| 0.192808924 | 0.847108615 | NA |
| NA | NA | NA |
| -1.099734575 | 0.271447786 | 0.377820056 |
| -2.114551621 | 0.034468181 | 0.067024438 |
| -1.298423208 | 0.194141949 | 0.288492815 |
| 1.273618284 | 0.202798729 | 0.299081181 |
| 0.085801041 | 0.931624579 | 0.953451495 |
| -2.041617694 | 0.041189469 | 0.078163184 |
| -1.205308312 | 0.228084302 | 0.328459242 |
| -1.015556051 | 0.309840842 | 0.419841722 |
| -1.03750473 | 0.299500693 | 0.408417452 |
| -5.757666297 | 8.53E-09 | 5.24E-08 |

| | | | |
|---|---|---|---|
| | -0.852161634 | 0.394124387 | 0.506127619 |
| | 4.425071938 | 9.64E-06 | 3.94E-05 |
| | -1.862254517 | 0.062567223 | 0.11197323 |
| | -1.072913017 | 0.283310145 | 0.390272588 |
| | -6.404461726 | 1.51E-10 | 1.12E-09 |
| NA | NA | NA | |
| | -3.947349951 | 7.90E-05 | 0.000278616 |
| | -3.839198322 | 0.000123437 | 0.000419957 |
| | -5.572307142 | 2.51E-08 | 1.46E-07 |
| | -4.975003452 | 6.52E-07 | 3.14E-06 |
| | 5.358678632 | 8.38E-08 | 4.57E-07 |
| | -4.703894236 | 2.55E-06 | 1.13E-05 |
| | 10.33880905 | 4.70E-25 | 1.20E-23 |
| | 4.190092766 | 2.79E-05 | 0.000106053 |
| NA | NA | NA | |
| NA | NA | NA | |
| NA | NA | NA | |
| NA | NA | NA | |
| | 2.57347355 | 0.010068334 | 0.022928797 |
| | 12.3532391 | 4.68E-35 | 2.04E-33 |
| | 0.063708882 | 0.949202033 | NA |
| | 0.422806406 | 0.672436513 | NA |
| NA | NA | NA | |
| | -0.038305156 | 0.96944438 | 0.97875092 |
| | -2.477857317 | 0.013217399 | 0.029160996 |
| | -0.601043389 | 0.547811088 | 0.6512949 |
| | -0.7404859 | 0.459005214 | 0.568419775 |
| | 1.399464083 | 0.161673862 | 0.248004949 |
| | 0.851061984 | 0.394734922 | 0.50675566 |
| | -1.339006879 | 0.180568433 | 0.271857317 |
| | -1.387799478 | 0.165198114 | 0.252621505 |
| | -0.481888585 | 0.629885096 | 0.722343059 |
| | -2.326134112 | 0.020011397 | 0.041803144 |
| NA | NA | NA | |
| | 0.387089649 | 0.69868984 | 0.778281844 |
| | -0.065388772 | 0.947864464 | NA |
| | -2.307810504 | 0.021009678 | 0.043586594 |
| | -0.424055658 | 0.671525223 | 0.755749361 |
| | -2.899975519 | 0.003731918 | 0.009433439 |
| | -2.743899 | 0.006071421 | 0.014609467 |
| | -3.675480512 | 0.000237402 | 0.000768816 |
| | -0.449610158 | 0.652991562 | 0.741073625 |
| | 0.063708285 | 0.949202508 | NA |
| | 2.841960465 | 0.004483705 | 0.011148022 |
| | -0.568966805 | 0.569378668 | 0.670882081 |
| | -1.074480138 | 0.282607542 | 0.389592619 |
| | -4.3710441 | 1.24E-05 | 4.96E-05 |

| | | |
|---|---|---|
| -0.438225132 | 0.661223092 | 0.747872299 |
| 0.192809521 | 0.847108147 | NA |
| 0.83118638 | 0.405868348 | 0.517702937 |
| 0.192809521 | 0.847108147 | NA |
| 0.086687957 | 0.93091955 | NA |
| 2.415505485 | 0.015713386 | 0.033956315 |
| -0.065389369 | 0.947863988 | NA |
| 2.603092949 | 0.00923869 | 0.021220925 |
| 1.568699895 | 0.116717883 | 0.189409272 |
| -0.065390563 | 0.947863038 | NA |
| 0.320071032 | 0.748914485 | NA |
| 1.880416021 | 0.060051402 | 0.108151287 |
| 8.395904 | 4.62E-17 | 6.47E-16 |
| 8.644794359 | 5.39E-18 | 8.10E-17 |
| -0.311473824 | 0.755440439 | 0.82272437 |
| 0.642027383 | 0.520855404 | 0.626087055 |
| 0.192810118 | 0.847107679 | NA |
| 5.314597628 | 1.07E-07 | 5.74E-07 |
| -2.34764836 | 0.018892348 | 0.039760823 |
| 5.342904767 | 9.15E-08 | 4.96E-07 |
| 0.085936513 | 0.931516886 | NA |
| 14.02573283 | 1.08E-44 | 7.18E-43 |
| -1.320458124 | 0.186682109 | 0.279379464 |
| -0.378704733 | 0.704907139 | 0.783508401 |
| -3.378555419 | 0.000728677 | 0.002156906 |
| -1.935283563 | 0.052955508 | 0.097246983 |
| -2.867333064 | 0.004139471 | 0.010371468 |
| -1.820243135 | 0.068721987 | 0.121348242 |
| 0.603310697 | 0.546302018 | 0.649871181 |
| -1.507027698 | 0.131803558 | 0.209662729 |
| 3.859968996 | 0.000113401 | 0.000388295 |
| 0.796852491 | 0.425536706 | 0.537043501 |
| 2.230734404 | 0.025698727 | 0.051915992 |
| 11.65697784 | 2.11E-31 | 7.89E-30 |
| 6.68360029 | 2.33E-11 | 1.89E-10 |
| 0.442430623 | 0.658177623 | 0.745757449 |
| -0.948933939 | 0.342654212 | 0.453978235 |
| 4.466776733 | 7.94E-06 | 3.29E-05 |
| 9.410512926 | 4.94E-21 | 9.48E-20 |
| NA | NA | NA |
| -0.763213339 | 0.445336175 | 0.555179918 |
| -2.127534747 | 0.033375677 | 0.065198877 |
| 0.063708882 | 0.949202033 | NA |
| NA | NA | NA |
| -1.109298718 | 0.267301338 | 0.373127683 |
| 6.864499589 | 6.67E-12 | 5.67E-11 |
| -7.313293176 | 2.61E-13 | 2.57E-12 |

| | | |
|---|---|---|
| 1.5279027 | 0.126536698 | 0.202922447 |
| 0.192808924 | 0.847108615 | NA |
| 1.608124489 | 0.107807917 | 0.177155473 |
| -0.698735294 | 0.484717473 | 0.593130205 |
| 0.91439462 | 0.360509528 | 0.472640128 |
| -3.849286299 | 0.000118462 | 0.000404463 |
| -0.374758994 | 0.707839713 | NA |
| NA | NA | NA |
| 0.063708285 | 0.949202508 | NA |
| NA | NA | NA |
| -0.089395238 | 0.928767808 | NA |
| 2.851719516 | 0.004348345 | 0.010838387 |
| 13.57697068 | 5.48E-42 | 3.21E-40 |
| 4.875237418 | 1.09E-06 | 5.07E-06 |
| -2.186608194 | 0.028771139 | 0.057258615 |
| -0.065388772 | 0.947864464 | NA |
| 0.554809719 | 0.579024818 | 0.679765212 |
| -2.230095819 | 0.025741082 | 0.051972194 |
| 1.499332281 | 0.133787452 | 0.212305834 |
| -0.892815352 | 0.371956064 | 0.484747489 |
| 0.248208232 | 0.803973298 | 0.861128008 |
| -1.261709167 | 0.207053457 | 0.303958567 |
| 0.08624007 | 0.931275578 | NA |
| 0.063707194 | 0.949203376 | NA |
| -6.529596376 | 6.59E-11 | 5.11E-10 |
| -0.741553777 | 0.458357738 | 0.56789893 |
| -0.728369429 | 0.466387472 | 0.575681942 |
| -1.737420904 | 0.082312904 | 0.141503571 |
| 1.682778243 | 0.092418028 | 0.156077407 |
| -7.142225688 | 9.18E-13 | 8.56E-12 |
| 2.177119267 | 0.02947167 | 0.058476092 |
| 2.183639954 | 0.028988715 | 0.057622899 |
| 8.360144657 | 6.26E-17 | 8.66E-16 |
| -5.653319462 | 1.57E-08 | 9.40E-08 |
| 0.697407749 | 0.485547654 | 0.593843201 |
| -0.20193895 | 0.839964451 | 0.887775957 |
| 0.192809521 | 0.847108147 | NA |
| 0.132821301 | 0.894334708 | NA |
| -0.955763955 | 0.339191503 | NA |
| -5.505760134 | 3.68E-08 | 2.10E-07 |
| -0.814847062 | 0.415159859 | 0.526816287 |
| NA | NA | NA |
| NA | NA | NA |
| 3.863420885 | 0.00011181 | 0.000383371 |
| -0.643332263 | 0.520008526 | 0.625279276 |
| -1.309753431 | 0.190279262 | 0.283865176 |
| -1.700514026 | 0.08903428 | 0.151250715 |

| | | | |
|---|---|---|---|
| NA | NA | NA | |
| | 10.03212541 | 1.10E-23 | 2.59E-22 |
| | -7.999210664 | 1.25E-15 | 1.55E-14 |
| | 0.407334021 | 0.683762679 | NA |
| | -1.155667091 | 0.247817354 | 0.351039523 |
| | 1.311033132 | 0.18984657 | 0.283304099 |
| | 0.192810118 | 0.847107679 | NA |
| | 1.096105855 | 0.273032453 | 0.379730082 |
| | -0.285369886 | 0.775360776 | NA |
| | -0.890052755 | 0.37343756 | 0.486241384 |
| | -0.398874231 | 0.689985878 | NA |
| | -0.065389966 | 0.947863513 | NA |
| | -5.929902493 | 3.03E-09 | 1.96E-08 |
| | 2.131462699 | 0.033051042 | 0.064647027 |
| | 0.722859837 | 0.469766001 | 0.578767621 |
| | 0.06371127 | 0.949200131 | NA |
| | -0.329990486 | 0.741407151 | 0.81178973 |
| | 5.330939528 | 9.77E-08 | 5.27E-07 |
| | 3.494606111 | 0.000474762 | 0.001453613 |
| | -1.636802781 | 0.101671685 | 0.168845692 |
| | 0.063708285 | 0.949202508 | NA |
| | 0.54024686 | 0.589026801 | NA |
| | -8.237681984 | 1.76E-16 | 2.34E-15 |
| | 1.683494951 | 0.092279316 | 0.155898675 |
| NA | NA | NA | |
| | 9.409464203 | 4.99E-21 | 9.56E-20 |
| | 6.594303948 | 4.27E-11 | 3.38E-10 |
| | 2.136122124 | 0.032669461 | 0.063990897 |
| | -0.769762939 | 0.441440528 | 0.551470174 |
| | -4.003358548 | 6.24E-05 | 0.000223621 |
| | 0.719114092 | 0.472070625 | 0.580977862 |
| | -0.244816433 | 0.806598569 | 0.862985349 |
| | 0.773473111 | 0.439242436 | 0.549492112 |
| | -1.66105206 | 0.096702991 | NA |
| | -0.065389369 | 0.947863988 | NA |
| | -2.847140365 | 0.004411391 | 0.010985684 |
| | -1.519396067 | 0.128662831 | 0.205725736 |
| | -2.596558194 | 0.009416295 | 0.021589253 |
| | -7.375510536 | 1.64E-13 | 1.64E-12 |
| | -3.374745003 | 0.000738841 | 0.002184408 |
| | -0.065389369 | 0.947863988 | NA |
| | -4.509973207 | 6.48E-06 | 2.71E-05 |
| | 0.400746861 | 0.688606506 | 0.770247166 |
| | 2.122388066 | 0.033805164 | 0.065919542 |
| | 0.069489539 | 0.944599959 | 0.9623678 |
| | -4.64086736 | 3.47E-06 | 1.51E-05 |
| | 0.198308907 | 0.842803383 | NA |

| | | | |
|---|---|---|---|
| | 7.734458715 | 1.04E-14 | 1.18E-13 |
| | 3.438612583 | 0.000584703 | 0.001760722 |
| | -0.065388772 | 0.947864464 | NA |
| | -0.502749728 | 0.615140243 | 0.710487686 |
| | 2.879608853 | 0.003981688 | 0.01001524 |
| | -0.582841589 | 0.559999945 | 0.662435021 |
| | 0.241486509 | 0.809178072 | NA |
| | 2.050974419 | 0.040269438 | 0.076665357 |
| | 9.266552973 | 1.92E-20 | 3.52E-19 |
| | -3.159431331 | 0.001580773 | 0.004360454 |
| | -0.627110934 | 0.53058652 | 0.635404077 |
| | -0.065389369 | 0.947863988 | NA |
| | -1.504187454 | 0.132533107 | 0.210622455 |
| | -5.569906959 | 2.55E-08 | 1.48E-07 |
| | -2.167187188 | 0.030220584 | 0.059725323 |
| | -7.313665849 | 2.60E-13 | 2.56E-12 |
| NA | NA | NA | |
| | -3.266980064 | 0.001087013 | 0.003108058 |
| | -3.266654214 | 0.001088265 | 0.003111282 |
| | -2.70939244 | 0.006740656 | 0.016042466 |
| | 6.419611552 | 1.37E-10 | 1.02E-09 |
| | 6.731606639 | 1.68E-11 | 1.38E-10 |
| | 6.467962413 | 9.93E-11 | 7.56E-10 |
| | 3.904670394 | 9.44E-05 | 0.000328102 |
| | -2.27732178 | 0.022767017 | 0.046809466 |
| | -3.336566347 | 0.000848202 | 0.002478779 |
| | 0.068907162 | 0.945063518 | 0.962561133 |
| | -0.377336731 | 0.705923379 | 0.784254868 |
| | 0.288217989 | 0.77317988 | NA |
| | -6.984859538 | 2.85E-12 | 2.51E-11 |
| | 5.128857997 | 2.92E-07 | 1.48E-06 |
| | 0.447103929 | 0.654800022 | 0.742721918 |
| | 1.639973505 | 0.101010676 | 0.167926223 |
| NA | NA | NA | |
| | 0.274065198 | 0.784034518 | 0.845915082 |
| | -1.094428896 | 0.273766917 | 0.380434484 |
| | -5.355408942 | 8.54E-08 | 4.64E-07 |
| | 3.182886444 | 0.001458148 | 0.00405212 |
| | 7.43724143 | 1.03E-13 | 1.06E-12 |
| | 0.192810118 | 0.847107679 | NA |
| NA | NA | NA | |
| | 0.174341926 | 0.861596751 | NA |
| | -0.209937071 | 0.833716788 | 0.883485657 |
| | -2.645218783 | 0.008163811 | 0.018969569 |
| | -3.47908922 | 0.000503121 | 0.001534436 |
| | 1.052192513 | 0.292711234 | 0.400621292 |
| | 2.732341676 | 0.006288589 | 0.015071593 |

| | | |
|---|---|---|
| 4.549626994 | 5.37E-06 | 2.28E-05 |
| 0.192810118 | 0.847107679 | NA |
| -0.492384597 | 0.622447486 | NA |
| 0.317642674 | 0.750756 | NA |
| 2.580893827 | 0.009854488 | 0.022486812 |
| 0.879227913 | 0.379277713 | NA |
| 2.230222441 | 0.025732679 | 0.051959419 |
| 6.87196298 | 6.33E-12 | 5.39E-11 |
| 4.254826118 | 2.09E-05 | 8.11E-05 |
| 0.772335153 | 0.43991595 | 0.550121339 |
| 0.192809521 | 0.847108147 | NA |
| NA | NA | NA |
| NA | NA | NA |
| -0.327055164 | 0.743626162 | 0.813432516 |
| NA | NA | NA |
| 0.987702503 | 0.323298368 | 0.434221837 |
| NA | NA | NA |
| 0.192809521 | 0.847108147 | NA |
| -0.417149038 | 0.676569398 | NA |
| NA | NA | NA |
| NA | NA | NA |
| -1.553531398 | 0.120296234 | NA |
| NA | NA | NA |
| 0.114028449 | 0.909215242 | NA |
| NA | NA | NA |
| -3.218260125 | 0.001289708 | 0.003623458 |
| -4.984937378 | 6.20E-07 | 3.00E-06 |
| NA | NA | NA |
| -4.335176953 | 1.46E-05 | 5.78E-05 |
| -3.465742164 | 0.00052877 | 0.001605814 |
| NA | NA | NA |
| -3.473300323 | 0.0005141 | 0.001564679 |
| -3.329007855 | 0.000871559 | 0.002541668 |
| -3.363592216 | 0.000769351 | 0.002265247 |
| -0.108033632 | 0.913969014 | NA |
| -2.279893784 | 0.022613989 | 0.046529235 |
| NA | NA | NA |
| -1.947368003 | 0.051490631 | 0.094946994 |
| NA | NA | NA |
| 25.68601692 | 1.68E-145 | 6.55E-143 |
| 13.04201549 | 7.06E-39 | 3.72E-37 |
| NA | NA | NA |
| NA | NA | NA |
| NA | NA | NA |
| 0.431335892 | 0.666224157 | NA |
| -0.278347918 | 0.780745296 | NA |
| 3.84304292 | 0.000121518 | 0.000413936 |

| | | | |
|---|---|---|---|
| | -0.095258147 | 0.924109786 | NA |
| | 0.467878124 | 0.639871748 | 0.730489845 |
| | 2.505204508 | 0.012238061 | 0.027235589 |
| | -0.065389369 | 0.947863988 | NA |
| NA | NA | NA | |
| | -1.790349206 | 0.07339779 | 0.128220735 |
| | -0.247506811 | 0.804516021 | 0.861551191 |
| NA | NA | NA | |
| | 0.063708285 | 0.949202508 | NA |
| NA | NA | NA | |
| | 0.496473495 | 0.619560383 | 0.714043484 |
| | 6.726649348 | 1.74E-11 | 1.42E-10 |
| | -0.396779915 | 0.691529763 | 0.772343491 |
| | 0.46878741 | 0.6392216 | 0.729947381 |
| | 0.063708882 | 0.949202033 | NA |
| NA | NA | NA | |
| | 0.901680788 | 0.367226461 | 0.479682618 |
| | -0.341761119 | 0.732530672 | 0.805024707 |
| | 0.06353712 | 0.949338802 | NA |
| NA | NA | NA | |
| | 0.235102644 | 0.814129066 | 0.869279521 |
| | -1.215168362 | 0.224301885 | 0.324112468 |
| | -0.19448762 | 0.845794089 | NA |
| NA | NA | NA | |
| | -2.164254501 | 0.030444823 | 0.060114262 |
| | 3.35220057 | 0.000801719 | 0.002351973 |
| | 1.378359389 | 0.168092355 | 0.256202147 |
| | 0.06371127 | 0.949200131 | NA |
| NA | NA | NA | |
| NA | NA | NA | |
| | -0.118746078 | 0.90547653 | NA |
| NA | NA | NA | |
| | -0.367633621 | 0.713146442 | NA |
| | 2.235778893 | 0.025366258 | 0.05135626 |
| | 0.953043981 | 0.340567788 | 0.452089806 |
| | 0.241335017 | 0.809295473 | 0.865299688 |
| NA | NA | NA | |
| NA | NA | NA | |
| | -0.199519519 | 0.841856376 | 0.889239142 |
| | 7.638872309 | 2.19E-14 | 2.41E-13 |
| | -7.81278891 | 5.59E-15 | 6.52E-14 |
| | -6.810118703 | 9.75E-12 | 8.17E-11 |
| NA | NA | NA | |
| | -0.186028495 | 0.852422416 | NA |
| NA | NA | NA | |
| NA | NA | NA | |
| NA | NA | NA | |

| | | |
|---|---|---|
| 0.071152621 | 0.943276289 | NA |
| NA | NA | NA |
| NA | NA | NA |
| 0.063708285 | 0.949202508 | NA |
| NA | NA | NA |
| NA | NA | NA |
| 0.77906077 | 0.435943918 | 0.546403418 |
| -1.081416717 | 0.279511789 | 0.386507932 |
| -8.943332226 | 3.78E-19 | 6.30E-18 |
| 0.577675556 | 0.563483186 | 0.665570949 |
| -7.660835071 | 1.85E-14 | 2.05E-13 |
| 0.469318727 | 0.638841832 | 0.729680161 |
| -1.897841775 | 0.057716928 | 0.104662372 |
| -3.162087185 | 0.001566427 | 0.004323738 |
| NA | NA | NA |
| 2.445810455 | 0.014452696 | 0.031505755 |
| 1.68859358 | 0.091297342 | 0.154426804 |
| 0.333547261 | 0.738721221 | NA |
| 12.07856323 | 1.37E-33 | 5.65E-32 |
| 0.577694389 | 0.563470468 | NA |
| NA | NA | NA |
| NA | NA | NA |
| 3.436910396 | 0.00058839 | 0.001770547 |
| NA | NA | NA |
| NA | NA | NA |
| NA | NA | NA |
| NA | NA | NA |
| NA | NA | NA |
| NA | NA | NA |
| NA | NA | NA |
| -0.117287908 | 0.906631907 | NA |
| -3.575031041 | 0.000350186 | 0.001100233 |
| -0.065388772 | 0.947864464 | NA |
| -3.705279012 | 0.000211158 | 0.000690707 |
| -5.808283998 | 6.31E-09 | 3.94E-08 |
| -1.170605202 | 0.241757506 | 0.344294371 |
| 0.271397367 | 0.786085427 | NA |
| 0.402281244 | 0.687477056 | NA |
| 0.192808924 | 0.847108615 | NA |
| NA | NA | NA |
| 0.199526594 | 0.841850842 | NA |
| 0.063708285 | 0.949202508 | NA |
| NA | NA | NA |
| NA | NA | NA |
| NA | NA | NA |
| NA | NA | NA |
| -0.065389369 | 0.947863988 | NA |

| | | |
|---|---|---|
| NA | NA | NA |
| NA | NA | NA |
| NA | NA | NA |
| 0.46968931 | 0.638577006 | 0.729479893 |
| NA | NA | NA |
| NA | NA | NA |
| NA | NA | NA |
| 0.063708882 | 0.949202033 | NA |
| -0.093108474 | 0.925817385 | NA |
| 0.10696831 | 0.91481412 | NA |
| -0.194488217 | 0.845793621 | NA |
| 0.063708285 | 0.949202508 | NA |
| NA | NA | NA |
| NA | NA | NA |
| 9.17104253 | 4.68E-20 | 8.38E-19 |
| NA | NA | NA |
| 0.111043683 | 0.911581706 | NA |
| -0.065390563 | 0.947863038 | NA |
| 3.045375975 | 0.002323896 | 0.006161369 |
| 6.609438875 | 3.86E-11 | 3.06E-10 |
| -1.301262946 | 0.193168466 | 0.287341818 |
| 1.212463205 | 0.225335119 | 0.325398837 |
| 5.894504918 | 3.76E-09 | 2.41E-08 |
| -0.699973249 | 0.483944011 | 0.592444419 |
| -0.065390563 | 0.947863038 | NA |
| 0.469878781 | 0.638441625 | NA |
| NA | NA | NA |
| 2.555870906 | 0.010592241 | 0.024008312 |
| 1.812905788 | 0.069846353 | 0.123029854 |
| 6.620397513 | 3.58E-11 | 2.85E-10 |
| -1.017297964 | 0.309011705 | 0.418910469 |
| -6.948820819 | 3.68E-12 | 3.22E-11 |
| 0.192810118 | 0.847107679 | NA |
| NA | NA | NA |
| NA | NA | NA |
| -0.065389966 | 0.947863513 | NA |
| NA | NA | NA |
| 0.063708285 | 0.949202508 | NA |
| 0.77176961 | 0.440250894 | NA |
| NA | NA | NA |
| -6.450791728 | 1.11E-10 | 8.43E-10 |
| -3.949772935 | 7.82E-05 | 0.000276005 |
| -2.302307208 | 0.021317854 | 0.044156395 |
| 0.754549488 | 0.450519341 | 0.560434328 |
| 0.655482322 | 0.512157265 | 0.617827189 |
| -2.524399539 | 0.011589612 | 0.025988891 |
| 5.230780261 | 1.69E-07 | 8.83E-07 |

| | | |
|---|---|---|
| -0.799095251 | 0.424235183 | 0.535806168 |
| NA | NA | NA |
| 0.458102228 | 0.646878998 | NA |
| 0.192809521 | 0.847108147 | NA |
| 11.72655986 | 9.32E-32 | 3.53E-30 |
| -0.844648632 | 0.398307047 | 0.510216243 |
| 3.935379762 | 8.31E-05 | 0.000291479 |
| -10.09830826 | 5.62E-24 | 1.34E-22 |
| 0.468305165 | 0.639566375 | NA |
| 0.192808924 | 0.847108615 | NA |
| NA | NA | NA |
| -0.065388772 | 0.947864464 | NA |
| 0.769162033 | 0.441797126 | 0.551774369 |
| -3.213545772 | 0.001311069 | 0.003677625 |
| 0.539910978 | 0.589258427 | NA |
| 2.535829451 | 0.011218135 | 0.025264525 |
| -2.778472426 | 0.005461515 | 0.013310781 |
| -4.233240479 | 2.30E-05 | 8.86E-05 |
| -4.161503989 | 3.16E-05 | 0.000119024 |
| 0.440028241 | 0.659916654 | NA |
| 8.803411711 | 1.33E-18 | 2.12E-17 |
| 14.65338356 | 1.28E-48 | 9.58E-47 |
| 7.617229921 | 2.59E-14 | 2.83E-13 |
| -0.848448861 | 0.396188041 | 0.508125982 |
| 0.250299177 | 0.802355993 | NA |
| 0.434843042 | 0.66367636 | NA |
| -0.869624126 | 0.384505848 | 0.496744113 |
| NA | NA | NA |
| 5.149130745 | 2.62E-07 | 1.33E-06 |
| 7.217951024 | 5.28E-13 | 5.05E-12 |
| -2.169362676 | 0.03005516 | 0.059436401 |
| 2.87967239 | 0.003980886 | 0.010014229 |
| 1.964649174 | 0.049454856 | 0.09172694 |
| -5.272282342 | 1.35E-07 | 7.13E-07 |
| -1.317053075 | 0.187820835 | 0.280766573 |
| -2.973103868 | 0.002948047 | 0.007636304 |
| 3.366475377 | 0.000761354 | 0.002244307 |
| 0.72113027 | 0.470829369 | 0.579808847 |
| 1.649038968 | 0.099139652 | 0.165431032 |
| -2.913607799 | 0.003572785 | 0.009065977 |
| 0.122976706 | 0.902125543 | NA |
| 2.596491001 | 0.009418137 | 0.021591498 |
| -0.041499695 | 0.966897536 | 0.977534078 |
| 1.126782332 | 0.259834523 | 0.364820671 |
| 0.657023521 | 0.511165793 | NA |
| 9.774237035 | 1.45E-22 | 3.12E-21 |
| -0.849522693 | 0.395590508 | 0.507567686 |

| | | |
|---|---|---|
| -4.24057335 | 2.23E-05 | 8.60E-05 |
| -3.667217109 | 0.000245205 | 0.000791523 |
| -0.065388772 | 0.947864464 | NA |
| -3.885156509 | 0.000102264 | 0.000352863 |
| 0.192810118 | 0.847107679 | NA |
| 0.455520056 | 0.648735142 | NA |
| -2.728558755 | 0.006361176 | 0.015227642 |
| 0.093974254 | 0.925129609 | NA |
| -0.077191413 | 0.938471273 | NA |
| -2.682155586 | 0.007314943 | 0.0172177 |
| -1.131693486 | 0.257763321 | 0.362461569 |
| NA | NA | NA |
| -0.058161962 | 0.953619619 | 0.96824189 |
| 1.119718191 | 0.26283387 | 0.36820573 |
| -0.135419517 | 0.892280194 | NA |
| 0.458742089 | 0.646419386 | NA |
| 0.836936377 | 0.402628332 | NA |
| -6.438414774 | 1.21E-10 | 9.10E-10 |
| 0.109802879 | 0.912565707 | 0.939670876 |
| 0.063708285 | 0.949202508 | NA |
| -1.316775641 | 0.18791384 | 0.280888828 |
| 4.793694339 | 1.64E-06 | 7.46E-06 |
| 3.260992756 | 0.001110229 | 0.003168646 |
| -2.983054439 | 0.002853872 | 0.007413078 |
| -2.364958246 | 0.018032104 | 0.038247298 |
| 8.263196817 | 1.42E-16 | 1.90E-15 |
| 0.291029976 | 0.771028394 | NA |
| -2.843076034 | 0.004468041 | 0.011112387 |
| NA | NA | NA |
| 5.119534872 | 3.06E-07 | 1.55E-06 |
| 1.514482983 | 0.129903387 | 0.207259164 |
| -0.430087283 | 0.667132151 | 0.752447627 |
| NA | NA | NA |
| 5.529402678 | 3.21E-08 | 1.85E-07 |
| NA | NA | NA |
| 1.114794453 | 0.264938514 | NA |
| 0.347709792 | 0.728058142 | 0.801454707 |
| NA | NA | NA |
| 0.063708324 | 0.949202477 | NA |
| -1.39066184 | 0.164327994 | 0.251475281 |
| 0.192810118 | 0.847107679 | NA |
| 0.694575578 | 0.487321322 | 0.595414072 |
| 14.03361233 | 9.71E-45 | 6.45E-43 |
| 3.545349269 | 0.000392093 | 0.001219748 |
| 13.89129241 | 7.15E-44 | 4.60E-42 |
| -0.061442056 | 0.95100716 | 0.966569586 |
| NA | NA | NA |

| | | |
|---|---|---|
| -1.912864362 | 0.055765419 | 0.101668616 |
| -0.589778541 | 0.555339131 | 0.658025869 |
| 0.459986758 | 0.645525725 | NA |
| 1.521898634 | 0.128034481 | 0.2049436 |
| -0.280778859 | 0.77888002 | 0.841805212 |
| 5.06938676 | 3.99E-07 | 1.98E-06 |
| 7.104088495 | 1.21E-12 | 1.11E-11 |
| 0.066394788 | 0.94706352 | NA |
| -0.279594256 | 0.779788815 | 0.842623939 |
| NA | NA | NA |
| 2.365394838 | 0.018010858 | 0.038207625 |
| -2.120382139 | 0.033973832 | 0.066186567 |
| 20.58185453 | 3.99E-94 | 8.69E-92 |
| 0.80239704 | 0.422323325 | 0.534089636 |
| NA | NA | NA |
| 0.421867315 | 0.673121869 | NA |
| -5.702661275 | 1.18E-08 | 7.15E-08 |
| -2.366607399 | 0.017951965 | 0.038102067 |
| -1.621310096 | 0.104951151 | 0.173348773 |
| -3.204094453 | 0.00135488 | 0.003789546 |
| -3.975642869 | 7.02E-05 | 0.000249409 |
| -1.414302869 | 0.157272995 | 0.24277994 |
| NA | NA | NA |
| 0.192809521 | 0.847108147 | NA |
| 1.299297485 | 0.193841857 | 0.288132412 |
| -4.913668335 | 8.94E-07 | 4.22E-06 |
| -2.684888387 | 0.007255403 | 0.017093631 |
| -3.04934307 | 0.002293424 | 0.0060913 |
| -1.970675625 | 0.04876099 | 0.090587663 |
| -3.213029229 | 0.001313429 | 0.003682663 |
| 3.173932834 | 0.001503885 | 0.004164886 |
| 1.978818984 | 0.04783639 | 0.089108049 |
| -0.06620969 | 0.947210882 | NA |
| 1.994210816 | 0.046129021 | 0.08633239 |
| NA | NA | NA |
| NA | NA | NA |
| NA | NA | NA |
| NA | NA | NA |
| -0.065390563 | 0.947863038 | NA |
| -5.611284341 | 2.01E-08 | 1.19E-07 |
| -10.50548344 | 8.15E-26 | 2.21E-24 |
| -2.996192864 | 0.002733735 | 0.007128386 |
| -2.169359589 | 0.030055394 | 0.059436401 |
| -1.45891862 | 0.144587508 | 0.226458001 |
| -4.150456797 | 3.32E-05 | 0.000124498 |
| -4.128437974 | 3.65E-05 | 0.000136161 |
| 1.735528039 | 0.08264732 | 0.142010188 |

| | | |
|---|---|---|
| 0.192809521 | 0.847108147 | NA |
| NA | NA | NA |
| NA | NA | NA |
| 0.192810118 | 0.847107679 | NA |
| -1.337591508 | 0.181029634 | 0.272420594 |
| -2.973385022 | 0.002945347 | 0.007630892 |
| 0.063708285 | 0.949202508 | NA |
| -0.058894338 | 0.953036268 | 0.967963715 |
| NA | NA | NA |
| -0.682670919 | 0.49481481 | 0.601920549 |
| 0.192809521 | 0.847108147 | NA |
| -4.785352207 | 1.71E-06 | 7.76E-06 |
| -4.880827509 | 1.06E-06 | 4.94E-06 |
| -3.686359621 | 0.000227485 | 0.000739762 |
| 0.655616954 | 0.512070614 | NA |
| NA | NA | NA |
| NA | NA | NA |
| 4.448991658 | 8.63E-06 | 3.55E-05 |
| 0.6148803 | 0.538633774 | NA |
| 6.485468967 | 8.85E-11 | 6.76E-10 |
| 1.145405147 | 0.252041367 | 0.356028035 |
| 5.779098651 | 7.51E-09 | 4.64E-08 |
| 0.292967733 | 0.769546815 | 0.833986699 |
| -6.905323359 | 5.01E-12 | 4.33E-11 |
| 0.192809521 | 0.847108147 | NA |
| -1.905652077 | 0.056695361 | 0.103101197 |
| -4.148829428 | 3.34E-05 | 0.000125349 |
| -2.362762461 | 0.018139292 | 0.038411716 |
| -2.735911718 | 0.006220771 | 0.014932953 |
| -0.791395235 | 0.42871339 | 0.539882076 |
| -1.675654538 | 0.093805879 | 0.15797357 |
| -2.299460658 | 0.021478795 | 0.044441947 |
| NA | NA | NA |
| -1.705575904 | 0.088087069 | 0.149906916 |
| -0.350039566 | 0.726309004 | 0.800013835 |
| -0.806961801 | 0.419688491 | 0.531404438 |
| 0.192808924 | 0.847108615 | NA |
| 0.062883132 | 0.949859567 | NA |
| 0.3474247 | 0.728272279 | NA |
| NA | NA | NA |
| -0.065388772 | 0.947864464 | NA |
| -0.891101577 | 0.372874679 | 0.485691779 |
| -1.068079495 | 0.285484655 | 0.39261983 |
| -1.739890135 | 0.081878311 | 0.140901625 |
| -2.300546065 | 0.021417303 | 0.044333037 |
| 0.146485633 | 0.883538031 | 0.918488585 |
| -0.065388772 | 0.947864464 | NA |

| | | |
|---|---|---|
| 0.150649701 | 0.880252053 | 0.915863364 |
| -4.551006696 | 5.34E-06 | 2.26E-05 |
| 0.769723836 | 0.441463728 | NA |
| 7.916710847 | 2.44E-15 | 2.94E-14 |
| NA | NA | NA |
| NA | NA | NA |
| NA | NA | NA |
| NA | NA | NA |
| NA | NA | NA |
| NA | NA | NA |
| NA | NA | NA |
| NA | NA | NA |
| NA | NA | NA |
| NA | NA | NA |
| -1.606077927 | 0.108256789 | 0.177718071 |
| -0.233579906 | 0.815311129 | 0.870133354 |
| 3.249781552 | 0.001154937 | 0.003283888 |
| -0.028463982 | 0.977292095 | 0.984331461 |
| -2.083662413 | 0.037190885 | 0.071535223 |
| NA | NA | NA |
| 1.462946648 | 0.143481978 | 0.225078586 |
| 6.122409122 | 9.22E-10 | 6.30E-09 |
| NA | NA | NA |
| 1.864004821 | 0.062321027 | NA |
| 4.250309422 | 2.13E-05 | 8.26E-05 |
| NA | NA | NA |
| NA | NA | NA |
| 0.814764194 | 0.4152073 | 0.526849778 |
| NA | NA | NA |
| 9.398804788 | 5.52E-21 | 1.05E-19 |
| -1.797190702 | 0.07230535 | 0.126648209 |
| 3.966085027 | 7.31E-05 | 0.00025904 |
| NA | NA | NA |
| 0.192809521 | 0.847108147 | NA |
| 0.736832226 | 0.461224383 | 0.570631521 |
| -0.338681409 | 0.734849748 | 0.806618981 |
| 0.37145882 | 0.710295822 | NA |
| -6.373552125 | 1.85E-10 | 1.36E-09 |
| -0.400181831 | 0.689022596 | NA |
| -3.061576541 | 0.002201747 | 0.005874094 |
| 1.108933117 | 0.267459036 | 0.373326995 |
| NA | NA | NA |
| NA | NA | NA |
| NA | NA | NA |
| -0.27541979 | 0.782993742 | 0.845120024 |
| 1.324573693 | 0.185312586 | 0.277746948 |
| -2.379292908 | 0.017345887 | 0.036994508 |

| | | | |
|---|---|---|---|
| | 0.192810118 | 0.847107679 | NA |
| | -3.456374613 | 0.000547494 | 0.001657447 |
| | 1.124370756 | 0.260855773 | NA |
| | 1.817076917 | 0.069205337 | 0.122040092 |
| | 0.88480103 | 0.376263954 | 0.489012242 |
| | 2.306429788 | 0.021086629 | 0.043731736 |
| | 0.812635333 | 0.416427168 | 0.528049648 |
| | -0.738451326 | 0.460240235 | 0.569639177 |
| | -0.09784576 | 0.92205477 | 0.946443752 |
| | 0.116665749 | 0.907124933 | NA |
| | -0.065389369 | 0.947863988 | NA |
| | -0.065389369 | 0.947863988 | NA |
| NA | NA | NA | |
| | -0.066204891 | 0.947214703 | NA |
| NA | NA | NA | |
| | 7.069661002 | 1.55E-12 | 1.41E-11 |
| | 0.110719647 | 0.911838666 | NA |
| | 0.763721356 | 0.445033311 | 0.554933681 |
| | 1.215517637 | 0.224168726 | 0.323997201 |
| | 5.295895452 | 1.18E-07 | 6.32E-07 |
| NA | NA | NA | |
| | 0.192810118 | 0.847107679 | NA |
| | 1.115770943 | 0.264520194 | 0.370112386 |
| | -3.55052646 | 0.000384461 | 0.001197495 |
| NA | NA | NA | |
| | -11.08959831 | 1.41E-28 | 4.52E-27 |
| | 2.515420044 | 0.011889064 | 0.026560449 |
| | 0.003680494 | 0.997063397 | 0.997999007 |
| | 1.191069344 | 0.233626363 | 0.334898796 |
| NA | NA | NA | |
| | 6.413012602 | 1.43E-10 | 1.06E-09 |
| | 12.07578435 | 1.42E-33 | 5.83E-32 |
| | 9.212615734 | 3.18E-20 | 5.74E-19 |
| | 1.115666058 | 0.264565105 | NA |
| | -0.424555055 | 0.671161063 | NA |
| | -2.274941676 | 0.022909429 | 0.047063608 |
| | 0.192808924 | 0.847108615 | NA |
| | 2.474443829 | 0.01334438 | 0.029411494 |
| | -0.159599638 | 0.873196464 | 0.911734837 |
| | 2.475964128 | 0.013287693 | 0.029295424 |
| | -3.822846561 | 0.00013192 | 0.000446572 |
| | -2.019290964 | 0.043456986 | 0.081987516 |
| | -5.542135965 | 2.99E-08 | 1.72E-07 |
| | 0.42534918 | 0.670582147 | 0.755056436 |
| NA | NA | NA | |
| | 2.108845504 | 0.034957919 | 0.067850242 |
| | 0.744977876 | 0.456285094 | 0.566003366 |

| | | |
|---|---|---|
| -0.400782883 | 0.688579982 | NA |
| NA | NA | NA |
| 1.572994828 | 0.115720016 | 0.188009347 |
| NA | NA | NA |
| 1.841063895 | 0.065612196 | 0.116630443 |
| -0.260088399 | 0.794795586 | 0.854029229 |
| 0.142334167 | 0.88681606 | 0.920617394 |
| -1.241022352 | 0.214597492 | 0.312853176 |
| 3.683012074 | 0.000230494 | 0.000748551 |
| 0.063708882 | 0.949202033 | NA |
| NA | NA | NA |
| 0.470601958 | 0.637925009 | 0.729032188 |
| 0.144777852 | 0.884886269 | 0.919262195 |
| 0.510624327 | 0.609614138 | 0.705980316 |
| 1.456118376 | 0.145359898 | 0.227468475 |
| 0.027674154 | 0.977922038 | 0.984677254 |
| -3.82179277 | 0.000132485 | 0.000448243 |
| -2.940499534 | 0.003276835 | 0.008390709 |
| -6.209942147 | 5.30E-10 | 3.71E-09 |
| 0.192810118 | 0.847107679 | NA |
| 0.06353712 | 0.949338802 | NA |
| -2.762622392 | 0.005733906 | 0.013892137 |
| -4.526660652 | 5.99E-06 | 2.52E-05 |
| -3.463025973 | 0.000534137 | 0.001620932 |
| 1.730614856 | 0.08352048 | 0.143225378 |
| 0.570124333 | 0.568593372 | 0.670209188 |
| -2.354075678 | 0.01856883 | 0.039191991 |
| -0.448631297 | 0.653697653 | NA |
| -0.08339265 | 0.933539332 | 0.954825494 |
| 1.335706047 | 0.181645372 | NA |
| -1.280884623 | 0.200234195 | 0.295999403 |
| -3.385226147 | 0.000711196 | 0.002108403 |
| -0.442837503 | 0.657883275 | NA |
| 3.641072413 | 0.000271505 | 0.000869802 |
| NA | NA | NA |
| 0.192810118 | 0.847107679 | NA |
| 0.114097037 | 0.909160871 | NA |
| 3.220999615 | 0.001277443 | 0.003592413 |
| NA | NA | NA |
| 3.504197937 | 0.000457985 | 0.001405685 |
| NA | NA | NA |
| 9.414451769 | 4.76E-21 | 9.16E-20 |
| -0.065390563 | 0.947863038 | NA |
| 0.06371127 | 0.949200131 | NA |
| 0.423407673 | 0.671997848 | NA |
| -0.065389369 | 0.947863988 | NA |
| -3.975120714 | 7.03E-05 | 0.000249922 |

| | | |
|---|---|---|
| NA | NA | NA |
| NA | NA | NA |
| NA | NA | NA |
| -0.065388772 | 0.947864464 | NA |
| -1.998127509 | 0.045702838 | 0.085650048 |
| 0.588605266 | 0.556126101 | NA |
| 0.695169058 | 0.486949361 | 0.595075799 |
| -3.012038965 | 0.002594993 | 0.006801327 |
| -0.605340902 | 0.54495251 | 0.648820878 |
| 0.183516435 | 0.854392825 | 0.898775002 |
| -1.82989555 | 0.067265559 | 0.11912948 |
| -1.789218947 | 0.073579561 | 0.128484472 |
| -2.551307343 | 0.010731964 | 0.024292011 |
| NA | NA | NA |
| NA | NA | NA |
| -0.065389369 | 0.947863988 | NA |
| 0.138888167 | 0.889538522 | NA |
| 1.068854563 | 0.285135209 | NA |
| NA | NA | NA |
| 0.063708882 | 0.949202033 | NA |
| 4.241530865 | 2.22E-05 | 8.57E-05 |
| -3.263299103 | 0.001101232 | 0.003145839 |
| -2.803537366 | 0.005054537 | 0.012426586 |
| 0.343329054 | 0.731350925 | NA |
| 0.063708285 | 0.949202508 | NA |
| 3.250354809 | 0.001152611 | 0.003278765 |
| -0.065389369 | 0.947863988 | NA |
| -0.592631174 | 0.553428012 | 0.656538373 |
| NA | NA | NA |
| 0.192810118 | 0.847107679 | NA |
| 2.704108328 | 0.006848796 | 0.016267352 |
| 1.267145162 | 0.20510339 | 0.301653737 |
| -0.880670405 | 0.378496239 | 0.491122907 |
| -0.100530638 | 0.91992306 | 0.944991964 |
| -7.022068257 | 2.19E-12 | 1.95E-11 |
| 0.727808438 | 0.466730858 | NA |
| -6.492235349 | 8.46E-11 | 6.48E-10 |
| 0.659380047 | 0.509651751 | 0.615605798 |
| -0.091442676 | 0.927140853 | NA |
| -4.533780604 | 5.79E-06 | 2.44E-05 |
| -3.628765155 | 0.00028478 | 0.000908943 |
| -4.184475602 | 2.86E-05 | 0.000108496 |
| 0.063708285 | 0.949202508 | NA |
| 0.199709364 | 0.84170789 | NA |
| NA | NA | NA |
| NA | NA | NA |
| NA | NA | NA |

| | | |
|---|---|---|
| 1.016748715 | 0.309272985 | 0.419140333 |
| 0.063708285 | 0.949202508 | NA |
| NA | NA | NA |
| -5.21338002 | 1.85E-07 | 9.63E-07 |
| -2.324772132 | 0.020084148 | 0.041930622 |
| NA | NA | NA |
| NA | NA | NA |
| 4.509274689 | 6.50E-06 | 2.72E-05 |
| -0.129185557 | 0.897210824 | 0.928328177 |
| 4.128523075 | 3.65E-05 | 0.000136131 |
| 2.105637425 | 0.035235859 | 0.068310244 |
| 11.11126671 | 1.11E-28 | 3.58E-27 |
| 3.351925274 | 0.000802517 | 0.002354037 |
| -0.847903774 | 0.396491562 | 0.508400596 |
| -0.26176171 | 0.793505161 | NA |
| -5.603209081 | 2.10E-08 | 1.24E-07 |
| 0.759407915 | 0.44760858 | 0.557506109 |
| 5.192773521 | 2.07E-07 | 1.07E-06 |
| -0.426105969 | 0.670030631 | 0.754764477 |
| 6.564943626 | 5.21E-11 | 4.08E-10 |
| 0.980669908 | 0.326755548 | 0.437784572 |
| -3.984067452 | 6.77E-05 | 0.000241308 |
| NA | NA | NA |
| 0.298719414 | 0.765154143 | 0.830700854 |
| -1.364711656 | 0.172343695 | 0.261598707 |
| 2.043131383 | 0.041039434 | 0.077913913 |
| 0.147168237 | 0.88299923 | 0.918063917 |
| -0.741313891 | 0.458503141 | 0.568022848 |
| -0.834815109 | 0.403821818 | 0.515486541 |
| -0.869343158 | 0.384659464 | 0.496853461 |
| 0.894392948 | 0.371111684 | 0.483871723 |
| 6.509710147 | 7.53E-11 | 5.80E-10 |
| 3.906393951 | 9.37E-05 | 0.000326043 |
| 2.466210595 | 0.013655102 | 0.030015563 |
| 0.063708882 | 0.949202033 | NA |
| 19.13391986 | 1.32E-81 | 2.14E-79 |
| 0.063708882 | 0.949202033 | NA |
| 0.192808924 | 0.847108615 | NA |
| 12.71248737 | 5.04E-37 | 2.41E-35 |
| 2.576772025 | 0.00997277 | 0.02273434 |
| 12.8437678 | 9.32E-38 | 4.64E-36 |
| 2.450792329 | 0.014254217 | 0.0311329 |
| 0.192809521 | 0.847108147 | NA |
| NA | NA | NA |
| 1.717313778 | 0.085921856 | 0.146690134 |
| 0.461233948 | 0.644630768 | 0.734014563 |
| 0.762885685 | 0.445531574 | 0.55536132 |

| | | |
|---|---|---|
| -2.731395735 | 0.006306669 | 0.015110169 |
| -1.672304343 | 0.094464324 | 0.158915392 |
| 3.096594947 | 0.001957572 | 0.005285084 |
| -0.511574511 | 0.608948828 | 0.70548434 |
| 18.02941894 | 1.14E-72 | 1.56E-70 |
| 1.139734885 | 0.254396769 | 0.358674654 |
| 3.466484764 | 0.000527312 | 0.001602471 |
| 1.752333939 | 0.079716404 | 0.137682969 |
| -3.483001273 | 0.000495826 | 0.001513848 |
| -2.546696453 | 0.010874798 | 0.024559794 |
| 2.019410662 | 0.043444554 | 0.081970239 |
| 1.906705963 | 0.056558674 | 0.102906645 |
| NA | NA | NA |
| -3.118576115 | 0.001817272 | 0.004939869 |
| 4.560649811 | 5.10E-06 | 2.17E-05 |
| -2.453616982 | 0.014142753 | 0.03093346 |
| -0.361313244 | 0.717865294 | 0.793752189 |
| -3.224217657 | 0.001263174 | 0.003554895 |
| -2.660082834 | 0.007812144 | 0.018238779 |
| -1.202812263 | 0.229048978 | 0.329528657 |
| 9.470013648 | 2.80E-21 | 5.49E-20 |
| 20.03237183 | 2.88E-89 | 5.67E-87 |
| 6.035805342 | 1.58E-09 | 1.05E-08 |
| 0.756695698 | 0.449232192 | 0.559194439 |
| -0.726690377 | 0.46741565 | 0.57675197 |
| 7.419872494 | 1.17E-13 | 1.20E-12 |
| 0.468336718 | 0.639543814 | 0.730251482 |
| NA | NA | NA |
| 2.245084392 | 0.024762715 | 0.05030115 |
| -0.065389369 | 0.947863988 | NA |
| 0.7707622 | 0.440847896 | 0.550949812 |
| NA | NA | NA |
| 9.095592022 | 9.41E-20 | 1.65E-18 |
| 13.49047292 | 1.78E-41 | 1.02E-39 |
| 13.13340769 | 2.12E-39 | 1.13E-37 |
| 0.822358057 | 0.410873143 | 0.522675075 |
| 1.507213222 | 0.131756012 | NA |
| 1.707522269 | 0.087725022 | 0.149391571 |
| 0.192810118 | 0.847107679 | NA |
| 12.78579958 | 1.97E-37 | 9.62E-36 |
| 3.042514849 | 0.002346102 | 0.006213411 |
| 1.967565699 | 0.04911803 | 0.091196816 |
| NA | NA | NA |
| 0.904067179 | 0.365959781 | NA |
| NA | NA | NA |
| NA | NA | NA |
| 5.504696888 | 3.70E-08 | 2.11E-07 |

| | | |
|---|---|---|
| 18.54110484 | 9.62E-77 | 1.44E-74 |
| 6.80905527 | 9.82E-12 | 8.23E-11 |
| 0.917517704 | 0.35887142 | 0.470808305 |
| 11.22693897 | 3.01E-29 | 9.97E-28 |
| -2.621445895 | 0.008755766 | 0.020215554 |
| 2.262315278 | 0.023677931 | 0.048392146 |
| 8.40560157 | 4.26E-17 | 5.98E-16 |
| 7.416276714 | 1.20E-13 | 1.23E-12 |
| -0.497593732 | 0.618770423 | 0.71339584 |
| NA | NA | NA |
| 0.765535268 | 0.443952874 | 0.553917573 |
| 1.754969154 | 0.079264584 | 0.136987678 |
| -2.680605505 | 0.007348909 | 0.017292771 |
| 2.429004103 | 0.015140361 | 0.032856185 |
| 2.161468181 | 0.030659192 | 0.06048708 |
| -0.990504 | 0.321927835 | 0.432938626 |
| -0.202312801 | 0.839672193 | 0.887691966 |
| -0.129230368 | 0.897175367 | 0.928328177 |
| 2.256002247 | 0.02407049 | 0.049042277 |
| -3.943635455 | 8.03E-05 | 0.000282373 |
| -5.270753966 | 1.36E-07 | 7.19E-07 |
| 0.386443387 | 0.699168323 | 0.778685731 |
| -0.834426366 | 0.404040765 | 0.515713429 |
| 2.49503608 | 0.012594432 | 0.027944435 |
| 0.46895895 | 0.639098978 | NA |
| 1.104086509 | 0.269555613 | NA |
| 13.97274868 | 2.29E-44 | 1.51E-42 |
| -0.05067781 | 0.959582259 | 0.97223389 |
| 16.88465385 | 5.84E-64 | 6.46E-62 |
| 0.349291888 | 0.726870188 | NA |
| 0.381892277 | 0.702541269 | 0.781711846 |
| 0.021429435 | 0.982903094 | 0.987955362 |
| 0.451011515 | 0.651981249 | 0.740445501 |
| 9.368316663 | 7.37E-21 | 1.39E-19 |
| 2.533607524 | 0.01128951 | 0.025402412 |
| 0.545373801 | 0.585496455 | 0.685645002 |
| 0.862335197 | 0.38850309 | 0.500707087 |
| NA | NA | NA |
| -0.082329643 | 0.934384583 | NA |
| NA | NA | NA |
| NA | NA | NA |
| NA | NA | NA |
| 0.192809521 | 0.847108147 | NA |
| -0.065389369 | 0.947863988 | NA |
| NA | NA | NA |
| 0.192809521 | 0.847108147 | NA |
| NA | NA | NA |

| | | | |
|---|---|---|---|
| NA | NA | NA | |
| | -0.065389369 | 0.947863988 | NA |
| NA | NA | NA | |
| | -0.085802466 | 0.931623446 | NA |
| NA | NA | NA | |
| NA | NA | NA | |
| | 0.330704598 | 0.740867629 | NA |
| NA | NA | NA | |
| | 0.350236135 | 0.72616149 | NA |
| | 3.121750456 | 0.001797793 | 0.004893296 |
| | -4.387049969 | 1.15E-05 | 4.64E-05 |
| | 12.25736283 | 1.53E-34 | 6.55E-33 |
| | 11.57364153 | 5.61E-31 | 2.05E-29 |
| | 11.32565651 | 9.79E-30 | 3.32E-28 |
| NA | NA | NA | |
| | 9.926830879 | 3.18E-23 | 7.23E-22 |
| | 12.31462368 | 7.56E-35 | 3.27E-33 |
| NA | NA | NA | |
| | 11.1895909 | 4.59E-29 | 1.51E-27 |
| | 7.475012192 | 7.72E-14 | 8.03E-13 |
| | 0.559171841 | 0.576044449 | NA |
| | 19.17529148 | 5.95E-82 | 9.74E-80 |
| | 15.68816397 | 1.82E-55 | 1.61E-53 |
| | 9.401666223 | 5.37E-21 | 1.02E-19 |
| | 3.867532387 | 0.000109942 | 0.000377535 |
| | 10.12875171 | 4.12E-24 | 9.90E-23 |
| | 18.61527899 | 2.42E-77 | 3.69E-75 |
| | 11.84526683 | 2.28E-32 | 8.86E-31 |
| | 20.29601714 | 1.39E-91 | 2.91E-89 |
| | 20.1892249 | 1.22E-90 | 2.50E-88 |
| | 4.97340071 | 6.58E-07 | 3.17E-06 |
| | 11.39598808 | 4.38E-30 | 1.51E-28 |
| | 4.083500822 | 4.44E-05 | 0.000162694 |
| | 1.830063501 | 0.067240444 | NA |
| | 2.104513665 | 0.035333664 | NA |
| NA | NA | NA | |
| | 0.402281244 | 0.687477056 | NA |
| | 0.192809521 | 0.847108147 | NA |
| NA | NA | NA | |
| NA | NA | NA | |
| NA | NA | NA | |
| NA | NA | NA | |
| NA | NA | NA | |
| | 0.192810118 | 0.847107679 | NA |
| NA | NA | NA | |
| NA | NA | NA | |
| | 2.696312828 | 0.007011179 | 0.016595821 |

| | | |
|---|---|---|
| NA | NA | NA |
| 0.192809521 | 0.847108147 | NA |
| NA | NA | NA |
| NA | NA | NA |
| NA | NA | NA |
| -0.194487022 | 0.845794556 | NA |
| 0.126093806 | 0.89965767 | NA |
| -8.894688528 | 5.86E-19 | 9.65E-18 |
| 2.368006033 | 0.017884244 | 0.037977656 |
| -0.595517235 | 0.551497782 | 0.654776088 |
| -2.553870657 | 0.010653282 | 0.024129188 |
| 0.758625678 | 0.448076508 | NA |
| -1.978732923 | 0.047846084 | 0.089119474 |
| 10.46929203 | 1.20E-25 | 3.19E-24 |
| -0.198702818 | 0.84249522 | 0.889699407 |
| -2.10567202 | 0.035232852 | 0.068309704 |
| -1.147192805 | 0.251301945 | NA |
| -0.065389369 | 0.947863988 | NA |
| 3.423890054 | 0.000617316 | 0.001850691 |
| NA | NA | NA |
| 0.192808327 | 0.847109082 | NA |
| -1.168802676 | 0.242483142 | 0.345113951 |
| -4.946436686 | 7.56E-07 | 3.61E-06 |
| -3.025309149 | 0.002483791 | 0.006537254 |
| -4.053218537 | 5.05E-05 | 0.000184002 |
| -0.065388772 | 0.947864464 | NA |
| -0.58777919 | 0.556680514 | 0.659291622 |
| -0.065389369 | 0.947863988 | NA |
| 0.3474247 | 0.728272279 | NA |
| -0.428847474 | 0.668034228 | NA |
| 0.418180567 | 0.675815103 | NA |
| NA | NA | NA |
| -3.244530769 | 0.001176443 | 0.003339727 |
| -2.089584092 | 0.036655177 | 0.070673169 |
| -2.738931876 | 0.006163914 | 0.014809251 |
| -0.599032432 | 0.549151257 | NA |
| -0.789505861 | 0.429816405 | 0.540944521 |
| NA | NA | NA |
| -1.875051132 | 0.06078569 | 0.109198723 |
| -0.649853132 | 0.515787094 | 0.621128264 |
| -0.065388772 | 0.947864464 | NA |
| -0.581954436 | 0.560597372 | 0.66283844 |
| -2.46783145 | 0.013593431 | 0.029893127 |
| -2.546229776 | 0.010889348 | 0.024585362 |
| -0.276975133 | 0.7817992 | 0.844304008 |
| -1.794036992 | 0.072807265 | 0.127340216 |
| -1.840399255 | 0.065709643 | 0.116778782 |

| | | |
|---|---|---|
| -2.249926591 | 0.024453606 | 0.04973379 |
| -2.883983223 | 0.003926797 | 0.009891086 |
| -0.672485226 | 0.501274843 | 0.608174505 |
| -2.257531 | 0.023974915 | 0.048895308 |
| -1.519595028 | 0.128612788 | 0.205671997 |
| -2.114906579 | 0.03443791 | 0.066981187 |
| -1.049643186 | 0.293882194 | 0.401982147 |
| -0.133089392 | 0.894122685 | 0.926166645 |
| -5.088959861 | 3.60E-07 | 1.80E-06 |
| -1.25786249 | 0.208441491 | 0.305533528 |
| -0.877154571 | 0.380402687 | 0.492932246 |
| 0.032447715 | 0.974115012 | NA |
| 0.601847366 | 0.547275743 | NA |
| 0.11481598 | 0.908590983 | NA |
| 0.202714101 | 0.839358501 | NA |
| -1.853699746 | 0.063782128 | 0.11379761 |
| NA | NA | NA |
| -2.23249039 | 0.02558257 | 0.051731436 |
| -8.72042289 | 2.77E-18 | 4.28E-17 |
| 0.065061479 | 0.948125051 | NA |
| NA | NA | NA |
| -1.298006705 | 0.194285031 | 0.28863689 |
| -1.650834121 | 0.098772451 | 0.164906246 |
| 0.599429216 | 0.548886699 | NA |
| -0.441605038 | 0.658775038 | NA |
| 7.158714053 | 8.14E-13 | 7.64E-12 |
| -2.26734241 | 0.023369317 | 0.047866939 |
| 0.426694144 | 0.669602117 | 0.754410306 |
| -3.62714426 | 0.000286573 | 0.000914215 |
| -1.805229811 | 0.071038728 | 0.124848993 |
| -2.733833909 | 0.006260161 | 0.015017428 |
| -0.638168161 | 0.52336422 | 0.62858956 |
| -0.065389369 | 0.947863988 | NA |
| 0.063708285 | 0.949202508 | NA |
| -3.386992941 | 0.000706632 | 0.002095616 |
| -5.484361759 | 4.15E-08 | 2.36E-07 |
| -2.231778402 | 0.025629612 | 0.051801453 |
| -2.853333336 | 0.004326321 | 0.010788234 |
| -1.256861955 | 0.208803627 | 0.305930601 |
| -6.998325238 | 2.59E-12 | 2.29E-11 |
| -0.406786604 | 0.684164727 | NA |
| -2.008705527 | 0.044568372 | 0.083818946 |
| NA | NA | NA |
| -6.399484969 | 1.56E-10 | 1.15E-09 |
| 1.087444295 | 0.276840504 | 0.383704404 |
| -0.22899168 | 0.818875383 | 0.872672152 |
| -3.159615325 | 0.001579776 | 0.004358182 |

| | | |
|---|---|---|
| -1.123226064 | 0.261341496 | 0.366524179 |
| 1.835290024 | 0.066462728 | 0.117983127 |
| -9.523391545 | 1.68E-21 | 3.36E-20 |
| -4.567100884 | 4.95E-06 | 2.10E-05 |
| 0.135908579 | 0.891893554 | NA |
| -4.790551645 | 1.66E-06 | 7.57E-06 |
| -0.717156001 | 0.473277845 | 0.582091517 |
| 5.499331099 | 3.81E-08 | 2.17E-07 |
| -0.612651246 | 0.540106966 | 0.644339038 |
| -1.850407506 | 0.064254838 | 0.114469651 |
| 9.724407769 | 2.37E-22 | 5.02E-21 |
| 4.2780485 | 1.89E-05 | 7.36E-05 |
| 0.56820978 | 0.569892532 | 0.67136066 |
| 2.883732804 | 0.00392992 | 0.009896962 |
| -7.044892111 | 1.86E-12 | 1.67E-11 |
| -7.007540438 | 2.43E-12 | 2.15E-11 |
| -0.092497988 | 0.92630239 | NA |
| -1.080892714 | 0.27974484 | 0.386709931 |
| -2.771357955 | 0.005582302 | 0.013580039 |
| 0.801046755 | 0.423104579 | 0.53481 |
| 0.143209478 | 0.886124745 | NA |
| NA | NA | NA |
| 5.133735497 | 2.84E-07 | 1.44E-06 |
| -0.065389966 | 0.947863513 | NA |
| -0.842484148 | 0.399517009 | 0.51134745 |
| 1.475049606 | 0.140199207 | 0.22070868 |
| 0.778502676 | 0.43627273 | NA |
| -0.369567434 | 0.71170482 | 0.788848943 |
| NA | NA | NA |
| 4.16563156 | 3.10E-05 | 0.000117166 |
| 0.881253291 | 0.378180741 | 0.490815304 |
| NA | NA | NA |
| -3.905226479 | 9.41E-05 | 0.00032747 |
| 1.07929842 | 0.280454717 | 0.387405164 |
| -3.023764342 | 0.002496508 | 0.006565651 |
| 0.9226712 | 0.356178579 | 0.468085153 |
| -0.065388772 | 0.947864464 | NA |
| NA | NA | NA |
| -1.765930444 | 0.077407519 | 0.134260287 |
| 0.192809521 | 0.847108147 | NA |
| NA | NA | NA |
| 0.063708285 | 0.949202508 | NA |
| 0.741156961 | 0.458598276 | NA |
| NA | NA | NA |
| 0.192810118 | 0.847107679 | NA |
| -2.660129471 | 0.007811062 | 0.018237955 |
| NA | NA | NA |

| | | |
|---|---|---|
| 4.829735145 | 1.37E-06 | 6.28E-06 |
| 3.460440438 | 0.000539293 | 0.001633804 |
| -2.854842887 | 0.004305812 | 0.010740929 |
| 2.715664198 | 0.006614297 | 0.015768726 |
| 0.345975663 | 0.729361005 | 0.802492679 |
| -6.094545676 | 1.10E-09 | 7.45E-09 |
| -3.068688534 | 0.002150006 | 0.005753732 |
| NA | NA | NA |
| NA | NA | NA |
| 0.351808794 | 0.724981663 | 0.79917204 |
| -0.331014947 | 0.740633195 | 0.811186666 |
| 0.063708381 | 0.949202431 | NA |
| 0.064436689 | 0.948622517 | 0.965006231 |
| 5.636512299 | 1.74E-08 | 1.03E-07 |
| 1.41611046 | 0.156743166 | 0.242147972 |
| 0.192809521 | 0.847108147 | NA |
| -2.066871913 | 0.038746225 | 0.074116776 |
| -2.86558472 | 0.004162399 | 0.010423696 |
| -2.927129249 | 0.003421067 | 0.008723441 |
| -2.293448175 | 0.021822216 | 0.045056001 |
| -2.98850212 | 0.002803486 | 0.007294308 |
| -0.09551596 | 0.923905014 | NA |
| 3.152350525 | 0.001619617 | 0.004457286 |
| 0.063708285 | 0.949202508 | NA |
| NA | NA | NA |
| 4.117592881 | 3.83E-05 | 0.000142198 |
| -8.039124548 | 9.05E-16 | 1.13E-14 |
| 1.696878624 | 0.08971961 | 0.152249617 |
| -0.065389369 | 0.947863988 | NA |
| -0.994461836 | 0.319998077 | 0.430713694 |
| -4.164723797 | 3.12E-05 | 0.000117598 |
| 9.093029746 | 9.63E-20 | 1.68E-18 |
| -4.488418903 | 7.18E-06 | 2.98E-05 |
| -4.654150593 | 3.25E-06 | 1.42E-05 |
| NA | NA | NA |
| -0.92607973 | 0.354404547 | NA |
| 0.063708285 | 0.949202508 | NA |
| NA | NA | NA |
| 0.874590596 | 0.381796706 | 0.494324608 |
| -8.386673836 | 5.00E-17 | 6.96E-16 |
| 0.260606205 | 0.794396204 | NA |
| 3.52704419 | 0.000420227 | 0.001299033 |
| 0.743752411 | 0.457026275 | 0.566698091 |
| -0.088104255 | 0.929793814 | NA |
| -1.37729764 | 0.168420245 | 0.256613265 |
| 0.163921873 | 0.869792648 | 0.909373729 |
| 5.103782201 | 3.33E-07 | 1.67E-06 |

| | | |
|---|---|---|
| 12.16409922 | 4.83E-34 | 2.03E-32 |
| -1.172673821 | 0.240926633 | 0.343298956 |
| -0.083614731 | 0.933362754 | 0.954768603 |
| 1.411536211 | 0.158086566 | 0.243681904 |
| 0.232009435 | 0.816530685 | 0.870877921 |
| 2.379033057 | 0.017358119 | 0.037014289 |
| -0.017731448 | 0.985853093 | 0.989966417 |
| -0.085607355 | 0.931778551 | NA |
| 0.06353712 | 0.949338802 | NA |
| 0.19280706 | 0.847110074 | NA |
| 0.063708285 | 0.949202508 | NA |
| 0.063707194 | 0.949203376 | NA |
| 3.432224651 | 0.000598651 | 0.001800342 |
| 2.926514849 | 0.003427831 | 0.00873802 |
| 0.192809521 | 0.847108147 | NA |
| NA | NA | NA |
| -0.065388772 | 0.947864464 | NA |
| 0.192808924 | 0.847108615 | NA |
| NA | NA | NA |
| 0.693117146 | 0.488236038 | NA |
| 0.11084824 | 0.911736691 | 0.939246509 |
| -0.13165686 | 0.895255709 | NA |
| NA | NA | NA |
| 11.26670294 | 1.92E-29 | 6.42E-28 |
| 0.618586993 | 0.536188473 | NA |
| 0.657199822 | 0.51105244 | 0.616821419 |
| 4.166794281 | 3.09E-05 | 0.000116633 |
| NA | NA | NA |
| NA | NA | NA |
| 0.391387478 | 0.695510847 | 0.77578594 |
| -5.303928829 | 1.13E-07 | 6.06E-07 |
| 0.630040909 | 0.528667819 | NA |
| 0.327217779 | 0.743503175 | NA |
| NA | NA | NA |
| NA | NA | NA |
| -1.340163001 | 0.180192357 | 0.271389055 |
| -0.442074526 | 0.658435277 | NA |
| NA | NA | NA |
| NA | NA | NA |
| -2.604318127 | 0.009205726 | 0.021152973 |
| NA | NA | NA |
| NA | NA | NA |
| NA | NA | NA |
| -0.165935185 | 0.868207958 | 0.908193129 |
| -2.179425709 | 0.029300058 | 0.058163253 |
| -1.790982889 | 0.073296041 | 0.128051923 |
| 5.81836769 | 5.94E-09 | 3.72E-08 |

| | | |
|---|---|---|
| -1.639282919 | 0.101154352 | 0.168109251 |
| -6.322188008 | 2.58E-10 | 1.87E-09 |
| -8.802126034 | 1.34E-18 | 2.14E-17 |
| -2.378711194 | 0.017373282 | 0.037043465 |
| NA | NA | NA |
| 0.110280842 | 0.91218665 | NA |
| -0.571681003 | 0.567538107 | 0.669312044 |
| -2.739168754 | 0.006159475 | 0.014801426 |
| NA | NA | NA |
| NA | NA | NA |
| NA | NA | NA |
| NA | NA | NA |
| 0.192808327 | 0.847109082 | NA |
| 4.199650278 | 2.67E-05 | 0.000101907 |
| 0.192810118 | 0.847107679 | NA |
| -0.188985072 | 0.850104517 | 0.895581298 |
| -1.032113879 | 0.302018773 | 0.411223951 |
| -0.028186801 | 0.977513165 | 0.984435282 |
| -1.212821901 | 0.22519792 | 0.325219475 |
| NA | NA | NA |
| 0.448188091 | 0.654017456 | 0.742002399 |
| 0.392544237 | 0.694656132 | NA |
| 0.063708075 | 0.949202675 | NA |
| 0.063976859 | 0.948988653 | NA |
| 0.268106797 | 0.788617114 | 0.849650756 |
| -0.060226654 | 0.951975118 | NA |
| -1.190285553 | 0.233934177 | 0.335205659 |
| -0.685347739 | 0.493124514 | 0.600335923 |
| 0.123751295 | 0.901512197 | 0.931431119 |
| -1.80988674 | 0.070313353 | 0.123756643 |
| -0.963842786 | 0.335124752 | 0.446373436 |
| -1.06894664 | 0.285093714 | 0.392197006 |
| 0.645638082 | 0.518513769 | NA |
| -0.891318152 | 0.372758513 | 0.485591006 |
| -3.032895028 | 0.002422198 | 0.006391948 |
| -1.159517409 | 0.246245345 | 0.349287146 |
| NA | NA | NA |
| 0.063708882 | 0.949202033 | NA |
| -3.223755882 | 0.001265212 | 0.003559831 |
| 0.063708882 | 0.949202033 | NA |
| -3.402797527 | 0.000666997 | 0.001986782 |
| NA | NA | NA |
| NA | NA | NA |
| -3.57869183 | 0.000345318 | 0.00108639 |
| -0.104891963 | 0.916461536 | 0.942674129 |
| -0.116408872 | 0.907328504 | NA |
| -6.003465538 | 1.93E-09 | 1.28E-08 |

| | | |
|---|---|---|
| -3.234590068 | 0.001218175 | 0.003440248 |
| 2.203651039 | 0.027548895 | 0.055159494 |
| -4.100616678 | 4.12E-05 | 0.000152006 |
| -0.065389369 | 0.947863988 | NA |
| 0.063708882 | 0.949202033 | NA |
| 3.516130151 | 0.000437886 | 0.00134979 |
| -0.19448762 | 0.845794089 | NA |
| -3.880977719 | 0.000104037 | 0.000358488 |
| 0.568413893 | 0.569753959 | 0.671229483 |
| 0.758452535 | 0.448180118 | NA |
| 0.262595821 | 0.792862121 | 0.852682758 |
| -3.889829627 | 0.000100315 | 0.000346614 |
| -4.372271292 | 1.23E-05 | 4.93E-05 |
| 1.664215461 | 0.096069379 | 0.161145729 |
| -0.441279053 | 0.659010989 | 0.746192368 |
| 0.305691466 | 0.759839571 | 0.826149362 |
| 0.097281592 | 0.922502774 | NA |
| 0.3474247 | 0.728272279 | NA |
| 0.063708882 | 0.949202033 | NA |
| NA | NA | NA |
| NA | NA | NA |
| -3.242502685 | 0.001184848 | 0.003360539 |
| 0.144069302 | 0.885445745 | 0.919690721 |
| 2.073578083 | 0.038118512 | 0.073066807 |
| 0.192810118 | 0.847107679 | NA |
| -2.032886241 | 0.042064017 | 0.079653681 |
| -3.260791248 | 0.001111018 | 0.003170537 |
| 0.372887057 | 0.709232505 | 0.78691986 |
| -0.661105009 | 0.508544973 | NA |
| 0.109002014 | 0.913200893 | NA |
| -3.325282902 | 0.000883289 | 0.002574073 |
| 2.40461412 | 0.016189551 | 0.034867015 |
| NA | NA | NA |
| 3.199177185 | 0.001378204 | 0.003848333 |
| NA | NA | NA |
| 0.963130212 | 0.335482182 | NA |
| 12.78074358 | 2.10E-37 | 1.02E-35 |
| NA | NA | NA |
| -2.433961193 | 0.014934596 | 0.03244904 |
| -2.168724346 | 0.030103618 | 0.059519045 |
| -0.286881733 | 0.774202878 | 0.837654937 |
| -0.846850582 | 0.39707841 | 0.508928906 |
| -1.250076781 | 0.211271501 | 0.309012193 |
| 1.534775342 | 0.124839022 | 0.200624123 |
| NA | NA | NA |
| -1.834707301 | 0.06654907 | 0.118094521 |
| 0.192808924 | 0.847108615 | NA |

| | | | |
|---|---|---|---|
| | 0.728431169 | 0.466349689 | NA |
| | 1.955456763 | 0.050529182 | 0.093456397 |
| | 7.154370602 | 8.41E-13 | 7.87E-12 |
| | 1.552243921 | 0.120603871 | 0.194755799 |
| | -0.098382029 | 0.921628944 | 0.9462217 |
| | -5.155188979 | 2.53E-07 | 1.29E-06 |
| NA | NA | NA | |
| NA | NA | NA | |
| NA | NA | NA | |
| | -1.4740027 | 0.140480864 | 0.221021814 |
| | 2.531721518 | 0.01135041 | 0.025514212 |
| | -4.85836148 | 1.18E-06 | 5.49E-06 |
| | 3.929787191 | 8.50E-05 | 0.000297925 |
| | 2.30335211 | 0.02125904 | 0.044041861 |
| | -0.095748031 | 0.923720693 | 0.947630605 |
| | -0.405122408 | 0.685387536 | NA |
| | 5.048475976 | 4.45E-07 | 2.20E-06 |
| NA | NA | NA | |
| | -1.302289383 | 0.19281748 | 0.286882335 |
| NA | NA | NA | |
| | -1.870499664 | 0.061414464 | 0.110185938 |
| | -0.065389966 | 0.947863513 | NA |
| | -1.832908582 | 0.066816167 | 0.11846771 |
| | -1.731039217 | 0.08344477 | 0.143115157 |
| | 1.631365911 | 0.102813133 | 0.17037955 |
| | 1.107423987 | 0.26811066 | NA |
| NA | NA | NA | |
| | -0.180607203 | 0.856675901 | 0.900498959 |
| | 0.597417971 | 0.550228358 | NA |
| | -3.241605531 | 0.001188584 | 0.003370371 |
| | 1.128025059 | 0.259309336 | 0.364308076 |
| | 0.0615769 | 0.950899773 | NA |
| | -6.027010164 | 1.67E-09 | 1.11E-08 |
| | -2.268500582 | 0.023298714 | 0.047734043 |
| | 0.789996817 | 0.429529627 | 0.540637965 |
| NA | NA | NA | |
| | -1.447243646 | 0.147828696 | 0.230582182 |
| | -1.077605316 | 0.281209929 | 0.388202593 |
| | -0.060421008 | 0.951820327 | 0.967121165 |
| | 10.81941691 | 2.79E-27 | 8.21E-26 |
| | -0.065389966 | 0.947863513 | NA |
| | 0.063708882 | 0.949202033 | NA |
| | -0.020624959 | 0.983544831 | 0.988441625 |
| | -0.941181699 | 0.346611754 | 0.45815865 |
| | 0.344383536 | 0.730557869 | 0.803350472 |
| | -0.956347988 | 0.338896453 | 0.450266905 |
| | -0.958766842 | 0.337676217 | 0.448812431 |

| | | | |
|---|---|---|---|
| NA | NA | NA | |
| | -7.058923879 | 1.68E-12 | 1.51E-11 |
| NA | NA | NA | |
| NA | NA | NA | |
| | 0.192808327 | 0.847109082 | NA |
| NA | NA | NA | |
| NA | NA | NA | |
| NA | NA | NA | |
| | -0.266255164 | 0.790042701 | NA |
| | -6.86822061 | 6.50E-12 | 5.53E-11 |
| | -3.460794347 | 0.000538584 | 0.001632053 |
| | -4.155424517 | 3.25E-05 | 0.000122025 |
| | -3.460527702 | 0.000539118 | 0.001633473 |
| | -2.75412223 | 0.005884979 | 0.014210015 |
| | -0.065388772 | 0.947864464 | NA |
| | -0.225963728 | 0.821229632 | 0.874585273 |
| | 0.474004791 | 0.635496491 | NA |
| | -5.431509832 | 5.59E-08 | 3.11E-07 |
| | 0.212393462 | 0.831800086 | 0.882313691 |
| | -2.486394899 | 0.012904468 | 0.028551121 |
| | 0.063708882 | 0.949202033 | NA |
| | -4.228016187 | 2.36E-05 | 9.06E-05 |
| | -3.56509771 | 0.000363721 | 0.001138276 |
| | 1.446607944 | 0.14800676 | 0.230816773 |
| | -3.743442852 | 0.000181516 | 0.000600336 |
| NA | NA | NA | |
| NA | NA | NA | |
| | -2.408382636 | 0.016023377 | 0.034550752 |
| | -4.926460595 | 8.37E-07 | 3.97E-06 |
| | 0.980831088 | 0.326676045 | 0.437771785 |
| | -0.306585392 | 0.759158974 | 0.825732189 |
| | 0.192809521 | 0.847108147 | NA |
| | -0.065389369 | 0.947863988 | NA |
| NA | NA | NA | |
| NA | NA | NA | |
| | 0.450794256 | 0.652137842 | 0.74047406 |
| NA | NA | NA | |
| NA | NA | NA | |
| | -0.745826122 | 0.455772458 | 0.565479559 |
| | -3.064045854 | 0.002183654 | 0.00583131 |
| | -0.065389966 | 0.947863513 | NA |
| | -2.17067324 | 0.029955881 | 0.059264343 |
| | -0.075437792 | 0.939866391 | 0.959102078 |
| | 0.063708882 | 0.949202033 | NA |
| NA | NA | NA | |
| | 0.192809521 | 0.847108147 | NA |
| | -3.05037701 | 0.002285543 | 0.006072944 |

| | | | |
|---|---|---|---|
| | -1.851544117 | 0.064091314 | 0.114202719 |
| | 0.126968641 | 0.898965218 | 0.929681 |
| | -0.2768658 | 0.781883154 | NA |
| | 4.87602155 | 1.08E-06 | 5.05E-06 |
| | 2.897500273 | 0.003761494 | 0.009501483 |
| | -2.363559742 | 0.018100308 | 0.038347625 |
| | -3.496360725 | 0.000471651 | 0.001444794 |
| | -0.192788356 | 0.847124723 | 0.893193977 |
| | -1.642368327 | 0.100513695 | 0.167199959 |
| | 1.938321624 | 0.052583996 | 0.096699538 |
| | -0.065390563 | 0.947863038 | NA |
| | 7.003649373 | 2.49E-12 | 2.21E-11 |
| | -7.357899324 | 1.87E-13 | 1.86E-12 |
| | -3.710559606 | 0.000206802 | 0.000677696 |
| | 2.186516025 | 0.028777874 | NA |
| NA | NA | NA | |
| | 2.112624109 | 0.034632954 | 0.067303014 |
| | -2.021432202 | 0.043235044 | 0.081599544 |
| | 5.518502498 | 3.42E-08 | 1.96E-07 |
| | 0.539715995 | 0.589392907 | 0.688918673 |
| | -0.225508224 | 0.821583928 | 0.87485092 |
| | 0.063708285 | 0.949202508 | NA |
| | -2.609581905 | 0.009065294 | 0.020860928 |
| | -3.246890576 | 0.001166732 | 0.003315168 |
| | 1.415417459 | 0.156946134 | 0.242401686 |
| | -2.886188069 | 0.003899391 | 0.009827988 |
| | 4.613009344 | 3.97E-06 | 1.71E-05 |
| | -5.226549032 | 1.73E-07 | 9.01E-07 |
| | 0.355639023 | 0.722110915 | NA |
| | -4.755221597 | 1.98E-06 | 8.94E-06 |
| | 2.390473573 | 0.01682666 | 0.0360561 |
| | 5.039856991 | 4.66E-07 | 2.29E-06 |
| | 10.85665989 | 1.85E-27 | 5.56E-26 |
| | -0.253514925 | 0.79987033 | NA |
| | 5.519176919 | 3.41E-08 | 1.95E-07 |
| | 1.480402819 | 0.138765778 | 0.218791019 |
| | -0.675607458 | 0.499289905 | 0.606425967 |
| | -0.085607355 | 0.931778551 | NA |
| | 11.43471635 | 2.80E-30 | 9.75E-29 |
| | 0.664959976 | 0.506076093 | NA |
| NA | NA | NA | |
| | 3.409079765 | 0.000651824 | 0.001945523 |
| | -3.535713862 | 0.000406675 | 0.001261347 |
| NA | NA | NA | |
| | 2.213097537 | 0.026890922 | 0.053975924 |
| | -0.128306536 | 0.897906393 | 0.928894277 |
| | 0.491999988 | 0.622719352 | NA |

| | | |
|---|---|---|
| -3.302053998 | 0.000959796 | 0.002776663 |
| -3.929735339 | 8.50E-05 | 0.000297947 |
| -5.837630231 | 5.29E-09 | 3.33E-08 |
| -2.381018125 | 0.017264862 | 0.036867824 |
| -0.949705941 | 0.34226169 | 0.453606381 |
| -4.338163186 | 1.44E-05 | 5.71E-05 |
| -2.049778432 | 0.040386057 | 0.076848336 |
| 0.263094147 | 0.792478015 | 0.852416189 |
| 0.063708285 | 0.949202508 | NA |
| 14.69676572 | 6.76E-49 | 5.11E-47 |
| 0.655832193 | 0.5119321 | NA |
| NA | NA | NA |
| NA | NA | NA |
| -5.187129565 | 2.14E-07 | 1.10E-06 |
| -4.11298553 | 3.91E-05 | 0.000144767 |
| 1.162590568 | 0.244995659 | 0.347869376 |
| -2.692343189 | 0.007095189 | 0.016773006 |
| 5.204360981 | 1.95E-07 | 1.01E-06 |
| -0.065389369 | 0.947863988 | NA |
| -5.706211981 | 1.16E-08 | 7.01E-08 |
| NA | NA | NA |
| 0.364584686 | 0.715421448 | 0.791854552 |
| 0.192810118 | 0.847107679 | NA |
| NA | NA | NA |
| NA | NA | NA |
| NA | NA | NA |
| NA | NA | NA |
| 6.239501181 | 4.39E-10 | 3.10E-09 |
| 1.477621607 | 0.139509088 | 0.21978301 |
| -0.039339092 | 0.96862004 | 0.978471208 |
| 9.920440179 | 3.39E-23 | 7.68E-22 |
| 3.049983878 | 0.002288537 | 0.006079608 |
| 3.284713286 | 0.001020862 | 0.002937707 |
| 0.063708882 | 0.949202033 | NA |
| -0.066872027 | 0.946683582 | NA |
| 3.383337363 | 0.000716106 | 0.002121702 |
| -4.823599496 | 1.41E-06 | 6.47E-06 |
| 5.581360522 | 2.39E-08 | 1.40E-07 |
| 0.344972708 | 0.730114889 | 0.803004467 |
| -0.922827795 | 0.356096954 | 0.468002455 |
| -0.091925728 | 0.92675705 | 0.94982921 |
| NA | NA | NA |
| 0.353828976 | 0.723467056 | NA |
| -5.973307693 | 2.32E-09 | 1.52E-08 |
| 0.253475679 | 0.799900653 | NA |
| -0.194488217 | 0.845793621 | NA |
| -0.718937111 | 0.472179669 | NA |

| | | | |
|---|---|---|---|
| | 0.645177404 | 0.518812227 | NA |
| | 5.816564213 | 6.01E-09 | 3.76E-08 |
| | 2.687792176 | 0.007192615 | 0.016961548 |
| | 6.353862215 | 2.10E-10 | 1.54E-09 |
| | 0.359012128 | 0.719586017 | NA |
| | -2.860024099 | 0.004236088 | 0.010585991 |
| | -0.065389369 | 0.947863988 | NA |
| | 3.14770993 | 0.001645549 | 0.004522188 |
| | -0.676184978 | 0.498923209 | NA |
| NA | NA | NA | |
| | 1.560466985 | 0.118649566 | 0.192058374 |
| NA | NA | NA | |
| | 4.667182254 | 3.05E-06 | 1.34E-05 |
| | -1.13509315 | 0.256336291 | 0.360982186 |
| | -4.152605036 | 3.29E-05 | 0.000123464 |
| | 0.063707688 | 0.949202983 | NA |
| | 0.492978977 | 0.622027442 | 0.7162272 |
| | 5.472628664 | 4.43E-08 | 2.51E-07 |
| | 0.063710673 | 0.949200606 | NA |
| | -0.570561997 | 0.568296586 | 0.669985566 |
| | 0.767016294 | 0.443071828 | 0.553094003 |
| | -3.008898934 | 0.002621963 | 0.006864826 |
| | 1.951442533 | 0.051004426 | 0.094205598 |
| | 6.317517898 | 2.66E-10 | 1.92E-09 |
| | 3.989997031 | 6.61E-05 | 0.000235656 |
| | -6.445435978 | 1.15E-10 | 8.70E-10 |
| NA | NA | NA | |
| | 1.046044397 | 0.295540535 | 0.403869314 |
| NA | NA | NA | |
| | 8.752375172 | 2.09E-18 | 3.27E-17 |
| | -0.270034818 | 0.787133468 | 0.848307888 |
| | 9.024311349 | 1.81E-19 | 3.10E-18 |
| | 0.085566393 | 0.931811115 | NA |
| NA | NA | NA | |
| | 0.023488224 | 0.981260832 | 0.987018109 |
| | 0.841008352 | 0.400343256 | 0.512012247 |
| | -6.83357687 | 8.28E-12 | 6.97E-11 |
| | -4.750546627 | 2.03E-06 | 9.13E-06 |
| | -5.674021162 | 1.39E-08 | 8.38E-08 |
| | -6.446709258 | 1.14E-10 | 8.63E-10 |
| NA | NA | NA | |
| | -0.223581637 | 0.823082853 | 0.876037072 |
| | 1.011908699 | 0.31158171 | 0.421711445 |
| | 3.397497015 | 0.000680053 | 0.002023025 |
| | -1.924302991 | 0.054316616 | 0.099396468 |
| | -2.388651489 | 0.016910335 | 0.036207501 |
| | -0.644577888 | 0.519200768 | 0.624518008 |

| | | |
|---|---|---|
| 3.626571747 | 0.000287209 | 0.000916126 |
| -10.47122826 | 1.17E-25 | 3.13E-24 |
| 3.97639445 | 7.00E-05 | 0.000248729 |
| -1.292948429 | 0.196028896 | 0.29079614 |
| NA | NA | NA |
| 1.732306399 | 0.083219023 | 0.142845436 |
| 1.979849854 | 0.047720403 | 0.088938323 |
| 0.488367962 | 0.625289231 | NA |
| 1.866478754 | 0.061974414 | NA |
| -0.065389966 | 0.947863513 | NA |
| NA | NA | NA |
| -1.275088893 | 0.20227778 | 0.298473905 |
| -0.099472165 | 0.920763387 | NA |
| 6.751103208 | 1.47E-11 | 1.21E-10 |
| 0.192809521 | 0.847108147 | NA |
| 0.15296915 | 0.878422604 | 0.91476425 |
| NA | NA | NA |
| NA | NA | NA |
| 1.388446525 | 0.165001118 | NA |
| 0.505685026 | 0.613077782 | NA |
| 0.397214236 | 0.691209485 | NA |
| NA | NA | NA |
| -0.106264656 | 0.915372373 | 0.941747318 |
| 4.646288912 | 3.38E-06 | 1.47E-05 |
| 0.149379997 | 0.881253795 | 0.916668938 |
| 3.036529902 | 0.002393184 | 0.006324716 |
| 0.733223725 | 0.463421987 | NA |
| 0.750298742 | 0.4530748 | NA |
| 0.113296314 | 0.909795639 | NA |
| 3.921282515 | 8.81E-05 | 0.000308036 |
| 7.139880314 | 9.34E-13 | 8.70E-12 |
| -1.203487939 | 0.228787556 | 0.329282532 |
| 5.009388797 | 5.46E-07 | 2.66E-06 |
| 2.864981717 | 0.004170333 | 0.010441476 |
| 0.30922356 | 0.757151473 | NA |
| 0.192809521 | 0.847108147 | NA |
| 1.232132229 | 0.217899694 | 0.316726442 |
| -1.54891204 | 0.121402866 | 0.195856509 |
| -0.57762661 | 0.563516237 | 0.665572453 |
| 0.192810118 | 0.847107679 | NA |
| 2.871365543 | 0.004087026 | 0.010252382 |
| 0.101009225 | 0.919543137 | 0.944834348 |
| 9.539726348 | 1.43E-21 | 2.88E-20 |
| -0.704490843 | 0.481127151 | 0.589803731 |
| 1.162634447 | 0.244977849 | 0.347869376 |
| -0.323047031 | 0.746659624 | 0.815823747 |
| -4.124255272 | 3.72E-05 | 0.000138432 |

| | | |
|---|---|---|
| 4.432603794 | 9.31E-06 | 3.81E-05 |
| 4.941869771 | 7.74E-07 | 3.69E-06 |
| 0.815118158 | 0.415004679 | NA |
| NA | NA | NA |
| 0.192810118 | 0.847107679 | NA |
| 0.288217989 | 0.77317988 | NA |
| NA | NA | NA |
| NA | NA | NA |
| NA | NA | NA |
| 0.192809521 | 0.847108147 | NA |
| NA | NA | NA |
| NA | NA | NA |
| NA | NA | NA |
| NA | NA | NA |
| NA | NA | NA |
| NA | NA | NA |
| NA | NA | NA |
| NA | NA | NA |
| NA | NA | NA |
| NA | NA | NA |
| NA | NA | NA |
| 2.790759825 | 0.005258448 | 0.012865977 |
| 1.97293862 | 0.048502556 | 0.090140999 |
| -0.340941384 | 0.73314771 | NA |
| -0.601329453 | 0.547620576 | NA |
| -0.880590682 | 0.378539404 | 0.491127995 |
| -0.758991636 | 0.44785756 | NA |
| 4.379375413 | 1.19E-05 | 4.79E-05 |
| 1.065795714 | 0.286515999 | 0.393735338 |
| 0.661015167 | 0.508602587 | 0.614753852 |
| -2.294073575 | 0.021786273 | 0.045003748 |
| NA | NA | NA |
| -0.215561736 | 0.829329385 | 0.880738025 |
| 9.458027554 | 3.14E-21 | 6.15E-20 |
| -0.76193168 | 0.446100781 | NA |
| -5.49187518 | 3.98E-08 | 2.26E-07 |
| -0.973178928 | 0.330464382 | 0.441429962 |
| -1.646016583 | 0.099760345 | 0.166200841 |
| 5.542751748 | 2.98E-08 | 1.72E-07 |
| -0.115230853 | 0.908262145 | NA |
| 1.302520579 | 0.192738488 | 0.28680303 |
| -0.221725276 | 0.824527752 | 0.877050345 |
| 0.354904564 | 0.722661089 | 0.797293916 |
| -1.156303873 | 0.247556885 | 0.350730108 |
| -0.399739779 | 0.68934819 | 0.770766783 |
| -1.241820966 | 0.214302625 | 0.312568889 |
| NA | NA | NA |

| | | |
|---|---|---|
| NA | NA | NA |
| 4.409850757 | 1.03E-05 | 4.21E-05 |
| -2.932443236 | 0.003363064 | 0.008588663 |
| 0.721751773 | 0.470447105 | NA |
| -0.065389369 | 0.947863988 | NA |
| -2.333818834 | 0.01960521 | 0.041064302 |
| 3.146528443 | 0.001652212 | 0.004539502 |
| -4.094078056 | 4.24E-05 | 0.000155946 |
| NA | NA | NA |
| 9.070219106 | 1.19E-19 | 2.06E-18 |
| 0.139574703 | 0.888996029 | NA |
| NA | NA | NA |
| 0.192809521 | 0.847108147 | NA |
| -1.352892948 | 0.176089833 | 0.266428611 |
| -0.847127789 | 0.396923898 | 0.508809032 |
| 1.168786217 | 0.242489775 | 0.345113951 |
| -1.886145595 | 0.059275338 | 0.106945988 |
| -1.285811725 | 0.198508777 | 0.293847933 |
| 0.592062095 | 0.553809008 | 0.656937539 |
| -1.043901123 | 0.296531138 | 0.405052085 |
| NA | NA | NA |
| 4.631010356 | 3.64E-06 | 1.58E-05 |
| 4.554304012 | 5.26E-06 | 2.23E-05 |
| -0.779212879 | 0.435854325 | 0.546389894 |
| 1.074924723 | 0.282408432 | NA |
| -0.201202638 | 0.840540129 | 0.888196877 |
| -3.632057873 | 0.00028117 | 0.000898237 |
| 3.057720996 | 0.002230271 | 0.005942483 |
| -0.955556901 | 0.339296145 | 0.45072617 |
| -0.701063457 | 0.483263414 | 0.591698051 |
| NA | NA | NA |
| NA | NA | NA |
| 1.919934243 | 0.054866206 | 0.100255594 |
| 0.104655972 | 0.916648799 | 0.94278926 |
| NA | NA | NA |
| 2.543026292 | 0.010989696 | 0.024787969 |
| 2.581012646 | 0.009851097 | 0.022481124 |
| -1.612960653 | 0.106753056 | 0.175664256 |
| 0.664735604 | 0.506219617 | NA |
| -12.11980568 | 8.30E-34 | 3.45E-32 |
| -1.931764204 | 0.053388616 | 0.097934559 |
| -4.96474482 | 6.88E-07 | 3.30E-06 |
| 0.742441917 | 0.457819631 | 0.567485037 |
| NA | NA | NA |
| 0.088026469 | 0.929855638 | 0.95238099 |
| 0.128294427 | 0.897915976 | NA |
| 4.04062238 | 5.33E-05 | 0.000193607 |

| | | |
|---|---|---|
| 12.56090289 | 3.46E-36 | 1.59E-34 |
| -4.398410627 | 1.09E-05 | 4.42E-05 |
| 2.156413124 | 0.031051421 | 0.06117411 |
| -0.718776722 | 0.472278502 | 0.581205119 |
| 1.769294837 | 0.076844685 | 0.133404248 |
| 1.796334803 | 0.072441286 | 0.126815316 |
| 8.967375289 | 3.04E-19 | 5.12E-18 |
| 7.442113226 | 9.91E-14 | 1.02E-12 |
| -1.522087755 | 0.127987094 | 0.204907059 |
| 5.409053483 | 6.34E-08 | 3.50E-07 |
| -0.332884274 | 0.739221641 | 0.810070472 |
| -4.69339515 | 2.69E-06 | 1.19E-05 |
| 0.718375421 | 0.472525838 | 0.58142374 |
| 3.349180912 | 0.000810509 | 0.002374974 |
| NA | NA | NA |
| -0.765548184 | 0.443945186 | 0.553917573 |
| -3.521522682 | 0.000429076 | 0.0013241 |
| NA | NA | NA |
| -2.371518894 | 0.017715141 | 0.0376825 |
| NA | NA | NA |
| -0.767567709 | 0.442744053 | 0.55276754 |
| -2.439991111 | 0.014687623 | 0.031975933 |
| 0.769002963 | 0.441891552 | NA |
| NA | NA | NA |
| NA | NA | NA |
| NA | NA | NA |
| NA | NA | NA |
| 0.263135833 | 0.792445886 | NA |
| 4.774192359 | 1.80E-06 | 8.19E-06 |
| -4.090932007 | 4.30E-05 | 0.000157938 |
| -0.875259504 | 0.381432723 | 0.493985671 |
| -0.59566039 | 0.551402125 | 0.654724629 |
| -0.813564247 | 0.415894627 | 0.527481252 |
| NA | NA | NA |
| 2.062327953 | 0.039176523 | 0.074825516 |
| 2.326191053 | 0.020008361 | 0.04180029 |
| 8.528539138 | 1.48E-17 | 2.16E-16 |
| -6.299610125 | 2.98E-10 | 2.14E-09 |
| -2.175056421 | 0.029625889 | 0.058735524 |
| -1.379211394 | 0.167829586 | 0.255848379 |
| NA | NA | NA |
| -0.065388772 | 0.947864464 | NA |
| -2.535055695 | 0.011242945 | 0.025313566 |
| -1.681659096 | 0.092634964 | 0.156394453 |
| 0.288760435 | 0.772764711 | NA |
| 5.257103542 | 1.46E-07 | 7.71E-07 |
| 2.385881413 | 0.017038244 | 0.03643151 |

| | | | | | |
|---|---|---|---|---|---|
| | -0.079944323 | | 0.936281538 | | 0.956535791 |
| | -0.488049945 | | 0.625514465 | | 0.719025613 |
| | 8.5456621 | | 1.28E-17 | | 1.87E-16 |
| NA | | NA | | NA | |
| | -2.336193547 | | 0.019481156 | | 0.040846048 |
| | 2.747121011 | | 0.006012095 | | 0.014478271 |
| | 3.944143994 | | 8.01E-05 | | 0.000281853 |
| | 0.192808924 | | 0.847108615 | NA | |
| NA | | NA | | NA | |
| | -3.116476363 | | 0.001830264 | | 0.004970865 |
| | 1.289374662 | | 0.197267866 | | 0.292270314 |
| | -2.79815134 | | 0.005139602 | | 0.012609709 |
| NA | | NA | | NA | |
| | -0.309887642 | | 0.756646401 | | 0.823536088 |
| | -2.32103114 | | 0.020285163 | | 0.042318523 |
| | -2.361469282 | | 0.01820268 | | 0.038529647 |
| | -2.812759185 | | 0.004911842 | | 0.012115038 |
| | -9.417776432 | | 4.61E-21 | | 8.88E-20 |
| | -5.421880938 | | 5.90E-08 | | 3.27E-07 |
| | -2.390528354 | | 0.01682415 | | 0.036053808 |
| NA | | NA | | NA | |
| | -1.436791798 | | 0.150777162 | | 0.234494368 |
| | 0.078356692 | | 0.937544322 | NA | |
| NA | | NA | | NA | |
| | 2.449601202 | | 0.014301451 | | 0.031216999 |
| | 9.669341305 | | 4.07E-22 | | 8.46E-21 |
| | 0.062472152 | | 0.950186838 | | 0.965931954 |
| | -3.05536069 | | 0.0022479 | | 0.00598245 |
| | 8.370432891 | | 5.74E-17 | | 7.97E-16 |
| | 4.767721685 | | 1.86E-06 | | 8.43E-06 |
| | 0.044343137 | | 0.964630887 | NA | |
| | 1.021134396 | | 0.307190769 | | 0.416927934 |
| | 1.510896254 | | 0.130814886 | | 0.208407904 |
| | 1.286639244 | | 0.198220058 | | 0.293472617 |
| | -1.387312027 | | 0.165346637 | | 0.252786851 |
| NA | | NA | | NA | |
| | -1.217165412 | | 0.223541287 | | 0.323237343 |
| NA | | NA | | NA | |
| | -6.048672905 | | 1.46E-09 | | 9.77E-09 |
| | -4.105298663 | | 4.04E-05 | | 0.00014929 |
| | 2.533320545 | | 0.011298758 | | 0.025418651 |
| | -5.43910851 | | 5.35E-08 | | 2.99E-07 |
| NA | | NA | | NA | |
| | -0.584946727 | | 0.558583538 | | 0.660994041 |
| | -1.869345092 | | 0.061574818 | | 0.110426145 |
| NA | | NA | | NA | |
| NA | | NA | | NA | |

| | | | |
|---|---|---|---|
| | 4.43415164 | 9.24E-06 | 3.79E-05 |
| | -2.192012495 | 0.028378604 | 0.056580888 |
| | 5.118759888 | 3.08E-07 | 1.56E-06 |
| | 0.482923215 | 0.629150258 | NA |
| | 0.192809521 | 0.847108147 | NA |
| | 0.634792341 | 0.525563859 | 0.63074719 |
| | 0.237484388 | 0.81228103 | NA |
| | 0.192809521 | 0.847108147 | NA |
| | 5.357473302 | 8.44E-08 | 4.59E-07 |
| | -8.842792981 | 9.34E-19 | 1.51E-17 |
| NA | NA | NA | |
| | -1.698044613 | 0.089499342 | 0.151937638 |
| | -2.25759742 | 0.02397077 | 0.048890838 |
| | -3.119120217 | 0.001813919 | 0.004932363 |
| | -1.884734764 | 0.059465656 | 0.10725844 |
| NA | NA | NA | |
| | 3.496442373 | 0.000471506 | 0.001444529 |
| | 2.67542179 | 0.007463529 | 0.017537751 |
| | 0.934331503 | 0.350132919 | 0.461861763 |
| | 0.36465355 | 0.715370036 | 0.791832663 |
| | 9.131665314 | 6.75E-20 | 1.19E-18 |
| | 0.067953551 | 0.94582261 | NA |
| | 0.063708882 | 0.949202033 | NA |
| | -2.290121428 | 0.022014279 | 0.045418512 |
| | 0.93742571 | 0.348539621 | 0.460172313 |
| | 14.13771792 | 2.22E-45 | 1.51E-43 |
| | 1.887886766 | 0.059041154 | 0.106650101 |
| | 0.192809521 | 0.847108147 | NA |
| | 4.388776449 | 1.14E-05 | 4.60E-05 |
| | -0.686534092 | 0.492376373 | 0.599629282 |
| | 6.222954349 | 4.88E-10 | 3.43E-09 |
| NA | NA | NA | |
| NA | NA | NA | |
| NA | NA | NA | |
| NA | NA | NA | |
| | 0.293234498 | 0.769342917 | 0.833837932 |
| | 2.602533253 | 0.009253784 | 0.021253644 |
| NA | NA | NA | |
| | 0.946661641 | 0.343811223 | 0.455177981 |
| NA | NA | NA | |
| NA | NA | NA | |
| | 0.192809521 | 0.847108147 | NA |
| | -2.015234111 | 0.043880125 | 0.082678402 |
| | -2.734664282 | 0.006244393 | 0.014985345 |
| NA | NA | NA | |
| | -7.790910394 | 6.65E-15 | 7.71E-14 |
| NA | NA | NA | |

| | | |
|---|---|---|
| NA | NA | NA |
| NA | NA | NA |
| NA | NA | NA |
| NA | NA | NA |
| NA | NA | NA |
| -3.321481493 | 0.000895409 | 0.00260303 |
| -1.598772073 | 0.109871257 | 0.179955343 |
| 18.12045196 | 2.20E-73 | 3.06E-71 |
| NA | NA | NA |
| -2.3737973 | 0.017606213 | 0.037475396 |
| NA | NA | NA |
| 13.6274514 | 2.75E-42 | 1.64E-40 |
| 16.27090187 | 1.59E-59 | 1.57E-57 |
| NA | NA | NA |
| -0.650888701 | 0.515118336 | NA |
| 0.285955351 | 0.774912319 | NA |
| NA | NA | NA |
| NA | NA | NA |
| 0.459923268 | 0.645571298 | 0.734918342 |
| -0.065389369 | 0.947863988 | NA |
| NA | NA | NA |
| NA | NA | NA |
| NA | NA | NA |
| -1.717399777 | 0.085906152 | 0.146673328 |
| -1.102874215 | 0.270081793 | 0.376253536 |
| NA | NA | NA |
| NA | NA | NA |
| -0.550470294 | 0.581996846 | NA |
| 2.204276707 | 0.02750489 | 0.055080198 |
| 1.037719551 | 0.29940064 | 0.40830326 |
| -4.611552785 | 4.00E-06 | 1.72E-05 |
| -1.969072138 | 0.048944808 | 0.09090217 |
| -2.199286544 | 0.027857554 | 0.055679516 |
| 1.496204383 | 0.134600403 | 0.213298391 |
| 0.357350822 | 0.720829188 | 0.795869191 |
| 8.65834296 | 4.79E-18 | 7.23E-17 |
| 3.675428356 | 0.000237451 | 0.000768873 |
| -4.85721345 | 1.19E-06 | 5.52E-06 |
| 0.691236868 | 0.489416699 | 0.597158037 |
| -1.666422369 | 0.095629319 | 0.160558213 |
| NA | NA | NA |
| NA | NA | NA |
| -3.023339299 | 0.002500018 | 0.006573741 |
| 8.330184696 | 8.07E-17 | 1.11E-15 |
| 0.010787349 | 0.991393108 | 0.993974344 |
| -3.675641942 | 0.000237252 | 0.000768528 |
| 0.116822912 | 0.907000387 | 0.935708604 |

| | | |
|---|---|---|
| -0.071206596 | 0.943233332 | NA |
| NA | NA | NA |
| -0.450351353 | 0.652457116 | 0.740702178 |
| 1.130852159 | 0.258117322 | 0.362877812 |
| -3.175807157 | 0.001494202 | 0.004140362 |
| NA | NA | NA |
| -1.367093859 | 0.171595888 | 0.2606533 |
| 0.228255412 | 0.819447686 | NA |
| 2.382369985 | 0.017201605 | 0.036743145 |
| 7.800935071 | 6.15E-15 | 7.14E-14 |
| 0.063707135 | 0.949203424 | NA |
| 0.063707194 | 0.949203376 | NA |
| 0.192810118 | 0.847107679 | NA |
| 0.089906082 | 0.928361848 | NA |
| -0.109799636 | 0.91256828 | NA |
| -3.918840052 | 8.90E-05 | 0.000310783 |
| 8.183637895 | 2.75E-16 | 3.60E-15 |
| NA | NA | NA |
| 4.310304222 | 1.63E-05 | 6.42E-05 |
| -1.818620282 | 0.068969381 | 0.121724976 |
| 0.086966698 | 0.930697983 | 0.952804571 |
| NA | NA | NA |
| NA | NA | NA |
| NA | NA | NA |
| -1.964971241 | 0.049417566 | 0.091677156 |
| -3.32396283 | 0.00088748 | 0.002584182 |
| -2.830893042 | 0.004641824 | 0.01150459 |
| 3.289799714 | 0.001002587 | 0.002889557 |
| 0.192810118 | 0.847107679 | NA |
| NA | NA | NA |
| NA | NA | NA |
| 3.256481096 | 0.001128024 | 0.003214671 |
| 1.284023684 | 0.199133669 | 0.294593804 |
| -0.066588298 | 0.946909462 | NA |
| 3.508575841 | 0.000450513 | 0.001385469 |
| -3.721734093 | 0.000197859 | 0.000650949 |
| -0.120584507 | 0.904020141 | NA |
| -1.273535995 | 0.202827908 | 0.299091701 |
| 0.359966159 | 0.71887244 | 0.794472686 |
| 0.063708285 | 0.949202508 | NA |
| -2.202010225 | 0.027664586 | 0.055346862 |
| NA | NA | NA |
| -0.051296044 | 0.959089621 | 0.971824678 |
| 1.123159031 | 0.261369959 | 0.366524179 |
| 1.942250297 | 0.052106807 | 0.095945532 |
| -1.703754301 | 0.088426999 | 0.150354492 |
| -8.780388958 | 1.63E-18 | 2.58E-17 |

| | | |
|---|---|---|
| -0.858196575 | 0.390783927 | 0.502895717 |
| 5.40493433 | 6.48E-08 | 3.58E-07 |
| -0.065389369 | 0.947863988 | NA |
| 6.480540062 | 9.14E-11 | 6.98E-10 |
| 0.146112224 | 0.883832797 | 0.918663465 |
| 3.134852936 | 0.001719403 | 0.004702945 |
| 2.689588504 | 0.007154017 | 0.016883623 |
| 11.855835 | 2.01E-32 | 7.85E-31 |
| -1.897724128 | 0.057732432 | 0.104682902 |
| NA | NA | NA |
| 1.307476542 | 0.191050917 | NA |
| 2.451951725 | 0.014208372 | 0.031051738 |
| -0.194488217 | 0.845793621 | NA |
| -0.535620512 | 0.59222085 | NA |
| 0.192809521 | 0.847108147 | NA |
| NA | NA | NA |
| -0.270861779 | 0.786497342 | 0.847731826 |
| 10.35176074 | 4.11E-25 | 1.06E-23 |
| -0.701387799 | 0.483061034 | 0.591493035 |
| -4.018291089 | 5.86E-05 | 0.000211001 |
| 10.63439382 | 2.06E-26 | 5.78E-25 |
| 10.67630372 | 1.31E-26 | 3.73E-25 |
| -0.39872306 | 0.690097274 | NA |
| 0.433266762 | 0.664820982 | 0.750942936 |
| -0.99659805 | 0.318959653 | 0.429616322 |
| -1.199749355 | 0.230236699 | 0.330911546 |
| -1.036939658 | 0.29976398 | 0.408665165 |
| -4.304859594 | 1.67E-05 | 6.57E-05 |
| 0.141641557 | 0.88736314 | NA |
| 0.285391104 | 0.775344522 | 0.838744636 |
| 8.142906584 | 3.86E-16 | 4.98E-15 |
| 5.381240919 | 7.40E-08 | 4.05E-07 |
| 1.047766024 | 0.29474642 | 0.403033842 |
| 4.345265786 | 1.39E-05 | 5.54E-05 |
| -0.627117438 | 0.530582257 | 0.635404077 |
| NA | NA | NA |
| -1.803817937 | 0.071259853 | 0.125174426 |
| 3.851581865 | 0.000117357 | 0.000400908 |
| -4.8091304 | 1.52E-06 | 6.93E-06 |
| -0.065389966 | 0.947863513 | NA |
| 3.533168938 | 0.00041061 | 0.001271662 |
| -0.989580871 | 0.322379024 | 0.433405681 |
| 2.083235202 | 0.03722979 | 0.071604552 |
| 1.648849258 | 0.099178521 | 0.165484859 |
| 1.429867634 | 0.152755013 | 0.236996093 |
| -6.309635497 | 2.80E-10 | 2.01E-09 |
| 0.909378462 | 0.363150387 | 0.475574435 |

| | | |
|---|---|---|
| 0.149812249 | 0.880912745 | 0.916427515 |
| 8.316561285 | 9.06E-17 | 1.23E-15 |
| 13.60436387 | 3.77E-42 | 2.22E-40 |
| -0.517177751 | 0.605032082 | 0.701986418 |
| 0.768475892 | 0.442204508 | 0.552176545 |
| 8.926732214 | 4.39E-19 | 7.29E-18 |
| 0.963183056 | 0.335455667 | 0.446667483 |
| -3.42114937 | 0.000623571 | 0.001867876 |
| NA | NA | NA |
| 2.344622728 | 0.019046341 | 0.040044512 |
| 0.815842666 | 0.414590127 | 0.526194664 |
| -2.27482437 | 0.022916468 | 0.047074205 |
| NA | NA | NA |
| -0.065389966 | 0.947863513 | NA |
| -3.766187304 | 0.000165759 | 0.00055187 |
| NA | NA | NA |
| -1.57260417 | 0.115810502 | 0.188144146 |
| 1.730057277 | 0.083620042 | 0.143376466 |
| -4.050872222 | 5.10E-05 | 0.000185686 |
| -1.381707962 | 0.167061389 | 0.254956805 |
| -1.793374149 | 0.072913118 | 0.127481433 |
| -1.756863528 | 0.078941074 | 0.136560583 |
| 0.063708217 | 0.949202562 | NA |
| 0.184882082 | 0.853321525 | 0.898100991 |
| 0.04748312 | 0.962128183 | 0.973878536 |
| -5.900187057 | 3.63E-09 | 2.33E-08 |
| 0.268315239 | 0.788456677 | 0.849514469 |
| -0.287969481 | 0.7733701 | 0.837111505 |
| -0.19448762 | 0.845794089 | NA |
| 0.322751319 | 0.746883583 | 0.81599721 |
| -0.445735399 | 0.655788391 | NA |
| 1.834454241 | 0.066586594 | 0.118119239 |
| 0.063708285 | 0.949202508 | NA |
| -7.404402542 | 1.32E-13 | 1.33E-12 |
| -6.761972326 | 1.36E-11 | 1.13E-10 |
| -13.52723123 | 1.08E-41 | 6.23E-40 |
| -2.001777575 | 0.045308659 | 0.085013173 |
| NA | NA | NA |
| NA | NA | NA |
| -0.686990458 | 0.492088741 | 0.599419717 |
| NA | NA | NA |
| NA | NA | NA |
| NA | NA | NA |
| 2.671297056 | 0.007555875 | 0.017723132 |
| 0.784601554 | 0.432687218 | 0.543382724 |
| 4.097598006 | 4.17E-05 | 0.000153866 |
| 0.742021307 | 0.458074426 | 0.567617999 |

| | | |
|---|---|---|
| NA | NA | NA |
| 2.173885178 | 0.029713759 | 0.05889574 |
| 1.177611281 | 0.23895161 | 0.341032989 |
| -1.423041466 | 0.154724135 | 0.239575942 |
| -1.688083396 | 0.091395221 | 0.154548195 |
| -4.690584243 | 2.72E-06 | 1.20E-05 |
| 12.70550812 | 5.51E-37 | 2.63E-35 |
| 0.063708285 | 0.949202508 | NA |
| 5.222107373 | 1.77E-07 | 9.21E-07 |
| -4.405047268 | 1.06E-05 | 4.30E-05 |
| 7.881183748 | 3.24E-15 | 3.86E-14 |
| NA | NA | NA |
| -1.916970697 | 0.055241654 | 0.100868005 |
| -1.721318614 | 0.085193024 | 0.145644517 |
| 5.464987981 | 4.63E-08 | 2.61E-07 |
| -6.830796841 | 8.44E-12 | 7.10E-11 |
| 6.549225319 | 5.78E-11 | 4.51E-10 |
| 2.858787418 | 0.004252637 | 0.010624164 |
| 9.269058878 | 1.88E-20 | 3.44E-19 |
| 0.192810118 | 0.847107679 | NA |
| 0.192808924 | 0.847108615 | NA |
| -4.003595554 | 6.24E-05 | 0.000223461 |
| 0.192808882 | 0.847108647 | NA |
| -0.696020435 | 0.486416033 | 0.594598227 |
| -0.592025967 | 0.5538332 | NA |
| -0.152424098 | 0.878852452 | NA |
| 7.358397401 | 1.86E-13 | 1.86E-12 |
| NA | NA | NA |
| 3.063193421 | 0.002189885 | 0.005845453 |
| -3.999016225 | 6.36E-05 | 0.000227471 |
| -1.139114568 | 0.254655372 | 0.358978597 |
| -3.086115002 | 0.002027904 | 0.005459657 |
| -4.155124332 | 3.25E-05 | 0.000122148 |
| NA | NA | NA |
| NA | NA | NA |
| 1.637714376 | 0.10148129 | 0.168585432 |
| NA | NA | NA |
| 0.555291038 | 0.578695608 | 0.679492943 |
| 0.308902638 | 0.75739559 | 0.824279822 |
| NA | NA | NA |
| NA | NA | NA |
| -0.069867013 | 0.944299508 | NA |
| -0.689480206 | 0.490521125 | 0.597980897 |
| -0.432402088 | 0.665449201 | 0.751395926 |
| NA | NA | NA |
| -0.408051277 | 0.68323603 | 0.765814026 |
| -1.355053541 | 0.175400499 | 0.265594344 |

| | | |
|---|---|---|
| -4.515659248 | 6.31E-06 | 2.65E-05 |
| -5.315474847 | 1.06E-07 | 5.72E-07 |
| -2.831311722 | 0.004635752 | 0.011491816 |
| 0.65939213 | 0.509643995 | 0.615605798 |
| -9.849075853 | 6.92E-23 | 1.52E-21 |
| 2.082962656 | 0.037254628 | 0.071635809 |
| 0.388153401 | 0.697902516 | 0.777691211 |
| 0.063708285 | 0.949202508 | NA |
| 1.318028933 | 0.187493962 | 0.280344915 |
| -6.487023118 | 8.75E-11 | 6.70E-10 |
| 0.754633941 | 0.450468653 | 0.560399124 |
| 7.959784712 | 1.72E-15 | 2.11E-14 |
| 0.707884994 | 0.479016679 | NA |
| NA | NA | NA |
| -5.340236774 | 9.28E-08 | 5.02E-07 |
| -0.049860005 | 0.960233949 | 0.97262983 |
| -0.865884343 | 0.38655359 | 0.498791416 |
| NA | NA | NA |
| -0.679759996 | 0.49665644 | 0.603784094 |
| 4.015314547 | 5.94E-05 | 0.000213283 |
| 0.637836026 | 0.523580426 | NA |
| 1.961588123 | 0.049810456 | 0.092297581 |
| 3.614755984 | 0.000300631 | 0.000955528 |
| -0.958908143 | 0.337605022 | 0.448741633 |
| 2.376929222 | 0.017457438 | 0.037191221 |
| 4.187023574 | 2.83E-05 | 0.000107383 |
| -1.461678541 | 0.14382932 | 0.225496266 |
| 7.613054152 | 2.68E-14 | 2.92E-13 |
| 2.217888881 | 0.02656241 | 0.053393601 |
| 0.06371127 | 0.949200131 | NA |
| 0.680082002 | 0.49645254 | 0.603594845 |
| NA | NA | NA |
| NA | NA | NA |
| NA | NA | NA |
| -4.585619396 | 4.53E-06 | 1.94E-05 |
| -0.108111502 | 0.913907244 | NA |
| 0.819482724 | 0.412511055 | 0.524172591 |
| 1.315782961 | 0.1882469 | NA |
| -0.65195746 | 0.514428616 | 0.620004155 |
| NA | NA | NA |
| -4.47148355 | 7.77E-06 | 3.22E-05 |
| -7.513444223 | 5.76E-14 | 6.06E-13 |
| -0.065389369 | 0.947863988 | NA |
| -1.976811684 | 0.048062921 | 0.08947675 |
| -0.454372255 | 0.649560919 | NA |
| -0.069647865 | 0.944473938 | NA |
| 1.513596082 | 0.130128316 | 0.207551885 |

| | | | |
|---|---|---|---|
| | -2.237143298 | 0.025276976 | 0.051212782 |
| | -3.206744232 | 0.001342463 | 0.003759016 |
| NA | NA | NA | |
| | -0.338002966 | 0.735360953 | NA |
| NA | NA | NA | |
| | 0.19464298 | 0.845672453 | 0.891903027 |
| | 4.063708707 | 4.83E-05 | 0.000176434 |
| | 1.203970031 | 0.228601162 | 0.329033185 |
| | -0.624673971 | 0.53218506 | 0.636661852 |
| | -0.08277479 | 0.934030615 | NA |
| | 0.063708285 | 0.949202508 | NA |
| | -2.633913222 | 0.008440706 | 0.019563929 |
| | 4.244832056 | 2.19E-05 | 8.45E-05 |
| | -1.200657027 | 0.22988427 | 0.330480919 |
| | 4.617167676 | 3.89E-06 | 1.68E-05 |
| | -7.907835028 | 2.62E-15 | 3.15E-14 |
| | 3.793236563 | 0.000148696 | 0.000498843 |
| | -1.07918864 | 0.280503643 | 0.387451365 |
| | 23.93971684 | 1.18E-126 | 3.89E-124 |
| | 8.808356878 | 1.27E-18 | 2.03E-17 |
| | 4.560979602 | 5.09E-06 | 2.16E-05 |
| | 5.70809003 | 1.14E-08 | 6.94E-08 |
| NA | NA | NA | |
| NA | NA | NA | |
| | 0.273694368 | 0.784319507 | NA |
| | -0.171564928 | 0.863779582 | NA |
| NA | NA | NA | |
| | 0.47838917 | 0.63237324 | NA |
| | -0.284999511 | 0.775644516 | NA |
| NA | NA | NA | |
| NA | NA | NA | |
| NA | NA | NA | |
| | 0.063708882 | 0.949202033 | NA |
| NA | NA | NA | |
| NA | NA | NA | |
| | 0.063708285 | 0.949202508 | NA |
| NA | NA | NA | |
| NA | NA | NA | |
| NA | NA | NA | |
| | -0.840415586 | 0.400675413 | 0.51230617 |
| | -0.08089136 | 0.935528351 | NA |
| | -0.098267909 | 0.921719559 | 0.946254894 |
| | -3.951563193 | 7.76E-05 | 0.000274064 |
| | -6.668275373 | 2.59E-11 | 2.08E-10 |
| | 0.192809521 | 0.847108147 | NA |
| | 0.063708285 | 0.949202508 | NA |
| | -3.250526457 | 0.001151915 | 0.003277481 |

| | | | |
|---|---|---|---|
| | -3.808742212 | 0.000139675 | 0.000470916 |
| | -0.169221327 | 0.865622555 | 0.906246133 |
| | 5.055603532 | 4.29E-07 | 2.12E-06 |
| | 13.27168871 | 3.38E-40 | 1.86E-38 |
| | -5.32921252 | 9.86E-08 | 5.32E-07 |
| | 2.16090654 | 0.03070256 | 0.060558319 |
| | 0.914636391 | 0.360382548 | 0.472518343 |
| | -2.714320515 | 0.006641188 | 0.015828312 |
| NA | NA | NA | |
| | -0.71175924 | 0.476613877 | 0.585532804 |
| NA | NA | NA | |
| | 0.397864302 | 0.690730215 | 0.771691541 |
| | 0.493920604 | 0.621362253 | 0.715602998 |
| | -0.395773078 | 0.692272439 | 0.772931535 |
| | 11.33620667 | 8.68E-30 | 2.95E-28 |
| NA | NA | NA | |
| | -3.426871221 | 0.000610579 | 0.001831809 |
| | 0.415251941 | 0.677957479 | NA |
| | -0.266395207 | 0.789934856 | NA |
| NA | NA | NA | |
| | -0.065389369 | 0.947863988 | NA |
| | -0.31560811 | 0.752299981 | 0.82033931 |
| | 9.793811892 | 1.20E-22 | 2.59E-21 |
| | 1.0404737 | 0.298119877 | NA |
| NA | NA | NA | |
| | 0.063708381 | 0.949202431 | NA |
| | 0.504125697 | 0.61417305 | 0.709729984 |
| | 0.611180682 | 0.541079971 | 0.645179286 |
| NA | NA | NA | |
| | -0.065389966 | 0.947863513 | NA |
| | 0.330493631 | 0.741027004 | NA |
| NA | NA | NA | |
| | 0.192810118 | 0.847107679 | NA |
| NA | NA | NA | |
| NA | NA | NA | |
| | 0.066394788 | 0.94706352 | NA |
| | -3.478973822 | 0.000503338 | 0.00153491 |
| NA | NA | NA | |
| NA | NA | NA | |
| | 0.192809521 | 0.847108147 | NA |
| NA | NA | NA | |
| | 2.034919891 | 0.041858934 | 0.079331349 |
| | -0.065388772 | 0.947864464 | NA |
| | -0.465242594 | 0.641757739 | 0.732108664 |
| | 1.19793713 | 0.230941493 | 0.331714999 |
| | -3.748407654 | 0.000177961 | 0.000589201 |
| | -4.372127577 | 1.23E-05 | 4.94E-05 |

| | | |
|---|---|---|
| -1.205325568 | 0.228077643 | 0.328459242 |
| 3.73579642 | 0.000187122 | 0.00061749 |
| 3.067987541 | 0.002155056 | 0.005766013 |
| 7.066985063 | 1.58E-12 | 1.43E-11 |
| 2.032216901 | 0.042131703 | 0.079763748 |
| NA | NA | NA |
| 0.851201197 | 0.394657598 | 0.50670837 |
| -3.619235367 | 0.000295475 | 0.000940694 |
| -1.545909639 | 0.122126392 | 0.196909534 |
| -3.674548452 | 0.00023827 | 0.000771428 |
| 0.392220909 | 0.694894996 | 0.775271752 |
| 8.35172276 | 6.73E-17 | 9.27E-16 |
| 0.795854655 | 0.42611652 | 0.537531322 |
| -6.639984689 | 3.14E-11 | 2.51E-10 |
| 8.955350579 | 3.39E-19 | 5.69E-18 |
| -1.319045308 | 0.187153966 | 0.279936881 |
| 12.44050771 | 1.57E-35 | 7.04E-34 |
| -11.12562161 | 9.41E-29 | 3.06E-27 |
| NA | NA | NA |
| -0.400637512 | 0.688687023 | NA |
| -0.786988365 | 0.431288677 | 0.542197669 |
| 1.751062675 | 0.079935115 | 0.138003581 |
| 0.192636473 | 0.847243678 | NA |
| -3.941564886 | 8.10E-05 | 0.000284662 |
| 1.324857868 | 0.185218297 | 0.277633183 |
| -6.858599075 | 6.95E-12 | 5.90E-11 |
| -0.448024136 | 0.654135778 | NA |
| -4.309247459 | 1.64E-05 | 6.45E-05 |
| 1.018872526 | 0.308263488 | 0.418134456 |
| 2.809454502 | 0.004962553 | 0.012224457 |
| -2.814106222 | 0.004891306 | 0.012075096 |
| 0.257930883 | 0.79646025 | 0.855132594 |
| 0.438472069 | 0.661044113 | 0.747771236 |
| 12.39366634 | 2.83E-35 | 1.25E-33 |
| -3.457572949 | 0.000545065 | 0.001650492 |
| 17.82042618 | 4.91E-71 | 6.50E-69 |
| -1.403557939 | 0.160450528 | 0.246490929 |
| 2.605703884 | 0.009168569 | 0.021079203 |
| 13.7762996 | 3.54E-43 | 2.19E-41 |
| 6.947680733 | 3.71E-12 | 3.24E-11 |
| -4.036960742 | 5.41E-05 | 0.00019631 |
| -0.723188492 | 0.469564088 | 0.578661273 |
| 5.014261441 | 5.32E-07 | 2.59E-06 |
| -1.025995293 | 0.304893803 | 0.414441982 |
| 1.177123725 | 0.239146127 | 0.34120889 |
| -3.68383949 | 0.000229747 | 0.000746245 |
| -6.565345817 | 5.19E-11 | 4.07E-10 |

| | | |
|---|---|---|
| 0.870053162 | 0.384271353 | 0.496505882 |
| -8.509875068 | 1.74E-17 | 2.52E-16 |
| 2.309577114 | 0.020911578 | 0.043426267 |
| -1.731554041 | 0.083352995 | 0.143006749 |
| NA | NA | NA |
| 3.148529652 | 0.001640941 | 0.004510515 |
| -13.42723914 | 4.19E-41 | 2.36E-39 |
| 8.471178768 | 2.43E-17 | 3.47E-16 |
| -4.260576863 | 2.04E-05 | 7.92E-05 |
| -6.694146683 | 2.17E-11 | 1.76E-10 |
| 0.106705451 | 0.915022658 | 0.941503623 |
| -6.315139256 | 2.70E-10 | 1.95E-09 |
| NA | NA | NA |
| -0.501066031 | 0.616324652 | 0.711323848 |
| -0.298238945 | 0.765520799 | 0.830882685 |
| NA | NA | NA |
| 2.200689388 | 0.027758021 | 0.055507171 |
| -3.467667693 | 0.000524996 | 0.001595708 |
| 2.652062472 | 0.008000174 | 0.018629114 |
| 0.06371127 | 0.949200131 | NA |
| -1.091704071 | 0.274963195 | 0.381636493 |
| -0.569800105 | 0.568813284 | NA |
| 0.883378748 | 0.377031669 | 0.489781123 |
| -0.544015264 | 0.586430969 | 0.686386119 |
| -4.680209109 | 2.87E-06 | 1.26E-05 |
| -1.57650722 | 0.114908956 | 0.186934302 |
| -2.9521964 | 0.003155222 | 0.008110835 |
| -2.239687587 | 0.025111212 | 0.050918149 |
| -1.733474201 | 0.083011419 | 0.142528181 |
| 0.609378208 | 0.542273779 | 0.646307922 |
| -0.954918281 | 0.339619024 | 0.451083267 |
| -2.972999689 | 0.002949048 | 0.007638106 |
| -3.887174545 | 0.000101418 | 0.000350233 |
| 6.024695975 | 1.69E-09 | 1.13E-08 |
| 0.372969058 | 0.709171473 | NA |
| 6.535307364 | 6.35E-11 | 4.93E-10 |
| -6.544440177 | 5.97E-11 | 4.65E-10 |
| 3.255886471 | 0.001130389 | 0.003220677 |
| 0.544079247 | 0.586386941 | 0.686366682 |
| 0.063707194 | 0.949203376 | NA |
| 4.767204042 | 1.87E-06 | 8.44E-06 |
| 4.133873068 | 3.57E-05 | 0.000133257 |
| -1.923156626 | 0.054460383 | 0.099615856 |
| 3.319940444 | 0.000900367 | 0.002615619 |
| 0.199919974 | 0.841543169 | 0.889069117 |
| NA | NA | NA |
| -6.199867051 | 5.65E-10 | 3.94E-09 |

| | | |
|---|---|---|
| 0.949846159 | 0.342190428 | 0.453569696 |
| -0.810475688 | 0.417666832 | 0.529353426 |
| 0.053922418 | 0.956996976 | 0.970365225 |
| 1.650155812 | 0.098911072 | 0.165115655 |
| -2.512867485 | 0.011975431 | 0.026712863 |
| 9.096966445 | 9.29E-20 | 1.63E-18 |
| -6.403637247 | 1.52E-10 | 1.13E-09 |
| -2.449295436 | 0.014313599 | 0.03124079 |
| -6.913139741 | 4.74E-12 | 4.10E-11 |
| -1.37561033 | 0.168942307 | 0.257231337 |
| -3.142455111 | 0.001675374 | 0.004593057 |
| 3.122653968 | 0.001792283 | 0.004880954 |
| 1.277696747 | 0.201356373 | 0.297429921 |
| 0.370855435 | 0.71074521 | 0.788002862 |
| 0.30922356 | 0.757151473 | NA |
| NA | NA | NA |
| 5.15836239 | 2.49E-07 | 1.27E-06 |
| -0.62407434 | 0.532578765 | 0.636967053 |
| 1.954974985 | 0.050586023 | 0.093547598 |
| 3.778607258 | 0.000157708 | 0.000526535 |
| -0.431986881 | 0.665750948 | 0.75159202 |
| 0.063708217 | 0.949202562 | NA |
| 2.30700349 | 0.021054626 | 0.043676221 |
| 1.284833531 | 0.198850462 | 0.294266706 |
| 2.74376764 | 0.006073851 | 0.014613909 |
| -9.404524553 | 5.23E-21 | 9.99E-20 |
| -6.513989329 | 7.32E-11 | 5.65E-10 |
| -6.752065765 | 1.46E-11 | 1.20E-10 |
| 4.927510857 | 8.33E-07 | 3.95E-06 |
| 3.543247838 | 0.000395231 | 0.001227983 |
| 0.112893378 | 0.910115086 | NA |
| 3.205772753 | 0.001347003 | 0.003769621 |
| -3.920908884 | 8.82E-05 | 0.000308428 |
| 1.095582488 | 0.273261529 | 0.379837622 |
| 3.743285374 | 0.00018163 | 0.000600633 |
| 2.456557471 | 0.014027536 | 0.030737042 |
| 2.891560654 | 0.003833336 | 0.009673193 |
| 5.245921058 | 1.56E-07 | 8.17E-07 |
| 6.363974926 | 1.97E-10 | 1.44E-09 |
| 0.073065847 | 0.941753719 | 0.960401548 |
| -2.530088076 | 0.01140339 | 0.0256241 |
| 1.437530977 | 0.150567177 | 0.234226007 |
| 0.258484731 | 0.796032832 | 0.854880584 |
| 0.325838112 | 0.744546842 | 0.814226121 |
| 1.289777411 | 0.197127953 | 0.292097601 |
| 1.692639926 | 0.090524027 | 0.153356777 |
| -1.279509686 | 0.200717631 | 0.296626457 |

| | | | |
|---|---|---|---|
| | -6.075098886 | 1.24E-09 | 8.35E-09 |
| | 0.382721062 | 0.701926596 | NA |
| | 8.833888178 | 1.01E-18 | 1.63E-17 |
| | 7.00800803 | 2.42E-12 | 2.14E-11 |
| | 4.461532119 | 8.14E-06 | 3.37E-05 |
| | -0.194488217 | 0.845793621 | NA |
| NA | NA | NA | |
| | -3.349064471 | 0.000810849 | 0.002375694 |
| | -2.408916282 | 0.015999968 | 0.034512168 |
| | -0.342221111 | 0.732184499 | NA |
| | -6.609782694 | 3.85E-11 | 3.05E-10 |
| | -2.327291329 | 0.019949765 | 0.041695275 |
| | -2.740968415 | 0.00612584 | 0.014731914 |
| | 0.06371127 | 0.949200131 | NA |
| | 0.777793841 | 0.436690559 | 0.5470653 |
| | -6.194502875 | 5.85E-10 | 4.07E-09 |
| | 0.247088433 | 0.804839786 | 0.861824164 |
| | 0.192810118 | 0.847107679 | NA |
| | -0.572929911 | 0.566692151 | 0.668566391 |
| | -0.030057549 | 0.976021156 | 0.983293121 |
| | -1.196270299 | 0.231591095 | 0.332438212 |
| | -0.157444788 | 0.874894318 | 0.912726807 |
| | -1.29626132 | 0.19488547 | 0.289288537 |
| | -0.181367705 | 0.856078966 | 0.900097242 |
| | -0.779560967 | 0.435649339 | 0.546198304 |
| | -0.959760222 | 0.337175906 | 0.44847996 |
| | 0.192809521 | 0.847108147 | NA |
| | -0.634099932 | 0.526015607 | 0.63113804 |
| NA | NA | NA | |
| | 1.486472421 | 0.137154206 | 0.216744371 |
| | -1.646419341 | 0.099677454 | 0.166140153 |
| | 0.340790178 | 0.733261545 | 0.80538127 |
| | -2.944103445 | 0.003238917 | 0.008306363 |
| | -3.760217031 | 0.000169766 | 0.000563635 |
| | -2.570875227 | 0.010144187 | 0.023091033 |
| | 0.584538031 | 0.558858386 | NA |
| | 9.018116145 | 1.91E-19 | 3.26E-18 |
| | 2.919309402 | 0.003508078 | 0.00892259 |
| | 8.893857488 | 5.90E-19 | 9.71E-18 |
| | 0.063708285 | 0.949202508 | NA |
| NA | NA | NA | |
| | -1.342485165 | 0.179438737 | 0.27046765 |
| | -2.299730195 | 0.02146351 | 0.044421338 |
| | 6.688829061 | 2.25E-11 | 1.82E-10 |
| | 10.59998661 | 2.98E-26 | 8.25E-25 |
| | -5.849323173 | 4.94E-09 | 3.12E-08 |
| | -3.190726862 | 0.001419154 | 0.003951654 |

| | | | |
|---|---|---|---|
| NA | NA | NA | |
| | -3.017230442 | 0.002550958 | 0.006696432 |
| | 0.192810118 | 0.847107679 | NA |
| | 7.23497375 | 4.66E-13 | 4.48E-12 |
| | 0.216763006 | 0.828393043 | NA |
| | -0.951668539 | 0.34126511 | 0.452837072 |
| | 0.926941186 | 0.353957072 | 0.465899532 |
| | -1.600610022 | 0.109463321 | 0.179392845 |
| | 7.920516336 | 2.37E-15 | 2.86E-14 |
| | -2.256708992 | 0.024026265 | 0.048972101 |
| | 0.063708285 | 0.949202508 | NA |
| | -6.832940738 | 8.32E-12 | 7.00E-11 |
| | 1.091161337 | 0.275201897 | 0.381919687 |
| | -0.609987926 | 0.541869806 | 0.645918718 |
| | -2.189174305 | 0.028584173 | 0.056949921 |
| | -0.041286528 | 0.967067473 | 0.977534078 |
| NA | NA | NA | |
| | 3.623436525 | 0.000290715 | 0.000926449 |
| NA | NA | NA | |
| | -3.242678083 | 0.001184119 | 0.003358851 |
| | 5.275727344 | 1.32E-07 | 7.01E-07 |
| | 5.031271379 | 4.87E-07 | 2.39E-06 |
| | -1.670315212 | 0.094857015 | 0.159507631 |
| | 5.40753197 | 6.39E-08 | 3.53E-07 |
| | 2.507354001 | 0.012163882 | 0.027082545 |
| | 1.501326924 | 0.133271025 | 0.211635762 |
| | 6.013813923 | 1.81E-09 | 1.20E-08 |
| | 1.089135883 | 0.276093968 | 0.382881712 |
| | -4.535796091 | 5.74E-06 | 2.42E-05 |
| | 0.192809521 | 0.847108147 | NA |
| | 6.409747166 | 1.46E-10 | 1.08E-09 |
| | 11.81492888 | 3.27E-32 | 1.27E-30 |
| NA | NA | NA | |
| | -0.356656981 | 0.721348614 | 0.796324899 |
| | -1.723686515 | 0.084764451 | 0.144981153 |
| | 6.012907176 | 1.82E-09 | 1.21E-08 |
| | -5.541172533 | 3.00E-08 | 1.73E-07 |
| NA | NA | NA | |
| | 0.163701701 | 0.869965978 | 0.909386028 |
| | -1.306418075 | 0.191410422 | 0.285195718 |
| | 0.256240552 | 0.797765097 | 0.85618988 |
| | -0.619852085 | 0.535355174 | 0.63970688 |
| | -0.12865934 | 0.89762721 | 0.928682225 |
| NA | NA | NA | |
| | 3.181833045 | 0.001463462 | 0.00406463 |
| | -0.308339833 | 0.757823759 | 0.824602386 |
| NA | NA | NA | |

| | | |
|---|---|---|
| -0.530148946 | 0.596008665 | NA |
| -0.065389369 | 0.947863988 | NA |
| NA | NA | NA |
| -4.844440684 | 1.27E-06 | 5.86E-06 |
| -1.705478699 | 0.088105182 | 0.149926759 |
| 0.266884854 | 0.789557819 | NA |
| -1.863766835 | 0.062354455 | 0.111632359 |
| -4.227091773 | 2.37E-05 | 9.09E-05 |
| 0.392944364 | 0.694360574 | 0.774848202 |
| 3.366276329 | 0.000761903 | 0.002245431 |
| NA | NA | NA |
| 1.939913959 | 0.052390147 | 0.096371382 |
| 8.158217159 | 3.40E-16 | 4.41E-15 |
| 7.27315577 | 3.51E-13 | 3.41E-12 |
| 1.712262144 | 0.08684837 | 0.14808002 |
| 0.821845086 | 0.41116507 | 0.522940127 |
| -6.045296643 | 1.49E-09 | 9.96E-09 |
| 0.376253218 | 0.706728655 | 0.784875092 |
| -2.590999788 | 0.009569755 | 0.021890968 |
| 4.634011546 | 3.59E-06 | 1.56E-05 |
| NA | NA | NA |
| -0.585307899 | 0.558340704 | 0.660769165 |
| 3.895090812 | 9.82E-05 | 0.000339927 |
| 4.597884291 | 4.27E-06 | 1.84E-05 |
| 1.717955311 | 0.085804767 | 0.146530212 |
| -2.853998661 | 0.004317271 | 0.010768441 |
| 5.572348944 | 2.51E-08 | 1.46E-07 |
| 1.835075854 | 0.066494451 | 0.1180227 |
| -0.477954538 | 0.632682563 | 0.724397079 |
| 1.499514218 | 0.133740283 | 0.212257896 |
| -4.206277071 | 2.60E-05 | 9.93E-05 |
| 1.444625573 | 0.148563086 | 0.231525686 |
| 3.784151847 | 0.000154234 | 0.000515967 |
| 0.063708882 | 0.949202033 | NA |
| -5.844894936 | 5.07E-09 | 3.20E-08 |
| -1.619588822 | 0.105320632 | 0.173809944 |
| 8.009308336 | 1.15E-15 | 1.43E-14 |
| NA | NA | NA |
| -0.440996035 | 0.659215868 | 0.746310243 |
| 0.620445448 | 0.534964558 | 0.63937685 |
| -0.778091536 | 0.436515052 | 0.546922447 |
| 7.204107319 | 5.84E-13 | 5.57E-12 |
| -0.027375081 | 0.978160573 | NA |
| -2.176824266 | 0.029493682 | 0.058505853 |
| 0.186404113 | 0.852127867 | 0.897234342 |
| -0.423685686 | 0.671795056 | 0.755935544 |
| NA | NA | NA |

| | | |
|---|---|---|
| NA | NA | NA |
| -0.081248729 | 0.935244147 | 0.955944177 |
| NA | NA | NA |
| -0.065389369 | 0.947863988 | NA |
| 0.324874675 | 0.745275925 | NA |
| NA | NA | NA |
| -0.940892609 | 0.346759896 | 0.458306061 |
| 0.739286859 | 0.459732827 | NA |
| -0.74834564 | 0.454251702 | 0.564124044 |
| NA | NA | NA |
| NA | NA | NA |
| 6.897919439 | 5.28E-12 | 4.54E-11 |
| -0.510664631 | 0.609585911 | NA |
| -5.950685772 | 2.67E-09 | 1.74E-08 |
| -5.156867893 | 2.51E-07 | 1.28E-06 |
| -2.31403853 | 0.020665609 | 0.042986802 |
| 13.08094287 | 4.23E-39 | 2.24E-37 |
| -0.224614383 | 0.822279275 | NA |
| 0.098737635 | 0.921346586 | 0.945988394 |
| 10.61023588 | 2.67E-26 | 7.44E-25 |
| 4.395785444 | 1.10E-05 | 4.47E-05 |
| 0.67937269 | 0.496901749 | 0.604023641 |
| 1.966762911 | 0.04921055 | 0.09134827 |
| -0.243847656 | 0.80734881 | NA |
| -2.756395181 | 0.005844234 | 0.014126623 |
| 0.259780914 | 0.795032774 | NA |
| 1.643629955 | 0.100252663 | 0.166876647 |
| -0.725366062 | 0.468227488 | 0.57744058 |
| 0.940811962 | 0.34680123 | 0.45833649 |
| NA | NA | NA |
| 0.417163746 | 0.676558641 | 0.759718322 |
| 3.472051519 | 0.000516497 | 0.001571211 |
| -0.065389369 | 0.947863988 | NA |
| NA | NA | NA |
| -3.329278915 | 0.000870712 | 0.002539787 |
| NA | NA | NA |
| 0.312294397 | 0.754816798 | 0.822259844 |
| 0.7359102 | 0.461785349 | 0.571151829 |
| NA | NA | NA |
| -0.036481386 | 0.970898521 | 0.979876453 |
| -0.065388772 | 0.947864464 | NA |
| 2.750343225 | 0.005953287 | 0.014351418 |
| -0.172955366 | 0.862686511 | 0.904444435 |
| 0.935901965 | 0.349323665 | 0.460964326 |
| NA | NA | NA |
| -3.614374151 | 0.000301074 | 0.000956695 |
| -0.95302871 | 0.340575524 | 0.452089806 |

| | | |
|---|---|---|
| -3.026975565 | 0.002470139 | 0.006504059 |
| 0.410151397 | 0.681694892 | 0.764346299 |
| -0.404100119 | 0.686139097 | 0.76813944 |
| 3.118437833 | 0.001818125 | 0.004941651 |
| NA | NA | NA |
| 0.13263931 | 0.894478642 | 0.926458678 |
| 0.360534001 | 0.718447834 | 0.79418581 |
| -0.542633929 | 0.587381874 | 0.687242009 |
| -3.013754953 | 0.002580361 | 0.006767939 |
| -1.515331625 | 0.129688443 | 0.2070149 |
| -0.686179431 | 0.492599967 | 0.599843209 |
| NA | NA | NA |
| 0.660295365 | 0.509064304 | 0.615074316 |
| 0.448726544 | 0.653628934 | NA |
| NA | NA | NA |
| 3.503775611 | 0.000458712 | 0.001407743 |
| NA | NA | NA |
| NA | NA | NA |
| 0.392023906 | 0.69504055 | NA |
| NA | NA | NA |
| 6.819060208 | 9.16E-12 | 7.69E-11 |
| NA | NA | NA |
| NA | NA | NA |
| 10.59414908 | 3.17E-26 | 8.76E-25 |
| 4.253732365 | 2.10E-05 | 8.14E-05 |
| 0.063708285 | 0.949202508 | NA |
| 2.165980508 | 0.030312676 | 0.059888412 |
| -4.758279861 | 1.95E-06 | 8.81E-06 |
| NA | NA | NA |
| -2.542789435 | 0.010997148 | 0.024802542 |
| -3.084549075 | 0.00203861 | 0.005483172 |
| 0.192809521 | 0.847108147 | NA |
| -5.049218741 | 4.44E-07 | 2.19E-06 |
| 3.073613037 | 0.002114836 | 0.005665061 |
| 3.447387598 | 0.000566036 | 0.001709477 |
| NA | NA | NA |
| 0.192809521 | 0.847108147 | NA |
| 1.638628506 | 0.101290652 | 0.168313418 |
| 0.601187838 | 0.547714884 | 0.651211455 |
| 1.2087782 | 0.226748069 | 0.326925864 |
| -0.739873501 | 0.459376755 | 0.568767322 |
| 1.309430259 | 0.190388647 | 0.283994508 |
| NA | NA | NA |
| NA | NA | NA |
| NA | NA | NA |
| NA | NA | NA |
| NA | NA | NA |

| | | | |
|---|---|---|---|
| NA | NA | NA | |
| NA | NA | NA | |
| NA | NA | NA | |
| NA | NA | NA | |
| NA | NA | NA | |
| NA | NA | NA | |
| NA | NA | NA | |
| NA | NA | NA | |
| NA | NA | NA | |
| | -0.651951388 | 0.514432533 | 0.620004155 |
| | -0.765994174 | 0.443679769 | 0.553714829 |
| | -1.631058403 | 0.102877997 | 0.170453184 |
| | 0.295622597 | 0.767518314 | NA |
| NA | NA | NA | |
| | -0.194487022 | 0.845794556 | NA |
| | -4.868530311 | 1.12E-06 | 5.23E-06 |
| | 0.504006247 | 0.614256987 | 0.70979421 |
| | 0.063710673 | 0.949200606 | NA |
| | 4.438925783 | 9.04E-06 | 3.71E-05 |
| | 3.615367831 | 0.000299921 | 0.000953516 |
| | -0.943823759 | 0.345259717 | 0.456757438 |
| NA | NA | NA | |
| | -2.59230963 | 0.009533393 | 0.02182574 |
| NA | NA | NA | |
| | -3.522246051 | 0.000427907 | 0.001320817 |
| NA | NA | NA | |
| NA | NA | NA | |
| NA | NA | NA | |
| NA | NA | NA | |
| NA | NA | NA | |
| NA | NA | NA | |
| NA | NA | NA | |
| NA | NA | NA | |
| NA | NA | NA | |
| NA | NA | NA | |
| NA | NA | NA | |
| NA | NA | NA | |
| NA | NA | NA | |
| NA | NA | NA | |
| | -0.116243422 | 0.907459624 | NA |
| | 0.159490825 | 0.873282187 | NA |
| | -2.46308372 | 0.013774775 | 0.030241845 |
| NA | NA | NA | |
| | -2.403801021 | 0.016225602 | 0.034935641 |
| NA | NA | NA | |
| | -4.031502903 | 5.54E-05 | 0.000200524 |
| | -1.132605866 | 0.257379804 | 0.361983283 |

| | | |
|---|---|---|
| 0.951163436 | 0.341521417 | 0.453045055 |
| 3.544436918 | 0.000393453 | 0.001223217 |
| NA | NA | NA |
| NA | NA | NA |
| NA | NA | NA |
| -0.922898887 | 0.356059901 | 0.467983464 |
| -4.268509356 | 1.97E-05 | 7.66E-05 |
| -0.522735539 | 0.601158303 | 0.6987551 |
| -0.065388772 | 0.947864464 | NA |
| -5.09109072 | 3.56E-07 | 1.78E-06 |
| NA | NA | NA |
| -0.188683455 | 0.85034092 | NA |
| 0.585582452 | 0.558156144 | NA |
| 0.063708285 | 0.949202508 | NA |
| -0.36369102 | 0.71608875 | 0.792196125 |
| -0.959253529 | 0.33743104 | 0.44863979 |
| 3.119601442 | 0.001810959 | 0.004924848 |
| 6.072067057 | 1.26E-09 | 8.50E-09 |
| 0.586277721 | 0.5576889 | NA |
| NA | NA | NA |
| -6.872583696 | 6.30E-12 | 5.37E-11 |
| 0.069839618 | 0.944321313 | NA |
| -0.065389966 | 0.947863513 | NA |
| -0.134073176 | 0.893344711 | NA |
| -5.440420247 | 5.32E-08 | 2.97E-07 |
| -1.893440112 | 0.058299365 | 0.105512156 |
| 2.730089108 | 0.006331721 | 0.015165476 |
| NA | NA | NA |
| NA | NA | NA |
| NA | NA | NA |
| NA | NA | NA |
| 3.847589958 | 0.000119285 | 0.00040694 |
| -5.417247288 | 6.05E-08 | 3.36E-07 |
| 3.22105838 | 0.001277181 | 0.003592297 |
| NA | NA | NA |
| -4.811180995 | 1.50E-06 | 6.87E-06 |
| -0.611127141 | 0.541115413 | 0.645179286 |
| -2.157855692 | 0.030939053 | 0.060967131 |
| 0.008688479 | 0.993067684 | 0.995159569 |
| 4.862697513 | 1.16E-06 | 5.38E-06 |
| 0.842747146 | 0.399369874 | 0.511206207 |
| NA | NA | NA |
| 0.192808327 | 0.847109082 | NA |
| 0.213395169 | 0.831018748 | 0.881821234 |
| 9.097038764 | 9.28E-20 | 1.63E-18 |
| -0.083000608 | 0.933851056 | NA |
| 0.419243822 | 0.675037949 | NA |

| | | |
|---|---|---|
| NA | NA | NA |
| NA | NA | NA |
| 0.889224706 | 0.373882327 | 0.486649748 |
| -5.308674777 | 1.10E-07 | 5.92E-07 |
| -0.092527504 | 0.92627894 | NA |
| -4.00509814 | 6.20E-05 | 0.000222173 |
| -0.315357106 | 0.752490531 | 0.820494715 |
| 0.195316213 | 0.845145406 | NA |
| NA | NA | NA |
| NA | NA | NA |
| -9.453398908 | 3.28E-21 | 6.41E-20 |
| NA | NA | NA |
| -2.100672883 | 0.035669691 | 0.068990988 |
| NA | NA | NA |
| NA | NA | NA |
| 0.335713995 | 0.737086551 | 0.808505034 |
| 3.15005176 | 0.001632415 | 0.004488559 |
| 0.063708285 | 0.949202508 | NA |
| 8.369280699 | 5.80E-17 | 8.04E-16 |
| -0.686791342 | 0.492214227 | 0.599490151 |
| -1.129694644 | 0.258604915 | 0.363481638 |
| -1.008669266 | 0.313133276 | 0.423477734 |
| -2.583751685 | 0.009773213 | 0.022315592 |
| -2.295413417 | 0.021709443 | 0.04485985 |
| 1.532737103 | 0.125340643 | 0.201301014 |
| -5.987211696 | 2.13E-09 | 1.40E-08 |
| 0.063708882 | 0.949202033 | NA |
| 1.89129293 | 0.058585252 | 0.105968265 |
| -0.065389369 | 0.947863988 | NA |
| -4.725514022 | 2.30E-06 | 1.03E-05 |
| -0.312702278 | 0.754506865 | 0.821993768 |
| -2.214806132 | 0.026773374 | 0.05378311 |
| -1.487150012 | 0.136975195 | 0.216513365 |
| -1.875090049 | 0.060780336 | 0.109196944 |
| -0.726280526 | 0.467666815 | 0.576893018 |
| -0.065389369 | 0.947863988 | NA |
| 0.192809521 | 0.847108147 | NA |
| 1.621714712 | 0.104864447 | 0.173226879 |
| 2.753267572 | 0.005900365 | 0.014241672 |
| NA | NA | NA |
| 2.926732572 | 0.003425433 | 0.008733685 |
| 1.971463533 | 0.04867088 | 0.090433682 |
| 0.261359224 | 0.793815499 | NA |
| 5.494129928 | 3.93E-08 | 2.23E-07 |
| -1.517690555 | 0.129092426 | 0.206267694 |
| NA | NA | NA |
| 1.072097951 | 0.283676039 | 0.39066912 |

| | | |
|---|---|---|
| 0.200068104 | 0.841427318 | NA |
| NA | NA | NA |
| 9.689361928 | 3.35E-22 | 7.01E-21 |
| 0.836265343 | 0.403005645 | 0.514656974 |
| -3.12116365 | 0.001801379 | 0.004901991 |
| NA | NA | NA |
| NA | NA | NA |
| -2.352259951 | 0.018659729 | 0.039360613 |
| -1.121293967 | 0.262162753 | 0.367471227 |
| 10.42136299 | 1.98E-25 | 5.21E-24 |
| -1.202951757 | 0.22899499 | 0.329486353 |
| -0.761365163 | 0.446438989 | 0.55640512 |
| 11.84605754 | 2.26E-32 | 8.79E-31 |
| NA | NA | NA |
| -1.565712718 | 0.117415887 | 0.190332211 |
| 3.418379724 | 0.000629952 | 0.001885861 |
| 0.68569479 | 0.492905593 | NA |
| NA | NA | NA |
| NA | NA | NA |
| -0.065388772 | 0.947864464 | NA |
| NA | NA | NA |
| NA | NA | NA |
| -0.408950968 | 0.682575645 | NA |
| 7.954306708 | 1.80E-15 | 2.20E-14 |
| -1.267344742 | 0.205032049 | 0.301577962 |
| 0.668432743 | 0.503857401 | NA |
| NA | NA | NA |
| NA | NA | NA |
| -1.570682055 | 0.116256519 | 0.188733992 |
| -0.619126824 | 0.535832814 | 0.640185935 |
| -2.078916777 | 0.037625 | 0.07227026 |
| -2.0509876 | 0.040268154 | 0.076665357 |
| -3.778047985 | 0.000158062 | 0.000527577 |
| NA | NA | NA |
| 0.063708882 | 0.949202033 | NA |
| 11.4379167 | 2.70E-30 | 9.44E-29 |
| 9.103127514 | 8.78E-20 | 1.54E-18 |
| 6.353806611 | 2.10E-10 | 1.54E-09 |
| 2.519185008 | 0.011762683 | 0.026322751 |
| NA | NA | NA |
| 8.409410258 | 4.12E-17 | 5.80E-16 |
| 0.849146134 | 0.395799982 | 0.507732347 |
| NA | NA | NA |
| 1.080309095 | 0.280004561 | NA |
| NA | NA | NA |
| 1.559448079 | 0.118890364 | NA |
| 0.192810118 | 0.847107679 | NA |

| | | |
|---|---|---|
| NA | NA | NA |
| 1.718443654 | 0.085715724 | 0.146388139 |
| -0.616735402 | 0.537409266 | 0.641640619 |
| 2.046395901 | 0.040717435 | 0.077414159 |
| -0.981249641 | 0.32646965 | 0.437542051 |
| -2.608332068 | 0.009098464 | 0.020927638 |
| 0.207171563 | 0.835875875 | NA |
| -0.63125334 | 0.527874891 | NA |
| 1.819974114 | 0.068762947 | 0.121412004 |
| -10.86801089 | 1.64E-27 | 4.93E-26 |
| 10.77759148 | 4.39E-27 | 1.28E-25 |
| -5.69343236 | 1.25E-08 | 7.52E-08 |
| 5.503024029 | 3.73E-08 | 2.13E-07 |
| -1.443198096 | 0.148964676 | 0.232079287 |
| -0.503095161 | 0.614897369 | 0.710403007 |
| -0.449935319 | 0.65275708 | 0.740908294 |
| -0.169837186 | 0.865138181 | 0.906032769 |
| 0.192809521 | 0.847108147 | NA |
| 4.167546555 | 3.08E-05 | 0.000116274 |
| 4.490704621 | 7.10E-06 | 2.96E-05 |
| -0.016699998 | 0.986675949 | 0.990633735 |
| 3.125137232 | 0.001777221 | 0.004843625 |
| NA | NA | NA |
| 2.31937301 | 0.02037482 | 0.042473702 |
| NA | NA | NA |
| 1.778873945 | 0.075260431 | 0.131017428 |
| -2.496579318 | 0.012539762 | 0.027840185 |
| 2.280808296 | 0.022559793 | 0.04644445 |
| NA | NA | NA |
| -5.592258092 | 2.24E-08 | 1.31E-07 |
| -2.716483617 | 0.006597946 | 0.015738739 |
| NA | NA | NA |
| -6.322938286 | 2.57E-10 | 1.86E-09 |
| -2.048212551 | 0.040539177 | 0.077110389 |
| NA | NA | NA |
| -0.545192026 | 0.585621455 | NA |
| -0.119618938 | 0.904785017 | NA |
| -2.690555125 | 0.007133325 | 0.016844068 |
| 4.839541947 | 1.30E-06 | 6.00E-06 |
| -0.543759439 | 0.586607024 | 0.686523994 |
| NA | NA | NA |
| 2.297208792 | 0.021606862 | 0.044666316 |
| 2.388111986 | 0.016935181 | 0.036248296 |
| 0.20108039 | 0.840635716 | 0.888222887 |
| 6.743932771 | 1.54E-11 | 1.27E-10 |
| 1.925638273 | 0.054149556 | 0.09915061 |
| -2.937905571 | 0.003304377 | 0.008454313 |

| | | | |
|---|---|---|---|
| | 4.458204797 | 8.26E-06 | 3.41E-05 |
| | 8.384755199 | 5.08E-17 | 7.07E-16 |
| | -8.061140634 | 7.56E-16 | 9.54E-15 |
| | 4.964125265 | 6.90E-07 | 3.31E-06 |
| | 0.063708882 | 0.949202033 | NA |
| | -4.633700497 | 3.59E-06 | 1.56E-05 |
| | -2.462417816 | 0.01380038 | 0.030289679 |
| | -9.845286182 | 7.18E-23 | 1.58E-21 |
| | -2.846531292 | 0.004419839 | 0.011004532 |
| | -0.252872845 | 0.800366474 | 0.858282544 |
| | -2.778067906 | 0.005468319 | 0.013324768 |
| | -2.988216369 | 0.002806108 | 0.007299614 |
| | -0.928562806 | 0.353115705 | 0.464987695 |
| | -1.035029834 | 0.300654978 | 0.409679031 |
| | -0.763204098 | 0.445341685 | 0.555179918 |
| | -2.258262301 | 0.023929311 | 0.048823028 |
| | 10.94624971 | 6.93E-28 | 2.15E-26 |
| | 5.256651208 | 1.47E-07 | 7.73E-07 |
| | -1.455244965 | 0.145601457 | 0.227775279 |
| | 12.96176124 | 2.02E-38 | 1.04E-36 |
| | -3.417571814 | 0.000631824 | 0.001890663 |
| | -0.782754585 | 0.433771241 | 0.544470897 |
| | 0.063708882 | 0.949202033 | NA |
| | -6.458697632 | 1.06E-10 | 8.01E-10 |
| NA | NA | NA | |
| | -2.350390239 | 0.018753738 | 0.039518947 |
| | -0.07026533 | 0.943982477 | 0.962111992 |
| | 10.72224138 | 8.00E-27 | 2.29E-25 |
| | 0.138178936 | 0.890099002 | 0.923088184 |
| | 0.84434191 | 0.398478373 | 0.510357349 |
| | 3.370583824 | 0.000750091 | 0.002215051 |
| | 11.62777788 | 2.98E-31 | 1.10E-29 |
| | 2.052189986 | 0.040151202 | 0.076471206 |
| | 0.982959438 | 0.325627402 | 0.436717242 |
| | 2.592462997 | 0.009529143 | 0.021818007 |
| | 1.777520953 | 0.075482566 | 0.131349321 |
| | -1.390266487 | 0.164447971 | 0.251628115 |
| | 1.796715883 | 0.072380736 | 0.126753643 |
| | 1.300199155 | 0.19353272 | 0.287792537 |
| | 5.748291332 | 9.01E-09 | 5.54E-08 |
| | -0.643693269 | 0.519774355 | 0.625027726 |
| NA | NA | NA | |
| | -0.800515327 | 0.423412287 | 0.535063843 |
| | -0.550653701 | 0.581871089 | 0.68219885 |
| | 0.133506226 | 0.89379304 | NA |
| | 0.192810118 | 0.847107679 | NA |
| | -0.065388772 | 0.947864464 | NA |

| | | | |
|---|---|---|---|
| NA | NA | NA | |
| | 0.248664424 | 0.80362037 | NA |
| NA | NA | NA | |
| | -1.198434089 | 0.230748068 | 0.331513267 |
| NA | NA | NA | |
| | -0.065388772 | 0.947864464 | NA |
| | 1.926430826 | 0.054050601 | 0.099011018 |
| | 4.67026986 | 3.01E-06 | 1.32E-05 |
| | 6.613948933 | 3.74E-11 | 2.97E-10 |
| | -0.847809018 | 0.39654434 | 0.508400596 |
| | 0.293055392 | 0.769479812 | 0.833950193 |
| | 0.063708285 | 0.949202508 | NA |
| NA | NA | NA | |
| | 2.664180603 | 0.007717612 | 0.018048381 |
| | 7.084663585 | 1.39E-12 | 1.27E-11 |
| | -7.941239309 | 2.00E-15 | 2.43E-14 |
| | 2.54620734 | 0.010890048 | 0.024585362 |
| NA | NA | NA | |
| | -1.009271016 | 0.312844676 | 0.423247514 |
| | 11.69054784 | 1.42E-31 | 5.35E-30 |
| | 2.468379621 | 0.013572629 | 0.029852629 |
| | 3.628736323 | 0.000284812 | 0.000908943 |
| | 34.68491399 | 1.33E-263 | 2.56E-260 |
| | -1.302409691 | 0.192776371 | 0.286838222 |
| NA | NA | NA | |
| | -1.454185802 | 0.1458948 | 0.228148622 |
| | 4.037730978 | 5.40E-05 | 0.000195752 |
| | -0.065765347 | 0.947564646 | 0.96427637 |
| | -0.064942513 | 0.948219771 | 0.964700274 |
| | -0.683728865 | 0.494146393 | 0.601287517 |
| | 0.729835532 | 0.465490722 | 0.574826689 |
| NA | NA | NA | |
| | 1.330340853 | 0.183405993 | 0.275432769 |
| | -0.081381505 | 0.935138557 | NA |
| | -0.131670081 | 0.895245251 | NA |
| | -2.997615774 | 0.002721005 | 0.007097411 |
| | 3.096553259 | 0.001957847 | 0.005285257 |
| NA | NA | NA | |
| | -1.902709999 | 0.057078396 | 0.103699809 |
| | -1.452814453 | 0.146275278 | 0.228615061 |
| | 0.312829511 | 0.754410194 | NA |
| | -4.019702737 | 5.83E-05 | 0.000209952 |
| | 4.219339966 | 2.45E-05 | 9.38E-05 |
| | -3.713057803 | 0.00020477 | 0.000671831 |
| | -6.020613603 | 1.74E-09 | 1.15E-08 |
| | 11.44289432 | 2.55E-30 | 8.95E-29 |
| NA | NA | NA | |

| | | | |
|---|---|---|---|
| | 0.446134528 | 0.655500073 | NA |
| | 3.079475011 | 0.002073658 | 0.005564285 |
| | 4.594886862 | 4.33E-06 | 1.86E-05 |
| | 5.275036991 | 1.33E-07 | 7.04E-07 |
| NA | NA | NA | |
| | 3.707417279 | 0.000209384 | 0.00068553 |
| | -0.065389369 | 0.947863988 | NA |
| | 7.720650717 | 1.16E-14 | 1.31E-13 |
| | 0.295230597 | 0.76781773 | 0.832689678 |
| | 0.429819496 | 0.66732695 | 0.752531697 |
| | 0.508518839 | 0.611089531 | 0.707027638 |
| | -0.562496026 | 0.57377811 | 0.675017207 |
| | 7.63334476 | 2.29E-14 | 2.51E-13 |
| | -2.121038415 | 0.03391857 | 0.066104633 |
| | 5.41269451 | 6.21E-08 | 3.44E-07 |
| | 4.995805424 | 5.86E-07 | 2.84E-06 |
| | 2.691981516 | 0.007102888 | 0.01678645 |
| | 10.01160516 | 1.36E-23 | 3.16E-22 |
| | 4.673022241 | 2.97E-06 | 1.30E-05 |
| | 4.696188955 | 2.65E-06 | 1.17E-05 |
| | 5.409970733 | 6.30E-08 | 3.49E-07 |
| | 1.273625595 | 0.202796137 | 0.299081181 |
| | 4.003818355 | 6.23E-05 | 0.000223347 |
| | 10.55250571 | 4.95E-26 | 1.35E-24 |
| | 8.427630482 | 3.53E-17 | 4.98E-16 |
| | 0.354968218 | 0.722613401 | 0.797276445 |
| | 9.753114383 | 1.79E-22 | 3.82E-21 |
| | -0.157288573 | 0.875017426 | 0.912760415 |
| | 0.192810118 | 0.847107679 | NA |
| NA | NA | NA | |
| | -0.065389369 | 0.947863988 | NA |
| NA | NA | NA | |
| NA | NA | NA | |
| NA | NA | NA | |
| NA | NA | NA | |
| NA | NA | NA | |
| NA | NA | NA | |
| NA | NA | NA | |
| NA | NA | NA | |
| NA | NA | NA | |
| NA | NA | NA | |
| | 0.063708882 | 0.949202033 | NA |
| NA | NA | NA | |
| NA | NA | NA | |
| NA | NA | NA | |
| NA | NA | NA | |
| NA | NA | NA | |

| | | |
|---|---|---|
| NA | NA | NA |
| NA | NA | NA |
| 0.063708285 | 0.949202508 | NA |
| NA | NA | NA |
| NA | NA | NA |
| NA | NA | NA |
| NA | NA | NA |
| 0.06371127 | 0.949200131 | NA |
| NA | NA | NA |
| NA | NA | NA |
| NA | NA | NA |
| NA | NA | NA |
| NA | NA | NA |
| NA | NA | NA |
| 0.192809521 | 0.847108147 | NA |
| NA | NA | NA |
| NA | NA | NA |
| NA | NA | NA |
| NA | NA | NA |
| NA | NA | NA |
| NA | NA | NA |
| NA | NA | NA |
| NA | NA | NA |
| NA | NA | NA |
| NA | NA | NA |
| NA | NA | NA |
| 0.192809521 | 0.847108147 | NA |
| 0.192809521 | 0.847108147 | NA |
| NA | NA | NA |
| NA | NA | NA |
| NA | NA | NA |
| NA | NA | NA |
| NA | NA | NA |
| 0.063708882 | 0.949202033 | NA |
| NA | NA | NA |
| NA | NA | NA |
| NA | NA | NA |
| NA | NA | NA |
| NA | NA | NA |
| NA | NA | NA |
| NA | NA | NA |
| 0.063708882 | 0.949202033 | NA |
| NA | NA | NA |
| NA | NA | NA |
| 0.063708882 | 0.949202033 | NA |
| NA | NA | NA |
| 0.063708882 | 0.949202033 | NA |

| | | |
|---|---|---|
| 0.920759496 | 0.357176006 | 0.468977361 |
| 4.692396653 | 2.70E-06 | 1.19E-05 |
| -3.330164725 | 0.000867946 | 0.002532311 |
| 4.07264216 | 4.65E-05 | 0.000170072 |
| -3.892032555 | 9.94E-05 | 0.000343814 |
| -3.846490661 | 0.000119822 | 0.00040849 |
| -4.164449789 | 3.12E-05 | 0.000117717 |
| NA | NA | NA |
| 0.254117618 | 0.799404693 | NA |
| 0.459997179 | 0.645518245 | NA |
| -2.857198187 | 0.004273989 | 0.010670052 |
| -0.389630241 | 0.696809987 | 0.776681242 |
| 2.150160662 | 0.031542509 | 0.062014688 |
| 0.666536086 | 0.505068506 | 0.611545313 |
| 4.487720863 | 7.20E-06 | 2.99E-05 |
| -0.107463428 | 0.914421337 | 0.941194427 |
| -2.096793549 | 0.036011855 | 0.069577517 |
| 0.566452231 | 0.571086397 | 0.672482712 |
| -1.609621178 | 0.107480582 | 0.176652372 |
| 1.802804628 | 0.071418903 | 0.125385084 |
| -2.471928933 | 0.013438623 | 0.029589086 |
| 6.211784737 | 5.24E-10 | 3.67E-09 |
| -0.334771259 | 0.737797643 | 0.809143189 |
| -2.217934868 | 0.026559274 | 0.053391585 |
| 4.109799877 | 3.96E-05 | 0.000146634 |
| -2.338666009 | 0.019352726 | 0.040606817 |
| 0.033942362 | 0.972923112 | 0.98135263 |
| -0.5645024 | 0.572412267 | NA |
| 5.995355687 | 2.03E-09 | 1.34E-08 |
| -6.271321644 | 3.58E-10 | 2.55E-09 |
| 2.458506571 | 0.013951622 | 0.030592125 |
| 3.064914427 | 0.002177323 | 0.005817687 |
| 0.848897061 | 0.395938574 | 0.507832053 |
| NA | NA | NA |
| NA | NA | NA |
| NA | NA | NA |
| 0.835536566 | 0.403415668 | NA |
| 3.600760579 | 0.000317288 | 0.001004644 |
| NA | NA | NA |
| NA | NA | NA |
| NA | NA | NA |
| NA | NA | NA |
| 2.934982866 | 0.003335661 | 0.008526511 |
| 0.201524559 | 0.840288428 | NA |
| -2.656793961 | 0.007888764 | 0.0184005 |
| -8.327821132 | 8.23E-17 | 1.13E-15 |
| 0.192808924 | 0.847108615 | NA |

| | | | |
|---|---|---|---|
| NA | NA | NA | |
| | 4.220443091 | 2.44E-05 | 9.34E-05 |
| | 2.481789887 | 0.013072433 | 0.028886996 |
| | 5.292879209 | 1.20E-07 | 6.42E-07 |
| | 7.587212725 | 3.27E-14 | 3.52E-13 |
| | -0.83101726 | 0.405963879 | 0.517798404 |
| | -1.08245855 | 0.279048824 | 0.386059698 |
| | 7.689186934 | 1.48E-14 | 1.66E-13 |
| | 0.082800965 | 0.934009802 | 0.955111547 |
| | -0.491339605 | 0.623186273 | 0.717132656 |
| | -0.691555521 | 0.489216502 | 0.597030183 |
| | -2.832527434 | 0.00461816 | 0.011455015 |
| | -5.936926103 | 2.90E-09 | 1.88E-08 |
| | -3.806197052 | 0.00014112 | 0.00047521 |
| NA | NA | NA | |
| | 0.449852648 | 0.652816693 | 0.740942358 |
| | 3.666347489 | 0.00024604 | 0.000794014 |
| | -0.640506556 | 0.521843328 | 0.627123992 |
| | 0.022443652 | 0.98209406 | 0.987641213 |
| | 0.275053149 | 0.783275406 | 0.845314681 |
| | 7.387958265 | 1.49E-13 | 1.50E-12 |
| | 5.125790351 | 2.96E-07 | 1.50E-06 |
| | 0.063708882 | 0.949202033 | NA |
| | 8.684038709 | 3.82E-18 | 5.83E-17 |
| | 11.36048462 | 6.58E-30 | 2.25E-28 |
| NA | NA | NA | |
| NA | NA | NA | |
| | -1.127038988 | 0.259725997 | 0.364750138 |
| | 2.298703646 | 0.021521773 | 0.044523513 |
| | 2.549558827 | 0.010785931 | 0.024396517 |
| | -0.106971675 | 0.914811451 | 0.941363699 |
| NA | NA | NA | |
| | 0.752499578 | 0.451750684 | NA |
| | 10.33819126 | 4.73E-25 | 1.21E-23 |
| | 1.154233968 | 0.24840426 | 0.351671868 |
| | 2.86060059 | 0.004228394 | 0.010570985 |
| | 0.04335894 | 0.965415408 | 0.976574025 |
| | 3.082563833 | 0.002052257 | 0.005515138 |
| | -4.913685793 | 8.94E-07 | 4.22E-06 |
| | -0.64975718 | 0.515849082 | NA |
| | 1.25921254 | 0.207953573 | 0.304962645 |
| | -2.346815728 | 0.018934617 | 0.039843081 |
| | -1.115653097 | 0.264570655 | 0.370162297 |
| | -1.021350391 | 0.30708846 | 0.416834282 |
| | 0.067524971 | 0.946163784 | NA |
| NA | NA | NA | |
| | 4.031939265 | 5.53E-05 | 0.000200206 |

| | | |
|---|---|---|
| -2.195725481 | 0.028111598 | 0.056117217 |
| 0.219410159 | 0.826330557 | 0.87852373 |
| 0.122570042 | 0.902447578 | 0.932166676 |
| -3.694101855 | 0.000220665 | 0.000718986 |
| 7.749968688 | 9.19E-15 | 1.05E-13 |
| NA | NA | NA |
| 1.090724724 | 0.275394028 | 0.382059224 |
| NA | NA | NA |
| NA | NA | NA |
| 6.089207619 | 1.13E-09 | 7.69E-09 |
| -2.006076176 | 0.044848121 | 0.084262759 |
| NA | NA | NA |
| -0.065388772 | 0.947864464 | NA |
| 0.481843553 | 0.629917087 | 0.722343059 |
| 0.194173692 | 0.846039881 | NA |
| 3.353651081 | 0.000797529 | 0.002341051 |
| 1.098800092 | 0.271855272 | 0.378303077 |
| -2.412713768 | 0.015834249 | 0.034184169 |
| 10.09132364 | 6.04E-24 | 1.43E-22 |
| -5.12611268 | 2.96E-07 | 1.50E-06 |
| 1.413335115 | 0.157557214 | 0.24303057 |
| 2.195118884 | 0.028155071 | 0.056175482 |
| 4.732023037 | 2.22E-06 | 9.96E-06 |
| 2.547048292 | 0.01086384 | 0.024543904 |
| -0.137499223 | 0.890636207 | NA |
| 0.657856936 | 0.510630064 | 0.616400796 |
| 0.260037194 | 0.794835084 | NA |
| 9.134803467 | 6.55E-20 | 1.16E-18 |
| -6.289105572 | 3.19E-10 | 2.28E-09 |
| -3.31582135 | 0.000913742 | 0.002652014 |
| -0.824713999 | 0.409533982 | 0.521448549 |
| -0.924881342 | 0.355027633 | 0.466964879 |
| 1.386313233 | 0.165651278 | 0.253159817 |
| 1.623155807 | 0.104556102 | 0.172821716 |
| -0.08091295 | 0.935511181 | NA |
| 5.689229226 | 1.28E-08 | 7.70E-08 |
| 3.326383582 | 0.000879808 | 0.002564526 |
| NA | NA | NA |
| -3.364285619 | 0.000767421 | 0.002260094 |
| -4.26340035 | 2.01E-05 | 7.83E-05 |
| -1.10393345 | 0.269622007 | 0.375776845 |
| -0.56473513 | 0.572253936 | 0.673509006 |
| -2.223074878 | 0.026210751 | 0.052822472 |
| 0.019480701 | 0.984457633 | NA |
| 0.405492012 | 0.685115889 | NA |
| -0.086025746 | 0.931445951 | NA |
| -4.844789989 | 1.27E-06 | 5.86E-06 |

| | | |
|---|---|---|
| NA | NA | NA |
| 1.523214806 | 0.127704973 | 0.204507708 |
| 14.52013808 | 9.03E-48 | 6.59E-46 |
| 7.204991133 | 5.80E-13 | 5.53E-12 |
| NA | NA | NA |
| 0.063708882 | 0.949202033 | NA |
| 1.631943452 | 0.102691399 | 0.170245445 |
| NA | NA | NA |
| -2.762791319 | 0.005730939 | 0.013887638 |
| 3.785832758 | 0.000153195 | 0.000512903 |
| -0.811528895 | 0.417062006 | 0.528747506 |
| -1.613415321 | 0.106654307 | 0.175528973 |
| 0.192809521 | 0.847108147 | NA |
| -0.656427665 | 0.511548995 | NA |
| -4.12426207 | 3.72E-05 | 0.000138432 |
| 0.192809521 | 0.847108147 | NA |
| NA | NA | NA |
| NA | NA | NA |
| 0.127088718 | 0.898870181 | NA |
| NA | NA | NA |
| NA | NA | NA |
| 0.488367962 | 0.625289231 | NA |
| NA | NA | NA |
| -9.84917237 | 6.91E-23 | 1.52E-21 |
| NA | NA | NA |
| 2.358955429 | 0.018326457 | 0.038762144 |
| 10.79095568 | 3.80E-27 | 1.11E-25 |
| 0.646392946 | 0.518024907 | NA |
| NA | NA | NA |
| NA | NA | NA |
| NA | NA | NA |
| NA | NA | NA |
| 0.345444204 | 0.72976045 | 0.802781493 |
| 4.69235111 | 2.70E-06 | 1.19E-05 |
| NA | NA | NA |
| 0.444654634 | 0.656569359 | 0.744121849 |
| NA | NA | NA |
| 5.10661974 | 3.28E-07 | 1.65E-06 |
| 0.272593468 | 0.785165734 | 0.846770576 |
| 1.653400579 | 0.098249365 | 0.164186247 |
| 10.71791194 | 8.39E-27 | 2.40E-25 |
| 1.49081027 | 0.136011315 | 0.215207232 |
| 2.105476586 | 0.035249843 | 0.06832677 |
| NA | NA | NA |
| 1.37857338 | 0.168026329 | 0.256132704 |
| NA | NA | NA |
| NA | NA | NA |

| | | |
|---|---|---|
| -3.54315204 | 0.000395375 | 0.001228277 |
| -1.513668561 | 0.130109923 | 0.207535774 |
| -3.144956825 | 0.001661113 | 0.004559955 |
| -0.124196093 | 0.901160016 | NA |
| -0.890297509 | 0.373306158 | NA |
| 0.311351029 | 0.755533778 | 0.822754425 |
| 2.104501818 | 0.035334696 | 0.068459435 |
| 0.429236314 | 0.667751258 | NA |
| -0.065389966 | 0.947863513 | NA |
| 0.495961646 | 0.619921471 | NA |
| 16.34150312 | 5.00E-60 | 5.03E-58 |
| 0.063708285 | 0.949202508 | NA |
| 5.305635468 | 1.12E-07 | 6.01E-07 |
| -0.457124172 | 0.647581797 | 0.736470188 |
| 0.791338052 | 0.428746749 | NA |
| NA | NA | NA |
| -0.420147598 | 0.674377633 | 0.757989679 |
| NA | NA | NA |
| 0.192810118 | 0.847107679 | NA |
| -4.139505588 | 3.48E-05 | 0.000130241 |
| 0.764361136 | 0.444652062 | 0.554568794 |
| NA | NA | NA |
| NA | NA | NA |
| 1.221611045 | 0.22185475 | 0.321225452 |
| NA | NA | NA |
| 1.031655059 | 0.302233738 | NA |
| NA | NA | NA |
| NA | NA | NA |
| NA | NA | NA |
| NA | NA | NA |
| NA | NA | NA |
| NA | NA | NA |
| 1.211348352 | 0.225761925 | 0.325827191 |
| NA | NA | NA |
| NA | NA | NA |
| NA | NA | NA |
| NA | NA | NA |
| NA | NA | NA |
| -8.330404437 | 8.06E-17 | 1.10E-15 |
| -2.451648532 | 0.014220348 | 0.031069773 |
| 5.984437195 | 2.17E-09 | 1.43E-08 |
| 3.164663289 | 0.001552625 | 0.004288953 |
| -1.901636341 | 0.057218714 | 0.103901944 |
| NA | NA | NA |
| NA | NA | NA |
| NA | NA | NA |
| 3.301844028 | 0.000960515 | 0.002778422 |

| | | |
|---|---|---|
| 2.701340431 | 0.006906061 | 0.016381606 |
| 4.883642212 | 1.04E-06 | 4.88E-06 |
| 0.943221412 | 0.345567663 | 0.4570682 |
| 19.77142697 | 5.25E-87 | 9.73E-85 |
| NA | NA | NA |
| -4.640457456 | 3.48E-06 | 1.51E-05 |
| 1.081071648 | 0.279665244 | 0.38663469 |
| -7.212781811 | 5.48E-13 | 5.24E-12 |
| 0.426654185 | 0.669631225 | 0.754410306 |
| 6.873844839 | 6.25E-12 | 5.33E-11 |
| 1.91672507 | 0.055272869 | 0.100902917 |
| NA | NA | NA |
| NA | NA | NA |
| NA | NA | NA |
| 1.027404561 | 0.304230005 | 0.413717104 |
| 1.41516491 | 0.15702015 | 0.24250104 |
| 0.087854441 | 0.929992367 | 0.952443083 |
| 0.06371127 | 0.949200131 | NA |
| NA | NA | NA |
| 0.521817537 | 0.601797378 | 0.699368018 |
| NA | NA | NA |
| 1.014589884 | 0.310301363 | 0.420420219 |
| -0.216782542 | 0.828377818 | 0.879951482 |
| -0.065393498 | 0.947860701 | NA |
| 0.464569096 | 0.642240067 | 0.732491975 |
| NA | NA | NA |
| -0.074832904 | 0.940347661 | 0.959436734 |
| NA | NA | NA |
| -0.430615639 | 0.666747868 | 0.752183673 |
| -2.526582547 | 0.011517832 | 0.025846452 |
| -2.085771229 | 0.036999351 | 0.071237981 |
| -1.200886044 | 0.229795409 | 0.330391124 |
| -2.051031048 | 0.040263923 | 0.076665357 |
| 1.060596954 | 0.288873107 | NA |
| 2.027750818 | 0.042585686 | 0.080513613 |
| -0.382094002 | 0.702391641 | NA |
| -0.402974324 | 0.686967113 | NA |
| -0.065389369 | 0.947863988 | NA |
| 0.31685483 | 0.751353758 | NA |
| 0.3474247 | 0.728272279 | NA |
| 0.063708285 | 0.949202508 | NA |
| -4.715692989 | 2.41E-06 | 1.07E-05 |
| NA | NA | NA |
| NA | NA | NA |
| -2.653139038 | 0.007974701 | 0.018574984 |
| 4.827829591 | 1.38E-06 | 6.34E-06 |
| NA | NA | NA |

| | | |
|---|---|---|
| -1.670181231 | 0.094883512 | 0.159541469 |
| 0.132629791 | 0.894486171 | NA |
| 4.385061929 | 1.16E-05 | 4.68E-05 |
| -1.06368283 | 0.287472405 | 0.39487621 |
| NA | NA | NA |
| 6.71643144 | 1.86E-11 | 1.52E-10 |
| -0.387059178 | 0.698712398 | 0.778281844 |
| -1.463636107 | 0.143293402 | 0.224839132 |
| 2.566655334 | 0.010268462 | 0.023339957 |
| NA | NA | NA |
| 8.517860793 | 1.63E-17 | 2.36E-16 |
| NA | NA | NA |
| 1.070219787 | 0.28452039 | 0.391659537 |
| 2.255668543 | 0.024091397 | 0.049076884 |
| -0.065389369 | 0.947863988 | NA |
| NA | NA | NA |
| NA | NA | NA |
| 8.852176154 | 8.58E-19 | 1.40E-17 |
| 0.461323878 | 0.644566256 | 0.733974495 |
| NA | NA | NA |
| 0.063706581 | 0.949203865 | NA |
| 0.521295443 | 0.602160975 | NA |
| 8.779620642 | 1.64E-18 | 2.60E-17 |
| NA | NA | NA |
| NA | NA | NA |
| 0.107734654 | 0.91420618 | 0.941046355 |
| 2.378533375 | 0.017381663 | 0.037051868 |
| -1.78224891 | 0.07470865 | 0.130196468 |
| 0.354136188 | 0.723236823 | 0.797812136 |
| NA | NA | NA |
| NA | NA | NA |
| NA | NA | NA |
| NA | NA | NA |
| -3.696958706 | 0.000218198 | 0.00071178 |
| -1.79443769 | 0.072743336 | 0.127268951 |
| -0.572220676 | 0.567172482 | 0.668943889 |
| -0.276824054 | 0.78191521 | 0.844356431 |
| 0.250441544 | 0.802245906 | NA |
| -0.065388772 | 0.947864464 | NA |
| 1.239549193 | 0.215142183 | 0.313419949 |
| -5.474504243 | 4.39E-08 | 2.48E-07 |
| NA | NA | NA |
| -0.078634545 | 0.937323309 | NA |
| NA | NA | NA |
| 2.883760335 | 0.003929577 | 0.009896962 |
| 0.063707688 | 0.949202983 | NA |
| 0.132896263 | 0.894275422 | 0.926286523 |

| | | | |
|---|---|---|---|
| | -0.257722191 | 0.796621318 | NA |
| | -0.497518508 | 0.618823455 | NA |
| NA | NA | NA | |
| | -1.439695291 | 0.149953626 | 0.233387593 |
| | -0.681718911 | 0.495416707 | 0.602511063 |
| | 0.063708285 | 0.949202508 | NA |
| | 2.619372698 | 0.008809165 | 0.020321971 |
| | 0.324874675 | 0.745275925 | NA |
| NA | NA | NA | |
| | 1.846160049 | 0.06486897 | 0.115424065 |
| NA | NA | NA | |
| | 1.406990871 | 0.159430096 | 0.245299686 |
| | -1.817512774 | 0.069138634 | 0.121946305 |
| | 6.234260211 | 4.54E-10 | 3.20E-09 |
| | 0.276830269 | 0.781910438 | 0.844356431 |
| | -0.113891368 | 0.909323908 | NA |
| | 0.747219405 | 0.454931132 | 0.564771662 |
| | 0.192809521 | 0.847108147 | NA |
| NA | NA | NA | |
| | 6.371062495 | 1.88E-10 | 1.38E-09 |
| | -5.930772067 | 3.02E-09 | 1.95E-08 |
| | -1.596996297 | 0.110266533 | 0.180508261 |
| | -2.737392397 | 0.006192838 | 0.014874458 |
| | -0.267555363 | 0.789041594 | NA |
| | -0.053513074 | 0.957323114 | 0.9705848 |
| | 0.585931511 | 0.55792154 | 0.660398002 |
| | -1.87196366 | 0.061211631 | 0.109885041 |
| | -2.730388645 | 0.00632597 | 0.015154961 |
| | -1.784180424 | 0.074394352 | 0.129726253 |
| | 4.152342654 | 3.29E-05 | 0.000123587 |
| | 1.731682846 | 0.083330046 | 0.142977177 |
| | 8.059800945 | 7.64E-16 | 9.63E-15 |
| | 0.192810118 | 0.847107679 | NA |
| | -1.134984394 | 0.256381857 | 0.361026041 |
| | -0.797984359 | 0.424879567 | 0.536484667 |
| NA | NA | NA | |
| NA | NA | NA | |
| | -1.357989335 | 0.174467068 | 0.26438919 |
| | -3.55717092 | 0.00037487 | 0.001169805 |
| | -6.353319602 | 2.11E-10 | 1.54E-09 |
| NA | NA | NA | |
| | -0.152039653 | 0.879155661 | NA |
| NA | NA | NA | |
| | 0.063707194 | 0.949203376 | NA |
| | 1.076126472 | 0.281870698 | 0.388924869 |
| | 0.192810118 | 0.847107679 | NA |
| NA | NA | NA | |

| | | |
|---|---|---|
| NA | NA | NA |
| NA | NA | NA |
| NA | NA | NA |
| NA | NA | NA |
| 0.171383166 | 0.863922491 | 0.905433835 |
| NA | NA | NA |
| -0.24662222 | 0.805200606 | NA |
| -4.80749756 | 1.53E-06 | 6.99E-06 |
| NA | NA | NA |
| NA | NA | NA |
| NA | NA | NA |
| -0.358591543 | 0.719900674 | 0.795124608 |
| -0.950617958 | 0.341798349 | 0.453236495 |
| 6.892845877 | 5.47E-12 | 4.70E-11 |
| 0.192808924 | 0.847108615 | NA |
| NA | NA | NA |
| -1.178073692 | 0.238767229 | 0.34094037 |
| -0.214845067 | 0.829888115 | NA |
| NA | NA | NA |
| NA | NA | NA |
| NA | NA | NA |
| NA | NA | NA |
| NA | NA | NA |
| NA | NA | NA |
| NA | NA | NA |
| NA | NA | NA |
| 0.116778639 | 0.907035472 | 0.935708604 |
| NA | NA | NA |
| 1.705589532 | 0.08808453 | 0.149906916 |
| NA | NA | NA |
| -9.432730295 | 4.00E-21 | 7.75E-20 |
| -0.544284763 | 0.58624553 | 0.686265346 |
| NA | NA | NA |
| NA | NA | NA |
| 0.196692259 | 0.844068368 | 0.890970145 |
| 0.192809521 | 0.847108147 | NA |
| NA | NA | NA |
| 3.513888533 | 0.000441598 | 0.001360561 |
| 1.049462087 | 0.293965496 | NA |
| 15.53376486 | 2.05E-54 | 1.75E-52 |
| -1.047036857 | 0.29508258 | 0.403356938 |
| 7.720448265 | 1.16E-14 | 1.31E-13 |
| -2.240243647 | 0.025075109 | 0.050853183 |
| NA | NA | NA |
| 0.192808924 | 0.847108615 | NA |
| NA | NA | NA |
| 0.193341124 | 0.846691821 | NA |

| | | |
|---|---|---|
| 1.109470496 | 0.267227265 | 0.373056497 |
| 4.540377591 | 5.62E-06 | 2.37E-05 |
| -2.215136508 | 0.026750696 | 0.053741868 |
| -2.631607248 | 0.008498204 | 0.019688083 |
| 2.326214503 | 0.02000711 | 0.04180029 |
| 1.351623998 | 0.176495629 | 0.266849025 |
| NA | NA | NA |
| 0.074978317 | 0.940231964 | 0.959357795 |
| 0.548227804 | 0.583535493 | 0.683668479 |
| 0.126721802 | 0.899160589 | 0.929769047 |
| 0.139120144 | 0.88935521 | NA |
| -0.191904085 | 0.847817338 | 0.893736016 |
| -6.33464612 | 2.38E-10 | 1.73E-09 |
| -4.183672402 | 2.87E-05 | 0.000108781 |
| 0.03201598 | 0.974459307 | NA |
| NA | NA | NA |
| NA | NA | NA |
| -6.733152625 | 1.66E-11 | 1.36E-10 |
| 0.063708285 | 0.949202508 | NA |
| -2.132152956 | 0.032994274 | 0.06456467 |
| -0.629912003 | 0.528752159 | 0.633571093 |
| 0.809569848 | 0.418187443 | 0.529798639 |
| 0.369220905 | 0.711963075 | NA |
| -1.59064772 | 0.111688879 | 0.182454812 |
| -1.741009943 | 0.081681836 | 0.140592515 |
| NA | NA | NA |
| -0.73110957 | 0.46471223 | NA |
| 0.270830778 | 0.786521187 | NA |
| 0.416915303 | 0.67674036 | 0.759860543 |
| 0.063708882 | 0.949202033 | NA |
| -4.775733383 | 1.79E-06 | 8.13E-06 |
| 0.74041719 | 0.459046892 | 0.568443264 |
| 0.144670767 | 0.884970821 | 0.919311876 |
| -0.388104528 | 0.697938681 | 0.777696888 |
| 0.810423063 | 0.417697067 | 0.529364941 |
| NA | NA | NA |
| -2.218397826 | 0.026527719 | 0.053338751 |
| NA | NA | NA |
| NA | NA | NA |
| 5.220108373 | 1.79E-07 | 9.30E-07 |
| 3.765292948 | 0.000166354 | 0.000553555 |
| -0.065388772 | 0.947864464 | NA |
| 0.502263473 | 0.6154822 | 0.71071782 |
| NA | NA | NA |
| 1.568273947 | 0.116817213 | 0.189540218 |
| 2.27840539 | 0.022702436 | 0.046695863 |
| -9.434824482 | 3.92E-21 | 7.61E-20 |

| | | | |
|---|---|---|---|
| | 0.480493736 | 0.630876356 | NA |
| | 0.063708285 | 0.949202508 | NA |
| | 0.070101598 | 0.944112795 | NA |
| | -0.901456515 | 0.367345644 | 0.479780315 |
| | 8.718361431 | 2.82E-18 | 4.36E-17 |
| | -2.322800086 | 0.020189894 | 0.042137335 |
| | 1.774556219 | 0.075971192 | 0.132098571 |
| | 0.192809521 | 0.847108147 | NA |
| | 0.28738371 | 0.773818533 | 0.837383393 |
| | 0.257767112 | 0.796586648 | 0.855145128 |
| | 0.260371648 | NA | NA |
| | -2.344310078 | 0.019062316 | 0.040074733 |
| | 10.66200661 | 1.53E-26 | 4.34E-25 |
| | -2.925962769 | 0.00343392 | 0.008751759 |
| | -1.241167634 | 0.214543828 | 0.312818207 |
| | 4.576289558 | 4.73E-06 | 2.02E-05 |
| | 6.482421663 | 9.03E-11 | 6.90E-10 |
| NA | | NA | NA |
| NA | | NA | NA |
| | 0.192809521 | 0.847108147 | NA |
| NA | | NA | NA |
| NA | | NA | NA |
| NA | | NA | NA |
| | 0.107256042 | 0.914585857 | NA |
| | 2.946197309 | 0.003217071 | 0.008255412 |
| NA | | NA | NA |
| | 0.988423595 | 0.322945237 | 0.433817382 |
| | -0.065389369 | 0.947863988 | NA |
| | -2.621597955 | 0.008751861 | 0.020208402 |
| | 6.808793352 | 9.84E-12 | 8.24E-11 |
| | 0.361184658 | 0.71796141 | 0.793823399 |
| | -0.313806396 | 0.753668086 | 0.821294449 |
| | -2.761628494 | 0.005751388 | 0.01392371 |
| | -3.138729108 | 0.001696822 | 0.004646768 |
| | -0.74370082 | 0.457057493 | 0.566708726 |
| | -2.368769847 | 0.017847356 | 0.037913546 |
| | 0.288217989 | 0.77317988 | NA |
| | 0.159251661 | 0.873470604 | NA |
| NA | | NA | NA |
| | -5.294102681 | 1.20E-07 | 6.38E-07 |
| | -0.065389369 | 0.947863988 | NA |
| | 0.063708882 | 0.949202033 | NA |
| | -0.080297022 | 0.936001027 | 0.956436594 |
| | -2.090703386 | 0.036554661 | 0.070506525 |
| | 19.07219082 | 4.30E-81 | 6.90E-79 |
| NA | | NA | NA |
| | 0.43538885 | 0.663280202 | NA |

| | | |
|---|---|---|
| 22.60376191 | 3.98E-113 | 1.17E-110 |
| 0.063708882 | 0.949202033 | NA |
| 1.930807891 | 0.053506814 | 0.098129804 |
| 8.02727601 | 9.97E-16 | 1.24E-14 |
| NA | NA | NA |
| NA | NA | NA |
| NA | NA | NA |
| -3.685608953 | 0.000228156 | 0.000741753 |
| -1.524283644 | 0.127437871 | 0.204171408 |
| NA | NA | NA |
| -3.12843859 | 0.001757377 | 0.004797901 |
| -1.64307332 | 0.100367764 | 0.167001605 |
| -3.107121622 | 0.001889187 | 0.005115355 |
| 0.259257068 | 0.795436902 | 0.854460566 |
| 0.059694893 | 0.952398639 | 0.967567454 |
| -1.616432757 | 0.106000781 | 0.174681564 |
| 8.397153867 | 4.57E-17 | 6.41E-16 |
| -3.042054831 | 0.002349691 | 0.006221598 |
| NA | NA | NA |
| -7.531428542 | 5.02E-14 | 5.31E-13 |
| NA | NA | NA |
| NA | NA | NA |
| NA | NA | NA |
| -0.065388772 | 0.947864464 | NA |
| -0.065388772 | 0.947864464 | NA |
| 0.192809521 | 0.847108147 | NA |
| -2.548688752 | 0.010812876 | 0.024448625 |
| 0.06371127 | 0.949200131 | NA |
| -1.633088279 | 0.102450431 | 0.169913484 |
| NA | NA | NA |
| NA | NA | NA |
| NA | NA | NA |
| NA | NA | NA |
| 0.2961526 | 0.767113545 | NA |
| NA | NA | NA |
| NA | NA | NA |
| NA | NA | NA |
| 0.17794706 | 0.858764552 | 0.901895554 |
| -6.640018564 | 3.14E-11 | 2.51E-10 |
| 4.302151637 | 1.69E-05 | 6.65E-05 |
| 2.524045066 | 0.011601306 | 0.026008122 |
| 4.640914823 | 3.47E-06 | 1.51E-05 |
| NA | NA | NA |
| -0.477991453 | 0.632656288 | 0.724397079 |
| NA | NA | NA |
| NA | NA | NA |
| 3.902214826 | 9.53E-05 | 0.000331081 |

| | | | |
|---|---|---|---|
| NA | NA | NA | |
| | 6.426070627 | 1.31E-10 | 9.82E-10 |
| NA | NA | NA | |
| | -7.201326912 | 5.96E-13 | 5.67E-12 |
| | -5.845121781 | 5.06E-09 | 3.20E-08 |
| | -1.380320531 | 0.167487976 | 0.255483213 |
| | -6.963290077 | 3.32E-12 | 2.91E-11 |
| NA | NA | NA | |
| | -1.810162096 | 0.070270654 | NA |
| NA | NA | NA | |
| | -6.109320415 | 1.00E-09 | 6.82E-09 |
| | 3.031326636 | 0.002434817 | 0.006421862 |
| | -1.108005626 | 0.267859386 | 0.373781593 |
| NA | NA | NA | |
| | 0.192810118 | 0.847107679 | NA |
| NA | NA | NA | |
| | 11.35607167 | 6.92E-30 | 2.36E-28 |
| | 4.230626878 | 2.33E-05 | 8.96E-05 |
| NA | NA | NA | |
| | -0.787163638 | 0.43118608 | 0.542177614 |
| | -3.127228464 | 0.001764627 | 0.004814544 |
| NA | NA | NA | |
| | -1.902927053 | 0.057050064 | 0.103655859 |
| NA | NA | NA | |
| NA | NA | NA | |
| NA | NA | NA | |
| | -3.020627593 | 0.002522514 | 0.006628717 |
| | 6.341824871 | 2.27E-10 | 1.65E-09 |
| | -0.14496177 | 0.884741056 | 0.919225795 |
| | 0.522605653 | 0.601248706 | 0.698827727 |
| NA | NA | NA | |
| | 0.192810118 | 0.847107679 | NA |
| | 7.104201421 | 1.21E-12 | 1.11E-11 |
| | -1.332340428 | 0.182748348 | 0.274586268 |
| NA | NA | NA | |
| | -0.168724057 | 0.866013696 | 0.906508752 |
| | -3.165427854 | 0.001548551 | 0.00427817 |
| | 1.029400815 | 0.303291366 | NA |
| | -1.530487689 | 0.125896058 | 0.202037465 |
| NA | NA | NA | |
| | 0.018525639 | 0.985219524 | 0.989507396 |
| NA | NA | NA | |
| NA | NA | NA | |
| NA | NA | NA | |
| NA | NA | NA | |
| NA | NA | NA | |
| | -0.770523748 | 0.440989273 | 0.551098982 |

| | | | |
|---|---|---|---|
| NA | | NA | NA |
| NA | | NA | NA |
| | 0.063708882 | 0.949202033 | NA |
| | -0.065389369 | 0.947863988 | NA |
| | 2.544364864 | 0.010947666 | 0.024699846 |
| NA | | NA | NA |
| | 6.317219137 | 2.66E-10 | 1.92E-09 |
| | -4.574880698 | 4.76E-06 | 2.03E-05 |
| | 2.496041629 | 0.012558786 | 0.027872532 |
| | 1.301251962 | 0.193172224 | 0.287341818 |
| NA | | NA | NA |
| | 4.141125325 | 3.46E-05 | 0.000129383 |
| | 0.490794612 | 0.623571722 | 0.717411298 |
| | -0.40104687 | 0.688385616 | 0.770144563 |
| | 0.637178475 | 0.524008597 | 0.629152164 |
| | -0.091498571 | 0.927096441 | NA |
| | 1.083515992 | 0.278579456 | 0.385559513 |
| | 0.123240863 | 0.901916368 | NA |
| | -8.071349511 | 6.95E-16 | 8.82E-15 |
| | 6.84077003 | 7.88E-12 | 6.64E-11 |
| | -3.480128758 | 0.000501173 | 0.00152924 |
| | 13.60778249 | 3.60E-42 | 2.13E-40 |
| | 0.193472986 | 0.84658856 | NA |
| | 1.178329984 | 0.238665079 | 0.340852843 |
| | 8.918615741 | 4.72E-19 | 7.83E-18 |
| | 7.63876174 | 2.19E-14 | 2.41E-13 |
| | 0.555509634 | 0.578546122 | 0.679406084 |
| | 13.0028504 | 1.18E-38 | 6.12E-37 |
| | 0.399043737 | 0.689860978 | NA |
| NA | | NA | NA |
| | 0.608885372 | 0.542600421 | NA |
| NA | | NA | NA |
| | 0.392175023 | 0.694928897 | NA |
| | 0.202714101 | 0.839358501 | NA |
| | -0.740363458 | 0.459079486 | NA |
| NA | | NA | NA |
| | 0.192808924 | 0.847108615 | NA |
| | 2.55264523 | 0.010690833 | 0.024203288 |
| | 3.782556766 | 0.000155226 | 0.000518731 |
| | 6.091531311 | 1.12E-09 | 7.59E-09 |
| NA | | NA | NA |
| | -7.617575793 | 2.58E-14 | 2.82E-13 |
| | -0.954874111 | 0.339641363 | 0.451089002 |
| | -3.268284433 | 0.001082016 | 0.003096245 |
| | -0.216826093 | 0.828343877 | 0.879951482 |
| | 0.06371127 | 0.949200131 | NA |
| NA | | NA | NA |

| | | |
|---|---|---|
| 0.06371127 | 0.949200131 | NA |
| NA | NA | NA |
| 1.167908354 | 0.242843734 | NA |
| 2.308333718 | 0.020980582 | 0.043540667 |
| 8.896110338 | 5.78E-19 | 9.53E-18 |
| NA | NA | NA |
| NA | NA | NA |
| NA | NA | NA |
| 3.572963089 | 0.000352965 | 0.001108495 |
| 2.959637777 | 0.003080009 | 0.007938692 |
| -0.825052286 | 0.409341907 | 0.521257143 |
| 3.001614001 | 0.002685525 | 0.007018762 |
| 38.37185464 | 0 | 0 |
| 19.10431207 | 2.32E-81 | 3.75E-79 |
| 12.50028555 | 7.44E-36 | 3.37E-34 |
| 17.25958628 | 9.48E-67 | 1.12E-64 |
| 15.79408017 | 3.42E-56 | 3.09E-54 |
| 8.359242494 | 6.31E-17 | 8.72E-16 |
| 17.37262397 | 1.33E-67 | 1.62E-65 |
| 0.923148668 | 0.355929735 | NA |
| 21.67273677 | 3.71E-104 | 9.48E-102 |
| 16.10760501 | 2.26E-58 | 2.16E-56 |
| 2.489939774 | 0.012776474 | 0.028297952 |
| 21.47290994 | 2.79E-102 | 6.78E-100 |
| 1.193977467 | 0.232486782 | NA |
| NA | NA | NA |
| 0.137505335 | 0.890631376 | 0.923429159 |
| 0.192809521 | 0.847108147 | NA |
| 3.803896323 | 0.000142438 | 0.000479196 |
| -1.373893511 | 0.169474744 | 0.257869474 |
| 8.186526974 | 2.69E-16 | 3.53E-15 |
| -0.829559697 | 0.406787772 | 0.518585002 |
| 1.110255757 | 0.266888832 | 0.372676568 |
| -0.748365314 | 0.454239838 | NA |
| 0.193341124 | 0.846691821 | NA |
| 0.071932862 | 0.942655337 | NA |
| NA | NA | NA |
| 0.192810118 | 0.847107679 | NA |
| NA | NA | NA |
| 0.192809521 | 0.847108147 | NA |
| NA | NA | NA |
| NA | NA | NA |
| 0.063708882 | 0.949202033 | NA |
| NA | NA | NA |
| NA | NA | NA |
| NA | NA | NA |
| NA | NA | NA |

| | | | |
|---|---|---|---|
| | 0.192809521 | 0.847108147 | NA |
| NA | NA | NA | |
| | 10.65523276 | 1.65E-26 | 4.65E-25 |
| | 4.625824635 | 3.73E-06 | 1.62E-05 |
| | 1.352383218 | 0.176252755 | 0.266578431 |
| | 3.939157249 | 8.18E-05 | 0.000287291 |
| NA | NA | NA | |
| NA | NA | NA | |
| | 0.063708285 | 0.949202508 | NA |
| | 5.150517845 | 2.60E-07 | 1.33E-06 |
| | -5.971208668 | 2.36E-09 | 1.54E-08 |
| | -0.065389369 | 0.947863988 | NA |
| | -1.869475512 | 0.061556687 | 0.110409451 |
| | 0.017468833 | 0.986062597 | 0.990137072 |
| | -1.422651991 | 0.154837064 | 0.239691491 |
| | 0.288217989 | 0.77317988 | NA |
| | -2.987942819 | 0.002808621 | 0.007305016 |
| NA | NA | NA | |
| | -1.485558826 | 0.137395853 | 0.217013832 |
| | 9.028684893 | 1.74E-19 | 2.98E-18 |
| | -3.12561167 | 0.001774357 | 0.004837398 |
| | -0.065390563 | 0.947863038 | NA |
| | -4.93603826 | 7.97E-07 | 3.80E-06 |
| | -1.523331101 | 0.12767589 | 0.2044873 |
| | 4.105112997 | 4.04E-05 | 0.000149381 |
| | 3.928825055 | 8.54E-05 | 0.000298952 |
| | -0.065388772 | 0.947864464 | NA |
| | 0.663074844 | 0.507282621 | 0.61375116 |
| | 0.233942337 | 0.815029745 | 0.869907233 |
| | -1.412666245 | 0.157753881 | 0.24330394 |
| | 1.259252273 | 0.207939226 | 0.304962645 |
| NA | NA | NA | |
| | -1.223238674 | 0.221239562 | 0.320483028 |
| | -4.763923521 | 1.90E-06 | 8.57E-06 |
| NA | NA | NA | |
| | 0.063708882 | 0.949202033 | NA |
| NA | NA | NA | |
| | -4.370931201 | 1.24E-05 | 4.96E-05 |
| | -1.259006351 | 0.208028038 | 0.305029359 |
| NA | NA | NA | |
| NA | NA | NA | |
| | -2.689742342 | 0.007150721 | 0.016878776 |
| | -6.584382525 | 4.57E-11 | 3.60E-10 |
| | -4.612362514 | 3.98E-06 | 1.72E-05 |
| | 0.063708285 | 0.949202508 | NA |
| | 1.021396932 | 0.307066418 | NA |
| | -0.065388772 | 0.947864464 | NA |

| | | | |
|---|---|---|---|
| | -0.065389369 | 0.947863988 | NA |
| NA | NA | NA | |
| | 0.192809521 | 0.847108147 | NA |
| NA | NA | NA | |
| NA | NA | NA | |
| NA | NA | NA | |
| NA | NA | NA | |
| | -3.571458232 | 0.000354999 | 0.001114325 |
| | -1.325573722 | 0.184980935 | 0.277349515 |
| NA | NA | NA | |
| | 0.06371127 | 0.949200131 | NA |
| NA | NA | NA | |
| NA | NA | NA | |
| | 0.192809521 | 0.847108147 | NA |
| | 0.074728106 | 0.940431045 | NA |
| | 3.536718613 | 0.000405131 | 0.001257026 |
| | -1.635882449 | 0.101864192 | 0.169109287 |
| | -2.350504281 | 0.018747992 | 0.039510681 |
| | 2.94795529 | 0.003198833 | 0.008210295 |
| | -2.97742416 | 0.002906816 | 0.007539647 |
| | 5.411253648 | 6.26E-08 | 3.47E-07 |
| | 2.234669383 | 0.025439062 | 0.051478674 |
| | -1.975528027 | 0.048208257 | 0.089673951 |
| | -1.607265125 | 0.107996221 | 0.177383386 |
| | -2.6487297 | 0.008079492 | 0.018791088 |
| | -0.358724083 | 0.71980151 | 0.795085251 |
| | 2.398023098 | 0.016483826 | 0.035415464 |
| | -0.435829727 | 0.662960274 | 0.749295388 |
| | 2.691555961 | 0.007111957 | 0.01679995 |
| | 6.441218063 | 1.19E-10 | 8.93E-10 |
| | -1.084269349 | 0.278245391 | 0.385210542 |
| NA | NA | NA | |
| | 0.420026166 | 0.674466339 | 0.758026737 |
| | -1.027919991 | 0.303987465 | 0.413432202 |
| | -0.048145507 | 0.961600279 | NA |
| | 0.396993674 | 0.691372125 | 0.77227081 |
| NA | NA | NA | |
| | 0.644491585 | 0.519256713 | 0.624555287 |
| | 0.133243269 | 0.894000992 | 0.926078915 |
| | 0.369131931 | 0.712029389 | NA |
| NA | NA | NA | |
| NA | NA | NA | |
| | 0.501032788 | 0.616348047 | 0.711323848 |
| | 0.192809521 | 0.847108147 | NA |
| NA | NA | NA | |
| | -2.117081661 | 0.034252918 | 0.066668004 |
| | -2.004183806 | 0.045050375 | 0.084591964 |

| | | |
|---|---|---|
| 3.066369693 | 0.002166752 | 0.00579483 |
| 2.699189616 | 0.006950856 | 0.016475372 |
| -5.079112107 | 3.79E-07 | 1.89E-06 |
| 8.335141124 | 7.74E-17 | 1.06E-15 |
| -0.486727932 | 0.626451148 | 0.719830705 |
| NA | NA | NA |
| NA | NA | NA |
| -2.062948555 | 0.039117516 | 0.074718516 |
| 0.592239138 | 0.553690464 | NA |
| 1.113364079 | 0.265552096 | 0.371161992 |
| 1.905552278 | 0.056708319 | 0.10311727 |
| 2.36436017 | 0.018061244 | 0.038288885 |
| 0.063708882 | 0.949202033 | NA |
| -0.46812281 | 0.639696768 | 0.730390024 |
| -2.589073012 | 0.009623469 | 0.022007805 |
| 0.294730507 | 0.768199757 | 0.832995707 |
| -0.732490705 | 0.463869114 | NA |
| 1.471992948 | 0.141022782 | 0.22172112 |
| 0.060616027 | 0.951665009 | 0.967002605 |
| -1.84461602 | 0.065093415 | 0.115774048 |
| -2.93432495 | 0.00334274 | 0.008539375 |
| -2.425208875 | 0.015299581 | 0.033141369 |
| -0.531014359 | 0.595408829 | NA |
| -8.240550654 | 1.71E-16 | 2.29E-15 |
| -3.436299751 | 0.000589718 | 0.001774329 |
| -0.873957185 | 0.382141571 | 0.494612229 |
| -7.438599609 | 1.02E-13 | 1.05E-12 |
| -2.512435495 | 0.011990102 | 0.026740823 |
| -7.949792774 | 1.87E-15 | 2.27E-14 |
| 0.532170805 | 0.594607701 | NA |
| NA | NA | NA |
| 0.471829667 | 0.637048373 | NA |
| 2.2106482 | 0.027060208 | 0.0542635 |
| NA | NA | NA |
| -1.83174841 | 0.066988913 | 0.118765582 |
| 1.169960891 | 0.242016708 | 0.344597428 |
| NA | NA | NA |
| 0.450917721 | 0.652048851 | 0.740445501 |
| 22.25303087 | 1.05E-109 | 2.93E-107 |
| 3.648048305 | 0.00026424 | 0.000848266 |
| 8.062223047 | 7.49E-16 | 9.46E-15 |
| 0.148332437 | 0.882080417 | 0.917222917 |
| NA | NA | NA |
| NA | NA | NA |
| NA | NA | NA |
| 1.830044041 | 0.067243354 | 0.119115438 |
| -0.425104004 | 0.67076086 | 0.755128437 |

| | | |
|---|---|---|
| 0.884678516 | 0.376330046 | 0.489072744 |
| 13.80315857 | 2.44E-43 | 1.51E-41 |
| -2.405908372 | 0.01613231 | 0.0347557 |
| 0.322968501 | 0.746719097 | NA |
| NA | NA | NA |
| NA | NA | NA |
| 2.956430808 | 0.003112221 | 0.008012639 |
| 0.496567089 | 0.619494366 | 0.714000275 |
| -0.065388772 | 0.947864464 | NA |
| -6.100904141 | 1.05E-09 | 7.17E-09 |
| -0.334203924 | 0.738225683 | 0.809399836 |
| 4.407222277 | 1.05E-05 | 4.25E-05 |
| 2.466926651 | 0.013627827 | 0.029958239 |
| NA | NA | NA |
| 6.296127968 | 3.05E-10 | 2.19E-09 |
| -1.332489098 | 0.182699522 | 0.274553856 |
| -1.183658035 | 0.236548457 | 0.338389889 |
| NA | NA | NA |
| -2.635169818 | 0.00840952 | 0.019495256 |
| 0.134409133 | 0.89307906 | NA |
| -1.209985154 | 0.22628459 | 0.326355706 |
| 0.063707194 | 0.949203376 | NA |
| -4.935741253 | 7.98E-07 | 3.80E-06 |
| NA | NA | NA |
| 4.130602568 | 3.62E-05 | 0.000134946 |
| 4.459261055 | 8.22E-06 | 3.40E-05 |
| 7.478017457 | 7.55E-14 | 7.86E-13 |
| 0.192808924 | 0.847108615 | NA |
| 6.447295215 | 1.14E-10 | 8.60E-10 |
| 3.557097616 | 0.000374975 | 0.001169985 |
| 1.979276403 | 0.047784895 | 0.089033147 |
| 1.562478121 | 0.118175396 | 0.191381551 |
| NA | NA | NA |
| 0.519024983 | 0.60374332 | NA |
| 2.062117155 | 0.039196583 | 0.074846695 |
| -4.529982914 | 5.90E-06 | 2.48E-05 |
| -0.065390563 | 0.947863038 | NA |
| -2.166258737 | 0.030291421 | 0.059855867 |
| -1.591226474 | 0.111558617 | 0.182301507 |
| 4.897099009 | 9.73E-07 | 4.58E-06 |
| 2.195502439 | 0.028127576 | 0.056134048 |
| NA | NA | NA |
| 5.422036137 | 5.89E-08 | 3.27E-07 |
| 0.516288212 | 0.605653128 | NA |
| NA | NA | NA |
| NA | NA | NA |
| -1.269268054 | 0.204345476 | 0.300833271 |

| | | |
|---|---|---|
| 0.192636473 | 0.847243678 | NA |
| 0.192809521 | 0.847108147 | NA |
| -1.45718474 | 0.145065392 | 0.227092798 |
| 0.430987523 | 0.666477443 | 0.752086412 |
| 0.399043737 | 0.689860978 | NA |
| NA | NA | NA |
| -0.390519815 | 0.696152206 | 0.776293759 |
| NA | NA | NA |
| -7.654878196 | 1.93E-14 | 2.14E-13 |
| -0.065388772 | 0.947864464 | NA |
| 1.678014176 | 0.093344332 | 0.157376305 |
| 1.58020449 | 0.114060044 | 0.185753663 |
| 0.847050502 | 0.396966973 | 0.508823032 |
| NA | NA | NA |
| 1.353950464 | 0.175752182 | 0.266030305 |
| NA | NA | NA |
| -0.263865796 | 0.791883333 | NA |
| -0.068677197 | 0.945246569 | NA |
| -1.050063459 | 0.293688938 | 0.4018276 |
| 0.939046612 | 0.347706813 | 0.459290809 |
| -0.065390563 | 0.947863038 | NA |
| NA | NA | NA |
| 0.192636473 | 0.847243678 | NA |
| NA | NA | NA |
| NA | NA | NA |
| 6.295591012 | 3.06E-10 | 2.19E-09 |
| 0.182045332 | 0.855547151 | NA |
| 0.355086882 | 0.722524504 | NA |
| 3.195017275 | 0.001398225 | 0.003897436 |
| 6.593509199 | 4.30E-11 | 3.39E-10 |
| -1.463086927 | 0.143443595 | 0.225032477 |
| 3.534330766 | 0.000408809 | 0.001267025 |
| 21.10732529 | 6.81E-99 | 1.56E-96 |
| 33.98369576 | 3.88E-253 | 5.39E-250 |
| 34.37234537 | 6.53E-259 | 1.17E-255 |
| 0.330282312 | 0.741186656 | NA |
| -3.627510716 | 0.000286167 | 0.000913035 |
| -2.034539809 | 0.041897199 | 0.079397857 |
| 1.035938865 | 0.300230665 | 0.409234523 |
| 0.707973783 | 0.478961538 | 0.587695834 |
| 1.82788727 | 0.067566472 | 0.119603168 |
| 0.891987817 | 0.372399462 | 0.485199029 |
| NA | NA | NA |
| 0.320071032 | 0.748914485 | NA |
| 7.612748024 | 2.68E-14 | 2.92E-13 |
| NA | NA | NA |
| NA | NA | NA |

| | | | |
|---|---|---|---|
| NA | NA | NA | |
| 0.063708882 | 0.949202033 | NA | |
| NA | NA | NA | |
| 3.441260096 | 0.000579012 | | 0.001745473 |
| 0.063708882 | 0.949202033 | NA | |
| NA | NA | NA | |
| -0.194488217 | 0.845793621 | NA | |
| -0.065388772 | 0.947864464 | NA | |
| NA | NA | NA | |
| NA | NA | NA | |
| 0.707512168 | 0.479248254 | NA | |
| 0.06371127 | 0.949200131 | NA | |
| -0.065389966 | 0.947863513 | NA | |
| -0.868841972 | 0.384933572 | | 0.497079261 |
| NA | NA | NA | |
| NA | NA | NA | |
| NA | NA | NA | |
| NA | NA | NA | |
| NA | NA | NA | |
| NA | NA | NA | |
| NA | NA | NA | |
| NA | NA | NA | |
| 0.192809521 | 0.847108147 | NA | |
| NA | NA | NA | |
| 0.259481947 | 0.795263409 | | 0.854347531 |
| NA | NA | NA | |
| NA | NA | NA | |
| NA | NA | NA | |
| -3.868899542 | 0.000109328 | | 0.000375579 |
| 0.350807759 | 0.725732576 | | 0.799739244 |
| -1.997331707 | 0.045789161 | | 0.08577329 |
| 2.783878675 | 0.005371312 | | 0.013116482 |
| -0.084501006 | 0.932658103 | NA | |
| -0.065388772 | 0.947864464 | NA | |
| -0.358874267 | 0.719689151 | | 0.795009842 |
| -1.533185179 | 0.125230235 | | 0.201138436 |
| -3.689254957 | 0.000224912 | | 0.000731965 |
| -2.516431706 | 0.011854987 | | 0.026498512 |
| 5.936303031 | 2.92E-09 | | 1.89E-08 |
| 1.825281357 | 0.067958582 | | 0.120152808 |
| -0.398252416 | 0.690444132 | | 0.771509671 |
| 0.425838616 | 0.670225446 | | 0.75480377 |
| -1.665034902 | 0.095905793 | | 0.16090368 |
| -1.012536349 | 0.311281676 | | 0.421474648 |
| -5.694317359 | 1.24E-08 | | 7.49E-08 |
| -0.080519055 | 0.935824442 | | 0.956342048 |
| 1.169255333 | 0.242300774 | | 0.344923413 |

| | | |
|---|---|---|
| -2.393401474 | 0.016692964 | 0.035800265 |
| -0.649256893 | 0.516172343 | 0.621472594 |
| -1.207533795 | 0.227226638 | 0.327398568 |
| 0.207863816 | 0.835335303 | 0.884611625 |
| 1.502457384 | 0.132979026 | 0.211237229 |
| -1.304018492 | 0.192227277 | 0.286225387 |
| 4.288004102 | 1.80E-05 | 7.06E-05 |
| -0.290534031 | 0.77140772 | NA |
| -7.062992176 | 1.63E-12 | 1.47E-11 |
| NA | NA | NA |
| 1.041876984 | 0.297468716 | 0.406155464 |
| NA | NA | NA |
| -5.110623908 | 3.21E-07 | 1.62E-06 |
| -2.751399745 | 0.005934118 | 0.014313479 |
| 4.48306161 | 7.36E-06 | 3.05E-05 |
| 2.727989985 | 0.006372154 | 0.015252464 |
| NA | NA | NA |
| 5.458356921 | 4.81E-08 | 2.71E-07 |
| -5.900926543 | 3.61E-09 | 2.32E-08 |
| 12.92535613 | 3.24E-38 | 1.65E-36 |
| 0.381375239 | 0.702924832 | NA |
| 0.731113103 | 0.464710072 | NA |
| NA | NA | NA |
| NA | NA | NA |
| 1.833740032 | 0.066692593 | 0.118256986 |
| -1.483895572 | 0.137836627 | 0.217600174 |
| -0.615776131 | 0.538042283 | NA |
| -1.270550085 | 0.203888754 | 0.300302196 |
| 1.767867188 | 0.077083109 | 0.13378951 |
| 0.033304903 | 0.973431444 | 0.981588471 |
| -0.716336482 | 0.473783605 | NA |
| 3.922359829 | 8.77E-05 | 0.00030679 |
| -0.918841554 | 0.358178454 | 0.47007166 |
| -0.065389369 | 0.947863988 | NA |
| -0.070666592 | 0.94366311 | NA |
| -0.865852957 | 0.386570804 | 0.498791416 |
| 0.452474727 | 0.650927024 | 0.739636073 |
| 2.789975811 | 0.005271198 | 0.012894653 |
| 0.419866584 | 0.67458292 | 0.75808425 |
| NA | NA | NA |
| -5.082541956 | 3.72E-07 | 1.86E-06 |
| 2.230603732 | 0.025707389 | 0.051921012 |
| 3.707143657 | 0.00020961 | 0.000686181 |
| NA | NA | NA |
| NA | NA | NA |
| -4.420266233 | 9.86E-06 | 4.02E-05 |
| 1.177081763 | 0.239162873 | NA |

| | | |
|---|---|---|
| -2.129953983 | 0.033175413 | 0.064848151 |
| NA | NA | NA |
| -0.330051315 | 0.741361188 | 0.81178973 |
| 3.559897429 | 0.000371 | 0.001159027 |
| -5.008468049 | 5.49E-07 | 2.67E-06 |
| -1.545862371 | 0.12213781 | 0.196915258 |
| 0.366483087 | 0.71400463 | NA |
| 4.229190459 | 2.35E-05 | 9.01E-05 |
| 1.383602323 | 0.166480257 | 0.254162905 |
| 3.841216416 | 0.000122426 | 0.000416802 |
| NA | NA | NA |
| NA | NA | NA |
| NA | NA | NA |
| -0.065389369 | 0.947863988 | NA |
| NA | NA | NA |
| NA | NA | NA |
| NA | NA | NA |
| NA | NA | NA |
| NA | NA | NA |
| NA | NA | NA |
| NA | NA | NA |
| NA | NA | NA |
| NA | NA | NA |
| NA | NA | NA |
| NA | NA | NA |
| NA | NA | NA |
| NA | NA | NA |
| 3.529605057 | 0.00041618 | 0.001287639 |
| 7.677454941 | 1.62E-14 | 1.81E-13 |
| 2.956697323 | 0.003109532 | 0.008007365 |
| -0.421135647 | 0.673656033 | NA |
| -0.065388772 | 0.947864464 | NA |
| 6.167879312 | 6.92E-10 | 4.77E-09 |
| 8.367336145 | 5.89E-17 | 8.17E-16 |
| 2.763348975 | 0.005721156 | 0.013867958 |
| -0.065388772 | 0.947864464 | NA |
| 0.192810118 | 0.847107679 | NA |
| 0.324874675 | 0.745275925 | NA |
| 0.269654315 | 0.78742621 | 0.848513778 |
| 4.808588725 | 1.52E-06 | 6.95E-06 |
| -1.474841008 | 0.140255293 | 0.220777938 |
| 1.246387968 | 0.212621996 | 0.310497583 |
| 4.570646146 | 4.86E-06 | 2.07E-05 |
| 7.467780349 | 8.16E-14 | 8.46E-13 |
| -3.831728076 | 0.000127246 | 0.00043192 |
| 6.202138664 | 5.57E-10 | 3.89E-09 |
| NA | NA | NA |

| | | |
|---|---|---|
| -1.674633903 | 0.094006083 | 0.158226006 |
| 0.169696871 | 0.865248535 | 0.906081953 |
| -0.787674151 | 0.430887329 | 0.541856403 |
| 3.431850618 | 0.000599478 | 0.00180261 |
| -0.317851133 | 0.750597862 | 0.819000424 |
| NA | NA | NA |
| -0.221921377 | 0.824375089 | 0.876997021 |
| -0.098841965 | 0.921263748 | NA |
| -7.847042223 | 4.26E-15 | 5.00E-14 |
| -1.15485206 | 0.248151014 | 0.351392843 |
| 4.504238811 | 6.66E-06 | 2.78E-05 |
| -1.645144595 | 0.099939996 | 0.1664669 |
| 0.440801288 | 0.659356861 | 0.746436103 |
| -7.708699167 | 1.27E-14 | 1.43E-13 |
| 0.149280204 | 0.881332535 | 0.916673732 |
| 7.555775333 | 4.16E-14 | 4.45E-13 |
| 0.422053602 | 0.672985893 | 0.756799214 |
| 7.407813478 | 1.28E-13 | 1.30E-12 |
| 1.646465769 | 0.099667902 | 0.166135295 |
| -6.083202527 | 1.18E-09 | 7.97E-09 |
| -5.694526066 | 1.24E-08 | 7.48E-08 |
| -0.571395388 | 0.567731654 | NA |
| -1.99736295 | 0.04578577 | 0.08577329 |
| NA | NA | NA |
| 0.760211086 | 0.44712842 | NA |
| -0.233860868 | 0.815092994 | 0.869937644 |
| 0.192809521 | 0.847108147 | NA |
| -1.246859003 | 0.212449202 | 0.310353888 |
| 0.431915223 | 0.66580303 | 0.75159202 |
| -6.029583833 | 1.64E-09 | 1.09E-08 |
| -1.756049878 | 0.079079892 | 0.136715686 |
| -4.652116421 | 3.29E-06 | 1.43E-05 |
| -1.528086367 | 0.126491096 | 0.202862314 |
| -0.833341443 | 0.404652189 | 0.516414842 |
| -0.672899305 | 0.501011354 | 0.608000154 |
| NA | NA | NA |
| -1.007501411 | 0.31369388 | 0.424098402 |
| -1.680117777 | 0.092934403 | 0.156778919 |
| 5.166518954 | 2.38E-07 | 1.22E-06 |
| NA | NA | NA |
| 6.175414144 | 6.60E-10 | 4.56E-09 |
| 9.605025148 | 7.61E-22 | 1.56E-20 |
| 0.080739901 | 0.935648804 | 0.956240642 |
| 0.074663323 | 0.94048259 | 0.959496178 |
| 0.789063436 | 0.430074931 | 0.541133842 |
| -0.978520995 | 0.327816713 | 0.43883048 |
| -1.260190844 | 0.207600525 | 0.304621945 |

| | | | |
|---|---|---|---|
| NA | NA | NA | |
| | 0.841552126 | 0.400038696 | 0.511734122 |
| | -0.040615909 | 0.967602101 | NA |
| | 0.142597867 | 0.886607782 | 0.920547694 |
| | 0.610293539 | 0.541667376 | NA |
| | 0.164722896 | 0.869162094 | 0.908963283 |
| | -0.065389369 | 0.947863988 | NA |
| | -2.439804704 | 0.014695204 | 0.031984431 |
| | -0.416363976 | 0.677143685 | 0.760211098 |
| NA | NA | NA | |
| NA | NA | NA | |
| | 5.461197749 | 4.73E-08 | 2.67E-07 |
| | -0.153893668 | 0.877693577 | 0.914503526 |
| | 0.119400848 | 0.904957789 | 0.934219808 |
| | 0.780973676 | 0.434817977 | 0.545429186 |
| NA | NA | NA | |
| | 2.51619114 | 0.011863083 | 0.026511871 |
| NA | NA | NA | |
| | -0.018976556 | 0.984859807 | 0.989361701 |
| | 2.170762446 | 0.029949134 | 0.059259038 |
| | 1.057449336 | 0.290306565 | 0.397874026 |
| NA | NA | NA | |
| | -1.576534302 | 0.11490272 | 0.186934302 |
| | 0.207537456 | 0.835590144 | 0.88469202 |
| | 0.063708285 | 0.949202508 | NA |
| NA | NA | NA | |
| | 1.089081978 | 0.276117737 | 0.38289346 |
| | 4.276444913 | 1.90E-05 | 7.40E-05 |
| | 1.33271394 | 0.182625697 | 0.274492355 |
| | -0.194487022 | 0.845794556 | NA |
| NA | NA | NA | |
| | -0.847864453 | 0.396513463 | 0.508400596 |
| | 1.604275454 | 0.108653349 | 0.178298914 |
| | -3.30398694 | 0.000953203 | 0.002759184 |
| | -2.185318896 | 0.028865473 | 0.057418961 |
| NA | NA | NA | |
| | 0.552888392 | 0.580339837 | NA |
| | 0.517739789 | 0.604639833 | NA |
| | 2.656693824 | 0.007891107 | 0.018402536 |
| | -6.653999098 | 2.85E-11 | 2.29E-10 |
| | -3.055532745 | 0.002246611 | 0.00598029 |
| | -1.383367152 | 0.166552317 | 0.25425741 |
| | -3.749868556 | 0.000176927 | 0.000585934 |
| | 0.192809521 | 0.847108147 | NA |
| | 1.474214889 | 0.140423742 | 0.220973612 |
| | 3.016313709 | 0.002558684 | 0.006715259 |
| NA | NA | NA | |

| | | |
|---|---|---|
| -6.847464334 | 7.52E-12 | 6.35E-11 |
| -3.167772926 | 0.001536115 | 0.004247094 |
| 1.897108213 | 0.057813657 | 0.104792226 |
| -1.602384007 | 0.109070719 | 0.178866547 |
| -2.138137695 | 0.032505569 | 0.063717942 |
| -2.526439262 | 0.011522531 | 0.025852362 |
| -1.387682793 | 0.165233658 | 0.252644988 |
| NA | NA | NA |
| -0.530427234 | 0.595815748 | 0.694221589 |
| NA | NA | NA |
| 0.07890933 | 0.937104742 | NA |
| 1.871135472 | 0.061326306 | 0.110051437 |
| 4.881248668 | 1.05E-06 | 4.93E-06 |
| 8.625263654 | 6.39E-18 | 9.57E-17 |
| -1.410312946 | 0.158447297 | 0.244087633 |
| NA | NA | NA |
| 0.087301421 | 0.930431925 | NA |
| -1.342806111 | 0.179334764 | 0.270357715 |
| 0.063708882 | 0.949202033 | NA |
| -2.099685121 | 0.035756549 | 0.069142957 |
| -2.906494663 | 0.003655031 | 0.009255912 |
| -2.671842292 | 0.007543609 | 0.017696115 |
| -2.838332138 | 0.004534997 | 0.011268834 |
| -0.172211301 | 0.863271414 | 0.904920156 |
| NA | NA | NA |
| -5.58401735 | 2.35E-08 | 1.38E-07 |
| 0.084572383 | 0.932601356 | NA |
| 5.071274301 | 3.95E-07 | 1.97E-06 |
| -1.694160951 | 0.090234703 | 0.152937067 |
| 0.952439912 | 0.340873926 | NA |
| 1.340809056 | 0.179982455 | 0.271121862 |
| -2.848755225 | 0.004389063 | 0.010934434 |
| -2.328043526 | 0.019909793 | 0.041625636 |
| 1.87323215 | 0.061036335 | 0.109625391 |
| 3.585451756 | 0.000336495 | 0.001060364 |
| 0.063708285 | 0.949202508 | NA |
| -0.162548568 | 0.870873885 | 0.910031033 |
| NA | NA | NA |
| -0.045581667 | 0.963643682 | 0.975178691 |
| 7.346436621 | 2.04E-13 | 2.03E-12 |
| 0.192808924 | 0.847108615 | NA |
| -1.08848214 | 0.276382317 | 0.383217892 |
| 13.61160129 | 3.42E-42 | 2.02E-40 |
| 4.255286357 | 2.09E-05 | 8.09E-05 |
| 2.034439932 | 0.041907259 | 0.079410909 |
| NA | NA | NA |
| -2.632836616 | 0.008467507 | 0.019622414 |

| | | |
|---|---|---|
| -7.769943849 | 7.85E-15 | 9.03E-14 |
| 4.016152236 | 5.92E-05 | 0.000212636 |
| 0.685988085 | 0.492720622 | 0.599931759 |
| 4.026428702 | 5.66E-05 | 0.00020454 |
| 0.066587943 | 0.946909745 | NA |
| -1.383615389 | 0.166476254 | 0.254162905 |
| 0.743971965 | 0.456893436 | 0.566561442 |
| -3.263135538 | 0.001101868 | 0.003147296 |
| 0.503052363 | 0.614927457 | 0.710404256 |
| -1.354414399 | 0.175604206 | 0.265838471 |
| NA | NA | NA |
| NA | NA | NA |
| -4.113106535 | 3.90E-05 | 0.000144712 |
| 5.040783748 | 4.64E-07 | 2.28E-06 |
| -4.947078479 | 7.53E-07 | 3.60E-06 |
| -0.946213068 | 0.344039922 | 0.455360332 |
| -1.247504094 | 0.21221272 | 0.310098919 |
| 2.655200842 | 0.00792612 | 0.018475581 |
| -0.138746824 | 0.889650216 | NA |
| 0.168217457 | 0.866412208 | NA |
| -7.320114608 | 2.48E-13 | 2.44E-12 |
| -1.625591317 | 0.104036624 | 0.172104667 |
| -1.949483269 | 0.051237739 | 0.094550325 |
| 2.015210064 | 0.043882643 | 0.082678402 |
| 0.670376501 | 0.502617811 | 0.609314854 |
| -3.291928118 | 0.00099503 | 0.002869971 |
| -2.087280839 | 0.036862756 | 0.071029621 |
| -0.975092971 | 0.329514145 | 0.440373998 |
| NA | NA | NA |
| NA | NA | NA |
| -3.065431735 | 0.00217356 | 0.005810553 |
| -0.705202906 | 0.480683973 | 0.589375911 |
| -5.348216136 | 8.88E-08 | 4.82E-07 |
| 1.619143493 | 0.105416393 | 0.173910645 |
| -1.831140553 | 0.067079567 | 0.118901038 |
| -10.50062368 | 8.58E-26 | 2.32E-24 |
| -1.330506288 | 0.183351516 | 0.275384845 |
| -2.27427795 | 0.02294928 | 0.047122271 |
| -1.095919654 | 0.273113937 | 0.379801203 |
| -0.807648355 | 0.419293042 | 0.530984308 |
| 3.807729832 | 0.000140248 | 0.000472529 |
| 3.720591857 | 0.000198756 | 0.000653471 |
| -0.313195373 | 0.754132234 | NA |
| -5.219222399 | 1.80E-07 | 9.34E-07 |
| 2.080226492 | 0.037504763 | 0.072072494 |
| 0.192810118 | 0.847107679 | NA |
| -5.223716477 | 1.75E-07 | 9.14E-07 |

| | | |
|---|---|---|
| 1.549064427 | 0.121366233 | 0.195810031 |
| 9.048001211 | 1.46E-19 | 2.51E-18 |
| -0.077047762 | 0.938585549 | 0.958224754 |
| 2.983855772 | 0.002846409 | 0.007395227 |
| -3.641551195 | 0.000271 | 0.000868408 |
| NA | NA | NA |
| NA | NA | NA |
| -1.450363154 | 0.146957276 | 0.22946604 |
| NA | NA | NA |
| -1.508777573 | 0.131355634 | 0.209029914 |
| 3.672121623 | 0.000240545 | 0.000777686 |
| -0.390743479 | 0.695986857 | NA |
| 12.59288714 | 2.31E-36 | 1.07E-34 |
| 6.94669685 | 3.74E-12 | 3.26E-11 |
| NA | NA | NA |
| 2.171321853 | 0.029906852 | 0.059189943 |
| NA | NA | NA |
| 3.153234091 | 0.001614723 | 0.004445282 |
| 3.129182927 | 0.001752931 | 0.004786285 |
| 2.870071781 | 0.004103786 | 0.010291331 |
| 1.523111886 | 0.127730716 | 0.204535846 |
| 5.152977266 | 2.56E-07 | 1.31E-06 |
| 0.063708285 | 0.949202508 | NA |
| 7.374760462 | 1.65E-13 | 1.65E-12 |
| -0.712973349 | 0.475862249 | NA |
| -7.121162025 | 1.07E-12 | 9.91E-12 |
| 4.4509689 | 8.55E-06 | 3.52E-05 |
| -7.851963446 | 4.10E-15 | 4.82E-14 |
| 0.313364108 | 0.75400405 | 0.821553255 |
| NA | NA | NA |
| 1.134397705 | 0.256627761 | 0.361234449 |
| -0.225747667 | 0.821397682 | 0.874727025 |
| 1.795421265 | 0.072586607 | 0.127034175 |
| 7.36766234 | 1.74E-13 | 1.74E-12 |
| -0.270904471 | 0.786464507 | 0.847731826 |
| 1.509338269 | 0.13121236 | 0.208868314 |
| 7.502113175 | 6.28E-14 | 6.58E-13 |
| 8.365373653 | 5.99E-17 | 8.29E-16 |
| NA | NA | NA |
| 0.441263409 | 0.659022313 | 0.746192368 |
| -5.09186672 | 3.55E-07 | 1.78E-06 |
| 2.64193499 | 0.008243388 | 0.019136709 |
| 6.829372016 | 8.53E-12 | 7.17E-11 |
| -5.174877969 | 2.28E-07 | 1.17E-06 |
| -2.070146597 | 0.038438618 | 0.073614174 |
| NA | NA | NA |
| -0.40673691 | 0.684201229 | NA |

| | | |
|---|---|---|
| 2.074179699 | 0.038062625 | 0.072982036 |
| 1.371509164 | 0.170216289 | 0.258855133 |
| 6.469893604 | 9.81E-11 | 7.47E-10 |
| 7.442375077 | 9.89E-14 | 1.02E-12 |
| 5.233887974 | 1.66E-07 | 8.69E-07 |
| -1.164048699 | 0.244404277 | 0.347206938 |
| 4.48860886 | 7.17E-06 | 2.98E-05 |
| NA | NA | NA |
| -8.135046427 | 4.12E-16 | 5.30E-15 |
| 0.627607108 | 0.530261351 | 0.635126811 |
| 1.925630135 | 0.054150573 | 0.09915061 |
| NA | NA | NA |
| -0.842135686 | 0.399712008 | 0.511492411 |
| 3.177921437 | 0.001483349 | 0.00411302 |
| -6.762799645 | 1.35E-11 | 1.12E-10 |
| -1.839339805 | 0.065865222 | 0.11700543 |
| 3.992465946 | 6.54E-05 | 0.000233348 |
| 0.063708882 | 0.949202033 | NA |
| -1.164366828 | 0.244275385 | 0.347122339 |
| -0.118635159 | 0.905564409 | 0.93461476 |
| -2.022209456 | 0.043154718 | 0.081478659 |
| 2.139816112 | 0.032369631 | 0.063471355 |
| -0.950510282 | 0.341853032 | 0.453256775 |
| 1.781921129 | 0.074762094 | 0.13026768 |
| -1.77740063 | 0.075502347 | 0.131365477 |
| 2.614747677 | 0.008929339 | 0.020574553 |
| 0.421497639 | 0.673391735 | 0.757153548 |
| 1.653275539 | 0.098274799 | 0.164206823 |
| NA | NA | NA |
| 0.192809521 | 0.847108147 | NA |
| 9.717309306 | 2.54E-22 | 5.37E-21 |
| NA | NA | NA |
| 3.983088424 | 6.80E-05 | 0.000242167 |
| 21.65809058 | 5.10E-104 | 1.29E-101 |
| 10.96907158 | 5.38E-28 | 1.67E-26 |
| 1.24344391 | 0.213704298 | 0.311805222 |
| -0.675081959 | 0.499623695 | 0.606713618 |
| 0.169533173 | 0.865377281 | NA |
| 0.349693629 | 0.726568635 | NA |
| 7.518626283 | 5.54E-14 | 5.84E-13 |
| -0.406722179 | 0.684212049 | 0.766564952 |
| 1.910697183 | 0.056043507 | 0.102093812 |
| -5.013626117 | 5.34E-07 | 2.60E-06 |
| 0.346622043 | 0.728875284 | 0.80209929 |
| 0.053827445 | 0.957072644 | 0.970370165 |
| -1.098368703 | 0.272043523 | 0.378501911 |
| 0.487312321 | 0.626037018 | NA |

| | | |
|---|---|---|
| 0.532170805 | 0.594607701 | NA |
| -4.173471009 | 3.00E-05 | 0.000113478 |
| -1.206109186 | 0.227775393 | 0.328071015 |
| 10.35654066 | 3.91E-25 | 1.01E-23 |
| 0.935956448 | 0.349295612 | 0.460964326 |
| -3.855135633 | 0.000115665 | 0.00039529 |
| 0.287393196 | 0.77381127 | 0.837383393 |
| -8.525964057 | 1.52E-17 | 2.21E-16 |
| -1.376793994 | 0.168575948 | 0.25681412 |
| 3.358645017 | 0.000783256 | 0.002301882 |
| NA | NA | NA |
| -0.066872027 | 0.946683582 | NA |
| NA | NA | NA |
| NA | NA | NA |
| -1.229003166 | 0.219070619 | 0.317985247 |
| 9.889815969 | 4.61E-23 | 1.03E-21 |
| -0.608440781 | 0.542895171 | 0.646863718 |
| -2.625017175 | 0.00866446 | 0.020034312 |
| -1.987238549 | 0.046895968 | 0.087584396 |
| -0.089440543 | 0.928731804 | NA |
| 1.230418077 | 0.218540587 | 0.31739996 |
| 0.450908555 | 0.652055457 | 0.740445501 |
| 2.956389627 | 0.003112636 | 0.008012885 |
| -1.121085105 | 0.262251638 | 0.367575232 |
| -1.264714805 | 0.205973581 | 0.302731576 |
| 4.989891936 | 6.04E-07 | 2.92E-06 |
| 1.259439723 | 0.207871549 | 0.304930251 |
| -1.059024586 | 0.289588583 | 0.397085692 |
| NA | NA | NA |
| NA | NA | NA |
| 0.645210054 | 0.518791072 | NA |
| 8.862925914 | 7.79E-19 | 1.27E-17 |
| NA | NA | NA |
| NA | NA | NA |
| NA | NA | NA |
| 1.101358438 | 0.270740687 | NA |
| 0.573193348 | 0.566513788 | NA |
| -2.047877839 | 0.040571971 | 0.077149315 |
| NA | NA | NA |
| -1.779224797 | 0.075202915 | 0.130953733 |
| -0.582138223 | 0.560473581 | 0.662782938 |
| NA | NA | NA |
| 0.31685483 | 0.751353758 | NA |
| NA | NA | NA |
| 0.822627625 | 0.410719784 | 0.522613864 |
| 3.474197679 | 0.000512383 | 0.001560024 |
| -6.218046342 | 5.03E-10 | 3.53E-09 |

| | | | |
|---|---|---|---|
| | -2.31633352 | 0.020540065 | 0.042768149 |
| | 3.414141686 | 0.000639833 | 0.001912241 |
| | 1.887840045 | 0.059047427 | 0.106650101 |
| | -3.567817285 | 0.000359967 | 0.001128081 |
| | -4.404368513 | 1.06E-05 | 4.31E-05 |
| | -0.140380016 | 0.888359747 | NA |
| NA | NA | NA | |
| | 35.91321036 | 1.90E-282 | 4.76E-279 |
| | 3.892673726 | 9.91E-05 | 0.00034305 |
| NA | NA | NA | |
| | 0.33253772 | 0.739483262 | 0.810281751 |
| NA | NA | NA | |
| | -1.737510745 | 0.082297059 | 0.141486049 |
| | -1.344837248 | 0.178677799 | 0.269497156 |
| | 10.50367745 | 8.31E-26 | 2.25E-24 |
| | 1.410783385 | 0.158308495 | 0.243925786 |
| | 2.573808875 | 0.010058582 | 0.022910757 |
| | -0.624609509 | 0.532227377 | 0.636668479 |
| | -2.096852021 | 0.036006677 | 0.069572883 |
| NA | NA | NA | |
| | -1.061958963 | 0.288254316 | 0.39568366 |
| | 19.06582282 | 4.86E-81 | 7.74E-79 |
| | -1.246563175 | 0.212557711 | 0.310476162 |
| | 1.461445534 | 0.143893212 | 0.225539928 |
| NA | NA | NA | |
| | 0.293264375 | 0.769320082 | 0.833837932 |
| | -2.179690047 | 0.029280445 | 0.05813552 |
| NA | NA | NA | |
| | -0.398936726 | 0.689939827 | 0.771083849 |
| | 0.213508286 | 0.830930527 | 0.881765002 |
| | -3.309500039 | 0.000934628 | 0.002709803 |
| | -3.943587349 | 8.03E-05 | 0.00028239 |
| | -1.132396804 | 0.257467648 | 0.362086483 |
| | 1.675687807 | 0.093799359 | 0.15797357 |
| | 4.544912629 | 5.50E-06 | 2.33E-05 |
| | 3.756460604 | 0.000172333 | 0.000571552 |
| | 3.313313129 | 0.000921977 | 0.002674674 |
| | 7.5544069 | 4.21E-14 | 4.49E-13 |
| | 3.158897831 | 0.00158367 | 0.004367481 |
| | -0.340878959 | 0.733194705 | 0.80538127 |
| NA | NA | NA | |
| | 1.694301237 | 0.090208056 | 0.152902257 |
| | -0.096210188 | 0.923353639 | 0.947388464 |
| | -2.567110259 | 0.010255 | 0.023317828 |
| | -0.821977078 | 0.411089943 | 0.522897717 |
| NA | NA | NA | |
| NA | NA | NA | |

| | | |
|---|---|---|
| NA | NA | NA |
| 0.063708324 | 0.949202477 | NA |
| NA | NA | NA |
| NA | NA | NA |
| NA | NA | NA |
| NA | NA | NA |
| NA | NA | NA |
| NA | NA | NA |
| NA | NA | NA |
| NA | NA | NA |
| NA | NA | NA |
| NA | NA | NA |
| NA | NA | NA |
| NA | NA | NA |
| NA | NA | NA |
| NA | NA | NA |
| 0.30922356 | 0.757151473 | NA |
| NA | NA | NA |
| 0.192810118 | 0.847107679 | NA |
| NA | NA | NA |
| 0.192809521 | 0.847108147 | NA |
| NA | NA | NA |
| NA | NA | NA |
| NA | NA | NA |
| NA | NA | NA |
| NA | NA | NA |
| NA | NA | NA |
| NA | NA | NA |
| NA | NA | NA |
| -0.065389369 | 0.947863988 | NA |
| NA | NA | NA |
| NA | NA | NA |
| NA | NA | NA |
| 0.834880677 | 0.403784895 | NA |
| NA | NA | NA |
| NA | NA | NA |
| -0.065389369 | 0.947863988 | NA |
| NA | NA | NA |
| NA | NA | NA |
| NA | NA | NA |
| NA | NA | NA |
| NA | NA | NA |
| 0.192809521 | 0.847108147 | NA |
| 0.063708882 | 0.949202033 | NA |
| NA | NA | NA |
| NA | NA | NA |
| NA | NA | NA |

| | | |
|---|---|---|
| NA | NA | NA |
| 0.192809521 | 0.847108147 | NA |
| 0.192809521 | 0.847108147 | NA |
| -0.065388772 | 0.947864464 | NA |
| -3.512128792 | 0.000444533 | 0.001368592 |
| 0.81416122 | 0.415552598 | 0.527176792 |
| -3.856536888 | 0.000115005 | 0.000393247 |
| -0.341110661 | 0.733020276 | 0.80538127 |
| -2.715298694 | 0.006621602 | 0.015783134 |
| NA | NA | NA |
| 0.063708285 | 0.949202508 | NA |
| -0.872247103 | 0.383073588 | 0.495495422 |
| -0.729784614 | 0.465521851 | 0.574826689 |
| -0.7944764 | 0.426918143 | 0.538256654 |
| NA | NA | NA |
| 6.082593547 | 1.18E-09 | 7.99E-09 |
| -0.373408362 | 0.708844537 | 0.7866273 |
| 1.395333438 | 0.162915313 | 0.249710367 |
| -5.651061674 | 1.59E-08 | 9.51E-08 |
| -1.416225752 | 0.156709418 | 0.242113277 |
| NA | NA | NA |
| NA | NA | NA |
| NA | NA | NA |
| -0.886176355 | 0.375522503 | 0.488303731 |
| 2.200392149 | 0.027779084 | 0.055540417 |
| 4.222527494 | 2.42E-05 | 9.26E-05 |
| -1.085316434 | 0.27778153 | 0.384713507 |
| -0.120058966 | 0.904436438 | 0.933874182 |
| 3.181134138 | 0.001466997 | 0.004072189 |
| 2.52423864 | 0.011594919 | 0.025998461 |
| -0.089937715 | 0.92833671 | 0.950989594 |
| -2.629381844 | 0.008554025 | 0.01979908 |
| 0.458652326 | 0.646483854 | 0.735622625 |
| -0.065388772 | 0.947864464 | NA |
| 0.080463241 | 0.935868831 | 0.956348364 |
| 1.262361763 | 0.206818642 | 0.303750467 |
| 0.063710673 | 0.949200606 | NA |
| 0.108967543 | 0.913228234 | NA |
| 4.151588535 | 3.30E-05 | 0.00012394 |
| 7.007766261 | 2.42E-12 | 2.15E-11 |
| 3.916015959 | 9.00E-05 | 0.000314094 |
| 1.67314077 | 0.094299587 | 0.158666219 |
| NA | NA | NA |
| 3.923636804 | 8.72E-05 | 0.000305296 |
| 1.434096199 | 0.151544812 | 0.235527634 |
| 9.288062522 | 1.57E-20 | 2.89E-19 |
| 3.565897498 | 0.000362613 | 0.001135093 |

| | | |
|---|---|---|
| -1.322080871 | 0.186141223 | 0.278771902 |
| -0.864080065 | 0.387543909 | 0.499771724 |
| NA | NA | NA |
| 0.355606525 | 0.722135257 | 0.796953624 |
| -0.820451844 | 0.411958571 | 0.523630055 |
| 9.038355607 | 1.59E-19 | 2.74E-18 |
| 3.567499978 | 0.000360403 | 0.001129165 |
| 0.744063924 | 0.456837803 | 0.566520522 |
| -0.065388772 | 0.947864464 | NA |
| -5.211382388 | 1.87E-07 | 9.73E-07 |
| -2.867862373 | 0.004132553 | 0.01035517 |
| 0.026513166 | 0.978848032 | 0.985303879 |
| -2.113834409 | 0.034529413 | 0.067127861 |
| 0.208453878 | 0.834874591 | 0.884338173 |
| -2.772970347 | 0.005554718 | 0.013519506 |
| -0.692867538 | 0.488392683 | 0.596298952 |
| -4.438006341 | 9.08E-06 | 3.73E-05 |
| NA | NA | NA |
| -3.526865491 | 0.00042051 | 0.00129975 |
| -0.414165456 | 0.678752939 | 0.761744431 |
| -1.40322649 | 0.160549311 | 0.246597278 |
| 2.362209929 | 0.018166352 | 0.038456004 |
| 2.630113503 | 0.008535636 | 0.019763828 |
| 4.802122761 | 1.57E-06 | 7.16E-06 |
| -4.548621783 | 5.40E-06 | 2.29E-05 |
| 0.691148639 | 0.489472137 | NA |
| 1.581487408 | 0.113766636 | 0.185340985 |
| 0.532270167 | 0.594538892 | NA |
| -1.472414157 | 0.140909073 | 0.221584098 |
| 2.70341322 | 0.006863137 | 0.016298322 |
| 5.330044845 | 9.82E-08 | 5.30E-07 |
| 1.424739613 | 0.154232484 | 0.23890334 |
| 0.258557746 | 0.795976489 | NA |
| 0.577556033 | 0.563563898 | 0.665597337 |
| -1.177603492 | 0.238954717 | 0.341032989 |
| 0.769617562 | 0.441526785 | 0.551550403 |
| NA | NA | NA |
| 1.98741608 | 0.046876307 | 0.087560744 |
| NA | NA | NA |
| 1.528430949 | 0.126405576 | NA |
| -3.24351654 | 0.00118064 | 0.0033505 |
| -1.642659068 | 0.100453492 | 0.167122028 |
| 2.407879055 | 0.016045496 | 0.034592485 |
| -0.745946349 | 0.455699825 | 0.565456367 |
| NA | NA | NA |
| 1.240012867 | 0.214970635 | 0.313253923 |
| -7.330092075 | 2.30E-13 | 2.28E-12 |

| | | |
|---|---|---|
| -1.565179635 | 0.117540795 | 0.190497676 |
| 0.98802907 | 0.323138411 | 0.434053581 |
| 2.271537393 | 0.023114466 | 0.047407005 |
| 3.392879765 | 0.00069162 | 0.002054987 |
| 6.059325548 | 1.37E-09 | 9.16E-09 |
| 0.748241244 | 0.454314658 | NA |
| -6.01114124 | 1.84E-09 | 1.22E-08 |
| 2.269178732 | 0.023257459 | 0.047653422 |
| NA | NA | NA |
| NA | NA | NA |
| 0.23363542 | 0.815268027 | NA |
| 4.040663016 | 5.33E-05 | 0.000193601 |
| -0.261996244 | 0.793324338 | NA |
| 0.348928263 | 0.727143166 | NA |
| 7.586097829 | 3.30E-14 | 3.55E-13 |
| 0.643905338 | 0.51963682 | NA |
| -1.6167777 | 0.105926275 | 0.174583875 |
| -0.235903864 | 0.81350727 | 0.868837983 |
| -6.416105821 | 1.40E-10 | 1.04E-09 |
| -0.221759497 | 0.824501111 | 0.877050345 |
| -0.7961451 | 0.425947703 | 0.537345447 |
| 0.024434359 | 0.980506142 | 0.986459996 |
| -0.698134002 | 0.485093396 | 0.593445147 |
| NA | NA | NA |
| NA | NA | NA |
| 0.258749957 | 0.795828172 | NA |
| 3.58813536 | 0.000333051 | 0.001050041 |
| 0.726361072 | 0.467617449 | 0.576887217 |
| 0.65925493 | 0.509732079 | 0.615673113 |
| 0.688355653 | 0.491228843 | 0.598722034 |
| 1.719252321 | 0.085568438 | 0.14619645 |
| -8.635142456 | 5.87E-18 | 8.79E-17 |
| NA | NA | NA |
| 3.920666805 | 8.83E-05 | 0.000308609 |
| -0.047688231 | 0.961964714 | 0.973823966 |
| -1.596022898 | 0.110483681 | 0.180816434 |
| 1.27744352 | 0.201445709 | 0.297520207 |
| 0.611219138 | 0.541054516 | 0.645179286 |
| -3.517733614 | 0.000435249 | 0.001341991 |
| 1.628569385 | 0.10340421 | 0.171234361 |
| 2.934884398 | 0.00333672 | 0.008526605 |
| 7.06399501 | 1.62E-12 | 1.46E-11 |
| -0.057345603 | 0.954269894 | NA |
| -4.71005131 | 2.48E-06 | 1.10E-05 |
| -0.065389369 | 0.947863988 | NA |
| -1.411192174 | 0.158187957 | 0.243793151 |
| -3.379945571 | 0.000725002 | 0.002146534 |

| | | |
|---|---|---|
| -0.887264753 | 0.374936378 | 0.487793416 |
| -4.709243138 | 2.49E-06 | 1.10E-05 |
| -0.071880766 | 0.942696797 | NA |
| NA | NA | NA |
| NA | NA | NA |
| 0.063708882 | 0.949202033 | NA |
| -1.069690802 | 0.284758508 | 0.391922658 |
| -6.398456245 | 1.57E-10 | 1.16E-09 |
| -1.202035677 | 0.229349706 | 0.329845043 |
| -1.104398069 | 0.269420498 | 0.375541297 |
| 1.545401674 | 0.122249136 | 0.197082047 |
| -1.262633807 | 0.206720812 | 0.303687166 |
| 6.113480607 | 9.75E-10 | 6.65E-09 |
| -5.451417037 | 5.00E-08 | 2.81E-07 |
| -0.66451672 | 0.506359652 | 0.612812208 |
| -1.140584274 | 0.254042965 | 0.358236361 |
| -2.974422031 | 0.00293541 | 0.007606722 |
| -7.357940123 | 1.87E-13 | 1.86E-12 |
| NA | NA | NA |
| -1.162838582 | 0.244895 | 0.347785637 |
| NA | NA | NA |
| 3.048016036 | 0.002303576 | 0.006115019 |
| NA | NA | NA |
| NA | NA | NA |
| -7.994768071 | 1.30E-15 | 1.60E-14 |
| 8.919833649 | 4.67E-19 | 7.75E-18 |
| 23.58307349 | 5.75E-123 | 1.87E-120 |
| -3.507928926 | 0.00045161 | 0.001388269 |
| 0.931691921 | 0.351495763 | NA |
| NA | NA | NA |
| 3.816387517 | 0.00013542 | 0.000457492 |
| NA | NA | NA |
| -9.679233679 | 3.69E-22 | 7.71E-21 |
| NA | NA | NA |
| NA | NA | NA |
| -2.449037809 | 0.014323841 | 0.031260419 |
| -8.180426979 | 2.83E-16 | 3.69E-15 |
| 4.323744107 | 1.53E-05 | 6.07E-05 |
| NA | NA | NA |
| 5.463106777 | 4.68E-08 | 2.64E-07 |
| -8.745621218 | 2.22E-18 | 3.46E-17 |
| -2.066537798 | 0.038777727 | 0.07416494 |
| -0.065389966 | 0.947863513 | NA |
| NA | NA | NA |
| NA | NA | NA |
| 0.063708882 | 0.949202033 | NA |
| 0.119306565 | 0.905032483 | NA |

| | | | |
|---|---|---|---|
| | -0.065389369 | 0.947863988 | NA |
| | 0.768927303 | 0.441936468 | NA |
| | 1.479184544 | 0.139090999 | NA |
| | 0.063706581 | 0.949203865 | NA |
| NA | NA | NA | |
| | -0.110578491 | 0.911950604 | NA |
| | 1.495214244 | 0.134858537 | 0.213639822 |
| | 4.257136506 | 2.07E-05 | 8.03E-05 |
| | 0.405243743 | 0.685298354 | 0.767507173 |
| | 4.653701124 | 3.26E-06 | 1.42E-05 |
| | 0.236752982 | 0.812848429 | 0.868356645 |
| | 4.119230587 | 3.80E-05 | 0.000141233 |
| | 14.13532207 | 2.30E-45 | 1.56E-43 |
| | 0.392544237 | 0.694656132 | NA |
| | 12.59479984 | 2.26E-36 | 1.05E-34 |
| | -2.313382687 | 0.020701608 | 0.043043799 |
| | -2.104018853 | 0.035376802 | 0.068525102 |
| | -3.693285196 | 0.000221376 | 0.000721019 |
| | -1.75489586 | 0.079277123 | 0.136999888 |
| | 7.045709869 | 1.85E-12 | 1.66E-11 |
| | -7.412793962 | 1.24E-13 | 1.26E-12 |
| | 1.441819765 | 0.149353225 | 0.232597762 |
| | -1.56531799 | 0.117508367 | 0.190457452 |
| | 0.927692372 | 0.353567166 | 0.465459747 |
| NA | NA | NA | |
| | 0.106496541 | 0.9151884 | 0.941596746 |
| | 0.063708882 | 0.949202033 | NA |
| | -7.636195037 | 2.24E-14 | 2.46E-13 |
| NA | NA | NA | |
| | -0.065389369 | 0.947863988 | NA |
| NA | NA | NA | |
| | -3.543457323 | 0.000394917 | 0.001227313 |
| NA | NA | NA | |
| | 0.479334599 | 0.631700612 | 0.723570672 |
| | -1.042593136 | 0.29713677 | 0.405790776 |
| | 0.192810118 | 0.847107679 | NA |
| | 13.35980484 | 1.04E-40 | 5.79E-39 |
| | 11.5623312 | 6.39E-31 | 2.32E-29 |
| | 6.566418745 | 5.15E-11 | 4.05E-10 |
| | 12.91936143 | 3.50E-38 | 1.78E-36 |
| | 8.989228037 | 2.49E-19 | 4.22E-18 |
| | 0.57861044 | 0.562852059 | NA |
| | 3.580442818 | 0.000343012 | 0.001079542 |
| NA | NA | NA | |
| NA | NA | NA | |
| | -0.394205139 | 0.693429592 | 0.773912814 |
| NA | NA | NA | |

| | | |
|---|---|---|
| NA | NA | NA |
| 0.857971185 | 0.390908375 | 0.503004145 |
| 0.131056421 | 0.895730675 | 0.927486815 |
| 0.891516941 | 0.372651907 | 0.485477398 |
| 2.914533538 | 0.003562205 | 0.009043716 |
| -0.3029988 | 0.761890776 | 0.827912049 |
| -1.766702468 | 0.07727807 | 0.134082219 |
| -0.345886865 | 0.72942774 | NA |
| 2.563405703 | 0.010365085 | 0.023531799 |
| 4.055804012 | 5.00E-05 | 0.000182137 |
| 1.6239004 | 0.104397067 | 0.172604438 |
| 0.512454904 | 0.608332674 | NA |
| 21.46808614 | 3.10E-102 | 7.45E-100 |
| 0.39353732 | 0.693922665 | NA |
| 22.86902219 | 9.45E-116 | 2.92E-113 |
| 3.46101533 | 0.000538142 | 0.001631307 |
| 27.96007909 | 4.97E-172 | 2.65E-169 |
| 8.479151279 | 2.27E-17 | 3.26E-16 |
| 10.72861319 | 7.47E-27 | 2.15E-25 |
| 1.405664624 | 0.159823736 | NA |
| 19.93712877 | 1.94E-88 | 3.70E-86 |
| 13.00745644 | 1.11E-38 | 5.77E-37 |
| 15.53356125 | 2.06E-54 | 1.75E-52 |
| 8.167431035 | 3.15E-16 | 4.10E-15 |
| 11.02622133 | 2.86E-28 | 9.00E-27 |
| 4.752325393 | 2.01E-06 | 9.06E-06 |
| 1.436012484 | 0.150998789 | 0.234780699 |
| 0.166850382 | 0.867487775 | NA |
| -3.079695898 | 0.00207212 | 0.005561949 |
| -1.913492739 | 0.055685002 | 0.101536796 |
| 4.144098185 | 3.41E-05 | 0.000127792 |
| -2.364356951 | 0.018061401 | 0.038288885 |
| NA | NA | NA |
| -8.275408802 | 1.28E-16 | 1.72E-15 |
| -0.304287049 | 0.760909211 | 0.827060863 |
| 1.506660852 | 0.131897611 | 0.209764588 |
| 6.768966442 | 1.30E-11 | 1.08E-10 |
| 2.954469409 | 0.003132072 | 0.008057118 |
| 8.350575515 | 6.79E-17 | 9.36E-16 |
| 1.364977086 | 0.172260252 | 0.261487907 |
| 12.63931119 | 1.28E-36 | 6.03E-35 |
| 11.12537818 | 9.44E-29 | 3.06E-27 |
| 2.608918297 | 0.009082893 | 0.020893742 |
| 11.74033355 | 7.92E-32 | 3.02E-30 |
| NA | NA | NA |
| 8.680711915 | 3.93E-18 | 5.99E-17 |
| 10.41006809 | 2.23E-25 | 5.85E-24 |

| | | | |
|---|---|---|---|
| | 0.192809521 | 0.847108147 | NA |
| NA | NA | NA | |
| | 0.192810118 | 0.847107679 | NA |
| NA | NA | NA | |
| NA | NA | NA | |
| | 9.85715134 | 6.38E-23 | 1.41E-21 |
| | 0.380742228 | 0.703394531 | 0.78241779 |
| | 0.063707005 | 0.949203527 | NA |
| | 1.603733106 | 0.108772895 | 0.178471657 |
| NA | NA | NA | |
| | -0.065389369 | 0.947863988 | NA |
| | -1.730980821 | 0.083455185 | 0.143123212 |
| NA | NA | NA | |
| | -0.060855846 | 0.951474014 | 0.966847782 |
| | 6.512393398 | 7.40E-11 | 5.71E-10 |
| | -1.483239365 | 0.138010826 | 0.217820226 |
| | -0.958930491 | 0.337593763 | 0.448741633 |
| | -3.282065251 | 0.001030498 | 0.002962328 |
| | 0.170336933 | 0.864745168 | 0.9057443 |
| | 0.063707688 | 0.949202983 | NA |
| | -1.617030799 | 0.105871633 | 0.174523641 |
| | 2.878701459 | 0.003993161 | 0.010040063 |
| | -1.470243388 | 0.141495847 | 0.222367116 |
| | -2.508570152 | 0.01212209 | 0.027003907 |
| NA | NA | NA | |
| NA | NA | NA | |
| | 1.442459727 | 0.149172725 | 0.232360029 |
| | -0.94043352 | 0.346995235 | 0.458544467 |
| | 0.073892965 | 0.941095554 | NA |
| | 1.142484818 | 0.253252553 | 0.357297364 |
| | 0.702811032 | 0.482173521 | 0.590650416 |
| | -1.927281149 | 0.053944602 | 0.098845782 |
| | 0.265965772 | 0.79026557 | 0.850800664 |
| | -0.026743976 | 0.978663938 | 0.985158189 |
| | -6.16988899 | 6.83E-10 | 4.72E-09 |
| | -7.048780969 | 1.80E-12 | 1.62E-11 |
| | -3.031416074 | 0.002434096 | 0.006420636 |
| | -1.445895531 | 0.148206506 | 0.231085083 |
| | -2.314451429 | 0.020642973 | 0.042953995 |
| NA | NA | NA | |
| | 2.518923227 | 0.011771432 | 0.026337619 |
| | -0.538410701 | 0.59029354 | 0.689733087 |
| | -0.577713775 | 0.563457378 | NA |
| | -0.880855871 | 0.378395835 | 0.491043537 |
| NA | NA | NA | |
| | -1.559840135 | 0.118797664 | 0.1922608 |
| | -6.460458418 | 1.04E-10 | 7.92E-10 |

| | | | |
|---|---|---|---|
| | -0.61086643 | 0.541288011 | 0.645317405 |
| | -0.065388772 | 0.947864464 | NA |
| NA | NA | NA | |
| NA | NA | NA | |
| | -1.586993016 | 0.112514226 | 0.183587424 |
| | 1.241440033 | 0.214443238 | 0.312727071 |
| | 2.187029068 | 0.028740402 | 0.057206542 |
| NA | NA | NA | |
| | -0.288327136 | 0.773096337 | 0.836891329 |
| NA | NA | NA | |
| | 10.46163089 | 1.30E-25 | 3.45E-24 |
| | -1.164545481 | 0.244203023 | 0.34705621 |
| | 10.13832223 | 3.73E-24 | 9.04E-23 |
| | -0.780003748 | 0.435388669 | 0.545926179 |
| | 4.275932352 | 1.90E-05 | 7.42E-05 |
| NA | NA | NA | |
| NA | NA | NA | |
| NA | NA | NA | |
| | 0.695714648 | 0.48660755 | 0.594779603 |
| | -1.47971044 | 0.138950538 | 0.219013348 |
| | -0.560274337 | 0.575292329 | 0.676417089 |
| | 8.841803244 | 9.42E-19 | 1.52E-17 |
| NA | NA | NA | |
| | 1.953427545 | 0.050768955 | 0.093851325 |
| | 1.886466019 | 0.059232183 | 0.106891244 |
| | 0.864054964 | 0.387557698 | 0.499771724 |
| | 4.015782728 | 5.92E-05 | 0.000212921 |
| | -5.547618913 | 2.90E-08 | 1.67E-07 |
| NA | NA | NA | |
| | -2.954537896 | 0.003131377 | 0.00805622 |
| | -0.085607355 | 0.931778551 | NA |
| | 2.992492793 | 0.002767092 | 0.007209359 |
| | -0.065389369 | 0.947863988 | NA |
| | 4.016605933 | 5.90E-05 | 0.000212301 |
| | 1.280156944 | 0.200489945 | 0.296342466 |
| | -0.690771337 | 0.489709251 | 0.5974143 |
| | -0.371732752 | 0.710091837 | NA |
| | -10.31623331 | 5.95E-25 | 1.51E-23 |
| | -4.203826411 | 2.62E-05 | 0.000100242 |
| NA | NA | NA | |
| | -2.389192697 | 0.016885443 | 0.036169675 |
| | -0.940100699 | 0.34716591 | 0.458721572 |
| | 6.302023596 | 2.94E-10 | 2.11E-09 |
| | -1.878620177 | 0.060296374 | 0.108483197 |
| | 0.269754526 | 0.787349109 | 0.848467223 |
| | 0.320051528 | 0.748929269 | NA |
| NA | NA | NA | |

| | | |
|---|---|---|
| 0.063708882 | 0.949202033 | NA |
| NA | NA | NA |
| -0.736423874 | 0.46147278 | NA |
| NA | NA | NA |
| -0.532593541 | 0.594314974 | NA |
| -6.174371997 | 6.64E-10 | 4.59E-09 |
| -0.748687812 | 0.454045391 | 0.563951774 |
| -0.818185588 | 0.413251223 | 0.524846667 |
| -1.560328582 | 0.118682252 | 0.19209886 |
| NA | NA | NA |
| -0.482805576 | 0.629233791 | 0.721895074 |
| 2.851940119 | 0.004345329 | 0.010833025 |
| 7.823096229 | 5.15E-15 | 6.02E-14 |
| -1.031718904 | 0.302203819 | 0.411453525 |
| 0.063707688 | 0.949202983 | NA |
| -7.089826996 | 1.34E-12 | 1.23E-11 |
| 5.099706421 | 3.40E-07 | 1.71E-06 |
| 0.199601859 | 0.841791974 | NA |
| 5.77298643 | 7.79E-09 | 4.81E-08 |
| 3.07534139 | 0.002102617 | 0.005635951 |
| 1.536336628 | 0.124455841 | NA |
| -0.090765623 | 0.927678826 | NA |
| 0.192810118 | 0.847107679 | NA |
| -0.344849123 | 0.730207801 | 0.80301204 |
| NA | NA | NA |
| 0.192809521 | 0.847108147 | NA |
| -5.33138687 | 9.75E-08 | 5.26E-07 |
| 0.212524895 | 0.831697558 | 0.882273648 |
| -0.814513627 | 0.415350769 | 0.527003985 |
| 4.967810209 | 6.77E-07 | 3.26E-06 |
| -1.103486771 | 0.269815834 | 0.375966727 |
| 4.976110666 | 6.49E-07 | 3.13E-06 |
| 3.535052606 | 0.000407694 | 0.001264195 |
| 4.292078022 | 1.77E-05 | 6.93E-05 |
| -0.940391798 | 0.347016628 | NA |
| NA | NA | NA |
| 0.799275514 | 0.424130674 | 0.535701204 |
| NA | NA | NA |
| 0.0639902 | 0.94897803 | 0.965095079 |
| -0.802363647 | 0.422342635 | 0.534089636 |
| -2.739727478 | 0.006149015 | 0.014779128 |
| NA | NA | NA |
| -6.79716687 | 1.07E-11 | 8.92E-11 |
| -0.194488217 | 0.845793621 | NA |
| NA | NA | NA |
| NA | NA | NA |
| 9.395864332 | 5.67E-21 | 1.08E-19 |

| | | |
|---|---|---|
| -0.065388772 | 0.947864464 | NA |
| -0.144314877 | 0.885251831 | 0.919558191 |
| -0.065389369 | 0.947863988 | NA |
| 1.045120892 | 0.295967099 | NA |
| 26.74201063 | 1.53E-157 | 6.72E-155 |
| -2.09339221 | 0.036314157 | 0.070096657 |
| 0.826687394 | 0.408414268 | 0.520261038 |
| NA | NA | NA |
| 0.192810118 | 0.847107679 | NA |
| NA | NA | NA |
| NA | NA | NA |
| -4.273253034 | 1.93E-05 | 7.50E-05 |
| -0.32359698 | 0.746243175 | NA |
| -4.389439784 | 1.14E-05 | 4.59E-05 |
| -4.568974817 | 4.90E-06 | 2.09E-05 |
| 3.321544292 | 0.000895208 | 0.00260303 |
| -1.631453887 | 0.102794582 | 0.170360087 |
| NA | NA | NA |
| NA | NA | NA |
| -6.54827109 | 5.82E-11 | 4.54E-10 |
| -11.14974404 | 7.18E-29 | 2.35E-27 |
| -2.552688539 | 0.010689504 | 0.024202468 |
| -0.203790186 | 0.838517462 | 0.886984776 |
| -1.239530547 | 0.215149084 | 0.313419949 |
| 0.693721315 | 0.487856996 | 0.595923118 |
| -2.623286168 | 0.008708609 | 0.020123383 |
| NA | NA | NA |
| -0.459655539 | 0.645763488 | NA |
| NA | NA | NA |
| NA | NA | NA |
| NA | NA | NA |
| NA | NA | NA |
| NA | NA | NA |
| NA | NA | NA |
| NA | NA | NA |
| -8.064260151 | 7.37E-16 | 9.31E-15 |
| NA | NA | NA |
| 0.063708285 | 0.949202508 | NA |
| NA | NA | NA |
| 4.14414736 | 3.41E-05 | 0.000127784 |
| -4.401229117 | 1.08E-05 | 4.37E-05 |
| 1.742327466 | 0.08145116 | 0.140243691 |
| 0.192809521 | 0.847108147 | NA |
| 2.940508034 | 0.003276745 | 0.008390709 |
| -12.20149272 | 3.05E-34 | 1.29E-32 |
| 4.901374138 | 9.52E-07 | 4.49E-06 |
| -3.89217989 | 9.93E-05 | 0.000343652 |

| | | |
|---|---|---|
| -4.359133346 | 1.31E-05 | 5.22E-05 |
| -0.503004741 | 0.614960939 | 0.710404256 |
| -0.065388772 | 0.947864464 | NA |
| -0.909442659 | 0.363116512 | 0.475554955 |
| 0.257881913 | 0.796498044 | 0.855132594 |
| 7.128264379 | 1.02E-12 | 9.43E-12 |
| -3.617906518 | 0.000296996 | 0.000944935 |
| -0.726278189 | 0.467668247 | 0.576893018 |
| -5.452895361 | 4.96E-08 | 2.78E-07 |
| -0.619756334 | 0.535418221 | 0.639737942 |
| -2.606587461 | 0.009144947 | 0.021028756 |
| -2.174831848 | 0.02964272 | 0.058764238 |
| -0.118480828 | 0.905686685 | 0.934702418 |
| NA | NA | NA |
| 8.239769515 | 1.73E-16 | 2.30E-15 |
| 4.872520575 | 1.10E-06 | 5.14E-06 |
| 7.467518777 | 8.17E-14 | 8.47E-13 |
| -0.343141703 | 0.731491859 | 0.804200872 |
| -1.062544192 | 0.287988707 | 0.395433504 |
| -0.111311781 | 0.911369114 | NA |
| -0.355115243 | 0.722503258 | 0.797190061 |
| 0.424625959 | 0.671109366 | NA |
| 0.063708285 | 0.949202508 | NA |
| NA | NA | NA |
| NA | NA | NA |
| NA | NA | NA |
| NA | NA | NA |
| NA | NA | NA |
| NA | NA | NA |
| 3.306217193 | 0.000945648 | 0.002739215 |
| NA | NA | NA |
| 1.228494448 | 0.219261412 | 0.318225278 |
| NA | NA | NA |
| -5.809614174 | 6.26E-09 | 3.91E-08 |
| -0.339204534 | 0.734455654 | 0.806361305 |
| -5.863299175 | 4.54E-09 | 2.88E-08 |
| -1.901316472 | 0.057260573 | 0.103962871 |
| 3.466417827 | 0.000527443 | 0.001602561 |
| -0.687521086 | 0.491754418 | 0.599077161 |
| -1.238118275 | 0.215672209 | 0.314090614 |
| 1.167572438 | 0.242979273 | 0.34565338 |
| 1.297250147 | 0.194545131 | 0.288886132 |
| 0.767350095 | 0.442873392 | NA |
| -0.273877838 | 0.784178503 | NA |
| 3.226138509 | 0.001254726 | 0.003533108 |
| 0.307624806 | 0.758367842 | 0.825015086 |
| 0.063705462 | 0.949204756 | NA |

| | | |
|---|---|---|
| 0.16880313 | 0.865951496 | 0.906508752 |
| 0.092176936 | 0.926557462 | NA |
| 5.527120725 | 3.26E-08 | 1.87E-07 |
| 1.574124671 | 0.11545863 | 0.187682135 |
| NA | NA | NA |
| -3.051852566 | 0.002274338 | 0.006047665 |
| 2.599204284 | 0.009344015 | 0.021433348 |
| NA | NA | NA |
| 2.464666571 | 0.01371408 | 0.030129328 |
| -2.264189677 | 0.023562451 | 0.048183673 |
| -1.463582393 | 0.143308086 | 0.224848078 |
| -1.448777895 | 0.147399619 | 0.230056274 |
| -7.292622848 | 3.04E-13 | 2.98E-12 |
| -3.012930217 | 0.002587384 | 0.006784937 |
| -0.98074521 | 0.326718403 | 0.437784572 |
| 0.327806351 | 0.743058085 | 0.813065012 |
| 11.52211692 | 1.02E-30 | 3.65E-29 |
| 10.09246487 | 5.97E-24 | 1.42E-22 |
| 1.276259732 | 0.20186372 | 0.298038631 |
| -1.367251069 | 0.171546623 | 0.260604948 |
| 1.808031725 | 0.070601563 | 0.124194049 |
| NA | NA | NA |
| NA | NA | NA |
| 1.247257021 | 0.212303271 | 0.310213128 |
| NA | NA | NA |
| NA | NA | NA |
| NA | NA | NA |
| NA | NA | NA |
| NA | NA | NA |
| 1.128229219 | 0.259223127 | 0.364248295 |
| 4.491463843 | 7.07E-06 | 2.95E-05 |
| -2.963122134 | 0.003045357 | 0.007860708 |
| NA | NA | NA |
| 0.522182735 | 0.601543106 | 0.699137441 |
| -1.404357433 | 0.16021244 | 0.246215839 |
| NA | NA | NA |
| NA | NA | NA |
| NA | NA | NA |
| NA | NA | NA |
| -0.441735356 | 0.658680722 | 0.745974321 |
| 0.063710673 | 0.949200606 | NA |
| NA | NA | NA |
| 0.063708882 | 0.949202033 | NA |
| 0.548184902 | 0.583564948 | 0.683670988 |
| 0.800395101 | 0.423481918 | 0.535093149 |
| 0.363245836 | 0.716421251 | NA |
| 0.501994266 | 0.615671554 | 0.710805304 |

| | | |
|---|---|---|
| 0.864084541 | 0.387541451 | 0.499771724 |
| -0.090498911 | 0.927890759 | NA |
| 1.34941974 | 0.177202184 | 0.267710363 |
| 0.859984621 | 0.38979752 | 0.501910177 |
| 1.18524531 | 0.23592047 | 0.337645913 |
| 0.703806258 | 0.481553436 | 0.590152884 |
| 1.408129783 | 0.159092644 | 0.244870795 |
| 0.192810118 | 0.847107679 | NA |
| 0.063708882 | 0.949202033 | NA |
| -0.469896657 | 0.638428853 | NA |
| 5.337184483 | 9.44E-08 | 5.10E-07 |
| 0.107487194 | 0.914402484 | NA |
| 1.531357553 | 0.125681049 | NA |
| 6.691181826 | 2.21E-11 | 1.80E-10 |
| -0.039158654 | 0.968763898 | 0.978537545 |
| 2.156339026 | 0.031057202 | 0.061180684 |
| 3.661567995 | 0.000250676 | 0.000807936 |
| -0.228902048 | 0.818945048 | 0.872709237 |
| -0.19448762 | 0.845794089 | NA |
| -0.10782225 | 0.914136693 | NA |
| NA | NA | NA |
| 0.869606893 | 0.38451527 | 0.496744113 |
| -0.199357551 | 0.841983063 | 0.889271112 |
| 0.057638743 | 0.954036389 | 0.968586469 |
| 1.595820976 | 0.110528769 | 0.180865982 |
| -3.56313797 | 0.000366448 | 0.001146237 |
| 3.885525666 | 0.000102109 | 0.000352376 |
| -0.882517356 | 0.377497098 | 0.490156786 |
| 1.663126636 | 0.096287089 | 0.161456813 |
| -0.531099506 | 0.595349827 | 0.693872516 |
| 2.054786063 | 0.039899673 | 0.076055808 |
| 0.124255511 | 0.901112971 | NA |
| 3.749110016 | 0.000177463 | 0.000587631 |
| 0.012185398 | 0.9902777 | 0.993174293 |
| 1.4455195 | 0.14831202 | 0.231206393 |
| -1.756445227 | 0.079012416 | 0.136665108 |
| NA | NA | NA |
| 4.116801054 | 3.84E-05 | 0.000142603 |
| 0.063708285 | 0.949202508 | NA |
| 1.137585893 | 0.25529344 | 0.359678893 |
| NA | NA | NA |
| 5.107528668 | 3.26E-07 | 1.64E-06 |
| 0.37842362 | 0.705115924 | 0.783654307 |
| -1.976763783 | 0.048068337 | 0.089480179 |
| -1.02458706 | 0.305558073 | 0.415187255 |
| 2.336188355 | 0.019481427 | 0.040846048 |
| 0.192809521 | 0.847108147 | NA |

| | | | |
|---|---|---|---|
| | 3.513014539 | 0.000443053 | 0.001364541 |
| | -3.278153473 | 0.001044886 | 0.002999941 |
| | 1.344787091 | 0.178694 | 0.269505352 |
| | 0.291144164 | 0.770941064 | 0.835027688 |
| | -10.00565295 | 1.44E-23 | 3.35E-22 |
| | -3.86396992 | 0.000111559 | 0.000382667 |
| | -1.040355763 | 0.298174646 | NA |
| | 0.347664246 | 0.728092351 | NA |
| | -8.959030784 | 3.28E-19 | 5.51E-18 |
| | -6.464120811 | 1.02E-10 | 7.75E-10 |
| NA | NA | NA | |
| | -6.423600859 | 1.33E-10 | 9.96E-10 |
| NA | NA | NA | |
| | 0.139412004 | 0.889124588 | 0.922440041 |
| | -3.233758189 | 0.001221728 | 0.003449116 |
| | 5.979175595 | 2.24E-09 | 1.47E-08 |
| | -6.534997595 | 6.36E-11 | 4.94E-10 |
| NA | NA | NA | |
| | 0.216349054 | 0.828715675 | 0.88023566 |
| | 3.066706014 | 0.002164316 | 0.005789551 |
| | 0.27536091 | 0.783038973 | 0.8451324 |
| | 0.567771488 | 0.570190143 | 0.671553702 |
| | 5.148301987 | 2.63E-07 | 1.34E-06 |
| | 0.430914007 | 0.666530898 | 0.752108394 |
| | 0.485627925 | 0.627230994 | 0.720329973 |
| | 1.600974241 | 0.109382625 | 0.179284074 |
| | -7.104784726 | 1.21E-12 | 1.11E-11 |
| | 0.173680827 | 0.862116306 | 0.904239488 |
| | -0.404855097 | 0.685584027 | NA |
| | -4.178248223 | 2.94E-05 | 0.000111272 |
| | -0.067039719 | 0.946550083 | NA |
| | 0.192808924 | 0.847108615 | NA |
| | 4.968261012 | 6.76E-07 | 3.25E-06 |
| | -4.768486503 | 1.86E-06 | 8.40E-06 |
| | -1.291857706 | 0.196406427 | 0.291227894 |
| | 4.967387278 | 6.79E-07 | 3.26E-06 |
| NA | NA | NA | |
| | -1.987949076 | 0.046817322 | 0.08746362 |
| | 0.063708882 | 0.949202033 | NA |
| | -4.750341175 | 2.03E-06 | 9.14E-06 |
| NA | NA | NA | |
| | 2.754381327 | 0.005880321 | 0.014202879 |
| | 0.716553434 | 0.473649686 | 0.582491605 |
| | 3.494943112 | 0.000474163 | 0.001451957 |
| | 0.265841104 | 0.790361586 | 0.850834394 |
| | 1.328090862 | 0.184148092 | 0.276332539 |
| NA | NA | NA | |

| | | |
|---|---|---|
| 0.590715933 | 0.55471077 | 0.657623767 |
| -3.129719051 | 0.001749736 | 0.004778601 |
| -4.058923111 | 4.93E-05 | 0.0001798 |
| -0.100775111 | 0.919728985 | 0.944870156 |
| 8.50207668 | 1.86E-17 | 2.70E-16 |
| -6.339604632 | 2.30E-10 | 1.67E-09 |
| 1.148576704 | 0.250730568 | 0.354424019 |
| -0.389748721 | 0.696722366 | 0.776618161 |
| 8.221069434 | 2.02E-16 | 2.68E-15 |
| NA | NA | NA |
| 0.347422444 | 0.728273975 | 0.801649043 |
| 1.793746142 | 0.072853697 | 0.127412523 |
| -1.568040462 | 0.11687169 | 0.189597407 |
| 12.97227069 | 1.76E-38 | 9.07E-37 |
| 0.063708285 | 0.949202508 | NA |
| 0.195609901 | 0.84491551 | 0.891615802 |
| 6.425094685 | 1.32E-10 | 9.87E-10 |
| NA | NA | NA |
| 1.688109335 | 0.091390243 | 0.154548195 |
| 0.071152621 | 0.943276289 | NA |
| NA | NA | NA |
| 0.742892103 | 0.457547007 | 0.567203288 |
| 6.024599739 | 1.70E-09 | 1.13E-08 |
| 0.837946072 | 0.402060992 | 0.513815308 |
| -0.175156348 | 0.860956784 | 0.903279692 |
| -3.236297711 | 0.00121091 | 0.003423981 |
| 0.142790503 | 0.886455638 | NA |
| -8.397724251 | 4.55E-17 | 6.38E-16 |
| -6.877041795 | 6.11E-12 | 5.22E-11 |
| -5.444050504 | 5.21E-08 | 2.92E-07 |
| 0.385512486 | 0.699857757 | 0.779245537 |
| 1.148230177 | 0.250873555 | 0.354546049 |
| -9.02840029 | 1.74E-19 | 2.99E-18 |
| -4.125870751 | 3.69E-05 | 0.00013754 |
| NA | NA | NA |
| 4.971256758 | 6.65E-07 | 3.20E-06 |
| 7.430831683 | 1.08E-13 | 1.10E-12 |
| 0.780704172 | 0.434976506 | 0.545510067 |
| 1.414218581 | 0.157297734 | 0.24277994 |
| 7.432015713 | 1.07E-13 | 1.10E-12 |
| NA | NA | NA |
| NA | NA | NA |
| 5.507797007 | 3.63E-08 | 2.08E-07 |
| 1.252667609 | 0.210326699 | 0.307864168 |
| 7.075118756 | 1.49E-12 | 1.36E-11 |
| 0.251153833 | 0.801695176 | 0.859302427 |
| -2.701763443 | 0.006897282 | 0.016365433 |

| | | |
|---|---|---|
| -0.601261767 | 0.54766565 | 0.651211455 |
| NA | NA | NA |
| -0.55180594 | 0.581081317 | 0.681658583 |
| 1.138684967 | 0.254834575 | 0.359170528 |
| 0.354307124 | 0.723108729 | NA |
| 0.278704665 | 0.780471482 | 0.843161172 |
| -1.839358693 | 0.065862445 | 0.11700543 |
| 0.662221771 | 0.507829104 | 0.614174839 |
| -7.038265179 | 1.95E-12 | 1.75E-11 |
| -5.918384443 | 3.25E-09 | 2.10E-08 |
| -0.633728659 | 0.526257918 | 0.631368245 |
| -0.137425166 | 0.89069474 | NA |
| NA | NA | NA |
| NA | NA | NA |
| NA | NA | NA |
| NA | NA | NA |
| NA | NA | NA |
| NA | NA | NA |
| NA | NA | NA |
| NA | NA | NA |
| NA | NA | NA |
| NA | NA | NA |
| NA | NA | NA |
| NA | NA | NA |
| NA | NA | NA |
| NA | NA | NA |
| NA | NA | NA |
| NA | NA | NA |
| NA | NA | NA |
| NA | NA | NA |
| 0.063708882 | 0.949202033 | NA |
| NA | NA | NA |
| NA | NA | NA |
| 0.628856507 | 0.529443002 | NA |
| NA | NA | NA |
| NA | NA | NA |
| NA | NA | NA |
| NA | NA | NA |
| NA | NA | NA |
| NA | NA | NA |
| NA | NA | NA |
| 0.192809521 | 0.847108147 | NA |
| NA | NA | NA |
| NA | NA | NA |
| NA | NA | NA |
| NA | NA | NA |
| NA | NA | NA |

| | | |
|---|---|---|
| NA | NA | NA |
| NA | NA | NA |
| NA | NA | NA |
| 0.063708285 | 0.949202508 | NA |
| NA | NA | NA |
| NA | NA | NA |
| NA | NA | NA |
| NA | NA | NA |
| NA | NA | NA |
| NA | NA | NA |
| 0.192810118 | 0.847107679 | NA |
| NA | NA | NA |
| 2.540126168 | 0.011081249 | 0.024985466 |
| NA | NA | NA |
| NA | NA | NA |
| NA | NA | NA |
| NA | NA | NA |
| 0.063708882 | 0.949202033 | NA |
| 0.412143228 | 0.680234446 | NA |
| NA | NA | NA |
| 1.258970395 | 0.208041025 | 0.305029359 |
| NA | NA | NA |
| NA | NA | NA |
| NA | NA | NA |
| NA | NA | NA |
| NA | NA | NA |
| NA | NA | NA |
| NA | NA | NA |
| 0.210360696 | 0.833386166 | NA |
| NA | NA | NA |
| -0.433241652 | 0.664839222 | 0.750942936 |
| -1.093189277 | 0.274310704 | 0.381020917 |
| 0.223483479 | 0.82315924 | 0.87608112 |
| 1.286391696 | 0.198306395 | 0.293583076 |
| 1.631485821 | 0.102787849 | 0.170360087 |
| NA | NA | NA |
| 0.10424596 | 0.916974162 | 0.942935619 |
| 3.320282243 | 0.000899265 | 0.002613025 |
| 2.820107769 | 0.004800752 | 0.011866874 |
| NA | NA | NA |
| 1.438874938 | 0.150185959 | 0.233691071 |
| 5.043724937 | 4.57E-07 | 2.25E-06 |
| 4.393043441 | 1.12E-05 | 4.52E-05 |
| 0.141486415 | 0.887485691 | 0.921121531 |
| 7.864962554 | 3.69E-15 | 4.38E-14 |
| 2.765021772 | 0.0056919 | 0.013805063 |
| -1.380634741 | 0.167391296 | 0.255366864 |

| | | | |
|---|---|---|---|
| | 4.86187778 | 1.16E-06 | 5.40E-06 |
| | -0.85429224 | 0.392943086 | NA |
| | 1.761802986 | 0.07810259 | 0.135304253 |
| | 7.237191222 | 4.58E-13 | 4.41E-12 |
| | -2.913204226 | 0.003577406 | 0.009074943 |
| | -0.113289382 | 0.909801135 | 0.937750058 |
| | -1.011897845 | 0.3115869 | 0.421711445 |
| | -2.558508515 | 0.010512225 | 0.023837741 |
| | 0.071582775 | 0.942933948 | 0.961291819 |
| | -0.343007234 | 0.731593018 | NA |
| | 0.703279855 | 0.481881362 | 0.59039673 |
| | 2.050621054 | 0.040303864 | 0.076709431 |
| | 1.351086812 | 0.176667626 | 0.267076805 |
| | 9.920207706 | 3.40E-23 | 7.69E-22 |
| | -0.229605304 | 0.818398485 | 0.872349633 |
| | 0.118908045 | 0.905348208 | 0.934468687 |
| | 24.55793156 | 3.56E-133 | 1.27E-130 |
| | 5.011146799 | 5.41E-07 | 2.63E-06 |
| | 0.922465972 | 0.356285572 | NA |
| | 9.33437841 | 1.02E-20 | 1.90E-19 |
| | -5.785252305 | 7.24E-09 | 4.49E-08 |
| | -1.660610334 | 0.096791732 | 0.162118379 |
| | 6.241941227 | 4.32E-10 | 3.05E-09 |
| | -0.065388772 | 0.947864464 | NA |
| | 3.873993911 | 0.000107066 | 0.000368316 |
| | 9.637869908 | 5.53E-22 | 1.14E-20 |
| | 14.83681029 | 8.47E-50 | 6.52E-48 |
| | 11.84617092 | 2.25E-32 | 8.79E-31 |
| | -4.934834196 | 8.02E-07 | 3.82E-06 |
| | -1.74838159 | 0.080397974 | 0.13868789 |
| | 8.005877606 | 1.19E-15 | 1.47E-14 |
| | -1.887846 | 0.059046628 | 0.106650101 |
| | 4.36436235 | 1.27E-05 | 5.10E-05 |
| | 4.745300559 | 2.08E-06 | 9.35E-06 |
| | 0.481333108 | 0.63027977 | 0.722472186 |
| | 3.900628548 | 9.59E-05 | 0.000332843 |
| | -1.17713812 | 0.239140382 | 0.34120889 |
| NA | NA | | NA |
| | -5.984796134 | 2.17E-09 | 1.42E-08 |
| | 0.366148092 | 0.714254573 | 0.790994625 |
| | -11.28861531 | 1.49E-29 | 5.04E-28 |
| | 3.392208472 | 0.000693317 | 0.002059544 |
| | -1.936405811 | 0.052818018 | 0.097037214 |
| | 4.432793441 | 9.30E-06 | 3.81E-05 |
| | -0.456772668 | 0.647834453 | 0.736724051 |
| | -0.153606637 | 0.877919903 | 0.914615531 |
| | -0.116448024 | 0.907297476 | NA |

| | | |
|---|---|---|
| 0.327601119 | 0.743213275 | 0.813087473 |
| -0.484184941 | 0.628254623 | 0.721146623 |
| 3.520951318 | 0.000430002 | 0.001326545 |
| NA | NA | NA |
| 3.652711204 | 0.000259486 | 0.000834074 |
| 0.372802944 | 0.709295111 | 0.786952865 |
| 1.401285413 | 0.161128742 | 0.247305151 |
| NA | NA | NA |
| NA | NA | NA |
| NA | NA | NA |
| NA | NA | NA |
| -0.397113154 | 0.69128402 | 0.772206856 |
| NA | NA | NA |
| -2.037021943 | 0.041647843 | 0.078979128 |
| -1.939935549 | 0.052387523 | 0.096371382 |
| 1.085431985 | 0.277730372 | 0.384682155 |
| 0.063708882 | 0.949202033 | NA |
| -0.429560632 | 0.667515279 | 0.752676252 |
| 6.191086144 | 5.98E-10 | 4.15E-09 |
| 23.95591952 | 8.02E-127 | 2.68E-124 |
| 0.603712732 | 0.546034648 | 0.649614869 |
| 4.293690892 | 1.76E-05 | 6.89E-05 |
| -3.711235814 | 0.00020625 | 0.000676154 |
| 0.31685483 | 0.751353758 | NA |
| 3.907290261 | 9.33E-05 | 0.000324882 |
| 2.332562842 | 0.019671101 | 0.041188528 |
| NA | NA | NA |
| 16.0919481 | 2.91E-58 | 2.73E-56 |
| NA | NA | NA |
| -0.647795152 | 0.517117452 | 0.622371006 |
| 0.192809521 | 0.847108147 | NA |
| -0.83806552 | 0.401993907 | 0.513755807 |
| 0.852422192 | 0.393979807 | 0.505993872 |
| 0.376754054 | 0.706356388 | 0.784631447 |
| 0.570474576 | 0.568355861 | 0.670023888 |
| 1.071675522 | 0.283865799 | 0.39084599 |
| 1.800947761 | 0.071711114 | 0.125774591 |
| 4.402078838 | 1.07E-05 | 4.35E-05 |
| -2.604528265 | 0.009200083 | 0.021145829 |
| 2.457742001 | 0.013981358 | 0.030643904 |
| -0.075586801 | 0.939747837 | NA |
| 0.0731897 | 0.941655163 | NA |
| 0.3474247 | 0.728272279 | NA |
| -0.636222237 | 0.524631581 | 0.629749112 |
| -0.065392754 | 0.947861293 | NA |
| -5.587419887 | 2.30E-08 | 1.35E-07 |
| -0.112233322 | 0.910638409 | NA |

| | | |
|---|---|---|
| -6.202346143 | 5.56E-10 | 3.88E-09 |
| -0.471860688 | 0.63702623 | 0.728100569 |
| 3.153869853 | 0.001611209 | 0.004438539 |
| -10.24439784 | 1.25E-24 | 3.11E-23 |
| 0.192810118 | 0.847107679 | NA |
| 0.428708418 | 0.668135434 | 0.753145249 |
| 0.192809521 | 0.847108147 | NA |
| 0.145112203 | 0.884622284 | 0.919140546 |
| 0.379440295 | 0.704360933 | 0.783214311 |
| NA | NA | NA |
| 0.347768159 | 0.728014305 | 0.801454707 |
| 4.887225809 | 1.02E-06 | 4.79E-06 |
| 0.94637554 | 0.343957077 | 0.455298833 |
| -4.888157882 | 1.02E-06 | 4.77E-06 |
| NA | NA | NA |
| 2.168503951 | 0.030120364 | 0.059541361 |
| -4.051142617 | 5.10E-05 | 0.000185507 |
| -0.594580577 | 0.552123867 | 0.655332909 |
| -6.70964554 | 1.95E-11 | 1.59E-10 |
| -0.131995104 | 0.894988164 | 0.926842771 |
| 1.453926713 | 0.145966626 | 0.22818966 |
| -0.065390563 | 0.947863038 | NA |
| -3.350804778 | 0.000805771 | 0.002361921 |
| 4.970336083 | 6.68E-07 | 3.22E-06 |
| NA | NA | NA |
| 21.8935792 | 2.99E-106 | 8.14E-104 |
| -6.79090798 | 1.11E-11 | 9.30E-11 |
| 5.56673206 | 2.60E-08 | 1.51E-07 |
| 0.063707688 | 0.949202983 | NA |
| -2.555472156 | 0.010604384 | 0.02403366 |
| -2.904919332 | 0.003673478 | 0.009297921 |
| NA | NA | NA |
| -2.126336211 | 0.033475275 | 0.065357733 |
| -6.691840582 | 2.20E-11 | 1.79E-10 |
| -2.153877399 | 0.031249788 | 0.061506817 |
| -0.798460427 | 0.424603349 | NA |
| 0.892828696 | 0.371948916 | 0.484747489 |
| NA | NA | NA |
| NA | NA | NA |
| 0.133926464 | 0.893460724 | NA |
| 2.895447389 | 0.003786185 | 0.009559993 |
| -0.012461756 | 0.990057215 | 0.992992951 |
| -1.408172615 | 0.159079963 | 0.244866335 |
| -6.373426034 | 1.85E-10 | 1.36E-09 |
| 3.757385396 | 0.000171698 | 0.000569596 |
| NA | NA | NA |
| -0.608444793 | 0.542892511 | 0.646863718 |

| | | |
|---|---|---|
| 1.07765322 | 0.281188542 | 0.388202593 |
| -0.241416985 | 0.80923195 | NA |
| 0.158813663 | 0.873815684 | 0.912077004 |
| -0.89269632 | 0.372019821 | NA |
| 18.87328047 | 1.89E-79 | 2.94E-77 |
| 5.104942072 | 3.31E-07 | 1.66E-06 |
| -0.947031481 | 0.343622738 | 0.455024713 |
| 0.085595035 | 0.931788346 | 0.953580109 |
| 4.429018093 | 9.47E-06 | 3.87E-05 |
| -0.350596381 | 0.725891172 | NA |
| 5.201144251 | 1.98E-07 | 1.03E-06 |
| -2.370380189 | 0.017769802 | 0.037779508 |
| -0.25783658 | 0.796533032 | 0.855132594 |
| -2.42218208 | 0.015427618 | 0.033401374 |
| -0.008351218 | 0.993336769 | 0.995325034 |
| 1.360148174 | 0.173783039 | 0.263463929 |
| -1.661861446 | 0.096540559 | 0.161795124 |
| -1.504735943 | 0.132391978 | 0.210442578 |
| -2.56384681 | 0.010351922 | 0.023508312 |
| 6.203210429 | 5.53E-10 | 3.86E-09 |
| -0.569537726 | 0.568991276 | 0.670583466 |
| 8.662013316 | 4.64E-18 | 7.01E-17 |
| 5.202505511 | 1.97E-07 | 1.02E-06 |
| 0.204886216 | 0.837661025 | 0.886464377 |
| -0.030585516 | 0.975600093 | 0.983022838 |
| 8.739001098 | 2.35E-18 | 3.66E-17 |
| 2.443682412 | 0.014538218 | 0.031675801 |
| 3.441557131 | 0.000578376 | 0.001743768 |
| 0.192810118 | 0.847107679 | NA |
| 0.193341124 | 0.846691821 | NA |
| 1.506818546 | 0.131857174 | 0.209721362 |
| 7.608613189 | 2.77E-14 | 3.01E-13 |
| -0.187260194 | 0.851456631 | 0.896741542 |
| -0.022487264 | 0.982059271 | 0.987641213 |
| 3.758825772 | 0.000170713 | 0.000566477 |
| -0.275986503 | 0.782558433 | 0.844795893 |
| -8.772160702 | 1.75E-18 | 2.76E-17 |
| -1.803762522 | 0.071268543 | 0.125174426 |
| 5.27645241 | 1.32E-07 | 6.99E-07 |
| -8.259602297 | 1.46E-16 | 1.96E-15 |
| 0.06371127 | 0.949200131 | NA |
| -10.97264127 | 5.17E-28 | 1.61E-26 |
| -0.065390563 | 0.947863038 | NA |
| NA | NA | NA |
| NA | NA | NA |
| -2.665835935 | 0.007679716 | 0.017971512 |
| 1.246747052 | 0.21249026 | 0.310395752 |

| | | |
|---|---|---|
| NA | NA | NA |
| 2.277980487 | 0.02272774 | 0.04674407 |
| -6.322120967 | 2.58E-10 | 1.87E-09 |
| NA | NA | NA |
| 0.64393645 | 0.519616644 | NA |
| NA | NA | NA |
| 0.838901968 | 0.401524323 | 0.513234287 |
| 2.671840503 | 0.007543649 | 0.017696115 |
| -0.065389369 | 0.947863988 | NA |
| -0.890632662 | 0.373126271 | 0.485951243 |
| 0.192808327 | 0.847109082 | NA |
| -0.194487022 | 0.845794556 | NA |
| -0.065389966 | 0.947863513 | NA |
| -0.065388772 | 0.947864464 | NA |
| -0.422348711 | 0.672770508 | 0.756624992 |
| NA | NA | NA |
| 1.197582321 | 0.231079662 | 0.33187537 |
| -0.235094007 | 0.81413577 | NA |
| 0.398194802 | 0.690486597 | NA |
| -2.304943382 | 0.021169745 | 0.043885929 |
| -1.25612457 | 0.209070809 | 0.306258697 |
| 7.268392291 | 3.64E-13 | 3.53E-12 |
| 2.472479408 | 0.013417944 | 0.029551355 |
| -1.959133289 | 0.050097179 | 0.092732761 |
| -1.507535222 | 0.131673523 | 0.209469192 |
| 0.349532517 | 0.726689563 | NA |
| -5.889383216 | 3.88E-09 | 2.48E-08 |
| -0.548259067 | 0.583514029 | 0.683668479 |
| -2.237243964 | 0.025270399 | 0.051203602 |
| -3.784465011 | 0.00015404 | 0.000515387 |
| -3.740811058 | 0.000183427 | 0.000605937 |
| 0.895624776 | 0.370453199 | 0.483216612 |
| -0.43156189 | 0.666059862 | NA |
| -1.081765312 | 0.279356824 | 0.386421735 |
| 0.063707194 | 0.949203376 | NA |
| -3.615699441 | 0.000299538 | 0.000952659 |
| -6.000335635 | 1.97E-09 | 1.30E-08 |
| 0.68168998 | 0.495435005 | NA |
| 2.338592092 | 0.019356554 | 0.040611447 |
| 0.884377796 | 0.376492305 | 0.489156626 |
| 1.882786637 | 0.059729289 | 0.107664129 |
| 0.069889846 | 0.944281335 | NA |
| 0.059735049 | 0.952366657 | 0.967567454 |
| -0.836367666 | 0.402948097 | 0.51463368 |
| -3.44738251 | 0.000566047 | 0.001709477 |
| NA | NA | NA |
| 1.320097884 | 0.186802339 | 0.279511152 |

| | | | |
|---|---|---|---|
| | -0.502972515 | 0.614983596 | 0.710404256 |
| | 5.443249685 | 5.23E-08 | 2.93E-07 |
| | 4.624503593 | 3.75E-06 | 1.63E-05 |
| | -1.527283966 | 0.126690415 | 0.203147019 |
| | 3.5781656 | 0.000346014 | 0.001088179 |
| | 2.251481449 | 0.024355061 | 0.049573649 |
| | 0.875601541 | NA | NA |
| | 3.658588968 | 0.000253608 | 0.000816542 |
| | 2.902717262 | 0.003699405 | 0.009356922 |
| | 1.484429986 | 0.137694885 | 0.217431264 |
| | 7.778766733 | 7.32E-15 | 8.45E-14 |
| | 6.386997862 | 1.69E-10 | 1.25E-09 |
| | 1.945851239 | 0.051672612 | 0.095240462 |
| | -0.103283175 | 0.917738229 | NA |
| NA | NA | NA | |
| | -3.92992604 | 8.50E-05 | 0.000297795 |
| | 15.80812396 | 2.73E-56 | 2.48E-54 |
| | 10.08582277 | 6.38E-24 | 1.51E-22 |
| | 0.496346442 | 0.619650005 | 0.714113892 |
| | -0.872117561 | 0.383144247 | 0.495535601 |
| | -0.370914642 | 0.710701109 | 0.787988871 |
| NA | NA | NA | |
| NA | NA | NA | |
| | -1.68858621 | 0.091298755 | 0.154426804 |
| | 0.034259203 | 0.972670457 | 0.981176865 |
| | 4.375328253 | 1.21E-05 | 4.87E-05 |
| | 0.063708285 | 0.949202508 | NA |
| | 1.335926704 | 0.181573232 | NA |
| | 2.792175248 | 0.005235499 | 0.012816092 |
| NA | NA | NA | |
| | -0.220504209 | 0.825478495 | 0.877802988 |
| | -3.291769678 | 0.000995591 | 0.002871258 |
| | -3.160994454 | 0.001572315 | 0.004338556 |
| | 0.066394788 | 0.94706352 | NA |
| | 5.087158326 | 3.63E-07 | 1.82E-06 |
| | -0.484481084 | 0.628044486 | 0.720999564 |
| | 0.192810118 | 0.847107679 | NA |
| | -10.13369976 | 3.92E-24 | 9.44E-23 |
| NA | NA | NA | |
| | 1.707309038 | 0.087764627 | 0.149448848 |
| | 0.063708882 | 0.949202033 | NA |
| | -0.065388772 | 0.947864464 | NA |
| | 3.813174273 | 0.000137193 | 0.000463108 |
| | 5.731670509 | 9.94E-09 | 6.08E-08 |
| | -0.411745065 | 0.68052629 | 0.763289704 |
| | -0.364325069 | 0.715615282 | NA |
| | 0.165349919 | 0.86866857 | 0.908523078 |

| | | |
|---|---|---|
| NA | NA | NA |
| -5.669019446 | 1.44E-08 | 8.62E-08 |
| 0.399127724 | 0.689799095 | 0.770995417 |
| -1.240007721 | 0.214972538 | 0.313253923 |
| -5.958972332 | 2.54E-09 | 1.66E-08 |
| -2.660669697 | 0.007798542 | 0.018213819 |
| NA | NA | NA |
| NA | NA | NA |
| 2.496007621 | 0.01255999 | 0.027872733 |
| -6.546582582 | 5.89E-11 | 4.59E-10 |
| NA | NA | NA |
| NA | NA | NA |
| -0.065388772 | 0.947864464 | NA |
| 1.846790039 | 0.064777576 | 0.115302427 |
| 0.063708882 | 0.949202033 | NA |
| NA | NA | NA |
| -3.884464056 | 0.000102556 | 0.000353675 |
| NA | NA | NA |
| -0.065388772 | 0.947864464 | NA |
| -1.458413454 | 0.144726614 | 0.226661692 |
| -1.206071158 | 0.227790055 | 0.328073253 |
| -5.524234902 | 3.31E-08 | 1.90E-07 |
| 1.901948805 | 0.057177848 | 0.103842804 |
| 0.163563161 | 0.870075047 | 0.909460894 |
| 0.063708285 | 0.949202508 | NA |
| -0.284901308 | 0.775719754 | NA |
| 0.063708285 | 0.949202508 | NA |
| -4.4138584 | 1.02E-05 | 4.14E-05 |
| NA | NA | NA |
| 2.353571943 | 0.018594009 | 0.039238517 |
| 12.30311366 | 8.71E-35 | 3.76E-33 |
| 2.517274377 | 0.011826669 | 0.026439936 |
| -1.221366148 | 0.221947418 | 0.321322452 |
| NA | NA | NA |
| 3.782859406 | 0.000155037 | 0.00051817 |
| -0.617568165 | 0.536860035 | 0.641180355 |
| -5.053429537 | 4.34E-07 | 2.15E-06 |
| NA | NA | NA |
| 2.370084744 | 0.017784009 | 0.0378065 |
| 0.063708285 | 0.949202508 | NA |
| -0.176667287 | 0.859769738 | 0.902572518 |
| -3.700477295 | 0.000215194 | 0.000702991 |
| NA | NA | NA |
| 0.31685483 | 0.751353758 | NA |
| NA | NA | NA |
| NA | NA | NA |
| -2.378398145 | 0.01738804 | 0.037059148 |

| | | |
|---|---|---|
| NA | NA | NA |
| -10.48625779 | 9.99E-26 | 2.68E-24 |
| -4.380072554 | 1.19E-05 | 4.78E-05 |
| -2.776633575 | 0.005492506 | 0.013381101 |
| -7.897527228 | 2.84E-15 | 3.40E-14 |
| -9.505355765 | 1.99E-21 | 3.96E-20 |
| -0.626157548 | 0.531211609 | 0.635925803 |
| -0.434662724 | 0.663807259 | 0.750151054 |
| -4.870507419 | 1.11E-06 | 5.18E-06 |
| 1.85995467 | 0.06289194 | 0.11248172 |
| NA | NA | NA |
| 0.063976859 | 0.948988653 | NA |
| 0.063708882 | 0.949202033 | NA |
| NA | NA | NA |
| NA | NA | NA |
| -5.242649879 | 1.58E-07 | 8.31E-07 |
| 9.944638628 | 2.66E-23 | 6.09E-22 |
| 0.746412345 | 0.455418364 | 0.565175289 |
| NA | NA | NA |
| 1.669371948 | 0.095043689 | 0.159789375 |
| 1.842182725 | 0.065448426 | NA |
| 4.587366332 | 4.49E-06 | 1.93E-05 |
| -0.52873095 | 0.596992104 | 0.695268582 |
| 1.154682276 | 0.24822056 | 0.351451557 |
| 2.719769523 | 0.006532743 | 0.015601046 |
| 4.368103887 | 1.25E-05 | 5.02E-05 |
| 0.734923752 | 0.462385932 | NA |
| 1.00701925 | 0.313925523 | 0.42436573 |
| 1.126448476 | 0.259975738 | 0.364896132 |
| 4.504383218 | 6.66E-06 | 2.78E-05 |
| 0.315439194 | 0.752428212 | NA |
| -4.035773446 | 5.44E-05 | 0.00019722 |
| -0.065389966 | 0.947863513 | NA |
| 0.417281549 | 0.676472483 | NA |
| -0.339101735 | 0.734533092 | 0.806410951 |
| -0.065389369 | 0.947863988 | NA |
| -0.065388772 | 0.947864464 | NA |
| NA | NA | NA |
| -1.253115235 | 0.210163772 | 0.307661669 |
| NA | NA | NA |
| 1.872863228 | 0.061087274 | 0.109701139 |
| -2.525873224 | 0.011541112 | 0.02589173 |
| 3.562598932 | 0.000367201 | 0.001148021 |
| -1.75296766 | 0.079607558 | 0.137527784 |
| -8.710872218 | 3.02E-18 | 4.65E-17 |
| 0.08086109 | 0.935552424 | NA |
| -1.106256365 | 0.268615573 | 0.374604402 |

| | | |
|---|---|---|
| 4.893717997 | 9.89E-07 | 4.65E-06 |
| 0.439262522 | 0.660471327 | 0.747393522 |
| 6.659402597 | 2.75E-11 | 2.21E-10 |
| NA | NA | NA |
| 1.290965304 | 0.196715709 | 0.291573056 |
| 7.364214239 | 1.78E-13 | 1.78E-12 |
| 13.54590866 | 8.38E-42 | 4.84E-40 |
| 2.425895579 | 0.015270663 | 0.033090183 |
| 2.231637426 | 0.025638936 | 0.051812031 |
| 2.769811769 | 0.00560887 | 0.013628775 |
| 3.946307858 | 7.94E-05 | 0.000279634 |
| 5.208697028 | 1.90E-07 | 9.87E-07 |
| -3.214761772 | 0.001305528 | 0.003663382 |
| 17.27436603 | 7.34E-67 | 8.75E-65 |
| 6.504245086 | 7.81E-11 | 6.00E-10 |
| 1.631154324 | 0.10285776 | 0.170442221 |
| 8.666861909 | 4.44E-18 | 6.73E-17 |
| -7.023882434 | 2.16E-12 | 1.92E-11 |
| 0.063707194 | 0.949203376 | NA |
| -3.375832307 | 0.000735928 | 0.002176565 |
| -5.351717848 | 8.71E-08 | 4.74E-07 |
| -1.032159592 | 0.301997361 | 0.411217166 |
| 1.113660794 | 0.265424735 | 0.371087589 |
| 2.488625074 | 0.012823812 | 0.028390237 |
| -4.598984985 | 4.25E-06 | 1.83E-05 |
| 0.487053513 | 0.626220408 | 0.719649896 |
| 0.50690371 | 0.612222383 | 0.7079661 |
| -1.078852422 | 0.280653522 | 0.387615934 |
| -1.291114865 | 0.19666385 | 0.291545485 |
| -1.685370662 | 0.09191708 | 0.155336204 |
| 0.511795777 | 0.608793946 | 0.705337552 |
| NA | NA | NA |
| -1.074050315 | 0.282800131 | 0.389720161 |
| 0.736672232 | 0.461321697 | NA |
| 5.485852479 | 4.11E-08 | 2.34E-07 |
| -0.478101918 | 0.632577667 | 0.724343245 |
| 1.856732424 | 0.063349232 | 0.113138158 |
| 2.362456827 | 0.018154256 | 0.038436899 |
| 0.537182668 | 0.591141442 | 0.690439918 |
| -2.884504453 | 0.003920302 | 0.009876715 |
| -2.658201306 | 0.007855895 | 0.018332043 |
| NA | NA | NA |
| 5.552037934 | 2.82E-08 | 1.63E-07 |
| 4.565309687 | 4.99E-06 | 2.12E-05 |
| -5.359895875 | 8.33E-08 | 4.54E-07 |
| -0.065389369 | 0.947863988 | NA |
| 3.830234321 | 0.000128021 | 0.000434138 |

| | | |
|---|---|---|
| -2.341410394 | 0.019211037 | 0.040350109 |
| 2.523976569 | 0.011603566 | 0.02601086 |
| NA | NA | NA |
| NA | NA | NA |
| -2.889960716 | 0.0038529 | 0.009720601 |
| -0.433131957 | 0.664918907 | NA |
| 1.140254889 | 0.254180126 | 0.358409585 |
| 1.602697732 | 0.109001405 | 0.178764589 |
| 5.100622059 | 3.39E-07 | 1.70E-06 |
| 0.275131662 | 0.783215087 | 0.845286031 |
| -0.284057767 | 0.776366112 | 0.839566771 |
| 2.474962521 | 0.013325016 | 0.029372536 |
| -0.605665128 | 0.544737145 | NA |
| 0.063708882 | 0.949202033 | NA |
| 3.660982329 | 0.00025125 | 0.000809681 |
| 6.75058473 | 1.47E-11 | 1.21E-10 |
| 0.278769668 | 0.780421594 | 0.843143668 |
| 0.774468555 | 0.438653757 | 0.549002627 |
| -1.6323454 | 0.102606744 | 0.170138908 |
| 8.444193073 | 3.06E-17 | 4.34E-16 |
| 4.08142361 | 4.48E-05 | 0.000164035 |
| 9.345596528 | 9.14E-21 | 1.72E-19 |
| -2.701474993 | 0.006903267 | 0.016376531 |
| 14.27331895 | 3.21E-46 | 2.23E-44 |
| -1.321067739 | 0.186478779 | 0.279171232 |
| 3.94066168 | 8.13E-05 | 0.000285615 |
| -3.301466866 | 0.000961807 | 0.002781517 |
| 2.54838963 | 0.010822153 | 0.024460762 |
| 1.331768378 | 0.182936312 | 0.274843691 |
| -0.959622252 | 0.337245366 | 0.44847996 |
| -0.961624898 | 0.33623806 | 0.447542602 |
| 6.488926321 | 8.65E-11 | 6.62E-10 |
| -1.531876324 | 0.125552957 | 0.201600979 |
| -1.631521768 | 0.102780271 | 0.170358928 |
| 0.280302294 | 0.779245591 | 0.842127591 |
| NA | NA | NA |
| 8.845588804 | 9.10E-19 | 1.48E-17 |
| -7.989945516 | 1.35E-15 | 1.66E-14 |
| -0.502296585 | 0.615458911 | 0.71071782 |
| 6.306131298 | 2.86E-10 | 2.06E-09 |
| 0.730294403 | 0.465210248 | NA |
| NA | NA | NA |
| 1.3216229 | 0.186293754 | 0.278966955 |
| NA | NA | NA |
| 5.703870787 | 1.17E-08 | 7.10E-08 |
| 7.091738962 | 1.32E-12 | 1.21E-11 |
| -4.979625949 | 6.37E-07 | 3.07E-06 |

| | | |
|---|---|---|
| NA | NA | NA |
| -3.243415635 | 0.001181058 | 0.003351307 |
| 17.26031001 | 9.36E-67 | 1.11E-64 |
| -7.455978881 | 8.92E-14 | 9.21E-13 |
| -3.221302583 | 0.001276093 | 0.00358964 |
| 3.01948356 | 0.00253206 | 0.006651708 |
| NA | NA | NA |
| NA | NA | NA |
| NA | NA | NA |
| -2.673204359 | 0.007513047 | 0.01763755 |
| 0.192810118 | 0.847107679 | NA |
| NA | NA | NA |
| 0.482637744 | 0.629352975 | NA |
| -2.127035018 | 0.033417174 | 0.065249384 |
| 1.552498843 | 0.120542909 | 0.194682476 |
| NA | NA | NA |
| -0.298251825 | 0.76551097 | 0.830882685 |
| 3.901832709 | 9.55E-05 | 0.000331521 |
| 3.783675034 | 0.00015453 | 0.00051675 |
| -8.217761892 | 2.07E-16 | 2.75E-15 |
| -0.333502566 | 0.738754953 | 0.809710858 |
| -0.065388772 | 0.947864464 | NA |
| -3.066069402 | 0.00216893 | 0.005800033 |
| NA | NA | NA |
| NA | NA | NA |
| -0.069988098 | 0.944203133 | NA |
| -0.031830751 | 0.974607024 | 0.982417696 |
| -7.099026346 | 1.26E-12 | 1.15E-11 |
| -1.378179379 | 0.168147912 | 0.256255617 |
| -2.241982701 | 0.02496249 | 0.050649411 |
| 10.60132796 | 2.94E-26 | 8.15E-25 |
| -1.877327093 | 0.060473277 | 0.108770202 |
| -0.336105822 | 0.736791068 | NA |
| -0.09675114 | 0.922924026 | NA |
| -1.544975044 | 0.1223523 | 0.197197557 |
| 1.942807609 | 0.052039408 | 0.095845945 |
| 0.785480971 | 0.432171623 | 0.542953156 |
| -0.848255274 | 0.39629582 | 0.508238171 |
| -5.220712359 | 1.78E-07 | 9.27E-07 |
| 6.274378975 | 3.51E-10 | 2.50E-09 |
| 6.930098089 | 4.21E-12 | 3.65E-11 |
| -0.065389966 | 0.947863513 | NA |
| 0.063708285 | 0.949202508 | NA |
| 0.063708882 | 0.949202033 | NA |
| 0.063708285 | 0.949202508 | NA |
| 15.37863025 | 2.28E-53 | 1.89E-51 |
| 5.063259464 | 4.12E-07 | 2.05E-06 |

| | | |
|---|---|---|
| 0.926296566 | 0.35429188 | 0.466180833 |
| 3.051697621 | 0.002275512 | 0.006049503 |
| -0.041984389 | 0.966511143 | 0.977326946 |
| -0.689541161 | 0.49048278 | 0.59796624 |
| -0.829430552 | 0.406860819 | 0.518651709 |
| 0.192810118 | 0.847107679 | NA |
| 13.46397144 | 2.55E-41 | 1.45E-39 |
| 10.39545008 | 2.60E-25 | 6.77E-24 |
| 0.21765406 | 0.82769866 | NA |
| 1.408578446 | 0.158959856 | 0.244711557 |
| 0.396091757 | 0.692037339 | 0.77270351 |
| 3.009391945 | 0.002617712 | 0.006855129 |
| -2.964895336 | 0.003027859 | 0.007821996 |
| 0.769992532 | 0.441304322 | 0.551373876 |
| 1.846711045 | 0.06478903 | 0.115306415 |
| -2.112190244 | 0.034670135 | 0.067359581 |
| -1.588069395 | 0.112270648 | 0.183263386 |
| 33.5594064 | 6.56E-247 | 8.22E-244 |
| 7.323226927 | 2.42E-13 | 2.39E-12 |
| 7.679904737 | 1.59E-14 | 1.78E-13 |
| 0.403678851 | 0.686448893 | NA |
| 0.336583495 | 0.7364309 | 0.807938865 |
| -0.437672995 | 0.661623348 | 0.748189772 |
| -7.655363229 | 1.93E-14 | 2.13E-13 |
| 1.80681794 | 0.070790671 | NA |
| -2.802308324 | 0.005073836 | 0.012466684 |
| -3.259353891 | 0.001116663 | 0.003184247 |
| 0.192810118 | 0.847107679 | NA |
| -3.793079046 | 0.000148791 | 0.000499093 |
| -3.874122061 | 0.00010701 | 0.000368172 |
| 1.25782312 | 0.208455732 | 0.30553651 |
| 1.914955182 | 0.05549822 | 0.101240466 |
| 0.192636473 | 0.847243678 | NA |
| -2.24158211 | 0.024988393 | 0.05069375 |
| -7.777915216 | 7.37E-15 | 8.50E-14 |
| -0.353592364 | 0.723644398 | NA |
| NA | NA | NA |
| -0.534443156 | 0.593034968 | 0.6921499 |
| -0.065389369 | 0.947863988 | NA |
| 4.090201331 | 4.31E-05 | 0.000158389 |
| -1.059851593 | 0.289212121 | 0.396656404 |
| 5.301464409 | 1.15E-07 | 6.14E-07 |
| 2.178256221 | 0.029386967 | 0.058326522 |
| 1.177404755 | 0.239033993 | 0.341107232 |
| -0.591587096 | 0.554127117 | 0.657180845 |
| 5.382392735 | 7.35E-08 | 4.03E-07 |
| 36.95671473 | 5.68E-299 | 1.78E-295 |

| | | |
|---|---|---|
| -3.461652769 | 0.000536869 | 0.001628237 |
| -1.724406461 | 0.084634492 | 0.144818248 |
| 0.471852997 | 0.63703172 | 0.728100569 |
| -0.541325547 | 0.588283215 | 0.688007143 |
| NA | NA | NA |
| 0.967730989 | 0.333178737 | 0.444367938 |
| 1.123304732 | 0.261308095 | NA |
| 2.177779994 | 0.02942242 | 0.058392259 |
| -3.936702424 | 8.26E-05 | 0.000290041 |
| 1.27578858 | 0.202030266 | 0.298214162 |
| -2.619130438 | 0.008815423 | 0.020332662 |
| -8.929528925 | 4.28E-19 | 7.12E-18 |
| -1.471753823 | 0.141087367 | 0.2217948 |
| 0.182887879 | 0.854885994 | NA |
| -4.515111952 | 6.33E-06 | 2.65E-05 |
| -0.611427071 | 0.540916886 | 0.645090035 |
| -0.450560613 | 0.652306259 | 0.740598098 |
| 0.946684189 | 0.34379973 | 0.455177981 |
| 5.893443091 | 3.78E-09 | 2.42E-08 |
| 0.474041829 | 0.635470079 | 0.726890416 |
| -4.959869105 | 7.05E-07 | 3.39E-06 |
| 0.497237016 | 0.619021921 | 0.713606026 |
| -3.640302931 | 0.000272317 | 0.000872182 |
| -3.780096788 | 0.000156767 | 0.000523604 |
| 6.476127788 | 9.41E-11 | 7.17E-10 |
| NA | NA | NA |
| 5.287878851 | 1.24E-07 | 6.59E-07 |
| NA | NA | NA |
| NA | NA | NA |
| -1.918464152 | 0.055052183 | 0.100558721 |
| -1.52282781 | 0.12780179 | 0.204623477 |
| -1.82313543 | 0.068282882 | 0.120640963 |
| 9.445548066 | 3.54E-21 | 6.88E-20 |
| 1.454696183 | 0.14575339 | 0.227984461 |
| 0.03700942 | 0.970477495 | 0.979556887 |
| -0.676842923 | 0.498505621 | 0.605649728 |
| 6.045065896 | 1.49E-09 | 9.98E-09 |
| -3.534582615 | 0.00040842 | 0.001265975 |
| 7.224108375 | 5.04E-13 | 4.84E-12 |
| 2.565165195 | 0.010312669 | 0.023429801 |
| -0.042076019 | 0.966438097 | 0.977292565 |
| 0.083757469 | 0.933249264 | NA |
| -2.620319996 | 0.008784729 | 0.020273075 |
| 0.192810118 | 0.847107679 | NA |
| 0.790979677 | 0.428955851 | 0.540105888 |
| 2.924828831 | 0.003446457 | 0.008780135 |
| 0.063707019 | 0.949203516 | NA |

| | | |
|---|---|---|
| 9.734018919 | 2.16E-22 | 4.58E-21 |
| 8.577653844 | 9.68E-18 | 1.43E-16 |
| 0.030052136 | 0.976025473 | 0.983293121 |
| 2.383130393 | 0.017166113 | 0.036676722 |
| -0.237600362 | 0.812191069 | 0.867796596 |
| 1.085311424 | 0.277783748 | 0.384713507 |
| -2.26487966 | 0.023520065 | 0.048124518 |
| 1.833959912 | 0.066659945 | 0.118209882 |
| NA | NA | NA |
| -0.631623721 | 0.527632783 | 0.632562915 |
| NA | NA | NA |
| 0.58911503 | 0.555784111 | 0.658459616 |
| 4.890451995 | 1.01E-06 | 4.72E-06 |
| -2.002565458 | 0.045223951 | 0.084879685 |
| -1.798859261 | 0.072040946 | 0.126264609 |
| -1.496981375 | 0.134398104 | 0.213058747 |
| -0.928092956 | 0.353359353 | 0.465235107 |
| 3.741019803 | 0.000183275 | 0.000605594 |
| -0.502634867 | 0.615221011 | NA |
| -0.141925445 | 0.887138895 | 0.920876148 |
| -3.263337419 | 0.001101083 | 0.003145773 |
| -4.49130658 | 7.08E-06 | 2.95E-05 |
| 1.756375278 | 0.079024351 | 0.136666863 |
| -1.320098226 | 0.186802225 | 0.279511152 |
| 9.639677001 | 5.44E-22 | 1.12E-20 |
| NA | NA | NA |
| -3.616238345 | 0.000298915 | 0.0009508 |
| 1.229396059 | 0.218923347 | 0.317845208 |
| 2.584169994 | 0.009761367 | 0.022294645 |
| -2.340987268 | 0.019232823 | 0.040385701 |
| -2.501119876 | 0.012380127 | 0.027527278 |
| 0.063707664 | 0.949203002 | NA |
| 9.178043403 | 4.39E-20 | 7.87E-19 |
| 4.032654618 | 5.52E-05 | 0.00019968 |
| NA | NA | NA |
| 0.192809521 | 0.847108147 | NA |
| 0.063710673 | 0.949200606 | NA |
| -0.065389369 | 0.947863988 | NA |
| 5.159533448 | 2.48E-07 | 1.27E-06 |
| 2.39433848 | 0.016650375 | 0.035733421 |
| 8.881850069 | 6.58E-19 | 1.08E-17 |
| NA | NA | NA |
| NA | NA | NA |
| 1.544912722 | 0.122367376 | 0.197209157 |
| NA | NA | NA |
| 0.192810118 | 0.847107679 | NA |
| 1.853848929 | 0.063760776 | 0.113767624 |

| | | |
|---|---|---|
| -0.170933178 | 0.864276308 | 0.905510726 |
| 1.83558292 | 0.066419365 | 0.117922876 |
| -4.46215079 | 8.11E-06 | 3.36E-05 |
| 9.963620974 | 2.20E-23 | 5.05E-22 |
| -8.787971103 | 1.52E-18 | 2.42E-17 |
| -1.275150775 | 0.202255881 | 0.298459185 |
| 5.238741909 | 1.62E-07 | 8.47E-07 |
| -4.56861904 | 4.91E-06 | 2.09E-05 |
| -0.92211251 | 0.356469892 | 0.468345042 |
| NA | NA | NA |
| -3.496655567 | 0.00047113 | 0.001443552 |
| -7.183313373 | 6.80E-13 | 6.44E-12 |
| 0.66227209 | 0.50779686 | 0.614165519 |
| 4.166773589 | 3.09E-05 | 0.000116633 |
| 0.773821679 | 0.439036251 | 0.549272127 |
| 8.892274056 | 5.99E-19 | 9.84E-18 |
| 1.47715642 | 0.139633713 | NA |
| NA | NA | NA |
| NA | NA | NA |
| NA | NA | NA |
| NA | NA | NA |
| NA | NA | NA |
| NA | NA | NA |
| NA | NA | NA |
| NA | NA | NA |
| NA | NA | NA |
| -3.695375416 | 0.000219562 | 0.000715858 |
| 5.226373229 | 1.73E-07 | 9.02E-07 |
| -1.074070026 | 0.282791298 | 0.389720161 |
| -7.248768454 | 4.21E-13 | 4.06E-12 |
| 2.805136675 | 0.005029525 | 0.012368737 |
| -0.013962733 | 0.988859713 | 0.992152349 |
| -7.076483288 | 1.48E-12 | 1.35E-11 |
| -1.424028255 | 0.154438293 | 0.239203337 |
| -1.142459966 | 0.253262878 | 0.357297364 |
| -4.616846412 | 3.90E-06 | 1.68E-05 |
| 3.802162274 | 0.000143439 | 0.000482304 |
| 1.509883459 | 0.131073164 | 0.208699828 |
| -1.409322888 | 0.158739714 | 0.244432792 |
| -8.773043118 | 1.74E-18 | 2.74E-17 |
| -5.378081182 | 7.53E-08 | 4.12E-07 |
| 1.929163274 | 0.053710596 | 0.098453037 |
| -2.048539896 | 0.040507127 | 0.077055281 |
| -1.159367731 | 0.246306324 | 0.349334049 |
| NA | NA | NA |
| NA | NA | NA |
| -0.026096564 | 0.979180318 | 0.985519456 |

| | | |
|---|---|---|
| 0.368195837 | 0.712727214 | 0.789647537 |
| 0.192808924 | 0.847108615 | NA |
| -5.029314201 | 4.92E-07 | 2.41E-06 |
| -6.380530112 | 1.76E-10 | 1.30E-09 |
| NA | NA | NA |
| NA | NA | NA |
| -6.42275403 | 1.34E-10 | 1.00E-09 |
| 2.223998749 | 0.026148528 | 0.052714052 |
| 3.951864623 | 7.75E-05 | 0.000273796 |
| -5.745179937 | 9.18E-09 | 5.63E-08 |
| -1.580185136 | 0.114064475 | 0.185753663 |
| 0.107100764 | 0.914709041 | 0.941363699 |
| -3.118187769 | 0.001819668 | 0.004945309 |
| -3.181300195 | 0.001466156 | 0.004070759 |
| NA | NA | NA |
| -3.281656054 | 0.001031994 | 0.00296565 |
| -6.654435371 | 2.84E-11 | 2.28E-10 |
| -0.065388772 | 0.947864464 | NA |
| -0.152335907 | 0.878922005 | NA |
| 0.113242484 | 0.909838315 | NA |
| 2.270141626 | 0.023198991 | 0.047556982 |
| 1.133774534 | 0.256889136 | 0.361416607 |
| 0.503194275 | 0.61482769 | 0.710355293 |
| -1.255412454 | 0.209329071 | 0.306583185 |
| 8.707353487 | 3.11E-18 | 4.79E-17 |
| 4.446221171 | 8.74E-06 | 3.59E-05 |
| 0.192810118 | 0.847107679 | NA |
| 2.390921475 | 0.016806147 | 0.036018311 |
| -0.089580369 | 0.928620685 | NA |
| 10.56207408 | 4.47E-26 | 1.23E-24 |
| -0.087320677 | 0.930416619 | NA |
| 10.83269108 | 2.41E-27 | 7.18E-26 |
| -0.63629597 | 0.524583531 | 0.629721633 |
| -4.544555628 | 5.51E-06 | 2.33E-05 |
| 0.834923635 | 0.403760706 | 0.515434818 |
| 2.090465976 | 0.036575962 | 0.070542173 |
| NA | NA | NA |
| -1.079484597 | 0.280371757 | 0.387311941 |
| NA | NA | NA |
| NA | NA | NA |
| 0.459314837 | 0.646008095 | 0.735248391 |
| -0.046947059 | 0.962555422 | 0.974271592 |
| 1.157099991 | 0.24723151 | 0.350348452 |
| -0.994034953 | 0.320205851 | 0.430947008 |
| NA | NA | NA |
| -0.065390563 | 0.947863038 | NA |
| 1.811819422 | 0.070014103 | 0.123281956 |

| | | |
|---|---|---|
| 2.437971596 | 0.014769934 | 0.03213032 |
| -0.732940006 | 0.463595022 | 0.572998044 |
| -9.837927175 | 7.73E-23 | 1.69E-21 |
| 0.731443401 | 0.464508364 | 0.573928539 |
| -3.567083939 | 0.000360976 | 0.001130675 |
| NA | NA | NA |
| 0.284204169 | 0.776253921 | NA |
| 5.279451172 | 1.30E-07 | 6.88E-07 |
| NA | NA | NA |
| -5.019348406 | 5.18E-07 | 2.53E-06 |
| 0.192809521 | 0.847108147 | NA |
| NA | NA | NA |
| NA | NA | NA |
| NA | NA | NA |
| 0.063708285 | 0.949202508 | NA |
| -0.165450529 | 0.868589386 | NA |
| -7.367329205 | 1.74E-13 | 1.74E-12 |
| -0.065389369 | 0.947863988 | NA |
| 1.627570571 | 0.103615973 | 0.171547274 |
| 2.482048149 | 0.013062962 | 0.028871165 |
| 1.337079638 | 0.181196643 | 0.272622744 |
| NA | NA | NA |
| -0.273342966 | 0.78458959 | NA |
| -0.372835836 | 0.70927063 | NA |
| -8.052312449 | 8.12E-16 | 1.02E-14 |
| 0.605035772 | 0.545155228 | 0.648985422 |
| -0.065389369 | 0.947863988 | NA |
| NA | NA | NA |
| 0.192809521 | 0.847108147 | NA |
| -1.872876267 | 0.061085473 | 0.109701139 |
| -5.9073598 | 3.48E-09 | 2.24E-08 |
| -1.342294345 | 0.179500576 | 0.270542508 |
| 0.243837729 | 0.807356498 | 0.863484731 |
| -4.516139461 | 6.30E-06 | 2.64E-05 |
| -10.08052917 | 6.74E-24 | 1.59E-22 |
| 1.386416108 | 0.165619881 | 0.253127289 |
| NA | NA | NA |
| 0.086133117 | 0.931360598 | 0.953235203 |
| -0.760145297 | 0.447167739 | 0.55709564 |
| -1.169044026 | 0.242385894 | 0.345024962 |
| 9.15306662 | 5.53E-20 | 9.85E-19 |
| 3.50227539 | 0.000461303 | 0.001414827 |
| -5.889641423 | 3.87E-09 | 2.47E-08 |
| 8.076690765 | 6.65E-16 | 8.45E-15 |
| -5.026700645 | 4.99E-07 | 2.45E-06 |
| 3.486128088 | 0.000490066 | 0.001497358 |
| 2.094119162 | 0.036249367 | 0.069982387 |

| | | |
|---|---|---|
| 9.681149921 | 3.63E-22 | 7.58E-21 |
| 2.19614954 | 0.028081242 | 0.056068401 |
| NA | NA | NA |
| -1.082233112 | 0.279148959 | 0.386155546 |
| -2.609611031 | 0.009064522 | 0.020860928 |
| -0.846412637 | 0.397322591 | 0.509163653 |
| 0.210781401 | 0.833057853 | 0.88301169 |
| 0.633277537 | 0.52655242 | NA |
| -0.689147185 | 0.490730649 | 0.598207186 |
| NA | NA | NA |
| 0.063708285 | 0.949202508 | NA |
| NA | NA | NA |
| -0.065388772 | 0.947864464 | NA |
| 0.063708285 | 0.949202508 | NA |
| 0.570980893 | 0.568012595 | NA |
| -2.052868942 | 0.04008529 | 0.076363103 |
| NA | NA | NA |
| -1.545149693 | 0.122310059 | 0.197154868 |
| -0.907455899 | 0.36416576 | 0.47665476 |
| 0.245779731 | 0.805852748 | 0.86261362 |
| -1.888373054 | 0.058975886 | 0.106559332 |
| -3.120612813 | 0.001804752 | 0.004909034 |
| 2.467240823 | 0.013615875 | 0.029934594 |
| -1.093741375 | 0.274068423 | 0.380726641 |
| 1.275721303 | 0.202054055 | 0.29823169 |
| 2.331736256 | 0.01971457 | 0.041269189 |
| 0.024041965 | 0.980819135 | 0.986613469 |
| -1.706109023 | 0.087987783 | 0.149788084 |
| 7.416088949 | 1.21E-13 | 1.23E-12 |
| NA | NA | NA |
| -3.459162636 | 0.000541858 | 0.001641377 |
| -0.496010796 | 0.619886794 | 0.714288116 |
| -1.158831012 | 0.246525074 | 0.349558454 |
| NA | NA | NA |
| -0.628022001 | 0.529989528 | 0.634901784 |
| NA | NA | NA |
| 0.063708882 | 0.949202033 | NA |
| -0.551199668 | 0.581496808 | 0.681950914 |
| 0.066587943 | 0.946909745 | NA |
| 0.370774956 | 0.710805156 | NA |
| NA | NA | NA |
| 0.711628668 | 0.47669475 | NA |
| NA | NA | NA |
| -0.339213706 | 0.734448745 | NA |
| NA | NA | NA |
| -0.479131456 | 0.631845113 | NA |
| -0.065389369 | 0.947863988 | NA |

| | | |
|---|---|---|
| NA | NA | NA |
| 0.084099977 | 0.932976943 | NA |
| 0.192809521 | 0.847108147 | NA |
| 0.566468483 | 0.571075352 | NA |
| 0.110682929 | 0.911867783 | 0.939338654 |
| -0.966055945 | 0.334016188 | 0.445224474 |
| -0.333247644 | 0.738947357 | 0.809871807 |
| 3.100666245 | 0.001930858 | 0.005218591 |
| -6.004997181 | 1.91E-09 | 1.27E-08 |
| 5.34039758 | 9.27E-08 | 5.02E-07 |
| NA | NA | NA |
| -6.02099601 | 1.73E-09 | 1.15E-08 |
| -7.424537886 | 1.13E-13 | 1.16E-12 |
| 5.410362247 | 6.29E-08 | 3.48E-07 |
| NA | NA | NA |
| NA | NA | NA |
| -5.069536067 | 3.99E-07 | 1.98E-06 |
| -2.374985086 | 0.017549659 | 0.037371786 |
| 22.40966327 | 3.17E-111 | 9.01E-109 |
| 16.50563061 | 3.34E-61 | 3.46E-59 |
| -3.866687013 | 0.000110324 | 0.000378742 |
| 3.704432331 | 0.000211865 | 0.000692837 |
| NA | NA | NA |
| NA | NA | NA |
| NA | NA | NA |
| -0.663757805 | 0.506845336 | 0.613311035 |
| 0.255589302 | 0.79826798 | NA |
| -3.114549679 | 0.001842259 | 0.004999106 |
| 1.620687414 | 0.105084694 | 0.173534993 |
| 5.340953777 | 9.25E-08 | 5.01E-07 |
| 0.109240142 | 0.913012022 | 0.940091773 |
| -1.211439252 | 0.225727103 | 0.325827191 |
| NA | NA | NA |
| -5.259218954 | 1.45E-07 | 7.63E-07 |
| -1.523620633 | 0.127603506 | 0.204384447 |
| NA | NA | NA |
| -6.429754599 | 1.28E-10 | 9.60E-10 |
| 1.053740398 | 0.292001788 | 0.399825205 |
| -1.410334307 | 0.158440993 | 0.244087633 |
| -0.007337236 | 0.994145785 | 0.995856606 |
| -2.139934402 | 0.032360068 | 0.063462549 |
| 0.355766635 | 0.722015338 | NA |
| NA | NA | NA |
| NA | NA | NA |
| 0.192810118 | 0.847107679 | NA |
| 2.354125011 | 0.018566366 | 0.039190094 |
| 5.398396641 | 6.72E-08 | 3.71E-07 |

| | | |
|---|---|---|
| 2.953447069 | 0.003142465 | 0.008081362 |
| -2.258255729 | 0.023929721 | 0.048823028 |
| NA | NA | NA |
| 3.490402882 | 0.000482293 | 0.001474868 |
| -0.065388772 | 0.947864464 | NA |
| -1.177458363 | 0.239012607 | 0.34109616 |
| -2.060632139 | 0.039338148 | 0.075088374 |
| 1.663594445 | 0.096193503 | 0.161321499 |
| -0.657470405 | 0.510878494 | 0.616670945 |
| NA | NA | NA |
| 3.167974356 | 0.001535051 | 0.004244622 |
| NA | NA | NA |
| NA | NA | NA |
| 0.873696314 | 0.382283659 | 0.494612229 |
| -0.850857188 | 0.394848689 | 0.506875715 |
| 3.390662738 | 0.000697238 | 0.002070211 |
| -2.912499243 | 0.003585491 | 0.009090846 |
| 1.447765976 | 0.147682511 | 0.230411598 |
| -1.541778417 | 0.123127449 | 0.1982809 |
| 0.88617411 | 0.375523713 | 0.488303731 |
| -8.952518987 | 3.47E-19 | 5.84E-18 |
| 12.07268069 | 1.47E-33 | 6.04E-32 |
| NA | NA | NA |
| 5.858698801 | 4.67E-09 | 2.96E-08 |
| 17.1576694 | 5.51E-66 | 6.38E-64 |
| 7.103638863 | 1.22E-12 | 1.12E-11 |
| -2.650634131 | 0.008034082 | 0.018699374 |
| 3.239663865 | 0.001196707 | 0.003389947 |
| NA | NA | NA |
| -1.531849267 | 0.125559635 | 0.201600979 |
| 0.063708285 | 0.949202508 | NA |
| -3.465805262 | 0.000528646 | 0.001605814 |
| NA | NA | NA |
| 2.212469223 | 0.02693426 | 0.054049911 |
| -0.065389966 | 0.947863513 | NA |
| NA | NA | NA |
| -1.371478155 | 0.170225949 | 0.258855133 |
| 5.136706342 | 2.80E-07 | 1.42E-06 |
| NA | NA | NA |
| -0.116490634 | 0.907263709 | 0.935905482 |
| 1.999194536 | 0.045587309 | 0.085465535 |
| -0.256598516 | 0.797488721 | 0.855966668 |
| -7.62839618 | 2.38E-14 | 2.60E-13 |
| -1.412306307 | 0.15785979 | 0.24341941 |
| NA | NA | NA |
| NA | NA | NA |
| NA | NA | NA |

| | | |
|---|---|---|
| 0.063708285 | 0.949202508 | NA |
| -5.567659246 | 2.58E-08 | 1.50E-07 |
| NA | NA | NA |
| NA | NA | NA |
| NA | NA | NA |
| 3.58951604 | 0.000331292 | 0.00104507 |
| 0.06371127 | 0.949200131 | NA |
| 0.108685182 | 0.913452195 | NA |
| -3.792089738 | 0.000149385 | 0.000500952 |
| 1.408658504 | 0.158936171 | 0.244690143 |
| -3.463790295 | 0.000532622 | 0.001616725 |
| -1.595930342 | 0.110504346 | 0.18083843 |
| 0.433446722 | 0.664690264 | NA |
| -8.721073046 | 2.76E-18 | 4.26E-17 |
| 0.192809521 | 0.847108147 | NA |
| -1.127136349 | 0.259684837 | 0.364733263 |
| 3.292102473 | 0.000994414 | 0.002868523 |
| NA | NA | NA |
| NA | NA | NA |
| NA | NA | NA |
| -1.326513976 | 0.184669509 | NA |
| -3.784921529 | 0.000153757 | 0.000514717 |
| -9.100965111 | 8.95E-20 | 1.57E-18 |
| -5.782567013 | 7.36E-09 | 4.55E-08 |
| NA | NA | NA |
| NA | NA | NA |
| 2.918288193 | 0.003519589 | 0.008948229 |
| -1.560761718 | 0.118579983 | 0.191970569 |
| 2.982577266 | 0.002858325 | 0.007423104 |
| 2.858975326 | 0.004250118 | 0.010618932 |
| 0.128763521 | 0.897544772 | NA |
| 10.66468743 | 1.49E-26 | 4.22E-25 |
| -0.093570706 | 0.925450181 | 0.948729276 |
| 1.785758076 | 0.074138438 | 0.129316034 |
| -0.101556797 | 0.919108473 | 0.944504047 |
| 0.03956545 | 0.968439572 | 0.978407366 |
| -4.772467372 | 1.82E-06 | 8.25E-06 |
| -1.818210701 | 0.069031935 | 0.121801025 |
| 0.067524971 | 0.946163784 | NA |
| -4.998040107 | 5.79E-07 | 2.81E-06 |
| NA | NA | NA |
| NA | NA | NA |
| 6.059447438 | 1.37E-09 | 9.16E-09 |
| -0.065389369 | 0.947863988 | NA |
| -3.476953547 | 0.000507146 | 0.001545581 |
| -0.253336793 | 0.800007966 | 0.858008373 |
| -7.113872502 | 1.13E-12 | 1.04E-11 |

| | | | |
|---|---|---|---|
| | 0.324874675 | 0.745275925 | NA |
| | -3.428104386 | 0.000607812 | 0.001824791 |
| | 0.764613086 | 0.444501975 | NA |
| NA | NA | NA | |
| | -0.065388772 | 0.947864464 | NA |
| NA | NA | NA | |
| | -5.305639442 | 1.12E-07 | 6.01E-07 |
| NA | NA | NA | |
| NA | NA | NA | |
| | -0.994392758 | 0.320031692 | 0.430735778 |
| NA | NA | NA | |
| NA | NA | NA | |
| | -2.697135694 | 0.006993876 | 0.016563225 |
| | -4.57765743 | 4.70E-06 | 2.01E-05 |
| | -3.218865088 | 0.00128699 | 0.003616229 |
| | -1.647096924 | 0.099538125 | 0.165963181 |
| | 0.209569518 | 0.834003673 | NA |
| NA | NA | NA | |
| | -0.065389369 | 0.947863988 | NA |
| | -1.252566026 | 0.210363686 | 0.307900301 |
| NA | NA | NA | |
| | 0.063708285 | 0.949202508 | NA |
| | -3.415752094 | 0.000636061 | 0.001901877 |
| | 7.400847235 | 1.35E-13 | 1.37E-12 |
| | 0.72309948 | 0.469618769 | 0.578700167 |
| | 0.48856441 | 0.625150115 | 0.718755147 |
| | -2.863990497 | 0.004183406 | 0.010471064 |
| | -0.374157946 | 0.70828681 | NA |
| | -0.749478397 | 0.453568915 | 0.563499773 |
| | 3.961609224 | 7.44E-05 | 0.0002636 |
| | 0.787603371 | 0.430928742 | NA |
| | 2.330388502 | 0.019785627 | 0.041397161 |
| | 0.44149679 | 0.658853385 | NA |
| | -1.941361522 | 0.052214442 | 0.096111743 |
| | -3.323019095 | 0.000890488 | 0.002591132 |
| | 0.324874675 | 0.745275925 | NA |
| | -0.633617077 | 0.526330754 | 0.6313951 |
| | 0.785734044 | 0.432023314 | 0.542821322 |
| | -7.203357591 | 5.87E-13 | 5.60E-12 |
| NA | NA | NA | |
| | 0.117857225 | 0.906180786 | 0.93509668 |
| | 0.288217989 | 0.77317988 | NA |
| | 0.68637005 | 0.492479786 | 0.599726041 |
| NA | NA | NA | |
| NA | NA | NA | |
| | 0.904622757 | 0.365665275 | 0.478242337 |
| NA | NA | NA | |

| | | |
|---|---|---|
| -0.073164678 | 0.941675074 | NA |
| 0.849439784 | 0.395636624 | 0.507600835 |
| -0.265925921 | 0.790296262 | 0.850800664 |
| 1.282238304 | 0.199759063 | 0.29543662 |
| -2.28066624 | 0.022568204 | 0.046457945 |
| 0.458111809 | 0.646872115 | 0.735897148 |
| -0.636429045 | 0.524496815 | 0.629647735 |
| 0.323742634 | 0.74613289 | 0.815354999 |
| 3.16633011 | 0.001543755 | 0.004266805 |
| -0.916499807 | 0.359404809 | 0.471458643 |
| 9.393192526 | 5.82E-21 | 1.10E-19 |
| 4.536534481 | 5.72E-06 | 2.41E-05 |
| 12.61394689 | 1.77E-36 | 8.23E-35 |
| NA | NA | NA |
| NA | NA | NA |
| NA | NA | NA |
| -3.761948438 | 0.000168595 | 0.000560192 |
| -0.221882659 | 0.82440523 | 0.876997021 |
| -1.172776861 | 0.240885299 | 0.343270487 |
| 1.262364203 | 0.206817764 | 0.303750467 |
| -0.666215237 | 0.505273535 | 0.611734381 |
| 3.185562135 | 0.001444731 | 0.004017957 |
| 2.346431804 | 0.018954134 | 0.039877447 |
| 5.58326722 | 2.36E-08 | 1.38E-07 |
| -0.04100327 | 0.967293289 | 0.977647039 |
| 7.589464037 | 3.21E-14 | 3.47E-13 |
| 0.011624866 | 0.990724908 | 0.993543188 |
| 1.03884189 | 0.298878276 | 0.407768628 |
| -6.333453724 | 2.40E-10 | 1.74E-09 |
| -0.063448617 | 0.949409275 | 0.965361523 |
| NA | NA | NA |
| -3.148072269 | 0.001643511 | 0.004517082 |
| NA | NA | NA |
| NA | NA | NA |
| 0.063708285 | 0.949202508 | NA |
| -0.137819341 | 0.890383198 | NA |
| -1.080138084 | 0.280080695 | 0.387080769 |
| -0.321641413 | 0.747724366 | 0.816559376 |
| -1.848616876 | 0.064513154 | 0.114872608 |
| -1.095680457 | 0.273218638 | 0.379834705 |
| NA | NA | NA |
| -1.050518751 | 0.293479675 | NA |
| -0.104239282 | 0.916979461 | 0.942935619 |
| -2.318665527 | 0.020413179 | 0.042543037 |
| NA | NA | NA |
| 0.335915797 | 0.736934364 | NA |
| -1.208799157 | 0.226740016 | 0.326925864 |

| | | |
|---|---|---|
| 3.131069578 | 0.001741709 | 0.004759277 |
| -1.250659933 | 0.211058574 | 0.308724038 |
| NA | NA | NA |
| 0.063708285 | 0.949202508 | NA |
| NA | NA | NA |
| NA | NA | NA |
| -1.049956399 | 0.293738159 | 0.401851013 |
| -2.483264878 | 0.013018424 | 0.028785439 |
| 0.192810118 | 0.847107679 | NA |
| NA | NA | NA |
| -1.896783336 | 0.057856539 | 0.104854767 |
| 8.687394676 | 3.71E-18 | 5.67E-17 |
| 0.063708882 | 0.949202033 | NA |
| -4.327785407 | 1.51E-05 | 5.97E-05 |
| 2.914571856 | 0.003561768 | 0.009043523 |
| 2.213547842 | 0.026859899 | 0.053926627 |
| 0.110296913 | 0.912173905 | 0.939460677 |
| -0.998961355 | 0.317813412 | 0.428349023 |
| 0.792186403 | 0.428251996 | 0.539463888 |
| 1.067360532 | 0.285809064 | 0.39291486 |
| -0.869136223 | 0.384772626 | 0.496922699 |
| 1.649818969 | 0.098979968 | 0.165197613 |
| -2.622614628 | 0.008725791 | 0.020159364 |
| 0.220589495 | 0.825412082 | 0.877769656 |
| NA | NA | NA |
| -0.065389369 | 0.947863988 | NA |
| NA | NA | NA |
| NA | NA | NA |
| NA | NA | NA |
| NA | NA | NA |
| 0.637339191 | 0.523903928 | 0.629093825 |
| NA | NA | NA |
| -8.481851865 | 2.22E-17 | 3.19E-16 |
| -2.320533842 | 0.020312016 | 0.042357728 |
| NA | NA | NA |
| NA | NA | NA |
| NA | NA | NA |
| NA | NA | NA |
| NA | NA | NA |
| NA | NA | NA |
| 0.202714101 | 0.839358501 | NA |
| NA | NA | NA |
| 0.826366188 | 0.408596397 | 0.520440071 |
| -4.450702477 | 8.56E-06 | 3.53E-05 |
| -0.573000632 | 0.566644266 | 0.668541408 |
| -1.264679876 | 0.205986107 | 0.302732209 |
| -0.412915897 | 0.679668233 | 0.762498135 |

| | | | |
|---|---|---|---|
| | -3.057063346 | 0.00223517 | 0.005954903 |
| | -3.394727172 | 0.00068697 | 0.002042631 |
| | -3.720470223 | 0.000198852 | 0.0006537 |
| | 4.537037925 | 5.70E-06 | 2.41E-05 |
| | 0.199526594 | 0.841850842 | NA |
| NA | NA | NA | |
| NA | NA | NA | |
| NA | NA | NA | |
| NA | NA | NA | |
| | 0.192810118 | 0.847107679 | NA |
| | -8.777257984 | 1.68E-18 | 2.65E-17 |
| | -1.669270785 | 0.095063727 | 0.159812284 |
| NA | NA | NA | |
| NA | NA | NA | |
| NA | NA | NA | |
| | 0.199526594 | 0.841850842 | NA |
| | 0.080618145 | 0.935745636 | NA |
| | 3.063464792 | 0.002187899 | 0.0058414 |
| | 2.203516871 | 0.027558339 | 0.055173991 |
| NA | NA | NA | |
| NA | NA | NA | |
| | 1.887309612 | 0.059118694 | 0.106732853 |
| | 5.657419128 | 1.54E-08 | 9.19E-08 |
| | 7.195690607 | 6.21E-13 | 5.90E-12 |
| | 2.988883705 | 0.002799987 | 0.00728672 |
| | -4.413902777 | 1.02E-05 | 4.14E-05 |
| | 4.010251457 | 6.07E-05 | 0.00021769 |
| | -4.223702375 | 2.40E-05 | 9.21E-05 |
| NA | NA | NA | |
| | -2.629829113 | 0.00854278 | 0.019776709 |
| | 0.843519049 | 0.398938217 | 0.510789481 |
| NA | NA | NA | |
| | 1.405696126 | 0.159814378 | 0.245785184 |
| NA | NA | NA | |
| | 0.110856935 | 0.911729796 | 0.939246509 |
| | 9.950966755 | 2.50E-23 | 5.72E-22 |
| | 0.192810118 | 0.847107679 | NA |
| | 7.488197655 | 6.98E-14 | 7.28E-13 |
| | -2.417932826 | 0.015608958 | 0.033750192 |
| | -0.598406092 | 0.549569002 | 0.652950687 |
| | -4.066005218 | 4.78E-05 | 0.000174782 |
| | -5.046398766 | 4.50E-07 | 2.22E-06 |
| | -2.001272749 | 0.045363005 | 0.085102384 |
| | -1.094329528 | 0.273810479 | 0.380473896 |
| | -0.803124152 | 0.421902979 | 0.533722409 |
| | -2.706445095 | 0.006800783 | 0.016162513 |
| | 0.288217989 | 0.77317988 | NA |

| | | | |
|---|---|---|---|
| | -12.48780105 | 8.70E-36 | 3.92E-34 |
| | -1.181853555 | 0.237263813 | 0.339258112 |
| | -1.721683367 | 0.085126892 | 0.145551343 |
| | -1.234515765 | 0.217010776 | 0.315617648 |
| | 7.158518467 | 8.16E-13 | 7.65E-12 |
| NA | NA | NA | |
| | -0.933900672 | 0.350355132 | 0.462081829 |
| | -0.066715524 | 0.946808176 | 0.963710652 |
| | 5.415197123 | 6.12E-08 | 3.39E-07 |
| | 0.091838578 | 0.926826293 | NA |
| | -4.161467535 | 3.16E-05 | 0.000119024 |
| | 5.234814053 | 1.65E-07 | 8.65E-07 |
| | 0.063708285 | 0.949202508 | NA |
| | -1.390344929 | 0.164424161 | 0.251607064 |
| NA | NA | NA | |
| | -1.53376173 | 0.125088281 | 0.200960185 |
| | 7.633301187 | 2.29E-14 | 2.51E-13 |
| | -3.285635304 | 0.001017527 | 0.002928446 |
| | -1.360536964 | 0.173660064 | 0.263309389 |
| | 0.349291458 | 0.726870511 | NA |
| | -2.725830631 | 0.00641399 | 0.015341023 |
| | -2.004487904 | 0.045017821 | 0.084543523 |
| | 0.940655009 | 0.346881682 | 0.458418612 |
| | -0.212662582 | 0.831590154 | NA |
| NA | NA | NA | |
| NA | NA | NA | |
| | -1.551981592 | 0.120666628 | 0.194844572 |
| | -1.479118276 | 0.139108706 | 0.219221238 |
| | -2.051577205 | 0.040210769 | 0.076573004 |
| | -0.651134314 | 0.514959788 | 0.62036106 |
| | -2.189311173 | 0.028574231 | 0.056934644 |
| | 3.20786823 | 0.001337228 | 0.003744776 |
| NA | NA | NA | |
| | -9.888851663 | 4.65E-23 | 1.04E-21 |
| | 0.757173522 | 0.448945911 | 0.558893671 |
| NA | NA | NA | |
| | -3.398636752 | 0.000677226 | 0.002015333 |
| | 2.060154425 | 0.03938378 | 0.075164012 |
| | -3.204125266 | 0.001354735 | 0.003789546 |
| | -4.24933699 | 2.14E-05 | 8.29E-05 |
| | 0.563114066 | 0.573357214 | 0.674680601 |
| NA | NA | NA | |
| NA | NA | NA | |
| | 2.922746244 | 0.003469592 | 0.008832778 |
| | -3.185069938 | 0.00144719 | 0.00402435 |
| | 0.532385669 | 0.594458909 | 0.693253809 |
| NA | NA | NA | |

| | | |
|---|---|---|
| 0.350766081 | 0.725763846 | 0.799739244 |
| 0.842283045 | 0.399629539 | 0.511465326 |
| NA | NA | NA |
| 0.865683364 | 0.386663825 | 0.498876059 |
| 5.693721827 | 1.24E-08 | 7.51E-08 |
| 5.94282757 | 2.80E-09 | 1.82E-08 |
| -2.860423769 | 0.004230753 | 0.010575179 |
| NA | NA | NA |
| -0.099243818 | 0.920944685 | 0.945639218 |
| -6.019940255 | 1.74E-09 | 1.16E-08 |
| NA | NA | NA |
| -2.137953998 | 0.032520477 | 0.063737181 |
| -2.566075331 | 0.010285649 | 0.023372654 |
| -1.18050048 | 0.23780122 | 0.339832402 |
| -8.703020388 | 3.23E-18 | 4.96E-17 |
| 10.77491068 | 4.52E-27 | 1.31E-25 |
| 3.862122551 | 0.000112406 | 0.000385309 |
| -1.252076221 | 0.210542092 | 0.308143407 |
| 1.139758756 | 0.254386821 | NA |
| 6.545962035 | 5.91E-11 | 4.61E-10 |
| 2.754302439 | 0.005881739 | 0.014204933 |
| 2.517734517 | 0.011811231 | 0.0264125 |
| -0.065389369 | 0.947863988 | NA |
| 0.678633512 | 0.497370103 | 0.604446188 |
| 0.37795348 | 0.705465153 | 0.783877868 |
| 0.946068643 | 0.344113576 | NA |
| 0.453277928 | 0.650348624 | 0.739045948 |
| -11.44008711 | 2.64E-30 | 9.22E-29 |
| -0.742184701 | 0.457975436 | 0.567569184 |
| 4.074510729 | 4.61E-05 | 0.000168787 |
| -1.867391606 | 0.061846921 | 0.110826778 |
| 2.674212082 | 0.007490506 | 0.017589584 |
| 2.232568305 | 0.025577426 | 0.051725214 |
| NA | NA | NA |
| -0.625893054 | 0.531385091 | 0.635964848 |
| -0.065388772 | 0.947864464 | NA |
| -2.999448036 | 0.002704693 | 0.007060012 |
| -7.280265307 | 3.33E-13 | 3.25E-12 |
| NA | NA | NA |
| 0.925184613 | 0.354869886 | 0.466781925 |
| 0.369399539 | 0.711829941 | 0.78892594 |
| 0.510294292 | 0.609845301 | 0.7060654 |
| -1.623735362 | 0.1044323 | 0.172650897 |
| 10.91080155 | 1.02E-27 | 3.14E-26 |
| NA | NA | NA |
| -2.073660552 | 0.038110847 | 0.073057709 |
| -2.348928893 | 0.018827502 | 0.03963768 |

| | | |
|---|---|---|
| -1.214678773 | 0.224488631 | 0.324307425 |
| NA | NA | NA |
| NA | NA | NA |
| NA | NA | NA |
| -3.562998212 | 0.000366643 | 0.001146562 |
| -1.350886557 | 0.176731775 | 0.267141514 |
| 4.03785579 | 5.39E-05 | 0.000195677 |
| 0.192810118 | 0.847107679 | NA |
| 1.825527539 | 0.067921459 | 0.120104142 |
| 8.890037128 | 6.11E-19 | 1.00E-17 |
| 6.01574728 | 1.79E-09 | 1.19E-08 |
| 5.44403659 | 5.21E-08 | 2.92E-07 |
| -4.469884382 | 7.83E-06 | 3.24E-05 |
| 2.299597856 | 0.021471014 | 0.04442952 |
| NA | NA | NA |
| 6.719994304 | 1.82E-11 | 1.49E-10 |
| -1.198999481 | 0.230528147 | 0.33125435 |
| -2.35786051 | 0.018380598 | 0.038860239 |
| NA | NA | NA |
| 3.583652541 | 0.000338823 | 0.001067296 |
| -0.129769969 | 0.896748424 | 0.9280629 |
| 4.107330538 | 4.00E-05 | 0.000148114 |
| 1.21206756 | 0.22548652 | 0.325542343 |
| NA | NA | NA |
| 9.5998871 | 8.00E-22 | 1.64E-20 |
| 1.93058901 | 0.053533898 | NA |
| -2.086038077 | 0.036975174 | 0.071196908 |
| 9.30296812 | 1.37E-20 | 2.53E-19 |
| -1.675665606 | 0.09380371 | 0.15797357 |
| NA | NA | NA |
| -0.389759159 | 0.696714647 | 0.776618161 |
| -6.898561628 | 5.25E-12 | 4.53E-11 |
| -4.833099699 | 1.34E-06 | 6.19E-06 |
| -2.201139483 | 0.027726151 | 0.055452302 |
| 0.288217989 | 0.77317988 | NA |
| 4.66850953 | 3.03E-06 | 1.33E-05 |
| 3.933240487 | 8.38E-05 | 0.000293963 |
| -0.272596901 | 0.785163094 | 0.846770576 |
| -3.529107729 | 0.000416963 | 0.001289583 |
| -7.428771272 | 1.10E-13 | 1.12E-12 |
| 0.433991152 | 0.664294866 | 0.750566537 |
| 7.677255536 | 1.63E-14 | 1.81E-13 |
| NA | NA | NA |
| 0.333816862 | 0.738517758 | NA |
| NA | NA | NA |
| -2.026091535 | 0.042755405 | 0.080791771 |
| 0.192809521 | 0.847108147 | NA |

| | | | |
|---|---|---|---|
| | 3.847681128 | 0.000119241 | 0.000406844 |
| | -0.081634198 | 0.934937606 | 0.955826013 |
| | -5.36981368 | 7.88E-08 | 4.31E-07 |
| | -4.565527193 | 4.98E-06 | 2.12E-05 |
| | -1.979056099 | 0.047809691 | 0.089071571 |
| | -1.693220796 | 0.090413448 | 0.153198524 |
| | -1.754645416 | 0.079319978 | 0.137055022 |
| | -4.255633127 | 2.08E-05 | 8.08E-05 |
| NA | NA | NA | |
| | -3.554314028 | 0.000378966 | 0.001181556 |
| | -1.144119827 | 0.252573946 | 0.356606829 |
| | -2.813545744 | 0.004899841 | 0.012090204 |
| | 0.192810118 | 0.847107679 | NA |
| | -1.518120174 | 0.128984106 | 0.206120934 |
| | -2.374400058 | 0.017577494 | 0.037421509 |
| | -1.619695592 | 0.105297684 | 0.17378353 |
| | 3.194678701 | 0.001399866 | 0.003901419 |
| | 0.031509005 | 0.974863611 | 0.982499888 |
| | -1.073915888 | 0.282860382 | 0.389750929 |
| | 0.75112464 | 0.452577648 | 0.56249161 |
| | -7.376766249 | 1.62E-13 | 1.63E-12 |
| | -5.064831697 | 4.09E-07 | 2.03E-06 |
| NA | NA | NA | |
| | -4.409644387 | 1.04E-05 | 4.22E-05 |
| | -0.359313772 | 0.719360373 | 0.794808428 |
| NA | NA | NA | |
| NA | NA | NA | |
| | -4.670436446 | 3.01E-06 | 1.32E-05 |
| NA | NA | NA | |
| NA | NA | NA | |
| | 0.109442495 | 0.91285153 | NA |
| | -0.171176952 | 0.86408463 | 0.905433929 |
| | 0.063708285 | 0.949202508 | NA |
| NA | NA | NA | |
| | -1.953847649 | 0.050719237 | 0.093766342 |
| | -3.803608326 | 0.000142604 | 0.000479625 |
| | -1.630946186 | 0.102901675 | 0.170469848 |
| | -1.579308284 | 0.114265363 | 0.185994673 |
| | -0.065388772 | 0.947864464 | NA |
| | -1.53356796 | 0.125135976 | 0.201011004 |
| | -0.366409638 | 0.714059429 | 0.790871609 |
| | 5.575349177 | 2.47E-08 | 1.44E-07 |
| | -6.050455072 | 1.44E-09 | 9.67E-09 |
| | -0.6184351 | 0.536288566 | 0.640638706 |
| | 0.873968801 | 0.382135244 | 0.494612229 |
| | 0.816289696 | 0.414334466 | 0.525956864 |
| | -1.162393859 | 0.245075517 | 0.347943289 |

| | | |
|---|---|---|
| 3.534018166 | 0.000409293 | 0.001268053 |
| 5.158099498 | 2.49E-07 | 1.28E-06 |
| 1.181407052 | 0.237441058 | 0.339453375 |
| -3.597085368 | 0.000321803 | 0.001018039 |
| -0.920197734 | 0.357469438 | 0.469288788 |
| -0.182273018 | 0.855368473 | 0.899461113 |
| NA | NA | NA |
| -1.03938645 | 0.298625043 | 0.407467555 |
| -8.591890449 | 8.55E-18 | 1.27E-16 |
| -4.894844549 | 9.84E-07 | 4.63E-06 |
| 3.422036661 | 0.000621539 | 0.001862237 |
| -0.192231474 | 0.847560893 | 0.893557831 |
| -1.352528553 | 0.176206291 | 0.266537409 |
| 2.74897101 | 0.005978267 | 0.014407475 |
| -1.881084441 | 0.059960433 | 0.108018543 |
| 1.537944536 | 0.124062177 | 0.19955506 |
| -4.319437597 | 1.56E-05 | 6.18E-05 |
| -5.908509942 | 3.45E-09 | 2.22E-08 |
| NA | NA | NA |
| NA | NA | NA |
| -5.967380592 | 2.41E-09 | 1.58E-08 |
| -0.471105197 | 0.637565614 | 0.728654733 |
| 0.184612647 | 0.853532866 | 0.898134596 |
| -1.439604302 | 0.149979382 | 0.233413165 |
| 0.986092992 | 0.324087482 | 0.43509492 |
| -7.438788842 | 1.02E-13 | 1.04E-12 |
| 16.45333186 | 7.94E-61 | 8.14E-59 |
| 1.638331319 | 0.101352598 | 0.168405173 |
| -1.625424103 | 0.104072223 | 0.172133403 |
| -1.001087027 | 0.316784736 | 0.427314284 |
| 0.093460388 | 0.925537818 | NA |
| 6.09228414 | 1.11E-09 | 7.55E-09 |
| -0.081807613 | 0.934799702 | 0.955763103 |
| 7.620778193 | 2.52E-14 | 2.75E-13 |
| -5.588444514 | 2.29E-08 | 1.34E-07 |
| -3.626380199 | 0.000287422 | 0.000916689 |
| 1.079952223 | 0.280163456 | 0.387131021 |
| -0.296287231 | 0.767010737 | 0.83195869 |
| 0.79670864 | 0.425620265 | 0.537094794 |
| -2.399852818 | 0.016401665 | 0.035266187 |
| 3.714627628 | 0.000203503 | 0.000668199 |
| 6.930381096 | 4.20E-12 | 3.65E-11 |
| -3.238380412 | 0.001202104 | 0.00340408 |
| 0.417302998 | 0.676456796 | NA |
| -3.134100152 | 0.00172382 | 0.004713997 |
| 2.398765004 | 0.016450468 | 0.035352899 |
| 8.531894979 | 1.44E-17 | 2.10E-16 |

| | | |
|---|---|---|
| 6.425393172 | 1.32E-10 | 9.86E-10 |
| 3.211421576 | 0.0013208 | 0.003700014 |
| -1.47569741 | 0.140025142 | 0.220484983 |
| 7.34064959 | 2.13E-13 | 2.11E-12 |
| NA | NA | NA |
| NA | NA | NA |
| -0.534214697 | 0.593193002 | 0.692261322 |
| -1.273140647 | 0.202968137 | 0.299263223 |
| -6.010172026 | 1.85E-09 | 1.23E-08 |
| 0.065548599 | 0.947737213 | NA |
| 7.061671012 | 1.65E-12 | 1.49E-11 |
| 0.197502823 | 0.843434073 | 0.890540574 |
| 1.273810925 | 0.202730432 | 0.299053673 |
| -0.385385392 | 0.699951904 | NA |
| 2.132680031 | 0.032950983 | 0.064490039 |
| 0.063707688 | 0.949202983 | NA |
| -0.386591601 | 0.699058577 | 0.778598148 |
| NA | NA | NA |
| NA | NA | NA |
| -5.816543426 | 6.01E-09 | 3.76E-08 |
| -0.31498043 | 0.752776512 | 0.820644463 |
| NA | NA | NA |
| 0.263726495 | 0.791990678 | NA |
| NA | NA | NA |
| 5.182445679 | 2.19E-07 | 1.13E-06 |
| 1.30354742 | 0.192387938 | 0.286396461 |
| -4.422605344 | 9.75E-06 | 3.98E-05 |
| -0.999698569 | 0.317456405 | 0.427960026 |
| -2.402520939 | 0.016282502 | 0.035046095 |
| -0.865406649 | 0.386815633 | 0.499046198 |
| -2.151359647 | 0.031447824 | 0.061857684 |
| -2.311929677 | 0.020781559 | 0.043192097 |
| -1.495229399 | 0.134854583 | 0.213639822 |
| -2.330217178 | 0.019794676 | 0.041412632 |
| -3.149053471 | 0.001638002 | 0.004502932 |
| 0.143679422 | 0.885753621 | 0.91992454 |
| -0.27586639 | 0.782650689 | 0.844859048 |
| 3.300282836 | 0.000965874 | 0.002792013 |
| 2.949571388 | 0.00318215 | 0.008169989 |
| 0.192810118 | 0.847107679 | NA |
| -0.395112494 | 0.692759868 | 0.773337771 |
| 7.590418983 | 3.19E-14 | 3.45E-13 |
| 0.645000256 | 0.51892702 | 0.624248729 |
| 2.256218343 | 0.02405696 | 0.049022691 |
| 0.06353712 | 0.949338802 | NA |
| 0.063708285 | 0.949202508 | NA |
| NA | NA | NA |

| | | |
|---|---|---|
| 5.796724973 | 6.76E-09 | 4.21E-08 |
| 7.45102702 | 9.26E-14 | 9.56E-13 |
| NA | NA | NA |
| NA | NA | NA |
| NA | NA | NA |
| NA | NA | NA |
| NA | NA | NA |
| NA | NA | NA |
| NA | NA | NA |
| NA | NA | NA |
| NA | NA | NA |
| NA | NA | NA |
| NA | NA | NA |
| NA | NA | NA |
| NA | NA | NA |
| NA | NA | NA |
| NA | NA | NA |
| NA | NA | NA |
| NA | NA | NA |
| NA | NA | NA |
| 0.20947264 | 0.834079292 | 0.883819707 |
| NA | NA | NA |
| NA | NA | NA |
| NA | NA | NA |
| NA | NA | NA |
| -0.066204891 | 0.947214703 | NA |
| NA | NA | NA |
| 1.207879483 | 0.227093622 | 0.327303274 |
| -4.15552079 | 3.25E-05 | 0.000121997 |
| -0.54779588 | 0.583832068 | NA |
| NA | NA | NA |
| -1.276829516 | 0.201662443 | 0.297776588 |
| -7.501096185 | 6.33E-14 | 6.63E-13 |
| -3.080600709 | 0.002065835 | 0.005546862 |
| -0.477574066 | 0.632953393 | 0.72457459 |
| NA | NA | NA |
| NA | NA | NA |
| NA | NA | NA |
| -1.762054163 | 0.078060147 | 0.135261023 |
| NA | NA | NA |
| NA | NA | NA |
| -1.076854391 | 0.281545322 | 0.388547136 |
| 0.330493631 | 0.741027004 | NA |
| 4.88667843 | 1.03E-06 | 4.81E-06 |
| 0.264875225 | 0.791105585 | 0.851321799 |
| 2.453780837 | 0.014136311 | 0.030923952 |
| -2.317001386 | 0.020503656 | 0.042703155 |

| | | | |
|---|---|---|---|
| | 0.31685483 | 0.751353758 | NA |
| NA | NA | NA | |
| | -2.873664739 | 0.004057394 | 0.010186216 |
| | 0.063707135 | 0.949203424 | NA |
| NA | NA | NA | |
| NA | NA | NA | |
| | 1.262345256 | 0.206824579 | 0.303750467 |
| | 0.192809521 | 0.847108147 | NA |
| | 0.063708285 | 0.949202508 | NA |
| | 0.218708665 | 0.826876998 | 0.878879412 |
| | -7.913137891 | 2.51E-15 | 3.02E-14 |
| NA | NA | NA | |
| NA | NA | NA | |
| | 2.950959438 | 0.003167885 | 0.008139208 |
| NA | NA | NA | |
| | -4.515589346 | 6.31E-06 | 2.65E-05 |
| | 12.41071931 | 2.29E-35 | 1.02E-33 |
| | 0.659186783 | 0.509775833 | 0.615696251 |
| NA | NA | NA | |
| | -2.362675729 | 0.018143537 | 0.038417455 |
| | 7.604723016 | 2.86E-14 | 3.10E-13 |
| | 0.770100424 | 0.441240324 | 0.551357662 |
| | -5.263128659 | 1.42E-07 | 7.48E-07 |
| | -3.844166873 | 0.000120963 | 0.000412212 |
| | 1.1571557 | 0.247208753 | 0.350336038 |
| | 0.528395732 | 0.597224699 | NA |
| | -0.587291264 | 0.557008107 | 0.659535121 |
| | 0.422902325 | 0.672366526 | NA |
| | -2.414806755 | 0.01574356 | 0.034011851 |
| | -1.120347723 | 0.262565613 | 0.36795349 |
| | -0.183081568 | 0.854734018 | 0.899051195 |
| | 0.378417616 | 0.705120384 | 0.783654307 |
| | 0.809964231 | 0.417960734 | 0.529591452 |
| | -2.953555285 | 0.003141363 | 0.008079359 |
| | 6.977965728 | 2.99E-12 | 2.63E-11 |
| | 0.063708285 | 0.949202508 | NA |
| | 4.384606861 | 1.16E-05 | 4.69E-05 |
| | 0.972190919 | 0.330955577 | 0.441874332 |
| | -0.449710415 | 0.652919261 | 0.74102517 |
| | -4.440875535 | 8.96E-06 | 3.68E-05 |
| | 1.044886949 | 0.296075222 | 0.404495541 |
| | -6.694089656 | 2.17E-11 | 1.76E-10 |
| | -7.113752956 | 1.13E-12 | 1.04E-11 |
| | 0.063705652 | 0.949204604 | NA |
| | 1.593187158 | 0.111118211 | 0.181705188 |
| | 2.802324675 | 0.005073579 | 0.012466684 |
| | 0.259068872 | 0.795582102 | 0.8545346 |

| | | |
|---|---|---|
| -1.056502476 | 0.29073871 | 0.398335427 |
| 0.116685195 | 0.907109522 | NA |
| -2.101278102 | 0.035616561 | 0.068909525 |
| 6.094901831 | 1.10E-09 | 7.44E-09 |
| -1.269547337 | 0.204245919 | 0.300739769 |
| NA | NA | NA |
| NA | NA | NA |
| NA | NA | NA |
| -0.065388772 | 0.947864464 | NA |
| NA | NA | NA |
| -2.043826403 | 0.0409707 | 0.077801124 |
| NA | NA | NA |
| 6.277465337 | 3.44E-10 | 2.45E-09 |
| 2.772971958 | 0.005554691 | 0.013519506 |
| NA | NA | NA |
| NA | NA | NA |
| 0.698268611 | 0.485009226 | 0.593400182 |
| -0.400071365 | 0.689103954 | 0.77052812 |
| 5.580157215 | 2.40E-08 | 1.40E-07 |
| -1.655104783 | 0.097903245 | 0.163684338 |
| -2.123898701 | 0.033678615 | 0.065683009 |
| NA | NA | NA |
| 13.97896683 | 2.10E-44 | 1.38E-42 |
| 16.32078993 | 7.02E-60 | 7.03E-58 |
| 1.854036988 | 0.063733868 | 0.11372772 |
| 1.801564656 | 0.071613926 | 0.125674583 |
| 5.262981611 | 1.42E-07 | 7.49E-07 |
| -1.669103014 | 0.095096966 | 0.15984669 |
| 1.513060631 | 0.130264259 | 0.207742236 |
| -0.86182321 | 0.388784811 | 0.500992783 |
| 0.433702046 | 0.664504821 | 0.750702097 |
| 0.063708285 | 0.949202508 | NA |
| NA | NA | NA |
| 1.853399798 | 0.063825076 | 0.113858004 |
| 0.790457763 | 0.429260479 | 0.540407896 |
| 7.207955574 | 5.68E-13 | 5.42E-12 |
| 0.816589685 | 0.414162952 | 0.525791169 |
| 2.593801736 | 0.009492121 | 0.021741195 |
| 4.959191754 | 7.08E-07 | 3.40E-06 |
| NA | NA | NA |
| -5.880443389 | 4.09E-09 | 2.61E-08 |
| -3.795764004 | 0.000147189 | 0.000493987 |
| -0.644327516 | 0.519363076 | 0.624653202 |
| 0.670645128 | 0.502446627 | 0.6092516 |
| -2.957658957 | 0.003099849 | 0.007985716 |
| -2.806139078 | 0.005013905 | 0.012333959 |
| 3.270363067 | 0.001074095 | 0.003076042 |

| | | | |
|---|---|---|---|
| | -0.817873979 | 0.41342915 | 0.525019362 |
| | -1.307597011 | 0.191010031 | 0.284751702 |
| | 0.886371165 | 0.375417552 | 0.488216414 |
| | 7.180380324 | 6.95E-13 | 6.56E-12 |
| | 3.330988422 | 0.000865382 | 0.002525418 |
| | 24.19486418 | 2.52E-129 | 8.76E-127 |
| | -0.065389369 | 0.947863988 | NA |
| | -1.03734848 | 0.29957348 | 0.408472201 |
| | 4.306486716 | 1.66E-05 | 6.52E-05 |
| NA | NA | NA | |
| | -0.084501006 | 0.932658103 | NA |
| NA | NA | NA | |
| | 0.192809521 | 0.847108147 | NA |
| NA | NA | NA | |
| NA | NA | NA | |
| | 3.03712481 | 0.002388465 | 0.006314914 |
| NA | NA | NA | |
| NA | NA | NA | |
| NA | NA | NA | |
| | -1.125727464 | 0.260280895 | NA |
| NA | NA | NA | |
| | 1.659897815 | 0.096935011 | 0.162325773 |
| | 0.317642674 | 0.750756 | NA |
| | -0.321924872 | 0.74750961 | 0.816404485 |
| | 0.594824454 | 0.55196082 | NA |
| NA | NA | NA | |
| | -0.081937211 | 0.934696644 | NA |
| | 4.389582467 | 1.14E-05 | 4.59E-05 |
| NA | NA | NA | |
| | -5.426675427 | 5.74E-08 | 3.20E-07 |
| | -0.194488217 | 0.845793621 | NA |
| | -0.065390883 | 0.947862783 | NA |
| | 0.31685483 | 0.751353758 | NA |
| | 0.192809521 | 0.847108147 | NA |
| NA | NA | NA | |
| | 0.769934421 | 0.441338795 | NA |
| | 0.324874675 | 0.745275925 | NA |
| | 2.545337914 | 0.010917203 | 0.024644445 |
| NA | NA | NA | |
| | 5.129714549 | 2.90E-07 | 1.47E-06 |
| | 7.649232696 | 2.02E-14 | 2.23E-13 |
| NA | NA | NA | |
| | 12.77719901 | 2.20E-37 | 1.07E-35 |
| | 10.87145917 | 1.58E-27 | 4.76E-26 |
| | 7.630467791 | 2.34E-14 | 2.57E-13 |
| NA | NA | NA | |
| | 8.26153174 | 1.44E-16 | 1.93E-15 |

| | | |
|---|---|---|
| 0.908124008 | 0.363812708 | 0.476265232 |
| NA | NA | NA |
| 11.21360936 | 3.50E-29 | 1.16E-27 |
| NA | NA | NA |
| 0.005062419 | 0.995960791 | NA |
| 0.063707688 | 0.949202983 | NA |
| -2.379518805 | 0.017335258 | 0.036974991 |
| -9.189648848 | 3.94E-20 | 7.09E-19 |
| -0.171290604 | 0.863995268 | 0.905433929 |
| 1.614277322 | 0.106467287 | 0.175286206 |
| 5.67694005 | 1.37E-08 | 8.24E-08 |
| 0.178221166 | 0.858549288 | 0.901820835 |
| NA | NA | NA |
| 0.312099765 | 0.754964705 | 0.822385176 |
| 0.097933823 | 0.921984842 | NA |
| 0.063707688 | 0.949202983 | NA |
| NA | NA | NA |
| NA | NA | NA |
| NA | NA | NA |
| NA | NA | NA |
| -2.227875692 | 0.025888806 | 0.052249379 |
| -1.352960395 | 0.176068283 | 0.26641211 |
| 7.23861259 | 4.53E-13 | 4.37E-12 |
| 4.567754293 | 4.93E-06 | 2.10E-05 |
| -4.858026431 | 1.19E-06 | 5.50E-06 |
| NA | NA | NA |
| 1.916133087 | 0.055348158 | 0.101015248 |
| 1.7820561 | 0.074740084 | 0.130238394 |
| 1.771302311 | 0.076510443 | 0.132906961 |
| NA | NA | NA |
| -0.832811538 | 0.404951027 | 0.516743523 |
| NA | NA | NA |
| 0.415888655 | 0.677491482 | NA |
| NA | NA | NA |
| 0.720087889 | 0.471470884 | NA |
| 0.30922356 | 0.757151473 | NA |
| 0.192808924 | 0.847108615 | NA |
| NA | NA | NA |
| NA | NA | NA |
| NA | NA | NA |
| NA | NA | NA |
| -1.501505315 | 0.133224913 | NA |
| -0.175122499 | 0.86098338 | NA |
| NA | NA | NA |
| NA | NA | NA |
| -0.404824089 | 0.68560682 | NA |
| NA | NA | NA |

| | | |
|---|---|---|
| 0.117887787 | 0.90615657 | NA |
| NA | NA | NA |
| NA | NA | NA |
| -0.084880256 | 0.932356588 | NA |
| -0.34093621 | 0.733151604 | NA |
| -8.248213408 | 1.61E-16 | 2.15E-15 |
| -2.558371153 | 0.010516378 | 0.023845 |
| 3.938315859 | 8.21E-05 | 0.000288219 |
| -7.037714672 | 1.95E-12 | 1.75E-11 |
| -0.098104154 | 0.921849589 | 0.946310758 |
| -0.013051746 | 0.989586509 | 0.992640176 |
| -7.502744573 | 6.25E-14 | 6.55E-13 |
| -0.983448342 | 0.325386828 | 0.436423343 |
| NA | NA | NA |
| -0.754762836 | 0.450391296 | 0.560330739 |
| -4.784874844 | 1.71E-06 | 7.78E-06 |
| -1.079554349 | 0.280340679 | 0.387290386 |
| -1.268570731 | 0.204594209 | 0.30112426 |
| NA | NA | NA |
| 2.878839855 | 0.00399141 | 0.010036667 |
| -3.534804171 | 0.000408077 | 0.001265227 |
| -0.260289018 | 0.794640843 | 0.8540086 |
| -0.509676648 | 0.610278016 | 0.706443621 |
| 8.945218937 | 3.71E-19 | 6.21E-18 |
| -0.676543908 | 0.498695379 | 0.605821458 |
| -0.425309258 | 0.670611246 | 0.755056436 |
| -7.389673678 | 1.47E-13 | 1.48E-12 |
| -1.007750592 | 0.31357421 | 0.424012572 |
| NA | NA | NA |
| NA | NA | NA |
| 0.397333236 | 0.691121742 | 0.772060037 |
| NA | NA | NA |
| -3.360495256 | 0.000778029 | 0.002288373 |
| -2.364497741 | 0.018054537 | 0.038280819 |
| NA | NA | NA |
| NA | NA | NA |
| NA | NA | NA |
| -0.518166321 | 0.604342231 | 0.701543736 |
| -1.530839406 | 0.125809088 | 0.201949687 |
| -2.753423616 | 0.005897553 | 0.014236257 |
| -4.183623419 | 2.87E-05 | 0.000108788 |
| NA | NA | NA |
| -2.554250487 | 0.010641667 | 0.024105061 |
| -2.987054189 | 0.002816798 | 0.007324377 |
| -3.029037891 | 0.002453339 | 0.006463903 |
| -2.239205361 | 0.025142557 | 0.050973449 |
| 6.195818106 | 5.80E-10 | 4.04E-09 |

| | | |
|---|---|---|
| -2.745054847 | 0.006050079 | 0.01456231 |
| -3.913773262 | 9.09E-05 | 0.000316894 |
| -3.043077371 | 0.002341721 | 0.006203777 |
| -2.21473709 | 0.026778116 | 0.053788318 |
| -2.307850692 | 0.021007442 | 0.043585567 |
| NA | NA | NA |
| -1.889955355 | 0.058763931 | 0.106237718 |
| -2.083065173 | 0.037245283 | 0.071623344 |
| -8.873269667 | 7.10E-19 | 1.16E-17 |
| -5.538063823 | 3.06E-08 | 1.76E-07 |
| -4.262595118 | 2.02E-05 | 7.85E-05 |
| -3.388326646 | 0.000703205 | 0.002086688 |
| -1.946822063 | 0.051556071 | 0.095053658 |
| -3.116275978 | 0.001831508 | 0.004973166 |
| NA | NA | NA |
| NA | NA | NA |
| -5.865704682 | 4.47E-09 | 2.84E-08 |
| NA | NA | NA |
| -5.198708451 | 2.01E-07 | 1.04E-06 |
| NA | NA | NA |
| NA | NA | NA |
| -2.757628021 | 0.005822241 | 0.0140789 |
| NA | NA | NA |
| NA | NA | NA |
| -3.628270456 | 0.000285326 | 0.000910469 |
| -4.328567321 | 1.50E-05 | 5.95E-05 |
| NA | NA | NA |
| -3.91337256 | 9.10E-05 | 0.000317332 |
| NA | NA | NA |
| -1.818430853 | 0.068998306 | 0.121758856 |
| -1.210299238 | 0.22616409 | 0.326257113 |
| -1.422477928 | 0.154887555 | 0.239739997 |
| -1.200023357 | 0.23013027 | 0.330777572 |
| -1.414184741 | 0.157307667 | 0.2427803 |
| -5.354407169 | 8.58E-08 | 4.67E-07 |
| -2.876497458 | 0.004021154 | 0.010104482 |
| -2.513674292 | 0.011948072 | 0.026663717 |
| -3.180320469 | 0.001471123 | 0.004082284 |
| -1.414121735 | 0.157326162 | 0.242780805 |
| -3.513287011 | 0.000442599 | 0.001363368 |
| -1.810957514 | 0.070147429 | 0.123484119 |
| -2.322033378 | 0.020231138 | 0.042216373 |
| -1.887308617 | 0.059118828 | 0.106732853 |
| NA | NA | NA |
| NA | NA | NA |
| -1.715350908 | 0.08628091 | 0.147192692 |
| -2.007347495 | 0.044712675 | 0.084039999 |

| | | |
|---|---|---|
| -1.887148248 | 0.059140389 | 0.106756378 |
| -2.112446295 | 0.034648188 | 0.067322166 |
| -2.222586014 | 0.026243728 | 0.052867647 |
| -1.963598186 | 0.049576708 | 0.091905273 |
| -4.081293672 | 4.48E-05 | 0.000164103 |
| -2.265128863 | 0.023504773 | 0.048105026 |
| -2.247360073 | 0.024617025 | 0.050013322 |
| -2.369247018 | 0.017824344 | 0.037874214 |
| -0.566290851 | 0.571196078 | 0.672540753 |
| -3.477416185 | 0.000506271 | 0.00154348 |
| -2.737613599 | 0.006188674 | 0.014865884 |
| -1.653040279 | 0.098322666 | 0.164264874 |
| -0.22754324 | 0.82000135 | 0.873686097 |
| -0.065388772 | 0.947864464 | NA |
| -1.527023045 | 0.126755281 | 0.203233917 |
| -1.912759462 | 0.055778853 | 0.101685702 |
| -2.514188896 | 0.011930651 | 0.026636473 |
| -2.78405596 | 0.005368377 | 0.013110594 |
| NA | NA | NA |
| -2.36578935 | 0.017991678 | 0.038176645 |
| -3.22098469 | 0.00127751 | 0.003592413 |
| -3.010882577 | 0.002604896 | 0.006823426 |
| -3.408992816 | 0.000652032 | 0.001945912 |
| -1.940469016 | 0.052322716 | 0.096275644 |
| NA | NA | NA |
| NA | NA | NA |
| NA | NA | NA |
| -2.277850179 | 0.022735505 | 0.046753218 |
| -1.344309253 | 0.178848405 | 0.269705722 |
| -2.335638996 | 0.019510064 | 0.040895535 |
| NA | NA | NA |
| -3.462493342 | 0.000535195 | 0.001623553 |
| -1.143985703 | 0.252629566 | 0.35666524 |
| NA | NA | NA |
| -2.613020363 | 0.008974595 | 0.02067312 |
| -1.49732133 | 0.134309667 | 0.212945523 |
| NA | NA | NA |
| -1.299215146 | 0.193870105 | 0.288157288 |
| 0.066629133 | 0.946876953 | NA |
| -4.249040919 | 2.15E-05 | 8.30E-05 |
| -4.451851917 | 8.51E-06 | 3.51E-05 |
| 3.94899596 | 7.85E-05 | 0.000276746 |
| -0.734214198 | 0.462818202 | 0.572292243 |
| 0.680451859 | 0.496218394 | NA |
| -1.056481674 | 0.290748209 | NA |
| 1.824954921 | 0.068007832 | 0.1202229 |
| -1.882565393 | 0.05975929 | 0.107696092 |

| | | | |
|---|---|---|---|
| | -2.997762782 | 0.002719693 | 0.007095467 |
| | -1.135941778 | 0.255980932 | NA |
| | 1.178075569 | 0.238766481 | 0.34094037 |
| | -2.226326433 | 0.025992324 | 0.052428711 |
| | 0.990119195 | 0.322115862 | 0.433168217 |
| | 0.476986826 | 0.633371503 | 0.724887458 |
| | -7.543433882 | 4.58E-14 | 4.87E-13 |
| | 0.819321833 | 0.412602819 | 0.52423162 |
| | -2.836402005 | 0.004562498 | 0.011331546 |
| | 2.084017591 | 0.037158567 | 0.071484047 |
| | 0.423585206 | 0.671868346 | 0.755984028 |
| | 0.991899898 | 0.321246359 | 0.432161473 |
| | -4.707295106 | 2.51E-06 | 1.11E-05 |
| | -3.642739185 | 0.000269752 | 0.00086463 |
| | -1.497660978 | 0.134221355 | 0.212865463 |
| | -0.252412801 | 0.800722006 | 0.858479903 |
| | 4.123518654 | 3.73E-05 | 0.000138855 |
| | 4.874446948 | 1.09E-06 | 5.09E-06 |
| | 12.31946759 | 7.12E-35 | 3.09E-33 |
| | 0.289504371 | 0.772195435 | 0.836133057 |
| | 6.319851126 | 2.62E-10 | 1.89E-09 |
| | -2.689583565 | 0.007154123 | 0.016883623 |
| | -2.746249927 | 0.006028082 | 0.014513553 |
| | 3.036171757 | 0.002396028 | 0.006331566 |
| | -2.784493302 | 0.005361143 | 0.013094204 |
| | 4.848633892 | 1.24E-06 | 5.75E-06 |
| | 2.351306869 | 0.018707598 | 0.039438321 |
| | -2.016524858 | 0.043745122 | 0.082493814 |
| | -0.065389369 | 0.947863988 | NA |
| | -0.065389369 | 0.947863988 | NA |
| | -3.259947704 | 0.001114328 | 0.003178532 |
| | -3.969375032 | 7.21E-05 | 0.000255771 |
| | -1.035168677 | 0.300590144 | 0.409635287 |
| | -3.685258466 | 0.000228471 | 0.000742582 |
| | -1.198482758 | 0.230729131 | 0.331505089 |
| | 0.050384594 | 0.959815913 | 0.972324442 |
| | 0.819324908 | 0.412601065 | 0.52423162 |
| | 15.20842798 | 3.11E-52 | 2.51E-50 |
| | 0.785471321 | 0.432177279 | NA |
| NA | NA | NA | |
| NA | NA | NA | |
| | -1.783543901 | 0.074497808 | 0.129888558 |
| | 1.598996139 | 0.10982146 | NA |
| | 0.151987361 | 0.879196904 | 0.915136962 |
| NA | NA | NA | |
| | -4.892521047 | 9.96E-07 | 4.68E-06 |
| | -4.511370489 | 6.44E-06 | 2.70E-05 |

| | | | |
|---|---|---|---|
| | 4.145265789 | 3.39E-05 | 0.0001272 |
| | 2.130426281 | 0.033136436 | 0.064792203 |
| | 1.514946385 | 0.129785982 | 0.207137865 |
| | 0.482639927 | 0.629351424 | 0.721970125 |
| | 8.495638964 | 1.97E-17 | 2.84E-16 |
| | -1.852356321 | 0.063974673 | 0.114071599 |
| | -5.84924495 | 4.94E-09 | 3.12E-08 |
| | -0.573164806 | 0.566533112 | NA |
| | 21.0232052 | 4.02E-98 | 9.07E-96 |
| | 3.716076375 | 0.00020234 | 0.000664643 |
| | 6.043393136 | 1.51E-09 | 1.01E-08 |
| | 6.137924868 | 8.36E-10 | 5.74E-09 |
| | 6.883437716 | 5.84E-12 | 5.00E-11 |
| | -4.133191002 | 3.58E-05 | 0.000133613 |
| | -0.562813606 | 0.573561815 | 0.67488559 |
| | -0.501811511 | 0.615800116 | 0.710871857 |
| | 0.103716319 | 0.917394476 | 0.943284876 |
| | -1.00658278 | 0.314135313 | 0.424557607 |
| | -4.818548698 | 1.45E-06 | 6.63E-06 |
| | -1.857713204 | 0.063209752 | 0.112953531 |
| | 4.239695322 | 2.24E-05 | 8.63E-05 |
| | 0.766798211 | 0.4432015 | NA |
| | -2.029653252 | 0.042391798 | 0.080195503 |
| | 1.804496765 | 0.071153466 | 0.125015531 |
| | 1.756284846 | 0.079039784 | 0.136684108 |
| | 7.915774741 | 2.46E-15 | 2.96E-14 |
| | 1.867009614 | 0.061900245 | 0.110914392 |
| | 0.063708285 | 0.949202508 | NA |
| | -0.630381647 | 0.528444916 | 0.633324216 |
| NA | NA | NA | |
| | 3.505943499 | 0.000454992 | 0.001397527 |
| | 0.320051528 | 0.748929269 | NA |
| | 3.280781487 | 0.001035199 | 0.002974177 |
| | 3.58262861 | 0.000340154 | 0.001071355 |
| | -0.065388772 | 0.947864464 | NA |
| | 0.949806326 | 0.342210671 | 0.453569696 |
| | 2.323149295 | 0.020171134 | 0.042101691 |
| | 0.192810118 | 0.847107679 | NA |
| | 1.399409711 | 0.161690156 | 0.248014745 |
| NA | NA | NA | |
| | 0.192810118 | 0.847107679 | NA |
| | -1.279821451 | 0.200607938 | 0.296499362 |
| | 1.268450906 | 0.204636972 | 0.30112426 |
| NA | NA | NA | |
| NA | NA | NA | |
| NA | NA | NA | |
| NA | NA | NA | |

| | | |
|---|---|---|
| 0.232776481 | 0.815934977 | NA |
| 0.288217989 | 0.77317988 | NA |
| 0.841936315 | 0.399823602 | 0.511582903 |
| -0.851160791 | 0.39468004 | 0.506711192 |
| NA | NA | NA |
| NA | NA | NA |
| NA | NA | NA |
| NA | NA | NA |
| NA | NA | NA |
| NA | NA | NA |
| NA | NA | NA |
| NA | NA | NA |
| NA | NA | NA |
| NA | NA | NA |
| NA | NA | NA |
| NA | NA | NA |
| NA | NA | NA |
| NA | NA | NA |
| NA | NA | NA |
| NA | NA | NA |
| NA | NA | NA |
| NA | NA | NA |
| NA | NA | NA |
| NA | NA | NA |
| -0.425243395 | 0.670659253 | NA |
| NA | NA | NA |
| NA | NA | NA |
| NA | NA | NA |
| NA | NA | NA |
| NA | NA | NA |
| NA | NA | NA |
| NA | NA | NA |
| NA | NA | NA |
| NA | NA | NA |
| 1.339045914 | 0.180555726 | NA |
| NA | NA | NA |
| 0.063708882 | 0.949202033 | NA |
| NA | NA | NA |
| 0.192809521 | 0.847108147 | NA |
| -2.206284774 | 0.027364067 | 0.054824509 |
| NA | NA | NA |
| NA | NA | NA |
| NA | NA | NA |
| 0.192810118 | 0.847107679 | NA |
| NA | NA | NA |
| NA | NA | NA |
| NA | NA | NA |

| | | |
|---|---|---|
| NA | NA | NA |
| NA | NA | NA |
| 0.063708285 | 0.949202508 | NA |
| NA | NA | NA |
| NA | NA | NA |
| NA | NA | NA |
| NA | NA | NA |
| NA | NA | NA |
| NA | NA | NA |
| NA | NA | NA |
| NA | NA | NA |
| NA | NA | NA |
| 0.192809521 | 0.847108147 | NA |
| NA | NA | NA |
| NA | NA | NA |
| NA | NA | NA |
| NA | NA | NA |
| NA | NA | NA |
| 0.623891798 | 0.532698647 | NA |
| NA | NA | NA |
| NA | NA | NA |
| 0.926350717 | 0.354263747 | 0.466180833 |
| NA | NA | NA |
| NA | NA | NA |
| NA | NA | NA |
| 0.508204875 | 0.611309673 | NA |
| 0.063708882 | 0.949202033 | NA |
| NA | NA | NA |
| NA | NA | NA |
| NA | NA | NA |
| 0.464964985 | 0.641956532 | NA |
| NA | NA | NA |
| NA | NA | NA |
| NA | NA | NA |
| NA | NA | NA |
| NA | NA | NA |
| NA | NA | NA |
| NA | NA | NA |
| NA | NA | NA |
| NA | NA | NA |
| NA | NA | NA |
| NA | NA | NA |
| NA | NA | NA |
| NA | NA | NA |
| NA | NA | NA |
| NA | NA | NA |
| NA | NA | NA |

| | | |
|---|---|---|
| NA | NA | NA |
| NA | NA | NA |
| NA | NA | NA |
| NA | NA | NA |
| NA | NA | NA |
| 0.773849747 | 0.43901965 | 0.549272127 |
| NA | NA | NA |
| NA | NA | NA |
| NA | NA | NA |
| 0.192810118 | 0.847107679 | NA |
| NA | NA | NA |
| NA | NA | NA |
| NA | NA | NA |
| NA | NA | NA |
| NA | NA | NA |
| NA | NA | NA |
| NA | NA | NA |
| 0.299923392 | 0.764235591 | NA |
| NA | NA | NA |
| NA | NA | NA |
| NA | NA | NA |
| NA | NA | NA |
| 0.192809521 | 0.847108147 | NA |
| NA | NA | NA |
| NA | NA | NA |
| NA | NA | NA |
| NA | NA | NA |
| NA | NA | NA |
| NA | NA | NA |
| NA | NA | NA |
| NA | NA | NA |
| NA | NA | NA |
| NA | NA | NA |
| NA | NA | NA |
| 0.869147594 | 0.384766407 | NA |
| NA | NA | NA |
| NA | NA | NA |
| NA | NA | NA |
| NA | NA | NA |
| NA | NA | NA |
| NA | NA | NA |
| NA | NA | NA |
| NA | NA | NA |
| NA | NA | NA |
| 0.3474247 | 0.728272279 | NA |
| NA | NA | NA |
| NA | NA | NA |

| | | |
|---|---|---|
| NA | NA | NA |
| NA | NA | NA |
| NA | NA | NA |
| NA | NA | NA |
| 0.192810118 | 0.847107679 | NA |
| NA | NA | NA |
| NA | NA | NA |
| NA | NA | NA |
| NA | NA | NA |
| NA | NA | NA |
| NA | NA | NA |
| NA | NA | NA |
| NA | NA | NA |
| NA | NA | NA |
| 0.192810118 | 0.847107679 | NA |
| NA | NA | NA |
| NA | NA | NA |
| NA | NA | NA |
| NA | NA | NA |
| NA | NA | NA |
| NA | NA | NA |
| NA | NA | NA |
| NA | NA | NA |
| NA | NA | NA |
| NA | NA | NA |
| NA | NA | NA |
| NA | NA | NA |
| NA | NA | NA |
| NA | NA | NA |
| NA | NA | NA |
| NA | NA | NA |
| NA | NA | NA |
| NA | NA | NA |
| NA | NA | NA |
| NA | NA | NA |
| NA | NA | NA |
| 0.063708882 | 0.949202033 | NA |
| NA | NA | NA |
| NA | NA | NA |
| NA | NA | NA |
| NA | NA | NA |
| NA | NA | NA |
| NA | NA | NA |
| NA | NA | NA |
| 2.747111373 | 0.006012271 | 0.014478271 |
| NA | NA | NA |
| NA | NA | NA |

| | | |
|---|---|---|
| NA | NA | NA |
| 0.063708217 | 0.949202562 | NA |
| NA | NA | NA |
| NA | NA | NA |
| NA | NA | NA |
| NA | NA | NA |
| NA | NA | NA |
| NA | NA | NA |
| NA | NA | NA |
| NA | NA | NA |
| NA | NA | NA |
| NA | NA | NA |
| NA | NA | NA |
| NA | NA | NA |
| NA | NA | NA |
| 0.192809521 | 0.847108147 | NA |
| NA | NA | NA |
| NA | NA | NA |
| NA | NA | NA |
| NA | NA | NA |
| NA | NA | NA |
| 0.108897936 | 0.913283444 | NA |
| 0.192809521 | 0.847108147 | NA |
| NA | NA | NA |
| NA | NA | NA |
| 0.063708285 | 0.949202508 | NA |
| NA | NA | NA |
| NA | NA | NA |
| NA | NA | NA |
| NA | NA | NA |
| NA | NA | NA |
| NA | NA | NA |
| NA | NA | NA |
| NA | NA | NA |
| NA | NA | NA |
| NA | NA | NA |
| NA | NA | NA |
| NA | NA | NA |
| NA | NA | NA |
| 0.320051528 | 0.748929269 | NA |
| NA | NA | NA |
| NA | NA | NA |
| NA | NA | NA |
| NA | NA | NA |
| NA | NA | NA |
| NA | NA | NA |
| NA | NA | NA |

| | | |
|---|---|---|
| 0.192809521 | 0.847108147 | NA |
| NA | NA | NA |
| NA | NA | NA |
| NA | NA | NA |
| NA | NA | NA |
| -1.356588135 | 0.174912114 | 0.26493496 |
| NA | NA | NA |
| NA | NA | NA |
| NA | NA | NA |
| NA | NA | NA |
| NA | NA | NA |
| NA | NA | NA |
| NA | NA | NA |
| 3.471912438 | 0.000516765 | 0.001571834 |
| 0.063708882 | 0.949202033 | NA |
| NA | NA | NA |
| 1.40031373 | 0.161419391 | 0.247675312 |
| -1.651266023 | 0.098684267 | 0.164802989 |
| 1.84887895 | 0.064475294 | NA |
| NA | NA | NA |
| NA | NA | NA |
| -2.719086786 | 0.006546243 | 0.015630303 |
| -0.194488217 | 0.845793621 | NA |
| NA | NA | NA |
| -3.257010047 | 0.001125924 | 0.003209051 |
| 7.54104768 | 4.66E-14 | 4.95E-13 |
| -0.18707872 | 0.851598912 | NA |
| -0.065389369 | 0.947863988 | NA |
| -8.779458334 | 1.64E-18 | 2.60E-17 |
| 0.195978728 | 0.844626815 | 0.891460865 |
| -3.655908099 | 0.000256273 | 0.0008247 |
| -2.323945159 | 0.020128434 | 0.04201607 |
| -0.767788829 | 0.442612652 | 0.552631051 |
| -2.981539035 | 0.002868035 | 0.007444458 |
| -3.920750147 | 8.83E-05 | 0.000308545 |
| -0.26341647 | 0.792229597 | 0.852185604 |
| 2.428643371 | 0.015155432 | 0.032883188 |
| 0.192809521 | 0.847108147 | NA |
| -4.658746136 | 3.18E-06 | 1.39E-05 |
| -0.843839998 | 0.398758821 | 0.510585906 |
| 0.192809521 | 0.847108147 | NA |
| NA | NA | NA |
| -0.404547544 | 0.685810123 | 0.767874184 |
| NA | NA | NA |
| 3.179487788 | 0.001475356 | 0.004093123 |
| 3.32410256 | 0.000887036 | 0.002583703 |
| 0.489426317 | 0.624539908 | 0.718228068 |

| | | | |
|---|---|---|---|
| NA | NA | NA | |
| | -5.046061807 | 4.51E-07 | 2.22E-06 |
| | -0.728694219 | 0.46618873 | 0.575521771 |
| NA | NA | NA | |
| NA | NA | NA | |
| | -2.064927181 | 0.038929891 | 0.074405556 |
| | -4.676756231 | 2.91E-06 | 1.28E-05 |
| | 3.848888386 | 0.000118655 | 0.000405065 |
| | 7.182505004 | 6.84E-13 | 6.47E-12 |
| | 3.600312225 | 0.000317835 | 0.001006123 |
| | 1.562687772 | 0.118126052 | 0.19134708 |
| | 0.204082933 | 0.838288691 | 0.886906662 |
| | 0.063705791 | NA | NA |
| | 0.063708882 | 0.949202033 | NA |
| NA | NA | NA | |
| | 1.200956192 | 0.229768195 | 0.330370974 |
| | 0.192810118 | 0.847107679 | NA |
| | -0.652654421 | 0.513979092 | 0.619547156 |
| NA | NA | NA | |
| | 2.222291113 | 0.026263638 | 0.052899241 |
| | -0.561814389 | 0.574242488 | 0.675500022 |
| | 5.624199968 | 1.86E-08 | 1.10E-07 |
| | 0.063708285 | 0.949202508 | NA |
| NA | NA | NA | |
| | 0.8577719 | 0.391018429 | 0.503068172 |
| | 0.31685483 | 0.751353758 | NA |
| | 2.434543257 | 0.014910598 | 0.032405336 |
| NA | NA | NA | |
| | -0.544578953 | 0.586043133 | 0.686060504 |
| | -0.81863301 | 0.412995828 | 0.524575539 |
| | 0.065474733 | 0.947796023 | NA |
| NA | NA | NA | |
| NA | NA | NA | |
| | 1.686822013 | 0.091637581 | NA |
| | 10.81895003 | 2.80E-27 | 8.24E-26 |
| NA | NA | NA | |
| NA | NA | NA | |
| | 3.380742452 | 0.000722903 | 0.002140825 |
| | -0.897805582 | 0.369289208 | 0.481848975 |
| | 2.965426301 | 0.003022638 | 0.007810925 |
| NA | NA | NA | |
| | 0.665592484 | 0.505671612 | NA |
| | -10.44204912 | 1.59E-25 | 4.22E-24 |
| | 1.216723866 | 0.223709295 | 0.32344291 |
| | -3.126498809 | 0.001769012 | 0.004823878 |
| | 0.695686448 | 0.486625215 | 0.594779603 |
| | 0.520693425 | 0.602580355 | NA |

| | | |
|---|---|---|
| 2.343100684 | 0.019124221 | 0.040188 |
| 0.06371127 | 0.949200131 | NA |
| 0.192809521 | 0.847108147 | NA |
| 0.324874675 | 0.745275925 | NA |
| 0.145348608 | 0.884435638 | NA |
| 0.192809521 | 0.847108147 | NA |
| 4.872658587 | 1.10E-06 | 5.13E-06 |
| 1.704169104 | 0.0883495 | 0.150270996 |
| -0.234246307 | 0.814793767 | 0.869766632 |
| NA | NA | NA |
| 1.553526577 | 0.120297385 | 0.194348644 |
| -2.023564999 | 0.043014929 | 0.081233113 |
| -0.065388772 | 0.947864464 | NA |
| -4.654336532 | 3.25E-06 | 1.42E-05 |
| 0.92655586 | NA | NA |
| -9.274047527 | 1.79E-20 | 3.29E-19 |
| -0.564908486 | 0.572136012 | 0.673433547 |
| NA | NA | NA |
| NA | NA | NA |
| -6.561567436 | 5.32E-11 | 4.17E-10 |
| 0.350236135 | 0.72616149 | NA |
| 0.730051743 | 0.465358557 | 0.574716461 |
| -4.974375974 | 6.55E-07 | 3.15E-06 |
| -1.08503701 | 0.277905264 | 0.384839211 |
| 4.668900316 | 3.03E-06 | 1.33E-05 |
| 12.82316742 | 1.22E-37 | 6.03E-36 |
| -2.476927775 | 0.013251872 | 0.029226746 |
| -1.570178594 | 0.116373568 | 0.188899508 |
| -0.427933888 | 0.668699254 | 0.753705693 |
| 2.56673661 | 0.010266056 | 0.023337122 |
| NA | NA | NA |
| 0.160659931 | 0.87236125 | NA |
| -2.003285241 | 0.045146681 | 0.084760082 |
| -0.563488102 | 0.573102559 | NA |
| 0.192809521 | 0.847108147 | NA |
| 0.760939602 | 0.446693143 | 0.556642904 |
| NA | NA | NA |
| NA | NA | NA |
| 0.88705586 | 0.375048827 | 0.487888994 |
| NA | NA | NA |
| NA | NA | NA |
| 7.654020247 | 1.95E-14 | 2.15E-13 |
| 2.073529381 | 0.03812304 | 0.073069889 |
| -1.848901895 | 0.06447198 | 0.114823798 |
| -2.132852352 | 0.03293684 | 0.0644674 |
| -2.722939967 | 0.006470381 | 0.015462445 |
| 2.091288973 | 0.036502168 | 0.070419449 |

| | | | |
|---|---|---|---|
| | -1.579673054 | 0.11418176 | 0.185916291 |
| | 2.893275769 | 0.003812464 | 0.009623434 |
| | 0.130217092 | 0.896394672 | 0.927867278 |
| | -1.452939167 | 0.146240645 | 0.228575206 |
| NA | NA | NA | |
| | -7.725379742 | 1.12E-14 | 1.26E-13 |
| | 5.715321901 | 1.09E-08 | 6.67E-08 |
| | 8.176272286 | 2.93E-16 | 3.82E-15 |
| | 0.192809521 | 0.847108147 | NA |
| | -1.391914273 | 0.163948359 | 0.250986387 |
| | -2.693323638 | 0.007074356 | 0.016728496 |
| | -3.622981846 | 0.000291226 | 0.000927876 |
| | -0.989240779 | 0.322545351 | 0.433471614 |
| | 5.546960642 | 2.91E-08 | 1.68E-07 |
| | -0.359041854 | 0.719563779 | NA |
| | -1.721441472 | 0.085170744 | 0.145616375 |
| NA | NA | NA | |
| NA | NA | NA | |
| NA | NA | NA | |
| | 0.385949819 | 0.699533834 | 0.778954171 |
| | 1.376545777 | 0.168652725 | 0.256884178 |
| | 0.296827766 | 0.766598006 | 0.831619126 |
| | 1.836208968 | 0.066326757 | 0.11776681 |
| | -0.065388772 | 0.947864464 | NA |
| | 0.463636829 | 0.642907963 | NA |
| | 1.916114333 | 0.055350544 | 0.101015248 |
| | 1.184318627 | 0.236286958 | 0.338073772 |
| | 0.063707194 | 0.949203376 | NA |
| | 1.845551892 | 0.064957297 | 0.115564798 |
| | -1.77962635 | 0.075137131 | 0.130863973 |
| | -3.442544576 | 0.000576269 | 0.001738251 |
| | 0.192809521 | 0.847108147 | NA |
| | -7.057721518 | 1.69E-12 | 1.52E-11 |
| NA | NA | NA | |
| | 0.193076837 | 0.846898793 | NA |
| | 0.192810118 | 0.847107679 | NA |
| | 0.063708285 | 0.949202508 | NA |
| | 0.392779812 | 0.694482116 | NA |
| NA | NA | NA | |
| | 1.789889233 | 0.07347172 | 0.128323017 |
| NA | NA | NA | |
| NA | NA | NA | |
| | 0.192808924 | 0.847108615 | NA |
| | 0.71939902 | 0.471895101 | 0.580847523 |
| NA | NA | NA | |
| | 0.192810118 | 0.847107679 | NA |
| NA | NA | NA | |

| | | |
|---|---|---|
| NA | NA | NA |
| 0.833963177 | 0.404301734 | NA |
| 3.998971564 | 6.36E-05 | 0.000227481 |
| -1.5963774 | 0.110404559 | 0.180710575 |
| -0.949529596 | 0.342351327 | 0.453653116 |
| -1.447602835 | 0.147728158 | 0.230454085 |
| 5.035620864 | 4.76E-07 | 2.34E-06 |
| 0.836077291 | 0.403111423 | NA |
| 2.524937281 | 0.011571894 | 0.025956135 |
| 10.40025495 | 2.47E-25 | 6.45E-24 |
| 1.92371193 | 0.054390702 | 0.099495671 |
| -1.616260836 | 0.106037931 | 0.174705727 |
| NA | NA | NA |
| 9.374851536 | 6.93E-21 | 1.31E-19 |
| 1.782235587 | 0.074710822 | 0.130196468 |
| NA | NA | NA |
| NA | NA | NA |
| 0.426069196 | 0.670057426 | NA |
| NA | NA | NA |
| 4.500817549 | 6.77E-06 | 2.83E-05 |
| NA | NA | NA |
| NA | NA | NA |
| 0.024775659 | 0.980233907 | 0.986341897 |
| 0.192809521 | 0.847108147 | NA |
| 0.830305057 | 0.40636633 | 0.518226451 |
| -0.404244043 | 0.686033269 | NA |
| -2.982362279 | 0.002860333 | 0.007427548 |
| -1.373918396 | 0.169467018 | 0.257869474 |
| -4.239990365 | 2.24E-05 | 8.62E-05 |
| 0.915381309 | 0.359991487 | 0.472153999 |
| -3.374268787 | 0.000740121 | 0.002187932 |
| NA | NA | NA |
| 0.948548423 | 0.342850334 | 0.45417009 |
| -0.533777262 | 0.593495647 | 0.692485365 |
| 0.188144977 | 0.850763004 | NA |
| NA | NA | NA |
| -3.949349207 | 7.84E-05 | 0.000276416 |
| NA | NA | NA |
| NA | NA | NA |
| NA | NA | NA |
| 0.459712773 | 0.6457224 | 0.734990072 |
| -1.12003517 | 0.262698775 | 0.368057674 |
| 0.192809521 | 0.847108147 | NA |
| NA | NA | NA |
| -0.194487022 | 0.845794556 | NA |
| NA | NA | NA |
| -0.19448762 | 0.845794089 | NA |

| | | | |
|---|---|---|---|
| | 0.192810118 | 0.847107679 | NA |
| NA | NA | NA | |
| | 0.063708882 | 0.949202033 | NA |
| | 4.161455515 | 3.16E-05 | 0.000119024 |
| | 2.654438115 | 0.007944061 | 0.018508782 |
| | 0.063707688 | 0.949202983 | NA |
| | 3.244676476 | 0.001175841 | 0.003338776 |
| NA | NA | NA | |
| | 1.468680405 | 0.141919495 | 0.222990895 |
| | 5.081007551 | 3.75E-07 | 1.88E-06 |
| | 3.289787985 | 0.001002629 | 0.002889557 |
| | 1.962201968 | 0.049738975 | 0.092185601 |
| | 4.201325223 | 2.65E-05 | 0.000101279 |
| | 0.288217989 | 0.77317988 | NA |
| | 2.851093737 | 0.004356912 | 0.010857578 |
| | 2.23904684 | 0.025152869 | 0.050990224 |
| | 4.370732454 | 1.24E-05 | 4.96E-05 |
| | 3.450061043 | 0.00056046 | 0.001694036 |
| | 1.764964949 | 0.077569657 | 0.134486794 |
| NA | NA | NA | |
| | 1.936613698 | 0.052792582 | 0.096997602 |
| | 4.04390152 | 5.26E-05 | 0.000191056 |
| | 1.261411195 | 0.207160737 | 0.304047921 |
| | 0.063708882 | 0.949202033 | NA |
| | 2.764378015 | 0.005703143 | 0.013829651 |
| | 3.799459022 | 0.000145012 | 0.000487006 |
| | 0.920615456 | 0.35725123 | 0.469051525 |
| | 0.063708285 | 0.949202508 | NA |
| | 1.895522807 | 0.058023173 | 0.105133924 |
| NA | NA | NA | |
| | 0.771363487 | 0.440491511 | 0.550641888 |
| | 0.192810118 | 0.847107679 | NA |
| | 1.288942663 | 0.197418021 | NA |
| | 0.31685483 | 0.751353758 | NA |
| | -3.241480772 | 0.001189105 | 0.003371082 |
| NA | NA | NA | |
| | -0.065388772 | 0.947864464 | NA |
| | -0.153873292 | 0.877709643 | NA |
| | 1.385882852 | 0.165782678 | 0.253316234 |
| NA | NA | NA | |
| NA | NA | NA | |
| | 0.192808327 | 0.847109082 | NA |
| | 0.063976859 | 0.948988653 | NA |
| | 3.150906209 | 0.001627647 | 0.004476925 |
| | 3.014730419 | 0.002572078 | 0.006747626 |
| | 5.463440255 | 4.67E-08 | 2.64E-07 |
| | 2.117285167 | 0.034235653 | 0.066639583 |

| | | | |
|---|---|---|---|
| | 0.192810118 | 0.847107679 | NA |
| | 3.739333608 | 0.000184509 | 0.000609429 |
| NA | NA | NA | |
| NA | NA | NA | |
| NA | NA | NA | |
| NA | NA | NA | |
| NA | NA | NA | |
| | 1.064841084 | 0.28694785 | NA |
| NA | NA | NA | |
| NA | NA | NA | |
| NA | NA | NA | |
| NA | NA | NA | |
| | 0.451387854 | 0.651710035 | NA |
| NA | NA | NA | |
| NA | NA | NA | |
| NA | NA | NA | |
| NA | NA | NA | |
| | 0.063708285 | 0.949202508 | NA |
| NA | NA | NA | |
| | 0.192809521 | 0.847108147 | NA |
| NA | NA | NA | |
| | -0.127012659 | 0.898930379 | NA |
| NA | NA | NA | |
| NA | NA | NA | |
| | 0.192809521 | 0.847108147 | NA |
| NA | NA | NA | |
| | 0.192810118 | 0.847107679 | NA |
| NA | NA | NA | |
| | 1.131084417 | 0.258019562 | NA |
| NA | NA | NA | |
| NA | NA | NA | |
| | 0.063708285 | 0.949202508 | NA |
| NA | NA | NA | |
| NA | NA | NA | |
| NA | NA | NA | |
| NA | NA | NA | |
| NA | NA | NA | |
| | 0.192810118 | 0.847107679 | NA |
| NA | NA | NA | |
| NA | NA | NA | |
| | -0.065389369 | 0.947863988 | NA |
| | -1.675738259 | 0.093789472 | 0.15797357 |
| | -1.428504792 | 0.153146616 | 0.237471181 |
| | -0.777071823 | 0.437116397 | 0.547406985 |
| NA | NA | NA | |
| | 0.192810118 | 0.847107679 | NA |
| NA | NA | NA | |

| | | |
|---|---|---|
| 1.00401314 | 0.31537228 | NA |
| NA | NA | NA |
| 0.669835421 | 0.502962711 | 0.609614869 |
| NA | NA | NA |
| NA | NA | NA |
| NA | NA | NA |
| 0.124460124 | 0.900950971 | NA |
| 0.539910978 | 0.589258427 | NA |
| NA | NA | NA |
| NA | NA | NA |
| NA | NA | NA |
| NA | NA | NA |
| 0.192810118 | 0.847107679 | NA |
| NA | NA | NA |
| NA | NA | NA |
| NA | NA | NA |
| NA | NA | NA |
| 0.063708285 | 0.949202508 | NA |
| NA | NA | NA |
| 0.063710673 | 0.949200606 | NA |
| NA | NA | NA |
| NA | NA | NA |
| NA | NA | NA |
| NA | NA | NA |
| NA | NA | NA |
| NA | NA | NA |
| NA | NA | NA |
| NA | NA | NA |
| NA | NA | NA |
| NA | NA | NA |
| NA | NA | NA |
| 0.192810118 | 0.847107679 | NA |
| NA | NA | NA |
| NA | NA | NA |
| NA | NA | NA |
| NA | NA | NA |
| NA | NA | NA |
| NA | NA | NA |
| 0.320051528 | 0.748929269 | NA |
| 0.063708285 | 0.949202508 | NA |
| NA | NA | NA |
| 0.193341124 | 0.846691821 | NA |
| -3.142529149 | 0.00167495 | 0.004592902 |
| NA | NA | NA |
| 3.115455143 | 0.001836613 | 0.004985946 |
| 3.694484817 | 0.000220333 | 0.000718091 |
| 5.676387519 | 1.38E-08 | 8.27E-08 |

| | | |
|---|---|---|
| -1.32570132 | 0.18493865 | 0.277302726 |
| NA | NA | NA |
| 0.159177259 | 0.87352922 | NA |
| -1.087124455 | 0.276981811 | 0.383857745 |
| 4.868940418 | 1.12E-06 | 5.22E-06 |
| 0.31685483 | 0.751353758 | NA |
| 3.603236228 | 0.00031428 | 0.000995875 |
| 4.527273692 | 5.97E-06 | 2.51E-05 |
| 5.024149212 | 5.06E-07 | 2.47E-06 |
| 0.360644449 | 0.718365257 | 0.794129599 |
| 4.33755716 | 1.44E-05 | 5.72E-05 |
| 3.277627366 | 0.001046835 | 0.003005193 |
| 1.387612664 | 0.165255023 | 0.252662222 |
| -0.169087535 | 0.86572779 | NA |
| 6.195964358 | 5.79E-10 | 4.04E-09 |
| 3.405210749 | 0.00066113 | 0.001970479 |
| 2.691583649 | 0.007111366 | 0.01679995 |
| -0.065388772 | 0.947864464 | NA |
| 1.724634086 | 0.084593436 | 0.144757894 |
| 4.630547472 | 3.65E-06 | 1.58E-05 |
| 5.58751707 | 2.30E-08 | 1.35E-07 |
| NA | NA | NA |
| 2.349183439 | 0.018814635 | 0.039620588 |
| NA | NA | NA |
| 0.192808924 | 0.847108615 | NA |
| 1.187818705 | 0.234904843 | 0.336372252 |
| -0.065388772 | 0.947864464 | NA |
| 8.649449424 | 5.17E-18 | 7.78E-17 |
| 1.416220229 | 0.156711035 | 0.242113277 |
| 1.887313814 | 0.05911813 | 0.106732853 |
| 8.919699589 | 4.68E-19 | 7.76E-18 |
| NA | NA | NA |
| 9.335559512 | 1.00E-20 | 1.88E-19 |
| 6.858181345 | 6.97E-12 | 5.91E-11 |
| 0.192808327 | 0.847109082 | NA |
| -0.065388772 | 0.947864464 | NA |
| 3.030295982 | 0.002443142 | 0.006440426 |
| 6.445417835 | 1.15E-10 | 8.70E-10 |
| 0.192810118 | 0.847107679 | NA |
| 5.341778837 | 9.20E-08 | 4.99E-07 |
| -0.662325104 | 0.507762891 | NA |
| 0.363662767 | 0.71610985 | 0.792196125 |
| 0.46244344 | 0.643763354 | 0.733460638 |
| 4.05957096 | 4.92E-05 | 0.000179354 |
| 4.039919311 | 5.35E-05 | 0.000194132 |
| 0.350386091 | 0.726048964 | 0.799973833 |
| NA | NA | NA |

| | | | |
|---|---|---|---|
| | 1.333329685 | 0.182423636 | NA |
| | 0.17337293 | 0.862358302 | 0.904341725 |
| | 0.192809521 | 0.847108147 | NA |
| NA | NA | NA | |
| NA | NA | NA | |
| | 1.259913413 | 0.2077006 | NA |
| NA | NA | NA | |
| NA | NA | NA | |
| | 1.585873449 | 0.11276802 | 0.183941579 |
| NA | NA | NA | |
| | 8.198460376 | 2.43E-16 | 3.21E-15 |
| NA | NA | NA | |
| | -0.065389369 | 0.947863988 | NA |
| NA | NA | NA | |
| NA | NA | NA | |
| | 4.013833739 | 5.97E-05 | 0.000214534 |
| | 0.063708882 | 0.949202033 | NA |
| | -0.365338043 | 0.714859084 | NA |
| NA | NA | NA | |
| NA | NA | NA | |
| | -5.052152755 | 4.37E-07 | 2.16E-06 |
| | 1.911512859 | 0.055938706 | 0.101925152 |
| | -2.918919383 | 0.00351247 | 0.008931644 |
| | 0.27823453 | 0.780832329 | 0.843441792 |
| | 1.042268647 | 0.297287144 | 0.405951839 |
| | 12.62653606 | 1.51E-36 | 7.05E-35 |
| | -0.572386416 | 0.567060217 | NA |
| | -2.491793261 | 0.012709999 | 0.028175779 |
| | -1.235821936 | 0.216524759 | 0.315075561 |
| | -7.200150693 | 6.01E-13 | 5.72E-12 |
| | -1.431552938 | 0.152271807 | 0.23643693 |
| | -16.59575034 | 7.48E-62 | 7.87E-60 |
| | -2.122051412 | 0.033833421 | 0.065969504 |
| | -4.767677806 | 1.86E-06 | 8.43E-06 |
| | -3.06490441 | 0.002177396 | 0.005817687 |
| | 4.397589618 | 1.09E-05 | 4.43E-05 |
| NA | NA | NA | |
| | -3.930360475 | 8.48E-05 | 0.000297299 |
| | 0.192809521 | 0.847108147 | NA |
| | 0.120231281 | 0.904299939 | 0.93377176 |
| NA | NA | NA | |
| | 0.549761263 | 0.582483131 | NA |
| | 0.063708285 | 0.949202508 | NA |
| NA | NA | NA | |
| | 2.739009802 | 0.006162454 | 0.014807163 |
| NA | NA | NA | |
| NA | NA | NA | |

| | | | |
|---|---|---|---|
| NA | NA | NA | |
| 0.766089159 | 0.443623253 | | 0.553681184 |
| 0.192809521 | 0.847108147 | NA | |
| NA | NA | NA | |
| NA | NA | NA | |
| NA | NA | NA | |
| NA | NA | NA | |
| NA | NA | NA | |
| NA | NA | NA | |
| NA | NA | NA | |
| 0.192809521 | 0.847108147 | NA | |
| NA | NA | NA | |
| NA | NA | NA | |
| NA | NA | NA | |
| NA | NA | NA | |
| NA | NA | NA | |
| NA | NA | NA | |
| NA | NA | NA | |
| NA | NA | NA | |
| NA | NA | NA | |
| NA | NA | NA | |
| NA | NA | NA | |
| 0.192809521 | 0.847108147 | NA | |
| 0.063708882 | 0.949202033 | NA | |
| 0.063708882 | 0.949202033 | NA | |
| NA | NA | NA | |
| NA | NA | NA | |
| NA | NA | NA | |
| NA | NA | NA | |
| NA | NA | NA | |
| NA | NA | NA | |
| 5.965331129 | 2.44E-09 | | 1.60E-08 |
| -1.920996006 | 0.054732211 | | 0.100061883 |
| -0.199685219 | 0.841726774 | | 0.889206446 |
| -0.065465566 | 0.947803322 | NA | |
| NA | NA | NA | |
| 0.194173692 | 0.846039881 | NA | |
| -1.28523288 | 0.198710917 | | 0.294094977 |
| 0.724844389 | 0.4685475 | | 0.577721376 |
| -3.667526653 | 0.000244908 | | 0.00079077 |
| 0.065342075 | 0.947901643 | NA | |
| 0.161986728 | 0.871316306 | | 0.910303328 |
| NA | NA | NA | |
| -0.158216949 | 0.874285849 | | 0.912451647 |
| -0.065389369 | 0.947863988 | NA | |
| 0.06371127 | 0.949200131 | NA | |
| NA | NA | NA | |

| | | |
|---|---|---|
| 2.380915265 | 0.017269684 | 0.03687283 |
| NA | NA | NA |
| 0.110944962 | 0.911659991 | NA |
| -1.558067457 | 0.119217258 | 0.192845744 |
| 0.31685483 | 0.751353758 | NA |
| 0.018748926 | 0.985041398 | 0.989389494 |
| -0.065389369 | 0.947863988 | NA |
| -1.292029999 | 0.196346756 | 0.291164127 |
| NA | NA | NA |
| 10.02180362 | 1.22E-23 | 2.86E-22 |
| 3.838626704 | 0.000123724 | 0.000420821 |
| -1.846735006 | 0.064785556 | 0.115306415 |
| 1.002448698 | 0.316126932 | 0.426639898 |
| 0.192809521 | 0.847108147 | NA |
| NA | NA | NA |
| -0.248510038 | 0.803739804 | NA |
| NA | NA | NA |
| -0.400113636 | 0.689072821 | NA |
| -1.106679726 | 0.268432424 | NA |
| 0.812871805 | 0.416291561 | 0.527904437 |
| NA | NA | NA |
| -2.377027942 | 0.017452766 | 0.0371876 |
| NA | NA | NA |
| 0.871485317 | 0.383489219 | 0.495891118 |
| -1.525351105 | 0.127171547 | 0.203796903 |
| NA | NA | NA |
| 0.730192434 | 0.465272567 | 0.574645828 |
| NA | NA | NA |
| 4.17831534 | 2.94E-05 | 0.000111257 |
| 0.063708285 | 0.949202508 | NA |
| 0.725017099 | 0.468441541 | NA |
| -0.147854543 | 0.882457563 | NA |
| 6.212864653 | 5.20E-10 | 3.65E-09 |
| -0.413253735 | 0.679420722 | 0.762391315 |
| 0.063708285 | 0.949202508 | NA |
| NA | NA | NA |
| -1.12876639 | 0.258996394 | 0.363991002 |
| NA | NA | NA |
| 2.741003564 | 0.006125184 | 0.014731754 |
| -0.732499632 | 0.463863667 | 0.573301777 |
| -8.019395273 | 1.06E-15 | 1.32E-14 |
| 0.478403934 | 0.632362734 | NA |
| 3.98374026 | 6.78E-05 | 0.000241607 |
| 2.143041628 | 0.032109756 | 0.063025991 |
| 10.86212723 | 1.75E-27 | 5.25E-26 |
| 13.64156603 | 2.27E-42 | 1.36E-40 |
| 7.241851619 | 4.43E-13 | 4.27E-12 |

| | | |
|---|---|---|
| 4.259198034 | 2.05E-05 | 7.96E-05 |
| NA | NA | NA |
| NA | NA | NA |
| -0.061669412 | 0.950826099 | NA |
| NA | NA | NA |
| NA | NA | NA |
| 0.192810118 | 0.847107679 | NA |
| NA | NA | NA |
| NA | NA | NA |
| NA | NA | NA |
| NA | NA | NA |
| NA | NA | NA |
| NA | NA | NA |
| NA | NA | NA |
| NA | NA | NA |
| NA | NA | NA |
| NA | NA | NA |
| NA | NA | NA |
| NA | NA | NA |
| NA | NA | NA |
| NA | NA | NA |
| NA | NA | NA |
| NA | NA | NA |
| -0.355571338 | 0.722161612 | 0.796953624 |
| 1.113442603 | 0.265518386 | 0.371156328 |
| 7.893332037 | 2.94E-15 | 3.51E-14 |
| -0.987423966 | 0.32343484 | 0.434358518 |
| 4.786280396 | 1.70E-06 | 7.73E-06 |
| 12.04421694 | 2.08E-33 | 8.49E-32 |
| 0.279655305 | 0.779741972 | NA |
| 1.306156799 | 0.191499241 | 0.285311071 |
| -2.775720729 | 0.00550795 | 0.013416113 |
| 4.640511636 | 3.48E-06 | 1.51E-05 |
| 11.36781976 | 6.05E-30 | 2.08E-28 |
| 8.136522604 | 4.07E-16 | 5.23E-15 |
| 1.672277446 | 0.094469625 | 0.158915392 |
| -2.476593418 | 0.013264291 | 0.029251559 |
| 3.702522297 | 0.000213467 | 0.00069762 |
| -0.260365116 | 0.794582148 | NA |
| 0.192810118 | 0.847107679 | NA |
| -2.127092494 | 0.033412399 | 0.065249384 |
| -0.152593762 | 0.878718645 | 0.914943536 |
| -0.19448762 | 0.845794089 | NA |
| 0.192809521 | 0.847108147 | NA |
| 1.226539004 | 0.219995907 | 0.319087764 |
| -0.162050869 | 0.871265796 | 0.910303328 |
| 2.349860269 | 0.01878046 | 0.039558605 |

| | | | |
|---|---|---|---|
| | 0.444284664 | 0.656836788 | NA |
| NA | NA | NA | |
| | 1.431703745 | 0.152228624 | 0.23640516 |
| NA | NA | NA | |
| | -0.059529776 | 0.95253015 | 0.967567454 |
| | -5.355400086 | 8.54E-08 | 4.64E-07 |
| | -1.417061356 | 0.156464989 | 0.241807773 |
| | -0.424562197 | 0.671155856 | 0.755454021 |
| | 0.063708285 | 0.949202508 | NA |
| | -3.040755207 | 0.002359856 | 0.00624389 |
| | 8.055070799 | 7.94E-16 | 9.99E-15 |
| | 6.261395727 | 3.82E-10 | 2.71E-09 |
| | -0.418266049 | 0.67575261 | NA |
| NA | NA | NA | |
| | 0.113207509 | 0.909866043 | NA |
| NA | NA | NA | |
| NA | NA | NA | |
| | -1.026988365 | 0.304425943 | 0.413938578 |
| | 2.32823802 | 0.019899468 | 0.041607526 |
| | -2.24115377 | 0.025016116 | 0.050745878 |
| | 3.335452275 | 0.000851608 | 0.002488122 |
| NA | NA | NA | |
| | 0.374707949 | 0.70787768 | NA |
| | -2.833159623 | 0.004609036 | 0.01143465 |
| | -0.532177542 | 0.594603036 | 0.693357279 |
| | -0.244293732 | 0.807003337 | 0.863180784 |
| | 6.207485879 | 5.38E-10 | 3.77E-09 |
| | -0.060862961 | 0.951468347 | 0.966847782 |
| | 5.059527114 | 4.20E-07 | 2.08E-06 |
| | 0.192809521 | 0.847108147 | NA |
| | 1.444439221 | 0.148615465 | 0.231578478 |
| | -2.276205869 | 0.022833691 | 0.046931128 |
| | 0.215432239 | 0.829430337 | 0.880770495 |
| NA | NA | NA | |
| NA | NA | NA | |
| | -0.065389369 | 0.947863988 | NA |
| | 1.009131105 | 0.312911762 | NA |
| | 0.022401377 | 0.982127782 | 0.987641213 |
| | 0.717206392 | 0.473246756 | 0.582081882 |
| NA | NA | NA | |
| | 0.475706908 | 0.634283199 | 0.725764958 |
| | 9.082312641 | 1.06E-19 | 1.85E-18 |
| | 1.915025341 | 0.055489272 | 0.101231643 |
| | -1.579853269 | 0.114140473 | 0.185865334 |
| | 4.851755762 | 1.22E-06 | 5.67E-06 |
| | 4.205287586 | 2.61E-05 | 9.97E-05 |
| | 1.190437469 | 0.233874494 | 0.335169354 |

| | | | |
|---|---|---|---|
| | 1.123784554 | 0.261104435 | NA |
| | -1.414638134 | 0.157174622 | 0.242664742 |
| | -1.996945792 | 0.045831072 | 0.085845374 |
| | -7.441064429 | 9.99E-14 | 1.03E-12 |
| | 0.284003059 | 0.776408036 | NA |
| | 7.998988987 | 1.25E-15 | 1.55E-14 |
| | 0.307503436 | 0.758460208 | 0.825079708 |
| | -0.650521115 | 0.515355667 | NA |
| | -5.887801343 | 3.91E-09 | 2.50E-08 |
| | -0.130095276 | 0.896491048 | NA |
| | 1.796460821 | 0.072421258 | 0.126802428 |
| | 0.06371127 | 0.949200131 | NA |
| | 0.192809521 | 0.847108147 | NA |
| NA | NA | NA | |
| | 0.3474247 | 0.728272279 | NA |
| | 1.194365541 | 0.23233501 | 0.333295811 |
| | 0.248415817 | 0.803812697 | 0.861050969 |
| | 6.431715121 | 1.26E-10 | 9.48E-10 |
| NA | NA | NA | |
| | -6.494190588 | 8.35E-11 | 6.41E-10 |
| | -0.175325133 | 0.860824164 | NA |
| NA | NA | NA | |
| | -2.546936451 | 0.010867322 | 0.02454734 |
| NA | NA | NA | |
| | 0.192808924 | 0.847108615 | NA |
| | -0.950945528 | 0.341632029 | 0.453089945 |
| | -4.106023068 | 4.03E-05 | 0.000148867 |
| | -4.889733025 | 1.01E-06 | 4.74E-06 |
| | 1.275962825 | 0.201968661 | 0.298158394 |
| | 3.778182745 | 0.000157977 | 0.000527362 |
| | -2.412414147 | 0.015847269 | 0.034209327 |
| | 8.775992497 | 1.69E-18 | 2.68E-17 |
| | -2.107467711 | 0.035077058 | 0.068044544 |
| | 1.888539271 | 0.058953591 | 0.106526739 |
| NA | NA | NA | |
| | -0.289268777 | 0.772375704 | 0.836255899 |
| NA | NA | NA | |
| | 0.192809521 | 0.847108147 | NA |
| NA | NA | NA | |
| | 0.606948106 | 0.543885352 | 0.647766005 |
| | -0.065388772 | 0.947864464 | NA |
| | 2.315253749 | 0.020599049 | 0.042869727 |
| NA | NA | NA | |
| | 3.430352106 | 0.000602798 | 0.001811073 |
| | 0.642775035 | 0.520370084 | NA |
| | -0.085117327 | 0.932168115 | NA |
| NA | NA | NA | |

| | | | |
|---|---|---|---|
| | 3.904254485 | 9.45E-05 | 0.00032853 |
| NA | NA | NA | |
| | -7.869695007 | 3.56E-15 | 4.22E-14 |
| | -2.927587094 | 0.003416034 | 0.008713271 |
| | 2.879086378 | 0.003988291 | 0.010029832 |
| | 0.192810118 | 0.847107679 | NA |
| | 1.890560484 | 0.058683039 | 0.106137472 |
| | -5.047921886 | 4.47E-07 | 2.20E-06 |
| | -4.039334153 | 5.36E-05 | 0.000194504 |
| | -2.763494343 | 0.005718608 | 0.013863125 |
| | -0.004981403 | 0.996025432 | 0.997420147 |
| | 8.274359186 | 1.29E-16 | 1.74E-15 |
| | -4.02657409 | 5.66E-05 | 0.000204472 |
| | -1.512251921 | 0.130469787 | 0.207977249 |
| NA | NA | NA | |
| NA | NA | NA | |
| | 0.996388368 | 0.319061483 | 0.429730355 |
| | -2.615401064 | 0.008912274 | 0.020539012 |
| | -1.63417104 | 0.102222942 | 0.169621205 |
| | 0.043010352 | 0.965693282 | 0.976697222 |
| | 6.886508672 | 5.72E-12 | 4.90E-11 |
| | 0.951145639 | 0.341530449 | 0.453045055 |
| | -0.577975919 | 0.563280378 | NA |
| | -6.318915782 | 2.63E-10 | 1.90E-09 |
| | -4.980004948 | 6.36E-07 | 3.07E-06 |
| | -3.193658728 | 0.001404821 | 0.003913938 |
| | -4.42204015 | 9.78E-06 | 3.99E-05 |
| | -1.504558821 | 0.13243754 | 0.210497309 |
| | 1.906536899 | 0.056580583 | 0.102929864 |
| NA | NA | NA | |
| NA | NA | NA | |
| | 0.447779461 | 0.654312368 | 0.742236056 |
| | 0.895944108 | 0.370282615 | 0.483044449 |
| | -2.40388852 | 0.016221719 | 0.034930285 |
| NA | NA | NA | |
| | 0.353502264 | 0.723711931 | 0.798242173 |
| NA | NA | NA | |
| | 0.864438866 | 0.38734685 | 0.499602786 |
| | -3.195603001 | 0.00139539 | 0.003890319 |
| | 5.082482243 | 3.73E-07 | 1.86E-06 |
| | -3.27104843 | 0.001071496 | 0.003068948 |
| | -6.1012621 | 1.05E-09 | 7.16E-09 |
| | 8.080608852 | 6.44E-16 | 8.20E-15 |
| | 8.898603859 | 5.66E-19 | 9.33E-18 |
| | 2.009669964 | 0.044466131 | 0.08367067 |
| | 8.167000787 | 3.16E-16 | 4.11E-15 |
| | -0.297889894 | 0.765787201 | 0.830991661 |

| | | |
|---|---|---|
| 1.37929866 | 0.16780269 | 0.255822958 |
| 0.299390046 | 0.764642456 | 0.830217354 |
| 0.717326968 | 0.473172371 | 0.582018991 |
| 6.871865212 | 6.34E-12 | 5.40E-11 |
| -2.550109276 | 0.010768916 | 0.024366838 |
| 0.50456487 | 0.613864489 | 0.709515937 |
| -1.919202998 | 0.054958648 | 0.100409851 |
| 2.851757646 | 0.004347824 | 0.010838166 |
| 6.950632014 | 3.64E-12 | 3.18E-11 |
| -0.052608924 | 0.958043506 | 0.971013803 |
| -2.707010586 | 0.006789209 | 0.016141139 |
| 0.484816082 | 0.627806814 | 0.720825897 |
| NA | NA | NA |
| NA | NA | NA |
| NA | NA | NA |
| NA | NA | NA |
| NA | NA | NA |
| NA | NA | NA |
| NA | NA | NA |
| 10.11820843 | 4.59E-24 | 1.10E-22 |
| 0.295551365 | 0.76757272 | NA |
| NA | NA | NA |
| -0.194488217 | 0.845793621 | NA |
| 0.192810118 | 0.847107679 | NA |
| NA | NA | NA |
| NA | NA | NA |
| NA | NA | NA |
| 0.007782398 | 0.993790607 | 0.995620353 |
| 1.035745204 | 0.300321027 | 0.409335402 |
| 3.720964942 | 0.000198463 | 0.000652678 |
| 0.479809343 | 0.631362968 | NA |
| 13.46957809 | 2.36E-41 | 1.35E-39 |
| -10.2802347 | 8.65E-25 | 2.17E-23 |
| 1.578275959 | 0.114502226 | 0.18630896 |
| 0.304332541 | 0.760874556 | NA |
| 2.961801915 | 0.003058445 | 0.007891235 |
| -4.100404706 | 4.12E-05 | 0.000152123 |
| 7.08704624 | 1.37E-12 | 1.25E-11 |
| NA | NA | NA |
| 8.334704718 | 7.77E-17 | 1.07E-15 |
| 0.31685483 | 0.751353758 | NA |
| NA | NA | NA |
| -6.617869388 | 3.64E-11 | 2.90E-10 |
| 13.20980531 | 7.70E-40 | 4.20E-38 |
| -4.739795511 | 2.14E-06 | 9.59E-06 |
| 1.229633144 | 0.218834513 | 0.317753095 |
| 5.687052606 | 1.29E-08 | 7.79E-08 |

| | | |
|---|---|---|
| 1.297699347 | 0.194390668 | 0.28872528 |
| 6.960914599 | 3.38E-12 | 2.96E-11 |
| -2.351943793 | 0.018675597 | 0.039390763 |
| NA | NA | NA |
| 1.63624884 | 0.101787519 | 0.169015629 |
| 0.192636473 | 0.847243678 | NA |
| -3.710935198 | 0.000206495 | 0.00067678 |
| -2.528145808 | 0.011466672 | 0.025752427 |
| -1.138528004 | 0.254900072 | 0.359222386 |
| 0.193076837 | 0.846898793 | NA |
| -9.732740046 | 2.19E-22 | 4.63E-21 |
| NA | NA | NA |
| NA | NA | NA |
| 2.194347146 | 0.028210463 | 0.056272544 |
| 1.728486603 | 0.083901021 | 0.143808981 |
| -3.042530358 | 0.002345982 | 0.006213411 |
| 0.466236953 | 0.641045902 | 0.731522636 |
| -0.73702954 | 0.461104385 | 0.570511259 |
| -2.44107069 | 0.014643788 | 0.031886387 |
| 2.743043661 | 0.006087259 | 0.014644761 |
| -5.063402981 | 4.12E-07 | 2.05E-06 |
| 8.151689712 | 3.59E-16 | 4.64E-15 |
| 7.51614198 | 5.64E-14 | 5.94E-13 |
| -2.415018329 | 0.015734418 | 0.033995543 |
| -1.643593281 | 0.100260243 | 0.166878167 |
| -5.258366959 | 1.45E-07 | 7.67E-07 |
| -7.79505814 | 6.44E-15 | 7.47E-14 |
| NA | NA | NA |
| NA | NA | NA |
| -1.026887651 | 0.30447337 | 0.413980577 |
| NA | NA | NA |
| -0.231842717 | 0.816660178 | NA |
| 1.779429741 | 0.075169334 | 0.130904365 |
| -0.095146975 | 0.924198087 | NA |
| -4.882104085 | 1.05E-06 | 4.91E-06 |
| 16.31252086 | 8.04E-60 | 8.02E-58 |
| 1.360974216 | 0.173521838 | 0.263131686 |
| 1.928673931 | 0.053771355 | 0.098547923 |
| 0.941102253 | 0.346652461 | 0.458188261 |
| NA | NA | NA |
| 7.144938044 | 9.00E-13 | 8.40E-12 |
| -4.631228507 | 3.64E-06 | 1.58E-05 |
| -1.595790235 | 0.110535634 | 0.180865982 |
| 4.292926163 | 1.76E-05 | 6.91E-05 |
| -0.772857773 | 0.439606558 | 0.549837704 |
| 5.935559329 | 2.93E-09 | 1.90E-08 |
| -7.668183333 | 1.74E-14 | 1.94E-13 |

| | | |
|---|---|---|
| -0.083902417 | 0.933134017 | 0.95464183 |
| -1.27753116 | 0.201414787 | 0.297498649 |
| 1.952155074 | 0.050919796 | 0.094068078 |
| 1.075363263 | 0.282212123 | 0.389247263 |
| NA | NA | NA |
| -2.612883185 | 0.008978198 | 0.020679516 |
| -6.242639503 | 4.30E-10 | 3.04E-09 |
| NA | NA | NA |
| 7.677966282 | 1.62E-14 | 1.80E-13 |
| NA | NA | NA |
| 7.936026122 | 2.09E-15 | 2.53E-14 |
| 4.901190636 | 9.53E-07 | 4.49E-06 |
| -3.940413962 | 8.13E-05 | 0.00028583 |
| 9.861880742 | 6.09E-23 | 1.35E-21 |
| -0.856199866 | 0.391887237 | 0.503805562 |
| -0.065390563 | 0.947863038 | NA |
| 7.220063749 | 5.20E-13 | 4.98E-12 |
| -0.550156978 | 0.582211709 | 0.682468931 |
| -1.928981416 | 0.053733169 | 0.098487202 |
| 2.021840375 | 0.043192845 | 0.081544495 |
| -0.515518748 | 0.606190572 | 0.703069831 |
| NA | NA | NA |
| -2.949871928 | 0.003179057 | 0.008164559 |
| -0.240350281 | 0.810058718 | 0.865856844 |
| -0.065389369 | 0.947863988 | NA |
| 7.425957462 | 1.12E-13 | 1.14E-12 |
| NA | NA | NA |
| 9.593505717 | 8.51E-22 | 1.74E-20 |
| NA | NA | NA |
| -1.001532291 | 0.316569537 | NA |
| -4.710236478 | 2.47E-06 | 1.10E-05 |
| NA | NA | NA |
| NA | NA | NA |
| 0.60735075 | 0.543618165 | 0.647509407 |
| NA | NA | NA |
| -2.388250081 | 0.016928818 | 0.036240875 |
| 0.063708882 | 0.949202033 | NA |
| 19.35578719 | 1.82E-83 | 3.08E-81 |
| NA | NA | NA |
| 2.028736125 | 0.042485174 | 0.08035393 |
| 0.192809521 | 0.847108147 | NA |
| -0.122711815 | 0.902335308 | NA |
| -2.573570977 | 0.0100655 | 0.022924428 |
| -2.82238987 | 0.004766719 | 0.011789622 |
| NA | NA | NA |
| -8.064453181 | 7.36E-16 | 9.30E-15 |
| -5.741951406 | 9.36E-09 | 5.73E-08 |

| | | | |
|---|---|---|---|
| | -1.933634279 | 0.053158109 | 0.097539331 |
| | 1.391669752 | 0.164022426 | 0.251084418 |
| | -0.281170343 | 0.77857975 | NA |
| | 10.95094879 | 6.58E-28 | 2.04E-26 |
| | 3.332009182 | 0.000862214 | 0.002517054 |
| | -0.302016079 | 0.762639806 | 0.828600828 |
| | -1.940612727 | 0.052305269 | 0.096257694 |
| | -1.793004788 | 0.072972158 | 0.127566219 |
| | -0.065389369 | 0.947863988 | NA |
| NA | NA | NA | |
| | 7.368630121 | 1.72E-13 | 1.73E-12 |
| | 4.489158375 | 7.15E-06 | 2.98E-05 |
| | 0.267660793 | 0.788960432 | 0.849764603 |
| | 3.324079415 | 0.000887109 | 0.002583703 |
| | -2.150942833 | 0.031480712 | 0.061906103 |
| | -8.706794002 | 3.13E-18 | 4.81E-17 |
| | -8.910153443 | 5.10E-19 | 8.43E-18 |
| | -1.510152042 | 0.131004633 | 0.208617251 |
| | 2.125691247 | 0.033528976 | 0.065442161 |
| | -0.085181149 | 0.932117377 | NA |
| | 0.513923138 | 0.60730573 | 0.703841386 |
| | 1.252933526 | 0.2102299 | 0.307740475 |
| | -1.408959625 | 0.158847107 | 0.244583113 |
| | -0.065389966 | 0.947863513 | NA |
| | -1.786697557 | 0.073986384 | 0.129113797 |
| | -3.142549457 | 0.001674834 | 0.004592902 |
| | -4.043522154 | 5.27E-05 | 0.00019131 |
| | 11.8119005 | 3.39E-32 | 1.31E-30 |
| | 0.400757376 | 0.688598763 | 0.770247166 |
| | -0.527269941 | 0.598006145 | 0.696319957 |
| NA | NA | NA | |
| NA | NA | NA | |
| | -2.389085606 | 0.016890366 | 0.036174028 |
| NA | NA | NA | |
| | -0.065388772 | 0.947864464 | NA |
| NA | NA | NA | |
| | 5.651409474 | 1.59E-08 | 9.50E-08 |
| | 11.82707842 | 2.83E-32 | 1.10E-30 |
| NA | NA | NA | |
| NA | NA | NA | |
| | 14.8029824 | 1.40E-49 | 1.07E-47 |
| NA | NA | NA | |
| NA | NA | NA | |
| NA | NA | NA | |
| NA | NA | NA | |
| NA | NA | NA | |
| NA | NA | NA | |

| | | |
|---|---|---|
| NA | NA | NA |
| NA | NA | NA |
| NA | NA | NA |
| NA | NA | NA |
| 1.582581091 | 0.113516976 | 0.185030601 |
| NA | NA | NA |
| NA | NA | NA |
| -0.417820037 | 0.6760787 | 0.759356063 |
| 0.063708285 | 0.949202508 | NA |
| -0.013860733 | 0.988941089 | 0.992191577 |
| 0.011759556 | 0.990617448 | 0.993475227 |
| 0.088739328 | 0.929289076 | NA |
| -0.72490959 | 0.468507497 | 0.577716676 |
| 1.725131374 | 0.0845038 | 0.144638348 |
| 0.192810118 | 0.847107679 | NA |
| NA | NA | NA |
| 0.6162237 | 0.537746888 | 0.642013099 |
| 0.063710673 | 0.949200606 | NA |
| 0.418909496 | 0.675282276 | 0.758699855 |
| 0.078449344 | 0.937470623 | 0.957437967 |
| -0.034527984 | 0.972456127 | NA |
| -0.846507016 | 0.397269962 | 0.509122274 |
| -2.912821727 | 0.00358179 | 0.009084224 |
| -0.065389369 | 0.947863988 | NA |
| -3.673216376 | 0.000239517 | 0.000774861 |
| NA | NA | NA |
| -2.107588683 | 0.035066583 | 0.068029498 |
| NA | NA | NA |
| NA | NA | NA |
| NA | NA | NA |
| -1.590424901 | 0.111739061 | 0.182521961 |
| 1.789974498 | 0.073458011 | 0.128316981 |
| 0.932331401 | 0.351165285 | 0.462913452 |
| 0.059575917 | 0.9524934 | 0.967567454 |
| 0.612126103 | 0.540454328 | 0.644599821 |
| 0.059551862 | 0.952512559 | 0.967567454 |
| 0.414878215 | 0.678231058 | NA |
| -0.063490317 | 0.94937607 | 0.965361523 |
| -2.481949667 | 0.013066573 | 0.028876596 |
| 1.465787664 | 0.142706143 | 0.224142492 |
| 0.498667632 | 0.618013551 | 0.712785881 |
| NA | NA | NA |
| -5.543609782 | 2.96E-08 | 1.71E-07 |
| -0.841542637 | 0.40004401 | 0.511734122 |
| 2.791557998 | 0.005245496 | 0.012836797 |
| -0.065389369 | 0.947863988 | NA |
| 8.354356994 | 6.58E-17 | 9.08E-16 |

| | | | |
|---|---|---|---|
| | 1.349950563 | 0.177031841 | 0.267514307 |
| | -5.565926161 | 2.61E-08 | 1.52E-07 |
| | -3.320878864 | 0.000897345 | 0.002608051 |
| | 2.793123002 | 0.005220184 | 0.012785805 |
| | -2.91944343 | 0.00350657 | 0.00891966 |
| | -0.325554496 | 0.744761445 | 0.814296182 |
| | 10.01375858 | 1.33E-23 | 3.10E-22 |
| | 1.376167205 | 0.168769873 | 0.257015692 |
| | -0.552205347 | 0.580807671 | NA |
| | -2.797484837 | 0.005150218 | 0.01263204 |
| | -7.446311514 | 9.60E-14 | 9.90E-13 |
| NA | NA | NA | |
| NA | NA | NA | |
| | 0.192636473 | 0.847243678 | NA |
| | 2.699880627 | 0.006936436 | 0.01644275 |
| NA | NA | NA | |
| NA | NA | NA | |
| | 1.935329428 | 0.052949883 | 0.097243788 |
| | 1.18222139 | 0.237117868 | 0.339088169 |
| | -3.05945976 | 0.002217366 | 0.005909984 |
| NA | NA | NA | |
| NA | NA | NA | |
| | 0.337312816 | 0.735881099 | NA |
| NA | NA | NA | |
| NA | NA | NA | |
| NA | NA | NA | |
| | -0.203426292 | 0.838801851 | 0.887071588 |
| NA | NA | NA | |
| NA | NA | NA | |
| NA | NA | NA | |
| NA | NA | NA | |
| NA | NA | NA | |
| NA | NA | NA | |
| NA | NA | NA | |
| NA | NA | NA | |
| NA | NA | NA | |
| NA | NA | NA | |
| NA | NA | NA | |
| NA | NA | NA | |
| NA | NA | NA | |
| | -0.319878756 | 0.749060243 | NA |
| | 3.473452726 | 0.000513808 | 0.001564171 |
| | 1.287829961 | 0.19780516 | 0.292962317 |
| | -1.434892225 | 0.151317812 | 0.235232894 |
| | -0.258489623 | 0.796029056 | 0.854880584 |
| | 0.679472702 | 0.496838398 | 0.603975964 |
| | -2.202774345 | 0.027610657 | 0.055265474 |

| | | | |
|---|---|---|---|
| | -0.944349225 | 0.344991219 | NA |
| | -0.975090911 | 0.329515167 | 0.440373998 |
| | -1.462705186 | 0.143548066 | 0.225154036 |
| | -0.227082018 | 0.820359967 | 0.873966191 |
| | 3.211444142 | 0.001320696 | 0.003700014 |
| | 3.848485148 | 0.000118851 | 0.000405621 |
| | -0.777444009 | 0.436896856 | 0.547269363 |
| | -1.829896333 | 0.067265442 | 0.11912948 |
| | 3.27735625 | 0.001047841 | 0.003007391 |
| | -0.115304458 | 0.908203805 | NA |
| | 3.69245138 | 0.000222103 | 0.000723293 |
| | 0.083592589 | 0.933380359 | NA |
| | -0.292419756 | 0.769965705 | 0.834223945 |
| | 5.274781941 | 1.33E-07 | 7.05E-07 |
| | 6.403486912 | 1.52E-10 | 1.13E-09 |
| | -0.910575968 | 0.362518838 | 0.474921308 |
| | 2.07862382 | 0.03765194 | 0.072316456 |
| | -0.135122775 | 0.892514803 | NA |
| NA | NA | NA | |
| | -0.205737351 | 0.83699608 | 0.885985448 |
| | -0.138280787 | 0.89001851 | 0.923061378 |
| NA | NA | NA | |
| | 0.201685361 | 0.840162708 | 0.887948004 |
| | 1.833953659 | 0.066660873 | 0.118209882 |
| | -0.841732757 | 0.399937558 | 0.51167168 |
| | 4.005298947 | 6.19E-05 | 0.000222016 |
| | -0.736453883 | 0.461454523 | 0.570859819 |
| | 2.676709937 | 0.007434897 | 0.017478677 |
| NA | NA | NA | |
| | -0.16948829 | 0.865412583 | 0.906177918 |
| NA | NA | NA | |
| | 3.095703933 | 0.001963463 | 0.005299275 |
| | 3.419542928 | 0.000627264 | 0.001878266 |
| | 2.478676369 | 0.01318709 | 0.029102251 |
| NA | NA | NA | |
| | -0.153800995 | 0.87776665 | 0.914522945 |
| | -1.268444163 | 0.204639379 | 0.30112426 |
| | 8.574981948 | 9.91E-18 | 1.46E-16 |
| | -0.344835531 | 0.73021802 | 0.80301204 |
| | -0.196769892 | 0.844007613 | 0.890958228 |
| | -0.334234837 | 0.738202358 | 0.809399836 |
| | 2.035946408 | 0.041755737 | 0.079153749 |
| | 0.08210235 | 0.934565324 | 0.955601539 |
| | -4.898723103 | 9.65E-07 | 4.54E-06 |
| | -10.82790967 | 2.54E-27 | 7.53E-26 |
| NA | NA | NA | |
| | -2.364925738 | 0.018033687 | 0.038247298 |

| | | | |
|---|---|---|---|
| NA | NA | NA | |
| NA | NA | NA | |
| NA | NA | NA | |
| | 10.6619016 | 1.53E-26 | 4.34E-25 |
| | 4.783430556 | 1.72E-06 | 7.83E-06 |
| | 0.099377103 | 0.920838862 | NA |
| | -1.600857674 | 0.109408446 | 0.179314655 |
| | 0.07848539 | 0.937441951 | 0.957437967 |
| | -2.701041218 | 0.006912277 | 0.016394797 |
| | 4.558488001 | 5.15E-06 | 2.19E-05 |
| | 0.192808924 | 0.847108615 | NA |
| | 0.317642674 | 0.750756 | NA |
| | -0.121651289 | 0.903175193 | NA |
| NA | NA | NA | |
| NA | NA | NA | |
| | -2.523470698 | 0.011620274 | 0.026038986 |
| | -5.901306431 | 3.61E-09 | 2.32E-08 |
| | -0.521583176 | 0.601960579 | 0.699427783 |
| | 0.240040136 | 0.810299142 | NA |
| NA | NA | NA | |
| | 0.484178746 | 0.628259019 | 0.721146623 |
| | 2.827756165 | 0.00468755 | 0.011607545 |
| | 0.334104705 | 0.73830055 | 0.809425271 |
| | 1.770106343 | 0.076709427 | 0.133178675 |
| | 1.100567951 | 0.27108474 | 0.377419683 |
| | -0.498046947 | 0.618450952 | 0.713093207 |
| | -1.255729477 | 0.209214068 | 0.306432683 |
| | 0.261408382 | 0.793777593 | 0.853373987 |
| | 6.311865063 | 2.76E-10 | 1.99E-09 |
| | -4.338985125 | 1.43E-05 | 5.69E-05 |
| | 6.404424039 | 1.51E-10 | 1.12E-09 |
| | -4.116908982 | 3.84E-05 | 0.000142557 |
| | 0.264686584 | 0.791250914 | NA |
| | 2.999830734 | 0.002701297 | 0.007052129 |
| | -0.065389369 | 0.947863988 | NA |
| | 10.63505394 | 2.05E-26 | 5.74E-25 |
| | -2.873983499 | 0.004053301 | 0.010177983 |
| NA | NA | NA | |
| | 4.072177704 | 4.66E-05 | 0.000170362 |
| | -0.574741 | 0.565466474 | NA |
| | -5.037001773 | 4.73E-07 | 2.32E-06 |
| | 0.250441544 | 0.802245906 | NA |
| | -1.406702634 | 0.159515584 | 0.24540105 |
| | -3.05495617 | 0.002250934 | 0.005989887 |
| | -8.186000546 | 2.70E-16 | 3.54E-15 |
| | 0.979153347 | 0.327504216 | NA |
| | -0.544137814 | 0.58634664 | 0.686351607 |

| | | |
|---|---|---|
| -3.02029763 | 0.002525264 | 0.006635246 |
| -1.023196917 | 0.30621475 | 0.415851294 |
| -0.389051264 | 0.697238225 | 0.777054756 |
| 5.635808682 | 1.74E-08 | 1.04E-07 |
| -3.065707292 | 0.002171558 | 0.005805821 |
| NA | NA | NA |
| -3.188716565 | 0.001429059 | 0.003976582 |
| 5.608312811 | 2.04E-08 | 1.21E-07 |
| -2.206178251 | 0.027371521 | 0.054835055 |
| 10.46596981 | 1.24E-25 | 3.30E-24 |
| NA | NA | NA |
| 1.160228237 | 0.245955894 | 0.348995239 |
| 25.60521792 | 1.33E-144 | 5.14E-142 |
| 11.02057887 | 3.04E-28 | 9.56E-27 |
| 0.19482206 | 0.845532252 | 0.891903027 |
| 1.009408032 | 0.312778988 | 0.423181517 |
| 1.322464234 | 0.186013612 | 0.278597457 |
| NA | NA | NA |
| NA | NA | NA |
| 5.947747521 | 2.72E-09 | 1.77E-08 |
| -0.089918594 | 0.928351905 | 0.950989594 |
| 7.853776001 | 4.04E-15 | 4.76E-14 |
| -5.39258017 | 6.95E-08 | 3.82E-07 |
| NA | NA | NA |
| -1.202807757 | 0.229050722 | 0.329528657 |
| 0.265758864 | 0.790424926 | 0.850865985 |
| -0.11187799 | 0.910920148 | NA |
| 2.935279552 | 0.003332473 | 0.008519232 |
| 0.667589805 | 0.504395467 | 0.610932094 |
| -2.896902704 | 0.003768666 | 0.009518639 |
| -1.956555334 | 0.050399771 | 0.093244605 |
| 2.353992353 | 0.018572993 | 0.039197472 |
| NA | NA | NA |
| -7.018696554 | 2.24E-12 | 1.99E-11 |
| NA | NA | NA |
| NA | NA | NA |
| -0.936837832 | 0.348841982 | 0.460498645 |
| -3.7637046 | 0.000167415 | 0.00055664 |
| -5.447835662 | 5.10E-08 | 2.86E-07 |
| -3.293807891 | 0.0009884 | 0.002852723 |
| -0.065389369 | 0.947863988 | NA |
| -2.438775763 | 0.01473711 | 0.032064487 |
| -1.065611252 | 0.286599411 | 0.393828342 |
| -6.788427017 | 1.13E-11 | 9.46E-11 |
| -8.181058078 | 2.81E-16 | 3.68E-15 |
| NA | NA | NA |
| NA | NA | NA |

| | | |
|---|---|---|
| 0.192809521 | 0.847108147 | NA |
| NA | NA | NA |
| -0.137444972 | 0.890679086 | NA |
| 0.063710673 | 0.949200606 | NA |
| NA | NA | NA |
| 0.59111132 | 0.554445835 | 0.657433358 |
| 1.318474831 | 0.187344745 | NA |
| -1.561014055 | 0.118520435 | 0.191898989 |
| -3.351155721 | 0.00080475 | 0.002359759 |
| 2.251211758 | 0.024372129 | 0.049592259 |
| NA | NA | NA |
| -1.396337004 | 0.162613036 | 0.249277577 |
| 4.309841153 | 1.63E-05 | 6.43E-05 |
| -3.882162209 | 0.000103532 | 0.000356893 |
| -7.23380235 | 4.70E-13 | 4.52E-12 |
| -0.791299143 | 0.428769448 | 0.539903825 |
| -1.02051921 | 0.307482283 | 0.417255709 |
| NA | NA | NA |
| NA | NA | NA |
| -7.973881542 | 1.54E-15 | 1.88E-14 |
| -4.317261062 | 1.58E-05 | 6.24E-05 |
| -3.120519382 | 0.001805324 | 0.004910058 |
| -0.031765262 | 0.974659249 | 0.982430775 |
| NA | NA | NA |
| 0.169244544 | 0.865604294 | 0.906246133 |
| -1.92264284 | 0.05452492 | 0.09971933 |
| NA | NA | NA |
| -2.096553834 | 0.03603309 | 0.069602426 |
| 1.066212879 | 0.286327422 | 0.393497796 |
| -0.265672259 | 0.79049163 | NA |
| -0.065389369 | 0.947863988 | NA |
| 0.067149635 | 0.94646258 | 0.963437243 |
| -2.160783543 | 0.030712064 | 0.060569749 |
| -1.723930022 | 0.084720477 | 0.144925747 |
| NA | NA | NA |
| -0.129420327 | 0.897025064 | 0.928212715 |
| 0.281918511 | 0.778005996 | 0.841042156 |
| NA | NA | NA |
| NA | NA | NA |
| NA | NA | NA |
| NA | NA | NA |
| NA | NA | NA |
| NA | NA | NA |
| -4.67380318 | 2.96E-06 | 1.30E-05 |
| -6.318808241 | 2.64E-10 | 1.90E-09 |
| -0.559627202 | 0.575733747 | 0.676808927 |
| 9.719423055 | 2.49E-22 | 5.26E-21 |

| | | | |
|---|---|---|---|
| | -0.065388772 | 0.947864464 | NA |
| | 9.416669581 | 4.66E-21 | 8.97E-20 |
| | 1.381651417 | 0.167078758 | 0.254967767 |
| | -1.950282065 | 0.051142509 | 0.094388512 |
| | 1.691633492 | 0.090715877 | 0.153611483 |
| | -0.138467872 | 0.88987066 | 0.922946282 |
| | -0.312452571 | 0.754696603 | 0.822164692 |
| | 0.839482565 | 0.401198569 | 0.512870285 |
| NA | NA | NA | |
| | 0.4881809 | 0.625421712 | NA |
| NA | NA | NA | |
| NA | NA | NA | |
| | -3.736361959 | 0.000186702 | 0.000616185 |
| | 6.230605286 | 4.65E-10 | 3.27E-09 |
| | -0.531201247 | 0.595279329 | 0.693822659 |
| NA | NA | NA | |
| | -7.357799542 | 1.87E-13 | 1.87E-12 |
| | -1.447867436 | 0.147654128 | 0.230398773 |
| | -4.644314089 | 3.41E-06 | 1.49E-05 |
| | 1.350540205 | 0.176842766 | 0.267277002 |
| | -1.004437172 | 0.31516794 | 0.425696519 |
| | 1.810936479 | 0.070150685 | 0.123484119 |
| NA | NA | NA | |
| NA | NA | NA | |
| | 2.428403162 | 0.015165474 | 0.032902126 |
| | 0.181364731 | 0.8560813 | 0.900097242 |
| | -1.05637297 | 0.29079785 | 0.398394646 |
| | -5.225655314 | 1.74E-07 | 9.05E-07 |
| | 3.059897528 | 0.002214127 | 0.005901982 |
| NA | NA | NA | |
| NA | NA | NA | |
| | 10.12757742 | 4.17E-24 | 1.00E-22 |
| | 0.61750278 | 0.536903148 | 0.641180355 |
| | -5.323624642 | 1.02E-07 | 5.48E-07 |
| | -0.431120439 | 0.6663808 | 0.75206645 |
| | -1.692311734 | 0.090586552 | 0.153450283 |
| NA | NA | NA | |
| | -3.141122926 | 0.001683014 | 0.004611476 |
| | -1.904001623 | 0.056909973 | 0.103446377 |
| | -7.381010298 | 1.57E-13 | 1.58E-12 |
| | -4.869227966 | 1.12E-06 | 5.22E-06 |
| | 3.41839711 | 0.000629911 | 0.001885861 |
| | 0.772153965 | 0.440023243 | 0.550221401 |
| | -0.694765154 | 0.487202489 | 0.595356066 |
| | -5.593247812 | 2.23E-08 | 1.31E-07 |
| | -1.563337236 | 0.117973294 | 0.191124373 |
| | 0.432021709 | 0.665725635 | NA |

| | | | |
|---|---|---|---|
| | -0.591362058 | 0.554277857 | 0.657297331 |
| | -2.143982844 | 0.032034261 | 0.062887652 |
| | -2.831504154 | 0.004632963 | 0.011487179 |
| NA | NA | NA | |
| | -5.572196473 | 2.52E-08 | 1.46E-07 |
| | -0.063933316 | 0.949023324 | NA |
| | 0.063708882 | 0.949202033 | NA |
| | 0.678532746 | 0.497433968 | NA |
| | -0.503746398 | 0.614439599 | 0.709939674 |
| | 10.3243043 | 5.47E-25 | 1.39E-23 |
| | 0.582741386 | 0.560067408 | NA |
| | -10.05262146 | 8.95E-24 | 2.11E-22 |
| | -0.069466525 | 0.944618277 | 0.9623678 |
| | 4.286186096 | 1.82E-05 | 7.11E-05 |
| | -0.912164542 | 0.361682106 | 0.473890298 |
| | 0.063703882 | 0.949206013 | NA |
| | 5.72399557 | 1.04E-08 | 6.34E-08 |
| | 0.861342845 | 0.389049245 | 0.501179344 |
| | 0.295422184 | 0.767671389 | 0.832567047 |
| | -4.659013996 | 3.18E-06 | 1.39E-05 |
| | 0.063708285 | 0.949202508 | NA |
| | 2.956470177 | 0.003111823 | 0.008012441 |
| | -1.262276424 | 0.206849337 | 0.303765675 |
| | -0.599572524 | 0.548791163 | 0.652243248 |
| | -0.054993524 | 0.956143623 | 0.970056945 |
| | 3.364445335 | 0.000766977 | 0.002259318 |
| NA | NA | NA | |
| | -1.43903963 | 0.150139295 | 0.233632984 |
| | 2.844621265 | 0.004446426 | 0.011063025 |
| | 12.92032105 | 3.46E-38 | 1.76E-36 |
| | 5.281277502 | 1.28E-07 | 6.82E-07 |
| | 0.063707688 | 0.949202983 | NA |
| | 4.143648486 | 3.42E-05 | 0.000128024 |
| | -0.065389966 | 0.947863513 | NA |
| | 0.39561313 | 0.69239045 | 0.772994333 |
| | 1.239542972 | 0.215144485 | 0.313419949 |
| NA | NA | NA | |
| | -1.278828974 | 0.200957288 | 0.296928036 |
| | 0.641893286 | 0.520942474 | 0.626161646 |
| | 3.240294458 | 0.001194063 | 0.003383607 |
| NA | NA | NA | |
| | -0.365537731 | 0.714710047 | 0.791347097 |
| | -1.194905109 | 0.232124107 | 0.333031433 |
| NA | NA | NA | |
| | -3.667421474 | 0.000245009 | 0.000790993 |
| NA | NA | NA | |
| | -5.524349475 | 3.31E-08 | 1.90E-07 |

| | | |
|---|---|---|
| 5.00564466 | 5.57E-07 | 2.70E-06 |
| 4.683788425 | 2.82E-06 | 1.24E-05 |
| -1.599243063 | 0.109766605 | 0.179839559 |
| 0.445707694 | 0.655808406 | 0.743596123 |
| -5.426383371 | 5.75E-08 | 3.20E-07 |
| 0.063707019 | 0.949203516 | NA |
| 2.287624049 | 0.022159424 | 0.045699134 |
| -0.163808945 | 0.869881549 | 0.909373729 |
| 1.121953773 | 0.261882095 | 0.367139512 |
| 1.10149061 | 0.270683189 | 0.376902549 |
| 7.97302865 | 1.55E-15 | 1.90E-14 |
| -6.347575901 | 2.19E-10 | 1.60E-09 |
| 0.790416716 | 0.429284443 | 0.540410884 |
| -1.514543513 | 0.129888046 | 0.2072479 |
| 1.299019241 | 0.193937326 | 0.288240085 |
| 7.595435638 | 3.07E-14 | 3.32E-13 |
| -1.723120977 | 0.08486665 | 0.145146037 |
| 1.898740395 | 0.057598618 | 0.104493252 |
| 0.207660165 | 0.835494323 | 0.88469202 |
| NA | NA | NA |
| NA | NA | NA |
| 1.429442578 | 0.152877068 | 0.237141362 |
| NA | NA | NA |
| NA | NA | NA |
| 0.193472986 | 0.84658856 | NA |
| 7.997142478 | 1.27E-15 | 1.57E-14 |
| -1.010483509 | 0.312263696 | 0.422552864 |
| 7.468770031 | 8.09E-14 | 8.40E-13 |
| 2.101794138 | 0.035571313 | 0.06884859 |
| NA | NA | NA |
| -0.194488217 | 0.845793621 | NA |
| NA | NA | NA |
| 4.41551297 | 1.01E-05 | 4.11E-05 |
| 1.551861908 | 0.120695269 | 0.194878247 |
| 0.622402048 | 0.533677534 | 0.638067766 |
| 0.4906226 | 0.623693399 | 0.717487253 |
| 1.074382572 | 0.28265125 | 0.389600793 |
| -3.357686976 | 0.000785976 | 0.002309034 |
| -2.663934811 | 0.007723253 | 0.018058199 |
| -0.089656978 | 0.928559805 | NA |
| NA | NA | NA |
| -0.273655547 | 0.784349342 | 0.846218277 |
| -6.086380793 | 1.15E-09 | 7.81E-09 |
| 0.822956212 | 0.410532895 | 0.522428103 |
| 1.368433059 | 0.171176562 | 0.260174238 |
| -2.291406592 | 0.021939909 | 0.045276273 |
| 4.240510611 | 2.23E-05 | 8.60E-05 |

| | | |
|---|---|---|
| 19.49453004 | 1.22E-84 | 2.12E-82 |
| 11.24912381 | 2.34E-29 | 7.80E-28 |
| 9.52314945 | 1.68E-21 | 3.36E-20 |
| 2.912561662 | 0.003584774 | 0.009090846 |
| 1.973515723 | 0.048436835 | 0.090025542 |
| 1.716986051 | 0.085981721 | 0.146772316 |
| -1.971038838 | 0.048719434 | 0.090517178 |
| 0.100095563 | 0.920268458 | 0.945307978 |
| -2.147292758 | 0.031769982 | 0.062403096 |
| NA | NA | NA |
| 1.778663113 | 0.07529501 | 0.13106851 |
| 1.916674258 | 0.055279328 | 0.100907349 |
| -2.166212185 | 0.030294976 | 0.059858167 |
| 4.731014443 | 2.23E-06 | 1.00E-05 |
| -4.006805149 | 6.15E-05 | 0.000220732 |
| 0.869284194 | 0.384691706 | 0.496869467 |
| 1.240878413 | 0.214650669 | 0.312912419 |
| 1.063770545 | 0.287432658 | NA |
| 2.952033017 | 0.003156892 | 0.008113461 |
| -0.827673544 | 0.407855406 | 0.519654915 |
| 0.879179374 | 0.379304026 | 0.49187908 |
| 0.077233675 | 0.938437653 | NA |
| 0.901835859 | 0.367144068 | 0.479625053 |
| NA | NA | NA |
| 0.430982122 | 0.66648137 | 0.752086412 |
| 1.265353929 | 0.205744482 | NA |
| NA | NA | NA |
| NA | NA | NA |
| 3.601019591 | 0.000316972 | 0.00100377 |
| -1.731046511 | 0.083443469 | 0.143115157 |
| NA | NA | NA |
| NA | NA | NA |
| 22.81654275 | 3.14E-115 | 9.47E-113 |
| -3.938724049 | 8.19E-05 | 0.00028777 |
| 1.182001853 | 0.237204966 | 0.339193344 |
| 1.549713667 | 0.121210256 | NA |
| -4.095908133 | 4.21E-05 | 0.000154924 |
| -4.554218531 | 5.26E-06 | 2.23E-05 |
| NA | NA | NA |
| -0.640565018 | 0.521805334 | NA |
| NA | NA | NA |
| NA | NA | NA |
| -1.924615847 | 0.054277435 | 0.099332032 |
| 7.000310801 | 2.55E-12 | 2.26E-11 |
| 1.423709024 | 0.15453072 | 0.239320872 |
| -8.151623827 | 3.59E-16 | 4.64E-15 |
| 0.860897625 | 0.38929443 | 0.501417155 |

| | | |
|---|---|---|
| -1.643233783 | 0.100334573 | 0.166974817 |
| 0.849401506 | 0.395657915 | 0.507602133 |
| 2.309100781 | 0.020937989 | 0.043466689 |
| 10.01917999 | 1.26E-23 | 2.93E-22 |
| -0.396916963 | 0.691428695 | 0.772299533 |
| -0.065388772 | 0.947864464 | NA |
| 0.720933503 | 0.470950429 | 0.579884358 |
| 1.986280879 | 0.047002145 | 0.087763049 |
| -0.213727338 | 0.830759691 | 0.881621092 |
| -7.462100066 | 8.52E-14 | 8.81E-13 |
| 1.319598375 | 0.186969146 | 0.27971058 |
| -0.20303826 | 0.839105128 | 0.887279851 |
| -0.60846319 | 0.542880312 | 0.646863718 |
| -2.197659485 | 0.027973381 | 0.05586641 |
| NA | NA | NA |
| -0.639565503 | 0.522455116 | NA |
| 0.604811701 | 0.545304118 | NA |
| -4.708940515 | 2.49E-06 | 1.10E-05 |
| -0.26176171 | 0.793505161 | NA |
| -0.802028847 | 0.422536272 | 0.53425353 |
| -4.888486438 | 1.02E-06 | 4.77E-06 |
| -0.932868234 | 0.350888005 | 0.462613999 |
| -2.944382807 | 0.003235995 | 0.008300569 |
| 1.482071105 | 0.138321376 | 0.218214046 |
| 0.892330026 | 0.372216065 | NA |
| 0.187830656 | 0.851009404 | 0.896345959 |
| -1.080879839 | 0.279750568 | 0.386709931 |
| 2.395837032 | 0.016582461 | 0.03559988 |
| -0.774534826 | 0.438614582 | 0.548983596 |
| -2.371756874 | 0.017703736 | 0.037663768 |
| -1.496810048 | 0.134442691 | 0.213115931 |
| -0.380365779 | 0.703673914 | NA |
| 0.886936567 | 0.375113053 | 0.487921827 |
| -5.437925776 | 5.39E-08 | 3.01E-07 |
| NA | NA | NA |
| NA | NA | NA |
| 2.863441595 | 0.004190661 | 0.010487126 |
| -0.182581424 | 0.85512646 | 0.899343876 |
| -2.950299993 | 0.003174655 | 0.008154928 |
| -0.586443141 | 0.557577761 | 0.660084725 |
| -6.949268742 | 3.67E-12 | 3.21E-11 |
| 0.026935326 | 0.978511318 | NA |
| 4.396913724 | 1.10E-05 | 4.45E-05 |
| 1.916385141 | 0.055316091 | 0.100967093 |
| -2.179987094 | 0.029258418 | 0.058112856 |
| 0.991600705 | 0.321392345 | NA |
| NA | NA | NA |

| | | | |
|---|---|---|---|
| NA | NA | NA | |
| | 4.28197084 | 1.85E-05 | 7.24E-05 |
| NA | NA | NA | |
| NA | NA | NA | |
| | 0.063708285 | 0.949202508 | NA |
| NA | NA | NA | |
| | 4.592666859 | 4.38E-06 | 1.88E-05 |
| | 3.121308573 | 0.001800493 | 0.004900112 |
| | 0.606574427 | 0.544133376 | NA |
| | 0.707463819 | 0.47927829 | NA |
| | 6.213613928 | 5.18E-10 | 3.63E-09 |
| | -0.362763645 | 0.716781452 | 0.792728868 |
| NA | NA | NA | |
| NA | NA | NA | |
| | 4.092726118 | 4.26E-05 | 0.000156835 |
| | 3.151488018 | 0.001624408 | 0.004468506 |
| | -0.347068937 | 0.728539529 | NA |
| | -0.198521673 | 0.84263693 | NA |
| NA | NA | NA | |
| | 0.545060679 | 0.585711785 | 0.685865075 |
| | -1.638114673 | 0.101397775 | 0.168457872 |
| | 7.391389382 | 1.45E-13 | 1.47E-12 |
| | 2.517761842 | 0.011810315 | 0.0264125 |
| | 5.477932521 | 4.30E-08 | 2.44E-07 |
| | -3.371324026 | 0.000748078 | 0.00220989 |
| | 0.973407902 | 0.330350613 | 0.441325568 |
| | 1.617475777 | 0.10577562 | 0.174422806 |
| | 0.503092796 | 0.614899032 | NA |
| | 2.445233194 | 0.014475851 | 0.031548154 |
| | -0.291659048 | 0.770547324 | 0.834709578 |
| | 0.028232498 | 0.977476718 | 0.984435282 |
| | -0.31498039 | 0.752776543 | 0.820644463 |
| | -0.065389369 | 0.947863988 | NA |
| | 0.192810118 | 0.847107679 | NA |
| NA | NA | NA | |
| | 0.464760005 | 0.642103332 | 0.732469529 |
| | 8.605829689 | 7.58E-18 | 1.13E-16 |
| | 11.71430356 | 1.08E-31 | 4.07E-30 |
| NA | NA | NA | |
| | 8.181614909 | 2.80E-16 | 3.66E-15 |
| | 4.113549421 | 3.90E-05 | 0.000144477 |
| | -3.369795691 | 0.000752239 | 0.002220872 |
| NA | NA | NA | |
| | 5.778081772 | 7.56E-09 | 4.67E-08 |
| | 2.625102855 | 0.00866228 | 0.020031121 |
| | -4.279793284 | 1.87E-05 | 7.30E-05 |
| | -0.325674617 | 0.744670551 | NA |

| | | | |
|---|---|---|---|
| | 1.624711183 | 0.104224113 | 0.172341254 |
| | 0.9851748 | 0.324538217 | 0.43554758 |
| NA | | NA | NA |
| NA | | NA | NA |
| | 5.428590966 | 5.68E-08 | 3.16E-07 |
| | 6.183059626 | 6.29E-10 | 4.36E-09 |
| | 4.683676316 | 2.82E-06 | 1.24E-05 |
| | 3.104860831 | 0.001903686 | 0.005150157 |
| NA | | NA | NA |
| NA | | NA | NA |
| | -0.70587632 | 0.480265055 | 0.588948818 |
| | -1.34254228 | 0.179420231 | 0.27046765 |
| NA | | NA | NA |
| | 4.235735482 | 2.28E-05 | 8.77E-05 |
| | -2.398841082 | 0.016447051 | 0.035348591 |
| | 0.760262125 | 0.447097917 | 0.557036379 |
| | 1.288296128 | 0.1976429 | 0.292771079 |
| | 0.214289958 | 0.830320949 | 0.881304951 |
| NA | | NA | NA |
| | 0.063708882 | 0.949202033 | NA |
| | 3.707796474 | 0.000209071 | 0.000684684 |
| | 0.428880954 | 0.668009862 | 0.753064308 |
| NA | | NA | NA |
| | -5.026620527 | 4.99E-07 | 2.45E-06 |
| | -2.171150404 | 0.029919805 | 0.059210367 |
| NA | | NA | NA |
| | -2.385618496 | 0.017050428 | 0.036451335 |
| | 2.816889258 | 0.004849124 | 0.011980415 |
| | -4.304489988 | 1.67E-05 | 6.58E-05 |
| | -0.0653931 | 0.947861018 | NA |
| | -1.000136232 | 0.317244584 | 0.427766629 |
| | -0.196999848 | 0.843827657 | 0.890833347 |
| | -0.842221114 | 0.399664197 | 0.511476627 |
| | -1.851628096 | 0.064079246 | 0.114200103 |
| NA | | NA | NA |
| | -0.053991963 | 0.956941567 | 0.970365225 |
| | -5.780810046 | 7.43E-09 | 4.60E-08 |
| | 4.803780366 | 1.56E-06 | 7.11E-06 |
| NA | | NA | NA |
| NA | | NA | NA |
| | 2.175401471 | 0.029600044 | 0.058688935 |
| NA | | NA | NA |
| | 0.261213554 | 0.793927827 | NA |
| NA | | NA | NA |
| NA | | NA | NA |
| NA | | NA | NA |
| NA | | NA | NA |

| | | |
|---|---|---|
| -0.378901018 | 0.704761368 | 0.783495113 |
| NA | NA | NA |
| NA | NA | NA |
| 2.137406273 | 0.032564962 | NA |
| NA | NA | NA |
| 1.250650123 | 0.211062155 | 0.308724038 |
| NA | NA | NA |
| 0.320071032 | 0.748914485 | NA |
| 3.783262283 | 0.000154786 | 0.00051747 |
| NA | NA | NA |
| NA | NA | NA |
| 0.772742185 | 0.439674976 | 0.5498958 |
| 9.079666125 | 1.09E-19 | 1.90E-18 |
| NA | NA | NA |
| NA | NA | NA |
| 0.252900123 | 0.800345394 | 0.858282544 |
| 0.97263818 | 0.33073316 | 0.441695176 |
| NA | NA | NA |
| -0.254487446 | 0.799119002 | NA |
| NA | NA | NA |
| NA | NA | NA |
| -0.194488217 | 0.845793621 | NA |
| 2.67429481 | 0.007488659 | 0.017586895 |
| -1.490022744 | 0.136218256 | 0.215452951 |
| NA | NA | NA |
| -0.79357683 | 0.427441826 | 0.538741573 |
| -1.593185047 | 0.111118685 | 0.181705188 |
| NA | NA | NA |
| NA | NA | NA |
| -0.022262259 | 0.982238755 | 0.987641213 |
| 0.355613515 | 0.722130021 | 0.796953624 |
| NA | NA | NA |
| -0.065389369 | 0.947863988 | NA |
| NA | NA | NA |
| NA | NA | NA |
| NA | NA | NA |
| NA | NA | NA |
| 0.063708882 | 0.949202033 | NA |
| 0.400384553 | 0.6888733 | 0.770387495 |
| NA | NA | NA |
| NA | NA | NA |
| NA | NA | NA |
| NA | NA | NA |
| 0.192810118 | 0.847107679 | NA |
| -0.002011396 | 0.998395139 | 0.998874024 |
| NA | NA | NA |
| NA | NA | NA |

| | | | |
|---|---|---|---|
| | 1.122932219 | 0.261466283 | NA |
| NA | NA | NA | |
| | -0.065389369 | 0.947863988 | NA |
| NA | NA | NA | |
| | -0.055631092 | 0.955635695 | 0.969659537 |
| NA | NA | NA | |
| | 0.320071032 | 0.748914485 | NA |
| | 4.170996029 | 3.03E-05 | 0.000114648 |
| NA | NA | NA | |
| | 0.063708882 | 0.949202033 | NA |
| | -0.93967454 | 0.347384528 | 0.458913535 |
| NA | NA | NA | |
| | -0.194488217 | 0.845793621 | NA |
| | 2.203479138 | 0.027560996 | 0.055174896 |
| NA | NA | NA | |
| | 0.192636473 | 0.847243678 | NA |
| NA | NA | NA | |
| | 0.063708285 | 0.949202508 | NA |
| | 4.547655667 | 5.42E-06 | 2.30E-05 |
| NA | NA | NA | |
| | 0.063708285 | 0.949202508 | NA |
| | 1.320677876 | 0.186608794 | 0.279305036 |
| | 2.101325639 | 0.035612391 | 0.068906782 |
| NA | NA | NA | |
| | -0.065388772 | 0.947864464 | NA |
| NA | NA | NA | |
| | -0.065389369 | 0.947863988 | NA |
| NA | NA | NA | |
| | 0.06353712 | 0.949338802 | NA |
| NA | NA | NA | |
| | -0.19448762 | 0.845794089 | NA |
| NA | NA | NA | |
| | -0.11907437 | 0.905216436 | 0.934409741 |
| NA | NA | NA | |
| | 0.192809521 | 0.847108147 | NA |
| | 1.411280496 | 0.158161923 | 0.243768038 |
| NA | NA | NA | |
| | -0.144869288 | 0.884814075 | NA |
| NA | NA | NA | |
| NA | NA | NA | |
| | 2.309275308 | 0.020928309 | 0.043450197 |
| NA | NA | NA | |
| | -0.695967709 | 0.486449053 | NA |
| NA | NA | NA | |
| NA | NA | NA | |
| NA | NA | NA | |
| | 1.104568082 | 0.269346788 | 0.375480373 |

| | | | |
|---|---|---|---|
| | 0.31685483 | 0.751353758 | NA |
| | -0.569122525 | 0.569272993 | 0.67081221 |
| NA | NA | NA | |
| | 0.362026528 | 0.717332207 | NA |
| NA | NA | NA | |
| | -0.046312804 | 0.963060934 | 0.974736804 |
| NA | NA | NA | |
| | -0.065388772 | 0.947864464 | NA |
| NA | NA | NA | |
| | -0.24662222 | 0.805200606 | NA |
| NA | NA | NA | |
| NA | NA | NA | |
| NA | NA | NA | |
| | 0.142922848 | 0.886351114 | 0.920401941 |
| NA | NA | NA | |
| | 0.392603364 | 0.694612454 | NA |
| NA | NA | NA | |
| | -0.065389369 | 0.947863988 | NA |
| | -0.002304859 | 0.99816099 | 0.998679681 |
| NA | NA | NA | |
| NA | NA | NA | |
| NA | NA | NA | |
| NA | NA | NA | |
| NA | NA | NA | |
| | -0.069305451 | 0.944746487 | 0.962419527 |
| NA | NA | NA | |
| NA | NA | NA | |
| | 0.192636473 | 0.847243678 | NA |
| | 0.684750251 | 0.493501533 | 0.600678048 |
| NA | NA | NA | |
| NA | NA | NA | |
| | 0.282379522 | 0.777652516 | NA |
| NA | NA | NA | |
| | 2.548556738 | 0.010816969 | 0.024455671 |
| | 0.39390925 | 0.693648041 | NA |
| NA | NA | NA | |
| NA | NA | NA | |
| | 2.054641937 | 0.039913602 | 0.076074202 |
| NA | NA | NA | |
| | 0.192809521 | 0.847108147 | NA |
| NA | NA | NA | |
| | 0.803571615 | 0.421644422 | 0.533422283 |
| | 1.663775784 | 0.096157245 | 0.161271498 |
| | -0.180898617 | 0.856447154 | NA |
| NA | NA | NA | |
| NA | NA | NA | |
| | -0.000818239 | 0.99934714 | 0.999506869 |

| | | | |
|---|---|---|---|
| NA | NA | NA | |
| NA | NA | NA | |
| NA | NA | NA | |
| NA | NA | NA | |
| | 0.045252537 | 0.963906019 | 0.975323105 |
| NA | NA | NA | |
| NA | NA | NA | |
| NA | NA | NA | |
| NA | NA | NA | |
| | -0.45775249 | 0.647130273 | NA |
| | 2.44789951 | 0.014369173 | 0.031353883 |
| NA | NA | NA | |
| | 0.063708882 | 0.949202033 | NA |
| | -1.789108683 | 0.073597314 | 0.128506507 |
| NA | NA | NA | |
| | 0.192808924 | 0.847108615 | NA |
| NA | NA | NA | |
| NA | NA | NA | |
| | -0.41643793 | 0.677089578 | NA |
| | 2.350035986 | 0.018771596 | 0.039543263 |
| NA | NA | NA | |
| | 0.192809521 | 0.847108147 | NA |
| | 1.073896277 | 0.282869172 | 0.389750929 |
| | 0.063708882 | 0.949202033 | NA |
| | 3.005534068 | 0.002651149 | 0.006933263 |
| | 0.06371127 | 0.949200131 | NA |
| | 2.273705989 | 0.02298367 | 0.047181273 |
| | 0.063710673 | 0.949200606 | NA |
| | 2.644489483 | 0.008181425 | 0.019005203 |
| NA | NA | NA | |
| | 1.126671876 | 0.259881238 | 0.364824867 |
| | 0.150116531 | 0.880672678 | NA |
| | 0.06371127 | 0.949200131 | NA |
| | 2.67042305 | 0.007575573 | 0.017764343 |
| | 0.192809521 | 0.847108147 | NA |
| | 0.192810118 | 0.847107679 | NA |
| NA | NA | NA | |
| | 0.192808924 | 0.847108615 | NA |
| | 0.192809521 | 0.847108147 | NA |
| | -0.678189024 | 0.497651851 | 0.604729868 |
| | 1.535582352 | 0.124640845 | 0.20034423 |
| | 0.567341218 | 0.570482378 | NA |
| | -1.261929683 | 0.20697409 | 0.303863069 |
| | -0.19448762 | 0.845794089 | NA |
| NA | NA | NA | |
| | 0.679124382 | 0.497059055 | NA |
| | -1.411983797 | 0.157954732 | 0.24352368 |

| | | |
|---|---|---|
| -0.51620395 | 0.605711972 | 0.702644976 |
| -0.065388772 | 0.947864464 | NA |
| 0.063710673 | 0.949200606 | NA |
| 1.155624579 | 0.24783475 | 0.351044298 |
| 6.755708342 | 1.42E-11 | 1.17E-10 |
| -4.173231288 | 3.00E-05 | 0.00011358 |
| -0.122763408 | 0.902294451 | 0.932150576 |
| 9.86982389 | 5.63E-23 | 1.25E-21 |
| -2.84321183 | 0.004466138 | 0.011108757 |
| -0.947258121 | 0.343507265 | 0.454919939 |
| 1.193139184 | 0.232814868 | 0.333869379 |
| 2.75836046 | 0.00580921 | 0.014051462 |
| NA | NA | NA |
| -7.297816886 | 2.92E-13 | 2.87E-12 |
| NA | NA | NA |
| 5.602237294 | 2.12E-08 | 1.25E-07 |
| -2.727345347 | 0.006384618 | 0.015277914 |
| NA | NA | NA |
| -0.494650881 | 0.62084658 | NA |
| NA | NA | NA |
| NA | NA | NA |
| -0.152797042 | 0.87855833 | 0.914852727 |
| 0.378466068 | 0.705084396 | NA |
| -1.282199307 | 0.199772739 | 0.295439389 |
| NA | NA | NA |
| NA | NA | NA |
| NA | NA | NA |
| -5.889527084 | 3.87E-09 | 2.48E-08 |
| -0.065390563 | 0.947863038 | NA |
| -1.345178911 | 0.178567465 | 0.269346971 |
| 0.063708285 | 0.949202508 | NA |
| -4.959125185 | 7.08E-07 | 3.40E-06 |
| -10.23975442 | 1.32E-24 | 3.26E-23 |
| -3.54582697 | 0.000391383 | 0.001217842 |
| -1.15963957 | 0.246195583 | 0.349256147 |
| -0.323274023 | 0.746487726 | 0.815671532 |
| -2.852402702 | 0.004339009 | 0.010818349 |
| NA | NA | NA |
| 0.718628689 | 0.472369732 | 0.581260233 |
| 0.168610962 | 0.866102658 | 0.906517856 |
| NA | NA | NA |
| 1.19065336 | 0.233789695 | 0.335094546 |
| NA | NA | NA |
| 3.37175507 | 0.000746908 | 0.002206695 |
| 2.378491395 | 0.017383643 | 0.037052932 |
| -0.356853111 | 0.721201773 | 0.796245441 |
| 5.575106127 | 2.47E-08 | 1.44E-07 |

| | | | |
|---|---|---|---|
| NA | NA | NA | |
| | -0.065389966 | 0.947863513 | NA |
| | 9.677319341 | 3.76E-22 | 7.85E-21 |
| NA | NA | NA | |
| | 2.179748655 | 0.029276098 | 0.05813552 |
| | -7.518129397 | 5.56E-14 | 5.86E-13 |
| | -5.015745564 | 5.28E-07 | 2.58E-06 |
| | -3.566251122 | 0.000362124 | 0.001133989 |
| | 2.675004164 | 0.007472832 | 0.017553021 |
| | 1.626267525 | 0.103892757 | 0.171906902 |
| | -1.6395603 | 0.101096623 | 0.168046784 |
| | -6.263446961 | 3.77E-10 | 2.67E-09 |
| | -8.341226511 | 7.35E-17 | 1.01E-15 |
| NA | NA | NA | |
| NA | NA | NA | |
| | 0.202714101 | 0.839358501 | NA |
| NA | NA | NA | |
| | -6.703730862 | 2.03E-11 | 1.66E-10 |
| | 4.572631013 | 4.82E-06 | 2.05E-05 |
| | -7.85861941 | 3.88E-15 | 4.58E-14 |
| | 3.060976847 | 0.002206162 | 0.005884508 |
| | 0.486426165 | 0.626665043 | 0.71994428 |
| | 1.048277023 | 0.294510993 | 0.402776207 |
| | -4.27853341 | 1.88E-05 | 7.34E-05 |
| | -0.870770141 | 0.383879672 | 0.49620472 |
| | -3.284345197 | 0.001022196 | 0.002940871 |
| NA | NA | NA | |
| NA | NA | NA | |
| NA | NA | NA | |
| | -0.135468718 | 0.892241296 | NA |
| | -0.137402958 | 0.890712293 | NA |
| NA | NA | NA | |
| | 1.745122711 | 0.080963511 | 0.139557653 |
| NA | NA | NA | |
| | 0.101394995 | 0.919236909 | NA |
| | 2.497121796 | 0.012520594 | 0.027800096 |
| | -0.717768192 | 0.472900229 | NA |
| NA | NA | NA | |
| | 0.06371127 | 0.949200131 | NA |
| | 8.140122462 | 3.95E-16 | 5.09E-15 |
| | 6.512802105 | 7.38E-11 | 5.69E-10 |
| | -2.975555226 | 0.002924587 | 0.007580243 |
| | 0.989530451 | 0.322403679 | 0.433415548 |
| | 4.357993613 | 1.31E-05 | 5.24E-05 |
| NA | NA | NA | |
| | 0.192810118 | 0.847107679 | NA |
| | 5.637524389 | 1.73E-08 | 1.03E-07 |

| | | |
|---|---|---|
| 3.897649659 | 9.71E-05 | 0.000336729 |
| -2.370678438 | 0.017755471 | 0.037755453 |
| 2.2446734 | 0.024789107 | 0.050347935 |
| 4.953494831 | 7.29E-07 | 3.49E-06 |
| 0.330377319 | 0.741114877 | 0.811607691 |
| NA | NA | NA |
| 9.013490277 | 2.00E-19 | 3.40E-18 |
| -1.18788734 | 0.234877798 | 0.336365218 |
| -8.049844634 | 8.29E-16 | 1.04E-14 |
| -0.511318142 | 0.609128303 | 0.705629287 |
| -0.964840113 | 0.334624901 | 0.44584591 |
| -1.539863443 | 0.123593643 | 0.198929258 |
| 0.445388891 | 0.656038739 | 0.743621485 |
| 0.560886463 | 0.574874939 | 0.676021599 |
| 6.681042326 | 2.37E-11 | 1.92E-10 |
| 0.15693603 | 0.875295264 | 0.912954326 |
| -0.222952204 | 0.823572706 | NA |
| -3.372852315 | 0.000743938 | 0.002198699 |
| -0.604040863 | 0.545816475 | 0.649417044 |
| 6.798969477 | 1.05E-11 | 8.81E-11 |
| 2.64345268 | 0.008206524 | 0.019056432 |
| 7.404157311 | 1.32E-13 | 1.34E-12 |
| -13.42439942 | 4.35E-41 | 2.45E-39 |
| 8.654151887 | 4.97E-18 | 7.49E-17 |
| 6.543527226 | 6.01E-11 | 4.67E-10 |
| 1.509497093 | 0.131171798 | 0.208830307 |
| -3.398775048 | 0.000676884 | 0.002014554 |
| -1.591549428 | 0.111485981 | 0.18220684 |
| -0.108597177 | 0.913522 | NA |
| -6.235107577 | 4.51E-10 | 3.18E-09 |
| 0.565817987 | 0.571517518 | 0.672863757 |
| 2.118776995 | 0.034109318 | 0.066414331 |
| 2.279286596 | 0.022650034 | 0.046595739 |
| 1.579413705 | 0.114241196 | 0.185993049 |
| -0.638816061 | 0.522942599 | NA |
| 2.258762908 | 0.023898137 | 0.048778469 |
| 0.969460814 | 0.332315323 | 0.443473691 |
| 1.117515158 | 0.263774122 | 0.369295574 |
| NA | NA | NA |
| 0.052952561 | 0.957769706 | 0.970919132 |
| -1.158404303 | 0.246699084 | 0.34973256 |
| 3.203883427 | 0.001355874 | 0.003791478 |
| NA | NA | NA |
| NA | NA | NA |
| 1.094784064 | 0.27361125 | 0.380239275 |
| -5.879474434 | 4.12E-09 | 2.62E-08 |
| -6.529763484 | 6.59E-11 | 5.11E-10 |

| | | |
|---|---|---|
| NA | NA | NA |
| -0.203817875 | 0.838495824 | 0.886984776 |
| NA | NA | NA |
| 5.119402303 | 3.07E-07 | 1.55E-06 |
| -0.897213212 | 0.369605156 | 0.482236086 |
| -0.225391447 | 0.821674764 | NA |
| -1.261555356 | 0.207108829 | 0.303989562 |
| 7.712174575 | 1.24E-14 | 1.39E-13 |
| -0.065388772 | 0.947864464 | NA |
| 0.364296074 | 0.715636932 | 0.79202301 |
| 0.969075552 | 0.332507495 | NA |
| NA | NA | NA |
| -2.513692828 | 0.011947444 | 0.026663717 |
| -1.803117284 | 0.071369797 | 0.125307662 |
| 4.303481226 | 1.68E-05 | 6.61E-05 |
| -1.164580349 | 0.244188902 | 0.34705621 |
| 0.462670194 | 0.643600786 | NA |
| 0.463725222 | 0.642844624 | 0.732787126 |
| 0.111320837 | 0.911361933 | NA |
| 2.063938812 | 0.039023518 | 0.074550348 |
| NA | NA | NA |
| NA | NA | NA |
| 4.191939359 | 2.77E-05 | 0.000105258 |
| 4.248771053 | 2.15E-05 | 8.31E-05 |
| NA | NA | NA |
| NA | NA | NA |
| NA | NA | NA |
| NA | NA | NA |
| 0.06371127 | 0.949200131 | NA |
| -0.037735682 | 0.969898426 | 0.979090849 |
| 0.192810118 | 0.847107679 | NA |
| -1.095628541 | 0.273241366 | 0.379834705 |
| NA | NA | NA |
| NA | NA | NA |
| NA | NA | NA |
| NA | NA | NA |
| 0.063708285 | 0.949202508 | NA |
| NA | NA | NA |
| NA | NA | NA |
| NA | NA | NA |
| NA | NA | NA |
| NA | NA | NA |
| 0.255447283 | 0.798377656 | NA |
| NA | NA | NA |
| -7.731940279 | 1.06E-14 | 1.20E-13 |
| NA | NA | NA |
| NA | NA | NA |

| | | |
|---|---|---|
| -2.131831782 | 0.033020678 | 0.064606235 |
| 0.44212154 | 0.658401258 | 0.745869748 |
| 9.247301913 | 2.30E-20 | 4.18E-19 |
| 2.325305463 | 0.020055632 | 0.041881568 |
| NA | NA | NA |
| -0.681542516 | 0.495528274 | 0.602617467 |
| -2.0048172 | 0.044982593 | 0.084502725 |
| -1.116005338 | 0.26441985 | 0.369992667 |
| -5.399036218 | 6.70E-08 | 3.69E-07 |
| NA | NA | NA |
| -2.297801741 | 0.021573076 | 0.04460384 |
| -3.80201183 | 0.000143526 | 0.000482532 |
| NA | NA | NA |
| -0.730276057 | 0.46522146 | 0.574634417 |
| -1.049823494 | 0.293799271 | 0.401912613 |
| -0.248195385 | 0.803983237 | 0.861128008 |
| NA | NA | NA |
| -6.463945885 | 1.02E-10 | 7.75E-10 |
| 0.254229472 | 0.799318283 | 0.857378899 |
| 3.821676605 | 0.000132547 | 0.000448393 |
| -7.28104959 | 3.31E-13 | 3.23E-12 |
| -2.853320237 | 0.0043265 | 0.010788234 |
| -0.090182268 | 0.928142375 | NA |
| -0.303543754 | 0.761475508 | 0.827532649 |
| 3.319533441 | 0.00090168 | 0.002619131 |
| -1.742515262 | 0.081418323 | 0.140196796 |
| -0.842834639 | 0.399320932 | 0.511192785 |
| 0.350112445 | 0.726254312 | 0.800013835 |
| -1.184924697 | 0.236047222 | 0.337808002 |
| -3.886874464 | 0.000101543 | 0.000350569 |
| 1.862668635 | 0.062508901 | 0.111884854 |
| 0.21456982 | 0.830102726 | 0.881148059 |
| 0.373260976 | 0.708954218 | NA |
| 3.219435051 | 0.001284435 | 0.003609453 |
| 2.135522134 | 0.032718384 | 0.064074889 |
| 2.035168114 | 0.04183396 | 0.079290022 |
| -0.065388772 | 0.947864464 | NA |
| -0.41712997 | 0.676583345 | 0.759718322 |
| -3.965707974 | 7.32E-05 | 0.000259368 |
| 6.659245497 | 2.75E-11 | 2.21E-10 |
| 4.615180542 | 3.93E-06 | 1.70E-05 |
| -0.926632374 | 0.35411744 | 0.466037096 |
| NA | NA | NA |
| -0.637828911 | 0.523585057 | 0.628794453 |
| 4.033652483 | 5.49E-05 | 0.000198923 |
| -1.33959614 | 0.18037668 | 0.271633974 |
| 4.143166247 | 3.43E-05 | 0.000128274 |

| | | |
|---|---|---|
| 0.46559822 | 0.641503117 | 0.731914293 |
| 1.464061333 | 0.143177191 | 0.224699047 |
| 0.073138669 | 0.94169577 | 0.96038158 |
| 2.212644426 | 0.02692217 | 0.054034312 |
| -0.434205826 | 0.664138983 | 0.750424284 |
| -0.492479733 | 0.622380246 | 0.716559287 |
| 2.638607192 | 0.008324738 | 0.019311231 |
| 7.060263148 | 1.66E-12 | 1.50E-11 |
| -3.614535211 | 0.000300887 | 0.000956221 |
| -4.870085167 | 1.12E-06 | 5.19E-06 |
| 9.241687716 | 2.43E-20 | 4.39E-19 |
| 0.850532676 | 0.395029002 | 0.50701547 |
| -0.340731665 | 0.733305599 | 0.80538127 |
| -1.999765406 | 0.045525602 | 0.085375426 |
| -2.56817427 | 0.010223574 | 0.02325271 |
| -0.599597479 | 0.548774528 | 0.652243248 |
| 2.740416607 | 0.006136135 | 0.014752422 |
| 6.898937193 | 5.24E-12 | 4.52E-11 |
| -1.734735605 | 0.082787647 | 0.142202498 |
| -0.261867683 | 0.793423456 | NA |
| -2.182829763 | 0.029048349 | 0.057716232 |
| 3.911192827 | 9.18E-05 | 0.000320032 |
| 4.183686121 | 2.87E-05 | 0.000108781 |
| -0.065388772 | 0.947864464 | NA |
| -0.415400109 | 0.677849027 | NA |
| NA | NA | NA |
| 8.312323403 | 9.38E-17 | 1.28E-15 |
| 0.806990137 | 0.419672166 | 0.531404438 |
| 0.128059334 | 0.898102018 | 0.929028695 |
| NA | NA | NA |
| 3.954496134 | 7.67E-05 | 0.000271068 |
| 3.968123855 | 7.24E-05 | 0.000256899 |
| -1.313867902 | 0.188890662 | 0.282096257 |
| -0.25085512 | 0.801926124 | 0.859513162 |
| 0.462030739 | 0.644059277 | NA |
| -0.194487022 | 0.845794556 | NA |
| -2.949705899 | 0.003180766 | 0.008168109 |
| -1.277923519 | 0.201276395 | 0.297346878 |
| 0.682143544 | 0.495148189 | 0.602272289 |
| 0.965577944 | 0.334255416 | 0.44545556 |
| -1.723032034 | 0.084882732 | 0.145163623 |
| -2.365514539 | 0.018005036 | 0.038198513 |
| 0.125055207 | 0.900479845 | 0.930594984 |
| -0.72074428 | 0.471066863 | 0.579942137 |
| 0.061089587 | 0.951287862 | NA |
| 0.822208651 | 0.410958156 | 0.522756652 |
| 0.063708882 | 0.949202033 | NA |

| | | |
|---|---|---|
| 2.450236428 | 0.014276244 | 0.031172851 |
| -0.36489862 | 0.715187084 | 0.791705404 |
| 12.89579983 | 4.75E-38 | 2.40E-36 |
| 4.685308924 | 2.80E-06 | 1.23E-05 |
| NA | NA | NA |
| 4.807764683 | 1.53E-06 | 6.98E-06 |
| -2.313553456 | 0.020692229 | 0.043030846 |
| -7.202950766 | 5.89E-13 | 5.61E-12 |
| -0.390458298 | 0.696197687 | 0.77630989 |
| NA | NA | NA |
| 0.609502528 | 0.542191398 | 0.646240509 |
| 0.519102107 | 0.603689539 | NA |
| 1.396557233 | 0.162546758 | 0.249191239 |
| 1.659991676 | 0.096916127 | 0.162305009 |
| NA | NA | NA |
| 0.317642674 | 0.750756 | NA |
| NA | NA | NA |
| 3.168345239 | 0.001533093 | 0.004240147 |
| NA | NA | NA |
| -0.36618381 | 0.714227922 | 0.790994625 |
| 4.436723397 | 9.13E-06 | 3.75E-05 |
| -0.208161064 | 0.835103209 | 0.884468046 |
| 0.13677888 | 0.891205578 | 0.923833166 |
| -2.504552343 | 0.012260646 | 0.027283426 |
| -4.623655194 | 3.77E-06 | 1.63E-05 |
| 0.613093409 | 0.539814578 | 0.644113023 |
| 0.36877229 | 0.712297461 | 0.789304296 |
| 8.862883211 | 7.80E-19 | 1.27E-17 |
| 0.452385852 | 0.650991038 | 0.739675231 |
| -0.427288152 | 0.669169462 | 0.754046555 |
| 0.063708285 | 0.949202508 | NA |
| -4.99346962 | 5.93E-07 | 2.87E-06 |
| 2.073802101 | 0.038097694 | 0.073038088 |
| 4.164422508 | 3.12E-05 | 0.000117717 |
| -0.949808896 | 0.342209365 | NA |
| -7.320248146 | 2.48E-13 | 2.44E-12 |
| -1.024845546 | 0.305436072 | 0.415064058 |
| -0.046272667 | 0.963092925 | 0.974736804 |
| NA | NA | NA |
| NA | NA | NA |
| NA | NA | NA |
| NA | NA | NA |
| NA | NA | NA |
| 0.088043758 | 0.929841897 | NA |
| -2.669036243 | 0.007606924 | 0.0178245 |
| -0.116407041 | 0.907329955 | NA |
| 0.548715979 | 0.583200379 | NA |

| | | |
|---|---|---|
| 0.5366015 | 0.591542908 | NA |
| -0.008246469 | 0.993420344 | 0.995368929 |
| 1.857585194 | 0.063227943 | 0.112967289 |
| 3.130939892 | 0.001742478 | 0.004760859 |
| NA | NA | NA |
| NA | NA | NA |
| -4.09788403 | 4.17E-05 | 0.000153698 |
| NA | NA | NA |
| NA | NA | NA |
| -0.88324617 | 0.377103281 | 0.489824184 |
| 0.066394788 | 0.94706352 | NA |
| NA | NA | NA |
| -1.272902059 | 0.203052799 | 0.299370416 |
| NA | NA | NA |
| -0.065389369 | 0.947863988 | NA |
| -0.065389369 | 0.947863988 | NA |
| NA | NA | NA |
| -0.065389966 | 0.947863513 | NA |
| -0.97645637 | 0.328838359 | 0.439727755 |
| 5.019200029 | 5.19E-07 | 2.53E-06 |
| -2.317006777 | 0.020503363 | 0.042703155 |
| 1.073347357 | 0.283115295 | 0.390047106 |
| -0.098181347 | 0.921788294 | 0.946286646 |
| 3.175334851 | 0.001496637 | 0.004146028 |
| 0.063708882 | 0.949202033 | NA |
| NA | NA | NA |
| NA | NA | NA |
| NA | NA | NA |
| 1.226170778 | 0.220134415 | NA |
| 0.903568263 | 0.366224376 | 0.478748427 |
| NA | NA | NA |
| NA | NA | NA |
| NA | NA | NA |
| 0.063708285 | 0.949202508 | NA |
| -0.065388772 | 0.947864464 | NA |
| NA | NA | NA |
| NA | NA | NA |
| NA | NA | NA |
| NA | NA | NA |
| NA | NA | NA |
| NA | NA | NA |
| NA | NA | NA |
| NA | NA | NA |
| NA | NA | NA |
| NA | NA | NA |
| NA | NA | NA |
| NA | NA | NA |

| | | |
|---|---|---|
| NA | NA | NA |
| NA | NA | NA |
| NA | NA | NA |
| NA | NA | NA |
| NA | NA | NA |
| NA | NA | NA |
| NA | NA | NA |
| NA | NA | NA |
| NA | NA | NA |
| NA | NA | NA |
| NA | NA | NA |
| NA | NA | NA |
| NA | NA | NA |
| NA | NA | NA |
| NA | NA | NA |
| NA | NA | NA |
| NA | NA | NA |
| NA | NA | NA |
| 0.063708882 | 0.949202033 | NA |
| NA | NA | NA |
| NA | NA | NA |
| NA | NA | NA |
| NA | NA | NA |
| NA | NA | NA |
| NA | NA | NA |
| NA | NA | NA |
| NA | NA | NA |
| NA | NA | NA |
| NA | NA | NA |
| NA | NA | NA |
| NA | NA | NA |
| NA | NA | NA |
| NA | NA | NA |
| NA | NA | NA |
| -0.065388772 | 0.947864464 | NA |
| NA | NA | NA |
| NA | NA | NA |
| NA | NA | NA |
| -0.065388772 | 0.947864464 | NA |
| NA | NA | NA |
| NA | NA | NA |
| NA | NA | NA |
| NA | NA | NA |
| NA | NA | NA |
| NA | NA | NA |
| NA | NA | NA |
| NA | NA | NA |

| | | | |
|---|---|---|---|
| NA | NA | NA | |
| NA | NA | NA | |
| NA | NA | NA | |
| NA | NA | NA | |
| NA | NA | NA | |
| NA | NA | NA | |
| NA | NA | NA | |
| NA | NA | NA | |
| NA | NA | NA | |
| | 0.06353712 | 0.949338802 | NA |
| | -0.959674506 | 0.337219058 | 0.44847996 |
| | 2.114596118 | 0.034464385 | 0.067022263 |
| NA | NA | NA | |
| | -0.052297248 | 0.958291846 | NA |
| | -1.46256059 | 0.143587654 | 0.225199147 |
| NA | NA | NA | |
| NA | NA | NA | |
| | 0.552486533 | 0.580615059 | NA |
| NA | NA | NA | |
| NA | NA | NA | |
| | 2.244429075 | 0.024804808 | 0.050374388 |
| NA | NA | NA | |
| NA | NA | NA | |
| | 0.192636473 | 0.847243678 | NA |
| | 0.626854101 | 0.530754876 | NA |
| NA | NA | NA | |
| | 0.192809521 | 0.847108147 | NA |
| NA | NA | NA | |
| NA | NA | NA | |
| NA | NA | NA | |
| NA | NA | NA | |
| | 0.912556357 | 0.361475916 | NA |
| NA | NA | NA | |
| NA | NA | NA | |
| NA | NA | NA | |
| | 0.324874675 | 0.745275925 | NA |
| | 2.453283879 | 0.014155857 | 0.030953275 |
| NA | NA | NA | |
| | 1.479537238 | 0.138996787 | NA |
| NA | NA | NA | |
| NA | NA | NA | |
| | 0.06371127 | 0.949200131 | NA |
| NA | NA | NA | |
| NA | NA | NA | |
| NA | NA | NA | |
| | 0.063708882 | 0.949202033 | NA |
| NA | NA | NA | |

| | | | |
|---|---|---|---|
| NA | NA | NA | |
| | -0.29373198 | 0.768962715 | 0.833643674 |
| NA | NA | NA | |
| NA | NA | NA | |
| NA | NA | NA | |
| NA | NA | NA | |
| NA | NA | NA | |
| NA | NA | NA | |
| NA | NA | NA | |
| NA | NA | NA | |
| | 0.192809521 | 0.847108147 | NA |
| NA | NA | NA | |
| | 2.528192243 | 0.011465155 | 0.025751332 |
| NA | NA | NA | |
| NA | NA | NA | |
| | 1.442012911 | 0.149298731 | 0.232541832 |
| | -0.065388772 | 0.947864464 | NA |
| | 0.063708882 | 0.949202033 | NA |
| NA | NA | NA | |
| | 0.192809521 | 0.847108147 | NA |
| NA | NA | NA | |
| NA | NA | NA | |
| NA | NA | NA | |
| | 0.063708882 | 0.949202033 | NA |
| NA | NA | NA | |
| | 0.912764571 | 0.361366375 | NA |
| | 0.063707019 | 0.949203516 | NA |
| NA | NA | NA | |
| NA | NA | NA | |
| | 0.358414178 | 0.720033383 | NA |
| | -0.065389369 | 0.947863988 | NA |
| | -1.212260002 | 0.22541287 | 0.325473558 |
| NA | NA | NA | |
| NA | NA | NA | |
| NA | NA | NA | |
| NA | NA | NA | |
| NA | NA | NA | |
| | 0.063708285 | 0.949202508 | NA |
| NA | NA | NA | |
| NA | NA | NA | |
| | 0.623802519 | 0.532757285 | 0.637150119 |
| | 0.566294251 | 0.571193768 | 0.672540753 |
| NA | NA | NA | |
| | 1.132747984 | 0.257320101 | 0.361960331 |
| | 2.815178426 | 0.004875016 | 0.012037254 |
| | 6.280605257 | 3.37E-10 | 2.41E-09 |
| | -4.016748884 | 5.90E-05 | 0.000212234 |

| | | |
|---|---|---|
| NA | NA | NA |
| -7.025488735 | 2.13E-12 | 1.90E-11 |
| 0.461704574 | 0.644293191 | 0.733763698 |
| 6.419375923 | 1.37E-10 | 1.02E-09 |
| -2.221623009 | 0.026308794 | 0.052981664 |
| 3.492596701 | 0.000478348 | 0.00146352 |
| NA | NA | NA |
| 2.60368685 | 0.009222698 | 0.02118808 |
| -1.271347702 | 0.203604979 | 0.29995922 |
| 2.250196811 | 0.024436455 | 0.049711026 |
| -3.361018973 | 0.000776555 | 0.002284362 |
| NA | NA | NA |
| 3.206664231 | 0.001342836 | 0.003759641 |
| -0.065390563 | 0.947863038 | NA |
| 1.717121256 | 0.085957019 | 0.146740158 |
| 0.063707194 | 0.949203376 | NA |
| -0.050643 | 0.959609998 | 0.97223389 |
| NA | NA | NA |
| 0.157748723 | 0.874654805 | NA |
| NA | NA | NA |
| 4.91446731 | 8.90E-07 | 4.21E-06 |
| 0.063708882 | 0.949202033 | NA |
| -0.934505576 | 0.350043161 | 0.461792037 |
| -0.426092492 | 0.670040451 | NA |
| 0.063708882 | 0.949202033 | NA |
| 0.192809521 | 0.847108147 | NA |
| 0.192809521 | 0.847108147 | NA |
| 3.382858613 | 0.000717356 | 0.002125152 |
| 1.901591337 | 0.057224601 | 0.103905098 |
| 0.98344021 | 0.325390828 | 0.436423343 |
| -0.695222948 | 0.486915593 | 0.595063586 |
| 4.801970136 | 1.57E-06 | 7.16E-06 |
| -3.018105708 | 0.002543602 | 0.006679222 |
| NA | NA | NA |
| -0.345909826 | 0.729410484 | 0.802511842 |
| 0.063707135 | 0.949203424 | NA |
| 11.719452 | 1.01E-31 | 3.84E-30 |
| 2.065831775 | 0.038844367 | 0.074259108 |
| -0.627674152 | 0.530217421 | 0.635113996 |
| 4.090587022 | 4.30E-05 | 0.000158149 |
| 9.115954009 | 7.80E-20 | 1.38E-18 |
| 15.92062249 | 4.56E-57 | 4.18E-55 |
| 6.277407997 | 3.44E-10 | 2.45E-09 |
| -3.111146393 | 0.001863625 | 0.005050513 |
| 1.317202436 | 0.187770778 | 0.280725276 |
| 4.032630206 | 5.52E-05 | 0.00019968 |
| -1.303937748 | 0.192254809 | 0.286232326 |

| | | |
|---|---|---|
| 3.964635983 | 7.35E-05 | 0.000260499 |
| -3.243258721 | 0.001181709 | 0.003352773 |
| -2.317922979 | 0.020453507 | 0.042612892 |
| NA | NA | NA |
| 2.309991966 | 0.020888599 | 0.043382147 |
| -2.056616722 | 0.039723108 | 0.075753859 |
| NA | NA | NA |
| 0.981528665 | 0.326332106 | 0.437403794 |
| -1.506230943 | 0.132007899 | 0.209894404 |
| NA | NA | NA |
| NA | NA | NA |
| -0.164440503 | 0.869384381 | 0.9090818 |
| 2.210196134 | 0.027091553 | 0.054322004 |
| -1.086416495 | 0.277294767 | 0.3841851 |
| 1.281483023 | 0.200024059 | 0.295758637 |
| NA | NA | NA |
| -5.603940575 | 2.10E-08 | 1.23E-07 |
| -3.946743938 | 7.92E-05 | 0.000279243 |
| NA | NA | NA |
| -7.133899185 | 9.76E-13 | 9.07E-12 |
| -1.867772316 | 0.061793814 | 0.11074747 |
| NA | NA | NA |
| NA | NA | NA |
| NA | NA | NA |
| NA | NA | NA |
| NA | NA | NA |
| -0.483619032 | 0.628656265 | NA |
| NA | NA | NA |
| NA | NA | NA |
| -0.065388772 | 0.947864464 | NA |
| NA | NA | NA |
| NA | NA | NA |
| 0.192809521 | 0.847108147 | NA |
| NA | NA | NA |
| -0.19448762 | 0.845794089 | NA |
| 0.063976859 | 0.948988653 | NA |
| 0.192809521 | 0.847108147 | NA |
| 0.688994108 | 0.490826975 | 0.598295471 |
| NA | NA | NA |
| NA | NA | NA |
| NA | NA | NA |
| NA | NA | NA |
| -0.065388772 | 0.947864464 | NA |
| NA | NA | NA |
| NA | NA | NA |
| NA | NA | NA |
| NA | NA | NA |

| | | |
|---|---|---|
| NA | NA | NA |
| NA | NA | NA |
| NA | NA | NA |
| NA | NA | NA |
| 0.434937002 | 0.663608156 | NA |
| 0.083058248 | 0.933805225 | 0.954999687 |
| 1.246954404 | 0.212414217 | 0.310320892 |
| -1.447629785 | 0.147720616 | 0.230454085 |
| 1.688645308 | 0.091287422 | 0.154426804 |
| 1.646016967 | 0.099760266 | 0.166200841 |
| 13.54810349 | 8.13E-42 | 4.71E-40 |
| -2.184620555 | 0.02891668 | 0.05750711 |
| 0.692122749 | 0.488860246 | 0.596740891 |
| -0.0719378 | 0.942651408 | 0.96104292 |
| -3.833302042 | 0.000126435 | 0.000429455 |
| NA | NA | NA |
| 3.060320132 | 0.002211005 | 0.005895543 |
| 1.950940868 | 0.05106408 | 0.094264616 |
| 9.789345003 | 1.25E-22 | 2.70E-21 |
| 0.192810118 | 0.847107679 | NA |
| 4.109305383 | 3.97E-05 | 0.000146919 |
| 0.794231076 | 0.427060921 | 0.53839704 |
| 0.192809521 | 0.847108147 | NA |
| 0.192810118 | 0.847107679 | NA |
| 0.063708882 | 0.949202033 | NA |
| -0.065389966 | 0.947863513 | NA |
| 2.01533951 | 0.043869088 | 0.082665307 |
| 3.085575207 | 0.002031588 | 0.005467177 |
| NA | NA | NA |
| 8.403620087 | 4.33E-17 | 6.08E-16 |
| -0.750563311 | 0.452915508 | 0.562799741 |
| -0.065389966 | 0.947863513 | NA |
| 0.192809521 | 0.847108147 | NA |
| 0.192809521 | 0.847108147 | NA |
| -0.076259932 | 0.939212302 | NA |
| NA | NA | NA |
| NA | NA | NA |
| NA | NA | NA |
| 0.192810118 | 0.847107679 | NA |
| NA | NA | NA |
| NA | NA | NA |
| NA | NA | NA |
| NA | NA | NA |
| NA | NA | NA |
| 0.063708285 | 0.949202508 | NA |
| NA | NA | NA |
| NA | NA | NA |

| | | | |
|---|---|---|---|
| NA | NA | NA | |
| NA | NA | NA | |
| NA | NA | NA | |
| | -0.326463148 | 0.744073967 | NA |
| NA | NA | NA | |
| | 0.192808924 | 0.847108615 | NA |
| NA | NA | NA | |
| NA | NA | NA | |
| | 0.606574427 | 0.544133376 | NA |
| NA | NA | NA | |
| NA | NA | NA | |
| NA | NA | NA | |
| | -2.765084611 | 0.005690803 | 0.013803741 |
| | -1.189378754 | 0.234290657 | 0.335601188 |
| | -0.448340614 | 0.653907394 | 0.741982954 |
| | -0.270050578 | 0.787121343 | NA |
| | -0.086961293 | 0.930702279 | NA |
| | 6.208082651 | 5.36E-10 | 3.75E-09 |
| | -0.829782662 | 0.406661675 | 0.518503476 |
| | 1.335220914 | 0.181804054 | 0.273418324 |
| | 1.029847703 | 0.303081504 | 0.412424162 |
| | 0.192810118 | 0.847107679 | NA |
| | -0.065390563 | 0.947863038 | NA |
| | 4.586021715 | 4.52E-06 | 1.94E-05 |
| | 0.221683242 | 0.824560476 | 0.877050345 |
| | 3.667557649 | 0.000244878 | 0.00079077 |
| | 1.978651922 | 0.047855209 | 0.089129838 |
| | -1.120047001 | 0.262693734 | 0.368057674 |
| | -0.166825102 | 0.867507667 | 0.907733498 |
| | -1.881895887 | 0.059850153 | 0.10784316 |
| | -0.981622826 | 0.326285698 | 0.43736577 |
| NA | NA | NA | |
| | 9.413741748 | 4.79E-21 | 9.20E-20 |
| | 0.246242162 | 0.80549478 | 0.862313679 |
| | -3.71612599 | 0.000202301 | 0.0006646 |
| | -0.254192019 | 0.799347216 | NA |
| NA | NA | NA | |
| | 3.848608687 | 0.000118791 | 0.000405472 |
| NA | NA | NA | |
| NA | NA | NA | |
| NA | NA | NA | |
| | 0.063708882 | 0.949202033 | NA |
| | 0.192810118 | 0.847107679 | NA |
| | -0.119314322 | 0.905026337 | NA |
| | -1.208323043 | 0.226923028 | NA |
| | -0.266679173 | 0.789716191 | NA |
| NA | NA | NA | |

| | | | |
|---|---|---|---|
| | -0.345413057 | 0.729783863 | 0.802781493 |
| | 0.063708285 | 0.949202508 | NA |
| | 0.434937002 | 0.663608156 | NA |
| | 2.327459796 | 0.019940806 | 0.041680032 |
| | 0.691294139 | 0.489380714 | NA |
| NA | NA | NA | |
| | 1.066254814 | 0.28630847 | NA |
| | 1.95925292 | 0.050083174 | 0.092713694 |
| NA | NA | NA | |
| | 0.560367539 | 0.575228768 | NA |
| | 1.614624622 | 0.10639201 | NA |
| NA | NA | NA | |
| NA | NA | NA | |
| NA | NA | NA | |
| | 0.520971041 | 0.602386945 | NA |
| | 1.065686463 | 0.286565399 | NA |
| | 0.192810118 | 0.847107679 | NA |
| NA | NA | NA | |
| | 1.792635918 | 0.073031159 | NA |
| | 0.063708285 | 0.949202508 | NA |
| | 0.063708882 | 0.949202033 | NA |
| | 0.192636473 | 0.847243678 | NA |
| | 0.192810118 | 0.847107679 | NA |
| NA | NA | NA | |
| NA | NA | NA | |
| NA | NA | NA | |
| | 0.192810118 | 0.847107679 | NA |
| NA | NA | NA | |
| | 0.41775227 | 0.676128251 | NA |
| NA | NA | NA | |
| NA | NA | NA | |
| NA | NA | NA | |
| NA | NA | NA | |
| | 1.260000278 | 0.207669262 | 0.304704943 |
| NA | NA | NA | |
| | 0.063708285 | 0.949202508 | NA |
| | 0.651644573 | 0.514630486 | NA |
| NA | NA | NA | |
| NA | NA | NA | |
| NA | NA | NA | |
| NA | NA | NA | |
| NA | NA | NA | |
| NA | NA | NA | |
| NA | NA | NA | |
| | 0.330493631 | 0.741027004 | NA |
| NA | NA | NA | |
| NA | NA | NA | |

| | | |
|---|---|---|
| NA | NA | NA |
| 0.192809521 | 0.847108147 | NA |
| NA | NA | NA |
| -0.155707049 | 0.87626394 | NA |
| NA | NA | NA |
| 1.310599453 | 0.189993124 | NA |
| 0.879276873 | 0.379251173 | 0.491847505 |
| 0.192809521 | 0.847108147 | NA |
| 1.442593354 | 0.149135057 | NA |
| NA | NA | NA |
| NA | NA | NA |
| 0.201524559 | 0.840288428 | NA |
| NA | NA | NA |
| NA | NA | NA |
| 0.909127212 | 0.36328298 | NA |
| NA | NA | NA |
| NA | NA | NA |
| NA | NA | NA |
| 0.192809521 | 0.847108147 | NA |
| NA | NA | NA |
| 0.696238979 | 0.48627918 | 0.59445997 |
| NA | NA | NA |
| NA | NA | NA |
| NA | NA | NA |
| NA | NA | NA |
| NA | NA | NA |
| NA | NA | NA |
| NA | NA | NA |
| NA | NA | NA |
| NA | NA | NA |
| NA | NA | NA |
| NA | NA | NA |
| NA | NA | NA |
| NA | NA | NA |
| NA | NA | NA |
| NA | NA | NA |
| 0.192636473 | 0.847243678 | NA |
| 0.063707688 | 0.949202983 | NA |
| NA | NA | NA |
| NA | NA | NA |
| NA | NA | NA |
| NA | NA | NA |
| NA | NA | NA |
| 0.063708882 | 0.949202033 | NA |
| 0.492221807 | 0.62256255 | NA |
| 0.192809521 | 0.847108147 | NA |
| 0.362026528 | 0.717332207 | NA |

| | | | |
|---|---|---|---|
| NA | NA | NA | |
| NA | NA | NA | |
| 0.063708882 | 0.949202033 | NA | |
| 0.210360696 | 0.833386166 | NA | |
| 1.179382161 | 0.238246035 | NA | |
| 0.430491827 | 0.66683791 | NA | |
| 0.495101805 | 0.620528259 | NA | |
| -0.065389369 | 0.947863988 | NA | |
| NA | NA | NA | |
| 0.574475814 | 0.565645863 | NA | |
| 0.193472986 | 0.84658856 | NA | |
| 0.063708285 | 0.949202508 | NA | |
| 0.412143228 | 0.680234446 | NA | |
| NA | NA | NA | |
| NA | NA | NA | |
| 0.192810118 | 0.847107679 | NA | |
| 0.886937245 | 0.375112688 | NA | |
| NA | NA | NA | |
| 0.324874675 | 0.745275925 | NA | |
| NA | NA | NA | |
| 0.882822453 | 0.377332207 | NA | |
| NA | NA | NA | |
| NA | NA | NA | |
| NA | NA | NA | |
| 0.493471196 | 0.621679687 | NA | |
| 0.063708217 | 0.949202562 | NA | |
| NA | NA | NA | |
| NA | NA | NA | |
| NA | NA | NA | |
| 0.557632165 | 0.577095588 | | 0.67802772 |
| 0.446789653 | 0.655026942 | NA | |
| 0.188826074 | 0.850229135 | NA | |
| NA | NA | NA | |
| NA | NA | NA | |
| 0.063708381 | 0.949202431 | NA | |
| NA | NA | NA | |
| NA | NA | NA | |
| NA | NA | NA | |
| NA | NA | NA | |
| 0.346721347 | 0.728800672 | NA | |
| NA | NA | NA | |
| NA | NA | NA | |
| -0.354077849 | 0.723280542 | NA | |
| 0.063708882 | 0.949202033 | NA | |
| 0.330493631 | 0.741027004 | NA | |
| 1.516985559 | 0.12927033 | NA | |
| NA | NA | NA | |

| | | | |
|---|---|---|---|
| NA | NA | NA | |
| NA | NA | NA | |
| 0.192810118 | 0.847107679 | NA | |
| 0.60616447 | 0.544405544 | NA | |
| 0.085681794 | 0.931719375 | NA | |
| NA | NA | NA | |
| NA | NA | NA | |
| 0.192809521 | 0.847108147 | NA | |
| -0.194488217 | 0.845793621 | NA | |
| NA | NA | NA | |
| -0.153482596 | 0.878017714 | NA | |
| 1.655709582 | 0.097780646 | | 0.163516302 |
| NA | NA | NA | |
| NA | NA | NA | |
| 0.29141163 | 0.770736521 | NA | |
| 0.198201446 | 0.842887456 | NA | |
| 0.673351994 | 0.500723382 | | 0.607813106 |
| 1.667101366 | 0.095494251 | NA | |
| 6.067633903 | 1.30E-09 | | 8.72E-09 |
| -0.347044203 | 0.728558111 | NA | |
| -0.153608817 | 0.877918185 | NA | |
| NA | NA | NA | |
| 0.127184131 | 0.898794665 | NA | |
| 0.063708285 | 0.949202508 | NA | |
| 0.192809521 | 0.847108147 | NA | |
| 0.192809521 | 0.847108147 | NA | |
| 0.192810118 | 0.847107679 | NA | |
| NA | NA | NA | |
| 0.575305927 | 0.565084414 | NA | |
| 0.192809521 | 0.847108147 | NA | |
| -0.068677197 | 0.945246569 | NA | |
| 0.192809521 | 0.847108147 | NA | |
| 0.192808327 | 0.847109082 | NA | |
| -0.116388807 | 0.907344405 | NA | |
| NA | NA | NA | |
| 0.458102228 | 0.646878998 | NA | |
| 0.063707194 | 0.949203376 | NA | |
| 1.08113363 | 0.279637676 | NA | |
| 1.549713667 | 0.121210256 | NA | |
| 0.665173001 | 0.505939847 | NA | |
| 0.063708882 | 0.949202033 | NA | |
| 1.156789732 | 0.247358278 | NA | |
| 1.920562998 | 0.054786824 | NA | |
| 1.267939257 | 0.204819643 | NA | |
| 1.504693055 | 0.132403009 | NA | |
| 2.261089744 | 0.0237537 | | 0.048523229 |
| 0.330493631 | 0.741027004 | NA | |

| | | | |
|---|---|---|---|
| | 0.263490142 | 0.792172821 | NA |
| | 0.699535399 | 0.484217499 | NA |
| | 0.63826869 | 0.52329879 | NA |
| | 0.835997648 | 0.403156226 | 0.514820219 |
| | 1.450200557 | 0.1470026 | NA |
| | 2.47937289 | 0.013161363 | 0.029059492 |
| | 0.420088277 | 0.674420966 | NA |
| | 1.380809393 | 0.167337575 | 0.255316031 |
| | 0.192809521 | 0.847108147 | NA |
| | 1.889235007 | 0.058860346 | 0.10638897 |
| | 1.137178592 | 0.255463634 | 0.359854509 |
| | 1.750451114 | 0.080040503 | 0.138166468 |
| | 1.51616263 | 0.129478235 | 0.20672579 |
| | 1.163701249 | 0.244545103 | 0.347367569 |
| | 1.952501465 | 0.050878697 | 0.094026417 |
| | -0.112207441 | 0.910658929 | NA |
| | 0.738265222 | 0.460353296 | NA |
| | 1.414888274 | 0.157101257 | NA |
| | 0.192808924 | 0.847108615 | NA |
| | 1.28530734 | 0.198684906 | NA |
| | 0.362030575 | 0.717329183 | NA |
| | 0.692087054 | 0.488882661 | NA |
| | 1.500534097 | 0.133476108 | NA |
| | 2.237600373 | 0.025247127 | 0.051164731 |
| | 0.585582452 | 0.558156144 | NA |
| | 0.481049613 | 0.630481238 | NA |
| | 0.429236314 | 0.667751258 | NA |
| | 1.87130087 | 0.06130339 | NA |
| | 1.860450281 | 0.062821847 | NA |
| | 0.975406288 | 0.329358766 | NA |
| NA | NA | NA | |
| | 1.427099947 | 0.153551088 | NA |
| | 0.500009504 | 0.617068385 | NA |
| | 10.1943494 | 2.10E-24 | 5.15E-23 |
| | 0.982021316 | 0.326089347 | 0.437149395 |
| | 4.727662921 | 2.27E-06 | 1.02E-05 |
| | -0.065389369 | 0.947863988 | NA |
| | 0.394257461 | 0.693390966 | NA |
| NA | NA | NA | |
| NA | NA | NA | |
| | 0.063710673 | 0.949200606 | NA |
| | 4.466886578 | 7.94E-06 | 3.29E-05 |
| | 12.284457 | 1.10E-34 | 4.71E-33 |
| | 6.881994249 | 5.90E-12 | 5.05E-11 |
| | 12.8455278 | 9.11E-38 | 4.54E-36 |
| NA | NA | NA | |
| | 10.79497419 | 3.64E-27 | 1.06E-25 |

| | | | |
|---|---|---|---|
| | 3.292243918 | 0.000993914 | 0.002867411 |
| | 6.522153944 | 6.93E-11 | 5.36E-10 |
| | 2.649615609 | 0.00805834 | 0.018745376 |
| | 7.627628457 | 2.39E-14 | 2.62E-13 |
| NA | NA | NA | |
| | 2.516627585 | 0.011848399 | 0.026486151 |
| | 5.091299635 | 3.56E-07 | 1.78E-06 |
| NA | NA | NA | |
| | -0.993234485 | 0.320595696 | 0.43140209 |
| | 0.630180215 | 0.528576683 | 0.633451803 |
| | -0.065390563 | 0.947863038 | NA |
| | 2.066510597 | 0.038780293 | 0.07416494 |
| | -0.168311079 | 0.866338558 | 0.906654435 |
| | -1.023660783 | 0.305995525 | 0.415621228 |
| | 1.055318391 | 0.291279732 | 0.398969954 |
| | 5.170368098 | 2.34E-07 | 1.20E-06 |
| | 4.003059203 | 6.25E-05 | 0.000223872 |
| | 2.135474381 | 0.032722281 | 0.064075782 |
| NA | NA | NA | |
| | -2.602203724 | 0.009262681 | 0.021266273 |
| | -2.154468397 | 0.031203458 | 0.06143012 |
| | -2.171426727 | 0.02989893 | 0.059183094 |
| | -5.19470484 | 2.05E-07 | 1.06E-06 |
| | 0.063704484 | 0.949205534 | NA |
| | -2.739771618 | 0.006148189 | 0.014778562 |
| | -10.49470926 | 9.14E-26 | 2.46E-24 |
| | 0.469746194 | 0.63853636 | 0.729479893 |
| | 0.982347551 | 0.325928656 | 0.437056483 |
| | -4.155510726 | 3.25E-05 | 0.000121997 |
| | 2.213453761 | 0.026866378 | 0.053935309 |
| NA | NA | NA | |
| | -1.944540079 | 0.051830357 | 0.095510111 |
| | 4.204006008 | 2.62E-05 | 0.000100178 |
| | -0.087659735 | 0.930147124 | NA |
| NA | NA | NA | |
| | 0.192810118 | 0.847107679 | NA |
| | 5.543691034 | 2.96E-08 | 1.71E-07 |
| | -0.076441601 | 0.939067774 | 0.958560678 |
| | -0.065389369 | 0.947863988 | NA |
| | -1.52563608 | 0.127100521 | 0.203722211 |
| | -4.506342772 | 6.60E-06 | 2.76E-05 |
| | -3.355396306 | 0.000792514 | 0.002327695 |
| | -4.453186867 | 8.46E-06 | 3.49E-05 |
| | -4.193828425 | 2.74E-05 | 0.000104448 |
| | 1.592541686 | 0.111263044 | 0.181877874 |
| | -11.09181344 | 1.37E-28 | 4.42E-27 |
| NA | NA | NA | |

| | | |
|---|---|---|
| -1.625576484 | 0.104039781 | 0.172104667 |
| 1.727201236 | 0.084131529 | 0.144154721 |
| 0.986493196 | 0.323891153 | 0.434901322 |
| -7.87359536 | 3.45E-15 | 4.10E-14 |
| -8.584280034 | 9.14E-18 | 1.35E-16 |
| -1.565954679 | 0.117359228 | 0.190265008 |
| 2.033252419 | 0.042027028 | 0.079595679 |
| -0.102006902 | 0.918751197 | 0.944261988 |
| 1.38392567 | 0.166381216 | 0.254058188 |
| -0.065393256 | 0.947860894 | NA |
| 0.505685026 | 0.613077782 | NA |
| 20.49705936 | 2.29E-93 | 4.89E-91 |
| -4.155235697 | 3.25E-05 | 0.000122107 |
| 17.57835089 | 3.61E-69 | 4.49E-67 |
| 13.13461818 | 2.09E-39 | 1.12E-37 |
| 28.24608949 | 1.59E-175 | 8.65E-173 |
| 1.14273908 | 0.253146939 | 0.357203415 |
| 1.310493922 | 0.190028799 | 0.283525324 |
| 5.286592 | 1.25E-07 | 6.63E-07 |
| -1.004815957 | 0.314985479 | 0.425545773 |
| -0.717487202 | 0.47307353 | 0.581927014 |
| 1.385856473 | 0.165790734 | 0.253316234 |
| -0.667558263 | 0.504415607 | 0.610932094 |
| 3.688288996 | 0.000225767 | 0.000734558 |
| 0.063707135 | 0.949203424 | NA |
| 16.62623836 | 4.50E-62 | 4.75E-60 |
| 1.909823855 | 0.056155897 | 0.102261338 |
| 14.4055897 | 4.77E-47 | 3.44E-45 |
| 7.638970571 | 2.19E-14 | 2.41E-13 |
| -4.085984661 | 4.39E-05 | 0.000161081 |
| 2.12427917 | 0.033646806 | 0.065626086 |
| 6.776333452 | 1.23E-11 | 1.03E-10 |
| 7.66767284 | 1.75E-14 | 1.94E-13 |
| 9.886888991 | 4.75E-23 | 1.06E-21 |
| 3.45571496 | 0.000548835 | 0.001661307 |
| -0.872267425 | 0.383062504 | 0.495495422 |
| -1.378219029 | 0.168135674 | 0.256252567 |
| -0.403018635 | 0.686934516 | 0.768926744 |
| 5.151400379 | 2.59E-07 | 1.32E-06 |
| -3.230892658 | 0.001234043 | 0.003481928 |
| 2.410742782 | 0.015920072 | 0.03434279 |
| 0.074774392 | 0.940394217 | NA |
| 1.87796034 | 0.060386591 | 0.108622089 |
| 1.599866876 | 0.109628119 | NA |
| NA | NA | NA |
| 0.874941065 | 0.381605973 | 0.494133343 |
| 0.158269914 | 0.874244114 | NA |

| | | | |
|---|---|---|---|
| NA | NA | NA | |
| NA | NA | NA | |
| 0.591974618 | 0.553867585 | NA | |
| 0.063708882 | 0.949202033 | NA | |
| 0.192809521 | 0.847108147 | NA | |
| NA | NA | NA | |
| NA | NA | NA | |
| 3.769279482 | 0.000163719 | | 0.000545514 |
| 3.635572731 | 0.000277364 | | 0.000887436 |
| NA | NA | NA | |
| NA | NA | NA | |
| 13.16873387 | 1.33E-39 | | 7.16E-38 |
| 0.074438505 | 0.940661472 | | 0.959639561 |
| 0.873799527 | 0.382227438 | | 0.494612229 |
| 0.192809521 | 0.847108147 | NA | |
| 8.970305735 | 2.96E-19 | | 4.99E-18 |
| NA | NA | NA | |
| 15.64240089 | 3.74E-55 | | 3.26E-53 |
| NA | NA | NA | |
| -0.803692903 | 0.421574354 | | 0.533360598 |
| 1.893073673 | 0.058348073 | | 0.105592673 |
| -4.517511316 | 6.26E-06 | | 2.63E-05 |
| -4.102932425 | 4.08E-05 | | 0.000150648 |
| 0.06371127 | 0.949200131 | NA | |
| -1.731205438 | 0.08341513 | | 0.143093736 |
| 1.243225587 | 0.213784716 | | 0.311886195 |
| NA | NA | NA | |
| NA | NA | NA | |
| NA | NA | NA | |
| -2.987504415 | 0.002812652 | | 0.007314357 |
| 0.573477492 | 0.566321434 | NA | |
| 1.086836455 | 0.277109094 | | 0.383991619 |
| 9.40345575 | 5.28E-21 | | 1.01E-19 |
| -3.361505311 | 0.000775189 | | 0.002280824 |
| -4.411345672 | 1.03E-05 | | 4.19E-05 |
| -4.738486765 | 2.15E-06 | | 9.65E-06 |
| 0.038772977 | 0.96907139 | | 0.978683157 |
| -2.173986026 | 0.029706185 | | 0.058885389 |
| -0.065388772 | 0.947864464 | NA | |
| -2.506807809 | 0.012182693 | | 0.027122015 |
| -4.999644326 | 5.74E-07 | | 2.79E-06 |
| -0.811902837 | 0.416847386 | | 0.528528954 |
| -0.649778248 | 0.515835472 | | 0.621156637 |
| 0.730245115 | 0.46524037 | | 0.574634417 |
| 5.509415505 | 3.60E-08 | | 2.06E-07 |
| -0.065390563 | 0.947863038 | NA | |
| -1.130878617 | 0.258106185 | | 0.362877812 |

| | | |
|---|---|---|
| 1.064340765 | 0.287174357 | 0.394510108 |
| NA | NA | NA |
| 2.828069128 | 0.004682969 | 0.01159738 |
| -0.322197598 | 0.747303004 | 0.816277294 |
| -1.380737848 | 0.16735958 | 0.255334041 |
| 0.185139277 | 0.853119796 | 0.897964192 |
| 7.849232574 | 4.19E-15 | 4.92E-14 |
| -1.934300812 | 0.053076153 | 0.097439938 |
| -6.188625463 | 6.07E-10 | 4.21E-09 |
| NA | NA | NA |
| NA | NA | NA |
| 0.261213554 | 0.793927827 | NA |
| 1.55451167 | 0.120062415 | 0.194059222 |
| 0.192810118 | 0.847107679 | NA |
| NA | NA | NA |
| -2.233705829 | 0.025502435 | 0.051581895 |
| 7.8635534 | 3.73E-15 | 4.42E-14 |
| 11.06340986 | 1.89E-28 | 6.02E-27 |
| 16.56305007 | 1.29E-61 | 1.34E-59 |
| 17.7914922 | 8.23E-71 | 1.08E-68 |
| 11.76481431 | 5.93E-32 | 2.27E-30 |
| 10.65179321 | 1.71E-26 | 4.82E-25 |
| 3.271248811 | 0.001070737 | 0.003067125 |
| 0.795693766 | 0.426210051 | 0.537568031 |
| -2.4878741 | 0.012850922 | 0.028447738 |
| 1.634259645 | 0.102204343 | 0.169617737 |
| -5.809165205 | 6.28E-09 | 3.92E-08 |
| -2.509906574 | 0.012076311 | 0.026918698 |
| 0.330370322 | 0.741120163 | NA |
| NA | NA | NA |
| NA | NA | NA |
| 1.223695047 | 0.221067289 | 0.32038181 |
| -0.087448001 | 0.930315416 | NA |
| -1.703679883 | 0.088440908 | 0.150365163 |
| -5.689404697 | 1.27E-08 | 7.69E-08 |
| -2.388862085 | 0.016900645 | 0.036192946 |
| 10.75540909 | 5.59E-27 | 1.62E-25 |
| 1.578134371 | 0.114534744 | 0.186349755 |
| 0.364080175 | 0.715798141 | 0.792031198 |
| -1.244234891 | 0.213413126 | 0.311525663 |
| 6.046011306 | 1.48E-09 | 9.93E-09 |
| -4.457606421 | 8.29E-06 | 3.42E-05 |
| 0.148776779 | 0.881729774 | 0.916972588 |
| -0.087670103 | 0.930138882 | 0.952476219 |
| 12.3880855 | 3.03E-35 | 1.34E-33 |
| -0.491936272 | 0.622764395 | 0.716943833 |
| 0.483144243 | 0.628993322 | 0.721736639 |

| | | |
|---|---|---|
| 2.747303256 | 0.006008755 | 0.014472588 |
| 2.495033234 | 0.012594533 | 0.027944435 |
| 0.239557066 | 0.810673652 | 0.866366146 |
| 0.617178539 | 0.53711697 | 0.641352437 |
| -1.425148628 | 0.154114243 | 0.238749736 |
| -0.93617847 | 0.349181306 | 0.460849359 |
| -0.065389966 | 0.947863513 | NA |
| 0.093460388 | 0.925537818 | NA |
| 0.447456301 | 0.654545633 | NA |
| -0.180662283 | 0.856632665 | 0.900498959 |
| 0.063707688 | 0.949202983 | NA |
| NA | NA | NA |
| 1.261347917 | 0.207183524 | 0.304063536 |
| -0.065388772 | 0.947864464 | NA |
| -2.796294886 | 0.005169221 | 0.012672439 |
| -2.650869091 | 0.008028495 | 0.018688109 |
| 0.656821328 | 0.511295809 | 0.617055646 |
| NA | NA | NA |
| -1.024583179 | 0.305559905 | 0.415187255 |
| 0.431938626 | 0.66578602 | 0.75159202 |
| 3.018240275 | 0.002542472 | 0.006677658 |
| 2.683221508 | 0.007291668 | 0.017169374 |
| NA | NA | NA |
| -2.39889127 | 0.016444797 | 0.035346781 |
| NA | NA | NA |
| -0.110311035 | 0.912162706 | 0.939460677 |
| NA | NA | NA |
| 0.138470404 | 0.889868658 | 0.922946282 |
| -2.805359972 | 0.005026041 | 0.012361385 |
| 0.180310247 | 0.856909012 | 0.900654331 |
| 2.411767918 | 0.015875383 | 0.034267061 |
| NA | NA | NA |
| -0.130249707 | 0.896368869 | 0.927867278 |
| -0.728373825 | 0.466384782 | 0.575681942 |
| NA | NA | NA |
| 0.24880206 | 0.803513897 | NA |
| NA | NA | NA |
| 0.192636473 | 0.847243678 | NA |
| -2.034169673 | 0.041934491 | 0.079456496 |
| NA | NA | NA |
| 12.08393251 | 1.28E-33 | 5.30E-32 |
| 2.650453793 | 0.008038372 | 0.01870762 |
| 0.37985166 | 0.704055532 | 0.782979072 |
| 0.182771792 | 0.854977083 | NA |
| 1.140894897 | 0.253913663 | NA |
| 0.192808327 | 0.847109082 | NA |
| 9.420145726 | 4.50E-21 | 8.69E-20 |

| | | | |
|---|---|---|---|
| | -0.591828058 | 0.553965733 | NA |
| | 0.12494259 | 0.900569 | NA |
| | 0.88073983 | 0.378458654 | 0.491099596 |
| | -1.026808232 | 0.304510772 | 0.41400668 |
| | -4.859918625 | 1.17E-06 | 5.45E-06 |
| | -2.183139036 | 0.029025573 | 0.057687 |
| | -0.467034557 | 0.640475154 | 0.731045333 |
| | 3.438127995 | 0.000585751 | 0.001763664 |
| | 0.203595243 | 0.838669811 | NA |
| | 0.463414682 | 0.643067157 | NA |
| | 2.187069145 | 0.028737477 | 0.057205268 |
| NA | NA | NA | |
| | -1.388688299 | 0.164927554 | 0.252269413 |
| | 0.108289391 | 0.913766138 | NA |
| | -2.166615003 | 0.030264222 | 0.059806844 |
| | -0.084400959 | 0.932737644 | 0.954317497 |
| | -3.226764439 | 0.001251985 | 0.003526579 |
| NA | NA | NA | |
| NA | NA | NA | |
| | -2.234472652 | 0.02545199 | 0.051497127 |
| | -1.211196401 | 0.225820142 | 0.325892421 |
| | -5.215435507 | 1.83E-07 | 9.52E-07 |
| | -1.835085034 | 0.066493091 | 0.1180227 |
| NA | NA | NA | |
| | 5.083357255 | 3.71E-07 | 1.85E-06 |
| | 1.583720027 | 0.113257445 | 0.1846677 |
| | -7.189127614 | 6.52E-13 | 6.18E-12 |
| | 0.226954819 | 0.820458875 | 0.873987558 |
| | -2.758795222 | 0.005801488 | 0.014034139 |
| | -1.390695502 | 0.164317782 | 0.251475028 |
| | 3.879189279 | 0.000104805 | 0.000360935 |
| | 1.874113601 | 0.060914769 | 0.109414903 |
| | 3.889117269 | 0.000100609 | 0.000347585 |
| | 5.926804686 | 3.09E-09 | 2.00E-08 |
| | 3.195850796 | 0.001394192 | 0.003888203 |
| | 3.696598046 | 0.000218508 | 0.000712606 |
| | 7.070282869 | 1.55E-12 | 1.40E-11 |
| | 3.551546964 | 0.000382974 | 0.001193158 |
| | 5.106832446 | 3.28E-07 | 1.65E-06 |
| | -0.654211514 | 0.512975545 | 0.6185758 |
| | 0.863628405 | 0.387792055 | NA |
| | 0.340661817 | 0.733358186 | NA |
| | -5.015219008 | 5.30E-07 | 2.58E-06 |
| | -0.979265815 | 0.327448657 | 0.438501946 |
| | -1.706695659 | 0.087878635 | 0.149632804 |
| | 0.861991686 | 0.388692093 | 0.500899093 |
| | 5.397405097 | 6.76E-08 | 3.72E-07 |

| | | |
|---|---|---|
| -3.586367672 | 0.000335316 | 0.001056782 |
| NA | NA | NA |
| -1.413980973 | 0.157367489 | 0.242812739 |
| 0.266426158 | 0.789911022 | NA |
| -3.299880881 | 0.000967259 | 0.002795668 |
| 0.047595462 | 0.962038648 | 0.973827289 |
| -7.15265809 | 8.51E-13 | 7.96E-12 |
| NA | NA | NA |
| NA | NA | NA |
| -3.408245835 | 0.00065382 | 0.001950631 |
| 5.457189087 | 4.84E-08 | 2.72E-07 |
| NA | NA | NA |
| 0.349291458 | 0.726870511 | NA |
| -0.080149674 | 0.936118216 | 0.956456249 |
| NA | NA | NA |
| -0.69802661 | 0.485160553 | NA |
| NA | NA | NA |
| NA | NA | NA |
| NA | NA | NA |
| NA | NA | NA |
| 1.112149005 | 0.26607409 | 0.371767024 |
| 3.070871518 | 0.00213435 | 0.005714277 |
| -1.195287778 | 0.231974613 | 0.332886577 |
| -1.948070692 | 0.051406505 | 0.094812823 |
| 2.702126664 | 0.006889751 | 0.016353766 |
| -2.235792088 | 0.025365393 | 0.05135626 |
| -0.572475474 | 0.566999897 | 0.668834885 |
| -3.768664328 | 0.000164123 | 0.000546714 |
| NA | NA | NA |
| 0.465664828 | 0.641455432 | NA |
| 0.621700334 | 0.534138932 | 0.638558412 |
| -0.249012234 | 0.803351318 | 0.860635315 |
| 2.192812046 | 0.028320924 | 0.056470383 |
| -0.275633905 | 0.782829265 | 0.845015375 |
| NA | NA | NA |
| NA | NA | NA |
| 0.984659331 | 0.324791436 | 0.43578288 |
| 0.49606901 | 0.619845723 | NA |
| 0.467178938 | 0.640371861 | 0.730960767 |
| NA | NA | NA |
| 0.192810118 | 0.847107679 | NA |
| NA | NA | NA |
| NA | NA | NA |
| NA | NA | NA |
| NA | NA | NA |
| -0.06879015 | 0.945156659 | NA |
| NA | NA | NA |

| | | |
|---|---|---|
| -0.065389369 | 0.947863988 | NA |
| NA | NA | NA |
| NA | NA | NA |
| NA | NA | NA |
| 0.192636473 | 0.847243678 | NA |
| 0.192810118 | 0.847107679 | NA |
| NA | NA | NA |
| NA | NA | NA |
| 1.975681313 | 0.048190883 | 0.089661622 |
| NA | NA | NA |
| 0.063708285 | 0.949202508 | NA |
| 7.097901671 | 1.27E-12 | 1.16E-11 |
| NA | NA | NA |
| NA | NA | NA |
| -0.069981114 | 0.944208692 | 0.962218155 |
| -0.455224486 | 0.648947746 | 0.737856005 |
| -4.661642332 | 3.14E-06 | 1.37E-05 |
| 8.363068333 | 6.11E-17 | 8.45E-16 |
| -5.772743871 | 7.80E-09 | 4.81E-08 |
| -0.370496921 | 0.71101227 | 0.788264035 |
| -1.644129984 | 0.100149356 | 0.166726862 |
| 9.832691965 | 8.14E-23 | 1.78E-21 |
| -3.552448306 | 0.000381664 | 0.00118967 |
| 0.073938323 | 0.941059462 | 0.959886935 |
| -2.430460745 | 0.015079639 | 0.032735765 |
| 4.26237585 | 2.02E-05 | 7.86E-05 |
| -0.422435418 | 0.67270723 | 0.756621819 |
| -1.063717673 | 0.287456616 | 0.394876192 |
| 0.211624162 | 0.832400258 | 0.882651067 |
| 7.772253197 | 7.71E-15 | 8.87E-14 |
| 1.61928861 | 0.10538518 | 0.173874547 |
| 6.616906288 | 3.67E-11 | 2.91E-10 |
| 0.192808924 | 0.847108615 | NA |
| -0.065390563 | 0.947863038 | NA |
| 2.330803236 | 0.019763737 | 0.041358277 |
| -1.996367116 | 0.045893977 | 0.085950337 |
| -7.574156055 | 3.61E-14 | 3.88E-13 |
| 1.441361266 | 0.149482647 | 0.232741395 |
| -1.362102801 | 0.173165444 | 0.262654895 |
| -2.904285966 | 0.003680918 | 0.009314869 |
| 4.772906159 | 1.82E-06 | 8.23E-06 |
| -0.791291682 | 0.428773801 | 0.539903825 |
| 2.243274909 | 0.024879092 | 0.050500663 |
| -2.201247662 | 0.027718496 | 0.055441422 |
| -7.991674156 | 1.33E-15 | 1.64E-14 |
| 0.254539972 | 0.799078427 | 0.85719509 |
| 0.0507346 | 0.959537005 | 0.97223389 |

| | | |
|---|---|---|
| -4.433763422 | 9.26E-06 | 3.80E-05 |
| 11.68842856 | 1.46E-31 | 5.47E-30 |
| 0.192809521 | 0.847108147 | NA |
| -2.959944388 | 0.003076946 | 0.007931612 |
| NA | NA | NA |
| NA | NA | NA |
| NA | NA | NA |
| 15.75276169 | 6.57E-56 | 5.88E-54 |
| 1.233501431 | 0.217388743 | NA |
| 8.216507774 | 2.10E-16 | 2.77E-15 |
| -0.526531578 | 0.598518917 | 0.696607807 |
| -1.952212663 | 0.050912961 | 0.094068078 |
| 2.138465959 | 0.032478944 | 0.063670737 |
| -0.662673143 | 0.507539914 | 0.613884414 |
| 6.205084811 | 5.47E-10 | 3.82E-09 |
| 7.917640296 | 2.42E-15 | 2.92E-14 |
| 0.06439044 | 0.948659342 | 0.965006231 |
| -0.997914976 | 0.318320589 | 0.428917116 |
| -2.011003953 | 0.044325039 | 0.083417724 |
| 1.843708857 | 0.065225581 | 0.115984392 |
| 1.130733074 | 0.258167456 | 0.362907527 |
| 2.386744097 | 0.016998319 | 0.036358564 |
| NA | NA | NA |
| -1.222479656 | 0.221526292 | 0.32082411 |
| -2.627686829 | 0.008596762 | 0.019890641 |
| 10.6298511 | 2.16E-26 | 6.05E-25 |
| 4.277234648 | 1.89E-05 | 7.38E-05 |
| 11.1005953 | 1.25E-28 | 4.02E-27 |
| 0.42694442 | 0.669419811 | 0.754246134 |
| 20.67236112 | 6.14E-95 | 1.37E-92 |
| NA | NA | NA |
| 0.263490142 | 0.792172821 | NA |
| NA | NA | NA |
| NA | NA | NA |
| -1.935673918 | 0.052907651 | 0.097180487 |
| -1.765685547 | 0.077448619 | 0.134322265 |
| -0.065389966 | 0.947863513 | NA |
| -0.065390563 | 0.947863038 | NA |
| NA | NA | NA |
| -0.133811298 | 0.893551793 | 0.925728534 |
| -1.207746685 | 0.227144715 | 0.327328229 |
| 2.573876827 | 0.010056607 | 0.022908343 |
| -0.587618435 | 0.556788434 | 0.65936859 |
| 10.35537144 | 3.96E-25 | 1.02E-23 |
| -0.163880372 | 0.869825319 | 0.909373729 |
| 8.151397955 | 3.60E-16 | 4.65E-15 |
| -8.250898197 | 1.57E-16 | 2.10E-15 |

| | | |
|---|---|---|
| -0.98939273 | 0.32247103 | 0.433471614 |
| -1.551118611 | 0.120873262 | 0.195112374 |
| 3.951955952 | 7.75E-05 | 0.000273769 |
| 0.923357089 | 0.355821146 | 0.467738212 |
| -3.566490148 | 0.000361794 | 0.001133097 |
| -0.073466102 | 0.941435218 | 0.960194104 |
| -2.735995785 | 0.006219182 | 0.014930571 |
| -0.930977284 | 0.351865315 | 0.463633862 |
| 8.584790087 | 9.10E-18 | 1.35E-16 |
| 4.446587284 | 8.72E-06 | 3.59E-05 |
| -2.518363054 | 0.011790172 | 0.026372476 |
| 2.258520076 | 0.023913254 | 0.048805345 |
| NA | NA | NA |
| -2.694372072 | 0.007052139 | 0.016686996 |
| NA | NA | NA |
| 0.063707135 | 0.949203424 | NA |
| 4.060969466 | 4.89E-05 | 0.000178335 |
| 17.93363943 | 6.44E-72 | 8.62E-70 |
| NA | NA | NA |
| 1.384864206 | 0.166093994 | 0.253681515 |
| NA | NA | NA |
| NA | NA | NA |
| 1.308051121 | 0.190855971 | 0.284555929 |
| NA | NA | NA |
| -4.928726048 | 8.28E-07 | 3.93E-06 |
| -2.520372225 | 0.011723079 | 0.026243509 |
| 0.063708882 | 0.949202033 | NA |
| -7.364204744 | 1.78E-13 | 1.78E-12 |
| NA | NA | NA |
| 2.673801731 | 0.007499678 | 0.017609468 |
| -2.270880347 | 0.023154222 | 0.047472984 |
| 1.569092246 | 0.116626446 | 0.18927316 |
| -3.951882983 | 7.75E-05 | 0.000273796 |
| -0.20268874 | 0.839378324 | 0.887531238 |
| NA | NA | NA |
| 2.211916378 | 0.026972443 | 0.054117856 |
| 15.14825875 | 7.78E-52 | 6.24E-50 |
| NA | NA | NA |
| 0.218581057 | 0.82697641 | 0.87891043 |
| 1.462534411 | 0.143594822 | 0.225199147 |
| NA | NA | NA |
| -0.346939122 | 0.728637055 | 0.801872383 |
| 0.287633901 | 0.773626992 | 0.837248513 |
| 2.660028106 | 0.007813413 | 0.018240042 |
| 0.063708882 | 0.949202033 | NA |
| -0.107903854 | 0.91407196 | 0.940950901 |
| NA | NA | NA |

| | | | |
|---|---|---|---|
| NA | NA | NA | |
| NA | NA | NA | |
| | -0.183686164 | 0.854259664 | 0.898672693 |
| NA | NA | NA | |
| | 0.063708285 | 0.949202508 | NA |
| | 3.414502357 | 0.000638986 | 0.001909939 |
| NA | NA | NA | |
| NA | NA | NA | |
| NA | NA | NA | |
| | -1.486235164 | 0.137216929 | 0.216785946 |
| | 4.043818949 | 5.26E-05 | 0.000191096 |
| | -0.065388772 | 0.947864464 | NA |
| | -0.065388772 | 0.947864464 | NA |
| | -5.273451024 | 1.34E-07 | 7.09E-07 |
| NA | NA | NA | |
| NA | NA | NA | |
| | -0.065388772 | 0.947864464 | NA |
| | 3.571907208 | 0.000354391 | 0.001112695 |
| NA | NA | NA | |
| NA | NA | NA | |
| NA | NA | NA | |
| NA | NA | NA | |
| | -8.653014982 | 5.02E-18 | 7.55E-17 |
| | -0.319438564 | 0.749393971 | 0.817954435 |
| NA | NA | NA | |
| NA | NA | NA | |
| | -0.53537574 | 0.592390062 | NA |
| | 2.717653179 | 0.006574671 | 0.015693689 |
| | -0.065389369 | 0.947863988 | NA |
| NA | NA | NA | |
| NA | NA | NA | |
| NA | NA | NA | |
| NA | NA | NA | |
| | 0.788844532 | 0.430202878 | NA |
| | -6.972283034 | 3.12E-12 | 2.74E-11 |
| | 0.181114273 | 0.856277882 | 0.900266114 |
| NA | NA | NA | |
| NA | NA | NA | |
| NA | NA | NA | |
| NA | NA | NA | |
| | 1.818476607 | 0.068991319 | 0.121755109 |
| | -2.813959096 | 0.004893545 | 0.012078241 |
| | -1.177861405 | 0.238851864 | 0.340998506 |
| | 2.502136996 | 0.012344615 | 0.027458074 |
| | 0.995008903 | 0.319731934 | 0.430424908 |
| NA | NA | NA | |
| | 4.02583954 | 5.68E-05 | 0.000204964 |

| | | | |
|---|---|---|---|
| NA | NA | NA | |
| NA | NA | NA | |
| | 0.245705299 | 0.80591037 | NA |
| NA | NA | NA | |
| | 0.085232335 | 0.932076684 | NA |
| | -4.387732996 | 1.15E-05 | 4.62E-05 |
| | -0.706265494 | 0.480023048 | 0.588680886 |
| NA | NA | NA | |
| NA | NA | NA | |
| NA | NA | NA | |
| NA | NA | NA | |
| NA | NA | NA | |
| | 0.529092132 | 0.596741539 | NA |
| NA | NA | NA | |
| | -2.115592504 | 0.03437948 | 0.066877934 |
| | -0.068752064 | 0.945186975 | 0.962606031 |
| | 0.070155456 | 0.944069928 | 0.962136413 |
| | 0.526843343 | 0.598302381 | 0.696502888 |
| | -5.757656671 | 8.53E-09 | 5.24E-08 |
| NA | NA | NA | |
| NA | NA | NA | |
| | -3.064950216 | 0.002177062 | 0.005817687 |
| NA | NA | NA | |
| | -2.218867871 | 0.026495715 | 0.053293776 |
| NA | NA | NA | |
| | 10.9869821 | 4.41E-28 | 1.38E-26 |
| | 6.860793846 | 6.85E-12 | 5.81E-11 |
| | -1.401590456 | 0.161037578 | 0.247210708 |
| | -3.973544036 | 7.08E-05 | 0.000251511 |
| | -1.203167706 | 0.228911429 | 0.329403994 |
| | 0.720944552 | 0.47094363 | 0.579884358 |
| | -5.851804662 | 4.86E-09 | 3.08E-08 |
| | -0.068186498 | 0.945637175 | NA |
| | -0.065389966 | 0.947863513 | NA |
| | -1.710786169 | 0.087120594 | 0.148478992 |
| | 0.192809521 | 0.847108147 | NA |
| | 1.362215058 | 0.173130025 | 0.262633002 |
| | -0.860973163 | 0.389252824 | 0.501389367 |
| NA | NA | NA | |
| NA | NA | NA | |
| NA | NA | NA | |
| | -0.134737963 | 0.892819056 | 0.925160877 |
| | 1.503167968 | 0.132795735 | 0.211010638 |
| NA | NA | NA | |
| | -3.083776356 | 0.002043912 | 0.005496252 |
| NA | NA | NA | |
| | 0.324874675 | 0.745275925 | NA |

| | | |
|---|---|---|
| -0.177479854 | 0.859131488 | NA |
| -4.745854999 | 2.08E-06 | 9.33E-06 |
| 5.204759269 | 1.94E-07 | 1.01E-06 |
| NA | NA | NA |
| 3.462720753 | 0.000534743 | 0.001622378 |
| NA | NA | NA |
| NA | NA | NA |
| NA | NA | NA |
| NA | NA | NA |
| NA | NA | NA |
| NA | NA | NA |
| NA | NA | NA |
| NA | NA | NA |
| NA | NA | NA |
| -0.383510901 | 0.70134099 | 0.780584507 |
| -9.029200443 | 1.73E-19 | 2.97E-18 |
| NA | NA | NA |
| NA | NA | NA |
| NA | NA | NA |
| NA | NA | NA |
| NA | NA | NA |
| NA | NA | NA |
| -1.857124274 | 0.063293475 | 0.113058902 |
| NA | NA | NA |
| -3.613197893 | 0.000302444 | 0.000960559 |
| NA | NA | NA |
| NA | NA | NA |
| 12.17498785 | 4.22E-34 | 1.78E-32 |
| NA | NA | NA |
| NA | NA | NA |
| NA | NA | NA |
| 4.229626924 | 2.34E-05 | 9.00E-05 |
| NA | NA | NA |
| NA | NA | NA |
| NA | NA | NA |
| NA | NA | NA |
| NA | NA | NA |
| NA | NA | NA |
| -6.784795857 | 1.16E-11 | 9.69E-11 |
| NA | NA | NA |
| NA | NA | NA |
| -0.065388772 | 0.947864464 | NA |
| NA | NA | NA |
| -0.740782906 | 0.458825081 | 0.568337298 |
| NA | NA | NA |
| 0.368045143 | 0.712839574 | 0.789730206 |
| 0.063708285 | 0.949202508 | NA |

| | | |
|---|---|---|
| 2.717337614 | 0.006580944 | 0.015707164 |
| -2.864156389 | 0.004181215 | 0.010466734 |
| -4.178690975 | 2.93E-05 | 0.00011109 |
| -1.626703228 | 0.103800143 | 0.171776369 |
| 0.063708285 | 0.949202508 | NA |
| NA | NA | NA |
| NA | NA | NA |
| 0.192810118 | 0.847107679 | NA |
| NA | NA | NA |
| 3.10773747 | 0.001885255 | 0.005106365 |
| NA | NA | NA |
| 0.249244798 | 0.803171429 | 0.860504999 |
| NA | NA | NA |
| -0.065389369 | 0.947863988 | NA |
| -7.773383407 | 7.64E-15 | 8.80E-14 |
| NA | NA | NA |
| -6.747660109 | 1.50E-11 | 1.24E-10 |
| 6.863108272 | 6.74E-12 | 5.72E-11 |
| -7.238271646 | 4.54E-13 | 4.38E-12 |
| -2.339005778 | 0.019335134 | 0.040583515 |
| -6.144229703 | 8.04E-10 | 5.52E-09 |
| -2.632568503 | 0.008474193 | 0.019636091 |
| 0.179607404 | 0.857460792 | 0.901093443 |
| 3.388210829 | 0.000703502 | 0.002087322 |
| -1.951307576 | 0.051020468 | 0.094211901 |
| -4.993723521 | 5.92E-07 | 2.87E-06 |
| 0.30825259 | 0.757890138 | NA |
| NA | NA | NA |
| 6.920566055 | 4.50E-12 | 3.90E-11 |
| -3.364948201 | 0.00076558 | 0.002255685 |
| -9.111702407 | 8.11E-20 | 1.43E-18 |
| 0.7265837 | 0.467481017 | 0.576752927 |
| 1.494558618 | 0.135029673 | 0.213897393 |
| -6.518152047 | 7.12E-11 | 5.50E-10 |
| -0.600130649 | 0.548419168 | 0.651893991 |
| 0.928591323 | 0.353100921 | 0.464987695 |
| 6.544390329 | 5.97E-11 | 4.65E-10 |
| 2.23198157 | 0.025616181 | 0.051786852 |
| 1.225286379 | 0.220467339 | 0.3196604 |
| 1.31912531 | 0.187127223 | 0.279925532 |
| 2.910514982 | 0.003608337 | 0.00914414 |
| 1.063571881 | 0.287522686 | 0.394923604 |
| -1.120200649 | 0.262628268 | 0.368020689 |
| -2.97200633 | 0.002958606 | 0.007661277 |
| -0.194488217 | 0.845793621 | NA |
| -1.127531244 | 0.259517938 | 0.36456022 |
| -3.280888975 | 0.001034805 | 0.002973386 |

| | | |
|---|---|---|
| 0.15988669 | 0.872970333 | 0.91161281 |
| -2.300594084 | 0.021414586 | 0.044333037 |
| -0.164233862 | 0.869547045 | 0.909213908 |
| -2.430254203 | 0.015088236 | 0.032751587 |
| 0.063708285 | 0.949202508 | NA |
| -3.886142882 | 0.00010185 | 0.000351578 |
| 1.563713153 | 0.117884947 | 0.191005971 |
| -2.550004855 | 0.010772142 | 0.024370189 |
| -1.384446197 | 0.166221872 | 0.253845848 |
| -2.854946366 | 0.00430441 | 0.010739571 |
| -13.85045512 | 1.26E-43 | 7.99E-42 |
| -2.629989492 | 0.008538751 | 0.01976921 |
| NA | NA | NA |
| -0.065388772 | 0.947864464 | NA |
| -8.187582358 | 2.67E-16 | 3.50E-15 |
| -1.59913958 | 0.109789592 | 0.179856894 |
| -1.970048199 | 0.048832847 | 0.090707693 |
| 0.063708285 | 0.949202508 | NA |
| -0.718547905 | 0.472419521 | NA |
| -0.283862434 | 0.776515806 | 0.839685084 |
| 0.123620316 | 0.901615906 | NA |
| -2.360955011 | 0.018227942 | 0.038579856 |
| NA | NA | NA |
| 0.192809521 | 0.847108147 | NA |
| NA | NA | NA |
| 4.465482386 | 7.99E-06 | 3.31E-05 |
| 0.259766706 | 0.795043734 | 0.854198553 |
| 1.240339501 | 0.214849848 | 0.313129829 |
| -0.506991238 | 0.612160966 | 0.707960495 |
| -4.576776533 | 4.72E-06 | 2.02E-05 |
| NA | NA | NA |
| -5.793673051 | 6.89E-09 | 4.28E-08 |
| 3.659461511 | 0.000252746 | 0.000814005 |
| -0.269087362 | 0.787862455 | NA |
| 0.81035466 | 0.417736368 | 0.529387945 |
| NA | NA | NA |
| -0.065389369 | 0.947863988 | NA |
| 6.713936671 | 1.89E-11 | 1.55E-10 |
| 3.532526948 | 0.000411608 | 0.001274438 |
| 1.041944207 | 0.297437547 | 0.406135061 |
| -2.12043193 | 0.033969636 | 0.066183545 |
| 5.424687724 | 5.81E-08 | 3.23E-07 |
| -0.746954734 | 0.455090885 | 0.564829913 |
| 0.168952532 | 0.865833978 | 0.906429565 |
| 8.847944944 | 8.91E-19 | 1.45E-17 |
| -0.861127294 | 0.389167938 | 0.501305825 |
| -0.396463033 | 0.691763474 | NA |

| | | | |
|---|---|---|---|
| NA | NA | NA | |
| | 10.09799996 | 5.64E-24 | 1.34E-22 |
| NA | NA | NA | |
| | -0.082851036 | 0.933969988 | NA |
| NA | NA | NA | |
| | 16.87460143 | 6.92E-64 | 7.63E-62 |
| | 0.985270995 | 0.324490976 | NA |
| | 3.761585681 | 0.00016884 | 0.000560857 |
| | 1.619348752 | 0.105372247 | 0.173874547 |
| | 15.67885624 | 2.11E-55 | 1.85E-53 |
| | 0.207251126 | 0.835813741 | NA |
| | 4.825433817 | 1.40E-06 | 6.41E-06 |
| | -0.283214589 | 0.777012346 | 0.840076844 |
| NA | NA | NA | |
| | -0.065388772 | 0.947864464 | NA |
| | 0.192810118 | 0.847107679 | NA |
| | 7.147079497 | 8.86E-13 | 8.28E-12 |
| | 0.683859622 | 0.494063813 | 0.601216259 |
| | 1.877031553 | 0.060513769 | 0.108834417 |
| | -0.454876046 | 0.649198417 | 0.73793989 |
| | -3.099402686 | 0.001939113 | 0.005237508 |
| | -2.918898211 | 0.003512709 | 0.008931644 |
| | 2.359937802 | 0.018277999 | 0.038672724 |
| NA | NA | NA | |
| | -0.065390563 | 0.947863038 | NA |
| | 2.911464099 | 0.003597392 | 0.009118251 |
| | 0.589947979 | 0.555225526 | 0.657953552 |
| | 0.792662231 | 0.427974641 | NA |
| NA | NA | NA | |
| | -3.941230974 | 8.11E-05 | 0.000285018 |
| | -2.346549181 | 0.018948165 | 0.03986824 |
| | -2.492929556 | 0.012669398 | 0.028098089 |
| | -8.623065268 | 6.52E-18 | 9.73E-17 |
| | 0.063707019 | 0.949203516 | NA |
| | 0.261213554 | 0.793927827 | NA |
| | 3.527317156 | 0.000419794 | 0.001297855 |
| | 11.18949692 | 4.59E-29 | 1.51E-27 |
| | 0.195654901 | 0.844880286 | 0.891615802 |
| NA | NA | NA | |
| | -5.662920478 | 1.49E-08 | 8.92E-08 |
| NA | NA | NA | |
| | 0.193341124 | 0.846691821 | NA |
| | -3.764319178 | 0.000167003 | 0.000555421 |
| | 5.222885854 | 1.76E-07 | 9.17E-07 |
| | -0.210810096 | 0.83303546 | 0.88301169 |
| | -0.134007205 | 0.893396877 | NA |
| | 0.192810118 | 0.847107679 | NA |

| | | | |
|---|---|---|---|
| | 0.108340983 | 0.913725214 | NA |
| | 9.100886699 | 8.96E-20 | 1.57E-18 |
| | 1.783585965 | 0.074490968 | 0.129885679 |
| | 2.027329418 | 0.042628735 | 0.080588915 |
| | -0.651547727 | 0.514692979 | 0.620198597 |
| | -0.229364346 | 0.818585745 | 0.872474547 |
| | 1.383868212 | 0.166398812 | 0.254069558 |
| | 0.791625772 | 0.428578915 | 0.539767045 |
| NA | NA | NA | |
| | 8.443781636 | 3.07E-17 | 4.35E-16 |
| | -8.87153228 | 7.21E-19 | 1.18E-17 |
| | 0.349291458 | 0.726870511 | NA |
| NA | NA | NA | |
| | -3.831327077 | 0.000127454 | 0.000432566 |
| | 0.063708285 | 0.949202508 | NA |
| | 1.404666183 | 0.160120566 | 0.246119981 |
| NA | NA | NA | |
| | -0.041321282 | 0.967039767 | 0.977534078 |
| | -4.829688799 | 1.37E-06 | 6.28E-06 |
| | -0.067464376 | 0.946212021 | 0.963221364 |
| | 4.873154903 | 1.10E-06 | 5.12E-06 |
| | -0.335575119 | 0.737191289 | 0.808549054 |
| | -2.393625475 | 0.016682774 | 0.035790677 |
| | -0.677008638 | 0.498400474 | 0.60558077 |
| | -3.543504447 | 0.000394847 | 0.001227246 |
| NA | NA | NA | |
| | -3.346800179 | 0.000817501 | 0.002394343 |
| | -4.559181875 | 5.14E-06 | 2.18E-05 |
| NA | NA | NA | |
| | -0.139743304 | 0.888862811 | NA |
| | -2.008049663 | 0.044638015 | 0.083937308 |
| NA | NA | NA | |
| NA | NA | NA | |
| NA | NA | NA | |
| | -8.265519107 | 1.39E-16 | 1.87E-15 |
| NA | NA | NA | |
| NA | NA | NA | |
| | 0.063708285 | 0.949202508 | NA |
| NA | NA | NA | |
| | 0.642063157 | 0.520832176 | 0.626087055 |
| | -2.420389637 | 0.015503885 | 0.033534586 |
| | -7.290875935 | 3.08E-13 | 3.01E-12 |
| NA | NA | NA | |
| NA | NA | NA | |
| | 0.311104013 | 0.75572155 | 0.822923101 |
| | -3.599215633 | 0.000319178 | 0.001009992 |
| | -3.000686746 | 0.002693715 | 0.007037855 |

| | | |
|---|---|---|
| -5.673819961 | 1.40E-08 | 8.39E-08 |
| -3.767620998 | 0.000164811 | 0.00054893 |
| -1.06925372 | 0.284955358 | 0.392102048 |
| NA | NA | NA |
| NA | NA | NA |
| -5.006539715 | 5.54E-07 | 2.69E-06 |
| -0.077964336 | 0.937856422 | 0.957714751 |
| -3.711301484 | 0.000206196 | 0.000676067 |
| -2.044416853 | 0.040912384 | 0.077713969 |
| NA | NA | NA |
| NA | NA | NA |
| -8.154409118 | 3.51E-16 | 4.54E-15 |
| 1.41712329 | 0.156446884 | 0.241794722 |
| 0.192805732 | 0.847111115 | NA |
| -3.606342398 | 0.000310543 | 0.000984534 |
| -0.377366912 | 0.705900953 | 0.784254868 |
| 2.312904996 | 0.020727863 | 0.04309481 |
| 0.534841024 | 0.592759793 | 0.691884798 |
| -0.056647996 | 0.9548256 | 0.969073591 |
| -1.19602558 | 0.231686577 | 0.332546676 |
| 3.153782082 | 0.001611694 | 0.004439386 |
| -2.774019778 | 0.005536831 | 0.013482526 |
| -0.998487424 | 0.31804306 | 0.428612383 |
| 1.584080401 | 0.113175424 | 0.1845587 |
| NA | NA | NA |
| -10.9148025 | 9.79E-28 | 3.00E-26 |
| NA | NA | NA |
| 4.27842216 | 1.88E-05 | 7.35E-05 |
| 1.054980153 | 0.291434402 | 0.399135646 |
| -2.004565333 | 0.045009536 | 0.084537329 |
| -6.528800353 | 6.63E-11 | 5.14E-10 |
| NA | NA | NA |
| NA | NA | NA |
| NA | NA | NA |
| NA | NA | NA |
| NA | NA | NA |
| -1.546758835 | 0.12192141 | 0.196629696 |
| -2.245780086 | 0.024718097 | 0.050214591 |
| -1.418671283 | 0.155994873 | 0.241173337 |
| 0.192804347 | 0.847112199 | NA |
| 7.132085515 | 9.89E-13 | 9.18E-12 |
| -0.065388772 | 0.947864464 | NA |
| -0.244470635 | 0.806866342 | 0.863144907 |
| 1.659448785 | 0.097025393 | 0.16244452 |
| -0.211392129 | 0.832581298 | 0.88275094 |
| -9.57829835 | 9.87E-22 | 2.00E-20 |
| -5.258318685 | 1.45E-07 | 7.67E-07 |

| | | |
|---|---|---|
| 6.129981819 | 8.79E-10 | 6.01E-09 |
| -2.316146814 | 0.020550254 | 0.042782405 |
| NA | NA | NA |
| 0.249739732 | 0.80278863 | 0.860179769 |
| 0.404860484 | 0.685580066 | NA |
| -4.483871308 | 7.33E-06 | 3.04E-05 |
| 1.564195094 | 0.117771759 | 0.190834928 |
| 4.583516551 | 4.57E-06 | 1.96E-05 |
| 0.073771891 | 0.941191894 | NA |
| 0.657557132 | 0.510822748 | NA |
| -2.22567491 | 0.026035965 | 0.052512505 |
| 2.669111311 | 0.007605224 | 0.017823854 |
| NA | NA | NA |
| -0.909075809 | 0.363310111 | 0.475708939 |
| NA | NA | NA |
| 2.42714881 | 0.015218012 | 0.032990373 |
| 0.442063591 | 0.65844319 | NA |
| 5.95171053 | 2.65E-09 | 1.73E-08 |
| NA | NA | NA |
| -5.156219016 | 2.52E-07 | 1.29E-06 |
| -3.440346582 | 0.00058097 | 0.001750361 |
| -1.251659583 | 0.210693934 | 0.308320208 |
| -0.632094946 | 0.527324838 | 0.632375464 |
| -1.551629516 | 0.120750897 | 0.194942914 |
| -4.425647908 | 9.62E-06 | 3.93E-05 |
| 2.348105919 | 0.018869155 | 0.03971869 |
| -0.065389966 | 0.947863513 | NA |
| -5.024588405 | 5.05E-07 | 2.47E-06 |
| -2.546968583 | 0.010866322 | 0.024547295 |
| NA | NA | NA |
| NA | NA | NA |
| -1.627113601 | 0.103712973 | 0.171643461 |
| -1.018676489 | 0.308356578 | 0.418206088 |
| NA | NA | NA |
| 0.449351605 | 0.653178037 | 0.741251644 |
| 1.349413698 | 0.177204123 | 0.267710363 |
| NA | NA | NA |
| 2.759131522 | 0.00579552 | 0.014021059 |
| -2.76876924 | 0.005626847 | 0.013664499 |
| -0.321304183 | 0.747979887 | 0.816802782 |
| 1.304834962 | 0.191949051 | 0.285828113 |
| NA | NA | NA |
| 0.57257538 | 0.566932234 | 0.668786587 |
| -1.318250385 | 0.187419844 | 0.280267576 |
| -0.914764898 | 0.360315066 | 0.472454613 |
| NA | NA | NA |
| 12.80870797 | 1.47E-37 | 7.24E-36 |

| | | | |
|---|---|---|---|
| | -2.054623324 | 0.039915401 | 0.076074202 |
| NA | NA | NA | |
| | -1.58346413 | 0.113315716 | 0.184738642 |
| | -2.806846949 | 0.005002901 | 0.012311729 |
| | 0.180356889 | 0.856872397 | NA |
| | 0.083728242 | 0.933272502 | NA |
| | 1.258207598 | 0.208316688 | 0.305368474 |
| NA | NA | NA | |
| | 0.569376575 | 0.569100611 | 0.670676079 |
| | -6.309158859 | 2.81E-10 | 2.02E-09 |
| | -0.156739198 | 0.875450393 | NA |
| | -1.4009996 | 0.161214193 | 0.247421132 |
| | -3.581228948 | 0.000341982 | 0.00107657 |
| NA | NA | NA | |
| | -0.065388772 | 0.947864464 | NA |
| NA | NA | NA | |
| | -1.707858127 | 0.08766267 | 0.149305705 |
| | -0.78215661 | 0.434122541 | 0.544829884 |
| | -3.515872495 | 0.000438311 | 0.001350768 |
| | -0.980453537 | 0.326862294 | 0.437880712 |
| | -3.0448454 | 0.002327999 | 0.006170037 |
| | 2.269192063 | 0.023256649 | 0.047653422 |
| | 1.007605604 | 0.313643837 | 0.424053652 |
| NA | NA | NA | |
| | -0.066204891 | 0.947214703 | NA |
| NA | NA | NA | |
| | 0.756464912 | 0.449370502 | NA |
| | 4.336415763 | 1.45E-05 | 5.75E-05 |
| | -0.134056335 | 0.893358027 | NA |
| | 3.667943746 | 0.000244509 | 0.000789787 |
| | -0.516716605 | 0.605354003 | NA |
| | 0.31685483 | 0.751353758 | NA |
| | -1.272739691 | 0.203110428 | 0.299437744 |
| | 1.595130997 | 0.110682945 | 0.181083347 |
| | -2.519989579 | 0.011735831 | 0.026267356 |
| | 6.284943277 | 3.28E-10 | 2.34E-09 |
| | 0.063708381 | 0.949202431 | NA |
| | -0.837284042 | 0.402432927 | 0.514185614 |
| | 1.775691384 | 0.075783798 | 0.131809358 |
| | 6.538438774 | 6.22E-11 | 4.83E-10 |
| | 0.288217989 | 0.77317988 | NA |
| | -0.67749802 | 0.498090026 | 0.605232941 |
| | -7.511927405 | 5.83E-14 | 6.12E-13 |
| NA | NA | NA | |
| | 2.481748839 | 0.013073939 | 0.028887773 |
| | -0.577736339 | 0.563442141 | 0.665570949 |
| | 5.448789604 | 5.07E-08 | 2.84E-07 |

| | | |
|---|---|---|
| -2.762495953 | 0.005736127 | 0.013896174 |
| NA | NA | NA |
| NA | NA | NA |
| 0.769967139 | 0.441319386 | 0.551373876 |
| -2.162760253 | 0.030559624 | 0.060309658 |
| NA | NA | NA |
| -5.825508754 | 5.69E-09 | 3.57E-08 |
| 0.339478625 | 0.734249197 | NA |
| 0.169439718 | 0.865450784 | 0.90618001 |
| 0.333816862 | 0.738517758 | NA |
| 7.786305544 | 6.90E-15 | 7.98E-14 |
| 9.505693624 | 1.99E-21 | 3.95E-20 |
| -1.236553159 | 0.216253019 | 0.314808251 |
| 0.968517467 | 0.332786 | 0.443962988 |
| 0.471546284 | 0.637250676 | NA |
| 6.157840895 | 7.37E-10 | 5.08E-09 |
| -0.741380604 | 0.458462701 | 0.568000862 |
| -0.597564677 | 0.550130439 | 0.653462647 |
| 1.182732844 | 0.236915043 | 0.33883684 |
| -1.868151748 | 0.061740922 | 0.110676451 |
| NA | NA | NA |
| NA | NA | NA |
| 0.326218213 | 0.744259263 | 0.81394719 |
| -1.240159644 | 0.214916352 | 0.313208517 |
| -0.713036615 | 0.4758231 | 0.584704806 |
| -2.264313015 | 0.02355487 | 0.048176041 |
| -0.367056719 | 0.713576711 | 0.790441895 |
| -10.05225662 | 8.98E-24 | 2.12E-22 |
| -1.593132762 | 0.111130411 | 0.181708531 |
| -1.573575859 | 0.115585538 | 0.187827437 |
| 0.367466416 | 0.713271138 | NA |
| 0.437384145 | 0.661832781 | NA |
| -0.065389966 | 0.947863513 | NA |
| -11.31816401 | 1.07E-29 | 3.61E-28 |
| -0.456865133 | 0.647767987 | NA |
| NA | NA | NA |
| NA | NA | NA |
| NA | NA | NA |
| 0.192809521 | 0.847108147 | NA |
| NA | NA | NA |
| -10.10364003 | 5.32E-24 | 1.27E-22 |
| -4.34631757 | 1.38E-05 | 5.51E-05 |
| 0.063708285 | 0.949202508 | NA |
| NA | NA | NA |
| NA | NA | NA |
| NA | NA | NA |
| -1.806944703 | 0.070770902 | 0.124456943 |

| | | |
|---|---|---|
| -1.631776434 | 0.102726592 | 0.170292509 |
| 4.77328841 | 1.81E-06 | 8.22E-06 |
| 2.308963903 | 0.020945584 | 0.043475658 |
| 0.981494358 | 0.326349015 | 0.437403794 |
| -1.346017141 | 0.178296989 | 0.269003835 |
| 0.129896632 | 0.896648209 | 0.928014583 |
| 1.866351423 | 0.061992215 | 0.111055335 |
| NA | NA | NA |
| NA | NA | NA |
| 3.141176305 | 0.001682707 | 0.00461114 |
| -0.661265862 | 0.50844183 | 0.614589222 |
| 0.192809521 | 0.847108147 | NA |
| -7.741618187 | 9.82E-15 | 1.12E-13 |
| -3.7114896 | 0.000206043 | 0.000675653 |
| -2.070416115 | 0.038413394 | 0.0735757 |
| NA | NA | NA |
| NA | NA | NA |
| 7.041814643 | 1.90E-12 | 1.70E-11 |
| NA | NA | NA |
| 0.368282397 | 0.712662676 | NA |
| -5.453907732 | 4.93E-08 | 2.77E-07 |
| -7.327563208 | 2.34E-13 | 2.32E-12 |
| NA | NA | NA |
| -7.597181571 | 3.03E-14 | 3.28E-13 |
| -3.700233089 | 0.000215402 | 0.000703484 |
| -3.52452952 | 0.000424236 | 0.001310876 |
| 1.18414457 | 0.23635584 | 0.338152996 |
| -1.211408543 | 0.225738867 | 0.325827191 |
| -2.0740903 | 0.038070925 | 0.07299236 |
| NA | NA | NA |
| -1.857672272 | 0.063215568 | 0.112955859 |
| 0.063708285 | 0.949202508 | NA |
| 5.177562039 | 2.25E-07 | 1.16E-06 |
| 6.63991105 | 3.14E-11 | 2.51E-10 |
| 1.441693244 | 0.14938893 | 0.232638893 |
| 9.073806751 | 1.15E-19 | 2.00E-18 |
| -0.034766067 | 0.972266279 | 0.981006368 |
| NA | NA | NA |
| 0.736993322 | 0.461126411 | NA |
| -3.774993704 | 0.000160012 | 0.000533657 |
| 1.751600435 | 0.079842538 | 0.137853261 |
| 1.194187243 | 0.232404732 | 0.333338516 |
| 7.852629547 | 4.07E-15 | 4.79E-14 |
| -2.059530785 | 0.039443419 | 0.075260617 |
| 1.164064853 | 0.244397731 | 0.347206938 |
| 0.070244835 | 0.943998789 | 0.962111992 |
| -2.848131478 | 0.004397675 | 0.010953708 |

| | | |
|---|---|---|
| -0.985158014 | 0.324546461 | 0.43554758 |
| -0.532096844 | 0.594658923 | 0.693390145 |
| 2.632847918 | 0.008467225 | 0.019622414 |
| 1.841947862 | 0.065482777 | 0.116408658 |
| NA | NA | NA |
| 0.100847189 | 0.919671766 | 0.944850154 |
| -2.579046152 | 0.009907355 | 0.022599207 |
| -2.292352211 | 0.021885327 | 0.045178535 |
| 2.663474561 | 0.007733826 | 0.018081231 |
| NA | NA | NA |
| 0.963836645 | 0.335127831 | 0.446373436 |
| 2.792453718 | 0.005230995 | 0.012807572 |
| NA | NA | NA |
| -4.081260135 | 4.48E-05 | 0.000164103 |
| 0.552478249 | 0.580620733 | 0.681273999 |
| -2.696291942 | 0.007011618 | 0.016595821 |
| NA | NA | NA |
| 0.297978939 | 0.765719238 | 0.830957767 |
| 9.303057882 | 1.36E-20 | 2.53E-19 |
| 0.171173802 | 0.864087106 | 0.905433929 |
| -0.131437814 | 0.895428977 | 0.927212779 |
| -2.038805405 | 0.041469453 | 0.078664676 |
| 1.130378101 | 0.258316938 | 0.363097263 |
| -0.537644923 | 0.59082221 | NA |
| 0.798884131 | 0.424357602 | 0.535906699 |
| 5.456855836 | 4.85E-08 | 2.73E-07 |
| 1.355582157 | 0.175232152 | 0.265355483 |
| 6.300641948 | 2.96E-10 | 2.13E-09 |
| 0.192810118 | 0.847107679 | NA |
| NA | NA | NA |
| NA | NA | NA |
| 0.063708285 | 0.949202508 | NA |
| -0.065389369 | 0.947863988 | NA |
| 1.946873549 | 0.051549896 | 0.095049275 |
| -1.677414416 | 0.093461472 | 0.157510143 |
| 0.069134878 | 0.944882259 | NA |
| NA | NA | NA |
| 0.192810118 | 0.847107679 | NA |
| 9.398788098 | 5.52E-21 | 1.05E-19 |
| 0.148685811 | 0.881801558 | 0.91700914 |
| NA | NA | NA |
| -0.690413085 | 0.489934451 | 0.597585853 |
| NA | NA | NA |
| -0.688324148 | 0.491248678 | 0.598722034 |
| 0.298704597 | 0.765165449 | NA |
| NA | NA | NA |
| 0.363594285 | 0.716160995 | NA |

| | | |
|---|---|---|
| 1.036963856 | 0.299752702 | 0.408665165 |
| -0.065389369 | 0.947863988 | NA |
| NA | NA | NA |
| -1.41004814 | 0.158525469 | 0.24414795 |
| 0.192809521 | 0.847108147 | NA |
| NA | NA | NA |
| 0.320071032 | 0.748914485 | NA |
| NA | NA | NA |
| 1.137660313 | 0.255262351 | NA |
| NA | NA | NA |
| 1.64164832 | 0.100662908 | 0.167425914 |
| NA | NA | NA |
| -0.708433394 | 0.478676161 | 0.587445708 |
| NA | NA | NA |
| -5.436590941 | 5.43E-08 | 3.03E-07 |
| 1.419525287 | 0.15574593 | 0.240874977 |
| -11.85649481 | 1.99E-32 | 7.80E-31 |
| NA | NA | NA |
| NA | NA | NA |
| 0.192810118 | 0.847107679 | NA |
| NA | NA | NA |
| -2.32836138 | 0.019892923 | 0.041597314 |
| -0.818755018 | 0.4129262 | 0.524540336 |
| 1.278597721 | 0.201038753 | NA |
| -1.104518033 | 0.269368486 | 0.375489708 |
| 1.399611058 | 0.161629821 | 0.247964306 |
| -11.3563918 | 6.89E-30 | 2.35E-28 |
| 0.063708285 | 0.949202508 | NA |
| -0.752344318 | 0.451844023 | 0.561775079 |
| -3.724100552 | 0.000196013 | 0.000645383 |
| 5.829663764 | 5.55E-09 | 3.49E-08 |
| -1.052905451 | 0.292384327 | 0.40027481 |
| 4.924055753 | 8.48E-07 | 4.01E-06 |
| NA | NA | NA |
| 10.39904081 | 2.50E-25 | 6.53E-24 |
| 2.372473915 | 0.017669411 | 0.03759481 |
| 0.56814851 | 0.569934131 | 0.67136066 |
| -0.081017037 | 0.935428404 | NA |
| -0.404899649 | 0.685551276 | 0.767653042 |
| 6.863812222 | 6.70E-12 | 5.70E-11 |
| -1.059327682 | 0.289450572 | NA |
| -2.109294915 | 0.034919133 | 0.067790732 |
| 1.614738406 | 0.106367357 | 0.175144723 |
| -0.778158834 | 0.436475382 | 0.546905227 |
| -0.890585587 | 0.373151534 | 0.485951243 |
| -1.148331072 | 0.250831917 | 0.35450722 |
| -0.700967245 | 0.483323457 | 0.59174262 |

| | | |
|---|---|---|
| 1.805610161 | 0.070979254 | 0.124760881 |
| NA | NA | NA |
| NA | NA | NA |
| -0.47304923 | 0.636178057 | 0.72746753 |
| -1.793412622 | 0.072906971 | 0.127481433 |
| NA | NA | NA |
| NA | NA | NA |
| -0.979654229 | 0.327256827 | 0.438291956 |
| NA | NA | NA |
| -0.065389369 | 0.947863988 | NA |
| 4.752634816 | 2.01E-06 | 9.05E-06 |
| 0.063708324 | 0.949202477 | NA |
| -1.979578196 | 0.047750945 | 0.08898862 |
| NA | NA | NA |
| -6.00431372 | 1.92E-09 | 1.27E-08 |
| -8.031408069 | 9.64E-16 | 1.20E-14 |
| -1.75252432 | 0.079683692 | 0.137635967 |
| -11.55834739 | 6.70E-31 | 2.42E-29 |
| 0.573785523 | 0.566112947 | NA |
| 3.587621403 | 0.000333708 | 0.001051847 |
| 0.728414513 | 0.466359882 | 0.575681942 |
| -3.137920993 | 0.001701507 | 0.00465756 |
| -3.031243501 | 0.002435487 | 0.006422764 |
| NA | NA | NA |
| 4.249319622 | 2.14E-05 | 8.29E-05 |
| -0.87371103 | 0.382275643 | 0.494612229 |
| -0.488535584 | 0.625170527 | 0.718755147 |
| -0.793065447 | 0.427739693 | 0.538994738 |
| -1.370658898 | 0.170481313 | 0.259196482 |
| -1.800544121 | 0.071774763 | 0.125859768 |
| -1.519027362 | 0.128755609 | 0.205860932 |
| -0.926278708 | 0.354301158 | 0.466180833 |
| 0.800211679 | 0.423588164 | 0.535151007 |
| 1.582451466 | 0.113546544 | 0.185042645 |
| 1.075640309 | 0.282088153 | 0.389103189 |
| -3.89580235 | 9.79E-05 | 0.000339024 |
| NA | NA | NA |
| 0.264686584 | 0.791250914 | NA |
| -0.322383666 | 0.747162057 | 0.816265823 |
| 1.751978666 | 0.079777476 | 0.137759934 |
| 0.733083893 | 0.463507263 | 0.57294616 |
| 3.684516745 | 0.000229137 | 0.000744554 |
| NA | NA | NA |
| 0.164468186 | 0.86936259 | 0.9090818 |
| 0.580676152 | 0.561458737 | 0.663643379 |
| 0.896428753 | 0.370023816 | 0.482731998 |
| 1.351922791 | 0.176400016 | 0.266720575 |

| | | | |
|---|---|---|---|
| | 0.126314312 | 0.899483128 | 0.929948892 |
| | 5.864569546 | 4.50E-09 | 2.86E-08 |
| | 0.192809521 | 0.847108147 | NA |
| | 0.192810118 | 0.847107679 | NA |
| | 1.476411618 | 0.139833426 | 0.220210812 |
| NA | NA | NA | |
| NA | NA | NA | |
| | 1.741361568 | 0.08162022 | 0.140505785 |
| | 1.067862413 | 0.28558258 | 0.392711348 |
| | 3.636216605 | 0.000276672 | 0.000885335 |
| | 3.823008431 | 0.000131833 | 0.000446339 |
| | 2.629504911 | 0.00855093 | 0.019793745 |
| | 22.67826301 | 7.34E-114 | 2.19E-111 |
| | 7.970483774 | 1.58E-15 | 1.93E-14 |
| | 4.31734077 | 1.58E-05 | 6.24E-05 |
| | 1.483327864 | 0.137987323 | 0.217796865 |
| | 27.73331708 | 2.77E-169 | 1.44E-166 |
| | 2.597723273 | 0.009384409 | 0.021518116 |
| | 0.063708882 | 0.949202033 | NA |
| | 15.28095806 | 1.02E-52 | 8.32E-51 |
| | 10.82390667 | 2.65E-27 | 7.85E-26 |
| | 0.430180397 | 0.667064421 | NA |
| | 13.16146415 | 1.46E-39 | 7.87E-38 |
| | 7.99726962 | 1.27E-15 | 1.57E-14 |
| | 1.731809806 | 0.083307432 | NA |
| NA | NA | NA | |
| | 0.063708285 | 0.949202508 | NA |
| NA | NA | NA | |
| NA | NA | NA | |
| NA | NA | NA | |
| NA | NA | NA | |
| | 0.288217989 | 0.77317988 | NA |
| | 0.192810118 | 0.847107679 | NA |
| NA | NA | NA | |
| | 0.296893239 | 0.766548018 | NA |
| NA | NA | NA | |
| | 0.347873358 | 0.727935294 | NA |
| NA | NA | NA | |
| | -0.334497779 | 0.738003966 | 0.809227637 |
| | 0.603111262 | 0.546434675 | NA |
| | 2.325910521 | 0.020023325 | 0.041824569 |
| | -0.196966541 | 0.843853722 | 0.890833347 |
| | 1.065323088 | 0.286729748 | 0.393964187 |
| | -0.34808645 | 0.72777526 | 0.801276272 |
| | -0.949216525 | 0.342510499 | 0.45384001 |
| | -0.445845625 | 0.655708762 | 0.743516822 |
| | -0.178472664 | 0.858351787 | 0.901726905 |

| | | | |
|---|---|---|---|
| NA | | NA | NA |
| | 0.192808924 | 0.847108615 | NA |
| | 18.44387226 | 5.84E-76 | 8.60E-74 |
| | -0.056480103 | 0.954959346 | 0.969169705 |
| | 0.653136897 | 0.513668027 | 0.619212041 |
| | 3.375238105 | 0.000737519 | 0.002180797 |
| | 0.028316423 | 0.977409782 | 0.984410384 |
| NA | | NA | NA |
| | 2.119283433 | 0.034066521 | 0.066355537 |
| | 4.397095901 | 1.10E-05 | 4.44E-05 |
| | 0.470170712 | 0.638233057 | 0.729284338 |
| | -0.470222732 | 0.638195895 | 0.729275167 |
| | 0.710611883 | 0.477324777 | 0.586147239 |
| | -0.252709177 | 0.800492956 | 0.858381402 |
| | 3.128362072 | 0.001757835 | 0.004798627 |
| | 0.504213587 | 0.614111293 | 0.709691383 |
| | 0.633626382 | 0.526324679 | 0.6313951 |
| | -0.22700384 | 0.820420756 | 0.873984147 |
| NA | | NA | NA |
| | 3.082080972 | 0.002055589 | 0.005522314 |
| | -6.254832884 | 3.98E-10 | 2.82E-09 |
| | -0.110572025 | 0.911955732 | 0.939351502 |
| | 3.797786091 | 0.000145994 | 0.000490107 |
| | -1.661462394 | 0.096620616 | 0.161890605 |
| | -2.099429441 | 0.035779062 | 0.069165115 |
| | 30.28649409 | 1.73E-201 | 1.27E-198 |
| | 6.830559246 | 8.46E-12 | 7.11E-11 |
| NA | | NA | NA |
| | 2.888522781 | 0.00387056 | 0.009764172 |
| | 0.199526594 | 0.841850842 | NA |
| NA | | NA | NA |
| | 0.892567568 | 0.372088794 | 0.484895221 |
| | -2.649203437 | 0.008068175 | 0.01876651 |
| | 4.50904691 | 6.51E-06 | 2.73E-05 |
| | -1.682946408 | 0.092385467 | 0.156032944 |
| | 1.358022227 | 0.174456631 | 0.26438919 |
| | 0.680800715 | 0.495997598 | 0.60310031 |
| | 4.221577943 | 2.43E-05 | 9.29E-05 |
| | 2.38984177 | 0.016855633 | 0.036112 |
| | 4.05540378 | 5.00E-05 | 0.000182423 |
| | 0.69361968 | 0.487920749 | 0.595961062 |
| | -1.641184468 | 0.100759129 | 0.16754142 |
| NA | | NA | NA |
| | 2.584694608 | 0.009746528 | 0.022264817 |
| | 0.730684119 | 0.464972118 | 0.574444823 |
| | 1.481855602 | 0.138378721 | 0.21826651 |
| | -1.193186835 | 0.23279621 | 0.33386175 |

| | | |
|---|---|---|
| -13.11911366 | 2.56E-39 | 1.36E-37 |
| 0.829695487 | 0.406710974 | 0.518539918 |
| 2.152332539 | 0.031371173 | 0.061726315 |
| -1.039425664 | 0.298606814 | 0.407467555 |
| 2.312837692 | 0.020731565 | 0.043098925 |
| 3.203208736 | 0.001359054 | 0.003799523 |
| 0.761065298 | 0.446618067 | 0.556577058 |
| -3.819300423 | 0.000133831 | 0.000452429 |
| -10.70692636 | 9.44E-27 | 2.70E-25 |
| -0.976554851 | 0.32878958 | 0.43968676 |
| -3.842152813 | 0.00012196 | 0.000415327 |
| -7.352833372 | 1.94E-13 | 1.93E-12 |
| 0.738667636 | 0.460108842 | 0.56952969 |
| 1.44455436 | 0.1485831 | 0.231542461 |
| 1.32570662 | 0.184936894 | 0.277302726 |
| -3.990563883 | 6.59E-05 | 0.000235127 |
| -4.522140341 | 6.12E-06 | 2.57E-05 |
| 0.063708285 | 0.949202508 | NA |
| -6.066360857 | 1.31E-09 | 8.79E-09 |
| 0.021670302 | 0.982710953 | 0.98785177 |
| 2.898018418 | 0.003755285 | 0.009487715 |
| 6.176576845 | 6.55E-10 | 4.53E-09 |
| -0.464664328 | 0.642171858 | 0.732480933 |
| 7.269856819 | 3.60E-13 | 3.49E-12 |
| -0.12647554 | 0.89935551 | 0.929893771 |
| 12.6550862 | 1.05E-36 | 4.95E-35 |
| 0.988936179 | 0.322694368 | 0.433583323 |
| -0.333485077 | 0.738768153 | 0.809710858 |
| -0.065389369 | 0.947863988 | NA |
| -2.376987306 | 0.017454689 | 0.037188531 |
| 5.650387676 | 1.60E-08 | 9.55E-08 |
| 0.612086895 | 0.540480268 | NA |
| 12.95492287 | 2.20E-38 | 1.13E-36 |
| 1.235986832 | 0.216463458 | 0.315022987 |
| 0.207171563 | 0.835875875 | NA |
| 0.845498453 | 0.397832596 | NA |
| 0.203595243 | 0.838669811 | NA |
| NA | NA | NA |
| 1.11394879 | 0.265301157 | NA |
| 0.75383326 | 0.45094935 | NA |
| 8.610630031 | 7.27E-18 | 1.08E-16 |
| 0.229429265 | 0.818535292 | NA |
| 5.007179962 | 5.52E-07 | 2.68E-06 |
| 3.289513047 | 0.001003609 | 0.002892049 |
| 10.81171769 | 3.03E-27 | 8.90E-26 |
| -0.374364544 | 0.708133118 | 0.786046918 |
| 12.6638703 | 9.38E-37 | 4.44E-35 |

| | | | |
|---|---|---|---|
| | 2.783659674 | 0.00537494 | 0.013124061 |
| | 1.718483873 | 0.085708393 | 0.146385608 |
| | -1.91350456 | 0.05568349 | 0.101536796 |
| | 9.20131602 | 3.54E-20 | 6.37E-19 |
| | 0.884931354 | 0.376193656 | 0.488949052 |
| | -6.458759492 | 1.06E-10 | 8.00E-10 |
| | -4.82547615 | 1.40E-06 | 6.41E-06 |
| | -1.633990354 | 0.102260876 | 0.16964487 |
| | -0.326800303 | 0.74381893 | 0.813588175 |
| | 16.20680344 | 4.51E-59 | 4.36E-57 |
| | 18.71980673 | 3.41E-78 | 5.27E-76 |
| NA | NA | NA | |
| | 21.06410464 | 1.70E-98 | 3.86E-96 |
| NA | NA | NA | |
| NA | NA | NA | |
| | 1.522997072 | 0.127759438 | 0.204568752 |
| | 6.353807498 | 2.10E-10 | 1.54E-09 |
| | 0.485130667 | 0.627583659 | 0.720635792 |
| | -3.581560971 | 0.000341547 | 0.00107557 |
| | 2.858222298 | 0.004260218 | 0.01064098 |
| | 13.80633925 | 2.33E-43 | 1.45E-41 |
| | 4.16874081 | 3.06E-05 | 0.000115684 |
| | -0.624319376 | 0.532417862 | 0.636835472 |
| | 0.498187659 | 0.61835178 | 0.713044552 |
| | 19.4902642 | 1.33E-84 | 2.29E-82 |
| | 18.31053546 | 6.82E-75 | 9.81E-73 |
| | -6.820908823 | 9.05E-12 | 7.59E-11 |
| | 3.571262746 | 0.000355264 | 0.001115018 |
| | 0.776802589 | 0.437275249 | 0.547578521 |
| | 7.028763804 | 2.08E-12 | 1.86E-11 |
| | 2.564277431 | 0.010339087 | 0.023483424 |
| | 0.31685483 | 0.751353758 | NA |
| | 2.588058432 | 0.009651861 | 0.022070717 |
| | -1.346152909 | 0.178253208 | 0.268970208 |
| | -8.091942566 | 5.87E-16 | 7.48E-15 |
| | -1.648183425 | 0.099315037 | 0.165668469 |
| | 10.52735249 | 6.46E-26 | 1.76E-24 |
| | 7.438665823 | 1.02E-13 | 1.05E-12 |
| | 10.841783 | 2.18E-27 | 6.52E-26 |
| NA | NA | NA | |
| | 1.772660601 | 0.076284963 | 0.132561277 |
| | 1.696143298 | 0.089858746 | 0.152438534 |
| | 1.677546302 | 0.093435703 | 0.157487923 |
| | 6.669736994 | 2.56E-11 | 2.07E-10 |
| NA | NA | NA | |
| | 2.369713044 | 0.017801896 | 0.037838099 |
| | 4.372400839 | 1.23E-05 | 4.93E-05 |

| | | |
|---|---|---|
| 2.749586058 | 0.005967059 | 0.014383233 |
| 5.345159582 | 9.03E-08 | 4.90E-07 |
| 3.682793819 | 0.000230692 | 0.000749022 |
| -4.268427389 | 1.97E-05 | 7.66E-05 |
| -4.927716164 | 8.32E-07 | 3.95E-06 |
| NA | NA | NA |
| -1.523217367 | 0.127704333 | 0.204507708 |
| -1.246428983 | 0.212606946 | 0.31049372 |
| NA | NA | NA |
| 8.276839064 | 1.26E-16 | 1.71E-15 |
| 9.788917117 | 1.26E-22 | 2.71E-21 |
| 0.581841494 | 0.560673452 | 0.662871688 |
| -2.535290879 | 0.011235399 | 0.025301128 |
| -0.340313773 | 0.733620246 | NA |
| NA | NA | NA |
| -6.210112882 | 5.29E-10 | 3.71E-09 |
| 4.589436789 | 4.44E-06 | 1.91E-05 |
| -1.976235658 | 0.048128093 | 0.089572581 |
| -1.809449295 | 0.070381231 | 0.123858694 |
| -0.469622766 | 0.638624556 | NA |
| -4.582430916 | 4.60E-06 | 1.97E-05 |
| -4.911596455 | 9.03E-07 | 4.26E-06 |
| 3.369591825 | 0.000752796 | 0.002221991 |
| 0.611119297 | 0.541120605 | 0.645179286 |
| 1.262248988 | 0.206859206 | 0.303765675 |
| 3.267737593 | 0.001084108 | 0.003101169 |
| -0.278273851 | 0.780802148 | 0.843441792 |
| 3.462899258 | 0.000534388 | 0.001621499 |
| 0.217787977 | 0.827594313 | NA |
| 0.742002798 | 0.45808564 | 0.567617999 |
| -3.152062463 | 0.001621216 | 0.004461195 |
| -7.409769262 | 1.27E-13 | 1.29E-12 |
| -0.503889364 | 0.614339125 | NA |
| -7.314537149 | 2.58E-13 | 2.55E-12 |
| 18.52435483 | 1.31E-76 | 1.96E-74 |
| -5.055698491 | 4.29E-07 | 2.12E-06 |
| 2.768022767 | 0.005639752 | 0.013690523 |
| NA | NA | NA |
| 3.791991904 | 0.000149444 | 0.000501082 |
| -0.078317958 | 0.937575133 | 0.957505634 |
| NA | NA | NA |
| 1.234723878 | 0.216933286 | 0.315550611 |
| 0.094589793 | 0.924640658 | 0.948190794 |
| -11.95361887 | 6.22E-33 | 2.51E-31 |
| 1.298979326 | 0.193951024 | 0.288243328 |
| -5.727824585 | 1.02E-08 | 6.21E-08 |
| 4.514701868 | 6.34E-06 | 2.66E-05 |

| | | |
|---|---|---|
| 9.093870587 | 9.56E-20 | 1.67E-18 |
| 4.457394082 | 8.30E-06 | 3.42E-05 |
| -6.442748216 | 1.17E-10 | 8.85E-10 |
| 3.190991189 | 0.001417856 | 0.003948919 |
| NA | NA | NA |
| 10.40365075 | 2.39E-25 | 6.24E-24 |
| NA | NA | NA |
| 5.865354695 | 4.48E-09 | 2.85E-08 |
| -5.027687446 | 4.96E-07 | 2.43E-06 |
| NA | NA | NA |
| 8.95114857 | 3.52E-19 | 5.90E-18 |
| -0.46307077 | 0.643313641 | 0.733148497 |
| -2.067257699 | 0.038709878 | 0.074058568 |
| -1.646531182 | 0.099654446 | 0.166129082 |
| NA | NA | NA |
| -0.292074227 | 0.770229872 | NA |
| 1.047451722 | 0.294891288 | 0.403186155 |
| 2.153934182 | 0.031245334 | 0.06150481 |
| 0.790584544 | 0.429186469 | 0.540341901 |
| 1.8951325 | 0.05807485 | 0.105204711 |
| 0.065474733 | 0.947796023 | NA |
| -1.391879283 | 0.163958956 | NA |
| 2.338393487 | 0.019366845 | 0.040626227 |
| 0.192810118 | 0.847107679 | NA |
| 0.420880312 | 0.673842483 | 0.757553559 |
| 0.606717611 | 0.544038333 | 0.647886548 |
| NA | NA | NA |
| -2.795351314 | 0.005184334 | 0.012705756 |
| -1.579347207 | 0.11425644 | 0.185994673 |
| -0.149332459 | 0.881291304 | 0.916668938 |
| -1.490285117 | 0.136149284 | 0.215371079 |
| -0.561640282 | 0.574361129 | 0.675607832 |
| -0.683478458 | 0.494304558 | 0.601436944 |
| -3.312034004 | 0.000926203 | 0.00268631 |
| -8.796760562 | 1.41E-18 | 2.24E-17 |
| -0.484541443 | 0.62800166 | 0.720983467 |
| 1.786509269 | 0.074016838 | 0.129148933 |
| 5.724276416 | 1.04E-08 | 6.33E-08 |
| -5.142862423 | 2.71E-07 | 1.38E-06 |
| -0.907897233 | 0.363932519 | 0.476374384 |
| -4.081869906 | 4.47E-05 | 0.000163744 |
| -3.901698902 | 9.55E-05 | 0.00033165 |
| 4.214452778 | 2.50E-05 | 9.58E-05 |
| 0.063708285 | 0.949202508 | NA |
| 0.475769381 | 0.634238686 | 0.725747203 |
| -0.822469004 | 0.410810021 | 0.522647902 |
| 3.276072912 | 0.001052614 | 0.003019705 |

| | | |
|---|---|---|
| 0.063708882 | 0.949202033 | NA |
| 0.490620235 | 0.623695072 | 0.717487253 |
| 1.627345586 | 0.103663721 | 0.171595989 |
| -4.381455164 | 1.18E-05 | 4.75E-05 |
| -0.086025746 | 0.931445951 | NA |
| -0.113951911 | 0.909275914 | 0.937401711 |
| 2.934704085 | 0.003338659 | 0.008529819 |
| 2.012047755 | 0.044214903 | 0.083248009 |
| 6.286142776 | 3.25E-10 | 2.33E-09 |
| -8.367287128 | 5.90E-17 | 8.17E-16 |
| 0.762500877 | 0.44576112 | NA |
| 7.058244411 | 1.69E-12 | 1.52E-11 |
| 0.874732936 | 0.381719235 | 0.494254433 |
| 1.5820352 | 0.113641535 | 0.185173336 |
| NA | NA | NA |
| 1.603691329 | 0.108782108 | 0.178471657 |
| 1.906887536 | 0.056535153 | 0.102877126 |
| NA | NA | NA |
| NA | NA | NA |
| NA | NA | NA |
| NA | NA | NA |
| NA | NA | NA |
| 0.192809521 | 0.847108147 | NA |
| NA | NA | NA |
| NA | NA | NA |
| 2.004430093 | 0.045024008 | 0.084548798 |
| NA | NA | NA |
| 0.192809521 | 0.847108147 | NA |
| NA | NA | NA |
| NA | NA | NA |
| 0.551758995 | 0.581113485 | NA |
| 0.3474247 | 0.728272279 | NA |
| 0.192809521 | 0.847108147 | NA |
| 1.638154235 | 0.101389524 | NA |
| 0.884977946 | 0.376168526 | 0.488949052 |
| NA | NA | NA |
| 1.444838006 | 0.148503393 | 0.231447069 |
| -7.218523581 | 5.26E-13 | 5.03E-12 |
| 0.94002377 | 0.347205368 | 0.458749491 |
| -3.024800095 | 0.002487975 | 0.006546202 |
| -2.086147472 | 0.036965267 | 0.071183307 |
| -0.868408634 | 0.38517067 | 0.497334117 |
| 9.585179396 | 9.23E-22 | 1.88E-20 |
| 4.194131173 | 2.74E-05 | 0.00010434 |
| 0.192809521 | 0.847108147 | NA |
| 5.251525933 | 1.51E-07 | 7.93E-07 |
| 5.746587138 | 9.11E-09 | 5.59E-08 |

| | | |
|---|---|---|
| -0.168635581 | 0.866083292 | 0.906517856 |
| 6.919245992 | 4.54E-12 | 3.93E-11 |
| 6.64815313 | 2.97E-11 | 2.38E-10 |
| 9.523318289 | 1.68E-21 | 3.36E-20 |
| -3.085390851 | 0.002032848 | 0.005468851 |
| 4.250063597 | 2.14E-05 | 8.27E-05 |
| 0.229804117 | 0.818243985 | 0.872296391 |
| -5.096962823 | 3.45E-07 | 1.73E-06 |
| NA | NA | NA |
| 2.648483378 | 0.008085383 | 0.018801294 |
| 4.705609718 | 2.53E-06 | 1.12E-05 |
| 17.7217687 | 2.85E-70 | 3.66E-68 |
| 4.133044983 | 3.58E-05 | 0.000133678 |
| -8.215319428 | 2.12E-16 | 2.80E-15 |
| -10.31117036 | 6.27E-25 | 1.59E-23 |
| 0.104674246 | 0.916634298 | 0.94278926 |
| NA | NA | NA |
| -5.484257415 | 4.15E-08 | 2.36E-07 |
| NA | NA | NA |
| -6.31004274 | 2.79E-10 | 2.01E-09 |
| -2.36741478 | 0.017912845 | 0.038022261 |
| 0.09451447 | 0.924700488 | 0.948190794 |
| 7.997561022 | 1.27E-15 | 1.57E-14 |
| 0.192809521 | 0.847108147 | NA |
| 8.093088738 | 5.82E-16 | 7.41E-15 |
| -2.786790061 | 0.005323295 | 0.013006841 |
| -3.101196611 | 0.001927403 | 0.005210378 |
| 2.318593933 | 0.020417064 | 0.042544049 |
| 7.845619301 | 4.31E-15 | 5.06E-14 |
| -4.566949812 | 4.95E-06 | 2.11E-05 |
| 1.511036405 | 0.130779176 | 0.20837754 |
| -3.907798099 | 9.31E-05 | 0.000324335 |
| 4.40720008 | 1.05E-05 | 4.25E-05 |
| -2.159381702 | 0.030820566 | 0.060757563 |
| NA | NA | NA |
| -0.708667909 | 0.478530584 | 0.587365486 |
| 3.30445216 | 0.000951622 | 0.002755246 |
| 2.303463176 | 0.021252797 | 0.044032572 |
| 2.247747992 | 0.024592264 | 0.049971129 |
| 1.329588058 | 0.183654035 | 0.275739937 |
| 7.385997994 | 1.51E-13 | 1.52E-12 |
| 1.722333791 | 0.08500907 | 0.145359818 |
| NA | NA | NA |
| NA | NA | NA |
| 2.455134141 | 0.014083203 | 0.030837421 |
| -1.658875231 | 0.097140937 | 0.162627092 |
| -0.463323876 | 0.643132234 | 0.733011644 |

| | | |
|---|---|---|
| -0.63913499 | 0.522735119 | 0.627954504 |
| -2.513757655 | 0.011945249 | 0.02666217 |
| -0.514918826 | 0.606609744 | 0.703393028 |
| -0.137825397 | 0.890378412 | 0.923312001 |
| -3.044345462 | 0.002331872 | 0.006178992 |
| 0.949566862 | 0.342332383 | 0.453652031 |
| -0.003507326 | 0.997201564 | 0.997999007 |
| -2.562990875 | 0.010377478 | 0.023557796 |
| 5.913249093 | 3.35E-09 | 2.16E-08 |
| -1.371711138 | 0.17015338 | 0.258776224 |
| -3.902626515 | 9.52E-05 | 0.000330656 |
| 0.366109976 | 0.714283014 | 0.790994625 |
| NA | NA | NA |
| -0.777413595 | 0.436914794 | 0.547269363 |
| NA | NA | NA |
| 0.669774747 | 0.503001394 | 0.609632234 |
| -2.258185955 | 0.023934069 | 0.048827919 |
| 1.926259359 | 0.054071997 | 0.099042963 |
| 0.063708285 | 0.949202508 | NA |
| NA | NA | NA |
| -4.102615587 | 4.09E-05 | 0.000150832 |
| NA | NA | NA |
| NA | NA | NA |
| -0.875121393 | 0.381507858 | 0.494057413 |
| 1.422046974 | 0.155012615 | 0.239874235 |
| 3.37112139 | 0.000748629 | 0.002211255 |
| 8.414836046 | 3.93E-17 | 5.55E-16 |
| NA | NA | NA |
| 4.86188569 | 1.16E-06 | 5.40E-06 |
| -0.968236004 | 0.332926517 | 0.444126778 |
| -2.139685716 | 0.032380174 | 0.063487056 |
| -1.461952918 | 0.143754112 | 0.225420713 |
| -0.125316225 | 0.900273208 | NA |
| NA | NA | NA |
| -7.761809772 | 8.37E-15 | 9.61E-14 |
| 1.018655759 | 0.308366423 | 0.418206088 |
| 4.301968631 | 1.69E-05 | 6.65E-05 |
| -1.964202552 | 0.049506607 | 0.091809319 |
| -2.087471813 | 0.036845506 | 0.07100185 |
| -1.997618586 | 0.045758027 | 0.085734217 |
| -3.518643441 | 0.000433759 | 0.001337892 |
| -3.981059273 | 6.86E-05 | 0.000244137 |
| 7.937788411 | 2.06E-15 | 2.50E-14 |
| NA | NA | NA |
| 2.273558235 | 0.022992561 | 0.047195654 |
| NA | NA | NA |
| -0.811577066 | 0.417034356 | NA |

| | | |
|---|---|---|
| -7.298943507 | 2.90E-13 | 2.85E-12 |
| -4.74534258 | 2.08E-06 | 9.35E-06 |
| -1.575680639 | 0.115099422 | 0.18720765 |
| 2.994565977 | 0.002748356 | 0.007164273 |
| 0.192808924 | 0.847108615 | NA |
| 6.719387332 | 1.82E-11 | 1.49E-10 |
| 0.043976661 | 0.964923008 | 0.976154833 |
| -0.26033994 | 0.794601567 | 0.854003058 |
| 0.350882206 | 0.725676721 | 0.799739244 |
| -1.990007449 | 0.046590115 | 0.087097661 |
| -3.338334275 | 0.000842823 | 0.002463894 |
| NA | NA | NA |
| 0.235444205 | 0.813863979 | 0.869070623 |
| -0.07389278 | 0.941095701 | 0.959886935 |
| 1.389263386 | 0.164752673 | 0.25201732 |
| -6.602652393 | 4.04E-11 | 3.20E-10 |
| -0.820790652 | 0.411765524 | 0.52343784 |
| -5.626138708 | 1.84E-08 | 1.09E-07 |
| -9.410393329 | 4.94E-21 | 9.49E-20 |
| -3.245004443 | 0.001174488 | 0.00333569 |
| -1.855263213 | 0.063558649 | 0.113447406 |
| -1.697594888 | 0.089584248 | 0.15204053 |
| -4.13964772 | 3.48E-05 | 0.00013018 |
| NA | NA | NA |
| NA | NA | NA |
| 1.286212665 | 0.198368852 | 0.293658172 |
| NA | NA | NA |
| 1.098930115 | 0.27179855 | 0.378245174 |
| NA | NA | NA |
| 2.220296313 | 0.02639866 | 0.053129136 |
| 2.191094351 | 0.028444965 | 0.056704164 |
| -5.525671023 | 3.28E-08 | 1.89E-07 |
| -10.02714376 | 1.16E-23 | 2.71E-22 |
| -1.68104024 | 0.092755099 | 0.156541037 |
| -5.483946277 | 4.16E-08 | 2.36E-07 |
| -5.0607334 | 4.18E-07 | 2.07E-06 |
| -6.408602044 | 1.47E-10 | 1.09E-09 |
| -3.143337437 | 0.001670332 | 0.004582747 |
| 0.063708285 | 0.949202508 | NA |
| NA | NA | NA |
| -2.408113267 | 0.016035205 | 0.034573278 |
| -0.949879521 | 0.342173474 | 0.453569696 |
| 1.727637085 | 0.08405331 | 0.144030557 |
| 1.210326122 | 0.226153778 | 0.326257113 |
| -14.26332394 | 3.70E-46 | 2.57E-44 |
| -1.533816689 | 0.125074756 | 0.200951354 |
| -5.124621462 | 2.98E-07 | 1.51E-06 |

| | | |
|---|---|---|
| -4.670990414 | 3.00E-06 | 1.32E-05 |
| 0.147699503 | 0.882579924 | 0.9177042 |
| 0.959495954 | 0.337308957 | 0.44853866 |
| -2.457401627 | 0.013994613 | 0.030670271 |
| -3.760572199 | 0.000169525 | 0.00056291 |
| -0.065389369 | 0.947863988 | NA |
| -0.46146574 | 0.644464496 | 0.733892007 |
| 2.411287031 | 0.015896333 | 0.034300449 |
| -4.567102083 | 4.95E-06 | 2.10E-05 |
| -4.685840175 | 2.79E-06 | 1.23E-05 |
| 1.128496211 | 0.259110416 | 0.364130802 |
| 8.63231025 | 6.01E-18 | 9.00E-17 |
| 7.938849761 | 2.04E-15 | 2.48E-14 |
| 0.112446825 | 0.91046913 | 0.938322655 |
| NA | NA | NA |
| NA | NA | NA |
| 0.442351314 | 0.658235004 | 0.745773228 |
| -7.382694287 | 1.55E-13 | 1.56E-12 |
| NA | NA | NA |
| 3.646874921 | 0.000265449 | 0.000851492 |
| 1.648807115 | 0.099187157 | 0.165488237 |
| 3.568367544 | 0.000359212 | 0.001125856 |
| 10.5952785 | 3.13E-26 | 8.67E-25 |
| -0.048636072 | 0.961209322 | 0.973530124 |
| 0.063708285 | 0.949202508 | NA |
| 1.300126787 | 0.193557518 | 0.287799529 |
| -2.533040394 | 0.011307792 | 0.025436689 |
| 0.841831159 | 0.399882468 | 0.51163207 |
| -3.50514385 | 0.000456361 | 0.001401387 |
| NA | NA | NA |
| 4.594405745 | 4.34E-06 | 1.87E-05 |
| -7.007971304 | 2.42E-12 | 2.14E-11 |
| NA | NA | NA |
| 1.531471551 | 0.125652892 | 0.201737774 |
| 0.351028164 | 0.725567219 | NA |
| NA | NA | NA |
| -2.662146616 | 0.007764405 | 0.018140862 |
| 2.03153645 | 0.042200606 | 0.079870031 |
| 3.282517095 | 0.001028847 | 0.002958306 |
| 0.855785524 | 0.392116424 | 0.50404037 |
| 3.752867786 | 0.000174823 | 0.000579119 |
| -2.779800879 | 0.005439224 | 0.013265495 |
| 0.063708882 | 0.949202033 | NA |
| -1.32849766 | 0.184013756 | 0.27614751 |
| -0.188156586 | 0.850753904 | 0.896152269 |
| 1.805784431 | 0.070952018 | 0.124731635 |
| 7.292440361 | 3.04E-13 | 2.98E-12 |

| | | |
|---|---|---|
| NA | NA | NA |
| -0.743661907 | 0.45708104 | 0.566709849 |
| -0.510358349 | 0.609800431 | 0.7060654 |
| 7.182679943 | 6.84E-13 | 6.47E-12 |
| 0.063708285 | 0.949202508 | NA |
| -3.820615772 | 0.000133119 | 0.000450266 |
| -0.660590897 | 0.508874708 | 0.61496398 |
| 1.000027927 | 0.317296993 | 0.427791206 |
| -3.426988595 | 0.000610315 | 0.001831237 |
| -0.639613085 | 0.522424173 | 0.627701472 |
| -4.25353234 | 2.10E-05 | 8.15E-05 |
| -0.434405845 | 0.663993754 | NA |
| NA | NA | NA |
| -4.368220765 | 1.25E-05 | 5.02E-05 |
| NA | NA | NA |
| NA | NA | NA |
| 0.06371127 | 0.949200131 | NA |
| 0.063708882 | 0.949202033 | NA |
| NA | NA | NA |
| 0.192810118 | 0.847107679 | NA |
| 2.124621008 | 0.033618249 | 0.065580608 |
| -0.066649018 | 0.946861122 | NA |
| 3.177368319 | 0.001486182 | 0.004120417 |
| 1.08256714 | 0.279000599 | 0.386031181 |
| -1.028268324 | 0.303823627 | 0.413276745 |
| -2.198853404 | 0.027888348 | 0.055727714 |
| -0.523957897 | 0.600307824 | 0.698090728 |
| -1.146881989 | 0.251430398 | 0.355232719 |
| 0.291366177 | 0.77077128 | NA |
| 0.063710673 | 0.949200606 | NA |
| -3.197562191 | 0.001385945 | 0.00386779 |
| -10.41468488 | 2.13E-25 | 5.58E-24 |
| 1.42845151 | 0.153161942 | 0.237480233 |
| 2.61616926 | 0.008892247 | 0.020502297 |
| -2.273368008 | 0.023004012 | 0.047215287 |
| 3.172816152 | 0.001509681 | 0.00417955 |
| -5.220811423 | 1.78E-07 | 9.27E-07 |
| 1.091873731 | 0.274888605 | 0.381611857 |
| 0.363716865 | 0.716069448 | NA |
| -8.176280558 | 2.93E-16 | 3.82E-15 |
| -3.377053401 | 0.000732668 | 0.002168206 |
| 1.157250513 | 0.247170025 | 0.350320823 |
| -1.795388164 | 0.072591877 | 0.127034516 |
| -2.064079502 | 0.039010179 | 0.074536243 |
| 5.022431083 | 5.10E-07 | 2.49E-06 |
| 1.211392393 | 0.225745053 | 0.325827191 |
| 1.407585123 | 0.159253955 | 0.245075503 |

| | | | |
|---|---|---|---|
| | -0.57995714 | 0.561943521 | 0.664059597 |
| | -2.593336804 | 0.009504964 | 0.021766628 |
| | 0.063708285 | 0.949202508 | NA |
| | 7.08210658 | 1.42E-12 | 1.29E-11 |
| | 1.893008589 | 0.058356727 | 0.105600137 |
| | 3.08578147 | 0.00203018 | 0.005464021 |
| NA | NA | NA | |
| | -0.05499807 | 0.956140001 | 0.970056945 |
| | -1.401628696 | 0.161026153 | 0.24720833 |
| | -1.641387164 | 0.100717073 | 0.167504872 |
| NA | NA | NA | |
| NA | NA | NA | |
| | -11.2412609 | 2.56E-29 | 8.50E-28 |
| | 0.033605595 | 0.97319166 | 0.981474724 |
| | -0.158974005 | 0.873689354 | 0.912000568 |
| NA | NA | NA | |
| | -0.754205101 | 0.450726075 | 0.560663635 |
| | -3.815593103 | 0.000135856 | 0.000458904 |
| | 5.402004761 | 6.59E-08 | 3.64E-07 |
| | 0.436271948 | 0.662639432 | 0.749034231 |
| | 8.522888518 | 1.56E-17 | 2.26E-16 |
| | -4.214349443 | 2.50E-05 | 9.59E-05 |
| | -0.19448762 | 0.845794089 | NA |
| | 2.205167822 | 0.02744232 | 0.054972491 |
| | -7.2798985 | 3.34E-13 | 3.26E-12 |
| | 0.192809521 | 0.847108147 | NA |
| | 0.336410497 | 0.736561335 | NA |
| | 0.719298335 | 0.471957122 | 0.580866733 |
| | -0.118378285 | 0.905767931 | 0.934747724 |
| | -0.604346624 | 0.545613215 | 0.64935162 |
| | -5.069142994 | 4.00E-07 | 1.99E-06 |
| | -1.099689187 | 0.271467568 | 0.37782658 |
| | -5.788325965 | 7.11E-09 | 4.41E-08 |
| | -1.677720272 | 0.09340172 | 0.157451849 |
| | 4.204400182 | 2.62E-05 | 0.000100019 |
| | -0.557895695 | 0.576915611 | 0.677943556 |
| | -1.218371431 | 0.223082857 | 0.322664759 |
| | 0.756799832 | 0.449169794 | NA |
| | 8.838166029 | 9.73E-19 | 1.57E-17 |
| | 0.195400213 | 0.84507965 | 0.891713824 |
| | 0.473367423 | 0.635951067 | 0.727307649 |
| | 0.324874675 | 0.745275925 | NA |
| | 0.374599278 | 0.70795851 | NA |
| | -1.900127738 | 0.057416358 | 0.104200366 |
| | -0.461128115 | 0.644706691 | NA |
| NA | NA | NA | |
| | 4.661674214 | 3.14E-06 | 1.37E-05 |

| | | |
|---|---|---|
| -1.148988469 | 0.250560736 | 0.354203954 |
| -6.559546793 | 5.40E-11 | 4.22E-10 |
| 4.996914634 | 5.83E-07 | 2.82E-06 |
| -0.102531454 | 0.918334847 | NA |
| -1.387851265 | 0.16518234 | 0.252612817 |
| -0.778699991 | 0.436156461 | 0.546567359 |
| NA | NA | NA |
| 0.192809521 | 0.847108147 | NA |
| NA | NA | NA |
| 2.829879005 | 0.004656561 | 0.011537688 |
| 0.152951128 | 0.878436816 | 0.91476425 |
| NA | NA | NA |
| -1.422004271 | 0.155025011 | 0.239878587 |
| 1.559129403 | 0.118965756 | 0.192483055 |
| -0.598495438 | 0.549509402 | 0.652910867 |
| -1.765323081 | 0.077509483 | 0.134409197 |
| -3.900811349 | 9.59E-05 | 0.000332638 |
| -0.286955474 | 0.774146413 | 0.837654937 |
| -0.194488217 | 0.845793621 | NA |
| -0.734235176 | 0.462805418 | 0.572292243 |
| 4.872936326 | 1.10E-06 | 5.13E-06 |
| 5.174120327 | 2.29E-07 | 1.18E-06 |
| 10.23686962 | 1.36E-24 | 3.36E-23 |
| NA | NA | NA |
| 9.218256981 | 3.02E-20 | 5.45E-19 |
| 6.878361327 | 6.05E-12 | 5.18E-11 |
| -0.065390563 | 0.947863038 | NA |
| -5.981757444 | 2.21E-09 | 1.45E-08 |
| -9.065679773 | 1.24E-19 | 2.15E-18 |
| 3.694255937 | 0.000220532 | 0.000718644 |
| -0.723839665 | 0.469164175 | 0.57833929 |
| 14.8248962 | 1.01E-49 | 7.77E-48 |
| 0.192810118 | 0.847107679 | NA |
| -6.619625884 | 3.60E-11 | 2.87E-10 |
| 0.259705945 | 0.795090607 | 0.854198553 |
| -1.088398226 | 0.276419344 | 0.383248002 |
| 0.391022247 | 0.695780792 | 0.775914155 |
| -1.808462332 | 0.070534574 | 0.124084931 |
| 1.069208789 | 0.284975599 | 0.392102048 |
| 0.423991594 | 0.671571943 | 0.755752439 |
| 5.450879381 | 5.01E-08 | 2.81E-07 |
| 0.192809521 | 0.847108147 | NA |
| 1.276305617 | 0.201847505 | 0.298032271 |
| -0.150714291 | 0.880201099 | NA |
| -0.065389369 | 0.947863988 | NA |
| NA | NA | NA |
| 1.487321272 | 0.136929979 | 0.216455562 |

| | | |
|---|---|---|
| -0.194487022 | 0.845794556 | NA |
| NA | NA | NA |
| NA | NA | NA |
| -0.371491365 | 0.710271587 | 0.787664873 |
| NA | NA | NA |
| 4.63323008 | 3.60E-06 | 1.56E-05 |
| NA | NA | NA |
| NA | NA | NA |
| NA | NA | NA |
| NA | NA | NA |
| 1.167564371 | 0.242982529 | NA |
| NA | NA | NA |
| 5.397355176 | 6.76E-08 | 3.72E-07 |
| NA | NA | NA |
| 1.667574888 | 0.095400147 | 0.16021643 |
| NA | NA | NA |
| -0.502689565 | 0.615182547 | NA |
| NA | NA | NA |
| 0.230473895 | 0.817723546 | NA |
| 1.589207008 | 0.112013665 | 0.182901822 |
| 0.104456909 | NA | NA |
| -0.977268961 | 0.328436015 | 0.439471478 |
| -4.133715978 | 3.57E-05 | 0.000133328 |
| 0.324874675 | 0.745275925 | NA |
| NA | NA | NA |
| -0.097727738 | 0.922148489 | NA |
| 0.346807984 | 0.728735578 | NA |
| -0.279068303 | 0.780192407 | NA |
| 6.10656151 | 1.02E-09 | 6.94E-09 |
| 3.255525235 | 0.001131828 | 0.00322441 |
| 0.969523586 | 0.332284018 | 0.443458756 |
| 2.339093184 | 0.019330611 | 0.040577424 |
| -3.841948614 | 0.000122061 | 0.000415616 |
| 1.312859777 | 0.189230203 | 0.282519056 |
| 9.171618373 | 4.66E-20 | 8.34E-19 |
| -4.832299355 | 1.35E-06 | 6.21E-06 |
| 0.438541297 | 0.660993941 | 0.747748275 |
| 12.21032866 | 2.74E-34 | 1.16E-32 |
| 1.52145094 | 0.128146714 | 0.205103513 |
| -0.951476086 | 0.341362752 | 0.452894609 |
| 5.451002445 | 5.01E-08 | 2.81E-07 |
| 4.322215761 | 1.54E-05 | 6.11E-05 |
| 1.445099526 | 0.148429932 | 0.231346983 |
| 1.200382376 | 0.22999087 | 0.33060389 |
| 0.618883426 | 0.535993158 | 0.64034694 |
| -1.114382227 | 0.265115245 | 0.370711196 |
| NA | NA | NA |

| | | |
|---|---|---|
| 5.634021245 | 1.76E-08 | 1.05E-07 |
| 1.012400127 | 0.311346778 | 0.421539993 |
| -0.737375883 | 0.460893797 | 0.570307086 |
| 0.192810118 | 0.847107679 | NA |
| 6.718492782 | 1.84E-11 | 1.50E-10 |
| 4.863901957 | 1.15E-06 | 5.35E-06 |
| 1.104754221 | 0.269266102 | 0.375388801 |
| -9.96320543 | 2.21E-23 | 5.07E-22 |
| 0.412128299 | 0.680245388 | 0.763043025 |
| -1.282594184 | 0.199634289 | 0.29526953 |
| -0.870860039 | 0.383830578 | 0.496166884 |
| NA | NA | NA |
| 1.280178307 | 0.200482433 | 0.296342466 |
| NA | NA | NA |
| 3.442305874 | 0.000576778 | 0.001739576 |
| 1.395193769 | 0.162957416 | 0.249759605 |
| NA | NA | NA |
| -2.766554074 | 0.005665219 | 0.013749678 |
| 0.415075566 | 0.678086586 | NA |
| NA | NA | NA |
| NA | NA | NA |
| NA | NA | NA |
| NA | NA | NA |
| NA | NA | NA |
| NA | NA | NA |
| NA | NA | NA |
| NA | NA | NA |
| 0.100944874 | 0.919594221 | 0.94484805 |
| NA | NA | NA |
| 0.192808924 | 0.847108615 | NA |
| 0.30922356 | 0.757151473 | NA |
| 0.414944476 | 0.67818255 | 0.761172558 |
| NA | NA | NA |
| 0.214124678 | 0.830449832 | 0.881400519 |
| -0.364419319 | 0.715544912 | NA |
| 0.08251396 | 0.934238018 | 0.95530589 |
| 0.109227158 | 0.91302232 | NA |
| -0.385650291 | 0.699755682 | NA |
| -1.618566564 | 0.105540554 | 0.174081059 |
| 0.490924191 | 0.623480067 | 0.717338822 |
| -3.931475998 | 8.44E-05 | 0.000295964 |
| 0.459757722 | 0.645690133 | 0.734986767 |
| 4.861041244 | 1.17E-06 | 5.42E-06 |
| 6.135897899 | 8.47E-10 | 5.80E-09 |
| -1.115340291 | 0.264704626 | 0.370308338 |
| 7.681153587 | 1.58E-14 | 1.76E-13 |
| 0.341471887 | 0.732748366 | 0.805193239 |

| | | | |
|---|---|---|---|
| | 12.7643212 | 2.59E-37 | 1.26E-35 |
| | 0.192808924 | 0.847108615 | NA |
| | -0.10795104 | 0.91403453 | NA |
| | 0.252471443 | 0.800676685 | 0.858479903 |
| | -0.071591321 | 0.942927147 | NA |
| | 19.03165736 | 9.33E-81 | 1.47E-78 |
| | -8.065276398 | 7.31E-16 | 9.25E-15 |
| | -5.089940597 | 3.58E-07 | 1.79E-06 |
| | -1.08117193 | 0.279620642 | 0.38659438 |
| | -0.065389369 | 0.947863988 | NA |
| NA | | NA | NA |
| | -0.245949289 | 0.805721488 | NA |
| NA | | NA | NA |
| NA | | NA | NA |
| NA | | NA | NA |
| NA | | NA | NA |
| | 9.630421045 | 5.95E-22 | 1.23E-20 |
| | 0.662508019 | 0.507645697 | NA |
| | 1.677122294 | 0.09351857 | 0.15758515 |
| | 0.192810118 | 0.847107679 | NA |
| | -2.336760918 | 0.019451619 | 0.04079722 |
| | 0.304123487 | 0.761033814 | 0.827124462 |
| | -0.869176065 | 0.384750837 | 0.496920199 |
| | -4.252057824 | 2.12E-05 | 8.20E-05 |
| | -1.460454845 | 0.144165109 | 0.225923659 |
| | 1.348892545 | 0.177371483 | 0.26793049 |
| | -1.951916747 | 0.05094809 | 0.094112966 |
| NA | | NA | NA |
| | -1.868708453 | 0.061663387 | 0.110569137 |
| | -0.040999366 | 0.967296401 | 0.977647039 |
| | -1.619789546 | 0.105277493 | 0.173761664 |
| | 0.61247514 | 0.540223441 | 0.644416562 |
| | 4.720629201 | 2.35E-06 | 1.05E-05 |
| | -1.354699647 | 0.17551327 | 0.265732952 |
| | -2.133472667 | 0.032885972 | 0.064377903 |
| | -0.745764332 | 0.45580979 | NA |
| | 2.104916504 | 0.035298577 | 0.068401281 |
| | -1.98692304 | 0.046930926 | 0.087636607 |
| | -0.882010193 | 0.377771295 | 0.490411053 |
| | -2.464175025 | 0.013732903 | 0.030163376 |
| | 0.40592812 | 0.684795416 | NA |
| | -6.523903259 | 6.85E-11 | 5.30E-10 |
| | -3.703731236 | 0.000212451 | 0.000694665 |
| | -4.389615713 | 1.14E-05 | 4.59E-05 |
| NA | | NA | NA |
| | 1.850565575 | 0.064232076 | 0.114445392 |
| | 2.396246194 | 0.016563961 | 0.035563213 |

| | | | |
|---|---|---|---|
| | 1.15132115 | 0.249600139 | 0.353065357 |
| | -5.11479753 | 3.14E-07 | 1.59E-06 |
| | 7.613640308 | 2.66E-14 | 2.90E-13 |
| | 2.160768916 | 0.030713194 | 0.060569749 |
| | 1.677355452 | 0.093472995 | 0.157518957 |
| | 5.868834558 | 4.39E-09 | 2.79E-08 |
| | -0.349439332 | 0.726759509 | 0.800404388 |
| | -5.814289326 | 6.09E-09 | 3.81E-08 |
| | -2.799543471 | 0.005117493 | 0.012560388 |
| | 1.313188008 | 0.189119604 | NA |
| | -2.964752469 | 0.003029266 | 0.007824821 |
| | -2.260792791 | 0.023772091 | 0.048552872 |
| | 1.641314868 | 0.100732071 | 0.167518686 |
| | 0.721111472 | 0.470840933 | 0.579808847 |
| | -2.274070705 | 0.022961736 | 0.047143979 |
| | -3.060668272 | 0.002208436 | 0.00588932 |
| | -0.065389369 | 0.947863988 | NA |
| | 0.477768774 | 0.632814789 | 0.724482192 |
| | 0.851587782 | 0.394442923 | NA |
| | -8.448416946 | 2.95E-17 | 4.19E-16 |
| | 0.105649443 | 0.915860494 | 0.942210776 |
| | 1.456650494 | 0.145212882 | 0.227295256 |
| NA | NA | NA | |
| | 7.689258951 | 1.48E-14 | 1.66E-13 |
| | 0.215481986 | 0.829391555 | 0.880766679 |
| | 6.114881998 | 9.66E-10 | 6.59E-09 |
| | 3.286136023 | 0.001015719 | 0.002923917 |
| | 0.192809521 | 0.847108147 | NA |
| | 0.083963038 | 0.933085819 | NA |
| | -0.194487022 | 0.845794556 | NA |
| | 2.672531549 | 0.00752813 | 0.017668117 |
| | 4.226341982 | 2.38E-05 | 9.12E-05 |
| | 1.13633853 | 0.255814911 | 0.36032905 |
| NA | NA | NA | |
| | -2.925241393 | 0.003441891 | 0.008771181 |
| | -10.56509065 | 4.33E-26 | 1.19E-24 |
| NA | NA | NA | |
| NA | NA | NA | |
| NA | NA | NA | |
| | -2.873237684 | 0.004062883 | 0.010197951 |
| NA | NA | NA | |
| | 0.407425151 | 0.683695758 | 0.766151039 |
| NA | NA | NA | |
| | -7.095922926 | 1.28E-12 | 1.18E-11 |
| NA | NA | NA | |
| | -0.024042028 | 0.980819085 | 0.986613469 |
| | -1.255739145 | 0.209210562 | 0.306432683 |

| | | |
|---|---|---|
| 2.857810985 | 0.004265744 | 0.010652656 |
| 7.972616457 | 1.55E-15 | 1.90E-14 |
| 0.770017514 | 0.441289503 | 0.551373876 |
| -0.50424194 | 0.614091372 | 0.709691383 |
| NA | NA | NA |
| 3.413388045 | 0.000641605 | 0.001917309 |
| NA | NA | NA |
| 1.118173847 | 0.263492752 | 0.368984199 |
| -6.657338416 | 2.79E-11 | 2.24E-10 |
| NA | NA | NA |
| NA | NA | NA |
| 1.776578742 | 0.075637576 | 0.131573322 |
| -0.07734905 | 0.938345871 | 0.958058216 |
| NA | NA | NA |
| NA | NA | NA |
| 7.235052217 | 4.65E-13 | 4.48E-12 |
| -3.958200945 | 7.55E-05 | 0.000267087 |
| 3.761743002 | 0.000168733 | 0.000560578 |
| 7.944804682 | 1.94E-15 | 2.37E-14 |
| 11.48151164 | 1.63E-30 | 5.78E-29 |
| 0.709429196 | 0.478058176 | 0.586846304 |
| -3.035734373 | 0.002399507 | 0.006340087 |
| 0.409712733 | 0.682016689 | NA |
| 13.89577989 | 6.72E-44 | 4.35E-42 |
| 1.299568174 | 0.193749013 | NA |
| NA | NA | NA |
| 0.394978309 | 0.692858896 | NA |
| 3.600035648 | 0.000318174 | 0.001007067 |
| 1.875331758 | 0.060747097 | 0.10914506 |
| 0.063708285 | 0.949202508 | NA |
| -0.402177683 | 0.687553265 | 0.7694817 |
| 5.70579592 | 1.16E-08 | 7.03E-08 |
| 1.280214639 | 0.200469659 | NA |
| 10.24515008 | 1.24E-24 | 3.09E-23 |
| NA | NA | NA |
| NA | NA | NA |
| 2.555298134 | 0.010609688 | 0.024042178 |
| -5.806145668 | 6.39E-09 | 3.99E-08 |
| -3.506108124 | 0.00045471 | 0.001397005 |
| 4.904923616 | 9.35E-07 | 4.41E-06 |
| 0.60819849 | 0.543055836 | 0.647024352 |
| 0.087569491 | 0.930218852 | 0.952503424 |
| -0.340668234 | 0.733353355 | 0.80539838 |
| -0.653200477 | 0.513627043 | 0.619212041 |
| 0.694513428 | 0.487360283 | 0.59543261 |
| 6.998324915 | 2.59E-12 | 2.29E-11 |
| -3.056519386 | 0.00223923 | 0.005962545 |

| | | | |
|---|---|---|---|
| | 3.389336881 | 0.000700619 | 0.002079754 |
| | 0.985073845 | 0.3245878 | NA |
| | 1.226139828 | 0.22014606 | 0.319268549 |
| | -5.309508273 | 1.10E-07 | 5.89E-07 |
| | 0.063707194 | 0.949203376 | NA |
| | 0.419238142 | 0.675042099 | 0.758532155 |
| | 2.796370154 | 0.005168017 | 0.012670729 |
| | 0.199526594 | 0.841850842 | NA |
| | -8.414124294 | 3.96E-17 | 5.58E-16 |
| | 0.06353712 | 0.949338802 | NA |
| | -1.06374222 | 0.287445493 | NA |
| | -0.691136427 | 0.48947981 | 0.59720593 |
| | -6.780864814 | 1.19E-11 | 9.95E-11 |
| | -0.317887001 | 0.750570653 | NA |
| | -0.506522449 | 0.612489934 | 0.70817734 |
| | -2.624176859 | 0.008685867 | 0.020078944 |
| NA | NA | NA | |
| | -0.908683373 | 0.363517283 | 0.475930411 |
| | -0.065389369 | 0.947863988 | NA |
| | -1.241154856 | 0.214548548 | 0.312818207 |
| | 3.873306494 | 0.000107369 | 0.000369255 |
| | 0.735972527 | 0.461747417 | 0.571137471 |
| NA | NA | NA | |
| | 0.192808327 | 0.847109082 | NA |
| | 3.957732902 | 7.57E-05 | 0.000267535 |
| | -1.662771449 | 0.096358194 | 0.161554397 |
| | -0.418948543 | 0.675253739 | 0.758699855 |
| | 5.475026743 | 4.37E-08 | 2.48E-07 |
| | -7.588849328 | 3.23E-14 | 3.49E-13 |
| | 0.342276862 | 0.732142547 | 0.804739502 |
| | 0.512454904 | 0.608332674 | NA |
| | 2.797906318 | 0.005143503 | 0.012618041 |
| | -6.559886576 | 5.38E-11 | 4.22E-10 |
| | 0.192810118 | 0.847107679 | NA |
| NA | NA | NA | |
| | -2.182814082 | 0.029049504 | 0.057716232 |
| | -6.114660065 | 9.68E-10 | 6.60E-09 |
| NA | NA | NA | |
| | 0.129253303 | 0.89715722 | NA |
| | 8.370126125 | 5.76E-17 | 7.99E-16 |
| | 0.178648807 | 0.858213468 | 0.901619438 |
| | -0.076654729 | 0.938898219 | 0.95842669 |
| NA | NA | NA | |
| | -2.696340196 | 0.007010603 | 0.016595821 |
| | -1.66216284 | 0.096480129 | 0.161704675 |
| | 0.063708285 | 0.949202508 | NA |
| | 0.333073926 | 0.739078481 | NA |

| | | |
|---|---|---|
| NA | NA | NA |
| 0.258866089 | 0.795738564 | 0.8546379 |
| 1.00101818 | 0.316818019 | 0.42732958 |
| -4.029113745 | 5.60E-05 | 0.000202422 |
| NA | NA | NA |
| -0.085117327 | 0.932168115 | NA |
| NA | NA | NA |
| -3.897593588 | 9.72E-05 | 0.00033676 |
| 5.175154543 | 2.28E-07 | 1.17E-06 |
| -5.90820881 | 3.46E-09 | 2.23E-08 |
| 0.385282141 | 0.700028391 | 0.77939141 |
| 12.687198 | 6.96E-37 | 3.31E-35 |
| 11.09595061 | 1.31E-28 | 4.23E-27 |
| 1.253727602 | 0.209941032 | NA |
| -1.909724304 | 0.05616872 | 0.102277249 |
| 2.525577127 | 0.011550842 | 0.025911237 |
| -2.963418501 | 0.003042426 | 0.007853953 |
| NA | NA | NA |
| -1.42157583 | 0.155149426 | 0.240041422 |
| NA | NA | NA |
| 0.478608033 | 0.632217502 | NA |
| -1.964225706 | 0.049503923 | 0.091809319 |
| 0.113007313 | 0.910024758 | 0.937941926 |
| 0.320071032 | 0.748914485 | NA |
| 0.063708285 | 0.949202508 | NA |
| -0.748145396 | 0.454372462 | 0.564218025 |
| 0.051296625 | 0.959089158 | 0.971824678 |
| -2.593896938 | 0.009489493 | 0.021739155 |
| -7.050195326 | 1.79E-12 | 1.61E-11 |
| NA | NA | NA |
| -2.259758496 | 0.023836244 | 0.048664044 |
| -6.748630536 | 1.49E-11 | 1.23E-10 |
| -1.64622749 | 0.099716932 | 0.166161694 |
| 3.451689333 | 0.000557089 | 0.001685271 |
| -1.677679045 | 0.093409772 | 0.157454819 |
| -1.084488706 | 0.278148172 | 0.385132972 |
| 0.063708285 | 0.949202508 | NA |
| -0.065390563 | 0.947863038 | NA |
| 7.147763456 | 8.82E-13 | 8.24E-12 |
| -2.599225996 | 0.009343424 | 0.021433348 |
| NA | NA | NA |
| -4.510264194 | 6.47E-06 | 2.71E-05 |
| 5.576123721 | 2.46E-08 | 1.43E-07 |
| -0.813796877 | 0.415761324 | 0.527365629 |
| 0.063710673 | 0.949200606 | NA |
| -3.40310967 | 0.000666235 | 0.001985223 |
| -5.118025153 | 3.09E-07 | 1.56E-06 |

| | | |
|---|---|---|
| -0.20574224 | 0.836992262 | 0.885985448 |
| -8.338239914 | 7.54E-17 | 1.04E-15 |
| -1.635751367 | 0.101891634 | 0.169143627 |
| 4.33233362 | 1.48E-05 | 5.85E-05 |
| -0.019632086 | 0.984336868 | 0.989039051 |
| 5.501824299 | 3.76E-08 | 2.15E-07 |
| 6.822245718 | 8.96E-12 | 7.53E-11 |
| NA | NA | NA |
| -7.98046723 | 1.46E-15 | 1.79E-14 |
| -4.414407641 | 1.01E-05 | 4.13E-05 |
| 6.19376773 | 5.87E-10 | 4.09E-09 |
| -0.658367827 | 0.510301804 | 0.616153119 |
| 3.168553197 | 0.001531997 | 0.004237583 |
| -0.972107658 | 0.330996993 | 0.441882486 |
| -1.461397734 | 0.143906322 | 0.225546352 |
| -3.994171166 | 6.49E-05 | 0.000231775 |
| 7.209977307 | 5.60E-13 | 5.35E-12 |
| NA | NA | NA |
| -4.756356924 | 1.97E-06 | 8.89E-06 |
| -1.267473529 | 0.204986023 | 0.301545704 |
| -0.597473503 | 0.550191293 | 0.653497022 |
| -1.7596184 | 0.07847253 | 0.135825145 |
| -0.071942718 | 0.942647494 | 0.96104292 |
| 0.43684664 | 0.662222572 | 0.748698269 |
| -0.065389369 | 0.947863988 | NA |
| 2.710260551 | 0.006723037 | 0.016006622 |
| -0.704207714 | 0.481303428 | NA |
| -1.851569012 | 0.064087736 | 0.114202719 |
| 4.000679474 | 6.32E-05 | 0.000226039 |
| -0.065389369 | 0.947863988 | NA |
| -1.114357394 | 0.265125894 | 0.370711196 |
| -7.132895792 | 9.83E-13 | 9.13E-12 |
| -3.173571096 | 0.00150576 | 0.004169618 |
| -7.017324849 | 2.26E-12 | 2.01E-11 |
| NA | NA | NA |
| -0.19448762 | 0.845794089 | NA |
| -8.071281309 | 6.96E-16 | 8.83E-15 |
| 5.985557259 | 2.16E-09 | 1.42E-08 |
| -1.095638242 | 0.273237119 | 0.379834705 |
| 5.035618174 | 4.76E-07 | 2.34E-06 |
| -2.80999659 | 0.004954202 | 0.01220749 |
| NA | NA | NA |
| NA | NA | NA |
| -3.056551707 | 0.002238989 | 0.005962537 |
| -0.205570965 | 0.837126059 | 0.886048091 |
| -3.364335871 | 0.000767281 | 0.002259949 |
| -0.81323537 | 0.416083124 | 0.527693585 |

| | | | |
|---|---|---|---|
| NA | NA | NA | |
| | 3.594069306 | 0.000325553 | 0.001028344 |
| | 1.9416971 | 0.05217378 | 0.096058089 |
| | 0.296678867 | 0.766711693 | 0.831670365 |
| | -0.005634266 | 0.99550453 | 0.997002158 |
| | -1.620367081 | 0.105153447 | 0.173614167 |
| | 1.332999874 | 0.182531845 | 0.27438336 |
| | -0.295990919 | 0.767237016 | NA |
| | -2.165371512 | 0.030359246 | 0.059961487 |
| | -8.45602666 | 2.77E-17 | 3.94E-16 |
| | 12.7876889 | 1.92E-37 | 9.41E-36 |
| | 2.907498047 | 0.003643326 | 0.009229071 |
| | 4.642321871 | 3.45E-06 | 1.50E-05 |
| | -5.105684443 | 3.30E-07 | 1.66E-06 |
| | -0.065388772 | 0.947864464 | NA |
| | 3.063394177 | 0.002188416 | 0.005842156 |
| | -1.126893131 | 0.259787668 | 0.364775347 |
| | -0.946907499 | 0.343685916 | 0.45507518 |
| NA | NA | NA | |
| NA | NA | NA | |
| | 0.127730455 | 0.898362288 | NA |
| | 0.95261206 | 0.340786665 | NA |
| | 1.138746232 | 0.254809014 | 0.359154725 |
| NA | NA | NA | |
| | 4.519500014 | 6.20E-06 | 2.60E-05 |
| NA | NA | NA | |
| | 0.063708882 | 0.949202033 | NA |
| | 1.091837025 | 0.274904742 | 0.381613093 |
| NA | NA | NA | |
| NA | NA | NA | |
| NA | NA | NA | |
| | 2.709234049 | 0.006743875 | 0.01604555 |
| NA | NA | NA | |
| | 0.192809521 | 0.847108147 | NA |
| | 0.954311532 | 0.339925972 | NA |
| | 0.192810118 | 0.847107679 | NA |
| | 2.808065082 | 0.004984015 | 0.01227451 |
| | 0.192810118 | 0.847107679 | NA |
| | -0.094412664 | 0.924781356 | NA |
| | 0.376406755 | 0.706614525 | NA |
| | 0.192809521 | 0.847108147 | NA |
| NA | NA | NA | |
| | 0.248626819 | 0.803649461 | NA |
| | 0.125773595 | 0.899911144 | NA |
| NA | NA | NA | |
| NA | NA | NA | |
| | -0.194488217 | 0.845793621 | NA |

| | | |
|---|---|---|
| NA | NA | NA |
| NA | NA | NA |
| 0.14503292 | 0.88468488 | NA |
| 0.063708882 | 0.949202033 | NA |
| 10.17631307 | 2.53E-24 | 6.18E-23 |
| 2.862468027 | 0.004203557 | 0.010515193 |
| NA | NA | NA |
| 11.06005491 | 1.96E-28 | 6.23E-27 |
| -4.348817743 | 1.37E-05 | 5.45E-05 |
| -3.033818535 | 0.002414796 | 0.006375776 |
| 1.080027649 | 0.280129868 | 0.387127352 |
| -11.03735713 | 2.52E-28 | 7.99E-27 |
| 4.065739137 | 4.79E-05 | 0.000174956 |
| 3.166052549 | 0.001545229 | 0.004270407 |
| -3.817611553 | 0.00013475 | 0.000455413 |
| -0.073164678 | 0.941675074 | NA |
| NA | NA | NA |
| -0.065389369 | 0.947863988 | NA |
| 4.339605857 | 1.43E-05 | 5.67E-05 |
| -0.314530209 | 0.753118376 | NA |
| -2.125453811 | 0.033548764 | 0.065475678 |
| 0.034413875 | 0.972547119 | 0.981097375 |
| -0.090977841 | 0.927510199 | NA |
| 4.274748542 | 1.91E-05 | 7.45E-05 |
| -5.841752935 | 5.17E-09 | 3.26E-08 |
| NA | NA | NA |
| 4.030778678 | 5.56E-05 | 0.00020111 |
| -2.235012709 | 0.025416514 | 0.051437205 |
| -3.784819588 | 0.00015382 | 0.00051479 |
| 6.077013045 | 1.22E-09 | 8.26E-09 |
| 0.035514391 | 0.971669571 | 0.980483344 |
| 7.16917989 | 7.54E-13 | 7.09E-12 |
| 1.436883483 | 0.150751104 | 0.234482981 |
| -10.85293957 | 1.93E-27 | 5.79E-26 |
| -4.348920268 | 1.37E-05 | 5.45E-05 |
| -0.134581655 | 0.892942647 | 0.925212338 |
| -2.635989239 | 0.008389239 | 0.019453646 |
| -3.269622664 | 0.00107691 | 0.003083398 |
| 0.063708285 | 0.949202508 | NA |
| 6.045324701 | 1.49E-09 | 9.96E-09 |
| NA | NA | NA |
| 0.063708285 | 0.949202508 | NA |
| NA | NA | NA |
| -5.773334829 | 7.77E-09 | 4.80E-08 |
| -0.065389966 | 0.947863513 | NA |
| NA | NA | NA |
| -2.13172641 | 0.033029344 | 0.06461309 |

| | | |
|---|---|---|
| 1.038592886 | 0.298994116 | 0.407882206 |
| 0.222749121 | 0.823730769 | 0.876428529 |
| NA | NA | NA |
| -4.183678643 | 2.87E-05 | 0.000108781 |
| 1.912503407 | 0.055811656 | 0.101738093 |
| -7.417626437 | 1.19E-13 | 1.22E-12 |
| 1.607351922 | 0.10797719 | 0.177363767 |
| 0.549300253 | 0.582799414 | NA |
| -1.070129238 | 0.28456114 | 0.39169409 |
| 4.131939939 | 3.60E-05 | 0.000134203 |
| -0.689883204 | 0.490267639 | 0.597875713 |
| -0.347168772 | 0.72846453 | 0.801741332 |
| 12.16001879 | 5.07E-34 | 2.13E-32 |
| NA | NA | NA |
| 0.310805976 | 0.755948126 | NA |
| NA | NA | NA |
| 4.704639549 | 2.54E-06 | 1.13E-05 |
| 2.648500365 | 0.008084976 | 0.018801294 |
| -4.802879502 | 1.56E-06 | 7.14E-06 |
| -1.767216604 | 0.07719196 | 0.133951384 |
| 0.06371127 | 0.949200131 | NA |
| -2.767942018 | 0.005641149 | 0.013692587 |
| -0.704507044 | 0.481117065 | 0.589803731 |
| NA | NA | NA |
| NA | NA | NA |
| 1.951399615 | 0.051009527 | 0.094205598 |
| 1.61566673 | 0.106166388 | 0.174871327 |
| 6.239214941 | 4.40E-10 | 3.10E-09 |
| -0.608523439 | 0.542840365 | 0.646863718 |
| 1.052434103 | 0.292600429 | 0.400513439 |
| 1.180157807 | 0.237937457 | 0.339988271 |
| 4.104880717 | 4.05E-05 | 0.000149494 |
| 5.241770869 | 1.59E-07 | 8.35E-07 |
| NA | NA | NA |
| 1.078618158 | 0.280757983 | NA |
| -3.174664982 | 0.001500096 | 0.004154852 |
| NA | NA | NA |
| NA | NA | NA |
| NA | NA | NA |
| NA | NA | NA |
| NA | NA | NA |
| 1.610670669 | 0.107251522 | 0.176356957 |
| 2.262631488 | 0.023658415 | 0.048356209 |
| 0.109730744 | 0.912622918 | NA |
| -0.416691046 | 0.676904404 | NA |
| 0.063708882 | 0.949202033 | NA |
| NA | NA | NA |

| | | |
|---|---|---|
| -1.263699661 | 0.206337847 | 0.303213545 |
| 2.903799243 | 0.003686645 | 0.009328419 |
| NA | NA | NA |
| NA | NA | NA |
| -0.672017365 | 0.501572642 | 0.608430901 |
| 3.976831086 | 6.98E-05 | 0.000248308 |
| 31.77060102 | 1.65E-221 | 1.59E-218 |
| 0.170475022 | 0.864636576 | 0.90566846 |
| -2.375330088 | 0.017533263 | 0.037340046 |
| -1.491026234 | 0.135954608 | 0.21515831 |
| 6.583052144 | 4.61E-11 | 3.63E-10 |
| 1.237048981 | 0.216068899 | 0.314576812 |
| -3.304220174 | 0.00095241 | 0.002757208 |
| 5.135185885 | 2.82E-07 | 1.43E-06 |
| NA | NA | NA |
| -0.785382363 | 0.432229418 | 0.542971259 |
| -0.470475982 | 0.63801499 | 0.72910173 |
| 5.792323966 | 6.94E-09 | 4.31E-08 |
| 0.712816401 | 0.475959375 | NA |
| -1.12430264 | 0.260884659 | 0.365987166 |
| NA | NA | NA |
| -0.111157831 | 0.91149119 | NA |
| -0.986653686 | 0.323812442 | 0.43481896 |
| -0.353109436 | 0.724006399 | 0.798496592 |
| -2.345493549 | 0.019001907 | 0.039962064 |
| 4.531535945 | 5.86E-06 | 2.47E-05 |
| NA | NA | NA |
| -0.809719443 | 0.418101441 | NA |
| 6.191703614 | 5.95E-10 | 4.14E-09 |
| 0.231282445 | 0.817095384 | NA |
| -2.391076579 | 0.016799049 | 0.036006181 |
| 3.165887248 | 0.001546107 | 0.004272363 |
| NA | NA | NA |
| 0.203227347 | 0.838957339 | 0.887198538 |
| 3.13354802 | 0.001727067 | 0.004721328 |
| -7.945489853 | 1.93E-15 | 2.35E-14 |
| -3.075936431 | 0.002098426 | 0.005625921 |
| -1.446727093 | 0.147973373 | 0.230779085 |
| -7.492976709 | 6.73E-14 | 7.03E-13 |
| -6.93126433 | 4.17E-12 | 3.62E-11 |
| -1.121756335 | 0.261966057 | 0.367216084 |
| 9.384902737 | 6.30E-21 | 1.19E-19 |
| -5.441849131 | 5.27E-08 | 2.95E-07 |
| -0.005512668 | 0.99560155 | 0.99703556 |
| 1.36138113 | 0.173393276 | NA |
| 0.637735386 | 0.523645947 | NA |
| -0.750767027 | 0.452792876 | 0.56267529 |

| | | |
|---|---|---|
| 2.61147846 | 0.009015167 | 0.020757025 |
| NA | NA | NA |
| NA | NA | NA |
| -0.065389966 | 0.947863513 | NA |
| 0.278475486 | 0.78064738 | NA |
| 2.266324542 | 0.02343152 | 0.047982569 |
| 1.726848956 | 0.084194794 | 0.144253247 |
| 0.422448409 | 0.67269775 | 0.756621819 |
| -12.13557819 | 6.84E-34 | 2.85E-32 |
| 5.934126819 | 2.95E-09 | 1.91E-08 |
| 2.045359754 | 0.040819404 | 0.077566784 |
| 1.06906543 | 0.285040188 | NA |
| NA | NA | NA |
| -1.436839693 | 0.150763549 | 0.234487767 |
| -1.058475271 | 0.289838819 | 0.397341748 |
| -4.708269107 | 2.50E-06 | 1.11E-05 |
| 0.776323596 | 0.437557944 | 0.5478503 |
| NA | NA | NA |
| 14.62642111 | 1.91E-48 | 1.42E-46 |
| 0.382624269 | 0.701998373 | 0.781246689 |
| -3.042059038 | 0.002349658 | 0.006221598 |
| -0.004593578 | 0.996334868 | 0.997521005 |
| -2.084583775 | 0.037107098 | 0.071412477 |
| NA | NA | NA |
| -1.174571113 | 0.240166346 | 0.342371776 |
| 0.800040877 | 0.423687114 | NA |
| -5.084892054 | 3.68E-07 | 1.84E-06 |
| -3.225549819 | 0.00125731 | 0.003539586 |
| NA | NA | NA |
| 0.003556709 | 0.997162162 | 0.997999007 |
| 2.168715117 | 0.030104319 | 0.059519045 |
| -2.364916359 | 0.018034143 | 0.038247298 |
| 0.063708882 | 0.949202033 | NA |
| 1.256896958 | 0.20879095 | 0.305930601 |
| -0.266788362 | 0.789632115 | 0.850378274 |
| 8.132541401 | 4.20E-16 | 5.40E-15 |
| NA | NA | NA |
| 1.830796843 | 0.067130872 | 0.118975127 |
| -3.836633162 | 0.000124733 | 0.000423847 |
| -0.014094295 | 0.988754752 | 0.992123931 |
| 12.06688638 | 1.58E-33 | 6.46E-32 |
| 1.268885571 | 0.204481879 | NA |
| -3.019586491 | 0.0025312 | 0.006650146 |
| 0.809927949 | 0.417981588 | 0.529591452 |
| 1.827348214 | 0.06764743 | 0.119712612 |
| NA | NA | NA |
| 0.797485044 | 0.425169386 | 0.536715237 |

| | | |
|---|---|---|
| 1.178223437 | 0.238707541 | 0.340894035 |
| -1.929999516 | 0.053606898 | 0.098284548 |
| 5.871599213 | 4.32E-09 | 2.75E-08 |
| 4.188537404 | 2.81E-05 | 0.000106702 |
| 1.154881421 | 0.248138988 | 0.351392843 |
| 7.391248066 | 1.45E-13 | 1.47E-12 |
| NA | NA | NA |
| 0.764550643 | 0.444539169 | 0.554455621 |
| NA | NA | NA |
| 0.32175779 | 0.747636194 | 0.81649871 |
| -7.701982686 | 1.34E-14 | 1.51E-13 |
| 0.162125509 | 0.871207019 | NA |
| NA | NA | NA |
| 1.440095401 | 0.14984041 | NA |
| 0.763325779 | 0.445269132 | NA |
| 0.192810118 | 0.847107679 | NA |
| NA | NA | NA |
| 0.921506064 | 0.356786277 | 0.468637726 |
| 0.793658564 | 0.427394229 | 0.538735839 |
| 1.799474189 | 0.071943703 | 0.126120668 |
| -1.694995255 | 0.090076321 | 0.152720335 |
| 0.192810118 | 0.847107679 | NA |
| 0.065061479 | 0.948125051 | NA |
| 2.252725656 | 0.024276452 | 0.049422057 |
| 5.625967831 | 1.84E-08 | 1.09E-07 |
| -4.408195961 | 1.04E-05 | 4.24E-05 |
| -0.574651507 | 0.56552701 | 0.667537895 |
| -0.132250864 | 0.894785871 | 0.926700196 |
| NA | NA | NA |
| 3.284503642 | 0.001021622 | 0.002939555 |
| 5.223028804 | 1.76E-07 | 9.17E-07 |
| -0.80116965 | 0.423033439 | 0.534763321 |
| -4.568802511 | 4.91E-06 | 2.09E-05 |
| 0.080138309 | 0.936127255 | 0.956456249 |
| 8.562792789 | 1.10E-17 | 1.62E-16 |
| NA | NA | NA |
| 0.3474247 | 0.728272279 | NA |
| 4.738115937 | 2.16E-06 | 9.67E-06 |
| 2.029221098 | 0.042435776 | 0.080272632 |
| 5.298564312 | 1.17E-07 | 6.23E-07 |
| 0.112531359 | 0.910402107 | NA |
| -6.575585759 | 4.85E-11 | 3.81E-10 |
| -0.143809698 | 0.885650745 | NA |
| 0.697184493 | 0.485687342 | 0.593968544 |
| 6.435343945 | 1.23E-10 | 9.27E-10 |
| 0.069839618 | 0.944321313 | NA |
| 3.601282084 | 0.000316652 | 0.001003011 |

| | | |
|---|---|---|
| -9.615350122 | 6.89E-22 | 1.41E-20 |
| NA | NA | NA |
| -0.156369882 | 0.875741476 | 0.913325381 |
| NA | NA | NA |
| 0.192810118 | 0.847107679 | NA |
| 1.033056328 | 0.301577539 | 0.410667853 |
| 0.128342884 | 0.897877629 | NA |
| 0.192809521 | 0.847108147 | NA |
| -5.557112782 | 2.74E-08 | 1.59E-07 |
| 0.968017793 | 0.333035483 | 0.444224788 |
| -4.106732998 | 4.01E-05 | 0.000148476 |
| -0.829167392 | 0.407009694 | 0.518815066 |
| -0.44133666 | 0.65896929 | 0.746192368 |
| NA | NA | NA |
| 38.35316205 | 0 | 0 |
| -0.065389369 | 0.947863988 | NA |
| 34.08340838 | 1.30E-254 | 1.91E-251 |
| 34.1916393 | 3.22E-256 | 5.04E-253 |
| 0.854321333 | 0.39292697 | 0.504874849 |
| -1.268125572 | 0.204753111 | 0.301238486 |
| 0.31070316 | 0.756026295 | 0.823075906 |
| 0.19944167 | 0.841917267 | 0.889239142 |
| -0.557436916 | 0.577228949 | 0.678120745 |
| NA | NA | NA |
| 0.063708882 | 0.949202033 | NA |
| NA | NA | NA |
| 1.669871978 | 0.094944696 | NA |
| NA | NA | NA |
| 1.451274734 | 0.146703374 | 0.229156426 |
| NA | NA | NA |
| 0.254408903 | 0.799179674 | NA |
| 0.192808924 | 0.847108615 | NA |
| NA | NA | NA |
| 2.23163667 | 0.025638986 | 0.051812031 |
| NA | NA | NA |
| NA | NA | NA |
| NA | NA | NA |
| NA | NA | NA |
| NA | NA | NA |
| NA | NA | NA |
| NA | NA | NA |
| NA | NA | NA |
| 0.063708285 | 0.949202508 | NA |
| 0.06371127 | 0.949200131 | NA |
| 0.063708285 | 0.949202508 | NA |
| NA | NA | NA |
| NA | NA | NA |

| | | |
|---|---|---|
| 0.202714101 | 0.839358501 | NA |
| NA | NA | NA |
| NA | NA | NA |
| 0.392023906 | 0.69504055 | NA |
| 2.077568616 | 0.037749109 | 0.072480838 |
| NA | NA | NA |
| 14.34457922 | 1.15E-46 | 8.15E-45 |
| 3.552018179 | 0.000382288 | 0.00119132 |
| -6.001804928 | 1.95E-09 | 1.29E-08 |
| 0.08878279 | 0.929254535 | NA |
| NA | NA | NA |
| 3.127876805 | 0.00176074 | 0.004805508 |
| -2.121769318 | 0.033857114 | 0.066010438 |
| -1.945923133 | 0.051663974 | 0.095231553 |
| 0.61357602 | 0.539495534 | NA |
| 19.95659146 | 1.31E-88 | 2.55E-86 |
| 2.104160659 | 0.035364435 | 0.068506448 |
| -0.859752007 | 0.38992576 | 0.502036197 |
| -5.166525245 | 2.38E-07 | 1.22E-06 |
| -3.35944329 | 0.000780997 | 0.002296294 |
| 3.815100143 | 0.000136128 | 0.000459697 |
| 4.794241744 | 1.63E-06 | 7.44E-06 |
| 0.347735698 | 0.728038685 | 0.801454707 |
| -1.056596651 | 0.29069571 | 0.398298315 |
| 6.137785446 | 8.37E-10 | 5.74E-09 |
| NA | NA | NA |
| 0.410127234 | 0.681712616 | 0.764346299 |
| NA | NA | NA |
| -0.315741509 | 0.752198717 | 0.82031191 |
| 0.063707688 | 0.949202983 | NA |
| -6.148233916 | 7.84E-10 | 5.39E-09 |
| NA | NA | NA |
| NA | NA | NA |
| NA | NA | NA |
| -0.245123876 | 0.806360516 | NA |
| 0.996313628 | 0.319097784 | 0.429756123 |
| 0.063708882 | 0.949202033 | NA |
| -1.227866778 | 0.219496983 | 0.318511768 |
| -0.114978004 | 0.908462557 | NA |
| 0.288838667 | 0.772704841 | 0.836503705 |
| 11.44139545 | 2.60E-30 | 9.09E-29 |
| 1.834508094 | 0.066578607 | 0.118113441 |
| -0.761333547 | 0.446457868 | 0.55640512 |
| 6.758715329 | 1.39E-11 | 1.15E-10 |
| 1.718616094 | 0.085684299 | 0.146354443 |
| NA | NA | NA |
| 2.439166127 | 0.014721199 | 0.032032653 |

| | | |
|---|---|---|
| 3.145031078 | 0.001660692 | 0.004559298 |
| 0.207251126 | 0.835813741 | NA |
| 8.189774485 | 2.62E-16 | 3.44E-15 |
| -0.467390233 | 0.640220709 | 0.730830787 |
| -0.838844916 | 0.401556342 | 0.513249004 |
| NA | NA | NA |
| 2.619731266 | 0.008799908 | 0.020302489 |
| NA | NA | NA |
| NA | NA | NA |
| NA | NA | NA |
| 1.803760202 | 0.071268907 | 0.125174426 |
| -1.31601095 | 0.188170367 | 0.281205104 |
| NA | NA | NA |
| 0.423407673 | 0.671997848 | NA |
| -10.89036321 | 1.28E-27 | 3.90E-26 |
| -1.400870488 | 0.161252805 | 0.247465219 |
| -4.084507953 | 4.42E-05 | 0.000162061 |
| -3.976082346 | 7.01E-05 | 0.000248985 |
| NA | NA | NA |
| NA | NA | NA |
| -1.985417605 | 0.047098029 | 0.087912644 |
| NA | NA | NA |
| NA | NA | NA |
| NA | NA | NA |
| -4.194035032 | 2.74E-05 | 0.000104369 |
| -1.392792566 | 0.163682527 | 0.250625415 |
| -0.408255041 | 0.683086442 | 0.765763682 |
| 0.786184683 | 0.431759298 | 0.542584532 |
| -1.61497765 | 0.106315535 | 0.175070911 |
| 0.064913412 | 0.948242942 | 0.964700274 |
| -1.959894666 | 0.050008103 | 0.092602118 |
| NA | NA | NA |
| -1.982552258 | 0.047417465 | 0.088436974 |
| 1.068947709 | 0.285093233 | 0.392197006 |
| 0.06353712 | 0.949338802 | NA |
| NA | NA | NA |
| 8.824175025 | 1.10E-18 | 1.77E-17 |
| -2.508850105 | 0.012112487 | 0.026988358 |
| -0.208506596 | 0.834833433 | NA |
| NA | NA | NA |
| -0.434703939 | 0.663777339 | 0.750151054 |
| -0.176253917 | 0.860094464 | 0.902761959 |
| -2.708860885 | 0.006751464 | 0.016062081 |
| -3.37614277 | 0.000735098 | 0.002174624 |
| -1.199316668 | 0.230404837 | 0.331115178 |
| NA | NA | NA |
| -2.420681958 | 0.015491424 | 0.033513427 |

| | | | |
|---|---|---|---|
| | 5.927802369 | 3.07E-09 | 1.99E-08 |
| | -5.189019288 | 2.11E-07 | 1.09E-06 |
| | -4.159849212 | 3.18E-05 | 0.000119829 |
| | -3.815283011 | 0.000136027 | 0.000459419 |
| | -3.231907509 | 0.001229669 | 0.003470358 |
| | -4.008011308 | 6.12E-05 | 0.000219639 |
| | -1.329376792 | 0.183723691 | 0.27581143 |
| | -1.305257473 | 0.19180519 | 0.285681874 |
| | 2.130925718 | 0.033095262 | 0.064721807 |
| | 0.749512927 | 0.45354811 | 0.563499773 |
| | 7.474867517 | 7.73E-14 | 8.04E-13 |
| | 11.87167438 | 1.66E-32 | 6.56E-31 |
| | 0.792001307 | 0.428359914 | 0.539572676 |
| | 0.862502742 | 0.388410925 | 0.50063986 |
| | 13.71375409 | 8.40E-43 | 5.12E-41 |
| | 0.373551896 | 0.708737729 | 0.786613398 |
| | 5.357054692 | 8.46E-08 | 4.60E-07 |
| | 0.662785285 | 0.507468079 | 0.613884414 |
| | 5.896550514 | 3.71E-09 | 2.38E-08 |
| | 7.877325332 | 3.34E-15 | 3.98E-14 |
| | 1.290299963 | 0.19694653 | 0.29186333 |
| | -0.356584737 | 0.721402705 | 0.796324899 |
| | 0.860490324 | 0.389518815 | 0.501628727 |
| | 0.192809521 | 0.847108147 | NA |
| | 0.192810118 | 0.847107679 | NA |
| | -0.041287255 | 0.967066893 | 0.977534078 |
| | -0.591721503 | 0.554037095 | 0.657114505 |
| | -1.475860667 | 0.1399813 | 0.220429818 |
| | -3.643240282 | 0.000269227 | 0.000863169 |
| | -0.913821563 | 0.360810614 | 0.47295573 |
| | 0.063708285 | 0.949202508 | NA |
| | 1.251629909 | 0.210704751 | 0.308320208 |
| | 0.31379056 | 0.753680115 | NA |
| | 0.330493631 | 0.741027004 | NA |
| | -9.050746855 | 1.42E-19 | 2.45E-18 |
| NA | NA | NA | |
| | 3.84659552 | 0.00011977 | 0.000408371 |
| | 2.209170108 | 0.027162811 | 0.054447438 |
| NA | NA | NA | |
| | -3.675493198 | 0.00023739 | 0.000768816 |
| NA | NA | NA | |
| NA | NA | NA | |
| NA | NA | NA | |
| NA | NA | NA | |
| NA | NA | NA | |
| | 3.262685988 | 0.001103617 | 0.003151933 |
| NA | NA | NA | |

| | | |
|---|---|---|
| 1.622179476 | 0.104764925 | NA |
| NA | NA | NA |
| 1.314434283 | 0.1887001 | 0.281845298 |
| 2.812761551 | 0.004911806 | 0.012115038 |
| NA | NA | NA |
| 0.063708285 | 0.949202508 | NA |
| 3.617997999 | 0.000296891 | 0.000944721 |
| -1.084865042 | 0.277981433 | 0.384923394 |
| -0.483577755 | 0.628685565 | 0.721437727 |
| 3.235662534 | 0.001213608 | 0.003430059 |
| -0.859454032 | 0.39009007 | 0.502209364 |
| -0.065388772 | 0.947864464 | NA |
| 0.063708285 | 0.949202508 | NA |
| -0.065389369 | 0.947863988 | NA |
| -3.807962295 | 0.000140117 | 0.000472149 |
| 0.106285468 | 0.915355861 | NA |
| 0.799774053 | 0.423841719 | NA |
| 6.769998809 | 1.29E-11 | 1.07E-10 |
| NA | NA | NA |
| -0.065388772 | 0.947864464 | NA |
| 6.761294086 | 1.37E-11 | 1.13E-10 |
| 0.611706953 | 0.540731661 | NA |
| -0.065389369 | 0.947863988 | NA |
| 0.458578086 | 0.646537176 | 0.735649856 |
| -1.530170202 | 0.125974605 | 0.202150555 |
| NA | NA | NA |
| NA | NA | NA |
| NA | NA | NA |
| 0.971562825 | 0.331268084 | 0.442197224 |
| 4.955961093 | 7.20E-07 | 3.45E-06 |
| NA | NA | NA |
| -6.688524393 | 2.25E-11 | 1.83E-10 |
| NA | NA | NA |
| NA | NA | NA |
| NA | NA | NA |
| NA | NA | NA |
| NA | NA | NA |
| NA | NA | NA |
| -1.528206059 | 0.126461385 | 0.202853657 |
| NA | NA | NA |
| NA | NA | NA |
| NA | NA | NA |
| 9.879497577 | 5.11E-23 | 1.14E-21 |
| -4.135735671 | 3.54E-05 | 0.0001322 |
| -4.31699384 | 1.58E-05 | 6.25E-05 |
| 1.623061985 | 0.104576155 | 0.172843447 |
| NA | NA | NA |

| | | |
|---|---|---|
| 0.902721222 | 0.36667387 | 0.479135838 |
| -1.303622975 | 0.192362163 | 0.286375123 |
| 2.623094924 | 0.008713499 | 0.020132824 |
| 5.541267929 | 3.00E-08 | 1.73E-07 |
| -0.21248704 | 0.831727087 | 0.882273648 |
| -2.199764325 | 0.02782362 | 0.055616133 |
| -0.065389966 | 0.947863513 | NA |
| 4.200176799 | 2.67E-05 | 0.000101701 |
| -0.800364554 | 0.423499612 | 0.535093149 |
| 6.51774324 | 7.14E-11 | 5.52E-10 |
| 0.888539388 | 0.374250679 | 0.487027893 |
| -0.065389369 | 0.947863988 | NA |
| 5.76617872 | 8.11E-09 | 4.99E-08 |
| 0.256647848 | 0.797450634 | NA |
| NA | NA | NA |
| 2.169532816 | 0.030042256 | 0.059424503 |
| 0.36953505 | 0.711728953 | 0.788848943 |
| 0.535828027 | 0.59207741 | 0.69131397 |
| -0.885815314 | 0.375717056 | 0.488504385 |
| 3.580767298 | 0.000342587 | 0.001078338 |
| -1.037360927 | 0.299567681 | 0.408472201 |
| 1.18309531 | 0.236771376 | 0.338650717 |
| -0.202641834 | 0.839414989 | 0.887532513 |
| 3.14080561 | 0.001684838 | 0.00461597 |
| 1.445953077 | 0.148190363 | 0.231074308 |
| NA | NA | NA |
| 0.063708882 | 0.949202033 | NA |
| NA | NA | NA |
| -1.732533217 | 0.083178668 | 0.142785959 |
| -1.806621224 | 0.070821358 | 0.124519429 |
| 4.576358983 | 4.73E-06 | 2.02E-05 |
| 4.033182048 | 5.50E-05 | 0.000199293 |
| NA | NA | NA |
| -3.431147862 | 0.000601033 | 0.001806419 |
| 6.370534744 | 1.88E-10 | 1.38E-09 |
| 3.340305632 | 0.000836862 | 0.00244704 |
| NA | NA | NA |
| NA | NA | NA |
| NA | NA | NA |
| NA | NA | NA |
| 0.192809521 | 0.847108147 | NA |
| NA | NA | NA |
| NA | NA | NA |
| NA | NA | NA |
| NA | NA | NA |
| NA | NA | NA |
| 0.434937002 | 0.663608156 | NA |

| | | | |
|---|---|---|---|
| | 0.192808327 | 0.847109082 | NA |
| NA | NA | NA | |
| NA | NA | NA | |
| | 0.192809521 | 0.847108147 | NA |
| NA | NA | NA | |
| NA | NA | NA | |
| | 0.192810118 | 0.847107679 | NA |
| NA | NA | NA | |
| NA | NA | NA | |
| | -0.167858774 | 0.866694383 | NA |
| NA | NA | NA | |
| NA | NA | NA | |
| NA | NA | NA | |
| NA | NA | NA | |
| | -0.065388772 | 0.947864464 | NA |
| | 0.192809521 | 0.847108147 | NA |
| | 0.063708285 | 0.949202508 | NA |
| | -0.065388772 | 0.947864464 | NA |
| | 0.827216118 | 0.408114578 | 0.519958662 |
| | 0.192810118 | 0.847107679 | NA |
| | 3.619119429 | 0.000295607 | 0.000940876 |
| | 1.728310854 | 0.083932508 | 0.143853101 |
| | 1.625397992 | 0.104077783 | 0.172133403 |
| | 0.863402282 | 0.387916325 | 0.500131123 |
| | -9.351944482 | 8.61E-21 | 1.62E-19 |
| | -0.166910016 | 0.867440852 | NA |
| | -2.899885358 | 0.003732992 | 0.009434748 |
| | -1.1641743 | 0.244353383 | 0.347193754 |
| | -0.194487022 | 0.845794556 | NA |
| NA | NA | NA | |
| NA | NA | NA | |
| NA | NA | NA | |
| NA | NA | NA | |
| NA | NA | NA | |
| | 0.898702094 | 0.368811362 | 0.481325845 |
| NA | NA | NA | |
| NA | NA | NA | |
| | 1.849694349 | 0.064357615 | 0.114628273 |
| | 6.427266346 | 1.30E-10 | 9.74E-10 |
| | 1.897242008 | 0.057796005 | 0.104767816 |
| NA | NA | NA | |
| | 0.063708882 | 0.949202033 | NA |
| | 12.48053891 | 9.53E-36 | 4.28E-34 |
| | -3.443202059 | 0.00057487 | 0.001734449 |
| | -2.248323944 | 0.024555541 | 0.049908664 |
| | 0.665842554 | 0.505511742 | 0.611963576 |
| | 7.825103807 | 5.07E-15 | 5.94E-14 |

| | | |
|---|---|---|
| 1.053181664 | 0.292257741 | 0.40015378 |
| 5.944710561 | 2.77E-09 | 1.80E-08 |
| 0.188804083 | 0.850246371 | NA |
| -0.669445525 | 0.50321132 | 0.609853352 |
| -4.200605342 | 2.66E-05 | 0.00010154 |
| NA | NA | NA |
| NA | NA | NA |
| 0.766402426 | 0.44343689 | 0.553494507 |
| 0.873764912 | 0.382246292 | 0.494612229 |
| -2.152702404 | 0.031342074 | 0.061673909 |
| 0.063708882 | 0.949202033 | NA |
| NA | NA | NA |
| -8.214960177 | 2.12E-16 | 2.81E-15 |
| 0.651041748 | 0.515019538 | 0.62036106 |
| NA | NA | NA |
| -2.476207367 | 0.013278643 | 0.02928063 |
| NA | NA | NA |
| NA | NA | NA |
| 0.192809521 | 0.847108147 | NA |
| 0.283513734 | 0.776783055 | NA |
| 6.914882598 | 4.68E-12 | 4.05E-11 |
| 0.549064639 | 0.582961091 | NA |
| NA | NA | NA |
| -8.866358771 | 7.56E-19 | 1.23E-17 |
| NA | NA | NA |
| -7.860122156 | 3.84E-15 | 4.53E-14 |
| NA | NA | NA |
| -2.781295676 | 0.00541424 | 0.013212125 |
| -1.369822192 | 0.170742413 | 0.259576965 |
| NA | NA | NA |
| -0.474199877 | 0.635357381 | NA |
| 0.743213702 | 0.457352309 | NA |
| 1.379986388 | 0.167590836 | 0.255562241 |
| NA | NA | NA |
| 5.043468091 | 4.57E-07 | 2.25E-06 |
| NA | NA | NA |
| -0.065389369 | 0.947863988 | NA |
| 12.03648138 | 2.28E-33 | 9.30E-32 |
| NA | NA | NA |
| NA | NA | NA |
| NA | NA | NA |
| NA | NA | NA |
| -0.140388254 | 0.888353238 | 0.921830834 |
| 2.377987325 | 0.017407424 | 0.037097304 |
| 0.063708285 | 0.949202508 | NA |
| 0.192810118 | 0.847107679 | NA |
| 13.92371272 | 4.55E-44 | 2.95E-42 |

| | | | |
|---|---|---|---|
| | -0.39457502 | 0.693156552 | 0.773655214 |
| | -1.078506805 | 0.280807647 | 0.387785677 |
| | -0.651354813 | 0.514817473 | 0.620288888 |
| | -1.398771826 | 0.161881418 | 0.248232058 |
| | -1.374042055 | 0.169428627 | 0.257862005 |
| | -2.421089358 | 0.015474073 | 0.033487467 |
| | 0.219114645 | 0.826560743 | 0.878681286 |
| | -0.666376446 | 0.505170514 | 0.611639238 |
| | -0.94796937 | 0.343145047 | 0.454512385 |
| | 0.152967987 | 0.878423521 | 0.91476425 |
| | -1.439774791 | 0.149931126 | 0.233383145 |
| | -0.065389369 | 0.947863988 | NA |
| | 0.595243338 | 0.551680825 | 0.654900216 |
| | 0.617475246 | 0.536921304 | 0.641180355 |
| | 0.20241502 | 0.839592287 | 0.887644982 |
| | 0.287005967 | 0.774107752 | NA |
| | -0.531338483 | 0.595184243 | 0.693744137 |
| | 0.576346657 | 0.564380892 | 0.666436464 |
| | 0.315235826 | 0.752582606 | 0.820540254 |
| | 0.442790952 | 0.657916948 | NA |
| | 0.192809521 | 0.847108147 | NA |
| NA | NA | NA | |
| | -0.236655879 | 0.812923766 | 0.868390573 |
| | 4.375221142 | 1.21E-05 | 4.87E-05 |
| NA | NA | NA | |
| | 2.427426948 | 0.015206349 | 0.032973656 |
| NA | NA | NA | |
| | 0.089617125 | 0.928591475 | 0.951164046 |
| | 0.391397493 | 0.695503445 | NA |
| NA | NA | NA | |
| NA | NA | NA | |
| NA | NA | NA | |
| NA | NA | NA | |
| NA | NA | NA | |
| NA | NA | NA | |
| | -5.456587955 | 4.85E-08 | 2.73E-07 |
| | -0.510452385 | 0.609734565 | NA |
| NA | NA | NA | |
| | -0.194487022 | 0.845794556 | NA |
| NA | NA | NA | |
| NA | NA | NA | |
| | 0.741156961 | 0.458598276 | NA |
| | 7.443422623 | 9.81E-14 | 1.01E-12 |
| NA | NA | NA | |
| | 3.193621761 | 0.001405001 | 0.003913986 |
| | 0.063708882 | 0.949202033 | NA |
| | -2.772436391 | 0.005563839 | 0.013540388 |

| | | |
|---|---|---|
| -1.121036825 | 0.262272188 | NA |
| -1.375663413 | 0.168925864 | 0.257221948 |
| 5.007296334 | 5.52E-07 | 2.68E-06 |
| -5.144217064 | 2.69E-07 | 1.37E-06 |
| NA | NA | NA |
| 12.25263381 | 1.63E-34 | 6.93E-33 |
| -0.242944486 | 0.808048403 | 0.864187811 |
| 6.847446849 | 7.52E-12 | 6.35E-11 |
| 3.35321028 | 0.0007988 | 0.002344233 |
| NA | NA | NA |
| -1.826482389 | 0.067777632 | 0.119892164 |
| -1.068923037 | 0.285104351 | 0.392197006 |
| -0.653732708 | 0.513284027 | 0.618917969 |
| -0.274933198 | 0.783367561 | 0.845341239 |
| 0.469348916 | 0.638820256 | 0.729680161 |
| 3.765384149 | 0.000166293 | 0.000553426 |
| -2.45486472 | 0.014093761 | 0.030852444 |
| NA | NA | NA |
| 0.192808924 | 0.847108615 | NA |
| 0.505492167 | 0.613213199 | 0.708895056 |
| NA | NA | NA |
| 10.68189455 | 1.24E-26 | 3.52E-25 |
| NA | NA | NA |
| -1.168515704 | 0.242598809 | 0.345227274 |
| -6.94032617 | 3.91E-12 | 3.41E-11 |
| 0.06353712 | 0.949338802 | NA |
| -0.19733955 | 0.843561831 | NA |
| NA | NA | NA |
| NA | NA | NA |
| NA | NA | NA |
| NA | NA | NA |
| NA | NA | NA |
| NA | NA | NA |
| 0.324874675 | 0.745275925 | NA |
| NA | NA | NA |
| NA | NA | NA |
| NA | NA | NA |
| NA | NA | NA |
| NA | NA | NA |
| 0.192810118 | 0.847107679 | NA |
| NA | NA | NA |
| NA | NA | NA |
| NA | NA | NA |
| NA | NA | NA |
| NA | NA | NA |
| NA | NA | NA |
| NA | NA | NA |

| | | | |
|---|---|---|---|
| NA | NA | NA | |
| NA | NA | NA | |
| NA | NA | NA | |
| NA | NA | NA | |
| NA | NA | NA | |
| NA | NA | NA | |
| NA | NA | NA | |
| NA | NA | NA | |
| NA | NA | NA | |
| NA | NA | NA | |
| NA | NA | NA | |
| NA | NA | NA | |
| NA | NA | NA | |
| NA | NA | NA | |
| NA | NA | NA | |
| NA | NA | NA | |
| NA | NA | NA | |
| NA | NA | NA | |
| NA | NA | NA | |
| | 0.192809521 | 0.847108147 | NA |
| NA | NA | NA | |
| | -0.065389966 | 0.947863513 | NA |
| | 0.063708882 | 0.949202033 | NA |
| | -3.053174422 | 0.002264343 | 0.006023648 |
| | 1.067717894 | 0.285647784 | 0.392779434 |
| | -0.003118465 | 0.997511829 | 0.998189785 |
| | 1.114365403 | 0.265122459 | 0.370711196 |
| | 2.257344016 | 0.023986587 | 0.048913599 |
| | -0.065389966 | 0.947863513 | NA |
| | 1.265584618 | 0.205661836 | 0.302338936 |
| | -1.66587647 | 0.095738021 | 0.160676054 |
| | 4.860834473 | 1.17E-06 | 5.43E-06 |
| | 0.053942904 | 0.956980654 | 0.970365225 |
| | 1.924736974 | 0.054262272 | 0.099311544 |
| | -1.058250794 | 0.28994112 | 0.397460224 |
| | 3.219557706 | 0.001283885 | 0.003608719 |
| | -4.408315762 | 1.04E-05 | 4.24E-05 |
| NA | NA | NA | |
| | 0.063708882 | 0.949202033 | NA |
| | 1.035428794 | 0.300468704 | NA |
| | -0.815269576 | 0.414918019 | 0.526536099 |
| | 2.71291665 | 0.006669388 | 0.015887959 |
| | -0.369046137 | 0.712093336 | 0.789182917 |
| | -0.178097007 | 0.858646793 | 0.901847561 |
| | -3.422847175 | 0.000619689 | 0.00185736 |
| | 4.517230343 | 6.27E-06 | 2.63E-05 |
| | 0.063707194 | 0.949203376 | NA |

| | | |
|---|---|---|
| 2.142911425 | 0.032120211 | 0.06304155 |
| NA | NA | NA |
| NA | NA | NA |
| 14.06199449 | 6.50E-45 | 4.34E-43 |
| NA | NA | NA |
| 14.39542979 | 5.53E-47 | 3.98E-45 |
| -1.146923571 | 0.25141321 | 0.355228487 |
| 2.697688553 | 0.006982273 | 0.01653731 |
| 0.063705218 | 0.949204949 | NA |
| NA | NA | NA |
| 0.905026497 | 0.36545135 | 0.478031041 |
| 4.393738891 | 1.11E-05 | 4.51E-05 |
| -3.508345371 | 0.000450903 | 0.0013865 |
| 4.776253051 | 1.79E-06 | 8.11E-06 |
| -1.615614166 | 0.106177759 | 0.174878549 |
| 4.559783691 | 5.12E-06 | 2.17E-05 |
| -0.306443178 | 0.759267238 | 0.825774933 |
| -0.817217658 | 0.413804053 | 0.525425715 |
| -0.670611163 | 0.502468269 | 0.6092516 |
| -0.26481177 | 0.791154469 | 0.851321799 |
| -3.291680542 | 0.000995907 | 0.002871837 |
| -9.084122371 | 1.05E-19 | 1.82E-18 |
| -0.082051403 | 0.934605838 | NA |
| -0.144574828 | 0.885046573 | NA |
| 0.241681374 | 0.809027063 | 0.865086611 |
| NA | NA | NA |
| NA | NA | NA |
| 11.95202665 | 6.34E-33 | 2.55E-31 |
| 8.007591758 | 1.17E-15 | 1.45E-14 |
| -1.405882655 | 0.159758973 | 0.245730172 |
| -5.804543066 | 6.45E-09 | 4.02E-08 |
| -1.001903596 | 0.316390156 | 0.426913509 |
| NA | NA | NA |
| 0.673961563 | 0.500335751 | NA |
| 0.140777587 | 0.888045649 | 0.921588068 |
| 6.042410388 | 1.52E-09 | 1.01E-08 |
| -1.189439224 | 0.234266874 | 0.335586324 |
| -6.58635625 | 4.51E-11 | 3.55E-10 |
| 15.60638626 | 6.59E-55 | 5.72E-53 |
| 5.669520707 | 1.43E-08 | 8.59E-08 |
| NA | NA | NA |
| NA | NA | NA |
| NA | NA | NA |
| 0.192809521 | 0.847108147 | NA |
| -2.999308462 | 0.002705932 | 0.007062511 |
| -3.917244762 | 8.96E-05 | 0.000312672 |
| -0.754046931 | 0.450821042 | 0.560735256 |

| | | |
|---|---|---|
| NA | NA | NA |
| 1.582533965 | 0.113527725 | 0.185033751 |
| -1.74445835 | 0.081079198 | 0.139728196 |
| 5.163985481 | 2.42E-07 | 1.24E-06 |
| 10.99467601 | 4.05E-28 | 1.27E-26 |
| 7.663890026 | 1.80E-14 | 2.00E-13 |
| 1.576152254 | 0.114990719 | 0.187043001 |
| 0.396782033 | 0.691528201 | 0.772343491 |
| -5.506171159 | 3.67E-08 | 2.10E-07 |
| -1.387875524 | 0.165174952 | 0.252612817 |
| 0.959937228 | 0.337086808 | 0.448410013 |
| 1.520470887 | 0.12839267 | 0.20542499 |
| -0.065388772 | 0.947864464 | NA |
| -12.01773515 | 2.87E-33 | 1.17E-31 |
| -3.873473615 | 0.000107295 | 0.000369052 |
| 4.366931414 | 1.26E-05 | 5.04E-05 |
| 2.576623029 | 0.009977069 | 0.022741649 |
| -2.091269758 | 0.036503889 | 0.070419449 |
| -5.864214951 | 4.51E-09 | 2.87E-08 |
| NA | NA | NA |
| 0.086111037 | 0.93137815 | NA |
| 5.514095727 | 3.51E-08 | 2.01E-07 |
| 5.050485515 | 4.41E-07 | 2.18E-06 |
| 2.935415622 | 0.003331012 | 0.008516367 |
| 0.192809521 | 0.847108147 | NA |
| 0.06371127 | 0.949200131 | NA |
| 2.881616719 | 0.003956407 | 0.009957653 |
| NA | NA | NA |
| 1.885149162 | 0.059409702 | 0.107172962 |
| -4.026505024 | 5.66E-05 | 0.000204503 |
| NA | NA | NA |
| -1.739855038 | 0.081884475 | 0.140902545 |
| -1.994359889 | 0.046112739 | 0.086308371 |
| -0.124597859 | 0.900841923 | NA |
| 5.704760389 | 1.17E-08 | 7.07E-08 |
| -4.785332007 | 1.71E-06 | 7.76E-06 |
| -0.065389369 | 0.947863988 | NA |
| -2.24102952 | 0.025024163 | 0.050758087 |
| -2.272397881 | 0.02306249 | 0.047315911 |
| -0.466842721 | 0.640612408 | 0.731135314 |
| 0.233175768 | 0.815624921 | 0.870282709 |
| 0.088036847 | 0.92984739 | NA |
| -6.267883906 | 3.66E-10 | 2.60E-09 |
| -7.274630696 | 3.47E-13 | 3.38E-12 |
| -0.61714017 | 0.537142275 | 0.641352437 |
| 21.61308774 | 1.35E-103 | 3.39E-101 |
| 5.112932273 | 3.17E-07 | 1.60E-06 |

| | | |
|---|---|---|
| 2.834729257 | 0.004586454 | 0.011384266 |
| NA | NA | NA |
| 26.62219123 | 3.76E-156 | 1.62E-153 |
| -0.8246111 | 0.409592418 | 0.521449453 |
| 1.646277734 | 0.099706591 | 0.166161694 |
| -4.229323245 | 2.34E-05 | 9.01E-05 |
| -0.411519023 | 0.680691995 | 0.763407144 |
| -0.065388772 | 0.947864464 | NA |
| 3.759529931 | 0.000170233 | 0.000565035 |
| -0.211239413 | 0.832700457 | 0.882819791 |
| 6.264220245 | 3.75E-10 | 2.66E-09 |
| 1.826398933 | 0.067790192 | 0.119905909 |
| NA | NA | NA |
| NA | NA | NA |
| -0.27061107 | 0.786690181 | NA |
| -1.785620254 | 0.074160765 | 0.129345966 |
| 0.170576577 | 0.864556717 | 0.905640126 |
| -3.317059395 | 0.000909703 | 0.002641515 |
| -0.843893247 | 0.398729062 | 0.510573921 |
| 1.25920983 | 0.207954552 | 0.304962645 |
| 0.130336314 | 0.896300351 | NA |
| NA | NA | NA |
| -2.186115204 | 0.028807178 | 0.057316666 |
| -0.531771433 | 0.594884311 | 0.693556025 |
| -0.103254845 | 0.917760713 | NA |
| 1.813448547 | 0.069762668 | 0.122899745 |
| -2.772358954 | 0.005565163 | 0.013542294 |
| 0.192810118 | 0.847107679 | NA |
| 0.540520051 | 0.588838437 | 0.68842056 |
| 1.061851755 | 0.28830299 | 0.395691388 |
| -0.194848135 | 0.845511838 | 0.891903027 |
| -0.194487022 | 0.845794556 | NA |
| NA | NA | NA |
| 5.050157316 | 4.41E-07 | 2.18E-06 |
| -4.024774987 | 5.70E-05 | 0.000205834 |
| -4.584314746 | 4.55E-06 | 1.95E-05 |
| -2.447509763 | 0.014384723 | 0.031378269 |
| -1.811228034 | 0.07010556 | 0.123434312 |
| 1.666159496 | 0.095681651 | 0.160624529 |
| 2.973917025 | 0.002940246 | 0.007618463 |
| 6.931455202 | 4.17E-12 | 3.62E-11 |
| -0.726647726 | 0.467441783 | 0.576752927 |
| 1.725986926 | 0.084349765 | 0.144459433 |
| -5.53335492 | 3.14E-08 | 1.81E-07 |
| 0.075654317 | 0.939694121 | 0.958993584 |
| 0.262571712 | 0.792880706 | NA |
| -1.722769422 | 0.08493023 | 0.14523493 |

| | | | |
|---|---|---|---|
| | 1.379565735 | 0.167720394 | 0.255728646 |
| | -4.928924659 | 8.27E-07 | 3.93E-06 |
| | 0.97512531 | 0.329498105 | 0.440373998 |
| | -3.991558155 | 6.56E-05 | 0.00023421 |
| | 4.225896382 | 2.38E-05 | 9.13E-05 |
| | 7.125130429 | 1.04E-12 | 9.65E-12 |
| | 12.6266384 | 1.51E-36 | 7.05E-35 |
| | -1.890490616 | 0.058692374 | 0.106139936 |
| | -0.065388772 | 0.947864464 | NA |
| NA | NA | NA | |
| | 1.591516406 | 0.111493406 | 0.18220684 |
| | 0.042685736 | 0.965952051 | 0.976879993 |
| | 3.008978387 | 0.002621278 | 0.006863749 |
| | -2.025684296 | 0.042797147 | 0.080852335 |
| NA | NA | NA | |
| | -6.065409235 | 1.32E-09 | 8.84E-09 |
| | 0.192808327 | 0.847109082 | NA |
| | 0.332640584 | 0.739405604 | NA |
| | 3.6331849 | 0.000279944 | 0.000894664 |
| NA | NA | NA | |
| NA | NA | NA | |
| | -2.678241665 | 0.00740098 | 0.01740548 |
| | -3.870781461 | 0.000108487 | 0.000372794 |
| | 12.83691603 | 1.02E-37 | 5.06E-36 |
| | 3.323202239 | 0.000889904 | 0.002590034 |
| | -2.440782403 | 0.014655482 | 0.031909075 |
| NA | NA | NA | |
| | 2.840522877 | 0.004503964 | 0.011195057 |
| | -0.127070744 | 0.898884407 | 0.929637084 |
| | -5.50918599 | 3.60E-08 | 2.06E-07 |
| | -3.257735511 | 0.00112305 | 0.003201589 |
| | 0.066361664 | 0.947089891 | 0.963928201 |
| | 1.960378216 | 0.0499516 | 0.092511177 |
| | -4.399638647 | 1.08E-05 | 4.40E-05 |
| | -1.800704339 | 0.071749493 | 0.125824271 |
| | -1.67994659 | 0.092967708 | 0.156815266 |
| | 1.851617829 | 0.064080721 | 0.114200103 |
| | -7.499191478 | 6.42E-14 | 6.72E-13 |
| NA | NA | NA | |
| | -0.194488217 | 0.845793621 | NA |
| | -0.065388772 | 0.947864464 | NA |
| | 3.529944334 | 0.000415647 | 0.001286148 |
| | -2.601779562 | 0.009274144 | 0.021288686 |
| | -0.063404143 | 0.949444688 | 0.965361523 |
| | 0.217705071 | 0.827658912 | 0.879374419 |
| NA | NA | NA | |
| | 0.31685483 | 0.751353758 | NA |

| | | |
|---|---|---|
| -1.479882527 | 0.1389046 | 0.218968518 |
| -2.284103746 | 0.022365434 | 0.046078428 |
| -1.210541006 | 0.226071366 | 0.326179749 |
| -0.402792444 | 0.68710092 | 0.769078617 |
| -2.287313421 | 0.022177536 | 0.045732717 |
| 2.160730512 | 0.030716163 | 0.06057083 |
| -5.094007325 | 3.51E-07 | 1.76E-06 |
| -1.310901757 | 0.189890957 | 0.283336552 |
| 0.320071032 | 0.748914485 | NA |
| -3.452813136 | 0.000554773 | 0.001678672 |
| 0.60616447 | 0.544405544 | NA |
| NA | NA | NA |
| 0.927572321 | 0.353629461 | NA |
| 2.389092557 | 0.016890047 | 0.036174028 |
| -1.13423426 | 0.256696297 | 0.361290435 |
| -4.346593788 | 1.38E-05 | 5.51E-05 |
| NA | NA | NA |
| -1.998395891 | 0.045673757 | 0.085608367 |
| 1.172605862 | 0.240953897 | 0.343298956 |
| -7.181649252 | 6.89E-13 | 6.51E-12 |
| 1.477745057 | 0.13947603 | 0.219744761 |
| 0.844519001 | 0.39837945 | 0.510267981 |
| 6.076548231 | 1.23E-09 | 8.28E-09 |
| -4.293181323 | 1.76E-05 | 6.90E-05 |
| -2.83000718 | 0.004654696 | 0.011534208 |
| -2.96860428 | 0.002991556 | 0.007739394 |
| -1.817176023 | 0.069190165 | 0.122028596 |
| -4.422824497 | 9.74E-06 | 3.98E-05 |
| NA | NA | NA |
| -0.636549829 | 0.524418114 | 0.629583452 |
| -2.111415222 | 0.034736638 | 0.067462604 |
| 1.557276472 | 0.119404859 | 0.193081182 |
| 4.953605708 | 7.29E-07 | 3.49E-06 |
| NA | NA | NA |
| -2.278934723 | 0.022670946 | 0.046634926 |
| -0.401184247 | 0.688284478 | 0.770144563 |
| -3.603843523 | 0.000313546 | 0.000993676 |
| -0.422952104 | 0.672330207 | 0.756367705 |
| 2.706458965 | 0.006800499 | 0.016162513 |
| -3.490568541 | 0.000481994 | 0.001474134 |
| 7.29010191 | 3.10E-13 | 3.03E-12 |
| 0.23819727 | 0.811728094 | 0.867418956 |
| 6.028828494 | 1.65E-09 | 1.10E-08 |
| NA | NA | NA |
| 0.192809521 | 0.847108147 | NA |
| NA | NA | NA |
| 0.348406474 | 0.727534941 | NA |

| | | |
|---|---|---|
| 0.33754632 | 0.735705101 | NA |
| 1.610544245 | 0.107279094 | 0.176390707 |
| 0.063708285 | 0.949202508 | NA |
| -0.444606678 | 0.65660402 | NA |
| -3.747348181 | 0.000178714 | 0.000591538 |
| -0.065388772 | 0.947864464 | NA |
| NA | NA | NA |
| NA | NA | NA |
| -1.245296684 | 0.213022714 | 0.311010181 |
| NA | NA | NA |
| -1.612773242 | 0.106793781 | 0.175696618 |
| -2.983990138 | 0.002845159 | 0.007392748 |
| -2.10442075 | 0.035341761 | 0.068467823 |
| -2.335674776 | 0.019508198 | 0.040895535 |
| -0.123337669 | 0.901839713 | NA |
| 1.393753987 | 0.163391907 | NA |
| NA | NA | NA |
| NA | NA | NA |
| 0.063708882 | 0.949202033 | NA |
| -4.161521585 | 3.16E-05 | 0.000119024 |
| 4.320770806 | 1.55E-05 | 6.15E-05 |
| 6.426531286 | 1.31E-10 | 9.79E-10 |
| -0.985815404 | 0.324223705 | 0.435254457 |
| NA | NA | NA |
| -2.207195335 | 0.027300416 | 0.054701361 |
| NA | NA | NA |
| 3.830710596 | 0.000127774 | 0.000433533 |
| -0.057504079 | 0.954143657 | 0.968616796 |
| 1.268462739 | 0.204632749 | 0.30112426 |
| 9.094518678 | 9.50E-20 | 1.66E-18 |
| NA | NA | NA |
| 5.582296919 | 2.37E-08 | 1.39E-07 |
| 1.113058473 | 0.265683317 | 0.371324665 |
| NA | NA | NA |
| 4.271217137 | 1.94E-05 | 7.57E-05 |
| 3.955657966 | 7.63E-05 | 0.000269792 |
| -2.568619303 | 0.010210456 | 0.023227094 |
| 5.042575759 | 4.59E-07 | 2.26E-06 |
| -1.880228904 | 0.060076888 | 0.108189402 |
| 3.89360737 | 9.88E-05 | 0.000341919 |
| -1.887397756 | 0.059106847 | 0.106732853 |
| NA | NA | NA |
| 2.407264951 | 0.016072505 | 0.03463878 |
| -1.691440954 | 0.090752616 | 0.153648402 |
| NA | NA | NA |
| 0.792340026 | 0.428162439 | 0.539405368 |
| -1.149452302 | 0.250369524 | 0.353993628 |

| | | |
|---|---|---|
| 1.407579988 | 0.159255477 | 0.245075503 |
| 0.517615751 | 0.60472639 | 0.701742573 |
| 0.245087801 | 0.806388449 | 0.862932143 |
| 5.701105841 | 1.19E-08 | 7.21E-08 |
| -0.194626645 | 0.845685242 | 0.891903027 |
| -8.276252612 | 1.27E-16 | 1.71E-15 |
| 3.606736627 | 0.000310072 | 0.000983165 |
| 0.192810118 | 0.847107679 | NA |
| NA | NA | NA |
| -3.522397563 | 0.000427662 | 0.001320388 |
| 0.192809521 | 0.847108147 | NA |
| -2.479720654 | 0.013148535 | 0.029034656 |
| -2.009579595 | 0.044475703 | 0.0836761 |
| NA | NA | NA |
| 1.634152836 | 0.102226763 | 0.169621205 |
| 7.647771622 | 2.04E-14 | 2.26E-13 |
| 8.312860584 | 9.34E-17 | 1.27E-15 |
| 1.976390616 | 0.048110554 | NA |
| 0.482226932 | 0.629644747 | 0.722207314 |
| 0.192810118 | 0.847107679 | NA |
| 1.750227323 | 0.080079096 | 0.138214024 |
| 0.400687315 | 0.688650351 | NA |
| 0.409647917 | 0.682064241 | NA |
| NA | NA | NA |
| NA | NA | NA |
| 0.192809521 | 0.847108147 | NA |
| NA | NA | NA |
| NA | NA | NA |
| NA | NA | NA |
| NA | NA | NA |
| NA | NA | NA |
| NA | NA | NA |
| NA | NA | NA |
| NA | NA | NA |
| NA | NA | NA |
| NA | NA | NA |
| 0.063708882 | 0.949202033 | NA |
| NA | NA | NA |
| NA | NA | NA |
| NA | NA | NA |
| NA | NA | NA |
| NA | NA | NA |
| NA | NA | NA |
| NA | NA | NA |
| NA | NA | NA |
| NA | NA | NA |
| NA | NA | NA |

| | | | |
|---|---|---|---|
| NA | NA | NA | |
| NA | NA | NA | |
| NA | NA | NA | |
| NA | NA | NA | |
| NA | NA | NA | |
| NA | NA | NA | |
| NA | NA | NA | |
| NA | NA | NA | |
| NA | NA | NA | |
| NA | NA | NA | |
| NA | NA | NA | |
| NA | NA | NA | |
| NA | NA | NA | |
| 0.496627515 | 0.619451746 | NA | |
| 0.807954747 | 0.419116633 | | 0.530787762 |
| 0.198308907 | 0.842803383 | NA | |
| NA | NA | NA | |
| NA | NA | NA | |
| NA | NA | NA | |
| 0.31685483 | 0.751353758 | NA | |
| NA | NA | NA | |
| NA | NA | NA | |
| 0.063710673 | 0.949200606 | NA | |
| 0.372342228 | 0.709638061 | NA | |
| NA | NA | NA | |
| -1.163703394 | 0.244544233 | | 0.347367569 |
| NA | NA | NA | |
| NA | NA | NA | |
| 3.53878335 | 0.000401976 | | 0.001248009 |
| NA | NA | NA | |
| NA | NA | NA | |
| -0.19448762 | 0.845794089 | NA | |
| 0.987252195 | 0.32351902 | NA | |
| 0.192809521 | 0.847108147 | NA | |
| NA | NA | NA | |
| NA | NA | NA | |
| NA | NA | NA | |
| NA | NA | NA | |
| NA | NA | NA | |
| NA | NA | NA | |
| 0.192808924 | 0.847108615 | NA | |
| NA | NA | NA | |
| 2.020989707 | 0.043280831 | | 0.081673642 |
| NA | NA | NA | |
| 0.192810118 | 0.847107679 | NA | |
| NA | NA | NA | |
| 4.987305659 | 6.12E-07 | | 2.96E-06 |

| | | |
|---|---|---|
| 3.089049534 | 0.00200798 | 0.005410672 |
| 6.509349567 | 7.55E-11 | 5.81E-10 |
| 0.702775947 | 0.482195389 | 0.590650416 |
| 2.654876417 | 0.007933747 | 0.018489915 |
| -2.354562147 | 0.018544542 | 0.039153935 |
| 1.236086115 | 0.216426555 | 0.315005913 |
| 1.006921308 | 0.313972591 | 0.424406435 |
| -0.12482576 | 0.900661493 | 0.930744279 |
| 1.536374522 | 0.124446552 | 0.200070473 |
| 1.724727304 | 0.084576628 | 0.144739026 |
| -0.851781659 | 0.394335288 | 0.506372474 |
| 1.361356238 | 0.173401139 | 0.262964587 |
| NA | NA | NA |
| 0.192809521 | 0.847108147 | NA |
| 2.386660477 | 0.017002185 | 0.036363727 |
| 1.980139734 | 0.04768783 | 0.088884234 |
| -0.841702833 | 0.399954312 | 0.51167168 |
| -0.177797834 | 0.858881748 | 0.901942948 |
| NA | NA | NA |
| 3.212224729 | 0.001317113 | 0.003691338 |
| NA | NA | NA |
| 0.192810118 | 0.847107679 | NA |
| 2.101620991 | 0.03558649 | 0.068867314 |
| 4.132745697 | 3.58E-05 | 0.000133833 |
| 1.100977427 | 0.270906482 | 0.377192481 |
| 4.655267409 | 3.24E-06 | 1.41E-05 |
| 1.237211615 | 0.216008531 | 0.31452524 |
| 2.221926404 | 0.02628828 | 0.052944612 |
| NA | NA | NA |
| 0.358787056 | 0.719754397 | NA |
| NA | NA | NA |
| 0.013173804 | 0.989489129 | 0.992582274 |
| -1.562483251 | 0.118174188 | 0.191381551 |
| 10.53247814 | 6.12E-26 | 1.67E-24 |
| NA | NA | NA |
| 0.063708285 | 0.949202508 | NA |
| NA | NA | NA |
| NA | NA | NA |
| NA | NA | NA |
| NA | NA | NA |
| NA | NA | NA |
| NA | NA | NA |
| NA | NA | NA |
| 0.146625656 | 0.883427502 | NA |
| -1.171801765 | 0.241276654 | 0.343719673 |
| NA | NA | NA |
| NA | NA | NA |

| | | |
|---|---|---|
| NA | NA | NA |
| NA | NA | NA |
| NA | NA | NA |
| 0.254385151 | 0.799198022 | NA |
| -0.117394102 | 0.906547758 | NA |
| 0.110117865 | 0.9123159 | 0.939490948 |
| 3.510541744 | 0.000447195 | 0.001375603 |
| -1.34874432 | 0.177419105 | 0.267986252 |
| 0.063705421 | 0.949204788 | NA |
| 7.82026121 | 5.27E-15 | 6.16E-14 |
| 0.347315421 | 0.728354366 | NA |
| 5.448194992 | 5.09E-08 | 2.85E-07 |
| -1.341995488 | 0.179597459 | 0.270672229 |
| 9.191787996 | 3.86E-20 | 6.95E-19 |
| -0.134099399 | 0.893323975 | NA |
| 0.728279181 | 0.466442704 | 0.575721726 |
| -0.903504568 | 0.366258165 | 0.478767593 |
| 0.795810759 | 0.426142037 | 0.537536421 |
| -2.368686009 | 0.017851401 | 0.037917564 |
| -0.033593877 | 0.973201005 | 0.981474724 |
| 17.03739453 | 4.34E-65 | 4.89E-63 |
| -0.920990591 | 0.357055339 | 0.468892715 |
| 0.574391398 | 0.565702973 | NA |
| -4.337256752 | 1.44E-05 | 5.73E-05 |
| -4.437782468 | 9.09E-06 | 3.73E-05 |
| -1.253243008 | 0.210117283 | 0.307629597 |
| -3.826259692 | 0.000130105 | 0.000440846 |
| 0.976750928 | 0.328692476 | 0.439650118 |
| -1.761334691 | 0.078181772 | 0.135406155 |
| -1.36978993 | 0.170752486 | 0.259576965 |
| -0.78191798 | 0.434262778 | 0.544923915 |
| 0.31685483 | 0.751353758 | NA |
| NA | NA | NA |
| NA | NA | NA |
| NA | NA | NA |
| NA | NA | NA |
| NA | NA | NA |
| NA | NA | NA |
| 1.632376392 | 0.102600219 | 0.170138908 |
| 2.599108787 | 0.009346615 | 0.021437347 |
| NA | NA | NA |
| NA | NA | NA |
| NA | NA | NA |
| 0.584372778 | 0.558969537 | NA |
| 6.058077877 | 1.38E-09 | 9.23E-09 |
| -3.238296884 | 0.001202456 | 0.003404692 |
| 5.09752361 | 3.44E-07 | 1.73E-06 |

| | | |
|---|---|---|
| NA | NA | NA |
| -1.029540971 | 0.303225538 | 0.412530449 |
| -4.186196542 | 2.84E-05 | 0.000107755 |
| -0.696680612 | 0.486002695 | 0.594296128 |
| 1.895662676 | 0.058004664 | 0.105107995 |
| 5.655022099 | 1.56E-08 | 9.31E-08 |
| -0.065390563 | 0.947863038 | NA |
| 0.192809521 | 0.847108147 | NA |
| 0.033757007 | 0.97307092 | 0.981462166 |
| NA | NA | NA |
| -3.1031923 | 0.001914452 | 0.005177605 |
| NA | NA | NA |
| -5.620847505 | 1.90E-08 | 1.12E-07 |
| 1.710514912 | 0.087170698 | 0.148535174 |
| 0.379042486 | 0.704656315 | 0.783438349 |
| -2.148323724 | 0.031688047 | 0.06226659 |
| -0.693595879 | 0.487935679 | 0.595961062 |
| -0.491682397 | 0.622943884 | 0.716952613 |
| 0.622785303 | 0.533425617 | 0.637797038 |
| -2.881858323 | 0.003953374 | 0.009952403 |
| -12.80469938 | 1.54E-37 | 7.60E-36 |
| 2.99150044 | 0.002776101 | 0.007231327 |
| -7.507482121 | 6.03E-14 | 6.32E-13 |
| -0.198462909 | 0.842682902 | 0.889860062 |
| NA | NA | NA |
| -3.64502459 | 0.000267366 | 0.000857312 |
| NA | NA | NA |
| -0.755496696 | 0.449951014 | 0.559866468 |
| 2.116063348 | 0.03433942 | 0.066806861 |
| 3.848195853 | 0.000118991 | 0.000406045 |
| 3.226259962 | 0.001254194 | 0.003532006 |
| 2.29966964 | 0.021466943 | 0.04442477 |
| -7.499180591 | 6.42E-14 | 6.72E-13 |
| 5.396341203 | 6.80E-08 | 3.74E-07 |
| -2.586212149 | 0.009703718 | 0.022181194 |
| 3.968215509 | 7.24E-05 | 0.000256836 |
| -0.069728076 | 0.944410095 | 0.962295441 |
| 3.654709531 | 0.000257473 | 0.000828349 |
| -0.065389369 | 0.947863988 | NA |
| 0.816792963 | 0.414046754 | 0.525670315 |
| NA | NA | NA |
| -1.625361532 | 0.104085548 | 0.17213487 |
| -0.429899739 | 0.667268575 | 0.752531697 |
| 11.25450579 | 2.20E-29 | 7.34E-28 |
| -1.214732246 | 0.22446823 | 0.324296669 |
| 1.991919431 | 0.046379899 | 0.086737064 |
| -0.630650088 | 0.528269341 | 0.633235075 |

| | | | |
|---|---|---|---|
| | -1.389534283 | 0.164670343 | 0.251922174 |
| | -0.125866448 | 0.899837642 | 0.930161732 |
| | 6.833931822 | 8.26E-12 | 6.95E-11 |
| | 3.96831769 | 7.24E-05 | 0.000256799 |
| | 1.693525477 | 0.09035549 | 0.15312105 |
| | -2.970330245 | 0.002974798 | 0.007700134 |
| | 4.112514913 | 3.91E-05 | 0.000145019 |
| | 0.192809521 | 0.847108147 | NA |
| | 10.06086563 | 8.23E-24 | 1.94E-22 |
| | -2.372741001 | 0.01765664 | 0.037574027 |
| | -4.717115199 | 2.39E-06 | 1.07E-05 |
| | -1.976112505 | 0.048142036 | 0.089590719 |
| | -2.275347628 | 0.022885084 | 0.047021315 |
| | -0.80867463 | 0.418702325 | 0.530343565 |
| | 7.914028192 | 2.49E-15 | 3.00E-14 |
| | 1.743246602 | 0.081290548 | 0.140005672 |
| | -1.404225088 | 0.160251833 | 0.246261259 |
| | -2.29575058 | 0.021690147 | 0.044823676 |
| | -0.65103161 | 0.515026082 | 0.62036106 |
| | 3.892986784 | 9.90E-05 | 0.0003427 |
| | 0.840265038 | 0.4007598 | 0.512335553 |
| NA | NA | NA | |
| | 1.122975206 | 0.261448025 | 0.366592575 |
| | 0.192810118 | 0.847107679 | NA |
| NA | NA | NA | |
| NA | NA | NA | |
| NA | NA | NA | |
| | -0.307673647 | 0.758330674 | 0.825015086 |
| | 1.741505095 | 0.081595081 | 0.14047217 |
| | -0.740975162 | 0.458708501 | 0.568249135 |
| | -0.065389369 | 0.947863988 | NA |
| | -3.654161933 | 0.000258023 | 0.000829906 |
| | 0.904637188 | 0.365657627 | 0.478242337 |
| | -1.820428345 | 0.068693799 | 0.121315585 |
| | -7.795351852 | 6.42E-15 | 7.46E-14 |
| | -6.132477523 | 8.65E-10 | 5.93E-09 |
| | -2.338506532 | 0.019360987 | 0.040617342 |
| NA | NA | NA | |
| | -2.02166977 | 0.043210479 | 0.081571256 |
| | 1.939649969 | 0.052422244 | 0.096423336 |
| | -5.399692193 | 6.68E-08 | 3.68E-07 |
| | 0.445532538 | 0.655934951 | 0.743604177 |
| NA | NA | NA | |
| | 6.986675388 | 2.81E-12 | 2.48E-11 |
| | 2.778535477 | 0.005460455 | 0.013309494 |
| | -9.010649465 | 2.05E-19 | 3.49E-18 |
| NA | NA | NA | |

| | | |
|---|---|---|
| -0.940374083 | 0.347025711 | 0.45856053 |
| -2.249631314 | 0.024472359 | 0.049767886 |
| 0.085451449 | 0.931902492 | 0.953657933 |
| 2.771248675 | 0.005584176 | 0.013583278 |
| -10.38103721 | 3.02E-25 | 7.86E-24 |
| 2.607342665 | 0.009124799 | 0.020984356 |
| -1.75689845 | 0.07893512 | 0.136559722 |
| -0.287768709 | 0.773523791 | 0.837173023 |
| -1.008993293 | 0.312977851 | 0.42329917 |
| -5.265991634 | 1.39E-07 | 7.37E-07 |
| NA | NA | NA |
| -3.198977025 | 0.001379162 | 0.003850576 |
| -3.884152145 | 0.000102688 | 0.000354081 |
| 0.268798119 | 0.788085041 | 0.849187161 |
| -4.628388121 | 3.69E-06 | 1.60E-05 |
| -1.007704148 | 0.313596513 | 0.424012572 |
| -2.327684033 | 0.019928888 | 0.041658598 |
| -1.566619626 | 0.117203628 | 0.190049676 |
| 0.221386987 | 0.824791124 | 0.877258396 |
| -0.364031899 | 0.71583419 | 0.792031198 |
| -1.676390551 | 0.093661719 | 0.15781574 |
| -1.655507623 | 0.097821571 | 0.16356964 |
| -1.517357164 | 0.129176533 | 0.206375733 |
| -3.193207212 | 0.001407019 | 0.003919174 |
| 1.937542202 | 0.0526791 | 0.096824634 |
| -6.745375432 | 1.53E-11 | 1.26E-10 |
| -4.642107212 | 3.45E-06 | 1.50E-05 |
| -0.976959072 | 0.328589415 | 0.439590315 |
| 1.061964771 | 0.288251678 | 0.39568366 |
| 0.439525322 | 0.660280938 | 0.747211858 |
| 1.776864729 | 0.075590499 | 0.131500569 |
| 4.745037094 | 2.08E-06 | 9.36E-06 |
| 0.063710673 | 0.949200606 | NA |
| -5.538362911 | 3.05E-08 | 1.76E-07 |
| -4.538650226 | 5.66E-06 | 2.39E-05 |
| 2.221369239 | 0.026325963 | 0.053003447 |
| 1.887718209 | 0.059063791 | 0.106671959 |
| -0.959137621 | 0.33748942 | 0.448683284 |
| 0.380302023 | 0.703721234 | 0.782718658 |
| 0.192808327 | 0.847109082 | NA |
| -10.68363048 | 1.21E-26 | 3.45E-25 |
| 1.570940012 | 0.116196584 | 0.188661163 |
| 0.328928313 | 0.742209873 | NA |
| -2.201059003 | 0.027731847 | 0.055459263 |
| 0.517119224 | 0.605072935 | 0.702001277 |
| NA | NA | NA |
| NA | NA | NA |

| | | |
|---|---|---|
| 6.310436623 | 2.78E-10 | 2.00E-09 |
| 1.723946137 | 0.084717567 | 0.144925747 |
| -0.04154663 | 0.96686012 | 0.977534078 |
| NA | NA | NA |
| 0.336169089 | 0.736743361 | 0.808199427 |
| -0.119594143 | 0.90480466 | 0.934177315 |
| 0.289455699 | 0.772232677 | 0.836137211 |
| NA | NA | NA |
| -0.786082379 | 0.431819227 | 0.542592131 |
| 0.192810118 | 0.847107679 | NA |
| 2.479940835 | 0.013140418 | 0.029019294 |
| -7.48943003 | 6.92E-14 | 7.22E-13 |
| NA | NA | NA |
| NA | NA | NA |
| -4.631919955 | 3.62E-06 | 1.57E-05 |
| 14.56521614 | 4.68E-48 | 3.43E-46 |
| 8.389303709 | 4.89E-17 | 6.82E-16 |
| NA | NA | NA |
| NA | NA | NA |
| 0.845053327 | 0.398081066 | 0.509952867 |
| -1.735652653 | 0.08262527 | 0.141982048 |
| 5.658385221 | 1.53E-08 | 9.14E-08 |
| 0.192809521 | 0.847108147 | NA |
| 0.063708882 | 0.949202033 | NA |
| NA | NA | NA |
| 1.995924137 | 0.045942181 | 0.086012775 |
| NA | NA | NA |
| NA | NA | NA |
| -2.71666176 | 0.006594396 | 0.015731771 |
| NA | NA | NA |
| 1.829953337 | 0.067256917 | 0.11912948 |
| 8.763503392 | 1.89E-18 | 2.97E-17 |
| -1.190886723 | 0.233698057 | 0.334982383 |
| -1.404092024 | 0.160291448 | 0.246291894 |
| 5.831333007 | 5.50E-09 | 3.46E-08 |
| 0.324699329 | 0.745408643 | 0.814776992 |
| -6.54958016 | 5.77E-11 | 4.51E-10 |
| 0.389795043 | 0.69668811 | 0.776618161 |
| 0.07655416 | 0.938978227 | NA |
| 3.671548282 | 0.000241086 | 0.000779232 |
| 0.342233342 | 0.732175295 | 0.804740159 |
| 8.573976883 | 1.00E-17 | 1.47E-16 |
| 0.381805449 | 0.702605676 | 0.78171547 |
| -0.795012405 | 0.426606286 | 0.538013571 |
| 2.560732808 | 0.010445165 | 0.023702855 |
| NA | NA | NA |
| -0.465586825 | 0.641511276 | 0.731914293 |

| | | |
|---|---|---|
| 0.391582251 | 0.695366904 | 0.77565995 |
| -0.065419841 | 0.947839727 | 0.964443791 |
| -4.212897046 | 2.52E-05 | 9.64E-05 |
| -5.057311984 | 4.25E-07 | 2.11E-06 |
| 9.935945412 | 2.90E-23 | 6.62E-22 |
| 6.643398589 | 3.07E-11 | 2.45E-10 |
| 1.26625409 | 0.205422128 | 0.302045225 |
| 6.668595061 | 2.58E-11 | 2.08E-10 |
| 5.507315838 | 3.64E-08 | 2.08E-07 |
| 1.6559222 | 0.097737575 | 0.163490165 |
| 4.912546306 | 8.99E-07 | 4.25E-06 |
| NA | NA | NA |
| 0.130254918 | 0.896364747 | 0.927867278 |
| 2.155751952 | 0.031103039 | 0.061256517 |
| NA | NA | NA |
| -0.056785925 | 0.954715726 | NA |
| NA | NA | NA |
| 0.31685483 | 0.751353758 | NA |
| NA | NA | NA |
| -0.580700249 | 0.561442494 | 0.663643379 |
| -2.94419735 | 0.003237935 | 0.008304693 |
| -5.654053831 | 1.57E-08 | 9.36E-08 |
| -0.15541299 | 0.876495744 | 0.913769356 |
| -0.108029621 | 0.913972196 | 0.940915844 |
| 0.523181448 | 0.600847991 | 0.698491719 |
| 0.551758995 | 0.581113485 | NA |
| 0.514467835 | 0.606924943 | 0.703562955 |
| 1.591067613 | 0.11159436 | 0.182336108 |
| 0.428649523 | 0.6681783 | NA |
| 0.192809521 | 0.847108147 | NA |
| -0.065389369 | 0.947863988 | NA |
| 0.00837238 | 0.993319885 | 0.995325034 |
| -0.480517227 | 0.630859657 | 0.722933796 |
| -2.08568457 | 0.037007205 | 0.071247623 |
| 0.459110011 | 0.646155168 | 0.735382342 |
| 0.192810118 | 0.847107679 | NA |
| 4.232518905 | 2.31E-05 | 8.89E-05 |
| 3.860396826 | 0.000113203 | 0.000387669 |
| -0.065389369 | 0.947863988 | NA |
| 8.761090336 | 1.93E-18 | 3.04E-17 |
| 0.248092209 | 0.804063063 | NA |
| 3.277103707 | 0.001048778 | 0.003009392 |
| 1.142927505 | 0.253068692 | 0.35714419 |
| -3.983371734 | 6.79E-05 | 0.000241947 |
| 3.920755111 | 8.83E-05 | 0.000308545 |
| 4.664526873 | 3.09E-06 | 1.35E-05 |
| -0.835961742 | 0.403176426 | 0.514820219 |

| | | |
|---|---|---|
| 3.666231229 | 0.000246151 | 0.000794272 |
| -4.810042621 | 1.51E-06 | 6.90E-06 |
| -0.23236855 | 0.816251774 | 0.870654645 |
| 2.911841823 | 0.003593045 | 0.009108155 |
| 10.72307949 | 7.93E-27 | 2.28E-25 |
| NA | NA | NA |
| -1.008979786 | 0.312984329 | 0.42329917 |
| -2.874381358 | 0.004048198 | 0.010167209 |
| NA | NA | NA |
| NA | NA | NA |
| -0.901056284 | 0.367558395 | NA |
| -0.780678809 | 0.434991427 | 0.545510067 |
| 1.087893851 | 0.276641971 | 0.383471703 |
| 1.967204945 | 0.049159588 | 0.091267207 |
| -5.750149792 | 8.92E-09 | 5.48E-08 |
| -5.272614302 | 1.34E-07 | 7.12E-07 |
| -3.316150504 | 0.000912667 | 0.002649199 |
| NA | NA | NA |
| 7.883869213 | 3.17E-15 | 3.78E-14 |
| 2.689389907 | 0.007158275 | 0.016890237 |
| NA | NA | NA |
| 0.192810118 | 0.847107679 | NA |
| -1.553787595 | 0.120235091 | 0.194285624 |
| 0.805701504 | 0.420414982 | NA |
| 5.478211786 | 4.30E-08 | 2.44E-07 |
| NA | NA | NA |
| 0.770790197 | 0.440831299 | 0.550949812 |
| -0.631956102 | 0.527415562 | 0.632423662 |
| NA | NA | NA |
| 8.43516494 | 3.31E-17 | 4.68E-16 |
| 0.481625447 | 0.630072046 | 0.722432584 |
| -1.439771276 | 0.14993212 | 0.233383145 |
| NA | NA | NA |
| 0.824227714 | 0.409810183 | NA |
| 0.833140522 | 0.404765482 | NA |
| -2.262859889 | 0.023644328 | 0.048331362 |
| 5.163545516 | 2.42E-07 | 1.24E-06 |
| 0.191055948 | 0.84848176 | 0.894210461 |
| 3.727892588 | 0.000193088 | 0.000636254 |
| -0.281991858 | 0.777949753 | 0.841017679 |
| 3.307461336 | 0.000941457 | 0.002728656 |
| -1.214404048 | 0.22459347 | 0.324440155 |
| -1.168486016 | 0.242610777 | 0.345227274 |
| -0.194487022 | 0.845794556 | NA |
| 2.310027778 | 0.020886616 | 0.043381629 |
| -0.065389966 | 0.947863513 | NA |
| -3.316736191 | 0.000910756 | 0.002644266 |

| | | |
|---|---|---|
| 6.459068624 | 1.05E-10 | 7.99E-10 |
| 0.192308348 | 0.84750068 | NA |
| 0.969430092 | 0.332330644 | 0.443473691 |
| 1.380184427 | 0.167529868 | 0.25551597 |
| NA | NA | NA |
| NA | NA | NA |
| -0.904999746 | 0.365465522 | 0.478031041 |
| -1.461446761 | 0.143892876 | 0.225539928 |
| -1.917418371 | 0.055184802 | 0.100786253 |
| -4.50324905 | 6.69E-06 | 2.80E-05 |
| 1.065335225 | 0.286724258 | 0.393964187 |
| 1.116923564 | 0.264027008 | 0.36956694 |
| -1.996632881 | 0.045865078 | 0.085902642 |
| -1.52022792 | 0.128453703 | 0.205496368 |
| -3.645346485 | 0.000267032 | 0.000856349 |
| -2.184442481 | 0.02892975 | 0.057528532 |
| -0.118424305 | 0.905731468 | NA |
| -0.065389369 | 0.947863988 | NA |
| 7.854243773 | 4.02E-15 | 4.74E-14 |
| 7.496189584 | 6.57E-14 | 6.87E-13 |
| NA | NA | NA |
| 0.748745322 | 0.454010721 | 0.563936695 |
| 6.328392228 | 2.48E-10 | 1.79E-09 |
| 2.065918416 | 0.038836184 | NA |
| 0.192808924 | 0.847108615 | NA |
| -9.686264264 | 3.45E-22 | 7.21E-21 |
| -4.09106557 | 4.29E-05 | 0.00015787 |
| -1.828484385 | 0.067476887 | 0.11948684 |
| -2.427636688 | 0.015197559 | 0.032963163 |
| -1.161306089 | 0.245517444 | 0.348511406 |
| -2.640832473 | 0.00827026 | 0.019195532 |
| 0.547221566 | 0.584226521 | NA |
| -4.879251793 | 1.06E-06 | 4.98E-06 |
| -2.036799834 | 0.041670105 | 0.079011675 |
| -3.85907839 | 0.000113815 | 0.0003895 |
| 0.723043489 | 0.469653166 | 0.578714062 |
| 0.428435268 | 0.668334252 | 0.753328215 |
| 0.063708183 | 0.949202589 | NA |
| 1.801684933 | 0.07159499 | 0.125650161 |
| 0.004915889 | 0.996077704 | 0.997432586 |
| -3.041223727 | 0.002356187 | 0.006236819 |
| 0.13400901 | 0.89339545 | 0.925643187 |
| -3.132377687 | 0.001733967 | 0.004739155 |
| -2.621994049 | 0.008741696 | 0.020188655 |
| 0.063708882 | 0.949202033 | NA |
| -0.065389369 | 0.947863988 | NA |
| -5.8215546 | 5.83E-09 | 3.65E-08 |

| | | |
|---|---|---|
| -2.261104179 | 0.023752807 | 0.048523229 |
| 0.897379603 | 0.369516393 | NA |
| -3.014510277 | 0.002573945 | 0.006751817 |
| -0.568508817 | 0.569689521 | 0.671185159 |
| -2.416131286 | 0.015686404 | 0.033903005 |
| -5.727381733 | 1.02E-08 | 6.22E-08 |
| NA | NA | NA |
| NA | NA | NA |
| -3.690581839 | 0.000223742 | 0.000728346 |
| 16.65361508 | 2.85E-62 | 3.02E-60 |
| 1.283648602 | 0.199264936 | 0.294758072 |
| -1.433593751 | 0.151688228 | 0.235706254 |
| 0.278584686 | 0.780563566 | 0.843224258 |
| 1.826774058 | 0.067733748 | 0.119839946 |
| -3.376254934 | 0.000734798 | 0.002174065 |
| NA | NA | NA |
| -1.959522876 | 0.050051584 | 0.092662066 |
| 1.303223884 | 0.192498338 | 0.286543765 |
| 8.498370924 | 1.92E-17 | 2.78E-16 |
| 0.609947997 | 0.541896256 | NA |
| -1.49592932 | 0.134672075 | 0.213398457 |
| 2.773344103 | 0.005548342 | 0.013507927 |
| 0.609208432 | 0.542386292 | 0.646411241 |
| -1.676226854 | 0.093693766 | 0.157859112 |
| 0.192810118 | 0.847107679 | NA |
| 1.005156177 | 0.314821653 | 0.425393326 |
| 0.445600191 | 0.655886073 | 0.743604177 |
| NA | NA | NA |
| 3.562200334 | 0.00036776 | 0.001149622 |
| -0.486197984 | 0.6268268 | 0.719981236 |
| 15.64929818 | 3.36E-55 | 2.94E-53 |
| 26.96283279 | 4.03E-160 | 1.87E-157 |
| -1.231175749 | 0.218257139 | 0.317098688 |
| 4.993923175 | 5.92E-07 | 2.87E-06 |
| -1.505393287 | 0.132222994 | 0.210209701 |
| -0.01152082 | 0.990807919 | 0.993586627 |
| 7.326927149 | 2.35E-13 | 2.33E-12 |
| 0.995249646 | 0.319614861 | 0.430313591 |
| 3.9951813 | 6.46E-05 | 0.00023092 |
| -2.697827865 | 0.006979352 | 0.016531956 |
| -0.46191414 | 0.644142894 | 0.733759461 |
| 10.13772261 | 3.76E-24 | 9.09E-23 |
| NA | NA | NA |
| 38.03818661 | 0 | 0 |
| 2.444410056 | 0.014508925 | 0.03161473 |
| -0.065388772 | 0.947864464 | NA |
| -0.065388772 | 0.947864464 | NA |

| | | |
|---|---|---|
| -0.717632883 | 0.472983677 | 0.581927014 |
| -0.695444115 | 0.486777022 | 0.594923285 |
| 2.046111244 | 0.040745427 | 0.077457352 |
| 4.316221167 | 1.59E-05 | 6.27E-05 |
| 0.170555432 | 0.864573344 | 0.905640126 |
| 10.30986791 | 6.36E-25 | 1.61E-23 |
| -0.637094993 | 0.52406297 | NA |
| -1.223382297 | 0.221185337 | 0.320430221 |
| NA | NA | NA |
| 2.208444415 | 0.027213309 | 0.054539924 |
| 8.943915539 | 3.76E-19 | 6.27E-18 |
| NA | NA | NA |
| 7.167049911 | 7.66E-13 | 7.20E-12 |
| 6.229987128 | 4.66E-10 | 3.28E-09 |
| 7.719902316 | 1.16E-14 | 1.31E-13 |
| 24.03408632 | 1.22E-127 | 4.20E-125 |
| 36.96765387 | 3.79E-299 | 1.36E-295 |
| 27.69469189 | 8.09E-169 | 4.13E-166 |
| 1.906127227 | 0.056633701 | 0.103011521 |
| 30.97078478 | 1.33E-210 | 1.19E-207 |
| 2.355970064 | 0.018474406 | 0.039025605 |
| 2.798812565 | 0.00512909 | 0.012586385 |
| 0.342385066 | 0.732061126 | NA |
| 14.7286896 | 4.22E-49 | 3.20E-47 |
| 13.0177394 | 9.70E-39 | 5.07E-37 |
| 0.787379661 | 0.431059649 | NA |
| 10.72975232 | 7.38E-27 | 2.13E-25 |
| 1.712055455 | 0.086886449 | 0.148134856 |
| 12.75668724 | 2.86E-37 | 1.38E-35 |
| 1.247281968 | 0.212294127 | NA |
| 3.298282218 | 0.000972783 | 0.002810661 |
| NA | NA | NA |
| NA | NA | NA |
| -1.345857957 | 0.17834833 | 0.269065077 |
| 2.326227353 | 0.020006425 | 0.04180029 |
| NA | NA | NA |
| NA | NA | NA |
| NA | NA | NA |
| NA | NA | NA |
| NA | NA | NA |
| 2.397010724 | 0.01652944 | 0.035501277 |
| 7.611145149 | 2.72E-14 | 2.96E-13 |
| NA | NA | NA |
| 11.69554017 | 1.34E-31 | 5.06E-30 |
| -3.895922067 | 9.78E-05 | 0.000338904 |
| 0.114922714 | 0.908506382 | 0.936917092 |
| -5.369564168 | 7.89E-08 | 4.31E-07 |

| | | | |
|---|---|---|---|
| | -0.065390563 | 0.947863038 | NA |
| | -1.262185302 | 0.206882116 | 0.303781495 |
| | -3.020813431 | 0.002520966 | 0.006625421 |
| NA | NA | NA | |
| | -2.096705079 | 0.036019691 | 0.069587284 |
| | -4.544672426 | 5.50E-06 | 2.33E-05 |
| | -0.065389369 | 0.947863988 | NA |
| | 2.541071201 | 0.011051341 | 0.024920276 |
| | 3.850443955 | 0.000117904 | 0.000402665 |
| | -1.259790073 | 0.207745102 | 0.304798353 |
| | -0.065388772 | 0.947864464 | NA |
| | 1.135329364 | 0.256237343 | 0.360863147 |
| | 29.70199389 | 7.24E-194 | 4.77E-191 |
| | 0.648576833 | 0.516611934 | NA |
| | 0.063710673 | 0.949200606 | NA |
| NA | NA | NA | |
| | 0.192809521 | 0.847108147 | NA |
| | 0.350236135 | 0.72616149 | NA |
| | 13.65241036 | 1.95E-42 | 1.18E-40 |
| | 2.216759389 | 0.026639538 | 0.053531442 |
| | 3.101245274 | 0.001927086 | 0.005210084 |
| | 1.389337232 | 0.164730227 | 0.251998386 |
| NA | NA | NA | |
| NA | NA | NA | |
| | 4.841992936 | 1.29E-06 | 5.93E-06 |
| | 11.43742765 | 2.72E-30 | 9.48E-29 |
| NA | NA | NA | |
| NA | NA | NA | |
| | 13.94963224 | 3.16E-44 | 2.07E-42 |
| | -0.658533027 | 0.510195682 | 0.616128348 |
| | -4.991861642 | 5.98E-07 | 2.89E-06 |
| | -1.027649018 | 0.304114958 | 0.413583124 |
| | 3.206386684 | 0.001344132 | 0.003762428 |
| | 8.710135682 | 3.04E-18 | 4.68E-17 |
| NA | NA | NA | |
| NA | NA | NA | |
| | 1.597463842 | 0.110162351 | 0.180396706 |
| | 3.669233116 | 0.000243279 | 0.000786017 |
| | 3.885933701 | 0.000101937 | 0.000351833 |
| | 1.336324443 | 0.181443251 | 0.272928159 |
| | 4.956196115 | 7.19E-07 | 3.45E-06 |
| | -4.397600073 | 1.09E-05 | 4.43E-05 |
| | 1.989876865 | 0.046604501 | 0.087118049 |
| | 5.846020606 | 5.03E-09 | 3.18E-08 |
| | -3.259341244 | 0.001116713 | 0.003184247 |
| | -0.547729864 | 0.583877403 | 0.683941005 |
| | 0.195323091 | 0.845140021 | 0.891739934 |

| | | |
|---|---|---|
| NA | NA | NA |
| 16.67543137 | 1.98E-62 | 2.12E-60 |
| -0.114616953 | 0.908748742 | 0.937089812 |
| -1.497370438 | 0.134296896 | 0.212945523 |
| 1.60576607 | 0.108325318 | NA |
| -2.22727747 | 0.025928735 | 0.052325747 |
| 2.762700768 | 0.005732529 | 0.013890146 |
| -0.894632705 | 0.370983463 | 0.483782228 |
| 3.235495412 | 0.001214318 | 0.00343168 |
| 0.192808924 | 0.847108615 | NA |
| 0.142545149 | 0.88664942 | 0.920547694 |
| 6.399644466 | 1.56E-10 | 1.15E-09 |
| 1.681412122 | 0.092682893 | 0.156450823 |
| 0.364757961 | 0.715292089 | 0.791781398 |
| 2.513598373 | 0.011950644 | 0.026667079 |
| 1.672259948 | 0.094473074 | 0.158915392 |
| -1.543009986 | 0.122828353 | 0.197850162 |
| -1.078366389 | 0.28087028 | 0.387850773 |
| -0.399311012 | 0.689664054 | 0.770972974 |
| -1.92539701 | 0.054179709 | 0.099189447 |
| 12.91530192 | 3.69E-38 | 1.87E-36 |
| 0.136605807 | 0.891342386 | NA |
| 10.01062968 | 1.37E-23 | 3.19E-22 |
| 9.393416877 | 5.81E-21 | 1.10E-19 |
| 0.907509674 | 0.364137336 | NA |
| 10.13104117 | 4.02E-24 | 9.68E-23 |
| 0.559541996 | 0.575791879 | 0.676818713 |
| NA | NA | NA |
| 4.870695494 | 1.11E-06 | 5.18E-06 |
| 0.157838271 | 0.874584241 | NA |
| -9.272679811 | 1.82E-20 | 3.33E-19 |
| 2.004550285 | 0.045011146 | 0.084537329 |
| NA | NA | NA |
| 0.193437579 | 0.846616288 | 0.892733104 |
| -1.165660123 | 0.243751889 | 0.346535828 |
| -1.328890219 | 0.183884191 | 0.276019269 |
| 30.62448371 | 5.78E-206 | 4.99E-203 |
| 1.331163248 | 0.183135301 | 0.275093116 |
| 0.373693917 | 0.708632051 | 0.78655275 |
| 16.09873125 | 2.60E-58 | 2.47E-56 |
| -0.862774586 | 0.388261415 | 0.500472923 |
| 10.82654055 | 2.58E-27 | 7.63E-26 |
| 0.063708285 | 0.949202508 | NA |
| 9.499945438 | 2.10E-21 | 4.16E-20 |
| 31.66033624 | 5.47E-220 | 5.07E-217 |
| 12.79642333 | 1.72E-37 | 8.43E-36 |
| 0.815452803 | 0.41481317 | 0.526429734 |

| | | |
|---|---|---|
| 20.48167057 | 3.14E-93 | 6.65E-91 |
| 16.16361756 | 9.11E-59 | 8.73E-57 |
| 1.385919001 | 0.165771638 | 0.253316234 |
| 7.953544532 | 1.81E-15 | 2.21E-14 |
| NA | NA | NA |
| NA | NA | NA |
| 2.143041503 | 0.032109766 | 0.063025991 |
| 5.053488227 | 4.34E-07 | 2.15E-06 |
| -0.440348884 | 0.65968444 | 0.746739397 |
| 5.25149684 | 1.51E-07 | 7.93E-07 |
| -0.065389966 | 0.947863513 | NA |
| NA | NA | NA |
| -0.065389966 | 0.947863513 | NA |
| -0.065389966 | 0.947863513 | NA |
| 0.464282325 | 0.642445486 | NA |
| -0.332657325 | 0.739392966 | NA |
| -6.370485302 | 1.88E-10 | 1.38E-09 |
| 1.359809181 | 0.173890316 | 0.2636106 |
| 8.174468674 | 2.97E-16 | 3.87E-15 |
| -0.427775596 | 0.668814506 | 0.753801652 |
| -0.179265537 | 0.857729206 | 0.901261997 |
| 0.46651466 | 0.640847157 | NA |
| 0.275770074 | 0.78272467 | NA |
| 0.192636473 | 0.847243678 | NA |
| 5.40319195 | 6.55E-08 | 3.62E-07 |
| 0.264249229 | 0.791587879 | 0.851695927 |
| -0.959672216 | 0.337220211 | 0.44847996 |
| 1.725392451 | 0.084456771 | 0.144583337 |
| 2.732332599 | 0.006288762 | 0.015071593 |
| 0.582525562 | 0.560212729 | 0.66257101 |
| 13.29690214 | 2.41E-40 | 1.34E-38 |
| 0.06371127 | 0.949200131 | NA |
| 5.199406686 | 2.00E-07 | 1.03E-06 |
| -2.572134896 | 0.01010735 | 0.023013461 |
| -0.829087168 | 0.407055084 | 0.518846502 |
| -6.669787222 | 2.56E-11 | 2.07E-10 |
| 5.023339338 | 5.08E-07 | 2.48E-06 |
| 5.118566862 | 3.08E-07 | 1.56E-06 |
| NA | NA | NA |
| 0.854532845 | 0.392809819 | 0.504750232 |
| -0.826761357 | 0.408372337 | 0.5202341 |
| 2.9891158 | 0.002797861 | 0.007281944 |
| NA | NA | NA |
| 1.888761814 | 0.058923752 | 0.106488195 |
| -0.237562846 | 0.81222017 | 0.867796596 |
| -0.38599979 | 0.699496825 | 0.778954171 |
| 10.25286935 | 1.15E-24 | 2.86E-23 |

| | | | |
|---|---|---|---|
| | 25.73800837 | 4.39E-146 | 1.74E-143 |
| | 36.82012087 | 8.80E-297 | 2.45E-293 |
| | 30.53475238 | 9.01E-205 | 7.52E-202 |
| | 9.824281794 | 8.85E-23 | 1.93E-21 |
| | 21.8696224 | 5.06E-106 | 1.35E-103 |
| | 10.80417277 | 3.29E-27 | 9.64E-26 |
| | 6.359194559 | 2.03E-10 | 1.49E-09 |
| NA | NA | NA | |
| | 3.684401301 | 0.000229241 | 0.000744795 |
| | 7.532494192 | 4.98E-14 | 5.27E-13 |
| | 6.323368191 | 2.56E-10 | 1.85E-09 |
| | 0.942165188 | 0.346108074 | NA |
| | 19.27520779 | 8.68E-83 | 1.45E-80 |
| | 6.8755437 | 6.18E-12 | 5.27E-11 |
| | 0.324874675 | 0.745275925 | NA |
| | 7.849961235 | 4.16E-15 | 4.89E-14 |
| | -0.243390232 | 0.80770311 | NA |
| | 0.192810118 | 0.847107679 | NA |
| | 0.193472986 | 0.84658856 | NA |
| | 5.316808614 | 1.06E-07 | 5.68E-07 |
| | 2.620005008 | 0.008792848 | 0.020288069 |
| | 0.27349457 | 0.784473066 | NA |
| NA | NA | NA | |
| | 0.751944626 | 0.45208436 | NA |
| | -0.065388772 | 0.947864464 | NA |
| NA | NA | NA | |
| | 2.959983206 | 0.003076558 | 0.00793143 |
| | 0.882822453 | 0.377332207 | NA |
| | 0.095291971 | 0.92408292 | NA |
| | 1.560491467 | 0.118643785 | 0.192058374 |
| | 0.073671169 | 0.94127204 | NA |
| NA | NA | NA | |
| NA | NA | NA | |
| NA | NA | NA | |
| NA | NA | NA | |
| NA | NA | NA | |
| | -0.065389966 | 0.947863513 | NA |
| | -0.327938983 | 0.742957798 | NA |
| | 0.473490159 | 0.635863521 | 0.727240755 |
| | -1.944022094 | 0.051892787 | 0.095604038 |
| | -0.065388772 | 0.947864464 | NA |
| NA | NA | NA | |
| NA | NA | NA | |
| NA | NA | NA | |
| NA | NA | NA | |
| NA | NA | NA | |
| NA | NA | NA | |

| | | | |
|---|---|---|---|
| | 0.063708882 | 0.949202033 | NA |
| NA | NA | NA | |
| | 0.39390925 | 0.693648041 | NA |
| | 0.063708882 | 0.949202033 | NA |
| NA | NA | NA | |
| NA | NA | NA | |
| | 0.192809521 | 0.847108147 | NA |
| NA | NA | NA | |
| NA | NA | NA | |
| | 2.531960709 | 0.01134267 | 0.025499105 |
| | 0.362026528 | 0.717332207 | NA |
| | 0.501493897 | 0.616023572 | NA |
| NA | NA | NA | |
| NA | NA | NA | |
| NA | NA | NA | |
| NA | NA | NA | |
| NA | NA | NA | |
| NA | NA | NA | |
| NA | NA | NA | |
| | 0.092644471 | 0.926186013 | NA |
| | 0.192809521 | 0.847108147 | NA |
| NA | NA | NA | |
| | 15.53169697 | 2.12E-54 | 1.80E-52 |
| NA | NA | NA | |
| | 4.15972786 | 3.19E-05 | 0.000119856 |
| | 3.121054727 | 0.001802045 | 0.004902739 |
| | -1.38020504 | 0.167523523 | 0.25551597 |
| | 1.482148419 | 0.138300808 | 0.21819535 |
| | 0.192809521 | 0.847108147 | NA |
| | 0.345178988 | 0.729959814 | 0.802869189 |
| | 4.887920288 | 1.02E-06 | 4.78E-06 |
| | 3.485404705 | 0.000491393 | 0.001501046 |
| | 1.870215945 | 0.061453836 | 0.110240774 |
| NA | NA | NA | |
| | -2.083165825 | 0.037236111 | 0.071611207 |
| | -2.357907565 | 0.018378269 | 0.038858596 |
| | -3.982984796 | 6.81E-05 | 0.000242238 |
| | -3.856021952 | 0.000115247 | 0.000393968 |
| | -1.697775365 | 0.089550167 | 0.15201361 |
| | -1.452425759 | 0.146383258 | 0.228726696 |
| | -6.678475343 | 2.41E-11 | 1.95E-10 |
| | 0.192809521 | 0.847108147 | NA |
| | 0.192808327 | 0.847109082 | NA |
| | 0.063707135 | 0.949203424 | NA |
| | 0.192809521 | 0.847108147 | NA |
| | 2.458304354 | 0.013959481 | 0.030606677 |
| | 0.92883463 | 0.352974795 | NA |

| | | | |
|---|---|---|---|
| | 5.761341018 | 8.34E-09 | 5.14E-08 |
| | 7.837874544 | 4.58E-15 | 5.37E-14 |
| | 8.162650031 | 3.28E-16 | 4.26E-15 |
| | 0.192810118 | 0.847107679 | NA |
| | 9.407367364 | 5.09E-21 | 9.73E-20 |
| | 0.639341327 | 0.522600909 | NA |
| | 0.06371127 | 0.949200131 | NA |
| | 0.192809521 | 0.847108147 | NA |
| NA | NA | NA | |
| | 4.222570419 | 2.42E-05 | 9.26E-05 |
| | 2.796472841 | 0.005166375 | 0.012667944 |
| NA | NA | NA | |
| | 0.063708882 | 0.949202033 | NA |
| | 20.27056048 | 2.34E-91 | 4.84E-89 |
| | 4.844742227 | 1.27E-06 | 5.86E-06 |
| NA | NA | NA | |
| NA | NA | NA | |
| | 0.063976859 | 0.948988653 | NA |
| | -0.065388772 | 0.947864464 | NA |
| NA | NA | NA | |
| NA | NA | NA | |
| NA | NA | NA | |
| NA | NA | NA | |
| NA | NA | NA | |
| NA | NA | NA | |
| NA | NA | NA | |
| NA | NA | NA | |
| NA | NA | NA | |
| NA | NA | NA | |
| NA | NA | NA | |
| | 0.192808924 | 0.847108615 | NA |
| | 2.456068678 | 0.014046631 | 0.030773494 |
| | -0.404440584 | 0.685888761 | 0.767927881 |
| | 1.286787001 | 0.198168539 | 0.293431054 |
| | 0.063708882 | 0.949202033 | NA |
| NA | NA | NA | |
| NA | NA | NA | |
| NA | NA | NA | |
| | 0.192809521 | 0.847108147 | NA |
| NA | NA | NA | |
| | 0.192809521 | 0.847108147 | NA |
| NA | NA | NA | |
| | 0.192810118 | 0.847107679 | NA |
| NA | NA | NA | |
| | -1.853125456 | 0.063864379 | 0.113919998 |
| | 0.343222146 | 0.731431346 | NA |
| NA | NA | NA | |

| | | |
|---|---|---|
| 0.192636473 | 0.847243678 | NA |
| NA | NA | NA |
| -0.065389369 | 0.947863988 | NA |
| NA | NA | NA |
| NA | NA | NA |
| 0.407334021 | 0.683762679 | NA |
| NA | NA | NA |
| -0.108690324 | 0.913448116 | NA |
| NA | NA | NA |
| NA | NA | NA |
| NA | NA | NA |
| NA | NA | NA |
| NA | NA | NA |
| NA | NA | NA |
| NA | NA | NA |
| NA | NA | NA |
| 0.20929253 | 0.834219883 | 0.88383169 |
| NA | NA | NA |
| -0.065388772 | 0.947864464 | NA |
| NA | NA | NA |
| -0.065388772 | 0.947864464 | NA |
| -0.065389369 | 0.947863988 | NA |
| 0.192810118 | 0.847107679 | NA |
| NA | NA | NA |
| NA | NA | NA |
| NA | NA | NA |
| NA | NA | NA |
| NA | NA | NA |
| 0.063710673 | 0.949200606 | NA |
| NA | NA | NA |
| NA | NA | NA |
| NA | NA | NA |
| NA | NA | NA |
| -0.065389369 | 0.947863988 | NA |
| NA | NA | NA |
| NA | NA | NA |
| NA | NA | NA |
| NA | NA | NA |
| NA | NA | NA |
| NA | NA | NA |
| NA | NA | NA |
| 7.199882027 | 6.03E-13 | 5.73E-12 |
| 0.429610934 | 0.667478682 | 0.752668893 |
| 1.127346767 | 0.259595897 | 0.364649267 |
| NA | NA | NA |
| NA | NA | NA |
| 0.204950389 | 0.837610886 | 0.886448796 |

| | | | |
|---|---|---|---|
| NA | NA | NA | |
| NA | NA | NA | |
| | 0.633748541 | 0.526244941 | NA |
| NA | NA | NA | |
| NA | NA | NA | |
| NA | NA | NA | |
| NA | NA | NA | |
| NA | NA | NA | |
| NA | NA | NA | |
| | 0.304966188 | 0.760391905 | NA |
| | 0.314020606 | 0.753505389 | NA |
| NA | NA | NA | |
| | 2.569037685 | 0.010198137 | 0.023201178 |
| | -0.257164985 | 0.797051413 | NA |
| | -0.912607089 | 0.361449225 | 0.473693601 |
| | 6.87264367 | 6.30E-12 | 5.37E-11 |
| | -2.938214661 | 0.003301084 | 0.008447615 |
| | 0.063708882 | 0.949202033 | NA |
| | -2.716698115 | 0.006593672 | 0.015731542 |
| | 0.448205756 | 0.654004709 | 0.742002399 |
| | 2.643215701 | 0.00821227 | 0.019066238 |
| | -1.320428461 | 0.186692006 | 0.279379464 |
| | 2.231837793 | 0.025625686 | 0.051797699 |
| | -4.76746273 | 1.87E-06 | 8.43E-06 |
| | 0.461925311 | 0.644134882 | 0.733759461 |
| | 6.878172093 | 6.06E-12 | 5.18E-11 |
| | 0.192809521 | 0.847108147 | NA |
| | 2.56947052 | 0.010185406 | 0.023178535 |
| | 6.278662084 | 3.41E-10 | 2.43E-09 |
| | 3.271502374 | 0.001069777 | 0.003064746 |
| | -0.30323504 | 0.761710748 | NA |
| | -3.244429822 | 0.00117686 | 0.003340532 |
| | -5.457821441 | 4.82E-08 | 2.71E-07 |
| | -1.23807683 | 0.215687574 | 0.314094716 |
| NA | NA | NA | |
| | -2.322559931 | 0.020202805 | 0.042160765 |
| | 0.063708882 | 0.949202033 | NA |
| | -9.058095086 | 1.33E-19 | 2.29E-18 |
| | 9.568246618 | 1.09E-21 | 2.20E-20 |
| | 9.499776124 | 2.10E-21 | 4.16E-20 |
| | 5.229169277 | 1.70E-07 | 8.90E-07 |
| NA | NA | NA | |
| | 1.048934206 | 0.294208401 | 0.402384366 |
| | 0.063710673 | 0.949200606 | NA |
| | 2.514160631 | 0.011931608 | 0.026636473 |
| | 0.895253618 | 0.370651528 | 0.483424926 |
| NA | NA | NA | |

| | | | |
|---|---|---|---|
| | 2.293720401 | 0.021806564 | 0.045034513 |
| | -2.031461655 | 0.042208186 | 0.079878337 |
| | -0.424657418 | 0.671086429 | 0.755409846 |
| | -0.667631192 | 0.504369042 | 0.610932094 |
| | -2.781268014 | 0.005414701 | 0.013212125 |
| | -0.826448782 | 0.408549561 | 0.520406896 |
| | -6.428674095 | 1.29E-10 | 9.67E-10 |
| | 9.790842402 | 1.23E-22 | 2.67E-21 |
| | 3.712067419 | 0.000205573 | 0.000674289 |
| | 3.107480803 | 0.001886892 | 0.005110249 |
| | -1.551286535 | 0.120833032 | 0.195062934 |
| | -0.352560849 | 0.724417693 | 0.798739036 |
| NA | NA | NA | |
| NA | NA | NA | |
| | -0.373430582 | 0.708828002 | 0.7866273 |
| | -8.951407698 | 3.51E-19 | 5.89E-18 |
| | 0.310971365 | 0.755822391 | NA |
| | 2.585340051 | 0.0097283 | 0.022229263 |
| | -0.509975227 | 0.610068818 | NA |
| | -4.11720158 | 3.84E-05 | 0.000142413 |
| | -5.091599911 | 3.55E-07 | 1.78E-06 |
| | 4.176861757 | 2.96E-05 | 0.000111885 |
| | -1.739375979 | 0.081968651 | 0.140998923 |
| | 3.322126303 | 0.000893342 | 0.002598229 |
| | 3.887450298 | 0.000101303 | 0.000349884 |
| | 2.567018312 | 0.01025772 | 0.023321893 |
| | 4.104044913 | 4.06E-05 | 0.000149969 |
| | -0.821594685 | 0.411307615 | 0.523057264 |
| NA | NA | NA | |
| | 4.099438844 | 4.14E-05 | 0.000152718 |
| | 4.609084151 | 4.04E-06 | 1.74E-05 |
| | -7.580252099 | 3.45E-14 | 3.71E-13 |
| | 3.779723829 | 0.000157002 | 0.000524249 |
| | 0.938788214 | 0.347839491 | NA |
| | -6.470968491 | 9.74E-11 | 7.42E-10 |
| | -2.817237142 | 0.004843875 | 0.011968626 |
| | -3.473348148 | 0.000514008 | 0.00156459 |
| | -0.699824971 | 0.484036618 | 0.592499832 |
| | -4.609105647 | 4.04E-06 | 1.74E-05 |
| | 1.700723404 | 0.088994938 | 0.151194142 |
| | 0.139672038 | 0.88891912 | 0.922303332 |
| | -4.344616899 | 1.40E-05 | 5.55E-05 |
| | -0.091952038 | 0.926736146 | NA |
| | 1.308672445 | 0.19064533 | 0.284292678 |
| | -8.189764845 | 2.62E-16 | 3.44E-15 |
| | -0.153945642 | 0.877652596 | 0.914503526 |
| | -0.177156917 | 0.859385136 | 0.902244451 |

| | | |
|---|---|---|
| 2.043427319 | 0.041010155 | 0.07787014 |
| -2.147382448 | 0.031762847 | 0.062393977 |
| 2.22507353 | 0.026076302 | 0.052585389 |
| -1.508813018 | 0.131346573 | 0.209028785 |
| 7.777972432 | 7.37E-15 | 8.50E-14 |
| -5.074943064 | 3.88E-07 | 1.93E-06 |
| 1.714747847 | 0.086391467 | 0.14735117 |
| -1.756391543 | 0.079021576 | 0.136666863 |
| -0.938033182 | 0.348227358 | 0.459881326 |
| 0.866098106 | 0.386436364 | 0.498685409 |
| 4.616894546 | 3.90E-06 | 1.68E-05 |
| 0.20780639 | 0.835380143 | 0.884611625 |
| 3.112932141 | 0.001852386 | 0.005023862 |
| 6.857804569 | 6.99E-12 | 5.93E-11 |
| -3.715325853 | 0.000202942 | 0.000666444 |
| -4.676308374 | 2.92E-06 | 1.28E-05 |
| 3.524171409 | 0.000424809 | 0.001312389 |
| 0.440507249 | 0.659569763 | NA |
| -1.326746852 | 0.184592437 | 0.276866533 |
| -1.53572571 | 0.124605667 | 0.200300548 |
| 1.177960584 | 0.23881232 | 0.340965854 |
| NA | NA | NA |
| -8.208823108 | 2.23E-16 | 2.95E-15 |
| -2.067789664 | 0.038659806 | 0.073979736 |
| 0.707692337 | 0.479136337 | 0.587881496 |
| 0.879159459 | 0.379314822 | 0.49187908 |
| 2.910453044 | 0.003609052 | 0.009145026 |
| 2.482624247 | 0.013041858 | 0.028832158 |
| -0.137221532 | 0.890855692 | NA |
| 2.325755149 | 0.020031617 | 0.041838398 |
| 3.533647699 | 0.000409867 | 0.001269518 |
| 9.153824624 | 5.50E-20 | 9.79E-19 |
| 4.213009707 | 2.52E-05 | 9.64E-05 |
| 0.889857917 | 0.373542183 | 0.486282875 |
| -2.314549086 | 0.020637622 | 0.042946432 |
| 0.82345518 | 0.410249195 | 0.522093622 |
| -2.11356259 | 0.034552644 | 0.06716259 |
| -1.556758349 | 0.119527869 | 0.193242641 |
| -0.177410595 | 0.859185886 | 0.902110941 |
| -3.548828425 | 0.000386949 | 0.001204644 |
| -1.458003689 | 0.144839526 | 0.226810144 |
| -1.705573419 | 0.088087532 | 0.149906916 |
| -0.03254787 | 0.974035142 | NA |
| 0.643485027 | 0.519909427 | NA |
| 0.141414929 | 0.887542161 | 0.921141946 |
| 0.493024204 | 0.621995485 | 0.716223351 |
| -0.065388772 | 0.947864464 | NA |

| | | | |
|---|---|---|---|
| NA | NA | NA | |
| -4.413739893 | 1.02E-05 | | 4.14E-05 |
| 0.529656814 | 0.596349895 | NA | |
| NA | NA | NA | |
| NA | NA | NA | |
| NA | NA | NA | |
| NA | NA | NA | |
| NA | NA | NA | |
| 8.064360566 | 7.36E-16 | | 9.31E-15 |
| 7.790798878 | 6.66E-15 | | 7.72E-14 |
| 6.379097175 | 1.78E-10 | | 1.31E-09 |
| 3.180896692 | 0.0014682 | | 0.004074625 |
| 16.5196844 | 2.65E-61 | | 2.75E-59 |
| 10.25682094 | 1.10E-24 | | 2.75E-23 |
| 0.76405603 | 0.444833854 | | 0.55474024 |
| -0.570791517 | 0.568140974 | | 0.669833659 |
| 3.769030446 | 0.000163883 | | 0.000545985 |
| 1.695487633 | 0.089982954 | | 0.152593045 |
| 0.247600382 | 0.804443615 | | 0.861510512 |
| NA | NA | NA | |
| 0.266348516 | 0.789970812 | | 0.850624918 |
| NA | NA | NA | |
| NA | NA | NA | |
| -2.314079584 | 0.020663357 | | 0.042985691 |
| -0.400637512 | 0.688687023 | NA | |
| NA | NA | NA | |
| 0.192809521 | 0.847108147 | NA | |
| 0.107096977 | 0.914712045 | NA | |
| 5.415571581 | 6.11E-08 | | 3.39E-07 |
| -3.721102986 | 0.000198355 | | 0.000652407 |
| -0.924555921 | 0.35519695 | | 0.467138486 |
| NA | NA | NA | |
| 0.063708285 | 0.949202508 | NA | |
| 7.007676268 | 2.42E-12 | | 2.15E-11 |
| NA | NA | NA | |
| NA | NA | NA | |
| -3.114744892 | 0.001841041 | | 0.00499634 |
| -5.135242061 | 2.82E-07 | | 1.43E-06 |
| 3.294715091 | 0.000985215 | | 0.002844283 |
| -2.258334088 | 0.023924839 | | 0.048821026 |
| -0.257309657 | 0.796939738 | | 0.855487483 |
| -5.441103859 | 5.30E-08 | | 2.96E-07 |
| -0.644952496 | 0.518957971 | | 0.624255959 |
| -2.016327308 | 0.043765761 | | 0.0825043 |
| -2.536671144 | 0.011191203 | | 0.025212943 |
| -1.895008882 | 0.058091225 | | 0.105219145 |
| -2.090747393 | 0.036550714 | | 0.070504344 |

| | | |
|---|---|---|
| -2.667651098 | 0.007638353 | 0.017886424 |
| -3.105669504 | 0.001898488 | 0.005137205 |
| -3.574541624 | 0.000350842 | 0.001101967 |
| -2.789682354 | 0.005275978 | 0.012903822 |
| 4.950231614 | 7.41E-07 | 3.55E-06 |
| 0.154290749 | 0.877380492 | NA |
| -1.107593145 | 0.268037565 | 0.373925998 |
| 8.296436116 | 1.07E-16 | 1.45E-15 |
| 4.010608251 | 6.06E-05 | 0.000217424 |
| 0.405422617 | 0.685166889 | NA |
| 6.41666467 | 1.39E-10 | 1.04E-09 |
| 0.192808924 | 0.847108615 | NA |
| 3.237221907 | 0.001206995 | 0.003414211 |
| -0.074014514 | 0.940998836 | NA |
| NA | NA | NA |
| 0.192806584 | 0.847110447 | NA |
| NA | NA | NA |
| NA | NA | NA |
| NA | NA | NA |
| -0.678377203 | 0.49753256 | NA |
| 1.541515749 | 0.123191313 | 0.198348962 |
| NA | NA | NA |
| NA | NA | NA |
| NA | NA | NA |
| NA | NA | NA |
| NA | NA | NA |
| -0.271694065 | 0.785857265 | 0.847297323 |
| NA | NA | NA |
| NA | NA | NA |
| -0.110197645 | 0.91225263 | 0.939490948 |
| 0.192809521 | 0.847108147 | NA |
| NA | NA | NA |
| NA | NA | NA |
| -0.297050656 | 0.766427835 | 0.831506598 |
| -7.782872203 | 7.09E-15 | 8.20E-14 |
| 0.396434461 | 0.691784548 | 0.772524638 |
| 1.484067123 | 0.137791114 | 0.217555768 |
| 0.639325586 | 0.522611147 | 0.627865845 |
| NA | NA | NA |
| 3.997210541 | 6.41E-05 | 0.000229048 |
| 1.139623775 | 0.254443076 | 0.358719736 |
| 2.70463184 | 0.006838013 | 0.016243567 |
| 1.947751735 | 0.051444676 | 0.094869244 |
| 1.206995972 | 0.227433695 | 0.327635418 |
| 4.944507747 | 7.63E-07 | 3.64E-06 |
| 4.44542037 | 8.77E-06 | 3.61E-05 |
| -0.338679105 | 0.734851483 | 0.806618981 |

| | | |
|---|---|---|
| 3.035332907 | 0.002402703 | 0.006347194 |
| -8.241619155 | 1.70E-16 | 2.27E-15 |
| -5.072407806 | 3.93E-07 | 1.96E-06 |
| -2.675232556 | 0.007467743 | 0.017546006 |
| -2.486567886 | 0.012898196 | 0.028539767 |
| -0.939693709 | 0.347374693 | NA |
| 2.793832709 | 0.005208742 | 0.012760576 |
| 2.306863614 | 0.021062425 | 0.043688778 |
| -2.45492998 | 0.014091203 | 0.030849542 |
| -1.338244344 | 0.180816798 | 0.272165762 |
| 5.592459692 | 2.24E-08 | 1.31E-07 |
| -0.585260109 | 0.558372833 | 0.660775945 |
| 10.5894593 | 3.34E-26 | 9.19E-25 |
| -8.915967944 | 4.84E-19 | 8.01E-18 |
| -3.630613122 | 0.000282749 | 0.000902935 |
| 0.063707135 | 0.949203424 | NA |
| 6.70590339 | 2.00E-11 | 1.63E-10 |
| -0.432298884 | 0.665524198 | 0.751412815 |
| -1.016236348 | 0.309516852 | 0.419448121 |
| 1.013444468 | 0.310847908 | 0.421023979 |
| 9.021786915 | 1.85E-19 | 3.16E-18 |
| 4.467784378 | 7.90E-06 | 3.27E-05 |
| NA | NA | NA |
| 10.44349988 | 1.57E-25 | 4.16E-24 |
| -4.466544164 | 7.95E-06 | 3.29E-05 |
| -8.453206748 | 2.83E-17 | 4.03E-16 |
| NA | NA | NA |
| 0.317642674 | 0.750756 | NA |
| -1.243916659 | 0.213530237 | 0.311664478 |
| -0.980682166 | 0.326749501 | 0.437784572 |
| -0.78785943 | 0.430778934 | 0.541774531 |
| 3.976246856 | 7.00E-05 | 0.000248848 |
| 3.78312201 | 0.000154873 | 0.000517693 |
| -1.631115119 | 0.102866031 | 0.170444642 |
| 3.199197798 | 0.001378106 | 0.003848333 |
| -3.640401858 | 0.000272213 | 0.000871959 |
| -0.808924256 | 0.418558716 | 0.530215328 |
| -0.244009704 | 0.807223304 | 0.863379169 |
| -3.61800653 | 0.000296881 | 0.000944721 |
| -0.717000876 | 0.473373556 | 0.582180627 |
| 6.549343294 | 5.78E-11 | 4.51E-10 |
| 1.12095264 | 0.262308022 | NA |
| 0.753948264 | 0.450880288 | 0.560758935 |
| 0.091648006 | 0.926977708 | 0.949977565 |
| 0.192809521 | 0.847108147 | NA |
| 3.411409012 | 0.000646281 | 0.001930589 |
| -1.347295193 | 0.17788518 | 0.268528198 |

| | | |
|---|---|---|
| -0.32618468 | 0.744284632 | NA |
| 6.022059104 | 1.72E-09 | 1.14E-08 |
| 5.026401097 | 5.00E-07 | 2.45E-06 |
| -0.724112286 | 0.468996802 | 0.5782184 |
| -0.39540263 | 0.692545768 | NA |
| -10.3552974 | 3.96E-25 | 1.02E-23 |
| 1.071529463 | 0.28393143 | 0.390913295 |
| 12.52648093 | 5.35E-36 | 2.45E-34 |
| -7.739056596 | 1.00E-14 | 1.14E-13 |
| -0.77308985 | 0.439469208 | 0.549720853 |
| 4.329150966 | 1.50E-05 | 5.93E-05 |
| 1.134390263 | 0.256630882 | 0.361234449 |
| 5.791127525 | 6.99E-09 | 4.34E-08 |
| 14.05860358 | 6.82E-45 | 4.54E-43 |
| NA | NA | NA |
| 6.023671463 | 1.71E-09 | 1.13E-08 |
| 6.897435235 | 5.29E-12 | 4.56E-11 |
| 21.46354537 | 3.41E-102 | 8.14E-100 |
| 9.887956546 | 4.70E-23 | 1.05E-21 |
| NA | NA | NA |
| 0.063708285 | 0.949202508 | NA |
| NA | NA | NA |
| -0.784135433 | 0.432960646 | 0.543698825 |
| 1.904597718 | 0.056832383 | 0.103327853 |
| -1.950041107 | 0.05117122 | 0.094434537 |
| 0.464301778 | 0.64243155 | 0.732504476 |
| 0.324874675 | 0.745275925 | NA |
| 1.432428062 | 0.152021352 | 0.236150589 |
| NA | NA | NA |
| NA | NA | NA |
| NA | NA | NA |
| NA | NA | NA |
| NA | NA | NA |
| NA | NA | NA |
| -0.50055633 | 0.616683403 | 0.711612446 |
| -0.239185104 | 0.810962053 | NA |
| NA | NA | NA |
| -3.412605027 | 0.000643451 | 0.001922366 |
| -6.343218339 | 2.25E-10 | 1.64E-09 |
| -1.41160377 | 0.158066662 | 0.243681236 |
| NA | NA | NA |
| -3.305766404 | 0.00094717 | 0.002742991 |
| -2.87854128 | 0.00399519 | 0.010044155 |
| NA | NA | NA |
| 0.27946001 | 0.779891823 | 0.842680437 |
| -0.49293485 | 0.622058622 | 0.716230153 |
| -7.273333801 | 3.51E-13 | 3.41E-12 |

| | | | |
|---|---|---|---|
| | -1.642873354 | 0.100409139 | 0.167059343 |
| NA | | NA | NA |
| NA | | NA | NA |
| | 7.344348937 | 2.07E-13 | 2.06E-12 |
| NA | | NA | NA |
| NA | | NA | NA |
| NA | | NA | NA |
| | -0.044219141 | 0.964729725 | 0.976061817 |
| NA | | NA | NA |
| | 4.983347197 | 6.25E-07 | 3.02E-06 |
| | 1.173577496 | 0.240564299 | 0.342900023 |
| | 6.698983761 | 2.10E-11 | 1.71E-10 |
| | 5.772599124 | 7.81E-09 | 4.81E-08 |
| | -0.080005474 | 0.936232902 | 0.956525146 |
| | 8.480394865 | 2.24E-17 | 3.23E-16 |
| | -4.550328326 | 5.36E-06 | 2.27E-05 |
| | 0.806725739 | 0.419824512 | 0.53152289 |
| | 0.078934652 | 0.937084601 | NA |
| | 4.87233192 | 1.10E-06 | 5.14E-06 |
| | 0.154374174 | 0.877314716 | 0.914318497 |
| | -6.990795299 | 2.73E-12 | 2.41E-11 |
| | -3.048075728 | 0.002303119 | 0.006114453 |
| | 0.541864379 | 0.587911937 | 0.687701458 |
| | 0.854844035 | 0.392637497 | 0.504603521 |
| | -4.405875498 | 1.05E-05 | 4.28E-05 |
| | -0.184909881 | 0.853299721 | 0.898100991 |
| NA | | NA | NA |
| | -0.065388772 | 0.947864464 | NA |
| | 0.494685841 | 0.620821897 | 0.7151343 |
| | 6.596392325 | 4.21E-11 | 3.33E-10 |
| | 0.665586517 | 0.505675426 | NA |
| | -2.473051859 | 0.01339647 | 0.029514448 |
| | 0.063708882 | 0.949202033 | NA |
| | 1.794135345 | 0.072791569 | 0.127337132 |
| | -8.6251815 | 6.40E-18 | 9.57E-17 |
| | 0.074985411 | 0.94022632 | NA |
| | -0.140297568 | 0.888424886 | 0.921866964 |
| | -1.956942227 | 0.050354261 | 0.093167294 |
| | 1.549998491 | 0.121141878 | 0.1954892 |
| | 1.440376868 | 0.149760805 | 0.233145475 |
| NA | | NA | NA |
| | -5.730661185 | 1.00E-08 | 6.11E-08 |
| | 2.899091194 | 0.00374246 | 0.009457222 |
| | 0.566261108 | 0.571216295 | 0.672540753 |
| | 15.69180729 | 1.72E-55 | 1.52E-53 |
| | 2.603319555 | 0.009232585 | 0.021208848 |
| NA | | NA | NA |

| | | |
|---|---|---|
| 2.241674839 | 0.024982395 | 0.05068569 |
| 1.744114371 | 0.081139149 | 0.13981226 |
| 0.19280579 | 0.847111069 | NA |
| NA | NA | NA |
| -1.35325281 | 0.175974879 | 0.266319078 |
| 10.63574997 | 2.03E-26 | 5.71E-25 |
| -0.48385667 | 0.628487593 | 0.721342831 |
| 2.782082224 | 0.005401135 | 0.013184164 |
| 11.46790221 | 1.91E-30 | 6.73E-29 |
| 8.639021903 | 5.67E-18 | 8.51E-17 |
| 0.025944178 | 0.979301863 | 0.985602157 |
| 0.486137332 | 0.626869799 | 0.719981236 |
| 0.204454056 | 0.837998692 | 0.886671764 |
| 4.245384558 | 2.18E-05 | 8.43E-05 |
| 0.063708285 | 0.949202508 | NA |
| 0.372248965 | 0.709707492 | 0.787305701 |
| -2.833233311 | 0.004607974 | 0.011433147 |
| 2.481251064 | 0.013092212 | 0.028923043 |
| 8.119496626 | 4.68E-16 | 6.00E-15 |
| -0.355520504 | 0.722199687 | 0.796960505 |
| -0.330582033 | 0.740960219 | 0.811473826 |
| -0.578524168 | 0.562910286 | 0.665045051 |
| 5.423736028 | 5.84E-08 | 3.24E-07 |
| NA | NA | NA |
| 0.751892977 | 0.452115422 | 0.562056671 |
| 1.323705141 | 0.185600991 | 0.2780793 |
| 1.061898974 | 0.288281551 | 0.39568366 |
| -1.552882399 | 0.120451232 | 0.194554452 |
| NA | NA | NA |
| 6.464835568 | 1.01E-10 | 7.71E-10 |
| 9.247294178 | 2.30E-20 | 4.18E-19 |
| 0.063708285 | 0.949202508 | NA |
| 0.714669237 | 0.474813451 | 0.583779383 |
| 0.063707194 | 0.949203376 | NA |
| -1.931967534 | 0.053363513 | 0.097895686 |
| 0.305831315 | 0.759733084 | NA |
| -1.036655115 | 0.299896617 | 0.408801456 |
| -2.015842253 | 0.043816474 | 0.082572379 |
| 6.342940513 | 2.25E-10 | 1.64E-09 |
| 21.83964815 | 9.75E-106 | 2.54E-103 |
| NA | NA | NA |
| -0.376105115 | 0.706838752 | 0.784923423 |
| 1.486367356 | 0.137181979 | 0.216771777 |
| 0.501731772 | 0.615856212 | 0.710887336 |
| -1.686126808 | 0.091771378 | 0.155121393 |
| 1.636529051 | 0.101728911 | 0.168929519 |
| 6.331169714 | 2.43E-10 | 1.76E-09 |

| | | | |
|---|---|---|---|
| | 1.327102045 | 0.184474929 | 0.276723449 |
| | 1.77197611 | 0.076398523 | 0.132730966 |
| | 0.38834224 | 0.697762782 | NA |
| | 3.685014759 | 0.000228689 | 0.000743196 |
| | 13.33264218 | 1.50E-40 | 8.30E-39 |
| | 0.063976859 | 0.948988653 | NA |
| | -0.811442668 | 0.417111504 | 0.528756695 |
| | 3.278179504 | 0.001044789 | 0.002999941 |
| NA | NA | NA | |
| NA | NA | NA | |
| NA | NA | NA | |
| | -1.615756707 | 0.106146925 | 0.174850776 |
| | 0.063708882 | 0.949202033 | NA |
| | -1.13261366 | 0.25737653 | 0.361983283 |
| | -0.469812967 | 0.63848865 | 0.729479893 |
| | 5.420320657 | 5.95E-08 | 3.30E-07 |
| | 7.078131753 | 1.46E-12 | 1.33E-11 |
| | -1.717814501 | 0.085830456 | 0.146564081 |
| | -0.065389966 | 0.947863513 | NA |
| | -9.477738783 | 2.60E-21 | 5.12E-20 |
| | -6.701906314 | 2.06E-11 | 1.68E-10 |
| | -3.55488296 | 0.000378147 | 0.001179393 |
| | 20.60937529 | 2.26E-94 | 4.96E-92 |
| | 30.04088696 | 2.87E-198 | 2.00E-195 |
| | 0.199526594 | 0.841850842 | NA |
| | 4.846270575 | 1.26E-06 | 5.82E-06 |
| | 1.149592161 | 0.250311888 | 0.353939696 |
| | -1.01688153 | 0.309209791 | 0.419077381 |
| | 2.012897003 | 0.044125466 | 0.083085866 |
| | -4.944939718 | 7.62E-07 | 3.64E-06 |
| | 3.651815877 | 0.000260393 | 0.000836666 |
| NA | NA | NA | |
| | -1.243873389 | 0.213546165 | 0.311665335 |
| | -2.46307887 | 0.013774961 | 0.030241845 |
| | -1.111187232 | 0.266487766 | 0.372261903 |
| | -0.804630207 | 0.421033106 | 0.532810487 |
| | -0.153037115 | 0.878369007 | 0.91476425 |
| | 13.84841298 | 1.30E-43 | 8.20E-42 |
| | 0.199526594 | 0.841850842 | NA |
| | -5.359234338 | 8.36E-08 | 4.55E-07 |
| | -0.122450714 | 0.902542076 | 0.932187347 |
| NA | NA | NA | |
| NA | NA | NA | |
| | 1.224413077 | 0.220796439 | 0.320081945 |
| | -0.782018858 | 0.434203491 | 0.544876836 |
| | 2.724706917 | 0.006435859 | 0.015390231 |
| | 0.862282168 | 0.388532263 | 0.500718904 |

| | | |
|---|---|---|
| 1.160918571 | 0.245675016 | 0.348697998 |
| -4.039638448 | 5.35E-05 | 0.00019428 |
| 0.34715139 | 0.728477587 | 0.801741332 |
| -0.562739465 | 0.573612307 | 0.67488559 |
| -0.459833982 | 0.64563539 | NA |
| -2.731550324 | 0.006303711 | 0.015104527 |
| -5.015795755 | 5.28E-07 | 2.58E-06 |
| 0.295789065 | 0.767391174 | 0.832335273 |
| -0.112920952 | 0.910093225 | NA |
| 0.471339641 | 0.637398212 | NA |
| -5.278549714 | 1.30E-07 | 6.91E-07 |
| 3.23486814 | 0.001216989 | 0.003437287 |
| -3.307265279 | 0.000942116 | 0.002730002 |
| -0.50530786 | 0.613342623 | NA |
| NA | NA | NA |
| NA | NA | NA |
| NA | NA | NA |
| NA | NA | NA |
| 3.011506427 | 0.002599549 | 0.006811129 |
| NA | NA | NA |
| -0.214515747 | 0.830144889 | 0.881155446 |
| NA | NA | NA |
| 0.693407433 | 0.488053899 | 0.596047284 |
| -2.127375371 | 0.033388907 | 0.06521963 |
| -0.693134602 | 0.488225084 | 0.596227257 |
| -6.032849993 | 1.61E-09 | 1.07E-08 |
| -3.702813936 | 0.000213221 | 0.000696909 |
| 3.676101035 | 0.000236826 | 0.000767346 |
| 0.945112768 | 0.344601308 | 0.455982804 |
| 0.192810118 | 0.847107679 | NA |
| 0.651149009 | 0.514950302 | NA |
| 9.015761549 | 1.96E-19 | 3.33E-18 |
| -1.882560682 | 0.059759929 | 0.107696092 |
| -0.04821078 | 0.961548259 | 0.97360651 |
| 0.397439157 | 0.691043645 | 0.77200725 |
| -3.562667769 | 0.000367105 | 0.001147863 |
| -0.295406379 | 0.767683461 | NA |
| -1.679023203 | 0.093147522 | 0.157097397 |
| 1.152839853 | 0.248976123 | 0.352361889 |
| -0.065390563 | 0.947863038 | NA |
| -2.878203979 | 0.003999464 | 0.010052882 |
| -0.984285092 | 0.324975358 | 0.436006282 |
| -5.851186676 | 4.88E-09 | 3.09E-08 |
| 0.176259751 | 0.860089881 | 0.902761959 |
| -9.597560274 | 8.19E-22 | 1.67E-20 |
| -1.23908905 | 0.215312522 | 0.313639785 |
| 0.113507484 | 0.90962823 | 0.937649065 |

| | | |
|---|---|---|
| NA | NA | NA |
| -2.2633757 | 0.023612539 | 0.048278211 |
| 0.063710673 | 0.949200606 | NA |
| -0.868082871 | 0.385348968 | NA |
| -3.61948851 | 0.000295186 | 0.000939894 |
| -0.733116597 | 0.463487318 | NA |
| -0.918954467 | 0.358119389 | NA |
| -0.065389369 | 0.947863988 | NA |
| -0.19448762 | 0.845794089 | NA |
| -0.515309936 | 0.606336457 | NA |
| -0.360212113 | 0.718688517 | 0.794311546 |
| 1.96148709 | 0.04982223 | 0.092312562 |
| -0.519150221 | 0.603655989 | NA |
| -0.542535736 | 0.587449497 | 0.687281237 |
| -5.576203042 | 2.46E-08 | 1.43E-07 |
| NA | NA | NA |
| 5.116329962 | 3.12E-07 | 1.57E-06 |
| NA | NA | NA |
| 1.419401048 | 0.155782127 | 0.240907951 |
| 0.063707019 | 0.949203516 | NA |
| NA | NA | NA |
| 0.222540612 | 0.823893062 | NA |
| 4.672520417 | 2.98E-06 | 1.31E-05 |
| 0.742848106 | 0.457573647 | 0.56720822 |
| 1.976062405 | 0.04814771 | 0.089594616 |
| NA | NA | NA |
| 0.884273736 | 0.376548463 | 0.489204195 |
| -4.915912896 | 8.84E-07 | 4.18E-06 |
| -0.065389369 | 0.947863988 | NA |
| 7.902487445 | 2.73E-15 | 3.28E-14 |
| -0.462566415 | 0.643675186 | 0.733393569 |
| 6.265124941 | 3.73E-10 | 2.65E-09 |
| -0.975883146 | 0.329122375 | 0.440005849 |
| 0.619782129 | 0.535401236 | NA |
| 1.525513939 | 0.127130959 | 0.203744904 |
| -4.374624575 | 1.22E-05 | 4.89E-05 |
| -0.065389369 | 0.947863988 | NA |
| 0.495984725 | 0.619905188 | NA |
| 6.382980891 | 1.74E-10 | 1.28E-09 |
| -3.703161505 | 0.000212929 | 0.000696045 |
| -0.951935595 | 0.341129646 | 0.452681319 |
| -0.07741303 | 0.938294976 | NA |
| 0.67190392 | 0.501644865 | 0.608459535 |
| -1.801044678 | 0.071695838 | 0.125774591 |
| 0.761014352 | 0.446648495 | NA |
| -1.057780118 | 0.290155699 | NA |
| -0.102170225 | 0.918621561 | NA |

| | | |
|---|---|---|
| NA | NA | NA |
| 1.544664089 | 0.122427536 | 0.197268007 |
| -6.242815784 | 4.30E-10 | 3.04E-09 |
| -2.403400241 | 0.016243398 | 0.034970949 |
| 2.709715734 | 0.00673409 | 0.016028363 |
| -0.392062979 | 0.695011681 | 0.775367372 |
| -2.393418972 | 0.016692168 | 0.035800265 |
| 0.063708882 | 0.949202033 | NA |
| 1.2198656 | 0.222515828 | 0.321977754 |
| -7.71779337 | 1.18E-14 | 1.34E-13 |
| NA | NA | NA |
| -0.943916445 | 0.345212348 | 0.456718911 |
| -1.357640087 | 0.174577916 | 0.264523606 |
| -3.041496835 | 0.00235405 | 0.006232482 |
| -0.194487022 | 0.845794556 | NA |
| -0.560794796 | 0.574937435 | NA |
| -1.268487967 | 0.204623745 | 0.30112426 |
| NA | NA | NA |
| -1.229988565 | 0.218701387 | 0.317615069 |
| 0.063707194 | 0.949203376 | NA |
| NA | NA | NA |
| NA | NA | NA |
| NA | NA | NA |
| NA | NA | NA |
| 1.215009748 | 0.224362374 | 0.324178762 |
| NA | NA | NA |
| 0.319226478 | 0.74955478 | NA |
| -0.734785262 | 0.462470284 | 0.571918538 |
| 2.527859038 | 0.011476042 | 0.025766534 |
| -2.770515858 | 0.005596757 | 0.013608591 |
| -6.933448247 | 4.11E-12 | 3.57E-11 |
| 0.066394788 | 0.94706352 | NA |
| -1.00375551 | 0.315496474 | 0.42600619 |
| NA | NA | NA |
| 1.988539559 | 0.046752048 | 0.087361237 |
| NA | NA | NA |
| 0.317642674 | 0.750756 | NA |
| 0.192808327 | 0.847109082 | NA |
| 9.427618297 | 4.20E-21 | 8.12E-20 |
| 1.69764693 | 0.08957442 | 0.152034158 |
| 3.625775142 | 0.000288096 | 0.000918721 |
| 0.417818162 | 0.676080071 | NA |
| 3.822479556 | 0.000132116 | 0.000447117 |
| NA | NA | NA |
| NA | NA | NA |
| NA | NA | NA |
| 2.815188906 | 0.004874857 | 0.012037254 |

| | | |
|---|---|---|
| 12.14153812 | 6.36E-34 | 2.66E-32 |
| -0.222206819 | 0.824152886 | NA |
| 2.120769369 | 0.033941216 | 0.066138468 |
| -3.692160175 | 0.000222357 | 0.000724028 |
| 0.926021215 | 0.354434955 | NA |
| 4.177032062 | 2.95E-05 | 0.000111835 |
| 1.578769931 | 0.114388838 | 0.186148665 |
| -0.44327886 | 0.657564045 | 0.745147955 |
| -2.68776307 | 0.007193242 | 0.016961548 |
| -1.956381503 | 0.050420229 | 0.093275561 |
| -4.730336734 | 2.24E-06 | 1.00E-05 |
| -1.189048737 | 0.234420489 | 0.335767945 |
| NA | NA | NA |
| -0.589123079 | 0.555778712 | 0.658459616 |
| -0.486192262 | 0.626830857 | 0.719981236 |
| 2.298634894 | 0.02152568 | 0.044527916 |
| 1.427824943 | 0.153342251 | 0.237700907 |
| 2.534601751 | 0.011257523 | 0.025341829 |
| NA | NA | NA |
| 0.47635149 | 0.633823988 | 0.725372156 |
| 0.02680838 | 0.97861257 | 0.985146099 |
| NA | NA | NA |
| 0.063708882 | 0.949202033 | NA |
| 1.772889342 | 0.076247045 | 0.132513785 |
| -3.330901318 | 0.000865653 | 0.002525914 |
| 0.356809846 | 0.721234165 | 0.796246081 |
| -4.586515024 | 4.51E-06 | 1.93E-05 |
| 0.127022074 | 0.898922928 | NA |
| NA | NA | NA |
| 3.782314646 | 0.000155377 | 0.000519167 |
| -1.894963562 | 0.058097229 | 0.105220647 |
| -1.60226512 | 0.109096995 | 0.178897916 |
| NA | NA | NA |
| NA | NA | NA |
| 1.24173558 | 0.214334138 | 0.312596636 |
| 0.860222051 | 0.389666652 | 0.501793296 |
| -1.840265038 | 0.065729336 | 0.116805486 |
| NA | NA | NA |
| 0.192810118 | 0.847107679 | NA |
| 7.508097218 | 6.00E-14 | 6.30E-13 |
| NA | NA | NA |
| 0.866287441 | 0.386332552 | 0.498643401 |
| NA | NA | NA |
| NA | NA | NA |
| NA | NA | NA |
| -0.469686433 | 0.638579063 | 0.729479893 |
| -2.601672389 | 0.009277043 | 0.021293386 |

| | | | |
|---|---|---|---|
| | 6.343515393 | 2.25E-10 | 1.64E-09 |
| | -1.594945082 | 0.110724517 | 0.18113952 |
| | 1.875661669 | 0.060701752 | 0.109094914 |
| | -4.42979327 | 9.43E-06 | 3.86E-05 |
| NA | NA | NA | |
| | -0.413305167 | 0.679383044 | 0.762383215 |
| | -0.936873818 | 0.348823469 | 0.460498493 |
| | 1.861386677 | 0.062689591 | 0.112168165 |
| | 3.109130406 | 0.001876389 | 0.005083452 |
| | -5.110070918 | 3.22E-07 | 1.62E-06 |
| | -1.511743997 | 0.130599001 | 0.208143457 |
| | -5.467246271 | 4.57E-08 | 2.58E-07 |
| | -1.732063325 | 0.083262288 | 0.142900101 |
| | 0.192809521 | 0.847108147 | NA |
| NA | NA | NA | |
| | -2.311830906 | 0.020787004 | 0.043197274 |
| NA | NA | NA | |
| | 0.192808924 | 0.847108615 | NA |
| | 0.86134197 | 0.389049726 | 0.501179344 |
| | 7.692734223 | 1.44E-14 | 1.62E-13 |
| | -1.126632843 | 0.259897747 | 0.364827582 |
| | -6.120838656 | 9.31E-10 | 6.36E-09 |
| | 1.15297551 | 0.248920436 | 0.352302998 |
| | -3.342431616 | 0.000830478 | 0.002430076 |
| | 7.019899209 | 2.22E-12 | 1.98E-11 |
| | 0.063710673 | 0.949200606 | NA |
| | 0.951562462 | 0.341318927 | 0.452884475 |
| | 1.964673402 | 0.04945205 | 0.09172694 |
| | 0.351505941 | 0.725208816 | 0.799294437 |
| | -6.278933853 | 3.41E-10 | 2.43E-09 |
| | -1.016972133 | 0.309166686 | 0.419064343 |
| | -2.45151988 | 0.014225433 | 0.031078169 |
| | -0.732053217 | 0.464136087 | NA |
| NA | NA | NA | |
| | 1.446254757 | 0.14810576 | 0.230956774 |
| | 0.192810118 | 0.847107679 | NA |
| | 2.302361333 | 0.021314804 | 0.044153731 |
| | -1.706069153 | 0.087995205 | 0.149790532 |
| | 1.87068812 | 0.061388322 | 0.110146933 |
| | -1.455965478 | 0.145402163 | 0.227520389 |
| | -11.5186817 | 1.06E-30 | 3.79E-29 |
| | 0.255070733 | 0.798668469 | NA |
| | -4.343029851 | 1.41E-05 | 5.59E-05 |
| | -2.108176801 | 0.035015699 | 0.067941314 |
| | 0.380012449 | 0.703936174 | NA |
| NA | NA | NA | |
| | 2.795219185 | 0.005186454 | 0.012709706 |

| | | |
|---|---|---|
| 0.469498505 | 0.638713353 | 0.729600001 |
| 3.21799872 | 0.001290884 | 0.003626163 |
| 3.415932232 | 0.000635641 | 0.001901074 |
| -0.02200401 | 0.982444757 | 0.987732659 |
| 2.099511563 | 0.03577183 | 0.069161818 |
| 4.04441374 | 5.25E-05 | 0.000190666 |
| -8.576147162 | 9.81E-18 | 1.45E-16 |
| -1.834671976 | 0.066554307 | 0.118095442 |
| 7.369837692 | 1.71E-13 | 1.71E-12 |
| -2.913961684 | 0.003568737 | 0.009058461 |
| 3.324492642 | 0.000885796 | 0.002581079 |
| 0.063708882 | 0.949202033 | NA |
| 0.057813281 | 0.953897359 | 0.968484602 |
| -0.197831904 | 0.843176584 | 0.890343819 |
| NA | NA | NA |
| 6.631056216 | 3.33E-11 | 2.66E-10 |
| 2.25732472 | 0.023987792 | 0.048913599 |
| NA | NA | NA |
| -4.90103935 | 9.53E-07 | 4.49E-06 |
| NA | NA | NA |
| 0.192809521 | 0.847108147 | NA |
| NA | NA | NA |
| NA | NA | NA |
| -0.065388772 | 0.947864464 | NA |
| NA | NA | NA |
| NA | NA | NA |
| 1.342480947 | 0.179440104 | 0.27046765 |
| -0.402262441 | 0.687490893 | 0.769465177 |
| 3.179451561 | NA | NA |
| -2.950184769 | 0.003175839 | 0.008157133 |
| -0.657420525 | NA | NA |
| -0.077892384 | 0.937913658 | 0.957734124 |
| NA | NA | NA |
| 1.696486588 | 0.089793768 | 0.1523548 |
| 0.742327819 | 0.457888741 | 0.567514493 |
| -0.032530442 | 0.97404904 | 0.982052988 |
| -8.394408842 | 4.68E-17 | 6.55E-16 |
| -0.06220183 | 0.950402105 | 0.96611155 |
| -6.848881451 | 7.44E-12 | 6.29E-11 |
| 4.599155339 | 4.24E-06 | 1.83E-05 |
| -0.886599047 | 0.375294807 | 0.488132873 |
| 8.697989118 | 3.38E-18 | 5.18E-17 |
| 2.774427774 | 0.005529891 | 0.013466937 |
| NA | NA | NA |
| 0.083888369 | 0.933145187 | NA |
| -3.979791638 | 6.90E-05 | 0.000245341 |
| -0.065389369 | 0.947863988 | NA |

| | | |
|---|---|---|
| NA | NA | NA |
| -2.918253244 | 0.003519983 | 0.008948323 |
| -2.24937151 | 0.02448887 | 0.049797417 |
| 1.353196832 | 0.175992757 | 0.266326673 |
| -2.900655015 | 0.003723836 | 0.009414911 |
| -10.31140367 | 6.26E-25 | 1.58E-23 |
| NA | NA | NA |
| -1.788788744 | 0.073648844 | 0.128581467 |
| -1.315809613 | 0.188237952 | 0.28128931 |
| -8.058169846 | 7.74E-16 | 9.75E-15 |
| NA | NA | NA |
| NA | NA | NA |
| -0.479664296 | 0.63146612 | 0.723368283 |
| -0.642975602 | 0.520239931 | 0.625527476 |
| 3.537592117 | 0.000403793 | 0.001253031 |
| 4.470400005 | 7.81E-06 | 3.24E-05 |
| 5.120167647 | 3.05E-07 | 1.55E-06 |
| 3.808379044 | 0.000139881 | 0.000471481 |
| -2.898708774 | 0.003747028 | 0.009467807 |
| 0.591415815 | 0.554241846 | NA |
| 1.665703018 | 0.095772581 | 0.160723278 |
| -2.290216302 | 0.022008781 | 0.045410913 |
| 0.192810118 | 0.847107679 | NA |
| 2.285242226 | 0.022298628 | 0.045952138 |
| -4.185877461 | 2.84E-05 | 0.000107877 |
| 16.26271619 | 1.82E-59 | 1.79E-57 |
| 0.063708882 | 0.949202033 | NA |
| -0.748987885 | 0.453864508 | 0.563811039 |
| -2.173420173 | 0.029748708 | 0.058941673 |
| -0.844100811 | 0.398613074 | 0.510451512 |
| -7.983623448 | 1.42E-15 | 1.75E-14 |
| -4.104264089 | 4.06E-05 | 0.000149849 |
| -2.8311047 | 0.004638753 | 0.011498118 |
| -4.499779927 | 6.80E-06 | 2.84E-05 |
| -4.630131316 | 3.65E-06 | 1.58E-05 |
| NA | NA | NA |
| NA | NA | NA |
| NA | NA | NA |
| -1.112166586 | 0.266066532 | 0.371767024 |
| NA | NA | NA |
| -4.471572749 | 7.76E-06 | 3.22E-05 |
| -0.378872981 | 0.704782189 | 0.783495113 |
| NA | NA | NA |
| -0.294491425 | 0.768382415 | NA |
| -5.025075278 | 5.03E-07 | 2.46E-06 |
| NA | NA | NA |
| 1.012237681 | 0.311424424 | 0.421576707 |

| | | | |
|---|---|---|---|
| | 0.479703601 | 0.631438167 | 0.723368283 |
| NA | NA | NA | |
| | 0.959619212 | 0.337246896 | 0.44847996 |
| | 1.285247886 | 0.198705675 | 0.294094977 |
| | -1.352822718 | 0.176112273 | 0.266446458 |
| | 3.694667484 | 0.000220175 | 0.000717668 |
| | -0.007862362 | 0.993726807 | 0.995596285 |
| | 4.208156139 | 2.57E-05 | 9.85E-05 |
| | 0.949977652 | 0.342123608 | 0.453567474 |
| | 1.756145033 | 0.079063647 | 0.136706486 |
| | 0.063708882 | 0.949202033 | NA |
| | -0.232632967 | 0.816046426 | 0.870472705 |
| | 0.735878466 | 0.461804663 | 0.571151829 |
| | -1.655911671 | 0.097739708 | 0.163490165 |
| | -0.495236682 | 0.62043306 | NA |
| | -0.115933728 | 0.907705065 | NA |
| | 0.063708285 | 0.949202508 | NA |
| | -0.569851183 | 0.568778637 | 0.670364425 |
| | -0.065388772 | 0.947864464 | NA |
| | 0.192809521 | 0.847108147 | NA |
| | -0.266314409 | 0.789997077 | 0.850624918 |
| | 5.002888436 | 5.65E-07 | 2.74E-06 |
| NA | NA | NA | |
| | -4.396320392 | 1.10E-05 | 4.46E-05 |
| | -9.095362591 | 9.43E-20 | 1.65E-18 |
| | 9.51722115 | 1.78E-21 | 3.55E-20 |
| NA | NA | NA | |
| | 1.061231132 | 0.288584875 | NA |
| | 2.117519744 | 0.034215761 | 0.066611224 |
| | 0.063708882 | 0.949202033 | NA |
| | 4.099432883 | 4.14E-05 | 0.000152718 |
| NA | NA | NA | |
| | -0.970856409 | 0.331619788 | 0.442595888 |
| | -6.479910017 | 9.18E-11 | 7.01E-10 |
| NA | NA | NA | |
| | -3.493502317 | 0.000476729 | 0.0014591 |
| | -5.217149784 | 1.82E-07 | 9.44E-07 |
| | -1.139540428 | 0.254477816 | 0.358748507 |
| | 2.747703417 | 0.006001427 | 0.01445772 |
| | 2.068036591 | 0.038636582 | 0.073957912 |
| | 2.069770968 | 0.038473797 | 0.073674481 |
| NA | NA | NA | |
| NA | NA | NA | |
| NA | NA | NA | |
| | 1.452586236 | 0.14633867 | 0.228699857 |
| | -7.020579068 | 2.21E-12 | 1.97E-11 |
| | 2.672301425 | 0.007533295 | 0.017676794 |

| | | |
|---|---|---|
| -2.658175992 | 0.007856485 | 0.018332043 |
| 0.382395287 | 0.702168186 | 0.781364766 |
| 1.761427157 | 0.078166132 | 0.135388436 |
| -0.239977021 | 0.810348071 | NA |
| 0.526972529 | 0.598212665 | NA |
| -3.008072399 | 0.002629105 | 0.006880646 |
| -2.003218636 | 0.045153826 | 0.084767139 |
| -1.51799911 | 0.129014623 | 0.206156538 |
| 5.542077146 | 2.99E-08 | 1.72E-07 |
| -2.511660075 | 0.012016477 | 0.026794871 |
| NA | NA | NA |
| NA | NA | NA |
| 2.275743487 | 0.022861367 | 0.046980297 |
| 0.414674468 | 0.678380225 | NA |
| 0.356059827 | 0.721795762 | NA |
| 0.764318687 | 0.444677352 | 0.554572702 |
| 0.769198835 | 0.441775282 | 0.551774369 |
| 0.435523881 | 0.663182208 | NA |
| -4.003025142 | 6.25E-05 | 0.000223873 |
| 0.112958056 | 0.910063809 | 0.937943553 |
| 8.793897946 | 1.44E-18 | 2.30E-17 |
| 6.45763862 | 1.06E-10 | 8.06E-10 |
| -0.261252665 | 0.793897667 | 0.853466417 |
| -1.114698148 | 0.264979795 | 0.370589723 |
| 9.269588286 | 1.87E-20 | 3.43E-19 |
| -3.510828317 | 0.000446713 | 0.001374459 |
| 2.037959701 | 0.041553963 | 0.078819013 |
| -0.260495077 | 0.794481912 | NA |
| -3.619137486 | 0.000295587 | 0.000940876 |
| 1.011888541 | 0.311591349 | 0.421711445 |
| NA | NA | NA |
| NA | NA | NA |
| NA | NA | NA |
| -0.065388772 | 0.947864464 | NA |
| -2.261706513 | 0.023715542 | 0.0484611 |
| -4.850626892 | 1.23E-06 | 5.70E-06 |
| -0.858462937 | 0.390636888 | 0.502758191 |
| -0.092470838 | 0.92632396 | NA |
| 2.304354939 | 0.021202727 | 0.043943385 |
| -4.115969015 | 3.86E-05 | 0.000143076 |
| -0.886943273 | 0.375109442 | 0.487921827 |
| -2.369932674 | 0.017791325 | 0.037818841 |
| 0.656771918 | 0.511327584 | 0.617064241 |
| 9.515541381 | 1.81E-21 | 3.61E-20 |
| 13.89052995 | 7.23E-44 | 4.64E-42 |
| NA | NA | NA |
| 3.044534847 | 0.002330404 | 0.006175756 |

| | | | |
|---|---|---|---|
| | -1.805582309 | 0.070983608 | 0.124760881 |
| | -1.212299863 | 0.225397616 | 0.325470309 |
| NA | NA | NA | |
| | -6.606256351 | 3.94E-11 | 3.12E-10 |
| | -5.073479842 | 3.91E-07 | 1.95E-06 |
| | -7.17333417 | 7.32E-13 | 6.90E-12 |
| | 0.699172068 | 0.484444504 | 0.592912128 |
| | 4.78861843 | 1.68E-06 | 7.64E-06 |
| | 10.17240301 | 2.63E-24 | 6.41E-23 |
| | -1.927094557 | 0.053967847 | 0.098881137 |
| | 0.137741649 | 0.890444602 | 0.923312001 |
| | 3.422956548 | 0.00061944 | 0.001856836 |
| | -1.71911984 | 0.085592553 | 0.14621769 |
| | -0.755189902 | 0.450135047 | 0.560067616 |
| | 5.904651934 | 3.53E-09 | 2.27E-08 |
| | 0.361436363 | 0.717773269 | 0.793687359 |
| | 9.045867616 | 1.48E-19 | 2.56E-18 |
| | -0.349653447 | 0.726598795 | 0.800262599 |
| | 1.96519387 | 0.049391803 | 0.09165074 |
| | 1.681183669 | 0.092727245 | 0.156515135 |
| | 3.112467094 | 0.001855307 | 0.005030694 |
| | 1.771098529 | 0.076544319 | 0.132947352 |
| | 2.367606046 | 0.017903588 | 0.038012284 |
| | -1.414410348 | 0.157241454 | 0.24275295 |
| | -0.817203557 | 0.41381211 | 0.525425715 |
| | 3.822492505 | 0.000132109 | 0.000447117 |
| | 0.192809521 | 0.847108147 | NA |
| | 0.524190099 | 0.600146327 | 0.69796778 |
| | -5.279049339 | 1.30E-07 | 6.89E-07 |
| | -1.515790597 | 0.12957231 | 0.206862797 |
| | -4.132246453 | 3.59E-05 | 0.000134064 |
| | -6.156554722 | 7.43E-10 | 5.12E-09 |
| | -2.66142131 | 0.007781153 | 0.018176598 |
| | -0.49927101 | 0.617588475 | 0.712368141 |
| | -5.499505834 | 3.81E-08 | 2.17E-07 |
| | -2.715518759 | 0.006617203 | 0.015774151 |
| | -6.568617687 | 5.08E-11 | 3.99E-10 |
| NA | NA | NA | |
| | -5.990202291 | 2.10E-09 | 1.38E-08 |
| NA | NA | NA | |
| | -4.814198671 | 1.48E-06 | 6.77E-06 |
| | -3.70030436 | 0.000215341 | 0.000703378 |
| | -5.938154936 | 2.88E-09 | 1.87E-08 |
| | -6.796679695 | 1.07E-11 | 8.95E-11 |
| | -3.437318513 | 0.000587504 | 0.001768519 |
| | -2.73691605 | 0.006201812 | 0.014894583 |
| | -6.512099733 | 7.41E-11 | 5.72E-10 |

| | | |
|---|---|---|
| -4.396185414 | 1.10E-05 | 4.46E-05 |
| -5.496928584 | 3.86E-08 | 2.20E-07 |
| -4.70043439 | 2.60E-06 | 1.15E-05 |
| 0.262133444 | 0.793218563 | 0.852956164 |
| 0.288217989 | 0.77317988 | NA |
| -2.084061882 | 0.037154539 | 0.071482574 |
| -5.21786257 | 1.81E-07 | 9.41E-07 |
| -6.048141069 | 1.47E-09 | 9.80E-09 |
| -1.900669525 | 0.057345313 | 0.104094073 |
| -4.253783089 | 2.10E-05 | 8.14E-05 |
| -4.987275535 | 6.12E-07 | 2.96E-06 |
| -5.812392708 | 6.16E-09 | 3.85E-08 |
| -4.714109196 | 2.43E-06 | 1.08E-05 |
| -3.636374682 | 0.000276502 | 0.000885132 |
| -4.534524336 | 5.77E-06 | 2.43E-05 |
| -3.521877561 | 0.000428502 | 0.001322491 |
| -6.153274591 | 7.59E-10 | 5.22E-09 |
| 0.651653512 | 0.514624718 | 0.620176057 |
| -2.803306786 | 0.005058153 | 0.012434253 |
| -4.612580525 | 3.98E-06 | 1.72E-05 |
| -4.948282364 | 7.49E-07 | 3.58E-06 |
| -5.592480194 | 2.24E-08 | 1.31E-07 |
| -5.077223724 | 3.83E-07 | 1.91E-06 |
| -6.51790966 | 7.13E-11 | 5.51E-10 |
| -5.946789898 | 2.73E-09 | 1.78E-08 |
| -5.977775522 | 2.26E-09 | 1.48E-08 |
| -5.426153494 | 5.76E-08 | 3.20E-07 |
| -4.882448612 | 1.05E-06 | 4.91E-06 |
| -3.770499309 | 0.000162921 | 0.000542999 |
| -6.561678515 | 5.32E-11 | 4.17E-10 |
| 0.063708285 | 0.949202508 | NA |
| -0.065550486 | 0.947735711 | 0.964380228 |
| 6.074056625 | 1.25E-09 | 8.40E-09 |
| 0.708173319 | 0.478837633 | 0.587572603 |
| 1.346070364 | 0.178279825 | 0.268994154 |
| -0.687691221 | 0.491647249 | 0.599016518 |
| 0.192809521 | 0.847108147 | NA |
| -4.488461202 | 7.17E-06 | 2.98E-05 |
| -2.8440283 | 0.004454709 | 0.011081432 |
| -2.614811639 | 0.008927667 | 0.020572594 |
| -3.916823334 | 8.97E-05 | 0.000313175 |
| -0.655079749 | 0.51241641 | 0.61805044 |
| -2.878309225 | 0.00399813 | 0.010050537 |
| -2.135248522 | 0.032740715 | 0.064103575 |
| -3.444413493 | 0.0005723 | 0.001727528 |
| 0.820216876 | 0.412092483 | 0.523747073 |
| -4.830111757 | 1.36E-06 | 6.27E-06 |

| | | | |
|---|---|---|---|
| | -4.270553635 | 1.95E-05 | 7.59E-05 |
| | -5.534882902 | 3.11E-08 | 1.79E-07 |
| | -2.243669646 | 0.024853665 | 0.050456758 |
| | -4.983578751 | 6.24E-07 | 3.02E-06 |
| | -3.322710911 | 0.000891472 | 0.002593695 |
| | -0.872125074 | 0.383140149 | 0.495535601 |
| | -1.18640879 | 0.235460902 | 0.337065277 |
| | -3.642276432 | 0.000270238 | 0.000866075 |
| | -2.881223212 | 0.00396135 | 0.009969092 |
| | -3.301156778 | 0.000962871 | 0.00278395 |
| | -5.320105729 | 1.04E-07 | 5.58E-07 |
| | 2.515497857 | 0.01188644 | 0.026556957 |
| | -1.119463682 | 0.262942376 | 0.368295897 |
| | -5.778191049 | 7.55E-09 | 4.67E-08 |
| NA | NA | NA | |
| | 0.864448891 | 0.387341345 | 0.499602786 |
| | 7.577014676 | 3.54E-14 | 3.80E-13 |
| | 1.263686376 | 0.206342617 | NA |
| | -3.212270307 | 0.001316904 | 0.003691166 |
| | -3.760159663 | 0.000169805 | 0.000563689 |
| | 6.507333423 | 7.65E-11 | 5.89E-10 |
| | 2.738640554 | 0.006169378 | 0.014820956 |
| NA | NA | NA | |
| NA | NA | NA | |
| | -4.510236885 | 6.48E-06 | 2.71E-05 |
| | -0.465409836 | 0.641637992 | 0.732005421 |
| | -0.632041996 | 0.527359436 | 0.632386656 |
| | 0.451782898 | 0.651425391 | 0.740034382 |
| | -0.923757789 | 0.355612437 | 0.467512963 |
| | 2.906542011 | 0.003654478 | 0.009255448 |
| | 2.087150489 | 0.036874533 | 0.071046846 |
| | -2.736383679 | 0.006211855 | 0.014917273 |
| | 9.687248395 | 3.42E-22 | 7.15E-21 |
| NA | NA | NA | |
| | 1.021966504 | 0.306796754 | 0.416551275 |
| | 0.058973092 | 0.95297354 | 0.967939281 |
| NA | NA | NA | |
| NA | NA | NA | |
| | -0.563712198 | 0.572950015 | 0.674233139 |
| | -1.234715767 | 0.216936306 | 0.315550611 |
| NA | NA | NA | |
| | 0.063708285 | 0.949202508 | NA |
| | -3.343836868 | 0.000826283 | 0.002418366 |
| | -1.086734653 | 0.277154095 | 0.384032717 |
| | -0.065389966 | 0.947863513 | NA |
| NA | NA | NA | |
| | 0.638176588 | 0.523358736 | 0.62858956 |

| | | |
|---|---|---|
| -2.479034334 | 0.013173862 | 0.029080322 |
| 0.192809521 | 0.847108147 | NA |
| 6.039243601 | 1.55E-09 | 1.03E-08 |
| -7.220380231 | 5.18E-13 | 4.97E-12 |
| -6.233206767 | 4.57E-10 | 3.22E-09 |
| -0.115834957 | 0.907783345 | NA |
| NA | NA | NA |
| 0.192809521 | 0.847108147 | NA |
| NA | NA | NA |
| 3.880272584 | 0.000104339 | 0.000359479 |
| -0.510763898 | 0.609516391 | NA |
| -2.349644658 | 0.018791341 | 0.039578194 |
| 1.855764496 | 0.063487133 | 0.11335209 |
| 1.643187632 | 0.100344118 | 0.166974817 |
| 5.863770431 | 4.52E-09 | 2.87E-08 |
| 0.958938716 | 0.337589619 | 0.448741633 |
| 1.365963112 | 0.171950542 | 0.261144473 |
| NA | NA | NA |
| 1.695802102 | 0.089923364 | 0.152522993 |
| 1.205120786 | 0.228156676 | 0.328506766 |
| 0.498489532 | 0.618139045 | 0.71289777 |
| -1.328657697 | 0.183960927 | 0.276101338 |
| 8.65348319 | 5.00E-18 | 7.53E-17 |
| -7.781908476 | 7.14E-15 | 8.26E-14 |
| 0.063708285 | 0.949202508 | NA |
| NA | NA | NA |
| 0.018150349 | 0.985518912 | 0.989720316 |
| 2.397128921 | 0.016524109 | 0.035492872 |
| 1.327648587 | 0.184294226 | 0.276502425 |
| 0.879325719 | 0.379224695 | 0.49183865 |
| 4.920731322 | 8.62E-07 | 4.08E-06 |
| NA | NA | NA |
| -3.731201361 | 0.000190569 | 0.000628533 |
| NA | NA | NA |
| NA | NA | NA |
| -0.065388772 | 0.947864464 | NA |
| 0.063707801 | 0.949202893 | NA |
| -0.054257635 | 0.956729902 | 0.970365225 |
| NA | NA | NA |
| NA | NA | NA |
| NA | NA | NA |
| NA | NA | NA |
| NA | NA | NA |
| NA | NA | NA |
| -0.065388772 | 0.947864464 | NA |
| NA | NA | NA |
| -0.061951903 | 0.950601134 | NA |

| | | | |
|---|---|---|---|
| | 0.217940737 | 0.827475286 | 0.879216641 |
| | 8.214662333 | 2.13E-16 | 2.81E-15 |
| | -0.156627431 | 0.875538483 | 0.913151724 |
| | -3.754468654 | 0.00017371 | 0.000575583 |
| NA | NA | NA | |
| | -3.762442442 | 0.000168262 | 0.000559309 |
| NA | NA | NA | |
| | -9.474363557 | 2.68E-21 | 5.28E-20 |
| | 0.147799886 | 0.882500699 | NA |
| | -1.081489782 | 0.279479304 | 0.386484363 |
| NA | NA | NA | |
| | 0.150510984 | 0.880361486 | NA |
| | -3.307551534 | 0.000941154 | 0.002728093 |
| | 1.814793348 | 0.069555673 | 0.122558133 |
| | 4.615684496 | 3.92E-06 | 1.69E-05 |
| | -1.547639081 | 0.121709217 | 0.196320408 |
| | 0.437253324 | 0.661927641 | NA |
| | -0.283472213 | 0.776814879 | 0.839899629 |
| | 0.523905855 | 0.600344022 | 0.698095411 |
| | 4.14283385 | 3.43E-05 | 0.000128441 |
| | 4.201159466 | 2.66E-05 | 0.000101322 |
| | 4.14586589 | 3.39E-05 | 0.000126886 |
| | 7.984544132 | 1.41E-15 | 1.73E-14 |
| | 3.998091322 | 6.39E-05 | 0.000228231 |
| | 4.660331931 | 3.16E-06 | 1.38E-05 |
| | 2.677876199 | 0.00740906 | 0.017421209 |
| | -1.067232391 | 0.285866909 | 0.3929728 |
| NA | NA | NA | |
| | 1.668328662 | 0.095250501 | 0.159997318 |
| | -0.027638927 | 0.977950134 | 0.984677254 |
| NA | NA | NA | |
| NA | NA | NA | |
| | -3.546375645 | 0.000390569 | 0.00121546 |
| | 2.164236486 | 0.030446204 | 0.060114262 |
| NA | NA | NA | |
| NA | NA | NA | |
| NA | NA | NA | |
| | -1.275357151 | 0.202182858 | 0.298369019 |
| | -0.263704351 | 0.792007742 | 0.852020191 |
| NA | NA | NA | |
| | -0.771843327 | 0.440207226 | 0.550423962 |
| | 9.35044607 | 8.73E-21 | 1.64E-19 |
| | -4.238174954 | 2.25E-05 | 8.69E-05 |
| | -2.990047536 | 0.00278934 | 0.007261278 |
| | 11.51067754 | 1.17E-30 | 4.15E-29 |
| NA | NA | NA | |
| NA | NA | NA | |

| | | |
|---|---|---|
| 0.06371127 | 0.949200131 | NA |
| -6.343213747 | 2.25E-10 | 1.64E-09 |
| NA | NA | NA |
| NA | NA | NA |
| 2.315936721 | 0.020561724 | 0.042802724 |
| NA | NA | NA |
| NA | NA | NA |
| 2.623938955 | 0.008691936 | 0.02009042 |
| NA | NA | NA |
| 7.406929584 | 1.29E-13 | 1.31E-12 |
| 1.152321952 | 0.2491888 | 0.352563205 |
| 0.330493631 | 0.741027004 | NA |
| NA | NA | NA |
| -0.797867047 | 0.424947648 | 0.536489439 |
| 10.91793412 | 9.46E-28 | 2.91E-26 |
| 0.884412251 | 0.376473712 | 0.489156626 |
| 3.899156628 | 9.65E-05 | 0.000334826 |
| 0.774338268 | 0.438730779 | 0.549016668 |
| -2.530975678 | 0.011374574 | 0.025566234 |
| -2.350499424 | 0.018748237 | 0.039510681 |
| -1.699916499 | 0.089146633 | 0.151400477 |
| 2.236748087 | 0.025302809 | 0.051259101 |
| -1.809304289 | 0.070403743 | 0.123872184 |
| 0.337316516 | 0.735878311 | 0.807498208 |
| -3.896378939 | 9.76E-05 | 0.000338312 |
| NA | NA | NA |
| -0.429756499 | 0.66737278 | NA |
| NA | NA | NA |
| NA | NA | NA |
| 0.658401513 | 0.510280164 | 0.616153119 |
| -1.376352753 | 0.168712448 | NA |
| NA | NA | NA |
| -0.600266556 | 0.548328604 | NA |
| NA | NA | NA |
| NA | NA | NA |
| -0.065388772 | 0.947864464 | NA |
| NA | NA | NA |
| NA | NA | NA |
| 5.007494775 | 5.51E-07 | 2.68E-06 |
| -1.734390144 | 0.082848883 | 0.142268629 |
| NA | NA | NA |
| 2.744566865 | 0.006059081 | 0.014582577 |
| -0.614226233 | 0.53906584 | 0.643342296 |
| 6.487570553 | 8.72E-11 | 6.68E-10 |
| 3.277022062 | 0.001049082 | 0.003009917 |
| 1.647877499 | 0.099377812 | 0.165751091 |
| -1.743742422 | 0.081204015 | 0.1399144 |

| | | |
|---|---|---|
| 1.817820427 | 0.069091583 | 0.121880494 |
| 3.85762685 | 0.000114493 | 0.000391605 |
| 2.281699324 | 0.022507098 | 0.046343589 |
| 1.224273016 | 0.220849253 | 0.320121427 |
| 1.272685443 | 0.203129685 | NA |
| 1.000528885 | 0.317054626 | 0.427587643 |
| NA | NA | NA |
| -0.065389369 | 0.947863988 | NA |
| -5.869815237 | 4.36E-09 | 2.78E-08 |
| -4.396404299 | 1.10E-05 | 4.46E-05 |
| -4.051869188 | 5.08E-05 | 0.000184959 |
| NA | NA | NA |
| 1.934921933 | 0.052999876 | 0.09731418 |
| NA | NA | NA |
| 5.746138571 | 9.13E-09 | 5.60E-08 |
| -2.185533404 | 0.028849759 | 0.057392265 |
| 3.811620309 | 0.000138059 | 0.000465779 |
| -4.269186805 | 1.96E-05 | 7.63E-05 |
| NA | NA | NA |
| NA | NA | NA |
| NA | NA | NA |
| 21.88886887 | 3.32E-106 | 8.93E-104 |
| 10.97469633 | 5.06E-28 | 1.58E-26 |
| 12.72068724 | 4.54E-37 | 2.18E-35 |
| 19.79556827 | 3.25E-87 | 6.07E-85 |
| 37.36951346 | 1.22E-305 | 5.10E-302 |
| 2.499982784 | 0.012419934 | 0.027605981 |
| 11.86099041 | 1.89E-32 | 7.42E-31 |
| 9.855074097 | 6.52E-23 | 1.44E-21 |
| 7.64648537 | 2.07E-14 | 2.28E-13 |
| 2.720849932 | 0.006511431 | 0.015554603 |
| 0.063708285 | 0.949202508 | NA |
| -2.167466978 | 0.030199265 | 0.059687903 |
| 8.30886299 | 9.66E-17 | 1.31E-15 |
| 5.532067235 | 3.16E-08 | 1.82E-07 |
| 0.063976859 | 0.948988653 | NA |
| 12.98584432 | 1.47E-38 | 7.63E-37 |
| 0.063708285 | 0.949202508 | NA |
| 7.299924258 | 2.88E-13 | 2.83E-12 |
| 5.639614529 | 1.70E-08 | 1.01E-07 |
| -8.078722427 | 6.54E-16 | 8.32E-15 |
| -3.672858517 | 0.000239852 | 0.000775646 |
| -3.277179999 | 0.001048495 | 0.003008924 |
| -6.618663056 | 3.62E-11 | 2.88E-10 |
| -0.816264671 | 0.414348776 | NA |
| 4.84853792 | 1.24E-06 | 5.75E-06 |
| -2.354430716 | 0.018551102 | 0.03916448 |

| | | | |
|---|---|---|---|
| NA | NA | NA | |
| | -2.376325486 | 0.017486031 | 0.037245796 |
| | -1.07260459 | 0.283448564 | 0.390420293 |
| | 5.156370735 | 2.52E-07 | 1.29E-06 |
| | -0.112011778 | 0.910814068 | 0.938593548 |
| | 3.075706127 | 0.002100047 | 0.005629665 |
| | -6.905753205 | 4.99E-12 | 4.32E-11 |
| NA | NA | NA | |
| | -2.8154904 | 0.004870285 | 0.012029133 |
| | 0.192810118 | 0.847107679 | NA |
| | 3.91886267 | 8.90E-05 | 0.000310783 |
| | 3.8388538 | 0.00012361 | 0.000420489 |
| | 3.489118971 | 0.000484615 | 0.001481608 |
| | 0.073164869 | 0.941674922 | NA |
| | 0.067524971 | 0.946163784 | NA |
| | -2.006115054 | 0.044843974 | 0.084261292 |
| | 1.401673379 | 0.161012804 | 0.247202998 |
| | 0.463279173 | 0.643164272 | 0.733011644 |
| | -2.921374628 | 0.003484905 | 0.008869057 |
| | -0.60127755 | 0.54765514 | 0.651211455 |
| | -2.348292783 | 0.01885969 | 0.039702106 |
| | -2.532347522 | 0.011330164 | 0.025480143 |
| | -8.454855889 | 2.79E-17 | 3.98E-16 |
| NA | NA | NA | |
| | 0.721576001 | 0.470555199 | 0.579654344 |
| | -2.352814406 | 0.018631931 | 0.039305287 |
| | 0.9801119 | 0.327030886 | 0.438083118 |
| | -2.499072755 | 0.012451874 | 0.027667147 |
| | 4.081101219 | 4.48E-05 | 0.000164191 |
| | 0.419090334 | 0.675150115 | 0.758619473 |
| | 0.324874675 | 0.745275925 | NA |
| | -0.035827768 | 0.971419692 | 0.980293047 |
| | -1.297858316 | 0.194336026 | 0.28867284 |
| | -0.506941921 | 0.612195571 | 0.7079661 |
| | -2.679607458 | 0.007370854 | 0.017337889 |
| | -5.050172072 | 4.41E-07 | 2.18E-06 |
| | 2.754191778 | 0.005883728 | 0.014208366 |
| NA | NA | NA | |
| NA | NA | NA | |
| NA | NA | NA | |
| NA | NA | NA | |
| NA | NA | NA | |
| | 4.872805747 | 1.10E-06 | 5.13E-06 |
| | -5.579898127 | 2.41E-08 | 1.41E-07 |
| NA | NA | NA | |
| | 1.550230657 | 0.121086164 | 0.195421155 |
| | -4.64310782 | 3.43E-06 | 1.49E-05 |

| | | |
|---|---|---|
| -1.451799734 | 0.146557298 | 0.22897005 |
| 0.80375467 | 0.421538674 | 0.533360598 |
| -1.188476786 | 0.23464562 | 0.336051947 |
| 5.800288371 | 6.62E-09 | 4.12E-08 |
| 5.055696784 | 4.29E-07 | 2.12E-06 |
| NA | NA | NA |
| NA | NA | NA |
| -1.62370416 | 0.104438963 | 0.172650897 |
| 0.192809521 | 0.847108147 | NA |
| -0.858129408 | 0.39082101 | 0.502917582 |
| 0.192809521 | 0.847108147 | NA |
| 5.66667478 | 1.46E-08 | 8.73E-08 |
| -1.915807364 | 0.05538962 | 0.101071823 |
| -2.701964004 | 0.006893122 | 0.016358666 |
| -1.725735807 | 0.084394953 | 0.144508967 |
| 0.963763028 | 0.335164747 | 0.446376911 |
| -3.272117211 | 0.001067453 | 0.00305877 |
| NA | NA | NA |
| 0.034621686 | 0.972381409 | NA |
| NA | NA | NA |
| 0.759738886 | 0.44741068 | 0.557342822 |
| 7.363889691 | 1.79E-13 | 1.79E-12 |
| -4.886647605 | 1.03E-06 | 4.81E-06 |
| -0.621925752 | 0.533990691 | NA |
| -2.01464496 | 0.043941863 | 0.082777514 |
| 12.62457181 | 1.55E-36 | 7.22E-35 |
| NA | NA | NA |
| NA | NA | NA |
| 0.599262451 | 0.548997883 | 0.652396003 |
| -6.117942405 | 9.48E-10 | 6.47E-09 |
| -0.794457108 | 0.42692937 | NA |
| 3.867791658 | 0.000109825 | 0.000377238 |
| -2.257149877 | 0.023998711 | 0.048931878 |
| -0.726576298 | 0.467485552 | 0.576752927 |
| -1.965329699 | 0.04937609 | 0.091628377 |
| 13.76435342 | 4.18E-43 | 2.57E-41 |
| -0.979889197 | 0.327140816 | 0.438206926 |
| NA | NA | NA |
| -5.405539126 | 6.46E-08 | 3.57E-07 |
| -3.090965785 | 0.001995066 | 0.00537993 |
| 0.063707688 | 0.949202983 | NA |
| 1.231935666 | 0.217973117 | 0.316814768 |
| 0.657957291 | 0.510565575 | 0.616382402 |
| 3.846911058 | 0.000119616 | 0.000407902 |
| -1.893955316 | 0.058230941 | 0.105403562 |
| 7.89656855 | 2.87E-15 | 3.43E-14 |
| 2.564120372 | 0.010343766 | 0.023491922 |

| | | |
|---|---|---|
| -1.026488235 | 0.304661506 | 0.414168885 |
| -10.50015552 | 8.62E-26 | 2.33E-24 |
| NA | NA | NA |
| -3.411177377 | 0.00064683 | 0.001931999 |
| 1.257614486 | 0.208531212 | 0.305611349 |
| 2.53756058 | 0.011162805 | 0.025155758 |
| NA | NA | NA |
| 0.41775227 | 0.676128251 | NA |
| NA | NA | NA |
| -0.128634206 | 0.897647098 | NA |
| NA | NA | NA |
| -2.587753719 | 0.009660402 | 0.022088231 |
| -2.69349981 | 0.007070619 | 0.016722818 |
| NA | NA | NA |
| 0.553010584 | 0.580256163 | 0.68100585 |
| 0.063708882 | 0.949202033 | NA |
| 0.197532278 | 0.843411025 | 0.890540574 |
| -3.441762281 | 0.000577938 | 0.001742656 |
| 0.063708882 | 0.949202033 | NA |
| 3.855948042 | 0.000115282 | 0.000394034 |
| -0.173317817 | 0.86240162 | 0.904349262 |
| 2.349135017 | 0.018817082 | 0.039622408 |
| -0.470537502 | 0.637971047 | NA |
| 9.805298278 | 1.07E-22 | 2.32E-21 |
| -8.110893961 | 5.02E-16 | 6.42E-15 |
| NA | NA | NA |
| NA | NA | NA |
| 5.962344807 | 2.49E-09 | 1.62E-08 |
| NA | NA | NA |
| -0.065389369 | 0.947863988 | NA |
| 3.437095835 | 0.000587988 | 0.00176976 |
| -3.230129568 | 0.001237341 | 0.003490044 |
| 2.086574143 | 0.036926647 | 0.071125353 |
| -5.660014256 | 1.51E-08 | 9.06E-08 |
| 0.901596125 | 0.36727145 | 0.479716349 |
| 3.558548247 | 0.00037291 | 0.00116456 |
| -2.726478335 | 0.006401415 | 0.015316646 |
| 2.976905912 | 0.002911734 | 0.007550694 |
| 1.359270064 | 0.174061026 | 0.263837431 |
| 1.855660726 | 0.063501932 | 0.113370425 |
| 1.606788268 | 0.108100823 | 0.177520246 |
| 1.574392959 | 0.11539663 | 0.187617906 |
| -2.25709317 | 0.024002254 | 0.048935115 |
| NA | NA | NA |
| -0.725373232 | 0.46822309 | 0.57744058 |
| 0.902740192 | 0.3666638 | 0.479135838 |
| -3.697292367 | 0.000217911 | 0.000710939 |

| | | | |
|---|---|---|---|
| NA | | NA | NA |
| NA | | NA | NA |
| | 10.68370075 | 1.21E-26 | 3.45E-25 |
| | 7.060372124 | 1.66E-12 | 1.50E-11 |
| | -0.230231377 | 0.817911981 | 0.872016735 |
| | -0.21765212 | 0.827700172 | 0.879380929 |
| | -0.241276371 | 0.809340923 | 0.86531132 |
| | 0.640336857 | 0.521953624 | 0.627226425 |
| | -6.389831318 | 1.66E-10 | 1.23E-09 |
| | -0.086207166 | 0.931301734 | NA |
| NA | | NA | NA |
| NA | | NA | NA |
| | 0.488947501 | 0.624878865 | 0.718584857 |
| | 4.226278449 | 2.38E-05 | 9.12E-05 |
| | -4.181430549 | 2.90E-05 | 0.000109809 |
| | 2.016195602 | 0.043779526 | 0.082515176 |
| NA | | NA | NA |
| | -1.100115415 | 0.271281838 | 0.377610078 |
| NA | | NA | NA |
| | 1.784808411 | 0.074292399 | 0.129566523 |
| | -1.069302912 | 0.284933199 | 0.392102048 |
| | 3.153319127 | 0.001614252 | 0.004444476 |
| | -2.49092379 | 0.012741145 | 0.028232193 |
| | -3.34265343 | 0.000829815 | 0.002428418 |
| | -2.717271225 | 0.006582264 | 0.015707467 |
| | -2.868821445 | 0.004120043 | 0.010326926 |
| | -5.020309082 | 5.16E-07 | 2.52E-06 |
| NA | | NA | NA |
| | 3.235763634 | 0.001213178 | 0.003429619 |
| | -6.563229437 | 5.27E-11 | 4.13E-10 |
| | -0.991041952 | 0.321665096 | 0.432678275 |
| NA | | NA | NA |
| | -1.517192345 | 0.129218128 | 0.206429012 |
| | 1.806417095 | 0.070853213 | 0.124566687 |
| | -0.932270024 | 0.351196995 | 0.462913452 |
| | -1.464380957 | 0.143089888 | 0.224604283 |
| | 1.026776646 | 0.304525649 | 0.41400668 |
| | -2.622177761 | 0.008736985 | 0.020179637 |
| | 0.06371127 | 0.949200131 | NA |
| | -1.581584905 | 0.113744362 | 0.18531676 |
| | 0.580686673 | 0.561451645 | NA |
| | 6.667154614 | 2.61E-11 | 2.10E-10 |
| | -0.140527327 | 0.888243363 | 0.921755032 |
| | -2.188868855 | 0.028606373 | 0.056989615 |
| NA | | NA | NA |
| NA | | NA | NA |
| | -4.053025633 | 5.06E-05 | 0.000184127 |

| | | |
|---|---|---|
| NA | NA | NA |
| 0.324874675 | 0.745275925 | NA |
| 0.47751133 | 0.632998055 | 0.724592578 |
| -0.102796458 | 0.918124517 | NA |
| 3.555522309 | 0.000377229 | 0.001176725 |
| -1.366332838 | 0.171834518 | 0.260984102 |
| 4.866459425 | 1.14E-06 | 5.29E-06 |
| 2.064535047 | 0.038967014 | 0.074470823 |
| -1.737170745 | 0.082357037 | 0.141559994 |
| 3.417698347 | 0.000631531 | 0.001890136 |
| -1.475323055 | 0.140125711 | 0.220615583 |
| -2.212571052 | 0.026927233 | 0.05404014 |
| 2.910882306 | 0.003604097 | 0.009134322 |
| 0.82995872 | 0.406562123 | 0.518402952 |
| 1.977265886 | 0.048011584 | 0.089401126 |
| 13.24032537 | 5.13E-40 | 2.82E-38 |
| 1.965952327 | 0.049304118 | 0.091508384 |
| 0.064320904 | 0.948714709 | 0.965010012 |
| -1.815000734 | 0.069523797 | 0.122522047 |
| -1.773517159 | 0.076143051 | 0.132351428 |
| -2.159740364 | 0.030792775 | 0.060712339 |
| NA | NA | NA |
| 0.063708882 | 0.949202033 | NA |
| NA | NA | NA |
| NA | NA | NA |
| 0.91967498 | 0.357742631 | 0.469598178 |
| NA | NA | NA |
| 1.19655782 | 0.231478949 | 0.332328103 |
| -2.39288017 | 0.0167167 | 0.035844469 |
| 1.628505334 | 0.103417779 | 0.171245502 |
| -4.17696277 | 2.95E-05 | 0.000111852 |
| -4.068258677 | 4.74E-05 | 0.000173176 |
| -0.359139984 | 0.719490371 | NA |
| -0.065389369 | 0.947863988 | NA |
| -0.22626737 | 0.820993475 | 0.874370976 |
| 4.429800393 | 9.43E-06 | 3.86E-05 |
| NA | NA | NA |
| NA | NA | NA |
| -1.059148818 | 0.28953201 | 0.39702987 |
| -2.477879728 | 0.013216569 | 0.029160996 |
| -1.490978323 | 0.135967186 | 0.215164612 |
| -0.842717904 | 0.399386231 | 0.511206207 |
| 2.884390912 | 0.003921716 | 0.009879283 |
| 1.612099156 | 0.106940364 | 0.17591465 |
| 10.54654627 | 5.27E-26 | 1.44E-24 |
| -0.904210444 | 0.365883823 | 0.478436302 |
| -3.952371614 | 7.74E-05 | 0.000273332 |

| | | |
|---|---|---|
| NA | NA | NA |
| NA | NA | NA |
| NA | NA | NA |
| -0.359456676 | 0.719253482 | 0.794760492 |
| 0.248314386 | 0.803891169 | NA |
| 1.220837915 | 0.222147395 | 0.321518983 |
| -1.155855002 | 0.247740471 | 0.350950478 |
| NA | NA | NA |
| -3.1156207 | 0.001835582 | 0.004983688 |
| -5.955966891 | 2.59E-09 | 1.69E-08 |
| 0.083043937 | 0.933816603 | NA |
| NA | NA | NA |
| 2.49368756 | 0.012642377 | 0.028040647 |
| -0.102437417 | 0.918409485 | 0.944172892 |
| 0.063708285 | 0.949202508 | NA |
| 5.198620236 | 2.01E-07 | 1.04E-06 |
| -4.492607595 | 7.04E-06 | 2.93E-05 |
| -5.386669405 | 7.18E-08 | 3.94E-07 |
| -0.884927382 | 0.376195799 | 0.488949052 |
| NA | NA | NA |
| NA | NA | NA |
| NA | NA | NA |
| NA | NA | NA |
| 1.13420951 | 0.256706676 | 0.361290435 |
| NA | NA | NA |
| 2.363197533 | 0.01811801 | 0.038369894 |
| 0.362026528 | 0.717332207 | NA |
| NA | NA | NA |
| 1.61608215 | 0.106076553 | 0.174757857 |
| 3.366446912 | 0.000761432 | 0.002244307 |
| -2.248474653 | 0.02454594 | 0.0498932 |
| 7.905356459 | 2.67E-15 | 3.21E-14 |
| -0.85681393 | 0.391547726 | 0.503567957 |
| 4.684350241 | 2.81E-06 | 1.24E-05 |
| 2.084326307 | 0.037130496 | 0.071452012 |
| NA | NA | NA |
| 5.040646314 | 4.64E-07 | 2.28E-06 |
| 1.565716879 | 0.117414913 | 0.190332211 |
| -2.707752971 | 0.006774042 | 0.01610814 |
| -1.068035194 | 0.285504637 | 0.392625738 |
| -2.186562937 | 0.028774446 | 0.057258969 |
| 20.14012863 | 3.28E-90 | 6.63E-88 |
| -1.68906254 | 0.091207446 | 0.154314071 |
| 5.622102787 | 1.89E-08 | 1.12E-07 |
| 18.25376658 | 1.93E-74 | 2.76E-72 |
| 0.875801998 | 0.381137683 | 0.49370575 |
| -2.199887121 | 0.027814905 | 0.055603152 |

| | | |
|---|---|---|
| 1.232554117 | 0.217742164 | 0.316589391 |
| 0.356723368 | 0.72129891 | NA |
| -2.789901144 | 0.005272414 | 0.012896366 |
| 0.409586829 | 0.68210906 | NA |
| -0.627187723 | 0.53053619 | 0.635404077 |
| -2.512733198 | 0.01197999 | 0.026720651 |
| -1.211107897 | 0.225854055 | 0.325922572 |
| -0.510383729 | 0.609782654 | 0.7060654 |
| 2.900817376 | 0.003721907 | 0.009410985 |
| NA | NA | NA |
| -0.30070743 | 0.763637603 | 0.829352835 |
| 3.369752635 | 0.000752357 | 0.002220957 |
| -7.059469314 | 1.67E-12 | 1.51E-11 |
| -0.06148492 | 0.950973024 | 0.966569586 |
| 2.258434325 | 0.023918595 | 0.048812265 |
| 3.912211286 | 9.15E-05 | 0.000318774 |
| 1.900136621 | 0.057415193 | 0.104200366 |
| 11.32763344 | 9.58E-30 | 3.25E-28 |
| 8.882048152 | 6.56E-19 | 1.07E-17 |
| -2.77099726 | 0.00558849 | 0.013591129 |
| -6.431028095 | 1.27E-10 | 9.52E-10 |
| 0.36238013 | 0.717067986 | 0.792975687 |
| 8.071219695 | 6.96E-16 | 8.83E-15 |
| 1.871318663 | 0.061300925 | 0.110021666 |
| 7.992274761 | 1.32E-15 | 1.63E-14 |
| 13.85454831 | 1.19E-43 | 7.57E-42 |
| -0.065387578 | 0.947865415 | NA |
| 6.461352893 | 1.04E-10 | 7.88E-10 |
| -0.682759594 | 0.494758766 | 0.601895115 |
| NA | NA | NA |
| -8.724637969 | 2.67E-18 | 4.14E-17 |
| -4.281958813 | 1.85E-05 | 7.24E-05 |
| 0.830265902 | 0.406388462 | 0.518226451 |
| 8.306513387 | 9.86E-17 | 1.34E-15 |
| 1.910467339 | 0.056073068 | 0.102140229 |
| 2.044605635 | 0.040893753 | 0.077684476 |
| 1.0731911 | 0.283185383 | NA |
| -4.900582067 | 9.56E-07 | 4.50E-06 |
| 3.1374512 | 0.001704236 | 0.00466452 |
| 3.548489268 | 0.000387448 | 0.001206046 |
| -0.065389369 | 0.947863988 | NA |
| -3.381786569 | 0.000720161 | 0.00213321 |
| -5.620711695 | 1.90E-08 | 1.13E-07 |
| -3.331147158 | 0.000864889 | 0.002524273 |
| -1.899028023 | 0.057560792 | 0.104439769 |
| -2.512295918 | 0.011994846 | 0.02674902 |
| -2.858082649 | 0.004262094 | 0.010644602 |

| | | |
|---|---|---|
| -2.691307789 | 0.00711725 | 0.016809281 |
| 1.568700824 | 0.116717666 | NA |
| 1.655826724 | 0.097756914 | 0.163505216 |
| -0.902741186 | 0.366663272 | 0.479135838 |
| -7.834376084 | 4.71E-15 | 5.52E-14 |
| -1.111610366 | 0.266305714 | 0.372028354 |
| 0.939300272 | 0.347576599 | 0.459143038 |
| -5.715205401 | 1.10E-08 | 6.67E-08 |
| NA | NA | NA |
| 3.1017947 | 0.001923513 | 0.005200987 |
| -5.332012213 | 9.71E-08 | 5.24E-07 |
| 0.618577126 | 0.536194974 | 0.640557474 |
| -0.51695255 | 0.605189283 | 0.702103719 |
| -2.98508637 | 0.002834983 | 0.007369365 |
| -10.59195622 | 3.25E-26 | 8.96E-25 |
| 7.6557204 | 1.92E-14 | 2.13E-13 |
| 0.169533719 | 0.865376852 | 0.906177918 |
| 4.158614536 | 3.20E-05 | 0.000120388 |
| -3.392980135 | 0.000691367 | 0.002054482 |
| 5.294724058 | 1.19E-07 | 6.36E-07 |
| -4.243341083 | 2.20E-05 | 8.50E-05 |
| 2.268051691 | 0.023326056 | 0.047786151 |
| -5.869563209 | 4.37E-09 | 2.78E-08 |
| -7.278970031 | 3.36E-13 | 3.28E-12 |
| -2.045460565 | 0.040809473 | 0.077553801 |
| 1.431690363 | 0.152232455 | 0.23640516 |
| -8.248125865 | 1.61E-16 | 2.15E-15 |
| NA | NA | NA |
| -3.244591588 | 0.001176192 | 0.003339392 |
| 2.936912128 | 0.00331498 | 0.00847711 |
| 5.166257505 | 2.39E-07 | 1.22E-06 |
| 1.537794471 | 0.124098876 | 0.199588438 |
| -0.152636341 | 0.878685065 | NA |
| -0.804095716 | 0.421341698 | 0.533147096 |
| 0.063708882 | 0.949202033 | NA |
| 2.369575087 | 0.017808539 | 0.037849004 |
| 5.313639547 | 1.07E-07 | 5.77E-07 |
| NA | NA | NA |
| 9.988235547 | 1.72E-23 | 3.97E-22 |
| -1.135762097 | 0.256056144 | 0.360628251 |
| -0.123981305 | 0.901330078 | 0.931281397 |
| 10.50183464 | 8.47E-26 | 2.29E-24 |
| 9.966521516 | 2.14E-23 | 4.91E-22 |
| -1.570719561 | 0.116247804 | 0.188732083 |
| 4.317503424 | 1.58E-05 | 6.24E-05 |
| -8.89448356 | 5.87E-19 | 9.66E-18 |
| -2.667314691 | 0.007646004 | 0.017897642 |

| | | |
|---|---|---|
| 2.509415138 | 0.012093127 | 0.026951382 |
| 6.171827851 | 6.75E-10 | 4.66E-09 |
| -3.323330551 | 0.000889494 | 0.002589445 |
| 1.654870077 | 0.097950855 | 0.163742063 |
| 7.088928002 | 1.35E-12 | 1.24E-11 |
| 9.432850085 | 3.99E-21 | 7.74E-20 |
| -1.141014279 | 0.253863981 | 0.358044483 |
| NA | NA | NA |
| 5.799286467 | 6.66E-09 | 4.14E-08 |
| 1.918724385 | 0.055019224 | 0.100505851 |
| 4.408516821 | 1.04E-05 | 4.23E-05 |
| 0.851544208 | 0.394467117 | NA |
| -0.886883087 | 0.375141849 | NA |
| -0.231354217 | 0.81703963 | 0.871267477 |
| -4.021861839 | 5.77E-05 | 0.000208246 |
| 1.710757098 | 0.087125962 | 0.148478992 |
| -1.150993719 | 0.249734819 | 0.353215942 |
| 0.232923641 | 0.8158207 | 0.870373642 |
| NA | NA | NA |
| -0.155509691 | 0.876419515 | 0.913732184 |
| NA | NA | NA |
| 4.772167173 | 1.82E-06 | 8.26E-06 |
| NA | NA | NA |
| NA | NA | NA |
| NA | NA | NA |
| 0.2854296 | 0.775315033 | 0.838744636 |
| 8.673290355 | 4.20E-18 | 6.37E-17 |
| 5.424201474 | 5.82E-08 | 3.24E-07 |
| -1.797757607 | 0.072215428 | 0.126518691 |
| 5.557511898 | 2.74E-08 | 1.59E-07 |
| -1.875476606 | 0.060727185 | 0.109123792 |
| 6.151711176 | 7.67E-10 | 5.27E-09 |
| 0.066587943 | 0.946909745 | NA |
| 8.063569208 | 7.41E-16 | 9.36E-15 |
| -0.661457163 | 0.508319178 | 0.614500315 |
| -0.724888222 | 0.468520607 | 0.577716676 |
| 6.248153437 | 4.15E-10 | 2.94E-09 |
| NA | NA | NA |
| 0.612679103 | 0.540088542 | 0.644339038 |
| 6.079250528 | 1.21E-09 | 8.16E-09 |
| 2.42719257 | 0.015216177 | 0.032989251 |
| -8.387765328 | 4.95E-17 | 6.90E-16 |
| -0.625163586 | 0.531863699 | 0.636446354 |
| 0.24151761 | 0.80915397 | NA |
| 0.192809521 | 0.847108147 | NA |
| 2.244087693 | 0.02482676 | 0.050410789 |
| 4.313418406 | 1.61E-05 | 6.34E-05 |

| | | |
|---|---|---|
| -1.4094011 | 0.158716599 | 0.244412235 |
| 0.950936332 | 0.341636697 | 0.453089945 |
| -4.724997003 | 2.30E-06 | 1.03E-05 |
| -0.460624208 | 0.64506824 | 0.734445872 |
| 14.12578826 | 2.63E-45 | 1.78E-43 |
| NA | NA | NA |
| 1.271004899 | 0.203726906 | 0.300081472 |
| 5.766758975 | 8.08E-09 | 4.98E-08 |
| -0.153866079 | 0.877715331 | NA |
| -5.00389796 | 5.62E-07 | 2.73E-06 |
| -2.640167495 | 0.008286506 | 0.019231456 |
| 0.324874675 | 0.745275925 | NA |
| 0.095974993 | 0.923540434 | NA |
| -0.423001414 | 0.67229423 | 0.756361226 |
| 5.725037202 | 1.03E-08 | 6.31E-08 |
| -1.879314203 | 0.060201604 | 0.10833643 |
| -2.073267149 | 0.038147424 | 0.073111028 |
| -7.117134796 | 1.10E-12 | 1.02E-11 |
| NA | NA | NA |
| -1.990420595 | 0.046544623 | 0.087019115 |
| 3.001491655 | 0.002686604 | 0.00702085 |
| -5.615608117 | 1.96E-08 | 1.16E-07 |
| -7.556415805 | 4.14E-14 | 4.43E-13 |
| 0.764601659 | 0.444508781 | 0.554445347 |
| -6.545335569 | 5.94E-11 | 4.63E-10 |
| -1.674977085 | 0.093938728 | 0.158165368 |
| 1.39321332 | 0.163555293 | 0.250491892 |
| NA | NA | NA |
| 1.121205293 | 0.262200487 | NA |
| 0.37784138 | 0.705548431 | 0.783907903 |
| NA | NA | NA |
| -0.065389369 | 0.947863988 | NA |
| -0.454170717 | 0.649705958 | 0.738468319 |
| NA | NA | NA |
| 17.13505369 | 8.13E-66 | 9.33E-64 |
| 3.823082715 | 0.000131794 | 0.000446265 |
| 3.763158384 | 0.000167781 | 0.000557784 |
| -3.800016186 | 0.000144687 | 0.000486108 |
| NA | NA | NA |
| -3.86064922 | 0.000113086 | 0.000387428 |
| -7.157423976 | 8.22E-13 | 7.70E-12 |
| -3.206517597 | 0.001343521 | 0.003761137 |
| 0.532946538 | 0.594070589 | 0.692962384 |
| 4.71429561 | 2.43E-06 | 1.08E-05 |
| 6.437747899 | 1.21E-10 | 9.13E-10 |
| -0.525687131 | 0.599105607 | 0.697048916 |
| -3.375232705 | 0.000737533 | 0.002180797 |

| | | |
|---|---|---|
| -0.973768498 | 0.330171498 | 0.441156921 |
| 3.549980334 | 0.00038526 | 0.001199534 |
| -3.032096341 | 0.002428617 | 0.006406859 |
| 19.27451657 | 8.79E-83 | 1.46E-80 |
| 2.493970509 | 0.012632304 | 0.028020788 |
| -2.868734923 | 0.00412117 | 0.010328716 |
| -7.162480869 | 7.92E-13 | 7.44E-12 |
| -2.680153989 | 0.00735883 | 0.017314487 |
| -4.234184616 | 2.29E-05 | 8.83E-05 |
| -0.767098787 | 0.443022783 | 0.553060362 |
| NA | NA | NA |
| NA | NA | NA |
| -7.451922277 | 9.20E-14 | 9.50E-13 |
| -2.967149897 | 0.003005744 | 0.007772083 |
| 0.192809521 | 0.847108147 | NA |
| -0.129584536 | 0.896895138 | 0.928155013 |
| -0.785086026 | 0.432403132 | 0.543134962 |
| -0.242188321 | 0.808634244 | 0.864777403 |
| -4.227110346 | 2.37E-05 | 9.09E-05 |
| -0.653188742 | 0.513634607 | 0.619212041 |
| -11.02607516 | 2.86E-28 | 9.01E-27 |
| 3.832182685 | 0.000127011 | 0.000431181 |
| 7.148315783 | 8.78E-13 | 8.21E-12 |
| -2.275377405 | 0.0228833 | 0.047021315 |
| 1.126322 | 0.260029249 | 0.364930311 |
| 1.062561584 | 0.287980816 | 0.395433504 |
| NA | NA | NA |
| NA | NA | NA |
| NA | NA | NA |
| -0.065389369 | 0.947863988 | NA |
| NA | NA | NA |
| NA | NA | NA |
| NA | NA | NA |
| 0.569342914 | 0.569123449 | 0.670676079 |
| 0.192808924 | 0.847108615 | NA |
| 0.509124143 | 0.61066521 | 0.706720465 |
| -6.254894024 | 3.98E-10 | 2.82E-09 |
| -4.292788264 | 1.76E-05 | 6.91E-05 |
| -7.762023019 | 8.36E-15 | 9.60E-14 |
| -0.065390563 | 0.947863038 | NA |
| 11.06651049 | 1.82E-28 | 5.82E-27 |
| 0.838082557 | 0.401984339 | 0.513755807 |
| 1.514234293 | 0.129966427 | 0.207346527 |
| 0.582110183 | 0.560492467 | 0.662782938 |
| 3.066586259 | 0.002165183 | 0.005791252 |
| 1.724208053 | 0.084670291 | 0.1448696 |
| 2.068099073 | 0.038630708 | 0.073952323 |

| | | |
|---|---|---|
| -0.103330262 | 0.917700859 | 0.943522386 |
| 2.428728886 | 0.015151858 | 0.032878284 |
| 3.764948627 | 0.000166583 | 0.000554171 |
| -1.428378763 | 0.153182868 | 0.237497968 |
| -0.51422098 | 0.607097501 | 0.703730397 |
| NA | NA | NA |
| -0.978191349 | 0.327979694 | 0.439001698 |
| 5.901683182 | 3.60E-09 | 2.31E-08 |
| 3.914782286 | 9.05E-05 | 0.00031566 |
| 1.553332781 | 0.120343653 | 0.194410845 |
| 7.220817764 | 5.17E-13 | 4.96E-12 |
| -1.838842817 | 0.065938309 | 0.117118639 |
| -7.303446055 | 2.80E-13 | 2.76E-12 |
| 0.928786081 | 0.352999959 | 0.464908654 |
| 3.264317121 | 0.001097283 | 0.003135273 |
| 5.16187141 | 2.44E-07 | 1.25E-06 |
| 2.003990896 | 0.045071036 | 0.084624411 |
| -0.557688958 | 0.577056799 | 0.678013974 |
| 3.870842014 | 0.00010846 | 0.000372752 |
| -1.350907279 | 0.176725136 | 0.267141514 |
| -0.600150609 | 0.548405867 | 0.651893991 |
| 5.284366901 | 1.26E-07 | 6.71E-07 |
| 3.957805152 | 7.56E-05 | 0.000267492 |
| -1.311449736 | 0.189705865 | 0.283127887 |
| 2.94359792 | 0.003244212 | 0.008318236 |
| 3.507528238 | 0.00045229 | 0.001389911 |
| 7.213522132 | 5.45E-13 | 5.21E-12 |
| 3.406410905 | 0.00065823 | 0.001962772 |
| 6.434518983 | 1.24E-10 | 9.31E-10 |
| 15.5670584 | 1.22E-54 | 1.05E-52 |
| NA | NA | NA |
| -5.653033592 | 1.58E-08 | 9.41E-08 |
| 0.645714789 | 0.518464081 | 0.623781771 |
| NA | NA | NA |
| 2.962031773 | 0.003056162 | 0.007886159 |
| -0.065390563 | 0.947863038 | NA |
| 4.529521751 | 5.91E-06 | 2.49E-05 |
| -0.345769024 | 0.729516306 | NA |
| -0.604718467 | 0.545366076 | 0.649128008 |
| -0.254439702 | 0.799155882 | NA |
| NA | NA | NA |
| -0.952715926 | 0.340734022 | 0.452276214 |
| 0.192810118 | 0.847107679 | NA |
| 6.514700462 | 7.28E-11 | 5.62E-10 |
| 2.116672454 | 0.034287656 | 0.066720053 |
| 5.080818106 | 3.76E-07 | 1.88E-06 |
| 1.856339843 | 0.063405133 | 0.11321376 |

| | | |
|---|---|---|
| -1.32259364 | 0.18597055 | 0.27854963 |
| -3.187401107 | 0.001435575 | 0.003993826 |
| -1.059365226 | 0.28943348 | 0.396916502 |
| -0.92839891 | 0.353200684 | 0.465050664 |
| 5.709449045 | 1.13E-08 | 6.89E-08 |
| 2.712644572 | 0.006674866 | 0.015899495 |
| 6.866925148 | 6.56E-12 | 5.58E-11 |
| -5.645882004 | 1.64E-08 | 9.80E-08 |
| -9.519331297 | 1.74E-21 | 3.48E-20 |
| -1.232269846 | 0.2178483 | 0.316706502 |
| -5.845683097 | 5.04E-09 | 3.19E-08 |
| 2.953561369 | 0.003141301 | 0.008079359 |
| 6.424847403 | 1.32E-10 | 9.88E-10 |
| 0.499631003 | 0.617334923 | 0.712167264 |
| 2.943503367 | 0.003245203 | 0.008319138 |
| 2.160293054 | 0.03074999 | 0.06063276 |
| 4.728678028 | 2.26E-06 | 1.01E-05 |
| -1.636107092 | 0.101817176 | 0.16905366 |
| 3.290012347 | 0.00100183 | 0.002887919 |
| 1.747041327 | 0.08063017 | 0.139030942 |
| 0.192809521 | 0.847108147 | NA |
| 4.266222184 | 1.99E-05 | 7.73E-05 |
| 9.251766528 | 2.21E-20 | 4.02E-19 |
| 4.771080214 | 1.83E-06 | 8.30E-06 |
| 8.675567506 | 4.11E-18 | 6.26E-17 |
| 17.06854645 | 2.54E-65 | 2.88E-63 |
| -4.13808219 | 3.50E-05 | 0.000130993 |
| -1.134918468 | 0.256409481 | 0.361026386 |
| 2.756650849 | 0.005839667 | 0.01411831 |
| 0.179944158 | 0.857196408 | NA |
| 0.929072274 | 0.352851632 | 0.464737758 |
| 10.09046095 | 6.09E-24 | 1.44E-22 |
| 0.063708285 | 0.949202508 | NA |
| -0.01888271 | 0.984934673 | 0.989361701 |
| 0.93710151 | 0.348706345 | NA |
| 5.322833186 | 1.02E-07 | 5.50E-07 |
| 15.54530008 | 1.71E-54 | 1.47E-52 |
| 8.888094817 | 6.22E-19 | 1.02E-17 |
| 3.557244551 | 0.000374765 | 0.001169769 |
| 0.925927504 | 0.354483657 | NA |
| 2.970099426 | 0.002977034 | 0.00770501 |
| 2.198803764 | 0.027891879 | 0.055730321 |
| NA | NA | NA |
| 0.153562679 | 0.877954566 | NA |
| 1.919255901 | 0.054951956 | NA |
| 3.835696358 | 0.000125209 | 0.000425408 |
| NA | NA | NA |

| | | |
|---|---|---|
| 7.180400063 | 6.95E-13 | 6.56E-12 |
| 15.39864793 | 1.67E-53 | 1.39E-51 |
| 1.77596279 | 0.07573905 | 0.131740683 |
| 11.97039081 | 5.08E-33 | 2.05E-31 |
| 1.187118246 | 0.235180982 | NA |
| 15.59620947 | 7.72E-55 | 6.69E-53 |
| 1.87931266 | 0.060201814 | 0.10833643 |
| 12.03874143 | 2.22E-33 | 9.06E-32 |
| 6.548502975 | 5.81E-11 | 4.53E-10 |
| 21.52641561 | 8.81E-103 | 2.16E-100 |
| 6.839846809 | 7.93E-12 | 6.68E-11 |
| 9.16228206 | 5.08E-20 | 9.08E-19 |
| 2.679965841 | 0.007362967 | 0.017320965 |
| 19.32309096 | 3.43E-83 | 5.77E-81 |
| 8.259388815 | 1.46E-16 | 1.96E-15 |
| 1.54510279 | 0.122321402 | 0.197160454 |
| 3.344159126 | 0.000825324 | 0.002415841 |
| 18.87554167 | 1.81E-79 | 2.84E-77 |
| NA | NA | NA |
| 2.406179472 | 0.016120343 | 0.034732908 |
| 1.875680465 | 0.06069917 | 0.109094914 |
| 1.463275111 | 0.143392115 | NA |
| 1.707864718 | 0.087661447 | NA |
| NA | NA | NA |
| NA | NA | NA |
| NA | NA | NA |
| 0.439122991 | 0.660572421 | NA |
| 0.192809521 | 0.847108147 | NA |
| 0.790322522 | 0.429339437 | 0.540425754 |
| 2.789627018 | 0.005276879 | 0.012904767 |
| 0.063708882 | 0.949202033 | NA |
| NA | NA | NA |
| -0.276706088 | 0.782005797 | NA |
| 2.343992677 | 0.019078546 | 0.040098749 |
| 0.192808924 | 0.847108615 | NA |
| 7.275012724 | 3.46E-13 | 3.37E-12 |
| 0.394170842 | 0.693454911 | NA |
| NA | NA | NA |
| -0.262205383 | 0.793163103 | 0.85293317 |
| NA | NA | NA |
| 0.192809521 | 0.847108147 | NA |
| 0.063708882 | 0.949202033 | NA |
| NA | NA | NA |
| NA | NA | NA |
| 0.063710673 | 0.949200606 | NA |
| 0.063708882 | 0.949202033 | NA |
| NA | NA | NA |

| | | | |
|---|---|---|---|
| NA | NA | NA | |
| NA | NA | NA | |
| NA | NA | NA | |
| -0.065390563 | 0.947863038 | NA | |
| NA | NA | NA | |
| NA | NA | NA | |
| 0.743191973 | 0.457365462 | NA | |
| 0.063708285 | 0.949202508 | NA | |
| 0.06371127 | 0.949200131 | NA | |
| NA | NA | NA | |
| 25.46201331 | 5.20E-143 | | 1.97E-140 |
| 5.545832231 | 2.93E-08 | | 1.69E-07 |
| 9.88590742 | 4.79E-23 | | 1.07E-21 |
| -0.522847597 | 0.601080314 | | 0.698696896 |
| 0.790338202 | 0.429330282 | | 0.540425754 |
| 5.026608086 | 4.99E-07 | | 2.45E-06 |
| 0.913689227 | 0.360880166 | | 0.473022128 |
| 0.192809521 | 0.847108147 | NA | |
| 1.048112435 | 0.294586808 | NA | |
| 0.317642674 | 0.750756 | NA | |
| 0.758506128 | 0.448148046 | NA | |
| NA | NA | NA | |
| -0.624814949 | 0.532092518 | | 0.636611565 |
| 2.222745428 | 0.026232971 | | 0.052857703 |
| 16.09238664 | 2.88E-58 | | 2.72E-56 |
| 1.511666097 | 0.130618827 | | 0.208161801 |
| 1.90367148 | 0.056952983 | | 0.103509524 |
| 0.192809521 | 0.847108147 | NA | |
| NA | NA | NA | |
| NA | NA | NA | |
| 0.192808924 | 0.847108615 | NA | |
| 0.192809521 | 0.847108147 | NA | |
| NA | NA | NA | |
| NA | NA | NA | |
| NA | NA | NA | |
| NA | NA | NA | |
| NA | NA | NA | |
| NA | NA | NA | |
| NA | NA | NA | |
| 0.192809521 | 0.847108147 | NA | |
| NA | NA | NA | |
| NA | NA | NA | |
| NA | NA | NA | |
| NA | NA | NA | |
| NA | NA | NA | |
| NA | NA | NA | |
| 0.063708882 | 0.949202033 | NA | |

| | | | |
|---|---|---|---|
| NA | NA | NA | |
| NA | NA | NA | |
| NA | NA | NA | |
| NA | NA | NA | |
| NA | NA | NA | |
| NA | NA | NA | |
| NA | NA | NA | |
| NA | NA | NA | |
| | -0.065388772 | 0.947864464 | NA |
| NA | NA | NA | |
| NA | NA | NA | |
| NA | NA | NA | |
| NA | NA | NA | |
| NA | NA | NA | |
| NA | NA | NA | |
| NA | NA | NA | |
| | 0.192810118 | 0.847107679 | NA |
| NA | NA | NA | |
| NA | NA | NA | |
| | -0.065389369 | 0.947863988 | NA |
| NA | NA | NA | |
| NA | NA | NA | |
| | -0.065389369 | 0.947863988 | NA |
| NA | NA | NA | |
| NA | NA | NA | |
| NA | NA | NA | |
| NA | NA | NA | |
| NA | NA | NA | |
| | 2.092508941 | 0.036393013 | 0.070238038 |
| NA | NA | NA | |
| NA | NA | NA | |
| | 1.326085893 | 0.184811248 | 0.277161507 |
| NA | NA | NA | |
| | 1.285241229 | 0.198708 | 0.294094977 |
| NA | NA | NA | |
| | -4.291081998 | 1.78E-05 | 6.96E-05 |
| | -2.312524253 | 0.02074881 | 0.043127614 |
| | -6.593609315 | 4.29E-11 | 3.39E-10 |
| | -1.609689136 | 0.107465738 | 0.176639573 |
| | -0.527452944 | 0.597879086 | 0.696209256 |
| | -0.86280614 | 0.388244063 | 0.500472923 |
| NA | NA | NA | |
| | 4.137555921 | 3.51E-05 | 0.000131274 |
| | -6.087825208 | 1.14E-09 | 7.75E-09 |
| | 0.784338518 | 0.432841504 | NA |
| | 0.293663067 | 0.769015379 | 0.83366337 |
| | 2.487764944 | 0.012854866 | 0.028453953 |

| | | |
|---|---|---|
| NA | NA | NA |
| NA | NA | NA |
| 0.926058237 | 0.354415716 | 0.466282543 |
| 2.250551914 | 0.024413932 | 0.049669245 |
| -1.019517794 | 0.30795721 | 0.417809581 |
| NA | NA | NA |
| NA | NA | NA |
| NA | NA | NA |
| 12.07604204 | 1.41E-33 | 5.82E-32 |
| 0.063708285 | 0.949202508 | NA |
| -0.740622866 | 0.45892214 | 0.56838903 |
| -0.097016482 | 0.922713306 | 0.946809086 |
| 0.2852366 | 0.775462882 | 0.83880017 |
| 1.078783948 | 0.280684052 | 0.387636384 |
| 8.589984012 | 8.70E-18 | 1.29E-16 |
| NA | NA | NA |
| 2.447513567 | 0.014384571 | 0.031378269 |
| NA | NA | NA |
| NA | NA | NA |
| 11.35648122 | 6.89E-30 | 2.35E-28 |
| -1.176363006 | 0.239449848 | 0.341544888 |
| NA | NA | NA |
| 4.164378976 | 3.12E-05 | 0.000117722 |
| -1.74997134 | 0.080123259 | 0.138280713 |
| -0.057445467 | 0.954190346 | 0.968624908 |
| -0.998262701 | 0.318151989 | 0.428736098 |
| -4.897584453 | 9.70E-07 | 4.57E-06 |
| 0.063708882 | 0.949202033 | NA |
| 2.0854238 | 0.037030848 | 0.071287658 |
| 1.222851229 | 0.221385892 | 0.320657883 |
| 6.320970794 | 2.60E-10 | 1.88E-09 |
| -1.394439843 | 0.163184824 | 0.250031595 |
| -8.393631653 | 4.71E-17 | 6.58E-16 |
| NA | NA | NA |
| -1.68526012 | 0.091938396 | 0.155361738 |
| -1.544672145 | 0.122425586 | 0.197268007 |
| 2.477162216 | 0.01324317 | 0.029210128 |
| -0.50547684 | 0.613223962 | 0.708895056 |
| 0.64238193 | 0.520625229 | 0.625909976 |
| -1.471838693 | 0.141064442 | 0.221772689 |
| -1.394806462 | 0.16307421 | 0.249923308 |
| 0.349291458 | 0.726870511 | NA |
| -1.448484146 | 0.147481697 | 0.230155675 |
| 2.280205311 | 0.022595514 | 0.046498868 |
| 3.252081923 | 0.00114563 | 0.00326113 |
| NA | NA | NA |
| NA | NA | NA |

| | | |
|---|---|---|
| 3.133937966 | 0.001724773 | 0.004716088 |
| -1.992270437 | 0.046341394 | 0.08667153 |
| -2.017744374 | 0.043617891 | 0.08227248 |
| -5.468721828 | 4.53E-08 | 2.56E-07 |
| -0.598565274 | 0.549462819 | 0.652886511 |
| 0.3474247 | 0.728272279 | NA |
| 10.25642406 | 1.11E-24 | 2.76E-23 |
| 1.301417162 | 0.193115702 | 0.287291889 |
| 0.063976859 | 0.948988653 | NA |
| -4.36857156 | 1.25E-05 | 5.01E-05 |
| -0.319966246 | 0.748993918 | NA |
| 0.67904871 | 0.497106999 | 0.604214451 |
| 11.94451859 | 6.94E-33 | 2.79E-31 |
| 9.446446317 | 3.51E-21 | 6.83E-20 |
| 0.726352074 | 0.467622963 | NA |
| 1.987275207 | 0.046891908 | 0.087583348 |
| 0.694589752 | 0.487312436 | NA |
| 3.631602145 | 0.000281667 | 0.00089971 |
| NA | NA | NA |
| -0.194488217 | 0.845793621 | NA |
| 14.64122661 | 1.53E-48 | 1.14E-46 |
| NA | NA | NA |
| 4.750036468 | 2.03E-06 | 9.15E-06 |
| NA | NA | NA |
| NA | NA | NA |
| 2.214106133 | 0.026821479 | 0.053862453 |
| -0.0889968 | 0.929084453 | NA |
| -2.653459513 | 0.007967133 | 0.018559082 |
| NA | NA | NA |
| 0.192810118 | 0.847107679 | NA |
| -9.749508631 | 1.85E-22 | 3.96E-21 |
| 0.192810118 | 0.847107679 | NA |
| 0.420303405 | 0.674263823 | 0.757895792 |
| 0.06371127 | 0.949200131 | NA |
| -4.440509186 | 8.97E-06 | 3.69E-05 |
| 0.436478987 | 0.662489241 | 0.74889828 |
| 3.347367812 | 0.000815829 | 0.002389725 |
| NA | NA | NA |
| -5.729646411 | 1.01E-08 | 6.15E-08 |
| 0.063708217 | 0.949202562 | NA |
| NA | NA | NA |
| -0.065388772 | 0.947864464 | NA |
| 15.58115755 | 9.78E-55 | 8.44E-53 |
| 1.375407525 | 0.169005137 | 0.257295699 |
| -0.066404945 | 0.947055434 | NA |
| 0.06353712 | 0.949338802 | NA |
| 0.668958613 | 0.503521879 | 0.610056278 |

| | | | |
|---|---|---|---|
| | 0.192810118 | 0.847107679 | NA |
| | -5.410466472 | 6.29E-08 | 3.48E-07 |
| | 0.431780725 | 0.66590079 | 0.751668475 |
| | -5.055601493 | 4.29E-07 | 2.12E-06 |
| | -4.12152775 | 3.76E-05 | 0.000139957 |
| | -3.650954839 | 0.000261267 | 0.000839153 |
| | 2.396249757 | 0.016563799 | 0.035563213 |
| | 1.764257185 | 0.077688689 | 0.134654656 |
| | 0.016205305 | 0.987070603 | 0.990950473 |
| | 2.236198527 | 0.02533877 | 0.05131721 |
| | -1.102240788 | 0.270357004 | NA |
| | 0.888845123 | 0.374086321 | 0.486839319 |
| | 11.79695563 | 4.05E-32 | 1.56E-30 |
| | 3.47653079 | 0.000507946 | 0.001547831 |
| | 2.335293059 | 0.019528117 | 0.040913096 |
| | 6.99425081 | 2.67E-12 | 2.35E-11 |
| | 26.60098557 | 6.61E-156 | 2.81E-153 |
| | 5.805126046 | 6.43E-09 | 4.01E-08 |
| | 5.606963011 | 2.06E-08 | 1.21E-07 |
| | 0.382354918 | 0.702198124 | 0.781364766 |
| | -4.657696378 | 3.20E-06 | 1.40E-05 |
| | 3.313784288 | 0.000920425 | 0.00267048 |
| | -4.276266992 | 1.90E-05 | 7.41E-05 |
| | -0.937567222 | 0.348466863 | 0.460149067 |
| | 7.123595266 | 1.05E-12 | 9.75E-12 |
| NA | NA | NA | |
| NA | NA | NA | |
| | 2.666066212 | 0.007674458 | 0.017960886 |
| | -1.170592534 | 0.2417626 | 0.344294371 |
| NA | NA | NA | |
| | -0.05357599 | 0.957272986 | NA |
| NA | NA | NA | |
| NA | NA | NA | |
| | 1.31211239 | 0.189482216 | 0.282861565 |
| | -0.539032945 | 0.589864122 | NA |
| | -5.840868193 | 5.19E-09 | 3.27E-08 |
| NA | NA | NA | |
| | -0.079583924 | 0.936568181 | 0.956750534 |
| | 1.925104867 | 0.05421624 | 0.099241962 |
| | 2.587061768 | 0.009679824 | 0.022130617 |
| | -1.074465297 | 0.282614191 | 0.389592619 |
| | -1.691587219 | 0.090724705 | 0.153611538 |
| | 2.700469054 | 0.006924178 | 0.016419912 |
| | 2.053897805 | 0.039985584 | 0.076184758 |
| | 1.259742012 | 0.207762445 | 0.304805931 |
| | 3.476998351 | 0.000507061 | 0.001545511 |
| | -4.755070576 | 1.98E-06 | 8.94E-06 |

| | | |
|---|---|---|
| 0.063708882 | 0.949202033 | NA |
| 2.425926566 | 0.01526936 | 0.033090183 |
| NA | NA | NA |
| 0.333816862 | 0.738517758 | NA |
| NA | NA | NA |
| 0.106410254 | 0.915256858 | NA |
| 7.595017604 | 3.08E-14 | 3.33E-13 |
| NA | NA | NA |
| NA | NA | NA |
| 0.06353712 | 0.949338802 | NA |
| 0.6552338 | 0.512317236 | 0.6179606 |
| 0.063710673 | 0.949200606 | NA |
| 0.519067346 | 0.603713779 | NA |
| NA | NA | NA |
| 0.063707688 | 0.949202983 | NA |
| 0.192809521 | 0.847108147 | NA |
| -0.19448762 | 0.845794089 | NA |
| 0.847666164 | 0.396623915 | NA |
| NA | NA | NA |
| -0.093562861 | 0.925456413 | 0.948729276 |
| -1.168551915 | 0.242584211 | 0.345227274 |
| 0.416388637 | 0.677125642 | 0.760211098 |
| 4.30200866 | 1.69E-05 | 6.65E-05 |
| 10.8992991 | 1.16E-27 | 3.55E-26 |
| 1.502613937 | 0.132938627 | 0.211186459 |
| NA | NA | NA |
| 14.25661853 | 4.08E-46 | 2.82E-44 |
| 0.98929691 | 0.322517895 | 0.433471614 |
| 6.692741816 | 2.19E-11 | 1.78E-10 |
| 24.99226648 | 7.42E-138 | 2.69E-135 |
| 32.55644679 | 1.70E-232 | 1.85E-229 |
| 11.5621455 | 6.41E-31 | 2.32E-29 |
| 27.22488753 | 3.30E-163 | 1.59E-160 |
| 16.01918479 | 9.39E-58 | 8.70E-56 |
| 32.1367138 | 1.35E-226 | 1.41E-223 |
| -4.744516148 | 2.09E-06 | 9.38E-06 |
| NA | NA | NA |
| NA | NA | NA |
| -5.207694812 | 1.91E-07 | 9.91E-07 |
| -1.735100641 | 0.082722981 | 0.142110927 |
| 4.547513387 | 5.43E-06 | 2.30E-05 |
| NA | NA | NA |
| NA | NA | NA |
| NA | NA | NA |
| NA | NA | NA |
| NA | NA | NA |
| -0.402401852 | 0.687388307 | NA |

| | | |
|---|---|---|
| 0.320069237 | 0.748915845 | 0.817575175 |
| 3.492337642 | 0.000478813 | 0.001464583 |
| 3.680606477 | 0.00023268 | 0.000754401 |
| 0.914715989 | 0.360340748 | NA |
| 13.77447313 | 3.63E-43 | 2.24E-41 |
| NA | NA | NA |
| -0.308436951 | 0.757749869 | 0.824588925 |
| -2.084124058 | 0.037148884 | 0.071481901 |
| -2.408747315 | 0.016007377 | 0.034522199 |
| 1.00426977 | 0.3152486 | 0.425717425 |
| -1.074836477 | 0.282447946 | 0.389470697 |
| 1.499631287 | 0.133709938 | 0.212224751 |
| -6.090481766 | 1.13E-09 | 7.63E-09 |
| -2.267771124 | 0.02334316 | 0.047817277 |
| -1.295243854 | 0.195236121 | 0.289774674 |
| 0.281048768 | 0.778672995 | 0.841617802 |
| 10.40663865 | 2.31E-25 | 6.06E-24 |
| NA | NA | NA |
| 2.539039944 | 0.011115713 | 0.025060917 |
| -2.086576152 | 0.036926465 | 0.071125353 |
| 4.505972685 | 6.61E-06 | 2.76E-05 |
| 2.554589489 | 0.01063131 | 0.02408596 |
| 4.653978911 | 3.26E-06 | 1.42E-05 |
| NA | NA | NA |
| -1.739381567 | 0.081967668 | 0.140998923 |
| -0.158456872 | 0.874096803 | 0.912294344 |
| 0.474679205 | 0.635015642 | 0.726403799 |
| 9.410114058 | 4.96E-21 | 9.51E-20 |
| -1.650191624 | 0.098903749 | 0.165114443 |
| 5.991968598 | 2.07E-09 | 1.37E-08 |
| NA | NA | NA |
| -0.106745897 | 0.91499057 | NA |
| -0.170896557 | 0.864305104 | 0.905510726 |
| 0.654535552 | 0.51276683 | 0.618413499 |
| -1.345793721 | 0.178369051 | 0.26908012 |
| -6.307393669 | 2.84E-10 | 2.04E-09 |
| 0.024430878 | 0.980508919 | 0.986459996 |
| 0.310799452 | 0.755953086 | 0.823032003 |
| NA | NA | NA |
| 7.164750023 | 7.79E-13 | 7.32E-12 |
| NA | NA | NA |
| 4.064210505 | 4.82E-05 | 0.000176081 |
| -0.661784171 | 0.508109552 | 0.614335914 |
| -1.430307751 | 0.152628711 | 0.236858865 |
| 2.716061612 | 0.006606362 | 0.01575281 |
| 1.068115624 | 0.285468359 | 0.39261983 |
| NA | NA | NA |

| | | |
|---|---|---|
| 1.59093016 | 0.111625294 | 0.182374746 |
| 0.192810118 | 0.847107679 | NA |
| 0.192810118 | 0.847107679 | NA |
| 0.464282325 | 0.642445486 | NA |
| 0.063708882 | 0.949202033 | NA |
| 0.120600738 | 0.904007284 | NA |
| NA | NA | NA |
| 0.063708882 | 0.949202033 | NA |
| -0.247174184 | 0.804773424 | NA |
| 0.063708285 | 0.949202508 | NA |
| 0.063708882 | 0.949202033 | NA |
| 0.063708882 | 0.949202033 | NA |
| 0.241893141 | 0.808862965 | NA |
| -0.36608595 | 0.714300941 | NA |
| 0.430718809 | 0.666672841 | 0.752183673 |
| 4.169697048 | 3.05E-05 | 0.000115252 |
| 5.077304148 | 3.83E-07 | 1.91E-06 |
| 6.701690474 | 2.06E-11 | 1.68E-10 |
| 3.681440785 | 0.00023192 | 0.000752424 |
| 1.080844593 | 0.279766249 | 0.38671025 |
| -4.724648471 | 2.31E-06 | 1.03E-05 |
| 6.915059457 | 4.68E-12 | 4.05E-11 |
| -1.149086367 | 0.250520369 | 0.354186898 |
| 4.57505125 | 4.76E-06 | 2.03E-05 |
| 0.032155181 | 0.974348298 | 0.98223601 |
| 0.728135341 | 0.466530742 | 0.575801995 |
| NA | NA | NA |
| 4.453037488 | 8.47E-06 | 3.49E-05 |
| 7.322416153 | 2.44E-13 | 2.41E-12 |
| 3.662926464 | 0.00024935 | 0.000803765 |
| -0.264293185 | 0.791554012 | 0.851695927 |
| 0.518107346 | 0.604383375 | 0.701558961 |
| -0.620834495 | 0.534708524 | 0.639117251 |
| 25.41318263 | 1.80E-142 | 6.74E-140 |
| 0.063708882 | 0.949202033 | NA |
| -3.461345094 | 0.000537483 | 0.001629639 |
| -0.035010348 | 0.972071489 | 0.980849366 |
| 1.284491474 | 0.198970045 | 0.294416242 |
| -0.607781844 | 0.543332173 | 0.64732278 |
| -7.746391619 | 9.45E-15 | 1.08E-13 |
| -0.194488217 | 0.845793621 | NA |
| -1.647708886 | 0.099412424 | 0.165786401 |
| 2.111901729 | 0.034694879 | 0.067391962 |
| -3.705683563 | 0.000210821 | 0.000689695 |
| 6.220757764 | 4.95E-10 | 3.48E-09 |
| 0.744908877 | 0.456326809 | 0.566027059 |
| 2.758129158 | 0.005813322 | 0.01406005 |

| | | |
|---|---|---|
| 3.345571702 | 0.000821131 | 0.002404131 |
| 1.460052888 | 0.144275539 | 0.226054256 |
| NA | NA | NA |
| 8.759040318 | 1.97E-18 | 3.09E-17 |
| 0.887294107 | 0.374920577 | 0.487793416 |
| 1.990641125 | 0.046520356 | 0.086980242 |
| -2.599680602 | 0.009331056 | 0.021407547 |
| -4.245327561 | 2.18E-05 | 8.43E-05 |
| 8.757025164 | 2.00E-18 | 3.14E-17 |
| 0.221341313 | 0.824826684 | 0.87725894 |
| 0.391132133 | 0.69569957 | 0.775914155 |
| -1.175959455 | 0.239611078 | 0.341735914 |
| -3.401835787 | 0.000669348 | 0.00199355 |
| -3.711073794 | 0.000206382 | 0.000676498 |
| 1.301044039 | 0.193243381 | 0.287430582 |
| -0.401495214 | 0.688055561 | NA |
| 1.677761684 | 0.093393632 | 0.157448819 |
| 0.888954805 | 0.37402737 | 0.486787909 |
| 0.29222879 | 0.7701117 | NA |
| 5.743602415 | 9.27E-09 | 5.68E-08 |
| NA | NA | NA |
| -4.163746785 | 3.13E-05 | 0.000118031 |
| 1.61413194 | 0.106498811 | 0.175326574 |
| -2.140896143 | 0.032282414 | 0.063320181 |
| -0.226808238 | 0.820572858 | 0.874071777 |
| NA | NA | NA |
| -0.646039117 | 0.518254023 | 0.623588991 |
| -6.944832023 | 3.79E-12 | 3.30E-11 |
| -3.053090968 | 0.002264973 | 0.006024683 |
| 0.288217989 | 0.77317988 | NA |
| -0.065389369 | 0.947863988 | NA |
| NA | NA | NA |
| 6.643912073 | 3.05E-11 | 2.45E-10 |
| 1.138166832 | 0.255050826 | 0.359394368 |
| 5.969796449 | 2.38E-09 | 1.55E-08 |
| 4.779282887 | 1.76E-06 | 7.99E-06 |
| 0.515613324 | 0.606124502 | 0.703058357 |
| -0.34714027 | 0.728485942 | 0.801741332 |
| -2.914967853 | 0.003557252 | 0.009033889 |
| NA | NA | NA |
| 0.76336899 | 0.445243368 | 0.555140299 |
| 0.048272567 | 0.961499017 | 0.97360651 |
| 2.783179676 | 0.005382899 | 0.013142212 |
| 2.448565735 | 0.014342626 | 0.031298685 |
| -0.245039149 | 0.806426119 | 0.862932143 |
| 1.809380306 | 0.070391941 | 0.123868832 |
| 4.017265132 | 5.89E-05 | 0.00021183 |

| | | |
|---|---|---|
| 5.827061891 | 5.64E-09 | 3.54E-08 |
| -2.860639683 | 0.004227873 | 0.010570737 |
| -8.703530602 | 3.22E-18 | 4.94E-17 |
| NA | NA | NA |
| 1.196494875 | 0.231503497 | 0.332331529 |
| -3.871446597 | 0.000108191 | 0.000371931 |
| -2.03224524 | 0.042128835 | 0.079763748 |
| 0.801147457 | 0.423046285 | 0.534763321 |
| 0.192810118 | 0.847107679 | NA |
| -0.575023347 | 0.565275507 | 0.667335438 |
| -0.665310064 | 0.505852195 | 0.612316493 |
| -0.637915847 | 0.523528462 | 0.628756652 |
| 0.771804637 | 0.440230145 | 0.550425123 |
| NA | NA | NA |
| -0.065389369 | 0.947863988 | NA |
| -0.194487022 | 0.845794556 | NA |
| -0.267888177 | 0.788785394 | 0.849733788 |
| -0.003868143 | 0.996913676 | 0.997950301 |
| -7.318382833 | 2.51E-13 | 2.48E-12 |
| 3.589369772 | 0.000331478 | 0.001045477 |
| -1.309338725 | 0.190419637 | 0.284006884 |
| 5.636172812 | 1.74E-08 | 1.03E-07 |
| -2.500043623 | 0.012417801 | 0.027603692 |
| -4.027531942 | 5.64E-05 | 0.0002037 |
| 0.713810773 | 0.475344197 | 0.584231046 |
| -0.125999325 | 0.899732459 | 0.930091415 |
| NA | NA | NA |
| 0.626136468 | 0.531225434 | 0.635925803 |
| 0.107308715 | 0.914544071 | 0.941282047 |
| 0.942697137 | 0.345835838 | 0.457326238 |
| -0.792477314 | 0.428082414 | 0.539358846 |
| 0.596150774 | 0.551074507 | 0.654397709 |
| -4.29920119 | 1.71E-05 | 6.73E-05 |
| 2.023904686 | 0.04297996 | 0.081173198 |
| 0.845822306 | 0.39765188 | 0.509499773 |
| 2.04799659 | 0.040560334 | 0.077138907 |
| 1.126512304 | 0.259948736 | 0.364878693 |
| -0.344041529 | 0.730815055 | NA |
| 4.567982513 | 4.92E-06 | 2.10E-05 |
| 0.936747705 | 0.348888351 | 0.460535567 |
| 2.496442762 | 0.012544591 | 0.027848436 |
| -9.900070841 | 4.16E-23 | 9.35E-22 |
| 0.946885336 | 0.343697211 | 0.45507518 |
| 0.385251561 | 0.700051046 | 0.77939141 |
| 3.687028366 | 0.000226888 | 0.000737917 |
| -3.918939081 | 8.89E-05 | 0.000310742 |
| 5.713868395 | 1.10E-08 | 6.72E-08 |

| | | |
|---|---|---|
| 1.66864638 | 0.095187482 | 0.159934394 |
| 2.831360353 | 0.004635047 | 0.011491207 |
| 0.179925889 | 0.857210751 | 0.900906327 |
| 1.387060884 | 0.165423198 | 0.252888454 |
| -0.106175676 | 0.915442969 | NA |
| 0.969652943 | 0.332219514 | NA |
| 13.39667523 | 6.32E-41 | 3.53E-39 |
| -10.92942139 | 8.34E-28 | 2.58E-26 |
| -0.430709765 | 0.666679418 | 0.752183673 |
| -0.655801817 | 0.511951647 | 0.617642927 |
| -0.222915886 | 0.823600972 | 0.87632768 |
| -3.602836323 | 0.000314764 | 0.000997283 |
| 0.769118301 | 0.441823085 | NA |
| -1.693764477 | 0.090310047 | 0.153054403 |
| -2.595857284 | 0.009435525 | 0.021621458 |
| 2.474000786 | 0.01336094 | 0.029443946 |
| 1.068727682 | 0.285192394 | 0.392282553 |
| 1.375561609 | 0.168957399 | 0.257238668 |
| -2.807851028 | 0.004987329 | 0.012278238 |
| 2.210940022 | 0.027039991 | 0.054227303 |
| 8.950152801 | 3.55E-19 | 5.95E-18 |
| 0.4001112 | 0.689074616 | NA |
| 0.063708882 | 0.949202033 | NA |
| -3.07470442 | 0.002107113 | 0.005646791 |
| 3.274450022 | 0.001058679 | 0.003035364 |
| -5.588154211 | 2.29E-08 | 1.34E-07 |
| -1.725613853 | 0.084416905 | 0.144524976 |
| -1.549350639 | 0.121297453 | 0.195724294 |
| -3.401761001 | 0.000669532 | 0.001993858 |
| -0.065389369 | 0.947863988 | NA |
| 1.465592459 | 0.142759347 | 0.224211988 |
| 0.758569886 | 0.448109893 | 0.558074963 |
| 4.717229237 | 2.39E-06 | 1.06E-05 |
| 0.233416754 | 0.815437805 | 0.870214774 |
| NA | NA | NA |
| 0.442952555 | 0.657800053 | NA |
| -1.081658859 | 0.27940414 | 0.386444473 |
| -8.624212356 | 6.45E-18 | 9.64E-17 |
| -5.965385326 | 2.44E-09 | 1.59E-08 |
| -1.512303597 | 0.130456647 | 0.207977249 |
| 0.233342711 | 0.815495295 | 0.87021859 |
| 1.746785141 | 0.080674615 | 0.139088415 |
| 0.49617555 | 0.61977056 | 0.714187059 |
| -0.629054948 | 0.529313083 | NA |
| 0.185655208 | 0.852715156 | 0.897689282 |
| NA | NA | NA |
| NA | NA | NA |

| | | |
|---|---|---|
| 2.482143244 | 0.013059477 | 0.028868559 |
| 9.76883132 | 1.53E-22 | 3.28E-21 |
| 0.192809521 | 0.847108147 | NA |
| -0.032441776 | 0.974119748 | 0.982084718 |
| -0.565345354 | 0.571838886 | 0.673147125 |
| NA | NA | NA |
| 2.446266215 | 0.014434438 | 0.031468859 |
| 2.08634534 | 0.036947352 | 0.071154284 |
| NA | NA | NA |
| NA | NA | NA |
| 0.216730434 | 0.828418429 | NA |
| -0.019192081 | 0.984687875 | 0.989312307 |
| NA | NA | NA |
| -0.194487022 | 0.845794556 | NA |
| NA | NA | NA |
| NA | NA | NA |
| NA | NA | NA |
| NA | NA | NA |
| 6.505700744 | 7.73E-11 | 5.95E-10 |
| -0.990035244 | 0.322156893 | 0.43320012 |
| 0.317642674 | 0.750756 | NA |
| 4.3260237 | 1.52E-05 | 6.01E-05 |
| NA | NA | NA |
| -7.699785234 | 1.36E-14 | 1.53E-13 |
| 0.433700853 | 0.664505687 | NA |
| 4.665141162 | 3.08E-06 | 1.35E-05 |
| 1.091055555 | 0.275248438 | 0.381963098 |
| 1.774434412 | 0.075991322 | 0.132124395 |
| 3.808697513 | 0.000139701 | 0.000470937 |
| 2.355455985 | 0.018499989 | 0.039075672 |
| -1.556974317 | 0.119476583 | 0.193172203 |
| -3.074103405 | 0.002111362 | 0.005656969 |
| -0.065389369 | 0.947863988 | NA |
| -0.333501592 | 0.738755688 | 0.809710858 |
| -7.459921587 | 8.66E-14 | 8.95E-13 |
| -0.304769202 | 0.76054194 | 0.826768114 |
| 1.678557147 | 0.093238384 | 0.157240046 |
| -4.741227399 | 2.12E-06 | 9.53E-06 |
| 1.731845966 | 0.083300992 | 0.142945974 |
| 0.177878249 | 0.858818593 | 0.901914466 |
| NA | NA | NA |
| -5.706208688 | 1.16E-08 | 7.01E-08 |
| 13.61775899 | 3.14E-42 | 1.86E-40 |
| 4.588059836 | 4.47E-06 | 1.92E-05 |
| -1.96383997 | 0.049548653 | 0.091873679 |
| 3.849688904 | 0.000118268 | 0.000403854 |
| 0.187508527 | 0.851261938 | 0.896574219 |

| | | |
|---|---|---|
| -0.74776783 | 0.45460021 | 0.564416828 |
| NA | NA | NA |
| 0.063708285 | 0.949202508 | NA |
| 6.844218746 | 7.69E-12 | 6.49E-11 |
| 0.069717895 | 0.944418199 | 0.962295441 |
| 17.25824951 | 9.70E-67 | 1.14E-64 |
| 1.46413021 | 0.143158374 | 0.224683603 |
| 5.847107587 | 5.00E-09 | 3.16E-08 |
| 3.697350404 | 0.000217862 | 0.000710869 |
| -5.471603072 | 4.46E-08 | 2.52E-07 |
| -1.074335758 | 0.282672224 | 0.389608247 |
| -1.479903699 | 0.138898948 | 0.218968518 |
| 0.128005965 | 0.898144253 | 0.929028695 |
| 0.976663425 | 0.328735808 | NA |
| 3.272931797 | 0.001064382 | 0.003050667 |
| -4.652332585 | 3.28E-06 | 1.43E-05 |
| -4.367182533 | 1.26E-05 | 5.04E-05 |
| -1.829575827 | 0.067313391 | 0.119205758 |
| -2.381988492 | 0.017219436 | 0.036778093 |
| NA | NA | NA |
| -5.60230319 | 2.12E-08 | 1.25E-07 |
| 0.320506792 | 0.748584182 | 0.81724875 |
| 0.06371127 | 0.949200131 | NA |
| 9.32692689 | 1.09E-20 | 2.03E-19 |
| 0.985310966 | 0.324471348 | 0.435516829 |
| 0.371239867 | 0.710458882 | 0.787755197 |
| 0.040161647 | 0.967964254 | 0.978164047 |
| NA | NA | NA |
| 0.063710673 | 0.949200606 | NA |
| -0.109366048 | 0.912912162 | NA |
| 1.755486471 | 0.079176133 | 0.136844262 |
| NA | NA | NA |
| NA | NA | NA |
| -9.474148552 | 2.69E-21 | 5.29E-20 |
| -2.688471072 | 0.007178005 | 0.016928812 |
| 0.972468879 | 0.33081734 | 0.441780604 |
| -4.900671172 | 9.55E-07 | 4.50E-06 |
| 1.035191361 | 0.300579553 | 0.409635287 |
| -0.065389369 | 0.947863988 | NA |
| -0.301648723 | 0.762919862 | 0.828778444 |
| -0.58439353 | 0.558955579 | 0.661277976 |
| -0.590264045 | 0.555013641 | 0.65782704 |
| 2.806145116 | 0.005013811 | 0.012333959 |
| -4.175075425 | 2.98E-05 | 0.000112732 |
| -2.399893748 | 0.016399831 | 0.035265273 |
| -0.528809872 | 0.596937349 | NA |
| -2.093263285 | 0.036325658 | 0.07011345 |

| | | |
|---|---|---|
| -4.100739871 | 4.12E-05 | 0.00015197 |
| 1.07723389 | 0.281375789 | 0.388355996 |
| -0.188629437 | 0.850383259 | 0.895799545 |
| 1.410768112 | 0.158313 | 0.243925786 |
| 0.524311123 | 0.600062162 | 0.697902324 |
| NA | NA | NA |
| -0.800420469 | 0.423467226 | 0.535093149 |
| 5.618362982 | 1.93E-08 | 1.14E-07 |
| -0.604146749 | 0.545746081 | 0.649406859 |
| -2.670536671 | 0.00757301 | 0.017759996 |
| 3.973198573 | 7.09E-05 | 0.000251841 |
| 0.295079439 | 0.767933196 | 0.832742739 |
| 2.414191998 | 0.01577015 | 0.034060476 |
| 2.515996977 | 0.01186962 | 0.026521745 |
| NA | NA | NA |
| -3.660907639 | 0.000251323 | 0.000809812 |
| 1.882991482 | 0.059701522 | 0.107629581 |
| -2.740886231 | 0.006127372 | 0.014734184 |
| -0.078508538 | 0.937423539 | NA |
| NA | NA | NA |
| -2.235331462 | 0.025395596 | 0.05139963 |
| NA | NA | NA |
| -0.478697365 | 0.632153941 | 0.723957409 |
| NA | NA | NA |
| 8.67355867 | 4.19E-18 | 6.36E-17 |
| NA | NA | NA |
| NA | NA | NA |
| NA | NA | NA |
| -1.601903144 | 0.109177029 | 0.179005701 |
| -0.590128778 | 0.555104316 | 0.657872208 |
| 0.063708285 | 0.949202508 | NA |
| 1.590374902 | 0.111750324 | 0.182521961 |
| 0.206933198 | 0.83606203 | 0.885071587 |
| 1.271835083 | 0.203431721 | 0.299787811 |
| 0.192810118 | 0.847107679 | NA |
| 1.176652151 | 0.239334373 | 0.34143855 |
| 4.159252818 | 3.19E-05 | 0.000120088 |
| -2.426190041 | 0.015258278 | 0.033069071 |
| 1.705283656 | 0.088141534 | 0.149978423 |
| 1.117253711 | 0.263885861 | 0.369390029 |
| 0.260848991 | 0.794208963 | 0.853680426 |
| 0.063708285 | 0.949202508 | NA |
| 0.338267486 | 0.735161623 | NA |
| 5.879096242 | 4.13E-09 | 2.63E-08 |
| NA | NA | NA |
| 0.270954455 | 0.786426062 | 0.847731826 |
| -0.065389369 | 0.947863988 | NA |

| | | | |
|---|---|---|---|
| | -0.065390563 | 0.947863038 | NA |
| | 0.063708882 | 0.949202033 | NA |
| | 8.6015061 | 7.87E-18 | 1.17E-16 |
| | -0.291666853 | 0.770541355 | NA |
| NA | NA | NA | |
| | -2.583578517 | 0.009778121 | 0.022324762 |
| | -1.688008581 | 0.091409582 | 0.154562037 |
| NA | NA | NA | |
| | -2.185054652 | 0.02888484 | 0.057448354 |
| | -2.403895484 | 0.01622141 | 0.034930285 |
| | -4.359984528 | 1.30E-05 | 5.20E-05 |
| | -3.587918023 | 0.000333329 | 0.001050784 |
| | 1.298309801 | 0.1941809 | 0.288516438 |
| | -3.622312867 | 0.000291981 | 0.000930043 |
| NA | NA | NA | |
| NA | NA | NA | |
| | 2.002024751 | 0.04528207 | 0.084969653 |
| | 2.432179909 | 0.01500825 | 0.032592097 |
| | 0.192809521 | 0.847108147 | NA |
| | 0.46697527 | 0.640517572 | 0.731060411 |
| NA | NA | NA | |
| | -0.814474447 | 0.415373205 | 0.527003985 |
| | 7.196990011 | 6.16E-13 | 5.85E-12 |
| | 0.528839849 | 0.596916552 | 0.695245291 |
| NA | NA | NA | |
| | -0.171540934 | 0.863798447 | 0.905394686 |
| | 2.235392175 | 0.025391613 | 0.05139963 |
| | 4.721803213 | 2.34E-06 | 1.04E-05 |
| | 2.548456233 | 0.010820086 | 0.024460055 |
| | 0.567341218 | 0.570482378 | NA |
| | 0.441203915 | 0.659065379 | 0.746207375 |
| | 5.56887345 | 2.56E-08 | 1.49E-07 |
| | -0.684114593 | 0.493902808 | 0.601108007 |
| NA | NA | NA | |
| | -0.065388772 | 0.947864464 | NA |
| | -2.18587222 | 0.028824955 | 0.057347479 |
| | 0.06371127 | 0.949200131 | NA |
| | 0.514005889 | 0.607247873 | 0.703806921 |
| | -0.837651802 | 0.40222629 | 0.513974068 |
| | 0.053432556 | 0.957387266 | 0.970610526 |
| | -2.777993341 | 0.005469574 | 0.013326529 |
| | 1.044623135 | 0.296197183 | NA |
| | -3.511808322 | 0.000445069 | 0.001370074 |
| | 3.241332749 | 0.001189722 | 0.003372452 |
| | -1.974378373 | 0.048338735 | 0.089869906 |
| NA | NA | NA | |
| | 0.883522501 | 0.376954031 | NA |

| | | |
|---|---|---|
| 2.809522714 | 0.004961502 | 0.01222307 |
| -0.461743395 | 0.644265347 | NA |
| 1.164268798 | 0.244315097 | 0.347159062 |
| 0.529689019 | 0.596327562 | 0.694720916 |
| 0.279600948 | 0.77978368 | NA |
| 2.47442619 | 0.013345039 | 0.029411494 |
| 2.433636456 | 0.014948 | 0.032475344 |
| -5.345656016 | 9.01E-08 | 4.89E-07 |
| 0.326654803 | 0.743928988 | 0.813657093 |
| -3.10235623 | 0.001919868 | 0.005191691 |
| 0.192809521 | 0.847108147 | NA |
| -3.794056972 | 0.000148206 | 0.000497264 |
| -0.69627182 | 0.486258617 | 0.59445997 |
| -1.055006387 | 0.291422404 | 0.399135646 |
| NA | NA | NA |
| -0.445116002 | 0.656235925 | NA |
| 0.575805444 | 0.564746693 | 0.666774044 |
| 0.19909715 | 0.842186751 | NA |
| 0.063708882 | 0.949202033 | NA |
| NA | NA | NA |
| NA | NA | NA |
| -1.488728676 | 0.136558831 | 0.215923407 |
| NA | NA | NA |
| NA | NA | NA |
| NA | NA | NA |
| NA | NA | NA |
| NA | NA | NA |
| NA | NA | NA |
| NA | NA | NA |
| 3.313970145 | 0.000919813 | 0.002669325 |
| 0.192809521 | 0.847108147 | NA |
| 0.892338181 | 0.372211695 | 0.485030129 |
| -1.615481506 | 0.106206462 | 0.174902806 |
| NA | NA | NA |
| 0.070392489 | 0.943881269 | 0.962100838 |
| -0.031482401 | 0.974884827 | 0.982499888 |
| -0.674476373 | 0.500008502 | 0.60700421 |
| -3.060196773 | 0.002211916 | 0.005897344 |
| 0.968552829 | 0.332768349 | 0.443962988 |
| 7.991350001 | 1.33E-15 | 1.64E-14 |
| -0.525055227 | 0.5995448 | 0.697397822 |
| 0.192810118 | 0.847107679 | NA |
| 7.33670787 | 2.19E-13 | 2.17E-12 |
| -0.340281252 | 0.733644735 | 0.805638431 |
| 1.622965615 | 0.104596756 | 0.172866081 |
| -1.14414037 | 0.252565428 | 0.356606829 |
| 0.085267118 | 0.932049033 | 0.9537689 |

| | | | |
|---|---|---|---|
| | 1.680389084 | 0.092881638 | 0.15673346 |
| | 0.81073628 | 0.417517134 | 0.529190493 |
| | 0.675150066 | 0.499580428 | 0.60669051 |
| | 2.096647965 | 0.03602475 | 0.069591687 |
| | -0.067700047 | 0.946024412 | 0.963108722 |
| | 1.195265685 | 0.231983242 | 0.332886577 |
| | -1.775085358 | 0.075883794 | 0.131955771 |
| | -0.14342724 | 0.885952771 | 0.920026463 |
| | -0.076664994 | 0.938890053 | 0.95842669 |
| | 0.640038797 | 0.522147377 | 0.627429133 |
| | 0.454148002 | 0.649722306 | 0.738468319 |
| | 0.500800337 | 0.616511648 | 0.711447052 |
| | 0.52570951 | 0.599090055 | NA |
| | -0.065388772 | 0.947864464 | NA |
| | 0.190138984 | 0.849200223 | 0.894892286 |
| | -1.152758226 | 0.249009635 | 0.352389391 |
| | 0.2793968 | 0.779940326 | 0.842696467 |
| | 0.267256654 | 0.789271559 | 0.850041343 |
| | 0.635777181 | 0.524921663 | NA |
| | 0.625902172 | 0.53137911 | 0.635964848 |
| | 0.200304827 | 0.841242188 | NA |
| | 0.736035405 | 0.461709152 | 0.571118363 |
| | 1.473573163 | 0.140596552 | 0.221162122 |
| | 0.714407095 | 0.474975486 | 0.583921238 |
| | 1.878143405 | 0.06036155 | 0.108592654 |
| | 0.110525229 | 0.911992843 | 0.939351502 |
| | 0.208683593 | 0.834695249 | 0.884223044 |
| | -1.58090808 | 0.113899057 | 0.185532564 |
| | 0.00059529 | 0.999525027 | 0.999600945 |
| | 0.821415085 | 0.411409874 | 0.523118575 |
| | 2.034102797 | 0.041941232 | 0.079457239 |
| | -0.713811261 | 0.475343895 | 0.584231046 |
| NA | NA | NA | |
| NA | NA | NA | |
| | 1.071754632 | 0.283830255 | 0.390838493 |
| NA | NA | NA | |
| | 0.685103038 | 0.493278904 | NA |
| | 0.684383417 | 0.493733085 | NA |
| NA | NA | NA | |
| NA | NA | NA | |
| | 0.063708882 | 0.949202033 | NA |
| | 0.063708882 | 0.949202033 | NA |
| NA | NA | NA | |
| NA | NA | NA | |
| NA | NA | NA | |
| | 0.192810118 | 0.847107679 | NA |
| NA | NA | NA | |

| | | | |
|---|---|---|---|
| | 0.063708285 | 0.949202508 | NA |
| NA | NA | NA | |
| | -0.065389369 | 0.947863988 | NA |
| | 0.152441452 | 0.878838765 | NA |
| | -0.450480661 | 0.652363895 | NA |
| | 0.334970895 | 0.737647041 | NA |
| NA | NA | NA | |
| NA | NA | NA | |
| NA | NA | NA | |
| | 0.330493631 | 0.741027004 | NA |
| | 0.192810118 | 0.847107679 | NA |
| NA | NA | NA | |
| | 0.260606205 | 0.794396204 | NA |
| NA | NA | NA | |
| | -0.194488217 | 0.845793621 | NA |
| | 0.181127434 | 0.856267551 | NA |
| | 0.063708285 | 0.949202508 | NA |
| | -0.065388772 | 0.947864464 | NA |
| | 0.083031301 | 0.933826651 | NA |
| | -0.802453231 | 0.422290832 | 0.534078092 |
| | -0.441924238 | 0.65854403 | 0.745974321 |
| | -0.952439881 | 0.340873942 | 0.452413954 |
| | -0.086734923 | 0.930882216 | NA |
| | 0.406611392 | 0.684293429 | NA |
| | 1.449847491 | 0.147101053 | 0.229618918 |
| | 1.725085757 | 0.084512019 | 0.144638348 |
| | 0.15204725 | 0.879149669 | 0.915125851 |
| NA | NA | NA | |
| | -0.811978113 | 0.416804191 | 0.528500957 |
| | 0.87753679 | 0.380195145 | 0.492765352 |
| | 0.481056525 | 0.630476326 | 0.722654276 |
| | 0.143428926 | 0.885951439 | 0.920026463 |
| | 1.04324848 | 0.296833225 | 0.405420475 |
| | -0.720331406 | 0.471320972 | 0.580220851 |
| | -0.263114001 | 0.792462713 | NA |
| | 0.03935127 | 0.968610331 | 0.978471208 |
| | -1.41897775 | 0.155905503 | 0.241077083 |
| | 0.34770012 | 0.728065407 | 0.801454707 |
| | -0.139187764 | 0.889301777 | 0.922547392 |
| | -0.91343252 | 0.361015108 | NA |
| | 0.145461423 | 0.884346572 | NA |
| | 0.490967037 | 0.623449762 | 0.717337821 |
| | 0.778045711 | 0.436542066 | NA |
| | -0.088767216 | 0.929266913 | NA |
| | 0.533095477 | 0.59396749 | 0.692939002 |
| | -0.575389014 | 0.565028232 | 0.66707498 |
| | 0.175685864 | 0.860540742 | NA |

| | | | |
|---|---|---|---|
| | -0.464716518 | 0.642134478 | 0.732471676 |
| | 1.115163298 | 0.264780451 | 0.370373012 |
| | -0.04075424 | 0.967491821 | 0.977805066 |
| | 0.642697837 | 0.520420185 | 0.625714152 |
| | 0.886390002 | 0.375407405 | 0.488216414 |
| | -0.38465124 | 0.700495826 | NA |
| | 0.06977913 | 0.944369458 | NA |
| | -0.526877623 | 0.598278574 | 0.696502888 |
| | 0.753928257 | 0.450892302 | 0.560758935 |
| | 0.981258669 | 0.326465199 | NA |
| | 0.512602511 | 0.608229397 | 0.704781343 |
| | -0.328236389 | 0.742732936 | 0.81288174 |
| | -0.299503107 | 0.764556202 | NA |
| NA | NA | NA | |
| | 0.094497879 | 0.924713667 | NA |
| | 0.152985974 | 0.878409336 | 0.91476425 |
| NA | NA | NA | |
| NA | NA | NA | |
| | 0.693454599 | 0.488024308 | NA |
| NA | NA | NA | |
| NA | NA | NA | |
| | -0.084882241 | 0.932355011 | 0.954004003 |
| | 5.607525827 | 2.05E-08 | 1.21E-07 |
| | 1.534355593 | 0.124942196 | 0.20076415 |
| | 0.657565876 | 0.510817127 | NA |
| | 1.786418694 | 0.074031491 | 0.129165497 |
| | 3.945267038 | 7.97E-05 | 0.000280772 |
| NA | NA | NA | |
| | 1.072719728 | 0.283396886 | 0.390370594 |
| | 6.122870907 | 9.19E-10 | 6.29E-09 |
| | -1.779158318 | 0.07521381 | 0.130963594 |
| | -1.602726363 | 0.108995081 | 0.178764589 |
| | 0.063708882 | 0.949202033 | NA |
| | -0.051689497 | 0.958776106 | 0.971585665 |
| NA | NA | NA | |
| | 8.078169084 | 6.57E-16 | 8.35E-15 |
| | 3.857437478 | 0.000114582 | 0.000391855 |
| | -3.584565234 | 0.00033764 | 0.001063839 |
| | 3.229723501 | 0.0012391 | 0.003493823 |
| | 0.441759817 | 0.658663019 | NA |
| NA | NA | NA | |
| | 12.852425 | 8.34E-38 | 4.16E-36 |
| | 1.465873293 | 0.142682809 | 0.224119906 |
| | 10.11138292 | 4.92E-24 | 1.18E-22 |
| | -1.908539743 | 0.056321492 | 0.102525595 |
| | -2.773293704 | 0.005549201 | 0.013508705 |
| | 0.073164869 | 0.941674922 | NA |

| | | |
|---|---|---|
| 0.063708285 | 0.949202508 | NA |
| -7.286864151 | 3.17E-13 | 3.10E-12 |
| -2.939154671 | 0.003291088 | 0.008423758 |
| -1.974484062 | 0.048326728 | 0.089860932 |
| 4.550138717 | 5.36E-06 | 2.27E-05 |
| 0.711367476 | 0.476856548 | 0.585658459 |
| 0.412985851 | 0.67961698 | 0.762496861 |
| -0.985436505 | 0.324409706 | 0.435466464 |
| NA | NA | NA |
| 1.044009022 | 0.296481215 | 0.405005994 |
| 0.608830485 | 0.542636805 | 0.646679009 |
| -11.19406903 | 4.36E-29 | 1.44E-27 |
| -1.075212942 | 0.282279403 | 0.389302631 |
| -3.124599114 | 0.001780475 | 0.004850908 |
| -2.214511449 | 0.026793616 | 0.053815135 |
| -0.70346831 | 0.481763949 | 0.590353062 |
| 1.394597907 | 0.163137127 | 0.249989119 |
| -1.106359015 | 0.268571158 | 0.374566004 |
| -1.491621172 | 0.135798485 | 0.214952006 |
| NA | NA | NA |
| -2.26456476 | 0.023539402 | 0.048152274 |
| 0.42435554 | 0.67130654 | 0.755582678 |
| 0.334654281 | 0.737885893 | 0.809182303 |
| 5.525465854 | 3.29E-08 | 1.89E-07 |
| 15.29435077 | 8.34E-53 | 6.80E-51 |
| -0.973698505 | 0.33020626 | 0.441179817 |
| NA | NA | NA |
| -1.908900468 | 0.056274933 | 0.102455744 |
| -2.683871882 | 0.007277499 | 0.017140848 |
| -3.786881299 | 0.00015255 | 0.000510881 |
| 0.783164855 | 0.43353031 | 0.54425036 |
| 2.624957711 | 0.008665973 | 0.02003596 |
| -2.845546832 | 0.004433524 | 0.011034215 |
| 4.963368875 | 6.93E-07 | 3.33E-06 |
| 0.112909235 | 0.910102515 | NA |
| -2.996275078 | 0.002732998 | 0.007127207 |
| 0.192808327 | 0.847109082 | NA |
| -0.434405845 | 0.663993754 | NA |
| -7.24295591 | 4.39E-13 | 4.24E-12 |
| -0.445045986 | 0.656286522 | 0.743859498 |
| -2.128063954 | 0.033331781 | 0.065123292 |
| -1.031191831 | 0.302450871 | 0.41170031 |
| -2.453607424 | 0.014143129 | 0.03093346 |
| 1.987110426 | 0.046910161 | 0.087604367 |
| -1.393447263 | 0.163484582 | 0.250405142 |
| -2.280539367 | 0.022575719 | 0.046469592 |
| -0.494809058 | 0.62073491 | NA |

| | | |
|---|---|---|
| -0.18262736 | 0.855090415 | 0.899343876 |
| 0.488041164 | 0.625520685 | 0.719025613 |
| -1.917427602 | 0.05518363 | 0.100786253 |
| -1.980465371 | 0.047651261 | 0.088829303 |
| 0.582822218 | 0.560012987 | 0.662435021 |
| 0.439412421 | 0.660362728 | NA |
| 0.304473105 | 0.76076748 | NA |
| -2.153921723 | 0.031246311 | 0.06150481 |
| -0.965531758 | 0.334278537 | 0.44545556 |
| -0.610063567 | 0.5418197 | 0.64588975 |
| -1.613010773 | 0.106742167 | 0.175657886 |
| -0.966160015 | 0.333964117 | 0.445178775 |
| -2.392854441 | 0.016717872 | 0.035844469 |
| 0.469332844 | 0.638831742 | NA |
| NA | NA | NA |
| NA | NA | NA |
| -0.065389369 | 0.947863988 | NA |
| NA | NA | NA |
| 2.108539371 | 0.034984361 | 0.067891034 |
| NA | NA | NA |
| NA | NA | NA |
| 0.093300397 | 0.925664916 | 0.948904159 |
| -6.541821081 | 6.08E-11 | 4.73E-10 |
| 0.063708882 | 0.949202033 | NA |
| -1.701858027 | 0.088781986 | 0.150863075 |
| NA | NA | NA |
| -3.214636522 | 0.001306098 | 0.00366457 |
| 0.063403744 | 0.949445006 | 0.965361523 |
| 0.063708882 | 0.949202033 | NA |
| NA | NA | NA |
| NA | NA | NA |
| 0.469883289 | 0.638438404 | NA |
| 0.192808924 | 0.847108615 | NA |
| 3.189777842 | 0.001423822 | 0.003963771 |
| -0.251742468 | 0.801240129 | NA |
| -1.464425285 | 0.143077784 | 0.224604283 |
| 0.966340458 | 0.333873848 | 0.445082148 |
| -2.938347937 | 0.003299665 | 0.008444848 |
| -1.289116749 | 0.197357502 | NA |
| NA | NA | NA |
| -1.794105297 | 0.072796364 | 0.127337132 |
| 1.133584257 | 0.25696898 | 0.361507057 |
| 0.711608577 | 0.476707194 | 0.585561225 |
| -4.094757945 | 4.23E-05 | 0.000155581 |
| -2.022881253 | 0.043085392 | 0.081360042 |
| NA | NA | NA |
| 0.192810118 | 0.847107679 | NA |

| | | |
|---|---|---|
| 0.502636833 | 0.615219628 | NA |
| 2.319199439 | 0.020384225 | 0.042489769 |
| 3.909880478 | 9.23E-05 | 0.000321686 |
| -3.736564028 | 0.000186552 | 0.000615934 |
| 0.20478375 | 0.837741084 | 0.886484775 |
| -0.065388772 | 0.947864464 | NA |
| 2.346288434 | 0.018961428 | 0.039889438 |
| 1.019413594 | 0.308006655 | 0.417854015 |
| 0.732313838 | 0.463977035 | 0.573370503 |
| NA | NA | NA |
| -1.404171796 | 0.160267698 | 0.246270519 |
| -4.279655358 | 1.87E-05 | 7.31E-05 |
| 1.514719625 | 0.129843422 | 0.207216326 |
| NA | NA | NA |
| NA | NA | NA |
| -0.088144457 | 0.929761862 | NA |
| -2.752231979 | 0.005919058 | 0.014284035 |
| 0.927515932 | 0.353658724 | NA |
| NA | NA | NA |
| 0.067649016 | 0.946065036 | NA |
| -0.095670104 | 0.923782585 | NA |
| 6.337737262 | 2.33E-10 | 1.69E-09 |
| 1.777554713 | 0.075477017 | 0.131349321 |
| 0.571222683 | 0.567848704 | NA |
| NA | NA | NA |
| -0.065465566 | 0.947803322 | NA |
| -8.559691983 | 1.13E-17 | 1.66E-16 |
| -3.000334524 | 0.002696832 | 0.007043167 |
| 0.324874675 | 0.745275925 | NA |
| -2.260634732 | 0.023781885 | 0.048560988 |
| 0.192809521 | 0.847108147 | NA |
| -0.12625663 | 0.899528786 | 0.929953838 |
| -1.330309143 | 0.183416436 | 0.275432769 |
| 0.925229821 | 0.354846375 | 0.466775529 |
| -1.299998668 | 0.193601426 | 0.287843403 |
| -1.482396066 | 0.13823494 | 0.218118928 |
| 0.099349618 | 0.920860684 | NA |
| -5.395613179 | 6.83E-08 | 3.76E-07 |
| -2.621197263 | 0.008762155 | 0.020226573 |
| -5.231457408 | 1.68E-07 | 8.80E-07 |
| -0.831712674 | 0.405571145 | NA |
| 0.063708285 | 0.949202508 | NA |
| NA | NA | NA |
| 1.341808659 | 0.179658044 | 0.270727025 |
| -0.081614138 | 0.934953558 | NA |
| -2.680824636 | 0.007344099 | 0.017283076 |
| 0.089879251 | 0.928383169 | 0.950989594 |

| | | |
|---|---|---|
| 0.030199843 | 0.975907674 | 0.983253596 |
| -4.322570706 | 1.54E-05 | 6.10E-05 |
| NA | NA | NA |
| NA | NA | NA |
| 0.462626734 | 0.643631943 | 0.733377682 |
| -0.501791203 | 0.615814402 | 0.710871857 |
| 1.711671831 | 0.086957163 | 0.148235223 |
| -0.959238159 | 0.337438781 | 0.44863979 |
| -1.231593073 | 0.21810113 | 0.316927215 |
| -0.233960009 | 0.815016025 | 0.869907233 |
| 1.113399422 | 0.265536923 | 0.371161511 |
| 0.098356239 | 0.921649422 | 0.9462217 |
| 0.669277636 | 0.50331839 | 0.609927829 |
| -0.133064404 | 0.894142446 | NA |
| 0.063708882 | 0.949202033 | NA |
| 0.214588724 | 0.830087986 | NA |
| 16.65871341 | 2.62E-62 | 2.79E-60 |
| -1.141796615 | 0.25353857 | 0.357625833 |
| -0.660496077 | 0.508935535 | 0.614978104 |
| -1.004352046 | 0.315208956 | 0.425696519 |
| NA | NA | NA |
| 2.879919731 | 0.003977764 | 0.010007382 |
| -1.16702621 | 0.243199785 | 0.345888438 |
| -3.807415282 | 0.000140427 | 0.000473066 |
| 3.117988476 | 0.001820899 | 0.004948117 |
| 3.507770083 | 0.000451879 | 0.001388819 |
| -1.07885189 | 0.280653758 | 0.387615934 |
| 1.464967953 | 0.142929661 | 0.22445131 |
| 0.793237608 | 0.4276394 | 0.538926373 |
| -2.549695406 | 0.010781707 | 0.024389167 |
| 0.418577516 | 0.675524924 | NA |
| -1.733635434 | 0.082982789 | 0.142488798 |
| 0.155737998 | 0.876239544 | 0.91365448 |
| -0.371148017 | 0.710527289 | NA |
| NA | NA | NA |
| -0.58892792 | 0.555909627 | 0.658577148 |
| -4.805808936 | 1.54E-06 | 7.04E-06 |
| NA | NA | NA |
| NA | NA | NA |
| 1.406278297 | 0.159641502 | 0.245579669 |
| 3.960218996 | 7.49E-05 | 0.00026499 |
| NA | NA | NA |
| NA | NA | NA |
| 0.346721347 | 0.728800672 | NA |
| 0.210645135 | 0.83316419 | NA |
| NA | NA | NA |
| 0.192810118 | 0.847107679 | NA |

| | | |
|---|---|---|
| 11.91498121 | 9.89E-33 | 3.94E-31 |
| 15.76989974 | 5.01E-56 | 4.50E-54 |
| 1.719494343 | 0.085524397 | 0.146148968 |
| 1.242167882 | 0.214174627 | NA |
| NA | NA | NA |
| NA | NA | NA |
| -0.112238445 | 0.910634347 | NA |
| NA | NA | NA |
| -1.209839243 | 0.226340585 | 0.326416615 |
| NA | NA | NA |
| -5.843048187 | 5.13E-09 | 3.23E-08 |
| NA | NA | NA |
| -2.984036233 | 0.002844731 | 0.007392402 |
| -2.819124807 | 0.004815479 | 0.011900814 |
| -3.108966157 | 0.001877432 | 0.005085728 |
| 2.545207815 | 0.010921272 | 0.024647212 |
| 0.063708285 | 0.949202508 | NA |
| -0.160198751 | 0.872724513 | 0.911432159 |
| 2.107353281 | 0.035086968 | 0.06805322 |
| 0.749529294 | 0.453538249 | 0.563499773 |
| 8.774329618 | 1.72E-18 | 2.71E-17 |
| 0.433717073 | 0.664493907 | 0.750702097 |
| 0.103742285 | 0.917373869 | NA |
| NA | NA | NA |
| NA | NA | NA |
| -6.28536955 | 3.27E-10 | 2.34E-09 |
| NA | NA | NA |
| -2.354610228 | 0.018542143 | 0.039153379 |
| 2.285381289 | 0.02229048 | 0.045942912 |
| NA | NA | NA |
| -2.131736096 | 0.033028548 | 0.06461309 |
| -7.139995023 | 9.33E-13 | 8.69E-12 |
| 0.477014928 | 0.633351491 | NA |
| 11.60033239 | 4.10E-31 | 1.50E-29 |
| 3.291619141 | 0.000996124 | 0.002872133 |
| NA | NA | NA |
| 0.06371127 | 0.949200131 | NA |
| -0.16468604 | 0.869191105 | NA |
| 1.711757314 | 0.086941401 | 0.14821845 |
| 0.472647675 | 0.636464563 | 0.727658677 |
| 0.063708285 | 0.949202508 | NA |
| -0.504758353 | 0.61372857 | NA |
| -10.79777624 | 3.53E-27 | 1.03E-25 |
| 1.33937308 | 0.180449248 | 0.271726909 |
| -2.378599237 | 0.017378558 | 0.037048405 |
| 2.25603955 | 0.024068154 | 0.04904151 |
| 1.501556865 | 0.13321159 | 0.211566377 |

| | | |
|---|---|---|
| -0.4171588 | 0.676562258 | 0.759718322 |
| 0.491900869 | 0.622789424 | NA |
| 0.497275607 | 0.618994711 | NA |
| -5.768551175 | 8.00E-09 | 4.93E-08 |
| -1.597116295 | 0.110239787 | 0.180476281 |
| -1.398987006 | 0.16181688 | 0.248163501 |
| 2.499032708 | 0.012453282 | 0.027667818 |
| -1.472888466 | 0.140781113 | 0.221410697 |
| 4.352503007 | 1.35E-05 | 5.37E-05 |
| 8.029197267 | 9.81E-16 | 1.23E-14 |
| -1.935124493 | 0.052975021 | 0.097275678 |
| -1.621713698 | 0.104864664 | 0.173226879 |
| -1.442479254 | 0.14916722 | 0.232360029 |
| 1.575211206 | 0.115207701 | 0.187347243 |
| -4.336455057 | 1.45E-05 | 5.75E-05 |
| NA | NA | NA |
| 0.854811836 | 0.392655324 | 0.504603521 |
| 0.720832113 | 0.471012815 | 0.57990412 |
| -4.587709397 | 4.48E-06 | 1.92E-05 |
| 0.288217989 | 0.77317988 | NA |
| -1.124959987 | 0.26060599 | 0.365637216 |
| -2.067502343 | 0.038686844 | 0.074025816 |
| 8.459039542 | 2.70E-17 | 3.84E-16 |
| NA | NA | NA |
| 0.06353712 | 0.949338802 | NA |
| 1.116135476 | 0.264364149 | 0.369935408 |
| -2.815424532 | 0.004871284 | 0.012030412 |
| -6.365966061 | 1.94E-10 | 1.42E-09 |
| -11.17944765 | 5.14E-29 | 1.69E-27 |
| -1.952719685 | 0.05085282 | 0.093985533 |
| -5.15575468 | 2.53E-07 | 1.29E-06 |
| -1.49702636 | 0.134386399 | 0.213053685 |
| -0.121871325 | 0.903000925 | NA |
| -2.39271872 | 0.016724057 | 0.035854659 |
| -2.752133098 | 0.005920845 | 0.014286971 |
| -0.19448762 | 0.845794089 | NA |
| -0.327938983 | 0.742957798 | NA |
| -0.268711546 | 0.788151666 | NA |
| 0.320071032 | 0.748914485 | NA |
| 0.082721219 | 0.934073213 | NA |
| -1.743347363 | 0.081272957 | 0.140004274 |
| -3.061741734 | 0.002200532 | 0.005871996 |
| 1.243908813 | 0.213533125 | 0.311664478 |
| -0.065388772 | 0.947864464 | NA |
| NA | NA | NA |
| 0.192809521 | 0.847108147 | NA |
| 0.192809521 | 0.847108147 | NA |

| | | |
|---|---|---|
| NA | NA | NA |
| -0.065389369 | 0.947863988 | NA |
| -0.083774515 | 0.933235711 | NA |
| -3.891130271 | 9.98E-05 | 0.000344809 |
| -0.786999777 | 0.431281997 | 0.542197669 |
| -1.720580744 | 0.08532693 | 0.145843558 |
| -0.788080469 | 0.430649639 | 0.541699052 |
| -0.930070779 | 0.352334439 | 0.464178694 |
| 0.193341124 | 0.846691821 | NA |
| -0.170666104 | 0.864486317 | 0.905640126 |
| -1.295290375 | 0.195220078 | 0.289768046 |
| 0.192810118 | 0.847107679 | NA |
| 1.22265002 | 0.221461912 | 0.32074943 |
| 0.135499875 | 0.892216663 | 0.924689788 |
| 3.182262447 | 0.001461293 | 0.004059605 |
| -3.565988772 | 0.000362487 | 0.00113484 |
| -7.16961999 | 7.52E-13 | 7.07E-12 |
| -4.718034654 | 2.38E-06 | 1.06E-05 |
| 1.53318054 | 0.125231378 | 0.201138436 |
| 6.690788143 | 2.22E-11 | 1.80E-10 |
| -2.755697612 | 0.005856711 | 0.014152683 |
| -0.722866856 | 0.469761688 | 0.578767621 |
| -2.993639453 | 0.002756715 | 0.007185161 |
| -1.140805544 | 0.253950853 | 0.358146825 |
| 1.393092811 | 0.163591727 | 0.250532362 |
| -0.31782979 | 0.750614052 | 0.819000424 |
| 6.568008761 | 5.10E-11 | 4.00E-10 |
| -2.125814695 | 0.033518692 | 0.065432293 |
| 0.408237743 | 0.683099141 | 0.765763682 |
| 11.57334734 | 5.62E-31 | 2.05E-29 |
| 0.334239139 | 0.738199112 | NA |
| NA | NA | NA |
| 9.954615616 | 2.41E-23 | 5.52E-22 |
| NA | NA | NA |
| NA | NA | NA |
| NA | NA | NA |
| 7.546453246 | 4.47E-14 | 4.76E-13 |
| -4.678895549 | 2.88E-06 | 1.27E-05 |
| 0.083297288 | 0.933615156 | NA |
| NA | NA | NA |
| 0.760783913 | 0.446786144 | 0.55673108 |
| -0.387740434 | 0.69820813 | 0.777893238 |
| 1.545412023 | 0.122246634 | NA |
| NA | NA | NA |
| 0.679929793 | 0.496548916 | NA |
| 4.295650198 | 1.74E-05 | 6.83E-05 |
| 7.069883829 | 1.55E-12 | 1.41E-11 |

| | | |
|---|---|---|
| 5.423306948 | 5.85E-08 | 3.25E-07 |
| 0.871608371 | 0.383422063 | 0.49584367 |
| NA | NA | NA |
| NA | NA | NA |
| 4.311760304 | 1.62E-05 | 6.38E-05 |
| 1.006738763 | 0.314060329 | 0.424502108 |
| 9.155998662 | 5.39E-20 | 9.61E-19 |
| 0.542505501 | 0.587470319 | 0.687281237 |
| 0.173166019 | 0.862520933 | 0.904360714 |
| -2.059700004 | 0.039427229 | 0.075241197 |
| 2.721571544 | 0.006497232 | 0.015523646 |
| -3.040365723 | 0.00236291 | 0.00625131 |
| 1.045253934 | 0.295905622 | NA |
| 1.510535594 | 0.130906814 | 0.208527815 |
| 0.529464799 | 0.596483057 | 0.69480505 |
| -1.612824471 | 0.106782648 | 0.175689849 |
| -9.030988758 | 1.70E-19 | 2.92E-18 |
| -0.231315153 | 0.817069976 | 0.871267477 |
| 0.008840624 | 0.992946295 | 0.99509312 |
| 3.561585252 | 0.000368622 | 0.001152031 |
| NA | NA | NA |
| 1.073502835 | 0.283045567 | 0.389972508 |
| NA | NA | NA |
| NA | NA | NA |
| -3.993871822 | 6.50E-05 | 0.000232034 |
| -0.173383911 | 0.862349671 | NA |
| 3.079639833 | 0.002072511 | 0.0055624 |
| -0.617532658 | 0.536883447 | 0.641180355 |
| -1.061902041 | 0.288280159 | 0.39568366 |
| 3.720803274 | 0.00019859 | 0.00065301 |
| 0.229319955 | 0.818620245 | 0.872474547 |
| 0.830548529 | 0.406228722 | 0.518083405 |
| NA | NA | NA |
| 1.35444075 | 0.175595804 | 0.265838471 |
| -0.653012293 | 0.513748353 | NA |
| -1.195597635 | 0.231853615 | 0.332757797 |
| -5.515485784 | 3.48E-08 | 1.99E-07 |
| 0.190079725 | 0.849246658 | 0.894903542 |
| -9.260091673 | 2.04E-20 | 3.72E-19 |
| -0.430090383 | 0.667129895 | 0.752447627 |
| 4.617708548 | 3.88E-06 | 1.68E-05 |
| 1.842096646 | 0.065461014 | 0.116378236 |
| 3.244728448 | 0.001175627 | 0.003338545 |
| 1.085698294 | 0.277612496 | 0.384540164 |
| 4.504999355 | 6.64E-06 | 2.78E-05 |
| -0.999000861 | 0.317794274 | NA |
| 0.192809521 | 0.847108147 | NA |

| | | |
|---|---|---|
| 5.075557086 | 3.86E-07 | 1.93E-06 |
| NA | NA | NA |
| 0.063708882 | 0.949202033 | NA |
| 0.885555966 | 0.375856849 | 0.488635378 |
| -3.449495416 | 0.000561635 | 0.001696974 |
| 0.433779648 | 0.664448462 | 0.750702097 |
| -0.786356225 | 0.431658822 | 0.542499513 |
| NA | NA | NA |
| NA | NA | NA |
| 0.192809521 | 0.847108147 | NA |
| 4.225923845 | 2.38E-05 | 9.13E-05 |
| 1.390860249 | 0.164267809 | 0.251413921 |
| -1.422690067 | 0.154826021 | 0.239689221 |
| 2.01613908 | 0.043785435 | 0.082520098 |
| -1.227365819 | 0.219685128 | 0.318692404 |
| 0.500416448 | 0.616781874 | 0.711637657 |
| 0.200068104 | 0.841427318 | NA |
| 0.063707019 | 0.949203516 | NA |
| -3.693464117 | 0.00022122 | 0.000720605 |
| -4.447881682 | 8.67E-06 | 3.57E-05 |
| -3.319409912 | 0.000902079 | 0.002619986 |
| 0.482149038 | 0.629700076 | 0.722237681 |
| -0.246436355 | 0.805344467 | NA |
| -1.301535974 | 0.193075059 | 0.287248498 |
| NA | NA | NA |
| -1.870025784 | 0.061480237 | 0.110280231 |
| -1.860835763 | 0.062767373 | 0.112275234 |
| 1.940797883 | 0.052282798 | NA |
| -0.913023783 | 0.361230033 | 0.473431131 |
| 0.192809521 | 0.847108147 | NA |
| 3.237413518 | 0.001206185 | 0.003413319 |
| 0.801261826 | 0.422980086 | 0.53473365 |
| -1.98973905 | 0.046619689 | 0.08713993 |
| 5.438431284 | 5.38E-08 | 3.00E-07 |
| -0.237077515 | 0.812596656 | 0.86812473 |
| 0.199470624 | 0.841894621 | 0.889239142 |
| 0.022514063 | 0.982037894 | 0.987641213 |
| 1.731817004 | 0.08330615 | 0.142945974 |
| -3.57023571 | 0.00035666 | 0.001119118 |
| NA | NA | NA |
| -3.337824728 | 0.00084437 | 0.002468128 |
| -4.93203246 | 8.14E-07 | 3.87E-06 |
| -1.971880045 | 0.048623302 | 0.090351985 |
| -0.946161658 | 0.344066139 | 0.455370953 |
| 3.58814189 | 0.000333043 | 0.001050041 |
| 6.95872432 | 3.43E-12 | 3.01E-11 |
| 5.021223898 | 5.13E-07 | 2.51E-06 |

| | | |
|---|---|---|
| -6.677100816 | 2.44E-11 | 1.97E-10 |
| 3.950007106 | 7.81E-05 | 0.000275774 |
| 3.081702688 | 0.002058203 | 0.005528742 |
| 0.964459681 | 0.334815513 | 0.44605239 |
| 0.28814782 | 0.773233589 | 0.837003708 |
| -5.738532808 | 9.55E-09 | 5.85E-08 |
| 0.194845313 | 0.845514048 | 0.891903027 |
| 5.390104769 | 7.04E-08 | 3.87E-07 |
| -1.632010219 | 0.102677333 | 0.170244678 |
| 2.216936134 | 0.026627456 | 0.053511459 |
| NA | NA | NA |
| 1.160739455 | 0.245747871 | 0.348739609 |
| 4.710528853 | 2.47E-06 | 1.10E-05 |
| NA | NA | NA |
| -7.507618419 | 6.02E-14 | 6.32E-13 |
| -2.492876105 | 0.012671305 | 0.028099829 |
| NA | NA | NA |
| 0.232842114 | 0.81588401 | 0.870373642 |
| -2.081751557 | 0.037365169 | 0.07183181 |
| -1.142081804 | 0.253420018 | 0.357478757 |
| -0.360216209 | 0.718685454 | 0.794311546 |
| 2.643300853 | 0.008210205 | 0.019063212 |
| 0.719307466 | 0.471951497 | 0.580866733 |
| -1.668738576 | 0.0951692 | 0.159925149 |
| 0.209054699 | 0.834405538 | 0.883953554 |
| 1.830496813 | 0.067175683 | 0.119036546 |
| 4.205672777 | 2.60E-05 | 9.95E-05 |
| -0.974426708 | 0.329844714 | NA |
| -0.605256834 | 0.545008358 | 0.648856507 |
| -0.065387578 | 0.947865415 | NA |
| 2.837509232 | 0.004546703 | 0.01129568 |
| 5.060153743 | 4.19E-07 | 2.08E-06 |
| 4.20144829 | 2.65E-05 | 0.00010124 |
| 6.750391422 | 1.47E-11 | 1.21E-10 |
| 1.336569483 | 0.181363206 | 0.272840548 |
| 1.904171387 | 0.056887867 | 0.103413705 |
| 2.792954273 | 0.005222907 | 0.012789021 |
| -0.723749416 | 0.469219589 | 0.578379115 |
| -6.477196252 | 9.34E-11 | 7.13E-10 |
| -4.502703485 | 6.71E-06 | 2.80E-05 |
| -1.362236891 | 0.173123137 | 0.262633002 |
| 6.33073922 | 2.44E-10 | 1.77E-09 |
| -0.263497069 | 0.792167482 | 0.85215541 |
| -0.194487022 | 0.845794556 | NA |
| -1.617865121 | 0.105691668 | 0.174307337 |
| 4.971299653 | 6.65E-07 | 3.20E-06 |
| 5.992027988 | 2.07E-09 | 1.37E-08 |

| | | | |
|---|---|---|---|
| NA | NA | NA | |
| | 2.801277963 | 0.005090066 | 0.012499199 |
| | -0.880244694 | 0.378726766 | 0.491320149 |
| | -0.551713307 | 0.581144792 | 0.681697166 |
| | -0.500404132 | 0.616790545 | 0.711637657 |
| NA | NA | NA | |
| | 4.762123899 | 1.92E-06 | 8.65E-06 |
| | -0.06259634 | 0.950087944 | 0.96590988 |
| | -0.441106645 | 0.659135793 | 0.746253343 |
| | 1.557158494 | 0.11943286 | 0.193113985 |
| | -8.723692877 | 2.69E-18 | 4.17E-17 |
| | 0.063707688 | 0.949202983 | NA |
| | -1.294002706 | 0.195664486 | 0.290341599 |
| | 1.000445914 | 0.317094759 | 0.427587643 |
| | 0.116263957 | 0.90744335 | NA |
| | 2.016325978 | 0.0437659 | 0.0825043 |
| | -0.981115792 | 0.326535644 | 0.437607065 |
| | 0.370551357 | 0.710971718 | NA |
| | -2.427290902 | 0.015212053 | 0.032983167 |
| | -1.15242454 | 0.249146661 | 0.352523513 |
| | -0.345886865 | 0.72942774 | NA |
| | -1.109924608 | 0.267031515 | 0.372834207 |
| | -5.464560777 | 4.64E-08 | 2.62E-07 |
| | -2.281006781 | 0.022548046 | 0.046424083 |
| | -1.05887459 | 0.289656898 | 0.397137769 |
| | -1.061667472 | 0.288386671 | 0.395762837 |
| | -0.483989111 | 0.628393597 | 0.721268021 |
| | -0.788073025 | 0.430653993 | 0.541699052 |
| | 1.71568981 | 0.086218831 | 0.147136929 |
| | 2.490886277 | 0.01274249 | 0.028232674 |
| | 2.99032988 | 0.002786763 | 0.007255324 |
| | -0.321119074 | 0.748120156 | 0.816856769 |
| | -8.223100396 | 1.98E-16 | 2.63E-15 |
| | 0.063708285 | 0.949202508 | NA |
| | -0.065389369 | 0.947863988 | NA |
| | 4.541897545 | 5.58E-06 | 2.36E-05 |
| | -7.334927386 | 2.22E-13 | 2.20E-12 |
| | 8.567241378 | 1.06E-17 | 1.56E-16 |
| | 1.109552222 | 0.267192029 | 0.373037511 |
| | 0.641730895 | 0.521047926 | NA |
| | 2.285030453 | 0.022311042 | 0.045973935 |
| | 2.708625018 | 0.006756265 | 0.01606892 |
| NA | NA | NA | |
| NA | NA | NA | |
| | 1.185612509 | 0.23577536 | 0.337476827 |
| | 3.599940727 | 0.00031829 | 0.001007307 |
| | -0.278125479 | 0.780916037 | 0.843495811 |

| | | |
|---|---|---|
| -3.27150054 | 0.001069784 | 0.003064746 |
| 9.296192066 | 1.46E-20 | 2.69E-19 |
| 0.063708285 | 0.949202508 | NA |
| -9.721655543 | 2.44E-22 | 5.15E-21 |
| -2.415012881 | 0.015734653 | 0.033995543 |
| 13.72898225 | 6.81E-43 | 4.17E-41 |
| 0.713583834 | 0.475484557 | NA |
| NA | NA | NA |
| -0.065389966 | 0.947863513 | NA |
| 3.189736917 | 0.001424024 | 0.003963891 |
| NA | NA | NA |
| 2.351448 | 0.018700503 | 0.039426684 |
| NA | NA | NA |
| NA | NA | NA |
| 1.178409857 | 0.23863325 | 0.340826833 |
| -3.268507257 | 0.001081164 | 0.00309484 |
| 0.740860609 | 0.458777961 | 0.568307056 |
| 4.019756292 | 5.83E-05 | 0.000209935 |
| 5.761549329 | 8.33E-09 | 5.13E-08 |
| -0.097240929 | 0.922535066 | 0.946703813 |
| -5.438283064 | 5.38E-08 | 3.00E-07 |
| NA | NA | NA |
| 0.529525531 | 0.596440938 | 0.694788322 |
| 0.670495628 | 0.502541893 | 0.609295678 |
| 11.47728146 | 1.72E-30 | 6.06E-29 |
| -4.185256156 | 2.85E-05 | 0.000108156 |
| 7.034946265 | 1.99E-12 | 1.78E-11 |
| NA | NA | NA |
| -6.468838169 | 9.88E-11 | 7.52E-10 |
| -4.025535526 | 5.68E-05 | 0.000205199 |
| -1.86398759 | 0.062323447 | 0.111584827 |
| -4.131123456 | 3.61E-05 | 0.00013466 |
| -0.065698092 | 0.947618191 | NA |
| -1.540880767 | 0.123345808 | 0.198580547 |
| 1.572240197 | 0.115894856 | 0.18825675 |
| 0.192810118 | 0.847107679 | NA |
| -1.321210351 | 0.186431236 | 0.279122732 |
| -3.122738177 | 0.001791771 | 0.004880089 |
| -0.29031584 | 0.771574622 | 0.835605433 |
| -0.113970581 | 0.909261114 | 0.937401711 |
| -1.903114729 | 0.057025576 | 0.103626411 |
| 1.225934256 | 0.220223416 | 0.319362231 |
| NA | NA | NA |
| 0.594672557 | 0.55206237 | 0.65529099 |
| 0.192808924 | 0.847108615 | NA |
| 0.192809521 | 0.847108147 | NA |
| -0.457371616 | 0.647403962 | 0.736301398 |

| | | | |
|---|---|---|---|
| | -0.086690433 | 0.930917581 | NA |
| | 0.98379832 | 0.325214685 | 0.436257226 |
| | 0.253270362 | 0.800059297 | 0.858026659 |
| | -0.878112183 | 0.379882842 | 0.492489238 |
| | -2.532065396 | 0.011339284 | 0.025493782 |
| | -1.797752474 | 0.072216242 | 0.126518691 |
| | 0.699368003 | 0.484322079 | NA |
| | -0.261984328 | 0.793333524 | 0.853043136 |
| NA | NA | NA | |
| NA | NA | NA | |
| | 2.676137599 | 0.007447606 | 0.017505266 |
| | 1.084033981 | 0.278349732 | 0.385284179 |
| | 0.277850054 | 0.781127465 | 0.843687773 |
| | 8.532613977 | 1.43E-17 | 2.09E-16 |
| | 0.605915694 | 0.544570737 | 0.648428027 |
| | 0.369559967 | 0.711710384 | 0.788848943 |
| | 0.063708285 | 0.949202508 | NA |
| | -0.147040168 | 0.883100316 | NA |
| | 3.58134368 | 0.000341832 | 0.001076232 |
| | 1.30281081 | 0.192639359 | 0.286719579 |
| | 2.355980814 | 0.018473872 | 0.039025605 |
| NA | NA | NA | |
| NA | NA | NA | |
| | 9.30150153 | 1.38E-20 | 2.56E-19 |
| | 0.063708285 | 0.949202508 | NA |
| | 2.661265431 | 0.007784756 | 0.018183319 |
| | 0.522895836 | 0.601046743 | 0.69869032 |
| | 1.662709755 | 0.096370549 | 0.161564289 |
| | 4.071705852 | 4.67E-05 | 0.000170683 |
| | -0.314216589 | 0.753356544 | 0.821079931 |
| NA | NA | NA | |
| | 2.484437408 | 0.012975632 | 0.02870096 |
| | 0.625476504 | 0.531658364 | 0.63626148 |
| | 9.414012001 | 4.78E-21 | 9.19E-20 |
| | 0.489545416 | 0.62445561 | 0.718197119 |
| NA | NA | NA | |
| | 0.317642674 | 0.750756 | NA |
| | 15.71210226 | 1.25E-55 | 1.11E-53 |
| | 2.607389495 | 0.009123551 | 0.020983414 |
| | -0.065388772 | 0.947864464 | NA |
| NA | NA | NA | |
| | 0.628856507 | 0.529443002 | NA |
| | -7.187592335 | 6.59E-13 | 6.25E-12 |
| | -5.929507101 | 3.04E-09 | 1.97E-08 |
| | -1.586564349 | 0.112611347 | 0.183721941 |
| | -2.993541884 | 0.002757597 | 0.007186116 |
| | -0.065388772 | 0.947864464 | NA |

| | | |
|---|---|---|
| 1.191438407 | 0.233481523 | 0.334748683 |
| 0.559535733 | 0.575796151 | 0.676818713 |
| -0.681464471 | 0.495577641 | 0.602648222 |
| NA | NA | NA |
| -3.743508587 | 0.000181468 | 0.000600258 |
| NA | NA | NA |
| 2.631931096 | 0.008490108 | 0.019671147 |
| 2.460931516 | 0.013857681 | 0.030404783 |
| NA | NA | NA |
| 0.372987218 | 0.709157957 | 0.786912319 |
| 0.409851401 | 0.681914957 | NA |
| 1.055357029 | 0.291262068 | 0.398969954 |
| -0.065388772 | 0.947864464 | NA |
| 0.542891996 | 0.587204167 | NA |
| -0.19448762 | 0.845794089 | NA |
| 0.063707688 | 0.949202983 | NA |
| 0.3474247 | 0.728272279 | NA |
| 0.392785496 | 0.694477918 | NA |
| -0.923657026 | 0.355664913 | 0.467557394 |
| NA | NA | NA |
| NA | NA | NA |
| NA | NA | NA |
| 0.3474247 | 0.728272279 | NA |
| NA | NA | NA |
| 0.192809521 | 0.847108147 | NA |
| NA | NA | NA |
| NA | NA | NA |
| NA | NA | NA |
| NA | NA | NA |
| NA | NA | NA |
| NA | NA | NA |
| 5.240166354 | 1.60E-07 | 8.41E-07 |
| NA | NA | NA |
| NA | NA | NA |
| NA | NA | NA |
| 0.192810118 | 0.847107679 | NA |
| NA | NA | NA |
| NA | NA | NA |
| NA | NA | NA |
| NA | NA | NA |
| NA | NA | NA |
| NA | NA | NA |
| NA | NA | NA |
| NA | NA | NA |
| NA | NA | NA |
| NA | NA | NA |
| NA | NA | NA |

| | | | |
|---|---|---|---|
| NA | NA | NA | |
| NA | NA | NA | |
| NA | NA | NA | |
| NA | NA | NA | |
| NA | NA | NA | |
| NA | NA | NA | |
| NA | NA | NA | |
| NA | NA | NA | |
| | -0.065388772 | 0.947864464 | NA |
| NA | NA | NA | |
| NA | NA | NA | |
| NA | NA | NA | |
| | 1.729885188 | 0.08365079 | 0.14340954 |
| | 0.683420325 | 0.494341281 | 0.601436944 |
| | -0.40041953 | 0.688847542 | 0.770387495 |
| | 5.592431415 | 2.24E-08 | 1.31E-07 |
| | 0.022331178 | 0.982183779 | 0.987641213 |
| | -3.830533996 | 0.000127865 | 0.000433727 |
| | -10.55606781 | 4.76E-26 | 1.30E-24 |
| | -2.368040077 | 0.017882599 | 0.037977384 |
| NA | NA | NA | |
| | -4.159753863 | 3.19E-05 | 0.000119856 |
| | 0.916270685 | 0.35952494 | 0.471591512 |
| | 4.380064298 | 1.19E-05 | 4.78E-05 |
| | -0.894359792 | 0.371129418 | 0.483871723 |
| | -0.065389369 | 0.947863988 | NA |
| | -4.082668557 | 4.45E-05 | 0.000163206 |
| | -0.658158366 | 0.510436375 | 0.616285874 |
| NA | NA | NA | |
| | 0.490234361 | 0.623968069 | 0.717768315 |
| | -0.327938983 | 0.742957798 | NA |
| | 0.063708882 | 0.949202033 | NA |
| | 12.87381165 | 6.32E-38 | 3.19E-36 |
| | 5.689220644 | 1.28E-08 | 7.70E-08 |
| | 0.693038022 | 0.488285691 | 0.59626241 |
| | -0.038328536 | 0.969425739 | 0.97875092 |
| NA | NA | NA | |
| NA | NA | NA | |
| | 0.039570633 | 0.96843544 | 0.978407366 |
| | 0.063707135 | 0.949203424 | NA |
| | -3.590773456 | 0.000329698 | 0.001040386 |
| NA | NA | NA | |
| | -2.309467569 | 0.020917649 | 0.043433028 |
| | -0.083185833 | 0.933703777 | NA |
| | 0.147680043 | 0.882595282 | NA |
| | -1.204593227 | 0.228360373 | 0.328743322 |
| NA | NA | NA | |

| | | | |
|---|---|---|---|
| | 0.192808327 | 0.847109082 | NA |
| | -0.297239495 | 0.766283671 | 0.831422268 |
| | 0.209372286 | 0.834157626 | 0.88383169 |
| | 7.35770022 | 1.87E-13 | 1.87E-12 |
| | 4.116099585 | 3.85E-05 | 0.000143016 |
| | -2.156295594 | 0.031060591 | 0.061182545 |
| NA | NA | NA | |
| | -5.771505191 | 7.86E-09 | 4.84E-08 |
| | -1.194942055 | 0.23210967 | 0.333029811 |
| | 4.804727731 | 1.55E-06 | 7.08E-06 |
| | -2.670142578 | 0.007581904 | 0.017777524 |
| NA | NA | NA | |
| | 0.192809521 | 0.847108147 | NA |
| NA | NA | NA | |
| NA | NA | NA | |
| NA | NA | NA | |
| | 0.192809521 | 0.847108147 | NA |
| | 0.999347622 | 0.317626324 | 0.428142974 |
| NA | NA | NA | |
| | 0.192810118 | 0.847107679 | NA |
| NA | NA | NA | |
| NA | NA | NA | |
| | 6.308551606 | 2.82E-10 | 2.03E-09 |
| | -7.262764018 | 3.79E-13 | 3.67E-12 |
| | -0.383519505 | 0.701334611 | NA |
| NA | NA | NA | |
| NA | NA | NA | |
| | 1.607069194 | 0.10803919 | 0.177430676 |
| NA | NA | NA | |
| | -2.243707263 | 0.024851242 | 0.050456408 |
| NA | NA | NA | |
| | 0.052855765 | 0.957846829 | 0.970919132 |
| | 5.267039479 | 1.39E-07 | 7.33E-07 |
| | 6.357531784 | 2.05E-10 | 1.50E-09 |
| | -0.08862269 | 0.929381775 | NA |
| | -0.065389369 | 0.947863988 | NA |
| | -5.731368023 | 9.96E-09 | 6.09E-08 |
| | -0.116874969 | 0.906959134 | NA |
| | 1.165734731 | 0.243721713 | 0.346512609 |
| | -0.937496782 | 0.348503078 | 0.460156728 |
| | 8.57931234 | 9.54E-18 | 1.41E-16 |
| NA | NA | NA | |
| | 1.422092875 | 0.154999291 | 0.239868446 |
| NA | NA | NA | |
| NA | NA | NA | |
| | 0.234942868 | 0.814253077 | NA |
| | 3.105838043 | 0.001897406 | 0.005134833 |

| | | | |
|---|---|---|---|
| | -0.065389369 | 0.947863988 | NA |
| | -0.276053389 | 0.78250706 | 0.844795893 |
| NA | NA | NA | |
| NA | NA | NA | |
| | -0.065389369 | 0.947863988 | NA |
| NA | NA | NA | |
| NA | NA | NA | |
| | 0.063708285 | 0.949202508 | NA |
| NA | NA | NA | |
| | 1.071111485 | 0.284119305 | NA |
| NA | NA | NA | |
| | 2.615830723 | 0.008901067 | 0.020518854 |
| | 1.314481811 | 0.188684115 | 0.281838241 |
| NA | NA | NA | |
| NA | NA | NA | |
| | 0.192808924 | 0.847108615 | NA |
| | 0.063708882 | 0.949202033 | NA |
| | -1.376713958 | 0.168600702 | 0.256820567 |
| | -5.896410306 | 3.71E-09 | 2.38E-08 |
| | 2.311821872 | 0.020787502 | 0.043197274 |
| | -3.902512409 | 9.52E-05 | 0.000330738 |
| | -1.937244459 | 0.052715468 | 0.096884365 |
| | 0.683930005 | 0.494019366 | 0.601191401 |
| NA | NA | NA | |
| | 0.288217989 | 0.77317988 | NA |
| | 1.73907274 | 0.082021969 | 0.141080943 |
| | -4.405339644 | 1.06E-05 | 4.29E-05 |
| | -2.304702109 | 0.021183263 | 0.043910316 |
| | 1.307324856 | 0.191102406 | 0.284868879 |
| | 0.86824854 | 0.385258286 | 0.497403812 |
| NA | NA | NA | |
| | -3.228163742 | 0.001245876 | 0.003510953 |
| | 4.351306148 | 1.35E-05 | 5.40E-05 |
| | -2.277069484 | 0.022782077 | 0.046832735 |
| | -1.167405221 | 0.243046764 | 0.345729739 |
| | 0.065474733 | 0.947796023 | NA |
| | 0.376155756 | 0.706801106 | 0.784916441 |
| | -0.5186898 | 0.603977076 | 0.701184889 |
| NA | NA | NA | |
| | 1.088053293 | 0.276571581 | 0.383413971 |
| | 6.651432497 | 2.90E-11 | 2.33E-10 |
| NA | NA | NA | |
| NA | NA | NA | |
| | 0.465075116 | 0.641877665 | NA |
| | -0.065388772 | 0.947864464 | NA |
| | 0.119783369 | 0.904654757 | NA |
| | 0.192809521 | 0.847108147 | NA |

| | | | |
|---|---|---|---|
| NA | NA | NA | |
| | 0.063708882 | 0.949202033 | NA |
| NA | NA | NA | |
| | -0.194488217 | 0.845793621 | NA |
| NA | NA | NA | |
| | -0.461815704 | 0.644213489 | 0.733763698 |
| | 14.69316141 | 7.13E-49 | 5.36E-47 |
| | 2.941718406 | 0.003263966 | 0.008362889 |
| | 16.35331905 | 4.12E-60 | 4.16E-58 |
| | 0.89447649 | 0.371067003 | 0.48386596 |
| | -1.816408976 | 0.069307659 | 0.122192767 |
| | -1.297857169 | 0.19433642 | 0.28867284 |
| NA | NA | NA | |
| NA | NA | NA | |
| | -0.265412129 | 0.790691994 | NA |
| | -0.860677857 | 0.389415492 | 0.501547276 |
| | -1.002792795 | 0.315960846 | 0.426495172 |
| | -9.383807219 | 6.36E-21 | 1.20E-19 |
| NA | NA | NA | |
| | 0.672747343 | 0.501108043 | 0.60804413 |
| | 6.856075444 | 7.08E-12 | 5.99E-11 |
| | -0.328756312 | 0.742339887 | 0.812522624 |
| | 8.003125109 | 1.21E-15 | 1.50E-14 |
| | 0.366404893 | 0.714062969 | NA |
| | 4.340389774 | 1.42E-05 | 5.65E-05 |
| | 2.980598202 | 0.00287686 | 0.007465043 |
| NA | NA | NA | |
| NA | NA | NA | |
| NA | NA | NA | |
| | 3.333959996 | 0.00085619 | 0.002500925 |
| NA | NA | NA | |
| | -6.142946056 | 8.10E-10 | 5.56E-09 |
| | -0.094467073 | 0.924738138 | NA |
| NA | NA | NA | |
| NA | NA | NA | |
| | 0.273802548 | 0.784236365 | NA |
| | -5.629554736 | 1.81E-08 | 1.07E-07 |
| NA | NA | NA | |
| | -2.785636072 | 0.005342281 | 0.013050683 |
| | -0.055887455 | 0.955431465 | 0.969491611 |
| | -1.177750633 | 0.238896035 | 0.341027019 |
| | -4.803551055 | 1.56E-06 | 7.11E-06 |
| | -9.448369672 | 3.44E-21 | 6.71E-20 |
| | 0.357279321 | 0.720882709 | NA |
| NA | NA | NA | |
| | 0.192810118 | 0.847107679 | NA |
| NA | NA | NA | |

| | | | |
|---|---|---|---|
| NA | NA | NA | |
| | -5.233317055 | 1.66E-07 | 8.71E-07 |
| | -1.983332916 | 0.047330255 | 0.088290079 |
| | -0.065389369 | 0.947863988 | NA |
| NA | NA | NA | |
| | 0.943632727 | 0.345357362 | 0.456857926 |
| | -5.362075394 | 8.23E-08 | 4.49E-07 |
| NA | NA | NA | |
| NA | NA | NA | |
| | 2.760962325 | 0.005763132 | 0.013949444 |
| NA | NA | NA | |
| NA | NA | NA | |
| | 0.292917527 | 0.769585192 | 0.833988915 |
| | 5.087722294 | 3.62E-07 | 1.81E-06 |
| | -0.180602959 | 0.856679233 | 0.900498959 |
| | 14.17724611 | 1.27E-45 | 8.69E-44 |
| | 2.402805667 | 0.016269831 | 0.035021833 |
| | -3.055390824 | 0.002247674 | 0.00598245 |
| | 0.604279677 | 0.545657716 | 0.64935162 |
| | 0.192810118 | 0.847107679 | NA |
| | 2.924942639 | 0.003445197 | 0.008777818 |
| NA | NA | NA | |
| | -8.615152662 | 6.98E-18 | 1.04E-16 |
| | -3.012065542 | 0.002594766 | 0.006801327 |
| | -0.972368648 | 0.330867184 | 0.441803446 |
| | -7.369103385 | 1.72E-13 | 1.72E-12 |
| | -0.194487022 | 0.845794556 | NA |
| | 0.063708882 | 0.949202033 | NA |
| | -0.800913802 | 0.423181549 | 0.534880279 |
| | -0.942581226 | 0.345895146 | 0.457380501 |
| | -7.399288231 | 1.37E-13 | 1.38E-12 |
| | 0.285290064 | 0.775421925 | NA |
| NA | NA | NA | |
| NA | NA | NA | |
| | -2.335471415 | 0.019518807 | 0.040903864 |
| | 0.850720439 | 0.394924667 | 0.506947249 |
| | 2.761554605 | 0.00575269 | 0.013925515 |
| | 1.821633424 | 0.068510626 | 0.121017711 |
| | -2.760873087 | 0.005764707 | 0.013951907 |
| | -2.62336791 | 0.00870652 | 0.020120413 |
| | 1.689609961 | 0.091102599 | 0.154167626 |
| | -0.768220767 | 0.442356038 | 0.552338203 |
| | -0.163024502 | 0.870499143 | 0.909753384 |
| | 0.206020247 | 0.836775096 | NA |
| | 1.601950216 | 0.109166619 | 0.179000358 |
| | 3.681876551 | 0.000231524 | 0.000751236 |
| | 0.008011651 | 0.993607695 | 0.995516797 |

| | | |
|---|---|---|
| 7.88457294 | 3.16E-15 | 3.76E-14 |
| 1.772306792 | 0.076343644 | 0.132654038 |
| 0.469883289 | 0.638438404 | NA |
| 5.621551466 | 1.89E-08 | 1.12E-07 |
| NA | NA | NA |
| NA | NA | NA |
| NA | NA | NA |
| 4.171962988 | 3.02E-05 | 0.000114197 |
| 10.08880929 | 6.19E-24 | 1.47E-22 |
| 0.574475814 | 0.565645863 | NA |
| NA | NA | NA |
| NA | NA | NA |
| NA | NA | NA |
| NA | NA | NA |
| 7.999086284 | 1.25E-15 | 1.55E-14 |
| 0.176310914 | 0.860049688 | NA |
| 6.348057729 | 2.18E-10 | 1.59E-09 |
| NA | NA | NA |
| -0.291668459 | 0.770540127 | 0.834709578 |
| -7.800935389 | 6.14E-15 | 7.14E-14 |
| NA | NA | NA |
| -4.590899408 | 4.41E-06 | 1.90E-05 |
| 8.481956147 | 2.21E-17 | 3.19E-16 |
| 1.352105014 | 0.176341724 | 0.266664653 |
| NA | NA | NA |
| 0.114021913 | 0.909220422 | NA |
| 1.340026997 | 0.180236568 | 0.271439308 |
| 3.554861419 | 0.000378178 | 0.001179393 |
| 25.90968074 | 5.18E-148 | 2.13E-145 |
| 5.549872617 | 2.86E-08 | 1.65E-07 |
| 0.118850735 | 0.905393613 | NA |
| -7.46624104 | 8.25E-14 | 8.55E-13 |
| -2.057300834 | 0.039657297 | 0.07563988 |
| -0.065388772 | 0.947864464 | NA |
| NA | NA | NA |
| -5.494572915 | 3.92E-08 | 2.23E-07 |
| NA | NA | NA |
| 0.726814147 | 0.467339815 | 0.576686826 |
| NA | NA | NA |
| 0.109770915 | 0.912591058 | NA |
| -4.366737176 | 1.26E-05 | 5.05E-05 |
| 4.784841679 | 1.71E-06 | 7.78E-06 |
| -0.697386217 | 0.485561125 | 0.593843201 |
| NA | NA | NA |
| NA | NA | NA |
| -3.238159318 | 0.001203036 | 0.003405685 |
| NA | NA | NA |

| | | |
|---|---|---|
| 0.063707019 | 0.949203516 | NA |
| -5.823377335 | 5.77E-09 | 3.62E-08 |
| 0.063708882 | 0.949202033 | NA |
| -8.021413201 | 1.05E-15 | 1.30E-14 |
| -3.350694731 | 0.000806091 | 0.002362583 |
| -1.037953267 | 0.299291813 | 0.408221572 |
| 5.619511776 | 1.91E-08 | 1.13E-07 |
| 0.611173619 | 0.541084646 | 0.645179286 |
| 0.324874675 | 0.745275925 | NA |
| NA | NA | NA |
| -3.427193891 | 0.000609853 | 0.001830292 |
| -0.637708397 | 0.523663518 | NA |
| -12.50084265 | 7.39E-36 | 3.36E-34 |
| -0.418596995 | 0.675510686 | 0.758922415 |
| -3.9186726 | 8.90E-05 | 0.000310912 |
| -0.085589846 | 0.931792471 | NA |
| -0.279673671 | 0.77972788 | 0.842612418 |
| -1.81114408 | 0.070118552 | 0.123448502 |
| 0.90211756 | 0.366994422 | 0.479454584 |
| -0.663682116 | 0.506893788 | 0.613340013 |
| -0.065388772 | 0.947864464 | NA |
| 0.063710673 | 0.949200606 | NA |
| NA | NA | NA |
| NA | NA | NA |
| NA | NA | NA |
| NA | NA | NA |
| NA | NA | NA |
| NA | NA | NA |
| 0.015465591 | 0.987660735 | 0.991372849 |
| -2.351706275 | 0.018687525 | 0.03940928 |
| 4.276135859 | 1.90E-05 | 7.41E-05 |
| -0.802055846 | 0.422520655 | 0.53425353 |
| 0.192808924 | 0.847108615 | NA |
| -0.065389966 | 0.947863513 | NA |
| 0.513772177 | 0.607411283 | 0.703931124 |
| -0.900464939 | 0.367872875 | 0.480351509 |
| -1.069941934 | 0.284645446 | 0.391788591 |
| -1.431206885 | 0.152370931 | 0.236532151 |
| -0.900116071 | 0.368058484 | 0.480568798 |
| -1.096269056 | 0.272961047 | 0.379651868 |
| -0.065389369 | 0.947863988 | NA |
| 3.920956335 | 8.82E-05 | 0.00030841 |
| 0.063707019 | 0.949203516 | NA |
| 0.950498949 | 0.341858788 | NA |
| -0.122581364 | 0.902438612 | NA |
| 3.064486755 | 0.002180438 | 0.005823964 |
| -0.828434776 | 0.407424318 | 0.519264254 |

| | | |
|---|---|---|
| -1.76631271 | 0.0773434 | 0.134176969 |
| -1.271598124 | 0.203515944 | 0.29989427 |
| -0.426979743 | 0.669394082 | 0.754246134 |
| -1.076103022 | 0.281881184 | 0.388924869 |
| -0.921804607 | 0.356630504 | 0.468457702 |
| 4.260672607 | 2.04E-05 | 7.92E-05 |
| 5.593148458 | 2.23E-08 | 1.31E-07 |
| 4.529921774 | 5.90E-06 | NA |
| 2.663979976 | 0.007722216 | 0.018057461 |
| NA | NA | NA |
| NA | NA | NA |
| 6.222894768 | 4.88E-10 | 3.43E-09 |
| 1.402761088 | 0.160688094 | 0.246763565 |
| -1.968239464 | 0.049040491 | 0.091066363 |
| -0.457924181 | 0.647006913 | 0.73595015 |
| -4.952195437 | 7.34E-07 | 3.51E-06 |
| -1.588032341 | 0.112279026 | 0.183263386 |
| 0.481476035 | 0.630178208 | 0.722448347 |
| -1.506647199 | 0.131901112 | 0.209764588 |
| -0.084501006 | 0.932658103 | NA |
| 2.221468089 | 0.026319274 | 0.052994242 |
| -0.071511503 | 0.94299067 | NA |
| -5.893667429 | 3.78E-09 | 2.42E-08 |
| 2.786515284 | 0.005327811 | 0.013016603 |
| 1.853526804 | 0.063806888 | 0.113833671 |
| -0.925571779 | 0.354668565 | 0.466566152 |
| -1.664954961 | 0.095921742 | 0.160913791 |
| -5.090476611 | 3.57E-07 | 1.79E-06 |
| 0.3968255 | 0.691496145 | NA |
| 1.054861741 | 0.291488562 | 0.399166141 |
| 2.604824933 | 0.009192121 | 0.02112947 |
| 3.266982976 | 0.001087002 | 0.003108058 |
| 2.147837734 | 0.031726648 | 0.062327761 |
| 4.61641838 | 3.90E-06 | 1.69E-05 |
| 4.667581116 | 3.05E-06 | 1.34E-05 |
| -0.071805711 | 0.942756528 | NA |
| -2.172297611 | 0.029833222 | 0.059090349 |
| -0.065389966 | 0.947863513 | NA |
| 0.925667841 | 0.354618625 | 0.466524976 |
| 5.164016972 | 2.42E-07 | 1.24E-06 |
| -2.453213189 | 0.01415864 | 0.030956565 |
| 0.746001618 | 0.455666438 | NA |
| -0.194488217 | 0.845793621 | NA |
| -3.293204987 | 0.000990522 | 0.002857957 |
| -2.566728955 | 0.010266283 | 0.023337122 |
| 1.405606048 | 0.159841139 | 0.245811238 |
| -0.33463828 | 0.737897965 | 0.809182303 |

| | | |
|---|---|---|
| -2.339799674 | 0.019294086 | 0.040507546 |
| 0.063708217 | 0.949202562 | NA |
| 6.447936214 | 1.13E-10 | 8.57E-10 |
| 1.333260263 | 0.182446409 | 0.274272289 |
| NA | NA | NA |
| -5.39025766 | 7.04E-08 | 3.86E-07 |
| 0.124377168 | 0.90101665 | NA |
| -5.935089 | 2.94E-09 | 1.90E-08 |
| -1.747855112 | 0.080489119 | 0.138825982 |
| -1.734845748 | 0.082768131 | 0.142178732 |
| 0.192636473 | 0.847243678 | NA |
| -0.065392853 | 0.947861215 | NA |
| 3.755206847 | 0.000173198 | 0.000574041 |
| 0.192810118 | 0.847107679 | NA |
| 0.948797561 | 0.342723583 | 0.454026213 |
| NA | NA | NA |
| NA | NA | NA |
| -2.274584695 | 0.022930855 | 0.047099894 |
| NA | NA | NA |
| -5.144692975 | 2.68E-07 | 1.37E-06 |
| NA | NA | NA |
| -0.117713201 | 0.906294907 | NA |
| -0.441183901 | 0.659079867 | NA |
| -0.03873263 | 0.969103558 | 0.978683157 |
| 2.91289526 | 0.003580947 | 0.009083006 |
| -0.349861133 | 0.726442917 | NA |
| 0.143783857 | 0.885671151 | NA |
| 0.14218332 | 0.886935207 | 0.920702896 |
| NA | NA | NA |
| NA | NA | NA |
| 0.192810118 | 0.847107679 | NA |
| -1.028609777 | 0.303663082 | 0.413080812 |
| 0.537600802 | 0.590852677 | NA |
| -0.038910397 | 0.968961827 | 0.978683157 |
| 4.398686659 | 1.09E-05 | 4.41E-05 |
| -0.906799997 | 0.364512571 | 0.476978717 |
| NA | NA | NA |
| 0.202110911 | 0.839830018 | 0.887695518 |
| -3.412939177 | 0.000642663 | 0.00192024 |
| -3.052798378 | 0.002267182 | 0.006029278 |
| NA | NA | NA |
| 0.63431576 | 0.525874774 | NA |
| NA | NA | NA |
| -3.533927815 | 0.000409433 | 0.00126833 |
| -1.777534302 | 0.075480372 | 0.131349321 |
| -1.787155189 | 0.07391241 | 0.129002692 |
| -9.647370807 | 5.04E-22 | 1.05E-20 |

| | | |
|---|---|---|
| 1.2830165 | 0.199486295 | 0.295068075 |
| -0.201445455 | 0.840350276 | NA |
| -0.824648593 | 0.409571125 | 0.521449453 |
| -0.071030282 | 0.943373655 | 0.961700915 |
| 10.89450356 | 1.22E-27 | 3.74E-26 |
| NA | NA | NA |
| 2.501659492 | 0.012361275 | 0.027490246 |
| 0.106908374 | 0.91486167 | 0.941376674 |
| 0.967551277 | 0.333268521 | 0.44444089 |
| -7.533262791 | 4.95E-14 | 5.24E-13 |
| -1.87234426 | 0.061158991 | 0.109806294 |
| NA | NA | NA |
| -3.415097683 | 0.000637591 | 0.001906225 |
| NA | NA | NA |
| -0.917696611 | 0.358777722 | 0.470759404 |
| -0.87352141 | 0.382378942 | 0.494690108 |
| NA | NA | NA |
| -0.065390563 | 0.947863038 | NA |
| NA | NA | NA |
| NA | NA | NA |
| -0.065389369 | 0.947863988 | NA |
| -3.727791472 | 0.000193165 | 0.000636425 |
| NA | NA | NA |
| 0.837286347 | 0.402431632 | 0.514185614 |
| NA | NA | NA |
| NA | NA | NA |
| 0.061918218 | 0.95062796 | 0.966301894 |
| 0.437743209 | 0.661572443 | NA |
| NA | NA | NA |
| NA | NA | NA |
| NA | NA | NA |
| NA | NA | NA |
| NA | NA | NA |
| NA | NA | NA |
| NA | NA | NA |
| NA | NA | NA |
| 0.073834402 | 0.941142153 | NA |
| 0.358454124 | 0.720003494 | 0.795203082 |
| -0.108326299 | 0.913736862 | 0.940760711 |
| -0.555438903 | 0.57859449 | 0.679406084 |
| -0.065389369 | 0.947863988 | NA |
| -0.065388772 | 0.947864464 | NA |
| -0.781361078 | 0.434590156 | 0.545252713 |
| 0.06353712 | 0.949338802 | NA |
| NA | NA | NA |
| 1.553295484 | 0.120352559 | 0.194412684 |
| -0.314504187 | 0.753138137 | 0.820967144 |

| | | |
|---|---|---|
| -1.984226306 | 0.047230617 | 0.088112858 |
| NA | NA | NA |
| NA | NA | NA |
| -0.029534936 | 0.976437956 | 0.983629494 |
| -4.01250905 | 6.01E-05 | 0.000215711 |
| 1.085203861 | 0.277831374 | 0.384758177 |
| 5.078773627 | 3.80E-07 | 1.90E-06 |
| 0.494982108 | 0.62061275 | 0.714960056 |
| NA | NA | NA |
| 0.063708285 | 0.949202508 | NA |
| NA | NA | NA |
| -0.065389369 | 0.947863988 | NA |
| -0.065389369 | 0.947863988 | NA |
| -0.42727035 | 0.669182426 | 0.754046555 |
| 11.14723592 | 7.39E-29 | 2.41E-27 |
| -0.199310203 | 0.842020099 | 0.889272703 |
| 1.622225876 | 0.104754993 | 0.173081885 |
| NA | NA | NA |
| 1.763931499 | 0.077743513 | 0.134738918 |
| 2.8874314 | 0.003884013 | 0.009792188 |
| -1.508601273 | 0.131400709 | 0.209088349 |
| 1.177713163 | 0.238910977 | 0.3410289 |
| 1.512105437 | 0.130507042 | 0.208010141 |
| 0.163518424 | 0.870110268 | 0.909460894 |
| 1.181691964 | 0.237327948 | 0.339330432 |
| -3.457185899 | 0.000545848 | 0.001652665 |
| NA | NA | NA |
| NA | NA | NA |
| 2.802873607 | 0.005064951 | 0.012449743 |
| -5.45808864 | 4.81E-08 | 2.71E-07 |
| -1.453310585 | 0.14613754 | 0.228428316 |
| 1.661483518 | 0.096616376 | 0.161890605 |
| -1.103904599 | 0.269634524 | 0.375776845 |
| 1.143407772 | 0.252869327 | 0.356903082 |
| 3.229559185 | 0.001239812 | 0.003495438 |
| -1.362335668 | 0.173091976 | 0.262623032 |
| -0.407450154 | 0.683677397 | 0.766151039 |
| -0.520108627 | 0.602987866 | 0.700198853 |
| -1.656784823 | 0.097562987 | 0.163235398 |
| -5.773826916 | 7.75E-09 | 4.79E-08 |
| -2.036584013 | 0.041691746 | 0.079044418 |
| 0.645018148 | 0.518915425 | 0.624248729 |
| NA | NA | NA |
| 0.063708882 | 0.949202033 | NA |
| -6.528421782 | 6.65E-11 | 5.15E-10 |
| 0.476235897 | 0.633906328 | 0.72543322 |
| -0.890547061 | 0.373172211 | 0.485952887 |

| | | |
|---|---|---|
| -0.932506118 | 0.351075026 | NA |
| 0.755701788 | 0.449828012 | 0.559783206 |
| NA | NA | NA |
| 0.704718944 | 0.48098516 | NA |
| 1.699046515 | 0.08931042 | 0.151647772 |
| NA | NA | NA |
| NA | NA | NA |
| 1.569641476 | 0.116498544 | 0.189090107 |
| -0.194487022 | 0.845794556 | NA |
| NA | NA | NA |
| -0.967468699 | 0.333309782 | 0.444472234 |
| 2.558914821 | 0.010499947 | 0.023814214 |
| 0.851544208 | 0.394467117 | NA |
| 0.192809521 | 0.847108147 | NA |
| 0.063708285 | 0.949202508 | NA |
| 7.189681901 | 6.49E-13 | 6.16E-12 |
| -2.416805972 | 0.015657361 | 0.033843155 |
| NA | NA | NA |
| 0.192810118 | 0.847107679 | NA |
| -2.257628058 | 0.023968858 | 0.048890838 |
| 2.135636165 | 0.032709081 | 0.064061683 |
| 9.082300189 | 1.06E-19 | 1.85E-18 |
| 3.036722944 | 0.002391652 | 0.006321335 |
| -3.472466764 | 0.000515699 | 0.001568973 |
| NA | NA | NA |
| NA | NA | NA |
| 0.259163815 | 0.795508849 | 0.85450118 |
| NA | NA | NA |
| -5.777493336 | 7.58E-09 | 4.69E-08 |
| NA | NA | NA |
| -6.894922942 | 5.39E-12 | 4.64E-11 |
| -2.667575403 | 0.007640074 | 0.017887107 |
| NA | NA | NA |
| 0.179350163 | 0.857662761 | 0.901239777 |
| -3.008258303 | 0.002627497 | 0.006877157 |
| 2.844340615 | 0.004450345 | 0.011071675 |
| 0.924164659 | 0.355400593 | 0.467308092 |
| -0.133306047 | 0.893951345 | 0.926065813 |
| 0.362222328 | 0.717185896 | NA |
| 1.601242689 | 0.109323177 | 0.179210107 |
| NA | NA | NA |
| NA | NA | NA |
| 0.427399909 | 0.669088073 | NA |
| NA | NA | NA |
| NA | NA | NA |
| NA | NA | NA |
| 1.921550256 | 0.054662373 | 0.099948805 |

| | | | |
|---|---|---|---|
| NA | NA | NA | |
| | -0.065388772 | 0.947864464 | NA |
| | -0.32347968 | 0.746331994 | NA |
| NA | NA | NA | |
| NA | NA | NA | |
| | -0.526220124 | 0.598735274 | 0.696715198 |
| | 0.192808327 | 0.847109082 | NA |
| | 2.342892715 | 0.019134884 | 0.040207031 |
| NA | NA | NA | |
| NA | NA | NA | |
| NA | NA | NA | |
| NA | NA | NA | |
| NA | NA | NA | |
| | 1.450165553 | 0.147012359 | 0.229523412 |
| NA | NA | NA | |
| | 3.387539654 | 0.000705225 | 0.00209194 |
| | 0.06371127 | 0.949200131 | NA |
| | 0.192810118 | 0.847107679 | NA |
| | -5.86409642 | 4.52E-09 | 2.87E-08 |
| | 0.147719372 | 0.882564243 | NA |
| NA | NA | NA | |
| NA | NA | NA | |
| NA | NA | NA | |
| NA | NA | NA | |
| | -0.065388772 | 0.947864464 | NA |
| | 0.192810118 | 0.847107679 | NA |
| NA | NA | NA | |
| NA | NA | NA | |
| NA | NA | NA | |
| | -0.065389369 | 0.947863988 | NA |
| | -0.089835293 | 0.928418102 | NA |
| | 5.400533719 | 6.64E-08 | 3.67E-07 |
| | 6.291361264 | 3.15E-10 | 2.25E-09 |
| | -1.393433813 | 0.163488647 | 0.250405142 |
| | 0.259034991 | 0.795608243 | 0.8545346 |
| NA | NA | NA | |
| | 0.192810118 | 0.847107679 | NA |
| | -0.119978459 | 0.904500212 | 0.933901506 |
| | 2.142115827 | 0.032184163 | 0.063157161 |
| | 7.024018909 | 2.16E-12 | 1.92E-11 |
| | 6.637747605 | 3.19E-11 | 2.54E-10 |
| | -0.87008114 | 0.384256064 | 0.496505882 |
| | 0.063708882 | 0.949202033 | NA |
| | 2.68581403 | 0.007235335 | 0.01704956 |
| | -2.516273797 | 0.011860301 | 0.026508021 |
| | 0.020279251 | 0.983820608 | NA |
| | 0.173059158 | 0.862604927 | 0.904410897 |

| | | |
|---|---|---|
| 19.3740123 | 1.28E-83 | 2.18E-81 |
| 21.11057308 | 6.36E-99 | 1.47E-96 |
| 0.192808924 | 0.847108615 | NA |
| 2.798247657 | 0.00513807 | 0.012607185 |
| 3.215516436 | 0.0013021 | 0.003654173 |
| -0.625973289 | 0.531332462 | 0.635964848 |
| 0.251464041 | 0.80145536 | NA |
| 9.300082511 | 1.40E-20 | 2.60E-19 |
| 0.541645045 | 0.588063054 | NA |
| 0.482734547 | 0.62928423 | NA |
| 0.003776778 | 0.996986574 | 0.997983362 |
| 6.704936838 | 2.01E-11 | 1.64E-10 |
| 6.577521664 | 4.78E-11 | 3.77E-10 |
| -0.521325597 | 0.602139973 | NA |
| 4.337165932 | 1.44E-05 | 5.73E-05 |
| 0.856713532 | 0.391603223 | NA |
| 5.165819346 | 2.39E-07 | 1.23E-06 |
| 0.240534544 | 0.809915887 | 0.865778119 |
| 0.298311409 | 0.765465497 | 0.830882685 |
| 2.360173286 | 0.0182664 | 0.038654718 |
| -0.065388772 | 0.947864464 | NA |
| 0.192809521 | 0.847108147 | NA |
| -0.065389369 | 0.947863988 | NA |
| NA | NA | NA |
| NA | NA | NA |
| NA | NA | NA |
| NA | NA | NA |
| -0.065389369 | 0.947863988 | NA |
| NA | NA | NA |
| -0.497213845 | 0.619038259 | 0.713606026 |
| NA | NA | NA |
| NA | NA | NA |
| 0.192810118 | 0.847107679 | NA |
| NA | NA | NA |
| NA | NA | NA |
| 0.192809521 | 0.847108147 | NA |
| NA | NA | NA |
| NA | NA | NA |
| NA | NA | NA |
| NA | NA | NA |
| -0.065388772 | 0.947864464 | NA |
| NA | NA | NA |
| NA | NA | NA |
| 0.192810118 | 0.847107679 | NA |
| 0.063710673 | 0.949200606 | NA |
| NA | NA | NA |
| NA | NA | NA |

| | | | |
|---|---|---|---|
| NA | NA | NA | |
| -0.065389369 | 0.947863988 | NA | |
| NA | NA | NA | |
| NA | NA | NA | |
| NA | NA | NA | |
| 0.063708882 | 0.949202033 | NA | |
| NA | NA | NA | |
| NA | NA | NA | |
| NA | NA | NA | |
| NA | NA | NA | |
| NA | NA | NA | |
| NA | NA | NA | |
| NA | NA | NA | |
| NA | NA | NA | |
| NA | NA | NA | |
| NA | NA | NA | |
| 0.192808924 | 0.847108615 | NA | |
| NA | NA | NA | |
| NA | NA | NA | |
| NA | NA | NA | |
| NA | NA | NA | |
| NA | NA | NA | |
| NA | NA | NA | |
| NA | NA | NA | |
| NA | NA | NA | |
| -0.246436355 | 0.805344467 | NA | |
| NA | NA | NA | |
| NA | NA | NA | |
| NA | NA | NA | |
| NA | NA | NA | |
| NA | NA | NA | |
| NA | NA | NA | |
| 0.192808924 | 0.847108615 | NA | |
| NA | NA | NA | |
| NA | NA | NA | |
| NA | NA | NA | |
| NA | NA | NA | |
| NA | NA | NA | |
| NA | NA | NA | |
| NA | NA | NA | |
| 0.063708285 | 0.949202508 | NA | |
| NA | NA | NA | |
| 0.642340129 | 0.520652364 | | 0.625909976 |
| NA | NA | NA | |
| NA | NA | NA | |
| NA | NA | NA | |
| NA | NA | NA | |

| | | |
|---|---|---|
| NA | NA | NA |
| NA | NA | NA |
| NA | NA | NA |
| NA | NA | NA |
| NA | NA | NA |
| NA | NA | NA |
| NA | NA | NA |
| NA | NA | NA |
| NA | NA | NA |
| NA | NA | NA |
| NA | NA | NA |
| NA | NA | NA |
| NA | NA | NA |
| NA | NA | NA |
| NA | NA | NA |
| NA | NA | NA |
| NA | NA | NA |
| NA | NA | NA |
| NA | NA | NA |
| NA | NA | NA |
| NA | NA | NA |
| NA | NA | NA |
| NA | NA | NA |
| NA | NA | NA |
| NA | NA | NA |
| NA | NA | NA |
| NA | NA | NA |
| NA | NA | NA |
| NA | NA | NA |
| NA | NA | NA |
| NA | NA | NA |
| NA | NA | NA |
| NA | NA | NA |
| -0.065388772 | 0.947864464 | NA |
| NA | NA | NA |
| NA | NA | NA |
| NA | NA | NA |
| NA | NA | NA |
| NA | NA | NA |
| NA | NA | NA |
| NA | NA | NA |
| NA | NA | NA |
| 0.192809521 | 0.847108147 | NA |
| NA | NA | NA |
| NA | NA | NA |
| NA | NA | NA |
| NA | NA | NA |

| | | |
|---|---|---|
| NA | NA | NA |
| -0.333116482 | 0.739046358 | NA |
| NA | NA | NA |
| NA | NA | NA |
| NA | NA | NA |
| NA | NA | NA |
| NA | NA | NA |
| NA | NA | NA |
| NA | NA | NA |
| NA | NA | NA |
| -0.065389369 | 0.947863988 | NA |
| NA | NA | NA |
| NA | NA | NA |
| NA | NA | NA |
| NA | NA | NA |
| NA | NA | NA |
| NA | NA | NA |
| NA | NA | NA |
| NA | NA | NA |
| 0.732984173 | 0.463568083 | 0.572993042 |
| NA | NA | NA |
| NA | NA | NA |
| NA | NA | NA |
| NA | NA | NA |
| NA | NA | NA |
| NA | NA | NA |
| NA | NA | NA |
| 1.616764629 | 0.105929097 | 0.174583875 |
| NA | NA | NA |
| NA | NA | NA |
| -0.257602004 | 0.796714083 | NA |
| NA | NA | NA |
| 0.488699428 | 0.625054509 | 0.718720799 |
| NA | NA | NA |
| NA | NA | NA |
| NA | NA | NA |
| NA | NA | NA |
| NA | NA | NA |
| NA | NA | NA |
| NA | NA | NA |
| NA | NA | NA |
| NA | NA | NA |
| NA | NA | NA |
| NA | NA | NA |
| NA | NA | NA |
| NA | NA | NA |
| NA | NA | NA |

| | | |
|---|---|---|
| NA | NA | NA |
| NA | NA | NA |
| NA | NA | NA |
| NA | NA | NA |
| NA | NA | NA |
| NA | NA | NA |
| NA | NA | NA |
| NA | NA | NA |
| NA | NA | NA |
| NA | NA | NA |
| NA | NA | NA |
| NA | NA | NA |
| NA | NA | NA |
| NA | NA | NA |
| NA | NA | NA |
| NA | NA | NA |
| NA | NA | NA |
| NA | NA | NA |
| NA | NA | NA |
| NA | NA | NA |
| NA | NA | NA |
| NA | NA | NA |
| NA | NA | NA |
| NA | NA | NA |
| NA | NA | NA |
| NA | NA | NA |
| NA | NA | NA |
| 0.192809521 | 0.847108147 | NA |
| NA | NA | NA |
| NA | NA | NA |
| 0.192809521 | 0.847108147 | NA |
| NA | NA | NA |
| NA | NA | NA |
| NA | NA | NA |
| NA | NA | NA |
| NA | NA | NA |
| NA | NA | NA |
| NA | NA | NA |
| NA | NA | NA |
| 0.192810118 | 0.847107679 | NA |
| NA | NA | NA |
| NA | NA | NA |
| NA | NA | NA |
| NA | NA | NA |
| NA | NA | NA |
| NA | NA | NA |
| NA | NA | NA |

| | | |
|---|---|---|
| NA | NA | NA |
| NA | NA | NA |
| NA | NA | NA |
| NA | NA | NA |
| NA | NA | NA |
| NA | NA | NA |
| NA | NA | NA |
| NA | NA | NA |
| NA | NA | NA |
| 0.063708882 | 0.949202033 | NA |
| NA | NA | NA |
| NA | NA | NA |
| -0.065389369 | 0.947863988 | NA |
| -0.08180493 | 0.934801835 | NA |
| NA | NA | NA |
| NA | NA | NA |
| NA | NA | NA |
| NA | NA | NA |
| NA | NA | NA |
| NA | NA | NA |
| NA | NA | NA |
| NA | NA | NA |
| NA | NA | NA |
| NA | NA | NA |
| NA | NA | NA |
| NA | NA | NA |
| NA | NA | NA |
| -0.065388772 | 0.947864464 | NA |
| NA | NA | NA |
| NA | NA | NA |
| NA | NA | NA |
| NA | NA | NA |
| NA | NA | NA |
| NA | NA | NA |
| NA | NA | NA |
| NA | NA | NA |
| NA | NA | NA |
| 0.063708882 | 0.949202033 | NA |
| 0.192808924 | 0.847108615 | NA |
| NA | NA | NA |
| -0.065388772 | 0.947864464 | NA |
| -0.065388772 | 0.947864464 | NA |
| NA | NA | NA |
| NA | NA | NA |
| NA | NA | NA |
| NA | NA | NA |
| NA | NA | NA |

| | | | |
|---|---|---|---|
| 0.063708882 | 0.949202033 | NA | |
| NA | NA | NA | |
| NA | NA | NA | |
| 0.192809521 | 0.847108147 | NA | |
| NA | NA | NA | |
| -0.065388772 | 0.947864464 | NA | |
| -0.065388772 | 0.947864464 | NA | |
| NA | NA | NA | |
| NA | NA | NA | |
| NA | NA | NA | |
| NA | NA | NA | |
| NA | NA | NA | |
| NA | NA | NA | |
| -0.065389966 | 0.947863513 | NA | |
| NA | NA | NA | |
| 0.063708882 | 0.949202033 | NA | |
| NA | NA | NA | |
| NA | NA | NA | |
| NA | NA | NA | |
| -0.065389369 | 0.947863988 | NA | |
| NA | NA | NA | |
| NA | NA | NA | |
| NA | NA | NA | |
| -0.065389369 | 0.947863988 | NA | |
| NA | NA | NA | |
| NA | NA | NA | |
| -0.422467629 | 0.672683723 | NA | |
| NA | NA | NA | |
| 0.850608084 | 0.394987097 | NA | |
| NA | NA | NA | |
| -0.065389369 | 0.947863988 | NA | |
| 0.607025786 | 0.5438338 | NA | |
| 0.588263471 | 0.55635546 | NA | |
| NA | NA | NA | |
| -0.065388772 | 0.947864464 | NA | |
| 0.192810118 | 0.847107679 | NA | |
| 0.494915508 | 0.620659764 | NA | |
| 0.192809521 | 0.847108147 | NA | |
| 0.401175821 | 0.688290681 | NA | |
| 0.063708882 | 0.949202033 | NA | |
| -0.281363158 | 0.778431873 | | 0.841393522 |
| NA | NA | NA | |
| 0.192809521 | 0.847108147 | NA | |
| -0.533822724 | 0.59346419 | | 0.692480941 |
| NA | NA | NA | |
| 1.670512457 | 0.094818017 | | 0.159463479 |
| 0.3474247 | 0.728272279 | NA | |

| | | |
|---|---|---|
| NA | NA | NA |
| NA | NA | NA |
| 0.063708285 | 0.949202508 | NA |
| NA | NA | NA |
| 0.735003875 | 0.462337134 | NA |
| 0.258473652 | 0.796041381 | NA |
| 1.520124124 | 0.128479783 | 0.205524954 |
| 1.297168901 | 0.194573078 | 0.288893354 |
| 1.607377554 | 0.107971571 | 0.177363767 |
| 2.161017188 | 0.030694012 | 0.06054623 |
| 1.511436177 | 0.130677358 | 0.20824182 |
| 0.111714442 | 0.911049828 | 0.938661361 |
| -1.199409705 | 0.230368676 | 0.33108222 |
| -1.64457029 | 0.100058458 | 0.166586617 |
| -9.188790695 | 3.97E-20 | 7.14E-19 |
| -1.203091103 | 0.228941068 | 0.329427705 |
| 1.674632545 | 0.09400635 | 0.158226006 |
| 1.509571713 | 0.131152744 | 0.208813255 |
| -2.179700364 | 0.02927968 | 0.05813552 |
| -3.297477088 | 0.000975576 | 0.002818407 |
| 0.192809521 | 0.847108147 | NA |
| -0.157416187 | 0.874916857 | 0.912726807 |
| -0.660436318 | 0.508973872 | 0.614994739 |
| 4.162913468 | 3.14E-05 | 0.000118409 |
| 0.167903063 | 0.866659541 | 0.906914516 |
| 17.40451283 | 7.63E-68 | 9.36E-66 |
| -2.056051862 | 0.039777518 | 0.075836384 |
| -5.192802992 | 2.07E-07 | 1.07E-06 |
| 9.744560201 | 1.95E-22 | 4.14E-21 |
| -1.773707604 | 0.076111528 | 0.132305823 |
| NA | NA | NA |
| 7.118241266 | 1.09E-12 | 1.01E-11 |
| -2.906450087 | 0.003655552 | 0.009256294 |
| 1.429547871 | 0.152846826 | 0.237109145 |
| -3.414514178 | 0.000638959 | 0.001909939 |
| -4.154154215 | 3.26E-05 | 0.000122649 |
| 6.054296814 | 1.41E-09 | 9.45E-09 |
| -5.813753616 | 6.11E-09 | 3.82E-08 |
| -1.485271216 | 0.137471994 | 0.217106694 |
| 0.147674387 | 0.882599746 | NA |
| 0.130374271 | 0.896270321 | 0.927867278 |
| -0.375529707 | 0.707266558 | 0.785259135 |
| 2.868454345 | 0.004124827 | 0.010336847 |
| -0.394795184 | 0.692994049 | NA |
| 2.609789612 | 0.009059792 | 0.020852101 |
| 0.014256198 | 0.988625585 | 0.99211364 |
| 3.550299527 | 0.000384793 | 0.001198298 |

| | | | |
|---|---|---|---|
| | 4.785364892 | 1.71E-06 | 7.76E-06 |
| | -0.757445104 | 0.448783243 | 0.558746745 |
| | -0.688034474 | 0.491431072 | 0.598856852 |
| NA | NA | NA | |
| | -1.157586602 | 0.247032779 | 0.350183071 |
| | -0.562741012 | 0.573611253 | 0.67488559 |
| | -5.333842331 | 9.62E-08 | 5.19E-07 |
| | 4.358478009 | 1.31E-05 | 5.23E-05 |
| | -6.375246222 | 1.83E-10 | 1.35E-09 |
| | -1.783055792 | 0.074577222 | 0.129990798 |
| | 0.759109113 | 0.447787288 | NA |
| | 11.26598275 | 1.93E-29 | 6.46E-28 |
| NA | NA | NA | |
| | -2.090400029 | 0.03658188 | 0.070548152 |
| | -1.710736201 | 0.087129822 | 0.148478992 |
| | -0.33637565 | 0.736587609 | 0.808063984 |
| | -1.337888414 | 0.180932815 | 0.272291267 |
| | -2.771191847 | 0.005585151 | 0.013584329 |
| | -1.215844329 | 0.224044228 | 0.323889745 |
| | 3.133744293 | 0.001725912 | 0.004718686 |
| | -1.915197936 | 0.055467266 | 0.101206128 |
| | -2.6481392 | 0.008093619 | 0.018813455 |
| | -0.851437978 | 0.394526103 | 0.506585175 |
| | -4.491406626 | 7.08E-06 | 2.95E-05 |
| | -0.165730714 | 0.868368874 | 0.908285538 |
| | 2.674896782 | 0.007475226 | 0.017556996 |
| | 0.004668153 | 0.996275366 | 0.997521005 |
| NA | NA | NA | |
| | 0.192809521 | 0.847108147 | NA |
| NA | NA | NA | |
| NA | NA | NA | |
| | -5.893397423 | 3.78E-09 | 2.42E-08 |
| | 0.807772331 | 0.419221656 | NA |
| | 1.223473595 | 0.221150872 | 0.320410183 |
| | -0.909503208 | 0.363084565 | NA |
| | -3.380252593 | 0.000724192 | 0.002144391 |
| | -5.703224647 | 1.18E-08 | 7.13E-08 |
| | -2.659394515 | 0.007828124 | 0.018269272 |
| NA | NA | NA | |
| | 3.147253119 | 0.001648122 | 0.004528763 |
| | -1.431469873 | 0.152295595 | 0.236455108 |
| NA | NA | NA | |
| NA | NA | NA | |
| | 6.329415447 | 2.46E-10 | 1.78E-09 |
| | -0.160365872 | 0.872592872 | 0.911332704 |
| | 0.063708285 | 0.949202508 | NA |
| NA | NA | NA | |

| | | |
|---|---|---|
| 1.892979205 | 0.058360635 | 0.105600137 |
| 2.748396241 | 0.005988759 | 0.014431369 |
| NA | NA | NA |
| 0.083034503 | 0.933824105 | 0.954999687 |
| 0.645888259 | 0.518351724 | 0.623676569 |
| 0.770830602 | 0.440807346 | 0.550949812 |
| 2.9135241 | 0.003573743 | 0.009067489 |
| NA | NA | NA |
| -1.30857852 | 0.190677161 | 0.284323206 |
| 3.184302664 | 0.001451032 | 0.004033515 |
| 0.995809131 | 0.319342893 | 0.430016818 |
| 5.673620645 | 1.40E-08 | 8.40E-08 |
| 0.863807576 | 0.387693605 | 0.499921227 |
| 0.09010428 | 0.928204348 | 0.950923167 |
| NA | NA | NA |
| -2.249940847 | 0.024452701 | 0.04973379 |
| -0.665695993 | 0.505605436 | NA |
| -1.993116261 | 0.046248718 | 0.08652319 |
| 3.428513894 | 0.000606895 | 0.001822507 |
| -0.610890808 | 0.54127187 | 0.645317405 |
| 4.366746559 | 1.26E-05 | 5.05E-05 |
| NA | NA | NA |
| NA | NA | NA |
| NA | NA | NA |
| NA | NA | NA |
| NA | NA | NA |
| NA | NA | NA |
| NA | NA | NA |
| NA | NA | NA |
| NA | NA | NA |
| NA | NA | NA |
| NA | NA | NA |
| -6.586912523 | 4.49E-11 | 3.54E-10 |
| NA | NA | NA |
| -1.332968267 | 0.182542218 | 0.27438336 |
| -3.226712493 | 0.001252212 | 0.003526822 |
| -3.770909671 | 0.000162654 | 0.000542178 |
| -2.966377331 | 0.003013305 | 0.007789222 |
| NA | NA | NA |
| NA | NA | NA |
| -1.147265385 | 0.251271956 | 0.355048947 |
| NA | NA | NA |
| NA | NA | NA |
| -0.719870522 | 0.471604719 | NA |
| -2.419169029 | 0.01555601 | 0.033638611 |
| NA | NA | NA |
| NA | NA | NA |

| | | | |
|---|---|---|---|
| NA | NA | NA | |
| | 0.192808327 | 0.847109082 | NA |
| NA | NA | NA | |
| | -0.065388772 | 0.947864464 | NA |
| NA | NA | NA | |
| | -0.784979838 | 0.43246539 | 0.543185905 |
| NA | NA | NA | |
| | 0.397517277 | 0.690986048 | NA |
| | 0.975859248 | 0.329134219 | 0.440005849 |
| | 3.624173113 | 0.000289887 | 0.000924081 |
| | 4.947604719 | 7.51E-07 | 3.59E-06 |
| | 0.192808924 | 0.847108615 | NA |
| | 3.653334077 | 0.000258857 | 0.00083226 |
| | 1.037904549 | 0.299314495 | 0.408230265 |
| | 1.806667913 | 0.070814074 | 0.124515368 |
| | 4.463315094 | 8.07E-06 | 3.34E-05 |
| | 0.13519054 | 0.892461226 | NA |
| | -0.669047741 | 0.503465024 | 0.610016921 |
| | -4.162934646 | 3.14E-05 | 0.000118409 |
| | -1.134031242 | 0.256781443 | 0.361324462 |
| | -6.356999627 | 2.06E-10 | 1.51E-09 |
| | -4.187815645 | 2.82E-05 | 0.000107025 |
| | -8.755350445 | 2.03E-18 | 3.19E-17 |
| | 0.154676764 | 0.877076149 | 0.914145992 |
| | 15.50763789 | 3.08E-54 | 2.60E-52 |
| NA | NA | NA | |
| | 0.363833705 | 0.715982192 | 0.792136745 |
| | 6.776317265 | 1.23E-11 | 1.03E-10 |
| | 0.96146628 | 0.336317772 | 0.447577669 |
| | 1.455160945 | 0.14562471 | NA |
| | -0.112224728 | 0.910645223 | NA |
| NA | NA | NA | |
| | 1.819351666 | 0.068857796 | 0.121570898 |
| NA | NA | NA | |
| NA | NA | NA | |
| | -0.297974314 | 0.765722768 | 0.830957767 |
| | -3.431477267 | 0.000600303 | 0.00180466 |
| | -1.633250994 | 0.102416219 | 0.169867998 |
| NA | NA | NA | |
| | 6.970643712 | 3.15E-12 | 2.77E-11 |
| | -0.168718242 | 0.866018269 | 0.906508752 |
| | 9.599264868 | 8.05E-22 | 1.65E-20 |
| | 0.21198725 | 0.832116981 | NA |
| NA | NA | NA | |
| | 0.637672984 | 0.523686575 | 0.628886196 |
| NA | NA | NA | |
| NA | NA | NA | |

| | | | |
|---|---|---|---|
| NA | NA | NA | |
| NA | NA | NA | |
| NA | NA | NA | |
| NA | NA | NA | |
| | -0.194487022 | 0.845794556 | NA |
| | 8.183769379 | 2.75E-16 | 3.60E-15 |
| | 0.167650371 | 0.866858341 | NA |
| | -4.121459329 | 3.76E-05 | 0.000139978 |
| | -2.707613114 | 0.006776897 | 0.016113398 |
| | -0.59151041 | 0.554178482 | 0.657210623 |
| | -0.194487022 | 0.845794556 | NA |
| | -0.246230714 | 0.805503641 | 0.862313679 |
| | 3.114313713 | 0.001843733 | 0.005002563 |
| | 1.225001518 | 0.220574649 | 0.319778943 |
| | -6.409905856 | 1.46E-10 | 1.08E-09 |
| | -2.39454666 | 0.016640926 | 0.035716205 |
| | 1.225130984 | 0.220525873 | 0.319726749 |
| NA | NA | NA | |
| | 0.468004293 | 0.63978152 | 0.730420153 |
| | -2.151616347 | 0.031427584 | 0.061827589 |
| NA | NA | NA | |
| NA | NA | NA | |
| | 0.560366206 | 0.575229677 | NA |
| | -1.033332937 | 0.301448119 | 0.410580966 |
| NA | NA | NA | |
| | -0.520492986 | 0.602720015 | NA |
| | -0.607451256 | 0.543551482 | 0.647474254 |
| | -4.599486057 | 4.24E-06 | 1.82E-05 |
| | -0.207009306 | 0.83600259 | NA |
| | -0.417684833 | 0.676177563 | 0.759433025 |
| | 0.065749872 | 0.947576966 | 0.96427637 |
| | -0.332422935 | 0.739569922 | 0.810341234 |
| | 4.239576281 | 2.24E-05 | 8.63E-05 |
| | -0.17566996 | 0.860553237 | 0.903129875 |
| | 0.108343039 | 0.913723583 | NA |
| | 3.979376973 | 6.91E-05 | 0.000245734 |
| NA | NA | NA | |
| | 0.434937002 | 0.663608156 | NA |
| NA | NA | NA | |
| NA | NA | NA | |
| NA | NA | NA | |
| NA | NA | NA | |
| NA | NA | NA | |
| | 7.001917014 | 2.52E-12 | 2.23E-11 |
| | 0.164587098 | 0.869268987 | NA |
| | 6.636307593 | 3.22E-11 | 2.57E-10 |
| NA | NA | NA | |

| | | |
|---|---|---|
| -2.107373267 | 0.035085237 | 0.06805322 |
| -0.065389966 | 0.947863513 | NA |
| 2.022631997 | 0.043111103 | 0.081402452 |
| -1.012624765 | 0.311239426 | 0.42144024 |
| -2.189759974 | 0.028541649 | 0.056887838 |
| -1.405438752 | 0.15989085 | 0.245836531 |
| 0.353496336 | 0.723716375 | NA |
| 3.652643057 | 0.000259555 | 0.000834188 |
| 1.15946216 | 0.246267852 | 0.349299277 |
| 0.06371127 | 0.949200131 | NA |
| NA | NA | NA |
| -3.238149152 | 0.001203079 | 0.003405685 |
| -0.065388772 | 0.947864464 | NA |
| NA | NA | NA |
| -1.195084307 | 0.232054093 | NA |
| -2.147159535 | 0.031780583 | 0.06241902 |
| 0.063708882 | 0.949202033 | NA |
| -3.389444997 | 0.000700343 | 0.002079181 |
| -7.464885679 | 8.34E-14 | 8.63E-13 |
| 0.192809521 | 0.847108147 | NA |
| -0.430929341 | 0.666519749 | NA |
| 4.40812317 | 1.04E-05 | 4.24E-05 |
| 0.192636473 | 0.847243678 | NA |
| -1.001922523 | 0.316381014 | 0.426913509 |
| 0.741493448 | 0.458394303 | NA |
| NA | NA | NA |
| NA | NA | NA |
| -4.587740161 | 4.48E-06 | 1.92E-05 |
| 0.274591726 | 0.783629924 | 0.845545201 |
| NA | NA | NA |
| 0.06371127 | 0.949200131 | NA |
| 3.84306894 | 0.000121505 | 0.000413936 |
| 0.278377556 | 0.780722547 | NA |
| 0.192810118 | 0.847107679 | NA |
| NA | NA | NA |
| NA | NA | NA |
| 1.367312334 | 0.171527427 | 0.260596753 |
| -0.079057133 | 0.93698718 | 0.95706138 |
| NA | NA | NA |
| -0.723864147 | 0.469149143 | 0.57833929 |
| 0.422262608 | 0.672833347 | 0.756661665 |
| 1.542246278 | 0.123013758 | 0.198123311 |
| NA | NA | NA |
| NA | NA | NA |
| 0.085566393 | 0.931811115 | NA |
| 1.963671146 | 0.049568241 | 0.091896383 |
| 0.267107038 | 0.789386749 | NA |

| | | | |
|---|---|---|---|
| | -1.077586981 | 0.281218115 | 0.388202593 |
| NA | NA | NA | |
| | -2.729540649 | 0.006342263 | 0.015186726 |
| | -2.702570693 | 0.006880555 | 0.016336587 |
| NA | NA | NA | |
| | -0.136057793 | 0.891775593 | 0.924309209 |
| | -0.257332193 | 0.796922343 | NA |
| | -0.165083893 | 0.86887795 | NA |
| | -4.568125069 | 4.92E-06 | 2.10E-05 |
| | 1.630991344 | 0.102892146 | 0.170465344 |
| | -0.065389369 | 0.947863988 | NA |
| | 0.192809521 | 0.847108147 | NA |
| | -0.154527899 | 0.877193515 | NA |
| | -8.486510666 | 2.13E-17 | 3.07E-16 |
| | 6.757998689 | 1.40E-11 | 1.16E-10 |
| NA | NA | NA | |
| NA | NA | NA | |
| NA | NA | NA | |
| | 0.3474247 | 0.728272279 | NA |
| | 0.356317 | 0.72160318 | 0.796442701 |
| NA | NA | NA | |
| NA | NA | NA | |
| | 0.063708285 | 0.949202508 | NA |
| NA | NA | NA | |
| NA | NA | NA | |
| NA | NA | NA | |
| NA | NA | NA | |
| NA | NA | NA | |
| NA | NA | NA | |
| | 0.192810118 | 0.847107679 | NA |
| NA | NA | NA | |
| NA | NA | NA | |
| NA | NA | NA | |
| NA | NA | NA | |
| | 0.192808924 | 0.847108615 | NA |
| NA | NA | NA | |
| | 0.063708882 | 0.949202033 | NA |
| NA | NA | NA | |
| NA | NA | NA | |
| NA | NA | NA | |
| | -0.060419584 | 0.951821462 | NA |
| | 0.192636473 | 0.847243678 | NA |
| NA | NA | NA | |
| NA | NA | NA | |
| NA | NA | NA | |
| NA | NA | NA | |
| NA | NA | NA | |

| | | | |
|---|---|---|---|
| NA | | NA | NA |
| NA | | NA | NA |
| | 0.192809521 | 0.847108147 | NA |
| NA | | NA | NA |
| | 0.192809521 | 0.847108147 | NA |
| NA | | NA | NA |
| | -0.065388772 | 0.947864464 | NA |
| NA | | NA | NA |
| NA | | NA | NA |
| NA | | NA | NA |
| NA | | NA | NA |
| NA | | NA | NA |
| NA | | NA | NA |
| NA | | NA | NA |
| | -0.065388772 | 0.947864464 | NA |
| NA | | NA | NA |
| NA | | NA | NA |
| | 0.192809521 | 0.847108147 | NA |
| NA | | NA | NA |
| NA | | NA | NA |
| | 0.192809521 | 0.847108147 | NA |
| NA | | NA | NA |
| NA | | NA | NA |
| | -7.914915433 | 2.47E-15 | 2.98E-14 |
| | 0.677223252 | 0.498264317 | NA |
| | -1.801042993 | 0.071696104 | 0.125774591 |
| | -8.497493923 | 1.94E-17 | 2.80E-16 |
| | -0.065392899 | 0.947861177 | NA |
| | 0.76009436 | 0.447198184 | 0.55710584 |
| | 2.751510038 | 0.00593212 | 0.01431031 |
| NA | | NA | NA |
| | 2.106058458 | 0.035199275 | 0.068249891 |
| NA | | NA | NA |
| | -2.86382077 | 0.004185648 | 0.010475629 |
| | 1.674285035 | 0.094074594 | 0.158330224 |
| | -0.203710406 | 0.83857981 | NA |
| | -1.629691153 | 0.103166788 | 0.170869883 |
| | -3.219504482 | 0.001284124 | 0.003608984 |
| | 3.116329396 | 0.001831176 | 0.004972804 |
| | 0.27182906 | 0.78575346 | NA |
| | -6.290302295 | 3.17E-10 | 2.27E-09 |
| | -0.362982774 | 0.716617752 | 0.792652894 |
| | 2.667520744 | 0.007641317 | 0.017888344 |
| | -2.220291244 | 0.026399004 | 0.053129136 |
| | -0.065389369 | 0.947863988 | NA |
| NA | | NA | NA |
| | 0.063708285 | 0.949202508 | NA |

| | | |
|---|---|---|
| 0.063708882 | 0.949202033 | NA |
| 0.371292192 | 0.710419913 | 0.787755197 |
| -3.909056692 | 9.27E-05 | 0.000322695 |
| 2.655257751 | 0.007924783 | 0.018474185 |
| 0.160827401 | 0.872229344 | 0.911067081 |
| -2.99543266 | 0.002740558 | 0.007145434 |
| -4.426078536 | 9.60E-06 | 3.92E-05 |
| 2.811256784 | 0.004934838 | 0.012164566 |
| 0.060896036 | 0.951442007 | 0.966847782 |
| 3.575561109 | 0.000349478 | 0.00109837 |
| -0.554316153 | 0.579362497 | NA |
| -0.989307172 | 0.322512876 | 0.433471614 |
| 0.063708285 | 0.949202508 | NA |
| -0.870733628 | 0.383899613 | 0.49620487 |
| -2.26340077 | 0.023610995 | 0.048278211 |
| -0.126157982 | 0.899606871 | NA |
| -0.065389369 | 0.947863988 | NA |
| -1.338888308 | 0.180607037 | 0.271899081 |
| NA | NA | NA |
| -0.946931711 | 0.343673578 | NA |
| NA | NA | NA |
| 0.94586908 | 0.344215366 | NA |
| 0.462367622 | 0.643817714 | NA |
| 0.063708285 | 0.949202508 | NA |
| 1.12887166 | 0.258951977 | 0.363949014 |
| 0.656427522 | 0.511549087 | NA |
| -1.175874395 | 0.239645072 | 0.341764922 |
| -0.050278734 | 0.95990027 | NA |
| -4.235895702 | 2.28E-05 | 8.77E-05 |
| 0.544705219 | 0.585956275 | 0.686043419 |
| -4.469393451 | 7.84E-06 | 3.25E-05 |
| -1.067622737 | 0.285690723 | 0.392815056 |
| 0.651872154 | 0.51448365 | 0.620035907 |
| -0.081507465 | 0.935038389 | 0.955850965 |
| 1.501322422 | 0.133272188 | 0.211635762 |
| -1.217054154 | 0.223583612 | 0.323279869 |
| NA | NA | NA |
| 0.333816862 | 0.738517758 | NA |
| 3.51056001 | 0.000447164 | 0.001375603 |
| 0.063708324 | 0.949202477 | NA |
| 3.143371423 | 0.001670138 | 0.004582717 |
| -1.302674984 | 0.192685746 | 0.286754517 |
| 0.825185898 | 0.409266058 | 0.521213465 |
| 0.115491663 | 0.908055428 | NA |
| 0.75739392 | 0.448813897 | 0.558757118 |
| -5.134461367 | 2.83E-07 | 1.44E-06 |
| 0.374707949 | 0.70787768 | NA |

| | | | |
|---|---|---|---|
| | 0.58653885 | 0.557513462 | NA |
| | 0.192808327 | 0.847109082 | NA |
| NA | NA | NA | |
| | 3.946657527 | 7.92E-05 | 0.000279305 |
| NA | NA | NA | |
| | -0.065388772 | 0.947864464 | NA |
| | 12.15225942 | 5.58E-34 | 2.34E-32 |
| | -0.078634545 | 0.937323309 | NA |
| | 0.482461722 | 0.629477984 | 0.722082215 |
| NA | NA | NA | |
| | 0.192809521 | 0.847108147 | NA |
| | -0.34726064 | 0.728395518 | 0.801741332 |
| | -2.602391976 | 0.009257597 | 0.021260452 |
| | -1.232740884 | 0.217672453 | NA |
| NA | NA | NA | |
| | -2.660344705 | 0.007806072 | 0.018229704 |
| | 2.670094852 | 0.007582982 | 0.017778386 |
| NA | NA | NA | |
| | 0.854188412 | 0.393000603 | 0.504943539 |
| NA | NA | NA | |
| | 6.068911655 | 1.29E-09 | 8.66E-09 |
| | -3.106108884 | 0.001895669 | 0.005131242 |
| | -4.982929344 | 6.26E-07 | 3.03E-06 |
| | 5.02590008 | 5.01E-07 | 2.45E-06 |
| | 0.584205771 | 0.559081879 | 0.661396136 |
| | 4.64342685 | 3.43E-06 | 1.49E-05 |
| | 0.192806163 | 0.847110777 | NA |
| | -4.56248677 | 5.06E-06 | 2.15E-05 |
| | 2.957443461 | 0.003102016 | 0.007990477 |
| | -1.322654672 | 0.185950244 | 0.278535883 |
| | -5.238679705 | 1.62E-07 | 8.48E-07 |
| | -0.084089386 | 0.932985363 | 0.954531929 |
| | 5.283487067 | 1.27E-07 | 6.74E-07 |
| | 0.571242974 | 0.567834952 | NA |
| | 0.066629133 | 0.946876953 | NA |
| | 6.065253053 | 1.32E-09 | 8.85E-09 |
| | 0.691311531 | 0.489369787 | 0.597129906 |
| | 6.55703104 | 5.49E-11 | 4.29E-10 |
| | 2.723973416 | 0.00645017 | 0.015421506 |
| | 5.892072619 | 3.81E-09 | 2.44E-08 |
| | -2.204957666 | 0.027457065 | 0.054993226 |
| | 1.09077602 | 0.275371451 | 0.382049078 |
| | 4.731331071 | 2.23E-06 | 9.99E-06 |
| | -0.483607633 | 0.628664357 | 0.721437727 |
| | 1.912251478 | 0.055843946 | 0.101774718 |
| | -0.065389966 | 0.947863513 | NA |
| | 3.352254005 | 0.000801564 | 0.002351794 |

| | | |
|---|---|---|
| 10.88232326 | 1.40E-27 | 4.25E-26 |
| NA | NA | NA |
| 4.864960204 | 1.14E-06 | 5.32E-06 |
| 0.368087159 | 0.712808245 | NA |
| 0.539589566 | 0.589480114 | 0.688988431 |
| 8.721325686 | 2.75E-18 | 4.25E-17 |
| 4.649930323 | 3.32E-06 | 1.45E-05 |
| 0.192808924 | 0.847108615 | NA |
| -0.352643015 | 0.724356085 | 0.79871943 |
| -3.355348546 | 0.00079265 | 0.002327824 |
| -10.8843313 | 1.37E-27 | 4.16E-26 |
| -0.892932482 | 0.371893331 | 0.484716209 |
| 4.903959047 | 9.39E-07 | 4.43E-06 |
| NA | NA | NA |
| 3.247507896 | 0.001164204 | 0.003308736 |
| 0.948904415 | 0.342669229 | 0.453978235 |
| -0.133755924 | 0.893595581 | NA |
| -2.608949399 | 0.009082067 | 0.020893742 |
| -2.026279887 | 0.042736111 | 0.080767507 |
| -8.092873809 | 5.83E-16 | 7.42E-15 |
| -7.869776228 | 3.55E-15 | 4.22E-14 |
| -1.619306457 | 0.105381342 | 0.173874547 |
| NA | NA | NA |
| -11.27145103 | 1.82E-29 | 6.09E-28 |
| NA | NA | NA |
| -0.630593332 | 0.52830646 | 0.633249243 |
| -4.109787643 | 3.96E-05 | 0.000146634 |
| -0.194488217 | 0.845793621 | NA |
| NA | NA | NA |
| 0.557725519 | 0.577031829 | 0.678013974 |
| 0.31685483 | 0.751353758 | NA |
| -1.035419058 | 0.300473249 | 0.409520579 |
| -1.368374112 | 0.171195003 | 0.260186467 |
| 13.93359445 | 3.96E-44 | 2.58E-42 |
| -4.337304842 | 1.44E-05 | 5.73E-05 |
| -0.340137567 | 0.733752932 | 0.805695805 |
| 1.634416364 | 0.102171455 | 0.169574399 |
| 1.109451543 | 0.267235437 | 0.373056497 |
| -0.306716189 | 0.759059406 | 0.825659769 |
| 5.143431976 | 2.70E-07 | 1.37E-06 |
| 2.221571912 | 0.02631225 | 0.052984362 |
| -3.321491931 | 0.000895376 | 0.00260303 |
| -4.967775583 | 6.77E-07 | 3.26E-06 |
| 0.195448797 | 0.845041619 | 0.891711287 |
| 2.855720303 | 0.004293932 | 0.010714698 |
| 0.796326775 | 0.425842127 | 0.537293499 |
| 4.716344268 | 2.40E-06 | 1.07E-05 |

| | | |
|---|---|---|
| -2.998609749 | 0.002712145 | 0.007076512 |
| NA | NA | NA |
| -2.545204253 | 0.010921383 | 0.024647212 |
| -5.660580433 | 1.51E-08 | 9.03E-08 |
| -1.895148178 | 0.058072773 | 0.105204711 |
| -4.731650463 | 2.23E-06 | 9.97E-06 |
| 0.180327349 | 0.856895586 | 0.900654331 |
| -0.71748588 | 0.473074346 | 0.581927014 |
| 0.125680433 | 0.899984892 | 0.930237109 |
| -2.061268355 | 0.039277445 | 0.074983942 |
| 1.367569976 | 0.17144672 | 0.260505761 |
| 0.907118887 | 0.364343931 | NA |
| NA | NA | NA |
| -0.272606637 | 0.78515561 | 0.846770576 |
| 3.911290065 | 9.18E-05 | 0.000319948 |
| -0.078161535 | 0.937699559 | NA |
| -1.086042915 | 0.277460006 | 0.38435021 |
| 8.036874705 | 9.22E-16 | 1.15E-14 |
| -2.13119476 | 0.033073101 | 0.064683521 |
| -4.485953068 | 7.26E-06 | 3.02E-05 |
| -9.167071919 | 4.86E-20 | 8.69E-19 |
| -1.207524954 | 0.227230041 | 0.327398568 |
| -2.300491817 | 0.021420372 | 0.044335725 |
| -9.865227455 | 5.89E-23 | 1.31E-21 |
| 0.55746352 | 0.577210777 | 0.678120745 |
| -2.407639329 | 0.016056034 | 0.034609244 |
| -2.667801653 | 0.007634932 | 0.017880084 |
| NA | NA | NA |
| 9.218950559 | 3.00E-20 | 5.42E-19 |
| -0.521311256 | 0.602149961 | 0.699582878 |
| 1.007947356 | 0.313479736 | 0.423923377 |
| 2.998630722 | 0.002711958 | 0.007076512 |
| 1.187293836 | 0.235111739 | 0.336603948 |
| -2.471981901 | 0.013436632 | 0.029587305 |
| 3.361012286 | 0.000776574 | 0.002284362 |
| -0.448294437 | 0.653940715 | 0.741982954 |
| -0.705409709 | 0.480555304 | 0.589247012 |
| 0.451700888 | 0.651484478 | 0.740037865 |
| -0.237426105 | 0.81232624 | 0.867872878 |
| 0.143644605 | 0.885781116 | 0.91992454 |
| -3.949066781 | 7.85E-05 | 0.000276704 |
| -0.065389369 | 0.947863988 | NA |
| 1.367692808 | 0.171408252 | 0.260463122 |
| -2.862338852 | 0.00420527 | 0.010518429 |
| NA | NA | NA |
| -0.227748707 | 0.819841605 | 0.873553078 |
| 1.305931672 | 0.191575795 | 0.285391153 |

| | | |
|---|---|---|
| 1.310939529 | 0.189878195 | 0.283334399 |
| NA | NA | NA |
| 1.340847555 | 0.179969953 | 0.271119345 |
| -0.335099411 | 0.737550098 | 0.808907148 |
| 35.77836624 | 2.40E-280 | 5.45E-277 |
| 6.435586826 | 1.23E-10 | 9.25E-10 |
| 0.39390925 | 0.693648041 | NA |
| NA | NA | NA |
| -0.065389369 | 0.947863988 | NA |
| 0.893848342 | 0.37140304 | 0.484178026 |
| -3.061570955 | 0.002201788 | 0.005874094 |
| 1.415694398 | 0.156864999 | 0.242321232 |
| 0.116376109 | 0.907354469 | NA |
| 1.730136139 | 0.083605955 | 0.143362133 |
| -1.259053471 | 0.208011019 | 0.305027582 |
| -1.047006081 | 0.295096774 | 0.403356938 |
| NA | NA | NA |
| 10.50067321 | 8.58E-26 | 2.32E-24 |
| NA | NA | NA |
| 1.665946528 | 0.095724065 | 0.160663405 |
| 10.20500045 | 1.88E-24 | 4.64E-23 |
| 2.363239254 | 0.01811597 | 0.038369894 |
| 4.729782954 | 2.25E-06 | 1.01E-05 |
| 11.25705617 | 2.14E-29 | 7.14E-28 |
| NA | NA | NA |
| -0.032752901 | 0.973871638 | 0.981992793 |
| -1.145836916 | 0.251862637 | 0.35580324 |
| -0.431223377 | 0.666305957 | 0.752024084 |
| 8.51971142 | 1.60E-17 | 2.32E-16 |
| 6.282536422 | 3.33E-10 | 2.38E-09 |
| 0.192809521 | 0.847108147 | NA |
| -3.230319655 | 0.001236519 | 0.003488118 |
| 0.065647654 | 0.947658348 | NA |
| 0.833469778 | 0.404579836 | NA |
| NA | NA | NA |
| -2.81163684 | 0.004929012 | 0.012152597 |
| -4.061602162 | 4.87E-05 | 0.000177878 |
| NA | NA | NA |
| -2.297873535 | 0.021568988 | 0.044599072 |
| -2.336500978 | 0.019465147 | 0.040822172 |
| -0.995328362 | 0.319576588 | 0.430285207 |
| -0.705679155 | 0.480387687 | 0.589070341 |
| -0.502708316 | 0.615169362 | 0.710487686 |
| NA | NA | NA |
| 0.350236135 | 0.72616149 | NA |
| 9.513804337 | 1.84E-21 | 3.66E-20 |
| 1.894203351 | 0.058198023 | 0.105359216 |

| | | |
|---|---|---|
| -0.317651505 | 0.750749301 | 0.819112288 |
| 2.781403497 | 0.005412442 | 0.013209187 |
| -6.188409329 | 6.08E-10 | 4.22E-09 |
| -2.104909176 | 0.035299215 | 0.068401281 |
| 3.24913507 | 0.001157565 | 0.003290987 |
| -0.051830944 | 0.958663399 | 0.971510785 |
| -6.163493865 | 7.12E-10 | 4.91E-09 |
| -1.783406335 | 0.074520183 | 0.12990947 |
| 0.401244323 | 0.688240251 | NA |
| 2.354598774 | 0.018542715 | 0.039153379 |
| NA | NA | NA |
| NA | NA | NA |
| 11.89651727 | 1.23E-32 | 4.90E-31 |
| 0.145451085 | 0.884354734 | 0.918978042 |
| -5.592891072 | 2.23E-08 | 1.31E-07 |
| 8.782313203 | 1.60E-18 | 2.54E-17 |
| 2.34514114 | 0.019019878 | 0.039995594 |
| -2.248721838 | 0.024530199 | 0.049869305 |
| -2.266286306 | 0.023433859 | 0.047983434 |
| 3.814805489 | 0.00013629 | 0.000460184 |
| 0.606724062 | 0.544034051 | 0.647886548 |
| NA | NA | NA |
| -1.40053638 | 0.161352757 | 0.247588249 |
| NA | NA | NA |
| 11.75456831 | 6.69E-32 | 2.56E-30 |
| 0.674973418 | 0.499692653 | 0.606767919 |
| -0.100877409 | 0.919647777 | 0.944850154 |
| NA | NA | NA |
| 5.007135794 | 5.52E-07 | 2.69E-06 |
| 4.954856626 | 7.24E-07 | 3.47E-06 |
| 8.202079497 | 2.36E-16 | 3.11E-15 |
| -1.80420085 | 0.071199826 | 0.125088205 |
| 4.499898156 | 6.80E-06 | 2.84E-05 |
| 2.513224927 | 0.011963303 | 0.02669044 |
| 3.760629862 | 0.000169486 | 0.000562855 |
| -2.272150443 | 0.023077425 | 0.047342673 |
| -3.78967275 | 0.000150846 | 0.000505512 |
| 10.42931967 | 1.82E-25 | 4.82E-24 |
| 2.471393379 | 0.013458768 | 0.029623018 |
| -5.349770236 | 8.81E-08 | 4.78E-07 |
| -3.190893647 | 0.001418335 | 0.003949813 |
| -6.436851781 | 1.22E-10 | 9.18E-10 |
| 1.540679056 | 0.123394917 | 0.198634945 |
| -2.120552317 | 0.033959494 | 0.066168936 |
| -0.267768579 | 0.788877458 | 0.849748366 |
| NA | NA | NA |
| -1.084039861 | 0.278347126 | 0.385284179 |

| | | |
|---|---|---|
| -0.952627966 | 0.340778603 | 0.452311402 |
| -4.427253755 | 9.54E-06 | 3.90E-05 |
| NA | NA | NA |
| NA | NA | NA |
| -0.065389369 | 0.947863988 | NA |
| 1.914029774 | 0.055616352 | 0.101433786 |
| 0.155168724 | 0.876688304 | 0.913893976 |
| 1.025222 | 0.30525845 | 0.414867731 |
| 5.387838096 | 7.13E-08 | 3.91E-07 |
| NA | NA | NA |
| -0.065389369 | 0.947863988 | NA |
| NA | NA | NA |
| 8.800238788 | 1.37E-18 | 2.18E-17 |
| -0.065389369 | 0.947863988 | NA |
| 8.263344852 | 1.42E-16 | 1.90E-15 |
| 0.240040136 | 0.810299142 | NA |
| 1.611102137 | 0.107157463 | 0.176225446 |
| NA | NA | NA |
| 0.063708285 | 0.949202508 | NA |
| -0.065389369 | 0.947863988 | NA |
| NA | NA | NA |
| NA | NA | NA |
| 1.604136841 | 0.108683893 | 0.178337343 |
| 5.556014045 | 2.76E-08 | 1.60E-07 |
| NA | NA | NA |
| -1.229893174 | 0.218737111 | 0.317648519 |
| 0.678328437 | 0.497563472 | 0.604651828 |
| -0.277253631 | 0.781585361 | 0.844145916 |
| NA | NA | NA |
| -1.40485403 | 0.160064689 | 0.246049202 |
| NA | NA | NA |
| -1.071319662 | 0.284025722 | NA |
| 0.66268962 | 0.507529359 | 0.613884414 |
| -1.090045589 | 0.275693062 | 0.382389303 |
| -2.05788375 | 0.039601294 | 0.075544577 |
| 1.619836558 | 0.105267391 | 0.173756449 |
| -2.693396802 | 0.007072804 | 0.016726406 |
| -5.301350705 | 1.15E-07 | 6.14E-07 |
| 1.184540579 | 0.236199143 | 0.337986768 |
| 3.322365547 | 0.000892577 | 0.002596606 |
| 2.417816736 | 0.015613938 | 0.033758044 |
| -0.866240083 | 0.386358517 | 0.498643401 |
| 3.184282966 | 0.001451131 | 0.004033515 |
| -1.042327601 | 0.29725982 | 0.405936674 |
| 2.070907685 | 0.038367424 | 0.073493275 |
| 1.256631026 | 0.208887275 | 0.306025665 |
| -1.653850285 | 0.098157936 | 0.164055364 |

| | | |
|---|---|---|
| 0.266426158 | 0.789911022 | NA |
| 3.135714517 | 0.00171436 | 0.004689665 |
| -0.204770888 | 0.837751133 | 0.886484775 |
| 2.471018308 | 0.013472892 | 0.029646285 |
| NA | NA | NA |
| NA | NA | NA |
| 8.92232418 | 4.57E-19 | 7.58E-18 |
| 0.155095401 | 0.876746108 | 0.91391617 |
| 2.956748075 | 0.00310902 | 0.008006871 |
| -3.361765941 | 0.000774457 | 0.002278941 |
| -0.089744118 | 0.928490556 | NA |
| -1.770634077 | 0.076621571 | 0.133053832 |
| -5.006638998 | 5.54E-07 | 2.69E-06 |
| -2.158231606 | 0.030909829 | 0.060919136 |
| 0.192809521 | 0.847108147 | NA |
| -1.253129865 | 0.210158449 | 0.307661669 |
| -0.284820328 | 0.775781797 | 0.839092213 |
| -1.839852443 | 0.065789904 | 0.116896521 |
| NA | NA | NA |
| 0.192809521 | 0.847108147 | NA |
| 0.192810118 | 0.847107679 | NA |
| 0.192809521 | 0.847108147 | NA |
| -1.306924211 | 0.191238452 | 0.285024322 |
| 4.876141444 | 1.08E-06 | 5.05E-06 |
| -1.699958475 | 0.089138736 | 0.151397338 |
| 2.110535494 | 0.034812257 | 0.067593732 |
| -1.338808604 | 0.180632989 | NA |
| NA | NA | NA |
| NA | NA | NA |
| NA | NA | NA |
| NA | NA | NA |
| -3.901826783 | 9.55E-05 | 0.000331521 |
| -7.276728137 | 3.42E-13 | 3.33E-12 |
| 0.893594179 | 0.371539062 | 0.484320158 |
| -2.411520055 | 0.015886178 | 0.034286405 |
| 0.192809521 | 0.847108147 | NA |
| NA | NA | NA |
| 0.230088074 | 0.818023332 | NA |
| 0.192809521 | 0.847108147 | NA |
| -2.491011541 | 0.012737998 | 0.02822772 |
| -1.675959054 | 0.093746213 | 0.157926216 |
| 1.034634007 | 0.300839866 | 0.409886334 |
| NA | NA | NA |
| 0.93548434 | 0.34953875 | 0.461199521 |
| 0.820751681 | 0.411787727 | 0.52343948 |
| -0.376619152 | 0.706456653 | 0.784673176 |
| -0.723266832 | 0.469515966 | 0.578630459 |

| | | |
|---|---|---|
| 0.191672495 | 0.847998752 | 0.893889609 |
| 0.279300781 | 0.780014007 | 0.842739699 |
| -8.830661343 | 1.04E-18 | 1.68E-17 |
| 2.90202503 | 0.003707589 | 0.009375728 |
| 1.148479486 | 0.250770677 | 0.354460698 |
| 9.594589026 | 8.43E-22 | 1.72E-20 |
| -1.061073254 | 0.288656611 | 0.396048708 |
| -5.257466265 | 1.46E-07 | 7.70E-07 |
| 0.557591829 | 0.577123137 | NA |
| 0.491225214 | 0.623267168 | 0.717192773 |
| 5.055866573 | 4.28E-07 | 2.12E-06 |
| -8.943827829 | 3.76E-19 | 6.27E-18 |
| -2.648387121 | 0.008087685 | 0.018804901 |
| -4.372561035 | 1.23E-05 | 4.93E-05 |
| NA | NA | NA |
| -2.480516884 | 0.013119205 | 0.028980117 |
| 30.30294192 | 1.05E-201 | 7.95E-199 |
| 5.243009735 | 1.58E-07 | 8.29E-07 |
| 0.333816862 | 0.738517758 | NA |
| -5.992598486 | 2.07E-09 | 1.36E-08 |
| -0.136052899 | 0.891779463 | NA |
| -2.374560876 | 0.017569839 | 0.037408393 |
| 3.041172294 | 0.002356589 | 0.006237225 |
| -5.088621147 | 3.61E-07 | 1.81E-06 |
| -0.374913036 | 0.707725144 | 0.785628896 |
| 3.400179813 | 0.000673416 | 0.002004947 |
| -1.976230198 | 0.048128711 | 0.089572581 |
| -1.009024546 | 0.312962862 | 0.42329917 |
| -3.246756091 | 0.001167284 | 0.003315982 |
| -2.104806855 | 0.035308124 | 0.068413249 |
| 4.251064327 | 2.13E-05 | 8.24E-05 |
| NA | NA | NA |
| NA | NA | NA |
| -0.065388772 | 0.947864464 | NA |
| 7.697331059 | 1.39E-14 | 1.56E-13 |
| -6.449351306 | 1.12E-10 | 8.50E-10 |
| -1.761438442 | 0.078164223 | 0.135388436 |
| 0.139431824 | 0.889108926 | NA |
| -1.751719043 | 0.079822131 | 0.137827534 |
| NA | NA | NA |
| 0.063708882 | 0.949202033 | NA |
| 7.008533032 | 2.41E-12 | 2.14E-11 |
| NA | NA | NA |
| 4.748131695 | 2.05E-06 | 9.23E-06 |
| 1.168075159 | 0.242776449 | NA |
| 0.402019378 | 0.687669765 | NA |
| 3.136768327 | 0.00170821 | 0.004674886 |

| | | |
|---|---|---|
| 1.270343509 | 0.203962296 | 0.300357491 |
| 0.063708882 | 0.949202033 | NA |
| -0.058503744 | 0.95334738 | 0.968122563 |
| 2.1101299 | 0.034847168 | 0.06765627 |
| 6.382546861 | 1.74E-10 | 1.29E-09 |
| -1.341555812 | 0.179740062 | 0.270821921 |
| -8.515240649 | 1.66E-17 | 2.41E-16 |
| -0.562539275 | 0.573748653 | 0.675014279 |
| 3.79074082 | 0.000150199 | 0.000503478 |
| NA | NA | NA |
| 0.657269384 | 0.511007719 | NA |
| -2.707910298 | 0.006770832 | 0.016102036 |
| 0.112979468 | 0.910046833 | NA |
| NA | NA | NA |
| NA | NA | NA |
| 0.063708285 | 0.949202508 | NA |
| -1.209520883 | 0.226462793 | 0.326556268 |
| 1.306845371 | 0.191265233 | 0.285043564 |
| -1.691619555 | 0.090718535 | 0.153611483 |
| -0.651576441 | 0.51467445 | 0.620198597 |
| 2.194874667 | 0.028172589 | 0.056205955 |
| -0.745413684 | 0.456021674 | 0.565760719 |
| -0.43604092 | 0.66280704 | 0.749156027 |
| 3.015695787 | 0.002563904 | 0.006727593 |
| -1.057563772 | 0.290254366 | 0.397846054 |
| -1.291030119 | 0.196693234 | 0.291557008 |
| -0.065390563 | 0.947863038 | NA |
| NA | NA | NA |
| NA | NA | NA |
| NA | NA | NA |
| 0.192809521 | 0.847108147 | NA |
| NA | NA | NA |
| -0.065389369 | 0.947863988 | NA |
| NA | NA | NA |
| 3.098745306 | 0.00194342 | 0.005248576 |
| NA | NA | NA |
| NA | NA | NA |
| 3.48478904 | 0.000492525 | 0.00150432 |
| -1.883748036 | 0.059599065 | 0.10748358 |
| -0.065389369 | 0.947863988 | NA |
| -1.300644351 | 0.193380219 | 0.287599933 |
| -9.420675154 | 4.48E-21 | 8.66E-20 |
| -0.614178336 | 0.539097487 | 0.64334939 |
| 0.703262246 | 0.481892334 | 0.59039673 |
| -0.926891757 | 0.353982737 | 0.465908814 |
| 0.324874675 | 0.745275925 | NA |
| -6.363571844 | 1.97E-10 | 1.45E-09 |

| | | | |
|---|---|---|---|
| | -3.153388925 | 0.001613866 | 0.00444402 |
| | 0.192809521 | 0.847108147 | NA |
| NA | NA | NA | |
| | -0.26176171 | 0.793505161 | NA |
| | -6.966390308 | 3.25E-12 | 2.85E-11 |
| | -5.994298747 | 2.04E-09 | 1.35E-08 |
| NA | NA | NA | |
| | 1.264957022 | 0.205886735 | NA |
| | 0.934717371 | 0.349933972 | NA |
| | -2.472504115 | 0.013417017 | 0.029551355 |
| | 0.063708285 | 0.949202508 | NA |
| | -9.855714416 | 6.48E-23 | 1.43E-21 |
| | 0.375794244 | 0.707069867 | NA |
| | -1.265433103 | 0.205716115 | 0.302388675 |
| | 1.328615847 | 0.183974741 | 0.276105515 |
| | 3.809847107 | 0.000139053 | 0.000468879 |
| | 0.192808327 | 0.847109082 | NA |
| | 0.607000381 | 0.543850659 | NA |
| NA | NA | NA | |
| | 4.344288757 | 1.40E-05 | 5.56E-05 |
| | -5.687240029 | 1.29E-08 | 7.78E-08 |
| | -1.313773493 | 0.18892244 | 0.282126883 |
| NA | NA | NA | |
| | 0.346721347 | 0.728800672 | NA |
| NA | NA | NA | |
| NA | NA | NA | |
| | -4.535164983 | 5.76E-06 | 2.43E-05 |
| NA | NA | NA | |
| | 0.422675052 | 0.67253236 | 0.756561122 |
| NA | NA | NA | |
| | -3.828739809 | 0.000128801 | 0.000436664 |
| | -1.856965267 | 0.063316096 | 0.113087047 |
| NA | NA | NA | |
| | 1.194622141 | 0.232234695 | 0.333170998 |
| | 0.192809521 | 0.847108147 | NA |
| | 2.693665897 | 0.007067097 | 0.016716068 |
| NA | NA | NA | |
| | -0.101901075 | 0.918835197 | 0.944261988 |
| NA | NA | NA | |
| | -8.010101407 | 1.15E-15 | 1.42E-14 |
| NA | NA | NA | |
| | -7.958156616 | 1.75E-15 | 2.13E-14 |
| | 0.192808924 | 0.847108615 | NA |
| | -3.506189761 | 0.000454571 | 0.001396748 |
| NA | NA | NA | |
| NA | NA | NA | |
| | -2.74084007 | 0.006128233 | 0.014734838 |

| | | |
|---|---|---|
| -3.15451069 | 0.001607675 | 0.004430266 |
| -1.738139572 | 0.082186224 | 0.141314912 |
| NA | NA | NA |
| 0.419243822 | 0.675037949 | NA |
| -2.793067279 | 0.005221083 | 0.012785805 |
| 0.063707194 | 0.949203376 | NA |
| 0.52093466 | 0.60241229 | 0.699822687 |
| 3.674068618 | 0.000238719 | 0.000772779 |
| 1.400550852 | 0.161348427 | NA |
| NA | NA | NA |
| NA | NA | NA |
| NA | NA | NA |
| -2.005445248 | 0.044915468 | 0.08438296 |
| -4.558988777 | 5.14E-06 | 2.18E-05 |
| -9.340861695 | 9.56E-21 | 1.79E-19 |
| 0.211741396 | 0.832308791 | 0.882628862 |
| 0.35866217 | 0.719847832 | 0.795101329 |
| 0.324874675 | 0.745275925 | NA |
| NA | NA | NA |
| -0.194488217 | 0.845793621 | NA |
| 1.989494047 | 0.046646698 | 0.087177395 |
| 9.715617472 | 2.59E-22 | 5.45E-21 |
| NA | NA | NA |
| -0.645068991 | 0.518882478 | 0.624248729 |
| 1.518608773 | 0.128861002 | 0.205950506 |
| 0.546752176 | 0.584549003 | 0.684631582 |
| -0.065389369 | 0.947863988 | NA |
| 10.73124651 | 7.26E-27 | 2.09E-25 |
| 1.169320608 | 0.242274483 | 0.344905603 |
| 2.114770148 | 0.034449542 | 0.066998605 |
| 2.066406267 | 0.038790135 | 0.074172427 |
| 2.345485124 | 0.019002337 | 0.039962064 |
| 0.300455256 | 0.763829921 | 0.829515052 |
| 1.11267815 | 0.265846683 | 0.3715115 |
| NA | NA | NA |
| -0.207930023 | 0.835283606 | 0.884611625 |
| -0.379555652 | 0.704275286 | 0.783153868 |
| NA | NA | NA |
| -1.982228928 | 0.047453624 | 0.08848724 |
| -12.52467545 | 5.47E-36 | 2.49E-34 |
| 2.007346592 | 0.044712771 | 0.084039999 |
| -1.474227139 | 0.140420445 | 0.220973612 |
| 8.589457875 | 8.74E-18 | 1.29E-16 |
| 0.687796684 | 0.491580825 | 0.598987971 |
| -3.657967894 | 0.000254223 | 0.000818418 |
| NA | NA | NA |
| -1.781824088 | 0.074777923 | 0.13028619 |

| | | | |
|---|---|---|---|
| | 1.058212665 | 0.289958498 | NA |
| NA | NA | NA | |
| | -7.63699204 | 2.22E-14 | 2.44E-13 |
| | -2.718540324 | 0.006557066 | 0.015654652 |
| | -2.704023185 | 0.006850551 | 0.016269977 |
| NA | NA | NA | |
| | 0.390336045 | 0.696288073 | NA |
| | -0.065388772 | 0.947864464 | NA |
| | 0.873850623 | 0.382199608 | 0.494612229 |
| | 0.192809521 | 0.847108147 | NA |
| | 11.54936586 | 7.44E-31 | 2.69E-29 |
| | -0.699079027 | 0.484502645 | 0.592954293 |
| NA | NA | NA | |
| | 8.344879219 | 7.13E-17 | 9.81E-16 |
| | 3.025928818 | 0.002478706 | 0.006525244 |
| | -5.037385743 | 4.72E-07 | 2.32E-06 |
| | -3.203503076 | 0.001357666 | 0.003796065 |
| | -2.97730233 | 0.002907971 | 0.007541862 |
| | 0.330493631 | 0.741027004 | NA |
| | -1.313313467 | 0.189077343 | 0.282324517 |
| NA | NA | NA | |
| | 0.311043314 | 0.755767693 | NA |
| | -0.515555861 | 0.606164645 | 0.703069831 |
| | -0.152669717 | 0.878658743 | 0.914919225 |
| | 6.027582744 | 1.66E-09 | 1.11E-08 |
| | 1.337171135 | 0.181166782 | 0.272594202 |
| | 0.063707135 | 0.949203424 | NA |
| | 4.192736391 | 2.76E-05 | 0.00010492 |
| | -0.102031653 | 0.91873155 | 0.944261988 |
| | -0.903241027 | 0.366397988 | 0.47887534 |
| NA | NA | NA | |
| | -0.065388772 | 0.947864464 | NA |
| | 0.126648093 | 0.899218931 | 0.929790963 |
| NA | NA | NA | |
| NA | NA | NA | |
| | 0.432311411 | 0.665515095 | 0.751412815 |
| NA | NA | NA | |
| NA | NA | NA | |
| | 0.192809521 | 0.847108147 | NA |
| NA | NA | NA | |
| NA | NA | NA | |
| | -1.134572045 | 0.256554672 | 0.361167797 |
| | 0.385718498 | 0.699705162 | 0.77911029 |
| NA | NA | NA | |
| NA | NA | NA | |
| NA | NA | NA | |
| | -7.512308876 | 5.81E-14 | 6.11E-13 |

| | | | |
|---|---|---|---|
| | 7.113126964 | 1.13E-12 | 1.05E-11 |
| | 0.192808327 | 0.847109082 | NA |
| | 19.50449418 | 1.01E-84 | 1.77E-82 |
| | -1.883621279 | 0.059616221 | 0.107500606 |
| NA | NA | NA | |
| | 3.350850523 | 0.000805638 | 0.002361808 |
| | 0.276032492 | 0.78252311 | 0.844795893 |
| | -7.116833679 | 1.10E-12 | 1.02E-11 |
| | 1.977164976 | 0.048022985 | 0.089415705 |
| | -0.065388772 | 0.947864464 | NA |
| | 0.350236135 | 0.72616149 | NA |
| | 1.012257719 | 0.311414845 | 0.421576707 |
| | 0.324874675 | 0.745275925 | NA |
| | 0.200068104 | 0.841427318 | NA |
| | 0.192810118 | 0.847107679 | NA |
| | 1.090117667 | 0.275661315 | 0.382366457 |
| | -3.020809989 | 0.002520995 | 0.006625421 |
| | 0.063708285 | 0.949202508 | NA |
| NA | NA | NA | |
| NA | NA | NA | |
| NA | NA | NA | |
| NA | NA | NA | |
| | -1.062646237 | 0.28794241 | 0.395433504 |
| | 0.192808924 | 0.847108615 | NA |
| NA | NA | NA | |
| | -0.194488217 | 0.845793621 | NA |
| | 0.415945136 | 0.67745015 | 0.760521047 |
| | -0.300832947 | 0.763541883 | 0.829310191 |
| | -3.70502431 | 0.00021137 | 0.000691311 |
| | -1.165331404 | 0.243884872 | 0.346665816 |
| | 0.620995906 | 0.534602317 | 0.639020821 |
| | 1.246216123 | 0.212685061 | 0.310564329 |
| NA | NA | NA | |
| | -0.432363921 | 0.665476936 | NA |
| | -0.675956899 | 0.49906801 | 0.606215292 |
| | 0.633564087 | 0.526365345 | 0.631406334 |
| | -0.967696307 | 0.333196063 | 0.444367938 |
| NA | NA | NA | |
| | -0.329972517 | 0.741420729 | 0.81178973 |
| | 1.63719053 | 0.101590665 | 0.168726977 |
| | -0.871112626 | 0.383692661 | 0.496091084 |
| | -2.227092053 | 0.025941122 | 0.052342305 |
| | -0.040613745 | 0.967603826 | 0.977878781 |
| | 0.219648545 | 0.826144881 | NA |
| | 0.875748311 | 0.381166875 | 0.493718013 |
| | -1.521986514 | 0.12801246 | 0.204934562 |
| NA | NA | NA | |

| | | | |
|---|---|---|---|
| | 9.800499425 | 1.12E-22 | 2.43E-21 |
| | 2.165802847 | 0.030326256 | 0.059910512 |
| | 2.184131383 | 0.028952595 | 0.057564815 |
| | -6.601528146 | 4.07E-11 | 3.22E-10 |
| | -1.680136128 | 0.092930833 | 0.156778919 |
| | -1.740909969 | 0.081699361 | 0.140603342 |
| | -0.266893662 | 0.789551037 | NA |
| NA | NA | NA | |
| | 0.627002606 | 0.530657527 | NA |
| | -4.303560863 | 1.68E-05 | 6.61E-05 |
| | -2.380996712 | 0.017265866 | 0.036867824 |
| | -0.23627423 | 0.813219882 | 0.868605174 |
| NA | NA | NA | |
| | 4.879581474 | 1.06E-06 | 4.97E-06 |
| | 0.192808924 | 0.847108615 | NA |
| | -1.031042112 | 0.302521072 | 0.411772463 |
| | -0.140364386 | 0.888372095 | NA |
| NA | NA | NA | |
| | -0.4790804 | 0.631881433 | 0.723711552 |
| NA | NA | NA | |
| NA | NA | NA | |
| NA | NA | NA | |
| NA | NA | NA | |
| NA | NA | NA | |
| NA | NA | NA | |
| | -0.831726932 | 0.405563095 | 0.517419046 |
| NA | NA | NA | |
| NA | NA | NA | |
| NA | NA | NA | |
| NA | NA | NA | |
| NA | NA | NA | |
| | 0.231912591 | 0.816605905 | NA |
| NA | NA | NA | |
| NA | NA | NA | |
| | 1.126712181 | 0.259864191 | 0.364821397 |
| NA | NA | NA | |
| | 0.515244935 | 0.606381873 | NA |
| | 5.554955761 | 2.78E-08 | 1.61E-07 |
| | -0.540417797 | 0.588908937 | 0.688449428 |
| | -0.62833665 | 0.529783427 | 0.634715642 |
| | 8.042662248 | 8.79E-16 | 1.10E-14 |
| | -4.837607134 | 1.31E-06 | 6.06E-06 |
| | -0.862835491 | 0.388227923 | 0.500472923 |
| | 0.259708306 | 0.795088785 | 0.854198553 |
| | -1.494402488 | 0.135070452 | 0.213934912 |
| | -4.017551309 | 5.88E-05 | 0.000211604 |
| | 5.187435846 | 2.13E-07 | 1.10E-06 |

| | | |
|---|---|---|
| 0.089763161 | 0.928475423 | NA |
| NA | NA | NA |
| NA | NA | NA |
| -0.727652119 | 0.466826567 | 0.5761387 |
| 0.161196102 | 0.871938951 | NA |
| 4.646240283 | 3.38E-06 | 1.47E-05 |
| 0.223926529 | 0.822814472 | 0.875788665 |
| 0.192808924 | 0.847108615 | NA |
| 1.516613968 | 0.129364177 | 0.206589323 |
| 9.128910927 | 6.92E-20 | 1.22E-18 |
| 0.400996812 | 0.688422471 | 0.770144563 |
| 1.986078575 | 0.0470246 | 0.087798256 |
| 13.20509052 | 8.20E-40 | 4.46E-38 |
| 0.518946187 | 0.603798268 | 0.701009817 |
| 3.452240039 | 0.000555953 | 0.001682038 |
| -1.383025524 | 0.16665704 | 0.254401763 |
| 5.429080621 | 5.66E-08 | 3.16E-07 |
| 0.573590952 | 0.566244636 | 0.6681959 |
| 10.63903187 | 1.96E-26 | 5.52E-25 |
| -0.295318752 | 0.767750393 | NA |
| 1.813328603 | 0.069781154 | 0.12292366 |
| -2.364159132 | 0.018071049 | 0.038306093 |
| -3.410259757 | 0.00064901 | 0.001937587 |
| 1.01392465 | 0.310618708 | 0.420757678 |
| -0.384928838 | 0.700290139 | 0.779622924 |
| -1.618206143 | 0.10561818 | 0.174197617 |
| -0.614943953 | 0.538591735 | 0.642991087 |
| 0.192810118 | 0.847107679 | NA |
| NA | NA | NA |
| -0.93160066 | 0.351542942 | 0.463306647 |
| 0.192810118 | 0.847107679 | NA |
| NA | NA | NA |
| NA | NA | NA |
| NA | NA | NA |
| 0.192808924 | 0.847108615 | NA |
| NA | NA | NA |
| -0.065389369 | 0.947863988 | NA |
| NA | NA | NA |
| 0.288217989 | 0.77317988 | NA |
| NA | NA | NA |
| NA | NA | NA |
| -0.112124554 | 0.910724649 | NA |
| -1.413560211 | 0.157491071 | 0.242943517 |
| -0.401231405 | 0.688249761 | NA |
| 5.592941032 | 2.23E-08 | 1.31E-07 |
| 0.436137907 | 0.662736676 | 0.749110323 |
| -0.42236546 | 0.672758285 | 0.756624992 |

| | | |
|---|---|---|
| 9.51408937 | 1.83E-21 | 3.65E-20 |
| -0.065389369 | 0.947863988 | NA |
| 0.134677539 | 0.892866833 | 0.925172082 |
| -1.756683555 | 0.078971762 | 0.13660423 |
| 7.257960322 | 3.93E-13 | 3.80E-12 |
| 10.03135725 | 1.11E-23 | 2.60E-22 |
| NA | NA | NA |
| 8.022989894 | 1.03E-15 | 1.29E-14 |
| 5.122231091 | 3.02E-07 | 1.53E-06 |
| -7.582135517 | 3.40E-14 | 3.66E-13 |
| -0.194693086 | 0.845633225 | 0.891903027 |
| 0.803570124 | 0.421645283 | NA |
| 10.76777595 | 4.89E-27 | 1.42E-25 |
| -4.595644907 | 4.31E-06 | 1.86E-05 |
| 1.635706635 | 0.101901 | 0.169147956 |
| 21.5386542 | 6.76E-103 | 1.68E-100 |
| NA | NA | NA |
| 1.85810247 | 0.063154464 | 0.112877473 |
| -5.482349757 | 4.20E-08 | 2.38E-07 |
| 9.403935744 | 5.26E-21 | 1.00E-19 |
| -0.595934966 | 0.551218673 | 0.654537852 |
| 0.866103874 | 0.386433201 | 0.498685409 |
| -3.689452737 | 0.000224737 | 0.000731491 |
| -1.033150974 | 0.301533252 | 0.410629885 |
| -4.48638126 | 7.24E-06 | 3.01E-05 |
| -0.094138167 | 0.924999402 | NA |
| 0.040181727 | 0.967948246 | 0.978164047 |
| 4.547494069 | 5.43E-06 | 2.30E-05 |
| NA | NA | NA |
| -0.162438652 | 0.870960435 | 0.91008348 |
| 2.136901673 | 0.03260599 | 0.06388977 |
| 3.306183725 | 0.000945761 | 0.002739225 |
| 2.236351638 | 0.025328746 | 0.051305213 |
| 0.987911566 | 0.32319596 | 0.434107587 |
| 0.193341124 | 0.846691821 | NA |
| 0.351609739 | 0.725130961 | 0.799279018 |
| 1.012298797 | 0.31139521 | 0.421576707 |
| -0.10721583 | 0.914617757 | 0.941319181 |
| 0.368700675 | 0.712350846 | 0.789328508 |
| -0.075786787 | 0.939588728 | NA |
| NA | NA | NA |
| NA | NA | NA |
| -0.34093621 | 0.733151604 | NA |
| NA | NA | NA |
| -0.33656973 | 0.736441278 | 0.807938865 |
| 0.286888673 | 0.774197564 | NA |
| 0.906825061 | 0.364499315 | 0.476978717 |

| | | |
|---|---|---|
| -0.587754482 | 0.5566971 | 0.659291622 |
| -0.744270082 | 0.456713097 | 0.566393937 |
| -0.034551256 | 0.97243757 | 0.981060556 |
| -0.767204694 | 0.442959822 | 0.553009345 |
| 0.11016455 | 0.912278876 | 0.939490948 |
| -1.431334304 | 0.152334427 | 0.236490151 |
| -2.379812315 | 0.017321458 | 0.036955003 |
| -9.557544356 | 1.21E-21 | 2.44E-20 |
| -0.935931039 | 0.349308695 | 0.460964326 |
| 0.160262267 | 0.872674481 | NA |
| -2.788654402 | 0.005292751 | 0.012938525 |
| -4.611561045 | 4.00E-06 | 1.72E-05 |
| -0.607434219 | 0.543562785 | 0.647474254 |
| -3.956100987 | 7.62E-05 | 0.00026933 |
| -0.087373182 | 0.930374885 | NA |
| 7.404806629 | 1.31E-13 | 1.33E-12 |
| 2.226376483 | 0.025988974 | 0.052426179 |
| NA | NA | NA |
| 0.192808924 | 0.847108615 | NA |
| NA | NA | NA |
| 7.263225587 | 3.78E-13 | 3.66E-12 |
| 0.199136792 | 0.842155742 | NA |
| 6.655822446 | 2.82E-11 | 2.26E-10 |
| 1.050734028 | 0.293380763 | 0.401449841 |
| 3.472765827 | 0.000515125 | 0.001567417 |
| -0.065389369 | 0.947863988 | NA |
| 2.213709935 | 0.026848739 | 0.053908546 |
| 0.066587943 | 0.946909745 | NA |
| -0.541582106 | 0.588106421 | 0.687832519 |
| -1.524374058 | 0.127415297 | 0.204148309 |
| 1.371832463 | 0.170115599 | 0.258750209 |
| NA | NA | NA |
| 0.399197784 | 0.689747476 | 0.770995417 |
| -2.173525413 | 0.029740795 | 0.058935326 |
| -2.888409158 | 0.003871959 | 0.009766716 |
| -0.766076661 | 0.443630689 | 0.553681184 |
| -5.903156431 | 3.57E-09 | 2.29E-08 |
| -5.185001974 | 2.16E-07 | 1.11E-06 |
| 0.367622201 | 0.713154958 | 0.79004464 |
| -1.246454925 | 0.212597428 | 0.31049372 |
| -0.145274321 | 0.884494288 | NA |
| 0.791524567 | 0.428637946 | 0.539814228 |
| -0.292428698 | 0.769958869 | 0.834223945 |
| -1.910294475 | 0.056095309 | 0.102173306 |
| 0.192810118 | 0.847107679 | NA |
| -1.194283869 | 0.232366945 | 0.333321161 |
| NA | NA | NA |

| | | |
|---|---|---|
| 2.224489689 | 0.026115515 | 0.052655981 |
| 0.014256754 | 0.988625142 | 0.99211364 |
| 4.146167235 | 3.38E-05 | 0.000126776 |
| NA | NA | NA |
| 9.246115333 | 2.33E-20 | 4.22E-19 |
| -2.334887159 | 0.019549316 | 0.040950656 |
| -1.985369878 | 0.047103335 | 0.087912644 |
| 9.024710471 | 1.80E-19 | 3.09E-18 |
| 2.009492218 | 0.044484959 | 0.083687225 |
| 0.444113916 | 0.656960226 | NA |
| 2.988297511 | 0.002805363 | 0.007298435 |
| 0.178899295 | 0.858016776 | 0.90148146 |
| 1.243683309 | 0.213616141 | 0.311718594 |
| -2.136110815 | 0.032670382 | 0.063990897 |
| -1.772219279 | 0.076358165 | 0.132670058 |
| 2.061102874 | 0.039293227 | 0.075008348 |
| 1.21000546 | 0.226276798 | 0.326355706 |
| NA | NA | NA |
| -3.145325169 | 0.001659023 | 0.004556216 |
| -2.454680475 | 0.014100986 | 0.030865561 |
| -0.015943348 | 0.987279588 | 0.991080777 |
| -2.520339023 | 0.011724185 | 0.026243637 |
| -1.357284886 | 0.174690708 | 0.26463163 |
| -1.028014244 | 0.303943128 | 0.413394364 |
| -1.266854524 | 0.205207312 | 0.301782551 |
| 1.52920729 | 0.126213064 | 0.202481284 |
| -5.556518509 | 2.75E-08 | 1.60E-07 |
| 4.023024207 | 5.75E-05 | 0.000207251 |
| 2.425374059 | 0.015292621 | 0.033129159 |
| 1.526235102 | 0.126951324 | NA |
| NA | NA | NA |
| 0.063707688 | 0.949202983 | NA |
| 3.784036199 | 0.000154305 | 0.000516138 |
| 0.067057933 | 0.946535583 | NA |
| 0.455119702 | 0.649023125 | NA |
| -1.249037826 | 0.211651239 | 0.309441035 |
| -5.5923447 | 2.24E-08 | 1.31E-07 |
| 0.357667975 | 0.720591802 | 0.795655058 |
| NA | NA | NA |
| -6.014357047 | 1.81E-09 | 1.20E-08 |
| NA | NA | NA |
| 0.192810118 | 0.847107679 | NA |
| 4.261467621 | 2.03E-05 | 7.89E-05 |
| 1.124084537 | 0.260977165 | 0.36609642 |
| -3.823729562 | 0.000131448 | 0.000445156 |
| -3.22764828 | 0.001248123 | 0.003516889 |
| -1.895287823 | 0.058054281 | 0.105182674 |

| | | | |
|---|---|---|---|
| NA | NA | NA | |
| | 0.137767028 | 0.890424543 | 0.923312001 |
| | -0.816238356 | 0.414363823 | 0.525966149 |
| | 0.330493631 | 0.741027004 | NA |
| | 3.982450193 | 6.82E-05 | 0.000242749 |
| | 7.165665932 | 7.74E-13 | 7.27E-12 |
| | 0.729277469 | 0.465831952 | 0.575138053 |
| | 0.000864469 | 0.999310253 | 0.999506869 |
| | -0.856188541 | 0.3918935 | 0.503805562 |
| NA | NA | NA | |
| | 4.772343462 | 1.82E-06 | 8.25E-06 |
| | 5.95878059 | 2.54E-09 | 1.66E-08 |
| | -0.658514719 | 0.510207442 | 0.616128348 |
| NA | NA | NA | |
| | -1.482174715 | 0.138293813 | 0.21819535 |
| | -10.26318353 | 1.03E-24 | 2.58E-23 |
| | -2.484848329 | 0.012960664 | 0.028670387 |
| | -1.083655665 | 0.2785175 | 0.38549508 |
| | 1.535083262 | 0.124763377 | 0.200528307 |
| | -2.450913041 | 0.014249437 | 0.031125178 |
| | -1.97686247 | 0.048057178 | 0.089472714 |
| | 22.28633348 | 5.01E-110 | 1.41E-107 |
| | 15.09055239 | 1.87E-51 | 1.48E-49 |
| | 1.077455258 | 0.281276929 | 0.388262369 |
| | -2.204702953 | 0.027474946 | 0.055024635 |
| | 7.555614908 | 4.17E-14 | 4.46E-13 |
| | 1.552452529 | 0.120553983 | 0.194687799 |
| | 2.665762486 | 0.007681394 | 0.017973711 |
| | -0.187995005 | 0.850880567 | 0.896247973 |
| | 2.054265399 | 0.039950012 | 0.076128572 |
| | -0.606634582 | 0.544093445 | 0.647921353 |
| | -3.537761256 | 0.000403535 | 0.001252384 |
| | 0.891280732 | 0.372778583 | 0.485591878 |
| | 0.083247481 | 0.93365476 | 0.954904533 |
| | -1.107477424 | 0.268087568 | 0.373969946 |
| | 0.330493631 | 0.741027004 | NA |
| NA | NA | NA | |
| NA | NA | NA | |
| | -0.439030571 | 0.660639385 | 0.747541893 |
| | 0.801687684 | 0.422733642 | 0.534476085 |
| | -6.090648287 | 1.12E-09 | 7.62E-09 |
| | 1.307264951 | 0.191122743 | 0.284868879 |
| | 0.869093301 | 0.3847961 | 0.496927375 |
| | 0.063708285 | 0.949202508 | NA |
| | 17.86460184 | 2.23E-71 | 2.96E-69 |
| | 2.631545402 | 0.008499751 | 0.019689844 |
| | -3.959056459 | 7.52E-05 | 0.00026617 |

| | | |
|---|---|---|
| 0.676602468 | 0.498658213 | NA |
| NA | NA | NA |
| 0.063708285 | 0.949202508 | NA |
| 8.662115715 | 4.63E-18 | 7.01E-17 |
| 2.318269202 | 0.020434695 | 0.042577243 |
| NA | NA | NA |
| NA | NA | NA |
| NA | NA | NA |
| -3.777423774 | 0.000158459 | 0.000528689 |
| NA | NA | NA |
| -3.048453841 | 0.002300222 | 0.006107412 |
| NA | NA | NA |
| NA | NA | NA |
| -4.642584171 | 3.44E-06 | 1.50E-05 |
| 0.25947095 | 0.795271893 | NA |
| -1.144288811 | 0.252503882 | 0.35656222 |
| -0.721108481 | 0.470842774 | 0.579808847 |
| 0.294386595 | 0.768462509 | 0.833244525 |
| -4.18146432 | 2.90E-05 | 0.000109809 |
| -5.344632356 | 9.06E-08 | 4.91E-07 |
| 1.201776947 | 0.22944996 | 0.329970266 |
| 1.180059182 | 0.237976677 | 0.340005493 |
| 0.091074266 | 0.927433581 | NA |
| -3.836678629 | 0.000124709 | 0.000423826 |
| -0.360283618 | 0.718635049 | 0.794311546 |
| 1.180215838 | 0.237914381 | 0.339974707 |
| 1.392722045 | 0.163703859 | 0.250642745 |
| 0.09817548 | 0.921792953 | NA |
| NA | NA | NA |
| -0.19448762 | 0.845794089 | NA |
| NA | NA | NA |
| NA | NA | NA |
| NA | NA | NA |
| NA | NA | NA |
| NA | NA | NA |
| NA | NA | NA |
| 0.859995651 | 0.38979144 | 0.501910177 |
| 14.22208048 | 6.68E-46 | 4.60E-44 |
| 1.059947085 | 0.289168673 | NA |
| NA | NA | NA |
| 0.063708882 | 0.949202033 | NA |
| -2.532208271 | 0.011334664 | 0.025485685 |
| -3.714510777 | 0.000203597 | 0.00066842 |
| 0.527013427 | 0.598184264 | 0.696462561 |
| -4.356342062 | 1.32E-05 | 5.28E-05 |
| -0.019515821 | 0.984429616 | 0.989092537 |
| -0.065389966 | 0.947863513 | NA |

| | | |
|---|---|---|
| 14.5374492 | 7.02E-48 | 5.13E-46 |
| -0.319138505 | 0.749621486 | 0.818167087 |
| 0.063704039 | 0.949205889 | NA |
| -6.033921592 | 1.60E-09 | 1.07E-08 |
| NA | NA | NA |
| -0.065389369 | 0.947863988 | NA |
| NA | NA | NA |
| -0.065389966 | 0.947863513 | NA |
| -0.065193062 | 0.948020285 | NA |
| NA | NA | NA |
| 0.225080218 | 0.82191687 | 0.875093762 |
| 0.132881669 | 0.894286964 | NA |
| -9.976962474 | 1.92E-23 | 4.44E-22 |
| 1.168516641 | 0.242598431 | NA |
| -5.270556422 | 1.36E-07 | 7.20E-07 |
| -2.088462611 | 0.036756125 | 0.070856886 |
| NA | NA | NA |
| NA | NA | NA |
| NA | NA | NA |
| -0.065389369 | 0.947863988 | NA |
| 1.281108535 | 0.200155546 | 0.295900616 |
| 0.083076052 | 0.933791068 | NA |
| 3.275836977 | 0.001053494 | 0.003021882 |
| -5.373300193 | 7.73E-08 | 4.23E-07 |
| 0.192810118 | 0.847107679 | NA |
| -3.341646397 | 0.000832831 | 0.002436675 |
| 1.925837157 | 0.05412471 | 0.099132263 |
| 3.1424872 | 0.00167519 | 0.004593057 |
| 0.192809521 | 0.847108147 | NA |
| 0.474218714 | 0.63534395 | NA |
| NA | NA | NA |
| -0.309902647 | 0.75663499 | 0.823536088 |
| NA | NA | NA |
| -2.007707352 | 0.044674399 | 0.083993105 |
| -0.233392093 | 0.815456953 | 0.870214774 |
| 4.417274692 | 1.00E-05 | 4.08E-05 |
| -0.491512266 | 0.623064179 | 0.71705809 |
| -0.145982194 | 0.883935445 | 0.918732007 |
| -0.475102586 | 0.634713855 | 0.72613549 |
| -0.220098995 | 0.825794057 | NA |
| -0.084061579 | 0.933007472 | NA |
| NA | NA | NA |
| 0.591485267 | 0.554195324 | NA |
| NA | NA | NA |
| -1.830830007 | 0.06712592 | 0.118974776 |
| 8.846102499 | 9.06E-19 | 1.47E-17 |
| 0.964330714 | 0.334880147 | 0.446114752 |

| | | | |
|---|---|---|---|
| | 1.470401412 | 0.141453069 | 0.222313999 |
| | 5.009990962 | 5.44E-07 | 2.65E-06 |
| | 11.48248233 | 1.62E-30 | 5.72E-29 |
| | 7.860634347 | 3.82E-15 | 4.52E-14 |
| | -0.320803545 | 0.748359273 | 0.817072888 |
| | -4.854198715 | 1.21E-06 | 5.60E-06 |
| | -3.078238784 | 0.00208228 | 0.005585662 |
| | 4.009677977 | 6.08E-05 | 0.000218188 |
| | -3.518515487 | 0.000433968 | 0.001338372 |
| | 4.4367954 | 9.13E-06 | 3.75E-05 |
| | -1.868480922 | 0.061695067 | 0.110610094 |
| NA | NA | NA | |
| | -3.609333413 | 0.000306985 | 0.000974487 |
| | -0.065389369 | 0.947863988 | NA |
| | 0.445267379 | 0.656126539 | NA |
| | 0.335915797 | 0.736934364 | NA |
| | 0.192809521 | 0.847108147 | NA |
| NA | NA | NA | |
| NA | NA | NA | |
| | 0.066629133 | 0.946876953 | NA |
| NA | NA | NA | |
| NA | NA | NA | |
| | 0.192810118 | 0.847107679 | NA |
| | 0.443344846 | 0.657516323 | NA |
| | 7.724774464 | 1.12E-14 | 1.27E-13 |
| | 0.192810118 | 0.847107679 | NA |
| | -3.230043918 | 0.001237712 | 0.003490697 |
| | -2.916287832 | 0.003542236 | 0.008998493 |
| | 9.352480245 | 8.56E-21 | 1.61E-19 |
| | 4.018847806 | 5.85E-05 | 0.000210564 |
| | -0.787816699 | 0.430803932 | 0.541778748 |
| | -0.158531921 | 0.874037669 | NA |
| | 0.063710673 | 0.949200606 | NA |
| | 0.532985808 | 0.594043405 | 0.692962384 |
| | 2.74913132 | 0.005975344 | 0.014401816 |
| | -0.440431271 | 0.65962478 | 0.746705633 |
| | 1.688111065 | 0.091389911 | 0.154548195 |
| NA | NA | NA | |
| | 0.063708882 | 0.949202033 | NA |
| | 0.063708285 | 0.949202508 | NA |
| | 0.278377556 | 0.780722547 | NA |
| NA | NA | NA | |
| NA | NA | NA | |
| NA | NA | NA | |
| | 0.192810118 | 0.847107679 | NA |
| NA | NA | NA | |
| | 0.192809521 | 0.847108147 | NA |

| | | |
|---|---|---|
| 0.354627914 | 0.722868361 | NA |
| NA | NA | NA |
| NA | NA | NA |
| NA | NA | NA |
| NA | NA | NA |
| 0.774425936 | 0.438678951 | 0.549006708 |
| 4.743253557 | 2.10E-06 | 9.44E-06 |
| -0.07030088 | 0.943954182 | NA |
| -0.374936811 | 0.707707462 | 0.785628896 |
| 4.622346335 | 3.79E-06 | 1.64E-05 |
| 1.062121908 | 0.288180346 | 0.395674934 |
| -6.400019226 | 1.55E-10 | 1.15E-09 |
| 4.435290622 | 9.19E-06 | 3.77E-05 |
| -1.357988638 | 0.17446729 | 0.26438919 |
| -0.425083103 | 0.670776095 | 0.755128437 |
| 1.219998911 | 0.222465288 | 0.321941845 |
| -0.678890994 | 0.497206933 | 0.604277229 |
| 8.699422619 | 3.34E-18 | 5.12E-17 |
| 2.455414876 | 0.014072208 | 0.030818738 |
| 1.47879328 | 0.139195572 | 0.219330512 |
| 0.698397506 | 0.484928635 | 0.593330583 |
| 0.656006459 | 0.511819969 | 0.617541973 |
| 0.419732535 | 0.674680855 | 0.758160268 |
| -12.32388662 | 6.74E-35 | 2.93E-33 |
| 1.492940274 | 0.135452822 | 0.214499818 |
| 12.29638672 | 9.47E-35 | 4.07E-33 |
| NA | NA | NA |
| -3.195776876 | 0.001394549 | 0.003888766 |
| NA | NA | NA |
| NA | NA | NA |
| 1.623949968 | 0.104386487 | 0.172598346 |
| -3.392131848 | 0.000693511 | 0.002059875 |
| -0.359075353 | 0.719538719 | 0.7949353 |
| -3.203968731 | 0.001355472 | 0.003790778 |
| 4.018237034 | 5.86E-05 | 0.000211019 |
| 0.571523693 | 0.567644704 | 0.669377231 |
| NA | NA | NA |
| 1.513455761 | 0.13016393 | 0.207595462 |
| 3.905535303 | 9.40E-05 | 0.000327153 |
| 1.668865984 | 0.095143942 | 0.159904175 |
| NA | NA | NA |
| -1.009908525 | 0.312539118 | NA |
| -0.964694213 | 0.334697995 | 0.445919562 |
| 0.059551652 | 0.952512726 | NA |
| -0.415317532 | 0.677909468 | 0.760934302 |
| 0.063708324 | 0.949202477 | NA |
| NA | NA | NA |

| | | |
|---|---|---|
| NA | NA | NA |
| NA | NA | NA |
| 0.463670001 | 0.642884193 | 0.732792538 |
| 3.36407107 | 0.000768018 | 0.002261586 |
| 9.773288604 | 1.47E-22 | 3.14E-21 |
| NA | NA | NA |
| NA | NA | NA |
| NA | NA | NA |
| 0.160672331 | 0.872351483 | NA |
| 1.234928344 | 0.216857174 | NA |
| 1.323710353 | 0.185599259 | 0.2780793 |
| 1.223782759 | 0.22103419 | 0.32035239 |
| -0.730384246 | 0.465155346 | 0.574634417 |
| 8.896287915 | 5.77E-19 | 9.52E-18 |
| 0.386983605 | 0.698768345 | 0.778309527 |
| NA | NA | NA |
| NA | NA | NA |
| -3.254103139 | 0.00113751 | 0.00323912 |
| 1.234146192 | 0.217148433 | 0.315762812 |
| 0.607673981 | 0.543403723 | 0.64737721 |
| -0.903035946 | 0.366506818 | 0.478992567 |
| 3.200285927 | 0.001372913 | 0.003835698 |
| 0.192806378 | 0.847110608 | NA |
| 2.01633506 | 0.043764951 | 0.0825043 |
| 18.65202186 | 1.22E-77 | 1.87E-75 |
| 10.20467552 | 1.89E-24 | 4.65E-23 |
| NA | NA | NA |
| NA | NA | NA |
| NA | NA | NA |
| NA | NA | NA |
| 0.16646194 | 0.867793433 | 0.907882847 |
| 0.063708882 | 0.949202033 | NA |
| -5.608082749 | 2.05E-08 | 1.21E-07 |
| -5.755891321 | 8.62E-09 | 5.30E-08 |
| -2.966381123 | 0.003013268 | 0.007789222 |
| 0.430520204 | 0.666817273 | 0.752228066 |
| -1.072984769 | 0.28327795 | 0.390249716 |
| 0.063707019 | 0.949203516 | NA |
| 11.07140427 | 1.73E-28 | 5.53E-27 |
| 2.915271262 | 0.003553795 | 0.009026026 |
| -2.215292848 | 0.02673997 | 0.053724632 |
| 15.03533453 | 4.31E-51 | 3.37E-49 |
| 0.192809521 | 0.847108147 | NA |
| 5.251679422 | 1.51E-07 | 7.93E-07 |
| -0.528028381 | 0.597479637 | NA |
| NA | NA | NA |
| 0.551005291 | 0.581630047 | 0.682010685 |

| | | |
|---|---|---|
| NA | NA | NA |
| 5.818008619 | 5.96E-09 | 3.73E-08 |
| -0.107994175 | 0.914000313 | 0.940915844 |
| -1.126716589 | 0.259862327 | 0.364821397 |
| 11.92107496 | 9.19E-33 | 3.66E-31 |
| 1.64042877 | 0.100916047 | 0.167791196 |
| -1.198242777 | 0.230822516 | 0.331563133 |
| -1.019055703 | 0.308176522 | 0.418039148 |
| 1.917109875 | 0.055223974 | 0.100843079 |
| -0.065388772 | 0.947864464 | NA |
| 7.860567431 | 3.82E-15 | 4.52E-14 |
| -0.544603854 | 0.586026003 | 0.686060504 |
| 0.147426332 | 0.882795523 | NA |
| NA | NA | NA |
| 0.618074094 | 0.536526498 | 0.640892347 |
| 0.329549476 | 0.741740404 | NA |
| 0.301023421 | 0.763396635 | 0.829224397 |
| -2.102249179 | 0.035531454 | 0.0687874 |
| -0.031590036 | 0.974798989 | 0.982499888 |
| -0.160444919 | 0.872530608 | 0.911332704 |
| -2.83548057 | 0.00457568 | 0.011359776 |
| 2.23026005 | 0.025730184 | 0.051958572 |
| 0.063708285 | 0.949202508 | NA |
| -4.408562739 | 1.04E-05 | 4.23E-05 |
| NA | NA | NA |
| -0.065390563 | 0.947863038 | NA |
| 0.063708285 | 0.949202508 | NA |
| 2.581507972 | 0.009836972 | 0.022450935 |
| -7.336537619 | 2.19E-13 | 2.18E-12 |
| -0.11870859 | 0.905506231 | 0.934593252 |
| -1.376217147 | 0.168754415 | 0.257007789 |
| -0.194487022 | 0.845794556 | NA |
| 0.063707019 | 0.949203516 | NA |
| 1.499852292 | 0.133652668 | 0.212159201 |
| -0.065389369 | 0.947863988 | NA |
| NA | NA | NA |
| -0.472024919 | 0.636909002 | 0.72807053 |
| 0.063708285 | 0.949202508 | NA |
| 0.424546952 | 0.671166971 | NA |
| NA | NA | NA |
| NA | NA | NA |
| NA | NA | NA |
| 0.083873413 | 0.933157078 | NA |
| NA | NA | NA |
| -1.091791361 | 0.274924817 | 0.381619797 |
| -4.543441565 | 5.53E-06 | 2.34E-05 |
| 0.762624927 | 0.445687114 | 0.555527537 |

| | | |
|---|---|---|
| NA | NA | NA |
| NA | NA | NA |
| 11.53083903 | 9.22E-31 | 3.31E-29 |
| 0.30922356 | 0.757151473 | NA |
| 2.693992837 | 0.007060168 | 0.016704415 |
| -0.381344205 | 0.702947856 | 0.781990438 |
| -0.876778054 | 0.380607202 | 0.493095148 |
| 0.997061979 | 0.318734426 | 0.429336061 |
| 1.779655683 | 0.075132327 | 0.130863973 |
| 0.593801626 | 0.552644801 | 0.655795732 |
| 2.46238393 | 0.013801684 | 0.030289885 |
| 1.45697658 | 0.145122845 | 0.227168531 |
| 1.672182196 | 0.0944884 | 0.158930492 |
| 0.192808924 | 0.847108615 | NA |
| 1.151365586 | 0.249581866 | 0.35305946 |
| 3.475857319 | 0.000509223 | 0.001551157 |
| NA | NA | NA |
| 0.224679734 | 0.822228431 | NA |
| NA | NA | NA |
| 0.31685483 | 0.751353758 | NA |
| NA | NA | NA |
| NA | NA | NA |
| NA | NA | NA |
| NA | NA | NA |
| -3.945730035 | 7.96E-05 | 0.00028027 |
| 0.679091087 | 0.49708015 | NA |
| NA | NA | NA |
| NA | NA | NA |
| 1.52378485 | 0.127562465 | 0.204344864 |
| NA | NA | NA |
| 1.096176245 | 0.273001653 | NA |
| -1.492392396 | 0.135596307 | 0.214686291 |
| 4.243317663 | 2.20E-05 | 8.50E-05 |
| 3.136112937 | 0.001712033 | 0.00468381 |
| 3.445987403 | 0.000568977 | 0.001717912 |
| -0.66513017 | 0.505967239 | 0.612403794 |
| 8.054157884 | 8.00E-16 | 1.01E-14 |
| 1.709384011 | 0.087379841 | 0.148864512 |
| -0.396557873 | 0.691693523 | 0.772491918 |
| 2.456386924 | 0.014034196 | 0.030748943 |
| -1.641233199 | 0.100749017 | 0.167535735 |
| NA | NA | NA |
| 2.17220953 | 0.029839862 | 0.059090349 |
| -0.257921001 | 0.796467876 | 0.855132594 |
| 2.066560091 | 0.038775625 | 0.07416494 |
| 4.017170387 | 5.89E-05 | 0.000211885 |
| NA | NA | NA |

| | | |
|---|---|---|
| NA | NA | NA |
| 0.687027015 | 0.492065704 | NA |
| 1.33508552 | 0.181848358 | 0.273455356 |
| 1.175186724 | 0.23992002 | 0.342080542 |
| 19.95256998 | 1.42E-88 | 2.74E-86 |
| 6.770595345 | 1.28E-11 | 1.06E-10 |
| 2.465947038 | 0.013665154 | 0.030032382 |
| NA | NA | NA |
| NA | NA | NA |
| -0.258462176 | 0.796050237 | NA |
| -2.265542754 | 0.023479394 | 0.048060944 |
| 3.121812651 | 0.001797413 | 0.004892794 |
| NA | NA | NA |
| 0.084306951 | 0.932812386 | NA |
| -4.119428508 | 3.80E-05 | 0.000141154 |
| NA | NA | NA |
| NA | NA | NA |
| 0.57881588 | 0.562713415 | 0.664843841 |
| 0.225694319 | 0.821439177 | 0.874733997 |
| -3.632447968 | 0.000280745 | 0.000896994 |
| 3.803730216 | 0.000142533 | 0.000479453 |
| -2.912186067 | 0.003589088 | 0.009099044 |
| NA | NA | NA |
| -0.093544986 | 0.925470613 | NA |
| 0.354377823 | 0.723055752 | NA |
| -0.418535768 | 0.675555441 | 0.758938631 |
| -0.26008719 | 0.794796519 | 0.854029229 |
| -0.910528953 | 0.36254362 | NA |
| -0.113458458 | 0.909667096 | NA |
| -0.883634044 | 0.376893795 | 0.48962743 |
| 0.305216104 | 0.760201568 | 0.826471149 |
| -0.33189951 | 0.739965141 | 0.810738782 |
| 19.57255122 | 2.65E-85 | 4.74E-83 |
| NA | NA | NA |
| 1.803317701 | 0.071338334 | 0.125274621 |
| -0.223333585 | 0.82327589 | 0.876152474 |
| 3.410641416 | 0.000648103 | 0.001935108 |
| NA | NA | NA |
| -0.065389369 | 0.947863988 | NA |
| -4.194786467 | 2.73E-05 | 0.000104071 |
| -6.30027546 | 2.97E-10 | 2.13E-09 |
| 0.461190708 | 0.644661788 | NA |
| 0.409676637 | 0.68204317 | NA |
| NA | NA | NA |
| -0.244824041 | 0.806592679 | 0.862985349 |
| -1.519428846 | 0.128654586 | 0.205725693 |
| -0.293248386 | 0.769332302 | NA |

| | | |
|---|---|---|
| -1.868067837 | 0.061752616 | 0.110689486 |
| -0.261996244 | 0.793324338 | NA |
| -1.029746391 | 0.303129072 | 0.412466468 |
| -1.088180772 | 0.276515311 | 0.383359823 |
| -4.375112343 | 1.21E-05 | 4.88E-05 |
| NA | NA | NA |
| -3.75923402 | 0.000170434 | 0.000565629 |
| -1.796444879 | 0.072423792 | 0.126802428 |
| 0.079823642 | 0.936377521 | NA |
| -0.003242593 | 0.99741279 | 0.998130583 |
| 0.293994676 | 0.768761972 | 0.833533124 |
| NA | NA | NA |
| -0.500861726 | 0.616468441 | 0.711429991 |
| -2.393107123 | 0.016706363 | 0.03582593 |
| NA | NA | NA |
| -0.315150557 | 0.752647344 | 0.820575094 |
| 0.543910651 | 0.586502959 | 0.686438281 |
| -0.71596948 | 0.474010195 | 0.58287768 |
| NA | NA | NA |
| 0.480673712 | 0.630748418 | 0.722876964 |
| 0.330754714 | 0.74082977 | NA |
| NA | NA | NA |
| -1.826065162 | 0.067840447 | 0.119969365 |
| 0.740564271 | 0.458957678 | 0.56838903 |
| -0.065388772 | 0.947864464 | NA |
| 1.835311275 | 0.066459581 | 0.117983127 |
| NA | NA | NA |
| -1.004017712 | 0.315370077 | 0.425858492 |
| -0.531414595 | 0.59513151 | 0.693715344 |
| 0.336191489 | 0.736726471 | NA |
| -0.317887001 | 0.750570653 | NA |
| -1.439459125 | 0.150020483 | 0.233462614 |
| -0.326211617 | 0.744264253 | NA |
| -2.095988472 | 0.036083214 | 0.069684361 |
| -0.327796706 | 0.743065378 | 0.813065012 |
| 0.14764561 | 0.882622458 | NA |
| -0.327938983 | 0.742957798 | NA |
| -0.334087465 | 0.738313559 | 0.809425271 |
| NA | NA | NA |
| 0.729765016 | 0.465533832 | 0.574826689 |
| -1.600400716 | 0.109509716 | 0.17945713 |
| NA | NA | NA |
| 0.483767003 | 0.628551236 | 0.721382797 |
| -0.799965765 | 0.423730633 | 0.535303979 |
| -3.100101057 | 0.001934546 | 0.005227431 |
| 0.280721894 | 0.778923714 | 0.841816087 |
| 0.192809521 | 0.847108147 | NA |

| | | |
|---|---|---|
| -0.744760477 | 0.456416532 | 0.566082245 |
| 0.040758167 | 0.967488689 | NA |
| -1.500719725 | 0.133428069 | 0.211869858 |
| -1.547761705 | 0.12167968 | 0.196290435 |
| NA | NA | NA |
| 0.073671169 | 0.94127204 | NA |
| -0.707462183 | 0.479279306 | 0.587970447 |
| -0.408009864 | 0.683266433 | 0.765814026 |
| -1.410079633 | 0.15851617 | 0.24414795 |
| -0.172361364 | 0.863153444 | 0.904834389 |
| -1.842547502 | 0.065395105 | 0.116269319 |
| -2.84777664 | 0.004402581 | 0.010964837 |
| -1.514560335 | 0.129883784 | 0.2072479 |
| -1.176599519 | 0.23935539 | 0.341449071 |
| -3.780009137 | 0.000156823 | 0.000523718 |
| -3.764111721 | 0.000167142 | 0.000555808 |
| 0.455971406 | 0.64841054 | NA |
| -1.719215927 | 0.085575062 | 0.146197788 |
| -2.382479249 | 0.017196501 | 0.036738513 |
| -0.008657854 | 0.993092118 | 0.995159569 |
| 0.192808924 | 0.847108615 | NA |
| 1.241885273 | 0.214278894 | 0.31255249 |
| -2.136175817 | 0.032665086 | 0.063990538 |
| -2.699950609 | 0.006934977 | 0.01644275 |
| -0.194487022 | 0.845794556 | NA |
| -1.834386849 | 0.06659659 | 0.118125158 |
| 0.065061479 | 0.948125051 | NA |
| -1.584073871 | 0.11317691 | 0.1845587 |
| 0.479797063 | 0.631371702 | 0.723336973 |
| -0.551198895 | 0.581497338 | 0.681950914 |
| -1.24791727 | 0.212061356 | 0.309913921 |
| -0.724194053 | 0.468946609 | 0.578184997 |
| 0.194624965 | 0.845686558 | 0.891903027 |
| -1.857107787 | 0.063295821 | 0.113058902 |
| 0.751844202 | 0.452144757 | 0.562065223 |
| -3.282885448 | 0.001027504 | 0.002955122 |
| -0.54718499 | 0.584251647 | 0.68434736 |
| -0.55033745 | 0.582087942 | 0.682387766 |
| -7.980425692 | 1.46E-15 | 1.79E-14 |
| -4.671802571 | 2.99E-06 | 1.31E-05 |
| -1.863098916 | 0.062448351 | 0.111784468 |
| -7.318114709 | 2.51E-13 | 2.48E-12 |
| -0.180854724 | 0.856481608 | 0.900404664 |
| NA | NA | NA |
| 2.693903602 | 0.007062059 | 0.01670573 |
| NA | NA | NA |
| NA | NA | NA |

| | | | |
|---|---|---|---|
| NA | NA | NA | |
| | -5.358295538 | 8.40E-08 | 4.57E-07 |
| | -0.065389369 | 0.947863988 | NA |
| | -1.994084292 | 0.046142844 | 0.086351804 |
| | -0.966740447 | 0.333673802 | 0.444862856 |
| | 0.31685483 | 0.751353758 | NA |
| | -8.140057786 | 3.95E-16 | 5.09E-15 |
| | 8.050251134 | 8.26E-16 | 1.04E-14 |
| | -7.506668614 | 6.07E-14 | 6.36E-13 |
| | 0.800255258 | 0.42356292 | 0.535146125 |
| | -10.14372003 | 3.53E-24 | 8.56E-23 |
| | -0.265598544 | 0.790548407 | NA |
| | -2.453075295 | 0.014164069 | 0.03096573 |
| | 7.190655922 | 6.45E-13 | 6.12E-12 |
| | 3.891160298 | 9.98E-05 | 0.000344809 |
| NA | NA | NA | |
| | 1.988463955 | 0.046760402 | 0.087370324 |
| | -0.882169414 | 0.3776852 | 0.490333755 |
| | -1.223961889 | 0.220966605 | 0.320272982 |
| | -0.065389369 | 0.947863988 | NA |
| | 0.193076837 | 0.846898793 | NA |
| NA | NA | NA | |
| | -3.86505166 | 0.000111066 | 0.000381027 |
| | 0.063708285 | 0.949202508 | NA |
| | -0.729170249 | 0.465897527 | NA |
| | 1.936740054 | 0.052777127 | 0.096976324 |
| | -8.460997956 | 2.65E-17 | 3.78E-16 |
| | 3.323167549 | 0.000890014 | 0.002590055 |
| | 16.78609672 | 3.08E-63 | 3.33E-61 |
| | -2.63941857 | 0.008304837 | 0.019272212 |
| | -0.322307878 | 0.747219466 | 0.816277294 |
| | -2.359376111 | 0.018305692 | 0.038721497 |
| NA | NA | NA | |
| | 5.790627342 | 7.01E-09 | 4.35E-08 |
| | 1.067360978 | 0.285808862 | 0.39291486 |
| NA | NA | NA | |
| | -0.662844878 | 0.507429908 | 0.613870641 |
| | 0.436679322 | 0.662343927 | 0.748767829 |
| | -0.893978339 | 0.371333481 | 0.484112559 |
| | 0.193348586 | 0.846685978 | NA |
| | -0.19448762 | 0.845794089 | NA |
| | 0.401175821 | 0.688290681 | NA |
| | 0.597487559 | 0.550181911 | NA |
| | -4.038400613 | 5.38E-05 | 0.000195251 |
| | 5.235530986 | 1.65E-07 | 8.61E-07 |
| | -3.167519575 | 0.001537454 | 0.004250327 |
| | -8.311985155 | 9.41E-17 | 1.28E-15 |

| | | | |
|---|---|---|---|
| | -1.321750761 | 0.18625116 | 0.278919859 |
| | -0.768778768 | 0.442024655 | 0.551979501 |
| | -2.626825697 | 0.008618547 | 0.019935517 |
| | 3.081080771 | 0.002062507 | 0.005539115 |
| | -2.490238715 | 0.012765732 | 0.028279164 |
| | -4.123443945 | 3.73E-05 | 0.00013888 |
| | -4.174563541 | 2.99E-05 | 0.000112969 |
| NA | NA | NA | |
| | 0.148846112 | 0.881675064 | 0.916953789 |
| | 3.000128324 | 0.002698659 | 0.007047202 |
| | 1.023902074 | 0.305881531 | 0.41551149 |
| | 1.130793398 | 0.258142059 | 0.362892207 |
| | 1.348559312 | 0.177478557 | 0.268043702 |
| | 21.86344613 | 5.79E-106 | 1.53E-103 |
| | 3.082299435 | 0.002054081 | 0.005518854 |
| | -0.708596387 | 0.478574979 | 0.587365486 |
| | -3.482032311 | 0.000497624 | 0.001518966 |
| | -0.840315131 | 0.40073172 | 0.512325822 |
| | 0.946267288 | 0.344012273 | 0.455347816 |
| | -2.20254557 | 0.027626794 | 0.05528893 |
| | -1.690943486 | 0.090847597 | 0.153798806 |
| | -1.818637022 | 0.068966826 | 0.121724976 |
| | -3.416556311 | 0.000634185 | 0.001897152 |
| | -0.08329421 | 0.933617604 | NA |
| NA | NA | NA | |
| | 3.164461426 | 0.001553703 | 0.004291456 |
| | -1.882716215 | 0.059738837 | 0.107673586 |
| | -3.63859296 | 0.000274132 | 0.000877768 |
| | 1.140723144 | 0.253985152 | 0.358175016 |
| | -0.787256786 | 0.431131561 | 0.542136296 |
| | -2.569542155 | 0.0101833 | 0.02317585 |
| NA | NA | NA | |
| | 3.836924698 | 0.000124585 | 0.000423517 |
| | -2.913426969 | 0.003574854 | 0.00906939 |
| NA | NA | NA | |
| | 5.076959121 | 3.84E-07 | 1.91E-06 |
| | -5.933344912 | 2.97E-09 | 1.92E-08 |
| | 4.71527207 | 2.41E-06 | 1.07E-05 |
| NA | NA | NA | |
| | -5.110980717 | 3.20E-07 | 1.62E-06 |
| NA | NA | NA | |
| | -3.940627443 | 8.13E-05 | 0.000285616 |
| | -5.059164229 | 4.21E-07 | 2.09E-06 |
| | -4.991290896 | 6.00E-07 | 2.90E-06 |
| | -1.196535038 | 0.231487834 | 0.332328103 |
| | 0.192809521 | 0.847108147 | NA |
| NA | NA | NA | |

| | | | |
|---|---|---|---|
| NA | NA | NA | |
| NA | NA | NA | |
| NA | NA | NA | |
| | 2.192813488 | 0.02832082 | 0.056470383 |
| | -0.065388772 | 0.947864464 | NA |
| NA | NA | NA | |
| | -0.479139149 | 0.63183964 | NA |
| NA | NA | NA | |
| | 0.153498734 | 0.878004989 | NA |
| | 1.766166947 | 0.077367845 | 0.134210073 |
| | -0.592645408 | 0.553418484 | 0.656538373 |
| NA | NA | NA | |
| NA | NA | NA | |
| NA | NA | NA | |
| NA | NA | NA | |
| NA | NA | NA | |
| NA | NA | NA | |
| NA | NA | NA | |
| | -1.481377031 | 0.138506132 | 0.218436676 |
| NA | NA | NA | |
| | 0.156084431 | 0.87596647 | 0.913407797 |
| | -1.43251891 | 0.15199537 | 0.236125963 |
| | 0.507328657 | 0.611924235 | 0.707784824 |
| | -1.241022247 | 0.21459753 | 0.312853176 |
| NA | NA | NA | |
| | 3.151494557 | 0.001624372 | 0.004468506 |
| | 2.522722356 | 0.01164503 | 0.02608045 |
| | 0.063708882 | 0.949202033 | NA |
| NA | NA | NA | |
| NA | NA | NA | |
| | -0.364878516 | 0.715202092 | NA |
| | 0.083904682 | 0.933132217 | NA |
| NA | NA | NA | |
| | 0.063708285 | 0.949202508 | NA |
| NA | NA | NA | |
| NA | NA | NA | |
| NA | NA | NA | |
| | -0.065388772 | 0.947864464 | NA |
| | 0.083462637 | 0.933483685 | NA |
| NA | NA | NA | |
| NA | NA | NA | |
| | 0.733295115 | 0.463378454 | 0.572815226 |
| NA | NA | NA | |
| | 0.597605122 | 0.550103446 | 0.653461594 |
| NA | NA | NA | |
| | -3.385895147 | 0.000709465 | 0.002103768 |
| | 8.706564581 | 3.13E-18 | 4.81E-17 |

| | | |
|---|---|---|
| 6.452709403 | 1.10E-10 | 8.32E-10 |
| 1.567263739 | 0.117053057 | NA |
| 0.320051528 | 0.748929269 | NA |
| 1.043150056 | 0.2968788 | NA |
| 2.879347997 | 0.003984983 | 0.010022522 |
| NA | NA | NA |
| -1.191385777 | 0.233502174 | 0.334757453 |
| -6.369852973 | 1.89E-10 | 1.39E-09 |
| 0.68274821 | 0.494765961 | 0.601895115 |
| 0.165641978 | 0.86843871 | 0.908320627 |
| -1.888144181 | 0.059006597 | 0.106599432 |
| -1.112654896 | 0.265856674 | NA |
| NA | NA | NA |
| 0.670372004 | 0.502620677 | NA |
| -0.091377393 | 0.927192725 | NA |
| 4.247218571 | 2.16E-05 | 8.37E-05 |
| NA | NA | NA |
| -1.82829152 | 0.067505812 | 0.119518972 |
| 5.970422517 | 2.37E-09 | 1.55E-08 |
| NA | NA | NA |
| -0.719885234 | 0.47159566 | NA |
| 0.11486469 | 0.908552373 | NA |
| 0.010629578 | 0.991518984 | 0.994060729 |
| 0.192810118 | 0.847107679 | NA |
| 0.127480965 | 0.898559738 | 0.929416514 |
| -8.687198806 | 3.71E-18 | 5.67E-17 |
| NA | NA | NA |
| NA | NA | NA |
| -3.368966089 | 0.000754507 | 0.002225925 |
| NA | NA | NA |
| -2.273238168 | 0.023011831 | 0.047227463 |
| NA | NA | NA |
| 0.478281401 | 0.632449933 | NA |
| 3.26065421 | 0.001111555 | 0.003171708 |
| 1.056295347 | 0.290833301 | NA |
| -0.377919924 | 0.705490081 | 0.783877868 |
| NA | NA | NA |
| NA | NA | NA |
| NA | NA | NA |
| NA | NA | NA |
| 4.637075953 | 3.53E-06 | 1.54E-05 |
| 2.221176794 | 0.026338989 | 0.053025408 |
| NA | NA | NA |
| NA | NA | NA |
| 2.364880387 | 0.018035895 | 0.038247772 |
| 0.063708285 | 0.949202508 | NA |
| NA | NA | NA |

| | | |
|---|---|---|
| 0.607612833 | 0.543444287 | 0.647394722 |
| NA | NA | NA |
| 0.510308718 | 0.609835196 | 0.7060654 |
| -4.036075639 | 5.44E-05 | 0.000196995 |
| 0.877712097 | 0.380099978 | 0.492693032 |
| 1.087029805 | 0.277023638 | 0.383894456 |
| 0.192809521 | 0.847108147 | NA |
| NA | NA | NA |
| -0.19448762 | 0.845794089 | NA |
| NA | NA | NA |
| NA | NA | NA |
| NA | NA | NA |
| -0.065388772 | 0.947864464 | NA |
| NA | NA | NA |
| NA | NA | NA |
| NA | NA | NA |
| NA | NA | NA |
| 0.113778407 | 0.909413456 | NA |
| NA | NA | NA |
| 0.039925844 | 0.968152247 | 0.978275029 |
| -0.065389966 | 0.947863513 | NA |
| 0.063708285 | 0.949202508 | NA |
| -1.060851217 | 0.288757522 | 0.396163165 |
| -4.01439695 | 5.96E-05 | 0.000214053 |
| 0.331074127 | 0.740588494 | 0.811186666 |
| -3.851000416 | 0.000117636 | 0.000401806 |
| -3.202734164 | 0.001361296 | 0.003804515 |
| 0.063708285 | 0.949202508 | NA |
| -1.907250515 | 0.056488155 | 0.102799078 |
| -6.20335735 | 5.53E-10 | 3.86E-09 |
| 0.324874675 | 0.745275925 | NA |
| NA | NA | NA |
| 0.107506976 | 0.914386791 | NA |
| -0.033336484 | 0.97340626 | 0.981588471 |
| NA | NA | NA |
| 0.192810118 | 0.847107679 | NA |
| NA | NA | NA |
| 0.345240757 | 0.72991338 | NA |
| -3.673590247 | 0.000239166 | 0.000774028 |
| NA | NA | NA |
| 2.014954616 | 0.043909405 | 0.082722595 |
| 2.873141001 | 0.004064127 | 0.010199027 |
| 0.192810118 | 0.847107679 | NA |
| 11.41585955 | 3.48E-30 | 1.21E-28 |
| NA | NA | NA |
| -0.065389369 | 0.947863988 | NA |
| 0.092267435 | 0.926485561 | 0.949628731 |

| | | |
|---|---|---|
| -2.327039828 | 0.019963146 | 0.041719757 |
| -1.405800325 | 0.159783425 | 0.245752682 |
| 0.933064776 | 0.350786524 | 0.462504567 |
| NA | NA | NA |
| 0.341383248 | 0.732815085 | NA |
| 3.671096998 | 0.000241512 | 0.000780508 |
| 9.50506157 | 2.00E-21 | 3.96E-20 |
| 7.073971084 | 1.51E-12 | 1.37E-11 |
| NA | NA | NA |
| 7.903374742 | 2.71E-15 | 3.26E-14 |
| 0.192809521 | 0.847108147 | NA |
| 1.87806376 | 0.060372443 | NA |
| 7.940197861 | 2.02E-15 | 2.45E-14 |
| NA | NA | NA |
| NA | NA | NA |
| NA | NA | NA |
| 2.742181354 | 0.006103263 | 0.014680442 |
| NA | NA | NA |
| 0.761044571 | 0.446630446 | NA |
| -0.657595072 | 0.510798362 | NA |
| -0.128191109 | 0.897997736 | NA |
| -1.070851703 | 0.284236115 | 0.391311257 |
| NA | NA | NA |
| NA | NA | NA |
| -5.276444801 | 1.32E-07 | 6.99E-07 |
| -0.708415481 | 0.478687282 | 0.587445708 |
| NA | NA | NA |
| 0.371978693 | 0.709908713 | NA |
| 0.063710673 | 0.949200606 | NA |
| 1.523656195 | 0.127594617 | 0.204383289 |
| -3.332541564 | 0.000860566 | 0.002512536 |
| NA | NA | NA |
| 2.426838695 | 0.015231026 | 0.033015726 |
| -1.24142562 | 0.21444856 | 0.312727071 |
| 0.846310298 | 0.397379664 | 0.509184652 |
| NA | NA | NA |
| NA | NA | NA |
| NA | NA | NA |
| NA | NA | NA |
| -1.391414757 | 0.164099692 | 0.251187335 |
| -0.246503929 | 0.805292163 | NA |
| NA | NA | NA |
| -1.095133647 | 0.27345809 | 0.380047528 |
| -2.574542227 | 0.010037283 | 0.022866406 |
| 3.351502683 | 0.000803743 | 0.002357356 |
| -4.6859193 | 2.79E-06 | 1.23E-05 |
| -1.463824031 | 0.143242035 | 0.224772624 |

| | | | |
|---|---|---|---|
| | -1.125852784 | 0.260227838 | 0.365168065 |
| | 3.578542882 | 0.000345515 | 0.001086873 |
| | -0.113081513 | 0.909965931 | NA |
| | -0.065389369 | 0.947863988 | NA |
| | -2.354245481 | 0.018560349 | 0.039180699 |
| | 4.113496324 | 3.90E-05 | 0.000144489 |
| NA | NA | NA | |
| | 1.84086295 | 0.065641645 | 0.11666622 |
| | 0.063707019 | 0.949203516 | NA |
| NA | NA | NA | |
| | 0.346199816 | 0.729192553 | 0.802342606 |
| | -0.105695914 | 0.915823622 | NA |
| | 1.512774059 | 0.13033706 | NA |
| | 10.4809926 | 1.06E-25 | 2.83E-24 |
| | 0.897833944 | 0.369274085 | 0.481848975 |
| | 0.163231756 | 0.870335964 | 0.909620827 |
| | 4.457814426 | 8.28E-06 | 3.42E-05 |
| | 0.650390817 | 0.515439807 | 0.620769782 |
| NA | NA | NA | |
| | 1.830428891 | 0.067185831 | 0.119038816 |
| | -0.065389369 | 0.947863988 | NA |
| | 0.216968119 | 0.828233191 | 0.87990988 |
| | 0.192809521 | 0.847108147 | NA |
| NA | NA | NA | |
| NA | NA | NA | |
| NA | NA | NA | |
| NA | NA | NA | |
| NA | NA | NA | |
| | 0.063707688 | 0.949202983 | NA |
| | -0.514729703 | 0.606741914 | 0.703509155 |
| | -0.400816848 | 0.688554973 | NA |
| | 0.139620454 | 0.888959879 | NA |
| NA | NA | NA | |
| NA | NA | NA | |
| NA | NA | NA | |
| | -1.060261033 | 0.289025862 | 0.39645535 |
| | -1.791126138 | 0.073273056 | 0.128020702 |
| NA | NA | NA | |
| | -3.74831114 | 0.000178029 | 0.00058935 |
| | -5.139993646 | 2.75E-07 | 1.40E-06 |
| | 0.659812309 | 0.509374283 | 0.615359733 |
| | 0.175315086 | 0.860832058 | NA |
| NA | NA | NA | |
| NA | NA | NA | |
| NA | NA | NA | |
| | 2.414676681 | 0.015749183 | 0.034021062 |
| | 0.192810118 | 0.847107679 | NA |

| | | |
|---|---|---|
| 3.145109084 | 0.001660249 | 0.004559082 |
| 3.657622839 | 0.000254565 | 0.000819415 |
| NA | NA | NA |
| 1.872547906 | 0.061130841 | 0.109764605 |
| -0.422536534 | 0.67263344 | 0.756606822 |
| 4.161546919 | 3.16E-05 | 0.000119024 |
| NA | NA | NA |
| NA | NA | NA |
| -0.15190362 | 0.879262953 | 0.915167653 |
| -0.400443486 | 0.6888299 | NA |
| NA | NA | NA |
| 0.946053944 | 0.344121073 | 0.455419577 |
| -2.849571415 | 0.004377818 | 0.010908589 |
| NA | NA | NA |
| -2.460291407 | 0.013882425 | 0.030451064 |
| 0.442411743 | 0.658191283 | 0.745757449 |
| -1.784294441 | 0.074375833 | 0.129702996 |
| -0.711199352 | 0.47696071 | 0.585757646 |
| -0.065389369 | 0.947863988 | NA |
| -0.398359952 | 0.690364874 | NA |
| NA | NA | NA |
| NA | NA | NA |
| NA | NA | NA |
| 4.78896848 | 1.68E-06 | 7.63E-06 |
| NA | NA | NA |
| -1.135551982 | 0.256144115 | NA |
| 1.514562427 | 0.129883253 | 0.2072479 |
| -0.355366594 | 0.722314972 | NA |
| 0.063708285 | 0.949202508 | NA |
| 18.58355209 | 4.37E-77 | 6.58E-75 |
| 1.472129714 | 0.140985853 | 0.221676983 |
| 10.36349999 | 3.63E-25 | 9.42E-24 |
| NA | NA | NA |
| 0.18411517 | 0.853923106 | 0.898469685 |
| -1.574453919 | 0.115382547 | 0.187607194 |
| NA | NA | NA |
| -0.029593502 | 0.976391248 | 0.983622029 |
| NA | NA | NA |
| -5.758281931 | 8.50E-09 | 5.23E-08 |
| -4.035219923 | 5.46E-05 | 0.000197657 |
| -2.825076361 | 0.004726935 | 0.011697003 |
| 4.838213074 | 1.31E-06 | 6.04E-06 |
| -0.338299766 | 0.7351373 | 0.806826564 |
| 0.808232553 | 0.418956721 | 0.53061209 |
| -4.050849238 | 5.10E-05 | 0.000185686 |
| -0.512223267 | 0.608494761 | 0.705023555 |
| 2.154604608 | 0.031192788 | 0.061418777 |

| | | | |
|---|---|---|---|
| 0.192808924 | 0.847108615 | NA | |
| NA | NA | NA | |
| NA | NA | NA | |
| 0.860931579 | 0.389275728 | NA | |
| -7.342774851 | 2.09E-13 | | 2.08E-12 |
| -5.243176777 | 1.58E-07 | | 8.29E-07 |
| -5.698775464 | 1.21E-08 | | 7.30E-08 |
| -2.644876553 | 0.008172072 | | 0.01898524 |
| -1.700453355 | 0.089045683 | | 0.15125982 |
| 0.339334084 | 0.734358069 | NA | |
| 0.063708882 | 0.949202033 | NA | |
| 0.132139668 | 0.89487382 | NA | |
| -0.432777777 | 0.66517622 | NA | |
| 4.691868723 | 2.71E-06 | | 1.20E-05 |
| 0.487076727 | 0.626203959 | NA | |
| 1.094254316 | 0.273843455 | | 0.380498595 |
| 0.983889626 | 0.325169785 | | 0.436220373 |
| -0.333010162 | 0.739126612 | | 0.810032799 |
| 0.163876323 | 0.869828507 | NA | |
| NA | NA | NA | |
| 1.622466486 | 0.104703503 | | 0.173019653 |
| -0.065389369 | 0.947863988 | NA | |
| 0.063708285 | 0.949202508 | NA | |
| -0.354758218 | 0.722770732 | | 0.797379735 |
| 0.156143159 | 0.875920179 | | 0.913407797 |
| 2.425816358 | 0.015273997 | | 0.033094542 |
| -1.940130644 | 0.052363815 | | 0.096344184 |
| 0.39390925 | 0.693648041 | NA | |
| -0.087493249 | 0.930279452 | NA | |
| -1.271394645 | 0.203588287 | | 0.29995922 |
| 2.231290259 | 0.025661909 | | 0.051849981 |
| 0.429052701 | 0.667884873 | | 0.752957317 |
| -0.792603179 | 0.428009056 | | 0.539293565 |
| 0.063707019 | 0.949203516 | NA | |
| 3.340613585 | 0.000835935 | | 0.002444899 |
| 1.892608596 | 0.05840994 | | 0.105681711 |
| -2.107976092 | 0.035033058 | | 0.067969726 |
| NA | NA | NA | |
| -1.598943144 | 0.109833236 | | 0.179904843 |
| -0.390480795 | 0.696181054 | NA | |
| 7.120947252 | 1.07E-12 | | 9.92E-12 |
| 7.435157202 | 1.04E-13 | | 1.07E-12 |
| 1.273589812 | 0.202808825 | | 0.299081181 |
| 0.771371737 | 0.440486623 | | 0.550641888 |
| -0.732159162 | 0.464071427 | | 0.573445291 |
| NA | NA | NA | |
| -2.802782649 | 0.00506638 | | 0.012452032 |

| | | |
|---|---|---|
| -3.899082643 | 9.66E-05 | 0.000334882 |
| NA | NA | NA |
| -10.60201234 | 2.92E-26 | 8.10E-25 |
| 0.063708285 | 0.949202508 | NA |
| -0.590913579 | 0.554578327 | 0.657497893 |
| -2.345682813 | 0.018992262 | 0.03994759 |
| -0.806329654 | 0.420052796 | NA |
| 1.054902744 | 0.291469807 | 0.399162295 |
| NA | NA | NA |
| 4.424720492 | 9.66E-06 | 3.95E-05 |
| 1.156379432 | 0.247525991 | 0.35070619 |
| -0.125764484 | 0.899918356 | 0.93020675 |
| 0.997278072 | 0.318629554 | 0.429240998 |
| -1.339173848 | 0.180514084 | 0.271791838 |
| 4.928820796 | 8.27E-07 | 3.93E-06 |
| -1.697702027 | 0.089564015 | 0.152026807 |
| -0.595306266 | 0.551638768 | 0.654881349 |
| NA | NA | NA |
| -0.248822858 | 0.803497809 | 0.86075541 |
| -3.910886187 | 9.20E-05 | 0.000320394 |
| NA | NA | NA |
| 0.919638788 | 0.35776155 | 0.469598385 |
| -3.127835644 | 0.001760986 | 0.004805657 |
| -3.291073771 | 0.000998057 | 0.002877376 |
| 0.192809521 | 0.847108147 | NA |
| NA | NA | NA |
| 0.063976859 | 0.948988653 | NA |
| NA | NA | NA |
| -0.32338754 | 0.746401764 | NA |
| 1.591996407 | 0.111385512 | 0.182066176 |
| -0.404178092 | 0.686081763 | NA |
| -1.105205238 | 0.269070668 | 0.375137237 |
| 0.063710673 | 0.949200606 | NA |
| -4.835358651 | 1.33E-06 | 6.12E-06 |
| -4.401286817 | 1.08E-05 | 4.37E-05 |
| 1.094109661 | 0.273906886 | 0.380562629 |
| NA | NA | NA |
| NA | NA | NA |
| NA | NA | NA |
| 5.364614421 | 8.11E-08 | 4.43E-07 |
| -1.88979064 | 0.058785966 | 0.106269878 |
| 2.498196969 | 0.012482681 | 0.027725754 |
| -1.990735846 | 0.046509936 | 0.086967256 |
| -2.982116942 | 0.002862626 | 0.007432732 |
| -2.430951605 | 0.015059226 | 0.032697122 |
| -3.283625386 | 0.00102481 | 0.002948053 |
| -0.405444011 | 0.685151166 | 0.767376664 |

| | | |
|---|---|---|
| NA | NA | NA |
| -2.970717906 | 0.002971045 | 0.007691898 |
| -0.692574274 | 0.48857676 | 0.596453021 |
| NA | NA | NA |
| NA | NA | NA |
| 0.48402001 | 0.628371668 | NA |
| 0.231370566 | 0.81702693 | NA |
| NA | NA | NA |
| NA | NA | NA |
| -4.227233072 | 2.37E-05 | 9.09E-05 |
| 3.598545577 | 0.000320002 | 0.001012469 |
| NA | NA | NA |
| 2.151263335 | 0.03145542 | 0.061867765 |
| -1.103069422 | 0.269997019 | 0.376156372 |
| NA | NA | NA |
| -0.19448762 | 0.845794089 | NA |
| NA | NA | NA |
| 1.794746978 | 0.072694022 | 0.127195482 |
| -0.638308828 | 0.523272667 | NA |
| -0.596423078 | 0.550892627 | 0.654243806 |
| -0.208503457 | 0.834835883 | 0.884334596 |
| NA | NA | NA |
| 0.207251126 | 0.835813741 | NA |
| NA | NA | NA |
| -0.358067529 | 0.720292778 | 0.795452382 |
| 1.314482441 | 0.188683903 | 0.281838241 |
| -1.09818981 | 0.272121615 | 0.378589518 |
| -0.548906267 | 0.583069777 | 0.683282762 |
| -0.502041731 | 0.615638167 | 0.710805304 |
| -2.085382548 | 0.03703459 | 0.071289378 |
| -3.447337724 | 0.00056614 | 0.001709554 |
| -2.961346394 | 0.003062972 | 0.007901288 |
| -0.307491907 | 0.758468982 | NA |
| -1.296624356 | 0.194760468 | 0.289137278 |
| 0.192808924 | 0.847108615 | NA |
| -2.719878801 | 0.006530585 | 0.015597379 |
| NA | NA | NA |
| -1.076325943 | 0.28178151 | 0.388830211 |
| -1.76183844 | 0.078096598 | 0.135304253 |
| -0.899076378 | 0.368611979 | 0.481140897 |
| -0.551091463 | 0.581570977 | 0.681973369 |
| -2.779713023 | 0.005440696 | 0.013267791 |
| 1.643886668 | 0.100199615 | 0.166799438 |
| 0.192810118 | 0.847107679 | NA |
| NA | NA | NA |
| 8.131491647 | 4.24E-16 | 5.44E-15 |
| NA | NA | NA |

| | | |
|---|---|---|
| -0.782226071 | 0.434081726 | 0.544814369 |
| 0.222689079 | 0.823777501 | NA |
| 2.224762341 | 0.026097197 | 0.052623285 |
| 5.593684021 | 2.22E-08 | 1.31E-07 |
| NA | NA | NA |
| 3.574552573 | 0.000350827 | 0.001101967 |
| 5.095702048 | 3.47E-07 | 1.74E-06 |
| -4.173927652 | 2.99E-05 | 0.000113268 |
| -1.938697117 | 0.05253823 | 0.096629576 |
| NA | NA | NA |
| NA | NA | NA |
| -2.108942604 | 0.034949536 | 0.067839231 |
| 0.570060066 | 0.568636959 | NA |
| 0.06371127 | 0.949200131 | NA |
| NA | NA | NA |
| 0.192809521 | 0.847108147 | NA |
| NA | NA | NA |
| 0.856137555 | 0.391921698 | 0.503815936 |
| -0.824412419 | 0.409705261 | 0.521533981 |
| 5.364003803 | 8.14E-08 | 4.44E-07 |
| 6.708639984 | 1.96E-11 | 1.60E-10 |
| 5.203520862 | 1.96E-07 | 1.01E-06 |
| 0.063708285 | 0.949202508 | NA |
| 1.729948705 | 0.08363944 | 0.143399903 |
| NA | NA | NA |
| 2.161307085 | 0.030671626 | 0.060506841 |
| -0.400135106 | 0.689057008 | 0.770510048 |
| 0.532170805 | 0.594607701 | NA |
| -0.16681616 | 0.867514703 | 0.907733498 |
| 7.692007099 | 1.45E-14 | 1.63E-13 |
| -0.155666632 | 0.8762958 | 0.913675076 |
| -1.492640108 | 0.135531419 | 0.214597129 |
| 5.393272291 | 6.92E-08 | 3.80E-07 |
| -0.276164091 | 0.782422037 | 0.844795893 |
| NA | NA | NA |
| 0.674748282 | 0.499835704 | NA |
| -0.065389369 | 0.947863988 | NA |
| -1.414225993 | 0.157295558 | 0.24277994 |
| 3.703337441 | 0.000212782 | 0.000695653 |
| -1.694408398 | 0.090187705 | 0.152878115 |
| -1.465511446 | 0.142781432 | 0.224232604 |
| -3.755502173 | 0.000172994 | 0.000573529 |
| 3.874808223 | 0.000106709 | 0.000367238 |
| 0.137791119 | 0.890405503 | NA |
| -0.194488217 | 0.845793621 | NA |
| NA | NA | NA |
| -0.083108651 | 0.933765147 | NA |

| | | | |
|---|---|---|---|
| | -0.088144457 | 0.929761862 | NA |
| | 0.386018564 | 0.699482921 | NA |
| NA | NA | NA | |
| NA | NA | NA | |
| NA | NA | NA | |
| | 0.090025187 | 0.9282672 | NA |
| | 1.019680032 | 0.307880235 | 0.41772779 |
| NA | NA | NA | |
| NA | NA | NA | |
| NA | NA | NA | |
| NA | NA | NA | |
| | -0.841378325 | 0.400136024 | 0.511825668 |
| | -4.235837498 | 2.28E-05 | 8.77E-05 |
| | -1.266887182 | 0.205195633 | NA |
| | 2.266043194 | 0.023448739 | 0.048006047 |
| NA | NA | NA | |
| NA | NA | NA | |
| NA | NA | NA | |
| | -5.024821264 | 5.04E-07 | 2.47E-06 |
| | -6.670596869 | 2.55E-11 | 2.06E-10 |
| NA | NA | NA | |
| | 0.063708285 | 0.949202508 | NA |
| | -0.065388772 | 0.947864464 | NA |
| | 5.292448661 | 1.21E-07 | 6.43E-07 |
| NA | NA | NA | |
| | -0.259755806 | 0.795052143 | NA |
| NA | NA | NA | |
| NA | NA | NA | |
| NA | NA | NA | |
| | -0.982339271 | 0.325932734 | 0.437056483 |
| | 0.176420747 | 0.859963406 | 0.902737966 |
| NA | NA | NA | |
| | 0.626083979 | 0.53125986 | 0.6359366 |
| NA | NA | NA | |
| | 0.255670233 | 0.798205482 | 0.856515614 |
| | -4.264406276 | 2.00E-05 | 7.79E-05 |
| | 1.211026436 | 0.225885273 | 0.325948829 |
| NA | NA | NA | |
| | -3.138641432 | 0.00169733 | 0.00464765 |
| | -3.341601663 | 0.000832965 | 0.002436783 |
| | -0.152338572 | 0.878919904 | NA |
| | -2.724848784 | 0.006433094 | 0.01538509 |
| | -0.446519282 | 0.655222187 | 0.743066075 |
| | -1.190560125 | 0.233826313 | 0.335127842 |
| | -1.438581744 | 0.150269062 | 0.233791311 |
| | -1.113448982 | 0.265515648 | 0.371156328 |
| | 1.312947183 | 0.189200746 | 0.282491928 |

| | | |
|---|---|---|
| 0.180277149 | 0.856934994 | 0.900654331 |
| 0.490623886 | 0.623692489 | NA |
| NA | NA | NA |
| NA | NA | NA |
| NA | NA | NA |
| -0.065389966 | 0.947863513 | NA |
| -0.19448762 | 0.845794089 | NA |
| 0.68955258 | 0.490475597 | NA |
| -0.875099315 | 0.38151987 | NA |
| -3.570030408 | 0.00035694 | 0.001119855 |
| -0.877350027 | 0.380296547 | 0.492845738 |
| -5.974538831 | 2.31E-09 | 1.51E-08 |
| -1.258357998 | 0.208262315 | 0.30530665 |
| -5.759010103 | 8.46E-09 | 5.21E-08 |
| -3.078236821 | 0.002082294 | 0.005585662 |
| 1.352009738 | 0.1763722 | 0.266694627 |
| 0.097648006 | 0.922211803 | 0.946488496 |
| -2.294009403 | 0.021789959 | 0.045007647 |
| NA | NA | NA |
| 0.192809521 | 0.847108147 | NA |
| NA | NA | NA |
| NA | NA | NA |
| NA | NA | NA |
| -0.089632422 | 0.928579319 | NA |
| NA | NA | NA |
| NA | NA | NA |
| NA | NA | NA |
| NA | NA | NA |
| NA | NA | NA |
| NA | NA | NA |
| NA | NA | NA |
| NA | NA | NA |
| NA | NA | NA |
| -3.571936467 | 0.000354351 | 0.001112695 |
| -4.254468793 | 2.10E-05 | 8.12E-05 |
| -4.226167064 | 2.38E-05 | 9.12E-05 |
| -1.267133192 | 0.205107669 | 0.301653737 |
| -8.458199112 | 2.72E-17 | 3.87E-16 |
| 0.31287131 | 0.754378436 | NA |
| -5.59416477 | 2.22E-08 | 1.30E-07 |
| -1.880997474 | 0.059972262 | 0.108024897 |
| NA | NA | NA |
| -0.138840954 | 0.889575831 | NA |
| 0.06353712 | 0.949338802 | NA |
| NA | NA | NA |
| NA | NA | NA |
| NA | NA | NA |

| | | | |
|---|---|---|---|
| NA | | NA | NA |
| | 0.85831628 | 0.390717842 | 0.502836526 |
| NA | | NA | NA |
| NA | | NA | NA |
| | 0.507823256 | 0.611577299 | 0.70742383 |
| NA | | NA | NA |
| NA | | NA | NA |
| | 0.063708285 | 0.949202508 | NA |
| NA | | NA | NA |
| NA | | NA | NA |
| | 0.372978763 | 0.70916425 | 0.786912319 |
| NA | | NA | NA |
| NA | | NA | NA |
| | -0.065389369 | 0.947863988 | NA |
| NA | | NA | NA |
| NA | | NA | NA |
| | 0.194173692 | 0.846039881 | NA |
| NA | | NA | NA |
| | 0.763297205 | 0.445286169 | NA |
| | 2.285422443 | 0.022288069 | 0.045941726 |
| NA | | NA | NA |
| NA | | NA | NA |
| | 1.184063358 | 0.236387983 | 0.338179654 |
| NA | | NA | NA |
| NA | | NA | NA |
| NA | | NA | NA |
| NA | | NA | NA |
| | -2.194820572 | 0.028176471 | 0.056209219 |
| | 0.192809521 | 0.847108147 | NA |
| | 7.313033728 | 2.61E-13 | 2.57E-12 |
| NA | | NA | NA |
| NA | | NA | NA |
| NA | | NA | NA |
| NA | | NA | NA |
| | 2.302198891 | 0.021323958 | 0.044165385 |
| | 0.448294001 | 0.653941029 | 0.741982954 |
| | 0.390204786 | 0.696385123 | NA |
| | 0.063708882 | 0.949202033 | NA |
| | 3.877401157 | 0.000105578 | 0.000363497 |
| | 1.220506081 | 0.222273086 | NA |
| | 0.31685483 | 0.751353758 | NA |
| | 1.896916069 | 0.057839016 | 0.104830599 |
| | 7.253004117 | 4.08E-13 | 3.94E-12 |
| | 0.535596484 | 0.592237459 | NA |
| | -0.065389369 | 0.947863988 | NA |
| NA | | NA | NA |
| | 8.693231002 | 3.52E-18 | 5.39E-17 |

| | | |
|---|---|---|
| 3.65146527 | 0.000260748 | 0.000837594 |
| 4.382217371 | 1.17E-05 | 4.73E-05 |
| -1.408868764 | 0.158873977 | 0.244609439 |
| NA | NA | NA |
| 0.389547452 | 0.696871215 | 0.7767149 |
| -0.065389369 | 0.947863988 | NA |
| 0.05051531 | 0.95971175 | 0.972258271 |
| 1.04724852 | 0.294984973 | 0.403292215 |
| -0.933440574 | 0.35059254 | 0.462307203 |
| -1.151380644 | 0.249575673 | 0.35305946 |
| 3.54491988 | 0.000392732 | 0.001221433 |
| -0.517599391 | 0.604737806 | 0.701742573 |
| 0.782855531 | 0.433711954 | 0.544451086 |
| 0.517920936 | 0.604513434 | 0.701677391 |
| -2.095982775 | 0.036083719 | 0.069684361 |
| -5.400911567 | 6.63E-08 | 3.66E-07 |
| -1.48900666 | 0.136485615 | 0.215834909 |
| -6.088779056 | 1.14E-09 | 7.71E-09 |
| NA | NA | NA |
| -0.065393533 | 0.947860673 | NA |
| NA | NA | NA |
| 4.295908661 | 1.74E-05 | 6.83E-05 |
| 1.481569163 | 0.138454969 | NA |
| -2.027218361 | 0.042640086 | 0.080604287 |
| 0.063707019 | 0.949203516 | NA |
| -3.743223637 | 0.000181674 | 0.000600701 |
| -0.065390563 | 0.947863038 | NA |
| 0.279356032 | 0.77997161 | NA |
| -3.182127427 | 0.001461975 | 0.004060951 |
| -0.986699646 | 0.323789904 | 0.434812023 |
| -2.444775486 | 0.014494234 | 0.031585467 |
| -2.76843236 | 0.005632668 | 0.013674653 |
| -4.517755366 | 6.25E-06 | 2.62E-05 |
| 0.123528044 | 0.901688969 | NA |
| -3.76658422 | 0.000165496 | 0.000551067 |
| -2.344643656 | 0.019045272 | 0.040044512 |
| -3.921766288 | 8.79E-05 | 0.000307461 |
| -3.322303774 | 0.000892774 | 0.002596879 |
| -3.205645148 | 0.001347601 | 0.003770871 |
| NA | NA | NA |
| 0.746959827 | 0.455087811 | NA |
| NA | NA | NA |
| 0.058718257 | 0.953176518 | 0.9680276 |
| NA | NA | NA |
| 1.712725544 | 0.086763044 | 0.147951237 |
| -0.215913321 | 0.829055313 | NA |
| -1.197030475 | 0.231294677 | 0.332127001 |

| | | |
|---|---|---|
| -0.014147969 | 0.988711931 | 0.992120736 |
| -0.55294476 | 0.580301236 | 0.681026817 |
| 0.425361177 | 0.670573403 | 0.755056436 |
| -0.585475517 | 0.558228024 | 0.66066705 |
| -0.368778751 | 0.712292645 | 0.789304296 |
| 1.43144609 | 0.152302407 | 0.236455108 |
| 0.084601341 | 0.932578333 | NA |
| NA | NA | NA |
| -2.90292995 | 0.003696893 | 0.009351515 |
| 0.163830799 | 0.869864346 | 0.909373729 |
| 4.147381696 | 3.36E-05 | 0.000126125 |
| -0.119993129 | 0.904488591 | NA |
| -2.586051629 | 0.009708239 | 0.022185448 |
| -2.384549316 | 0.017100056 | 0.036554309 |
| -1.858798772 | 0.063055667 | 0.112725036 |
| NA | NA | NA |
| -1.70186407 | 0.088780853 | 0.150863075 |
| NA | NA | NA |
| 0.249820475 | 0.802726185 | 0.860179769 |
| 9.975349979 | 1.95E-23 | 4.51E-22 |
| 0.430180397 | 0.667064421 | NA |
| -1.004547238 | 0.315114914 | 0.425651716 |
| -1.669136586 | 0.095090314 | 0.159846243 |
| -0.366884479 | 0.713705189 | NA |
| 0.192810118 | 0.847107679 | NA |
| -5.141199542 | 2.73E-07 | 1.39E-06 |
| NA | NA | NA |
| 2.909470005 | 0.003620421 | 0.009172907 |
| -0.135135571 | 0.892504687 | NA |
| 0.407418784 | 0.683700433 | NA |
| -0.604081634 | 0.545789369 | 0.649415664 |
| NA | NA | NA |
| NA | NA | NA |
| 3.211915247 | 0.001318533 | 0.003694903 |
| NA | NA | NA |
| 0.324874675 | 0.745275925 | NA |
| NA | NA | NA |
| -3.840612752 | 0.000122728 | 0.000417658 |
| 3.000659645 | 0.002693955 | 0.007037855 |
| -0.315856147 | 0.752111699 | 0.820276942 |
| NA | NA | NA |
| NA | NA | NA |
| 0.876945629 | 0.380516171 | 0.493042822 |
| 0.281528634 | 0.778304969 | NA |
| -0.202081667 | 0.839852879 | 0.887695518 |
| 3.056966785 | 0.002235891 | 0.005955849 |
| -4.38727044 | 1.15E-05 | 4.63E-05 |

| | | |
|---|---|---|
| -0.661816056 | 0.508089115 | 0.614335914 |
| 3.310250905 | 0.000932124 | 0.002702857 |
| 2.371445669 | 0.017718652 | 0.037683563 |
| 0.637330268 | 0.523909739 | 0.629093825 |
| -0.152968676 | 0.878422977 | 0.91476425 |
| -0.328439436 | 0.742579429 | 0.812749273 |
| -1.628083356 | 0.103507212 | 0.171382253 |
| 0.481443612 | 0.630201247 | 0.722448347 |
| 0.063707135 | 0.949203424 | NA |
| 0.398354886 | 0.690368607 | NA |
| -3.502713956 | 0.000460544 | 0.001412846 |
| -3.620458024 | 0.000294082 | 0.000936617 |
| -3.755258981 | 0.000173162 | 0.000573997 |
| 1.248110769 | 0.211990495 | 0.309846545 |
| 0.3474247 | 0.728272279 | NA |
| -4.937599114 | 7.91E-07 | 3.77E-06 |
| -4.634644221 | 3.58E-06 | 1.55E-05 |
| -0.080655098 | 0.935716247 | NA |
| 1.98438054 | 0.047213434 | 0.088098423 |
| 3.481977564 | 0.000497725 | 0.001519091 |
| -7.902776588 | 2.73E-15 | 3.27E-14 |
| -0.675877088 | 0.499118685 | 0.606247426 |
| 0.063708285 | 0.949202508 | NA |
| 0.194173692 | 0.846039881 | NA |
| -3.475814707 | 0.000509304 | 0.001551214 |
| -3.940946166 | 8.12E-05 | 0.000285317 |
| -2.41377472 | 0.015788221 | 0.034094956 |
| -2.227956954 | 0.025883386 | 0.052242653 |
| -3.371909629 | 0.000746489 | 0.002205717 |
| NA | NA | NA |
| 0.330493631 | 0.741027004 | NA |
| 0.192810118 | 0.847107679 | NA |
| 1.245076848 | 0.213103504 | 0.311109985 |
| 0.192809521 | 0.847108147 | NA |
| NA | NA | NA |
| 9.927203334 | 3.17E-23 | 7.21E-22 |
| 0.146896081 | 0.883214045 | 0.91821098 |
| 0.07400941 | 0.941002898 | NA |
| 13.41757555 | 4.77E-41 | 2.67E-39 |
| NA | NA | NA |
| 0.234680396 | 0.814456805 | 0.869592365 |
| NA | NA | NA |
| -1.515124792 | 0.129740804 | 0.207078965 |
| 2.586153446 | 0.009705371 | 0.022182946 |
| -2.685420532 | 0.00724386 | 0.017068042 |
| -0.31173911 | 0.755238803 | 0.822582388 |
| NA | NA | NA |

| | | |
|---|---|---|
| 0.539265769 | 0.589703485 | 0.689185144 |
| -0.212583892 | 0.831651536 | 0.882268287 |
| 0.083757469 | 0.933249264 | NA |
| -0.452326745 | 0.651033611 | NA |
| -4.189857554 | 2.79E-05 | 0.000106131 |
| -0.243847656 | 0.80734881 | NA |
| -0.065389966 | 0.947863513 | NA |
| -3.045551338 | 0.002322541 | 0.006158831 |
| 0.921791146 | 0.356637526 | NA |
| 0.461761044 | 0.64425269 | NA |
| -1.309384444 | 0.190404158 | 0.28400072 |
| -5.881098011 | 4.08E-09 | 2.60E-08 |
| -1.99593427 | 0.045941078 | 0.086012775 |
| NA | NA | NA |
| NA | NA | NA |
| NA | NA | NA |
| 2.625701411 | 0.008647063 | 0.01999963 |
| 8.646003367 | 5.33E-18 | 8.02E-17 |
| 2.654588933 | 0.007940511 | 0.018502232 |
| -0.796643797 | 0.425657934 | 0.537115249 |
| 5.376572365 | 7.59E-08 | 4.16E-07 |
| -0.743431304 | 0.457220598 | 0.566826725 |
| -0.742356375 | 0.457871443 | 0.567514493 |
| -2.400863639 | 0.01635643 | 0.035190085 |
| -0.352671519 | 0.724334714 | 0.79871943 |
| -1.516174063 | 0.129475345 | 0.20672579 |
| -0.992838168 | 0.320788826 | 0.431638766 |
| -0.009484645 | 0.992432461 | 0.994697654 |
| -0.050119931 | 0.960026817 | 0.972498734 |
| NA | NA | NA |
| 0.063707194 | 0.949203376 | NA |
| 1.490483407 | 0.136097176 | 0.215315866 |
| 0.06371127 | 0.949200131 | NA |
| -3.81862641 | 0.000134197 | 0.000453605 |
| -9.022294498 | 1.84E-19 | 3.15E-18 |
| 9.160321032 | 5.17E-20 | 9.24E-19 |
| 12.58605838 | 2.52E-36 | 1.16E-34 |
| 0.192809521 | 0.847108147 | NA |
| 3.324160156 | 0.000886852 | 0.002583557 |
| -9.509190427 | 1.92E-21 | 3.82E-20 |
| -3.261145224 | 0.001109632 | 0.003167304 |
| 7.897651377 | 2.84E-15 | 3.40E-14 |
| 1.274812739 | 0.202375531 | 0.298582938 |
| 0.192810118 | 0.847107679 | NA |
| 0.686742704 | 0.492244881 | 0.599498315 |
| NA | NA | NA |
| NA | NA | NA |

| | | | |
|---|---|---|---|
| | 0.063708285 | 0.949202508 | NA |
| NA | NA | NA | |
| NA | NA | NA | |
| NA | NA | NA | |
| NA | NA | NA | |
| NA | NA | NA | |
| | 0.67021965 | 0.502717779 | 0.609377016 |
| | 0.192809521 | 0.847108147 | NA |
| NA | NA | NA | |
| | 0.192809521 | 0.847108147 | NA |
| | 0.192809521 | 0.847108147 | NA |
| | 0.192809521 | 0.847108147 | NA |
| NA | NA | NA | |
| NA | NA | NA | |
| | -0.065388772 | 0.947864464 | NA |
| | 2.067390959 | 0.03869733 | 0.07404022 |
| NA | NA | NA | |
| | -0.546331727 | 0.584837931 | 0.684905882 |
| | 5.322943287 | 1.02E-07 | 5.50E-07 |
| | -0.138882492 | 0.889543006 | 0.922722688 |
| | -0.771223616 | 0.440574399 | 0.550717998 |
| | 0.063708882 | 0.949202033 | NA |
| | -0.410404521 | 0.68150923 | NA |
| NA | NA | NA | |
| | 0.442093262 | 0.65842172 | NA |
| | 0.106510417 | 0.915177391 | 0.941596746 |
| | -2.082719199 | 0.037276826 | 0.071672989 |
| | -5.39798271 | 6.74E-08 | 3.71E-07 |
| | 0.192809521 | 0.847108147 | NA |
| | 0.192810118 | 0.847107679 | NA |
| | -1.321012817 | 0.186497091 | 0.279171232 |
| | 1.046018901 | 0.295552306 | 0.403869314 |
| | 0.516221395 | 0.605699789 | 0.702644976 |
| | -1.144353269 | 0.25247716 | 0.356550627 |
| | -1.066188848 | 0.286338283 | NA |
| | 0.203595243 | 0.838669811 | NA |
| | 2.024131246 | 0.042956649 | 0.081135296 |
| | 2.887957805 | 0.003877519 | 0.0097768 |
| | 1.756218108 | 0.079051174 | 0.136694361 |
| | -1.074579706 | 0.282562942 | 0.389586341 |
| | -0.315442982 | 0.752425336 | NA |
| | 0.263457145 | 0.79219825 | NA |
| NA | NA | NA | |
| | 3.089180627 | 0.002007094 | 0.005408867 |
| | -7.032682757 | 2.03E-12 | 1.81E-11 |
| NA | NA | NA | |
| | -0.884439319 | 0.376459106 | 0.489156626 |

| | | | |
|---|---|---|---|
| NA | NA | NA | |
| NA | NA | NA | |
| | 3.050294768 | 0.002286169 | 0.006073963 |
| NA | NA | NA | |
| NA | NA | NA | |
| NA | NA | NA | |
| | 0.31685483 | 0.751353758 | NA |
| | -1.20065752 | 0.229884079 | 0.330480919 |
| | 3.859696219 | 0.000113528 | 0.000388622 |
| | 0.272458936 | 0.785269162 | 0.846794021 |
| | -1.369071121 | 0.170977044 | 0.259886769 |
| | 0.063708381 | 0.949202431 | NA |
| | 2.116955264 | 0.034263645 | 0.066683697 |
| | 0.324874675 | 0.745275925 | NA |
| | 0.279572028 | 0.77980587 | 0.842623939 |
| | 0.533617541 | 0.59360617 | NA |
| | 0.192809521 | 0.847108147 | NA |
| | 0.542125411 | 0.587732115 | NA |
| | 0.192809521 | 0.847108147 | NA |
| | 0.063708217 | 0.949202562 | NA |
| | 0.352770413 | 0.724260567 | NA |
| | 0.414990094 | 0.678149155 | NA |
| NA | NA | NA | |
| NA | NA | NA | |
| | -0.457076399 | 0.647616133 | NA |
| NA | NA | NA | |
| | 0.436933212 | 0.662159785 | NA |
| | 0.573867431 | 0.566057514 | 0.668038077 |
| | 0.317407179 | 0.75093466 | 0.819243105 |
| | -0.468069461 | 0.639734917 | NA |
| NA | NA | NA | |
| | 0.212191773 | 0.831957425 | 0.882443188 |
| NA | NA | NA | |
| | -0.898889131 | 0.368711718 | 0.481220894 |
| | 0.063708285 | 0.949202508 | NA |
| NA | NA | NA | |
| | 0.192809521 | 0.847108147 | NA |
| NA | NA | NA | |
| | -2.176625541 | 0.029508518 | 0.058530644 |
| | -5.497063993 | 3.86E-08 | 2.20E-07 |
| | -2.899868696 | 0.00373319 | 0.009434748 |
| | 1.167590246 | 0.242972087 | 0.34565338 |
| | 7.51956112 | 5.50E-14 | 5.80E-13 |
| | 1.002842034 | 0.315937084 | 0.426486098 |
| | 0.446789653 | 0.655026942 | NA |
| | -1.547619446 | 0.121713947 | 0.196320408 |
| | 4.050036759 | 5.12E-05 | 0.000186251 |

| | | |
|---|---|---|
| NA | NA | NA |
| 5.611597843 | 2.00E-08 | 1.18E-07 |
| 1.119908946 | 0.262752566 | 0.368112432 |
| NA | NA | NA |
| 0.198201446 | 0.842887456 | NA |
| -2.200093006 | 0.027800297 | 0.055578389 |
| -3.542164776 | 0.000396857 | 0.001232578 |
| NA | NA | NA |
| 0.58653885 | 0.557513462 | NA |
| 0.418180567 | 0.675815103 | NA |
| 0.714281511 | 0.475053122 | 0.583987999 |
| 3.903480605 | 9.48E-05 | 0.000329537 |
| 0.535423386 | 0.592357122 | 0.691511533 |
| 1.281317436 | 0.200082191 | 0.295827116 |
| -2.075475579 | 0.03794248 | 0.072807442 |
| NA | NA | NA |
| NA | NA | NA |
| -1.338509438 | 0.180730426 | 0.272057234 |
| NA | NA | NA |
| NA | NA | NA |
| 0.192809521 | 0.847108147 | NA |
| NA | NA | NA |
| NA | NA | NA |
| 10.86608541 | 1.67E-27 | 5.03E-26 |
| -0.403307618 | 0.686721938 | NA |
| NA | NA | NA |
| 0.086946268 | 0.930714222 | NA |
| NA | NA | NA |
| -2.484060182 | 0.012989385 | 0.028728843 |
| -5.349142634 | 8.84E-08 | 4.80E-07 |
| -0.431715065 | 0.665948517 | 0.75168845 |
| NA | NA | NA |
| 3.003573911 | 0.002668287 | 0.006976625 |
| 0.523290525 | 0.600772093 | 0.698468271 |
| -2.229952358 | 0.025750606 | 0.051987229 |
| -2.05235781 | 0.040134902 | 0.076445977 |
| -1.943827637 | 0.05191624 | 0.095640206 |
| -0.06620969 | 0.947210882 | NA |
| NA | NA | NA |
| NA | NA | NA |
| 8.661852118 | 4.64E-18 | 7.02E-17 |
| 2.179594331 | 0.029287545 | 0.058143025 |
| NA | NA | NA |
| 0.195316213 | 0.845145406 | NA |
| 0.063708882 | 0.949202033 | NA |
| -5.286918619 | 1.24E-07 | 6.62E-07 |
| -2.342003372 | 0.019180542 | 0.040292821 |

| | | |
|---|---|---|
| -2.335119292 | 0.01953719 | 0.040928679 |
| -3.261644492 | 0.00110768 | 0.003162453 |
| NA | NA | NA |
| 1.033215793 | 0.301502924 | 0.410610925 |
| -0.469657952 | 0.638599414 | NA |
| NA | NA | NA |
| -0.065389369 | 0.947863988 | NA |
| NA | NA | NA |
| -2.339735861 | 0.019297382 | 0.04051107 |
| -2.230874664 | 0.025689432 | 0.051901404 |
| -2.886130613 | 0.003900103 | 0.009828793 |
| 3.511489384 | 0.000445603 | 0.001371382 |
| -0.194488217 | 0.845793621 | NA |
| 3.303870603 | 0.000953598 | 0.00276001 |
| -0.090348778 | 0.928010059 | 0.950801956 |
| 7.680176883 | 1.59E-14 | 1.77E-13 |
| 0.065152415 | 0.948052648 | NA |
| 0.144240189 | 0.885310806 | NA |
| -5.559301069 | 2.71E-08 | 1.57E-07 |
| 8.805596576 | 1.30E-18 | 2.08E-17 |
| -1.541782672 | 0.123126414 | 0.1982809 |
| -0.100664057 | 0.919817145 | 0.944921943 |
| -0.144847491 | 0.884831285 | NA |
| 0.349291458 | 0.726870511 | NA |
| -5.663138257 | 1.49E-08 | 8.91E-08 |
| NA | NA | NA |
| NA | NA | NA |
| NA | NA | NA |
| -1.742952777 | 0.081341864 | 0.140074775 |
| -3.425636667 | 0.00061336 | 0.001839493 |
| 0.192810118 | 0.847107679 | NA |
| NA | NA | NA |
| 0.192810118 | 0.847107679 | NA |
| 3.696045515 | 0.000218984 | 0.000714065 |
| -2.400212135 | 0.016385573 | 0.035242821 |
| NA | NA | NA |
| NA | NA | NA |
| 1.342918535 | 0.179298354 | 0.270319106 |
| 0.397517277 | 0.690986048 | NA |
| 0.346721347 | 0.728800672 | NA |
| NA | NA | NA |
| NA | NA | NA |
| NA | NA | NA |
| NA | NA | NA |
| NA | NA | NA |
| -0.092432603 | 0.926354337 | NA |
| NA | NA | NA |

| | | | |
|---|---|---|---|
| NA | NA | NA | |
| NA | NA | NA | |
| NA | NA | NA | |
| NA | NA | NA | |
| NA | NA | NA | |
| NA | NA | NA | |
| NA | NA | NA | |
| NA | NA | NA | |
| NA | NA | NA | |
| | -1.495434913 | 0.134800974 | 0.213589187 |
| | -5.231057352 | 1.69E-07 | 8.81E-07 |
| | -9.867871101 | 5.74E-23 | 1.27E-21 |
| | -4.710791976 | 2.47E-06 | 1.10E-05 |
| | -2.172997542 | 0.029780502 | 0.058995328 |
| | -2.711063787 | 0.006706772 | 0.015970936 |
| NA | NA | NA | |
| | -2.133160987 | 0.032911522 | 0.064422883 |
| NA | NA | NA | |
| | -2.76176437 | 0.005748995 | 0.013919264 |
| NA | NA | NA | |
| NA | NA | NA | |
| NA | NA | NA | |
| NA | NA | NA | |
| NA | NA | NA | |
| | -1.183499113 | 0.236611398 | 0.338460584 |
| | -2.515117577 | 0.011899269 | 0.026578503 |
| | 2.787393854 | 0.005313386 | 0.012985164 |
| | 7.462973434 | 8.46E-14 | 8.75E-13 |
| NA | NA | NA | |
| | 4.570495483 | 4.87E-06 | 2.07E-05 |
| | -7.606582112 | 2.81E-14 | 3.06E-13 |
| | -4.738591159 | 2.15E-06 | 9.65E-06 |
| | -0.095387118 | 0.924007347 | NA |
| NA | NA | NA | |
| | 0.063708882 | 0.949202033 | NA |
| NA | NA | NA | |
| | -1.081197705 | 0.279609179 | 0.38659438 |
| NA | NA | NA | |
| | -0.065388772 | 0.947864464 | NA |
| NA | NA | NA | |
| | -4.576771124 | 4.72E-06 | 2.02E-05 |
| | -3.129897995 | 0.00174867 | 0.004776213 |
| | 0.063708285 | 0.949202508 | NA |
| | 0.112375181 | 0.910525933 | NA |
| NA | NA | NA | |
| | 0.063708285 | 0.949202508 | NA |
| | 0.063708882 | 0.949202033 | NA |

| | | | |
|---|---|---|---|
| NA | NA | NA | |
| NA | NA | NA | |
| NA | NA | NA | |
| NA | NA | NA | |
| | 0.063708882 | 0.949202033 | NA |
| NA | NA | NA | |
| | 1.269326677 | 0.204324576 | 0.300820194 |
| | 0.224192989 | 0.822607138 | 0.875642454 |
| NA | NA | NA | |
| NA | NA | NA | |
| | 0.661850706 | 0.508066906 | NA |
| | 0.192809521 | 0.847108147 | NA |
| | 0.052592482 | 0.958056607 | 0.971013803 |
| | 2.341006846 | 0.019231814 | 0.040385701 |
| NA | NA | NA | |
| | -2.119106427 | 0.034081474 | 0.066373327 |
| NA | NA | NA | |
| | 0.486371394 | 0.626703869 | NA |
| | 0.506610187 | 0.612428359 | 0.708160123 |
| | 0.483110698 | 0.629017139 | NA |
| | 8.882733467 | 6.52E-19 | 1.07E-17 |
| | 0.842890443 | 0.399289719 | 0.511187237 |
| | 0.589882582 | 0.555269372 | 0.657974359 |
| | 0.063708285 | 0.949202508 | NA |
| | -1.003631445 | 0.315556292 | 0.42605295 |
| | 0.225443949 | 0.821633925 | 0.874866939 |
| | -0.510009911 | 0.610044518 | 0.706230715 |
| | 0.240803097 | 0.809707729 | 0.865592571 |
| | -7.59770916 | 3.01E-14 | 3.27E-13 |
| | -2.939867178 | 0.00328353 | 0.008406991 |
| | -5.818355431 | 5.94E-09 | 3.72E-08 |
| NA | NA | NA | |
| | -0.067996262 | 0.945788609 | NA |
| | -6.215662199 | 5.11E-10 | 3.59E-09 |
| | -0.51762314 | 0.604721234 | 0.701742573 |
| | 0.063708882 | 0.949202033 | NA |
| | -0.19448762 | 0.845794089 | NA |
| | -0.065390563 | 0.947863038 | NA |
| | -0.743132013 | 0.45740176 | NA |
| | 1.000949022 | 0.316851455 | 0.427351649 |
| | 0.959446408 | 0.337333906 | NA |
| | -6.849043071 | 7.43E-12 | 6.28E-11 |
| NA | NA | NA | |
| | -4.662098208 | 3.13E-06 | 1.37E-05 |
| NA | NA | NA | |
| | 0.289058872 | 0.772536326 | 0.836393626 |
| | 0.407334021 | 0.683762679 | NA |

| | | |
|---|---|---|
| 2.454090172 | 0.014124156 | 0.030902764 |
| 0.23363542 | 0.815268027 | NA |
| NA | NA | NA |
| NA | NA | NA |
| NA | NA | NA |
| 0.889987609 | 0.373472539 | 0.486242791 |
| NA | NA | NA |
| -1.197811616 | 0.230990364 | 0.331766156 |
| 6.784106524 | 1.17E-11 | 9.74E-11 |
| -0.535662957 | 0.592191509 | 0.691414939 |
| -0.168414104 | 0.866257513 | NA |
| -6.371336904 | 1.87E-10 | 1.38E-09 |
| -10.74810988 | 6.05E-27 | 1.75E-25 |
| -0.3958386 | 0.692224099 | NA |
| -8.161998156 | 3.30E-16 | 4.28E-15 |
| 3.387782391 | 0.000704601 | 0.002090337 |
| NA | NA | NA |
| -0.207626023 | 0.835520984 | NA |
| 0.063708882 | 0.949202033 | NA |
| NA | NA | NA |
| NA | NA | NA |
| NA | NA | NA |
| 1.434051253 | 0.151557637 | 0.235532576 |
| -0.48856609 | 0.625148925 | 0.718755147 |
| -0.590997875 | 0.554521845 | 0.65746207 |
| -0.145218395 | 0.884538443 | NA |
| 1.311627948 | 0.189645698 | 0.283071849 |
| 6.052573653 | 1.43E-09 | 9.55E-09 |
| 1.644971552 | 0.099975678 | 0.166515252 |
| 4.413230295 | 1.02E-05 | 4.15E-05 |
| -4.49282195 | 7.03E-06 | 2.93E-05 |
| -7.187356135 | 6.61E-13 | 6.26E-12 |
| 0.909206422 | 0.363241175 | 0.475643558 |
| 0.277841331 | 0.781134162 | NA |
| 0.06371127 | 0.949200131 | NA |
| NA | NA | NA |
| NA | NA | NA |
| -2.489447586 | 0.012794178 | 0.028332149 |
| -0.15973075 | 0.873093177 | 0.911703055 |
| NA | NA | NA |
| 18.09741762 | 3.34E-73 | 4.57E-71 |
| -4.11362146 | 3.89E-05 | 0.000144454 |
| 2.883005062 | 0.003939011 | 0.009918858 |
| 0.605015803 | 0.545168497 | 0.648985422 |
| -5.074466965 | 3.89E-07 | 1.94E-06 |
| -0.838814786 | 0.401573252 | NA |
| 0.063708285 | 0.949202508 | NA |

| | | |
|---|---|---|
| 4.524150473 | 6.06E-06 | 2.55E-05 |
| 1.646516151 | 0.099657538 | 0.166129082 |
| -1.87176541 | 0.061239066 | 0.109926407 |
| -0.90830629 | 0.36371642 | 0.476166222 |
| -1.782576513 | 0.074655266 | 0.130117771 |
| -0.094510242 | 0.924703847 | 0.948190794 |
| 6.743479072 | 1.55E-11 | 1.27E-10 |
| -5.019906752 | 5.17E-07 | 2.52E-06 |
| -0.391302273 | 0.695573819 | 0.775821607 |
| NA | NA | NA |
| NA | NA | NA |
| NA | NA | NA |
| -0.756090548 | 0.449594909 | NA |
| -2.087895431 | 0.036807268 | 0.070933626 |
| NA | NA | NA |
| -0.723301644 | 0.469494583 | 0.578630459 |
| -1.986047814 | 0.047028015 | 0.087798256 |
| -0.086413875 | 0.931137418 | 0.953147782 |
| -3.154089972 | 0.001609994 | 0.004435681 |
| -1.597315212 | 0.110195462 | 0.180427318 |
| -0.465130708 | 0.641837856 | NA |
| NA | NA | NA |
| NA | NA | NA |
| 0.13888062 | 0.889544485 | 0.922722688 |
| NA | NA | NA |
| NA | NA | NA |
| NA | NA | NA |
| -0.298865708 | 0.765042513 | 0.830615689 |
| -0.065390563 | 0.947863038 | NA |
| -0.953580068 | 0.340296249 | 0.4518389 |
| 5.319803129 | 1.04E-07 | 5.59E-07 |
| 0.289915183 | 0.771881126 | 0.835865042 |
| -0.344259417 | 0.730651201 | 0.803396163 |
| 0.875010488 | 0.381568199 | 0.494109991 |
| NA | NA | NA |
| NA | NA | NA |
| -8.395836427 | 4.63E-17 | 6.47E-16 |
| NA | NA | NA |
| NA | NA | NA |
| -1.620026979 | 0.105226482 | 0.173700378 |
| -0.750804466 | 0.452770341 | 0.562675222 |
| 1.960872033 | 0.049893952 | 0.092438611 |
| 5.29124475 | 1.21E-07 | 6.48E-07 |
| -0.701369681 | 0.483072338 | 0.591493035 |
| -3.037203903 | 0.002387839 | 0.006313924 |
| 0.192809521 | 0.847108147 | NA |
| NA | NA | NA |

| | | | |
|---|---|---|---|
| | 0.063708882 | 0.949202033 | NA |
| | 2.169539387 | 0.030041757 | NA |
| | 8.330824908 | 8.03E-17 | 1.10E-15 |
| NA | NA | NA | |
| NA | NA | NA | |
| NA | NA | NA | |
| NA | NA | NA | |
| | 7.99856654 | 1.26E-15 | 1.56E-14 |
| | 1.124956346 | 0.260607533 | NA |
| | 3.800677849 | 0.000144301 | 0.000484877 |
| | -8.379160999 | 5.33E-17 | 7.41E-16 |
| | 5.435753369 | 5.46E-08 | 3.04E-07 |
| | 4.379039971 | 1.19E-05 | 4.80E-05 |
| | 1.883496282 | 0.059633143 | 0.107521796 |
| | 6.511905085 | 7.42E-11 | 5.72E-10 |
| | 0.372516892 | 0.709508038 | NA |
| NA | NA | NA | |
| NA | NA | NA | |
| | -4.821584725 | 1.42E-06 | 6.53E-06 |
| | 0.739744372 | 0.459455119 | 0.568836208 |
| | 7.56918462 | 3.76E-14 | 4.03E-13 |
| | 1.269126562 | 0.204395928 | 0.300888773 |
| | 6.816262846 | 9.34E-12 | 7.84E-11 |
| | 8.988582317 | 2.50E-19 | 4.24E-18 |
| | 2.406980901 | 0.016085011 | 0.034662749 |
| | 6.062318766 | 1.34E-09 | 9.00E-09 |
| | 3.708962713 | 0.00020811 | 0.000681716 |
| | -2.792335557 | 0.005232906 | 0.012810997 |
| | 6.042999734 | 1.51E-09 | 1.01E-08 |
| | 7.754815846 | 8.85E-15 | 1.01E-13 |
| NA | NA | NA | |
| | 11.5484592 | 7.52E-31 | 2.71E-29 |
| | -11.06195192 | 1.92E-28 | 6.11E-27 |
| | 2.095542506 | 0.036122794 | 0.069754169 |
| | 1.617056889 | 0.105866002 | 0.174523641 |
| | 3.178388404 | 0.001480962 | 0.004107766 |
| | 1.090814572 | 0.275354483 | 0.382049078 |
| | -0.065389369 | 0.947863988 | NA |
| | 4.919826109 | 8.66E-07 | 4.10E-06 |
| | -5.091771641 | 3.55E-07 | 1.78E-06 |
| | -1.336756529 | 0.181302123 | 0.27276505 |
| | -3.237501877 | 0.001205811 | 0.003412648 |
| | -6.728135105 | 1.72E-11 | 1.41E-10 |
| | -1.890271768 | 0.058721623 | 0.106176567 |
| | -0.376248026 | 0.706732514 | 0.784875092 |
| | -0.802321 | 0.422367298 | 0.53409384 |
| | -1.603071365 | 0.108918899 | 0.178664398 |

| | | |
|---|---|---|
| 4.585246833 | 4.53E-06 | 1.94E-05 |
| -0.992658012 | 0.320876643 | 0.431710513 |
| 0.375587794 | 0.707223367 | 0.785246013 |
| 0.401946135 | 0.687723669 | 0.76956918 |
| 8.113472235 | 4.92E-16 | 6.29E-15 |
| 6.763084909 | 1.35E-11 | 1.12E-10 |
| 1.578877546 | 0.114364147 | 0.186120586 |
| 5.261823046 | 1.43E-07 | 7.53E-07 |
| -5.461539159 | 4.72E-08 | 2.66E-07 |
| 0.582242746 | 0.560403184 | 0.662739981 |
| 0.192810118 | 0.847107679 | NA |
| 1.001455203 | 0.316606787 | 0.427136737 |
| -0.065388772 | 0.947864464 | NA |
| 1.668663518 | 0.095184083 | 0.159934394 |
| 4.439539591 | 9.02E-06 | 3.70E-05 |
| 4.337971578 | 1.44E-05 | 5.71E-05 |
| 2.736323096 | 0.006212999 | 0.014918589 |
| 2.415496275 | 0.015713783 | 0.033956315 |
| NA | NA | NA |
| NA | NA | NA |
| 7.054828106 | 1.73E-12 | 1.56E-11 |
| -2.86606009 | 0.004156153 | 0.010410139 |
| -0.169820482 | 0.865151319 | 0.906032769 |
| -0.444413328 | 0.656743779 | NA |
| 0.412959025 | 0.679636634 | 0.762496861 |
| 0.124648059 | 0.90080218 | 0.930851232 |
| -4.201878571 | 2.65E-05 | 0.000101063 |
| 6.387491067 | 1.69E-10 | 1.25E-09 |
| 3.086980879 | 0.002022006 | 0.005445467 |
| -0.77034168 | 0.441097238 | 0.551206384 |
| -0.514646524 | 0.606800048 | 0.703509155 |
| NA | NA | NA |
| -0.021025076 | 0.983225653 | 0.988200212 |
| 0.697597588 | 0.48542889 | 0.593768515 |
| -0.282557441 | 0.777516108 | 0.840585185 |
| 3.902499682 | 9.52E-05 | 0.000330738 |
| -2.999820147 | 0.002701391 | 0.007052129 |
| -0.472667309 | 0.636450553 | 0.727658677 |
| -8.196464593 | 2.48E-16 | 3.26E-15 |
| -1.167606978 | 0.242965334 | 0.34565338 |
| -1.212950122 | 0.225148891 | 0.325167431 |
| -1.492977604 | 0.135443049 | 0.214497914 |
| -0.266395207 | 0.789934856 | NA |
| -0.813006899 | 0.416214103 | 0.527832953 |
| 3.149245511 | 0.001636926 | 0.004500468 |
| -0.070610168 | 0.943708018 | NA |
| -3.052923544 | 0.002266237 | 0.006027405 |

| | | |
|---|---|---|
| -4.880556155 | 1.06E-06 | 4.95E-06 |
| NA | NA | NA |
| NA | NA | NA |
| NA | NA | NA |
| -7.675976645 | 1.64E-14 | 1.83E-13 |
| 0.378762978 | 0.704863882 | 0.783495113 |
| -0.586284151 | 0.55768458 | 0.660179958 |
| NA | NA | NA |
| -1.55754762 | 0.119340524 | 0.193002087 |
| -0.552649836 | 0.580503209 | 0.681231906 |
| -1.761016255 | 0.078235652 | 0.135480723 |
| 4.537676346 | 5.69E-06 | 2.40E-05 |
| -1.307264855 | 0.191122776 | 0.284868879 |
| -4.712556362 | 2.45E-06 | 1.09E-05 |
| NA | NA | NA |
| 10.82912926 | 2.51E-27 | 7.44E-26 |
| 0.353799914 | 0.723488838 | 0.798031272 |
| 3.879499099 | 0.000104672 | 0.000360543 |
| 0.064556667 | 0.948526988 | 0.964950029 |
| 0.34140522 | 0.732798546 | NA |
| 7.224635225 | 5.02E-13 | 4.83E-12 |
| -3.697633861 | 0.000217618 | 0.000710261 |
| 7.194888094 | 6.25E-13 | 5.93E-12 |
| 4.132706991 | 3.59E-05 | 0.000133835 |
| 1.805695803 | 0.070965869 | 0.124747222 |
| 2.688799093 | 0.007170956 | 0.016915756 |
| NA | NA | NA |
| -3.62932056 | 0.000284168 | 0.00090712 |
| 1.450126664 | 0.147023201 | 0.229526022 |
| 2.072055695 | 0.038260247 | 0.073299196 |
| 0.392764722 | 0.694493263 | 0.774961721 |
| 4.451110705 | 8.54E-06 | 3.52E-05 |
| 0.926133868 | 0.354376415 | 0.466255344 |
| -3.576158105 | 0.000348681 | 0.001096142 |
| 8.185291381 | 2.72E-16 | 3.56E-15 |
| 0.068009815 | 0.945777821 | 0.962975181 |
| -0.357972434 | 0.720363943 | NA |
| 0.713327604 | 0.47564306 | 0.584512264 |
| 2.920719195 | 0.003492244 | 0.008885928 |
| 9.701560289 | 2.97E-22 | 6.23E-21 |
| 0.495355602 | 0.620349128 | 0.71475115 |
| 10.14529523 | 3.48E-24 | 8.43E-23 |
| 0.153088314 | 0.878328631 | 0.91476425 |
| -4.710613667 | 2.47E-06 | 1.10E-05 |
| 11.07078306 | 1.74E-28 | 5.56E-27 |
| -8.065990942 | 7.26E-16 | 9.20E-15 |
| -1.433862042 | 0.151611636 | 0.235601865 |

| | | |
|---|---|---|
| 1.069208025 | 0.284975943 | 0.392102048 |
| NA | NA | NA |
| 5.601648342 | 2.12E-08 | 1.25E-07 |
| 17.74102892 | 2.02E-70 | 2.61E-68 |
| 1.443820304 | 0.148789529 | 0.231835279 |
| -4.112675435 | 3.91E-05 | 0.00014494 |
| 0.458102228 | 0.646878998 | NA |
| 0.192809521 | 0.847108147 | NA |
| 7.000108391 | 2.56E-12 | 2.26E-11 |
| -0.84088214 | 0.400413965 | 0.512076515 |
| 1.106226207 | 0.268628622 | 0.374604402 |
| -2.808001902 | 0.004984993 | 0.012274902 |
| -2.087981249 | 0.036799526 | 0.070929627 |
| 3.328956266 | 0.000871721 | 0.002541842 |
| -1.902421389 | 0.057116087 | 0.103753223 |
| 0.3474247 | 0.728272279 | NA |
| 8.439692349 | 3.18E-17 | 4.50E-16 |
| -0.296985435 | 0.766477629 | 0.831524577 |
| 0.405406182 | 0.685178967 | NA |
| 0.880152639 | 0.378776626 | 0.491359366 |
| NA | NA | NA |
| 4.603332125 | 4.16E-06 | 1.79E-05 |
| 8.653857582 | 4.98E-18 | 7.51E-17 |
| 3.946405752 | 7.93E-05 | 0.000279559 |
| 1.438141232 | 0.150393986 | 0.233971127 |
| 12.35684352 | 4.47E-35 | 1.96E-33 |
| 4.075756025 | 4.59E-05 | 0.00016791 |
| -3.542449942 | 0.000396429 | 0.001231399 |
| NA | NA | NA |
| 0.346471969 | 0.728988047 | 0.802188113 |
| -0.079714773 | 0.93646411 | 0.956683265 |
| -0.221960535 | 0.824344606 | 0.876997021 |
| 2.095981191 | 0.036083859 | 0.069684361 |
| 5.196140427 | 2.03E-07 | 1.05E-06 |
| 0.054528412 | 0.956514173 | 0.970275572 |
| 0.192809521 | 0.847108147 | NA |
| -0.703311897 | 0.481861398 | 0.59039673 |
| -0.710247196 | 0.477550858 | 0.586367328 |
| 3.141823735 | 0.001678991 | 0.004601461 |
| 1.519603548 | 0.128610646 | 0.205671997 |
| 5.944499742 | 2.77E-09 | 1.80E-08 |
| -0.22202802 | 0.824292071 | 0.876988461 |
| 9.624629506 | 6.29E-22 | 1.30E-20 |
| NA | NA | NA |
| NA | NA | NA |
| 4.89302014 | 9.93E-07 | 4.67E-06 |
| -1.020452718 | 0.307513803 | NA |

| | | | |
|---|---|---|---|
| NA | NA | NA | |
| | 7.386019847 | 1.51E-13 | 1.52E-12 |
| | -0.240039146 | 0.81029991 | 0.866003704 |
| | 14.51355967 | 9.94E-48 | 7.23E-46 |
| NA | NA | NA | |
| | 4.019337024 | 5.84E-05 | 0.000210206 |
| | 6.79564804 | 1.08E-11 | 9.01E-11 |
| | 6.69046569 | 2.22E-11 | 1.80E-10 |
| NA | NA | NA | |
| NA | NA | NA | |
| | 0.063708285 | 0.949202508 | NA |
| | -0.350046437 | 0.726303848 | 0.800013835 |
| NA | NA | NA | |
| NA | NA | NA | |
| NA | NA | NA | |
| NA | NA | NA | |
| | -0.106781367 | 0.91496243 | NA |
| NA | NA | NA | |
| NA | NA | NA | |
| | 4.457184206 | 8.30E-06 | 3.43E-05 |
| | -0.10595854 | 0.915615248 | NA |
| NA | NA | NA | |
| | -1.290318233 | 0.196940189 | 0.29186333 |
| NA | NA | NA | |
| NA | NA | NA | |
| NA | NA | NA | |
| NA | NA | NA | |
| NA | NA | NA | |
| | -0.409544633 | 0.68214002 | NA |
| NA | NA | NA | |
| NA | NA | NA | |
| | 0.324874675 | 0.745275925 | NA |
| | -0.065388772 | 0.947864464 | NA |
| | 1.199331642 | 0.230399016 | NA |
| NA | NA | NA | |
| NA | NA | NA | |
| | 0.46503342 | 0.641907524 | NA |
| NA | NA | NA | |
| NA | NA | NA | |
| NA | NA | NA | |
| | 0.218327194 | 0.827174188 | NA |
| NA | NA | NA | |
| | 0.192810118 | 0.847107679 | NA |
| NA | NA | NA | |
| NA | NA | NA | |
| NA | NA | NA | |
| | 3.051714476 | 0.002275384 | 0.006049503 |

| | | | |
|---|---|---|---|
| NA | NA | NA | |
| -0.065390638 | 0.947862978 | NA | |
| NA | NA | NA | |
| NA | NA | NA | |
| 6.14263137 | 8.12E-10 | | 5.57E-09 |
| NA | NA | NA | |
| -0.078080006 | 0.937764412 | NA | |
| NA | NA | NA | |
| NA | NA | NA | |
| NA | NA | NA | |
| 0.063708381 | 0.949202431 | NA | |
| 0.063706666 | 0.949203797 | NA | |
| 0.063707801 | 0.949202893 | NA | |
| NA | NA | NA | |
| NA | NA | NA | |
| 0.063707378 | 0.94920323 | NA | |
| -0.194491094 | 0.845791368 | NA | |
| 1.770886944 | 0.076579504 | | 0.132990008 |
| NA | NA | NA | |
| 1.634192762 | 0.102218382 | NA | |
| NA | NA | NA | |
| 1.338063092 | 0.180875871 | | 0.272238307 |
| 0.063707378 | 0.94920323 | NA | |
| -0.214718195 | 0.829987036 | NA | |
| 0.063708381 | 0.949202431 | NA | |
| NA | NA | NA | |
| 0.063706305 | 0.949204085 | | 0.965285753 |
| NA | NA | NA | |
| 0.063708381 | 0.949202431 | NA | |
| -0.0653916 | 0.947862212 | NA | |
| NA | NA | NA | |
| NA | NA | NA | |
| NA | NA | NA | |
| 0.364128192 | 0.715762286 | NA | |
| NA | NA | NA | |
| NA | NA | NA | |
| NA | NA | NA | |
| NA | NA | NA | |
| 0.192810118 | 0.847107679 | NA | |
| 3.083195227 | 0.002047908 | | 0.005505814 |
| 0.063708882 | 0.949202033 | NA | |
| 0.081165102 | 0.935310653 | NA | |
| NA | NA | NA | |
| NA | NA | NA | |
| NA | NA | NA | |
| NA | NA | NA | |
| 0.063705937 | NA | NA | |

| | | |
|---|---|---|
| 0.317642674 | 0.750756 | NA |
| -0.065395473 | NA | NA |
| 2.441339266 | 0.014632901 | 0.031868224 |
| NA | NA | NA |
| 0.063708882 | 0.949202033 | NA |
| 0.192810118 | 0.847107679 | NA |
| -0.053895408 | NA | NA |
| NA | NA | NA |
| NA | NA | NA |
| 0.193341124 | 0.846691821 | NA |
| 0.063707135 | 0.949203424 | NA |
| -0.200509253 | 0.841082322 | NA |
| -0.392418978 | 0.694748666 | 0.775212157 |
| 0.649568718 | 0.515970845 | 0.621259873 |
| -0.108325859 | 0.913737211 | NA |
| NA | NA | NA |
| 11.97060948 | 5.07E-33 | 2.05E-31 |
| NA | NA | NA |
| 13.73794253 | 6.02E-43 | 3.69E-41 |
| NA | NA | NA |
| NA | NA | NA |
| NA | NA | NA |
| NA | NA | NA |
| NA | NA | NA |
| NA | NA | NA |
| NA | NA | NA |
| NA | NA | NA |
| NA | NA | NA |
| NA | NA | NA |
| NA | NA | NA |
| NA | NA | NA |
| 0.063708285 | 0.949202508 | NA |
| NA | NA | NA |
| NA | NA | NA |
| NA | NA | NA |
| NA | NA | NA |
| NA | NA | NA |
| NA | NA | NA |
| NA | NA | NA |
| 0.063708882 | 0.949202033 | NA |
| NA | NA | NA |
| NA | NA | NA |
| NA | NA | NA |
| NA | NA | NA |
| NA | NA | NA |
| NA | NA | NA |
| NA | NA | NA |

| | | | |
|---|---|---|---|
| NA | NA | NA | |
| NA | NA | NA | |
| NA | NA | NA | |
| NA | NA | NA | |
| NA | NA | NA | |
| NA | NA | NA | |
| | 0.063708882 | 0.949202033 | NA |
| NA | NA | NA | |
| NA | NA | NA | |
| | 0.192810118 | 0.847107679 | NA |
| NA | NA | NA | |
| NA | NA | NA | |
| NA | NA | NA | |
| | -4.551681707 | 5.32E-06 | 2.26E-05 |
| NA | NA | NA | |
| | -0.290816386 | 0.771191752 | NA |
| | -1.190115401 | 0.234001038 | 0.33528227 |
| | 0.425977619 | 0.670124154 | 0.754767924 |
| | 0.192809521 | 0.847108147 | NA |
| | -2.076958231 | 0.037805414 | 0.072577813 |
| | -0.194488217 | 0.845793621 | NA |
| | -1.938400221 | 0.052574414 | 0.096689021 |
| | -2.180104381 | 0.029249725 | 0.0581002 |
| | -5.039284935 | 4.67E-07 | 2.30E-06 |
| | -4.289781467 | 1.79E-05 | 7.00E-05 |
| | 0.063708882 | 0.949202033 | NA |
| | 1.55805278 | 0.119220737 | 0.192845744 |
| | 7.323838092 | 2.41E-13 | 2.38E-12 |
| | -0.030660925 | 0.975539954 | 0.983001814 |
| | -0.895149364 | 0.370707249 | 0.483472407 |
| | -6.971829817 | 3.13E-12 | 2.75E-11 |
| | -0.698904301 | 0.484611839 | 0.593058932 |
| | -1.149032805 | 0.250542454 | 0.354198115 |
| | -1.018555589 | 0.308413997 | 0.418247947 |
| | -2.255051844 | 0.024130075 | 0.049151674 |
| | -0.065389966 | 0.947863513 | NA |
| | -0.355187271 | 0.7224493 | 0.797165667 |
| | 4.05967703 | 4.91E-05 | 0.000179299 |
| NA | NA | NA | |
| | 0.151011519 | 0.879966629 | NA |
| | 0.151049228 | 0.879936883 | 0.915748097 |
| | 0.353428733 | 0.723767048 | 0.798267789 |
| | -0.871468539 | 0.383498377 | 0.495891118 |
| NA | NA | NA | |
| | -6.709390813 | 1.95E-11 | 1.60E-10 |
| | -1.743163135 | 0.081305123 | 0.140021138 |
| | -3.471248446 | 0.000518044 | 0.001575344 |

| | | |
|---|---|---|
| 7.28193348 | 3.29E-13 | 3.21E-12 |
| 0.074728106 | 0.940431045 | NA |
| -0.273377057 | 0.784563387 | 0.846376243 |
| 1.908798554 | 0.056288084 | 0.102472233 |
| 5.899402294 | 3.65E-09 | 2.34E-08 |
| -0.869528269 | 0.384558253 | 0.496774002 |
| 17.07118153 | 2.43E-65 | 2.77E-63 |
| 0.078918318 | 0.937097593 | 0.9571351 |
| -1.526684205 | 0.126839557 | 0.203356015 |
| 3.672900266 | 0.000239813 | 0.00077562 |
| 0.532972763 | 0.594052435 | 0.692962384 |
| 5.880034905 | 4.10E-09 | 2.62E-08 |
| 1.170214528 | 0.241914648 | 0.344491303 |
| -2.518713279 | 0.011778453 | 0.026348616 |
| NA | NA | NA |
| 5.409772376 | 6.31E-08 | 3.49E-07 |
| -2.648142473 | 0.008093541 | 0.018813455 |
| 4.351579362 | 1.35E-05 | 5.39E-05 |
| -1.486317836 | 0.137195071 | 0.216778779 |
| 0.192809521 | 0.847108147 | NA |
| 11.99942401 | 3.58E-33 | 1.45E-31 |
| 1.314926386 | 0.188534644 | 0.281665402 |
| -4.776663224 | 1.78E-06 | 8.09E-06 |
| NA | NA | NA |
| -0.980705242 | 0.326738118 | 0.437784572 |
| -0.235697132 | 0.813667695 | 0.868972241 |
| -4.372690157 | 1.23E-05 | 4.93E-05 |
| 10.10963003 | 5.01E-24 | 1.20E-22 |
| -1.868366361 | 0.061711022 | 0.110630776 |
| -3.056839006 | 0.002236844 | 0.005957459 |
| -0.216588385 | 0.828529138 | 0.880074874 |
| 3.649042328 | 0.00026322 | 0.000845099 |
| -0.105046034 | 0.916339281 | 0.942621766 |
| 1.812035195 | 0.069980758 | 0.123240581 |
| 7.565688694 | 3.86E-14 | 4.14E-13 |
| -1.442997219 | 0.149021255 | 0.232152984 |
| NA | NA | NA |
| -0.906735552 | 0.364546658 | 0.476978717 |
| NA | NA | NA |
| -0.583401735 | 0.55962289 | NA |
| -0.065388772 | 0.947864464 | NA |
| -2.747317093 | 0.006008501 | 0.014472588 |
| -1.410054758 | 0.158523515 | 0.24414795 |
| -3.723790185 | 0.000196254 | 0.000646007 |
| 10.20083864 | 1.97E-24 | 4.83E-23 |
| 19.04465166 | 7.28E-81 | 1.15E-78 |
| 5.212221539 | 1.87E-07 | 9.68E-07 |

| | | |
|---|---|---|
| 0.330493631 | 0.741027004 | NA |
| 6.783092505 | 1.18E-11 | 9.80E-11 |
| -0.457773033 | 0.647115512 | 0.736006783 |
| 4.06282615 | 4.85E-05 | 0.000177051 |
| 1.545963552 | 0.122113371 | 0.196901222 |
| -2.264464565 | 0.023545557 | 0.048160929 |
| 12.40858159 | 2.35E-35 | 1.04E-33 |
| 9.126920465 | 7.05E-20 | 1.25E-18 |
| 12.30876942 | 8.13E-35 | 3.51E-33 |
| NA | NA | NA |
| 0.196664498 | 0.844090094 | 0.890970145 |
| -4.039643319 | 5.35E-05 | 0.00019428 |
| 0.431838883 | 0.665858517 | NA |
| -2.428363169 | 0.015167147 | 0.032902903 |
| 0.358148839 | 0.720231932 | NA |
| 2.024518742 | 0.042916806 | 0.081066157 |
| 0.692477402 | 0.488637573 | NA |
| -0.504550711 | 0.613874436 | 0.709515937 |
| 0.116151747 | 0.907532278 | 0.936066795 |
| -6.770213941 | 1.29E-11 | 1.07E-10 |
| 9.077191694 | 1.11E-19 | 1.94E-18 |
| 5.590653529 | 2.26E-08 | 1.33E-07 |
| -3.783882638 | 0.000154401 | 0.000516388 |
| -1.18141805 | 0.237436691 | 0.339453375 |
| -3.384233381 | 0.000713773 | 0.00211529 |
| NA | NA | NA |
| 7.751183394 | 9.10E-15 | 1.04E-13 |
| -0.685874816 | 0.492792052 | 0.599989545 |
| 0.317642674 | 0.750756 | NA |
| -0.849897089 | 0.395382304 | 0.507357095 |
| 8.341325034 | 7.35E-17 | 1.01E-15 |
| -0.061221387 | 0.951182897 | 0.966669695 |
| 6.756554194 | 1.41E-11 | 1.17E-10 |
| -0.748473058 | 0.45417487 | 0.564056614 |
| -11.24763114 | 2.38E-29 | 7.92E-28 |
| -0.157737158 | 0.874663919 | 0.912678041 |
| 0.3688776 | 0.71221896 | 0.789287194 |
| -0.61798807 | 0.536583203 | 0.640929495 |
| 3.121063093 | 0.001801994 | 0.004902739 |
| 0.624171667 | 0.532514852 | 0.636921047 |
| -3.465764215 | 0.000528727 | 0.001605814 |
| 3.176338671 | 0.001491467 | 0.004133698 |
| -0.167981879 | 0.866597535 | 0.906887545 |
| 2.118851364 | 0.03410303 | 0.066407254 |
| NA | NA | NA |
| -0.067927152 | 0.945843625 | 0.963003007 |
| 0.065548599 | 0.947737213 | NA |

| | | |
|---|---|---|
| 2.710940309 | 0.00670927 | 0.015975365 |
| NA | NA | NA |
| NA | NA | NA |
| 0.567011215 | 0.570706562 | 0.672067055 |
| 2.387430728 | 0.0169666 | 0.036303128 |
| 0.520205987 | 0.602920013 | 0.700152544 |
| 0.357843478 | 0.720460451 | 0.795567349 |
| 2.665453561 | 0.007688455 | 0.017986917 |
| 5.203175684 | 1.96E-07 | 1.01E-06 |
| 20.10573615 | 6.57E-90 | 1.32E-87 |
| NA | NA | NA |
| NA | NA | NA |
| NA | NA | NA |
| 1.29454637 | 0.195476763 | 0.290097438 |
| 0.161530855 | 0.871675312 | NA |
| NA | NA | NA |
| NA | NA | NA |
| NA | NA | NA |
| 3.608736905 | 0.000307691 | 0.000976337 |
| 0.010439643 | 0.991670521 | 0.994172834 |
| 0.496220235 | 0.619739036 | 0.714183613 |
| 3.655739126 | 0.000256442 | 0.000825137 |
| NA | NA | NA |
| -0.065388772 | 0.947864464 | NA |
| NA | NA | NA |
| 0.350190254 | 0.72619592 | 0.800013835 |
| NA | NA | NA |
| 0.192809521 | 0.847108147 | NA |
| NA | NA | NA |
| NA | NA | NA |
| NA | NA | NA |
| 2.187340252 | 0.028717694 | 0.057174983 |
| -2.070138645 | 0.038439363 | 0.073614174 |
| 0.776743153 | 0.437310322 | 0.547595045 |
| 13.48467377 | 1.93E-41 | 1.10E-39 |
| 0.844885305 | 0.39817488 | NA |
| -2.593845366 | 0.009490916 | 0.021740426 |
| -6.655482777 | 2.82E-11 | 2.27E-10 |
| 5.287782372 | 1.24E-07 | 6.59E-07 |
| -0.526081497 | 0.598831584 | 0.696762484 |
| 3.114062309 | 0.001845305 | 0.005006285 |
| -1.594279639 | 0.110873416 | 0.181359404 |
| -0.914003031 | 0.360715253 | 0.472855493 |
| 0.063708285 | 0.949202508 | NA |
| -2.565013655 | 0.010317174 | 0.023437909 |
| 0.438989472 | 0.660669165 | 0.747541893 |
| -4.751691531 | 2.02E-06 | 9.08E-06 |

| | | | |
|---|---|---|---|
| | 3.326097273 | 0.000880712 | 0.002566863 |
| | -1.144264743 | 0.25251386 | 0.35656222 |
| | -0.974332153 | 0.329891645 | 0.440830061 |
| | 6.150067116 | 7.75E-10 | 5.33E-09 |
| | -1.052884105 | 0.292394112 | 0.40027481 |
| | 4.954812757 | 7.24E-07 | 3.47E-06 |
| | 2.098316724 | 0.035877176 | 0.069338717 |
| | 0.063707019 | 0.949203516 | NA |
| | -2.523012262 | 0.011635434 | 0.026065956 |
| | 2.538174491 | 0.011143241 | 0.025116193 |
| NA | NA | NA | |
| NA | NA | NA | |
| | 0.105006467 | 0.916370677 | NA |
| | -0.065389966 | 0.947863513 | NA |
| NA | NA | NA | |
| | -3.523836771 | 0.000425346 | 0.001313886 |
| | 1.52657651 | 0.126866352 | 0.203385947 |
| | -0.065389369 | 0.947863988 | NA |
| | -5.217713668 | 1.81E-07 | 9.41E-07 |
| NA | NA | NA | |
| NA | NA | NA | |
| | 1.420707162 | 0.15540191 | 0.240402336 |
| | -1.092291546 | 0.274704975 | 0.381399242 |
| | -2.475134801 | 0.01331859 | 0.029360956 |
| | 2.38798396 | 0.016941081 | 0.036257825 |
| NA | NA | NA | |
| | -3.235245245 | 0.001215383 | 0.003433913 |
| NA | NA | NA | |
| | -0.633363853 | 0.526496064 | 0.631472351 |
| | 0.192636473 | 0.847243678 | NA |
| | 4.126970139 | 3.68E-05 | 0.000136951 |
| | 2.86282085 | 0.004198879 | 0.010504542 |
| | 9.295051704 | 1.47E-20 | 2.72E-19 |
| | -0.000557517 | 0.999555166 | 0.999600945 |
| | -1.247962403 | 0.212044826 | 0.309907859 |
| | 0.373044889 | 0.709115035 | NA |
| | 0.192809521 | 0.847108147 | NA |
| | 0.081590913 | 0.934972027 | NA |
| NA | NA | NA | |
| | -0.849827589 | 0.395420949 | 0.507376139 |
| | -0.671402323 | 0.501964266 | 0.608817443 |
| | 1.354750071 | 0.175497199 | 0.265724693 |
| | -5.871677127 | 4.31E-09 | 2.75E-08 |
| NA | NA | NA | |
| | -0.293248386 | 0.769332302 | NA |
| NA | NA | NA | |
| NA | NA | NA | |

| | | | |
|---|---|---|---|
| NA | NA | NA | |
| NA | NA | NA | |
| | 0.192808327 | 0.847109082 | NA |
| NA | NA | NA | |
| NA | NA | NA | |
| NA | NA | NA | |
| NA | NA | NA | |
| | -0.425238312 | 0.670662958 | NA |
| | -0.065388772 | 0.947864464 | NA |
| NA | NA | NA | |
| NA | NA | NA | |
| | -1.51006194 | 0.13102762 | 0.208640583 |
| NA | NA | NA | |
| | 0.192809521 | 0.847108147 | NA |
| | -1.403387088 | 0.160501441 | 0.246538881 |
| | 0.34424867 | 0.730659283 | NA |
| | 0.192808327 | 0.847109082 | NA |
| NA | NA | NA | |
| | -0.098431204 | 0.921589898 | NA |
| NA | NA | NA | |
| NA | NA | NA | |
| NA | NA | NA | |
| | -1.51866406 | 0.128847078 | 0.205950506 |
| | -2.69085695 | 0.007126875 | 0.016830425 |
| NA | NA | NA | |
| NA | NA | NA | |
| | -2.470631048 | 0.013487489 | 0.029673187 |
| | 1.400050163 | 0.161498297 | 0.247781193 |
| NA | NA | NA | |
| NA | NA | NA | |
| | 2.363596136 | 0.01809853 | 0.038347625 |
| NA | NA | NA | |
| NA | NA | NA | |
| | 0.204079259 | 0.838291562 | 0.886906662 |
| NA | NA | NA | |
| NA | NA | NA | |
| | -1.500576696 | 0.133465083 | 0.211915187 |
| | 1.658360255 | 0.097244774 | 0.162757384 |
| | -1.410258979 | 0.158463226 | 0.244097148 |
| NA | NA | NA | |
| NA | NA | NA | |
| NA | NA | NA | |
| NA | NA | NA | |
| | 18.38978298 | 1.59E-75 | 2.29E-73 |
| | 0.661898197 | 0.508036468 | 0.614314308 |
| | 3.015425439 | 0.00256619 | 0.006732887 |
| | -3.861684007 | 0.000112608 | 0.000385896 |

| | | |
|---|---|---|
| -3.5911619 | 0.000329207 | 0.001038968 |
| -2.708803293 | 0.006752636 | 0.016063342 |
| -2.350038676 | 0.01877146 | 0.039543263 |
| -2.383302026 | 0.017158111 | 0.036665885 |
| NA | NA | NA |
| 5.069426474 | 3.99E-07 | 1.98E-06 |
| NA | NA | NA |
| 5.055687856 | 4.29E-07 | 2.12E-06 |
| -1.189967558 | 0.234059143 | 0.335346329 |
| 6.843525251 | 7.73E-12 | 6.51E-11 |
| NA | NA | NA |
| 5.46111867 | 4.73E-08 | 2.67E-07 |
| 4.672954378 | 2.97E-06 | 1.31E-05 |
| -2.430482118 | 0.01507875 | 0.032735765 |
| NA | NA | NA |
| -0.065388772 | 0.947864464 | NA |
| 0.063708882 | 0.949202033 | NA |
| 8.492550098 | 2.02E-17 | 2.92E-16 |
| -0.249361872 | 0.803080875 | 0.860456073 |
| NA | NA | NA |
| 0.950654811 | 0.341779635 | 0.453236495 |
| -1.14017029 | 0.254215363 | 0.358439079 |
| -2.28719606 | 0.022184382 | 0.045743066 |
| NA | NA | NA |
| -3.879486928 | 0.000104677 | 0.000360543 |
| -3.164100512 | 0.001555631 | 0.004296307 |
| -3.933270843 | 8.38E-05 | 0.000293963 |
| 0.192809521 | 0.847108147 | NA |
| -0.194487022 | 0.845794556 | NA |
| 3.216990861 | 0.001295427 | 0.003636669 |
| -1.203386085 | 0.22882695 | 0.329320294 |
| -0.674909751 | 0.499733105 | 0.606787601 |
| NA | NA | NA |
| NA | NA | NA |
| 0.690750738 | 0.489722198 | 0.5974143 |
| -0.80480131 | 0.420934346 | 0.532739377 |
| -1.713175891 | 0.086680186 | 0.147833541 |
| 2.462086047 | 0.013813152 | 0.03030974 |
| 0.233243305 | 0.815572481 | 0.870263853 |
| -0.291311767 | 0.770812888 | NA |
| -3.932410782 | 8.41E-05 | 0.000294939 |
| -2.186543146 | 0.028775892 | 0.057258969 |
| NA | NA | NA |
| 7.122190851 | 1.06E-12 | 9.84E-12 |
| 3.06487317 | 0.002177623 | 0.005817687 |
| 0.201524559 | 0.840288428 | NA |
| -0.138342326 | 0.889969876 | NA |

| | | |
|---|---|---|
| NA | NA | NA |
| 0.380374799 | 0.703667219 | NA |
| -0.464329352 | 0.642411798 | 0.732504476 |
| -10.60898923 | 2.71E-26 | 7.54E-25 |
| 0.782079808 | 0.434167673 | 0.544859205 |
| 0.198308907 | 0.842803383 | NA |
| 1.937073251 | 0.05273639 | 0.0969157 |
| NA | NA | NA |
| NA | NA | NA |
| NA | NA | NA |
| 0.764947655 | 0.444302715 | NA |
| -0.065389369 | 0.947863988 | NA |
| -3.224419248 | 0.001262285 | 0.003553192 |
| 5.081809995 | 3.74E-07 | 1.87E-06 |
| 7.18679156 | 6.63E-13 | 6.28E-12 |
| -2.179676598 | 0.029281442 | 0.05813552 |
| 2.98855872 | 0.002802966 | 0.007293715 |
| NA | NA | NA |
| 0.40089638 | 0.688496415 | NA |
| 4.321426218 | 1.55E-05 | 6.13E-05 |
| -1.302714205 | 0.19267235 | 0.286751631 |
| NA | NA | NA |
| NA | NA | NA |
| NA | NA | NA |
| -1.538458588 | 0.123936527 | 0.199417026 |
| -2.524664649 | 0.011580874 | 0.02597395 |
| NA | NA | NA |
| 1.360654475 | 0.173622908 | 0.263269 |
| NA | NA | NA |
| 1.566096305 | 0.117326073 | 0.190223579 |
| -1.476853738 | 0.139714848 | 0.220065612 |
| 1.817001698 | 0.069216853 | 0.122049865 |
| NA | NA | NA |
| NA | NA | NA |
| 9.898480644 | 4.23E-23 | 9.50E-22 |
| NA | NA | NA |
| 3.522764974 | 0.00042707 | 0.001318722 |
| -2.452491913 | 0.014187057 | 0.031007862 |
| 0.907299205 | 0.364248594 | 0.476738251 |
| -2.360070396 | 0.018271467 | 0.038662172 |
| 1.703745603 | 0.088428624 | 0.150354492 |
| 0.439968425 | 0.659959976 | NA |
| 0.670409335 | 0.502596886 | 0.609314854 |
| -2.207799173 | 0.027258276 | 0.054625672 |
| -1.339304295 | 0.180471631 | 0.271744265 |
| 8.391050739 | 4.82E-17 | 6.73E-16 |
| 2.185277961 | 0.028868472 | 0.057420364 |

| | | | |
|---|---|---|---|
| | 3.067607608 | 0.002157798 | 0.005772731 |
| | 0.063708285 | 0.949202508 | NA |
| | 0.952261401 | 0.340964428 | 0.452486063 |
| | -0.973132311 | 0.330487547 | 0.441429962 |
| | -3.590332332 | 0.000330257 | 0.001042017 |
| | -4.450327187 | 8.57E-06 | 3.53E-05 |
| | -3.807151103 | 0.000140577 | 0.000473508 |
| | -2.934950799 | 0.003336006 | 0.008526522 |
| NA | NA | NA | |
| | -3.006704843 | 0.002640961 | 0.006909506 |
| | -1.347495118 | 0.177820825 | 0.268449394 |
| | 0.201524559 | 0.840288428 | NA |
| | -3.393720907 | 0.000689499 | 0.002049663 |
| | -2.871117724 | 0.004090232 | 0.010258367 |
| | -3.289351278 | 0.001004186 | 0.002893378 |
| | -4.3878819 | 1.14E-05 | 4.62E-05 |
| | -3.600629644 | 0.000317447 | 0.001005023 |
| | -2.48034135 | 0.013125666 | 0.02899183 |
| | -3.428890467 | 0.000606054 | 0.001820417 |
| NA | NA | NA | |
| | -1.575397312 | 0.115164765 | 0.187289736 |
| | -2.932831697 | 0.003358859 | 0.008578801 |
| | -0.065389369 | 0.947863988 | NA |
| | -0.135125937 | 0.892512303 | NA |
| | -2.696468586 | 0.007007901 | 0.016591729 |
| | -4.337325082 | 1.44E-05 | 5.73E-05 |
| | -3.123879999 | 0.001784832 | 0.00486225 |
| | -0.076155158 | 0.939295658 | 0.958701546 |
| | -0.527497123 | 0.597848414 | 0.696209256 |
| | 5.978787699 | 2.25E-09 | 1.47E-08 |
| | -1.470599998 | 0.141399324 | 0.22225729 |
| | -1.622270263 | 0.104745493 | 0.173077614 |
| NA | NA | NA | |
| | 0.301304004 | 0.763182688 | NA |
| NA | NA | NA | |
| NA | NA | NA | |
| | 6.529539668 | 6.60E-11 | 5.11E-10 |
| | 0.539910978 | 0.589258427 | NA |
| | 0.313098098 | 0.754206134 | 0.821737672 |
| | 2.727838 | 0.006375091 | 0.015256575 |
| | 16.23774846 | 2.73E-59 | 2.65E-57 |
| | 17.7571232 | 1.52E-70 | 1.98E-68 |
| | 9.856226858 | 6.44E-23 | 1.42E-21 |
| | -1.263041426 | 0.206574293 | 0.303507545 |
| | -1.22641423 | 0.220042834 | 0.319137335 |
| | 0.025006472 | 0.980049801 | 0.98623594 |
| | 0.31685483 | 0.751353758 | NA |

| | | |
|---|---|---|
| 4.225551593 | 2.38E-05 | 9.14E-05 |
| -1.130397413 | 0.258308804 | NA |
| -0.526487476 | 0.598549551 | 0.696607807 |
| 16.58073538 | 9.60E-62 | 1.01E-59 |
| 0.063708882 | 0.949202033 | NA |
| -3.852848825 | 0.000116751 | 0.000398893 |
| -3.85797442 | 0.000114331 | 0.000391102 |
| -5.659280495 | 1.52E-08 | 9.09E-08 |
| 1.660679418 | 0.096777849 | 0.162105974 |
| -0.081284791 | 0.935215469 | 0.955944177 |
| -1.033713271 | 0.301270227 | 0.41040567 |
| 2.746069567 | 0.006031397 | 0.014520138 |
| -2.235369264 | 0.025393116 | 0.05139963 |
| -1.518262242 | 0.128948302 | 0.206076874 |
| -2.223565244 | 0.026177709 | 0.052768628 |
| -0.709626531 | 0.477935763 | 0.58672481 |
| 0.464251966 | 0.642467234 | 0.732504476 |
| 3.605996237 | 0.000310958 | 0.000985723 |
| 1.507767751 | 0.131613979 | 0.209387776 |
| 8.474716081 | 2.36E-17 | 3.38E-16 |
| 0.794630998 | 0.426828182 | 0.538239175 |
| 0.710878934 | 0.477159261 | 0.585972736 |
| -0.624815487 | 0.532092165 | 0.636611565 |
| -1.476579852 | 0.139788296 | 0.220167445 |
| -1.327669638 | 0.184287268 | 0.276502425 |
| -3.566124267 | 0.0003623 | 0.001134396 |
| NA | NA | NA |
| 3.847116512 | 0.000119516 | 0.000407615 |
| 0.158952856 | 0.873706017 | 0.912000568 |
| NA | NA | NA |
| -1.726712273 | 0.08421935 | 0.144285444 |
| -3.633737706 | 0.000279345 | 0.00089309 |
| 9.619922353 | 6.59E-22 | 1.35E-20 |
| NA | NA | NA |
| 0.317642674 | 0.750756 | NA |
| 1.38059223 | 0.167404374 | NA |
| NA | NA | NA |
| 1.386958939 | 0.165454284 | 0.252920528 |
| NA | NA | NA |
| 6.236291552 | 4.48E-10 | 3.16E-09 |
| 9.395793363 | 5.68E-21 | 1.08E-19 |
| 7.06197955 | 1.64E-12 | 1.48E-11 |
| 7.142597704 | 9.16E-13 | 8.54E-12 |
| NA | NA | NA |
| 0.701537709 | 0.48296751 | 0.591445273 |
| 1.273952778 | 0.202680152 | 0.298997125 |
| 1.802704269 | 0.071434671 | 0.125403971 |

| | | |
|---|---|---|
| -0.183044458 | 0.854763136 | 0.899051195 |
| 15.09482028 | 1.75E-51 | 1.40E-49 |
| -0.135353125 | 0.892332683 | NA |
| 0.782804572 | 0.433741882 | 0.54446135 |
| -1.791325018 | 0.073241154 | 0.127973897 |
| 2.249090705 | 0.024506726 | 0.049829679 |
| 2.130189554 | 0.033155967 | 0.064815203 |
| 0.192809521 | 0.847108147 | NA |
| -8.404764731 | 4.29E-17 | 6.02E-16 |
| 0.31685483 | 0.751353758 | NA |
| 2.394795476 | 0.016629638 | 0.035695039 |
| NA | NA | NA |
| 1.039844415 | 0.29841219 | 0.407265928 |
| 2.946823498 | 0.003210564 | 0.008239559 |
| -1.002549091 | 0.316078469 | 0.426607931 |
| 3.994232549 | 6.49E-05 | 0.000231748 |
| -0.065388772 | 0.947864464 | NA |
| 0.063707194 | 0.949203376 | NA |
| -5.68916675 | 1.28E-08 | 7.70E-08 |
| -1.263499935 | 0.20640957 | 0.303283331 |
| 0.70841574 | 0.478687121 | NA |
| 3.876630996 | 0.000105913 | 0.000364549 |
| -0.696360749 | 0.486202938 | 0.594424829 |
| 0.083031301 | 0.933826651 | NA |
| -5.021438234 | 5.13E-07 | 2.51E-06 |
| 13.01059324 | 1.07E-38 | 5.55E-37 |
| -0.400633811 | 0.688689749 | NA |
| 0.192809521 | 0.847108147 | NA |
| -0.175120827 | 0.860984694 | 0.903279692 |
| NA | NA | NA |
| -0.701471953 | 0.483008532 | NA |
| NA | NA | NA |
| -11.74254828 | 7.71E-32 | 2.94E-30 |
| -5.312791182 | 1.08E-07 | 5.79E-07 |
| 0.192810118 | 0.847107679 | NA |
| NA | NA | NA |
| -2.326916521 | 0.019969709 | 0.041729989 |
| -0.648587753 | 0.516604874 | 0.621843801 |
| -0.00100176 | 0.999200712 | 0.999506869 |
| 0.519235259 | 0.603596694 | NA |
| NA | NA | NA |
| 0.350608161 | 0.725882333 | 0.799825444 |
| 0.063708285 | 0.949202508 | NA |
| -4.110106817 | 3.95E-05 | 0.000146475 |
| 0.37015191 | 0.711269306 | NA |
| 1.022524167 | 0.30653288 | 0.41621558 |
| NA | NA | NA |

| | | |
|---|---|---|
| NA | NA | NA |
| 0.331542811 | 0.74023451 | 0.810944091 |
| -5.958439897 | 2.55E-09 | 1.66E-08 |
| -0.720301121 | 0.471339615 | 0.580220851 |
| 1.583655446 | 0.113272149 | 0.184679643 |
| 6.735558272 | 1.63E-11 | 1.34E-10 |
| -6.010618786 | 1.85E-09 | 1.22E-08 |
| 0.452671454 | 0.650785338 | NA |
| -5.773396205 | 7.77E-09 | 4.80E-08 |
| 1.087453168 | 0.276836585 | 0.383704404 |
| -4.852594968 | 1.22E-06 | 5.64E-06 |
| -7.17808737 | 7.07E-13 | 6.66E-12 |
| -3.498380617 | 0.000468093 | 0.00143481 |
| -0.761964354 | 0.446081279 | NA |
| -0.097031555 | 0.922701335 | 0.946809086 |
| 6.560847697 | 5.35E-11 | 4.19E-10 |
| 0.10440906 | 0.916844733 | 0.942913291 |
| -11.35493854 | 7.01E-30 | 2.38E-28 |
| -3.730939741 | 0.000190767 | 0.000629103 |
| -1.845444337 | 0.064972929 | 0.115581384 |
| -1.709182114 | 0.087417221 | 0.148907924 |
| -4.062211428 | 4.86E-05 | 0.00017744 |
| -0.455141269 | 0.64900761 | 0.737890551 |
| -9.453912305 | 3.26E-21 | 6.38E-20 |
| 5.39961042 | 6.68E-08 | 3.68E-07 |
| NA | NA | NA |
| -0.289905935 | 0.771888201 | NA |
| -4.458830112 | 8.24E-06 | 3.40E-05 |
| NA | NA | NA |
| -2.41453395 | 0.015755355 | 0.034031457 |
| -1.0501608 | 0.293644189 | 0.401788337 |
| 0.929178596 | 0.352796538 | 0.464689646 |
| NA | NA | NA |
| 4.859519006 | 1.18E-06 | 5.46E-06 |
| -0.065388772 | 0.947864464 | NA |
| 2.63072465 | 0.008520304 | 0.019730151 |
| -5.374187646 | 7.69E-08 | 4.21E-07 |
| NA | NA | NA |
| 4.932394421 | 8.12E-07 | 3.86E-06 |
| 2.062191207 | 0.039189536 | 0.074838947 |
| NA | NA | NA |
| 0.953897993 | 0.34013528 | 0.451657206 |
| 10.13374848 | 3.91E-24 | 9.44E-23 |
| 4.13575422 | 3.54E-05 | 0.0001322 |
| 0.31685483 | 0.751353758 | NA |
| 1.061549356 | 0.288440315 | 0.395804859 |
| -0.408075028 | 0.683218593 | 0.765814026 |

| | | |
|---|---|---|
| 0.14549979 | 0.884316282 | 0.918978042 |
| -3.080936713 | 0.002063505 | 0.005541201 |
| NA | NA | NA |
| NA | NA | NA |
| NA | NA | NA |
| NA | NA | NA |
| -1.770213668 | 0.076691553 | 0.13315688 |
| 10.1201813 | 4.50E-24 | 1.08E-22 |
| NA | NA | NA |
| 7.619795401 | 2.54E-14 | 2.77E-13 |
| -9.454435952 | 3.25E-21 | 6.36E-20 |
| NA | NA | NA |
| NA | NA | NA |
| 2.24742005 | 0.024613195 | 0.0500096 |
| 4.5272099 | 5.98E-06 | 2.51E-05 |
| 3.074128447 | 0.002111185 | 0.005656969 |
| NA | NA | NA |
| 0.253941913 | 0.799540434 | NA |
| NA | NA | NA |
| NA | NA | NA |
| 0.803706584 | 0.421566451 | 0.533360598 |
| 0.063708285 | 0.949202508 | NA |
| NA | NA | NA |
| NA | NA | NA |
| NA | NA | NA |
| 0.063707688 | 0.949202983 | NA |
| NA | NA | NA |
| NA | NA | NA |
| 0.561628487 | 0.574369167 | NA |
| NA | NA | NA |
| NA | NA | NA |
| -0.109163436 | 0.913072861 | 0.940115743 |
| NA | NA | NA |
| 0.192810118 | 0.847107679 | NA |
| NA | NA | NA |
| 0.192810118 | 0.847107679 | NA |
| NA | NA | NA |
| 0.202714101 | 0.839358501 | NA |
| 0.063707194 | 0.949203376 | NA |
| 0.063708285 | 0.949202508 | NA |
| NA | NA | NA |
| 0.445923379 | 0.655652594 | NA |
| -2.308279969 | 0.02098357 | 0.043543256 |
| -3.620066847 | 0.000294527 | 0.000937915 |
| NA | NA | NA |
| -0.194488217 | 0.845793621 | NA |
| 0.063708285 | 0.949202508 | NA |

| | | |
|---|---|---|
| 2.314142726 | 0.020659894 | 0.042982059 |
| NA | NA | NA |
| NA | NA | NA |
| NA | NA | NA |
| -4.091703953 | 4.28E-05 | 0.000157459 |
| 0.192809521 | 0.847108147 | NA |
| NA | NA | NA |
| NA | NA | NA |
| 0.063707688 | 0.949202983 | NA |
| 5.525643131 | 3.28E-08 | 1.89E-07 |
| 1.092929186 | 0.274424893 | 0.381137225 |
| -2.672315035 | 0.007532989 | 0.017676794 |
| 4.050054042 | 5.12E-05 | 0.000186251 |
| 0.089105343 | 0.928998191 | NA |
| -0.663634182 | 0.506924474 | NA |
| 0.541035083 | 0.588483401 | 0.688112657 |
| 3.304769017 | 0.000950547 | 0.002752452 |
| -1.163061175 | 0.244804683 | 0.347696829 |
| 0.882838214 | 0.37732369 | 0.490033309 |
| NA | NA | NA |
| -1.4261734 | 0.153818296 | 0.238365029 |
| NA | NA | NA |
| 14.90409359 | 3.10E-50 | 2.40E-48 |
| -0.099637994 | 0.920631729 | 0.945536345 |
| 2.220251375 | 0.026401709 | 0.053130308 |
| 0.09515006 | 0.924195636 | NA |
| -0.529295358 | 0.596600575 | NA |
| -9.144080723 | 6.01E-20 | 1.07E-18 |
| 5.896856755 | 3.70E-09 | 2.37E-08 |
| NA | NA | NA |
| NA | NA | NA |
| NA | NA | NA |
| 2.467724948 | 0.013597476 | 0.029899395 |
| 0.435692301 | 0.663059992 | NA |
| 1.868487715 | 0.061694121 | 0.110610094 |
| 2.045006891 | 0.040854179 | 0.077615187 |
| -3.450719922 | 0.000559093 | 0.00169011 |
| 2.734602096 | 0.006245572 | 0.014986739 |
| 0.150933285 | 0.880028343 | 0.91577116 |
| 1.171095082 | 0.24156056 | 0.344062938 |
| -0.417940816 | 0.675990389 | 0.759290946 |
| 8.159765951 | 3.36E-16 | 4.35E-15 |
| -1.082539691 | 0.279012789 | 0.386031181 |
| -1.166445792 | 0.243434253 | 0.3461629 |
| 0.531016354 | 0.595407446 | 0.693907361 |
| NA | NA | NA |
| 0.088923592 | 0.929142634 | NA |

| | | |
|---|---|---|
| 2.03206972 | 0.042146598 | 0.079785914 |
| -4.613542371 | 3.96E-06 | 1.71E-05 |
| 0.787044536 | 0.431255796 | NA |
| -0.160289051 | 0.872653383 | NA |
| NA | NA | NA |
| 0.192810118 | 0.847107679 | NA |
| 11.53770783 | 8.52E-31 | 3.06E-29 |
| 0.086931452 | 0.930725999 | 0.952804571 |
| -3.840450915 | 0.000122809 | 0.000417876 |
| 0.063707688 | 0.949202983 | NA |
| NA | NA | NA |
| NA | NA | NA |
| 0.192810118 | 0.847107679 | NA |
| NA | NA | NA |
| 0.466009984 | 0.641208354 | 0.731648665 |
| 2.291770489 | 0.02191889 | 0.045236628 |
| -2.969634136 | 0.002981546 | 0.007715891 |
| 0.473507012 | 0.6358515 | 0.727240755 |
| -2.857288024 | 0.004272779 | 0.01066916 |
| 1.147785135 | 0.251057275 | 0.35478566 |
| -0.440879843 | 0.659299987 | NA |
| NA | NA | NA |
| NA | NA | NA |
| -2.198624803 | 0.027904612 | 0.055746862 |
| NA | NA | NA |
| -1.920749146 | 0.05476334 | 0.100104171 |
| -2.463205865 | 0.013770083 | 0.030236439 |
| -2.430072656 | 0.015095796 | 0.032765156 |
| -2.32077048 | 0.020299235 | 0.04234082 |
| -1.426222513 | 0.153804123 | 0.238365029 |
| 2.062247516 | 0.039184177 | 0.074834423 |
| -0.450870193 | 0.652083108 | 0.740445501 |
| NA | NA | NA |
| 5.397834047 | 6.75E-08 | 3.72E-07 |
| -1.536996885 | 0.124294072 | 0.199863846 |
| -1.229326779 | 0.218949311 | 0.317864466 |
| -0.885622931 | 0.375820751 | 0.488613827 |
| NA | NA | NA |
| NA | NA | NA |
| NA | NA | NA |
| NA | NA | NA |
| NA | NA | NA |
| -5.693756344 | 1.24E-08 | 7.51E-08 |
| 0.063708285 | 0.949202508 | NA |
| -0.856538357 | 0.391700066 | 0.503712117 |
| -0.194294914 | 0.845944968 | NA |
| 0.192809521 | 0.847108147 | NA |

| | | | |
|---|---|---|---|
| | -2.127154161 | 0.033407276 | 0.065245328 |
| | 0.063708285 | 0.949202508 | NA |
| NA | NA | NA | |
| | -9.929080124 | 3.11E-23 | 7.08E-22 |
| | -2.091967449 | 0.036441428 | 0.070326055 |
| | 0.282742491 | 0.777374242 | NA |
| | 2.008283127 | 0.044613214 | 0.083896975 |
| | -0.359697346 | 0.719073477 | 0.794596668 |
| NA | NA | NA | |
| | 8.170914971 | 3.06E-16 | 3.99E-15 |
| | 14.06901888 | 5.89E-45 | 3.95E-43 |
| | 3.027518611 | 0.002465705 | 0.006493891 |
| | 8.023317615 | 1.03E-15 | 1.28E-14 |
| | 1.942973342 | 0.052019379 | 0.095816106 |
| | 2.664894016 | 0.007701259 | 0.018013505 |
| NA | NA | NA | |
| | 0.320051528 | 0.748929269 | NA |
| NA | NA | NA | |
| | 0.325791008 | 0.744582483 | 0.814229523 |
| | 7.035022645 | 1.99E-12 | 1.78E-11 |
| | -3.076126862 | 0.002097086 | 0.005623534 |
| | -1.204145176 | 0.228533471 | 0.328954672 |
| | -3.698111655 | 0.000217209 | 0.000709018 |
| | -1.637177157 | 0.101593459 | 0.168726977 |
| | 0.813494337 | 0.415934692 | NA |
| NA | NA | NA | |
| | 8.772421982 | 1.75E-18 | 2.75E-17 |
| | -3.379233333 | 0.000726883 | 0.002151848 |
| | -3.635391268 | 0.000277559 | 0.000887948 |
| | 0.145926516 | 0.883979399 | NA |
| NA | NA | NA | |
| | -0.065389369 | 0.947863988 | NA |
| | 0.010236726 | 0.991832417 | 0.994295314 |
| | -0.194488217 | 0.845793621 | NA |
| | 6.017614264 | 1.77E-09 | 1.17E-08 |
| | -4.019460149 | 5.83E-05 | 0.000210138 |
| | 0.471953149 | 0.63696023 | 0.728095838 |
| | 6.463919514 | 1.02E-10 | 7.75E-10 |
| | -1.044956573 | 0.296043041 | 0.404473653 |
| NA | NA | NA | |
| | 3.928436171 | 8.55E-05 | 0.000299394 |
| | 3.466465477 | 0.000527349 | 0.001602471 |
| | 0.063707688 | 0.949202983 | NA |
| | -1.839782587 | 0.065800163 | 0.116906451 |
| | -5.901572488 | 3.60E-09 | 2.31E-08 |
| NA | NA | NA | |
| | -0.065388772 | 0.947864464 | NA |

| | | |
|---|---|---|
| -3.983119334 | 6.80E-05 | 0.000242167 |
| 0.319804015 | 0.749116904 | NA |
| 0.063708285 | 0.949202508 | NA |
| -1.965006589 | 0.049413475 | 0.091677156 |
| -1.213747837 | 0.224844031 | 0.32476462 |
| -0.880712302 | 0.378473557 | NA |
| 0.798655575 | 0.424490153 | 0.536038595 |
| 0.427528097 | 0.668994724 | 0.753936873 |
| NA | NA | NA |
| -0.093316276 | 0.925652302 | NA |
| 5.193430596 | 2.06E-07 | 1.07E-06 |
| 3.824713146 | 0.000130924 | 0.000443562 |
| -0.203807531 | 0.838503908 | 0.886984776 |
| 0.748822834 | 0.453963995 | 0.56390664 |
| 0.063708285 | 0.949202508 | NA |
| NA | NA | NA |
| -2.92016646 | 0.003498445 | 0.0089008 |
| 0.063708285 | 0.949202508 | NA |
| -1.689578768 | 0.091108571 | 0.154167626 |
| -2.579211511 | 0.009902613 | 0.022590449 |
| -1.489140473 | 0.136450382 | 0.215792826 |
| 11.11334936 | 1.08E-28 | 3.50E-27 |
| -0.077075932 | 0.93856314 | 0.958224754 |
| 0.192809521 | 0.847108147 | NA |
| 0.260149108 | 0.794748759 | 0.854029229 |
| -2.567663417 | 0.010238652 | 0.023284886 |
| -0.145211686 | 0.88454374 | NA |
| 0.893224598 | 0.371736908 | 0.484537562 |
| NA | NA | NA |
| 0.192809521 | 0.847108147 | NA |
| -3.500015756 | 0.000465231 | 0.00142635 |
| -3.569098112 | 0.000358212 | 0.001123284 |
| NA | NA | NA |
| 0.063707135 | 0.949203424 | NA |
| 0.063708285 | 0.949202508 | NA |
| NA | NA | NA |
| 0.3474247 | 0.728272279 | NA |
| NA | NA | NA |
| -4.087952167 | 4.35E-05 | 0.000159839 |
| 4.020818392 | 5.80E-05 | 0.000209051 |
| NA | NA | NA |
| NA | NA | NA |
| 0.09817548 | 0.921792953 | NA |
| 0.192809521 | 0.847108147 | NA |
| NA | NA | NA |
| NA | NA | NA |
| NA | NA | NA |

| | | | |
|---|---|---|---|
| NA | NA | NA | |
| | -0.074559618 | 0.940565105 | NA |
| | 0.075392854 | 0.939902145 | NA |
| | -0.194488217 | 0.845793621 | NA |
| NA | NA | NA | |
| | -3.827146564 | 0.000129637 | 0.000439321 |
| | -2.785281822 | 0.005348122 | 0.013063676 |
| | 0.18437553 | 0.853718865 | 0.898292551 |
| NA | NA | NA | |
| | 0.084628933 | 0.932556396 | NA |
| | 5.2798597 | 1.29E-07 | 6.87E-07 |
| | 0.333154828 | 0.739017415 | NA |
| | 0.88788908 | 0.374600419 | 0.48743234 |
| | 0.60547614 | 0.544862674 | 0.648744778 |
| | 5.410276369 | 6.29E-08 | 3.48E-07 |
| | 0.004194858 | 0.996652998 | 0.997729254 |
| | 1.142151581 | 0.253391018 | 0.357457994 |
| | -0.234346405 | 0.814716063 | 0.869752857 |
| | 0.853183502 | 0.393557551 | 0.505477499 |
| | -0.097977687 | 0.921950012 | 0.946375032 |
| NA | NA | NA | |
| | 0.635150169 | 0.52533048 | 0.630497334 |
| | 0.639570364 | 0.522451955 | 0.62770472 |
| | -7.122509396 | 1.06E-12 | 9.82E-12 |
| | -2.928462475 | 0.003406429 | 0.008690544 |
| | -3.501253095 | 0.000463076 | 0.001419918 |
| | -2.976879992 | 0.00291198 | 0.007550694 |
| NA | NA | NA | |
| | -3.227013659 | 0.001250895 | 0.003523905 |
| | -3.431188583 | 0.000600943 | 0.001806364 |
| | -2.593962901 | 0.009487672 | 0.021736974 |
| | 1.478951124 | 0.139153378 | 0.21927783 |
| | -1.860985632 | 0.062746206 | 0.112245392 |
| | -2.420745193 | 0.01548873 | 0.033510494 |
| | -2.611664208 | 0.009010271 | 0.020747661 |
| | -3.242328367 | 0.001185573 | 0.003362214 |
| | -3.035471669 | 0.002401598 | 0.006344943 |
| | -2.266611675 | 0.023413958 | 0.047954453 |
| | -1.29230865 | 0.196250278 | 0.291078807 |
| | -2.232321992 | 0.025593689 | 0.051749741 |
| NA | NA | NA | |
| NA | NA | NA | |
| | 1.183299357 | 0.236690528 | 0.338554426 |
| NA | NA | NA | |
| NA | NA | NA | |
| NA | NA | NA | |
| NA | NA | NA | |

| | | | |
|---|---|---|---|
| NA | | NA | NA |
| | 2.069280628 | 0.03851976 | 0.073756855 |
| NA | | NA | NA |
| | -0.524911886 | 0.599644446 | 0.697481319 |
| | 7.209544214 | 5.61E-13 | 5.36E-12 |
| | 1.638629106 | 0.101290527 | 0.168313418 |
| | 2.47118485 | 0.013466619 | 0.029635086 |
| | 1.412279601 | 0.15786765 | 0.24341941 |
| NA | | NA | NA |
| | 11.18348357 | 4.91E-29 | 1.61E-27 |
| | -3.250498495 | 0.001152029 | 0.003277481 |
| | -1.686173551 | 0.091762377 | 0.155116653 |
| | -0.78542657 | 0.432203508 | 0.542965961 |
| | -0.525319456 | 0.599361134 | 0.697248987 |
| | 0.600428062 | 0.54822099 | NA |
| | 10.60405899 | 2.85E-26 | 7.94E-25 |
| | 0.494554414 | 0.620914687 | 0.715176229 |
| | -0.065388772 | 0.947864464 | NA |
| NA | | NA | NA |
| | 0.658091078 | 0.510479608 | 0.616308344 |
| NA | | NA | NA |
| NA | | NA | NA |
| NA | | NA | NA |
| | 0.063708882 | 0.949202033 | NA |
| NA | | NA | NA |
| | -0.507832681 | 0.611570689 | 0.70742383 |
| NA | | NA | NA |
| | -1.661416284 | 0.09662987 | 0.161890605 |
| | 0.398641462 | 0.690157406 | NA |
| | 0.192809521 | 0.847108147 | NA |
| | -0.111973425 | 0.910844478 | 0.938593548 |
| NA | | NA | NA |
| NA | | NA | NA |
| NA | | NA | NA |
| | -0.194487022 | 0.845794556 | NA |
| | 0.185687078 | 0.852690162 | 0.897689282 |
| | 0.213302193 | 0.831091263 | NA |
| | -6.367537339 | 1.92E-10 | 1.41E-09 |
| | -6.352186183 | 2.12E-10 | 1.55E-09 |
| | -0.595603357 | 0.551440233 | 0.654738819 |
| | 0.06371127 | 0.949200131 | NA |
| | -0.606507644 | 0.544177708 | NA |
| NA | | NA | NA |
| | -1.174834518 | 0.240060927 | 0.342240987 |
| | 2.030859224 | 0.042269277 | 0.079987904 |
| | 0.39633286 | 0.691859489 | 0.772539392 |
| | 0.997302282 | 0.318617806 | 0.429240998 |

| | | | |
|---|---|---|---|
| | 1.350540627 | 0.176842631 | 0.267277002 |
| | -1.61180045 | 0.10700537 | 0.17599845 |
| | 1.747401997 | 0.080567631 | 0.13894225 |
| | 6.422904697 | 1.34E-10 | 1.00E-09 |
| | 1.167138545 | 0.243154424 | 0.345843575 |
| | 2.162085542 | 0.030611583 | 0.060397913 |
| NA | NA | NA | |
| | -6.270390966 | 3.60E-10 | 2.56E-09 |
| | 0.067524971 | 0.946163784 | NA |
| | -0.779190273 | 0.435867639 | 0.546389894 |
| | -2.609507807 | 0.009067258 | 0.020863528 |
| | -3.205257694 | 0.001349416 | 0.003775529 |
| | -2.566201792 | 0.010281899 | 0.023368377 |
| | -6.471776956 | 9.69E-11 | 7.38E-10 |
| | 5.237238342 | 1.63E-07 | 8.54E-07 |
| | -1.222038423 | 0.221693097 | 0.321028535 |
| | 6.260712221 | 3.83E-10 | 2.72E-09 |
| | -1.648340699 | 0.099282778 | 0.165636734 |
| | -0.340590088 | 0.733412192 | NA |
| | -0.48766376 | 0.625788025 | 0.719266854 |
| NA | NA | NA | |
| | -0.065388772 | 0.947864464 | NA |
| | -0.194488217 | 0.845793621 | NA |
| NA | NA | NA | |
| | 7.090094934 | 1.34E-12 | 1.23E-11 |
| | 0.192809521 | 0.847108147 | NA |
| NA | NA | NA | |
| | 0.063708285 | 0.949202508 | NA |
| | 0.78181312 | 0.43432441 | 0.544946612 |
| | 0.192809521 | 0.847108147 | NA |
| | -0.130721153 | 0.895995898 | 0.927684688 |
| | -2.399183587 | 0.016431674 | 0.035324642 |
| | -3.045363732 | 0.002323991 | 0.006161369 |
| | -0.45057111 | 0.652298693 | 0.740598098 |
| | 0.274339353 | 0.783823845 | NA |
| | -7.585496716 | 3.31E-14 | 3.57E-13 |
| | -1.210493662 | 0.226089521 | 0.326187142 |
| | 5.5151476 | 3.48E-08 | 2.00E-07 |
| NA | NA | NA | |
| | -3.480201651 | 0.000501037 | 0.00152901 |
| | -0.065388772 | 0.947864464 | NA |
| | 0.192808924 | 0.847108615 | NA |
| NA | NA | NA | |
| NA | NA | NA | |
| | -0.065388772 | 0.947864464 | NA |
| | -1.623497684 | 0.104483058 | 0.172712386 |
| NA | NA | NA | |

| | | |
|---|---|---|
| NA | NA | NA |
| NA | NA | NA |
| 0.22330737 | 0.823296292 | 0.876152474 |
| 1.803376846 | 0.071329051 | 0.125271271 |
| -4.626375304 | 3.72E-06 | 1.61E-05 |
| -2.92876332 | 0.003403134 | 0.008684794 |
| NA | NA | NA |
| -2.162230317 | 0.030600428 | 0.060380663 |
| -1.506466303 | 0.131947511 | 0.209825045 |
| -1.582230247 | 0.113597018 | 0.185112849 |
| -3.635146446 | 0.000277823 | 0.000888678 |
| NA | NA | NA |
| 1.68010729 | 0.092936443 | 0.156778919 |
| NA | NA | NA |
| -0.865839301 | 0.386578293 | 0.498791416 |
| -0.857611803 | 0.391106855 | 0.503104358 |
| 6.07014619 | 1.28E-09 | 8.60E-09 |
| 0.644177528 | 0.519460321 | 0.624740142 |
| -1.087175001 | 0.276959477 | 0.383848045 |
| -6.103436027 | 1.04E-09 | 7.07E-09 |
| 0.063708285 | 0.949202508 | NA |
| -0.444022315 | 0.657026451 | 0.744572466 |
| -0.146194223 | 0.883768066 | 0.918634331 |
| NA | NA | NA |
| NA | NA | NA |
| NA | NA | NA |
| NA | NA | NA |
| 0.063708285 | 0.949202508 | NA |
| -0.194487022 | 0.845794556 | NA |
| -4.781719809 | 1.74E-06 | 7.90E-06 |
| -2.592270564 | 0.009534475 | 0.021826223 |
| NA | NA | NA |
| NA | NA | NA |
| -2.11667931 | 0.034287074 | 0.066720053 |
| -6.059736891 | 1.36E-09 | 9.15E-09 |
| NA | NA | NA |
| 0.445694569 | 0.655817889 | NA |
| -1.613403755 | 0.106656818 | 0.175528973 |
| -0.065389966 | 0.947863513 | NA |
| NA | NA | NA |
| 0.192809521 | 0.847108147 | NA |
| 0.425818606 | 0.670240028 | 0.75480377 |
| 0.259680703 | 0.795110079 | NA |
| 0.063708285 | 0.949202508 | NA |
| NA | NA | NA |
| NA | NA | NA |
| NA | NA | NA |

| | | |
|---|---|---|
| NA | NA | NA |
| NA | NA | NA |
| 8.776632332 | 1.68E-18 | 2.66E-17 |
| 4.018300366 | 5.86E-05 | 0.000211001 |
| -1.755712414 | 0.079137526 | 0.136786981 |
| -1.760622094 | 0.078302387 | 0.135577528 |
| -2.824543061 | 0.004734809 | 0.011715328 |
| -2.811031688 | 0.004938292 | 0.012170683 |
| -3.483431032 | 0.000495031 | 0.001511604 |
| -1.975613057 | 0.048198619 | 0.089669349 |
| 5.402700574 | 6.56E-08 | 3.63E-07 |
| NA | NA | NA |
| -1.100198861 | 0.271245487 | 0.377594814 |
| -0.144277416 | 0.885281411 | 0.919558191 |
| -0.095380872 | 0.924012309 | 0.947831158 |
| 7.547010251 | 4.45E-14 | 4.74E-13 |
| -0.413044917 | 0.679573705 | 0.762494613 |
| 5.120691721 | 3.04E-07 | 1.54E-06 |
| 0.088720201 | 0.929304278 | NA |
| 0.973157739 | 0.330474911 | 0.441429962 |
| -1.20789576 | 0.227087361 | 0.327303274 |
| NA | NA | NA |
| -0.745931852 | 0.455708583 | 0.565456367 |
| 0.424748057 | 0.671020347 | NA |
| 0.346034622 | 0.729316695 | NA |
| 1.655028711 | 0.097918674 | 0.163699199 |
| -0.065388772 | 0.947864464 | NA |
| 0.063708285 | 0.949202508 | NA |
| 0.518786686 | 0.603909503 | NA |
| -6.55181356 | 5.68E-11 | 4.44E-10 |
| -2.074663489 | 0.038017733 | 0.072912717 |
| -0.065389369 | 0.947863988 | NA |
| NA | NA | NA |
| -0.065388772 | 0.947864464 | NA |
| 0.192808924 | 0.847108615 | NA |
| NA | NA | NA |
| NA | NA | NA |
| 0.324874675 | 0.745275925 | NA |
| 7.358931352 | 1.85E-13 | 1.85E-12 |
| -0.486166679 | 0.626848994 | NA |
| 0.063708285 | 0.949202508 | NA |
| -7.311991802 | 2.63E-13 | 2.59E-12 |
| -0.291563986 | 0.770620015 | NA |
| -0.065390563 | 0.947863038 | NA |
| NA | NA | NA |
| NA | NA | NA |
| NA | NA | NA |

| | | |
|---|---|---|
| NA | NA | NA |
| 0.976943754 | 0.328596999 | 0.439590315 |
| 1.22108351 | 0.222054402 | 0.321458744 |
| -9.505230036 | 2.00E-21 | 3.96E-20 |
| NA | NA | NA |
| -2.372654564 | 0.017660772 | 0.037579625 |
| -0.711631359 | 0.476693083 | 0.585561225 |
| NA | NA | NA |
| 0.394559684 | 0.693167872 | 0.773655214 |
| 1.422153533 | 0.154981685 | 0.23985603 |
| 8.420197804 | 3.76E-17 | 5.30E-16 |
| 0.157291025 | 0.875015493 | NA |
| 0.247409252 | 0.804591515 | 0.861595175 |
| 3.126680677 | 0.001767918 | 0.00482142 |
| -1.300585005 | 0.193400542 | 0.28761307 |
| 0.756002165 | 0.449647897 | NA |
| -0.502923934 | 0.615017753 | 0.710410926 |
| 0.192810118 | 0.847107679 | NA |
| NA | NA | NA |
| -0.562136657 | 0.574022914 | 0.675273466 |
| 1.485917197 | 0.137301024 | 0.216891426 |
| 0.192809521 | 0.847108147 | NA |
| 0.247247612 | 0.8047166 | NA |
| 0.101922176 | 0.918818447 | 0.944261988 |
| NA | NA | NA |
| NA | NA | NA |
| -0.341233278 | 0.732927973 | NA |
| 1.818351446 | 0.069010434 | 0.121771672 |
| 1.429281128 | 0.152923448 | 0.237189723 |
| -0.065388772 | 0.947864464 | NA |
| 0.483655215 | 0.628630582 | NA |
| 2.076537544 | 0.037844263 | 0.072641249 |
| NA | NA | NA |
| NA | NA | NA |
| -1.749616901 | 0.080184441 | 0.138372893 |
| NA | NA | NA |
| NA | NA | NA |
| 0.192810118 | 0.847107679 | NA |
| -1.6121431 | 0.106930804 | 0.175910484 |
| 0.711416577 | 0.47682613 | 0.585649837 |
| -5.69952974 | 1.20E-08 | 7.28E-08 |
| -2.399697532 | 0.016408624 | 0.035278119 |
| NA | NA | NA |
| -0.365072848 | 0.715057028 | 0.791626229 |
| -0.19448762 | 0.845794089 | NA |
| 0.960124197 | 0.336992712 | 0.448308667 |
| -1.841015059 | 0.065619352 | 0.11663488 |

| | | | |
|---|---|---|---|
| | 0.320071032 | 0.748914485 | NA |
| | -0.065389966 | 0.947863513 | NA |
| | 0.366404893 | 0.714062969 | NA |
| NA | NA | NA | |
| | 0.192808924 | 0.847108615 | NA |
| NA | NA | NA | |
| NA | NA | NA | |
| NA | NA | NA | |
| NA | NA | NA | |
| NA | NA | NA | |
| | 0.336191489 | 0.736726471 | NA |
| | 0.444284664 | 0.656836788 | NA |
| NA | NA | NA | |
| | -1.678348443 | 0.093279096 | 0.157287509 |
| | -3.601243311 | 0.000316699 | 0.001003034 |
| | 2.380443072 | 0.017291833 | 0.036916971 |
| | -1.804758641 | 0.071112459 | 0.124952253 |
| | -1.646255663 | 0.099711134 | 0.166161694 |
| | 0.192808327 | 0.847109082 | NA |
| | 10.13169657 | 4.00E-24 | 9.63E-23 |
| | 19.67048893 | 3.86E-86 | 7.05E-84 |
| | 29.30362117 | 9.33E-189 | 5.84E-186 |
| NA | NA | NA | |
| | 30.47748917 | 5.18E-204 | 4.05E-201 |
| | 26.38275461 | 2.16E-153 | 9.02E-151 |
| | 26.95642088 | 4.80E-160 | 2.18E-157 |
| | 1.99756521 | 0.045763818 | 0.08573865 |
| NA | NA | NA | |
| | -0.19448762 | 0.845794089 | NA |
| | -0.990780882 | 0.321792586 | 0.432803248 |
| | -0.065388772 | 0.947864464 | NA |
| | 5.836350458 | 5.34E-09 | 3.36E-08 |
| NA | NA | NA | |
| | 6.881223622 | 5.93E-12 | 5.08E-11 |
| NA | NA | NA | |
| | 0.605753232 | 0.54467863 | NA |
| | -1.693290332 | 0.090400217 | 0.153186476 |
| NA | NA | NA | |
| | -0.337178757 | 0.73598215 | 0.807541347 |
| | 0.961066447 | 0.336518759 | NA |
| | -0.122855467 | 0.902221551 | NA |
| | 2.106615508 | 0.035150921 | 0.068171979 |
| | 1.346845092 | 0.178030128 | 0.268714594 |
| | 0.454962383 | 0.649136302 | 0.737939656 |
| | 0.192809521 | 0.847108147 | NA |
| | -1.422971559 | 0.1547444 | 0.239592499 |
| | -1.715589444 | 0.086237212 | 0.147158264 |

| | | |
|---|---|---|
| -0.566007657 | 0.571388575 | NA |
| NA | NA | NA |
| -0.065388772 | 0.947864464 | NA |
| -3.375934546 | 0.000735654 | 0.002176013 |
| NA | NA | NA |
| -0.429854739 | 0.667301312 | NA |
| 0.240691091 | 0.809794544 | NA |
| -0.089440543 | 0.928731804 | NA |
| -1.76093728 | 0.078249019 | 0.135494497 |
| 0.063708285 | 0.949202508 | NA |
| 0.192810118 | 0.847107679 | NA |
| 1.52093386 | 0.128276436 | 0.205278383 |
| -3.801888619 | 0.000143597 | 0.000482707 |
| NA | NA | NA |
| 0.133986612 | 0.893413161 | NA |
| -3.959801384 | 7.50E-05 | 0.000265379 |
| -3.416527114 | 0.000634253 | 0.001897152 |
| -3.994700548 | 6.48E-05 | 0.000231323 |
| -2.609374735 | 0.009070785 | 0.020869726 |
| 5.01362141 | 5.34E-07 | 2.60E-06 |
| NA | NA | NA |
| 0.063708882 | 0.949202033 | NA |
| 8.475883308 | 2.33E-17 | 3.35E-16 |
| -0.590203065 | 0.555054518 | 0.657844338 |
| 2.616281095 | 0.008889334 | 0.02049747 |
| 0.79749674 | 0.425162595 | 0.536715237 |
| -2.013258798 | 0.04408741 | 0.083032948 |
| 0.934038764 | 0.350283898 | 0.46203657 |
| -0.366699587 | 0.713843115 | 0.790689452 |
| -0.19448762 | 0.845794089 | NA |
| 0.459856884 | 0.64561895 | NA |
| 0.093230399 | 0.925720525 | NA |
| NA | NA | NA |
| -0.367784807 | 0.713033699 | NA |
| NA | NA | NA |
| -0.068186498 | 0.945637175 | NA |
| 0.502672469 | 0.615194569 | NA |
| -0.065391638 | 0.947862182 | NA |
| -0.207182747 | 0.835867141 | 0.8849027 |
| -4.955579359 | 7.21E-07 | 3.46E-06 |
| NA | NA | NA |
| 0.192809521 | 0.847108147 | NA |
| -0.54696364 | 0.584403712 | 0.684493445 |
| 6.040920705 | 1.53E-09 | 1.02E-08 |
| 1.902315676 | 0.057129897 | 0.103763249 |
| NA | NA | NA |
| 4.493632319 | 7.00E-06 | 2.92E-05 |

| | | |
|---|---|---|
| 2.759246161 | 0.005793488 | 0.014017496 |
| 8.429559575 | 3.47E-17 | 4.90E-16 |
| 2.835232362 | 0.004579237 | 0.011367479 |
| -3.714304089 | 0.000203764 | 0.000668792 |
| NA | NA | NA |
| -1.046344548 | 0.295401985 | 0.403730025 |
| NA | NA | NA |
| 1.663906854 | 0.096131045 | 0.161238361 |
| 7.384684321 | 1.53E-13 | 1.54E-12 |
| 1.462469795 | 0.143612516 | 0.225212785 |
| NA | NA | NA |
| 1.387344559 | 0.165336722 | NA |
| NA | NA | NA |
| NA | NA | NA |
| -5.171208503 | 2.33E-07 | 1.19E-06 |
| 3.534282349 | 0.000408884 | 0.0012671 |
| -0.534432672 | 0.593042219 | 0.6921499 |
| 9.871896007 | 5.51E-23 | 1.23E-21 |
| 2.341542281 | 0.019204251 | 0.040339241 |
| NA | NA | NA |
| 3.610248736 | 0.000305904 | 0.000971178 |
| 0.192810118 | 0.847107679 | NA |
| 1.642469662 | 0.100492708 | 0.167176159 |
| 4.119316086 | 3.80E-05 | 0.000141202 |
| 0.870896357 | 0.383810746 | 0.496166884 |
| 0.192810118 | 0.847107679 | NA |
| -7.897804744 | 2.84E-15 | 3.40E-14 |
| 0.063708285 | 0.949202508 | NA |
| 2.859010083 | 0.004249653 | 0.010618829 |
| -9.70183226 | 2.96E-22 | 6.22E-21 |
| -2.408404078 | 0.016022436 | 0.034550752 |
| NA | NA | NA |
| NA | NA | NA |
| NA | NA | NA |
| 0.339112524 | 0.734524964 | NA |
| NA | NA | NA |
| -0.065389369 | 0.947863988 | NA |
| NA | NA | NA |
| NA | NA | NA |
| NA | NA | NA |
| NA | NA | NA |
| -5.714120807 | 1.10E-08 | 6.71E-08 |
| -2.476012914 | 0.013285877 | 0.029294002 |
| -2.387949131 | 0.016942687 | 0.036258161 |
| 0.520454598 | 0.602746764 | 0.700048792 |
| 0.06371127 | 0.949200131 | NA |
| -3.755160713 | 0.00017323 | 0.00057407 |

| | | | |
|---|---|---|---|
| | -6.611078391 | 3.82E-11 | 3.03E-10 |
| | 0.504232133 | 0.614098263 | NA |
| | -0.167230228 | 0.867188901 | NA |
| NA | NA | NA | |
| | 4.179600577 | 2.92E-05 | 0.000110663 |
| | 2.199003174 | 0.027877697 | 0.055710878 |
| NA | NA | NA | |
| NA | NA | NA | |
| NA | NA | NA | |
| | 0.109580682 | 0.912741932 | NA |
| | -0.563339863 | 0.573203479 | NA |
| | 1.248116844 | 0.211988271 | NA |
| | 0.192809521 | 0.847108147 | NA |
| | 0.59205019 | 0.55381698 | 0.656937539 |
| | -0.259226734 | 0.795460305 | NA |
| | 28.80318688 | 1.96E-182 | 1.14E-179 |
| | 16.06458479 | 4.52E-58 | 4.24E-56 |
| | -0.950515868 | 0.341850195 | 0.453256775 |
| | 0.0850295 | 0.932237938 | NA |
| NA | NA | NA | |
| NA | NA | NA | |
| | 0.320071032 | 0.748914485 | NA |
| | 0.403695729 | 0.686436481 | NA |
| | 0.063708285 | 0.949202508 | NA |
| | 0.239566349 | 0.810666455 | NA |
| | -0.244297613 | 0.807000332 | 0.863180784 |
| | 0.0614441 | 0.951005532 | 0.966569586 |
| | 0.063976859 | 0.948988653 | NA |
| | 0.359810036 | 0.718989197 | 0.794538614 |
| | -3.35892296 | 0.000782469 | 0.002300081 |
| | -1.74422109 | 0.081120545 | 0.139789828 |
| | -3.608676817 | 0.000307763 | 0.000976337 |
| | -1.538280397 | 0.123980071 | 0.199462121 |
| | -5.949107652 | 2.70E-09 | 1.76E-08 |
| | -4.383061158 | 1.17E-05 | 4.72E-05 |
| | -1.513989652 | 0.130028464 | 0.207432279 |
| | -0.001890636 | 0.998491491 | 0.998930494 |
| | -3.532352564 | 0.00041188 | 0.001275121 |
| | -0.92922336 | 0.352773343 | 0.464683548 |
| | -3.30028043 | 0.000965883 | 0.002792013 |
| | -5.718515228 | 1.07E-08 | 6.55E-08 |
| | -2.828913687 | 0.004670629 | 0.01156911 |
| | -2.966048222 | 0.003016532 | 0.007795952 |
| | 0.414611089 | 0.678426629 | NA |
| | -6.132356863 | 8.66E-10 | 5.93E-09 |
| | -2.983005887 | 0.002854325 | 0.007413485 |
| | 4.056736939 | 4.98E-05 | 0.000181438 |

| | | | |
|---|---|---|---|
| | 0.745829636 | 0.455770335 | NA |
| | 0.192810118 | 0.847107679 | NA |
| | -1.491207401 | 0.135907051 | 0.215096648 |
| | -3.225789952 | 0.001256255 | 0.003537015 |
| | -1.529108815 | 0.12623747 | 0.202507459 |
| | -2.637535653 | 0.008351085 | 0.019366965 |
| | 0.192809521 | 0.847108147 | NA |
| | -1.347746534 | 0.17773992 | 0.268357473 |
| | -0.801335489 | 0.422937451 | 0.534706758 |
| NA | NA | NA | |
| NA | NA | NA | |
| NA | NA | NA | |
| NA | NA | NA | |
| NA | NA | NA | |
| NA | NA | NA | |
| | 2.450029467 | 0.014284452 | 0.031188053 |
| | -0.065389369 | 0.947863988 | NA |
| | -0.941786932 | 0.346301737 | 0.457869763 |
| NA | NA | NA | |
| | 5.433179597 | 5.54E-08 | 3.09E-07 |
| NA | NA | NA | |
| | 0.309559594 | 0.756895887 | 0.823771809 |
| NA | NA | NA | |
| | -0.1891838 | 0.849948764 | 0.895454902 |
| | -2.732751291 | 0.006280774 | 0.015059658 |
| | -4.416588302 | 1.00E-05 | 4.09E-05 |
| | -0.915582762 | 0.359885776 | 0.472040087 |
| | -2.943500954 | 0.003245228 | 0.008319138 |
| NA | NA | NA | |
| | -0.182552457 | 0.855149191 | 0.899343876 |
| | -1.477287529 | 0.13959858 | 0.219896315 |
| | -3.340052329 | 0.000837626 | 0.002448987 |
| | -3.701260631 | 0.000214531 | 0.000701007 |
| | -2.081658938 | 0.037373634 | 0.071842566 |
| | -2.001355006 | 0.045354146 | 0.085092142 |
| | -0.065389369 | 0.947863988 | NA |
| | 1.150543981 | 0.249919891 | 0.353457728 |
| NA | NA | NA | |
| | 5.020716788 | 5.15E-07 | 2.51E-06 |
| NA | NA | NA | |
| | -0.445628302 | 0.655865763 | 0.743604177 |
| | -0.121448525 | 0.903335784 | 0.932894439 |
| NA | NA | NA | |
| | -0.065388772 | 0.947864464 | NA |
| | -0.496571502 | 0.619491254 | 0.714000275 |
| | 0.083999021 | 0.93305721 | NA |
| NA | NA | NA | |

| | | |
|---|---|---|
| -0.065389369 | 0.947863988 | NA |
| -0.40248659 | 0.687325956 | NA |
| -0.065389369 | 0.947863988 | NA |
| NA | NA | NA |
| NA | NA | NA |
| NA | NA | NA |
| -0.74670204 | 0.455243438 | 0.564991237 |
| -5.207949295 | 1.91E-07 | 9.90E-07 |
| -3.197019788 | 0.001388554 | 0.003873343 |
| -3.907487878 | 9.33E-05 | 0.000324661 |
| -3.754306244 | 0.000173822 | 0.00057588 |
| -3.740927456 | 0.000183342 | 0.000605737 |
| -4.587916113 | 4.48E-06 | 1.92E-05 |
| 2.330913881 | 0.019757901 | 0.041349521 |
| -2.876903378 | 0.004015985 | 0.010093394 |
| -4.892040279 | 9.98E-07 | 4.69E-06 |
| 0.063708285 | 0.949202508 | NA |
| -2.652685959 | 0.007985413 | 0.018596472 |
| NA | NA | NA |
| -0.194488217 | 0.845793621 | NA |
| NA | NA | NA |
| 0.986996015 | 0.323644592 | NA |
| 1.15978314 | 0.24613711 | 0.349192987 |
| -2.986511898 | 0.002821799 | 0.007336618 |
| -2.482055086 | 0.013062708 | 0.028871165 |
| -0.556798069 | 0.577665403 | 0.678568474 |
| 0.063708882 | 0.949202033 | NA |
| 0.355082911 | 0.722527479 | NA |
| NA | NA | NA |
| 7.428219985 | 1.10E-13 | 1.13E-12 |
| -0.96235605 | 0.335870783 | 0.447125077 |
| 0.133652375 | 0.893677466 | NA |
| 3.451502014 | 0.000557476 | 0.001686237 |
| -2.593532242 | 0.009499563 | 0.021756251 |
| -0.961860203 | 0.336119832 | 0.447432824 |
| -0.480112653 | 0.631147292 | 0.723135435 |
| 0.045697564 | 0.963551305 | 0.975161296 |
| -4.929961655 | 8.22E-07 | 3.91E-06 |
| -4.649408928 | 3.33E-06 | 1.45E-05 |
| 0.557780506 | 0.576994276 | 0.678004165 |
| 0.159021377 | 0.873652032 | NA |
| -0.620244491 | 0.535096834 | NA |
| NA | NA | NA |
| NA | NA | NA |
| -3.789353614 | 0.00015104 | 0.000506027 |
| 1.073518634 | 0.283038483 | NA |
| 3.479899662 | 0.000501602 | 0.001530175 |

| | | |
|---|---|---|
| -5.384015479 | 7.28E-08 | 3.99E-07 |
| 2.260681404 | 0.023778993 | 0.048560988 |
| -2.141200354 | 0.032257884 | 0.063277026 |
| NA | NA | NA |
| NA | NA | NA |
| NA | NA | NA |
| NA | NA | NA |
| NA | NA | NA |
| NA | NA | NA |
| -1.597168035 | 0.110228256 | 0.180469207 |
| -0.065389369 | 0.947863988 | NA |
| NA | NA | NA |
| NA | NA | NA |
| 0.063708285 | 0.949202508 | NA |
| 2.903526413 | 0.003689858 | 0.009335607 |
| NA | NA | NA |
| -2.499702344 | 0.012429769 | 0.027625389 |
| 4.443877458 | 8.84E-06 | 3.63E-05 |
| -0.122428429 | 0.902559725 | NA |
| 1.323291766 | 0.185738369 | 0.278235165 |
| 18.47822921 | 3.09E-76 | 4.58E-74 |
| 2.845556115 | 0.004433395 | 0.011034215 |
| -0.902369425 | 0.366860658 | 0.479304847 |
| -0.113649484 | 0.909515659 | 0.937576869 |
| -0.002596581 | 0.99792823 | 0.998486713 |
| 8.353937915 | 6.60E-17 | 9.11E-16 |
| NA | NA | NA |
| NA | NA | NA |
| -0.372007838 | 0.709887014 | 0.787374187 |
| -3.60760527 | 0.000309036 | 0.000980129 |
| -4.191156337 | 2.78E-05 | 0.00010559 |
| 1.497358769 | 0.134299931 | 0.212945523 |
| NA | NA | NA |
| 0.192808924 | 0.847108615 | NA |
| 0.063708381 | 0.949202431 | NA |
| -0.600311313 | 0.548298781 | 0.651812799 |
| -1.086192201 | 0.277393967 | 0.384279998 |
| -0.194488217 | 0.845793621 | NA |
| NA | NA | NA |
| -0.582668692 | 0.560116353 | 0.662495684 |
| NA | NA | NA |
| NA | NA | NA |
| NA | NA | NA |
| NA | NA | NA |
| -2.941835417 | 0.003262733 | 0.008360586 |
| NA | NA | NA |
| NA | NA | NA |

| | | |
|---|---|---|
| -0.352627123 | 0.724368001 | 0.79871943 |
| -3.444845277 | 0.000571386 | 0.001724979 |
| NA | NA | NA |
| -0.615620012 | 0.53814534 | NA |
| 8.708626629 | 3.08E-18 | 4.73E-17 |
| 9.379529493 | 6.63E-21 | 1.25E-19 |
| 1.040352563 | 0.298176132 | 0.407010339 |
| 2.903251544 | 0.003693099 | 0.009342861 |
| -0.561325357 | 0.574575758 | 0.675733272 |
| NA | NA | NA |
| NA | NA | NA |
| NA | NA | NA |
| -3.127962025 | 0.001760229 | 0.004804639 |
| 2.282504931 | 0.022459547 | 0.046253287 |
| NA | NA | NA |
| 0.334843442 | 0.737743189 | NA |
| -0.26176171 | 0.793505161 | NA |
| 1.284864531 | 0.198839627 | NA |
| NA | NA | NA |
| NA | NA | NA |
| -0.537264281 | 0.591085075 | 0.690412964 |
| -3.814688342 | 0.000136355 | 0.00046034 |
| NA | NA | NA |
| 0.192809521 | 0.847108147 | NA |
| -0.109179307 | 0.913060273 | NA |
| -2.460368115 | 0.013879457 | 0.030447223 |
| NA | NA | NA |
| 0.173384133 | 0.862349496 | 0.904341725 |
| 0.526341422 | 0.598651008 | 0.69664953 |
| 0.063708285 | 0.949202508 | NA |
| 1.013398668 | 0.310869775 | 0.421030814 |
| -3.905505413 | 9.40E-05 | 0.000327153 |
| 2.344020231 | 0.019077137 | 0.040098749 |
| 0.091294499 | 0.927258589 | NA |
| -4.027227491 | 5.64E-05 | 0.000203934 |
| -0.065389369 | 0.947863988 | NA |
| NA | NA | NA |
| -2.049233941 | 0.040439245 | 0.076943694 |
| -1.886286694 | 0.059256331 | 0.106919404 |
| NA | NA | NA |
| NA | NA | NA |
| NA | NA | NA |
| 0.65959498 | 0.509513776 | 0.615498544 |
| NA | NA | NA |
| -2.283332672 | 0.02241078 | 0.046164252 |
| NA | NA | NA |
| -0.065389369 | 0.947863988 | NA |

| | | |
|---|---|---|
| NA | NA | NA |
| 2.383269955 | 0.017159606 | 0.036665949 |
| 0.439913262 | 0.65999993 | 0.746927623 |
| 0.06353712 | 0.949338802 | NA |
| 2.866267686 | 0.004153429 | 0.010404356 |
| 0.505685026 | 0.613077782 | NA |
| 13.62244091 | 2.95E-42 | 1.76E-40 |
| 0.068036266 | 0.945756765 | 0.962975181 |
| -0.588686483 | 0.556071607 | 0.658706688 |
| NA | NA | NA |
| -0.004615731 | 0.996317192 | 0.997521005 |
| 0.375794244 | 0.707069867 | NA |
| 1.876657953 | 0.060564989 | 0.108903856 |
| 1.607373757 | 0.107972403 | 0.177363767 |
| 0.974839893 | 0.329639685 | 0.440516888 |
| -4.278215711 | 1.88E-05 | 7.35E-05 |
| 0.702993727 | 0.482059659 | 0.590562075 |
| 0.156937051 | 0.875294459 | NA |
| -0.019662487 | 0.984312616 | 0.989039051 |
| 0.445158964 | 0.65620488 | NA |
| NA | NA | NA |
| 3.078531142 | 0.002080238 | 0.005581344 |
| NA | NA | NA |
| 2.527423544 | 0.011490284 | 0.025793884 |
| -0.551114833 | 0.581554958 | 0.681973369 |
| NA | NA | NA |
| NA | NA | NA |
| 9.839702104 | 7.59E-23 | 1.66E-21 |
| 3.427134745 | 0.000609986 | 0.001830471 |
| 0.192810118 | 0.847107679 | NA |
| NA | NA | NA |
| -2.926548465 | 0.003427461 | 0.008737966 |
| -2.494975577 | 0.01259658 | 0.027946499 |
| -0.065388772 | 0.947864464 | NA |
| -1.52474428 | 0.127322893 | 0.204026377 |
| -0.065388772 | 0.947864464 | NA |
| -0.413085216 | 0.679544181 | NA |
| 5.585966431 | 2.32E-08 | 1.36E-07 |
| -1.497860814 | 0.134169417 | 0.212817522 |
| -2.559847611 | 0.010471807 | 0.02375685 |
| -1.330469363 | 0.183363674 | 0.275386581 |
| 1.480601691 | 0.138712745 | 0.218721179 |
| -1.10869456 | 0.267561969 | 0.373449846 |
| -1.435171752 | 0.151238161 | 0.23513828 |
| 1.086682896 | 0.277176976 | 0.384043161 |
| -0.065389369 | 0.947863988 | NA |
| 1.279644003 | 0.200670367 | 0.296574119 |

| | | |
|---|---|---|
| NA | NA | NA |
| -4.793670229 | 1.64E-06 | 7.46E-06 |
| -0.329807528 | 0.741545399 | NA |
| NA | NA | NA |
| NA | NA | NA |
| 0.709002589 | 0.47832287 | 0.587142438 |
| 0.063708285 | 0.949202508 | NA |
| NA | NA | NA |
| -7.838548888 | 4.56E-15 | 5.35E-14 |
| 0.901704157 | 0.367214044 | NA |
| NA | NA | NA |
| 2.733113685 | 0.006273867 | 0.015045981 |
| NA | NA | NA |
| NA | NA | NA |
| 0.038754172 | 0.969086383 | 0.978683157 |
| -2.510676072 | 0.012050021 | 0.02686249 |
| 0.514127472 | 0.607162871 | 0.703773579 |
| NA | NA | NA |
| -1.454119636 | 0.14591314 | 0.22815593 |
| 1.706433219 | 0.08792745 | 0.14970574 |
| 0.591293281 | 0.554323931 | 0.657320825 |
| -8.490009204 | 2.07E-17 | 2.98E-16 |
| NA | NA | NA |
| NA | NA | NA |
| -5.491927326 | 3.98E-08 | 2.26E-07 |
| NA | NA | NA |
| NA | NA | NA |
| 0.126725397 | 0.899157744 | NA |
| NA | NA | NA |
| -1.067590928 | 0.285705078 | 0.392815056 |
| -1.677425341 | 0.093459337 | 0.157510143 |
| NA | NA | NA |
| -3.297077709 | 0.000976965 | 0.002822092 |
| 0.192810118 | 0.847107679 | NA |
| 1.898753764 | 0.057596859 | 0.104493252 |
| 1.314721202 | 0.188603618 | 0.28175163 |
| 5.351855632 | 8.71E-08 | 4.73E-07 |
| 0.192809521 | 0.847108147 | NA |
| 2.301158119 | 0.021382695 | 0.04427971 |
| -1.483961784 | 0.137819059 | 0.217586165 |
| 0.261522889 | 0.793689301 | 0.853352371 |
| 7.624948772 | 2.44E-14 | 2.67E-13 |
| -0.318989547 | 0.749734439 | 0.818254688 |
| 0.840623009 | 0.400559165 | 0.512209864 |
| 0.620426846 | 0.534976803 | 0.63937685 |
| 0.199526594 | 0.841850842 | NA |
| 0.192809521 | 0.847108147 | NA |

| | | |
|---|---|---|
| 0.06371127 | 0.949200131 | NA |
| -0.436928286 | 0.662163357 | 0.748698269 |
| NA | NA | NA |
| NA | NA | NA |
| -0.065389369 | 0.947863988 | NA |
| NA | NA | NA |
| 0.126823811 | 0.899079849 | NA |
| 1.531953199 | 0.125533984 | 0.20159859 |
| 0.320071032 | 0.748914485 | NA |
| -4.594905189 | 4.33E-06 | 1.86E-05 |
| NA | NA | NA |
| 0.063708285 | 0.949202508 | NA |
| 0.330493631 | 0.741027004 | NA |
| NA | NA | NA |
| NA | NA | NA |
| NA | NA | NA |
| NA | NA | NA |
| 1.490868145 | 0.135996116 | 0.215196788 |
| NA | NA | NA |
| NA | NA | NA |
| NA | NA | NA |
| 5.620266008 | 1.91E-08 | 1.13E-07 |
| NA | NA | NA |
| 0.819201519 | 0.412671448 | NA |
| 9.525276404 | 1.65E-21 | 3.30E-20 |
| -4.773272121 | 1.81E-06 | 8.22E-06 |
| -1.396992076 | 0.162415954 | 0.249021214 |
| -3.936920955 | 8.25E-05 | 0.000289818 |
| NA | NA | NA |
| 1.434650769 | 0.15138664 | NA |
| 2.277357285 | 0.022764899 | 0.046808955 |
| 0.063708285 | 0.949202508 | NA |
| -2.747729219 | 0.006000955 | 0.01445772 |
| 1.144822577 | 0.252282661 | 0.356336268 |
| 6.115035619 | 9.65E-10 | 6.59E-09 |
| -5.638608787 | 1.71E-08 | 1.02E-07 |
| 0.794482089 | 0.426914832 | 0.538256654 |
| -0.644804116 | 0.519054134 | NA |
| NA | NA | NA |
| NA | NA | NA |
| NA | NA | NA |
| NA | NA | NA |
| NA | NA | NA |
| -0.822429382 | 0.410832563 | 0.522650015 |
| 2.621164763 | 0.00876299 | 0.020226636 |
| -0.032179953 | 0.974328543 | 0.98223601 |
| 2.009093708 | 0.044527197 | 0.083754095 |

| | | | |
|---|---|---|---|
| | -5.299878781 | 1.16E-07 | 6.19E-07 |
| | 0.211897103 | 0.832187311 | 0.882574822 |
| | -5.392392912 | 6.95E-08 | 3.82E-07 |
| | 0.192809521 | 0.847108147 | NA |
| NA | NA | NA | |
| NA | NA | NA | |
| | -1.449032081 | 0.147328624 | 0.229959806 |
| | 0.015001202 | 0.988031221 | 0.991676216 |
| | -0.336925975 | 0.736172704 | 0.807715021 |
| | -7.694529178 | 1.42E-14 | 1.60E-13 |
| | 0.414807055 | 0.678283155 | NA |
| NA | NA | NA | |
| NA | NA | NA | |
| NA | NA | NA | |
| NA | NA | NA | |
| | 0.063708285 | 0.949202508 | NA |
| NA | NA | NA | |
| | -1.504425112 | 0.132471942 | 0.210538619 |
| | -0.194488217 | 0.845793621 | NA |
| | 0.483711736 | 0.628590463 | 0.721394741 |
| | -2.413702949 | 0.015791331 | 0.034097396 |
| | 0.583695982 | 0.559424871 | NA |
| NA | NA | NA | |
| | -4.536055146 | 5.73E-06 | 2.42E-05 |
| | 3.061827481 | 0.002199902 | 0.00587094 |
| | -0.942816879 | 0.345774577 | 0.457269385 |
| | -0.108194776 | 0.913841189 | NA |
| | -0.760638164 | 0.446873218 | 0.55681186 |
| | -0.157712673 | 0.874683214 | 0.912678041 |
| | -3.404304359 | 0.000663328 | 0.001976795 |
| | -1.422859633 | 0.15477685 | 0.239627919 |
| | 0.590584177 | 0.554799069 | 0.657654335 |
| | -1.169855504 | 0.242059123 | 0.344638216 |
| | -0.065389369 | 0.947863988 | NA |
| NA | NA | NA | |
| | 0.063708285 | 0.949202508 | NA |
| | -0.065389966 | 0.947863513 | NA |
| NA | NA | NA | |
| | 1.687616975 | 0.091484778 | 0.154657842 |
| NA | NA | NA | |
| | -6.966337711 | 3.25E-12 | 2.85E-11 |
| NA | NA | NA | |
| NA | NA | NA | |
| NA | NA | NA | |
| NA | NA | NA | |
| NA | NA | NA | |
| | -0.154598604 | 0.877137771 | 0.914172152 |

| | | |
|---|---|---|
| NA | NA | NA |
| NA | NA | NA |
| -0.441236305 | 0.659041933 | NA |
| NA | NA | NA |
| NA | NA | NA |
| NA | NA | NA |
| NA | NA | NA |
| 0.063708882 | 0.949202033 | NA |
| NA | NA | NA |
| 0.310025982 | 0.756541198 | 0.823493202 |
| NA | NA | NA |
| NA | NA | NA |
| -0.264588435 | 0.79132653 | NA |
| NA | NA | NA |
| NA | NA | NA |
| NA | NA | NA |
| NA | NA | NA |
| NA | NA | NA |
| NA | NA | NA |
| -4.294038725 | 1.75E-05 | 6.88E-05 |
| -0.765143007 | 0.444186393 | 0.554136618 |
| 0.194173692 | 0.846039881 | NA |
| NA | NA | NA |
| NA | NA | NA |
| NA | NA | NA |
| NA | NA | NA |
| NA | NA | NA |
| 0.523787264 | 0.600426512 | NA |
| 0.192809521 | 0.847108147 | NA |
| NA | NA | NA |
| -0.394795184 | 0.692994049 | NA |
| 0.063708285 | 0.949202508 | NA |
| 0.065152415 | 0.948052648 | NA |
| 2.788819393 | 0.005290056 | 0.012933199 |
| NA | NA | NA |
| NA | NA | NA |
| 0.656099977 | 0.511759799 | 0.617526287 |
| NA | NA | NA |
| NA | NA | NA |
| -2.239440207 | 0.025127288 | 0.050946619 |
| -3.113864133 | 0.001846545 | 0.005009106 |
| -2.529481711 | 0.011423113 | 0.02566381 |
| 9.639074899 | 5.47E-22 | 1.13E-20 |
| 0.063708882 | 0.949202033 | NA |
| 0.282742491 | 0.777374242 | NA |
| -0.159242165 | 0.873478085 | 0.911838698 |
| NA | NA | NA |

| | | | |
|---|---|---|---|
| | -1.564526026 | 0.117694086 | 0.190721415 |
| | 0.774531381 | 0.438616618 | 0.548983596 |
| | -3.113112389 | 0.001851255 | 0.005021339 |
| NA | NA | NA | |
| | 2.331641815 | 0.019719542 | 0.041276145 |
| | -5.33296356 | 9.66E-08 | 5.22E-07 |
| | 0.063708882 | 0.949202033 | NA |
| | -3.592415616 | 0.000327627 | 0.001034372 |
| | 1.909621803 | 0.056181926 | 0.102293853 |
| NA | NA | NA | |
| | -0.102148683 | 0.918638659 | 0.944261988 |
| | 1.960709163 | 0.04991296 | 0.092466981 |
| | -3.984346776 | 6.77E-05 | 0.000241059 |
| NA | NA | NA | |
| | -5.939737096 | 2.85E-09 | 1.85E-08 |
| | -3.540657963 | 0.000399131 | 0.001239484 |
| | 0.178173284 | 0.85858689 | 0.901822487 |
| | 0.125186614 | 0.900375815 | 0.930525895 |
| | 1.084426698 | 0.278175652 | 0.385149716 |
| NA | NA | NA | |
| | 0.063708217 | 0.949202562 | NA |
| NA | NA | NA | |
| | -3.156115801 | 0.001598854 | 0.004406927 |
| NA | NA | NA | |
| | 0.176534198 | 0.859874285 | NA |
| NA | NA | NA | |
| | 0.067782637 | 0.945958666 | NA |
| | 3.384985426 | 0.00071182 | 0.002110003 |
| | 0.31685483 | 0.751353758 | NA |
| | 0.192809521 | 0.847108147 | NA |
| NA | NA | NA | |
| NA | NA | NA | |
| | 0.549428521 | 0.582711405 | 0.682894769 |
| | 3.673940779 | 0.000238838 | 0.000773066 |
| | 0.274555299 | 0.783657913 | 0.845545201 |
| | -1.960409331 | 0.049947966 | 0.092511177 |
| | -1.973580478 | 0.048429465 | 0.090018529 |
| | 0.473112592 | 0.636132854 | 0.727449074 |
| | -0.706370544 | 0.479957734 | 0.588638108 |
| | -1.934064849 | 0.053105155 | 0.097464587 |
| | 3.563714206 | 0.000365644 | 0.001143866 |
| | -0.702323144 | 0.482477663 | 0.590967258 |
| NA | NA | NA | |
| | 1.202257964 | 0.229263598 | 0.3297591 |
| NA | NA | NA | |
| | -0.065389369 | 0.947863988 | NA |
| | 0.063710673 | 0.949200606 | NA |

| | | | |
|---|---|---|---|
| | -1.853045234 | 0.063875875 | 0.113932385 |
| NA | NA | NA | |
| | 0.192810118 | 0.847107679 | NA |
| | 2.368756601 | 0.017847995 | 0.037913546 |
| | 1.660774542 | 0.096758736 | 0.162084805 |
| | -1.284162267 | 0.199085185 | 0.294544401 |
| NA | NA | NA | |
| NA | NA | NA | |
| | -6.201403781 | 5.60E-10 | 3.90E-09 |
| | -4.476252592 | 7.60E-06 | 3.15E-05 |
| NA | NA | NA | |
| | -2.472325561 | 0.013423721 | 0.029561475 |
| | -1.916925484 | 0.055247399 | 0.100871136 |
| NA | NA | NA | |
| NA | NA | NA | |
| NA | NA | NA | |
| | -1.306514834 | 0.191377538 | 0.285168664 |
| | 1.230964752 | 0.218336048 | 0.31717651 |
| NA | NA | NA | |
| | -0.352172322 | 0.724709033 | 0.798989874 |
| | -0.065389369 | 0.947863988 | NA |
| NA | NA | NA | |
| NA | NA | NA | |
| NA | NA | NA | |
| | 0.076268466 | 0.939205513 | NA |
| NA | NA | NA | |
| NA | NA | NA | |
| NA | NA | NA | |
| NA | NA | NA | |
| NA | NA | NA | |
| | 0.08710423 | 0.930588663 | NA |
| NA | NA | NA | |
| | 0.063708882 | 0.949202033 | NA |
| | -0.065390563 | 0.947863038 | NA |
| NA | NA | NA | |
| NA | NA | NA | |
| | 0.192810118 | 0.847107679 | NA |
| NA | NA | NA | |
| | 0.063708882 | 0.949202033 | NA |
| NA | NA | NA | |
| | -1.197333246 | 0.231176691 | 0.331976625 |
| | 27.31308646 | 2.97E-164 | 1.46E-161 |
| | 3.173033973 | 0.001508549 | 0.004176878 |
| | 1.309070167 | 0.190510584 | 0.284125598 |
| | -0.936949721 | 0.348784421 | 0.460471231 |
| NA | NA | NA | |
| | 0.096682309 | 0.92297869 | NA |

| | | |
|---|---|---|
| -2.780798832 | 0.005422533 | 0.013229944 |
| 0.382405941 | 0.702160285 | NA |
| NA | NA | NA |
| -2.69993762 | 0.006935248 | 0.01644275 |
| -0.808705013 | 0.418684844 | 0.530343565 |
| -2.198493271 | 0.027913974 | 0.055760622 |
| 3.286421905 | 0.001014689 | 0.002921287 |
| -0.284430306 | 0.776080637 | NA |
| 3.130525164 | 0.00174494 | 0.004767066 |
| 2.045640719 | 0.040791732 | 0.077531859 |
| 14.57831053 | 3.86E-48 | 2.84E-46 |
| 0.562742821 | 0.573610021 | NA |
| -0.877315793 | 0.380315137 | NA |
| 1.885020954 | 0.059427009 | 0.107196457 |
| 0.059654288 | 0.952430979 | 0.967567454 |
| 0.063708285 | 0.949202508 | NA |
| 4.735923073 | 2.18E-06 | 9.77E-06 |
| 3.929533989 | 8.51E-05 | 0.000298155 |
| -1.487518307 | 0.136877972 | 0.216387017 |
| 0.525436181 | 0.599280007 | 0.697187013 |
| -0.311818627 | 0.755178367 | NA |
| NA | NA | NA |
| 1.880895465 | 0.05998614 | 0.108041526 |
| -1.097866691 | 0.272262705 | 0.378743706 |
| -0.430085243 | 0.667133634 | NA |
| 4.496107947 | 6.92E-06 | 2.89E-05 |
| -1.764960828 | 0.077570349 | 0.134486794 |
| NA | NA | NA |
| 2.013111747 | 0.044102875 | 0.083055824 |
| -1.324835885 | 0.18522559 | 0.277633183 |
| -0.857008379 | 0.391440254 | 0.503455606 |
| 0.175389712 | 0.860773424 | NA |
| 0.069839618 | 0.944321313 | NA |
| 4.00096769 | 6.31E-05 | 0.000225796 |
| NA | NA | NA |
| -0.918755673 | 0.358223383 | 0.470105976 |
| -1.941143175 | 0.052240913 | NA |
| 7.405637138 | 1.31E-13 | 1.32E-12 |
| 3.742386965 | 0.000182281 | 0.000602626 |
| 0.901713636 | 0.367209008 | 0.479682618 |
| -5.172591732 | 2.31E-07 | 1.19E-06 |
| 0.673257039 | 0.500783779 | 0.607856942 |
| -0.043061639 | 0.965652399 | 0.976695338 |
| -4.145999113 | 3.38E-05 | 0.000126837 |
| 0.039755131 | 0.968288348 | 0.978373056 |
| -1.761297683 | 0.078188032 | 0.135407628 |
| -3.943022112 | 8.05E-05 | 0.000282977 |

| | | |
|---|---|---|
| -1.181071944 | 0.237574144 | 0.339599938 |
| -2.028001453 | 0.0425601 | 0.080477396 |
| -4.1899471 | 2.79E-05 | 0.000106105 |
| 0.660110666 | 0.509182815 | 0.61518781 |
| 4.423353515 | 9.72E-06 | 3.97E-05 |
| 0.603852813 | 0.545941503 | 0.649534928 |
| 0.087200399 | 0.930512223 | 0.952702619 |
| 1.324421833 | 0.185362988 | 0.277789222 |
| -6.195162864 | 5.82E-10 | 4.05E-09 |
| -10.57384184 | 3.94E-26 | 1.08E-24 |
| 1.696129336 | 0.089861389 | 0.152438534 |
| 0.063707688 | 0.949202983 | NA |
| NA | NA | NA |
| NA | NA | NA |
| NA | NA | NA |
| NA | NA | NA |
| NA | NA | NA |
| NA | NA | NA |
| NA | NA | NA |
| 0.192810118 | 0.847107679 | NA |
| -0.930651449 | 0.352033892 | NA |
| NA | NA | NA |
| NA | NA | NA |
| 0.192809521 | 0.847108147 | NA |
| 0.06371127 | 0.949200131 | NA |
| 0.063707194 | 0.949203376 | NA |
| 0.631370519 | 0.527798288 | 0.632731028 |
| 0.063708882 | 0.949202033 | NA |
| NA | NA | NA |
| 0.100792124 | 0.91971548 | NA |
| NA | NA | NA |
| NA | NA | NA |
| NA | NA | NA |
| NA | NA | NA |
| NA | NA | NA |
| NA | NA | NA |
| -3.591199665 | 0.000329159 | 0.001038948 |
| NA | NA | NA |
| 0.220877413 | 0.825187885 | NA |
| NA | NA | NA |
| NA | NA | NA |
| -0.065389966 | 0.947863513 | NA |
| 0.220381707 | 0.825573891 | NA |
| NA | NA | NA |
| NA | NA | NA |
| 0.290157392 | 0.771695831 | 0.835700542 |
| -0.050403657 | 0.959800722 | NA |

| | | |
|---|---|---|
| -5.892093665 | 3.81E-09 | 2.44E-08 |
| -4.769891792 | 1.84E-06 | 8.34E-06 |
| -2.797689643 | 0.005146954 | 0.012625271 |
| 7.545028841 | 4.52E-14 | 4.81E-13 |
| 2.532946742 | 0.011310814 | 0.025441199 |
| -0.982260621 | 0.325971469 | 0.437069849 |
| -2.509124794 | 0.012103072 | 0.026971144 |
| 1.292296937 | 0.196254332 | 0.291078807 |
| -8.227440552 | 1.91E-16 | 2.54E-15 |
| -1.733107515 | 0.083076561 | 0.142630243 |
| 5.54840914 | 2.88E-08 | 1.67E-07 |
| 6.239099095 | 4.40E-10 | 3.11E-09 |
| -1.545248469 | 0.122286175 | 0.197129062 |
| -1.480771023 | 0.138667601 | 0.218677548 |
| 2.701657607 | 0.006899477 | 0.016369091 |
| -1.275407699 | 0.202164975 | NA |
| 9.264499119 | 1.96E-20 | 3.58E-19 |
| -3.53471787 | 0.000408211 | 0.001265484 |
| -2.97326178 | 0.00294653 | 0.007633166 |
| NA | NA | NA |
| NA | NA | NA |
| 1.554505157 | 0.120063968 | 0.194059222 |
| -2.961207063 | 0.003064358 | 0.007903335 |
| -1.975219416 | 0.048243253 | 0.089725712 |
| -1.024050532 | 0.305811408 | 0.415438781 |
| 16.79149703 | 2.82E-63 | 3.05E-61 |
| 3.422052891 | 0.000621502 | 0.001862237 |
| 2.457786857 | 0.013979612 | 0.03064276 |
| 1.250758464 | 0.211022613 | 0.308720323 |
| 0.30564061 | 0.759878297 | 0.82615558 |
| -0.271053891 | 0.786349584 | 0.847718584 |
| -0.887367535 | 0.374881056 | 0.487772149 |
| -1.530670288 | 0.125850901 | 0.20200385 |
| 14.33898784 | 1.25E-46 | 8.80E-45 |
| 7.072025869 | 1.53E-12 | 1.39E-11 |
| -5.829378326 | 5.56E-09 | 3.49E-08 |
| 4.568135403 | 4.92E-06 | 2.10E-05 |
| -0.109259919 | 0.912996336 | NA |
| 1.171849922 | 0.241257316 | 0.343711686 |
| -1.158808199 | 0.246534375 | 0.349558454 |
| 0.943603739 | 0.345372181 | 0.456857926 |
| -3.457720916 | 0.000544765 | 0.001649786 |
| 0.844503319 | 0.39838821 | 0.510267981 |
| 1.363893999 | 0.17260093 | 0.261973275 |
| -2.453980545 | 0.014128462 | 0.030909485 |
| 2.340990922 | 0.019232635 | 0.040385701 |
| -9.13338341 | 6.64E-20 | 1.18E-18 |

| | | |
|---|---|---|
| -1.467909493 | 0.142128809 | 0.223291746 |
| NA | NA | NA |
| NA | NA | NA |
| 2.913963411 | 0.003568717 | 0.009058461 |
| 0.787030114 | 0.431264238 | 0.542197669 |
| 0.38478338 | 0.700397914 | NA |
| NA | NA | NA |
| NA | NA | NA |
| NA | NA | NA |
| NA | NA | NA |
| -0.065388772 | 0.947864464 | NA |
| NA | NA | NA |
| 0.06371127 | 0.949200131 | NA |
| -0.602743662 | 0.546679231 | 0.650282168 |
| -0.651977816 | 0.514415484 | 0.620004155 |
| 15.34777615 | 3.67E-53 | 3.01E-51 |
| 1.796356994 | 0.072437759 | 0.126815316 |
| 2.667912107 | 0.007632422 | 0.01787588 |
| -0.068037904 | 0.945755461 | 0.962975181 |
| 0.269113449 | 0.787842381 | NA |
| 1.203688432 | 0.228710025 | NA |
| -0.149985659 | 0.88077593 | NA |
| -2.29794101 | 0.021565147 | 0.044598498 |
| 0.357652404 | 0.720603456 | 0.795655058 |
| 6.617965544 | 3.64E-11 | 2.90E-10 |
| NA | NA | NA |
| NA | NA | NA |
| NA | NA | NA |
| 7.904210464 | 2.70E-15 | 3.24E-14 |
| -0.19448762 | 0.845794089 | NA |
| 1.847135223 | 0.064727545 | 0.11522341 |
| 1.11268784 | 0.26584252 | 0.3715115 |
| 0.194539923 | 0.845753139 | 0.891935665 |
| -0.056647589 | 0.954825925 | 0.969073591 |
| -3.293785115 | 0.00098848 | 0.002852723 |
| -0.926732129 | 0.354065632 | 0.465993415 |
| NA | NA | NA |
| NA | NA | NA |
| 1.517548341 | 0.129128298 | 0.206311841 |
| -5.735842446 | 9.70E-09 | 5.94E-08 |
| 7.283724521 | 3.25E-13 | 3.17E-12 |
| 1.946407691 | 0.051605787 | 0.09513831 |
| 3.466244291 | 0.000527784 | 0.001603401 |
| -3.441972582 | 0.000577489 | 0.001741511 |
| 2.653055617 | 0.007976673 | 0.018577846 |
| NA | NA | NA |
| 2.755196061 | 0.005865697 | 0.014171661 |

| | | | |
|---|---|---|---|
| | -0.255610734 | 0.798251429 | NA |
| | -0.584502282 | 0.55888243 | 0.661241566 |
| | 4.282968768 | 1.84E-05 | 7.21E-05 |
| NA | NA | NA | |
| | 2.724619809 | 0.006437557 | 0.015392821 |
| | 0.532539948 | 0.594352082 | 0.693193821 |
| NA | NA | NA | |
| | 3.389201589 | 0.000700965 | 0.002080534 |
| | 0.086449216 | 0.931109325 | NA |
| NA | NA | NA | |
| | -0.75769156 | 0.448635653 | 0.558618565 |
| | -1.363634805 | 0.172682532 | 0.262065348 |
| | 0.478281401 | 0.632449933 | NA |
| | -0.065389369 | 0.947863988 | NA |
| | 0.699111915 | 0.484482093 | NA |
| | 0.063708285 | 0.949202508 | NA |
| | 7.719679827 | 1.17E-14 | 1.32E-13 |
| | 5.981799289 | 2.21E-09 | 1.45E-08 |
| | 11.70055939 | 1.27E-31 | 4.78E-30 |
| | 1.62751654 | 0.103627439 | 0.171547274 |
| | 15.07372392 | 2.41E-51 | 1.90E-49 |
| | -5.227282131 | 1.72E-07 | 8.98E-07 |
| | 3.577458309 | 0.000346952 | 0.001090842 |
| | -0.042092355 | 0.966425075 | NA |
| | 1.621856017 | 0.104834181 | 0.17320129 |
| | 0.441641294 | 0.658748798 | 0.746017664 |
| | -0.857484498 | 0.391177178 | 0.503153488 |
| NA | NA | NA | |
| | -0.19448762 | 0.845794089 | NA |
| | -1.674738571 | 0.093985536 | 0.158214473 |
| | 0.683976363 | 0.493990092 | NA |
| | -0.272960672 | 0.784883447 | 0.846612052 |
| | 1.318093899 | 0.187472217 | 0.280329147 |
| | -0.065390563 | 0.947863038 | NA |
| | 5.785989562 | 7.21E-09 | 4.47E-08 |
| | 3.064956541 | 0.002177016 | 0.005817687 |
| | 15.08340104 | 2.08E-51 | 1.65E-49 |
| | -0.099266104 | 0.92092699 | NA |
| | 1.716530393 | 0.08606501 | NA |
| | -1.756080966 | 0.079074585 | 0.136715686 |
| | 4.728580691 | 2.26E-06 | 1.01E-05 |
| | -6.102079521 | 1.05E-09 | 7.12E-09 |
| | -0.376979401 | 0.706188913 | 0.784480228 |
| | -3.178056776 | 0.001482657 | 0.004111556 |
| NA | NA | NA | |
| | -0.163245942 | 0.870324794 | NA |
| | -0.730325886 | 0.465191009 | 0.574634417 |

| | | |
|---|---|---|
| -1.26166626 | 0.207068902 | 0.303958567 |
| NA | NA | NA |
| -0.375664616 | 0.707166246 | 0.785217423 |
| -3.099449152 | 0.001938809 | 0.005237252 |
| -0.265266273 | 0.790804344 | 0.851164385 |
| 0.54090861 | 0.588570576 | 0.688182441 |
| -1.729798479 | 0.083666286 | 0.143426282 |
| 6.622476156 | 3.53E-11 | 2.81E-10 |
| NA | NA | NA |
| NA | NA | NA |
| 0.192809521 | 0.847108147 | NA |
| NA | NA | NA |
| NA | NA | NA |
| -1.958373341 | 0.050186221 | 0.092883844 |
| -0.024823727 | 0.980195565 | 0.986341897 |
| 5.661485319 | 1.50E-08 | 8.99E-08 |
| -2.10227432 | 0.035529253 | 0.0687874 |
| 0.202578279 | 0.839464669 | 0.887547548 |
| 4.267669659 | 1.98E-05 | 7.68E-05 |
| 1.455167197 | 0.14562298 | 0.227794712 |
| -0.236017191 | 0.813419331 | 0.868781133 |
| -7.378229152 | 1.60E-13 | 1.61E-12 |
| -2.61326018 | 0.0089683 | 0.02066052 |
| -1.402698158 | 0.160706867 | 0.246763565 |
| NA | NA | NA |
| NA | NA | NA |
| NA | NA | NA |
| 0.063708285 | 0.949202508 | NA |
| NA | NA | NA |
| -6.942133228 | 3.86E-12 | 3.36E-11 |
| -1.265564814 | 0.20566893 | NA |
| -1.053120278 | 0.29228587 | 0.400170405 |
| -0.065389966 | 0.947863513 | NA |
| 0.375550276 | 0.707251263 | NA |
| -0.065390705 | 0.947862925 | NA |
| 0.063708285 | 0.949202508 | NA |
| 1.122711391 | 0.261560088 | 0.366716827 |
| NA | NA | NA |
| 0.625040226 | 0.531944657 | 0.636512801 |
| 0.932026693 | 0.351322731 | 0.463040806 |
| -0.398620713 | 0.690172697 | 0.771293443 |
| -2.813100258 | 0.004906635 | 0.012104869 |
| -3.369218653 | 0.000753816 | 0.002224477 |
| -0.941547744 | 0.346424234 | 0.457983341 |
| -5.791055462 | 6.99E-09 | 4.34E-08 |
| -2.182960333 | 0.029038731 | 0.05770857 |
| -3.443935529 | 0.000573312 | 0.001730167 |

| | | | |
|---|---|---|---|
| | -0.065388772 | 0.947864464 | NA |
| NA | NA | NA | |
| | 0.192808924 | 0.847108615 | NA |
| NA | NA | NA | |
| | -0.672880657 | 0.501023219 | 0.608000154 |
| NA | NA | NA | |
| NA | NA | NA | |
| NA | NA | NA | |
| | 1.17868454 | 0.238523814 | NA |
| NA | NA | NA | |
| | -0.08569216 | 0.931711135 | NA |
| NA | NA | NA | |
| | 0.063708217 | 0.949202562 | NA |
| NA | NA | NA | |
| | 2.43960115 | 0.014703486 | 0.031996892 |
| | -1.205201186 | 0.228125645 | 0.328480981 |
| | 0.390039422 | 0.696507396 | 0.776551453 |
| NA | NA | NA | |
| | 0.192810118 | 0.847107679 | NA |
| NA | NA | NA | |
| NA | NA | NA | |
| | 0.723797733 | 0.469189921 | NA |
| NA | NA | NA | |
| | 0.063708882 | 0.949202033 | NA |
| | 0.86098243 | 0.38924772 | NA |
| | 0.192809521 | 0.847108147 | NA |
| | -0.065390563 | 0.947863038 | NA |
| NA | NA | NA | |
| NA | NA | NA | |
| | 5.423444669 | 5.85E-08 | 3.25E-07 |
| | 7.02826495 | 2.09E-12 | 1.87E-11 |
| | 0.933076844 | 0.350780294 | 0.462504567 |
| | 0.192810118 | 0.847107679 | NA |
| | 2.697919081 | 0.00697744 | 0.0165297 |
| NA | NA | NA | |
| | -0.307349979 | 0.758576998 | NA |
| | 2.017796746 | 0.043612434 | 0.082268387 |
| | 0.336185442 | 0.73673103 | NA |
| | -7.115140759 | 1.12E-12 | 1.03E-11 |
| | -2.66015697 | 0.007810424 | 0.018237955 |
| | -6.70618468 | 2.00E-11 | 1.63E-10 |
| NA | NA | NA | |
| | 2.836222317 | 0.004565066 | 0.011336462 |
| NA | NA | NA | |
| | 3.727106858 | 0.00019369 | 0.000638072 |
| NA | NA | NA | |
| | -0.065388772 | 0.947864464 | NA |

| | | | |
|---|---|---|---|
| | -0.065388772 | 0.947864464 | NA |
| NA | NA | NA | |
| NA | NA | NA | |
| | -0.065389369 | 0.947863988 | NA |
| NA | NA | NA | |
| NA | NA | NA | |
| NA | NA | NA | |
| NA | NA | NA | |
| | 0.06353712 | 0.949338802 | NA |
| NA | NA | NA | |
| NA | NA | NA | |
| NA | NA | NA | |
| | -6.413019384 | 1.43E-10 | 1.06E-09 |
| | 0.192810118 | 0.847107679 | NA |
| | 0.424625959 | 0.671109366 | NA |
| NA | NA | NA | |
| | 0.085850022 | 0.931585641 | NA |
| NA | NA | NA | |
| NA | NA | NA | |
| | 4.778904921 | 1.76E-06 | 8.00E-06 |
| NA | NA | NA | |
| NA | NA | NA | |
| | -0.065388772 | 0.947864464 | NA |
| NA | NA | NA | |
| NA | NA | NA | |
| NA | NA | NA | |
| | 7.266334197 | 3.69E-13 | 3.58E-12 |
| | 0.464422869 | 0.642344809 | 0.732504476 |
| | 0.145820112 | 0.884063398 | 0.918826843 |
| | 0.194351939 | 0.845900319 | NA |
| NA | NA | NA | |
| | 3.68608269 | 0.000227732 | 0.00074047 |
| NA | NA | NA | |
| | 0.192808327 | 0.847109082 | NA |
| | -0.463717484 | 0.642850169 | 0.732787126 |
| NA | NA | NA | |
| | -0.065389369 | 0.947863988 | NA |
| | -4.274423909 | 1.92E-05 | 7.46E-05 |
| | -2.264756059 | 0.023527653 | 0.048132175 |
| | -0.429844676 | 0.667308632 | 0.752531697 |
| | -0.698554038 | 0.484830776 | 0.593239847 |
| | -5.85169608 | 4.87E-09 | 3.08E-08 |
| | -1.531915014 | 0.125543408 | 0.201600789 |
| | -0.065388772 | 0.947864464 | NA |
| | 0.192810118 | 0.847107679 | NA |
| | -9.42381745 | 4.35E-21 | 8.41E-20 |
| | -10.11385947 | 4.80E-24 | 1.15E-22 |

| | | |
|---|---|---|
| 5.246416284 | 1.55E-07 | 8.15E-07 |
| -5.814330552 | 6.09E-09 | 3.81E-08 |
| -0.570284562 | 0.56848471 | 0.670112663 |
| NA | NA | NA |
| -0.740601409 | 0.458935153 | 0.56838903 |
| 0.298008385 | 0.765696764 | 0.830957767 |
| -5.258136352 | 1.46E-07 | 7.67E-07 |
| NA | NA | NA |
| -3.112709038 | 0.001853787 | 0.005027116 |
| 0.687674432 | 0.491657824 | 0.599016518 |
| 0.537082633 | 0.591210537 | NA |
| -0.850303451 | 0.395156399 | 0.507140679 |
| NA | NA | NA |
| 0.310999995 | 0.755800625 | 0.822973404 |
| 9.402687952 | 5.32E-21 | 1.01E-19 |
| NA | NA | NA |
| -3.817467701 | 0.000134828 | 0.000455609 |
| -0.92727077 | 0.353785967 | 0.465698803 |
| -0.550651959 | 0.581872283 | 0.68219885 |
| -1.299723189 | 0.19369586 | 0.287949597 |
| -0.662897446 | 0.507396238 | NA |
| 0.111863074 | 0.910931975 | 0.938628671 |
| NA | NA | NA |
| -0.065389369 | 0.947863988 | NA |
| 0.486164455 | 0.626850571 | 0.719981236 |
| 2.508427538 | 0.012126984 | 0.027012406 |
| 1.365475029 | 0.172103796 | 0.261313802 |
| NA | NA | NA |
| -10.5129736 | 7.53E-26 | 2.04E-24 |
| 4.659672111 | 3.17E-06 | 1.39E-05 |
| 0.818919959 | 0.412832082 | 0.524447394 |
| NA | NA | NA |
| -0.610570477 | 0.541483973 | 0.645520282 |
| 0.434555644 | 0.663884997 | NA |
| 0.192805732 | 0.847111115 | NA |
| NA | NA | NA |
| 0.923101766 | 0.355954174 | 0.467888509 |
| 1.86447319 | 0.062255283 | 0.111478734 |
| 1.256426345 | 0.208961435 | NA |
| -3.450765206 | 0.000559 | 0.00169003 |
| -1.324521711 | 0.185329837 | 0.277756172 |
| 13.43080601 | 3.99E-41 | 2.26E-39 |
| NA | NA | NA |
| 1.082397904 | 0.279075759 | 0.386075624 |
| 5.453145735 | 4.95E-08 | 2.78E-07 |
| -3.985252624 | 6.74E-05 | 0.00024021 |
| 1.498179718 | 0.134086564 | 0.212713062 |

| | | |
|---|---|---|
| -2.07996072 | 0.037529136 | 0.072108257 |
| -0.696481095 | 0.486127593 | 0.594361745 |
| 0.192810118 | 0.847107679 | NA |
| NA | NA | NA |
| -0.437908155 | 0.661452864 | NA |
| 0.192808924 | 0.847108615 | NA |
| 0.063708882 | 0.949202033 | NA |
| NA | NA | NA |
| NA | NA | NA |
| NA | NA | NA |
| NA | NA | NA |
| NA | NA | NA |
| NA | NA | NA |
| NA | NA | NA |
| NA | NA | NA |
| NA | NA | NA |
| NA | NA | NA |
| NA | NA | NA |
| NA | NA | NA |
| NA | NA | NA |
| NA | NA | NA |
| NA | NA | NA |
| NA | NA | NA |
| NA | NA | NA |
| NA | NA | NA |
| NA | NA | NA |
| NA | NA | NA |
| NA | NA | NA |
| NA | NA | NA |
| NA | NA | NA |
| NA | NA | NA |
| NA | NA | NA |
| NA | NA | NA |
| NA | NA | NA |
| NA | NA | NA |
| NA | NA | NA |
| NA | NA | NA |
| NA | NA | NA |
| NA | NA | NA |
| NA | NA | NA |
| NA | NA | NA |
| NA | NA | NA |
| NA | NA | NA |
| NA | NA | NA |
| NA | NA | NA |
| -0.401495214 | 0.688055561 | NA |
| NA | NA | NA |

| | | | |
|---|---|---|---|
| NA | NA | NA | |
| NA | NA | NA | |
| NA | NA | NA | |
| NA | NA | NA | |
| NA | NA | NA | |
| NA | NA | NA | |
| | -0.090190523 | 0.928135815 | 0.950891877 |
| NA | NA | NA | |
| NA | NA | NA | |
| | -0.065390563 | 0.947863038 | NA |
| | 2.257041066 | 0.024005509 | 0.048937766 |
| | 9.106094673 | 8.54E-20 | 1.50E-18 |
| NA | NA | NA | |
| | 4.380001238 | 1.19E-05 | 4.78E-05 |
| | 6.404301732 | 1.51E-10 | 1.12E-09 |
| | 1.557281372 | 0.119403696 | 0.193081182 |
| | 3.233019002 | 0.001224894 | 0.003457273 |
| | 2.336225256 | 0.019479505 | 0.040846048 |
| | -3.282469769 | 0.00102902 | 0.002958463 |
| | 3.085429375 | 0.002032585 | 0.005468731 |
| | -1.665381048 | 0.095836758 | 0.160809416 |
| | -0.912347273 | 0.361585937 | 0.47382315 |
| | 4.903325119 | 9.42E-07 | 4.44E-06 |
| | 3.03232693 | 0.002426762 | 0.006402641 |
| | 1.926606754 | 0.054028656 | 0.098978063 |
| | -0.061563817 | 0.950910192 | NA |
| | 1.451145961 | 0.146739221 | 0.229197048 |
| | 0.192809521 | 0.847108147 | NA |
| NA | NA | NA | |
| NA | NA | NA | |
| NA | NA | NA | |
| NA | NA | NA | |
| | -1.610396061 | 0.10731142 | 0.176432268 |
| | 0.614403175 | 0.538948938 | 0.643264123 |
| | 0.110822682 | 0.911756958 | NA |
| | -4.942776867 | 7.70E-07 | 3.67E-06 |
| | -0.350575654 | 0.725906724 | NA |
| | 0.063708075 | 0.949202675 | NA |
| NA | NA | NA | |
| | 0.209074615 | 0.834389991 | NA |
| | 2.427497775 | 0.01520338 | 0.032970075 |
| | 16.85650545 | 9.40E-64 | 1.03E-61 |
| | 0.437240455 | 0.661936974 | NA |
| | -7.536464336 | 4.83E-14 | 5.12E-13 |
| | 4.483680366 | 7.34E-06 | 3.05E-05 |
| | 2.580633268 | 0.009861928 | 0.022499687 |
| NA | NA | NA | |

| | | |
|---|---|---|
| NA | NA | NA |
| NA | NA | NA |
| NA | NA | NA |
| NA | NA | NA |
| 0.552361462 | 0.580700729 | 0.681335923 |
| NA | NA | NA |
| 4.72030678 | 2.35E-06 | 1.05E-05 |
| 11.38991642 | 4.69E-30 | 1.62E-28 |
| 1.418253855 | 0.156116664 | 0.241329057 |
| 8.997107539 | 2.32E-19 | 3.93E-18 |
| 1.271305629 | 0.203619941 | 0.29995922 |
| 0.192810118 | 0.847107679 | NA |
| NA | NA | NA |
| NA | NA | NA |
| 0.063708285 | 0.949202508 | NA |
| NA | NA | NA |
| 5.766804237 | 8.08E-09 | 4.98E-08 |
| -6.244796337 | 4.24E-10 | 3.00E-09 |
| -5.066963301 | 4.04E-07 | 2.01E-06 |
| NA | NA | NA |
| NA | NA | NA |
| NA | NA | NA |
| 11.28509719 | 1.55E-29 | 5.24E-28 |
| NA | NA | NA |
| NA | NA | NA |
| NA | NA | NA |
| 0.944819981 | 0.344750789 | 0.456156486 |
| 0.194173692 | 0.846039881 | NA |
| NA | NA | NA |
| 0.202714101 | 0.839358501 | NA |
| NA | NA | NA |
| -0.815940619 | 0.414534099 | 0.526155603 |
| NA | NA | NA |
| 0.068827122 | 0.945127229 | 0.962586854 |
| 2.548192933 | 0.010828257 | 0.02447014 |
| 0.393204471 | 0.694168468 | 0.774668364 |
| -5.098445269 | 3.42E-07 | 1.72E-06 |
| 1.29371686 | 0.195763239 | 0.290453698 |
| -0.065389369 | 0.947863988 | NA |
| 6.603617694 | 4.01E-11 | 3.18E-10 |
| -0.862365247 | 0.388486558 | 0.500707087 |
| 0.090100512 | 0.928207342 | NA |
| -7.863198778 | 3.74E-15 | 4.43E-14 |
| NA | NA | NA |
| NA | NA | NA |
| NA | NA | NA |
| 1.417800611 | 0.156248986 | NA |

| | | | |
|---|---|---|---|
| -0.413347704 | 0.679351884 | NA | |
| 0.06371127 | 0.949200131 | NA | |
| NA | NA | NA | |
| NA | NA | NA | |
| -5.520828057 | 3.37E-08 | | 1.94E-07 |
| NA | NA | NA | |
| -3.497051901 | 0.00047043 | | 0.001441585 |
| 0.404637056 | 0.685744316 | NA | |
| -0.931072118 | 0.351816261 | | 0.46359363 |
| NA | NA | NA | |
| NA | NA | NA | |
| NA | NA | NA | |
| NA | NA | NA | |
| NA | NA | NA | |
| NA | NA | NA | |
| 0.192810118 | 0.847107679 | NA | |
| -0.931389278 | 0.351652235 | | 0.463414286 |
| NA | NA | NA | |
| NA | NA | NA | |
| NA | NA | NA | |
| NA | NA | NA | |
| NA | NA | NA | |
| NA | NA | NA | |
| -0.218947773 | 0.826690732 | | 0.878756063 |
| -0.065388772 | 0.947864464 | NA | |
| 0.286060748 | 0.774831595 | NA | |
| NA | NA | NA | |
| NA | NA | NA | |
| NA | NA | NA | |
| NA | NA | NA | |
| -0.065389966 | 0.947863513 | NA | |
| -3.043659295 | 0.002337197 | | 0.006192446 |
| -1.79780424 | 0.072208035 | | 0.126518691 |
| 0.90143184 | 0.367358759 | | 0.479780315 |
| -1.68045186 | 0.092869433 | | 0.15672343 |
| -0.594181257 | 0.552390887 | | 0.655525503 |
| NA | NA | NA | |
| 0.192809521 | 0.847108147 | NA | |
| NA | NA | NA | |
| NA | NA | NA | |
| NA | NA | NA | |
| -0.089304777 | 0.928839697 | NA | |
| -0.117297005 | 0.906624698 | | 0.935439062 |
| NA | NA | NA | |
| NA | NA | NA | |
| NA | NA | NA | |
| NA | NA | NA | |

| | | | |
|---|---|---|---|
| | 0.063708285 | 0.949202508 | NA |
| | -9.937566384 | 2.86E-23 | 6.52E-22 |
| NA | NA | NA | |
| | 0.31685483 | 0.751353758 | NA |
| NA | NA | NA | |
| | -3.398417278 | 0.00067777 | 0.002016471 |
| NA | NA | NA | |
| | 0.426578866 | 0.669686093 | NA |
| | -3.112403731 | 0.001855705 | 0.005031229 |
| | -3.777885621 | 0.000158165 | 0.000527781 |
| | -0.152154004 | 0.879065471 | NA |
| | -0.065388772 | 0.947864464 | NA |
| NA | NA | NA | |
| | 0.199526594 | 0.841850842 | NA |
| NA | NA | NA | |
| | 2.807294826 | 0.004995949 | 0.012298251 |
| NA | NA | NA | |
| | -3.142342116 | 0.001676021 | 0.004594327 |
| | 0.346366587 | 0.729067233 | NA |
| | 0.400148123 | 0.689047422 | NA |
| NA | NA | NA | |
| | 2.301025917 | 0.021390166 | 0.044291517 |
| | -4.097364702 | 4.18E-05 | 0.000153975 |
| | -0.93227342 | 0.351195241 | 0.462913452 |
| | -2.146604154 | 0.031824809 | 0.062496075 |
| | -1.409634155 | 0.158647736 | 0.244321223 |
| NA | NA | NA | |
| | 0.192809521 | 0.847108147 | NA |
| | -1.035381399 | 0.300490829 | NA |
| NA | NA | NA | |
| NA | NA | NA | |
| | -0.06129694 | 0.951122728 | 0.966647794 |
| NA | NA | NA | |
| | -0.85664256 | 0.391642457 | 0.503663911 |
| | -4.27605125 | 1.90E-05 | 7.41E-05 |
| NA | NA | NA | |
| | -3.439232322 | 0.000583366 | 0.001756908 |
| | 0.126214393 | 0.899562218 | 0.929953838 |
| | 0.089333795 | 0.928816637 | NA |
| | 0.063708285 | 0.949202508 | NA |
| | -4.874239604 | 1.09E-06 | 5.10E-06 |
| | -1.392561932 | 0.163752301 | NA |
| | -5.10845089 | 3.25E-07 | 1.64E-06 |
| NA | NA | NA | |
| NA | NA | NA | |
| | -0.418147731 | 0.675839109 | 0.759155091 |
| | 4.881690684 | 1.05E-06 | 4.92E-06 |

| | | | |
|---|---|---|---|
| | 1.066247534 | 0.28631176 | NA |
| | -0.351781707 | 0.725001979 | 0.79917204 |
| NA | NA | NA | |
| | -1.704247831 | 0.088334797 | 0.150256199 |
| | -1.792412446 | 0.073066923 | 0.127714041 |
| | -0.065389966 | 0.947863513 | NA |
| | 1.696693611 | 0.089754601 | 0.152298669 |
| | -0.229264413 | 0.818663411 | 0.872483402 |
| NA | NA | NA | |
| | 0.452010036 | 0.651261753 | NA |
| | 0.192809521 | 0.847108147 | NA |
| | -0.340253345 | 0.733665748 | NA |
| NA | NA | NA | |
| NA | NA | NA | |
| | 2.201895717 | 0.027672676 | 0.055358621 |
| | 0.063710673 | 0.949200606 | NA |
| | -0.553526569 | 0.579902893 | 0.680686991 |
| | -0.493242131 | 0.621841512 | NA |
| NA | NA | NA | |
| | -0.065388772 | 0.947864464 | NA |
| | -5.984377331 | 2.17E-09 | 1.43E-08 |
| | -0.689929102 | 0.490238773 | 0.59786964 |
| | 0.624798222 | 0.532103498 | 0.636611565 |
| NA | NA | NA | |
| | -8.216100331 | 2.10E-16 | 2.78E-15 |
| NA | NA | NA | |
| | -0.189267568 | 0.849883113 | 0.895423425 |
| | -0.70759028 | 0.479199731 | 0.587930458 |
| NA | NA | NA | |
| | 4.25833089 | 2.06E-05 | 7.99E-05 |
| | 0.063708285 | 0.949202508 | NA |
| | -0.771526684 | 0.440394812 | NA |
| NA | NA | NA | |
| NA | NA | NA | |
| | 1.631722471 | 0.102737964 | 0.170300082 |
| | 0.62015204 | 0.535157693 | 0.639520077 |
| | -1.516635336 | 0.129358779 | 0.206589323 |
| | 3.040854223 | 0.00235908 | 0.006242497 |
| | 13.12974375 | 2.22E-39 | 1.19E-37 |
| | 8.102268387 | 5.39E-16 | 6.89E-15 |
| | 1.157522766 | 0.247058844 | 0.350183071 |
| | 10.33370315 | 4.96E-25 | 1.27E-23 |
| | 3.090280931 | 0.001999673 | 0.00539177 |
| | -8.883598296 | 6.47E-19 | 1.06E-17 |
| | 0.272452501 | 0.785274109 | 0.846794021 |
| NA | NA | NA | |
| | 0.063708285 | 0.949202508 | NA |

| | | | |
|---|---|---|---|
| | 0.921830473 | 0.35661701 | 0.468457702 |
| | 0.102390497 | 0.918446727 | 0.944172892 |
| | 0.857690928 | 0.391063151 | 0.503073992 |
| | -4.945746668 | 7.59E-07 | 3.62E-06 |
| | -3.158148309 | 0.001587748 | 0.004377762 |
| | -0.863465245 | 0.38788172 | 0.500112268 |
| | 7.08713606 | 1.37E-12 | 1.25E-11 |
| | 7.41988488 | 1.17E-13 | 1.20E-12 |
| | 1.305835821 | 0.191608396 | 0.285422731 |
| | 1.355489189 | 0.175261751 | NA |
| NA | NA | NA | |
| | 1.31711798 | 0.187799081 | NA |
| | 0.529147884 | 0.596702867 | 0.695028747 |
| | -0.08959232 | 0.928611188 | NA |
| | 0.044304008 | 0.964662077 | NA |
| | 1.579332992 | 0.114259698 | 0.185994673 |
| | 0.063708285 | 0.949202508 | NA |
| | 0.192809521 | 0.847108147 | NA |
| | 0.404860484 | 0.685580066 | NA |
| | 5.165203783 | 2.40E-07 | 1.23E-06 |
| | 3.564403376 | 0.000364685 | 0.001141007 |
| | 0.326972305 | 0.743688832 | NA |
| | -0.379380385 | 0.704405414 | 0.783228978 |
| | -0.588872514 | 0.555946797 | 0.658590011 |
| | 0.108030776 | 0.91397128 | 0.940915844 |
| | -0.065389369 | 0.947863988 | NA |
| | 0.192636473 | 0.847243678 | NA |
| NA | NA | NA | |
| NA | NA | NA | |
| | -0.068007011 | 0.945780053 | NA |
| NA | NA | NA | |
| NA | NA | NA | |
| NA | NA | NA | |
| NA | NA | NA | |
| NA | NA | NA | |
| NA | NA | NA | |
| NA | NA | NA | |
| NA | NA | NA | |
| NA | NA | NA | |
| | 0.063710673 | 0.949200606 | NA |
| NA | NA | NA | |
| NA | NA | NA | |
| NA | NA | NA | |
| NA | NA | NA | |
| | 0.063708882 | 0.949202033 | NA |
| NA | NA | NA | |
| NA | NA | NA | |

| | | | |
|---|---|---|---|
| NA | NA | NA | |
| NA | NA | NA | |
| | 0.491674205 | 0.622949677 | NA |
| | 0.195065941 | 0.845341326 | NA |
| | 0.19280564 | 0.847111186 | NA |
| | 5.421597637 | 5.91E-08 | 3.28E-07 |
| | 4.937693676 | 7.91E-07 | 3.77E-06 |
| | 3.230859542 | 0.001234186 | 0.003481928 |
| | 0.365202845 | 0.714959995 | NA |
| | 0.192808924 | 0.847108615 | NA |
| | 0.142836242 | 0.886419514 | 0.920434782 |
| | -0.065388772 | 0.947864464 | NA |
| NA | NA | NA | |
| | 2.331857095 | 0.01970821 | 0.041259326 |
| NA | NA | NA | |
| NA | NA | NA | |
| | 0.192809521 | 0.847108147 | NA |
| | 3.564411315 | 0.000364674 | 0.001141007 |
| | 3.884638984 | 0.000102482 | 0.000353518 |
| | 0.621634464 | 0.534182254 | 0.638579702 |
| | 1.227492897 | 0.21963739 | 0.318662934 |
| NA | NA | NA | |
| | 0.063707688 | 0.949202983 | NA |
| | -1.70259268 | 0.088644321 | 0.150659833 |
| NA | NA | NA | |
| NA | NA | NA | |
| | 0.096073881 | 0.923461895 | NA |
| NA | NA | NA | |
| NA | NA | NA | |
| NA | NA | NA | |
| NA | NA | NA | |
| | 0.094496708 | 0.924714598 | NA |
| NA | NA | NA | |
| | -0.065389966 | 0.947863513 | NA |
| | -0.202116247 | 0.839825847 | 0.887695518 |
| NA | NA | NA | |
| | -1.319639147 | 0.186955526 | 0.279706923 |
| | -0.065388772 | 0.947864464 | NA |
| | 0.108967543 | 0.913228234 | NA |
| | 0.520693425 | 0.602580355 | NA |
| | 2.002493142 | 0.04523172 | 0.084887902 |
| | 1.3050841 | 0.191864212 | 0.285735779 |
| NA | NA | NA | |
| NA | NA | NA | |
| | -0.787919791 | 0.430743624 | 0.541757344 |
| NA | NA | NA | |
| NA | NA | NA | |

| | | |
|---|---|---|
| NA | NA | NA |
| NA | NA | NA |
| NA | NA | NA |
| NA | NA | NA |
| 0.909220436 | 0.363233779 | 0.475643558 |
| 0.276666592 | 0.782036127 | NA |
| NA | NA | NA |
| NA | NA | NA |
| NA | NA | NA |
| NA | NA | NA |
| NA | NA | NA |
| NA | NA | NA |
| 0.333737388 | 0.738577734 | NA |
| NA | NA | NA |
| NA | NA | NA |
| 0.192810118 | 0.847107679 | NA |
| 0.170919246 | 0.864287263 | 0.905510726 |
| NA | NA | NA |
| NA | NA | NA |
| -0.485343327 | 0.627432826 | 0.720495648 |
| -1.297456992 | 0.194473994 | NA |
| 6.396720052 | 1.59E-10 | 1.17E-09 |
| 0.630032714 | 0.528673181 | 0.633515948 |
| 1.934159454 | 0.053093526 | 0.097457535 |
| 1.73578206 | 0.082602377 | 0.141952455 |
| -0.496601941 | 0.619469784 | 0.714000275 |
| 3.999266509 | 6.35E-05 | 0.000227263 |
| 1.655723825 | 0.09777776 | 0.163516302 |
| 0.540493505 | 0.588856739 | 0.68842056 |
| 2.927512307 | 0.003416855 | 0.008714478 |
| 0.089955405 | 0.928322653 | NA |
| 1.376760059 | 0.168586443 | 0.256814477 |
| 0.150266234 | 0.880554572 | NA |
| 8.843406367 | 9.28E-19 | 1.50E-17 |
| 1.589515022 | 0.111944165 | 0.182812192 |
| -0.995360016 | 0.319561198 | 0.430285207 |
| 0.192809521 | 0.847108147 | NA |
| -1.330144722 | 0.183470593 | 0.275481041 |
| 2.872645216 | 0.00407051 | 0.010212997 |
| NA | NA | NA |
| 0.081330041 | 0.935179484 | NA |
| NA | NA | NA |
| NA | NA | NA |
| -1.75878201 | 0.078614541 | 0.136061539 |
| NA | NA | NA |
| -0.16856906 | 0.866135619 | 0.906517856 |
| -3.220496196 | 0.001279689 | 0.003598137 |

| | | |
|---|---|---|
| -0.86161272 | 0.388900669 | 0.501115623 |
| -0.065389369 | 0.947863988 | NA |
| -0.734036679 | 0.462926384 | 0.572397736 |
| NA | NA | NA |
| NA | NA | NA |
| NA | NA | NA |
| -0.065388772 | 0.947864464 | NA |
| NA | NA | NA |
| 0.330195015 | 0.741252613 | NA |
| NA | NA | NA |
| -0.318404719 | 0.750177959 | 0.818642432 |
| 0.802904108 | 0.422030161 | 0.533856317 |
| 0.846566093 | 0.39723702 | 0.509106124 |
| 0.658841697 | 0.509997428 | NA |
| -1.228786737 | 0.219151776 | 0.318084601 |
| NA | NA | NA |
| NA | NA | NA |
| NA | NA | NA |
| 0.788468838 | 0.430422519 | 0.541489528 |
| -2.295979999 | 0.021677025 | 0.044801291 |
| -1.154782557 | 0.248179482 | 0.351413273 |
| -0.342297516 | 0.732127005 | 0.804739502 |
| -0.186247819 | 0.852250426 | 0.897275518 |
| -0.594728973 | 0.552024652 | 0.655277295 |
| -1.313285481 | 0.189086769 | NA |
| NA | NA | NA |
| NA | NA | NA |
| -0.151561305 | 0.879532955 | NA |
| NA | NA | NA |
| 0.054919824 | 0.956202338 | 0.970061485 |
| NA | NA | NA |
| -2.515307389 | 0.011892864 | 0.026566568 |
| -0.065389369 | 0.947863988 | NA |
| -0.349881552 | 0.726427592 | 0.800109247 |
| -3.489258394 | 0.000484363 | 0.001481016 |
| -0.577667873 | 0.563488373 | 0.665570949 |
| 0.192809521 | 0.847108147 | NA |
| -2.076803852 | 0.037819667 | 0.072599605 |
| 0.355229143 | 0.722417934 | NA |
| 0.063708882 | 0.949202033 | NA |
| NA | NA | NA |
| 6.220724254 | 4.95E-10 | 3.48E-09 |
| -1.937922928 | 0.052632626 | 0.096767639 |
| 2.320526417 | 0.020312418 | 0.042357728 |
| -2.351260479 | 0.018709931 | 0.039439917 |
| -0.065389966 | 0.947863513 | NA |
| 0.978576716 | 0.327789169 | NA |

| | | |
|---|---|---|
| -5.915911574 | 3.30E-09 | 2.13E-08 |
| -3.044934228 | 0.002327312 | 0.006169521 |
| -1.127071606 | 0.259712207 | 0.364750138 |
| NA | NA | NA |
| NA | NA | NA |
| NA | NA | NA |
| NA | NA | NA |
| NA | NA | NA |
| NA | NA | NA |
| NA | NA | NA |
| NA | NA | NA |
| -0.359414184 | 0.719285266 | 0.794760526 |
| 1.680089537 | 0.092939896 | 0.156778919 |
| NA | NA | NA |
| NA | NA | NA |
| -2.702065136 | 0.006891026 | 0.016355242 |
| 3.479362749 | 0.000502608 | 0.001533058 |
| 4.965862664 | 6.84E-07 | 3.29E-06 |
| 0.106213312 | 0.915413109 | NA |
| -1.486245367 | 0.137214232 | 0.216785946 |
| NA | NA | NA |
| -0.878110551 | 0.379883728 | 0.492489238 |
| -3.722009542 | 0.000197644 | 0.00065041 |
| 0.107064812 | 0.914737563 | NA |
| 0.388634216 | 0.697546752 | NA |
| NA | NA | NA |
| -1.327626647 | 0.184301477 | 0.276502425 |
| NA | NA | NA |
| 1.500086979 | 0.133591873 | 0.212089597 |
| NA | NA | NA |
| NA | NA | NA |
| -5.11367158 | 3.16E-07 | 1.59E-06 |
| 0.192810118 | 0.847107679 | NA |
| 6.502853405 | 7.88E-11 | 6.05E-10 |
| 0.085936513 | 0.931516886 | NA |
| -0.479784001 | 0.63138099 | 0.723336973 |
| NA | NA | NA |
| NA | NA | NA |
| 0.377372407 | 0.70589687 | NA |
| 0.192809521 | 0.847108147 | NA |
| 1.750355609 | 0.080056971 | 0.138185365 |
| NA | NA | NA |
| -0.306403759 | 0.759297247 | 0.825774933 |
| NA | NA | NA |
| -0.716403748 | 0.473742081 | 0.582576609 |
| -5.073706947 | 3.90E-07 | 1.94E-06 |
| -8.811717708 | 1.23E-18 | 1.98E-17 |

| | | | |
|---|---|---|---|
| | -4.094531745 | 4.23E-05 | 0.00015571 |
| | 5.583001032 | 2.36E-08 | 1.38E-07 |
| | 4.958975675 | 7.09E-07 | 3.40E-06 |
| | -0.065390563 | 0.947863038 | NA |
| | 0.558818742 | 0.57628543 | 0.677312114 |
| | -0.065388772 | 0.947864464 | NA |
| NA | NA | NA | |
| NA | NA | NA | |
| | 0.317642674 | 0.750756 | NA |
| | 0.475883562 | 0.634157334 | NA |
| | -0.065388772 | 0.947864464 | NA |
| | 0.3474247 | 0.728272279 | NA |
| | -0.247836019 | 0.804261285 | 0.861386453 |
| | 3.897558081 | 9.72E-05 | 0.000336763 |
| NA | NA | NA | |
| | -7.683413512 | 1.55E-14 | 1.73E-13 |
| | -0.901043401 | 0.367565244 | NA |
| | 0.924429143 | 0.355262927 | 0.467176162 |
| | -3.705981357 | 0.000210574 | 0.000688975 |
| | -1.207836506 | 0.227110157 | 0.327303274 |
| | -1.635454315 | 0.101953844 | 0.169224451 |
| | 0.063708882 | 0.949202033 | NA |
| | 4.100676505 | 4.12E-05 | 0.000151989 |
| | -2.196663365 | 0.028044497 | 0.055999502 |
| | 0.259680703 | 0.795110079 | NA |
| | 0.192810118 | 0.847107679 | NA |
| | 2.423627651 | 0.015366352 | 0.033277365 |
| | 0.348544514 | 0.727431289 | NA |
| NA | NA | NA | |
| | 0.063708285 | 0.949202508 | NA |
| NA | NA | NA | |
| | 6.180734018 | 6.38E-10 | 4.42E-09 |
| NA | NA | NA | |
| NA | NA | NA | |
| | -7.382751763 | 1.55E-13 | 1.56E-12 |
| NA | NA | NA | |
| | -3.408234364 | 0.000653847 | 0.001950631 |
| | 0.318798974 | 0.749878956 | 0.818376729 |
| | 1.000470063 | 0.317083078 | 0.427587643 |
| | 0.063705652 | 0.949204604 | NA |
| | -2.546868863 | 0.010869427 | 0.024549879 |
| | 0.063710673 | 0.949200606 | NA |
| | -11.2717158 | 1.81E-29 | 6.08E-28 |
| | -4.773479824 | 1.81E-06 | 8.21E-06 |
| | 0.510343377 | 0.609810919 | 0.7060654 |
| NA | NA | NA | |
| | 0.06371127 | 0.949200131 | NA |

| | | | |
|---|---|---|---|
| | -0.310945917 | 0.755841736 | NA |
| NA | NA | NA | |
| | 0.06353712 | 0.949338802 | NA |
| | 0.914387634 | 0.360513197 | 0.472640128 |
| NA | NA | NA | |
| | 1.445483048 | 0.148322251 | 0.231207943 |
| | -0.247713765 | 0.804355881 | NA |
| NA | NA | NA | |
| | 0.063708882 | 0.949202033 | NA |
| NA | NA | NA | |
| | 3.593106017 | 0.00032676 | 0.001032024 |
| | -5.562213698 | 2.66E-08 | 1.55E-07 |
| | -0.065389369 | 0.947863988 | NA |
| NA | NA | NA | |
| | -0.752624107 | 0.451675827 | 0.561593858 |
| | -2.292133746 | 0.021897927 | 0.045197089 |
| | -0.362463536 | 0.717005668 | NA |
| | 0.478436883 | 0.632339288 | 0.724136541 |
| | -1.452550091 | 0.146348712 | 0.228701271 |
| | 0.211922406 | 0.83216757 | 0.882574822 |
| | -1.189742356 | 0.234147672 | 0.335453966 |
| | -5.327032585 | 9.98E-08 | 5.38E-07 |
| | 1.930118094 | 0.053592207 | 0.098264811 |
| | -5.31874713 | 1.04E-07 | 5.62E-07 |
| NA | NA | NA | |
| | -0.065388772 | 0.947864464 | NA |
| | -3.385844401 | 0.000709596 | 0.002103908 |
| | 0.192636473 | 0.847243678 | NA |
| | 0.543537101 | 0.586760053 | 0.68664286 |
| | -1.674731849 | 0.093986856 | 0.158214473 |
| | 0.628794729 | 0.529483451 | NA |
| | 0.660901196 | 0.508675679 | NA |
| | 2.794450664 | 0.005198797 | 0.01273746 |
| | 3.904524564 | 9.44E-05 | 0.000328254 |
| | 0.569905467 | 0.568741816 | 0.670352593 |
| | 1.420241675 | 0.155537335 | 0.240582097 |
| | 1.100531272 | 0.271100712 | 0.377420926 |
| | -2.146520178 | 0.031831501 | 0.062499409 |
| | 0.611152837 | 0.541098403 | NA |
| | 0.591789901 | 0.553991287 | 0.65711316 |
| | -0.436741026 | 0.662299172 | NA |
| NA | NA | NA | |
| | -1.428621028 | 0.153113187 | 0.237447907 |
| NA | NA | NA | |
| | 2.25354549 | 0.024224776 | 0.049324505 |
| | 4.618907814 | 3.86E-06 | 1.67E-05 |
| NA | NA | NA | |

| | | |
|---|---|---|
| -1.084647136 | 0.27807797 | NA |
| NA | NA | NA |
| 1.506906016 | 0.131834749 | 0.20969902 |
| NA | NA | NA |
| 1.014226325 | 0.31047477 | NA |
| -1.997971653 | 0.045719733 | 0.085673054 |
| -1.02927352 | 0.303351163 | 0.412678928 |
| -3.462386225 | 0.000535408 | 0.001624002 |
| 0.192809521 | 0.847108147 | NA |
| 5.73541419 | 9.73E-09 | 5.95E-08 |
| -2.422940875 | 0.015395432 | 0.033334573 |
| 3.388635974 | 0.000702412 | 0.002084583 |
| -0.565378816 | 0.57181613 | NA |
| 0.882647987 | 0.377426493 | 0.490115959 |
| -2.71333185 | 0.006661036 | 0.015871084 |
| -0.609410961 | 0.542252075 | NA |
| -1.034057635 | 0.30110922 | 0.410208664 |
| NA | NA | NA |
| -5.142690464 | 2.71E-07 | 1.38E-06 |
| 0.895794232 | 0.370362671 | 0.483123705 |
| -10.97304561 | 5.15E-28 | 1.60E-26 |
| NA | NA | NA |
| NA | NA | NA |
| -2.149933745 | 0.031560456 | 0.0620451 |
| -0.397877662 | 0.690720367 | 0.771691541 |
| 4.062366265 | 4.86E-05 | 0.000177349 |
| 0.192809521 | 0.847108147 | NA |
| 0.657335837 | 0.510964998 | 0.616745619 |
| 0.215113513 | 0.829678819 | 0.880884871 |
| 0.063710673 | 0.949200606 | NA |
| -1.457638874 | 0.144940109 | 0.226939254 |
| -2.217161144 | 0.026612082 | 0.053489152 |
| 0.328517213 | 0.742520631 | NA |
| 0.293118036 | 0.769431931 | NA |
| 9.058125396 | 1.33E-19 | 2.29E-18 |
| 4.804309159 | 1.55E-06 | 7.09E-06 |
| -0.963550621 | 0.335271274 | 0.446493215 |
| 3.017819269 | 0.002546007 | 0.006684136 |
| -0.861378455 | 0.389029638 | 0.501179344 |
| -3.681109869 | 0.000232221 | 0.000753303 |
| -2.400075188 | 0.016391705 | 0.035250826 |
| -3.615570147 | 0.000299687 | 0.000952893 |
| NA | NA | NA |
| -2.304155227 | 0.021213932 | 0.043957866 |
| 5.308711705 | 1.10E-07 | 5.92E-07 |
| 0.427445894 | 0.669054586 | 0.753970388 |
| 1.640200771 | 0.100963429 | 0.167858826 |

| | | | |
|---|---|---|---|
| | -5.496809913 | 3.87E-08 | 2.20E-07 |
| NA | NA | NA | |
| | 0.330834869 | 0.740769221 | NA |
| | 0.192636473 | 0.847243678 | NA |
| | -1.506369064 | 0.131972458 | 0.209851383 |
| | -1.462744434 | 0.143537323 | NA |
| | 1.16551019 | 0.243812538 | 0.34658268 |
| | -1.858772678 | 0.063059367 | 0.112725036 |
| | -1.667215313 | 0.095471599 | 0.160325671 |
| | -4.981374875 | 6.31E-07 | 3.05E-06 |
| | 1.416765759 | 0.156551424 | 0.241896545 |
| | -2.121535725 | 0.033876745 | 0.066033403 |
| | -3.725921316 | 0.000194603 | 0.00064091 |
| | -1.739574583 | 0.081933745 | 0.140958255 |
| NA | NA | NA | |
| NA | NA | NA | |
| | 0.063708285 | 0.949202508 | NA |
| NA | NA | NA | |
| | -1.204734106 | 0.228305965 | 0.328702808 |
| | 2.842601027 | 0.004474705 | 0.011126749 |
| | 2.310070106 | 0.020884273 | 0.043380362 |
| NA | NA | NA | |
| | -1.030356774 | 0.302842557 | 0.412121416 |
| | 3.721905196 | 0.000197725 | 0.000650593 |
| | -1.82771299 | 0.067592638 | 0.119641024 |
| | -6.28308177 | 3.32E-10 | 2.37E-09 |
| | -1.479981552 | 0.13887817 | 0.218954434 |
| | 0.012895035 | 0.989711536 | 0.992715062 |
| | -1.593964685 | 0.110943945 | 0.181462912 |
| | 3.10003066 | 0.001935006 | 0.005228109 |
| | -0.378195734 | 0.705285194 | 0.783789389 |
| NA | NA | NA | |
| | 9.715443069 | 2.59E-22 | 5.45E-21 |
| | 0.192809521 | 0.847108147 | NA |
| | 0.404104969 | 0.686135531 | 0.76813944 |
| | 1.723626512 | 0.084775289 | NA |
| | 0.259680703 | 0.795110079 | NA |
| | 0.453458983 | 0.650218272 | 0.738931367 |
| NA | NA | NA | |
| | 1.291756526 | 0.196441475 | NA |
| NA | NA | NA | |
| NA | NA | NA | |
| NA | NA | NA | |
| NA | NA | NA | |
| NA | NA | NA | |
| NA | NA | NA | |
| | 0.346721347 | 0.728800672 | NA |

| | | | |
|---|---|---|---|
| | 0.06371127 | 0.949200131 | NA |
| NA | NA | NA | |
| NA | NA | NA | |
| NA | NA | NA | |
| NA | NA | NA | |
| NA | NA | NA | |
| | -0.081530475 | 0.93502009 | NA |
| | 2.016205687 | 0.043778472 | 0.082515176 |
| | -0.031509845 | 0.97486294 | 0.982499888 |
| NA | NA | NA | |
| NA | NA | NA | |
| NA | NA | NA | |
| NA | NA | NA | |
| NA | NA | NA | |
| | -0.053338834 | 0.957461939 | 0.970646915 |
| | 0.063708285 | 0.949202508 | NA |
| | -0.116970025 | 0.906883807 | 0.935629269 |
| | 0.592239138 | 0.553690464 | NA |
| | 0.063708882 | 0.949202033 | NA |
| NA | NA | NA | |
| | 0.761764507 | 0.446200568 | 0.556139837 |
| | 2.969226928 | 0.0029855 | 0.007724527 |
| | 0.582462895 | 0.560254927 | NA |
| | 2.664754814 | 0.007704447 | 0.018019278 |
| | -0.574716203 | 0.565483247 | 0.667517718 |
| | 1.066363591 | 0.286259314 | 0.393425797 |
| | 1.788440281 | 0.073705002 | 0.128658638 |
| | 1.464392428 | 0.143086756 | 0.224604283 |
| | -6.686493664 | 2.29E-11 | 1.85E-10 |
| | -1.516404271 | 0.12941716 | 0.206654642 |
| NA | NA | NA | |
| NA | NA | NA | |
| | 0.06371127 | 0.949200131 | NA |
| | -0.284430306 | 0.776080637 | NA |
| | 0.192809521 | 0.847108147 | NA |
| NA | NA | NA | |
| | 4.585553925 | 4.53E-06 | 1.94E-05 |
| NA | NA | NA | |
| NA | NA | NA | |
| NA | NA | NA | |
| NA | NA | NA | |
| | 0.874040356 | 0.382096277 | NA |
| | 0.330493631 | 0.741027004 | NA |
| NA | NA | NA | |
| NA | NA | NA | |
| NA | NA | NA | |
| NA | NA | NA | |

| | | | |
|---|---|---|---|
| | -1.297417581 | 0.194487547 | 0.288834893 |
| NA | | NA | NA |
| NA | | NA | NA |
| | 4.551710278 | 5.32E-06 | 2.26E-05 |
| | 0.244604511 | 0.806762671 | 0.863107782 |
| | -4.860336421 | 1.17E-06 | 5.44E-06 |
| | -1.007735731 | 0.313581347 | 0.424012572 |
| | 7.980265256 | 1.46E-15 | 1.79E-14 |
| | 0.701518846 | 0.482979278 | 0.591445273 |
| | -0.093104523 | 0.925820524 | NA |
| | 1.262674837 | 0.206706061 | 0.303683319 |
| | 1.755724377 | 0.079135482 | 0.136786981 |
| | 0.203489613 | 0.838752363 | 0.887071588 |
| | -9.845623746 | 7.16E-23 | 1.57E-21 |
| | 3.594698572 | 0.000324767 | 0.001026121 |
| | 0.052712437 | 0.957961029 | 0.970995568 |
| | 6.460018057 | 1.05E-10 | 7.94E-10 |
| | 6.694303681 | 2.17E-11 | 1.76E-10 |
| | 2.981433534 | 0.002869023 | 0.007446252 |
| | -1.797032287 | 0.072330494 | 0.126683386 |
| | -3.464696015 | 0.000530831 | 0.001611486 |
| | 0.080258587 | 0.936031595 | 0.956436594 |
| | 0.06353712 | 0.949338802 | NA |
| | -0.283744511 | 0.776606181 | 0.839740503 |
| | -0.872199493 | 0.383099556 | NA |
| | -4.592902022 | 4.37E-06 | 1.88E-05 |
| | -1.124843358 | 0.260655418 | 0.365686066 |
| NA | | NA | NA |
| | -2.929532318 | 0.003394725 | 0.008664216 |
| | 2.528449838 | 0.011456746 | 0.025734752 |
| | 0.639903849 | 0.522235112 | 0.627504433 |
| | -0.189843629 | 0.849431666 | 0.895060815 |
| | -2.890648489 | 0.003844479 | 0.009700333 |
| | -1.989380463 | 0.046659224 | 0.087194295 |
| | 1.476447899 | 0.139823692 | 0.220209338 |
| | 7.066409976 | 1.59E-12 | 1.44E-11 |
| NA | | NA | NA |
| | 6.42525924 | 1.32E-10 | 9.86E-10 |
| | -0.332566299 | 0.739461686 | 0.810281751 |
| | 0.004399172 | 0.99648998 | 0.99760596 |
| | 0.653123999 | 0.513676342 | 0.619212041 |
| | -0.65433548 | 0.512895692 | 0.618509307 |
| | 5.070339688 | 3.97E-07 | 1.98E-06 |
| | -0.27233687 | 0.785363009 | 0.846800901 |
| | 0.580391837 | 0.561650409 | 0.663838586 |
| | -1.825358602 | 0.067946932 | 0.120140697 |
| NA | | NA | NA |

| | | |
|---|---|---|
| -0.065388772 | 0.947864464 | NA |
| -2.683085791 | 0.007294627 | 0.017174727 |
| -2.45916409 | 0.013926095 | 0.030541502 |
| -1.934532408 | 0.053047701 | 0.097394848 |
| -0.01417729 | 0.988688538 | 0.992120736 |
| -0.194488217 | 0.845793621 | NA |
| 0.192806037 | 0.847110875 | NA |
| 0.554793199 | 0.57903612 | 0.679765212 |
| -2.692122144 | 0.007099894 | 0.016780958 |
| 0.06371127 | 0.949200131 | NA |
| NA | NA | NA |
| -0.03079784 | 0.975430763 | 0.982970932 |
| -0.483127015 | 0.629005553 | NA |
| -2.335610226 | 0.019511565 | 0.040895535 |
| 4.101884194 | 4.10E-05 | 0.000151287 |
| -0.864551592 | 0.387284952 | 0.499574435 |
| 5.573945875 | 2.49E-08 | 1.45E-07 |
| -0.269710986 | 0.787382608 | NA |
| -0.285831906 | 0.77500687 | NA |
| 4.999735609 | 5.74E-07 | 2.79E-06 |
| 0.457995885 | 0.646955397 | 0.735946801 |
| NA | NA | NA |
| NA | NA | NA |
| 2.407342361 | 0.016069098 | 0.03463442 |
| NA | NA | NA |
| 3.998147146 | 6.38E-05 | 0.000228209 |
| 2.298964085 | 0.021506979 | 0.044496585 |
| 0.604294642 | 0.545647768 | 0.64935162 |
| 1.26959192 | 0.204230029 | 0.300734064 |
| -0.481505556 | 0.630157232 | 0.722448347 |
| NA | NA | NA |
| 2.770168082 | 0.005602737 | 0.013619162 |
| -1.950832314 | 0.051076996 | 0.094281506 |
| 1.748087237 | 0.080448923 | 0.138766214 |
| 0.192808327 | 0.847109082 | NA |
| -0.655282486 | 0.512285895 | 0.617952576 |
| 0.414549357 | 0.678471828 | NA |
| -5.313615355 | 1.07E-07 | 5.77E-07 |
| -1.538260663 | 0.123984894 | 0.199462121 |
| 2.59104504 | 0.009568497 | 0.02189009 |
| 1.346172181 | 0.178246994 | 0.268970208 |
| 0.382131398 | 0.702363903 | NA |
| 1.095760786 | 0.273183473 | 0.379834593 |
| -0.054289161 | 0.956704785 | 0.970365225 |
| -0.366168581 | 0.714239285 | NA |
| -1.687807952 | 0.091448101 | 0.154616722 |
| 0.192809521 | 0.847108147 | NA |

| | | |
|---|---|---|
| -0.882189432 | 0.377674376 | 0.490333755 |
| -10.88057117 | 1.43E-27 | 4.32E-26 |
| -1.643210932 | 0.100339299 | 0.166974817 |
| 2.662491492 | 0.007756453 | 0.018127359 |
| 4.716586989 | 2.40E-06 | 1.07E-05 |
| -0.141739323 | 0.887285913 | 0.92095237 |
| 0.601665332 | 0.547396931 | 0.651021184 |
| -0.549715568 | 0.582514477 | 0.682727915 |
| 0.372430761 | 0.709572154 | NA |
| 5.854878202 | 4.77E-09 | 3.02E-08 |
| NA | NA | NA |
| 1.107416511 | 0.268113891 | 0.373969946 |
| NA | NA | NA |
| 0.484585969 | 0.627970068 | 0.720980267 |
| -2.598566689 | 0.009361386 | 0.021469259 |
| 5.712017554 | 1.12E-08 | 6.79E-08 |
| 9.586847478 | 9.08E-22 | 1.85E-20 |
| 4.277598237 | 1.89E-05 | 7.37E-05 |
| -0.065387578 | 0.947865415 | NA |
| -0.26514923 | 0.790894504 | 0.851164385 |
| 6.925265668 | 4.35E-12 | 3.77E-11 |
| NA | NA | NA |
| NA | NA | NA |
| 0.063708882 | 0.949202033 | NA |
| -1.717523556 | 0.085883554 | 0.146644748 |
| -1.649868431 | 0.098969849 | 0.165191739 |
| 1.457262116 | 0.14504404 | 0.227074508 |
| NA | NA | NA |
| 0.607077795 | 0.543799285 | NA |
| NA | NA | NA |
| 0.063708882 | 0.949202033 | NA |
| 0.200068104 | 0.841427318 | NA |
| -0.070610168 | 0.943708018 | NA |
| 0.063708285 | 0.949202508 | NA |
| NA | NA | NA |
| NA | NA | NA |
| NA | NA | NA |
| NA | NA | NA |
| NA | NA | NA |
| NA | NA | NA |
| NA | NA | NA |
| NA | NA | NA |
| NA | NA | NA |
| 0.228023028 | 0.819628338 | NA |
| 0.886009733 | 0.375612282 | NA |
| 0.557591829 | 0.577123137 | NA |
| -0.085070848 | 0.932205066 | NA |

| | | |
|---|---|---|
| NA | NA | NA |
| NA | NA | NA |
| 0.707780907 | 0.479081325 | NA |
| NA | NA | NA |
| 0.359307553 | 0.719365024 | NA |
| 0.20927576 | 0.834232974 | NA |
| 8.448246028 | 2.96E-17 | 4.20E-16 |
| 0.573955798 | 0.565997713 | 0.667998998 |
| 0.31685483 | 0.751353758 | NA |
| -0.066588298 | 0.946909462 | NA |
| 2.597751069 | 0.009383649 | 0.021518116 |
| 2.432970249 | 0.014975531 | 0.032526688 |
| 7.41103625 | 1.25E-13 | 1.27E-12 |
| -4.003633326 | 6.24E-05 | 0.000223457 |
| -2.56943621 | 0.010186414 | 0.023178723 |
| 0.192810118 | 0.847107679 | NA |
| 2.584373972 | 0.009755595 | 0.022283496 |
| -0.08180493 | 0.934801835 | NA |
| 2.83150665 | 0.004632927 | 0.011487179 |
| 6.200974257 | 5.61E-10 | 3.91E-09 |
| 0.192809521 | 0.847108147 | NA |
| -1.455312635 | 0.14558273 | 0.227760219 |
| -2.067867837 | 0.038652453 | 0.073971319 |
| -1.359511941 | 0.173984421 | 0.263737286 |
| 2.668032397 | 0.00762969 | 0.017871153 |
| 0.228496319 | 0.819260418 | 0.873008142 |
| -2.76997692 | 0.005606026 | 0.013625834 |
| 0.063708217 | 0.949202562 | NA |
| -0.149344364 | 0.88128191 | NA |
| -0.790825616 | 0.42904576 | 0.54019192 |
| 0.192809521 | 0.847108147 | NA |
| NA | NA | NA |
| -3.68235191 | 0.000231092 | 0.000750128 |
| 0.076845225 | 0.938746672 | 0.95835016 |
| NA | NA | NA |
| -0.175025327 | 0.861059733 | 0.903320555 |
| 0.474004791 | 0.635496491 | NA |
| 0.75403458 | 0.450828458 | 0.560735256 |
| -0.140591195 | 0.888192904 | NA |
| -5.799918627 | 6.63E-09 | 4.13E-08 |
| 1.591520872 | 0.111492402 | 0.18220684 |
| -3.700186884 | 0.000215441 | 0.00070352 |
| -2.656371967 | 0.007898644 | 0.018418395 |
| 4.067500897 | 4.75E-05 | 0.000173715 |
| 0.067524971 | 0.946163784 | NA |
| 0.218645955 | 0.826925852 | 0.878894016 |
| 0.384116604 | 0.700892026 | 0.780154196 |

| | | |
|---|---|---|
| 0.978224801 | 0.327963153 | 0.439001698 |
| 0.430632967 | 0.666735267 | 0.752183673 |
| -1.660837558 | 0.096746076 | 0.162074445 |
| NA | NA | NA |
| 0.520506838 | 0.602710363 | 0.700048792 |
| -0.464560378 | 0.642246312 | NA |
| -9.175498078 | 4.49E-20 | 8.05E-19 |
| 0.691837275 | 0.489039525 | 0.596872407 |
| 0.070876291 | 0.943496213 | NA |
| 2.751503842 | 0.005932232 | 0.01431031 |
| 0.939692182 | 0.347375476 | 0.458913535 |
| -2.183957608 | 0.028965363 | 0.057585627 |
| -1.330238645 | 0.183439655 | 0.275451111 |
| -5.70770122 | 1.15E-08 | 6.96E-08 |
| 5.550637739 | 2.85E-08 | 1.65E-07 |
| NA | NA | NA |
| 0.537633555 | 0.590830059 | 0.690211723 |
| 0.095864378 | 0.923628286 | NA |
| NA | NA | NA |
| 0.063708285 | 0.949202508 | NA |
| NA | NA | NA |
| NA | NA | NA |
| 0.064192091 | 0.948817275 | 0.965010012 |
| NA | NA | NA |
| NA | NA | NA |
| NA | NA | NA |
| NA | NA | NA |
| NA | NA | NA |
| 0.063708285 | 0.949202508 | NA |
| NA | NA | NA |
| NA | NA | NA |
| NA | NA | NA |
| NA | NA | NA |
| NA | NA | NA |
| -1.13723734 | 0.255439081 | 0.359840174 |
| NA | NA | NA |
| NA | NA | NA |
| NA | NA | NA |
| NA | NA | NA |
| 0.192809521 | 0.847108147 | NA |
| NA | NA | NA |
| NA | NA | NA |
| 0.192810118 | 0.847107679 | NA |
| NA | NA | NA |
| NA | NA | NA |
| 0.182771792 | 0.854977083 | NA |
| 5.393827232 | 6.90E-08 | 3.79E-07 |

| | | |
|---|---|---|
| 10.31812953 | 5.83E-25 | 1.48E-23 |
| NA | NA | NA |
| 7.156995128 | 8.25E-13 | 7.72E-12 |
| 1.774056842 | 0.076053749 | 0.13222375 |
| -4.876679455 | 1.08E-06 | 5.04E-06 |
| 0.99384909 | 0.320296343 | 0.431045619 |
| 0.258749957 | 0.795828172 | NA |
| 0.856294637 | 0.391834827 | 0.503805562 |
| -1.604589557 | 0.108584161 | 0.17819706 |
| -1.4331236 | 0.151822518 | 0.235900281 |
| 6.660806345 | 2.72E-11 | 2.19E-10 |
| -3.451166834 | 0.000558168 | 0.001688129 |
| -1.590368086 | 0.11175186 | 0.182521961 |
| 0.006322892 | 0.994955096 | 0.996547661 |
| 0.192808924 | 0.847108615 | NA |
| 4.223996427 | 2.40E-05 | 9.20E-05 |
| NA | NA | NA |
| 1.337896961 | 0.180930028 | 0.272291267 |
| -0.028541492 | 0.977230276 | 0.984308805 |
| 1.808856812 | 0.070473251 | 0.123985765 |
| -1.235994648 | 0.216460553 | 0.315022987 |
| -3.241163516 | 0.001190429 | 0.003374072 |
| 0.063708285 | 0.949202508 | NA |
| NA | NA | NA |
| NA | NA | NA |
| NA | NA | NA |
| 0.741580157 | 0.45834175 | 0.56789893 |
| -0.340839397 | 0.73322449 | 0.80538127 |
| -1.134737746 | 0.256485217 | 0.36109033 |
| 5.644815648 | 1.65E-08 | 9.85E-08 |
| 0.995208786 | 0.31963473 | 0.430317195 |
| -1.68033199 | 0.09289274 | 0.156741627 |
| 0.089783839 | 0.928458991 | NA |
| 0.192805721 | 0.847111123 | NA |
| -1.95122524 | 0.051030258 | 0.094223027 |
| -2.427593786 | 0.015199356 | 0.032964205 |
| NA | NA | NA |
| 0.481816583 | 0.629936248 | 0.722343059 |
| -0.081720839 | 0.934868707 | 0.955794614 |
| 5.603026591 | 2.11E-08 | 1.24E-07 |
| 0.399690438 | 0.689384536 | 0.77077299 |
| -2.544926578 | 0.010930072 | 0.024662372 |
| 0.457600159 | 0.647239729 | NA |
| -0.06663263 | 0.946874169 | NA |
| 0.063707688 | 0.949202983 | NA |
| -0.054638789 | 0.956426236 | 0.97022569 |
| NA | NA | NA |

| | | |
|---|---|---|
| -0.520475012 | 0.60273254 | 0.700048792 |
| -1.902366069 | 0.057123314 | 0.10375882 |
| NA | NA | NA |
| -1.223068895 | 0.221303676 | 0.32055735 |
| 4.145988085 | 3.38E-05 | 0.000126837 |
| 5.037163893 | 4.72E-07 | 2.32E-06 |
| NA | NA | NA |
| 6.473539488 | 9.57E-11 | 7.30E-10 |
| 5.044873993 | 4.54E-07 | 2.24E-06 |
| -0.975771655 | 0.329177634 | 0.440040386 |
| -0.829615186 | 0.406756388 | 0.518571405 |
| -0.176148097 | 0.860177596 | 0.902811356 |
| -0.065390563 | 0.947863038 | NA |
| 3.064447439 | 0.002180725 | 0.005824108 |
| 0.11749107 | 0.90647092 | NA |
| -0.669122886 | 0.503417092 | 0.609988372 |
| -1.798239257 | 0.072139101 | 0.126418938 |
| -2.50974686 | 0.012081774 | 0.026928477 |
| 3.127397323 | 0.001763614 | 0.004812304 |
| -4.331003133 | 1.48E-05 | 5.89E-05 |
| -6.295191221 | 3.07E-10 | 2.20E-09 |
| -0.429425445 | 0.66761364 | 0.752753251 |
| 2.37408861 | 0.017592328 | 0.037449904 |
| 1.92675054 | 0.054010726 | 0.098952457 |
| -0.98857658 | 0.32287035 | 0.433763343 |
| -4.1515066 | 3.30E-05 | 0.000123965 |
| 3.552996755 | 0.000380869 | 0.00118734 |
| NA | NA | NA |
| 2.033031907 | 0.0420493 | 0.079631835 |
| 4.183872528 | 2.87E-05 | 0.000108735 |
| 0.572867198 | 0.566734616 | 0.668584976 |
| -5.767396409 | 8.05E-09 | 4.96E-08 |
| 13.10440783 | 3.11E-39 | 1.65E-37 |
| NA | NA | NA |
| -6.351849951 | 2.13E-10 | 1.55E-09 |
| 3.215690389 | 0.001301311 | 0.003652778 |
| 0.947282003 | 0.343495099 | 0.454919939 |
| 3.373892563 | 0.000741133 | 0.002190666 |
| 14.36549003 | 8.52E-47 | 6.08E-45 |
| 0.92628411 | 0.354298351 | 0.466180833 |
| -7.172662822 | 7.36E-13 | 6.92E-12 |
| 2.849199827 | 0.004382934 | 0.010920251 |
| 4.771856818 | 1.83E-06 | 8.27E-06 |
| 9.675385806 | 3.84E-22 | 8.00E-21 |
| 0.999016519 | 0.317786689 | NA |
| 6.371521663 | 1.87E-10 | 1.38E-09 |
| 2.385777202 | 0.017043073 | 0.036438721 |

| | | |
|---|---|---|
| 2.286271046 | 0.022238407 | 0.045846909 |
| -1.143827064 | 0.252695364 | 0.356717893 |
| 4.005544165 | 6.19E-05 | 0.000221849 |
| 2.308101747 | 0.020993478 | 0.043560205 |
| -2.662321867 | 0.007760363 | 0.018134805 |
| 3.673224701 | 0.000239509 | 0.000774861 |
| -0.500100935 | 0.617004008 | 0.711851132 |
| 0.480305969 | 0.631009845 | NA |
| -6.265965536 | 3.71E-10 | 2.63E-09 |
| -4.359366295 | 1.30E-05 | 5.21E-05 |
| 1.870286704 | 0.061444015 | 0.110231056 |
| 0.684055676 | 0.493940009 | 0.601124054 |
| -4.338546013 | 1.43E-05 | 5.70E-05 |
| 0.192810118 | 0.847107679 | NA |
| -1.580329841 | 0.114031349 | 0.185723886 |
| 2.838680922 | 0.004530043 | 0.011257643 |
| NA | NA | NA |
| 0.192809521 | 0.847108147 | NA |
| -7.442148557 | 9.91E-14 | 1.02E-12 |
| -0.360806339 | 0.718244223 | 0.794030869 |
| 8.450385725 | 2.90E-17 | 4.13E-16 |
| 0.266588972 | 0.789785647 | NA |
| -6.510314531 | 7.50E-11 | 5.78E-10 |
| 2.672165836 | 0.007536339 | 0.017682281 |
| NA | NA | NA |
| NA | NA | NA |
| -0.42564448 | 0.670366923 | 0.75490548 |
| -0.135733036 | 0.89203233 | 0.924537028 |
| NA | NA | NA |
| -3.607835304 | 0.000308762 | 0.000979384 |
| -3.235730933 | 0.001213317 | 0.003429624 |
| -2.428095293 | 0.015178355 | 0.032924362 |
| -2.280415185 | 0.022583076 | 0.046480913 |
| 0.509194355 | 0.610616 | 0.706700863 |
| -2.560429535 | 0.010454286 | 0.023721402 |
| NA | NA | NA |
| -0.306181719 | 0.759466291 | 0.825922891 |
| -0.920223178 | 0.357456145 | NA |
| -8.65229515 | 5.05E-18 | 7.60E-17 |
| 0.532747666 | 0.594208265 | 0.693090679 |
| NA | NA | NA |
| NA | NA | NA |
| 5.879374316 | 4.12E-09 | 2.62E-08 |
| 3.129336001 | 0.001752018 | 0.004784314 |
| 4.20092947 | 2.66E-05 | 0.00010141 |
| 2.432320824 | 0.015002412 | 0.032582245 |
| NA | NA | NA |

| | | | |
|---|---|---|---|
| | 0.063708882 | 0.949202033 | NA |
| | 0.447435669 | 0.654560526 | 0.742483911 |
| | 1.525746112 | 0.127073106 | 0.203704358 |
| | 0.192808924 | 0.847108615 | NA |
| | -0.775424394 | 0.438088925 | 0.548405391 |
| | -3.812658983 | 0.00013748 | 0.000463888 |
| | 0.063707688 | 0.949202983 | NA |
| | 0.669094949 | 0.503434912 | NA |
| | -1.875449581 | 0.0607309 | 0.109123792 |
| | -0.651120693 | 0.51496858 | 0.62036106 |
| | -0.517679295 | 0.604682047 | 0.701742573 |
| NA | NA | NA | |
| | 7.980235979 | 1.46E-15 | 1.79E-14 |
| | 2.955522142 | 0.003121403 | 0.008034627 |
| | -2.763244059 | 0.005722996 | 0.013870082 |
| | 0.063708882 | 0.949202033 | NA |
| | -0.321333631 | 0.747957573 | NA |
| | -1.094002641 | 0.27395382 | 0.380588562 |
| | -0.621590628 | 0.534211085 | 0.63858367 |
| | 0.192809521 | 0.847108147 | NA |
| | 11.42434661 | 3.16E-30 | 1.10E-28 |
| | 0.832584683 | 0.405079 | 0.516854125 |
| | 10.49014969 | 9.59E-26 | 2.58E-24 |
| | -1.933739754 | 0.053145133 | 0.097526394 |
| | -9.321393859 | 1.15E-20 | 2.14E-19 |
| | -0.223035392 | 0.823507961 | 0.876303217 |
| | -5.47671441 | 4.33E-08 | 2.46E-07 |
| | -5.503701549 | 3.72E-08 | 2.12E-07 |
| | 5.830078248 | 5.54E-09 | 3.48E-08 |
| | -3.812912282 | 0.000137339 | 0.000463475 |
| | -2.187302974 | 0.028720414 | 0.05717585 |
| NA | NA | NA | |
| | -0.582667425 | 0.560117206 | 0.662495684 |
| | 2.723640941 | 0.006456666 | 0.015435563 |
| | 1.673046113 | 0.094318218 | 0.1586869 |
| | -1.734427531 | 0.082842254 | 0.142267006 |
| | -0.493908251 | 0.621370977 | 0.715602998 |
| | 0.063708285 | 0.949202508 | NA |
| | -4.218558155 | 2.46E-05 | 9.41E-05 |
| NA | NA | NA | |
| | 1.901661446 | 0.057215429 | 0.103901944 |
| | -1.207719779 | 0.227155068 | 0.327328229 |
| | -0.311522177 | 0.755403686 | 0.82272014 |
| NA | NA | NA | |
| | 5.340437143 | 9.27E-08 | 5.02E-07 |
| | 9.044885035 | 1.50E-19 | 2.58E-18 |
| | 15.2076762 | 3.15E-52 | 2.53E-50 |

| | | | |
|---|---|---|---|
| | 0.254385151 | 0.799198022 | NA |
| | 1.52379609 | 0.127559657 | 0.204344864 |
| | 4.165997965 | 3.10E-05 | 0.000116995 |
| | 2.881116671 | 0.003962689 | 0.00997146 |
| | -3.545454598 | 0.000391937 | 0.001219412 |
| | -2.670028949 | 0.007584471 | 0.01778021 |
| | -2.590402072 | 0.009586389 | 0.021927015 |
| | 5.970772803 | 2.36E-09 | 1.55E-08 |
| | 1.095621944 | 0.273244255 | 0.379834705 |
| | -0.394611612 | 0.693129542 | 0.773655214 |
| | -0.085589846 | 0.931792471 | NA |
| | 0.259390675 | 0.795333824 | 0.854386507 |
| NA | NA | NA | |
| NA | NA | NA | |
| | 4.751995299 | 2.01E-06 | 9.07E-06 |
| | -2.354882335 | 0.018528572 | 0.039130117 |
| | 0.063708285 | 0.949202508 | NA |
| | 2.672678734 | 0.007524828 | 0.01766355 |
| | 0.085821552 | 0.931608273 | NA |
| | -0.997111649 | 0.318710319 | 0.429326694 |
| | -2.706470786 | 0.006800257 | 0.016162513 |
| | -5.254219199 | 1.49E-07 | 7.83E-07 |
| | -1.865482325 | 0.062113827 | 0.111265233 |
| | 13.02666831 | 8.63E-39 | 4.52E-37 |
| | 3.220035894 | 0.001281745 | 0.003603514 |
| | -0.136809004 | 0.891181766 | 0.923833166 |
| | 8.443002429 | 3.09E-17 | 4.38E-16 |
| | -1.962565737 | 0.049696655 | 0.092113987 |
| | 1.631962352 | 0.102687417 | 0.170245445 |
| | 0.338781041 | 0.734774684 | 0.806605427 |
| | -5.337885229 | 9.40E-08 | 5.08E-07 |
| | 0.708266819 | 0.478779579 | 0.587530169 |
| | 3.049060842 | 0.00229558 | 0.006096378 |
| | 0.564990688 | 0.572080098 | 0.673399401 |
| | -3.654416839 | 0.000257767 | 0.000829188 |
| | -0.351528522 | 0.725191879 | 0.799294437 |
| | -2.725827279 | 0.006414055 | 0.015341023 |
| | -1.041447833 | 0.297667753 | 0.40639175 |
| | 0.208012043 | 0.835219564 | NA |
| | -0.550009202 | 0.582313062 | 0.682550303 |
| | -0.095312389 | 0.924066703 | NA |
| | 11.50293373 | 1.28E-30 | 4.53E-29 |
| | -0.08180493 | 0.934801835 | NA |
| | 1.566952675 | 0.117125754 | NA |
| NA | NA | NA | |
| | 14.45426814 | 2.36E-47 | 1.71E-45 |
| | 6.192550962 | 5.92E-10 | 4.11E-09 |

| | | | |
|---|---|---|---|
| | -0.065389369 | 0.947863988 | NA |
| | -0.209299993 | 0.834214057 | NA |
| NA | NA | NA | |
| | 1.714929466 | 0.08635816 | 0.147304398 |
| | 0.192809521 | 0.847108147 | NA |
| | 0.192809521 | 0.847108147 | NA |
| | -7.939098766 | 2.04E-15 | 2.47E-14 |
| | -0.521764939 | 0.601834004 | 0.69937811 |
| | 0.77957957 | 0.435638385 | 0.546198304 |
| | -2.125071367 | 0.033580658 | 0.065526533 |
| | -1.940526777 | 0.052315703 | 0.096269817 |
| NA | NA | NA | |
| | -0.631623839 | 0.527632705 | 0.632562915 |
| | -2.675035573 | 0.007472132 | 0.017553021 |
| | -1.486465723 | 0.137155977 | 0.216744371 |
| | 1.02367565 | 0.3059885 | 0.415621228 |
| | 0.390287866 | 0.696323695 | 0.776415809 |
| NA | NA | NA | |
| | -1.429965183 | 0.152727012 | 0.236967339 |
| | -1.275419184 | 0.202160912 | 0.298369019 |
| | -0.45225328 | 0.651086529 | 0.739716573 |
| | 4.761197846 | 1.92E-06 | 8.68E-06 |
| | 0.419711894 | 0.674695935 | NA |
| | 2.105036926 | 0.035288094 | 0.068390321 |
| | 1.092528738 | 0.274600766 | 0.381322565 |
| | -1.330737182 | 0.183275504 | 0.275287198 |
| | -1.474567634 | 0.140328822 | 0.2208659 |
| | 0.063708285 | 0.949202508 | NA |
| | 1.525550068 | 0.127121955 | 0.203743519 |
| | 8.42632701 | 3.57E-17 | 5.04E-16 |
| | -1.039557958 | 0.298545318 | 0.407425403 |
| | 7.107840923 | 1.18E-12 | 1.08E-11 |
| NA | NA | NA | |
| NA | NA | NA | |
| NA | NA | NA | |
| NA | NA | NA | |
| NA | NA | NA | |
| | 2.264888326 | 0.023519533 | 0.048124518 |
| | 1.95702517 | 0.050344509 | 0.093156137 |
| | 4.047157592 | 5.18E-05 | 0.000188528 |
| | -0.535047184 | 0.59261723 | 0.691750654 |
| | 1.516731216 | 0.12933456 | 0.206588642 |
| | 0.021267023 | 0.98303265 | 0.988045907 |
| | 6.962140105 | 3.35E-12 | 2.94E-11 |
| NA | NA | NA | |
| NA | NA | NA | |
| | 1.715103385 | 0.086326274 | 0.147260044 |

| | | | |
|---|---|---|---|
| | 9.307997914 | 1.30E-20 | 2.42E-19 |
| | 0.83626212 | 0.403007458 | 0.514656974 |
| | -2.086818071 | 0.036904583 | 0.071097197 |
| | -0.856344994 | 0.391806981 | 0.503797839 |
| | 1.743254534 | 0.081289163 | 0.140005672 |
| NA | NA | NA | |
| | 13.14832947 | 1.74E-39 | 9.34E-38 |
| | 3.52863573 | 0.000417708 | 0.001291725 |
| | 0.231088104 | 0.817246357 | 0.871381312 |
| | 1.386647049 | 0.165549414 | 0.253035041 |
| | 1.157533838 | 0.247054323 | 0.350183071 |
| | 17.93637285 | 6.13E-72 | 8.30E-70 |
| NA | NA | NA | |
| | 4.103848475 | 4.06E-05 | 0.000150074 |
| | -0.207598942 | 0.835542131 | 0.88469202 |
| | 0.98641137 | 0.323931288 | 0.434908552 |
| | 0.717657184 | 0.472968689 | 0.581927014 |
| | 0.177688626 | 0.858967518 | 0.901957335 |
| NA | NA | NA | |
| | 13.70603506 | 9.34E-43 | 5.68E-41 |
| | -3.01650737 | 0.00255705 | 0.00671172 |
| | -1.911761639 | 0.055906774 | 0.101874385 |
| | 11.86536021 | 1.79E-32 | 7.05E-31 |
| NA | NA | NA | |
| | -0.0653931 | 0.947861018 | NA |
| | 0.621890441 | 0.534013911 | 0.638439444 |
| | 0.924671615 | 0.355136749 | 0.467083854 |
| | 5.274220033 | 1.33E-07 | 7.07E-07 |
| | -2.0316189 | 0.042192252 | 0.079860259 |
| | 2.833424668 | 0.004605216 | 0.011427437 |
| | -1.017430877 | 0.3089485 | NA |
| | 9.09721258 | 9.27E-20 | 1.63E-18 |
| | 5.366199492 | 8.04E-08 | 4.39E-07 |
| NA | NA | NA | |
| | 1.766472049 | 0.077316687 | 0.134139923 |
| | 2.942970607 | 0.003250793 | 0.008331696 |
| | -1.587919728 | 0.112304492 | 0.183281048 |
| | 0.387947028 | 0.698055235 | 0.777757513 |
| | 1.512405906 | 0.130430633 | 0.207967815 |
| | -2.063975127 | 0.039020075 | 0.07454946 |
| | 0.192809521 | 0.847108147 | NA |
| | 4.123265944 | 3.74E-05 | 0.000138966 |
| | 3.623911852 | 0.000290181 | 0.000924897 |
| | 1.796786024 | 0.072369596 | 0.126743002 |
| | 10.09633031 | 5.73E-24 | 1.36E-22 |
| | -0.194487022 | 0.845794556 | NA |
| NA | NA | NA | |

| | | |
|---|---|---|
| 4.850794416 | 1.23E-06 | 5.69E-06 |
| -0.928470759 | 0.353163429 | 0.465026073 |
| -2.056508252 | 0.039733552 | 0.075768002 |
| NA | NA | NA |
| -0.34093621 | 0.733151604 | NA |
| 0.514051386 | 0.607216064 | 0.703802644 |
| NA | NA | NA |
| -0.512898333 | 0.60802244 | 0.704574151 |
| -2.356194846 | 0.01846323 | 0.039011573 |
| 1.807057964 | 0.070753242 | 0.124434629 |
| -2.227110789 | 0.02593987 | 0.052342305 |
| 0.66905202 | 0.503462295 | NA |
| NA | NA | NA |
| 1.52080032 | 0.128309954 | NA |
| 2.007886273 | 0.044655378 | 0.083963651 |
| 15.3538695 | 3.34E-53 | 2.75E-51 |
| -2.487290405 | 0.012872028 | 0.02848942 |
| 3.361948326 | 0.000773946 | 0.002277972 |
| 2.056041653 | 0.039778501 | 0.075836384 |
| -0.065388772 | 0.947864464 | NA |
| NA | NA | NA |
| -0.310452393 | 0.756216957 | 0.823247671 |
| 0.819291114 | 0.412620341 | 0.52423162 |
| 6.887776401 | 5.67E-12 | 4.86E-11 |
| 5.89211245 | 3.81E-09 | 2.44E-08 |
| 1.977275683 | 0.048010477 | 0.089401126 |
| 9.487551809 | 2.37E-21 | 4.66E-20 |
| 1.684943982 | 0.09199938 | 0.155447402 |
| 2.462783724 | 0.013786305 | 0.030261441 |
| NA | NA | NA |
| 1.347118226 | 0.177942159 | 0.268598012 |
| -4.728621998 | 2.26E-06 | 1.01E-05 |
| 1.127662728 | 0.259462384 | 0.364502637 |
| -0.016820296 | 0.986579979 | 0.990577113 |
| -11.02887286 | 2.77E-28 | 8.75E-27 |
| -0.445079144 | 0.65626256 | NA |
| NA | NA | NA |
| -9.068238948 | 1.21E-19 | 2.10E-18 |
| 0.612920554 | 0.539928872 | 0.644218689 |
| 5.732429097 | 9.90E-09 | 6.05E-08 |
| 3.805039317 | 0.000141782 | 0.000477309 |
| 1.491373138 | 0.135863557 | 0.215041408 |
| -2.119088894 | 0.034082955 | 0.066373327 |
| 7.633220851 | 2.29E-14 | 2.51E-13 |
| -3.76725535 | 0.000165052 | 0.000549661 |
| -1.084049593 | 0.278342811 | 0.385284179 |
| 4.082876698 | 4.45E-05 | 0.000163108 |

| | | |
|---|---|---|
| 3.012855947 | 0.002588017 | 0.006785887 |
| 4.249551184 | 2.14E-05 | 8.29E-05 |
| NA | NA | NA |
| 11.14295348 | 7.75E-29 | 2.52E-27 |
| NA | NA | NA |
| 10.31640284 | 5.94E-25 | 1.51E-23 |
| 0.063708882 | 0.949202033 | NA |
| NA | NA | NA |
| NA | NA | NA |
| -0.664300792 | 0.506497815 | 0.61294978 |
| -2.177009348 | 0.02947987 | 0.058487726 |
| NA | NA | NA |
| -0.751623045 | 0.452277781 | 0.562174745 |
| -1.611755099 | 0.107015242 | 0.176003122 |
| -0.876081867 | 0.38098553 | NA |
| -1.549989873 | 0.121143947 | 0.1954892 |
| -2.742793137 | 0.006091905 | 0.01465453 |
| -0.065388772 | 0.947864464 | NA |
| -3.197373933 | 0.00138685 | 0.003869453 |
| NA | NA | NA |
| 0.826252214 | 0.408661034 | 0.520495913 |
| -0.066649018 | 0.946861122 | NA |
| 0.689420916 | 0.490558425 | NA |
| 4.269888136 | 1.96E-05 | 7.61E-05 |
| 2.816752845 | 0.004851184 | 0.011984321 |
| 2.667177793 | 0.00764912 | 0.01790326 |
| -0.095289497 | 0.924084885 | NA |
| -5.311612351 | 1.09E-07 | 5.83E-07 |
| NA | NA | NA |
| 2.90599256 | 0.003660902 | 0.009267965 |
| 5.785490315 | 7.23E-09 | 4.48E-08 |
| NA | NA | NA |
| 6.837321757 | 8.07E-12 | 6.79E-11 |
| 5.67930219 | 1.35E-08 | 8.14E-08 |
| 4.571579402 | 4.84E-06 | 2.06E-05 |
| 2.256587108 | 0.024033887 | 0.048983648 |
| 9.361282259 | 7.88E-21 | 1.49E-19 |
| 5.288647458 | 1.23E-07 | 6.56E-07 |
| -0.284901878 | 0.775719317 | NA |
| 8.237063122 | 1.76E-16 | 2.35E-15 |
| -0.152279169 | 0.878966754 | NA |
| -0.747904087 | 0.454518012 | 0.564342767 |
| 4.315795685 | 1.59E-05 | 6.28E-05 |
| NA | NA | NA |
| NA | NA | NA |
| 7.193004001 | 6.34E-13 | 6.01E-12 |
| 4.259472849 | 2.05E-05 | 7.95E-05 |

| | | |
|---|---|---|
| 13.70023589 | 1.01E-42 | 6.13E-41 |
| -0.390102963 | 0.696460412 | 0.776533659 |
| -1.778952955 | 0.075247475 | 0.131003986 |
| NA | NA | NA |
| -4.663020286 | 3.12E-06 | 1.36E-05 |
| NA | NA | NA |
| 1.457259957 | 0.145044636 | 0.227074508 |
| 2.11864926 | 0.034120119 | 0.066430195 |
| -0.115678435 | 0.907907398 | 0.936376537 |
| 5.784062638 | 7.29E-09 | 4.52E-08 |
| -0.948921068 | 0.342660759 | 0.453978235 |
| NA | NA | NA |
| NA | NA | NA |
| NA | NA | NA |
| 0.293750308 | 0.768948709 | NA |
| -0.28799555 | 0.773350145 | NA |
| -3.463273183 | 0.000533646 | 0.001619639 |
| 15.01017919 | 6.30E-51 | 4.90E-49 |
| 11.87054836 | 1.68E-32 | 6.64E-31 |
| 8.800426983 | 1.36E-18 | 2.17E-17 |
| -7.433160304 | 1.06E-13 | 1.09E-12 |
| -1.844780902 | 0.065069417 | 0.115739589 |
| 0.192808327 | 0.847109082 | NA |
| NA | NA | NA |
| 0.130867119 | 0.895880426 | NA |
| 0.192809521 | 0.847108147 | NA |
| NA | NA | NA |
| NA | NA | NA |
| -1.434104422 | 0.151542466 | NA |
| -0.523871913 | 0.600367631 | 0.698095411 |
| 1.046414787 | 0.295369568 | 0.403707765 |
| 5.565159172 | 2.62E-08 | 1.52E-07 |
| NA | NA | NA |
| -0.713376853 | 0.475612592 | 0.58450352 |
| -0.056267507 | 0.955128703 | 0.969278444 |
| NA | NA | NA |
| NA | NA | NA |
| 0.192808327 | 0.847109082 | NA |
| NA | NA | NA |
| 10.35951306 | 3.79E-25 | 9.81E-24 |
| NA | NA | NA |
| -0.236209016 | 0.813270484 | NA |
| -3.789927635 | 0.000150691 | 0.000505062 |
| -7.179438174 | 7.00E-13 | 6.60E-12 |
| 0.333197539 | 0.738985176 | NA |
| 2.787977191 | 0.005303828 | 0.012964337 |
| 7.061610598 | 1.65E-12 | 1.49E-11 |

| | | | |
|---|---|---|---|
| | 0.579653016 | 0.562148633 | NA |
| | 0.307371449 | 0.758560657 | 0.825153118 |
| NA | NA | NA | |
| | 0.106996834 | 0.914791491 | 0.941363699 |
| | 1.081529109 | 0.279461819 | 0.386481537 |
| | 0.0921369 | 0.926589271 | 0.949696141 |
| | 1.57311253 | 0.115692764 | 0.187989476 |
| | -0.333775502 | 0.738548971 | 0.809594209 |
| | 8.756013567 | 2.02E-18 | 3.17E-17 |
| | -3.069944471 | 0.002140986 | 0.005730205 |
| NA | NA | NA | |
| NA | NA | NA | |
| | 0.293563678 | 0.769091334 | 0.83366337 |
| | -2.967931514 | 0.002998111 | 0.007753577 |
| | -3.299579205 | 0.000968299 | 0.002798352 |
| | 0.722872786 | 0.469758045 | 0.578767621 |
| | 3.286742737 | 0.001013534 | 0.002918632 |
| | -7.673473704 | 1.67E-14 | 1.86E-13 |
| | -5.302534789 | 1.14E-07 | 6.11E-07 |
| | -0.065389369 | 0.947863988 | NA |
| | 2.934910536 | 0.003336439 | 0.008526605 |
| | 0.57105734 | 0.567960775 | 0.669652749 |
| | -0.821573176 | 0.411319861 | 0.523057264 |
| | 1.229172523 | 0.219007128 | 0.317911525 |
| | -0.520961681 | 0.602393465 | 0.699822687 |
| | 0.194173692 | 0.846039881 | NA |
| NA | NA | NA | |
| | 0.278377556 | 0.780722547 | NA |
| | 6.94813626 | 3.70E-12 | 3.23E-11 |
| | -0.218135449 | 0.827323578 | 0.879092765 |
| | -0.074439931 | 0.940660337 | NA |
| | 11.09016338 | 1.40E-28 | 4.50E-27 |
| | -3.734852012 | 0.000187826 | 0.00061973 |
| | 3.394923896 | 0.000686477 | 0.002041406 |
| | 1.272704704 | 0.203122848 | 0.299438417 |
| NA | NA | NA | |
| | 0.74004568 | 0.459272277 | 0.568666094 |
| | 6.62447458 | 3.48E-11 | 2.78E-10 |
| | 1.691688954 | 0.090705296 | 0.153611483 |
| | -1.618941639 | 0.105459821 | 0.173970824 |
| NA | NA | NA | |
| | -0.199148371 | 0.842146684 | 0.889368867 |
| NA | NA | NA | |
| | 0.972285419 | 0.330908575 | 0.441835146 |
| | 4.407576035 | 1.05E-05 | 4.25E-05 |
| | 1.260818308 | 0.207374316 | 0.3043257 |
| | 9.088095652 | 1.01E-19 | 1.76E-18 |

| | | | |
|---|---|---|---|
| | 0.118920028 | 0.905338714 | 0.934468687 |
| | 4.805641576 | 1.54E-06 | 7.05E-06 |
| NA | NA | NA | |
| | -0.962480324 | 0.335808382 | 0.447065784 |
| | 3.03434489 | 0.002410587 | 0.006367348 |
| | 0.362026528 | 0.717332207 | NA |
| | 0.087763409 | 0.930064721 | NA |
| | -1.318886455 | 0.187207075 | 0.279993491 |
| | 0.309884575 | 0.756648733 | NA |
| | 0.06371127 | 0.949200131 | NA |
| | 0.065042077 | 0.948140499 | 0.964674472 |
| | -1.702656781 | 0.088632317 | 0.150652407 |
| NA | NA | NA | |
| | -9.50176085 | 2.06E-21 | 4.09E-20 |
| | 0.418367716 | 0.675678287 | NA |
| | -0.043916012 | 0.964971352 | 0.976164287 |
| NA | NA | NA | |
| | 0.455742734 | 0.648574988 | 0.73749918 |
| | 0.873614177 | 0.382328403 | NA |
| NA | NA | NA | |
| | 0.192809521 | 0.847108147 | NA |
| | -0.880351906 | 0.378668702 | 0.491270285 |
| | -0.5498125 | 0.582447984 | NA |
| NA | NA | NA | |
| | 4.562033039 | 5.07E-06 | 2.15E-05 |
| NA | NA | NA | |
| | -0.699309066 | 0.484358903 | 0.592836349 |
| | 2.202713004 | 0.027614983 | 0.055269713 |
| | 0.363202125 | 0.7164539 | NA |
| | 0.192636473 | 0.847243678 | NA |
| | -0.099490974 | 0.920748454 | 0.945538501 |
| | 0.063708882 | 0.949202033 | NA |
| | -6.261293403 | 3.82E-10 | 2.71E-09 |
| | -0.949930218 | 0.342147711 | NA |
| | -3.175694973 | 0.00149478 | 0.004141505 |
| | 0.85528876 | 0.392391308 | 0.504341915 |
| | -5.233487419 | 1.66E-07 | 8.70E-07 |
| | -2.252140596 | 0.024313389 | 0.049492853 |
| | -7.196600557 | 6.17E-13 | 5.86E-12 |
| | 9.53045715 | 1.57E-21 | 3.15E-20 |
| NA | NA | NA | |
| | -1.606609808 | 0.10813999 | 0.177561266 |
| | 7.958835587 | 1.74E-15 | 2.12E-14 |
| | 1.08937796 | 0.275987247 | NA |
| | 1.323654348 | 0.185617867 | 0.278087939 |
| | -2.791929593 | 0.005239476 | 0.012823318 |
| | -0.87057978 | 0.383983641 | 0.496287852 |

| | | | |
|---|---|---|---|
| | -3.4277115 | 0.000608692 | 0.001827025 |
| | 0.063708882 | 0.949202033 | NA |
| | 0.320071032 | 0.748914485 | NA |
| | 9.257127276 | 2.10E-20 | 3.83E-19 |
| NA | NA | NA | |
| NA | NA | NA | |
| NA | NA | NA | |
| | -2.198129911 | 0.02793985 | 0.055803897 |
| | 1.134157726 | 0.256728393 | 0.361290435 |
| | 0.192809521 | 0.847108147 | NA |
| | -0.194487022 | 0.845794556 | NA |
| | 4.210206129 | 2.55E-05 | 9.76E-05 |
| | 5.502620773 | 3.74E-08 | 2.14E-07 |
| | 2.686723343 | 0.007215669 | 0.017006421 |
| | -2.958479565 | 0.003091607 | 0.007966124 |
| | -0.779065542 | 0.435941107 | 0.546403418 |
| | 7.546476033 | 4.47E-14 | 4.76E-13 |
| | 2.066263439 | 0.038803612 | 0.074192529 |
| | 1.227455524 | 0.219651428 | 0.318662934 |
| | -1.786589523 | 0.074003856 | 0.129135285 |
| | 2.470951751 | 0.0134754 | 0.029649196 |
| | -1.047763285 | 0.294747682 | 0.403033842 |
| NA | NA | NA | |
| | 1.265561274 | 0.205670198 | 0.302338936 |
| | -2.270780742 | 0.023160254 | 0.047481462 |
| | -5.367732831 | 7.97E-08 | 4.35E-07 |
| | -0.166703449 | 0.867603392 | 0.907750403 |
| NA | NA | NA | |
| NA | NA | NA | |
| | 12.44398423 | 1.51E-35 | 6.75E-34 |
| | -1.20036858 | 0.229996226 | 0.33060389 |
| | 0.526516802 | 0.598529181 | NA |
| | 6.40527719 | 1.50E-10 | 1.12E-09 |
| NA | NA | NA | |
| NA | NA | NA | |
| | 3.67586578 | 0.000237044 | 0.000767954 |
| | 6.070938425 | 1.27E-09 | 8.56E-09 |
| | 0.672111808 | 0.50151252 | 0.608387456 |
| | 0.928693485 | 0.353047958 | 0.464947405 |
| | -1.237347543 | 0.215958085 | 0.314470351 |
| | 0.193076837 | 0.846898793 | NA |
| | -2.616368729 | 0.008887053 | 0.020494097 |
| NA | NA | NA | |
| | -1.587535031 | 0.112391519 | 0.183411118 |
| | 0.838924886 | 0.401511462 | 0.513234287 |
| NA | NA | NA | |
| | 0.450772038 | 0.652153856 | NA |

| | | |
|---|---|---|
| NA | NA | NA |
| 1.162431334 | 0.245060302 | 0.347941423 |
| 13.2714776 | 3.39E-40 | 1.86E-38 |
| NA | NA | NA |
| 0.993313676 | 0.320557115 | 0.431373365 |
| 1.638187418 | 0.101382603 | 0.168443848 |
| 1.120701974 | 0.262414742 | 0.367783246 |
| 0.063708381 | 0.949202431 | NA |
| 6.427820327 | 1.29E-10 | 9.71E-10 |
| NA | NA | NA |
| 1.257753366 | 0.208480966 | 0.305555602 |
| NA | NA | NA |
| NA | NA | NA |
| NA | NA | NA |
| NA | NA | NA |
| NA | NA | NA |
| NA | NA | NA |
| NA | NA | NA |
| 0.277051294 | 0.78174072 | 0.844277279 |
| NA | NA | NA |
| NA | NA | NA |
| NA | NA | NA |
| -0.673001063 | 0.500946615 | 0.607971377 |
| NA | NA | NA |
| NA | NA | NA |
| NA | NA | NA |
| NA | NA | NA |
| NA | NA | NA |
| NA | NA | NA |
| NA | NA | NA |
| NA | NA | NA |
| NA | NA | NA |
| NA | NA | NA |
| NA | NA | NA |
| NA | NA | NA |
| NA | NA | NA |
| NA | NA | NA |
| NA | NA | NA |
| NA | NA | NA |
| NA | NA | NA |
| NA | NA | NA |
| NA | NA | NA |
| NA | NA | NA |
| 0.210980867 | 0.832902201 | 0.882927856 |
| NA | NA | NA |
| NA | NA | NA |
| NA | NA | NA |

| | | | |
|---|---|---|---|
| NA | NA | NA | |
| NA | NA | NA | |
| NA | NA | NA | |
| NA | NA | NA | |
| NA | NA | NA | |
| NA | NA | NA | |
| -0.320138271 | 0.748863515 | NA | |
| -0.065388772 | 0.947864464 | NA | |
| NA | NA | NA | |
| NA | NA | NA | |
| -0.065388772 | 0.947864464 | NA | |
| -0.065389369 | 0.947863988 | NA | |
| 0.192809521 | 0.847108147 | NA | |
| NA | NA | NA | |
| -0.422590064 | 0.672594377 | | 0.756596884 |
| NA | NA | NA | |
| NA | NA | NA | |
| 0.063708882 | 0.949202033 | NA | |
| NA | NA | NA | |
| 0.211051298 | 0.832847242 | NA | |
| -0.065389369 | 0.947863988 | NA | |
| 11.50723375 | 1.21E-30 | | 4.31E-29 |
| -0.481459215 | 0.63019016 | | 0.722448347 |
| -4.61812496 | 3.87E-06 | | 1.67E-05 |
| 15.7323465 | 9.08E-56 | | 8.09E-54 |
| 3.551979099 | 0.000382345 | | 0.001191348 |
| 0.128520898 | 0.897736761 | NA | |
| NA | NA | NA | |
| NA | NA | NA | |
| -0.194487022 | 0.845794556 | NA | |
| -0.19448762 | 0.845794089 | NA | |
| 0.199662929 | 0.841744208 | | 0.889206446 |
| NA | NA | NA | |
| 0.169533173 | 0.865377281 | NA | |
| 0.063708285 | 0.949202508 | NA | |
| NA | NA | NA | |
| 2.231921736 | 0.025620136 | | 0.051790664 |
| NA | NA | NA | |
| 6.346329319 | 2.21E-10 | | 1.61E-09 |
| NA | NA | NA | |
| -0.045578239 | 0.963646414 | | 0.975178691 |
| 0.283708011 | 0.776634155 | | 0.839740503 |
| NA | NA | NA | |
| 0.063707019 | 0.949203516 | NA | |
| NA | NA | NA | |
| NA | NA | NA | |
| 0.604904124 | 0.545242703 | NA | |

| | | |
|---|---|---|
| NA | NA | NA |
| 0.554970686 | 0.578914711 | 0.679718324 |
| 0.063708882 | 0.949202033 | NA |
| 4.092280912 | 4.27E-05 | 0.000157091 |
| -0.137282977 | 0.890807125 | 0.923573124 |
| NA | NA | NA |
| 2.269597937 | 0.023231989 | 0.047612927 |
| NA | NA | NA |
| 0.963281073 | 0.335406489 | 0.44662576 |
| NA | NA | NA |
| NA | NA | NA |
| NA | NA | NA |
| -1.474030817 | 0.140473294 | NA |
| NA | NA | NA |
| NA | NA | NA |
| NA | NA | NA |
| -1.381196787 | 0.167218463 | 0.255165404 |
| 0.192810118 | 0.847107679 | NA |
| -0.19448762 | 0.845794089 | NA |
| NA | NA | NA |
| 4.008026492 | 6.12E-05 | 0.000219639 |
| 1.406811297 | 0.159483352 | 0.245366543 |
| 0.261643719 | 0.793596135 | 0.853288855 |
| 0.378842653 | 0.704804711 | 0.783495113 |
| NA | NA | NA |
| NA | NA | NA |
| -0.112053594 | 0.910780912 | NA |
| 0.192809521 | 0.847108147 | NA |
| 0.063708285 | 0.949202508 | NA |
| 0.3474247 | 0.728272279 | NA |
| 1.945692492 | 0.051691689 | 0.095268609 |
| 0.192809521 | 0.847108147 | NA |
| 0.613697194 | 0.539415443 | NA |
| 0.46156349 | 0.644394381 | 0.733845551 |
| NA | NA | NA |
| NA | NA | NA |
| NA | NA | NA |
| 0.192809521 | 0.847108147 | NA |
| NA | NA | NA |
| NA | NA | NA |
| 0.754308161 | 0.450664203 | NA |
| 1.611979051 | 0.106966499 | 0.175946077 |
| 1.499258755 | 0.133806518 | 0.212322629 |
| -0.127131979 | 0.898835941 | 0.929625371 |
| 5.095717032 | 3.47E-07 | 1.74E-06 |
| 0.431089098 | 0.666403588 | 0.75206645 |
| -0.733400127 | 0.463314422 | 0.572764359 |

| | | | |
|---|---|---|---|
| NA | NA | NA | |
| | 0.265575578 | 0.790566096 | 0.850981352 |
| | -0.385867407 | 0.699594871 | NA |
| | 1.116619033 | 0.26415725 | 0.369687222 |
| NA | NA | NA | |
| | -0.028571822 | 0.977206085 | 0.984308805 |
| | 0.440511299 | 0.659566831 | NA |
| NA | NA | NA | |
| | -0.945580223 | 0.344362735 | 0.455691212 |
| | -0.290349728 | 0.771548699 | 0.835605433 |
| | 0.63278982 | 0.526870905 | 0.631891651 |
| NA | NA | NA | |
| | -0.96021501 | 0.336947014 | 0.4482717 |
| NA | NA | NA | |
| | -1.543494947 | 0.122710733 | 0.19769887 |
| | 0.721228142 | 0.47076916 | 0.579803772 |
| | 3.654063744 | 0.000258122 | 0.000830117 |
| | -0.19448762 | 0.845794089 | NA |
| | 0.649213435 | 0.516200429 | 0.621476515 |
| | 0.78518249 | 0.43234658 | NA |
| NA | NA | NA | |
| | 1.953335794 | 0.050779819 | 0.093864476 |
| | 1.514515206 | 0.12989522 | NA |
| | 10.72411519 | 7.84E-27 | 2.25E-25 |
| | 3.597027344 | 0.000321875 | 0.001018138 |
| | -0.338181339 | 0.735226538 | 0.806889123 |
| | -2.338959096 | 0.019337551 | 0.040585183 |
| | -1.39300972 | 0.163616851 | 0.25054018 |
| | -3.997044551 | 6.41E-05 | 0.000229176 |
| | -4.61249882 | 3.98E-06 | 1.72E-05 |
| | -0.372100854 | 0.709817761 | 0.787358234 |
| | 7.894009681 | 2.93E-15 | 3.49E-14 |
| | 7.572554385 | 3.66E-14 | 3.93E-13 |
| | 1.588311481 | 0.112215923 | 0.183196314 |
| | 0.181048714 | 0.85632934 | 0.9002824 |
| | 0.904494665 | 0.365733162 | 0.478306131 |
| | -0.52325054 | 0.600799915 | 0.698468271 |
| | 0.063708882 | 0.949202033 | NA |
| | 15.4707027 | 5.47E-54 | 4.61E-52 |
| | 0.690525635 | 0.489863695 | 0.597528668 |
| | -0.628424548 | 0.52972586 | NA |
| | 7.320165642 | 2.48E-13 | 2.44E-12 |
| | 0.314932707 | 0.752812747 | 0.820648189 |
| | 8.301484288 | 1.03E-16 | 1.39E-15 |
| | 8.988726098 | 2.50E-19 | 4.24E-18 |
| | 0.081528689 | 0.935021511 | NA |
| | -4.13823229 | 3.50E-05 | 0.000130927 |

| | | |
|---|---|---|
| -3.791331718 | 0.000149842 | 0.000502349 |
| -5.641346665 | 1.69E-08 | 1.00E-07 |
| -0.065388772 | 0.947864464 | NA |
| -1.779603265 | 0.075140911 | 0.130863973 |
| 5.312946134 | 1.08E-07 | 5.79E-07 |
| 0.614542434 | 0.538856941 | 0.64318499 |
| -6.861892085 | 6.80E-12 | 5.77E-11 |
| 4.072735854 | 4.65E-05 | 0.000170029 |
| -0.802818499 | 0.422079648 | 0.533891934 |
| -0.21845203 | 0.82707693 | 0.878930414 |
| 4.556840371 | 5.19E-06 | 2.20E-05 |
| NA | NA | NA |
| 1.443215904 | 0.148959661 | NA |
| 1.266706878 | 0.20526012 | 0.301842477 |
| -1.685391553 | 0.091913052 | 0.155336204 |
| -0.695669713 | 0.486635697 | 0.594779603 |
| 0.418381697 | 0.675668067 | 0.758997025 |
| NA | NA | NA |
| 2.356527719 | 0.018446691 | 0.038983508 |
| NA | NA | NA |
| -4.565554235 | 4.98E-06 | 2.12E-05 |
| 1.349072581 | 0.177313654 | 0.267859302 |
| 3.827549294 | 0.000129425 | 0.000438662 |
| NA | NA | NA |
| -4.836781846 | 1.32E-06 | 6.08E-06 |
| -3.999748107 | 6.34E-05 | 0.000226833 |
| -1.321482174 | 0.186340642 | 0.279003785 |
| 0.587059011 | 0.557164075 | 0.65968859 |
| 12.39602706 | 2.75E-35 | 1.21E-33 |
| -7.626151956 | 2.42E-14 | 2.65E-13 |
| -1.886789114 | 0.059188696 | 0.10682817 |
| 0.063707688 | 0.949202983 | NA |
| 0.192810118 | 0.847107679 | NA |
| 0.143409731 | 0.885966598 | NA |
| 1.485470805 | 0.137419152 | 0.217036936 |
| 1.818859834 | 0.068932817 | 0.121677603 |
| -2.439768743 | 0.014696667 | 0.031984833 |
| -0.065389369 | 0.947863988 | NA |
| 3.50536252 | 0.000455986 | 0.001400408 |
| -1.641753457 | 0.100641108 | 0.16740078 |
| NA | NA | NA |
| -2.621275523 | 0.008760143 | 0.020223795 |
| 1.534829899 | 0.124825616 | 0.20061546 |
| 0.343843062 | 0.730964314 | NA |
| -9.975078537 | 1.96E-23 | 4.52E-22 |
| 9.627514967 | 6.12E-22 | 1.26E-20 |
| NA | NA | NA |

| | | |
|---|---|---|
| -4.184053011 | 2.86E-05 | 0.000108665 |
| -0.411529099 | 0.680684609 | 0.763407144 |
| -0.583417507 | 0.559612275 | NA |
| 1.093301436 | 0.274261473 | NA |
| 0.382160883 | 0.702342035 | NA |
| 0.063708285 | 0.949202508 | NA |
| 0.063707019 | 0.949203516 | NA |
| NA | NA | NA |
| NA | NA | NA |
| 11.04955584 | 2.20E-28 | 7.00E-27 |
| 0.063708882 | 0.949202033 | NA |
| 3.89118422 | 9.98E-05 | 0.000344809 |
| -1.78877835 | 0.073650519 | 0.128581467 |
| 0.258749957 | 0.795828172 | NA |
| -1.718950478 | 0.08562339 | 0.146260386 |
| 4.836117906 | 1.32E-06 | 6.10E-06 |
| -3.253917259 | 0.001138254 | 0.003240872 |
| -1.15881548 | 0.246531406 | 0.349558454 |
| NA | NA | NA |
| 3.798637632 | 0.000145494 | 0.000488557 |
| 0.458140068 | 0.646851814 | NA |
| 1.538002079 | 0.124048106 | 0.199545251 |
| -1.456286548 | 0.145313423 | 0.227409964 |
| 1.249620111 | 0.211438353 | 0.309165906 |
| -2.836200103 | 0.004565383 | 0.011336462 |
| 2.934843035 | 0.003337165 | 0.008526871 |
| -3.114765476 | 0.001840912 | 0.00499634 |
| NA | NA | NA |
| NA | NA | NA |
| 0.063710673 | 0.949200606 | NA |
| 1.303960015 | 0.192247216 | 0.286232326 |
| 4.5946091 | 4.34E-06 | 1.86E-05 |
| 0.215318343 | 0.829519129 | 0.880790048 |
| 8.030436739 | 9.71E-16 | 1.21E-14 |
| -9.610763333 | 7.20E-22 | 1.48E-20 |
| NA | NA | NA |
| 2.532424776 | 0.011327667 | 0.025476818 |
| -6.341641139 | 2.27E-10 | 1.65E-09 |
| 6.977485172 | 3.01E-12 | 2.64E-11 |
| 0.584055045 | 0.559183279 | NA |
| 0.063708882 | 0.949202033 | NA |
| 0.192810118 | 0.847107679 | NA |
| 4.749215165 | 2.04E-06 | 9.18E-06 |
| 0.338911016 | 0.734676766 | 0.806533309 |
| 2.130390783 | 0.033139364 | 0.064792867 |
| 2.091440128 | 0.036488628 | 0.070400861 |
| 1.588311242 | 0.112215977 | 0.183196314 |

| | | |
|---|---|---|
| 1.626130245 | 0.103921951 | 0.171943842 |
| 4.122902305 | 3.74E-05 | 0.000139165 |
| -3.037019952 | 0.002389296 | 0.006316444 |
| -2.393696779 | 0.016679532 | 0.035786788 |
| -2.108250465 | 0.03500933 | 0.067934222 |
| 1.473270198 | 0.140678193 | 0.221262736 |
| 0.151834059 | 0.879317818 | NA |
| -2.895503703 | 0.003785506 | 0.009559242 |
| NA | NA | NA |
| -2.538944063 | 0.01111876 | 0.025065528 |
| -7.828064253 | 4.95E-15 | 5.80E-14 |
| 0.506583979 | 0.612446752 | 0.708160123 |
| -3.399580783 | 0.000674892 | 0.002009105 |
| -5.060920271 | 4.17E-07 | 2.07E-06 |
| 4.309277648 | 1.64E-05 | 6.45E-05 |
| -9.253249215 | 2.18E-20 | 3.96E-19 |
| -2.333500848 | 0.019621874 | 0.041095766 |
| -8.584324337 | 9.14E-18 | 1.35E-16 |
| 0.456359148 | 0.648131737 | NA |
| -2.970325656 | 0.002974842 | 0.007700134 |
| -3.898676785 | 9.67E-05 | 0.000335397 |
| NA | NA | NA |
| 1.33396559 | 0.182215134 | 0.273973978 |
| NA | NA | NA |
| 0.266884854 | 0.789557819 | NA |
| 0.506363264 | 0.612601658 | NA |
| -3.494442469 | 0.000475053 | 0.001454327 |
| NA | NA | NA |
| -0.065389369 | 0.947863988 | NA |
| -2.402064446 | 0.016302836 | 0.03508081 |
| 3.273980403 | 0.00106044 | 0.003039717 |
| -1.45927889 | 0.144488363 | 0.226316879 |
| 2.026280491 | 0.042736049 | 0.080767507 |
| 2.163512447 | 0.030501787 | 0.060214507 |
| -1.055314439 | 0.29128154 | 0.398969954 |
| -0.554792706 | 0.579036456 | NA |
| NA | NA | NA |
| NA | NA | NA |
| -2.913666342 | 0.003572115 | 0.009065196 |
| 0.066587943 | 0.946909745 | NA |
| NA | NA | NA |
| -1.172758229 | 0.240892773 | 0.343270487 |
| -2.696817702 | 0.007000558 | 0.016575917 |
| 0.155504531 | 0.876423582 | 0.913732184 |
| NA | NA | NA |
| 0.273180073 | 0.784714797 | 0.846503098 |
| 0.149845203 | 0.880886745 | 0.916427515 |

| | | |
|---|---|---|
| -2.064458938 | 0.038974223 | 0.074478914 |
| 1.8875138 | 0.059091252 | 0.106713855 |
| 0.963324301 | 0.335384802 | 0.446620642 |
| -6.102198435 | 1.05E-09 | 7.12E-09 |
| NA | NA | NA |
| -0.259706918 | 0.795089856 | 0.854198553 |
| NA | NA | NA |
| 0.688180852 | 0.4913389 | 0.598778193 |
| -1.414117541 | 0.157327394 | 0.242780805 |
| -1.864869231 | 0.062199736 | 0.111411149 |
| -0.441751007 | 0.658669395 | 0.745974321 |
| -3.963619079 | 7.38E-05 | 0.000261501 |
| 13.3550482 | 1.11E-40 | 6.16E-39 |
| -2.045325296 | 0.040822798 | 0.077567346 |
| -7.687560102 | 1.50E-14 | 1.68E-13 |
| -1.415593886 | 0.156894442 | 0.242336803 |
| -8.570138642 | 1.03E-17 | 1.52E-16 |
| -1.910988153 | 0.056006103 | 0.102040527 |
| -2.034106668 | 0.041940842 | 0.079457239 |
| 7.639194081 | 2.19E-14 | 2.41E-13 |
| -3.396300629 | 0.000683033 | 0.002031406 |
| 0.211830481 | 0.832239288 | 0.88259255 |
| 5.262272927 | 1.42E-07 | 7.52E-07 |
| -6.555126489 | 5.56E-11 | 4.35E-10 |
| 0.070233856 | 0.944007528 | 0.962111992 |
| -6.491085679 | 8.52E-11 | 6.53E-10 |
| 0.531701888 | 0.594932484 | 0.693579883 |
| -4.051997232 | 5.08E-05 | 0.000184884 |
| 4.345404559 | 1.39E-05 | 5.53E-05 |
| -2.586120301 | 0.009706305 | 0.022183054 |
| 0.063708882 | 0.949202033 | NA |
| 4.771202502 | 1.83E-06 | 8.29E-06 |
| -3.381335516 | 0.000721344 | 0.002136462 |
| 0.873963914 | 0.382137906 | 0.494612229 |
| 0.063710673 | 0.949200606 | NA |
| -1.25750555 | 0.208570631 | 0.305651223 |
| 3.808173513 | 0.000139997 | 0.000471809 |
| 0.004704759 | 0.996246159 | 0.997521005 |
| -3.931844688 | 8.43E-05 | 0.000295551 |
| NA | NA | NA |
| NA | NA | NA |
| 4.24147045 | 2.22E-05 | 8.57E-05 |
| -0.065388772 | 0.947864464 | NA |
| -2.168540303 | 0.030117602 | 0.059540602 |
| -6.928174395 | 4.26E-12 | 3.70E-11 |
| 2.019909029 | 0.043392825 | 0.081878809 |
| -1.248355101 | 0.211901044 | 0.30973389 |

| | | |
|---|---|---|
| 7.587355274 | 3.27E-14 | 3.52E-13 |
| 2.300551533 | 0.021416993 | 0.044333037 |
| 1.249963013 | 0.211313059 | 0.30903686 |
| NA | NA | NA |
| -1.023008686 | 0.306303739 | 0.415949576 |
| 2.836847215 | 0.004556141 | 0.011318004 |
| 1.031221199 | 0.302437102 | 0.41170031 |
| 3.095288828 | 0.001966213 | 0.005306126 |
| -0.065390563 | 0.947863038 | NA |
| 6.045622216 | 1.49E-09 | 9.95E-09 |
| -0.784897134 | 0.432513883 | 0.543192297 |
| -0.244402481 | 0.80691912 | 0.863164476 |
| 2.234468002 | 0.025452296 | 0.051497127 |
| 8.943251659 | 3.78E-19 | 6.30E-18 |
| -2.98944939 | 0.002794807 | 0.007274754 |
| -4.759238396 | 1.94E-06 | 8.77E-06 |
| NA | NA | NA |
| 4.11049048 | 3.95E-05 | 0.000146253 |
| -4.966201765 | 6.83E-07 | 3.28E-06 |
| 1.207178588 | 0.227363375 | 0.327552968 |
| -0.420905861 | 0.673823825 | 0.757553559 |
| 2.683234314 | 0.007291388 | 0.017169374 |
| -4.136941462 | 3.52E-05 | 0.000131586 |
| -1.599617792 | 0.109683399 | 0.179718217 |
| 0.194442267 | 0.845829597 | 0.891966605 |
| 6.943783524 | 3.82E-12 | 3.33E-11 |
| -0.348802323 | 0.727237719 | 0.800790115 |
| -0.977539882 | 0.328301943 | 0.439315565 |
| NA | NA | NA |
| 0.063708285 | 0.949202508 | NA |
| -0.065389369 | 0.947863988 | NA |
| -4.538014799 | 5.68E-06 | 2.40E-05 |
| 2.549268562 | 0.010794913 | 0.024414628 |
| 22.85373595 | 1.34E-115 | 4.10E-113 |
| 12.81535362 | 1.35E-37 | 6.65E-36 |
| 3.279608274 | 0.001039513 | 0.002985544 |
| 4.859761607 | 1.18E-06 | 5.45E-06 |
| -4.439798833 | 9.00E-06 | 3.70E-05 |
| NA | NA | NA |
| 4.752258476 | 2.01E-06 | 9.06E-06 |
| -0.455263861 | 0.648919422 | 0.737856005 |
| -0.242561458 | 0.808345142 | NA |
| NA | NA | NA |
| NA | NA | NA |
| NA | NA | NA |
| NA | NA | NA |
| NA | NA | NA |

| | | |
|---|---|---|
| NA | NA | NA |
| NA | NA | NA |
| NA | NA | NA |
| NA | NA | NA |
| NA | NA | NA |
| NA | NA | NA |
| 4.115184652 | 3.87E-05 | 0.000143521 |
| NA | NA | NA |
| 11.56962779 | 5.87E-31 | 2.14E-29 |
| 0.06371127 | 0.949200131 | NA |
| 0.847807573 | 0.396545145 | NA |
| -1.681965685 | 0.092575494 | 0.156311699 |
| -1.826666551 | 0.06774992 | 0.119860087 |
| NA | NA | NA |
| -1.700925869 | 0.088956908 | 0.151139792 |
| 1.609904719 | 0.107418659 | 0.176574926 |
| -2.682720684 | 0.007302595 | 0.01719187 |
| -2.656706269 | 0.007890816 | 0.018402536 |
| 0.152204966 | 0.879025277 | 0.915034422 |
| -5.192498749 | 2.07E-07 | 1.07E-06 |
| -0.514574595 | 0.606850321 | 0.703509155 |
| 0.418577516 | 0.675524924 | NA |
| 0.063707019 | 0.949203516 | NA |
| 2.347915765 | 0.01887879 | 0.039735631 |
| NA | NA | NA |
| -2.801577023 | 0.00508535 | 0.012491296 |
| -3.031220621 | 0.002435672 | 0.006422764 |
| 1.155588063 | 0.247849693 | 0.351045598 |
| 2.283065423 | 0.022426515 | 0.046192863 |
| 0.083430915 | 0.933508908 | 0.954825494 |
| -1.917130837 | 0.055221312 | 0.100843079 |
| -2.798943881 | 0.005127005 | 0.012582501 |
| -2.188128771 | 0.028660224 | 0.057083266 |
| 1.868819329 | 0.061647955 | 0.110549384 |
| -0.919730497 | 0.357713612 | 0.469584712 |
| 2.38733489 | 0.016971024 | 0.036309491 |
| -0.752265085 | 0.45189166 | 0.5618064 |
| -1.65878293 | 0.097159541 | 0.162636482 |
| NA | NA | NA |
| -10.92147765 | 9.10E-28 | 2.81E-26 |
| -4.349993784 | 1.36E-05 | 5.43E-05 |
| -3.345149034 | 0.000822384 | 0.002407516 |
| 1.533673224 | 0.125110064 | 0.20098228 |
| -2.793751915 | 0.005210043 | 0.012762515 |
| -3.224265123 | 0.001262964 | 0.003554705 |
| -3.524509705 | 0.000424267 | 0.001310876 |
| -3.701203376 | 0.000214579 | 0.000701073 |

| | | |
|---|---|---|
| 0.541114772 | 0.588428477 | 0.688112657 |
| 0.973498677 | 0.330305518 | 0.441288877 |
| 0.82391514 | 0.409987777 | 0.521813997 |
| 0.384771524 | 0.700406698 | 0.779683329 |
| 1.021818171 | 0.306866968 | 0.416596539 |
| NA | NA | NA |
| 1.861423294 | 0.062684424 | 0.112166938 |
| -4.39968654 | 1.08E-05 | 4.40E-05 |
| -1.279474748 | 0.200729926 | 0.296627114 |
| -0.683372089 | 0.494371752 | NA |
| 0.861537083 | 0.388942306 | 0.501118337 |
| 0.311731823 | 0.755244341 | 0.822582388 |
| 3.867707204 | 0.000109863 | 0.000377316 |
| NA | NA | NA |
| NA | NA | NA |
| -0.628151054 | 0.529904991 | NA |
| NA | NA | NA |
| NA | NA | NA |
| 0.573193348 | 0.566513788 | NA |
| 0.41383631 | 0.67899399 | NA |
| -0.271500407 | 0.786006187 | NA |
| -1.862615051 | 0.062516445 | 0.111890356 |
| -7.178707346 | 7.04E-13 | 6.63E-12 |
| -2.550378392 | 0.010760606 | 0.024352438 |
| 1.879613618 | 0.060160756 | 0.108291122 |
| -4.984460933 | 6.21E-07 | 3.00E-06 |
| -3.891658346 | 9.96E-05 | 0.000344297 |
| NA | NA | NA |
| NA | NA | NA |
| 0.107615456 | 0.914300736 | NA |
| 0.065061479 | 0.948125051 | NA |
| 0.063708285 | 0.949202508 | NA |
| NA | NA | NA |
| NA | NA | NA |
| 6.081924198 | 1.19E-09 | 8.02E-09 |
| 1.70575112 | 0.088054427 | 0.149881151 |
| 3.512130499 | 0.00044453 | 0.001368592 |
| 0.063708882 | 0.949202033 | NA |
| -9.784978934 | 1.31E-22 | 2.82E-21 |
| 0.06371127 | 0.949200131 | NA |
| -2.955366723 | 0.003122976 | 0.00803785 |
| NA | NA | NA |
| -4.797117527 | 1.61E-06 | 7.34E-06 |
| 4.430729979 | 9.39E-06 | 3.85E-05 |
| -2.129370663 | 0.033223606 | 0.064932213 |
| -4.353154805 | 1.34E-05 | 5.35E-05 |
| -4.039670878 | 5.35E-05 | 0.00019428 |

| | | |
|---|---|---|
| 4.862789232 | 1.16E-06 | 5.38E-06 |
| 0.052900642 | 0.957811073 | 0.970919132 |
| -0.889123952 | 0.373936467 | 0.486694908 |
| 0.742132813 | 0.458006871 | 0.567576591 |
| 8.100427599 | 5.48E-16 | 6.99E-15 |
| 1.196918968 | 0.23133814 | 0.332170356 |
| -7.995072806 | 1.29E-15 | 1.60E-14 |
| 5.68360561 | 1.32E-08 | 7.94E-08 |
| 5.984938913 | 2.16E-09 | 1.42E-08 |
| 1.438697378 | 0.150236282 | 0.233754842 |
| -0.194487022 | 0.845794556 | NA |
| NA | NA | NA |
| 0.063707019 | 0.949203516 | NA |
| 8.560952501 | 1.12E-17 | 1.64E-16 |
| -1.648203113 | 0.099310998 | 0.165668469 |
| -0.153438855 | 0.878052206 | 0.914706235 |
| 2.763912112 | 0.005711292 | 0.013848071 |
| -0.177037332 | 0.859479066 | NA |
| -3.233516408 | 0.001222763 | 0.003451648 |
| -2.10227103 | 0.035529541 | 0.0687874 |
| -0.399027732 | 0.68987277 | 0.771043333 |
| -13.59684272 | 4.18E-42 | 2.46E-40 |
| -7.01060107 | 2.37E-12 | 2.11E-11 |
| NA | NA | NA |
| 0.180884848 | 0.856457962 | NA |
| 0.564443398 | 0.572452411 | NA |
| 0.970306462 | 0.331893757 | NA |
| 0.103599789 | 0.917486956 | NA |
| 0.748045911 | 0.454432466 | NA |
| 0.937306719 | 0.348600808 | NA |
| 0.811459128 | 0.417102055 | NA |
| -0.065388772 | 0.947864464 | NA |
| -2.079714435 | 0.037551733 | 0.072146138 |
| -2.306771286 | 0.021067574 | 0.043695838 |
| NA | NA | NA |
| NA | NA | NA |
| NA | NA | NA |
| NA | NA | NA |
| NA | NA | NA |
| NA | NA | NA |
| NA | NA | NA |
| NA | NA | NA |
| NA | NA | NA |
| NA | NA | NA |
| NA | NA | NA |
| NA | NA | NA |
| NA | NA | NA |

| | | | |
|---|---|---|---|
| NA | NA | NA | |
| NA | NA | NA | |
| NA | NA | NA | |
| NA | NA | NA | |
| NA | NA | NA | |
| NA | NA | NA | |
| NA | NA | NA | |
| NA | NA | NA | |
| NA | NA | NA | |
| NA | NA | NA | |
| NA | NA | NA | |
| NA | NA | NA | |
| | 0.447021132 | 0.654859802 | 0.742756065 |
| NA | NA | NA | |
| NA | NA | NA | |
| NA | NA | NA | |
| NA | NA | NA | |
| NA | NA | NA | |
| NA | NA | NA | |
| NA | NA | NA | |
| NA | NA | NA | |
| NA | NA | NA | |
| NA | NA | NA | |
| NA | NA | NA | |
| NA | NA | NA | |
| NA | NA | NA | |
| NA | NA | NA | |
| NA | NA | NA | |
| NA | NA | NA | |
| | -0.314450344 | 0.753179025 | NA |
| NA | NA | NA | |
| NA | NA | NA | |
| | 0.083462637 | 0.933483685 | NA |
| | 0.192809521 | 0.847108147 | NA |
| | 3.049592072 | 0.002291524 | 0.006086898 |
| NA | NA | NA | |
| | -0.573431144 | 0.566352808 | 0.668285329 |
| NA | NA | NA | |
| NA | NA | NA | |
| | 0.160053664 | 0.872838801 | 0.911513484 |
| NA | NA | NA | |
| | 2.033811867 | 0.041970568 | 0.07950078 |
| | -4.726552406 | 2.28E-06 | 1.02E-05 |
| NA | NA | NA | |
| | -3.001345682 | 0.002687892 | 0.007023483 |
| | -1.190384125 | 0.23389545 | 0.335169354 |

| | | | |
|---|---|---|---|
| | -2.691854255 | 0.007105599 | 0.016789686 |
| | -2.032009395 | 0.042152705 | 0.079791439 |
| NA | NA | NA | |
| NA | NA | NA | |
| | 0.192636473 | 0.847243678 | NA |
| | 0.562742821 | 0.573610021 | NA |
| NA | NA | NA | |
| NA | NA | NA | |
| NA | NA | NA | |
| NA | NA | NA | |
| NA | NA | NA | |
| NA | NA | NA | |
| | 2.380121847 | 0.017306914 | 0.036933418 |
| | -1.30251246 | 0.192741261 | 0.28680303 |
| NA | NA | NA | |
| | 0.955164376 | 0.339494578 | 0.450965838 |
| | -8.118217636 | 4.73E-16 | 6.06E-15 |
| | -1.220889375 | 0.222127908 | 0.32150937 |
| | -4.532721334 | 5.82E-06 | 2.45E-05 |
| | -4.496197721 | 6.92E-06 | 2.89E-05 |
| | -5.328271387 | 9.92E-08 | 5.35E-07 |
| | -4.742471452 | 2.11E-06 | 9.48E-06 |
| | -4.934036571 | 8.05E-07 | 3.83E-06 |
| | -2.66904184 | 0.007606797 | 0.0178245 |
| | 0.158644592 | 0.873948894 | NA |
| | -3.088016145 | 0.002014975 | 0.005427769 |
| | -3.022017425 | 0.002510961 | 0.006600436 |
| | 0.192810118 | 0.847107679 | NA |
| | 0.063708882 | 0.949202033 | NA |
| | -1.05436075 | 0.291717785 | NA |
| | 0.192809521 | 0.847108147 | NA |
| | 0.063708882 | 0.949202033 | NA |
| | 0.412473636 | 0.6799923 | NA |
| | 1.33480944 | 0.181938723 | NA |
| | 0.775726423 | 0.437910535 | NA |
| | 0.320071032 | 0.748914485 | NA |
| | 0.476098803 | 0.63400399 | NA |
| NA | NA | NA | |
| NA | NA | NA | |
| NA | NA | NA | |
| | 0.352770413 | 0.724260567 | NA |
| | 0.501957952 | 0.615697099 | NA |
| NA | NA | NA | |
| NA | NA | NA | |
| NA | NA | NA | |
| | -0.065389966 | 0.947863513 | NA |
| NA | NA | NA | |

| | | | |
|---|---|---|---|
| | -4.874107296 | 1.09E-06 | 5.10E-06 |
| | -8.775778363 | 1.70E-18 | 2.68E-17 |
| | -0.065389369 | 0.947863988 | NA |
| | -4.025943844 | 5.67E-05 | 0.000204902 |
| NA | NA | NA | |
| NA | NA | NA | |
| | 3.306771328 | 0.000943779 | 0.002734119 |
| | 0.303025451 | 0.761870466 | 0.827912049 |
| | 0.689572775 | 0.490462893 | 0.59796624 |
| | 2.727930572 | 0.006373302 | 0.015253753 |
| | 0.618531285 | 0.536225182 | NA |
| | 0.599378619 | 0.548920432 | 0.652334935 |
| NA | NA | NA | |
| NA | NA | NA | |
| NA | NA | NA | |
| NA | NA | NA | |
| NA | NA | NA | |
| | -4.674597868 | 2.95E-06 | 1.30E-05 |
| | 4.56738629 | 4.94E-06 | 2.10E-05 |
| | 0.192810118 | 0.847107679 | NA |
| | -8.342197442 | 7.29E-17 | 1.00E-15 |
| NA | NA | NA | |
| | 0.192810118 | 0.847107679 | NA |
| NA | NA | NA | |
| | 0.063708285 | 0.949202508 | NA |
| | -2.195319652 | 0.028140676 | 0.056151238 |
| | -2.182893521 | 0.029043652 | 0.057713768 |
| NA | NA | NA | |
| | -6.08944145 | 1.13E-09 | 7.68E-09 |
| | -0.19448762 | 0.845794089 | NA |
| NA | NA | NA | |
| | -3.371012507 | 0.000748925 | 0.002211868 |
| NA | NA | NA | |
| | -0.065388772 | 0.947864464 | NA |
| | -1.020674029 | 0.307408903 | 0.417178749 |
| | -3.213414537 | 0.001311668 | 0.003678393 |
| | -2.533938377 | 0.011278856 | 0.025383005 |
| | 0.179338328 | 0.857672053 | 0.901239777 |
| | -4.037574238 | 5.40E-05 | 0.000195855 |
| | 0.406656195 | 0.684260518 | NA |
| | -1.457732519 | 0.144914285 | 0.226913015 |
| | 2.77368173 | 0.005542587 | 0.01349523 |
| | 3.323240422 | 0.000889782 | 0.00258998 |
| | 5.738601642 | 9.55E-09 | 5.85E-08 |
| | 0.192810118 | 0.847107679 | NA |
| NA | NA | NA | |
| NA | NA | NA | |

| | | |
|---|---|---|
| -0.441839617 | 0.658605268 | 0.745974321 |
| 0.131019124 | 0.895760179 | NA |
| 7.761811727 | 8.37E-15 | 9.61E-14 |
| 3.323392677 | 0.000889296 | 0.002589169 |
| NA | NA | NA |
| -0.914869509 | 0.360260139 | 0.472432089 |
| 19.85977974 | 9.07E-88 | 1.72E-85 |
| -3.198623007 | 0.001380856 | 0.003854017 |
| -2.447493726 | 0.014385363 | 0.031378269 |
| -0.457854949 | 0.647056655 | 0.735973284 |
| 0.094996979 | 0.924317226 | NA |
| 0.661415068 | 0.508346165 | 0.614503261 |
| 0.384684592 | 0.700471112 | 0.779720357 |
| -4.810683062 | 1.50E-06 | 6.88E-06 |
| -0.930314955 | 0.352208037 | 0.464036592 |
| 6.009287434 | 1.86E-09 | 1.23E-08 |
| -0.065389369 | 0.947863988 | NA |
| 2.990355021 | 0.002786534 | 0.007255324 |
| 6.207095521 | 5.40E-10 | 3.77E-09 |
| -2.856812418 | 0.004279186 | 0.010680897 |
| -5.220804425 | 1.78E-07 | 9.27E-07 |
| NA | NA | NA |
| 0.662727242 | 0.507505259 | 0.613884414 |
| -1.490104255 | 0.136196826 | 0.215432668 |
| -3.726890057 | 0.000193857 | 0.000638537 |
| -2.288492411 | 0.022108862 | 0.045606132 |
| NA | NA | NA |
| NA | NA | NA |
| NA | NA | NA |
| 0.192808924 | 0.847108615 | NA |
| NA | NA | NA |
| 1.24479641 | 0.213206596 | 0.311242336 |
| 0.128001637 | 0.898147677 | 0.929028695 |
| 1.112107803 | 0.266091802 | 0.371771019 |
| -0.371997034 | 0.709895057 | 0.787374187 |
| -2.950605353 | 0.003171519 | 0.008147707 |
| -0.836387367 | 0.402937017 | 0.51463368 |
| 3.71653686 | 0.000201972 | 0.000663695 |
| NA | NA | NA |
| 0.955074608 | 0.339539969 | 0.451002198 |
| 2.048130809 | 0.040547184 | 0.077119758 |
| 3.352580116 | 0.000800621 | 0.002349301 |
| -0.070610168 | 0.943708018 | NA |
| -1.756925781 | 0.078930461 | 0.136559722 |
| -2.235957105 | 0.025354581 | 0.051345078 |
| 0.192809521 | 0.847108147 | NA |
| NA | NA | NA |

| | | |
|---|---|---|
| 0.307291868 | 0.758621225 | NA |
| NA | NA | NA |
| 0.507275069 | 0.611961829 | NA |
| NA | NA | NA |
| -1.81603537 | 0.069364947 | 0.122285154 |
| NA | NA | NA |
| -0.240361051 | 0.81005037 | 0.865856844 |
| 0.192809521 | 0.847108147 | NA |
| -3.817438709 | 0.000134844 | 0.000455609 |
| -1.592689746 | 0.111229809 | 0.18183542 |
| -9.912657824 | 3.67E-23 | 8.27E-22 |
| -0.693012808 | 0.488301514 | 0.59626241 |
| -1.178720194 | 0.238509612 | 0.340689127 |
| 2.498213358 | 0.012482104 | 0.027725754 |
| 0.501640755 | 0.615920246 | 0.710928464 |
| -2.592878034 | 0.009517652 | 0.02179369 |
| -0.146464819 | 0.883554461 | 0.918488585 |
| -0.224518725 | 0.822353699 | 0.875448324 |
| -0.452962969 | 0.650575407 | 0.739270098 |
| -0.40577039 | 0.684911317 | 0.767179334 |
| 0.54774891 | 0.583864323 | 0.683941005 |
| -0.11953074 | 0.904854888 | 0.934190638 |
| -0.764798614 | 0.444391473 | 0.554354278 |
| 0.723912369 | 0.469119536 | 0.57833929 |
| -0.907180575 | 0.364311314 | 0.476795409 |
| NA | NA | NA |
| -2.964594394 | 0.003030822 | 0.007828035 |
| 0.97908207 | 0.32753943 | 0.438600039 |
| -3.758150328 | 0.000171174 | 0.000567933 |
| -0.719874151 | 0.471602485 | 0.580515894 |
| 0.80580983 | 0.420352509 | 0.532056804 |
| NA | NA | NA |
| 0.192808924 | 0.847108615 | NA |
| -0.32878292 | 0.742319774 | 0.812522624 |
| -3.239692455 | 0.001196587 | 0.003389947 |
| NA | NA | NA |
| 9.306555917 | 1.32E-20 | 2.45E-19 |
| -3.904843484 | 9.43E-05 | 0.000327913 |
| -2.577486828 | 0.009952167 | 0.022695221 |
| -5.00276423 | 5.65E-07 | 2.74E-06 |
| -0.363076496 | 0.716547742 | 0.792610481 |
| -6.286136347 | 3.25E-10 | 2.33E-09 |
| 5.847314613 | 5.00E-09 | 3.16E-08 |
| -1.710493478 | 0.087174658 | 0.148535174 |
| NA | NA | NA |
| 0.79304714 | 0.427750358 | 0.538994738 |
| -1.009167267 | 0.312894422 | 0.423269058 |

| | | |
|---|---|---|
| 0.387227713 | 0.698587636 | 0.778212138 |
| -5.500953606 | 3.78E-08 | 2.16E-07 |
| -1.062558567 | 0.287982185 | 0.395433504 |
| -1.199027312 | 0.230517326 | 0.33125435 |
| -0.563833946 | 0.572867147 | 0.674167318 |
| 2.981636427 | 0.002867123 | 0.007442863 |
| 0.851410418 | 0.394541407 | 0.506585175 |
| 4.596240556 | 4.30E-06 | 1.85E-05 |
| -6.134384417 | 8.55E-10 | 5.86E-09 |
| NA | NA | NA |
| NA | NA | NA |
| 10.300424 | 7.02E-25 | 1.77E-23 |
| 0.185293973 | 0.852998465 | 0.897903875 |
| -2.218382987 | 0.02652873 | 0.053338751 |
| -0.479216631 | 0.631784524 | 0.723633672 |
| 3.640076415 | 0.000272557 | 0.000872838 |
| -2.36923467 | 0.01782494 | 0.037874214 |
| -4.201276964 | 2.65E-05 | 0.000101285 |
| 0.428699616 | 0.66814184 | 0.753145249 |
| NA | NA | NA |
| -2.282817452 | 0.022441124 | 0.046219149 |
| -2.554938252 | 0.010620664 | 0.024064018 |
| -1.897986429 | 0.05769787 | 0.104635392 |
| NA | NA | NA |
| 0.192808327 | 0.847109082 | NA |
| NA | NA | NA |
| -5.47660992 | 4.34E-08 | 2.46E-07 |
| NA | NA | NA |
| -2.668848978 | 0.007611166 | 0.017832771 |
| -0.19448762 | 0.845794089 | NA |
| 0.192810118 | 0.847107679 | NA |
| -0.633977435 | 0.526095549 | 0.6312037 |
| NA | NA | NA |
| -4.963123428 | 6.94E-07 | 3.33E-06 |
| 0.447714123 | 0.654359528 | NA |
| 0.003525598 | 0.997186986 | 0.997999007 |
| -0.387956216 | 0.698048435 | 0.777757513 |
| -0.065389369 | 0.947863988 | NA |
| -3.107278853 | 0.001888182 | 0.005113188 |
| NA | NA | NA |
| -5.114446828 | 3.15E-07 | 1.59E-06 |
| -3.625049687 | 0.000288906 | 0.000921069 |
| 0.127616757 | 0.898452269 | NA |
| -8.309007006 | 9.65E-17 | 1.31E-15 |
| NA | NA | NA |
| 0.281521372 | 0.778310538 | NA |
| 0.077354991 | 0.938341146 | NA |

| | | |
|---|---|---|
| -2.999846658 | 0.002701156 | 0.007052129 |
| 0.237504222 | 0.812265645 | NA |
| NA | NA | NA |
| 0.433370698 | 0.664745484 | NA |
| -3.741176272 | 0.000183161 | 0.000605297 |
| NA | NA | NA |
| 1.18891335 | 0.234473766 | 0.335825038 |
| 0.497915394 | 0.618543677 | NA |
| -3.175314032 | 0.001496744 | 0.004146028 |
| 0.330704598 | 0.740867629 | NA |
| NA | NA | NA |
| 0.330493631 | 0.741027004 | NA |
| -1.333540465 | 0.182354506 | 0.274167064 |
| 0.192810118 | 0.847107679 | NA |
| NA | NA | NA |
| NA | NA | NA |
| -6.288973985 | 3.20E-10 | 2.28E-09 |
| NA | NA | NA |
| NA | NA | NA |
| NA | NA | NA |
| -3.421647524 | 0.000622429 | 0.001864681 |
| -4.218136611 | 2.46E-05 | 9.43E-05 |
| -0.142066076 | 0.887027813 | NA |
| 0.192808924 | 0.847108615 | NA |
| -3.153404524 | 0.00161378 | 0.00444402 |
| 0.063976859 | 0.948988653 | NA |
| NA | NA | NA |
| -0.707294332 | 0.479383587 | 0.588069554 |
| 0.216845651 | 0.828328634 | 0.879951482 |
| 0.177197828 | 0.859353002 | 0.902244451 |
| -0.880604431 | 0.378531959 | 0.491127995 |
| -0.232745463 | 0.815959064 | 0.870416611 |
| -1.801124134 | 0.071683317 | 0.125769902 |
| 0.786461753 | 0.431597017 | 0.542499513 |
| -1.091619027 | 0.275000589 | 0.381661478 |
| 0.063708285 | 0.949202508 | NA |
| -3.810067302 | 0.000138929 | 0.000468525 |
| -3.205788714 | 0.001346928 | 0.003769621 |
| 2.620024407 | 0.008792347 | 0.020288069 |
| -2.827725318 | 0.004688001 | 0.011607545 |
| -4.811877895 | 1.50E-06 | 6.85E-06 |
| 0.394666331 | 0.693089154 | 0.773655214 |
| -0.794154428 | 0.427105535 | 0.538426167 |
| -5.399872465 | 6.67E-08 | 3.68E-07 |
| -3.197056542 | 0.001388377 | 0.003873281 |
| -8.740051293 | 2.33E-18 | 3.63E-17 |
| -4.372078179 | 1.23E-05 | 4.94E-05 |

| | | | |
|---|---|---|---|
| | -0.779144328 | 0.4358947 | 0.546396452 |
| | -1.232171505 | 0.217885025 | 0.316723514 |
| | 4.886359049 | 1.03E-06 | 4.81E-06 |
| | 0.216157829 | 0.828864724 | NA |
| NA | NA | NA | |
| NA | NA | NA | |
| | -1.454457832 | 0.145819416 | 0.228059235 |
| | -1.009142451 | 0.312906321 | NA |
| | 0.135367314 | 0.892321466 | NA |
| NA | NA | NA | |
| | -1.095140172 | 0.273455232 | 0.380047528 |
| | 2.807909024 | 0.004986431 | 0.012277235 |
| | 1.01140195 | 0.311824088 | NA |
| NA | NA | NA | |
| NA | NA | NA | |
| | 0.192808327 | 0.847109082 | NA |
| | 0.600571846 | 0.548125195 | NA |
| | -0.065389369 | 0.947863988 | NA |
| | 7.789641011 | 6.72E-15 | 7.78E-14 |
| | 1.232236723 | 0.217860669 | 0.316706502 |
| | -0.160394474 | 0.872570342 | 0.911332704 |
| | -3.706102483 | 0.000210473 | 0.000688736 |
| | -7.741157451 | 9.85E-15 | 1.12E-13 |
| | 5.791670513 | 6.97E-09 | 4.33E-08 |
| | -2.373774556 | 0.017607297 | 0.037475396 |
| | -0.639268071 | 0.522648556 | 0.62788065 |
| | 1.771830096 | 0.076422765 | 0.132763868 |
| | -6.772972367 | 1.26E-11 | 1.05E-10 |
| | 0.793943134 | 0.427228537 | 0.538554104 |
| | 0.949796058 | 0.342215889 | 0.453569696 |
| | -2.937453812 | 0.003309195 | 0.008464741 |
| | -11.0324894 | 2.66E-28 | 8.42E-27 |
| | -1.464856387 | 0.142960104 | 0.224480869 |
| | 0.192810118 | 0.847107679 | NA |
| NA | NA | NA | |
| | 0.08054679 | 0.935802384 | NA |
| | -2.38752791 | 0.016962115 | 0.036296634 |
| | -1.703320453 | 0.088508115 | 0.150469206 |
| | 12.34009499 | 5.51E-35 | 2.40E-33 |
| NA | NA | NA | |
| | 2.71600972 | 0.006607398 | 0.015753779 |
| | -3.425462007 | 0.000613755 | 0.001840235 |
| | -3.90934869 | 9.25E-05 | 0.00032235 |
| | -4.979870753 | 6.36E-07 | 3.07E-06 |
| | -2.802202225 | 0.005075505 | 0.012469561 |
| NA | NA | NA | |
| | 4.384205885 | 1.16E-05 | 4.69E-05 |

| | | |
|---|---|---|
| -0.22964189 | 0.818370053 | 0.872349633 |
| -2.844621274 | 0.004446426 | 0.011063025 |
| -2.42474186 | 0.015319275 | 0.033181158 |
| 2.548986704 | 0.010803642 | 0.024429954 |
| 10.91558706 | 9.71E-28 | 2.98E-26 |
| -1.914054382 | 0.055613207 | 0.101433786 |
| 1.935781455 | 0.052894473 | 0.097163413 |
| -3.989108026 | 6.63E-05 | 0.000236474 |
| 0.558006623 | 0.576839862 | 0.677886367 |
| 2.189312865 | 0.028574108 | 0.056934644 |
| -5.171054122 | 2.33E-07 | 1.20E-06 |
| 0.335614502 | 0.737161587 | 0.808549054 |
| 3.679324037 | 0.000233853 | 0.000758008 |
| -8.295752608 | 1.08E-16 | 1.46E-15 |
| -0.968741125 | 0.332674368 | 0.443861377 |
| 0.78898062 | 0.430123333 | NA |
| -1.73107741 | 0.083437959 | 0.143115157 |
| NA | NA | NA |
| -1.178606065 | 0.238555075 | 0.340734623 |
| -1.270369092 | 0.203953188 | 0.300357491 |
| -3.288068416 | 0.001008773 | 0.002905926 |
| -3.088369261 | 0.002012582 | 0.005421907 |
| 5.023947394 | 5.06E-07 | 2.48E-06 |
| 5.424271871 | 5.82E-08 | 3.23E-07 |
| 5.169964638 | 2.34E-07 | 1.20E-06 |
| 4.079508256 | 4.51E-05 | 0.000165296 |
| -1.405987582 | 0.159727812 | 0.245697342 |
| 0.505988286 | 0.612864873 | 0.708545394 |
| 4.399286475 | 1.09E-05 | 4.40E-05 |
| 0.711493528 | 0.47677846 | 0.585620024 |
| -4.975041755 | 6.52E-07 | 3.14E-06 |
| 0.409945893 | 0.681845639 | 0.764461202 |
| -1.533821083 | 0.125073675 | 0.200951354 |
| NA | NA | NA |
| NA | NA | NA |
| 0.70318223 | 0.481942192 | NA |
| -1.234999203 | 0.216830801 | 0.315465877 |
| -2.865753616 | 0.004160179 | 0.010419179 |
| 0.638250618 | 0.523310552 | 0.628585427 |
| -0.090656667 | 0.927765403 | NA |
| 1.361686956 | 0.1732967 | 0.262838053 |
| 1.249050468 | 0.211646616 | 0.309441035 |
| -0.744808489 | 0.456387502 | 0.566074291 |
| 4.191632819 | 2.77E-05 | 0.000105384 |
| 5.113479527 | 3.16E-07 | 1.60E-06 |
| -0.264816067 | 0.791151159 | 0.851321799 |
| 3.669998821 | 0.000242552 | 0.000783768 |

| | | | |
|---|---|---|---|
| NA | NA | NA | |
| | 3.993605557 | 6.51E-05 | 0.000232262 |
| | -0.027909959 | 0.977733965 | 0.984618031 |
| | 1.732247115 | 0.083229574 | 0.142853749 |
| | 0.157577741 | 0.874789545 | 0.912726807 |
| | -0.166703998 | 0.86760296 | 0.907750403 |
| | 0.26145272 | 0.793743406 | 0.853373886 |
| | 0.340366742 | 0.733580361 | NA |
| | 1.431101995 | 0.152400986 | 0.236564135 |
| | 3.647660001 | 0.00026464 | 0.000849113 |
| NA | NA | NA | |
| NA | NA | NA | |
| | 2.059637949 | 0.039433166 | 0.075246789 |
| NA | NA | NA | |
| | 0.063708285 | 0.949202508 | NA |
| NA | NA | NA | |
| | 0.634517975 | 0.52574284 | 0.630871244 |
| | 0.171730458 | 0.863649439 | 0.905278507 |
| | -2.446942928 | 0.014407365 | 0.031420785 |
| | -2.77946961 | 0.005444775 | 0.013275152 |
| NA | NA | NA | |
| | -0.922395626 | 0.35632225 | 0.468224786 |
| | 0.063708285 | 0.949202508 | NA |
| | 0.251134278 | 0.801710295 | NA |
| | -2.371734033 | 0.01770483 | 0.037663768 |
| | -0.83464151 | 0.403919583 | 0.515585046 |
| | 0.77311807 | 0.439452508 | 0.549720853 |
| | -2.229105789 | 0.025806867 | 0.052092409 |
| | -3.11771592 | 0.001822584 | 0.00495162 |
| | 4.097933362 | 4.17E-05 | 0.000153688 |
| NA | NA | NA | |
| NA | NA | NA | |
| | -4.600663749 | 4.21E-06 | 1.81E-05 |
| | -4.316658699 | 1.58E-05 | 6.25E-05 |
| NA | NA | NA | |
| | -4.885447767 | 1.03E-06 | 4.83E-06 |
| | -2.754733495 | 0.005873996 | 0.014190341 |
| | -3.079494843 | 0.00207352 | 0.005564285 |
| | -3.919917659 | 8.86E-05 | 0.000309526 |
| NA | NA | NA | |
| | 0.748649248 | 0.454068641 | 0.563952666 |
| | 3.125302877 | 0.001776221 | 0.004841952 |
| | 0.192810118 | 0.847107679 | NA |
| | 0.771535995 | 0.440389296 | NA |
| | 3.488375205 | 0.000485966 | 0.001485192 |
| | 0.480374117 | 0.630961396 | 0.722988635 |
| NA | NA | NA | |

| | | |
|---|---|---|
| NA | NA | NA |
| 4.720007346 | 2.36E-06 | 1.05E-05 |
| -0.881576633 | 0.378005796 | 0.490639135 |
| 0.519212488 | 0.603612572 | NA |
| 0.192810118 | 0.847107679 | NA |
| 3.860606508 | 0.000113106 | 0.000387442 |
| -0.005604395 | 0.995528363 | 0.997002158 |
| 4.507677772 | 6.55E-06 | 2.74E-05 |
| 0.192809521 | 0.847108147 | NA |
| 1.860743064 | 0.062780469 | 0.112290635 |
| 3.351238489 | 0.00080451 | 0.00235933 |
| NA | NA | NA |
| -3.294274098 | 0.000986762 | 0.002848421 |
| -1.33519416 | 0.181812808 | 0.273418324 |
| -1.219770572 | 0.22255186 | 0.322011276 |
| 0.038791606 | 0.969056537 | 0.978683157 |
| NA | NA | NA |
| -3.096463507 | 0.00195844 | 0.005286288 |
| 2.242754827 | 0.024912629 | 0.050560537 |
| NA | NA | NA |
| 0.986452142 | 0.323911289 | 0.43490503 |
| 3.00666978 | 0.002641265 | 0.00690958 |
| 1.293078042 | 0.195984069 | 0.290746873 |
| -3.476346163 | 0.000508296 | 0.001548709 |
| -12.7151341 | 4.87E-37 | 2.34E-35 |
| -1.10363781 | 0.269750283 | 0.375917241 |
| 0.855120484 | 0.392484451 | NA |
| -0.065389369 | 0.947863988 | NA |
| 0.063707688 | 0.949202983 | NA |
| 1.980905485 | 0.047601874 | 0.088743848 |
| NA | NA | NA |
| -5.900863068 | 3.62E-09 | 2.32E-08 |
| 0.085936513 | 0.931516886 | NA |
| -6.996695922 | 2.62E-12 | 2.31E-11 |
| -4.162994187 | 3.14E-05 | 0.000118403 |
| NA | NA | NA |
| 1.336337073 | 0.181439124 | 0.272928159 |
| -0.072414063 | 0.942272394 | NA |
| 7.407035686 | 1.29E-13 | 1.31E-12 |
| -0.521627983 | 0.601929376 | 0.699427783 |
| -2.391798078 | 0.016766064 | 0.035941638 |
| 0.063976859 | 0.948988653 | NA |
| -2.271655271 | 0.02310734 | 0.047400157 |
| 4.810632953 | 1.50E-06 | 6.88E-06 |
| 18.19789297 | 5.36E-74 | 7.58E-72 |
| 2.445780946 | 0.014453879 | 0.031505755 |
| -6.854166789 | 7.17E-12 | 6.07E-11 |

| | | |
|---|---|---|
| NA | NA | NA |
| -0.97243854 | 0.330832427 | 0.441780604 |
| 0.867221788 | 0.3858205 | 0.498070407 |
| 5.83679454 | 5.32E-09 | 3.35E-08 |
| -0.065465566 | 0.947803322 | NA |
| -1.880995054 | 0.059972592 | 0.108024897 |
| -0.065389369 | 0.947863988 | NA |
| NA | NA | NA |
| 0.379716159 | 0.704156124 | 0.783056148 |
| 0.771736921 | 0.440270259 | 0.550447781 |
| -3.891265706 | 9.97E-05 | 0.000344807 |
| -0.164035261 | 0.869703386 | NA |
| 0.43763768 | 0.661648952 | NA |
| -0.14237986 | 0.886779969 | 0.920617394 |
| -2.651818504 | 0.008005956 | 0.018640845 |
| 2.906035762 | 0.003660396 | 0.009267622 |
| -8.271734455 | 1.32E-16 | 1.77E-15 |
| 0.192810118 | 0.847107679 | NA |
| NA | NA | NA |
| 0.192810118 | 0.847107679 | NA |
| NA | NA | NA |
| NA | NA | NA |
| 1.499627501 | 0.13371092 | 0.212224751 |
| 0.805895591 | 0.420303054 | 0.53202111 |
| NA | NA | NA |
| -0.712294584 | 0.476282377 | 0.585182991 |
| 0.158173066 | 0.874320427 | 0.912451647 |
| 1.324929405 | 0.185194567 | 0.277633183 |
| -1.528172023 | 0.126469833 | 0.202854208 |
| 4.586360288 | 4.51E-06 | 1.93E-05 |
| 7.037948162 | 1.95E-12 | 1.75E-11 |
| 0.374168786 | 0.708278746 | NA |
| -6.473012178 | 9.61E-11 | 7.32E-10 |
| 0.519217738 | 0.603608911 | 0.70082248 |
| -0.414176804 | 0.678744628 | NA |
| -0.944534566 | 0.344896547 | 0.456325223 |
| 0.821280277 | 0.41148664 | 0.523141899 |
| 0.320381251 | 0.748679337 | NA |
| -0.022216966 | 0.982274884 | 0.987641213 |
| 5.13076658 | 2.89E-07 | 1.47E-06 |
| 1.518815461 | 0.128808953 | 0.205919914 |
| 3.091016541 | 0.001994725 | 0.00537959 |
| 0.920854734 | 0.357126274 | 0.468936661 |
| NA | NA | NA |
| -4.857057919 | 1.19E-06 | 5.52E-06 |
| NA | NA | NA |
| -0.0214534 | 0.982883976 | 0.987955362 |

| | | | |
|---|---|---|---|
| NA | NA | NA | |
| | 3.901086173 | 9.58E-05 | 0.000332306 |
| | 0.141191177 | 0.887718916 | NA |
| | 2.353120106 | 0.01861662 | 0.03928292 |
| | 5.826866667 | 5.65E-09 | 3.54E-08 |
| | -4.247760585 | 2.16E-05 | 8.35E-05 |
| | 0.567341218 | 0.570482378 | NA |
| | -2.411305649 | 0.015895521 | 0.034300449 |
| | 4.923462644 | 8.50E-07 | 4.03E-06 |
| | -4.91213519 | 9.01E-07 | 4.25E-06 |
| | 0.283730789 | 0.776616698 | NA |
| NA | NA | NA | |
| | -3.560592002 | 0.00037002 | 0.001156109 |
| | -2.195713196 | 0.028112478 | 0.056117217 |
| | -9.744902278 | 1.94E-22 | 4.13E-21 |
| | -0.486446735 | 0.626650462 | 0.71994428 |
| NA | NA | NA | |
| | -1.023497512 | 0.306072676 | 0.415681751 |
| NA | NA | NA | |
| | -0.11705545 | 0.906816112 | 0.935597992 |
| | -1.559670173 | 0.118837844 | 0.192313392 |
| NA | NA | NA | |
| | -0.809590919 | 0.418175329 | 0.529798639 |
| | -0.086753899 | 0.930867133 | NA |
| | -0.737383856 | 0.46088895 | 0.570307086 |
| | 0.192809521 | 0.847108147 | NA |
| | 3.146403229 | 0.001652919 | 0.004540948 |
| | -1.353848413 | 0.175784744 | 0.266063503 |
| | -3.969108649 | 7.21E-05 | 0.000256021 |
| | -3.121885919 | 0.001796966 | 0.004892109 |
| | 2.421474512 | 0.015457685 | 0.033460681 |
| | 5.240676095 | 1.60E-07 | 8.39E-07 |
| | 0.638310724 | 0.523271432 | 0.628568602 |
| | 2.568431268 | 0.010215997 | 0.023237587 |
| | 2.304136492 | 0.021214983 | 0.043957866 |
| | 0.063708882 | 0.949202033 | NA |
| | 11.65264579 | 2.22E-31 | 8.29E-30 |
| | 0.560705658 | 0.574998209 | NA |
| | 9.590801407 | 8.74E-22 | 1.78E-20 |
| NA | NA | NA | |
| NA | NA | NA | |
| | 0.537466014 | 0.590945755 | 0.690282445 |
| NA | NA | NA | |
| | 4.381184112 | 1.18E-05 | 4.75E-05 |
| | -0.363249875 | 0.716418233 | NA |
| | -0.34241502 | 0.732038587 | NA |
| NA | NA | NA | |

| | | |
|---|---|---|
| -0.066606123 | 0.946895271 | NA |
| NA | NA | NA |
| -0.065388772 | 0.947864464 | NA |
| 0.192809521 | 0.847108147 | NA |
| 0.218376629 | 0.827135673 | 0.878930414 |
| 0.31287131 | 0.754378436 | NA |
| -0.878179128 | 0.379846517 | 0.492489238 |
| NA | NA | NA |
| -2.863055844 | 0.004195766 | 0.010497803 |
| -3.845010198 | 0.000120547 | 0.000410909 |
| 1.594487248 | 0.110826945 | 0.181295237 |
| 0.192810118 | 0.847107679 | NA |
| -1.275388274 | 0.202171847 | 0.298369019 |
| NA | NA | NA |
| NA | NA | NA |
| -0.311892942 | 0.755121887 | 0.822520598 |
| NA | NA | NA |
| 2.049021557 | 0.040460007 | 0.076971496 |
| 0.083580704 | 0.933389809 | NA |
| -1.512272581 | 0.130464533 | 0.207977249 |
| 0.307639637 | 0.758356556 | 0.825015086 |
| NA | NA | NA |
| 0.30922356 | 0.757151473 | NA |
| -0.0909659 | 0.927519687 | 0.950455161 |
| 0.192808924 | 0.847108615 | NA |
| -0.437457048 | 0.661779919 | NA |
| -1.38008075 | 0.167561784 | 0.255547993 |
| -1.920185716 | 0.054834445 | 0.100204874 |
| -0.26445735 | 0.791427525 | NA |
| -2.222396475 | 0.026256523 | 0.052889166 |
| -2.234083626 | 0.025477571 | 0.051539934 |
| -4.151847242 | 3.30E-05 | 0.000123837 |
| -2.447435341 | 0.014387694 | 0.031380618 |
| -4.534965522 | 5.76E-06 | 2.43E-05 |
| 0.127933515 | 0.898201588 | NA |
| 0.083031301 | 0.933826651 | NA |
| 0.063708882 | 0.949202033 | NA |
| -1.171547326 | 0.241378847 | 0.343845685 |
| -1.931303288 | 0.053445557 | 0.098031826 |
| 0.31685483 | 0.751353758 | NA |
| -0.475661884 | 0.63431528 | NA |
| -2.688834317 | 0.0071702 | 0.016915756 |
| -0.594427129 | 0.552226468 | 0.655361457 |
| 0.19960581 | 0.841788883 | 0.889216111 |
| -0.88324492 | 0.377103956 | 0.489824184 |
| 7.296483461 | 2.95E-13 | 2.89E-12 |
| 0.063708882 | 0.949202033 | NA |

| | | |
|---|---|---|
| 11.85850131 | 1.94E-32 | 7.63E-31 |
| 8.68524645 | 3.78E-18 | 5.77E-17 |
| 3.175849761 | 0.001493983 | 0.004140212 |
| NA | NA | NA |
| 0.19712443 | 0.843730166 | 0.890778052 |
| NA | NA | NA |
| NA | NA | NA |
| NA | NA | NA |
| NA | NA | NA |
| -0.378764938 | 0.704862426 | 0.783495113 |
| 0.469754231 | 0.638530617 | 0.729479893 |
| -0.823990372 | 0.409945029 | 0.521786121 |
| -1.164984278 | 0.244025357 | 0.346845808 |
| 0.427004991 | 0.669375693 | NA |
| 0.39783287 | 0.690753386 | NA |
| 0.333816862 | 0.738517758 | NA |
| 0.763204481 | 0.445341457 | 0.555179918 |
| 3.181600402 | 0.001464637 | 0.004066993 |
| -2.125047189 | 0.033582676 | 0.065526533 |
| NA | NA | NA |
| NA | NA | NA |
| -1.941458681 | 0.052202667 | 0.096104203 |
| NA | NA | NA |
| 0.192810118 | 0.847107679 | NA |
| -0.531525878 | 0.595054414 | 0.69368972 |
| 0.192808327 | 0.847109082 | NA |
| -4.87807856 | 1.07E-06 | 5.01E-06 |
| -3.364952158 | 0.000765569 | 0.002255685 |
| -4.040307628 | 5.34E-05 | 0.000193839 |
| -1.050881827 | 0.293312868 | 0.401378879 |
| NA | NA | NA |
| -3.856329378 | 0.000115102 | 0.000393527 |
| 0.27517744 | 0.783179919 | NA |
| -4.537222074 | 5.70E-06 | 2.41E-05 |
| -3.237228485 | 0.001206967 | 0.003414211 |
| -3.358508811 | 0.000783642 | 0.00230272 |
| -4.830190901 | 1.36E-06 | 6.27E-06 |
| -5.610992287 | 2.01E-08 | 1.19E-07 |
| NA | NA | NA |
| 0.094001993 | 0.925107574 | NA |
| NA | NA | NA |
| 3.630854362 | 0.000282485 | 0.000902206 |
| 0.965743331 | 0.334172631 | 0.445409285 |
| 0.952402739 | 0.340892771 | 0.452414955 |
| -6.076667689 | 1.23E-09 | 8.27E-09 |
| -1.616327521 | 0.10602352 | 0.174693483 |
| -1.879963104 | 0.060113107 | 0.108236251 |

| | | |
|---|---|---|
| 12.61940787 | 1.65E-36 | 7.69E-35 |
| 4.573253076 | 4.80E-06 | 2.05E-05 |
| -5.841276247 | 5.18E-09 | 3.26E-08 |
| -0.252525274 | 0.800635081 | 0.858479903 |
| 1.205138698 | 0.228149763 | NA |
| -3.136318518 | 0.001710833 | 0.00468116 |
| -0.235500446 | 0.813820333 | 0.869061092 |
| 0.585582452 | 0.558156144 | NA |
| -1.731866222 | 0.083297384 | 0.142945974 |
| 6.985465038 | 2.84E-12 | 2.50E-11 |
| 0.08739807 | 0.930355103 | 0.952580718 |
| 0.692454754 | 0.488651791 | 0.596515525 |
| -0.379112222 | 0.704604531 | 0.783415574 |
| -0.237694866 | 0.812117766 | 0.867761267 |
| -0.327061973 | 0.743621013 | 0.813432516 |
| 0.786122223 | 0.431795886 | 0.542590041 |
| NA | NA | NA |
| 2.23847323 | 0.025190211 | 0.05106179 |
| NA | NA | NA |
| NA | NA | NA |
| -2.014100674 | 0.043998965 | 0.082872608 |
| NA | NA | NA |
| NA | NA | NA |
| NA | NA | NA |
| -7.217332794 | 5.30E-13 | 5.07E-12 |
| NA | NA | NA |
| -2.628727886 | 0.008570491 | 0.019833523 |
| -2.265715347 | 0.023468817 | 0.048043224 |
| -3.193657462 | 0.001404827 | 0.003913938 |
| -2.804183536 | 0.005044418 | 0.012402926 |
| -3.030468067 | 0.00244175 | 0.006437435 |
| -1.852915698 | 0.063894443 | 0.113957382 |
| NA | NA | NA |
| -1.519826312 | 0.128554634 | 0.20561841 |
| -11.09237412 | 1.37E-28 | 4.40E-27 |
| -4.835834699 | 1.33E-06 | 6.11E-06 |
| NA | NA | NA |
| -5.917240965 | 3.27E-09 | 2.11E-08 |
| NA | NA | NA |
| -5.748255685 | 9.02E-09 | 5.54E-08 |
| -4.670543267 | 3.00E-06 | 1.32E-05 |
| -4.403277701 | 1.07E-05 | 4.33E-05 |
| -4.965610842 | 6.85E-07 | 3.29E-06 |
| -4.329195362 | 1.50E-05 | 5.93E-05 |
| -3.905203836 | 9.41E-05 | 0.00032747 |
| NA | NA | NA |
| -4.585281793 | 4.53E-06 | 1.94E-05 |

| | | |
|---|---|---|
| -4.31465581 | 1.60E-05 | 6.30E-05 |
| -5.644345688 | 1.66E-08 | 9.88E-08 |
| -4.256804487 | 2.07E-05 | 8.04E-05 |
| -4.823777824 | 1.41E-06 | 6.47E-06 |
| -5.850732563 | 4.89E-09 | 3.10E-08 |
| -4.542441501 | 5.56E-06 | 2.35E-05 |
| -4.382191299 | 1.17E-05 | 4.73E-05 |
| -1.700076675 | 0.089116504 | NA |
| -3.647784602 | 0.000264511 | 0.00084881 |
| -4.934570228 | 8.03E-07 | 3.82E-06 |
| -4.346557944 | 1.38E-05 | 5.51E-05 |
| -5.073869235 | 3.90E-07 | 1.94E-06 |
| -4.016642088 | 5.90E-05 | 0.000212299 |
| -4.310763796 | 1.63E-05 | 6.41E-05 |
| -5.068613014 | 4.01E-07 | 1.99E-06 |
| NA | NA | NA |
| 0.192808924 | 0.847108615 | NA |
| -4.99389382 | 5.92E-07 | 2.87E-06 |
| 1.758728475 | 0.078623639 | 0.136067875 |
| 0.148149482 | 0.882224799 | NA |
| -1.423141677 | 0.154695089 | 0.239545785 |
| -2.709295814 | 0.006742619 | 0.016044088 |
| -0.232961213 | 0.815791525 | 0.870373642 |
| NA | NA | NA |
| -0.065388772 | 0.947864464 | NA |
| NA | NA | NA |
| -0.414241194 | 0.678697476 | NA |
| -0.065389369 | 0.947863988 | NA |
| -0.065389369 | 0.947863988 | NA |
| 0.192810118 | 0.847107679 | NA |
| NA | NA | NA |
| 0.103483409 | 0.917579316 | NA |
| NA | NA | NA |
| NA | NA | NA |
| NA | NA | NA |
| -3.16322163 | 0.001560335 | 0.004308349 |
| -2.12432634 | 0.033642865 | 0.065623512 |
| -2.451675253 | 0.014219293 | 0.031069773 |
| NA | NA | NA |
| -1.739782419 | 0.081897231 | 0.140914806 |
| -2.218000171 | 0.026554821 | 0.053386921 |
| -3.333378252 | 0.000857982 | 0.002505576 |
| -3.529411847 | 0.000416484 | 0.001288419 |
| -2.921952233 | 0.003478449 | 0.008853526 |
| -3.497751814 | 0.000469198 | 0.001437984 |
| -1.82835564 | 0.067496194 | 0.119512574 |
| -0.670624543 | 0.502459743 | 0.6092516 |

| | | |
|---|---|---|
| -2.990537863 | 0.002784866 | 0.007251893 |
| 2.714224281 | 0.006643117 | 0.015831403 |
| -1.904484768 | 0.056847079 | 0.103347064 |
| -1.92371656 | 0.054390121 | 0.099495671 |
| -0.149450039 | 0.88119853 | NA |
| 0.532605304 | 0.59430683 | 0.693173344 |
| NA | NA | NA |
| 0.063708882 | 0.949202033 | NA |
| 6.097975725 | 1.07E-09 | 7.30E-09 |
| -4.728086845 | 2.27E-06 | 1.01E-05 |
| -0.194487022 | 0.845794556 | NA |
| -9.912600696 | 3.67E-23 | 8.27E-22 |
| NA | NA | NA |
| -2.936961052 | 0.003314457 | 0.008476639 |
| -0.652604121 | 0.514011528 | NA |
| 1.554158775 | 0.120146548 | 0.194155085 |
| 1.512342056 | 0.130446867 | 0.207977249 |
| 8.131541174 | 4.24E-16 | 5.44E-15 |
| -1.305148507 | 0.191842284 | 0.285720122 |
| 2.984985103 | 0.002835921 | 0.00737104 |
| NA | NA | NA |
| NA | NA | NA |
| NA | NA | NA |
| 1.778466629 | 0.075327248 | NA |
| 3.213394578 | 0.001311759 | 0.003678393 |
| NA | NA | NA |
| -5.056642455 | 4.27E-07 | 2.11E-06 |
| NA | NA | NA |
| 2.970929139 | 0.002969003 | 0.007687405 |
| -2.464862781 | 0.013706573 | 0.030115478 |
| -0.633255717 | 0.526566666 | NA |
| 0.119245221 | 0.905081081 | NA |
| NA | NA | NA |
| 4.3479006 | 1.37E-05 | 5.48E-05 |
| -0.2581788 | 0.796268919 | NA |
| 0.407744301 | 0.683461409 | NA |
| 1.557824765 | 0.119274794 | 0.192908251 |
| -2.761862753 | 0.005747263 | 0.013916417 |
| -2.192911441 | 0.02831376 | 0.056465096 |
| -0.571245806 | 0.567833032 | 0.669565214 |
| -3.642831413 | 0.000269655 | 0.000864431 |
| NA | NA | NA |
| -0.040107356 | 0.968007537 | 0.978168295 |
| NA | NA | NA |
| 0.849000564 | 0.395880979 | 0.507810224 |
| 0.063708285 | 0.949202508 | NA |
| NA | NA | NA |

| | | |
|---|---|---|
| NA | NA | NA |
| NA | NA | NA |
| NA | NA | NA |
| NA | NA | NA |
| NA | NA | NA |
| 2.712598708 | 0.00667579 | 0.015900182 |
| -0.526211817 | 0.598741045 | NA |
| 0.063708882 | 0.949202033 | NA |
| -0.918695184 | 0.35825503 | 0.470122858 |
| 0.063708217 | 0.949202562 | NA |
| -0.097429433 | 0.922385372 | 0.946627818 |
| NA | NA | NA |
| NA | NA | NA |
| NA | NA | NA |
| NA | NA | NA |
| NA | NA | NA |
| NA | NA | NA |
| NA | NA | NA |
| 0.192810118 | 0.847107679 | NA |
| -2.942278552 | 0.003258067 | 0.008349485 |
| 2.1958701 | 0.028101242 | 0.05610386 |
| NA | NA | NA |
| -0.069275053 | 0.944770683 | 0.962419527 |
| -5.595041172 | 2.21E-08 | 1.30E-07 |
| -5.286071257 | 1.25E-07 | 6.65E-07 |
| -2.125782717 | 0.033521356 | 0.06543239 |
| -4.312429017 | 1.61E-05 | 6.36E-05 |
| -3.511689156 | 0.000445268 | 0.00137052 |
| -1.21022819 | 0.226191344 | 0.326277624 |
| -0.992691143 | 0.320860492 | 0.431710513 |
| -3.804173404 | 0.000142278 | 0.000478724 |
| -3.038473797 | 0.002377798 | 0.006289367 |
| -3.35864177 | 0.000783265 | 0.002301882 |
| -4.021491163 | 5.78E-05 | 0.000208514 |
| -2.886940226 | 0.003890081 | 0.0098065 |
| -3.367708765 | 0.000757956 | 0.002235377 |
| -3.033973983 | 0.002413552 | 0.006373836 |
| -1.298175927 | 0.194226888 | 0.288567639 |
| NA | NA | NA |
| NA | NA | NA |
| NA | NA | NA |
| NA | NA | NA |
| NA | NA | NA |
| NA | NA | NA |
| NA | NA | NA |
| NA | NA | NA |
| NA | NA | NA |

| | | |
|---|---|---|
| NA | NA | NA |
| NA | NA | NA |
| -0.646723912 | 0.517810644 | 0.623145364 |
| NA | NA | NA |
| -2.421025662 | 0.015476785 | 0.03349044 |
| NA | NA | NA |
| 0.063708882 | 0.949202033 | NA |
| NA | NA | NA |
| 0.063708285 | 0.949202508 | NA |
| -0.194487022 | 0.845794556 | NA |
| -2.120953351 | 0.033925728 | 0.066113437 |
| NA | NA | NA |
| NA | NA | NA |
| NA | NA | NA |
| NA | NA | NA |
| -0.246231387 | 0.80550312 | 0.862313679 |
| NA | NA | NA |
| NA | NA | NA |
| NA | NA | NA |
| NA | NA | NA |
| NA | NA | NA |
| -0.065389369 | 0.947863988 | NA |
| NA | NA | NA |
| NA | NA | NA |
| -0.343244076 | 0.731414848 | NA |
| NA | NA | NA |
| 6.068824489 | 1.29E-09 | 8.66E-09 |
| 0.07404034 | 0.940978287 | 0.959884522 |
| 0.492449326 | 0.622401736 | 0.716559287 |
| 4.472265353 | 7.74E-06 | 3.21E-05 |
| -5.275612803 | 1.32E-07 | 7.02E-07 |
| -5.325875676 | 1.00E-07 | 5.41E-07 |
| -2.460561947 | 0.013871962 | 0.030433448 |
| 0.287787545 | 0.773509372 | 0.837173023 |
| 7.233204999 | 4.72E-13 | 4.53E-12 |
| 0.421248522 | 0.673573616 | NA |
| -0.065389369 | 0.947863988 | NA |
| NA | NA | NA |
| 2.06787868 | 0.038651433 | 0.073971319 |
| 8.677917571 | 4.03E-18 | 6.14E-17 |
| NA | NA | NA |
| NA | NA | NA |
| NA | NA | NA |
| 8.816366542 | 1.18E-18 | 1.90E-17 |
| 0.063708285 | 0.949202508 | NA |
| 2.242023012 | 0.024959885 | 0.050648231 |
| NA | NA | NA |

| | | |
|---|---|---|
| -0.75084239 | 0.452747515 | 0.562674791 |
| 0.340755359 | 0.73328776 | NA |
| 0.748177951 | 0.454352828 | 0.564218025 |
| 0.063708882 | 0.949202033 | NA |
| NA | NA | NA |
| NA | NA | NA |
| -7.034443581 | 2.00E-12 | 1.79E-11 |
| -0.376708387 | 0.706390329 | 0.784634328 |
| -0.236439132 | 0.813091934 | NA |
| -0.442233627 | 0.658320155 | 0.745835942 |
| -0.350201701 | 0.72618733 | 0.800013835 |
| -0.065389966 | 0.947863513 | NA |
| NA | NA | NA |
| 0.192809521 | 0.847108147 | NA |
| -0.861551999 | 0.388934095 | NA |
| -2.692273233 | 0.007096678 | 0.016774941 |
| 6.981678451 | 2.92E-12 | 2.57E-11 |
| -1.270357975 | 0.203957146 | 0.300357491 |
| 9.644954954 | 5.16E-22 | 1.07E-20 |
| 9.968219111 | 2.10E-23 | 4.83E-22 |
| 7.60503609 | 2.85E-14 | 3.09E-13 |
| 0.192810118 | 0.847107679 | NA |
| 0.577744589 | 0.563436571 | 0.665570949 |
| 5.241492672 | 1.59E-07 | 8.36E-07 |
| -1.234253299 | 0.217108532 | 0.315723133 |
| 0.843449737 | 0.398976965 | 0.510812964 |
| -3.636218804 | 0.000276669 | 0.000885335 |
| 6.853439621 | 7.21E-12 | 6.10E-11 |
| -1.786140419 | 0.074076525 | 0.129235061 |
| -0.849890736 | 0.395385837 | 0.507357095 |
| 5.789388567 | 7.06E-09 | 4.38E-08 |
| 0.921448582 | 0.356816275 | 0.468650517 |
| -1.80222818 | 0.071509513 | 0.125508948 |
| -1.633966072 | 0.102265975 | 0.16964487 |
| 5.750523868 | 8.90E-09 | 5.47E-08 |
| 0.063708285 | 0.949202508 | NA |
| -1.231627079 | 0.218088421 | 0.316927147 |
| -6.049219435 | 1.46E-09 | 9.74E-09 |
| -3.420384585 | 0.000625327 | 0.001872912 |
| -2.294653708 | 0.021752978 | 0.044942388 |
| -3.415892866 | 0.000635733 | 0.001901121 |
| -3.268542463 | 0.00108103 | 0.003094838 |
| -3.028316706 | 0.002459202 | 0.006478668 |
| NA | NA | NA |
| -0.472611223 | 0.636490574 | 0.727658677 |
| -1.922962936 | 0.054484705 | 0.099653063 |
| 0.308399393 | 0.757778444 | 0.824588925 |

| | | | |
|---|---|---|---|
| | -0.194488217 | 0.845793621 | NA |
| | -1.373996174 | 0.16944287 | 0.257868003 |
| | -2.662718404 | 0.007751225 | 0.018116833 |
| | -0.264227999 | 0.791604237 | 0.851695927 |
| | -1.061753594 | 0.288347562 | 0.395730863 |
| | -2.277233795 | 0.022772268 | 0.046816416 |
| | -1.796635057 | 0.072393575 | 0.126767258 |
| | 0.320071032 | 0.748914485 | NA |
| | 0.469883289 | 0.638438404 | NA |
| NA | NA | NA | |
| | 9.619448941 | 6.62E-22 | 1.36E-20 |
| | 0.129968454 | 0.896591385 | NA |
| | 1.560617595 | 0.118614005 | NA |
| NA | NA | NA | |
| NA | NA | NA | |
| | 4.010308074 | 6.06E-05 | 0.000217669 |
| NA | NA | NA | |
| | 13.05675252 | 5.82E-39 | 3.07E-37 |
| | 4.60370186 | 4.15E-06 | 1.79E-05 |
| | 5.575699132 | 2.47E-08 | 1.44E-07 |
| | 0.256910418 | 0.797247928 | NA |
| NA | NA | NA | |
| NA | NA | NA | |
| NA | NA | NA | |
| NA | NA | NA | |
| | 0.192809521 | 0.847108147 | NA |
| | 4.117176808 | 3.84E-05 | 0.000142413 |
| | -0.961465727 | 0.33631805 | 0.447577669 |
| | -2.840698779 | 0.004501481 | 0.011189996 |
| | -0.978915763 | 0.327621603 | 0.438663139 |
| | -1.966502073 | 0.049240643 | 0.091397352 |
| | 0.063708882 | 0.949202033 | NA |
| | -2.687269959 | 0.00720387 | 0.016981812 |
| | -2.644916627 | 0.008171105 | 0.018984754 |
| | -1.85992392 | 0.062896291 | 0.11248172 |
| | -2.506640286 | 0.012188468 | 0.027132458 |
| | -1.357541619 | 0.174609179 | 0.264540146 |
| | -0.947045212 | 0.343615741 | 0.455024713 |
| | -2.792093802 | 0.005236817 | 0.012818065 |
| | -0.747312072 | 0.454875207 | 0.564730243 |
| | 0.192810118 | 0.847107679 | NA |
| | -5.94133701 | 2.83E-09 | 1.84E-08 |
| | -3.398967544 | 0.000676407 | 0.002013376 |
| | -2.719568676 | 0.006536712 | 0.015609035 |
| | -2.688727431 | 0.007172496 | 0.016917411 |
| NA | NA | NA | |
| | 0.192810118 | 0.847107679 | NA |

| | | |
|---|---|---|
| 0.02006644 | 0.983990372 | 0.988810503 |
| NA | NA | NA |
| 1.162614917 | 0.244985776 | 0.347869376 |
| 0.40223952 | 0.68750776 | 0.769465177 |
| 2.497424702 | 0.012509903 | 0.027778821 |
| 0.950304458 | 0.341957574 | 0.45337137 |
| -8.792092343 | 1.47E-18 | 2.33E-17 |
| -0.395627214 | 0.692380058 | 0.772994333 |
| -3.901397201 | 9.56E-05 | 0.000331971 |
| -2.71766454 | 0.006574446 | 0.015693689 |
| -2.684632218 | 0.007260966 | 0.017105127 |
| NA | NA | NA |
| -3.024962477 | 0.00248664 | 0.006543376 |
| -0.437005293 | 0.662107509 | 0.748669631 |
| 0.680206194 | 0.496373911 | 0.603528563 |
| -3.372653678 | 0.000744475 | 0.002200026 |
| -0.666599132 | 0.505028224 | 0.611526121 |
| -2.370987639 | 0.017740624 | 0.037727088 |
| -1.095514971 | 0.27329109 | 0.379857618 |
| -2.499429606 | 0.012439341 | 0.027641753 |
| -2.062058242 | 0.039202191 | 0.074851693 |
| -1.959618985 | 0.050040341 | 0.092648105 |
| 0.679256196 | 0.496975546 | 0.604084009 |
| 0.148800385 | 0.881711147 | NA |
| 0.063706061 | 0.949204279 | NA |
| 1.251609573 | 0.210712165 | 0.308320208 |
| 0.326366578 | 0.744147021 | NA |
| 0.058662938 | 0.95322058 | NA |
| -0.365555649 | 0.714696675 | 0.791347097 |
| 0.314787856 | 0.752922733 | NA |
| 0.192810118 | 0.847107679 | NA |
| NA | NA | NA |
| NA | NA | NA |
| -0.194488217 | 0.845793621 | NA |
| -0.711770282 | 0.476607038 | 0.585532804 |
| -4.917001875 | 8.79E-07 | 4.16E-06 |
| -3.56528236 | 0.000363465 | 0.001137617 |
| -3.98833475 | 6.65E-05 | 0.000237178 |
| -3.840904384 | 0.000122582 | 0.000417218 |
| -3.011411585 | 0.002600361 | 0.006812543 |
| -4.46118838 | 8.15E-06 | 3.37E-05 |
| -4.104812838 | 4.05E-05 | 0.000149516 |
| -3.863695908 | 0.000111684 | 0.000382992 |
| 0.124974262 | 0.900543927 | NA |
| NA | NA | NA |
| -0.139136637 | 0.889342178 | NA |
| 0.212137591 | 0.831999694 | 0.882450627 |

| | | |
|---|---|---|
| 1.937723146 | 0.052657008 | 0.096792067 |
| -0.121971237 | 0.902921798 | NA |
| -3.195597391 | 0.001395417 | 0.003890319 |
| 0.549448997 | 0.582697357 | 0.682894769 |
| NA | NA | NA |
| NA | NA | NA |
| NA | NA | NA |
| NA | NA | NA |
| NA | NA | NA |
| NA | NA | NA |
| -0.749496725 | 0.453557872 | NA |
| -2.285617825 | 0.022276626 | 0.045921921 |
| -9.250897651 | 2.23E-20 | 4.05E-19 |
| -0.069435456 | 0.944643007 | 0.9623678 |
| -7.68089575 | 1.58E-14 | 1.77E-13 |
| -6.120889649 | 9.31E-10 | 6.36E-09 |
| -2.709307832 | 0.006742375 | 0.016044088 |
| 0.063707019 | 0.949203516 | NA |
| -3.169266784 | 0.00152824 | 0.004228594 |
| 2.918991249 | 0.003511661 | 0.008930794 |
| NA | NA | NA |
| 5.696107937 | 1.23E-08 | 7.41E-08 |
| 0.173195055 | 0.862498111 | 0.904360714 |
| -0.194488217 | 0.845793621 | NA |
| NA | NA | NA |
| NA | NA | NA |
| -0.194487022 | 0.845794556 | NA |
| -0.083368646 | 0.933558418 | NA |
| 1.08928984 | 0.276026092 | 0.382830003 |
| -0.000121217 | 0.999903283 | 0.999903283 |
| 0.458011323 | 0.646944307 | NA |
| 0.314433745 | 0.75319163 | 0.820989701 |
| 0.876924926 | 0.380527416 | 0.493042822 |
| 0.593290179 | 0.55298697 | NA |
| 2.799597091 | 0.005116643 | 0.012559534 |
| NA | NA | NA |
| 5.122686028 | 3.01E-07 | 1.53E-06 |
| 7.419128723 | 1.18E-13 | 1.20E-12 |
| NA | NA | NA |
| 9.422739139 | 4.39E-21 | 8.49E-20 |
| 2.419751665 | 0.01553111 | 0.033587668 |
| -0.097676483 | 0.92218919 | 0.946488496 |
| 1.834818661 | 0.066532563 | 0.118073599 |
| 0.063708882 | 0.949202033 | NA |
| NA | NA | NA |
| -0.251508616 | 0.801420902 | 0.859075675 |
| -2.050732225 | 0.040293031 | 0.076694644 |

| | | |
|---|---|---|
| -0.244546224 | 0.806807807 | 0.863119178 |
| -0.648834438 | 0.516445395 | 0.621681731 |
| -2.514444522 | 0.011922006 | 0.026622161 |
| NA | NA | NA |
| -1.725729946 | 0.084396008 | 0.144508967 |
| -1.858094528 | 0.063155592 | 0.112877473 |
| 0.192810118 | 0.847107679 | NA |
| -0.065388772 | 0.947864464 | NA |
| NA | NA | NA |
| -4.496284195 | 6.92E-06 | 2.89E-05 |
| 0.870820257 | 0.383852303 | NA |
| -8.751177653 | 2.11E-18 | 3.30E-17 |
| 0.658725026 | 0.51007236 | 0.616024664 |
| 0.791864318 | 0.428439795 | 0.539646138 |
| 0.350136777 | 0.726236052 | NA |
| -1.008994565 | 0.312977241 | 0.42329917 |
| -0.085135148 | 0.932153948 | NA |
| NA | NA | NA |
| -6.857272954 | 7.02E-12 | 5.95E-11 |
| -0.50723349 | 0.611990999 | 0.707829338 |
| 4.84895705 | 1.24E-06 | 5.74E-06 |
| -0.399280396 | 0.68968661 | 0.770972974 |
| -0.650534068 | 0.515347303 | 0.62068824 |
| NA | NA | NA |
| 0.192809521 | 0.847108147 | NA |
| -6.493997058 | 8.36E-11 | 6.42E-10 |
| -0.425746318 | 0.670292708 | NA |
| 0.063708882 | 0.949202033 | NA |
| -4.57553741 | 4.75E-06 | 2.03E-05 |
| NA | NA | NA |
| 0.12219623 | 0.902743612 | NA |
| -0.14480593 | 0.8848641 | 0.919262195 |
| NA | NA | NA |
| 0.771582332 | 0.440361842 | 0.550534784 |
| -4.029690009 | 5.59E-05 | 0.000201985 |
| -3.465311602 | 0.000529617 | 0.001608178 |
| 0.295548531 | 0.767574884 | NA |
| -1.31157353 | 0.189664069 | 0.283082388 |
| -1.26248242 | 0.206775248 | 0.303749308 |
| -1.813612223 | 0.069737447 | 0.122863962 |
| -1.719916996 | 0.085447531 | 0.146029748 |
| -1.978526677 | 0.047869322 | 0.089149489 |
| -2.455023108 | 0.014087553 | 0.030844249 |
| -8.155513693 | 3.48E-16 | 4.50E-15 |
| 0.30922356 | 0.757151473 | NA |
| NA | NA | NA |
| NA | NA | NA |

| | | |
|---|---|---|
| NA | NA | NA |
| -0.065389369 | 0.947863988 | NA |
| 1.213481996 | 0.224945594 | 0.324892568 |
| 1.31670544 | 0.187937379 | 0.280907238 |
| -2.38034606 | 0.017296386 | 0.036923074 |
| -0.194487022 | 0.845794556 | NA |
| -0.530813074 | 0.59554832 | 0.694006911 |
| NA | NA | NA |
| 0.192810118 | 0.847107679 | NA |
| NA | NA | NA |
| 1.071673005 | 0.28386693 | 0.39084599 |
| -0.08355242 | 0.933412298 | NA |
| NA | NA | NA |
| -0.194487022 | 0.845794556 | NA |
| NA | NA | NA |
| NA | NA | NA |
| NA | NA | NA |
| NA | NA | NA |
| NA | NA | NA |
| -2.329354619 | 0.019840287 | 0.041497651 |
| 0.192808924 | 0.847108615 | NA |
| 0.541750273 | 0.587990552 | NA |
| NA | NA | NA |
| -0.749023512 | 0.453843034 | NA |
| 0.063708882 | 0.949202033 | NA |
| 0.192809521 | 0.847108147 | NA |
| NA | NA | NA |
| 13.56118956 | 6.80E-42 | 3.97E-40 |
| NA | NA | NA |
| NA | NA | NA |
| -0.626711336 | 0.530848472 | 0.635565735 |
| 4.343446225 | 1.40E-05 | 5.58E-05 |
| 3.16056556 | 0.001574632 | 0.00434447 |
| -0.757834823 | 0.448549873 | 0.558539541 |
| -0.979604254 | 0.327281504 | 0.438301555 |
| 0.960974623 | 0.336564928 | 0.447869612 |
| -1.920185884 | 0.054834424 | 0.100204874 |
| -2.264828298 | 0.023523218 | 0.048127035 |
| 0.06371127 | 0.949200131 | NA |
| 0.297749066 | 0.765894692 | 0.831072274 |
| 0.285346237 | 0.775378893 | 0.838745567 |
| NA | NA | NA |
| -0.065388772 | 0.947864464 | NA |
| 0.416706452 | 0.676893135 | 0.759997989 |
| 0.371268648 | 0.710437447 | 0.787755197 |
| -0.270947552 | 0.786431372 | 0.847731826 |
| -0.703812028 | 0.481549842 | 0.590152884 |

| | | |
|---|---|---|
| -5.70423412 | 1.17E-08 | 7.09E-08 |
| -0.065389369 | 0.947863988 | NA |
| -3.359946273 | 0.000779576 | 0.002292656 |
| -3.152022998 | 0.001621435 | 0.004461308 |
| 0.588188569 | 0.556405729 | 0.6590401 |
| 1.609236303 | 0.107564681 | 0.176778987 |
| 3.096755479 | 0.001956512 | 0.005282871 |
| 2.398072797 | 0.016481589 | 0.035413699 |
| NA | NA | NA |
| -0.048236218 | 0.961527986 | 0.97360651 |
| 4.559138891 | 5.14E-06 | 2.18E-05 |
| 1.922529282 | 0.054539193 | 0.099737293 |
| 0.080618145 | 0.935745636 | NA |
| NA | NA | NA |
| 2.055932769 | 0.039788997 | 0.075850616 |
| NA | NA | NA |
| 10.49926003 | 8.71E-26 | 2.35E-24 |
| 0.351417408 | 0.725275225 | NA |
| 2.82103008 | 0.004786971 | 0.011835035 |
| 0.941682775 | 0.346355077 | 0.457916098 |
| -1.702123353 | 0.088732247 | 0.150799032 |
| 12.37981082 | 3.36E-35 | 1.48E-33 |
| NA | NA | NA |
| 0.194173692 | 0.846039881 | NA |
| NA | NA | NA |
| NA | NA | NA |
| -0.067039719 | 0.946550083 | NA |
| NA | NA | NA |
| 0.284003059 | 0.776408036 | NA |
| NA | NA | NA |
| -0.216430406 | 0.828652267 | NA |
| NA | NA | NA |
| NA | NA | NA |
| NA | NA | NA |
| NA | NA | NA |
| NA | NA | NA |
| NA | NA | NA |
| NA | NA | NA |
| NA | NA | NA |
| 1.074951902 | 0.282396263 | 0.38943299 |
| -2.55015447 | 0.01076752 | 0.024365882 |
| -1.869354555 | 0.061573503 | 0.110426145 |
| -1.296693436 | 0.194736689 | 0.289119124 |
| -0.436698532 | 0.662329994 | 0.748767829 |
| -0.053949159 | 0.95697567 | 0.970365225 |
| -2.54999831 | 0.010772344 | 0.024370189 |
| 0.192808924 | 0.847108615 | NA |

| | | |
|---|---|---|
| 0.192809521 | 0.847108147 | NA |
| 3.5961405 | 0.000322973 | 0.001021097 |
| NA | NA | NA |
| NA | NA | NA |
| NA | NA | NA |
| -0.690598094 | 0.489818146 | 0.597502227 |
| -0.194488217 | 0.845793621 | NA |
| NA | NA | NA |
| NA | NA | NA |
| NA | NA | NA |
| NA | NA | NA |
| NA | NA | NA |
| -5.86443618 | 4.51E-09 | 2.86E-08 |
| 1.41424338 | 0.157290455 | 0.24277994 |
| -0.774171665 | 0.438829282 | 0.549112479 |
| NA | NA | NA |
| NA | NA | NA |
| -0.696333282 | 0.486220135 | NA |
| NA | NA | NA |
| -0.989775902 | 0.322283666 | 0.433324031 |
| -4.845831717 | 1.26E-06 | 5.83E-06 |
| 0.083031301 | 0.933826651 | NA |
| 5.459041018 | 4.79E-08 | 2.70E-07 |
| 0.31685483 | 0.751353758 | NA |
| 0.26081761 | 0.794233164 | 0.853680426 |
| 0.384427341 | 0.70066174 | NA |
| -0.065389369 | 0.947863988 | NA |
| NA | NA | NA |
| -0.487465365 | 0.625928581 | 0.719395371 |
| -5.256535378 | 1.47E-07 | 7.73E-07 |
| 0.191800466 | 0.847898506 | NA |
| 0.063707801 | 0.949202893 | NA |
| 3.437981784 | 0.000586067 | 0.001764405 |
| -0.065389966 | 0.947863513 | NA |
| 0.58653885 | 0.557513462 | NA |
| 0.063710673 | 0.949200606 | NA |
| NA | NA | NA |
| 0.06371127 | 0.949200131 | NA |
| 1.179326175 | 0.238268319 | 0.340383313 |
| NA | NA | NA |
| 3.391430846 | 0.000695287 | 0.002064907 |
| NA | NA | NA |
| NA | NA | NA |
| NA | NA | NA |
| NA | NA | NA |
| 0.192808924 | 0.847108615 | NA |
| 2.561397478 | 0.010425201 | 0.02366184 |

| | | |
|---|---|---|
| 0.794489971 | 0.426910245 | NA |
| NA | NA | NA |
| NA | NA | NA |
| 0.49606901 | 0.619845723 | NA |
| NA | NA | NA |
| NA | NA | NA |
| 1.456726699 | 0.145191837 | NA |
| 0.063708882 | 0.949202033 | NA |
| NA | NA | NA |
| 0.063708882 | 0.949202033 | NA |
| 0.192809521 | 0.847108147 | NA |
| 0.063708285 | 0.949202508 | NA |
| 0.063708882 | 0.949202033 | NA |
| NA | NA | NA |
| NA | NA | NA |
| -0.19448762 | 0.845794089 | NA |
| NA | NA | NA |
| NA | NA | NA |
| 0.3474247 | 0.728272279 | NA |
| 0.063708882 | 0.949202033 | NA |
| NA | NA | NA |
| NA | NA | NA |
| -0.065389369 | 0.947863988 | NA |
| -0.214831281 | 0.829898863 | 0.881006385 |
| 0.192810118 | 0.847107679 | NA |
| NA | NA | NA |
| NA | NA | NA |
| NA | NA | NA |
| 0.063708285 | 0.949202508 | NA |
| NA | NA | NA |
| NA | NA | NA |
| 0.670192256 | 0.50273524 | NA |
| NA | NA | NA |
| 0.063708285 | 0.949202508 | NA |
| NA | NA | NA |
| NA | NA | NA |
| NA | NA | NA |
| NA | NA | NA |
| NA | NA | NA |
| NA | NA | NA |
| NA | NA | NA |
| 2.282023856 | 0.022487932 | NA |
| -0.345246398 | 0.72990914 | NA |
| 1.077843761 | 0.281103487 | 0.38812998 |
| NA | NA | NA |
| 0.063976859 | 0.948988653 | NA |
| NA | NA | NA |

| | | | |
|---|---|---|---|
| NA | NA | NA | |
| NA | NA | NA | |
| NA | NA | NA | |
| NA | NA | NA | |
| NA | NA | NA | |
| NA | NA | NA | |
| NA | NA | NA | |
| NA | NA | NA | |
| NA | NA | NA | |
| NA | NA | NA | |
| NA | NA | NA | |
| NA | NA | NA | |
| | -1.470636823 | 0.14138936 | 0.222255585 |
| NA | NA | NA | |
| NA | NA | NA | |
| NA | NA | NA | |
| | 2.645502987 | 0.008156957 | 0.018955401 |
| | 0.338098497 | 0.735288963 | 0.806922253 |
| | 3.18900619 | 0.001427628 | 0.003973041 |
| | -1.460168604 | 0.144243742 | 0.226018584 |
| | 1.43082554 | 0.152480221 | 0.236657775 |
| | 1.300118323 | 0.193560419 | 0.287799529 |
| | 1.958635867 | 0.050155446 | 0.092833751 |
| | 9.926117119 | 3.20E-23 | 7.27E-22 |
| | -2.766158497 | 0.005672096 | 0.013762365 |
| NA | NA | NA | |
| | 0.538513287 | 0.590222735 | 0.68969538 |
| | -5.049248692 | 4.44E-07 | 2.19E-06 |
| | 0.840649422 | 0.400544363 | 0.512209864 |
| | 0.063708882 | 0.949202033 | NA |
| NA | NA | NA | |
| | -1.831551932 | 0.067018204 | 0.118808548 |
| | -3.430263703 | 0.000602995 | 0.001811445 |
| | 0.269116717 | 0.787839867 | NA |
| NA | NA | NA | |
| | 1.997951118 | 0.04572196 | 0.085673054 |
| | 6.857146817 | 7.02E-12 | 5.95E-11 |
| | 2.89795896 | 0.003755997 | 0.009488556 |
| | 1.239600884 | 0.215123054 | 0.313419949 |
| NA | NA | NA | |
| | 0.365193648 | 0.714966859 | 0.791592671 |
| | 2.754422842 | 0.005879575 | 0.014202448 |
| | 0.063708285 | 0.949202508 | NA |
| | 1.138422062 | 0.254944286 | NA |
| | -0.331626924 | 0.740170987 | NA |
| NA | NA | NA | |
| | 0.446273681 | 0.655399565 | 0.74320369 |

| | | |
|---|---|---|
| -2.457170172 | 0.014003634 | 0.030687353 |
| 0.111878367 | 0.91091985 | NA |
| 2.733081672 | 0.006274477 | 0.015046001 |
| 1.375072726 | 0.169108899 | 0.257422353 |
| -0.547827651 | 0.58381025 | 0.683926358 |
| 5.700316915 | 1.20E-08 | 7.24E-08 |
| -0.065388772 | 0.947864464 | NA |
| NA | NA | NA |
| NA | NA | NA |
| NA | NA | NA |
| 5.784228933 | 7.28E-09 | 4.51E-08 |
| 0.282341587 | 0.7776816 | NA |
| 6.209361749 | 5.32E-10 | 3.73E-09 |
| NA | NA | NA |
| -2.172123102 | 0.029846379 | 0.059097766 |
| -0.794712635 | 0.426780681 | 0.538206391 |
| 0.420430988 | 0.674170635 | NA |
| 5.99853528 | 1.99E-09 | 1.31E-08 |
| 2.513414888 | 0.011956863 | 0.026678576 |
| 0.102120801 | 0.91866079 | 0.944261988 |
| 1.243806412 | 0.21357082 | 0.311683145 |
| 7.757936715 | 8.63E-15 | 9.88E-14 |
| -3.859231757 | 0.000113744 | 0.000389309 |
| 0.819790929 | 0.412335303 | 0.52400247 |
| 0.184733446 | 0.853438111 | 0.898134596 |
| 0.574243826 | 0.565802816 | NA |
| -0.113822182 | 0.909378754 | NA |
| 1.184368207 | 0.23626734 | 0.338065027 |
| NA | NA | NA |
| -0.438179533 | 0.661256144 | 0.747875888 |
| 0.681310767 | 0.495674872 | NA |
| 6.5070193 | 7.67E-11 | 5.90E-10 |
| -0.906922261 | 0.364447908 | 0.476949239 |
| 4.736156871 | 2.18E-06 | 9.76E-06 |
| -0.065390563 | 0.947863038 | NA |
| 1.646255601 | 0.099711146 | 0.166161694 |
| -6.627498945 | 3.41E-11 | 2.72E-10 |
| -0.965558045 | 0.334265378 | 0.44545556 |
| 5.171519271 | 2.32E-07 | 1.19E-06 |
| 1.218171771 | 0.223158704 | 0.322740069 |
| -1.288582237 | 0.197543362 | 0.292643842 |
| 3.280358732 | 0.001036752 | 0.002978004 |
| 3.970194241 | 7.18E-05 | 0.000254929 |
| NA | NA | NA |
| -1.71270444 | 0.086766928 | 0.147951237 |
| 0.192810118 | 0.847107679 | NA |
| -1.32062133 | 0.186627657 | 0.279316566 |

| | | |
|---|---|---|
| 4.842260372 | 1.28E-06 | 5.93E-06 |
| 4.643880653 | 3.42E-06 | 1.49E-05 |
| 0.192809521 | 0.847108147 | NA |
| 0.063708882 | 0.949202033 | NA |
| 12.67777637 | 7.85E-37 | 3.72E-35 |
| 5.003998742 | 5.62E-07 | 2.73E-06 |
| 17.31257418 | 3.78E-67 | 4.57E-65 |
| 16.16503909 | 8.90E-59 | 8.57E-57 |
| 2.952779722 | 0.003149266 | 0.00809802 |
| 3.863776021 | 0.000111648 | 0.000382919 |
| NA | NA | NA |
| 7.639363288 | 2.18E-14 | 2.40E-13 |
| NA | NA | NA |
| -0.915112173 | 0.360132745 | 0.472289773 |
| 2.455916477 | 0.014052581 | 0.030781142 |
| -0.256787853 | 0.797342548 | 0.855846476 |
| 5.780965034 | 7.43E-09 | 4.60E-08 |
| -0.793306103 | 0.427599502 | NA |
| NA | NA | NA |
| NA | NA | NA |
| -0.85552897 | 0.392258374 | 0.504196944 |
| 0.192809521 | 0.847108147 | NA |
| -1.240353765 | 0.214844574 | 0.313129829 |
| -1.620540783 | 0.105116161 | 0.173575505 |
| -2.365581801 | 0.018001766 | 0.038194812 |
| -2.218514287 | 0.026519787 | 0.053333622 |
| -0.977736671 | 0.32820458 | 0.439208759 |
| -6.491437383 | 8.50E-11 | 6.51E-10 |
| -1.644624754 | 0.100047219 | 0.166578985 |
| -3.764347248 | 0.000166985 | 0.000555421 |
| 0.399583814 | 0.68946308 | 0.770826375 |
| -1.521169365 | 0.128217341 | 0.205196934 |
| -5.628574005 | 1.82E-08 | 1.08E-07 |
| 2.45328284 | 0.014155898 | 0.030953275 |
| NA | NA | NA |
| NA | NA | NA |
| NA | NA | NA |
| NA | NA | NA |
| NA | NA | NA |
| NA | NA | NA |
| NA | NA | NA |
| NA | NA | NA |
| NA | NA | NA |
| NA | NA | NA |
| NA | NA | NA |
| NA | NA | NA |
| NA | NA | NA |

| | | | |
|---|---|---|---|
| NA | NA | NA | |
| NA | NA | NA | |
| NA | NA | NA | |
| NA | NA | NA | |
| NA | NA | NA | |
| NA | NA | NA | |
| NA | NA | NA | |
| NA | NA | NA | |
| NA | NA | NA | |
| NA | NA | NA | |
| NA | NA | NA | |
| NA | NA | NA | |
| NA | NA | NA | |
| NA | NA | NA | |
| NA | NA | NA | |
| NA | NA | NA | |
| NA | NA | NA | |
| NA | NA | NA | |
| NA | NA | NA | |
| NA | NA | NA | |
| NA | NA | NA | |
| NA | NA | NA | |
| NA | NA | NA | |
| NA | NA | NA | |
| NA | NA | NA | |
| NA | NA | NA | |
| NA | NA | NA | |
| NA | NA | NA | |
| NA | NA | NA | |
| NA | NA | NA | |
| NA | NA | NA | |
| NA | NA | NA | |
| NA | NA | NA | |
| NA | NA | NA | |
| NA | NA | NA | |
| NA | NA | NA | |
| | 0.154248174 | 0.877414059 | 0.914383958 |
| NA | NA | NA | |
| | 10.26217822 | 1.04E-24 | 2.61E-23 |
| | 8.388292935 | 4.93E-17 | 6.88E-16 |
| | 4.378832411 | 1.19E-05 | 4.80E-05 |
| | 0.544789195 | 0.585898511 | 0.686019541 |
| | 2.517800441 | 0.011809021 | 0.026412277 |
| | -7.374597702 | 1.65E-13 | 1.65E-12 |
| | -0.092058339 | 0.926651689 | NA |
| | 0.517688767 | 0.604675437 | 0.701742573 |
| | 0.288217989 | 0.77317988 | NA |

| | | |
|---|---|---|
| 15.97005416 | 2.07E-57 | 1.91E-55 |
| 2.231433957 | 0.025652398 | 0.051834949 |
| 0.244888461 | 0.806542796 | NA |
| 0.934344607 | 0.350126162 | NA |
| 0.382131398 | 0.702363903 | NA |
| NA | NA | NA |
| 4.255382874 | 2.09E-05 | 8.09E-05 |
| 1.958402749 | 0.050182772 | NA |
| 0.649691104 | 0.515891771 | 0.621194547 |
| -0.087792604 | 0.930041517 | 0.952454448 |
| 0.619922598 | 0.535308747 | NA |
| 1.825205579 | 0.067970012 | 0.120164529 |
| 12.94710033 | 2.44E-38 | 1.25E-36 |
| 0.886525444 | 0.375334449 | 0.488159068 |
| -2.862059913 | 0.004208973 | 0.010525587 |
| -2.163816222 | 0.030478457 | 0.060173195 |
| 0.715402589 | 0.474360313 | 0.583250903 |
| 0.590371437 | 0.554941655 | 0.657772867 |
| NA | NA | NA |
| 0.198847804 | 0.8423818 | NA |
| -5.057806533 | 4.24E-07 | 2.10E-06 |
| -0.065389369 | 0.947863988 | NA |
| 9.244455815 | 2.36E-20 | 4.28E-19 |
| 1.827586766 | 0.067611594 | 0.119666114 |
| -0.065389369 | 0.947863988 | NA |
| NA | NA | NA |
| 0.332406488 | 0.73958234 | NA |
| 0.3474247 | 0.728272279 | NA |
| -4.994238813 | 5.91E-07 | 2.86E-06 |
| -0.536435204 | 0.591657806 | 0.690888495 |
| -2.38330573 | 0.017157938 | 0.036665885 |
| 0.183146922 | 0.85468274 | 0.899042193 |
| -0.38486375 | 0.700338364 | 0.779641934 |
| 0.549581891 | 0.582606182 | NA |
| 1.045679967 | 0.295708815 | 0.404039066 |
| -0.194280944 | 0.845955906 | NA |
| 0.820097499 | 0.412160528 | NA |
| NA | NA | NA |
| -6.814825123 | 9.44E-12 | 7.91E-11 |
| -3.709828552 | 0.0002074 | 0.000679567 |
| -0.175130081 | 0.860977423 | 0.903279692 |
| 1.535754637 | 0.124598569 | NA |
| 7.222928992 | 5.09E-13 | 4.88E-12 |
| -1.223603195 | 0.221101954 | 0.32038467 |
| -2.136825764 | 0.032612166 | 0.06389687 |
| -0.317887001 | 0.750570653 | NA |
| -1.331625982 | 0.182983123 | 0.274897519 |

| | | |
|---|---|---|
| -2.046101937 | 0.040746342 | 0.077457352 |
| NA | NA | NA |
| NA | NA | NA |
| NA | NA | NA |
| NA | NA | NA |
| -3.728370957 | 0.000192722 | 0.000635131 |
| 0.192810118 | 0.847107679 | NA |
| NA | NA | NA |
| 0.161652528 | 0.87157949 | 0.91050228 |
| 0.324874675 | 0.745275925 | NA |
| 0.06371127 | 0.949200131 | NA |
| -0.540166396 | 0.589082285 | 0.688619914 |
| 1.297208892 | 0.194559322 | 0.288890065 |
| 1.950977456 | 0.051059728 | 0.094263533 |
| 0.68751942 | 0.491755467 | 0.599077161 |
| -2.604403843 | 0.009203424 | 0.021151566 |
| NA | NA | NA |
| -5.758872267 | 8.47E-09 | 5.21E-08 |
| 1.743540488 | 0.081239248 | 0.139965473 |
| 0.696204536 | 0.486300747 | NA |
| 0.497440171 | 0.618878684 | 0.713487792 |
| -0.713861767 | 0.475312661 | NA |
| 3.843970413 | 0.00012106 | 0.000412486 |
| 1.725635284 | 0.084413047 | 0.144524976 |
| -0.228105934 | 0.819563887 | NA |
| 0.192810118 | 0.847107679 | NA |
| NA | NA | NA |
| 0.286888673 | 0.774197564 | NA |
| 0.253725823 | 0.799707383 | 0.857722749 |
| 2.194259721 | 0.028216743 | 0.056280587 |
| -0.445492259 | 0.655964052 | 0.743604177 |
| NA | NA | NA |
| NA | NA | NA |
| 4.537005206 | 5.71E-06 | 2.41E-05 |
| 12.70173662 | 5.78E-37 | 2.75E-35 |
| NA | NA | NA |
| 2.576765626 | 0.009972955 | 0.02273434 |
| 0.230088074 | 0.818023332 | NA |
| 14.12942608 | 2.50E-45 | 1.69E-43 |
| -1.490315301 | 0.136141351 | 0.215371079 |
| 5.296883343 | 1.18E-07 | 6.29E-07 |
| -1.236457471 | 0.216288565 | 0.314841685 |
| -2.157412468 | 0.030973541 | 0.061030286 |
| -2.458910201 | 0.013935947 | 0.030560431 |
| 1.306042506 | 0.191538103 | 0.285351986 |
| -0.086594846 | 0.930993563 | NA |
| 1.181189922 | 0.237527284 | 0.339557251 |

| | | | |
|---|---|---|---|
| | 7.75608692 | 8.76E-15 | 1.00E-13 |
| | 0.789184849 | 0.430003976 | 0.541103697 |
| | 0.063708285 | 0.949202508 | NA |
| | 0.063708882 | 0.949202033 | NA |
| NA | | NA | NA |
| NA | | NA | NA |
| | 1.554284207 | 0.120116639 | 0.194119284 |
| NA | | NA | NA |
| | 0.796752486 | 0.425594795 | 0.537089731 |
| | -1.665959621 | 0.095721457 | 0.160663405 |
| | -3.09184654 | 0.001989157 | 0.00536515 |
| | -3.40675534 | 0.0006574 | 0.001960764 |
| | -5.819119834 | 5.92E-09 | 3.71E-08 |
| | 1.220963873 | 0.222099699 | 0.32148713 |
| | 0.333816862 | 0.738517758 | NA |
| | -2.064267488 | 0.038992362 | 0.074507887 |
| | -1.072237898 | 0.283613192 | 0.390604061 |
| | 0.172977003 | 0.862669504 | NA |
| | 0.340743572 | 0.733296634 | 0.80538127 |
| | -1.013167436 | 0.310980192 | 0.42115757 |
| | -0.075714474 | 0.93964626 | NA |
| | 7.469156844 | 8.07E-14 | 8.38E-13 |
| | -6.59611605 | 4.22E-11 | 3.34E-10 |
| | -3.103207426 | 0.001914354 | 0.005177605 |
| | -3.239949176 | 0.00119551 | 0.003387324 |
| NA | | NA | NA |
| NA | | NA | NA |
| NA | | NA | NA |
| NA | | NA | NA |
| | 2.626869194 | 0.008617445 | 0.019934811 |
| | 5.473417049 | 4.41E-08 | 2.50E-07 |
| NA | | NA | NA |
| | 0.063708882 | 0.949202033 | NA |
| | 6.679841055 | 2.39E-11 | 1.93E-10 |
| NA | | NA | NA |
| | -1.152864086 | 0.248966175 | NA |
| | 2.101962051 | 0.0355566 | 0.068825435 |
| NA | | NA | NA |
| | 11.36258675 | 6.42E-30 | 2.20E-28 |
| | -6.409805593 | 1.46E-10 | 1.08E-09 |
| | -2.037349874 | 0.041614993 | 0.078928793 |
| NA | | NA | NA |
| | -0.162304809 | 0.871065828 | 0.910155612 |
| NA | | NA | NA |
| | -0.89443346 | 0.371090017 | 0.48387076 |
| | -1.809532974 | 0.070368242 | 0.123844544 |
| | 5.376744617 | 7.58E-08 | 4.15E-07 |

| | | | |
|---|---|---|---|
| | -0.498342367 | 0.618242751 | 0.712951673 |
| | -1.410972473 | 0.158252731 | 0.243862947 |
| | -4.186169463 | 2.84E-05 | 0.000107755 |
| | -8.673400667 | 4.19E-18 | 6.37E-17 |
| | -2.583492552 | 0.009780558 | 0.022328289 |
| | -0.32752371 | 0.743271813 | 0.813115973 |
| | -0.574887917 | 0.565367102 | 0.667412091 |
| | 0.192809521 | 0.847108147 | NA |
| NA | NA | NA | |
| | 1.202036795 | 0.229349273 | 0.329845043 |
| NA | NA | NA | |
| | 11.29876452 | 1.33E-29 | 4.50E-28 |
| | 2.791002265 | 0.00525451 | 0.012857601 |
| | -1.607804048 | 0.107878102 | 0.177247531 |
| | -2.75996872 | 0.00578069 | 0.013989235 |
| | -2.838031483 | 0.004539271 | 0.011278334 |
| | 1.012075422 | 0.311501993 | 0.421658909 |
| | -2.610373351 | 0.009044345 | 0.020820378 |
| | 2.382401894 | 0.017200114 | 0.036743097 |
| NA | NA | NA | |
| NA | NA | NA | |
| NA | NA | NA | |
| | 0.063708217 | 0.949202562 | NA |
| | 14.36934101 | 8.06E-47 | 5.78E-45 |
| | 29.95742485 | 3.52E-197 | 2.38E-194 |
| | 27.33658321 | 1.56E-164 | 7.81E-162 |
| | 29.24394985 | 5.36E-188 | 3.27E-185 |
| | 18.39401568 | 1.47E-75 | 2.15E-73 |
| | 20.56995072 | 5.10E-94 | 1.10E-91 |
| | 1.984294315 | 0.04722304 | 0.088109779 |
| | 2.534851243 | 0.011249509 | 0.025326066 |
| | -7.396757291 | 1.40E-13 | 1.41E-12 |
| | 1.777662063 | 0.075459374 | 0.131345489 |
| | 5.775292923 | 7.68E-09 | 4.75E-08 |
| | -1.101963081 | 0.270477722 | 0.376700278 |
| | 1.591120708 | 0.111582413 | 0.182328489 |
| | 3.821916468 | 0.000132419 | 0.000448078 |
| | -0.065388772 | 0.947864464 | NA |
| | -0.788384383 | 0.430471903 | 0.541524437 |
| | 0.192805606 | 0.847111213 | NA |
| | -4.927021091 | 8.35E-07 | 3.96E-06 |
| | -1.848846166 | 0.064480029 | 0.114829963 |
| NA | NA | NA | |
| NA | NA | NA | |
| | 1.155658973 | 0.247820676 | NA |
| | 0.542125411 | 0.587732115 | NA |
| | 0.063708285 | 0.949202508 | NA |

| | | | |
|---|---|---|---|
| NA | NA | NA | |
| | -0.065389369 | 0.947863988 | NA |
| | 0.067524971 | 0.946163784 | NA |
| | 0.063707194 | 0.949203376 | NA |
| NA | NA | NA | |
| | -0.454917311 | 0.649168729 | 0.737939656 |
| | 0.559844426 | 0.575585558 | 0.676666503 |
| | -0.360240641 | 0.718667185 | 0.794311546 |
| | 0.063708285 | 0.949202508 | NA |
| | 1.085195417 | 0.277835113 | NA |
| NA | NA | NA | |
| | -0.065389369 | 0.947863988 | NA |
| | -5.717082589 | 1.08E-08 | 6.60E-08 |
| | -2.639051676 | 0.008313831 | 0.019289505 |
| NA | NA | NA | |
| NA | NA | NA | |
| | -0.990901202 | 0.321733826 | 0.432747469 |
| | 0.149666725 | 0.881027562 | NA |
| NA | NA | NA | |
| | -3.430986224 | 0.000601391 | 0.001807061 |
| | 0.192809521 | 0.847108147 | NA |
| | -0.038658176 | 0.969162919 | 0.978703618 |
| | -2.537065939 | 0.01117859 | 0.025186794 |
| | -2.43981246 | 0.014694888 | 0.031984431 |
| NA | NA | NA | |
| | -4.245413441 | 2.18E-05 | 8.43E-05 |
| NA | NA | NA | |
| | -0.364315114 | 0.715622715 | 0.79202301 |
| | -0.045256736 | 0.963902672 | NA |
| | -4.712336702 | 2.45E-06 | 1.09E-05 |
| | -0.065389369 | 0.947863988 | NA |
| | -8.94961662 | 3.57E-19 | 5.98E-18 |
| | -2.750722469 | 0.0059464 | 0.014337577 |
| | -2.042714106 | 0.041080748 | 0.077986433 |
| | -3.709343809 | 0.000207797 | 0.00068078 |
| | 1.554352044 | 0.120100466 | 0.194105679 |
| | -5.476570958 | 4.34E-08 | 2.46E-07 |
| | 1.346360882 | 0.17818616 | 0.268901458 |
| | -2.251228586 | 0.024371063 | 0.049592259 |
| | -4.37808619 | 1.20E-05 | 4.81E-05 |
| | 0.065375237 | 0.94787524 | 0.964443791 |
| | 0.275770074 | 0.78272467 | NA |
| | -1.369079194 | 0.170974521 | 0.259886769 |
| | -0.086460871 | 0.93110006 | NA |
| | 1.341213736 | 0.179851068 | NA |
| | 5.376449433 | 7.60E-08 | 4.16E-07 |
| | 0.330493631 | 0.741027004 | NA |

| | | |
|---|---|---|
| -6.957105878 | 3.47E-12 | 3.04E-11 |
| -5.259314264 | 1.45E-07 | 7.63E-07 |
| NA | NA | NA |
| NA | NA | NA |
| 1.010273882 | 0.312364091 | 0.422665866 |
| 4.871070916 | 1.11E-06 | 5.17E-06 |
| NA | NA | NA |
| -0.802762265 | 0.422112156 | 0.533906073 |
| -0.000955359 | 0.999237734 | 0.999506869 |
| -3.033442729 | 0.002417806 | 0.006381702 |
| -0.978776006 | 0.327690668 | 0.438685212 |
| NA | NA | NA |
| -4.285188677 | 1.83E-05 | 7.14E-05 |
| 7.957866597 | 1.75E-15 | 2.14E-14 |
| 0.451696571 | 0.651487589 | 0.740037865 |
| NA | NA | NA |
| NA | NA | NA |
| NA | NA | NA |
| 4.731197004 | 2.23E-06 | 9.99E-06 |
| -1.639750318 | 0.101057092 | 0.16799223 |
| -1.20659131 | 0.227589575 | 0.32782224 |
| -1.773752973 | 0.07610402 | 0.13230196 |
| -2.717268115 | 0.006582326 | 0.015707467 |
| -0.6139683 | 0.539236275 | 0.643484337 |
| -0.559954587 | 0.575510414 | 0.676609942 |
| 8.123901768 | 4.51E-16 | 5.79E-15 |
| -1.8522786 | 0.063985827 | 0.114071599 |
| 6.682720851 | 2.35E-11 | 1.90E-10 |
| -0.648132256 | 0.516899413 | 0.622168421 |
| 0.513936281 | 0.60729654 | NA |
| -6.122658064 | 9.20E-10 | 6.29E-09 |
| 0.660636546 | 0.508845426 | 0.61496398 |
| 2.945100411 | 0.003228499 | 0.008283038 |
| -1.817069745 | 0.069206435 | 0.122040092 |
| -4.824434417 | 1.40E-06 | 6.45E-06 |
| -4.655485789 | 3.23E-06 | 1.41E-05 |
| 4.911370702 | 9.04E-07 | 4.27E-06 |
| -1.006592456 | 0.314130661 | 0.424557607 |
| 8.695274227 | 3.46E-18 | 5.30E-17 |
| -0.065388772 | 0.947864464 | NA |
| 0.796969681 | 0.42546864 | NA |
| NA | NA | NA |
| NA | NA | NA |
| NA | NA | NA |
| NA | NA | NA |
| 0.192809521 | 0.847108147 | NA |
| -0.511315257 | 0.609130324 | 0.705629287 |

| | | | |
|---|---|---|---|
| NA | NA | NA | |
| | -0.065389369 | 0.947863988 | NA |
| NA | NA | NA | |
| NA | NA | NA | |
| NA | NA | NA | |
| | 0.994722525 | 0.319871236 | NA |
| NA | NA | NA | |
| NA | NA | NA | |
| NA | NA | NA | |
| NA | NA | NA | |
| NA | NA | NA | |
| NA | NA | NA | |
| NA | NA | NA | |
| NA | NA | NA | |
| | -0.065389369 | 0.947863988 | NA |
| NA | NA | NA | |
| | 1.386861109 | 0.165484119 | 0.252950687 |
| | -0.425195301 | 0.670694309 | 0.755104289 |
| | 9.187562027 | 4.02E-20 | 7.22E-19 |
| | -0.085070848 | 0.932205066 | NA |
| NA | NA | NA | |
| | -0.957603651 | 0.338262657 | 0.449520272 |
| | -0.281458206 | 0.77835898 | 0.841351065 |
| | -0.131578231 | 0.895317905 | 0.92713612 |
| | -1.608561439 | 0.107712272 | 0.177009924 |
| | -1.385845423 | 0.165794109 | 0.253316234 |
| | -3.591774486 | 0.000328434 | 0.001036789 |
| | 0.646423783 | 0.518004942 | NA |
| | -3.406099218 | 0.000658982 | 0.001964681 |
| | -0.870416104 | 0.384073049 | 0.496352149 |
| | -2.505530179 | 0.012226796 | 0.027212939 |
| | -0.58249422 | 0.560233834 | 0.66257101 |
| | -0.66284407 | 0.507430426 | 0.613870641 |
| | 2.909113229 | 0.003624556 | 0.009182452 |
| | 0.647234009 | 0.517480503 | 0.622778008 |
| | 1.160611065 | 0.245800104 | 0.348793956 |
| | -3.029689304 | 0.002448054 | 0.006451337 |
| | 1.107820504 | 0.267939343 | 0.373828981 |
| | 5.394386727 | 6.88E-08 | 3.78E-07 |
| | 3.350936983 | 0.000805386 | 0.002361347 |
| | -2.176151653 | 0.029543923 | 0.058591584 |
| | -0.020735277 | 0.983456828 | 0.988392869 |
| | 8.842785186 | 9.34E-19 | 1.51E-17 |
| | 2.432139748 | 0.015009915 | 0.032592883 |
| | -1.661464948 | 0.096620103 | 0.161890605 |
| | -0.348226632 | 0.727669989 | 0.801195611 |
| | -0.487215384 | 0.626105704 | 0.719565902 |

| | | |
|---|---|---|
| 5.900502291 | 3.62E-09 | 2.33E-08 |
| NA | NA | NA |
| -0.065389966 | 0.947863513 | NA |
| 0.192810118 | 0.847107679 | NA |
| -0.26119582 | 0.793941501 | 0.853476884 |
| 0.192809521 | 0.847108147 | NA |
| -3.924807423 | 8.68E-05 | 0.000303858 |
| -0.099213538 | 0.920968726 | 0.945639218 |
| -0.285446359 | 0.775302195 | 0.838744636 |
| NA | NA | NA |
| -0.246241876 | 0.805495001 | 0.862313679 |
| -4.978872823 | 6.40E-07 | 3.09E-06 |
| 7.52178481 | 5.40E-14 | 5.70E-13 |
| 3.569829925 | 0.000357213 | 0.001120431 |
| 1.60610325 | 0.108251226 | 0.177718071 |
| 2.379540355 | 0.017334245 | 0.036974991 |
| -0.434565366 | 0.663877939 | 0.750197057 |
| -1.341008301 | 0.179917757 | 0.271057028 |
| 0.063708285 | 0.949202508 | NA |
| NA | NA | NA |
| 0.991623965 | 0.321380994 | 0.432319358 |
| 3.275549035 | 0.001054568 | 0.003024271 |
| -0.463725207 | 0.642844635 | 0.732787126 |
| -4.29050539 | 1.78E-05 | 6.98E-05 |
| 2.297380224 | 0.021597089 | 0.044649801 |
| 1.614059393 | 0.106514544 | 0.175340943 |
| 0.704250117 | 0.481277025 | NA |
| 1.228076474 | 0.219418261 | 0.318432006 |
| -1.430238079 | 0.1526487 | 0.236875199 |
| -0.576040124 | 0.564588061 | 0.666649645 |
| -1.934126587 | 0.053097566 | 0.097457804 |
| NA | NA | NA |
| -2.244663073 | 0.02478977 | 0.050347935 |
| -1.673576152 | 0.094213929 | 0.158543407 |
| -1.403933472 | 0.160338661 | 0.246349315 |
| NA | NA | NA |
| -2.638682171 | 0.008322897 | 0.019308751 |
| NA | NA | NA |
| 0.689825237 | 0.490304096 | 0.597891041 |
| 0.379151304 | 0.704575511 | NA |
| -6.985788174 | 2.83E-12 | 2.49E-11 |
| -4.849279221 | 1.24E-06 | 5.73E-06 |
| -2.256486964 | 0.024040151 | 0.048992426 |
| -2.044030656 | 0.040950519 | 0.077774602 |
| 0.198902723 | 0.842338839 | NA |
| -0.372885151 | 0.709233924 | 0.78691986 |
| NA | NA | NA |

| | | |
|---|---|---|
| NA | NA | NA |
| -0.44550836 | 0.65595242 | 0.743604177 |
| NA | NA | NA |
| NA | NA | NA |
| -4.169832907 | 3.05E-05 | 0.0001152 |
| NA | NA | NA |
| NA | NA | NA |
| 0.750347499 | 0.453045442 | NA |
| NA | NA | NA |
| NA | NA | NA |
| 1.615562433 | 0.106188952 | 0.174885476 |
| -4.416661975 | 1.00E-05 | 4.09E-05 |
| 7.916951671 | 2.43E-15 | 2.94E-14 |
| 4.253405802 | 2.11E-05 | 8.15E-05 |
| -0.04189097 | 0.966585615 | 0.977362768 |
| -1.498085066 | 0.134111151 | 0.212738583 |
| 1.485210439 | 0.137488088 | 0.217118412 |
| -2.992888271 | 0.002763509 | 0.007200773 |
| NA | NA | NA |
| -0.114269125 | 0.909024457 | NA |
| -6.445982412 | 1.15E-10 | 8.67E-10 |
| -1.992760883 | 0.046287637 | 0.086583929 |
| NA | NA | NA |
| 0.69390084 | 0.487744397 | 0.595814654 |
| 0.192808924 | 0.847108615 | NA |
| -0.069867013 | 0.944299508 | NA |
| -3.510863267 | 0.000446654 | 0.001374447 |
| -2.084711566 | 0.03709549 | 0.071395626 |
| 3.681002934 | 0.000232318 | 0.000753424 |
| NA | NA | NA |
| -10.33342492 | 4.98E-25 | 1.27E-23 |
| NA | NA | NA |
| 0.501144437 | 0.616269475 | NA |
| 0.969211738 | 0.332439556 | 0.443595377 |
| 0.721605064 | 0.470537325 | 0.579654344 |
| -5.900021544 | 3.63E-09 | 2.33E-08 |
| 0.192810118 | 0.847107679 | NA |
| -1.332323218 | 0.182754001 | 0.274586268 |
| -0.785224104 | 0.432322185 | NA |
| 0.192809521 | 0.847108147 | NA |
| -2.05615231 | 0.039767837 | 0.075827605 |
| 0.024046649 | 0.980815399 | 0.986613469 |
| -3.966077559 | 7.31E-05 | 0.00025904 |
| 1.912985776 | 0.055749873 | 0.101647679 |
| 0.75920712 | 0.447728667 | NA |
| NA | NA | NA |
| NA | NA | NA |

| | | | |
|---|---|---|---|
| | 0.117454658 | 0.906499773 | NA |
| | -5.929294695 | 3.04E-09 | 1.97E-08 |
| | 17.75499175 | 1.58E-70 | 2.05E-68 |
| | 0.063708285 | 0.949202508 | NA |
| | 0.588519472 | 0.556183668 | 0.658808253 |
| | 2.857209302 | 0.004273839 | 0.010670052 |
| | 1.047545292 | 0.294848154 | 0.403149203 |
| | 0.262131689 | 0.793219916 | NA |
| | 0.863664192 | 0.38777239 | 0.49999706 |
| | -3.61225014 | 0.000303552 | 0.000963955 |
| NA | NA | NA | |
| | 7.675939755 | 1.64E-14 | 1.83E-13 |
| | 5.558805213 | 2.72E-08 | 1.58E-07 |
| | 0.137069169 | 0.890976122 | NA |
| | -0.469141039 | 0.638968826 | NA |
| | -0.34355403 | 0.731181701 | 0.803895194 |
| | 0.662735349 | 0.507500066 | NA |
| NA | NA | NA | |
| | 4.0837369 | 4.43E-05 | 0.000162577 |
| | -2.693936048 | 0.007061371 | 0.016705683 |
| | -7.065886268 | 1.60E-12 | 1.45E-11 |
| | 0.348953938 | 0.727123891 | 0.800735227 |
| | -1.944497964 | 0.051835431 | 0.095512429 |
| | -2.054349165 | 0.039941909 | 0.076118927 |
| | 3.234504658 | 0.001218539 | 0.003440889 |
| | 2.945343951 | 0.003225958 | 0.008277369 |
| | 4.625875172 | 3.73E-06 | 1.62E-05 |
| NA | NA | NA | |
| | 0.288217989 | 0.77317988 | NA |
| | 0.003023799 | 0.997587361 | 0.998225459 |
| | 7.899898591 | 2.79E-15 | 3.34E-14 |
| | -1.393513972 | 0.163464423 | 0.250398684 |
| | -0.574116977 | 0.565888646 | 0.667901768 |
| | -6.207161179 | 5.40E-10 | 3.77E-09 |
| | 0.334541364 | 0.737971083 | 0.809227031 |
| | -0.19448762 | 0.845794089 | NA |
| | -0.555613639 | 0.578475005 | 0.679361394 |
| | 1.632788012 | 0.102513589 | 0.170006966 |
| | 6.476646842 | 9.38E-11 | 7.15E-10 |
| | -0.573221112 | 0.566494992 | 0.668396797 |
| | 0.06371127 | 0.949200131 | NA |
| | -1.045634708 | 0.295729718 | NA |
| | 0.50850374 | 0.611100117 | 0.707027638 |
| | -1.68313428 | 0.0923491 | 0.155982048 |
| | 7.516670762 | 5.62E-14 | 5.92E-13 |
| | -0.491724714 | 0.622913965 | 0.716952613 |
| | 0.538386688 | 0.590310115 | 0.689733087 |

| | | |
|---|---|---|
| 1.948220629 | 0.051388569 | 0.094786728 |
| 0.192808924 | 0.847108615 | NA |
| 0.840576031 | 0.400585491 | 0.512217361 |
| NA | NA | NA |
| 0.108117913 | 0.913902159 | NA |
| 2.007354244 | 0.044711957 | 0.084039999 |
| 0.192810118 | 0.847107679 | NA |
| 51.54598104 | 0 | 0 |
| 6.73114373 | 1.68E-11 | 1.38E-10 |
| 17.58148778 | 3.41E-69 | 4.27E-67 |
| -2.886334511 | 0.003897577 | 0.009824405 |
| -0.065389369 | 0.947863988 | NA |
| -0.065390563 | 0.947863038 | NA |
| 0.063708285 | 0.949202508 | NA |
| 0.063708285 | 0.949202508 | NA |
| -3.277518187 | 0.00104724 | 0.003006011 |
| -0.920447336 | 0.357339042 | 0.469142209 |
| 0.192809521 | 0.847108147 | NA |
| 0.192809521 | 0.847108147 | NA |
| 3.217981732 | 0.00129096 | 0.003626163 |
| 2.691308593 | 0.007117233 | 0.016809281 |
| 5.051429393 | 4.39E-07 | 2.17E-06 |
| 3.335204992 | 0.000852365 | 0.002490045 |
| -7.252896775 | 4.08E-13 | 3.94E-12 |
| -1.512238268 | 0.130473259 | NA |
| -2.90595273 | 0.003661368 | 0.009268207 |
| 3.722792665 | 0.000197031 | 0.00064848 |
| 0.063708285 | 0.949202508 | NA |
| 3.405661185 | 0.00066004 | 0.001967466 |
| NA | NA | NA |
| 0.627018068 | 0.530647392 | 0.635416171 |
| -2.972805989 | 0.002950909 | 0.007642137 |
| 0.756174386 | 0.449544647 | 0.559527724 |
| -2.364533466 | 0.018052796 | 0.03828037 |
| 0.821659083 | 0.411270953 | 0.523048216 |
| 0.372466765 | 0.709545352 | 0.78716072 |
| 0.389142073 | 0.697171052 | NA |
| -0.194487022 | 0.845794556 | NA |
| 1.510221116 | 0.130987012 | 0.208613172 |
| -3.368941799 | 0.000754574 | 0.002225925 |
| -3.512546046 | 0.000443835 | 0.001366781 |
| 0.704098274 | 0.481371575 | 0.590045569 |
| -0.427493158 | 0.669020168 | NA |
| 0.192808327 | 0.847109082 | NA |
| -2.826586821 | 0.004704699 | 0.011644281 |
| 1.047882092 | 0.294692934 | 0.403003013 |
| -0.515821717 | 0.605978934 | 0.702922083 |

| | | |
|---|---|---|
| 0.45452444 | 0.649451405 | NA |
| 6.559586858 | 5.40E-11 | 4.22E-10 |
| 0.835588536 | 0.40338642 | 0.515035814 |
| 10.12365739 | 4.34E-24 | 1.04E-22 |
| -0.300694564 | 0.763647414 | 0.829352835 |
| -0.068186498 | 0.945637175 | NA |
| -0.498137556 | 0.618387092 | 0.713052421 |
| 2.510929563 | 0.012041372 | 0.026845599 |
| -0.241481996 | 0.809181568 | 0.86521486 |
| -5.579448266 | 2.41E-08 | 1.41E-07 |
| 2.750784301 | 0.005945278 | 0.014336252 |
| -6.445050514 | 1.16E-10 | 8.72E-10 |
| 0.113354812 | 0.909749263 | NA |
| -0.739059938 | 0.459870602 | NA |
| -1.852326404 | 0.063978966 | 0.114071599 |
| 5.028492019 | 4.94E-07 | 2.42E-06 |
| NA | NA | NA |
| NA | NA | NA |
| -0.065388772 | 0.947864464 | NA |
| -2.452878488 | 0.01417182 | 0.030977265 |
| 1.999415415 | 0.045563425 | 0.085427156 |
| -0.988150604 | 0.323078896 | NA |
| 0.236872192 | 0.812755944 | NA |
| 0.746065521 | 0.455627836 | NA |
| 0.192809521 | 0.847108147 | NA |
| 0.145419387 | 0.884379759 | 0.918978042 |
| 0.192810118 | 0.847107679 | NA |
| 0.067649016 | 0.946065036 | NA |
| 13.0270369 | 8.59E-39 | 4.51E-37 |
| -1.477553402 | 0.139527355 | 0.219797954 |
| -5.79253245 | 6.93E-09 | 4.31E-08 |
| -3.914051026 | 9.08E-05 | 0.000316574 |
| -1.947825015 | 0.051435905 | 0.094860057 |
| -4.947421196 | 7.52E-07 | 3.59E-06 |
| -2.467269358 | 0.01361479 | 0.029934594 |
| 0.063708882 | 0.949202033 | NA |
| 0.063708285 | 0.949202508 | NA |
| NA | NA | NA |
| 2.450647895 | 0.014259937 | 0.03113996 |
| -2.310948868 | 0.020835679 | 0.043290201 |
| -0.985033156 | 0.324607785 | 0.435583169 |
| 1.897408096 | 0.057774098 | 0.104750864 |
| -0.884613817 | 0.376364952 | 0.489092713 |
| -0.475295804 | 0.63457615 | 0.726066969 |
| -0.587441327 | 0.556907345 | 0.659451043 |
| NA | NA | NA |
| -0.095258147 | 0.924109786 | NA |

| | | | |
|---|---|---|---|
| | -0.473139498 | 0.636113659 | 0.727449074 |
| | 1.248888388 | 0.211705899 | 0.309502871 |
| | 4.286883727 | 1.81E-05 | 7.09E-05 |
| NA | NA | NA | |
| | 1.645760342 | 0.09981311 | 0.166266614 |
| | 0.284683307 | 0.775886781 | 0.8391499 |
| | 2.733148243 | 0.006273209 | 0.015045981 |
| | -11.79026301 | 4.38E-32 | 1.68E-30 |
| | -2.618928557 | 0.008820642 | 0.020342824 |
| | -9.007826949 | 2.10E-19 | 3.57E-18 |
| | -0.903955429 | 0.366019036 | 0.478554977 |
| | -0.477781621 | 0.632805644 | NA |
| | 0.192809521 | 0.847108147 | NA |
| | -8.31557677 | 9.13E-17 | 1.24E-15 |
| | 1.203350752 | 0.228840617 | 0.329321028 |
| | -5.223620746 | 1.75E-07 | 9.14E-07 |
| | 5.387686478 | 7.14E-08 | 3.92E-07 |
| | -2.966091638 | 0.003016106 | 0.007795656 |
| | -0.344861777 | 0.730198288 | 0.80301204 |
| | -4.356422463 | 1.32E-05 | 5.28E-05 |
| | -7.934473951 | 2.11E-15 | 2.56E-14 |
| | -0.532417968 | 0.594436544 | 0.693253809 |
| | 5.085878278 | 3.66E-07 | 1.83E-06 |
| | -0.224124926 | 0.822660097 | 0.875661588 |
| | -1.620407526 | 0.105144764 | 0.173611284 |
| | 0.198308907 | 0.842803383 | NA |
| | 0.192809521 | 0.847108147 | NA |
| | -0.626555672 | 0.530950533 | 0.635657524 |
| NA | NA | NA | |
| | -1.654708197 | 0.097983703 | 0.163786036 |
| | -2.248231136 | 0.024561455 | 0.049916631 |
| | -1.679331028 | 0.093087547 | 0.157006827 |
| | -2.523725224 | 0.011611865 | 0.026022472 |
| | -1.794079916 | 0.072800414 | 0.127337132 |
| NA | NA | NA | |
| NA | NA | NA | |
| NA | NA | NA | |
| NA | NA | NA | |
| | 0.192809521 | 0.847108147 | NA |
| | -0.065388772 | 0.947864464 | NA |
| NA | NA | NA | |
| NA | NA | NA | |
| | -0.841079153 | 0.400303593 | 0.511987682 |
| | -1.249685079 | 0.21141461 | NA |
| NA | NA | NA | |
| NA | NA | NA | |
| | 0.389257077 | 0.697085985 | 0.776919682 |

| | | |
|---|---|---|
| 0.192809521 | 0.847108147 | NA |
| NA | NA | NA |
| NA | NA | NA |
| 0.94176285 | 0.346314069 | NA |
| NA | NA | NA |
| NA | NA | NA |
| NA | NA | NA |
| 9.070815545 | 1.18E-19 | 2.05E-18 |
| 1.783499809 | 0.074504979 | 0.129892013 |
| 2.635741383 | 0.008395369 | 0.019466057 |
| NA | NA | NA |
| 0.063705575 | 0.949204665 | NA |
| -1.795672791 | 0.072546572 | 0.126972987 |
| -0.8907702 | 0.373052465 | 0.48589807 |
| 1.057130215 | 0.290452163 | 0.398051777 |
| -0.687787622 | 0.491586532 | 0.598987971 |
| -1.847128073 | 0.064728581 | 0.11522341 |
| 2.575783411 | 0.010001327 | 0.022788617 |
| 12.80363089 | 1.56E-37 | 7.69E-36 |
| -7.220389742 | 5.18E-13 | 4.97E-12 |
| -2.822110108 | 0.004770879 | 0.011797581 |
| -2.780096737 | 0.005434271 | 0.013254706 |
| -3.492360424 | 0.000478772 | 0.001464583 |
| 6.711531683 | 1.93E-11 | 1.57E-10 |
| 0.265188363 | 0.790864359 | 0.851164385 |
| -0.363819397 | 0.715992877 | 0.792136745 |
| NA | NA | NA |
| 0.541192316 | 0.588375033 | 0.688082375 |
| -9.647271614 | 5.05E-22 | 1.05E-20 |
| 0.48990887 | 0.624198386 | 0.717967263 |
| 19.49977748 | 1.10E-84 | 1.93E-82 |
| 0.192809521 | 0.847108147 | NA |
| 13.83605616 | 1.54E-43 | 9.69E-42 |
| 3.7286597 | 0.000192501 | 0.000634487 |
| 1.412986582 | 0.15765967 | 0.243173621 |
| 9.784373055 | 1.31E-22 | 2.83E-21 |
| 17.28581463 | 6.02E-67 | 7.21E-65 |
| 0.192809521 | 0.847108147 | NA |
| NA | NA | NA |
| 0.30922356 | 0.757151473 | NA |
| 12.86349487 | 7.22E-38 | 3.63E-36 |
| 17.36359298 | 1.56E-67 | 1.89E-65 |
| 2.876478639 | 0.004021394 | 0.010104482 |
| -9.183086778 | 4.19E-20 | 7.52E-19 |
| -6.604220245 | 4.00E-11 | 3.17E-10 |
| 1.932226655 | 0.053331537 | 0.097844196 |
| -3.465281224 | 0.000529677 | 0.001608178 |

| | | |
|---|---|---|
| 7.554709312 | 4.20E-14 | 4.48E-13 |
| NA | NA | NA |
| -9.553408776 | 1.25E-21 | 2.53E-20 |
| -1.222241485 | 0.22161632 | 0.320935923 |
| 16.81099278 | 2.03E-63 | 2.22E-61 |
| 1.998665243 | 0.045644586 | 0.085560098 |
| 1.017507411 | 0.308912109 | 0.418809993 |
| 7.595765185 | 3.06E-14 | 3.31E-13 |
| -0.2146206 | 0.830063132 | 0.881143398 |
| 1.7997767 | 0.071895904 | 0.126054531 |
| -0.958424317 | 0.337848838 | 0.449018022 |
| NA | NA | NA |
| 0.535123262 | 0.592564625 | NA |
| -0.577366056 | 0.5636922 | 0.665717456 |
| -1.579651835 | 0.114186622 | 0.185916291 |
| 1.844017956 | 0.065180523 | 0.115912505 |
| -1.549185529 | 0.121337127 | 0.195775691 |
| -2.710202095 | 0.006724222 | 0.016007921 |
| 0.40109202 | 0.688352376 | 0.770144563 |
| -1.973759164 | 0.048409134 | 0.089987421 |
| 0.114428953 | 0.908897763 | 0.937166262 |
| 2.121341081 | 0.03389311 | 0.066060157 |
| -3.8605634 | 0.000113126 | 0.000387458 |
| -1.5984662 | 0.109939262 | 0.180054945 |
| NA | NA | NA |
| -2.750489334 | 0.005950633 | 0.014346402 |
| 2.223126806 | 0.026207251 | 0.05281967 |
| -0.12640191 | 0.89941379 | 0.929915617 |
| 1.581297776 | 0.113809968 | 0.185399512 |
| -0.336878923 | 0.736208175 | NA |
| -0.202153971 | 0.839796356 | 0.887695518 |
| -7.478864098 | 7.50E-14 | 7.81E-13 |
| NA | NA | NA |
| NA | NA | NA |
| -0.126642825 | 0.8992231 | NA |
| -1.699210008 | 0.089279621 | 0.151605761 |
| 0.690257537 | 0.490032246 | 0.597676012 |
| 0.460938956 | 0.644842402 | 0.734222141 |
| -2.591529732 | 0.009555028 | 0.021865229 |
| 6.672676422 | 2.51E-11 | 2.03E-10 |
| -1.353170465 | 0.176001179 | 0.266326673 |
| NA | NA | NA |
| -2.387227918 | 0.016975963 | 0.036316954 |
| 0.400325712 | 0.688916632 | 0.770387495 |
| -2.036293933 | 0.041720849 | 0.079093603 |
| -2.96791451 | 0.002998277 | 0.007753577 |
| 1.666478844 | 0.095618078 | 0.160550111 |

| | | |
|---|---|---|
| 0.272399335 | 0.785314984 | 0.846794021 |
| -0.344348066 | 0.730584541 | NA |
| 3.595173007 | 0.000324176 | 0.001024511 |
| -4.192637531 | 2.76E-05 | 0.00010495 |
| 2.922442399 | 0.003472979 | 0.008840502 |
| NA | NA | NA |
| -3.96294925 | 7.40E-05 | 0.000262199 |
| -3.068191088 | 0.002153589 | 0.005762703 |
| -4.281809796 | 1.85E-05 | 7.24E-05 |
| 1.227703516 | 0.219558286 | 0.318582255 |
| 1.325444961 | 0.185023613 | 0.277396887 |
| -3.65222444 | 0.000259979 | 0.000835443 |
| -2.698532871 | 0.006964586 | 0.016503227 |
| -0.196421698 | 0.844280114 | NA |
| 12.11417199 | 8.89E-34 | 3.69E-32 |
| NA | NA | NA |
| NA | NA | NA |
| 10.42442338 | 1.92E-25 | 5.05E-24 |
| -12.35856633 | 4.38E-35 | 1.92E-33 |
| -2.821865694 | 0.004774517 | 0.011805409 |
| -5.123249897 | 3.00E-07 | 1.52E-06 |
| 3.502826865 | 0.000460349 | 0.00141242 |
| -4.949345604 | 7.45E-07 | 3.56E-06 |
| -2.920997258 | 0.003489129 | 0.008878904 |
| NA | NA | NA |
| 0.035800017 | 0.97144182 | 0.980293047 |
| 0.414990094 | 0.678149155 | NA |
| 6.701591049 | 2.06E-11 | 1.68E-10 |
| 0.679957662 | 0.496531269 | 0.603661243 |
| -0.325064265 | 0.745132434 | 0.814510648 |
| -0.065389369 | 0.947863988 | NA |
| -0.510747408 | 0.609527939 | 0.705959202 |
| -2.801475277 | 0.005086954 | 0.01249401 |
| -2.446867099 | 0.014410396 | 0.031424658 |
| -5.826987622 | 5.64E-09 | 3.54E-08 |
| -3.833673046 | 0.000126244 | 0.000428866 |
| 2.802576956 | 0.005069612 | 0.012458752 |
| 1.208751611 | 0.226758287 | 0.326925864 |
| -1.499967292 | 0.133622875 | 0.212125361 |
| 2.96085631 | 0.00306785 | 0.007911425 |
| 1.038064419 | 0.299240065 | 0.408195478 |
| NA | NA | NA |
| -6.261456613 | 3.81E-10 | 2.71E-09 |
| 0.460564545 | 0.645111054 | 0.734461209 |
| -0.90117702 | 0.367494207 | 0.479932171 |
| -2.594085209 | 0.009484298 | 0.021731232 |
| -9.052866735 | 1.39E-19 | 2.41E-18 |

| | | | |
|---|---|---|---|
| | -2.860411479 | 0.004230917 | 0.010575179 |
| | -3.653832059 | 0.000258355 | 0.00083076 |
| | 1.894519426 | 0.058156098 | 0.105298549 |
| | 1.134914965 | 0.256410949 | 0.361026386 |
| | -1.461823385 | 0.143789614 | 0.225462259 |
| | 0.434484364 | 0.663936747 | 0.750229641 |
| | -1.184616387 | 0.236169155 | 0.337963178 |
| | -1.918824209 | 0.055006585 | 0.100490097 |
| | 0.865280339 | 0.386884939 | NA |
| | -0.047648398 | 0.96199646 | 0.973823966 |
| | -2.522126903 | 0.011664762 | 0.026122304 |
| | 2.758041818 | 0.005814876 | 0.014062449 |
| | -0.656241848 | 0.511668527 | 0.617445918 |
| | -1.120965114 | 0.262302712 | NA |
| | -1.21135305 | 0.225760125 | 0.325827191 |
| | 2.110954311 | 0.034776239 | 0.067529035 |
| | -1.276042212 | 0.201940598 | 0.29813455 |
| | -3.065070732 | 0.002176185 | 0.00581695 |
| NA | NA | NA | |
| | -3.228498156 | 0.001244421 | 0.003507641 |
| | 0.488426707 | 0.625247629 | 0.718810773 |
| | -0.682639032 | 0.494834964 | 0.601920549 |
| | -1.403393199 | 0.16049962 | 0.246538881 |
| | 4.429923847 | 9.43E-06 | 3.86E-05 |
| | 9.879180116 | 5.13E-23 | 1.14E-21 |
| | 2.39627755 | 0.016562543 | 0.035563213 |
| | 1.546108908 | 0.122078268 | 0.196857302 |
| | -0.520804885 | 0.6025027 | NA |
| | -2.698045931 | 0.006974782 | 0.016525822 |
| | -3.118787516 | 0.001815969 | 0.004936863 |
| NA | NA | NA | |
| | 3.105858105 | 0.001897277 | 0.005134833 |
| | -0.116270297 | 0.907438325 | NA |
| | -0.113293121 | 0.909798171 | NA |
| | 0.907742315 | 0.364014382 | NA |
| | -9.879151682 | 5.13E-23 | 1.14E-21 |
| | -5.051329018 | 4.39E-07 | 2.17E-06 |
| | 1.102449325 | 0.270266378 | 0.376468776 |
| | 0.889934522 | 0.373501046 | 0.486254612 |
| | -7.009505338 | 2.39E-12 | 2.12E-11 |
| | -0.553037125 | 0.580237989 | 0.68100585 |
| | -0.065388772 | 0.947864464 | NA |
| NA | NA | NA | |
| | -3.185026727 | 0.001447406 | 0.004024504 |
| NA | NA | NA | |
| | -2.820104751 | 0.004800798 | 0.011866874 |
| | -1.890079474 | 0.058747332 | 0.10621538 |

| | | | |
|---|---|---|---|
| | -0.977978469 | 0.328084973 | 0.439072175 |
| | 1.667983395 | 0.095319023 | 0.160099925 |
| | -5.578125564 | 2.43E-08 | 1.42E-07 |
| | -1.283999542 | 0.199142116 | 0.294593804 |
| NA | NA | NA | |
| | 6.078673258 | 1.21E-09 | 8.18E-09 |
| | 4.794783369 | 1.63E-06 | 7.42E-06 |
| | 0.346721347 | 0.728800672 | NA |
| | 1.985504401 | 0.047088381 | 0.087904399 |
| | -4.819525352 | 1.44E-06 | 6.60E-06 |
| | -3.929430834 | 8.51E-05 | 0.000298241 |
| | 8.058269569 | 7.74E-16 | 9.74E-15 |
| | -3.023261611 | 0.00250066 | 0.006574738 |
| | 2.584114427 | 0.00976294 | 0.022296203 |
| | 0.254852059 | 0.798837365 | 0.857083426 |
| | -0.821314759 | 0.411467003 | 0.523141899 |
| | 6.554837557 | 5.57E-11 | 4.35E-10 |
| NA | NA | NA | |
| | 0.063708285 | 0.949202508 | NA |
| | 0.774632593 | 0.438556792 | 0.548963618 |
| | 0.597443085 | 0.550211595 | 0.653497022 |
| | -3.744753012 | 0.000180571 | 0.000597528 |
| | -1.960226111 | 0.049969368 | 0.092537236 |
| | 8.475065178 | 2.35E-17 | 3.37E-16 |
| | 1.707661775 | 0.087699119 | 0.14935762 |
| | -0.528809872 | 0.596937349 | NA |
| | 0.499329681 | 0.617547149 | 0.712368141 |
| | 0.209060125 | 0.834401302 | 0.883953554 |
| | 1.551740607 | 0.120724302 | 0.194912552 |
| | -1.33577632 | 0.181622395 | 0.273181212 |
| NA | NA | NA | |
| | 7.787890776 | 6.81E-15 | 7.89E-14 |
| | -0.456417817 | 0.648089556 | 0.736980673 |
| | 7.676720081 | 1.63E-14 | 1.82E-13 |
| | 0.192809521 | 0.847108147 | NA |
| | -12.30156037 | 8.88E-35 | 3.83E-33 |
| | -0.924505067 | 0.355223415 | NA |
| | 2.050953052 | 0.040271519 | 0.076665357 |
| | -0.316642621 | 0.751514792 | 0.819768815 |
| | -7.472562321 | 7.86E-14 | 8.18E-13 |
| NA | NA | NA | |
| | 1.272136484 | 0.20332463 | 0.299665794 |
| NA | NA | NA | |
| | 1.119848225 | 0.262778445 | NA |
| | 1.326529529 | 0.184664361 | 0.276957814 |
| NA | NA | NA | |
| NA | NA | NA | |

| | | |
|---|---|---|
| -0.194488217 | 0.845793621 | NA |
| NA | NA | NA |
| -0.01905078 | 0.984800596 | 0.989346134 |
| -0.065389369 | 0.947863988 | NA |
| NA | NA | NA |
| 1.004973488 | 0.314909616 | NA |
| 0.286314326 | 0.774637387 | NA |
| 0.063708285 | 0.949202508 | NA |
| NA | NA | NA |
| 0.192809521 | 0.847108147 | NA |
| NA | NA | NA |
| -0.698178435 | 0.485065611 | 0.593440161 |
| NA | NA | NA |
| -1.272135508 | 0.203324977 | 0.299665794 |
| 0.359401919 | 0.719294439 | NA |
| 1.001076952 | 0.316789607 | 0.427314284 |
| 0.48081089 | 0.63065091 | 0.722798309 |
| -5.227688189 | 1.72E-07 | 8.97E-07 |
| -5.143198257 | 2.70E-07 | 1.38E-06 |
| -1.282186998 | 0.199777056 | NA |
| 0.568091894 | 0.569972572 | NA |
| -2.195329187 | 0.028139993 | 0.056151238 |
| -4.015673217 | 5.93E-05 | 0.00021299 |
| 0.081405154 | 0.93511975 | NA |
| 5.885013423 | 3.98E-09 | 2.54E-08 |
| 4.670050737 | 3.01E-06 | 1.32E-05 |
| NA | NA | NA |
| 0.063708285 | 0.949202508 | NA |
| -3.833202361 | 0.000126486 | 0.000429571 |
| -0.706398212 | 0.479940532 | 0.588638108 |
| -2.559133362 | 0.010493348 | 0.023801404 |
| -0.482274616 | 0.629610877 | 0.72220156 |
| NA | NA | NA |
| -2.489674597 | 0.01278601 | 0.028316566 |
| -0.06663263 | 0.946874169 | NA |
| -1.426807646 | 0.153635347 | 0.23812575 |
| 3.396849961 | 0.000681663 | 0.002027574 |
| 4.932076729 | 8.14E-07 | 3.87E-06 |
| 4.108691956 | 3.98E-05 | 0.000147288 |
| 3.865140676 | 0.000111025 | 0.00038094 |
| NA | NA | NA |
| 0.504645611 | 0.613807768 | 0.709504408 |
| -1.974426431 | 0.048333275 | 0.08986643 |
| -2.0396336 | 0.041386833 | 0.078525806 |
| -4.054124986 | 5.03E-05 | 0.000183343 |
| 2.646488761 | 0.008133221 | 0.018901998 |
| -0.703758378 | 0.481583258 | 0.590160536 |

| | | | |
|---|---|---|---|
| | 0.477154938 | 0.633251796 | NA |
| | -3.551162679 | 0.000383533 | 0.001194753 |
| | -1.734513602 | 0.082826994 | 0.142250561 |
| | -0.994599838 | 0.319930926 | 0.430646469 |
| | 3.538119733 | 0.000402987 | 0.00125084 |
| | 5.307374513 | 1.11E-07 | 5.96E-07 |
| | 0.048264384 | 0.961505539 | 0.97360651 |
| NA | NA | NA | |
| | 3.458685487 | 0.000542818 | 0.001644088 |
| | -6.378001183 | 1.79E-10 | 1.32E-09 |
| | -2.135394841 | 0.032728772 | NA |
| | 2.487102255 | 0.012878838 | 0.028501972 |
| | 1.735407544 | 0.082668645 | 0.14203708 |
| | -2.875050281 | 0.004039632 | 0.01014773 |
| | -2.659779621 | 0.00781918 | 0.018251801 |
| | -2.997344086 | 0.002723431 | 0.007102999 |
| | -1.5246204 | 0.127353806 | 0.204062849 |
| | 10.55753671 | 4.69E-26 | 1.29E-24 |
| | 2.627605448 | 0.008598819 | 0.01989356 |
| | 6.43608294 | 1.23E-10 | 9.23E-10 |
| | 5.456884689 | 4.85E-08 | 2.73E-07 |
| | -0.154695625 | 0.877061279 | 0.914145992 |
| | -3.061899005 | 0.002199376 | 0.005870163 |
| | 1.430928863 | 0.152450604 | 0.23662648 |
| | 1.459914616 | 0.144313542 | 0.226089092 |
| | -0.065388772 | 0.947864464 | NA |
| | 4.682219221 | 2.84E-06 | 1.25E-05 |
| | -0.203758046 | 0.838542579 | 0.886984776 |
| | -0.223174312 | 0.823399844 | 0.876225419 |
| | 4.518158772 | 6.24E-06 | 2.62E-05 |
| | -0.204619041 | 0.837869777 | 0.88657284 |
| | -2.80103483 | 0.005093902 | 0.012507393 |
| | -2.056671647 | 0.039717821 | 0.075749548 |
| NA | NA | NA | |
| | -0.957056684 | 0.338538645 | 0.449857671 |
| | 0.508884898 | 0.610832906 | 0.706836528 |
| | -4.517692922 | 6.25E-06 | 2.62E-05 |
| | -3.012957768 | 0.002587149 | 0.006784937 |
| | -2.706323557 | 0.006803272 | 0.016166895 |
| | -2.806206189 | 0.00501286 | 0.012333815 |
| | -4.697496451 | 2.63E-06 | 1.17E-05 |
| | -1.063236552 | 0.28767469 | 0.395089026 |
| | -6.106215721 | 1.02E-09 | 6.95E-09 |
| NA | NA | NA | |
| | -3.558365141 | 0.00037317 | 0.001165226 |
| | -4.227722607 | 2.36E-05 | 9.07E-05 |
| | 0.063710673 | 0.949200606 | NA |

| | | |
|---|---|---|
| -2.751227625 | 0.005937237 | 0.014319623 |
| -3.532860494 | 0.000411089 | 0.001272989 |
| -3.236557374 | 0.001209809 | 0.003421254 |
| -5.148365571 | 2.63E-07 | 1.34E-06 |
| NA | NA | NA |
| -4.410242865 | 1.03E-05 | 4.21E-05 |
| -3.228509562 | 0.001244371 | 0.003507641 |
| -4.358621918 | 1.31E-05 | 5.23E-05 |
| NA | NA | NA |
| -3.557174437 | 0.000374865 | 0.001169805 |
| -3.96826465 | 7.24E-05 | 0.00025682 |
| -4.612687816 | 3.97E-06 | 1.72E-05 |
| -5.775329616 | 7.68E-09 | 4.75E-08 |
| -4.834259682 | 1.34E-06 | 6.15E-06 |
| -4.672595626 | 2.97E-06 | 1.31E-05 |
| -6.38646586 | 1.70E-10 | 1.25E-09 |
| -4.767345542 | 1.87E-06 | 8.44E-06 |
| -5.996056006 | 2.02E-09 | 1.33E-08 |
| -3.615651321 | 0.000299593 | 0.000952715 |
| -5.120332622 | 3.05E-07 | 1.54E-06 |
| -5.099639778 | 3.40E-07 | 1.71E-06 |
| -3.741465762 | 0.00018295 | 0.00060468 |
| -0.065390563 | 0.947863038 | NA |
| 0.06371127 | 0.949200131 | NA |
| NA | NA | NA |
| NA | NA | NA |
| NA | NA | NA |
| 1.328770243 | 0.183923782 | 0.276062142 |
| NA | NA | NA |
| 0.192810118 | 0.847107679 | NA |
| -1.754702859 | 0.079310147 | 0.137047496 |
| NA | NA | NA |
| 0.162733451 | 0.870728308 | 0.909954889 |
| -2.017350332 | 0.043658967 | 0.082343752 |
| -5.668225657 | 1.44E-08 | 8.65E-08 |
| 5.558006118 | 2.73E-08 | 1.58E-07 |
| 2.519097851 | 0.011765595 | 0.026326914 |
| -1.121756351 | 0.26196605 | 0.367216084 |
| -1.014409701 | 0.310387297 | 0.420491126 |
| 9.428260459 | 4.17E-21 | 8.08E-20 |
| -3.425465565 | 0.000613747 | 0.001840235 |
| -3.217793731 | 0.001291807 | 0.003628133 |
| -1.827179673 | 0.067672759 | 0.119748969 |
| NA | NA | NA |
| -0.70686962 | 0.479647505 | 0.588364471 |
| NA | NA | NA |
| NA | NA | NA |

| | | |
|---|---|---|
| -0.543417685 | 0.586842252 | 0.686674843 |
| NA | NA | NA |
| -0.858639848 | 0.390539246 | 0.502710071 |
| 5.746422534 | 9.12E-09 | 5.59E-08 |
| -1.875890933 | 0.060670258 | 0.109058915 |
| -2.740222422 | 0.006139762 | 0.014759723 |
| 0.308721429 | 0.757533441 | 0.824394002 |
| -0.226210213 | 0.821037928 | NA |
| 3.002655356 | 0.002676353 | 0.006996253 |
| 14.60247068 | 2.71E-48 | 2.01E-46 |
| 1.875524419 | 0.060720613 | 0.109120976 |
| -2.729561889 | 0.006341854 | 0.015186726 |
| 0.129455473 | 0.896997256 | 0.928212715 |
| 0.426620258 | 0.66965594 | 0.754410306 |
| -3.688034237 | 0.000225993 | 0.000735184 |
| 0.063708882 | 0.949202033 | NA |
| NA | NA | NA |
| NA | NA | NA |
| 0.192810118 | 0.847107679 | NA |
| -4.676670915 | 2.92E-06 | 1.28E-05 |
| 0.730023733 | 0.465375678 | 0.574716461 |
| 0.601662532 | 0.547398795 | 0.651021184 |
| -1.602815941 | 0.108975296 | 0.178745195 |
| NA | NA | NA |
| -1.141144849 | 0.25380965 | 0.357988028 |
| 0.430233564 | 0.667025749 | 0.752395425 |
| -0.99619982 | 0.319153067 | 0.42980745 |
| -11.92924606 | 8.33E-33 | 3.33E-31 |
| 1.502326229 | 0.133012878 | 0.211264188 |
| -0.622798224 | 0.533417126 | 0.637797038 |
| -1.725116504 | 0.084506479 | 0.144638348 |
| -1.116685264 | 0.264128921 | 0.36968535 |
| -3.461325083 | 0.000537523 | 0.001629639 |
| NA | NA | NA |
| 0.405682574 | 0.684975848 | 0.767214636 |
| 5.36520435 | 8.09E-08 | 4.41E-07 |
| -1.268802139 | 0.204511642 | 0.301024781 |
| NA | NA | NA |
| -4.036668563 | 5.42E-05 | 0.000196527 |
| 0.06371127 | 0.949200131 | NA |
| 0.534257812 | 0.593163176 | NA |
| -0.43418633 | 0.664153139 | NA |
| 0.622921976 | 0.533335796 | 0.637750572 |
| 1.201374527 | 0.229605953 | 0.330175629 |
| NA | NA | NA |
| 5.804920282 | 6.44E-09 | 4.01E-08 |
| -1.005328793 | 0.314738555 | 0.425304002 |

| | | |
|---|---|---|
| NA | NA | NA |
| -1.095765398 | 0.273181455 | 0.379834593 |
| 0.063708882 | 0.949202033 | NA |
| -5.226430211 | 1.73E-07 | 9.02E-07 |
| 0.192636473 | 0.847243678 | NA |
| -0.122609988 | 0.902415945 | 0.932166676 |
| 0.481275713 | 0.630320556 | NA |
| 6.617459841 | 3.65E-11 | 2.90E-10 |
| 0.51253107 | 0.608279381 | 0.704806634 |
| 6.230435707 | 4.65E-10 | 3.27E-09 |
| NA | NA | NA |
| -0.543234204 | 0.586968559 | 0.686790531 |
| -1.297305742 | 0.194526009 | 0.288874875 |
| -4.873136859 | 1.10E-06 | 5.12E-06 |
| NA | NA | NA |
| 0.432801713 | 0.665158829 | 0.751203604 |
| -0.151511208 | 0.87957247 | NA |
| 8.878362766 | 6.79E-19 | 1.11E-17 |
| 0.535208666 | 0.592505575 | 0.691652576 |
| 0.825043038 | 0.409347157 | 0.521257143 |
| 2.209250401 | 0.027157229 | 0.054440609 |
| -0.674855156 | 0.499767794 | 0.606800285 |
| 6.770468062 | 1.28E-11 | 1.07E-10 |
| 1.051118972 | 0.293203951 | 0.401251772 |
| 5.494237122 | 3.92E-08 | 2.23E-07 |
| 18.14952081 | 1.30E-73 | 1.81E-71 |
| NA | NA | NA |
| -0.109314316 | 0.912953192 | NA |
| NA | NA | NA |
| 0.457969203 | 0.646974567 | 0.735946801 |
| 0.426022964 | 0.670091113 | 0.754767924 |
| -1.219145675 | 0.222788907 | 0.322316995 |
| 3.841170877 | 0.000122449 | 0.000416822 |
| -5.511949137 | 3.55E-08 | 2.03E-07 |
| 3.329577995 | 0.000869777 | 0.002537356 |
| NA | NA | NA |
| 3.431506268 | 0.000600239 | 0.00180466 |
| -3.76213014 | 0.000168472 | 0.000559934 |
| -1.044410449 | 0.296295531 | 0.404774432 |
| -2.769739902 | 0.005610107 | 0.013630459 |
| -1.837244474 | 0.066173813 | 0.117520259 |
| -2.188533073 | 0.028630795 | 0.057033728 |
| 0.063705357 | 0.949204839 | NA |
| -5.115561218 | 3.13E-07 | 1.58E-06 |
| 0.150224509 | 0.88058749 | NA |
| NA | NA | NA |
| -0.670477716 | 0.502553307 | 0.609295678 |

| | | |
|---|---|---|
| -4.023492297 | 5.73E-05 | 0.000206869 |
| 3.296265239 | 0.000979795 | 0.002829613 |
| 2.511393579 | 0.012025554 | 0.026812722 |
| 4.144811556 | 3.40E-05 | 0.000127433 |
| 13.83404545 | 1.59E-43 | 9.94E-42 |
| NA | NA | NA |
| -0.065389369 | 0.947863988 | NA |
| NA | NA | NA |
| 5.73738723 | 9.61E-09 | 5.89E-08 |
| 3.442741705 | 0.000575849 | 0.001737194 |
| 13.86598109 | 1.02E-43 | 6.49E-42 |
| 23.28920923 | 5.70E-120 | 1.83E-117 |
| 0.063708285 | 0.949202508 | NA |
| 0.192810118 | 0.847107679 | NA |
| -0.073980037 | 0.94102627 | NA |
| 2.363220725 | 0.018116876 | 0.038369894 |
| -2.366198399 | 0.017971811 | 0.038140955 |
| -4.12653764 | 3.68E-05 | 0.000137168 |
| 2.179933542 | 0.029262388 | 0.058116129 |
| -0.069025188 | 0.944969571 | 0.962531718 |
| 1.726595845 | 0.084240273 | 0.144297986 |
| 2.284314786 | 0.022353038 | 0.046056678 |
| NA | NA | NA |
| 1.405211288 | 0.159958458 | 0.245916112 |
| 7.53159419 | 5.01E-14 | 5.31E-13 |
| 5.506892018 | 3.65E-08 | 2.09E-07 |
| -0.254192019 | 0.799347216 | NA |
| 7.125005485 | 1.04E-12 | 9.65E-12 |
| -0.961587514 | 0.336256846 | 0.447543808 |
| NA | NA | NA |
| -0.652681164 | 0.513961848 | NA |
| 1.691648738 | 0.090712968 | 0.153611483 |
| 0.509666859 | 0.610284874 | 0.706443621 |
| -3.198704459 | 0.001380466 | 0.003853358 |
| -2.439963529 | 0.014688745 | 0.031975933 |
| 5.228425266 | 1.71E-07 | 8.93E-07 |
| NA | NA | NA |
| 0.234484403 | 0.814608941 | 0.869717701 |
| -7.46934102 | 8.06E-14 | 8.37E-13 |
| -2.205096459 | 0.027447327 | 0.05497812 |
| -0.732379976 | 0.463936677 | 0.573363701 |
| 0.371911183 | 0.709958978 | 0.787410193 |
| 2.730065363 | 0.006332177 | 0.015165476 |
| -1.728079321 | 0.083974004 | 0.143914367 |
| -0.753379671 | 0.451221796 | 0.561140833 |
| 0.495420334 | 0.620303445 | 0.714735314 |
| -1.563663779 | 0.117896548 | 0.191012402 |

| | | |
|---|---|---|
| -0.430667756 | 0.666709967 | 0.752183673 |
| 0.063708285 | 0.949202508 | NA |
| -0.332878877 | 0.739225715 | 0.810070472 |
| -1.25155218 | 0.210733089 | 0.3083328 |
| 0.804561603 | 0.421072707 | NA |
| 6.093045539 | 1.11E-09 | 7.52E-09 |
| -5.306106327 | 1.12E-07 | 6.00E-07 |
| 6.636991103 | 3.20E-11 | 2.56E-10 |
| 2.567244889 | 0.010251019 | 0.023310893 |
| 0.10230301 | 0.918516166 | 0.944205497 |
| -2.968355331 | 0.00299398 | 0.007744442 |
| 4.832385054 | 1.35E-06 | 6.21E-06 |
| 3.682126843 | 0.000231296 | 0.000750596 |
| 2.051859566 | 0.040183312 | 0.07652654 |
| 2.813556446 | 0.004899678 | 0.012090204 |
| 0.202714101 | 0.839358501 | NA |
| NA | NA | NA |
| NA | NA | NA |
| -1.847873055 | 0.064620709 | 0.115055937 |
| -0.240664196 | 0.809815391 | NA |
| 0.116350212 | 0.907374992 | 0.935974996 |
| -2.435003832 | 0.014891632 | 0.032372551 |
| -0.280297679 | 0.779249131 | NA |
| -0.173607518 | 0.862173923 | 0.904262028 |
| NA | NA | NA |
| 0.063708285 | 0.949202508 | NA |
| NA | NA | NA |
| NA | NA | NA |
| -0.158526475 | 0.87404196 | 0.912275146 |
| NA | NA | NA |
| 1.167879619 | 0.242855326 | 0.345535972 |
| 0.351903664 | 0.724910511 | 0.79917204 |
| 2.734160978 | 0.006253946 | 0.015005394 |
| 6.590848185 | 4.37E-11 | 3.45E-10 |
| NA | NA | NA |
| NA | NA | NA |
| 0.063708882 | 0.949202033 | NA |
| -0.604834169 | 0.545289188 | 0.649097985 |
| 1.834534366 | 0.066574711 | 0.118113441 |
| 0.551341405 | 0.58139966 | 0.681932216 |
| 0.437567139 | 0.661700096 | 0.748242757 |
| NA | NA | NA |
| NA | NA | NA |
| NA | NA | NA |
| 1.234891019 | 0.216871066 | 0.31550612 |
| 0.059043162 | 0.95291773 | 0.967921872 |
| -0.319604259 | 0.749268345 | 0.817888647 |

| | | |
|---|---|---|
| 0.364066175 | 0.715808594 | 0.792031198 |
| NA | NA | NA |
| -2.259759114 | 0.023836205 | 0.048664044 |
| 0.550569255 | 0.58192899 | 0.682233378 |
| 0.063708285 | 0.949202508 | NA |
| 1.138600702 | 0.254869736 | 0.359199858 |
| 0.220005973 | 0.825866503 | 0.878103673 |
| 0.192810118 | 0.847107679 | NA |
| -1.759620743 | 0.078472132 | 0.135825145 |
| 1.077349383 | 0.281324208 | 0.388306216 |
| 0.522041206 | 0.601641641 | 0.699219495 |
| -0.236622612 | 0.812949577 | 0.868390573 |
| -1.69823014 | 0.089464334 | 0.151888509 |
| NA | NA | NA |
| 0.417818162 | 0.676080071 | NA |
| 2.49165397 | 0.012714984 | 0.028184206 |
| -3.495906973 | 0.000472453 | 0.001446899 |
| 0.324874675 | 0.745275925 | NA |
| 1.925703855 | 0.054141362 | 0.09915061 |
| -0.065389369 | 0.947863988 | NA |
| 0.651431127 | 0.514768223 | 0.620259405 |
| -6.764653694 | 1.34E-11 | 1.11E-10 |
| -3.962473697 | 7.42E-05 | 0.000262685 |
| -1.105380058 | 0.268994941 | 0.375094338 |
| 7.746852046 | 9.42E-15 | 1.07E-13 |
| -3.937124252 | 8.25E-05 | 0.000289613 |
| 0.192809521 | 0.847108147 | NA |
| 4.693498835 | 2.69E-06 | 1.19E-05 |
| -5.979001901 | 2.25E-09 | 1.47E-08 |
| NA | NA | NA |
| -2.292935385 | 0.021851725 | 0.04511289 |
| NA | NA | NA |
| 2.141747961 | 0.032213769 | 0.063207238 |
| -0.341061096 | 0.733057588 | 0.80538127 |
| 4.488984289 | 7.16E-06 | 2.98E-05 |
| 5.032490878 | 4.84E-07 | 2.38E-06 |
| -0.341976562 | 0.732368531 | 0.804881859 |
| 0.417230204 | 0.676510035 | NA |
| 6.937319084 | 4.00E-12 | 3.48E-11 |
| 0.086306247 | 0.931222973 | NA |
| -1.747175569 | 0.080606888 | 0.139000373 |
| NA | NA | NA |
| 8.245069904 | 1.65E-16 | 2.20E-15 |
| -3.109722862 | 0.00187263 | 0.005073817 |
| NA | NA | NA |
| 11.65166418 | 2.25E-31 | 8.37E-30 |
| 2.356218733 | 0.018462043 | 0.039011573 |

| | | | |
|---|---|---|---|
| | 0.192810118 | 0.847107679 | NA |
| | 2.984422799 | 0.002841139 | 0.007383834 |
| | 1.474298003 | 0.140401373 | 0.220966195 |
| | -0.693097398 | 0.48824843 | NA |
| | -3.496220366 | 0.000471899 | 0.001445378 |
| NA | NA | NA | |
| | 0.090789508 | 0.927659847 | 0.950559871 |
| | -3.059923366 | 0.002213936 | 0.005901982 |
| | 0.324874675 | 0.745275925 | NA |
| | 0.18638912 | 0.852139624 | 0.897234342 |
| | -3.530138283 | 0.000415343 | 0.001285364 |
| | 0.087156496 | 0.930547119 | NA |
| | 0.356033948 | 0.721815142 | NA |
| | 2.154482359 | 0.031202364 | 0.06143012 |
| | -0.016074329 | 0.987175093 | 0.991015625 |
| | 7.68801416 | 1.49E-14 | 1.68E-13 |
| | 9.633883637 | 5.75E-22 | 1.19E-20 |
| | -1.324424494 | 0.185362104 | NA |
| NA | NA | NA | |
| | 1.107836606 | 0.267932388 | 0.373828981 |
| | -1.10166022 | 0.270609417 | 0.376841756 |
| | 2.864153188 | 0.004181257 | 0.010466734 |
| | 0.213302193 | 0.831091263 | NA |
| | 1.137958853 | 0.255137664 | 0.359496495 |
| | -0.902652285 | 0.366710468 | 0.479158645 |
| | 3.664179076 | 0.000248133 | 0.000800151 |
| | -2.611230642 | 0.009021703 | 0.020770163 |
| | -3.613860937 | 0.000301671 | 0.00095847 |
| | 0.351284813 | 0.725374687 | 0.799442051 |
| | -4.122400441 | 3.75E-05 | 0.000139448 |
| | -0.552187811 | 0.580819684 | 0.68144355 |
| | 11.76062526 | 6.23E-32 | 2.39E-30 |
| NA | NA | NA | |
| NA | NA | NA | |
| | -2.637613762 | 0.008349162 | 0.019364299 |
| | 0.307956229 | 0.758115639 | 0.824848269 |
| NA | NA | NA | |
| NA | NA | NA | |
| | 0.192810118 | 0.847107679 | NA |
| NA | NA | NA | |
| NA | NA | NA | |
| NA | NA | NA | |
| NA | NA | NA | |
| | -0.491688119 | 0.622939838 | 0.716952613 |
| | 1.512512646 | 0.130403498 | 0.207937794 |
| | -5.309757467 | 1.10E-07 | 5.88E-07 |
| | -3.366944485 | 0.00076006 | 0.002241317 |

| | | |
|---|---|---|
| NA | NA | NA |
| -4.489190027 | 7.15E-06 | 2.98E-05 |
| NA | NA | NA |
| 0.177500643 | 0.85911516 | 0.902074522 |
| -3.162168058 | 0.001565992 | 0.004323014 |
| 0.405965056 | 0.684768276 | 0.767085122 |
| 2.316798292 | 0.020514722 | 0.042722647 |
| -6.572322846 | 4.95E-11 | 3.89E-10 |
| -4.655901871 | 3.23E-06 | 1.41E-05 |
| 8.797937159 | 1.39E-18 | 2.22E-17 |
| -1.327592357 | 0.184312811 | 0.276502425 |
| 0.072175577 | 0.942462181 | 0.960959104 |
| -1.291319591 | 0.19659288 | 0.291460036 |
| -0.011088672 | 0.991152701 | 0.993812927 |
| NA | NA | NA |
| 0.06371127 | 0.949200131 | NA |
| NA | NA | NA |
| 0.192810118 | 0.847107679 | NA |
| -0.985243151 | 0.324504649 | 0.435538172 |
| 0.009590238 | 0.992348215 | 0.994653044 |
| 0.063708285 | 0.949202508 | NA |
| NA | NA | NA |
| -0.414774944 | 0.678306664 | 0.761277724 |
| -2.434113011 | 0.014928333 | 0.032438249 |
| -2.57136569 | 0.01012983 | 0.02306045 |
| -6.276365538 | 3.47E-10 | 2.47E-09 |
| 1.981282717 | 0.047559577 | 0.0886716 |
| -0.499657899 | 0.617315982 | 0.712167264 |
| 0.599907394 | 0.548567954 | 0.652039885 |
| 0.013959426 | 0.988862351 | 0.992152349 |
| -1.021362399 | 0.307082773 | 0.416834282 |
| -3.663842688 | 0.000248459 | 0.0008011 |
| -1.220966998 | 0.222098515 | 0.32148713 |
| -1.571990076 | 0.115952853 | 0.188338735 |
| -3.922593459 | 8.76E-05 | 0.000306536 |
| -2.490590271 | 0.012753109 | 0.028253702 |
| 6.250695612 | 4.09E-10 | 2.89E-09 |
| NA | NA | NA |
| NA | NA | NA |
| -1.415024525 | 0.157061305 | 0.242535119 |
| 3.895370931 | 9.80E-05 | 0.000339582 |
| 0.192809521 | 0.847108147 | NA |
| -0.138154752 | 0.890118115 | 0.923088184 |
| -1.242503989 | 0.21405067 | 0.312237792 |
| 4.665296989 | 3.08E-06 | 1.35E-05 |
| -4.863239249 | 1.15E-06 | 5.37E-06 |
| NA | NA | NA |

| | | | |
|---|---|---|---|
| | -2.564535075 | 0.010331414 | 0.023468126 |
| | 3.02404669 | 0.002494179 | 0.006561147 |
| | -0.938065837 | 0.348210577 | 0.459881326 |
| | 2.615403496 | 0.00891221 | 0.020539012 |
| | -5.822951001 | 5.78E-09 | 3.63E-08 |
| | -0.814077129 | 0.415600767 | NA |
| | 0.192211027 | 0.847576909 | 0.893557831 |
| | -3.60742098 | 0.000309256 | 0.000980701 |
| | 7.568829616 | 3.77E-14 | 4.04E-13 |
| NA | NA | NA | |
| | 1.137580116 | 0.255295854 | 0.359678893 |
| | -0.20198799 | 0.839926114 | NA |
| | -4.850252072 | 1.23E-06 | 5.71E-06 |
| NA | NA | NA | |
| | 1.881153607 | 0.059951027 | 0.108009371 |
| | -8.167597001 | 3.15E-16 | 4.09E-15 |
| | -2.769823244 | 0.005608672 | 0.013628775 |
| | -2.961351186 | 0.003062925 | 0.007901288 |
| | 6.203351004 | 5.53E-10 | 3.86E-09 |
| | -5.886090481 | 3.95E-09 | 2.53E-08 |
| | 1.323353208 | 0.185717945 | 0.278221221 |
| | 0.357968732 | 0.720366714 | NA |
| NA | NA | NA | |
| | 6.123361339 | 9.16E-10 | 6.27E-09 |
| | 0.145094608 | 0.884636175 | NA |
| | -3.070439205 | 0.002137442 | 0.005721944 |
| | -4.137130162 | 3.52E-05 | 0.000131498 |
| | -1.086218512 | 0.277382329 | 0.384279998 |
| | 6.4004767 | 1.55E-10 | 1.15E-09 |
| | 0.372737018 | 0.709344182 | NA |
| | 0.799900459 | 0.423768472 | 0.535324763 |
| NA | NA | NA | |
| NA | NA | NA | |
| NA | NA | NA | |
| | 0.063708285 | 0.949202508 | NA |
| | -0.065388772 | 0.947864464 | NA |
| | 1.99509468 | 0.046032554 | 0.086171179 |
| | 19.8550537 | 9.97E-88 | 1.88E-85 |
| | 2.454263133 | 0.014117364 | 0.030896005 |
| | 7.139130528 | 9.39E-13 | 8.74E-12 |
| | 6.321569674 | 2.59E-10 | 1.87E-09 |
| | 0.481655612 | 0.630050613 | 0.722432584 |
| | -0.369592759 | 0.711685947 | 0.788848943 |
| | 1.074529789 | 0.282585301 | 0.389592619 |
| | 1.25950324 | 0.207848621 | 0.304914487 |
| | 5.67055017 | 1.42E-08 | 8.54E-08 |
| NA | NA | NA | |

| | | |
|---|---|---|
| 2.337782739 | 0.019398521 | 0.040689264 |
| 1.191354651 | 0.233514388 | 0.334757453 |
| 7.095055018 | 1.29E-12 | 1.18E-11 |
| 0.192809521 | 0.847108147 | NA |
| 0.192809521 | 0.847108147 | NA |
| NA | NA | NA |
| -0.065389369 | 0.947863988 | NA |
| NA | NA | NA |
| NA | NA | NA |
| NA | NA | NA |
| NA | NA | NA |
| NA | NA | NA |
| NA | NA | NA |
| NA | NA | NA |
| 0.063708285 | 0.949202508 | NA |
| -1.307113668 | 0.19117411 | NA |
| NA | NA | NA |
| NA | NA | NA |
| NA | NA | NA |
| 0.93458563 | 0.350001888 | NA |
| NA | NA | NA |
| NA | NA | NA |
| NA | NA | NA |
| NA | NA | NA |
| 0.600376828 | 0.548255127 | NA |
| 0.192810118 | 0.847107679 | NA |
| 0.785124835 | 0.43238038 | 0.54313364 |
| -0.247794566 | 0.80429336 | 0.861386453 |
| 0.64887295 | 0.5164205 | 0.621681731 |
| -0.650050853 | 0.515659374 | 0.621004337 |
| 1.040696414 | 0.298016468 | 0.406814583 |
| -3.575047826 | 0.000350164 | 0.001100233 |
| -2.762141851 | 0.005742352 | 0.013907216 |
| -6.728298132 | 1.72E-11 | 1.41E-10 |
| 0.778024318 | 0.436554677 | 0.546922447 |
| 0.192810118 | 0.847107679 | NA |
| NA | NA | NA |
| NA | NA | NA |
| NA | NA | NA |
| NA | NA | NA |
| NA | NA | NA |
| 3.18442549 | 0.001450416 | 0.004032425 |
| -5.425114001 | 5.79E-08 | 3.22E-07 |
| 0.063707194 | 0.949203376 | NA |
| NA | NA | NA |
| 6.300563719 | 2.97E-10 | 2.13E-09 |
| -0.065389369 | 0.947863988 | NA |

| | | |
|---|---|---|
| 1.684932938 | 0.092001511 | 0.155447402 |
| NA | NA | NA |
| 0.192809521 | 0.847108147 | NA |
| 7.136146629 | 9.60E-13 | 8.92E-12 |
| 0.592665016 | 0.553405358 | 0.656538373 |
| 3.162250437 | 0.001565549 | 0.004322268 |
| 2.082062135 | 0.037336795 | 0.071782777 |
| 2.264220463 | 0.023560559 | 0.048183673 |
| -0.065388772 | 0.947864464 | NA |
| -0.031958901 | 0.974504826 | 0.982354241 |
| NA | NA | NA |
| NA | NA | NA |
| 0.192809521 | 0.847108147 | NA |
| NA | NA | NA |
| NA | NA | NA |
| NA | NA | NA |
| NA | NA | NA |
| NA | NA | NA |
| -4.711503855 | 2.46E-06 | 1.09E-05 |
| -1.082760053 | 0.278914941 | 0.385959806 |
| -0.164898527 | 0.869023852 | 0.908856684 |
| 0.063708285 | 0.949202508 | NA |
| 0.273514743 | 0.784457561 | 0.846298554 |
| -2.756018939 | 0.005850961 | 0.014140568 |
| 0.197559764 | 0.843389518 | NA |
| -3.154646625 | 0.001606926 | 0.00442869 |
| -3.28337866 | 0.001025708 | 0.002950295 |
| 7.722921493 | 1.14E-14 | 1.29E-13 |
| 0.31685483 | 0.751353758 | NA |
| NA | NA | NA |
| 9.198360053 | 3.63E-20 | 6.54E-19 |
| -1.014521285 | 0.310334078 | 0.420441785 |
| 1.353585502 | 0.175868655 | 0.266174412 |
| -3.364921958 | 0.000765653 | 0.002255685 |
| -7.470350434 | 8.00E-14 | 8.31E-13 |
| 0.847286729 | 0.396835322 | 0.508721542 |
| -2.582421629 | 0.009810964 | 0.022395662 |
| 6.100359474 | 1.06E-09 | 7.20E-09 |
| 0.273112031 | 0.784767099 | NA |
| 0.145139656 | 0.884600609 | 0.919140546 |
| NA | NA | NA |
| -0.065388772 | 0.947864464 | NA |
| NA | NA | NA |
| NA | NA | NA |
| NA | NA | NA |
| NA | NA | NA |
| 0.192809521 | 0.847108147 | NA |

| | | |
|---|---|---|
| -1.675814117 | 0.093774608 | 0.157963419 |
| NA | NA | NA |
| 0.451685082 | 0.651495867 | NA |
| 1.418665166 | 0.155996657 | 0.241173337 |
| 0.346721347 | 0.728800672 | NA |
| 0.14024788 | 0.888464143 | NA |
| 2.47927787 | 0.01316487 | 0.029063036 |
| -1.018236215 | 0.308565713 | 0.418431021 |
| 2.454403673 | 0.014111847 | 0.030886632 |
| 1.172990788 | 0.2407995 | 0.343176649 |
| -1.431648177 | 0.152244534 | 0.23640925 |
| -0.983735247 | 0.325245705 | 0.436275455 |
| 3.664267414 | 0.000248048 | 0.000799978 |
| NA | NA | NA |
| 0.01002183 | 0.99200387 | 0.99440575 |
| 0.131109294 | 0.895688849 | NA |
| 0.800821424 | 0.423235034 | 0.534893861 |
| -1.496699287 | 0.134471522 | 0.213126187 |
| -0.185829348 | 0.852578589 | NA |
| NA | NA | NA |
| NA | NA | NA |
| 14.22727807 | 6.21E-46 | 4.28E-44 |
| NA | NA | NA |
| 1.766126737 | 0.077374589 | 0.134212471 |
| -0.238029565 | 0.811858164 | 0.867520912 |
| -0.116616534 | 0.907163935 | NA |
| 0.303852939 | 0.761239933 | NA |
| 4.223579648 | 2.40E-05 | 9.22E-05 |
| NA | NA | NA |
| 0.063708882 | 0.949202033 | NA |
| 1.588050697 | 0.112274876 | 0.183263386 |
| 1.920440908 | 0.054802231 | 0.10016063 |
| -2.183880025 | 0.028971065 | 0.057592388 |
| -0.905768978 | 0.365058144 | 0.4775481 |
| 0.510600558 | 0.609630785 | 0.705980316 |
| 2.21419421 | 0.026815422 | 0.053854611 |
| 1.961974359 | 0.04976547 | 0.092227877 |
| 2.472580529 | 0.013414149 | 0.029550796 |
| -2.259515008 | 0.023851368 | 0.04869095 |
| NA | NA | NA |
| 0.91097766 | 0.362307145 | NA |
| NA | NA | NA |
| NA | NA | NA |
| 0.369227111 | 0.711958449 | NA |
| NA | NA | NA |
| 0.385991334 | 0.699503087 | 0.778954171 |
| NA | NA | NA |

| | | | |
|---|---|---|---|
| NA | NA | NA | |
| | -10.37437516 | 3.24E-25 | 8.41E-24 |
| | -2.874384594 | 0.004048157 | 0.010167209 |
| NA | NA | NA | |
| | -2.805978582 | 0.005016403 | 0.012338891 |
| | -0.785682844 | 0.432053317 | 0.54283177 |
| | -1.496511573 | 0.134520394 | 0.2131851 |
| | -0.723383556 | 0.469444271 | 0.578617282 |
| | -1.299651505 | 0.193720438 | 0.287969033 |
| | 0.063708285 | 0.949202508 | NA |
| | 0.067649016 | 0.946065036 | NA |
| | -1.167344594 | 0.243071237 | 0.345744903 |
| | -2.668108774 | 0.007627956 | 0.017868763 |
| | -0.674783432 | 0.499813368 | 0.606826183 |
| | -3.802896484 | 0.000143014 | 0.000480941 |
| | -1.70299054 | 0.088569839 | 0.150563914 |
| | -1.566834047 | 0.117153487 | 0.189980679 |
| | -1.061069598 | 0.288658273 | 0.396048708 |
| | -4.37155685 | 1.23E-05 | 4.95E-05 |
| | -2.931909639 | 0.003368848 | 0.008601678 |
| | 0.192809521 | 0.847108147 | NA |
| | -2.2744157 | 0.022941004 | 0.047115458 |
| | 0.195888478 | 0.844697456 | 0.891460865 |
| NA | NA | NA | |
| | -5.64559776 | 1.65E-08 | 9.81E-08 |
| | 1.182259508 | 0.237102747 | NA |
| | 4.952328677 | 7.33E-07 | 3.51E-06 |
| | 0.397517277 | 0.690986048 | NA |
| NA | NA | NA | |
| NA | NA | NA | |
| | -1.143443481 | 0.252854508 | 0.356902292 |
| | -1.637622129 | 0.101500544 | 0.168606226 |
| | 0.201808052 | 0.840066787 | NA |
| | 0.286967122 | 0.774137495 | NA |
| | 1.709956565 | 0.087273905 | NA |
| | 0.063708285 | 0.949202508 | NA |
| | 7.587814699 | 3.25E-14 | 3.51E-13 |
| NA | NA | NA | |
| | -0.069988098 | 0.944203133 | NA |
| | -0.194487022 | 0.845794556 | NA |
| NA | NA | NA | |
| NA | NA | NA | |
| NA | NA | NA | |
| | 0.192809521 | 0.847108147 | NA |
| | -0.55290661 | 0.580327361 | NA |
| | 0.120270825 | 0.904268614 | NA |
| NA | NA | NA | |

| | | | |
|---|---|---|---|
| NA | NA | NA | |
| | 0.192809521 | 0.847108147 | NA |
| NA | NA | NA | |
| | 16.25240069 | 2.15E-59 | 2.10E-57 |
| | 30.12951216 | 1.99E-199 | 1.42E-196 |
| | 0.195316213 | 0.845145406 | NA |
| | 22.46840308 | 8.46E-112 | 2.43E-109 |
| | 33.686523 | 9.11E-249 | 1.20E-245 |
| | 0.095261918 | 0.92410679 | 0.947850385 |
| | -0.947533762 | 0.343366861 | 0.454782121 |
| | -1.018766169 | 0.308313991 | 0.418180296 |
| | 0.258232509 | 0.79622747 | 0.855052925 |
| NA | NA | NA | |
| | -0.065389369 | 0.947863988 | NA |
| | 7.33270607 | 2.26E-13 | 2.24E-12 |
| NA | NA | NA | |
| | 7.794820612 | 6.45E-15 | 7.48E-14 |
| | 2.943321766 | 0.003247107 | 0.008323103 |
| NA | NA | NA | |
| NA | NA | NA | |
| NA | NA | NA | |
| | 8.573606978 | 1.00E-17 | 1.47E-16 |
| | -1.250701689 | 0.211043334 | 0.308724038 |
| | -4.055328252 | 5.01E-05 | 0.000182455 |
| | -1.191463809 | 0.233471556 | 0.334748683 |
| | 5.319879909 | 1.04E-07 | 5.58E-07 |
| NA | NA | NA | |
| NA | NA | NA | |
| | 2.507755207 | 0.012150081 | 0.027059037 |
| | -7.140235068 | 9.32E-13 | 8.68E-12 |
| | -3.647007075 | 0.000265313 | 0.000851163 |
| | -3.101156201 | 0.001927666 | 0.005210526 |
| | -4.003652711 | 6.24E-05 | 0.000223457 |
| | -3.21707997 | 0.001295025 | 0.003635948 |
| | -1.292056616 | 0.196337539 | 0.291164127 |
| | 2.358014665 | 0.018372967 | 0.038850669 |
| | 1.04705781 | 0.295072916 | 0.403356938 |
| | 3.811196834 | 0.000138296 | 0.000466515 |
| | 3.755496403 | 0.000172998 | 0.000573529 |
| | -9.140217459 | 6.23E-20 | 1.11E-18 |
| | 1.665342621 | 0.09584442 | 0.160811491 |
| | 0.074738084 | 0.940423106 | 0.959474599 |
| | 0.589459594 | 0.55555301 | NA |
| NA | NA | NA | |
| NA | NA | NA | |
| | 0.415507666 | 0.677770304 | NA |
| | 0.697614937 | 0.485418038 | NA |

| | | | |
|---|---|---|---|
| NA | NA | NA | |
| 2.388423004 | 0.016920853 | | 0.036226923 |
| -0.72685042 | 0.467317592 | NA | |
| NA | NA | NA | |
| 0.455033541 | 0.649085108 | | 0.737939656 |
| -4.499014617 | 6.83E-06 | | 2.85E-05 |
| 0.06371127 | 0.949200131 | NA | |
| NA | NA | NA | |
| -3.2887768 | 0.001006238 | | 0.002898956 |
| -2.79308299 | 0.00522083 | | 0.012785805 |
| -2.6356586 | 0.008397417 | | 0.019469003 |
| -2.526513777 | 0.011520087 | | 0.025849196 |
| NA | NA | NA | |
| -0.065388772 | 0.947864464 | NA | |
| -4.344058546 | 1.40E-05 | | 5.56E-05 |
| NA | NA | NA | |
| NA | NA | NA | |
| 0.599429216 | 0.548886699 | NA | |
| -4.302647715 | 1.69E-05 | | 6.63E-05 |
| NA | NA | NA | |
| 4.515390704 | 6.32E-06 | | 2.65E-05 |
| 4.086504778 | 4.38E-05 | | 0.000160744 |
| -0.076278806 | 0.939197287 | NA | |
| 2.795080924 | 0.005188673 | | 0.012713899 |
| -1.984216316 | 0.04723173 | | 0.088112858 |
| -1.54158182 | 0.123175246 | | 0.198345111 |
| 0.02904496 | 0.976828733 | | 0.983983547 |
| -4.136873175 | 3.52E-05 | | 0.000131606 |
| -1.000499407 | 0.317068884 | | 0.427587643 |
| NA | NA | NA | |
| 0.320051528 | 0.748929269 | NA | |
| -1.695434246 | 0.089993074 | | 0.152599867 |
| -1.715389538 | 0.086273832 | | 0.147192692 |
| -0.29606985 | 0.767176739 | NA | |
| -5.313728207 | 1.07E-07 | | 5.77E-07 |
| 1.265345171 | 0.205747621 | NA | |
| -3.665527506 | 0.000246829 | | 0.000796357 |
| -5.289201438 | 1.23E-07 | | 6.55E-07 |
| -1.118819143 | 0.263217303 | | 0.368660349 |
| NA | NA | NA | |
| -2.676596244 | 0.00743742 | | 0.017482966 |
| -3.719288847 | 0.000199784 | | 0.000656592 |
| 0.063708882 | 0.949202033 | NA | |
| -7.605450468 | 2.84E-14 | | 3.08E-13 |
| NA | NA | NA | |
| NA | NA | NA | |
| NA | NA | NA | |

| | | |
|---|---|---|
| 0.063708285 | 0.949202508 | NA |
| -5.22107573 | 1.78E-07 | 9.26E-07 |
| NA | NA | NA |
| 2.16779761 | 0.030174089 | 0.059642853 |
| -0.291563986 | 0.770620015 | NA |
| 0.083757469 | 0.933249264 | NA |
| 5.500360902 | 3.79E-08 | 2.16E-07 |
| 1.187806667 | 0.234909587 | 0.336372252 |
| 0.778734239 | 0.436136282 | 0.546567359 |
| 3.874408544 | 0.000106884 | 0.00036779 |
| -1.382361891 | 0.166860611 | 0.25468145 |
| -0.194487022 | 0.845794556 | NA |
| NA | NA | NA |
| -2.313595168 | 0.020689939 | 0.043030259 |
| -7.284235558 | 3.23E-13 | 3.16E-12 |
| -0.408186022 | 0.683137109 | 0.765771949 |
| 15.33226907 | 4.65E-53 | 3.81E-51 |
| NA | NA | NA |
| 0.074728106 | 0.940431045 | NA |
| 7.030358103 | 2.06E-12 | 1.84E-11 |
| NA | NA | NA |
| 1.511144248 | 0.130751703 | 0.208347029 |
| 1.567572339 | 0.116980972 | 0.189737799 |
| 10.82911568 | 2.51E-27 | 7.44E-26 |
| NA | NA | NA |
| -1.599219561 | 0.109771825 | 0.179839559 |
| 5.109877531 | 3.22E-07 | 1.62E-06 |
| 10.44566156 | 1.53E-25 | 4.08E-24 |
| NA | NA | NA |
| NA | NA | NA |
| 10.8414756 | 2.19E-27 | 6.54E-26 |
| 6.311205273 | 2.77E-10 | 1.99E-09 |
| -3.274265466 | 0.00105937 | 0.003036999 |
| NA | NA | NA |
| NA | NA | NA |
| 0.494646441 | 0.620849714 | 0.7151343 |
| -0.84923097 | 0.395752784 | 0.50769782 |
| -0.792394461 | 0.428130708 | 0.539392542 |
| 0.085057209 | 0.93221591 | 0.953900667 |
| 0.192810118 | 0.847107679 | NA |
| NA | NA | NA |
| -0.437911515 | 0.661450428 | 0.74806182 |
| 0.063708882 | 0.949202033 | NA |
| 0.478032609 | 0.632626996 | NA |
| NA | NA | NA |
| -1.536776074 | 0.124348154 | 0.199937965 |
| -3.419992831 | 0.000626228 | 0.001875387 |

| | | | |
|---|---|---|---|
| | -0.612177861 | 0.540420088 | 0.644589697 |
| | -0.118963346 | 0.905304395 | NA |
| NA | NA | NA | |
| | 0.063708285 | 0.949202508 | NA |
| | 0.413585237 | 0.679177887 | NA |
| | -2.101559331 | 0.035591896 | 0.06887245 |
| NA | NA | NA | |
| | -0.341145808 | 0.732993818 | NA |
| | -6.566029885 | 5.17E-11 | 4.06E-10 |
| | 2.991041224 | 0.00278028 | 0.007241457 |
| | -4.6937491 | 2.68E-06 | 1.19E-05 |
| | -0.994664166 | 0.319899628 | 0.430627498 |
| NA | NA | NA | |
| NA | NA | NA | |
| | 0.773807488 | 0.439044644 | 0.549272127 |
| | 1.134804583 | 0.256457205 | 0.361071203 |
| | 0.257873434 | 0.796504588 | 0.855132594 |
| | -1.013891592 | 0.310634484 | 0.420757678 |
| | 1.311670083 | 0.189631475 | 0.283067499 |
| | 0.717733306 | 0.472921743 | NA |
| NA | NA | NA | |
| | 0.063708882 | 0.949202033 | NA |
| NA | NA | NA | |
| NA | NA | NA | |
| NA | NA | NA | |
| NA | NA | NA | |
| NA | NA | NA | |
| NA | NA | NA | |
| NA | NA | NA | |
| | -0.065389369 | 0.947863988 | NA |
| NA | NA | NA | |
| NA | NA | NA | |
| NA | NA | NA | |
| | -7.286599481 | 3.18E-13 | 3.11E-12 |
| | -3.713386512 | 0.000204504 | 0.000671134 |
| | -4.689807977 | 2.73E-06 | 1.21E-05 |
| | -3.336563055 | 0.000848212 | 0.002478779 |
| | -1.844111992 | 0.065166821 | 0.115896371 |
| | -0.0822624 | 0.934438053 | NA |
| | -0.07082102 | 0.943540202 | NA |
| | -0.508784213 | 0.610903486 | 0.706865489 |
| NA | NA | NA | |
| | 0.694051146 | 0.487650135 | 0.59572858 |
| NA | NA | NA | |
| | 2.319933763 | 0.020344461 | 0.04241748 |
| | -5.45482698 | 4.90E-08 | 2.75E-07 |
| NA | NA | NA | |

| | | |
|---|---|---|
| 2.029675004 | 0.042389585 | 0.080195503 |
| -0.245931509 | 0.805735252 | 0.862524735 |
| -0.194488217 | 0.845793621 | NA |
| NA | NA | NA |
| NA | NA | NA |
| -0.065388772 | 0.947864464 | NA |
| -3.158069838 | 0.001588175 | 0.004378458 |
| -3.440874369 | 0.000579838 | 0.001747332 |
| NA | NA | NA |
| NA | NA | NA |
| NA | NA | NA |
| 1.41938604 | 0.1557865 | 0.240907951 |
| -0.84553712 | 0.397811017 | 0.50963301 |
| -4.925427154 | 8.42E-07 | 3.99E-06 |
| NA | NA | NA |
| -1.482028857 | 0.138332617 | 0.218218025 |
| -2.5375098 | 0.011164424 | 0.025157142 |
| -1.697356671 | 0.089629249 | 0.152106591 |
| NA | NA | NA |
| 1.249648053 | 0.211428141 | 0.309165906 |
| NA | NA | NA |
| 0.063708882 | 0.949202033 | NA |
| -0.694666619 | 0.487264252 | 0.595373407 |
| -0.737477851 | 0.460831808 | 0.570286767 |
| 0.380057477 | 0.70390275 | 0.782878731 |
| NA | NA | NA |
| -1.785921493 | 0.07411197 | 0.129285875 |
| -3.521175679 | 0.000429638 | 0.00132567 |
| -1.634164884 | 0.102224234 | 0.169621205 |
| -0.194488217 | 0.845793621 | NA |
| NA | NA | NA |
| NA | NA | NA |
| NA | NA | NA |
| NA | NA | NA |
| NA | NA | NA |
| -7.387735409 | 1.49E-13 | 1.51E-12 |
| 0.60917251 | 0.542410099 | NA |
| NA | NA | NA |
| NA | NA | NA |
| -4.484779291 | 7.30E-06 | 3.03E-05 |
| -2.485633562 | 0.012932105 | 0.028609739 |
| -0.321219895 | 0.748043756 | NA |
| NA | NA | NA |
| NA | NA | NA |
| NA | NA | NA |
| -4.761250869 | 1.92E-06 | 8.68E-06 |
| -1.23182143 | 0.218015796 | 0.316847985 |

| | | |
|---|---|---|
| NA | NA | NA |
| NA | NA | NA |
| 0.229694374 | 0.818329267 | 0.872349633 |
| -1.846389273 | 0.064835704 | 0.115381277 |
| -1.608041303 | 0.107826133 | 0.177173775 |
| -2.252722751 | 0.024276636 | 0.049422057 |
| 3.858980007 | 0.000113861 | 0.000389603 |
| -1.101765242 | 0.270563745 | 0.376799117 |
| 3.862044988 | 0.000112442 | 0.000385379 |
| 0.87763004 | 0.380144522 | 0.492725254 |
| -0.831253862 | 0.405830233 | 0.517680701 |
| 0.386046698 | 0.699462085 | NA |
| NA | NA | NA |
| 4.282917005 | 1.84E-05 | 7.21E-05 |
| -0.93464373 | 0.349971935 | 0.461722408 |
| -0.338500414 | 0.734986115 | 0.806731384 |
| -6.493469758 | 8.39E-11 | 6.44E-10 |
| -1.277422416 | 0.201453155 | 0.297520207 |
| -4.346148353 | 1.39E-05 | 5.52E-05 |
| NA | NA | NA |
| 0.31685483 | 0.751353758 | NA |
| NA | NA | NA |
| 0.159608838 | 0.873189216 | 0.911734837 |
| -1.520308667 | 0.128433417 | 0.205477048 |
| -5.026452659 | 5.00E-07 | 2.45E-06 |
| -8.556230387 | 1.17E-17 | 1.71E-16 |
| NA | NA | NA |
| 3.981027836 | 6.86E-05 | 0.000244137 |
| 1.128025489 | 0.259309155 | 0.364308076 |
| 0.535596484 | 0.592237459 | NA |
| 0.063708882 | 0.949202033 | NA |
| 8.32253638 | 8.61E-17 | 1.18E-15 |
| -0.704162733 | 0.481331436 | 0.590025263 |
| -6.417910507 | 1.38E-10 | 1.03E-09 |
| -0.209525922 | 0.834037702 | NA |
| -1.376336381 | 0.168717514 | 0.256967226 |
| 13.20187138 | 8.56E-40 | 4.65E-38 |
| 1.254646295 | 0.20960719 | 0.306972556 |
| -0.552520717 | 0.580591644 | 0.681271804 |
| -1.567336615 | 0.117036031 | 0.189814803 |
| NA | NA | NA |
| 0.063708882 | 0.949202033 | NA |
| 19.51709415 | 7.86E-85 | 1.39E-82 |
| 0.151516495 | 0.8795683 | 0.915447401 |
| -1.957100918 | 0.050335604 | 0.093146546 |
| -5.510987353 | 3.57E-08 | 2.04E-07 |
| 0.786767084 | 0.431418225 | 0.542333292 |

| | | |
|---|---|---|
| -2.043837203 | 0.040969633 | 0.077801124 |
| -8.353449726 | 6.63E-17 | 9.14E-16 |
| 0.063707688 | 0.949202983 | NA |
| -6.493144391 | 8.41E-11 | 6.45E-10 |
| -0.65988856 | 0.509325346 | 0.615330313 |
| -0.444225638 | 0.656879458 | NA |
| 0.118143671 | 0.905953821 | NA |
| 0.121488894 | 0.903303811 | 0.932894439 |
| NA | NA | NA |
| 0.601220849 | 0.5476929 | 0.651211455 |
| NA | NA | NA |
| NA | NA | NA |
| NA | NA | NA |
| NA | NA | NA |
| 0.063708285 | 0.949202508 | NA |
| -2.961203167 | 0.003064397 | 0.007903335 |
| NA | NA | NA |
| -0.065389369 | 0.947863988 | NA |
| 4.836070923 | 1.32E-06 | 6.10E-06 |
| NA | NA | NA |
| -2.173796064 | 0.029720454 | 0.058904345 |
| -0.531889487 | 0.594802539 | 0.693492992 |
| 5.802247416 | 6.54E-09 | 4.08E-08 |
| -1.320745604 | 0.186586203 | 0.279287923 |
| -0.478797714 | 0.632082543 | 0.723908764 |
| 6.104705431 | 1.03E-09 | 7.02E-09 |
| 0.063708882 | 0.949202033 | NA |
| 0.286881144 | 0.774203329 | 0.837654937 |
| 0.382131398 | 0.702363903 | NA |
| 6.227418171 | 4.74E-10 | 3.34E-09 |
| 0.192809521 | 0.847108147 | NA |
| NA | NA | NA |
| -10.57315241 | 3.97E-26 | 1.09E-24 |
| NA | NA | NA |
| NA | NA | NA |
| -1.551049374 | 0.120889852 | 0.195116908 |
| -2.011837581 | 0.044237061 | 0.083283463 |
| 0.063705268 | 0.94920491 | NA |
| 3.037699149 | 0.002383918 | 0.006304223 |
| 0.882570134 | 0.377468571 | 0.490145172 |
| 0.003603222 | 0.997125051 | 0.997999007 |
| 0.330493631 | 0.741027004 | NA |
| 0.407024242 | 0.683990184 | 0.766350685 |
| -0.579975479 | 0.561931154 | 0.664059597 |
| -3.634890557 | 0.000278099 | 0.000889333 |
| -0.065389369 | 0.947863988 | NA |
| -2.733142919 | 0.00627331 | 0.015045981 |

| | | |
|---|---|---|
| 5.42600779 | 5.76E-08 | 3.21E-07 |
| -2.846188325 | 0.004424602 | 0.0110142 |
| -2.683556359 | 0.00728437 | 0.017155417 |
| -2.162653321 | 0.030567854 | 0.060321143 |
| -2.218385675 | 0.026528547 | 0.053338751 |
| -3.428076177 | 0.000607875 | 0.001824791 |
| -1.974942108 | 0.048274718 | 0.089770891 |
| -1.659744465 | 0.09696587 | 0.162366586 |
| 13.65839821 | 1.80E-42 | 1.09E-40 |
| -0.999638788 | 0.317485345 | 0.42797599 |
| -0.777406389 | 0.436919044 | 0.547269363 |
| -0.419015372 | 0.675204898 | NA |
| 0.897070464 | 0.369681318 | 0.482310315 |
| 0.149860164 | 0.880874941 | NA |
| -1.237178259 | 0.216020911 | 0.31452524 |
| -0.065388772 | 0.947864464 | NA |
| NA | NA | NA |
| -5.299738089 | 1.16E-07 | 6.19E-07 |
| NA | NA | NA |
| -0.19448762 | 0.845794089 | NA |
| 3.474315708 | 0.000512158 | 0.001559527 |
| 0.192809521 | 0.847108147 | NA |
| -0.768987022 | 0.441901015 | 0.551852637 |
| 4.575392333 | 4.75E-06 | 2.03E-05 |
| NA | NA | NA |
| -2.226419969 | 0.025986064 | 0.052424533 |
| NA | NA | NA |
| NA | NA | NA |
| -1.527273697 | 0.126692967 | 0.203147019 |
| NA | NA | NA |
| -1.897350926 | 0.057781638 | 0.104752265 |
| NA | NA | NA |
| 0.063708882 | 0.949202033 | NA |
| -1.843034926 | 0.065323911 | 0.116150991 |
| 0.31685483 | 0.751353758 | NA |
| -5.180818971 | 2.21E-07 | 1.14E-06 |
| NA | NA | NA |
| -0.081538659 | 0.935013582 | 0.955850965 |
| 6.677959588 | 2.42E-11 | 1.96E-10 |
| 9.673863639 | 3.89E-22 | 8.11E-21 |
| 0.274339353 | 0.783823845 | NA |
| 0.784504912 | 0.432743901 | NA |
| 1.68997532 | 0.091032676 | 0.154060033 |
| 0.633411433 | 0.526465001 | 0.631465352 |
| 0.192810118 | 0.847107679 | NA |
| 2.599738566 | 0.009329481 | 0.021405894 |
| NA | NA | NA |

| | | |
|---|---|---|
| 6.755260695 | 1.43E-11 | 1.18E-10 |
| -0.992565548 | 0.320921721 | 0.431747954 |
| NA | NA | NA |
| NA | NA | NA |
| 2.238147678 | 0.025211426 | 0.051096518 |
| 1.674860137 | 0.093961677 | 0.158193365 |
| 0.192809521 | 0.847108147 | NA |
| NA | NA | NA |
| 0.898339184 | 0.369004748 | 0.481553121 |
| 21.8107004 | 1.84E-105 | 4.74E-103 |
| 0.041398385 | 0.9669783 | 0.977534078 |
| 1.324859153 | 0.185217871 | 0.277633183 |
| 0.193472986 | 0.84658856 | NA |
| 2.313527412 | 0.020693659 | 0.043030846 |
| -0.130341238 | 0.896296455 | 0.927867278 |
| 1.653126152 | 0.098305192 | 0.164246643 |
| -1.385715536 | 0.165833782 | 0.253361385 |
| 3.786764604 | 0.000152622 | 0.000511053 |
| -2.997655955 | 0.002720646 | 0.007097215 |
| -4.24048369 | 2.23E-05 | 8.60E-05 |
| 0.372190613 | 0.709750934 | NA |
| -0.13496936 | 0.8926361 | 0.925009587 |
| -0.042900054 | 0.965781206 | 0.97674668 |
| NA | NA | NA |
| -0.784069112 | 0.432999558 | 0.543720411 |
| NA | NA | NA |
| -0.345549558 | 0.72968126 | 0.802739185 |
| 0.535450952 | 0.592338065 | 0.691511533 |
| 0.739557738 | 0.459568393 | 0.568948308 |
| -1.603501882 | 0.108823894 | 0.178520256 |
| NA | NA | NA |
| NA | NA | NA |
| -0.121069055 | 0.90363634 | NA |
| -4.407204409 | 1.05E-05 | 4.25E-05 |
| -4.050556569 | 5.11E-05 | 0.000185891 |
| -0.065390563 | 0.947863038 | NA |
| 2.220008894 | 0.026418164 | 0.053151967 |
| 4.754869503 | 1.99E-06 | 8.95E-06 |
| 0.192809521 | 0.847108147 | NA |
| NA | NA | NA |
| 2.37321499 | 0.017633996 | 0.037529031 |
| 1.240447417 | 0.214809952 | 0.313126382 |
| 0.192808924 | 0.847108615 | NA |
| -1.407548497 | 0.159264807 | 0.245075503 |
| 2.656794243 | 0.007888757 | 0.0184005 |
| 2.130539691 | 0.033127083 | 0.064778975 |
| NA | NA | NA |

| | | |
|---|---|---|
| NA | NA | NA |
| -6.902046381 | 5.13E-12 | 4.42E-11 |
| NA | NA | NA |
| -4.381307438 | 1.18E-05 | 4.75E-05 |
| 29.60227293 | 1.40E-192 | 8.97E-190 |
| 0.098001612 | 0.921931014 | NA |
| 1.272225222 | 0.203293109 | 0.299657436 |
| -2.209423012 | 0.027145232 | 0.054420917 |
| -1.146660621 | 0.251521912 | 0.355341957 |
| -3.729259516 | 0.000192043 | 0.000633062 |
| NA | NA | NA |
| -0.868237387 | 0.385264391 | 0.497403812 |
| -0.038085406 | 0.969619588 | 0.978848846 |
| -2.73627069 | 0.006213989 | 0.014919534 |
| -1.367778779 | 0.171381332 | 0.260438027 |
| 7.803921849 | 6.00E-15 | 6.99E-14 |
| -5.197560712 | 2.02E-07 | 1.04E-06 |
| 2.882576599 | 0.003944372 | 0.009931359 |
| 4.740521508 | 2.13E-06 | 9.56E-06 |
| 0.192805387 | 0.847111385 | NA |
| -0.568593824 | 0.569631817 | NA |
| 6.289066879 | 3.19E-10 | 2.28E-09 |
| 31.99948607 | 1.11E-224 | 1.11E-221 |
| 10.80351698 | 3.31E-27 | 9.70E-26 |
| 0.748954178 | 0.453884824 | NA |
| -0.778690485 | 0.436162062 | 0.546567359 |
| 1.261553615 | 0.207109456 | NA |
| 0.075823031 | 0.939559893 | NA |
| -2.381042681 | 0.017263712 | 0.036867824 |
| 7.03644018 | 1.97E-12 | 1.77E-11 |
| 3.447493676 | 0.000565814 | 0.001709186 |
| 0.581938136 | 0.560608352 | 0.66283844 |
| -0.065389369 | 0.947863988 | NA |
| 5.300643227 | 1.15E-07 | 6.17E-07 |
| -1.021790591 | 0.306880023 | 0.416596539 |
| -2.201430927 | 0.027705532 | 0.05541992 |
| 1.079963774 | 0.280158312 | 0.387131021 |
| -0.19448762 | 0.845794089 | NA |
| -2.453345497 | 0.014153433 | 0.030953275 |
| -4.916582898 | 8.81E-07 | 4.17E-06 |
| -2.544108621 | 0.010955701 | 0.024715745 |
| -0.105003578 | 0.91637297 | 0.942621766 |
| NA | NA | NA |
| 0.06353712 | 0.949338802 | NA |
| 1.627193649 | 0.103695976 | 0.17162668 |
| -7.717476433 | 1.19E-14 | 1.34E-13 |
| -3.617338038 | 0.000297648 | 0.000946891 |

| | | |
|---|---|---|
| -4.671854048 | 2.98E-06 | 1.31E-05 |
| -4.12658907 | 3.68E-05 | 0.000137158 |
| -0.976914908 | 0.32861128 | 0.439590315 |
| -3.02751206 | 0.002465759 | 0.006493891 |
| -4.960442893 | 7.03E-07 | 3.38E-06 |
| -6.572702883 | 4.94E-11 | 3.89E-10 |
| 1.7468137 | 0.080669659 | 0.139088415 |
| 0.59948346 | 0.548850537 | 0.652282842 |
| -0.166048868 | 0.868118492 | 0.908154724 |
| 1.541506813 | 0.123193486 | 0.198348962 |
| NA | NA | NA |
| NA | NA | NA |
| NA | NA | NA |
| 0.823931015 | 0.409978756 | NA |
| 9.437192367 | 3.83E-21 | 7.45E-20 |
| NA | NA | NA |
| 1.992465441 | 0.046320013 | 0.086638016 |
| 3.00846344 | 0.002625724 | 0.006873235 |
| NA | NA | NA |
| NA | NA | NA |
| 2.478721464 | 0.013185423 | 0.029102251 |
| NA | NA | NA |
| NA | NA | NA |
| NA | NA | NA |
| NA | NA | NA |
| NA | NA | NA |
| NA | NA | NA |
| NA | NA | NA |
| NA | NA | NA |
| NA | NA | NA |
| 0.215104274 | 0.829686022 | NA |
| 3.5520941 | 0.000382178 | 0.001191124 |
| 0.969861752 | 0.332115407 | NA |
| 1.906423154 | 0.056595327 | 0.102949203 |
| 0.409607882 | 0.682093614 | NA |
| NA | NA | NA |
| NA | NA | NA |
| NA | NA | NA |
| NA | NA | NA |
| NA | NA | NA |
| NA | NA | NA |
| NA | NA | NA |
| NA | NA | NA |
| 5.413960261 | 6.16E-08 | 3.42E-07 |
| 3.09206117 | 0.001987719 | 0.00536185 |
| 4.944515034 | 7.63E-07 | 3.64E-06 |
| NA | NA | NA |

| | | |
|---|---|---|
| 3.569273074 | 0.000357973 | 0.001122675 |
| 2.068906587 | 0.038554853 | 0.073818403 |
| 0.390965141 | 0.695823003 | NA |
| 1.300972563 | 0.193267846 | 0.28744989 |
| 3.877090171 | 0.000105713 | 0.000363912 |
| NA | NA | NA |
| 2.856700887 | 0.00428069 | 0.010683585 |
| 0.996034934 | 0.319233172 | NA |
| NA | NA | NA |
| 0.192810118 | 0.847107679 | NA |
| NA | NA | NA |
| 0.192810118 | 0.847107679 | NA |
| NA | NA | NA |
| 0.192810118 | 0.847107679 | NA |
| -0.13388232 | 0.893495631 | NA |
| NA | NA | NA |
| NA | NA | NA |
| 3.247531122 | 0.001164109 | 0.003308736 |
| 0.192809521 | 0.847108147 | NA |
| 0.063708285 | 0.949202508 | NA |
| NA | NA | NA |
| NA | NA | NA |
| NA | NA | NA |
| NA | NA | NA |
| NA | NA | NA |
| NA | NA | NA |
| NA | NA | NA |
| -1.01515606 | 0.310031442 | 0.420077248 |
| NA | NA | NA |
| NA | NA | NA |
| -0.247713765 | 0.804355881 | NA |
| NA | NA | NA |
| NA | NA | NA |
| 0.192809521 | 0.847108147 | NA |
| 3.022988757 | 0.002502915 | 0.006579978 |
| 2.3659135 | 0.017985646 | 0.038167081 |
| 0.730766367 | 0.46492187 | NA |
| 1.770091311 | 0.076711931 | NA |
| 1.973609482 | 0.048426165 | NA |
| 1.131937225 | 0.257660827 | NA |
| 2.450989476 | 0.014246412 | 0.031121285 |
| 0.418180567 | 0.675815103 | NA |
| 8.480028523 | 2.25E-17 | 3.23E-16 |
| 1.046946512 | 0.295124248 | NA |
| 0.320071032 | 0.748914485 | NA |
| 0.653209897 | 0.513620971 | NA |
| -1.144730805 | 0.252320686 | 0.356369867 |

| | | | |
|---|---|---|---|
| | -0.065390563 | 0.947863038 | NA |
| | -0.091793473 | 0.92686213 | NA |
| | 1.346720935 | 0.178070127 | NA |
| | -1.372511267 | 0.169904334 | 0.258444575 |
| | 6.412364599 | 1.43E-10 | 1.07E-09 |
| | 2.31269561 | 0.020739381 | 0.043111593 |
| NA | NA | NA | |
| | -5.350345476 | 8.78E-08 | 4.77E-07 |
| | -0.378144912 | 0.705322946 | 0.783796543 |
| NA | NA | NA | |
| | 0.250305407 | 0.802351175 | NA |
| | 0.092721794 | 0.926124583 | NA |
| NA | NA | NA | |
| NA | NA | NA | |
| | 0.192810118 | 0.847107679 | NA |
| | 0.362026528 | 0.717332207 | NA |
| | 0.515191469 | 0.60641923 | NA |
| | 0.192809521 | 0.847108147 | NA |
| | 0.192809521 | 0.847108147 | NA |
| NA | NA | NA | |
| | 0.192809521 | 0.847108147 | NA |
| | -1.510897692 | 0.130814519 | 0.208407904 |
| | -0.469822703 | 0.638481693 | NA |
| | 3.444133579 | 0.000572893 | 0.001729109 |
| | 0.805590537 | 0.420478984 | 0.532189976 |
| | -1.308774497 | 0.190610748 | 0.284258045 |
| | -7.171186346 | 7.44E-13 | 7.00E-12 |
| NA | NA | NA | |
| | 7.340544508 | 2.13E-13 | 2.11E-12 |
| | 1.218434195 | 0.223059017 | 0.322651826 |
| | -0.762963129 | 0.445485385 | 0.555331401 |
| | -0.56909368 | 0.569292567 | 0.67081221 |
| | 0.661690315 | 0.508169713 | 0.614378975 |
| | -0.149467117 | 0.881185054 | 0.916668938 |
| | 0.063708882 | 0.949202033 | NA |
| NA | NA | NA | |
| NA | NA | NA | |
| | -0.278690983 | 0.780481983 | NA |
| NA | NA | NA | |
| | 0.192810118 | 0.847107679 | NA |
| | 3.91225766 | 9.14E-05 | 0.000318757 |
| | -6.191342762 | 5.97E-10 | 4.14E-09 |
| | 1.080818769 | 0.279777738 | NA |
| | 0.063705652 | 0.949204604 | NA |
| | 0.852614732 | 0.39387299 | NA |
| | 0.216157829 | 0.828864724 | NA |
| | 0.731530621 | 0.464455108 | NA |

| | | | |
|---|---|---|---|
| | 0.3474247 | 0.728272279 | NA |
| | 0.065474733 | 0.947796023 | NA |
| | 0.288217989 | 0.77317988 | NA |
| NA | NA | NA | |
| | -0.065388772 | 0.947864464 | NA |
| NA | NA | NA | |
| NA | NA | NA | |
| NA | NA | NA | |
| | 1.165179492 | 0.243946345 | NA |
| NA | NA | NA | |
| NA | NA | NA | |
| NA | NA | NA | |
| NA | NA | NA | |
| | 0.192810118 | 0.847107679 | NA |
| NA | NA | NA | |
| | 0.192810118 | 0.847107679 | NA |
| NA | NA | NA | |
| | 8.304673579 | 1.00E-16 | 1.36E-15 |
| | -0.453564372 | 0.650142402 | NA |
| | 0.31950278 | 0.749345283 | 0.817936961 |
| | 0.19131874 | 0.84827588 | NA |
| NA | NA | NA | |
| | -8.910372054 | 5.09E-19 | 8.41E-18 |
| | -1.238991331 | 0.215348709 | 0.313674242 |
| NA | NA | NA | |
| | 6.423329752 | 1.33E-10 | 9.98E-10 |
| | 1.639355322 | 0.101139281 | 0.168095365 |
| | 0.789133163 | 0.430034181 | 0.54110977 |
| | -0.013509352 | 0.989221424 | 0.992393276 |
| | 0.408021144 | 0.683258152 | NA |
| NA | NA | NA | |
| | -2.216384997 | 0.026665146 | 0.053578599 |
| NA | NA | NA | |
| | -1.372791782 | 0.169817086 | 0.25832756 |
| | 8.522040001 | 1.57E-17 | 2.28E-16 |
| | -0.065465566 | 0.947803322 | NA |
| | 0.135447386 | 0.89225816 | NA |
| | -0.411934926 | 0.68038712 | 0.763167807 |
| | -2.917106467 | 0.003532952 | 0.008977642 |
| | -2.952671388 | 0.003150371 | 0.00810003 |
| | 0.190791771 | 0.848688736 | 0.894390934 |
| | -1.10336444 | 0.269868933 | 0.376019783 |
| | -2.021616767 | 0.043215959 | 0.081571256 |
| | 3.235373894 | 0.001214835 | 0.003432753 |
| | 0.546704422 | 0.584581816 | 0.684637977 |
| | 0.386244748 | 0.699315416 | 0.778814899 |
| | -0.380752274 | 0.703387076 | 0.78241779 |

| | | |
|---|---|---|
| NA | NA | NA |
| -5.314448596 | 1.07E-07 | 5.75E-07 |
| -2.154922809 | 0.031167875 | 0.061374551 |
| -6.41104963 | 1.45E-10 | 1.08E-09 |
| -1.961890274 | 0.04977526 | 0.092239192 |
| 4.892740124 | 9.94E-07 | 4.68E-06 |
| 0.491891779 | 0.622795851 | 0.71694707 |
| -0.625378015 | 0.531722988 | 0.636308395 |
| -5.453188545 | 4.95E-08 | 2.78E-07 |
| -0.756241057 | 0.449504681 | 0.559505801 |
| 9.57856491 | 9.84E-22 | 2.00E-20 |
| 0.192809521 | 0.847108147 | NA |
| 1.886791466 | 0.059188379 | 0.10682817 |
| -1.752139851 | 0.079749763 | 0.137731082 |
| -4.393421144 | 1.12E-05 | 4.51E-05 |
| 10.4439166 | 1.56E-25 | 4.15E-24 |
| 8.80398722 | 1.32E-18 | 2.11E-17 |
| 12.23867837 | 1.93E-34 | 8.22E-33 |
| 11.48682654 | 1.54E-30 | 5.45E-29 |
| 0.82692882 | 0.408277407 | NA |
| 7.899859681 | 2.79E-15 | 3.34E-14 |
| 3.659872178 | 0.000252341 | 0.000812883 |
| 7.651839973 | 1.98E-14 | 2.19E-13 |
| 7.32669511 | 2.36E-13 | 2.33E-12 |
| 8.1624662 | 3.28E-16 | 4.26E-15 |
| 8.468959021 | 2.48E-17 | 3.54E-16 |
| 9.025954889 | 1.78E-19 | 3.05E-18 |
| 3.070392809 | 0.002137774 | 0.005722221 |
| 4.095311923 | 4.22E-05 | 0.000155255 |
| NA | NA | NA |
| NA | NA | NA |
| 0.192810118 | 0.847107679 | NA |
| 0.06371127 | 0.949200131 | NA |
| -0.065389369 | 0.947863988 | NA |
| 15.10990949 | 1.39E-51 | 1.11E-49 |
| -0.070819713 | 0.943541243 | 0.961832586 |
| 4.230637971 | 2.33E-05 | 8.96E-05 |
| 2.957198607 | 0.003104481 | 0.007996003 |
| 2.328864475 | 0.019866246 | 0.041545004 |
| 0.842194237 | 0.399679239 | 0.511476627 |
| 0.230844652 | 0.81743549 | 0.871545848 |
| 0.192809521 | 0.847108147 | NA |
| 2.375812954 | 0.017510337 | 0.037294395 |
| -0.193657701 | 0.846443915 | NA |
| 1.166775852 | 0.243300901 | 0.34599293 |
| -2.981229042 | 0.00287094 | 0.007450453 |
| -4.892045056 | 9.98E-07 | 4.69E-06 |

| | | |
|---|---|---|
| NA | NA | NA |
| 2.689947081 | 0.007146335 | 0.016871606 |
| -2.931535069 | 0.003372913 | 0.00861118 |
| 0.389881979 | 0.69662382 | 0.776618161 |
| -0.65103566 | 0.515023467 | 0.62036106 |
| 0.087119222 | 0.930576746 | NA |
| -0.065389369 | 0.947863988 | NA |
| 0.52144551 | 0.602056455 | 0.699506711 |
| NA | NA | NA |
| 0.201524559 | 0.840288428 | NA |
| 0.459229361 | 0.646069468 | NA |
| 0.133888364 | 0.893490852 | NA |
| 4.127365063 | 3.67E-05 | 0.000136757 |
| -0.231461875 | 0.816956001 | 0.871257294 |
| -1.834980646 | 0.066508557 | 0.118039367 |
| 6.924443198 | 4.38E-12 | 3.79E-11 |
| 0.360540156 | 0.718443233 | NA |
| 11.36707785 | 6.10E-30 | 2.09E-28 |
| 0.547887896 | 0.583768881 | NA |
| NA | NA | NA |
| NA | NA | NA |
| 0.207332115 | 0.835750495 | 0.884816635 |
| -0.972924715 | 0.33059072 | 0.441528506 |
| NA | NA | NA |
| -0.236202653 | 0.813275421 | NA |
| NA | NA | NA |
| -0.811910751 | 0.416842845 | NA |
| 1.215513411 | 0.224170337 | 0.323997201 |
| NA | NA | NA |
| 0.074096566 | 0.940933548 | 0.959878004 |
| 0.488305736 | 0.625333299 | 0.718876245 |
| 0.192808924 | 0.847108615 | NA |
| 1.127140481 | 0.259683091 | 0.364733263 |
| -1.092536 | 0.274597576 | 0.381322565 |
| -0.12809874 | 0.898070833 | NA |
| -0.049954438 | 0.960158696 | 0.972592965 |
| -0.024642803 | 0.980339878 | 0.986381755 |
| 0.099702691 | 0.920580364 | 0.945536345 |
| -0.517329572 | 0.604926114 | 0.701928545 |
| 0.063708075 | 0.949202675 | NA |
| NA | NA | NA |
| NA | NA | NA |
| NA | NA | NA |
| NA | NA | NA |
| NA | NA | NA |
| NA | NA | NA |
| NA | NA | NA |

| | | |
|---|---|---|
| NA | NA | NA |
| NA | NA | NA |
| NA | NA | NA |
| NA | NA | NA |
| NA | NA | NA |
| NA | NA | NA |
| NA | NA | NA |
| NA | NA | NA |
| NA | NA | NA |
| NA | NA | NA |
| NA | NA | NA |
| 0.281693361 | 0.778178645 | 0.841192463 |
| 1.434263998 | 0.15149694 | 0.235482109 |
| 0.706442235 | 0.479913164 | 0.588638108 |
| NA | NA | NA |
| -0.065389966 | 0.947863513 | NA |
| 2.872068874 | 0.004077941 | 0.010230616 |
| -0.660545512 | 0.508903822 | 0.614969472 |
| 9.462525004 | 3.01E-21 | 5.89E-20 |
| -0.712496139 | 0.476157601 | 0.585078261 |
| -2.288102466 | 0.022131555 | 0.04564542 |
| -1.012762638 | 0.311173551 | 0.421373835 |
| -0.225470211 | 0.821613497 | NA |
| -0.065389966 | 0.947863513 | NA |
| 4.865092945 | 1.14E-06 | 5.32E-06 |
| 1.815680404 | 0.069419412 | 0.122363935 |
| 0.931371082 | 0.351661644 | 0.463414286 |
| NA | NA | NA |
| 1.655903399 | 0.097741383 | 0.163490165 |
| -1.787875453 | 0.073796104 | 0.128808681 |
| 0.796569066 | 0.425701349 | NA |
| -0.074970427 | 0.940238242 | NA |
| 0.894986801 | 0.370794144 | 0.483560539 |
| -0.934378948 | 0.350108453 | 0.461853831 |
| 3.327510977 | 0.000876255 | 0.002554767 |
| -5.547264296 | 2.90E-08 | 1.68E-07 |
| -6.580803686 | 4.68E-11 | 3.69E-10 |
| 2.44177368 | 0.014615306 | 0.031832675 |
| -7.437696876 | 1.02E-13 | 1.05E-12 |
| 0.192810118 | 0.847107679 | NA |
| -0.157762142 | 0.874644231 | NA |
| -10.16095925 | 2.96E-24 | 7.20E-23 |
| 1.930986343 | 0.053484741 | 0.09809651 |
| 17.03120611 | 4.82E-65 | 5.41E-63 |
| NA | NA | NA |
| 1.877974234 | 0.06038469 | 0.108622089 |
| 6.591279776 | 4.36E-11 | 3.44E-10 |

| | | |
|---|---|---|
| -2.393421138 | 0.016692069 | 0.035800265 |
| 0.800839841 | 0.423224371 | 0.534893861 |
| -1.617176295 | 0.105840232 | 0.174494862 |
| NA | NA | NA |
| 3.884851952 | 0.000102392 | 0.000353257 |
| 0.892330026 | 0.372216065 | NA |
| -5.390267743 | 7.04E-08 | 3.86E-07 |
| 0.063708285 | 0.949202508 | NA |
| -8.328722642 | 8.17E-17 | 1.12E-15 |
| 0.895374287 | 0.370587041 | 0.483366006 |
| -0.461742756 | 0.644265806 | 0.733763698 |
| -5.636289147 | 1.74E-08 | 1.03E-07 |
| -1.210611112 | 0.226044483 | 0.326159764 |
| NA | NA | NA |
| -0.488148261 | 0.625444829 | NA |
| 0.975947197 | 0.329090632 | 0.439994579 |
| 4.904190655 | 9.38E-07 | 4.43E-06 |
| 2.872816434 | 0.004068304 | 0.010208487 |
| 1.859406734 | 0.062969508 | 0.11259657 |
| 12.09889756 | 1.07E-33 | 4.44E-32 |
| 0.05624838 | 0.95514394 | 0.969278444 |
| -0.073323935 | 0.941548346 | NA |
| 0.192808924 | 0.847108615 | NA |
| NA | NA | NA |
| NA | NA | NA |
| NA | NA | NA |
| NA | NA | NA |
| NA | NA | NA |
| NA | NA | NA |
| NA | NA | NA |
| NA | NA | NA |
| NA | NA | NA |
| NA | NA | NA |
| NA | NA | NA |
| NA | NA | NA |
| NA | NA | NA |
| NA | NA | NA |
| NA | NA | NA |
| NA | NA | NA |
| NA | NA | NA |
| NA | NA | NA |
| 0.063708882 | 0.949202033 | NA |
| 1.894939162 | 0.058100462 | 0.105220647 |
| NA | NA | NA |
| 29.1342977 | 1.32E-186 | 7.87E-184 |
| -1.479747165 | 0.138940733 | 0.219011686 |
| -2.65953787 | 0.007824793 | 0.018263202 |

| | | |
|---|---|---|
| NA | NA | NA |
| -4.935170761 | 8.01E-07 | 3.81E-06 |
| -2.647954891 | 0.008098033 | 0.018821967 |
| 0.063708285 | 0.949202508 | NA |
| NA | NA | NA |
| NA | NA | NA |
| NA | NA | NA |
| NA | NA | NA |
| NA | NA | NA |
| NA | NA | NA |
| NA | NA | NA |
| NA | NA | NA |
| -0.755151256 | 0.450158232 | 0.560068623 |
| NA | NA | NA |
| 0.063708381 | 0.949202431 | NA |
| 0.063707135 | 0.949203424 | NA |
| NA | NA | NA |
| 0.192808327 | 0.847109082 | NA |
| NA | NA | NA |
| NA | NA | NA |
| NA | NA | NA |
| -0.065388772 | 0.947864464 | NA |
| 0.066394788 | 0.94706352 | NA |
| NA | NA | NA |
| NA | NA | NA |
| 0.063708285 | 0.949202508 | NA |
| NA | NA | NA |
| NA | NA | NA |
| NA | NA | NA |
| -0.065388772 | 0.947864464 | NA |
| 0.192809521 | 0.847108147 | NA |
| NA | NA | NA |
| 3.804489876 | 0.000142097 | 0.000478177 |
| -0.903256399 | 0.366389831 | 0.47887534 |
| -0.139723594 | 0.888878384 | 0.922299298 |
| -1.867670935 | 0.061807952 | 0.110764878 |
| 1.432875071 | 0.151893543 | 0.235995988 |
| NA | NA | NA |
| 6.177643794 | 6.51E-10 | 4.50E-09 |
| -6.482120648 | 9.04E-11 | 6.91E-10 |
| NA | NA | NA |
| 0.19288673 | 0.847047678 | NA |
| NA | NA | NA |
| NA | NA | NA |
| -0.065389966 | 0.947863513 | NA |
| -5.187670713 | 2.13E-07 | 1.10E-06 |
| 2.451777313 | 0.01421526 | 0.03106408 |

| | | | |
|---|---|---|---|
| | -1.379357949 | 0.167784418 | 0.255810682 |
| | -5.172232008 | 2.31E-07 | 1.19E-06 |
| NA | NA | NA | |
| | -4.261989194 | 2.03E-05 | 7.87E-05 |
| | -0.508109467 | 0.611376577 | 0.707282109 |
| | 0.771371662 | 0.440486667 | NA |
| | 0.192810118 | 0.847107679 | NA |
| | 0.747096955 | 0.455005038 | 0.564779375 |
| | -1.557869639 | 0.119264154 | 0.192903507 |
| | 0.515255232 | 0.606374678 | 0.70321819 |
| | 1.187676426 | 0.234960915 | 0.336426503 |
| | 4.831050154 | 1.36E-06 | 6.25E-06 |
| | -2.669562858 | 0.007595006 | 0.017801572 |
| | 0.063708882 | 0.949202033 | NA |
| | -0.450062889 | 0.652665095 | 0.740892999 |
| | 1.208188757 | 0.226974665 | 0.327200131 |
| NA | NA | NA | |
| | -2.534507557 | 0.01126055 | 0.025346363 |
| NA | NA | NA | |
| NA | NA | NA | |
| | -0.350304817 | 0.72610995 | 0.800005813 |
| NA | NA | NA | |
| | 0.192809521 | 0.847108147 | NA |
| | 0.063705652 | 0.949204604 | NA |
| | 0.192810118 | 0.847107679 | NA |
| NA | NA | NA | |
| NA | NA | NA | |
| | 0.192809521 | 0.847108147 | NA |
| NA | NA | NA | |
| | -0.065388772 | 0.947864464 | NA |
| | 4.532082778 | 5.84E-06 | 2.46E-05 |
| | 1.331448299 | 0.183041546 | 0.274968785 |
| | 8.327768409 | 8.24E-17 | 1.13E-15 |
| | -0.274946187 | 0.783357582 | 0.845341239 |
| | 0.345341328 | 0.72983778 | 0.802805522 |
| | 6.84419689 | 7.69E-12 | 6.49E-11 |
| | -3.7300405 | 0.000191449 | 0.000631187 |
| | -1.699640596 | 0.089198549 | 0.15147837 |
| | 0.184022962 | 0.853995441 | 0.898508024 |
| | 1.147732128 | 0.251079164 | 0.35479656 |
| | -1.35760929 | 0.174587694 | 0.264523606 |
| | 7.767249639 | 8.02E-15 | 9.22E-14 |
| NA | NA | NA | |
| | 1.574236357 | 0.115432817 | 0.18766455 |
| | 0.063708882 | 0.949202033 | NA |
| | 2.944543904 | 0.003234311 | 0.008297099 |
| | 0.192810118 | 0.847107679 | NA |

| | | | |
|---|---|---|---|
| NA | NA | NA | |
| NA | NA | NA | |
| 1.71606087 | 0.086150902 | NA | |
| NA | NA | NA | |
| NA | NA | NA | |
| NA | NA | NA | |
| NA | NA | NA | |
| 0.063707664 | 0.949203002 | NA | |
| NA | NA | NA | |
| NA | NA | NA | |
| NA | NA | NA | |
| 0.324874675 | 0.745275925 | NA | |
| NA | NA | NA | |
| NA | NA | NA | |
| NA | NA | NA | |
| NA | NA | NA | |
| NA | NA | NA | |
| NA | NA | NA | |
| NA | NA | NA | |
| 0.202714101 | 0.839358501 | NA | |
| NA | NA | NA | |
| NA | NA | NA | |
| NA | NA | NA | |
| NA | NA | NA | |
| 0.063708882 | 0.949202033 | NA | |
| NA | NA | NA | |
| NA | NA | NA | |
| NA | NA | NA | |
| 0.192810118 | 0.847107679 | NA | |
| 0.520009438 | 0.603056997 | NA | |
| NA | NA | NA | |
| NA | NA | NA | |
| NA | NA | NA | |
| NA | NA | NA | |
| NA | NA | NA | |
| NA | NA | NA | |
| 0.807254596 | 0.419519817 | NA | |
| NA | NA | NA | |
| 0.201524559 | 0.840288428 | NA | |
| 0.063708285 | 0.949202508 | NA | |
| NA | NA | NA | |
| NA | NA | NA | |
| NA | NA | NA | |
| NA | NA | NA | |
| 0.209074615 | 0.834389991 | NA | |
| 3.047092867 | 0.002310663 | | 0.006131881 |
| NA | NA | NA | |

| | | |
|---|---|---|
| 0.324874675 | 0.745275925 | NA |
| NA | NA | NA |
| NA | NA | NA |
| 0.063707688 | 0.949202983 | NA |
| NA | NA | NA |
| NA | NA | NA |
| NA | NA | NA |
| NA | NA | NA |
| NA | NA | NA |
| 0.491836603 | 0.622834859 | 0.716952613 |
| 0.192809521 | 0.847108147 | NA |
| NA | NA | NA |
| NA | NA | NA |
| NA | NA | NA |
| NA | NA | NA |
| NA | NA | NA |
| NA | NA | NA |
| NA | NA | NA |
| NA | NA | NA |
| NA | NA | NA |
| 0.362026528 | 0.717332207 | NA |
| NA | NA | NA |
| NA | NA | NA |
| NA | NA | NA |
| 0.192809521 | 0.847108147 | NA |
| NA | NA | NA |
| NA | NA | NA |
| NA | NA | NA |
| NA | NA | NA |
| NA | NA | NA |
| NA | NA | NA |
| NA | NA | NA |
| NA | NA | NA |
| 0.43565198 | 0.663089251 | NA |
| 0.192808924 | 0.847108615 | NA |
| NA | NA | NA |
| NA | NA | NA |
| 0.063708882 | 0.949202033 | NA |
| 0.192809521 | 0.847108147 | NA |
| 0.31685483 | 0.751353758 | NA |
| NA | NA | NA |
| 0.192810118 | 0.847107679 | NA |
| 0.063708285 | 0.949202508 | NA |
| NA | NA | NA |
| NA | NA | NA |
| NA | NA | NA |
| NA | NA | NA |

| | | |
|---|---|---|
| 0.192810118 | 0.847107679 | NA |
| NA | NA | NA |
| NA | NA | NA |
| NA | NA | NA |
| NA | NA | NA |
| NA | NA | NA |
| NA | NA | NA |
| NA | NA | NA |
| NA | NA | NA |
| 0.063708882 | 0.949202033 | NA |
| NA | NA | NA |
| 0.192809521 | 0.847108147 | NA |
| -0.073551523 | 0.941367246 | NA |
| NA | NA | NA |
| NA | NA | NA |
| 0.192809521 | 0.847108147 | NA |
| NA | NA | NA |
| 0.063708285 | 0.949202508 | NA |
| NA | NA | NA |
| NA | NA | NA |
| NA | NA | NA |
| NA | NA | NA |
| NA | NA | NA |
| NA | NA | NA |
| NA | NA | NA |
| NA | NA | NA |
| NA | NA | NA |
| 0.366404893 | 0.714062969 | NA |
| NA | NA | NA |
| NA | NA | NA |
| NA | NA | NA |
| 0.86704156 | 0.385919239 | 0.498146488 |
| NA | NA | NA |
| NA | NA | NA |
| 0.192810118 | 0.847107679 | NA |
| NA | NA | NA |
| -0.065390563 | 0.947863038 | NA |
| NA | NA | NA |
| NA | NA | NA |
| 0.192810118 | 0.847107679 | NA |
| NA | NA | NA |
| NA | NA | NA |
| NA | NA | NA |
| NA | NA | NA |
| NA | NA | NA |
| NA | NA | NA |
| NA | NA | NA |

| | | | |
|---|---|---|---|
| 0.209569518 | 0.834003673 | NA | |
| 0.938770658 | 0.347848507 | | 0.459453727 |
| 0.192809521 | 0.847108147 | NA | |
| 0.624027198 | 0.532609723 | NA | |
| 0.624027198 | 0.532609723 | NA | |
| NA | NA | NA | |
| 0.3474247 | 0.728272279 | NA | |
| NA | NA | NA | |
| NA | NA | NA | |
| 1.04007691 | 0.29830417 | | 0.407162906 |
| NA | NA | NA | |
| 0.324874675 | 0.745275925 | NA | |
| NA | NA | NA | |
| NA | NA | NA | |
| NA | NA | NA | |
| NA | NA | NA | |
| NA | NA | NA | |
| 0.192809521 | 0.847108147 | NA | |
| -0.065389369 | 0.947863988 | NA | |
| 0.415622235 | 0.677686453 | | 0.760752206 |
| NA | NA | NA | |
| 0.586277721 | 0.5576889 | NA | |
| -0.065389369 | 0.947863988 | NA | |
| NA | NA | NA | |
| NA | NA | NA | |
| NA | NA | NA | |
| 0.320071032 | 0.748914485 | NA | |
| NA | NA | NA | |
| 0.192809521 | 0.847108147 | NA | |
| 0.192810118 | 0.847107679 | NA | |
| NA | NA | NA | |
| NA | NA | NA | |
| 1.173589171 | 0.240559621 | | 0.342900023 |
| NA | NA | NA | |
| NA | NA | NA | |
| NA | NA | NA | |
| 0.132797302 | 0.894353688 | NA | |
| -0.065388772 | 0.947864464 | NA | |
| 0.192810118 | 0.847107679 | NA | |
| -0.366675135 | 0.713861356 | NA | |
| NA | NA | NA | |
| -0.065389966 | 0.947863513 | NA | |
| 0.859408802 | 0.390115015 | | 0.502215647 |
| 0.192810118 | 0.847107679 | NA | |
| NA | NA | NA | |
| 1.107193916 | 0.268210097 | NA | |
| 0.192809521 | 0.847108147 | NA | |

| | | |
|---|---|---|
| NA | NA | NA |
| -1.0958773 | 0.273132474 | 0.37980588 |
| NA | NA | NA |
| NA | NA | NA |
| 0.063708285 | 0.949202508 | NA |
| NA | NA | NA |
| -0.065389369 | 0.947863988 | NA |
| NA | NA | NA |
| NA | NA | NA |
| 0.45412515 | 0.649738752 | NA |
| NA | NA | NA |
| 0.485694983 | 0.627183442 | NA |
| NA | NA | NA |
| NA | NA | NA |
| 3.71324996 | 0.000204615 | 0.000671408 |
| NA | NA | NA |
| 0.214588724 | 0.830087986 | NA |
| 0.366404893 | 0.714062969 | NA |
| NA | NA | NA |
| 0.669094949 | 0.503434912 | NA |
| NA | NA | NA |
| 0.504062275 | 0.614217616 | NA |
| 0.214588724 | 0.830087986 | NA |
| NA | NA | NA |
| 0.063708882 | 0.949202033 | NA |
| 0.422902325 | 0.672366526 | NA |
| 0.512266506 | 0.608464503 | NA |
| 5.617514762 | NA | NA |
| -1.361585166 | 0.173328839 | 0.26287087 |
| 0.490188816 | 0.624000294 | 0.717772397 |
| 0.038596909 | 0.969211767 | 0.97871346 |
| -0.065389966 | 0.947863513 | NA |
| NA | NA | NA |
| -0.065389369 | 0.947863988 | NA |
| NA | NA | NA |
| 3.24678493 | 0.001167166 | 0.003315982 |
| 0.636068883 | 0.524731526 | 0.629838877 |
| -0.070666592 | 0.94366311 | NA |
| -4.657541247 | 3.20E-06 | 1.40E-05 |
| 1.970127136 | 0.048823802 | 0.090697622 |
| 6.827483096 | 8.64E-12 | 7.26E-11 |
| 1.7676568 | 0.077118296 | 0.133832833 |
| 9.559007281 | 1.19E-21 | 2.41E-20 |
| 9.671585645 | 3.98E-22 | 8.28E-21 |
| -6.44947607 | 1.12E-10 | 8.49E-10 |
| -2.681788218 | 0.00732298 | 0.017234997 |
| -1.090298921 | 0.27558149 | 0.382276918 |

| | | |
|---|---|---|
| 2.298590136 | 0.021528224 | 0.044529498 |
| -2.931126435 | 0.003377353 | 0.008620757 |
| 1.786771156 | 0.073974483 | 0.12910203 |
| 3.12243983 | 0.001793588 | 0.004883975 |
| -0.21275998 | 0.83151418 | 0.882197352 |
| -0.427566352 | 0.668966868 | 0.753936873 |
| 7.963610367 | 1.67E-15 | 2.04E-14 |
| 0.231912591 | 0.816605905 | NA |
| -5.952493262 | 2.64E-09 | 1.72E-08 |
| -4.170087472 | 3.04E-05 | 0.000115089 |
| 4.714536923 | 2.42E-06 | 1.08E-05 |
| 0.402011926 | 0.687675249 | NA |
| 7.057043654 | 1.70E-12 | 1.53E-11 |
| -0.47745916 | 0.633035196 | 0.724601955 |
| 2.078369628 | 0.037675328 | 0.072355824 |
| 5.246851229 | 1.55E-07 | 8.13E-07 |
| 5.163205756 | 2.43E-07 | 1.24E-06 |
| 12.09737613 | 1.09E-33 | 4.52E-32 |
| 0.110843048 | 0.911740808 | 0.939246509 |
| 4.698695856 | 2.62E-06 | 1.16E-05 |
| 2.14889028 | 0.031643099 | 0.062183148 |
| 14.36889905 | 8.11E-47 | 5.80E-45 |
| -2.158254674 | 0.030908037 | 0.060919136 |
| -2.157991666 | 0.03092848 | 0.060951094 |
| 6.77350696 | 1.26E-11 | 1.04E-10 |
| -1.327581078 | 0.184316539 | 0.276502425 |
| 3.810689375 | 0.00013858 | 0.00046741 |
| 1.221565372 | 0.22187203 | 0.32123189 |
| -1.749510477 | 0.080202819 | 0.138372893 |
| -0.953375405 | 0.340399899 | 0.451928577 |
| 0.092484056 | 0.926313458 | NA |
| -0.065389966 | 0.947863513 | NA |
| -1.938063119 | 0.052615522 | 0.096743299 |
| 5.656485 | 1.55E-08 | 9.24E-08 |
| 0.330491737 | 0.741028435 | NA |
| 4.949978844 | 7.42E-07 | 3.55E-06 |
| -4.536575495 | 5.72E-06 | 2.41E-05 |
| -0.343252922 | 0.731408194 | NA |
| -0.095015892 | 0.924302204 | 0.948011972 |
| NA | NA | NA |
| NA | NA | NA |
| NA | NA | NA |
| NA | NA | NA |
| NA | NA | NA |
| NA | NA | NA |
| 0.437808672 | 0.661524984 | 0.748112337 |
| NA | NA | NA |

| | | |
|---|---|---|
| NA | NA | NA |
| 0.037051094 | 0.970444267 | 0.979556887 |
| -9.50991312 | 1.91E-21 | 3.80E-20 |
| -5.222768217 | 1.76E-07 | 9.18E-07 |
| NA | NA | NA |
| -0.096293488 | 0.923287483 | 0.947359421 |
| -0.19448762 | 0.845794089 | NA |
| NA | NA | NA |
| NA | NA | NA |
| NA | NA | NA |
| NA | NA | NA |
| 1.739617714 | 0.081926167 | NA |
| NA | NA | NA |
| NA | NA | NA |
| NA | NA | NA |
| NA | NA | NA |
| -0.880480878 | 0.378598859 | NA |
| -7.897262542 | 2.85E-15 | 3.41E-14 |
| NA | NA | NA |
| NA | NA | NA |
| -0.259914583 | 0.794929662 | NA |
| NA | NA | NA |
| -1.035712023 | 0.300336512 | NA |
| -0.809990636 | 0.417945558 | 0.529591452 |
| 0.819245844 | 0.412646164 | 0.524237817 |
| -2.67367292 | 0.007502559 | 0.01761458 |
| NA | NA | NA |
| 1.010583714 | 0.312215714 | 0.422510776 |
| 2.706700597 | 0.006795551 | 0.016154682 |
| 10.33288173 | 5.00E-25 | 1.28E-23 |
| 6.13657348 | 8.43E-10 | 5.78E-09 |
| 0.881179089 | 0.378220895 | NA |
| -3.16139902 | 0.001570132 | 0.004333012 |
| 1.744947321 | 0.080994039 | 0.139600661 |
| 12.86005549 | 7.55E-38 | 3.78E-36 |
| 10.17370111 | 2.60E-24 | 6.33E-23 |
| NA | NA | NA |
| 5.426500369 | 5.75E-08 | 3.20E-07 |
| 14.36227037 | 8.93E-47 | 6.35E-45 |
| -0.113405679 | 0.909708937 | 0.937693642 |
| -0.19448762 | 0.845794089 | NA |
| 0.373259565 | 0.708955267 | NA |
| 3.086227014 | 0.00202714 | 0.005458188 |
| 0.827831252 | 0.407766073 | 0.519567543 |
| NA | NA | NA |
| -2.099602484 | 0.035763824 | 0.069151682 |
| -0.095892535 | 0.923605923 | 0.94756963 |

| | | |
|---|---|---|
| NA | NA | NA |
| 0.256512829 | 0.797554876 | 0.856000967 |
| -1.653278916 | 0.098274112 | 0.164206823 |
| 1.644637917 | 0.100044503 | 0.166578985 |
| 4.313436763 | 1.61E-05 | 6.34E-05 |
| -4.929893025 | 8.23E-07 | 3.91E-06 |
| -4.237885551 | 2.26E-05 | 8.70E-05 |
| -8.501209717 | 1.88E-17 | 2.71E-16 |
| -4.10094903 | 4.11E-05 | 0.000151868 |
| -6.342761112 | 2.26E-10 | 1.64E-09 |
| 10.17414496 | 2.59E-24 | 6.30E-23 |
| 10.4289761 | 1.83E-25 | 4.83E-24 |
| 2.398104422 | 0.016480166 | 0.035413682 |
| -1.36531224 | 0.172154933 | 0.261348654 |
| -2.813996001 | 0.004892984 | 0.012078046 |
| 1.009227696 | 0.312865447 | 0.423252737 |
| -1.126947833 | 0.259764538 | 0.364763332 |
| 1.584047393 | 0.113182934 | 0.1845587 |
| 0.722297656 | 0.470111494 | 0.579164773 |
| NA | NA | NA |
| 0.643624233 | 0.519819132 | NA |
| 1.380311981 | 0.167490608 | NA |
| -1.063393667 | 0.287603463 | 0.395012878 |
| NA | NA | NA |
| NA | NA | NA |
| NA | NA | NA |
| 0.320071032 | 0.748914485 | NA |
| NA | NA | NA |
| -0.960706874 | 0.336699574 | 0.447966318 |
| -2.751142371 | 0.005938783 | 0.01432197 |
| NA | NA | NA |
| 5.039135113 | 4.68E-07 | 2.30E-06 |
| 1.00313105 | 0.315797635 | 0.426320846 |
| 1.46353883 | 0.143319996 | 0.224852671 |
| NA | NA | NA |
| 0.65600308 | 0.511822143 | 0.617541973 |
| -1.2363046 | 0.216345362 | 0.314906048 |
| 8.987371605 | 2.53E-19 | 4.29E-18 |
| 0.063708285 | 0.949202508 | NA |
| -3.765387857 | 0.000166291 | 0.000553426 |
| -2.388716496 | 0.016907344 | 0.036204193 |
| -0.169802615 | 0.86516537 | 0.906032769 |
| 0.192810118 | 0.847107679 | NA |
| 0.778439551 | 0.43630993 | 0.546725284 |
| 0.301093136 | 0.763343475 | 0.829202637 |
| 2.011128365 | 0.0443119 | 0.083399267 |
| 0.065474733 | 0.947796023 | NA |

| | | |
|---|---|---|
| NA | NA | NA |
| -0.708629015 | 0.478554726 | 0.587365486 |
| 0.136939754 | 0.891078416 | 0.923816126 |
| -6.361572332 | 2.00E-10 | 1.46E-09 |
| 2.22544867 | 0.026051133 | 0.052538866 |
| NA | NA | NA |
| NA | NA | NA |
| 3.282801471 | 0.00102781 | 0.002955663 |
| -5.179847133 | 2.22E-07 | 1.14E-06 |
| 5.934928347 | 2.94E-09 | 1.91E-08 |
| -7.737693264 | 1.01E-14 | 1.15E-13 |
| -0.245061552 | 0.806408772 | 0.862932143 |
| -4.003552098 | 6.24E-05 | 0.00022347 |
| -4.887222953 | 1.02E-06 | 4.79E-06 |
| -2.441848334 | 0.014612284 | 0.031828863 |
| -0.019071077 | 0.984784405 | 0.989346134 |
| -1.752854537 | 0.079626979 | 0.137547501 |
| -2.36394242 | 0.018081623 | 0.038325262 |
| 0.192810118 | 0.847107679 | NA |
| -0.783463652 | 0.433354891 | 0.544057426 |
| -0.261996244 | 0.793324338 | NA |
| -3.222729993 | 0.001269752 | 0.003572203 |
| -0.613668579 | 0.539434356 | 0.643690023 |
| -0.133886437 | 0.893492375 | 0.925705292 |
| 7.390346189 | 1.46E-13 | 1.48E-12 |
| -1.155319039 | 0.247959802 | 0.351181681 |
| NA | NA | NA |
| 2.746458102 | 0.006024258 | 0.014505741 |
| 0.120499022 | 0.904087854 | 0.933591278 |
| 0.322918585 | 0.746756901 | 0.815894418 |
| NA | NA | NA |
| 0.06371127 | 0.949200131 | NA |
| 0.133374594 | 0.893897137 | NA |
| -0.837006715 | 0.402588793 | 0.514332253 |
| -1.427963511 | 0.153302361 | 0.237653791 |
| 0.544963286 | 0.585778769 | 0.685911423 |
| -0.501116202 | 0.616289345 | 0.711321696 |
| -2.975039665 | 0.002929507 | 0.007592209 |
| 4.814096563 | 1.48E-06 | 6.77E-06 |
| 0.420483624 | 0.67413219 | NA |
| 1.060804002 | 0.288778983 | 0.396170893 |
| 0.602713248 | 0.546699467 | 0.650282168 |
| 5.138230909 | 2.77E-07 | 1.41E-06 |
| -5.117155899 | 3.10E-07 | 1.57E-06 |
| -4.489204082 | 7.15E-06 | 2.98E-05 |
| -1.464833235 | 0.142966422 | 0.224480869 |
| -1.05660502 | 0.290691888 | 0.398298315 |

| | | |
|---|---|---|
| -0.011399288 | 0.990904881 | 0.993644052 |
| 0.483125894 | 0.62900635 | NA |
| 0.435619695 | 0.663112679 | 0.749433799 |
| 6.168247442 | 6.91E-10 | 4.76E-09 |
| 1.059865295 | 0.289205886 | 0.396656404 |
| NA | NA | NA |
| 3.005887978 | 0.002648065 | 0.006925922 |
| 10.17756617 | 2.50E-24 | 6.10E-23 |
| 3.359710567 | 0.000780242 | 0.002294343 |
| -4.453250114 | 8.46E-06 | 3.49E-05 |
| 15.4582455 | 6.64E-54 | 5.58E-52 |
| 0.584478561 | 0.558898385 | 0.661241566 |
| -5.478657819 | 4.29E-08 | 2.43E-07 |
| 1.355877597 | 0.175138117 | 0.265245178 |
| 1.686010776 | 0.091793724 | 0.155148687 |
| 0.453881387 | 0.649914201 | NA |
| 1.305612084 | 0.191684509 | 0.285519118 |
| NA | NA | NA |
| 1.850213132 | 0.064282837 | 0.114511381 |
| NA | NA | NA |
| -4.095127142 | 4.22E-05 | 0.000155356 |
| 4.279263175 | 1.88E-05 | 7.32E-05 |
| 10.00971878 | 1.38E-23 | 3.22E-22 |
| 7.215716451 | 5.37E-13 | 5.13E-12 |
| -0.470004627 | 0.638351712 | 0.729386622 |
| -1.025798395 | 0.304986623 | 0.414520806 |
| -4.512621222 | 6.40E-06 | 2.68E-05 |
| -4.588714804 | 4.46E-06 | 1.91E-05 |
| 1.695622046 | 0.08995748 | 0.152570519 |
| -0.358388139 | 0.720052866 | 0.795222522 |
| 10.82095126 | 2.74E-27 | 8.09E-26 |
| 0.440003287 | 0.659934727 | 0.746906902 |
| 5.715009021 | 1.10E-08 | 6.68E-08 |
| -6.238255546 | 4.42E-10 | 3.12E-09 |
| 0.1208205 | 0.903833212 | NA |
| 1.176007169 | 0.239592011 | 0.341728192 |
| NA | NA | NA |
| -5.192497193 | 2.07E-07 | 1.07E-06 |
| -3.916646198 | 8.98E-05 | 0.000313337 |
| 0.429780538 | 0.667355291 | NA |
| 5.655728237 | 1.55E-08 | 9.27E-08 |
| 7.562327737 | 3.96E-14 | 4.24E-13 |
| NA | NA | NA |
| 7.277317593 | 3.41E-13 | 3.32E-12 |
| NA | NA | NA |
| 0.74041697 | 0.459047025 | NA |
| -0.215901234 | 0.829064735 | 0.880569053 |

| | | | |
|---|---|---|---|
| NA | NA | NA | |
| | 2.762368139 | 0.005738374 | 0.01390027 |
| | 10.48356611 | 1.03E-25 | 2.76E-24 |
| | 1.805135286 | 0.071053515 | 0.124857447 |
| | -2.586778366 | 0.009687788 | 0.022146803 |
| | 0.306134864 | 0.759501964 | 0.8259258 |
| | -3.44034043 | 0.000580983 | 0.001750361 |
| | -2.836557522 | 0.004560276 | 0.011327152 |
| | -0.769129736 | 0.441816297 | 0.551774369 |
| | 5.943803663 | 2.78E-09 | 1.81E-08 |
| | -1.530982287 | 0.12577377 | 0.201905944 |
| | 1.384038702 | 0.166346604 | 0.254020835 |
| | 0.810402751 | 0.417708737 | NA |
| NA | NA | NA | |
| | 0.659736718 | 0.5094228 | NA |
| | 0.31685483 | 0.751353758 | NA |
| | -0.316421392 | 0.751682682 | 0.819916221 |
| | -2.811126049 | 0.004936844 | 0.012168312 |
| | -2.441099955 | 0.014642601 | 0.031886387 |
| | -0.407314233 | 0.68377721 | 0.76618066 |
| NA | NA | NA | |
| | 0.192809521 | 0.847108147 | NA |
| NA | NA | NA | |
| NA | NA | NA | |
| NA | NA | NA | |
| NA | NA | NA | |
| | 0.192809521 | 0.847108147 | NA |
| NA | NA | NA | |
| NA | NA | NA | |
| NA | NA | NA | |
| NA | NA | NA | |
| NA | NA | NA | |
| NA | NA | NA | |
| NA | NA | NA | |
| NA | NA | NA | |
| NA | NA | NA | |
| NA | NA | NA | |
| NA | NA | NA | |
| | 0.063708285 | 0.949202508 | NA |
| NA | NA | NA | |
| NA | NA | NA | |
| NA | NA | NA | |
| | -0.382074673 | 0.702405977 | NA |
| | 0.731285159 | 0.464604993 | 0.574019596 |
| | 0.063708285 | 0.949202508 | NA |
| NA | NA | NA | |
| NA | NA | NA | |

| | | |
|---|---|---|
| NA | NA | NA |
| 0.846464029 | 0.397293932 | NA |
| -0.065389966 | 0.947863513 | NA |
| -1.126099678 | 0.26012333 | 0.365041879 |
| -5.066589211 | 4.05E-07 | 2.01E-06 |
| 1.020786361 | 0.307355668 | 0.41712912 |
| 12.52503272 | 5.45E-36 | 2.49E-34 |
| 0.202714101 | 0.839358501 | NA |
| 0.26385454 | 0.791892006 | 0.851932301 |
| -3.278320181 | 0.001044269 | 0.002998858 |
| -1.324286314 | 0.185407974 | 0.277840004 |
| -1.571562031 | 0.116052158 | 0.188463344 |
| NA | NA | NA |
| 1.343832824 | 0.179002453 | 0.269921766 |
| 0.458312316 | 0.646728077 | 0.735833618 |
| -7.687668408 | 1.50E-14 | 1.68E-13 |
| 0.06371127 | 0.949200131 | NA |
| 5.037163944 | 4.72E-07 | 2.32E-06 |
| 0.130054343 | 0.896523432 | 0.92792381 |
| -1.083033468 | 0.278793568 | 0.385834513 |
| 0.691668603 | 0.489145468 | 0.596972601 |
| -0.231394185 | 0.817008582 | 0.871267477 |
| 1.105327134 | 0.269017865 | 0.375105406 |
| 11.64423192 | 2.46E-31 | 9.10E-30 |
| -0.171357946 | 0.86394232 | 0.905433835 |
| 1.886413583 | 0.059239244 | 0.106896278 |
| -0.462844206 | 0.643476042 | 0.733289696 |
| -5.991478259 | 2.08E-09 | 1.37E-08 |
| -1.215394569 | 0.224215638 | 0.324030335 |
| 1.503664397 | 0.132667801 | 0.210823126 |
| 3.007503531 | 0.002634031 | 0.006892816 |
| -0.065389369 | 0.947863988 | NA |
| -5.827279833 | 5.63E-09 | 3.54E-08 |
| -3.944662633 | 7.99E-05 | 0.000281363 |
| 3.301189952 | 0.000962757 | 0.002783942 |
| 2.949034121 | 0.003187688 | 0.008183367 |
| -5.868188659 | 4.41E-09 | 2.80E-08 |
| -0.065390563 | 0.947863038 | NA |
| 7.587789057 | 3.25E-14 | 3.51E-13 |
| 6.436521208 | 1.22E-10 | 9.20E-10 |
| NA | NA | NA |
| -1.691657382 | 0.090711319 | 0.153611483 |
| NA | NA | NA |
| NA | NA | NA |
| NA | NA | NA |
| -8.63827938 | 5.71E-18 | 8.56E-17 |
| NA | NA | NA |

| | | | |
|---|---|---|---|
| NA | NA | NA | |
| -2.344054326 | 0.019075393 | | 0.040098749 |
| 3.923167236 | 8.74E-05 | | 0.000305849 |
| -0.374081537 | 0.708343655 | NA | |
| 0.192808327 | 0.847109082 | NA | |
| NA | NA | NA | |
| NA | NA | NA | |
| NA | NA | NA | |
| -1.137001576 | 0.255537627 | NA | |
| -1.074718904 | 0.282500598 | | 0.38952184 |
| 0.358817782 | 0.719731409 | NA | |
| NA | NA | NA | |
| -1.920669208 | 0.054773424 | | 0.100115292 |
| -3.377057125 | 0.000732658 | | 0.002168206 |
| -0.184538757 | 0.853590826 | NA | |
| NA | NA | NA | |
| NA | NA | NA | |
| -0.113643083 | 0.909520733 | | 0.937576869 |
| 0.321914724 | 0.747517299 | | 0.816404485 |
| 1.929738024 | 0.053639306 | | 0.09833313 |
| 0.342610156 | 0.731891761 | | 0.804569844 |
| 1.079573291 | 0.28033224 | | 0.387290386 |
| 6.770943025 | 1.28E-11 | | 1.06E-10 |
| -0.126137042 | 0.899623446 | NA | |
| 4.835405556 | 1.33E-06 | | 6.12E-06 |
| -0.203495122 | 0.838748058 | | 0.887071588 |
| -1.277040865 | 0.20158782 | | 0.297701525 |
| NA | NA | NA | |
| NA | NA | NA | |
| 4.991840519 | 5.98E-07 | | 2.89E-06 |
| 0.192809521 | 0.847108147 | NA | |
| -4.097422857 | 4.18E-05 | | 0.000153959 |
| -4.529500405 | 5.91E-06 | | 2.49E-05 |
| 0.063976859 | 0.948988653 | NA | |
| -3.179057404 | 0.001477548 | | 0.004098751 |
| -2.530161247 | 0.011401012 | | 0.025621056 |
| -0.466205129 | 0.641068679 | | 0.731522636 |
| NA | NA | NA | |
| 3.090015551 | 0.00200146 | | 0.005395428 |
| -3.799972288 | 0.000144712 | | 0.000486129 |
| 1.940093281 | 0.052368354 | | 0.096345455 |
| -2.280334455 | 0.02258786 | | 0.046486937 |
| 6.928271572 | 4.26E-12 | | 3.70E-11 |
| NA | NA | NA | |
| NA | NA | NA | |
| 2.230635632 | 0.025705275 | | 0.051921012 |
| 0.192809521 | 0.847108147 | NA | |

| | | |
|---|---|---|
| NA | NA | NA |
| 0.57645995 | 0.564304333 | 0.666384092 |
| 0.059974452 | 0.952175983 | 0.967443265 |
| -1.929722322 | 0.053641253 | 0.09833313 |
| -0.19448762 | 0.845794089 | NA |
| 3.663378039 | 0.000248911 | 0.000802452 |
| -2.465409893 | 0.013685659 | 0.030074807 |
| -1.987762805 | 0.046837929 | 0.087495587 |
| -0.152918869 | 0.878462256 | NA |
| -2.815974008 | 0.00486296 | 0.012012226 |
| NA | NA | NA |
| NA | NA | NA |
| -1.982533144 | 0.047419602 | 0.088436974 |
| -0.702174556 | 0.482570311 | 0.591051815 |
| NA | NA | NA |
| NA | NA | NA |
| -5.342445532 | 9.17E-08 | 4.97E-07 |
| NA | NA | NA |
| NA | NA | NA |
| -2.281991782 | 0.022489825 | 0.046311833 |
| -6.491950864 | 8.47E-11 | 6.49E-10 |
| 2.211193097 | 0.027022468 | 0.054205191 |
| 5.046195071 | 4.51E-07 | 2.22E-06 |
| 2.065868887 | 0.038840861 | 0.074258078 |
| NA | NA | NA |
| NA | NA | NA |
| -4.677871306 | 2.90E-06 | 1.28E-05 |
| 1.459906262 | 0.144315838 | 0.226089092 |
| 8.963603995 | 3.14E-19 | 5.29E-18 |
| 1.502371112 | 0.133001293 | 0.211259192 |
| -4.18439552 | 2.86E-05 | 0.000108517 |
| -3.214467903 | 0.001306865 | 0.003666312 |
| -0.764714106 | 0.444441805 | 0.554389433 |
| 1.793516903 | 0.07289031 | 0.12746765 |
| -2.33045766 | 0.019781975 | 0.041392981 |
| -0.912141988 | 0.361693978 | 0.473890298 |
| -0.656389328 | 0.511573655 | NA |
| 1.426184458 | 0.153815105 | 0.238365029 |
| 1.05591772 | 0.291005807 | 0.398657729 |
| 11.07157141 | 1.72E-28 | 5.52E-27 |
| -1.870789278 | 0.061374294 | 0.110129657 |
| -3.502985231 | 0.000460075 | 0.001411754 |
| -2.916016782 | 0.003545314 | 0.009005401 |
| -0.194488217 | 0.845793621 | NA |
| NA | NA | NA |
| NA | NA | NA |
| NA | NA | NA |

| | | |
|---|---|---|
| NA | NA | NA |
| NA | NA | NA |
| 3.471506702 | 0.000517546 | 0.00157402 |
| -1.429268078 | 0.152927197 | 0.237189723 |
| 0.486592398 | 0.626547212 | 0.719874993 |
| 2.591451819 | 0.009557192 | 0.021866227 |
| -3.361953101 | 0.000773933 | 0.002277972 |
| NA | NA | NA |
| 3.629572745 | 0.000283891 | 0.00090635 |
| 0.063708285 | 0.949202508 | NA |
| 1.705164568 | 0.088163736 | 0.150006003 |
| -2.233919859 | 0.025488347 | 0.051557566 |
| 0.87428904 | 0.381960866 | 0.494464985 |
| -7.355129265 | 1.91E-13 | 1.90E-12 |
| -0.065388772 | 0.947864464 | NA |
| NA | NA | NA |
| 6.382716216 | 1.74E-10 | 1.28E-09 |
| -0.065388772 | 0.947864464 | NA |
| 2.940702289 | 0.003274691 | 0.008386935 |
| -2.548254253 | 0.010826353 | 0.024468048 |
| -6.654377192 | 2.85E-11 | 2.28E-10 |
| 3.203078283 | 0.00135967 | 0.00380082 |
| 0.128819396 | 0.897500558 | NA |
| -9.124059561 | 7.24E-20 | 1.28E-18 |
| 1.996327232 | 0.045898316 | 0.085952031 |
| -1.278597095 | 0.201038974 | 0.297013665 |
| 2.82304629 | 0.00475697 | 0.011766673 |
| -2.42678524 | 0.01523327 | 0.033017731 |
| -1.258948898 | 0.20804879 | 0.305029359 |
| 3.139513732 | 0.001692285 | 0.004635357 |
| -1.658280272 | 0.097260909 | 0.162773506 |
| -3.235162818 | 0.001215734 | 0.003434517 |
| 0.192810118 | 0.847107679 | NA |
| 0.433746343 | 0.664472649 | NA |
| 3.488791447 | 0.00048521 | 0.001483243 |
| -0.482034755 | 0.629781257 | 0.722283545 |
| 9.777533532 | 1.41E-22 | 3.02E-21 |
| -2.865224599 | 0.004167136 | 0.010434514 |
| -5.443696396 | 5.22E-08 | 2.92E-07 |
| 5.83658165 | 5.33E-09 | 3.35E-08 |
| 0.527446933 | 0.597883259 | 0.696209256 |
| 2.088349052 | 0.03676636 | 0.070871158 |
| 0.633679511 | 0.526289999 | NA |
| -0.499262551 | 0.617594433 | 0.712368141 |
| 2.401525895 | 0.016326853 | 0.035129472 |
| -0.319010171 | 0.7497188 | NA |
| 6.300974674 | 2.96E-10 | 2.12E-09 |

| | | |
|---|---|---|
| 5.376322602 | 7.60E-08 | 4.16E-07 |
| 0.670295693 | 0.502669312 | 0.609347776 |
| -3.449750475 | 0.000561105 | 0.001695576 |
| 0.304945663 | 0.760407537 | 0.826659167 |
| 9.149839231 | 5.70E-20 | 1.01E-18 |
| -9.841772428 | 7.44E-23 | 1.63E-21 |
| NA | NA | NA |
| -1.412604887 | 0.157771931 | 0.243316786 |
| -8.728529992 | 2.58E-18 | 4.01E-17 |
| -9.404035132 | 5.25E-21 | 1.00E-19 |
| 2.679482569 | 0.007373604 | 0.017342728 |
| 2.716102855 | 0.006605539 | 0.015752348 |
| 2.963547293 | 0.003041153 | 0.007851477 |
| -2.458080974 | 0.013968168 | 0.030620357 |
| NA | NA | NA |
| -0.105249994 | 0.916177442 | 0.942498104 |
| -2.82948487 | 0.0046623 | 0.011550466 |
| -2.379911009 | 0.017316819 | 0.036948256 |
| 2.503652328 | 0.012291876 | 0.027345627 |
| 2.928643357 | 0.003404448 | 0.00868726 |
| -3.688299423 | 0.000225758 | 0.000734558 |
| 0.432839518 | 0.665131362 | NA |
| 1.518942105 | 0.128777069 | 0.205882093 |
| 1.907554563 | 0.056448813 | 0.102734951 |
| -0.327757142 | 0.743095294 | 0.813065012 |
| NA | NA | NA |
| 2.952303088 | 0.003154132 | 0.008108866 |
| NA | NA | NA |
| 1.55860369 | 0.119090209 | 0.19265951 |
| -0.065390563 | 0.947863038 | NA |
| 5.080821422 | 3.76E-07 | 1.88E-06 |
| 0.585529994 | 0.558191404 | 0.660654949 |
| -1.691835494 | 0.090677344 | 0.153593688 |
| -2.380232311 | 0.017301727 | 0.036928645 |
| -0.660611561 | 0.508861453 | 0.61496398 |
| 0.859734332 | 0.389935505 | 0.502036197 |
| 0.573815287 | 0.566092803 | NA |
| -3.978552418 | 6.93E-05 | 0.000246553 |
| -7.958339943 | 1.74E-15 | 2.13E-14 |
| 1.180873196 | 0.237653101 | 0.339659516 |
| 1.902970409 | 0.057044407 | 0.103653103 |
| 1.006497203 | 0.314176456 | 0.424590287 |
| 1.578717772 | 0.114400806 | 0.186156039 |
| -0.559964592 | 0.575503589 | 0.676609942 |
| 3.71576225 | 0.000202592 | 0.000665382 |
| NA | NA | NA |
| -2.21209897 | 0.026959827 | 0.05409688 |

| | | | |
|---|---|---|---|
| | 0.869147594 | 0.384766407 | NA |
| | 0.636346094 | 0.524550868 | NA |
| | 0.192809521 | 0.847108147 | NA |
| | -0.065389966 | 0.947863513 | NA |
| NA | NA | NA | |
| NA | NA | NA | |
| | 3.829675545 | 0.000128312 | 0.000435066 |
| | 1.660264282 | 0.096861298 | 0.162224041 |
| NA | NA | NA | |
| | 1.949136943 | 0.051279073 | 0.094619624 |
| | -3.227196795 | 0.001250095 | 0.003522047 |
| NA | NA | NA | |
| | 2.827300853 | 0.00469422 | 0.011621794 |
| NA | NA | NA | |
| | -0.43418633 | 0.664153139 | NA |
| | -1.580572971 | 0.11397571 | 0.185645345 |
| | -7.720290676 | 1.16E-14 | 1.31E-13 |
| | -0.671259238 | 0.502055397 | 0.608882706 |
| | 0.094569685 | 0.92465663 | 0.948190794 |
| | -6.806447839 | 1.00E-11 | 8.37E-11 |
| | 0.192809521 | 0.847108147 | NA |
| | 9.537022517 | 1.47E-21 | 2.96E-20 |
| | 3.516041874 | 0.000438032 | 0.001350072 |
| | 0.427262843 | 0.669187893 | NA |
| | 0.063708285 | 0.949202508 | NA |
| | 0.101472726 | 0.919175207 | 0.944533847 |
| | -7.178928232 | 7.03E-13 | 6.63E-12 |
| | 0.001825002 | 0.99854386 | NA |
| | 0.080765253 | 0.935628642 | 0.956240642 |
| | -1.413993632 | 0.157363772 | 0.242812739 |
| | -0.203642565 | 0.838632828 | 0.887042752 |
| | 3.609274403 | 0.000307055 | 0.000974585 |
| | -2.96432341 | 0.003033493 | 0.007834124 |
| NA | NA | NA | |
| | -0.887174598 | 0.374984907 | 0.487831196 |
| | -3.143806938 | 0.001667654 | 0.004576906 |
| | -6.328439624 | 2.48E-10 | 1.79E-09 |
| | -3.561762127 | 0.000368374 | 0.001151399 |
| NA | NA | NA | |
| | -3.543379668 | 0.000395034 | 0.001227522 |
| | -4.063366863 | 4.84E-05 | 0.000176667 |
| | -4.346430043 | 1.38E-05 | 5.51E-05 |
| | 5.611981916 | 2.00E-08 | 1.18E-07 |
| | 4.458647475 | 8.25E-06 | 3.41E-05 |
| | 5.179191519 | 2.23E-07 | 1.15E-06 |
| | -4.119801027 | 3.79E-05 | 0.000140947 |
| | -1.452490781 | 0.146365191 | 0.228712743 |

| | | |
|---|---|---|
| NA | NA | NA |
| 1.394121106 | 0.163281037 | 0.250163699 |
| -0.319991796 | 0.74897455 | 0.817603601 |
| 7.536269307 | 4.84E-14 | 5.13E-13 |
| 0.165830371 | 0.868290444 | 0.908241457 |
| -0.0861316 | 0.931361804 | 0.953235203 |
| -0.738937569 | 0.459944908 | 0.569358113 |
| NA | NA | NA |
| -3.588941108 | 0.000332024 | 0.001047065 |
| NA | NA | NA |
| -1.08855102 | 0.276351926 | 0.383196982 |
| 2.423905757 | 0.01535459 | 0.03325477 |
| NA | NA | NA |
| 10.76237013 | 5.18E-27 | 1.50E-25 |
| -6.881479137 | 5.92E-12 | 5.07E-11 |
| -3.353492006 | 0.000797987 | 0.002342122 |
| -2.682225472 | 0.007313415 | 0.017215722 |
| 0.063708285 | 0.949202508 | NA |
| -0.320762404 | 0.748390452 | 0.817072888 |
| -6.95407736 | 3.55E-12 | 3.10E-11 |
| -3.134578216 | 0.001721014 | 0.004706837 |
| -0.828174508 | 0.407571678 | 0.519425617 |
| -0.270481185 | 0.78679009 | 0.848010849 |
| 11.36774698 | 6.05E-30 | 2.08E-28 |
| -4.460644628 | 8.17E-06 | 3.38E-05 |
| 0.087681318 | 0.930129969 | NA |
| -0.87015003 | 0.38421842 | 0.496505882 |
| 3.960827204 | 7.47E-05 | 0.000264353 |
| 6.198038396 | 5.72E-10 | 3.98E-09 |
| 24.39487331 | 1.94E-131 | 6.83E-129 |
| -0.738634414 | 0.460129021 | 0.56952969 |
| NA | NA | NA |
| 2.294792948 | 0.021744993 | 0.0449296 |
| -1.571394023 | 0.116091153 | 0.18851444 |
| 9.655513193 | 4.66E-22 | 9.68E-21 |
| 0.82798911 | 0.407676667 | 0.519480068 |
| 0.06353712 | 0.949338802 | NA |
| NA | NA | NA |
| 0.210360696 | 0.833386166 | NA |
| NA | NA | NA |
| 3.801746978 | 0.000143679 | 0.000482919 |
| 0.063708285 | 0.949202508 | NA |
| 8.97093787 | 2.94E-19 | 4.97E-18 |
| -1.963922085 | 0.049539128 | 0.091862823 |
| NA | NA | NA |
| -2.980218651 | 0.002880427 | 0.007473524 |
| 2.296004302 | 0.021675635 | 0.044801291 |

| | | | |
|---|---|---|---|
| | 2.502066795 | 0.012347063 | 0.027461079 |
| NA | NA | NA | |
| | 3.892672111 | 9.91E-05 | 0.00034305 |
| | 0.783698459 | 0.433217067 | NA |
| | 1.620204731 | 0.105188306 | 0.173648812 |
| | 23.97259981 | 5.37E-127 | 1.82E-124 |
| | 1.204669334 | 0.228330979 | 0.328719913 |
| NA | NA | NA | |
| | 0.192810118 | 0.847107679 | NA |
| | 1.092429042 | 0.274644563 | 0.381336519 |
| | 1.156229779 | 0.247587182 | 0.350753179 |
| | 0.978786343 | 0.327685559 | 0.438685212 |
| | -3.417960627 | 0.000630922 | 0.001888542 |
| | 0.824473285 | 0.409670689 | 0.521516497 |
| | -0.594520628 | 0.55216395 | 0.655336424 |
| | -0.788839597 | 0.430205763 | 0.541244044 |
| | -1.788668136 | 0.073668278 | 0.1286035 |
| | 17.00819228 | 7.14E-65 | 7.98E-63 |
| | -1.06193306 | 0.288266075 | 0.39568366 |
| NA | NA | NA | |
| | 13.64440819 | 2.18E-42 | 1.31E-40 |
| | 11.43560348 | 2.78E-30 | 9.66E-29 |
| | 4.082733224 | 4.45E-05 | 0.000163185 |
| | 0.112228331 | 0.910642366 | NA |
| | 5.598057202 | 2.17E-08 | 1.28E-07 |
| | 0.254794962 | 0.798881466 | NA |
| NA | NA | NA | |
| | -0.368070077 | 0.712820982 | NA |
| | 4.916019066 | 8.83E-07 | 4.18E-06 |
| | 7.771488011 | 7.76E-15 | 8.92E-14 |
| | 8.775724117 | 1.70E-18 | 2.68E-17 |
| NA | NA | NA | |
| NA | NA | NA | |
| NA | NA | NA | |
| NA | NA | NA | |
| NA | NA | NA | |
| NA | NA | NA | |
| NA | NA | NA | |
| NA | NA | NA | |
| NA | NA | NA | |
| NA | NA | NA | |
| | 0.063708285 | 0.949202508 | NA |
| NA | NA | NA | |
| NA | NA | NA | |
| NA | NA | NA | |
| NA | NA | NA | |
| | 0.585257542 | 0.558374559 | NA |

| | | |
|---|---|---|
| NA | NA | NA |
| NA | NA | NA |
| 0.063708882 | 0.949202033 | NA |
| NA | NA | NA |
| NA | NA | NA |
| NA | NA | NA |
| NA | NA | NA |
| 1.539552044 | 0.123669583 | 0.199038686 |
| -0.787638601 | 0.430908129 | NA |
| -4.949926994 | 7.42E-07 | 3.55E-06 |
| 0.915514995 | 0.359921334 | NA |
| 1.509052815 | 0.131285286 | 0.208945658 |
| -8.115527069 | 4.84E-16 | 6.18E-15 |
| 7.865391196 | 3.68E-15 | 4.36E-14 |
| 19.62883299 | 8.77E-86 | 1.58E-83 |
| 14.30331399 | 2.09E-46 | 1.46E-44 |
| NA | NA | NA |
| 10.35430011 | 4.00E-25 | 1.03E-23 |
| 8.600928819 | 7.91E-18 | 1.17E-16 |
| 6.797438922 | 1.06E-11 | 8.91E-11 |
| 4.052173078 | 5.07E-05 | 0.000184772 |
| 5.337616846 | 9.42E-08 | 5.09E-07 |
| -0.73026656 | 0.465227264 | 0.574634417 |
| 0.112701884 | 0.910266908 | NA |
| -0.065388772 | 0.947864464 | NA |
| NA | NA | NA |
| NA | NA | NA |
| 18.39206463 | 1.52E-75 | 2.21E-73 |
| 1.229508749 | 0.21888112 | 0.317802333 |
| -1.496700876 | 0.134471108 | 0.213126187 |
| -2.966459263 | 0.003012503 | 0.007788755 |
| NA | NA | NA |
| -7.416919507 | 1.20E-13 | 1.22E-12 |
| -0.541991766 | 0.587824179 | 0.687630938 |
| 7.402164884 | 1.34E-13 | 1.36E-12 |
| -3.393279768 | 0.000690611 | 0.002052724 |
| -0.598124853 | 0.549756629 | 0.653111605 |
| 3.536412365 | 0.000405601 | 0.001258173 |
| 2.171318304 | 0.02990712 | 0.059189943 |
| -2.766297392 | 0.00566968 | 0.013757838 |
| -4.726835804 | 2.28E-06 | 1.02E-05 |
| -3.082721664 | 0.002051169 | 0.005512806 |
| 11.22659415 | 3.02E-29 | 1.00E-27 |
| -4.715983268 | 2.41E-06 | 1.07E-05 |
| -1.846184834 | 0.064865372 | 0.115424065 |
| -1.334528621 | 0.182030674 | 0.27371307 |
| -2.691743862 | 0.007107951 | 0.016793659 |

| | | |
|---|---|---|
| NA | NA | NA |
| -4.770826264 | 1.83E-06 | 8.31E-06 |
| 1.529837886 | 0.126056861 | 0.202269584 |
| -1.960501347 | 0.049937221 | 0.092499343 |
| -2.131452791 | 0.033051858 | 0.064647027 |
| 4.369550505 | 1.25E-05 | 4.99E-05 |
| 1.839940342 | 0.065776996 | 0.116881884 |
| NA | NA | NA |
| 0.057540044 | 0.954115009 | 0.968616796 |
| 4.877229738 | 1.08E-06 | 5.03E-06 |
| -2.434835634 | 0.014898556 | 0.032381977 |
| -0.818247796 | 0.413215709 | 0.524828192 |
| -2.615635321 | 0.008906162 | 0.020528708 |
| 0.192810118 | 0.847107679 | NA |
| NA | NA | NA |
| 4.202483668 | 2.64E-05 | 0.000100808 |
| -1.845652047 | 0.064942744 | 0.11554712 |
| -2.254752103 | 0.024148893 | 0.049186003 |
| 19.71159279 | 1.71E-86 | 3.16E-84 |
| 8.285918967 | 1.17E-16 | 1.58E-15 |
| NA | NA | NA |
| NA | NA | NA |
| 0.324874675 | 0.745275925 | NA |
| 12.60622959 | 1.95E-36 | 9.06E-35 |
| 0.824602694 | 0.409597191 | 0.521449453 |
| 17.65935987 | 8.62E-70 | 1.10E-67 |
| -0.324120467 | 0.745846832 | 0.815113575 |
| 9.281591629 | 1.67E-20 | 3.07E-19 |
| 3.044854678 | 0.002327928 | 0.006170037 |
| 0.479046804 | 0.631905333 | NA |
| 12.48251936 | 9.30E-36 | 4.19E-34 |
| -9.850147378 | 6.84E-23 | 1.51E-21 |
| 0.895620865 | 0.370455288 | NA |
| -0.579987442 | 0.561923087 | NA |
| -3.769595835 | 0.000163512 | 0.000544895 |
| NA | NA | NA |
| -2.363776023 | 0.018089745 | 0.038336217 |
| -0.970908847 | 0.331593672 | 0.442586536 |
| -3.368751267 | 0.000755095 | 0.002227202 |
| -3.949450297 | 7.83E-05 | 0.000276339 |
| 0.398085629 | 0.690567067 | 0.771578149 |
| -1.877003052 | 0.060517676 | 0.108834417 |
| 0.063708882 | 0.949202033 | NA |
| -0.551569233 | 0.581243521 | 0.681781026 |
| 1.69074815 | 0.090884914 | 0.153841171 |
| 0.845759974 | 0.397686658 | 0.509499773 |
| NA | NA | NA |

| | | |
|---|---|---|
| 1.04339632 | 0.296764777 | 0.405349106 |
| NA | NA | NA |
| 0.195732583 | 0.84481948 | NA |
| 0.651545954 | 0.514694123 | NA |
| -0.245233438 | 0.806275687 | 0.862932143 |
| -2.060547456 | 0.039346234 | 0.075098081 |
| -0.29143851 | 0.770715966 | 0.834820003 |
| 0.192810118 | 0.847107679 | NA |
| 1.87994279 | 0.060115875 | 0.108236251 |
| -0.273082755 | 0.784789603 | 0.846547309 |
| -1.262129453 | 0.206902209 | 0.303793177 |
| -0.77437228 | 0.438710671 | 0.549016668 |
| 0.000902803 | 0.999279667 | 0.999506869 |
| NA | NA | NA |
| -2.978089245 | 0.002900515 | 0.007524085 |
| 0.330493631 | 0.741027004 | NA |
| -1.198380301 | 0.230768998 | 0.331524308 |
| 0.067492576 | 0.946189572 | NA |
| NA | NA | NA |
| 0.192809521 | 0.847108147 | NA |
| 13.8146061 | 2.08E-43 | 1.30E-41 |
| 4.132106152 | 3.59E-05 | 0.000134126 |
| -1.951408088 | 0.05100852 | 0.094205598 |
| -0.821272761 | 0.411490919 | 0.523141899 |
| -5.926464878 | 3.10E-09 | 2.00E-08 |
| NA | NA | NA |
| -3.048902328 | 0.002296791 | 0.006098949 |
| 0.063708882 | 0.949202033 | NA |
| 0.063707688 | 0.949202983 | NA |
| NA | NA | NA |
| 0.983271546 | 0.32547381 | NA |
| -0.16645039 | 0.867802521 | 0.907882847 |
| 1.590707951 | 0.111675317 | 0.182444565 |
| -1.177031525 | 0.239182924 | 0.341241939 |
| 1.800978999 | 0.07170619 | 0.125774591 |
| 2.198660476 | 0.027902073 | 0.05574624 |
| 0.127870358 | 0.898251569 | NA |
| -7.619108058 | 2.55E-14 | 2.79E-13 |
| -1.567587832 | 0.116977354 | 0.189737799 |
| 2.149250255 | 0.031614567 | 0.062136837 |
| NA | NA | NA |
| 0.192810118 | 0.847107679 | NA |
| 3.737559286 | 0.000185815 | 0.000613664 |
| 11.22705002 | 3.00E-29 | 9.97E-28 |
| 0.104051517 | 0.917128465 | 0.943050098 |
| 1.644782657 | 0.10001464 | 0.166569062 |
| 3.046607755 | 0.002314395 | 0.006140482 |

| | | |
|---|---|---|
| -1.783154289 | 0.074561191 | 0.129971907 |
| 0.157492763 | 0.874856511 | 0.912726807 |
| -3.609234586 | 0.000307102 | 0.000974611 |
| 1.604749064 | 0.10854904 | 0.178151102 |
| 0.245042084 | 0.806423846 | 0.862932143 |
| -1.396713462 | 0.162499754 | 0.249134439 |
| 0.452341166 | 0.651023224 | 0.739678225 |
| 0.066587943 | 0.946909745 | NA |
| -1.727974543 | 0.083992788 | 0.143936704 |
| 4.221035451 | 2.43E-05 | 9.32E-05 |
| 3.558579416 | 0.000372866 | 0.00116456 |
| NA | NA | NA |
| 0.042400557 | 0.966179384 | NA |
| NA | NA | NA |
| 6.478500796 | 9.26E-11 | 7.07E-10 |
| 3.575617886 | 0.000349402 | 0.00109827 |
| 0.364249469 | 0.71567173 | NA |
| 0.808351629 | 0.418888189 | 0.530552139 |
| 6.244099333 | 4.26E-10 | 3.01E-09 |
| 0.822239084 | 0.410940839 | NA |
| -0.219334649 | 0.826389373 | 0.878547656 |
| 5.497443996 | 3.85E-08 | 2.20E-07 |
| NA | NA | NA |
| 0.317642674 | 0.750756 | NA |
| 4.689398718 | 2.74E-06 | 1.21E-05 |
| 0.192808924 | 0.847108615 | NA |
| 1.337216079 | 0.181152115 | 0.272588519 |
| NA | NA | NA |
| 5.346958453 | 8.94E-08 | 4.86E-07 |
| 1.647552413 | 0.099444552 | 0.16581821 |
| NA | NA | NA |
| 0.31685483 | 0.751353758 | NA |
| 15.96757107 | 2.15E-57 | 1.98E-55 |
| 0.192808924 | 0.847108615 | NA |
| 12.14895504 | 5.81E-34 | 2.43E-32 |
| 10.37632151 | 3.18E-25 | 8.25E-24 |
| 0.275770074 | 0.78272467 | NA |
| 10.54568177 | 5.32E-26 | 1.45E-24 |
| 3.741616519 | 0.00018284 | 0.000604397 |
| 1.090945552 | 0.275296842 | 0.382009089 |
| 3.130218809 | 0.001746761 | 0.00477152 |
| 5.469230225 | 4.52E-08 | 2.56E-07 |
| 12.5631193 | 3.37E-36 | 1.55E-34 |
| 7.007608741 | 2.42E-12 | 2.15E-11 |
| 0.192809521 | 0.847108147 | NA |
| 3.047101038 | 0.0023106 | 0.006131881 |
| -2.322000293 | 0.02023292 | 0.042216571 |

| | | | |
|---|---|---|---|
| | 5.553604766 | 2.80E-08 | 1.62E-07 |
| | 8.36539893 | 5.99E-17 | 8.29E-16 |
| | 12.18585191 | 3.70E-34 | 1.56E-32 |
| | 6.779077075 | 1.21E-11 | 1.01E-10 |
| | 8.126683725 | 4.41E-16 | 5.66E-15 |
| | 9.764295983 | 1.60E-22 | 3.43E-21 |
| NA | NA | NA | |
| | 0.192809521 | 0.847108147 | NA |
| | 5.417852369 | 6.03E-08 | 3.35E-07 |
| | 6.044096492 | 1.50E-09 | 1.00E-08 |
| | 0.192810118 | 0.847107679 | NA |
| | 0.063708285 | 0.949202508 | NA |
| | 6.137283183 | 8.39E-10 | 5.76E-09 |
| NA | NA | NA | |
| NA | NA | NA | |
| | 2.254719731 | 0.024150926 | 0.049186142 |
| NA | NA | NA | |
| NA | NA | NA | |
| NA | NA | NA | |
| | 1.416929886 | 0.156503428 | NA |
| NA | NA | NA | |
| NA | NA | NA | |
| NA | NA | NA | |
| | 0.192808327 | 0.847109082 | NA |
| NA | NA | NA | |
| NA | NA | NA | |
| NA | NA | NA | |
| | 2.51470369 | 0.011913246 | 0.026607348 |
| | 6.264654942 | 3.74E-10 | 2.65E-09 |
| | 0.851663256 | 0.39440102 | NA |
| | 2.351897973 | 0.018677897 | NA |
| NA | NA | NA | |
| | 7.108131578 | 1.18E-12 | 1.08E-11 |
| | 1.129133959 | 0.258841328 | NA |
| NA | NA | NA | |
| NA | NA | NA | |
| NA | NA | NA | |
| | 0.063708285 | 0.949202508 | NA |
| NA | NA | NA | |
| | -0.065389369 | 0.947863988 | NA |
| NA | NA | NA | |
| NA | NA | NA | |
| | 0.429236314 | 0.667751258 | NA |
| NA | NA | NA | |
| NA | NA | NA | |
| | 2.822115274 | 0.004770803 | 0.011797581 |
| | 11.52525438 | 9.84E-31 | 3.52E-29 |

| | | |
|---|---|---|
| -0.573400105 | 0.566373819 | 0.668285329 |
| 0.125389792 | 0.90021497 | 0.930398079 |
| 6.395442102 | 1.60E-10 | 1.18E-09 |
| 6.875066283 | 6.20E-12 | 5.29E-11 |
| -0.156091774 | 0.875960682 | 0.913407797 |
| 0.262131689 | 0.793219916 | NA |
| NA | NA | NA |
| NA | NA | NA |
| NA | NA | NA |
| NA | NA | NA |
| NA | NA | NA |
| NA | NA | NA |
| NA | NA | NA |
| NA | NA | NA |
| NA | NA | NA |
| NA | NA | NA |
| NA | NA | NA |
| NA | NA | NA |
| 0.192809521 | 0.847108147 | NA |
| NA | NA | NA |
| NA | NA | NA |
| NA | NA | NA |
| NA | NA | NA |
| NA | NA | NA |
| NA | NA | NA |
| NA | NA | NA |
| NA | NA | NA |
| NA | NA | NA |
| NA | NA | NA |
| NA | NA | NA |
| 0.192810118 | 0.847107679 | NA |
| NA | NA | NA |
| NA | NA | NA |
| 0.192810118 | 0.847107679 | NA |
| NA | NA | NA |
| NA | NA | NA |
| NA | NA | NA |
| NA | NA | NA |
| NA | NA | NA |
| NA | NA | NA |
| NA | NA | NA |
| NA | NA | NA |
| -1.321514258 | 0.186329952 | 0.279003785 |
| NA | NA | NA |
| 0.063708285 | 0.949202508 | NA |
| NA | NA | NA |
| NA | NA | NA |

| | | |
|---|---|---|
| 0.755189241 | 0.450135443 | NA |
| -0.065389369 | 0.947863988 | NA |
| 10.47646902 | 1.11E-25 | 2.97E-24 |
| 0.628856507 | 0.529443002 | NA |
| 1.566279871 | 0.117283111 | NA |
| 0.790003936 | 0.42952547 | NA |
| -7.228201398 | 4.89E-13 | 4.70E-12 |
| -0.988468238 | 0.322923382 | 0.433811306 |
| -1.292340066 | 0.196239402 | 0.291078807 |
| 5.019713989 | 5.17E-07 | 2.53E-06 |
| -1.400721848 | 0.161297267 | 0.247518275 |
| 1.402857862 | 0.160659229 | 0.246735826 |
| -1.488010889 | 0.136748022 | 0.216195237 |
| 1.17942751 | 0.238227986 | 0.340345119 |
| 1.766758556 | 0.077268672 | 0.134075207 |
| 1.800869077 | 0.071723518 | 0.125787532 |
| 2.491791662 | 0.012710057 | 0.028175779 |
| 4.761993486 | 1.92E-06 | 8.65E-06 |
| 0.580106907 | 0.561842526 | 0.664031139 |
| 0.550659306 | 0.581867246 | 0.68219885 |
| NA | NA | NA |
| -8.20234274 | 2.36E-16 | 3.11E-15 |
| 0.890621594 | 0.37313221 | 0.485951243 |
| NA | NA | NA |
| 2.965249374 | 0.003024377 | 0.007813805 |
| 12.27604455 | 1.22E-34 | 5.22E-33 |
| 0.890025802 | 0.373452032 | 0.486241384 |
| -7.048192059 | 1.81E-12 | 1.63E-11 |
| -0.680260557 | 0.496339494 | NA |
| 0.320071032 | 0.748914485 | NA |
| -4.384540823 | 1.16E-05 | 4.69E-05 |
| -4.323093231 | 1.54E-05 | 6.09E-05 |
| 0.192810118 | 0.847107679 | NA |
| -2.33562672 | 0.019510705 | 0.040895535 |
| 0.192808327 | 0.847109082 | NA |
| 0.193076837 | 0.846898793 | NA |
| -0.065389369 | 0.947863988 | NA |
| NA | NA | NA |
| NA | NA | NA |
| 0.581203698 | 0.561103175 | 0.663317077 |
| 1.940832383 | 0.052278612 | 0.09621571 |
| 3.706128442 | 0.000210452 | 0.000688736 |
| 0.671977618 | 0.501597945 | 0.608432109 |
| 8.724255417 | 2.68E-18 | 4.15E-17 |
| 1.613775666 | 0.106576095 | 0.175430728 |
| -2.687692907 | 0.007194753 | 0.016963514 |
| 1.445843886 | 0.148220994 | 0.231093277 |

| | | |
|---|---|---|
| 1.536113085 | 0.124510648 | 0.200160662 |
| 0.78221469 | 0.434088413 | 0.544814369 |
| 0.585780725 | 0.558022878 | 0.660486719 |
| -3.072574897 | 0.002122206 | 0.005684196 |
| -0.13923111 | 0.889267526 | 0.922547392 |
| 7.234455324 | 4.67E-13 | 4.50E-12 |
| 1.519617805 | 0.12860706 | 0.205671997 |
| 15.06893534 | 2.59E-51 | 2.03E-49 |
| 0.177696608 | 0.858961249 | 0.901957335 |
| -5.259371613 | 1.45E-07 | 7.63E-07 |
| -0.335974611 | 0.736890012 | 0.808324875 |
| 4.310695098 | 1.63E-05 | 6.41E-05 |
| 3.68268651 | 0.000230789 | 0.000749241 |
| 5.836019046 | 5.35E-09 | 3.36E-08 |
| NA | NA | NA |
| -0.850533266 | 0.395028673 | 0.50701547 |
| 9.345677402 | 9.13E-21 | 1.72E-19 |
| -11.01923865 | 3.09E-28 | 9.69E-27 |
| -1.890486812 | 0.058692883 | 0.106139936 |
| -3.045728998 | 0.002321169 | 0.006156498 |
| -3.003061164 | 0.002672787 | 0.00698766 |
| NA | NA | NA |
| -0.775779287 | 0.437879316 | NA |
| 6.22391749 | 4.85E-10 | 3.41E-09 |
| 0.54203616 | 0.587793596 | NA |
| 0.348613926 | 0.727379171 | 0.800910639 |
| 0.063708285 | 0.949202508 | NA |
| NA | NA | NA |
| NA | NA | NA |
| -4.281094882 | 1.86E-05 | 7.27E-05 |
| -3.172683524 | 0.001510371 | 0.004180997 |
| 1.720799304 | 0.085287249 | 0.145785689 |
| 0.595265498 | 0.551666014 | NA |
| 4.473675286 | 7.69E-06 | 3.19E-05 |
| -0.717512085 | 0.473058182 | 0.581927014 |
| -1.646109351 | 0.099741248 | 0.166191148 |
| 8.619878021 | 6.70E-18 | 9.99E-17 |
| -3.093033265 | 0.001981219 | 0.005345469 |
| -1.35775174 | 0.174542473 | 0.264487111 |
| -0.711116682 | 0.477011933 | NA |
| NA | NA | NA |
| -2.969526907 | 0.002982587 | 0.007717787 |
| 0.06371127 | 0.949200131 | NA |
| NA | NA | NA |
| 1.492231343 | 0.135638508 | 0.214725942 |
| NA | NA | NA |
| 1.479561105 | 0.138990413 | 0.21904861 |

| | | |
|---|---|---|
| 0.192809521 | 0.847108147 | NA |
| 1.255746352 | 0.209207948 | NA |
| 4.932082664 | 8.14E-07 | 3.87E-06 |
| NA | NA | NA |
| 0.192808924 | 0.847108615 | NA |
| 1.114140076 | 0.265219098 | 0.370820804 |
| -1.525671767 | 0.127091629 | 0.203721004 |
| 5.850171092 | 4.91E-09 | 3.11E-08 |
| -2.429870931 | 0.015104201 | 0.032780555 |
| 1.830655313 | 0.067152007 | 0.119004159 |
| -6.482564861 | 9.02E-11 | 6.89E-10 |
| 5.030387893 | 4.89E-07 | 2.40E-06 |
| -0.536693088 | 0.591479631 | 0.690712661 |
| NA | NA | NA |
| -3.428139048 | 0.000607734 | 0.001824791 |
| -1.537898213 | 0.124073504 | 0.199560456 |
| 0.234307692 | 0.814746115 | 0.869752857 |
| -14.30019172 | 2.18E-46 | 1.53E-44 |
| -5.521435879 | 3.36E-08 | 1.93E-07 |
| 5.029352904 | 4.92E-07 | 2.41E-06 |
| -0.494906239 | 0.620666307 | 0.714988846 |
| NA | NA | NA |
| NA | NA | NA |
| -0.065389369 | 0.947863988 | NA |
| -3.780158362 | 0.000156729 | 0.000523544 |
| -0.194488217 | 0.845793621 | NA |
| NA | NA | NA |
| -0.876841035 | 0.380572987 | 0.493076343 |
| 0.063708075 | 0.949202675 | NA |
| NA | NA | NA |
| NA | NA | NA |
| NA | NA | NA |
| 2.526831132 | 0.011509683 | 0.025830482 |
| -1.8238105 | 0.068180726 | 0.120494498 |
| -2.086490166 | 0.036934245 | 0.071134515 |
| 0.340005152 | 0.733852648 | NA |
| 8.452209267 | 2.86E-17 | 4.07E-16 |
| -0.810162777 | 0.417846628 | 0.529500866 |
| 0.192809521 | 0.847108147 | NA |
| -0.596459564 | 0.550868259 | 0.654243806 |
| 0.850515438 | 0.395038581 | 0.50701547 |
| -3.440963592 | 0.000579647 | 0.001747176 |
| 2.406896202 | 0.016088742 | 0.034667804 |
| NA | NA | NA |
| 0.576872103 | 0.564025857 | NA |
| 0.192636473 | 0.847243678 | NA |
| -1.003613081 | 0.315565147 | 0.42605295 |

| | | |
|---|---|---|
| NA | NA | NA |
| 0.526710219 | 0.598394838 | NA |
| NA | NA | NA |
| NA | NA | NA |
| 0.320071032 | 0.748914485 | NA |
| -1.031008615 | 0.30253678 | 0.411772463 |
| 0.395416634 | 0.692535435 | 0.773121714 |
| -2.78287284 | 0.005387992 | 0.013153364 |
| -2.14279028 | 0.032129942 | 0.063055704 |
| -0.065389369 | 0.947863988 | NA |
| 1.347490685 | 0.177822252 | 0.268449394 |
| -0.984807493 | 0.324718639 | 0.435708563 |
| 0.018831835 | 0.984975257 | 0.989362763 |
| 1.903597515 | 0.056962623 | 0.103519527 |
| -0.52158626 | 0.601958432 | 0.699427783 |
| 5.149798009 | 2.61E-07 | 1.33E-06 |
| 1.960496217 | 0.04993782 | 0.092499343 |
| -0.438646848 | 0.660917446 | 0.747695533 |
| 3.698815076 | 0.000216608 | 0.000707241 |
| -2.765574734 | 0.005682258 | 0.013785686 |
| 2.405342779 | 0.016157303 | 0.034803553 |
| 4.574774555 | 4.77E-06 | 2.04E-05 |
| 0.063708882 | 0.949202033 | NA |
| -0.877390756 | 0.380274433 | 0.492842595 |
| -1.294237123 | 0.195583528 | 0.290238673 |
| 1.651230191 | 0.098691581 | 0.164804207 |
| -4.972721304 | 6.60E-07 | 3.18E-06 |
| 0.463408908 | 0.643071295 | NA |
| 1.246436354 | 0.212604242 | 0.31049372 |
| 0.063708882 | 0.949202033 | NA |
| -0.4870311 | 0.626236292 | 0.719649896 |
| 0.304173871 | 0.76099543 | 0.82711866 |
| 1.02189599 | 0.306830131 | 0.416573995 |
| 0.215634835 | 0.8292724 | 0.880714875 |
| -0.065389369 | 0.947863988 | NA |
| -0.57762451 | 0.563517656 | NA |
| 2.145240886 | 0.031933592 | 0.062694942 |
| -7.548523049 | 4.40E-14 | 4.69E-13 |
| 1.172617278 | 0.240949317 | 0.343298956 |
| 0.536740279 | 0.591447029 | 0.690706814 |
| -1.675295657 | 0.093876237 | 0.158070785 |
| -5.358195341 | 8.41E-08 | 4.58E-07 |
| 0.063708882 | 0.949202033 | NA |
| 1.422506377 | 0.154879302 | 0.239739997 |
| 0.063708882 | 0.949202033 | NA |
| -0.065389966 | 0.947863513 | NA |
| -1.450181784 | 0.147007833 | NA |

| | | | |
|---|---|---|---|
| NA | NA | NA | |
| | -2.262994954 | 0.023636 | 0.048318286 |
| | -5.570550645 | 2.54E-08 | 1.48E-07 |
| | 0.95961227 | 0.337250391 | NA |
| | 6.171245024 | 6.78E-10 | 4.68E-09 |
| | 2.518873654 | 0.011773089 | 0.026338973 |
| NA | NA | NA | |
| | -7.755023943 | 8.83E-15 | 1.01E-13 |
| | 0.160220051 | 0.872707735 | NA |
| NA | NA | NA | |
| NA | NA | NA | |
| | -2.041699632 | 0.041181335 | 0.078153675 |
| NA | NA | NA | |
| | -1.426066906 | 0.15384903 | 0.238397897 |
| | 0.066587943 | 0.946909745 | NA |
| | 1.925476066 | 0.054169827 | 0.09917861 |
| | -0.152080709 | 0.87912328 | NA |
| | 0.026212805 | 0.979087603 | 0.985465767 |
| | 12.41041654 | 2.29E-35 | 1.02E-33 |
| | 0.400687315 | 0.688650351 | NA |
| | 0.135288389 | 0.892383865 | 0.924824782 |
| | 0.27464153 | 0.783591656 | NA |
| | -0.065388772 | 0.947864464 | NA |
| | -7.570771468 | 3.71E-14 | 3.98E-13 |
| | -1.365424126 | 0.172119785 | 0.261322227 |
| | 0.464295084 | 0.642436346 | 0.732504476 |
| | -0.870122173 | 0.384233642 | 0.496505882 |
| NA | NA | NA | |
| | 0.824166665 | 0.409844865 | 0.52168516 |
| | 2.74549814 | 0.006041911 | 0.01454405 |
| NA | NA | NA | |
| NA | NA | NA | |
| NA | NA | NA | |
| NA | NA | NA | |
| NA | NA | NA | |
| NA | NA | NA | |
| NA | NA | NA | |
| NA | NA | NA | |
| NA | NA | NA | |
| | -0.156913854 | 0.875312741 | 0.912954326 |
| | -0.241174546 | 0.809419838 | 0.865358728 |
| | -7.592464022 | 3.14E-14 | 3.40E-13 |
| | -2.39675843 | 0.016540825 | 0.035522681 |
| | -6.219490004 | 4.99E-10 | 3.50E-09 |
| NA | NA | NA | |
| | 3.02956912 | 0.002449029 | 0.006453225 |
| | 5.379864042 | NA | NA |

| | | |
|---|---|---|
| NA | NA | NA |
| 2.079254245 | 0.037593988 | 0.072216234 |
| 0.075347333 | 0.939938362 | 0.959136418 |
| 1.6157924 | 0.106139205 | 0.174849566 |
| 7.729269005 | 1.08E-14 | 1.23E-13 |
| -2.671106894 | 0.007560157 | 0.017731514 |
| -0.153229871 | 0.878217002 | 0.91476425 |
| NA | NA | NA |
| 1.968087327 | 0.049057991 | 0.0910921 |
| 0.296726814 | 0.766675084 | 0.831666696 |
| 0.421130352 | 0.673659899 | NA |
| 0.459859446 | 0.645617111 | 0.73493707 |
| 0.884469345 | 0.376442903 | 0.489156626 |
| 5.335326167 | 9.54E-08 | 5.15E-07 |
| 15.07857744 | 2.24E-51 | 1.77E-49 |
| 6.422647485 | 1.34E-10 | 1.00E-09 |
| NA | NA | NA |
| 4.532307178 | 5.83E-06 | 2.46E-05 |
| 5.261505028 | 1.43E-07 | 7.54E-07 |
| 8.018038257 | 1.07E-15 | 1.34E-14 |
| 0.92883463 | 0.352974795 | NA |
| 8.059409882 | 7.67E-16 | 9.66E-15 |
| 8.31959699 | 8.83E-17 | 1.20E-15 |
| 1.866458 | 0.061977315 | 0.11103659 |
| -1.809327566 | 0.070400129 | 0.123872184 |
| 5.982435651 | 2.20E-09 | 1.44E-08 |
| -1.418925873 | 0.155920628 | 0.241085578 |
| -0.065389369 | 0.947863988 | NA |
| -6.120259447 | 9.34E-10 | 6.38E-09 |
| -2.10905593 | 0.034939754 | 0.067825504 |
| -2.983438525 | 0.002850293 | 0.007404549 |
| -0.361434111 | 0.717774952 | 0.793687359 |
| 1.657655805 | 0.097386959 | 0.162962668 |
| NA | NA | NA |
| NA | NA | NA |
| -0.099553712 | 0.920698644 | 0.945538501 |
| NA | NA | NA |
| 8.472912992 | 2.39E-17 | 3.43E-16 |
| 6.863800687 | 6.71E-12 | 5.70E-11 |
| 0.698028324 | 0.485159481 | NA |
| 0.192810118 | 0.847107679 | NA |
| -0.065389369 | 0.947863988 | NA |
| -4.280493092 | 1.86E-05 | 7.28E-05 |
| NA | NA | NA |
| -6.635198628 | 3.24E-11 | 2.58E-10 |
| -1.88205003 | 0.059829223 | 0.107813208 |
| 0.425809797 | 0.670246448 | 0.75480377 |

| | | |
|---|---|---|
| -3.789205005 | 0.00015113 | 0.000506262 |
| -0.398556785 | 0.690219809 | 0.771293443 |
| 3.968600806 | 7.23E-05 | 0.00025653 |
| 0.739111913 | 0.459839043 | NA |
| -3.888573112 | 0.000100835 | 0.000348317 |
| 2.551686347 | 0.010720298 | 0.024267799 |
| NA | NA | NA |
| -0.892109034 | 0.372334493 | 0.485164889 |
| 5.992485421 | 2.07E-09 | 1.36E-08 |
| 1.910131158 | 0.056116328 | 0.102197625 |
| -9.743978829 | 1.96E-22 | 4.16E-21 |
| -0.572326565 | 0.567100756 | 0.668890812 |
| NA | NA | NA |
| -3.153060756 | 0.001615682 | 0.004447434 |
| -3.536501212 | 0.000405465 | 0.001257905 |
| 2.333145946 | 0.019640487 | 0.041131307 |
| 5.397694405 | 6.75E-08 | 3.72E-07 |
| 0.52981657 | 0.596239115 | NA |
| 0.682095204 | 0.495178753 | 0.602280198 |
| 6.193730435 | 5.88E-10 | 4.09E-09 |
| 3.321867785 | 0.000894171 | 0.002600336 |
| -4.939519306 | 7.83E-07 | 3.73E-06 |
| NA | NA | NA |
| -1.374697911 | 0.169225119 | 0.257567938 |
| -2.045548317 | 0.040800831 | 0.077543265 |
| 3.383406674 | 0.000715925 | 0.002121417 |
| -2.71211545 | 0.00668553 | 0.015921867 |
| 10.2806944 | 8.61E-25 | 2.16E-23 |
| 0.004555477 | 0.996365267 | 0.997521005 |
| -3.089917775 | 0.002002119 | 0.005396042 |
| -2.888375074 | 0.003872378 | 0.00976679 |
| 5.677818023 | 1.36E-08 | 8.21E-08 |
| -0.011217083 | 0.99105025 | 0.993750011 |
| 1.824721507 | 0.068043066 | 0.120276691 |
| -0.795554495 | 0.426291024 | 0.537643069 |
| 1.665999654 | 0.095713483 | 0.160663405 |
| -0.823497602 | 0.41022508 | 0.522089478 |
| -0.207528107 | 0.835597445 | 0.88469202 |
| -3.203023203 | 0.00135993 | 0.003801123 |
| 6.301501716 | 2.95E-10 | 2.12E-09 |
| -5.273544234 | 1.34E-07 | 7.09E-07 |
| NA | NA | NA |
| -0.045420025 | 0.96377252 | 0.975227449 |
| NA | NA | NA |
| -0.065388772 | 0.947864464 | NA |
| NA | NA | NA |
| 0.31685483 | 0.751353758 | NA |

| | | | |
|---|---|---|---|
| NA | NA | NA | |
| NA | NA | NA | |
| 0.288217989 | 0.77317988 | NA | |
| NA | NA | NA | |
| NA | NA | NA | |
| NA | NA | NA | |
| NA | NA | NA | |
| NA | NA | NA | |
| NA | NA | NA | |
| NA | NA | NA | |
| NA | NA | NA | |
| NA | NA | NA | |
| NA | NA | NA | |
| NA | NA | NA | |
| NA | NA | NA | |
| NA | NA | NA | |
| 0.324874675 | 0.745275925 | NA | |
| 0.550056725 | 0.582280467 | NA | |
| NA | NA | NA | |
| 3.681060725 | 0.000232266 | | 0.000753351 |
| -0.841242415 | 0.400212143 | | 0.511896875 |
| 1.095260369 | 0.273402584 | NA | |
| 1.355028708 | 0.175408411 | NA | |
| 0.3474247 | 0.728272279 | NA | |
| -1.709531018 | 0.087352631 | | 0.148828287 |
| -2.771377883 | 0.005581961 | | 0.013580039 |
| -1.716376179 | 0.086093214 | | 0.146952614 |
| -1.90108303 | 0.057291138 | | 0.104010821 |
| -5.373335366 | 7.73E-08 | | 4.23E-07 |
| -1.505969565 | 0.132074987 | | 0.209987736 |
| NA | NA | NA | |
| -3.012213162 | 0.002593504 | | 0.006798849 |
| NA | NA | NA | |
| NA | NA | NA | |
| -0.065388772 | 0.947864464 | NA | |
| 0.192809521 | 0.847108147 | NA | |
| 0.063708882 | 0.949202033 | NA | |
| NA | NA | NA | |
| NA | NA | NA | |
| -0.065389966 | 0.947863513 | NA | |
| NA | NA | NA | |
| -0.038323076 | 0.969430093 | | 0.97875092 |
| -6.343119773 | 2.25E-10 | | 1.64E-09 |
| -2.544997248 | 0.01092786 | | 0.024659605 |
| NA | NA | NA | |
| -1.25460113 | 0.209623593 | | 0.306978618 |
| NA | NA | NA | |

| | | |
|---|---|---|
| -1.11350626 | 0.265491061 | 0.371156328 |
| 4.4304914 | 9.40E-06 | 3.85E-05 |
| -0.972127372 | 0.330987187 | 0.441882486 |
| -1.219259976 | 0.222745535 | 0.322272875 |
| -2.563762375 | 0.010354441 | 0.023511898 |
| -1.716018914 | 0.086158581 | 0.147045012 |
| -0.329772929 | 0.741571544 | 0.81185915 |
| -4.817448606 | 1.45E-06 | 6.66E-06 |
| -0.274632474 | 0.783598615 | 0.845545201 |
| -0.114261147 | 0.909030781 | NA |
| -0.578882407 | 0.562668522 | 0.664822182 |
| 0.06371127 | 0.949200131 | NA |
| NA | NA | NA |
| NA | NA | NA |
| NA | NA | NA |
| NA | NA | NA |
| NA | NA | NA |
| NA | NA | NA |
| 0.063708882 | 0.949202033 | NA |
| NA | NA | NA |
| NA | NA | NA |
| -0.19448762 | 0.845794089 | NA |
| 0.192810118 | 0.847107679 | NA |
| NA | NA | NA |
| NA | NA | NA |
| -0.209525922 | 0.834037702 | NA |
| 0.192810118 | 0.847107679 | NA |
| -0.41111373 | 0.680989142 | NA |
| NA | NA | NA |
| NA | NA | NA |
| -0.065389369 | 0.947863988 | NA |
| NA | NA | NA |
| -0.065389369 | 0.947863988 | NA |
| NA | NA | NA |
| NA | NA | NA |
| NA | NA | NA |
| 0.063708217 | 0.949202562 | NA |
| -0.06842302 | 0.945448898 | 0.962796937 |
| -0.115607359 | 0.90796373 | NA |
| NA | NA | NA |
| -0.065388772 | 0.947864464 | NA |
| NA | NA | NA |
| NA | NA | NA |
| -0.065388772 | 0.947864464 | NA |
| NA | NA | NA |
| NA | NA | NA |
| -0.586888631 | 0.557278506 | NA |

| | | | |
|---|---|---|---|
| 0.34371353 | 0.731061735 | NA | |
| NA | NA | NA | |
| NA | NA | NA | |
| NA | NA | NA | |
| NA | NA | NA | |
| NA | NA | NA | |
| NA | NA | NA | |
| NA | NA | NA | |
| 0.192810118 | 0.847107679 | NA | |
| -0.065388772 | 0.947864464 | NA | |
| NA | NA | NA | |
| NA | NA | NA | |
| NA | NA | NA | |
| NA | NA | NA | |
| NA | NA | NA | |
| NA | NA | NA | |
| NA | NA | NA | |
| NA | NA | NA | |
| NA | NA | NA | |
| 0.324874675 | 0.745275925 | NA | |
| 0.427961561 | 0.668679107 | NA | |
| NA | NA | NA | |
| NA | NA | NA | |
| NA | NA | NA | |
| -0.065388772 | 0.947864464 | NA | |
| NA | NA | NA | |
| NA | NA | NA | |
| NA | NA | NA | |
| NA | NA | NA | |
| NA | NA | NA | |
| 0.192810118 | 0.847107679 | NA | |
| NA | NA | NA | |
| NA | NA | NA | |
| NA | NA | NA | |
| NA | NA | NA | |
| NA | NA | NA | |
| NA | NA | NA | |
| 0.063708882 | 0.949202033 | NA | |
| NA | NA | NA | |
| 0.14266915 | 0.886551482 | | 0.920533627 |
| NA | NA | NA | |
| NA | NA | NA | |
| NA | NA | NA | |
| NA | NA | NA | |
| NA | NA | NA | |
| NA | NA | NA | |
| 1.163640671 | 0.244569662 | | 0.34738274 |

| | | | |
|---|---|---|---|
| NA | NA | NA | |
| NA | NA | NA | |
| NA | NA | NA | |
| NA | NA | NA | |
| NA | NA | NA | |
| NA | NA | NA | |
| NA | NA | NA | |
| NA | NA | NA | |
| NA | NA | NA | |
| NA | NA | NA | |
| NA | NA | NA | |
| NA | NA | NA | |
| NA | NA | NA | |
| NA | NA | NA | |
| NA | NA | NA | |
| NA | NA | NA | |
| NA | NA | NA | |
| NA | NA | NA | |
| NA | NA | NA | |
| NA | NA | NA | |
| NA | NA | NA | |
| | 0.321951285 | 0.7474896 | 0.816404485 |
| NA | NA | NA | |
| NA | NA | NA | |
| NA | NA | NA | |
| NA | NA | NA | |
| | -1.780633737 | 0.074972305 | 0.130615771 |
| NA | NA | NA | |
| NA | NA | NA | |
| NA | NA | NA | |
| NA | NA | NA | |
| NA | NA | NA | |
| NA | NA | NA | |
| NA | NA | NA | |
| NA | NA | NA | |
| NA | NA | NA | |
| NA | NA | NA | |
| NA | NA | NA | |
| NA | NA | NA | |
| | -0.065388772 | 0.947864464 | NA |
| NA | NA | NA | |
| NA | NA | NA | |
| NA | NA | NA | |
| | -2.469470995 | 0.013531299 | 0.029764339 |
| NA | NA | NA | |
| | -2.189516374 | 0.02855933 | 0.056914015 |
| | 0.797221795 | 0.42532223 | NA |

| | | | |
|---|---|---|---|
| | 0.606958589 | 0.543878394 | 0.647766005 |
| | -0.676441163 | 0.49876059 | 0.605871273 |
| | 0.001686687 | 0.998654219 | 0.999053361 |
| | 0.976061191 | 0.329034142 | 0.439942552 |
| | -0.71459807 | 0.474857437 | 0.583804787 |
| | 0.021805253 | 0.982603304 | 0.987817803 |
| NA | NA | NA | |
| | -2.261365139 | 0.023736656 | 0.048496326 |
| | -2.892250158 | 0.003824933 | 0.009652961 |
| | 9.896404133 | 4.32E-23 | 9.69E-22 |
| | -1.261979474 | 0.206956172 | 0.303854587 |
| | 0.515194925 | 0.606416815 | 0.703234474 |
| | 9.743228527 | 1.97E-22 | 4.19E-21 |
| | 15.81787394 | 2.34E-56 | 2.13E-54 |
| NA | NA | NA | |
| | 2.801304047 | 0.005089654 | 0.012499199 |
| | 0.192809521 | 0.847108147 | NA |
| | 9.831955039 | 8.20E-23 | 1.79E-21 |
| | 2.162947434 | 0.030545223 | 0.06028599 |
| | -1.282129203 | 0.199797326 | 0.295458293 |
| | 1.126324089 | 0.260028365 | 0.364930311 |
| | 4.522526216 | 6.11E-06 | 2.57E-05 |
| | 5.373509168 | 7.72E-08 | 4.22E-07 |
| | 3.25124846 | 0.001148994 | 0.003269962 |
| | -6.187770978 | 6.10E-10 | 4.23E-09 |
| NA | NA | NA | |
| NA | NA | NA | |
| NA | NA | NA | |
| | -0.118714671 | 0.905501413 | NA |
| | 1.573806452 | 0.115532202 | 0.187765146 |
| | 0.324874675 | 0.745275925 | NA |
| NA | NA | NA | |
| | -1.830469294 | 0.067179794 | 0.119036546 |
| | 2.848186498 | 0.004396915 | 0.010952904 |
| | 5.278724711 | 1.30E-07 | 6.91E-07 |
| | 1.002646943 | 0.316031238 | 0.426567185 |
| | 0.334243658 | 0.738195702 | NA |
| NA | NA | NA | |
| | -0.51065809 | 0.609590492 | 0.705980316 |
| | 0.335128147 | 0.737528421 | NA |
| NA | NA | NA | |
| NA | NA | NA | |
| NA | NA | NA | |
| NA | NA | NA | |
| NA | NA | NA | |
| NA | NA | NA | |
| NA | NA | NA | |

| | | | |
|---|---|---|---|
| NA | NA | NA | |
| -0.065388772 | 0.947864464 | NA | |
| 0.214588724 | 0.830087986 | NA | |
| NA | NA | NA | |
| NA | NA | NA | |
| NA | NA | NA | |
| NA | NA | NA | |
| -0.065389966 | 0.947863513 | NA | |
| NA | NA | NA | |
| NA | NA | NA | |
| NA | NA | NA | |
| NA | NA | NA | |
| NA | NA | NA | |
| NA | NA | NA | |
| NA | NA | NA | |
| 0.962989502 | 0.335552792 | | 0.44677304 |
| NA | NA | NA | |
| NA | NA | NA | |
| NA | NA | NA | |
| -0.065388772 | 0.947864464 | NA | |
| 0.063707135 | 0.949203424 | NA | |
| 0.083879523 | 0.933152221 | NA | |
| -0.256367733 | 0.797666901 | NA | |
| -1.349869018 | 0.177058001 | | 0.267537686 |
| NA | NA | NA | |
| -0.15223098 | 0.87900476 | | 0.915034422 |
| NA | NA | NA | |
| NA | NA | NA | |
| -1.582728997 | 0.113483246 | | 0.184987669 |
| -0.081381505 | 0.935138557 | NA | |
| NA | NA | NA | |
| NA | NA | NA | |
| NA | NA | NA | |
| NA | NA | NA | |
| NA | NA | NA | |
| NA | NA | NA | |
| NA | NA | NA | |
| NA | NA | NA | |
| NA | NA | NA | |
| NA | NA | NA | |
| NA | NA | NA | |
| NA | NA | NA | |
| NA | NA | NA | |
| NA | NA | NA | |
| NA | NA | NA | |
| NA | NA | NA | |
| NA | NA | NA | |

| | | |
|---|---|---|
| NA | NA | NA |
| NA | NA | NA |
| NA | NA | NA |
| NA | NA | NA |
| NA | NA | NA |
| NA | NA | NA |
| NA | NA | NA |
| NA | NA | NA |
| NA | NA | NA |
| NA | NA | NA |
| NA | NA | NA |
| NA | NA | NA |
| NA | NA | NA |
| NA | NA | NA |
| NA | NA | NA |
| NA | NA | NA |
| NA | NA | NA |
| NA | NA | NA |
| NA | NA | NA |
| NA | NA | NA |
| NA | NA | NA |
| NA | NA | NA |
| 1.329502804 | 0.183682141 | 0.275765595 |
| NA | NA | NA |
| NA | NA | NA |
| NA | NA | NA |
| NA | NA | NA |
| NA | NA | NA |
| NA | NA | NA |
| NA | NA | NA |
| NA | NA | NA |
| NA | NA | NA |
| NA | NA | NA |
| NA | NA | NA |
| NA | NA | NA |
| NA | NA | NA |
| NA | NA | NA |
| NA | NA | NA |
| NA | NA | NA |
| NA | NA | NA |
| NA | NA | NA |
| NA | NA | NA |
| NA | NA | NA |
| NA | NA | NA |
| NA | NA | NA |
| NA | NA | NA |
| NA | NA | NA |

| | | | |
|---|---|---|---|
| NA | NA | NA | |
| NA | NA | NA | |
| NA | NA | NA | |
| NA | NA | NA | |
| | 0.541356021 | 0.588262214 | NA |
| | 0.278377556 | 0.780722547 | NA |
| | 0.086114805 | 0.931375155 | 0.953235203 |
| | -1.68920177 | 0.09118077 | NA |
| NA | NA | NA | |
| | 9.301537161 | 1.38E-20 | 2.56E-19 |
| | 5.940341773 | 2.84E-09 | 1.85E-08 |
| | 1.483478739 | 0.137947261 | 0.217747364 |
| | 0.079846339 | 0.936359468 | NA |
| | 14.15700116 | 1.69E-45 | 1.16E-43 |
| | 2.011540116 | 0.044268437 | 0.083323732 |
| | -1.236612928 | 0.216230818 | 0.314794241 |
| | -3.62289417 | 0.000291325 | 0.000928073 |
| | -6.878812213 | 6.04E-12 | 5.16E-11 |
| | -10.87259565 | 1.56E-27 | 4.71E-26 |
| | 0.152273594 | 0.878971151 | 0.915034422 |
| | 5.369214914 | 7.91E-08 | 4.32E-07 |
| | 2.655046363 | 0.007929751 | 0.018482323 |
| NA | NA | NA | |
| | 13.84557354 | 1.35E-43 | 8.51E-42 |
| | -0.152229689 | 0.879005778 | 0.915034422 |
| | 0.153946836 | 0.877651655 | 0.914503526 |
| | 2.150624981 | 0.031505812 | 0.061947405 |
| | 0.364124391 | 0.715765124 | 0.792031198 |
| | 1.181046429 | 0.23758428 | 0.339599938 |
| | 1.045975471 | 0.295572358 | 0.403874665 |
| | -0.878607076 | 0.379614358 | 0.492242002 |
| NA | NA | NA | |
| | -0.78126144 | 0.434648744 | 0.545298886 |
| | 0.288217989 | 0.77317988 | NA |
| | -1.219022999 | 0.222835463 | 0.322365718 |
| | -2.535226064 | 0.011237478 | 0.025303533 |
| | 0.324874675 | 0.745275925 | NA |
| NA | NA | NA | |
| | 0.275361891 | 0.783038219 | NA |
| | -1.218677185 | 0.222966741 | 0.322536989 |
| | 0.009796506 | 0.992183644 | 0.994527917 |
| | 6.515466586 | 7.25E-11 | 5.60E-10 |
| | -2.367551098 | 0.017906247 | 0.038014705 |
| | -3.575019256 | 0.000350202 | 0.001100233 |
| | 5.652820581 | 1.58E-08 | 9.42E-08 |
| | -3.723990258 | 0.000196098 | 0.00064558 |
| | -9.766361887 | 1.57E-22 | 3.36E-21 |

| | | | |
|---|---|---|---|
| | -2.002807891 | 0.045197913 | 0.084837177 |
| | 5.141725574 | 2.72E-07 | 1.39E-06 |
| | -7.495932572 | 6.58E-14 | 6.88E-13 |
| | -7.116917734 | 1.10E-12 | 1.02E-11 |
| NA | NA | NA | |
| | -5.266871643 | 1.39E-07 | 7.34E-07 |
| | -1.770945838 | 0.076569709 | 0.132982224 |
| | -2.562745249 | 0.010384822 | 0.02357233 |
| | -2.959184458 | 0.003084544 | 0.007948743 |
| | -0.133422272 | 0.893859432 | 0.926008923 |
| | -1.137545145 | 0.255310463 | 0.359679231 |
| | 0.450424123 | 0.652404654 | 0.740676214 |
| NA | NA | NA | |
| NA | NA | NA | |
| NA | NA | NA | |
| | -0.328962065 | 0.742184362 | NA |
| NA | NA | NA | |
| NA | NA | NA | |
| NA | NA | NA | |
| NA | NA | NA | |
| NA | NA | NA | |
| NA | NA | NA | |
| NA | NA | NA | |
| NA | NA | NA | |
| NA | NA | NA | |
| NA | NA | NA | |
| | -0.065388772 | 0.947864464 | NA |
| NA | NA | NA | |
| | 0.192809521 | 0.847108147 | NA |
| NA | NA | NA | |
| NA | NA | NA | |
| | 0.192809521 | 0.847108147 | NA |
| NA | NA | NA | |
| | 0.193472986 | 0.84658856 | NA |
| NA | NA | NA | |
| NA | NA | NA | |
| NA | NA | NA | |
| | 2.219477356 | 0.026454266 | 0.053218958 |
| | 6.349157297 | 2.16E-10 | 1.58E-09 |
| | 6.259037477 | 3.87E-10 | 2.74E-09 |
| | 0.329532201 | 0.741753459 | 0.81202279 |
| | -7.860890777 | 3.81E-15 | 4.51E-14 |
| | 1.374718052 | 0.169218873 | 0.257567938 |
| | -2.236730894 | 0.025303934 | 0.051259101 |
| | -1.2661924 | 0.205444208 | 0.302059946 |
| | -6.417999596 | 1.38E-10 | 1.03E-09 |
| | 1.89184033 | 0.058512258 | 0.105843884 |

| | | |
|---|---|---|
| 0.461846805 | 0.644191183 | 0.733763698 |
| 0.566421646 | 0.571107183 | NA |
| -0.989739161 | 0.322301628 | 0.433324905 |
| -0.317576223 | 0.750806412 | 0.819138894 |
| -0.836398858 | 0.402930555 | 0.51463368 |
| -3.085803751 | 0.002030028 | 0.005464021 |
| -0.378099914 | 0.705356372 | 0.78379889 |
| -2.072618085 | 0.038207837 | 0.073209994 |
| -0.822589437 | 0.410741508 | 0.522613864 |
| 0.813759576 | 0.415782697 | 0.527366013 |
| 4.603504818 | 4.15E-06 | 1.79E-05 |
| 3.25547604 | 0.001132024 | 0.003224601 |
| 0.2153835 | 0.829468333 | 0.880773476 |
| -6.289791415 | 3.18E-10 | 2.27E-09 |
| NA | NA | NA |
| 0.870443924 | 0.384057851 | 0.496352149 |
| 2.000508867 | 0.045445343 | 0.085235805 |
| -1.546323352 | 0.122026495 | 0.196786494 |
| 13.93990203 | 3.62E-44 | 2.37E-42 |
| NA | NA | NA |
| 1.483570364 | 0.137922937 | 0.217722698 |
| -1.860993035 | 0.06274516 | 0.112245392 |
| 1.704648713 | 0.08825996 | 0.150149311 |
| 2.963869962 | 0.003037966 | 0.007844058 |
| 6.875734346 | 6.17E-12 | 5.27E-11 |
| 3.234974428 | 0.001216536 | 0.003436396 |
| 6.491265733 | 8.51E-11 | 6.52E-10 |
| -0.027754471 | 0.977857979 | 0.984663685 |
| 0.919077332 | 0.358055125 | 0.469934444 |
| -2.251295166 | 0.024366849 | 0.049589578 |
| 4.31545432 | 1.59E-05 | 6.28E-05 |
| 1.048978581 | 0.294187977 | 0.402378418 |
| -2.470567287 | 0.013489894 | 0.029675869 |
| 0.375464795 | 0.707314824 | 0.785277891 |
| NA | NA | NA |
| 0.099625208 | 0.92064188 | 0.945536345 |
| 0.403205955 | 0.686796719 | 0.768806882 |
| -0.48301594 | 0.629084418 | 0.721796149 |
| -0.647838573 | 0.517089364 | 0.622367127 |
| 0.3043814 | 0.760837337 | 0.827018654 |
| 5.969032893 | 2.39E-09 | 1.56E-08 |
| 5.624636586 | 1.86E-08 | 1.10E-07 |
| -1.90266096 | 0.057084799 | 0.103703914 |
| -2.313925938 | 0.020671785 | 0.042996076 |
| 8.144754677 | 3.80E-16 | 4.91E-15 |
| 3.142830559 | 0.001673227 | 0.004589181 |
| 10.00243353 | 1.49E-23 | 3.45E-22 |

| | | | |
|---|---|---|---|
| | -0.114002238 | 0.90923602 | NA |
| | 2.720814761 | 0.006512124 | 0.015554773 |
| | -0.814441636 | 0.415391994 | 0.527003985 |
| | 0.661957736 | 0.507998308 | 0.614314308 |
| | -2.059009233 | 0.039493354 | 0.075344409 |
| | 6.175068351 | 6.61E-10 | 4.57E-09 |
| | 0.255131163 | 0.798621796 | 0.85688887 |
| NA | NA | NA | |
| | -0.150628505 | 0.880268774 | NA |
| | 13.87930969 | 8.46E-44 | 5.40E-42 |
| | -0.570285809 | 0.568483865 | 0.670112663 |
| | 3.38471677 | 0.000712517 | 0.002111819 |
| | -2.045120562 | 0.040842974 | 0.07759979 |
| | 0.935313 | 0.349627017 | 0.461291669 |
| | 4.308281554 | 1.65E-05 | 6.48E-05 |
| | 8.782552321 | 1.60E-18 | 2.53E-17 |
| NA | NA | NA | |
| | -1.348693616 | 0.177435397 | 0.267994689 |
| | 1.662175665 | 0.096477558 | NA |
| | 0.192808924 | 0.847108615 | NA |
| | 0.252511758 | 0.800645527 | NA |
| NA | NA | NA | |
| NA | NA | NA | |
| NA | NA | NA | |
| NA | NA | NA | |
| NA | NA | NA | |
| NA | NA | NA | |
| NA | NA | NA | |
| NA | NA | NA | |
| NA | NA | NA | |
| NA | NA | NA | |
| NA | NA | NA | |
| NA | NA | NA | |
| NA | NA | NA | |
| NA | NA | NA | |
| NA | NA | NA | |
| NA | NA | NA | |
| NA | NA | NA | |
| NA | NA | NA | |
| NA | NA | NA | |
| NA | NA | NA | |
| NA | NA | NA | |
| NA | NA | NA | |
| NA | NA | NA | |
| NA | NA | NA | |
| NA | NA | NA | |
| NA | NA | NA | |

| | | |
|---|---|---|
| NA | NA | NA |
| NA | NA | NA |
| NA | NA | NA |
| NA | NA | NA |
| NA | NA | NA |
| NA | NA | NA |
| NA | NA | NA |
| NA | NA | NA |
| NA | NA | NA |
| NA | NA | NA |
| NA | NA | NA |
| NA | NA | NA |
| NA | NA | NA |
| NA | NA | NA |
| NA | NA | NA |
| NA | NA | NA |
| NA | NA | NA |
| NA | NA | NA |
| NA | NA | NA |
| NA | NA | NA |
| NA | NA | NA |
| NA | NA | NA |
| NA | NA | NA |
| NA | NA | NA |
| NA | NA | NA |
| NA | NA | NA |
| NA | NA | NA |
| NA | NA | NA |
| NA | NA | NA |
| NA | NA | NA |
| NA | NA | NA |
| NA | NA | NA |
| NA | NA | NA |
| NA | NA | NA |
| NA | NA | NA |
| NA | NA | NA |
| NA | NA | NA |
| NA | NA | NA |
| NA | NA | NA |
| NA | NA | NA |
| NA | NA | NA |
| NA | NA | NA |
| NA | NA | NA |
| NA | NA | NA |
| NA | NA | NA |
| NA | NA | NA |
| NA | NA | NA |

| | | |
|---|---|---|
| NA | NA | NA |
| NA | NA | NA |
| NA | NA | NA |
| NA | NA | NA |
| NA | NA | NA |
| NA | NA | NA |
| NA | NA | NA |
| NA | NA | NA |
| NA | NA | NA |
| NA | NA | NA |
| NA | NA | NA |
| NA | NA | NA |
| NA | NA | NA |
| NA | NA | NA |
| NA | NA | NA |
| NA | NA | NA |
| NA | NA | NA |
| NA | NA | NA |
| NA | NA | NA |
| NA | NA | NA |
| NA | NA | NA |
| NA | NA | NA |
| NA | NA | NA |
| NA | NA | NA |
| NA | NA | NA |
| NA | NA | NA |
| NA | NA | NA |
| NA | NA | NA |
| NA | NA | NA |
| NA | NA | NA |
| NA | NA | NA |
| NA | NA | NA |
| -0.464282671 | 0.642445237 | 0.732504476 |
| 0.192809521 | 0.847108147 | NA |
| NA | NA | NA |
| NA | NA | NA |
| -0.065388772 | 0.947864464 | NA |
| NA | NA | NA |
| NA | NA | NA |
| NA | NA | NA |
| NA | NA | NA |
| NA | NA | NA |
| NA | NA | NA |
| NA | NA | NA |
| NA | NA | NA |
| NA | NA | NA |
| NA | NA | NA |

| | | |
|---|---|---|
| NA | NA | NA |
| NA | NA | NA |
| 1.450280519 | 0.146980309 | 0.229487689 |
| NA | NA | NA |
| NA | NA | NA |
| NA | NA | NA |
| NA | NA | NA |
| NA | NA | NA |
| -0.364070101 | 0.715805663 | 0.792031198 |
| -1.306819152 | 0.191274139 | 0.285043564 |
| -2.497760479 | 0.012498061 | 0.027754987 |
| -1.151495578 | 0.249528413 | 0.35302375 |
| -0.399151056 | 0.689781904 | 0.770995417 |
| -3.170387608 | 0.001522357 | 0.00421278 |
| -2.369260192 | 0.017823709 | 0.037874214 |
| -0.752852141 | 0.45153877 | 0.561451337 |
| -1.071822024 | 0.283799979 | 0.390818303 |
| -0.044140931 | 0.964792066 | 0.976061817 |
| NA | NA | NA |
| NA | NA | NA |
| 0.192810118 | 0.847107679 | NA |
| 0.197358978 | 0.843546629 | NA |
| 5.905124702 | 3.52E-09 | 2.27E-08 |
| 1.407849101 | 0.159175758 | 0.244983657 |
| -0.065389369 | 0.947863988 | NA |
| 10.88707264 | 1.33E-27 | 4.04E-26 |
| 0.76587422 | 0.443751147 | 0.553776298 |
| -0.604131792 | 0.545756024 | 0.649406859 |
| -8.788004377 | 1.52E-18 | 2.42E-17 |
| -0.065390563 | 0.947863038 | NA |
| 0.000901414 | 0.999280776 | 0.999506869 |
| -7.111358121 | 1.15E-12 | 1.06E-11 |
| 1.586999229 | 0.112512819 | 0.183587424 |
| 5.928540745 | 3.06E-09 | 1.98E-08 |
| NA | NA | NA |
| -0.435985734 | 0.66284708 | NA |
| NA | NA | NA |
| NA | NA | NA |
| NA | NA | NA |
| 8.315740795 | 9.12E-17 | 1.24E-15 |
| 0.412281605 | 0.68013303 | 0.762951182 |
| NA | NA | NA |
| 6.539469541 | 6.17E-11 | 4.80E-10 |
| NA | NA | NA |
| -0.065389369 | 0.947863988 | NA |
| 7.536808089 | 4.82E-14 | 5.11E-13 |
| 1.644630714 | 0.100045989 | 0.166578985 |

| | | |
|---|---|---|
| 2.90267957 | 0.00369985 | 0.009357102 |
| 0.913604786 | 0.360924551 | 0.473055532 |
| NA | NA | NA |
| -6.880690389 | 5.96E-12 | 5.10E-11 |
| -0.780606311 | 0.435034079 | NA |
| -7.339874419 | 2.14E-13 | 2.12E-12 |
| 1.906667114 | 0.056563708 | 0.102906645 |
| 1.618655956 | 0.105521308 | 0.174060785 |
| NA | NA | NA |
| 0.445431117 | 0.656008229 | 0.743620578 |
| 0.889687559 | 0.373633676 | 0.486351392 |
| -0.459061074 | 0.646190309 | NA |
| -1.329075591 | 0.183823032 | 0.275944014 |
| -1.45041118 | 0.146943891 | 0.22946604 |
| -1.882959058 | 0.059705917 | 0.107629751 |
| 3.259990025 | 0.001114161 | 0.00317842 |
| 0.467378934 | 0.640228792 | 0.730830787 |
| 2.403314079 | 0.016247226 | 0.034976183 |
| 1.075337385 | 0.282223704 | 0.389247263 |
| -7.087723886 | 1.36E-12 | 1.25E-11 |
| -9.573401742 | 1.03E-21 | 2.10E-20 |
| -0.065389369 | 0.947863988 | NA |
| 7.509012996 | 5.96E-14 | 6.26E-13 |
| -2.12958305 | 0.033206052 | 0.064902973 |
| 0.711309457 | 0.476892493 | NA |
| -0.085070848 | 0.932205066 | NA |
| 6.749949299 | 1.48E-11 | 1.22E-10 |
| 0.163749906 | 0.869928028 | 0.909384336 |
| -1.538246073 | 0.12398846 | 0.199462121 |
| -3.830424011 | 0.000127923 | 0.000433862 |
| 6.709285794 | 1.96E-11 | 1.60E-10 |
| -0.906013022 | 0.36492896 | 0.477429013 |
| 1.670384027 | 0.094843408 | 0.159495465 |
| -6.867532349 | 6.53E-12 | 5.56E-11 |
| -0.054359762 | 0.956648537 | 0.970365225 |
| 13.41779959 | 4.76E-41 | 2.67E-39 |
| 5.639187473 | 1.71E-08 | 1.02E-07 |
| 5.230397536 | 1.69E-07 | 8.84E-07 |
| 16.41701115 | 1.45E-60 | 1.47E-58 |
| -4.443189121 | 8.86E-06 | 3.64E-05 |
| -2.732052328 | 0.006294114 | 0.015082976 |
| 1.736889566 | 0.082406666 | 0.141635572 |
| NA | NA | NA |
| NA | NA | NA |
| NA | NA | NA |
| 1.864478321 | 0.062254563 | 0.111478734 |
| 0.192809521 | 0.847108147 | NA |

| | | |
|---|---|---|
| -1.894807935 | 0.058117852 | 0.105244525 |
| NA | NA | NA |
| 0.063708285 | 0.949202508 | NA |
| NA | NA | NA |
| NA | NA | NA |
| -0.065390563 | 0.947863038 | NA |
| 0.063708285 | 0.949202508 | NA |
| NA | NA | NA |
| NA | NA | NA |
| NA | NA | NA |
| NA | NA | NA |
| NA | NA | NA |
| -0.074007842 | 0.941004146 | NA |
| -0.405141758 | 0.685373314 | 0.76755678 |
| -0.759654283 | 0.447461263 | 0.557378093 |
| 0.192810118 | 0.847107679 | NA |
| 0.099856068 | 0.920458594 | NA |
| 1.858076367 | 0.06315817 | 0.112877473 |
| 0.063708285 | 0.949202508 | NA |
| 0.063710673 | 0.949200606 | NA |
| 3.426024057 | 0.000612486 | 0.001837091 |
| -0.291664339 | 0.770543277 | 0.834709578 |
| NA | NA | NA |
| NA | NA | NA |
| -1.432516478 | 0.151996066 | 0.236125963 |
| 3.636331147 | 0.000276549 | 0.000885168 |
| 0.574210932 | 0.565825072 | 0.667858227 |
| -1.034536664 | 0.300885346 | 0.409925986 |
| 5.953547793 | 2.62E-09 | 1.71E-08 |
| 10.64497875 | 1.84E-26 | 5.18E-25 |
| -0.666788422 | 0.504907289 | 0.611468427 |
| -0.614851107 | 0.538653054 | 0.643033622 |
| NA | NA | NA |
| -7.051407369 | 1.77E-12 | 1.59E-11 |
| 2.131963168 | 0.033009875 | 0.064590147 |
| NA | NA | NA |
| -4.840637701 | 1.29E-06 | 5.97E-06 |
| 1.937719007 | 0.052657514 | 0.096792067 |
| -0.173185862 | 0.862505337 | 0.904360714 |
| 7.26966433 | 3.60E-13 | 3.50E-12 |
| 6.933241473 | 4.11E-12 | 3.58E-11 |
| 5.04375487 | 4.56E-07 | 2.25E-06 |
| 2.400189772 | 0.016386574 | 0.035242821 |
| 0.933464119 | 0.350580388 | 0.462307203 |
| -8.758468959 | 1.98E-18 | 3.10E-17 |
| 7.445503528 | 9.66E-14 | 9.96E-13 |
| 4.60844471 | 4.06E-06 | 1.75E-05 |

| | | |
|---|---|---|
| -10.26055944 | 1.06E-24 | 2.65E-23 |
| -5.921946311 | 3.18E-09 | 2.06E-08 |
| -1.759667961 | 0.078464121 | 0.135825145 |
| -0.124255563 | 0.90111293 | 0.931133907 |
| 1.418441655 | 0.156061861 | 0.241259242 |
| 1.948892796 | 0.051308229 | 0.094659465 |
| 2.230571756 | 0.025709509 | 0.051921012 |
| 0.166027941 | 0.868134961 | 0.908154724 |
| 0.869742025 | 0.3844414 | 0.496699956 |
| -0.271627722 | 0.785908282 | 0.847315829 |
| 0.293921856 | 0.768817617 | 0.833557349 |
| 0.305897334 | 0.759682816 | 0.826086579 |
| 7.69746271 | 1.39E-14 | 1.56E-13 |
| -1.479602762 | 0.138979289 | 0.219044872 |
| -0.133425676 | 0.89385674 | NA |
| NA | NA | NA |
| 1.509050055 | 0.131285992 | 0.208945658 |
| 5.038310963 | 4.70E-07 | 2.31E-06 |
| -7.563366057 | 3.93E-14 | 4.21E-13 |
| 0.030228658 | 0.975884693 | 0.983253596 |
| -1.764410017 | 0.077662973 | 0.134619405 |
| -3.053511088 | 0.002261804 | 0.006017532 |
| -0.300392132 | 0.763878065 | 0.829530248 |
| 5.736054227 | 9.69E-09 | 5.93E-08 |
| 5.290429262 | 1.22E-07 | 6.50E-07 |
| 7.849984867 | 4.16E-15 | 4.89E-14 |
| 1.637205755 | 0.101587485 | 0.168726977 |
| 13.51289332 | 1.31E-41 | 7.55E-40 |
| 13.18252782 | 1.11E-39 | 5.98E-38 |
| 4.280983507 | 1.86E-05 | 7.27E-05 |
| 1.969231862 | 0.048926472 | 0.090874858 |
| 6.791038965 | 1.11E-11 | 9.29E-11 |
| 8.064598341 | 7.35E-16 | 9.30E-15 |
| -5.018993271 | 5.19E-07 | 2.53E-06 |
| 0.509535225 | 0.610377114 | 0.70651772 |
| 0.192809521 | 0.847108147 | NA |
| 0.192809521 | 0.847108147 | NA |
| NA | NA | NA |
| -0.786372627 | 0.431649215 | 0.542499513 |
| -0.112274323 | 0.9106059 | NA |
| -0.371476019 | 0.710283014 | 0.787664873 |
| -1.508292924 | 0.131479573 | 0.209200542 |
| 2.238385864 | 0.025195903 | 0.051069192 |
| 0.552536774 | 0.580580647 | 0.681271804 |
| 0.372183238 | 0.709756425 | 0.787325089 |
| -2.087900758 | 0.036806788 | 0.070933626 |
| 5.654599191 | 1.56E-08 | 9.33E-08 |

| | | |
|---|---|---|
| 2.006750422 | 0.044776244 | 0.084146662 |
| 9.296866379 | 1.45E-20 | 2.67E-19 |
| -1.020255984 | 0.307607073 | 0.417379792 |
| 0.538990733 | 0.589893249 | 0.689374732 |
| -0.025202857 | 0.979893158 | 0.986157603 |
| -0.430241319 | 0.667020108 | 0.752395425 |
| -2.363576206 | 0.018099504 | 0.038347625 |
| 9.29483637 | 1.47E-20 | 2.72E-19 |
| 3.724680847 | 0.000195562 | 0.000643985 |
| 10.17552816 | 2.55E-24 | 6.22E-23 |
| -7.2741978 | 3.48E-13 | 3.39E-12 |
| 0.323066476 | 0.746644898 | NA |
| -1.127428447 | 0.259561377 | NA |
| NA | NA | NA |
| 6.504095814 | 7.82E-11 | 6.01E-10 |
| NA | NA | NA |
| -5.833168339 | 5.44E-09 | 3.42E-08 |
| -0.943059757 | 0.345650338 | 0.457129239 |
| -0.6897812 | 0.490331793 | 0.597895688 |
| 2.702432279 | 0.006883421 | 0.01634184 |
| 5.689712397 | 1.27E-08 | 7.68E-08 |
| -2.04579489 | 0.040776555 | 0.077508898 |
| NA | NA | NA |
| 3.721646067 | 0.000197928 | 0.00065109 |
| 1.208454264 | 0.226872578 | 0.327071802 |
| 2.046525655 | 0.040704681 | 0.077395789 |
| 0.225391268 | 0.821674904 | 0.874873355 |
| 0.379908941 | 0.70401301 | 0.782966571 |
| -6.886705277 | 5.71E-12 | 4.90E-11 |
| -3.168275896 | 0.001533459 | 0.00424069 |
| -0.921964254 | 0.356547221 | 0.468397468 |
| -2.939300862 | 0.003289536 | 0.008420647 |
| -0.690229326 | 0.490049984 | NA |
| -1.556664839 | 0.119550081 | NA |
| NA | NA | NA |
| -2.275088648 | 0.022900613 | 0.047049358 |
| NA | NA | NA |
| 0.59170981 | 0.554044927 | 0.657114505 |
| -1.153371356 | 0.248757993 | 0.352112909 |
| -4.687054192 | 2.77E-06 | 1.22E-05 |
| -0.526506607 | 0.598536262 | 0.696607807 |
| -3.964327153 | 7.36E-05 | 0.000260763 |
| NA | NA | NA |
| NA | NA | NA |
| -0.740312443 | 0.459110433 | 0.568493823 |
| -3.268477546 | 0.001081278 | 0.00309484 |
| -0.065389369 | 0.947863988 | NA |

| | | |
|---|---|---|
| -3.317605428 | 0.000907926 | 0.002636663 |
| -2.954535546 | 0.003131401 | 0.00805622 |
| 0.966583341 | 0.333752366 | 0.444943899 |
| 1.464630744 | 0.143021689 | 0.224553562 |
| 0.806051747 | 0.420213012 | 0.531934036 |
| -0.922054262 | 0.356500273 | 0.468360373 |
| -2.925030955 | 0.00344422 | 0.008776221 |
| 0.280409999 | 0.779162966 | NA |
| -1.192154261 | 0.233200764 | 0.33438446 |
| 2.350245326 | 0.018761041 | 0.03953101 |
| NA | NA | NA |
| -0.601466788 | 0.547529126 | 0.651114312 |
| 2.410896988 | 0.015913342 | 0.034331232 |
| -5.225428804 | 1.74E-07 | 9.06E-07 |
| 0.856413396 | 0.391769157 | 0.503775084 |
| 6.492626485 | 8.44E-11 | 6.47E-10 |
| 0.383737811 | 0.701172785 | NA |
| 2.397734572 | 0.016496815 | 0.035440328 |
| 2.768462796 | 0.005632142 | 0.013674653 |
| -0.065390563 | 0.947863038 | NA |
| 8.0700839 | 7.02E-16 | 8.90E-15 |
| -0.406493483 | 0.684380044 | NA |
| 0.472170971 | 0.636804758 | 0.727984613 |
| -2.64134708 | 0.008257708 | 0.019168174 |
| -3.836730862 | 0.000124683 | 0.000423793 |
| NA | NA | NA |
| -0.420813139 | 0.673891537 | 0.757553559 |
| 2.488721259 | 0.012820344 | 0.028385069 |
| 1.138251647 | 0.255015418 | 0.359364706 |
| 0.015429708 | 0.987689362 | 0.991372849 |
| 0.164028803 | 0.869708469 | 0.909344709 |
| 14.3120306 | 1.84E-46 | 1.29E-44 |
| 5.800531192 | 6.61E-09 | 4.12E-08 |
| 3.653303165 | 0.000258888 | 0.00083226 |
| -7.207438268 | 5.70E-13 | 5.44E-12 |
| -7.818312612 | 5.35E-15 | 6.25E-14 |
| -3.973092375 | 7.09E-05 | 0.000251882 |
| -2.833554952 | 0.004603339 | 0.011425045 |
| -0.278690983 | 0.780481983 | NA |
| -1.772855241 | 0.076252697 | 0.132514407 |
| -9.062868559 | 1.27E-19 | 2.20E-18 |
| -0.46385817 | 0.642749364 | 0.732772334 |
| 5.733248057 | 9.85E-09 | 6.03E-08 |
| NA | NA | NA |
| 1.228047081 | 0.219429295 | 0.318432006 |
| 6.236615245 | 4.47E-10 | 3.15E-09 |
| 2.136594147 | 0.032631016 | 0.063928799 |

| | | |
|---|---|---|
| 0.066394788 | 0.94706352 | NA |
| NA | NA | NA |
| -0.89159064 | 0.372612389 | 0.485451181 |
| -0.538121581 | 0.590493114 | 0.689914702 |
| 9.809798648 | 1.02E-22 | 2.22E-21 |
| -0.234016503 | 0.814972167 | 0.869907233 |
| -1.965554441 | 0.049350101 | 0.091586938 |
| 1.185056629 | 0.235995058 | NA |
| -0.181590241 | 0.855904308 | 0.89998676 |
| -0.527751105 | 0.597672098 | NA |
| -1.415023535 | 0.157061596 | 0.242535119 |
| -0.482772667 | 0.629257161 | 0.721895074 |
| NA | NA | NA |
| 0.192809521 | 0.847108147 | NA |
| NA | NA | NA |
| -1.681100903 | 0.092743317 | 0.156531708 |
| 2.188070444 | 0.028664471 | 0.057087183 |
| -4.362432153 | 1.29E-05 | 5.14E-05 |
| -0.43765866 | 0.661633741 | NA |
| -1.253645129 | 0.20997102 | 0.307433439 |
| -3.099791462 | 0.001936569 | 0.005231768 |
| 2.800410053 | 0.005103773 | 0.012530401 |
| 0.3183917 | 0.750187833 | 0.818642432 |
| 4.162431477 | 3.15E-05 | 0.000118642 |
| -1.799035242 | 0.072013106 | NA |
| -0.056999076 | 0.954545931 | NA |
| 7.018129293 | 2.25E-12 | 2.00E-11 |
| 0.320071032 | 0.748914485 | NA |
| 0.063710673 | 0.949200606 | NA |
| 2.272698845 | 0.023044334 | 0.047282539 |
| 0.261359224 | 0.793815499 | NA |
| -0.811238686 | 0.417228614 | 0.528878365 |
| -1.058871523 | 0.289658295 | 0.397137769 |
| -2.100891669 | 0.035650477 | 0.068959153 |
| -2.242164146 | 0.024950765 | 0.05063383 |
| 10.00047159 | 1.52E-23 | 3.52E-22 |
| -0.360829665 | 0.718226786 | 0.794030869 |
| 11.79617247 | 4.08E-32 | 1.58E-30 |
| NA | NA | NA |
| -11.64064573 | 2.56E-31 | 9.48E-30 |
| 4.216513402 | 2.48E-05 | 9.50E-05 |
| 6.607249657 | 3.92E-11 | 3.10E-10 |
| 2.527905172 | 0.011474534 | 0.02576546 |
| -1.622799774 | 0.104632214 | 0.172913266 |
| -5.395630818 | 6.83E-08 | 3.76E-07 |
| -1.543030042 | 0.122823487 | 0.197850162 |
| -0.447897963 | 0.654226839 | 0.742172669 |

| | | |
|---|---|---|
| 0.057106724 | 0.95446018 | 0.96882024 |
| NA | NA | NA |
| -1.350172217 | 0.176960748 | 0.267439169 |
| NA | NA | NA |
| NA | NA | NA |
| NA | NA | NA |
| NA | NA | NA |
| NA | NA | NA |
| NA | NA | NA |
| NA | NA | NA |
| NA | NA | NA |
| NA | NA | NA |
| NA | NA | NA |
| NA | NA | NA |
| NA | NA | NA |
| NA | NA | NA |
| NA | NA | NA |
| NA | NA | NA |
| NA | NA | NA |
| NA | NA | NA |
| NA | NA | NA |
| NA | NA | NA |
| NA | NA | NA |
| NA | NA | NA |
| NA | NA | NA |
| NA | NA | NA |
| NA | NA | NA |
| NA | NA | NA |
| NA | NA | NA |
| NA | NA | NA |
| NA | NA | NA |
| NA | NA | NA |
| NA | NA | NA |
| NA | NA | NA |
| NA | NA | NA |
| NA | NA | NA |
| NA | NA | NA |
| NA | NA | NA |
| NA | NA | NA |
| NA | NA | NA |
| NA | NA | NA |
| NA | NA | NA |
| NA | NA | NA |
| NA | NA | NA |
| NA | NA | NA |
| NA | NA | NA |
| NA | NA | NA |

| | | |
|---|---|---|
| NA | NA | NA |
| NA | NA | NA |
| NA | NA | NA |
| NA | NA | NA |
| NA | NA | NA |
| NA | NA | NA |
| NA | NA | NA |
| NA | NA | NA |
| NA | NA | NA |
| NA | NA | NA |
| NA | NA | NA |
| NA | NA | NA |
| NA | NA | NA |
| NA | NA | NA |
| NA | NA | NA |
| NA | NA | NA |
| NA | NA | NA |
| NA | NA | NA |
| NA | NA | NA |
| NA | NA | NA |
| NA | NA | NA |
| NA | NA | NA |
| NA | NA | NA |
| NA | NA | NA |
| NA | NA | NA |
| NA | NA | NA |
| NA | NA | NA |
| NA | NA | NA |
| NA | NA | NA |
| NA | NA | NA |
| NA | NA | NA |
| NA | NA | NA |
| NA | NA | NA |
| NA | NA | NA |
| NA | NA | NA |
| NA | NA | NA |
| NA | NA | NA |
| NA | NA | NA |
| NA | NA | NA |
| NA | NA | NA |
| NA | NA | NA |
| NA | NA | NA |
| NA | NA | NA |
| NA | NA | NA |
| NA | NA | NA |
| NA | NA | NA |
| NA | NA | NA |

| | | |
|---|---|---|
| NA | NA | NA |
| NA | NA | NA |
| NA | NA | NA |
| NA | NA | NA |
| NA | NA | NA |
| NA | NA | NA |
| NA | NA | NA |
| NA | NA | NA |
| NA | NA | NA |
| NA | NA | NA |
| NA | NA | NA |
| NA | NA | NA |
| -0.065388772 | 0.947864464 | NA |
| NA | NA | NA |
| NA | NA | NA |
| NA | NA | NA |
| NA | NA | NA |
| NA | NA | NA |
| NA | NA | NA |
| NA | NA | NA |
| NA | NA | NA |
| NA | NA | NA |
| NA | NA | NA |
| NA | NA | NA |
| NA | NA | NA |
| NA | NA | NA |
| NA | NA | NA |
| NA | NA | NA |
| NA | NA | NA |
| NA | NA | NA |
| 0.192809521 | 0.847108147 | NA |
| NA | NA | NA |
| NA | NA | NA |
| 0.063708285 | 0.949202508 | NA |
| NA | NA | NA |
| 5.436881299 | 5.42E-08 | 3.03E-07 |
| 1.900779225 | 0.057330936 | 0.104075525 |
| 1.081304829 | 0.27956154 | 0.386555373 |
| NA | NA | NA |
| 3.819415751 | 0.000133768 | 0.000452278 |
| -5.530556491 | 3.19E-08 | 1.84E-07 |
| 0.406165476 | 0.684621019 | 0.766988815 |
| -1.303022451 | 0.192567097 | 0.28662907 |
| 0.158000685 | 0.874456259 | 0.912517309 |
| 3.894597945 | 9.84E-05 | 0.000340572 |
| 0.192809521 | 0.847108147 | NA |
| 7.801073149 | 6.14E-15 | 7.14E-14 |

| | | |
|---|---|---|
| NA | NA | NA |
| 5.345838836 | 9.00E-08 | 4.88E-07 |
| 0.074728106 | 0.940431045 | NA |
| NA | NA | NA |
| -0.91240966 | 0.361553106 | 0.473804934 |
| 2.074773832 | 0.038007501 | 0.072906837 |
| 0.236328127 | 0.813178063 | NA |
| -2.413760456 | 0.015788839 | 0.034094956 |
| -3.071495378 | 0.002129895 | 0.005703569 |
| 3.199929416 | 0.001374612 | 0.003840017 |
| 0.063707194 | 0.949203376 | NA |
| 8.263407857 | 1.42E-16 | 1.90E-15 |
| NA | NA | NA |
| NA | NA | NA |
| 8.279749617 | 1.23E-16 | 1.67E-15 |
| -8.562436963 | 1.11E-17 | 1.62E-16 |
| -7.686802921 | 1.51E-14 | 1.69E-13 |
| 0.109795436 | 0.91257161 | NA |
| 1.745935483 | 0.080822163 | 0.139323605 |
| 3.633652353 | 0.000279437 | 0.000893272 |
| 5.597365069 | 2.18E-08 | 1.28E-07 |
| 1.024223325 | 0.305729804 | 0.41539556 |
| NA | NA | NA |
| 0.063710673 | 0.949200606 | NA |
| 0.302523223 | 0.762253233 | 0.828234 |
| -0.776935738 | 0.437196685 | NA |
| 1.404486185 | 0.160174123 | 0.246172068 |
| 1.114941917 | 0.264875313 | 0.370485 |
| -1.633955781 | 0.102268136 | 0.16964487 |
| -0.495319937 | 0.6203743 | 0.71475115 |
| 2.555285814 | 0.010610064 | 0.024042178 |
| -0.543440211 | 0.586826746 | 0.686674843 |
| 0.075619579 | 0.93972176 | 0.958993584 |
| 0.449259926 | 0.653244163 | NA |
| -1.159802754 | 0.246129123 | 0.349192987 |
| 0.346341506 | 0.72908608 | 0.802260719 |
| 1.558898613 | 0.119020379 | 0.192558986 |
| 0.063707688 | 0.949202983 | NA |
| -2.008836811 | 0.044554443 | 0.083799046 |
| NA | NA | NA |
| NA | NA | NA |
| -0.596294947 | 0.550978206 | 0.654314394 |
| -2.82323657 | 0.004754148 | 0.011761173 |
| -2.987926754 | 0.002808769 | 0.007305016 |
| -3.012625031 | 0.002589987 | 0.00679034 |
| -2.906975491 | 0.003649418 | 0.009243567 |
| -2.547904964 | 0.010837199 | 0.024488137 |

| | | | |
|---|---|---|---|
| NA | NA | NA | |
| NA | NA | NA | |
| 0.063708217 | 0.949202562 | NA | |
| NA | NA | NA | |
| -0.135858696 | 0.891932989 | NA | |
| -2.602261752 | 0.009261114 | | 0.021264625 |
| NA | NA | NA | |
| NA | NA | NA | |
| -1.668443542 | 0.095227711 | | 0.159969962 |
| -0.978038482 | 0.328055292 | | 0.439055927 |
| NA | NA | NA | |
| 9.278509854 | 1.72E-20 | | 3.16E-19 |
| 3.435412619 | 0.000591652 | | 0.001779935 |
| 1.365845637 | 0.171987419 | | 0.261162443 |
| NA | NA | NA | |
| NA | NA | NA | |
| NA | NA | NA | |
| 0.063708285 | 0.949202508 | NA | |
| NA | NA | NA | |
| NA | NA | NA | |
| NA | NA | NA | |
| NA | NA | NA | |
| NA | NA | NA | |
| 0.31685483 | 0.751353758 | NA | |
| NA | NA | NA | |
| NA | NA | NA | |
| NA | NA | NA | |
| NA | NA | NA | |
| NA | NA | NA | |
| NA | NA | NA | |
| NA | NA | NA | |
| NA | NA | NA | |
| NA | NA | NA | |
| 0.157748723 | 0.874654805 | NA | |
| NA | NA | NA | |
| NA | NA | NA | |
| NA | NA | NA | |
| NA | NA | NA | |
| NA | NA | NA | |
| NA | NA | NA | |
| NA | NA | NA | |
| NA | NA | NA | |
| NA | NA | NA | |
| NA | NA | NA | |
| NA | NA | NA | |
| NA | NA | NA | |
| NA | NA | NA | |

| | | |
|---|---|---|
| NA | NA | NA |
| NA | NA | NA |
| NA | NA | NA |
| NA | NA | NA |
| -0.584872266 | 0.558633608 | 0.66102204 |
| 1.136303829 | 0.255829429 | 0.360329222 |
| 1.133771889 | 0.256890246 | 0.361416607 |
| 0.063370556 | 0.949471433 | 0.965361523 |
| -0.410407781 | 0.681506839 | 0.764218278 |
| -4.057731501 | 4.96E-05 | 0.000180693 |
| -2.319643316 | 0.020360181 | 0.04244672 |
| -3.35919644 | 0.000781695 | 0.002298076 |
| -1.763904435 | 0.07774807 | 0.134738918 |
| NA | NA | NA |
| -0.194488217 | 0.845793621 | NA |
| 0.692031765 | 0.488917381 | 0.596752428 |
| -0.093858287 | 0.92522173 | NA |
| 0.063708285 | 0.949202508 | NA |
| NA | NA | NA |
| 0.423407673 | 0.671997848 | NA |
| NA | NA | NA |
| NA | NA | NA |
| NA | NA | NA |
| 0.847730019 | 0.396588344 | 0.50843097 |
| NA | NA | NA |
| NA | NA | NA |
| 0.063708217 | 0.949202562 | NA |
| NA | NA | NA |
| -0.065389369 | 0.947863988 | NA |
| -4.309265582 | 1.64E-05 | 6.45E-05 |
| 0.063708882 | 0.949202033 | NA |
| NA | NA | NA |
| -2.113180986 | 0.03458528 | 0.067220808 |
| 0.069134878 | 0.944882259 | NA |
| 0.065474733 | 0.947796023 | NA |
| NA | NA | NA |
| -0.065389369 | 0.947863988 | NA |
| NA | NA | NA |
| NA | NA | NA |
| NA | NA | NA |
| NA | NA | NA |
| NA | NA | NA |
| NA | NA | NA |
| -0.19448762 | 0.845794089 | NA |
| NA | NA | NA |
| NA | NA | NA |
| NA | NA | NA |

| | | | |
|---|---|---|---|
| NA | NA | NA | |
| NA | NA | NA | |
| -0.394437355 | 0.693258169 | NA | |
| NA | NA | NA | |
| 3.523504725 | 0.00042588 | | 0.001315208 |
| -2.148090362 | 0.031706578 | | 0.062298111 |
| -1.855473221 | 0.06352868 | | 0.113410089 |
| -1.215265025 | 0.224265027 | | 0.324077918 |
| -0.439979742 | 0.65995178 | | 0.746906902 |
| 0.117568571 | 0.906409509 | NA | |
| NA | NA | NA | |
| NA | NA | NA | |
| -0.18719682 | 0.851506318 | | 0.896756138 |
| -1.923716698 | 0.054390104 | | 0.099495671 |
| NA | NA | NA | |
| 3.943301606 | 8.04E-05 | | 0.000282687 |
| 0.063708285 | 0.949202508 | NA | |
| 0.500009504 | 0.617068385 | NA | |
| 0.192809521 | 0.847108147 | NA | |
| 0.192809521 | 0.847108147 | NA | |
| NA | NA | NA | |
| NA | NA | NA | |
| NA | NA | NA | |
| 0.04211269 | 0.966408864 | | 0.977292565 |
| -0.52995667 | 0.596141973 | | 0.694537032 |
| 1.033283966 | 0.301471029 | | 0.410589827 |
| NA | NA | NA | |
| 0.571199605 | 0.567864346 | | 0.669570594 |
| NA | NA | NA | |
| NA | NA | NA | |
| NA | NA | NA | |
| NA | NA | NA | |
| NA | NA | NA | |
| NA | NA | NA | |
| 0.063708882 | 0.949202033 | NA | |
| 0.691336351 | 0.489354193 | | 0.597129906 |
| -0.411888514 | 0.68042114 | NA | |
| 0.684308144 | 0.493780605 | | 0.600988502 |
| -1.453524534 | 0.146078173 | | 0.228349779 |
| 0.018137789 | 0.985528932 | | 0.989720316 |
| -0.338343496 | 0.73510435 | | 0.806825778 |
| -1.856621196 | 0.063365066 | | 0.113158362 |
| NA | NA | NA | |
| NA | NA | NA | |
| NA | NA | NA | |
| -0.194488217 | 0.845793621 | NA | |
| -0.153173428 | 0.878261512 | NA | |

| | | | |
|---|---|---|---|
| NA | | NA | NA |
| NA | | NA | NA |
| NA | | NA | NA |
| | 2.099477144 | 0.035774861 | 0.069162336 |
| NA | | NA | NA |
| NA | | NA | NA |
| NA | | NA | NA |
| NA | | NA | NA |
| | -0.19448762 | 0.845794089 | NA |
| NA | | NA | NA |
| | -0.400637512 | 0.688687023 | NA |
| | 3.407967288 | 0.000654487 | 0.001952308 |
| | -0.058845308 | 0.953075321 | 0.967964101 |
| NA | | NA | NA |
| | 3.349418802 | 0.000809813 | 0.002373213 |
| | 0.192809521 | 0.847108147 | NA |
| | 10.78176846 | 4.20E-27 | 1.22E-25 |
| | 3.115348334 | 0.001837278 | 0.004987211 |
| | 2.255868998 | 0.024078837 | 0.049055289 |
| | 0.409317738 | 0.6823065 | NA |
| | 3.984713936 | 6.76E-05 | 0.000240721 |
| | -3.272233183 | 0.001067015 | 0.003057865 |
| | 0.083772312 | 0.933237462 | NA |
| | -1.092521274 | 0.274604045 | 0.381322565 |
| | -1.49442176 | 0.135065418 | 0.213934912 |
| | 2.044223596 | 0.040931463 | 0.07774431 |
| | 1.215385349 | 0.224219153 | 0.324030335 |
| | 2.099104719 | 0.035807671 | 0.069215074 |
| | 0.477232998 | 0.633196216 | 0.724719983 |
| | 1.153634091 | 0.248650215 | 0.351980256 |
| NA | | NA | NA |
| | -2.298508747 | 0.021532851 | 0.044535387 |
| | 0.080277591 | 0.936016481 | NA |
| NA | | NA | NA |
| | -0.342590475 | 0.731906568 | NA |
| | 1.362215229 | 0.173129971 | 0.262633002 |
| | 6.89115416 | 5.53E-12 | 4.76E-11 |
| | 6.48064922 | 9.13E-11 | 6.98E-10 |
| | 4.9227746 | 8.53E-07 | 4.04E-06 |
| | 7.915138871 | 2.47E-15 | 2.98E-14 |
| | 0.07690946 | 0.938695572 | NA |
| | 0.743915032 | 0.456927881 | NA |
| NA | | NA | NA |
| | 0.192810118 | 0.847107679 | NA |
| | 0.441759817 | 0.658663019 | NA |
| | 0.192809521 | 0.847108147 | NA |
| | 0.288217989 | 0.77317988 | NA |

| | | | |
|---|---|---|---|
| | 0.368108096 | 0.712792634 | NA |
| | 0.192809521 | 0.847108147 | NA |
| | 3.258787423 | 0.001118895 | 0.003190105 |
| NA | NA | NA | |
| NA | NA | NA | |
| | 0.069254174 | 0.944787302 | NA |
| | 0.192810118 | 0.847107679 | NA |
| NA | NA | NA | |
| NA | NA | NA | |
| NA | NA | NA | |
| | 0.548764119 | 0.583167337 | 0.683365096 |
| | 6.776109966 | 1.23E-11 | 1.03E-10 |
| | 1.234691647 | 0.216945286 | 0.315550611 |
| NA | NA | NA | |
| | -1.101548451 | 0.270658029 | 0.376888483 |
| | 0.590610467 | 0.55478145 | 0.657654335 |
| NA | NA | NA | |
| | -1.208072022 | 0.22701956 | 0.327246003 |
| NA | NA | NA | |
| NA | NA | NA | |
| NA | NA | NA | |
| NA | NA | NA | |
| | -3.995003953 | 6.47E-05 | 0.00023106 |
| | -10.54970198 | 5.10E-26 | 1.39E-24 |
| | 0.918035421 | 0.358600322 | 0.470551301 |
| | -9.336276441 | 9.98E-21 | 1.87E-19 |
| | -9.987545398 | 1.73E-23 | 4.00E-22 |
| | -3.934101952 | 8.35E-05 | 0.000292993 |
| | 0.009999061 | 0.992022037 | 0.99440575 |
| | -2.31024979 | 0.020874329 | 0.043363304 |
| | 0.410313778 | 0.681575786 | 0.764261353 |
| | -3.568579846 | 0.000358921 | 0.001125085 |
| | 0.915229625 | 0.360071095 | 0.472233666 |
| | 7.760533237 | 8.46E-15 | 9.70E-14 |
| | 3.416940876 | 0.00063329 | 0.001894723 |
| | 0.088322858 | 0.929620072 | NA |
| | 6.349485545 | 2.16E-10 | 1.58E-09 |
| | 10.65556437 | 1.64E-26 | 4.64E-25 |
| | 0.196015401 | 0.844598111 | 0.891460865 |
| | -4.491272307 | 7.08E-06 | 2.95E-05 |
| | -2.958080999 | 0.003095608 | 0.007975611 |
| | 0.421683449 | 0.673256087 | 0.757056091 |
| NA | NA | NA | |
| | -0.72131434 | 0.470716136 | 0.579766996 |
| | 3.987668277 | 6.67E-05 | 0.000237811 |
| | 13.45729198 | 2.79E-41 | 1.58E-39 |
| | 5.368208504 | 7.95E-08 | 4.34E-07 |

| | | |
|---|---|---|
| -0.090992946 | 0.927498196 | 0.950455161 |
| 0.703808242 | 0.4815522 | 0.590152884 |
| 9.007099529 | 2.12E-19 | 3.60E-18 |
| NA | NA | NA |
| 2.631174441 | 0.008509035 | 0.019705879 |
| -4.433460822 | 9.27E-06 | 3.80E-05 |
| 3.625140457 | 0.000288804 | 0.000920863 |
| -3.461323237 | 0.000537527 | 0.001629639 |
| -3.693500333 | 0.000221188 | 0.000720596 |
| -2.335544715 | 0.019514983 | 0.040899273 |
| -5.608577747 | 2.04E-08 | 1.20E-07 |
| 3.800794699 | 0.000144233 | 0.000484714 |
| -0.778997017 | 0.435981472 | 0.546423126 |
| 2.232198385 | 0.025601854 | 0.051762069 |
| 2.170805209 | 0.0299459 | 0.059257323 |
| -2.952043304 | 0.003156787 | 0.008113461 |
| -1.696240479 | 0.089840347 | 0.1524235 |
| 9.782836134 | 1.33E-22 | 2.87E-21 |
| -5.878408643 | 4.14E-09 | 2.64E-08 |
| 4.177608894 | 2.95E-05 | 0.000111569 |
| 6.097214918 | 1.08E-09 | 7.33E-09 |
| 7.05996521 | 1.67E-12 | 1.50E-11 |
| -0.30644596 | 0.75926512 | 0.825774933 |
| NA | NA | NA |
| 5.725452406 | 1.03E-08 | 6.29E-08 |
| -0.399715015 | 0.689366432 | NA |
| 5.101721499 | 3.37E-07 | 1.69E-06 |
| -1.251779794 | 0.210650115 | 0.308283481 |
| NA | NA | NA |
| NA | NA | NA |
| NA | NA | NA |
| NA | NA | NA |
| 8.079068425 | 6.53E-16 | 8.30E-15 |
| 6.382094396 | 1.75E-10 | 1.29E-09 |
| 2.936500384 | 0.003319384 | 0.008487505 |
| -6.579208256 | 4.73E-11 | 3.73E-10 |
| -1.041606457 | 0.297594174 | NA |
| 0.28793034 | 0.773400062 | 0.837111505 |
| -0.041485417 | 0.966908918 | 0.977534078 |
| 0.030710532 | 0.975500392 | 0.983001522 |
| -5.629710326 | 1.81E-08 | 1.07E-07 |
| 1.859081682 | 0.063015561 | 0.112662821 |
| 9.929292394 | 3.10E-23 | 7.07E-22 |
| 0.205072768 | 0.837515272 | 0.886385085 |
| NA | NA | NA |
| -0.183927391 | 0.854070416 | 0.89851137 |
| -0.955774147 | 0.339186353 | 0.450604236 |

| | | |
|---|---|---|
| 5.189986018 | 2.10E-07 | 1.09E-06 |
| 0.909699189 | 0.362981172 | 0.475440785 |
| -5.999406733 | 1.98E-09 | 1.31E-08 |
| -0.797882228 | 0.424938838 | 0.536489439 |
| -8.608293091 | 7.42E-18 | 1.10E-16 |
| -1.980407866 | 0.047657717 | 0.088834722 |
| 0.192809521 | 0.847108147 | NA |
| 0.311451612 | 0.755457323 | NA |
| 1.823563549 | 0.068218081 | 0.120552004 |
| 2.007179478 | 0.044730556 | 0.084067113 |
| 3.122363806 | 0.001794051 | 0.004884705 |
| 1.243672781 | 0.213620018 | 0.311718594 |
| 11.94184313 | 7.16E-33 | 2.87E-31 |
| NA | NA | NA |
| 8.191146282 | 2.59E-16 | 3.40E-15 |
| -0.772984313 | 0.439531665 | 0.549771504 |
| -0.356465241 | 0.721492178 | 0.796355304 |
| 7.553209171 | 4.25E-14 | 4.53E-13 |
| 6.207679698 | 5.38E-10 | 3.76E-09 |
| 12.49360041 | 8.09E-36 | 3.66E-34 |
| 0.063708882 | 0.949202033 | NA |
| 0.016276572 | 0.987013748 | 0.99093314 |
| 0.027401553 | 0.97813946 | 0.984788652 |
| -1.350126916 | 0.176975276 | 0.267444977 |
| 5.133665407 | 2.84E-07 | 1.44E-06 |
| 0.664587194 | 0.506314562 | 0.612787268 |
| 0.806871648 | 0.419740435 | 0.531443325 |
| 7.438487232 | 1.02E-13 | 1.05E-12 |
| 0.364892295 | 0.715191806 | 0.791705404 |
| -0.330912437 | 0.740710627 | 0.811235976 |
| 3.608687486 | 0.00030775 | 0.000976337 |
| -0.755716379 | 0.449819262 | 0.559783206 |
| -3.804793363 | 0.000141923 | 0.000477684 |
| -8.406396439 | 4.23E-17 | 5.95E-16 |
| -11.79357618 | 4.21E-32 | 1.62E-30 |
| 7.538101774 | 4.77E-14 | 5.06E-13 |
| 7.629458303 | 2.36E-14 | 2.58E-13 |
| -0.919339377 | 0.357918088 | 0.46977922 |
| NA | NA | NA |
| 3.595037067 | 0.000324345 | 0.001024917 |
| -8.803209895 | 1.33E-18 | 2.12E-17 |
| -0.941191597 | 0.346606682 | 0.45815865 |
| 1.79476933 | 0.072690459 | 0.127195482 |
| 4.579064635 | 4.67E-06 | 2.00E-05 |
| -3.150052035 | 0.001632414 | 0.004488559 |
| 4.380388053 | 1.18E-05 | 4.77E-05 |
| NA | NA | NA |

| | | |
|---|---|---|
| 0.382298295 | 0.70224012 | NA |
| -3.016283968 | 0.002558935 | 0.006715259 |
| 11.01504542 | 3.23E-28 | 1.01E-26 |
| 1.761778565 | 0.078106718 | 0.135304253 |
| 1.023496194 | 0.306073299 | 0.415681751 |
| -6.537813669 | 6.24E-11 | 4.85E-10 |
| -5.616596665 | 1.95E-08 | 1.15E-07 |
| -9.944188492 | 2.67E-23 | 6.11E-22 |
| -0.52021914 | 0.602910847 | 0.700152544 |
| -1.952403182 | 0.050890356 | 0.094041019 |
| NA | NA | NA |
| -0.775872953 | 0.437824004 | NA |
| -7.508122935 | 6.00E-14 | 6.30E-13 |
| -1.458227585 | 0.144777823 | 0.226727705 |
| -1.821133382 | 0.068586584 | 0.121143335 |
| NA | NA | NA |
| 0.523259813 | 0.600793463 | 0.698468271 |
| 1.888303026 | 0.058985281 | 0.106568615 |
| 10.44648672 | 1.52E-25 | 4.04E-24 |
| 1.401289285 | 0.161127584 | NA |
| 3.228241955 | 0.001245536 | 0.003510388 |
| 2.832404509 | 0.004619936 | 0.011458284 |
| 2.962148602 | 0.003055003 | 0.00788398 |
| 4.460286279 | 8.19E-06 | 3.38E-05 |
| 2.389761244 | 0.016859329 | 0.036116826 |
| 5.533308809 | 3.14E-08 | 1.81E-07 |
| 0.192810118 | 0.847107679 | NA |
| 2.386524567 | 0.017008471 | 0.036374063 |
| -4.572924126 | 4.81E-06 | 2.05E-05 |
| 0.063708882 | 0.949202033 | NA |
| 8.036132465 | 9.27E-16 | 1.16E-14 |
| 10.89606225 | 1.20E-27 | 3.68E-26 |
| -1.899675847 | 0.057475672 | 0.104300447 |
| -0.398585051 | 0.690198978 | 0.771293443 |
| 3.293751225 | 0.0009886 | 0.002852738 |
| NA | NA | NA |
| 0.192810118 | 0.847107679 | NA |
| NA | NA | NA |
| NA | NA | NA |
| NA | NA | NA |
| 0.195316213 | 0.845145406 | NA |
| 0.317642674 | 0.750756 | NA |
| 0.430180397 | 0.667064421 | NA |
| NA | NA | NA |
| NA | NA | NA |
| NA | NA | NA |
| NA | NA | NA |

| | | | |
|---|---|---|---|
| NA | NA | NA | |
| NA | NA | NA | |
| NA | NA | NA | |
| NA | NA | NA | |
| NA | NA | NA | |
| NA | NA | NA | |
| NA | NA | NA | |
| NA | NA | NA | |
| NA | NA | NA | |
| NA | NA | NA | |
| NA | NA | NA | |
| NA | NA | NA | |
| NA | NA | NA | |
| NA | NA | NA | |
| NA | NA | NA | |
| NA | NA | NA | |
| NA | NA | NA | |
| NA | NA | NA | |
| NA | NA | NA | |
| NA | NA | NA | |
| NA | NA | NA | |
| NA | NA | NA | |
| NA | NA | NA | |
| NA | NA | NA | |
| NA | NA | NA | |
| NA | NA | NA | |
| NA | NA | NA | |
| NA | NA | NA | |
| 0.192809521 | 0.847108147 | NA | |
| -0.407434563 | 0.683688846 | NA | |
| 0.31685483 | 0.751353758 | NA | |
| 0.063708285 | 0.949202508 | NA | |
| NA | NA | NA | |
| NA | NA | NA | |
| NA | NA | NA | |
| 0.063708882 | 0.949202033 | NA | |
| 0.837793356 | 0.402146771 | | 0.513898692 |
| 2.16939833 | 0.030052456 | | 0.059436401 |
| 6.317133643 | 2.66E-10 | | 1.92E-09 |
| 0.192809521 | 0.847108147 | NA | |
| 3.437037948 | 0.000588113 | | 0.001769926 |
| -3.4085531 | 0.000653084 | | 0.001948818 |
| -1.719469214 | 0.085528969 | | 0.146148968 |
| 0.192810118 | 0.847107679 | NA | |
| 4.840890238 | 1.29E-06 | | 5.96E-06 |
| 0.661984824 | 0.507980948 | | 0.614314308 |

| | | |
|---|---|---|
| NA | NA | NA |
| -0.337764856 | 0.735540396 | 0.807162791 |
| -3.651595694 | 0.000260616 | 0.000837276 |
| -0.065388772 | 0.947864464 | NA |
| -3.170498352 | 0.001521777 | 0.004211641 |
| -1.3945177 | 0.163161328 | 0.2500109 |
| -1.976305531 | 0.048120184 | 0.08957003 |
| -0.07925824 | 0.936827222 | 0.956959595 |
| -0.083821875 | 0.933198055 | NA |
| 0.751802774 | 0.452169673 | 0.562068282 |
| -1.066599683 | 0.286152644 | 0.393300789 |
| -5.15859519 | 2.49E-07 | 1.27E-06 |
| 0.256514144 | 0.79755386 | NA |
| NA | NA | NA |
| 0.276069368 | 0.782494788 | 0.844795893 |
| NA | NA | NA |
| NA | NA | NA |
| NA | NA | NA |
| NA | NA | NA |
| -0.502022678 | 0.615651569 | 0.710805304 |
| -7.512855429 | 5.79E-14 | 6.09E-13 |
| NA | NA | NA |
| NA | NA | NA |
| -2.570119517 | 0.010166343 | 0.023139363 |
| 0.43688466 | 0.662194997 | 0.748698269 |
| 5.41662812 | 6.07E-08 | 3.37E-07 |
| 0.418577516 | 0.675524924 | NA |
| 4.478900308 | 7.50E-06 | 3.11E-05 |
| -0.194819069 | 0.845534593 | 0.891903027 |
| -8.667386925 | 4.42E-18 | 6.71E-17 |
| NA | NA | NA |
| 1.512760581 | 0.130340485 | 0.207850557 |
| -7.141227058 | 9.25E-13 | 8.62E-12 |
| 0.154191892 | 0.877458435 | 0.914392132 |
| -4.465971711 | 7.97E-06 | 3.30E-05 |
| -1.892344683 | 0.058445071 | 0.105737631 |
| -8.247580095 | 1.62E-16 | 2.16E-15 |
| -8.125829038 | 4.44E-16 | 5.70E-15 |
| NA | NA | NA |
| 0.30061664 | 0.763706841 | NA |
| NA | NA | NA |
| 11.69545212 | 1.34E-31 | 5.06E-30 |
| 1.265671222 | 0.205630816 | 0.302316553 |
| 0.871396059 | 0.383537937 | 0.495916652 |
| -7.138641428 | 9.43E-13 | 8.77E-12 |
| -1.073936415 | 0.282851181 | 0.389750929 |
| -4.731168317 | 2.23E-06 | 9.99E-06 |

| | | |
|---|---|---|
| -3.761996532 | 0.000168562 | 0.000560159 |
| -2.17037346 | 0.029978566 | 0.059303209 |
| -0.313632148 | 0.75380044 | 0.821402918 |
| -0.923931377 | 0.355522046 | 0.467443232 |
| -2.289801442 | 0.02203283 | 0.045453039 |
| -2.149513341 | 0.03159373 | 0.062105635 |
| -4.416673981 | 1.00E-05 | 4.09E-05 |
| -3.241526238 | 0.001188915 | 0.003370927 |
| -1.793307573 | 0.072923757 | 0.127490511 |
| -1.65255974 | 0.098420497 | 0.16440637 |
| -4.275854185 | 1.90E-05 | 7.42E-05 |
| -1.520273324 | 0.128442296 | NA |
| 0.063708285 | 0.949202508 | NA |
| -0.207626023 | 0.835520984 | NA |
| -0.804761211 | 0.420957489 | 0.532741731 |
| 0.290939976 | 0.771097226 | NA |
| -1.752953027 | 0.07961007 | 0.137527784 |
| -0.789407167 | 0.429874068 | 0.540989891 |
| -0.51847402 | 0.604127582 | 0.701327089 |
| -2.520729895 | 0.011711171 | 0.026219196 |
| -1.731268186 | 0.083403943 | 0.143084352 |
| -1.963041174 | 0.04964139 | 0.092018366 |
| -0.909679901 | 0.362991347 | 0.475440785 |
| -3.085020339 | 0.002035382 | 0.00547508 |
| NA | NA | NA |
| -1.060346409 | 0.288987033 | 0.396434585 |
| -2.447824987 | 0.014372145 | 0.031357634 |
| -2.391726963 | 0.016769312 | 0.035945524 |
| -1.617261301 | 0.105821889 | 0.174476114 |
| -2.855714367 | 0.004294012 | 0.010714698 |
| -0.078323391 | 0.937570811 | NA |
| 0.063029682 | 0.949742869 | 0.965598278 |
| -0.065389966 | 0.947863513 | NA |
| -2.31636977 | 0.020538088 | 0.042767749 |
| -0.086309271 | 0.931220569 | NA |
| 0.151745575 | 0.87938761 | NA |
| -0.065388772 | 0.947864464 | NA |
| NA | NA | NA |
| NA | NA | NA |
| -0.065389369 | 0.947863988 | NA |
| -0.721411363 | 0.470656457 | 0.579722018 |
| -0.085181149 | 0.932117377 | NA |
| NA | NA | NA |
| NA | NA | NA |
| NA | NA | NA |
| -0.262490392 | 0.792943392 | NA |
| NA | NA | NA |

| | | |
|---|---|---|
| NA | NA | NA |
| -0.065389369 | 0.947863988 | NA |
| NA | NA | NA |
| 0.063707688 | 0.949202983 | NA |
| -1.542588589 | 0.122930629 | 0.198002164 |
| -0.7208871 | 0.47097898 | 0.579890987 |
| -0.98876858 | 0.32277638 | 0.433660376 |
| 0.26246791 | 0.792960723 | NA |
| 0.345524135 | 0.729700369 | NA |
| -0.305749404 | 0.759795454 | 0.826149362 |
| -1.391933047 | 0.163942673 | 0.250986387 |
| -0.680078118 | 0.496454999 | NA |
| 0.480575717 | 0.630818078 | 0.722923698 |
| 1.326973457 | 0.184517463 | 0.276770666 |
| 0.063707688 | 0.949202983 | NA |
| 0.195900794 | 0.844687816 | 0.891460865 |
| -0.065389369 | 0.947863988 | NA |
| 0.063708285 | 0.949202508 | NA |
| -1.232310985 | 0.217832938 | 0.316702976 |
| -0.178890713 | 0.858023515 | 0.90148146 |
| -1.446817011 | 0.147948181 | 0.230754173 |
| -1.428591621 | 0.153121644 | 0.237447907 |
| -0.246580038 | 0.805233255 | NA |
| -0.205097267 | 0.837496132 | NA |
| 0.078715558 | 0.93725887 | NA |
| -0.485977436 | 0.626983164 | NA |
| -1.405418486 | 0.159896873 | 0.245836531 |
| -3.757315875 | 0.000171746 | 0.000569678 |
| -1.65728525 | 0.097461818 | 0.163077031 |
| -0.191477271 | 0.848151686 | 0.894013168 |
| -2.374758038 | 0.017560457 | 0.037391599 |
| -1.13196805 | 0.257647867 | NA |
| 0.063708285 | 0.949202508 | NA |
| 0.108487005 | 0.913609388 | NA |
| -0.599626428 | 0.54875523 | NA |
| -1.952157827 | 0.050919469 | 0.094068078 |
| 0.189005896 | 0.850088196 | NA |
| -1.310129287 | 0.190152102 | 0.283692384 |
| -0.213350431 | 0.831053641 | NA |
| -1.81477195 | 0.069558963 | 0.122558133 |
| 1.110490827 | 0.266787579 | 0.372574327 |
| -1.317378799 | 0.187711684 | 0.280653691 |
| 0.06371127 | 0.949200131 | NA |
| 0.090532211 | 0.927864298 | NA |
| 0.104752356 | 0.916572316 | NA |
| -1.589966415 | 0.111842376 | 0.182657879 |
| -0.723210891 | 0.469550329 | NA |

| | | | |
|---|---|---|---|
| | -0.246580038 | 0.805233255 | NA |
| | -0.6197926 | 0.535394341 | NA |
| | -0.748404685 | 0.454216098 | NA |
| | -0.503902701 | 0.614329752 | 0.709845522 |
| NA | NA | NA | |
| NA | NA | NA | |
| | -0.220125339 | 0.825773541 | NA |
| | -0.065389369 | 0.947863988 | NA |
| | -0.632925885 | 0.526782043 | NA |
| NA | NA | NA | |
| NA | NA | NA | |
| NA | NA | NA | |
| | 0.063708882 | 0.949202033 | NA |
| NA | NA | NA | |
| NA | NA | NA | |
| NA | NA | NA | |
| NA | NA | NA | |
| | 0.067524971 | 0.946163784 | NA |
| NA | NA | NA | |
| NA | NA | NA | |
| NA | NA | NA | |
| NA | NA | NA | |
| | 0.115615398 | 0.907957359 | NA |
| | 0.474004791 | 0.635496491 | NA |
| NA | NA | NA | |
| NA | NA | NA | |
| NA | NA | NA | |
| | -0.105695914 | 0.915823622 | NA |
| | -0.316812758 | 0.751385683 | 0.819699427 |
| | -1.972009826 | 0.048608485 | 0.090331159 |
| | 0.292484042 | 0.769916559 | NA |
| | 0.063708285 | 0.949202508 | NA |
| | -0.11119828 | 0.911459116 | NA |
| | -1.277399234 | 0.201461335 | NA |
| | -0.159531316 | 0.873250288 | 0.911753001 |
| | 0.063707688 | 0.949202983 | NA |
| | -0.440691321 | 0.659436481 | NA |
| | -0.876425461 | 0.380798782 | NA |
| | -2.377852686 | 0.017413781 | 0.037107692 |
| | -0.923845999 | 0.355566502 | 0.467477127 |
| | -2.172260824 | 0.029835995 | 0.059090349 |
| | -2.315254241 | 0.020599022 | 0.042869727 |
| | -0.186052734 | 0.852403407 | NA |
| NA | NA | NA | |
| | -0.15215151 | 0.879067438 | NA |
| | -0.127331121 | 0.89867833 | 0.929500769 |
| NA | NA | NA | |

| | | |
|---|---|---|
| -0.313698781 | 0.753749826 | NA |
| -0.116787087 | 0.907028777 | NA |
| -0.34508181 | 0.730032868 | NA |
| -0.700535959 | 0.483592657 | NA |
| -0.083639162 | 0.933343329 | NA |
| -0.906447217 | 0.364699191 | NA |
| -0.410042961 | 0.681774433 | NA |
| -0.878290535 | 0.379786071 | 0.49243915 |
| 0.065474733 | 0.947796023 | NA |
| NA | NA | NA |
| 0.06371127 | 0.949200131 | NA |
| NA | NA | NA |
| -0.172926522 | 0.862709184 | 0.904444435 |
| -0.56273205 | 0.573617356 | NA |
| 0.192810118 | 0.847107679 | NA |
| -0.634138295 | 0.525990573 | 0.63113804 |
| NA | NA | NA |
| NA | NA | NA |
| NA | NA | NA |
| NA | NA | NA |
| NA | NA | NA |
| NA | NA | NA |
| -0.19448762 | 0.845794089 | NA |
| -0.749374949 | 0.453631245 | NA |
| NA | NA | NA |
| NA | NA | NA |
| -0.289905935 | 0.771888201 | NA |
| NA | NA | NA |
| -0.327305771 | 0.743436628 | NA |
| NA | NA | NA |
| NA | NA | NA |
| -0.065388772 | 0.947864464 | NA |
| 0.063708882 | 0.949202033 | NA |
| -0.194488217 | 0.845793621 | NA |
| -1.151037449 | 0.249716829 | 0.353210457 |
| -0.78894114 | 0.430146409 | NA |
| 0.251859234 | 0.801149871 | NA |
| -0.065465566 | 0.947803322 | NA |
| NA | NA | NA |
| -2.127050934 | 0.033415851 | 0.065249384 |
| -0.540208999 | 0.589052907 | NA |
| -0.627582218 | 0.53027766 | NA |
| NA | NA | NA |
| NA | NA | NA |
| -0.604769545 | 0.545332132 | NA |
| 0.194173692 | 0.846039881 | NA |
| -0.18544114 | 0.852883043 | 0.897828261 |

| | | | |
|---|---|---|---|
| | 0.415603364 | 0.677700265 | NA |
| | -0.68515021 | 0.493249139 | 0.600429236 |
| NA | NA | NA | |
| | -0.19448762 | 0.845794089 | NA |
| NA | NA | NA | |
| | 0.063708285 | 0.949202508 | NA |
| NA | NA | NA | |
| | -0.19448762 | 0.845794089 | NA |
| NA | NA | NA | |
| | 0.06371127 | 0.949200131 | NA |
| | -0.065389369 | 0.947863988 | NA |
| | 0.192809521 | 0.847108147 | NA |
| | 0.192808924 | 0.847108615 | NA |
| | 0.326726504 | 0.743874752 | NA |
| | -0.898999689 | 0.368652827 | 0.481169122 |
| | -0.080750035 | 0.935640744 | 0.956240642 |
| | -1.039972223 | 0.298352806 | 0.407207085 |
| | -4.269664175 | 1.96E-05 | 7.62E-05 |
| | -5.9242785 | 3.14E-09 | 2.03E-08 |
| NA | NA | NA | |
| NA | NA | NA | |
| | -8.60571114 | 7.58E-18 | 1.13E-16 |
| NA | NA | NA | |
| | 3.871158874 | 0.000108319 | 0.000372319 |
| | -4.719938936 | 2.36E-06 | 1.05E-05 |
| | -0.651172901 | 0.514934881 | 0.62036106 |
| | 1.12615559 | 0.260099667 | NA |
| | 0.478108317 | 0.632573113 | 0.724343245 |
| | 0.164557515 | 0.869292273 | 0.909061441 |
| | -0.08180493 | 0.934801835 | NA |
| | -1.738846939 | 0.08206169 | 0.141139565 |
| | 0.145324004 | 0.884455063 | NA |
| | -2.615856442 | 0.008900397 | 0.020518854 |
| | -0.61220578 | 0.540401618 | 0.644589697 |
| | -0.942260922 | 0.34605907 | NA |
| | -0.31290631 | 0.754351844 | 0.821860653 |
| | -0.804205995 | 0.421278017 | 0.533093467 |
| | -0.858553523 | 0.390586889 | 0.502719692 |
| | -3.819510121 | 0.000133717 | 0.000452166 |
| | -3.557809334 | 0.000373961 | 0.001167403 |
| | -2.506489422 | 0.012193671 | 0.027141625 |
| | -3.776017764 | 0.000159356 | 0.00053161 |
| | -2.074759366 | 0.038008842 | 0.072906837 |
| | -2.211113887 | 0.027027952 | 0.054209868 |
| | -1.596760255 | 0.110319159 | 0.1805826 |
| | -2.604350609 | 0.009204854 | 0.02115291 |
| | -3.201098932 | 0.001369045 | 0.003825318 |

| | | | |
|---|---|---|---|
| | -3.236902078 | 0.001208349 | 0.00341751 |
| | -0.465562245 | 0.641528873 | 0.731914293 |
| | -1.922501289 | 0.054542712 | 0.099737293 |
| | -1.186916045 | 0.235260738 | 0.336798002 |
| | 1.335407632 | 0.181742968 | 0.273346142 |
| | -0.539281754 | 0.589692457 | 0.689185144 |
| NA | NA | NA | |
| | -1.880080485 | 0.06009711 | 0.108218033 |
| | 1.389872837 | 0.164567495 | 0.251795611 |
| | -2.351873109 | 0.018679146 | 0.039394929 |
| | 1.668533517 | 0.095209864 | 0.159961263 |
| | -1.682556097 | 0.092461057 | 0.156129006 |
| | -3.596144421 | 0.000322969 | 0.001021097 |
| | -1.110681535 | 0.266705453 | 0.372524413 |
| | 4.719450187 | 2.36E-06 | 1.05E-05 |
| | 2.692652009 | 0.007088621 | 0.016759062 |
| | 0.063708882 | 0.949202033 | NA |
| | 2.68941464 | 0.007157745 | 0.016890237 |
| NA | NA | NA | |
| NA | NA | NA | |
| | 4.136724677 | 3.52E-05 | 0.000131671 |
| | -0.921379494 | 0.356852331 | 0.468650708 |
| | -0.170570519 | 0.86456148 | 0.905640126 |
| | 3.784865996 | 0.000153791 | 0.000514763 |
| | -2.213910993 | 0.026834902 | 0.053885087 |
| | -6.611931801 | 3.79E-11 | 3.01E-10 |
| | -7.530316897 | 5.06E-14 | 5.35E-13 |
| | -1.921017762 | 0.054729468 | 0.100061883 |
| | -2.197375705 | 0.027993625 | 0.05590238 |
| | 9.85091013 | 6.79E-23 | 1.50E-21 |
| NA | NA | NA | |
| | -1.537161223 | 0.124253833 | 0.199811979 |
| | -5.020193176 | 5.16E-07 | 2.52E-06 |
| | -0.057339345 | 0.954274879 | 0.968671434 |
| | 0.834294645 | 0.404114969 | 0.51578184 |
| | 0.063708324 | 0.949202477 | NA |
| | -1.507797613 | 0.131606333 | 0.209387776 |
| | -3.237247235 | 0.001206888 | 0.003414211 |
| | 4.766849107 | 1.87E-06 | 8.45E-06 |
| | 8.132991779 | 4.19E-16 | 5.38E-15 |
| | 1.434095375 | 0.151545048 | 0.235527634 |
| | 8.550554868 | 1.22E-17 | 1.79E-16 |
| NA | NA | NA | |
| | 12.74708043 | 3.24E-37 | 1.56E-35 |
| NA | NA | NA | |
| | 0.960520747 | 0.336793194 | NA |
| | 0.262131689 | 0.793219916 | NA |

| | | |
|---|---|---|
| 3.038715419 | 0.002375892 | 0.006284989 |
| 0.298298703 | 0.765475194 | 0.830882685 |
| 3.55791154 | 0.000373815 | 0.001167094 |
| 1.472247381 | 0.140954087 | 0.221640959 |
| -6.260233923 | 3.84E-10 | 2.72E-09 |
| -2.441863106 | 0.014611686 | 0.031828863 |
| -3.695190276 | 0.000219722 | 0.000716286 |
| -0.959453519 | 0.337330325 | 0.448543244 |
| -3.23582614 | 0.001212912 | 0.003429255 |
| -0.458974019 | 0.646252823 | 0.735420201 |
| 2.170664394 | 0.029956551 | 0.059264343 |
| -0.491445116 | 0.623111661 | 0.717079765 |
| NA | NA | NA |
| -0.065389369 | 0.947863988 | NA |
| -2.106281485 | 0.035179909 | 0.068222911 |
| -2.801688437 | 0.005083594 | 0.012488209 |
| 0.718685875 | 0.472334488 | 0.58124544 |
| 0.063708324 | 0.949202477 | NA |
| 7.642201169 | 2.14E-14 | 2.35E-13 |
| NA | NA | NA |
| NA | NA | NA |
| 8.841043762 | 9.48E-19 | 1.53E-17 |
| NA | NA | NA |
| NA | NA | NA |
| -0.065389966 | 0.947863513 | NA |
| NA | NA | NA |
| NA | NA | NA |
| NA | NA | NA |
| 0.528775736 | 0.596961031 | 0.695264744 |
| -2.521927786 | 0.011671367 | 0.026134756 |
| 0.063708882 | 0.949202033 | NA |
| -2.40776315 | 0.01605059 | 0.034600489 |
| 0.063708882 | 0.949202033 | NA |
| -0.065389369 | 0.947863988 | NA |
| NA | NA | NA |
| -0.243376547 | 0.807713711 | NA |
| -0.127847651 | 0.898269539 | NA |
| 0.192809521 | 0.847108147 | NA |
| 9.827035376 | 8.61E-23 | 1.88E-21 |
| NA | NA | NA |
| NA | NA | NA |
| 4.719519854 | 2.36E-06 | 1.05E-05 |
| NA | NA | NA |
| 4.378788158 | 1.19E-05 | 4.80E-05 |
| NA | NA | NA |
| 0.063708285 | 0.949202508 | NA |
| -0.065389966 | 0.947863513 | NA |

| | | |
|---|---|---|
| -5.611005185 | 2.01E-08 | 1.19E-07 |
| -3.01187083 | 0.002596431 | 0.006804031 |
| -0.704516729 | 0.481111035 | 0.589803731 |
| -1.089164114 | 0.276081521 | 0.382881712 |
| NA | NA | NA |
| -0.065388772 | 0.947864464 | NA |
| -3.262154391 | 0.001105689 | 0.00315749 |
| -1.807181214 | 0.070734029 | 0.124409581 |
| NA | NA | NA |
| NA | NA | NA |
| 2.184203133 | 0.028947325 | 0.057558909 |
| -0.065389369 | 0.947863988 | NA |
| NA | NA | NA |
| 5.329131182 | 9.87E-08 | 5.32E-07 |
| NA | NA | NA |
| NA | NA | NA |
| -4.24020044 | 2.23E-05 | 8.61E-05 |
| 0.063708285 | 0.949202508 | NA |
| -1.291543736 | 0.1965152 | 0.291362128 |
| NA | NA | NA |
| 12.95875677 | 2.10E-38 | 1.08E-36 |
| 0.54372453 | 0.58663105 | 0.686523994 |
| 0.201808052 | 0.840066787 | NA |
| 14.68516906 | 8.02E-49 | 6.01E-47 |
| NA | NA | NA |
| 5.04607209 | 4.51E-07 | 2.22E-06 |
| 0.508253258 | 0.611275745 | 0.707198147 |
| -1.806696417 | 0.070809627 | 0.124515368 |
| 0.087681318 | 0.930129969 | NA |
| NA | NA | NA |
| -3.44027881 | 0.000581115 | 0.001750549 |
| 1.413577602 | 0.157485962 | 0.242943517 |
| 2.69007365 | 0.007143625 | 0.016866799 |
| 2.423372119 | 0.015377166 | 0.033297904 |
| 1.904854403 | 0.056799 | 0.10327466 |
| -0.260876975 | 0.794187382 | NA |
| 9.320353361 | 1.16E-20 | 2.16E-19 |
| 15.40946516 | 1.41E-53 | 1.18E-51 |
| -2.338670174 | 0.01935251 | 0.040606817 |
| 5.227447938 | 1.72E-07 | 8.97E-07 |
| -1.761886932 | 0.078088403 | 0.135300618 |
| -8.401994809 | 4.39E-17 | 6.16E-16 |
| 7.002454799 | 2.52E-12 | 2.23E-11 |
| NA | NA | NA |
| -3.806223951 | 0.000141105 | 0.00047521 |
| 4.111291407 | 3.93E-05 | 0.000145768 |
| NA | NA | NA |

| | | | |
|---|---|---|---|
| | -2.250043827 | 0.024446163 | 0.049726735 |
| | -2.370549949 | 0.017761644 | 0.037765371 |
| | -3.649945345 | 0.000262296 | 0.000842349 |
| | -2.571493305 | 0.010126097 | 0.023054049 |
| | -3.751006102 | 0.000176126 | 0.000583359 |
| | -3.201837205 | 0.001365541 | 0.003815954 |
| | 5.570343464 | 2.54E-08 | 1.48E-07 |
| | 4.685291834 | 2.80E-06 | 1.23E-05 |
| | 0.749129704 | 0.453779033 | 0.563732837 |
| | -2.346035103 | 0.01897432 | 0.039913206 |
| | 0.811033298 | 0.417346549 | 0.52900107 |
| | 2.207257031 | 0.027296107 | 0.054697107 |
| | -0.451018178 | 0.651976448 | 0.740445501 |
| | -0.164690661 | 0.869187468 | NA |
| | -0.35458292 | 0.722902074 | 0.797489484 |
| | 11.83603338 | 2.54E-32 | 9.88E-31 |
| | -3.980446975 | 6.88E-05 | 0.0002447 |
| | 0.232309173 | 0.816297889 | NA |
| | -2.076088766 | 0.037885742 | 0.072715292 |
| | 2.202426802 | 0.027635174 | 0.055296859 |
| NA | NA | NA | |
| | 7.814666522 | 5.51E-15 | 6.43E-14 |
| | -3.792574434 | 0.000149094 | 0.000500042 |
| | 7.083246168 | 1.41E-12 | 1.28E-11 |
| | 6.367083634 | 1.93E-10 | 1.41E-09 |
| | -1.857563743 | 0.063230991 | 0.112967289 |
| NA | NA | NA | |
| | -1.174921618 | 0.240026076 | 0.342210793 |
| | 3.124932236 | 0.00177846 | 0.004845946 |
| | -1.24349178 | 0.213686668 | 0.311797674 |
| | 5.542868178 | 2.98E-08 | 1.72E-07 |
| | 0.481028325 | 0.630496368 | 0.722654276 |
| | -2.39487186 | 0.016626175 | 0.035690665 |
| | 3.103788041 | 0.001910602 | 0.005168308 |
| | -1.447357574 | 0.147796802 | 0.2305468 |
| | -9.155074683 | 5.43E-20 | 9.68E-19 |
| | 15.78906594 | 3.70E-56 | 3.33E-54 |
| | 10.19619108 | 2.06E-24 | 5.06E-23 |
| | 0.192809521 | 0.847108147 | NA |
| | -0.137505422 | 0.890631307 | 0.923429159 |
| | 3.000611149 | 0.002694384 | 0.007038242 |
| | 0.984020227 | 0.325105568 | 0.436157601 |
| | 2.150922306 | 0.031482333 | 0.061906103 |
| | 13.50604183 | 1.44E-41 | 8.27E-40 |
| | -2.222721753 | 0.026234568 | 0.052857703 |
| | -2.026117166 | 0.042752779 | 0.080791771 |
| | 0.10769217 | 0.914239881 | 0.941046355 |

| | | | |
|---|---|---|---|
| | 2.612783264 | 0.008980824 | 0.02068366 |
| NA | NA | NA | |
| | 1.249722953 | 0.21140077 | 0.309147071 |
| | -0.072655315 | 0.942080408 | 0.960656426 |
| | -0.966833425 | 0.333627312 | 0.444862856 |
| | 0.876247766 | 0.380895354 | 0.493417386 |
| | -0.301711329 | 0.762872131 | 0.828762563 |
| | -3.92183614 | 8.79E-05 | 0.000307415 |
| | 0.400334135 | 0.688910429 | 0.770387495 |
| | 4.340633205 | 1.42E-05 | 5.65E-05 |
| | -4.569033521 | 4.90E-06 | 2.09E-05 |
| NA | NA | NA | |
| | -4.793367173 | 1.64E-06 | 7.47E-06 |
| | -0.194487022 | 0.845794556 | NA |
| | -2.7301031 | 0.006331452 | 0.015165476 |
| NA | NA | NA | |
| NA | NA | NA | |
| | -1.117856367 | 0.263628343 | 0.369132772 |
| | -2.665731937 | 0.007682092 | 0.017973711 |
| | 7.724350886 | 1.12E-14 | 1.27E-13 |
| | -9.578226284 | 9.87E-22 | 2.00E-20 |
| NA | NA | NA | |
| | -1.817999645 | 0.069064187 | 0.121840753 |
| | -0.45894094 | 0.646276578 | 0.735420201 |
| | -3.106184801 | 0.001895182 | 0.00513048 |
| | -0.02086803 | 0.983350929 | NA |
| | 3.082483286 | 0.002052813 | 0.005516039 |
| | -4.256939863 | 2.07E-05 | 8.04E-05 |
| NA | NA | NA | |
| | 0.435259976 | 0.663373733 | NA |
| NA | NA | NA | |
| | -5.394896153 | 6.86E-08 | 3.77E-07 |
| | 8.138300215 | 4.01E-16 | 5.16E-15 |
| | -1.197439588 | 0.231135261 | 0.331936175 |
| | 3.255062356 | 0.001133675 | 0.003228567 |
| | 0.343034073 | 0.731572827 | NA |
| NA | NA | NA | |
| | 0.644615899 | 0.519176129 | NA |
| | -4.534192369 | 5.78E-06 | 2.44E-05 |
| NA | NA | NA | |
| | -0.259755806 | 0.795052143 | NA |
| | 1.673296682 | 0.094268905 | 0.158625257 |
| | 4.470392732 | 7.81E-06 | 3.24E-05 |
| | -2.787885256 | 0.005305333 | 0.01296675 |
| | 10.34180045 | 4.56E-25 | 1.17E-23 |
| | -1.890316908 | 0.058715589 | 0.106173327 |
| | -0.814130269 | 0.415570327 | 0.527176792 |

| | | | |
|---|---|---|---|
| | 4.381301162 | 1.18E-05 | 4.75E-05 |
| | 0.96092257 | 0.336591102 | 0.447869612 |
| | 0.552010305 | 0.580941293 | 0.68155428 |
| | 7.922566147 | 2.33E-15 | 2.81E-14 |
| | -1.217421088 | 0.223444043 | 0.323115397 |
| | 0.219083146 | 0.82658528 | 0.878681286 |
| | -0.777194156 | 0.43704423 | 0.547343994 |
| NA | NA | NA | |
| | -9.149047919 | 5.74E-20 | 1.02E-18 |
| | -0.661888636 | 0.508042596 | 0.614314308 |
| | -2.079435059 | 0.037577381 | 0.072189872 |
| | -1.10198265 | 0.270469215 | 0.376700278 |
| | 1.68768452 | 0.091471805 | 0.154646354 |
| | 0.970905983 | 0.331595098 | 0.442586536 |
| | 0.104964653 | 0.916403857 | NA |
| NA | NA | NA | |
| | 1.811934387 | 0.069996335 | 0.123259341 |
| | 0.192809521 | 0.847108147 | NA |
| | 0.192810118 | 0.847107679 | NA |
| | -0.19448762 | 0.845794089 | NA |
| NA | NA | NA | |
| NA | NA | NA | |
| NA | NA | NA | |
| NA | NA | NA | |
| NA | NA | NA | |
| NA | NA | NA | |
| NA | NA | NA | |
| NA | NA | NA | |
| NA | NA | NA | |
| NA | NA | NA | |
| NA | NA | NA | |
| NA | NA | NA | |
| NA | NA | NA | |
| NA | NA | NA | |
| NA | NA | NA | |
| NA | NA | NA | |
| NA | NA | NA | |
| NA | NA | NA | |
| NA | NA | NA | |
| NA | NA | NA | |
| NA | NA | NA | |
| NA | NA | NA | |
| NA | NA | NA | |
| NA | NA | NA | |
| NA | NA | NA | |
| NA | NA | NA | |

| | | |
|---|---|---|
| NA | NA | NA |
| NA | NA | NA |
| NA | NA | NA |
| 0.192636473 | 0.847243678 | NA |
| NA | NA | NA |
| NA | NA | NA |
| NA | NA | NA |
| NA | NA | NA |
| NA | NA | NA |
| NA | NA | NA |
| NA | NA | NA |
| NA | NA | NA |
| NA | NA | NA |
| NA | NA | NA |
| NA | NA | NA |
| NA | NA | NA |
| NA | NA | NA |
| NA | NA | NA |
| NA | NA | NA |
| 0.720295138 | 0.471343298 | NA |
| NA | NA | NA |
| NA | NA | NA |
| NA | NA | NA |
| NA | NA | NA |
| NA | NA | NA |
| NA | NA | NA |
| NA | NA | NA |
| NA | NA | NA |
| NA | NA | NA |
| 0.192809521 | 0.847108147 | NA |
| NA | NA | NA |
| NA | NA | NA |
| NA | NA | NA |
| NA | NA | NA |
| NA | NA | NA |
| NA | NA | NA |
| NA | NA | NA |
| 2.130310286 | 0.033146005 | 0.064800789 |
| 0.192809521 | 0.847108147 | NA |
| 0.192636473 | 0.847243678 | NA |
| NA | NA | NA |
| -0.79354766 | 0.427458813 | NA |
| 0.192810118 | 0.847107679 | NA |
| 7.625704389 | 2.43E-14 | 2.65E-13 |
| -0.742180109 | 0.457978219 | 0.567569184 |
| -8.462726392 | 2.61E-17 | 3.73E-16 |
| 0.192810118 | 0.847107679 | NA |

| | | |
|---|---|---|
| 21.17635925 | 1.58E-99 | 3.69E-97 |
| 0.063708285 | 0.949202508 | NA |
| 0.192809521 | 0.847108147 | NA |
| -1.25000457 | 0.211297878 | 0.309032715 |
| NA | NA | NA |
| 1.343453163 | 0.179125282 | 0.270090712 |
| -0.256020697 | 0.797934856 | NA |
| -2.327883692 | 0.01991828 | 0.041639903 |
| 0.063710673 | 0.949200606 | NA |
| 0.06371127 | 0.949200131 | NA |
| -1.42400415 | 0.154445271 | 0.239203337 |
| -0.032575794 | 0.974012873 | 0.982052988 |
| 7.150418456 | 8.65E-13 | 8.09E-12 |
| 0.335172531 | 0.737494942 | NA |
| -1.175184146 | 0.239921052 | 0.342080542 |
| 0.245195359 | 0.80630517 | 0.862932143 |
| -2.776050587 | 0.005502365 | 0.013403814 |
| 0.347873358 | 0.727935294 | NA |
| -0.561353041 | 0.57455689 | 0.675733272 |
| -0.985049802 | 0.324599609 | 0.435583169 |
| 5.022370436 | 5.10E-07 | 2.49E-06 |
| -1.223628088 | 0.221092559 | 0.32038467 |
| -0.065388772 | 0.947864464 | NA |
| 0.727489316 | 0.466926258 | 0.576233322 |
| 8.828371236 | 1.06E-18 | 1.71E-17 |
| 1.753157063 | 0.079575051 | 0.137486265 |
| NA | NA | NA |
| -8.272197481 | 1.32E-16 | 1.77E-15 |
| 0.325495478 | 0.744806105 | 0.814296182 |
| 1.145383788 | 0.252050211 | 0.356028035 |
| 4.246205122 | 2.17E-05 | 8.40E-05 |
| 0.733593419 | 0.463196573 | 0.572675239 |
| -5.303042366 | 1.14E-07 | 6.09E-07 |
| -0.065388772 | 0.947864464 | NA |
| NA | NA | NA |
| -2.548431205 | 0.010820863 | 0.024460055 |
| NA | NA | NA |
| NA | NA | NA |
| 2.343506951 | 0.019103406 | 0.040147629 |
| -2.478671095 | 0.013187285 | 0.029102251 |
| -0.374987847 | 0.707669505 | 0.785628896 |
| -0.058182314 | 0.953603408 | 0.96824189 |
| -0.462816181 | 0.643496132 | 0.733289696 |
| -1.663179682 | 0.096276473 | 0.161449828 |
| NA | NA | NA |
| -1.368095986 | 0.171282033 | 0.260302932 |
| -4.105089328 | 4.04E-05 | 0.000149381 |

| | | |
|---|---|---|
| 4.377728475 | 1.20E-05 | 4.82E-05 |
| 0.489847621 | 0.62424173 | 0.717984124 |
| 0.910272919 | 0.362678597 | 0.475103705 |
| 0.251739278 | 0.801242595 | 0.85892767 |
| -1.645919181 | 0.099780399 | 0.166223188 |
| NA | NA | NA |
| 1.596305668 | 0.110420565 | 0.180724956 |
| -0.09556214 | 0.923868335 | 0.947722313 |
| -0.44175393 | 0.65866728 | 0.745974321 |
| NA | NA | NA |
| -3.172607204 | 0.001510768 | 0.004181634 |
| -2.305410502 | 0.021143594 | 0.043846243 |
| -4.03214514 | 5.53E-05 | 0.000200059 |
| -2.478477126 | 0.013194457 | 0.029115513 |
| NA | NA | NA |
| NA | NA | NA |
| 0.551401162 | 0.581358705 | NA |
| 4.727347404 | 2.27E-06 | 1.02E-05 |
| 1.356937162 | 0.174801178 | 0.264782951 |
| NA | NA | NA |
| 1.118049181 | 0.263545989 | NA |
| NA | NA | NA |
| 0.072960039 | 0.941837917 | 0.960448281 |
| -0.675478152 | 0.499372028 | 0.606496281 |
| 8.959575812 | 3.26E-19 | 5.49E-18 |
| -3.673257438 | 0.000239478 | 0.000774861 |
| -0.316766897 | 0.751420484 | 0.819701666 |
| 3.083085249 | 0.002048665 | 0.005507257 |
| 2.066453817 | 0.038785649 | 0.074169516 |
| -1.752022604 | 0.079769921 | 0.137756391 |
| 4.819796049 | 1.44E-06 | 6.59E-06 |
| -3.28203684 | 0.001030601 | 0.002962328 |
| 2.66759012 | 0.00763974 | 0.017887107 |
| -2.187573523 | 0.028700682 | 0.057145659 |
| -2.08405741 | 0.037154945 | 0.071482574 |
| -2.195540872 | 0.028124822 | 0.056133028 |
| NA | NA | NA |
| -0.672998417 | 0.500948298 | 0.607971377 |
| 0.308293014 | 0.757859382 | 0.8246053 |
| -2.014290436 | 0.04397905 | 0.082841331 |
| -7.261269104 | 3.83E-13 | 3.71E-12 |
| -5.808260104 | 6.31E-09 | 3.94E-08 |
| -1.204435116 | 0.228421446 | 0.32881233 |
| 4.129717677 | 3.63E-05 | 0.000135446 |
| 4.233633864 | 2.30E-05 | 8.85E-05 |
| -0.300951847 | 0.763451214 | 0.829247695 |
| 0.732295875 | 0.463987996 | 0.573370503 |

| | | |
|---|---|---|
| 0.453754464 | 0.650005561 | 0.738756718 |
| 5.287851371 | 1.24E-07 | 6.59E-07 |
| 2.823227869 | 0.004754277 | 0.011761173 |
| 0.825015429 | 0.409362831 | 0.521257143 |
| 6.259364886 | 3.87E-10 | 2.74E-09 |
| 0.063708285 | 0.949202508 | NA |
| NA | NA | NA |
| -3.953496609 | 7.70E-05 | 0.000272127 |
| -2.91781128 | 0.003524976 | 0.008959195 |
| -0.489465132 | 0.624512434 | 0.718228068 |
| 0.148854979 | 0.881668067 | 0.916953789 |
| -0.065390563 | 0.947863038 | NA |
| 5.560822481 | 2.69E-08 | 1.56E-07 |
| NA | NA | NA |
| -4.767451451 | 1.87E-06 | 8.43E-06 |
| 0.33563868 | 0.737143352 | NA |
| 1.314967008 | 0.188520991 | 0.281664248 |
| -5.286231285 | 1.25E-07 | 6.64E-07 |
| 10.9290422 | 8.37E-28 | 2.58E-26 |
| 0.158070554 | 0.874401204 | 0.9124979 |
| -0.781148859 | 0.434714948 | 0.545354609 |
| 7.026090088 | 2.12E-12 | 1.90E-11 |
| NA | NA | NA |
| 4.31695259 | 1.58E-05 | 6.25E-05 |
| NA | NA | NA |
| -3.126947517 | 0.001766315 | 0.004817571 |
| -1.166112459 | 0.24356898 | 0.346334805 |
| -2.342133387 | 0.019173861 | 0.040282167 |
| -1.012860836 | 0.311126637 | 0.421333103 |
| NA | NA | NA |
| -2.873802721 | 0.004055622 | 0.010182789 |
| 0.45547931 | 0.648764449 | NA |
| 0.063708285 | 0.949202508 | NA |
| NA | NA | NA |
| 0.56761373 | 0.570297282 | 0.671648284 |
| 4.773533507 | 1.81E-06 | 8.21E-06 |
| 6.176641128 | 6.55E-10 | 4.53E-09 |
| -2.317562723 | 0.020473098 | 0.042650158 |
| -5.218524463 | 1.80E-07 | 9.38E-07 |
| -2.121614509 | 0.033870123 | 0.066025637 |
| 8.696452724 | 3.42E-18 | 5.25E-17 |
| -0.469046836 | 0.639036159 | 0.729835511 |
| 0.654774695 | 0.512612825 | 0.618257554 |
| -9.261804971 | 2.01E-20 | 3.67E-19 |
| -2.072682552 | 0.038201833 | 0.073204094 |
| 8.75031504 | 2.13E-18 | 3.32E-17 |
| 1.110623789 | 0.266730319 | 0.372535502 |

| | | | |
|---|---|---|---|
| | 0.641662779 | 0.521092162 | 0.626311491 |
| | -1.138999576 | 0.254703331 | 0.359025983 |
| | -2.560921946 | 0.010439481 | 0.023692103 |
| | -5.118001902 | 3.09E-07 | 1.56E-06 |
| | 16.71370836 | 1.04E-62 | 1.12E-60 |
| | 2.581929159 | 0.009824975 | 0.0224256 |
| | 3.531049369 | 0.000413914 | 0.001281103 |
| | -2.513195173 | 0.011964312 | 0.02669044 |
| | -0.065389966 | 0.947863513 | NA |
| | -4.139082368 | 3.49E-05 | 0.000130462 |
| | -2.479352813 | 0.013162104 | 0.029059492 |
| | -0.440122501 | 0.659848386 | 0.7468912 |
| | -5.678600889 | 1.36E-08 | 8.17E-08 |
| | 0.293599863 | 0.76906368 | 0.83366337 |
| | 2.895696652 | 0.003783179 | 0.009554331 |
| | -6.888042722 | 5.66E-12 | 4.86E-11 |
| NA | NA | NA | |
| | 11.87467169 | 1.60E-32 | 6.34E-31 |
| | 10.34361571 | 4.47E-25 | 1.15E-23 |
| | -0.763797526 | 0.444987911 | 0.554904714 |
| | -1.934173863 | 0.053091755 | 0.097457535 |
| | -3.096751103 | 0.001956541 | 0.005282871 |
| | -1.805182527 | 0.071046124 | 0.124853225 |
| | -2.050789303 | 0.04028747 | 0.07668989 |
| | -0.065389966 | 0.947863513 | NA |
| | 13.62060396 | 3.02E-42 | 1.80E-40 |
| | -5.169052257 | 2.35E-07 | 1.21E-06 |
| | -0.758601198 | 0.448091156 | NA |
| | 4.072378102 | 4.65E-05 | 0.00017024 |
| | 2.163180097 | 0.03052733 | 0.060255428 |
| | 0.563861906 | 0.572848117 | NA |
| | -3.158277978 | 0.001587041 | 0.004376297 |
| | 0.17985144 | 0.857269199 | 0.900929926 |
| | 5.041652728 | 4.62E-07 | 2.27E-06 |
| | 0.053993957 | 0.956939979 | 0.970365225 |
| | 5.427500437 | 5.71E-08 | 3.18E-07 |
| | -4.635996946 | 3.55E-06 | 1.54E-05 |
| | 0.063710673 | 0.949200606 | NA |
| | 0.630393336 | 0.52843727 | 0.633324216 |
| | 0.555702378 | 0.578414331 | 0.679322011 |
| | 4.330033986 | 1.49E-05 | 5.91E-05 |
| | 2.806966654 | 0.005001042 | 0.012309576 |
| | 2.085307953 | 0.037041356 | 0.071296919 |
| | 19.62919056 | 8.71E-86 | 1.58E-83 |
| | 4.449438699 | 8.61E-06 | 3.54E-05 |
| | -1.801458259 | 0.071630681 | 0.125695172 |
| | 1.123882157 | 0.261063022 | 0.366160788 |

| | | | |
|---|---|---|---|
| NA | NA | NA | |
| | 0.879475973 | 0.379143255 | 0.491783991 |
| | -0.100825882 | 0.919688681 | NA |
| | 0.111701246 | 0.911060292 | 0.938661361 |
| | 5.847833695 | 4.98E-09 | 3.15E-08 |
| | 5.171384767 | 2.32E-07 | 1.19E-06 |
| | 9.291265317 | 1.52E-20 | 2.81E-19 |
| | -5.471233965 | 4.47E-08 | 2.53E-07 |
| | 3.347555538 | 0.000815277 | 0.002388386 |
| NA | NA | NA | |
| NA | NA | NA | |
| NA | NA | NA | |
| NA | NA | NA | |
| NA | NA | NA | |
| NA | NA | NA | |
| | 30.51859168 | 1.48E-204 | 1.19E-201 |
| | 0.437321412 | 0.661878269 | NA |
| NA | NA | NA | |
| | 0.063708285 | 0.949202508 | NA |
| NA | NA | NA | |
| | 0.24166994 | 0.809035924 | NA |
| NA | NA | NA | |
| | -0.26176171 | 0.793505161 | NA |
| | 1.393552464 | 0.163452792 | 0.250396192 |
| | 0.343196468 | 0.731450662 | NA |
| | -0.676963092 | 0.498429372 | 0.605586485 |
| NA | NA | NA | |
| NA | NA | NA | |
| NA | NA | NA | |
| | -4.140088099 | 3.47E-05 | 0.00012995 |
| NA | NA | NA | |
| NA | NA | NA | |
| | 0.192809521 | 0.847108147 | NA |
| | 0.063708285 | 0.949202508 | NA |
| NA | NA | NA | |
| NA | NA | NA | |
| NA | NA | NA | |
| NA | NA | NA | |
| NA | NA | NA | |
| | 0.316744452 | 0.751437516 | NA |
| NA | NA | NA | |
| NA | NA | NA | |
| NA | NA | NA | |
| NA | NA | NA | |
| NA | NA | NA | |
| NA | NA | NA | |
| | 0.192809521 | 0.847108147 | NA |

| | | |
|---|---|---|
| 0.063708882 | 0.949202033 | NA |
| NA | NA | NA |
| 0.192809521 | 0.847108147 | NA |
| 0.288217989 | 0.77317988 | NA |
| NA | NA | NA |
| NA | NA | NA |
| NA | NA | NA |
| NA | NA | NA |
| NA | NA | NA |
| NA | NA | NA |
| -0.065389369 | 0.947863988 | NA |
| NA | NA | NA |
| NA | NA | NA |
| NA | NA | NA |
| NA | NA | NA |
| NA | NA | NA |
| NA | NA | NA |
| NA | NA | NA |
| NA | NA | NA |
| NA | NA | NA |
| 0.192810118 | 0.847107679 | NA |
| NA | NA | NA |
| NA | NA | NA |
| NA | NA | NA |
| NA | NA | NA |
| NA | NA | NA |
| -0.065389966 | 0.947863513 | NA |
| NA | NA | NA |
| NA | NA | NA |
| NA | NA | NA |
| NA | NA | NA |
| NA | NA | NA |
| NA | NA | NA |
| NA | NA | NA |
| NA | NA | NA |
| NA | NA | NA |
| NA | NA | NA |
| NA | NA | NA |
| -0.065389369 | 0.947863988 | NA |
| 0.063708285 | 0.949202508 | NA |
| NA | NA | NA |
| NA | NA | NA |
| NA | NA | NA |
| NA | NA | NA |
| NA | NA | NA |
| NA | NA | NA |
| NA | NA | NA |

| | | | |
|---|---|---|---|
| NA | NA | NA | |
| NA | NA | NA | |
| NA | NA | NA | |
| NA | NA | NA | |
| NA | NA | NA | |
| 0.192810118 | 0.847107679 | NA | |
| NA | NA | NA | |
| NA | NA | NA | |
| NA | NA | NA | |
| NA | NA | NA | |
| 0.063708882 | 0.949202033 | NA | |
| 1.120615874 | 0.262451405 | | 0.367814035 |
| 1.676168358 | 0.09370522 | | 0.157867784 |
| NA | NA | NA | |
| NA | NA | NA | |
| NA | NA | NA | |
| NA | NA | NA | |
| NA | NA | NA | |
| NA | NA | NA | |
| NA | NA | NA | |
| 1.528088009 | 0.126490688 | | 0.202862314 |
| NA | NA | NA | |
| -0.065389369 | 0.947863988 | NA | |
| NA | NA | NA | |
| -0.065389369 | 0.947863988 | NA | |
| NA | NA | NA | |
| NA | NA | NA | |
| NA | NA | NA | |
| 0.317642674 | 0.750756 | NA | |
| 0.192810118 | 0.847107679 | NA | |
| NA | NA | NA | |
| NA | NA | NA | |
| NA | NA | NA | |
| NA | NA | NA | |
| NA | NA | NA | |
| NA | NA | NA | |
| NA | NA | NA | |
| 0.063708882 | 0.949202033 | NA | |
| NA | NA | NA | |
| NA | NA | NA | |
| NA | NA | NA | |
| -0.065389369 | 0.947863988 | NA | |
| NA | NA | NA | |
| NA | NA | NA | |
| -0.431051048 | 0.666431253 | NA | |
| NA | NA | NA | |
| NA | NA | NA | |

| | | | |
|---|---|---|---|
| | 0.063708285 | 0.949202508 | NA |
| | 1.942243905 | 0.05210758 | 0.095945532 |
| | 7.55377343 | 4.23E-14 | 4.51E-13 |
| | 20.63920995 | 1.22E-94 | 2.70E-92 |
| NA | NA | NA | |
| | 4.159147256 | 3.19E-05 | 0.000120125 |
| NA | NA | NA | |
| | 1.251155314 | 0.210877818 | NA |
| | 11.46821541 | 1.91E-30 | 6.72E-29 |
| | 0.063708285 | 0.949202508 | NA |
| | 15.84057047 | 1.63E-56 | 1.49E-54 |
| NA | NA | NA | |
| | 8.929300542 | 4.29E-19 | 7.13E-18 |
| | 9.470222671 | 2.79E-21 | 5.49E-20 |
| | 28.31696442 | 2.14E-176 | 1.19E-173 |
| | -1.051870482 | 0.292858976 | 0.400801584 |
| | 17.29191382 | 5.41E-67 | 6.51E-65 |
| | 0.853850064 | 0.393188071 | 0.505132547 |
| NA | NA | NA | |
| NA | NA | NA | |
| | 0.063708285 | 0.949202508 | NA |
| | 0.192810118 | 0.847107679 | NA |
| NA | NA | NA | |
| | 0.330869843 | 0.740742802 | NA |
| | 0.755693089 | 0.449833228 | 0.559783206 |
| NA | NA | NA | |
| NA | NA | NA | |
| | 3.730496057 | 0.000191103 | 0.000630129 |
| NA | NA | NA | |
| NA | NA | NA | |
| NA | NA | NA | |
| NA | NA | NA | |
| NA | NA | NA | |
| NA | NA | NA | |
| NA | NA | NA | |
| | -0.210025412 | 0.833647839 | 0.883452308 |
| NA | NA | NA | |
| NA | NA | NA | |
| NA | NA | NA | |
| | 0.063708882 | 0.949202033 | NA |
| | 0.192810118 | 0.847107679 | NA |
| | 0.063708285 | 0.949202508 | NA |
| NA | NA | NA | |
| | 2.534170802 | 0.011271378 | 0.025368455 |
| NA | NA | NA | |
| | 0.192810118 | 0.847107679 | NA |
| | 0.192636473 | 0.847243678 | NA |

| | | |
|---|---|---|
| NA | NA | NA |
| NA | NA | NA |
| NA | NA | NA |
| NA | NA | NA |
| NA | NA | NA |
| -0.25287468 | 0.800365056 | NA |
| NA | NA | NA |
| 0.274205064 | 0.783927036 | NA |
| 1.539151133 | 0.123767404 | NA |
| 0.609232539 | 0.542370315 | NA |
| 0.223691394 | 0.822997443 | NA |
| -0.209901869 | 0.833744264 | NA |
| NA | NA | NA |
| NA | NA | NA |
| NA | NA | NA |
| -2.55992367 | 0.010469516 | 0.023753805 |
| -1.584046131 | 0.113183222 | 0.1845587 |
| 0.192809521 | 0.847108147 | NA |
| -9.240885802 | 2.44E-20 | 4.42E-19 |
| 0.063708285 | 0.949202508 | NA |
| -2.463520329 | 0.013758009 | 0.03021523 |
| -3.660065592 | 0.000252151 | 0.000812374 |
| 8.116075818 | 4.81E-16 | 6.16E-15 |
| 2.148030427 | 0.031711338 | 0.062302575 |
| -7.129752299 | 1.01E-12 | 9.34E-12 |
| 7.118710516 | 1.09E-12 | 1.01E-11 |
| -3.707626113 | 0.000209211 | 0.000685055 |
| 1.655117289 | 0.097900708 | 0.163684338 |
| -0.407444051 | 0.683681879 | 0.766151039 |
| 0.627580374 | 0.530278869 | 0.635126811 |
| 0.063708882 | 0.949202033 | NA |
| 1.948224389 | 0.05138812 | 0.094786728 |
| 5.185500239 | 2.15E-07 | 1.11E-06 |
| 7.504271344 | 6.18E-14 | 6.47E-13 |
| 4.128044873 | 3.66E-05 | 0.000136374 |
| 4.487364175 | 7.21E-06 | 3.00E-05 |
| -1.662515393 | 0.09640948 | 0.161597085 |
| -0.21841585 | 0.827105117 | 0.878930414 |
| 5.400236243 | 6.66E-08 | 3.67E-07 |
| 9.303115142 | 1.36E-20 | 2.53E-19 |
| 11.40329815 | 4.03E-30 | 1.39E-28 |
| -0.085296875 | 0.932025376 | NA |
| 6.074556462 | 1.24E-09 | 8.37E-09 |
| NA | NA | NA |
| 4.117914216 | 3.82E-05 | 0.000142021 |
| 0.192809521 | 0.847108147 | NA |
| NA | NA | NA |

| | | | |
|---|---|---|---|
| | 6.670117268 | 2.56E-11 | 2.06E-10 |
| | 3.667590789 | 0.000244847 | 0.00079077 |
| NA | NA | NA | |
| | 2.533335856 | 0.011298264 | 0.025418651 |
| | 10.02992462 | 1.13E-23 | 2.64E-22 |
| NA | NA | NA | |
| | 28.49956836 | 1.19E-178 | 6.75E-176 |
| | 8.136942443 | 4.05E-16 | 5.22E-15 |
| | 0.063710673 | 0.949200606 | NA |
| NA | NA | NA | |
| NA | NA | NA | |
| NA | NA | NA | |
| NA | NA | NA | |
| | 0.192809521 | 0.847108147 | NA |
| NA | NA | NA | |
| | 0.192810118 | 0.847107679 | NA |
| NA | NA | NA | |
| | -0.194488217 | 0.845793621 | NA |
| NA | NA | NA | |
| | -0.194487022 | 0.845794556 | NA |
| NA | NA | NA | |
| | -0.065388772 | 0.947864464 | NA |
| NA | NA | NA | |
| NA | NA | NA | |
| | -0.065389369 | 0.947863988 | NA |
| NA | NA | NA | |
| NA | NA | NA | |
| NA | NA | NA | |
| NA | NA | NA | |
| NA | NA | NA | |
| | -0.065388772 | 0.947864464 | NA |
| NA | NA | NA | |
| | 0.192810118 | 0.847107679 | NA |
| | 0.317642674 | 0.750756 | NA |
| NA | NA | NA | |
| | 2.136970011 | 0.032600431 | 0.063883879 |
| NA | NA | NA | |
| | 0.192809521 | 0.847108147 | NA |
| | 0.063708285 | 0.949202508 | NA |
| | -2.941641059 | 0.003264781 | 0.008364122 |
| | -0.038462482 | 0.969318945 | 0.978742716 |
| | 4.51891776 | 6.22E-06 | 2.61E-05 |
| NA | NA | NA | |
| | 5.447425856 | 5.11E-08 | 2.86E-07 |
| NA | NA | NA | |
| NA | NA | NA | |
| NA | NA | NA | |

| | | | |
|---|---|---|---|
| NA | NA | NA | |
| | 4.03726177 | 5.41E-05 | 0.000196087 |
| | -0.19448762 | 0.845794089 | NA |
| | 0.192810118 | 0.847107679 | NA |
| NA | NA | NA | |
| NA | NA | NA | |
| | 0.070531814 | 0.94377038 | NA |
| NA | NA | NA | |
| NA | NA | NA | |
| NA | NA | NA | |
| NA | NA | NA | |
| NA | NA | NA | |
| | 0.063708882 | 0.949202033 | NA |
| NA | NA | NA | |
| NA | NA | NA | |
| NA | NA | NA | |
| NA | NA | NA | |
| | 0.3474247 | 0.728272279 | NA |
| NA | NA | NA | |
| NA | NA | NA | |
| NA | NA | NA | |
| NA | NA | NA | |
| NA | NA | NA | |
| NA | NA | NA | |
| NA | NA | NA | |
| | 0.192809521 | 0.847108147 | NA |
| NA | NA | NA | |
| NA | NA | NA | |
| NA | NA | NA | |
| NA | NA | NA | |
| NA | NA | NA | |
| NA | NA | NA | |
| NA | NA | NA | |
| NA | NA | NA | |
| | 0.192809521 | 0.847108147 | NA |
| NA | NA | NA | |
| NA | NA | NA | |
| NA | NA | NA | |
| NA | NA | NA | |
| NA | NA | NA | |
| NA | NA | NA | |
| NA | NA | NA | |
| | 1.512196506 | 0.13048388 | 0.207986468 |
| | 5.873808488 | 4.26E-09 | 2.71E-08 |
| | 2.620497481 | 0.008780158 | 0.020264394 |
| | -7.110929842 | 1.15E-12 | 1.06E-11 |
| | 3.534190652 | 0.000409026 | 0.001267383 |

| | | | |
|---|---|---|---|
| | -1.230551564 | 0.21849063 | 0.317364234 |
| | 2.764647194 | 0.005698439 | 0.013819584 |
| | -0.821500094 | 0.41136147 | 0.523083601 |
| | 1.005541383 | 0.314636233 | 0.42518869 |
| | 0.438233803 | 0.661216807 | 0.747872299 |
| | 0.314251168 | 0.753330283 | 0.821079931 |
| | 5.88493961 | 3.98E-09 | 2.54E-08 |
| NA | NA | NA | |
| | -0.018502671 | 0.985237847 | 0.989507396 |
| | -0.090394335 | 0.927973858 | 0.950801956 |
| | 2.762005411 | 0.005744753 | 0.013911683 |
| | 5.380530795 | 7.43E-08 | 4.07E-07 |
| | 3.033748605 | 0.002415356 | 0.006376581 |
| | 1.08170367 | 0.279384222 | 0.386438278 |
| | 0.655796349 | 0.511955166 | 0.617642927 |
| | -5.531777363 | 3.17E-08 | 1.82E-07 |
| | -0.95692371 | 0.338605763 | 0.449904568 |
| | -1.777467157 | 0.07549141 | 0.131355578 |
| | -5.116551815 | 3.11E-07 | 1.57E-06 |
| | 2.400296433 | 0.0163818 | 0.035238608 |
| | 0.150266234 | 0.880554572 | NA |
| | 3.547311593 | 0.000389184 | 0.0012113 |
| NA | NA | NA | |
| | 4.371292628 | 1.24E-05 | 4.95E-05 |
| | 0.494053207 | 0.621268603 | 0.715550945 |
| | -0.175398803 | 0.860766281 | 0.903277713 |
| | -2.542409529 | 0.01100911 | 0.024827283 |
| | 5.074913615 | 3.88E-07 | 1.93E-06 |
| | -2.2922346 | 0.021892109 | 0.045188809 |
| | 8.499781373 | 1.90E-17 | 2.75E-16 |
| | 0.366049008 | 0.714328506 | 0.790994625 |
| NA | NA | NA | |
| | 0.192809521 | 0.847108147 | NA |
| NA | NA | NA | |
| | 0.466856248 | 0.64060273 | NA |
| | 6.899531089 | 5.22E-12 | 4.50E-11 |
| | 11.51435158 | 1.12E-30 | 3.98E-29 |
| | -1.797435751 | 0.072266469 | 0.126588966 |
| | 0.174495607 | 0.861475983 | 0.903643611 |
| | -1.759722324 | 0.078454899 | 0.135822805 |
| | -2.102313055 | 0.035525862 | 0.0687874 |
| | 0.063708285 | 0.949202508 | NA |
| | 0.620315299 | 0.535050224 | 0.639434068 |
| | 5.359272056 | 8.36E-08 | 4.55E-07 |
| NA | NA | NA | |
| | -5.578375391 | 2.43E-08 | 1.42E-07 |
| NA | NA | NA | |

| | | | |
|---|---|---|---|
| | 0.192809521 | 0.847108147 | NA |
| | 5.270489118 | 1.36E-07 | 7.20E-07 |
| | -2.316303604 | 0.020541698 | 0.042768149 |
| | -5.667645495 | 1.45E-08 | 8.68E-08 |
| | -5.649635885 | 1.61E-08 | 9.59E-08 |
| | 1.011387455 | 0.311831023 | 0.422013004 |
| | 1.65212986 | 0.098508081 | 0.164519735 |
| | 1.025026114 | 0.305350866 | 0.414970799 |
| | 0.098259736 | 0.921726049 | NA |
| | 10.4016431 | 2.44E-25 | 6.37E-24 |
| | -1.107884893 | 0.267911531 | 0.373828981 |
| | -3.076025229 | 0.002097801 | 0.005624848 |
| NA | NA | NA | |
| NA | NA | NA | |
| | 1.815368569 | 0.069467288 | 0.122439703 |
| | -3.038393585 | 0.002378431 | 0.006290377 |
| | 1.094615319 | 0.273685201 | NA |
| NA | NA | NA | |
| | -3.539510889 | 0.000400869 | 0.001244729 |
| | -0.194489411 | 0.845792686 | NA |
| NA | NA | NA | |
| | -0.101978268 | 0.918773925 | NA |
| | 0.192809521 | 0.847108147 | NA |
| | 6.037509882 | 1.57E-09 | 1.04E-08 |
| | 2.702131277 | 0.006889655 | 0.016353766 |
| | 6.54302358 | 6.03E-11 | 4.69E-10 |
| | 6.333110546 | 2.40E-10 | 1.74E-09 |
| | -4.031498124 | 5.54E-05 | 0.000200524 |
| | -0.135121132 | 0.892516102 | 0.924923531 |
| | 2.584953288 | 0.009739219 | 0.022250151 |
| | 10.68882953 | 1.15E-26 | 3.28E-25 |
| | 4.59042785 | 4.42E-06 | 1.90E-05 |
| | 3.439956306 | 0.000581808 | 0.001752426 |
| | 0.4387363 | 0.660852621 | 0.747655989 |
| NA | NA | NA | |
| | 0.192810118 | 0.847107679 | NA |
| NA | NA | NA | |
| | 0.112130324 | 0.910720074 | NA |
| | 3.194594253 | 0.001400276 | 0.003902126 |
| | -1.196641414 | 0.231446351 | 0.33230667 |
| | -0.747165307 | 0.454963783 | 0.564779375 |
| | 1.024076713 | 0.305799043 | 0.415438781 |
| | 2.50764019 | 0.012154036 | 0.027065437 |
| | -1.171065137 | 0.241572596 | 0.344062938 |
| | 7.465751753 | 8.28E-14 | 8.58E-13 |
| | 0.375984825 | 0.706928178 | 0.784987901 |
| | 1.982277169 | 0.047448228 | 0.088483769 |

| | | |
|---|---|---|
| 2.723083762 | 0.006467566 | 0.015458668 |
| NA | NA | NA |
| NA | NA | NA |
| 5.730796215 | 1.00E-08 | 6.11E-08 |
| NA | NA | NA |
| 1.454039395 | 0.145935384 | 0.228169322 |
| NA | NA | NA |
| 6.418166225 | 1.38E-10 | 1.03E-09 |
| 5.37510444 | 7.65E-08 | 4.19E-07 |
| 0.439999936 | 0.659937153 | 0.746906902 |
| NA | NA | NA |
| 0.342704238 | 0.731820974 | NA |
| 0.063708285 | 0.949202508 | NA |
| -4.633173527 | 3.60E-06 | 1.56E-05 |
| 14.34805166 | 1.10E-46 | 7.77E-45 |
| 0.809011243 | 0.418508679 | 0.530178775 |
| -7.776268302 | 7.47E-15 | 8.61E-14 |
| -0.631801162 | 0.527516814 | 0.632514772 |
| -0.325220757 | 0.745014 | 0.814452325 |
| -8.96605841 | 3.07E-19 | 5.18E-18 |
| 2.393774966 | 0.016675977 | 0.035782228 |
| -1.625513409 | 0.104053209 | 0.172115505 |
| 4.982248918 | 6.28E-07 | 3.04E-06 |
| 3.156503101 | 0.001596732 | 0.004401564 |
| -6.771872499 | 1.27E-11 | 1.06E-10 |
| -1.826314806 | 0.067802856 | 0.119919834 |
| NA | NA | NA |
| -0.131262054 | 0.895568009 | NA |
| NA | NA | NA |
| -0.19448762 | 0.845794089 | NA |
| -0.066588298 | 0.946909462 | NA |
| -0.218045791 | 0.827393434 | NA |
| 0.052431998 | 0.958184478 | 0.971104081 |
| 0.063708882 | 0.949202033 | NA |
| 0.192810118 | 0.847107679 | NA |
| -1.029698199 | 0.303151701 | 0.412474836 |
| 3.008699547 | 0.002623684 | 0.006868614 |
| 0.886026996 | 0.37560298 | 0.488381434 |
| 0.67246395 | 0.501288383 | 0.608174505 |
| 1.066956359 | 0.285991543 | 0.393109264 |
| 1.93603736 | 0.052863125 | 0.097112956 |
| 2.232972097 | 0.025550785 | 0.051675512 |
| -0.173778348 | 0.862039661 | 0.904196988 |
| -0.797030773 | 0.425433159 | 0.536939894 |
| 0.337023828 | 0.736098938 | NA |
| 5.711631891 | 1.12E-08 | 6.80E-08 |
| 1.180094347 | 0.237962693 | 0.340004921 |

| | | | |
|---|---|---|---|
| | 3.100574493 | 0.001931456 | 0.005219645 |
| | 9.496120904 | 2.18E-21 | 4.30E-20 |
| | 1.570948011 | 0.116194725 | 0.188661163 |
| | 1.571882484 | 0.115977807 | 0.188367044 |
| | 1.057614929 | 0.290231034 | NA |
| | 0.904158519 | 0.365911352 | 0.478439176 |
| | 0.350149461 | 0.726226533 | 0.800013835 |
| | 1.948831442 | 0.051315558 | 0.094666009 |
| | -2.701815804 | 0.006896195 | 0.016364408 |
| | 0.176900161 | 0.859586812 | 0.902418333 |
| | -3.578876062 | 0.000345075 | 0.001085761 |
| | -2.69189563 | 0.007104718 | 0.016789188 |
| | 0.192809521 | 0.847108147 | NA |
| | -1.771112068 | 0.076542068 | 0.132947352 |
| | 1.586204775 | 0.112692865 | 0.183842952 |
| | -0.635707775 | 0.524966908 | 0.630091191 |
| | 1.450637186 | 0.146880915 | 0.22940405 |
| NA | NA | NA | |
| | 3.671981844 | 0.000240677 | 0.000778011 |
| | 2.368335375 | 0.01786833 | 0.037950302 |
| NA | NA | NA | |
| | 11.93619492 | 7.67E-33 | 3.07E-31 |
| | 0.254419926 | 0.799171158 | 0.857257825 |
| | -0.322206147 | 0.747296528 | 0.816277294 |
| | -3.306987641 | 0.000943051 | 0.002732325 |
| | 0.978805513 | 0.327676085 | 0.438685212 |
| | 5.028951977 | 4.93E-07 | 2.42E-06 |
| | 4.016500534 | 5.91E-05 | 0.000212366 |
| | -2.768668213 | 0.005628592 | 0.01366741 |
| | 1.469592726 | 0.141672092 | 0.222616137 |
| | 3.005278245 | 0.00265338 | 0.006938372 |
| | 1.436084421 | 0.15097832 | 0.234763457 |
| | -0.480275594 | 0.631031441 | 0.723035796 |
| | 1.749501832 | 0.080204312 | 0.138372893 |
| | -0.194487022 | 0.845794556 | NA |
| | 1.002429409 | 0.316136244 | 0.426639898 |
| | -0.187788378 | 0.851042547 | NA |
| | 3.682241771 | 0.000231192 | 0.000750355 |
| | -1.609925832 | 0.107414049 | 0.176574926 |
| | -1.847797641 | 0.064631622 | 0.115067182 |
| NA | NA | NA | |
| | -9.647726842 | 5.03E-22 | 1.04E-20 |
| | -0.554437793 | 0.579279267 | 0.680018763 |
| | -2.661766058 | 0.007773188 | 0.018159688 |
| NA | NA | NA | |
| | -8.365569939 | 5.98E-17 | 8.29E-16 |
| | 0.537757695 | 0.590744342 | 0.690143798 |

| | | |
|---|---|---|
| -1.671075066 | 0.094706852 | 0.159287226 |
| 0.175518447 | 0.860672278 | 0.903216935 |
| 6.955969187 | 3.50E-12 | 3.06E-11 |
| -3.913609218 | 9.09E-05 | 0.000317066 |
| NA | NA | NA |
| -0.157217615 | 0.875073346 | 0.912780707 |
| -3.50790804 | 0.000451645 | 0.001388269 |
| 0.228140833 | 0.819536757 | NA |
| 1.102001551 | 0.270460998 | 0.376700278 |
| 0.325559456 | 0.744757692 | 0.814296182 |
| -1.852087271 | 0.064013292 | 0.114112435 |
| -1.704135328 | 0.088355808 | 0.150271514 |
| -3.250224358 | 0.00115314 | 0.003279524 |
| NA | NA | NA |
| 0.192810118 | 0.847107679 | NA |
| 2.473812235 | 0.013367993 | 0.029456895 |
| NA | NA | NA |
| NA | NA | NA |
| -0.21514613 | 0.82965339 | NA |
| NA | NA | NA |
| 0.345858301 | 0.729449208 | 0.802519171 |
| 6.3950042 | 1.61E-10 | 1.19E-09 |
| 17.76596619 | 1.30E-70 | 1.70E-68 |
| 15.44805292 | 7.78E-54 | 6.51E-52 |
| -3.467966868 | 0.000524412 | 0.001594126 |
| -0.483129601 | 0.629003717 | 0.721736639 |
| -6.168969474 | 6.87E-10 | 4.74E-09 |
| 3.613313942 | 0.000302308 | 0.000960251 |
| -0.065388772 | 0.947864464 | NA |
| 0.063708285 | 0.949202508 | NA |
| -2.499573566 | 0.012434288 | 0.027632978 |
| 1.425901574 | 0.153896755 | 0.238457087 |
| 10.44166438 | 1.60E-25 | 4.23E-24 |
| NA | NA | NA |
| -0.238453785 | 0.811529156 | 0.867243394 |
| 4.484287843 | 7.32E-06 | 3.04E-05 |
| 2.230594659 | 0.025707991 | 0.051921012 |
| 0.396456737 | 0.691768118 | 0.772524638 |
| -0.589326914 | 0.555641994 | 0.658353567 |
| 3.267704038 | 0.001084237 | 0.003101182 |
| 0.568662216 | 0.569585394 | 0.671094069 |
| 2.318604605 | 0.020416485 | 0.042544049 |
| 7.53140947 | 5.02E-14 | 5.31E-13 |
| -0.472929359 | 0.636263578 | 0.727532086 |
| 4.76883169 | 1.85E-06 | 8.38E-06 |
| NA | NA | NA |
| 10.27202753 | 9.42E-25 | 2.36E-23 |

| | | |
|---|---|---|
| -0.31240927 | 0.754729506 | NA |
| 11.14696683 | 7.41E-29 | 2.41E-27 |
| 7.055078914 | 1.73E-12 | 1.55E-11 |
| 14.85863788 | 6.12E-50 | 4.72E-48 |
| -1.582507798 | 0.113533694 | 0.185033751 |
| -1.789327972 | 0.073562012 | 0.128462789 |
| -1.037048203 | 0.299713393 | 0.408640714 |
| 0.688803359 | 0.490947022 | 0.598412659 |
| -1.470401053 | 0.141453166 | 0.222313999 |
| 2.993614209 | 0.002756943 | 0.007185161 |
| 3.804778663 | 0.000141931 | 0.000477684 |
| -3.989010236 | 6.63E-05 | 0.000236537 |
| -2.332294958 | 0.01968518 | 0.041214558 |
| -1.81518302 | 0.069495788 | 0.122481311 |
| -3.303438408 | 0.000955069 | 0.002763644 |
| -1.60641545 | 0.108182658 | 0.177619675 |
| 7.823062881 | 5.16E-15 | 6.02E-14 |
| 0.275476606 | 0.782950097 | 0.845109359 |
| 0.203461226 | 0.838774549 | 0.887071588 |
| -8.194140284 | 2.52E-16 | 3.32E-15 |
| -1.791737227 | 0.073175068 | 0.12787628 |
| -7.761735798 | 8.38E-15 | 9.61E-14 |
| 4.702234007 | 2.57E-06 | 1.14E-05 |
| -2.5575817 | 0.01054028 | 0.023897029 |
| -7.627021907 | 2.40E-14 | 2.63E-13 |
| 6.899063593 | 5.23E-12 | 4.51E-11 |
| 0.324874675 | 0.745275925 | NA |
| -0.194487022 | 0.845794556 | NA |
| -7.863799783 | 3.73E-15 | 4.41E-14 |
| 1.159889883 | 0.246093642 | 0.349170901 |
| -3.665417938 | 0.000246935 | 0.000796595 |
| -4.169003911 | 3.06E-05 | 0.000115568 |
| 1.764543525 | 0.077640514 | 0.134599118 |
| 2.102140527 | 0.035540968 | 0.068800497 |
| 14.29121721 | 2.48E-46 | 1.73E-44 |
| 0.073164869 | 0.941674922 | NA |
| -2.1160533 | 0.034340275 | 0.066806861 |
| -2.43704099 | 0.014808001 | 0.032201934 |
| -3.296893012 | 0.000977607 | 0.002823622 |
| 3.532001022 | 0.000412428 | 0.001276659 |
| NA | NA | NA |
| NA | NA | NA |
| 0.192809521 | 0.847108147 | NA |
| NA | NA | NA |
| 8.011066522 | 1.14E-15 | 1.41E-14 |
| 1.268334337 | 0.20467858 | 0.301164238 |
| 14.14389556 | 2.04E-45 | 1.39E-43 |

| | | | |
|---|---|---|---|
| NA | | NA | NA |
| | 3.309270301 | 0.000935395 | 0.002711713 |
| | 4.745470521 | 2.08E-06 | 9.35E-06 |
| | 1.455857071 | 0.145432135 | 0.227553063 |
| | 10.60011556 | 2.98E-26 | 8.25E-25 |
| | -0.973957744 | 0.33007752 | 0.441054897 |
| | 1.362110153 | 0.173163125 | 0.262654895 |
| | 0.549976246 | 0.582335666 | 0.682550303 |
| | 1.966837429 | 0.049201956 | 0.091339091 |
| | 0.192636473 | 0.847243678 | NA |
| | 19.37914215 | 1.16E-83 | 1.98E-81 |
| | 6.629558724 | 3.37E-11 | 2.68E-10 |
| | 2.285254269 | 0.022297923 | 0.045952138 |
| | 1.024181384 | 0.30574961 | 0.41539992 |
| | 0.45903447 | 0.646209413 | 0.73541064 |
| NA | | NA | NA |
| | 1.793371937 | 0.072913472 | 0.127481433 |
| | 3.085805789 | 0.002030014 | 0.005464021 |
| | 1.65569727 | 0.09778314 | 0.163516302 |
| | -2.779645415 | 0.005441828 | 0.013269261 |
| | 5.359807625 | 8.33E-08 | 4.54E-07 |
| | 2.923838145 | 0.003457445 | 0.008805438 |
| | 0.458153469 | 0.646842187 | 0.735896547 |
| NA | | NA | NA |
| | 1.31886519 | 0.187214185 | 0.279993491 |
| | -2.693261656 | 0.007075672 | 0.016730026 |
| | -0.96409296 | 0.334999322 | 0.446249762 |
| | -0.091293694 | 0.927259228 | NA |
| NA | | NA | NA |
| | -1.377599364 | 0.168327017 | 0.256497336 |
| | -1.633337101 | 0.102398118 | 0.169849231 |
| | 5.313661929 | 1.07E-07 | 5.77E-07 |
| | -1.155178421 | 0.248017369 | 0.351243338 |
| | -1.950433067 | 0.051124524 | 0.094362276 |
| | 2.813659014 | 0.004898115 | 0.012088329 |
| | -0.122669163 | 0.902369084 | 0.932162533 |
| | -0.026346821 | 0.97898071 | 0.985397803 |
| NA | | NA | NA |
| | -6.506062469 | 7.71E-11 | 5.93E-10 |
| | 4.313787076 | 1.60E-05 | 6.33E-05 |
| | -0.35654452 | 0.721432817 | 0.796324899 |
| | 7.800299891 | 6.18E-15 | 7.18E-14 |
| | -4.849463439 | 1.24E-06 | 5.73E-06 |
| | 3.736515008 | 0.000186588 | 0.000615973 |
| | -1.948916471 | 0.051305401 | 0.094659465 |
| | 4.512324978 | 6.41E-06 | 2.69E-05 |
| | -1.871980225 | 0.061209339 | 0.109885041 |

| | | |
|---|---|---|
| 0.192809521 | 0.847108147 | NA |
| 0.532304998 | 0.594514771 | 0.693286653 |
| 0.942504552 | 0.345934382 | 0.457408219 |
| -2.434263846 | 0.014922113 | 0.032427548 |
| -2.052558851 | 0.040115382 | 0.076414613 |
| -1.863197123 | 0.062434538 | 0.111767735 |
| -0.93756937 | 0.348465758 | 0.460149067 |
| 0.063707194 | 0.949203376 | NA |
| -0.199953569 | 0.841516895 | 0.889069117 |
| 12.09634078 | 1.10E-33 | 4.57E-32 |
| NA | NA | NA |
| 4.706898446 | 2.52E-06 | 1.12E-05 |
| NA | NA | NA |
| 16.4206671 | 1.36E-60 | 1.39E-58 |
| 12.7555783 | 2.90E-37 | 1.40E-35 |
| 5.183829778 | 2.17E-07 | 1.12E-06 |
| 0.787989449 | 0.430702877 | 0.541733318 |
| 4.840499991 | 1.30E-06 | 5.97E-06 |
| 7.697868675 | 1.38E-14 | 1.56E-13 |
| 5.115154209 | 3.13E-07 | 1.58E-06 |
| 1.421850908 | 0.155069537 | 0.239932652 |
| -1.883614818 | 0.059617096 | 0.107500606 |
| NA | NA | NA |
| -3.033448305 | 0.002417761 | 0.006381702 |
| -1.798678707 | 0.072069519 | 0.126305844 |
| -2.713470869 | 0.006658242 | 0.015865937 |
| NA | NA | NA |
| 10.83862718 | 2.26E-27 | 6.74E-26 |
| NA | NA | NA |
| 0.220980046 | 0.82510797 | 0.877483535 |
| NA | NA | NA |
| -2.684072138 | 0.007273141 | 0.017132197 |
| 7.159813744 | 8.08E-13 | 7.58E-12 |
| NA | NA | NA |
| 4.95238761 | 7.33E-07 | 3.51E-06 |
| NA | NA | NA |
| -0.065389369 | 0.947863988 | NA |
| -0.684862838 | 0.493430478 | 0.600620769 |
| -1.801254502 | 0.071662775 | 0.125742676 |
| NA | NA | NA |
| 0.186601545 | 0.851973055 | 0.897134437 |
| 1.871353767 | 0.061296063 | 0.110020828 |
| -0.910239857 | 0.362696029 | 0.475103705 |
| 1.069362904 | 0.284906176 | NA |
| -0.415360142 | 0.67787828 | NA |
| 4.166413263 | 3.09E-05 | 0.0001168 |
| -7.855027638 | 4.00E-15 | 4.71E-14 |

| | | | |
|---|---|---|---|
| NA | | NA | NA |
| | 7.210031404 | 5.59E-13 | 5.35E-12 |
| | -0.19441142 | 0.845853749 | 0.891966605 |
| | -0.617552362 | 0.536870455 | 0.641180355 |
| | 0.009257332 | 0.992613824 | 0.994839593 |
| | 3.672404604 | 0.000240279 | 0.000776925 |
| | 0.697141311 | 0.485714363 | NA |
| | 0.192810118 | 0.847107679 | NA |
| NA | | NA | NA |
| | -1.423579404 | 0.154568261 | 0.239364201 |
| | 1.290355963 | 0.196927094 | 0.29186333 |
| | 0.213320765 | 0.831076778 | 0.881845426 |
| NA | | NA | NA |
| | 0.629616275 | 0.528945673 | 0.633772628 |
| | -4.932154199 | 8.13E-07 | 3.87E-06 |
| | 4.233624119 | 2.30E-05 | 8.85E-05 |
| | -1.923883936 | 0.054369133 | 0.099485299 |
| | 2.546353191 | 0.010885499 | 0.024579525 |
| | -1.106513932 | 0.268504137 | 0.3744934 |
| | 4.710400521 | 2.47E-06 | 1.10E-05 |
| | 2.438667722 | 0.014741516 | 0.032071286 |
| | -6.686459591 | 2.29E-11 | 1.85E-10 |
| | 2.663251522 | 0.007738954 | 0.018091532 |
| | 0.253476181 | 0.799900265 | NA |
| | 0.434233791 | 0.664118677 | NA |
| | -2.866314179 | 0.004152819 | 0.010403869 |
| | -0.55023101 | 0.582160937 | 0.682441376 |
| | -0.857842188 | 0.390979611 | 0.50306006 |
| NA | | NA | NA |
| | -2.559558802 | 0.010480512 | 0.023774445 |
| | 2.1568905 | 0.031014197 | 0.061105586 |
| | -4.106261436 | 4.02E-05 | 0.000148757 |
| | -1.35278278 | 0.176125035 | 0.266449661 |
| | 4.889279144 | 1.01E-06 | 4.75E-06 |
| | -0.863289489 | 0.387978322 | 0.50018529 |
| | -0.333998007 | 0.738381063 | 0.809463823 |
| | -0.104347964 | 0.916893216 | 0.942924409 |
| | 0.047940351 | 0.961763781 | 0.973706566 |
| | 0.340757501 | 0.733286147 | 0.80538127 |
| | 0.192810118 | 0.847107679 | NA |
| | -0.527022864 | 0.598177711 | 0.696462561 |
| | 0.026878565 | 0.97855659 | 0.985129366 |
| | -2.114363042 | 0.034484272 | 0.067050518 |
| | 5.398235675 | 6.73E-08 | 3.71E-07 |
| | 0.053882219 | 0.957029003 | 0.970365225 |
| | -2.328935281 | 0.019862495 | 0.041540628 |
| | -4.623038444 | 3.78E-06 | 1.64E-05 |

| | | |
|---|---|---|
| 10.21580464 | 1.68E-24 | 4.16E-23 |
| -0.254439702 | 0.799155882 | NA |
| 2.880427095 | 0.003971368 | 0.009992294 |
| -1.735114139 | 0.08272059 | 0.142110927 |
| 4.717262827 | 2.39E-06 | 1.06E-05 |
| -1.09739979 | 0.272466665 | 0.378985309 |
| NA | NA | NA |
| 0.702194229 | 0.482558044 | NA |
| 1.100175215 | 0.271255787 | 0.377594814 |
| 0.06371127 | 0.949200131 | NA |
| 0.063707019 | 0.949203516 | NA |
| NA | NA | NA |
| -1.941408903 | 0.052208699 | 0.09610824 |
| -0.653426064 | 0.51348164 | 0.619126425 |
| 5.345426568 | 9.02E-08 | 4.89E-07 |
| 10.40487563 | 2.36E-25 | 6.17E-24 |
| NA | NA | NA |
| NA | NA | NA |
| 1.404513077 | 0.16016612 | 0.246172068 |
| -0.867463764 | 0.385687958 | 0.497924984 |
| 0.227325361 | 0.820170754 | 0.873829395 |
| 3.211843367 | 0.001318863 | 0.003695414 |
| NA | NA | NA |
| -1.268297156 | 0.204691852 | 0.301166063 |
| -4.382924129 | 1.17E-05 | 4.72E-05 |
| 2.464865262 | 0.013706478 | 0.030115478 |
| -1.485766188 | 0.137340977 | 0.216940846 |
| -1.568256462 | 0.116821292 | 0.189540218 |
| -4.115864371 | 3.86E-05 | 0.00014312 |
| -7.269220953 | 3.62E-13 | 3.51E-12 |
| -2.402275681 | 0.016293424 | 0.035064362 |
| NA | NA | NA |
| -2.762286253 | 0.005739813 | 0.013902412 |
| 3.071074059 | 0.002132902 | 0.005711013 |
| -0.544679856 | 0.585973721 | 0.686043419 |
| 7.86913742 | 3.57E-15 | 4.24E-14 |
| -5.140478472 | 2.74E-07 | 1.39E-06 |
| 1.075962805 | 0.281943891 | 0.388947068 |
| NA | NA | NA |
| NA | NA | NA |
| -7.18121445 | 6.91E-13 | 6.53E-12 |
| -5.487197359 | 4.08E-08 | 2.32E-07 |
| 1.177834888 | 0.238862437 | 0.340998506 |
| 1.472659043 | 0.140842996 | 0.221494105 |
| 0.192809521 | 0.847108147 | NA |
| NA | NA | NA |
| NA | NA | NA |

| | | |
|---|---|---|
| -1.275915679 | 0.201985328 | 0.298165414 |
| 9.240974829 | 2.44E-20 | 4.42E-19 |
| NA | NA | NA |
| -0.295149772 | 0.76787947 | 0.832720555 |
| 2.489293411 | 0.012799728 | 0.028341932 |
| -1.105237847 | 0.269056541 | 0.375137237 |
| -11.28304783 | 1.59E-29 | 5.35E-28 |
| NA | NA | NA |
| -1.005096332 | 0.314850466 | 0.425409294 |
| 0.26980614 | 0.787309399 | 0.84846096 |
| 2.919452674 | 0.003506466 | 0.00891966 |
| -2.074897419 | 0.037996043 | 0.072899046 |
| -4.604138854 | 4.14E-06 | 1.79E-05 |
| NA | NA | NA |
| 0.192810118 | 0.847107679 | NA |
| -0.065389369 | 0.947863988 | NA |
| -0.309061038 | 0.757275096 | NA |
| 0.464457116 | 0.642320277 | 0.732504476 |
| -0.065388772 | 0.947864464 | NA |
| 0.094001993 | 0.925107574 | NA |
| 0.063708882 | 0.949202033 | NA |
| 0.192808924 | 0.847108615 | NA |
| 0.320071032 | 0.748914485 | NA |
| -0.065389369 | 0.947863988 | NA |
| -0.065389369 | 0.947863988 | NA |
| NA | NA | NA |
| 0.210989877 | 0.83289517 | 0.882927856 |
| NA | NA | NA |
| -0.065389369 | 0.947863988 | NA |
| -0.689450171 | 0.49054002 | NA |
| -0.089787652 | 0.928455961 | NA |
| -0.547248969 | 0.584207698 | NA |
| 0.192809521 | 0.847108147 | NA |
| NA | NA | NA |
| 0.153357341 | 0.878116484 | 0.914735125 |
| 1.176382569 | 0.239442034 | 0.341544888 |
| 2.325531958 | 0.020043533 | 0.041859794 |
| 2.507606154 | 0.012155206 | 0.027065636 |
| 3.018315412 | 0.002541842 | 0.006676703 |
| -0.701711648 | 0.482859008 | 0.591376472 |
| -0.963201161 | 0.335446583 | NA |
| 4.77432812 | 1.80E-06 | 8.18E-06 |
| 5.314433854 | 1.07E-07 | 5.75E-07 |
| -2.600028795 | 0.009321594 | 0.021389759 |
| 6.035756473 | 1.58E-09 | 1.05E-08 |
| 3.577743404 | 0.000346573 | 0.001089791 |
| -3.778009398 | 0.000158087 | 0.000527589 |

| | | | |
|---|---|---|---|
| | -1.033625512 | 0.301311268 | 0.41043924 |
| | 10.05180071 | 9.02E-24 | 2.12E-22 |
| | 1.764882643 | 0.077583491 | NA |
| | 12.33636026 | 5.77E-35 | 2.51E-33 |
| | 12.42491779 | 1.91E-35 | 8.54E-34 |
| | 24.99633765 | 6.70E-138 | 2.47E-135 |
| | 8.69146935 | NA | NA |
| | 1.183098507 | 0.23677011 | NA |
| | -1.610765371 | 0.107230871 | 0.176334585 |
| | 4.029132693 | 5.60E-05 | 0.000202422 |
| | 5.995643234 | 2.03E-09 | 1.34E-08 |
| | 1.864481041 | 0.062254182 | 0.111478734 |
| NA | | NA | NA |
| | -1.930128662 | 0.053590898 | 0.098264811 |
| | 13.22815375 | 6.04E-40 | 3.31E-38 |
| | 0.192810118 | 0.847107679 | NA |
| | -2.913666986 | 0.003572107 | 0.009065196 |
| | -0.885455289 | 0.375911124 | 0.488680557 |
| | 0.619734973 | 0.535432287 | 0.639737942 |
| | 2.12760629 | 0.03336974 | 0.065192366 |
| | -0.065389369 | 0.947863988 | NA |
| | 3.863245881 | 0.00011189 | 0.000383593 |
| | 32.72654562 | 6.55E-235 | 7.45E-232 |
| | 2.496170908 | 0.01255421 | 0.027864846 |
| | 7.737121416 | 1.02E-14 | 1.16E-13 |
| | 4.890832671 | 1.00E-06 | 4.72E-06 |
| | -5.660142842 | 1.51E-08 | 9.05E-08 |
| | -2.771945976 | 0.005572228 | 0.013558168 |
| | 0.340582742 | 0.733417723 | 0.805433732 |
| | -2.900025986 | 0.003731317 | 0.009432873 |
| NA | | NA | NA |
| | 5.008279072 | 5.49E-07 | 2.67E-06 |
| | 0.593465283 | 0.55286981 | 0.656000538 |
| | -2.808043872 | 0.004984344 | 0.01227451 |
| | -3.679830443 | 0.000233389 | 0.000756603 |
| | 4.448035913 | 8.67E-06 | 3.56E-05 |
| | 3.080338484 | 0.002067655 | 0.005551153 |
| | 7.737364669 | 1.01E-14 | 1.15E-13 |
| NA | | NA | NA |
| | 8.145264981 | 3.78E-16 | 4.89E-15 |
| | -0.365155827 | 0.71499509 | 0.791592671 |
| | 2.639056831 | 0.008313704 | 0.019289505 |
| | 9.691724206 | 3.27E-22 | 6.85E-21 |
| | 4.094461887 | 4.23E-05 | 0.000155734 |
| | 6.65005279 | 2.93E-11 | 2.35E-10 |
| | -2.433274308 | 0.01496296 | 0.032502204 |
| | -1.223369133 | 0.221190306 | 0.320430221 |

| | | |
|---|---|---|
| 0.41407917 | 0.678816127 | NA |
| -0.539879776 | 0.589279946 | 0.688818804 |
| 0.749051995 | 0.453825867 | NA |
| 4.078856264 | 4.53E-05 | 0.000165736 |
| 2.789252468 | 0.005282986 | 0.01291844 |
| -0.400530513 | 0.688765814 | 0.770387495 |
| 1.764430455 | 0.077659534 | 0.134619405 |
| -2.137964038 | 0.032519662 | 0.063737181 |
| -7.665348078 | 1.78E-14 | 1.98E-13 |
| -1.417873971 | 0.156227563 | 0.241485574 |
| 0.704748494 | 0.480966767 | 0.589693763 |
| -0.132217826 | 0.894812002 | NA |
| -8.055973814 | 7.88E-16 | 9.92E-15 |
| 0.148145151 | 0.882228217 | NA |
| NA | NA | NA |
| NA | NA | NA |
| NA | NA | NA |
| 0.06371127 | 0.949200131 | NA |
| NA | NA | NA |
| NA | NA | NA |
| 0.192810118 | 0.847107679 | NA |
| NA | NA | NA |
| NA | NA | NA |
| 0.063708285 | 0.949202508 | NA |
| NA | NA | NA |
| -0.065389369 | 0.947863988 | NA |
| 4.483168739 | 7.35E-06 | 3.05E-05 |
| 7.823643544 | 5.13E-15 | 6.00E-14 |
| 0.063708285 | 0.949202508 | NA |
| 0.979655545 | 0.327256177 | 0.438291956 |
| 3.589503989 | 0.000331308 | 0.00104507 |
| 17.18837396 | 3.25E-66 | 3.78E-64 |
| -1.428186385 | 0.153238218 | 0.237569068 |
| 0.794553826 | 0.426873087 | 0.538256654 |
| -0.449971772 | 0.652730794 | 0.740908294 |
| NA | NA | NA |
| 10.89238532 | 1.25E-27 | 3.82E-26 |
| 0.447649523 | 0.654406156 | NA |
| NA | NA | NA |
| 0.192808327 | 0.847109082 | NA |
| -2.556830271 | 0.010563074 | 0.02394654 |
| NA | NA | NA |
| 0.642331332 | 0.520658075 | 0.625909976 |
| 0.398625966 | 0.690168826 | NA |
| 0.192636473 | 0.847243678 | NA |
| NA | NA | NA |
| -0.359102419 | 0.719518472 | 0.7949353 |

| | | |
|---|---|---|
| 0.999245203 | 0.317675923 | 0.428186772 |
| -0.772440935 | 0.439853317 | 0.550091362 |
| 2.25446978 | 0.02416663 | 0.049214119 |
| -4.000159754 | 6.33E-05 | 0.000226504 |
| 1.68348597 | 0.092281053 | 0.155898675 |
| 0.471316417 | 0.637414794 | NA |
| 6.758599624 | 1.39E-11 | 1.15E-10 |
| 1.510563835 | 0.130899614 | 0.208527815 |
| NA | NA | NA |
| 0.192809521 | 0.847108147 | NA |
| NA | NA | NA |
| -2.858748368 | 0.00425316 | 0.010624411 |
| 0.06371127 | 0.949200131 | NA |
| -0.678473523 | 0.497471506 | NA |
| NA | NA | NA |
| 0.063708882 | 0.949202033 | NA |
| 1.064868383 | 0.286935495 | 0.39422524 |
| -1.902667793 | 0.057083907 | NA |
| 0.192810118 | 0.847107679 | NA |
| NA | NA | NA |
| 0.199526594 | 0.841850842 | NA |
| -2.763235786 | 0.005723141 | 0.013870082 |
| 3.088462262 | 0.002011952 | 0.005420793 |
| 0.399043737 | 0.689860978 | NA |
| -0.976762127 | 0.32868693 | 0.439650118 |
| -1.915024819 | 0.055489339 | 0.101231643 |
| 8.304993662 | 9.98E-17 | 1.35E-15 |
| -1.527501007 | 0.126636477 | NA |
| NA | NA | NA |
| 0.063708882 | 0.949202033 | NA |
| 6.942222784 | 3.86E-12 | 3.36E-11 |
| 4.105824083 | 4.03E-05 | 0.000148973 |
| 6.183947701 | 6.25E-10 | 4.33E-09 |
| 2.575752299 | 0.010002227 | 0.022788617 |
| 0.063708882 | 0.949202033 | NA |
| 0.966799775 | 0.333644137 | 0.444862856 |
| 0.3474247 | 0.728272279 | NA |
| 0.257826504 | 0.796540808 | 0.855132594 |
| -1.069192719 | 0.284982839 | 0.392102048 |
| 0.192809521 | 0.847108147 | NA |
| NA | NA | NA |
| -5.782474362 | 7.36E-09 | 4.56E-08 |
| -0.065390563 | 0.947863038 | NA |
| -0.194487022 | 0.845794556 | NA |
| 6.643471799 | 3.06E-11 | 2.45E-10 |
| -1.352213833 | 0.17630692 | 0.266628131 |
| -0.653294717 | 0.513566297 | 0.619198672 |

| | | | |
|---|---|---|---|
| | 0.192809521 | 0.847108147 | NA |
| | 6.898070161 | 5.27E-12 | 4.54E-11 |
| | -3.633305342 | 0.000279814 | 0.00089436 |
| NA | NA | NA | |
| | -1.007059104 | 0.313906372 | 0.42436276 |
| NA | NA | NA | |
| | -1.195144935 | 0.232030408 | 0.332935171 |
| | 0.687352604 | 0.491860557 | 0.599176023 |
| | 1.235340815 | 0.21670369 | 0.315299271 |
| | 9.837375051 | 7.77E-23 | 1.70E-21 |
| | 4.021402296 | 5.79E-05 | 0.000208563 |
| | -0.207821709 | 0.835368181 | 0.884611625 |
| | 0.25696563 | 0.797205306 | NA |
| NA | NA | NA | |
| NA | NA | NA | |
| | -2.514543302 | 0.011918667 | 0.026617079 |
| NA | NA | NA | |
| | 3.946851916 | 7.92E-05 | 0.000279157 |
| | -0.797052834 | 0.425420347 | 0.536939894 |
| | 2.033291862 | 0.042023045 | 0.079594159 |
| | 1.706120772 | 0.087985596 | 0.149788084 |
| | 2.45554602 | 0.014067074 | 0.030810191 |
| | -3.295978583 | 0.000980795 | 0.002832175 |
| | -3.223935328 | 0.00126442 | 0.003558001 |
| | -2.117441593 | 0.034222387 | 0.066618942 |
| | 3.883991222 | 0.000102756 | 0.000354266 |
| | -4.936276917 | 7.96E-07 | 3.79E-06 |
| | 4.533975383 | 5.79E-06 | 2.44E-05 |
| | 0.192809521 | 0.847108147 | NA |
| | 0.3402497 | 0.733668493 | 0.805638431 |
| | -0.129006655 | 0.897352383 | 0.928436266 |
| | 1.038883539 | 0.298858903 | 0.407764423 |
| | 2.273707834 | 0.022983559 | 0.047181273 |
| | 7.061411219 | 1.65E-12 | 1.49E-11 |
| | -1.405492552 | 0.159874863 | 0.245832892 |
| | 4.02351961 | 5.73E-05 | 0.000206869 |
| | -0.846340413 | 0.397362869 | 0.509184652 |
| | 0.789178415 | 0.430007736 | 0.541103697 |
| | -0.09798922 | 0.921940853 | NA |
| NA | NA | NA | |
| | 1.919490431 | 0.054922296 | 0.100350761 |
| NA | NA | NA | |
| | 2.446597448 | 0.014421181 | 0.031445435 |
| | 0.192810118 | 0.847107679 | NA |
| NA | NA | NA | |
| | 0.063708882 | 0.949202033 | NA |
| | -1.585649085 | 0.112818935 | NA |

| | | | |
|---|---|---|---|
| | 0.091332088 | 0.927228722 | NA |
| | 1.802620874 | 0.071447776 | 0.125418181 |
| | 7.658154084 | 1.89E-14 | 2.09E-13 |
| NA | NA | NA | |
| | -1.481709142 | 0.138417704 | 0.218310972 |
| NA | NA | NA | |
| | -0.448324829 | 0.653918784 | NA |
| | 0.431281564 | 0.666263654 | 0.752010248 |
| | -7.025223776 | 2.14E-12 | 1.91E-11 |
| | -10.20435895 | 1.90E-24 | 4.66E-23 |
| | 2.467888868 | 0.013591251 | 0.029890959 |
| | -2.941391637 | 0.003267411 | 0.008369147 |
| | -3.138400497 | 0.001698726 | 0.004650964 |
| | 5.158909237 | 2.48E-07 | 1.27E-06 |
| | -11.59255618 | 4.50E-31 | 1.64E-29 |
| | -2.320005299 | 0.020340591 | 0.042412944 |
| NA | NA | NA | |
| | 0.860509106 | 0.389508466 | 0.501628727 |
| | -3.428517473 | 0.000606887 | 0.001822507 |
| | -4.019984345 | 5.82E-05 | 0.000209762 |
| | 0.192809521 | 0.847108147 | NA |
| | 0.76535671 | 0.444059164 | 0.554022573 |
| | -0.048374657 | 0.961417656 | 0.97360651 |
| | -0.672307651 | 0.50138786 | 0.60826571 |
| | 2.283811773 | 0.022382595 | 0.046109989 |
| | 0.504122235 | 0.614175483 | NA |
| | -0.214328636 | 0.830290789 | NA |
| NA | NA | NA | |
| NA | NA | NA | |
| NA | NA | NA | |
| | 1.169641002 | 0.24214547 | 0.344741545 |
| NA | NA | NA | |
| NA | NA | NA | |
| | 0.192810118 | 0.847107679 | NA |
| | -0.174895829 | 0.861161488 | 0.903351975 |
| NA | NA | NA | |
| NA | NA | NA | |
| NA | NA | NA | |
| NA | NA | NA | |
| | -8.180220151 | 2.83E-16 | 3.70E-15 |
| | 0.198308907 | 0.842803383 | NA |
| | -4.189452591 | 2.80E-05 | 0.000106305 |
| NA | NA | NA | |
| | -9.339611512 | 9.67E-21 | 1.81E-19 |
| | -2.125096659 | 0.033578548 | 0.065526533 |
| NA | NA | NA | |
| | 0.063708882 | 0.949202033 | NA |

| | | |
|---|---|---|
| NA | NA | NA |
| -1.227453786 | 0.219652081 | 0.318662934 |
| 1.739649495 | 0.081920583 | 0.140945297 |
| 4.1325238 | 3.59E-05 | 0.000133922 |
| -0.191210035 | 0.848361042 | 0.894158535 |
| NA | NA | NA |
| 6.910716908 | 4.82E-12 | 4.17E-11 |
| 2.220674555 | 0.026373012 | 0.053089633 |
| -2.02453952 | 0.04291467 | 0.081066157 |
| -2.481548568 | 0.013081288 | 0.02890146 |
| 12.70310536 | 5.68E-37 | 2.71E-35 |
| 7.013161357 | 2.33E-12 | 2.07E-11 |
| 0.099852033 | 0.920461798 | 0.945467778 |
| 12.52706049 | 5.31E-36 | 2.43E-34 |
| 4.753620639 | 2.00E-06 | 9.00E-06 |
| 0.150734289 | 0.880185324 | 0.915861096 |
| -0.780732325 | 0.434959944 | 0.545510067 |
| -3.161882187 | 0.00156753 | 0.004326306 |
| -2.47135988 | 0.013460029 | 0.029623188 |
| NA | NA | NA |
| 0.089186082 | 0.928934026 | NA |
| NA | NA | NA |
| -0.065388772 | 0.947864464 | NA |
| -3.310902651 | 0.000929956 | 0.002696882 |
| NA | NA | NA |
| -2.235868193 | 0.025360406 | 0.05135272 |
| -1.725826531 | 0.084378625 | 0.144498973 |
| -0.236421455 | 0.813105649 | 0.868520223 |
| -0.589116628 | 0.55578304 | NA |
| -8.02067299 | 1.05E-15 | 1.31E-14 |
| -3.820360593 | 0.000133257 | 0.000450671 |
| NA | NA | NA |
| NA | NA | NA |
| NA | NA | NA |
| -1.883132879 | 0.059682362 | 0.10760279 |
| NA | NA | NA |
| -0.848949604 | 0.395909335 | 0.507820573 |
| -0.395513609 | 0.69246388 | NA |
| 2.025947247 | 0.042770191 | 0.080813609 |
| 0.561009838 | 0.574790831 | NA |
| -0.433157105 | 0.664900639 | NA |
| -7.653378365 | 1.96E-14 | 2.16E-13 |
| -0.065389369 | 0.947863988 | NA |
| -2.716275263 | 0.0066021 | 0.015747148 |
| NA | NA | NA |
| NA | NA | NA |
| -0.255518765 | 0.798322452 | 0.856604406 |

| | | |
|---|---|---|
| 9.021588394 | 1.85E-19 | 3.17E-18 |
| 8.723447984 | 2.70E-18 | 4.18E-17 |
| 0.882873567 | 0.377304587 | NA |
| -0.065388772 | 0.947864464 | NA |
| 0.022256288 | 0.982243517 | 0.987641213 |
| NA | NA | NA |
| 1.430419904 | 0.152596539 | 0.236823622 |
| -0.398705074 | 0.690110529 | NA |
| 7.155095072 | 8.36E-13 | 7.83E-12 |
| 0.881838974 | 0.377863892 | 0.490480383 |
| -0.98923272 | 0.322549293 | 0.433471614 |
| -1.925681261 | 0.054144185 | 0.09915061 |
| -0.298339593 | 0.765443989 | 0.830882685 |
| -4.32948621 | 1.49E-05 | 5.92E-05 |
| -1.852336234 | 0.063977556 | 0.114071599 |
| -3.50880044 | 0.000450133 | 0.00138447 |
| 1.688771859 | 0.091263159 | 0.154397895 |
| NA | NA | NA |
| 1.365874879 | 0.171978239 | 0.261162443 |
| -0.293730496 | 0.768963849 | 0.833643674 |
| -5.748485613 | 9.00E-09 | 5.53E-08 |
| -0.381803681 | 0.702606988 | 0.78171547 |
| 4.06737592 | 4.75E-05 | 0.000173783 |
| 0.701507955 | 0.482986072 | 0.591445273 |
| 3.953740893 | 7.69E-05 | 0.000271887 |
| 1.692633632 | 0.090525225 | 0.153356777 |
| 0.192808327 | 0.847109082 | NA |
| 1.569209046 | 0.116599237 | 0.189241273 |
| NA | NA | NA |
| -0.19448762 | 0.845794089 | NA |
| 0.551801053 | 0.581084666 | 0.681658583 |
| -0.289316305 | 0.772339336 | NA |
| -5.062436378 | 4.14E-07 | 2.05E-06 |
| 0.055936354 | 0.95539251 | 0.969491386 |
| 3.123357748 | 0.001788003 | 0.004870357 |
| -4.029690836 | 5.59E-05 | 0.000201985 |
| -3.346741752 | 0.000817673 | 0.002394567 |
| -3.943717044 | 8.02E-05 | 0.000282316 |
| NA | NA | NA |
| -4.089892922 | 4.32E-05 | 0.000158554 |
| NA | NA | NA |
| 5.070729204 | 3.96E-07 | 1.97E-06 |
| 3.361900798 | 0.000774079 | 0.002278096 |
| 1.450504466 | 0.146917895 | 0.229447489 |
| -2.081074191 | 0.037427116 | 0.071934325 |
| 2.50883652 | 0.012112953 | 0.026988358 |
| -1.954810428 | 0.05060545 | 0.093569802 |

| | | |
|---|---|---|
| NA | NA | NA |
| 1.776989528 | 0.075569963 | 0.131473981 |
| NA | NA | NA |
| NA | NA | NA |
| NA | NA | NA |
| -1.108567667 | 0.267616733 | 0.373505455 |
| NA | NA | NA |
| -3.567684186 | 0.00036015 | 0.001128513 |
| -1.562177868 | 0.118246093 | 0.191479569 |
| NA | NA | NA |
| NA | NA | NA |
| NA | NA | NA |
| NA | NA | NA |
| NA | NA | NA |
| -3.831154323 | 0.000127543 | 0.000432811 |
| NA | NA | NA |
| NA | NA | NA |
| -0.358668089 | 0.719843403 | NA |
| 7.834172673 | 4.72E-15 | 5.53E-14 |
| -7.116278605 | 1.11E-12 | 1.02E-11 |
| 2.054097402 | 0.039966265 | 0.076153747 |
| 0.192810118 | 0.847107679 | NA |
| NA | NA | NA |
| NA | NA | NA |
| NA | NA | NA |
| 0.192808924 | 0.847108615 | NA |
| NA | NA | NA |
| NA | NA | NA |
| NA | NA | NA |
| -0.335273043 | 0.737419127 | NA |
| 0.063710673 | 0.949200606 | NA |
| NA | NA | NA |
| NA | NA | NA |
| NA | NA | NA |
| -0.092656501 | 0.926176455 | NA |
| NA | NA | NA |
| NA | NA | NA |
| NA | NA | NA |
| NA | NA | NA |
| NA | NA | NA |
| NA | NA | NA |
| NA | NA | NA |
| NA | NA | NA |
| 1.068900444 | 0.285114532 | 0.392197006 |
| NA | NA | NA |
| NA | NA | NA |
| NA | NA | NA |

| | | |
|---|---|---|
| NA | NA | NA |
| 0.870902766 | 0.383807246 | 0.496166884 |
| 1.683424333 | 0.092292976 | 0.155908294 |
| -0.953192235 | 0.34049268 | 0.452027781 |
| 1.25423537 | 0.209756467 | 0.307137263 |
| -0.425983056 | 0.670120193 | 0.754767924 |
| -1.76783802 | 0.077087987 | 0.13378951 |
| -0.155366482 | 0.876532407 | 0.913769518 |
| -0.358857789 | 0.719701479 | 0.795009842 |
| NA | NA | NA |
| 2.425503185 | 0.015287182 | 0.033120242 |
| -0.852180627 | 0.394113847 | 0.506127619 |
| -0.871842077 | 0.383294537 | 0.495704363 |
| -0.095291117 | 0.924083598 | NA |
| -0.484952229 | 0.627710233 | 0.720748068 |
| -0.311384038 | 0.755508687 | 0.822754425 |
| -0.065389369 | 0.947863988 | NA |
| -0.065389369 | 0.947863988 | NA |
| 2.271257236 | 0.02313141 | 0.047435703 |
| 2.416954478 | 0.015650974 | 0.033832273 |
| 10.87061508 | 1.59E-27 | 4.80E-26 |
| 2.680041756 | 0.007361298 | 0.017318665 |
| 3.502600667 | 0.00046074 | 0.001413274 |
| NA | NA | NA |
| NA | NA | NA |
| NA | NA | NA |
| NA | NA | NA |
| NA | NA | NA |
| NA | NA | NA |
| NA | NA | NA |
| NA | NA | NA |
| NA | NA | NA |
| 0.192808327 | 0.847109082 | NA |
| NA | NA | NA |
| 0.063708285 | 0.949202508 | NA |
| NA | NA | NA |
| NA | NA | NA |
| NA | NA | NA |
| NA | NA | NA |
| NA | NA | NA |
| NA | NA | NA |
| NA | NA | NA |
| NA | NA | NA |
| NA | NA | NA |
| NA | NA | NA |
| NA | NA | NA |
| NA | NA | NA |

| | | |
|---|---|---|
| NA | NA | NA |
| NA | NA | NA |
| NA | NA | NA |
| NA | NA | NA |
| NA | NA | NA |
| NA | NA | NA |
| NA | NA | NA |
| NA | NA | NA |
| NA | NA | NA |
| NA | NA | NA |
| NA | NA | NA |
| NA | NA | NA |
| NA | NA | NA |
| NA | NA | NA |
| NA | NA | NA |
| NA | NA | NA |
| NA | NA | NA |
| NA | NA | NA |
| NA | NA | NA |
| NA | NA | NA |
| NA | NA | NA |
| NA | NA | NA |
| NA | NA | NA |
| NA | NA | NA |
| NA | NA | NA |
| NA | NA | NA |
| NA | NA | NA |
| NA | NA | NA |
| NA | NA | NA |
| NA | NA | NA |
| NA | NA | NA |
| NA | NA | NA |
| 0.192636473 | 0.847243678 | NA |
| NA | NA | NA |
| 0.063976859 | 0.948988653 | NA |
| NA | NA | NA |
| 0.192809521 | 0.847108147 | NA |
| NA | NA | NA |
| 0.502444088 | 0.615355173 | 0.710636707 |
| 0.192809521 | 0.847108147 | NA |
| NA | NA | NA |
| 0.063708882 | 0.949202033 | NA |
| -0.438227614 | 0.661221293 | NA |
| -0.297503803 | 0.766081905 | NA |
| -0.19448762 | 0.845794089 | NA |
| NA | NA | NA |
| NA | NA | NA |

| | | |
|---|---|---|
| NA | NA | NA |
| NA | NA | NA |
| NA | NA | NA |
| -0.520728801 | 0.602555708 | 0.699931595 |
| 3.320832354 | 0.000897494 | 0.002608183 |
| 2.356166355 | 0.018464647 | 0.039011573 |
| -0.073200093 | 0.941646892 | 0.960370863 |
| 3.455393078 | 0.000549491 | 0.00166289 |
| 1.573940678 | 0.115501165 | 0.187726893 |
| 1.70931227 | 0.087393122 | 0.148877005 |
| NA | NA | NA |
| -0.960516644 | 0.336795259 | NA |
| -0.84782149 | 0.396537393 | 0.508400596 |
| 0.292513336 | 0.769894165 | 0.834218656 |
| 2.576948134 | 0.009967691 | 0.02272648 |
| -0.244947366 | 0.806497186 | 0.862971296 |
| 5.715342362 | 1.09E-08 | 6.67E-08 |
| -1.896356309 | 0.057912945 | 0.104949393 |
| 2.077253044 | 0.03777821 | 0.07253115 |
| 0.192808327 | 0.847109082 | NA |
| -0.402084624 | 0.687621748 | 0.769512083 |
| 0.306335375 | 0.759349307 | NA |
| 0.692958473 | 0.488335613 | 0.596274959 |
| NA | NA | NA |
| 12.63203075 | 1.41E-36 | 6.60E-35 |
| -2.250960131 | 0.024388063 | 0.049620648 |
| 3.480051987 | 0.000501316 | 0.001529492 |
| 0.611579667 | 0.540815894 | NA |
| 5.789633007 | 7.05E-09 | 4.38E-08 |
| -5.187281308 | 2.13E-07 | 1.10E-06 |
| NA | NA | NA |
| -3.026091138 | 0.002477376 | 0.006522428 |
| -0.477810028 | 0.632785423 | 0.72448171 |
| -1.930643886 | 0.053527107 | 0.098159827 |
| NA | NA | NA |
| 0.670075607 | 0.502809594 | 0.609458796 |
| 1.254282539 | 0.209739328 | 0.307130134 |
| -0.929700398 | 0.352526228 | 0.464406921 |
| -0.149350503 | 0.881277067 | 0.916668938 |
| 2.856896647 | 0.004278051 | 0.010679128 |
| NA | NA | NA |
| 6.862239507 | 6.78E-12 | 5.76E-11 |
| -4.316692124 | 1.58E-05 | 6.25E-05 |
| -0.66149375 | 0.508295722 | 0.614500315 |
| 0.150145829 | 0.880649563 | 0.916229885 |
| 3.712403559 | 0.0002053 | 0.000673482 |
| 2.095037441 | 0.036167664 | 0.069835427 |

| | | |
|---|---|---|
| 2.379594588 | 0.017331694 | 0.03697369 |
| 2.405548442 | 0.016148211 | 0.034786963 |
| 1.955194356 | 0.050560135 | 0.093506736 |
| NA | NA | NA |
| 1.425793494 | 0.15392796 | 0.238490673 |
| 0.933426326 | 0.350599893 | 0.462307203 |
| NA | NA | NA |
| 9.785392648 | 1.30E-22 | 2.81E-21 |
| -0.018891179 | 0.984927916 | 0.989361701 |
| 4.897272587 | 9.72E-07 | 4.57E-06 |
| 1.003326303 | 0.315703449 | 0.426216685 |
| 18.11297367 | 2.52E-73 | 3.48E-71 |
| -4.844677308 | 1.27E-06 | 5.86E-06 |
| NA | NA | NA |
| 0.345218368 | 0.729930211 | 0.802869189 |
| 7.723277547 | 1.13E-14 | 1.28E-13 |
| 0.192808924 | 0.847108615 | NA |
| -3.000341146 | 0.002696774 | 0.007043167 |
| -0.541610978 | 0.588086527 | 0.687832519 |
| 3.112008387 | 0.001858192 | 0.005037426 |
| NA | NA | NA |
| 0.267723987 | 0.788911785 | 0.849748772 |
| -0.702967375 | 0.482076082 | 0.590562075 |
| -2.378634295 | 0.017376906 | 0.037048037 |
| -7.476800208 | 7.62E-14 | 7.93E-13 |
| 0.46646864 | 0.64088009 | 0.731374123 |
| -7.334149474 | 2.23E-13 | 2.21E-12 |
| 1.521434756 | 0.128150772 | 0.205103513 |
| 0.853865431 | 0.393179556 | 0.505132547 |
| 0.086201974 | 0.931305862 | 0.953235203 |
| -2.181211361 | 0.029167787 | 0.057946639 |
| -5.440807498 | 5.30E-08 | 2.97E-07 |
| -4.315889483 | 1.59E-05 | 6.27E-05 |
| 0.50191714 | 0.615725808 | 0.710835154 |
| -7.926549659 | 2.25E-15 | 2.73E-14 |
| 0.192809521 | 0.847108147 | NA |
| -5.390929345 | 7.01E-08 | 3.85E-07 |
| 2.397605353 | 0.016502635 | 0.035449789 |
| NA | NA | NA |
| NA | NA | NA |
| -0.065389369 | 0.947863988 | NA |
| 7.60810382 | 2.78E-14 | 3.02E-13 |
| 3.87759385 | 0.000105495 | 0.000363259 |
| 0.84280967 | 0.399334899 | 0.511192785 |
| 2.042651156 | 0.041086983 | 0.077992355 |
| 4.881058182 | 1.06E-06 | 4.94E-06 |
| 1.666372308 | 0.095639283 | 0.160564173 |

| | | |
|---|---|---|
| 5.252903329 | 1.50E-07 | 7.88E-07 |
| 0.193341124 | 0.846691821 | NA |
| -0.367355645 | 0.713353752 | 0.790229892 |
| -2.335295977 | 0.019527964 | 0.040913096 |
| 14.5898242 | 3.26E-48 | 2.41E-46 |
| 1.203787783 | 0.228671613 | NA |
| -4.086896547 | 4.37E-05 | 0.00016052 |
| -3.369003511 | 0.000754405 | 0.002225925 |
| NA | NA | NA |
| NA | NA | NA |
| 0.486776342 | 0.626416838 | 0.719824325 |
| NA | NA | NA |
| NA | NA | NA |
| -0.6372451 | 0.523965205 | 0.629130245 |
| 2.032726237 | 0.042080189 | 0.079678276 |
| -1.346407221 | 0.178171223 | 0.268899537 |
| NA | NA | NA |
| 5.125093433 | 2.97E-07 | 1.51E-06 |
| 2.483906769 | 0.012994983 | 0.028737825 |
| 1.794419857 | 0.07274618 | 0.127268951 |
| 1.627535932 | 0.103623324 | 0.171547274 |
| 4.815592093 | 1.47E-06 | 6.72E-06 |
| 7.779784197 | 7.26E-15 | 8.39E-14 |
| NA | NA | NA |
| 0.444579266 | 0.656623834 | NA |
| 0.066587943 | 0.946909745 | NA |
| 1.935412381 | 0.052939711 | 0.09723224 |
| 4.972957968 | 6.59E-07 | 3.18E-06 |
| 0.855235818 | 0.392420611 | 0.504353679 |
| NA | NA | NA |
| 1.133783544 | 0.256885355 | 0.361416607 |
| -0.789596995 | 0.429763164 | 0.540904711 |
| 1.927435058 | 0.053925434 | 0.098817894 |
| 1.621222171 | 0.10497 | 0.173357026 |
| -9.318567786 | 1.18E-20 | 2.20E-19 |
| -2.669605506 | 0.007594041 | 0.017800978 |
| 1.206833748 | 0.227496176 | 0.327706566 |
| 0.901011432 | 0.367582241 | 0.480022095 |
| 2.62778067 | 0.008594391 | 0.019886993 |
| 10.63275323 | 2.10E-26 | 5.87E-25 |
| 1.229652466 | 0.218827274 | 0.317753095 |
| 0.041334469 | 0.967029254 | NA |
| 0.536926151 | 0.591318626 | 0.690589083 |
| -2.577374861 | 0.009955392 | 0.022700507 |
| 5.006344141 | 5.55E-07 | 2.69E-06 |
| NA | NA | NA |
| 0.8502644 | 0.395178104 | 0.507142532 |

| | | | |
|---|---|---|---|
| | 0.58653885 | 0.557513462 | NA |
| | -3.832782743 | 0.000126702 | 0.000430188 |
| | 0.267347788 | 0.789201396 | NA |
| | 3.756169217 | 0.000172534 | 0.000572142 |
| | 1.056976115 | 0.290522489 | 0.398126356 |
| | 6.655854616 | 2.82E-11 | 2.26E-10 |
| | 4.0192707 | 5.84E-05 | 0.000210217 |
| NA | NA | NA | |
| | 2.100111498 | 0.035719034 | 0.069075749 |
| | -0.065388772 | 0.947864464 | NA |
| NA | NA | NA | |
| NA | NA | NA | |
| NA | NA | NA | |
| | 0.192808924 | 0.847108615 | NA |
| | 6.044376194 | 1.50E-09 | 1.00E-08 |
| | 1.198276433 | 0.230809418 | 0.331563133 |
| | 0.194653292 | 0.84566438 | 0.891903027 |
| NA | NA | NA | |
| | 6.209260718 | 5.32E-10 | 3.73E-09 |
| | 2.629273816 | 0.008556743 | 0.01980354 |
| | 0.873931757 | 0.382155419 | 0.494612229 |
| | -1.241301109 | 0.214494535 | 0.312775894 |
| | -0.411483386 | 0.680718121 | NA |
| NA | NA | NA | |
| | 2.17590638 | 0.029562262 | 0.058623309 |
| NA | NA | NA | |
| | 2.151229674 | 0.031458076 | 0.061868126 |
| | -3.568625086 | 0.00035886 | 0.001125032 |
| | -3.830579292 | 0.000127842 | 0.000433706 |
| | 0.33526769 | 0.737423164 | NA |
| | 0.192808924 | 0.847108615 | NA |
| | -0.723359511 | 0.46945904 | 0.578617282 |
| | -4.861635173 | 1.16E-06 | 5.41E-06 |
| | 1.699994027 | 0.089132049 | 0.151396253 |
| NA | NA | NA | |
| | -5.544415986 | 2.95E-08 | 1.70E-07 |
| | -2.324936772 | 0.020075341 | 0.041915732 |
| NA | NA | NA | |
| | -1.592888065 | 0.111185304 | 0.181774537 |
| NA | NA | NA | |
| | -3.897848852 | 9.71E-05 | 0.000336499 |
| | 17.15159063 | 6.12E-66 | 7.05E-64 |
| | -1.248597728 | 0.211812244 | 0.309640256 |
| | -6.086984031 | 1.15E-09 | 7.79E-09 |
| | -2.172003381 | 0.029855408 | 0.059110968 |
| | -3.017966285 | 0.002544772 | 0.006681595 |
| | -1.897338771 | 0.057783241 | 0.104752265 |

| | | |
|---|---|---|
| 1.975092383 | 0.048257665 | 0.089745847 |
| -5.959756313 | 2.53E-09 | 1.65E-08 |
| 0.192808924 | 0.847108615 | NA |
| NA | NA | NA |
| NA | NA | NA |
| NA | NA | NA |
| 0.27464153 | 0.783591656 | NA |
| 0.324874675 | 0.745275925 | NA |
| 4.82025917 | 1.43E-06 | 6.58E-06 |
| 7.409770749 | 1.27E-13 | 1.29E-12 |
| -0.022040195 | 0.982415893 | 0.987732659 |
| 1.031648139 | 0.30223698 | 0.411453912 |
| -0.106131009 | 0.915478408 | NA |
| 0.464590973 | 0.642224398 | 0.732491975 |
| 0.192808327 | 0.847109082 | NA |
| 0.909623992 | 0.363020841 | 0.475454537 |
| -1.14963079 | 0.250295971 | 0.353939696 |
| 6.185894601 | 6.18E-10 | 4.28E-09 |
| 2.21310871 | 0.026890152 | 0.053975924 |
| 0.58653885 | 0.557513462 | NA |
| 1.246195807 | 0.212692518 | 0.310564329 |
| 0.058444712 | 0.953394401 | NA |
| NA | NA | NA |
| 2.269865128 | 0.023215768 | 0.047583579 |
| NA | NA | NA |
| NA | NA | NA |
| -0.08862269 | 0.929381775 | NA |
| 0.063708882 | 0.949202033 | NA |
| 1.777516236 | 0.075483342 | NA |
| NA | NA | NA |
| NA | NA | NA |
| NA | NA | NA |
| -0.065388772 | 0.947864464 | NA |
| -0.246580038 | 0.805233255 | NA |
| NA | NA | NA |
| NA | NA | NA |
| NA | NA | NA |
| NA | NA | NA |
| -2.841003836 | 0.004497177 | 0.011180408 |
| 0.063707688 | 0.949202983 | NA |
| 0.268017001 | 0.788686232 | NA |
| -0.646608656 | 0.517885254 | 0.623205188 |
| 1.900300916 | 0.057393641 | 0.104174245 |
| 1.791985503 | 0.073135288 | NA |
| 9.537256685 | 1.47E-21 | 2.95E-20 |
| -1.230431726 | 0.218535479 | 0.31739996 |
| 19.22034383 | 2.50E-82 | 4.12E-80 |

| | | |
|---|---|---|
| -4.831806556 | 1.35E-06 | 6.22E-06 |
| NA | NA | NA |
| 0.797015408 | 0.425442083 | NA |
| -0.776612617 | 0.437387355 | 0.547664107 |
| -2.573048109 | 0.010080719 | 0.022954913 |
| NA | NA | NA |
| -1.171411017 | 0.241433606 | 0.343904118 |
| -0.070651711 | 0.943674954 | 0.961929713 |
| 0.063708285 | 0.949202508 | NA |
| -1.933606978 | 0.053161469 | 0.097539331 |
| -0.92775146 | 0.353536508 | 0.465443867 |
| -4.667088809 | 3.05E-06 | 1.34E-05 |
| 1.214161185 | 0.22468618 | 0.324555348 |
| -6.757030422 | 1.41E-11 | 1.16E-10 |
| NA | NA | NA |
| -1.803206314 | 0.071355819 | 0.125291908 |
| -0.51125049 | 0.609175669 | 0.705649157 |
| -2.58528422 | 0.009729875 | 0.022230834 |
| 1.318512987 | 0.18733198 | 0.280152923 |
| NA | NA | NA |
| 1.36722866 | 0.171553645 | 0.260604948 |
| NA | NA | NA |
| 2.916624053 | 0.00353842 | 0.008990625 |
| -0.34376307 | 0.731024475 | 0.803757637 |
| -3.106893988 | 0.001890642 | 0.005118742 |
| 2.188464554 | 0.02863578 | 0.05703912 |
| 7.152240453 | 8.54E-13 | 7.98E-12 |
| 9.265680133 | 1.94E-20 | 3.54E-19 |
| 11.63603471 | 2.70E-31 | 9.99E-30 |
| -2.135453803 | 0.03272396 | 0.064075782 |
| 0.427649233 | 0.668906516 | NA |
| -4.801558795 | 1.57E-06 | 7.18E-06 |
| -2.788888232 | 0.005288931 | 0.012931714 |
| -4.400160609 | 1.08E-05 | 4.39E-05 |
| NA | NA | NA |
| -0.298015288 | 0.765691496 | 0.830957767 |
| 1.617570389 | 0.105755215 | 0.174400647 |
| NA | NA | NA |
| -0.119468513 | 0.904904184 | 0.934203 |
| -4.98536274 | 6.18E-07 | 2.99E-06 |
| -2.259270148 | 0.023866586 | 0.048718043 |
| 1.954943802 | 0.050589704 | 0.093547598 |
| -0.889821714 | 0.373561625 | 0.486282894 |
| -0.094276471 | 0.92488954 | NA |
| -0.065388772 | 0.947864464 | NA |
| -0.77565584 | 0.437952221 | 0.548261683 |
| -2.765233563 | 0.005688205 | 0.013798776 |

| | | |
|---|---|---|
| NA | NA | NA |
| -2.274364502 | 0.02294408 | 0.047115458 |
| -7.493290572 | 6.72E-14 | 7.01E-13 |
| 2.35849892 | 0.018349013 | 0.038803295 |
| -0.131783019 | 0.895155918 | NA |
| -1.875870911 | 0.060673008 | 0.109058915 |
| -0.065388772 | 0.947864464 | NA |
| 0.192810118 | 0.847107679 | NA |
| NA | NA | NA |
| 5.909045817 | 3.44E-09 | 2.22E-08 |
| NA | NA | NA |
| 0.572411904 | 0.567042953 | 0.668854152 |
| NA | NA | NA |
| NA | NA | NA |
| NA | NA | NA |
| 7.528937184 | 5.12E-14 | 5.40E-13 |
| 2.663027084 | 0.007744118 | 0.018101912 |
| -0.262989243 | 0.79255887 | 0.852466526 |
| 0.454918414 | 0.649167935 | 0.737939656 |
| 1.229253218 | 0.218976881 | 0.317886054 |
| -1.035067537 | 0.300637372 | 0.409677342 |
| -0.155609731 | 0.876340654 | NA |
| -9.338807029 | 9.74E-21 | 1.82E-19 |
| -0.433161742 | 0.664897271 | NA |
| -2.178512435 | 0.029367907 | 0.058293317 |
| -9.794499712 | 1.19E-22 | 2.58E-21 |
| 0.375794244 | 0.707069867 | NA |
| 0.497828975 | 0.618604592 | 0.713237502 |
| 10.35559934 | 3.95E-25 | 1.02E-23 |
| 0.063708285 | 0.949202508 | NA |
| -1.562467413 | 0.118177917 | 0.191381551 |
| 0.009154623 | 0.99269577 | 0.994881887 |
| -1.164083165 | 0.24439031 | 0.347206938 |
| -0.065390563 | 0.947863038 | NA |
| 0.192810118 | 0.847107679 | NA |
| 0.063708882 | 0.949202033 | NA |
| NA | NA | NA |
| NA | NA | NA |
| -1.89412952 | 0.05820782 | 0.105369331 |
| 10.18182871 | 2.39E-24 | 5.85E-23 |
| -0.326781079 | 0.743833471 | 0.813588175 |
| 7.206477159 | 5.74E-13 | 5.48E-12 |
| 4.924810081 | 8.44E-07 | 4.00E-06 |
| NA | NA | NA |
| -1.988078157 | 0.046803047 | 0.087443477 |
| -8.576517024 | 9.78E-18 | 1.44E-16 |
| -1.944403311 | 0.051846835 | 0.09552641 |

| | | | |
|---|---|---|---|
| NA | | NA | NA |
| | -5.303328661 | 1.14E-07 | 6.08E-07 |
| | -4.583870584 | 4.56E-06 | 1.95E-05 |
| NA | | NA | NA |
| NA | | NA | NA |
| NA | | NA | NA |
| | -0.171497468 | 0.863832621 | 0.905394686 |
| | -2.357641155 | 0.018391462 | 0.038879923 |
| | 0.192809521 | 0.847108147 | NA |
| | 0.315684384 | 0.752242081 | 0.82031191 |
| | -7.92144588 | 2.35E-15 | 2.84E-14 |
| | -2.665153882 | 0.00769531 | 0.018001272 |
| | -3.732647606 | 0.000189478 | 0.000625099 |
| NA | | NA | NA |
| | 0.320071032 | 0.748914485 | NA |
| | -4.880766799 | 1.06E-06 | 4.94E-06 |
| NA | | NA | NA |
| | 0.427894431 | 0.668727982 | NA |
| | -0.423768269 | 0.671734821 | 0.755901748 |
| | 10.04055597 | 1.01E-23 | 2.38E-22 |
| | 0.253476181 | 0.799900265 | NA |
| | 34.74514189 | 1.64E-264 | 3.42E-261 |
| | 0.612635989 | 0.540117056 | NA |
| NA | | NA | NA |
| | -7.414159762 | 1.22E-13 | 1.25E-12 |
| NA | | NA | NA |
| NA | | NA | NA |
| NA | | NA | NA |
| | -1.39830486 | 0.162021539 | 0.248431703 |
| | -0.09591505 | 0.923588041 | 0.94756963 |
| NA | | NA | NA |
| | 0.614555961 | 0.538848005 | NA |
| | -2.83061224 | 0.004645901 | 0.011513554 |
| | 4.014763325 | 5.95E-05 | 0.000213752 |
| | 0.039288331 | 0.96866051 | 0.978472601 |
| NA | | NA | NA |
| | 7.74369163 | 9.66E-15 | 1.10E-13 |
| NA | | NA | NA |
| NA | | NA | NA |
| | 0.192808924 | 0.847108615 | NA |
| NA | | NA | NA |
| NA | | NA | NA |
| NA | | NA | NA |
| NA | | NA | NA |
| | 2.402267442 | 0.016293791 | 0.035064362 |
| NA | | NA | NA |
| | -0.065389369 | 0.947863988 | NA |

| | | |
|---|---|---|
| -0.798969559 | 0.424308063 | 0.535871175 |
| 0.034407178 | 0.97255246 | 0.981097375 |
| 0.72361194 | 0.469304009 | NA |
| 0.872219498 | 0.383088645 | NA |
| 4.300424038 | 1.70E-05 | 6.69E-05 |
| -1.839046164 | 0.065908396 | 0.117073817 |
| NA | NA | NA |
| -5.044742715 | 4.54E-07 | 2.24E-06 |
| -7.469096159 | 8.07E-14 | 8.38E-13 |
| 1.910127289 | 0.056116826 | 0.102197625 |
| NA | NA | NA |
| NA | NA | NA |
| NA | NA | NA |
| NA | NA | NA |
| -3.063492625 | 0.002187696 | 0.0058414 |
| 8.734831924 | 2.44E-18 | 3.79E-17 |
| 1.306505025 | 0.191380871 | 0.285168664 |
| 0.535808449 | 0.592090942 | NA |
| NA | NA | NA |
| -5.953791428 | 2.62E-09 | 1.71E-08 |
| -4.628469393 | 3.68E-06 | 1.60E-05 |
| -3.86564544 | 0.000110796 | 0.000380205 |
| -0.089846921 | 0.928408862 | NA |
| 0.46192881 | 0.644132373 | 0.733759461 |
| 0.192808327 | 0.847109082 | NA |
| 3.071803309 | 0.002127699 | 0.005698299 |
| -2.113609591 | 0.034548626 | 0.067159996 |
| 0.06353712 | 0.949338802 | NA |
| NA | NA | NA |
| NA | NA | NA |
| -4.940077063 | 7.81E-07 | 3.72E-06 |
| -1.720849239 | 0.085278185 | 0.145780151 |
| -5.624263184 | 1.86E-08 | 1.10E-07 |
| 2.171833675 | 0.029868211 | 0.059131639 |
| NA | NA | NA |
| 3.195005625 | 0.001398281 | 0.003897436 |
| -1.510194756 | 0.130993736 | 0.208613172 |
| -7.198797066 | 6.07E-13 | 5.77E-12 |
| 2.596403365 | 0.00942054 | 0.021595029 |
| -5.482702193 | 4.19E-08 | 2.38E-07 |
| -0.342982867 | 0.731611349 | 0.804296911 |
| 0.980585313 | 0.32679728 | 0.437817048 |
| -0.066588298 | 0.946909462 | NA |
| 2.308960319 | 0.020945783 | 0.043475658 |
| 0.796436577 | 0.425778326 | 0.537240081 |
| 2.164445357 | 0.030430186 | 0.060092115 |
| -0.184645419 | 0.85350716 | 0.898134596 |

| | | | |
|---|---|---|---|
| | -2.523405636 | 0.011622425 | 0.026041473 |
| | 4.005994691 | 6.18E-05 | 0.000221458 |
| | -1.44056549 | 0.149707476 | 0.23307695 |
| NA | NA | NA | |
| | -2.914925701 | 0.003557732 | 0.009034193 |
| | 6.449518145 | 1.12E-10 | 8.49E-10 |
| | 6.138701007 | 8.32E-10 | 5.71E-09 |
| | 0.976641849 | 0.328746493 | 0.439675379 |
| | -5.580551582 | 2.40E-08 | 1.40E-07 |
| | 0.424036746 | 0.671539015 | 0.755749361 |
| | -3.324281509 | 0.000886467 | 0.002582733 |
| | -0.508963127 | 0.61077807 | 0.706818396 |
| | 1.2155157 | 0.224169464 | 0.323997201 |
| | 11.04904483 | 2.22E-28 | 7.02E-27 |
| | 0.50445392 | 0.613942436 | 0.709561766 |
| | 2.860237904 | 0.004233233 | 0.010579913 |
| | -1.798983807 | 0.072021242 | 0.126238914 |
| | -5.624877546 | 1.86E-08 | 1.10E-07 |
| | 3.66825158 | 0.000244215 | 0.000788939 |
| | -0.321667624 | 0.747704508 | NA |
| NA | NA | NA | |
| | 6.362768417 | 1.98E-10 | 1.45E-09 |
| | 9.918623746 | 3.45E-23 | 7.80E-22 |
| | 3.581537201 | 0.000341578 | 0.00107557 |
| | -0.376044707 | 0.70688366 | NA |
| | 1.430182435 | 0.152664665 | 0.236885288 |
| | -3.842524635 | 0.000121775 | 0.000414755 |
| | 0.56134669 | 0.574561218 | 0.675733272 |
| | -5.222990251 | 1.76E-07 | 9.17E-07 |
| | -0.170079713 | 0.864947448 | 0.905918262 |
| | 6.100047177 | 1.06E-09 | 7.21E-09 |
| | 3.002577984 | 0.002677034 | 0.006997301 |
| | -2.552869556 | 0.01068395 | 0.02419427 |
| NA | NA | NA | |
| | 8.539466611 | 1.35E-17 | 1.97E-16 |
| | -3.355024719 | 0.000793579 | 0.002330277 |
| | -5.996373861 | 2.02E-09 | 1.33E-08 |
| | 0.276238663 | 0.782364763 | 0.844795893 |
| | -4.670468438 | 3.01E-06 | 1.32E-05 |
| | 6.889978107 | 5.58E-12 | 4.79E-11 |
| | 2.204223598 | 0.027508623 | 0.055083267 |
| | 1.123025979 | 0.261426461 | 0.366582875 |
| | 2.420809558 | 0.015485988 | 0.033507458 |
| | 0.861577277 | 0.38892018 | 0.501115623 |
| | 2.164969346 | 0.030390033 | 0.060017557 |
| | -0.254755834 | 0.798911689 | 0.857126429 |
| | -6.172114163 | 6.74E-10 | 4.66E-09 |

| | | |
|---|---|---|
| 11.62994676 | 2.90E-31 | 1.07E-29 |
| 6.640908929 | 3.12E-11 | 2.49E-10 |
| 3.301758639 | 0.000960807 | 0.002778946 |
| 4.833678659 | 1.34E-06 | 6.17E-06 |
| 3.212884417 | 0.001314092 | 0.003684108 |
| -1.153930092 | 0.248528831 | 0.351828326 |
| 0.143049281 | 0.886251261 | NA |
| -2.111980984 | 0.03468808 | 0.06738884 |
| -12.47030433 | 1.08E-35 | 4.86E-34 |
| 2.634895132 | 0.008416328 | 0.019509233 |
| 10.30476235 | 6.71E-25 | 1.69E-23 |
| -6.639401328 | 3.15E-11 | 2.52E-10 |
| 2.07541489 | 0.037948099 | 0.072812643 |
| -10.80051287 | 3.42E-27 | 1.00E-25 |
| 0.288217989 | 0.77317988 | NA |
| 18.10089796 | 3.14E-73 | 4.31E-71 |
| 2.293445511 | 0.021822369 | 0.045056001 |
| -3.630246794 | 0.00028315 | 0.000904102 |
| -1.378520288 | 0.168042708 | 0.256142073 |
| -0.348866553 | 0.727189496 | 0.800772243 |
| -5.432402093 | 5.56E-08 | 3.10E-07 |
| 6.019401278 | 1.75E-09 | 1.16E-08 |
| NA | NA | NA |
| 1.090803088 | 0.275359538 | 0.382049078 |
| 1.117426089 | 0.263812185 | 0.36930755 |
| 0.194810053 | 0.845541652 | NA |
| 1.410428345 | 0.158413241 | 0.244065211 |
| -0.261996244 | 0.793324338 | NA |
| 0.063708882 | 0.949202033 | NA |
| 0.063708285 | 0.949202508 | NA |
| 4.816207004 | 1.46E-06 | 6.70E-06 |
| 9.434669132 | 3.92E-21 | 7.62E-20 |
| 1.31611734 | 0.188134662 | 0.281168533 |
| 1.573579406 | 0.115584717 | 0.187827437 |
| 2.279906015 | 0.022613263 | 0.046529235 |
| 2.811700161 | 0.004928042 | 0.012151403 |
| 0.623493125 | 0.532960518 | 0.637362722 |
| 0.024363649 | 0.980562543 | 0.986474295 |
| -4.558729164 | 5.15E-06 | 2.18E-05 |
| -2.591506848 | 0.009555664 | 0.021865229 |
| 3.010863395 | 0.00260506 | 0.006823426 |
| -6.145589979 | 7.97E-10 | 5.48E-09 |
| -1.796163725 | 0.072468482 | 0.126845182 |
| -1.28422649 | 0.199062719 | 0.294528572 |
| 27.02041451 | 8.51E-161 | 4.02E-158 |
| 7.975369196 | 1.52E-15 | 1.86E-14 |
| 4.321929509 | 1.55E-05 | 6.12E-05 |

| | | |
|---|---|---|
| 0.292629761 | 0.769805163 | 0.834158328 |
| -0.510209266 | 0.609904861 | 0.706101697 |
| 3.333555079 | 0.000857437 | 0.002504276 |
| -0.284539705 | 0.775996811 | 0.839232638 |
| -0.157390313 | 0.874937248 | 0.912726807 |
| -7.280146025 | 3.33E-13 | 3.25E-12 |
| -0.868082871 | 0.385348968 | NA |
| 20.0779603 | 1.15E-89 | 2.28E-87 |
| 2.198465434 | 0.027915955 | 0.055760622 |
| 5.785528163 | 7.23E-09 | 4.48E-08 |
| -8.008414007 | 1.16E-15 | 1.44E-14 |
| 1.601160927 | 0.109341281 | 0.179228045 |
| -2.408859354 | 0.016002464 | 0.034514577 |
| 2.291198848 | 0.021951916 | 0.045297316 |
| 0.191088183 | 0.848456506 | 0.894210461 |
| -0.515422856 | 0.606257564 | 0.703114949 |
| 9.042761961 | 1.53E-19 | 2.63E-18 |
| NA | NA | NA |
| NA | NA | NA |
| NA | NA | NA |
| NA | NA | NA |
| NA | NA | NA |
| 11.38992717 | 4.69E-30 | 1.62E-28 |
| -0.20878015 | 0.834619868 | NA |
| -1.31714008 | 0.187791674 | 0.280739749 |
| 9.962854686 | 2.22E-23 | 5.08E-22 |
| 7.587431939 | 3.26E-14 | 3.52E-13 |
| 0.316204602 | 0.751847215 | NA |
| NA | NA | NA |
| 3.143770603 | 0.001667861 | 0.004576973 |
| 1.466792031 | 0.142432638 | 0.223755032 |
| 3.936088545 | 8.28E-05 | 0.000290661 |
| 10.83090469 | 2.46E-27 | 7.31E-26 |
| -1.196008537 | 0.231693228 | 0.332546676 |
| 4.454742902 | 8.40E-06 | 3.46E-05 |
| 0.192810118 | 0.847107679 | NA |
| 1.160914293 | 0.245676756 | 0.348697998 |
| 4.208147407 | 2.57E-05 | 9.85E-05 |
| NA | NA | NA |
| 2.41149925 | 0.015887085 | 0.034286405 |
| 4.925637911 | 8.41E-07 | 3.99E-06 |
| 2.604183384 | 0.009209347 | 0.021159349 |
| -1.19532179 | 0.23196133 | 0.332886577 |
| 0.065727069 | 0.947595121 | 0.96427637 |
| -7.564639896 | 3.89E-14 | 4.17E-13 |
| -5.042778116 | 4.59E-07 | 2.26E-06 |
| -1.360243013 | 0.173753035 | 0.263434398 |

| | | | |
|---|---|---|---|
| | 3.275782684 | 0.001053696 | 0.003022117 |
| | 2.48686689 | 0.012887361 | 0.028518313 |
| | -4.161607131 | 3.16E-05 | 0.000119024 |
| NA | NA | NA | |
| | -0.362786808 | 0.716764147 | 0.792728868 |
| NA | NA | NA | |
| | 1.17328115 | 0.240683079 | 0.343049796 |
| NA | NA | NA | |
| NA | NA | NA | |
| | 0.25580338 | 0.798102664 | 0.856442001 |
| | 0.322270623 | 0.747247686 | 0.816277294 |
| | 3.366568651 | 0.000761096 | 0.002243845 |
| | 13.56022266 | 6.89E-42 | 4.01E-40 |
| | 7.326994091 | 2.35E-13 | 2.33E-12 |
| | -6.57775853 | 4.78E-11 | 3.76E-10 |
| | 2.606408798 | 0.009149719 | 0.021037797 |
| | 0.310819883 | 0.755937554 | 0.823032003 |
| | -0.571850658 | 0.567423154 | 0.669208008 |
| NA | NA | NA | |
| | 1.503081907 | 0.132817924 | 0.211010638 |
| NA | NA | NA | |
| NA | NA | NA | |
| | 0.192810118 | 0.847107679 | NA |
| | -3.569978782 | 0.00035701 | 0.001119935 |
| NA | NA | NA | |
| | -1.90608123 | 0.056639668 | 0.103014887 |
| | -5.335681128 | 9.52E-08 | 5.14E-07 |
| NA | NA | NA | |
| | -1.434461023 | 0.151440744 | 0.235409379 |
| | -1.384546185 | 0.166191276 | 0.25381461 |
| | 0.857698674 | 0.391058872 | 0.503073992 |
| | -1.128243021 | 0.2592173 | 0.364248295 |
| NA | NA | NA | |
| | -11.67587626 | 1.69E-31 | 6.33E-30 |
| | -0.44608138 | 0.655538463 | NA |
| NA | NA | NA | |
| | 0.493491931 | 0.62166504 | 0.71587578 |
| NA | NA | NA | |
| | -0.951070621 | 0.341568527 | 0.453071555 |
| | 0.260653038 | 0.794360084 | 0.853780184 |
| | 3.065917042 | 0.002170035 | 0.005802369 |
| | 6.853492273 | 7.21E-12 | 6.10E-11 |
| | -0.472833014 | 0.636332318 | 0.727577448 |
| | 0.924251853 | 0.355355204 | 0.467272958 |
| NA | NA | NA | |
| | -5.028910745 | 4.93E-07 | 2.42E-06 |
| | -0.111835848 | 0.910953563 | 0.938628671 |

| | | |
|---|---|---|
| -4.359687669 | 1.30E-05 | 5.21E-05 |
| 0.357856685 | 0.720450567 | 0.795567349 |
| 2.202497841 | 0.027630161 | 0.055291249 |
| 0.192636473 | 0.847243678 | NA |
| 1.874487181 | 0.060863307 | 0.109330313 |
| 1.177322084 | 0.239066976 | 0.34113485 |
| 4.449346277 | 8.61E-06 | 3.55E-05 |
| -0.065389369 | 0.947863988 | NA |
| 1.103265372 | 0.26991194 | 0.376058773 |
| 0.468021824 | 0.639768983 | 0.730420153 |
| NA | NA | NA |
| 0.667043548 | 0.504744317 | 0.611300637 |
| 1.431811083 | 0.152197895 | 0.23638082 |
| 0.79321419 | 0.427653041 | 0.538926373 |
| 3.744154789 | 0.000181002 | 0.000598795 |
| 5.437350816 | 5.41E-08 | 3.02E-07 |
| 3.14545687 | 0.001658276 | 0.004554665 |
| NA | NA | NA |
| 0.192808924 | 0.847108615 | NA |
| 4.660649687 | 3.15E-06 | 1.38E-05 |
| 1.760193016 | 0.078375086 | 0.135694017 |
| 0.262571712 | 0.792880706 | NA |
| NA | NA | NA |
| 6.537187559 | 6.27E-11 | 4.87E-10 |
| NA | NA | NA |
| NA | NA | NA |
| 0.092467785 | 0.926326385 | 0.949543346 |
| NA | NA | NA |
| 11.69108186 | 1.42E-31 | 5.32E-30 |
| NA | NA | NA |
| NA | NA | NA |
| 3.036935868 | 0.002389963 | 0.006317539 |
| -7.087581197 | 1.36E-12 | 1.25E-11 |
| -0.560661383 | 0.575028398 | 0.67617029 |
| 2.846665559 | 0.004417975 | 0.011000986 |
| 1.247214956 | 0.21231869 | 0.310217548 |
| -0.168434415 | 0.866241535 | 0.906590802 |
| 1.516678562 | 0.129347859 | 0.206589323 |
| NA | NA | NA |
| -4.934273894 | 8.04E-07 | 3.83E-06 |
| -3.667016986 | 0.000245396 | 0.000792041 |
| 0.192809521 | 0.847108147 | NA |
| 0.331522665 | 0.740249724 | 0.810944091 |
| -5.697594358 | 1.22E-08 | 7.35E-08 |
| 0.460324992 | 0.645282965 | 0.734623516 |
| 4.521031018 | 6.15E-06 | 2.59E-05 |
| 0.751341279 | 0.452447293 | 0.56235752 |

| | | |
|---|---|---|
| 5.83131384 | 5.50E-09 | 3.46E-08 |
| 10.04646641 | 9.52E-24 | 2.24E-22 |
| 17.67535413 | 6.49E-70 | 8.29E-68 |
| NA | NA | NA |
| NA | NA | NA |
| -4.926764343 | 8.36E-07 | 3.96E-06 |
| -9.984748533 | 1.78E-23 | 4.11E-22 |
| -0.132549473 | 0.894549695 | NA |
| -1.079824174 | 0.280220484 | 0.387188447 |
| 0.710478081 | 0.477407719 | 0.58622033 |
| 5.461840343 | 4.71E-08 | 2.66E-07 |
| -7.302488126 | 2.82E-13 | 2.77E-12 |
| -1.551093627 | 0.120879248 | 0.195112374 |
| -1.678519914 | 0.093245646 | 0.157241698 |
| NA | NA | NA |
| -1.725357876 | 0.084462998 | 0.144584108 |
| NA | NA | NA |
| -1.914133002 | 0.055603164 | 0.101424515 |
| 8.210644102 | 2.20E-16 | 2.90E-15 |
| 0.660673105 | 0.508821975 | 0.61496398 |
| -3.613657754 | 0.000301908 | 0.0009591 |
| -3.869002144 | 0.000109282 | 0.000375473 |
| -1.356186707 | 0.175039771 | 0.265112275 |
| -0.560349531 | 0.575241049 | 0.676388568 |
| 1.107788688 | 0.267953086 | 0.373828981 |
| 5.810201334 | 6.24E-09 | 3.90E-08 |
| 1.21571985 | 0.22409166 | 0.323939604 |
| -1.082896599 | 0.278854322 | 0.385897256 |
| NA | NA | NA |
| -2.563533959 | 0.010361256 | 0.02352524 |
| -2.472529065 | 0.01341608 | 0.029551355 |
| -0.289623741 | 0.772104102 | 0.83607033 |
| 0.137424442 | 0.890695312 | NA |
| 3.973145539 | 7.09E-05 | 0.000251861 |
| 2.217059706 | 0.026619012 | 0.053498785 |
| 1.807921442 | 0.070618728 | NA |
| NA | NA | NA |
| 0.132398728 | 0.894668921 | 0.926617416 |
| 2.119275583 | 0.034067184 | 0.066355537 |
| 4.100930141 | 4.11E-05 | 0.000151868 |
| 3.501572821 | 0.000462521 | 0.001418389 |
| 1.05685651 | 0.290577079 | 0.398179366 |
| 4.818107123 | 1.45E-06 | 6.64E-06 |
| NA | NA | NA |
| -5.81372165 | 6.11E-09 | 3.82E-08 |
| -0.089215702 | 0.928910487 | NA |
| 4.322050108 | 1.55E-05 | 6.12E-05 |

| | | |
|---|---|---|
| -2.753188696 | 0.005901787 | 0.01424373 |
| 3.64110022 | 0.000271475 | 0.000869802 |
| -0.250339914 | 0.802324492 | 0.859866497 |
| -0.087058824 | 0.930624754 | 0.952778861 |
| 4.696319235 | 2.65E-06 | 1.17E-05 |
| 1.18627691 | 0.235512962 | 0.337120521 |
| -1.460835258 | 0.144060657 | 0.225774107 |
| 1.578993941 | 0.114337447 | 0.186089232 |
| 1.097976157 | 0.272214901 | 0.378698253 |
| 9.495329543 | 2.20E-21 | 4.33E-20 |
| 0.869370411 | 0.384644562 | 0.496853461 |
| 5.458567111 | 4.80E-08 | 2.71E-07 |
| NA | NA | NA |
| 1.376082435 | 0.168796113 | 0.257040014 |
| -3.936646036 | 8.26E-05 | 0.000290048 |
| 0.331440975 | 0.740311419 | 0.810976183 |
| 6.874319406 | 6.23E-12 | 5.31E-11 |
| -11.15983651 | 6.41E-29 | 2.10E-27 |
| NA | NA | NA |
| -5.798030853 | 6.71E-09 | 4.17E-08 |
| -2.781465993 | 0.0054114 | 0.013207932 |
| -0.627885823 | 0.530078739 | 0.634978264 |
| 3.865702647 | 0.00011077 | 0.000380168 |
| 1.375342906 | 0.16902516 | 0.257310532 |
| 1.072248254 | 0.283608542 | 0.390604061 |
| 3.083679012 | 0.002044581 | 0.00549746 |
| 3.22997158 | 0.001238025 | 0.003491187 |
| -5.087169808 | 3.63E-07 | 1.82E-06 |
| -1.600814489 | 0.109418014 | NA |
| -0.750122955 | 0.453180656 | 0.563101261 |
| 1.296446535 | 0.194821689 | 0.289211012 |
| NA | NA | NA |
| -0.903909823 | 0.36604322 | 0.478561598 |
| 6.131978669 | 8.68E-10 | 5.94E-09 |
| -1.14298172 | 0.253046181 | 0.357132558 |
| 1.644728955 | 0.100025719 | 0.16657643 |
| 2.994576303 | 0.002748263 | 0.007164273 |
| -0.799603153 | 0.42394076 | 0.535515378 |
| NA | NA | NA |
| NA | NA | NA |
| NA | NA | NA |
| 4.434569258 | 9.23E-06 | 3.78E-05 |
| 0.381721344 | 0.702668067 | 0.781748676 |
| 0.063708285 | 0.949202508 | NA |
| -1.21787428 | 0.223271751 | 0.322884905 |
| NA | NA | NA |
| -0.96092996 | 0.336587385 | 0.447869612 |

| | | |
|---|---|---|
| 0.192636473 | 0.847243678 | NA |
| -1.84991653 | 0.064325581 | 0.114579369 |
| 2.394249055 | 0.016654436 | 0.035739071 |
| 1.358031038 | 0.174453836 | 0.26438919 |
| 1.192842216 | 0.232931173 | NA |
| -4.095406887 | 4.21E-05 | 0.000155214 |
| 0.55885036 | 0.57626385 | 0.677312114 |
| NA | NA | NA |
| -2.128979663 | 0.033255943 | 0.064984558 |
| NA | NA | NA |
| 0.063708285 | 0.949202508 | NA |
| 3.09009433 | 0.002000929 | 0.005394578 |
| NA | NA | NA |
| 2.260648897 | 0.023781007 | 0.048560988 |
| -0.159097118 | 0.873592358 | NA |
| 0.063708882 | 0.949202033 | NA |
| 1.059458429 | 0.289391052 | 0.396880063 |
| 0.592986066 | 0.553190477 | 0.656349908 |
| 10.28526075 | 8.21E-25 | 2.06E-23 |
| 4.374367287 | 1.22E-05 | 4.89E-05 |
| 2.884805369 | 0.003916557 | 0.009868273 |
| -0.19448762 | 0.845794089 | NA |
| NA | NA | NA |
| 4.433130852 | 9.29E-06 | 3.81E-05 |
| -0.218871611 | 0.826750062 | 0.878781809 |
| 0.53421942 | 0.593189735 | 0.692261322 |
| 0.06371127 | 0.949200131 | NA |
| 0.695379215 | 0.486817683 | NA |
| -0.246216359 | 0.805514752 | NA |
| NA | NA | NA |
| NA | NA | NA |
| NA | NA | NA |
| -1.475037443 | 0.140202477 | 0.22070868 |
| -3.893249765 | 9.89E-05 | 0.000342376 |
| 0.192809521 | 0.847108147 | NA |
| NA | NA | NA |
| -0.153595614 | 0.877928596 | 0.914615531 |
| 0.063708285 | 0.949202508 | NA |
| -4.451041122 | 8.55E-06 | 3.52E-05 |
| 1.698458293 | 0.089421298 | NA |
| 3.242964563 | 0.001182929 | 0.003355856 |
| 0.368302254 | 0.712647872 | 0.789622675 |
| -2.234188868 | 0.025470649 | 0.051530095 |
| NA | NA | NA |
| -1.37346206 | 0.169608748 | 0.258041999 |
| -0.571520501 | 0.567646868 | 0.669377231 |
| 5.976007766 | 2.29E-09 | 1.50E-08 |

| | | | |
|---|---|---|---|
| | 0.151011089 | 0.879966968 | 0.915748097 |
| | -5.67725907 | 1.37E-08 | 8.23E-08 |
| | 11.64512146 | 2.43E-31 | 9.02E-30 |
| | 12.23060806 | 2.13E-34 | 9.07E-33 |
| | -7.626930889 | 2.40E-14 | 2.63E-13 |
| | -0.255732822 | 0.798157149 NA | |
| NA | NA | NA | |
| | 0.717594946 | 0.473007075 | 0.581927014 |
| NA | NA | NA | |
| | 1.297486391 | 0.194463885 | 0.288816889 |
| | 0.725721257 | 0.468009668 | 0.577228835 |
| NA | NA | NA | |
| | -4.993946981 | 5.92E-07 | 2.87E-06 |
| NA | NA | NA | |
| | -2.695100901 | 0.007036732 | 0.016652113 |
| | 1.004337193 | 0.315216112 | 0.425696519 |
| | 0.063708285 | 0.949202508 NA | |
| | -3.251096053 | 0.00114961 | 0.003271344 |
| | -5.362634102 | 8.20E-08 | 4.47E-07 |
| NA | NA | NA | |
| | -3.400398353 | 0.000672877 | 0.002003584 |
| | -7.773962214 | 7.61E-15 | 8.76E-14 |
| | 0.318804313 | 0.749874908 NA | |
| | 2.357122151 | 0.018417188 | 0.038927732 |
| | -2.421368961 | 0.015462175 | 0.033467505 |
| | 0.063708285 | 0.949202508 NA | |
| | 0.259680703 | 0.795110079 NA | |
| | -0.065388772 | 0.947864464 NA | |
| NA | NA | NA | |
| | 2.449725782 | 0.014296505 | 0.031209386 |
| | 0.420019549 | 0.674471173 | 0.758026737 |
| | 2.566079265 | 0.010285532 | 0.023372654 |
| NA | NA | NA | |
| | 0.39201267 | 0.695048852 | 0.775374282 |
| | 0.904198718 | 0.36589004 | 0.478436302 |
| | 1.84866033 | 0.064506875 | 0.114869599 |
| | -4.289000722 | 1.79E-05 | 7.03E-05 |
| | -2.186921526 | 0.028748253 | 0.057217619 |
| NA | NA | NA | |
| | -4.363025665 | 1.28E-05 | 5.13E-05 |
| | -0.086309271 | 0.931220569 NA | |
| NA | NA | NA | |
| | -0.129182938 | 0.897212897 NA | |
| | -1.89913217 | 0.0575471 | 0.104422497 |
| NA | NA | NA | |
| | -2.295971261 | 0.021677525 | 0.044801291 |
| | 0.937367797 | 0.3485694 NA | |

| | | |
|---|---|---|
| NA | NA | NA |
| -0.161200786 | 0.871935262 | 0.910797914 |
| -2.928531723 | 0.003405671 | 0.008689494 |
| -9.022118972 | 1.84E-19 | 3.15E-18 |
| -0.090670487 | 0.927754422 | NA |
| 5.280612571 | 1.29E-07 | 6.84E-07 |
| -1.370787579 | 0.170441184 | 0.259166687 |
| -5.273288804 | 1.34E-07 | 7.10E-07 |
| -5.561428325 | 2.68E-08 | 1.55E-07 |
| 1.365302243 | 0.172158074 | 0.261348654 |
| 5.443543619 | 5.22E-08 | 2.92E-07 |
| 7.083915472 | 1.40E-12 | 1.28E-11 |
| 6.160062872 | 7.27E-10 | 5.01E-09 |
| 1.669056773 | 0.095106129 | 0.159851357 |
| 0.496728642 | 0.619380421 | 0.713967576 |
| 3.21268667 | 0.001314997 | 0.003686232 |
| -0.065389369 | 0.947863988 | NA |
| 1.792062287 | 0.073122989 | 0.127794192 |
| 2.600929382 | 0.009297159 | 0.021337602 |
| 0.791747237 | 0.428508073 | 0.53970498 |
| 9.888614008 | 4.66E-23 | 1.04E-21 |
| 34.29648673 | 8.85E-258 | 1.48E-254 |
| 2.199105059 | 0.027870453 | 0.055700849 |
| 8.394030917 | 4.70E-17 | 6.56E-16 |
| 7.094728018 | 1.30E-12 | 1.19E-11 |
| -0.37262036 | 0.709431017 | 0.787068763 |
| -0.40455113 | 0.685807487 | 0.767874184 |
| -3.56898776 | 0.000358363 | 0.001123616 |
| NA | NA | NA |
| -1.269412022 | 0.204294151 | 0.300793093 |
| -6.418048273 | 1.38E-10 | 1.03E-09 |
| -0.90384101 | 0.366079713 | 0.47858431 |
| 1.273721321 | 0.202762198 | 0.29906528 |
| 0.063708285 | 0.949202508 | NA |
| -3.731216433 | 0.000190557 | 0.000628533 |
| 5.344264699 | 9.08E-08 | 4.92E-07 |
| -2.471435423 | 0.013457185 | 0.02962214 |
| 5.809077745 | 6.28E-09 | 3.92E-08 |
| 10.63379208 | 2.08E-26 | 5.81E-25 |
| 0.253780977 | 0.799664771 | NA |
| 0.982623734 | 0.325792659 | 0.436915466 |
| 3.891169851 | 9.98E-05 | 0.000344809 |
| 1.309654435 | 0.190312765 | 0.283898236 |
| -0.932282165 | 0.351190722 | 0.462913452 |
| -0.246349338 | 0.80541182 | 0.862313679 |
| -0.584706041 | 0.558745392 | 0.661123057 |
| -1.605158642 | 0.108458897 | 0.178026505 |

| | | |
|---|---|---|
| 2.775261542 | 0.005515733 | 0.013433765 |
| 12.41641274 | 2.13E-35 | 9.48E-34 |
| -1.152646886 | 0.249055351 | 0.35243416 |
| -2.43662763 | 0.014824937 | 0.032233163 |
| 3.204230698 | 0.001354239 | 0.0037886 |
| 1.227070357 | 0.219796148 | 0.318816503 |
| 0.089304918 | 0.928839586 | NA |
| 0.822607596 | 0.410731178 | 0.522613864 |
| 1.775369144 | 0.075836955 | 0.131892648 |
| -12.76457284 | 2.59E-37 | 1.25E-35 |
| -0.183976404 | 0.854031966 | 0.898508684 |
| 1.403107604 | 0.160584754 | 0.246636583 |
| -2.163240955 | 0.030522652 | 0.060250944 |
| -0.997993182 | 0.318282665 | 0.428889104 |
| NA | NA | NA |
| NA | NA | NA |
| 1.258399515 | 0.208247307 | 0.30530253 |
| 3.006625131 | 0.002641653 | 0.006909873 |
| 5.639797008 | 1.70E-08 | 1.01E-07 |
| 0.84429988 | 0.398501853 | 0.510361307 |
| NA | NA | NA |
| NA | NA | NA |
| NA | NA | NA |
| 0.063708882 | 0.949202033 | NA |
| 1.850480071 | 0.064244388 | 0.114459181 |
| NA | NA | NA |
| NA | NA | NA |
| -0.194487022 | 0.845794556 | NA |
| -9.089367166 | 9.96E-20 | 1.74E-18 |
| 0.76997905 | 0.44131232 | NA |
| -4.374855637 | 1.22E-05 | 4.88E-05 |
| -2.257989744 | 0.023946299 | 0.048848888 |
| -4.557454065 | 5.18E-06 | 2.20E-05 |
| -0.909005053 | 0.363347459 | 0.475732954 |
| NA | NA | NA |
| 3.907743584 | 9.32E-05 | 0.000324363 |
| 0.192810118 | 0.847107679 | NA |
| NA | NA | NA |
| NA | NA | NA |
| -6.62541839 | 3.46E-11 | 2.76E-10 |
| -2.11611601 | 0.034334942 | 0.066806861 |
| 10.29511046 | 7.41E-25 | 1.86E-23 |
| 16.06311025 | 4.63E-58 | 4.32E-56 |
| -4.180626664 | 2.91E-05 | 0.000110182 |
| 0.199526594 | 0.841850842 | NA |
| 3.094434024 | 0.001971888 | 0.005320866 |
| 3.504331373 | 0.000457755 | 0.001405325 |

| | | | |
|---|---|---|---|
| | -0.065388772 | 0.947864464 | NA |
| | 0.624296961 | 0.532432579 | NA |
| NA | NA | NA | |
| | 0.31685483 | 0.751353758 | NA |
| | 9.449652692 | 3.40E-21 | 6.63E-20 |
| | -8.27468489 | 1.29E-16 | 1.73E-15 |
| | -7.80896394 | 5.77E-15 | 6.72E-14 |
| | -1.859804901 | 0.062913134 | 0.112503804 |
| NA | NA | NA | |
| | -0.329904232 | 0.741472325 | 0.81178973 |
| | -3.303044096 | 0.000956413 | 0.002767198 |
| | -2.026552497 | 0.042708199 | 0.080726946 |
| NA | NA | NA | |
| | 5.939906526 | 2.85E-09 | 1.85E-08 |
| | 4.520086586 | 6.18E-06 | 2.60E-05 |
| | -1.462829004 | 0.143514174 | 0.225114983 |
| NA | NA | NA | |
| | -0.194488217 | 0.845793621 | NA |
| NA | NA | NA | |
| | 0.06353712 | 0.949338802 | NA |
| NA | NA | NA | |
| NA | NA | NA | |
| NA | NA | NA | |
| | 0.320436416 | 0.748637524 | NA |
| NA | NA | NA | |
| | 0.211387677 | 0.832584772 | 0.88275094 |
| NA | NA | NA | |
| | 0.192808327 | 0.847109082 | NA |
| | -0.065389369 | 0.947863988 | NA |
| | 0.31685483 | 0.751353758 | NA |
| NA | NA | NA | |
| | 7.044077237 | 1.87E-12 | 1.68E-11 |
| | -0.671241451 | 0.502066726 | 0.608882706 |
| | 0.558801664 | 0.576297087 | 0.677312114 |
| | -3.142263651 | 0.00167647 | 0.004595055 |
| | -1.257436093 | 0.208595767 | 0.305670162 |
| | 3.649585316 | 0.000262664 | 0.000843423 |
| | 2.65603866 | 0.007906455 | 0.018433175 |
| | -2.723500426 | 0.006459413 | 0.015440656 |
| | -2.832769996 | 0.004614658 | 0.011447461 |
| | -6.346638684 | 2.20E-10 | 1.60E-09 |
| | 0.713566363 | 0.475495363 | 0.584388144 |
| | 8.313519791 | 9.29E-17 | 1.26E-15 |
| | 2.351637463 | 0.018690982 | 0.03940993 |
| | -0.601881916 | 0.547252743 | 0.65090934 |
| | -0.697532907 | 0.485469353 | 0.593788991 |
| | -2.243024688 | 0.024895223 | 0.050529308 |

| | | |
|---|---|---|
| 6.204432172 | 5.49E-10 | 3.83E-09 |
| 7.331983648 | 2.27E-13 | 2.25E-12 |
| -6.194648617 | 5.84E-10 | 4.07E-09 |
| -0.400385895 | 0.688872312 | 0.770387495 |
| 6.078132714 | 1.22E-09 | 8.21E-09 |
| -0.580941623 | 0.561279799 | 0.663494539 |
| 4.486866646 | 7.23E-06 | 3.00E-05 |
| -4.78013782 | 1.75E-06 | 7.96E-06 |
| -0.649285157 | 0.516154078 | NA |
| -0.090669501 | 0.927755206 | 0.950618667 |
| -0.76574748 | 0.44382657 | 0.553815197 |
| 0.495113468 | 0.620520028 | 0.714886141 |
| -1.647677573 | 0.099418853 | 0.165786401 |
| NA | NA | NA |
| -1.82826788 | 0.067509358 | 0.119518972 |
| NA | NA | NA |
| -1.31561812 | 0.188302249 | 0.281368592 |
| 0.162016246 | 0.871293061 | 0.910303328 |
| -5.467602761 | 4.56E-08 | 2.58E-07 |
| 2.482949029 | 0.013029973 | 0.028808429 |
| -1.689424011 | 0.091138203 | 0.154207342 |
| -0.11521152 | 0.908277467 | 0.936719613 |
| -0.86857097 | 0.385081838 | 0.497245069 |
| 0.921415649 | 0.356833462 | 0.468650517 |
| 1.82751882 | 0.067621799 | 0.119675715 |
| -1.436684973 | 0.150807527 | 0.23452702 |
| 3.477140573 | 0.000506792 | 0.001544879 |
| 0.833132741 | 0.40476987 | 0.516538689 |
| 1.52968825 | 0.126093913 | 0.202316067 |
| -1.221962414 | 0.221721841 | 0.321051584 |
| -0.378401527 | 0.705132334 | 0.783654307 |
| 2.536071229 | 0.011210393 | 0.02524936 |
| 1.923721548 | 0.054389496 | 0.099495671 |
| -0.469228914 | 0.63890602 | 0.729720177 |
| 18.61478367 | 2.44E-77 | 3.70E-75 |
| 5.059667213 | 4.20E-07 | 2.08E-06 |
| 8.737553339 | 2.38E-18 | 3.71E-17 |
| -0.065389369 | 0.947863988 | NA |
| 0.333816862 | 0.738517758 | NA |
| 7.901062243 | 2.77E-15 | 3.31E-14 |
| NA | NA | NA |
| -0.668687909 | 0.503694582 | 0.610206446 |
| 5.570085889 | 2.55E-08 | 1.48E-07 |
| 8.817536893 | 1.17E-18 | 1.88E-17 |
| NA | NA | NA |
| 0.201512299 | 0.840298014 | 0.888016014 |
| 1.49822446 | 0.134074943 | 0.212708108 |

| | | |
|---|---|---|
| 3.250091146 | 0.00115368 | 0.003280688 |
| -1.649743377 | 0.098995434 | 0.16521241 |
| 3.257051054 | 0.001125762 | 0.003208953 |
| 0.978947556 | 0.327605893 | 0.438663139 |
| 0.280283116 | 0.779260303 | NA |
| -7.548072189 | 4.42E-14 | 4.71E-13 |
| -2.522926725 | 0.011638265 | 0.026069963 |
| 4.385092271 | 1.16E-05 | 4.68E-05 |
| 0.063707135 | 0.949203424 | NA |
| 4.288165165 | 1.80E-05 | 7.05E-05 |
| 3.265129916 | 0.001094139 | 0.003126646 |
| 0.510446462 | 0.609738713 | NA |
| 4.967326185 | 6.79E-07 | 3.26E-06 |
| -0.786482818 | 0.431584682 | 0.542499513 |
| -5.906419791 | 3.50E-09 | 2.25E-08 |
| 2.35297185 | 0.018624044 | 0.039293584 |
| 4.566279133 | 4.96E-06 | 2.11E-05 |
| 2.45532359 | 0.014075782 | 0.030823869 |
| -1.710819706 | 0.087114401 | 0.148478992 |
| NA | NA | NA |
| -0.675858989 | 0.499130178 | NA |
| -1.726575295 | 0.084243966 | 0.144297986 |
| -0.144894373 | 0.884794269 | 0.919242925 |
| -0.298651177 | 0.765206213 | 0.830721353 |
| 3.455592953 | 0.000549084 | 0.001661858 |
| 4.33386988 | 1.47E-05 | 5.81E-05 |
| 0.063708285 | 0.949202508 | NA |
| 0.212600638 | 0.831638474 | 0.882268287 |
| NA | NA | NA |
| NA | NA | NA |
| -0.229449843 | 0.8185193 | 0.872441259 |
| 3.623412684 | 0.000290741 | 0.000926449 |
| -0.034662962 | 0.972348495 | 0.981049775 |
| -0.530848075 | 0.595524062 | 0.694006911 |
| 1.319790007 | 0.186905139 | 0.279648253 |
| 6.67822081 | 2.42E-11 | 1.95E-10 |
| 2.591819383 | 0.009546988 | 0.021852868 |
| -0.461754793 | 0.644257173 | 0.733763698 |
| 5.925247473 | 3.12E-09 | 2.02E-08 |
| 0.192809521 | 0.847108147 | NA |
| -0.065389966 | 0.947863513 | NA |
| -2.335423012 | 0.019521333 | 0.040905733 |
| -2.556270434 | 0.010580086 | 0.023982933 |
| -0.833577758 | 0.404518964 | 0.516271143 |
| NA | NA | NA |
| 0.234041249 | 0.814952956 | NA |
| -0.117338892 | 0.906591506 | 0.935439062 |

| | | | |
|---|---|---|---|
| NA | | NA | NA |
| | -0.917566275 | 0.358845981 | 0.470805209 |
| | 4.867535603 | 1.13E-06 | 5.26E-06 |
| | -2.59607491 | 0.00942955 | 0.021613704 |
| | 7.779678597 | 7.27E-15 | 8.39E-14 |
| | -1.239662612 | 0.215100211 | 0.313419949 |
| | -0.729540992 | 0.465670803 | 0.574967452 |
| | 5.946297677 | 2.74E-09 | 1.78E-08 |
| | 0.581922554 | 0.560618848 | 0.66283844 |
| | 1.630818537 | 0.102928615 | 0.170503192 |
| | 2.806896302 | 0.005002134 | 0.012311054 |
| | -1.416867039 | 0.156521805 | 0.241880643 |
| NA | | NA | NA |
| | -0.531934439 | 0.594771404 | 0.693488994 |
| | -6.076593828 | 1.23E-09 | 8.28E-09 |
| | -2.072413752 | 0.038226872 | 0.073240861 |
| | 1.861703325 | 0.06264492 | 0.112104264 |
| | -0.448076775 | 0.654097789 | 0.742059904 |
| | 1.292206821 | 0.19628553 | 0.291107828 |
| | -2.965359147 | 0.003023298 | 0.007811824 |
| NA | | NA | NA |
| | 0.192810118 | 0.847107679 | NA |
| | -3.126077167 | 0.001771551 | 0.004830274 |
| | 1.313505772 | 0.189012577 | 0.282244649 |
| NA | | NA | NA |
| | -3.917684756 | 8.94E-05 | 0.000312145 |
| NA | | NA | NA |
| NA | | NA | NA |
| NA | | NA | NA |
| NA | | NA | NA |
| | 6.059366558 | 1.37E-09 | 9.16E-09 |
| | -2.828852203 | 0.004671527 | 0.011570187 |
| | -4.188902414 | 2.80E-05 | 0.000106546 |
| | 1.425684328 | 0.153959482 | 0.238524749 |
| | 0.464226959 | 0.642485148 | 0.732504476 |
| | 9.716133417 | 2.57E-22 | 5.43E-21 |
| | 0.493833429 | 0.621423822 | 0.71563093 |
| | -0.425294998 | 0.670621639 | 0.755056436 |
| | 5.859904266 | 4.63E-09 | 2.94E-08 |
| | 0.362313734 | 0.717117596 | NA |
| | -4.258115177 | 2.06E-05 | 8.00E-05 |
| | -7.028004134 | 2.10E-12 | 1.87E-11 |
| | -1.047015155 | 0.295092589 | 0.403356938 |
| | 3.558680132 | 0.000372723 | 0.001164266 |
| | -0.235540993 | 0.813788866 | 0.869061092 |
| | -0.376354314 | 0.706653505 | 0.784856995 |
| | 8.445047792 | 3.04E-17 | 4.31E-16 |

| | | |
|---|---|---|
| NA | NA | NA |
| -1.401919478 | 0.160939293 | 0.247105294 |
| -2.565725098 | 0.010296039 | 0.023394141 |
| -3.136310956 | 0.001710877 | 0.00468116 |
| -2.732582168 | 0.006283999 | 0.015064506 |
| NA | NA | NA |
| 2.387078719 | 0.016982854 | 0.036328591 |
| 13.71825789 | 7.89E-43 | 4.82E-41 |
| -0.083474302 | 0.93347441 | 0.954825494 |
| NA | NA | NA |
| -2.146544633 | 0.031829552 | 0.062499409 |
| -0.065388772 | 0.947864464 | NA |
| -3.270326812 | 0.001074233 | 0.003076084 |
| -3.20941614 | 0.001330048 | 0.003725088 |
| 3.045987807 | 0.002319172 | 0.006151853 |
| 2.075481668 | 0.037941916 | 0.072807442 |
| NA | NA | NA |
| 1.016921754 | 0.309190654 | 0.419074136 |
| 1.432175963 | 0.152093469 | 0.236233294 |
| 0.054126437 | 0.956834429 | 0.970365225 |
| NA | NA | NA |
| 1.486178923 | 0.137231801 | 0.216795757 |
| -4.3013716 | 1.70E-05 | 6.67E-05 |
| -0.490965941 | 0.623450537 | 0.717337821 |
| -3.215518653 | 0.00130209 | 0.003654173 |
| 9.747434173 | 1.89E-22 | 4.03E-21 |
| 2.357150659 | 0.018415774 | 0.038927732 |
| 1.379846022 | 0.167634059 | 0.25561258 |
| NA | NA | NA |
| 0.314195157 | 0.753372821 | 0.821079931 |
| -5.687123629 | 1.29E-08 | 7.79E-08 |
| -0.879379456 | 0.379195568 | 0.491826358 |
| NA | NA | NA |
| NA | NA | NA |
| 0.433337604 | 0.664769523 | 0.750942936 |
| 0.356560558 | 0.721420809 | 0.796324899 |
| -1.80008418 | 0.071847347 | 0.125978221 |
| 3.032671227 | 0.002423995 | 0.006396016 |
| -0.526473164 | 0.598559493 | 0.696607807 |
| 0.520030127 | 0.603042577 | 0.700229899 |
| 0.066009206 | 0.947370496 | NA |
| -3.944539439 | 8.00E-05 | 0.000281457 |
| NA | NA | NA |
| 0.192809521 | 0.847108147 | NA |
| 2.329895463 | 0.019811677 | 0.041444736 |
| NA | NA | NA |
| -1.417611023 | 0.15630436 | 0.241589362 |

| | | |
|---|---|---|
| 4.895500746 | 9.81E-07 | 4.61E-06 |
| 0.957029243 | 0.338552495 | 0.449857671 |
| 0.870860786 | 0.38383017 | 0.496166884 |
| -0.154980323 | 0.87683683 | 0.913972675 |
| NA | NA | NA |
| -3.698490057 | 0.000216886 | 0.000708054 |
| 0.337252312 | 0.735926706 | 0.807515911 |
| -1.269094899 | 0.204407219 | 0.300888773 |
| -2.72223688 | 0.006484165 | 0.015493903 |
| -0.614697928 | 0.538754229 | 0.643093058 |
| 4.495447439 | 6.94E-06 | 2.90E-05 |
| NA | NA | NA |
| 1.801004108 | 0.071702233 | 0.125774591 |
| NA | NA | NA |
| -3.046790072 | 0.002312992 | 0.00613741 |
| 1.892185566 | 0.058466261 | 0.105768322 |
| -0.085401714 | 0.931942031 | NA |
| 1.021236574 | 0.307142368 | 0.416884847 |
| 12.48902013 | 8.57E-36 | 3.87E-34 |
| NA | NA | NA |
| -0.876407711 | 0.380808428 | 0.493330312 |
| -0.540701088 | 0.588713629 | 0.688317551 |
| 1.796227661 | 0.072458317 | 0.12683626 |
| -1.699188017 | 0.089283764 | NA |
| NA | NA | NA |
| -9.55741165 | 1.21E-21 | 2.44E-20 |
| NA | NA | NA |
| -0.890774685 | 0.373050059 | NA |
| 2.825777457 | 0.004716602 | 0.011672587 |
| 1.17013678 | 0.241945929 | 0.344516248 |
| 7.431209866 | 1.08E-13 | 1.10E-12 |
| -5.216244973 | 1.83E-07 | 9.48E-07 |
| -1.559339611 | 0.118916021 | 0.192415024 |
| -1.448572387 | 0.147457037 | 0.23013154 |
| 0.725128809 | 0.468373012 | 0.577591589 |
| -0.058406533 | 0.953424811 | 0.968161914 |
| 3.646073413 | 0.000266278 | 0.000854041 |
| 0.266557126 | 0.78981017 | 0.850533431 |
| -0.053953969 | 0.956971838 | 0.970365225 |
| 0.03363086 | 0.973171514 | 0.981474724 |
| 1.395434084 | 0.162884979 | 0.249679162 |
| 2.66827573 | 0.007624166 | 0.017861557 |
| 0.519461855 | 0.603438706 | 0.700657365 |
| 9.887856036 | 4.70E-23 | 1.05E-21 |
| 0.203156417 | 0.839012777 | 0.887219679 |
| 0.255890946 | 0.798035046 | 0.856406157 |
| NA | NA | NA |

| | | | |
|---|---|---|---|
| | 6.799951612 | 1.05E-11 | 8.76E-11 |
| | 0.271805119 | 0.785771869 | NA |
| | 2.583835377 | 0.009770842 | 0.022312214 |
| NA | NA | NA | |
| | -1.499072081 | 0.133854933 | 0.212372527 |
| | -3.111401331 | 0.001862017 | 0.005047247 |
| | -1.530535911 | 0.125884132 | 0.202031278 |
| | -0.373416534 | 0.708838456 | 0.7866273 |
| | -1.815759209 | 0.069407317 | 0.122351232 |
| | 1.157204405 | 0.247188858 | 0.350327678 |
| NA | NA | NA | |
| | -1.404948486 | 0.160036597 | 0.246021129 |
| | -0.371865199 | 0.709993217 | 0.787413276 |
| | -2.517519954 | 0.011818428 | 0.026426232 |
| | 4.151345103 | 3.31E-05 | 0.000124034 |
| | -0.310040089 | 0.756530471 | 0.823493202 |
| | 0.11124314 | 0.911423543 | NA |
| | -2.251393862 | 0.024360603 | 0.049580898 |
| | 0.087264587 | 0.930461202 | NA |
| NA | NA | NA | |
| | -1.743282585 | 0.081284266 | 0.140005672 |
| | -2.649631216 | 0.008057967 | 0.018745376 |
| | 1.262388521 | 0.206809018 | 0.303750467 |
| NA | NA | NA | |
| | 0.063707135 | 0.949203424 | NA |
| | 22.55325782 | 1.25E-112 | 3.63E-110 |
| | 0.063708285 | 0.949202508 | NA |
| NA | NA | NA | |
| | 1.823294175 | 0.068258848 | 0.120615529 |
| | 6.52393523 | 6.85E-11 | 5.30E-10 |
| | 0.199526594 | 0.841850842 | NA |
| | -2.508585141 | 0.012121575 | 0.027003907 |
| | 1.49182437 | 0.135745194 | 0.214881242 |
| NA | NA | NA | |
| | 0.686921102 | 0.492132449 | 0.599419717 |
| | -0.400763547 | 0.68859422 | NA |
| | -0.234385349 | 0.814685832 | 0.869752857 |
| NA | NA | NA | |
| | -1.495361004 | 0.134820252 | 0.21360621 |
| | 0.063976859 | 0.948988653 | NA |
| | -0.09562842 | 0.923815693 | NA |
| | -0.359223155 | 0.719428156 | NA |
| | 1.571637466 | 0.116034652 | 0.188447142 |
| | -6.69784598 | 2.12E-11 | 1.72E-10 |
| | 0.512956652 | 0.607981644 | 0.704559495 |
| | 2.177242295 | 0.029462494 | 0.05846252 |
| | -3.781284114 | 0.000156021 | 0.000521252 |

| | | |
|---|---|---|
| -3.871986922 | 0.000107952 | 0.000371158 |
| -0.053802717 | 0.957092345 | NA |
| -9.736566775 | 2.11E-22 | 4.47E-21 |
| 3.94451487 | 8.00E-05 | 0.000281457 |
| NA | NA | NA |
| 0.601956136 | 0.547203336 | NA |
| 0.337484188 | 0.73575193 | NA |
| 0.991187676 | 0.321593947 | NA |
| 10.42755394 | 1.86E-25 | 4.90E-24 |
| 0.703929402 | 0.48147674 | 0.590145576 |
| -0.521046483 | 0.602334391 | NA |
| 0.063708285 | 0.949202508 | NA |
| NA | NA | NA |
| 0.735881765 | 0.461802656 | NA |
| -3.234396358 | 0.001219002 | 0.003441806 |
| 2.2231584 | 0.026205121 | 0.052819631 |
| -1.256557414 | 0.208913944 | 0.306046823 |
| 4.034401398 | 5.47E-05 | 0.000198319 |
| 4.655913007 | 3.23E-06 | 1.41E-05 |
| -0.267855999 | 0.788810164 | 0.849733788 |
| 0.902975068 | 0.366539128 | NA |
| 3.081514316 | 0.002059506 | 0.005531648 |
| 1.363796027 | 0.172631771 | 0.262004198 |
| 17.45216491 | 3.31E-68 | 4.09E-66 |
| -1.698999382 | 0.0893193 | 0.151652563 |
| -1.211837465 | 0.225574604 | 0.325650729 |
| 3.256244864 | 0.001128963 | 0.00321698 |
| 1.399585564 | 0.161637459 | 0.247964306 |
| -2.384278495 | 0.017112646 | 0.036578099 |
| 2.11253779 | 0.034640348 | 0.067312159 |
| 13.85628084 | 1.17E-43 | 7.40E-42 |
| -0.41655889 | 0.677001084 | NA |
| 9.467646757 | 2.86E-21 | 5.61E-20 |
| 7.649303033 | 2.02E-14 | 2.23E-13 |
| 8.550907003 | 1.22E-17 | 1.79E-16 |
| 0.119219858 | 0.905101174 | 0.934329294 |
| -0.104471845 | 0.91679491 | NA |
| NA | NA | NA |
| -2.862288223 | 0.004205942 | 0.010519058 |
| 2.845455702 | 0.004434793 | 0.011036276 |
| 0.201808052 | 0.840066787 | NA |
| -2.799811101 | 0.005113252 | 0.012552442 |
| -0.249213939 | 0.803195298 | 0.860504999 |
| 0.151115321 | 0.879884747 | 0.915738678 |
| -1.593618453 | 0.111021519 | 0.181577928 |
| -0.537150994 | 0.591163319 | 0.690439918 |
| -2.041720148 | 0.041179299 | 0.078153675 |

| | | |
|---|---|---|
| -2.074798318 | 0.038005231 | 0.072906837 |
| 4.941925549 | 7.74E-07 | 3.69E-06 |
| NA | NA | NA |
| NA | NA | NA |
| -5.554661601 | 2.78E-08 | 1.61E-07 |
| NA | NA | NA |
| 0.141201091 | 0.887711084 | 0.921279062 |
| 2.413508773 | 0.015799747 | 0.034112627 |
| 5.787553999 | 7.14E-09 | 4.43E-08 |
| 1.33236389 | 0.182740642 | 0.274586268 |
| -0.77389821 | 0.438990988 | 0.549272127 |
| -5.346699868 | 8.96E-08 | 4.86E-07 |
| -5.321694873 | 1.03E-07 | 5.53E-07 |
| -1.123911813 | 0.261050439 | 0.366160788 |
| -1.53853315 | 0.12391831 | 0.199400533 |
| 0.520719819 | 0.602561966 | 0.699931595 |
| 8.623778393 | 6.48E-18 | 9.67E-17 |
| -0.10194104 | 0.918803474 | 0.944261988 |
| -2.728808798 | 0.006356355 | 0.015217557 |
| 3.243555252 | 0.001180479 | 0.003350424 |
| 0.192808924 | 0.847108615 | NA |
| -3.484059724 | 0.000493869 | 0.001508242 |
| -3.916631478 | 8.98E-05 | 0.000313337 |
| NA | NA | NA |
| 6.76527611 | 1.33E-11 | 1.10E-10 |
| -6.030053872 | 1.64E-09 | 1.09E-08 |
| 1.871193248 | 0.0613183 | 0.11004496 |
| -0.220299576 | 0.825637851 | 0.877897851 |
| NA | NA | NA |
| NA | NA | NA |
| NA | NA | NA |
| NA | NA | NA |
| NA | NA | NA |
| NA | NA | NA |
| NA | NA | NA |
| NA | NA | NA |
| NA | NA | NA |
| NA | NA | NA |
| NA | NA | NA |
| NA | NA | NA |
| NA | NA | NA |
| NA | NA | NA |
| NA | NA | NA |
| NA | NA | NA |
| NA | NA | NA |
| NA | NA | NA |
| NA | NA | NA |

| | | | |
|---|---|---|---|
| NA | | NA | NA |
| NA | | NA | NA |
| NA | | NA | NA |
| NA | | NA | NA |
| NA | | NA | NA |
| NA | | NA | NA |
| NA | | NA | NA |
| NA | | NA | NA |
| NA | | NA | NA |
| NA | | NA | NA |
| NA | | NA | NA |
| NA | | NA | NA |
| NA | | NA | NA |
| NA | | NA | NA |
| NA | | NA | NA |
| NA | | NA | NA |
| NA | | NA | NA |
| NA | | NA | NA |
| NA | | NA | NA |
| NA | | NA | NA |
| NA | | NA | NA |
| NA | | NA | NA |
| NA | | NA | NA |
| NA | | NA | NA |
| NA | | NA | NA |
| NA | | NA | NA |
| NA | | NA | NA |
| NA | | NA | NA |
| NA | | NA | NA |
| NA | | NA | NA |
| NA | | NA | NA |
| NA | | NA | NA |
| | 4.634481862 | 3.58E-06 | 1.55E-05 |
| | 2.471496863 | 0.013454873 | 0.029619655 |
| | -5.901398816 | 3.60E-09 | 2.32E-08 |
| | -3.267046388 | 0.001086759 | 0.003108041 |
| | -2.420110845 | 0.015515777 | 0.033557409 |
| | 1.20526639 | 0.22810048 | 0.328463642 |
| | 0.130372221 | 0.896271943 | 0.927867278 |
| | 0.126923236 | 0.899001156 | 0.929681 |
| | -2.51611928 | 0.011865502 | 0.02651491 |
| | -0.451392663 | 0.65170657 | 0.740219424 |
| | 3.387446605 | 0.000705464 | 0.002092401 |
| | 1.694698435 | 0.090132643 | 0.152805476 |
| | 0.498982296 | 0.617791856 | 0.712563023 |
| | -0.094313095 | 0.924860448 | 0.948273677 |
| | -0.456958642 | 0.647700773 | NA |

| | | | |
|---|---|---|---|
| 0.330493631 | 0.741027004 | NA | |
| NA | NA | NA | |
| -2.236205162 | 0.025338335 | | 0.05131721 |
| 1.017556539 | 0.30888875 | | 0.418801009 |
| -2.458248749 | 0.013961643 | | 0.030608735 |
| 2.464167449 | 0.013733194 | | 0.030163376 |
| -5.854242466 | 4.79E-09 | | 3.03E-08 |
| -2.074700019 | 0.038014346 | | 0.072911806 |
| -0.587436251 | 0.556910753 | | 0.659451043 |
| 1.845424201 | 0.064975855 | | 0.115581384 |
| 0.56707084 | 0.570666054 | | 0.67205097 |
| -1.34436549 | 0.178830228 | | 0.269694559 |
| 3.4263626 | 0.000611723 | | 0.001835023 |
| -3.240558196 | 0.001192959 | | 0.003380861 |
| -1.789915664 | 0.07346747 | | 0.128323017 |
| 2.25376481 | 0.024210968 | | 0.0493004 |
| -2.294181099 | 0.021780099 | | 0.044994707 |
| -2.057670989 | 0.039621727 | | 0.075577795 |
| -1.168854749 | 0.242462158 | | 0.345113893 |
| -0.092636176 | 0.926192603 | NA | |
| -0.090827669 | 0.927629524 | NA | |
| 2.494317648 | 0.012619955 | | 0.027995877 |
| 6.378525841 | 1.79E-10 | | 1.32E-09 |
| 0.122984315 | 0.902119519 | | 0.932020124 |
| -4.592662719 | 4.38E-06 | | 1.88E-05 |
| -0.604795447 | 0.54531492 | | 0.649097985 |
| -1.975563218 | 0.048204268 | | 0.089673194 |
| -0.316176975 | 0.751868183 | | 0.820082823 |
| -4.381685918 | 1.18E-05 | | 4.75E-05 |
| 1.719939776 | 0.085443389 | | 0.146029748 |
| -3.952469421 | 7.73E-05 | | 0.000273259 |
| NA | NA | NA | |
| -0.899883593 | 0.368182203 | | 0.480655116 |
| 4.75122315 | 2.02E-06 | | 9.10E-06 |
| -10.82346592 | 2.67E-27 | | 7.88E-26 |
| 0.092296884 | 0.926462164 | | 0.949628731 |
| 1.663015204 | 0.096309392 | | 0.161483393 |
| NA | NA | NA | |
| -1.486662501 | 0.137103971 | | 0.216703233 |
| -0.899649897 | 0.368306594 | | 0.48079243 |
| -0.822488553 | 0.410798899 | | 0.522647902 |
| 0.192808924 | 0.847108615 | NA | |
| 6.66264426 | 2.69E-11 | | 2.16E-10 |
| NA | NA | NA | |
| NA | NA | NA | |
| 9.776592808 | 1.42E-22 | | 3.05E-21 |
| 0.236872192 | 0.812755944 | NA | |

| | | | |
|---|---|---|---|
| | 0.974408458 | 0.329853772 | NA |
| | 0.267554472 | 0.789042279 | NA |
| NA | NA | NA | |
| | 0.113104051 | 0.909948063 | NA |
| | -0.209299993 | 0.834214057 | NA |
| | 1.35250136 | 0.176214984 | 0.266537409 |
| | -3.219910761 | 0.001282305 | 0.003604683 |
| | 6.414289078 | 1.41E-10 | 1.05E-09 |
| | 1.819065465 | 0.068901443 | 0.121630801 |
| | 0.324219936 | 0.74577153 | NA |
| | 0.149528542 | 0.881136589 | NA |
| | 0.088962902 | 0.929111393 | NA |
| NA | NA | NA | |
| | 0.063708285 | 0.949202508 | NA |
| | -0.78362449 | 0.433260481 | 0.543966184 |
| | -0.257465026 | 0.79681981 | NA |
| NA | NA | NA | |
| NA | NA | NA | |
| NA | NA | NA | |
| NA | NA | NA | |
| NA | NA | NA | |
| | -4.269833021 | 1.96E-05 | 7.61E-05 |
| | 1.341783311 | 0.179666265 | 0.270727025 |
| | -0.643749914 | 0.519737617 | 0.625013575 |
| | 4.315474732 | 1.59E-05 | 6.28E-05 |
| | -6.694697551 | 2.16E-11 | 1.76E-10 |
| | -0.236439132 | 0.813091934 | NA |
| | -1.052518195 | 0.292561867 | 0.400482556 |
| | -0.486374475 | 0.626701684 | 0.71995333 |
| | -0.065389369 | 0.947863988 | NA |
| | 4.027682467 | 5.63E-05 | 0.000203599 |
| | -8.474598522 | 2.36E-17 | 3.38E-16 |
| | -4.230364874 | 2.33E-05 | 8.97E-05 |
| NA | NA | NA | |
| | 16.92013841 | 3.20E-64 | 3.56E-62 |
| | 14.79814823 | 1.51E-49 | 1.15E-47 |
| | 16.2540532 | 2.09E-59 | 2.05E-57 |
| | -1.968969843 | 0.048956554 | 0.090917239 |
| NA | NA | NA | |
| NA | NA | NA | |
| | -5.23903071 | 1.61E-07 | 8.46E-07 |
| | 1.61460099 | 0.106397131 | 0.175182225 |
| | 7.758973856 | 8.56E-15 | 9.81E-14 |
| | 5.253532195 | 1.49E-07 | 7.85E-07 |
| NA | NA | NA | |
| | 7.035342119 | 1.99E-12 | 1.78E-11 |
| | 0.507275069 | 0.611961829 | NA |

| | | | |
|---|---|---|---|
| | 0.324874675 | 0.745275925 | NA |
| | 6.903587932 | 5.07E-12 | 4.38E-11 |
| | 6.069517945 | 1.28E-09 | 8.63E-09 |
| | 0.683861886 | 0.494062384 | NA |
| NA | NA | NA | |
| | 0.193076837 | 0.846898793 | NA |
| | 0.502735578 | 0.615150192 | 0.710487686 |
| | 8.948465699 | 3.60E-19 | 6.03E-18 |
| | -0.38626843 | 0.699297879 | NA |
| | 4.980527405 | 6.34E-07 | 3.06E-06 |
| | 12.91334471 | 3.78E-38 | 1.92E-36 |
| | 7.105168571 | 1.20E-12 | 1.10E-11 |
| | 0.063710673 | 0.949200606 | NA |
| | -0.586708279 | 0.557399646 | 0.659936292 |
| | -2.417175861 | 0.015641458 | 0.033814622 |
| | -0.104479422 | 0.916788898 | 0.942894609 |
| | 0.063708285 | 0.949202508 | NA |
| | -5.37021333 | 7.86E-08 | 4.30E-07 |
| | -9.926010036 | 3.21E-23 | 7.27E-22 |
| | -7.079051505 | 1.45E-12 | 1.32E-11 |
| | -5.445266951 | 5.17E-08 | 2.90E-07 |
| NA | NA | NA | |
| NA | NA | NA | |
| | -8.890541694 | 6.08E-19 | 9.99E-18 |
| | -1.21498014 | 0.224373666 | 0.324178762 |
| NA | NA | NA | |
| | -0.943096255 | 0.345631671 | 0.457128704 |
| | -0.115525838 | 0.908028342 | NA |
| NA | NA | NA | |
| NA | NA | NA | |
| NA | NA | NA | |
| | 0.605944583 | 0.544551553 | 0.648428027 |
| | -2.351033456 | 0.01872135 | 0.039460665 |
| | 1.957924655 | 0.050238854 | 0.092974383 |
| | 0.208385165 | 0.834928239 | 0.884357573 |
| NA | NA | NA | |
| | 7.596998486 | 3.03E-14 | 3.28E-13 |
| NA | NA | NA | |
| | -1.297835616 | 0.194343828 | 0.28867284 |
| | 1.952153091 | 0.050920032 | 0.094068078 |
| | 0.076123701 | 0.939320684 | 0.958701546 |
| | -0.68104154 | 0.495845206 | 0.602944302 |
| | 6.448364384 | 1.13E-10 | 8.55E-10 |
| | -0.082833243 | 0.933984136 | 0.955111547 |
| | -1.263873319 | 0.2062755 | 0.303139723 |
| | 3.904466217 | 9.44E-05 | 0.000328288 |
| | 10.22278283 | 1.57E-24 | 3.87E-23 |

| | | | |
|---|---|---|---|
| | 7.348947405 | 2.00E-13 | 1.99E-12 |
| | 0.323758732 | 0.746120702 | 0.815354999 |
| NA | NA | NA | |
| | 6.160838912 | 7.24E-10 | 4.99E-09 |
| | 0.432690543 | 0.665239601 | 0.751237686 |
| | 5.207669931 | 1.91E-07 | 9.91E-07 |
| | 2.050323613 | 0.040332862 | 0.076752948 |
| | 1.001565143 | 0.316553663 | 0.427111109 |
| | 1.114501288 | 0.265064192 | 0.370687045 |
| | 1.758610455 | 0.078643696 | 0.136093177 |
| | 5.932515031 | 2.98E-09 | 1.93E-08 |
| | 5.682990759 | 1.32E-08 | 7.97E-08 |
| | -0.186275398 | 0.852228799 | 0.897275518 |
| | -0.065389369 | 0.947863988 | NA |
| | -1.560859918 | 0.118556806 | 0.191945463 |
| | 5.517133782 | 3.45E-08 | 1.98E-07 |
| | 2.536239604 | 0.011205004 | 0.025241765 |
| | 2.812107843 | 0.0049218 | 0.012138403 |
| | -0.678965091 | 0.497159981 | 0.604249506 |
| NA | NA | NA | |
| | 0.59005312 | 0.555155037 | 0.657901169 |
| | -1.565458429 | 0.117475457 | 0.190416442 |
| | -0.065389369 | 0.947863988 | NA |
| | 0.548645277 | 0.583248907 | 0.683428686 |
| | -0.041399113 | 0.96697772 | 0.977534078 |
| | 4.75640946 | 1.97E-06 | 8.89E-06 |
| | 6.083136082 | 1.18E-09 | 7.97E-09 |
| | -0.170987335 | 0.864233724 | 0.905510726 |
| | -4.260545191 | 2.04E-05 | 7.92E-05 |
| | -3.075105132 | 0.002104284 | 0.005639814 |
| | -2.360487341 | 0.018250941 | 0.038625269 |
| | 0.133157103 | 0.894069135 | NA |
| | 1.225786106 | 0.220279177 | 0.319424587 |
| | 1.005013254 | 0.314890468 | 0.425440376 |
| | -4.087281967 | 4.36E-05 | 0.000160277 |
| | 0.192809521 | 0.847108147 | NA |
| | -7.27735897 | 3.40E-13 | 3.32E-12 |
| | -0.41363109 | 0.679144301 | 0.762149473 |
| NA | NA | NA | |
| | 1.18539449 | 0.235861509 | 0.337580831 |
| | -2.074230542 | 0.038057905 | 0.072981981 |
| | 0.192808327 | 0.847109082 | NA |
| | 0.436644557 | 0.662369143 | NA |
| | 0.248314386 | 0.803891169 | NA |
| | -1.886635705 | 0.059209341 | 0.106857726 |
| NA | NA | NA | |
| | 0.507275069 | 0.611961829 | NA |

| | | |
|---|---|---|
| -0.066588298 | 0.946909462 | NA |
| NA | NA | NA |
| NA | NA | NA |
| NA | NA | NA |
| NA | NA | NA |
| -0.065388772 | 0.947864464 | NA |
| NA | NA | NA |
| NA | NA | NA |
| NA | NA | NA |
| NA | NA | NA |
| NA | NA | NA |
| 0.192809521 | 0.847108147 | NA |
| NA | NA | NA |
| NA | NA | NA |
| NA | NA | NA |
| NA | NA | NA |
| NA | NA | NA |
| 0.192809521 | 0.847108147 | NA |
| NA | NA | NA |
| NA | NA | NA |
| NA | NA | NA |
| NA | NA | NA |
| NA | NA | NA |
| NA | NA | NA |
| NA | NA | NA |
| NA | NA | NA |
| 3.460874043 | 0.000538425 | 0.001631873 |
| 0.511144483 | 0.609249889 | 0.705702473 |
| 0.454468478 | 0.649491675 | NA |
| 0.146587814 | 0.883457373 | NA |
| NA | NA | NA |
| NA | NA | NA |
| NA | NA | NA |
| 0.240979646 | 0.809570892 | 0.865483254 |
| NA | NA | NA |
| NA | NA | NA |
| NA | NA | NA |
| 0.317642674 | 0.750756 | NA |
| 0.063710673 | 0.949200606 | NA |
| NA | NA | NA |
| NA | NA | NA |
| NA | NA | NA |
| NA | NA | NA |
| NA | NA | NA |
| NA | NA | NA |
| 0.192809521 | 0.847108147 | NA |
| NA | NA | NA |

| | | | |
|---|---|---|---|
| NA | | NA | NA |
| | 0.967762659 | 0.333162917 | 0.444367938 |
| | -1.85266406 | 0.063930524 | 0.114013611 |
| NA | | NA | NA |
| | 0.882771479 | 0.377359753 | 0.490054718 |
| | -0.587841457 | 0.556638715 | 0.659284864 |
| NA | | NA | NA |
| | -0.453520959 | 0.650173655 | 0.738914211 |
| | -3.847116782 | 0.000119516 | 0.000407615 |
| | -2.795401385 | 0.005183531 | 0.012705032 |
| | -0.058572198 | 0.953292855 | 0.96810647 |
| | -1.690490411 | 0.090934171 | 0.153903733 |
| | 1.403882484 | 0.160353846 | 0.246357524 |
| | 1.108144846 | 0.267799266 | 0.373718534 |
| | -5.105054789 | 3.31E-07 | 1.66E-06 |
| | 13.42856113 | 4.11E-41 | 2.32E-39 |
| NA | | NA | NA |
| | 2.011685327 | 0.044253118 | 0.083307427 |
| | 2.861760531 | 0.004212951 | 0.010534482 |
| | -3.329055583 | 0.00087141 | 0.002541528 |
| | 3.440925994 | 0.000579727 | 0.001747209 |
| NA | | NA | NA |
| | 0.175194766 | 0.860926597 | 0.903279692 |
| | 4.664552714 | 3.09E-06 | 1.35E-05 |
| | 2.554318886 | 0.010639577 | 0.024102507 |
| | -0.979677516 | 0.327245328 | 0.438291956 |
| | 12.38659549 | 3.09E-35 | 1.36E-33 |
| | 3.945214087 | 7.97E-05 | 0.000280795 |
| | 13.18560705 | 1.06E-39 | 5.75E-38 |
| | -1.830156295 | 0.067226571 | 0.119102567 |
| | -6.884469349 | 5.80E-12 | 4.97E-11 |
| | 0.488246986 | 0.625374908 | NA |
| | -0.175180252 | 0.860938001 | 0.903279692 |
| | 0.214084176 | 0.830481416 | 0.881400519 |
| | 1.836805876 | 0.066238558 | 0.117618552 |
| | 5.821937691 | 5.82E-09 | 3.65E-08 |
| NA | | NA | NA |
| | -0.406116161 | 0.684657252 | 0.766995077 |
| | 4.190485193 | 2.78E-05 | 0.000105886 |
| | 1.750840402 | 0.079973405 | 0.138060165 |
| | -6.498715388 | 8.10E-11 | 6.22E-10 |
| | 0.807535724 | 0.419357901 | 0.531039576 |
| | 6.313496749 | 2.73E-10 | 1.97E-09 |
| | 0.665897037 | 0.505476915 | 0.611951012 |
| | 0.423241377 | 0.672119161 | 0.756198256 |
| | 2.018240786 | 0.04356619 | 0.082187349 |
| | -1.984893025 | 0.047156375 | 0.088004883 |

| | | |
|---|---|---|
| 1.657714998 | 0.097375005 | 0.162953559 |
| -0.327681932 | 0.743152166 | 0.813087473 |
| 7.805952213 | 5.91E-15 | 6.88E-14 |
| 2.183151197 | 0.029024677 | 0.057687 |
| -0.325537711 | 0.744774146 | 0.814296182 |
| 1.394604206 | 0.163135226 | 0.249989119 |
| 6.852717372 | 7.25E-12 | 6.13E-11 |
| 0.317642674 | 0.750756 | NA |
| 3.59437198 | 0.000325175 | 0.001027279 |
| 15.36476815 | 2.82E-53 | 2.33E-51 |
| -0.906312501 | 0.364770471 | 0.477246609 |
| -1.16216847 | 0.24516704 | 0.348053485 |
| -2.770202283 | 0.005602149 | 0.013619055 |
| 3.251972122 | 0.001146073 | 0.003262019 |
| -1.515319867 | 0.12969142 | 0.2070149 |
| 1.889625438 | 0.058808073 | 0.106302163 |
| 0.462704079 | 0.643576494 | 0.733347886 |
| -0.273198492 | 0.784700639 | NA |
| 4.639689563 | 3.49E-06 | 1.52E-05 |
| 1.332630648 | 0.182653042 | 0.274506099 |
| 3.312210771 | 0.000925618 | 0.002684925 |
| -2.047915128 | 0.040568316 | 0.077148226 |
| NA | NA | NA |
| 0.290228139 | 0.771641711 | NA |
| 1.084258132 | 0.278250363 | 0.385210542 |
| 2.153010912 | 0.031317821 | 0.061635875 |
| -3.543580873 | 0.000394732 | 0.001227043 |
| -3.658909441 | 0.000253291 | 0.000815627 |
| -1.274243619 | 0.202577091 | 0.298862701 |
| -1.411558061 | 0.158080128 | 0.243681904 |
| -2.770211146 | 0.005601996 | 0.013619055 |
| -2.063501063 | 0.039065047 | 0.074623989 |
| 7.302747551 | 2.82E-13 | 2.77E-12 |
| -9.24676334 | 2.31E-20 | 4.20E-19 |
| -2.078298042 | 0.037681917 | 0.072362926 |
| -2.52999547 | 0.0114064 | 0.025628563 |
| -2.638183282 | 0.008335152 | 0.019333598 |
| -0.210473424 | 0.833298191 | 0.883229038 |
| 3.18092271 | 0.001468068 | 0.004074625 |
| NA | NA | NA |
| 11.633231 | 2.79E-31 | 1.03E-29 |
| 0.721459632 | 0.470626769 | 0.579713977 |
| 2.595866368 | 0.009435275 | 0.021621458 |
| 1.110583657 | 0.266747601 | NA |
| 0.063710673 | 0.949200606 | NA |
| 1.346398161 | 0.178174144 | 0.268899537 |
| -0.628117388 | 0.529927043 | 0.634857316 |

| | | |
|---|---|---|
| -1.102515974 | 0.270237418 | 0.376449386 |
| 1.436134539 | 0.150964062 | 0.234755869 |
| 1.352261923 | 0.176291541 | 0.266620982 |
| -0.382364267 | 0.702191191 | 0.781364766 |
| 1.695112145 | 0.090054149 | 0.152693086 |
| NA | NA | NA |
| -4.408647109 | 1.04E-05 | 4.23E-05 |
| NA | NA | NA |
| -0.433202016 | 0.664868014 | 0.750942936 |
| NA | NA | NA |
| NA | NA | NA |
| -2.293629834 | 0.02181177 | 0.045041548 |
| -2.273141315 | 0.023017665 | 0.047233854 |
| 3.023743689 | 0.002496679 | 0.006565651 |
| 2.881846288 | 0.003953525 | 0.009952403 |
| 0.785959964 | 0.431890943 | NA |
| -7.034122586 | 2.01E-12 | 1.79E-11 |
| -1.311348397 | 0.189740084 | 0.283162074 |
| NA | NA | NA |
| -0.261996244 | 0.793324338 | NA |
| 1.981728575 | 0.047509627 | 0.088585069 |
| NA | NA | NA |
| NA | NA | NA |
| 0.021657585 | 0.982721098 | 0.98785177 |
| 1.826624165 | 0.067756298 | 0.119862897 |
| -0.745239528 | 0.456126931 | 0.565863257 |
| 0.157375728 | 0.874948742 | 0.912726807 |
| -2.769218288 | 0.005619097 | 0.013647003 |
| 3.39284785 | 0.0006917 | 0.002054987 |
| NA | NA | NA |
| NA | NA | NA |
| NA | NA | NA |
| -9.840951529 | 7.50E-23 | 1.64E-21 |
| 0.084676542 | 0.932518546 | 0.95413233 |
| 0.209960823 | 0.83369825 | NA |
| -0.402057241 | 0.6876419 | 0.769512083 |
| -1.574087768 | 0.11546716 | 0.187683812 |
| 1.216099639 | 0.223946968 | 0.32376784 |
| -2.809657674 | 0.004959422 | 0.012219149 |
| -2.92728363 | 0.003419369 | 0.00872 |
| 5.624084748 | 1.86E-08 | 1.10E-07 |
| 0.192810118 | 0.847107679 | NA |
| -1.263644225 | 0.206357753 | 0.303224994 |
| 1.115421675 | 0.264669765 | 0.370280265 |
| -8.829041757 | 1.06E-18 | 1.70E-17 |
| -3.706663988 | 0.000210007 | 0.000687391 |
| -2.112940879 | 0.034605828 | 0.067255523 |

| | | |
|---|---|---|
| -10.00485156 | 1.45E-23 | 3.37E-22 |
| 1.530598557 | 0.125868639 | 0.202019367 |
| -4.933648039 | 8.07E-07 | 3.84E-06 |
| NA | NA | NA |
| NA | NA | NA |
| 2.115019411 | 0.034428293 | 0.066967685 |
| -2.152290849 | 0.031374454 | 0.061727919 |
| 0.780104018 | 0.435329652 | 0.545879524 |
| -5.206426807 | 1.93E-07 | 9.98E-07 |
| NA | NA | NA |
| 4.059105906 | 4.93E-05 | 0.000179686 |
| 2.039478757 | 0.041402269 | 0.07854914 |
| -6.811488223 | 9.66E-12 | 8.10E-11 |
| -1.765069784 | 0.077552038 | 0.134473676 |
| -5.610464925 | 2.02E-08 | 1.19E-07 |
| -0.021798894 | 0.982608376 | 0.987817803 |
| 0.591974618 | 0.553867585 | NA |
| 3.024563875 | 0.002489919 | 0.006550627 |
| 0.327819613 | 0.743048057 | 0.813065012 |
| -1.382424304 | 0.166841458 | 0.254667746 |
| -10.81538547 | 2.91E-27 | 8.56E-26 |
| 1.412551982 | 0.157787496 | 0.243325798 |
| 1.812438635 | 0.069918448 | 0.123139512 |
| 10.87948964 | 1.44E-27 | 4.37E-26 |
| 1.156933353 | 0.247299591 | 0.350425088 |
| 1.131460023 | 0.257861521 | 0.362558918 |
| 2.304225356 | 0.021209997 | 0.043954812 |
| 0.214588724 | 0.830087986 | NA |
| 2.576075969 | 0.009992869 | 0.022775588 |
| 2.152931282 | 0.03132408 | 0.061643346 |
| NA | NA | NA |
| 11.93074301 | 8.18E-33 | 3.27E-31 |
| NA | NA | NA |
| NA | NA | NA |
| -1.103567877 | 0.269780632 | 0.375938605 |
| NA | NA | NA |
| 0.192810118 | 0.847107679 | NA |
| NA | NA | NA |
| 1.156563035 | 0.247450932 | 0.35061969 |
| 4.421569125 | 9.80E-06 | 4.00E-05 |
| NA | NA | NA |
| NA | NA | NA |
| NA | NA | NA |
| -0.47588832 | 0.634153944 | NA |
| NA | NA | NA |
| NA | NA | NA |
| -5.716846011 | 1.09E-08 | 6.61E-08 |

| | | |
|---|---|---|
| -4.800727202 | 1.58E-06 | 7.21E-06 |
| -4.585161176 | 4.54E-06 | 1.94E-05 |
| -0.065389369 | 0.947863988 | NA |
| -0.323432628 | 0.746367623 | 0.815575902 |
| -1.749578401 | 0.080191089 | 0.138372893 |
| 6.464058521 | 1.02E-10 | 7.75E-10 |
| -0.759421827 | 0.447600261 | 0.557506109 |
| 3.197389116 | 0.001386777 | 0.003869453 |
| NA | NA | NA |
| 8.749687199 | 2.14E-18 | 3.34E-17 |
| 9.807170241 | 1.05E-22 | 2.28E-21 |
| 2.293952287 | 0.02179324 | 0.045010709 |
| -0.816287896 | 0.414335495 | 0.525956864 |
| 2.937738917 | 0.003306153 | 0.008457994 |
| NA | NA | NA |
| -0.065389369 | 0.947863988 | NA |
| -4.000068223 | 6.33E-05 | 0.000226559 |
| -1.924944911 | 0.05423625 | 0.099271177 |
| NA | NA | NA |
| NA | NA | NA |
| -5.082211183 | 3.73E-07 | 1.86E-06 |
| 2.190715561 | 0.028472382 | 0.056754299 |
| -1.878648872 | 0.060292453 | 0.108483197 |
| -1.459434166 | 0.144445648 | 0.226267903 |
| NA | NA | NA |
| NA | NA | NA |
| NA | NA | NA |
| NA | NA | NA |
| NA | NA | NA |
| 0.133832677 | 0.893534887 | NA |
| NA | NA | NA |
| NA | NA | NA |
| NA | NA | NA |
| -0.963760314 | 0.335166108 | 0.446376911 |
| NA | NA | NA |
| -0.065389369 | 0.947863988 | NA |
| -0.498412414 | 0.618193389 | 0.712927595 |
| NA | NA | NA |
| NA | NA | NA |
| NA | NA | NA |
| 0.192810118 | 0.847107679 | NA |
| NA | NA | NA |
| NA | NA | NA |
| NA | NA | NA |
| NA | NA | NA |
| NA | NA | NA |
| 0.3474247 | 0.728272279 | NA |

| | | | |
|---|---|---|---|
| | 0.788844532 | 0.430202878 | NA |
| | 1.450534576 | 0.146909504 | NA |
| | 0.320071032 | 0.748914485 | NA |
| | 2.305048318 | 0.021163867 | 0.04387738 |
| | 7.072206833 | 1.52E-12 | 1.39E-11 |
| | -0.527884342 | 0.597579613 | NA |
| | 0.06422648 | 0.948789893 | 0.965010012 |
| | 2.073208158 | 0.038152911 | 0.073115946 |
| | -4.924680529 | 8.45E-07 | 4.00E-06 |
| | -7.747837115 | 9.35E-15 | 1.07E-13 |
| | 0.324874675 | 0.745275925 | NA |
| NA | NA | NA | |
| NA | NA | NA | |
| | -4.770028697 | 1.84E-06 | 8.34E-06 |
| | 10.24532399 | 1.24E-24 | 3.09E-23 |
| | 3.928060488 | 8.56E-05 | 0.00029982 |
| | -2.464440194 | 0.013722746 | 0.03014572 |
| | 2.26992855 | 0.023211919 | 0.047579586 |
| | 8.639870308 | 5.63E-18 | 8.45E-17 |
| | 1.099497963 | 0.271550922 | 0.377921578 |
| | -0.746381654 | 0.455436899 | 0.565175289 |
| | 8.398662052 | 4.52E-17 | 6.33E-16 |
| | 5.02535429 | 5.03E-07 | 2.46E-06 |
| | -6.172049636 | 6.74E-10 | 4.66E-09 |
| | 0.094658805 | 0.92458584 | 0.948190794 |
| | 0.769418144 | 0.441645121 | 0.551670695 |
| | 11.37949689 | 5.29E-30 | 1.82E-28 |
| | -0.114429902 | 0.908897011 | NA |
| NA | NA | NA | |
| | 2.433434364 | 0.014956347 | 0.032490658 |
| NA | NA | NA | |
| NA | NA | NA | |
| | 0.063708882 | 0.949202033 | NA |
| | 0.192809521 | 0.847108147 | NA |
| NA | NA | NA | |
| | 0.085254203 | 0.932059299 | NA |
| | 0.06371127 | 0.949200131 | NA |
| | -0.104608774 | 0.916686252 | NA |
| NA | NA | NA | |
| NA | NA | NA | |
| | -0.19448762 | 0.845794089 | NA |
| NA | NA | NA | |
| | 0.063708882 | 0.949202033 | NA |
| NA | NA | NA | |
| NA | NA | NA | |
| NA | NA | NA | |
| NA | NA | NA | |

| | | |
|---|---|---|
| NA | NA | NA |
| NA | NA | NA |
| NA | NA | NA |
| NA | NA | NA |
| NA | NA | NA |
| NA | NA | NA |
| -0.065389369 | 0.947863988 | NA |
| NA | NA | NA |
| 0.063708882 | 0.949202033 | NA |
| NA | NA | NA |
| -0.065388772 | 0.947864464 | NA |
| 0.06371127 | 0.949200131 | NA |
| NA | NA | NA |
| NA | NA | NA |
| NA | NA | NA |
| NA | NA | NA |
| NA | NA | NA |
| NA | NA | NA |
| NA | NA | NA |
| NA | NA | NA |
| NA | NA | NA |
| NA | NA | NA |
| NA | NA | NA |
| NA | NA | NA |
| NA | NA | NA |
| NA | NA | NA |
| NA | NA | NA |
| NA | NA | NA |
| 18.22846963 | 3.07E-74 | 4.36E-72 |
| 5.242293278 | 1.59E-07 | 8.32E-07 |
| 1.8180959 | 0.069049477 | 0.121823388 |
| 0.770799995 | 0.44082549 | 0.550949812 |
| 0.08925156 | 0.92888199 | NA |
| -1.00655104 | 0.314150572 | NA |
| 0.111224404 | 0.9114384 | NA |
| 0.063707688 | 0.949202983 | NA |
| NA | NA | NA |
| NA | NA | NA |
| NA | NA | NA |
| NA | NA | NA |
| NA | NA | NA |
| NA | NA | NA |
| NA | NA | NA |
| NA | NA | NA |
| NA | NA | NA |
| NA | NA | NA |
| NA | NA | NA |

| | | |
|---|---|---|
| NA | NA | NA |
| NA | NA | NA |
| NA | NA | NA |
| NA | NA | NA |
| NA | NA | NA |
| NA | NA | NA |
| NA | NA | NA |
| NA | NA | NA |
| NA | NA | NA |
| 0.192809521 | 0.847108147 | NA |
| NA | NA | NA |
| NA | NA | NA |
| NA | NA | NA |
| 0.063707688 | 0.949202983 | NA |
| 1.187611475 | 0.234986515 | 0.336443912 |
| NA | NA | NA |
| NA | NA | NA |
| NA | NA | NA |
| NA | NA | NA |
| -0.065388772 | 0.947864464 | NA |
| NA | NA | NA |
| NA | NA | NA |
| NA | NA | NA |
| NA | NA | NA |
| 0.192809521 | 0.847108147 | NA |
| NA | NA | NA |
| NA | NA | NA |
| NA | NA | NA |
| NA | NA | NA |
| NA | NA | NA |
| 0.192809521 | 0.847108147 | NA |
| NA | NA | NA |
| 0.192636473 | 0.847243678 | NA |
| NA | NA | NA |
| 0.192809521 | 0.847108147 | NA |
| 0.063708381 | 0.949202431 | NA |
| NA | NA | NA |
| 0.192809521 | 0.847108147 | NA |
| NA | NA | NA |
| NA | NA | NA |
| 0.063708882 | 0.949202033 | NA |
| 0.063708381 | 0.949202431 | NA |
| NA | NA | NA |
| NA | NA | NA |
| 3.006946782 | 0.00263886 | 0.006904731 |
| -11.15494839 | 6.77E-29 | 2.22E-27 |
| -5.448687733 | 5.07E-08 | 2.85E-07 |

| | | |
|---|---|---|
| 0.017817424 | 0.985784505 | 0.989937258 |
| NA | NA | NA |
| NA | NA | NA |
| -0.324372546 | 0.745656002 | 0.814960834 |
| 0.231812047 | 0.816684 | NA |
| 1.999447525 | 0.045559954 | 0.085427047 |
| 4.944890536 | 7.62E-07 | 3.64E-06 |
| NA | NA | NA |
| NA | NA | NA |
| NA | NA | NA |
| 0.758752659 | 0.44800053 | 0.557966525 |
| 1.492820695 | 0.135484129 | 0.214535822 |
| 2.151452792 | 0.031440478 | 0.061848096 |
| -0.231210379 | 0.817151367 | 0.871317147 |
| -1.894774821 | 0.05812224 | 0.105244858 |
| NA | NA | NA |
| -0.745874274 | 0.455743367 | 0.565471495 |
| -1.373955556 | 0.16945548 | 0.257869474 |
| 0.068152451 | 0.945664278 | 0.962975181 |
| 0.173521862 | 0.862241245 | 0.904294744 |
| NA | NA | NA |
| -1.271556769 | 0.203530645 | 0.299898278 |
| -4.951562522 | 7.36E-07 | 3.52E-06 |
| NA | NA | NA |
| -0.1425123 | 0.886675364 | 0.920547694 |
| NA | NA | NA |
| 3.647873314 | 0.00026442 | 0.000848626 |
| -4.538573045 | 5.66E-06 | 2.39E-05 |
| NA | NA | NA |
| 0.19280579 | 0.847111069 | NA |
| -5.227476875 | 1.72E-07 | 8.97E-07 |
| NA | NA | NA |
| -2.649985037 | 0.008049534 | 0.018730113 |
| -1.587925051 | 0.112303288 | 0.183281048 |
| -4.120292976 | 3.78E-05 | 0.000140667 |
| 2.03538755 | 0.041811893 | 0.079254198 |
| 10.94200562 | 7.26E-28 | 2.25E-26 |
| 0.900856714 | 0.367664508 | 0.480104479 |
| 2.261919361 | 0.023702386 | 0.048438171 |
| NA | NA | NA |
| -5.036974892 | 4.73E-07 | 2.32E-06 |
| 0.063708285 | 0.949202508 | NA |
| -0.065388772 | 0.947864464 | NA |
| -3.865759563 | 0.000110744 | 0.000380132 |
| -0.595466225 | 0.551531869 | 0.6547855 |
| -4.397452166 | 1.10E-05 | 4.44E-05 |
| -3.634968421 | 0.000278015 | 0.000889178 |

| | | |
|---|---|---|
| NA | NA | NA |
| -1.492278141 | 0.135626245 | 0.214720108 |
| 1.052353155 | 0.292637553 | 0.40054235 |
| -1.42767888 | 0.153384308 | 0.237751376 |
| 2.498701711 | 0.012464918 | 0.027691213 |
| 0.847956514 | 0.396462189 | 0.508400596 |
| -3.853262971 | 0.000116554 | 0.000398273 |
| -0.752974783 | 0.451465067 | 0.561387582 |
| 3.250271037 | 0.001152951 | 0.003279359 |
| -8.558476858 | 1.14E-17 | 1.67E-16 |
| 0.553089131 | 0.580202379 | 0.68100585 |
| -5.155780217 | 2.53E-07 | 1.29E-06 |
| 2.904581046 | 0.00367745 | 0.009307034 |
| 2.362273408 | 0.018163241 | 0.038452671 |
| -1.589268048 | 0.111999889 | 0.18289126 |
| 1.668442971 | 0.095227824 | 0.159969962 |
| 3.579878973 | 0.000343753 | 0.001081738 |
| -0.546728638 | 0.584565176 | NA |
| 2.039319441 | 0.041418157 | 0.078573326 |
| 6.886064961 | 5.74E-12 | 4.92E-11 |
| NA | NA | NA |
| -3.409596757 | 0.00065059 | 0.001942071 |
| -0.743872176 | 0.45695381 | NA |
| 2.96043724 | 0.003072027 | 0.007920564 |
| 3.417557384 | 0.000631858 | 0.001890663 |
| -1.884324087 | 0.059521151 | 0.107350801 |
| 0.348508338 | 0.727458453 | NA |
| -0.065389369 | 0.947863988 | NA |
| 0.780536617 | 0.435075083 | 0.545587642 |
| -3.798306625 | 0.000145688 | 0.000489144 |
| -4.518379993 | 6.23E-06 | 2.62E-05 |
| -2.477326759 | 0.013237065 | 0.029199237 |
| 4.369128082 | 1.25E-05 | 5.00E-05 |
| 2.021608661 | 0.043216797 | 0.081571256 |
| -0.065389369 | 0.947863988 | NA |
| -1.345366238 | 0.178506993 | 0.269271984 |
| NA | NA | NA |
| 5.728676182 | 1.01E-08 | 6.18E-08 |
| -3.087757464 | 0.00201673 | 0.005431911 |
| -9.390155453 | 5.99E-21 | 1.14E-19 |
| 0.267238816 | 0.789285291 | 0.850041343 |
| -3.682987401 | 0.000230517 | 0.000748551 |
| NA | NA | NA |
| -0.360940375 | 0.718144021 | 0.793990231 |
| 2.578187295 | 0.009932015 | 0.022651329 |
| 8.169855712 | 3.09E-16 | 4.02E-15 |
| 8.968706388 | 3.00E-19 | 5.06E-18 |

| | | |
|---|---|---|
| 10.39143648 | 2.71E-25 | 7.06E-24 |
| 5.164288966 | 2.41E-07 | 1.24E-06 |
| 2.9437078 | 0.00324306 | 0.008316136 |
| 13.21575747 | 7.12E-40 | 3.89E-38 |
| -2.043207746 | 0.041031877 | 0.077905476 |
| -7.112929822 | 1.14E-12 | 1.05E-11 |
| -10.13422142 | 3.89E-24 | 9.41E-23 |
| 0.879492687 | 0.379134197 | 0.491783991 |
| -0.068734831 | 0.945200693 | NA |
| -1.732730781 | 0.083143531 | 0.14273543 |
| 17.59890863 | 2.51E-69 | 3.16E-67 |
| -0.065388772 | 0.947864464 | NA |
| 0.54199435 | 0.587822398 | 0.687630938 |
| NA | NA | NA |
| 1.543994043 | 0.122589776 | NA |
| -2.321670811 | 0.020250668 | 0.04225008 |
| -0.50522435 | 0.613401269 | 0.709067278 |
| -1.601740284 | 0.109213053 | 0.17904131 |
| -6.691850076 | 2.20E-11 | 1.79E-10 |
| NA | NA | NA |
| -2.315655058 | 0.02057711 | 0.04283119 |
| 0.063707135 | 0.949203424 | NA |
| -3.085545836 | 0.002031789 | 0.005467177 |
| -1.937888081 | 0.052636878 | 0.096768349 |
| -5.549886335 | 2.86E-08 | 1.65E-07 |
| -0.118256046 | 0.905864783 | 0.934809133 |
| 4.91682022 | 8.80E-07 | 4.16E-06 |
| 9.124252193 | 7.22E-20 | 1.28E-18 |
| 0.327635741 | 0.743187095 | 0.813087473 |
| -0.295652323 | 0.767495611 | 0.832412477 |
| 0.252413502 | 0.800721465 | 0.858479903 |
| 5.508830703 | 3.61E-08 | 2.07E-07 |
| 0.442110024 | 0.658409591 | 0.745869748 |
| 2.249018911 | 0.024511293 | 0.049834917 |
| 0.297924648 | 0.765760675 | NA |
| NA | NA | NA |
| 0.979127604 | 0.327516934 | NA |
| 5.532400232 | 3.16E-08 | 1.82E-07 |
| 3.046529538 | 0.002314997 | 0.006141429 |
| -0.24662222 | 0.805200606 | NA |
| 1.650927417 | 0.098753397 | 0.164885432 |
| -0.315442982 | 0.752425336 | NA |
| NA | NA | NA |
| -0.693495488 | 0.487998656 | 0.596008899 |
| 0.063708882 | 0.949202033 | NA |
| 0.063708285 | 0.949202508 | NA |
| -2.159068173 | 0.030844878 | 0.060800701 |

| | | |
|---|---|---|
| 2.63121774 | 0.008507951 | 0.019705192 |
| 0.205628504 | 0.83708111 | 0.886037983 |
| 0.890045163 | 0.373441636 | 0.486241384 |
| 6.792186966 | 1.10E-11 | 9.22E-11 |
| 9.496397633 | 2.17E-21 | 4.29E-20 |
| 1.160799043 | 0.245723632 | 0.348724982 |
| NA | NA | NA |
| 5.321073746 | 1.03E-07 | 5.55E-07 |
| -1.662597037 | 0.096393125 | 0.161580492 |
| 0.01721168 | 0.986267744 | 0.990303339 |
| 0.192809521 | 0.847108147 | NA |
| -5.495168133 | 3.90E-08 | 2.22E-07 |
| NA | NA | NA |
| NA | NA | NA |
| 1.742247146 | 0.081465208 | 0.14025823 |
| -0.778056822 | 0.436535516 | 0.546922447 |
| 0.063708285 | 0.949202508 | NA |
| -4.410049829 | 1.03E-05 | 4.21E-05 |
| 0.434440473 | 0.663968613 | NA |
| 0.729212924 | 0.465871427 | 0.57515842 |
| -6.521789343 | 6.95E-11 | 5.37E-10 |
| -0.933473016 | 0.350575797 | 0.462307203 |
| NA | NA | NA |
| 0.260008687 | 0.794857073 | NA |
| 1.975989076 | 0.048156014 | 0.089603408 |
| -1.38769258 | 0.165230677 | 0.252644988 |
| 0.590559841 | 0.554815379 | 0.657654335 |
| -0.699828471 | 0.484034432 | 0.592499832 |
| -0.13945826 | 0.889088038 | 0.922440041 |
| -0.065388772 | 0.947864464 | NA |
| 4.764465753 | 1.89E-06 | 8.55E-06 |
| 2.648326166 | 0.008089144 | 0.018806546 |
| -0.048077979 | 0.961654096 | 0.973674286 |
| 2.524135228 | 0.01159833 | 0.026003781 |
| -0.107032463 | 0.914763226 | 0.941363699 |
| -4.12581154 | 3.69E-05 | 0.00013754 |
| -9.747936532 | 1.88E-22 | 4.01E-21 |
| 0.67460242 | 0.499928395 | 0.606936399 |
| 3.261411204 | 0.001108591 | 0.003164695 |
| 5.856627324 | 4.72E-09 | 2.99E-08 |
| -0.210123171 | 0.833571541 | 0.883443946 |
| -0.6851714 | 0.493235769 | 0.600429236 |
| -0.002835505 | 0.997737597 | 0.998335881 |
| 8.228730533 | 1.89E-16 | 2.52E-15 |
| -0.409318902 | 0.682305646 | NA |
| -1.251115748 | 0.210892251 | 0.30854764 |
| 7.67315821 | 1.68E-14 | 1.87E-13 |

| | | |
|---|---|---|
| 2.371483423 | 0.017716841 | 0.037682915 |
| NA | NA | NA |
| NA | NA | NA |
| 0.771371662 | 0.440486667 | NA |
| 3.998851799 | 6.37E-05 | 0.000227564 |
| 0.191391878 | 0.848218583 | 0.894046032 |
| -2.304399148 | 0.021200248 | 0.043941885 |
| -0.112318627 | 0.910570773 | NA |
| -1.05878521 | 0.289697611 | 0.397169918 |
| -8.387905307 | 4.95E-17 | 6.90E-16 |
| -0.655616043 | 0.512071201 | 0.61775314 |
| 3.127001328 | 0.001765991 | 0.004817215 |
| 1.030514282 | 0.302768651 | 0.412043244 |
| 5.584896949 | 2.34E-08 | 1.37E-07 |
| NA | NA | NA |
| 5.09148929 | 3.55E-07 | 1.78E-06 |
| 0.775954344 | 0.437775944 | NA |
| NA | NA | NA |
| -10.68864314 | 1.15E-26 | 3.28E-25 |
| NA | NA | NA |
| 0.46888587 | 0.639151217 | 0.729933611 |
| NA | NA | NA |
| -3.14505429 | 0.00166056 | 0.004559298 |
| 0.145502495 | 0.884314146 | 0.918978042 |
| 0.192809521 | 0.847108147 | NA |
| 1.516598833 | 0.129368 | 0.206589323 |
| 7.858406448 | 3.89E-15 | 4.59E-14 |
| -1.193771842 | 0.232567229 | 0.333552472 |
| -2.631444483 | 0.008502276 | 0.01969387 |
| 1.159312864 | 0.24632868 | 0.349345961 |
| NA | NA | NA |
| -1.443437851 | 0.148897168 | 0.231988554 |
| 3.676594062 | 0.000236369 | 0.000765964 |
| 0.643807711 | 0.519700133 | 0.624998526 |
| 0.758267224 | 0.448291025 | 0.558272767 |
| 0.692710438 | 0.488491288 | 0.596377765 |
| -0.44916851 | 0.653310102 | NA |
| -11.19574966 | 4.28E-29 | 1.41E-27 |
| 2.991584242 | 0.00277534 | 0.007230095 |
| -0.446641606 | 0.65513385 | 0.743021888 |
| 0.192810118 | 0.847107679 | NA |
| 8.218751 | 2.06E-16 | 2.73E-15 |
| -7.878535393 | 3.31E-15 | 3.94E-14 |
| NA | NA | NA |
| 1.093268954 | 0.27427573 | 0.38099348 |
| -3.761256853 | 0.000169062 | 0.00056152 |
| -3.897246885 | 9.73E-05 | 0.000337149 |

| | | |
|---|---|---|
| -0.415093447 | 0.678073497 | 0.761084288 |
| 1.453948062 | 0.145960706 | 0.22818966 |
| -0.708396394 | 0.478699131 | NA |
| 3.187505312 | 0.001435058 | 0.003992831 |
| -2.211759277 | 0.026983302 | 0.054135304 |
| -1.854369559 | 0.063686307 | 0.113650952 |
| -2.545266531 | 0.010919436 | 0.024647212 |
| 0.063708324 | 0.949202477 | NA |
| 8.775366833 | 1.70E-18 | 2.69E-17 |
| -2.580732366 | 0.009859098 | 0.02249528 |
| -1.599000594 | 0.109820471 | 0.179895706 |
| 0.99585868 | 0.319318814 | 0.430007528 |
| -2.220184877 | 0.02640622 | 0.053135115 |
| 1.119524625 | 0.262916391 | 0.368280109 |
| 6.741262674 | 1.57E-11 | 1.29E-10 |
| NA | NA | NA |
| -3.513275614 | 0.000442618 | 0.001363368 |
| 2.503318937 | 0.012303462 | 0.027368969 |
| 0.353203278 | 0.723936051 | NA |
| NA | NA | NA |
| -6.885444749 | 5.76E-12 | 4.94E-11 |
| -1.538902078 | 0.123828204 | 0.199277259 |
| 0.192810118 | 0.847107679 | NA |
| -3.488648867 | 0.000485468 | 0.001483853 |
| 15.01444467 | 5.91E-51 | 4.60E-49 |
| -0.776097149 | 0.437691627 | 0.547962858 |
| 1.810917761 | 0.070153583 | 0.123484119 |
| 1.131872982 | 0.257687839 | 0.362375786 |
| -2.437222483 | 0.01480057 | 0.032188571 |
| 7.153143645 | 8.48E-13 | 7.94E-12 |
| 4.448931858 | 8.63E-06 | 3.55E-05 |
| -2.523921103 | 0.011605397 | 0.026012635 |
| -3.018155845 | 0.002543181 | 0.006678818 |
| -0.109757947 | 0.912601343 | NA |
| -3.974215646 | 7.06E-05 | 0.000250838 |
| -0.11056269 | 0.911963135 | 0.939351502 |
| 5.48813657 | 4.06E-08 | 2.31E-07 |
| 7.903356134 | 2.71E-15 | 3.26E-14 |
| 1.912392943 | 0.055825813 | 0.101756488 |
| 1.629673802 | 0.103170457 | 0.170869883 |
| 5.136125801 | 2.80E-07 | 1.43E-06 |
| -0.62685192 | 0.530756306 | 0.635485786 |
| 1.123182203 | 0.261360119 | 0.366524179 |
| 0.290961341 | 0.771080886 | 0.835142994 |
| -0.864120146 | 0.387521893 | 0.499771724 |
| -4.715564307 | 2.41E-06 | 1.07E-05 |
| -1.64987882 | 0.098967724 | 0.165191739 |

| | | |
|---|---|---|
| 0.189661132 | 0.84957468 | 0.895173827 |
| 0.063708285 | 0.949202508 | NA |
| 0.063707135 | 0.949203424 | NA |
| 9.75180091 | 1.81E-22 | 3.87E-21 |
| 1.597833009 | 0.110080146 | 0.180273885 |
| 0.950611711 | 0.341801522 | 0.453236495 |
| -7.686752668 | 1.51E-14 | 1.69E-13 |
| 4.230545225 | 2.33E-05 | 8.96E-05 |
| -2.933475829 | 0.003351898 | 0.008561894 |
| 0.800728516 | 0.42328883 | 0.534934839 |
| -0.699643329 | 0.484150076 | 0.592609732 |
| 0.965202465 | 0.334443413 | 0.445627821 |
| -2.039856772 | 0.041364593 | 0.078489559 |
| 9.527466399 | 1.61E-21 | 3.24E-20 |
| -2.386319233 | 0.017017971 | 0.036391271 |
| -4.949137804 | 7.45E-07 | 3.56E-06 |
| 3.63412659 | 0.000278924 | 0.000891858 |
| 6.921588775 | 4.47E-12 | 3.87E-11 |
| -0.318218677 | 0.750319066 | 0.818749944 |
| -0.033382653 | 0.973369443 | 0.981588471 |
| 0.747101167 | 0.455002496 | 0.564779375 |
| 0.260207735 | 0.794703538 | 0.854029229 |
| 0.246563337 | 0.805246181 | 0.862222447 |
| -2.770717326 | 0.005593296 | 0.013601496 |
| 0.284796675 | 0.77579992 | 0.839092213 |
| -2.812031186 | 0.004922973 | 0.0121401 |
| -5.455019908 | 4.90E-08 | 2.75E-07 |
| -5.23573447 | 1.64E-07 | 8.61E-07 |
| 1.416825874 | 0.156533843 | 0.241884312 |
| -0.267950475 | 0.78873744 | NA |
| 11.53281107 | 9.02E-31 | 3.24E-29 |
| NA | NA | NA |
| 4.829642638 | 1.37E-06 | 6.29E-06 |
| -0.145403309 | 0.884392452 | 0.918978042 |
| 2.338806325 | 0.019345459 | 0.040598377 |
| 19.99159715 | 6.52E-89 | 1.27E-86 |
| 0.063707688 | 0.949202983 | NA |
| 6.544908694 | 5.95E-11 | 4.64E-10 |
| 8.326127026 | 8.35E-17 | 1.14E-15 |
| NA | NA | NA |
| -0.19448762 | 0.845794089 | NA |
| 0.320071032 | 0.748914485 | NA |
| -5.883144006 | 4.03E-09 | 2.57E-08 |
| 4.293666173 | 1.76E-05 | 6.89E-05 |
| -1.031671499 | 0.302226033 | 0.411453912 |
| -9.288618389 | 1.56E-20 | 2.88E-19 |
| 1.823140182 | 0.068282162 | 0.120640963 |

| | | |
|---|---|---|
| NA | NA | NA |
| -1.080170286 | 0.280066358 | 0.387080769 |
| -1.629558988 | 0.103194738 | 0.170898789 |
| -0.707508993 | 0.479250226 | 0.587963589 |
| 8.878306499 | 6.79E-19 | 1.11E-17 |
| 0.874560789 | 0.381812931 | 0.494324608 |
| 5.513217797 | 3.52E-08 | 2.02E-07 |
| -0.990716408 | 0.321824077 | 0.432822344 |
| 2.483885934 | 0.012995743 | 0.028737825 |
| 5.49513122 | 3.90E-08 | 2.22E-07 |
| -1.088022886 | 0.276585005 | 0.383413971 |
| -4.106156053 | 4.02E-05 | 0.000148803 |
| 4.070867301 | 4.68E-05 | 0.000171273 |
| -6.395123621 | 1.60E-10 | 1.19E-09 |
| 0.094398948 | 0.924792251 | 0.948242598 |
| -6.274251433 | 3.51E-10 | 2.50E-09 |
| NA | NA | NA |
| 0.063708285 | 0.949202508 | NA |
| -0.494701951 | 0.620810524 | NA |
| 0.065548599 | 0.947737213 | NA |
| 0.063708285 | 0.949202508 | NA |
| 1.287496262 | 0.197921372 | 0.293107249 |
| -0.479902362 | 0.631296821 | NA |
| NA | NA | NA |
| -3.951049117 | 7.78E-05 | 0.000274615 |
| -0.396345974 | 0.691849816 | 0.772539392 |
| -2.769258475 | 0.005618404 | 0.013647003 |
| -0.381471592 | 0.702853347 | 0.781920053 |
| 10.85159835 | 1.96E-27 | 5.87E-26 |
| 1.47461516 | 0.140316037 | 0.220859666 |
| -0.444101165 | 0.656969444 | 0.744541574 |
| 6.949971678 | 3.65E-12 | 3.19E-11 |
| 4.723356774 | 2.32E-06 | 1.04E-05 |
| 3.184040229 | 0.001452348 | 0.004036451 |
| 1.613460323 | 0.106644537 | 0.175528973 |
| NA | NA | NA |
| -1.414875944 | 0.157104872 | 0.242586981 |
| NA | NA | NA |
| 1.516534991 | 0.129384129 | NA |
| 4.054953941 | 5.01E-05 | 0.000182721 |
| 0.656923336 | 0.511230213 | 0.617006231 |
| -4.175495359 | 2.97E-05 | 0.000112542 |
| NA | NA | NA |
| 2.437278245 | 0.014798288 | 0.032186404 |
| 5.083821898 | 3.70E-07 | 1.85E-06 |
| -0.065388772 | 0.947864464 | NA |
| 4.10841407 | 3.98E-05 | 0.000147443 |

| | | |
|---|---|---|
| 0.484341694 | 0.62814339 | 0.721080034 |
| -2.080810252 | 0.037451277 | 0.071975236 |
| -0.20215797 | 0.83979323 | 0.887695518 |
| NA | NA | NA |
| 1.658442311 | 0.097228223 | NA |
| 2.763515891 | 0.005718231 | 0.013863125 |
| -4.424177125 | 9.68E-06 | 3.96E-05 |
| -3.025352127 | 0.002483438 | 0.006537013 |
| -3.354665651 | 0.00079461 | 0.00233303 |
| -0.391087671 | 0.695732434 | 0.775914155 |
| -0.083858339 | 0.933169064 | 0.95464183 |
| 1.000077058 | 0.317273218 | 0.427782193 |
| 1.385462974 | 0.165910947 | 0.253448337 |
| 3.47290026 | 0.000514867 | 0.001566822 |
| 3.528175496 | 0.000418435 | 0.001293813 |
| NA | NA | NA |
| NA | NA | NA |
| 0.063708882 | 0.949202033 | NA |
| NA | NA | NA |
| -1.265164376 | 0.20581241 | 0.302512456 |
| -0.426224407 | 0.669944335 | 0.754701233 |
| 6.215671853 | 5.11E-10 | 3.59E-09 |
| -0.287157356 | 0.773991837 | 0.837534723 |
| 0.927550265 | 0.353640907 | 0.465532339 |
| -1.030967322 | 0.302556144 | 0.411776428 |
| -0.853602438 | 0.393325307 | 0.50525699 |
| 0.415429141 | 0.677827778 | 0.760876728 |
| -1.933727632 | 0.053146625 | 0.097526394 |
| 2.759936641 | 0.005781257 | 0.013989256 |
| NA | NA | NA |
| 0.321109739 | 0.74812723 | 0.816856769 |
| 2.367900506 | 0.017889346 | 0.037985267 |
| -0.476027656 | 0.634054675 | 0.725569813 |
| -2.066995755 | 0.038734554 | 0.074100114 |
| -1.399088506 | 0.161786444 | 0.248132028 |
| -0.16139596 | 0.871781548 | 0.910675354 |
| NA | NA | NA |
| -0.586220805 | 0.557727142 | 0.660199119 |
| NA | NA | NA |
| 1.464224251 | 0.143132686 | 0.224657373 |
| 4.089984514 | 4.31E-05 | 0.000158514 |
| 0.16189368 | 0.871389579 | 0.910341883 |
| -2.074211416 | 0.03805968 | 0.072981981 |
| -7.599387363 | 2.98E-14 | 3.23E-13 |
| -7.479269304 | 7.47E-14 | 7.79E-13 |
| -5.465926928 | 4.60E-08 | 2.60E-07 |
| 2.000486604 | 0.045447745 | 0.085235805 |

| | | | |
|---|---|---|---|
| | 2.550899196 | 0.01074454 | 0.024318277 |
| | -0.245667329 | 0.805939765 | 0.862632984 |
| | 1.28970973 | 0.19715146 | 0.292115139 |
| | 3.647999316 | 0.00026429 | 0.000848318 |
| NA | NA | NA | |
| | -2.175843016 | 0.029567001 | 0.058628063 |
| | -1.857171708 | 0.063286729 | 0.113058798 |
| | -0.548386832 | 0.583426316 | 0.683604564 |
| | -1.31422945 | 0.188769001 | 0.281931386 |
| | 0.192808327 | 0.847109082 | NA |
| | -2.09173877 | 0.03646189 | 0.070354696 |
| | 7.889573138 | 3.03E-15 | 3.62E-14 |
| | -5.282213397 | 1.28E-07 | 6.78E-07 |
| | 9.611630143 | 7.14E-22 | 1.47E-20 |
| | -0.630021026 | 0.528680828 | 0.633515948 |
| | 6.348102078 | 2.18E-10 | 1.59E-09 |
| | -2.139887044 | 0.032363896 | 0.063465084 |
| | -3.127121491 | 0.00176527 | 0.004815771 |
| | 5.385187088 | 7.24E-08 | 3.97E-07 |
| NA | NA | NA | |
| | 0.063708882 | 0.949202033 | NA |
| | 6.506127217 | 7.71E-11 | 5.93E-10 |
| NA | NA | NA | |
| | -2.869809353 | 0.004107194 | 0.010297812 |
| NA | NA | NA | |
| | 8.540518123 | 1.34E-17 | 1.95E-16 |
| | -0.213173958 | 0.83119128 | 0.881892151 |
| | 11.72898331 | 9.05E-32 | 3.44E-30 |
| | 5.369308883 | 7.90E-08 | 4.32E-07 |
| | 1.573064493 | 0.115703886 | 0.187995343 |
| | -4.575811637 | 4.74E-06 | 2.03E-05 |
| NA | NA | NA | |
| | 0.683441217 | 0.494328083 | 0.601436944 |
| | -3.050593762 | 0.002283894 | 0.006069851 |
| | -4.864725522 | 1.15E-06 | 5.33E-06 |
| NA | NA | NA | |
| | 2.480089063 | 0.013134957 | 0.029009792 |
| | -5.975375042 | 2.30E-09 | 1.50E-08 |
| | -1.964944256 | 0.04942069 | 0.091677156 |
| | -0.735833275 | 0.461832168 | 0.571157625 |
| | 0.050517196 | 0.959710246 | 0.972258271 |
| | -0.97310866 | 0.330499301 | 0.441429962 |
| | -0.802665073 | 0.422168345 | 0.533950161 |
| | 8.157475092 | 3.42E-16 | 4.43E-15 |
| | 8.942602438 | 3.80E-19 | 6.33E-18 |
| | -1.701109668 | 0.088922396 | 0.15109141 |
| NA | NA | NA | |

| | | | |
|---|---|---|---|
| | 0.368187532 | 0.712733407 | 0.789647537 |
| NA | NA | NA | |
| | -8.733324441 | 2.47E-18 | 3.84E-17 |
| | 0.141542698 | 0.887441232 | NA |
| | 0.284003059 | 0.776408036 | NA |
| | -0.259755806 | 0.795052143 | NA |
| | -6.182828831 | 6.30E-10 | 4.36E-09 |
| | 3.139796517 | 0.001690652 | 0.004631392 |
| | 1.79951543 | 0.071937185 | 0.126118074 |
| | -4.363431042 | 1.28E-05 | 5.12E-05 |
| NA | NA | NA | |
| NA | NA | NA | |
| NA | NA | NA | |
| | -0.065389369 | 0.947863988 | NA |
| | -3.475449784 | 0.000509997 | 0.001553137 |
| | -6.06393167 | 1.33E-09 | 8.92E-09 |
| | 0.369947695 | 0.711421463 | NA |
| | -1.791416395 | 0.0732265 | 0.127957225 |
| | 0.063710673 | 0.949200606 | NA |
| | 11.54778298 | 7.57E-31 | 2.73E-29 |
| | 1.376837304 | 0.168562555 | 0.256809346 |
| NA | NA | NA | |
| | -2.436459823 | 0.014831817 | 0.032245321 |
| | -3.550284612 | 0.000384815 | 0.001198298 |
| | 0.772326214 | 0.439921243 | 0.550121339 |
| | 6.64387798 | 3.06E-11 | 2.45E-10 |
| | -1.340613324 | 0.180046029 | 0.271201306 |
| | 0.231912591 | 0.816605905 | NA |
| | -9.903020068 | 4.04E-23 | 9.09E-22 |
| NA | NA | NA | |
| | -0.065389369 | 0.947863988 | NA |
| | 11.88164168 | 1.47E-32 | 5.84E-31 |
| | -0.19448762 | 0.845794089 | NA |
| | 2.700744107 | 0.006918455 | 0.016407895 |
| | 4.192829507 | 2.75E-05 | 0.000104893 |
| NA | NA | NA | |
| NA | NA | NA | |
| | -0.065389369 | 0.947863988 | NA |
| NA | NA | NA | |
| | -0.224327432 | 0.822502532 | 0.87556834 |
| | -2.235326922 | 0.025395894 | 0.05139963 |
| | 0.811587151 | 0.417028567 | 0.528731893 |
| | 2.654731292 | 0.007937161 | 0.018496148 |
| | 0.310067127 | 0.75650991 | 0.823493202 |
| NA | NA | NA | |
| | 2.924278732 | 0.003452554 | 0.008794773 |
| | 4.30542839 | 1.67E-05 | 6.55E-05 |

| | | |
|---|---|---|
| 10.92037823 | 9.21E-28 | 2.84E-26 |
| 3.356275852 | 0.000789997 | 0.002320576 |
| NA | NA | NA |
| 0.441738796 | 0.658678232 | 0.745974321 |
| -0.629460653 | 0.529047521 | 0.633864317 |
| NA | NA | NA |
| -1.568537785 | 0.116755679 | 0.189458324 |
| 5.324494072 | 1.01E-07 | 5.45E-07 |
| -3.346643787 | 0.000817962 | 0.002395133 |
| -3.033974829 | 0.002413546 | 0.006373836 |
| NA | NA | NA |
| -1.074424601 | 0.282632421 | 0.389596294 |
| NA | NA | NA |
| -2.380183951 | 0.017303998 | 0.036930343 |
| -1.585923057 | 0.112756765 | 0.183935207 |
| -2.132571627 | 0.032959883 | 0.064502414 |
| NA | NA | NA |
| -2.22039734 | 0.026391807 | 0.053123196 |
| 1.422254908 | 0.154952264 | 0.239825326 |
| 0.09086218 | 0.927602102 | NA |
| NA | NA | NA |
| 0.821870908 | 0.411150372 | 0.522940127 |
| 0.454502746 | 0.649467016 | NA |
| -0.352458214 | 0.724494651 | 0.798788702 |
| 0.193472986 | 0.84658856 | NA |
| -1.477839331 | 0.139450788 | 0.219718823 |
| -2.421167763 | 0.015470736 | 0.03348314 |
| -3.027103189 | 0.002469097 | 0.006501998 |
| -3.46879233 | 0.000522803 | 0.001589622 |
| 0.26284244 | 0.792672023 | 0.852514963 |
| -2.95139991 | 0.003163371 | 0.008128443 |
| -3.280354039 | 0.001036769 | 0.002978004 |
| -2.756955881 | 0.005834222 | 0.014106509 |
| -0.477081317 | 0.633304218 | NA |
| 0.714967331 | 0.47462923 | NA |
| 0.192809521 | 0.847108147 | NA |
| 0.063708285 | 0.949202508 | NA |
| -0.07484277 | 0.940339812 | NA |
| 2.960770705 | 0.003068703 | 0.007912809 |
| 1.405514133 | 0.15986845 | 0.245832892 |
| -0.775864825 | 0.437828803 | NA |
| -1.474066264 | 0.140463751 | 0.221010093 |
| 0.215610811 | 0.829291128 | NA |
| 0.456991567 | 0.647677107 | NA |
| -6.441259713 | 1.18E-10 | 8.93E-10 |
| -2.779296158 | 0.005447683 | 0.01328095 |
| -2.869061584 | 0.004116916 | 0.010320122 |

| | | |
|---|---|---|
| NA | NA | NA |
| 0.092716662 | 0.92612866 | NA |
| 0.330704598 | 0.740867629 | NA |
| 0.063708882 | 0.949202033 | NA |
| -1.142688992 | 0.253167742 | 0.357203415 |
| -2.452986571 | 0.014167563 | 0.030970663 |
| 4.574525156 | 4.77E-06 | 2.04E-05 |
| -2.34093428 | 0.019235553 | 0.040388045 |
| -3.358407325 | 0.00078393 | 0.002303295 |
| NA | NA | NA |
| -5.913205581 | 3.36E-09 | 2.16E-08 |
| NA | NA | NA |
| -3.544513464 | 0.000393338 | 0.001223014 |
| -1.363554383 | 0.172707857 | 0.262087891 |
| -10.34908514 | 4.23E-25 | 1.09E-23 |
| -0.267830262 | 0.788829976 | 0.849733788 |
| NA | NA | NA |
| 0.120420261 | 0.904150242 | NA |
| 0.192809521 | 0.847108147 | NA |
| -0.564194241 | 0.572621947 | 0.673910444 |
| -4.389654475 | 1.14E-05 | 4.59E-05 |
| -4.494286624 | 6.98E-06 | 2.91E-05 |
| -0.632747659 | 0.526898442 | 0.631894399 |
| -4.151805814 | 3.30E-05 | 0.000123841 |
| -0.226642106 | 0.820702048 | 0.874161389 |
| 4.660188131 | 3.16E-06 | 1.38E-05 |
| 0.22485065 | 0.822095462 | 0.875246678 |
| -1.856436124 | 0.06339142 | 0.113197349 |
| NA | NA | NA |
| 5.790112915 | 7.03E-09 | 4.37E-08 |
| -0.631686386 | 0.527591825 | 0.632562915 |
| 0.192810118 | 0.847107679 | NA |
| 0.487284587 | 0.626056669 | NA |
| -11.58153294 | 5.11E-31 | 1.87E-29 |
| -3.596014083 | 0.00032313 | 0.001021464 |
| -2.854900797 | 0.004305027 | 0.010740041 |
| -2.575801709 | 0.010000798 | 0.022788617 |
| -4.934100064 | 8.05E-07 | 3.83E-06 |
| NA | NA | NA |
| 1.581920126 | 0.113667806 | 0.185204087 |
| NA | NA | NA |
| NA | NA | NA |
| 0.063708285 | 0.949202508 | NA |
| NA | NA | NA |
| NA | NA | NA |
| NA | NA | NA |
| 0.366055258 | 0.714323843 | 0.790994625 |

| | | |
|---|---|---|
| NA | NA | NA |
| -0.335377545 | 0.737340305 | NA |
| -1.158562495 | 0.246634564 | 0.349660897 |
| -5.061852393 | 4.15E-07 | 2.06E-06 |
| -5.702239999 | 1.18E-08 | 7.17E-08 |
| -5.64238038 | 1.68E-08 | 9.99E-08 |
| 0.192810118 | 0.847107679 | NA |
| NA | NA | NA |
| -2.208771996 | 0.027190504 | 0.054498583 |
| -7.45610486 | 8.91E-14 | 9.21E-13 |
| -1.604944485 | 0.108506023 | 0.17809218 |
| 4.028737659 | 5.61E-05 | 0.000202717 |
| 0.087540923 | 0.930241558 | 0.952503424 |
| 0.461689452 | 0.644304036 | NA |
| 1.978942351 | 0.047822497 | 0.089088799 |
| 11.45762416 | 2.15E-30 | 7.57E-29 |
| -8.094764887 | 5.74E-16 | 7.32E-15 |
| 2.818846439 | 0.004819657 | 0.011909963 |
| -0.244796824 | 0.806613754 | 0.862985349 |
| -0.19448762 | 0.845794089 | NA |
| 0.485354229 | 0.627425094 | 0.720495648 |
| 2.923511306 | 0.003461077 | 0.008812894 |
| -0.658433085 | 0.510259881 | 0.616153119 |
| NA | NA | NA |
| NA | NA | NA |
| -4.704164367 | 2.55E-06 | 1.13E-05 |
| -3.656410556 | 0.000255772 | 0.000823192 |
| -0.696518024 | 0.486104474 | 0.594361745 |
| -1.180653589 | 0.237740366 | 0.339764837 |
| 0.192810118 | 0.847107679 | NA |
| 0.0569694 | 0.954569571 | 0.968891986 |
| NA | NA | NA |
| NA | NA | NA |
| NA | NA | NA |
| NA | NA | NA |
| NA | NA | NA |
| NA | NA | NA |
| NA | NA | NA |
| 0.565758212 | 0.571558157 | 0.672879952 |
| NA | NA | NA |
| NA | NA | NA |
| 0.39390925 | 0.693648041 | NA |
| 0.063708285 | 0.949202508 | NA |
| -7.79764543 | 6.31E-15 | 7.33E-14 |
| 0.450889308 | 0.65206933 | 0.740445501 |
| -0.914816489 | 0.360287977 | 0.472443842 |
| NA | NA | NA |

| | | | |
|---|---|---|---|
| | -0.241788955 | 0.808943698 | 0.865034423 |
| | -6.133388157 | 8.60E-10 | 5.89E-09 |
| | 0.961667149 | 0.336216829 | 0.447538143 |
| | -0.716560438 | 0.473645363 | NA |
| NA | NA | NA | |
| | -0.392333523 | 0.694811798 | 0.775213484 |
| | 0.922891391 | 0.356063807 | 0.467983464 |
| | 1.723945992 | 0.084717593 | 0.144925747 |
| NA | NA | NA | |
| | 0.063708882 | 0.949202033 | NA |
| | -1.726658848 | 0.08422895 | 0.144292015 |
| | 0.46328388 | 0.643160899 | 0.733011644 |
| | 0.696986663 | 0.485811141 | 0.594090915 |
| | 0.68969902 | 0.490383482 | 0.597929587 |
| | -2.675657819 | 0.007458275 | 0.017527051 |
| | -0.065388772 | 0.947864464 | NA |
| | 0.06353712 | 0.949338802 | NA |
| NA | NA | NA | |
| NA | NA | NA | |
| NA | NA | NA | |
| | 1.291537812 | 0.196517252 | NA |
| | 0.063708882 | 0.949202033 | NA |
| | 0.129420106 | 0.897025239 | NA |
| | -0.065389966 | 0.947863513 | NA |
| NA | NA | NA | |
| | 0.032594773 | 0.973997738 | 0.982052988 |
| | 2.826665772 | 0.004703539 | 0.011643713 |
| | 10.31267081 | 6.18E-25 | 1.56E-23 |
| | 0.669138799 | 0.503406942 | 0.609988372 |
| | 0.012858748 | 0.989740486 | 0.992715062 |
| | 7.062327745 | 1.64E-12 | 1.48E-11 |
| NA | NA | NA | |
| | -2.796633645 | 0.005163805 | 0.012662882 |
| | 12.06750119 | 1.57E-33 | 6.42E-32 |
| | 0.288217989 | 0.77317988 | NA |
| | 2.657390864 | 0.007874808 | 0.018373085 |
| NA | NA | NA | |
| | 7.952583522 | 1.83E-15 | 2.22E-14 |
| NA | NA | NA | |
| | -0.985422817 | 0.324416427 | 0.435466464 |
| | -0.178861929 | 0.858046117 | 0.90148146 |
| | 1.769235666 | 0.076854555 | 0.13341213 |
| | -6.864241393 | 6.68E-12 | 5.68E-11 |
| | -1.116637549 | 0.26414933 | NA |
| NA | NA | NA | |
| NA | NA | NA | |
| | 0.672809135 | 0.501068725 | 0.608025898 |

| | | | |
|---|---|---|---|
| NA | NA | NA | |
| NA | NA | NA | |
| | -4.490417502 | 7.11E-06 | 2.96E-05 |
| NA | NA | NA | |
| | -1.295183204 | 0.195257038 | 0.289788536 |
| | 0.192809521 | 0.847108147 | NA |
| | -0.337657865 | 0.735621031 | NA |
| | 0.819572082 | 0.412460094 | 0.524134445 |
| | 0.063707019 | 0.949203516 | NA |
| NA | NA | NA | |
| | -4.315389332 | 1.59E-05 | 6.28E-05 |
| | -7.108890006 | 1.17E-12 | 1.08E-11 |
| | 2.726233992 | 0.006406157 | 0.015326525 |
| | 0.201652944 | 0.840188052 | NA |
| | -2.477560854 | 0.013228385 | 0.029182661 |
| | -1.488034071 | 0.136741908 | 0.216195237 |
| | -1.152439255 | 0.249140618 | 0.352523513 |
| | -0.284366888 | 0.776129232 | NA |
| | -1.14392001 | 0.252656812 | 0.356683587 |
| NA | NA | NA | |
| NA | NA | NA | |
| | -0.264940255 | 0.791055487 | NA |
| | -1.371791939 | 0.170128217 | 0.258753678 |
| | 1.634076339 | 0.102242822 | 0.169636606 |
| | 0.226376714 | 0.820908437 | 0.874317611 |
| | -3.143930313 | 0.001666951 | 0.004575479 |
| | 1.83436811 | 0.06659937 | 0.118125158 |
| | -3.961103607 | 7.46E-05 | 0.000264122 |
| | 0.301212169 | 0.763252711 | NA |
| NA | NA | NA | |
| | -3.111148644 | 0.001863611 | 0.005050513 |
| | 4.318825422 | 1.57E-05 | 6.20E-05 |
| | 0.178287224 | 0.858497412 | 0.901804189 |
| | -0.883176123 | 0.37714112 | 0.489847036 |
| | 0.063707688 | 0.949202983 | NA |
| | -0.123208691 | 0.901941844 | NA |
| | 0.793475063 | 0.427501093 | 0.538789141 |
| | -1.812750314 | 0.06987034 | 0.123063445 |
| | 0.911815169 | 0.361866024 | 0.474090896 |
| | 3.806111478 | 0.000141169 | 0.00047531 |
| | -0.877748139 | 0.380080414 | 0.492693032 |
| NA | NA | NA | |
| NA | NA | NA | |
| | -0.552205347 | 0.580807671 | NA |
| | 1.513718162 | 0.130097336 | 0.207528923 |
| NA | NA | NA | |
| | 0.527730619 | 0.597686319 | NA |

| | | |
|---|---|---|
| NA | NA | NA |
| -0.1822812 | 0.855362052 | 0.899461113 |
| NA | NA | NA |
| NA | NA | NA |
| 13.80911635 | 2.25E-43 | 1.40E-41 |
| NA | NA | NA |
| NA | NA | NA |
| 0.836378009 | 0.40294228 | 0.51463368 |
| 1.763173556 | 0.077871222 | 0.134943 |
| -0.274907846 | 0.783387039 | NA |
| 2.497811095 | 0.012496276 | 0.027753487 |
| -0.754020821 | 0.45083672 | NA |
| -0.121445103 | 0.903338494 | 0.932894439 |
| 5.344483213 | 9.07E-08 | 4.92E-07 |
| 0.598253831 | 0.549670578 | 0.653040372 |
| 0.146118021 | 0.88382822 | NA |
| -3.165695845 | 0.001547125 | 0.004274703 |
| -0.148507036 | 0.881942634 | 0.917117745 |
| 0.867134784 | 0.385868164 | 0.498106248 |
| -5.687779027 | 1.29E-08 | 7.76E-08 |
| 4.415700484 | 1.01E-05 | 4.11E-05 |
| -3.230884156 | 0.001234079 | 0.003481928 |
| -0.75568285 | 0.449839369 | 0.559783206 |
| NA | NA | NA |
| -0.193918692 | 0.846239548 | NA |
| -2.156158444 | 0.031071295 | 0.061198813 |
| -0.445016048 | 0.656308157 | 0.743859498 |
| -7.555756326 | 4.16E-14 | 4.45E-13 |
| -2.826588147 | 0.004704679 | 0.011644281 |
| -2.538752181 | 0.011124859 | 0.02507702 |
| -0.788985122 | 0.430120702 | 0.541164229 |
| -0.466756733 | 0.640673935 | 0.731172195 |
| 2.514257334 | 0.011928336 | 0.026633921 |
| 0.706359422 | 0.479964648 | 0.588638108 |
| -1.202429657 | 0.229197105 | 0.329682405 |
| -0.086634921 | 0.930961707 | NA |
| NA | NA | NA |
| -0.085852801 | 0.931583432 | NA |
| 1.161464939 | 0.245452873 | 0.348439509 |
| 0.52202497 | 0.601652945 | NA |
| 0.393215746 | 0.694160141 | 0.774668364 |
| NA | NA | NA |
| NA | NA | NA |
| NA | NA | NA |
| 0.324874675 | 0.745275925 | NA |
| NA | NA | NA |
| NA | NA | NA |

| | | |
|---|---|---|
| NA | NA | NA |
| -0.034587876 | 0.97240837 | 0.981060556 |
| 4.204514906 | 2.62E-05 | 1.00E-04 |
| -6.13649758 | 8.44E-10 | 5.78E-09 |
| -0.517224185 | 0.604999671 | 0.701981354 |
| -0.256114586 | 0.797862359 | 0.856257551 |
| -4.237302821 | 2.26E-05 | 8.72E-05 |
| 0.998795381 | 0.317893824 | 0.428434332 |
| 3.813036947 | 0.00013727 | 0.000463303 |
| NA | NA | NA |
| NA | NA | NA |
| NA | NA | NA |
| -0.796231624 | 0.42589742 | 0.537309094 |
| 0.392345407 | 0.694803018 | 0.775213484 |
| 3.771688381 | 0.000162147 | 0.000540561 |
| -0.872312042 | 0.38303817 | 0.495495422 |
| 0.063708285 | 0.949202508 | NA |
| NA | NA | NA |
| NA | NA | NA |
| 0.027841302 | 0.977788724 | 0.98463356 |
| -3.303436848 | 0.000955075 | 0.002763644 |
| -3.146259523 | 0.001653732 | 0.004542681 |
| -1.143636448 | 0.25277444 | 0.356809397 |
| -1.02707714 | 0.304384142 | 0.413904225 |
| 8.192626854 | 2.56E-16 | 3.36E-15 |
| 0.066629133 | 0.946876953 | NA |
| 0.357743958 | 0.720534933 | NA |
| -0.409554422 | 0.682132837 | NA |
| 0.350755012 | 0.725772151 | 0.799739244 |
| NA | NA | NA |
| -0.55545237 | 0.57858528 | 0.679406084 |
| NA | NA | NA |
| NA | NA | NA |
| NA | NA | NA |
| 2.962872237 | 0.00304783 | 0.007866281 |
| NA | NA | NA |
| 7.734623234 | 1.04E-14 | 1.18E-13 |
| -1.482794749 | 0.138128952 | 0.217992917 |
| 1.11665669 | 0.264141143 | 0.36968535 |
| 0.086044261 | 0.931431232 | NA |
| 0.794459357 | 0.426928061 | 0.538256654 |
| NA | NA | NA |
| 0.06371127 | 0.949200131 | NA |
| 1.62526806 | 0.104105454 | 0.172156416 |
| NA | NA | NA |
| 5.338227327 | 9.39E-08 | 5.08E-07 |
| 1.816696547 | 0.069263591 | NA |

| | | |
|---|---|---|
| -0.672591177 | 0.501207417 | 0.608135229 |
| -0.065389369 | 0.947863988 | NA |
| 0.072278507 | 0.942380269 | 0.960923051 |
| 5.416769752 | 6.07E-08 | 3.36E-07 |
| -6.22672308 | 4.76E-10 | 3.35E-09 |
| 0.100756116 | 0.919744063 | NA |
| NA | NA | NA |
| NA | NA | NA |
| 0.12042439 | 0.904146972 | NA |
| -4.926592162 | 8.37E-07 | 3.97E-06 |
| 0.27316407 | 0.784727098 | NA |
| 0.06371127 | 0.949200131 | NA |
| 17.19419547 | 2.94E-66 | 3.43E-64 |
| 0.765813726 | 0.443787146 | 0.553793611 |
| 0.120685246 | 0.903940345 | NA |
| 1.649204627 | 0.09910572 | 0.165385438 |
| 0.578960509 | 0.56261582 | 0.664791294 |
| -0.341029662 | 0.733081252 | 0.80538127 |
| 3.824010547 | 0.000131298 | 0.000444709 |
| -2.526952797 | 0.011505697 | 0.025823851 |
| NA | NA | NA |
| 0.063708285 | 0.949202508 | NA |
| NA | NA | NA |
| NA | NA | NA |
| 0.063708285 | 0.949202508 | NA |
| -2.30124659 | 0.021377697 | 0.044273023 |
| -0.051489064 | 0.958935816 | NA |
| -0.065388772 | 0.947864464 | NA |
| -0.404583542 | 0.685783658 | NA |
| 0.014673194 | 0.988292905 | 0.991859319 |
| 0.581470102 | 0.560923663 | 0.663136184 |
| NA | NA | NA |
| -0.556760032 | 0.577691395 | 0.678568474 |
| -2.121738673 | 0.033859689 | 0.066010438 |
| 0.42432264 | 0.67133053 | 0.755582678 |
| NA | NA | NA |
| -0.124169457 | 0.901181105 | 0.931165912 |
| -2.80431096 | 0.005042425 | 0.012399243 |
| -3.086952658 | 0.002022198 | 0.005445467 |
| -0.448636742 | 0.653693724 | 0.741803234 |
| -1.235921073 | 0.216487903 | 0.315040245 |
| 0.063708285 | 0.949202508 | NA |
| 0.063708882 | 0.949202033 | NA |
| -1.266422147 | 0.205361986 | 0.301974534 |
| 6.993802622 | 2.68E-12 | 2.36E-11 |
| NA | NA | NA |
| 11.62266877 | 3.16E-31 | 1.16E-29 |

| | | |
|---|---|---|
| NA | NA | NA |
| 1.070782374 | 0.284267294 | 0.391332657 |
| NA | NA | NA |
| 0.027451007 | 0.978100016 | 0.984788552 |
| 0.192808924 | 0.847108615 | NA |
| -0.310839573 | 0.755922584 | 0.823032003 |
| NA | NA | NA |
| 5.693036434 | 1.25E-08 | 7.54E-08 |
| NA | NA | NA |
| NA | NA | NA |
| -0.78838242 | 0.430473051 | NA |
| 0.063710673 | 0.949200606 | NA |
| 1.675600211 | 0.093816528 | 0.157980872 |
| NA | NA | NA |
| NA | NA | NA |
| NA | NA | NA |
| NA | NA | NA |
| NA | NA | NA |
| -1.373635081 | 0.169555 | 0.257975908 |
| 0.192809521 | 0.847108147 | NA |
| -1.3851725 | 0.165999727 | 0.25355301 |
| -2.519724246 | 0.01174468 | 0.026284813 |
| 0.063708285 | 0.949202508 | NA |
| -0.671974219 | 0.501600109 | NA |
| 3.362725449 | 0.000771771 | 0.002272103 |
| 0.330026755 | 0.741379746 | 0.81178973 |
| 1.050017491 | 0.293710071 | 0.401834549 |
| NA | NA | NA |
| -0.676577348 | 0.498674155 | 0.605821458 |
| -2.538132561 | 0.011144576 | 0.025116941 |
| -4.032624387 | 5.52E-05 | 0.00019968 |
| -0.087688459 | 0.930124293 | 0.952476219 |
| -0.831509666 | 0.40568577 | 0.517522798 |
| -1.468314171 | 0.142018903 | 0.223133084 |
| 0.147058412 | 0.883085915 | 0.918115907 |
| -2.831874419 | 0.004627602 | 0.011476159 |
| 0.192809521 | 0.847108147 | NA |
| -4.393045585 | 1.12E-05 | 4.52E-05 |
| -8.32866451 | 8.18E-17 | 1.12E-15 |
| -0.11765063 | 0.906344487 | 0.935227048 |
| -1.519864634 | 0.128545001 | 0.205616141 |
| NA | NA | NA |
| 0.079230484 | 0.936849298 | 0.956959595 |
| 4.935145394 | 8.01E-07 | 3.81E-06 |
| -0.065389369 | 0.947863988 | NA |
| NA | NA | NA |
| 1.019356505 | 0.308033747 | 0.417868121 |

| | | | |
|---|---|---|---|
| | -0.066835228 | 0.946712878 | 0.96365284 |
| | -7.26270658 | 3.79E-13 | 3.67E-12 |
| | -4.021569699 | 5.78E-05 | 0.000208475 |
| | 0.192809521 | 0.847108147 | NA |
| | 5.99866587 | 1.99E-09 | 1.31E-08 |
| | -0.097376463 | 0.922427436 | 0.946632174 |
| | 2.077796559 | 0.037728101 | 0.072446058 |
| NA | NA | NA | |
| | 1.076021431 | 0.281917672 | 0.388947068 |
| | -1.999665786 | 0.045536365 | 0.085389213 |
| | 1.333317168 | 0.182427742 | 0.274260699 |
| | -0.797437 | 0.425197278 | 0.536723378 |
| NA | NA | NA | |
| | 1.15868349 | 0.246585223 | 0.349610747 |
| | 0.192808924 | 0.847108615 | NA |
| | 5.987652469 | 2.13E-09 | 1.40E-08 |
| | 1.202961526 | 0.228991209 | NA |
| | -0.210333116 | 0.833407691 | 0.883307694 |
| | -3.110505678 | 0.001867673 | 0.005060935 |
| | -0.766428753 | 0.44342123 | 0.553494507 |
| | 0.451554923 | 0.65158965 | 0.74012021 |
| | 0.364142666 | 0.715751478 | 0.792031198 |
| | -0.232185325 | 0.816394075 | 0.870769323 |
| | 1.108242679 | 0.267757023 | NA |
| | -1.579279695 | 0.114271917 | 0.185994673 |
| | 0.06371127 | 0.949200131 | NA |
| | -3.050544923 | 0.002284265 | 0.006070194 |
| | 0.192810118 | 0.847107679 | NA |
| | 1.768755636 | 0.076934663 | 0.133541929 |
| NA | NA | NA | |
| NA | NA | NA | |
| | 0.063708285 | 0.949202508 | NA |
| | -0.06879015 | 0.945156659 | NA |
| | 0.573433465 | 0.566351237 | NA |
| | 0.192809521 | 0.847108147 | NA |
| | 0.666005019 | 0.505407892 | NA |
| | -2.938176072 | 0.003301495 | 0.008447803 |
| | -2.297907771 | 0.021567039 | 0.044598726 |
| | -0.065389369 | 0.947863988 | NA |
| NA | NA | NA | |
| | 1.925104196 | 0.054216324 | 0.099241962 |
| | 2.705083739 | 0.006828717 | 0.01622428 |
| | 9.916829117 | 3.52E-23 | 7.94E-22 |
| | 0.94980935 | 0.342209134 | 0.453569696 |
| NA | NA | NA | |
| | -0.856236493 | 0.391866981 | 0.503805562 |
| | 8.03954276 | 9.02E-16 | 1.13E-14 |

| | | | |
|---|---|---|---|
| | 4.80528211 | 1.55E-06 | 7.06E-06 |
| | -0.286782923 | 0.774278539 | 0.837700101 |
| NA | NA | NA | |
| NA | NA | NA | |
| | 0.192808327 | 0.847109082 | NA |
| NA | NA | NA | |
| | -0.036494439 | 0.970888112 | 0.979876453 |
| | -0.480779996 | 0.630672869 | NA |
| | -0.113927401 | 0.909295344 | NA |
| NA | NA | NA | |
| NA | NA | NA | |
| | 1.025966162 | 0.304907534 | 0.414441982 |
| NA | NA | NA | |
| NA | NA | NA | |
| | 0.93641721 | 0.34905842 | 0.460735762 |
| | -0.049705062 | 0.960357422 | 0.972715534 |
| | -0.78493974 | 0.432488901 | 0.543188177 |
| | 0.948091536 | 0.343082857 | 0.454454059 |
| | 1.854371951 | 0.063685965 | 0.113650952 |
| | 0.192810118 | 0.847107679 | NA |
| | 0.192808924 | 0.847108615 | NA |
| | -0.187772727 | 0.851054816 | NA |
| | -4.294671864 | 1.75E-05 | 6.86E-05 |
| | 0.426539464 | 0.669714798 | NA |
| | -4.354046747 | 1.34E-05 | 5.33E-05 |
| | -1.876059145 | 0.060647159 | 0.109028111 |
| | -0.421209834 | 0.673601864 | 0.757355793 |
| NA | NA | NA | |
| | 1.736355936 | 0.082500918 | 0.141787832 |
| | -0.565455939 | 0.571763685 | 0.673090257 |
| | -0.611614916 | 0.540792567 | 0.644972504 |
| | 4.226783836 | 2.37E-05 | 9.10E-05 |
| | -1.069593406 | 0.284802365 | 0.391961466 |
| | -4.80450012 | 1.55E-06 | 7.08E-06 |
| | 0.06370649 | 0.949203937 | NA |
| NA | NA | NA | |
| NA | NA | NA | |
| | -1.286734405 | 0.198186877 | 0.293440848 |
| NA | NA | NA | |
| | 0.773412247 | 0.439278444 | NA |
| | 0.192810118 | 0.847107679 | NA |
| NA | NA | NA | |
| NA | NA | NA | |
| NA | NA | NA | |
| | 0.192636473 | 0.847243678 | NA |
| | 1.116216506 | 0.264329471 | 0.369907562 |
| | 1.464567129 | 0.143039056 | 0.224566742 |

| | | |
|---|---|---|
| -1.299886021 | 0.193640037 | 0.287883709 |
| 0.067830052 | 0.945920921 | 0.963042532 |
| -0.227070422 | 0.820368983 | 0.873966191 |
| -0.065390563 | 0.947863038 | NA |
| -3.5709142 | 0.000355737 | 0.001116363 |
| -2.87129179 | 0.00408798 | 0.010253746 |
| -1.566481918 | 0.117235838 | 0.190089592 |
| -6.598456939 | 4.15E-11 | 3.29E-10 |
| NA | NA | NA |
| 0.366672431 | 0.713863373 | 0.790689452 |
| -1.943445924 | 0.051962303 | 0.095718019 |
| 1.894306018 | 0.058184403 | 0.105342177 |
| -2.010883275 | 0.044337787 | 0.083435442 |
| -2.08679856 | 0.036906347 | 0.071097197 |
| -1.547429652 | 0.121759676 | 0.196381512 |
| -1.888691496 | 0.058933179 | 0.106497543 |
| -2.265284292 | 0.023495239 | 0.048089446 |
| 0.142610509 | 0.886597798 | NA |
| NA | NA | NA |
| NA | NA | NA |
| 1.20058235 | 0.229913251 | NA |
| NA | NA | NA |
| NA | NA | NA |
| 0.063707688 | 0.949202983 | NA |
| -1.708009874 | 0.08763451 | 0.1492679 |
| 0.753169868 | 0.451347844 | 0.5612697 |
| -9.86406358 | 5.96E-23 | 1.32E-21 |
| 0.837157277 | 0.402504169 | 0.514250387 |
| -0.08304876 | 0.933812769 | NA |
| -2.689876906 | 0.007147838 | 0.016873562 |
| NA | NA | NA |
| -2.863175469 | 0.004194182 | 0.01049489 |
| 0.192809521 | 0.847108147 | NA |
| 0.192808924 | 0.847108615 | NA |
| -1.953242563 | 0.05079086 | 0.093877952 |
| -1.537253441 | 0.124231258 | 0.199788512 |
| 0.325720476 | 0.744635851 | 0.814252309 |
| -4.348946704 | 1.37E-05 | 5.45E-05 |
| 0.192806724 | 0.847110337 | NA |
| 0.063705092 | 0.94920505 | NA |
| NA | NA | NA |
| 0.31685483 | 0.751353758 | NA |
| -3.142947348 | 0.001672559 | 0.004587853 |
| -0.737738344 | 0.460673466 | 0.570119004 |
| 0.069889846 | 0.944281335 | NA |
| 0.066009206 | 0.947370496 | NA |
| 7.024650918 | 2.15E-12 | 1.91E-11 |

| | | |
|---|---|---|
| 0.192810118 | 0.847107679 | NA |
| -1.014196095 | 0.310489191 | 0.420606401 |
| NA | NA | NA |
| 0.477403294 | 0.633074969 | 0.724614344 |
| 5.530606004 | 3.19E-08 | 1.84E-07 |
| 0.089238296 | 0.928892531 | NA |
| NA | NA | NA |
| NA | NA | NA |
| -8.24030012 | 1.72E-16 | 2.29E-15 |
| -4.453608318 | 8.44E-06 | 3.48E-05 |
| 3.449251427 | 0.000562143 | 0.001698303 |
| NA | NA | NA |
| 8.887026971 | 6.28E-19 | 1.03E-17 |
| 1.212079321 | 0.225482019 | 0.325542343 |
| 1.092451594 | 0.274634656 | 0.381336519 |
| 8.224225135 | 1.96E-16 | 2.61E-15 |
| 0.192810118 | 0.847107679 | NA |
| -1.867956748 | 0.0617681 | 0.110709312 |
| 0.370160581 | 0.711262846 | 0.788471989 |
| -0.325345108 | 0.744919895 | 0.814385017 |
| -0.058799738 | 0.953111617 | NA |
| -0.065389966 | 0.947863513 | NA |
| -5.197861578 | 2.02E-07 | 1.04E-06 |
| -0.331029324 | 0.740622335 | 0.811186666 |
| -3.595632172 | 0.000323605 | 0.001022835 |
| -0.079867593 | 0.936342564 | NA |
| -4.498047534 | 6.86E-06 | 2.86E-05 |
| NA | NA | NA |
| -0.314474997 | 0.753160303 | NA |
| -1.716016105 | 0.086159095 | 0.147045012 |
| NA | NA | NA |
| -0.432677562 | 0.665249033 | 0.751237686 |
| 1.946056165 | 0.051647993 | 0.095209108 |
| 0.192809521 | 0.847108147 | NA |
| -2.846338813 | 0.004422511 | 0.011010091 |
| 0.159338659 | 0.873402065 | 0.911838698 |
| 5.84510113 | 5.06E-09 | 3.20E-08 |
| -1.120164506 | 0.262643666 | 0.368021663 |
| 3.247993485 | 0.001162219 | 0.003303844 |
| 1.531295922 | 0.125696273 | 0.201794478 |
| -0.065388772 | 0.947864464 | NA |
| NA | NA | NA |
| NA | NA | NA |
| NA | NA | NA |
| NA | NA | NA |
| -6.768638193 | 1.30E-11 | 1.08E-10 |
| -2.697904805 | 0.006977739 | 0.0165297 |

| | | |
|---|---|---|
| -0.359932253 | 0.718897797 | 0.794472686 |
| -2.716756194 | 0.006592515 | 0.015730281 |
| -1.377676604 | 0.168303158 | 0.256476593 |
| 1.218263033 | 0.223124033 | NA |
| 1.898594087 | 0.057617867 | 0.104516664 |
| -0.19448762 | 0.845794089 | NA |
| NA | NA | NA |
| NA | NA | NA |
| 2.193159159 | 0.028295914 | 0.056434002 |
| -3.88168325 | 0.000103736 | 0.000357498 |
| 0.727236992 | 0.467080789 | 0.576395609 |
| 0.13203366 | 0.894957668 | NA |
| 0.068706839 | 0.945222974 | 0.962606031 |
| NA | NA | NA |
| -4.053585325 | 5.04E-05 | 0.00018374 |
| NA | NA | NA |
| 3.237702382 | 0.001204964 | 0.003410637 |
| NA | NA | NA |
| -1.543427461 | 0.122727095 | 0.197712505 |
| 0.737848407 | 0.460606574 | 0.570064406 |
| NA | NA | NA |
| NA | NA | NA |
| -0.327150278 | 0.743554226 | NA |
| NA | NA | NA |
| -6.194329932 | 5.85E-10 | 4.07E-09 |
| -0.844129853 | 0.398596847 | 0.510451512 |
| -0.057866476 | 0.953854986 | NA |
| -1.391366347 | 0.164114364 | 0.251194431 |
| -0.24975138 | 0.802779621 | 0.860179769 |
| 1.352552984 | 0.176198481 | 0.266537409 |
| -1.45644824 | 0.145268748 | 0.227354622 |
| 16.80556038 | 2.22E-63 | 2.42E-61 |
| 3.266576955 | 0.001088562 | 0.003111776 |
| -1.998942482 | 0.045614578 | 0.085510251 |
| -0.786166713 | 0.431769825 | 0.542584532 |
| 0.192809521 | 0.847108147 | NA |
| -1.261647743 | 0.207075569 | 0.303958567 |
| 0.487757162 | 0.625721858 | 0.719223829 |
| -2.081594723 | 0.037379504 | 0.071848332 |
| -3.025112231 | 0.002485409 | 0.006540825 |
| -4.737020131 | 2.17E-06 | 9.72E-06 |
| -0.946418428 | 0.34393521 | 0.455293965 |
| -5.075439571 | 3.87E-07 | 1.93E-06 |
| 3.199496237 | 0.00137668 | 0.003845363 |
| 26.87422952 | 4.40E-159 | 1.96E-156 |
| 7.431718581 | 1.07E-13 | 1.10E-12 |
| 0.869272464 | 0.38469812 | NA |

| | | |
|---|---|---|
| -1.858039597 | 0.063163391 | 0.112878744 |
| -1.35750177 | 0.174621831 | NA |
| NA | NA | NA |
| NA | NA | NA |
| NA | NA | NA |
| NA | NA | NA |
| -7.099049444 | 1.26E-12 | 1.15E-11 |
| 4.199838448 | 2.67E-05 | 0.000101838 |
| 1.009607493 | 0.312683379 | 0.423075029 |
| -0.274059473 | 0.784038918 | NA |
| 0.627046048 | 0.530629051 | 0.635416171 |
| 7.553048819 | 4.25E-14 | 4.53E-13 |
| -1.380049894 | 0.167571283 | 0.255547993 |
| -1.411384292 | 0.158131332 | 0.243735898 |
| 0.19734106 | 0.84356065 | 0.89063665 |
| -6.980518519 | 2.94E-12 | 2.59E-11 |
| -0.065389966 | 0.947863513 | NA |
| NA | NA | NA |
| NA | NA | NA |
| NA | NA | NA |
| 0.063708882 | 0.949202033 | NA |
| NA | NA | NA |
| 0.192809521 | 0.847108147 | NA |
| 8.985173152 | 2.58E-19 | 4.37E-18 |
| 0.192808327 | 0.847109082 | NA |
| 3.327275674 | 0.000876996 | 0.002556627 |
| 0.192809521 | 0.847108147 | NA |
| -0.06407439 | 0.948910994 | 0.965066116 |
| -1.83694411 | 0.066218147 | 0.11759065 |
| -1.4826726 | 0.138161418 | 0.218016662 |
| -2.59600109 | 0.009431577 | 0.021616369 |
| -0.065389369 | 0.947863988 | NA |
| NA | NA | NA |
| 5.671706186 | 1.41E-08 | 8.49E-08 |
| 2.049137314 | 0.04044869 | 0.076955815 |
| -3.237209975 | 0.001207046 | 0.003414211 |
| -2.422021961 | 0.015434418 | 0.033413205 |
| 0.265886584 | 0.790326558 | NA |
| 8.420691344 | 3.74E-17 | 5.28E-16 |
| -1.777565189 | 0.075475295 | 0.131349321 |
| -2.32980766 | 0.019816319 | 0.041450983 |
| 7.06880654 | 1.56E-12 | 1.42E-11 |
| 0.21765406 | 0.82769866 | NA |
| 5.230619599 | 1.69E-07 | 8.83E-07 |
| 4.891165934 | 1.00E-06 | 4.71E-06 |
| 1.207340291 | 0.22730112 | 0.32748213 |
| -0.240074959 | 0.810272147 | 0.866003704 |

| | | |
|---|---|---|
| 2.004562924 | 0.045009794 | 0.084537329 |
| -2.643585205 | 0.008203312 | 0.019052509 |
| 0.738992249 | 0.459911704 | 0.569345168 |
| 8.898988562 | 5.64E-19 | 9.31E-18 |
| -0.121048266 | 0.903652805 | 0.933180531 |
| -2.369067375 | 0.017833005 | 0.037888134 |
| -1.034830797 | 0.300747938 | 0.40978339 |
| 0.433864185 | 0.664387069 | NA |
| -3.827602306 | 0.000129398 | 0.000438627 |
| -2.778353115 | 0.005463521 | 0.013314373 |
| NA | NA | NA |
| -3.056951935 | 0.002236001 | 0.005955849 |
| -3.045569025 | 0.002322405 | 0.006158831 |
| 1.307882379 | 0.190913207 | 0.284624312 |
| -0.305730413 | 0.759809915 | NA |
| 1.272073284 | 0.203347083 | 0.299680727 |
| 0.3474247 | 0.728272279 | NA |
| -0.279063023 | 0.78019646 | NA |
| -8.818968609 | 1.16E-18 | 1.86E-17 |
| -3.056341698 | 0.002240558 | 0.005965446 |
| 33.18278056 | 1.91E-241 | 2.27E-238 |
| 2.912524682 | 0.003585199 | 0.009090846 |
| 1.996269391 | 0.045904608 | 0.085957383 |
| 2.553571782 | 0.01066243 | 0.024147722 |
| -0.109996848 | 0.912411875 | NA |
| 2.589729957 | 0.009605125 | 0.021967861 |
| 0.192809521 | 0.847108147 | NA |
| 0.226610234 | 0.820726834 | 0.874161389 |
| 1.540850439 | 0.12335319 | 0.198580547 |
| 0.333816862 | 0.738517758 | NA |
| 7.9372053 | 2.07E-15 | 2.51E-14 |
| -7.446858048 | 9.56E-14 | 9.86E-13 |
| -0.065389369 | 0.947863988 | NA |
| 5.970317985 | 2.37E-09 | 1.55E-08 |
| 1.308600794 | 0.190669612 | NA |
| NA | NA | NA |
| 2.399091542 | 0.016435805 | 0.035330488 |
| -6.414093413 | 1.42E-10 | 1.06E-09 |
| -0.941184615 | 0.34661026 | 0.45815865 |
| 6.428360193 | 1.29E-10 | 9.68E-10 |
| 0.41846943 | 0.675603933 | 0.758959044 |
| -3.268418364 | 0.001081504 | 0.003095134 |
| -0.69017164 | 0.490086256 | 0.59771276 |
| 3.684298221 | 0.000229334 | 0.000744999 |
| -6.98932051 | 2.76E-12 | 2.43E-11 |
| -3.697415009 | 0.000217806 | 0.000710781 |
| 3.561288206 | 0.00036904 | 0.001153192 |

| | | |
|---|---|---|
| 6.281586038 | 3.35E-10 | 2.39E-09 |
| -0.433229331 | 0.664848172 | 0.750942936 |
| 0.192809521 | 0.847108147 | NA |
| 8.845395053 | 9.12E-19 | 1.48E-17 |
| 3.41104484 | 0.000647144 | 0.001932708 |
| 1.238601722 | 0.21549303 | 0.313847931 |
| -2.57845804 | 0.009924235 | 0.022635648 |
| -0.568041293 | 0.570006929 | 0.671401103 |
| -1.220276199 | 0.222360189 | 0.321808357 |
| -0.065389369 | 0.947863988 | NA |
| NA | NA | NA |
| -6.694498307 | 2.16E-11 | 1.76E-10 |
| 2.591502849 | 0.009555775 | 0.021865229 |
| -0.420709237 | 0.673967415 | 0.757596641 |
| 1.210089201 | 0.226244667 | 0.326335735 |
| 7.167142768 | 7.66E-13 | 7.20E-12 |
| 5.576761769 | 2.45E-08 | 1.43E-07 |
| 1.338498998 | 0.180733827 | 0.272057234 |
| -1.864211103 | 0.062292065 | 0.111536618 |
| -2.173186167 | 0.029766308 | 0.058971878 |
| 4.046795428 | 5.19E-05 | 0.000188792 |
| -1.118361144 | 0.263412783 | 0.368892852 |
| -3.163309628 | 0.001559863 | 0.004307522 |
| NA | NA | NA |
| NA | NA | NA |
| NA | NA | NA |
| -0.982077628 | 0.326061606 | 0.437135619 |
| -6.264839191 | 3.73E-10 | 2.65E-09 |
| -2.27312382 | 0.023018719 | 0.047233854 |
| -2.113891203 | 0.034524561 | 0.067123641 |
| 4.570600494 | 4.86E-06 | 2.07E-05 |
| NA | NA | NA |
| 5.616201578 | 1.95E-08 | 1.15E-07 |
| 1.284477201 | 0.198975036 | 0.294416242 |
| 0.560456179 | 0.575168322 | NA |
| 0.291684799 | 0.770527633 | 0.834709578 |
| 0.063708882 | 0.949202033 | NA |
| -2.88171955 | 0.003955116 | 0.009955405 |
| -1.094079873 | 0.273919949 | 0.380562629 |
| 0.61657415 | 0.53751565 | NA |
| -4.259816197 | 2.05E-05 | 7.94E-05 |
| -0.692051651 | 0.488904893 | 0.596752428 |
| -1.401363648 | 0.161105357 | 0.247284423 |
| -0.509351618 | 0.610505783 | 0.706633977 |
| 0.31533402 | 0.752508057 | 0.820494715 |
| NA | NA | NA |
| 1.420285839 | 0.155524482 | 0.240577083 |

| | | |
|---|---|---|
| NA | NA | NA |
| 6.02993607 | 1.64E-09 | 1.09E-08 |
| 0.858715493 | 0.390497501 | 0.502682187 |
| 4.679953751 | 2.87E-06 | 1.26E-05 |
| 20.38041202 | 2.50E-92 | 5.25E-90 |
| 16.24213468 | 2.54E-59 | 2.47E-57 |
| -1.248538856 | 0.211833788 | 0.309653666 |
| 2.097301772 | 0.03596687 | 0.069501332 |
| 0.630984869 | 0.528050418 | 0.633002967 |
| -0.250491983 | 0.802206903 | 0.859777288 |
| -6.44978817 | 1.12E-10 | 8.48E-10 |
| NA | NA | NA |
| -0.065387578 | 0.947865415 | NA |
| 0.192809521 | 0.847108147 | NA |
| -0.194488217 | 0.845793621 | NA |
| NA | NA | NA |
| NA | NA | NA |
| NA | NA | NA |
| 0.192810118 | 0.847107679 | NA |
| 0.192809521 | 0.847108147 | NA |
| NA | NA | NA |
| NA | NA | NA |
| 0.192810118 | 0.847107679 | NA |
| 0.847134989 | 0.396919884 | NA |
| 5.129486638 | 2.91E-07 | 1.47E-06 |
| 0.645385546 | 0.518677368 | NA |
| -0.276044601 | 0.78251381 | NA |
| 0.192809521 | 0.847108147 | NA |
| 0.193341124 | 0.846691821 | NA |
| 0.063710673 | 0.949200606 | NA |
| 0.375794244 | 0.707069867 | NA |
| NA | NA | NA |
| 0.58653885 | 0.557513462 | NA |
| -0.127199333 | 0.898782634 | NA |
| 0.007172988 | 0.994276833 | 0.99594802 |
| -2.981744444 | 0.002866111 | 0.007441009 |
| 3.011854733 | 0.002596568 | 0.006804031 |
| 4.572129617 | 4.83E-06 | 2.06E-05 |
| 3.504343736 | 0.000457734 | 0.001405325 |
| 0.331497207 | 0.74026895 | NA |
| NA | NA | NA |
| 2.976783023 | 0.002912901 | 0.0075523 |
| -1.075963611 | 0.281943531 | 0.388947068 |
| NA | NA | NA |
| -3.959860044 | 7.50E-05 | 0.000265351 |
| NA | NA | NA |
| 1.50479014 | 0.13237804 | 0.210442578 |

| | | | |
|---|---|---|---|
| | -0.962779582 | 0.335658149 | 0.446889547 |
| | -1.058031537 | 0.290041065 | NA |
| | 1.164515951 | 0.244214983 | 0.34705621 |
| NA | NA | NA | |
| | -3.544739888 | 0.000393001 | 0.001222115 |
| | -0.665480763 | 0.505743044 | 0.612213976 |
| | 0.423401216 | 0.672002558 | 0.756101053 |
| | -2.012968086 | 0.044117987 | 0.083078033 |
| | 3.918711338 | 8.90E-05 | 0.000310906 |
| | -0.065389369 | 0.947863988 | NA |
| | -2.002419292 | 0.045239655 | 0.084896429 |
| | 1.621686777 | 0.104870431 | 0.173226879 |
| | 5.113770712 | 3.16E-07 | 1.59E-06 |
| | 4.596554378 | 4.30E-06 | 1.85E-05 |
| | -1.231291935 | 0.218213697 | 0.317053976 |
| | 2.459433532 | 0.013915646 | 0.030521261 |
| | -0.976912162 | 0.32861264 | 0.439590315 |
| | -1.402711834 | 0.160702787 | 0.246763565 |
| | 5.680094396 | 1.35E-08 | 8.10E-08 |
| | 3.556959049 | 0.000375173 | 0.001170456 |
| NA | NA | NA | |
| | -0.158061557 | 0.874408293 | NA |
| | 3.132153027 | 0.001735294 | 0.004742265 |
| | -0.123251129 | 0.901908239 | NA |
| | 0.373667563 | 0.708651661 | 0.78655275 |
| | -0.037320803 | 0.970229218 | 0.979385277 |
| | 0.414990094 | 0.678149155 | NA |
| | 0.553001254 | 0.580262551 | NA |
| | 1.134037649 | 0.256778756 | 0.361324462 |
| | 2.347538484 | 0.018897921 | 0.039769208 |
| | 8.408939347 | 4.14E-17 | 5.82E-16 |
| | 2.173546307 | 0.029739225 | 0.058935326 |
| | 1.481847815 | 0.138380793 | 0.21826651 |
| | -3.535435893 | 0.000407103 | 0.001262519 |
| | 0.06353712 | 0.949338802 | NA |
| | 1.850409683 | 0.064254524 | NA |
| NA | NA | NA | |
| | -0.685419712 | 0.493079109 | 0.600309845 |
| | -1.272218876 | 0.203295363 | 0.299657436 |
| | 0.063710673 | 0.949200606 | NA |
| | 3.837699576 | 0.000124192 | 0.000422298 |
| | -0.306112564 | 0.759518942 | NA |
| | 0.007676739 | 0.993874908 | 0.995625108 |
| | 5.340745393 | 9.26E-08 | 5.01E-07 |
| | 6.091109975 | 1.12E-09 | 7.60E-09 |
| | 14.09987816 | 3.80E-45 | 2.56E-43 |
| NA | NA | NA | |

| | | |
|---|---|---|
| 0.063708285 | 0.949202508 | NA |
| 1.480666797 | 0.138695386 | 0.218707586 |
| -0.450036109 | 0.652684404 | 0.740892999 |
| -0.103222606 | 0.917786299 | 0.943571472 |
| -0.933985161 | 0.350311547 | 0.462048692 |
| -2.367418866 | 0.017912647 | 0.038022261 |
| -2.894831179 | 0.003793625 | 0.009577813 |
| -7.623196032 | 2.47E-14 | 2.70E-13 |
| 0.063707688 | 0.949202983 | NA |
| NA | NA | NA |
| -2.011606059 | 0.04426148 | 0.083316902 |
| 0.745185016 | 0.45615988 | 0.565876086 |
| -6.605992621 | 3.95E-11 | 3.13E-10 |
| -1.473467865 | 0.140624923 | 0.221192849 |
| 1.654475017 | 0.098031035 | 0.163854211 |
| -0.241972629 | 0.808801372 | 0.864919181 |
| NA | NA | NA |
| NA | NA | NA |
| NA | NA | NA |
| 7.076233871 | 1.48E-12 | 1.35E-11 |
| NA | NA | NA |
| -11.75658634 | 6.53E-32 | 2.50E-30 |
| -1.23556304 | 0.21662103 | 0.315197325 |
| -7.324605578 | 2.40E-13 | 2.37E-12 |
| -4.680527325 | 2.86E-06 | 1.26E-05 |
| -0.065388772 | 0.947864464 | NA |
| 0.192810118 | 0.847107679 | NA |
| -1.568209815 | 0.116832175 | 0.189545588 |
| 5.455871246 | 4.87E-08 | 2.74E-07 |
| 0.828094657 | 0.407616895 | 0.519451844 |
| 1.160840518 | 0.245706762 | 0.348720813 |
| 1.190417824 | 0.233882211 | 0.335169354 |
| -1.664940247 | 0.095924678 | 0.160913791 |
| 0.063708882 | 0.949202033 | NA |
| -1.556259997 | 0.11964628 | 0.193421584 |
| -1.242652892 | 0.21399577 | 0.312175902 |
| 1.279288389 | 0.20079552 | 0.296706527 |
| 10.20512301 | 1.88E-24 | 4.64E-23 |
| 8.196994406 | 2.46E-16 | 3.25E-15 |
| 1.367312491 | 0.171527378 | 0.260596753 |
| 1.452340664 | 0.146406906 | 0.228749367 |
| NA | NA | NA |
| -9.268867192 | 1.88E-20 | 3.44E-19 |
| -5.265770014 | 1.40E-07 | 7.38E-07 |
| 3.435235494 | 0.000592039 | 0.001780884 |
| 0.192809521 | 0.847108147 | NA |
| 0.192810118 | 0.847107679 | NA |

| | | |
|---|---|---|
| -3.268096698 | 0.001082734 | 0.003097945 |
| 2.72591371 | 0.006412376 | 0.015339938 |
| NA | NA | NA |
| NA | NA | NA |
| NA | NA | NA |
| NA | NA | NA |
| 1.291086134 | 0.196673812 | 0.291545485 |
| NA | NA | NA |
| 0.203776651 | 0.83852804 | 0.886984776 |
| 0.141601692 | 0.88739463 | NA |
| 2.842681801 | 0.004473571 | 0.011125035 |
| 17.93476348 | 6.31E-72 | 8.50E-70 |
| 1.29025834 | 0.196960976 | 0.291867458 |
| -0.065389369 | 0.947863988 | NA |
| NA | NA | NA |
| 2.523734304 | 0.011611565 | 0.026022472 |
| NA | NA | NA |
| -0.163236505 | 0.870332225 | 0.909620827 |
| -0.818022827 | 0.413344153 | 0.524938056 |
| NA | NA | NA |
| 7.371371943 | 1.69E-13 | 1.69E-12 |
| -0.086460406 | 0.93110043 | 0.953147782 |
| -1.354285092 | 0.17564544 | 0.265884812 |
| NA | NA | NA |
| 0.192809521 | 0.847108147 | NA |
| -0.417619407 | 0.676225405 | NA |
| 0.063708217 | 0.949202562 | NA |
| -0.245687406 | 0.805924222 | 0.862632984 |
| 0.094623489 | 0.924613892 | 0.948190794 |
| 0.152439504 | 0.878840301 | 0.915032144 |
| 1.285146512 | 0.198741091 | 0.294122243 |
| -8.564256363 | 1.09E-17 | 1.60E-16 |
| -0.725778537 | 0.467974546 | 0.57721396 |
| -5.476565196 | 4.34E-08 | 2.46E-07 |
| 0.067515511 | 0.946171314 | 0.9632191 |
| 3.615114236 | 0.000300215 | 0.000954329 |
| 3.522259404 | 0.000427885 | 0.001320817 |
| -3.449902163 | 0.00056079 | 0.001694828 |
| 1.1083149 | 0.267725843 | 0.373636903 |
| 0.192808924 | 0.847108615 | NA |
| 0.725443308 | 0.468180112 | NA |
| -1.017143508 | 0.309085166 | 0.418976534 |
| -2.869943449 | 0.004105452 | 0.010294477 |
| 0.441537937 | 0.658823604 | 0.746068624 |
| -0.657903681 | 0.510600025 | 0.616394262 |
| 17.5692506 | 4.24E-69 | 5.25E-67 |
| 4.367651694 | 1.26E-05 | 5.03E-05 |

| | | |
|---|---|---|
| 0.526901487 | 0.598262001 | 0.696502888 |
| -5.456673283 | 4.85E-08 | 2.73E-07 |
| NA | NA | NA |
| 1.490503981 | 0.136091771 | 0.215315866 |
| -4.227604614 | 2.36E-05 | 9.07E-05 |
| -1.267543549 | 0.204961002 | 0.301526618 |
| 2.243641972 | 0.024855446 | 0.050456758 |
| 0.054890613 | 0.95622561 | 0.970061485 |
| -1.110024247 | 0.266988578 | 0.372795053 |
| -4.889616606 | 1.01E-06 | 4.74E-06 |
| 0.696628241 | 0.486035477 | 0.594307181 |
| -1.55946231 | 0.118886999 | 0.1923805 |
| 8.214798468 | 2.13E-16 | 2.81E-15 |
| 4.806083291 | 1.54E-06 | 7.03E-06 |
| 1.413850941 | 0.157405673 | 0.242856684 |
| 4.848692729 | 1.24E-06 | 5.75E-06 |
| 6.570240158 | 5.02E-11 | 3.95E-10 |
| -0.854626119 | 0.392758163 | 0.504709766 |
| 2.065146668 | 0.038909125 | 0.074371546 |
| 1.510306929 | 0.130965124 | 0.20859415 |
| 3.844727423 | 0.000120687 | 0.000411327 |
| -2.7560093 | 0.005851133 | 0.014140568 |
| -4.505307915 | 6.63E-06 | 2.77E-05 |
| -3.199406737 | 0.001377107 | 0.003846128 |
| -2.237506614 | 0.025253248 | 0.051172991 |
| 3.431010766 | 0.000601337 | 0.001807061 |
| 2.176903618 | 0.029487759 | 0.058498741 |
| 4.622157813 | 3.80E-06 | 1.64E-05 |
| 0.175755415 | 0.860486098 | 0.90309728 |
| NA | NA | NA |
| 0.925867628 | 0.354514777 | NA |
| 0.271196472 | 0.786239926 | 0.847636876 |
| 6.081169091 | 1.19E-09 | 8.06E-09 |
| -0.7655911 | 0.443919642 | NA |
| -3.951576746 | 7.76E-05 | 0.000274064 |
| NA | NA | NA |
| NA | NA | NA |
| -5.861030409 | 4.60E-09 | 2.92E-08 |
| -8.635258433 | 5.86E-18 | 8.78E-17 |
| 0.641492923 | 0.521202477 | 0.626414002 |
| 0.492221807 | 0.62256255 | NA |
| 0.333754673 | 0.738564689 | 0.809594209 |
| -6.183620159 | 6.26E-10 | 4.34E-09 |
| -2.103270532 | 0.035442128 | 0.068641014 |
| -1.879179736 | 0.060219956 | 0.108361286 |
| -5.714401896 | 1.10E-08 | 6.70E-08 |
| -3.478088176 | 0.000505004 | 0.001539803 |

| | | |
|---|---|---|
| 6.728454246 | 1.71E-11 | 1.41E-10 |
| -4.393525443 | 1.12E-05 | 4.51E-05 |
| -2.263224266 | 0.023621868 | 0.04829334 |
| -0.78182545 | 0.434317162 | 0.544946612 |
| -0.93985511 | 0.347291886 | 0.458839583 |
| -0.811442922 | 0.417111359 | 0.528756695 |
| -0.065389369 | 0.947863988 | NA |
| 0.076339117 | 0.939149306 | 0.958604809 |
| -4.161572109 | 3.16E-05 | 0.000119024 |
| -3.872982552 | 0.000107512 | 0.000369695 |
| -2.420485982 | 0.015499777 | 0.033528599 |
| 9.306783147 | 1.32E-20 | 2.45E-19 |
| 0.142347658 | 0.886805404 | NA |
| 12.86938843 | 6.69E-38 | 3.37E-36 |
| 3.696794639 | 0.000218339 | 0.000712148 |
| -0.873497995 | 0.382391699 | 0.494690108 |
| -2.998759031 | 0.002710816 | 0.007074521 |
| -2.359744147 | 0.018287543 | 0.038686376 |
| 2.753619172 | 0.005894031 | 0.0142305 |
| -1.223588069 | 0.221107663 | 0.32038467 |
| -0.604281016 | 0.545656825 | 0.64935162 |
| -3.960837221 | 7.47E-05 | 0.000264353 |
| -0.883053961 | 0.377207117 | 0.489907334 |
| -0.174895349 | 0.861161865 | 0.903351975 |
| 4.576532885 | 4.73E-06 | 2.02E-05 |
| 1.653683421 | 0.098191853 | 0.164101094 |
| 4.914977589 | 8.88E-07 | 4.20E-06 |
| -2.532241562 | 0.011333588 | 0.025485555 |
| -1.017281168 | 0.309019692 | 0.418910469 |
| 1.824159516 | 0.06812796 | 0.12041825 |
| -3.287431279 | 0.001011059 | 0.002912175 |
| -4.722749462 | 2.33E-06 | 1.04E-05 |
| -0.203337867 | 0.83887096 | NA |
| -0.291519396 | 0.770654112 | 0.83478913 |
| -1.703841864 | 0.088410634 | 0.150344329 |
| 1.8279131 | 0.067562595 | 0.119603168 |
| 0.847879291 | 0.396505199 | 0.508400596 |
| 10.70198967 | 9.96E-27 | 2.85E-25 |
| 1.486564625 | 0.137129837 | 0.216730429 |
| 1.117694544 | 0.263697474 | 0.369208915 |
| 0.840317148 | 0.40073059 | 0.512325822 |
| 1.182348596 | 0.237067411 | 0.339035384 |
| 0.795703043 | 0.426204658 | 0.537568031 |
| 7.523604925 | 5.33E-14 | 5.63E-13 |
| 1.938219681 | 0.052596427 | 0.096715292 |
| -4.6801757 | 2.87E-06 | 1.26E-05 |
| 0.192809521 | 0.847108147 | NA |

| | | |
|---|---|---|
| 0.59879903 | 0.549306909 | 0.652732242 |
| 2.363270527 | 0.018114441 | 0.038369894 |
| 8.949080001 | 3.58E-19 | 6.00E-18 |
| 0.568131365 | 0.569945772 | 0.67136066 |
| 10.41521929 | 2.11E-25 | 5.56E-24 |
| 1.491961186 | 0.135709321 | NA |
| 3.185898595 | 0.001443052 | 0.004013733 |
| 16.0204122 | 9.20E-58 | 8.56E-56 |
| 14.06387055 | 6.33E-45 | 4.24E-43 |
| -1.355632112 | 0.175216249 | 0.265347454 |
| -0.065389369 | 0.947863988 | NA |
| 0.192395457 | 0.847432451 | 0.893480802 |
| NA | NA | NA |
| NA | NA | NA |
| NA | NA | NA |
| NA | NA | NA |
| NA | NA | NA |
| NA | NA | NA |
| NA | NA | NA |
| NA | NA | NA |
| NA | NA | NA |
| NA | NA | NA |
| NA | NA | NA |
| NA | NA | NA |
| NA | NA | NA |
| NA | NA | NA |
| NA | NA | NA |
| NA | NA | NA |
| NA | NA | NA |
| 2.650049776 | 0.008047991 | 0.018728265 |
| NA | NA | NA |
| NA | NA | NA |
| NA | NA | NA |
| NA | NA | NA |
| NA | NA | NA |
| NA | NA | NA |
| NA | NA | NA |
| NA | NA | NA |
| NA | NA | NA |
| NA | NA | NA |
| NA | NA | NA |
| NA | NA | NA |
| NA | NA | NA |
| NA | NA | NA |
| NA | NA | NA |
| NA | NA | NA |
| NA | NA | NA |

| | | |
|---|---|---|
| NA | NA | NA |
| NA | NA | NA |
| NA | NA | NA |
| NA | NA | NA |
| NA | NA | NA |
| NA | NA | NA |
| NA | NA | NA |
| NA | NA | NA |
| NA | NA | NA |
| NA | NA | NA |
| NA | NA | NA |
| NA | NA | NA |
| NA | NA | NA |
| NA | NA | NA |
| NA | NA | NA |
| NA | NA | NA |
| NA | NA | NA |
| NA | NA | NA |
| NA | NA | NA |
| NA | NA | NA |
| NA | NA | NA |
| NA | NA | NA |
| NA | NA | NA |
| NA | NA | NA |
| NA | NA | NA |
| NA | NA | NA |
| NA | NA | NA |
| NA | NA | NA |
| NA | NA | NA |
| NA | NA | NA |
| NA | NA | NA |
| NA | NA | NA |
| NA | NA | NA |
| NA | NA | NA |
| NA | NA | NA |
| NA | NA | NA |
| NA | NA | NA |
| NA | NA | NA |
| NA | NA | NA |
| NA | NA | NA |
| NA | NA | NA |
| NA | NA | NA |
| NA | NA | NA |
| NA | NA | NA |
| NA | NA | NA |
| NA | NA | NA |
| NA | NA | NA |

| | | |
|---|---|---|
| NA | NA | NA |
| NA | NA | NA |
| NA | NA | NA |
| NA | NA | NA |
| NA | NA | NA |
| NA | NA | NA |
| NA | NA | NA |
| NA | NA | NA |
| NA | NA | NA |
| NA | NA | NA |
| NA | NA | NA |
| NA | NA | NA |
| NA | NA | NA |
| NA | NA | NA |
| NA | NA | NA |
| NA | NA | NA |
| NA | NA | NA |
| NA | NA | NA |
| NA | NA | NA |
| NA | NA | NA |
| NA | NA | NA |
| NA | NA | NA |
| NA | NA | NA |
| NA | NA | NA |
| NA | NA | NA |
| NA | NA | NA |
| 0.192809521 | 0.847108147 | NA |
| NA | NA | NA |
| NA | NA | NA |
| -0.065388772 | 0.947864464 | NA |
| NA | NA | NA |
| 0.192808924 | 0.847108615 | NA |
| NA | NA | NA |
| NA | NA | NA |
| NA | NA | NA |
| NA | NA | NA |
| NA | NA | NA |
| NA | NA | NA |
| NA | NA | NA |
| NA | NA | NA |
| NA | NA | NA |
| NA | NA | NA |
| NA | NA | NA |
| NA | NA | NA |
| NA | NA | NA |
| NA | NA | NA |
| NA | NA | NA |

| | | |
|---|---|---|
| NA | NA | NA |
| NA | NA | NA |
| NA | NA | NA |
| NA | NA | NA |
| NA | NA | NA |
| NA | NA | NA |
| NA | NA | NA |
| NA | NA | NA |
| NA | NA | NA |
| NA | NA | NA |
| NA | NA | NA |
| NA | NA | NA |
| NA | NA | NA |
| NA | NA | NA |
| NA | NA | NA |
| NA | NA | NA |
| NA | NA | NA |
| NA | NA | NA |
| NA | NA | NA |
| NA | NA | NA |
| NA | NA | NA |
| NA | NA | NA |
| NA | NA | NA |
| NA | NA | NA |
| NA | NA | NA |
| NA | NA | NA |
| NA | NA | NA |
| NA | NA | NA |
| NA | NA | NA |
| NA | NA | NA |
| NA | NA | NA |
| NA | NA | NA |
| NA | NA | NA |
| NA | NA | NA |
| NA | NA | NA |
| NA | NA | NA |
| NA | NA | NA |
| NA | NA | NA |
| NA | NA | NA |
| NA | NA | NA |
| NA | NA | NA |
| NA | NA | NA |
| NA | NA | NA |
| NA | NA | NA |
| NA | NA | NA |
| NA | NA | NA |
| NA | NA | NA |

| | | |
|---|---|---|
| NA | NA | NA |
| NA | NA | NA |
| NA | NA | NA |
| NA | NA | NA |
| NA | NA | NA |
| NA | NA | NA |
| NA | NA | NA |
| NA | NA | NA |
| NA | NA | NA |
| NA | NA | NA |
| NA | NA | NA |
| NA | NA | NA |
| NA | NA | NA |
| NA | NA | NA |
| NA | NA | NA |
| NA | NA | NA |
| NA | NA | NA |
| NA | NA | NA |
| NA | NA | NA |
| NA | NA | NA |
| NA | NA | NA |
| NA | NA | NA |
| NA | NA | NA |
| NA | NA | NA |
| NA | NA | NA |
| NA | NA | NA |
| NA | NA | NA |
| NA | NA | NA |
| NA | NA | NA |
| NA | NA | NA |
| NA | NA | NA |
| NA | NA | NA |
| NA | NA | NA |
| NA | NA | NA |
| NA | NA | NA |
| NA | NA | NA |
| NA | NA | NA |
| NA | NA | NA |
| NA | NA | NA |
| NA | NA | NA |
| NA | NA | NA |
| NA | NA | NA |
| NA | NA | NA |
| NA | NA | NA |
| NA | NA | NA |
| NA | NA | NA |
| NA | NA | NA |

| | | | |
|---|---|---|---|
| NA | | NA | NA |
| NA | | NA | NA |
| NA | | NA | NA |
| NA | | NA | NA |
| NA | | NA | NA |
| NA | | NA | NA |
| NA | | NA | NA |
| NA | | NA | NA |
| NA | | NA | NA |
| NA | | NA | NA |
| NA | | NA | NA |
| | 0.299483646 | 0.764571048 | 0.830206725 |
| NA | | NA | NA |
| | 0.691388291 | 0.48932156 | 0.597129279 |
| | 1.521918945 | 0.128029392 | 0.2049436 |
| | -2.543553675 | 0.01097312 | 0.02475281 |
| | -1.749523595 | 0.080200554 | 0.138372893 |
| | -1.079587973 | 0.280325699 | 0.387290386 |
| | -2.276082978 | 0.022841044 | 0.046942386 |
| | -1.473717187 | 0.140557753 | 0.221128886 |
| | 0.219408603 | 0.826331769 | 0.87852373 |
| | 0.60065477 | 0.54806995 | 0.651571713 |
| | 0.150890234 | 0.880062304 | 0.91577116 |
| | 1.312771719 | 0.189259882 | 0.282546514 |
| | -0.15925376 | 0.87346895 | 0.911838698 |
| | -0.507039504 | 0.612127101 | 0.70795404 |
| | 1.461686908 | 0.143827026 | 0.225496266 |
| | 0.475074593 | 0.634733807 | 0.72613549 |
| | -1.678181742 | 0.093311625 | 0.15733176 |
| | 0.192810118 | 0.847107679 | NA |
| | -0.694701594 | 0.487242329 | 0.595373407 |
| | -0.780991185 | 0.434807679 | 0.545429186 |
| | -1.049788636 | 0.293815301 | 0.401912613 |
| NA | | NA | NA |
| | -1.308420548 | 0.190730707 | 0.284386108 |
| NA | | NA | NA |
| | -1.797847773 | 0.072201134 | 0.126518691 |
| | -2.450657385 | 0.014259561 | 0.03113996 |
| | -0.244530817 | 0.806819737 | NA |
| | 0.063707135 | 0.949203424 | NA |
| | -0.175251911 | 0.860881696 | 0.903279692 |
| | -2.089972152 | 0.036620301 | 0.070616806 |
| | 3.030246748 | 0.00244354 | 0.00644073 |
| | -0.091806617 | 0.926851687 | 0.949887308 |
| | -3.213518885 | 0.001311192 | 0.003677625 |
| | 0.736664055 | 0.461326671 | 0.570729862 |
| | 0.205289855 | 0.83734567 | 0.88624306 |

| | | |
|---|---|---|
| NA | NA | NA |
| NA | NA | NA |
| NA | NA | NA |
| NA | NA | NA |
| NA | NA | NA |
| NA | NA | NA |
| NA | NA | NA |
| NA | NA | NA |
| NA | NA | NA |
| NA | NA | NA |
| NA | NA | NA |
| NA | NA | NA |
| NA | NA | NA |
| NA | NA | NA |
| NA | NA | NA |
| NA | NA | NA |
| NA | NA | NA |
| NA | NA | NA |
| NA | NA | NA |
| NA | NA | NA |
| NA | NA | NA |
| NA | NA | NA |
| NA | NA | NA |
| NA | NA | NA |
| NA | NA | NA |
| NA | NA | NA |
| NA | NA | NA |
| NA | NA | NA |
| NA | NA | NA |
| NA | NA | NA |
| NA | NA | NA |
| NA | NA | NA |
| NA | NA | NA |
| NA | NA | NA |
| NA | NA | NA |
| NA | NA | NA |
| NA | NA | NA |
| NA | NA | NA |
| NA | NA | NA |
| NA | NA | NA |
| NA | NA | NA |
| NA | NA | NA |
| NA | NA | NA |
| NA | NA | NA |
| NA | NA | NA |
| NA | NA | NA |
| NA | NA | NA |

| | | | |
|---|---|---|---|
| NA | | NA | NA |
| NA | | NA | NA |
| NA | | NA | NA |
| NA | | NA | NA |
| NA | | NA | NA |
| NA | | NA | NA |
| NA | | NA | NA |
| NA | | NA | NA |
| NA | | NA | NA |
| NA | | NA | NA |
| NA | | NA | NA |
| | 1.380534861 | 0.167422024 | 0.255398175 |
| | -0.065389369 | 0.947863988 | NA |
| | -0.065389966 | 0.947863513 | NA |
| | -2.331539631 | 0.019724923 | 0.041283955 |
| NA | | NA | NA |
| | -3.058332133 | 0.002225727 | 0.005931008 |
| | 0.669412208 | 0.503232567 | 0.609853352 |
| | 0.007696102 | 0.99385946 | 0.995625108 |
| NA | | NA | NA |
| | -3.858262967 | 0.000114196 | 0.000390694 |
| NA | | NA | NA |
| | -1.563157619 | 0.118015526 | 0.191180416 |
| | 0.60151322 | 0.547498209 | 0.65110848 |
| | 1.755717572 | 0.079136645 | 0.136786981 |
| | 1.460270354 | 0.144215786 | 0.225988927 |
| | 2.547710135 | 0.010843253 | 0.024499604 |
| NA | | NA | NA |
| | 0.063707194 | 0.949203376 | NA |
| | 0.136628688 | 0.891324299 | 0.923917969 |
| NA | | NA | NA |
| | 0.050214736 | 0.959951268 | NA |
| | -0.272683077 | 0.785096845 | 0.846770576 |
| | 0.063708285 | 0.949202508 | NA |
| | -0.321921986 | 0.747511797 | NA |
| | -2.180210553 | 0.029241858 | NA |
| | 1.115287488 | 0.264727246 | 0.370319284 |
| | 1.450371859 | 0.14695485 | 0.22946604 |
| | -1.989672705 | 0.046627001 | 0.087147091 |
| | 0.063708882 | 0.949202033 | NA |
| NA | | NA | NA |
| | 1.209624698 | 0.226422937 | NA |
| | 0.192809521 | 0.847108147 | NA |
| | -3.246053077 | 0.00117017 | 0.003323804 |
| | -1.31910146 | 0.187135195 | 0.279925532 |
| | -0.209442726 | 0.834102642 | 0.883819707 |
| | 2.558532684 | 0.010511494 | 0.023837741 |

| | | |
|---|---|---|
| 0.443124352 | 0.657675792 | 0.745240847 |
| 2.203074587 | 0.027589492 | 0.055227526 |
| -7.66563632 | 1.78E-14 | 1.98E-13 |
| -1.859300604 | 0.062984541 | 0.112615406 |
| 0.885025129 | 0.376143078 | 0.488949052 |
| 10.34720358 | 4.31E-25 | 1.11E-23 |
| -0.326261659 | 0.744226394 | 0.813946808 |
| NA | NA | NA |
| 3.744644075 | 0.00018065 | 0.000597708 |
| NA | NA | NA |
| -2.324675838 | 0.0200893 | 0.04193788 |
| -3.159380238 | 0.001581051 | 0.004360738 |
| -3.324043694 | 0.000887223 | 0.002583734 |
| -1.077901662 | 0.281077644 | 0.388115707 |
| 8.844873715 | 9.16E-19 | 1.48E-17 |
| 0.000550194 | 0.999561009 | 0.999600945 |
| -2.517395892 | 0.01182259 | 0.026433179 |
| -7.07659816 | 1.48E-12 | 1.35E-11 |
| 6.333177963 | 2.40E-10 | 1.74E-09 |
| -4.199117882 | 2.68E-05 | 0.000102131 |
| 2.129283242 | 0.033230833 | 0.064941268 |
| -3.286570042 | 0.001014155 | 0.002920087 |
| -0.842058129 | 0.399755417 | 0.511521808 |
| 0.192809521 | 0.847108147 | NA |
| 0.15376591 | 0.877794314 | NA |
| -1.20266315 | 0.2291067 | 0.329571304 |
| 0.379114205 | 0.704603058 | NA |
| -4.367196889 | 1.26E-05 | 5.04E-05 |
| NA | NA | NA |
| 1.200977305 | 0.229760005 | 0.330370974 |
| -0.614787546 | 0.538695035 | 0.643053068 |
| -0.19448762 | 0.845794089 | NA |
| 0.937484424 | 0.348509432 | 0.460156728 |
| -0.417397121 | 0.67638796 | 0.75960116 |
| 0.063708285 | 0.949202508 | NA |
| 0.468333306 | 0.639546254 | 0.730251482 |
| -2.528057176 | 0.011469567 | 0.025756618 |
| 13.89472191 | 6.82E-44 | 4.40E-42 |
| NA | NA | NA |
| -1.22724551 | 0.219730329 | 0.318739504 |
| -0.464513385 | 0.642279972 | NA |
| -8.795037643 | 1.43E-18 | 2.28E-17 |
| 4.51183737 | 6.43E-06 | 2.69E-05 |
| 0.541810539 | 0.587949031 | 0.687712708 |
| -0.097751656 | 0.922129496 | 0.946481641 |
| 6.699143836 | 2.10E-11 | 1.71E-10 |
| -6.29072383 | 3.16E-10 | 2.26E-09 |

| | | | |
|---|---|---|---|
| | -1.158910132 | 0.246492819 | 0.349558454 |
| | 2.623708456 | 0.00869782 | 0.020102164 |
| NA | NA | NA | |
| NA | NA | NA | |
| NA | NA | NA | |
| NA | NA | NA | |
| NA | NA | NA | |
| NA | NA | NA | |
| | 0.063708324 | 0.949202477 | NA |
| NA | NA | NA | |
| NA | NA | NA | |
| NA | NA | NA | |
| | -0.065388772 | 0.947864464 | NA |
| | 0.382131398 | 0.702363903 | NA |
| NA | NA | NA | |
| NA | NA | NA | |
| | -1.234257648 | 0.217106912 | 0.315723133 |
| NA | NA | NA | |
| | -0.119574487 | 0.904820231 | NA |
| | 0.864641419 | 0.387235632 | 0.49953656 |
| | -0.323542405 | 0.746284499 | NA |
| NA | NA | NA | |
| NA | NA | NA | |
| NA | NA | NA | |
| NA | NA | NA | |
| NA | NA | NA | |
| NA | NA | NA | |
| NA | NA | NA | |
| NA | NA | NA | |
| | 0.083076052 | 0.933791068 | NA |
| NA | NA | NA | |
| NA | NA | NA | |
| NA | NA | NA | |
| NA | NA | NA | |
| | 0.192809521 | 0.847108147 | NA |
| NA | NA | NA | |
| NA | NA | NA | |
| NA | NA | NA | |
| NA | NA | NA | |
| NA | NA | NA | |
| NA | NA | NA | |
| | -1.8175651 | 0.06913063 | 0.12194078 |
| | 0.063708882 | 0.949202033 | NA |
| | -5.166668463 | 2.38E-07 | 1.22E-06 |
| | -1.630274461 | 0.103043503 | 0.17068221 |
| | -0.576261731 | 0.564438286 | NA |
| | 1.822087031 | 0.068441781 | 0.120913169 |

| | | |
|---|---|---|
| 0.673286173 | 0.500765247 | NA |
| 0.063707688 | 0.949202983 | NA |
| 25.8022491 | 8.37E-147 | 3.38E-144 |
| -0.065389369 | 0.947863988 | NA |
| 0.116863935 | 0.906967878 | NA |
| -0.637128266 | 0.524041299 | 0.629161248 |
| -1.314962164 | 0.188522619 | 0.281664248 |
| -0.694052127 | 0.48764952 | 0.59572858 |
| -3.433519927 | 0.000595798 | 0.001791977 |
| 0.597695671 | 0.550043015 | 0.653420819 |
| 0.715739749 | 0.474152062 | 0.583023487 |
| -4.398725379 | 1.09E-05 | 4.41E-05 |
| -0.312630441 | 0.754561448 | NA |
| -1.104153906 | 0.269526381 | 0.375667965 |
| -9.923335287 | 3.30E-23 | 7.46E-22 |
| -4.519422318 | 6.20E-06 | 2.60E-05 |
| 0.74461608 | 0.456503844 | 0.566162482 |
| 0.192808924 | 0.847108615 | NA |
| -1.230884498 | 0.218366066 | 0.317201709 |
| -1.098418424 | 0.272021821 | 0.378492755 |
| 2.137648651 | 0.03254527 | 0.063780779 |
| -1.445771269 | 0.148241367 | 0.231110646 |
| NA | NA | NA |
| -0.065388772 | 0.947864464 | NA |
| 0.3535515 | 0.723675027 | NA |
| -0.065389369 | 0.947863988 | NA |
| NA | NA | NA |
| NA | NA | NA |
| NA | NA | NA |
| NA | NA | NA |
| NA | NA | NA |
| NA | NA | NA |
| NA | NA | NA |
| NA | NA | NA |
| NA | NA | NA |
| NA | NA | NA |
| NA | NA | NA |
| NA | NA | NA |
| NA | NA | NA |
| -1.454474618 | 0.145814765 | 0.228059235 |
| NA | NA | NA |
| -0.043102928 | 0.965619485 | 0.976695338 |
| NA | NA | NA |
| -0.065389369 | 0.947863988 | NA |
| 1.348350238 | 0.177545761 | NA |
| 0.88194999 | 0.377803852 | 0.490427882 |
| 0.836433916 | 0.402910839 | 0.51463368 |

| | | | |
|---|---|---|---|
| | 0.063708285 | 0.949202508 | NA |
| | 1.082682575 | 0.278949341 | 0.385986069 |
| | 0.870259882 | 0.384158398 | NA |
| | -7.759571583 | 8.52E-15 | 9.77E-14 |
| | -3.995436461 | 6.46E-05 | 0.000230705 |
| | 0.063708285 | 0.949202508 | NA |
| NA | NA | NA | |
| | 2.949651384 | 0.003181327 | 0.008168712 |
| | 0.362026528 | 0.717332207 | NA |
| | -3.98975834 | 6.61E-05 | 0.00023586 |
| | -2.119259787 | 0.034068518 | 0.066355537 |
| | 0.255447283 | 0.798377656 | NA |
| NA | NA | NA | |
| | 2.500675648 | 0.012395665 | 0.027556931 |
| NA | NA | NA | |
| | 1.876849747 | 0.06053869 | 0.108864388 |
| | -0.387444126 | 0.698427442 | 0.778068313 |
| NA | NA | NA | |
| | 3.498373703 | 0.000468105 | 0.00143481 |
| | -0.420803161 | 0.673898823 | 0.757553559 |
| | -0.194487022 | 0.845794556 | NA |
| | 0.372737018 | 0.709344182 | NA |
| | -1.439735524 | 0.149942239 | 0.233384381 |
| NA | NA | NA | |
| | 1.469954329 | 0.141574124 | 0.222476162 |
| | -6.205518576 | 5.45E-10 | 3.81E-09 |
| NA | NA | NA | |
| | 1.482729124 | 0.138146393 | 0.218006697 |
| NA | NA | NA | |
| NA | NA | NA | |
| NA | NA | NA | |
| NA | NA | NA | |
| NA | NA | NA | |
| | 0.282041257 | 0.777911875 | NA |
| NA | NA | NA | |
| NA | NA | NA | |
| | 0.078655732 | 0.937306457 | NA |
| NA | NA | NA | |
| NA | NA | NA | |
| NA | NA | NA | |
| NA | NA | NA | |
| NA | NA | NA | |
| | 0.244694817 | 0.806692741 | NA |
| NA | NA | NA | |
| | 0.288217989 | 0.77317988 | NA |
| | 0.192810118 | 0.847107679 | NA |
| | 0.192810118 | 0.847107679 | NA |

| | | |
|---|---|---|
| 1.192708519 | 0.232983548 | 0.334092135 |
| 2.068357969 | 0.038606374 | 0.073911393 |
| -2.73260244 | 0.006283613 | 0.015064506 |
| -0.698016556 | 0.48516684 | 0.593505988 |
| NA | NA | NA |
| -0.065389369 | 0.947863988 | NA |
| 1.388385538 | 0.165019678 | 0.2523949 |
| 0.30922356 | 0.757151473 | NA |
| NA | NA | NA |
| NA | NA | NA |
| NA | NA | NA |
| NA | NA | NA |
| -0.957611393 | 0.338258752 | 0.449520272 |
| NA | NA | NA |
| NA | NA | NA |
| 2.256863336 | 0.024016616 | 0.048956421 |
| NA | NA | NA |
| NA | NA | NA |
| 1.271320869 | 0.203614521 | 0.29995922 |
| 0.224517291 | 0.822354814 | 0.875448324 |
| 0.043085672 | 0.96563324 | 0.976695338 |
| -2.733953268 | 0.006257893 | 0.015013424 |
| 0.235445464 | 0.813863003 | NA |
| NA | NA | NA |
| NA | NA | NA |
| NA | NA | NA |
| NA | NA | NA |
| 0.63344967 | 0.526440038 | 0.631465352 |
| 0.192810118 | 0.847107679 | NA |
| -0.631173636 | 0.527926998 | NA |
| -0.362484691 | 0.716989861 | 0.792924324 |
| -5.904261117 | 3.54E-09 | 2.28E-08 |
| 0.776153964 | 0.437658084 | 0.547948271 |
| NA | NA | NA |
| 1.54434417 | 0.122504978 | 0.197380083 |
| 0.063707688 | 0.949202983 | NA |
| -1.343139283 | 0.179226877 | 0.270227621 |
| 0.797887632 | 0.424935701 | 0.536489439 |
| 0.063708882 | 0.949202033 | NA |
| NA | NA | NA |
| 1.447829323 | 0.14766479 | 0.230398773 |
| 0.425034791 | 0.670811313 | 0.75513412 |
| 0.044144311 | 0.964789372 | 0.976061817 |
| -0.025034913 | 0.980027116 | 0.98623594 |
| -2.200429887 | 0.027776409 | 0.055539505 |
| 6.669535531 | 2.57E-11 | 2.07E-10 |
| -4.463024166 | 8.08E-06 | 3.34E-05 |

| | | |
|---|---|---|
| -11.5672554 | 6.04E-31 | 2.19E-29 |
| -4.263596239 | 2.01E-05 | 7.82E-05 |
| NA | NA | NA |
| NA | NA | NA |
| 0.767248189 | 0.442933966 | NA |
| -6.364722457 | 1.96E-10 | 1.44E-09 |
| NA | NA | NA |
| 12.98227715 | 1.54E-38 | 7.98E-37 |
| NA | NA | NA |
| 2.569124668 | 0.010195577 | 0.023197463 |
| 1.437230518 | 0.150652504 | 0.234344179 |
| 2.931360426 | 0.00337481 | 0.008615144 |
| -1.993088936 | 0.046251709 | 0.08652319 |
| -1.744747257 | 0.081028873 | 0.139651085 |
| 1.906047898 | 0.056643992 | 0.103015266 |
| 1.117433919 | 0.263808839 | 0.36930755 |
| NA | NA | NA |
| 0.047986409 | 0.961727074 | 0.973706566 |
| NA | NA | NA |
| 0.063708882 | 0.949202033 | NA |
| -0.853725159 | 0.393257291 | 0.505195545 |
| -1.401466532 | 0.161074609 | 0.247252389 |
| 1.599638076 | 0.109678897 | 0.179718217 |
| -0.114291599 | 0.909006642 | 0.937201311 |
| NA | NA | NA |
| 0.138709691 | 0.88967956 | 0.922824557 |
| NA | NA | NA |
| 0.500430432 | 0.61677203 | 0.711637657 |
| -0.065388772 | 0.947864464 | NA |
| NA | NA | NA |
| 0.152347385 | 0.878912953 | 0.915034422 |
| NA | NA | NA |
| NA | NA | NA |
| NA | NA | NA |
| NA | NA | NA |
| NA | NA | NA |
| NA | NA | NA |
| NA | NA | NA |
| NA | NA | NA |
| NA | NA | NA |
| -5.137698379 | 2.78E-07 | 1.41E-06 |
| -1.56248583 | 0.118173582 | 0.191381551 |
| -0.065388772 | 0.947864464 | NA |
| 0.208330271 | 0.834971097 | 0.884365545 |
| NA | NA | NA |
| 0.353720202 | 0.723548581 | NA |
| NA | NA | NA |

| | | |
|---|---|---|
| NA | NA | NA |
| NA | NA | NA |
| NA | NA | NA |
| NA | NA | NA |
| NA | NA | NA |
| 0.192809521 | 0.847108147 | NA |
| NA | NA | NA |
| NA | NA | NA |
| NA | NA | NA |
| NA | NA | NA |
| NA | NA | NA |
| NA | NA | NA |
| NA | NA | NA |
| NA | NA | NA |
| NA | NA | NA |
| NA | NA | NA |
| NA | NA | NA |
| NA | NA | NA |
| NA | NA | NA |
| NA | NA | NA |
| NA | NA | NA |
| NA | NA | NA |
| NA | NA | NA |
| NA | NA | NA |
| NA | NA | NA |
| NA | NA | NA |
| NA | NA | NA |
| NA | NA | NA |
| NA | NA | NA |
| NA | NA | NA |
| NA | NA | NA |
| NA | NA | NA |
| NA | NA | NA |
| NA | NA | NA |
| NA | NA | NA |
| NA | NA | NA |
| NA | NA | NA |
| NA | NA | NA |
| NA | NA | NA |
| NA | NA | NA |
| NA | NA | NA |
| NA | NA | NA |
| NA | NA | NA |
| NA | NA | NA |
| NA | NA | NA |
| NA | NA | NA |
| NA | NA | NA |

| | | |
|---|---|---|
| NA | NA | NA |
| NA | NA | NA |
| NA | NA | NA |
| NA | NA | NA |
| NA | NA | NA |
| NA | NA | NA |
| NA | NA | NA |
| 1.500112892 | 0.133585162 | 0.212089597 |
| NA | NA | NA |
| NA | NA | NA |
| NA | NA | NA |
| NA | NA | NA |
| NA | NA | NA |
| NA | NA | NA |
| NA | NA | NA |
| NA | NA | NA |
| NA | NA | NA |
| NA | NA | NA |
| NA | NA | NA |
| NA | NA | NA |
| NA | NA | NA |
| NA | NA | NA |
| NA | NA | NA |
| NA | NA | NA |
| NA | NA | NA |
| NA | NA | NA |
| NA | NA | NA |
| NA | NA | NA |
| NA | NA | NA |
| NA | NA | NA |
| NA | NA | NA |
| NA | NA | NA |
| NA | NA | NA |
| NA | NA | NA |
| NA | NA | NA |
| NA | NA | NA |
| NA | NA | NA |
| NA | NA | NA |
| NA | NA | NA |
| NA | NA | NA |
| NA | NA | NA |
| NA | NA | NA |
| NA | NA | NA |
| NA | NA | NA |
| NA | NA | NA |
| NA | NA | NA |
| NA | NA | NA |

| | | |
|---|---|---|
| NA | NA | NA |
| 2.036787722 | 0.041671319 | 0.079011675 |
| NA | NA | NA |
| NA | NA | NA |
| NA | NA | NA |
| NA | NA | NA |
| NA | NA | NA |
| NA | NA | NA |
| NA | NA | NA |
| NA | NA | NA |
| NA | NA | NA |
| NA | NA | NA |
| NA | NA | NA |
| NA | NA | NA |
| NA | NA | NA |
| NA | NA | NA |
| NA | NA | NA |
| NA | NA | NA |
| NA | NA | NA |
| NA | NA | NA |
| NA | NA | NA |
| NA | NA | NA |
| NA | NA | NA |
| NA | NA | NA |
| 0.192809521 | 0.847108147 | NA |
| NA | NA | NA |
| 6.400038914 | 1.55E-10 | 1.15E-09 |
| 14.80095249 | 1.44E-49 | 1.10E-47 |
| -3.507950447 | 0.000451573 | 0.001388269 |
| NA | NA | NA |
| 1.785896724 | 0.074115981 | 0.129285875 |
| NA | NA | NA |
| 3.198873381 | 0.001379657 | 0.003851531 |
| 2.027832917 | 0.042577304 | 0.080503846 |
| 0.19280579 | 0.847111069 | NA |
| NA | NA | NA |
| 0.063708285 | 0.949202508 | NA |
| NA | NA | NA |
| 2.211096849 | 0.027029131 | 0.054209868 |
| -0.538630474 | 0.590141856 | 0.689633066 |
| -5.595604268 | 2.20E-08 | 1.29E-07 |
| 1.536565883 | 0.124399653 | 0.20000792 |
| 4.026363879 | 5.66E-05 | 0.000204566 |
| 1.178113032 | 0.238751547 | NA |
| 2.393531592 | 0.016687044 | 0.03579677 |
| NA | NA | NA |
| 0.304727353 | 0.760573815 | 0.826768114 |
| 2.400656131 | 0.016365707 | 0.035207018 |

| | | |
|---|---|---|
| 0.405767174 | 0.68491368 | 0.767179334 |
| 1.864519868 | 0.062248734 | 0.111478734 |
| -3.1528021 | 0.001617114 | 0.004450887 |
| -0.499164111 | 0.617663776 | NA |
| -5.907373174 | 3.48E-09 | 2.24E-08 |
| NA | NA | NA |
| 0.06353712 | 0.949338802 | NA |
| NA | NA | NA |
| NA | NA | NA |
| 5.662693888 | 1.49E-08 | 8.93E-08 |
| -0.342461268 | 0.732003788 | 0.804657655 |
| -2.065383792 | 0.038886701 | 0.074334361 |
| 2.445686291 | 0.014457674 | 0.031511283 |
| 0.015866161 | 0.987341166 | 0.991102843 |
| -1.719324266 | 0.085555344 | 0.146184057 |
| 1.061530413 | 0.288448919 | 0.395804859 |
| NA | NA | NA |
| NA | NA | NA |
| NA | NA | NA |
| NA | NA | NA |
| NA | NA | NA |
| NA | NA | NA |
| NA | NA | NA |
| NA | NA | NA |
| NA | NA | NA |
| NA | NA | NA |
| NA | NA | NA |
| NA | NA | NA |
| NA | NA | NA |
| NA | NA | NA |
| NA | NA | NA |
| NA | NA | NA |
| NA | NA | NA |
| NA | NA | NA |
| NA | NA | NA |
| NA | NA | NA |
| NA | NA | NA |
| NA | NA | NA |
| NA | NA | NA |
| NA | NA | NA |
| NA | NA | NA |
| NA | NA | NA |
| NA | NA | NA |
| NA | NA | NA |
| NA | NA | NA |
| NA | NA | NA |
| NA | NA | NA |

| | | |
|---|---|---|
| NA | NA | NA |
| NA | NA | NA |
| NA | NA | NA |
| NA | NA | NA |
| 0.063708882 | 0.949202033 | NA |
| NA | NA | NA |
| NA | NA | NA |
| NA | NA | NA |
| NA | NA | NA |
| 0.192809521 | 0.847108147 | NA |
| NA | NA | NA |
| 2.411232485 | 0.01589871 | 0.034302622 |
| 2.455939796 | 0.01405167 | 0.030781142 |
| -0.340066208 | 0.733806669 | 0.805719465 |
| NA | NA | NA |
| 0.063708285 | 0.949202508 | NA |
| 0.539910978 | 0.589258427 | NA |
| NA | NA | NA |
| 0.877212579 | 0.380371185 | 0.492916943 |
| 13.11384866 | 2.74E-39 | 1.46E-37 |
| -0.977093577 | 0.328522827 | 0.43956414 |
| 3.255301409 | 0.001132721 | 0.003226218 |
| -1.175389717 | 0.239838835 | 0.342021767 |
| 2.527021479 | 0.011503447 | 0.025821117 |
| 2.806236133 | 0.005012395 | 0.012333815 |
| 0.063708285 | 0.949202508 | NA |
| NA | NA | NA |
| NA | NA | NA |
| -0.513133831 | 0.607857708 | NA |
| NA | NA | NA |
| 0.514574445 | 0.606850426 | 0.703509155 |
| NA | NA | NA |
| NA | NA | NA |
| NA | NA | NA |
| NA | NA | NA |
| -0.065388772 | 0.947864464 | NA |
| NA | NA | NA |
| NA | NA | NA |
| -0.065389369 | 0.947863988 | NA |
| NA | NA | NA |
| -0.065388772 | 0.947864464 | NA |
| NA | NA | NA |
| NA | NA | NA |
| NA | NA | NA |
| NA | NA | NA |
| 1.23894458 | 0.215366023 | 0.313681207 |
| NA | NA | NA |

| | | |
|---|---|---|
| NA | NA | NA |
| NA | NA | NA |
| NA | NA | NA |
| NA | NA | NA |
| -11.4471582 | 2.43E-30 | 8.53E-29 |
| NA | NA | NA |
| NA | NA | NA |
| NA | NA | NA |
| 1.002189817 | 0.316251925 | 0.426773004 |
| NA | NA | NA |
| NA | NA | NA |
| NA | NA | NA |
| 0.421657796 | 0.673274814 | 0.757056091 |
| -0.065388772 | 0.947864464 | NA |
| -1.090540153 | 0.275475276 | 0.382150759 |
| 2.009588411 | 0.044474769 | 0.0836761 |
| 0.3474247 | 0.728272279 | NA |
| 1.499357647 | 0.133780875 | 0.212305834 |
| 5.569791188 | 2.55E-08 | 1.48E-07 |
| -6.699920617 | 2.09E-11 | 1.70E-10 |
| -1.690656494 | 0.090902428 | 0.153860412 |
| -1.72649857 | 0.084257756 | 0.144311731 |
| NA | NA | NA |
| -3.042540204 | 0.002345905 | 0.006213411 |
| NA | NA | NA |
| NA | NA | NA |
| NA | NA | NA |
| NA | NA | NA |
| NA | NA | NA |
| NA | NA | NA |
| -1.509419272 | 0.131191671 | 0.208848663 |
| NA | NA | NA |
| NA | NA | NA |
| -8.052368863 | 8.12E-16 | 1.02E-14 |
| NA | NA | NA |
| -4.447268501 | 8.70E-06 | 3.58E-05 |
| NA | NA | NA |
| NA | NA | NA |
| NA | NA | NA |
| NA | NA | NA |
| NA | NA | NA |
| NA | NA | NA |
| NA | NA | NA |
| NA | NA | NA |
| 0.192809521 | 0.847108147 | NA |
| NA | NA | NA |
| NA | NA | NA |

| | | | |
|---|---|---|---|
| | 0.320071032 | 0.748914485 | NA |
| | 0.233100121 | 0.815683661 | NA |
| | 0.07815334 | 0.937706077 | NA |
| NA | NA | NA | |
| | -0.449401087 | 0.653142347 | NA |
| | -0.065390563 | 0.947863038 | NA |
| | -7.662802595 | 1.82E-14 | 2.02E-13 |
| | 1.110460294 | 0.266800729 | 0.372574327 |
| | -7.732224536 | 1.06E-14 | 1.20E-13 |
| | -8.659723603 | 4.73E-18 | 7.14E-17 |
| | -3.745423752 | 0.00018009 | 0.000596012 |
| | 14.69631173 | 6.81E-49 | 5.13E-47 |
| | -3.143268582 | 0.001670725 | 0.004583323 |
| NA | NA | NA | |
| | 4.541630272 | 5.58E-06 | 2.36E-05 |
| | -1.445274168 | 0.148380891 | 0.231284948 |
| NA | NA | NA | |
| NA | NA | NA | |
| NA | NA | NA | |
| | -5.672304491 | 1.41E-08 | 8.46E-08 |
| | 21.20336175 | 8.89E-100 | 2.10E-97 |
| | -2.02969286 | 0.042387769 | 0.080195503 |
| | -2.332847394 | 0.019656156 | 0.041160678 |
| | -2.835979695 | 0.004568535 | 0.011343164 |
| | -5.746571741 | 9.11E-09 | 5.59E-08 |
| | -2.65417232 | 0.007950322 | 0.018521644 |
| | 0.149823351 | 0.880903986 | NA |
| | 1.504725372 | 0.132394697 | 0.210442578 |
| | 1.165539545 | 0.243800663 | 0.34658268 |
| NA | NA | NA | |
| | 0.159278972 | 0.873449087 | 0.911838698 |
| NA | NA | NA | |
| NA | NA | NA | |
| | 0.392329735 | 0.694814595 | NA |
| | 0.079638179 | 0.93652503 | NA |
| | 1.435086181 | 0.151262541 | 0.235161578 |
| NA | NA | NA | |
| | -1.123872931 | 0.261066936 | 0.366160788 |
| NA | NA | NA | |
| NA | NA | NA | |
| | -9.943141312 | 2.70E-23 | 6.17E-22 |
| NA | NA | NA | |
| | 0.063708882 | 0.949202033 | NA |
| NA | NA | NA | |
| | 0.663066018 | 0.507288274 | NA |
| NA | NA | NA | |
| | 2.744028142 | 0.006069033 | 0.014605125 |

| | | | |
|---|---|---|---|
| | 0.192810118 | 0.847107679 | NA |
| | 0.444307346 | 0.656820391 | NA |
| | 0.192809521 | 0.847108147 | NA |
| NA | NA | NA | |
| | -4.526372006 | 6.00E-06 | 2.52E-05 |
| | -3.083966773 | 0.002042604 | 0.005493325 |
| | 0.013187664 | 0.989478072 | 0.992582274 |
| | 6.415195036 | 1.41E-10 | 1.05E-09 |
| NA | NA | NA | |
| | 3.460856722 | 0.000538459 | 0.001631873 |
| NA | NA | NA | |
| | 0.514993604 | 0.606557489 | 0.70336502 |
| NA | NA | NA | |
| NA | NA | NA | |
| | -5.593634366 | 2.22E-08 | 1.31E-07 |
| | -0.960715289 | 0.336695342 | 0.447966318 |
| NA | NA | NA | |
| NA | NA | NA | |
| | 0.063708882 | 0.949202033 | NA |
| NA | NA | NA | |
| | 6.557711319 | 5.46E-11 | 4.28E-10 |
| | -3.176872221 | 0.001488726 | 0.004127015 |
| NA | NA | NA | |
| | -0.194488217 | 0.845793621 | NA |
| NA | NA | NA | |
| | 8.512009714 | 1.71E-17 | 2.48E-16 |
| NA | NA | NA | |
| NA | NA | NA | |
| NA | NA | NA | |
| NA | NA | NA | |
| NA | NA | NA | |
| NA | NA | NA | |
| | 0.192809521 | 0.847108147 | NA |
| | 2.721514654 | 0.00649835 | 0.015524836 |
| NA | NA | NA | |
| | -0.438248151 | 0.661206407 | 0.747872299 |
| | 0.063708285 | 0.949202508 | NA |
| NA | NA | NA | |
| | 2.704626002 | 0.006838133 | 0.016243567 |
| | 4.396073602 | 1.10E-05 | 4.46E-05 |
| | -1.230970182 | 0.218334017 | 0.31717651 |
| NA | NA | NA | |
| NA | NA | NA | |
| NA | NA | NA | |
| NA | NA | NA | |
| | -0.194487022 | 0.845794556 | NA |
| NA | NA | NA | |

| | | |
|---|---|---|
| -0.065388772 | 0.947864464 | NA |
| NA | NA | NA |
| -1.399111861 | 0.161779442 | 0.248132028 |
| NA | NA | NA |
| -1.89811618 | 0.057680779 | 0.104611978 |
| NA | NA | NA |
| NA | NA | NA |
| NA | NA | NA |
| 2.708759966 | 0.006753518 | 0.016063913 |
| -0.034137739 | 0.972767315 | 0.981235024 |
| NA | NA | NA |
| 1.109198648 | 0.267344496 | NA |
| -4.901010168 | 9.53E-07 | 4.49E-06 |
| -0.191190933 | 0.848376007 | NA |
| NA | NA | NA |
| -3.320054841 | 0.000899998 | 0.002614851 |
| -4.412180189 | 1.02E-05 | 4.17E-05 |
| -0.289905935 | 0.771888201 | NA |
| -2.619271146 | 0.008811788 | 0.020326149 |
| -1.447828262 | 0.147665087 | 0.230398773 |
| 0.192809521 | 0.847108147 | NA |
| NA | NA | NA |
| NA | NA | NA |
| -1.637524706 | 0.101520881 | NA |
| 3.422634744 | 0.000620174 | 0.00185859 |
| 0.063708285 | 0.949202508 | NA |
| NA | NA | NA |
| -0.048368086 | 0.961422893 | 0.97360651 |
| 0.063707135 | 0.949203424 | NA |
| -1.583057469 | 0.113408366 | 0.184877648 |
| NA | NA | NA |
| NA | NA | NA |
| NA | NA | NA |
| NA | NA | NA |
| -0.210022614 | 0.833650023 | 0.883452308 |
| 0.192809521 | 0.847108147 | NA |
| NA | NA | NA |
| NA | NA | NA |
| NA | NA | NA |
| NA | NA | NA |
| NA | NA | NA |
| NA | NA | NA |
| NA | NA | NA |
| NA | NA | NA |
| NA | NA | NA |
| NA | NA | NA |
| NA | NA | NA |

| | | |
|---|---|---|
| NA | NA | NA |
| NA | NA | NA |
| NA | NA | NA |
| NA | NA | NA |
| NA | NA | NA |
| NA | NA | NA |
| NA | NA | NA |
| NA | NA | NA |
| NA | NA | NA |
| NA | NA | NA |
| 0.192810118 | 0.847107679 | NA |
| NA | NA | NA |
| 2.276421992 | 0.022820765 | 0.046908413 |
| 0.902408945 | 0.366839671 | 0.479302446 |
| NA | NA | NA |
| NA | NA | NA |
| NA | NA | NA |
| -0.526370172 | 0.598631036 | 0.69664953 |
| -1.636053818 | 0.101828324 | 0.169060955 |
| -6.44869528 | 1.13E-10 | 8.53E-10 |
| -0.481344463 | 0.630271701 | 0.722472186 |
| -0.065389966 | 0.947863513 | NA |
| -5.340488834 | 9.27E-08 | 5.02E-07 |
| NA | NA | NA |
| -0.413260446 | 0.679415806 | NA |
| -2.053563497 | 0.040017958 | 0.076240637 |
| NA | NA | NA |
| -10.81136605 | 3.04E-27 | 8.92E-26 |
| NA | NA | NA |
| -3.138774986 | 0.001696557 | 0.004646549 |
| -1.381146472 | 0.167233929 | 0.255173448 |
| 1.82625254 | 0.067812231 | 0.11992794 |
| 1.025956541 | 0.30491207 | 0.414441982 |
| -0.575858808 | 0.56471062 | 0.666762905 |
| -0.899442074 | 0.368417237 | 0.480911783 |
| NA | NA | NA |
| -0.078508538 | 0.937423539 | NA |
| 1.570527689 | 0.116292398 | 0.188779994 |
| 0.520266663 | 0.602877728 | 0.700152544 |
| -4.65411965 | 3.25E-06 | 1.42E-05 |
| -0.1297439 | 0.896769049 | 0.9280629 |
| 0.682427482 | 0.494968683 | NA |
| -4.623544353 | 3.77E-06 | 1.63E-05 |
| 0.192810118 | 0.847107679 | NA |
| NA | NA | NA |
| NA | NA | NA |
| 0.192810118 | 0.847107679 | NA |

| | | | |
|---|---|---|---|
| NA | NA | NA | |
| NA | NA | NA | |
| NA | NA | NA | |
| | -2.349537034 | 0.018796774 | 0.039586307 |
| NA | NA | NA | |
| NA | NA | NA | |
| NA | NA | NA | |
| NA | NA | NA | |
| | 0.192809521 | 0.847108147 | NA |
| | 0.063708882 | 0.949202033 | NA |
| NA | NA | NA | |
| NA | NA | NA | |
| | 0.689937154 | 0.490233709 | NA |
| | -11.94789713 | 6.66E-33 | 2.68E-31 |
| | -1.553738009 | 0.120246923 | 0.1942922 |
| | -0.756819238 | 0.449158166 | 0.559130098 |
| NA | NA | NA | |
| | -0.065390563 | 0.947863038 | NA |
| NA | NA | NA | |
| NA | NA | NA | |
| NA | NA | NA | |
| | 2.644350281 | 0.008184791 | 0.019011257 |
| | 6.844239973 | 7.69E-12 | 6.49E-11 |
| | 0.192809521 | 0.847108147 | NA |
| | -3.340463465 | 0.000836387 | 0.002445936 |
| | 1.898578829 | 0.057619874 | 0.104516664 |
| NA | NA | NA | |
| | -0.685992481 | 0.49271785 | 0.599931759 |
| NA | NA | NA | |
| | -2.687582987 | 0.007197122 | 0.016967501 |
| | -0.345165728 | 0.729969782 | NA |
| NA | NA | NA | |
| NA | NA | NA | |
| | 0.124216473 | 0.90114388 | NA |
| NA | NA | NA | |
| | 0.956110998 | 0.339016159 | 0.450402041 |
| | -1.225708138 | 0.220308527 | 0.31944864 |
| NA | NA | NA | |
| NA | NA | NA | |
| | -0.065389966 | 0.947863513 | NA |
| | 0.192810118 | 0.847107679 | NA |
| | -4.2856167 | 1.82E-05 | 7.13E-05 |
| | -1.142771772 | 0.253133362 | 0.357203415 |
| | 0.560973803 | 0.574815396 | 0.675983338 |
| | -0.065389369 | 0.947863988 | NA |
| | 0.795459298 | 0.426346378 | NA |
| | 0.06371127 | 0.949200131 | NA |

| | | | |
|---|---|---|---|
| 0.063708285 | 0.949202508 | NA | |
| -3.1209234 | 0.001802849 | | 0.004904393 |
| 0.213921553 | 0.830608233 | | 0.881497734 |
| 0.278109782 | 0.780928087 | NA | |
| NA | NA | NA | |
| NA | NA | NA | |
| NA | NA | NA | |
| NA | NA | NA | |
| -0.065389369 | 0.947863988 | NA | |
| NA | NA | NA | |
| NA | NA | NA | |
| NA | NA | NA | |
| NA | NA | NA | |
| NA | NA | NA | |
| NA | NA | NA | |
| -3.406445983 | 0.000658146 | | 0.001962753 |
| NA | NA | NA | |
| NA | NA | NA | |
| NA | NA | NA | |
| NA | NA | NA | |
| NA | NA | NA | |
| NA | NA | NA | |
| NA | NA | NA | |
| -1.110593972 | 0.266743159 | | 0.372535502 |
| -1.646784993 | 0.099602247 | | 0.166059032 |
| NA | NA | NA | |
| NA | NA | NA | |
| NA | NA | NA | |
| NA | NA | NA | |
| NA | NA | NA | |
| NA | NA | NA | |
| NA | NA | NA | |
| NA | NA | NA | |
| NA | NA | NA | |
| NA | NA | NA | |
| -0.005859482 | 0.995324836 | | 0.996878102 |
| NA | NA | NA | |
| -1.60990154 | 0.107419353 | | 0.176574926 |
| 0.594418328 | 0.552232353 | NA | |
| 0.063708285 | 0.949202508 | NA | |
| NA | NA | NA | |
| NA | NA | NA | |
| 4.890962135 | 1.00E-06 | | 4.71E-06 |
| NA | NA | NA | |
| NA | NA | NA | |
| NA | NA | NA | |
| NA | NA | NA | |

| | | | |
|---|---|---|---|
| NA | NA | NA | |
| NA | NA | NA | |
| NA | NA | NA | |
| NA | NA | NA | |
| NA | NA | NA | |
| NA | NA | NA | |
| NA | NA | NA | |
| NA | NA | NA | |
| NA | NA | NA | |
| NA | NA | NA | |
| NA | NA | NA | |
| NA | NA | NA | |
| NA | NA | NA | |
| NA | NA | NA | |
| 0.063707194 | 0.949203376 | NA | |
| NA | NA | NA | |
| NA | NA | NA | |
| NA | NA | NA | |
| NA | NA | NA | |
| NA | NA | NA | |
| NA | NA | NA | |
| NA | NA | NA | |
| NA | NA | NA | |
| NA | NA | NA | |
| NA | NA | NA | |
| NA | NA | NA | |
| NA | NA | NA | |
| NA | NA | NA | |
| NA | NA | NA | |
| 1.993732672 | 0.046181277 | | 0.086417267 |
| NA | NA | NA | |
| NA | NA | NA | |
| 0.192808924 | 0.847108615 | NA | |
| 0.251985285 | 0.801052438 | | 0.858760601 |
| NA | NA | NA | |
| NA | NA | NA | |
| 5.174953585 | 2.28E-07 | | 1.17E-06 |
| NA | NA | NA | |
| 0.561300717 | 0.574592553 | NA | |
| NA | NA | NA | |
| NA | NA | NA | |
| -0.126804265 | 0.89909532 | | 0.929739966 |
| NA | NA | NA | |
| NA | NA | NA | |
| NA | NA | NA | |
| -1.432220607 | 0.152080696 | | 0.236228114 |
| -4.092655834 | 4.26E-05 | | 0.00015686 |

| | | | |
|---|---|---|---|
| | 0.664036541 | 0.506666924 | 0.613124788 |
| | -0.085782536 | 0.93163929 | NA |
| | -0.065389966 | 0.947863513 | NA |
| | -8.476768358 | 2.32E-17 | 3.32E-16 |
| | 2.083938172 | 0.037165792 | 0.07149245 |
| NA | NA | NA | |
| NA | NA | NA | |
| NA | NA | NA | |
| | 0.399135113 | 0.689793651 | NA |
| NA | NA | NA | |
| NA | NA | NA | |
| NA | NA | NA | |
| | -0.486289733 | 0.626761758 | 0.719981236 |
| NA | NA | NA | |
| NA | NA | NA | |
| NA | NA | NA | |
| | 1.321034363 | 0.186489907 | 0.279171232 |
| NA | NA | NA | |
| NA | NA | NA | |
| NA | NA | NA | |
| | 0.063708217 | 0.949202562 | NA |
| NA | NA | NA | |
| NA | NA | NA | |
| | 0.192809521 | 0.847108147 | NA |
| NA | NA | NA | |
| | -1.093603735 | 0.27412881 | NA |
| | 0.063708285 | 0.949202508 | NA |
| | -1.001473143 | 0.316598118 | 0.427136737 |
| | -2.100580669 | 0.035677792 | 0.069001325 |
| | -0.341589841 | 0.732659584 | 0.805131026 |
| | -0.537551735 | 0.590886559 | 0.690245511 |
| | -0.268045219 | 0.788664512 | 0.84966525 |
| | -1.325735081 | 0.184927463 | 0.277302726 |
| NA | NA | NA | |
| NA | NA | NA | |
| NA | NA | NA | |
| | -2.504405318 | 0.012265743 | 0.027292341 |
| | -0.857819708 | 0.390992025 | 0.50306006 |
| NA | NA | NA | |
| | 2.140093798 | 0.032347187 | 0.063442258 |
| | 0.686956185 | 0.49211034 | 0.599419717 |
| | 0.302832768 | 0.76201731 | 0.8280136 |
| | -6.256831359 | 3.93E-10 | 2.78E-09 |
| | -0.110062228 | 0.912360024 | 0.939497733 |
| | -0.250103763 | 0.802507106 | 0.860025384 |
| NA | NA | NA | |
| | 1.28748172 | 0.197926437 | 0.293107249 |

| | | |
|---|---|---|
| 0.192810118 | 0.847107679 | NA |
| -0.291563986 | 0.770620015 | NA |
| NA | NA | NA |
| -8.621867777 | 6.59E-18 | 9.83E-17 |
| NA | NA | NA |
| -0.992978589 | 0.320720389 | NA |
| -1.107424659 | 0.268110369 | 0.373969946 |
| NA | NA | NA |
| NA | NA | NA |
| -0.477679191 | 0.632878557 | 0.724522058 |
| NA | NA | NA |
| -1.463873634 | 0.143228479 | 0.224765443 |
| 1.879562898 | 0.060167674 | 0.108291122 |
| -1.370728588 | 0.17045958 | NA |
| 4.996979767 | 5.82E-07 | 2.82E-06 |
| -0.239185104 | 0.810962053 | NA |
| -1.348280493 | 0.177568184 | 0.268146705 |
| -10.30504011 | 6.69E-25 | 1.69E-23 |
| 2.071357276 | 0.03832542 | 0.073418435 |
| 1.293960561 | 0.195679044 | 0.290345988 |
| NA | NA | NA |
| NA | NA | NA |
| 0.797355013 | 0.425244879 | 0.536756395 |
| 1.451272277 | 0.146704058 | 0.229156426 |
| 0.063708285 | 0.949202508 | NA |
| NA | NA | NA |
| 1.177997437 | 0.238797628 | 0.340964326 |
| 4.670000975 | 3.01E-06 | 1.32E-05 |
| 0.930925522 | 0.351892091 | 0.463644736 |
| 3.892542013 | 9.92E-05 | 0.000343187 |
| 2.917723898 | 0.003525964 | 0.008960796 |
| NA | NA | NA |
| -0.832943243 | 0.40487674 | NA |
| NA | NA | NA |
| 2.126128549 | 0.033492557 | 0.065386375 |
| 1.683225782 | 0.092331392 | 0.155962662 |
| 0.451963815 | 0.651295051 | 0.739919895 |
| 0.094763301 | 0.924502837 | NA |
| 3.313900969 | 0.000920041 | 0.002669676 |
| 0.786400836 | 0.431632694 | 0.542499513 |
| 1.963705567 | 0.049564247 | 0.091895785 |
| 8.682896871 | 3.86E-18 | 5.88E-17 |
| -8.691992189 | 3.56E-18 | 5.45E-17 |
| 1.889139687 | 0.058873114 | 0.106404364 |
| -11.88924206 | 1.35E-32 | 5.34E-31 |
| 11.13543878 | 8.43E-29 | 2.74E-27 |
| 3.488054615 | 0.000486549 | 0.001486792 |

| | | |
|---|---|---|
| 7.530848742 | 5.04E-14 | 5.33E-13 |
| NA | NA | NA |
| 0.03111141 | 0.97518069 | 0.982758492 |
| -2.688791565 | 0.007171118 | 0.016915756 |
| -5.500256148 | 3.79E-08 | 2.16E-07 |
| 4.704847895 | 2.54E-06 | 1.13E-05 |
| -6.33882036 | 2.32E-10 | 1.68E-09 |
| -1.046050058 | 0.295537921 | 0.403869314 |
| 0.192809521 | 0.847108147 | NA |
| -13.88927586 | 7.36E-44 | 4.71E-42 |
| 4.448253835 | 8.66E-06 | 3.56E-05 |
| 0.426713653 | 0.669587905 | NA |
| -4.497681613 | 6.87E-06 | 2.87E-05 |
| 10.52172301 | 6.86E-26 | 1.86E-24 |
| 0.215706831 | 0.829216276 | 0.880692638 |
| 0.528504999 | 0.597148878 | NA |
| 5.089809651 | 3.58E-07 | 1.80E-06 |
| NA | NA | NA |
| -0.065389369 | 0.947863988 | NA |
| -3.253297828 | 0.001140739 | 0.003247576 |
| -0.270070768 | 0.787105811 | 0.848307888 |
| 0.429240411 | 0.667748277 | 0.752871143 |
| 0.063708882 | 0.949202033 | NA |
| 2.662179535 | 0.007763646 | 0.018140782 |
| 1.420954659 | 0.155329941 | 0.240305855 |
| 14.41389354 | 4.23E-47 | 3.06E-45 |
| 5.85466405 | 4.78E-09 | 3.03E-08 |
| NA | NA | NA |
| 7.56182758 | 3.97E-14 | 4.25E-13 |
| NA | NA | NA |
| 10.14743197 | 3.40E-24 | 8.26E-23 |
| NA | NA | NA |
| NA | NA | NA |
| -2.97934018 | 0.002888699 | 0.007494209 |
| -0.189303339 | 0.849855078 | 0.895423425 |
| -9.36102392 | 7.90E-21 | 1.49E-19 |
| 2.811598917 | 0.004929593 | 0.012152833 |
| -0.065389369 | 0.947863988 | NA |
| -5.156768315 | 2.51E-07 | 1.28E-06 |
| 1.936998338 | 0.052745547 | 0.096925411 |
| 0.380293508 | 0.703727554 | 0.782718658 |
| 6.178167278 | 6.48E-10 | 4.49E-09 |
| -0.530582122 | 0.595708387 | 0.694128808 |
| 4.245226491 | 2.18E-05 | 8.43E-05 |
| 10.45107233 | 1.45E-25 | 3.86E-24 |
| 0.192809521 | 0.847108147 | NA |
| NA | NA | NA |

| | | | | | |
|---|---|---|---|---|---|
| | -0.668478575 | | 0.503828153 | | 0.610338722 |
| NA | | NA | | NA | |
| NA | | NA | | NA | |
| NA | | NA | | NA | |
| NA | | NA | | NA | |
| NA | | NA | | NA | |
| NA | | NA | | NA | |
| NA | | NA | | NA | |
| | -0.231925918 | | 0.816595553 | | 0.870909996 |
| | 0.192809521 | | 0.847108147 | NA | |
| NA | | NA | | NA | |
| | 0.37145882 | | 0.710295822 | NA | |
| | 0.083873413 | | 0.933157078 | NA | |
| NA | | NA | | NA | |
| NA | | NA | | NA | |
| NA | | NA | | NA | |
| NA | | NA | | NA | |
| NA | | NA | | NA | |
| NA | | NA | | NA | |
| NA | | NA | | NA | |
| NA | | NA | | NA | |
| NA | | NA | | NA | |
| NA | | NA | | NA | |
| | 1.29183865 | | 0.196413027 | | 0.291227894 |
| NA | | NA | | NA | |
| NA | | NA | | NA | |
| NA | | NA | | NA | |
| NA | | NA | | NA | |
| | 0.063708882 | | 0.949202033 | NA | |
| | 0.06371127 | | 0.949200131 | NA | |
| NA | | NA | | NA | |
| | 1.07426793 | | 0.282702614 | | 0.389628679 |
| | 0.131421334 | | 0.895442013 | NA | |
| NA | | NA | | NA | |
| | 1.612885448 | | 0.106769397 | | 0.175679596 |
| | -1.122697615 | | 0.261565941 | | 0.366716827 |
| | 8.577938288 | | 9.66E-18 | | 1.43E-16 |
| | -0.194487022 | | 0.845794556 | NA | |
| | 0.116269467 | | 0.907438983 | NA | |
| | 2.656105487 | | 0.007904888 | | 0.018431239 |
| | 0.192810118 | | 0.847107679 | NA | |
| | -1.441448694 | | 0.149457962 | | 0.232726751 |
| | 0.815813312 | | 0.414606919 | | 0.526194664 |
| | -1.928651247 | | 0.053774172 | | 0.098547923 |
| | -0.914255648 | | 0.36058253 | | 0.472706265 |
| | -2.390280449 | | 0.016835512 | | 0.036071979 |
| | -0.690900051 | | 0.489628354 | | 0.597358048 |

| | | |
|---|---|---|
| 0.432087296 | 0.665677967 | 0.751552527 |
| 1.02290298 | 0.30635372 | 0.41599488 |
| 1.913571468 | 0.055674934 | 0.10153323 |
| 1.004558456 | 0.31510951 | 0.425651716 |
| -0.576451686 | 0.564309917 | 0.666384092 |
| 2.305100048 | 0.021160971 | 0.043876254 |
| 2.677313733 | 0.007421511 | 0.017448845 |
| NA | NA | NA |
| 0.969106748 | 0.332491931 | 0.443641614 |
| -0.184711198 | 0.853455562 | 0.898134596 |
| 0.066629133 | 0.946876953 | NA |
| -1.879569102 | 0.060166828 | 0.108291122 |
| NA | NA | NA |
| NA | NA | NA |
| NA | NA | NA |
| NA | NA | NA |
| 0.507813887 | 0.61158387 | 0.70742383 |
| NA | NA | NA |
| NA | NA | NA |
| 0.063708882 | 0.949202033 | NA |
| NA | NA | NA |
| NA | NA | NA |
| 0.452944158 | 0.650588953 | NA |
| NA | NA | NA |
| 0.192809521 | 0.847108147 | NA |
| NA | NA | NA |
| NA | NA | NA |
| 0.192810118 | 0.847107679 | NA |
| -0.065389369 | 0.947863988 | NA |
| NA | NA | NA |
| NA | NA | NA |
| 0.192809521 | 0.847108147 | NA |
| NA | NA | NA |
| 0.219648545 | 0.826144881 | NA |
| NA | NA | NA |
| -1.36316814 | 0.172829526 | 0.262240727 |
| NA | NA | NA |
| 0.218530872 | 0.827015507 | 0.878914663 |
| 5.480923862 | 4.23E-08 | 2.40E-07 |
| -0.352828809 | 0.724216784 | 0.798693431 |
| NA | NA | NA |
| -0.541383157 | 0.588243514 | NA |
| NA | NA | NA |
| 0.305716554 | 0.759820468 | 0.826149362 |
| 8.67504723 | 4.13E-18 | 6.29E-17 |
| -0.19448762 | 0.845794089 | NA |
| NA | NA | NA |

| | | | |
|---|---|---|---|
| | -0.365338043 | 0.714859084 | NA |
| | -0.19448762 | 0.845794089 | NA |
| NA | NA | NA | |
| NA | NA | NA | |
| NA | NA | NA | |
| | -0.274907846 | 0.783387039 | NA |
| NA | NA | NA | |
| | -0.19448762 | 0.845794089 | NA |
| | -0.194487022 | 0.845794556 | NA |
| NA | NA | NA | |
| | 0.192809521 | 0.847108147 | NA |
| NA | NA | NA | |
| NA | NA | NA | |
| NA | NA | NA | |
| NA | NA | NA | |
| | -7.240142104 | 4.48E-13 | 4.32E-12 |
| | -2.165508048 | 0.0303488 | 0.059950316 |
| NA | NA | NA | |
| NA | NA | NA | |
| NA | NA | NA | |
| NA | NA | NA | |
| | 0.192810118 | 0.847107679 | NA |
| | 0.760265545 | 0.447095873 | 0.557036379 |
| NA | NA | NA | |
| NA | NA | NA | |
| | 1.454103173 | 0.145917704 | 0.22815593 |
| | -0.221026306 | 0.82507195 | 0.877482512 |
| | 0.97641631 | 0.328858202 | 0.439730798 |
| | -0.313866998 | 0.753622057 | NA |
| | -0.151770323 | 0.87936809 | NA |
| | 1.529217638 | 0.1262105 | 0.202481284 |
| NA | NA | NA | |
| NA | NA | NA | |
| | 1.178968605 | 0.238410677 | 0.340567244 |
| | 2.363773784 | 0.018089855 | 0.038336217 |
| NA | NA | NA | |
| NA | NA | NA | |
| | -0.976096113 | 0.329016838 | NA |
| | -0.473508958 | 0.635850111 | 0.727240755 |
| NA | NA | NA | |
| | 0.063708882 | 0.949202033 | NA |
| | 5.215011191 | 1.84E-07 | 9.54E-07 |
| NA | NA | NA | |
| NA | NA | NA | |
| NA | NA | NA | |
| NA | NA | NA | |
| | -3.05125773 | 0.002278849 | 0.00605773 |

| | | |
|---|---|---|
| NA | NA | NA |
| NA | NA | NA |
| NA | NA | NA |
| NA | NA | NA |
| -4.232038741 | 2.32E-05 | 8.91E-05 |
| 0.573823318 | 0.566087368 | 0.668041812 |
| NA | NA | NA |
| 0.13704539 | 0.890994918 | NA |
| -3.773799941 | 0.00016078 | 0.000536075 |
| -1.702648152 | 0.088633933 | 0.150652407 |
| NA | NA | NA |
| 1.286347921 | 0.198321665 | NA |
| NA | NA | NA |
| NA | NA | NA |
| 2.699902883 | 0.006935972 | 0.01644275 |
| -1.080483427 | 0.279926963 | 0.386911032 |
| 2.60557087 | 0.00917213 | 0.021085452 |
| 0.063708285 | 0.949202508 | NA |
| NA | NA | NA |
| -0.329444278 | 0.741819905 | 0.812060014 |
| NA | NA | NA |
| 0.192808327 | 0.847109082 | NA |
| -0.593493424 | 0.552850983 | 0.656000538 |
| 12.26014604 | 1.48E-34 | 6.34E-33 |
| NA | NA | NA |
| -0.065388772 | 0.947864464 | NA |
| 0.656755757 | 0.511337976 | NA |
| 0.192810118 | 0.847107679 | NA |
| NA | NA | NA |
| NA | NA | NA |
| -0.138339369 | 0.889972213 | NA |
| NA | NA | NA |
| NA | NA | NA |
| NA | NA | NA |
| NA | NA | NA |
| NA | NA | NA |
| -6.840868042 | 7.87E-12 | 6.63E-11 |
| NA | NA | NA |
| -3.676873361 | 0.00023611 | 0.000765226 |
| -4.59448634 | 4.34E-06 | 1.87E-05 |
| -2.075651978 | 0.03792615 | 0.072787268 |
| -4.390390143 | 1.13E-05 | 4.57E-05 |
| NA | NA | NA |
| NA | NA | NA |
| NA | NA | NA |
| -0.085852801 | 0.931583432 | NA |
| NA | NA | NA |

| | | |
|---|---|---|
| NA | NA | NA |
| NA | NA | NA |
| 0.036465054 | 0.970911543 | 0.979876453 |
| NA | NA | NA |
| -3.737098223 | 0.000186156 | 0.000614708 |
| NA | NA | NA |
| NA | NA | NA |
| NA | NA | NA |
| -0.084501006 | 0.932658103 | NA |
| 0.063708882 | 0.949202033 | NA |
| NA | NA | NA |
| NA | NA | NA |
| NA | NA | NA |
| NA | NA | NA |
| NA | NA | NA |
| NA | NA | NA |
| NA | NA | NA |
| NA | NA | NA |
| NA | NA | NA |
| -0.835229481 | 0.403588514 | 0.515241278 |
| NA | NA | NA |
| NA | NA | NA |
| NA | NA | NA |
| NA | NA | NA |
| NA | NA | NA |
| 2.187965613 | 0.028672107 | 0.057097847 |
| NA | NA | NA |
| NA | NA | NA |
| NA | NA | NA |
| -0.065389369 | 0.947863988 | NA |
| NA | NA | NA |
| NA | NA | NA |
| 0.063708882 | 0.949202033 | NA |
| 0.404104215 | 0.686136085 | NA |
| NA | NA | NA |
| NA | NA | NA |
| NA | NA | NA |
| NA | NA | NA |
| NA | NA | NA |
| NA | NA | NA |
| NA | NA | NA |
| NA | NA | NA |
| NA | NA | NA |
| NA | NA | NA |
| NA | NA | NA |
| -0.150001763 | 0.880763224 | NA |
| NA | NA | NA |

| | | | |
|---|---|---|---|
| NA | NA | NA | |
| | 0.063708882 | 0.949202033 | NA |
| NA | NA | NA | |
| | 1.045584548 | 0.295752886 | NA |
| | -0.133869248 | 0.893505968 | NA |
| NA | NA | NA | |
| NA | NA | NA | |
| NA | NA | NA | |
| NA | NA | NA | |
| | -0.841086704 | 0.400299363 | NA |
| NA | NA | NA | |
| NA | NA | NA | |
| NA | NA | NA | |
| NA | NA | NA | |
| | -2.839718285 | 0.004515339 | 0.011222216 |
| | 0.997490347 | 0.318526557 | 0.429171541 |
| NA | NA | NA | |
| | -1.234673409 | 0.216952076 | 0.315550611 |
| NA | NA | NA | |
| | -0.52653154 | 0.598518944 | 0.696607807 |
| | 0.524866126 | 0.599676259 | 0.697485909 |
| NA | NA | NA | |
| NA | NA | NA | |
| | -0.065389369 | 0.947863988 | NA |
| NA | NA | NA | |
| | -0.065389369 | 0.947863988 | NA |
| NA | NA | NA | |
| | 0.130725433 | 0.895992512 | NA |
| | 2.380319317 | 0.017297642 | 0.036923074 |
| | 5.25143081 | 1.51E-07 | 7.94E-07 |
| | -2.308578513 | 0.020966981 | 0.043516049 |
| NA | NA | NA | |
| | 0.813157542 | 0.416127739 | NA |
| NA | NA | NA | |
| | 0.063708285 | 0.949202508 | NA |
| | 1.298509803 | 0.19411221 | 0.288465749 |
| | -4.975955842 | 6.49E-07 | 3.13E-06 |
| NA | NA | NA | |
| NA | NA | NA | |
| | -2.437818371 | 0.014776196 | 0.032141148 |
| NA | NA | NA | |
| | -4.095476788 | 4.21E-05 | 0.00015519 |
| NA | NA | NA | |
| | 0.192810118 | 0.847107679 | NA |
| | 0.253103899 | 0.800187926 | NA |
| | 0.192809521 | 0.847108147 | NA |
| | -3.010271565 | 0.002610142 | 0.006836021 |

| | | |
|---|---|---|
| 0.192809521 | 0.847108147 | NA |
| 0.407391913 | 0.683720166 | 0.766151039 |
| -1.898209839 | 0.057668445 | 0.104597187 |
| -3.125246716 | 0.00177656 | 0.004842349 |
| -0.194488217 | 0.845793621 | NA |
| 0.103610212 | 0.917478683 | NA |
| 0.410476279 | 0.681456601 | 0.76419618 |
| 0.192809521 | 0.847108147 | NA |
| 5.698788012 | 1.21E-08 | 7.30E-08 |
| -7.999806909 | 1.25E-15 | 1.54E-14 |
| -0.065388772 | 0.947864464 | NA |
| 0.063708285 | 0.949202508 | NA |
| 0.063707135 | 0.949203424 | NA |
| 0.063708882 | 0.949202033 | NA |
| 0.417530232 | 0.676290616 | 0.759525918 |
| -0.113885002 | 0.909328955 | NA |
| NA | NA | NA |
| NA | NA | NA |
| -3.182274281 | 0.001461234 | 0.004059605 |
| -5.295704702 | 1.19E-07 | 6.33E-07 |
| NA | NA | NA |
| 2.154959009 | 0.031165042 | 0.0613738 |
| 20.15525395 | 2.42E-90 | 4.92E-88 |
| NA | NA | NA |
| -0.58530233 | 0.558344449 | NA |
| NA | NA | NA |
| 7.313274156 | 2.61E-13 | 2.57E-12 |
| NA | NA | NA |
| 0.192808924 | 0.847108615 | NA |
| -0.069194723 | 0.944834623 | 0.962445492 |
| -0.065389369 | 0.947863988 | NA |
| 2.769873625 | 0.005607805 | 0.013628775 |
| 2.169324416 | 0.030058063 | 0.059436982 |
| -0.617251102 | 0.537069114 | 0.641326269 |
| 2.766323935 | 0.005669219 | 0.013757838 |
| NA | NA | NA |
| NA | NA | NA |
| NA | NA | NA |
| NA | NA | NA |
| 2.624165015 | 0.008686169 | 0.020078944 |
| NA | NA | NA |
| 8.899621466 | 5.60E-19 | 9.26E-18 |
| NA | NA | NA |
| 0.193472986 | 0.84658856 | NA |
| -0.594497832 | 0.552179192 | 0.655336424 |
| NA | NA | NA |
| 0.262551648 | 0.792896172 | 0.852682758 |

| | | |
|---|---|---|
| 1.455346488 | 0.145573363 | 0.227759799 |
| -6.554873112 | 5.57E-11 | 4.35E-10 |
| -1.567742308 | 0.116941284 | 0.189698012 |
| 0.192810118 | 0.847107679 | NA |
| 3.990761206 | 6.59E-05 | 0.000234965 |
| 0.192810118 | 0.847107679 | NA |
| -1.70474025 | 0.088242879 | 0.150130456 |
| NA | NA | NA |
| 2.325007813 | 0.020071542 | 0.041911296 |
| -0.065895875 | 0.947460725 | 0.964257215 |
| 0.282742491 | 0.777374242 | NA |
| -0.264388152 | 0.79148084 | NA |
| -2.311668467 | 0.02079596 | 0.043211264 |
| 0.099466044 | 0.920768247 | NA |
| 0.063708285 | 0.949202508 | NA |
| -2.600271497 | 0.009315003 | 0.021376596 |
| -4.667006104 | 3.06E-06 | 1.34E-05 |
| -1.605731931 | 0.108332822 | 0.177831226 |
| -2.351672476 | 0.018689223 | 0.039409541 |
| -0.314106036 | 0.753440505 | 0.821117945 |
| NA | NA | NA |
| 1.489694972 | 0.136304457 | 0.215575671 |
| NA | NA | NA |
| 2.678169457 | 0.007402576 | 0.017407598 |
| 0.075052761 | 0.940172733 | 0.959336466 |
| 0.701887728 | 0.482749183 | NA |
| NA | NA | NA |
| 2.434875591 | 0.014896911 | 0.032381213 |
| -2.643503064 | 0.008205302 | 0.019055364 |
| 6.353325257 | 2.11E-10 | 1.54E-09 |
| 4.005452982 | 6.19E-05 | 0.000221903 |
| -4.841063968 | 1.29E-06 | 5.96E-06 |
| 1.224316188 | 0.220832972 | 0.320116366 |
| 1.954279802 | 0.050668136 | 0.093678789 |
| 0.317642674 | 0.750756 | NA |
| -7.400390928 | 1.36E-13 | 1.37E-12 |
| -0.078196275 | 0.937671925 | 0.957565413 |
| NA | NA | NA |
| NA | NA | NA |
| NA | NA | NA |
| NA | NA | NA |
| 4.318988936 | 1.57E-05 | 6.20E-05 |
| -0.830751951 | 0.406113771 | 0.517963195 |
| 0.377495203 | 0.705805629 | NA |
| NA | NA | NA |
| -0.065389369 | 0.947863988 | NA |
| -0.297387515 | 0.766170675 | 0.831335702 |

| | | |
|---|---|---|
| 2.219998902 | 0.026418842 | 0.053151967 |
| NA | NA | NA |
| 16.10530268 | 2.34E-58 | 2.23E-56 |
| 2.521888985 | 0.011672654 | 0.026135301 |
| -1.029581315 | 0.30320659 | 0.412527094 |
| -8.325975959 | 8.36E-17 | 1.14E-15 |
| 2.356864798 | 0.018429956 | 0.03895143 |
| -9.343589026 | 9.31E-21 | 1.75E-19 |
| 7.852665076 | 4.07E-15 | 4.79E-14 |
| -0.70989928 | 0.477766597 | 0.586574676 |
| -1.017729956 | 0.308806307 | 0.418711911 |
| -0.906748749 | 0.364539678 | 0.476978717 |
| -2.266545288 | 0.023418018 | 0.047958843 |
| -4.892859111 | 9.94E-07 | 4.67E-06 |
| -0.482897811 | 0.629168296 | 0.721859298 |
| 0.063708882 | 0.949202033 | NA |
| 6.948279934 | 3.70E-12 | 3.23E-11 |
| 2.491071333 | 0.012735855 | 0.028225469 |
| NA | NA | NA |
| -1.465947557 | 0.142662574 | 0.224102186 |
| NA | NA | NA |
| NA | NA | NA |
| -0.818672684 | 0.412973185 | 0.524573399 |
| -9.409283375 | 5.00E-21 | 9.57E-20 |
| NA | NA | NA |
| NA | NA | NA |
| 4.741053683 | 2.13E-06 | 9.54E-06 |
| NA | NA | NA |
| 1.681463939 | 0.092672835 | 0.156444396 |
| 0.063707194 | 0.949203376 | NA |
| 6.689455502 | 2.24E-11 | 1.82E-10 |
| 0.063708882 | 0.949202033 | NA |
| 6.53909701 | 6.19E-11 | 4.81E-10 |
| NA | NA | NA |
| 2.172814975 | 0.029794246 | 0.059017883 |
| -3.53841636 | 0.000402535 | 0.00124959 |
| -0.066204891 | 0.947214703 | NA |
| -3.901351135 | 9.57E-05 | 0.000331989 |
| -4.878659581 | 1.07E-06 | 4.99E-06 |
| NA | NA | NA |
| NA | NA | NA |
| NA | NA | NA |
| NA | NA | NA |
| NA | NA | NA |
| 3.638253726 | 0.000274493 | 0.000878813 |
| NA | NA | NA |
| NA | NA | NA |

| | | | |
|---|---|---|---|
| NA | NA | NA | |
| NA | NA | NA | |
| NA | NA | NA | |
| NA | NA | NA | |
| NA | NA | NA | |
| NA | NA | NA | |
| NA | NA | NA | |
| NA | NA | NA | |
| NA | NA | NA | |
| NA | NA | NA | |
| NA | NA | NA | |
| -3.451097164 | 0.000558313 | | 0.001688361 |
| 0.192809521 | 0.847108147 | NA | |
| NA | NA | NA | |
| -0.057866476 | 0.953854986 | NA | |
| -0.137201159 | 0.890871795 | NA | |
| 0.672994286 | 0.500950926 | | 0.607971377 |
| -0.065389369 | 0.947863988 | NA | |
| -0.688174975 | 0.491342601 | | 0.598778193 |
| 0.164495614 | 0.869341 | NA | |
| -0.335999377 | 0.736871336 | NA | |
| 2.98577507 | 0.002828606 | | 0.007353553 |
| -2.108779021 | 0.03496366 | | 0.067856123 |
| -4.292178376 | 1.77E-05 | | 6.93E-05 |
| NA | NA | NA | |
| NA | NA | NA | |
| -1.887075243 | 0.059150206 | NA | |
| 0.103577121 | 0.917504944 | | 0.943359709 |
| -0.352171323 | 0.724709782 | NA | |
| -0.530192089 | 0.595978755 | | 0.694379195 |
| NA | NA | NA | |
| -0.153870969 | 0.877711474 | | 0.914503526 |
| NA | NA | NA | |
| 1.060243473 | 0.289033848 | | 0.39645535 |
| NA | NA | NA | |
| 0.20109316 | 0.84062573 | | 0.888222887 |
| -1.377286782 | 0.1684236 | | 0.256613265 |
| NA | NA | NA | |
| NA | NA | NA | |
| -0.879545826 | 0.379105397 | NA | |
| -0.800012678 | 0.423703452 | NA | |
| -0.396194401 | 0.691961622 | NA | |
| -2.885507707 | 0.003907829 | | 0.009847273 |
| NA | NA | NA | |
| -3.443411692 | 0.000574424 | | 0.001733314 |
| -2.059276313 | 0.039467776 | | 0.075301352 |
| 0.063708285 | 0.949202508 | NA | |

| | | |
|---|---|---|
| -2.105500564 | 0.035247758 | 0.06832677 |
| -4.016132526 | 5.92E-05 | 0.000212636 |
| -2.52909221 | 0.011435798 | 0.025690004 |
| -1.83458513 | 0.066567184 | 0.118109917 |
| NA | NA | NA |
| -3.51820178 | 0.000434482 | 0.001339791 |
| 0.21983428 | 0.826000221 | NA |
| -0.303189132 | 0.761745731 | NA |
| 0.192808327 | 0.847109082 | NA |
| NA | NA | NA |
| -0.064545442 | 0.948535925 | NA |
| 0.888251037 | 0.374405731 | 0.487204338 |
| -2.211298641 | 0.027015163 | 0.054194882 |
| NA | NA | NA |
| 0.063708882 | 0.949202033 | NA |
| -0.831565309 | 0.40565435 | 0.517509092 |
| 0.932249481 | 0.351207609 | 0.462913452 |
| -4.171274927 | 3.03E-05 | 0.000114526 |
| NA | NA | NA |
| 0.063707135 | 0.949203424 | NA |
| 0.769262845 | 0.441737289 | 0.551758289 |
| 4.324470724 | 1.53E-05 | 6.05E-05 |
| NA | NA | NA |
| NA | NA | NA |
| -0.194488217 | 0.845793621 | NA |
| NA | NA | NA |
| -8.504727559 | 1.82E-17 | 2.64E-16 |
| -1.413790452 | 0.157423438 | 0.242869123 |
| -1.298357191 | 0.194164622 | 0.28850938 |
| 5.841287493 | 5.18E-09 | 3.26E-08 |
| -2.449720437 | 0.014296717 | 0.031209386 |
| 1.398912008 | 0.161839371 | 0.248182788 |
| -1.723761193 | 0.084750963 | 0.14496799 |
| NA | NA | NA |
| -0.257863352 | 0.796512369 | 0.855132594 |
| 0.088208248 | 0.929711162 | NA |
| -1.738260791 | 0.082164872 | 0.141297612 |
| 1.575021048 | 0.115251586 | 0.187406432 |
| 0.063976859 | 0.948988653 | NA |
| NA | NA | NA |
| -3.765244089 | 0.000166386 | 0.000553589 |
| 0.916947119 | 0.359170351 | 0.471175781 |
| -8.000723815 | 1.24E-15 | 1.53E-14 |
| -0.783957777 | 0.433064886 | 0.543775162 |
| -0.806617957 | 0.419886625 | 0.531574641 |
| -2.103346644 | 0.035435479 | 0.068633447 |
| -3.965800741 | 7.31E-05 | 0.000259304 |

| | | |
|---|---|---|
| -1.851750849 | 0.064061608 | 0.114182302 |
| -3.664901837 | 0.000247433 | 0.0007981 |
| -0.209298498 | 0.834215224 | 0.88383169 |
| -0.065388772 | 0.947864464 | NA |
| NA | NA | NA |
| NA | NA | NA |
| NA | NA | NA |
| NA | NA | NA |
| 0.133211417 | 0.894026182 | NA |
| -2.021141267 | 0.043265144 | 0.081650196 |
| -5.161186298 | 2.45E-07 | 1.26E-06 |
| NA | NA | NA |
| -2.277843166 | 0.022735923 | 0.046753218 |
| NA | NA | NA |
| 4.462199947 | 8.11E-06 | 3.36E-05 |
| -1.149692729 | 0.25027045 | 0.35393352 |
| 8.94488248 | 3.72E-19 | 6.22E-18 |
| 16.29020216 | 1.16E-59 | 1.15E-57 |
| NA | NA | NA |
| 3.307258298 | 0.00094214 | 0.002730002 |
| 4.546907484 | 5.44E-06 | 2.30E-05 |
| 0.063708882 | 0.949202033 | NA |
| NA | NA | NA |
| 1.062061062 | 0.288207966 | 0.39568366 |
| NA | NA | NA |
| NA | NA | NA |
| NA | NA | NA |
| 1.595272447 | 0.110651324 | 0.181043447 |
| 0.780765887 | 0.434940201 | 0.545510067 |
| 0.192810118 | 0.847107679 | NA |
| 6.90420331 | 5.05E-12 | 4.36E-11 |
| 6.05573556 | 1.40E-09 | 9.36E-09 |
| 0.334970895 | 0.737647041 | NA |
| 0.409339218 | 0.682290739 | 0.76492597 |
| 0.192809521 | 0.847108147 | NA |
| NA | NA | NA |
| -0.093577099 | 0.925445102 | 0.948729276 |
| NA | NA | NA |
| NA | NA | NA |
| NA | NA | NA |
| NA | NA | NA |
| NA | NA | NA |
| -1.488768372 | 0.136548374 | 0.215920512 |
| 0.254560291 | 0.799062732 | 0.85719509 |
| 8.99978483 | 2.26E-19 | 3.84E-18 |
| 7.466414145 | 8.24E-14 | 8.54E-13 |
| NA | NA | NA |

| | | | |
|---|---|---|---|
| NA | NA | NA | |
| | 8.486442349 | 2.13E-17 | 3.07E-16 |
| | 0.063708285 | 0.949202508 | NA |
| NA | NA | NA | |
| NA | NA | NA | |
| | -3.058896653 | 0.002221538 | 0.005920474 |
| NA | NA | NA | |
| NA | NA | NA | |
| | -2.35989504 | 0.018280106 | 0.038673913 |
| | -0.709779368 | 0.477840965 | 0.586637206 |
| | 1.757775423 | 0.078785729 | 0.136320116 |
| NA | NA | NA | |
| NA | NA | NA | |
| NA | NA | NA | |
| NA | NA | NA | |
| | 0.192809521 | 0.847108147 | NA |
| | -2.363538082 | 0.018101366 | 0.038347625 |
| | 1.218342778 | 0.22309374 | 0.322664759 |
| | 0.502941955 | 0.615005083 | NA |
| | 0.356279174 | 0.721631504 | NA |
| | 0.211367843 | 0.832600247 | 0.88275094 |
| | 0.074728106 | 0.940431045 | NA |
| NA | NA | NA | |
| | 2.317363439 | 0.020483942 | 0.042669197 |
| NA | NA | NA | |
| NA | NA | NA | |
| | 9.520456095 | 1.72E-21 | 3.45E-20 |
| NA | NA | NA | |
| | 2.195682752 | 0.028114659 | 0.056117217 |
| NA | NA | NA | |
| NA | NA | NA | |
| NA | NA | NA | |
| NA | NA | NA | |
| | -0.065389966 | 0.947863513 | NA |
| | -2.191614044 | 0.028407387 | 0.056633763 |
| NA | NA | NA | |
| NA | NA | NA | |
| | 0.624656694 | 0.532196401 | 0.636661852 |
| | -5.305868157 | 1.12E-07 | 6.00E-07 |
| NA | NA | NA | |
| NA | NA | NA | |
| NA | NA | NA | |
| | 1.2433954 | 0.213722165 | 0.311813114 |
| NA | NA | NA | |
| | 0.917558122 | 0.358850251 | 0.470805209 |
| | 1.040735635 | 0.29799826 | 0.406811914 |
| NA | NA | NA | |

| | | |
|---|---|---|
| 6.306623816 | 2.85E-10 | 2.05E-09 |
| NA | NA | NA |
| 0.24166994 | 0.809035924 | NA |
| 1.393604965 | 0.163436928 | 0.250387215 |
| 0.176330302 | 0.860034457 | 0.902761959 |
| -1.097755704 | 0.272311179 | 0.378790086 |
| 0.401009486 | 0.68841314 | 0.770144563 |
| NA | NA | NA |
| NA | NA | NA |
| NA | NA | NA |
| NA | NA | NA |
| NA | NA | NA |
| NA | NA | NA |
| NA | NA | NA |
| 0.193341124 | 0.846691821 | NA |
| -0.491776089 | 0.622877641 | 0.716952613 |
| -0.12251278 | 0.902492925 | 0.932175047 |
| 0.055303391 | 0.955896761 | 0.969885116 |
| NA | NA | NA |
| 3.261871373 | 0.001106794 | 0.003160283 |
| NA | NA | NA |
| NA | NA | NA |
| NA | NA | NA |
| NA | NA | NA |
| NA | NA | NA |
| 0.3474247 | 0.728272279 | NA |
| -3.105111708 | 0.001902072 | 0.005146347 |
| 0.114540817 | 0.908809092 | 0.937113437 |
| 0.045472324 | 0.963730834 | 0.975224693 |
| -0.065390563 | 0.947863038 | NA |
| NA | NA | NA |
| 0.063708882 | 0.949202033 | NA |
| NA | NA | NA |
| NA | NA | NA |
| NA | NA | NA |
| NA | NA | NA |
| 0.945891739 | 0.344203808 | 0.455504986 |
| 0.063710673 | 0.949200606 | NA |
| NA | NA | NA |
| NA | NA | NA |
| NA | NA | NA |
| NA | NA | NA |
| 0.192809521 | 0.847108147 | NA |
| NA | NA | NA |
| NA | NA | NA |
| NA | NA | NA |
| NA | NA | NA |

| | | | |
|---|---|---|---|
| NA | NA | NA | |
| NA | NA | NA | |
| NA | NA | NA | |
| NA | NA | NA | |
| NA | NA | NA | |
| -0.065388772 | 0.947864464 | NA | |
| NA | NA | NA | |
| NA | NA | NA | |
| NA | NA | NA | |
| NA | NA | NA | |
| NA | NA | NA | |
| NA | NA | NA | |
| NA | NA | NA | |
| 0.115861689 | 0.907762159 | NA | |
| NA | NA | NA | |
| NA | NA | NA | |
| NA | NA | NA | |
| NA | NA | NA | |
| NA | NA | NA | |
| NA | NA | NA | |
| NA | NA | NA | |
| NA | NA | NA | |
| NA | NA | NA | |
| NA | NA | NA | |
| NA | NA | NA | |
| NA | NA | NA | |
| NA | NA | NA | |
| -0.526179247 | 0.598763672 | | 0.696715852 |
| 0.3474247 | 0.728272279 | NA | |
| NA | NA | NA | |
| -0.135388011 | 0.892305103 | NA | |
| -0.758114178 | 0.448382632 | | 0.558359069 |
| -5.11194343 | 3.19E-07 | | 1.61E-06 |
| NA | NA | NA | |
| NA | NA | NA | |
| -0.74698997 | 0.455069615 | | 0.564829913 |
| NA | NA | NA | |
| 0.234775509 | 0.814382977 | NA | |
| -0.065389369 | 0.947863988 | NA | |
| NA | NA | NA | |
| NA | NA | NA | |
| 2.622342645 | 0.008732758 | | 0.020171737 |
| -0.065388772 | 0.947864464 | NA | |
| NA | NA | NA | |
| NA | NA | NA | |
| NA | NA | NA | |
| 0.192808924 | 0.847108615 | NA | |

| | | | |
|---|---|---|---|
| NA | NA | NA | |
| 0.159645528 | 0.873160312 | NA | |
| 0.201808052 | 0.840066787 | NA | |
| -0.087005527 | 0.930667119 | NA | |
| -2.189693576 | 0.028546467 | | 0.056892912 |
| NA | NA | NA | |
| NA | NA | NA | |
| NA | NA | NA | |
| 0.063708882 | 0.949202033 | NA | |
| NA | NA | NA | |
| NA | NA | NA | |
| NA | NA | NA | |
| NA | NA | NA | |
| NA | NA | NA | |
| NA | NA | NA | |
| NA | NA | NA | |
| 0.192809521 | 0.847108147 | NA | |
| NA | NA | NA | |
| NA | NA | NA | |
| NA | NA | NA | |
| NA | NA | NA | |
| NA | NA | NA | |
| NA | NA | NA | |
| NA | NA | NA | |
| NA | NA | NA | |
| 0.063708882 | 0.949202033 | NA | |
| NA | NA | NA | |
| NA | NA | NA | |
| NA | NA | NA | |
| NA | NA | NA | |
| NA | NA | NA | |
| NA | NA | NA | |
| NA | NA | NA | |
| NA | NA | NA | |
| NA | NA | NA | |
| NA | NA | NA | |
| NA | NA | NA | |
| NA | NA | NA | |
| -0.065389369 | 0.947863988 | NA | |
| NA | NA | NA | |
| NA | NA | NA | |
| NA | NA | NA | |
| NA | NA | NA | |
| NA | NA | NA | |
| NA | NA | NA | |
| NA | NA | NA | |
| NA | NA | NA | |

| | | | |
|---|---|---|---|
| NA | NA | NA | |
| NA | NA | NA | |
| -0.065389369 | 0.947863988 | NA | |
| NA | NA | NA | |
| NA | NA | NA | |
| NA | NA | NA | |
| NA | NA | NA | |
| NA | NA | NA | |
| NA | NA | NA | |
| NA | NA | NA | |
| NA | NA | NA | |
| NA | NA | NA | |
| NA | NA | NA | |
| NA | NA | NA | |
| 0.709729927 | 0.47787163 | NA | |
| 0.243153646 | 0.807886375 | NA | |
| NA | NA | NA | |
| 0.192810118 | 0.847107679 | NA | |
| 0.329279929 | 0.741944113 | | 0.812160463 |
| NA | NA | NA | |
| NA | NA | NA | |
| NA | NA | NA | |
| NA | NA | NA | |
| NA | NA | NA | |
| NA | NA | NA | |
| NA | NA | NA | |
| NA | NA | NA | |
| NA | NA | NA | |
| NA | NA | NA | |
| NA | NA | NA | |
| 1.661423852 | 0.096628351 | | 0.161890605 |
| NA | NA | NA | |
| NA | NA | NA | |
| 0.230838446 | 0.817440312 | NA | |
| -0.065389369 | 0.947863988 | NA | |
| NA | NA | NA | |
| NA | NA | NA | |
| NA | NA | NA | |
| NA | NA | NA | |
| NA | NA | NA | |
| NA | NA | NA | |
| NA | NA | NA | |
| NA | NA | NA | |
| NA | NA | NA | |
| NA | NA | NA | |
| NA | NA | NA | |
| NA | NA | NA | |

| | | |
|---|---|---|
| NA | NA | NA |
| NA | NA | NA |
| NA | NA | NA |
| NA | NA | NA |
| -0.091700339 | 0.926936128 | NA |
| -10.10154712 | 5.44E-24 | 1.30E-22 |
| -1.518633724 | 0.128854717 | 0.205950506 |
| 0.39172146 | 0.695264032 | 0.775579763 |
| -0.9535168 | 0.340328288 | 0.451857471 |
| NA | NA | NA |
| 0.475048958 | 0.634752078 | 0.72613549 |
| 1.955813065 | 0.050487179 | 0.093385612 |
| 0.192808924 | 0.847108615 | NA |
| -2.602350648 | 0.009258713 | 0.021261064 |
| NA | NA | NA |
| -0.620128343 | 0.535173293 | 0.639520077 |
| -1.658426483 | 0.097231415 | 0.162745908 |
| 0.620056669 | 0.535220479 | NA |
| -1.620283747 | 0.105171339 | 0.173632254 |
| -3.516444453 | 0.000437368 | 0.001348359 |
| 3.11772543 | 0.001822525 | 0.00495162 |
| -7.936028596 | 2.09E-15 | 2.53E-14 |
| 1.441427506 | 0.149463944 | 0.232726751 |
| 0.796273528 | 0.425873069 | 0.537305454 |
| -0.065389369 | 0.947863988 | NA |
| 2.739554506 | 0.006152251 | 0.014785487 |
| 11.73717586 | 8.22E-32 | 3.13E-30 |
| NA | NA | NA |
| 1.668792071 | 0.095158595 | 0.159918063 |
| 12.53428197 | 4.85E-36 | 2.23E-34 |
| 2.65101066 | 0.008025131 | 0.018682015 |
| 0.252291482 | 0.800815772 | NA |
| 0.063708882 | 0.949202033 | NA |
| 0.508860203 | 0.610850217 | 0.706836528 |
| -5.037189384 | 4.72E-07 | 2.32E-06 |
| -3.596806178 | 0.000322148 | 0.001018875 |
| -0.768249684 | 0.442338861 | NA |
| 1.207832763 | 0.227111597 | 0.327303274 |
| 4.455736803 | 8.36E-06 | 3.45E-05 |
| -2.675033356 | 0.007472182 | 0.017553021 |
| -1.081576368 | 0.27944081 | 0.386473835 |
| -1.497330776 | 0.134307211 | 0.212945523 |
| NA | NA | NA |
| NA | NA | NA |
| 1.639391834 | 0.101131681 | 0.168093896 |
| -2.613594249 | 0.008959537 | 0.020642232 |
| -4.541172104 | 5.59E-06 | 2.36E-05 |

| | | | |
|---|---|---|---|
| NA | NA | NA | |
| NA | NA | NA | |
| NA | NA | NA | |
| | 0.063708882 | 0.949202033 | NA |
| | -4.407674025 | 1.04E-05 | 4.25E-05 |
| NA | NA | NA | |
| NA | NA | NA | |
| | 0.063708882 | 0.949202033 | NA |
| NA | NA | NA | |
| NA | NA | NA | |
| NA | NA | NA | |
| NA | NA | NA | |
| NA | NA | NA | |
| NA | NA | NA | |
| NA | NA | NA | |
| NA | NA | NA | |
| NA | NA | NA | |
| NA | NA | NA | |
| NA | NA | NA | |
| NA | NA | NA | |
| NA | NA | NA | |
| | 3.711827659 | 0.000205768 | 0.000674839 |
| NA | NA | NA | |
| | 0.199709364 | 0.84170789 | NA |
| | 0.763505735 | 0.445161843 | NA |
| | 0.063708882 | 0.949202033 | NA |
| NA | NA | NA | |
| | -8.007077829 | 1.17E-15 | 1.46E-14 |
| | -0.965211418 | 0.334438929 | 0.445627821 |
| NA | NA | NA | |
| NA | NA | NA | |
| | -1.690901214 | 0.090855671 | 0.153802073 |
| | 0.780748992 | 0.43495014 | 0.545510067 |
| | -0.128576576 | 0.897692702 | 0.928711595 |
| | -3.092581745 | 0.001984236 | 0.005353031 |
| | 0.459571 | 0.645824179 | 0.735072493 |
| | -0.407053895 | 0.683968405 | 0.766350685 |
| | -0.248377283 | 0.803842509 | 0.861050969 |
| | 1.209807145 | 0.226352904 | 0.326416615 |
| | -2.111358657 | 0.034741496 | 0.067466804 |
| | 2.552768595 | 0.010687047 | 0.024199095 |
| NA | NA | NA | |
| | -2.314352548 | 0.020648392 | 0.042961699 |
| | -1.606738832 | 0.108111671 | 0.177526414 |
| | -1.449904644 | 0.147085112 | 0.229608354 |
| | 4.685283987 | 2.80E-06 | 1.23E-05 |
| | -0.763369492 | 0.445243069 | 0.555140299 |

| | | | |
|---|---|---|---|
| | 0.192810118 | 0.847107679 | NA |
| NA | NA | NA | |
| NA | NA | NA | |
| | 0.192809521 | 0.847108147 | NA |
| | 0.192810118 | 0.847107679 | NA |
| | 0.063707688 | 0.949202983 | NA |
| NA | NA | NA | |
| NA | NA | NA | |
| NA | NA | NA | |
| NA | NA | NA | |
| NA | NA | NA | |
| NA | NA | NA | |
| | 0.06371127 | 0.949200131 | NA |
| | -0.188739282 | 0.850297162 | 0.895746547 |
| | -0.065388772 | 0.947864464 | NA |
| NA | NA | NA | |
| NA | NA | NA | |
| NA | NA | NA | |
| NA | NA | NA | |
| NA | NA | NA | |
| | -1.729265821 | 0.083761531 | 0.143579723 |
| | -0.817094056 | 0.413874679 | 0.525478504 |
| NA | NA | NA | |
| | -0.769853299 | 0.441386919 | 0.551430726 |
| | 4.401701537 | 1.07E-05 | 4.36E-05 |
| | -3.266243714 | 0.001089844 | 0.003115085 |
| | -1.852298789 | 0.063982929 | 0.114071599 |
| NA | NA | NA | |
| | -2.713002381 | 0.006667663 | 0.015885361 |
| | -3.028118268 | 0.002460818 | 0.006482242 |
| NA | NA | NA | |
| NA | NA | NA | |
| NA | NA | NA | |
| | -0.065388772 | 0.947864464 | NA |
| | -0.185258078 | 0.853026618 | 0.897903875 |
| | -1.995902849 | 0.045944499 | 0.086012775 |
| | -2.887960595 | 0.003877484 | 0.0097768 |
| | -3.168651223 | 0.001531481 | 0.004236622 |
| | -8.46537816 | 2.55E-17 | 3.65E-16 |
| NA | NA | NA | |
| | -1.493626879 | 0.13527317 | 0.214242435 |
| NA | NA | NA | |
| NA | NA | NA | |
| | -0.151378936 | 0.879676804 | NA |
| NA | NA | NA | |
| NA | NA | NA | |
| NA | NA | NA | |

| | | |
|---|---|---|
| NA | NA | NA |
| NA | NA | NA |
| NA | NA | NA |
| 0.063708882 | 0.949202033 | NA |
| NA | NA | NA |
| NA | NA | NA |
| -0.201521219 | 0.840291039 | 0.888016014 |
| -6.010021952 | 1.85E-09 | 1.23E-08 |
| NA | NA | NA |
| NA | NA | NA |
| NA | NA | NA |
| NA | NA | NA |
| NA | NA | NA |
| NA | NA | NA |
| NA | NA | NA |
| NA | NA | NA |
| NA | NA | NA |
| -0.6035019 | 0.546174852 | 0.649750786 |
| 0.192808924 | 0.847108615 | NA |
| NA | NA | NA |
| NA | NA | NA |
| NA | NA | NA |
| -2.695325695 | 0.007031986 | 0.016642455 |
| 0.073164869 | 0.941674922 | NA |
| NA | NA | NA |
| NA | NA | NA |
| NA | NA | NA |
| 0.284048971 | 0.776372852 | 0.839566771 |
| 2.379084671 | 0.017355689 | 0.03701226 |
| NA | NA | NA |
| NA | NA | NA |
| 1.576163455 | 0.114988138 | 0.187043001 |
| 0.325071335 | 0.745127083 | 0.814510648 |
| 1.125691953 | 0.260295931 | 0.365243141 |
| 0.192809521 | 0.847108147 | NA |
| -2.74782324 | 0.005999234 | 0.014455221 |
| 1.861075518 | 0.062733513 | 0.11223873 |
| 0.954449386 | 0.339856218 | 0.451344647 |
| NA | NA | NA |
| 0.220389076 | 0.825568152 | 0.877861033 |
| -0.97655373 | 0.328790135 | 0.43968676 |
| -0.065389369 | 0.947863988 | NA |
| 0.06371127 | 0.949200131 | NA |
| -0.33108846 | 0.740577668 | 0.811186666 |
| -3.601644375 | 0.000316211 | 0.001001741 |
| -0.116645796 | 0.907140746 | NA |
| -2.111812496 | 0.034702535 | 0.067401603 |

| | | |
|---|---|---|
| 4.649625327 | 3.33E-06 | 1.45E-05 |
| NA | NA | NA |
| NA | NA | NA |
| -0.989013856 | 0.322656363 | 0.433568946 |
| 0.975475788 | 0.329324306 | 0.440189436 |
| NA | NA | NA |
| 0.933706198 | 0.350455467 | NA |
| NA | NA | NA |
| NA | NA | NA |
| 0.063708674 | 0.949202198 | NA |
| 4.233654485 | 2.30E-05 | 8.85E-05 |
| 1.876200045 | 0.060627816 | 0.109008951 |
| 1.429703881 | 0.152802026 | 0.237054339 |
| NA | NA | NA |
| NA | NA | NA |
| NA | NA | NA |
| 5.680364841 | 1.34E-08 | 8.09E-08 |
| 0.792213106 | 0.428236428 | 0.539463888 |
| NA | NA | NA |
| -2.080026767 | 0.037523078 | 0.072102152 |
| 0.094020285 | 0.925093043 | 0.948473308 |
| NA | NA | NA |
| NA | NA | NA |
| 0.063708882 | 0.949202033 | NA |
| 0.607025786 | 0.5438338 | NA |
| 0.989052799 | 0.32263731 | 0.43356662 |
| NA | NA | NA |
| NA | NA | NA |
| 1.152977479 | 0.248919628 | 0.352302998 |
| 2.524625459 | 0.011582166 | 0.025974519 |
| NA | NA | NA |
| -0.667629147 | 0.504370347 | 0.610932094 |
| -0.240069239 | 0.810276581 | 0.866003704 |
| 6.950195124 | 3.65E-12 | 3.19E-11 |
| 1.288674673 | 0.197511211 | 0.292613531 |
| NA | NA | NA |
| NA | NA | NA |
| NA | NA | NA |
| 0.063708285 | 0.949202508 | NA |
| 0.063710673 | 0.949200606 | NA |
| NA | NA | NA |
| 0.743613333 | 0.457110435 | 0.566718222 |
| -2.239828816 | 0.025102038 | 0.050903672 |
| -0.881520489 | 0.378036169 | 0.490653115 |
| 0.192809521 | 0.847108147 | NA |
| 2.722960166 | 0.006469986 | 0.015462445 |
| -0.328138839 | 0.742806689 | 0.812897767 |

| | | |
|---|---|---|
| -1.643183399 | 0.100344994 | 0.166974817 |
| 0.192809521 | 0.847108147 | NA |
| 2.533789268 | 0.011283657 | 0.025391524 |
| NA | NA | NA |
| -1.473677162 | 0.140568535 | 0.221131948 |
| -0.131983182 | 0.894997594 | 0.926842771 |
| NA | NA | NA |
| -3.336007468 | 0.000849909 | 0.002483448 |
| 0.063708882 | 0.949202033 | NA |
| 0.482011499 | 0.629797777 | 0.722283545 |
| 0.102208618 | 0.918591087 | NA |
| NA | NA | NA |
| NA | NA | NA |
| 0.063708285 | 0.949202508 | NA |
| NA | NA | NA |
| -2.028834422 | 0.042475158 | 0.080341057 |
| -0.19448762 | 0.845794089 | NA |
| -8.861752592 | 7.88E-19 | 1.28E-17 |
| 0.873842198 | 0.382204196 | 0.494612229 |
| NA | NA | NA |
| -2.591498972 | 0.009555883 | 0.021865229 |
| -4.717930929 | 2.38E-06 | 1.06E-05 |
| -3.055891326 | 0.002243926 | 0.005973778 |
| -3.394403875 | 0.000687782 | 0.002044801 |
| 0.192809521 | 0.847108147 | NA |
| NA | NA | NA |
| 0.192809521 | 0.847108147 | NA |
| -0.065389369 | 0.947863988 | NA |
| 0.821025173 | 0.411631929 | 0.523294591 |
| -2.536162004 | 0.011207487 | 0.025245087 |
| -1.240344499 | 0.214848 | 0.313129829 |
| NA | NA | NA |
| -0.065389369 | 0.947863988 | NA |
| -0.065389966 | 0.947863513 | NA |
| 0.192810118 | 0.847107679 | NA |
| 0.027157819 | 0.978333859 | 0.984944755 |
| NA | NA | NA |
| NA | NA | NA |
| NA | NA | NA |
| 0.213245695 | 0.831135328 | 0.881870168 |
| NA | NA | NA |
| -0.45402129 | 0.649813504 | NA |
| NA | NA | NA |
| 0.26598224 | 0.790252887 | 0.850800664 |
| 0.039399741 | 0.968571686 | 0.978471208 |
| NA | NA | NA |
| NA | NA | NA |

| | | | |
|---|---|---|---|
| | -9.547758252 | 1.33E-21 | 2.67E-20 |
| | -0.825643518 | 0.409006341 | 0.520909211 |
| | -1.879560625 | 0.060167984 | 0.108291122 |
| | -1.746064573 | 0.080799732 | 0.13929453 |
| | -0.710180083 | 0.47759247 | 0.586389656 |
| | -2.288137973 | 0.022129487 | 0.045644917 |
| | -0.634716876 | 0.525613085 | 0.630776023 |
| | -0.53141414 | 0.595131825 | 0.693715344 |
| | -2.127248148 | 0.033399471 | 0.065235173 |
| | 0.586470883 | 0.557559123 | 0.660084725 |
| NA | NA | NA | |
| | -3.281864691 | 0.001031231 | 0.002963797 |
| | 0.567877067 | 0.570118445 | 0.671500855 |
| NA | NA | NA | |
| NA | NA | NA | |
| NA | NA | NA | |
| NA | NA | NA | |
| NA | NA | NA | |
| NA | NA | NA | |
| NA | NA | NA | |
| NA | NA | NA | |
| NA | NA | NA | |
| NA | NA | NA | |
| NA | NA | NA | |
| NA | NA | NA | |
| NA | NA | NA | |
| NA | NA | NA | |
| NA | NA | NA | |
| NA | NA | NA | |
| NA | NA | NA | |
| NA | NA | NA | |
| NA | NA | NA | |
| NA | NA | NA | |
| NA | NA | NA | |
| NA | NA | NA | |
| NA | NA | NA | |
| | -0.065388772 | 0.947864464 | NA |
| NA | NA | NA | |
| NA | NA | NA | |
| NA | NA | NA | |
| NA | NA | NA | |
| NA | NA | NA | |
| NA | NA | NA | |
| NA | NA | NA | |
| NA | NA | NA | |
| NA | NA | NA | |
| NA | NA | NA | |

| | | |
|---|---|---|
| NA | NA | NA |
| NA | NA | NA |
| NA | NA | NA |
| NA | NA | NA |
| NA | NA | NA |
| NA | NA | NA |
| NA | NA | NA |
| NA | NA | NA |
| NA | NA | NA |
| NA | NA | NA |
| NA | NA | NA |
| NA | NA | NA |
| NA | NA | NA |
| NA | NA | NA |
| NA | NA | NA |
| NA | NA | NA |
| 1.789566354 | 0.073523651 | 0.128404758 |
| -0.900006054 | 0.368117029 | 0.480620168 |
| -0.413109011 | 0.679526747 | 0.762476106 |
| -4.369754841 | 1.24E-05 | 4.99E-05 |
| NA | NA | NA |
| -0.935742017 | 0.349406032 | 0.46104871 |
| -2.260958183 | 0.023761847 | 0.048535908 |
| 2.489994045 | 0.012774524 | 0.028296135 |
| -2.177486534 | 0.029444285 | 0.058431021 |
| 0.908091077 | 0.363830104 | 0.476265232 |
| -1.1660601 | 0.243590147 | 0.346345228 |
| NA | NA | NA |
| NA | NA | NA |
| 0.26640786 | 0.789925113 | 0.850620615 |
| -0.555257089 | 0.578718825 | NA |
| 0.39390925 | 0.693648041 | NA |
| NA | NA | NA |
| 1.282421442 | 0.199694846 | NA |
| -6.764134063 | 1.34E-11 | 1.11E-10 |
| NA | NA | NA |
| -2.305089319 | 0.021161572 | 0.043876254 |
| -0.591073221 | 0.554471362 | 0.657433358 |
| NA | NA | NA |
| NA | NA | NA |
| -0.922494316 | 0.356270794 | 0.468181759 |
| -5.442055999 | 5.27E-08 | 2.95E-07 |
| 1.802312484 | 0.071496256 | 0.125494479 |
| 0.192808924 | 0.847108615 | NA |
| 3.117373984 | 0.001824699 | 0.004956291 |
| 2.309455755 | 0.020918304 | 0.043433028 |
| -2.147080799 | 0.03178685 | 0.06242643 |

| | | |
|---|---|---|
| 0.292877876 | 0.7696155 | 0.833988915 |
| -0.065465566 | 0.947803322 | NA |
| 1.033460206 | 0.301388584 | 0.410522217 |
| NA | NA | NA |
| 1.454308229 | 0.14586087 | 0.228109814 |
| 0.466595116 | 0.640789582 | NA |
| NA | NA | NA |
| NA | NA | NA |
| NA | NA | NA |
| -6.992080982 | 2.71E-12 | 2.39E-11 |
| NA | NA | NA |
| 0.013748753 | 0.989030428 | 0.992241437 |
| 0.146276352 | 0.883703234 | 0.918605089 |
| 0.536184175 | 0.59183127 | 0.691058811 |
| NA | NA | NA |
| 3.567115393 | 0.000360933 | 0.001130675 |
| 1.775119041 | 0.075878234 | 0.131955269 |
| 1.982793632 | 0.047390486 | 0.088395846 |
| 0.882156538 | 0.377692161 | 0.490333755 |
| NA | NA | NA |
| -8.117702759 | 4.75E-16 | 6.08E-15 |
| 5.12943926 | 2.91E-07 | 1.47E-06 |
| -5.449759928 | 5.04E-08 | 2.83E-07 |
| NA | NA | NA |
| 1.428589965 | 0.15312212 | 0.237447907 |
| -3.077760208 | 0.002085627 | 0.005593404 |
| -2.918029461 | 0.003522511 | 0.008953838 |
| 1.561954925 | 0.118298608 | 0.191552216 |
| -0.124887235 | 0.900612824 | NA |
| NA | NA | NA |
| 0.31685483 | 0.751353758 | NA |
| 0.192810118 | 0.847107679 | NA |
| -8.726342454 | 2.63E-18 | 4.08E-17 |
| 1.585825261 | 0.112778954 | 0.183947427 |
| NA | NA | NA |
| 0.192809521 | 0.847108147 | NA |
| 4.653640727 | 3.26E-06 | 1.42E-05 |
| 2.016488163 | 0.043748955 | 0.082494827 |
| 1.373346148 | 0.169644762 | 0.258081104 |
| -0.922296045 | 0.356374177 | 0.468243866 |
| -1.942690292 | 0.05205359 | 0.09586501 |
| 0.063708285 | 0.949202508 | NA |
| -1.441904661 | 0.149329271 | 0.232574927 |
| -1.456447415 | 0.145268976 | 0.227354622 |
| -5.243811909 | 1.57E-07 | 8.26E-07 |
| -0.264443293 | 0.791438356 | NA |
| 1.16285116 | 0.244889896 | 0.347785637 |

| | | |
|---|---|---|
| NA | NA | NA |
| -5.48992994 | 4.02E-08 | 2.29E-07 |
| NA | NA | NA |
| NA | NA | NA |
| NA | NA | NA |
| NA | NA | NA |
| 1.737940766 | 0.082221252 | 0.14136543 |
| 1.772952087 | 0.076236647 | 0.132504914 |
| -0.581566043 | 0.560859021 | NA |
| -2.101758413 | 0.035574444 | 0.068849326 |
| NA | NA | NA |
| NA | NA | NA |
| NA | NA | NA |
| NA | NA | NA |
| NA | NA | NA |
| NA | NA | NA |
| NA | NA | NA |
| NA | NA | NA |
| NA | NA | NA |
| NA | NA | NA |
| NA | NA | NA |
| NA | NA | NA |
| NA | NA | NA |
| NA | NA | NA |
| NA | NA | NA |
| NA | NA | NA |
| NA | NA | NA |
| NA | NA | NA |
| NA | NA | NA |
| -0.521056468 | 0.602327435 | NA |
| NA | NA | NA |
| NA | NA | NA |
| NA | NA | NA |
| -0.999813893 | 0.317400581 | 0.427907818 |
| NA | NA | NA |
| NA | NA | NA |
| NA | NA | NA |
| NA | NA | NA |
| NA | NA | NA |
| NA | NA | NA |
| NA | NA | NA |
| 0.192809521 | 0.847108147 | NA |
| NA | NA | NA |
| -1.231355712 | 0.218189854 | 0.317037736 |
| -0.065389369 | 0.947863988 | NA |
| 0.096365772 | 0.923230076 | NA |

| | | |
|---|---|---|
| 13.55172217 | 7.74E-42 | 4.49E-40 |
| NA | NA | NA |
| 5.476940681 | 4.33E-08 | 2.45E-07 |
| NA | NA | NA |
| 4.195709138 | 2.72E-05 | 0.000103664 |
| NA | NA | NA |
| 0.215610811 | 0.829291128 | NA |
| NA | NA | NA |
| -2.954546653 | 0.003131288 | 0.00805622 |
| 2.383529524 | 0.017147509 | 0.036649488 |
| -4.700742128 | 2.59E-06 | 1.15E-05 |
| -1.076327013 | 0.281781032 | 0.388830211 |
| NA | NA | NA |
| -2.780604203 | 0.005425784 | 0.013236587 |
| NA | NA | NA |
| 0.071962675 | 0.942631612 | NA |
| -1.497741279 | 0.134200483 | 0.21285331 |
| NA | NA | NA |
| 2.097488191 | 0.035950381 | 0.069474834 |
| NA | NA | NA |
| 0.524531508 | 0.599908911 | NA |
| -1.123659445 | 0.261157527 | NA |
| 1.95967504 | 0.050033784 | 0.092642819 |
| NA | NA | NA |
| -7.89516651 | 2.90E-15 | 3.46E-14 |
| -1.82188878 | 0.068471863 | 0.120957776 |
| 1.597338188 | 0.110190343 | 0.180427318 |
| NA | NA | NA |
| NA | NA | NA |
| 0.707094929 | 0.479507487 | NA |
| 0.192809521 | 0.847108147 | NA |
| NA | NA | NA |
| 0.991976128 | 0.321209171 | NA |
| NA | NA | NA |
| NA | NA | NA |
| -2.111951847 | 0.03469058 | 0.06738884 |
| 3.02396909 | 0.002494819 | 0.00656214 |
| 0.058195465 | 0.953592933 | 0.96824189 |
| -6.463150661 | 1.03E-10 | 7.79E-10 |
| 0.30922356 | 0.757151473 | NA |
| 0.192809521 | 0.847108147 | NA |
| NA | NA | NA |
| NA | NA | NA |
| NA | NA | NA |
| NA | NA | NA |
| 0.063708285 | 0.949202508 | NA |
| NA | NA | NA |

| | | | |
|---|---|---|---|
| NA | NA | NA | |
| NA | NA | NA | |
| | 4.085174962 | 4.40E-05 | 0.00016162 |
| | -6.893678389 | 5.44E-12 | 4.67E-11 |
| NA | NA | NA | |
| | -2.77923751 | 0.005448667 | 0.013282054 |
| | -0.623394761 | 0.53302514 | 0.637409548 |
| | 2.926162892 | 0.003431712 | 0.008747022 |
| | 1.516905623 | 0.129290514 | 0.206531467 |
| | 0.062498101 | 0.950166174 | 0.965931954 |
| | 0.488736396 | 0.625028333 | 0.718720799 |
| | 2.492055236 | 0.012700628 | 0.028162361 |
| NA | NA | NA | |
| NA | NA | NA | |
| | -7.144751229 | 9.02E-13 | 8.41E-12 |
| NA | NA | NA | |
| NA | NA | NA | |
| | 0.063708285 | 0.949202508 | NA |
| NA | NA | NA | |
| | 3.824456266 | 0.000131061 | 0.000443965 |
| | 1.288067169 | 0.197722583 | 0.292857343 |
| | -0.832074648 | 0.405366811 | 0.517194988 |
| | 0.115451102 | 0.908087576 | NA |
| | 3.5449702 | 0.000392657 | 0.001221352 |
| | 0.192636473 | 0.847243678 | NA |
| | 0.192810118 | 0.847107679 | NA |
| | -1.028054569 | 0.303924159 | 0.413391029 |
| | -6.1613891 | 7.21E-10 | 4.97E-09 |
| | -3.117383605 | 0.00182464 | 0.004956291 |
| | -0.873681043 | 0.382291977 | 0.494612229 |
| NA | NA | NA | |
| NA | NA | NA | |
| | 0.409647917 | 0.682064241 | NA |
| | -0.634608653 | 0.525683683 | 0.630830501 |
| | -0.315860628 | 0.752108297 | 0.820276942 |
| | -1.166960591 | 0.243226285 | 0.345906472 |
| | -1.866785156 | 0.061931596 | 0.110962624 |
| | -0.354107533 | 0.723258297 | 0.797812136 |
| | 0.799406489 | 0.424054749 | 0.535632336 |
| | -0.065389369 | 0.947863988 | NA |
| NA | NA | NA | |
| | 0.228140833 | 0.819536757 | NA |
| | 0.444308668 | 0.656819435 | NA |
| | -8.663595951 | 4.57E-18 | 6.93E-17 |
| | 2.388138665 | 0.016933951 | 0.036248296 |
| | -2.310847454 | 0.020841282 | 0.043298248 |
| NA | NA | NA | |

| | | |
|---|---|---|
| NA | NA | NA |
| NA | NA | NA |
| NA | NA | NA |
| -1.895095112 | 0.058079802 | 0.105206068 |
| NA | NA | NA |
| 5.058204016 | 4.23E-07 | 2.10E-06 |
| -2.349086607 | 0.018819529 | 0.039624227 |
| NA | NA | NA |
| NA | NA | NA |
| 0.063708285 | 0.949202508 | NA |
| 3.936629301 | 8.26E-05 | 0.000290048 |
| -1.202753782 | 0.229071615 | 0.329539773 |
| NA | NA | NA |
| -1.366949302 | 0.171641197 | 0.260706303 |
| -0.065388772 | 0.947864464 | NA |
| NA | NA | NA |
| NA | NA | NA |
| NA | NA | NA |
| NA | NA | NA |
| NA | NA | NA |
| NA | NA | NA |
| NA | NA | NA |
| 0.541035561 | 0.588483072 | 0.688112657 |
| 0.703259677 | 0.481893935 | 0.59039673 |
| NA | NA | NA |
| 1.740941593 | 0.081693817 | 0.140603342 |
| -0.099473034 | 0.920762697 | 0.945538501 |
| 0.96564651 | 0.334221094 | 0.445450159 |
| 0.765128802 | 0.444194851 | 0.554136618 |
| 1.19872068 | 0.230636574 | 0.331391128 |
| 0.192809521 | 0.847108147 | NA |
| 0.950961356 | 0.341623993 | 0.453089945 |
| -0.024627186 | 0.980352334 | 0.986381755 |
| NA | NA | NA |
| -1.607696682 | 0.107901626 | 0.177274545 |
| -0.530348433 | 0.595870372 | NA |
| -0.232870912 | 0.815861646 | 0.870373642 |
| -2.318926334 | 0.020399031 | 0.042517091 |
| 0.66673555 | 0.504941067 | 0.611479748 |
| 0.920327671 | 0.357401553 | NA |
| 0.540802538 | 0.588643693 | NA |
| NA | NA | NA |
| 1.827003007 | 0.067699317 | 0.119787496 |
| -1.474063165 | 0.140464585 | 0.221010093 |
| NA | NA | NA |
| -0.624560873 | 0.532259306 | 0.636676246 |
| 0.337632218 | 0.73564036 | NA |

| | | | |
|---|---|---|---|
| NA | NA | NA | |
| NA | NA | NA | |
| NA | NA | NA | |
| | -2.500678425 | 0.012395568 | 0.027556931 |
| NA | NA | NA | |
| | -4.538252034 | 5.67E-06 | 2.40E-05 |
| | 2.264954103 | 0.023515496 | 0.048123037 |
| | -3.838528553 | 0.000123774 | 0.000420932 |
| | 5.553708076 | 2.80E-08 | 1.62E-07 |
| | 4.86631137 | 1.14E-06 | 5.29E-06 |
| | 0.329264491 | 0.741955781 | NA |
| | 4.296250811 | 1.74E-05 | 6.82E-05 |
| | -2.279568324 | 0.022633303 | 0.046565147 |
| | 0.252113361 | 0.800953443 | NA |
| | -0.386735533 | 0.698952007 | NA |
| | 1.903712706 | 0.056947611 | 0.103507276 |
| | 0.305755909 | 0.759790501 | NA |
| NA | NA | NA | |
| | 0.192810118 | 0.847107679 | NA |
| NA | NA | NA | |
| | 0.192808327 | 0.847109082 | NA |
| | 2.09849185 | 0.035861719 | 0.069314195 |
| | 0.301954651 | 0.762686633 | 0.828600828 |
| | 3.720262146 | 0.000199016 | 0.000654153 |
| | 6.39699343 | 1.58E-10 | 1.17E-09 |
| | 10.19707039 | 2.04E-24 | 5.01E-23 |
| | 16.40470088 | 1.77E-60 | 1.79E-58 |
| | -1.057851638 | 0.290123086 | 0.397687889 |
| | 0.073100918 | 0.941725811 | NA |
| | 0.727738526 | 0.466773662 | NA |
| | -0.077360774 | 0.938336545 | 0.958058216 |
| | -8.252563923 | 1.55E-16 | 2.08E-15 |
| | 0.063976859 | 0.948988653 | NA |
| NA | NA | NA | |
| NA | NA | NA | |
| NA | NA | NA | |
| | 0.192808924 | 0.847108615 | NA |
| NA | NA | NA | |
| NA | NA | NA | |
| | 1.074932295 | 0.282405042 | 0.38943299 |
| NA | NA | NA | |
| | -5.110458005 | 3.21E-07 | 1.62E-06 |
| | 0.191800466 | 0.847898506 | NA |
| | 0.317642674 | 0.750756 | NA |
| | 1.993615504 | 0.04619409 | 0.08642832 |
| | -0.081879341 | 0.934742663 | 0.955743826 |
| | -2.928194222 | 0.00340937 | 0.00869716 |

| | | | |
|---|---|---|---|
| | -4.765664942 | 1.88E-06 | 8.50E-06 |
| | 0.063708882 | 0.949202033 | NA |
| | 5.340662801 | 9.26E-08 | 5.01E-07 |
| NA | NA | NA | |
| NA | NA | NA | |
| | 2.124802621 | 0.033603086 | 0.065561247 |
| | -0.075726246 | 0.939636895 | 0.958985178 |
| | 0.192810118 | 0.847107679 | NA |
| | -0.835301712 | 0.403547854 | 0.515215648 |
| | 0.199929773 | 0.841535505 | 0.889069117 |
| NA | NA | NA | |
| NA | NA | NA | |
| NA | NA | NA | |
| NA | NA | NA | |
| | 0.063708285 | 0.949202508 | NA |
| | -0.065389369 | 0.947863988 | NA |
| | 1.566876361 | 0.117143594 | 0.189976944 |
| NA | NA | NA | |
| | -0.3194719 | 0.749368696 | NA |
| | -0.194488217 | 0.845793621 | NA |
| | -1.797530519 | 0.072251438 | 0.126571492 |
| NA | NA | NA | |
| NA | NA | NA | |
| | 0.554168543 | 0.579463504 | 0.680203138 |
| | -1.259244976 | 0.207941861 | 0.304962645 |
| | -0.556799467 | 0.577664448 | 0.678568474 |
| | 0.407943893 | 0.683314867 | 0.765834018 |
| | 2.916552474 | 0.003539232 | 0.008991776 |
| | -0.564769901 | 0.572230282 | 0.673509006 |
| | 0.069451617 | 0.944630144 | 0.9623678 |
| | -0.465011877 | NA | NA |
| NA | NA | NA | |
| NA | NA | NA | |
| NA | NA | NA | |
| | -0.718499008 | 0.47244966 | NA |
| NA | NA | NA | |
| NA | NA | NA | |
| NA | NA | NA | |
| NA | NA | NA | |
| NA | NA | NA | |
| NA | NA | NA | |
| NA | NA | NA | |
| NA | NA | NA | |
| NA | NA | NA | |
| NA | NA | NA | |
| NA | NA | NA | |
| NA | NA | NA | |

| | | |
|---|---|---|
| NA | NA | NA |
| NA | NA | NA |
| NA | NA | NA |
| NA | NA | NA |
| NA | NA | NA |
| NA | NA | NA |
| NA | NA | NA |
| NA | NA | NA |
| NA | NA | NA |
| NA | NA | NA |
| NA | NA | NA |
| NA | NA | NA |
| NA | NA | NA |
| NA | NA | NA |
| NA | NA | NA |
| NA | NA | NA |
| NA | NA | NA |
| NA | NA | NA |
| NA | NA | NA |
| NA | NA | NA |
| NA | NA | NA |
| NA | NA | NA |
| NA | NA | NA |
| NA | NA | NA |
| NA | NA | NA |
| NA | NA | NA |
| NA | NA | NA |
| NA | NA | NA |
| NA | NA | NA |
| NA | NA | NA |
| NA | NA | NA |
| NA | NA | NA |
| NA | NA | NA |
| NA | NA | NA |
| NA | NA | NA |
| NA | NA | NA |
| NA | NA | NA |
| NA | NA | NA |
| NA | NA | NA |
| NA | NA | NA |
| NA | NA | NA |
| NA | NA | NA |
| NA | NA | NA |
| 0.192809521 | 0.847108147 | NA |
| NA | NA | NA |
| NA | NA | NA |
| NA | NA | NA |

| | | |
|---|---|---|
| NA | NA | NA |
| NA | NA | NA |
| NA | NA | NA |
| NA | NA | NA |
| NA | NA | NA |
| NA | NA | NA |
| NA | NA | NA |
| NA | NA | NA |
| NA | NA | NA |
| NA | NA | NA |
| NA | NA | NA |
| NA | NA | NA |
| NA | NA | NA |
| NA | NA | NA |
| NA | NA | NA |
| NA | NA | NA |
| NA | NA | NA |
| NA | NA | NA |
| NA | NA | NA |
| NA | NA | NA |
| NA | NA | NA |
| NA | NA | NA |
| NA | NA | NA |
| NA | NA | NA |
| NA | NA | NA |
| NA | NA | NA |
| NA | NA | NA |
| NA | NA | NA |
| NA | NA | NA |
| NA | NA | NA |
| NA | NA | NA |
| NA | NA | NA |
| 0.192809521 | 0.847108147 | NA |
| NA | NA | NA |
| NA | NA | NA |
| NA | NA | NA |
| NA | NA | NA |
| NA | NA | NA |
| NA | NA | NA |
| NA | NA | NA |
| NA | NA | NA |
| NA | NA | NA |
| NA | NA | NA |
| NA | NA | NA |
| NA | NA | NA |
| NA | NA | NA |
| NA | NA | NA |

| | | |
|---|---|---|
| 0.063708285 | 0.949202508 | NA |
| NA | NA | NA |
| NA | NA | NA |
| NA | NA | NA |
| NA | NA | NA |
| NA | NA | NA |
| NA | NA | NA |
| NA | NA | NA |
| NA | NA | NA |
| NA | NA | NA |
| NA | NA | NA |
| NA | NA | NA |
| NA | NA | NA |
| -0.08329421 | 0.933617604 | NA |
| NA | NA | NA |
| NA | NA | NA |
| NA | NA | NA |
| NA | NA | NA |
| NA | NA | NA |
| NA | NA | NA |
| NA | NA | NA |
| NA | NA | NA |
| NA | NA | NA |
| NA | NA | NA |
| NA | NA | NA |
| -0.433775282 | 0.664451633 | NA |
| NA | NA | NA |
| NA | NA | NA |
| NA | NA | NA |
| NA | NA | NA |
| NA | NA | NA |
| NA | NA | NA |
| NA | NA | NA |
| NA | NA | NA |
| NA | NA | NA |
| NA | NA | NA |
| NA | NA | NA |
| NA | NA | NA |
| NA | NA | NA |
| NA | NA | NA |
| NA | NA | NA |
| NA | NA | NA |
| NA | NA | NA |
| NA | NA | NA |
| NA | NA | NA |
| NA | NA | NA |
| NA | NA | NA |

| | | |
|---|---|---|
| NA | NA | NA |
| NA | NA | NA |
| NA | NA | NA |
| NA | NA | NA |
| NA | NA | NA |
| NA | NA | NA |
| NA | NA | NA |
| NA | NA | NA |
| NA | NA | NA |
| NA | NA | NA |
| NA | NA | NA |
| NA | NA | NA |
| NA | NA | NA |
| NA | NA | NA |
| NA | NA | NA |
| NA | NA | NA |
| NA | NA | NA |
| NA | NA | NA |
| NA | NA | NA |
| NA | NA | NA |
| NA | NA | NA |
| NA | NA | NA |
| NA | NA | NA |
| NA | NA | NA |
| NA | NA | NA |
| NA | NA | NA |
| NA | NA | NA |
| NA | NA | NA |
| NA | NA | NA |
| NA | NA | NA |
| NA | NA | NA |
| NA | NA | NA |
| NA | NA | NA |
| NA | NA | NA |
| NA | NA | NA |
| NA | NA | NA |
| NA | NA | NA |
| NA | NA | NA |
| NA | NA | NA |
| NA | NA | NA |
| NA | NA | NA |
| NA | NA | NA |
| NA | NA | NA |
| NA | NA | NA |
| NA | NA | NA |
| NA | NA | NA |
| NA | NA | NA |

| | | |
|---|---|---|
| NA | NA | NA |
| NA | NA | NA |
| NA | NA | NA |
| 0.063708285 | 0.949202508 | NA |
| NA | NA | NA |
| NA | NA | NA |
| NA | NA | NA |
| 1.916768016 | 0.05526741 | 0.100900312 |
| NA | NA | NA |
| NA | NA | NA |
| NA | NA | NA |
| NA | NA | NA |
| NA | NA | NA |
| NA | NA | NA |
| NA | NA | NA |
| NA | NA | NA |
| NA | NA | NA |
| NA | NA | NA |
| NA | NA | NA |
| NA | NA | NA |
| NA | NA | NA |
| NA | NA | NA |
| NA | NA | NA |
| NA | NA | NA |
| NA | NA | NA |
| NA | NA | NA |
| NA | NA | NA |
| NA | NA | NA |
| NA | NA | NA |
| NA | NA | NA |
| NA | NA | NA |
| NA | NA | NA |
| NA | NA | NA |
| NA | NA | NA |
| NA | NA | NA |
| NA | NA | NA |
| NA | NA | NA |
| NA | NA | NA |
| NA | NA | NA |
| NA | NA | NA |
| NA | NA | NA |
| NA | NA | NA |
| NA | NA | NA |
| NA | NA | NA |
| NA | NA | NA |
| NA | NA | NA |
| NA | NA | NA |

| | | |
|---|---|---|
| NA | NA | NA |
| NA | NA | NA |
| NA | NA | NA |
| NA | NA | NA |
| NA | NA | NA |
| NA | NA | NA |
| NA | NA | NA |
| NA | NA | NA |
| NA | NA | NA |
| NA | NA | NA |
| NA | NA | NA |
| NA | NA | NA |
| NA | NA | NA |
| NA | NA | NA |
| NA | NA | NA |
| NA | NA | NA |
| NA | NA | NA |
| NA | NA | NA |
| NA | NA | NA |
| NA | NA | NA |
| NA | NA | NA |
| NA | NA | NA |
| NA | NA | NA |
| NA | NA | NA |
| NA | NA | NA |
| NA | NA | NA |
| NA | NA | NA |
| NA | NA | NA |
| NA | NA | NA |
| NA | NA | NA |
| NA | NA | NA |
| NA | NA | NA |
| NA | NA | NA |
| NA | NA | NA |
| NA | NA | NA |
| NA | NA | NA |
| NA | NA | NA |
| NA | NA | NA |
| NA | NA | NA |
| NA | NA | NA |
| NA | NA | NA |
| NA | NA | NA |
| NA | NA | NA |
| NA | NA | NA |
| -0.065389369 | 0.947863988 | NA |
| NA | NA | NA |
| NA | NA | NA |

| | | |
|---|---|---|
| NA | NA | NA |
| NA | NA | NA |
| NA | NA | NA |
| NA | NA | NA |
| NA | NA | NA |
| NA | NA | NA |
| NA | NA | NA |
| NA | NA | NA |
| NA | NA | NA |
| NA | NA | NA |
| NA | NA | NA |
| NA | NA | NA |
| NA | NA | NA |
| NA | NA | NA |
| NA | NA | NA |
| NA | NA | NA |
| NA | NA | NA |
| NA | NA | NA |
| NA | NA | NA |
| NA | NA | NA |
| NA | NA | NA |
| NA | NA | NA |
| NA | NA | NA |
| NA | NA | NA |
| NA | NA | NA |
| NA | NA | NA |
| NA | NA | NA |
| NA | NA | NA |
| NA | NA | NA |
| NA | NA | NA |
| NA | NA | NA |
| NA | NA | NA |
| NA | NA | NA |
| NA | NA | NA |
| NA | NA | NA |
| NA | NA | NA |
| NA | NA | NA |
| NA | NA | NA |
| NA | NA | NA |
| NA | NA | NA |
| NA | NA | NA |
| NA | NA | NA |
| NA | NA | NA |
| NA | NA | NA |
| NA | NA | NA |
| NA | NA | NA |
| NA | NA | NA |

| | | |
|---|---|---|
| NA | NA | NA |
| NA | NA | NA |
| NA | NA | NA |
| NA | NA | NA |
| NA | NA | NA |
| NA | NA | NA |
| NA | NA | NA |
| NA | NA | NA |
| NA | NA | NA |
| NA | NA | NA |
| NA | NA | NA |
| NA | NA | NA |
| NA | NA | NA |
| NA | NA | NA |
| NA | NA | NA |
| NA | NA | NA |
| NA | NA | NA |
| NA | NA | NA |
| NA | NA | NA |
| NA | NA | NA |
| NA | NA | NA |
| NA | NA | NA |
| NA | NA | NA |
| NA | NA | NA |
| NA | NA | NA |
| NA | NA | NA |
| NA | NA | NA |
| NA | NA | NA |
| NA | NA | NA |
| NA | NA | NA |
| NA | NA | NA |
| NA | NA | NA |
| NA | NA | NA |
| NA | NA | NA |
| NA | NA | NA |
| NA | NA | NA |
| NA | NA | NA |
| NA | NA | NA |
| NA | NA | NA |
| NA | NA | NA |
| -0.065389369 | 0.947863988 | NA |
| NA | NA | NA |
| NA | NA | NA |
| NA | NA | NA |
| NA | NA | NA |
| NA | NA | NA |
| NA | NA | NA |

| | | |
|---|---|---|
| NA | NA | NA |
| NA | NA | NA |
| NA | NA | NA |
| NA | NA | NA |
| NA | NA | NA |
| NA | NA | NA |
| NA | NA | NA |
| NA | NA | NA |
| NA | NA | NA |
| NA | NA | NA |
| NA | NA | NA |
| NA | NA | NA |
| NA | NA | NA |
| NA | NA | NA |
| NA | NA | NA |
| NA | NA | NA |
| NA | NA | NA |
| NA | NA | NA |
| NA | NA | NA |
| NA | NA | NA |
| NA | NA | NA |
| NA | NA | NA |
| NA | NA | NA |
| NA | NA | NA |
| NA | NA | NA |
| NA | NA | NA |
| NA | NA | NA |
| NA | NA | NA |
| NA | NA | NA |
| NA | NA | NA |
| NA | NA | NA |
| NA | NA | NA |
| NA | NA | NA |
| NA | NA | NA |
| NA | NA | NA |
| NA | NA | NA |
| NA | NA | NA |
| NA | NA | NA |
| NA | NA | NA |
| NA | NA | NA |
| NA | NA | NA |
| NA | NA | NA |
| NA | NA | NA |
| NA | NA | NA |
| NA | NA | NA |
| NA | NA | NA |
| NA | NA | NA |

| | | |
|---|---|---|
| NA | NA | NA |
| NA | NA | NA |
| NA | NA | NA |
| NA | NA | NA |
| NA | NA | NA |
| NA | NA | NA |
| NA | NA | NA |
| NA | NA | NA |
| NA | NA | NA |
| NA | NA | NA |
| NA | NA | NA |
| NA | NA | NA |
| NA | NA | NA |
| NA | NA | NA |
| NA | NA | NA |
| NA | NA | NA |
| NA | NA | NA |
| NA | NA | NA |
| NA | NA | NA |
| NA | NA | NA |
| NA | NA | NA |
| NA | NA | NA |
| NA | NA | NA |
| NA | NA | NA |
| NA | NA | NA |
| NA | NA | NA |
| NA | NA | NA |
| NA | NA | NA |
| NA | NA | NA |
| NA | NA | NA |
| NA | NA | NA |
| NA | NA | NA |
| NA | NA | NA |
| NA | NA | NA |
| NA | NA | NA |
| NA | NA | NA |
| NA | NA | NA |
| NA | NA | NA |
| NA | NA | NA |
| NA | NA | NA |
| NA | NA | NA |
| NA | NA | NA |
| NA | NA | NA |
| NA | NA | NA |
| NA | NA | NA |
| NA | NA | NA |
| NA | NA | NA |

| | | |
|---|---|---|
| -0.065389369 | 0.947863988 | NA |
| NA | NA | NA |
| NA | NA | NA |
| NA | NA | NA |
| NA | NA | NA |
| NA | NA | NA |
| NA | NA | NA |
| NA | NA | NA |
| NA | NA | NA |
| NA | NA | NA |
| NA | NA | NA |
| NA | NA | NA |
| NA | NA | NA |
| NA | NA | NA |
| NA | NA | NA |
| NA | NA | NA |
| NA | NA | NA |
| NA | NA | NA |
| NA | NA | NA |
| NA | NA | NA |
| NA | NA | NA |
| NA | NA | NA |
| NA | NA | NA |
| NA | NA | NA |
| NA | NA | NA |
| NA | NA | NA |
| NA | NA | NA |
| NA | NA | NA |
| NA | NA | NA |
| NA | NA | NA |
| NA | NA | NA |
| NA | NA | NA |
| NA | NA | NA |
| NA | NA | NA |
| NA | NA | NA |
| NA | NA | NA |
| NA | NA | NA |
| NA | NA | NA |
| NA | NA | NA |
| NA | NA | NA |
| NA | NA | NA |
| NA | NA | NA |
| NA | NA | NA |
| NA | NA | NA |
| NA | NA | NA |
| NA | NA | NA |
| NA | NA | NA |

| | | |
|---|---|---|
| NA | NA | NA |
| NA | NA | NA |
| NA | NA | NA |
| NA | NA | NA |
| NA | NA | NA |
| NA | NA | NA |
| NA | NA | NA |
| NA | NA | NA |
| NA | NA | NA |
| NA | NA | NA |
| NA | NA | NA |
| NA | NA | NA |
| NA | NA | NA |
| NA | NA | NA |
| NA | NA | NA |
| NA | NA | NA |
| NA | NA | NA |
| NA | NA | NA |
| NA | NA | NA |
| NA | NA | NA |
| NA | NA | NA |
| NA | NA | NA |
| NA | NA | NA |
| NA | NA | NA |
| NA | NA | NA |
| NA | NA | NA |
| NA | NA | NA |
| NA | NA | NA |
| NA | NA | NA |
| NA | NA | NA |
| NA | NA | NA |
| NA | NA | NA |
| NA | NA | NA |
| NA | NA | NA |
| NA | NA | NA |
| NA | NA | NA |
| NA | NA | NA |
| NA | NA | NA |
| NA | NA | NA |
| NA | NA | NA |
| NA | NA | NA |
| NA | NA | NA |
| NA | NA | NA |
| NA | NA | NA |
| NA | NA | NA |
| NA | NA | NA |
| NA | NA | NA |

| | | |
|---|---|---|
| NA | NA | NA |
| NA | NA | NA |
| NA | NA | NA |
| NA | NA | NA |
| NA | NA | NA |
| NA | NA | NA |
| NA | NA | NA |
| NA | NA | NA |
| NA | NA | NA |
| NA | NA | NA |
| NA | NA | NA |
| NA | NA | NA |
| NA | NA | NA |
| NA | NA | NA |
| NA | NA | NA |
| NA | NA | NA |
| NA | NA | NA |
| NA | NA | NA |
| NA | NA | NA |
| NA | NA | NA |
| NA | NA | NA |
| NA | NA | NA |
| NA | NA | NA |
| NA | NA | NA |
| NA | NA | NA |
| NA | NA | NA |
| NA | NA | NA |
| NA | NA | NA |
| NA | NA | NA |
| NA | NA | NA |
| NA | NA | NA |
| NA | NA | NA |
| NA | NA | NA |
| NA | NA | NA |
| NA | NA | NA |
| NA | NA | NA |
| NA | NA | NA |
| NA | NA | NA |
| NA | NA | NA |
| NA | NA | NA |
| NA | NA | NA |
| NA | NA | NA |
| NA | NA | NA |
| NA | NA | NA |
| NA | NA | NA |
| NA | NA | NA |
| NA | NA | NA |

| | | |
|---|---|---|
| NA | NA | NA |
| NA | NA | NA |
| NA | NA | NA |
| NA | NA | NA |
| NA | NA | NA |
| NA | NA | NA |
| NA | NA | NA |
| NA | NA | NA |
| NA | NA | NA |
| NA | NA | NA |
| NA | NA | NA |
| NA | NA | NA |
| NA | NA | NA |
| NA | NA | NA |
| NA | NA | NA |
| NA | NA | NA |
| NA | NA | NA |
| NA | NA | NA |
| NA | NA | NA |
| NA | NA | NA |
| NA | NA | NA |
| NA | NA | NA |
| NA | NA | NA |
| NA | NA | NA |
| NA | NA | NA |
| NA | NA | NA |
| NA | NA | NA |
| NA | NA | NA |
| NA | NA | NA |
| NA | NA | NA |
| NA | NA | NA |
| NA | NA | NA |
| NA | NA | NA |
| NA | NA | NA |
| NA | NA | NA |
| NA | NA | NA |
| NA | NA | NA |
| NA | NA | NA |
| NA | NA | NA |
| NA | NA | NA |
| NA | NA | NA |
| NA | NA | NA |
| NA | NA | NA |
| NA | NA | NA |
| NA | NA | NA |
| NA | NA | NA |
| NA | NA | NA |

| | | |
|---|---|---|
| NA | NA | NA |
| NA | NA | NA |
| NA | NA | NA |
| NA | NA | NA |
| NA | NA | NA |
| NA | NA | NA |
| NA | NA | NA |
| NA | NA | NA |
| NA | NA | NA |
| NA | NA | NA |
| NA | NA | NA |
| NA | NA | NA |
| NA | NA | NA |
| NA | NA | NA |
| -0.256870426 | 0.797278801 | 0.855814752 |
| NA | NA | NA |
| NA | NA | NA |
| NA | NA | NA |
| NA | NA | NA |
| NA | NA | NA |
| NA | NA | NA |
| NA | NA | NA |
| NA | NA | NA |
| NA | NA | NA |
| NA | NA | NA |
| NA | NA | NA |
| NA | NA | NA |
| NA | NA | NA |
| NA | NA | NA |
| -0.065389369 | 0.947863988 | NA |
| NA | NA | NA |
| NA | NA | NA |
| NA | NA | NA |
| NA | NA | NA |
| NA | NA | NA |
| NA | NA | NA |
| NA | NA | NA |
| NA | NA | NA |
| NA | NA | NA |
| NA | NA | NA |
| NA | NA | NA |
| NA | NA | NA |
| NA | NA | NA |
| NA | NA | NA |
| NA | NA | NA |
| NA | NA | NA |
| NA | NA | NA |

| | | |
|---|---|---|
| NA | NA | NA |
| NA | NA | NA |
| NA | NA | NA |
| NA | NA | NA |
| NA | NA | NA |
| NA | NA | NA |
| NA | NA | NA |
| NA | NA | NA |
| NA | NA | NA |
| NA | NA | NA |
| NA | NA | NA |
| NA | NA | NA |
| NA | NA | NA |
| NA | NA | NA |
| NA | NA | NA |
| NA | NA | NA |
| NA | NA | NA |
| NA | NA | NA |
| -0.065389369 | 0.947863988 | NA |
| NA | NA | NA |
| NA | NA | NA |
| NA | NA | NA |
| NA | NA | NA |
| NA | NA | NA |
| NA | NA | NA |
| NA | NA | NA |
| NA | NA | NA |
| NA | NA | NA |
| NA | NA | NA |
| NA | NA | NA |
| NA | NA | NA |
| NA | NA | NA |
| NA | NA | NA |
| NA | NA | NA |
| NA | NA | NA |
| NA | NA | NA |
| NA | NA | NA |
| NA | NA | NA |
| NA | NA | NA |
| NA | NA | NA |
| NA | NA | NA |
| NA | NA | NA |
| NA | NA | NA |
| NA | NA | NA |
| NA | NA | NA |
| NA | NA | NA |
| NA | NA | NA |

| | | |
|---|---|---|
| NA | NA | NA |
| NA | NA | NA |
| NA | NA | NA |
| NA | NA | NA |
| NA | NA | NA |
| NA | NA | NA |
| NA | NA | NA |
| NA | NA | NA |
| NA | NA | NA |
| NA | NA | NA |
| NA | NA | NA |
| NA | NA | NA |
| NA | NA | NA |
| NA | NA | NA |
| NA | NA | NA |
| NA | NA | NA |
| NA | NA | NA |
| NA | NA | NA |
| NA | NA | NA |
| NA | NA | NA |
| NA | NA | NA |
| NA | NA | NA |
| NA | NA | NA |
| NA | NA | NA |
| NA | NA | NA |
| NA | NA | NA |
| NA | NA | NA |
| NA | NA | NA |
| NA | NA | NA |
| NA | NA | NA |
| NA | NA | NA |
| NA | NA | NA |
| NA | NA | NA |
| NA | NA | NA |
| NA | NA | NA |
| NA | NA | NA |
| NA | NA | NA |
| NA | NA | NA |
| NA | NA | NA |
| NA | NA | NA |
| NA | NA | NA |
| NA | NA | NA |
| NA | NA | NA |
| NA | NA | NA |
| NA | NA | NA |
| NA | NA | NA |
| NA | NA | NA |

| | | |
|---|---|---|
| NA | NA | NA |
| NA | NA | NA |
| NA | NA | NA |
| NA | NA | NA |
| NA | NA | NA |
| NA | NA | NA |
| NA | NA | NA |
| NA | NA | NA |
| NA | NA | NA |
| NA | NA | NA |
| NA | NA | NA |
| NA | NA | NA |
| NA | NA | NA |
| NA | NA | NA |
| NA | NA | NA |
| NA | NA | NA |
| NA | NA | NA |
| NA | NA | NA |
| NA | NA | NA |
| NA | NA | NA |
| NA | NA | NA |
| NA | NA | NA |
| NA | NA | NA |
| NA | NA | NA |
| NA | NA | NA |
| NA | NA | NA |
| NA | NA | NA |
| NA | NA | NA |
| NA | NA | NA |
| NA | NA | NA |
| NA | NA | NA |
| NA | NA | NA |
| NA | NA | NA |
| NA | NA | NA |
| NA | NA | NA |
| NA | NA | NA |
| NA | NA | NA |
| NA | NA | NA |
| NA | NA | NA |
| NA | NA | NA |
| NA | NA | NA |
| NA | NA | NA |
| NA | NA | NA |
| NA | NA | NA |
| NA | NA | NA |
| NA | NA | NA |
| NA | NA | NA |

| | | |
|---|---|---|
| NA | NA | NA |
| NA | NA | NA |
| NA | NA | NA |
| NA | NA | NA |
| NA | NA | NA |
| NA | NA | NA |
| NA | NA | NA |
| NA | NA | NA |
| NA | NA | NA |
| NA | NA | NA |
| NA | NA | NA |
| NA | NA | NA |
| NA | NA | NA |
| NA | NA | NA |
| NA | NA | NA |
| NA | NA | NA |
| NA | NA | NA |
| NA | NA | NA |
| NA | NA | NA |
| NA | NA | NA |
| NA | NA | NA |
| NA | NA | NA |
| NA | NA | NA |
| NA | NA | NA |
| NA | NA | NA |
| NA | NA | NA |
| NA | NA | NA |
| NA | NA | NA |
| NA | NA | NA |
| NA | NA | NA |
| NA | NA | NA |
| NA | NA | NA |
| NA | NA | NA |
| NA | NA | NA |
| NA | NA | NA |
| NA | NA | NA |
| NA | NA | NA |
| NA | NA | NA |
| NA | NA | NA |
| NA | NA | NA |
| NA | NA | NA |
| NA | NA | NA |
| NA | NA | NA |
| NA | NA | NA |
| NA | NA | NA |
| NA | NA | NA |
| NA | NA | NA |

| | | |
|---|---|---|
| NA | NA | NA |
| NA | NA | NA |
| NA | NA | NA |
| NA | NA | NA |
| NA | NA | NA |
| NA | NA | NA |
| NA | NA | NA |
| NA | NA | NA |
| NA | NA | NA |
| NA | NA | NA |
| NA | NA | NA |
| NA | NA | NA |
| NA | NA | NA |
| NA | NA | NA |
| NA | NA | NA |
| NA | NA | NA |
| NA | NA | NA |
| NA | NA | NA |
| NA | NA | NA |
| NA | NA | NA |
| NA | NA | NA |
| NA | NA | NA |
| NA | NA | NA |
| NA | NA | NA |
| NA | NA | NA |
| NA | NA | NA |
| NA | NA | NA |
| NA | NA | NA |
| NA | NA | NA |
| NA | NA | NA |
| NA | NA | NA |
| NA | NA | NA |
| NA | NA | NA |
| NA | NA | NA |
| NA | NA | NA |
| NA | NA | NA |
| NA | NA | NA |
| NA | NA | NA |
| NA | NA | NA |
| NA | NA | NA |
| NA | NA | NA |
| NA | NA | NA |
| NA | NA | NA |
| NA | NA | NA |
| NA | NA | NA |
| NA | NA | NA |
| NA | NA | NA |

| | | |
|---|---|---|
| NA | NA | NA |
| NA | NA | NA |
| NA | NA | NA |
| NA | NA | NA |
| NA | NA | NA |
| NA | NA | NA |
| NA | NA | NA |
| NA | NA | NA |
| NA | NA | NA |
| NA | NA | NA |
| NA | NA | NA |
| NA | NA | NA |
| NA | NA | NA |
| NA | NA | NA |
| NA | NA | NA |
| NA | NA | NA |
| NA | NA | NA |
| NA | NA | NA |
| NA | NA | NA |
| NA | NA | NA |
| NA | NA | NA |
| NA | NA | NA |
| NA | NA | NA |
| NA | NA | NA |
| NA | NA | NA |
| NA | NA | NA |
| NA | NA | NA |
| NA | NA | NA |
| NA | NA | NA |
| NA | NA | NA |
| NA | NA | NA |
| NA | NA | NA |
| NA | NA | NA |
| NA | NA | NA |
| NA | NA | NA |
| NA | NA | NA |
| NA | NA | NA |
| NA | NA | NA |
| NA | NA | NA |
| NA | NA | NA |
| NA | NA | NA |
| NA | NA | NA |
| NA | NA | NA |
| NA | NA | NA |
| NA | NA | NA |
| NA | NA | NA |
| NA | NA | NA |

| | | |
|---|---|---|
| NA | NA | NA |
| NA | NA | NA |
| NA | NA | NA |
| NA | NA | NA |
| NA | NA | NA |
| NA | NA | NA |
| NA | NA | NA |
| NA | NA | NA |
| NA | NA | NA |
| NA | NA | NA |
| NA | NA | NA |
| NA | NA | NA |
| NA | NA | NA |
| NA | NA | NA |
| NA | NA | NA |
| NA | NA | NA |
| NA | NA | NA |
| NA | NA | NA |
| NA | NA | NA |
| NA | NA | NA |
| NA | NA | NA |
| NA | NA | NA |
| NA | NA | NA |
| NA | NA | NA |
| NA | NA | NA |
| NA | NA | NA |
| NA | NA | NA |
| NA | NA | NA |
| NA | NA | NA |
| NA | NA | NA |
| NA | NA | NA |
| NA | NA | NA |
| NA | NA | NA |
| NA | NA | NA |
| NA | NA | NA |
| NA | NA | NA |
| NA | NA | NA |
| NA | NA | NA |
| NA | NA | NA |
| NA | NA | NA |
| NA | NA | NA |
| NA | NA | NA |
| NA | NA | NA |
| NA | NA | NA |
| NA | NA | NA |
| NA | NA | NA |
| NA | NA | NA |

| | | |
|---|---|---|
| NA | NA | NA |
| NA | NA | NA |
| NA | NA | NA |
| NA | NA | NA |
| NA | NA | NA |
| NA | NA | NA |
| NA | NA | NA |
| NA | NA | NA |
| NA | NA | NA |
| NA | NA | NA |
| NA | NA | NA |
| NA | NA | NA |
| NA | NA | NA |
| NA | NA | NA |
| NA | NA | NA |
| NA | NA | NA |
| NA | NA | NA |
| NA | NA | NA |
| NA | NA | NA |
| NA | NA | NA |
| NA | NA | NA |
| NA | NA | NA |
| NA | NA | NA |
| NA | NA | NA |
| NA | NA | NA |
| NA | NA | NA |
| NA | NA | NA |
| NA | NA | NA |
| NA | NA | NA |
| NA | NA | NA |
| NA | NA | NA |
| NA | NA | NA |
| NA | NA | NA |
| NA | NA | NA |
| NA | NA | NA |
| NA | NA | NA |
| NA | NA | NA |
| NA | NA | NA |
| NA | NA | NA |
| NA | NA | NA |
| NA | NA | NA |
| NA | NA | NA |
| NA | NA | NA |
| NA | NA | NA |
| NA | NA | NA |
| NA | NA | NA |
| NA | NA | NA |

| | | |
|---|---|---|
| NA | NA | NA |
| NA | NA | NA |
| NA | NA | NA |
| NA | NA | NA |
| NA | NA | NA |
| NA | NA | NA |
| NA | NA | NA |
| NA | NA | NA |
| NA | NA | NA |
| NA | NA | NA |
| NA | NA | NA |
| NA | NA | NA |
| NA | NA | NA |
| NA | NA | NA |
| NA | NA | NA |
| NA | NA | NA |
| NA | NA | NA |
| NA | NA | NA |
| NA | NA | NA |
| NA | NA | NA |
| NA | NA | NA |
| NA | NA | NA |
| NA | NA | NA |
| NA | NA | NA |
| NA | NA | NA |
| NA | NA | NA |
| NA | NA | NA |
| NA | NA | NA |
| NA | NA | NA |
| NA | NA | NA |
| NA | NA | NA |
| NA | NA | NA |
| NA | NA | NA |
| NA | NA | NA |
| NA | NA | NA |
| NA | NA | NA |
| NA | NA | NA |
| NA | NA | NA |
| NA | NA | NA |
| NA | NA | NA |
| NA | NA | NA |
| NA | NA | NA |
| NA | NA | NA |
| NA | NA | NA |
| NA | NA | NA |
| NA | NA | NA |
| NA | NA | NA |

| | | |
|---|---|---|
| NA | NA | NA |
| NA | NA | NA |
| NA | NA | NA |
| NA | NA | NA |
| NA | NA | NA |
| NA | NA | NA |
| NA | NA | NA |
| NA | NA | NA |
| NA | NA | NA |
| NA | NA | NA |
| NA | NA | NA |
| NA | NA | NA |
| NA | NA | NA |
| NA | NA | NA |
| NA | NA | NA |
| -0.065389369 | 0.947863988 | NA |
| NA | NA | NA |
| NA | NA | NA |
| NA | NA | NA |
| NA | NA | NA |
| NA | NA | NA |
| NA | NA | NA |
| NA | NA | NA |
| NA | NA | NA |
| NA | NA | NA |
| NA | NA | NA |
| NA | NA | NA |
| NA | NA | NA |
| NA | NA | NA |
| NA | NA | NA |
| NA | NA | NA |
| NA | NA | NA |
| NA | NA | NA |
| NA | NA | NA |
| NA | NA | NA |
| NA | NA | NA |
| NA | NA | NA |
| NA | NA | NA |
| NA | NA | NA |
| NA | NA | NA |
| NA | NA | NA |
| NA | NA | NA |
| NA | NA | NA |
| NA | NA | NA |
| NA | NA | NA |
| NA | NA | NA |
| NA | NA | NA |

| | | |
|---|---|---|
| NA | NA | NA |
| NA | NA | NA |
| NA | NA | NA |
| NA | NA | NA |
| NA | NA | NA |
| NA | NA | NA |
| NA | NA | NA |
| NA | NA | NA |
| NA | NA | NA |
| NA | NA | NA |
| NA | NA | NA |
| NA | NA | NA |
| NA | NA | NA |
| NA | NA | NA |
| NA | NA | NA |
| NA | NA | NA |
| NA | NA | NA |
| NA | NA | NA |
| NA | NA | NA |
| NA | NA | NA |
| NA | NA | NA |
| NA | NA | NA |
| NA | NA | NA |
| NA | NA | NA |
| NA | NA | NA |
| NA | NA | NA |
| NA | NA | NA |
| NA | NA | NA |
| NA | NA | NA |
| NA | NA | NA |
| NA | NA | NA |
| NA | NA | NA |
| NA | NA | NA |
| NA | NA | NA |
| NA | NA | NA |
| NA | NA | NA |
| NA | NA | NA |
| NA | NA | NA |
| NA | NA | NA |
| NA | NA | NA |
| NA | NA | NA |
| NA | NA | NA |
| NA | NA | NA |
| NA | NA | NA |
| NA | NA | NA |
| NA | NA | NA |
| NA | NA | NA |

| | | |
|---|---|---|
| NA | NA | NA |
| NA | NA | NA |
| NA | NA | NA |
| NA | NA | NA |
| NA | NA | NA |
| NA | NA | NA |
| NA | NA | NA |
| NA | NA | NA |
| NA | NA | NA |
| NA | NA | NA |
| NA | NA | NA |
| NA | NA | NA |
| NA | NA | NA |
| NA | NA | NA |
| NA | NA | NA |
| NA | NA | NA |
| NA | NA | NA |
| NA | NA | NA |
| NA | NA | NA |
| NA | NA | NA |
| NA | NA | NA |
| NA | NA | NA |
| NA | NA | NA |
| NA | NA | NA |
| NA | NA | NA |
| NA | NA | NA |
| NA | NA | NA |
| NA | NA | NA |
| NA | NA | NA |
| NA | NA | NA |
| NA | NA | NA |
| NA | NA | NA |
| NA | NA | NA |
| NA | NA | NA |
| NA | NA | NA |
| NA | NA | NA |
| NA | NA | NA |
| NA | NA | NA |
| NA | NA | NA |
| NA | NA | NA |
| NA | NA | NA |
| NA | NA | NA |
| NA | NA | NA |
| NA | NA | NA |
| NA | NA | NA |
| NA | NA | NA |
| NA | NA | NA |

| | | |
|---|---|---|
| NA | NA | NA |
| NA | NA | NA |
| NA | NA | NA |
| NA | NA | NA |
| NA | NA | NA |
| NA | NA | NA |
| NA | NA | NA |
| NA | NA | NA |
| NA | NA | NA |
| NA | NA | NA |
| NA | NA | NA |
| NA | NA | NA |
| NA | NA | NA |
| NA | NA | NA |
| NA | NA | NA |
| NA | NA | NA |
| NA | NA | NA |
| NA | NA | NA |
| NA | NA | NA |
| NA | NA | NA |
| NA | NA | NA |
| NA | NA | NA |
| NA | NA | NA |
| NA | NA | NA |
| NA | NA | NA |
| NA | NA | NA |
| NA | NA | NA |
| NA | NA | NA |
| NA | NA | NA |
| NA | NA | NA |
| NA | NA | NA |
| NA | NA | NA |
| NA | NA | NA |
| NA | NA | NA |
| NA | NA | NA |
| NA | NA | NA |
| NA | NA | NA |
| NA | NA | NA |
| NA | NA | NA |
| NA | NA | NA |
| NA | NA | NA |
| NA | NA | NA |
| NA | NA | NA |
| NA | NA | NA |
| NA | NA | NA |
| NA | NA | NA |
| NA | NA | NA |

| | | |
|---|---|---|
| NA | NA | NA |
| NA | NA | NA |
| NA | NA | NA |
| NA | NA | NA |
| NA | NA | NA |
| NA | NA | NA |
| NA | NA | NA |
| NA | NA | NA |
| NA | NA | NA |
| NA | NA | NA |
| NA | NA | NA |
| NA | NA | NA |
| NA | NA | NA |
| NA | NA | NA |
| NA | NA | NA |
| NA | NA | NA |
| NA | NA | NA |
| NA | NA | NA |
| NA | NA | NA |
| NA | NA | NA |
| NA | NA | NA |
| NA | NA | NA |
| NA | NA | NA |
| NA | NA | NA |
| NA | NA | NA |
| NA | NA | NA |
| NA | NA | NA |
| NA | NA | NA |
| NA | NA | NA |
| NA | NA | NA |
| NA | NA | NA |
| NA | NA | NA |
| NA | NA | NA |
| NA | NA | NA |
| NA | NA | NA |
| NA | NA | NA |
| NA | NA | NA |
| NA | NA | NA |
| NA | NA | NA |
| NA | NA | NA |
| NA | NA | NA |
| NA | NA | NA |
| NA | NA | NA |
| NA | NA | NA |
| NA | NA | NA |
| NA | NA | NA |
| NA | NA | NA |

| | | |
|---|---|---|
| NA | NA | NA |
| NA | NA | NA |
| NA | NA | NA |
| NA | NA | NA |
| NA | NA | NA |
| NA | NA | NA |
| NA | NA | NA |
| NA | NA | NA |
| NA | NA | NA |
| NA | NA | NA |
| NA | NA | NA |
| NA | NA | NA |
| NA | NA | NA |
| NA | NA | NA |
| NA | NA | NA |
| NA | NA | NA |
| NA | NA | NA |
| NA | NA | NA |
| NA | NA | NA |
| NA | NA | NA |
| NA | NA | NA |
| NA | NA | NA |
| NA | NA | NA |
| NA | NA | NA |
| NA | NA | NA |
| NA | NA | NA |
| NA | NA | NA |
| NA | NA | NA |
| NA | NA | NA |
| NA | NA | NA |
| NA | NA | NA |
| NA | NA | NA |
| NA | NA | NA |
| NA | NA | NA |
| NA | NA | NA |
| NA | NA | NA |
| NA | NA | NA |
| NA | NA | NA |
| NA | NA | NA |
| NA | NA | NA |
| NA | NA | NA |
| NA | NA | NA |
| NA | NA | NA |
| NA | NA | NA |
| NA | NA | NA |
| NA | NA | NA |
| NA | NA | NA |

| | | |
|---|---|---|
| NA | NA | NA |
| NA | NA | NA |
| NA | NA | NA |
| NA | NA | NA |
| NA | NA | NA |
| NA | NA | NA |
| NA | NA | NA |
| NA | NA | NA |
| NA | NA | NA |
| NA | NA | NA |
| NA | NA | NA |
| NA | NA | NA |
| NA | NA | NA |
| NA | NA | NA |
| NA | NA | NA |
| NA | NA | NA |
| NA | NA | NA |
| NA | NA | NA |
| NA | NA | NA |
| NA | NA | NA |
| NA | NA | NA |
| NA | NA | NA |
| NA | NA | NA |
| NA | NA | NA |
| NA | NA | NA |
| NA | NA | NA |
| NA | NA | NA |
| NA | NA | NA |
| NA | NA | NA |
| NA | NA | NA |
| NA | NA | NA |
| NA | NA | NA |
| NA | NA | NA |
| NA | NA | NA |
| NA | NA | NA |
| NA | NA | NA |
| NA | NA | NA |
| NA | NA | NA |
| NA | NA | NA |
| NA | NA | NA |
| NA | NA | NA |
| NA | NA | NA |
| NA | NA | NA |
| NA | NA | NA |
| NA | NA | NA |
| NA | NA | NA |
| NA | NA | NA |

| | | |
|---|---|---|
| NA | NA | NA |
| NA | NA | NA |
| NA | NA | NA |
| NA | NA | NA |
| NA | NA | NA |
| NA | NA | NA |
| NA | NA | NA |
| NA | NA | NA |
| NA | NA | NA |
| NA | NA | NA |
| NA | NA | NA |
| NA | NA | NA |
| NA | NA | NA |
| NA | NA | NA |
| NA | NA | NA |
| NA | NA | NA |
| NA | NA | NA |
| NA | NA | NA |
| NA | NA | NA |
| NA | NA | NA |
| NA | NA | NA |
| NA | NA | NA |
| NA | NA | NA |
| NA | NA | NA |
| NA | NA | NA |
| NA | NA | NA |
| NA | NA | NA |
| NA | NA | NA |
| NA | NA | NA |
| NA | NA | NA |
| NA | NA | NA |
| NA | NA | NA |
| NA | NA | NA |
| NA | NA | NA |
| NA | NA | NA |
| NA | NA | NA |
| NA | NA | NA |
| NA | NA | NA |
| 0.192809521 | 0.847108147 | NA |
| NA | NA | NA |
| NA | NA | NA |
| NA | NA | NA |
| NA | NA | NA |
| NA | NA | NA |
| NA | NA | NA |
| NA | NA | NA |
| NA | NA | NA |

| | | |
|---|---|---|
| NA | NA | NA |
| NA | NA | NA |
| NA | NA | NA |
| NA | NA | NA |
| NA | NA | NA |
| NA | NA | NA |
| NA | NA | NA |
| NA | NA | NA |
| NA | NA | NA |
| NA | NA | NA |
| NA | NA | NA |
| NA | NA | NA |
| NA | NA | NA |
| NA | NA | NA |
| NA | NA | NA |
| NA | NA | NA |
| NA | NA | NA |
| NA | NA | NA |
| NA | NA | NA |
| NA | NA | NA |
| NA | NA | NA |
| NA | NA | NA |
| NA | NA | NA |
| NA | NA | NA |
| NA | NA | NA |
| NA | NA | NA |
| 1.461592667 | 0.143852865 | 0.22550493 |
| 1.029840824 | 0.303084733 | NA |
| 0.066350308 | 0.947098931 | 0.963928201 |
| NA | NA | NA |
| NA | NA | NA |
